# The systemin signaling cascade as derived from phosphorylation time courses under stimulation by systemin and its inactive Thr17Ala (A17) analog

**DOI:** 10.1101/535070

**Authors:** Fatima Haj Ahmad, Xuna Wu, Annick Stintzi, Andreas Schaller, Waltraud X Schulze

## Abstract

Systemin is a small peptide with important functions in plant wound response signaling. Although transcriptional responses of systemin action are well described, the precise signaling cascades involved in its perception and signal transduction are poorly understood at the protein level. Here we use a phosphoproteomic profiling study involving stimulation time courses with systemin and its inactive analogon A17 to reconstruct a systemin-specific kinase/phosphatase signaling network. The time course analysis of systemin-induced phosphorylation patterns revealed early events at the plasma membrane, such as dephosphorylation of H^+^-ATPase, rapid phosphorylation of NADPH-oxidase and Ca^2+^-ATPase. Later responses involved transient phosphorylation of small GTPases and vesicle trafficking proteins, as well as transcription factors. Based on a correlation analysis of systemin-specific phosphorylation profiles, we predict substrate candidates for 44 systemin specific kinases and 9 phosphatases. Among the kinases are several systemin-specific receptor kinases as well as kinases with downstream signaling functions, such as MAP-kinases. A regulatory circuit for plasma membrane H^+^-ATPase was predicted and confirmed by in-vitro activity assays. In this regulatory model we propose that upon systemin treatment, H^+^-ATPase LHA1 is rapidly de-phosphorylated at its C-terminal regulatory residue T955 by phosphatase PLL5, resulting in the alkalization of the growth medium within two minutes of systemin treatment. We further propose that the H^+^-ATPase LHA1 is re-activated by MAP-Kinase MPK2 later in the systemin response. MPK2 was identified with increased phosphorylation at its activating TEY-motif at 15 minutes of treatment and the predicted interaction with LHA1 was confirmed by in-vitro kinase assays. Our data set provides a valuable resource of proteomic events involved in the systemin signaling cascade with a focus on predictions of substrates to systemin-specific kinases and phosphatases.

## Introduction

Almost 30 years ago, the quest for signaling molecules mediating systemic defense responses after local injury by insect herbivores culminated in the discovery of systemin as the first peptide with signaling function in plants (1, 2). The 18-amino-acid oligopeptide was established as an essential component of the wound signaling pathway that is responsible for the systemic regulation of defense gene expression (3, 4). Systemin was initially described as the long-sought hormonal signal that is released at the site of wounding, that travels through the vasculature and induces the defense response in distal, unwounded tissues (2, 5). However, this model had to be modified when it was shown that systemin rather acts locally at the site of wounding, where it induces and amplifies the production of jasmonates as systemic signals for defense gene activation in distal tissues (6, 7). Considering its paracrine immune-modulatory activity, systemin is thus better described as a plant cytokine, a phytocytokine (8) rather than a wound hormone. Systemin is synthesized as a larger precursor protein from which it is proteolytically released (4, 9). Whether or not precursor processing and systemin secretion are regulated processes still awaits an answer. On the other hand, if systemin is released passively simply as a result of tissue disruption, it could also be addressed as a damage-associated molecular pattern, a DAMP (8).

Local action of systemin is initiated by its interaction with a saturatable binding site at the cell surface (10, 11). Purification and characterization of the binding protein tentatively placed the receptor into the family of leucin-rich repeat receptor-like kinases (LRR-RLKs), but the protein initially declared as the systemin receptor later turned out to be the tomato ortholog of BRI1, the brassinosteroid receptor in Arabidopsis (12–15). Recent work identified the systemin receptor as SYR1, a LRR-RLK closely related to known pattern recognition receptors (16). The early cellular responses to systemin do in fact resemble those typically triggered by microbe-associated molecular patterns (MAMPs) including an increase in cytosolic calcium, extracellular alkalization, plasma membrane depolarization, an oxidative burst, and ethylene production (17–19). By still unknown mechanisms, these early signaling events translate into the activation of the octadecanoid pathway for jasmonate production (20–22). The locally produced jasmonate signal is then perceived in distal leaves resulting in the systemic activation of defense responses (23). In addition to the slow-moving jasmonate signal, rapidly propagating electrical and calcium signals have been linked to the activation of jasmonate signaling and activation of wound response gene expression in systemic tissues (24–26).

Despite almost 30 years of research, it is still unclear how early responses at the plasma membrane are activated by systemin, and how these early signaling events including the influx of calcium, plasma membrane depolarization, and the production of reactive oxygen species are linked to downstream events, like jasmonate production and defense gene activation. Since systemin is perceived by a receptor kinase, it is reasonable to assume that subsequent events are regulated by phosphorylation and can thus be captured by phosphoproteomics. Large scale phosphoproteomics was shown to be a powerful tool to identify global patterns and novel players in plant signaling networks. For example, time-courses studies of phosphorylation patterns in response to sucrose (27) have led to the identification of a novel kinase, SIRK1, regulating aquaporins (28). The cellular response to nitrate was well studied by large-scale phosphoproteomics to conclude about the cellular kinase network involved (29). Also for plant hormones, such as brassinolide, large-scale phosphoproteomics time-course studies revealed novel insights into the intracellular signaling network (30). Phosphoproteomics resulted in the identification of FERONIA as the receptor of rapid alkalization factor, and the plasma membrane H^+^-ATPase as a downstream target (31). In a similar approach, we used time-resolved phosphoproteomics in a systemin-responsive cell culture system to get further insight into the systemin signaling cascade and the sequence of early signaling events.

## Experimental procedures

### Experimental Design and statistical rationale

Batches of cell suspension cultures of *Solanum peruvianum* (200 ml) were subjected to stimulation with 10 nM systemin (sys) or 10 nM of the inactive Thr17Ala analog (A17). A third batch of cells was injected with equal volume of water as a control treatment for general handling. For each treatment, a batch of cells was harvested after 2, 5, 15 and 45 minutes (Figure 1A). At each time point, cells were harvested over a sieve and were immediately frozen in liquid nitrogen (32). Untreated cells were harvested as controls at time point 0. All treatments and time points were sampled from three independent replicates using independent batches of cells and both treatments were run at the same time. Data analysis was carried out jointly for all samples by averaging quantitative information for each time point and treatment across the three replicates. The same cell culture materials were used for phosphoproteome profiling as well as for activity assay. For phosphopeptide profiling, microsomal membranes were isolated from each sample, digested with trypsin, enriched for phosphopeptides and subjected to untargeted data-dependent acquisition by LC-MS/MS. All proteomic analyses were performed only on biological replicates. Statistical analysis were carried out (i) as pairwise t-tests between average ion intensities of systemin or A17 treatment at each time point when individual peptide profiles were studied, or (ii) as sum of square deviations when time profile patterns of phosphopeptides were compared between systemin and A17 treatments. Thereby, for each phosphopeptide identified under both treatments, the squared deviation from the respective consensus systemin-induced cluster (supplementary figure 1) was calculated and divided by the number of data points. (iii) Pairwise correlation analysis with a cutoff of r=0.8 was used when the similarity of patterns between different phosphopeptides was assessed.

**Figure 1:**
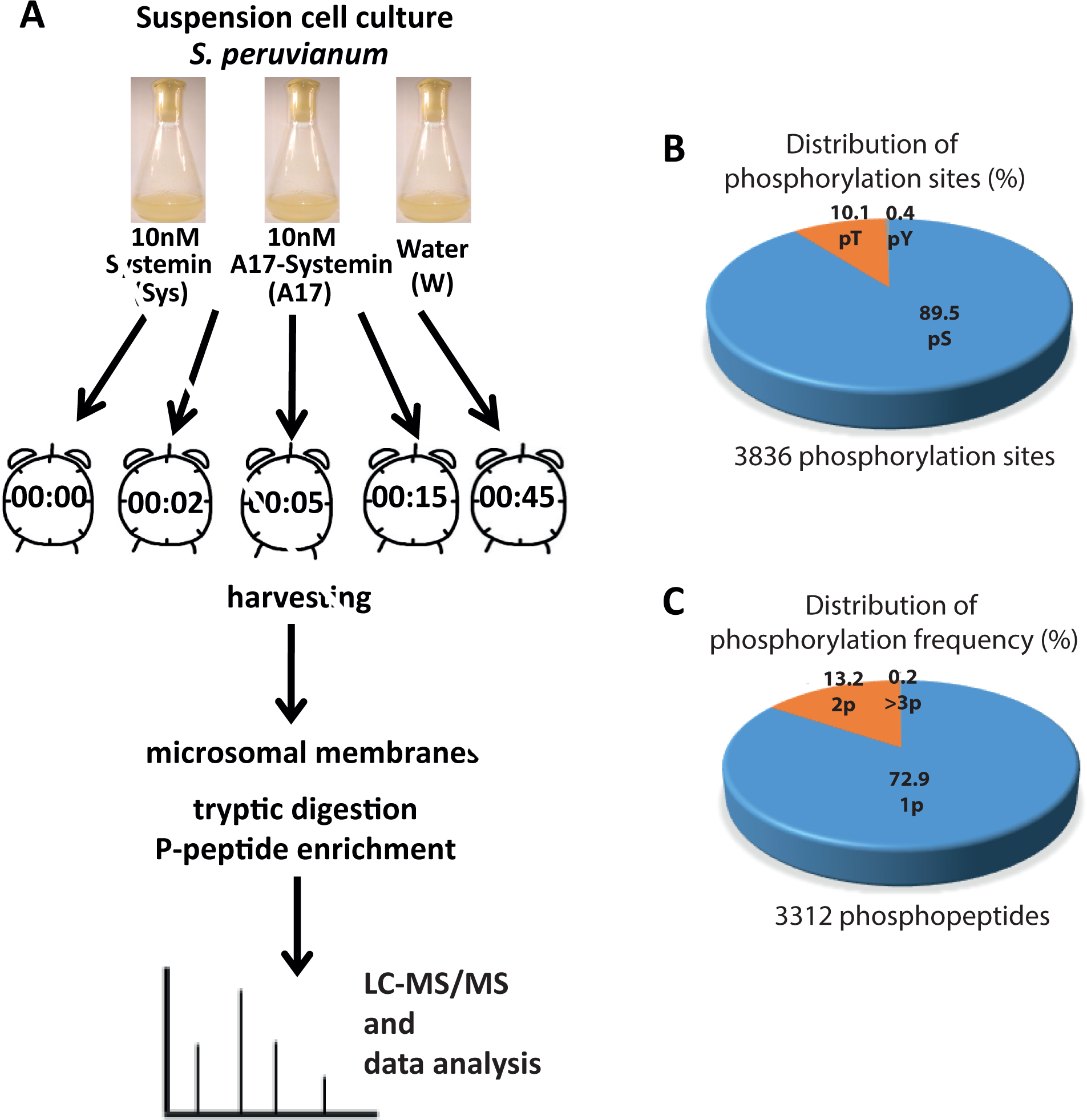
Analysis of the systemin signaling response. (**A**) Experimental design and analytical workflow. (**B**) Frequency distributions of 3312 identified phosphopeptides according to numbers of phosphorylation sites and phosphorylated amino acid as determined by MaxQuant. (**C**) Frequency of phosphosites per peptide as determined by MaxQuant.

### Cell suspension cultures and alkalization assay

The *Solanum peruvianum* cell suspension culture was kindly provided by Georg Felix (University of Tübingen). Cells were subcultured weekly by diluting 8 ml of the suspension in 70 ml fresh Murashige-Skoog medium containing Nitsch vitamins (33), 0.34 g/l KH_2_PO_4_, 5 mg/l 1-naphthaleneacetic acid and 2 mg/l benzylaminopurine and grown on a rotary shaker at 26 °C with 120 rpm. For the alkalization assay continuous measurements of extracellular pH was performed in 10 ml of cultured cells 6–8 days after subculture. Synthetic systemin and systemin-A17 peptides (Pepmic; Suzhou, China) were added from a 200-fold concentrated stock solution in water.

### Microsomal fraction extraction

The microsomal fraction was extracted according to (34) with minor modifications. A total of 1 g of harvested cells was homogenized in 10 ml extraction buffer (330 mM sucrose, 100 mM KCl, 1 mM EDTA, 50 mM Tris-MES pH 7.5 and fresh 5 mM DTT) in the presence of 1 % v/v proteinase inhibitor mixture (Serva) and 0.5 % of each phosphatase inhibitor cocktails 2 & 3 (Sigma-Aldrich, Germany) in a Dounce Homogenizer. At least 200 strokes were performed. The homogenate was filtered through four layers of miracloth and centrifuged for 15 min at 7500 × g at 4 °C. The supernatant was collected and centrifuged for 75 min at 48,000 × g at 4 °C. The pellet was washed with 5 ml of 100 mM Na_2_CO_3_ then again was centrifuged for 75 min at 48,000 x g at 4 °C. The microsomal membrane pellets used for LC-MS/MS were re-suspended in 100 µl UTU (6 M urea, 2 M thiourea and 10 mM Tris-HCl pH 8). Microsomal membrane pellets used for ATPase assay were re-suspended in 100 µl of re-suspension buffer (330 mM sucrose, 25 mM Tris-MES pH 7.5, 0.5 mM DTT). Protein concentrations were determined using Bradford assay (Roth, Germany) with BSA as protein standard. Samples were stored at −80 °C until further use.

### Protein preparation for LC-MS/MS

Microsomal proteins were predigested for three hours with endoproteinase Lys-C (0.5 µg/µl; Thermo Scientific) at 37 °C in 6 M urea 2 M thiourea, pH 8 (Tris-HCl). After 4-fold dilution with 10 mM Tris-HCl (pH 8), samples were digested with 4 µL Sequencing Grade Modified trypsin (0.5 µg/µl; GE Life Sciences) overnight at 37 °C. The reaction was stopped by addition of trifluoroacetic acid (TFA) to reach pH ⩽ 3 before further processing.

### Phosphopeptide enrichment

Phosphopeptides were enriched over titanium dioxide (TiO_2_, GL Sciences) with some modifications as described (35). TiO_2_ beads were equilibrated with 100 µl of 1 M glycolic acid, 80% acetonitrile and 5 % TFA. The digested peptides were mixed with the same volume of the equilibration solution and incubated for 20 minutes with 2 mg TiO_2_ beads with continuous shaking. After brief centrifugation, the supernatant was discarded and the beads were washed 3 times with 80 % acetonitrile and 1% TFA. Then the phosphopeptides were eluted with 240 µl of 1% ammonium hydroxide. Eluates were immediately acidified by adding 50 µl 10% formic acid to reach pH<3. Enriched phosphopeptides were desalted over C18-Stage tip (36) prior to mass spectrometric analysis. Only phosphopeptide enriched fractions were analyzed.

### LC-MS/MS analysis of peptides and phosphopeptides

Tryptic peptide mixtures were analyzed by LC/MS/MS using nanoflow Easy-nLC1000 (Thermo Scientific) as an HPLC-system and a Quadrupole-Orbitrap hybrid mass spectrometer (Q-Exactive Plus, Thermo Scientific) as a mass analyzer. Peptides were eluted from a 75 μm x 50 cm C18 analytical column (PepMan, Thermo Scientific) on a linear gradient running from 4 to 64% acetonitrile in 240 min and sprayed directly into the Q-Exactive mass spectrometer. Proteins were identified by MS/MS using information-dependent acquisition of fragmentation spectra of multiple charged peptides. Up to twelve data-dependent MS/MS spectra were acquired for each full-scan spectrum recorded at 70,000 full-width half-maximum resolution. Fragment spectra were acquired at a resolution of 35,000. Overall cycle time was approximately one second.

Protein identification and ion intensity quantitation was carried out by MaxQuant version 1.5.3.8 (37). Spectra were matched against the Tomato proteome (*Solanum lycopersicum*, ITAG2.4, 34725 entries) using Andromeda (38). Thereby, carbamidomethylation of cysteine was set as a fixed modification; oxidation of methionine, N-terminal protein acetylation as well as phosphorylation of serine, threonine and tyrosine were set as variable modifications. Up to two missed cleavages (trypsin) were allowed: Mass tolerance for the database search was set to 12 ppm on full scans and 20 ppm for fragment ions. Multiplicity was set to 1. For label-free quantitation, retention time matching between runs was chosen within a time window of two minutes. Peptide false discovery rate (FDR) and protein FDR were set to 0.01, while site FDR was set to 0.05. The score threshold for phosphorylation site localization was set to 50. Hits to contaminants (e.g. keratins) and reverse hits identified by MaxQuant were excluded from further analysis. For increasing quantitative coverage in label-free quantitation “match between runs” was selected in MaxQuant with a tolerance of 0.75 minutes within 20 minute time windows throughout the run. Raw files and identified spectra were submitted to ProteomeXChange and are available to the public under the accession number PXD010819. Representative annotated spectra of phosphopeptides are available as supplementary figure 2.

### Mass spectrometric data analysis and statistics

Reported ion intensity values were used for quantitative data analysis. cRacker (39) was used for label-free data analysis of phosphopeptide ion intensities based on the MaxQuant output (evidence.txt). All phosphopeptides and proteotypic non-phosphopeptides were used for quantitation. Within each sample, ion intensities of each peptide ions species (each m/z) were normalized against the total ion intensities in that sample (peptide ion intensity/total sum of ion intensities). Subsequently, each peptide ion species (i.e. each m/z value) was scaled against the average normalized intensities of that ion across all treatments. For each peptide, values from three biological replicates were averaged after normalization and scaling. In case of non-phosphopeptides, protein ion intensity sums were calculated from normalized scan scaled ion intensities of all proteotypic peptides. A list of quantified phosphopeptides including normalized ion intensities, standard deviation and number of spectra is available as Supplementary Table 1.

### H^+^-ATPase activity assay

Plasma membrane (PM) H^+^-ATPase activity was determined by resuspending 5 µg of microsomal proteins in 10µl resuspension buffer and 40 µl of reaction buffer containing 25 mM Tris-HCl pH 6.3, 50 mM KCl, 5 mM MgCl_2_, 5 mM CaCl_2_, 0.1 mM ouabain (Thermo Fisher Scientific, USA), in presence of 1 mM NaN_3_, 100 nM concanamycin A (Sigma-Aldrich, Germany), 0.02 % Brij58 (Sigma-Aldrich, Germany), 0.5 µg/µl BSA and 10 mM DTT. The reaction was initiated by the addition of 1 mM ATP, proceeded for 2 hours at room temperature, and stopped with 150 µl of stopping reagent (8% ascorbic acid, 2.1% HCl and 0.7% ammonium molybdate). After 5 min, 150 µl of 2% tri-sodium citrate and 2% acetic acid solution was added and incubated for 15 min at room temperature. The absorbance as a measure of inorganic phosphate was determined at 640 nm using plate reader (Tecan Spark). ATPase activity was measured with and without 400 µM orthovanadate (Na_3_VO_4_), and the difference between these two activities was attributed to the PM H^+^-ATPase.

### Recombinant protein expression

The whole coding sequence of MPK2 (Solyc08g014420.2.1) was amplified by PCR using the primers ATTA<u>CCATGG</u>CAGATGGTTCAGCTCCG (forward) and ATAT<u>CTCGAG</u>CATGTGCTGGTATTCGGGATT (reverse) with *S. lycopersicum* leaf cDNA as template. The PCR product was cloned into pCR2.1-TOPO vector (Thermo Scientific) and amplified in *E. coli*. The MPK2 insert was excised with *NcoI* and *XhoI* (underlined in the primer sequence) and ligated to pET21d (+) (Merck; Darmstadt, Germany) to produce 14420HispET21d, which was transformed into *E.coli* strain BL21 RIL (Agilent Technologies; Waldbronn, Germany) to be expressed as C-terminally 6xHis-tagged recombinant protein. MPK2 was expressed in unmodified form and as constitutively active mutant (CA) with two amino acid substitutions D216G/E220A (40).

The mutations were introduced using the following primers: ATTA<u>CCATGG</u>CAGATGGTTCAGCTCCG and 14420D216G/E220AR1 (GTCACAACATA<u>TGC</u>GGTCATAAA<u>GCC</u>AGTTTCAGAAG); or 14420D216G/E220AF2 (CTTCTGAAACT<u>GGC</u>TTTATGACC<u>GCA</u>TATGTTGTGAC) and (ATAT<u>CTCGAG</u>CATGTGCTGGTATTCGGGATT).

For PLL5 (Solyc06g076100.2.1), the catalytic domain (residues from 241-708) was amplified by PCR using the primers ATA<u>CCATGG</u>CACATCACCATCACCATCACTTCAGCAGTGAGTGTAGTTTG and AATT<u>CTCGAG</u>TTATGCACTGGATCTCCATATTC.

The PCR products were cloned into pET21d (+) as described above to produce His76100pET21d, which was transformed into *E.coli* strain BL21 RIL to be expressed as N-terminally 6xHis-tagged recombinant protein.

### Purification of His-tagged proteins

*E.coli* BL21 RIL cells were grown at 37 °C, 200 rpm to an A_600_ value of 0.8 and then induced with 1 mM isopropyl-β -d-thiogalactopyranoside (IPTG) for 4 h at 30 °C. Cells were pelleted by centrifugation and lysed in Bugbuster™ protein extraction reagent (Merck; Darmstadt, Germany) by continuous agitation for 20 min at room temperature. After centrifugation at 12,000 g for 20 min at 4 °C, the supernatants were collected. The supernatants were subjected to affinity purification on Ni-NTA agarose following the manufacturer’s instructions (Qiagen; Hilden, Germany).

### In-vitro kinase activity assay

Recombinant MPK2 (2 µg) was incubated with peptide substrate (GLDIETIQQSYTV; amount as indicated for each experiment) overnight in 50 µl of reaction buffer (20 mM HEPES pH 7.0, 10 mM MgCl_2_, 2 mM DTT and 0.1 mg/ml BSA) with 1 µM ATP at 30 °C. Kinase activity was measured as ATP consumption by addition of 50 µl of Kinase-Glo^®^ Plus Reagents (Promega; Fitchburg, WI), 10 min incubation at 30°C, and subsequent recording of the decrease in luminescence (Tecan Spark luminometer). To detect the phosphorylated peptides by mass spectrometry, the reaction was performed as described with 20 µg of peptide substrate. The reaction was stopped by addition of TFA until pH ≤ 3 and reaction products were purified over C18-Stage tips (36) prior to mass spectrometric analysis.

### In-vitro phosphatase activity assay

Recombinant PLL5 phosphatase (1 µg) was incubated with the phosphorylated peptide substrate (GLDIETIQQSYT(ph)V; amount as indicated for each experiment) in 50 µl of reaction buffer (20 mM Tris-HCl pH 7.5, 5 mM MgCl_2_, 1mM EGTA, 0.02 % β-mercaptoethanol and 0.1 mg/ml BSA) at room temperature for 30 minutes. Released phosphate was detected by addition of 150 µl of 8 % ascorbic acid, 2.1 % HCl and 0.7 % ammonium molybdate. After 5 min at room temperature the reaction was stopped by addition of 150 µl 2 % tri-sodium citrate and 2 % acetic acid (41). After incubation for 15 min at room temperature the reduced phoshomolybdenum complex was quantified at 640 nm using a plate reader (Tecan Spark) based on a standard curve of 0 - 0.1µmol K_2_HPO_4_. To detect the unphosphorylated peptide by mass spectrometry, the reaction was performed as above with 200 µg of phosphorylated peptide substrate. The reaction was stopped by addition of TFA to reach pH ⩽ 3 and reaction products were purified over C18-Stage tips (4) prior to mass spectrometric analysis.

## Results

### The inactive A17 systemin analogon allows definition of systemin specificity

Early cellular responses to systemin include a rapid influx of calcium and depolarization of the plasma membrane which are necessary and sufficient for systemin signaling and induction of the wound response (17–19, 42, 43). Systemin-induced changes in plasma membrane ion permeability can be assayed conveniently in tomato (*Solanum peruvianum*) cell cultures as an alkalization of the culture medium (18) (Figure 2A). Systemin-induced medium alkalization is a unique feature of this cell culture system that has been used by us and others to characterize systemin activity (44). This cell culture system was used here to analyze dynamic changes of protein phosphorylation in the systemin signaling network.

**Figure 2:**
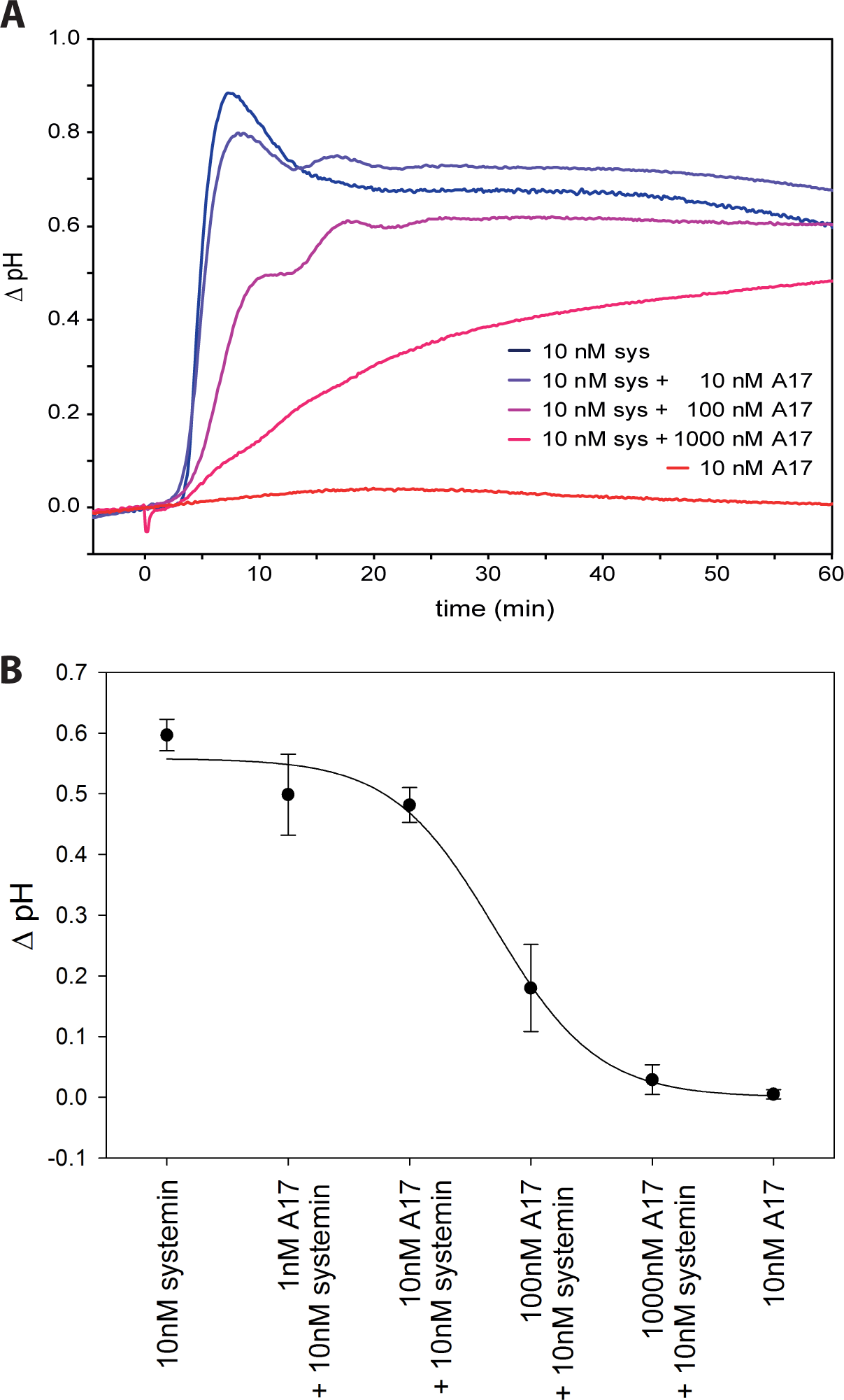
Alkalization response. (**A**) Time course of pH change after systemin treatment (blue) and A17 treatment (red) as well as competition of systemin response with increasing concentrations of A17 (blue to red). (**B**) Inhibition curve of A17. Data points show the mean with standard deviation from three independent experiments.

Systemin responses, including extracellular alkalization, the induction of ethylene biosynthesis, and expression of wound response genes are inhibited by A17, a systemin derivative in which the penultimate threonine residue is substituted by alanine (18, 45, 46). In our cell culture system, stimulation with 10 nM systemin resulted in a rapid alkalization of the medium from pH 5.5 to pH 6.4 within about 8 minutes (Figure 2A). A17 was completely inactive at this concentration. When A17 was added to the systemin treatment, the alkalization response was reduced with increasing concentrations of A17 (Figure 2B). To explain the antagonistic activity of A17, it was suggested that the peptide may interact with the binding site of the systemin perception system (45). Consistent with this proposition, binding to the putative receptor was shown to rely on the N-terminus of systemin, while the C-terminus, including Thr17, is required for activation (10). The receptor was recently identified as SYR1, and A17 was confirmed as a competitive systemin antagonist (16). By comparing phosphoproteomic responses triggered by A17 and by systemin, the data set can be filtered for systemin-specific phosphorylation changes. We therefore included the inactive A17 analog as a control in our experiments.

### Description of the phosphoproteomics data set

A total of 3312 phosphopeptide species were identified which differed in peptide sequence or in the position of their modification sites. Phosphorylation was observed most frequently on serines (89.5%), followed by threonines (10%) and at rather low frequency at tyrosines (<1%, Figure 1B). As described in a thorough meta-analysis of plant phosphorylation sites (47), this amino acid distribution of phosphorylation is not unexpected for plant tissues. Although tyrosine phosphorylation does occur, it is far less abundant in plants compared to animal signaling cascades. Most of these identified phosphopeptides were singly phosphorylated (Figure 1C). Quantitative information under at least one treatment condition was obtained for 2960 phosphopeptide species matching 1729 protein groups in at least one biological replicate. Thereby, the majority of phosphopeptides were identified at all time points.

### Phosphorylation time profiles

We studied the dynamics of protein phosphorylation in response to systemin and A17 treatment. As shown previously (48), no significant total protein abundance changes were observed within the time frame of minutes. Therefore, changes in phosphopeptide abundances observed here are considered to be attributed to changes in phosphorylation status rather than protein abundance changes.

Treatment of suspension cell cultures with systemin resulted in characteristic changes of phosphorylation patterns. K-means clustering was used to group these phosphorylation responses (Figure 3A; Supplementary Table 2). A high number of 827 phosphopeptides was rapidly dephosphorylated upon systemin treatment (cluster A). Three clusters were characterized by a transient increase in phosphorylation, with 465 phosphopeptides peaking at 2 minutes (cluster B), 501 at 5 minutes (cluster C), and 705 at 15 minutes after systemin treatment (cluster D). Further 1001 phosphopeptides were classified as not responsive (cluster F). The stimulation of suspension cells with A17 also resulted in characteristic changes in phosphorylation (Figure 3B). Interestingly, upon A17 stimulation no phosphopeptides displayed a transient increase in phosphorylation at 5 minutes, i.e. cluster C is missing. Instead, 1042 phosphopeptides showed an increase in phosphorylation at 45 minutes (cluster E). The missing cluster C and emergence of a “later” maximum in cluster E in A17 treatment suggests that the control treatment with A17 does trigger some cellular response, but weaker and delayed compared to systemin treatment. Stimulation of the suspension cells with water resulted in response groups very similar to the clusters observed upon A17 treatment (Supplementary Figure 1). Since the mock treatment with water and A17 showed highly similar responses, we did not further consider the water treatment for definition of systemin-specific responses.

**Figure 3:**
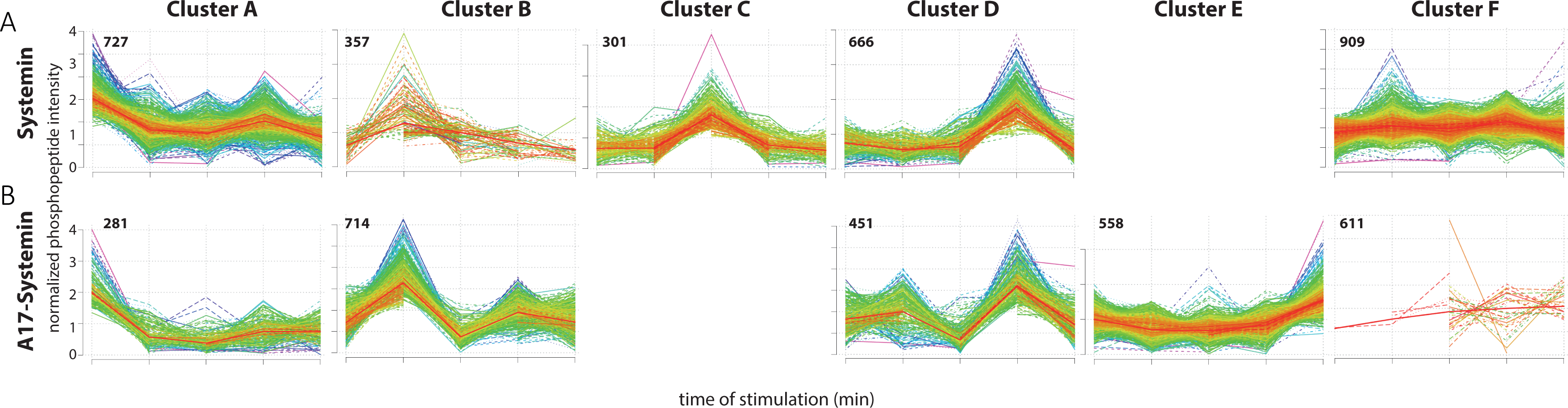
k-Means clustering of phosphorylation time courses. (**A**) Systemin treatment. (**B**) A17 treatment. (**C**) Water treatment. Numbers of identified phosphopeptides are indicated within each panel.

To substantiate the observed phosphorylation profiles as physiologically relevant, we analyzed the activity of the plasma membrane H^+^-ATPase (Figure 4) that has previously been implicated in the systemin-induced alkalization response (19). Two phosphopeptides of the C-terminal regulatory domain of plasma membrane ATPases Solyc07g017780 (AHA1) and Solyc03g113400 (LHA1) were identified with different time profiles under all three treatments. The peptides encompass the penultimate threonine residue, phosphorylation of which facilitates 14-3-3 protein binding and proton pump activation (49, 50). In agreement with the observed alkalization of the growth medium upon systemin treatment, the peptides were slightly dephosphorylated at 2 minutes of treatment (Figure 4A, B). Dephosphorylation of the regulatory threonine residue was associated indeed with a measureable decrease in H^+^-ATPase activity at 2 to 5 minutes of systemin treatment (Figure 4C). In contrast, A17 treatment induced a very transient increase in C-terminal phosphorylation and H^+^-ATPase activity at 2 minutes. We conclude that the time profiles presented in Figure 4 present a solid basis of quantitative measurements of phosphorylation status with corresponding effects on protein activity. At least in one example of the H^+^-ATPase, this relationship was verified by activity tests.

**Figure 4:**
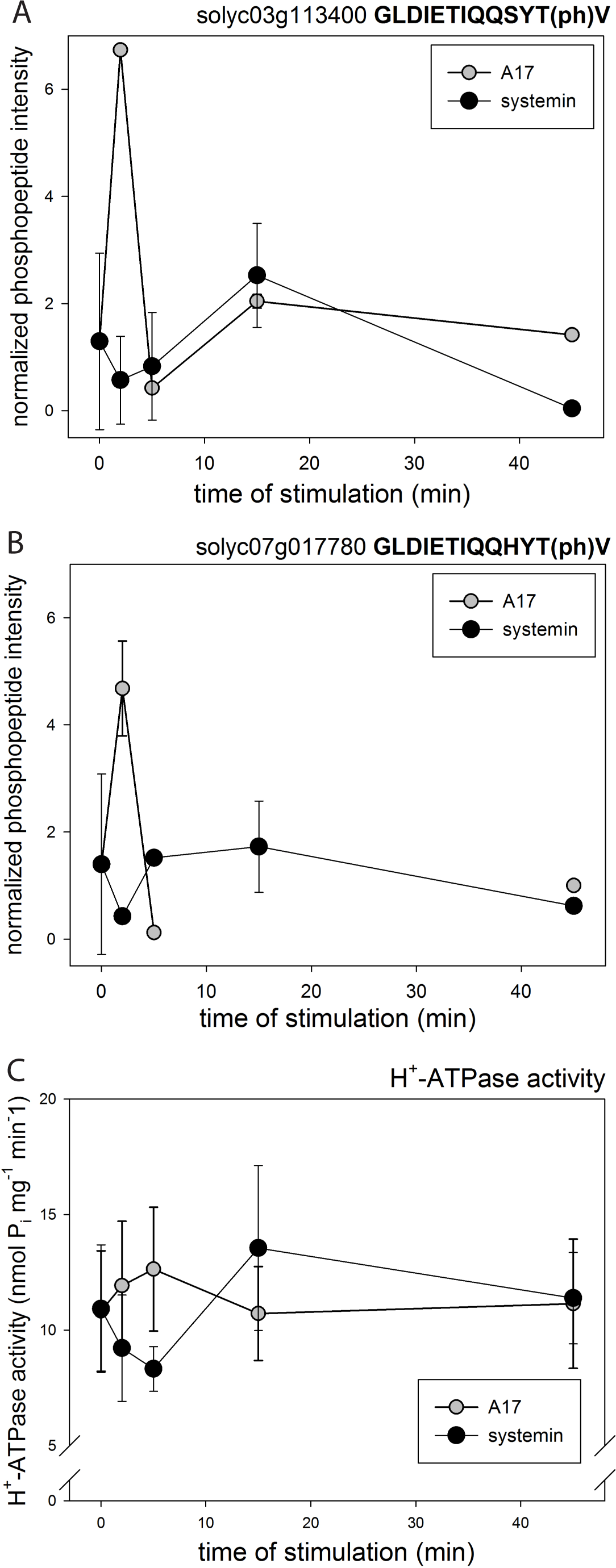
Phosphorylation of H^+^-ATPase and respiratory burst oxidase. (**A**) C-terminal activating phosphorylation site of H^+^-ATPase Solyc03g113400. (**B**) C-terminal activating phosphorylation site of H^+^-ATPase Solyc07g017780. (**C**) H^+^-ATPase activity under treatment with systemin, A17, and water. Black triangles: systemin treatment; white triangles: A17 treatment. White circles: water treatment. Mean values of three biological replicates are shown with standard deviation.

### Systemin-specific responses

To define precisely which of the phosphorylation events are indeed systemin-specific, we compared for each phosphopeptide the phosphorylation time profile induced by systemin with those induced by A17 (Figure 5A). For example, 649 out of 727 phosphopeptides in systemin-induced cluster A (de-phosphorylated after 2 min), were classified in a “later” cluster (B to F) upon A17 treatment, or were not identified at all (7 phosphopeptides). Thus, these proteins were considered systemin-specific. Only 71 phosphopeptides showed the same rapid de-phosphorylation kinetics (i.e. also classified as cluster A) when treated with A17. In contrast, 665 of the 1001 phosphopeptides that were clustered as non-responsive to systemin treatment (cluster F), upon A17 treatment showed a peak phosphorylation in an earlier cluster. A total of 161 proteins were found with equal classification upon A17 and systemin treatment, and 83 phosphopeptides were not identified in the A17 treatment. In total, 1515 phosphopeptides (45% of all identified phosphopeptides) were classified as systemin-specific.

**Figure 5:**
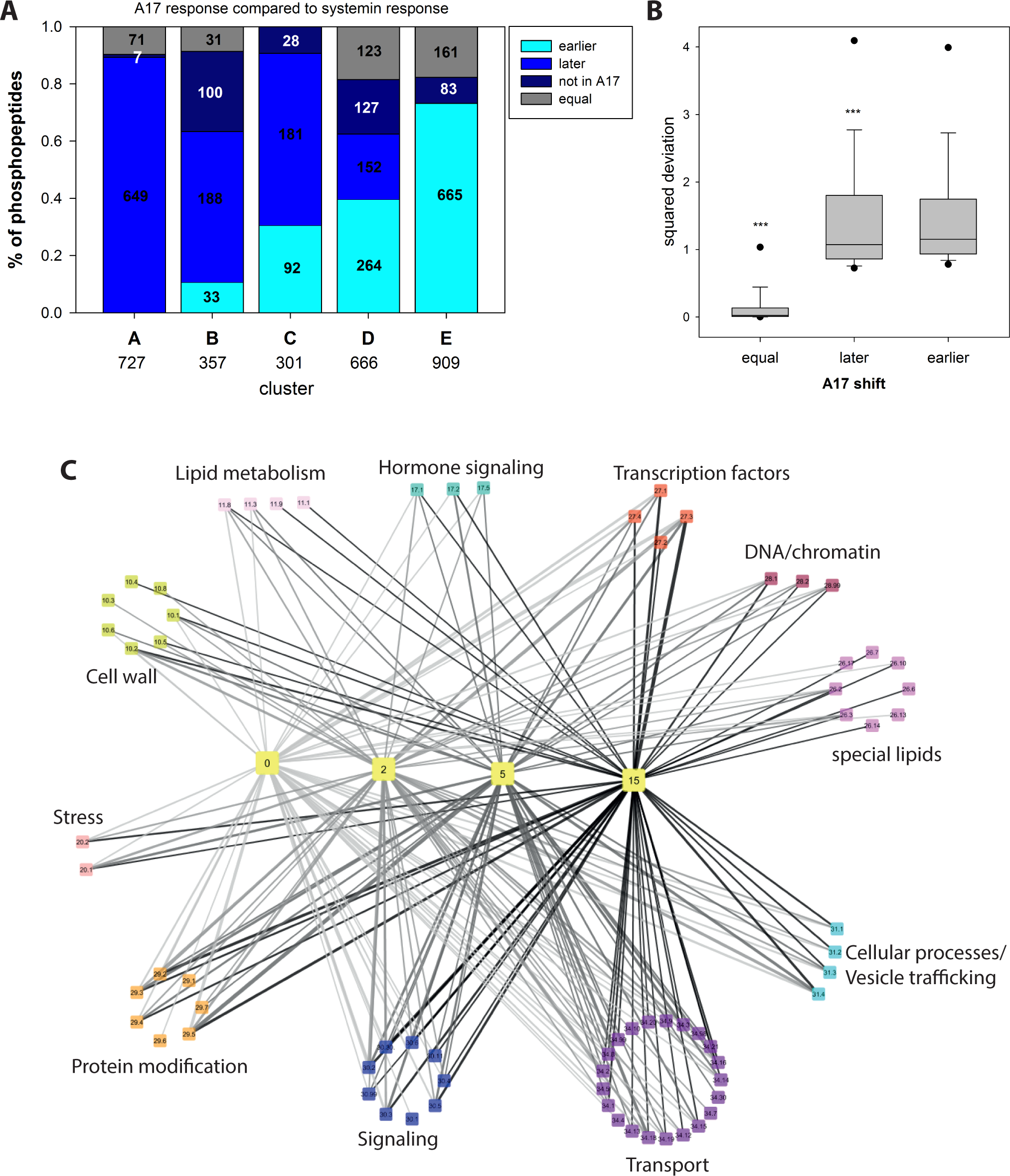
Over-representation analysis of functional groups in each cluster. (**A**) Comparison of systemin and A17 response. The proportion of phosphopeptides with a time-shifted response in A17 treatment compared to systemin treatment is displayed. “Late” peptides display a later phosphorylation maximum at A17 treatment compared to systemin treatment, or do not display any maximum at all. “Early” peptides have a phosphorylation maximum at earlier time points after A17 treatment compared to systemin stimulation, or they have a maximum in A17 treatment and remain unaltered under systemin treatment. Numbers indicate absolute numbers of classified phosphopeptides. (**B**) Boxplot of square deviations (χ2-values) of phosphopeptide classified as “later”, “earlier” or “equal” under A17 treatment relative to the respective consensus cluster profile under systemin treatment. (**C**) Network view of processes over-represented at different time points. Yellow colored “time”-nodes represent k-means clusters (figure 4) with maximum phosphorylation at respective time points. The “bin”-nodes represent the processes over-represented in the respective k-means clusters. Edges width represents the percentage of proteins of each bin within in each k-means cluster. Edge color indicates the processes involved at different time points. Bin code: 10 cell wall, 11 lipid metabolism, 17 hormone signaling, 20 stress responses, 26 special lipids, 27 RNA processes, 28 DNA processes, 29 protein processes, 30 signaling, 31 vesicle trafficking, 34 transport. Detailed listing in supplementary Table 3.

This classification was further substantiated by statistical analysis for which the average phosphopeptide ion intensities at each time point of the systemin-induced clusters A to F were used as an expected consensus profile. The response profiles of all the phosphopeptides observed upon the A17 induction were then compared against the expected model profiles of the systemin response clusters as described (51). The squared deviation from the respective consensus systemin-induced cluster (Supplementary Figure 1) was calculated for each phosphopeptide time profile identified also under A17 treatment and divided by the number of data points. Only phosphopeptides with a minimum of three quantification values were included in this statistical analysis. On average, phosphopeptides classified with a “later” or “earlier” response based on the k-means clustering had significantly higher squared deviations (p<0.05, Mann-Whitney Ranksum test) compared to phosphopeptides with equal time profile classification (Figure 5B). Phosphopeptides not identified in the A17 treatment were by definition considered systemin-specific (Supplementary Table 2). Using a threshold value for the squared deviations of 0.8, less than 5% of the phosphopeptides with “equal” time profiles (Figure 5A) under systemin and A17 treatment were classified as “systemin-specific”. We considered this an acceptable false classification rate and applied this threshold to all phosphopeptide profiles. As a result, 162 phosphopeptides in systemin-induced cluster A were considered systemin specific, as well as 266, 259, 474, and 672 phosphopeptides in systemin-induced clusters B, C, D and F, respectively. Based on this classification, most phosphopeptides with transient systemin-induced phosphorylation change displayed a delayed response upon A17 stimulation, if any.

To better understand the processes affected by systemin-specific phosphorylation at each time point, we carried out an over-representation analysis (Fisher Exact Test) of the systemin-specific phosphoproteins in clusters A to F (Figure 5C). Systemin-specific phosphopeptides in cluster A mainly related to transport proteins and signaling. Cluster B with transient increase in phosphorylation at 2 minutes showed highest enrichment of phosphopeptides matching proteins with functions in transport (Mapman bin 34; p-value <7.32E-17), cellular processes (bin 31; p-value <2.29E-07), signaling (bin 30; p-value <2.03E-05), and cell wall proteins (bin 10; p-value<1.51E-04).

Phosphopeptides of proteins in RNA processes (e.g. transcription factors) were mainly enriched in clusters C and D with transient increase in phosphorylation at later time points (5 minutes and 15 minutes, respectively). Vesicle transport proteins (bin 31.4) were highly abundant in clusters B and C, while transcription factors (bin 27.3) and protein post-translational modification (bin 29.4) were overrepresented in all clusters. The phosphopeptides not identified upon A17 treatment were particularly enriched for vesicle transport proteins (bin 31.4), cellulose synthases (bin 10.2), P- and V-ATPases (bin 34.1), and auxin-related processes (bin 17.2.3) (Supplementary Table 3).

### General insights into systemin-specific kinases and phosphatases

Among the systemin-specific phosphopeptides, 165 and 29 phosphopeptides matched 67 protein kinases and 17 phosphatases, respectively (Supplementary Table 4). Not all of these phosphopeptides met the requirement of at least four quantitative values across the time course and were therefore excluded from further analysis. The remaining phosphopeptides matched 56 kinases and 17 phosphatases, and their phosphorylation profiles were used to construct a correlation network of systemin-specific responses. Each systemin-specific phosphopeptide of a kinase or phosphatase was correlated against all other systemin-specific phosphopeptides of categories over-represented in the systemin-induced response clusters (Figure 5C). A stringent cutoff of r=0.85 was used to display only proteins that are highly correlated with systemin-specific kinases and phosphatases. Based on these criteria, phosphopeptides of 44 kinases and of 9 phosphatases showed high correlation with phosphopeptides of 207 putative substrate proteins (Figure 6A,B). We considered these correlating phosphopeptides as candidate substrates for respective kinases or phosphatases. In the displayed network, each node indicates a kinase or phosphatase with systemin-specific phosphorylation sites and each edge describes one correlation between the phosphorylation time profile of a kinase or phosphatase peptides and a potential substrate peptides. For each kinase/phosphatase displayed in the networks, node size reflects its degree, i.e. the number of correlating substrate phosphopeptides.

**Figure 6:**
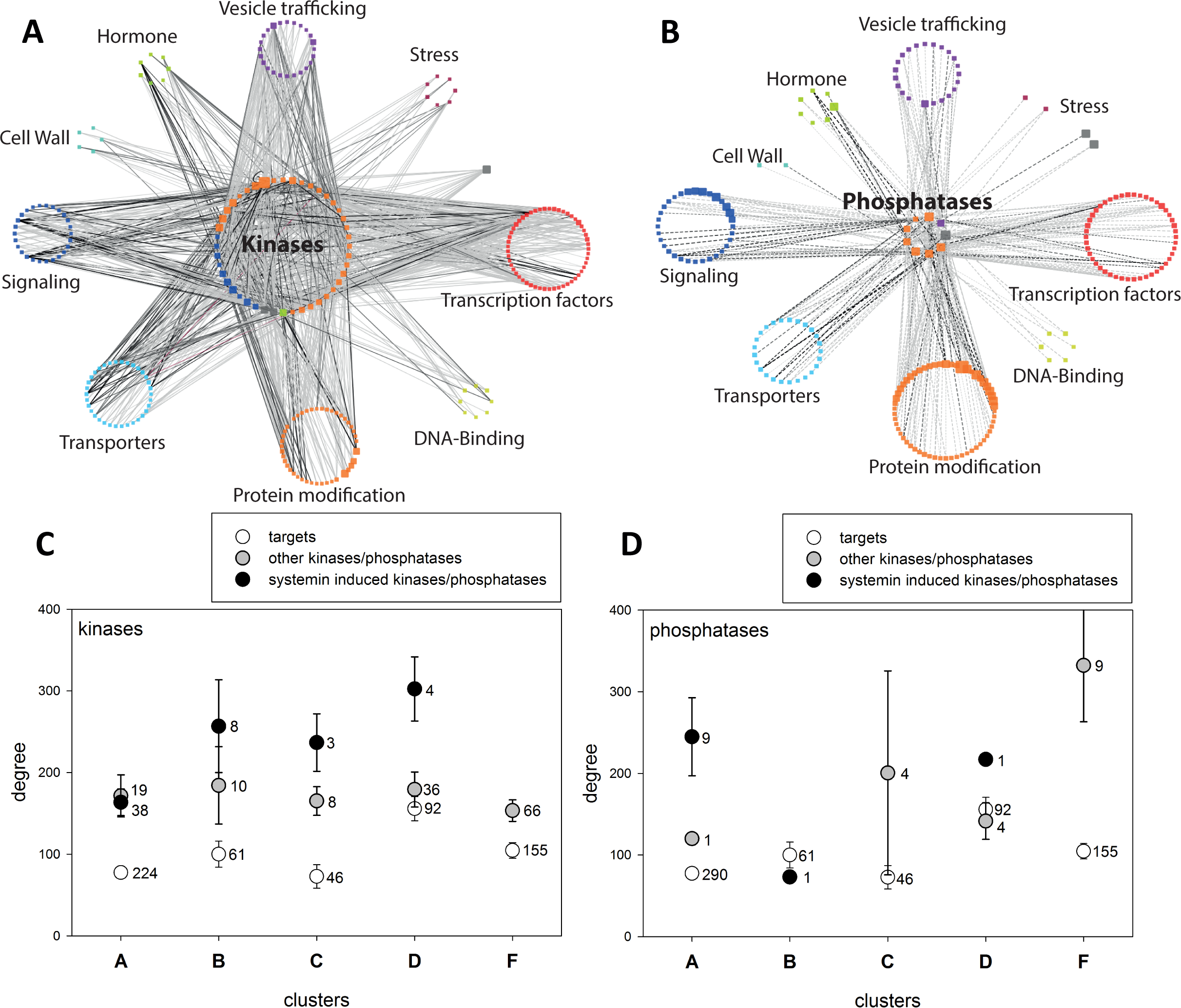
Correlation network of systemin specific kinases and phosphatases. (**A**) Kinase network. (**B**) Phosphatase network. Central nodes are the 57 kinases or 18 phosphatases, peripheral nodes represent individual proteins summarized under MapMan subbins. Node size is proportional to the degree, edge width is proportional to the correlation r-value (cutoff r=0.85). (**C**) Degree distribution of systemin-specific kinases, other kinases and their targets in different systemin response clusters. (**D**) Degree distribution of systemin-specific phosphatases, other phosphatases and their targets in different systemin response clusters. Values represent mean with standard error. For each class, the number of proteins is indicated.

Specifically, for the proteins in cluster A with fast dephosphorylation response, phosphopeptides from a MAP-triple kinase (Solyc11g033270), a protein phosphatase 1 regulatory subunit (Solyc02g070260) and a protein phosphatase 2C (Solyc10g005640) had highest numbers of systemin-specific correlation partners (highest degree, Table 1). Other kinases with high degree were an uncharacterized kinase family protein (Solyc02g076780), a calcium-dependent protein kinase (CDPK)-related kinase (Solyc10g078390), and a receptor like kinase (Solyc09g072810). Among proteins displaying transient phosphorylation at 2 minutes (cluster B) we found CBL-interacting protein kinase (Solyc04g076810), serine/threonine protein kinase (Solyc01g108920) and cell division protein kinase (Solyc07g063130) with highest numbers of interaction partners (Table 1). For proteins with transient phosphorylation at 5 minutes (cluster C), the protein kinases with highest degree were the ethylene receptor (Solyc05g055070), and two receptor kinases (Solyc09g083210 and Solyc12g010740). Among the proteins with transient phosphorylation at 15 minutes (cluster D) were a serine/threonine protein kinase (Solyc11g033270), the activating motif of a mitogen activated protein kinase MPK2 (Solyc08g014420), and a uncharacterized protein kinase (Solyc09g009090). MPK2 has previously been shown to be part of the systemin signaling network (52) thus corroborating our identification of systemin-specific kinase responses. Interestingly, these protein kinases with high degree often were identified with several phosphorylation sites which showed different response profiles during the systemin treatment.

**Table 1:**
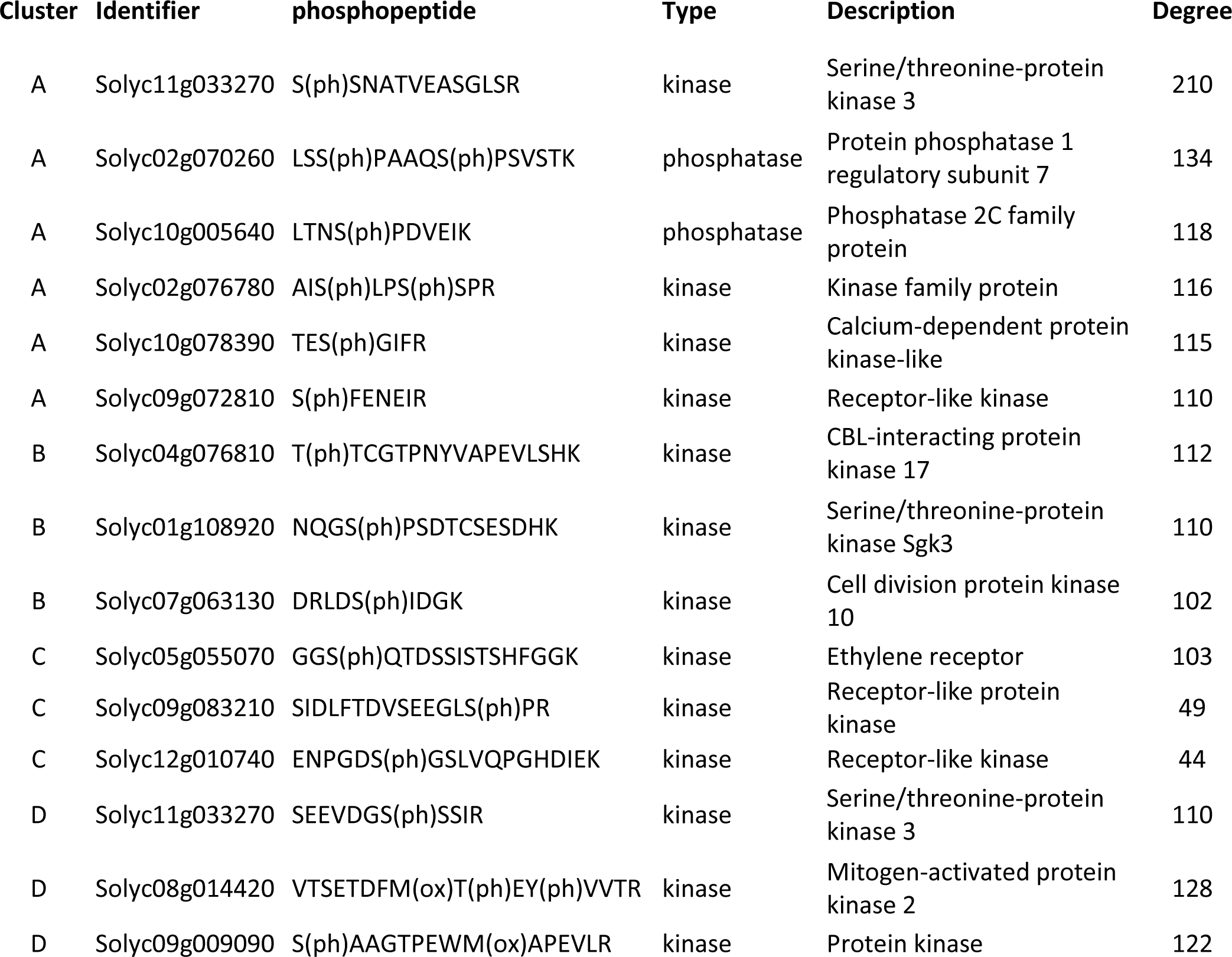
Phosphopeptides of kinases and phosphatases with highest numbers of interactions in the correlation network.

Within this kinase-substrate correlation network, we then displayed the degree distribution of phosphopeptides from systemin-specific kinases/phosphatases, other kinases/phosphatases and the putative targets for systemin-induced clusters A to F (Figure 6C, D). It becomes apparent that systemin-specific kinases in clusters B, C, and D (which showed a transient phosphorylation increase) showed a higher degree than kinases without systemin specificity, but kinases in generally had higher degree than their target proteins (Figure 6C). In contrast, for phosphatases, higher than average degree was observed for nine systemin-specific phosphatases in cluster A (rapid dephosphorylation response). In cluster F (no systemin-induced phosphorylation response), higher than average degree was observed for nine other phosphatases (those not linked to the systemin response). This observation supports the conclusion that the kinases identified as systemin-specific, are indeed responsible for the phosphorylation events that peak at 2 minutes (cluster B), 5 minutes (cluster C) and 15 minutes (cluster D) of stimulation, while systemin-specific phosphatases mediate rapid dephosphorylation responses (cluster A). A full list of kinase-target relationships proposed from the correlation analysis is available in Supplementary Table 5. Although our data set reveals correlative relationships between the phosphorylation sites that may be indicative of kinase/phosphatase-substrate interaction, it must be kept in mind that the data does not directly allow to derive the activity change of the respective proteins without further experiments. Selected candidate targets for systemin-specific kinases and phosphatases are addressed and discussed below.

### Connection to the MAP-Kinase signaling pathway

The MAP-triple kinase (MAPKKK) Solyc11g033270 (degree 210) was identified with two phosphorylation sites outside the kinase domain: S(ph)SNATVEASGLSR (Ser859) showed rapid dephosphorylation (cluster A) and SEEVDGS(ph)SSIR (Ser345) showed transient phosphorylation at 15 minutes (cluster D). The phosphorylation time profile of the rapidly dephosphorylated MAPKKK peptide S(ph)SNATVEASGLSR showed highest correlation with dephosphorylated peptides of an ubiquitin ligase (LEEGS(ph)SPEQR, Solyc05g007820), a YAK1 homologous protein kinase (TVYSY(ph)IQSR, Solyc03g097350) and a RNA recognition motif containing protein (SSDS(ph)QELTTTELK, Solyc02g062290), and a protein phosphatase 2C family protein (LTSS(ph)PEAEIK, Solyc06g065580). The finding that kinases and phosphatases correlate with the same phosphopeptide suggest kinase-phosphatase activation/inactivation loops upon systemin stimulation. At 15 minutes of systemin treatment, the MAPKKK phosphopeptide SEEVDGS(ph)SSIR of protein kinase Solyc11g033270 displayed a transient increase in normalized intensity. This kinase phosphopeptide showed highest correaltion with a transient phosphorylation of a calmodulin domain containing protein (DYS(ph)ASGYSSR, Solyc06g053530).

For Solyc08g014420, MAP-Kinase MPK2, which is the tomato homolog of Arabidopsis MPK6, several potential targets are proposed from the correlation network (Figure 6A), including protein kinase Solyc02g076780 (AIS(ph)LPS(ph)SPR), chromatin remodeling factor Solyc01g094800 (GGVVDS(ph)DDDLAGK), transcription initiation factor Solyc01g088370 (DGEAS(ph)DEEEEYEAK), kelch-repeat containing protein kinase Solyc11g071920 (LIHPIPPVSS(ph)PENS(ph)PER), and two phosphorylation sites of cell cycle-related protein Solyc10g078180 (ELEDGIELS(ph)AGNEK, ALENM(ox)DDAVGQS(ph)PQEVR). The majority of these peptides did contain the expected “(ph)SP” MAPK substrate motif. Interestingly, these corresponding proteins have already been confirmed as targets of MPK6 in Arabidopsis under various conditions (53). All of these proposed target phosphorylation sites were found in systemin response cluster D with high phosphorylation at 15 minutes of systemin treatment, suggesting activation of MAP-kinase signaling pathways after 15 minutes of systemin treatment. An unusual putative substrate peptide to MPK2 was found with the C-terminal peptide of H+-ATPase LHA1.

### Plasma membrane H^+^-ATPases

Phosphopeptides matching two different plasma membrane H^+^-ATPases (LHA1 Solyc03g113400; Solyc07g017780) showed systemin-induced correlation with phosphopeptides of distinct kinases and phosphatases (Figure 7A). Phosphopeptides of receptor kinases (Solyc02g079590, Solyc02g076780), MAP-kinase MPK2 (Solyc08g014420), a MAP-triple kinase (Solyc11g033270), were highly correlated to phosphopeptides encompassing the activating C-terminal phosphorylation site at T955 of LHA1 (GLDIETIQQSYT(ph)V) and T964 of H^+^-ATPase Solyc07g017780 (GLDIETIQQHYT(ph)V). We suggest that these kinases may directly phosphorylate the respective residues in the H^+^-ATPases. Thereby, kinases with highest correlation to the well-studied C-terminal activating phosphopeptide GLDIETIQQSYT(ph)V of LHA1 were receptor kinase Solyc02g079590 (r=0.99) and MPK2 (Solyc08g014420) (r=0.98). This C-terminal peptide also was highly correlated with a regulatory phosphorylation site in the N-terminus of the homolog of Arabidopsis POLTERGEIST-LIKE 5 phosphatase (PLL5, Solyc06g076100), a member of the PP2C clade C phosphatases (54, 55). Highest correlation with phosphopeptide GLDIETIQQHYT(ph)V of H^+^-ATPase Solyc07g017780 was found with receptor kinase Solyc11g011020 (r=0.94).

**Figure 7:**
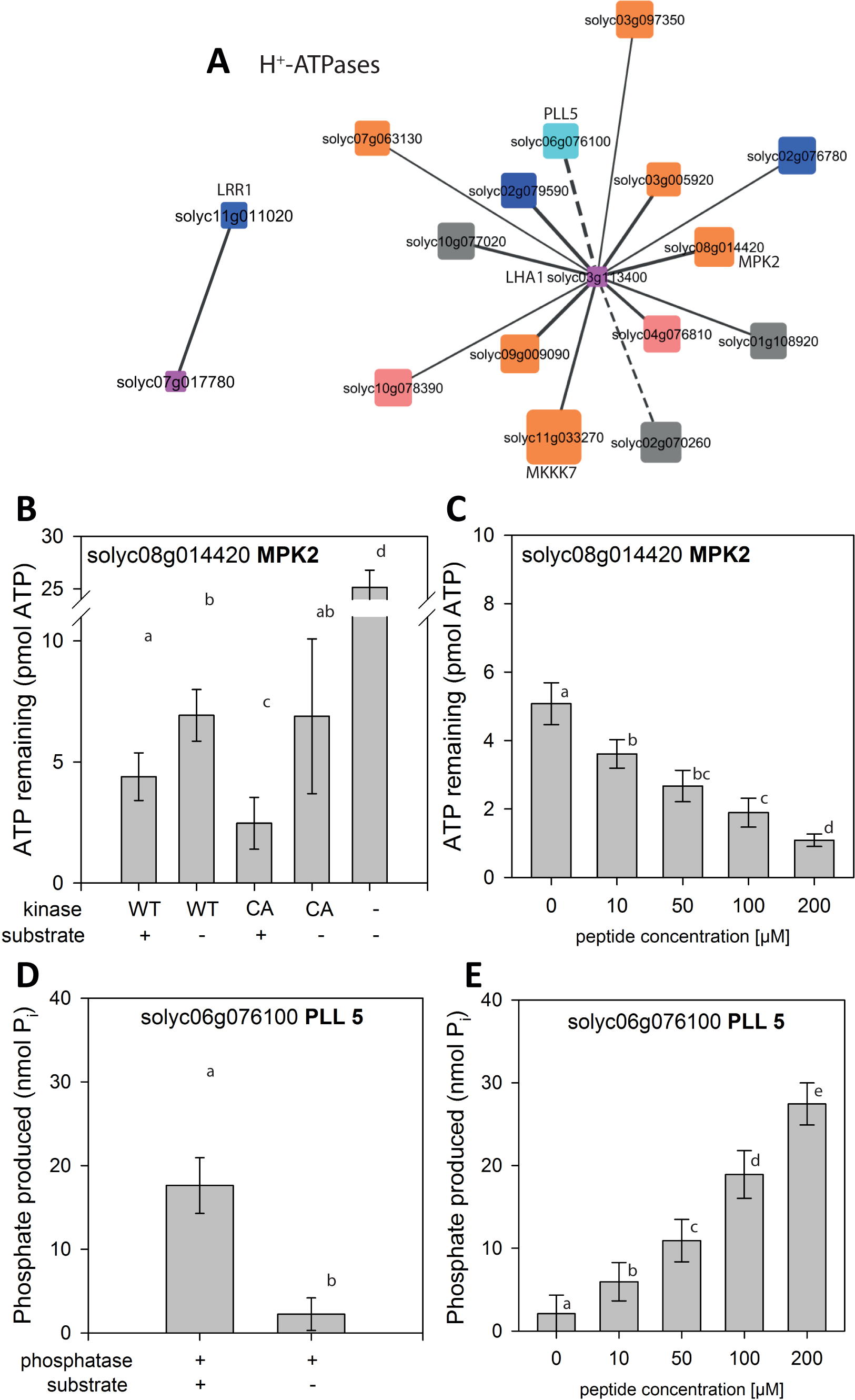
Kinase/phosphatase relationships with major targets. (**A**) H^+^-ATPases AHA1 and LHA1. Solid edges represent kinase-substrate relationships, dashed edges represent phosphatase-kinase relationships. Node size indicates the degree in the overall network (figure 7), Node color represents Mapman function. Blue: receptor kinase (30.2), cyan: Phosphatases, red: calcium-dependent protein kinases, orange: protein.posttranslational modification (29.4), gray: unclassified function. (35). (**B**) ATP remaining after kinase MPK2 reaction using H^+^-ATPase peptide GLDIETIQQSYTV as a substrate. (**C**) ATP remaining after MPK2 reaction with increasing concentrations of substrate peptide (**D**) Phosphate produced by Phosphatase PLL5 in the reaction with H^+^-ATPase phosphopeptide GLDIETIQQSY(pT)V as a substrate. (**E**) Phosphate produced by PLL5 reaction with increasing concentrations of substrate peptides. Data represent the mean with standard deviation. Activity assays were carried out on two independent protein isolations in two technical replicates.

In addition to the C-terminal phosphorylation site, another activating phosphorylation site (T(ph)LHGLQPPEASNLFNEK, (56)) and an inactivating phosphorylation site (ELS(ph)EIAEQAK, (31)) were identified for H^+^-ATPase Solyc07g017780. The number of data points identified for these two phosphorylation sites was too low to be included in the correlation analysis. Interestingly, the activating site was also identified at 15 minutes of systemin treatment, but phosphorylation at that site did not qualify as systemin-specific response based on the sum of squares deviation analysis. Interestingly, the phosphopeptide containing the inactivating serine residue was identified at 5 minutes of systemin treatment, the time point when systemin stimulation resulted in a minimum in measured H^+^-ATPase activity (Figure 4C).

The interaction of the C-terminal peptide of H^+^-ATPase LHA1 with candidate kinase MPK2 and candidate phosphatase PLL5 was investigated further by *in vitro* activity assays (Figure 8 B-E). Recombinant MPK2 was expressed either in wild-type form (WT) or as a constitutively active mutant (CA) with D216G and E220A amino acid substitutions (40). Kinase activity, assayed as ATP consumption, was observed only in presence of GLDIETIQQSYTV peptide substrate, and was higher for CA compared to WT (Figure 7B). Increasing the substrate peptide concentration from 10 µM to 200 µM resulted in increasing kinase activity (Figure 7C). Phosphorylation of the C-terminal LHA1 peptide substrate at T955 by recombinant MPK2 was confirmed by mass spectrometric analysis (supplementary figure 3A,C). Recombinant PLL5 was able to dephosphorylate synthetic phosphopeptide GLDIETIQQSY(pT)V *in vitro* (Figure 7D). PLL5 activity, assayed as release of inorganic phosphate, was observed only in presence of the phosphopeptide substrate, and increased with increasing substrate concentration (Figure 7E). De-phosphorylation of the LHA1-derived phosphopeptide substrate by PLL5 phosphatase was confirmed by mass spectrometric analysis (Supplementary Figure 3B,C). Interestingly PLL5 phosphatase was identified in systemin-induced cluster A (rapid dephosphorylation). Thus, the phosphatase may well be involved in dephosphorylation of LHA1 at early systemin stimulation.

MPK2, on the other hand, was identified in systemin-induced cluster D with maximum phosphorylation of peptide VTSETDFM(ox)T(ph)EY(ph)VVTR at 15 minutes. The identified phosphopeptide contained the typical MAP-Kinase activating motif (pT)E(pY). These data are fully consistent with a previous report showing that MPK2 is transiently activated in response to systemin with maximum activity between 15 and 30 minutes after treatment (44). MPK2 thus is a candidate kinase for re-phosphorylation (Figure 4A) with concomitant increase in H+-ATPase activity (Figure 4C) at later time points (15 min) of systemin treatment.

### Respiratory burst oxidases

Another protein with highly systemin-specific phosphorylation response was the respiratory burst oxidase. Phosphopeptides were identified for two isoforms of respiratory burst oxidases (NLSQM(ox)LS(ph)QK, Solyc03g117980; DVFSEPS(ph)QTGR, Solyc06g068680). Interestingly, Solyc03g117980 has previously been shown to be required for the systemic induction of wound-response genes (57) which is fully consistent with our classification of phosphorylation of Solyc03g117980 as systemin-specific. Potential interacting kinases include two receptor kinase (Solyc09g064270 and Solyc02g07000) and a CBL-interacting protein kinase (Solyc06g068450, r=0.90; Supplementary Figure 4A). Phosphorylation of the respiratory burst oxidase occurred at early time points with rapid dephosphorylation, and this response was not found with A17 treatment (Supplementary Tables 2 and 5)

### Processes at the cell wall

Among the target proteins with most significant systemin-induced phosphorylation responses were cellulose synthase proteins. Four phosphopeptides matching cellulose synthase like proteins (SSS(ph)RLNLSTR Solyc04g077470, SHS(ph)GLMR Solyc08g076320, SSS(ph)ESGLAELNK Solyc09g057640, GLIDSQSLSSS(ph)PVK Solyc09g075550) were identified as potential targets of systemin-specific kinases and phosphatases (Supplementary Figure 4B). Systemin-induced phosphorylation changes of cellulose synthase-like proteins were either rapid dephosphorylation (cluster A) or transient phosphorylation at 2 minutes (cluster B). The transient phosphorylation of an uncharacterized receptor kinase at 2 minutes (NNTS(ph)SVSPDSVTAK, solyc12g036330) showed strongest correlation with the transient phosphorylation of peptide S(ph)SEGDLTLLVDGKPK of cellulose synthase-like protein solyc04g077470, while dephosphorylation of peptide GQLPS(ph)GQVVAVK matching an uncharacterized receptor-like cytoplasmic kinase (Solyc05g024290) showed strongest correlation with dephosphorylation of peptide GLIDSQSLSSS(ph)PVK of cellulose-synthase-like protein (Solyc09g075550). These two peptides also showed a high correlation to a POLTERGEIST-LIKE 1 phosphatase (Solyc08g077150, peptide FVS(ph)PSQSLR; r=0.92).

### Vesicle trafficking and G-Protein signaling

In total, 52 kinases and 8 phosphatases were found with high correlations to phosphopeptides of vesicle trafficking proteins. While phosphopeptides of adaptins and syntaxins were rapidly dephosphorylated (cluster A), most phosphopeptides of vesicle-associated membrane proteins (VAMPs) were transiently phosphorylated at 2 minutes of systemin stimulation (cluster B). For example, phosphopeptide NS(ph)DDGYSSDSILR in cluster A matching a KEU-homolog (Solyc10g008490) showed strong correlation (r=0.99) to phosphopeptide VNS(ph)LVQLPR matching the catalytic subunit of a protein phosphatase (Solyc10g008490). Phosphopeptide VVYIS(ph)PHS(ph)SPGHSEDAFK of VAMP Solyc04g015130 with transient phosphorylation at 2 minutes showed high correlation with a protein phosphatase 2C Solyc07g053760 and protein kinase Solyc06g068920.

Several phosphopeptides of small GTPase proteins were also identified and these proteins may also have functions in vesicle budding and fusions. Most prominent members of this group included the ARF GTPase activator Solyc05g023750 and RAB GTPase activator Solyc12g009610. The phosphopeptide NS(ph)SELGSGLLSR of the ARF-GTPase activator was identified as being rapidly dephosphorylated (cluster A), while phosphopeptide TLS(ph)SLELPR of the RAB-GTPase activator Solyc12g009610 was transiently phosphorylated at 2 minutes of systemin treatment.

Phosphopeptide AELS(ph)VNR of a RAB-GTPase Solyc09g098170 was transiently phosphorylated at 5 minutes. Phosphopeptide NS(ph)SELGSGLLSR correlated highest with phosphopeptide ALIGEGS(ph)YGR of receptor kinase Solyc12g098980, phosphopeptide TLS(ph)SLELPR showed highest correlation with receptor kinase Solyc07g056270. Phosphorylation of the RAB-GTPase Solyc09g098170 correlated highest with receptor kinase Solyc07g055130 and the protein phosphatase 2C Solyc05g055790.

### Calcium-related responses

Four IQ-domain 13-containing proteins (FTLPAGAVS(ph)PR Solyc08g014280, EAS(ph)PKVT(ph)SPR Solyc01g108920, LSFPLTPSS(ph)TGSVK Solyc10g008490, FNSLS(ph)PR Solyc05g014760) were rapidly dephosphorylated upon systemin stimulation (cluster A). Phosphopeptides of CBL-interacting kinase (Solyc04g076810), CDPK (Solyc04g009800) and MAP-triple-kinases (Solyc12g088940, Solyc11g033270) as well as a phosphopeptide of phosphatases (Solyc10g008490 and Solyc01g067500) showed strong correlation with dephosphorylation of the IQ-domain 13-containing proteins. A phosphopeptide of calcium-ATPase Solyc02g064680 displayed a transient phosphorylation at 5 minutes (cluster C). Phosphopeptides of phosphatase 2C (solyc05g055790) and kinases Solyc07g055130 and Solyc01g094920 showed high correlation with the phosphorylation profile of the calcium ATPase.

### Transcription factors

Phosphopeptides of several transcription and splicing factor families (bin 27) were over-represented with similar abundance in all systemin-response clusters (supplementary Table 2). The highest number of identified phosphopeptides and highest degree for this functional protein group in the network (37) was observed for the RNA-processing factor SERRATE (Solyc01g009090) with matching phosphopeptides S(ph)PPFPPYK and NRS(ph)PPHHPPGR. Phosphopeptide S(ph)PPFPPYK was transiently phosphorylated at 2 minutes and showed best correlation to phosphorylation sites of kinases CTR1 (Solyc09g009090), a MAP-triple kinase (Solyc12g088940), and a CBL-interacting kinase (Solyc04g076810). Phosphopeptide NRS(ph)PPHHPPGR was transiently phosphorylated at 5 minutes of systemin treatment. Phosphorylation sites of receptor kinases Solyc02g023950 and Solyc09g072810, a CDK-related kinase Solyc10g078390 and a MAP-triple kinase Solyc11g033270 showed high correlation with that second SERRATE phosphorylation site.

### Interactions with other phytohormone signaling pathways

Among the specifically systemin-induced phosphorylation/dephosphorylation events were several phosphopeptides of proteins known from other hormone signaling pathways. For example, phosphopeptides of abscisic acid (ABA) response factors (ASSESSSEDEADS(ph)AEEAGSSK, Solyc04g081340; VGS(ph)PGNDFIAR, Solyc01g108080), auxin response factor (INS(ph)PSNLR, Solyc05g047460), EIN4 (GGT(ph)PELVDTR, Solyc05g055070) and PIN3 (PSNFEENCAPGGLVQS(ph)SPR, Solyc05g008060) were rapidly dephosphorylated upon systemin treatment (cluster A). Another phosphorylation site in ABA-response factor Solyc04g081340 (RSS(ph)ASATQVQAEEQAPR), SSS(ph)RLNLSTR of ACC-synthase (Solyc03g007070), and another phosphopeptide of EIN4 (RSS(ph)ASATQVQAEEQAPR) were transiently phosphorylated at 2 minutes of systemin treatment. A third phosphopeptide of EIN4 (GGS(ph)QTDSSISTSHFGGK) was transiently phosphorylated at 5 minutes of systemin treatment. Kinases with high correlations to the phosphorylation patterns in cluster A were CTR1 (Solyc09g009090), a receptor kinase (Solyc11g011020), and a CBL-interacting protein kinase (Solyc04g076810).

## Discussion

The aim of our work was to (i) identify early events induced by systemin and (ii) identify kinases and phosphatases involved in these signaling events. The maturation of the signaling peptide systemin is beginning to be understood (9), and the primary receptor of systemin was recently identified (16). While individual early responses to systemin perception, such as an increase in cytosolic calcium (17), ROS production (16, 58), the modulation of the H^+^-ATPase activity (19), or the activation of MAP-kinase signaling (52) have been identified, the proteome-wide effects of systemin treatment were not yet studied.

### The use of inactive analogon A17 to define systemin-specific responses

Antagonistic and inactive peptides—such as mutant peptide variants, chemically modified peptides or peptide mimetics, are important tools for the characterization of peptide-receptor interaction and to discover endogenous peptide function (59–61). For example, the dominant-negative effect produced by antagonistic peptides can be used for the characterization of peptide function when suitable mutants are not available, and in the case of genetic redundancy (62, 63). The use of inactive analoga in studying receptor signaling pathways provides an elegant way to define specific responses and rule-out common unspecific treatment responses. Comparing the responses to systemin and the antagonistic A17 analog, approximately half of the identified phosphopeptides could be identified as not specific to the systemin treatment and were excluded from the reconstruction of the systemin signaling network. Including the A17 analog thus allowed us to gain insight into systemin-specific protein signaling processes.

### Systemin-induced modulation of the plasma membrane H+-ATPase

Systemin-induced alkalization of the growth medium has been well described (18, 19, 60). Alkalization of the growth medium is a typical process connected with inhibition of the plasma membrane H^+^-ATPase, as it also occurs upon stimulation by flagellin and other microbe-associated molecular patterns (31, 64, 65). By pairwise correlation of phosphopeptide abundance profiles we identified the H^+^-ATPase LHA1 as putative target of 12 kinases and two phosphatases, while receptor like kinase Solyc11g011020 was the only kinase putatively involved in the phosphorylation of LHA1. These kinase-substrate predictions were confirmed by *in-vitro* kinase and phosphatase assays. MPK2 and PLL5 phosphatase were found to indeed act on the respective peptide of LHA1. While such verification can only be done for a limited number of kinases or phosphatases, the results obtained for MPK2 and PLL5 phosphatase suggest the general validity of the correlation analysis also for other candidate pairs. We thus consider the correlation analysis for prediction of kinase/phosphatase-substrate interactions as a powerful means to predict new regulatory circuits in the systemin signaling network. Arabidopsis homologs to 25 of the systemin-responsive kinases were experimentally characterized, as well as 49 Arabidospis homologs of the putative substrates. Among the systemin-induced predicted kinase-substrate relationships, seven kinase-substrate pairs were also experimentally identified in Arabidopsis (53).

We identified a novel direct connection of the MPK pathway with the plasma membrane H^+^-ATPase LHA1. MPK2 has previously been shown to play an essential but unknown role in the systemin signaling pathway (52). Our data suggest that MPK2 is responsible for re-phosphorylation of the H^+^-ATPase at later times of systemin stimulation. Consistent with the transient activation of MPK2 in response to systemin (44), phosphorylation of the typical MAPK activation motif (TEY) was observed at 15 minutes of systemin treatment (cluster D). Recombinant MPK2 phosphorylated the C-terminal peptide of LHA1 at the penultimate regulatory threonine residue (Figure 7) resulting in activation of the proton pump. MPK2-mediated activation of LHA1 is thus likely to be responsible for re-acidification of the extracellular space dampening the initial alkalization response.

While knowledge of specific interactions of phosphatases with their substrates is still limited in plants, several phosphatases have been implicated in the regulation of the plasma membrane proton pump. This includes a type 2A phosphatase activity partially purified from maize roots (66), an unidentified type 2C phosphatase in Arabidopsis seedlings and *Vicia faba* guard cells (67), and PP2C clade D phosphatases that are involved in proton pump regulation during auxin-induced acid growth of Arabidopsis hypocotyls (68). PP2C-D3 (alias PP2C38; At4g28400) also was shown to dephosphorylate BIK1 attenuating immune signaling through the FLS2 and EFR pattern recognition receptors (69). Likewise, EFR-mediated signaling also is downregulated by Arabidopsis PLL4 and PLL5 phosphatases, which belong to PP2C clade C (70). We show here that the tomato PLL5 homolog, is (i) involved in systemin signaling and (ii) dephosphorylates LHA1. The precise mechanisms of PP2C phosphatase regulation and signaling are still unclear (71). The identified phosphorylation site lies within the regulatory N-terminus (55), suggesting that PLL5 phosphatase may be activated by de-phosphorylation (consistent with its identification in systemin-induced cluster A). This phosphatase thus is a primary candidate for proteins involved in the downregulation of H^+^-ATPase activity and the resulting alkalization of the growth medium. It acts by dephosphorylation of activating threonine phosphorylation sites in the C-terminus of the H^+^-ATPase.

### Reconstruction of early vs. late responses from plasma membrane to the nucleus

The recently identified plasma membrane located LRR-RLK receptors for the systemin peptide (16) were not identified among the 3312 phosphopeptides from this work, while several other receptor kinases, among them also the phytosulfokine receptor PSKR2 (Solyc07g063000) or FERONIA (Solyc09g015830) were quantified in our experiments. There could be several reasons for missing these proteins in the phosphopeptide analysis: Firstly, a shotgun type phosphoproteome analysis was carried out depending on sufficient quantity and quality (peptide length, amino acid composition) of identified (phospho)peptides. There are examples of prominent conditional phosphorylation sites (e.g. Thr101 of NT1.1 (72)), which do not produce “findable” peptides by trypsin digestion. Secondly, some important receptor kinases may not have been enriched in the membrane purification protocol due to their strong embedding within the cell wall. This has in a previously published study using Arabidopsis cell cultures also been observed for the flagellin receptor FLS2, for which in that study also no phosphopeptides were identified upon a flg22-stimulation (73). Thirdly, considering the absence of phosphopeptides from recently identified systemin receptors, the earliest harvesting time point of 2 minutes may have been already too late to capture the immediate first systemin responses, i.e. receptor phosphorylation. Ligand-induced phosphorylation of ligand-perceiving receptor kinases is known to occur within seconds (74).

Nonetheless, our work identified systemin specific phosphorylation sites at 56 kinases and 17 phosphatases to be involved in systemin signaling, likely downstream of the primary systemin receptor SYR1 (16). Substrates for these kinases and phosphatases were identified by pairwise correlations. Correlation analysis was previously used to reconstruct topology of phosphorylation networks induced by external nutrient supply (75). Thereby, plasma membrane proteins were identified as signal induction layers within the network, while cytosolic and nuclear proteins were considered as an effector layer.

The identified phosphorylation patterns at 2 minutes may thus represent early responses of the systemin signaling cascade. This point of view is supported by the observation of rapid (de)phosphorylation changes for several cytosolic proteins such as a CBL-interacting kinase, MAP-triple kinase, and CDPKs already at 2 minutes (clusters A and B), while a large number of receptor kinases with systemin specific phosphorylation patterns peak only after 2 minutes or 5 minutes. Also observed very rapidly after systemin stimulation were plasma membrane-located proteins that are known to belong to the systemin signaling cascade, including the proton pump that is rapidly dephosphorylated after systemin treatment, and two RBOH isoforms also showing rapid dephosphorylation. In contrast, G-protein signaling or transcription factor phosphorylation was found at later times.

Later responses of the systemin signaling cascade included MAP-kinase signaling at 15 minutes of treatment. Here, phosphorylation of activating TEY motif particularly of MPK2 was observed. MPK2 was experimentally identified as a kinase re-phosphorylating the plasma membrane ATPase and seven additional putative substrates were also described for the Arabidopsis homolog MPK6. Thus, as suggested earlier (52), the MAP-Kinase signaling network is also activated upon systemin stimulation and here we present a range of putative substrate peptides for further study.

## Conclusions

Our study presents 56 systemin specific kinases and 17 systemin specific phosphatases. Putative substrates were identified by pairwise correlations of systemin induced phosphorylation time profiles. Thereby we identified novel candidates for the systemin signaling pathway. Two of these candidates, MPK2 and PLL5 were confirmed as kinase and phosphatase for reversible phosphorylation of the C-terminal regulatory site of the plasma membrane H^+^-ATPase. We highlighted several proteomic processes upon systemin stimulation ranging from plasma membrane processes to cytosolic and nuclear signaling, which will provide a valuable resource for further in-depth functional studies in future elucidation of systemin signaling cascades.

## Supporting information

Supplementary Material Description

## Acknowledgements

Fatima Haj Ahmad received a PhD fellowship from the German Academic Exchange Service (DAAD) Further support was provided by a grant from the Deutsche Forschungsgemeinschaft SFB 1101/D06 to Andreas Schaller. We thank Ursula Glück-Behrens for maintenance of the *Solanum peruvianum* cell culture. Denise Llanos Jara is acknowledged for excellent assistance during sample preparation and mass spectrometry.

## Data availability

The mass spectrometry proteomics data have been deposited to the ProteomeXchange Consortium via the PRIDE (76) partner repository (http://www.ebi.ac.uk/pride) with the dataset identifier PXD010819

