## Supplementary Material Description for "The systemin signaling cascade as derived from phosphorylation time courses under stimulation by systemin and its inactive Thr17Ala (A17) analog"

**Supplementary Figure 1:** Consensus cluster profiles upon systemin treatment, A17 treatment, and water treatment. The consensus cluster profiles were calculated from all phosphopeptide profiles within one k-means cluster and were used as model cluster profiles when calculating the best fit (sum of square deviation) for each phosphopeptide identified under A17 treatment. Averages with standard deviations are displayed.

**Supplementary Figure 2**: Representative annotated spectra of identified phosphopeptides under systemin and A17 treatment as exported from MaxQuant.

**Supplementary Figure 3**: Mass spectrometry analysis of in-vitro kinase and phosphatase assays. (**A**) Representative spectrum of kinase reaction with recombinant MPK2 resulting in phosphopeptide GLDIETIQQSY(pT)V. (**B**) Representative spectrum of phosphatase reaction with recombinant PLL5 phosphatase resulting in dephosphorylated peptide GLDIETIQQSYTV. (**C**) Summary of ion intensities of phosphorylated and dephosphorylated C-terminal peptide of H+-ATPase LHA1. Input peptides are shown as bars without patterns, reaction results are indicated as hatched bars. Averages with standard deviations of two biological replicates from two independent recombinant protein preparations are shown.

**Supplementary Figure 4:** Kinase and phosphatase relationship with proteins from respiratory burst oxidase and cell wall proteins. (**A**) Respiratory Burst Oxidases. (**B**) Cellulose synthase like proteins. Edge color represents the time cluster of this interaction, light color indicates earlier clusters (2min), dark color indicates later clusters (15min). Solid edges represent kinase-substrate relationships, dashed edges represent phosphatase-kinase relationships. Node size indicates the degree in the overall network (Figure 7), Node color represents MapMan function. Blue: receptor kinase (30.2), cyan: phosphatases, orange: protein.posttranslational modification (29.4), gray: unclassified function. (35)

**Supplementary Table 1**: list of normalized ion intensities of quantified phosphopeptides under systemin and A17 treatment. Quantitative values were obtained by cRracker analysis (38). Average normalized intensities, standard deviations and number of spectra are shown.

**Supplementary Table 2**: k-means clusters of identified phosphopeptides for systemin and A17 treatment.

**Supplementary Table 3**: Over-representation analysis for all clusters upon systemin stimulation and tabular overview of the abundance distribution of proteins in subcategories to statistically significant over-represented MapMan bins. Numbers represent the distribution of proteins in each subbin across clusters (in %).

**Supplementary Table 4**: List of kinases and phosphatases considered as systemin-specific based on their phosphorylation profiles under systemin stimulation in comparison with A17 and water treatment.

**Supplementary Table 5**: Correlation network of systemin specific kinases and phosphatases and their targets based on pairwise correlation analysis of phosphopeptide time profiles.
