## Supplementary figures and images for "The systemin signaling cascade as derived from phosphorylation time courses under stimulation by systemin and its inactive Thr17Ala (A17) analog"

### Supplementary Figure S1

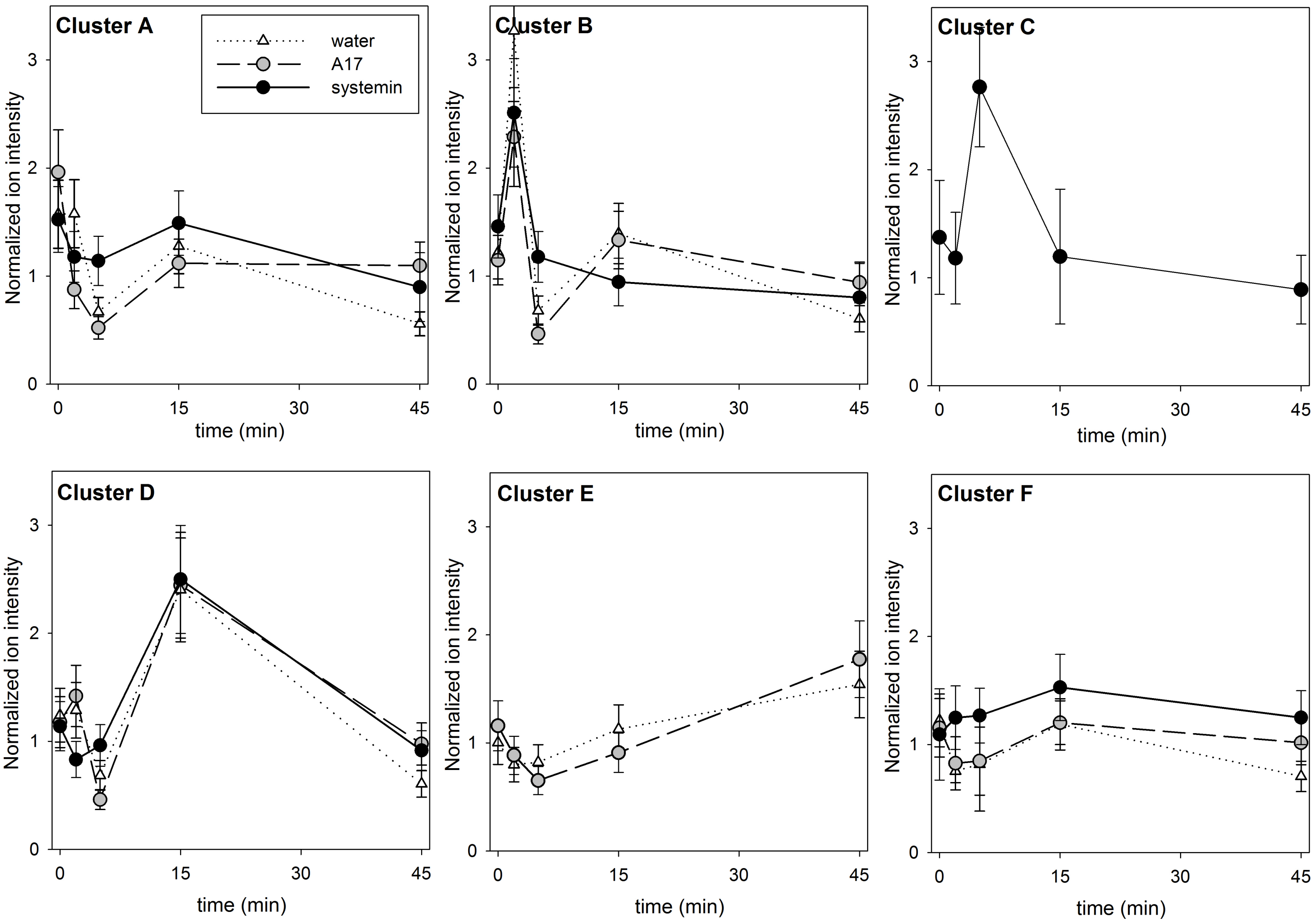

### Supplementary Figure S3

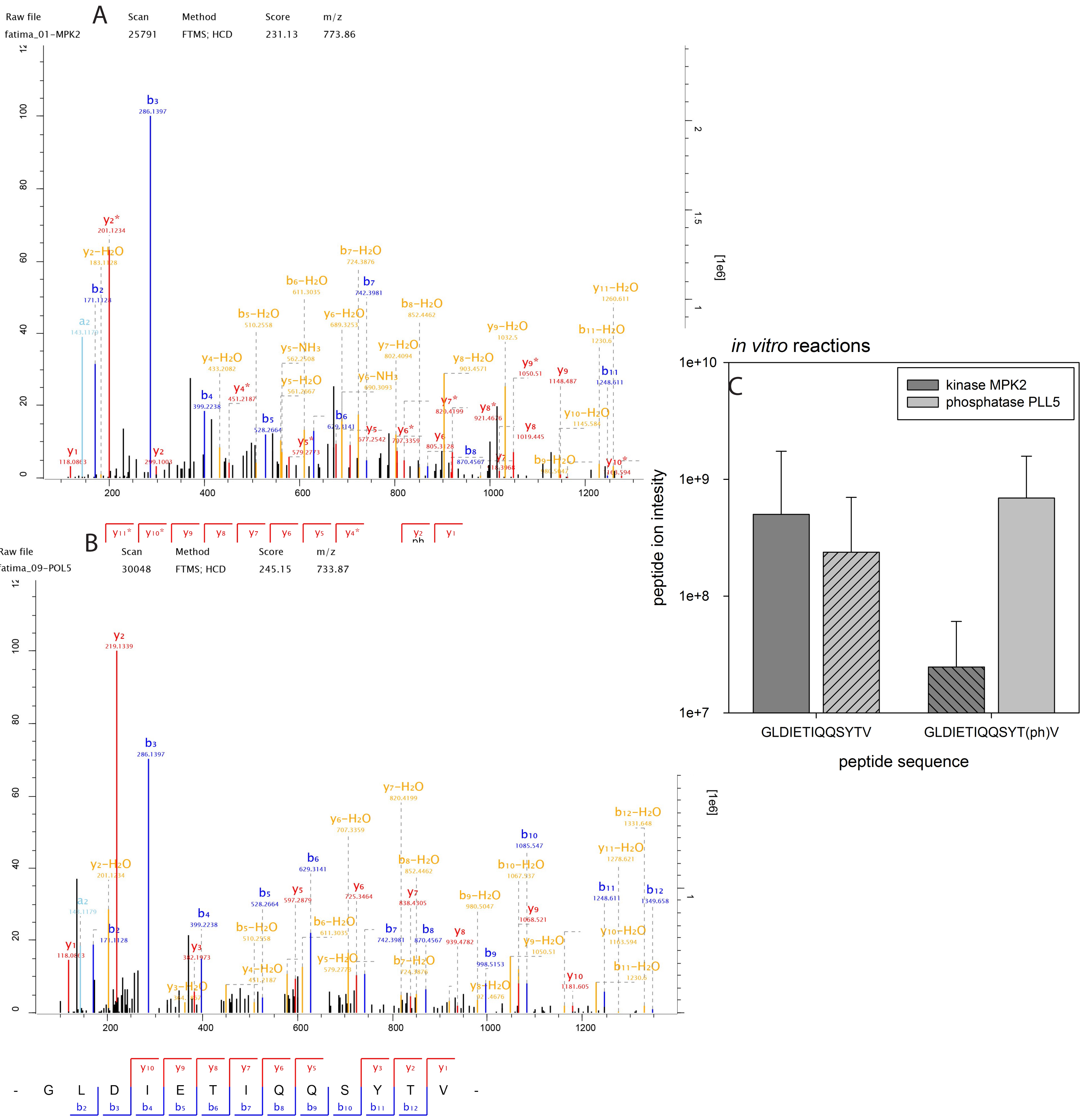

### Supplementary Figure S4

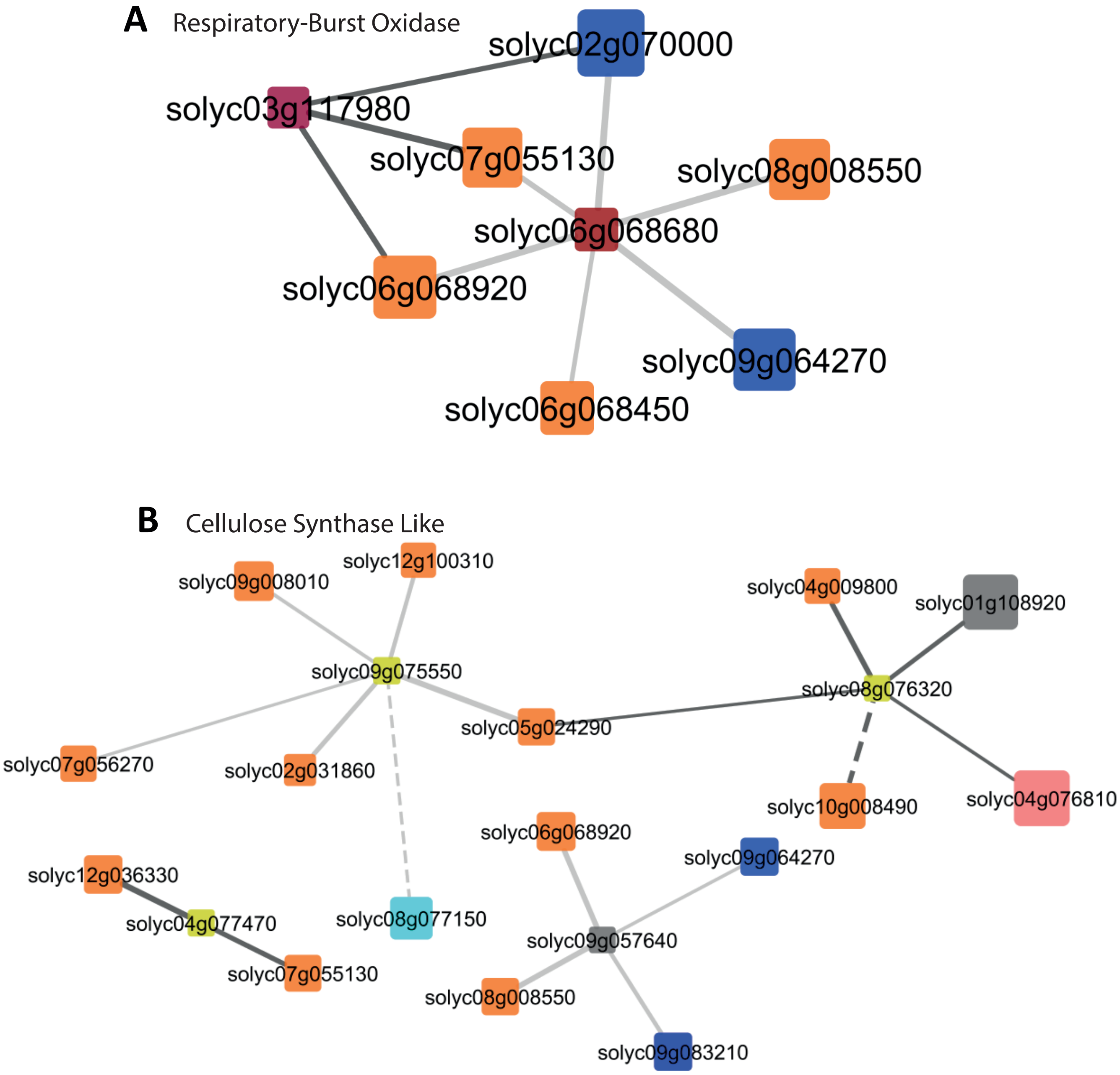
