## Supplementary Figure S2 for "The systemin signaling cascade as derived from phosphorylation time courses under stimulation by systemin and its inactive Thr17Ala (A17) analog"

| Raw file | Scan | Method | Score | m/z |
| --- | --- | --- | --- | --- |
| A17_00_1 | 12054 | FTMS; HCD | 44.32 | 706.3 |

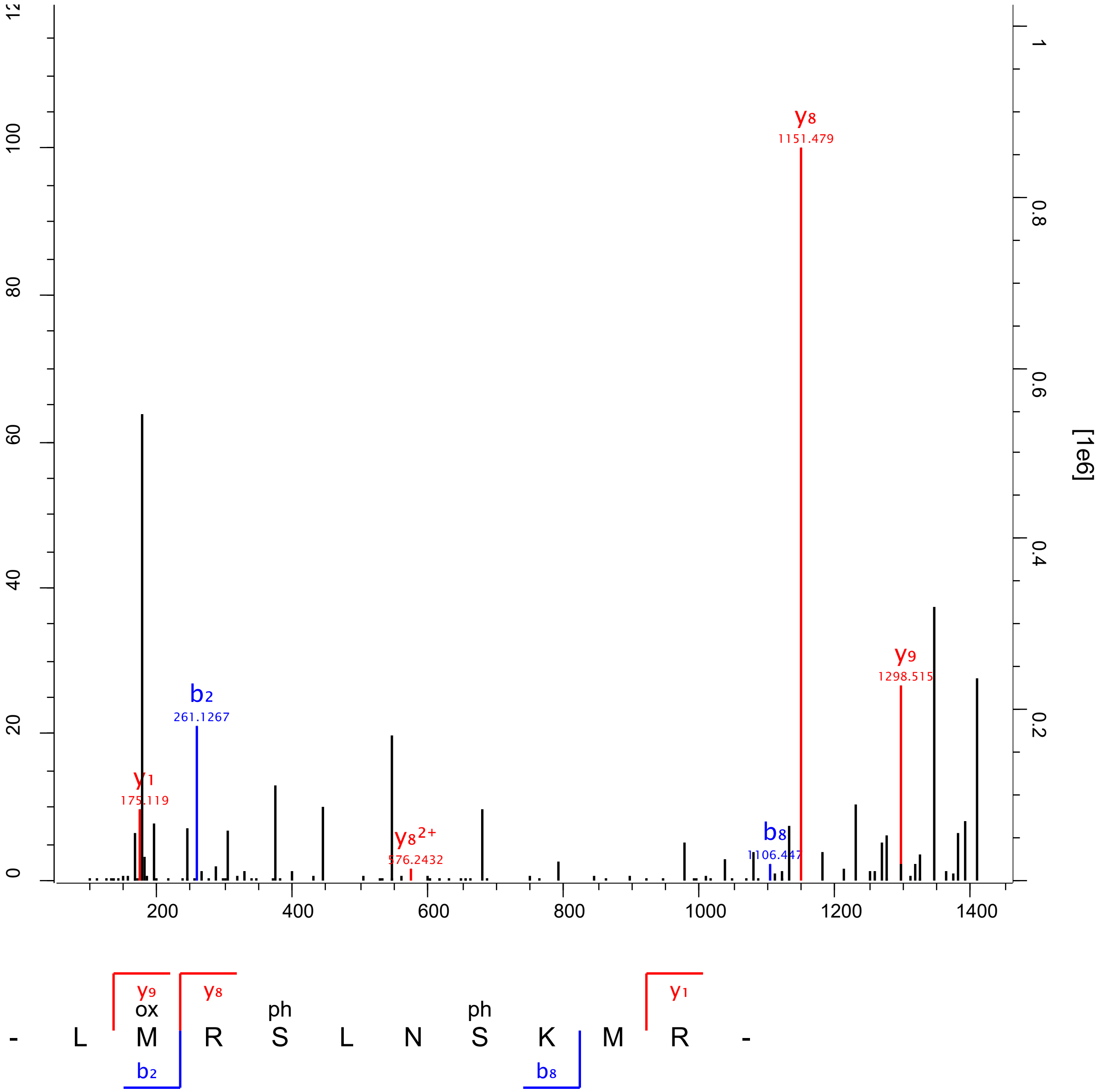

| Raw file | Scan | Method | Score | m/z |
| --- | --- | --- | --- | --- |
| A17_00_1 | 2027 | FTMS; HCD | 120.9 | 530.55 |

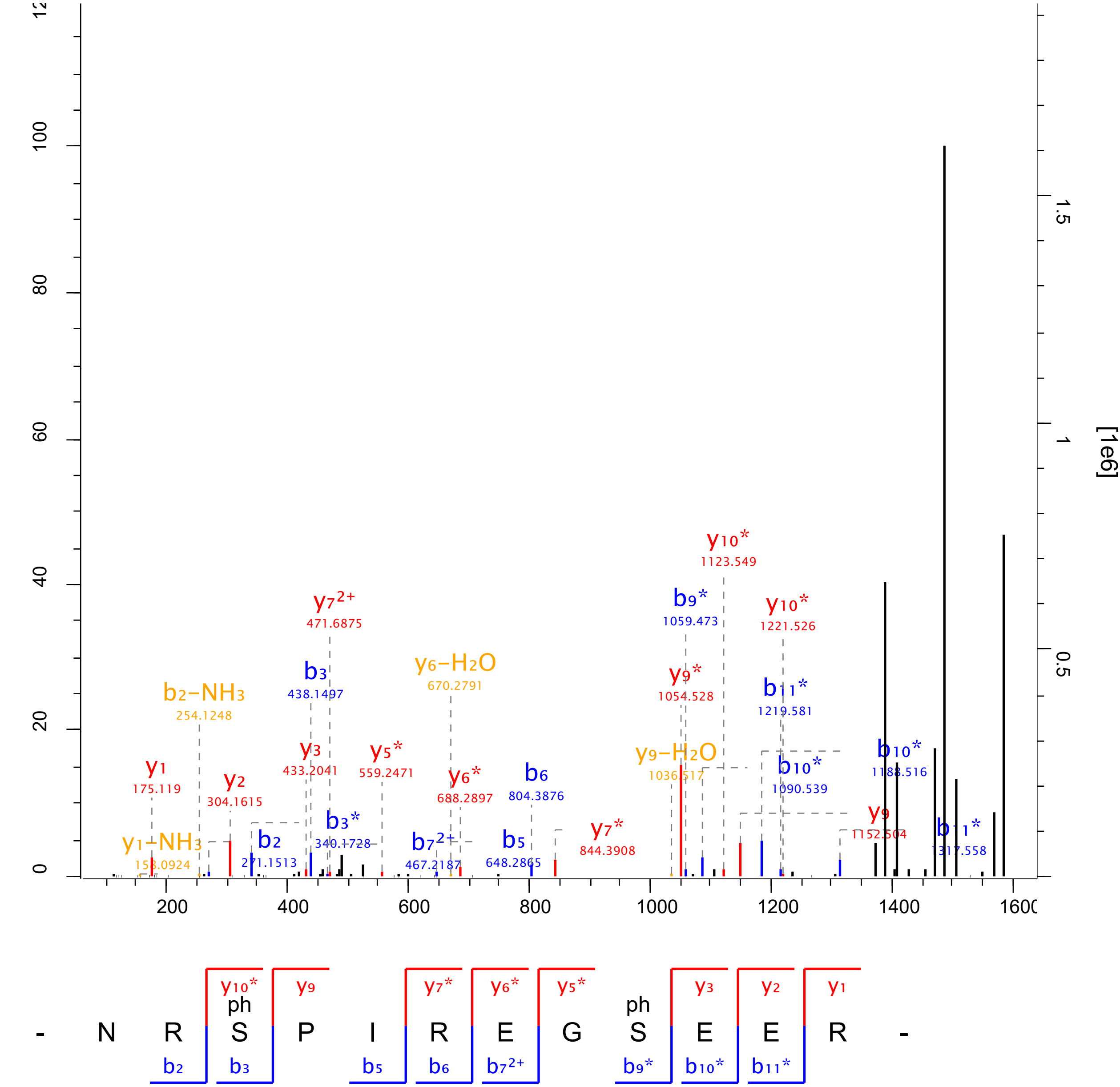

| Raw file | Scan | Method | Score | m/z |
|---|---|---|---|---|
| A17_00_1 | 26867 | FTMS; HCD | 51.29 | 684.38 |

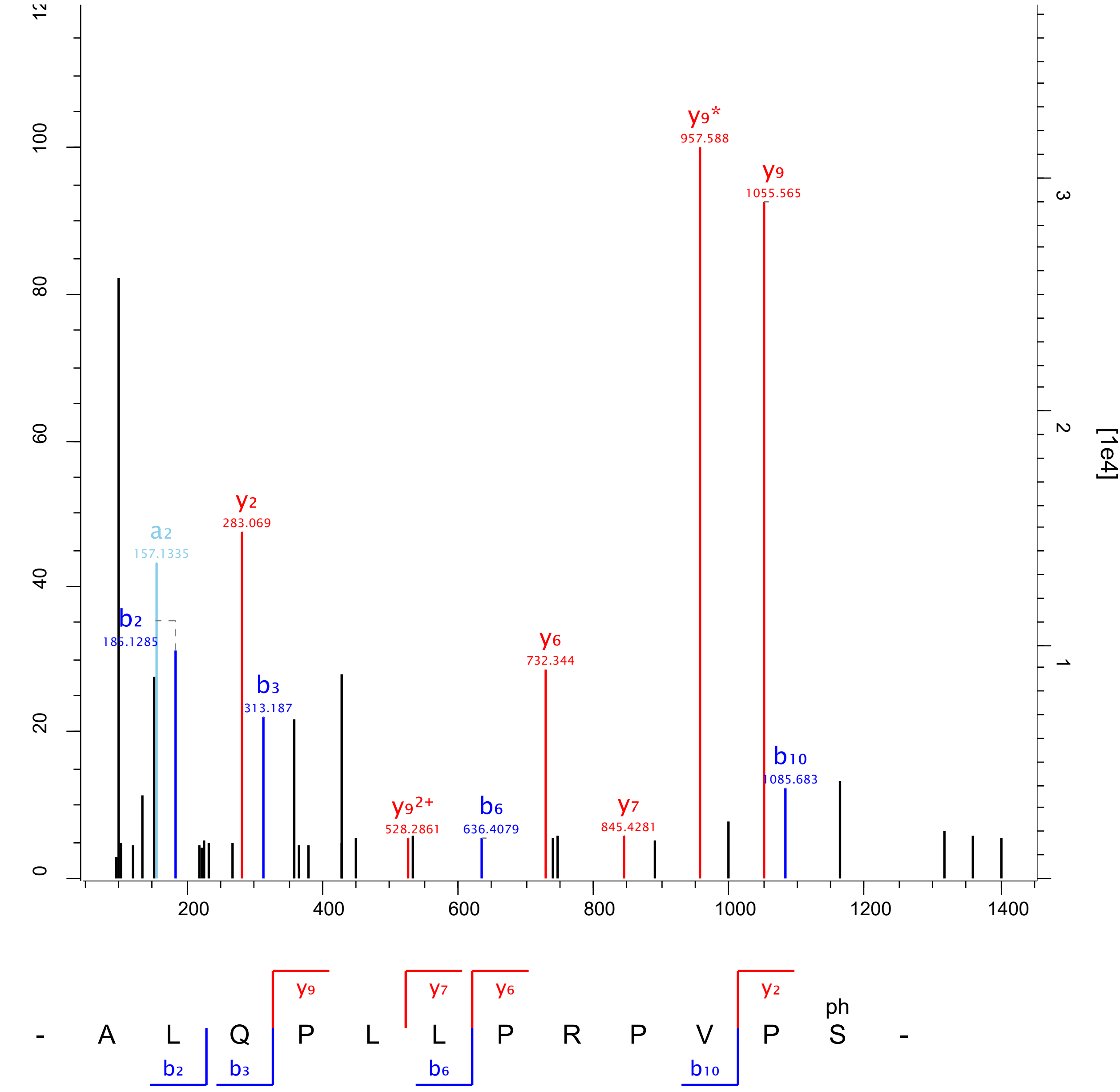

| Raw file | Scan | Method | Score | m/z |
|---|---|---|---|---|
| A17_00_1 | 33951 | FTMS; HCD | 50.78 | 773.7 |

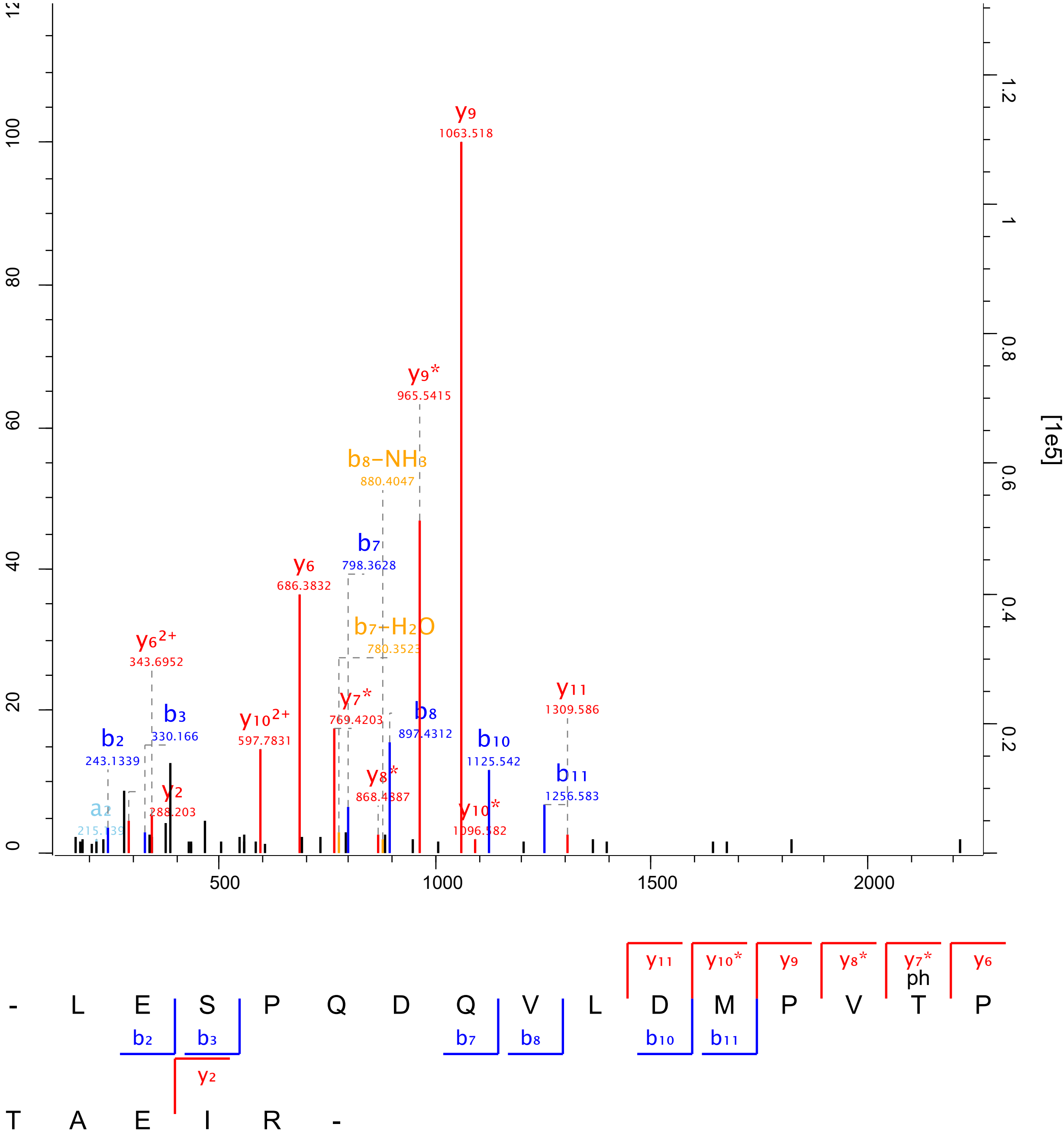

| Raw file | Scan | Method | Score | m/z |
|---|---|---|---|---|
| A17_00_1 | 38845 | FTMS; HCD | 74.03 | 922.92 |

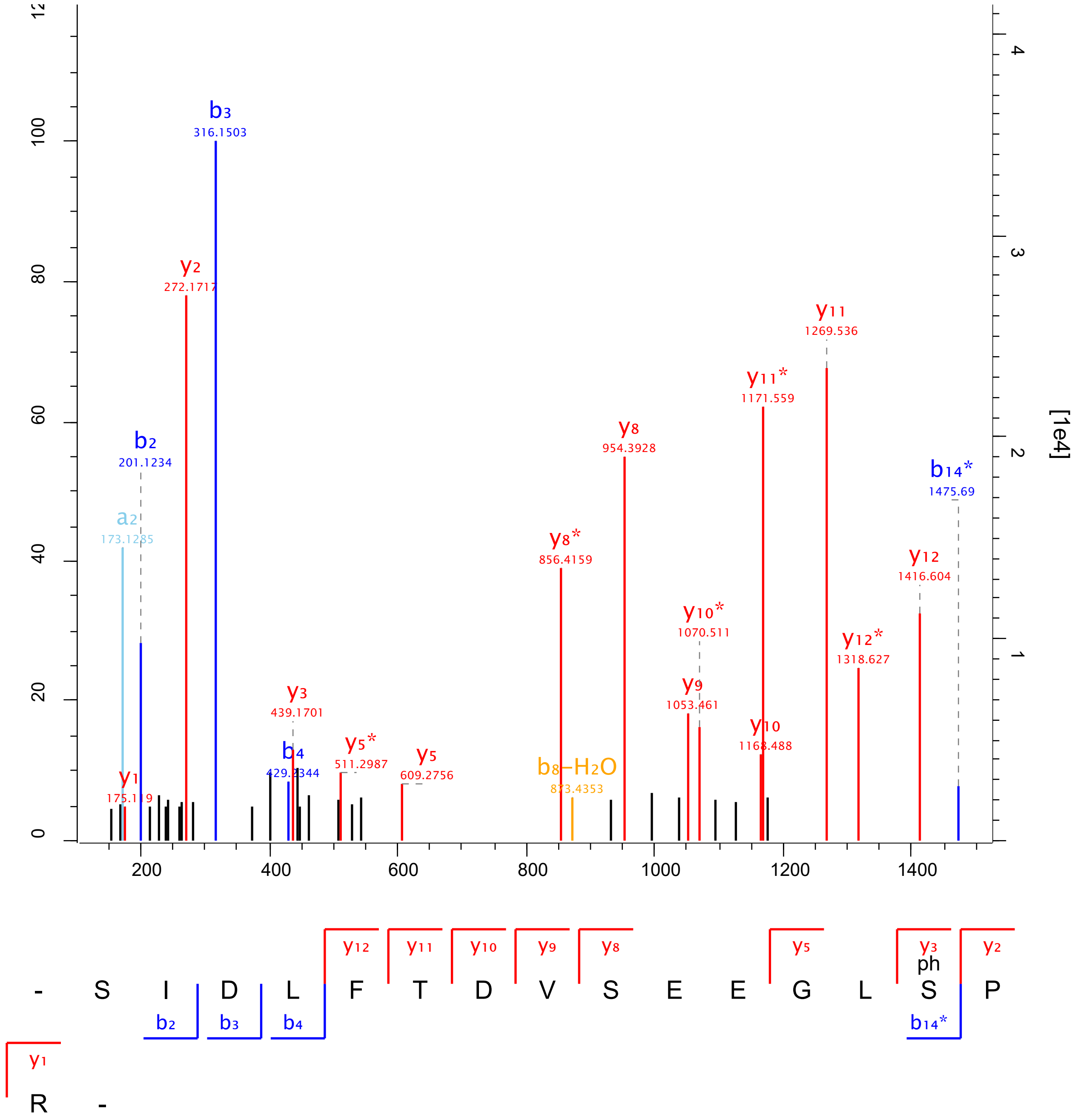

| Raw file | Scan | Method | Score | m/z |
| --- | --- | --- | --- | --- |
| A17_00_1 | 3964 | FTMS; HCD | 54.16 | 504.7 |

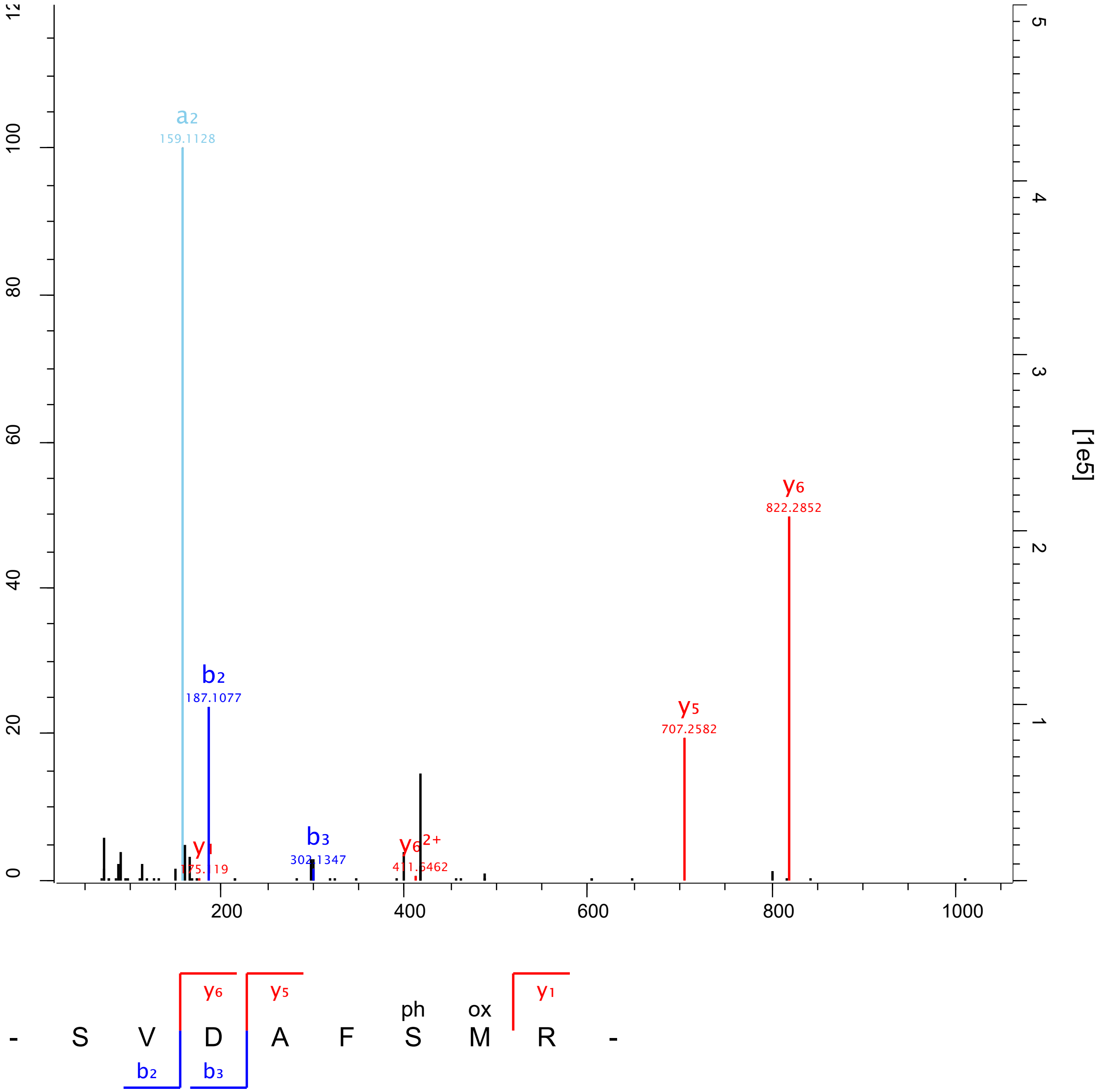

| Raw file | Scan | Method | Score | m/z |
|---|---|---|---|---|
| A17_00_1 | 43359 | FTMS; HCD | 80.73 | 881.7 |

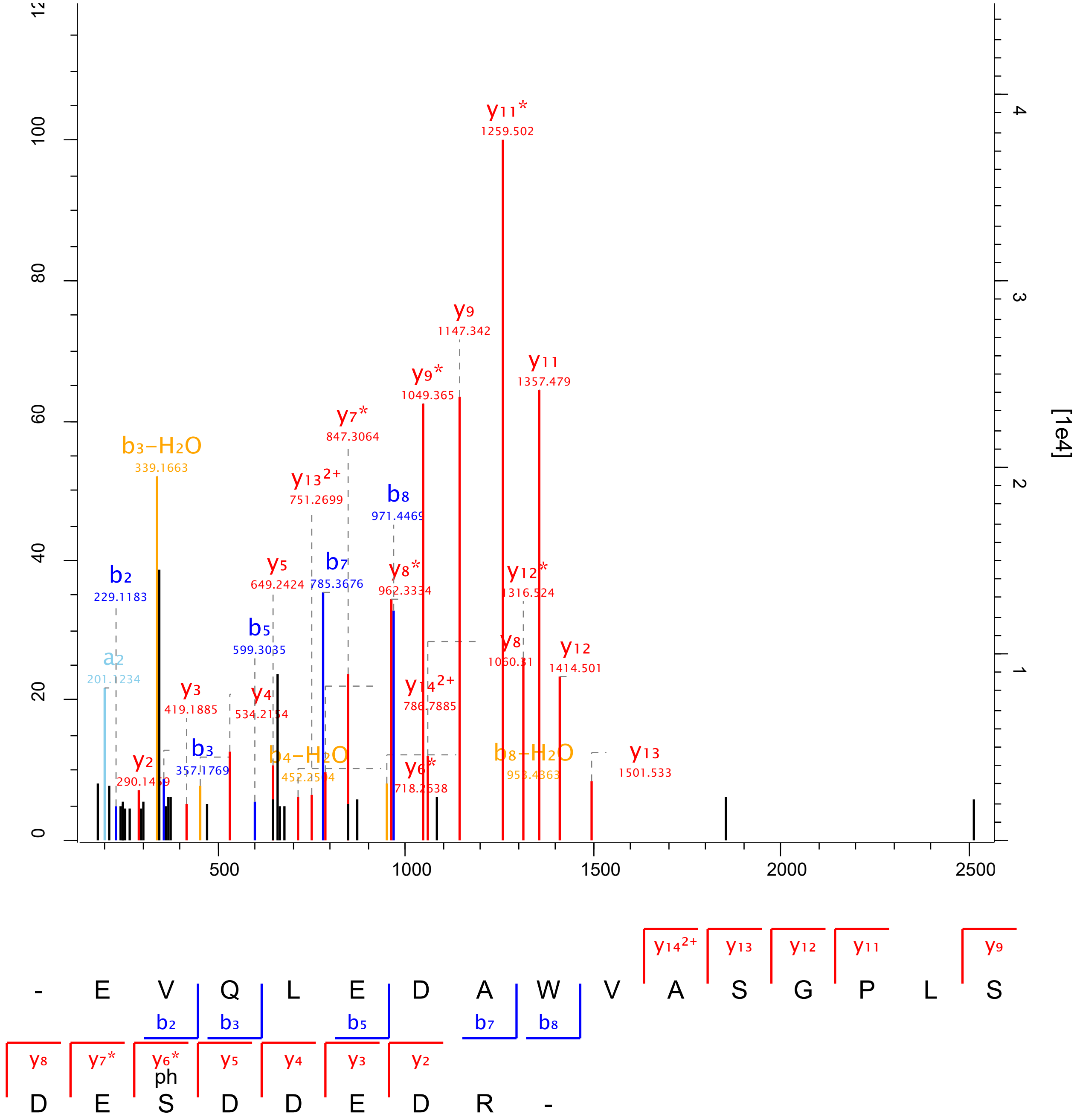

| Raw file | Scan | Method | Score | m/z |
|---|---|---|---|---|
| A17_00_1 | 6717 | FTMS; HCD | 43.12 | 503.75 |

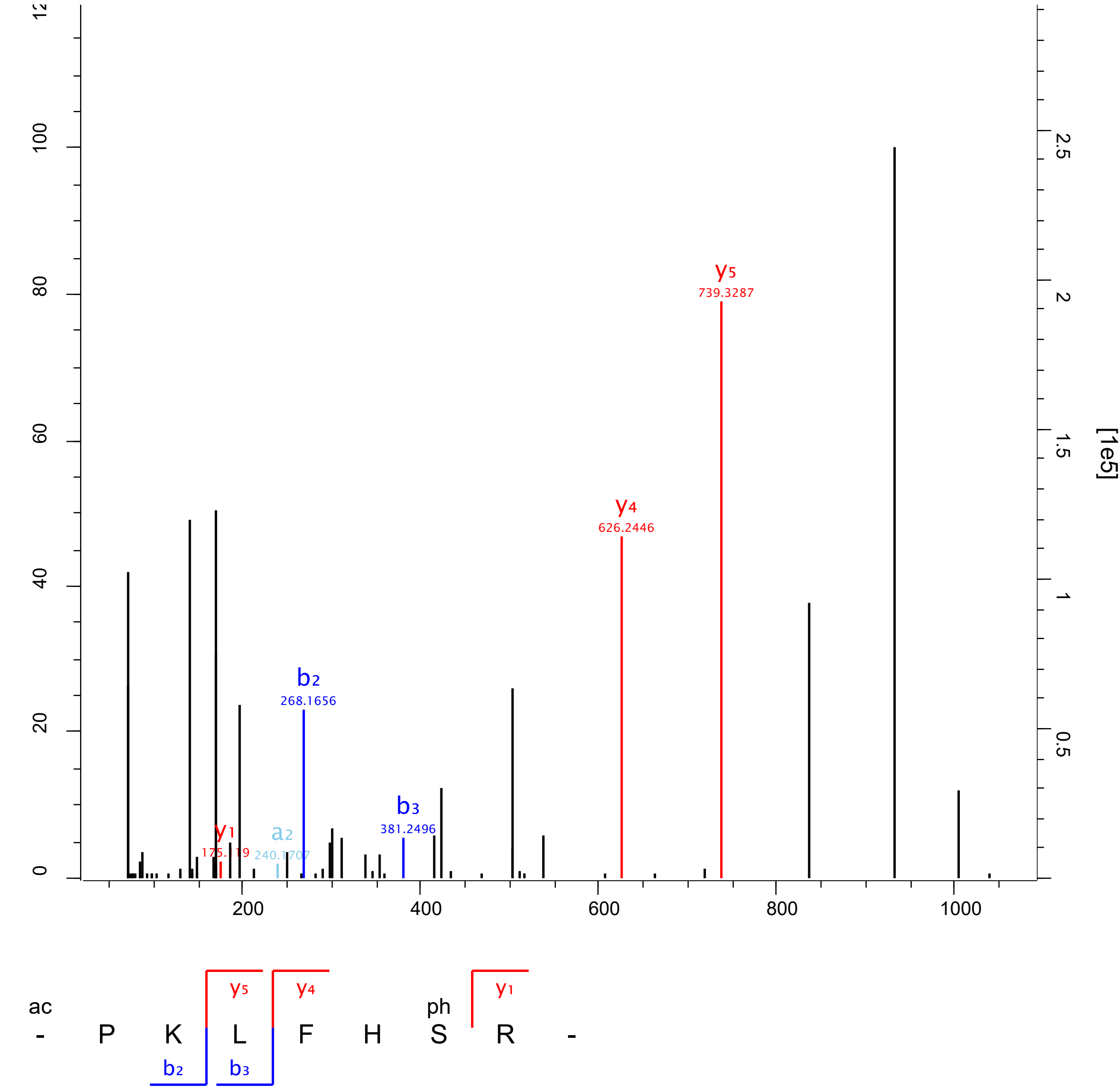

| Raw file | Scan | Method | Score | m/z |
|---|---|---|---|---|
| A17_00_2 | 10129 | FTMS; HCD | 58.98 | 663.8 |

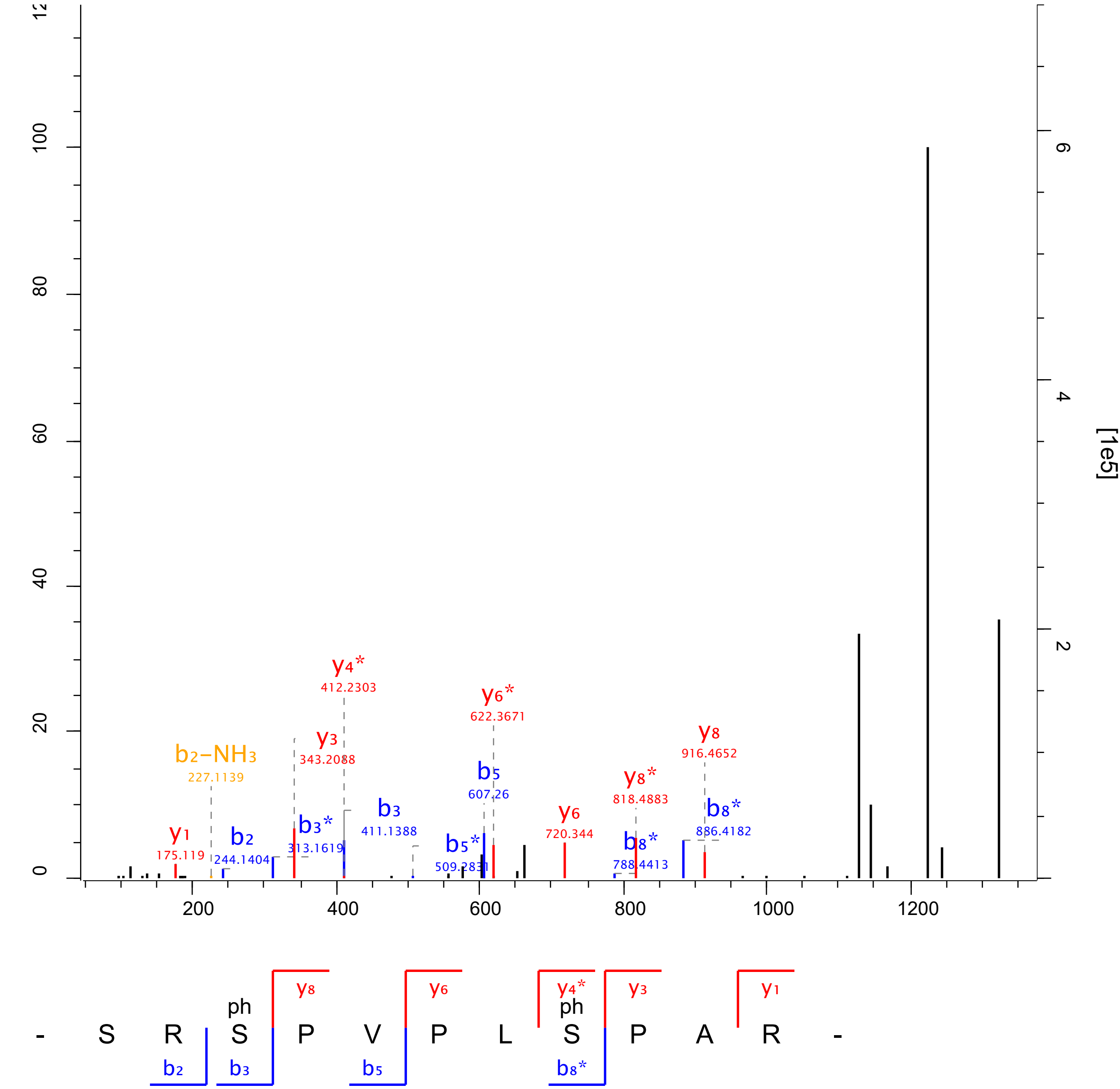

| Raw file | Scan | Method | Score | m/z |
|---|---|---|---|---|
| A17_00_2 | 10167 | FTMS; HCD | 88.19 | 679.8 |

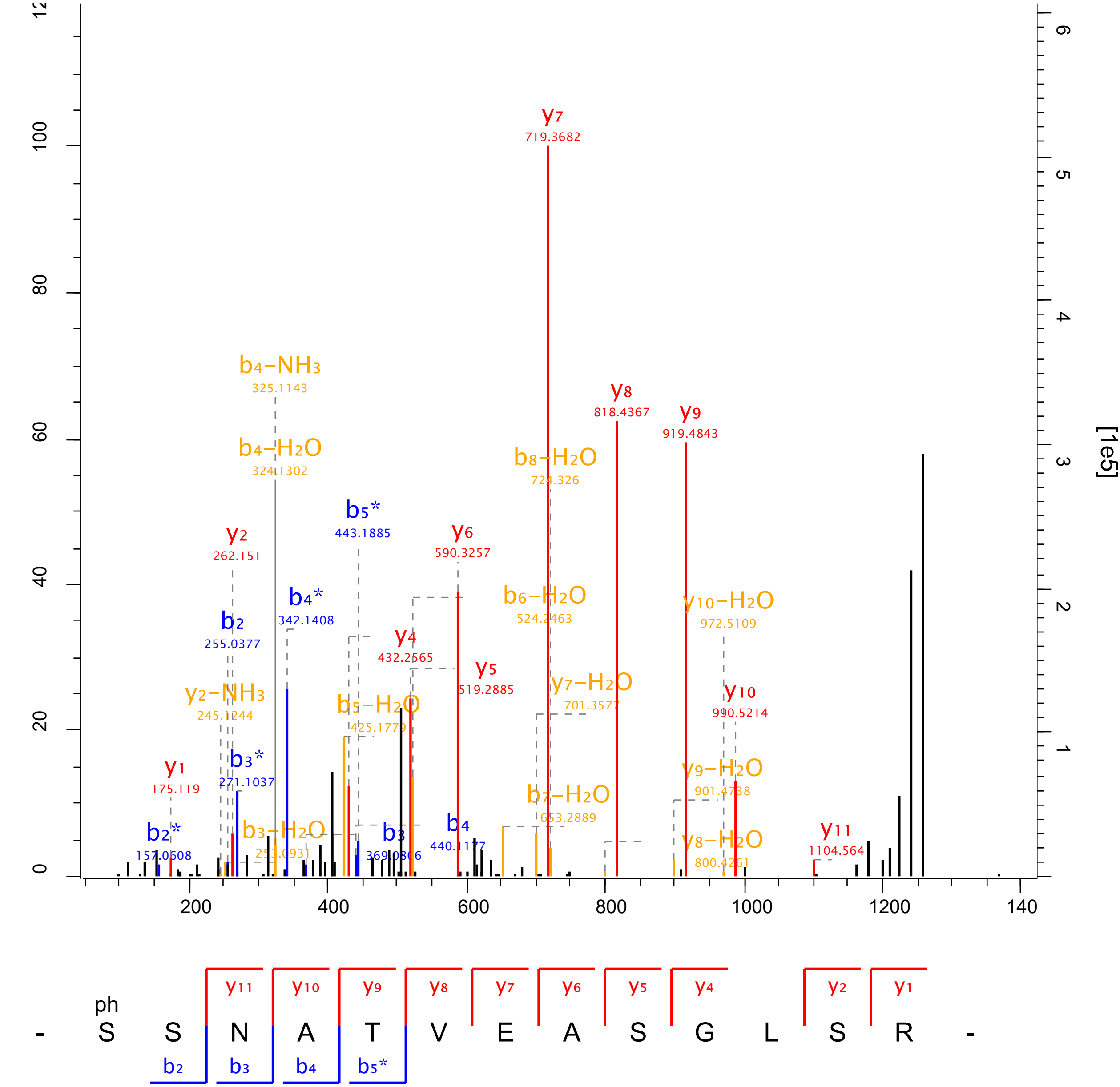

| Raw file | Scan | Method | Score | m/z |
| --- | --- | --- | --- | --- |
| A17_00_2 | 10259 | FTMS; HCD | 60.55 | 618.79 |

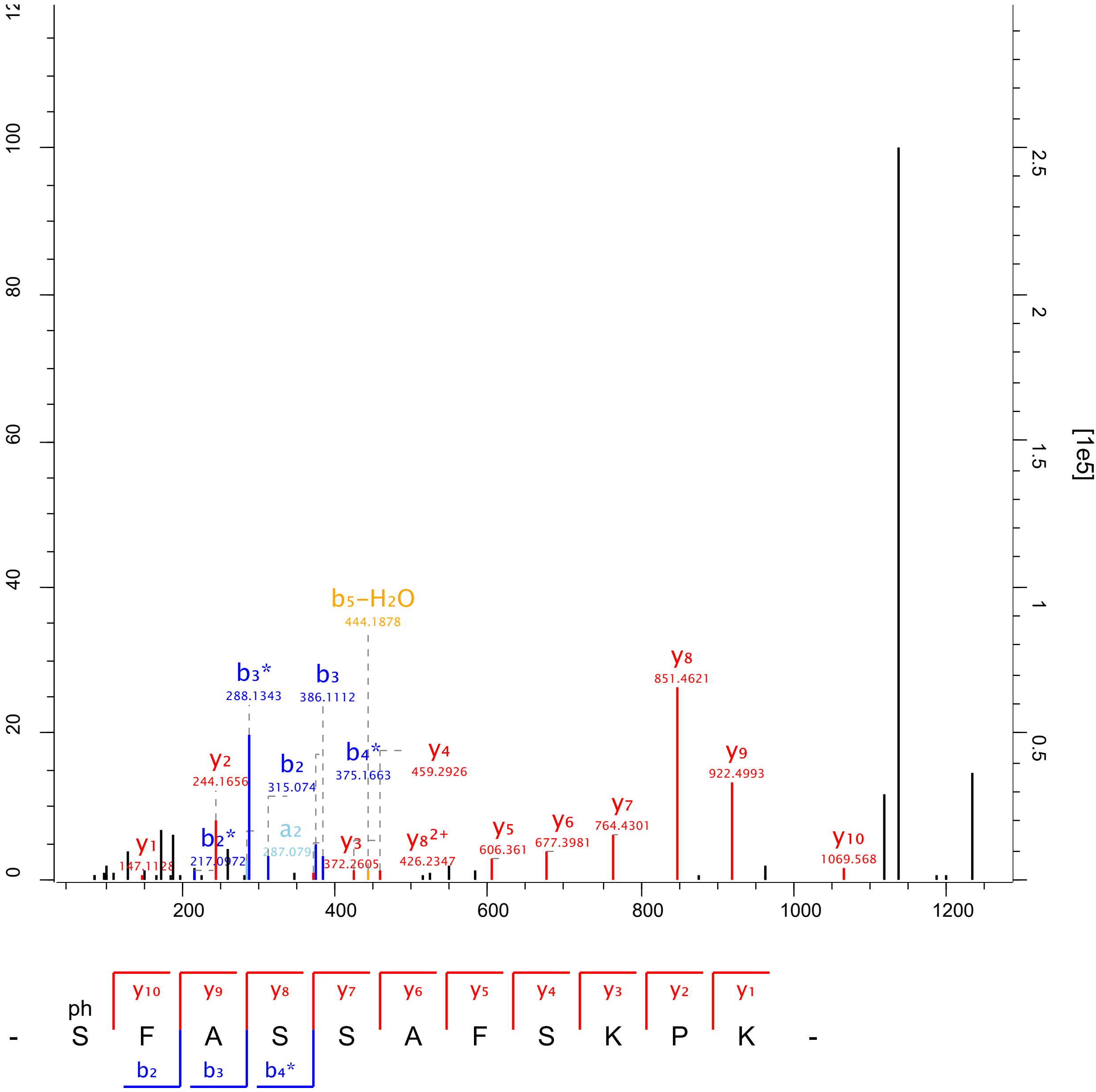

| Raw file | Scan | Method | Score | m/z |
|---|---|---|---|---|
| A17_00_2 | 10312 | FTMS; HCD | 190.96 | 684.84 |

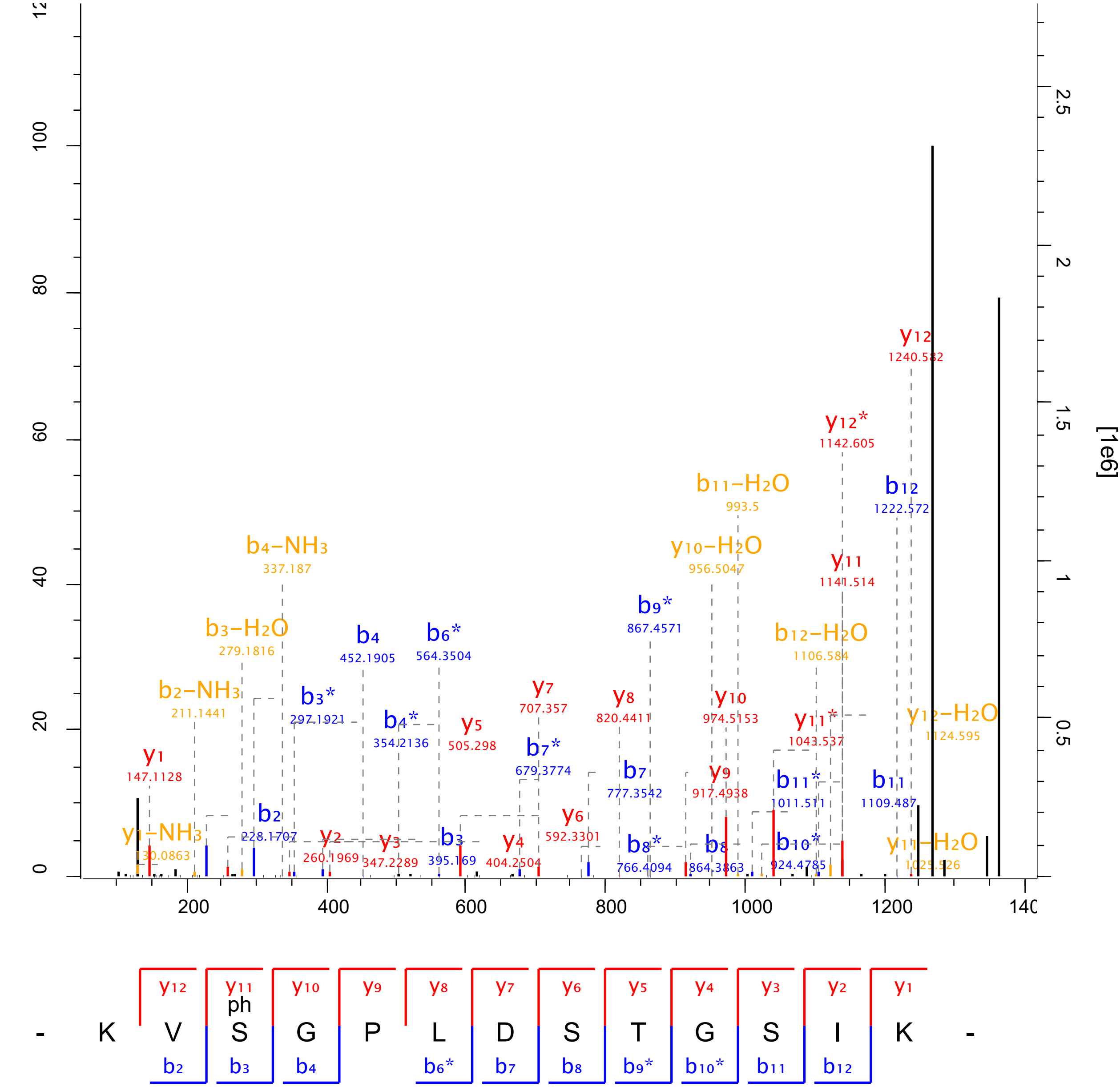

| Raw file | Scan | Method | Score | m/z |
|---|---|---|---|---|
| A17_00_2 | 10423 | FTMS; HCD | 52.25 | 602.75 |

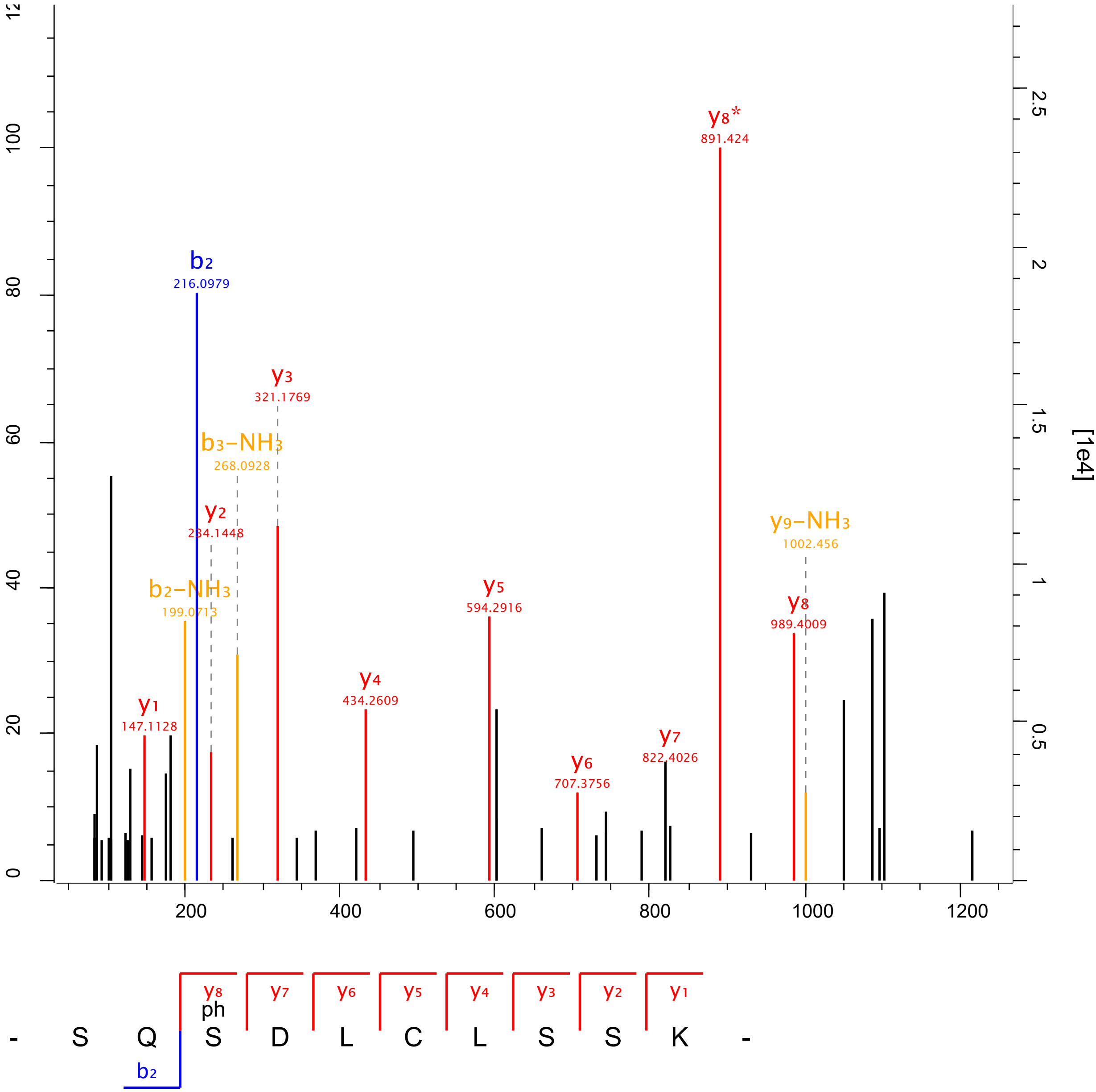

| Raw file | Scan | Method | Score | m/z |
| --- | --- | --- | --- | --- |
| A17_00_2 | 10432 | FTMS; HCD | 102.33 | 569.72 |

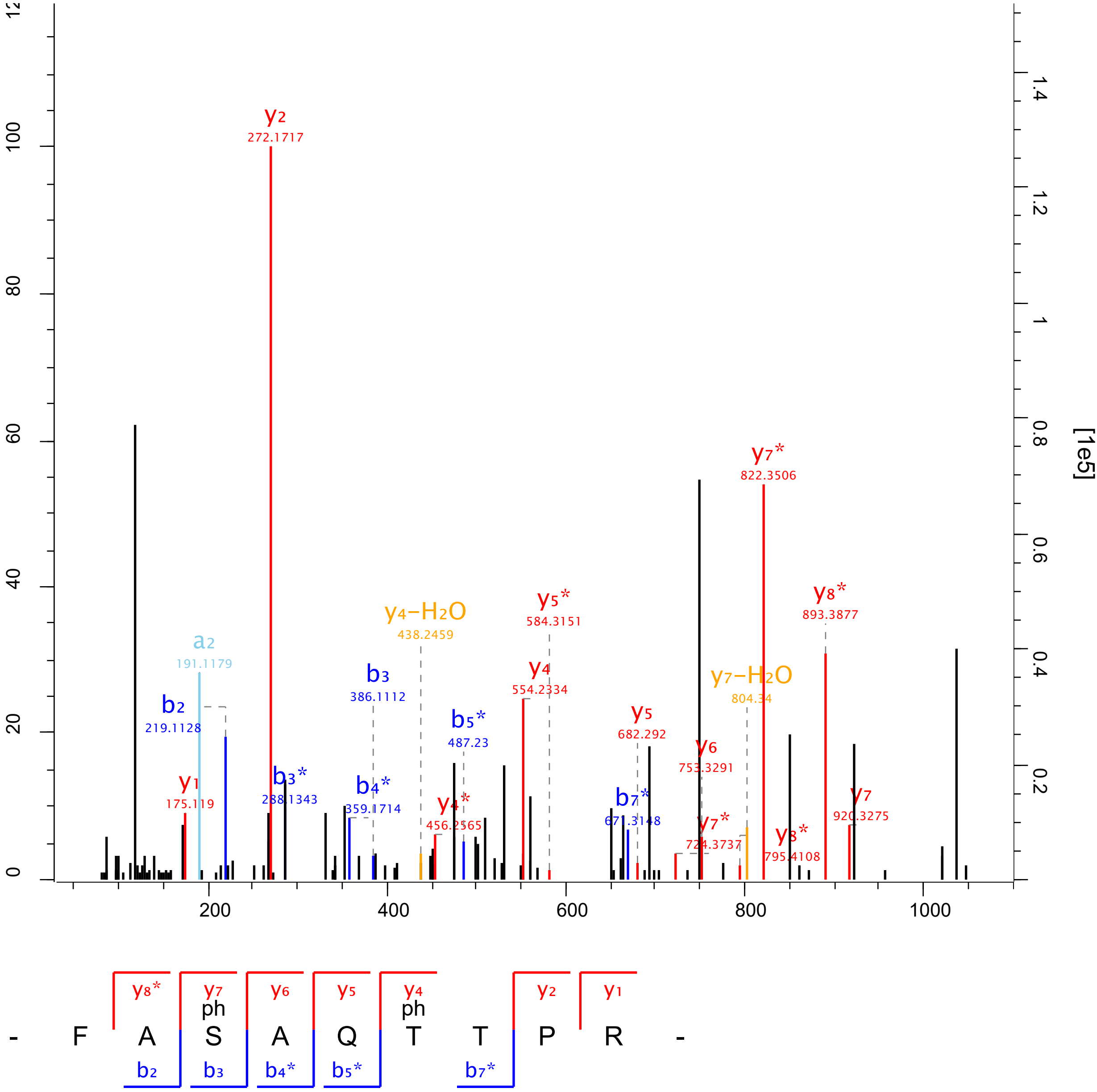

| Raw file | Scan | Method | Score | m/z |
|---|---|---|---|---|
| A17_00_2 | 10448 | FTMS; HCD | 45.61 | 671.78 |

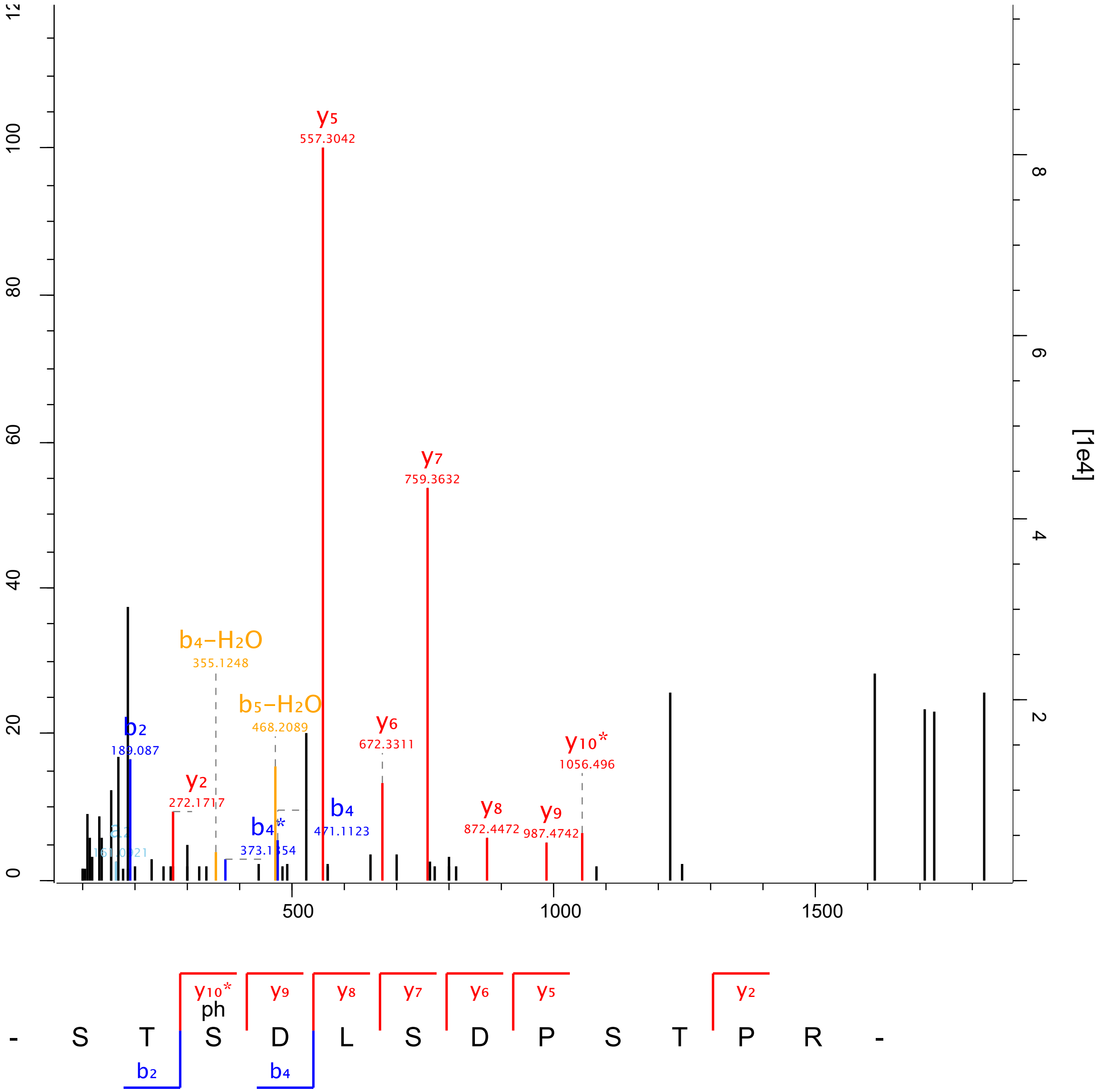

| Raw file | Scan | Method | Score | m/z |
|---|---|---|---|---|
| A17_00_2 | 10473 | FTMS; HCD | 80.32 | 744.31 |

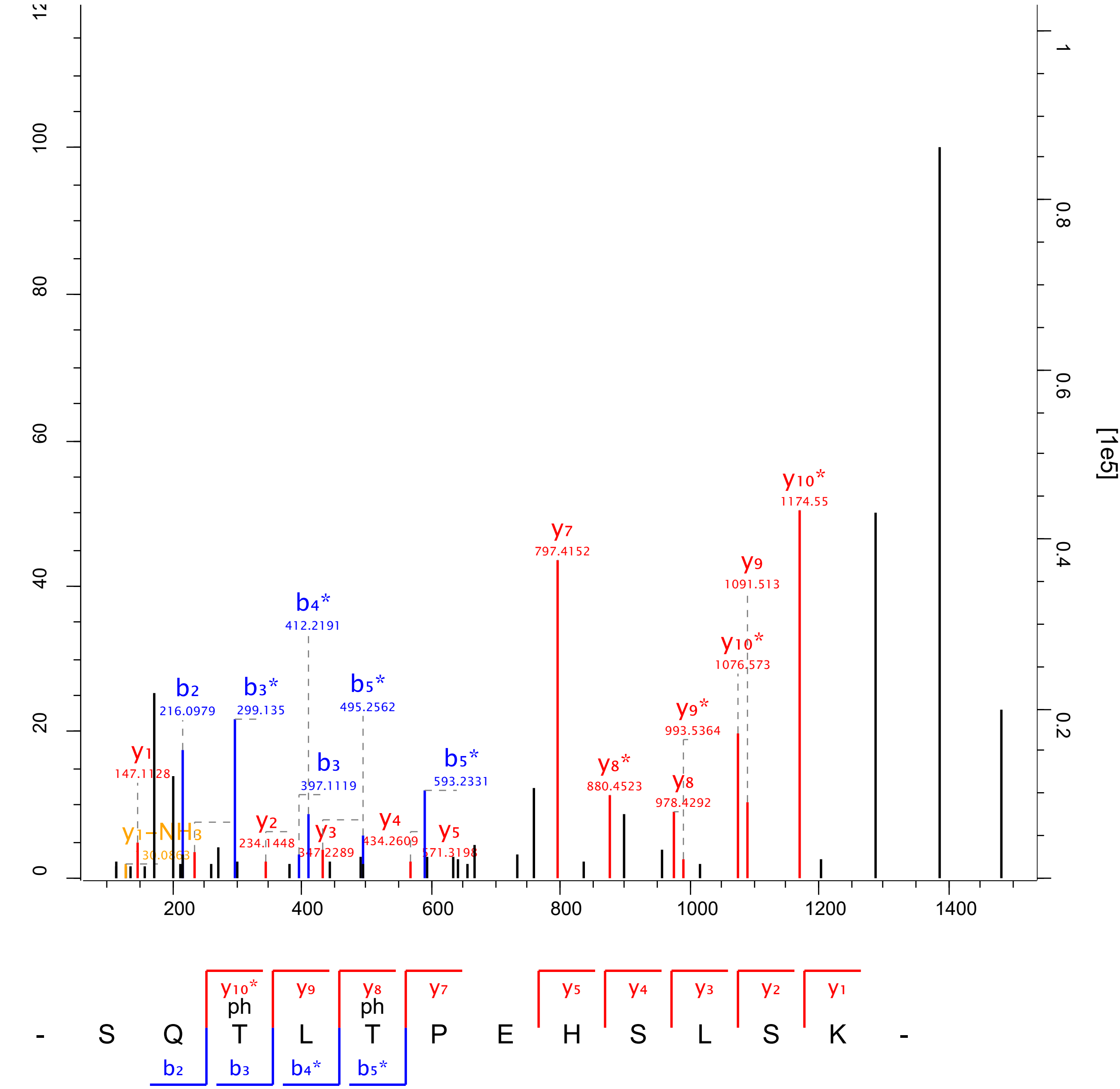

| Raw file | Scan | Method | Score | m/z |
|---|---|---|---|---|
| A17_00_2 | 10486 | FTMS; HCD | 55.04 | 564.24 |

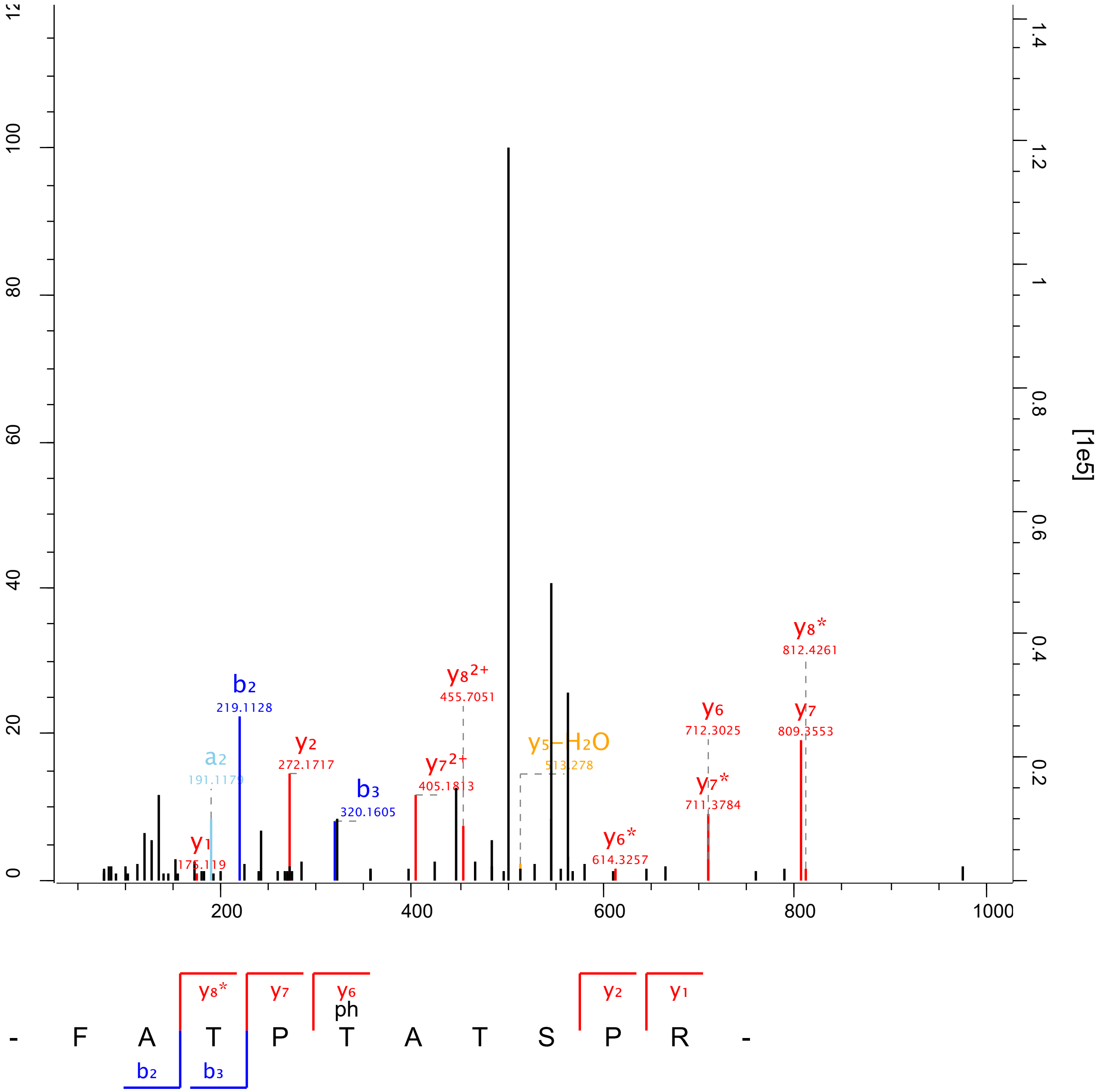

| Raw file | Scan | Method | Score | m/z |
|---|---|---|---|---|
| A17_00_2 | 10587 | FTMS; HCD | 47.33 | 751.97 |

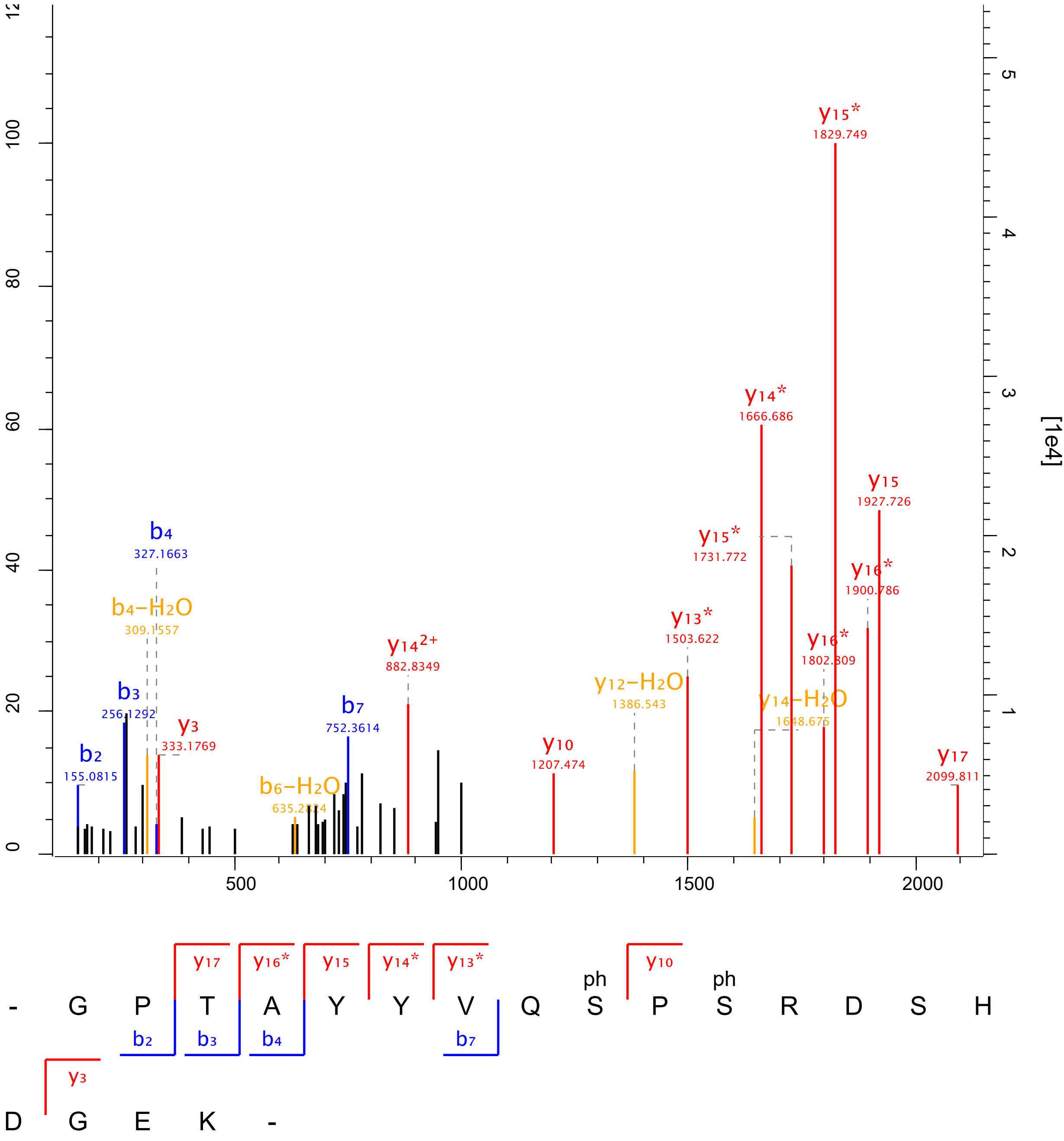

| Raw file | Scan | Method | Score | m/z |
|---|---|---|---|---|
| A17_00_2 | 10608 | FTMS; HCD | 41.7 | 729.76 |

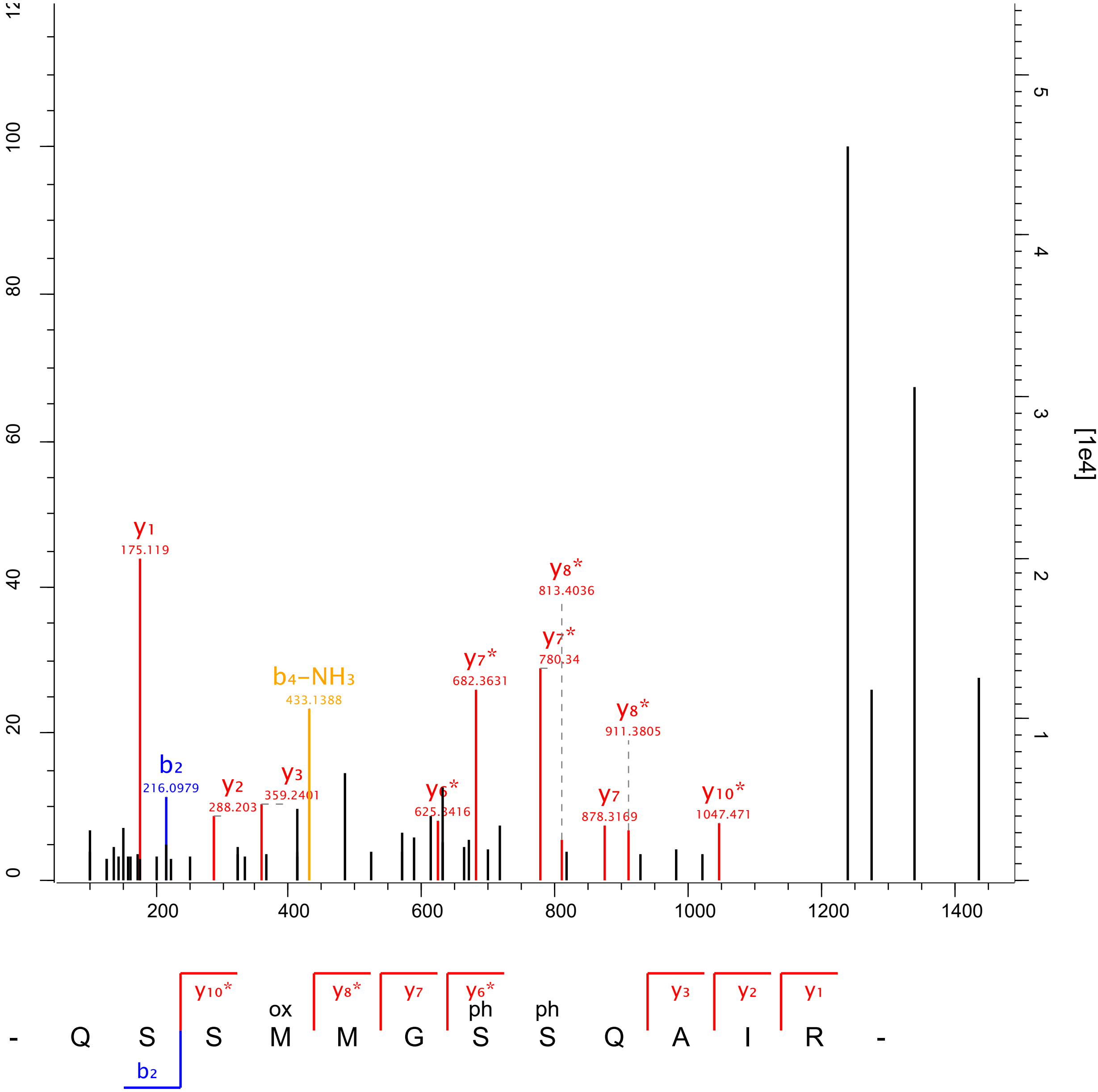

| Raw file | Scan | Method | Score | m/z |
|---|---|---|---|---|
| A17_00_2 | 10612 | FTMS; HCD | 146.78 | 671.58 |

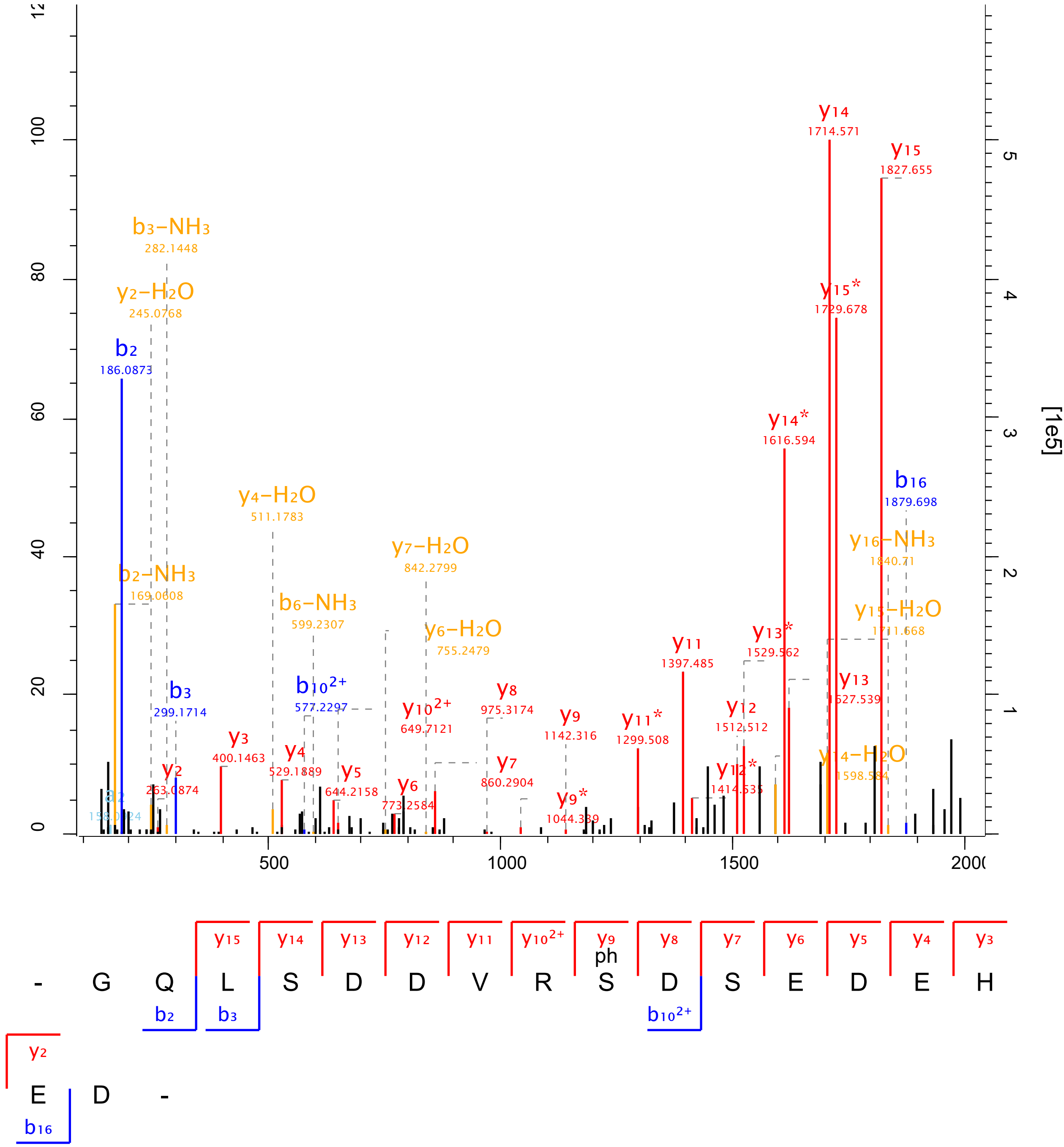

| Raw file | Scan | Method | Score | m/z |
| --- | --- | --- | --- | --- |
| A17_00_2 | 10631 | FTMS; HCD | 42.78 | 902.4 |

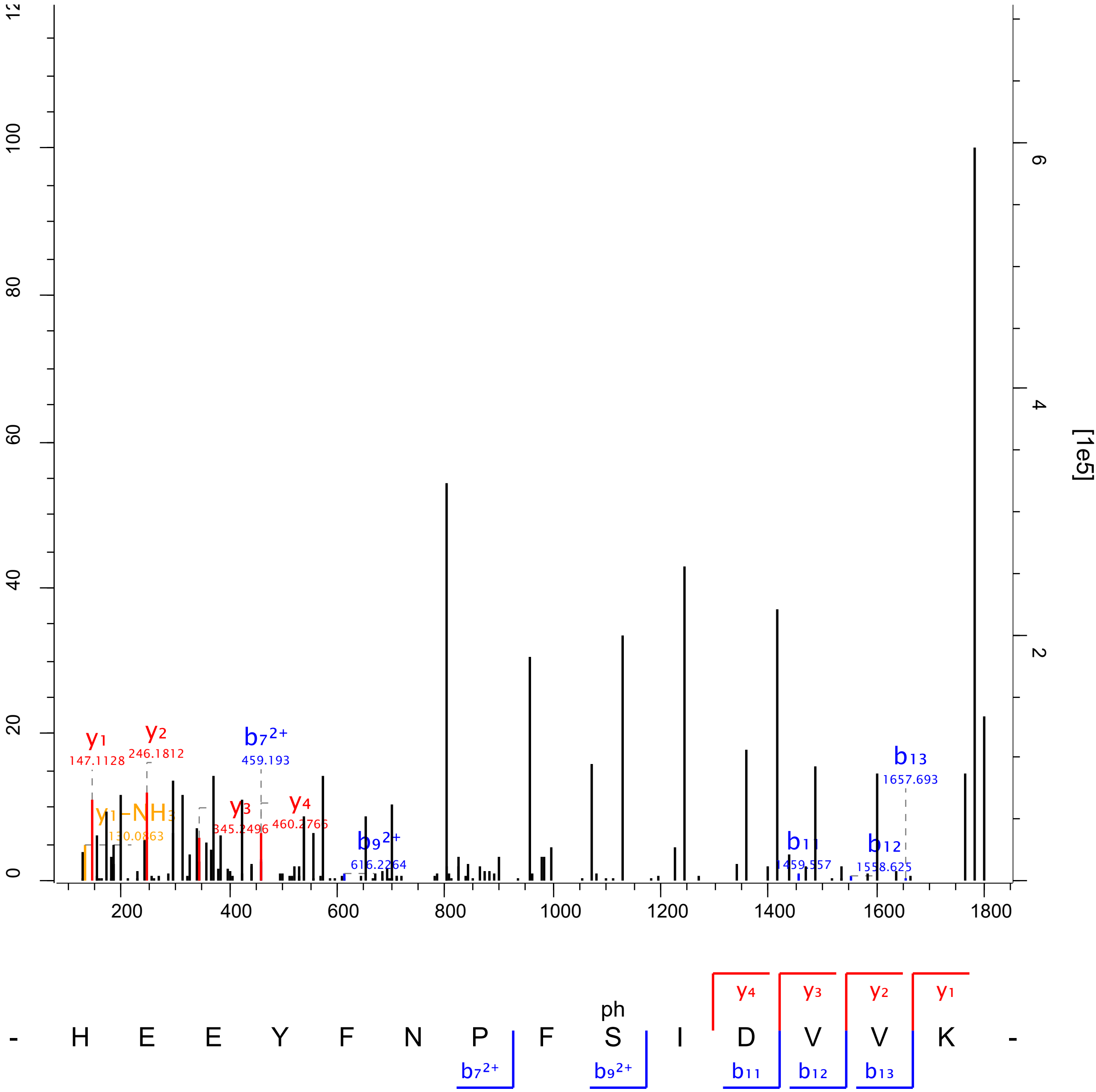

| Raw file | Scan | Method | Score | m/z |
|---|---|---|---|---|
| A17_00_2 | 10646 | FTMS; HCD | 117.52 | 503.73 |

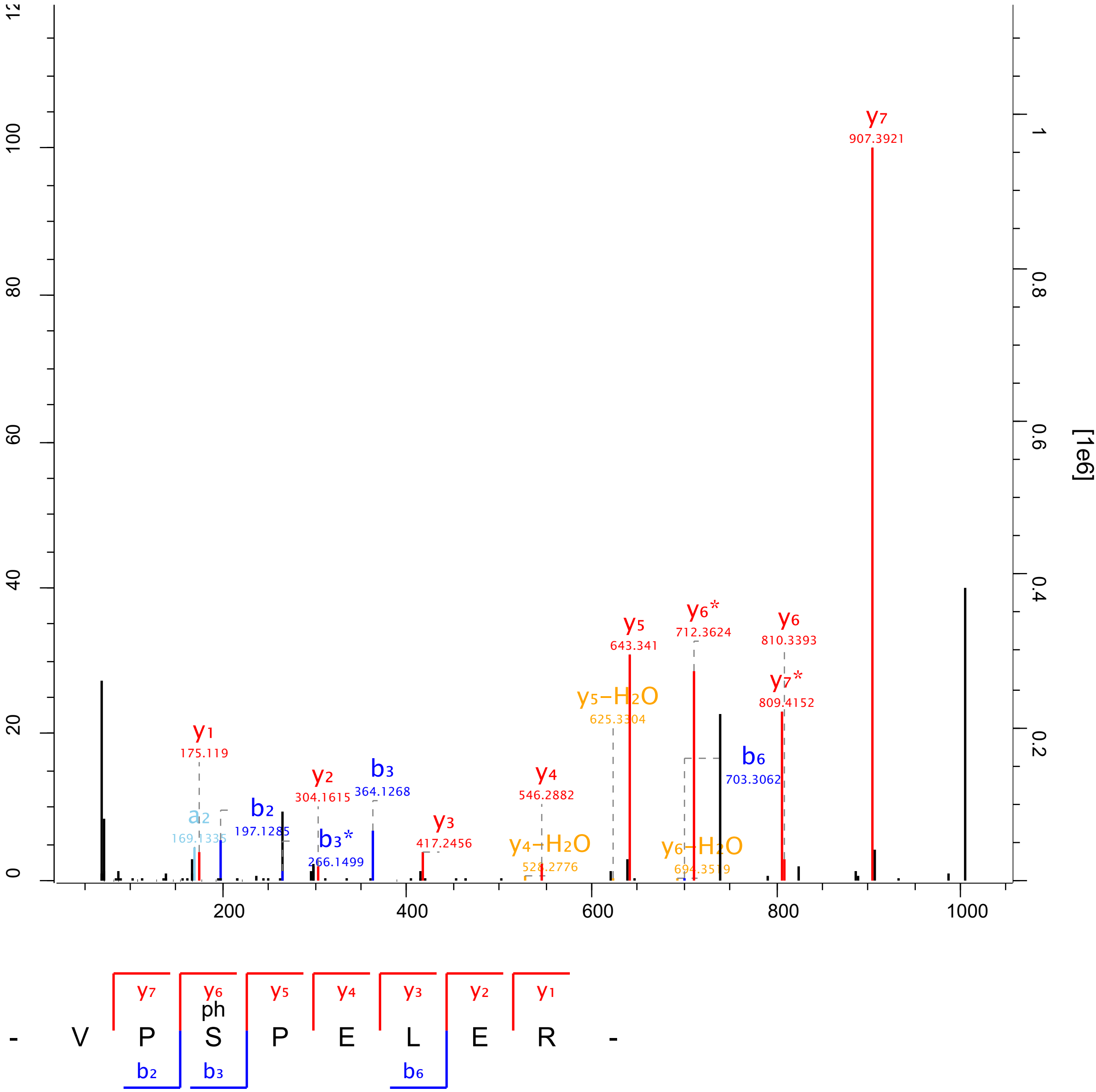

| Raw file | Scan | Method | Score | m/z |
|---|---|---|---|---|
| A17_00_2 | 10662 | FTMS; HCD | 50.09 | 555.28 |

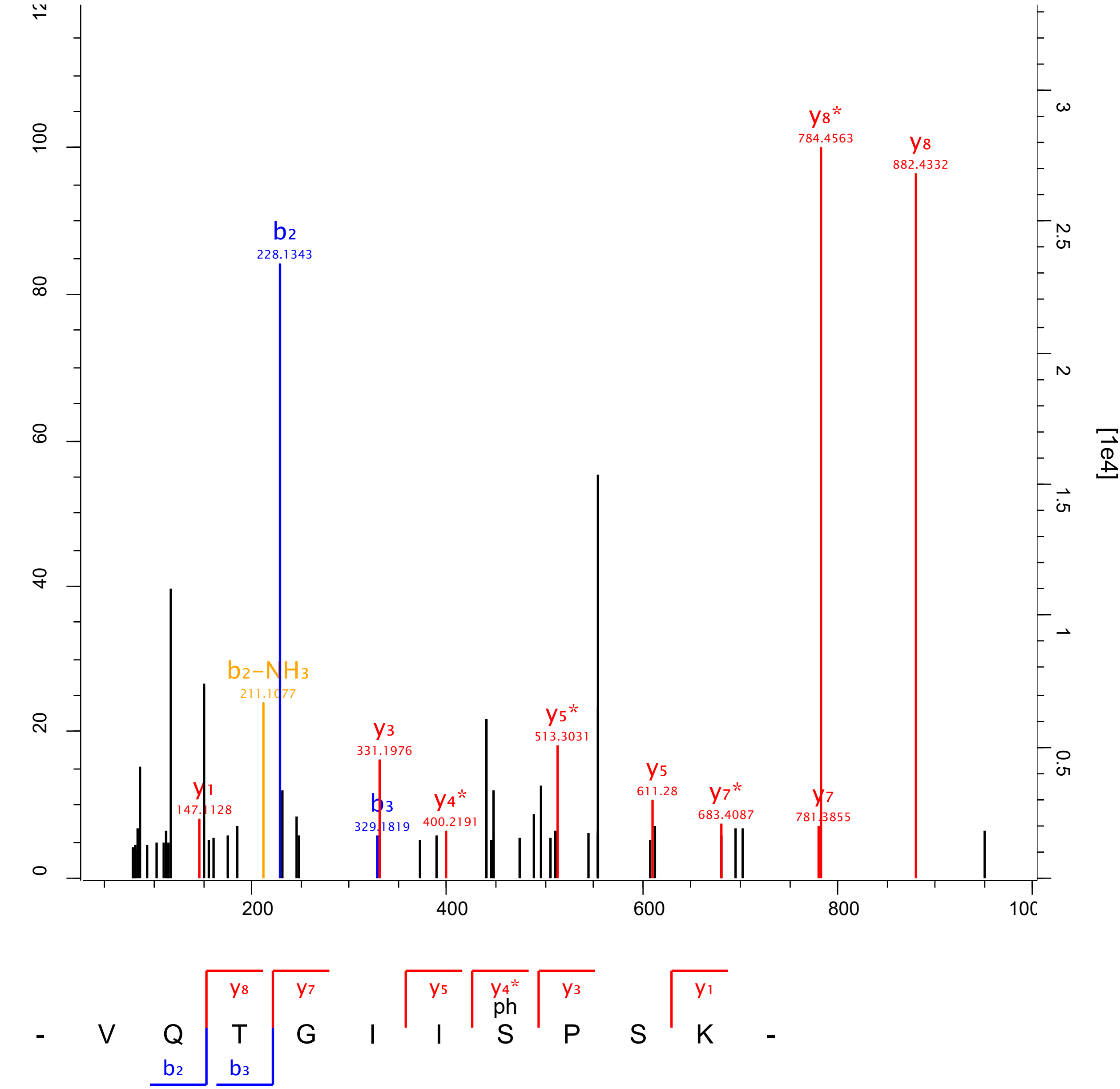

| Raw file | Scan | Method | Score | m/z |
|---|---|---|---|---|
| A17_00_2 | 10769 | FTMS; HCD | 73.5 | 460.75 |

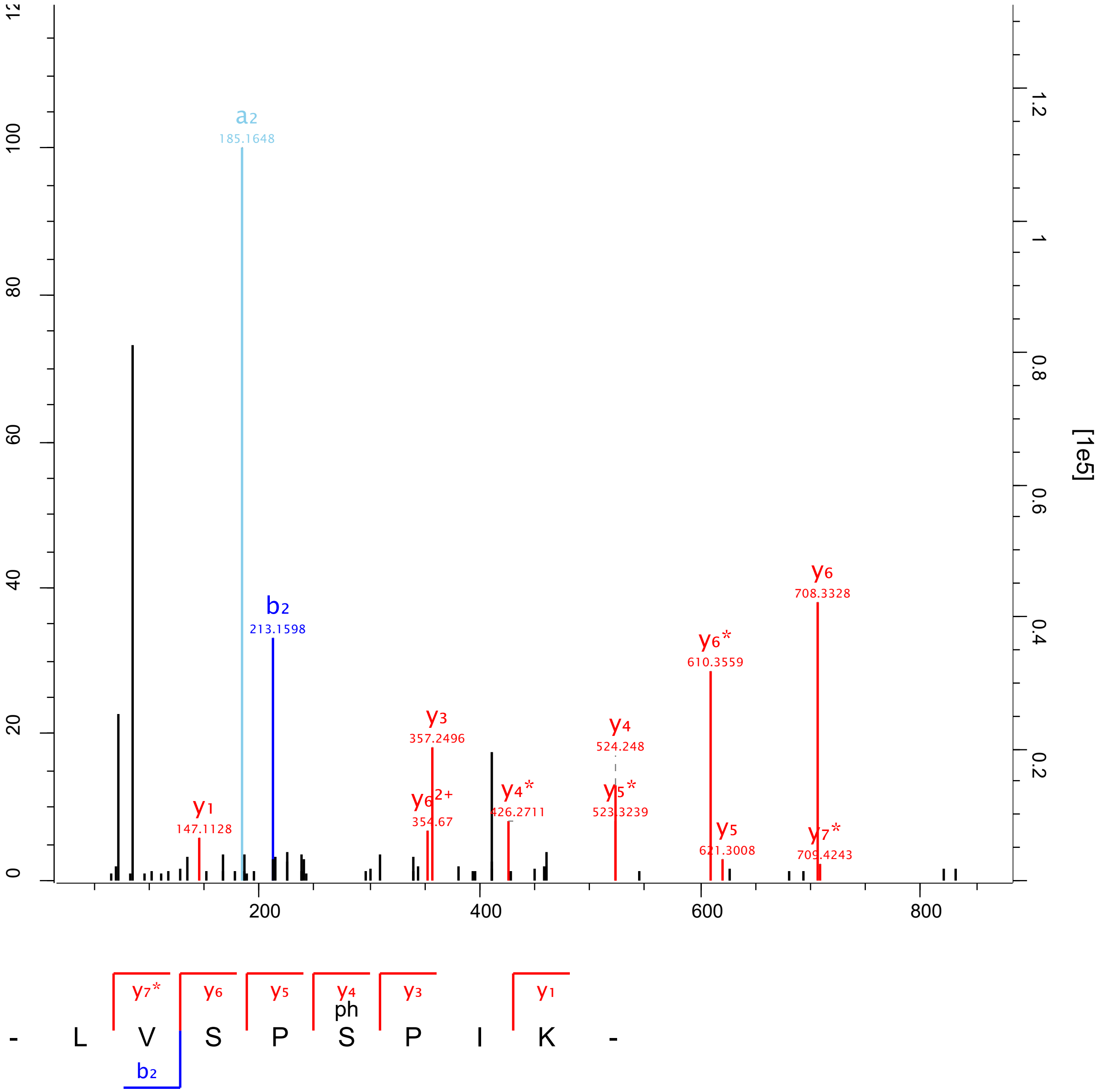

| Raw file | Scan | Method | Score | m/z |
|---|---|---|---|---|
| A17_00_2 | 10864 | FTMS; HCD | 149 | 562.6 |

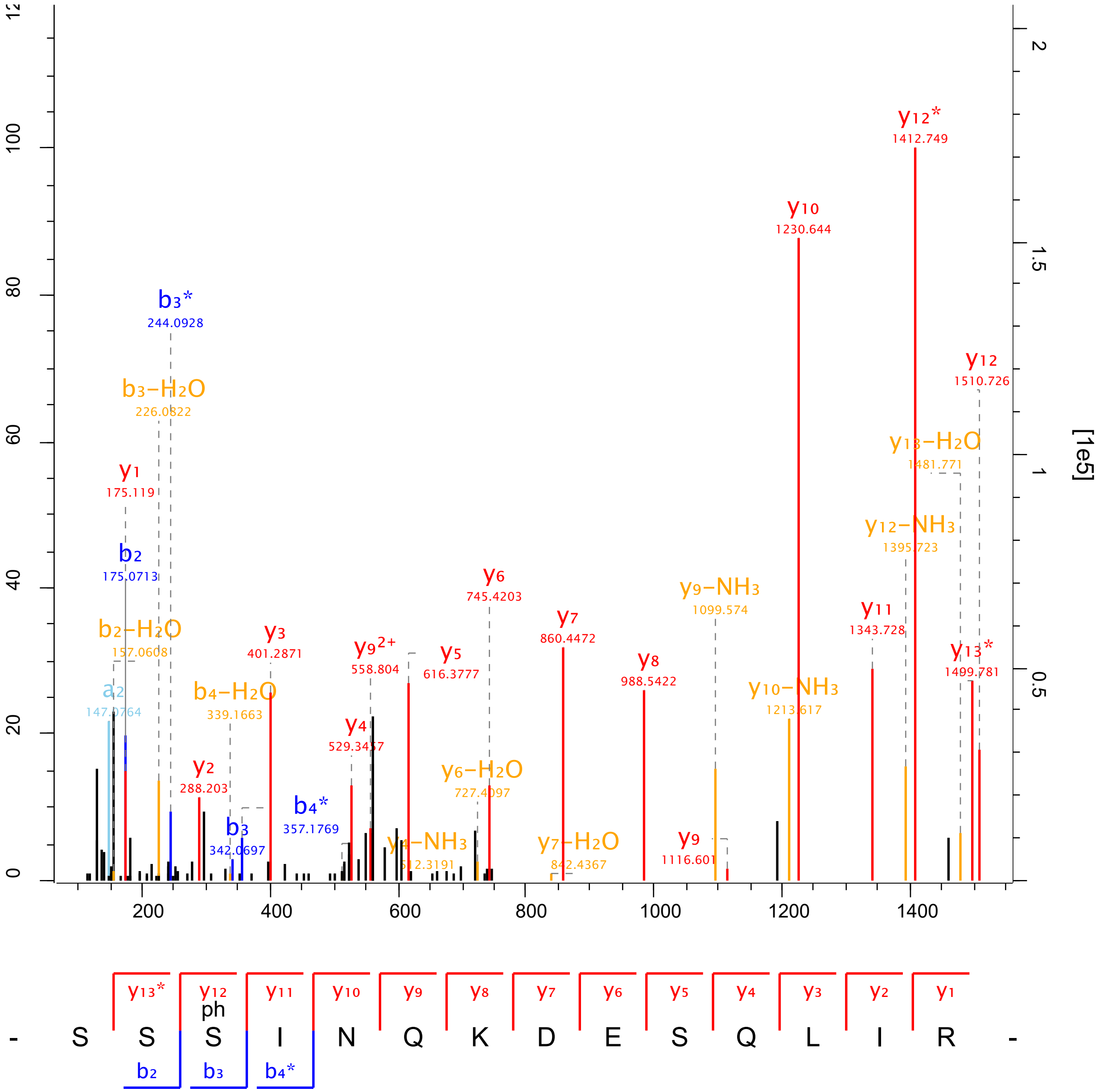

| Raw file | Scan | Method | Score | m/z |
|---|---|---|---|---|
| A17_00_2 | 10918 | FTMS; HCD | 184.98 | 585.73 |

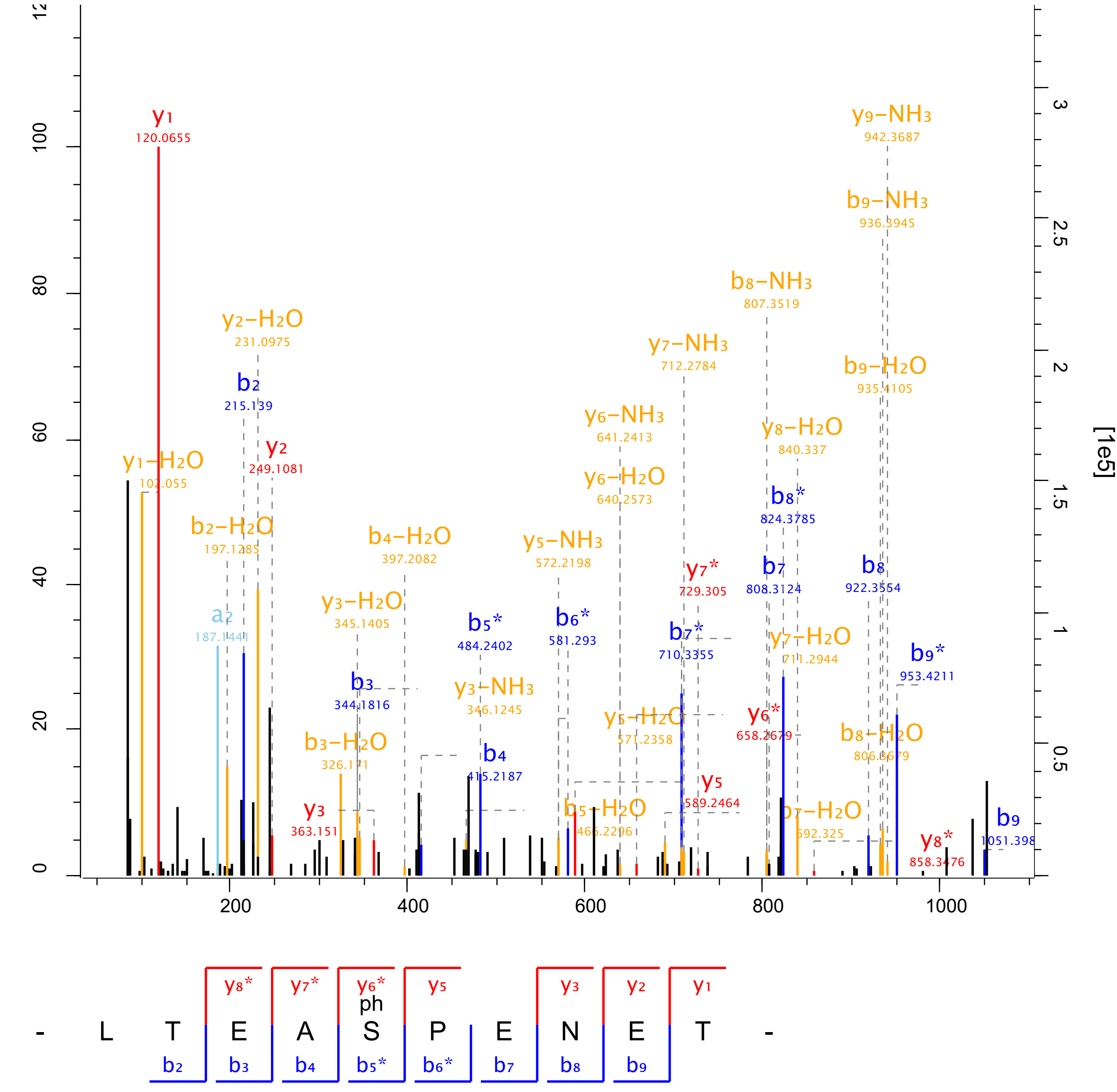

| Raw file | Scan | Method | Score | m/z |
|---|---|---|---|---|
| A17_00_2 | 10930 | FTMS; HCD | 206.67 | 600.28 |

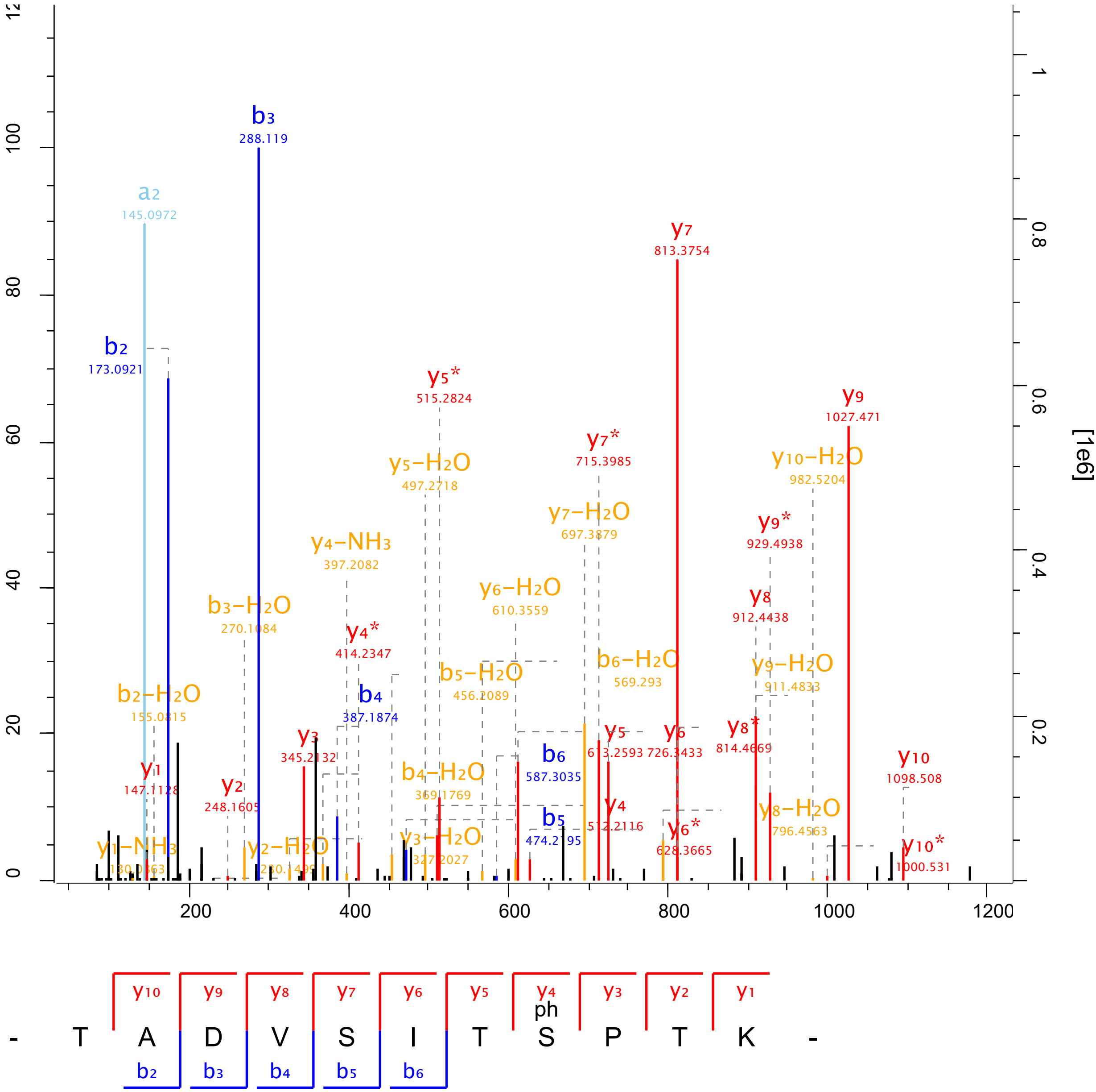

| Raw file | Scan | Method | Score | m/z |
|---|---|---|---|---|
| A17_00_2 | 10951 | FTMS; HCD | 90.71 | 667.76 |

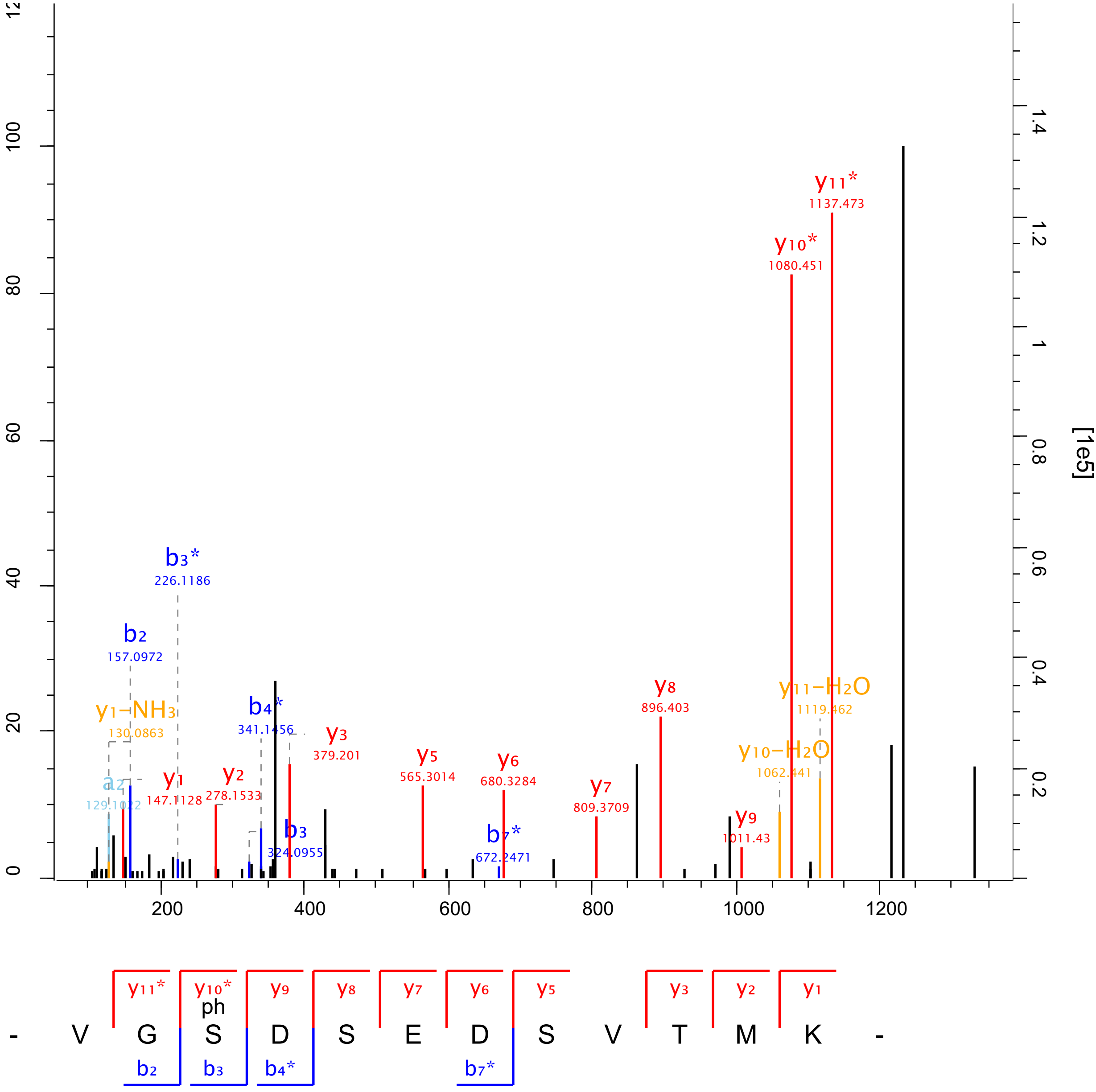

| Raw file | Scan | Method | Score | m/z |
|---|---|---|---|---|
| A17_00_2 | 10971 | FTMS; HCD | 62.09 | 572.25 |

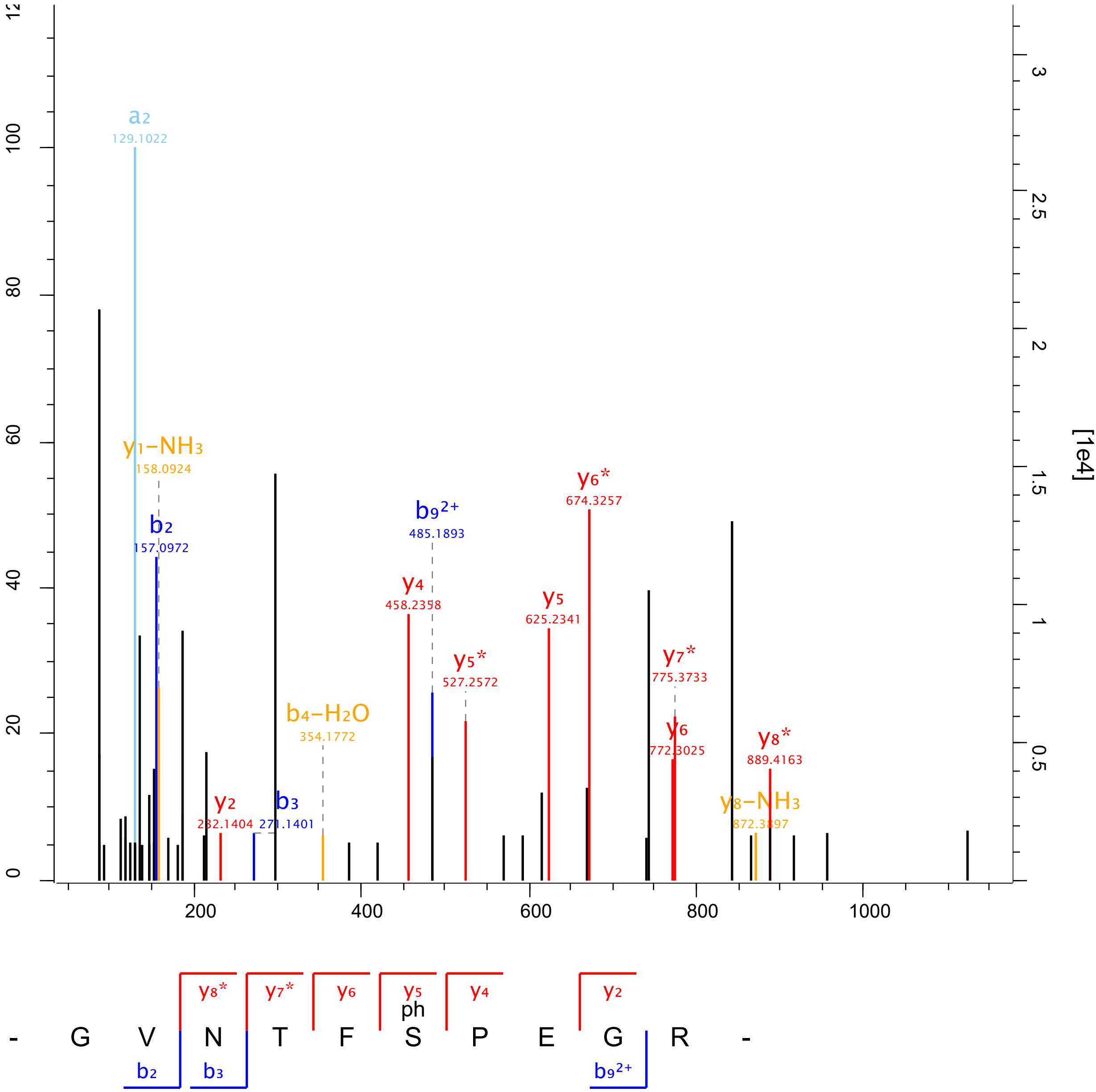

| Raw file | Scan | Method | Score | m/z |
|---|---|---|---|---|
| A17_00_2 | 11102 | FTMS; HCD | 190.19 | 681.78 |

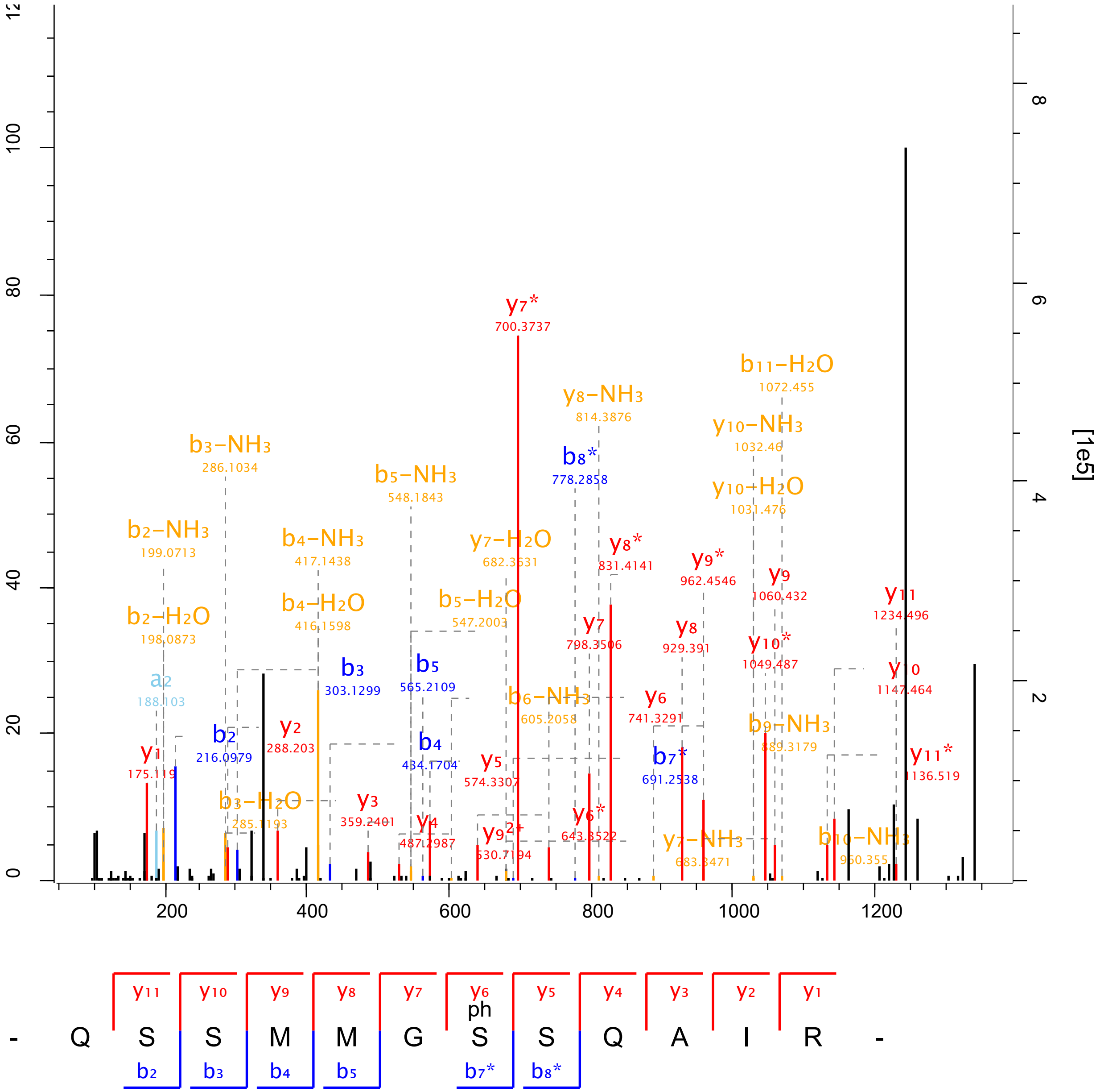

| Raw file | Scan | Method | Score | m/z |
| --- | --- | --- | --- | --- |
| A17_00_2 | 11387 | FTMS; HCD | 145.91 | 777.31 |

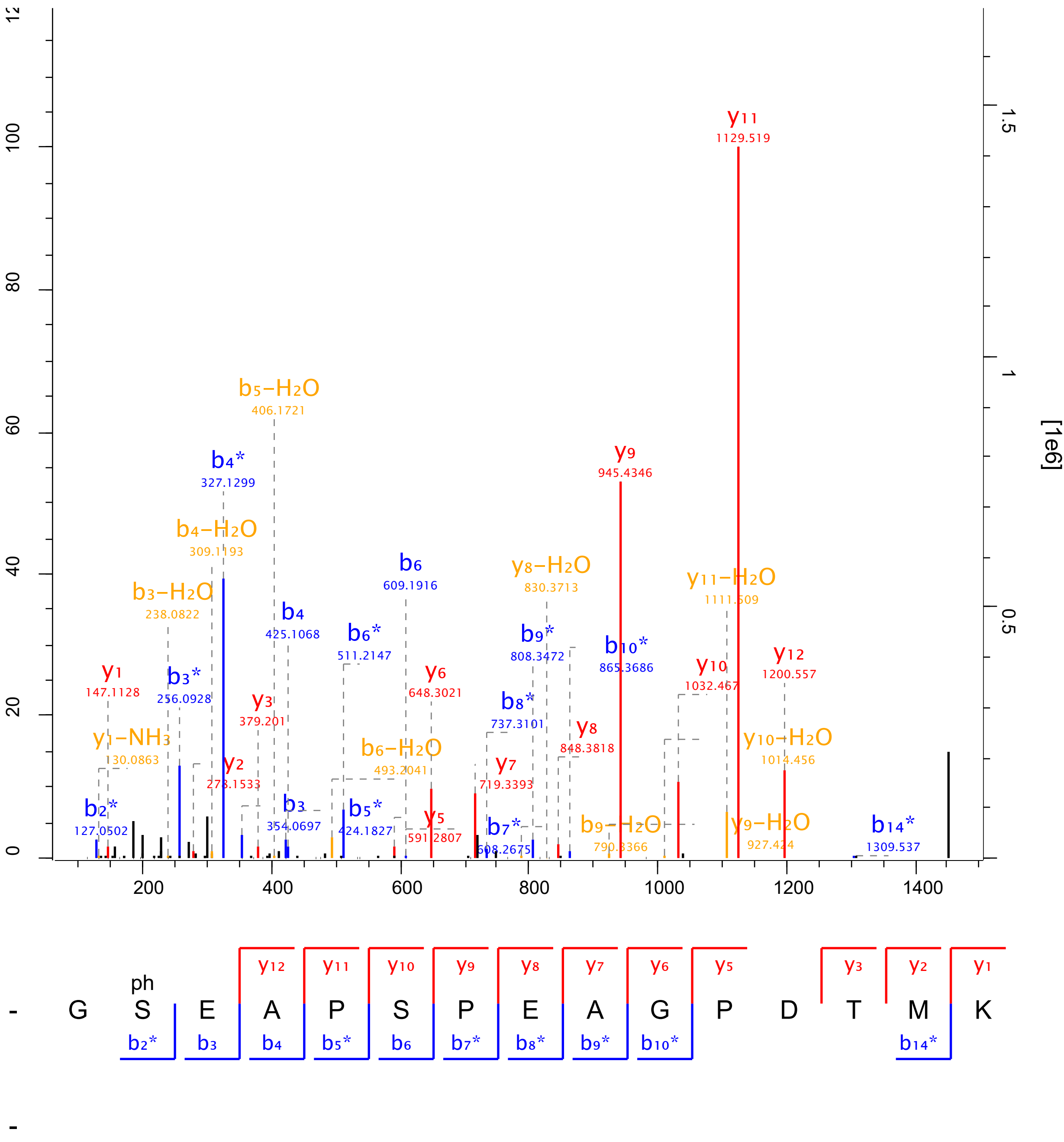

| Raw file | Scan | Method | Score | m/z |
| --- | --- | --- | --- | --- |
| A17_00_2 | 11488 | FTMS; HCD | 105.17 | 713.8 |

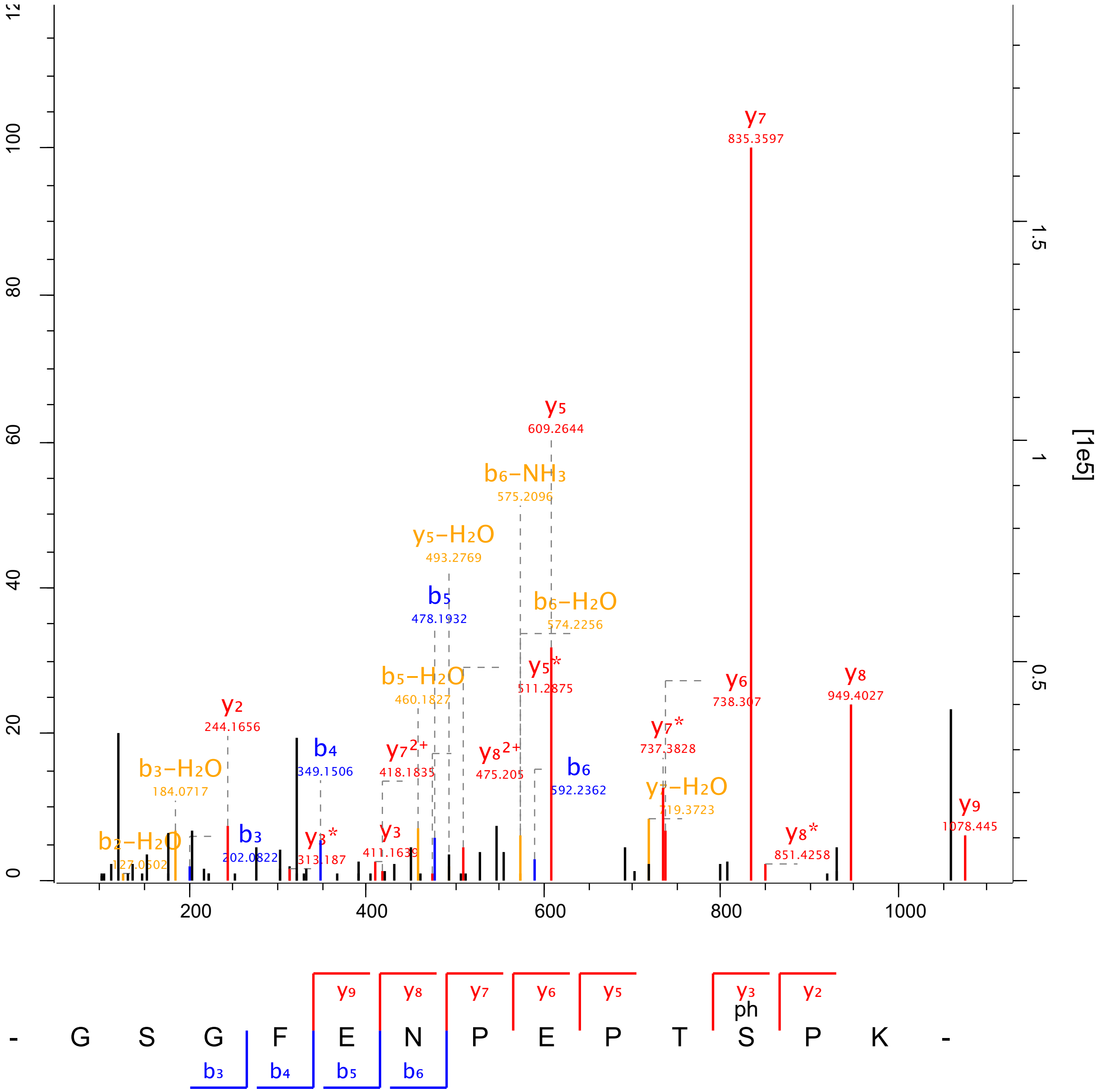

| Raw file | Scan | Method | Score | m/z |
|---|---|---|---|---|
| A17_00_2 | 11551 | FTMS; HCD | 77.74 | 546.22 |

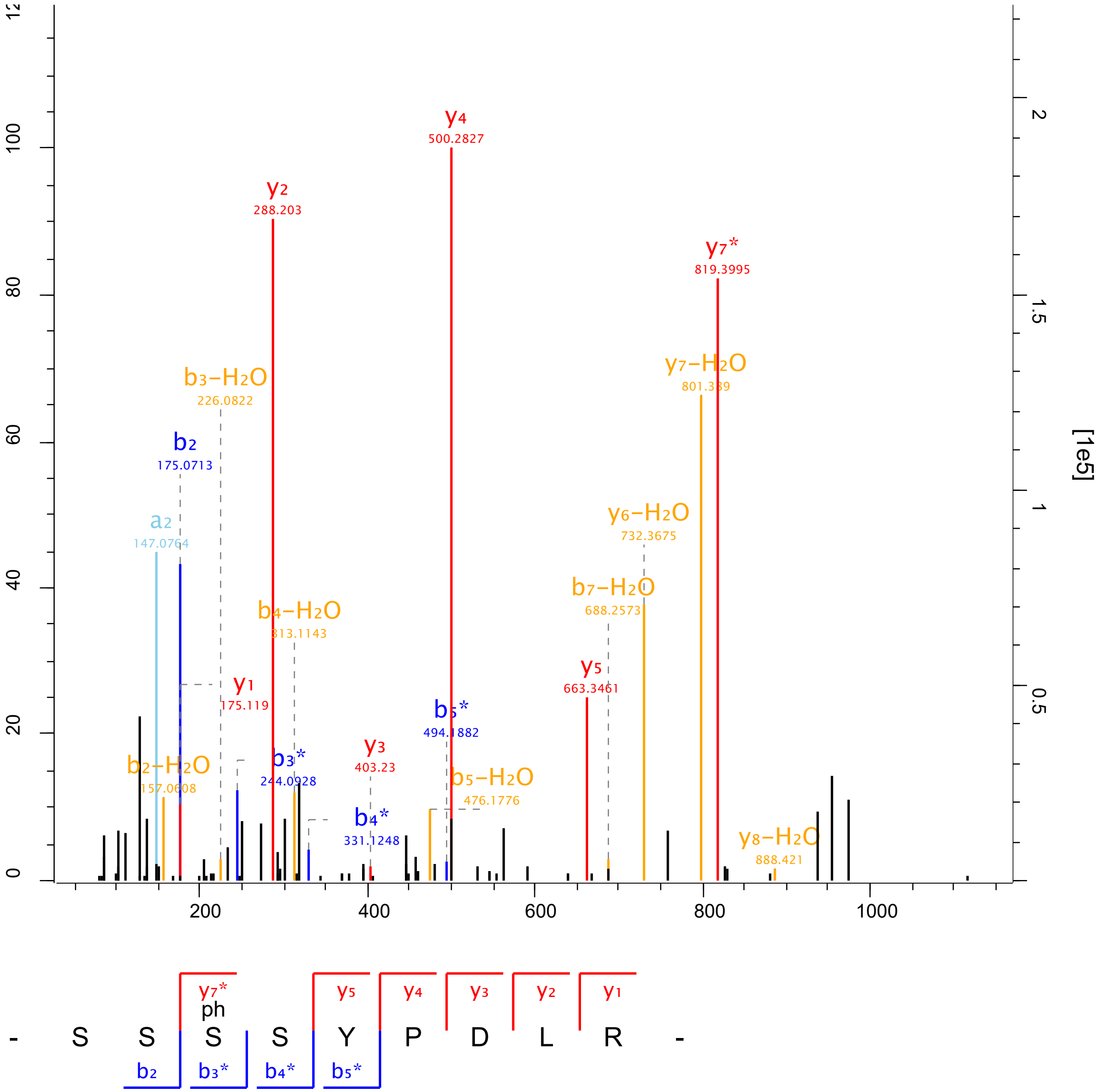

| Raw file | Scan | Method | Score | m/z |
|---|---|---|---|---|
| A17_00_2 | 11600 | FTMS; HCD | 56.12 | 687.75 |

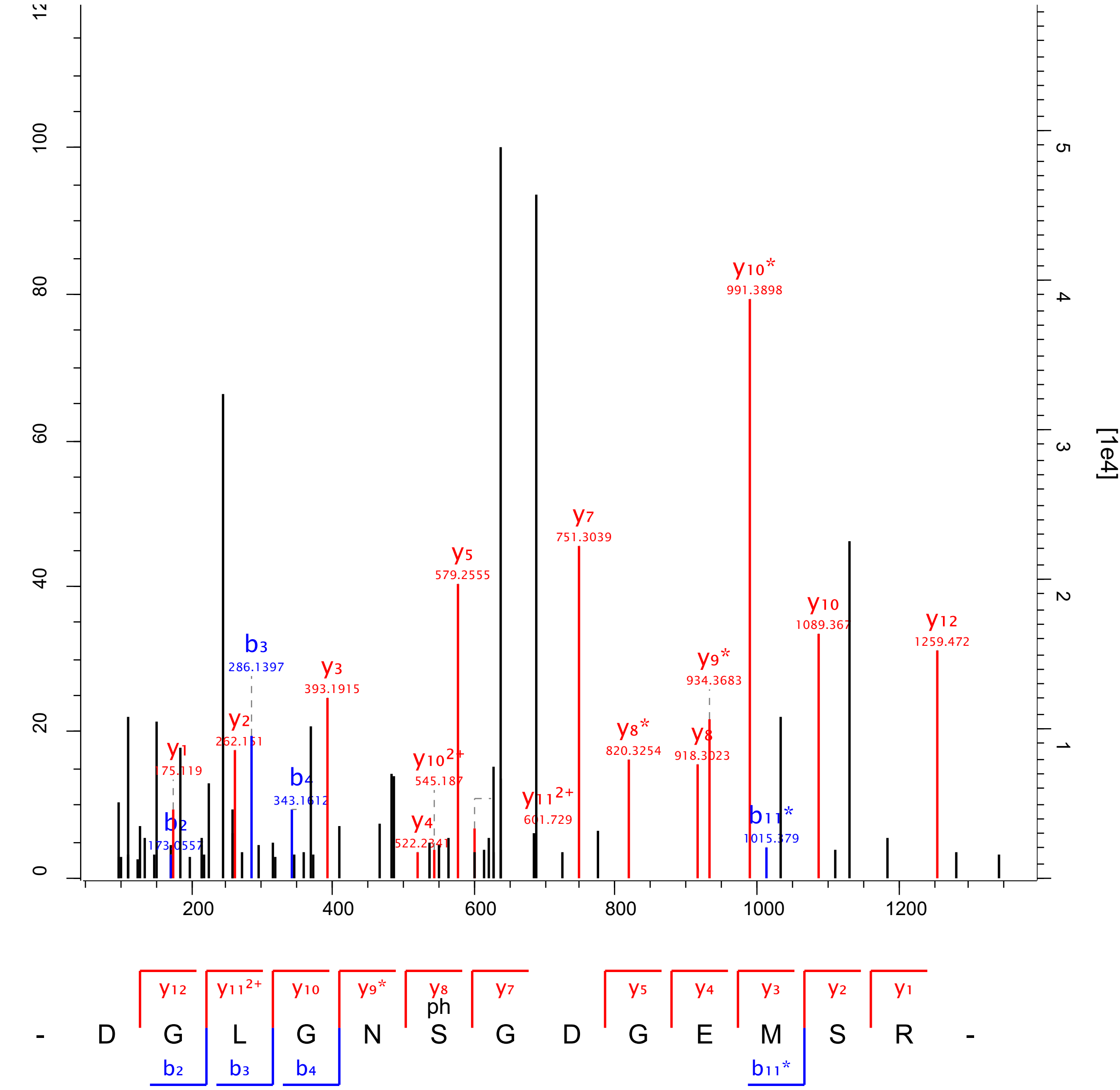

| Raw file | Scan | Method | Score | m/z |
|---|---|---|---|---|
| A17_00_2 | 11660 | FTMS; HCD | 68.97 | 589.76 |

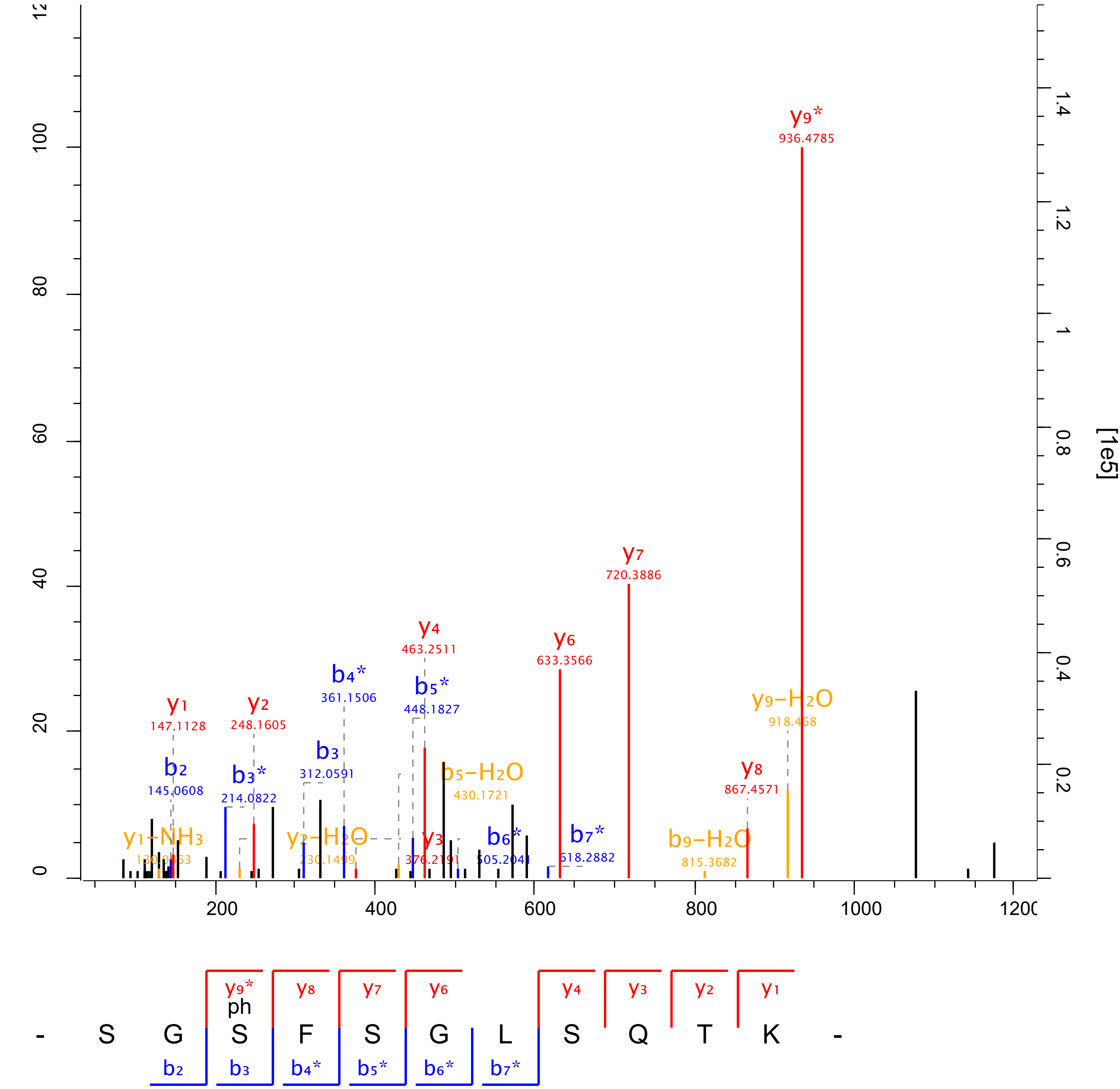

| Raw file | Scan | Method | Score | m/z |
| --- | --- | --- | --- | --- |
| A17_00_2 | 11664 | FTMS; HCD | 73.44 | 553.27 |

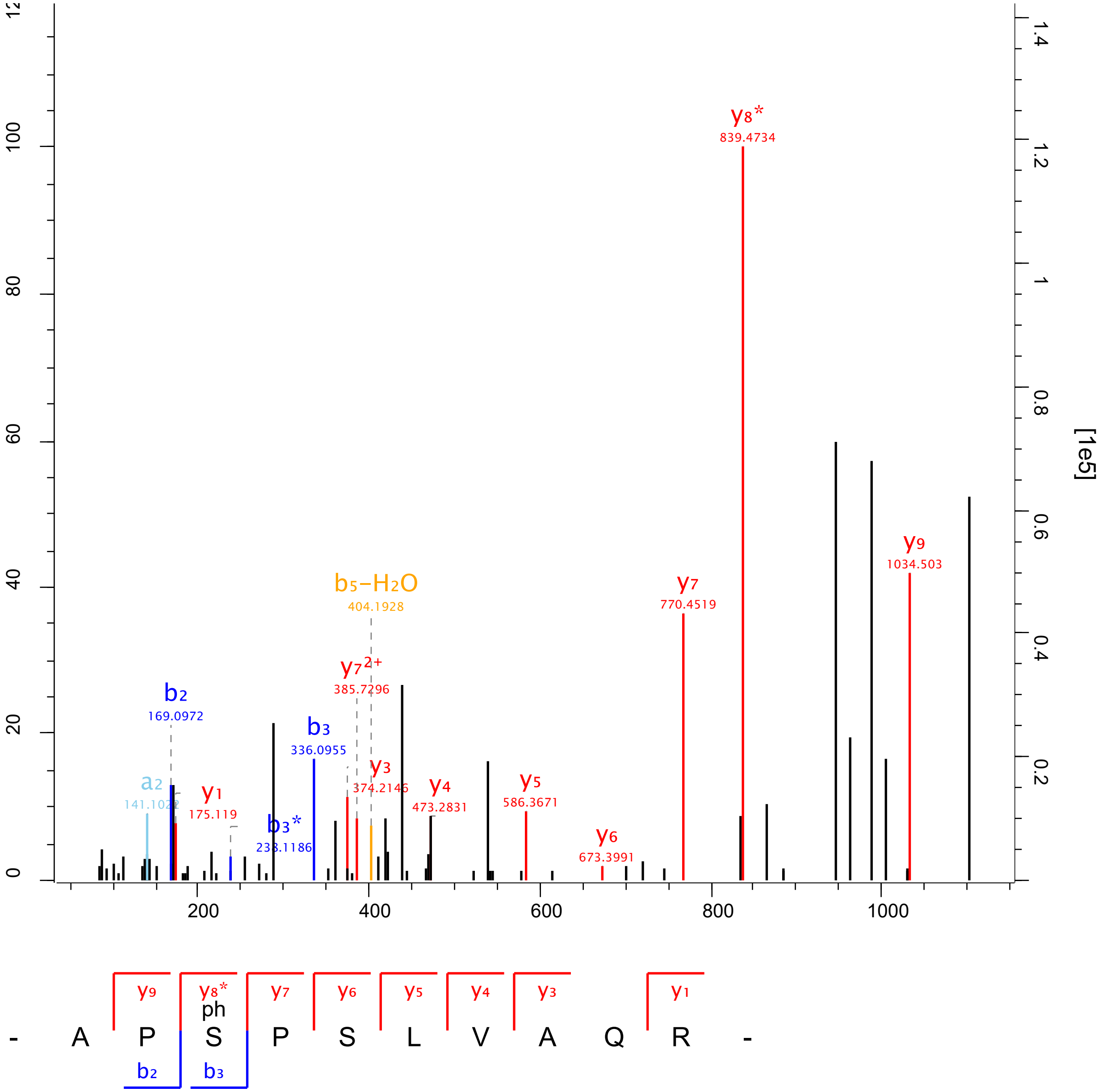

| Raw file | Scan | Method | Score | m/z |
|---|---|---|---|---|
| A17_00_2 | 11684 | FTMS; HCD | 73.5 | 448.72 |

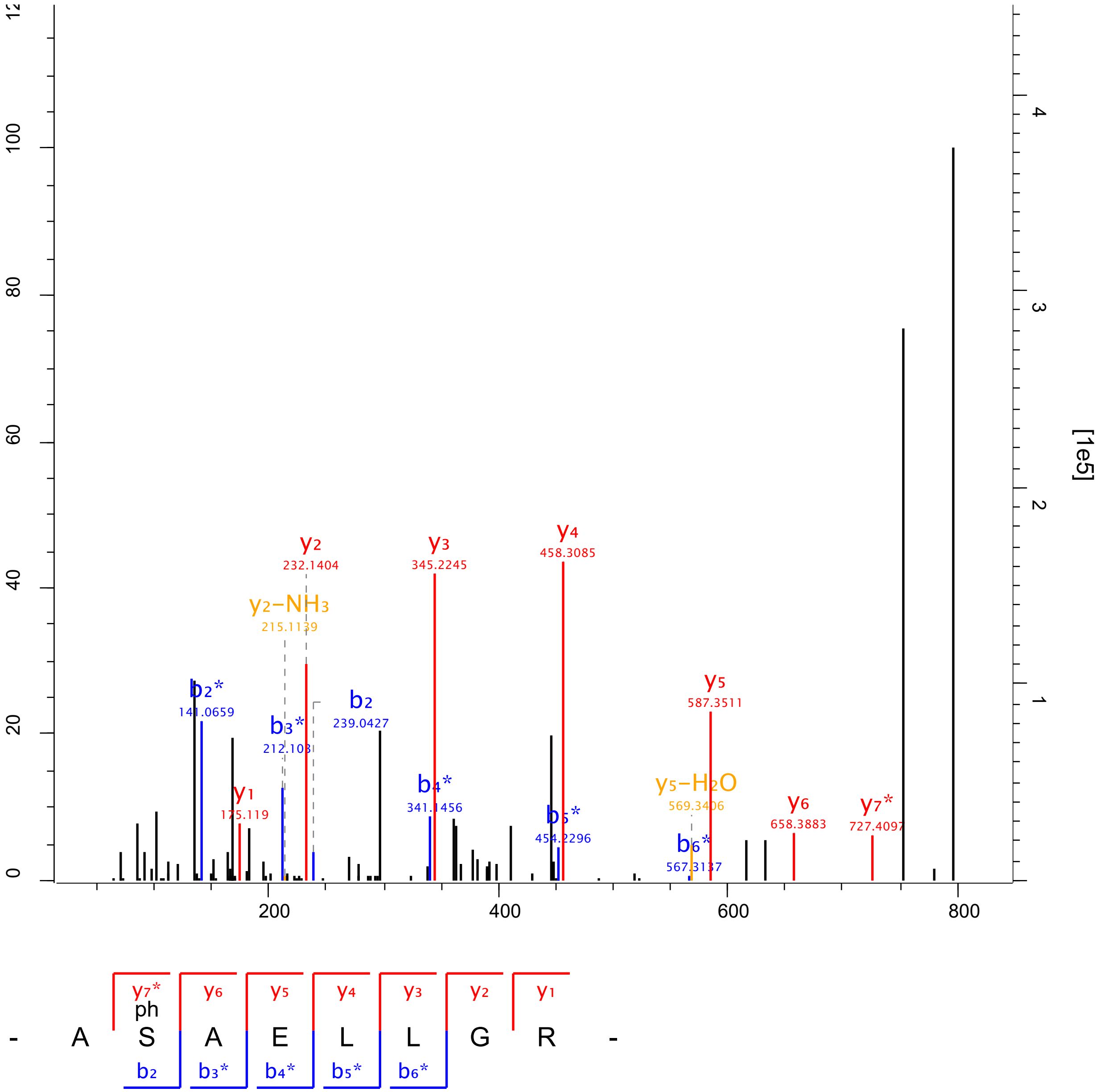

| Raw file | Scan | Method | Score | m/z |
| --- | --- | --- | --- | --- |
| A17_00_2 | 11702 | FTMS; HCD | 111.83 | 603.74 |

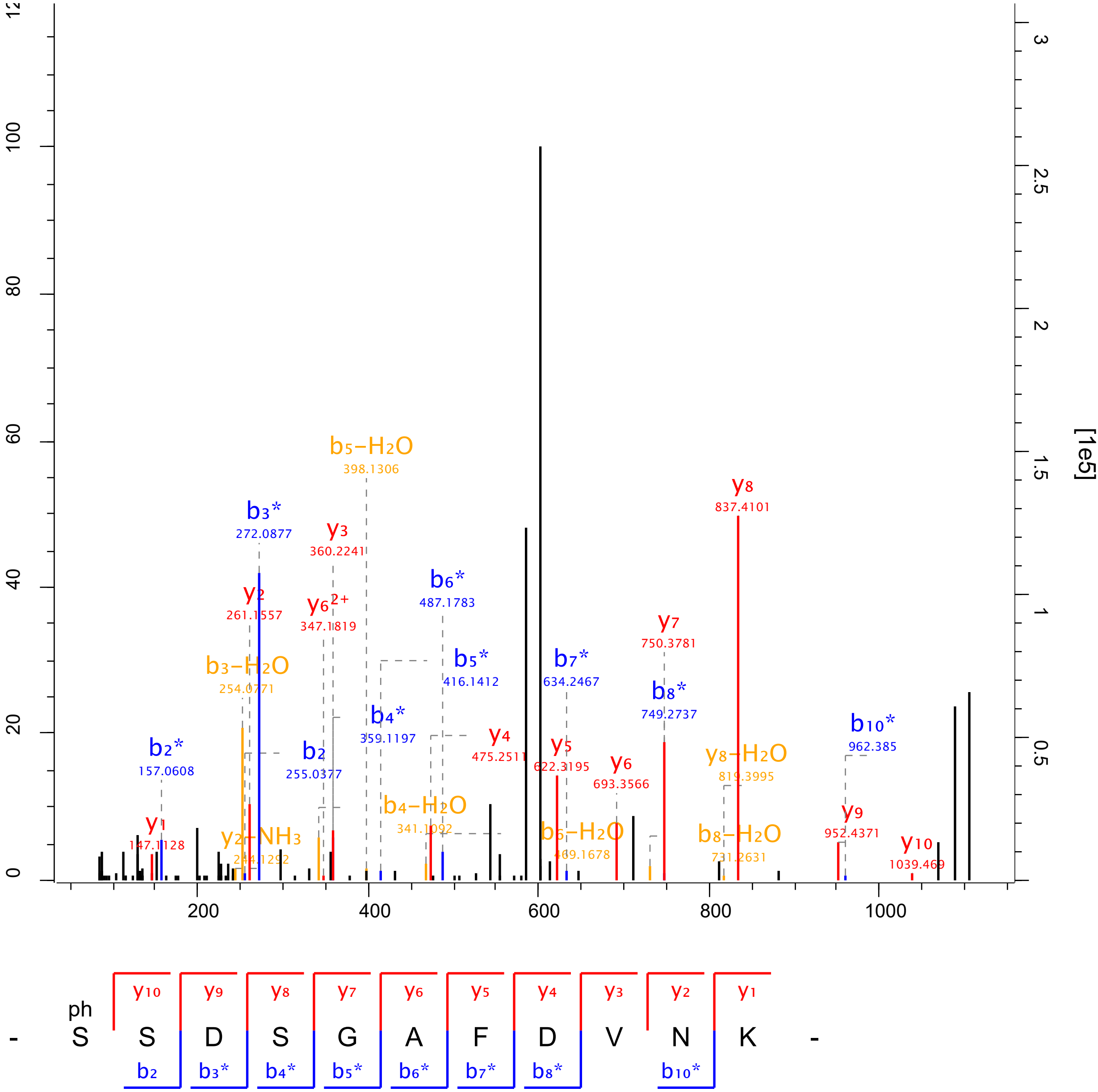

| Raw file | Scan | Method | Score | m/z |
|---|---|---|---|---|
| A17_00_2 | 11709 | FTMS; HCD | 95.27 | 547.22 |

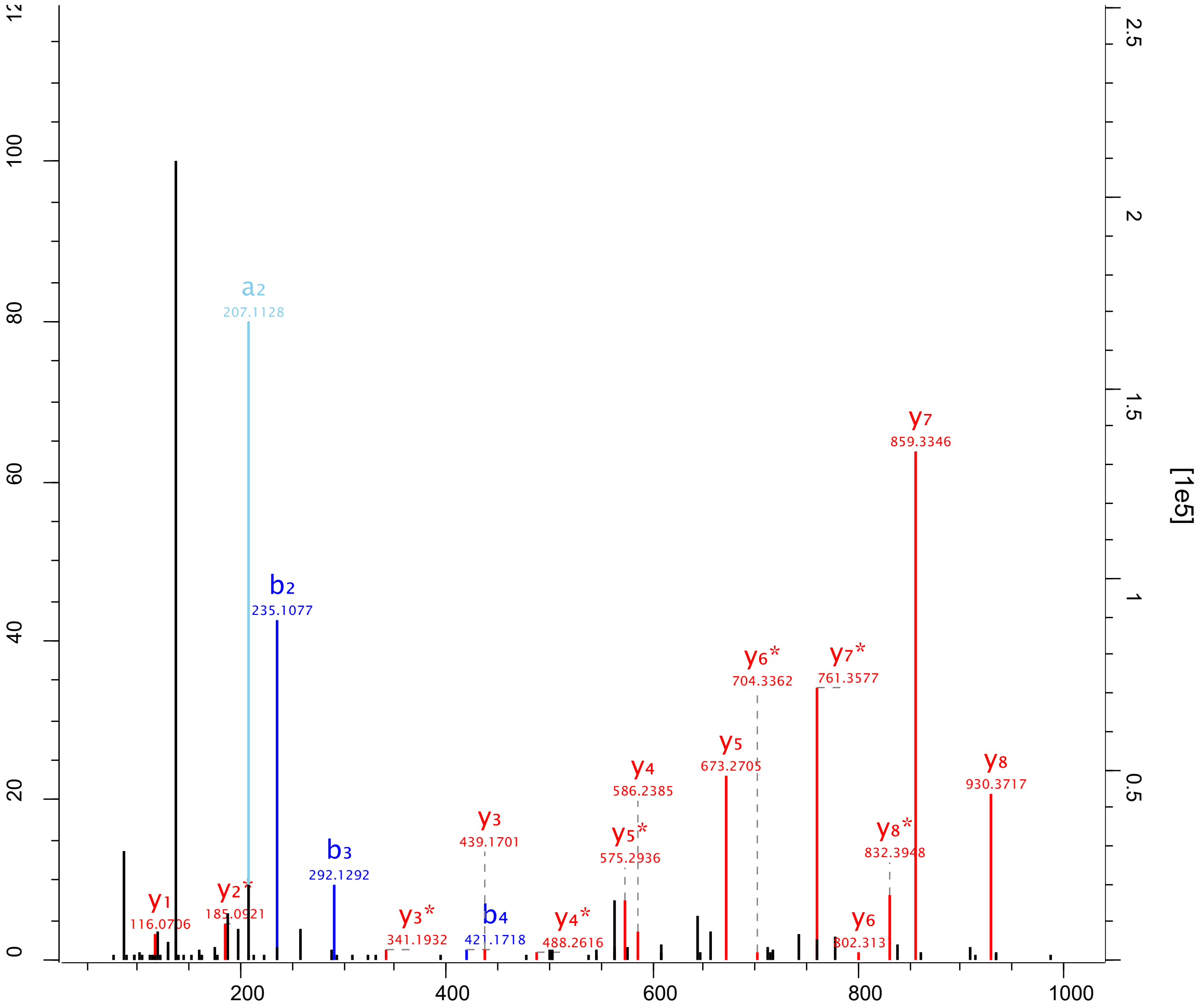


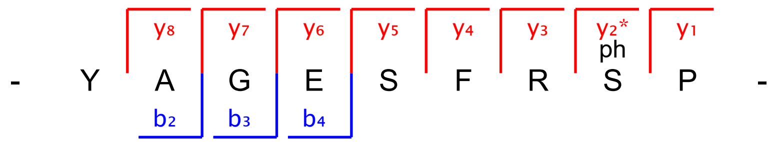

| Raw file | Scan | Method | Score | m/z |
| --- | --- | --- | --- | --- |
| A17_00_2 | 11728 | FTMS; HCD | 110.6 | 613.93 |

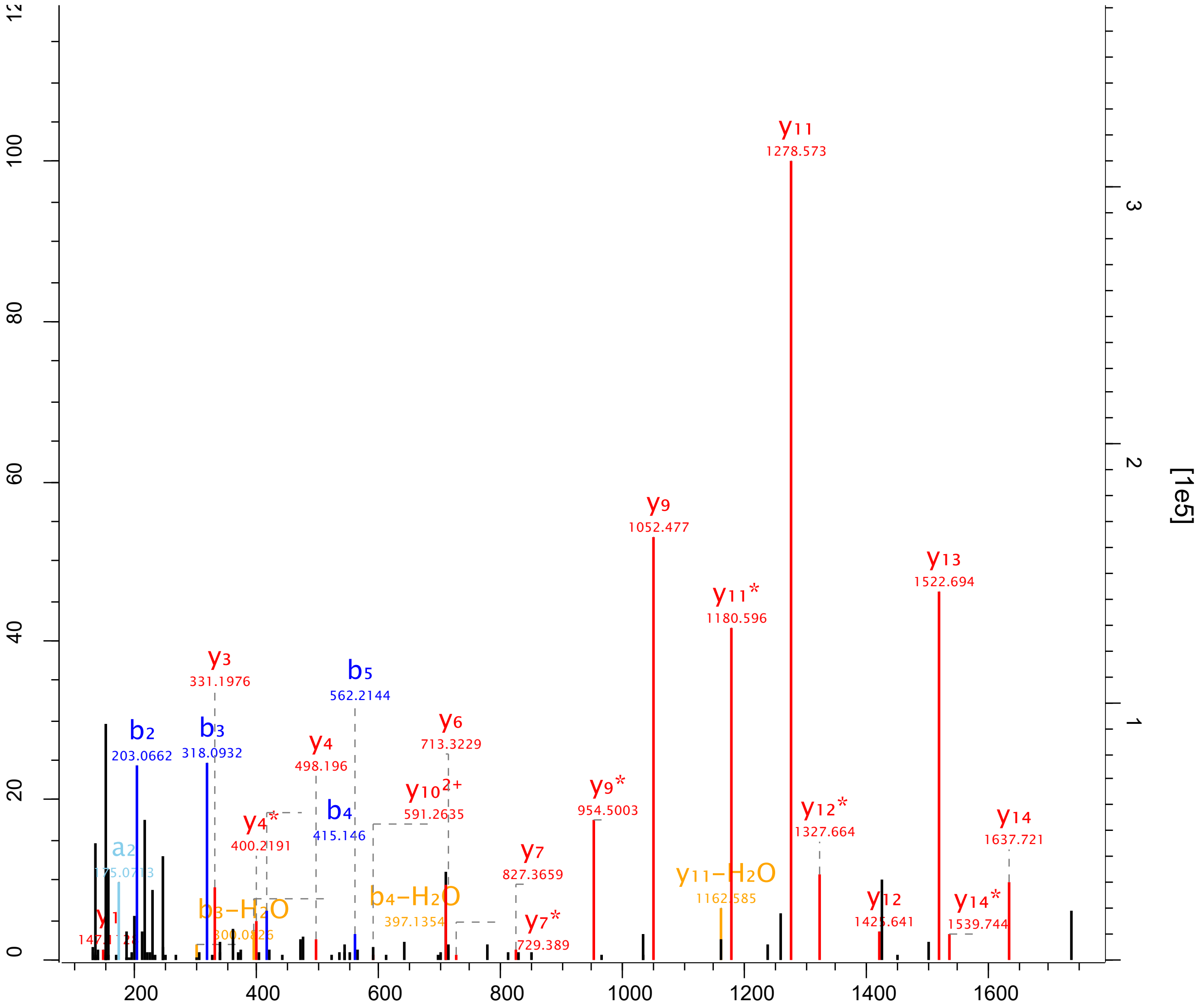


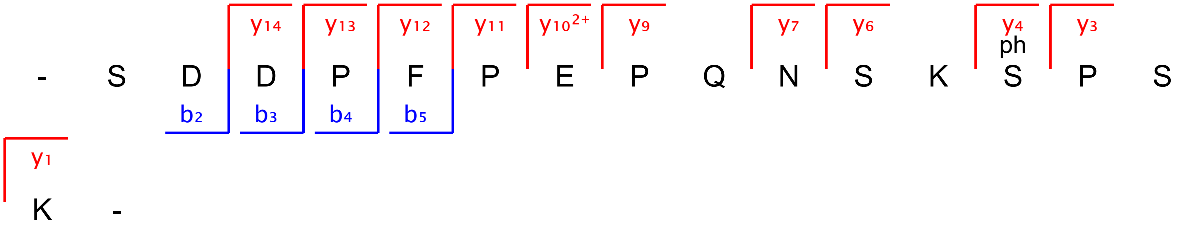

| Raw file | Scan | Method | Score | m/z |
|---|---|---|---|---|
| A17_00_2 | 11744 | FTMS; HCD | 101.3 | 640.59 |

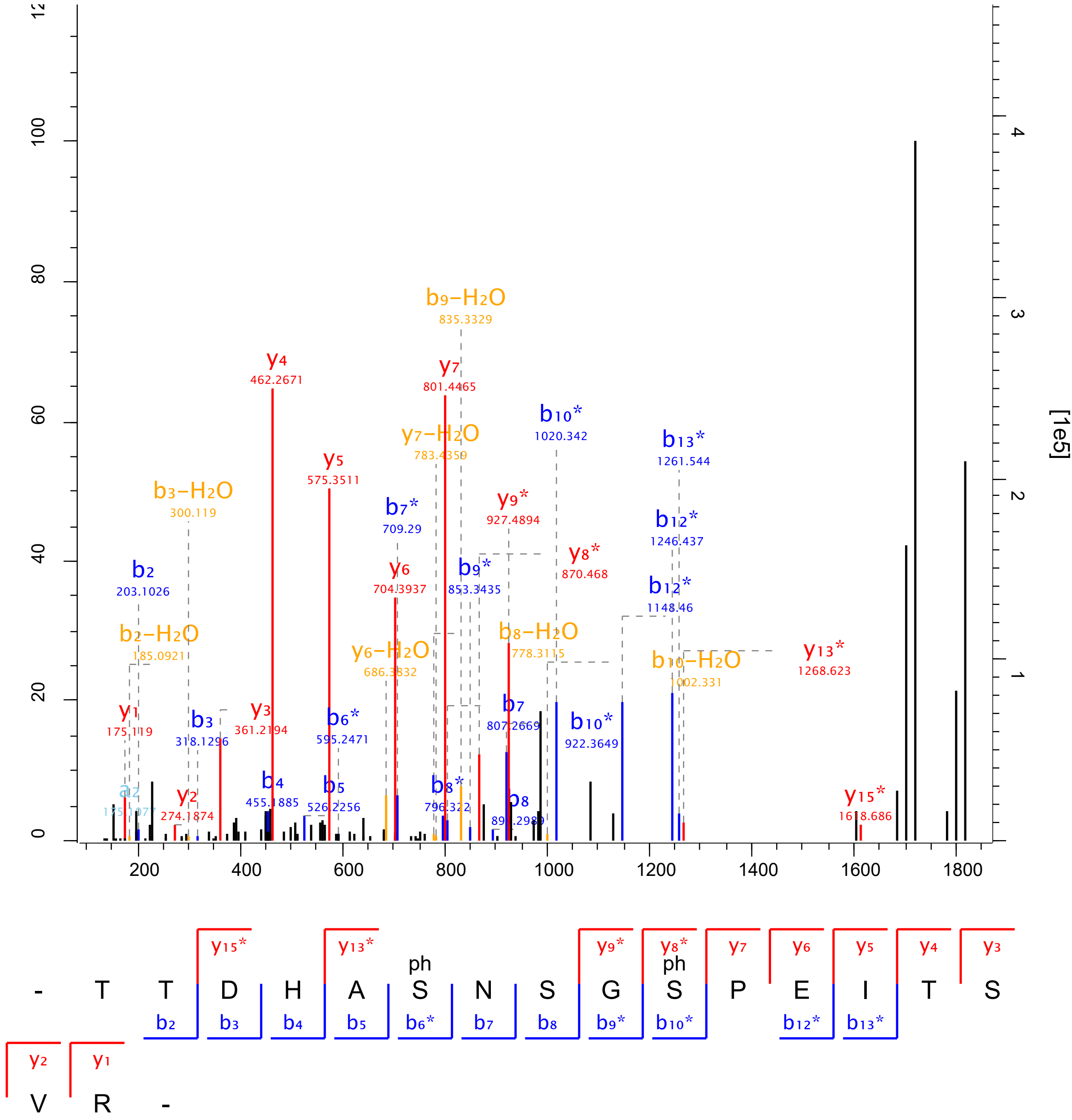

| Raw file | Scan | Method | Score | m/z |
| --- | --- | --- | --- | --- |
| A17_00_2 | 11809 | FTMS; HCD | 42.08 | 836.4 |

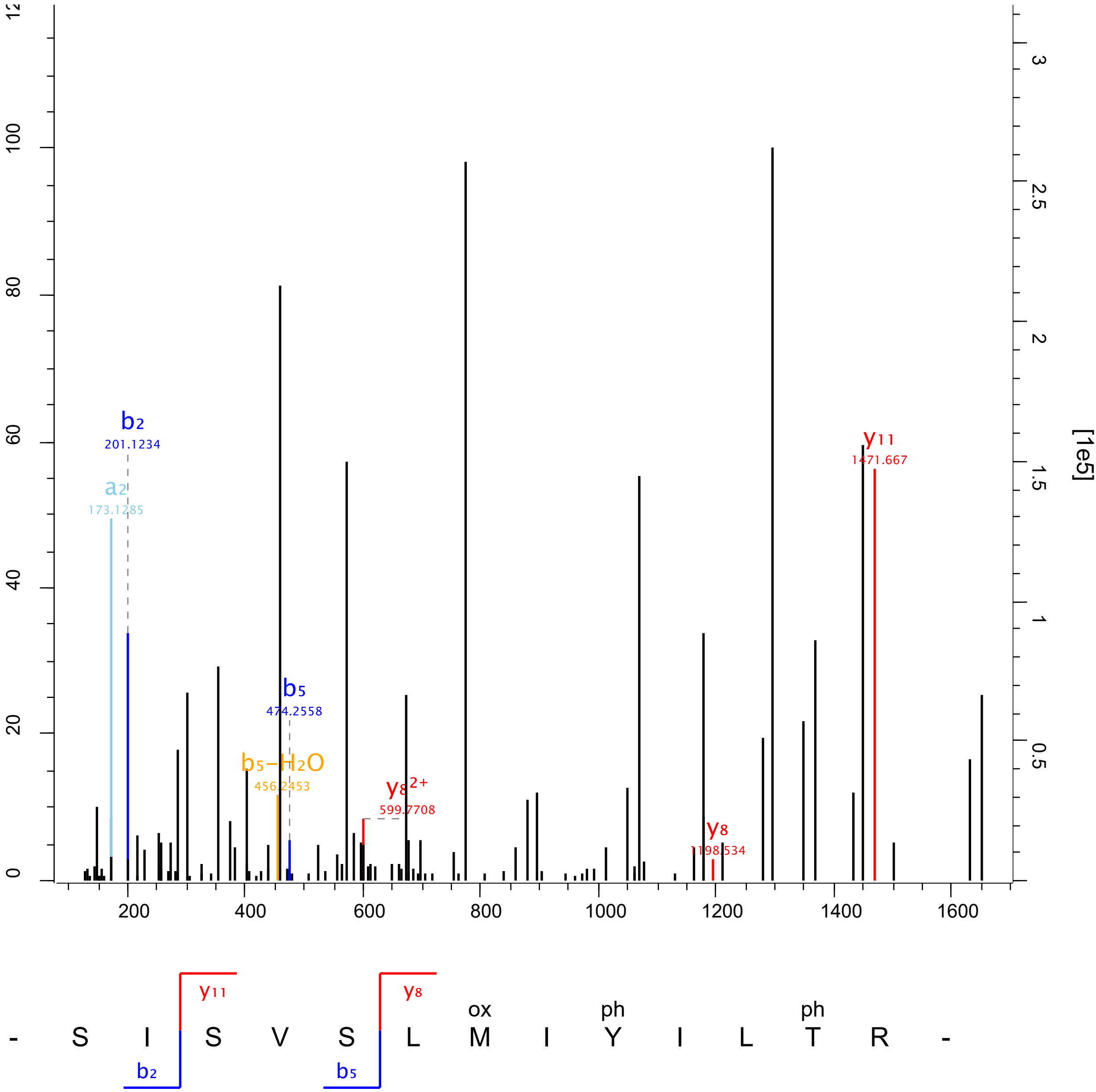

| Raw file | Scan | Method | Score | m/z |
|---|---|---|---|---|
| A17_00_2 | 12040 | FTMS; HCD | 115.57 | 619.71 |

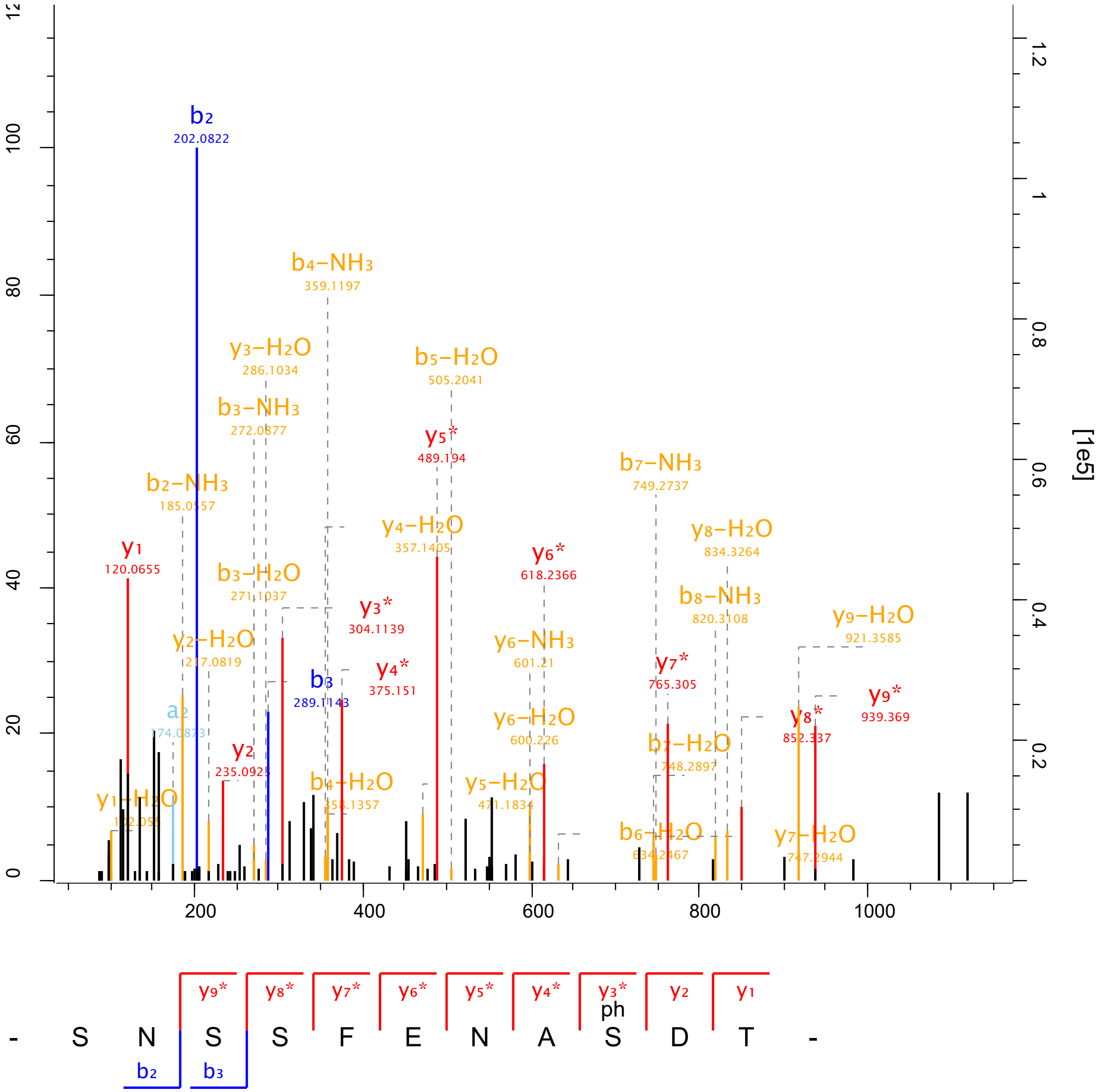

| Raw file | Scan | Method | Score | m/z |
|---|---|---|---|---|
| A17_00_2 | 12064 | FTMS; HCD | 237.87 | 679.25 |

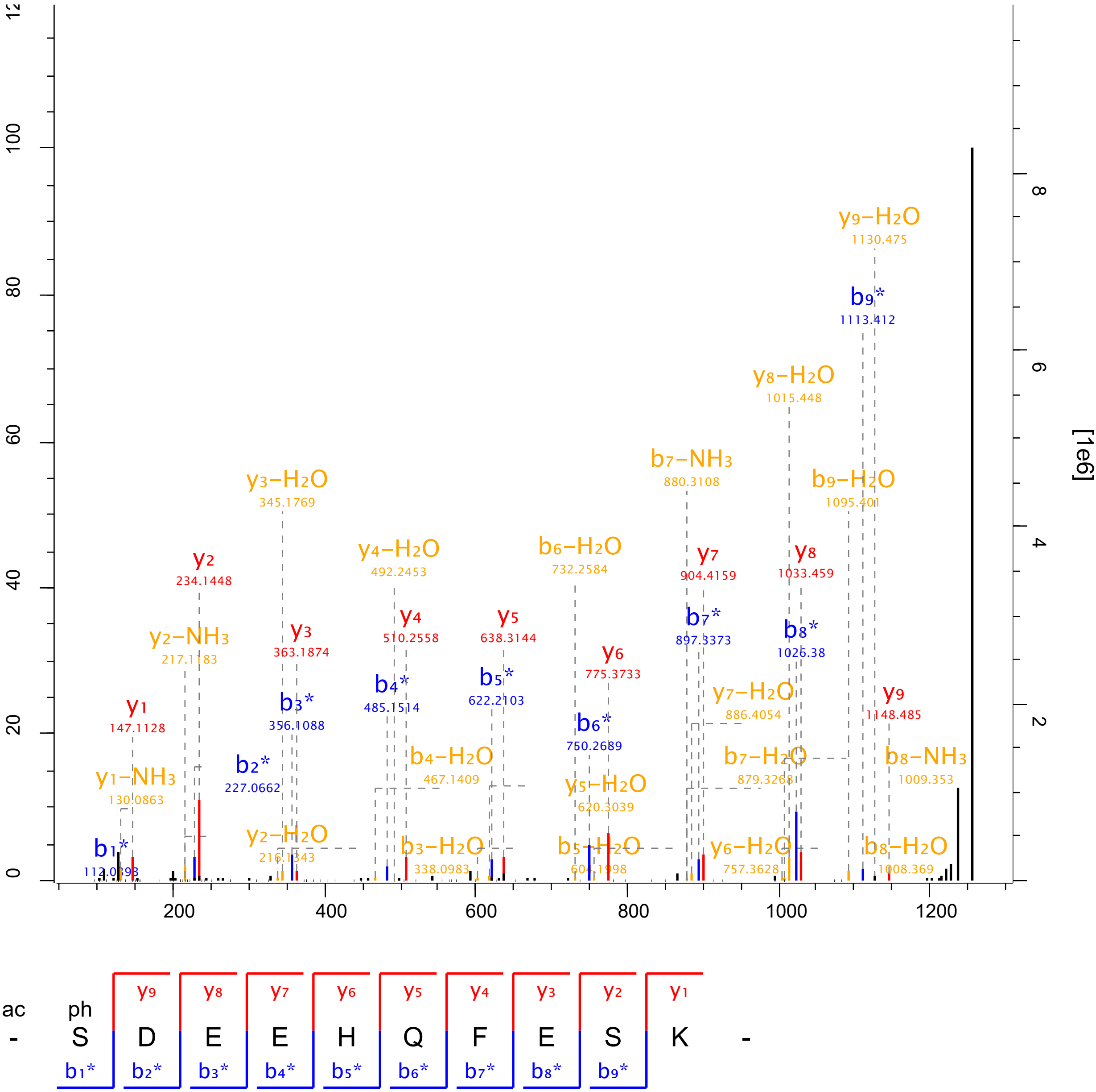

| Raw file | Scan | Method | Score | m/z |
| --- | --- | --- | --- | --- |
| A17_00_2 | 12074 | FTMS; HCD | 154.86 | 1002.9 |

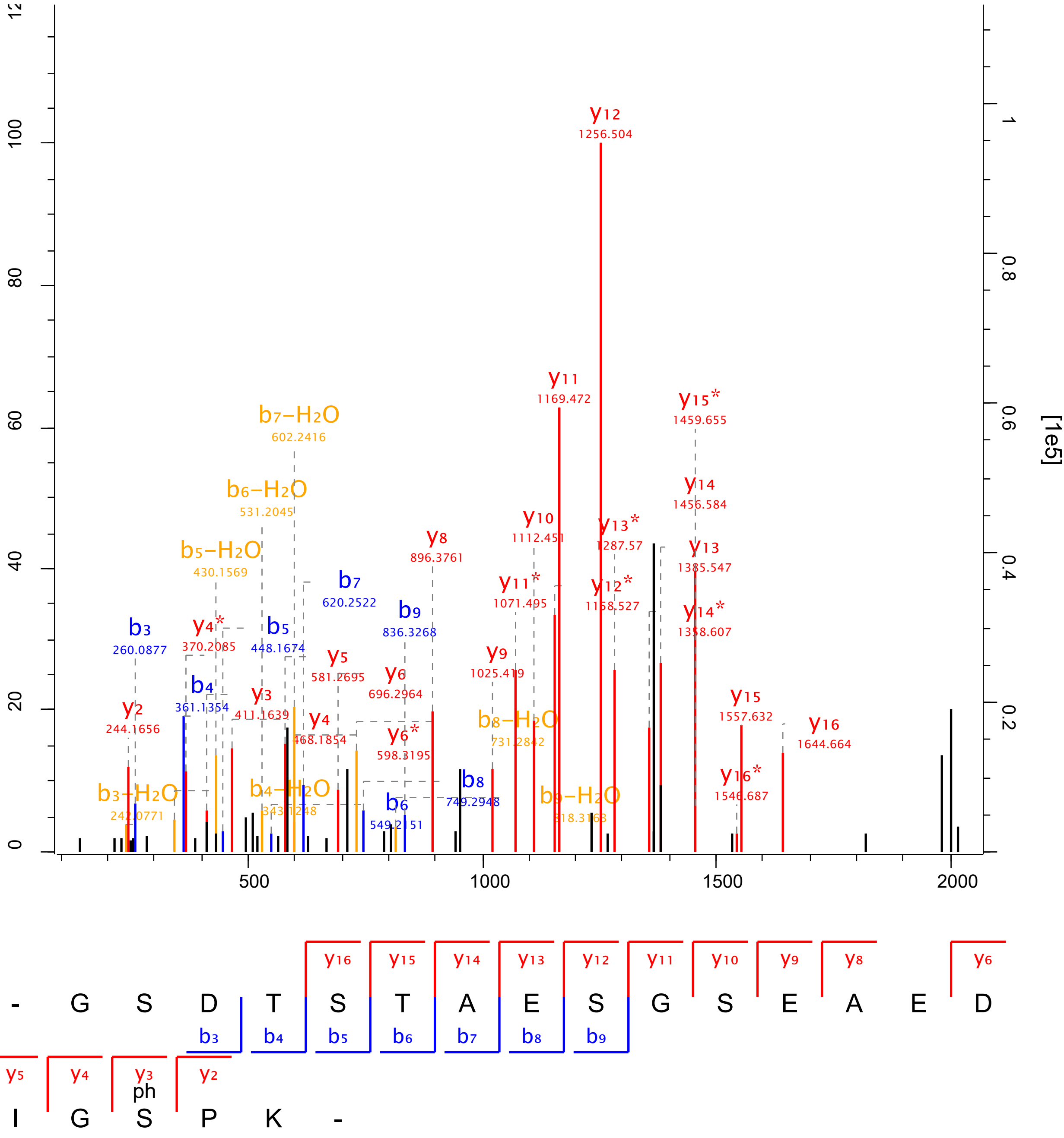

| Raw file | Scan | Method | Score | m/z |
|---|---|---|---|---|
| A17_00_2 | 12195 | FTMS; HCD | 46.25 | 711.82 |

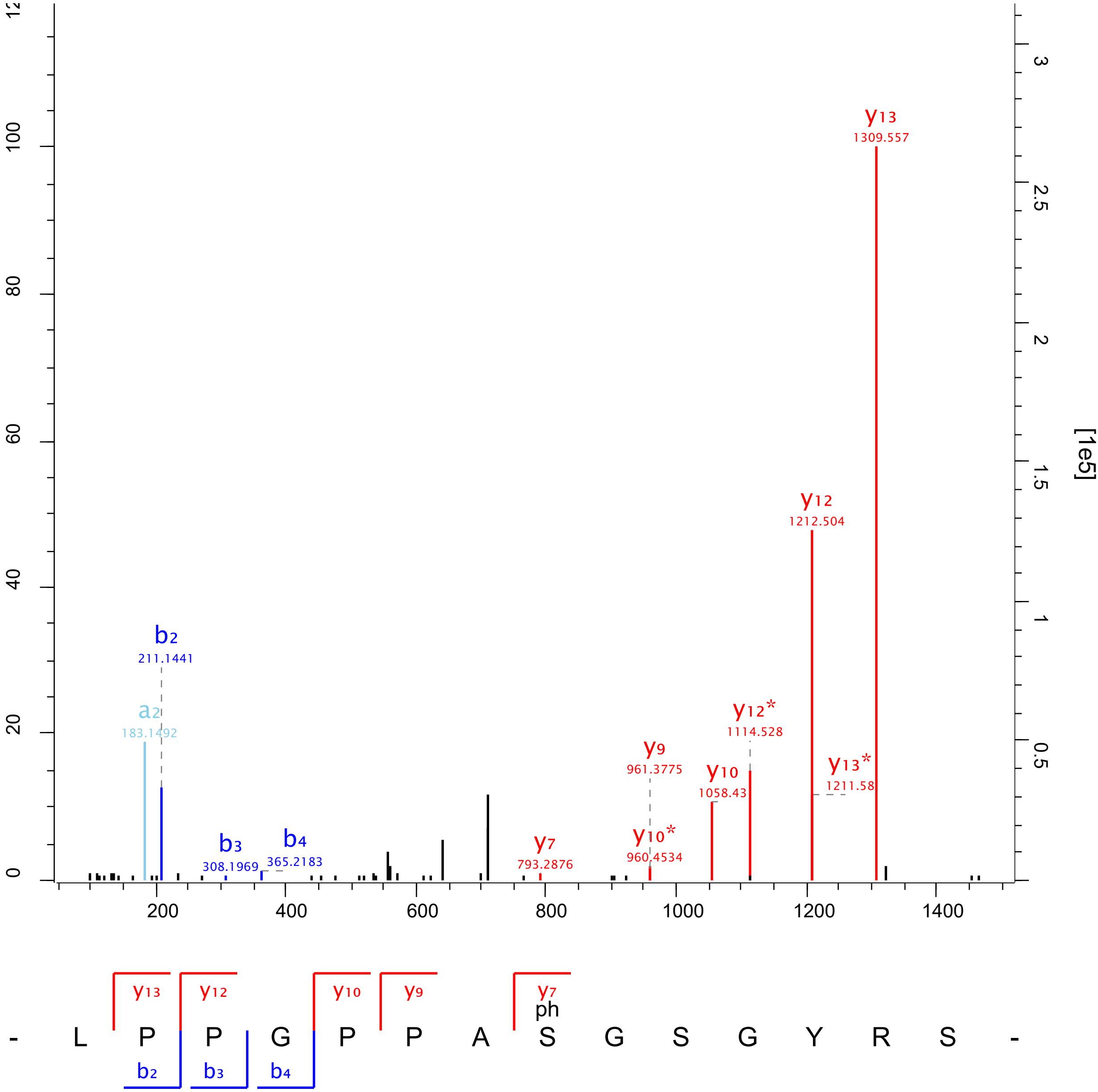

| Raw file | Scan | Method | Score | m/z |
| --- | --- | --- | --- | --- |
| A17_00_2 | 12740 | FTMS; HCD | 75.62 | 828 |

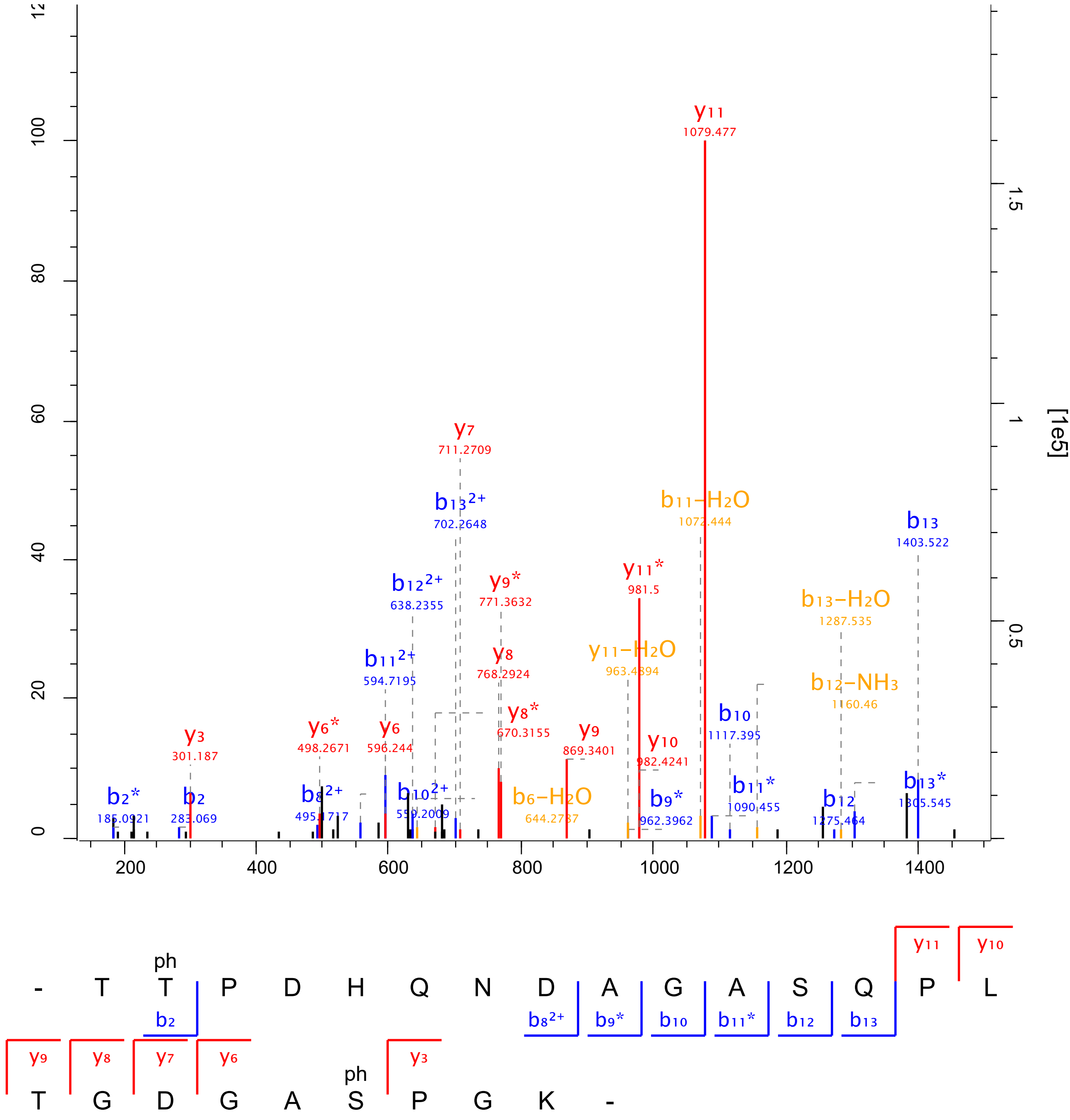

| Raw file | Scan | Method | Score | m/z |
|---|---|---|---|---|
| A17_00_2 | 12800 | FTMS; HCD | 65.18 | 573.73 |

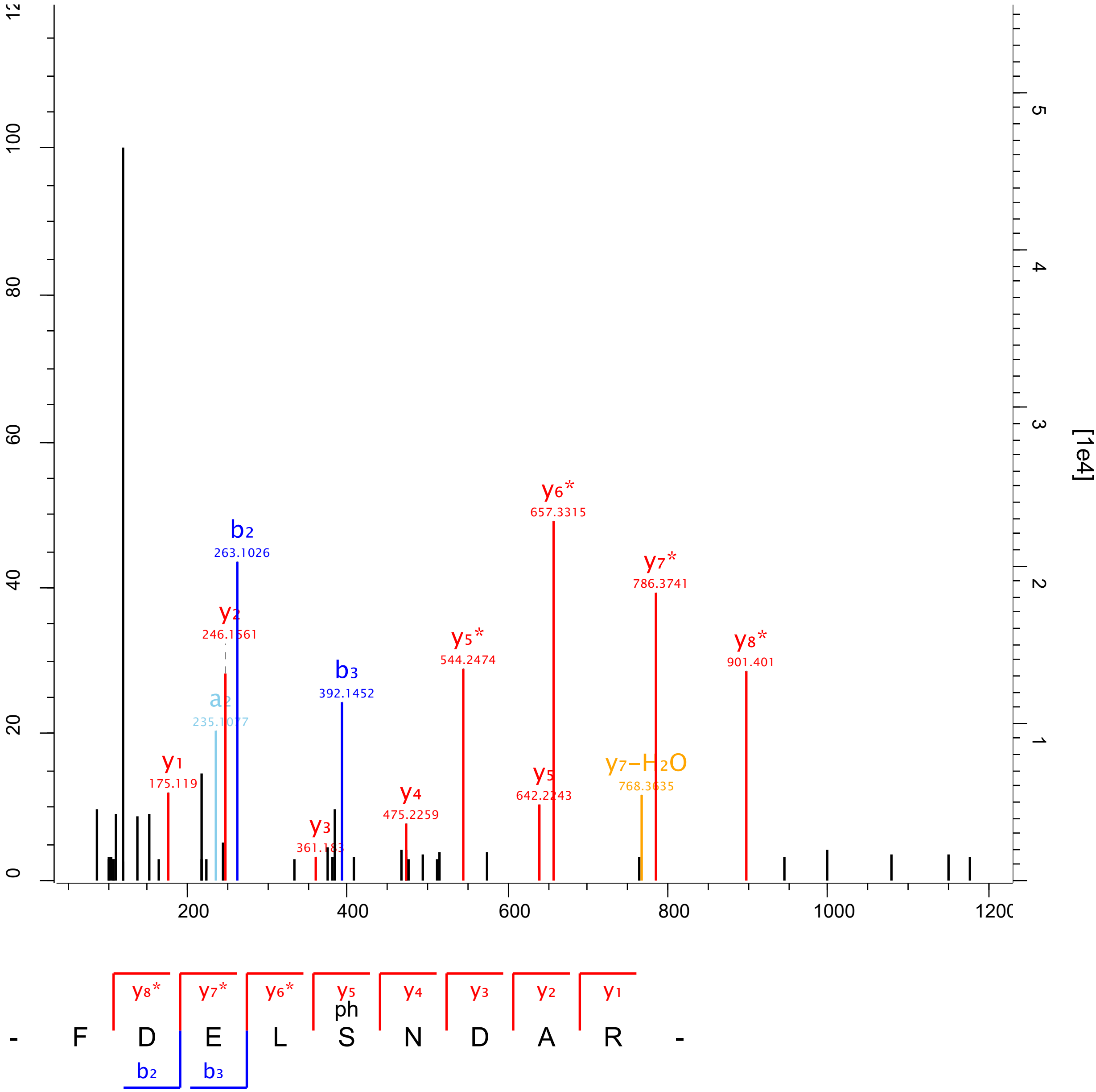

| Raw file | Scan | Method | Score | m/z |
|---|---|---|---|---|
| A17_00_2 | 12810 | FTMS; HCD | 321.17 | 886.35 |

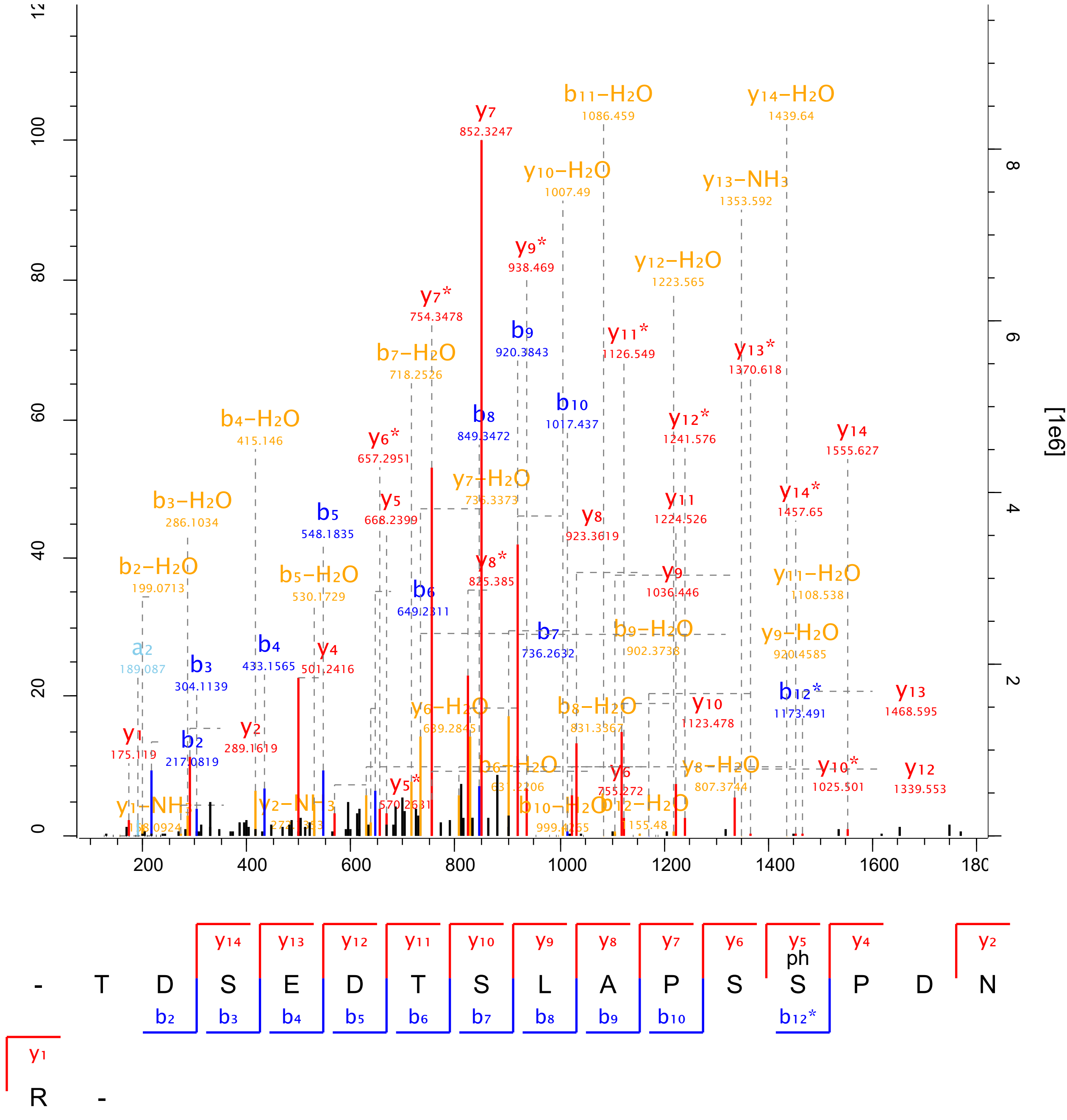

| Raw file | Scan | Method | Score | m/z |
|---|---|---|---|---|
| A17_00_2 | 13058 | FTMS; HCD | 140.3 | 734.28 |

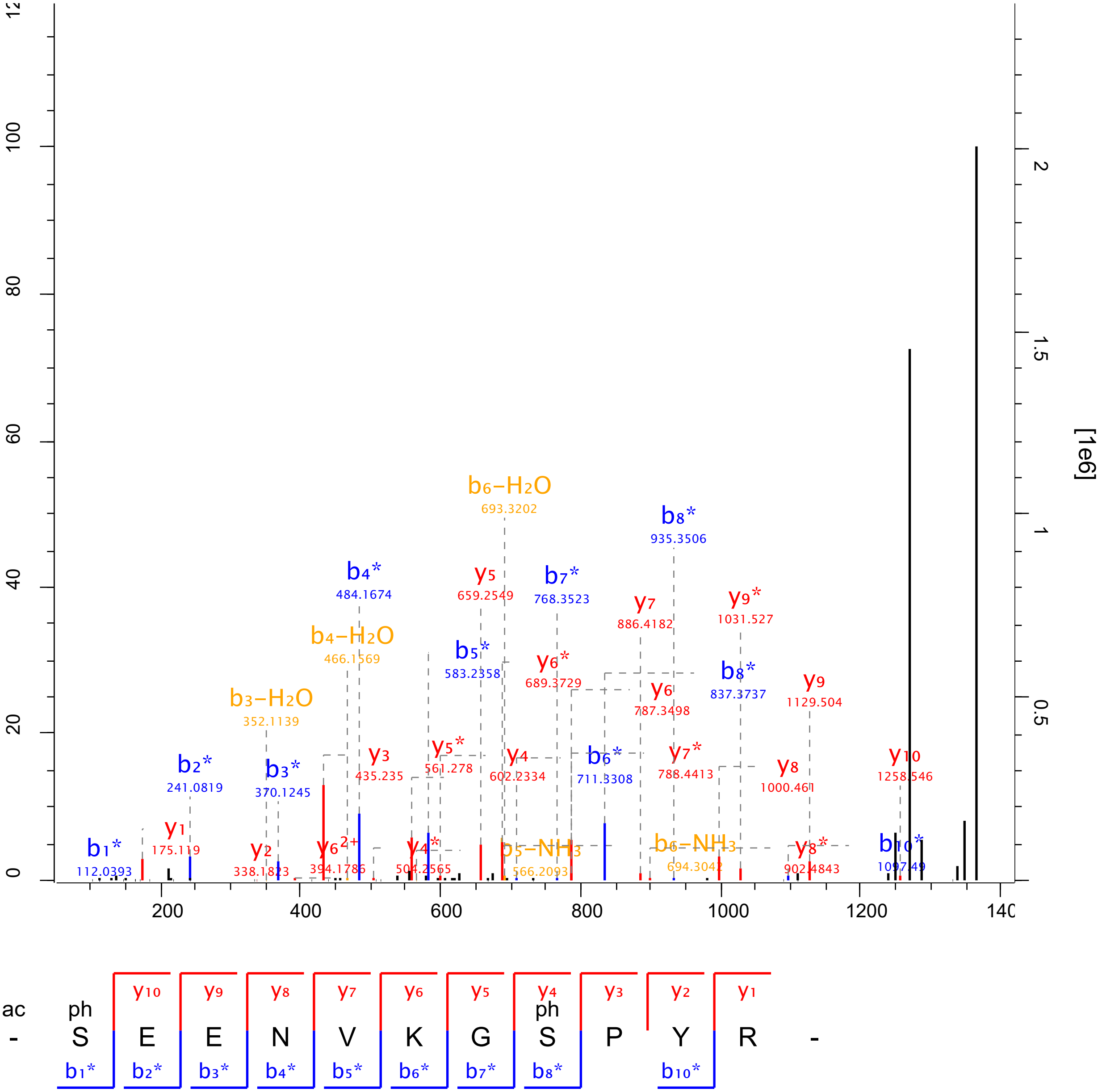

| Raw file | Scan | Method | Score | m/z |
| --- | --- | --- | --- | --- |
| A17_00_2 | 13153 | FTMS; HCD | 77.32 | 638.26 |

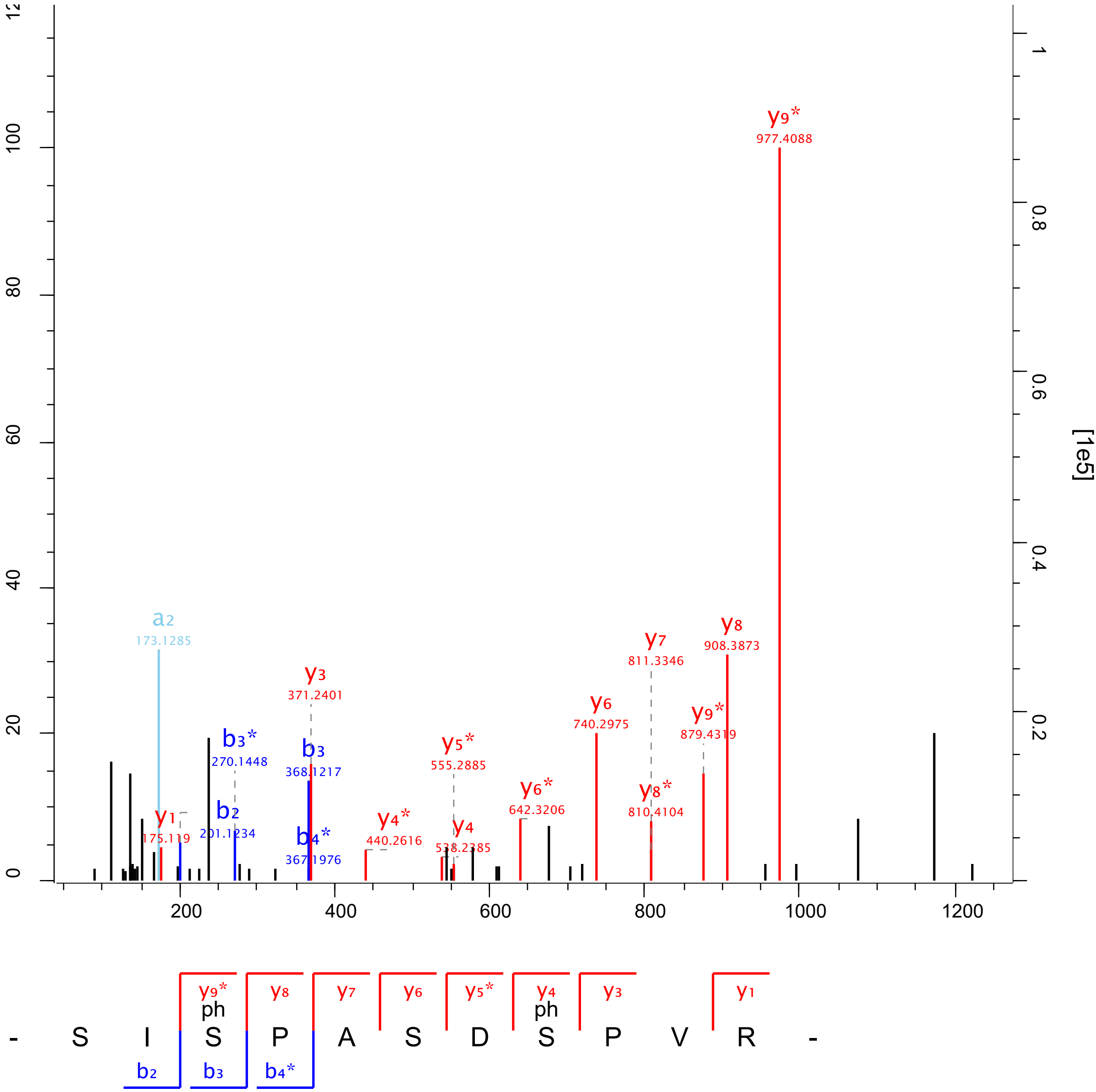

| Raw file | Scan | Method | Score | m/z |
|---|---|---|---|---|
| A17_00_2 | 13534 | FTMS; HCD | 306.92 | 703.31 |

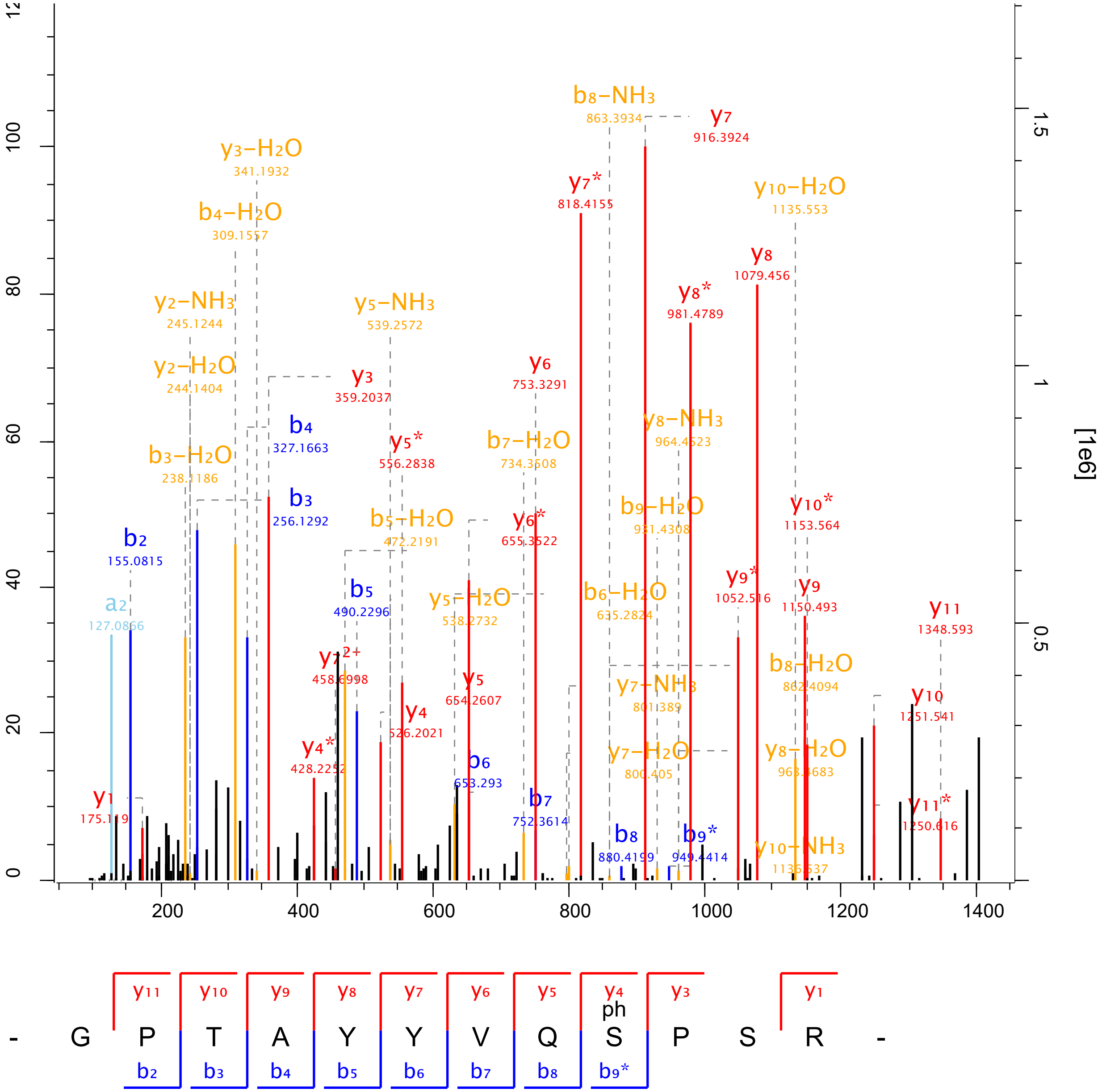

| Raw file | Scan | Method | Score | m/z |
|---|---|---|---|---|
| A17_00_2 | 13738 | FTMS; HCD | 101.5 | 828.87 |

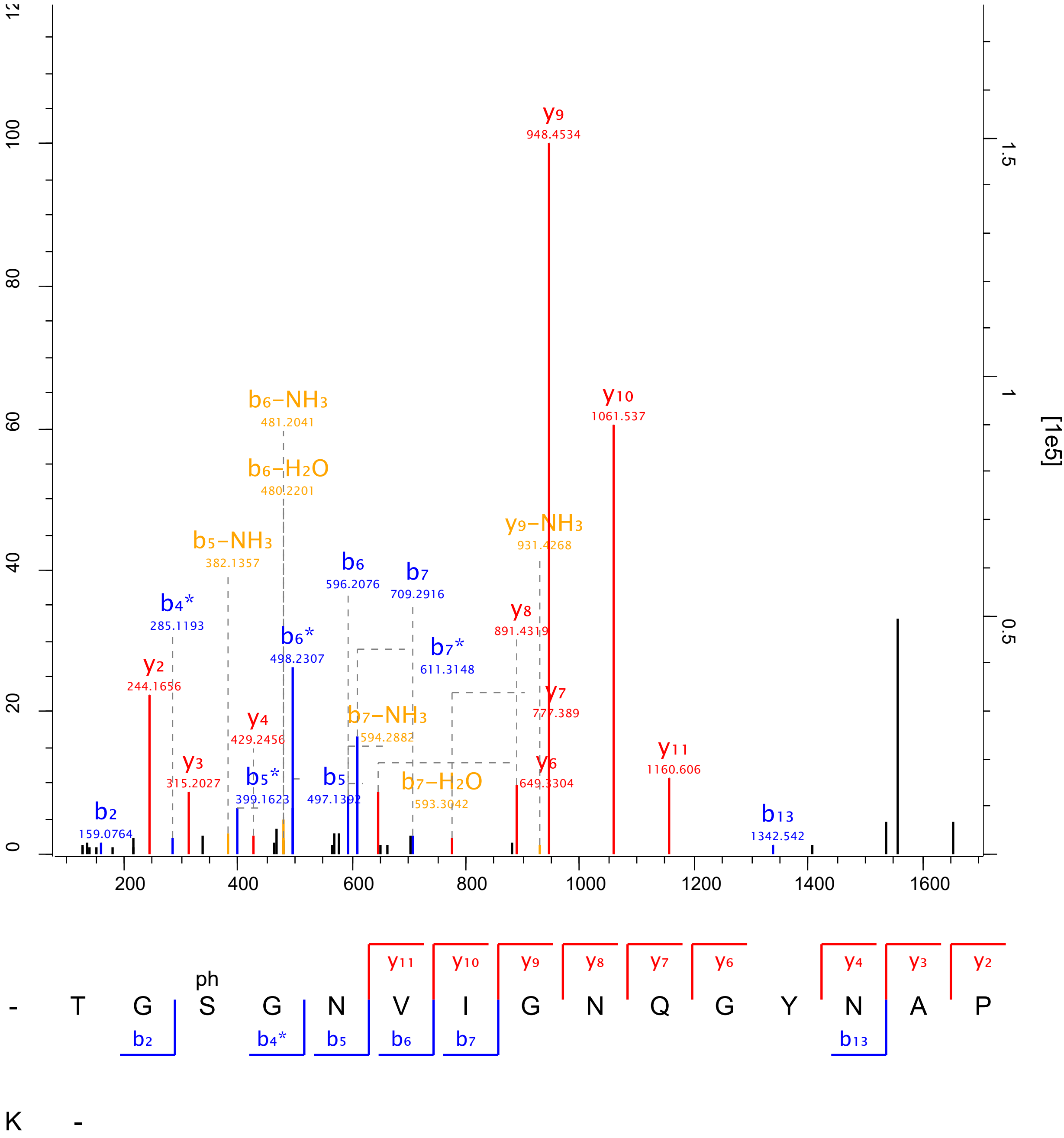

| Raw file | Scan | Method | Score | m/z |
| --- | --- | --- | --- | --- |
| A17_00_2 | 13759 | FTMS; HCD | 74.88 | 780.69 |

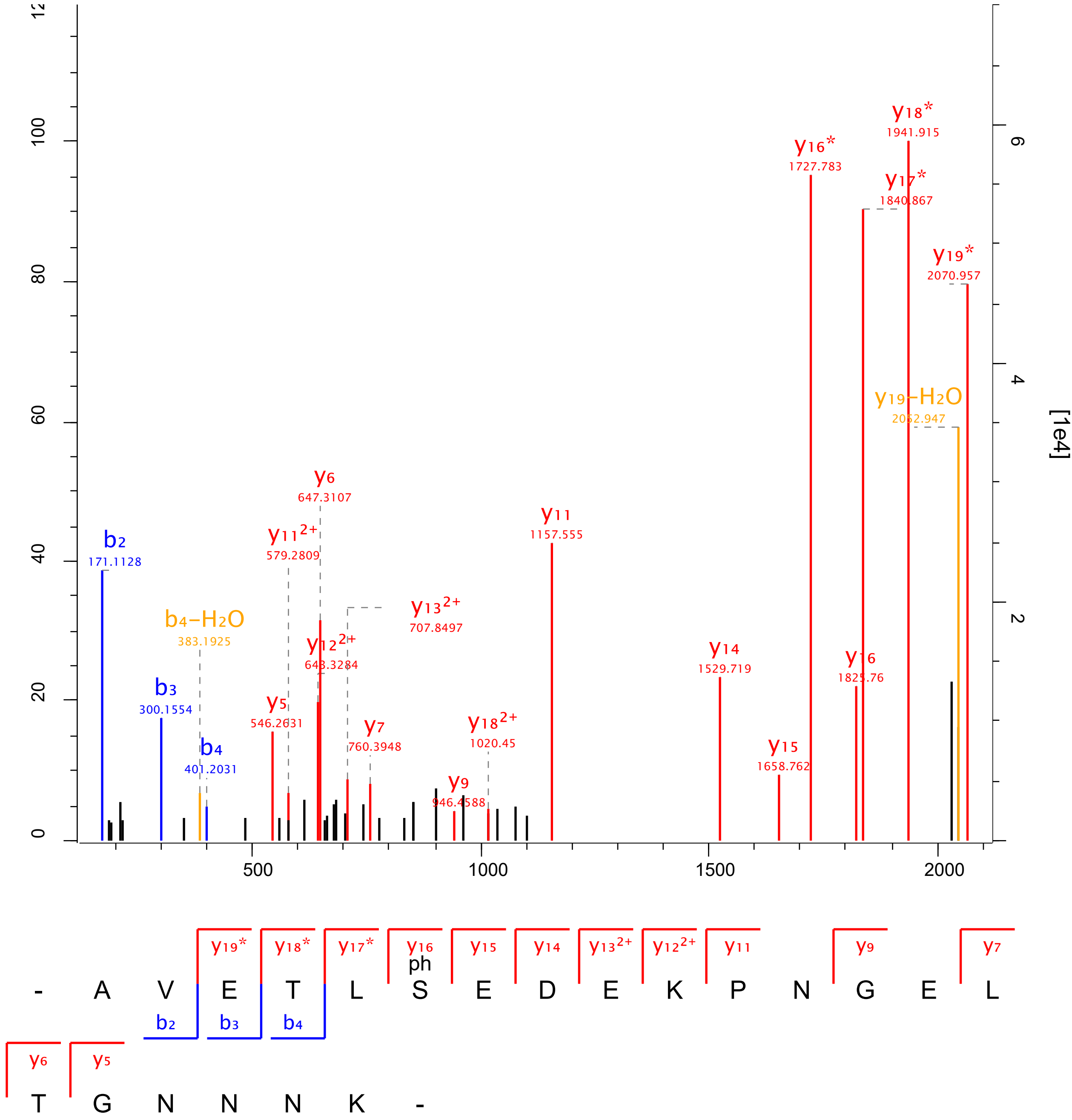

| Raw file | Scan | Method | Score | m/z |
|---|---|---|---|---|
| A17_00_2 | 13761 | FTMS; HCD | 41.45 | 886.71 |

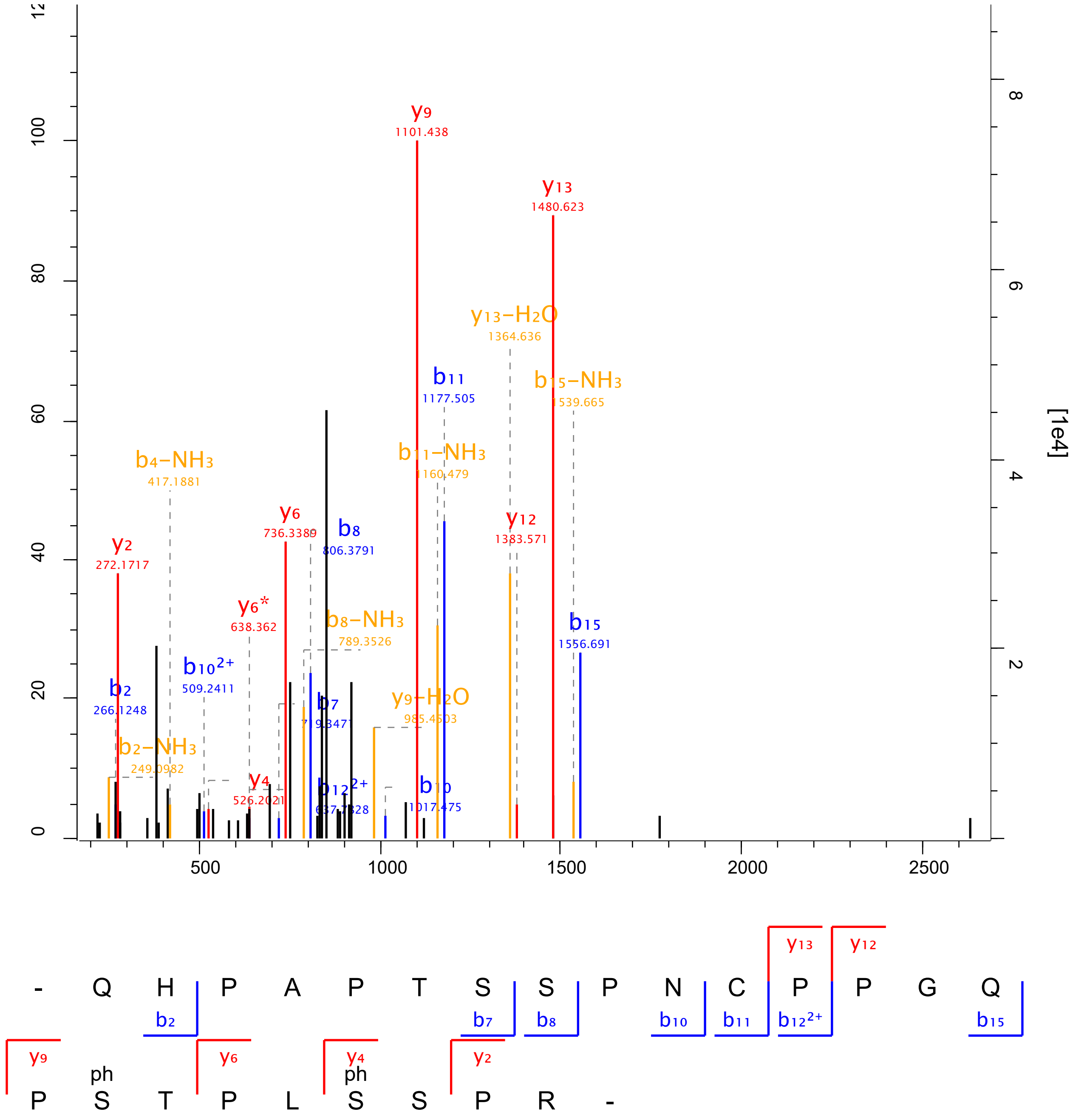

| Raw file | Scan | Method | Score | m/z |
| --- | --- | --- | --- | --- |
| A17_00_2 | 13880 | FTMS; HCD | 41.09 | 633.93 |

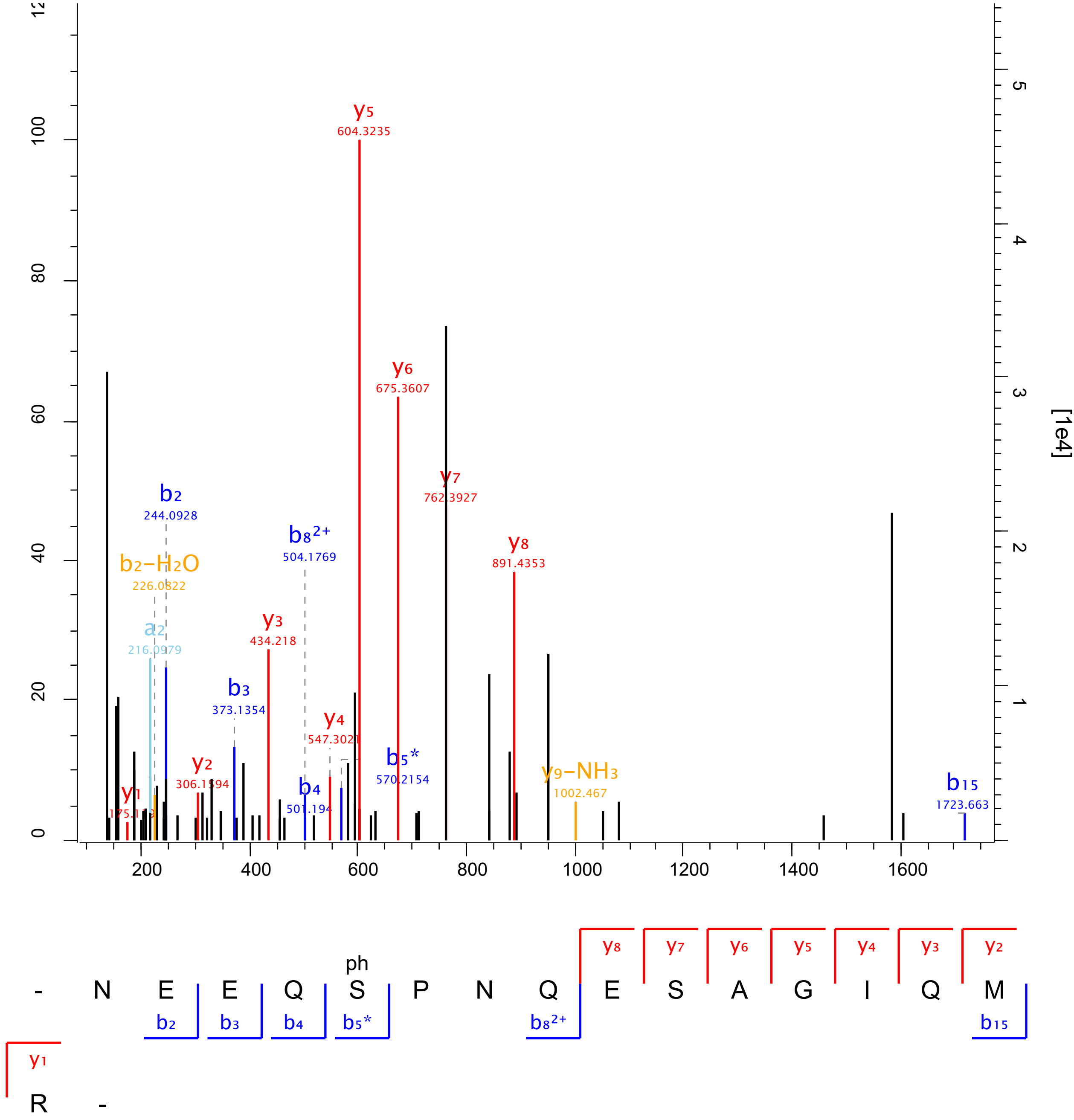

| Raw file | Scan | Method | Score | m/z |
| --- | --- | --- | --- | --- |
| A17_00_2 | 13951 | FTMS; HCD | 343.68 | 858.82 |

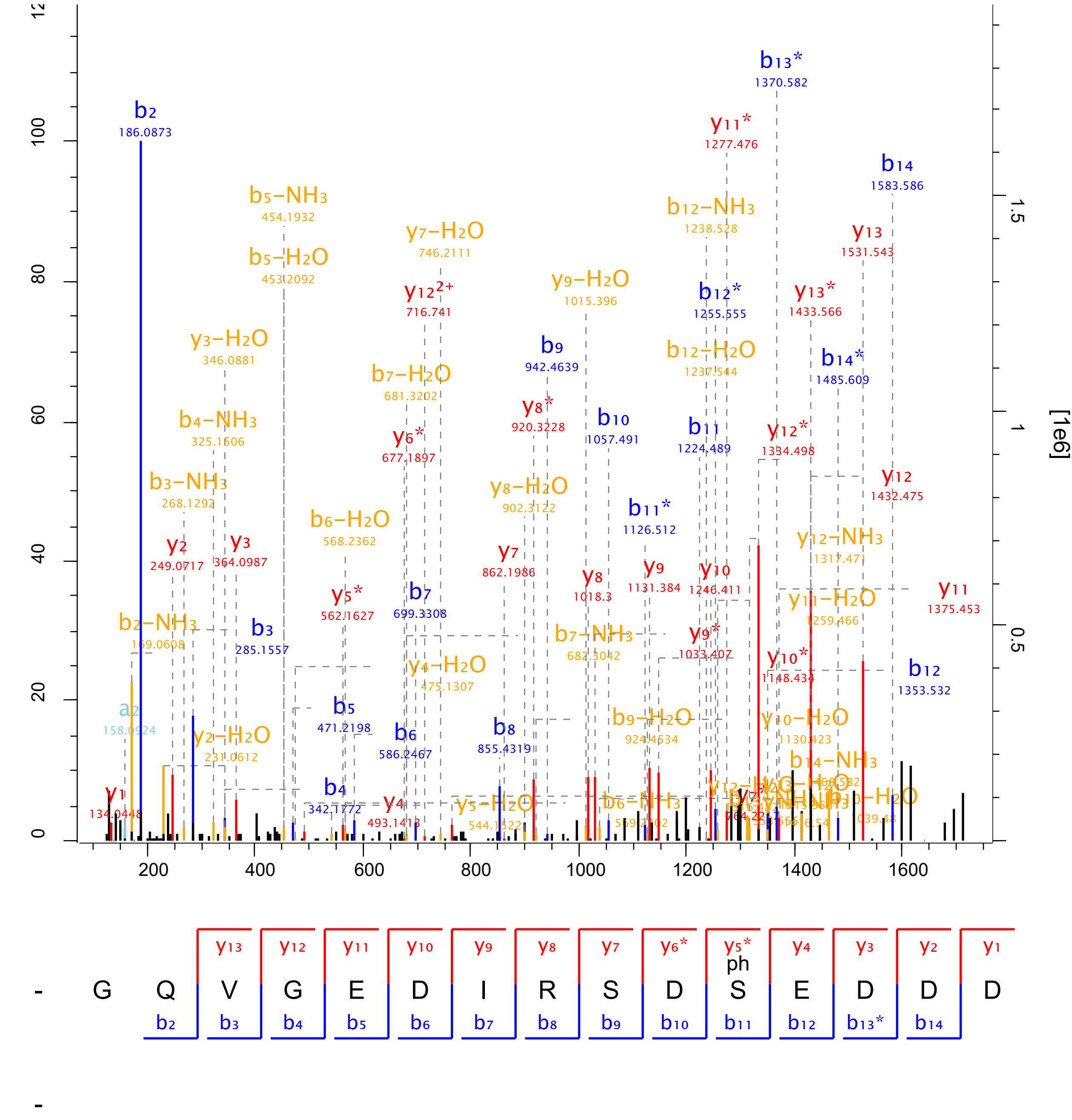

| Raw file | Scan | Method | Score | m/z |
|---|---|---|---|---|
| A17_00_2 | 13977 | FTMS; HCD | 337.91 | 998.36 |

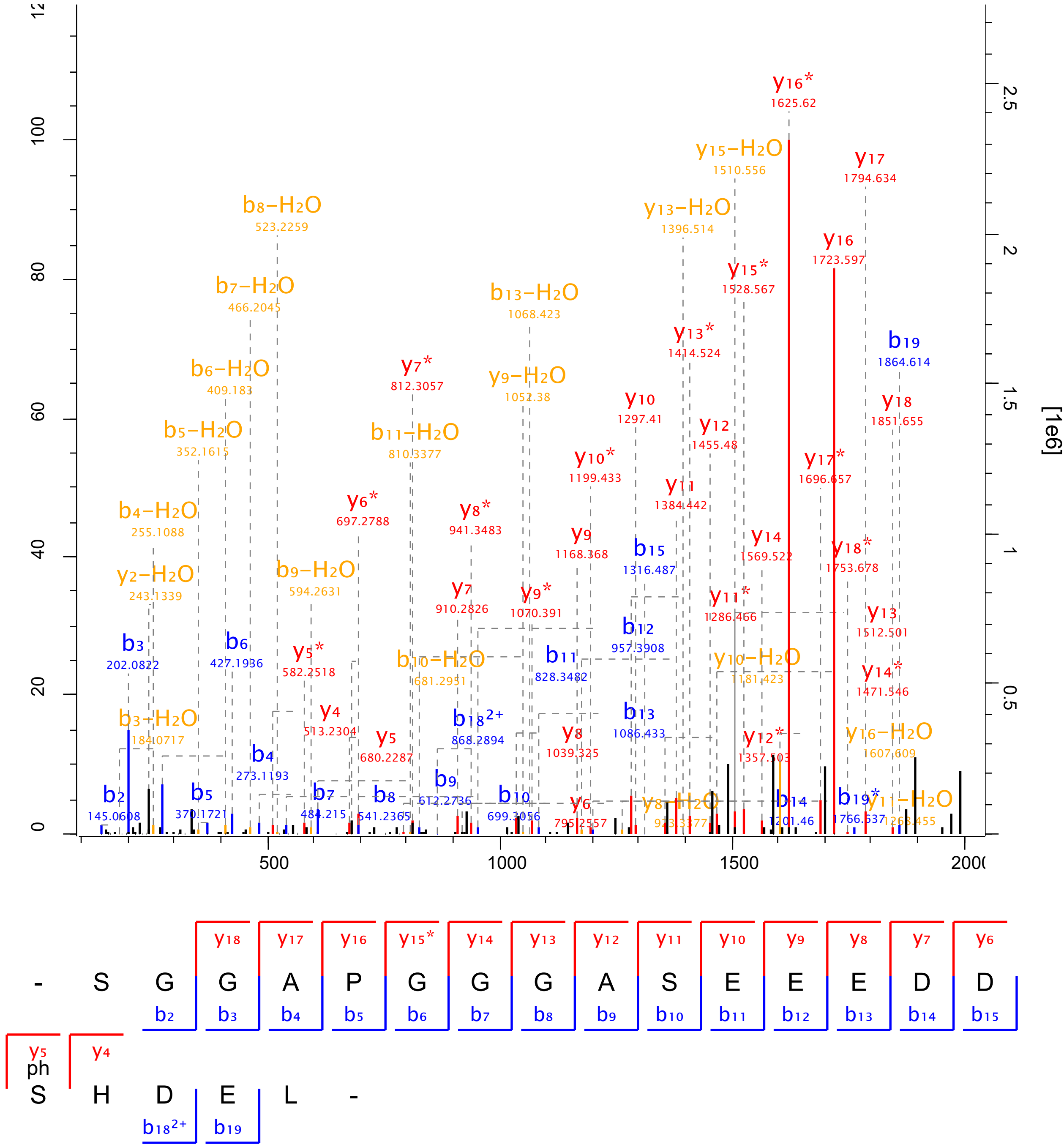

| Raw file | Scan | Method | Score | m/z |
|---|---|---|---|---|
| A17_00_2 | 13997 | FTMS; HCD | 48.57 | 524.21 |

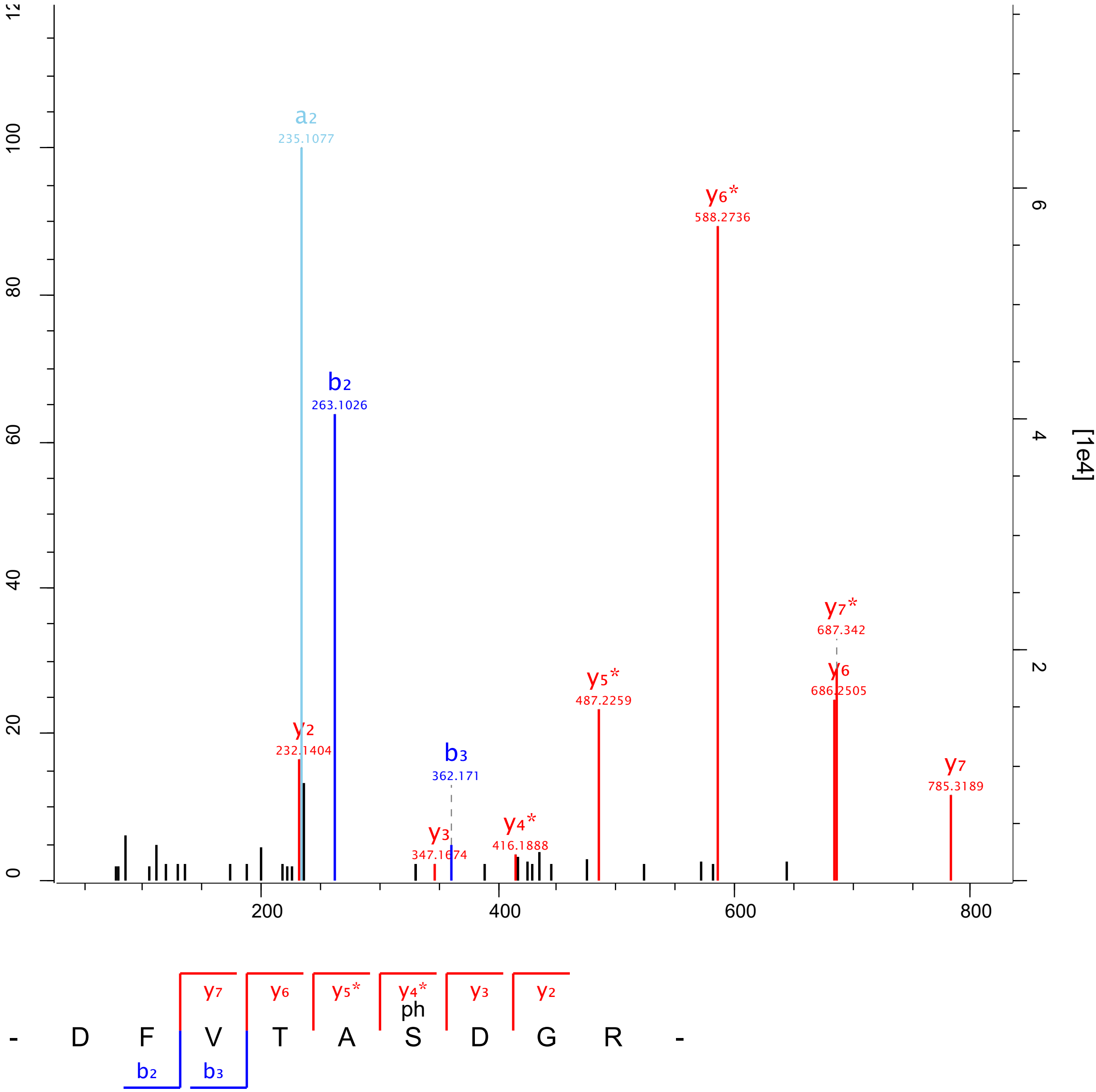

| Raw file | Scan | Method | Score | m/z |
| --- | --- | --- | --- | --- |
| A17_00_2 | 14010 | FTMS; HCD | 215.64 | 720.28 |

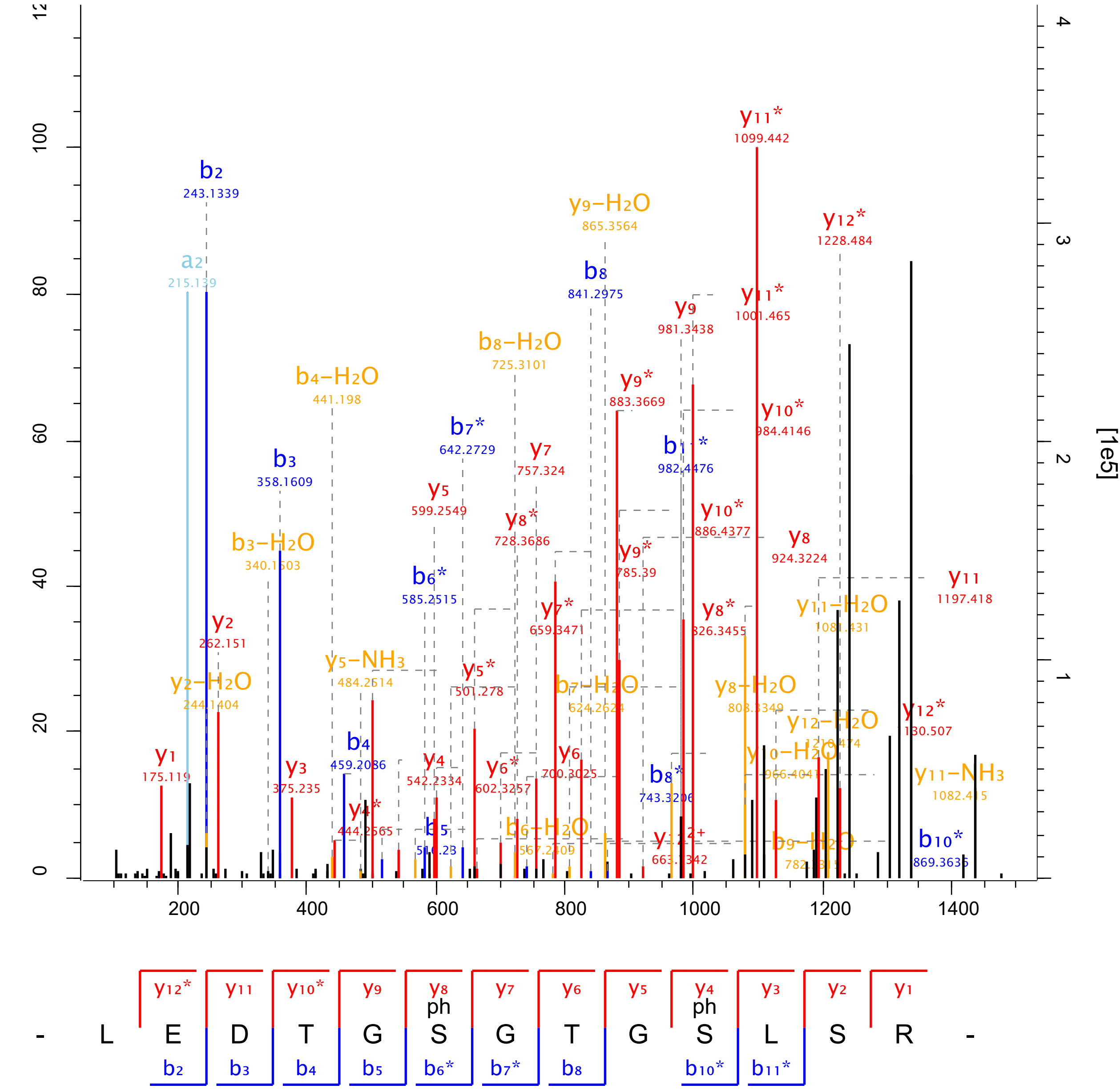

| Raw file | Scan | Method | Score | m/z |
| --- | --- | --- | --- | --- |
| A17_00_2 | 14067 | FTMS; HCD | 90.39 | 670.81 |

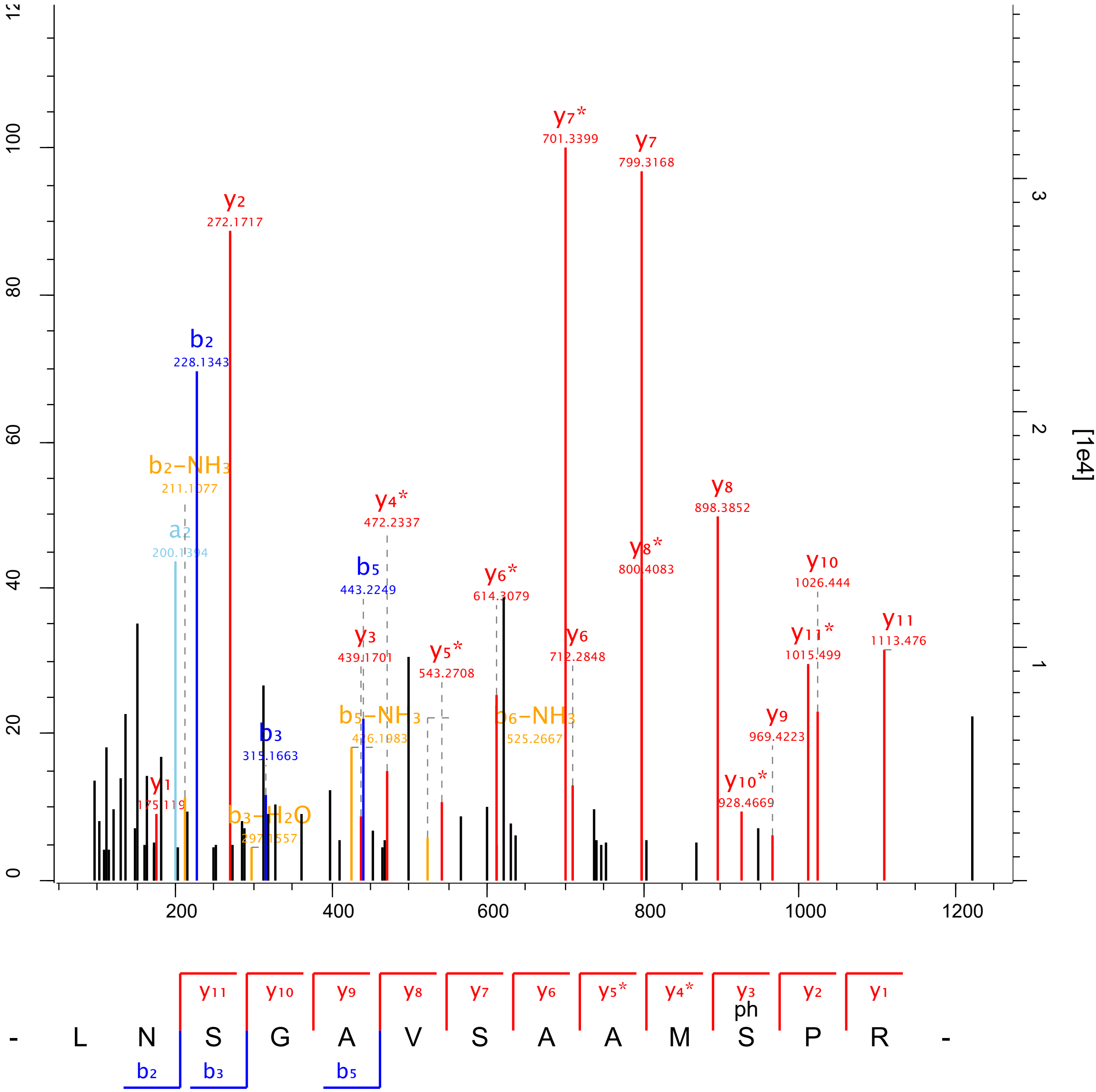

| Raw file | Scan | Method | Score | m/z |
|---|---|---|---|---|
| A17_00_2 | 14081 | FTMS; HCD | 188.19 | 651.28 |

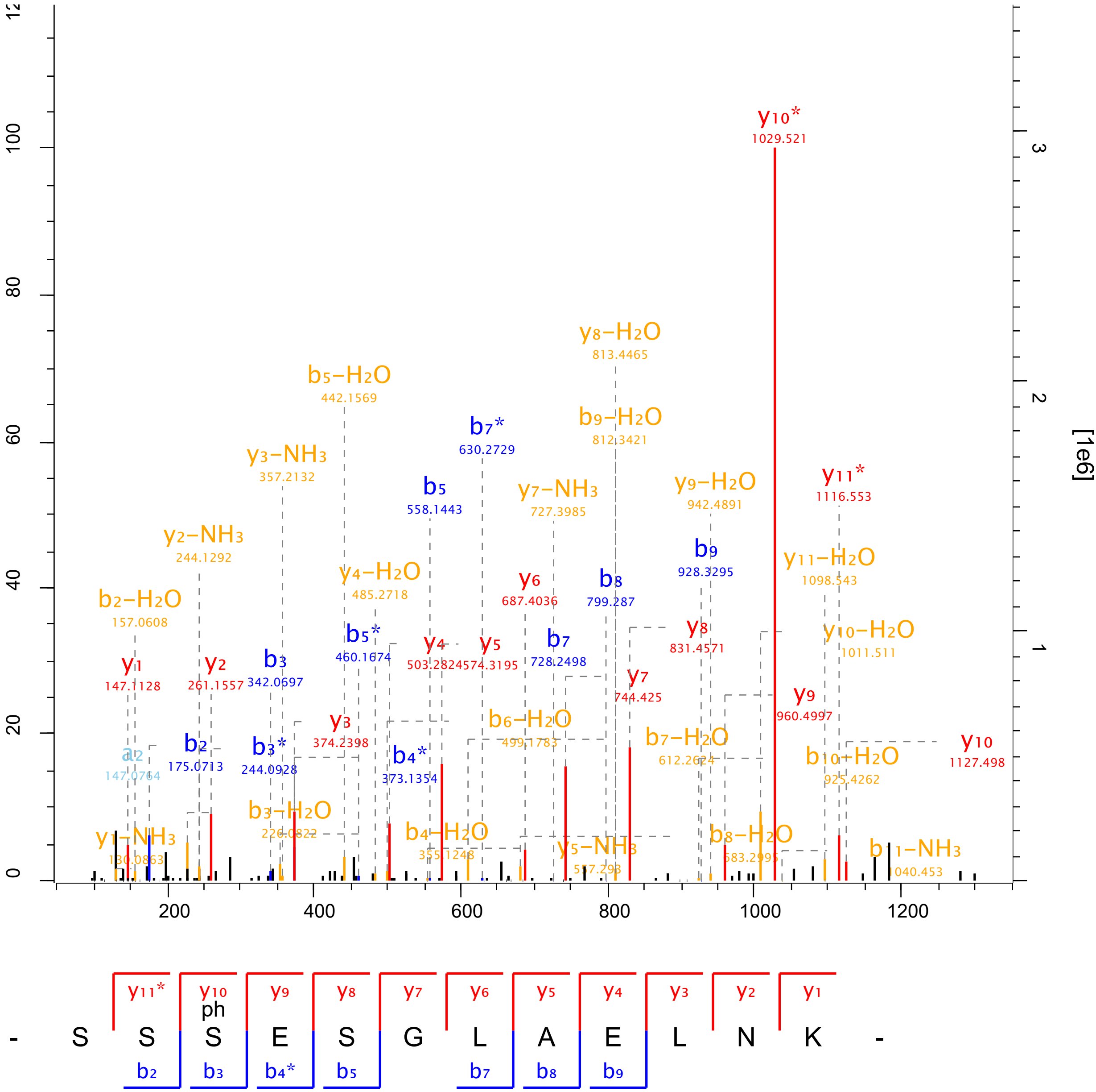

| Raw file | Scan | Method | Score | m/z |
|---|---|---|---|---|
| A17_00_2 | 14123 | FTMS; HCD | 54.52 | 795.96 |

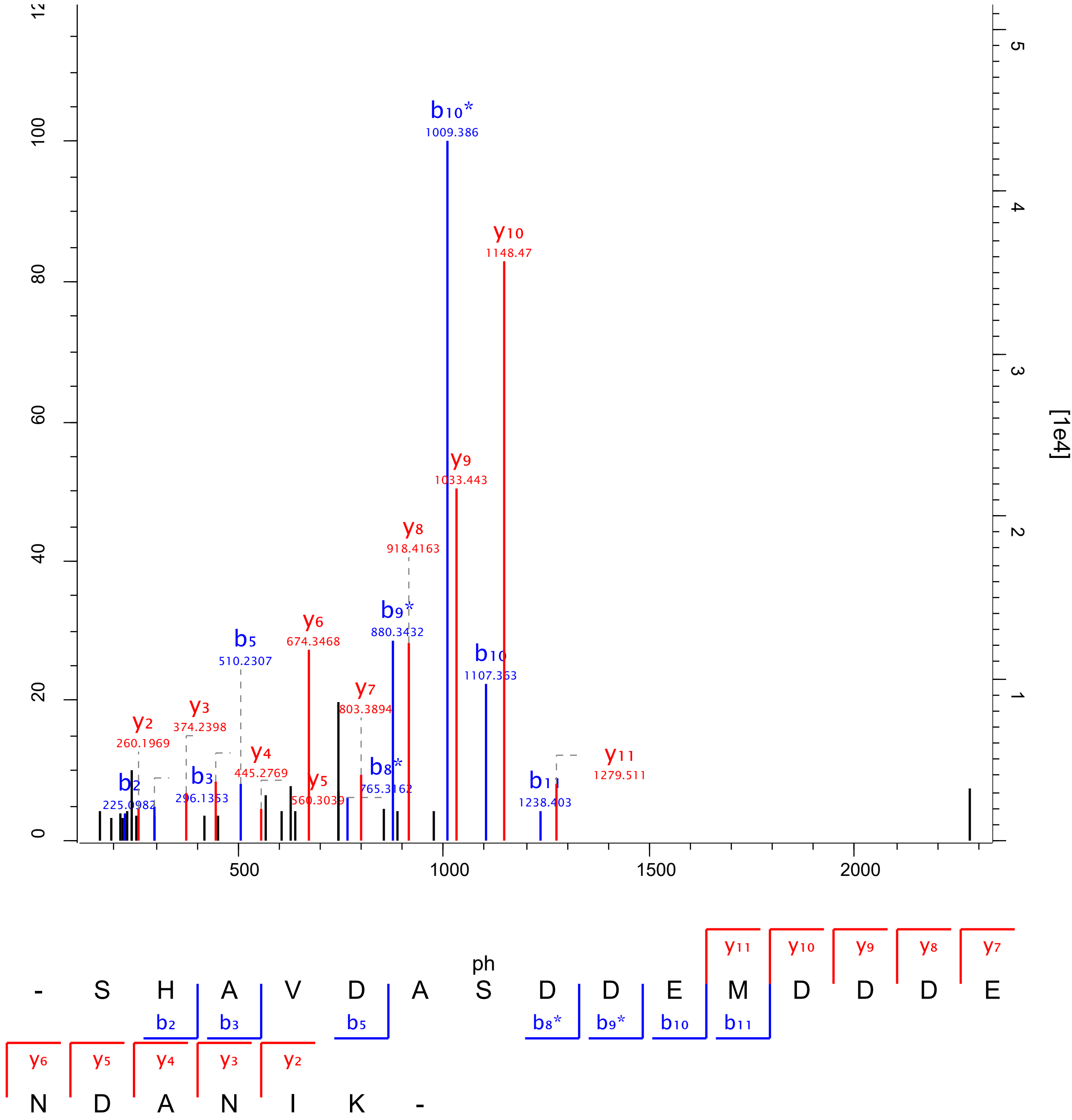

| Raw file | Scan | Method | Score | m/z |
| --- | --- | --- | --- | --- |
| A17_00_2 | 14124 | FTMS; HCD | 46.89 | 476.93 |

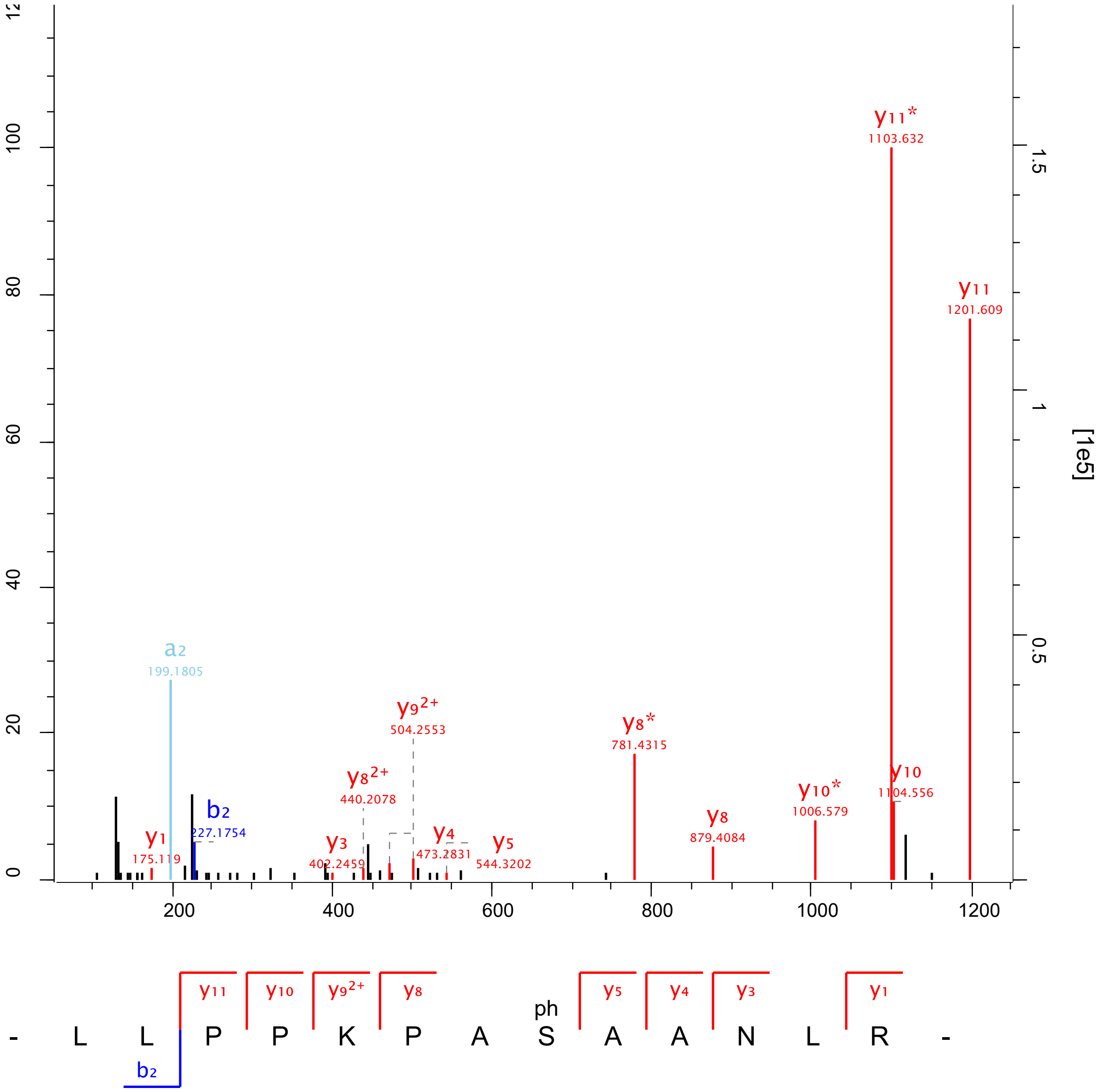

| Raw file | Scan | Method | Score | m/z |
|---|---|---|---|---|
| A17_00_2 | 14355 | FTMS; HCD | 155.09 | 606.26 |

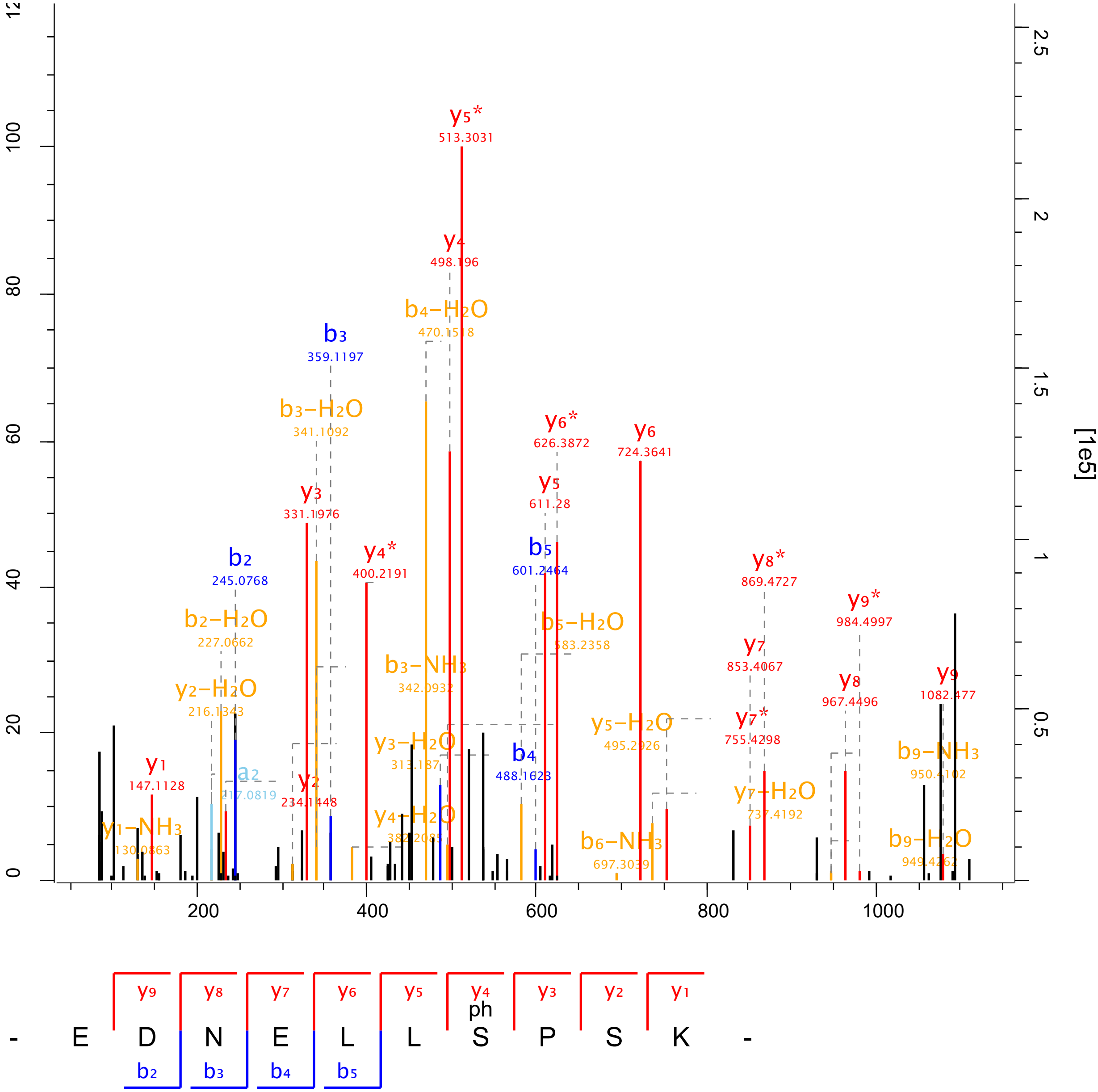

| Raw file | Scan | Method | Score | m/z |
|---|---|---|---|---|
| A17_00_2 | 14364 | FTMS; HCD | 41.4 | 568.26 |

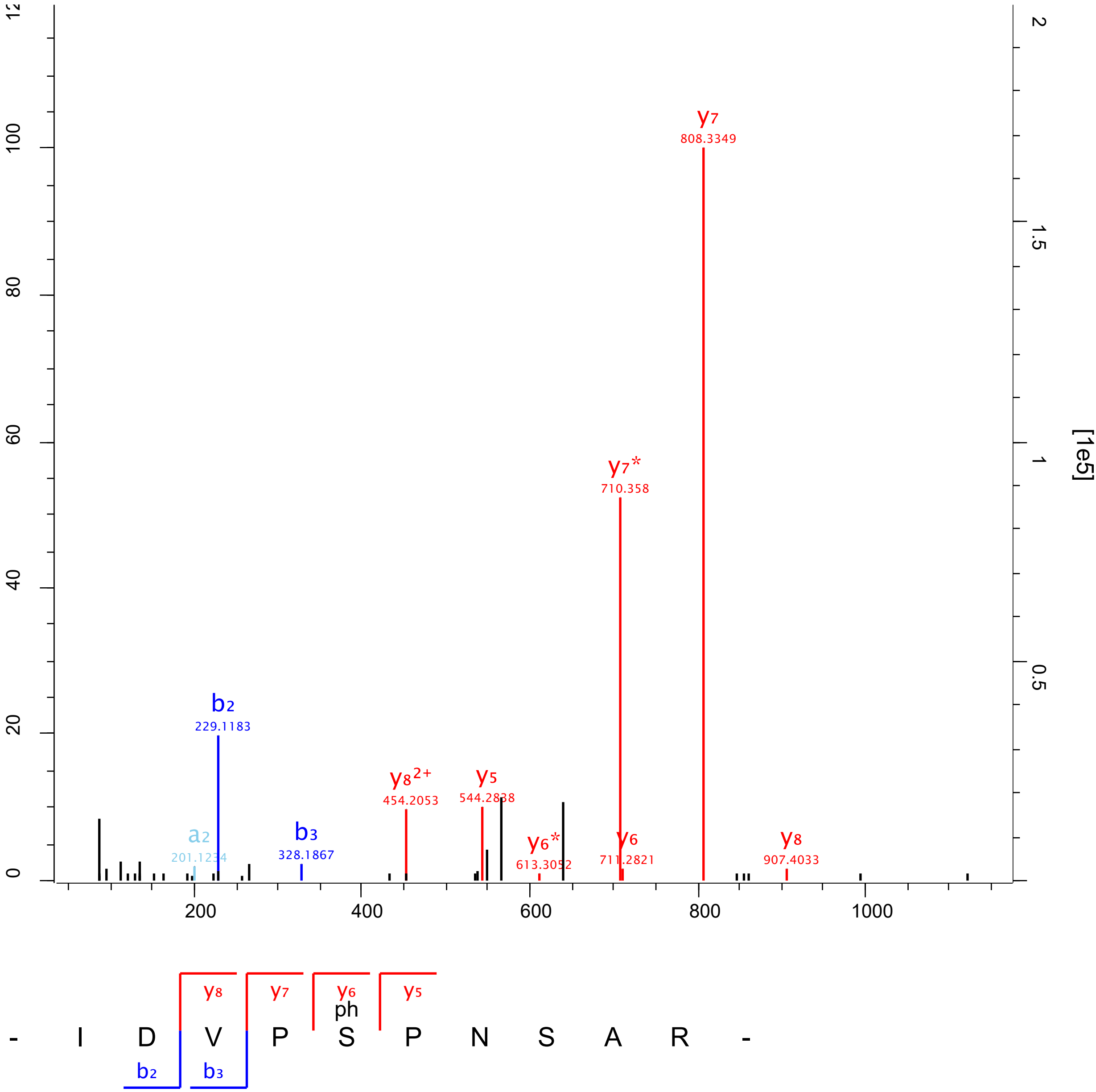

| Raw file | Scan | Method | Score | m/z |
|---|---|---|---|---|
| A17_00_2 | 14485 | FTMS; HCD | 77.75 | 693.98 |

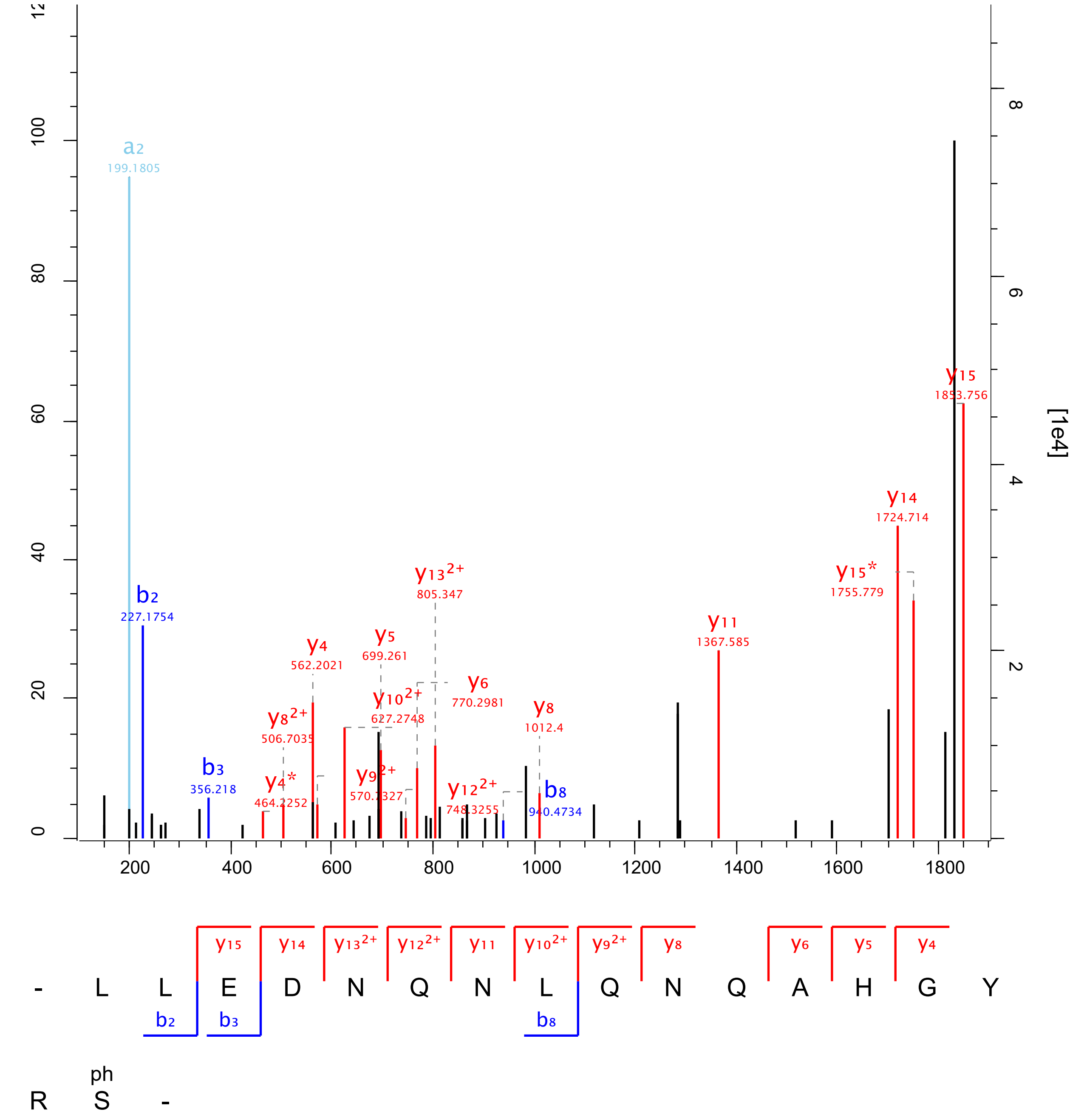

| Raw file | Scan | Method | Score | m/z |
|---|---|---|---|---|
| A17_00_2 | 14498 | FTMS; HCD | 142.62 | 630.28 |

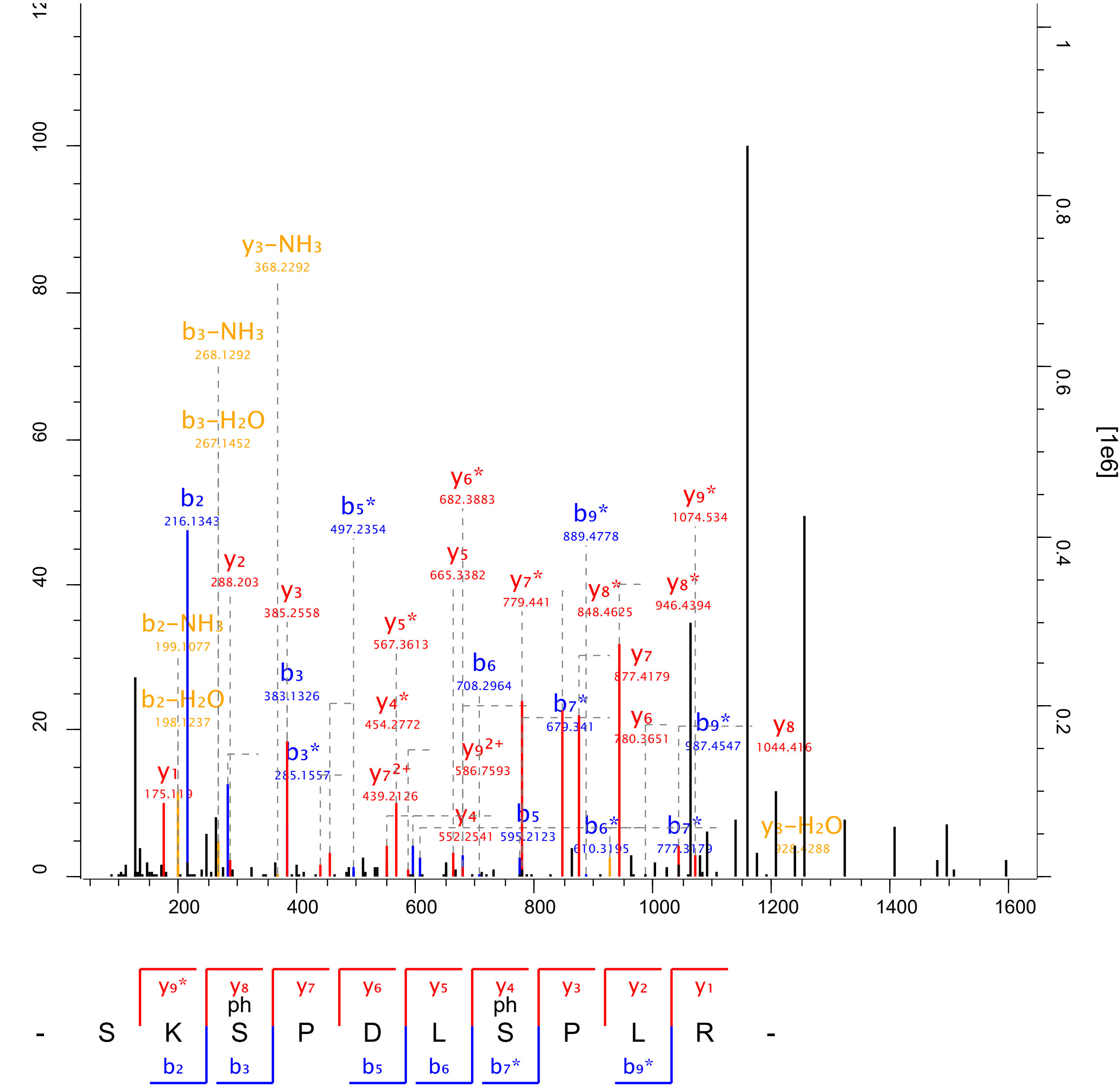

| Raw file | Scan | Method | Score | m/z |
|---|---|---|---|---|
| A17_00_2 | 14582 | FTMS; HCD | 40.94 | 599.74 |

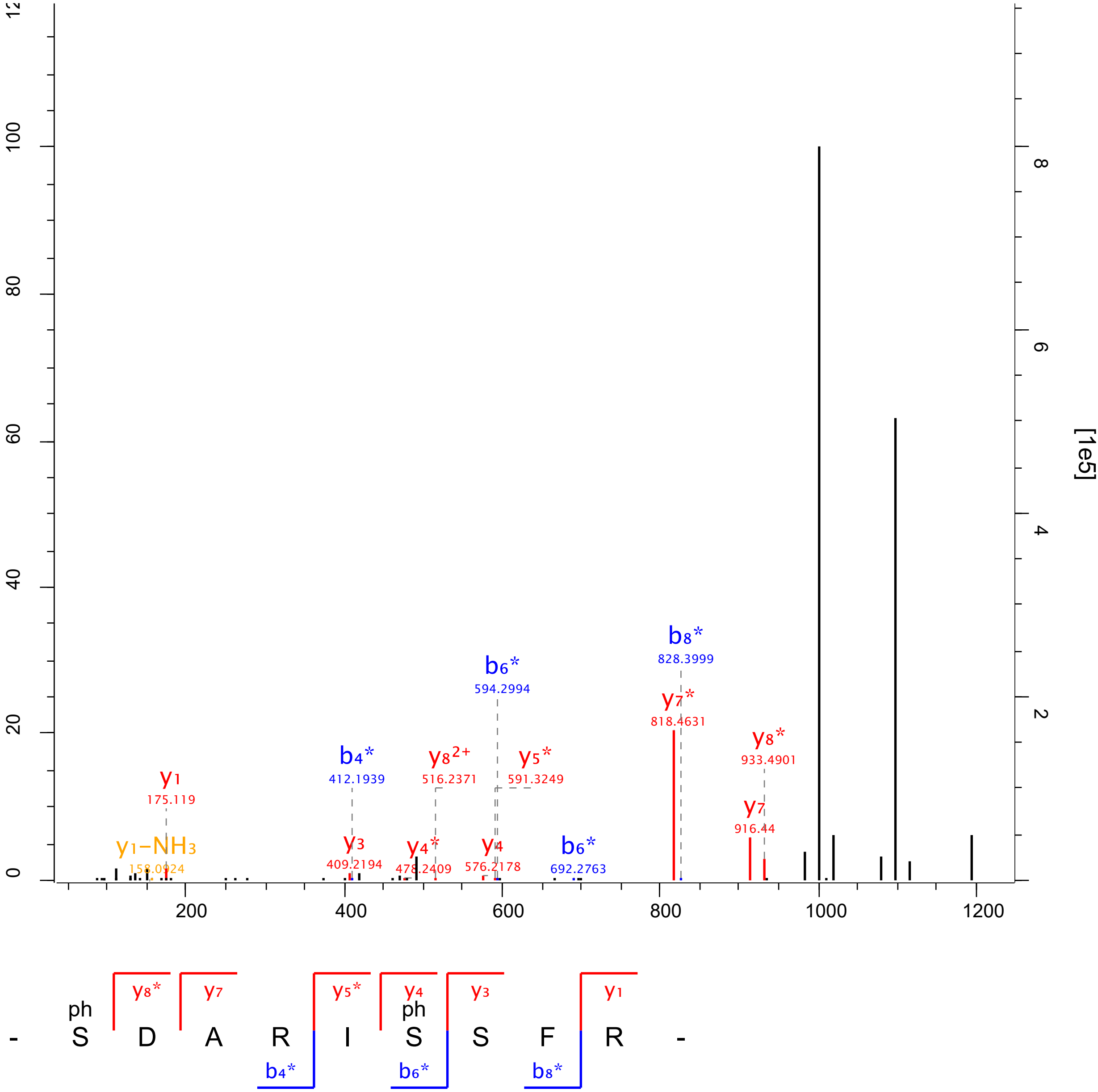

| Raw file | Scan | Method | Score | m/z |
|---|---|---|---|---|
| A17_00_2 | 14617 | FTMS; HCD | 136.02 | 485.23 |

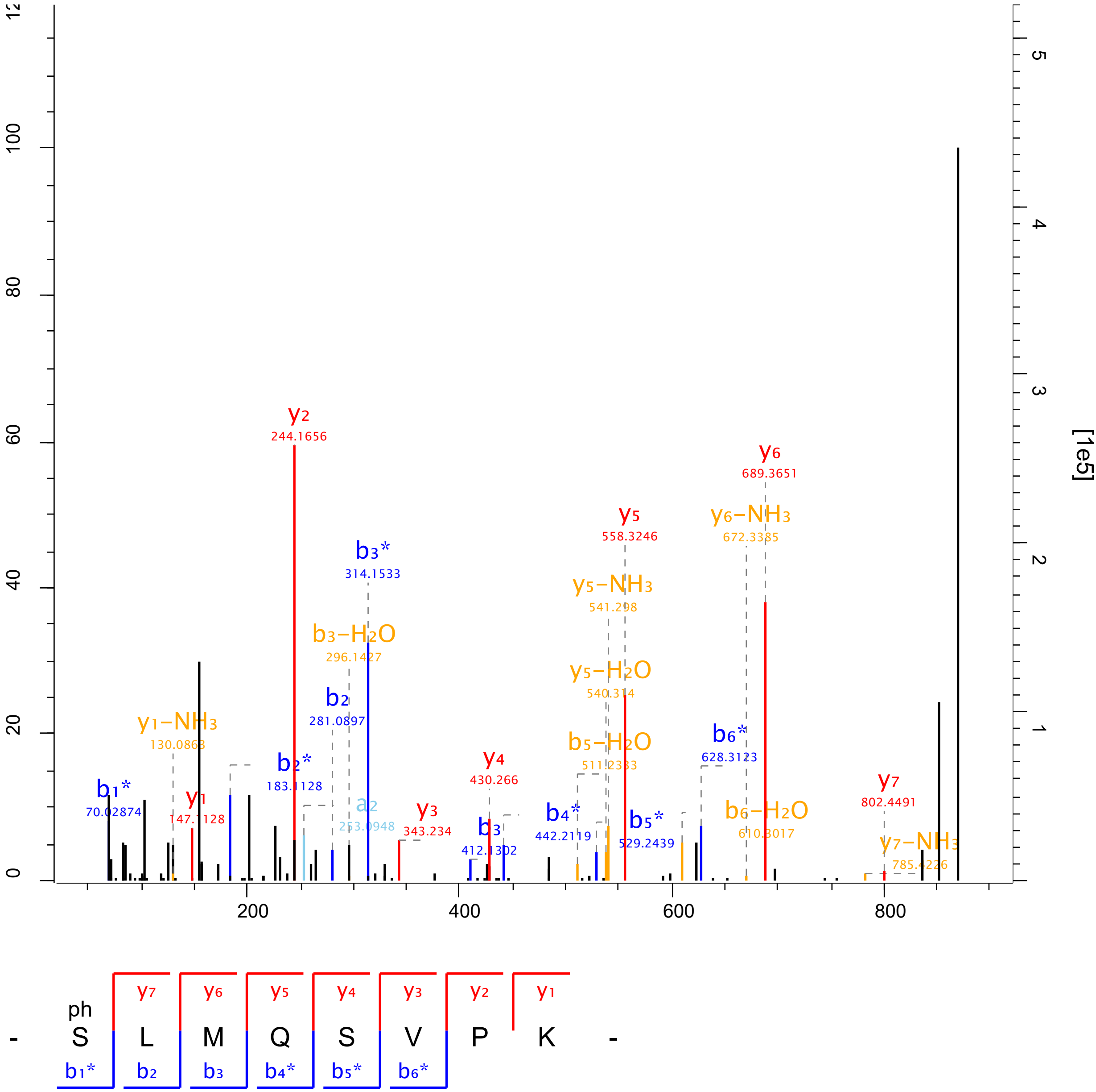

| Raw file | Scan | Method | Score | m/z |
|---|---|---|---|---|
| A17_00_2 | 14642 | FTMS; HCD | 162.81 | 866.32 |

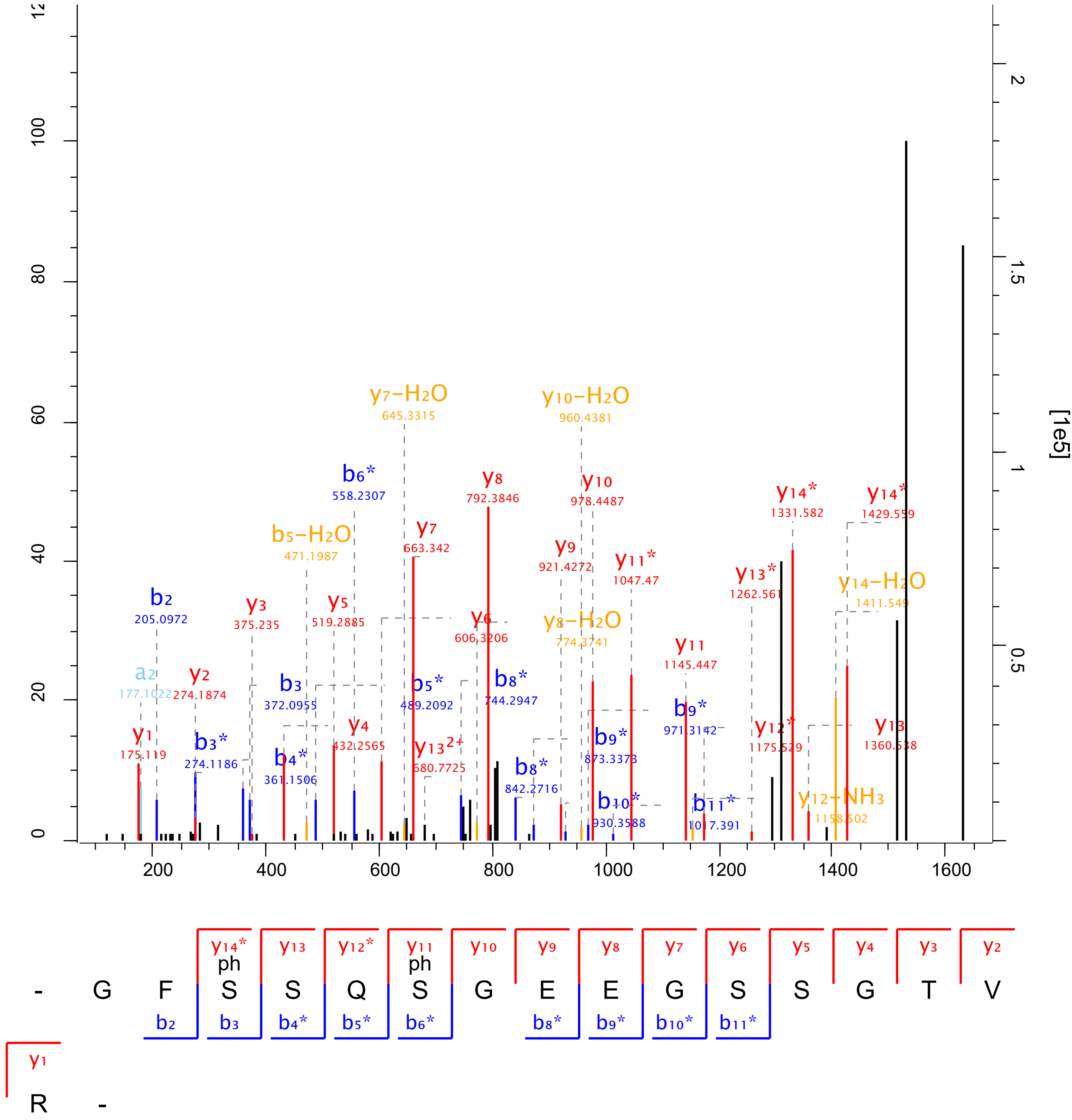

| Raw file | Scan | Method | Score | m/z |
|---|---|---|---|---|
| A17_00_2 | 1470 | FTMS; HCD | 41.58 | 333.91 |

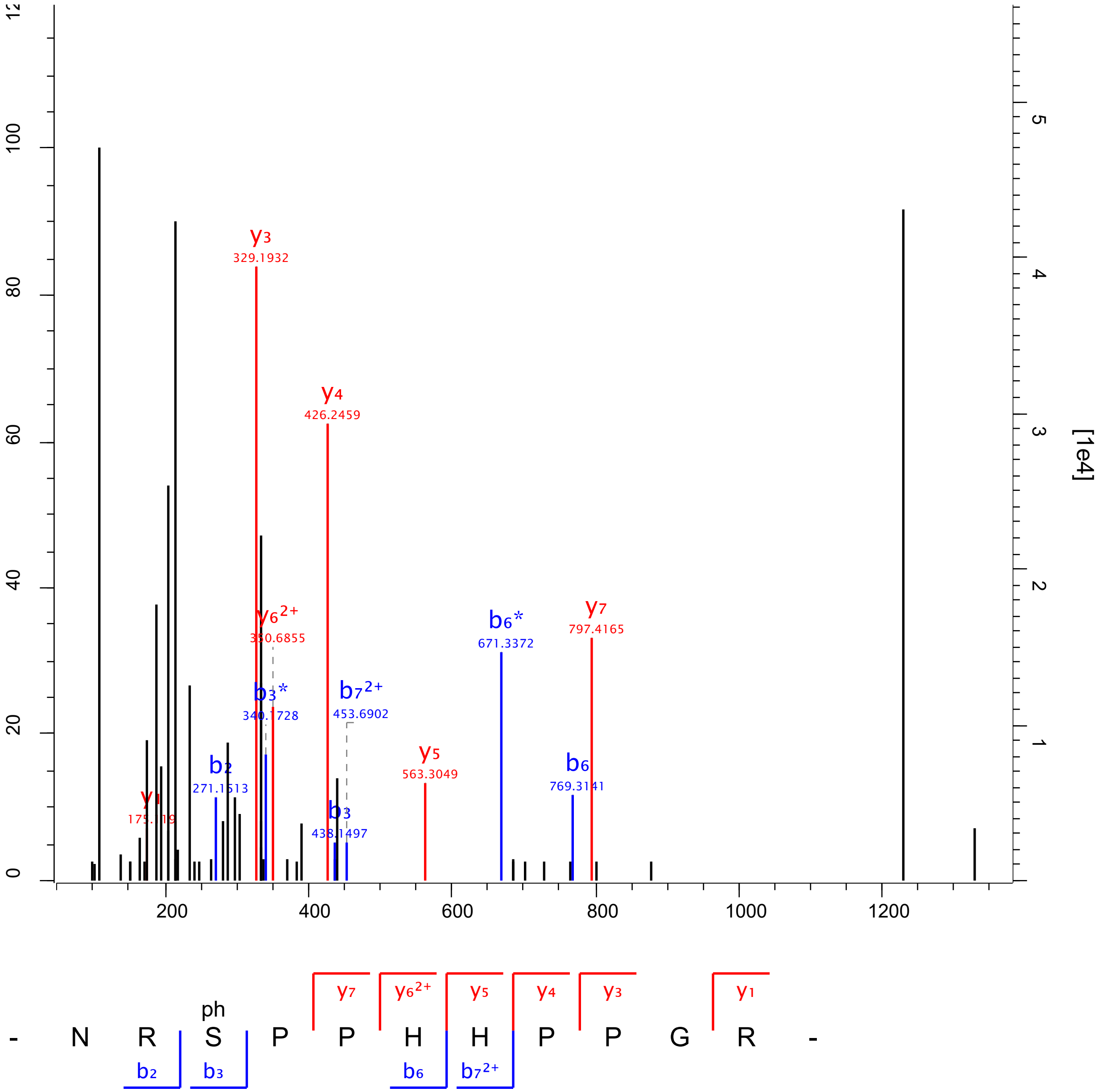

| Raw file | Scan | Method | Score | m/z |
| --- | --- | --- | --- | --- |
| A17_00_2 | 14730 | FTMS; HCD | 140.45 | 501.73 |

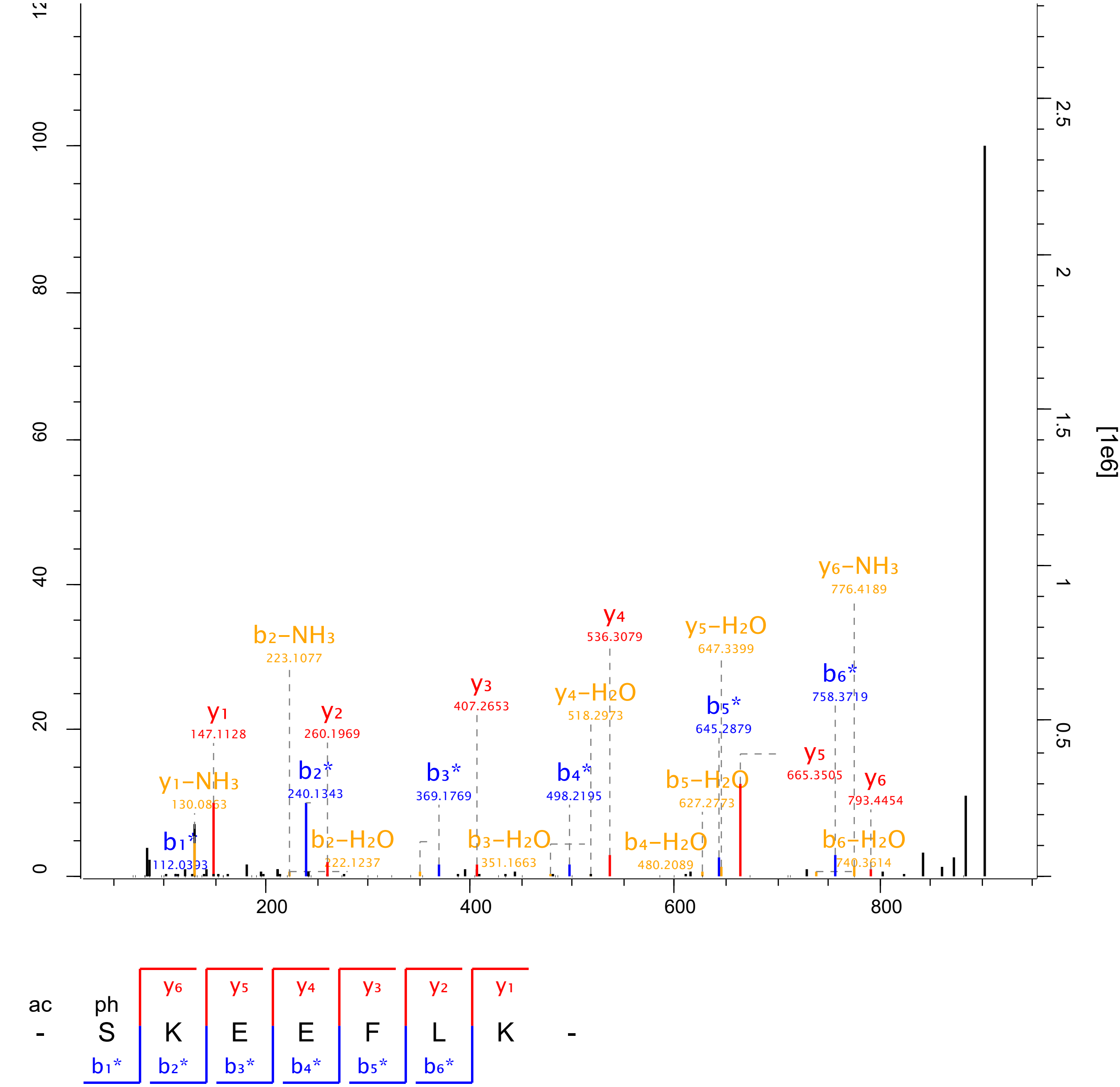

| Raw file | Scan | Method | Score | m/z |
|---|---|---|---|---|
| A17_00_2 | 1475 | FTMS; HCD | 58.71 | 577.21 |

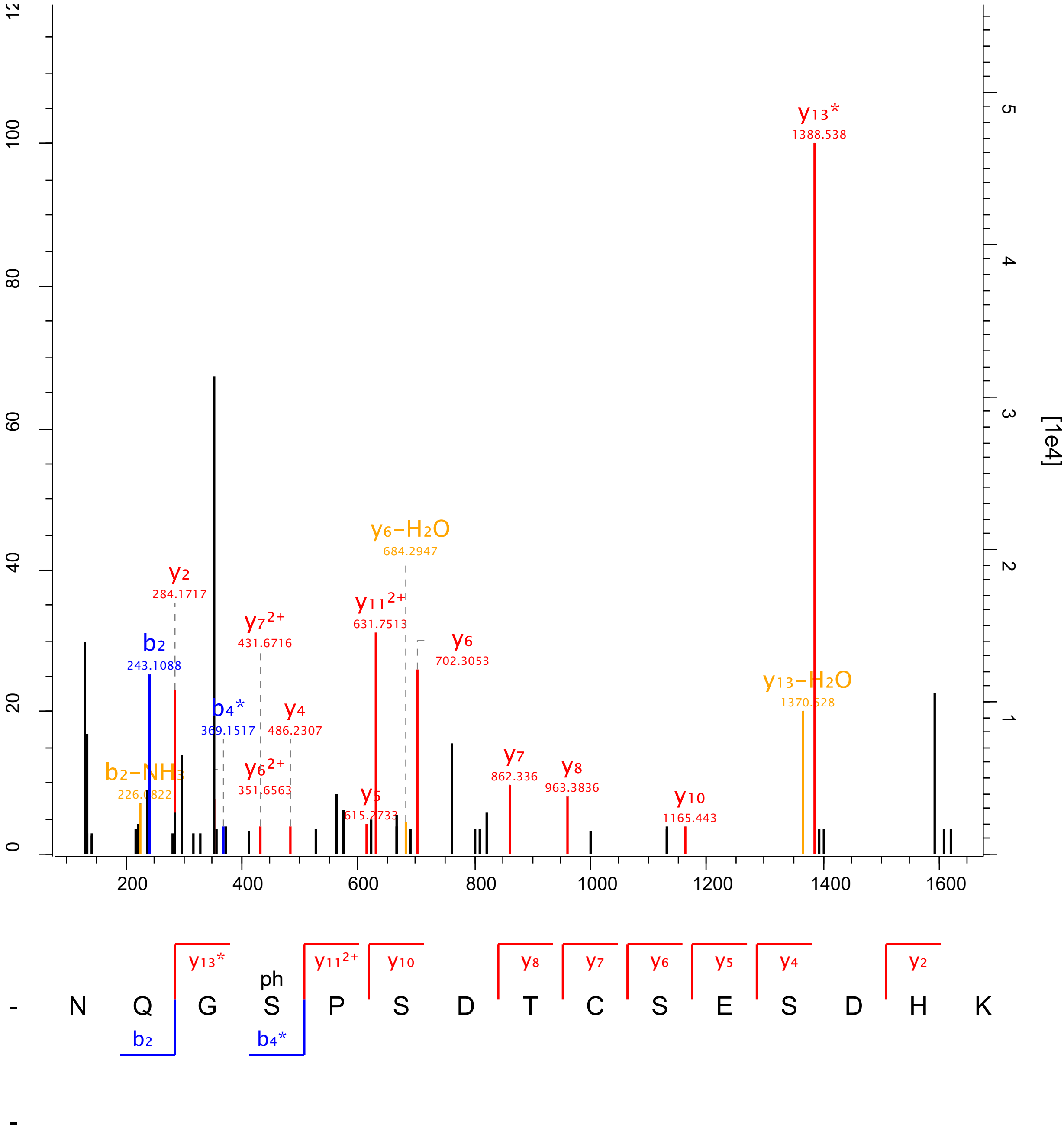

| Raw file | Scan | Method | Score | m/z |
| --- | --- | --- | --- | --- |
| A17_00_2 | 1492 | FTMS; HCD | 64.26 | 578.23 |

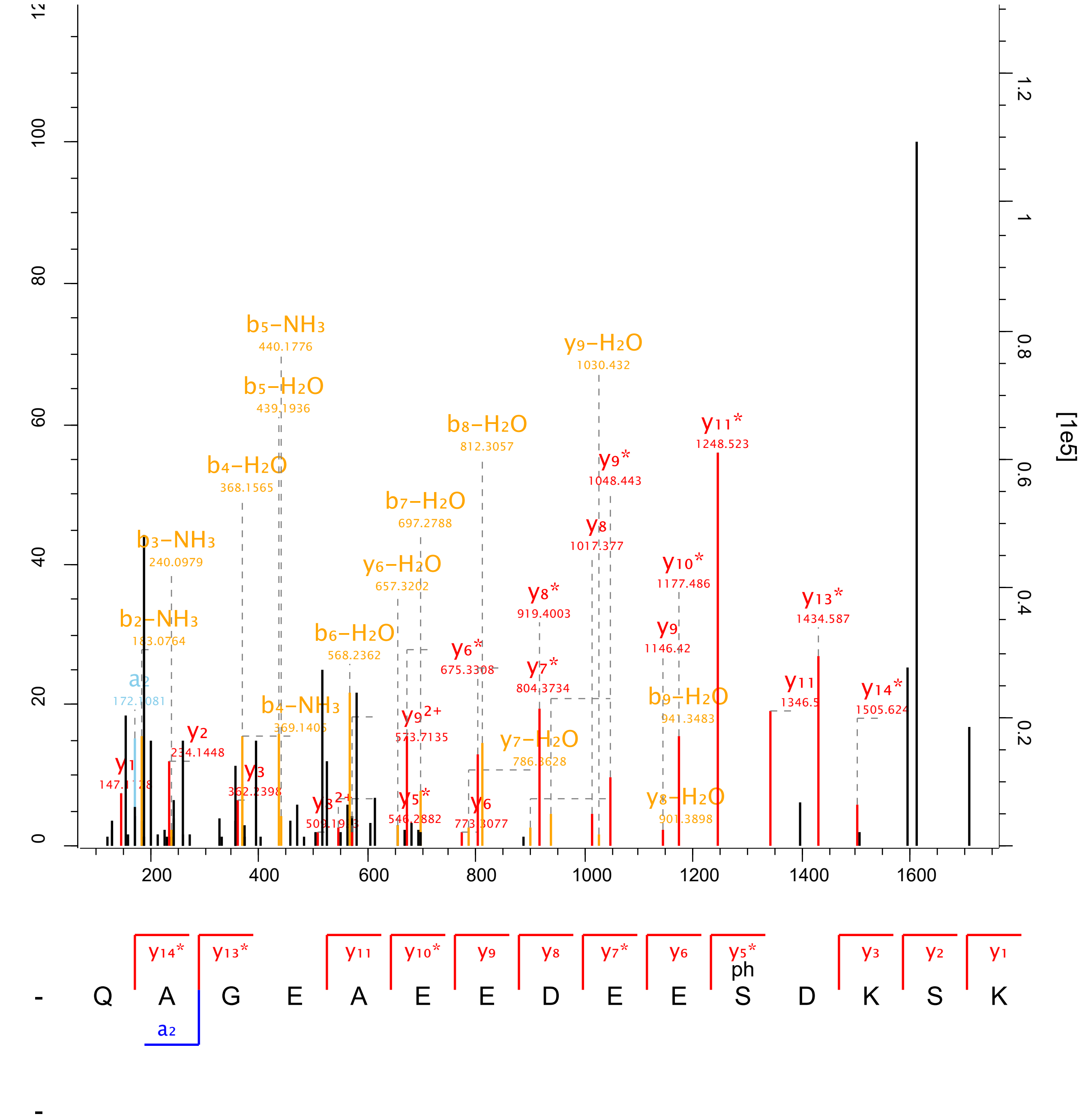

| Raw file | Scan | Method | Score | m/z |
|---|---|---|---|---|
| A17_00_2 | 1511 | FTMS; HCD | 41.4 | 353.53 |

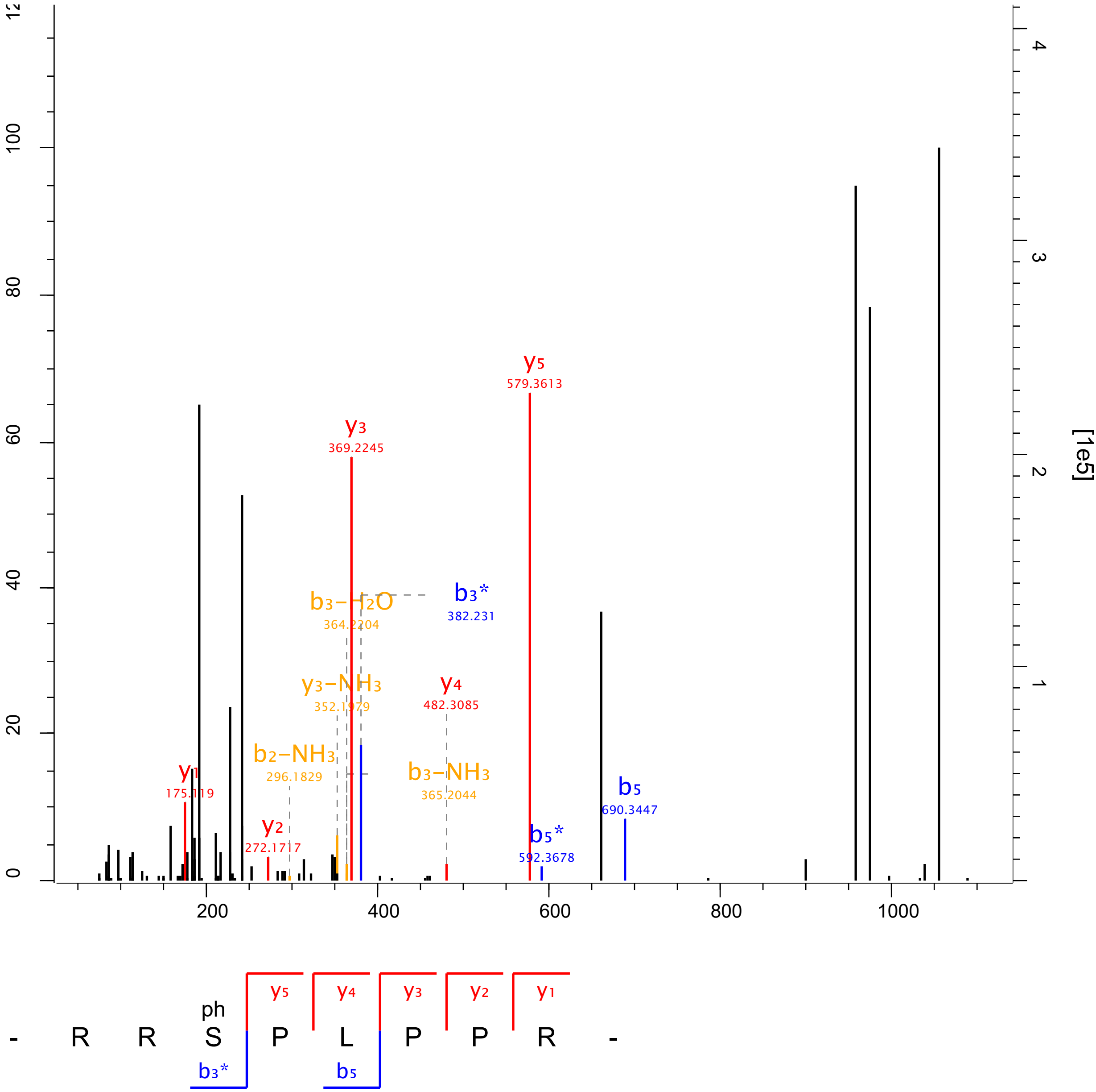

| Raw file | Scan | Method | Score | m/z |
|---|---|---|---|---|
| A17_00_2 | 15112 | FTMS; HCD | 154.19 | 578.74 |

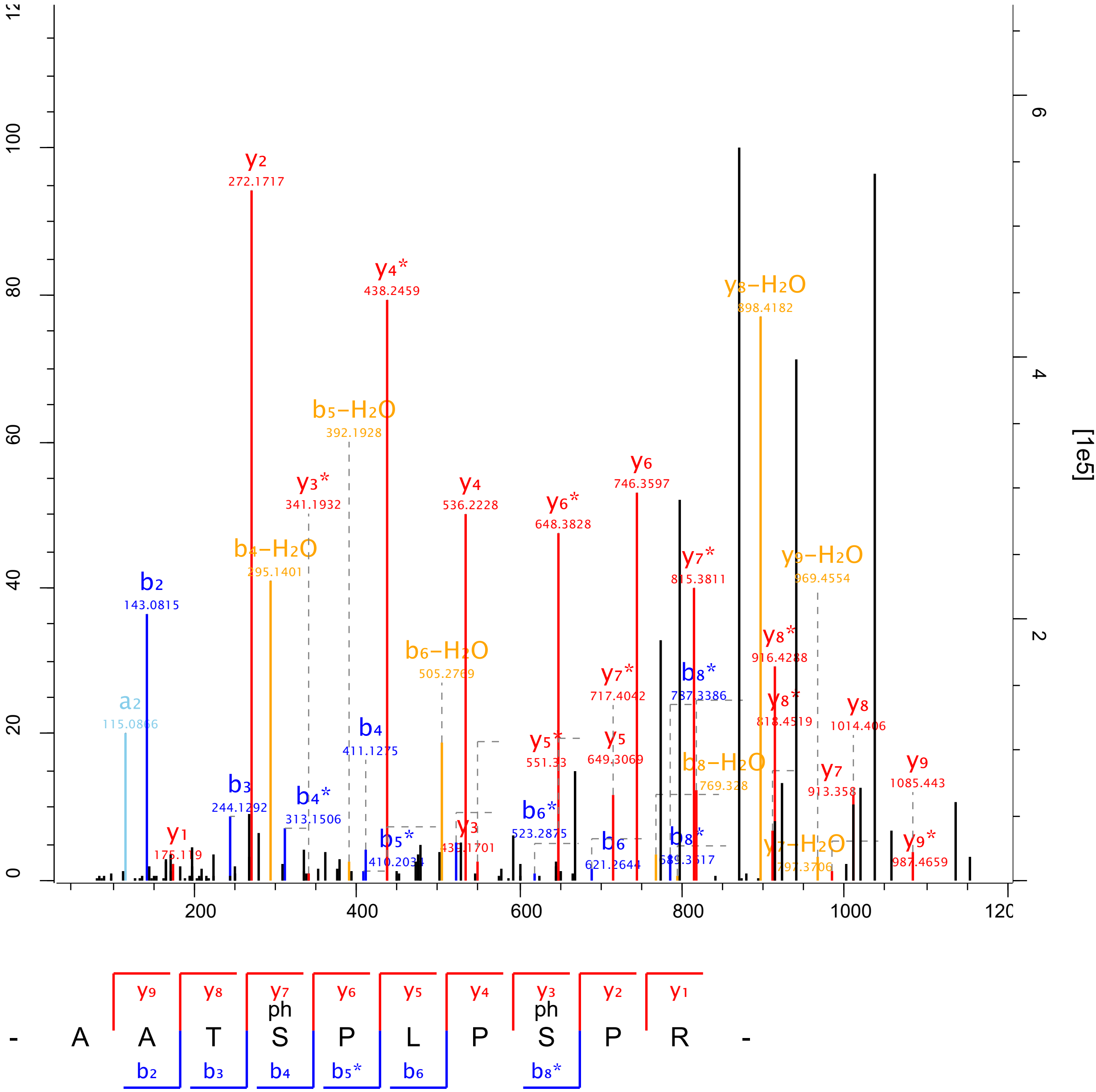

| Raw file | Scan | Method | Score | m/z |
| --- | --- | --- | --- | --- |
| A17_00_2 | 15145 | FTMS; HCD | 92.04 | 842.33 |

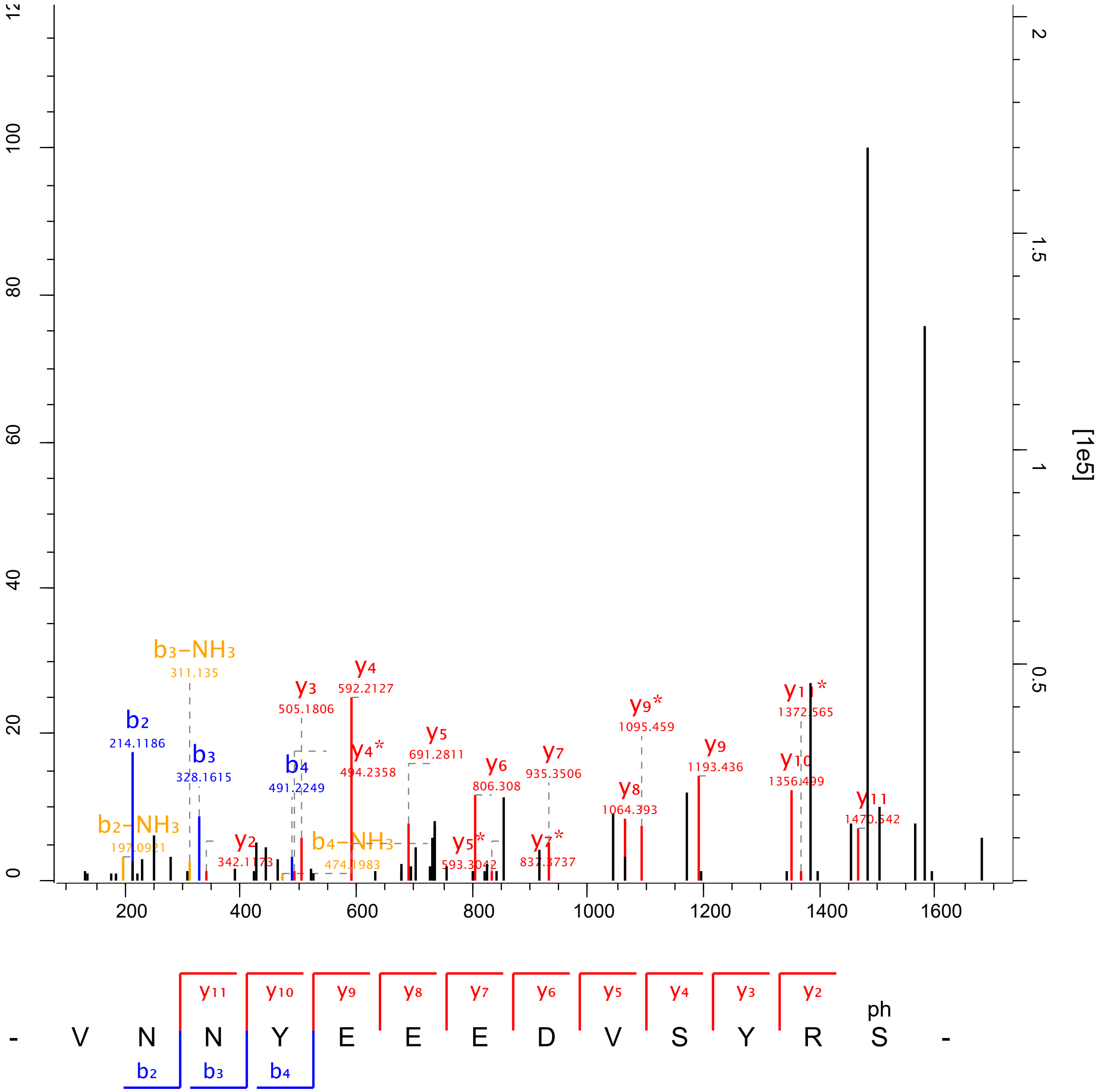

| Raw file | Scan | Method | Score | m/z |
|---|---|---|---|---|
| A17_00_2 | 15171 | FTMS; HCD | 118.76 | 598.72 |

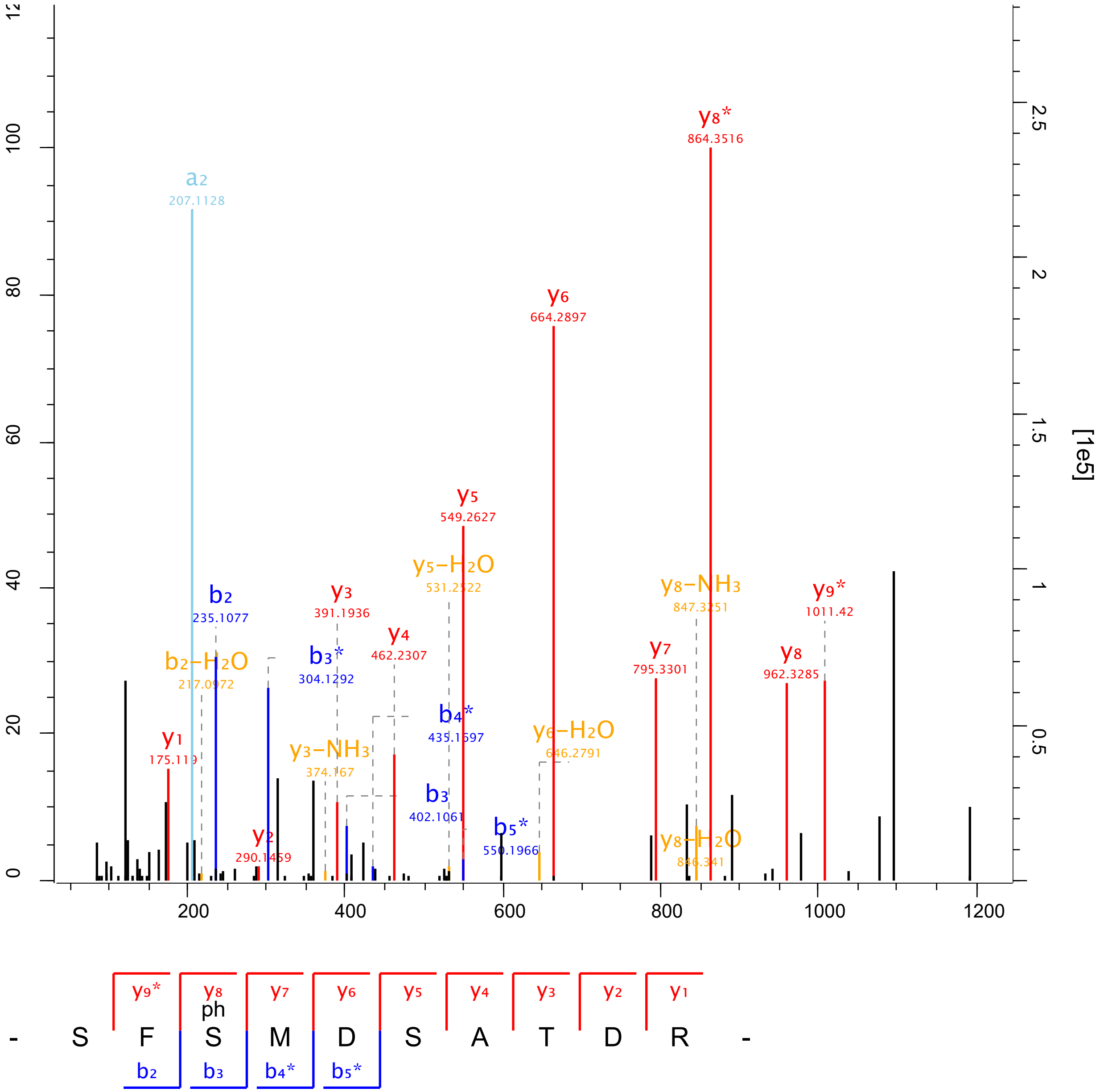

| Raw file | Scan | Method | Score | m/z |
|---|---|---|---|---|
| A17_00_2 | 15334 | FTMS; HCD | 43.21 | 611.25 |

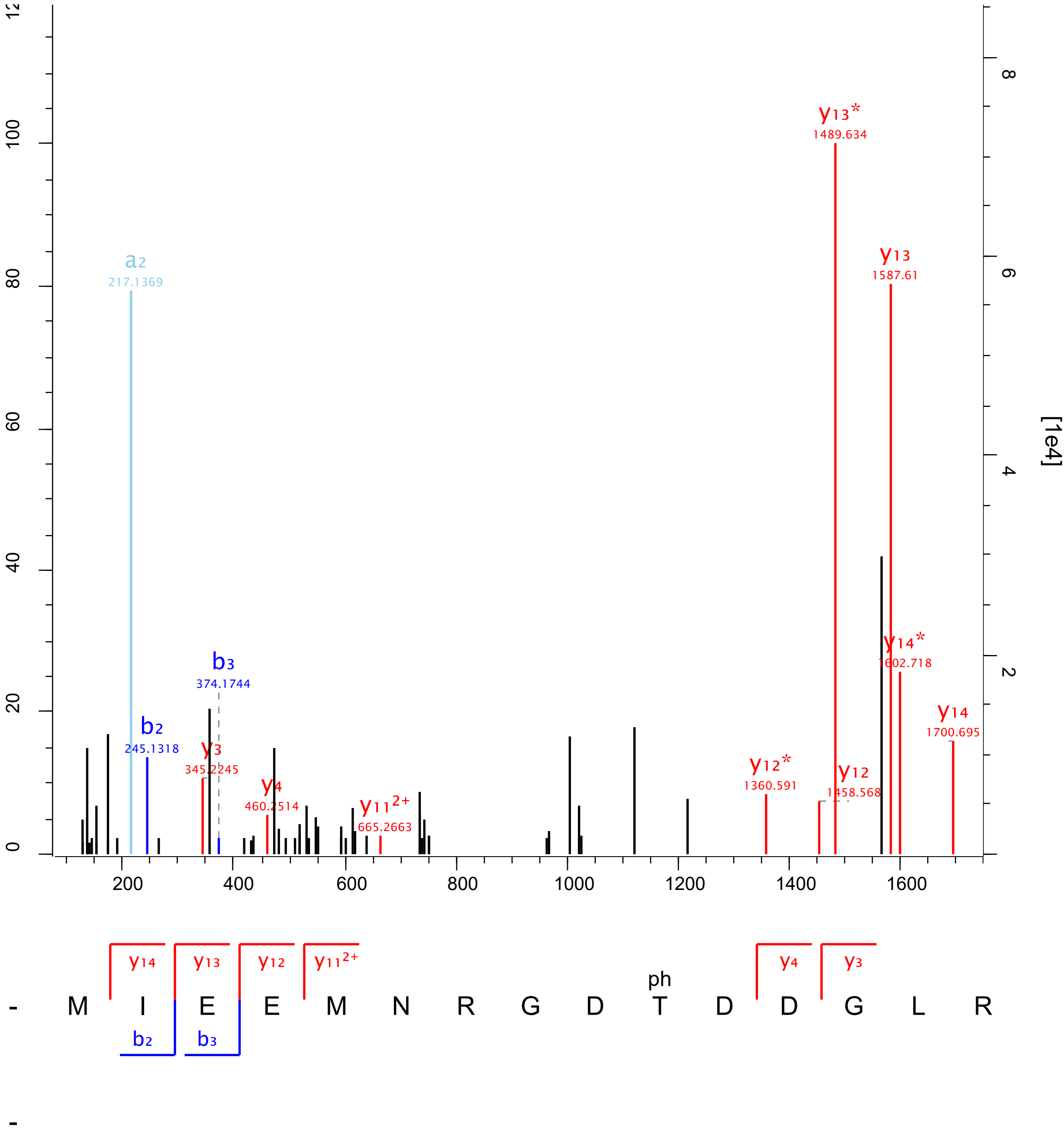

| Raw file | Scan | Method | Score | m/z |
| --- | --- | --- | --- | --- |
| A17_00_2 | 1535 | FTMS; HCD | 45.88 | 476.54 |

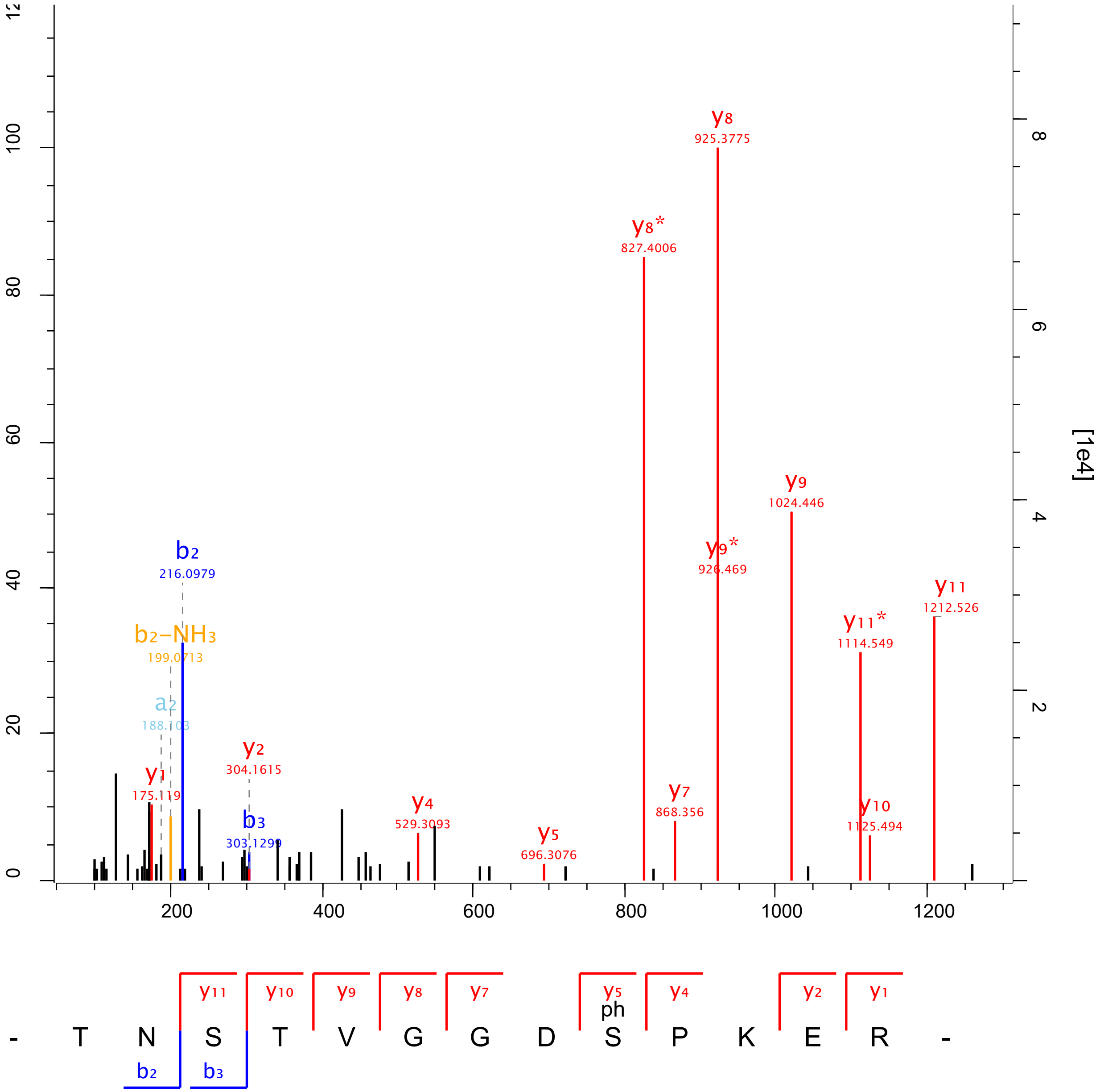

| Raw file | Scan | Method | Score | m/z |
|---|---|---|---|---|
| A17_00_2 | 15364 | FTMS; HCD | 159.06 | 958.86 |

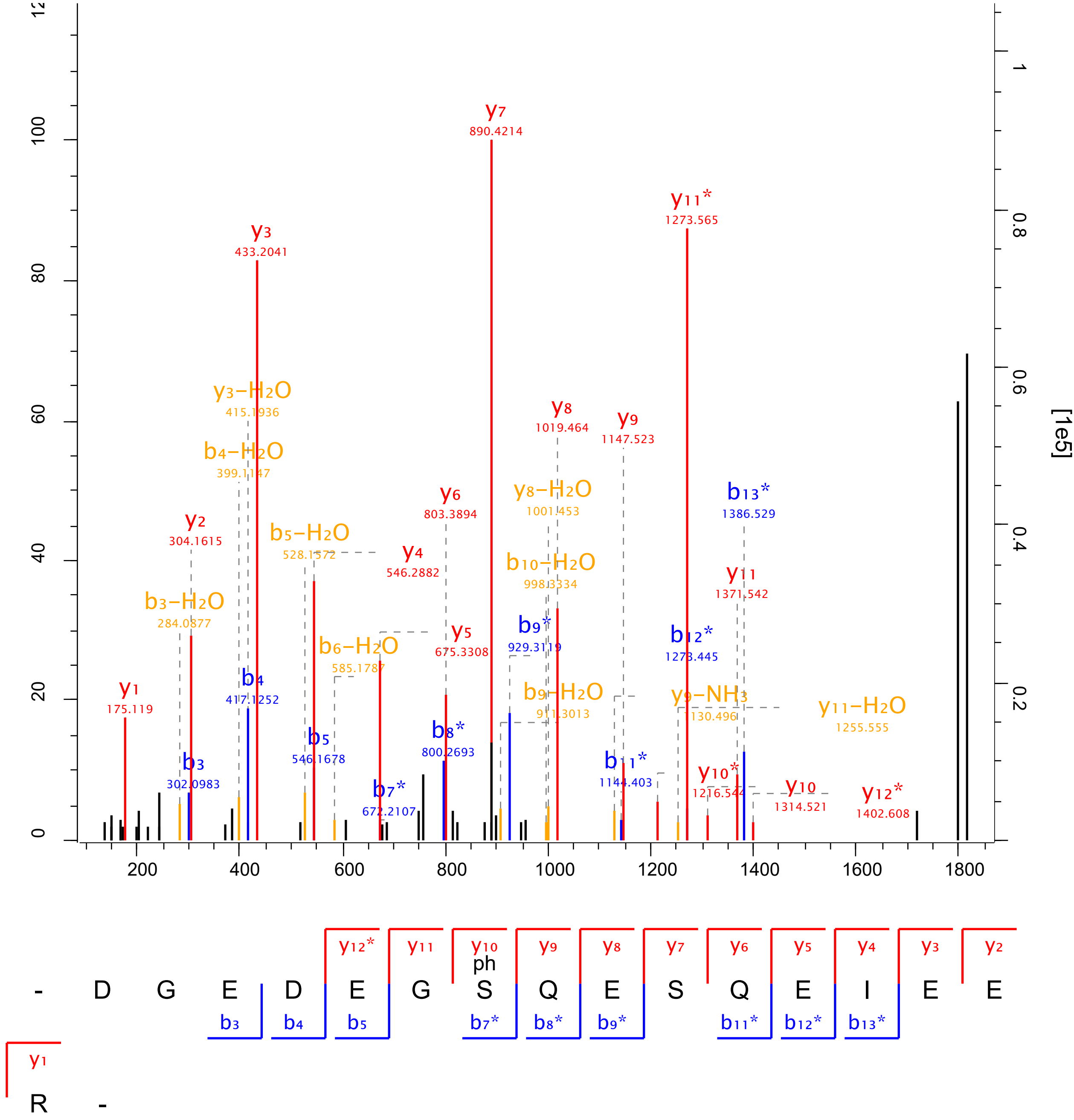

| Raw file | Scan | Method | Score | m/z |
|---|---|---|---|---|
| A17_00_2 | 15411 | FTMS; HCD | 57.35 | 469.17 |

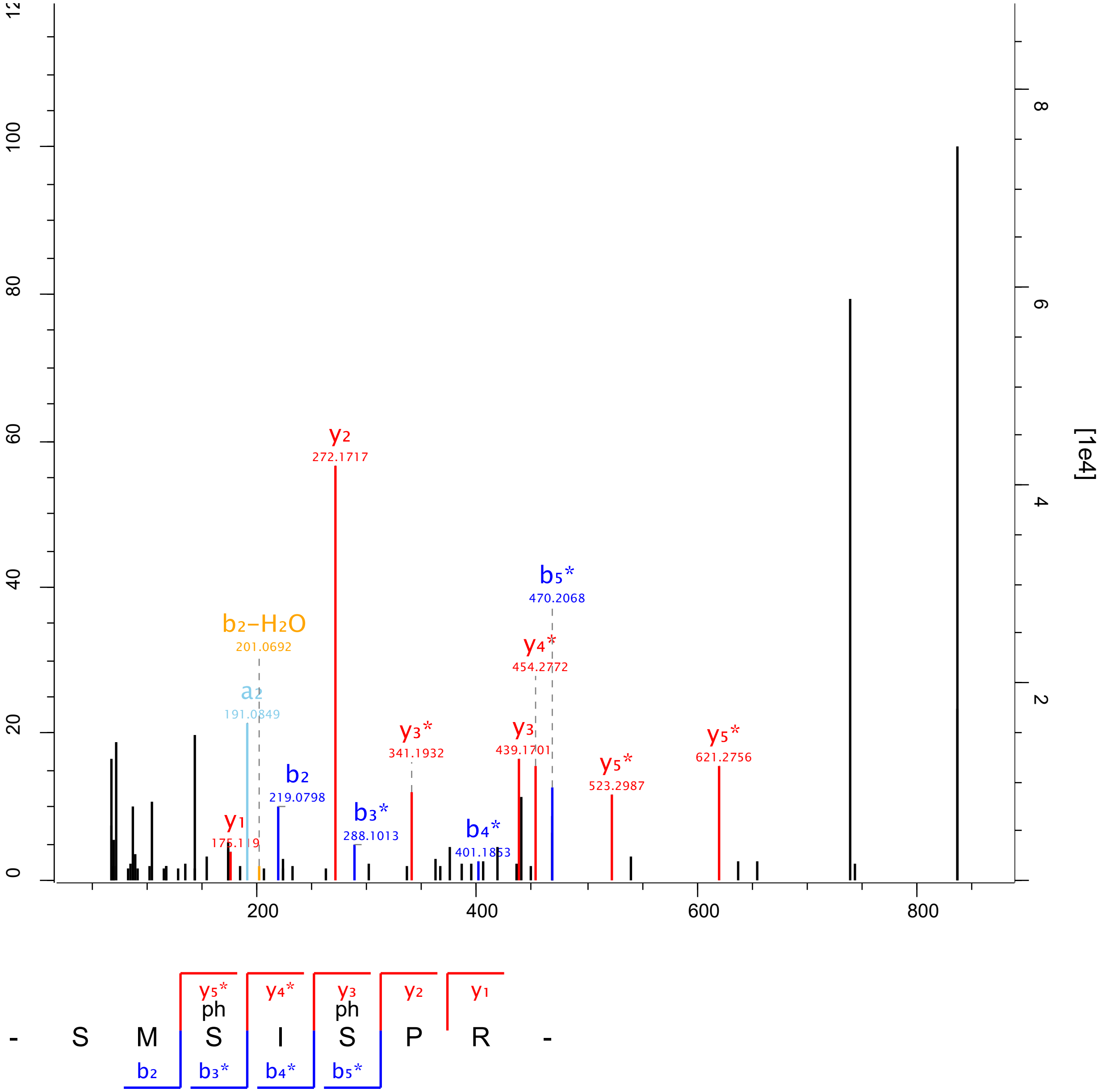

| Raw file | Scan | Method | Score | m/z |
|---|---|---|---|---|
| A17_00_2 | 15454 | FTMS; HCD | 52.97 | 745.31 |

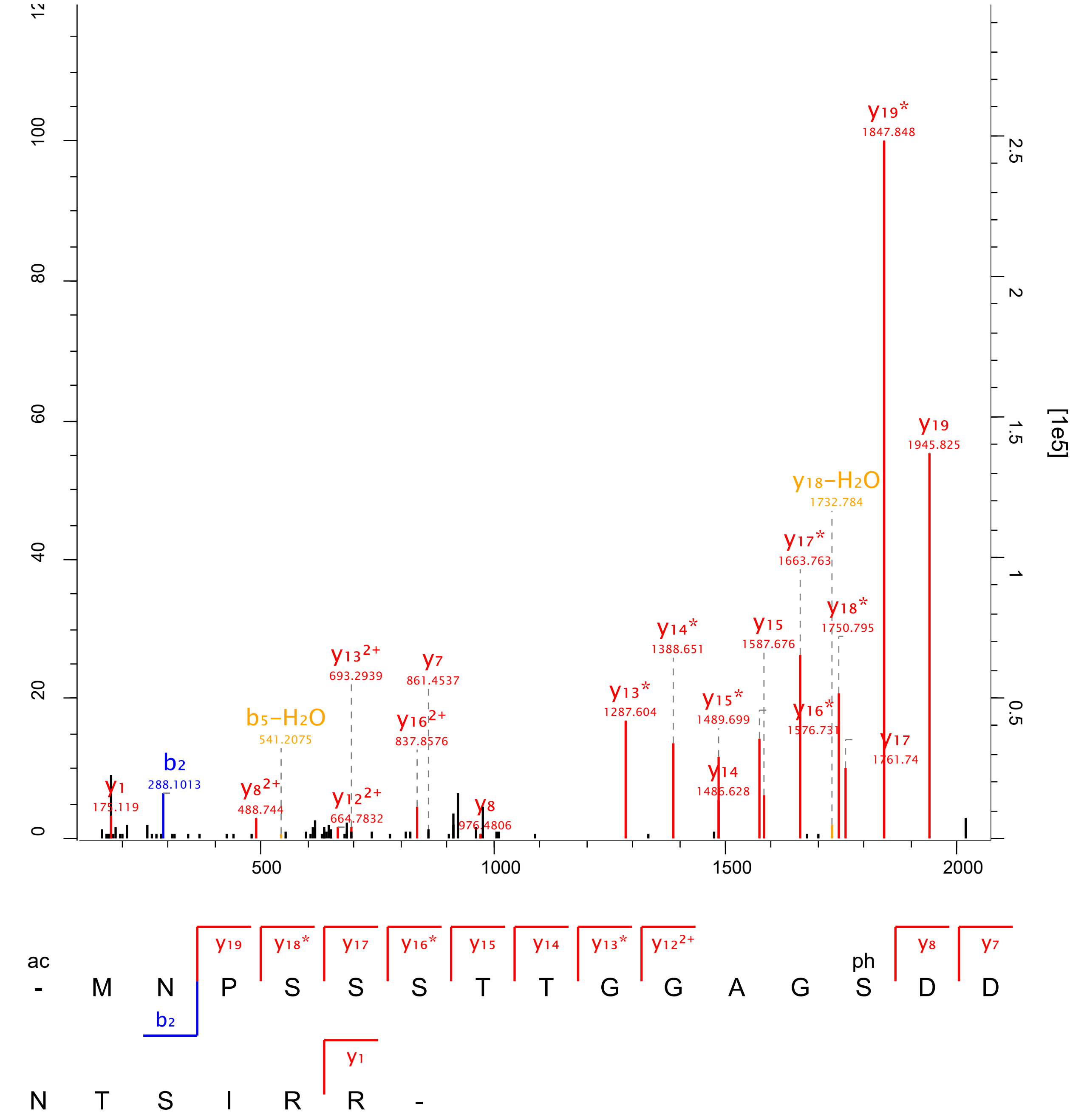

| Raw file | Scan | Method | Score | m/z |
| --- | --- | --- | --- | --- |
| A17_00_2 | 15570 | FTMS; HCD | 85.49 | 759.82 |

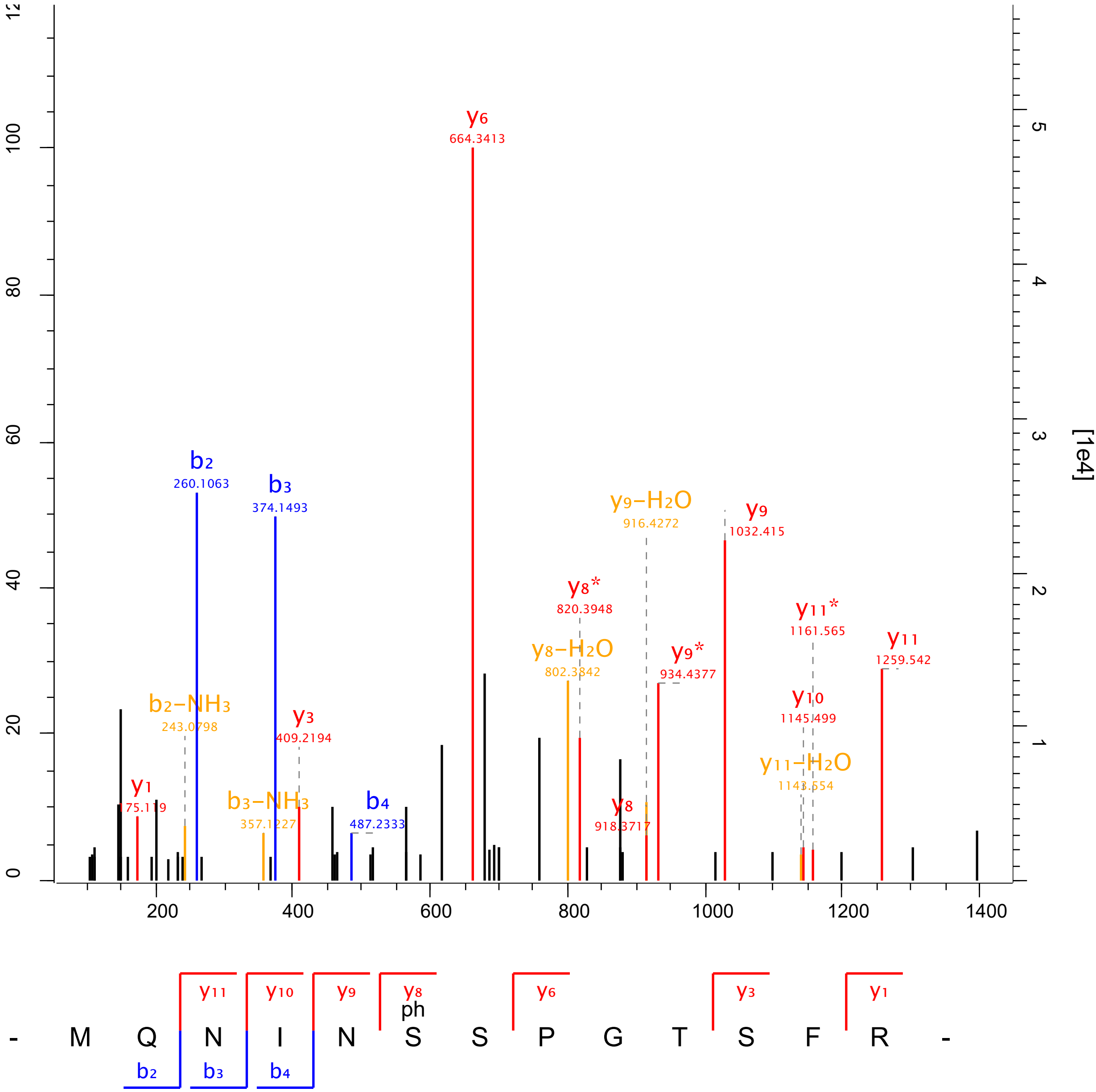

| Raw file | Scan | Method | Score | m/z |
|---|---|---|---|---|
| A17_00_2 | 15732 | FTMS; HCD | 45 | 736.36 |

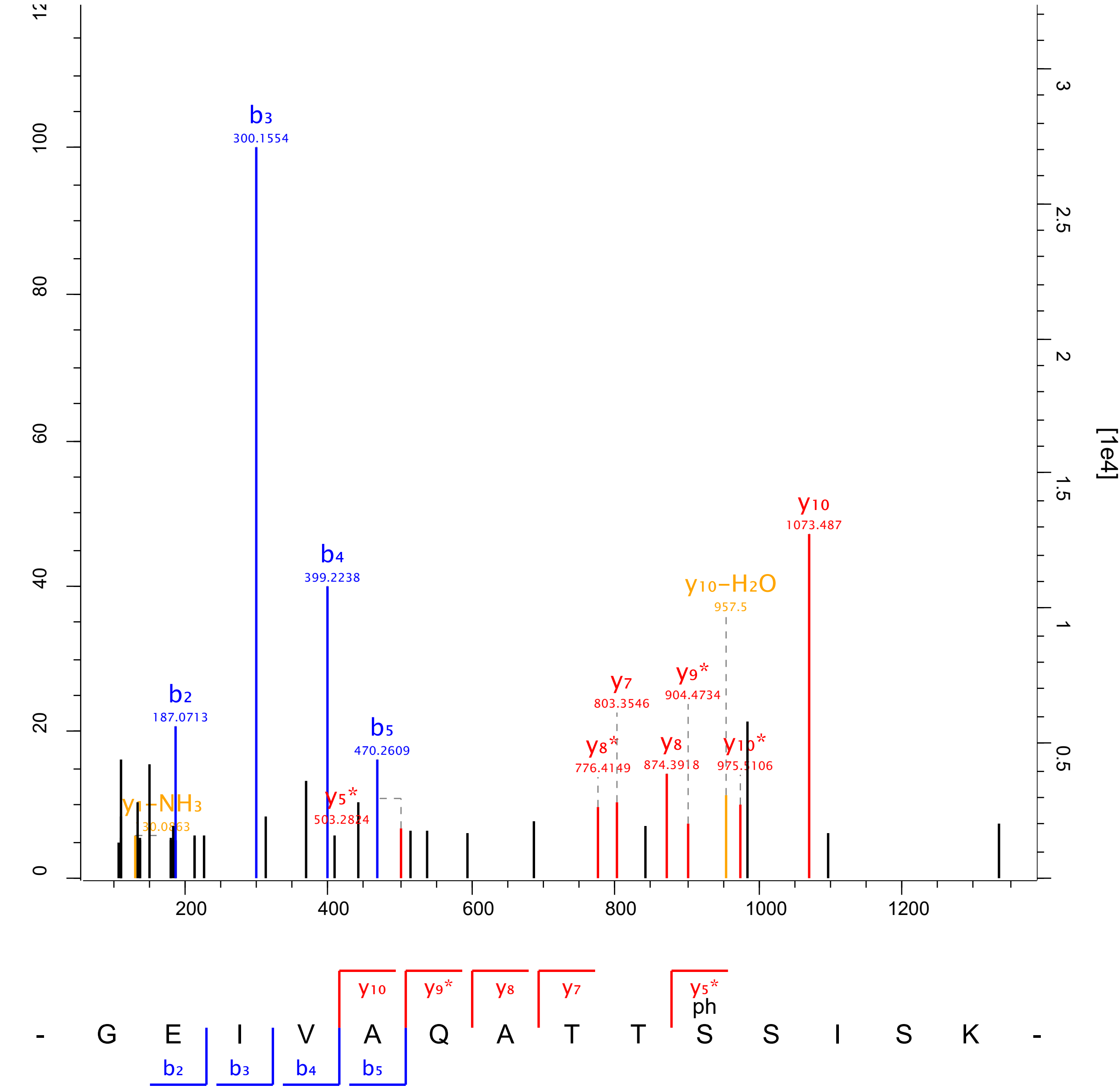

| Raw file | Scan | Method | Score | m/z |
| --- | --- | --- | --- | --- |
| A17_00_2 | 1585 | FTMS; HCD | 84.61 | 434.18 |

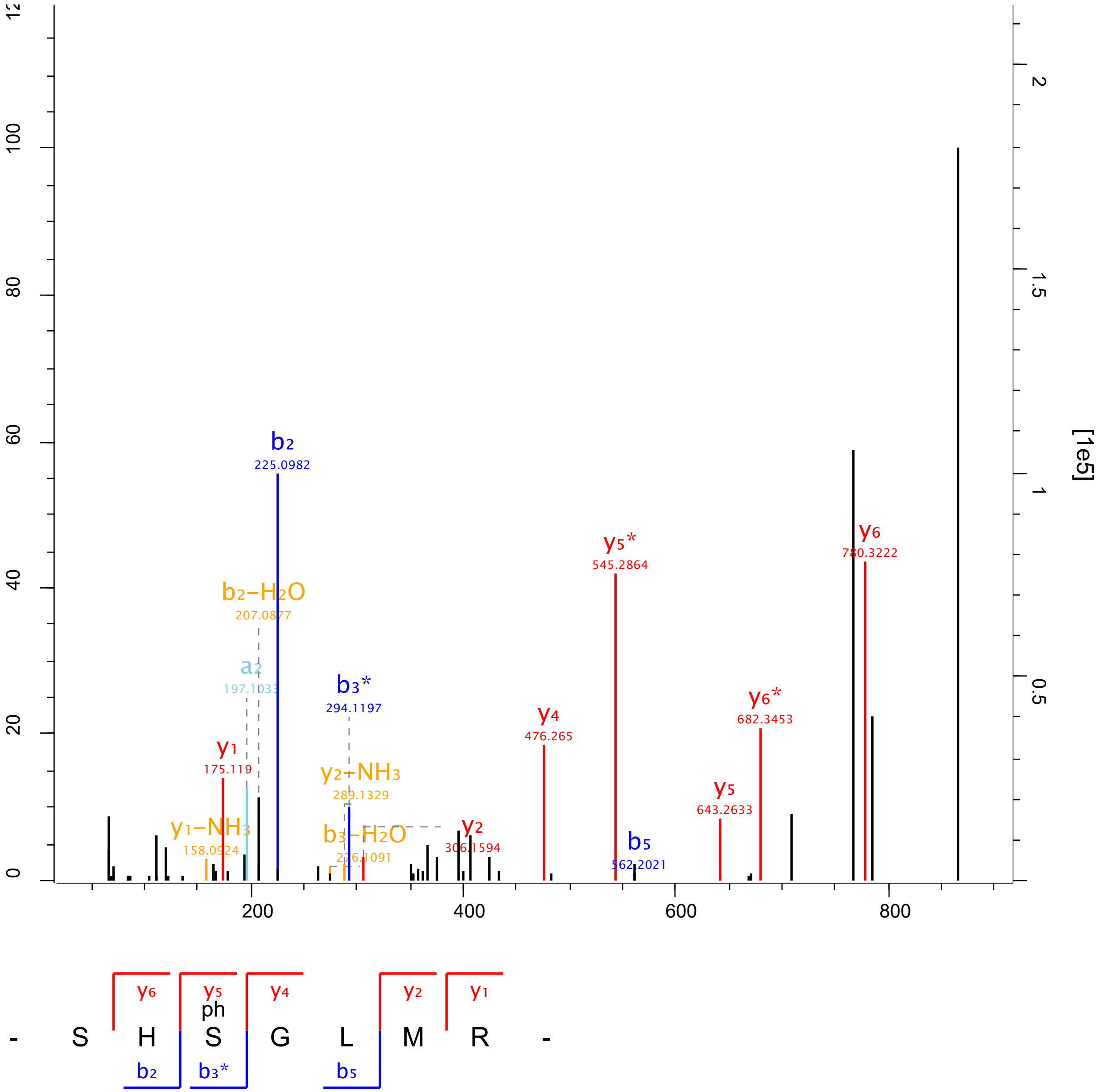

| Raw file | Scan | Method | Score | m/z |
| --- | --- | --- | --- | --- |
| A17_00_2 | 1587 | FTMS; HCD | 76.07 | 415.86 |

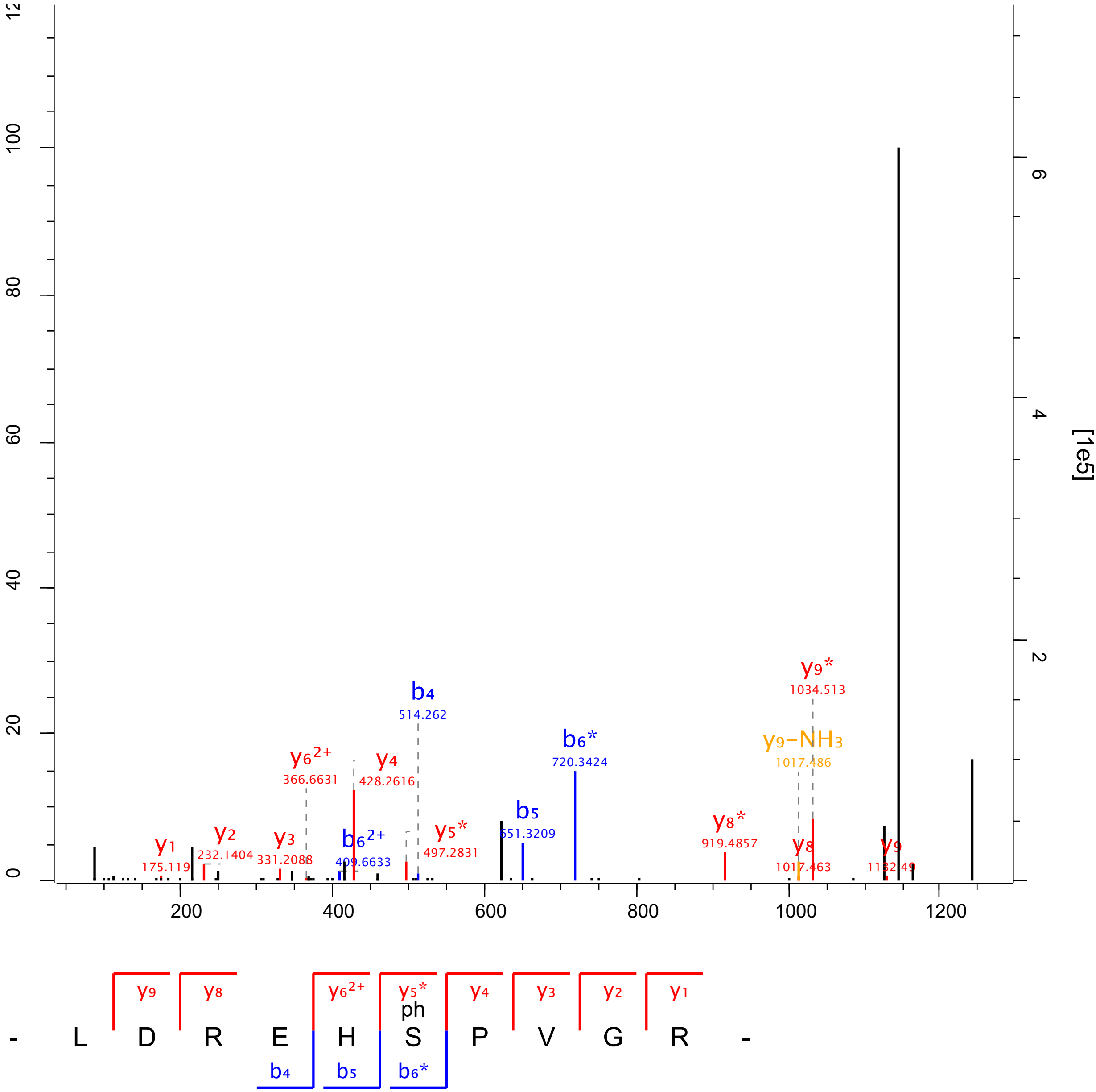

| Raw file | Scan | Method | Score | m/z |
|---|---|---|---|---|
| A17_00_2 | 15923 | FTMS; HCD | 111.86 | 482.73 |

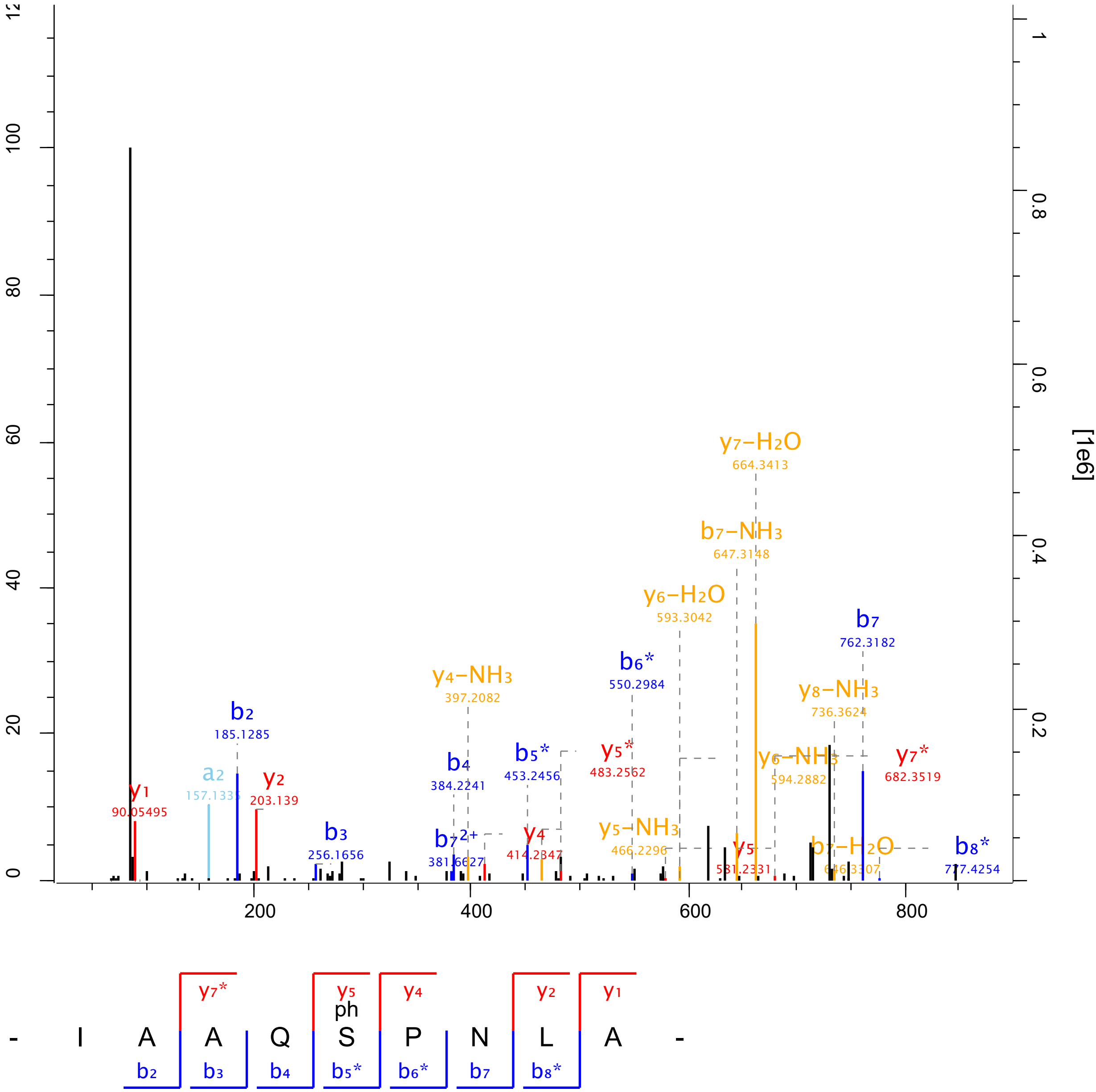

| Raw file | Scan | Method | Score | m/z |
|---|---|---|---|---|
| A17_00_2 | 15946 | FTMS; HCD | 46.89 | 619.59 |

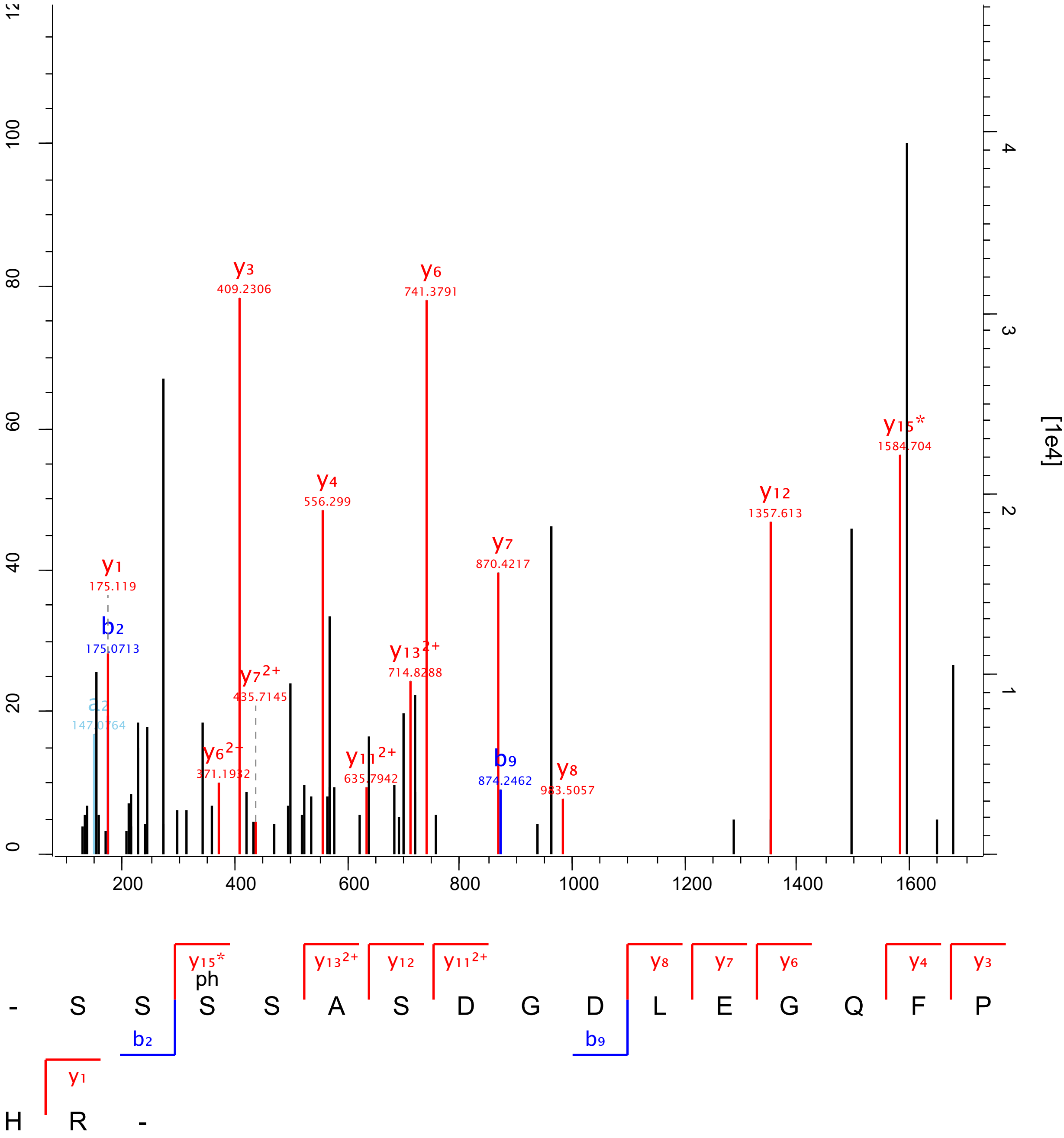

| Raw file | Scan | Method | Score | m/z |
|---|---|---|---|---|
| A17_00_2 | 15976 | FTMS; HCD | 43.84 | 673.97 |

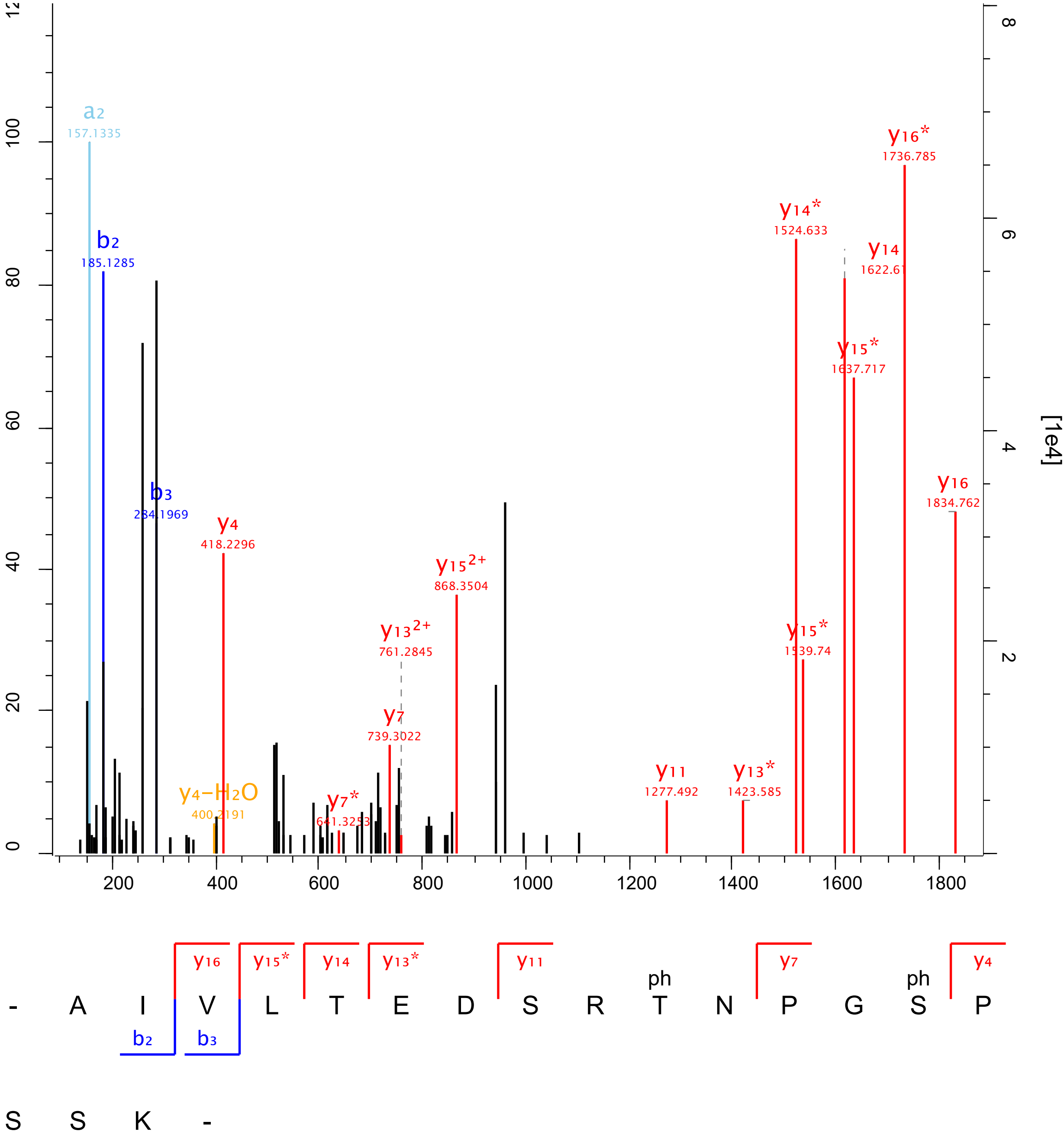

| Raw file | Scan | Method | Score | m/z |
| --- | --- | --- | --- | --- |
| A17_00_2 | 15977 | FTMS; HCD | 49.54 | 606.23 |

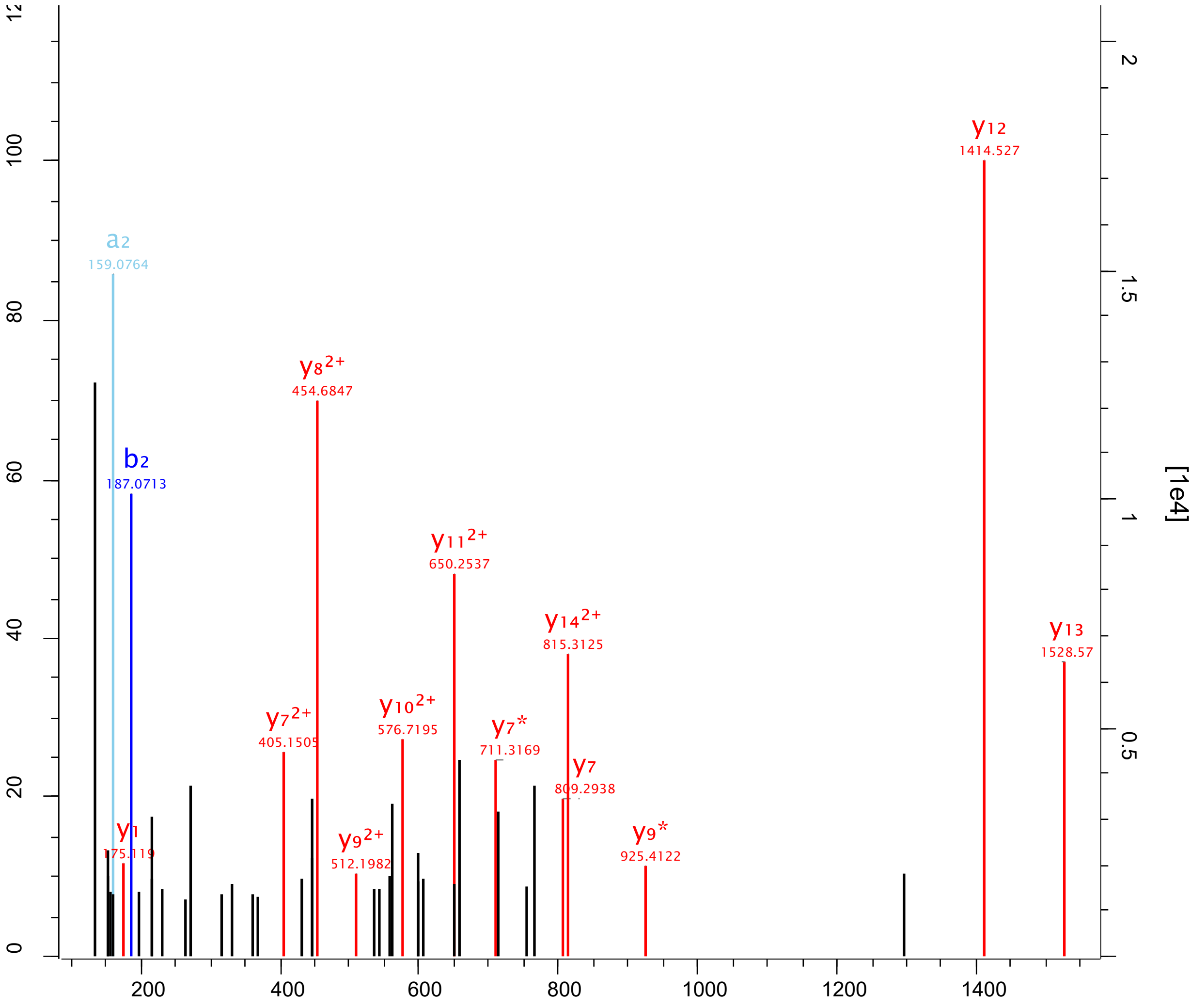


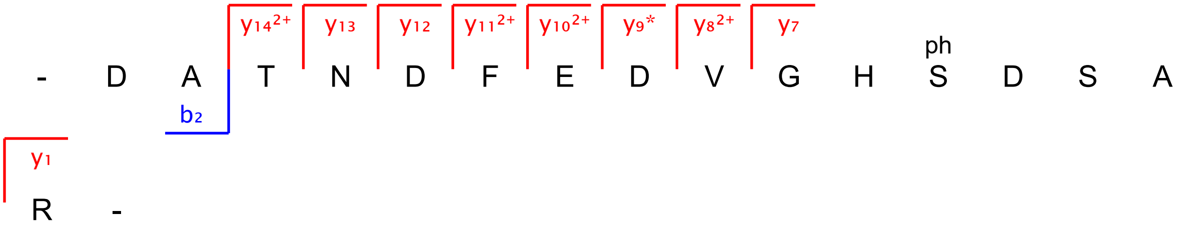

| Raw file | Scan | Method | Score | m/z |
|---|---|---|---|---|
| A17_00_2 | 16160 | FTMS; HCD | 132.81 | 727.82 |

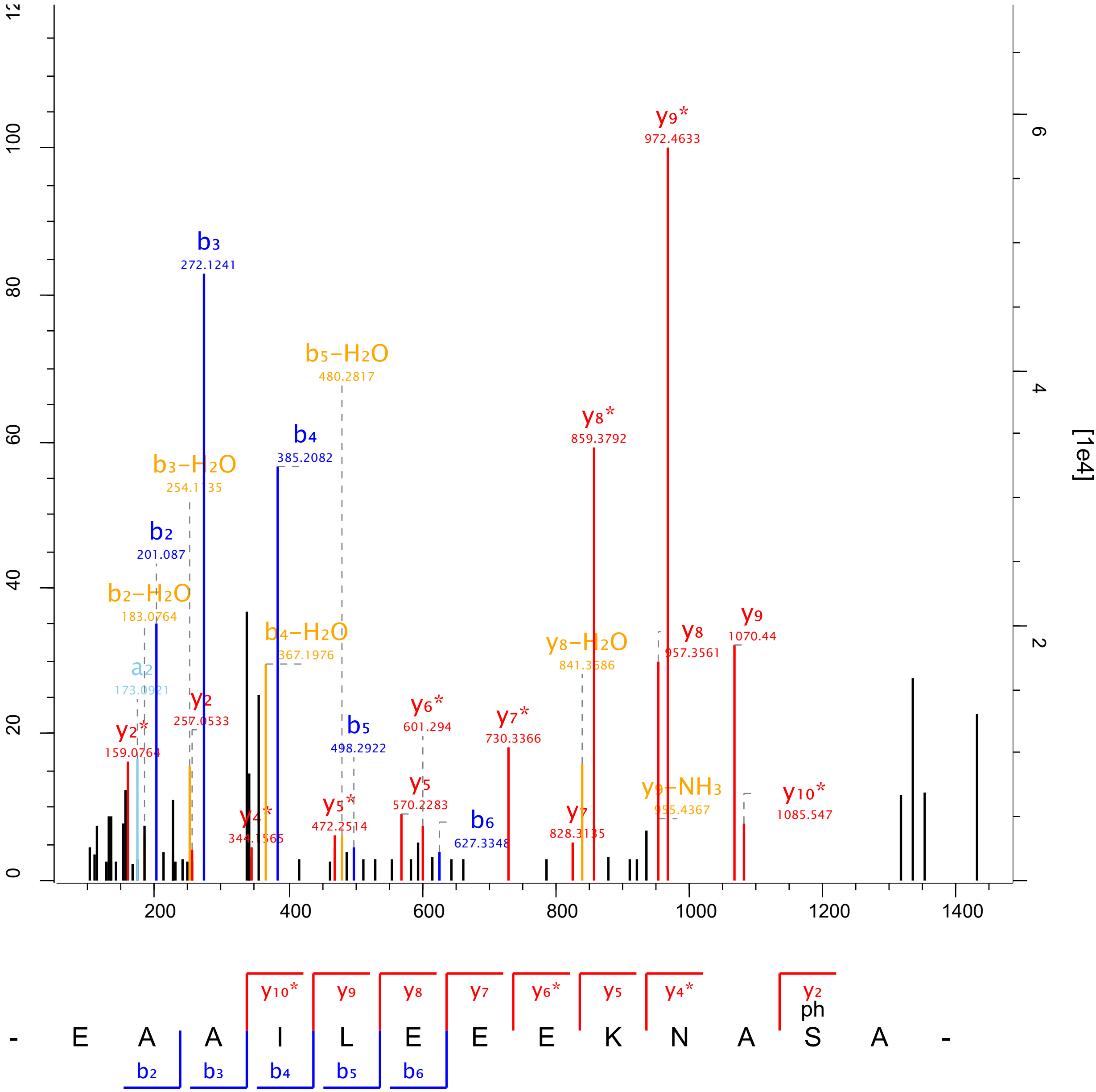

| Raw file | Scan | Method | Score | m/z |
|---|---|---|---|---|
| A17_00_2 | 16222 | FTMS; HCD | 68.97 | 590.93 |

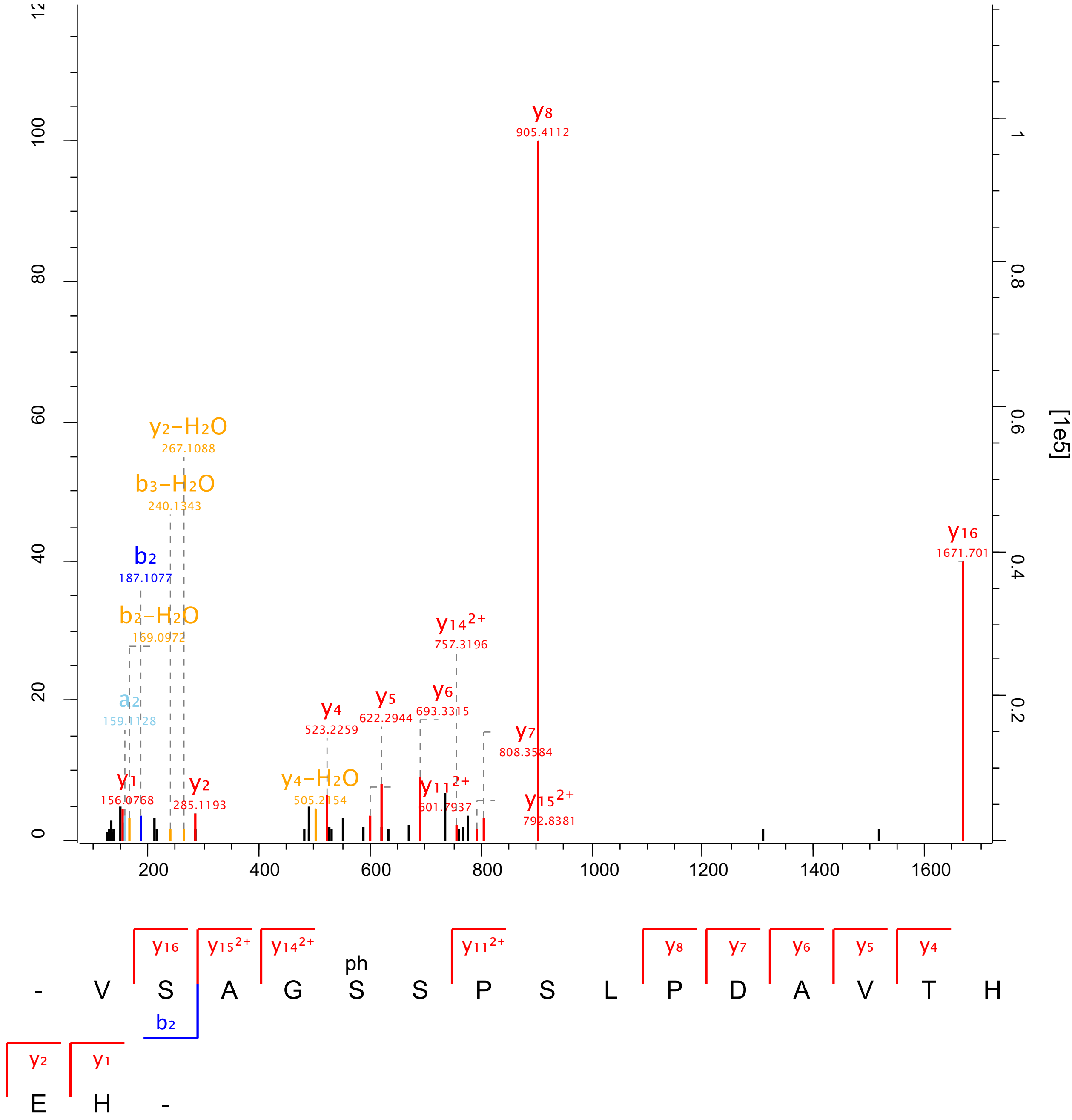

| Raw file | Scan | Method | Score | m/z |
|---|---|---|---|---|
| A17_00_2 | 1624 | FTMS; HCD | 114.46 | 611.24 |

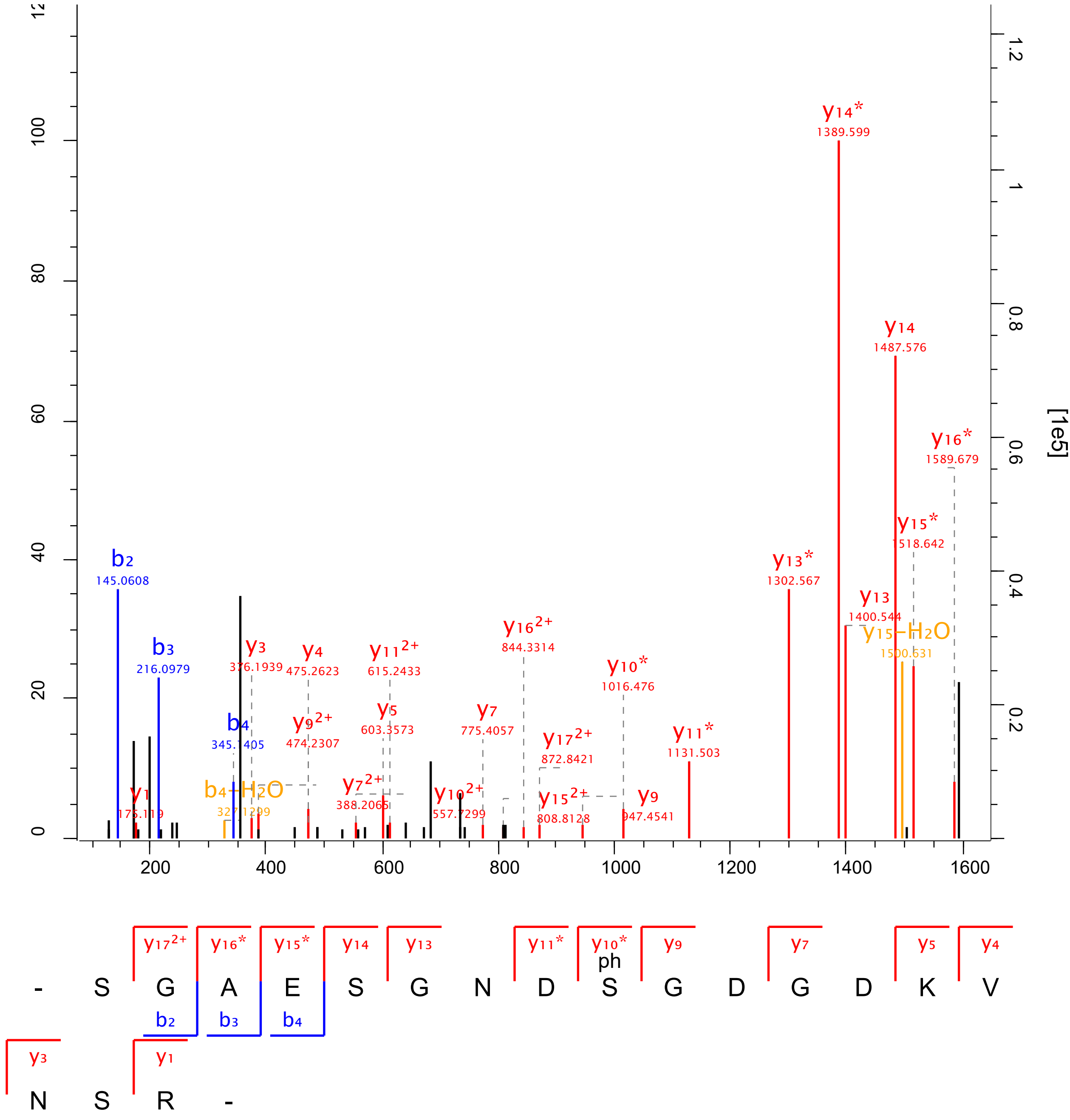

| Raw file | Scan | Method | Score | m/z |
|---|---|---|---|---|
| A17_00_2 | 16251 | FTMS; HCD | 113.73 | 698.24 |

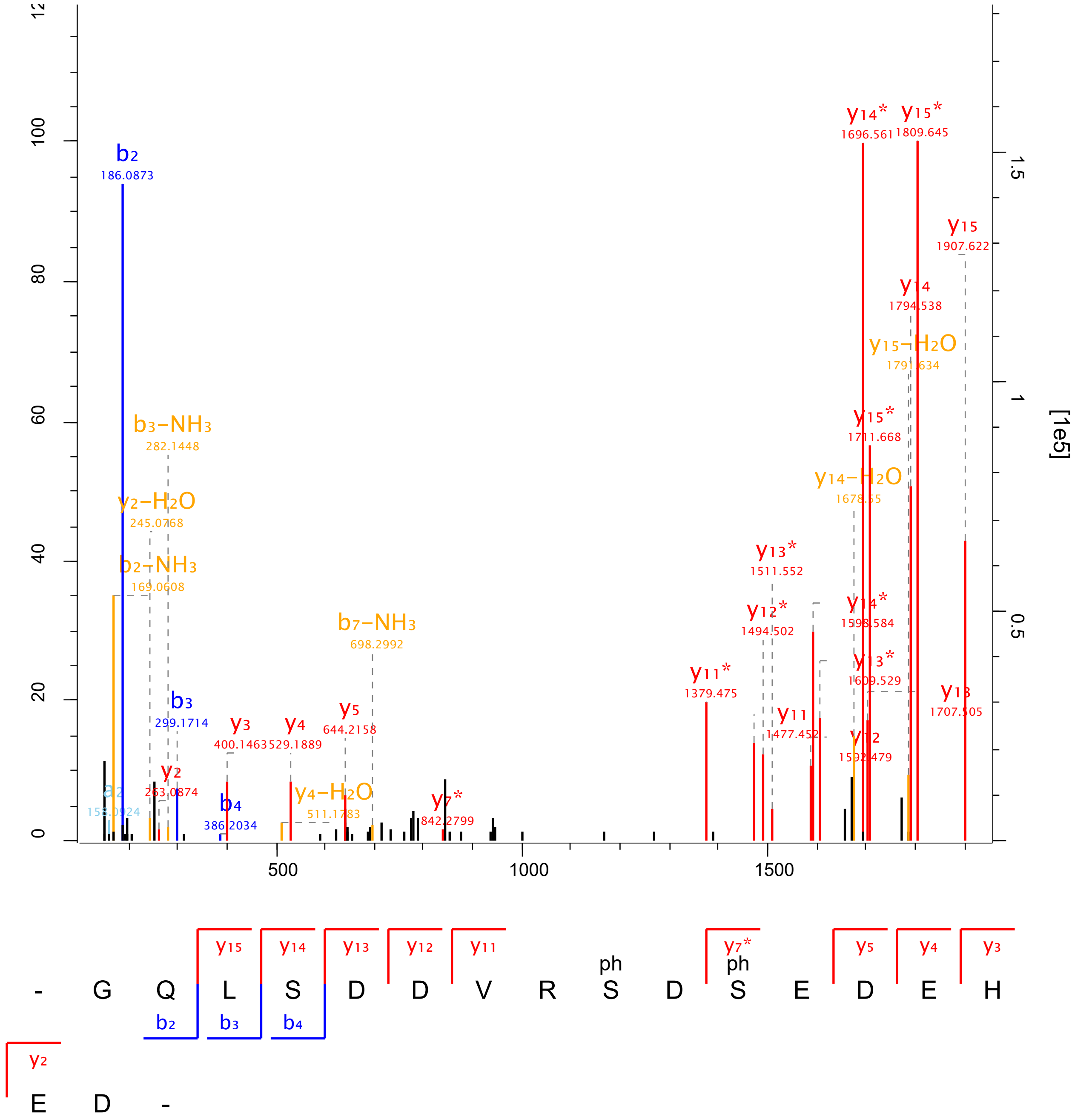

| Raw file | Scan | Method | Score | m/z |
|---|---|---|---|---|
| A17_00_2 | 16444 | FTMS; HCD | 93.38 | 569.94 |

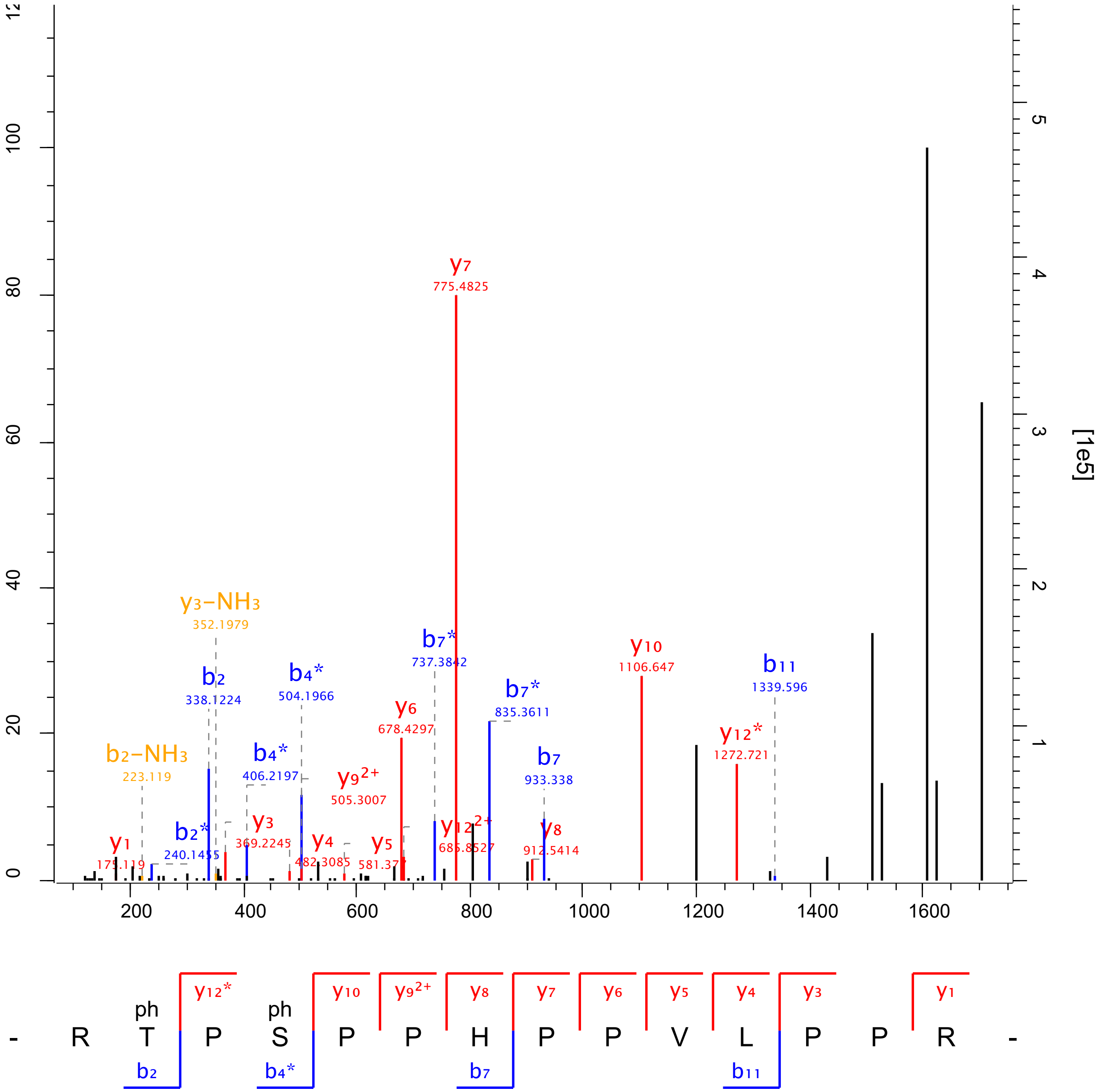

| Raw file | Scan | Method | Score | m/z |
| --- | --- | --- | --- | --- |
| A17_00_2 | 16702 | FTMS; HCD | 203.4 | 721.63 |

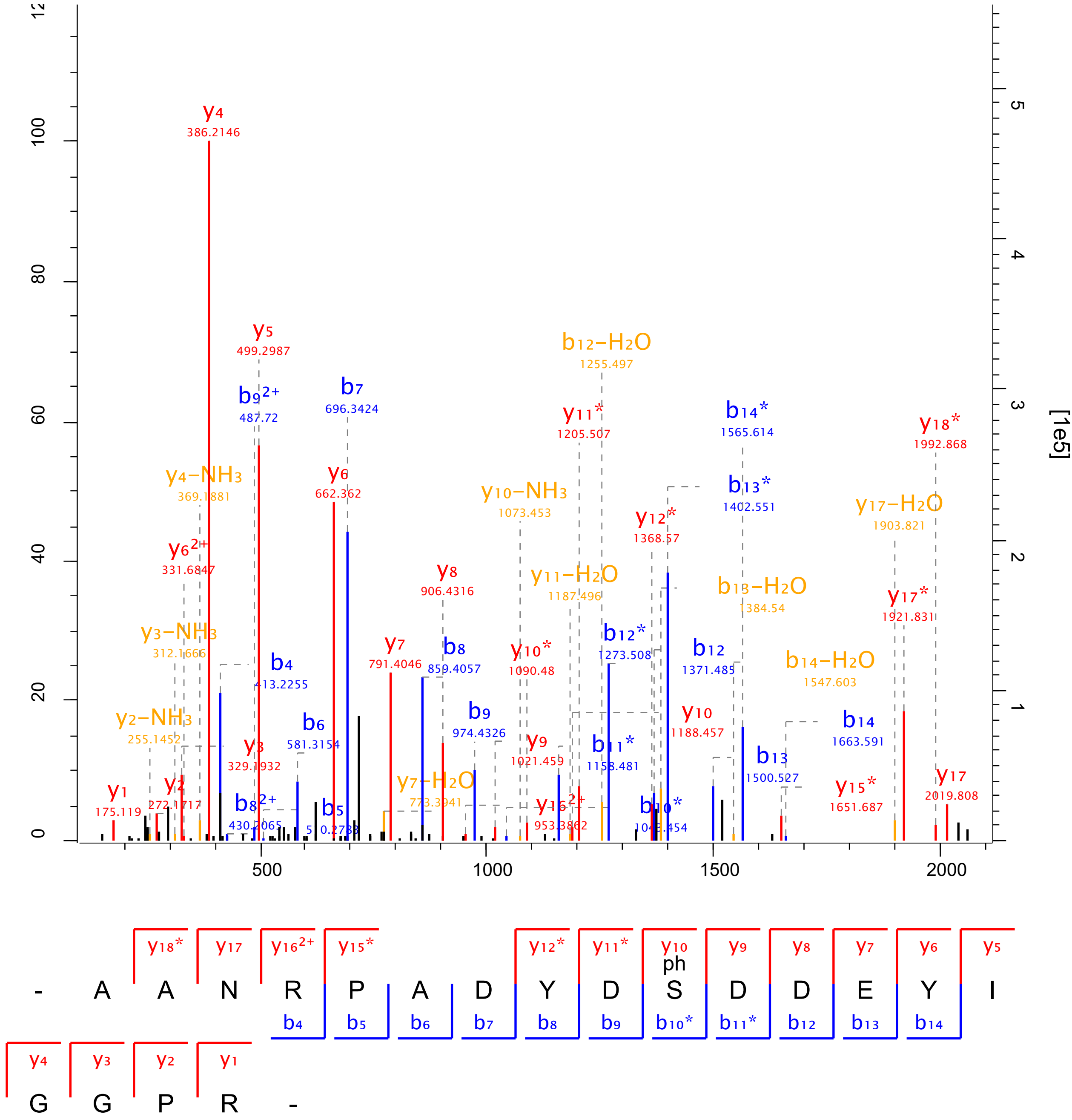

| Raw file | Scan | Method | Score | m/z |
| --- | --- | --- | --- | --- |
| A17_00_2 | 1676 | FTMS; HCD | 54.34 | 530.75 |

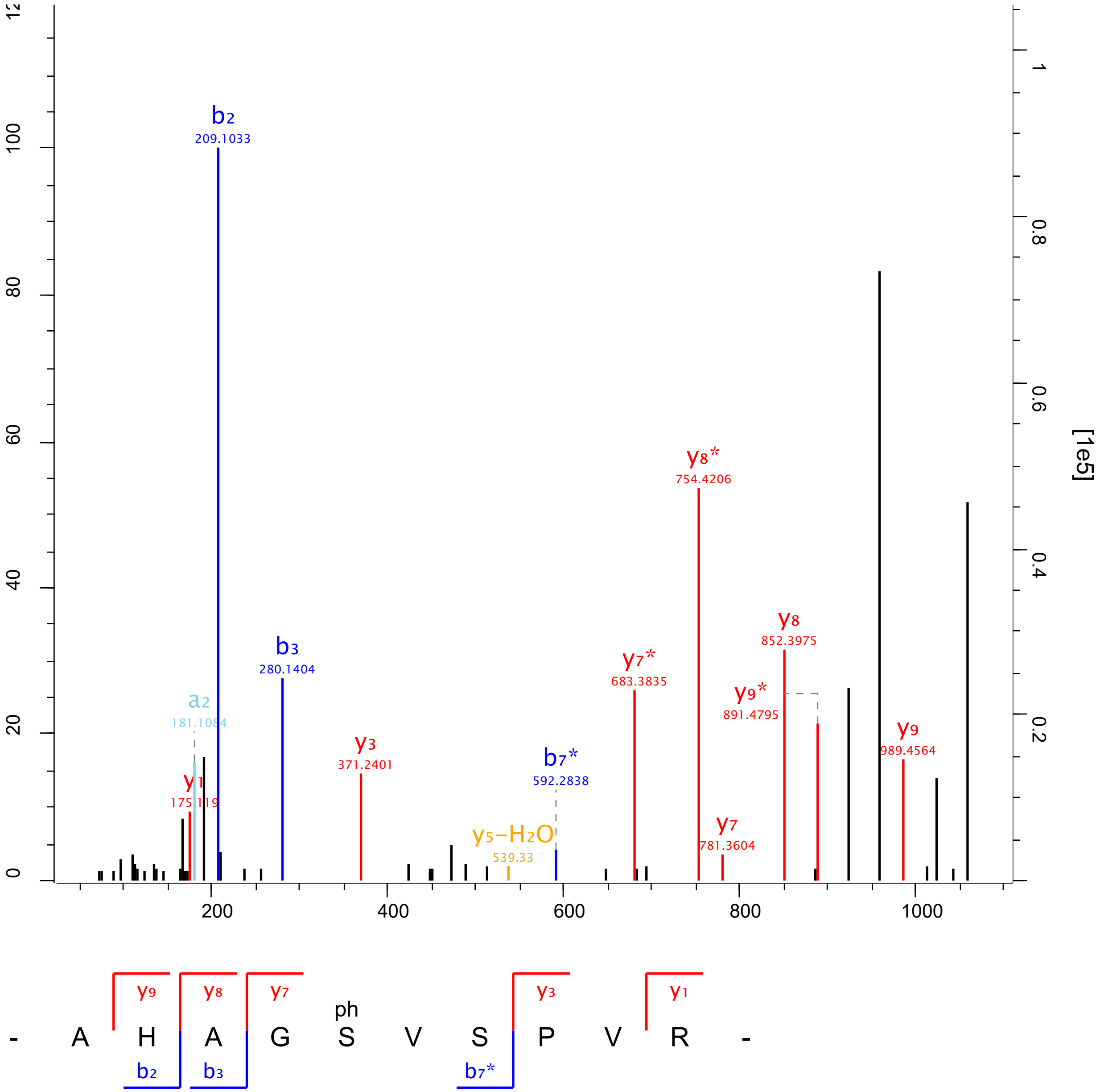

| Raw file | Scan | Method | Score | m/z |
|---|---|---|---|---|
| A17_00_2 | 16798 | FTMS; HCD | 106.82 | 584.23 |

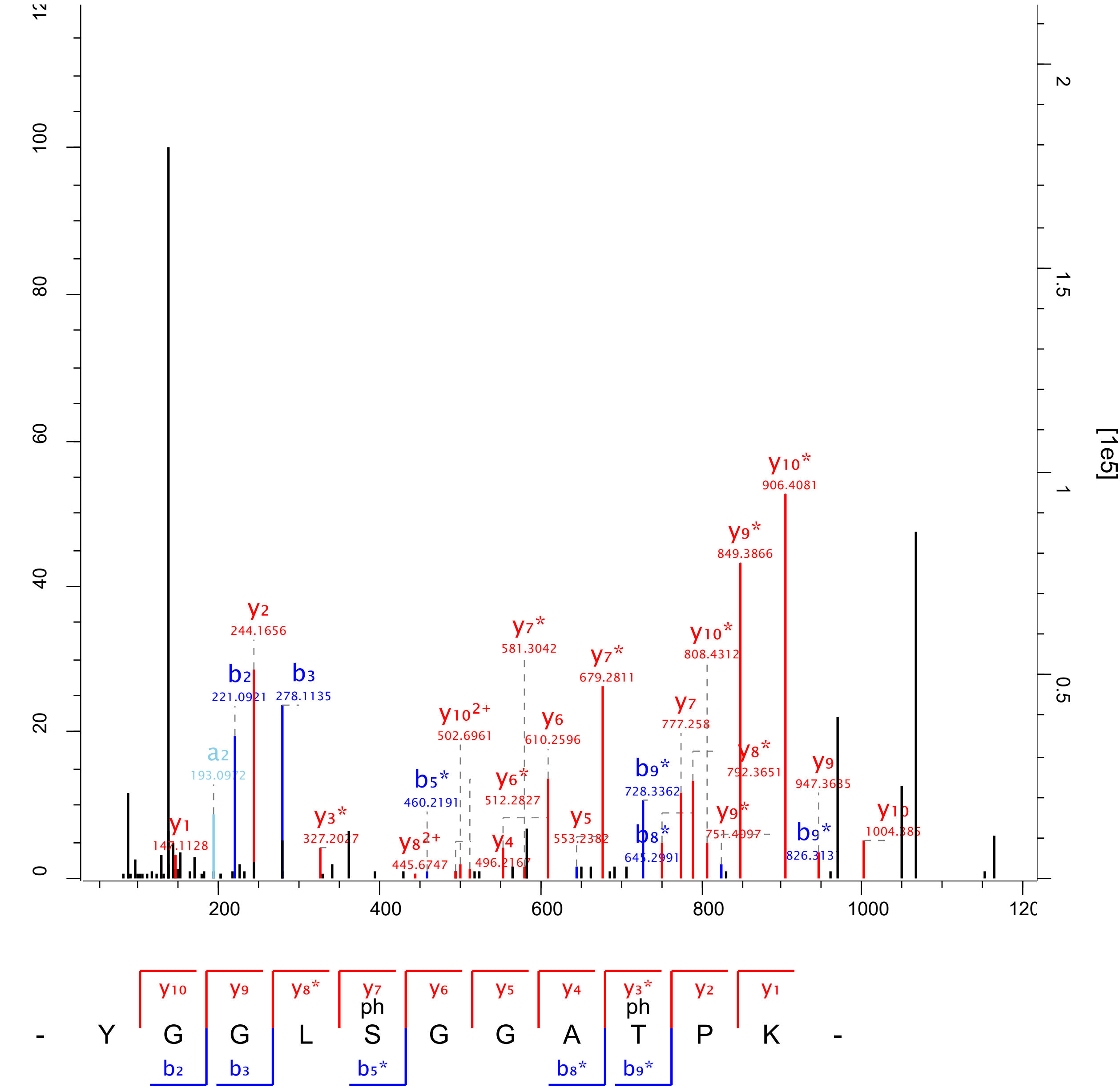

| Raw file | Scan | Method | Score | m/z |
|---|---|---|---|---|
| A17_00_2 | 16824 | FTMS; HCD | 76.59 | 735.83 |

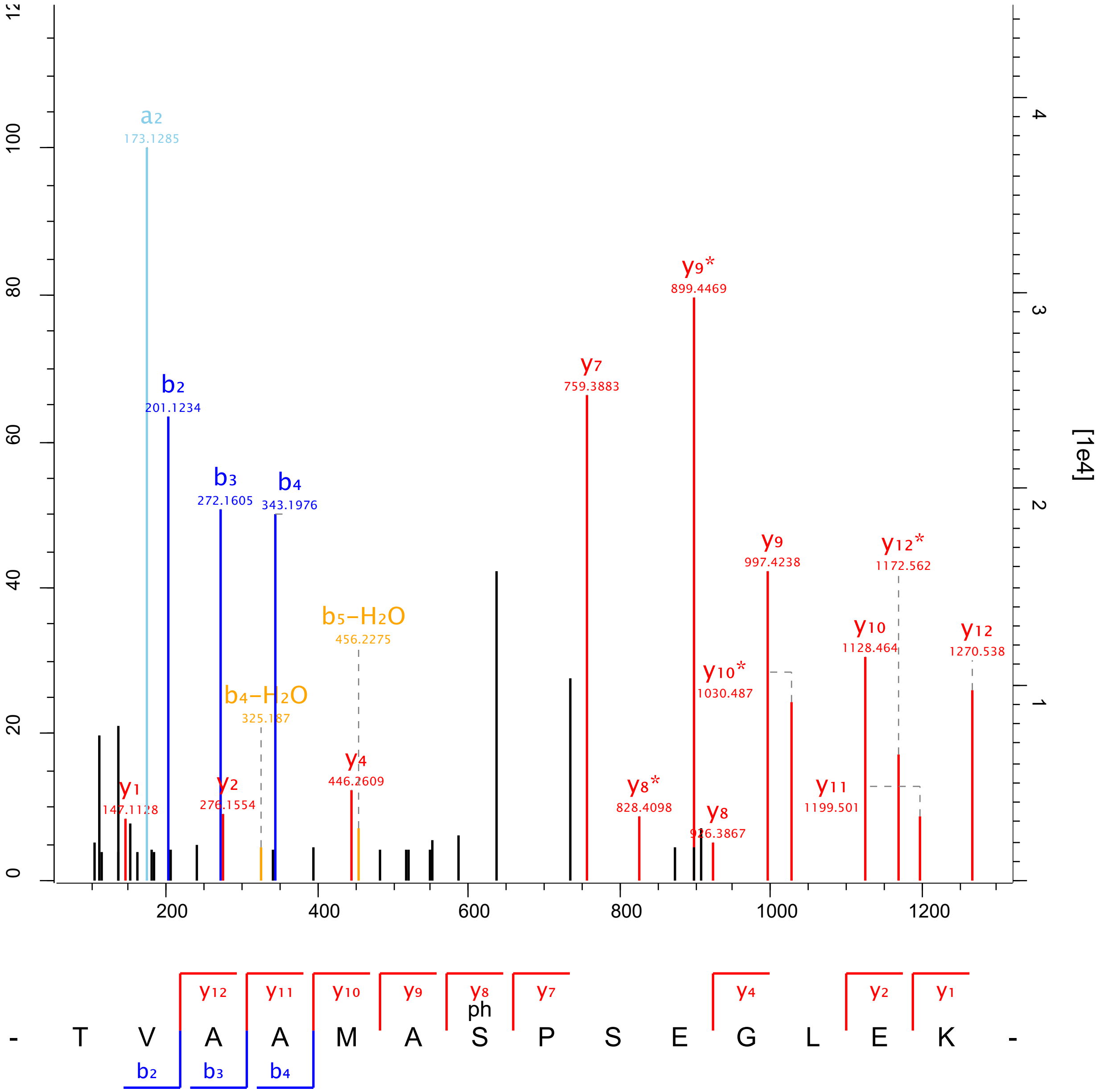

| Raw file | Scan | Method | Score | m/z |
| --- | --- | --- | --- | --- |
| A17_00_2 | 16985 | FTMS; HCD | 51.88 | 725.96 |

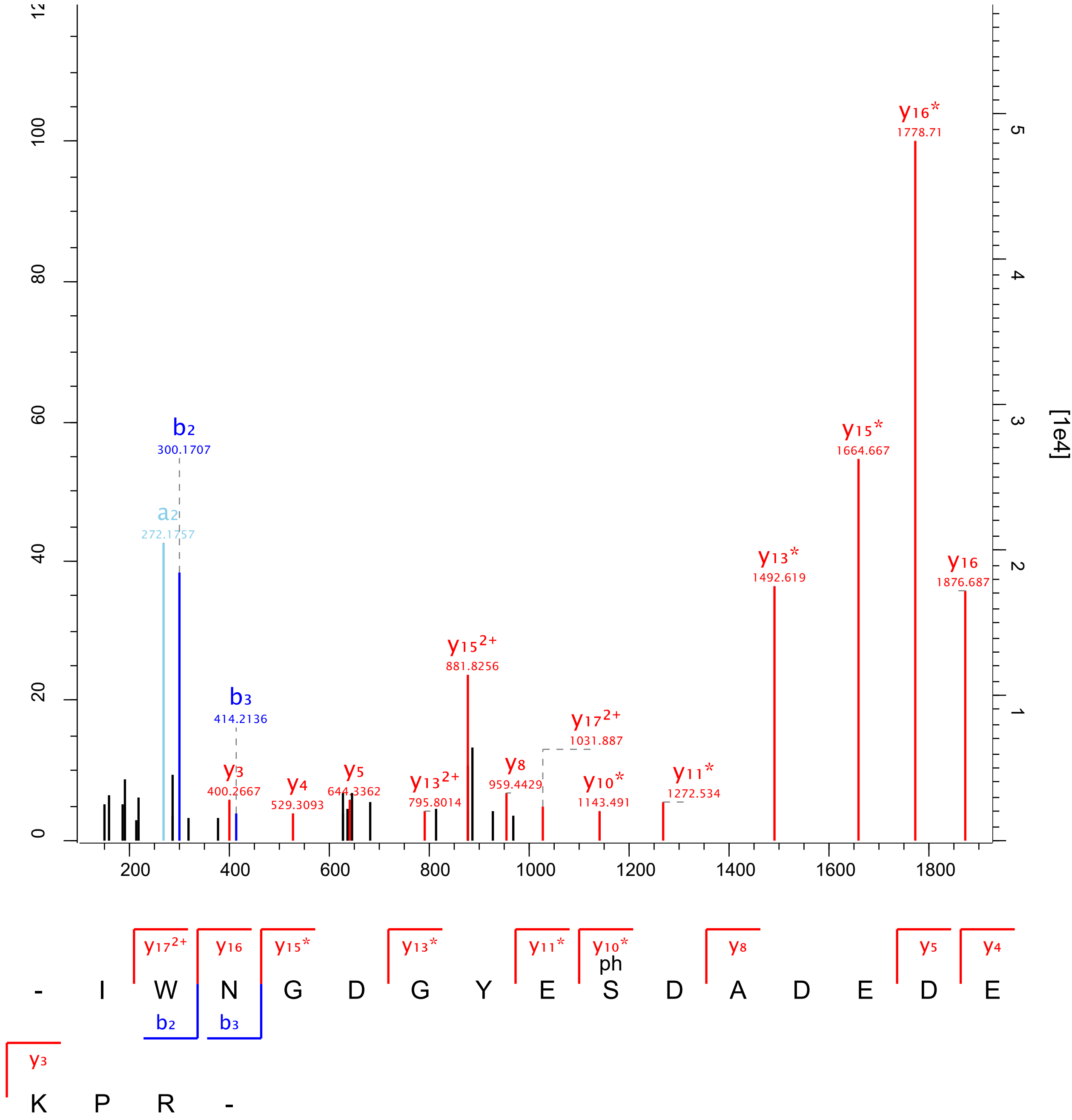

| Raw file | Scan | Method | Score | m/z |
|---|---|---|---|---|
| A17_00_2 | 17120 | FTMS; HCD | 69.03 | 536.67 |

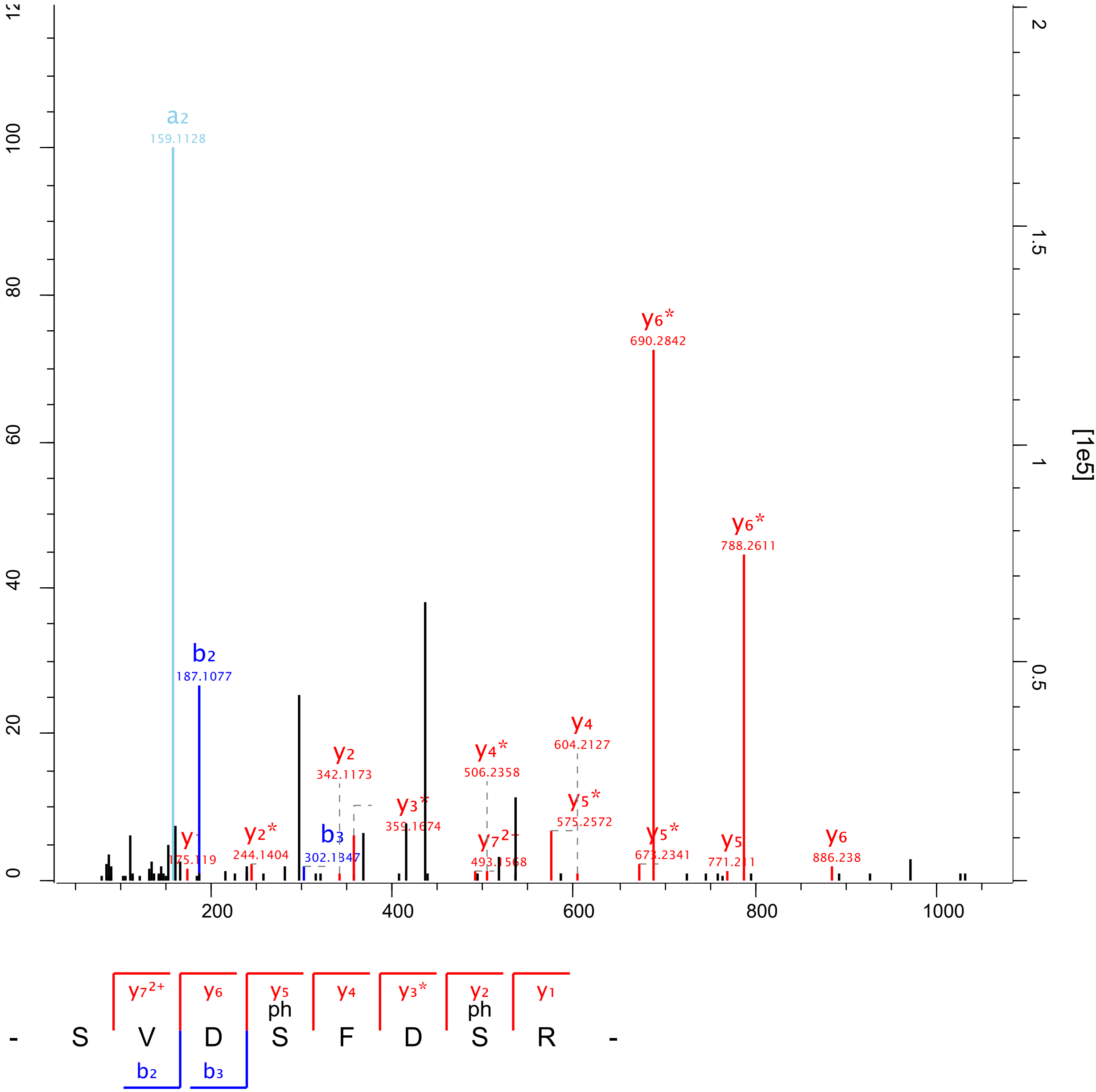

| Raw file | Scan | Method | Score | m/z |
|---|---|---|---|---|
| A17_00_2 | 1713 | FTMS; HCD | 47.77 | 434.19 |

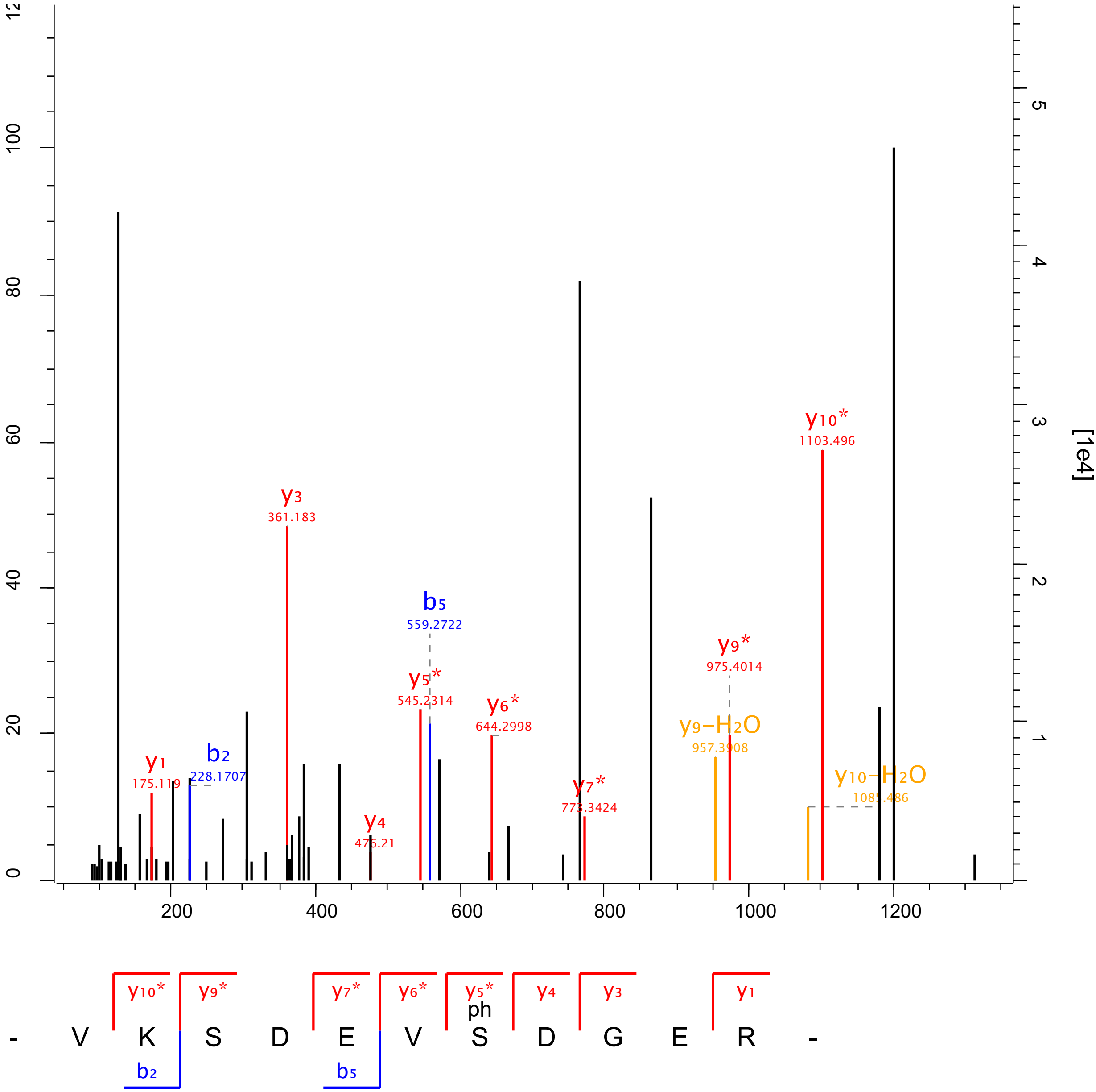

| Raw file | Scan | Method | Score | m/z |
| --- | --- | --- | --- | --- |
| A17_00_2 | 17210 | FTMS; HCD | 200.09 | 661.29 |

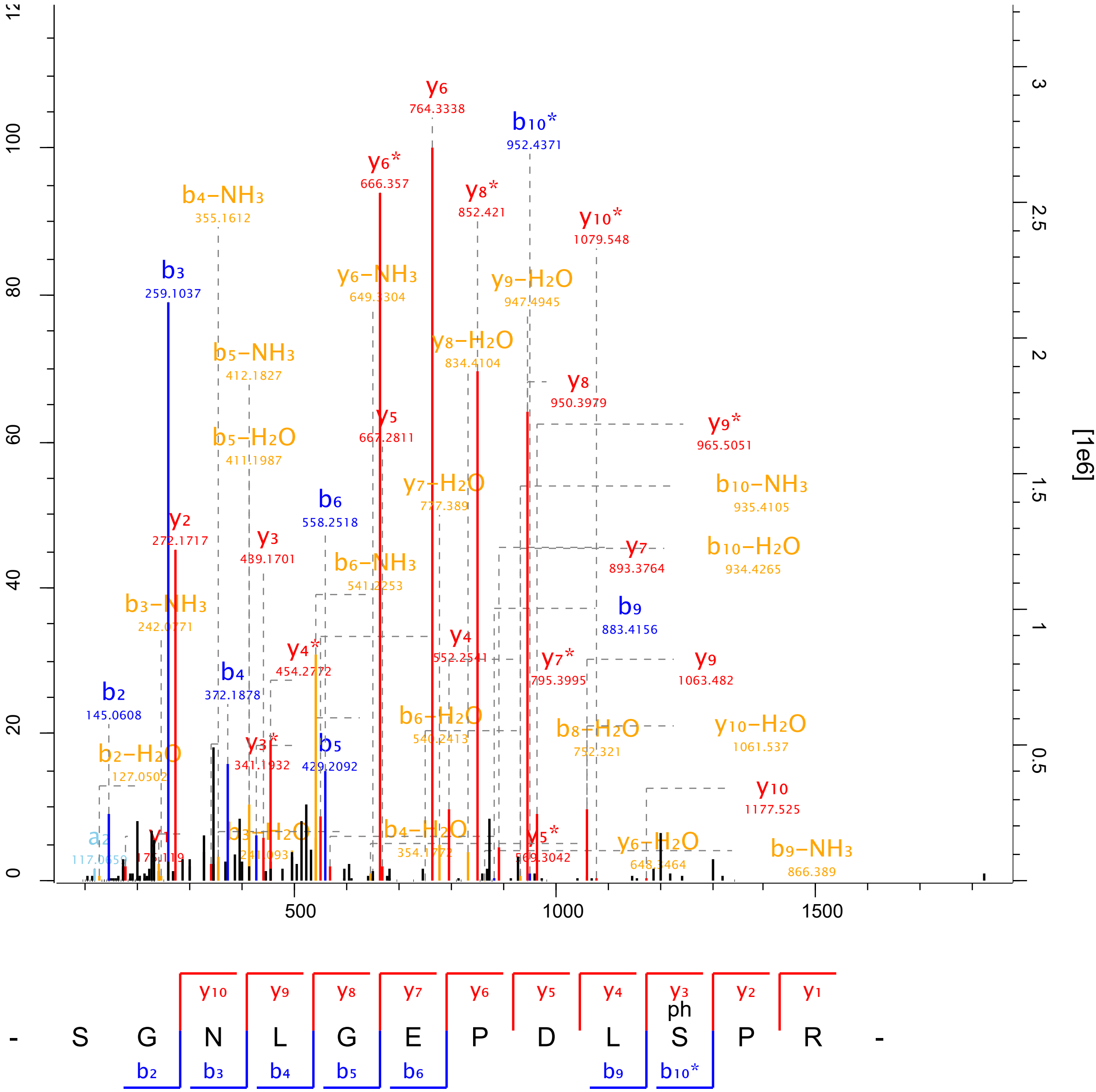

| Raw file | Scan | Method | Score | m/z |
| --- | --- | --- | --- | --- |
| A17_00_2 | 17221 | FTMS; HCD | 70.27 | 707.33 |

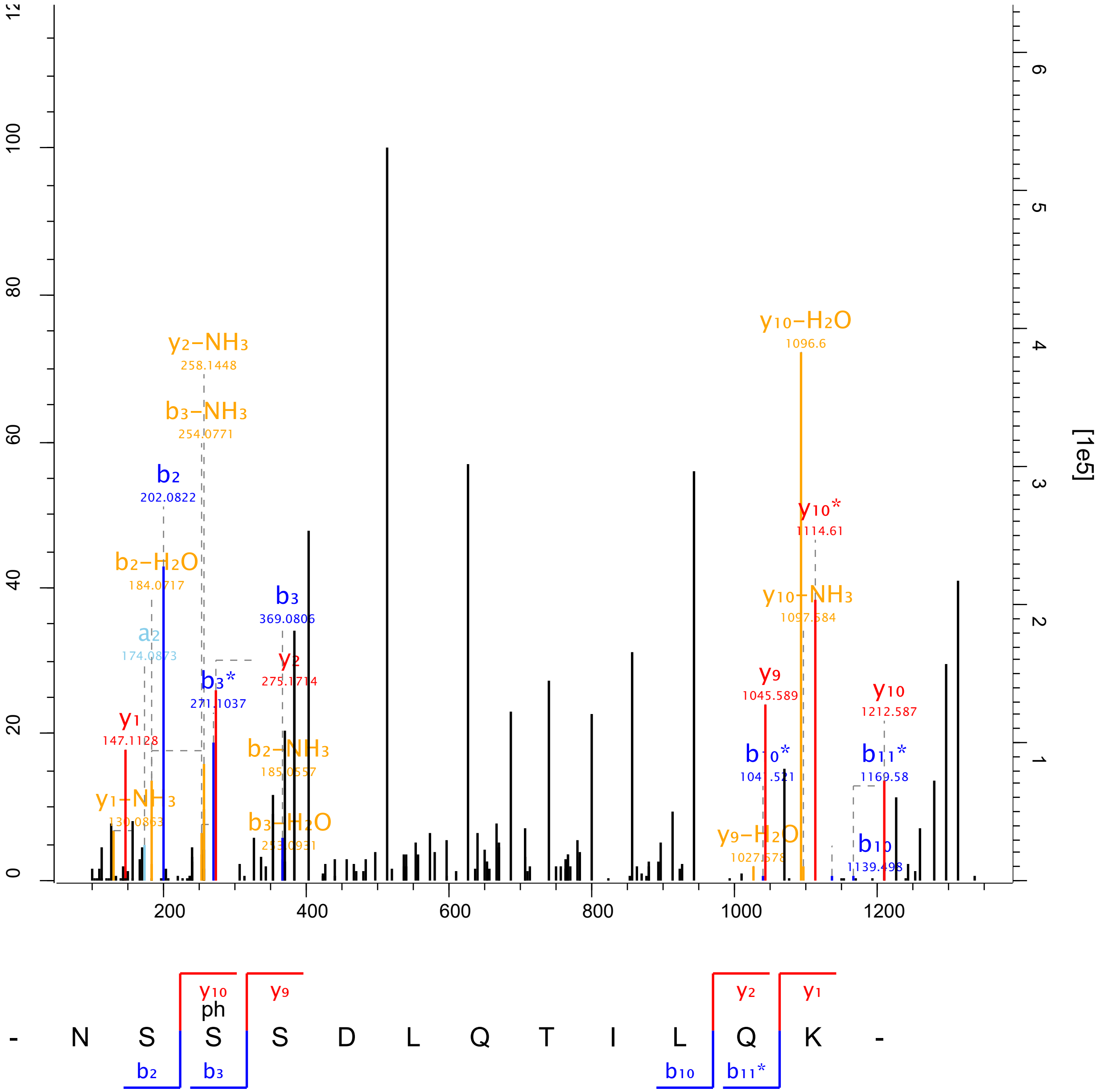

| Raw file | Scan | Method | Score | m/z |
|---|---|---|---|---|
| A17_00_2 | 17263 | FTMS; HCD | 133.86 | 900.38 |

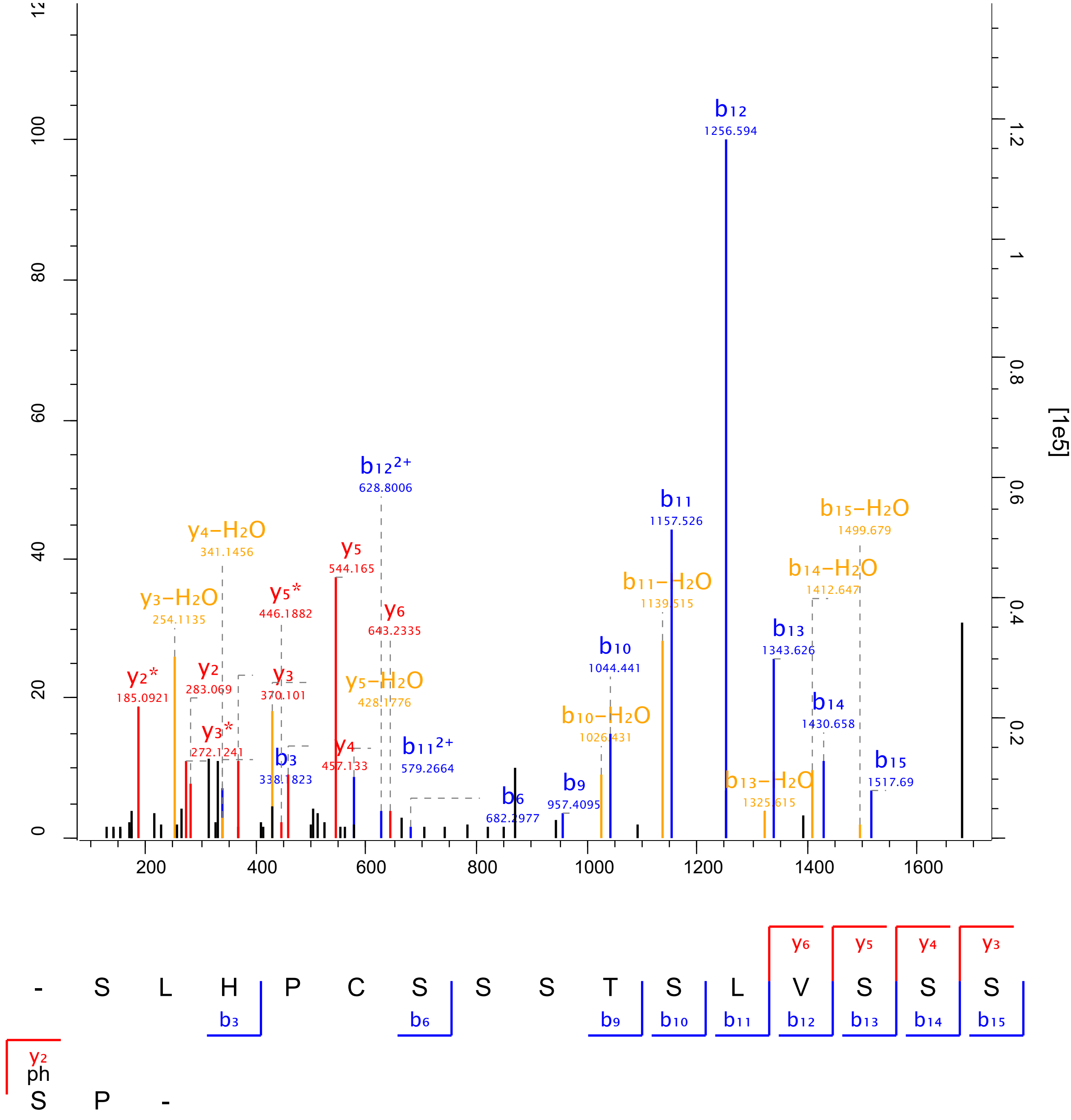

| Raw file | Scan | Method | Score | m/z |
|---|---|---|---|---|
| A17_00_2 | 17265 | FTMS; HCD | 43.59 | 628.78 |

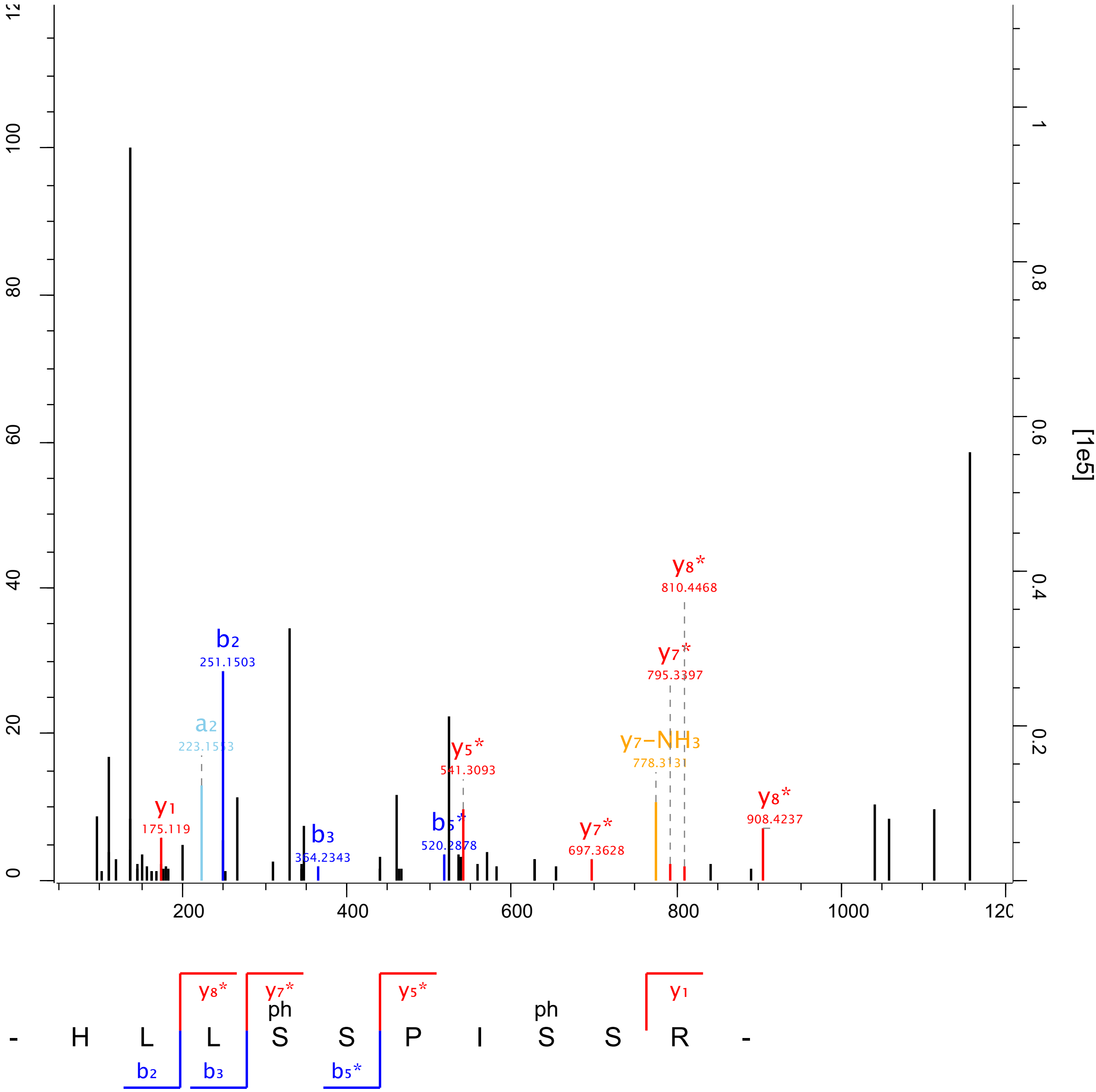

| Raw file | Scan | Method | Score | m/z |
| --- | --- | --- | --- | --- |
| A17_00_2 | 17328 | FTMS; HCD | 82.35 | 682.65 |

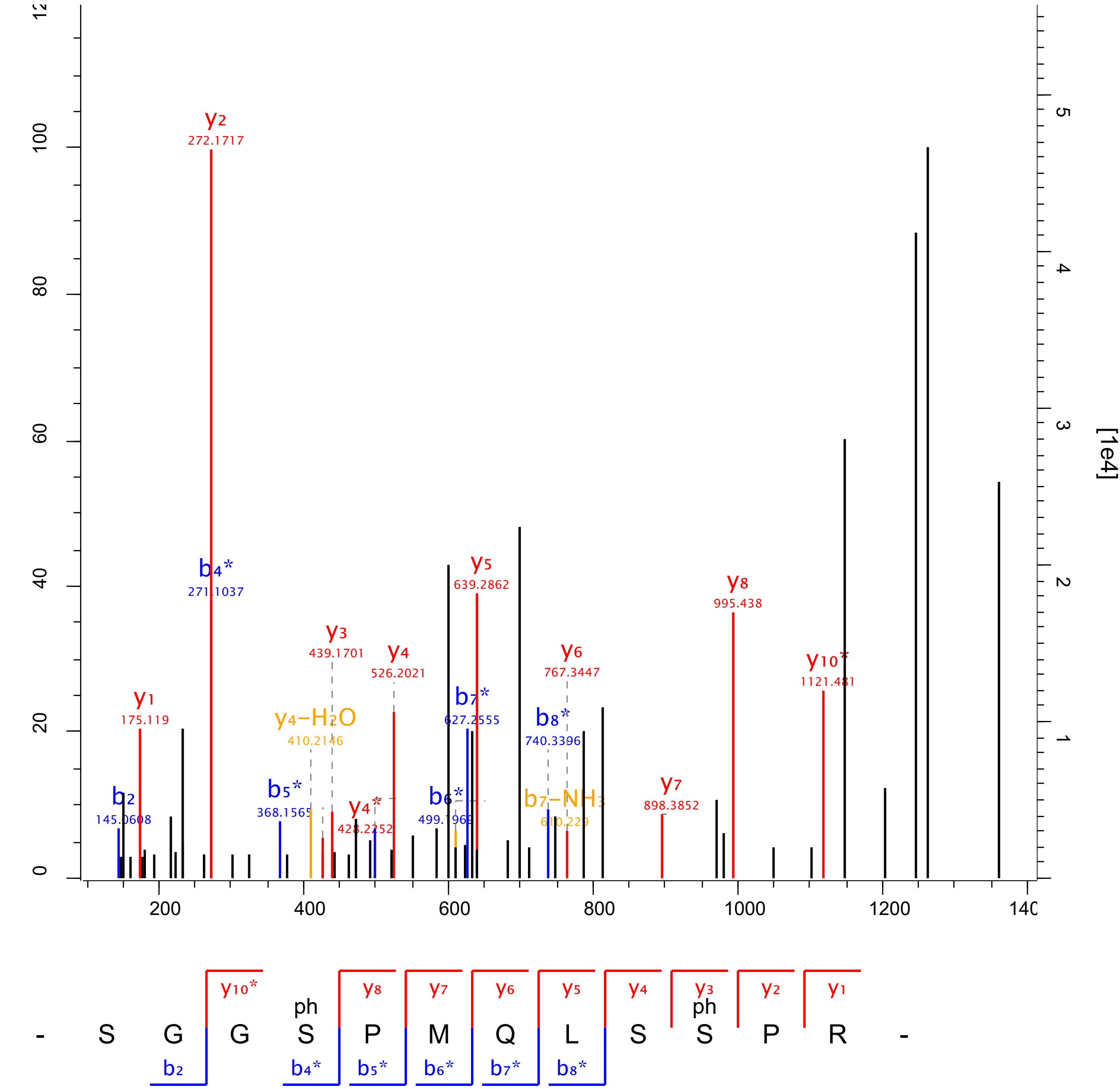

| Raw file | Scan | Method | Score | m/z |
|---|---|---|---|---|
| A17_00_2 | 17422 | FTMS; HCD | 102.69 | 858.8 |

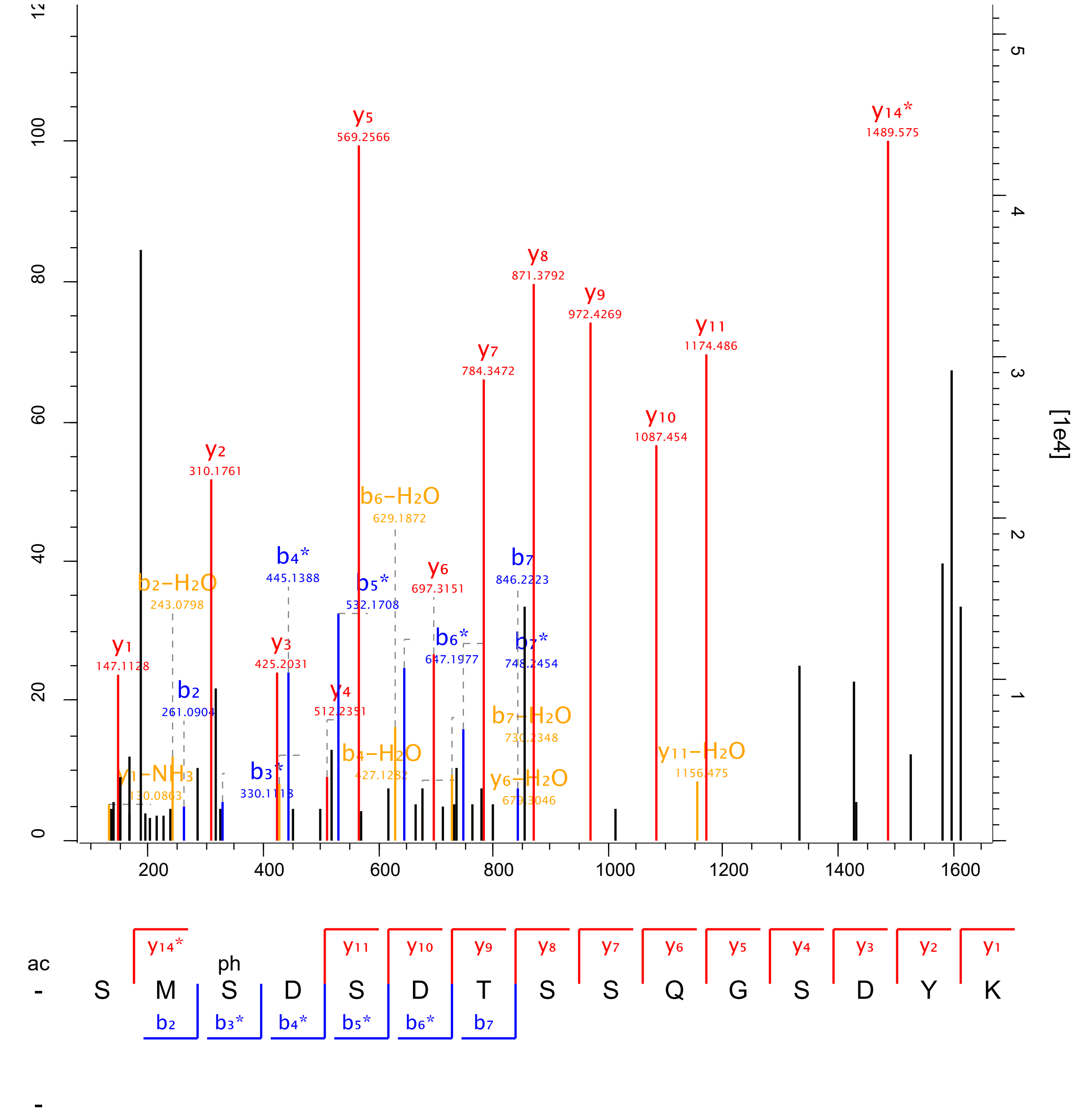

| Raw file | Scan | Method | Score | m/z |
| --- | --- | --- | --- | --- |
| A17_00_2 | 1749 | FTMS; HCD | 54.78 | 476.25 |

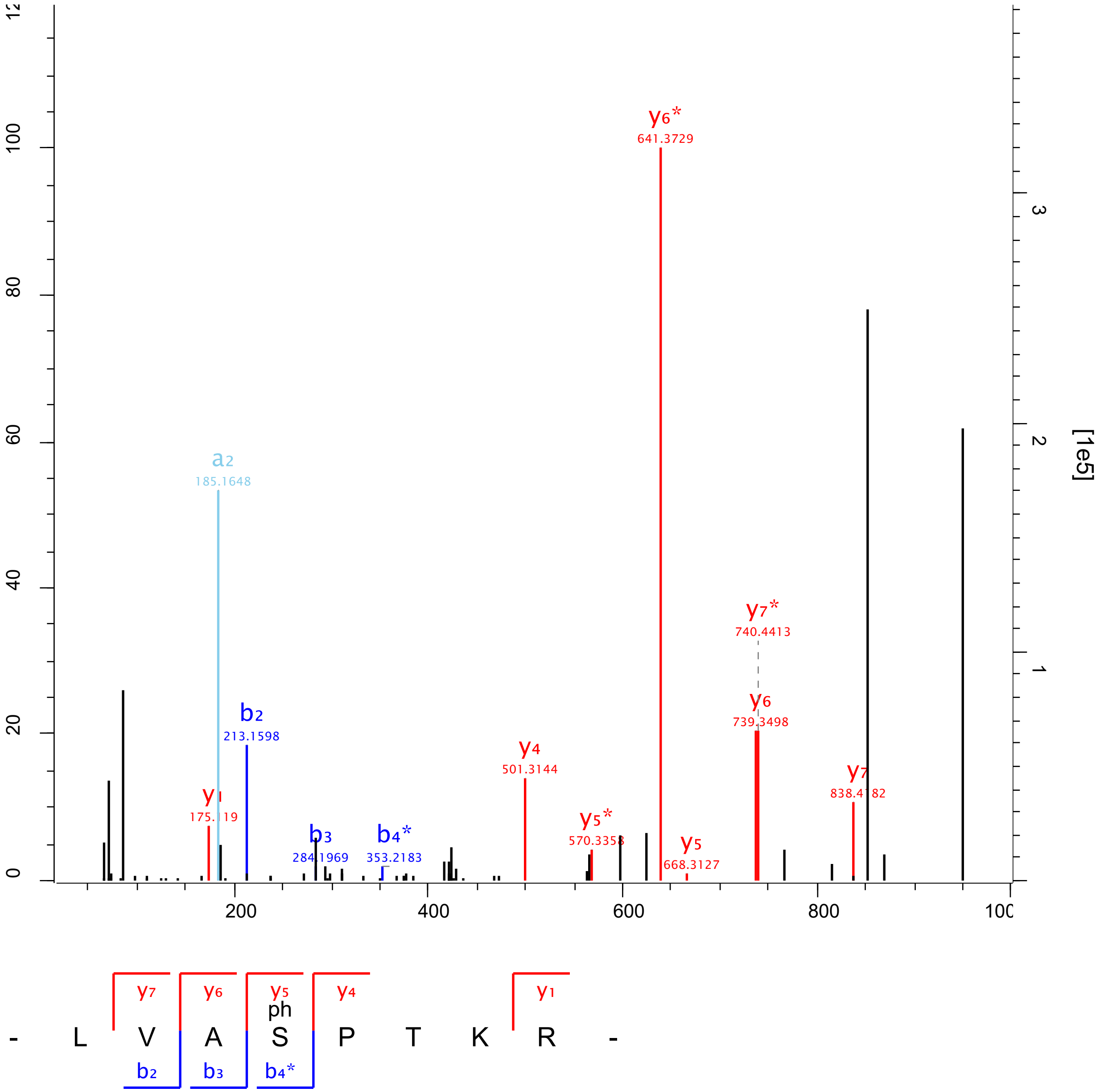

| Raw file | Scan | Method | Score | m/z |
|---|---|---|---|---|
| A17_00_2 | 17542 | FTMS; HCD | 78.46 | 817.3 |

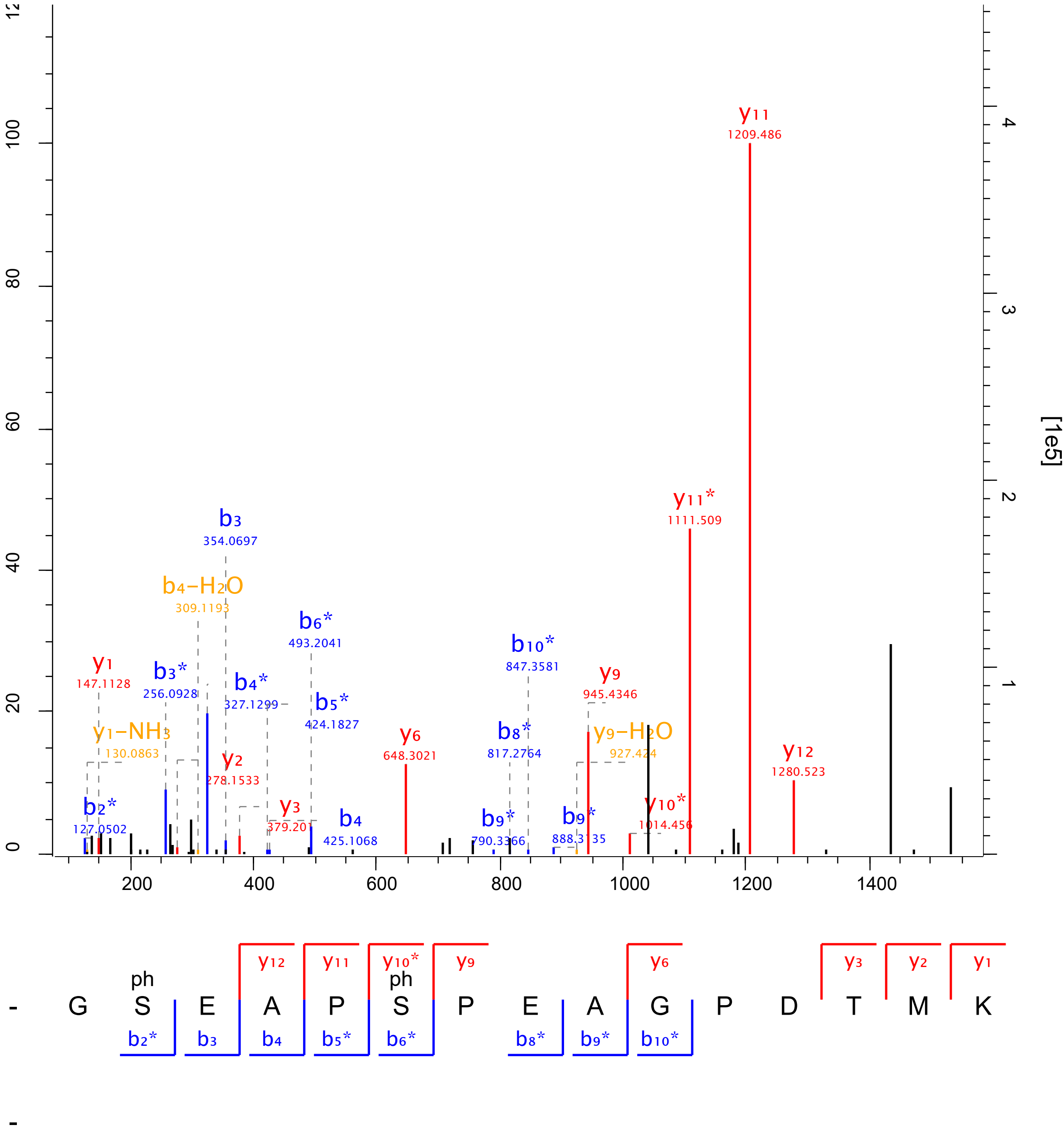

| Raw file | Scan | Method | Score | m/z |
| --- | --- | --- | --- | --- |
| A17_00_2 | 17576 | FTMS; HCD | 79.87 | 610.73 |

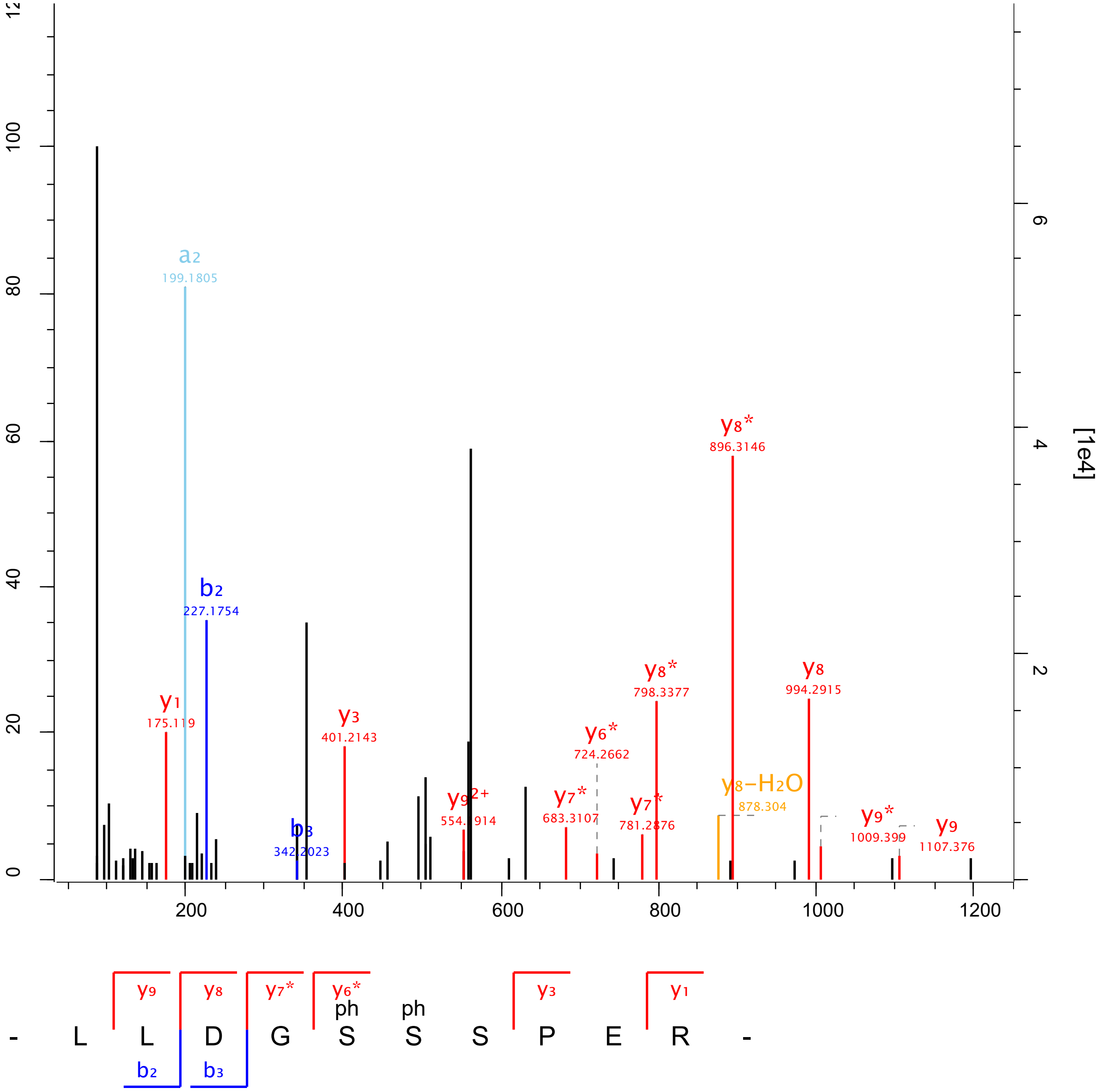

| Raw file | Scan | Method | Score | m/z |
|---|---|---|---|---|
| A17_00_2 | 17622 | FTMS; HCD | 151.95 | 580.27 |

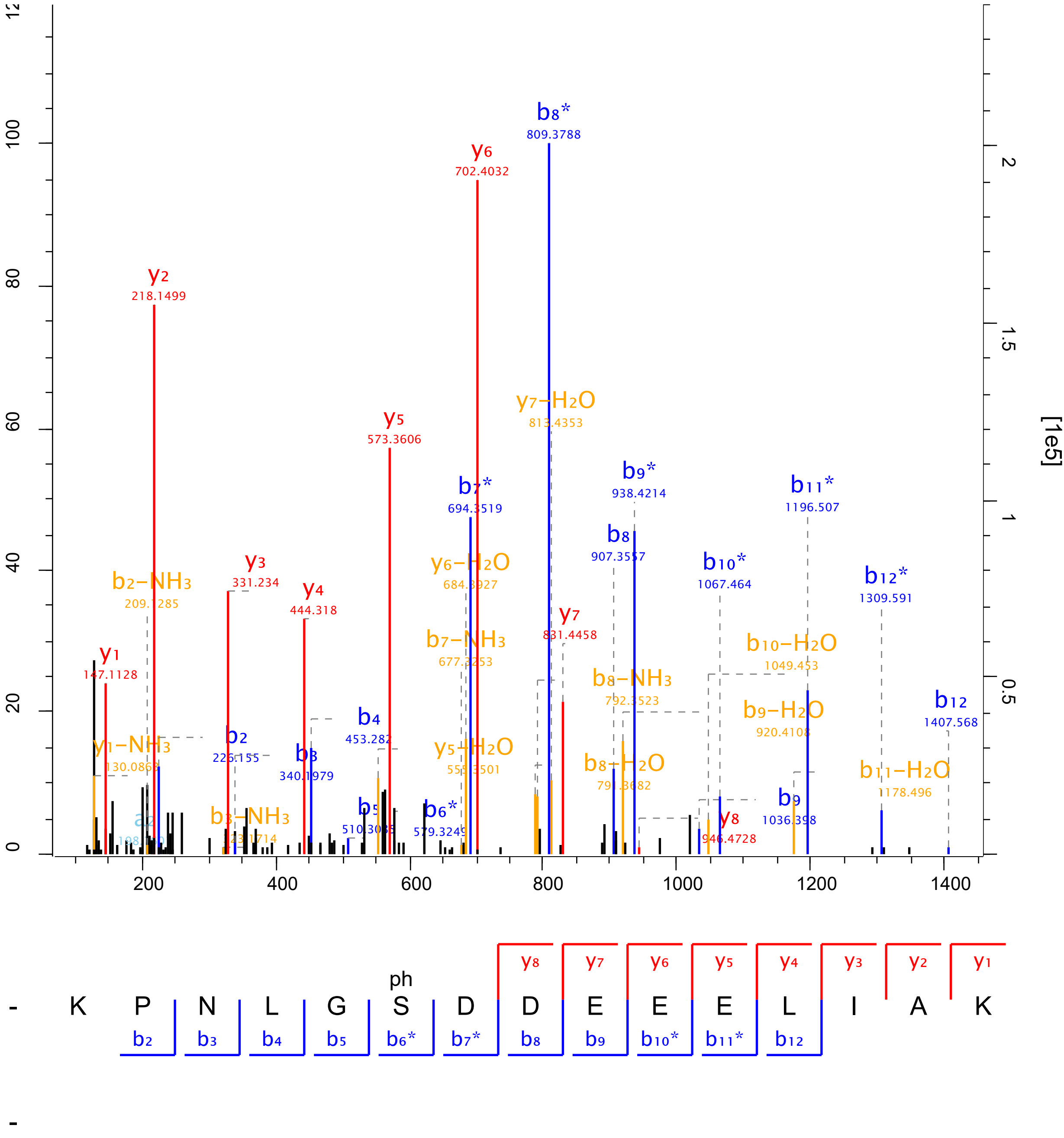

| Raw file | Scan | Method | Score | m/z |
| --- | --- | --- | --- | --- |
| A17_00_2 | 17642 | FTMS; HCD | 119.38 | 609.29 |

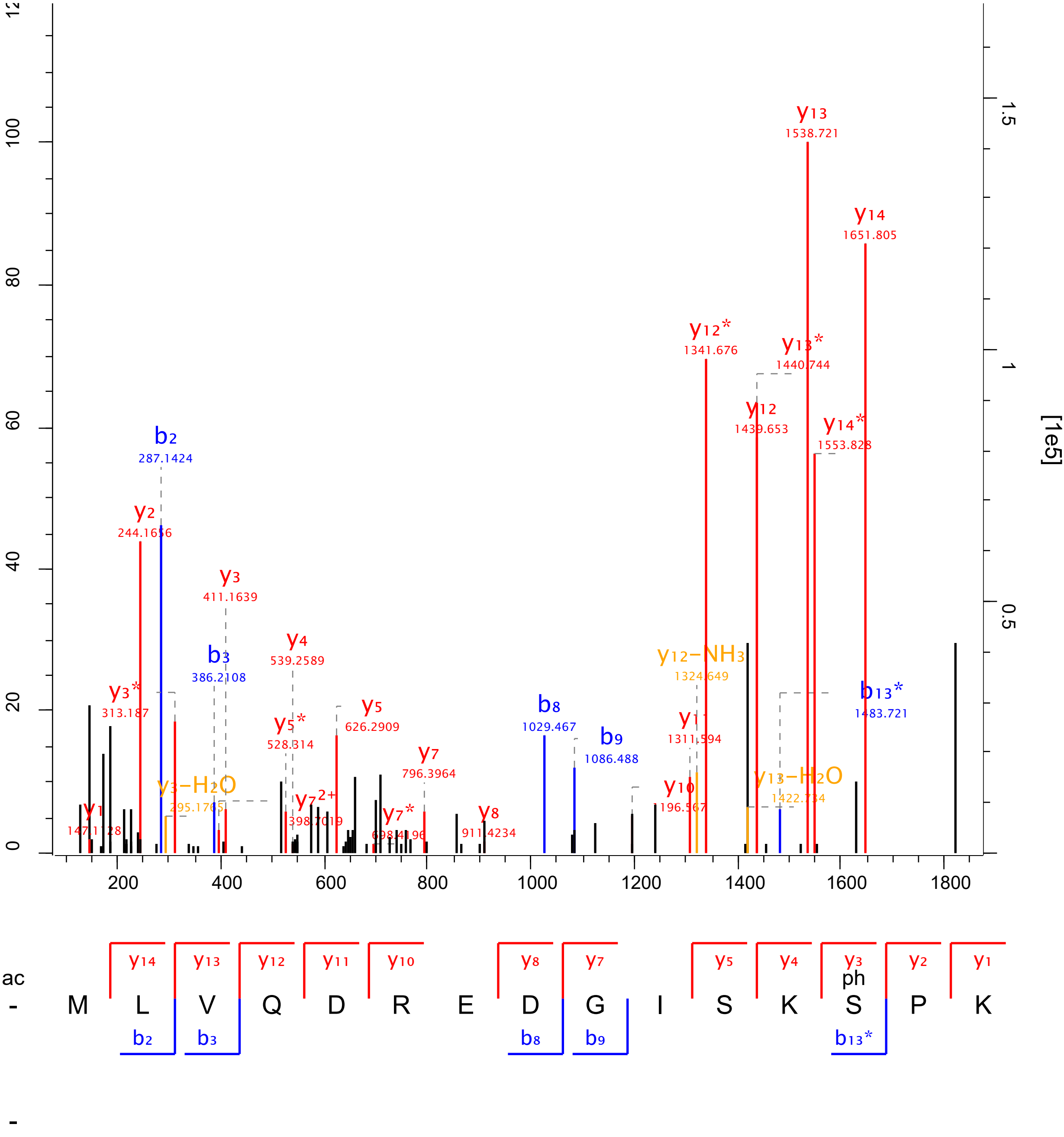

| Raw file | Scan | Method | Score | m/z |
| --- | --- | --- | --- | --- |
| A17_00_2 | 17705 | FTMS; HCD | 180.46 | 677.76 |

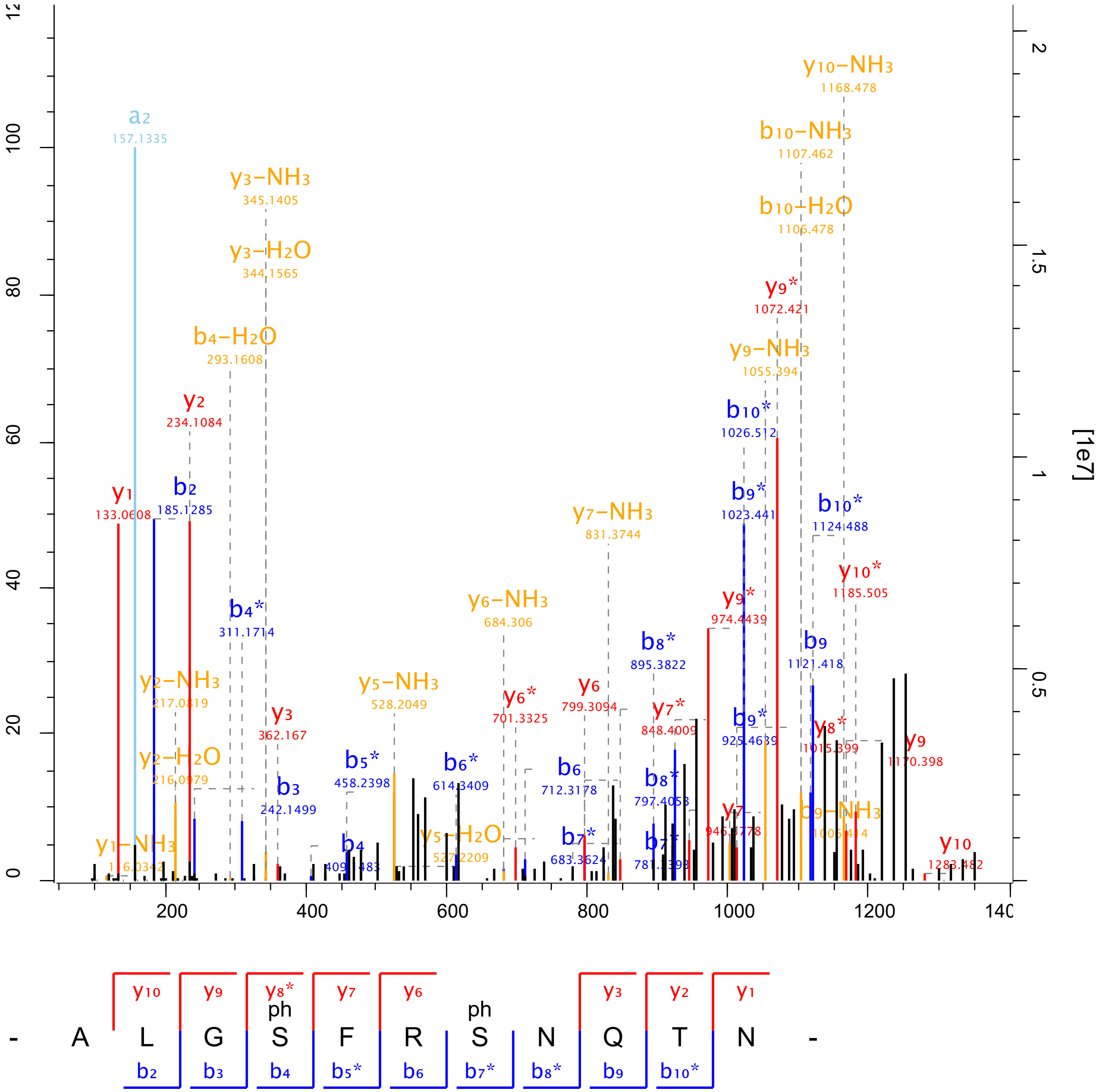

| Raw file | Scan | Method | Score | m/z |
|---|---|---|---|---|
| A17_00_2 | 17822 | FTMS; HCD | 168.67 | 668.77 |

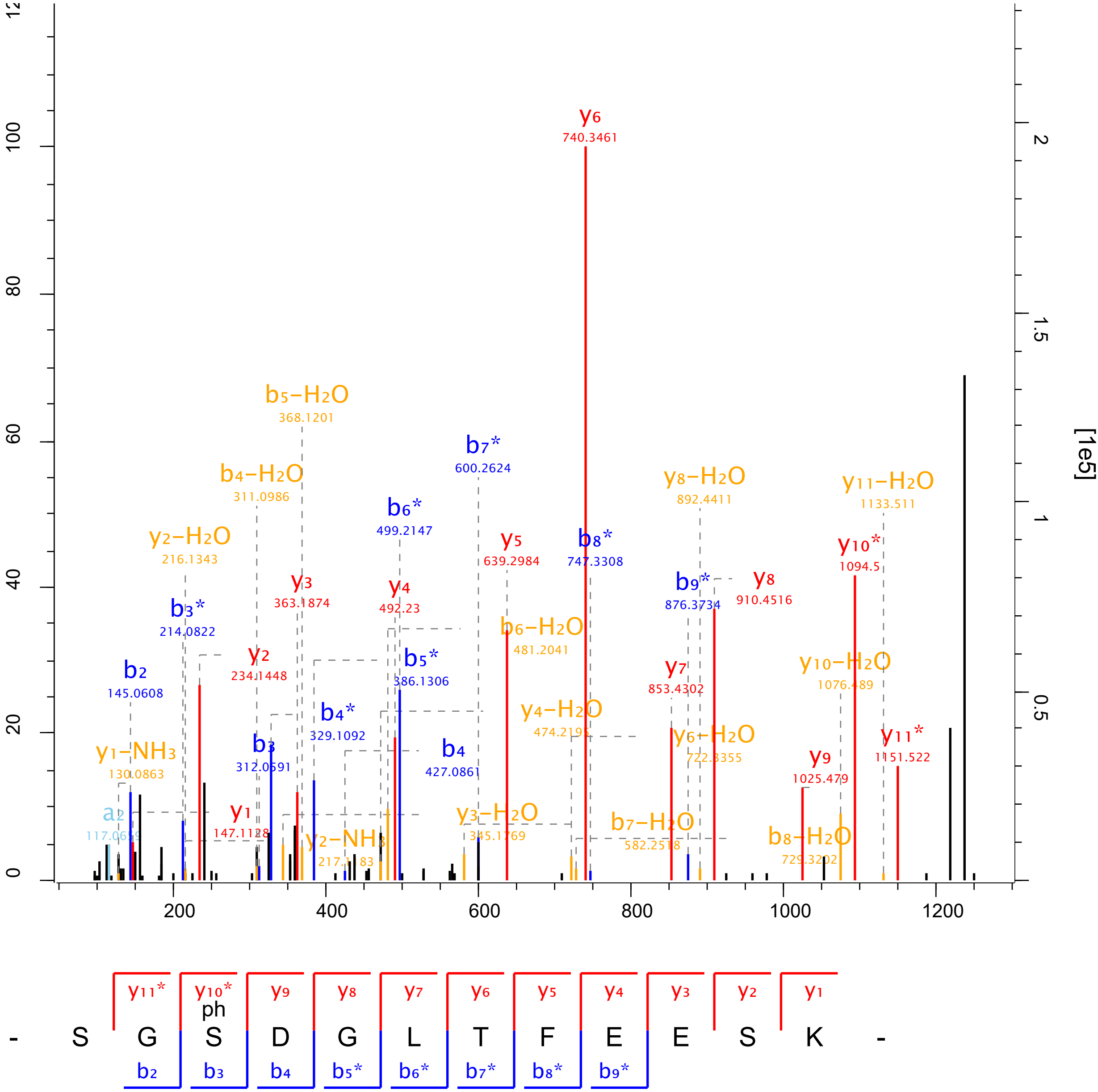

| Raw file | Scan | Method | Score | m/z |
|---|---|---|---|---|
| A17_00_2 | 17858 | FTMS; HCD | 114.7 | 550.71 |

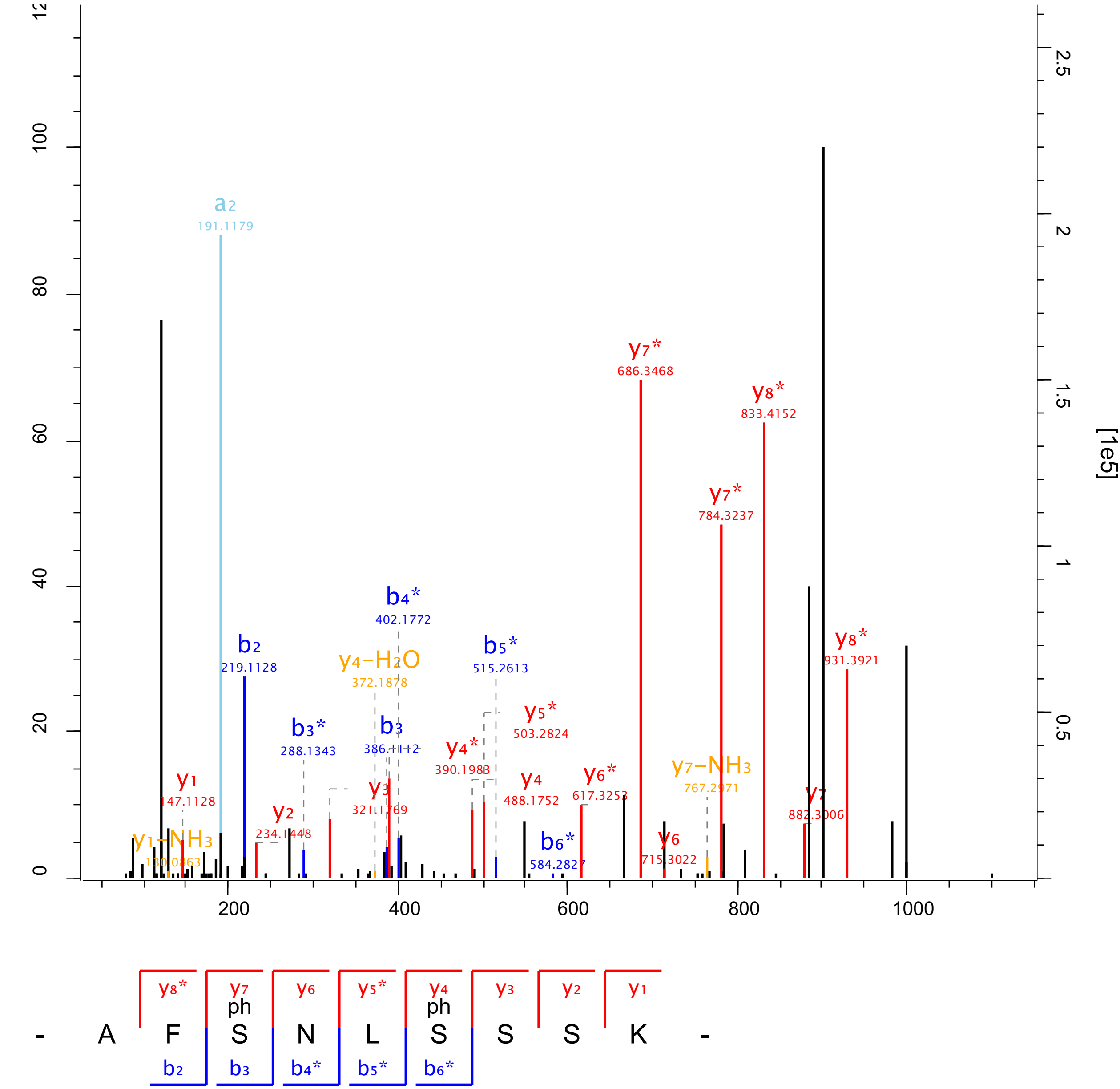

| Raw file | Scan | Method | Score | m/z |
|---|---|---|---|---|
| A17_00_2 | 17985 | FTMS; HCD | 61.24 | 596.28 |

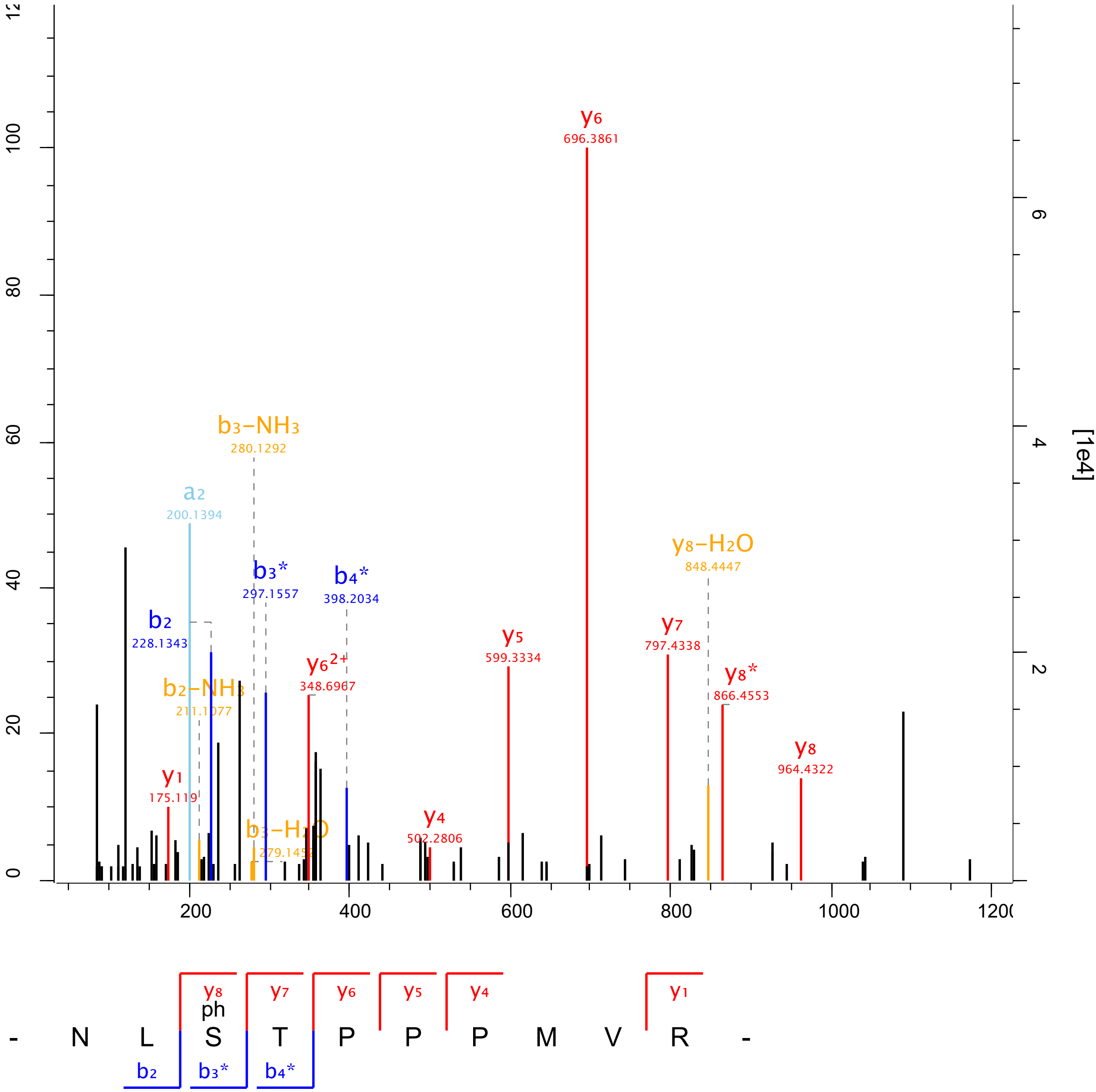

| Raw file | Scan | Method | Score | m/z |
|---|---|---|---|---|
| A17_00_2 | 18018 | FTMS; HCD | 148.72 | 706.3 |

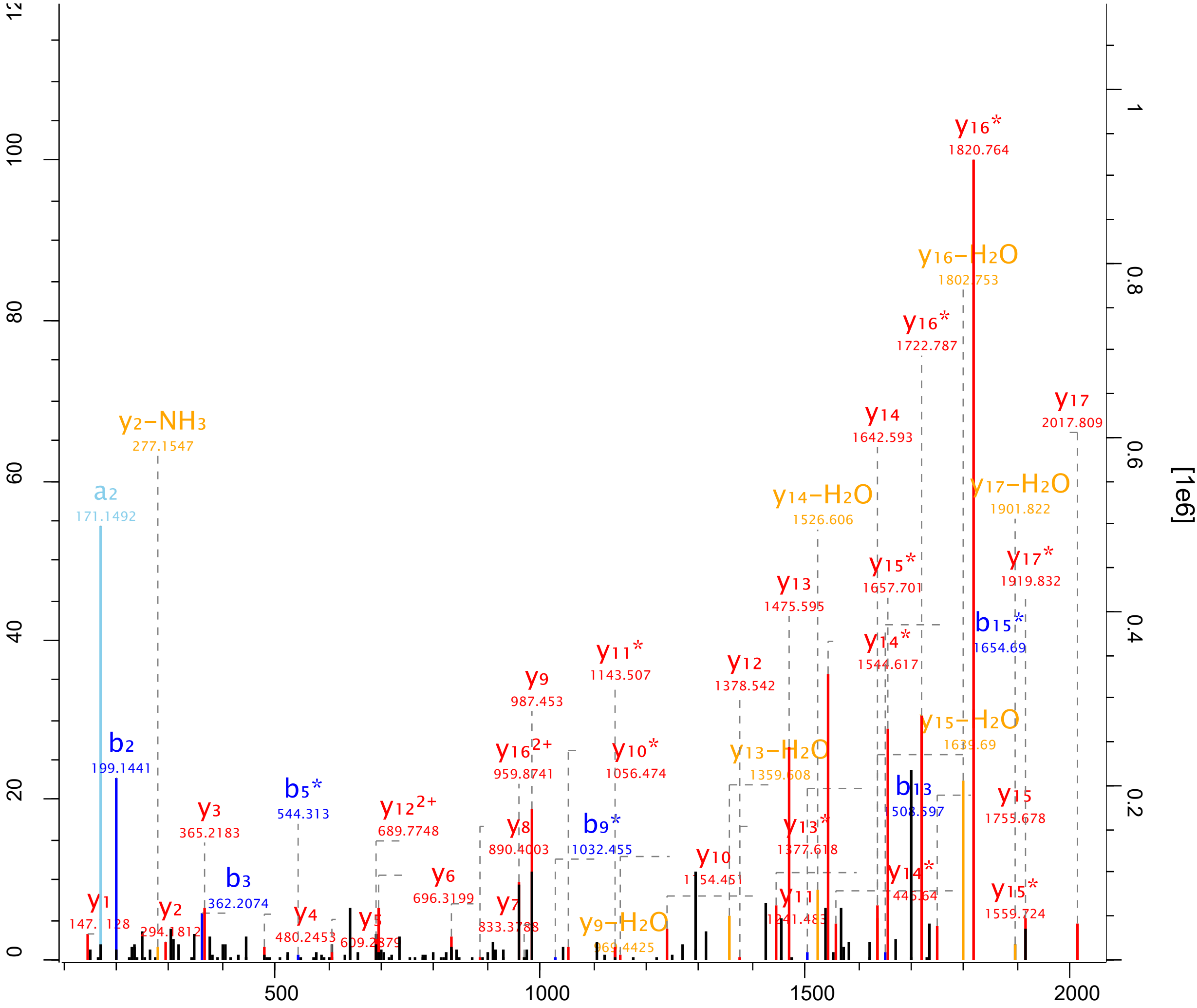


| | y17 | y16* | y15 | y14 | y13 | y12 | y11 | y10 | y9 | y8 | y7 | y6 | y5 | y4 |
|---|---|---|---|---|---|---|---|---|---|---|---|---|---|---|
| - V | V | Y | I | S (ph) | P | H | S | S (ph) | P | G | H | S | E | D |
| | b2 | b3 | | b5* | | | | b9* | | | | b13 | | b15* |

| y3 | y2 | y1 | |
|---|---|---|---|
| A | F | K | - |

| Raw file | Scan | Method | Score | m/z |
|---|---|---|---|---|
| A17_00_2 | 18093 | FTMS; HCD | 81.79 | 792.84 |

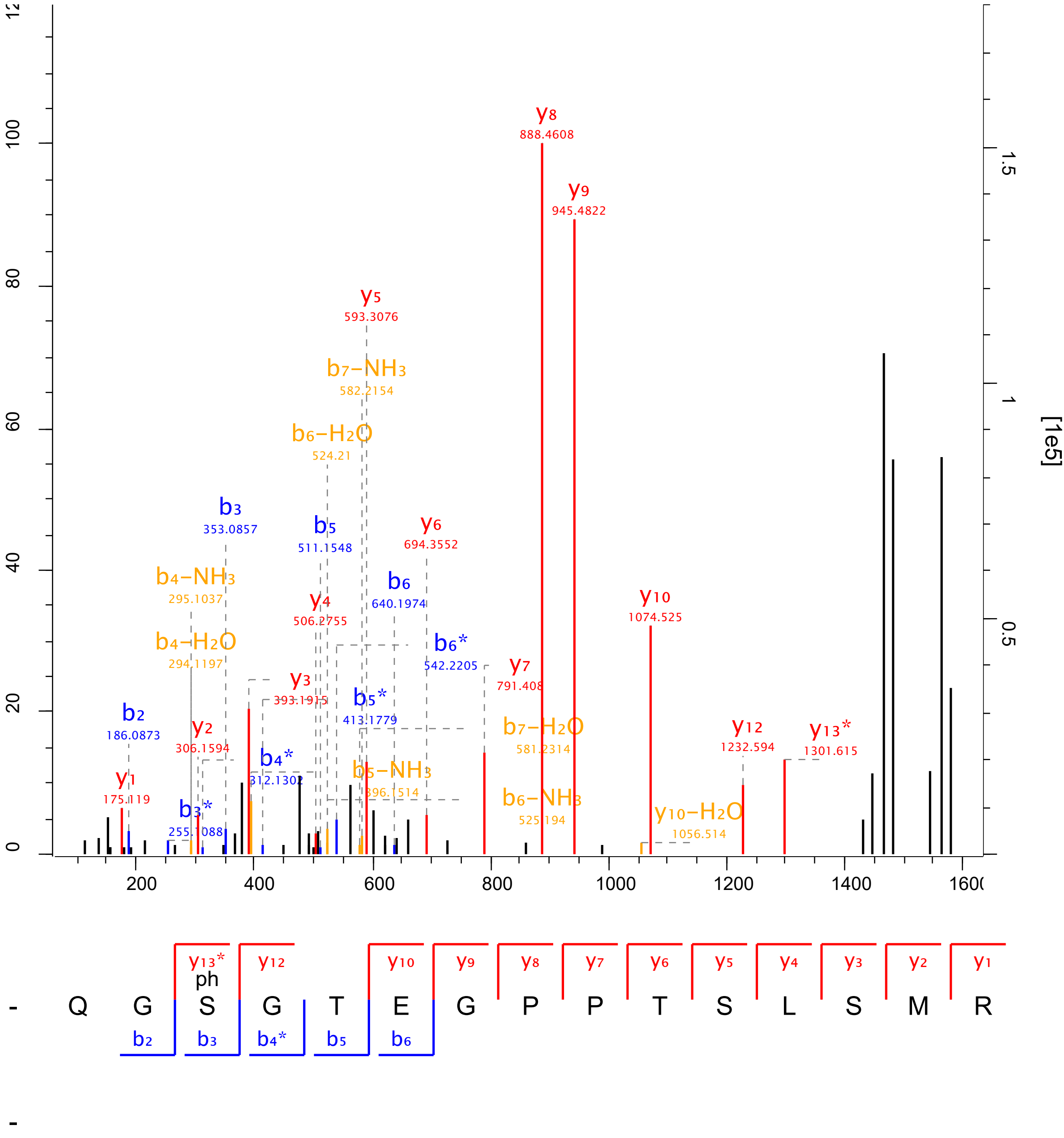

| Raw file | Scan | Method | Score | m/z |
| --- | --- | --- | --- | --- |
| A17_00_2 | 1817 | FTMS; HCD | 67.1 | 474.86 |

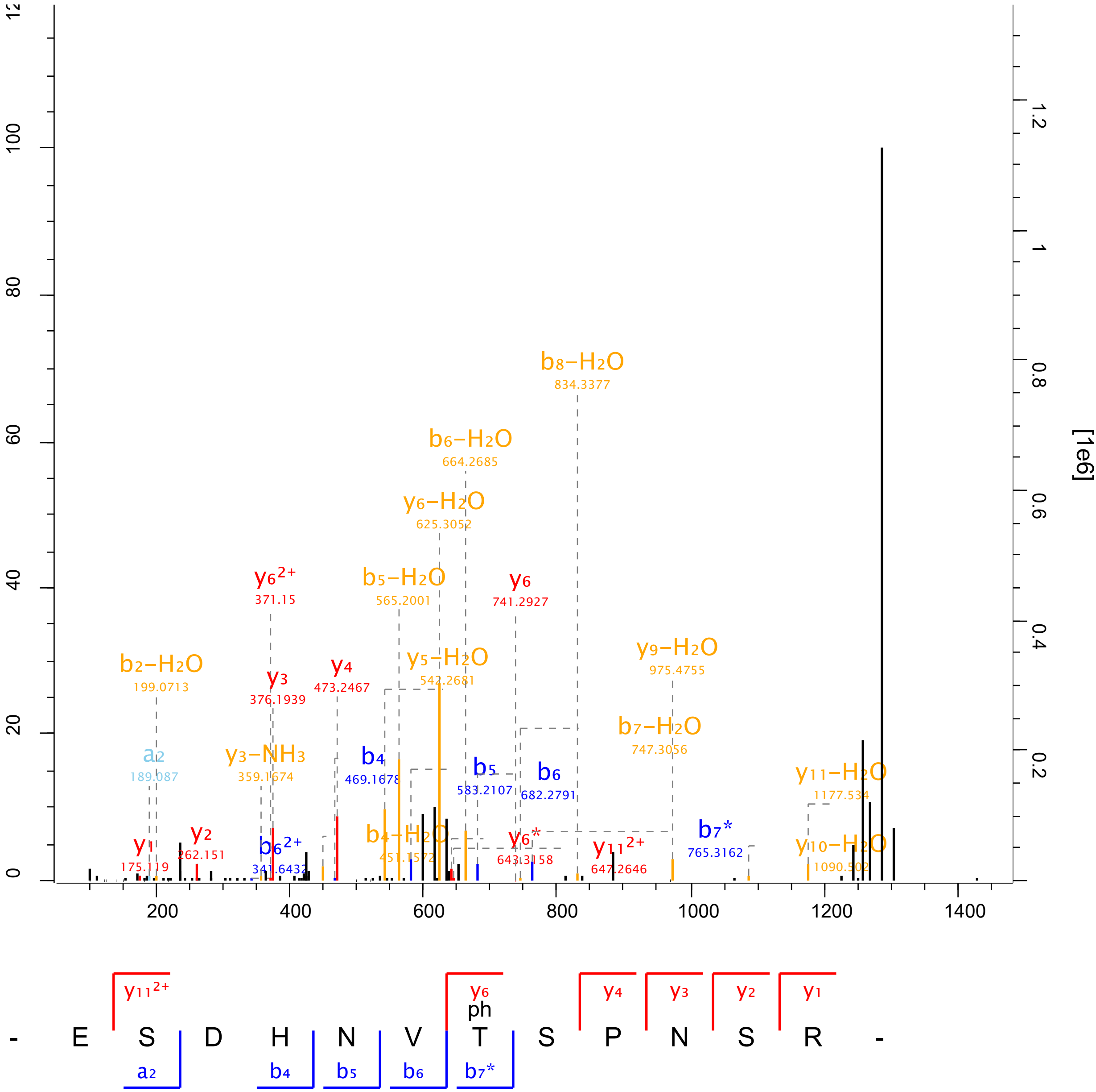

| Raw file | Scan | Method | Score | m/z |
|---|---|---|---|---|
| A17_00_2 | 18286 | FTMS; HCD | 92.39 | 565.59 |

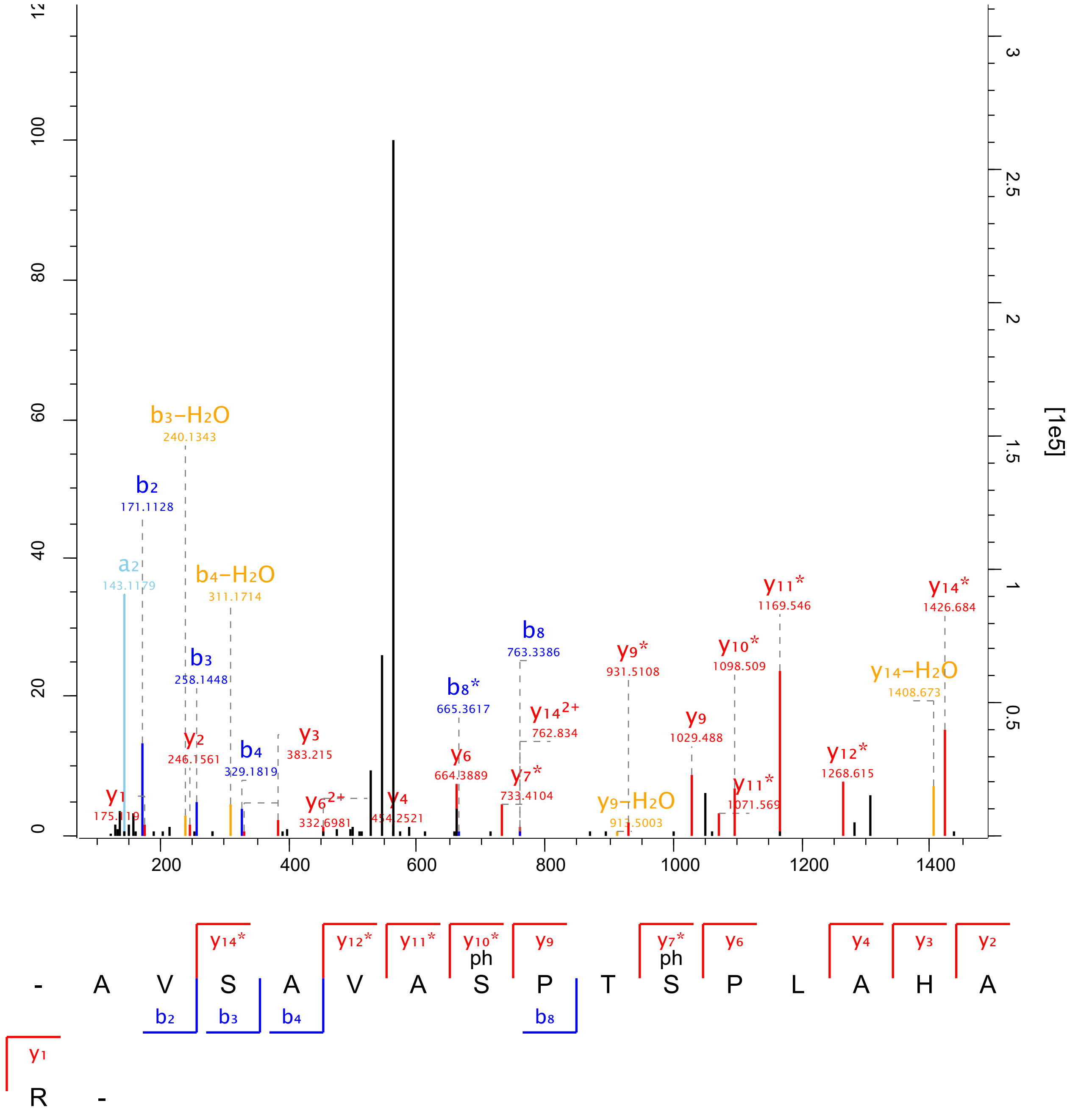

| Raw file | Scan | Method | Score | m/z |
| --- | --- | --- | --- | --- |
| A17_00_2 | 18310 | FTMS; HCD | 43.03 | 560.59 |

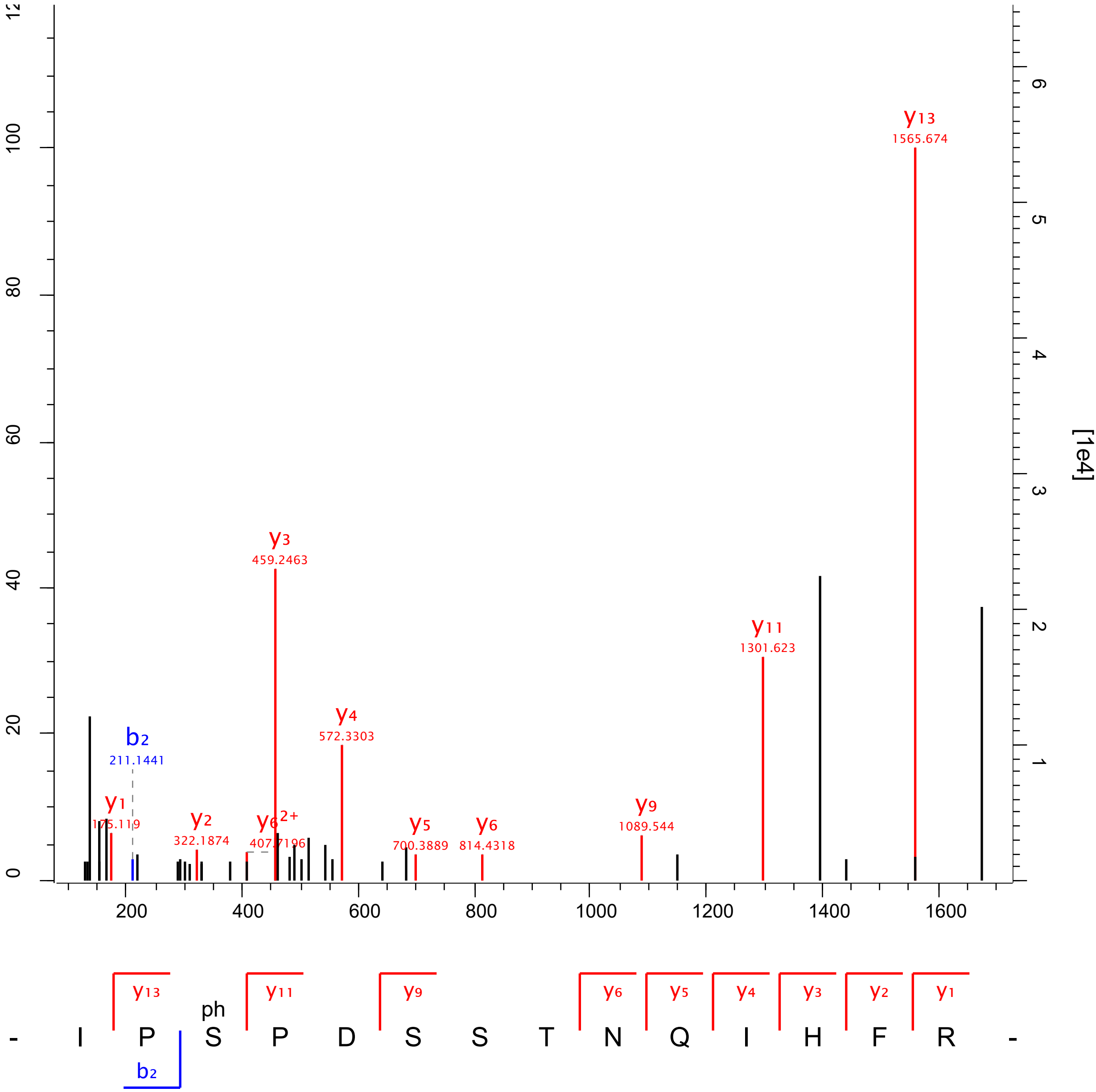

| Raw file | Scan | Method | Score | m/z |
|---|---|---|---|---|
| A17_00_2 | 1872 | FTMS; HCD | 130.47 | 598.75 |

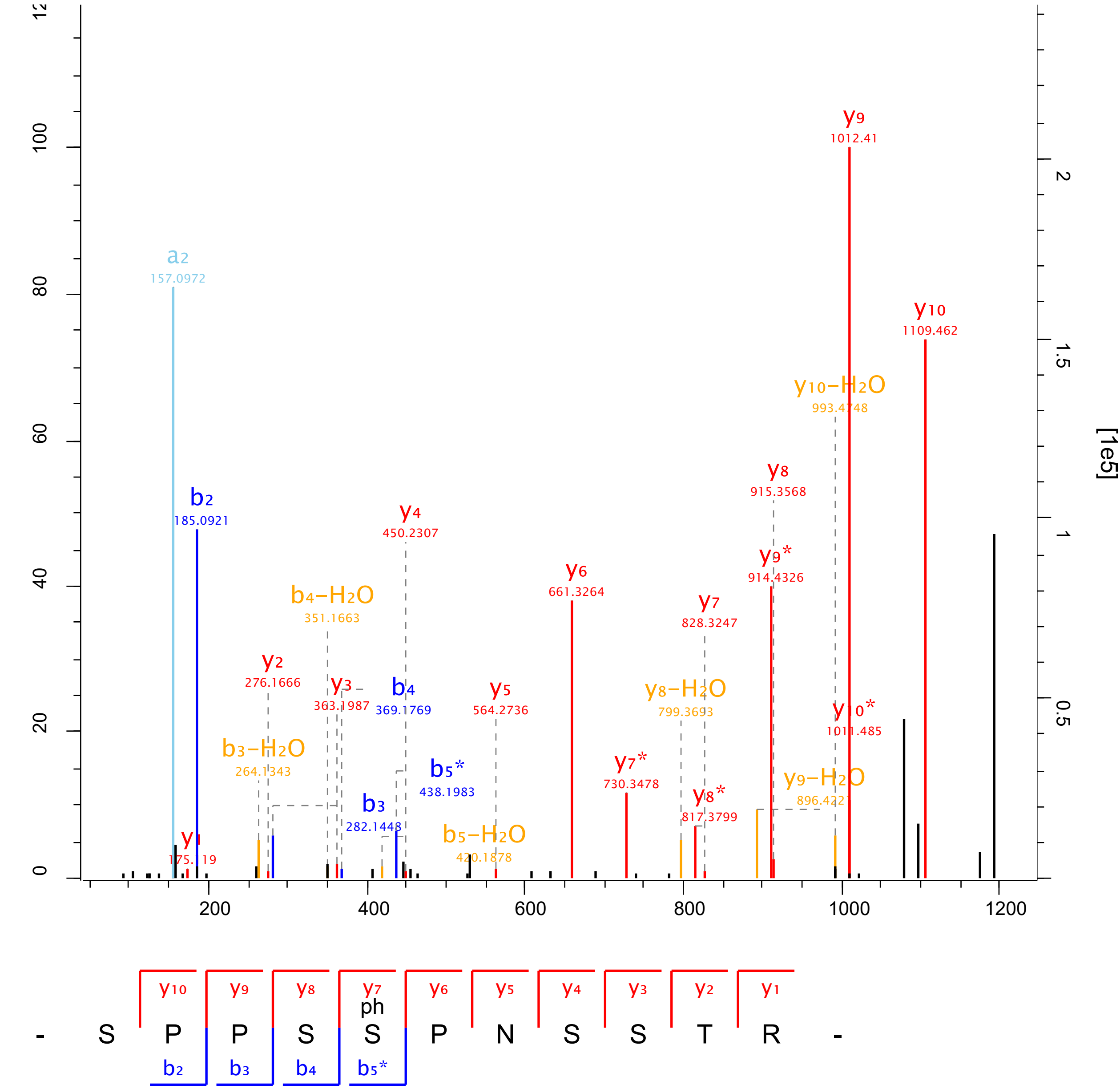

| Raw file | Scan | Method | Score | m/z |
| --- | --- | --- | --- | --- |
| A17_00_2 | 18875 | FTMS; HCD | 244.89 | 883.87 |

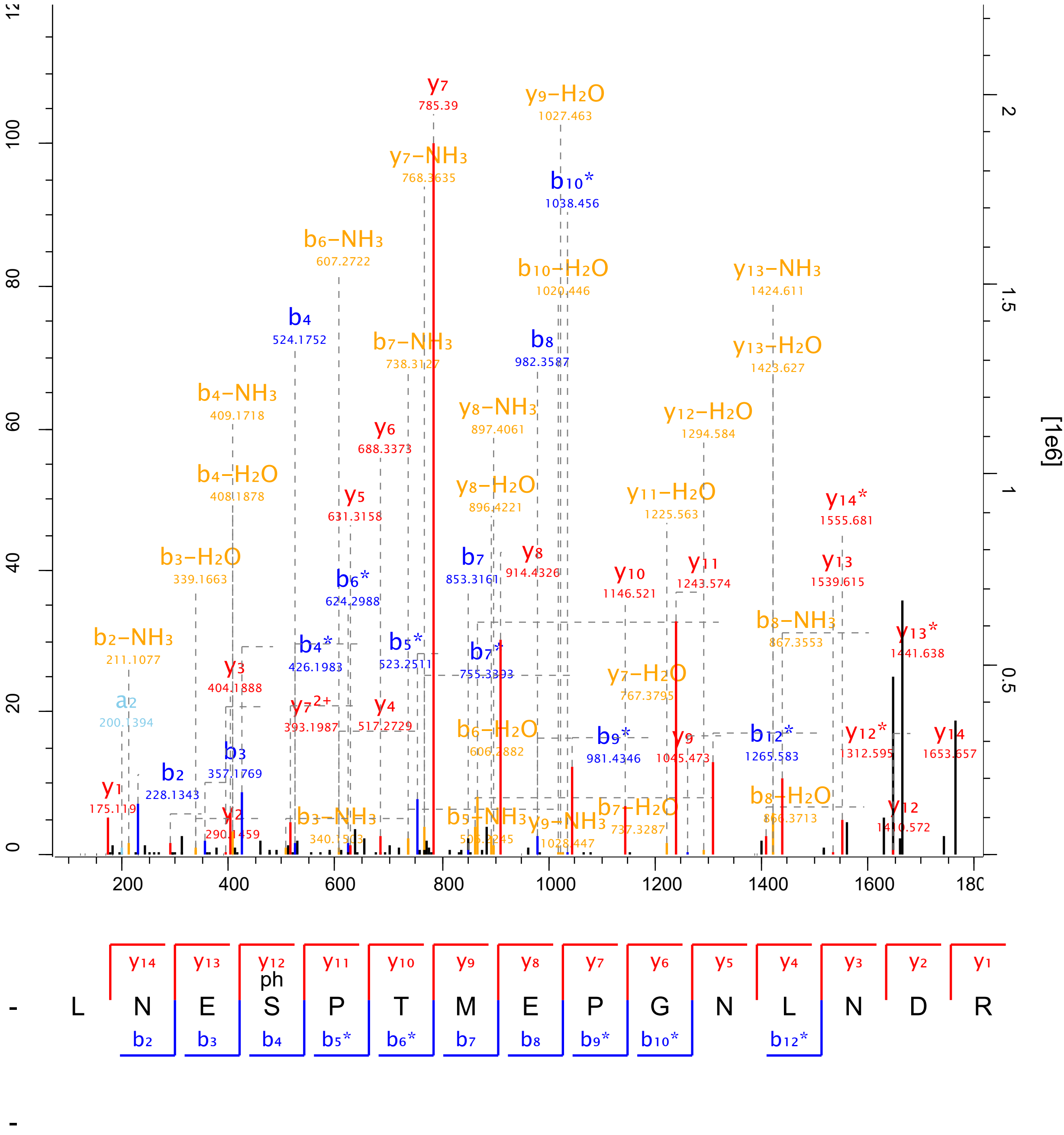

| Raw file | Scan | Method | Score | m/z |
|---|---|---|---|---|
| A17_00_2 | 1896 | FTMS; HCD | 49.59 | 497.21 |

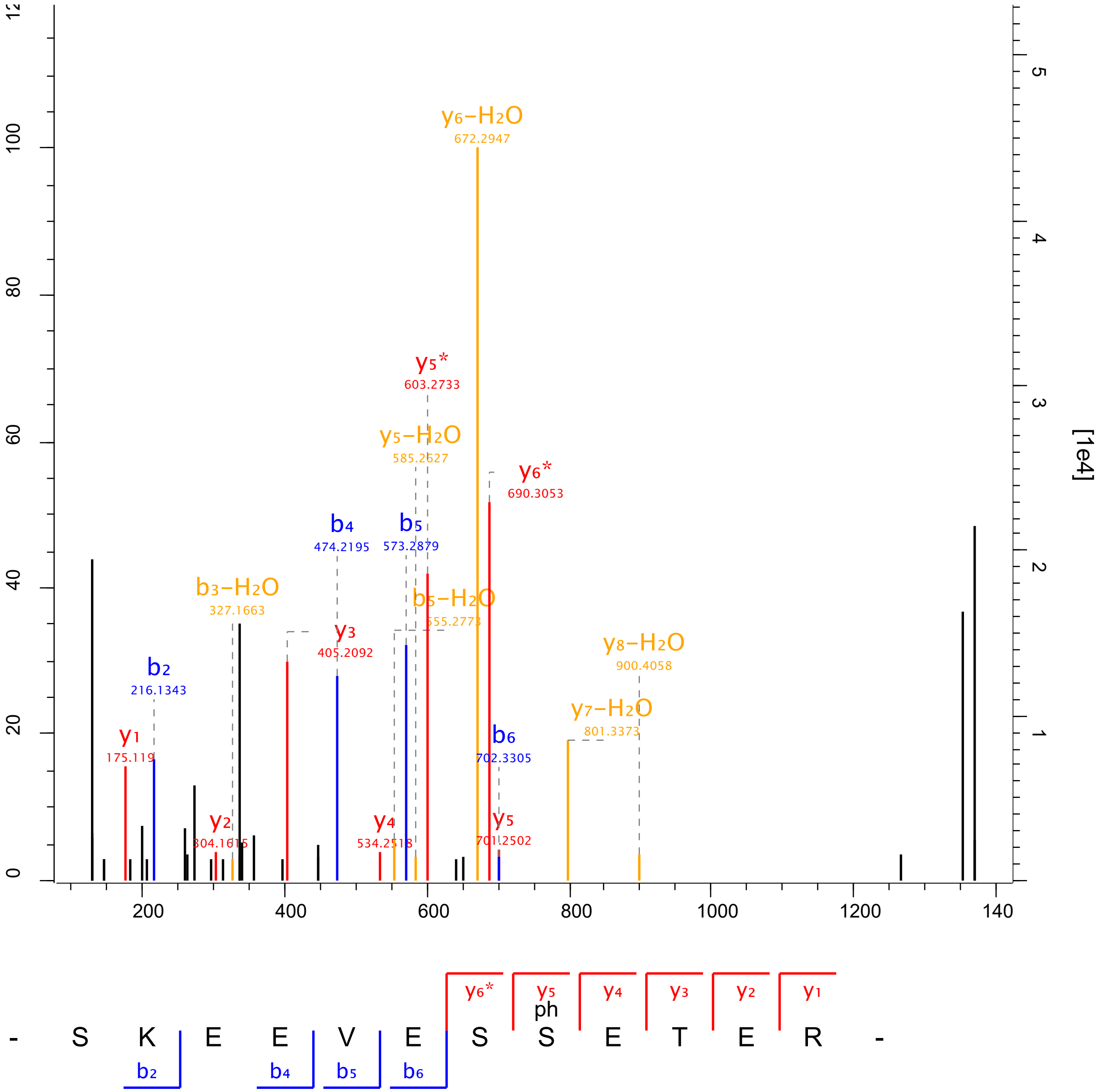

| Raw file | Scan | Method | Score | m/z |
| --- | --- | --- | --- | --- |
| A17_00_2 | 19243 | FTMS; HCD | 42.86 | 732.62 |

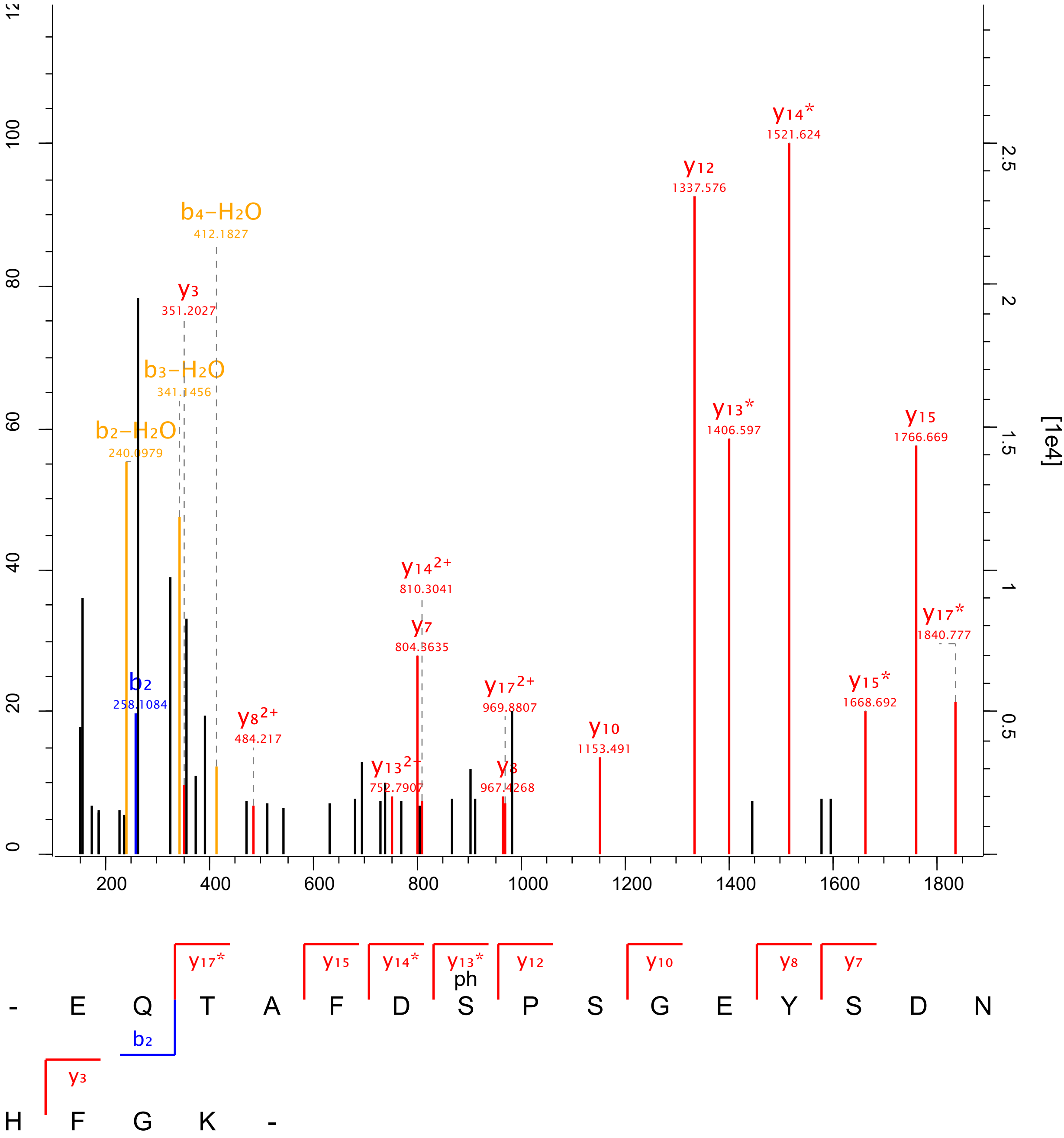

| Raw file | Scan | Method | Score | m/z |
|---|---|---|---|---|
| A17_00_2 | 19453 | FTMS; HCD | 52.17 | 768.32 |

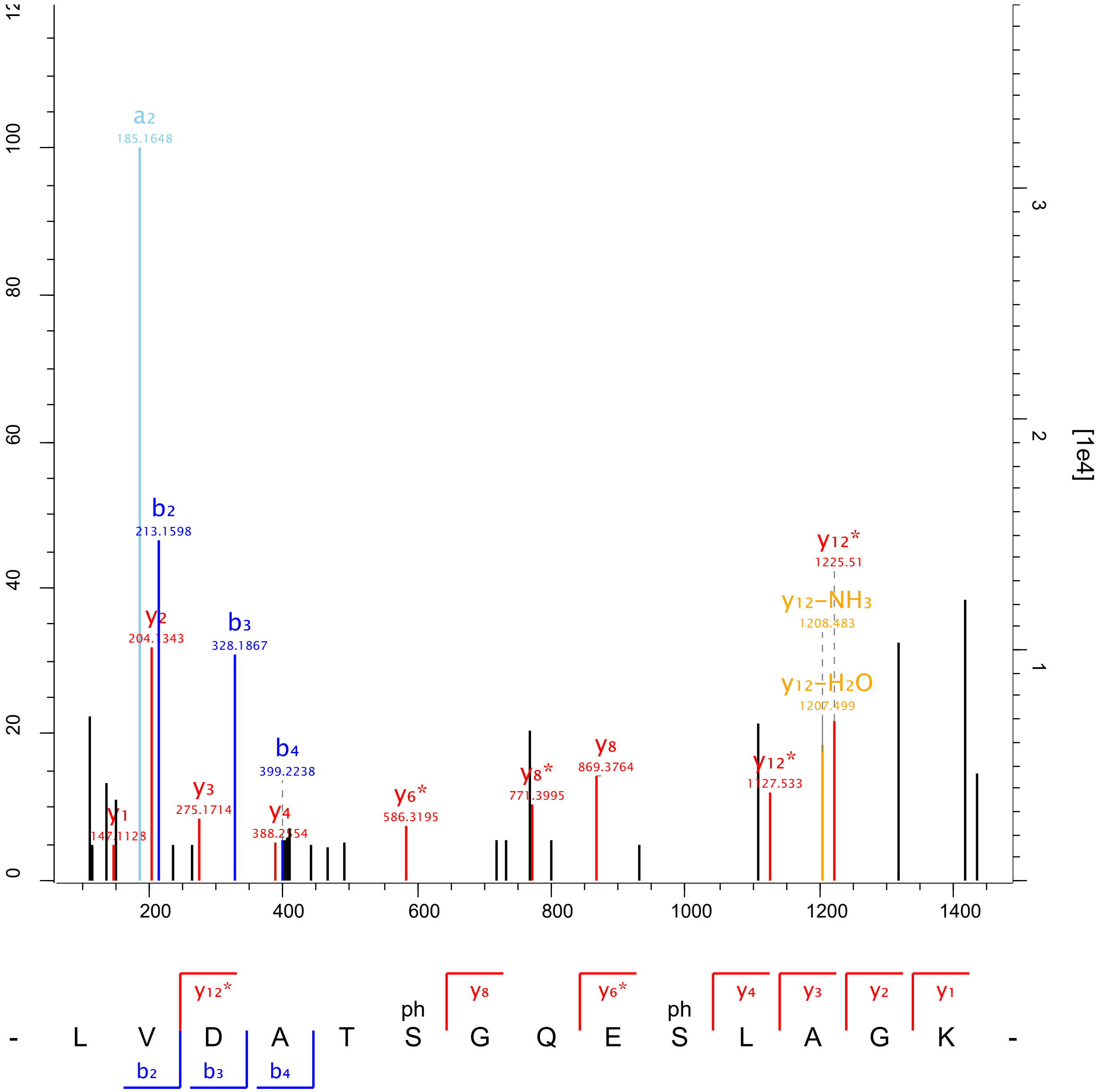

| Raw file | Scan | Method | Score | m/z |
| --- | --- | --- | --- | --- |
| A17_00_2 | 19599 | FTMS; HCD | 70.78 | 907.88 |

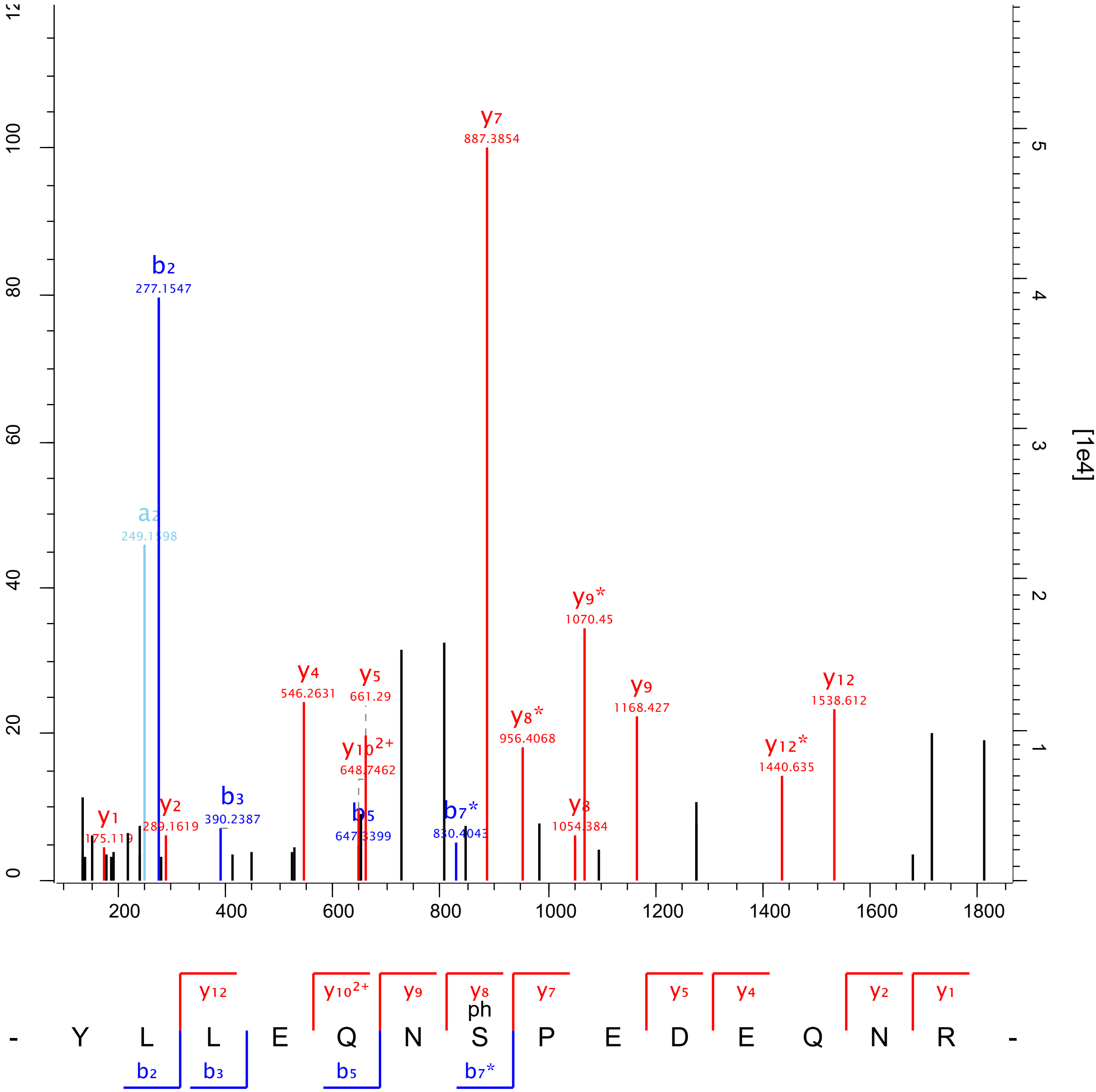

| Raw file | Scan | Method | Score | m/z |
|---|---|---|---|---|
| A17_00_2 | 1971 | FTMS; HCD | 141.65 | 592.75 |

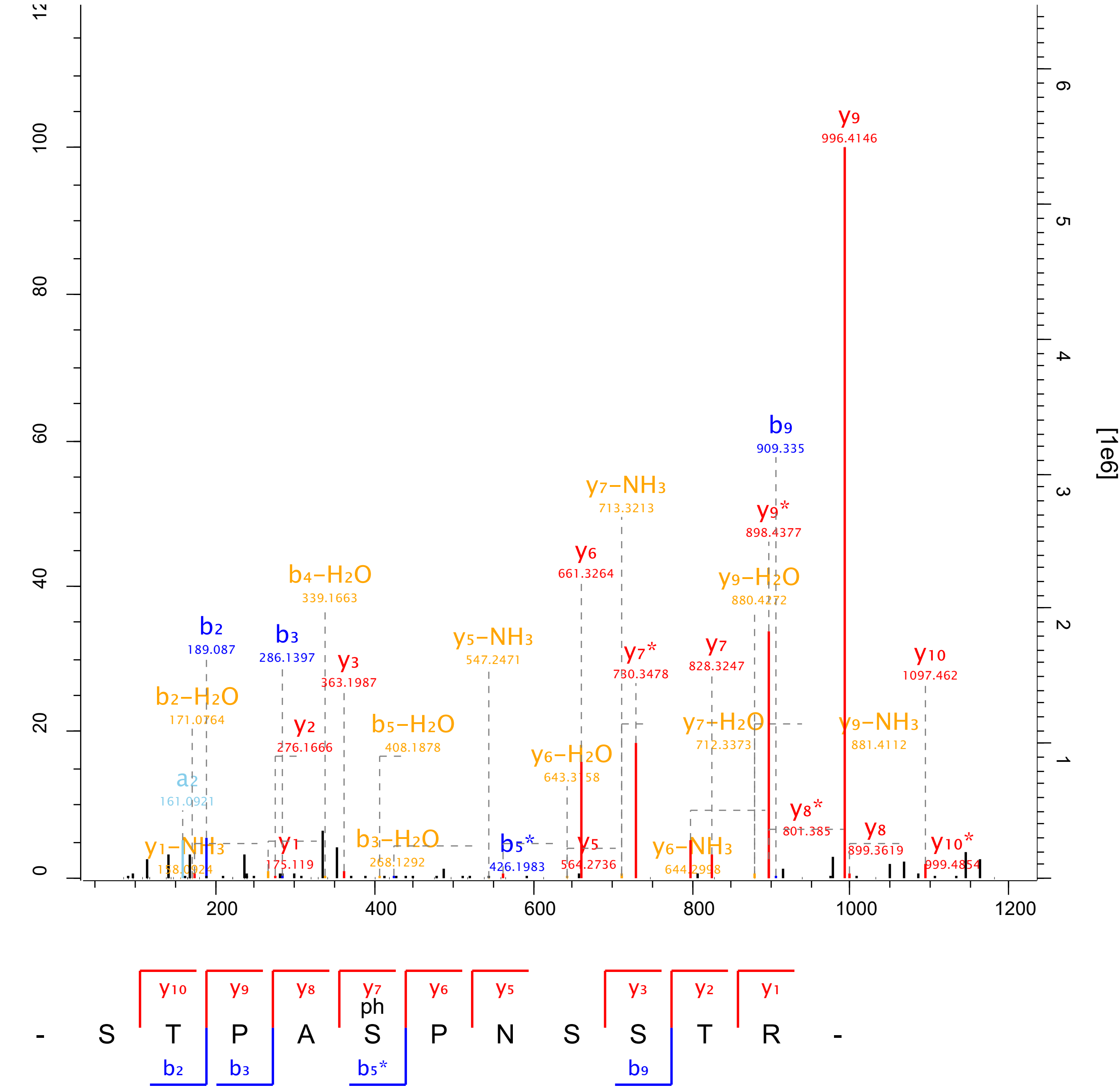

| Raw file | Scan | Method | Score | m/z |
| --- | --- | --- | --- | --- |
| A17_00_2 | 19877 | FTMS; HCD | 85.62 | 506.73 |

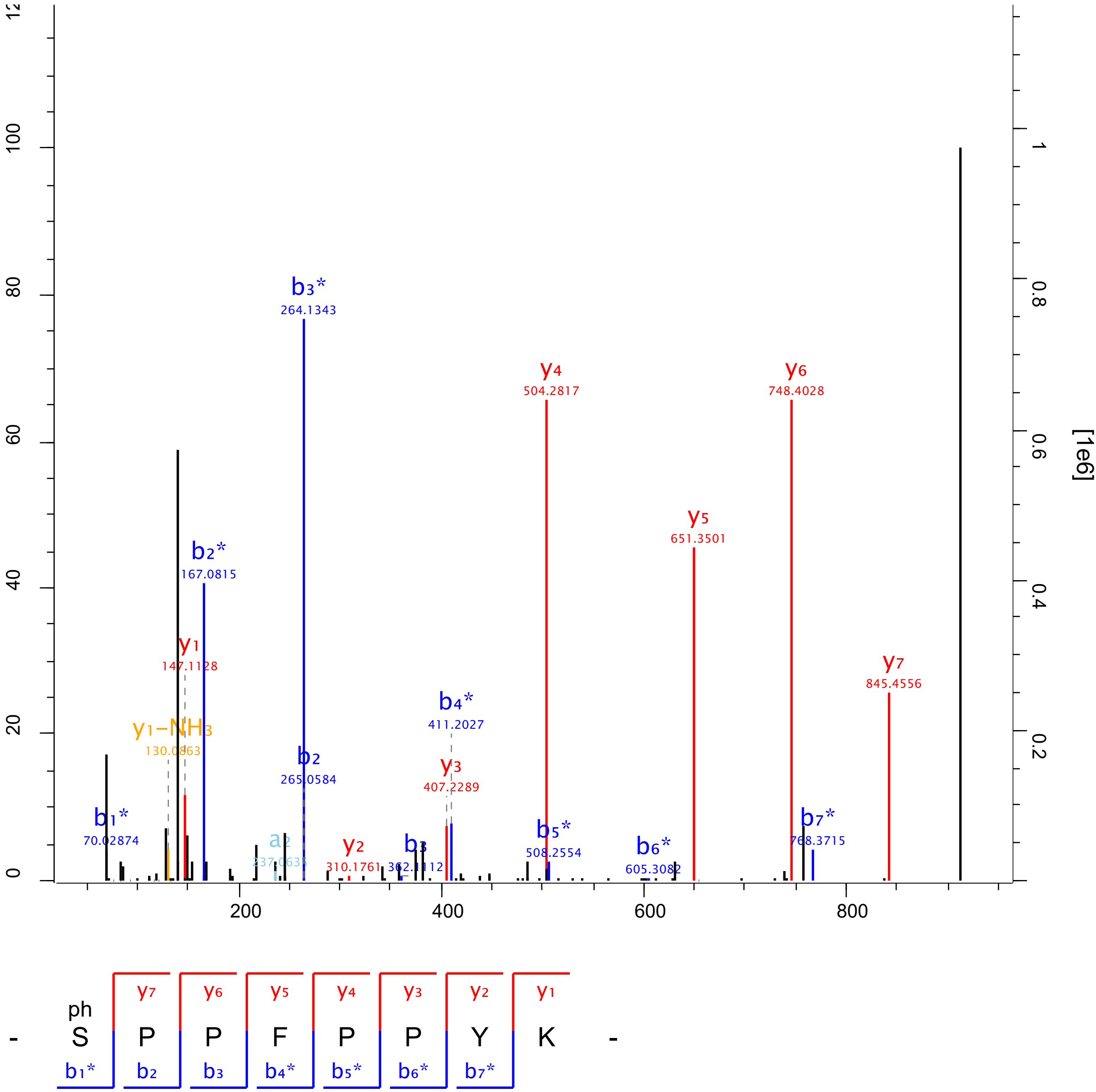

| Raw file | Scan | Method | Score | m/z |
|---|---|---|---|---|
| A17_00_2 | 20015 | FTMS; HCD | 42.12 | 652.3 |

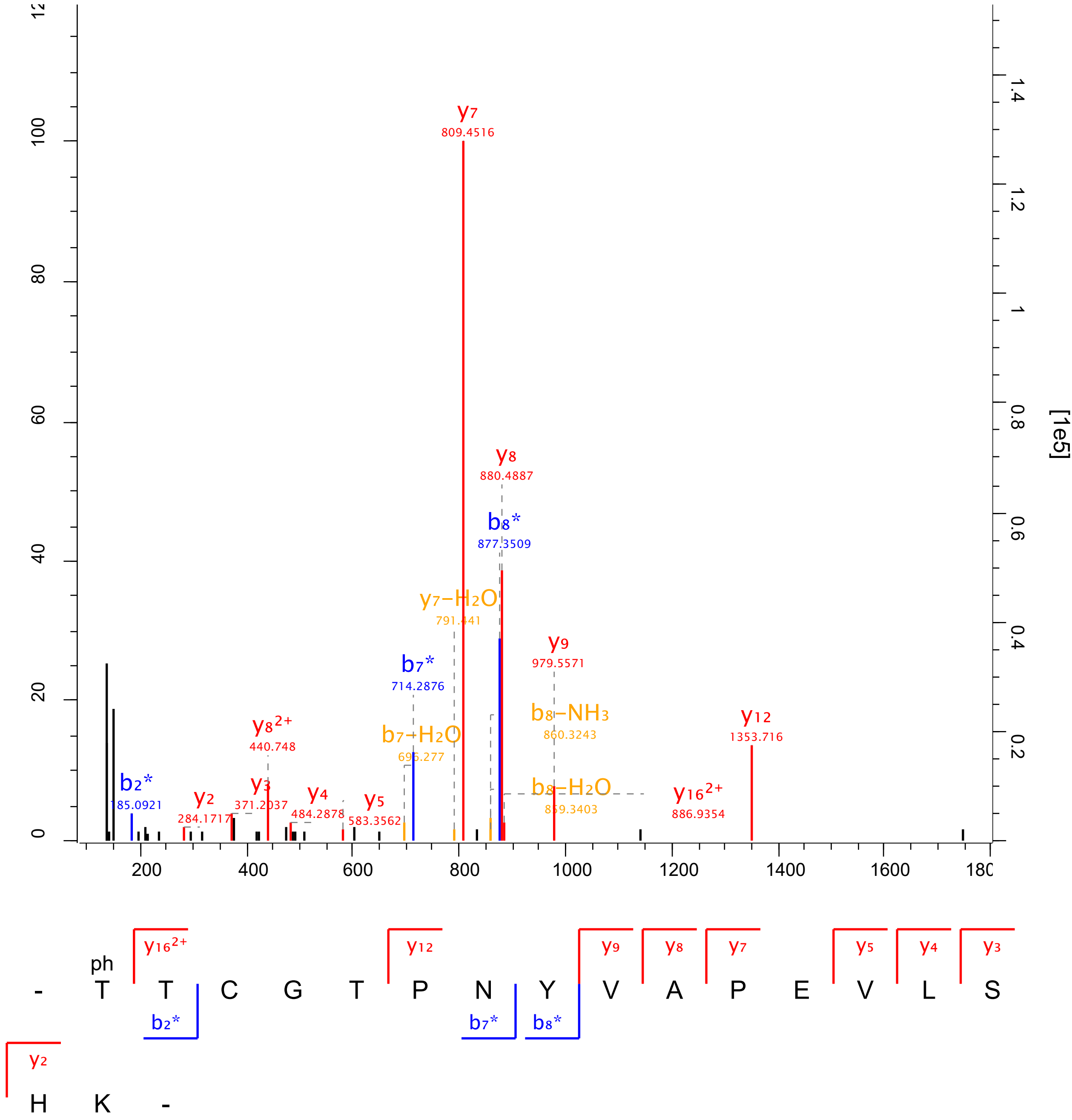

| Raw file | Scan | Method | Score | m/z |
|---|---|---|---|---|
| A17_00_2 | 2034 | FTMS; HCD | 102.65 | 645.78 |

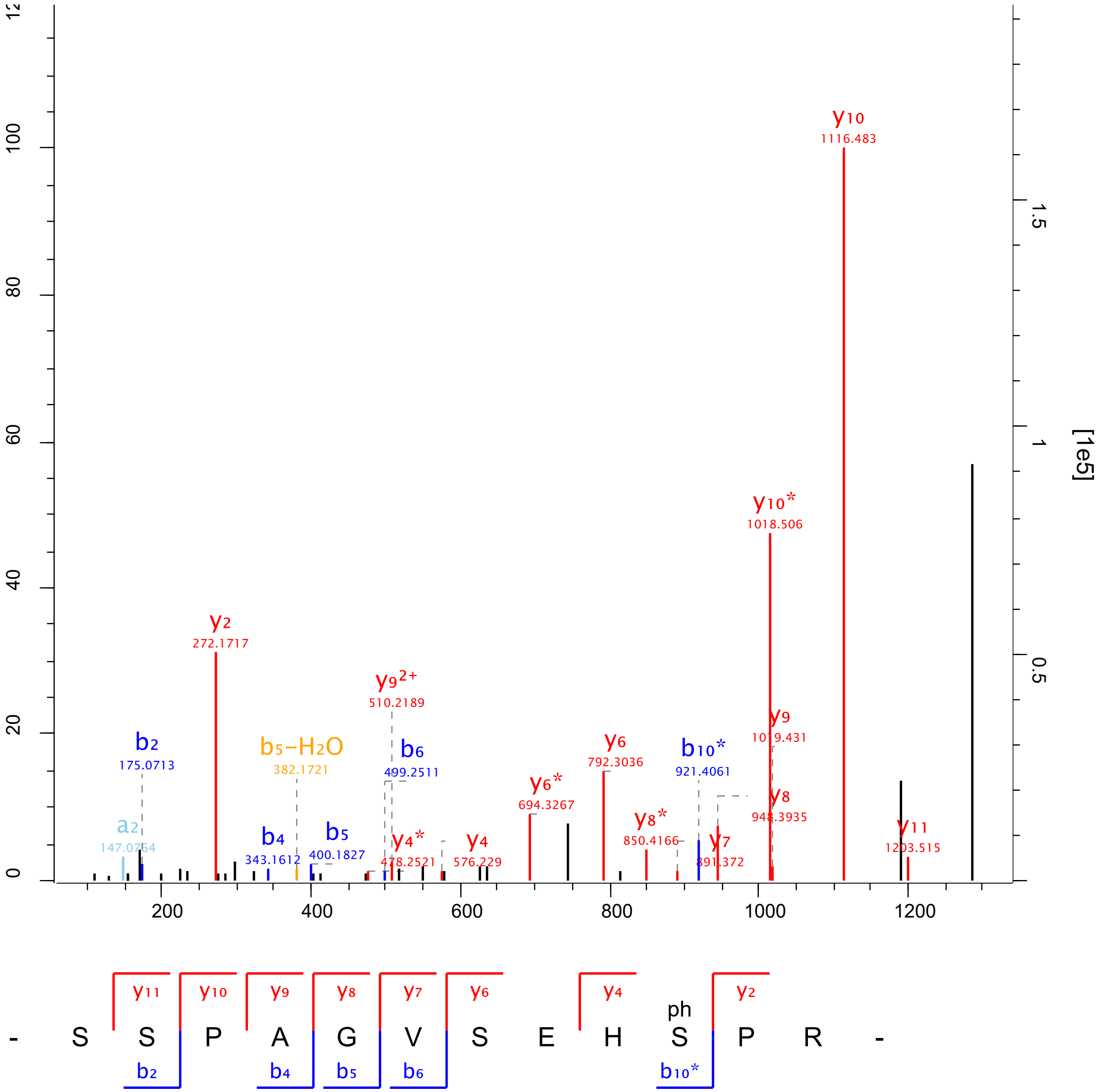

| Raw file | Scan | Method | Score | m/z |
|---|---|---|---|---|
| A17_00_2 | 2043 | FTMS; HCD | 102.39 | 428.53 |

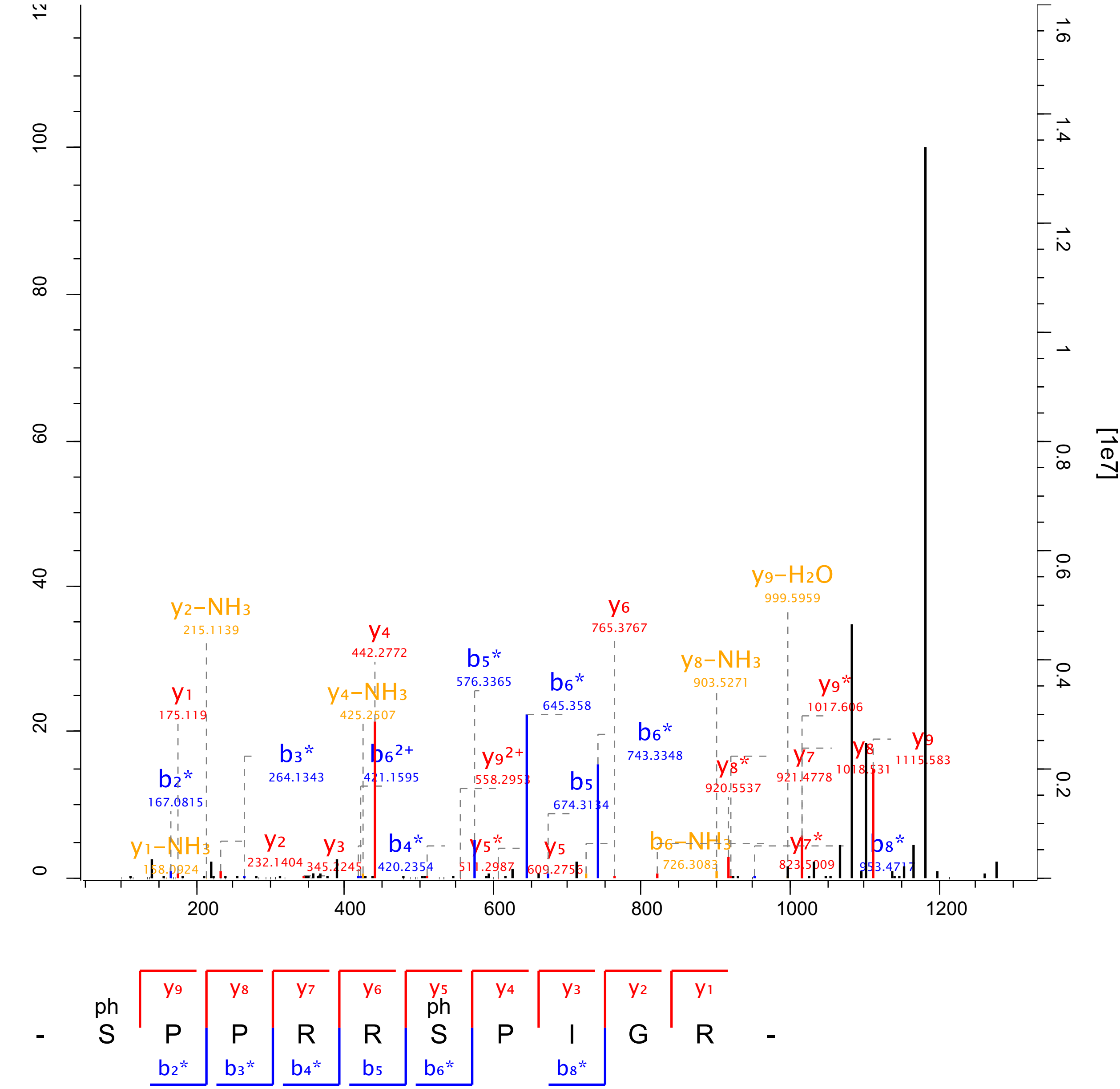

| Raw file | Scan | Method | Score | m/z |
|---|---|---|---|---|
| A17_00_2 | 20602 | FTMS; HCD | 111.55 | 807 |

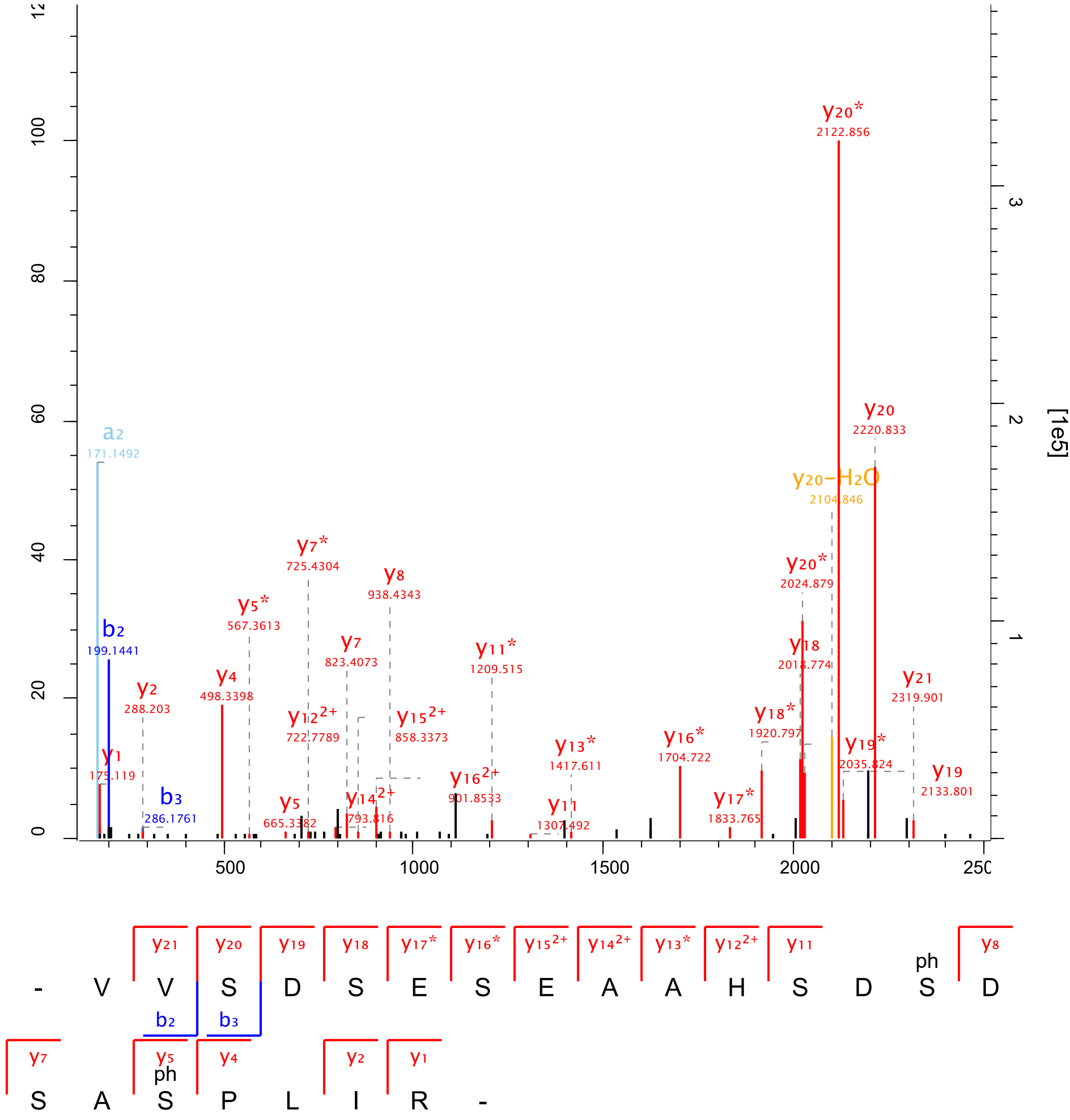

| Raw file | Scan | Method | Score | m/z |
|---|---|---|---|---|
| A17_00_2 | 20709 | FTMS; HCD | 55.66 | 627.74 |

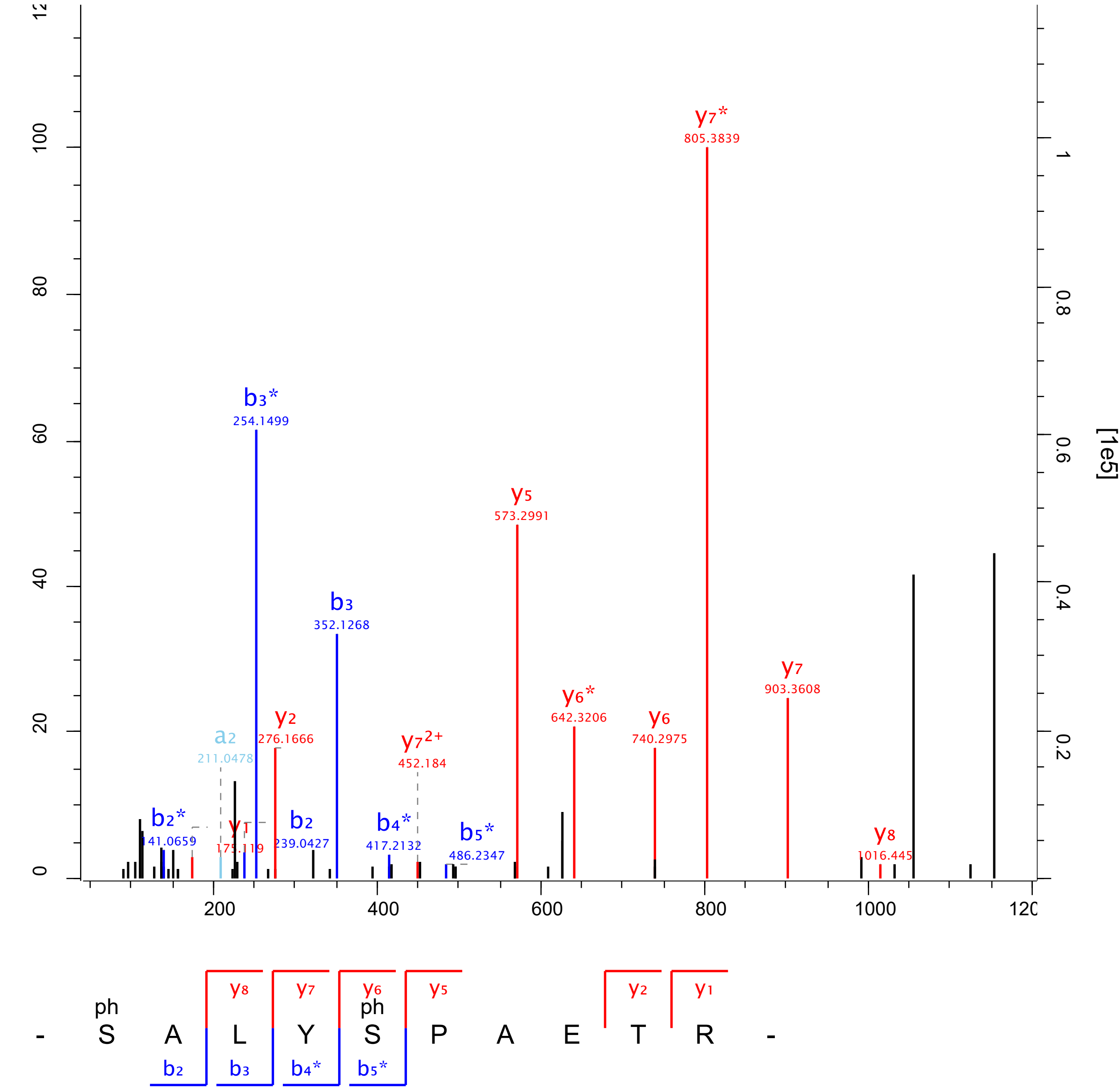

| Raw file | Scan | Method | Score | m/z |
|---|---|---|---|---|
| A17_00_2 | 20721 | FTMS; HCD | 156.02 | 599.28 |

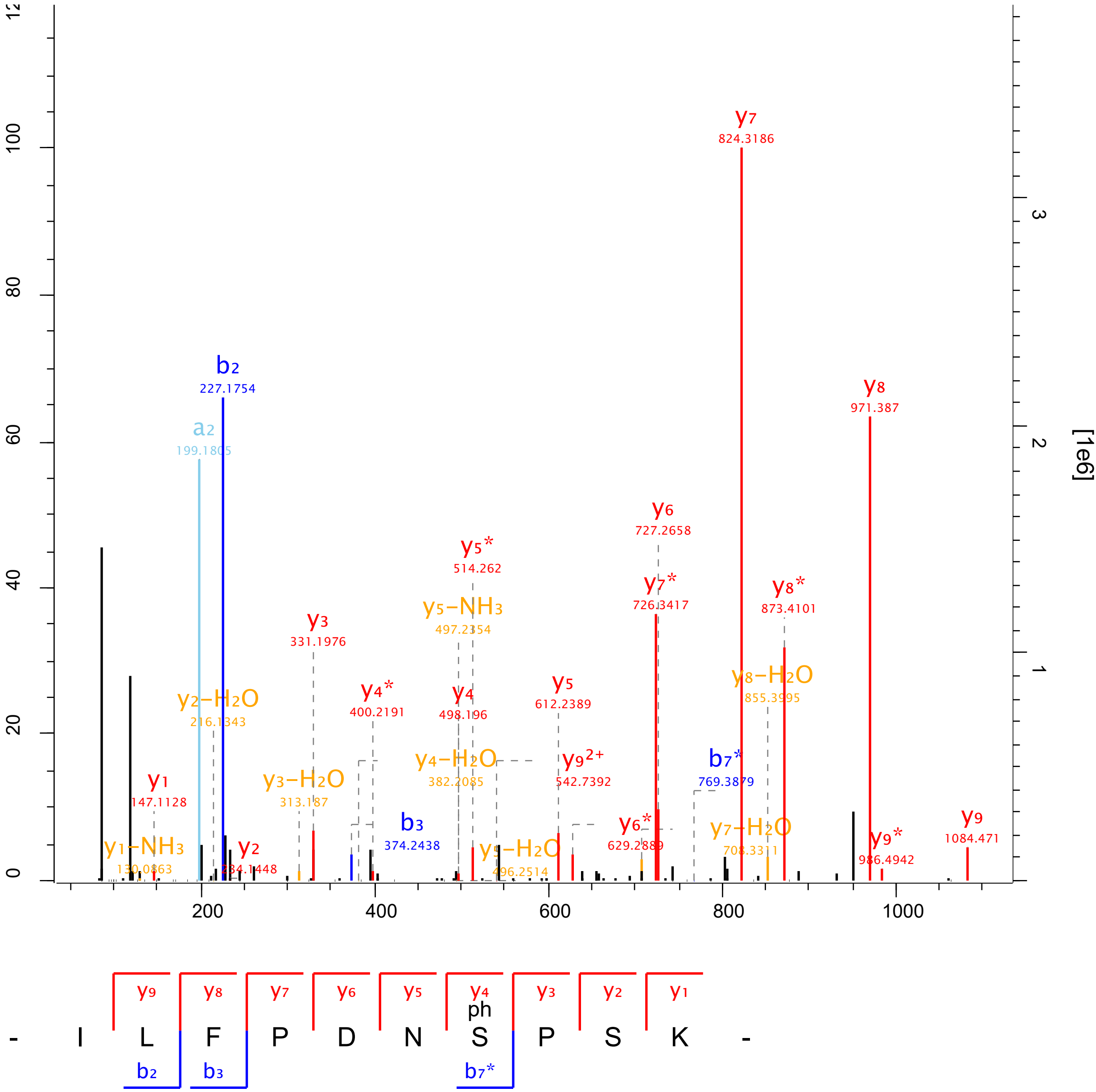

| Raw file | Scan | Method | Score | m/z |
|---|---|---|---|---|
| A17_00_2 | 2094 | FTMS; HCD | 48.9 | 410.2 |

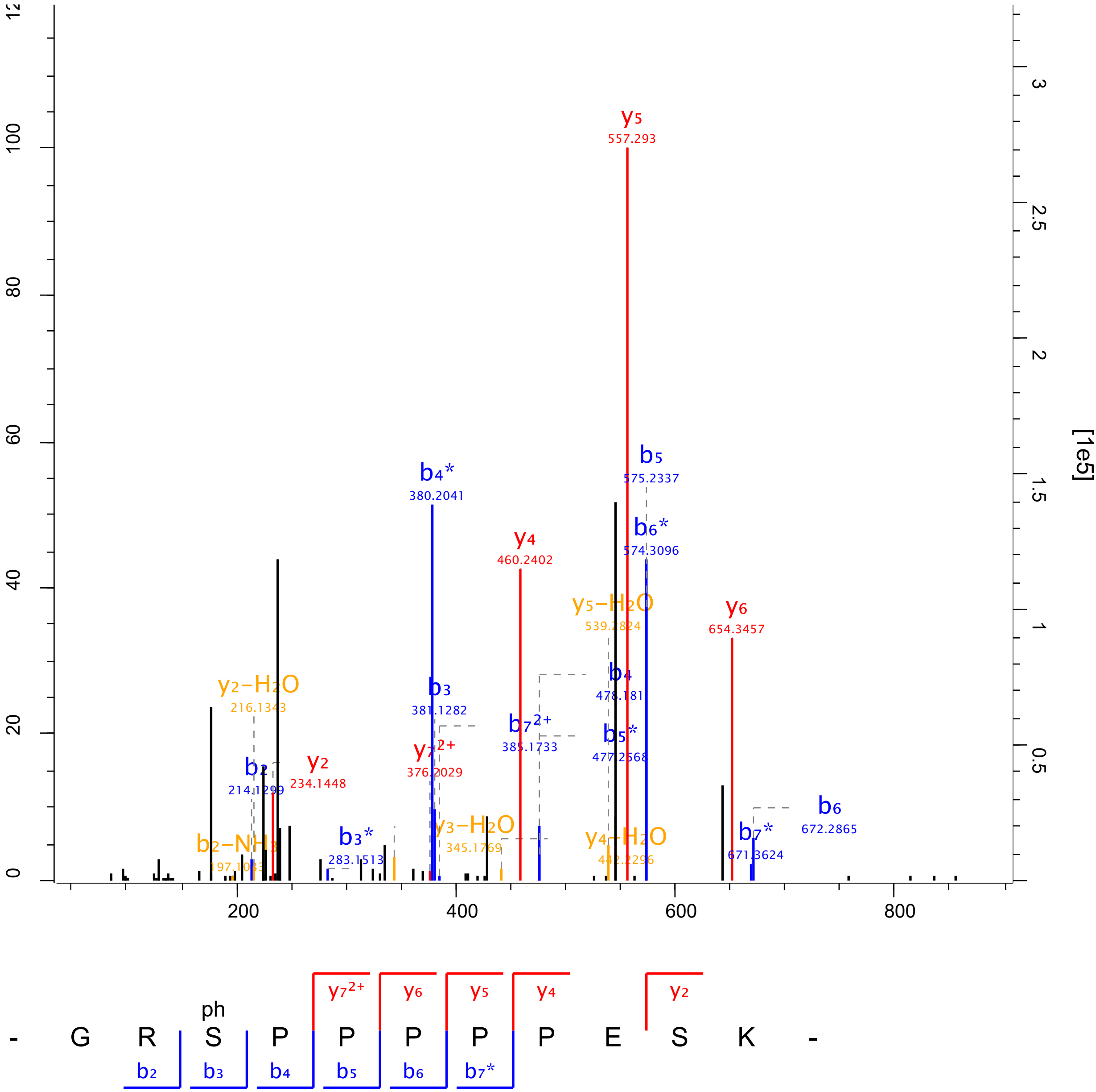

| Raw file | Scan | Method | Score | m/z |
| --- | --- | --- | --- | --- |
| A17_00_2 | 21198 | FTMS; HCD | 66.27 | 560.21 |

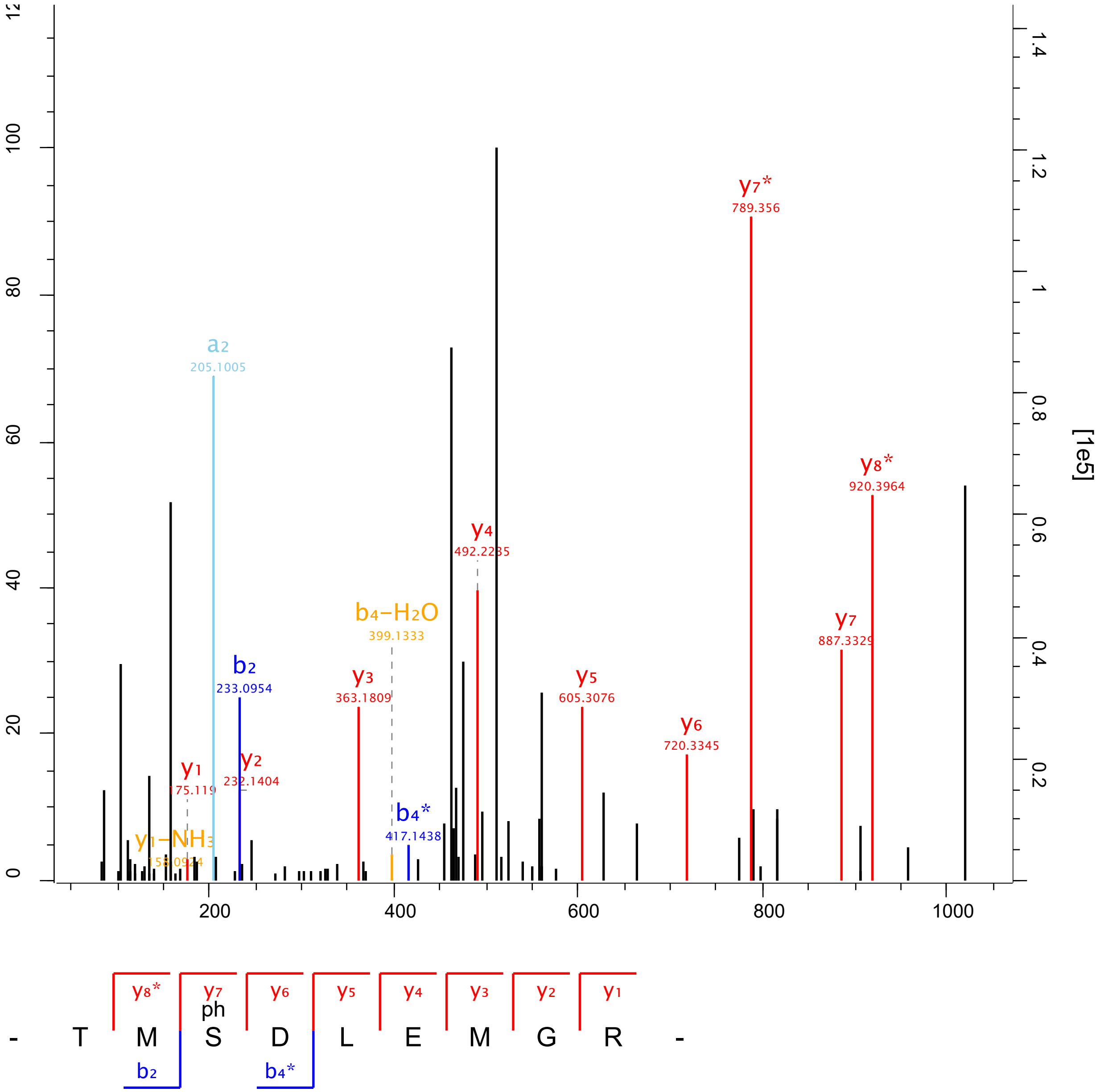

| Raw file | Scan | Method | Score | m/z |
|---|---|---|---|---|
| A17_00_2 | 21328 | FTMS; HCD | 214.11 | 1048.97 |

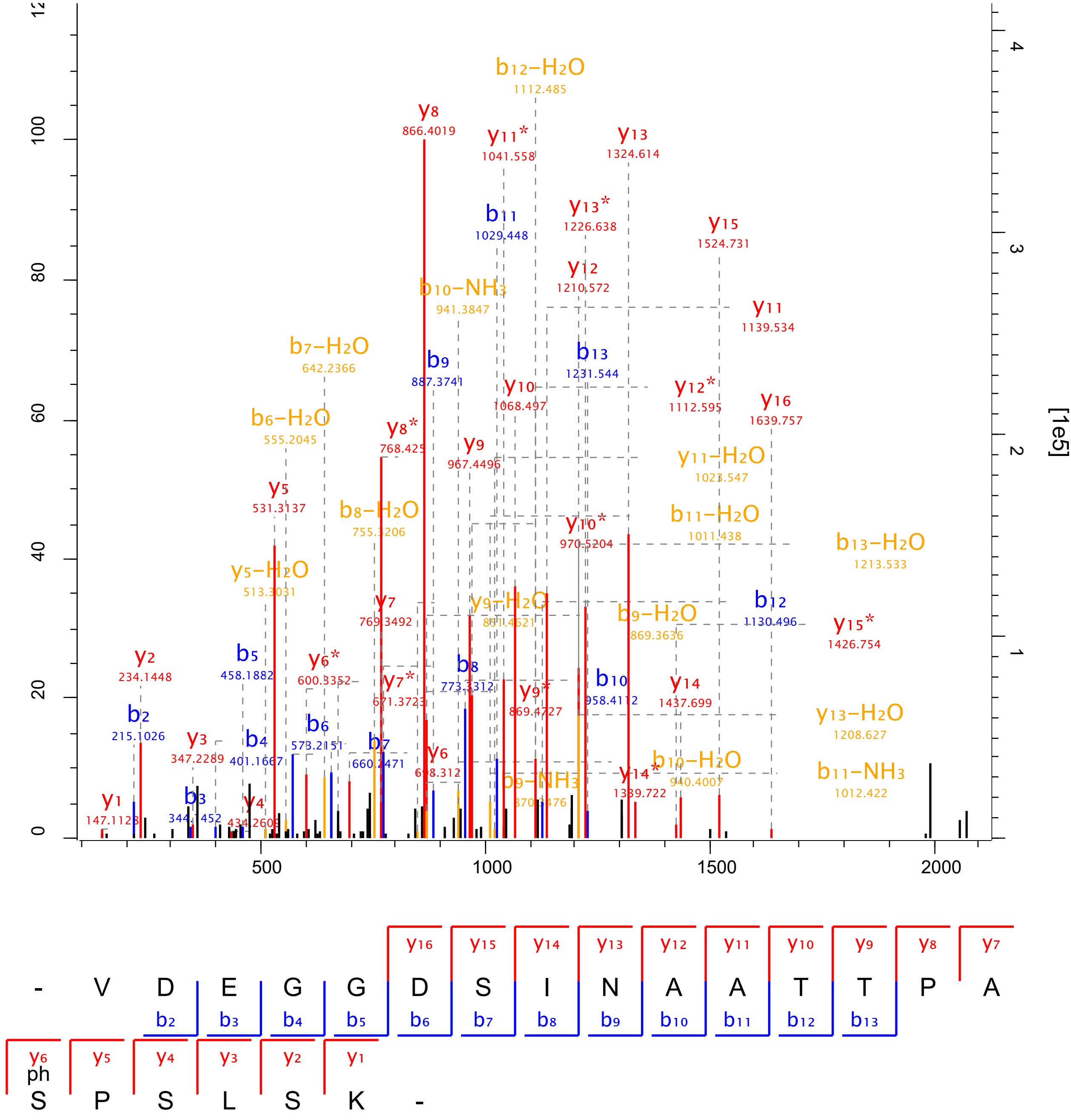

| Raw file | Scan | Method | Score | m/z |
| --- | --- | --- | --- | --- |
| A17_00_2 | 2133 | FTMS; HCD | 69.03 | 417.85 |

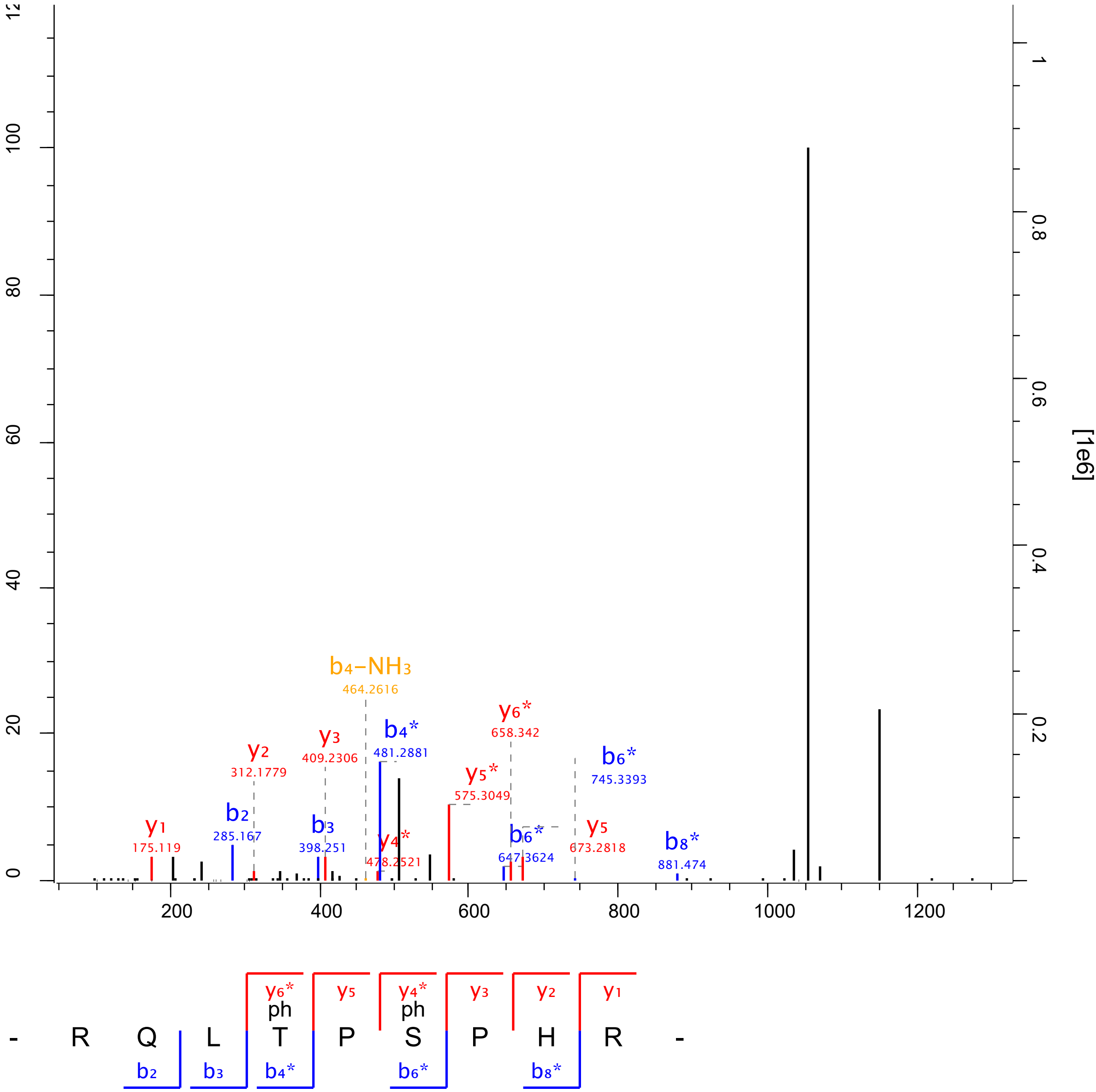

| Raw file | Scan | Method | Score | m/z |
|---|---|---|---|---|
| A17_00_2 | 21467 | FTMS; HCD | 241.76 | 708.32 |

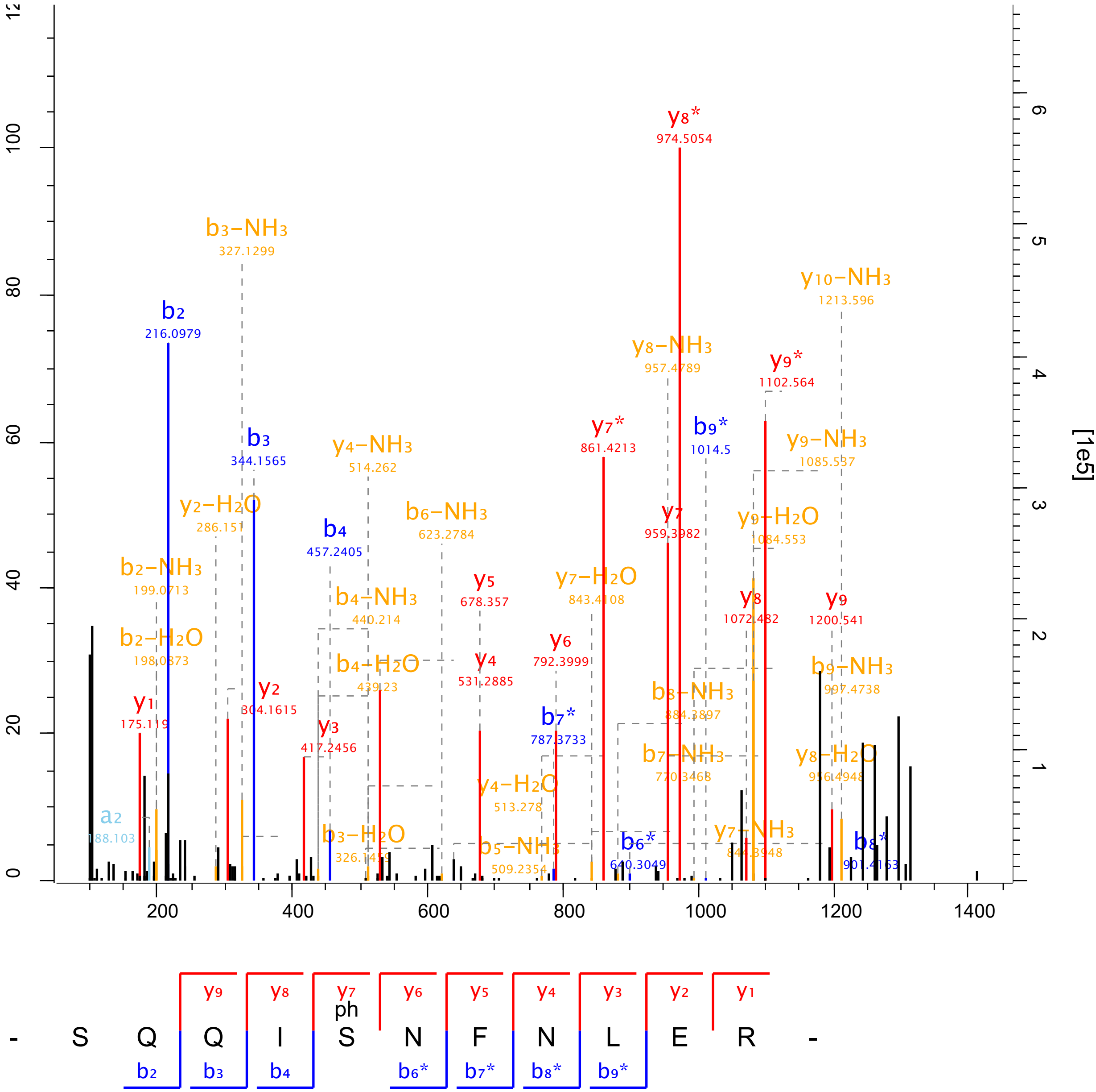

| Raw file | Scan | Method | Score | m/z |
|---|---|---|---|---|
| A17_00_2 | 21495 | FTMS; HCD | 65.86 | 926.34 |

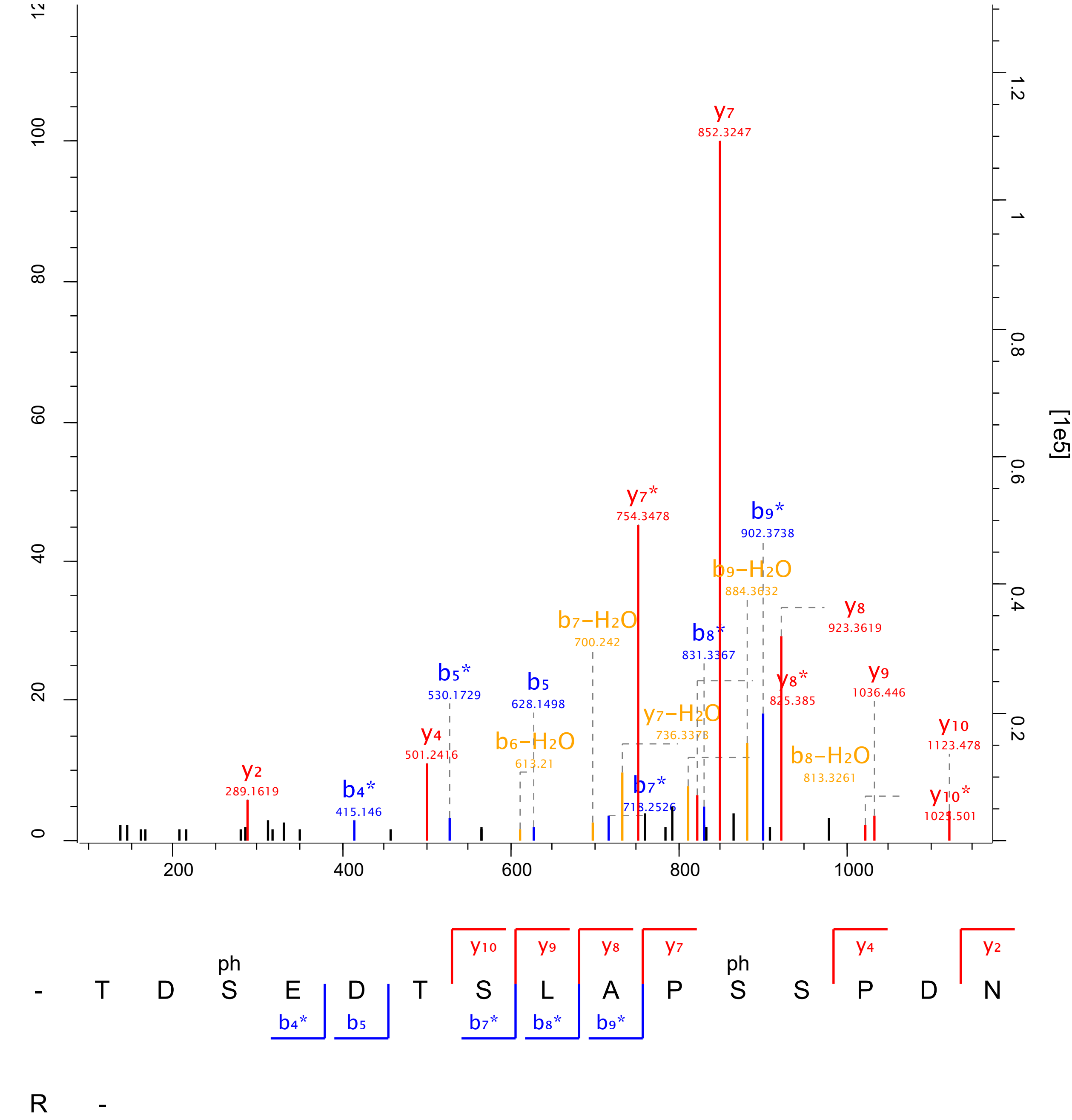

| Raw file | Scan | Method | Score | m/z |
|---|---|---|---|---|
| A17_00_2 | 21565 | FTMS; HCD | 123.14 | 501.61 |

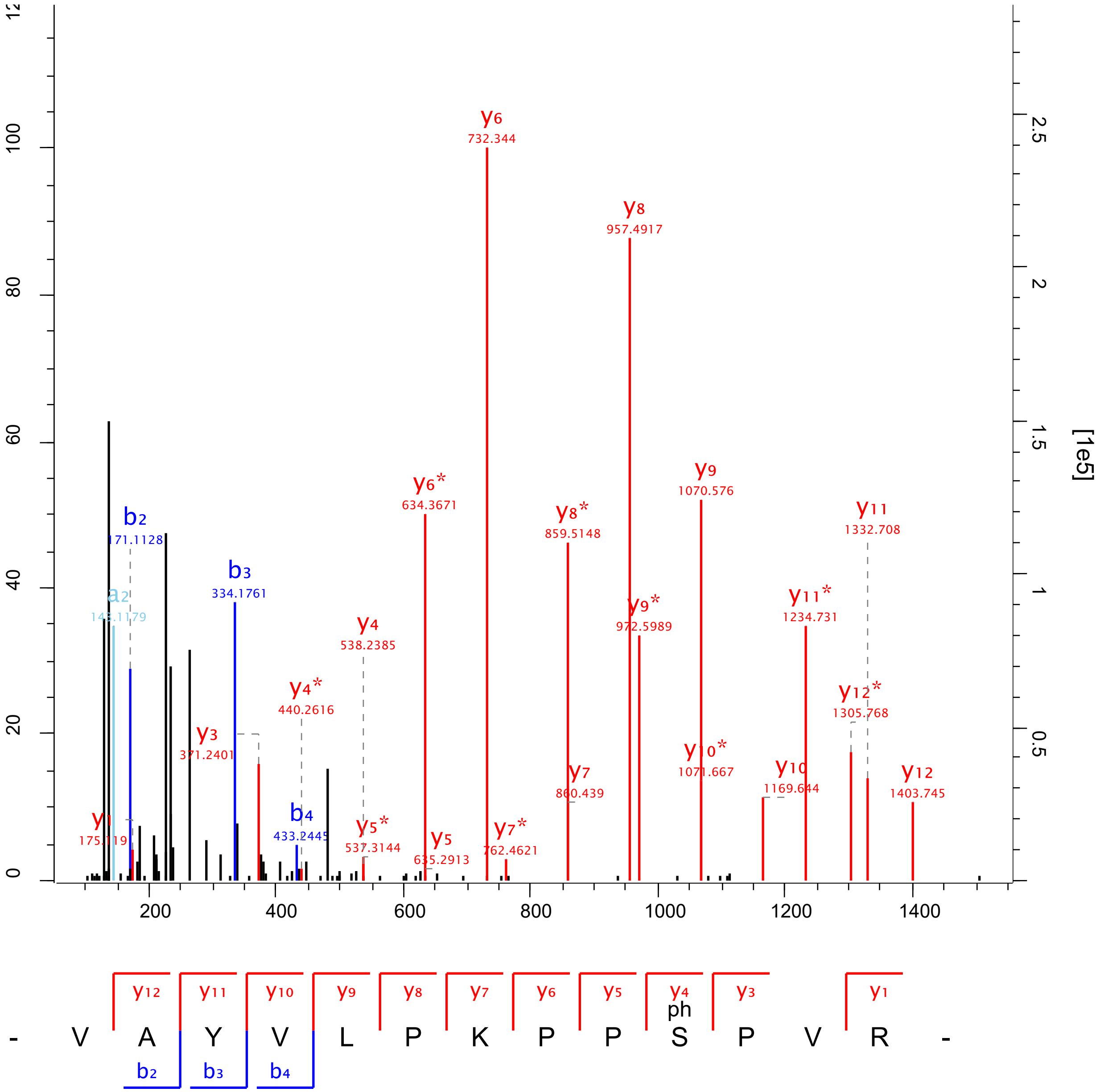

| Raw file | Scan | Method | Score | m/z |
| --- | --- | --- | --- | --- |
| A17_00_2 | 21613 | FTMS; HCD | 70.34 | 795.67 |

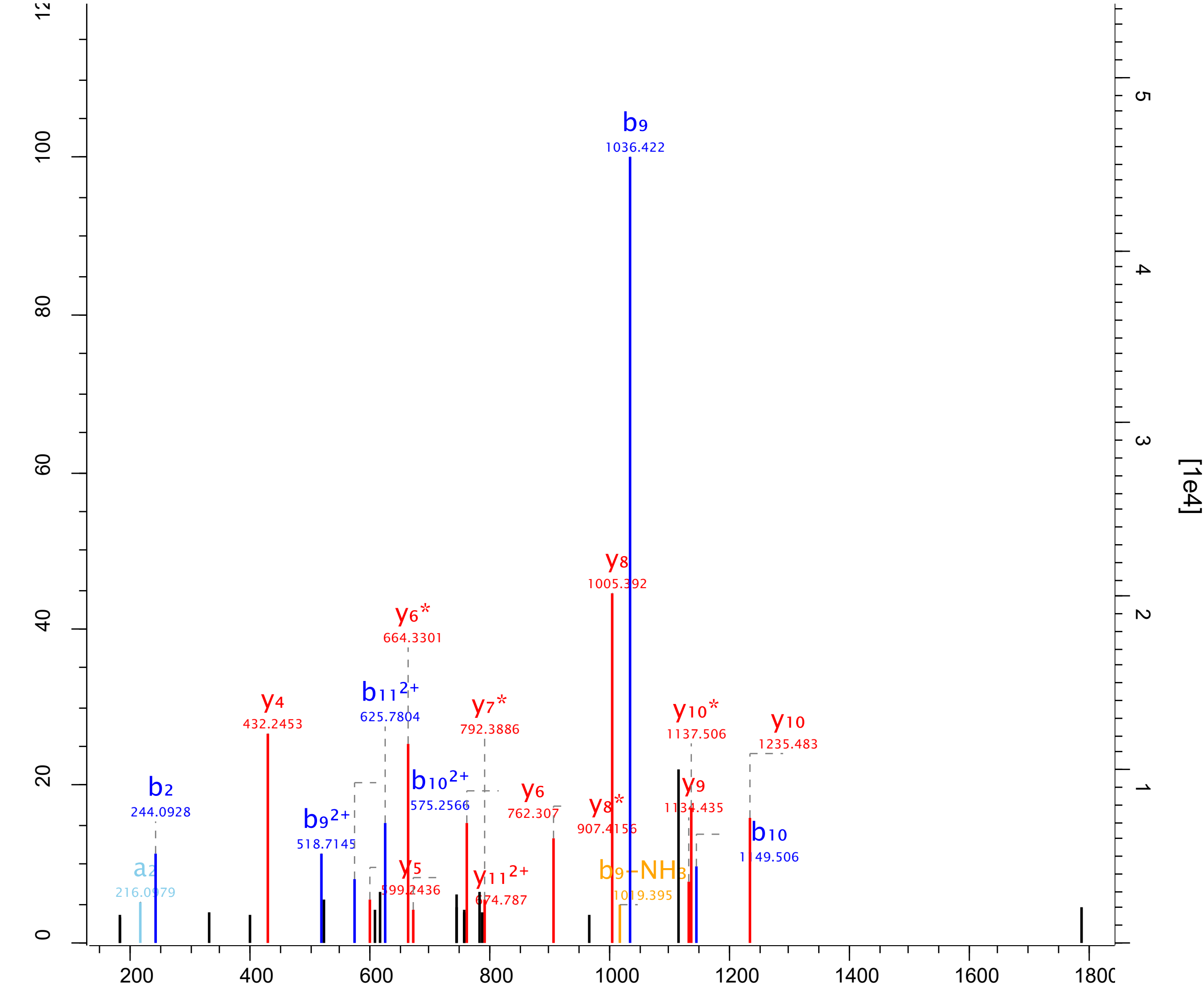


- N E D K F S D A E L T E D Q Y
S P T S K -

y11 2+, y10, y9, y8, y7*, y6, y5, y4

b2, b9, b10, b11 2+

ph

| Raw file | Scan | Method | Score | m/z |
| --- | --- | --- | --- | --- |
| A17_00_2 | 21730 | FTMS; HCD | 143.55 | 525.74 |

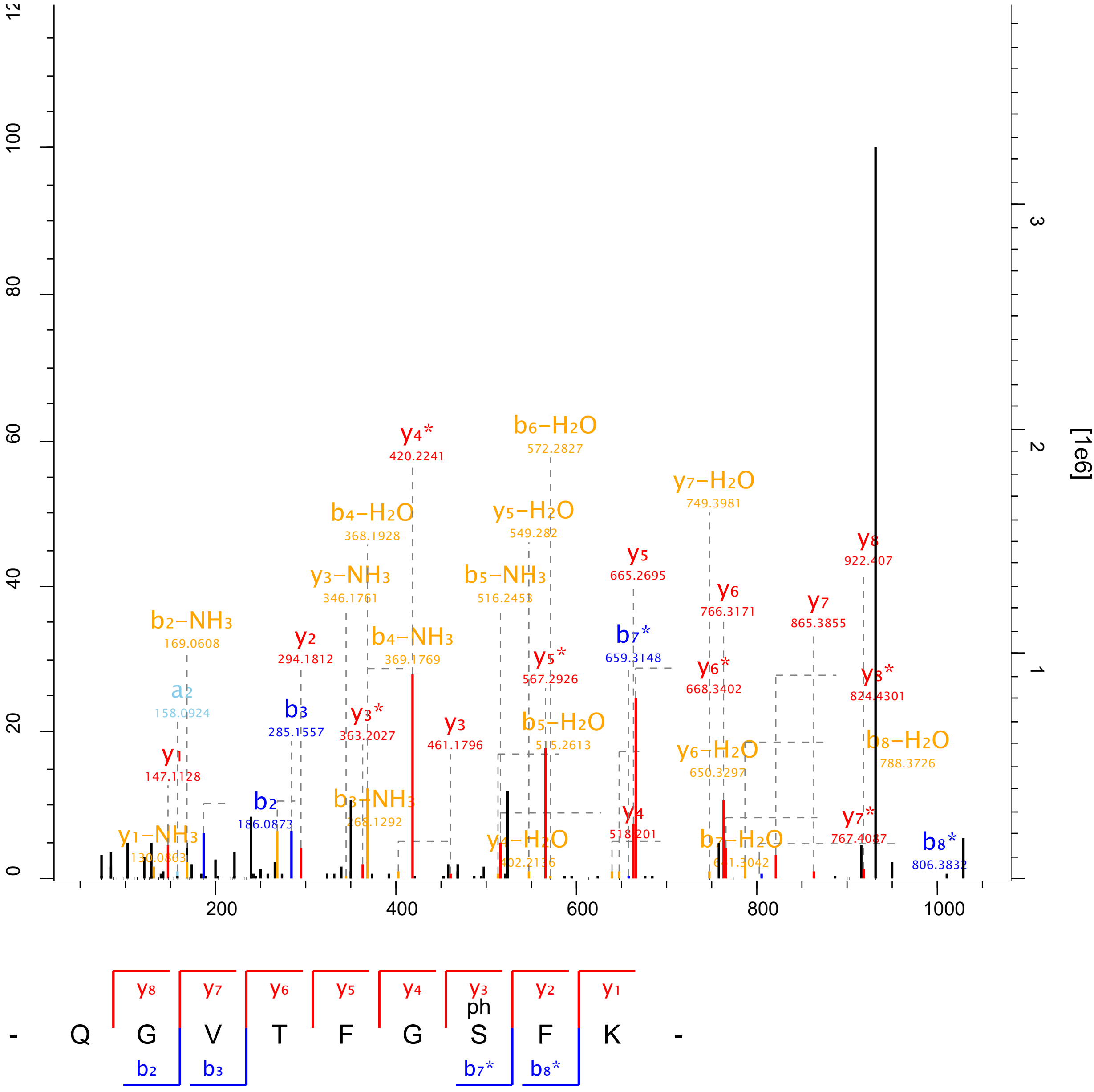

| Raw file | Scan | Method | Score | m/z |
| --- | --- | --- | --- | --- |
| A17_00_2 | 2194 | FTMS; HCD | 64.83 | 522.23 |

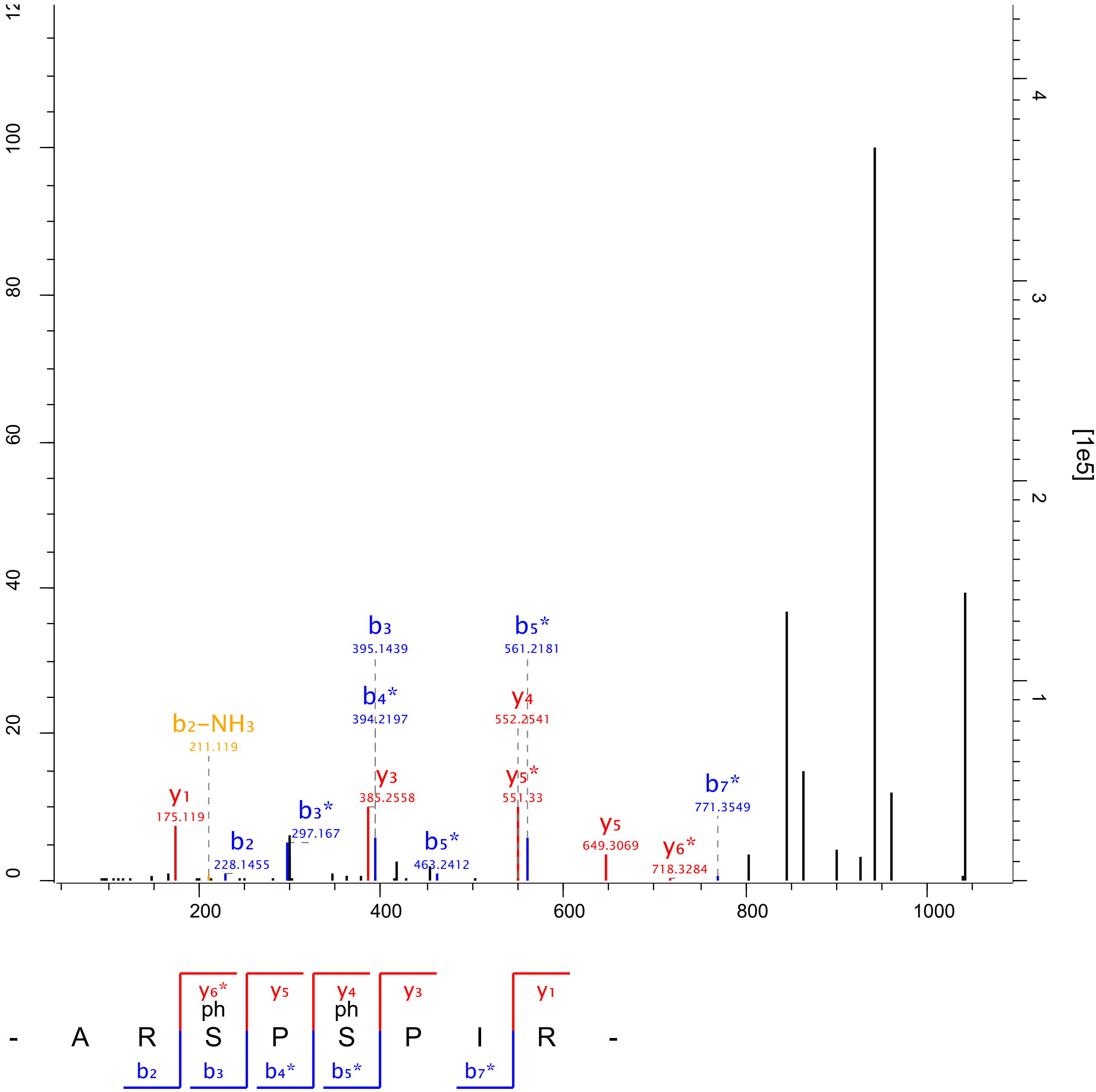

| Raw file | Scan | Method | Score | m/z |
|---|---|---|---|---|
| A17_00_2 | 22013 | FTMS; HCD | 51.77 | 640.3 |

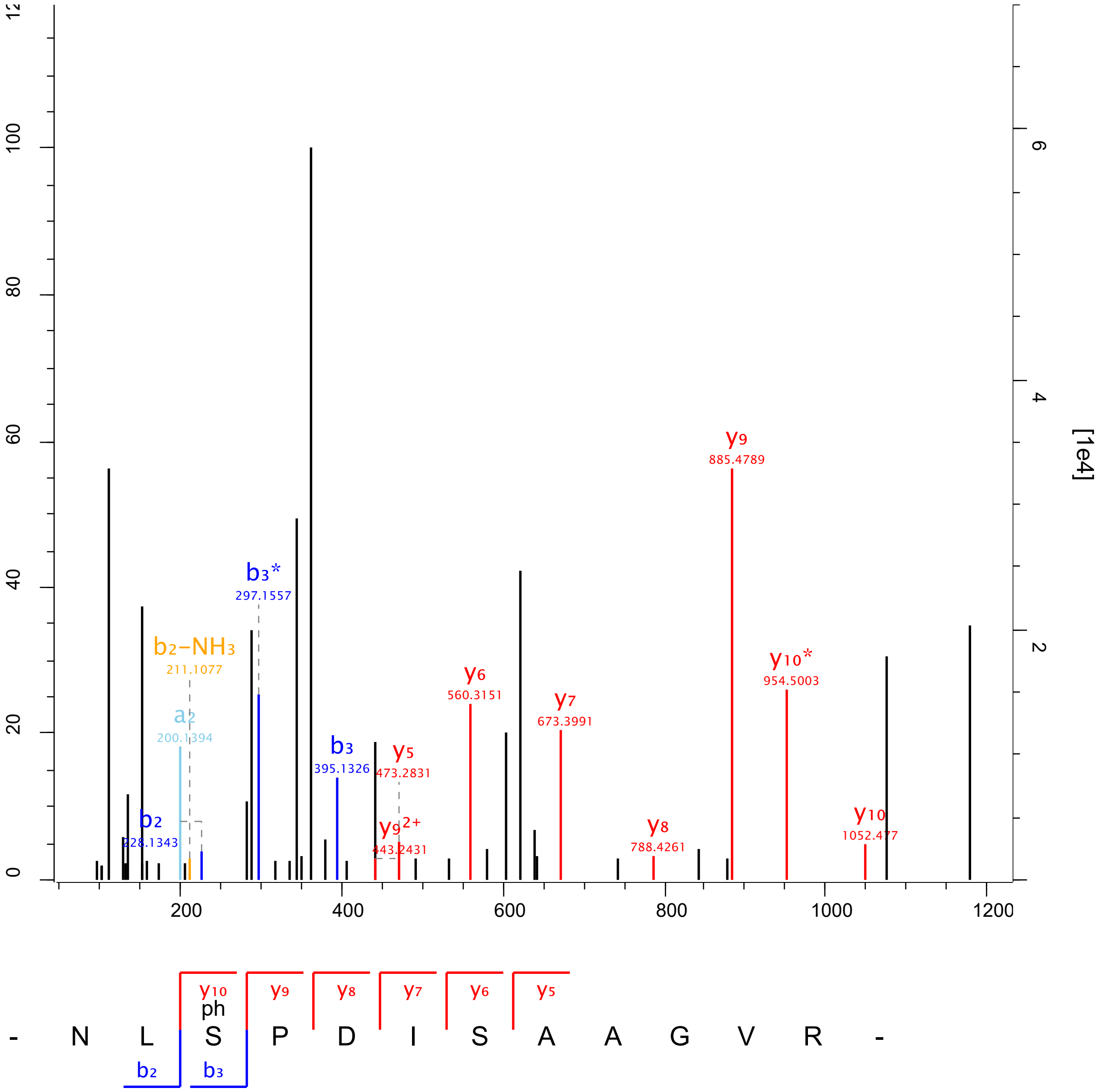

| Raw file | Scan | Method | Score | m/z |
|---|---|---|---|---|
| A17_00_2 | 2206 | FTMS; HCD | 77.75 | 480.89 |

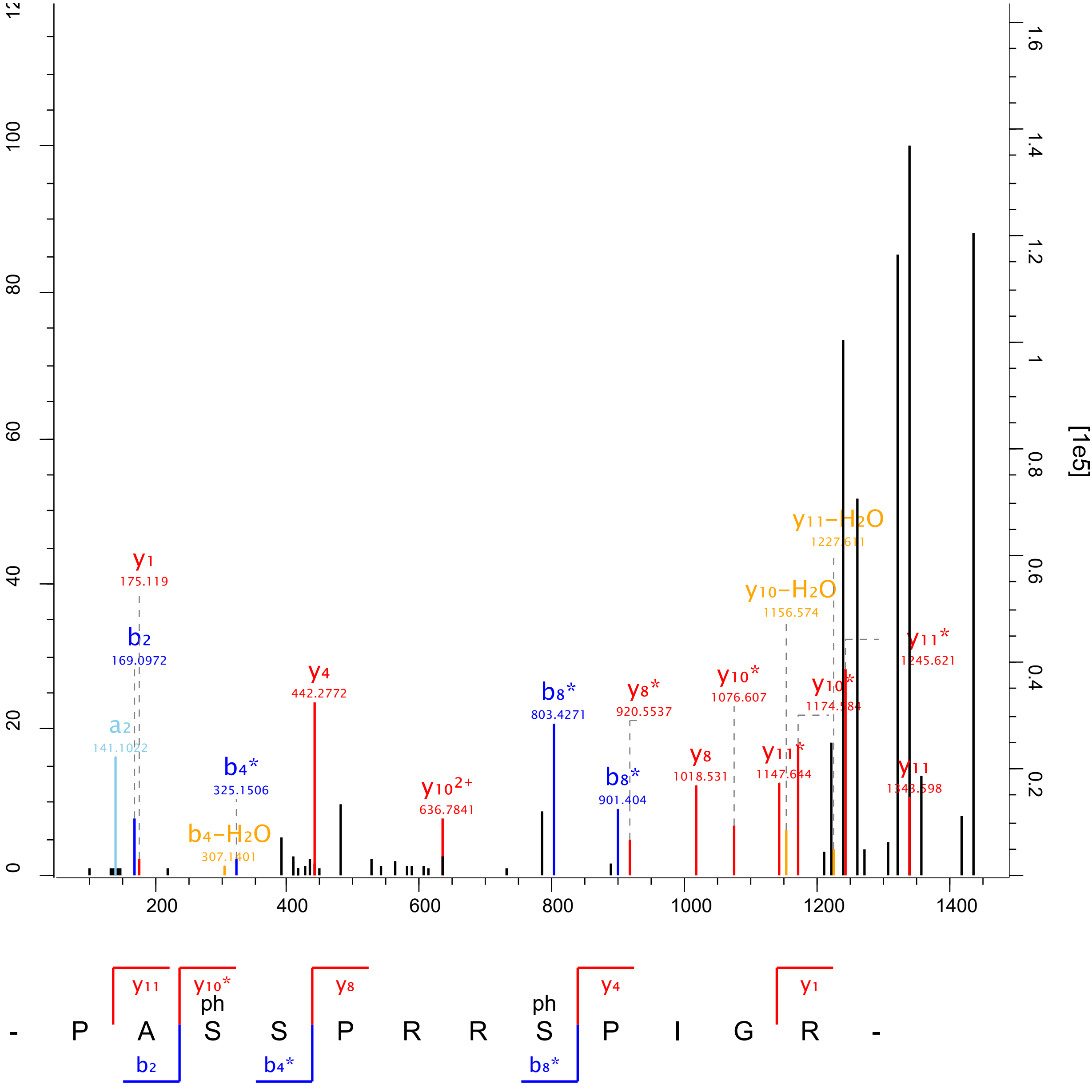

| Raw file | Scan | Method | Score | m/z |
| --- | --- | --- | --- | --- |
| A17_00_2 | 22118 | FTMS; HCD | 49.08 | 573.23 |

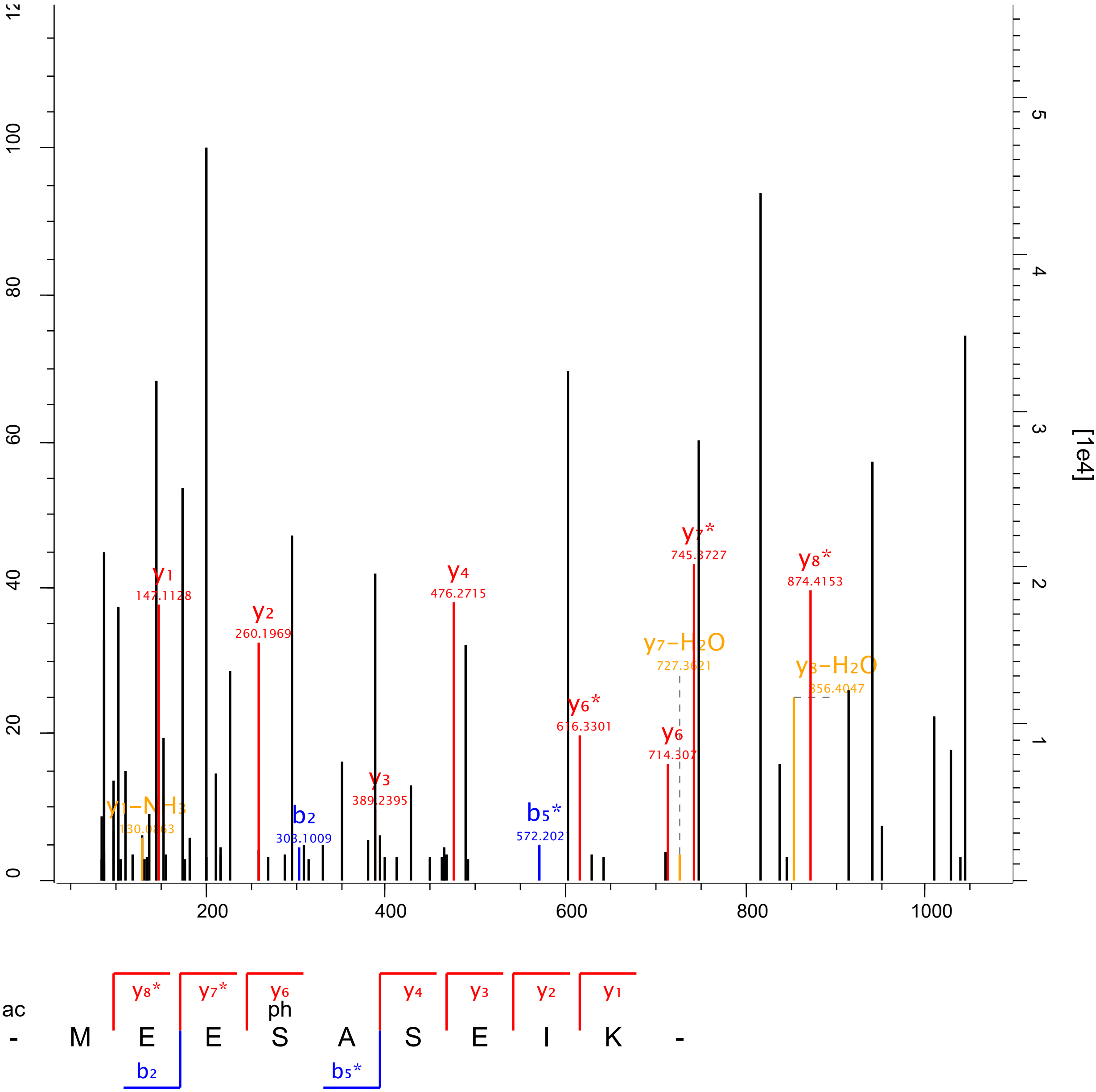

| Raw file | Scan | Method | Score | m/z |
|---|---|---|---|---|
| A17_00_2 | 22141 | FTMS; HCD | 57.53 | 636.31 |

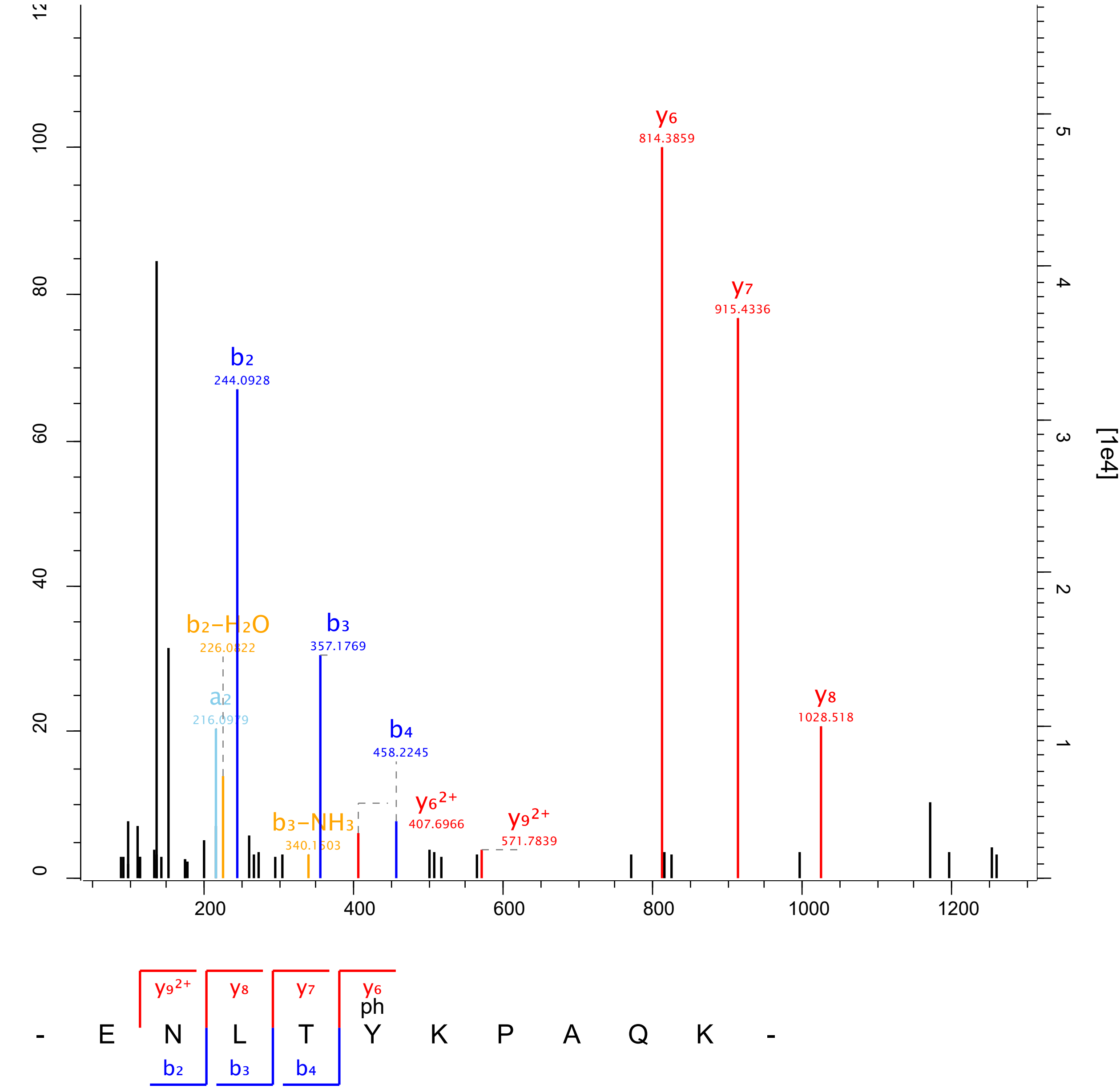

| Raw file | Scan | Method | Score | m/z |
| --- | --- | --- | --- | --- |
| A17_00_2 | 22194 | FTMS; HCD | 188.39 | 684.79 |

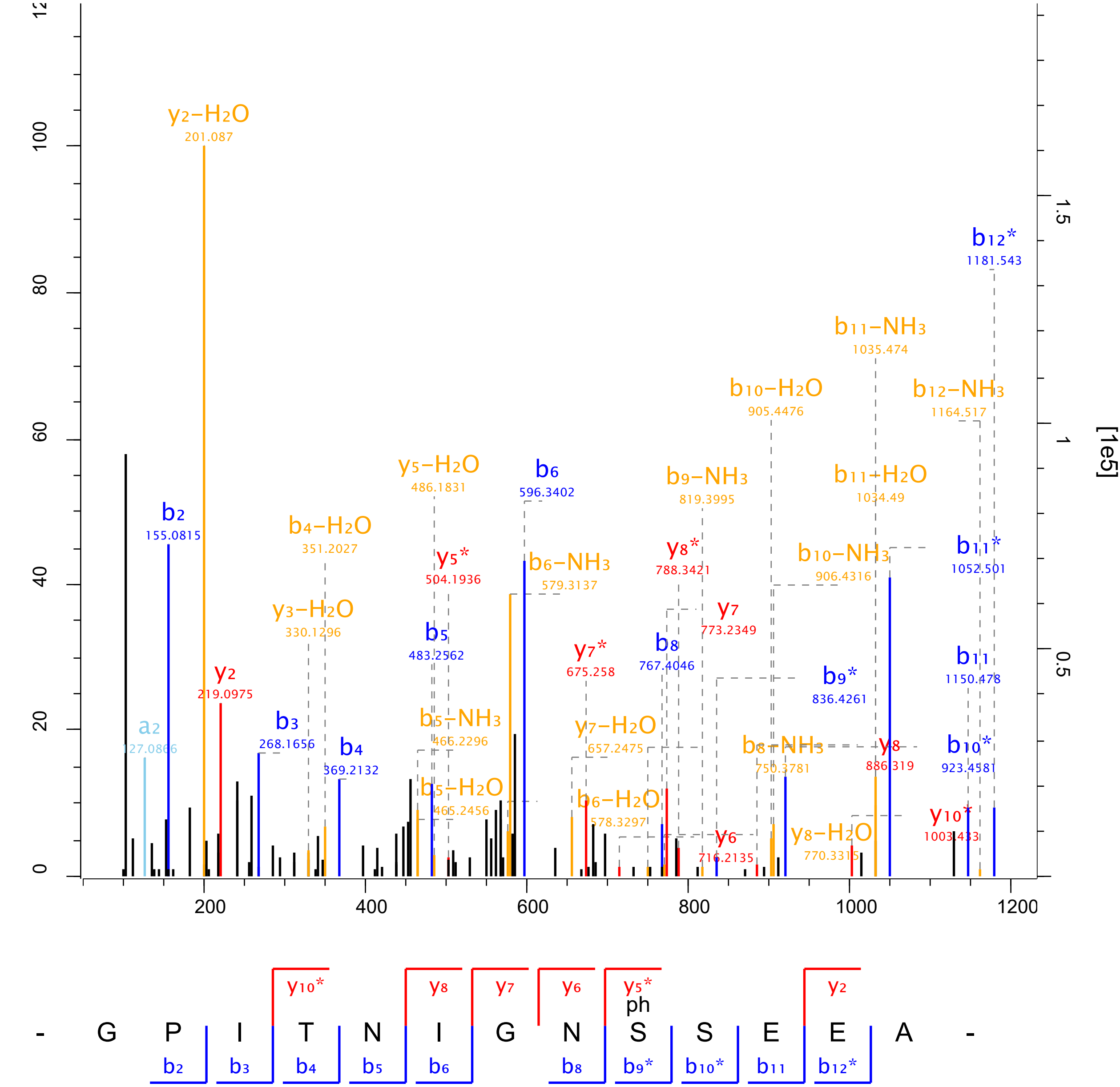

| Raw file | Scan | Method | Score | m/z |
|---|---|---|---|---|
| A17_00_2 | 22212 | FTMS; HCD | 87.18 | 496.71 |

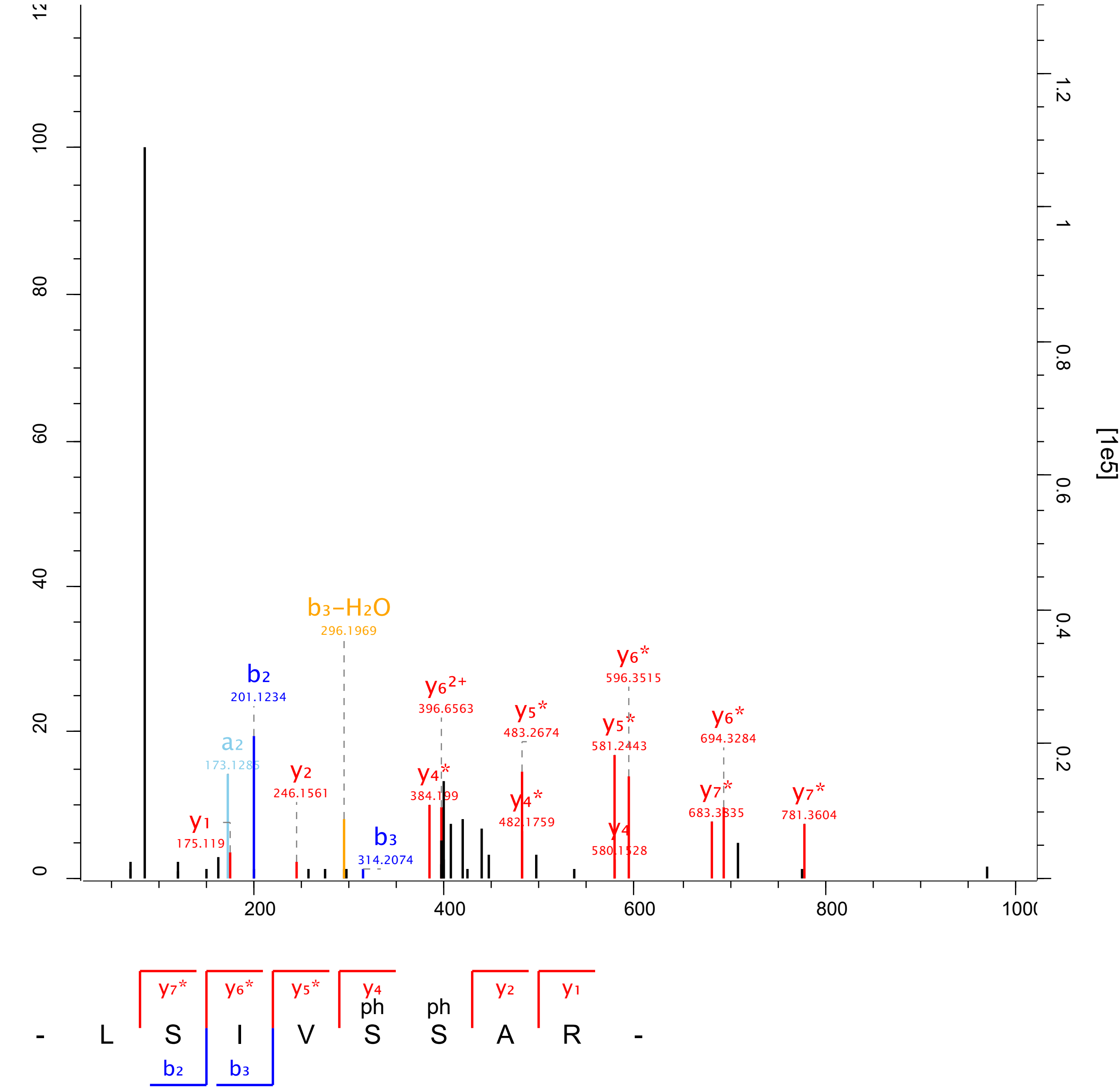

| Raw file | Scan | Method | Score | m/z |
| --- | --- | --- | --- | --- |
| A17_00_2 | 22553 | FTMS; HCD | 218.1 | 800.66 |

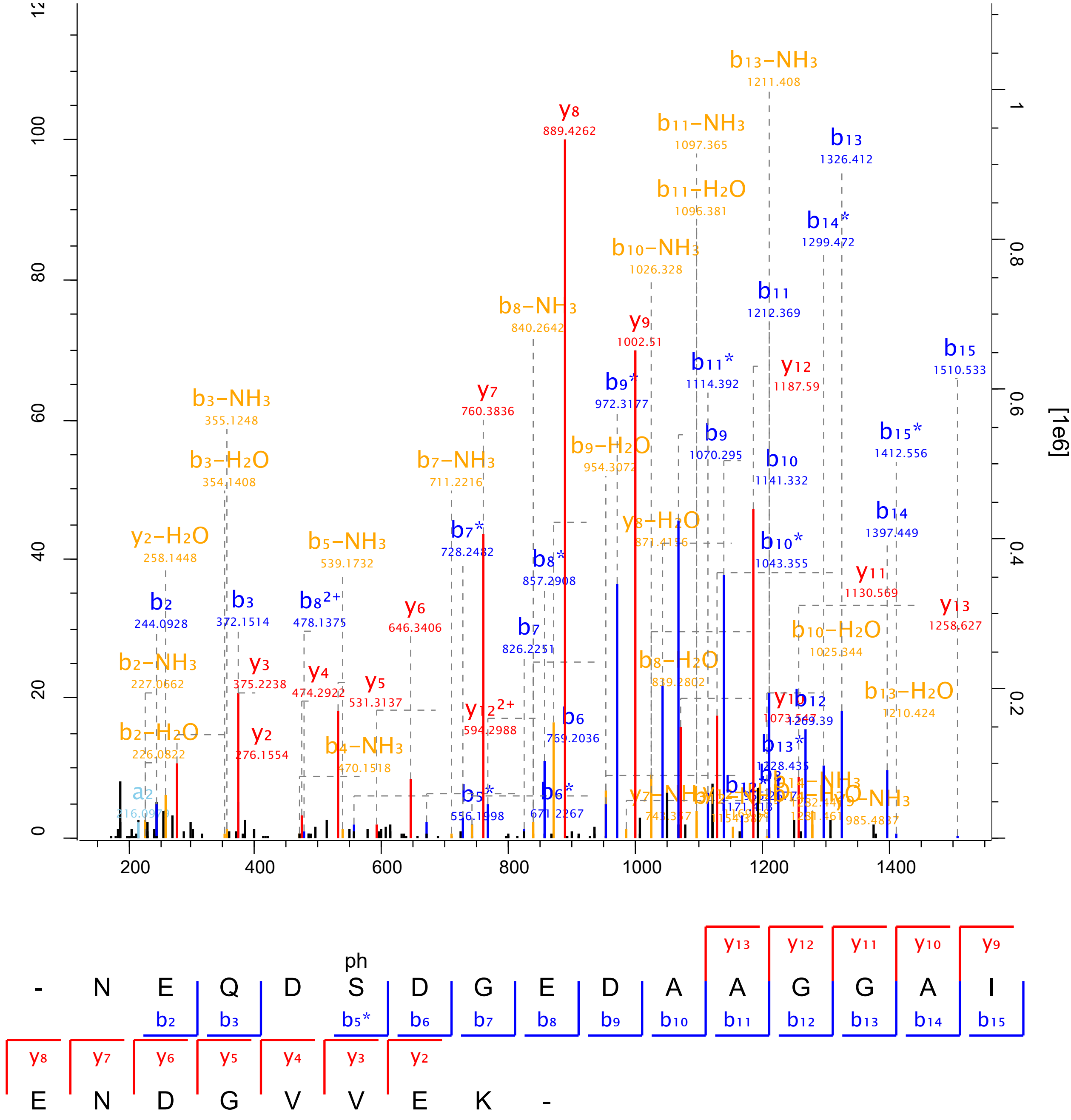

| Raw file | Scan | Method | Score | m/z |
| --- | --- | --- | --- | --- |
| A17_00_2 | 2265 | FTMS; HCD | 81.02 | 582.73 |

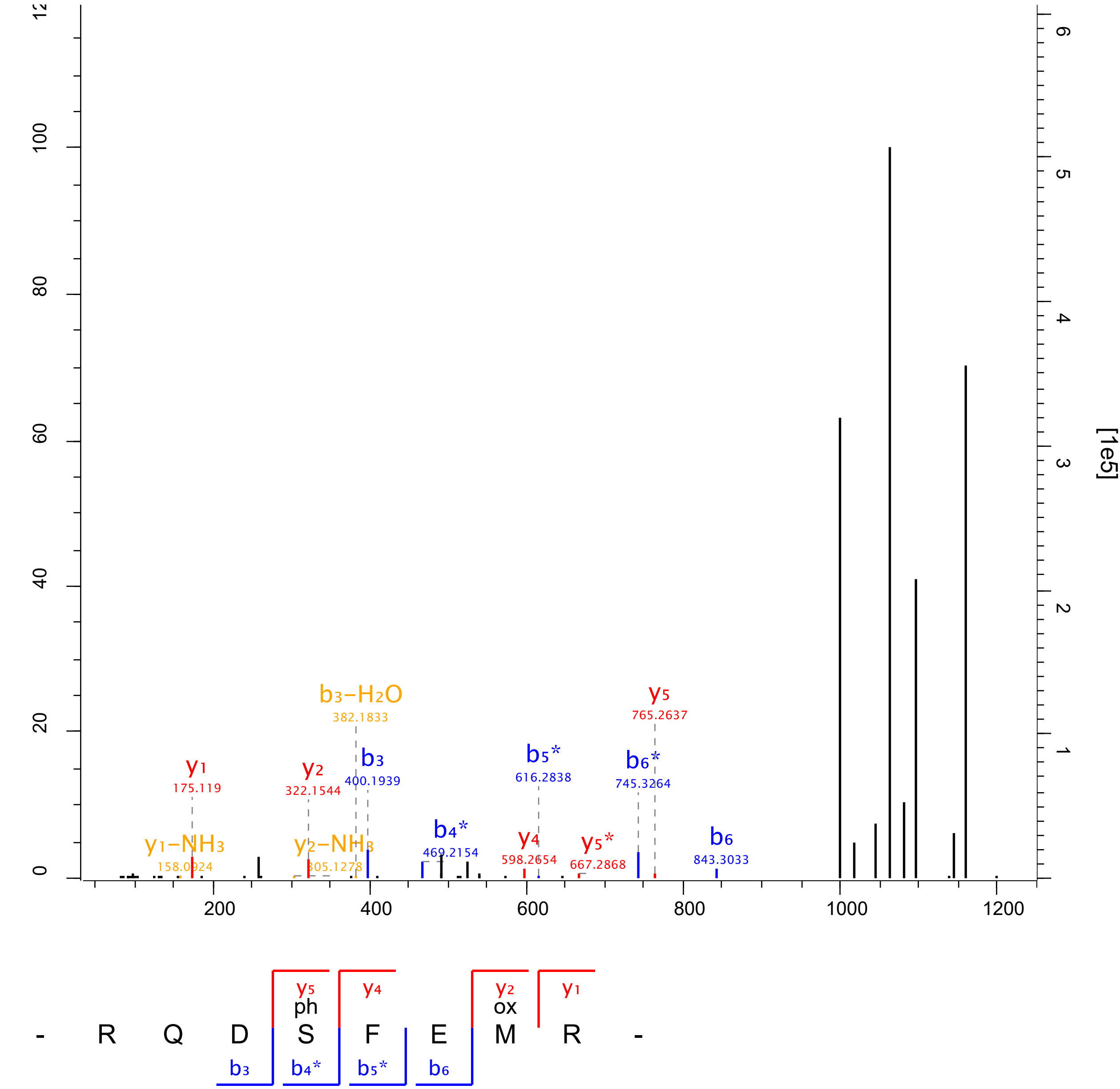

| Raw file | Scan | Method | Score | m/z |
| --- | --- | --- | --- | --- |
| A17_00_2 | 2268 | FTMS; HCD | 50.09 | 683.8 |

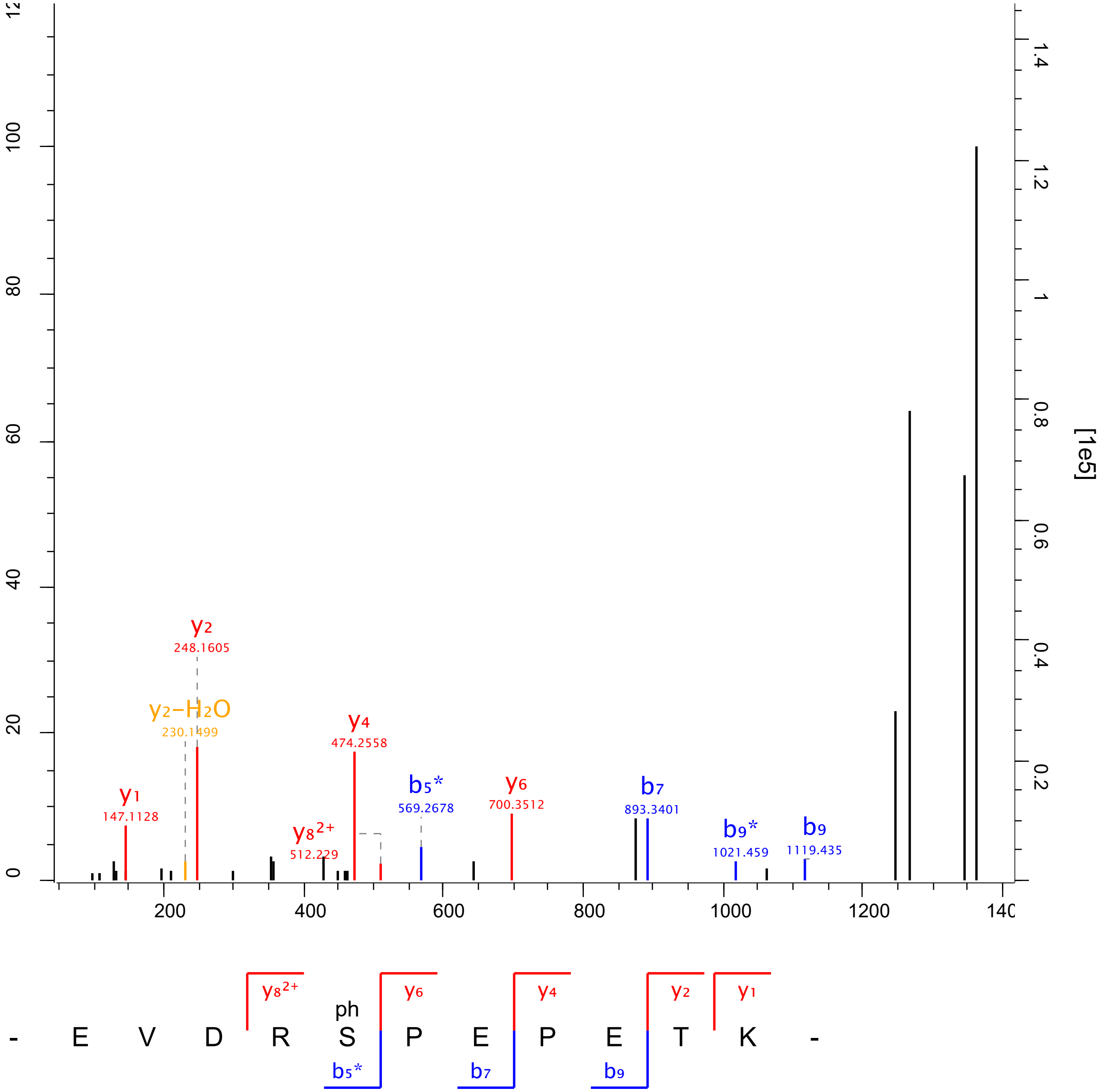

| Raw file | Scan | Method | Score | m/z |
|---|---|---|---|---|
| A17_00_2 | 22836 | FTMS; HCD | 111.12 | 672.8 |

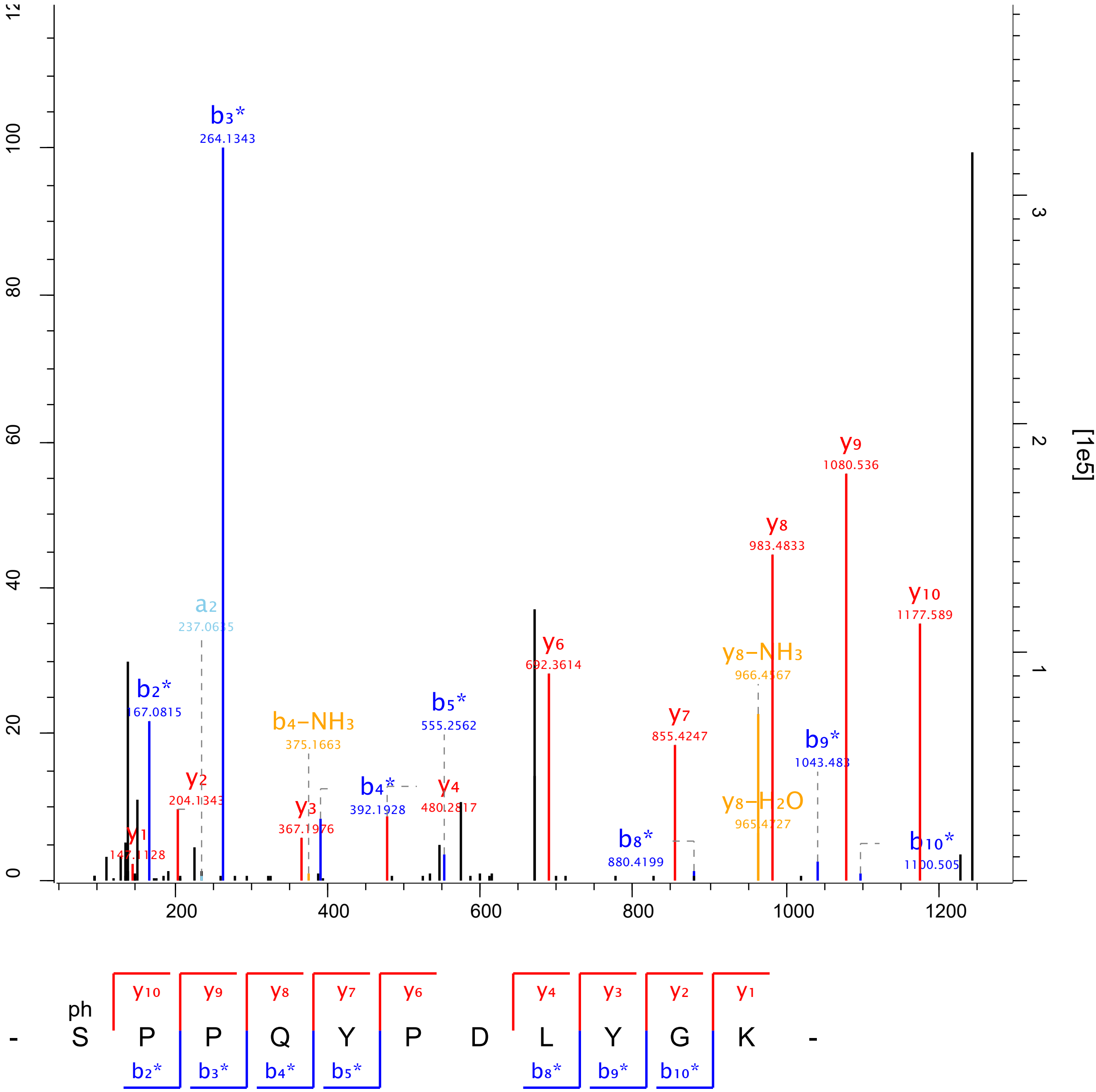

| Raw file | Scan | Method | Score | m/z |
|---|---|---|---|---|
| A17_00_2 | 23073 | FTMS; HCD | 207.21 | 752.82 |

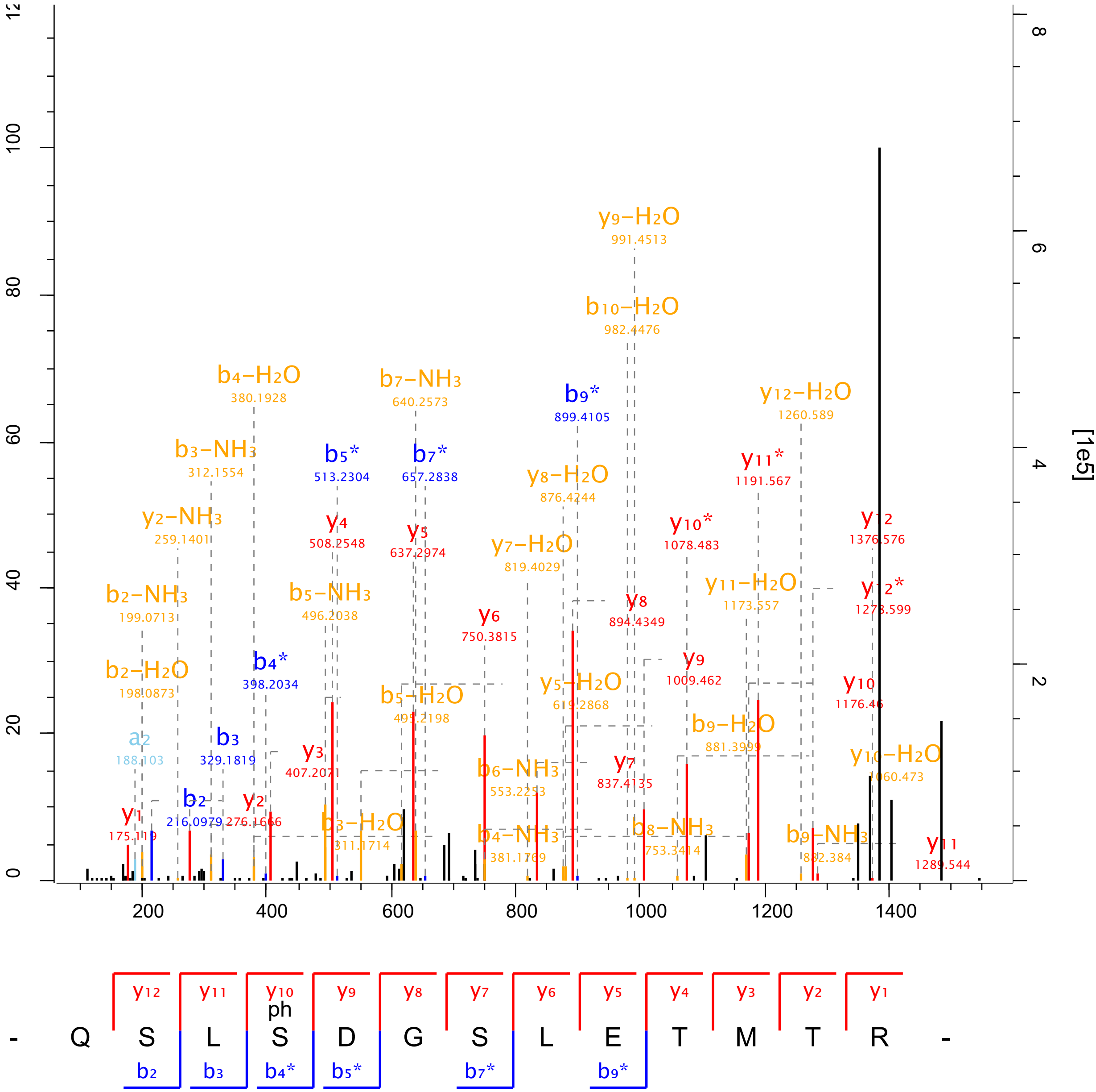

| Raw file | Scan | Method | Score | m/z |
|---|---|---|---|---|
| A17_00_2 | 23237 | FTMS; HCD | 92.86 | 867.07 |

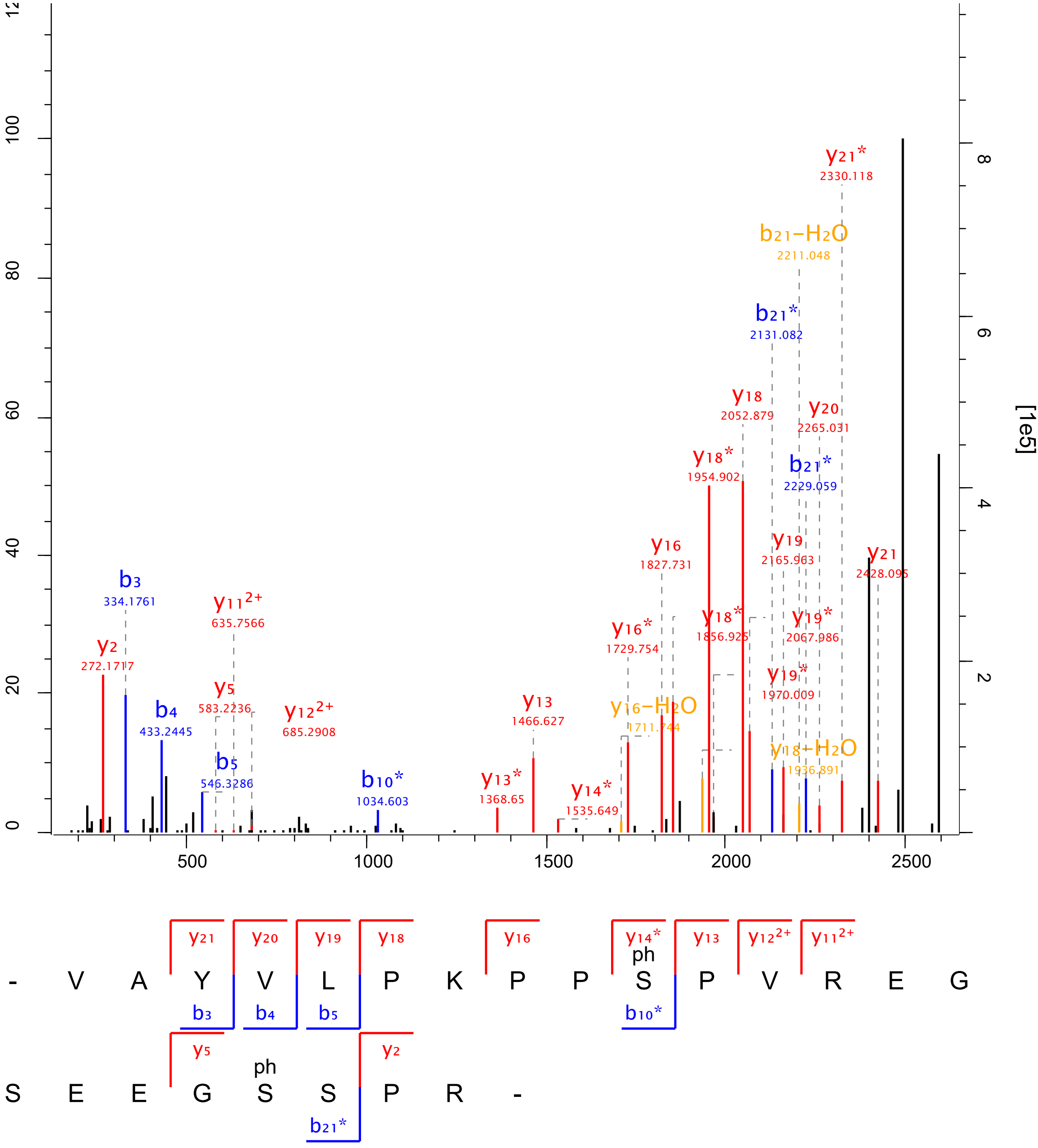

| Raw file | Scan | Method | Score | m/z |
|---|---|---|---|---|
| A17_00_2 | 23247 | FTMS; HCD | 145.88 | 851.85 |

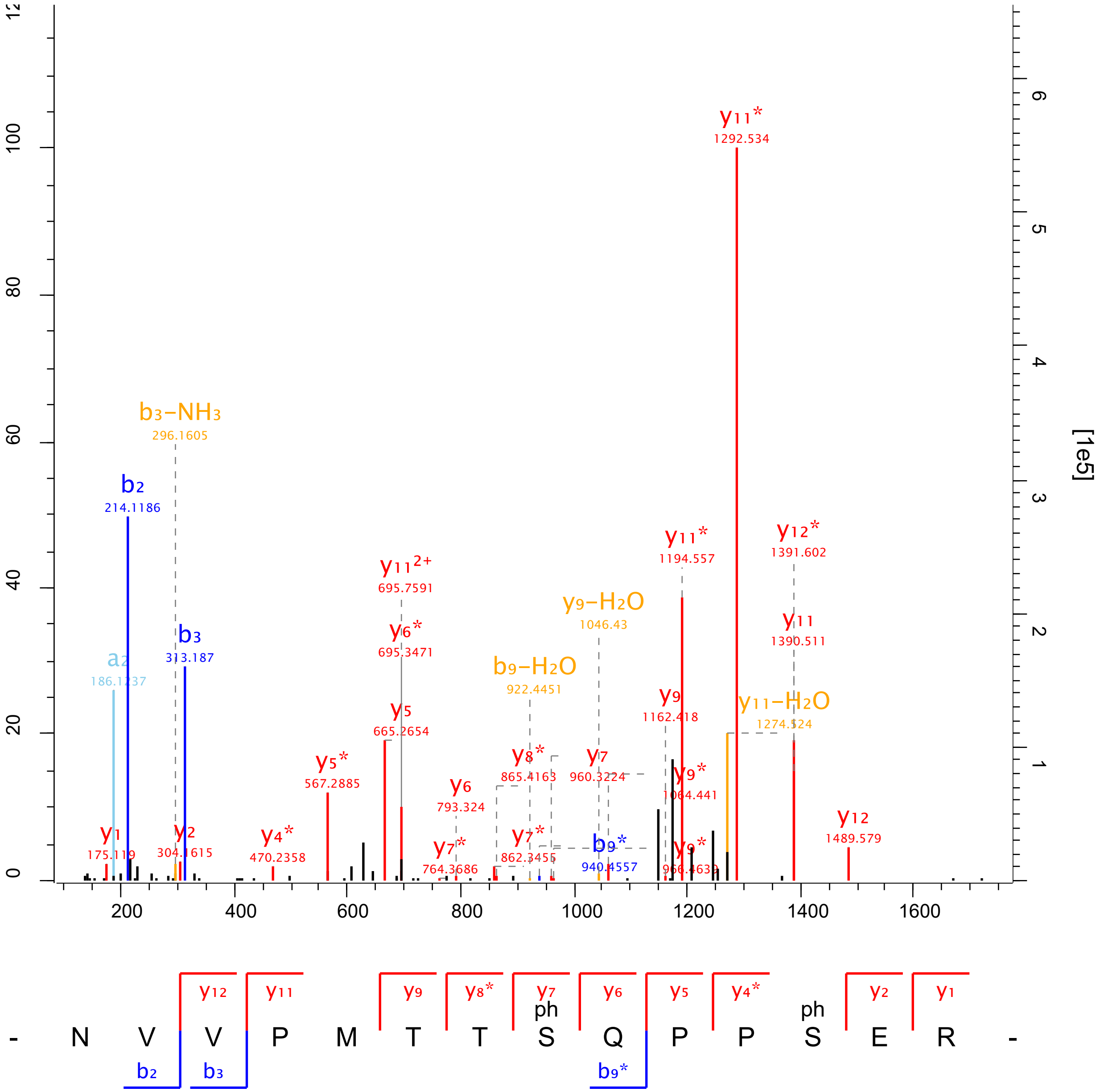

| Raw file | Scan | Method | Score | m/z |
|---|---|---|---|---|
| A17_00_2 | 23291 | FTMS; HCD | 44.24 | 618.63 |

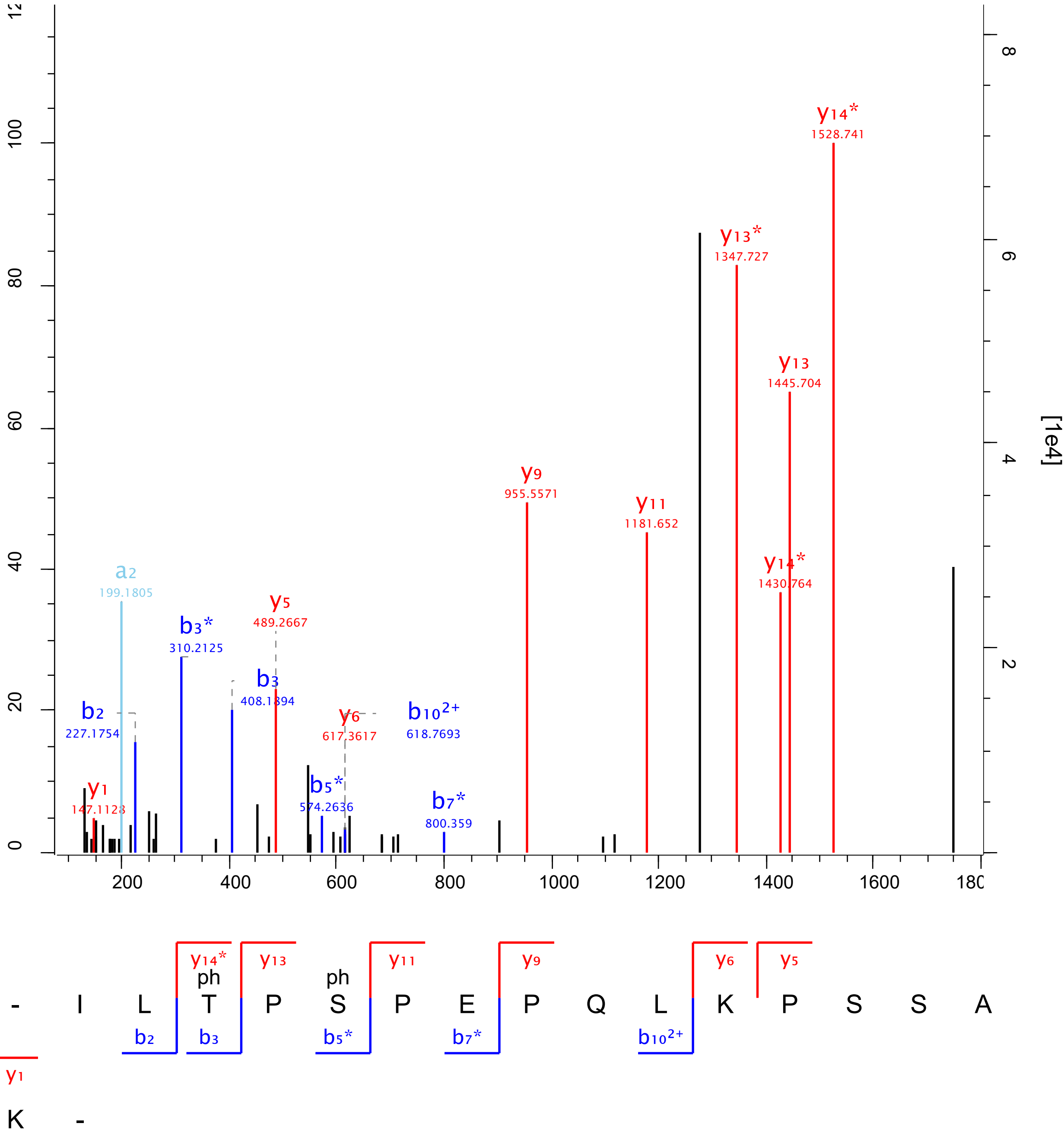

| Raw file | Scan | Method | Score | m/z |
|---|---|---|---|---|
| A17_00_2 | 23353 | FTMS; HCD | 78.67 | 809.83 |

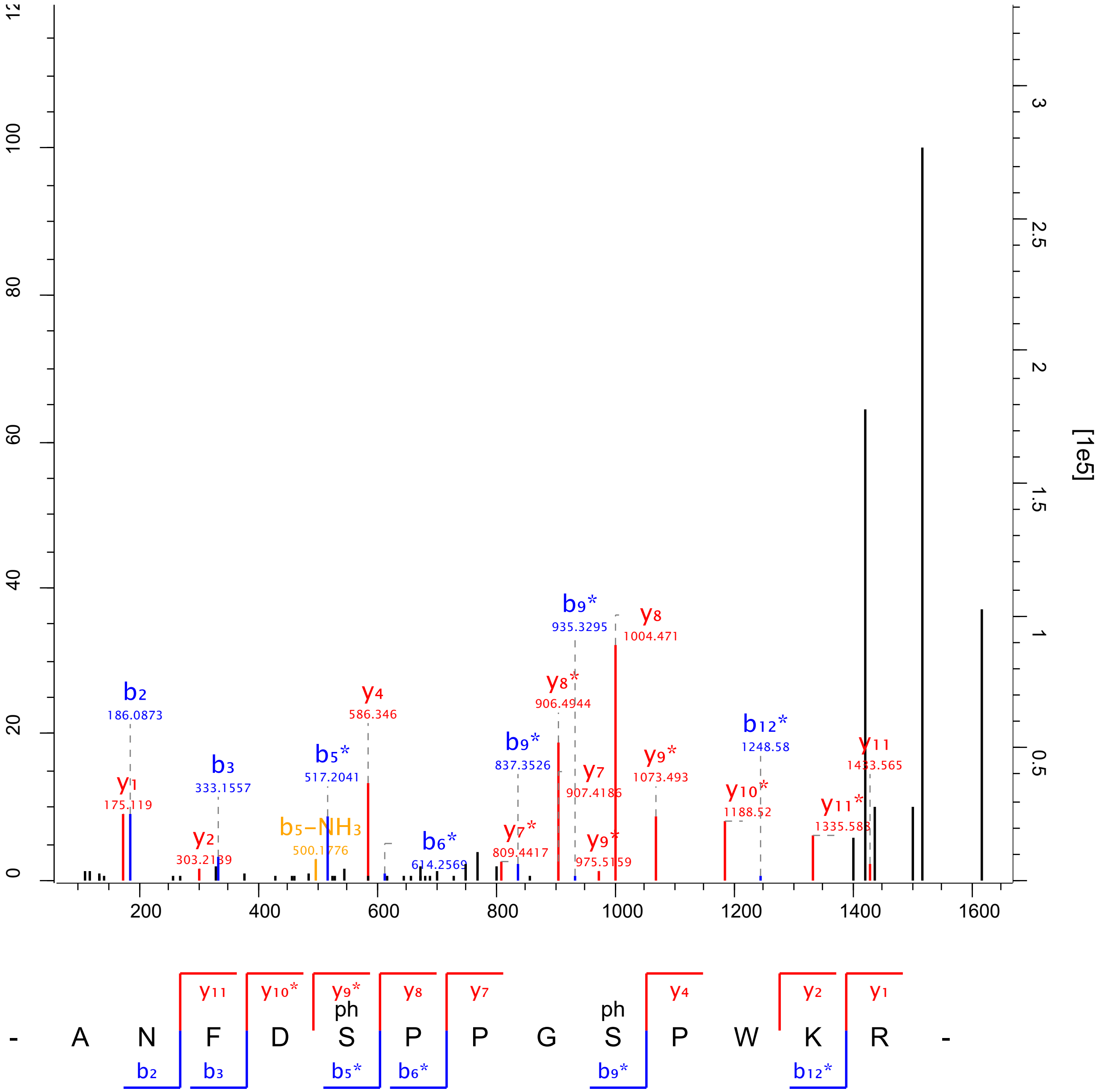

| Raw file | Scan | Method | Score | m/z |
|---|---|---|---|---|
| A17_00_2 | 23948 | FTMS; HCD | 141.24 | 1038.91 |

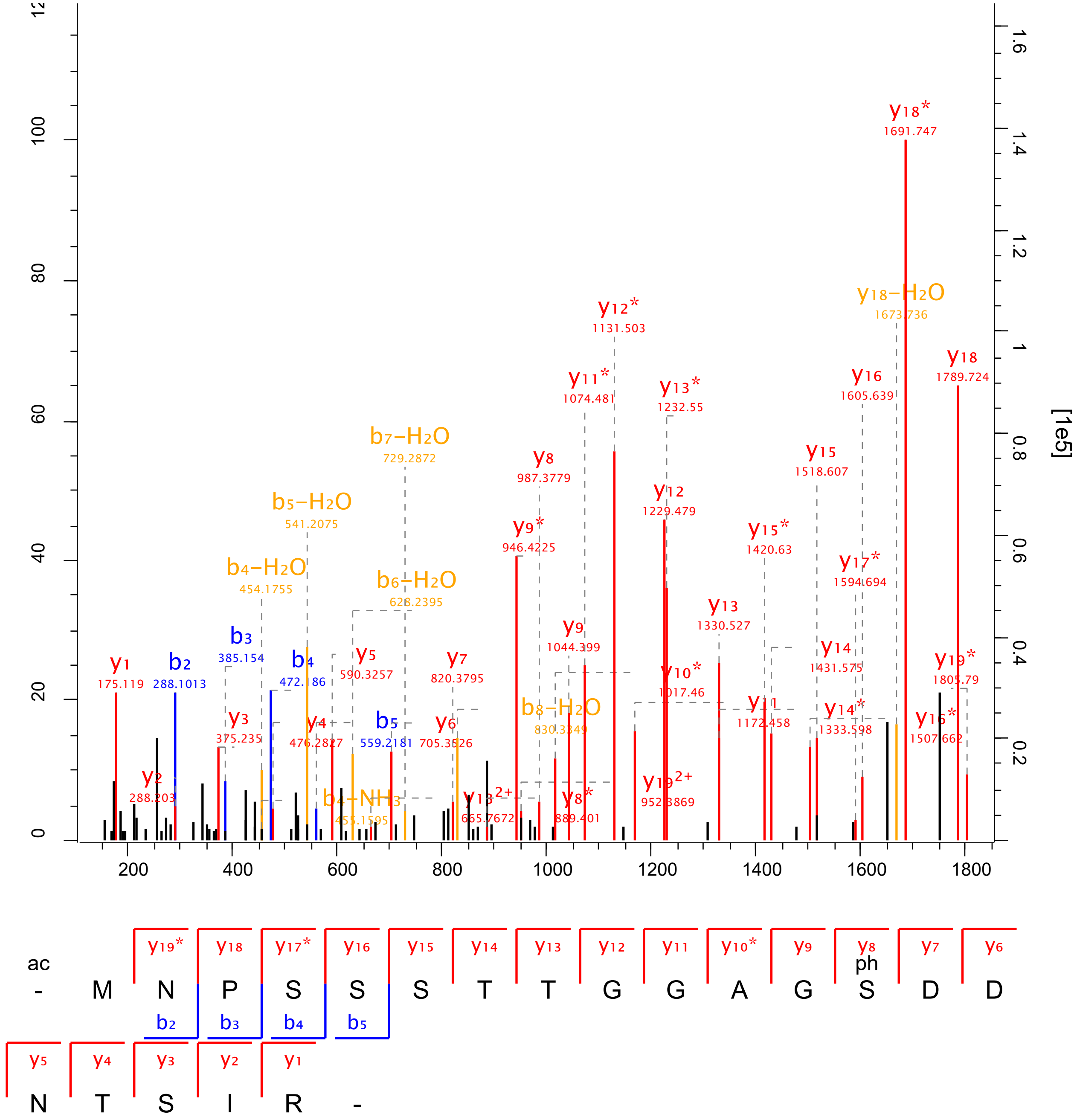

| Raw file | Scan | Method | Score | m/z |
|---|---|---|---|---|
| A17_00_2 | 23952 | FTMS; HCD | 138.28 | 834.33 |

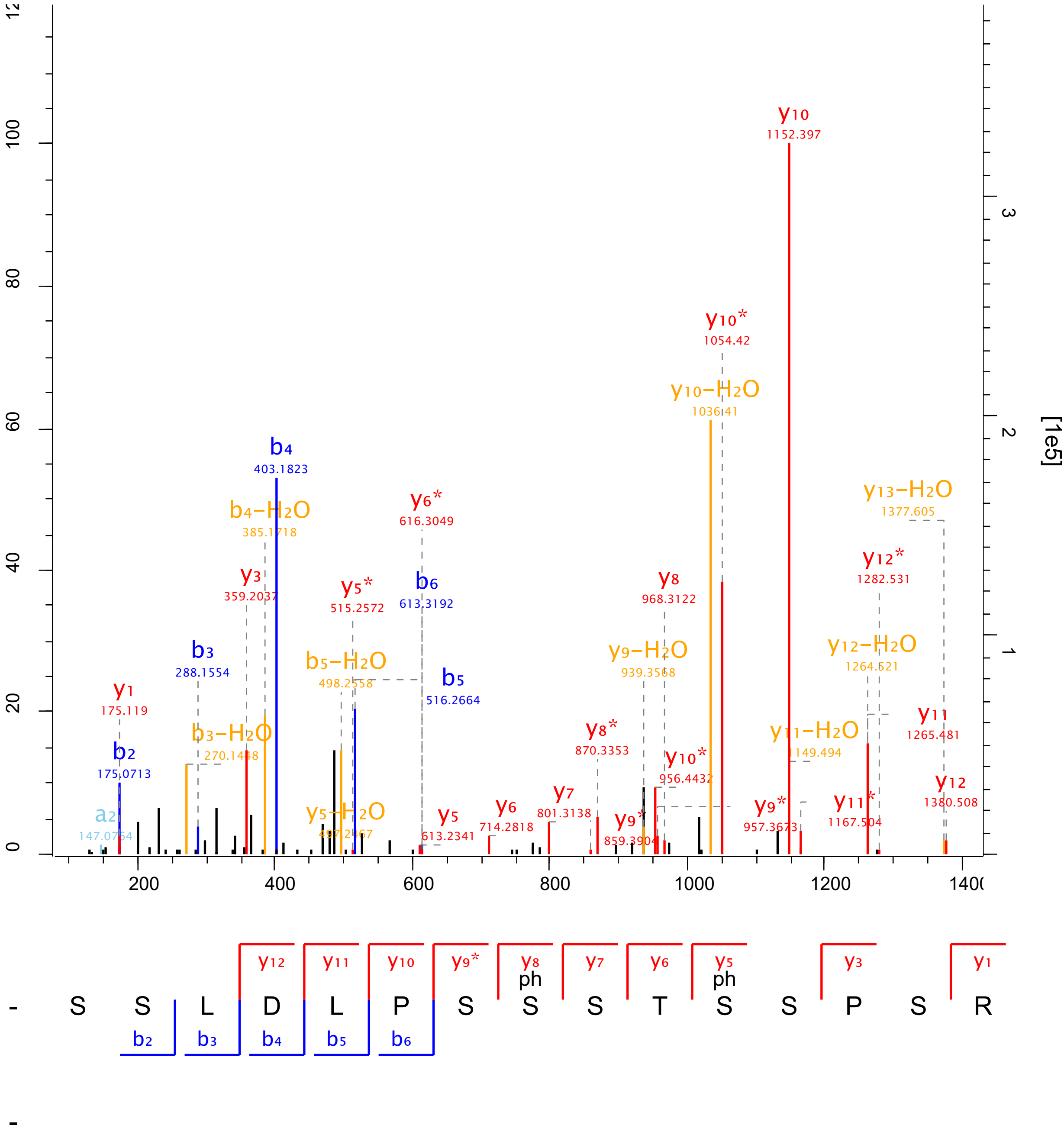

| Raw file | Scan | Method | Score | m/z |
| --- | --- | --- | --- | --- |
| A17_00_2 | 2408 | FTMS; HCD | 125.91 | 529.22 |

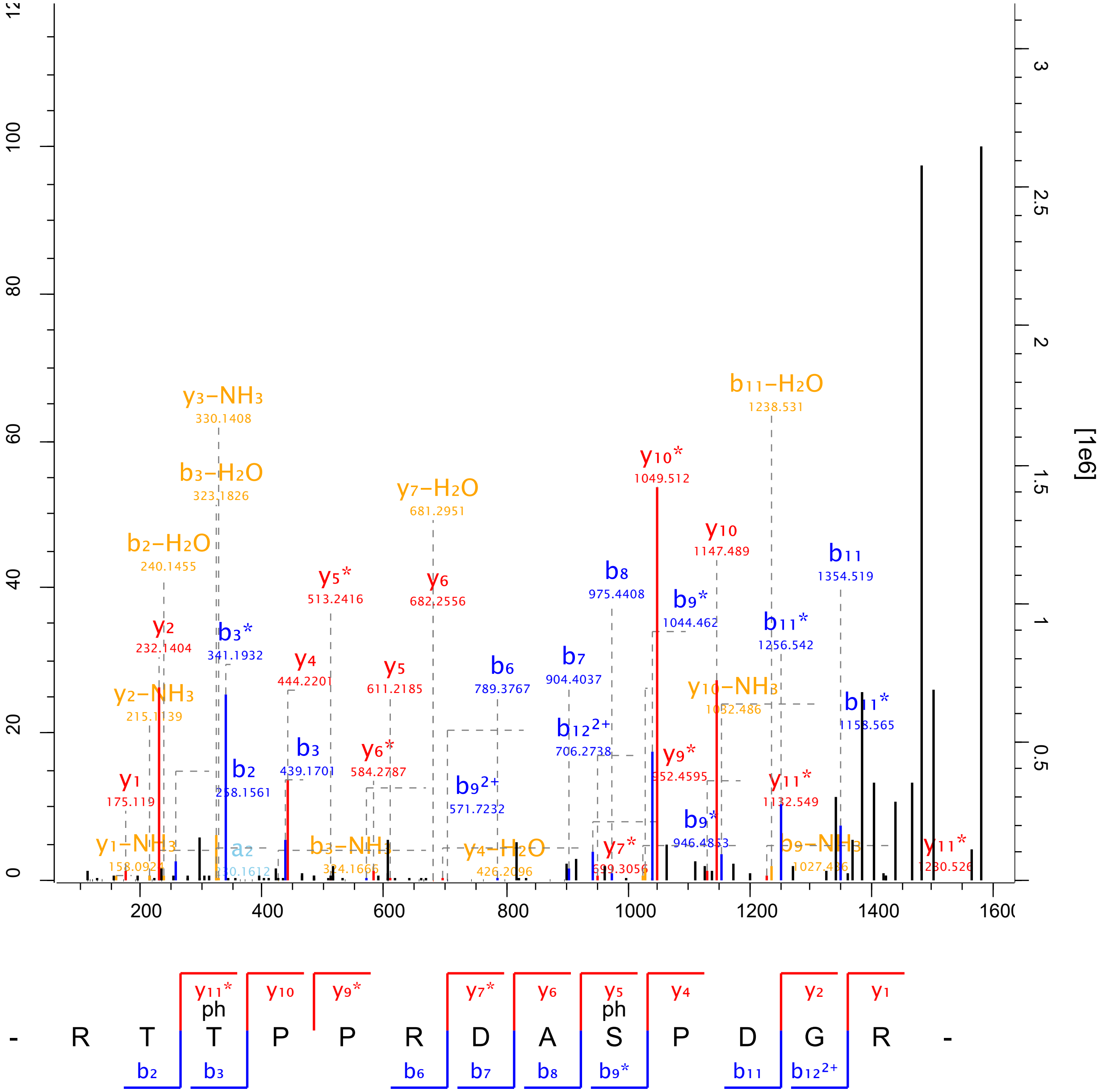

| Raw file | Scan | Method | Score | m/z |
|---|---|---|---|---|
| A17_00_2 | 24194 | FTMS; HCD | 64.52 | 805.35 |

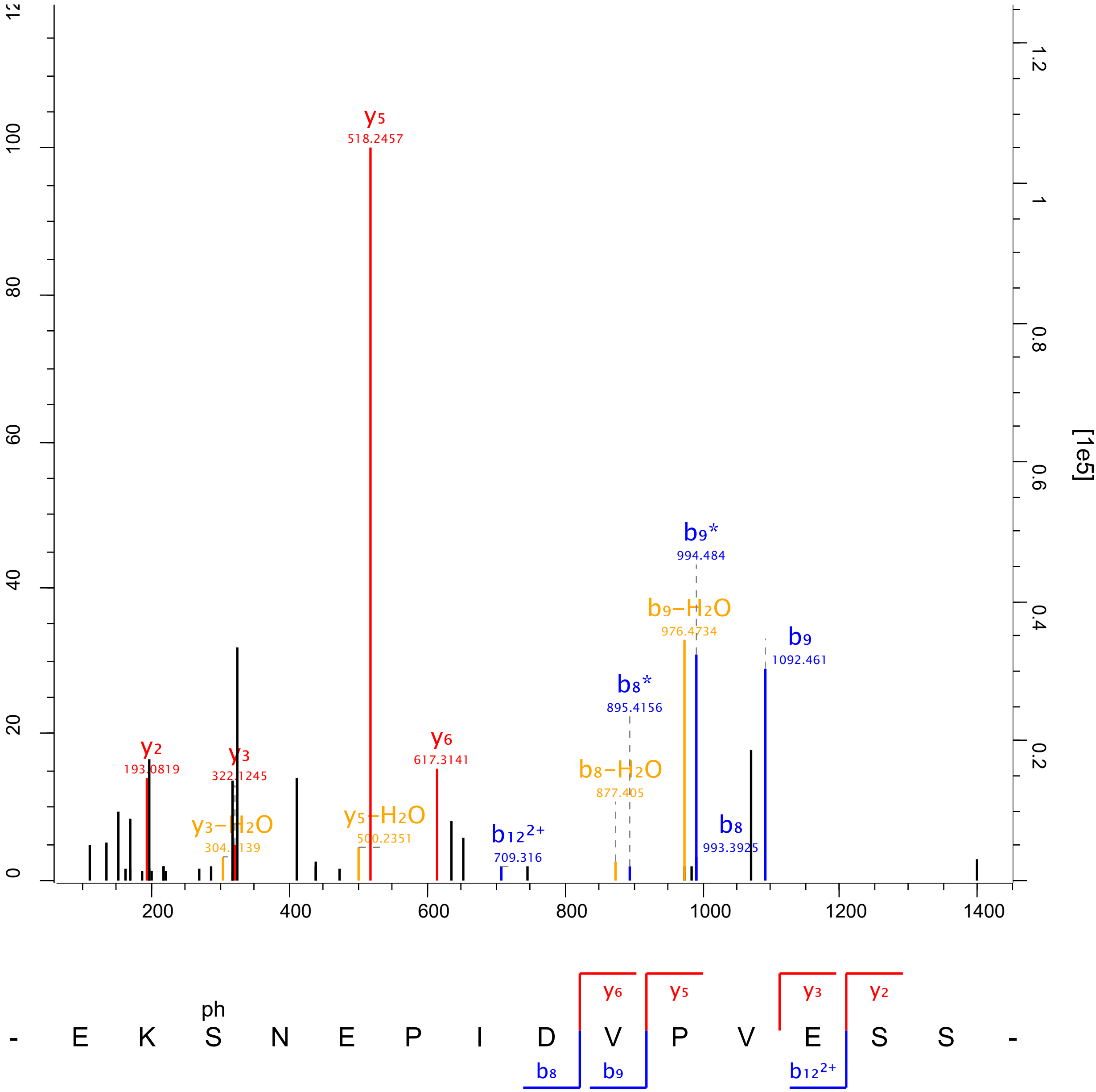

| Raw file | Scan | Method | Score | m/z |
| --- | --- | --- | --- | --- |
| A17_00_2 | 2481 | FTMS; HCD | 52.49 | 327.82 |

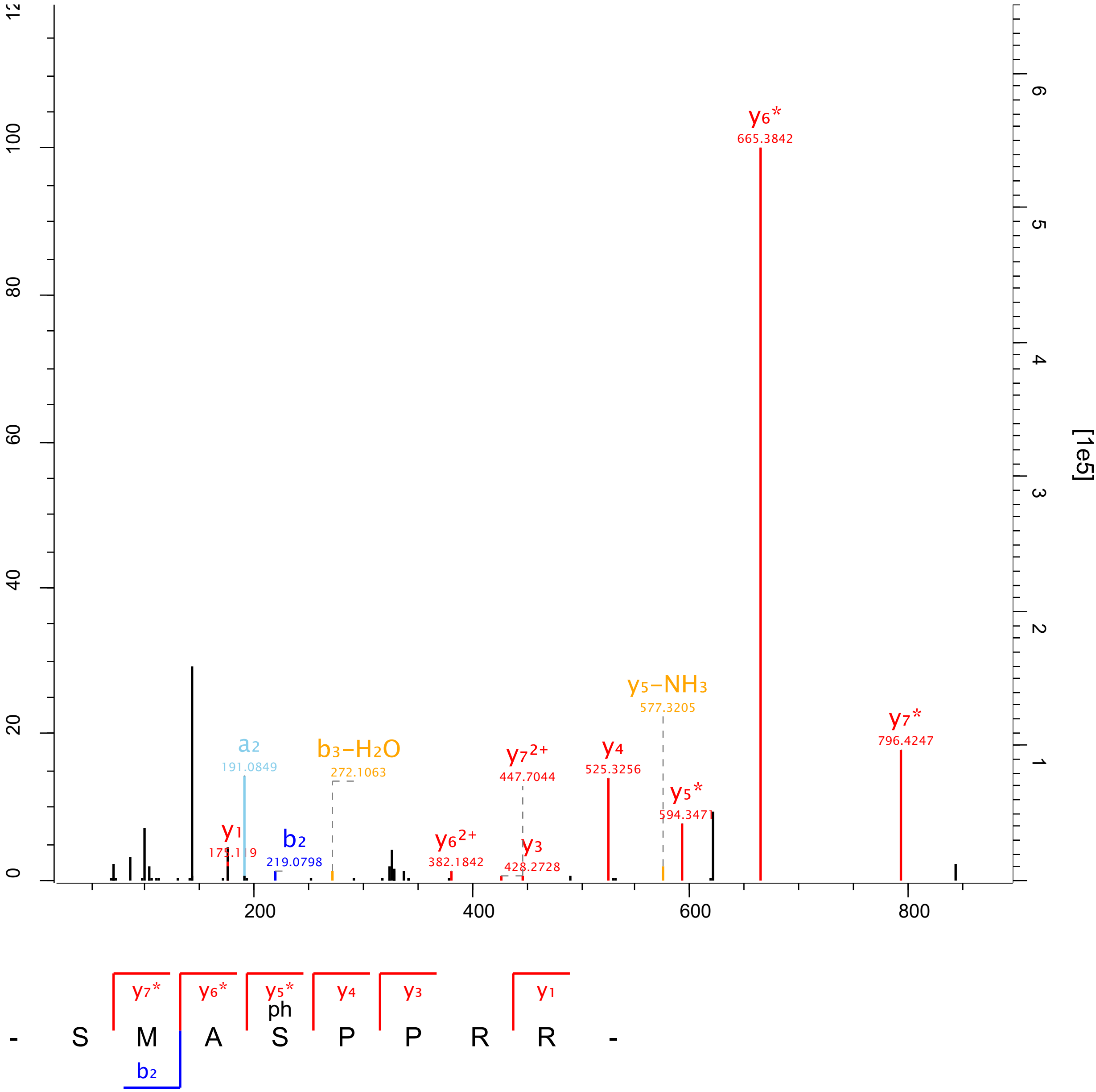

| Raw file | Scan | Method | Score | m/z |
|---|---|---|---|---|
| A17_00_2 | 24897 | FTMS; HCD | 180.09 | 850.86 |

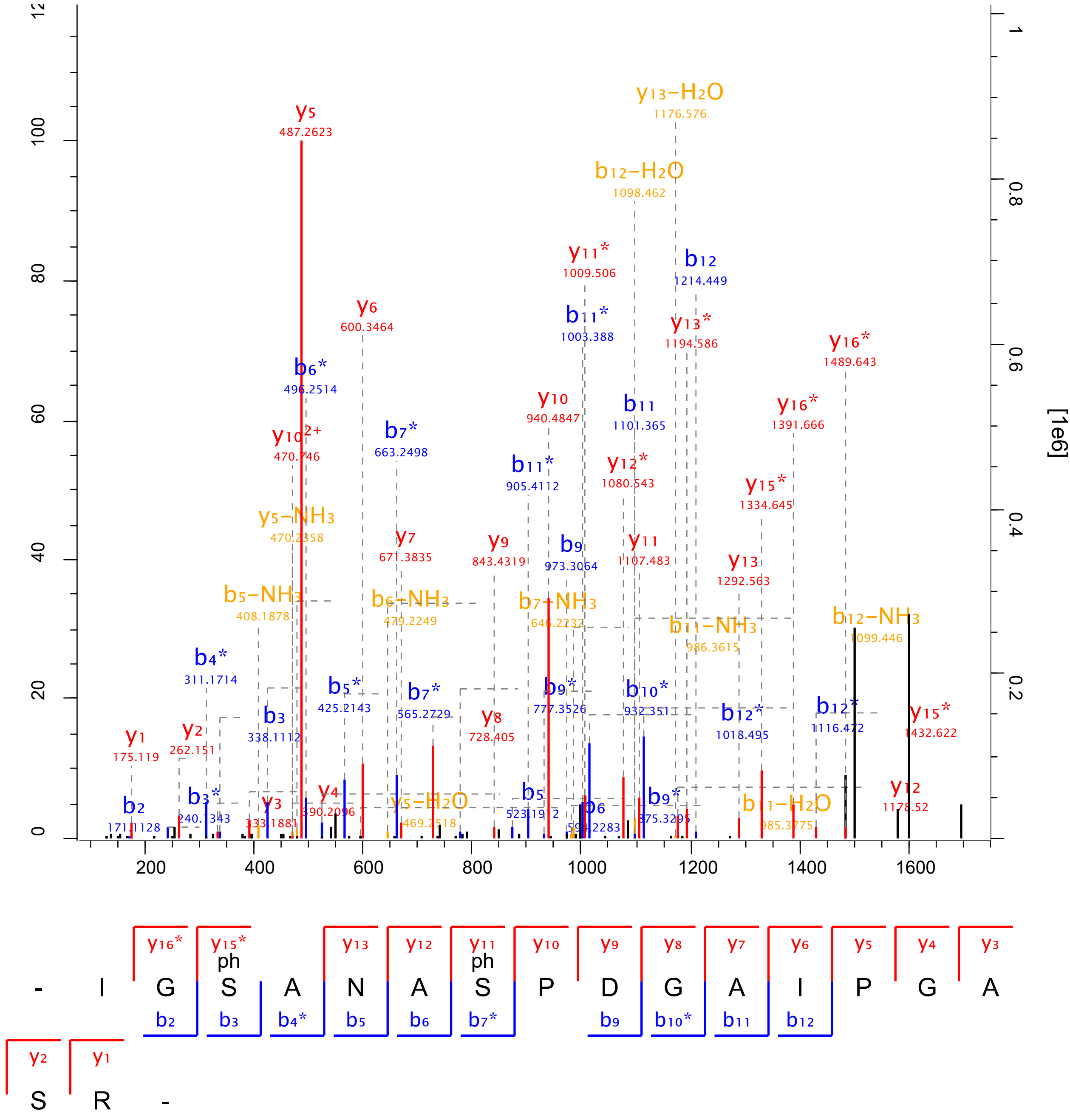

| Raw file | Scan | Method | Score | m/z |
| --- | --- | --- | --- | --- |
| A17_00_2 | 25341 | FTMS; HCD | 132.81 | 799.32 |

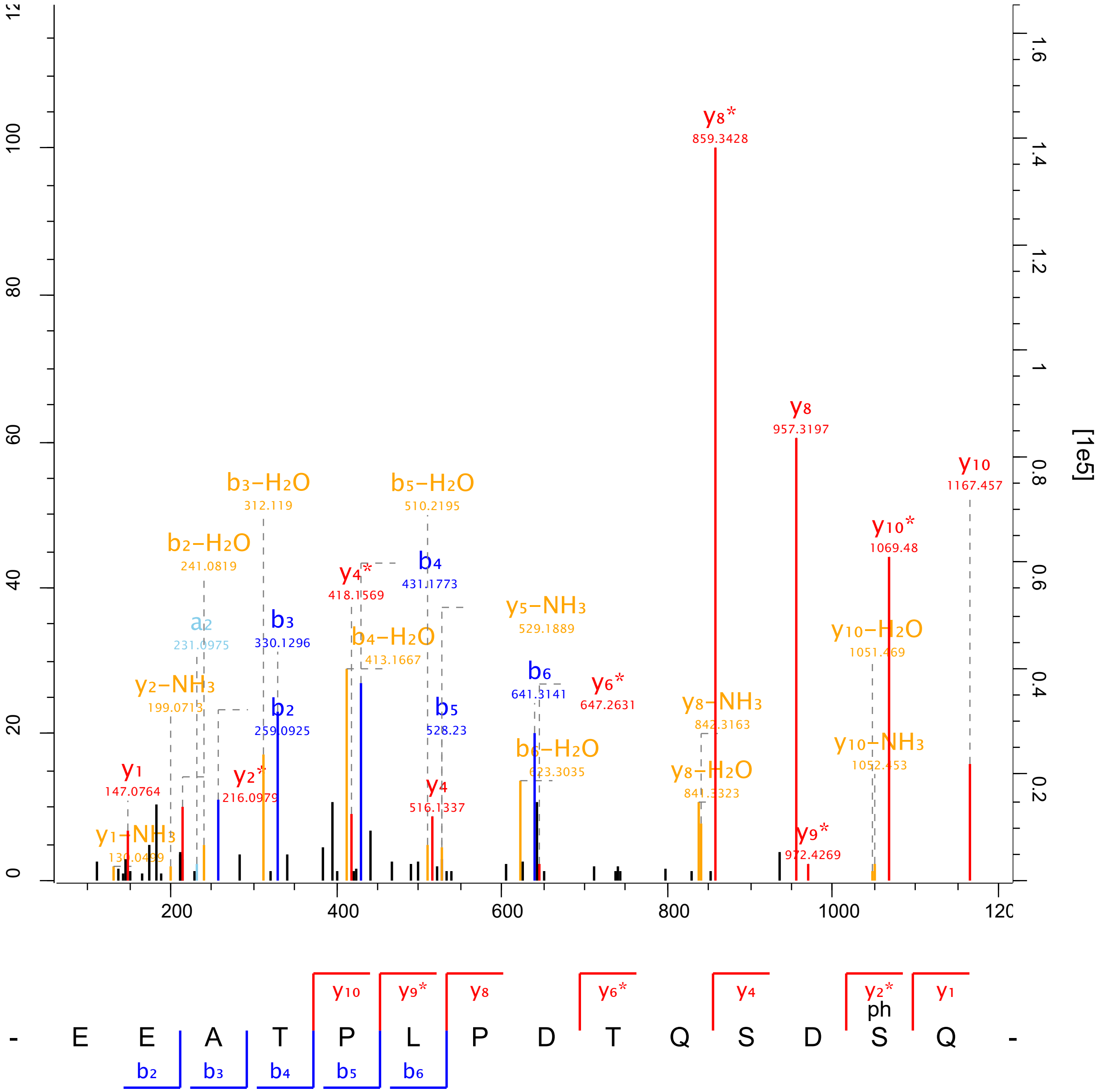

| Raw file | Scan | Method | Score | m/z |
| --- | --- | --- | --- | --- |
| A17_00_2 | 2559 | FTMS; HCD | 43.31 | 336.84 |

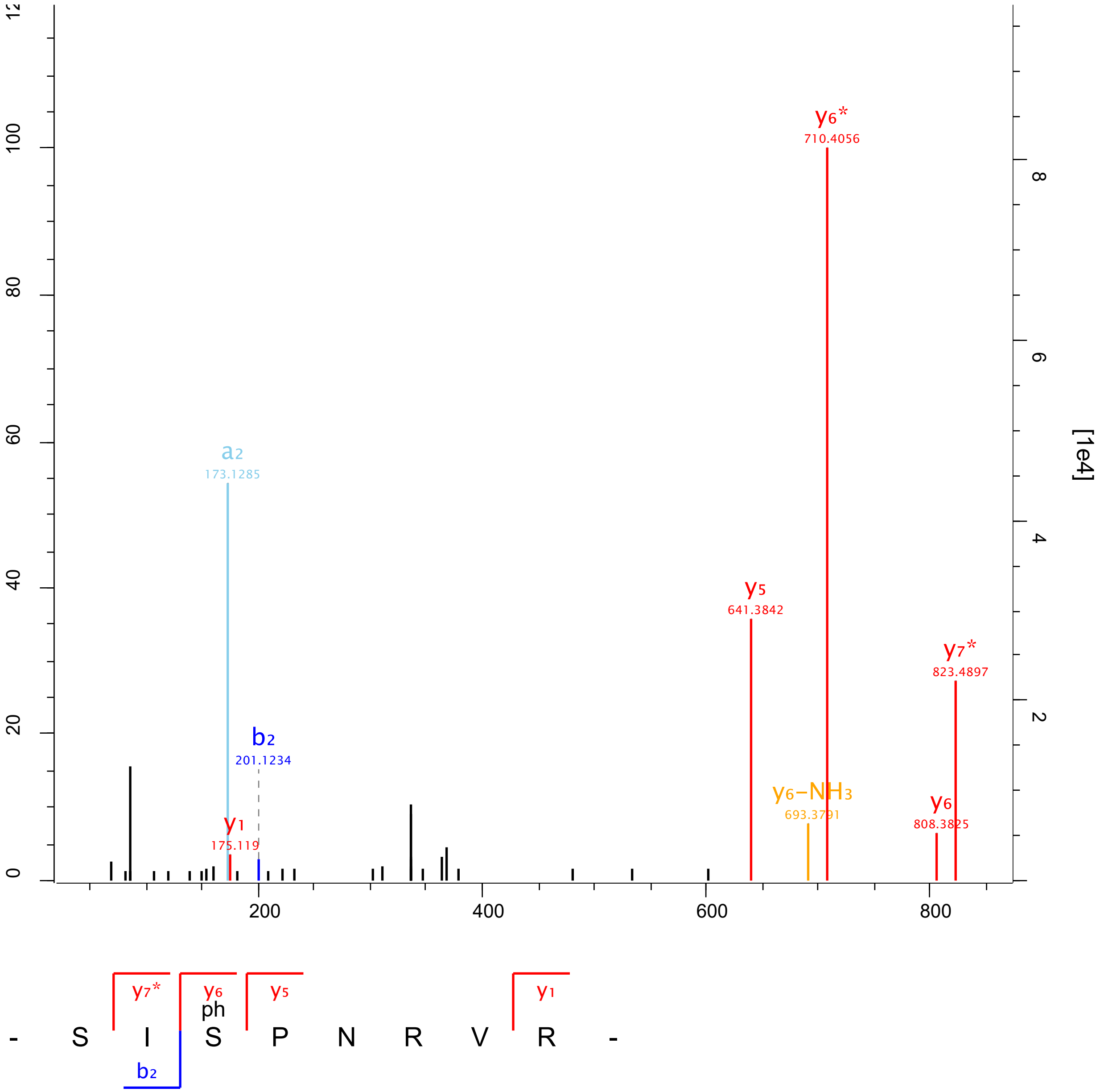

| Raw file | Scan | Method | Score | m/z |
|---|---|---|---|---|
| A17_00_2 | 25862 | FTMS; HCD | 91.08 | 655.78 |

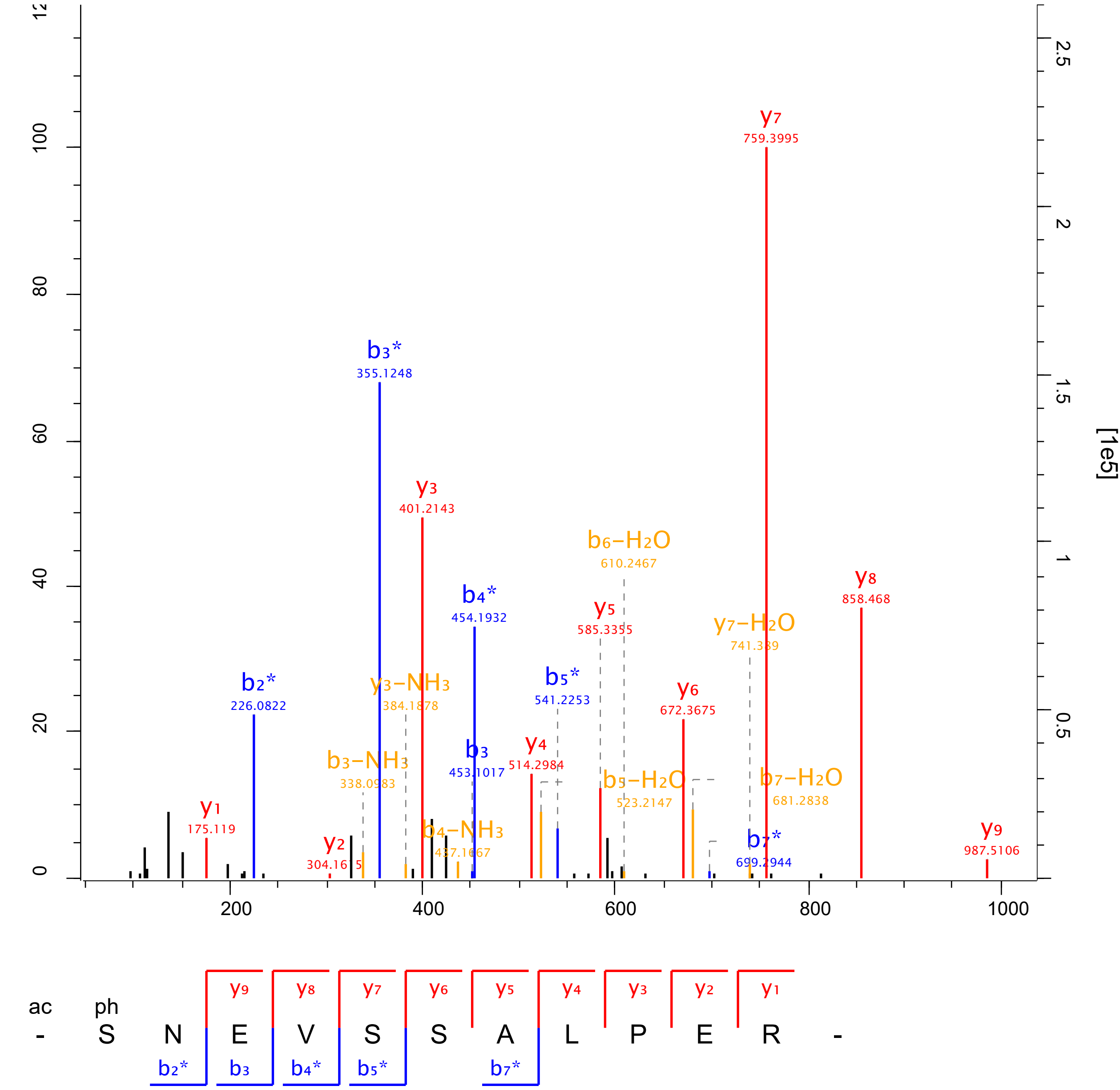

| Raw file | Scan | Method | Score | m/z |
| --- | --- | --- | --- | --- |
| A17_00_2 | 2611 | FTMS; HCD | 131.31 | 499.23 |

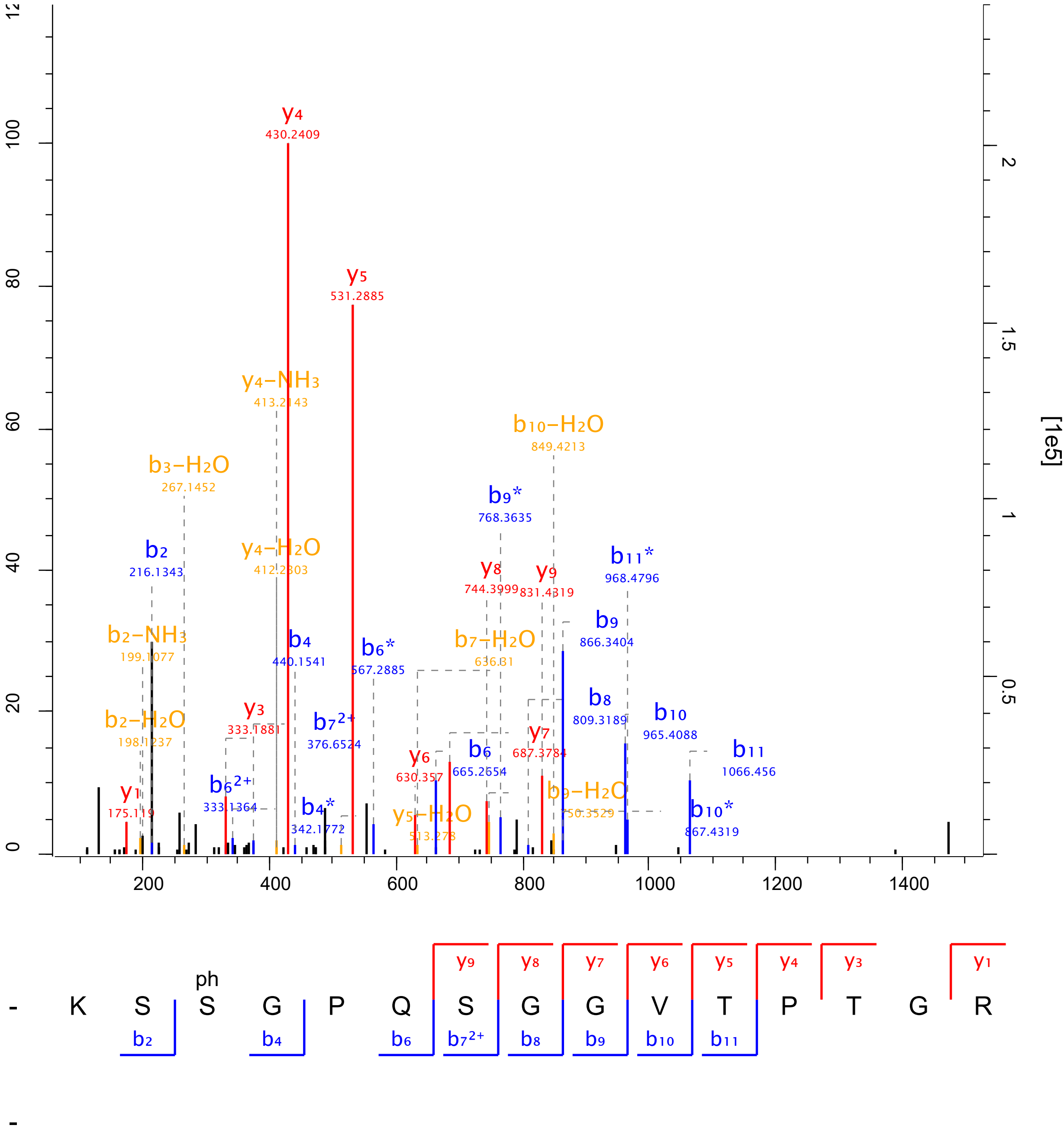

| Raw file | Scan | Method | Score | m/z |
| --- | --- | --- | --- | --- |
| A17_00_2 | 26493 | FTMS; HCD | 48.05 | 898.8 |

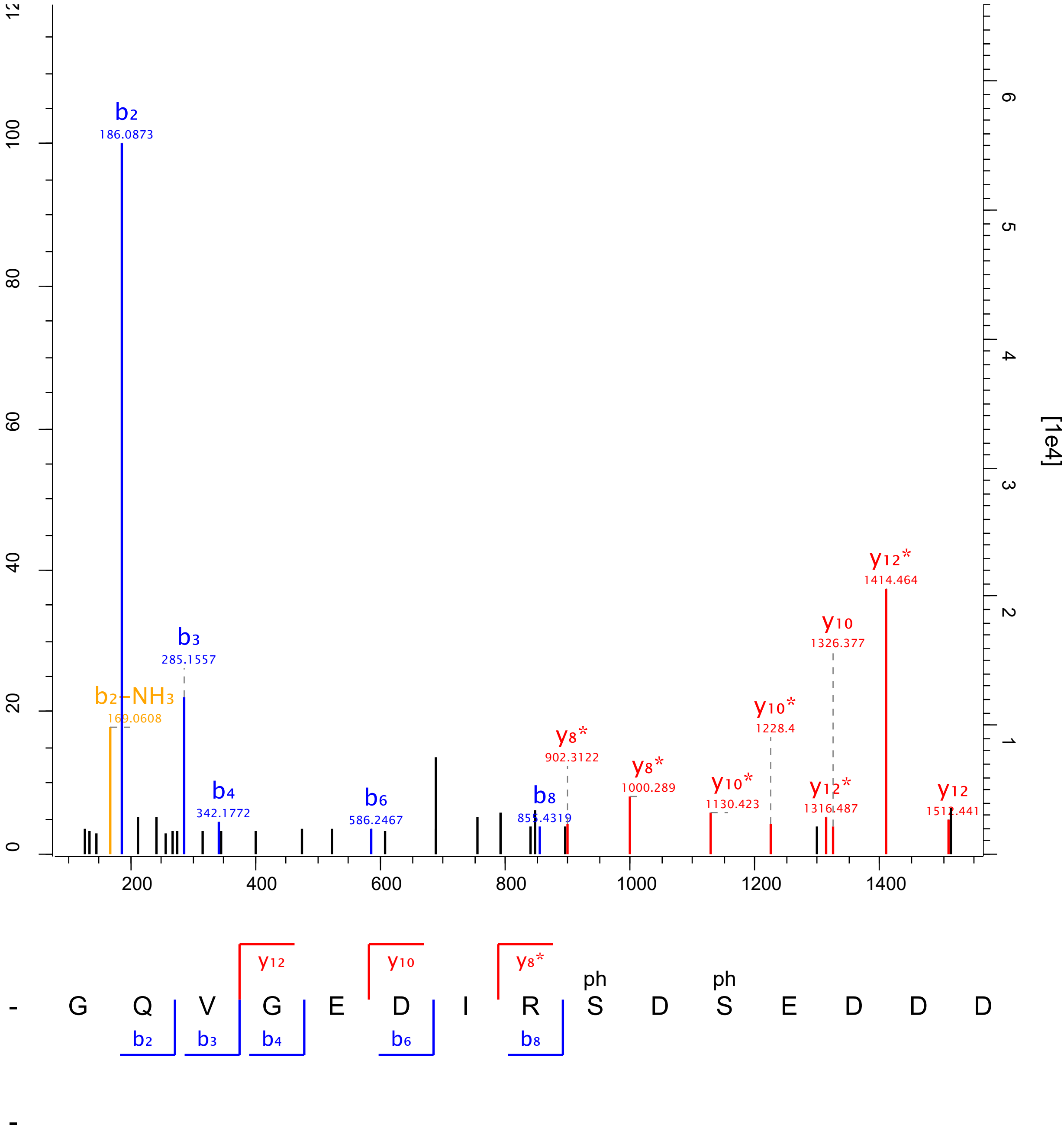

| Raw file | Scan | Method | Score | m/z |
|---|---|---|---|---|
| A17_00_2 | 26516 | FTMS; HCD | 291.51 | 722.8 |

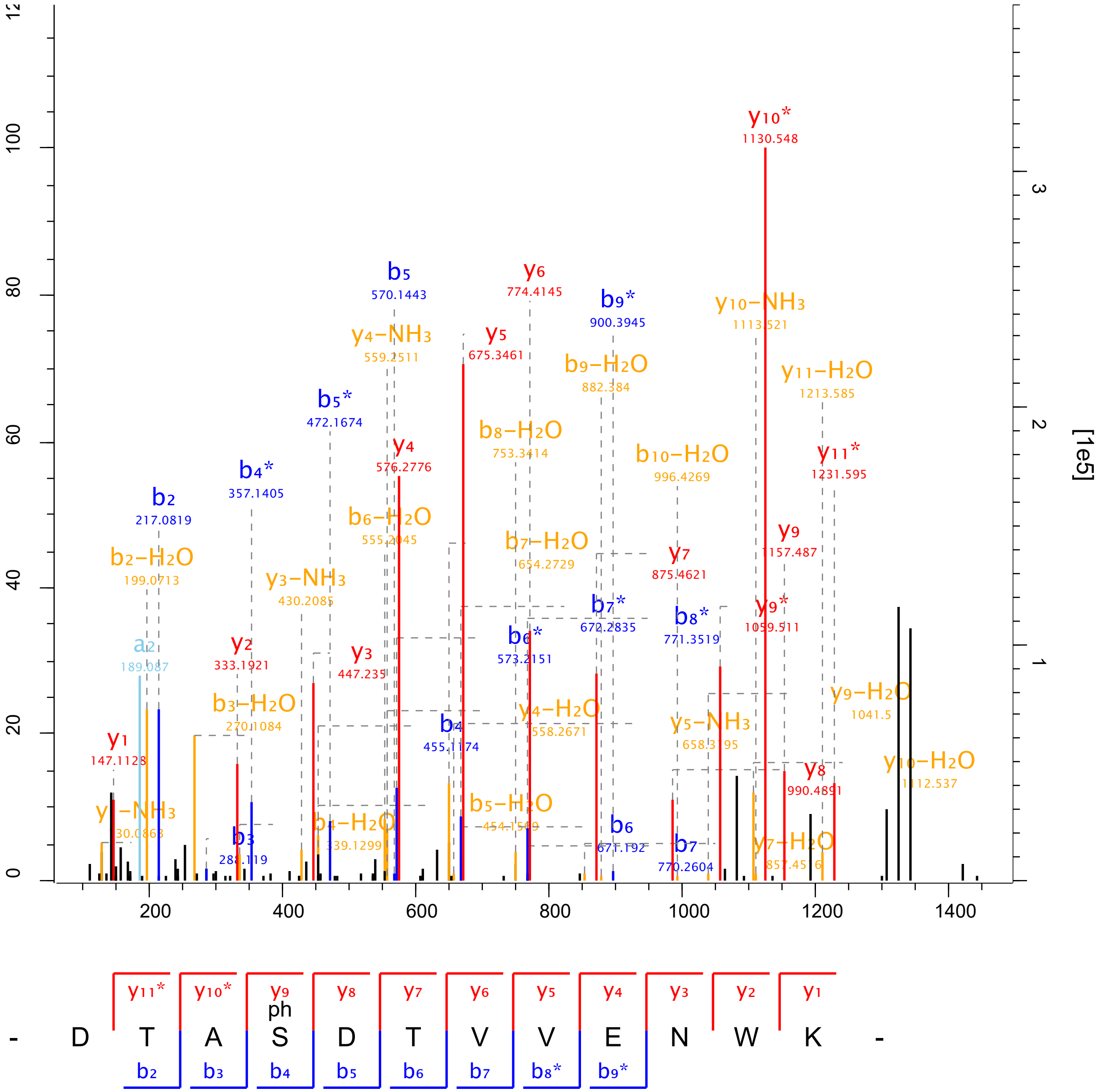

| Raw file | Scan | Method | Score | m/z |
|---|---|---|---|---|
| A17_00_2 | 2655 | FTMS; HCD | 46.35 | 398.85 |

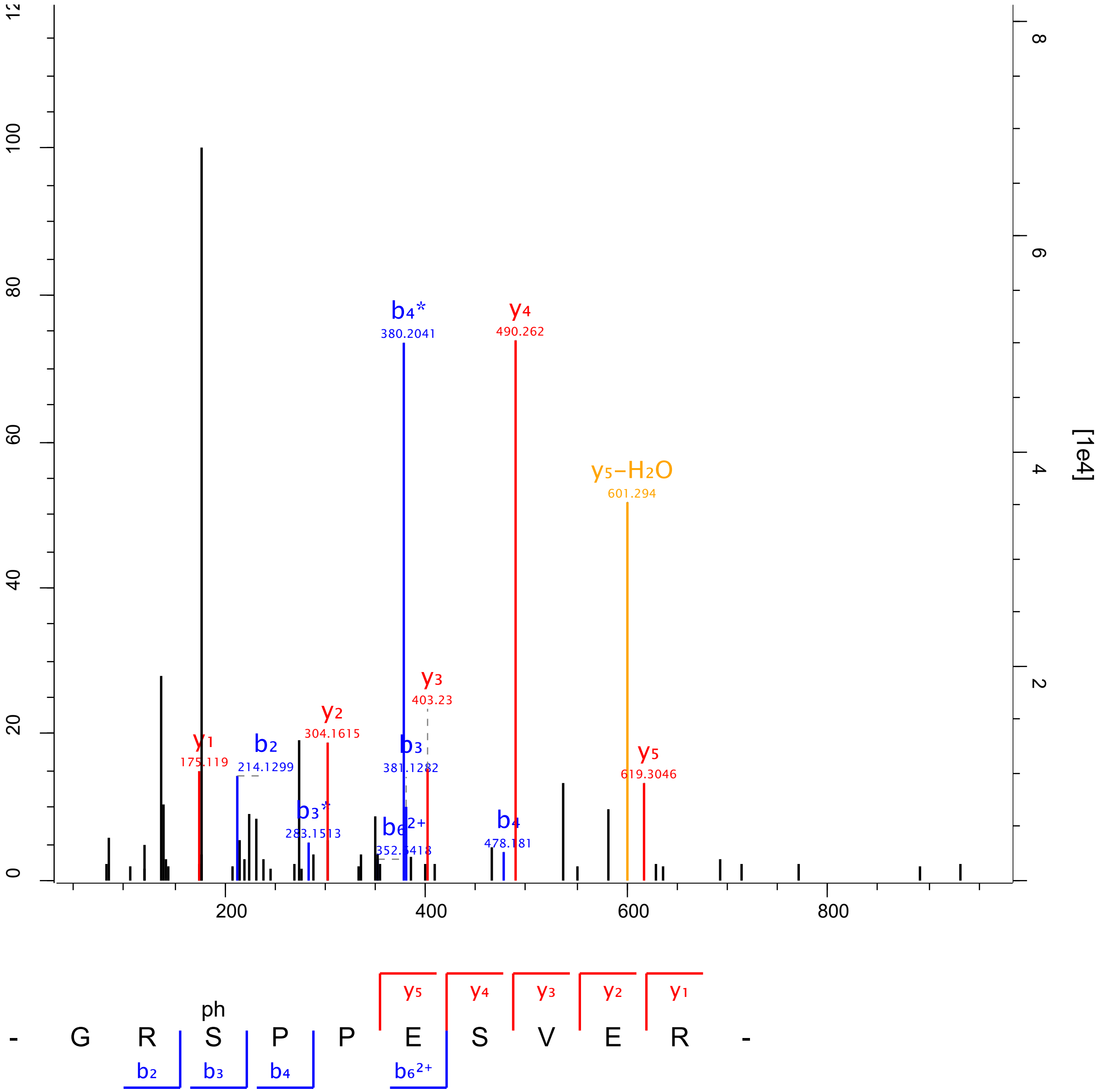

| Raw file | Scan | Method | Score | m/z |
| --- | --- | --- | --- | --- |
| A17_00_2 | 26609 | FTMS; HCD | 249.25 | 637.74 |

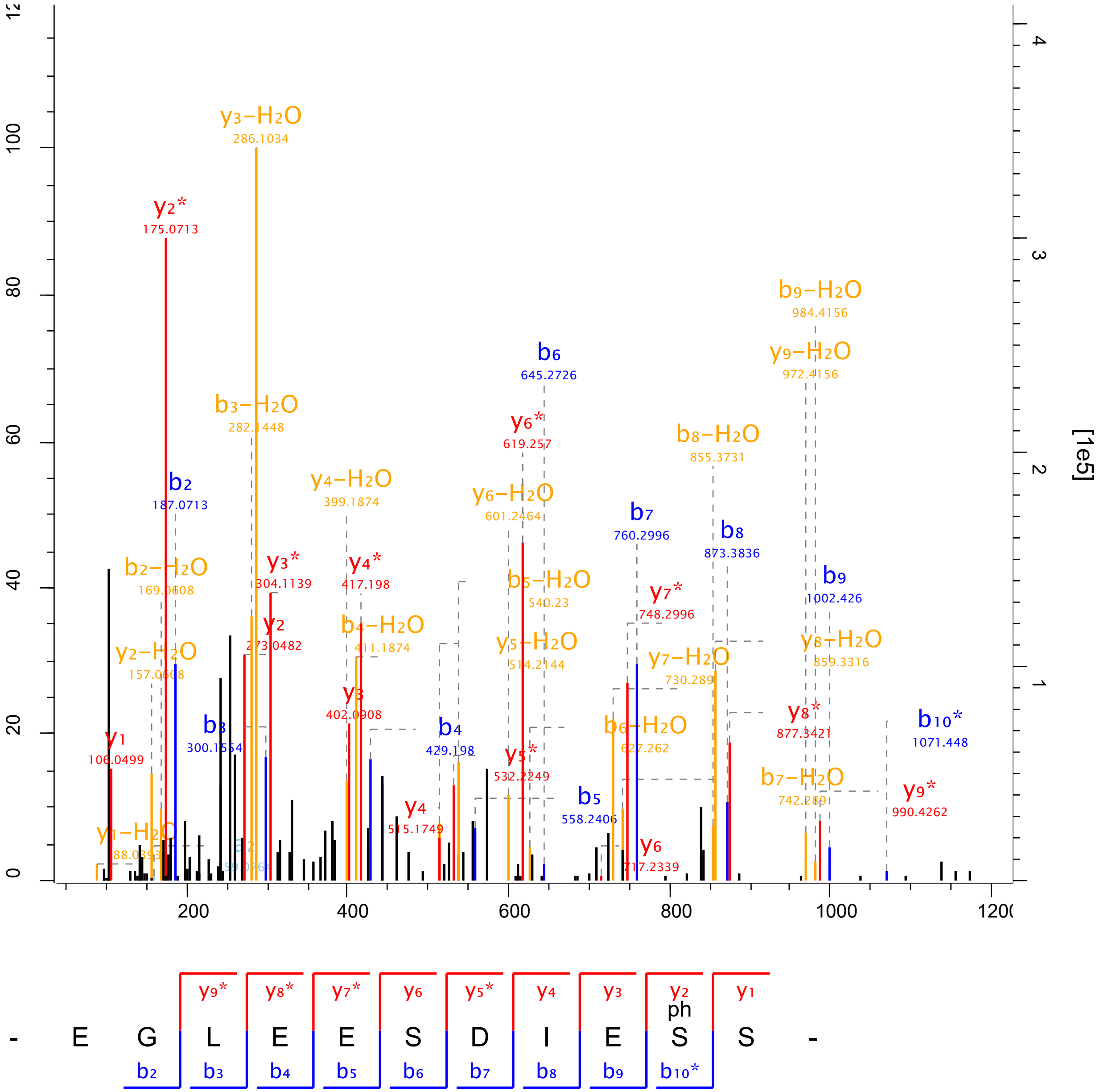

| Raw file | Scan | Method | Score | m/z |
| --- | --- | --- | --- | --- |
| A17_00_2 | 2667 | FTMS; HCD | 40.48 | 550.74 |

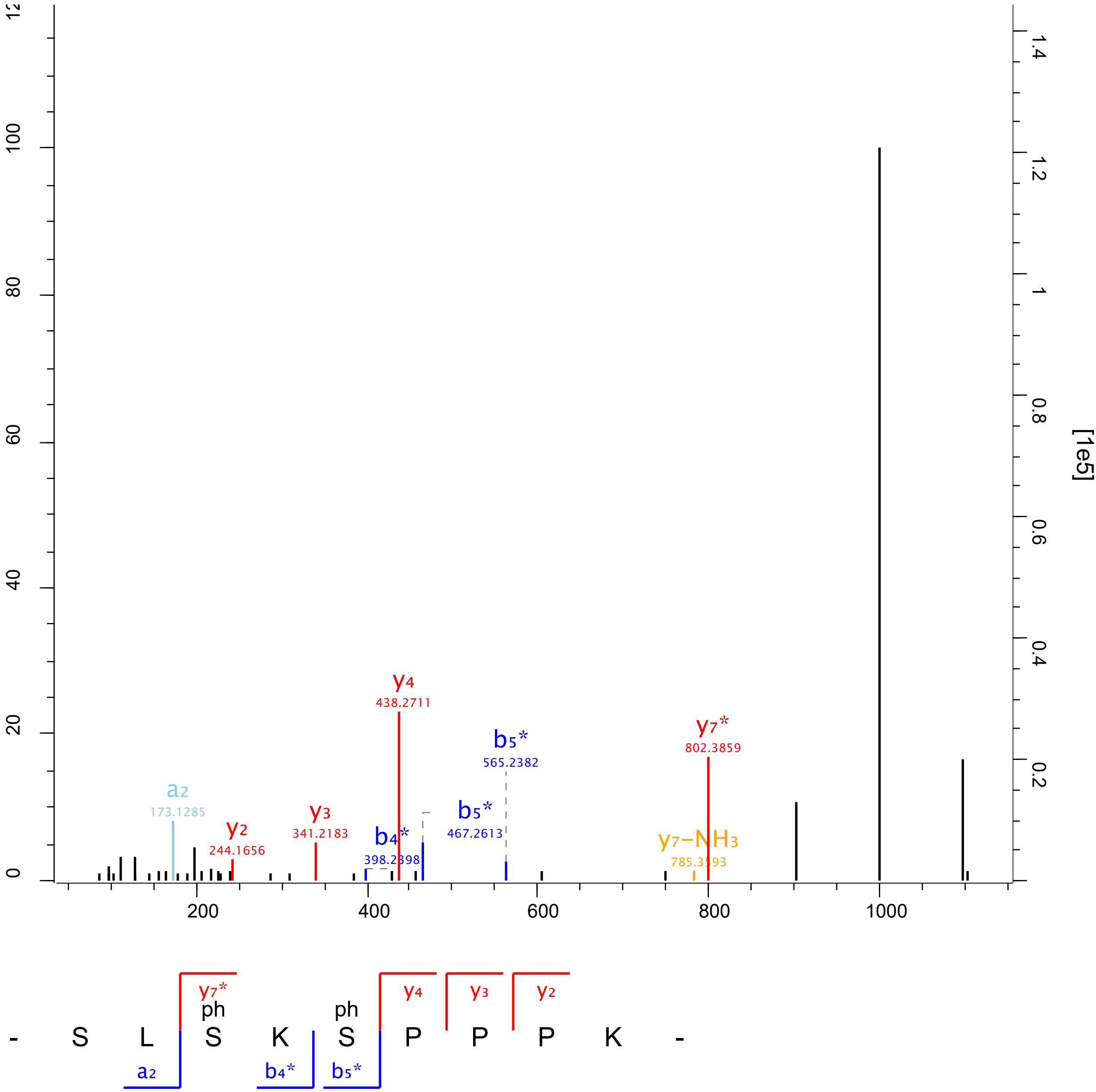

| Raw file | Scan | Method | Score | m/z |
| --- | --- | --- | --- | --- |
| A17_00_2 | 27067 | FTMS; HCD | 272.3 | 938.42 |

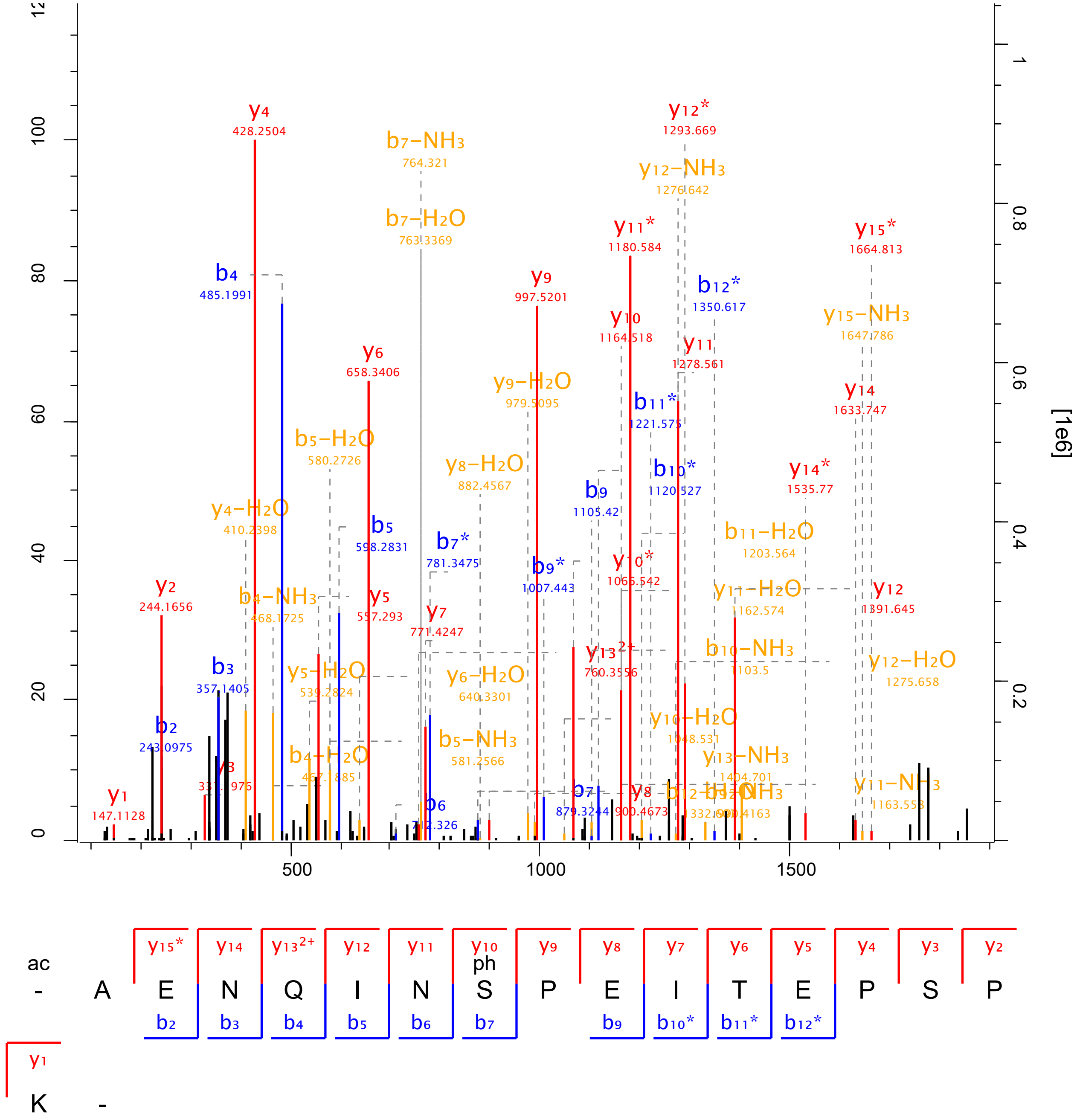

| Raw file | Scan | Method | Score | m/z |
| --- | --- | --- | --- | --- |
| A17_00_2 | 27766 | FTMS; HCD | 67.65 | 760.32 |

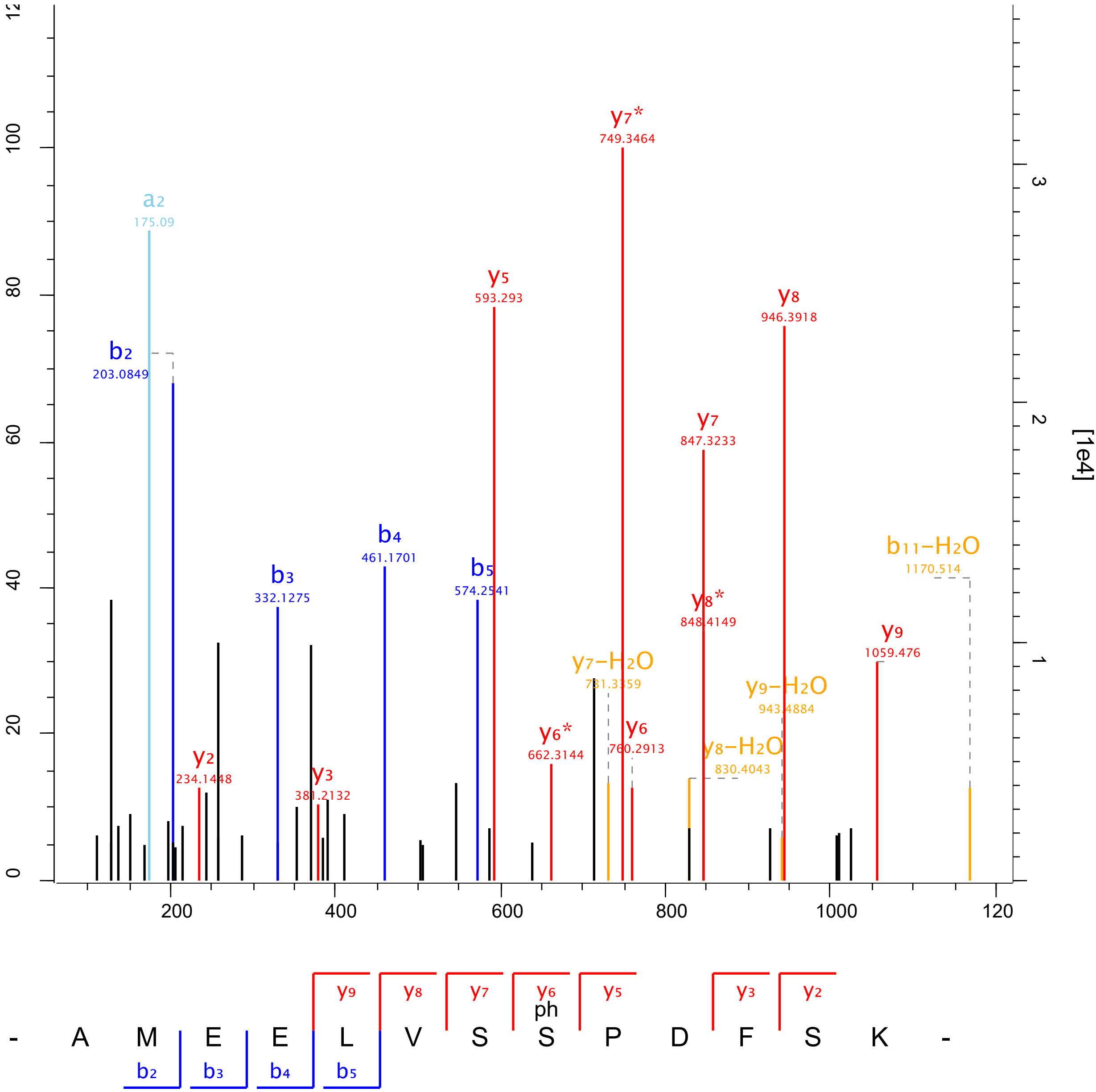

| Raw file | Scan | Method | Score | m/z |
|---|---|---|---|---|
| A17_00_2 | 27839 | FTMS; HCD | 79.68 | 806.85 |

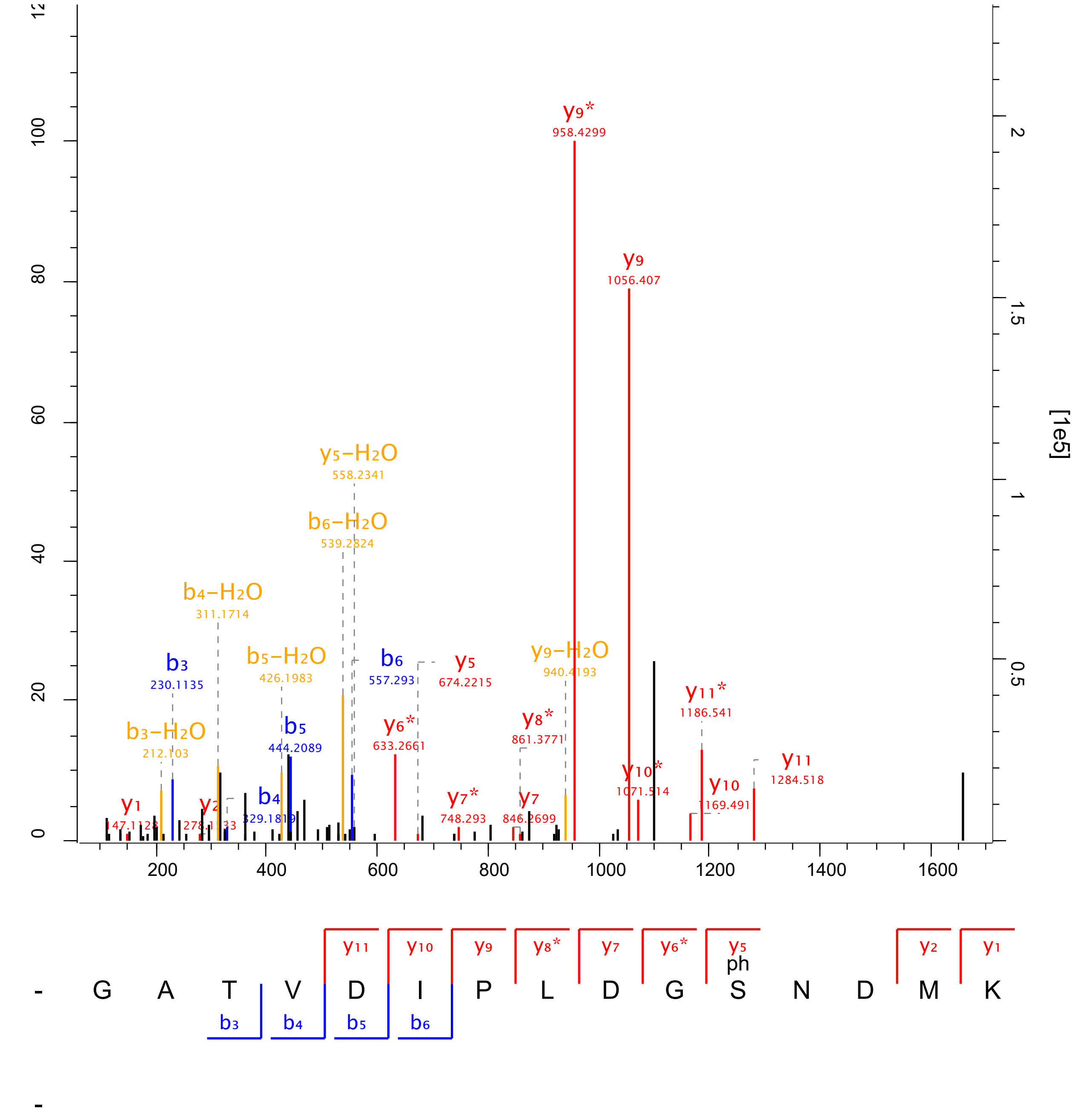

| Raw file | Scan | Method | Score | m/z |
|---|---|---|---|---|
| A17_00_2 | 2800 | FTMS; HCD | 113.43 | 590.27 |

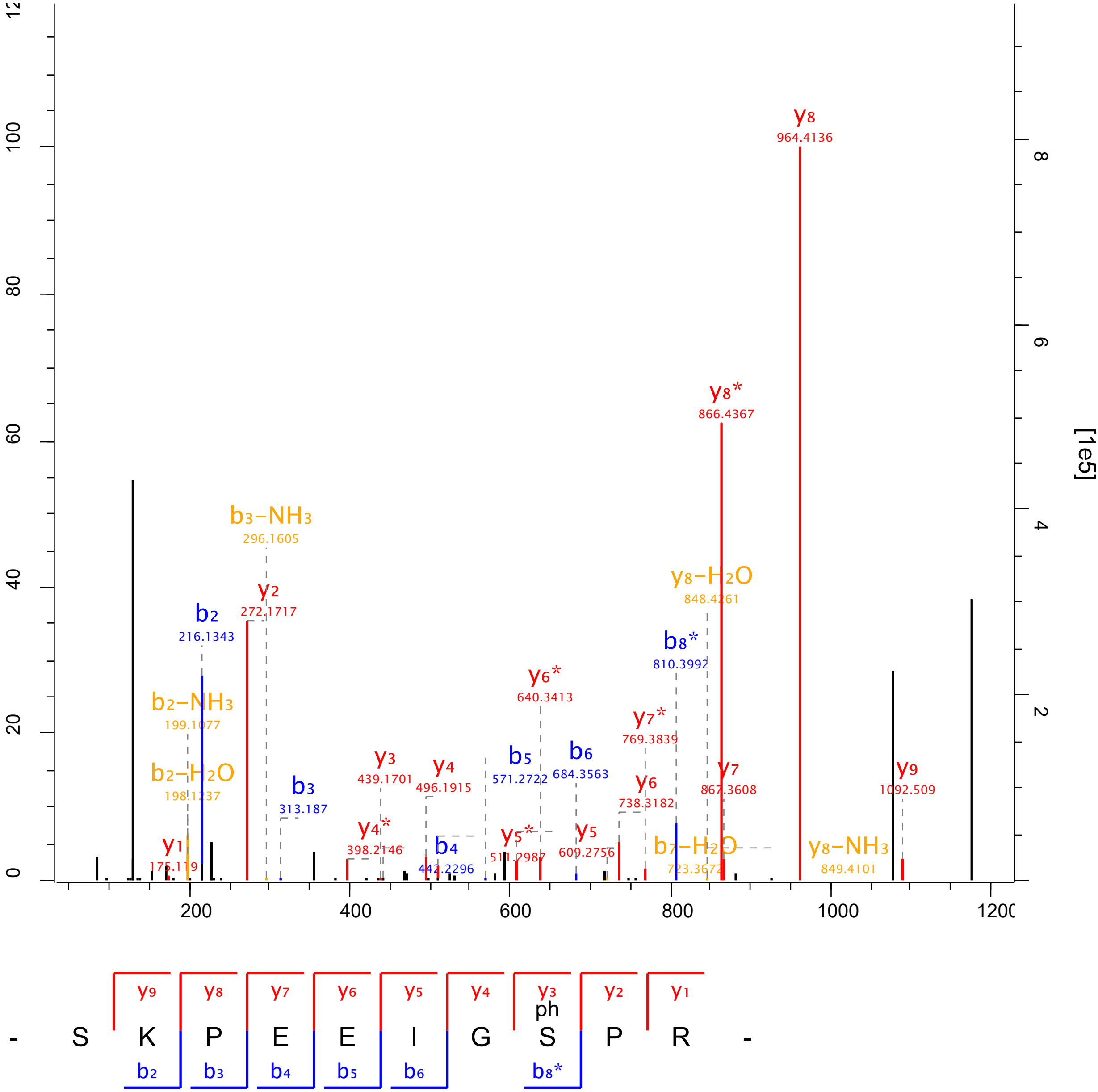

| Raw file | Scan | Method | Score | m/z |
| --- | --- | --- | --- | --- |
| A17_00_2 | 28163 | FTMS; HCD | 61.48 | 737.84 |

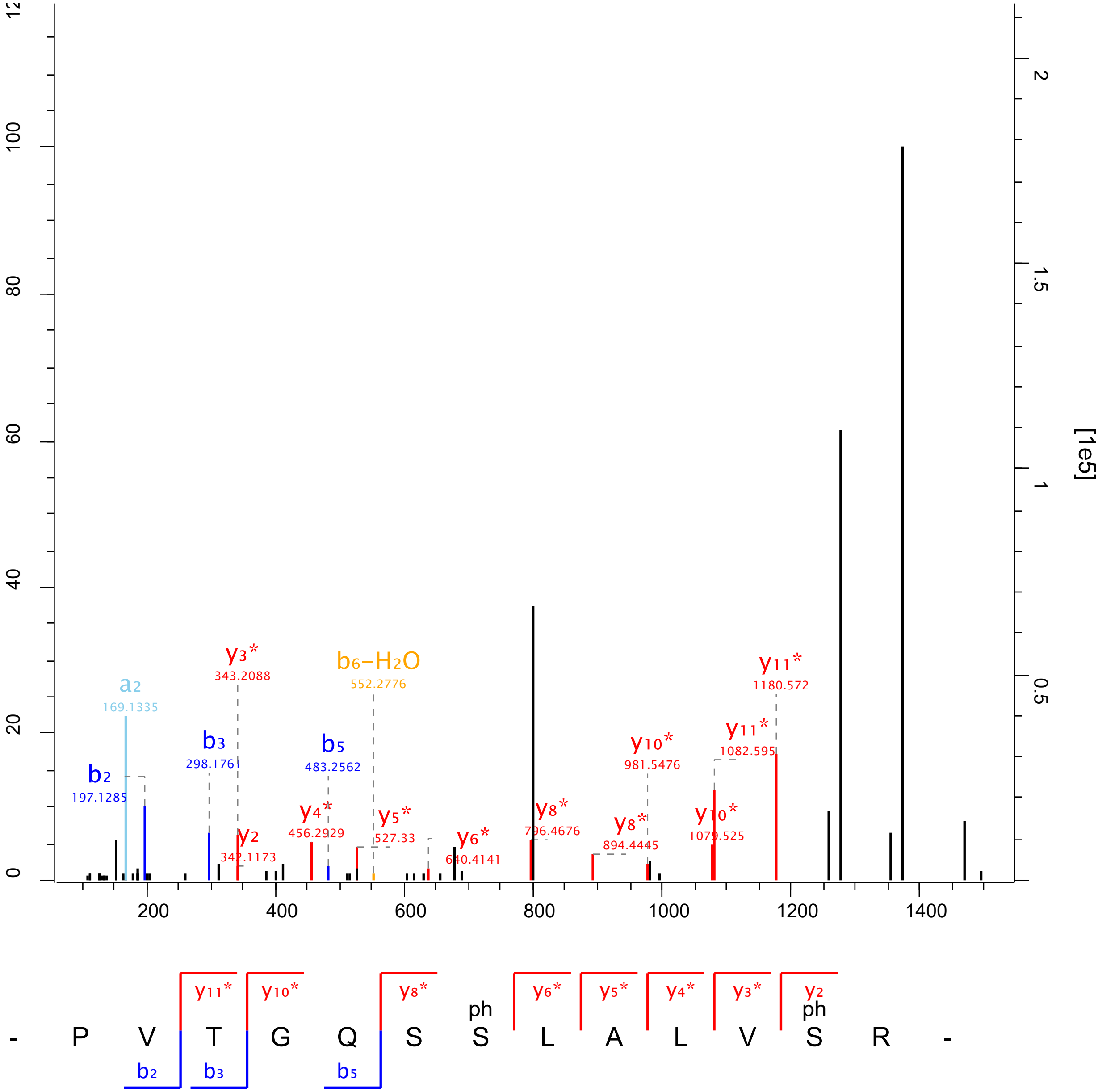

| Raw file | Scan | Method | Score | m/z |
|---|---|---|---|---|
| A17_00_2 | 2818 | FTMS; HCD | 73.14 | 737.81 |

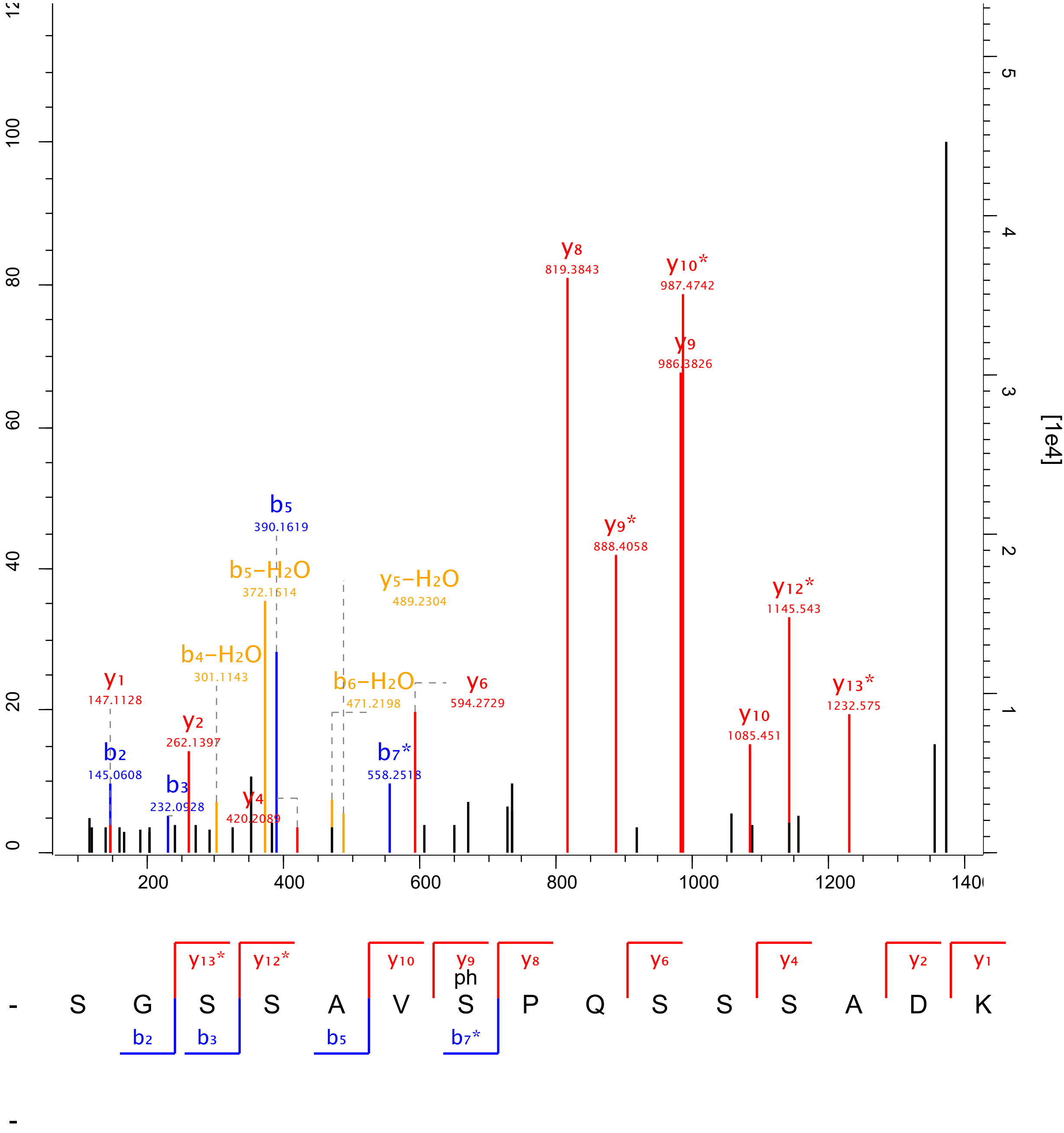

| Raw file | Scan | Method | Score | m/z |
| --- | --- | --- | --- | --- |
| A17_00_2 | 2834 | FTMS; HCD | 98.9 | 508.23 |

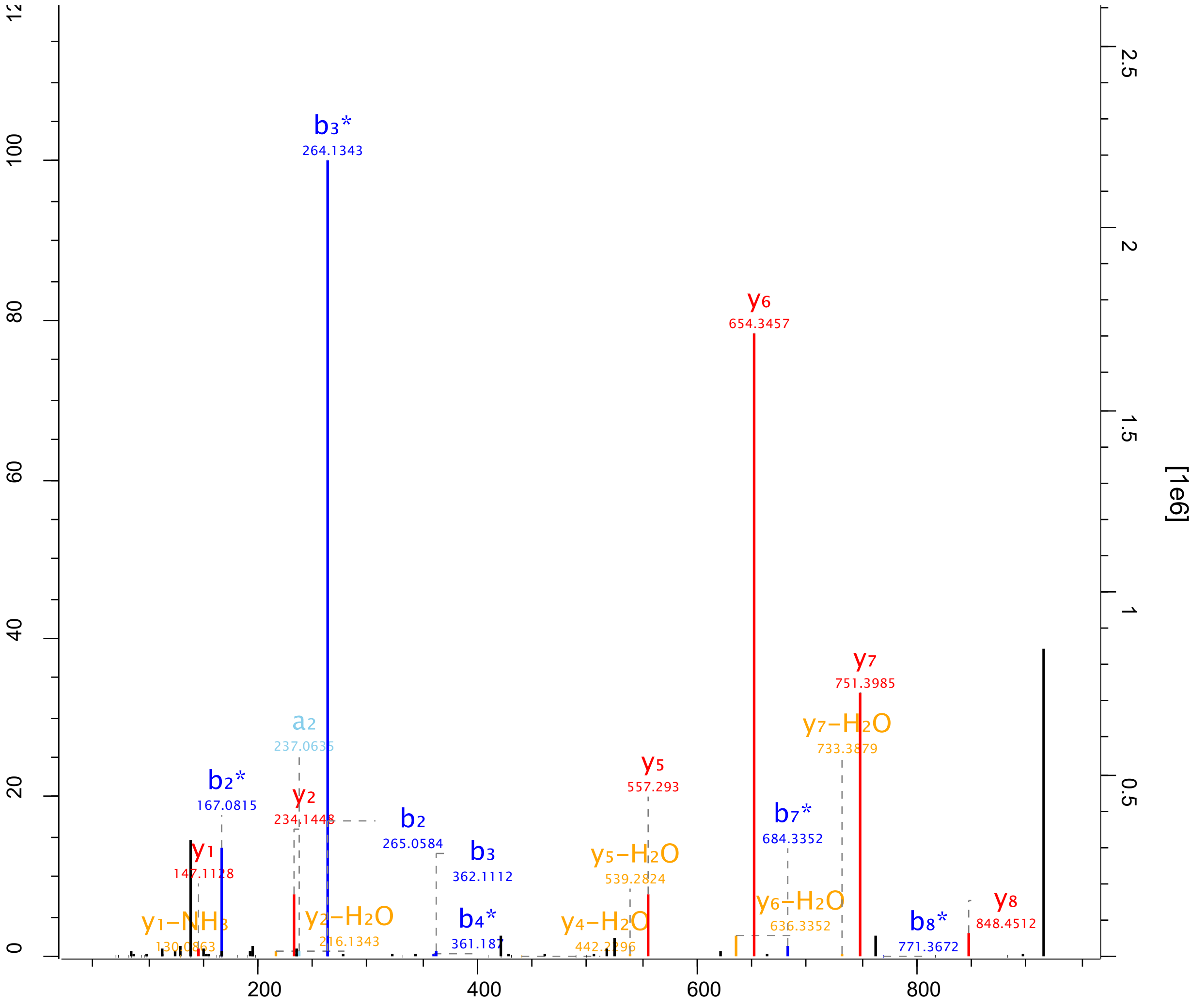


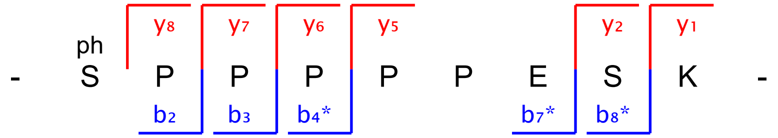

| Raw file | Scan | Method | Score | m/z |
| --- | --- | --- | --- | --- |
| A17_00_2 | 28787 | FTMS; HCD | 193.8 | 791.86 |

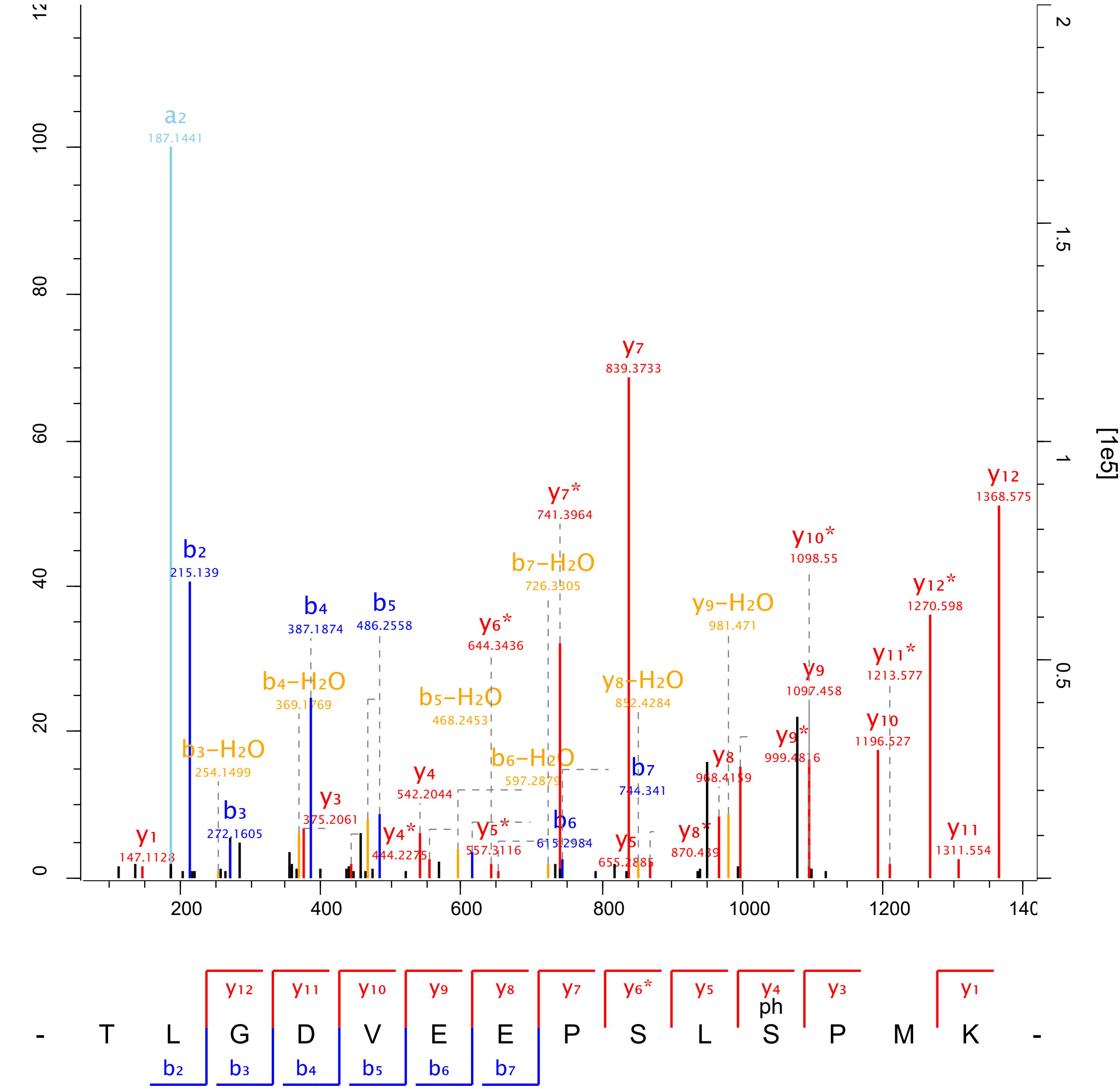

| Raw file | Scan | Method | Score | m/z |
| --- | --- | --- | --- | --- |
| A17_00_2 | 2946 | FTMS; HCD | 42.03 | 436.85 |

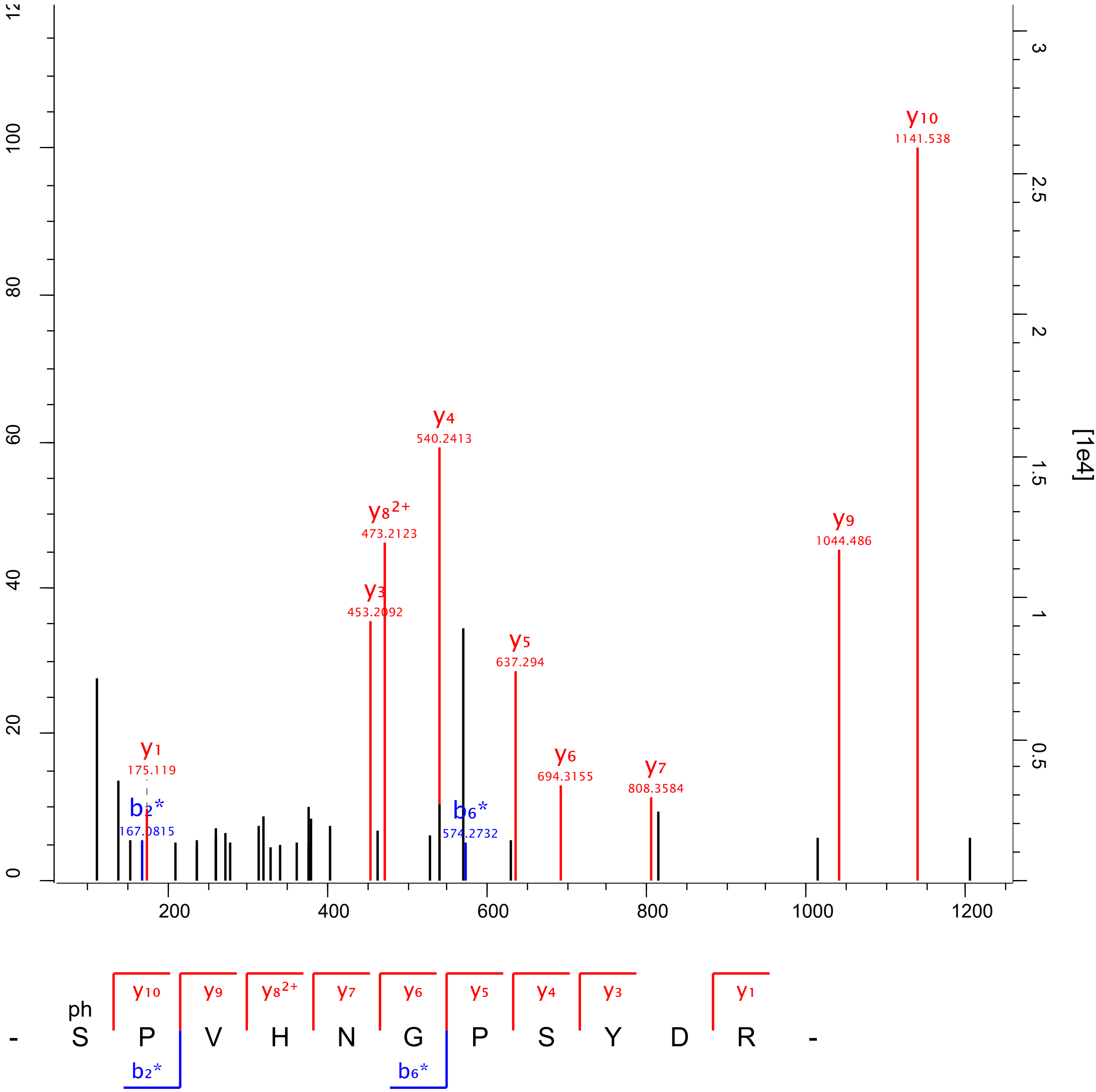

| Raw file | Scan | Method | Score | m/z |
| --- | --- | --- | --- | --- |
| A17_00_2 | 29759 | FTMS; HCD | 198.55 | 582.75 |

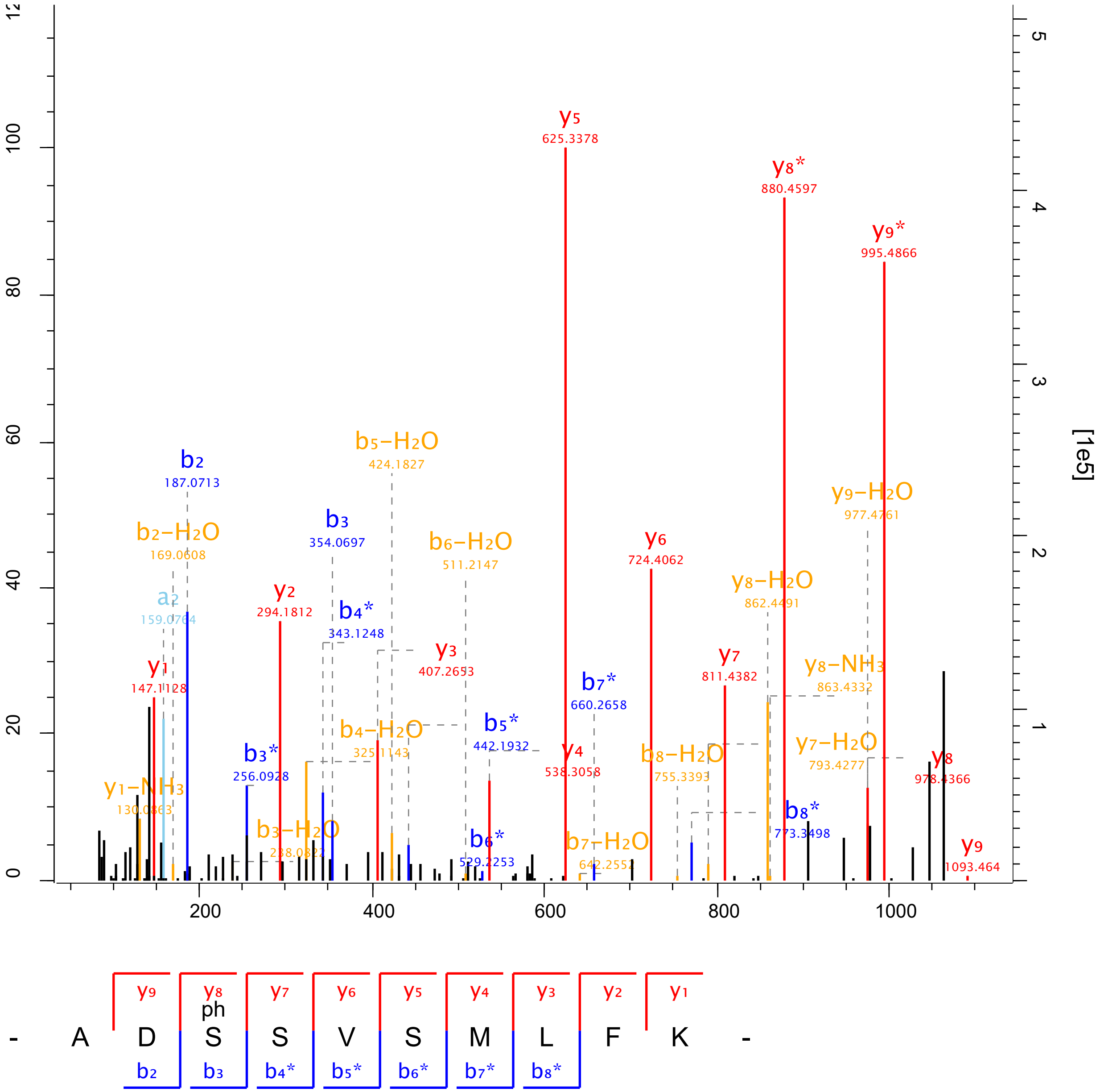

| Raw file | Scan | Method | Score | m/z |
| --- | --- | --- | --- | --- |
| A17_00_2 | 29899 | FTMS; HCD | 53.44 | 950.39 |

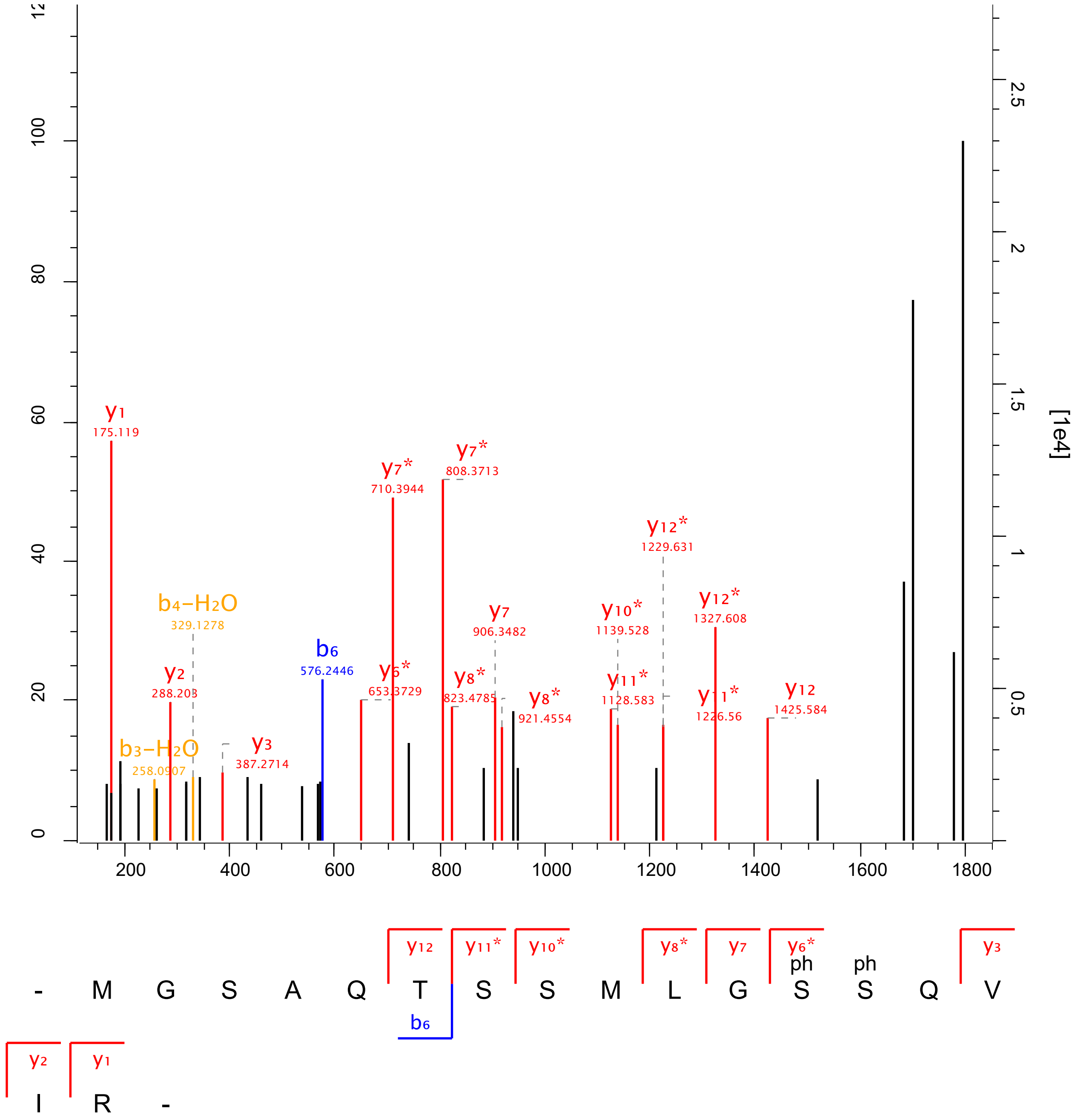

| Raw file | Scan | Method | Score | m/z |
| --- | --- | --- | --- | --- |
| A17_00_2 | 30014 | FTMS; HCD | 231.83 | 828.86 |

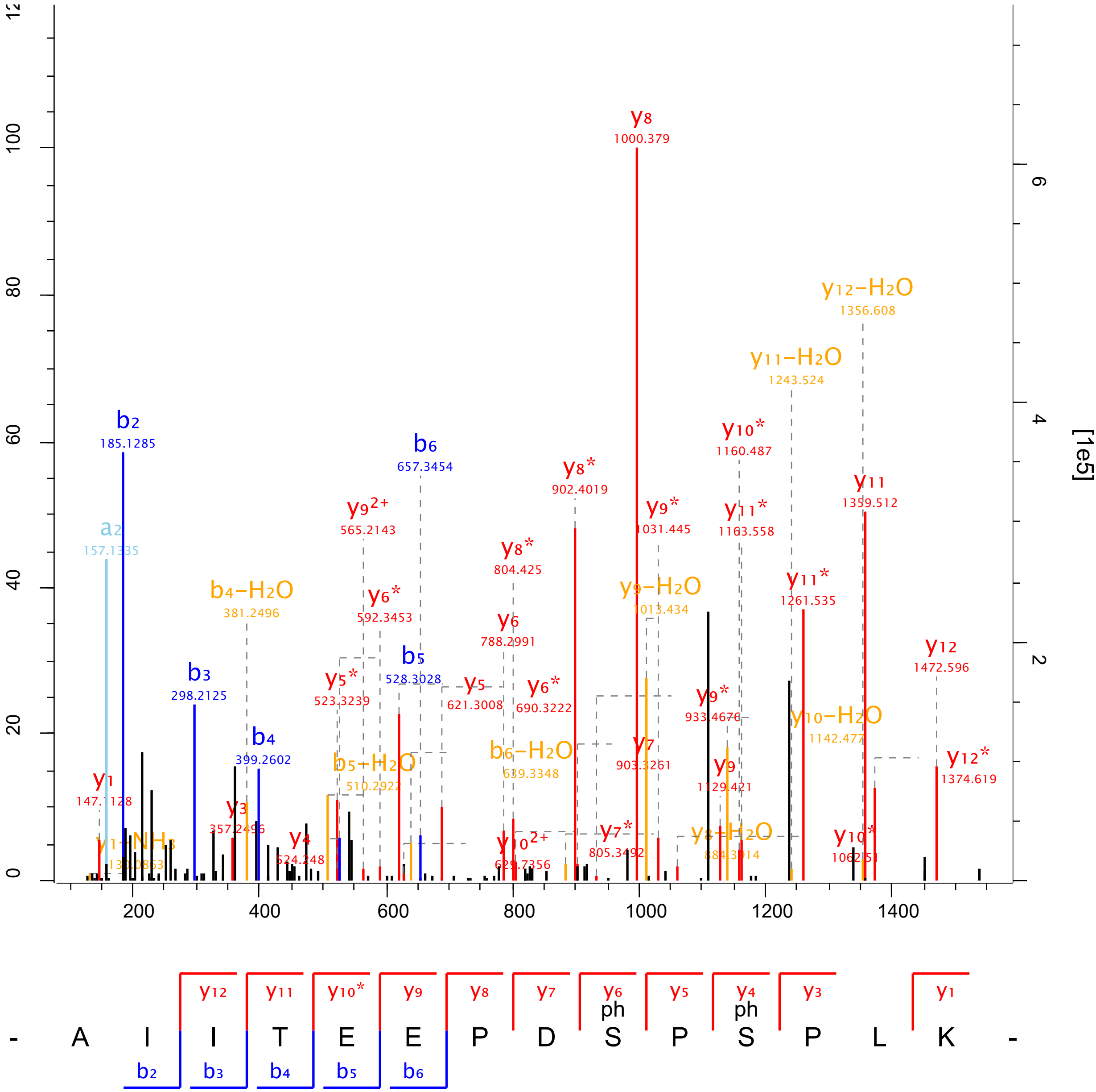

| Raw file | Scan | Method | Score | m/z |
|---|---|---|---|---|
| A17_00_2 | 30060 | FTMS; HCD | 68.24 | 715.28 |

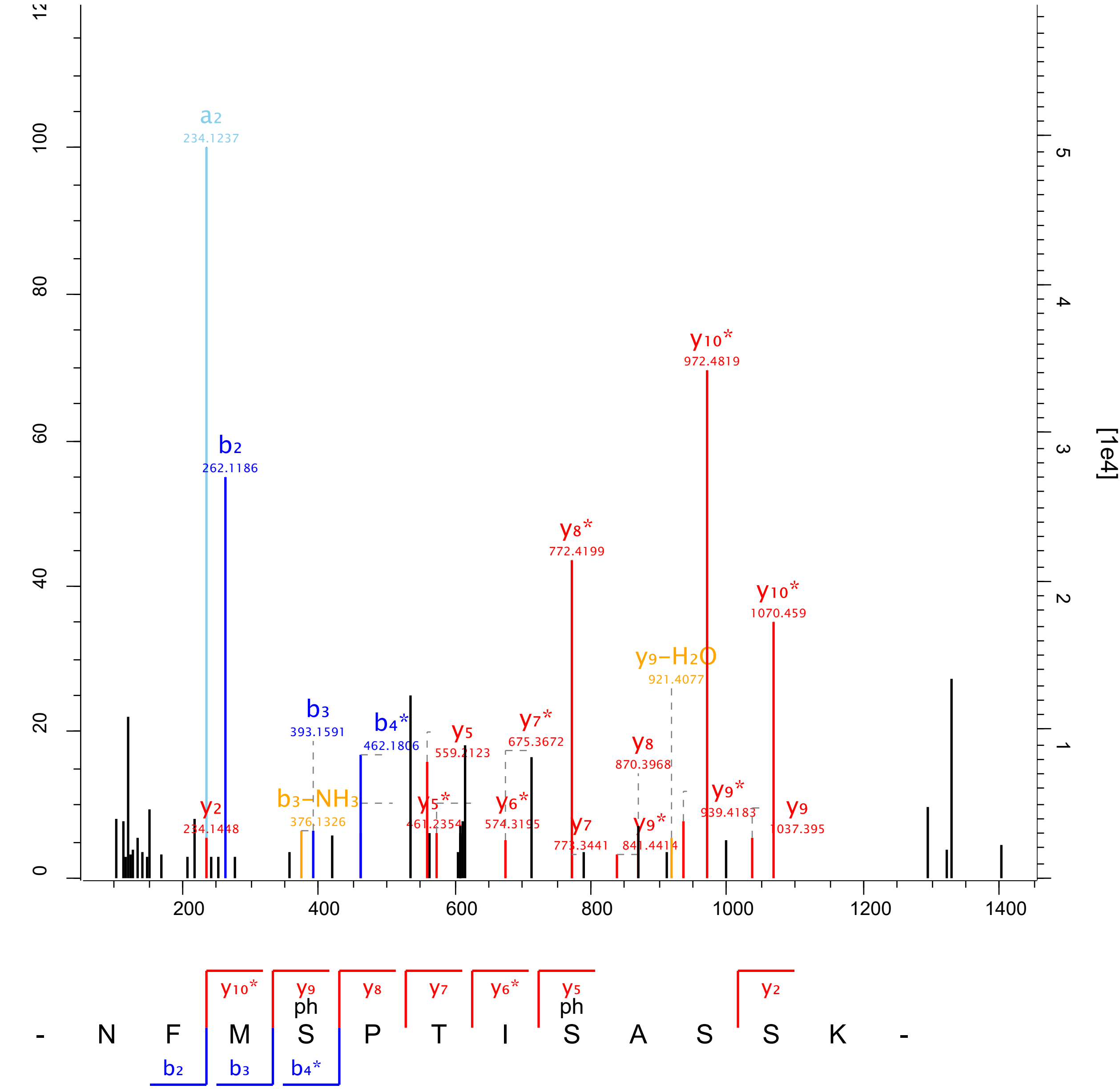

| Raw file | Scan | Method | Score | m/z |
|---|---|---|---|---|
| A17_00_2 | 3041 | FTMS; HCD | 130.36 | 632.57 |

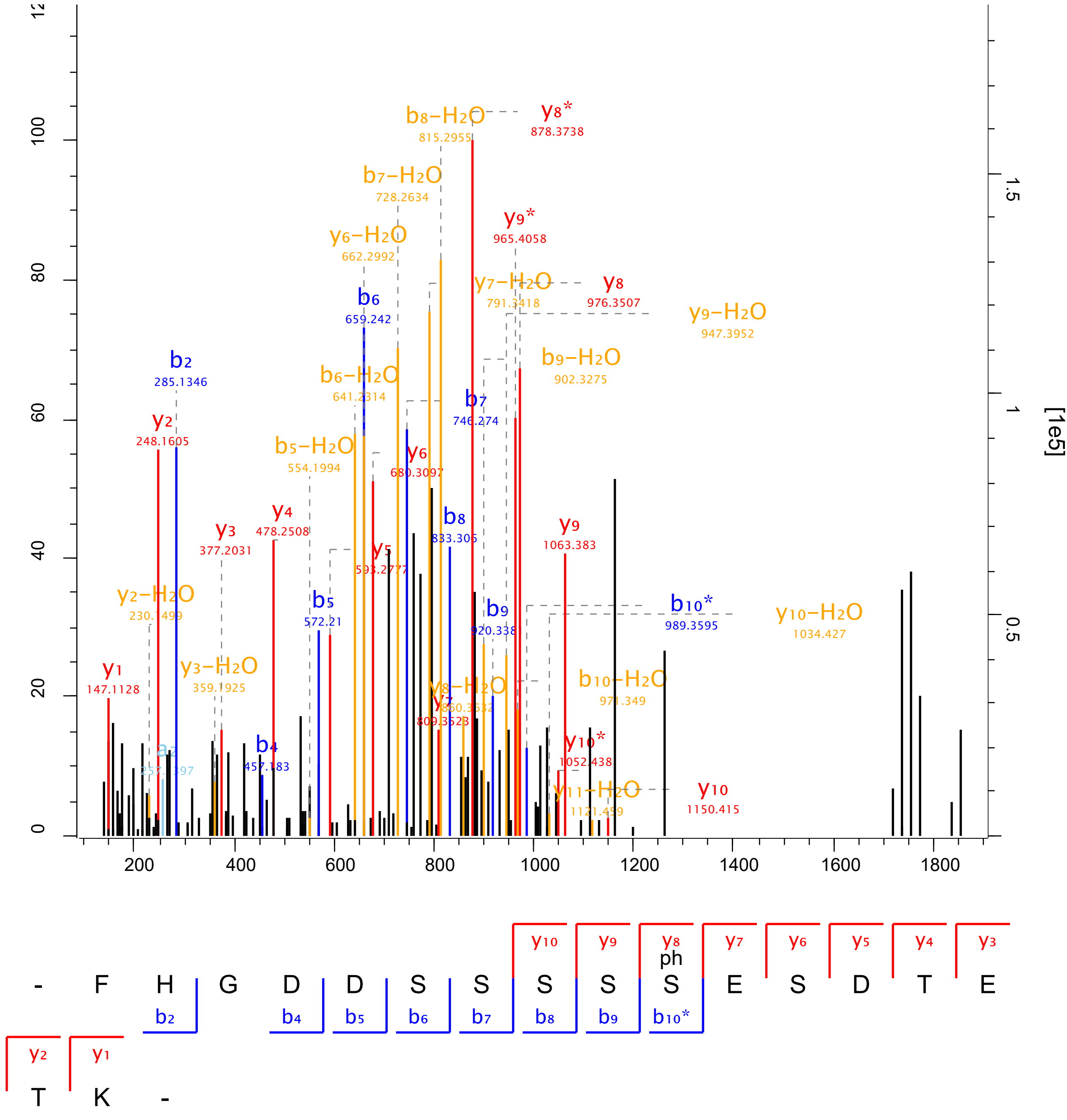

| Raw file | Scan | Method | Score | m/z |
|---|---|---|---|---|
| A17_00_2 | 3046 | FTMS; HCD | 44.54 | 558.76 |

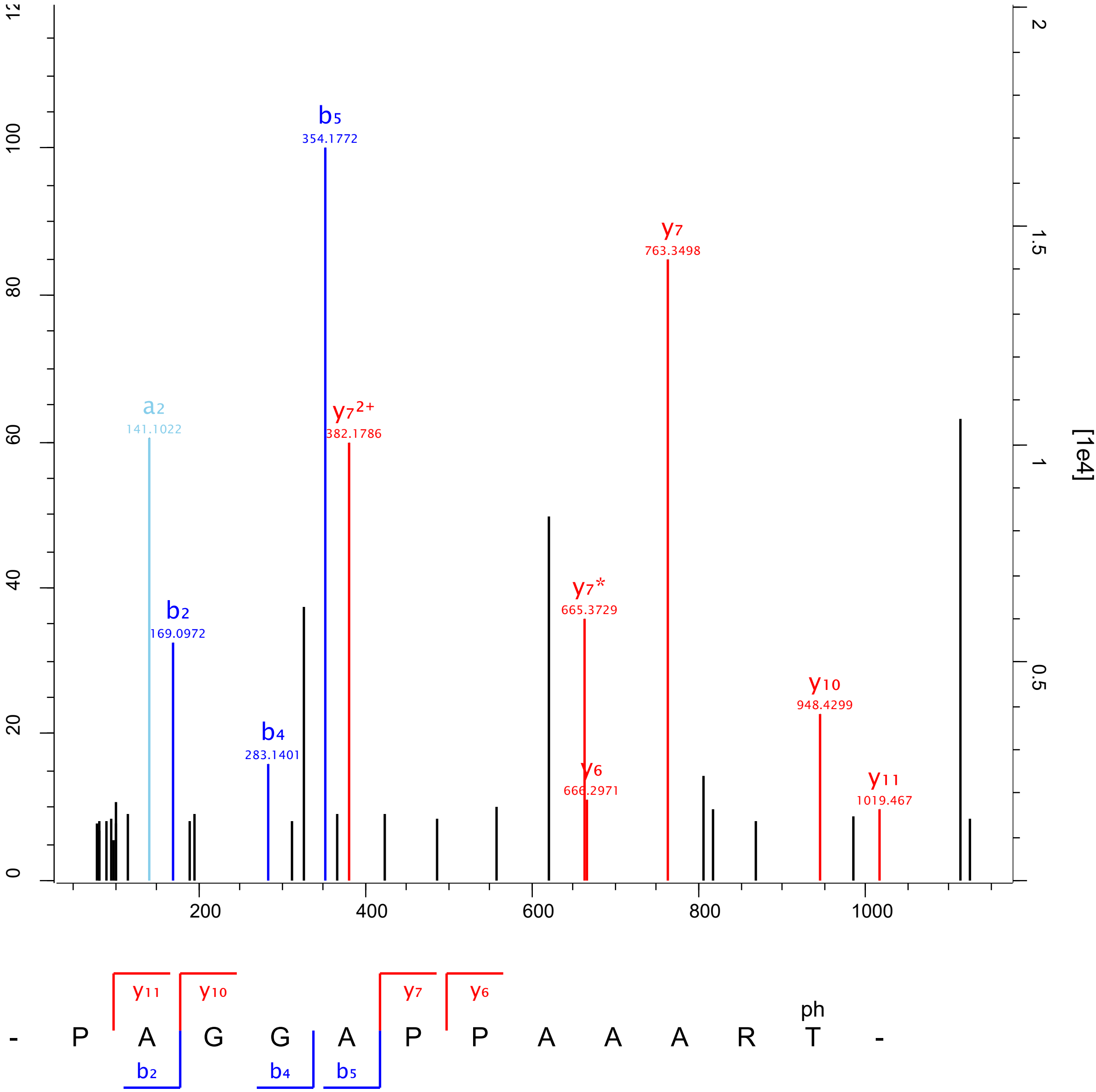

| Raw file | Scan | Method | Score | m/z |
|---|---|---|---|---|
| A17_00_2 | 30473 | FTMS; HCD | 118.55 | 717.75 |

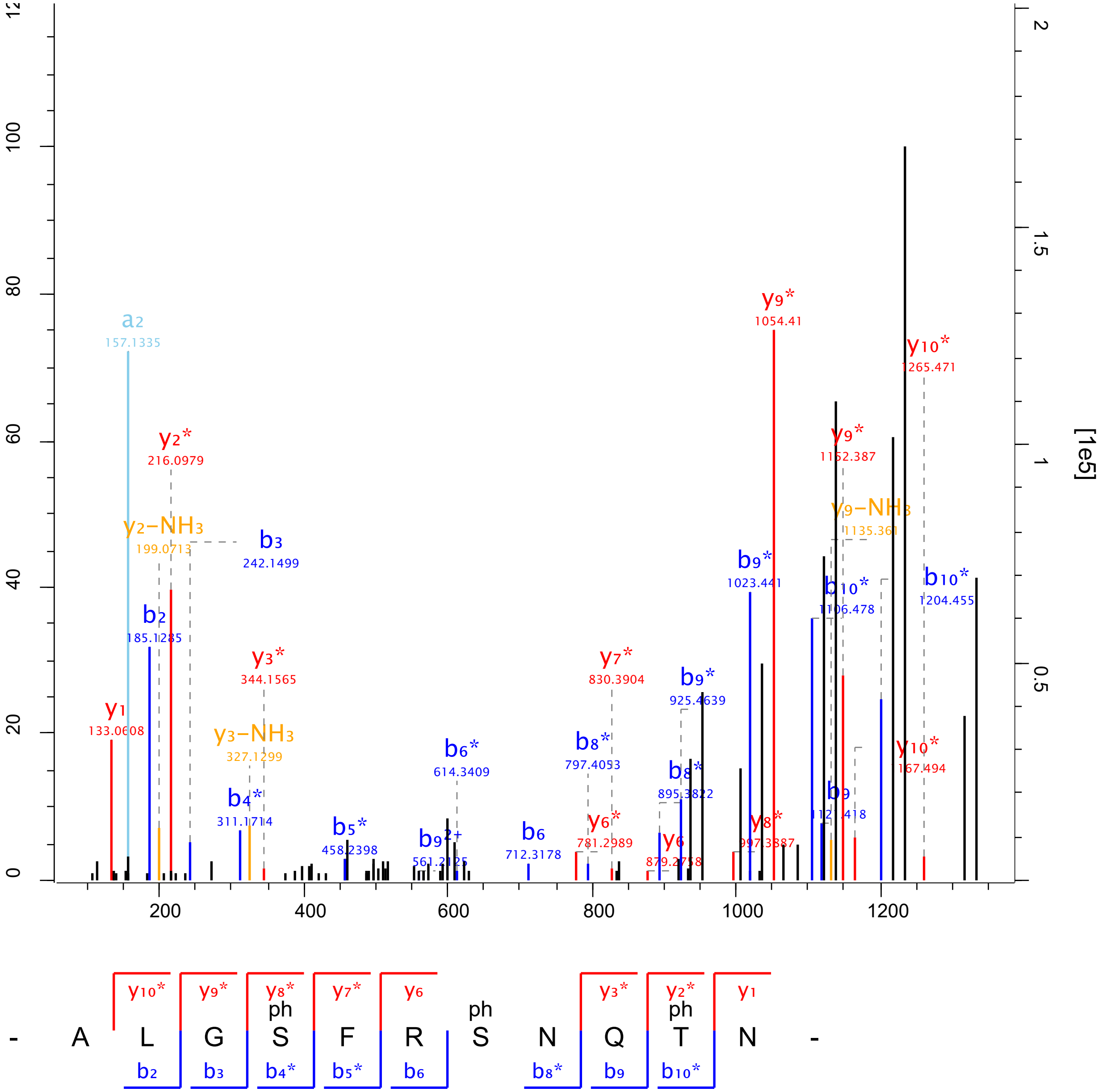

| Raw file | Scan | Method | Score | m/z |
|---|---|---|---|---|
| A17_00_2 | 3076 | FTMS; HCD | 53.95 | 580.2 |

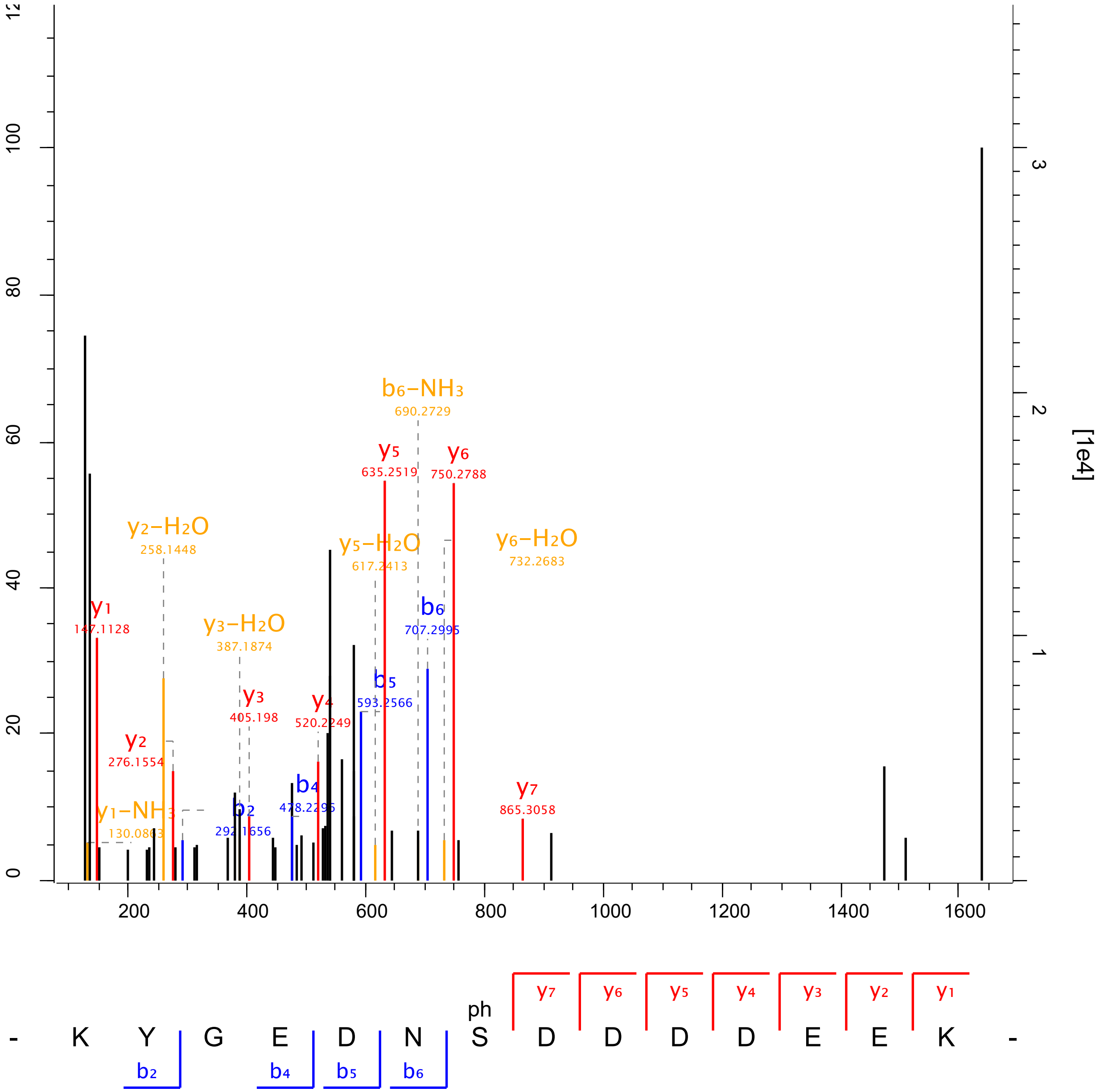

| Raw file | Scan | Method | Score | m/z |
|---|---|---|---|---|
| A17_00_2 | 31177 | FTMS; HCD | 117.48 | 701.34 |

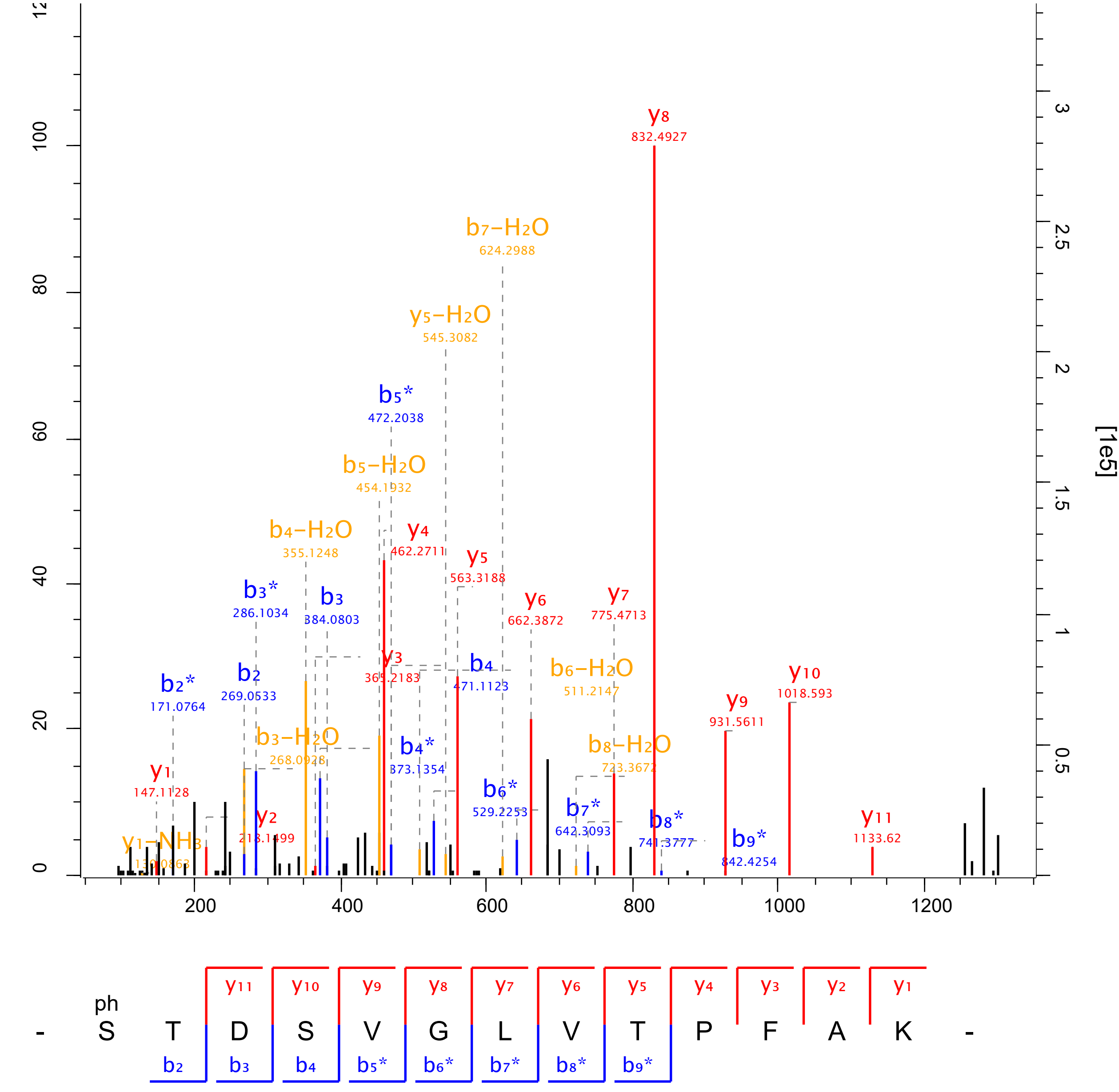

| Raw file | Scan | Method | Score | m/z |
| --- | --- | --- | --- | --- |
| A17_00_2 | 32123 | FTMS; HCD | 53.62 | 865.83 |

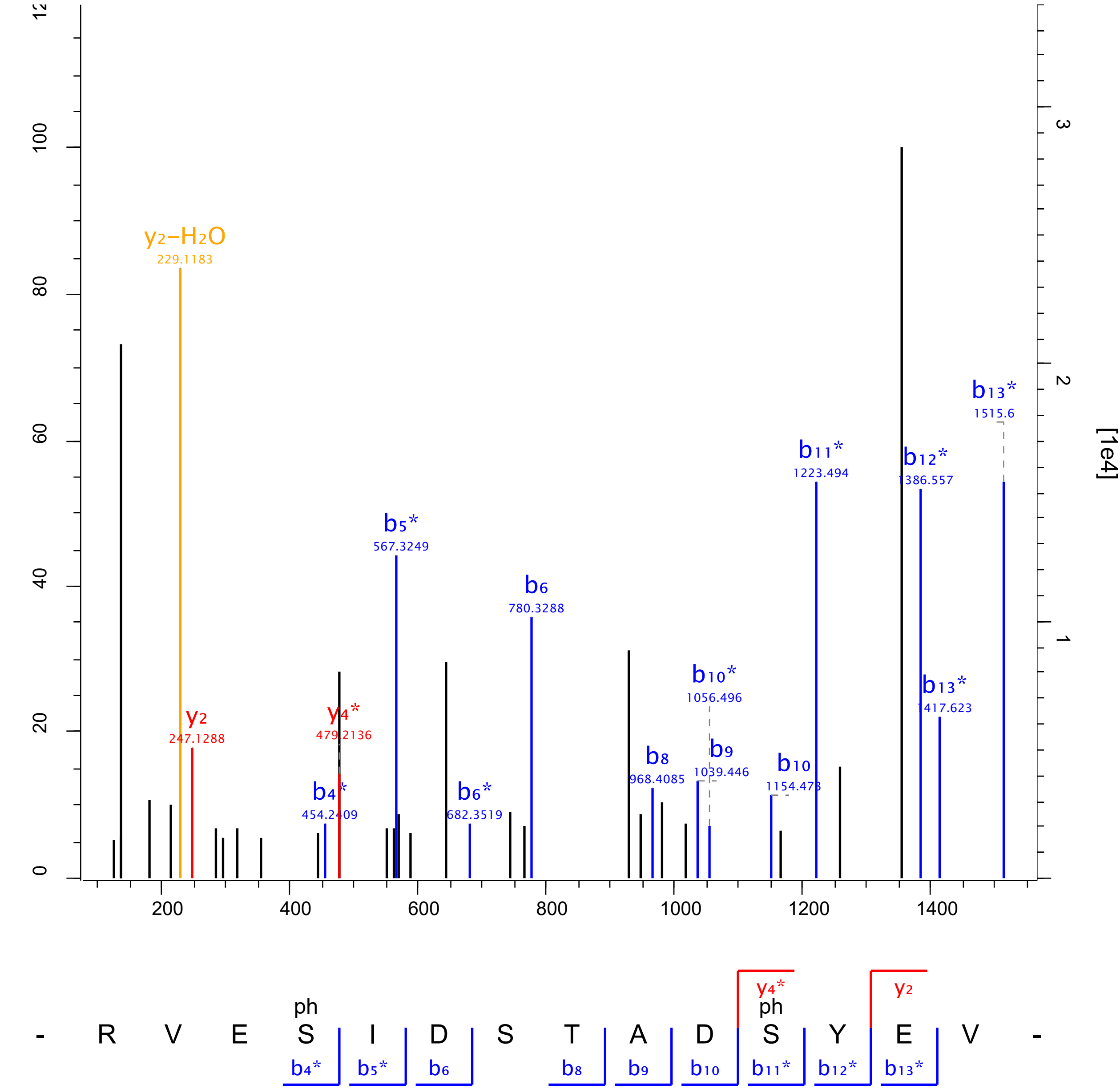

| Raw file | Scan | Method | Score | m/z |
| --- | --- | --- | --- | --- |
| A17_00_2 | 3220 | FTMS; HCD | 75.76 | 430.18 |

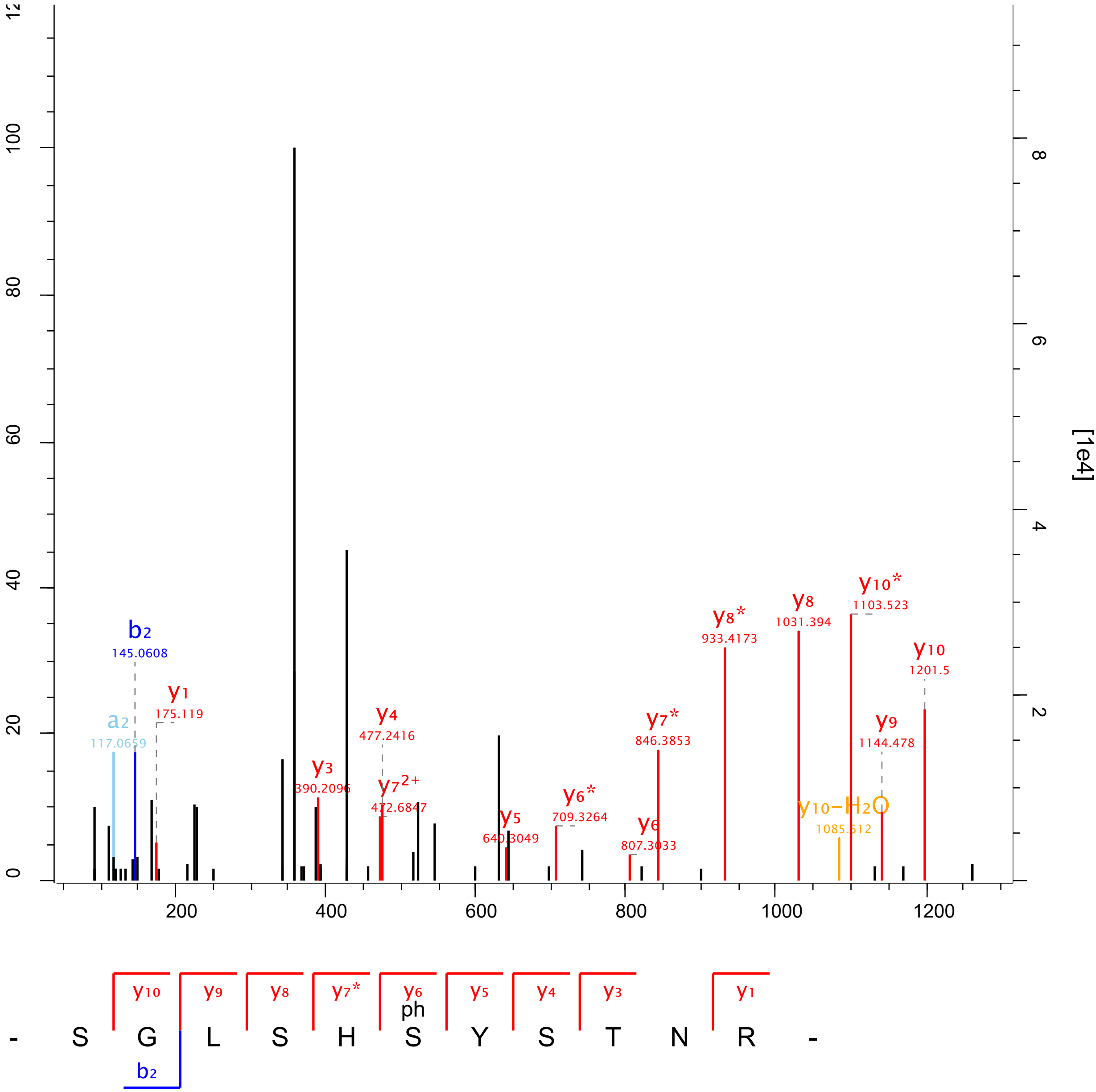

| Raw file | Scan | Method | Score | m/z |
|---|---|---|---|---|
| A17_00_2 | 32367 | FTMS; HCD | 65.48 | 689.24 |

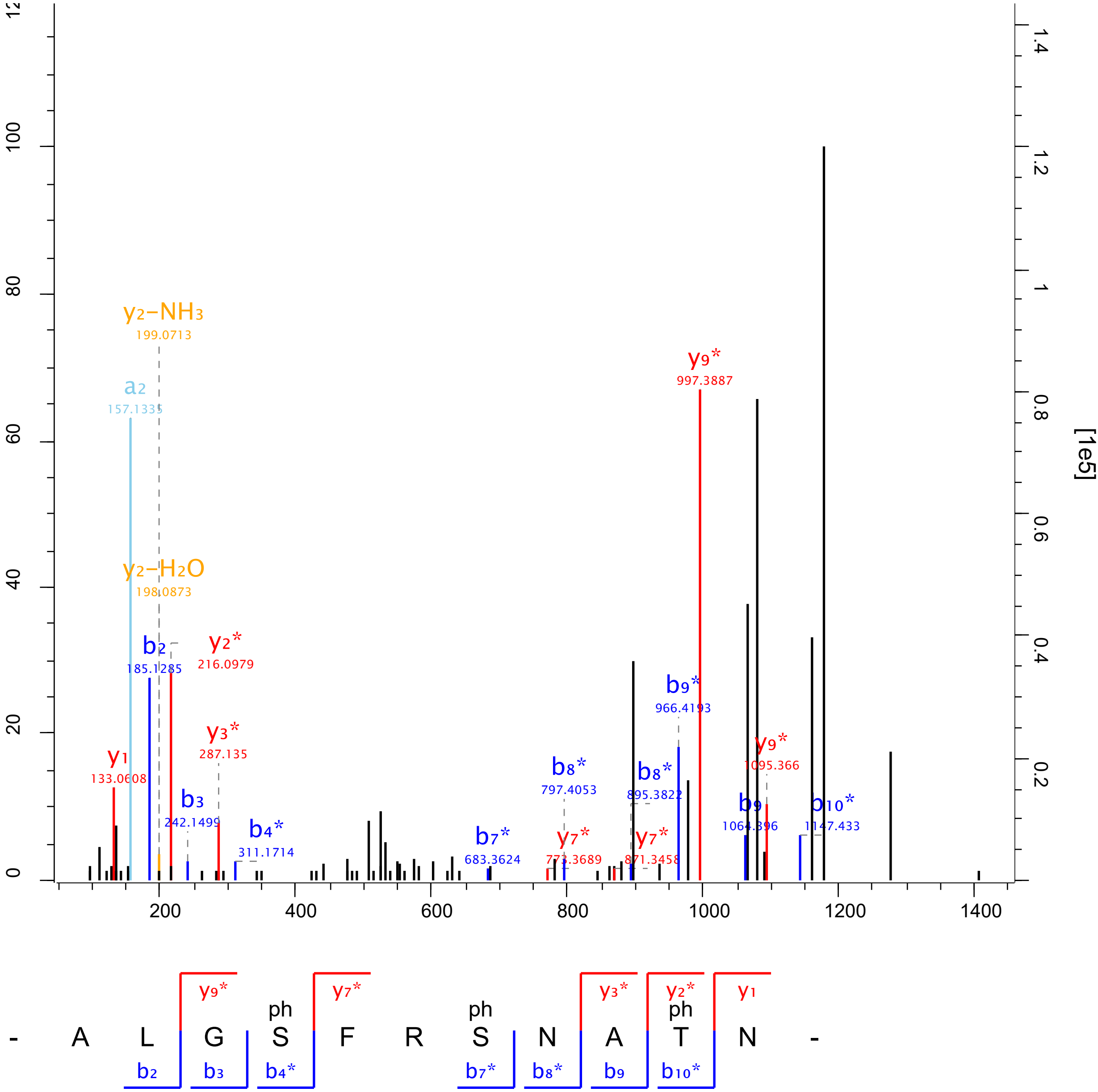

| Raw file | Scan | Method | Score | m/z |
|---|---|---|---|---|
| A17_00_2 | 3241 | FTMS; HCD | 57.16 | 563.24 |

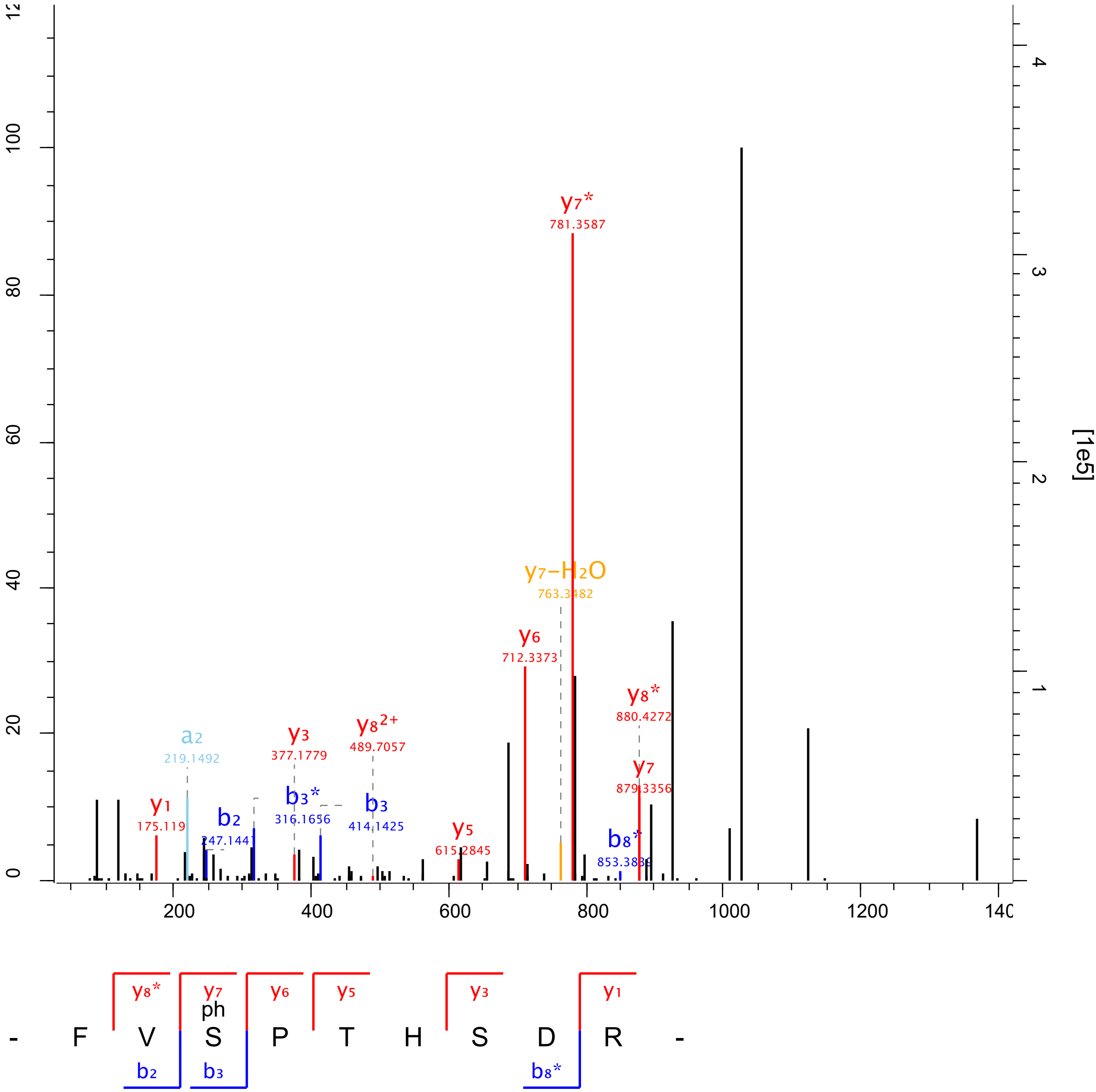

| Raw file | Scan | Method | Score | m/z |
|---|---|---|---|---|
| A17_00_2 | 32538 | FTMS; HCD | 78.34 | 535.18 |

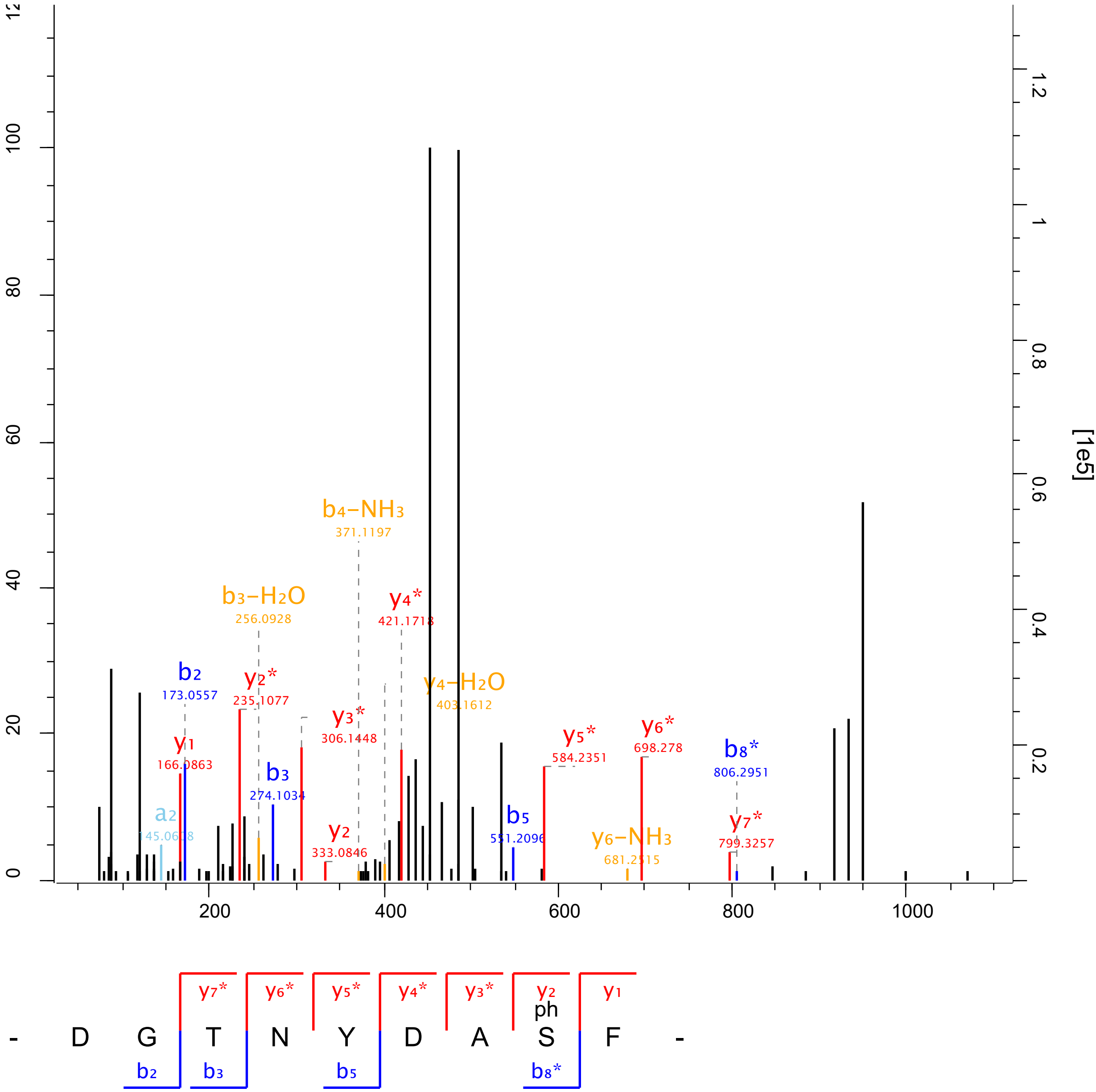

| Raw file | Scan | Method | Score | m/z |
|---|---|---|---|---|
| A17_00_2 | 33306 | FTMS; HCD | 98.94 | 686.81 |

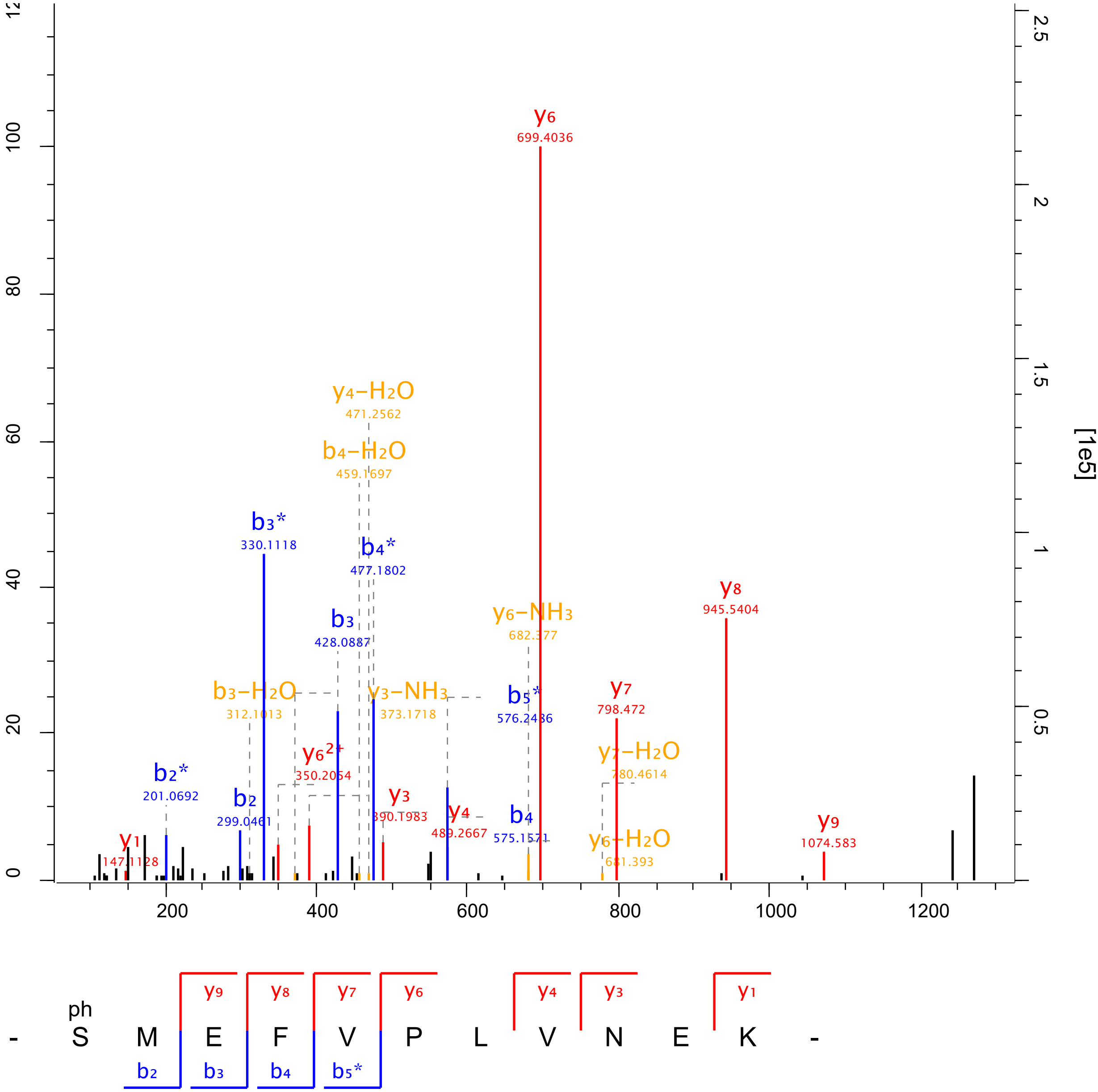

| Raw file | Scan | Method | Score | m/z |
|---|---|---|---|---|
| A17_00_2 | 33318 | FTMS; HCD | 63.19 | 648.76 |

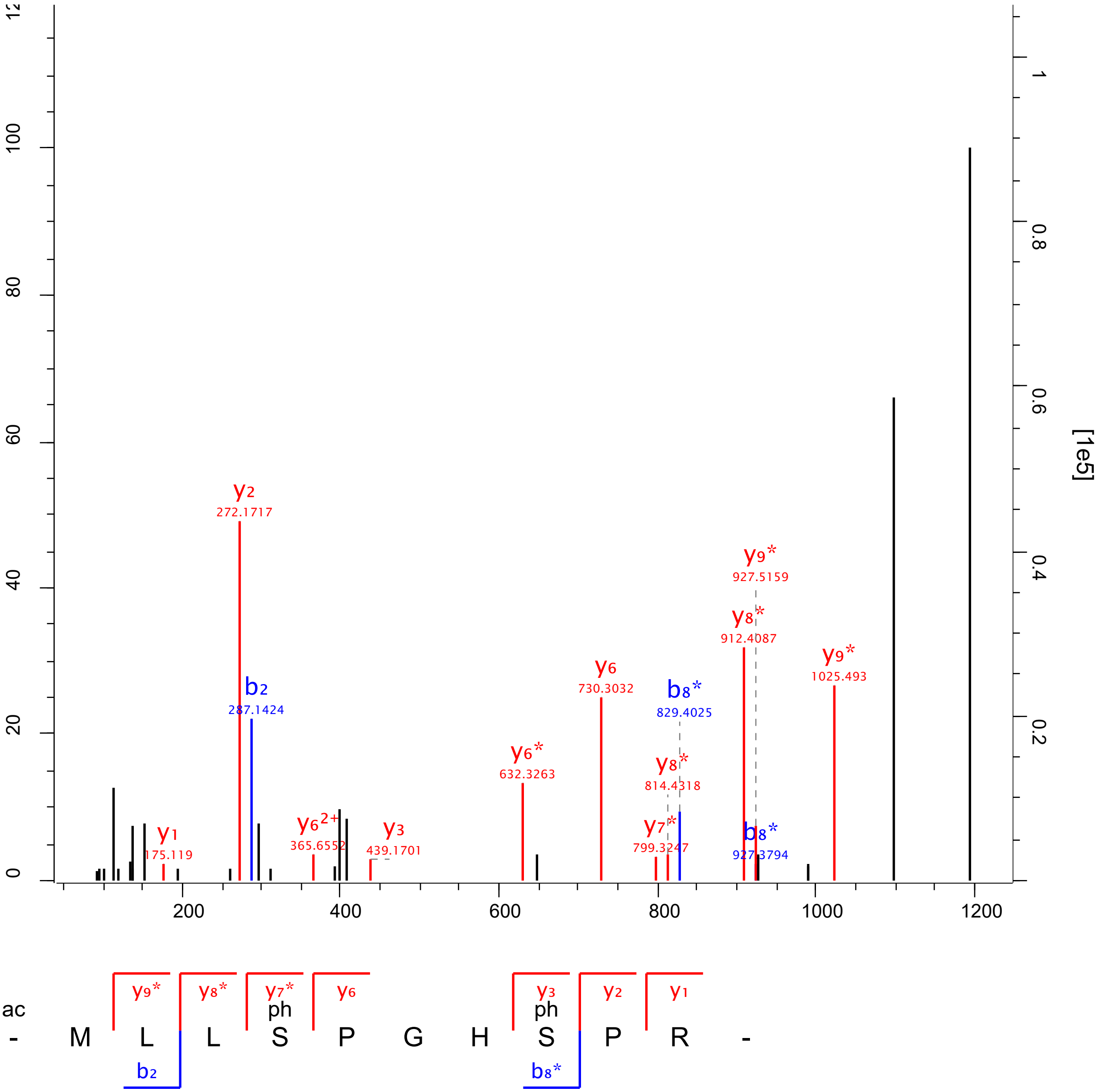

| Raw file | Scan | Method | Score | m/z |
|---|---|---|---|---|
| A17_00_2 | 3333 | FTMS; HCD | 86.9 | 469.21 |

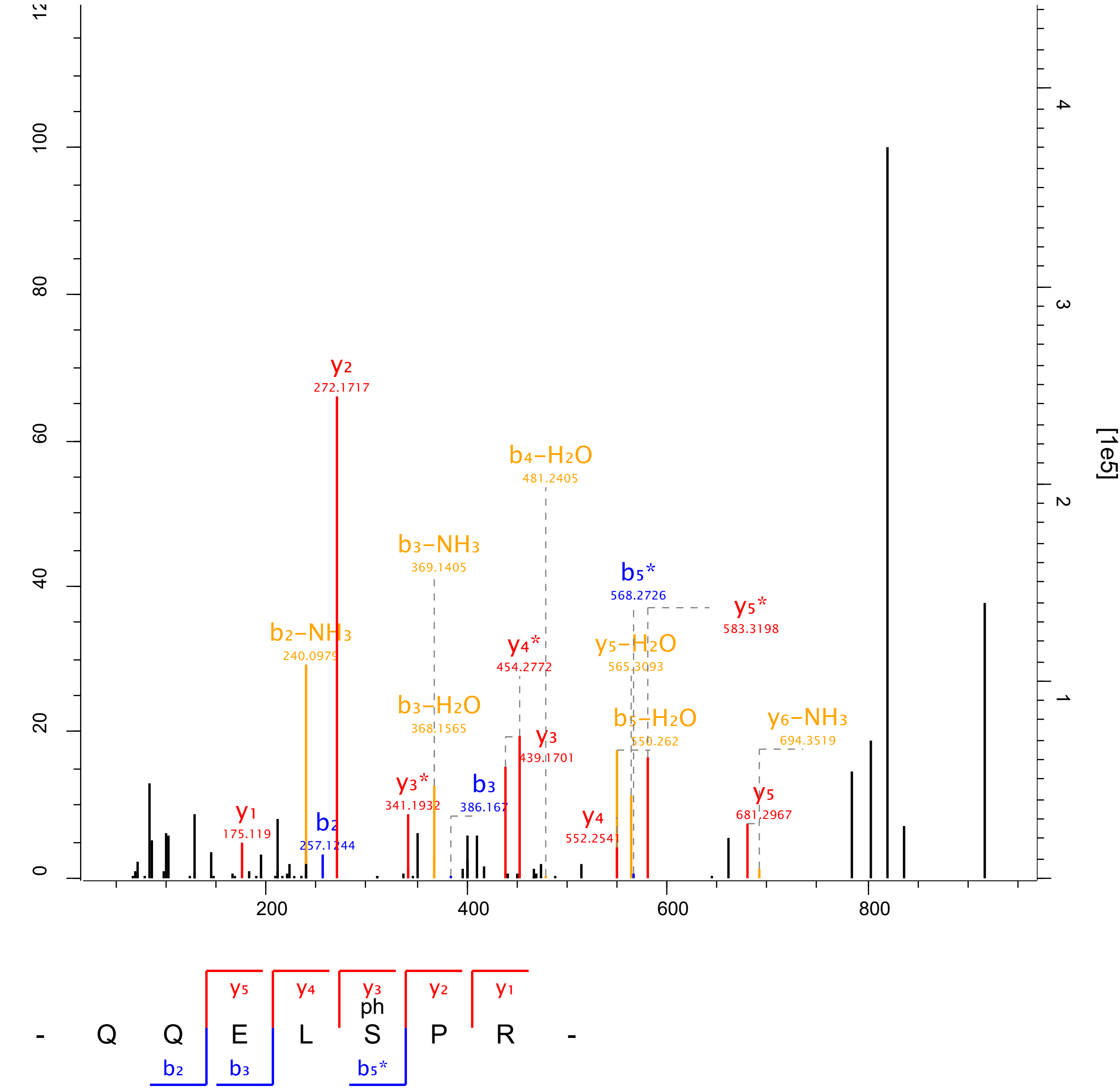

| Raw file | Scan | Method | Score | m/z |
|---|---|---|---|---|
| A17_00_2 | 33406 | FTMS; HCD | 392.07 | 1070.96 |

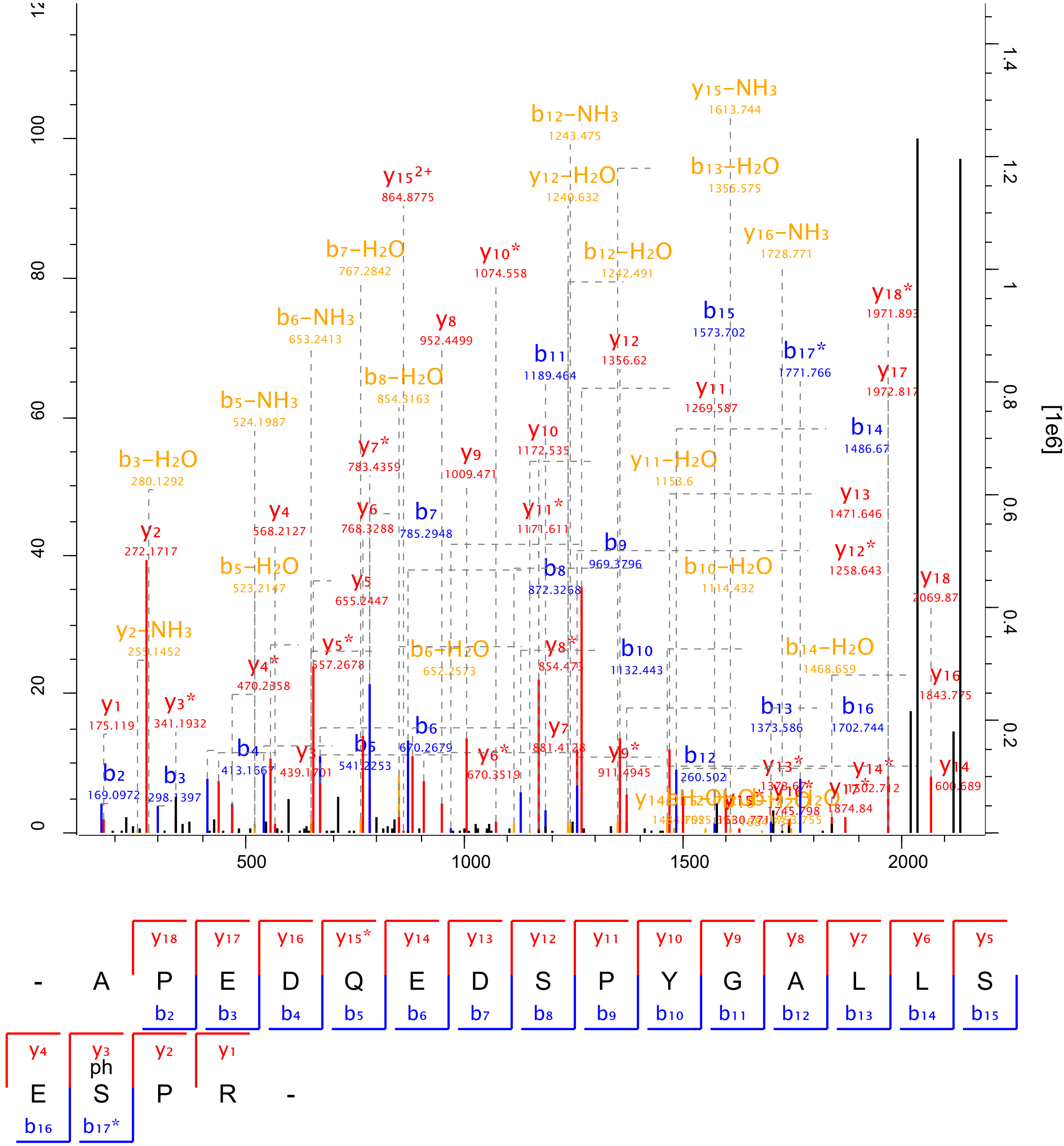

| Raw file | Scan | Method | Score | m/z |
|---|---|---|---|---|
| A17_00_2 | 33532 | FTMS; HCD | 59.34 | 738.95 |

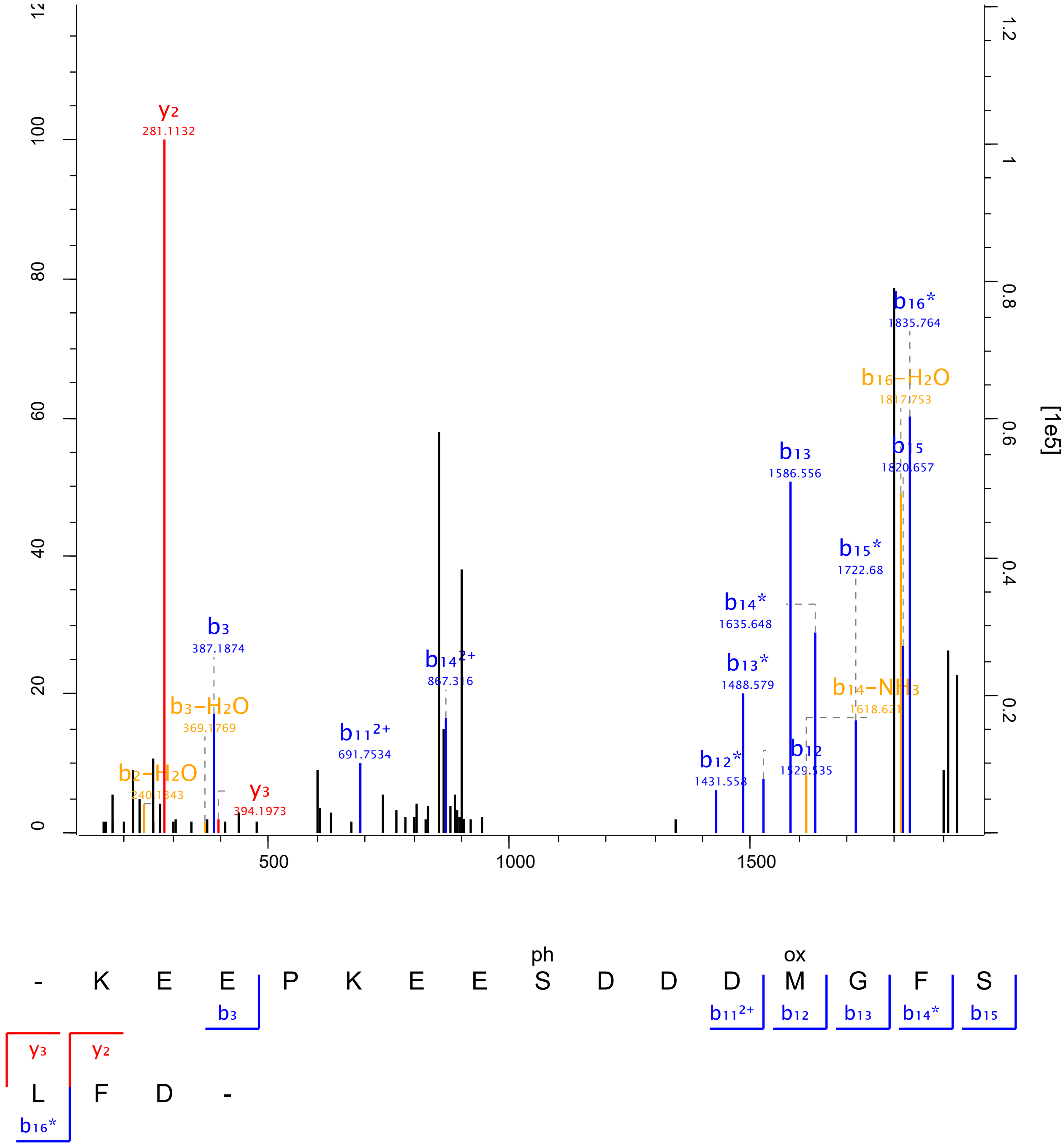

| Raw file | Scan | Method | Score | m/z |
| --- | --- | --- | --- | --- |
| A17_00_2 | 3357 | FTMS; HCD | 157.52 | 570.73 |

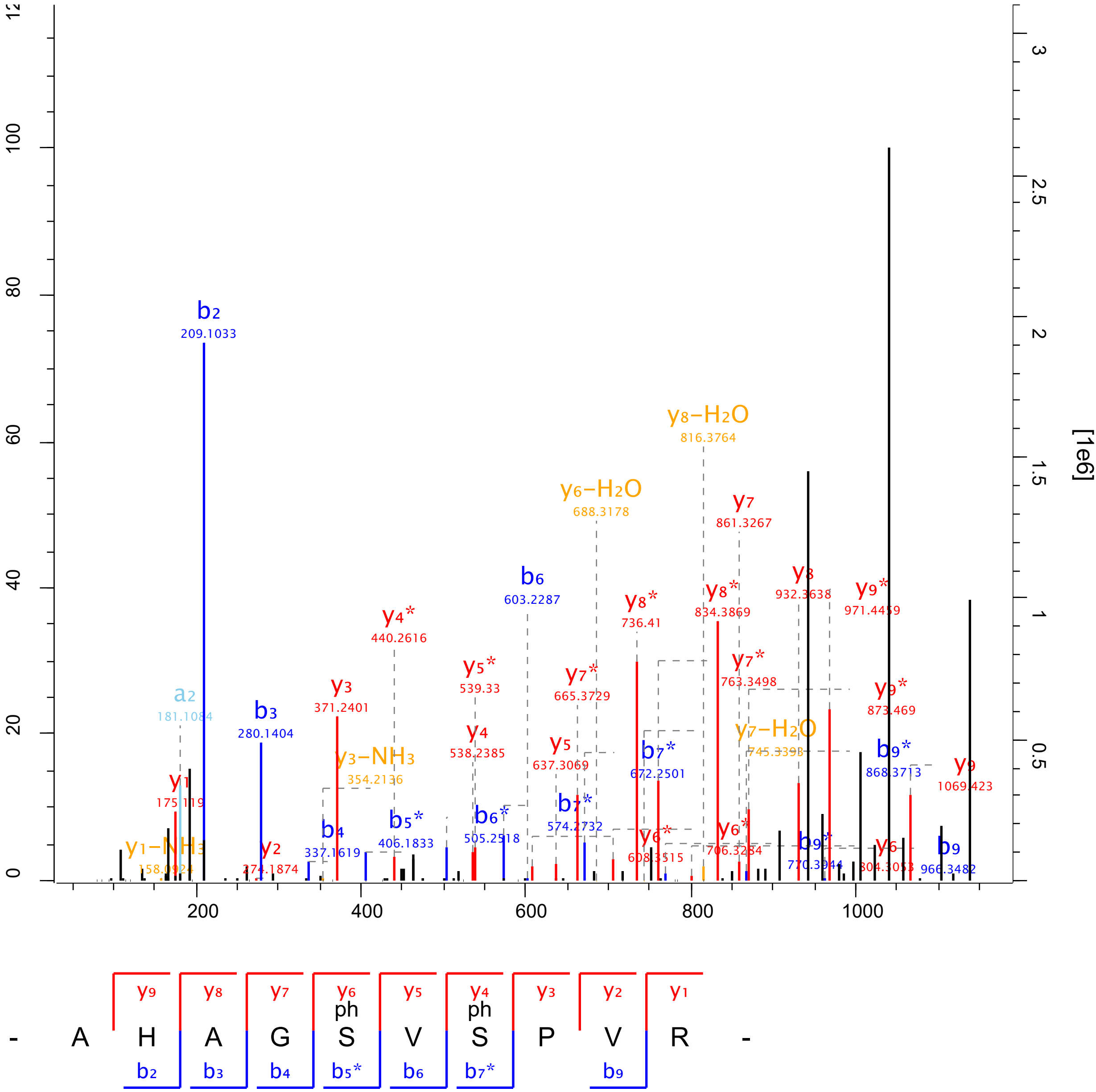

| Raw file | Scan | Method | Score | m/z |
|---|---|---|---|---|
| A17_00_2 | 3409 | FTMS; HCD | 63.71 | 557.61 |

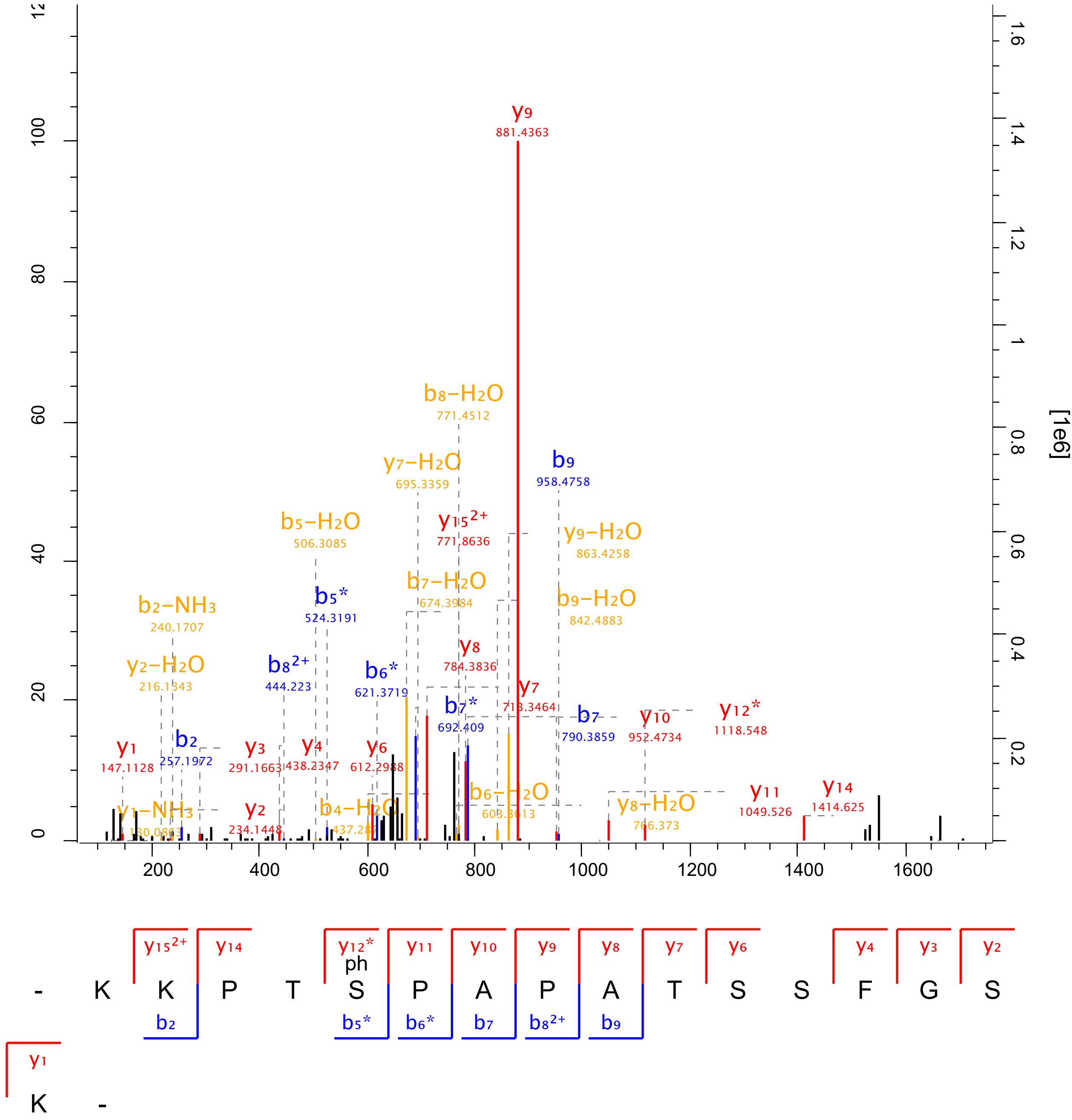

| Raw file | Scan | Method | Score | m/z |
| --- | --- | --- | --- | --- |
| A17_00_2 | 34618 | FTMS; HCD | 149.29 | 753.33 |

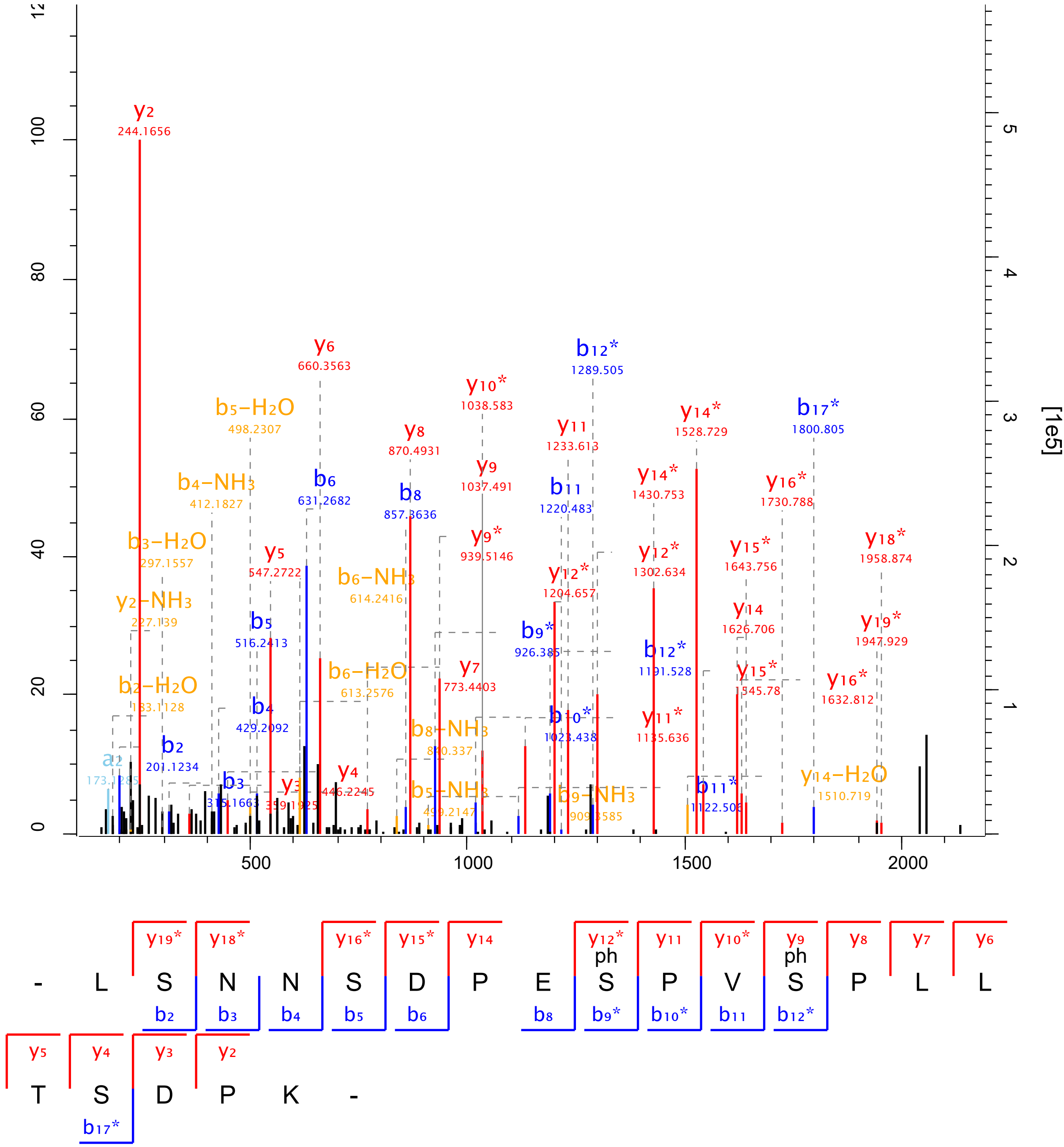

| Raw file | Scan | Method | Score | m/z |
|---|---|---|---|---|
| A17_00_2 | 3483 | FTMS; HCD | 57.41 | 470.54 |

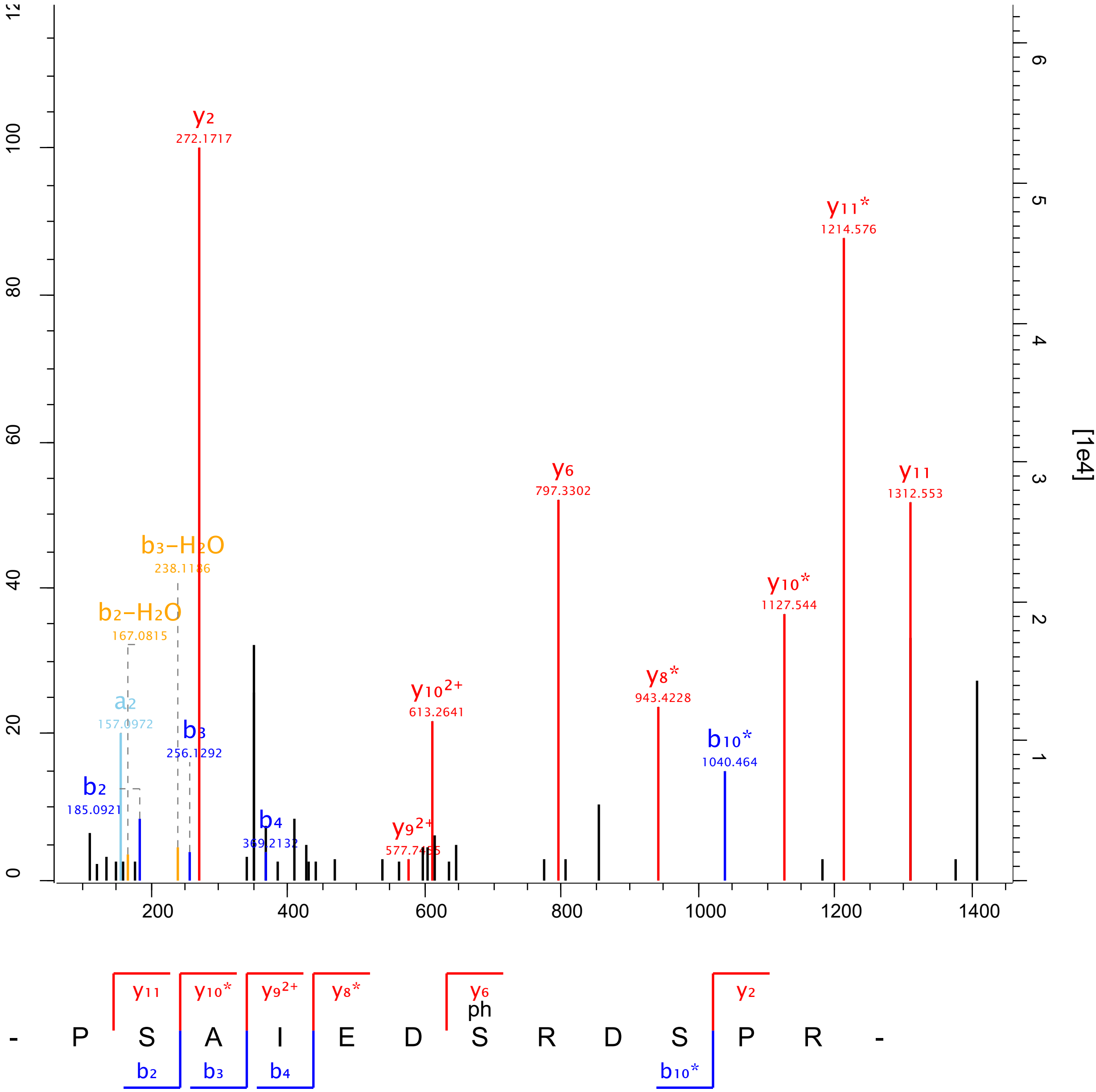

| Raw file | Scan | Method | Score | m/z |
| --- | --- | --- | --- | --- |
| A17_00_2 | 3575 | FTMS; HCD | 47.77 | 467.52 |

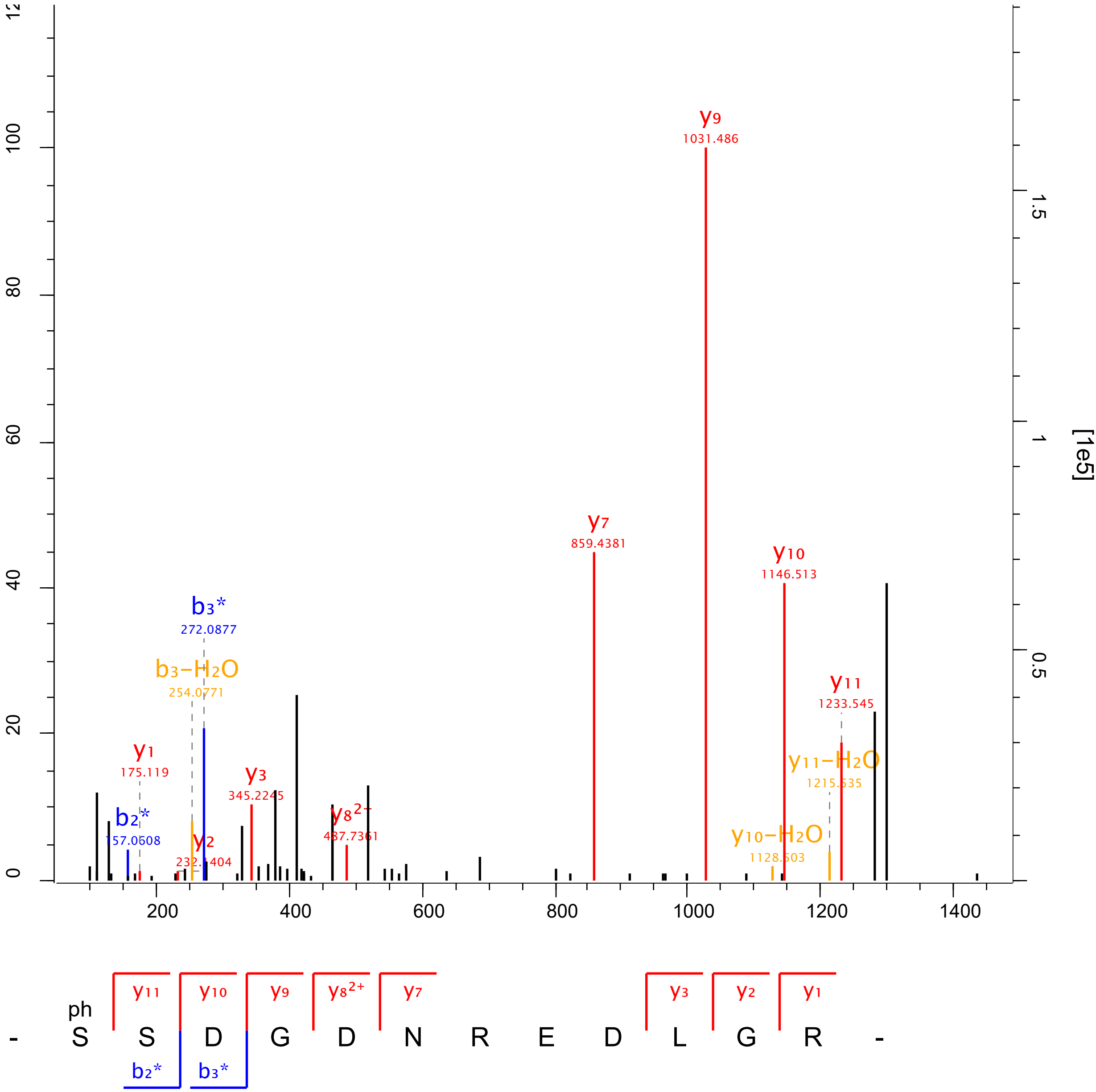

| Raw file | Scan | Method | Score | m/z |
|---|---|---|---|---|
| A17_00_2 | 3613 | FTMS; HCD | 88.43 | 685.76 |

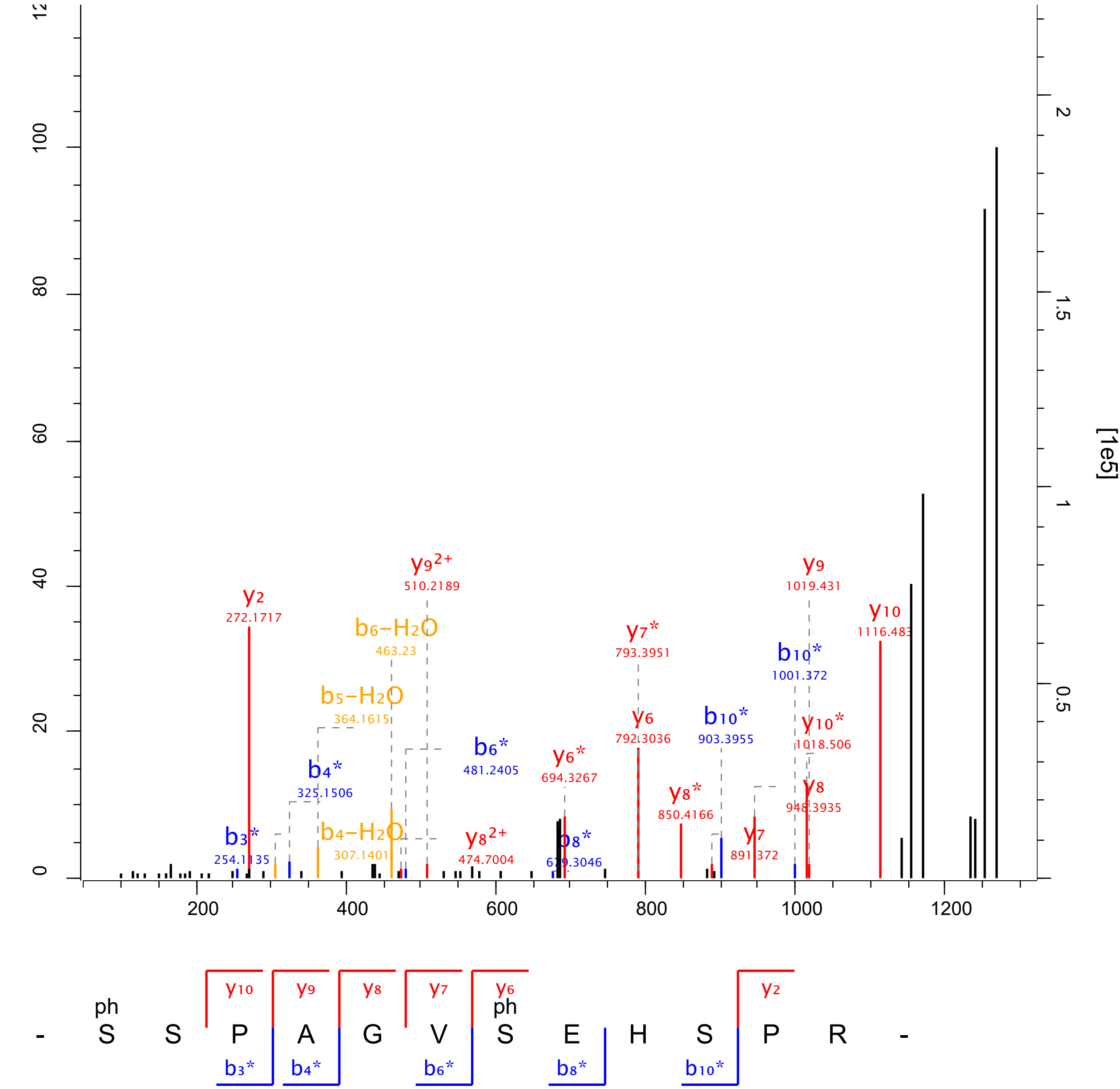

| Raw file | Scan | Method | Score | m/z |
|---|---|---|---|---|
| A17_00_2 | 3616 | FTMS; HCD | 68.54 | 583.23 |

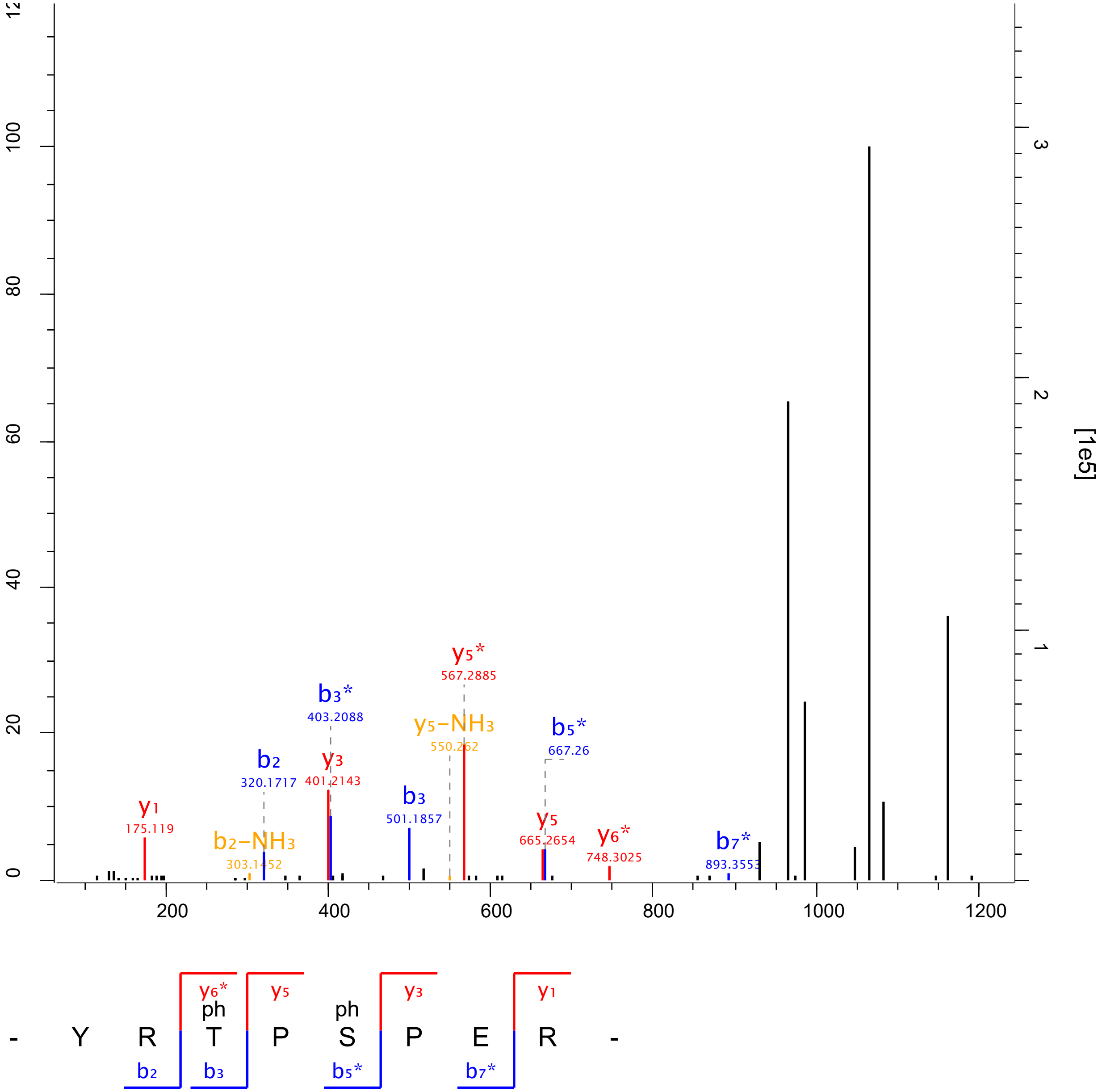

| Raw file | Scan | Method | Score | m/z |
|---|---|---|---|---|
| A17_00_2 | 3627 | FTMS; HCD | 91.36 | 482.23 |

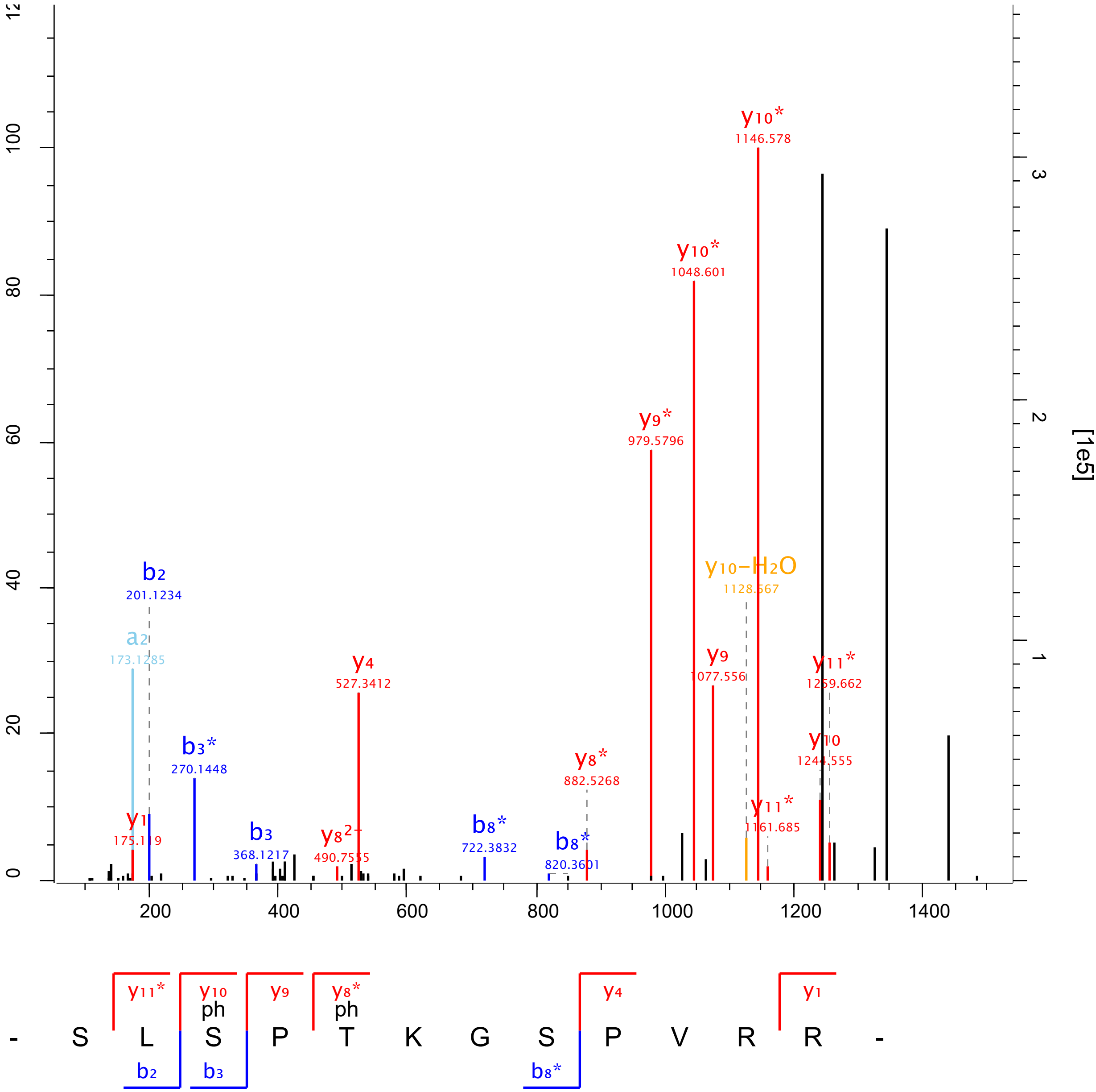

| Raw file | Scan | Method | Score | m/z |
| --- | --- | --- | --- | --- |
| A17_00_2 | 36472 | FTMS; HCD | 40.43 | 915.86 |

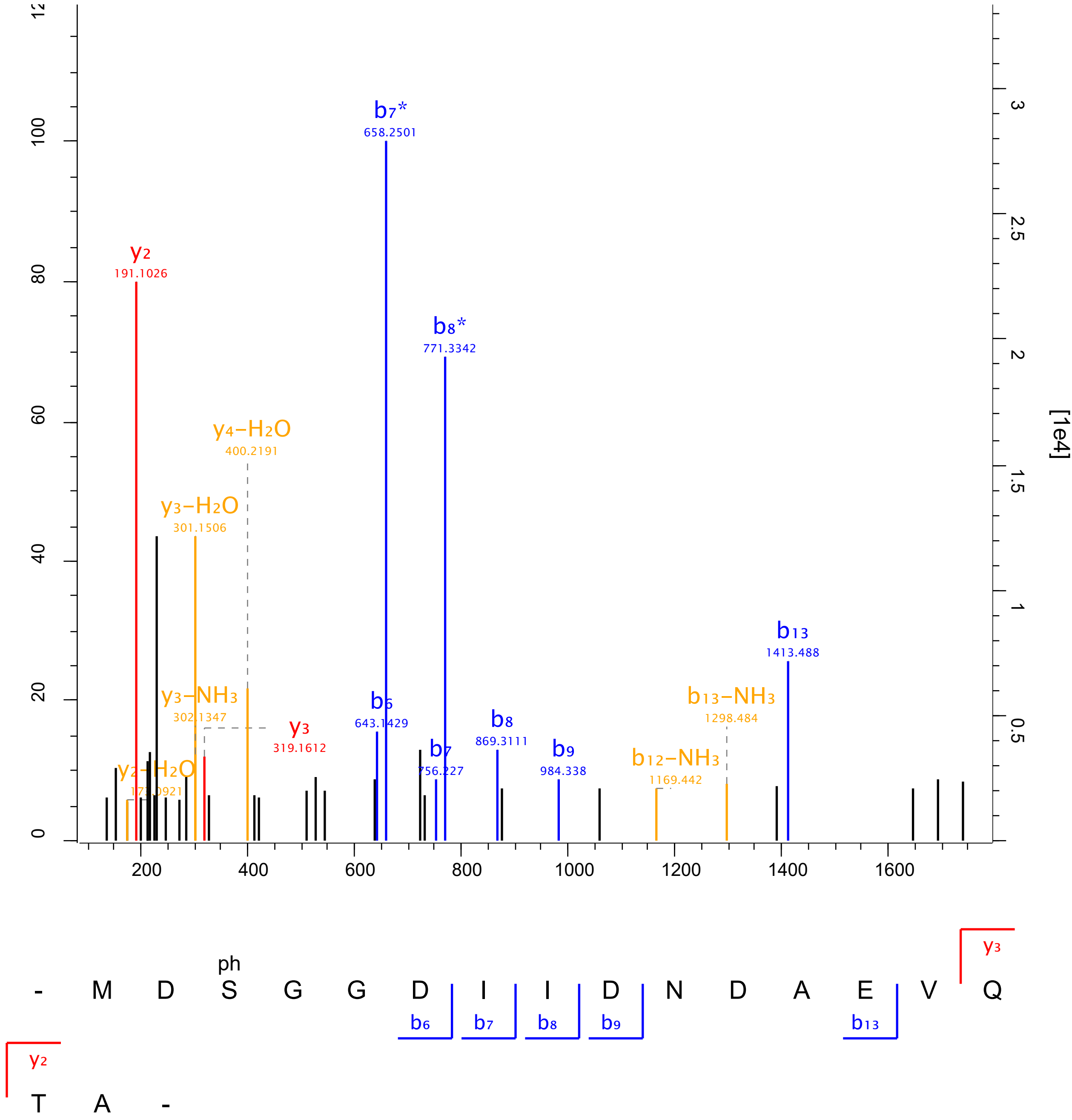

| Raw file | Scan | Method | Score | m/z |
|---|---|---|---|---|
| A17_00_2 | 37263 | FTMS; HCD | 248.67 | 1151.01 |

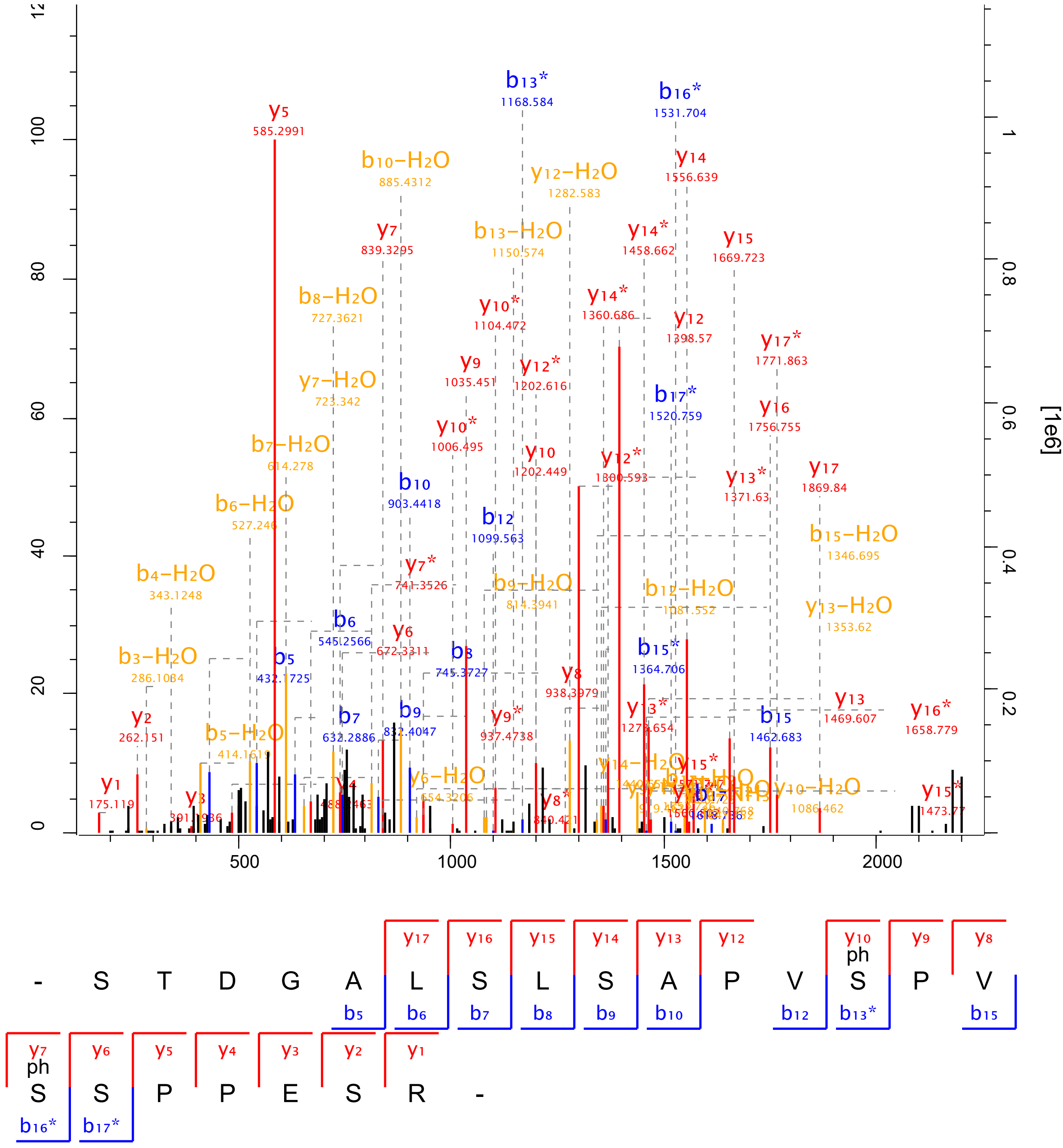

| Raw file | Scan | Method | Score | m/z |
| --- | --- | --- | --- | --- |
| A17_00_2 | 3809 | FTMS; HCD | 48.82 | 481.56 |

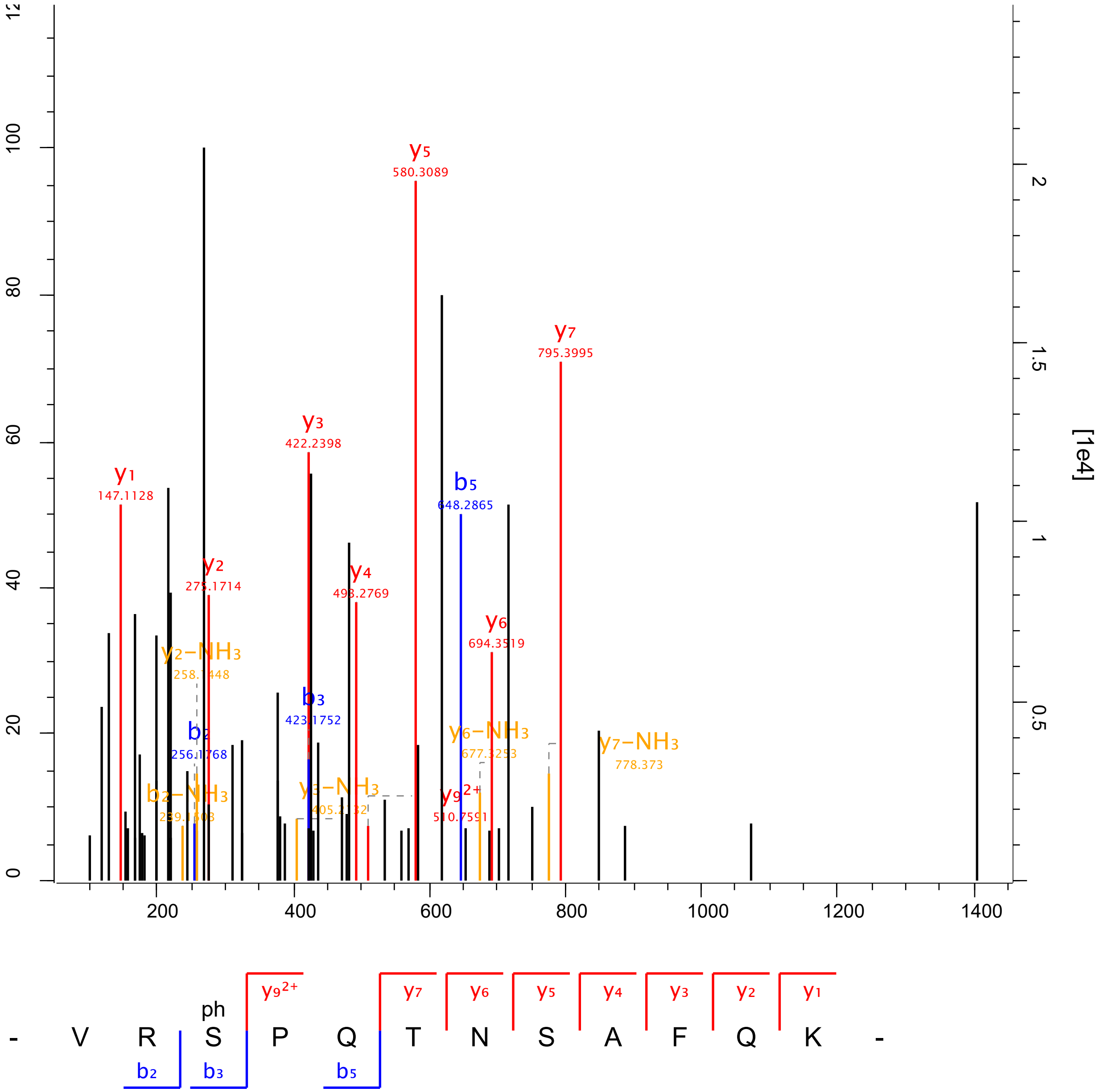

| Raw file | Scan | Method | Score | m/z |
| --- | --- | --- | --- | --- |
| A17_00_2 | 38174 | FTMS; HCD | 206.23 | 1016.42 |

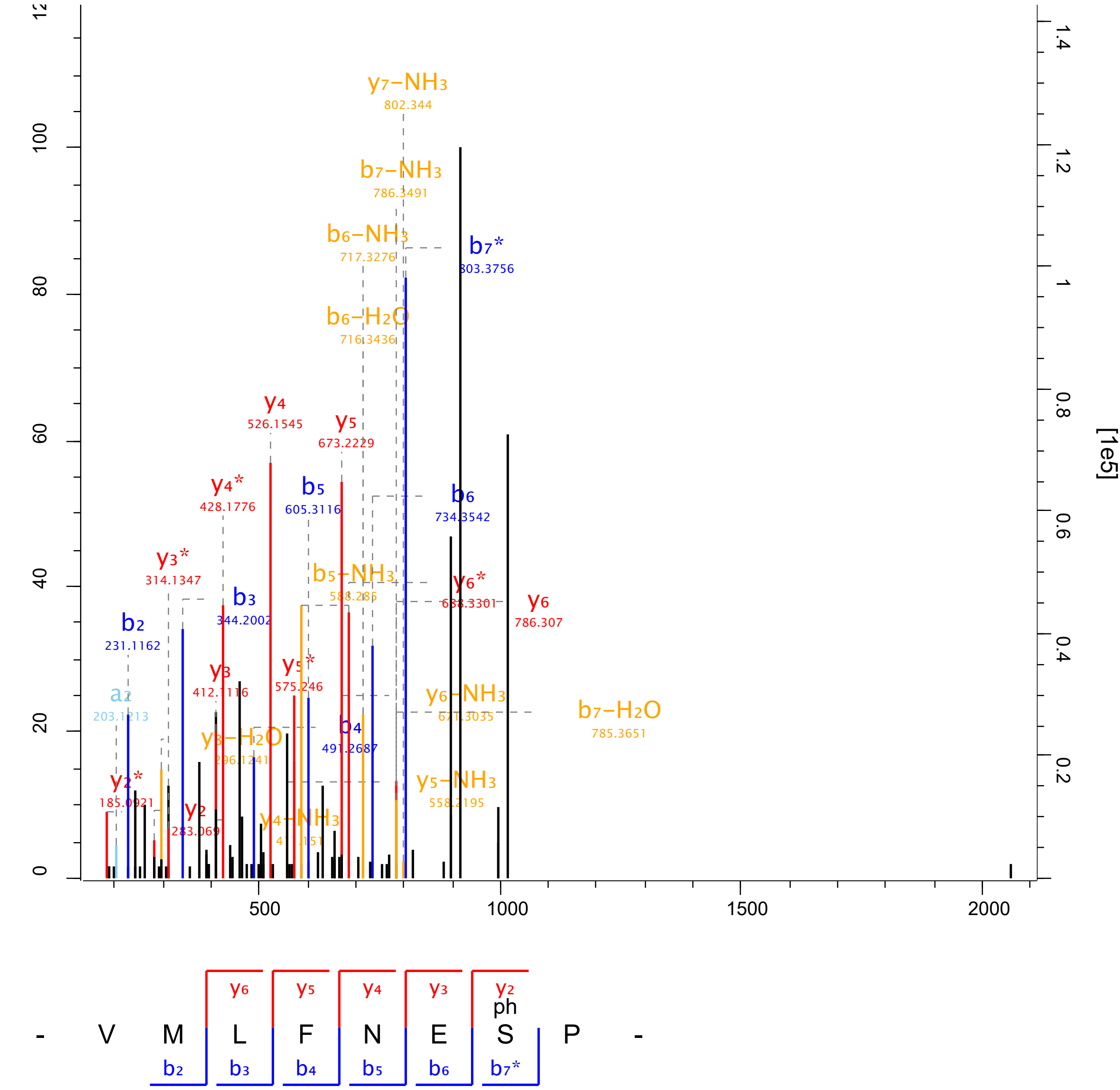

| Raw file | Scan | Method | Score | m/z |
|---|---|---|---|---|
| A17_00_2 | 3909 | FTMS; HCD | 42.08 | 617.23 |

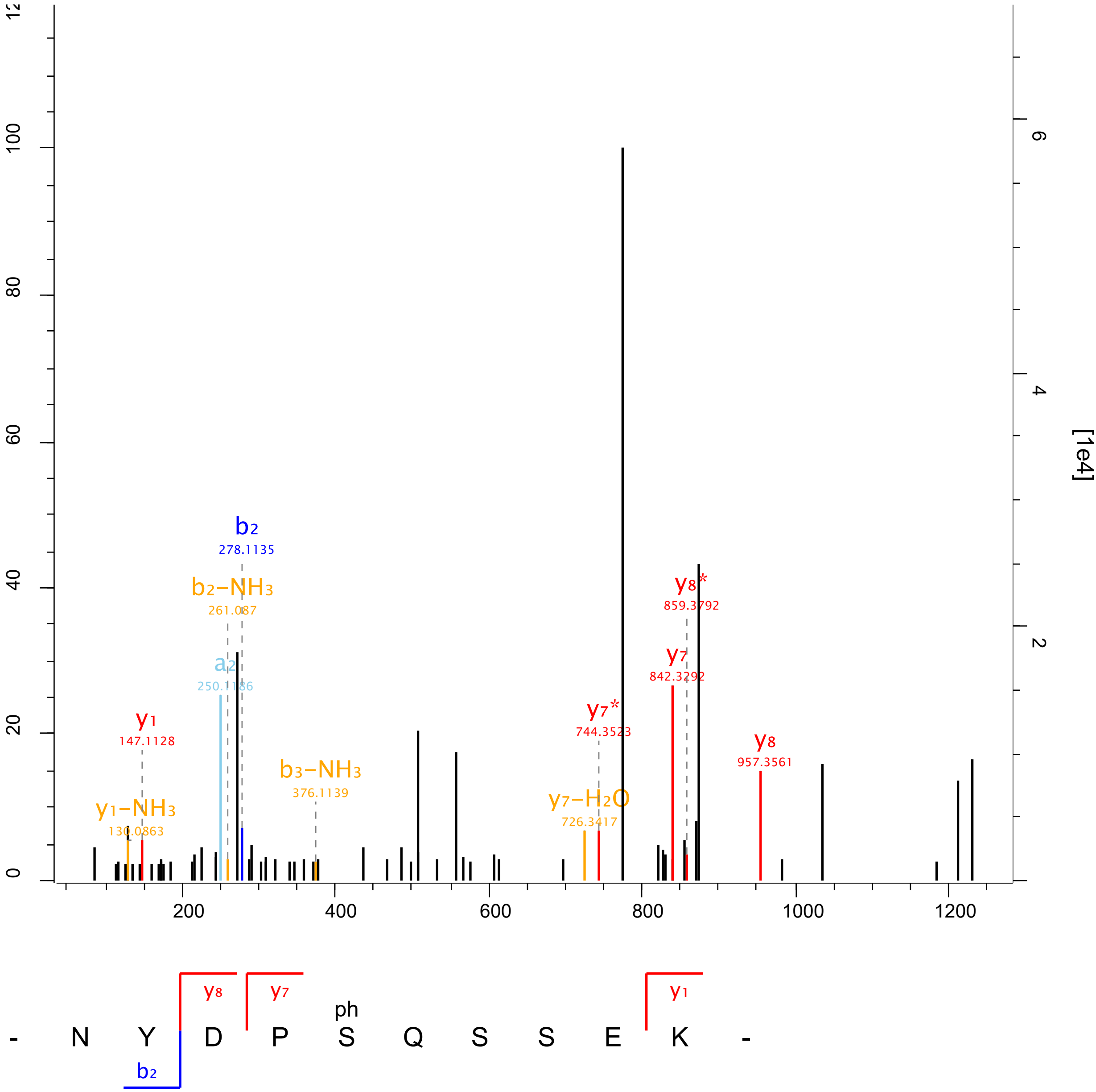

| Raw file | Scan | Method | Score | m/z |
| --- | --- | --- | --- | --- |
| A17_00_2 | 3968 | FTMS; HCD | 45 | 477.19 |

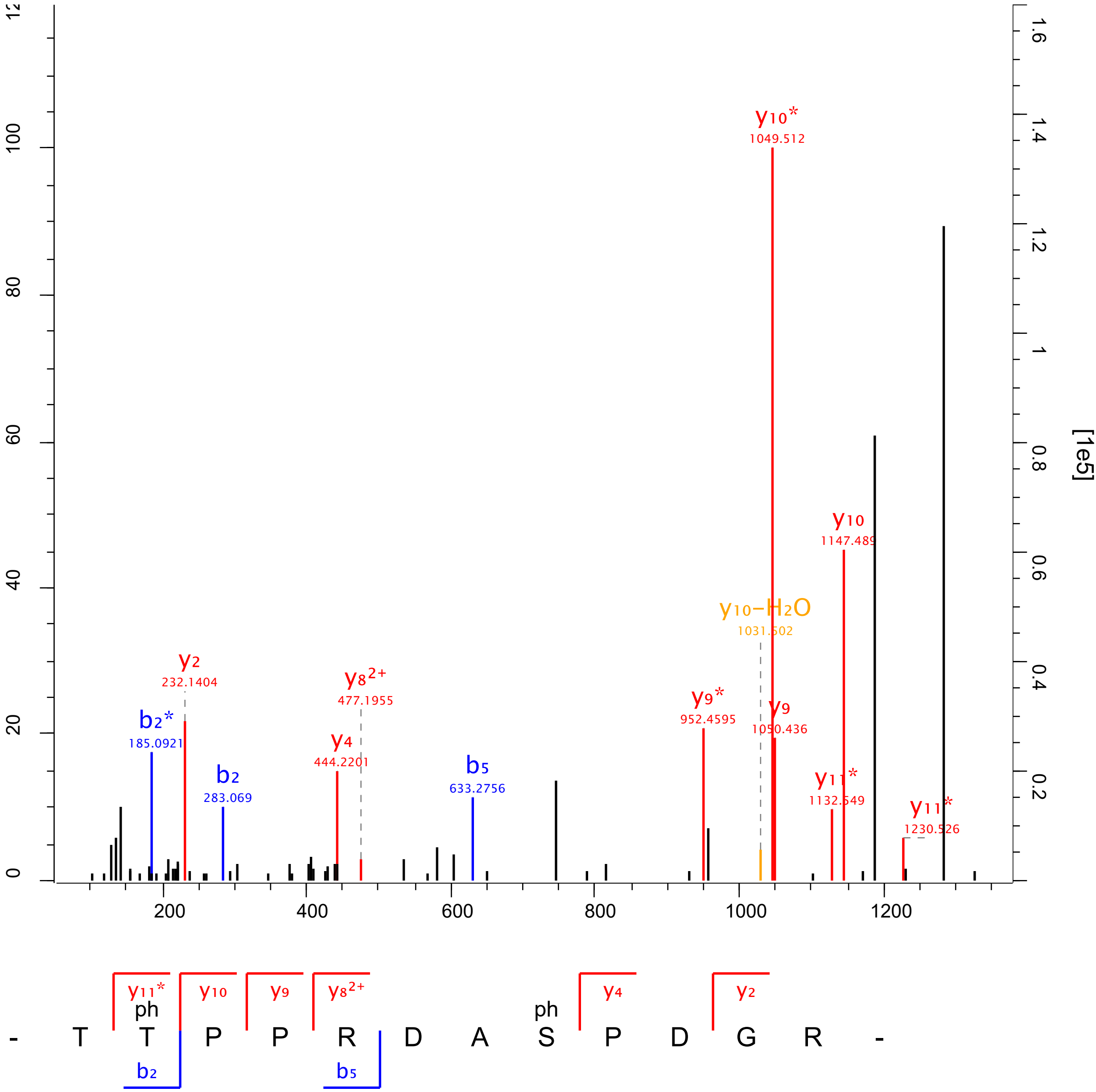

| Raw file | Scan | Method | Score | m/z |
|---|---|---|---|---|
| A17_00_2 | 4053 | FTMS; HCD | 45.36 | 620.25 |

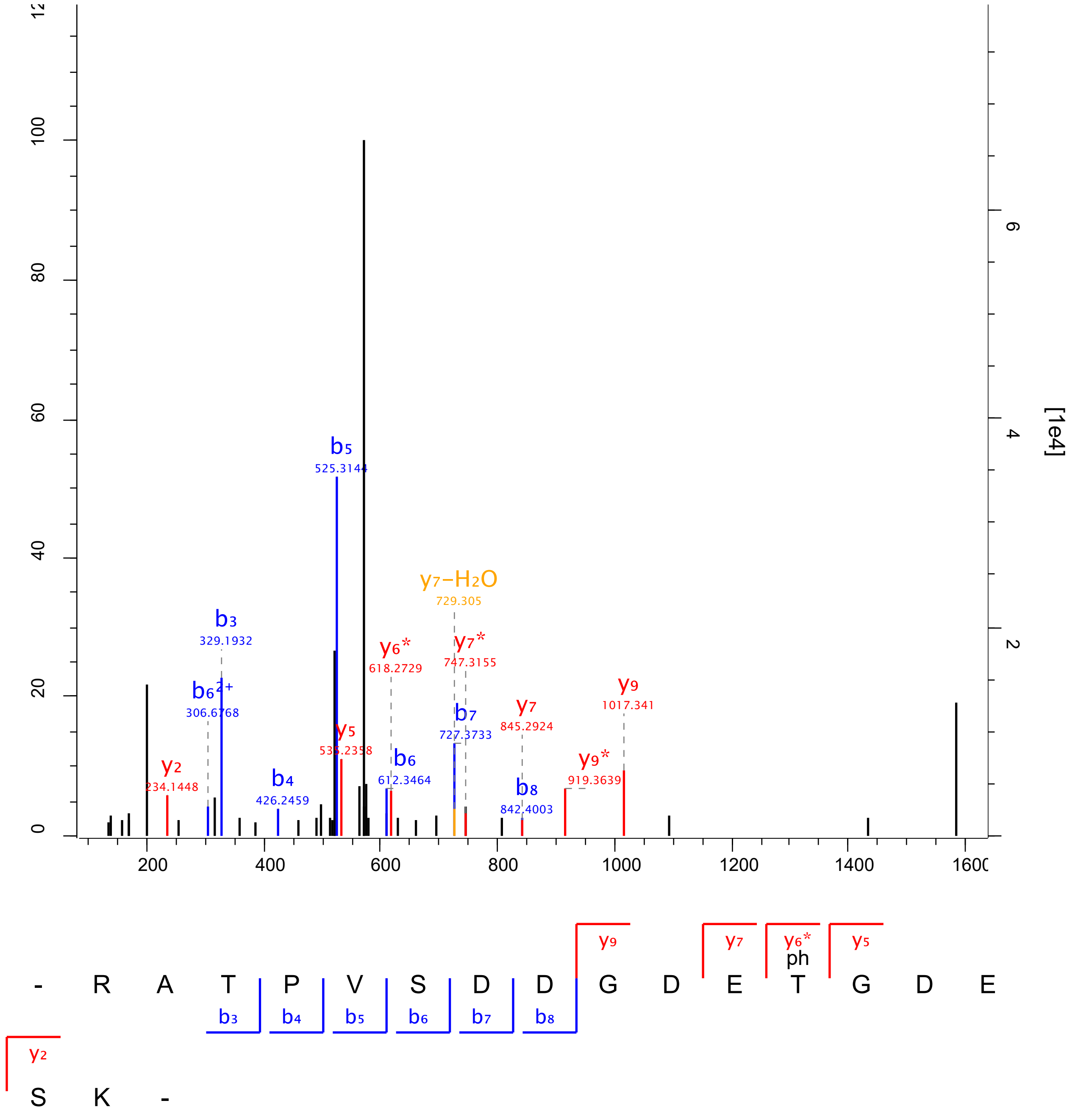

| Raw file | Scan | Method | Score | m/z |
|---|---|---|---|---|
| A17_00_2 | 4070 | FTMS; HCD | 92.77 | 672.3 |

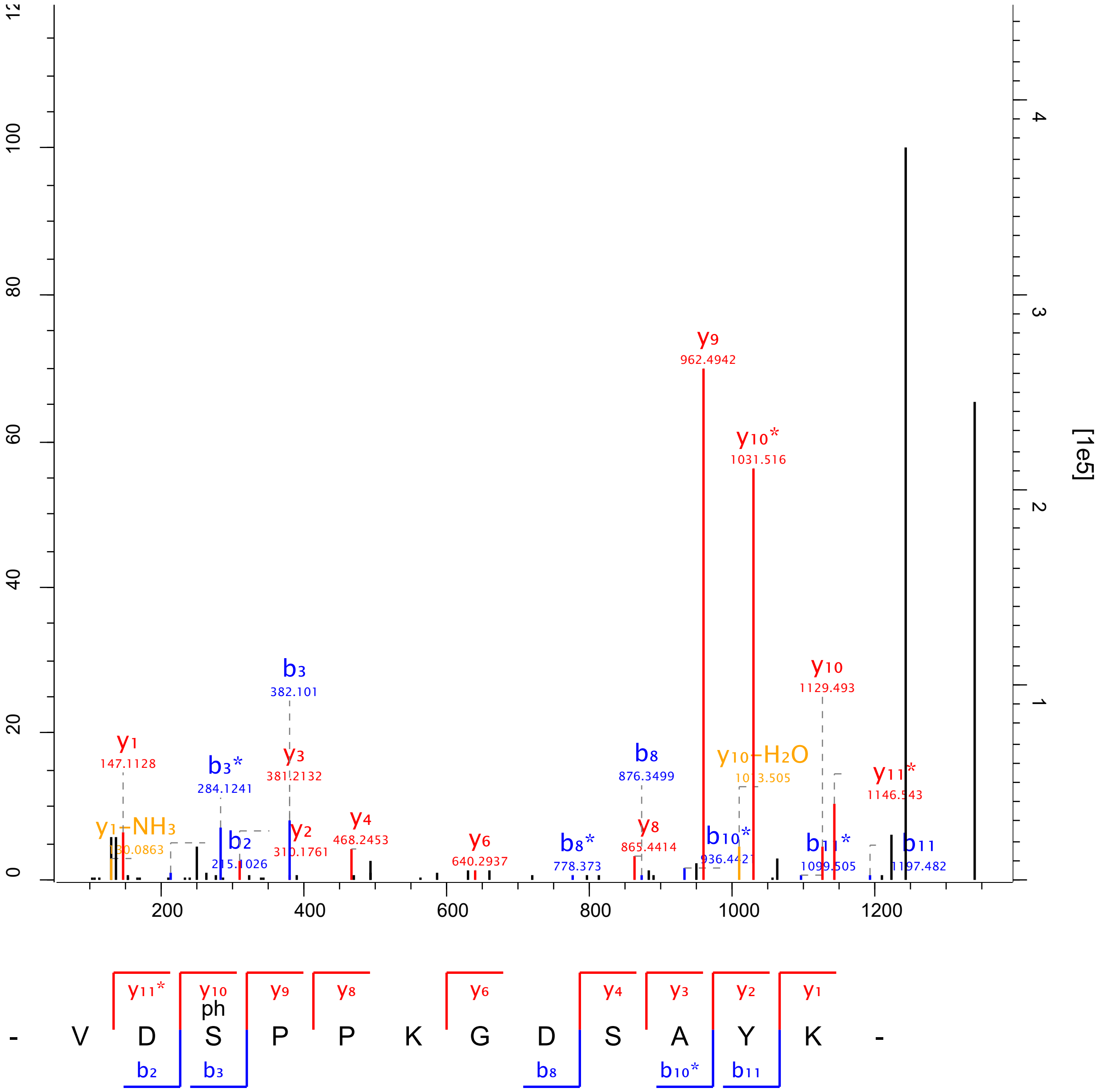

| Raw file | Scan | Method | Score | m/z |
| --- | --- | --- | --- | --- |
| A17_00_2 | 4157 | FTMS; HCD | 78.34 | 337.16 |

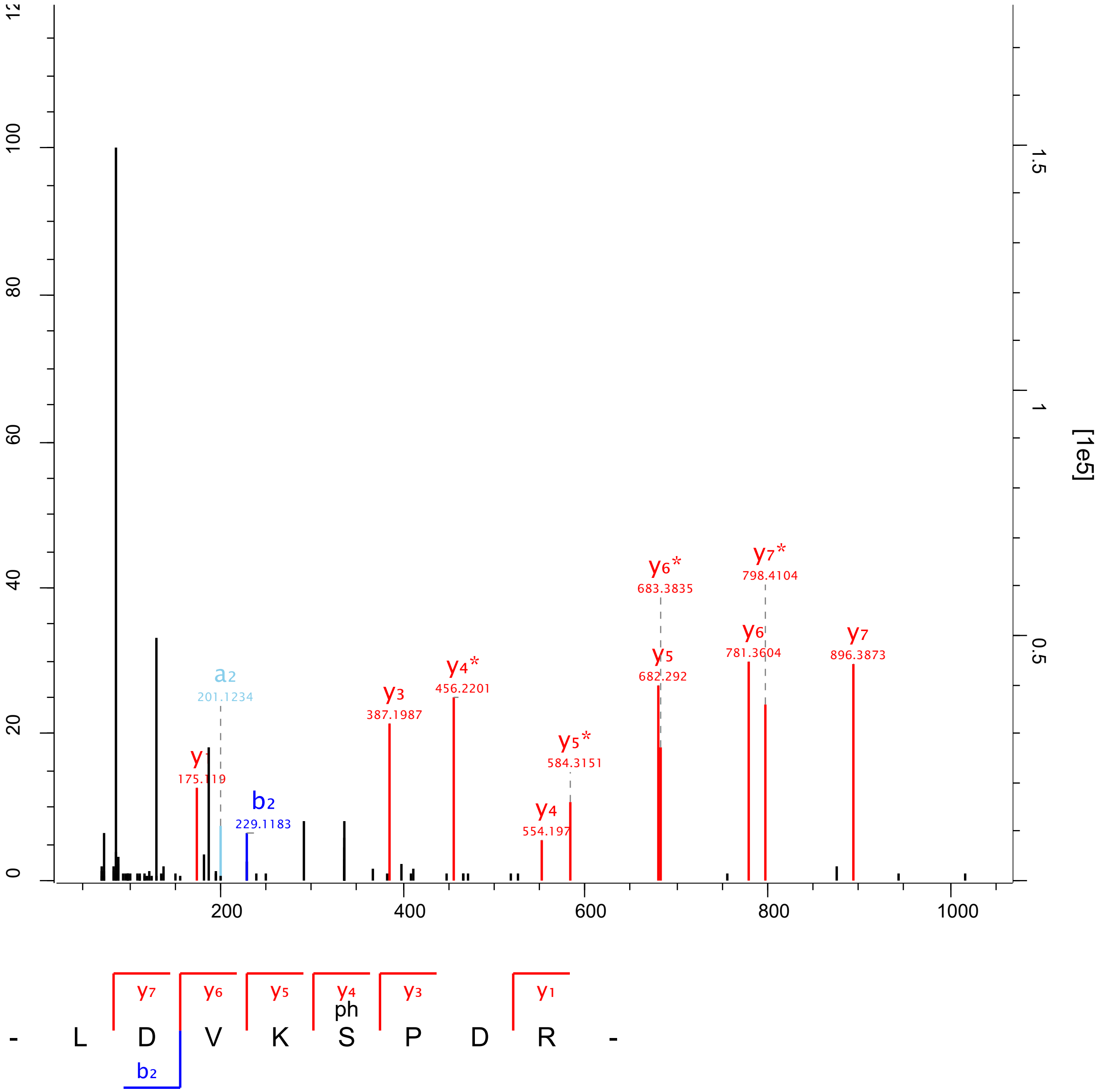

| Raw file | Scan | Method | Score | m/z |
|---|---|---|---|---|
| A17_00_2 | 42472 | FTMS; HCD | 83.4 | 989.39 |

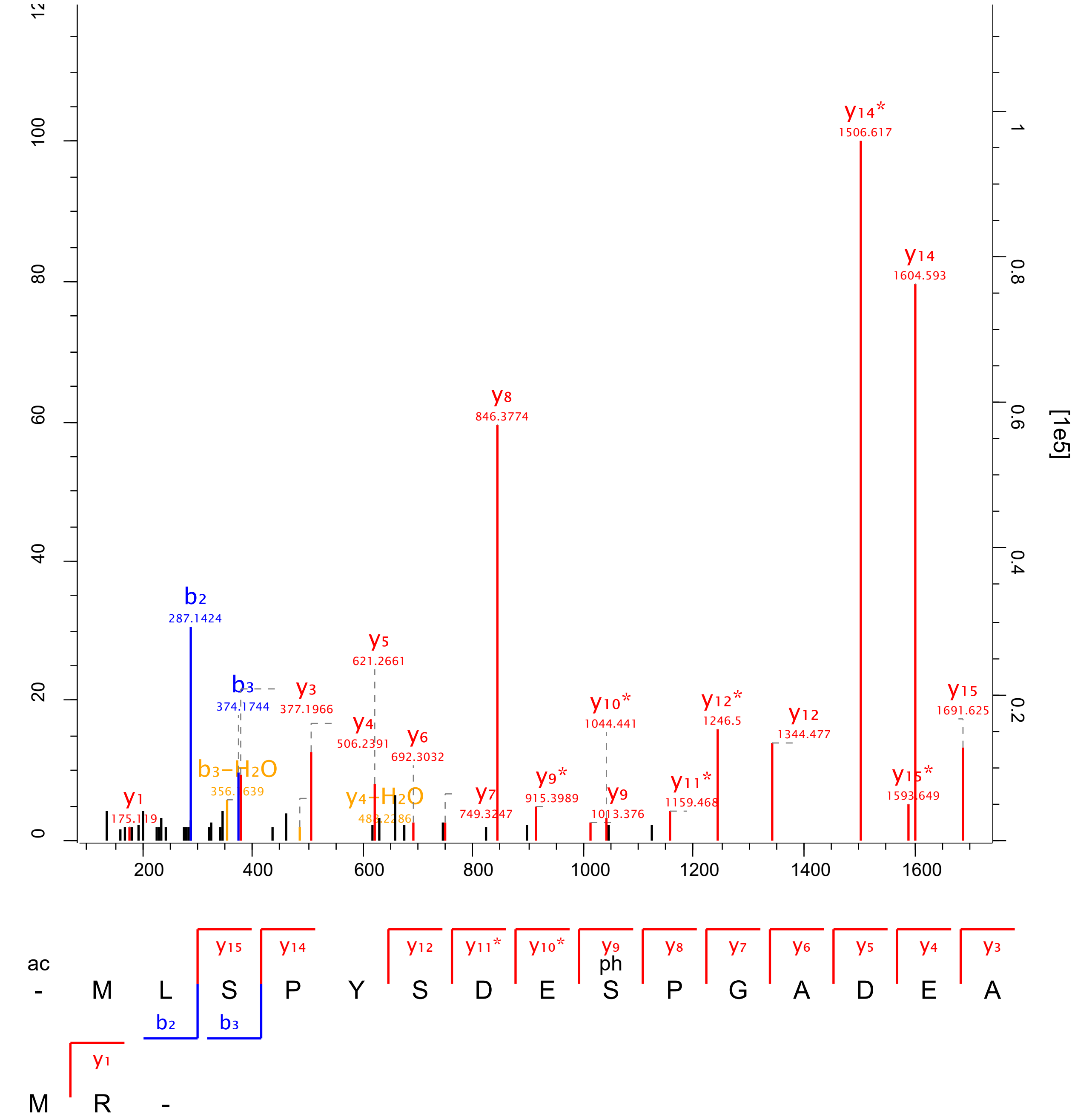

| Raw file | Scan | Method | Score | m/z |
|---|---|---|---|---|
| A17_00_2 | 4331 | FTMS; HCD | 68.36 | 553.23 |

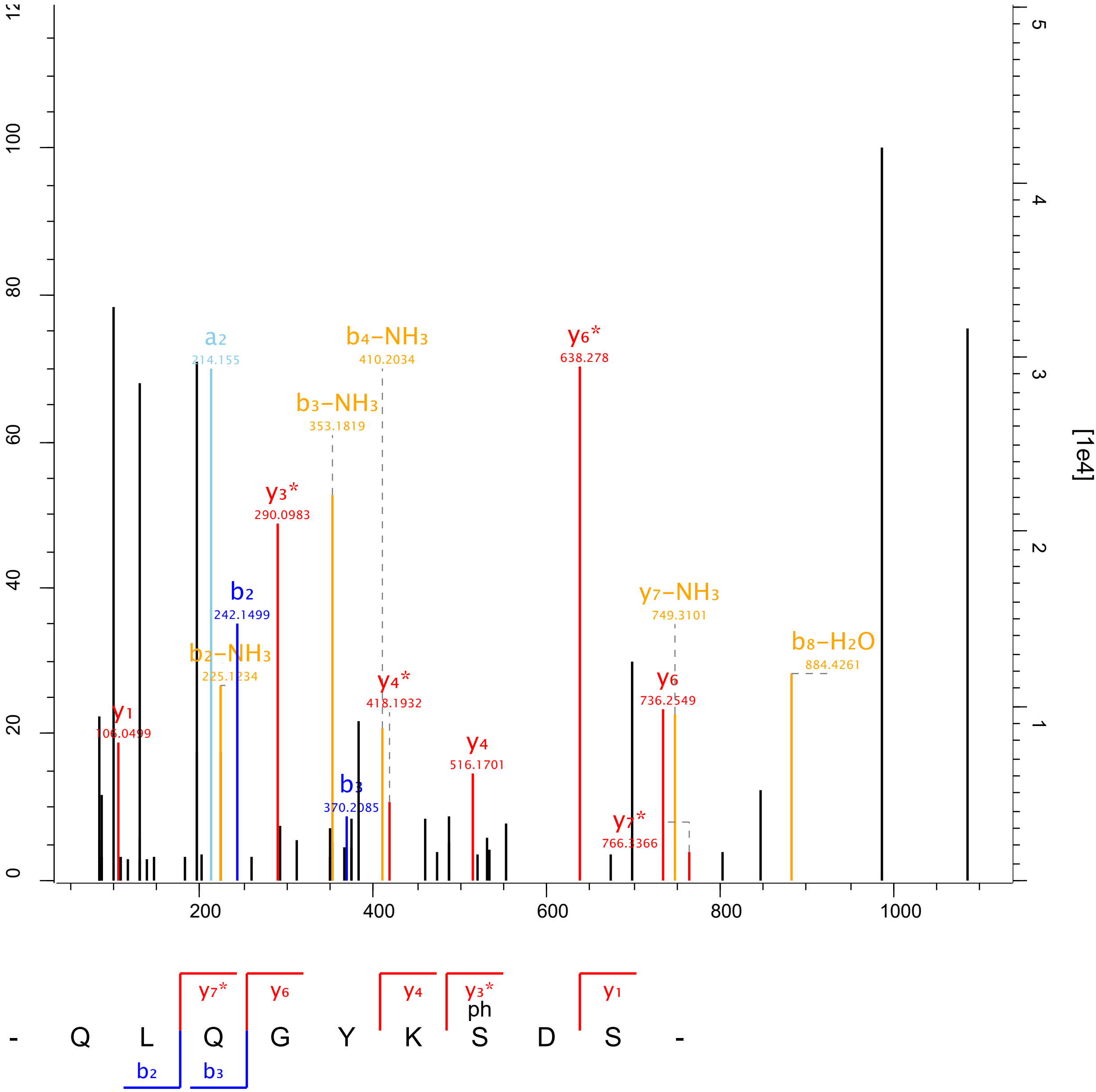

| Raw file | Scan | Method | Score | m/z |
|---|---|---|---|---|
| A17_00_2 | 4341 | FTMS; HCD | 75.76 | 649.78 |

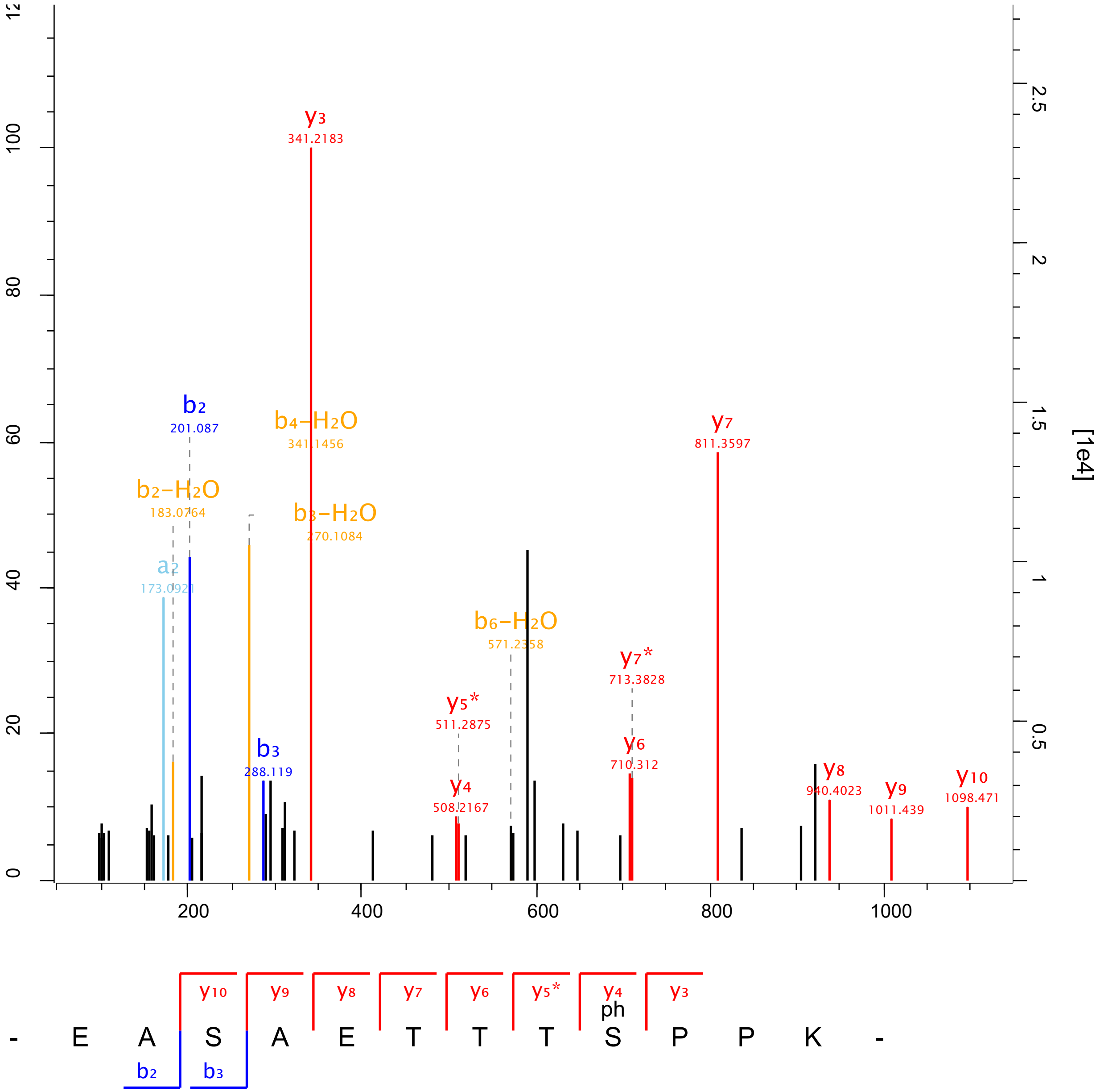

| Raw file | Scan | Method | Score | m/z |
|---|---|---|---|---|
| A17_00_2 | 4513 | FTMS; HCD | 125.31 | 788.33 |

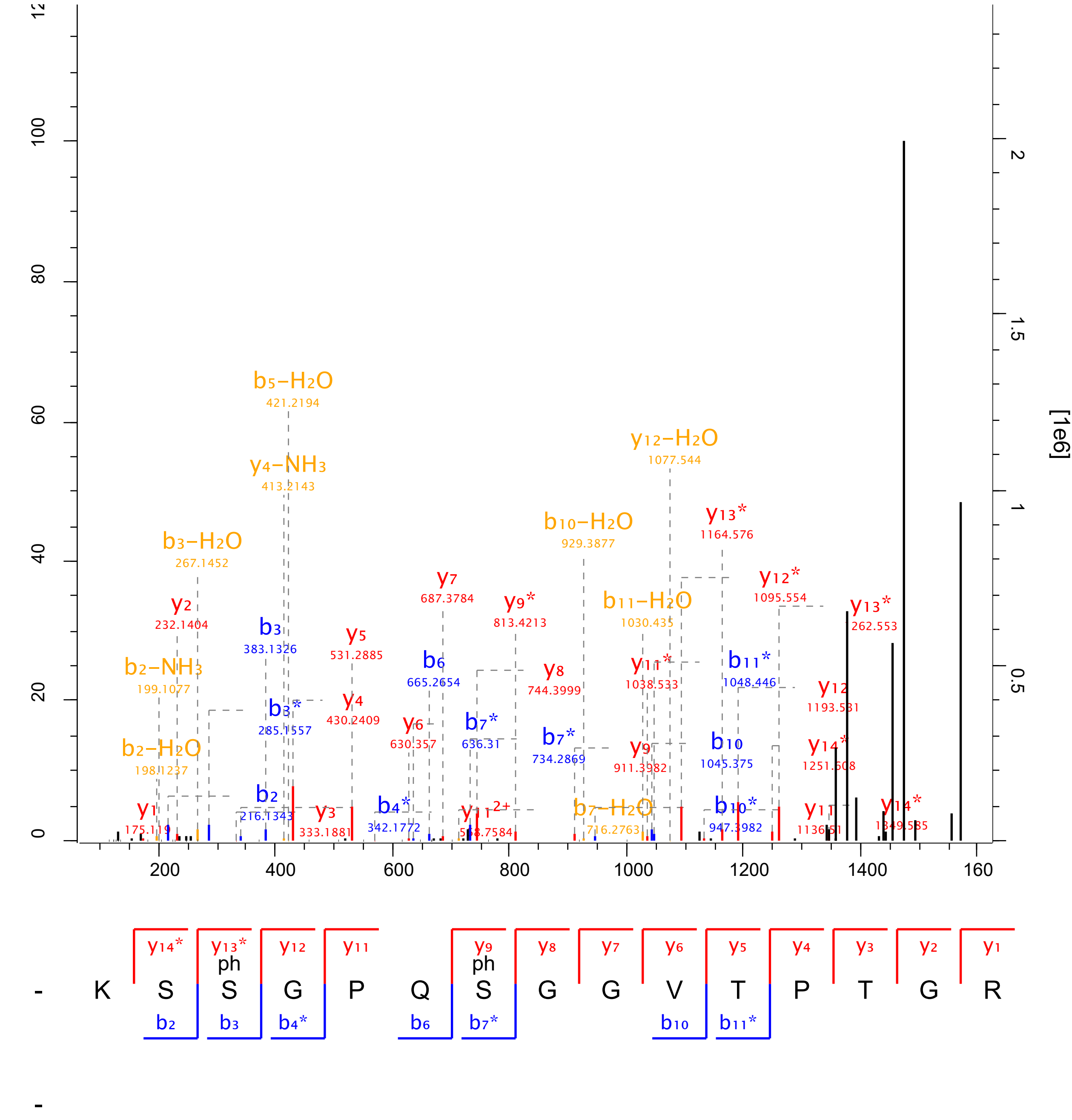

| Raw file | Scan | Method | Score | m/z |
| --- | --- | --- | --- | --- |
| A17_00_2 | 45304 | FTMS; HCD | 115.29 | 650.76 |

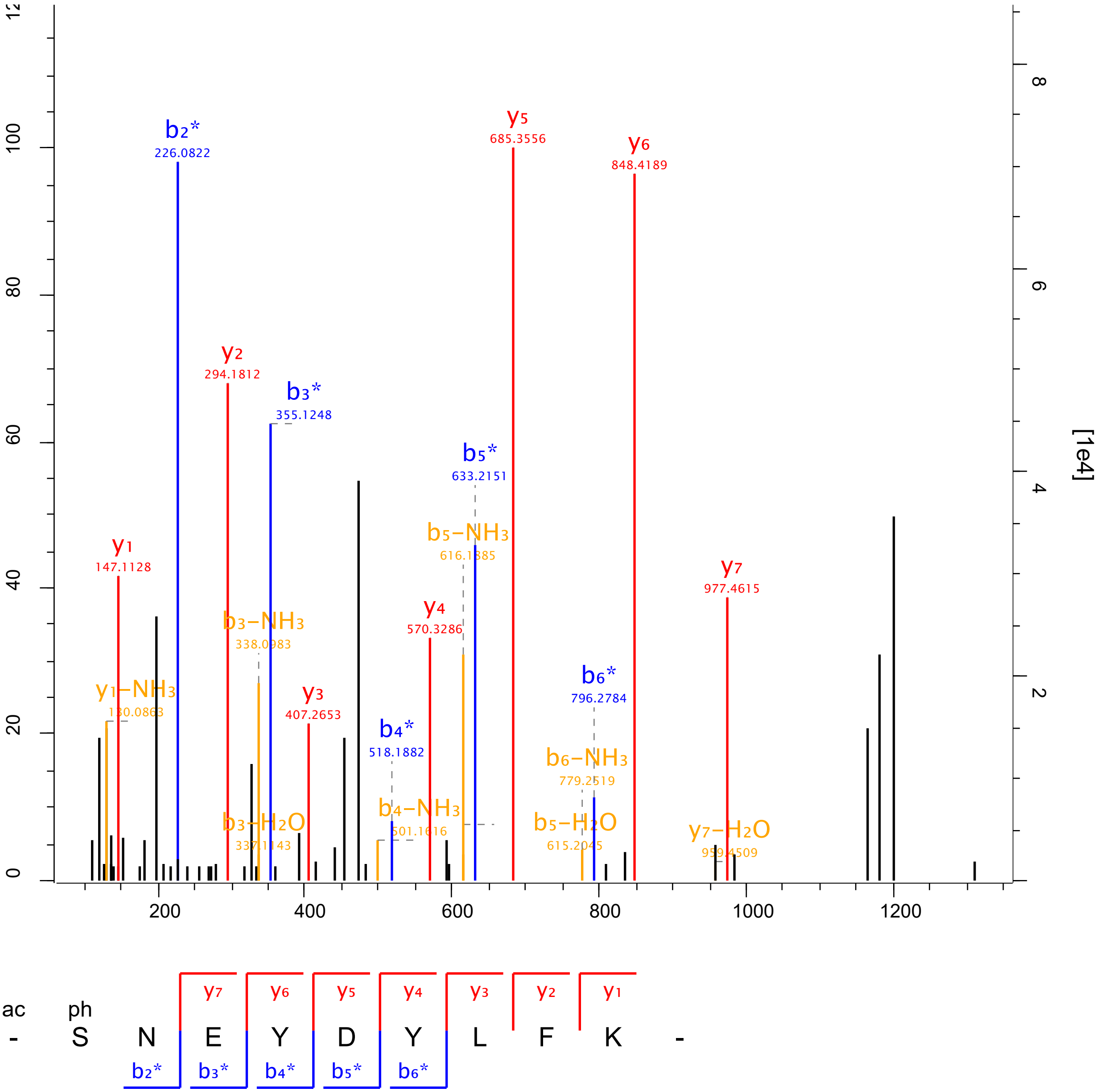

| Raw file | Scan | Method | Score | m/z |
|---|---|---|---|---|
| A17_00_2 | 4539 | FTMS; HCD | 66.27 | 561.23 |

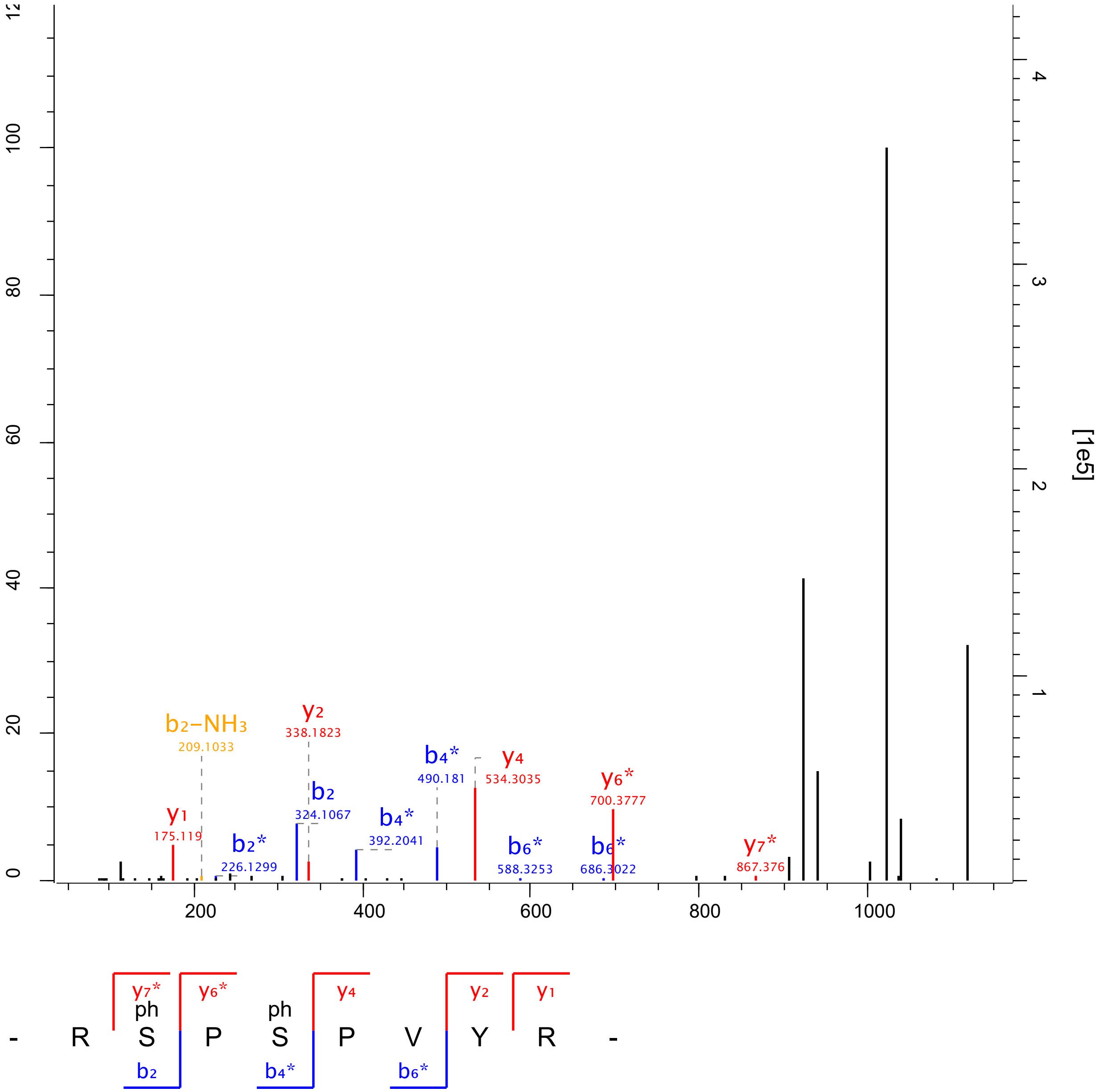

| Raw file | Scan | Method | Score | m/z |
|---|---|---|---|---|
| A17_00_2 | 4809 | FTMS; HCD | 49.59 | 671.76 |

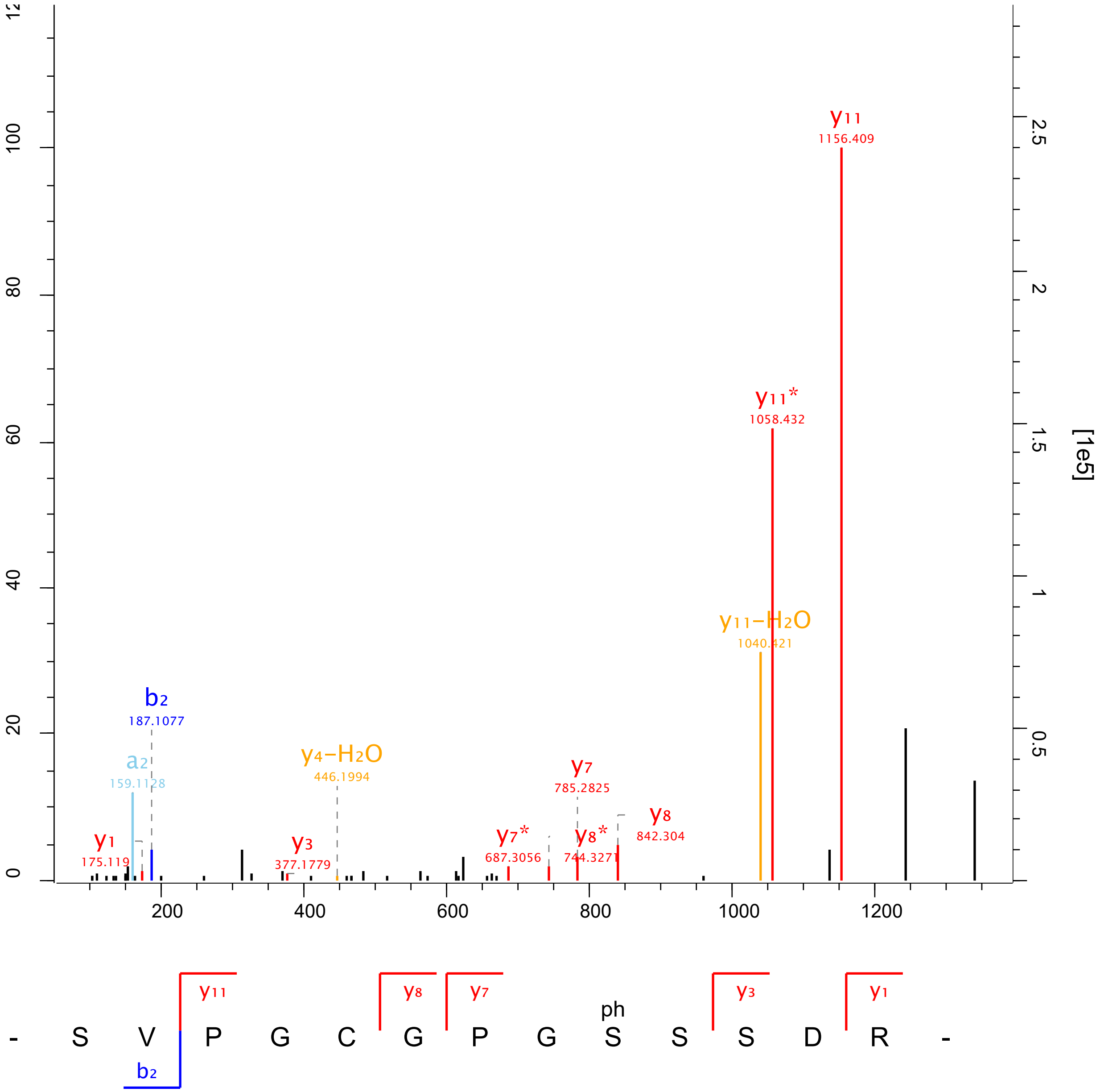

| Raw file | Scan | Method | Score | m/z |
|---|---|---|---|---|
| A17_00_2 | 48267 | FTMS; HCD | 88.39 | 893.82 |

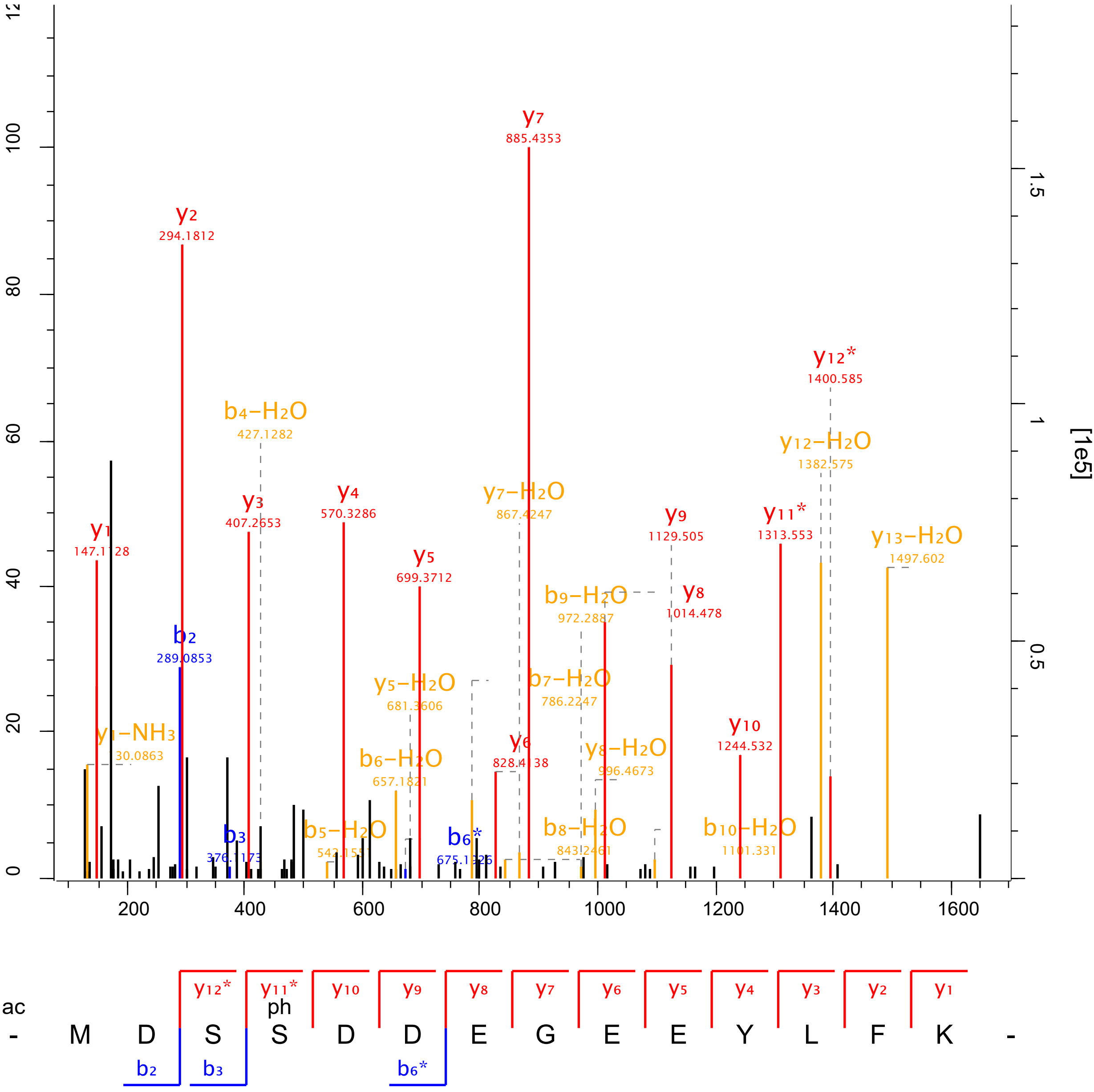

| Raw file | Scan | Method | Score | m/z |
|---|---|---|---|---|
| A17_00_2 | 4922 | FTMS; HCD | 217.48 | 515.24 |

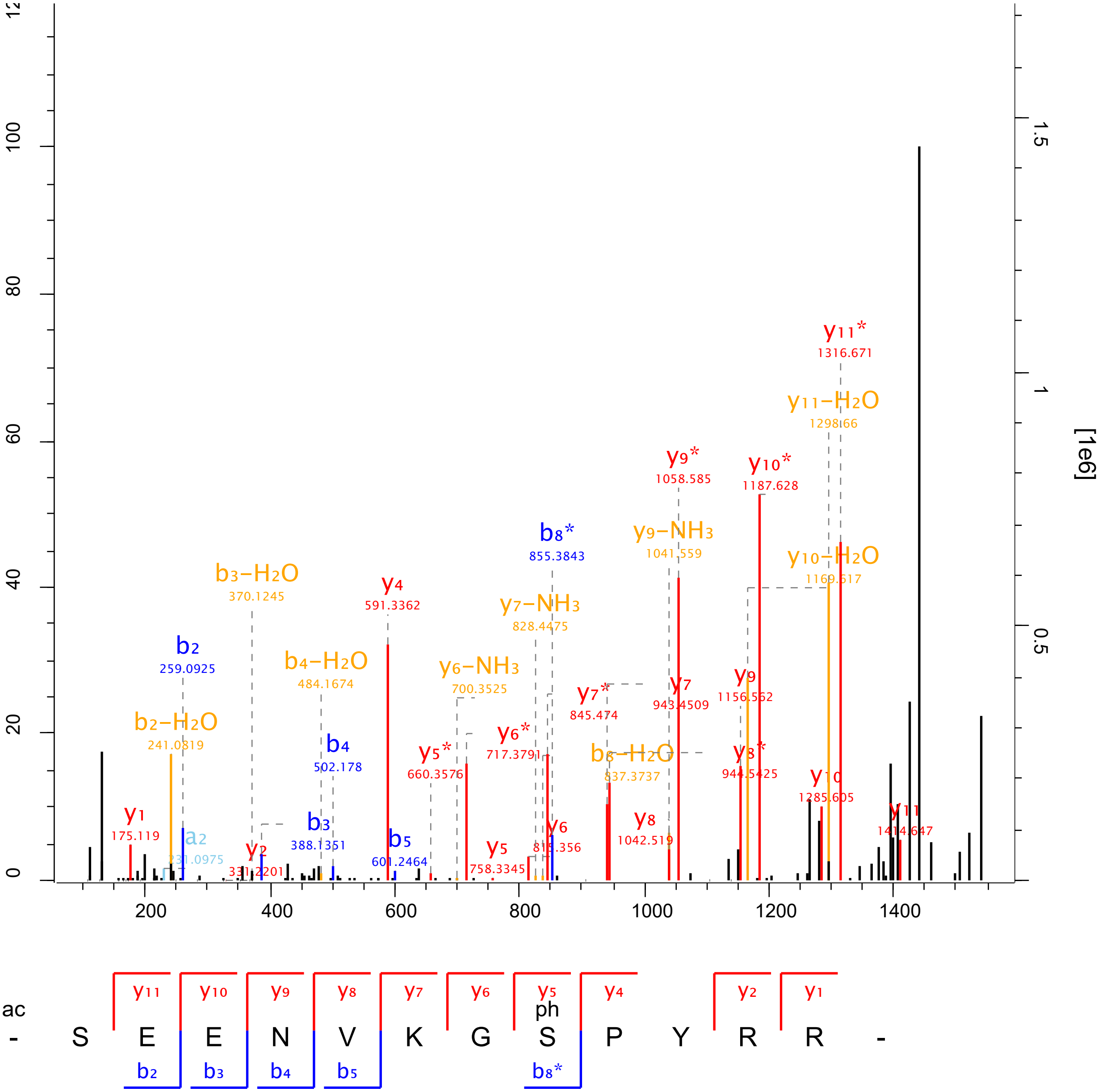

| Raw file | Scan | Method | Score | m/z |
|---|---|---|---|---|
| A17_00_2 | 4984 | FTMS; HCD | 55.31 | 493.22 |

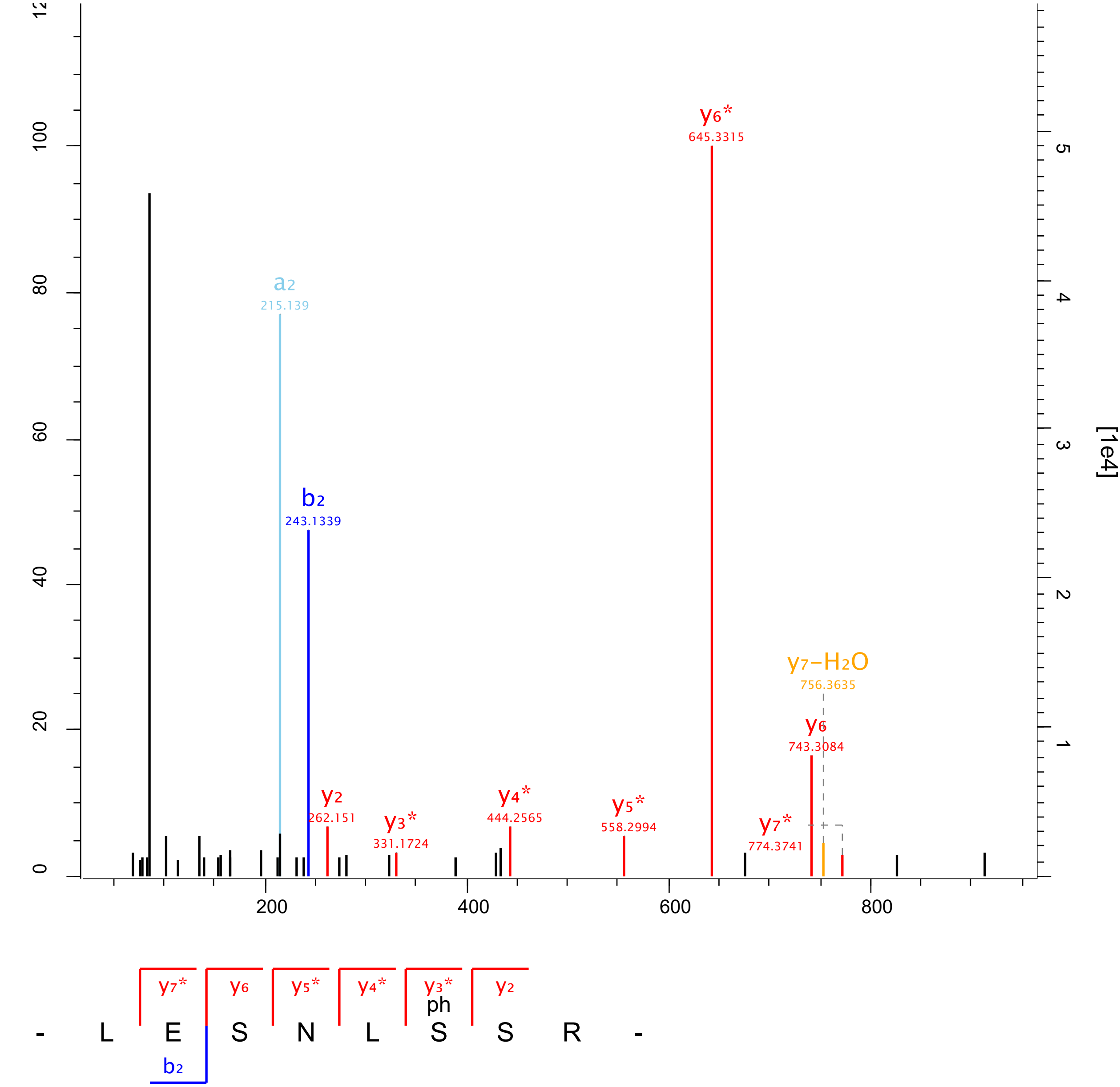

| Raw file | Scan | Method | Score | m/z |
|---|---|---|---|---|
| A17_00_2 | 5013 | FTMS; HCD | 56.41 | 588.22 |

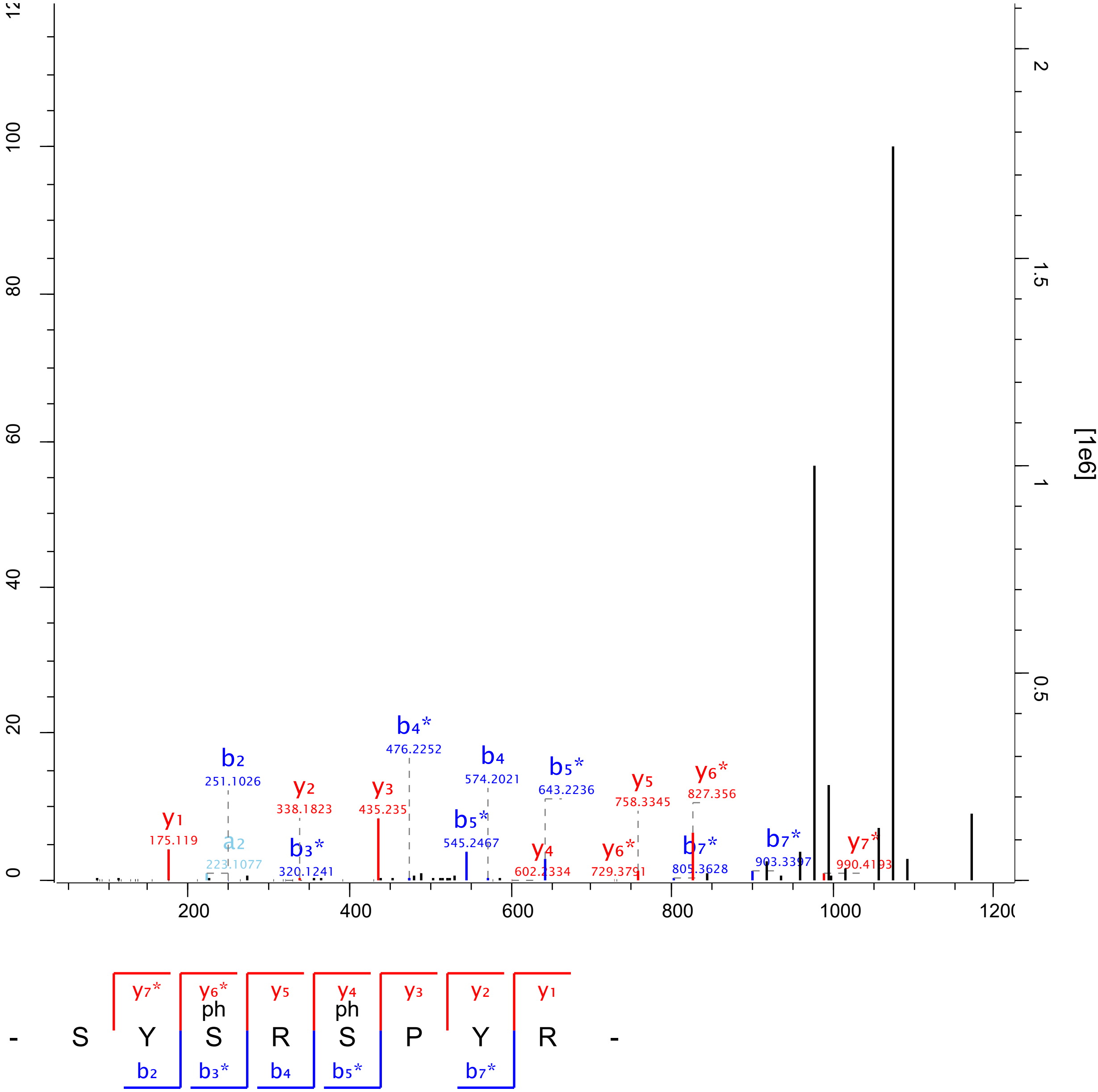

| Raw file | Scan | Method | Score | m/z |
|---|---|---|---|---|
| A17_00_2 | 5044 | FTMS; HCD | 63.62 | 555.76 |

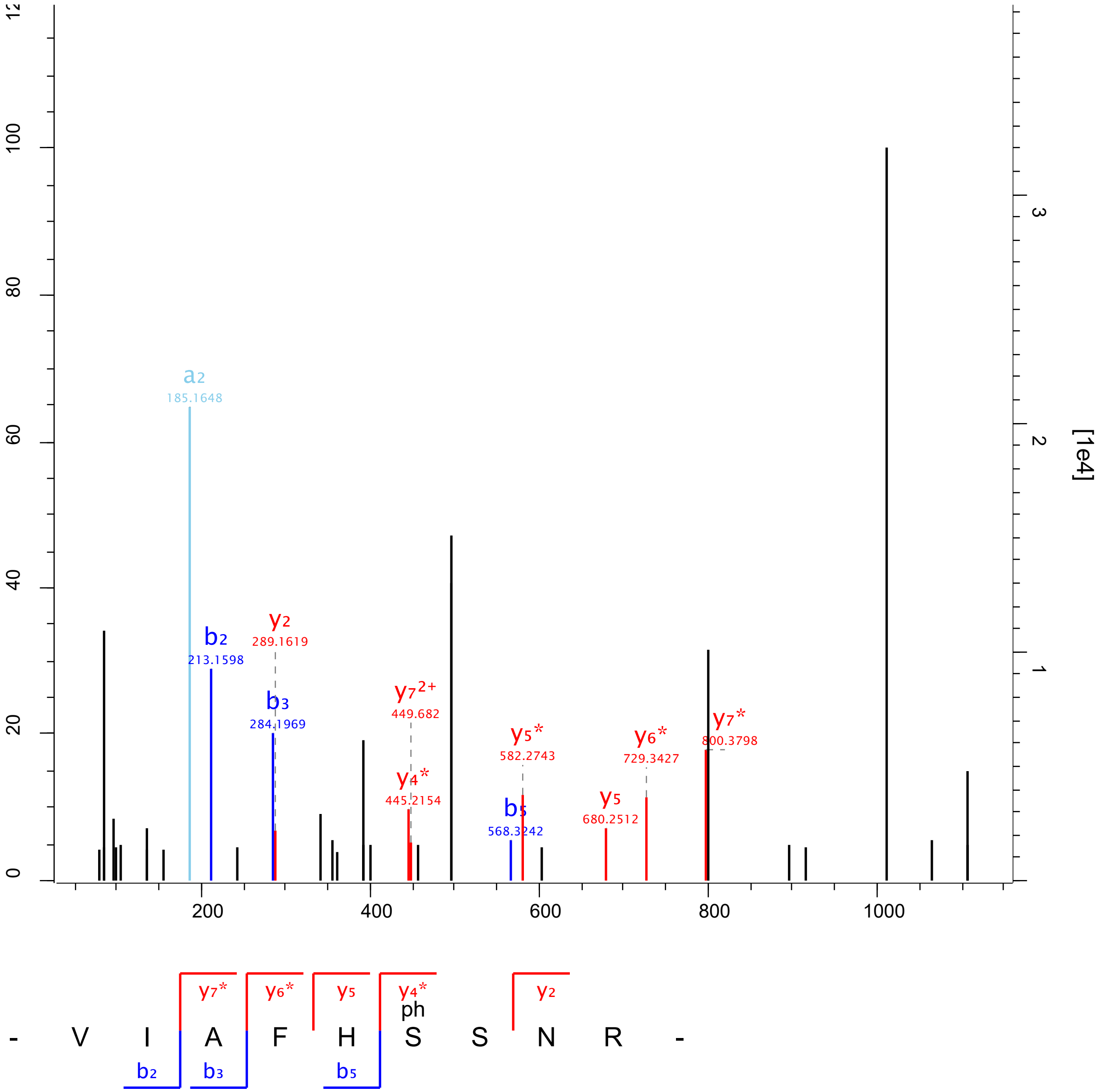

| Raw file | Scan | Method | Score | m/z |
|---|---|---|---|---|
| A17_00_2 | 5356 | FTMS; HCD | 75.38 | 729.29 |

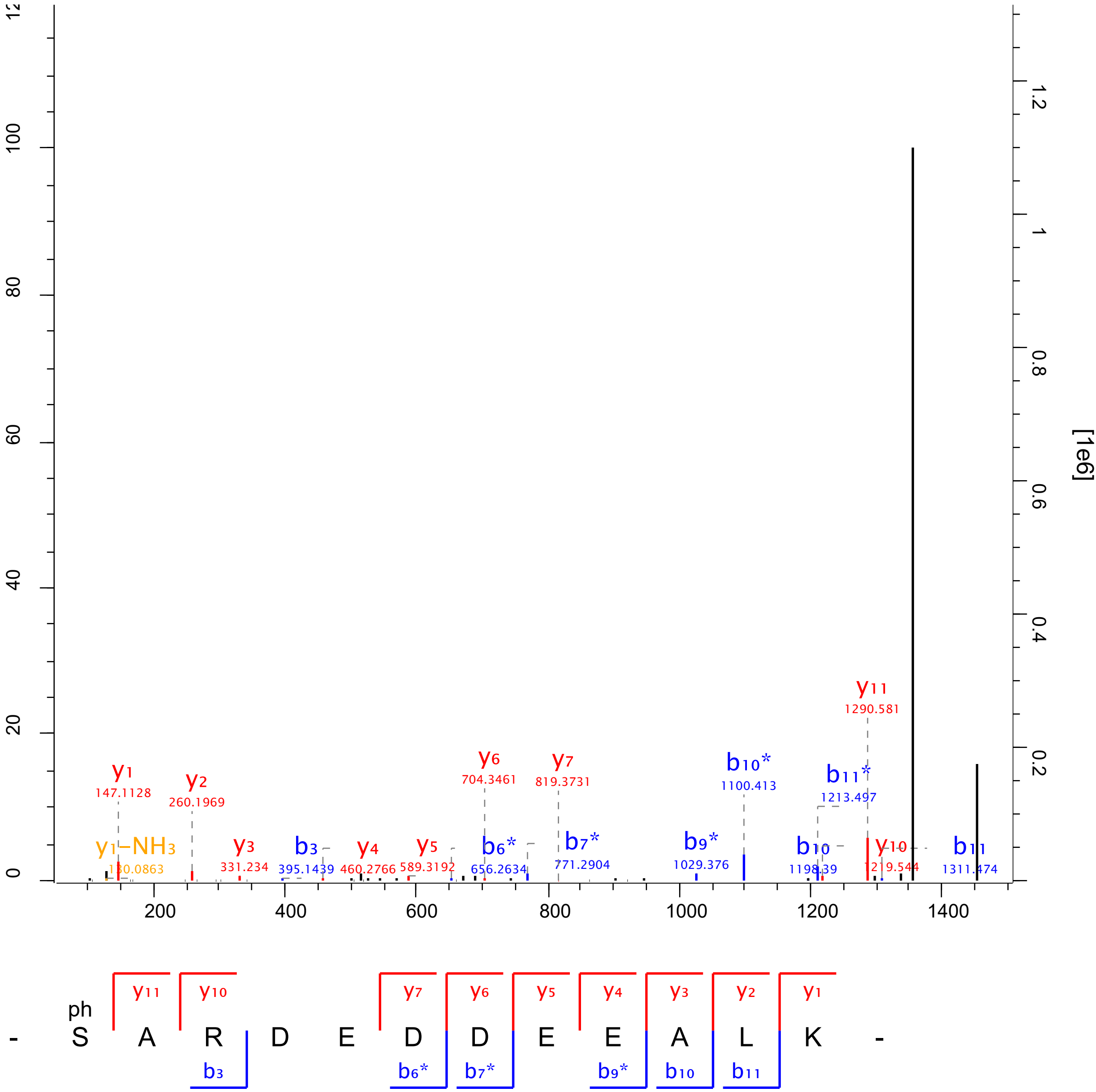

| Raw file | Scan | Method | Score | m/z |
| --- | --- | --- | --- | --- |
| A17_00_2 | 5362 | FTMS; HCD | 107.74 | 479.85 |

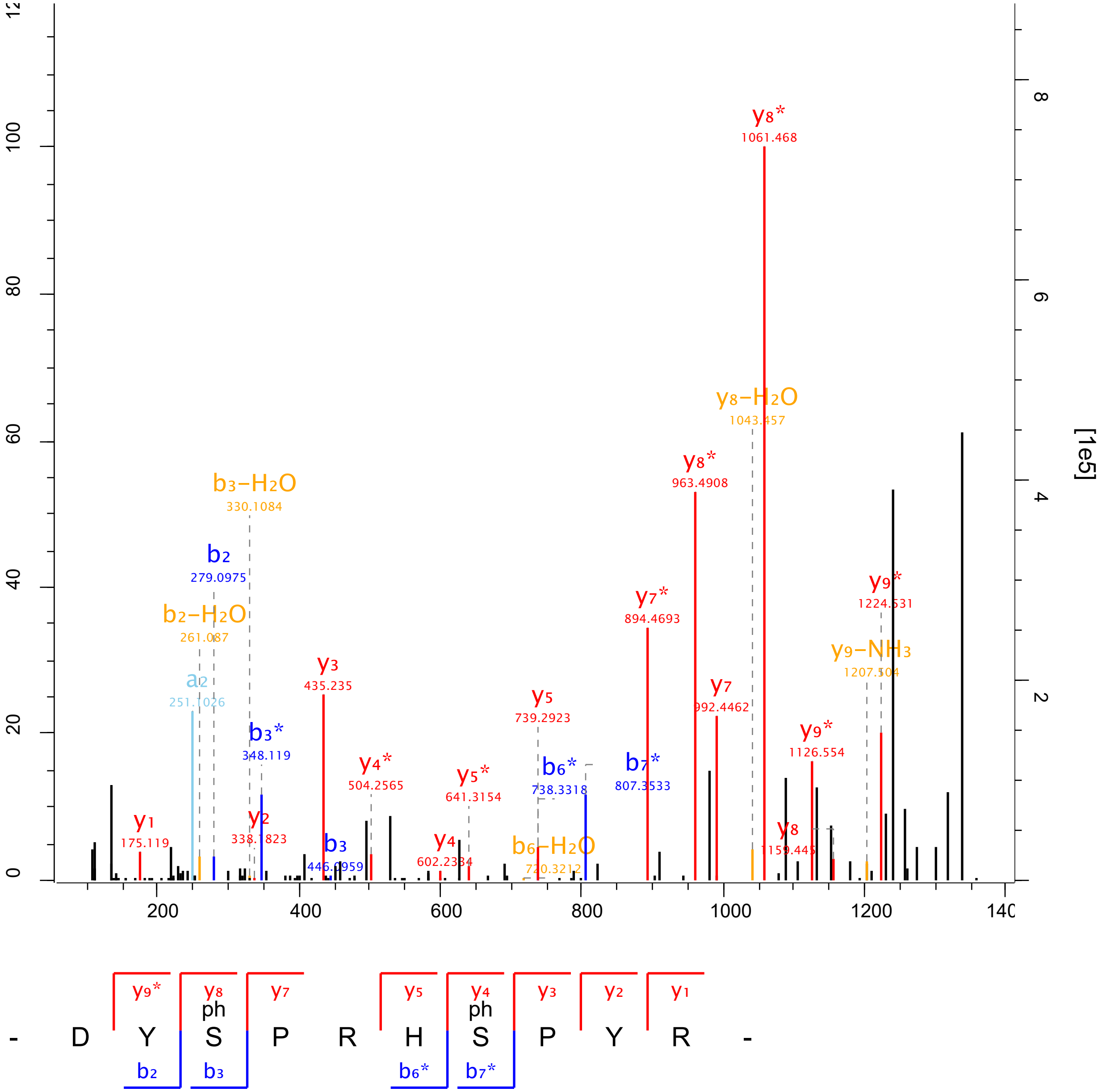

| Raw file | Scan | Method | Score | m/z |
|---|---|---|---|---|
| A17_00_2 | 5368 | FTMS; HCD | 131.4 | 497.56 |

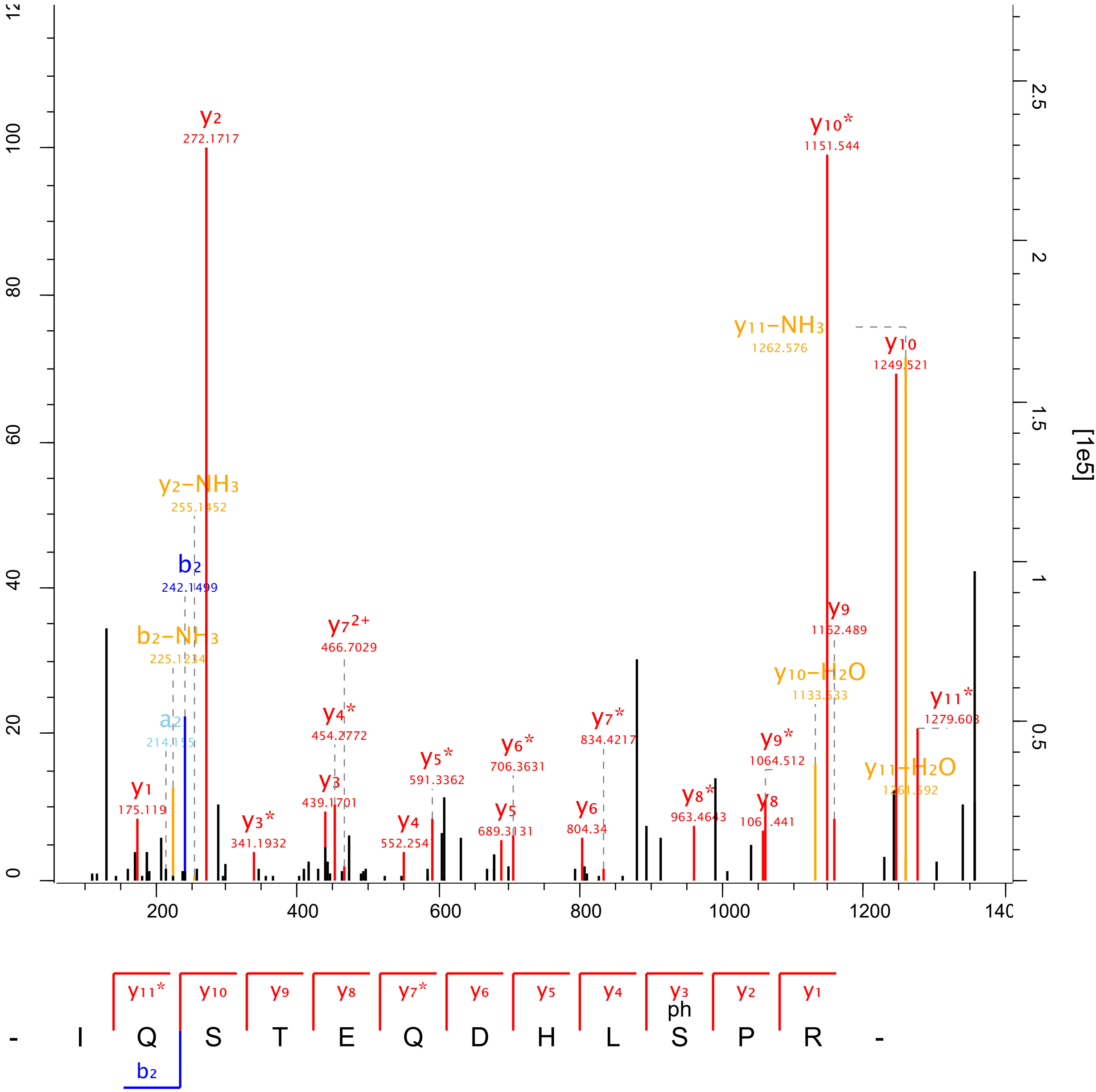

| Raw file | Scan | Method | Score | m/z |
|---|---|---|---|---|
| A17_00_2 | 5436 | FTMS; HCD | 248.78 | 771.86 |

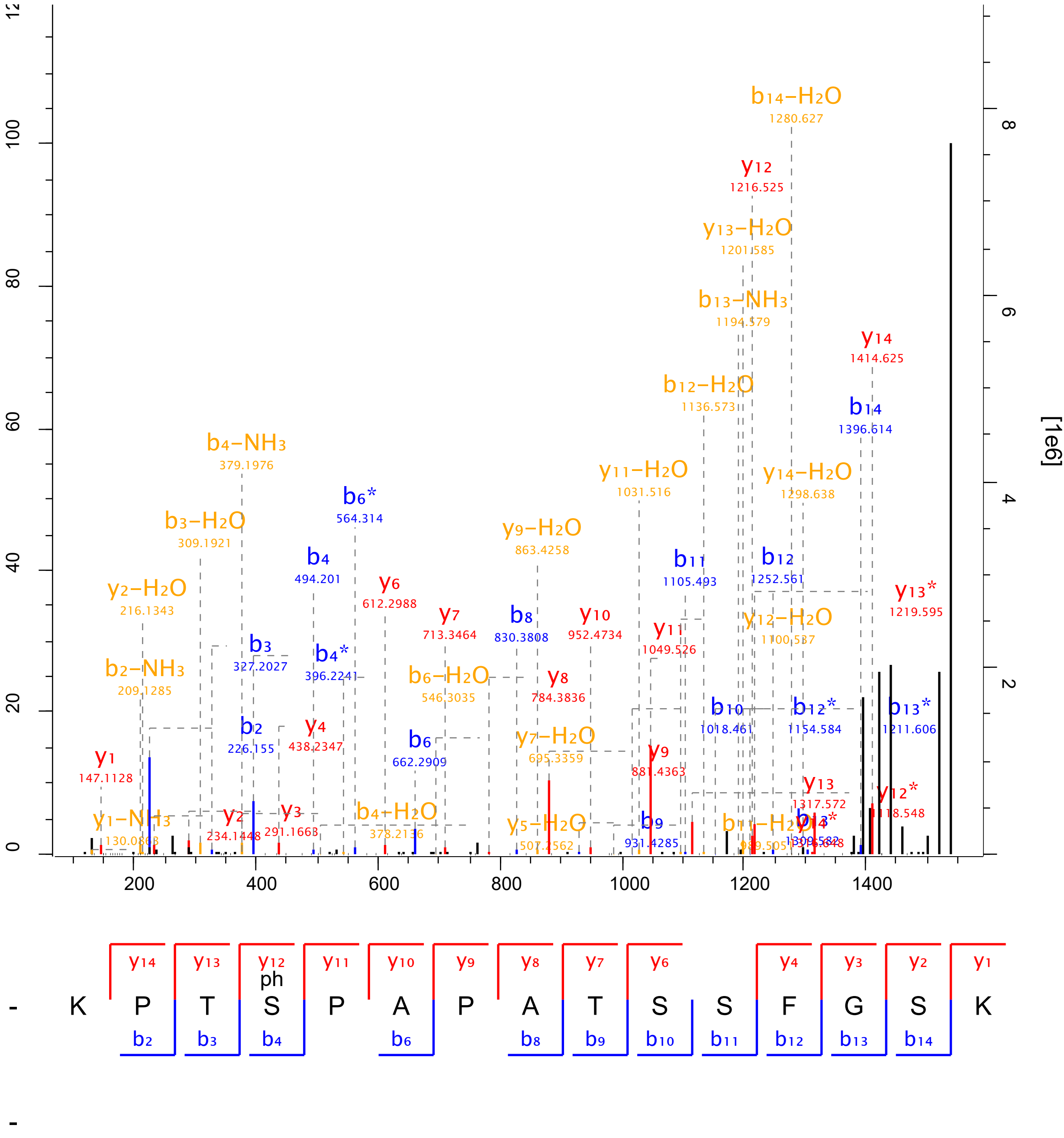

| Raw file | Scan | Method | Score | m/z |
|---|---|---|---|---|
| A17_00_2 | 5489 | FTMS; HCD | 87.57 | 493.25 |

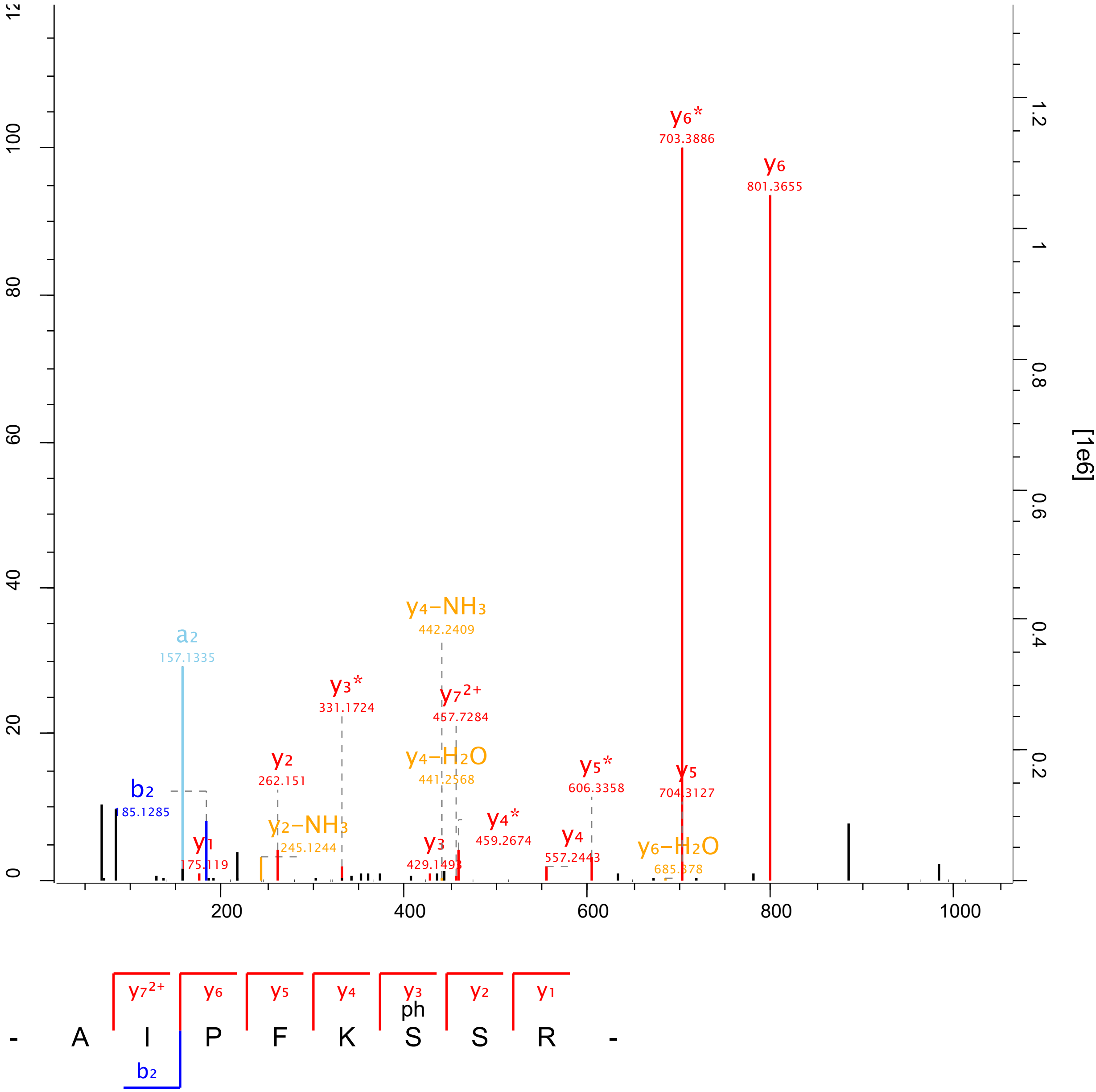

| Raw file | Scan | Method | Score | m/z |
|---|---|---|---|---|
| A17_00_2 | 5517 | FTMS; HCD | 116.86 | 512.54 |

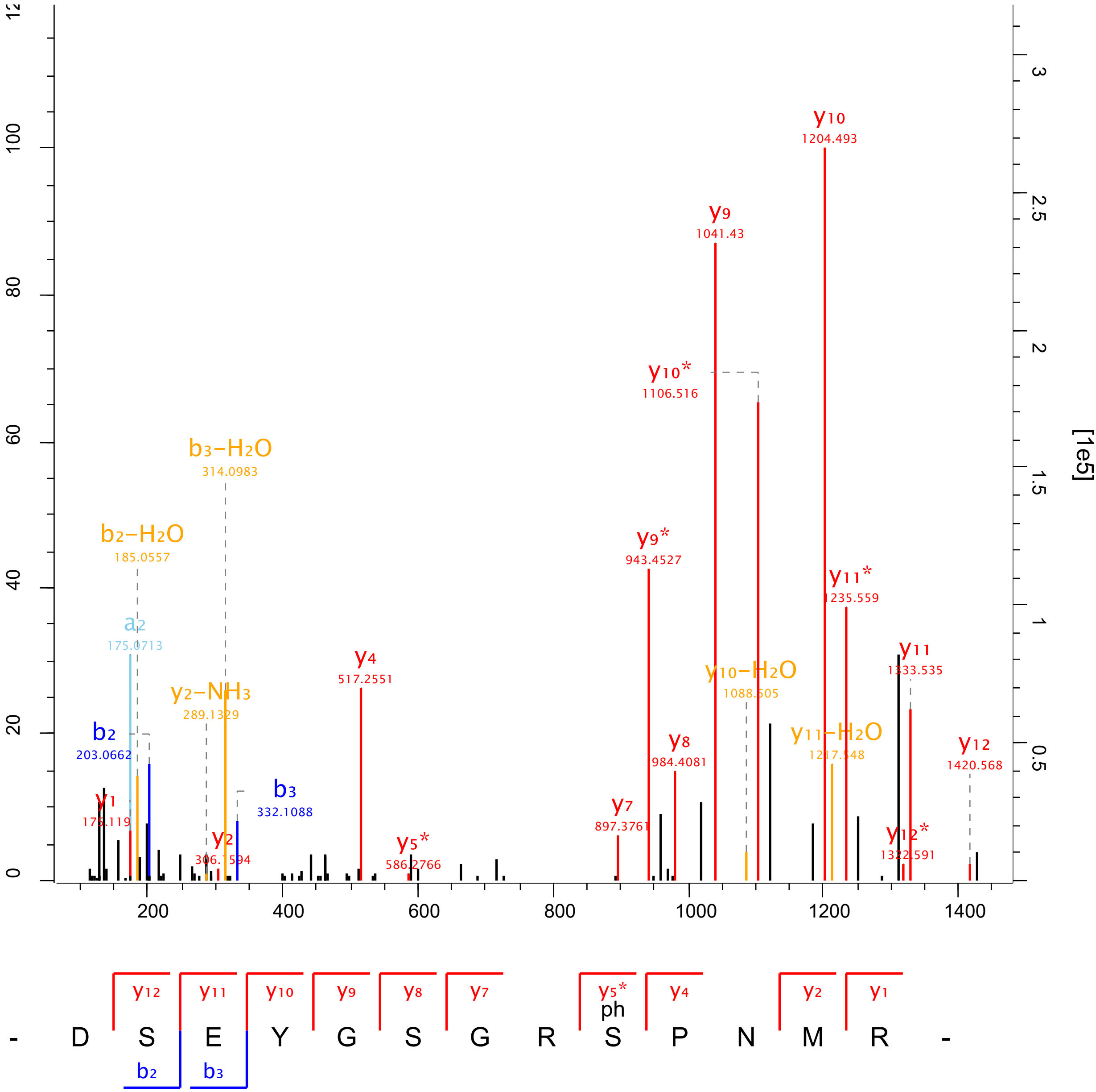

| Raw file | Scan | Method | Score | m/z |
|---|---|---|---|---|
| A17_00_2 | 5566 | FTMS; HCD | 73.93 | 748.29 |

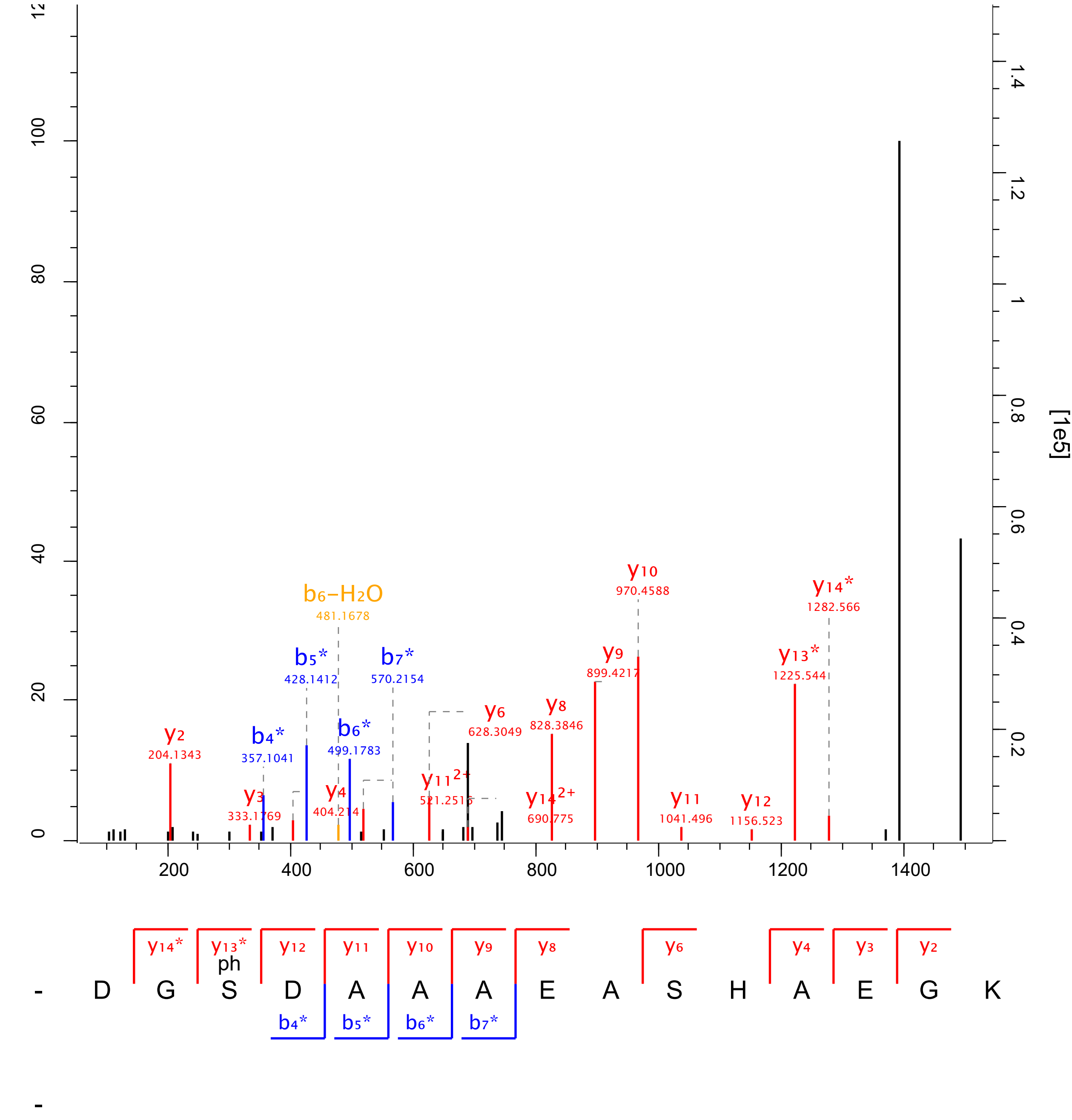

| Raw file | Scan | Method | Score | m/z |
|---|---|---|---|---|
| A17_00_2 | 5584 | FTMS; HCD | 150.46 | 500.75 |

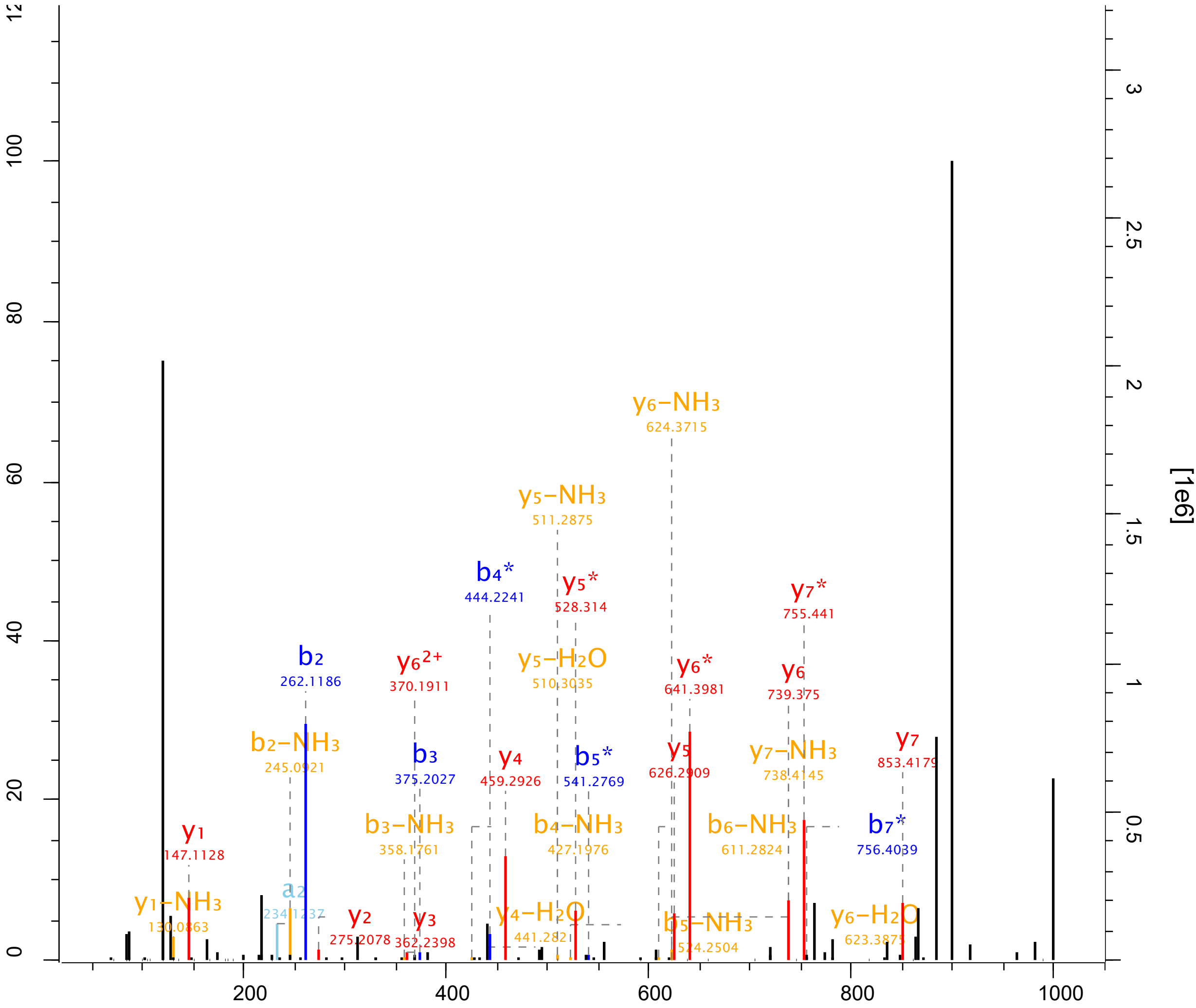


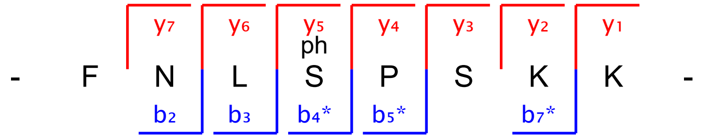

| Raw file | Scan | Method | Score | m/z |
|---|---|---|---|---|
| A17_00_2 | 5639 | FTMS; HCD | 94.49 | 775.31 |

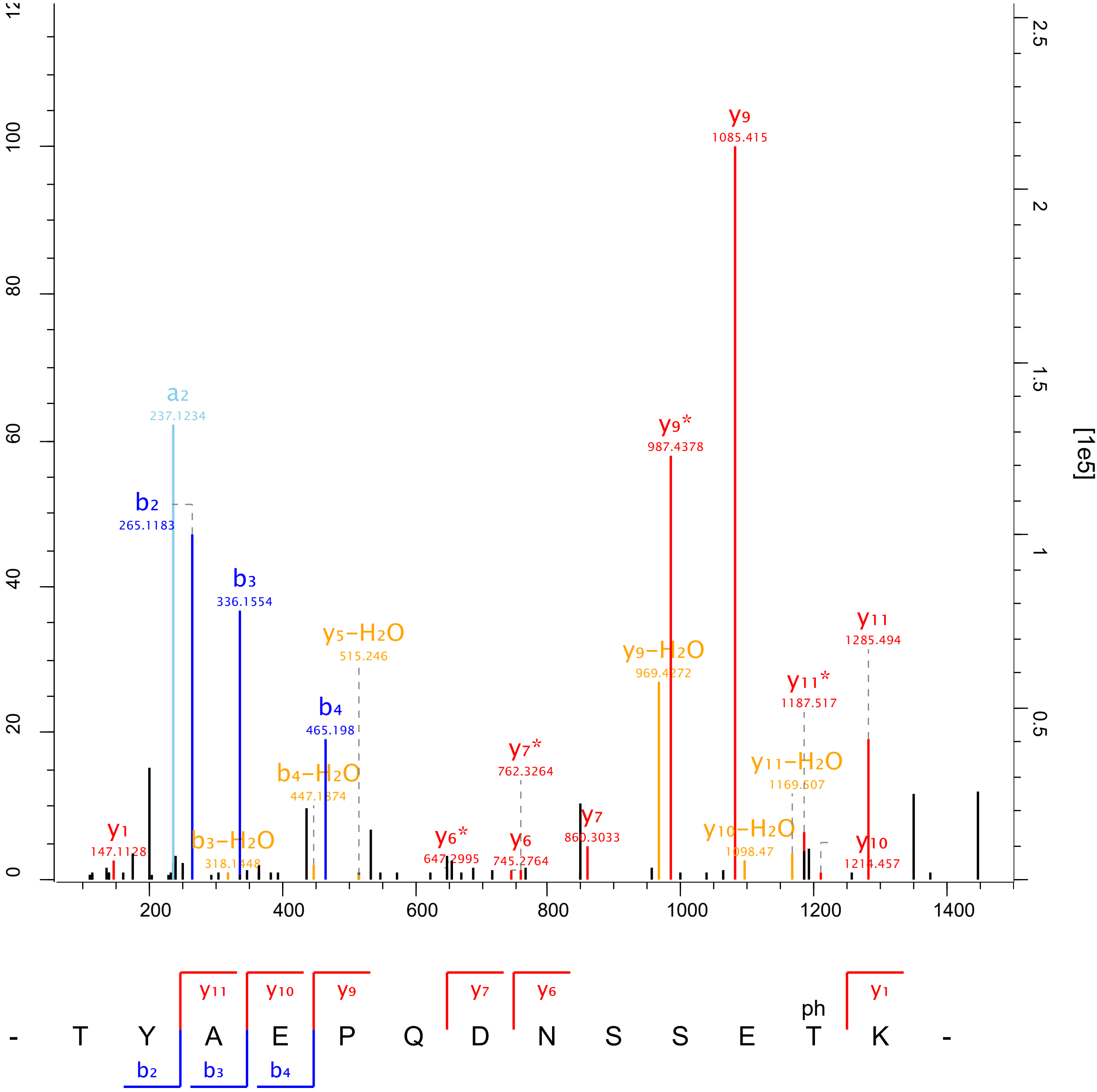

| Raw file | Scan | Method | Score | m/z |
| --- | --- | --- | --- | --- |
| A17_00_2 | 5682 | FTMS; HCD | 55.72 | 520.56 |

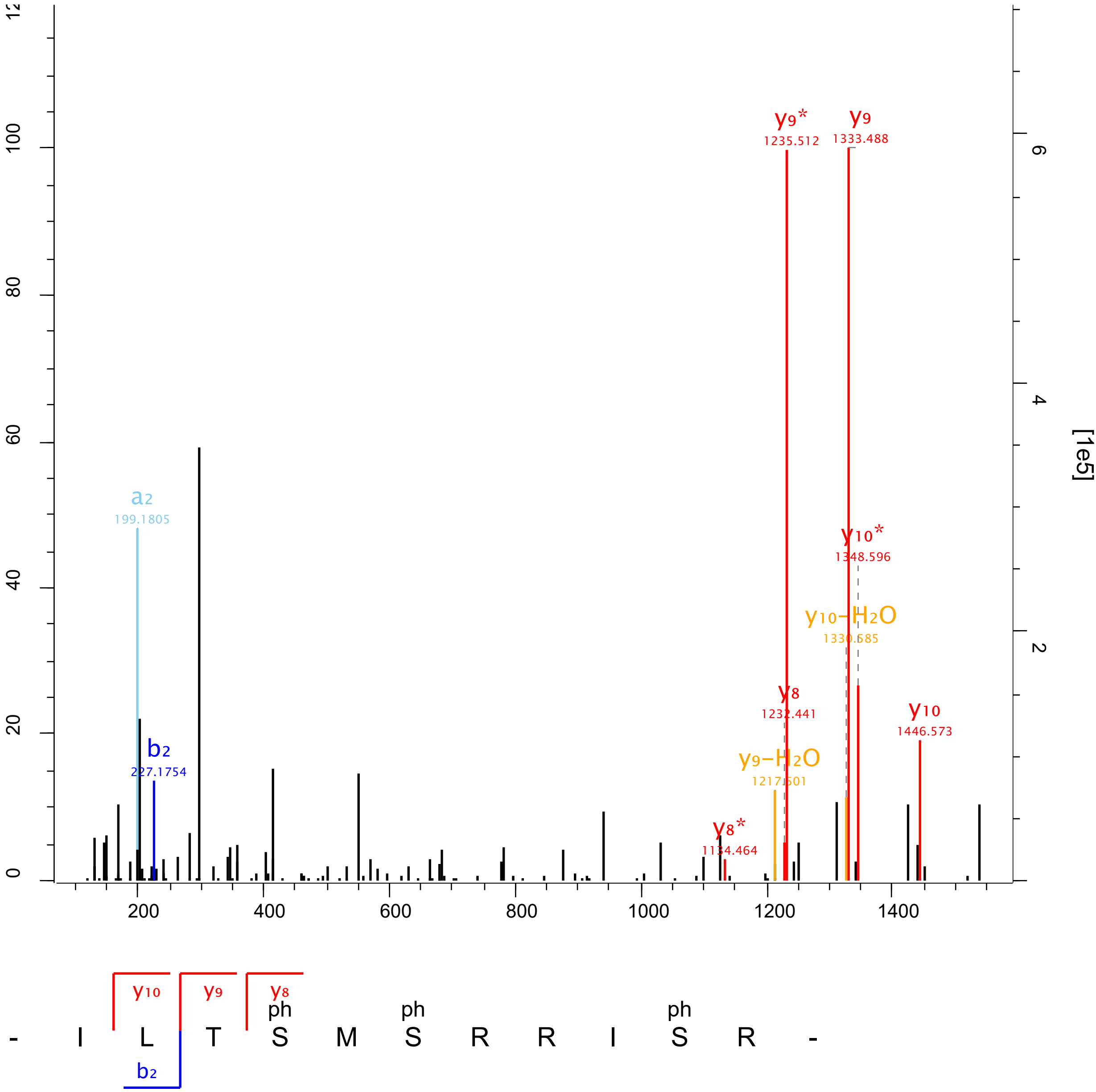

| Raw file | Scan | Method | Score | m/z |
|---|---|---|---|---|
| A17_00_2 | 5775 | FTMS; HCD | 218.95 | 920.85 |

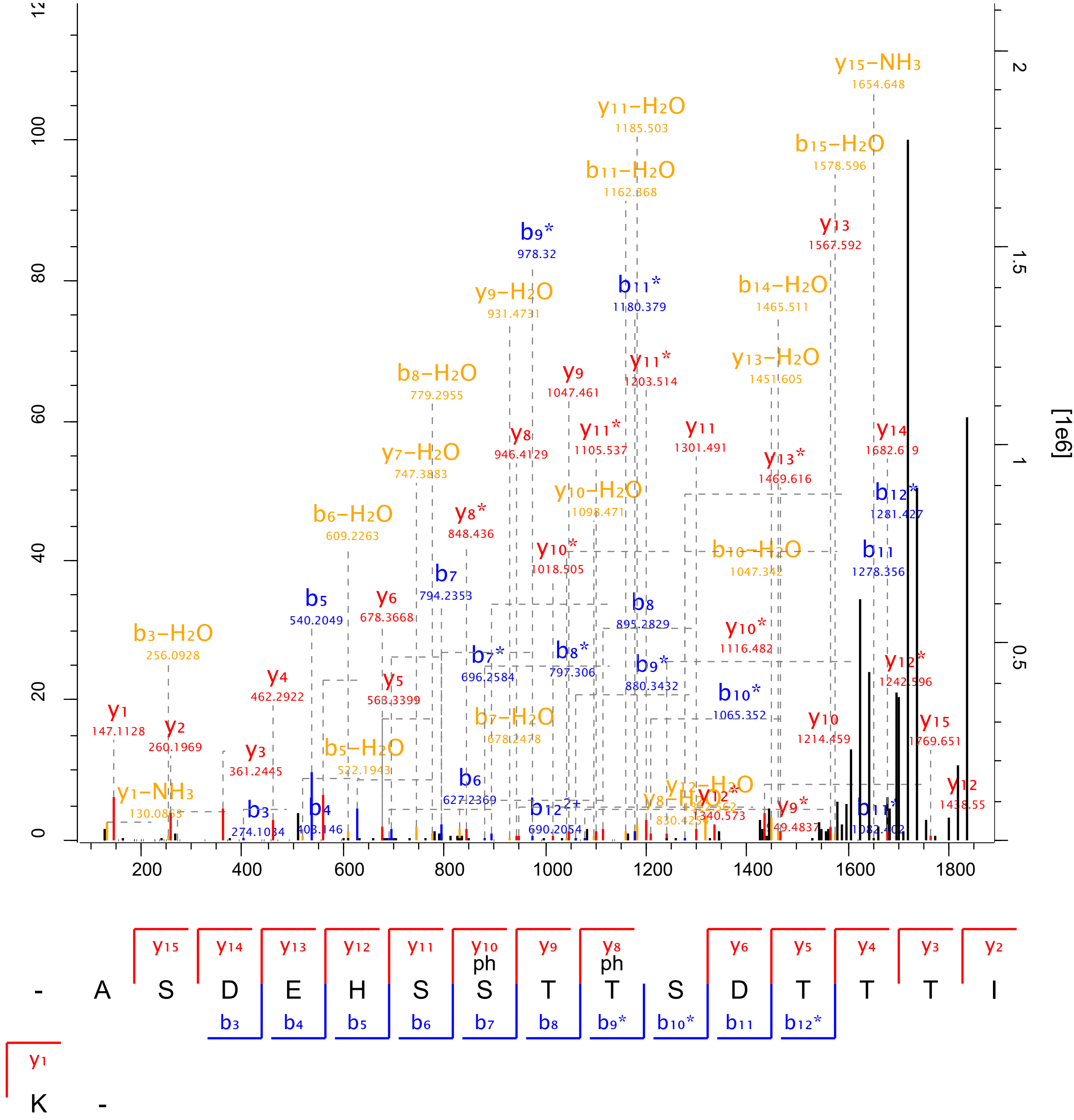

| Raw file | Scan | Method | Score | m/z |
|---|---|---|---|---|
| A17_00_2 | 5816 | FTMS; HCD | 133.25 | 689.78 |

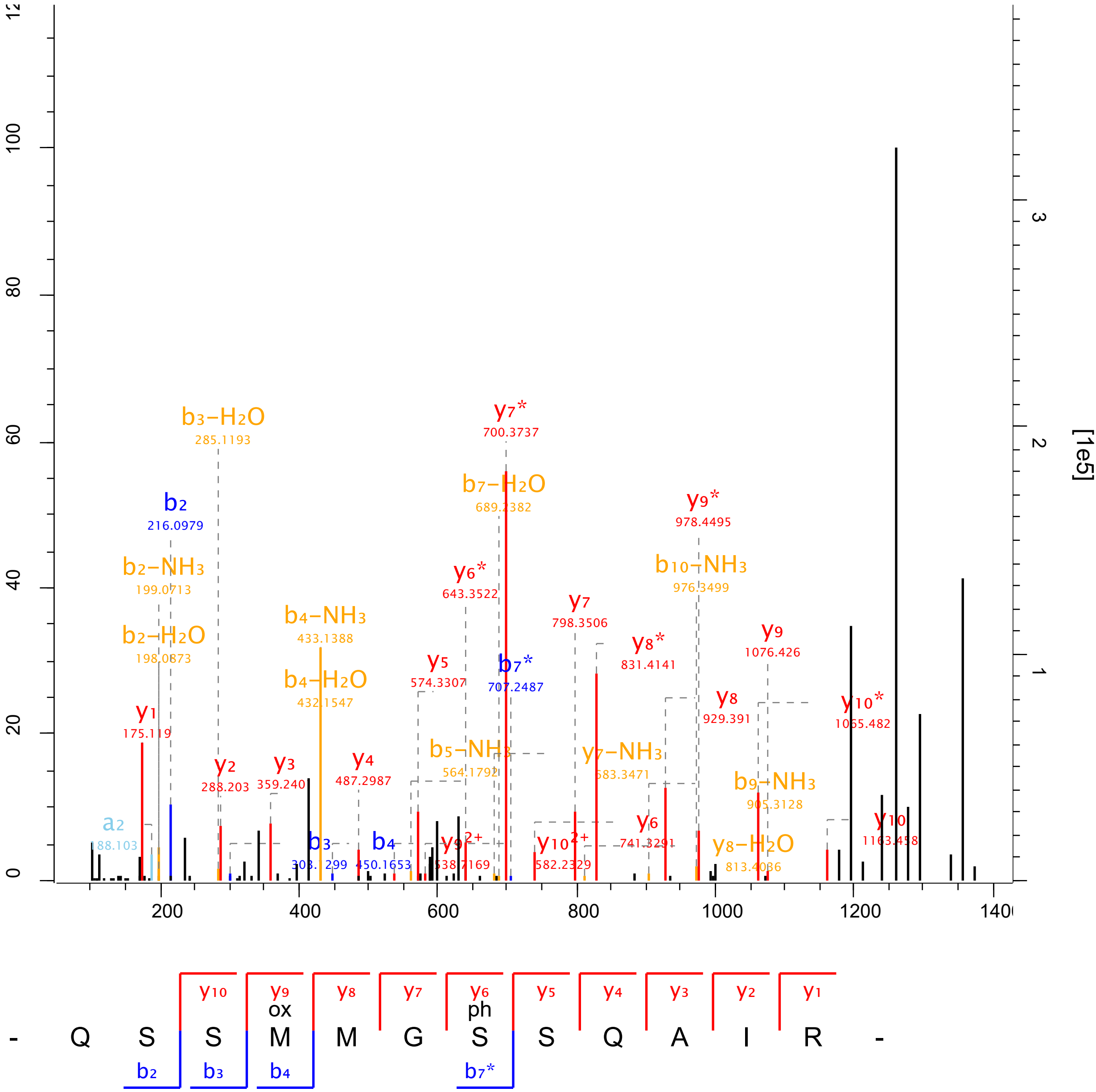

| Raw file | Scan | Method | Score | m/z |
| --- | --- | --- | --- | --- |
| A17_00_2 | 5905 | FTMS; HCD | 100.11 | 636.28 |

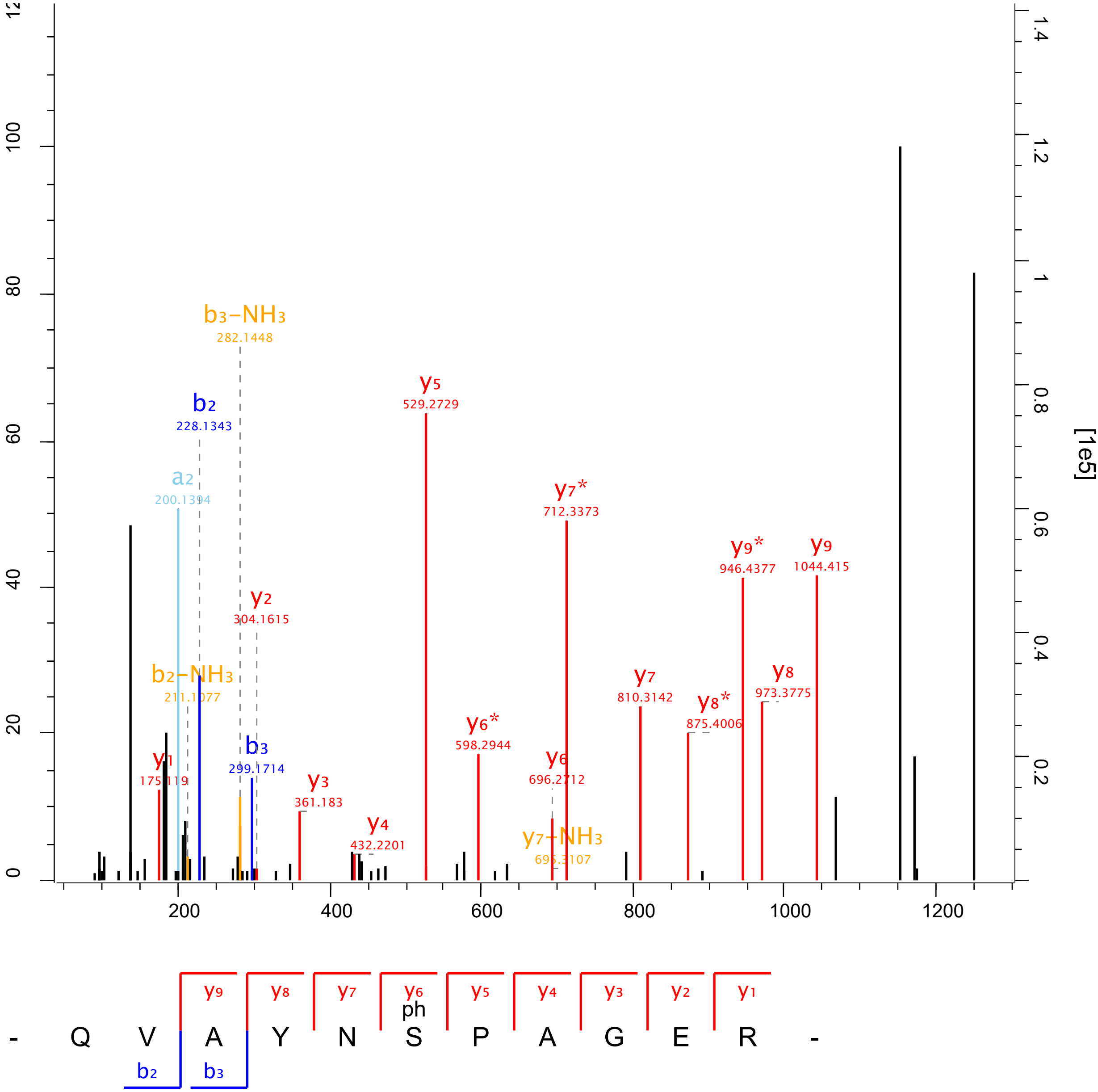

| Raw file | Scan | Method | Score | m/z |
|---|---|---|---|---|
| A17_00_2 | 5949 | FTMS; HCD | 171.71 | 504.88 |

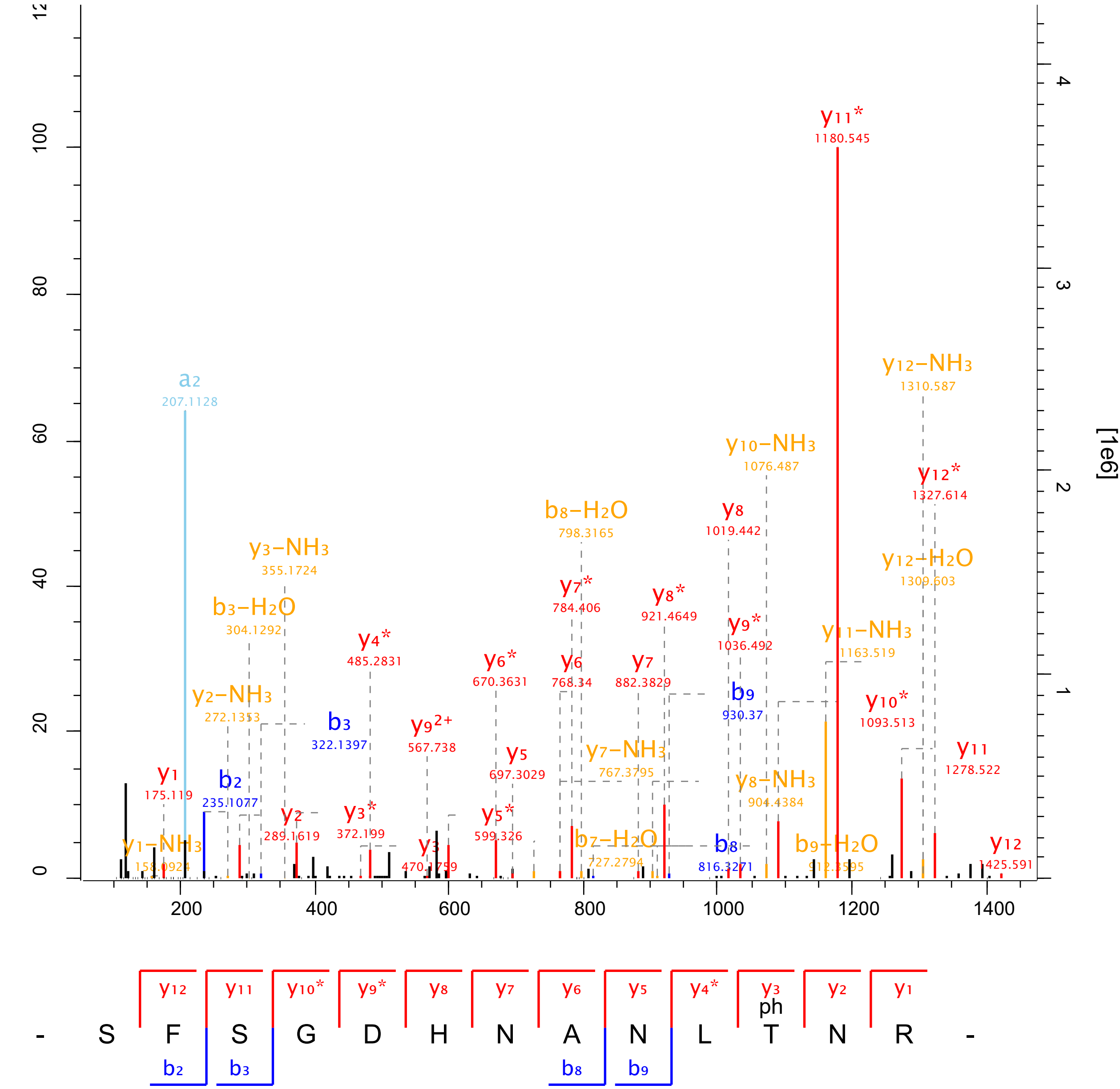

| Raw file | Scan | Method | Score | m/z |
|---|---|---|---|---|
| A17_00_2 | 6037 | FTMS; HCD | 400.69 | 1070.39 |

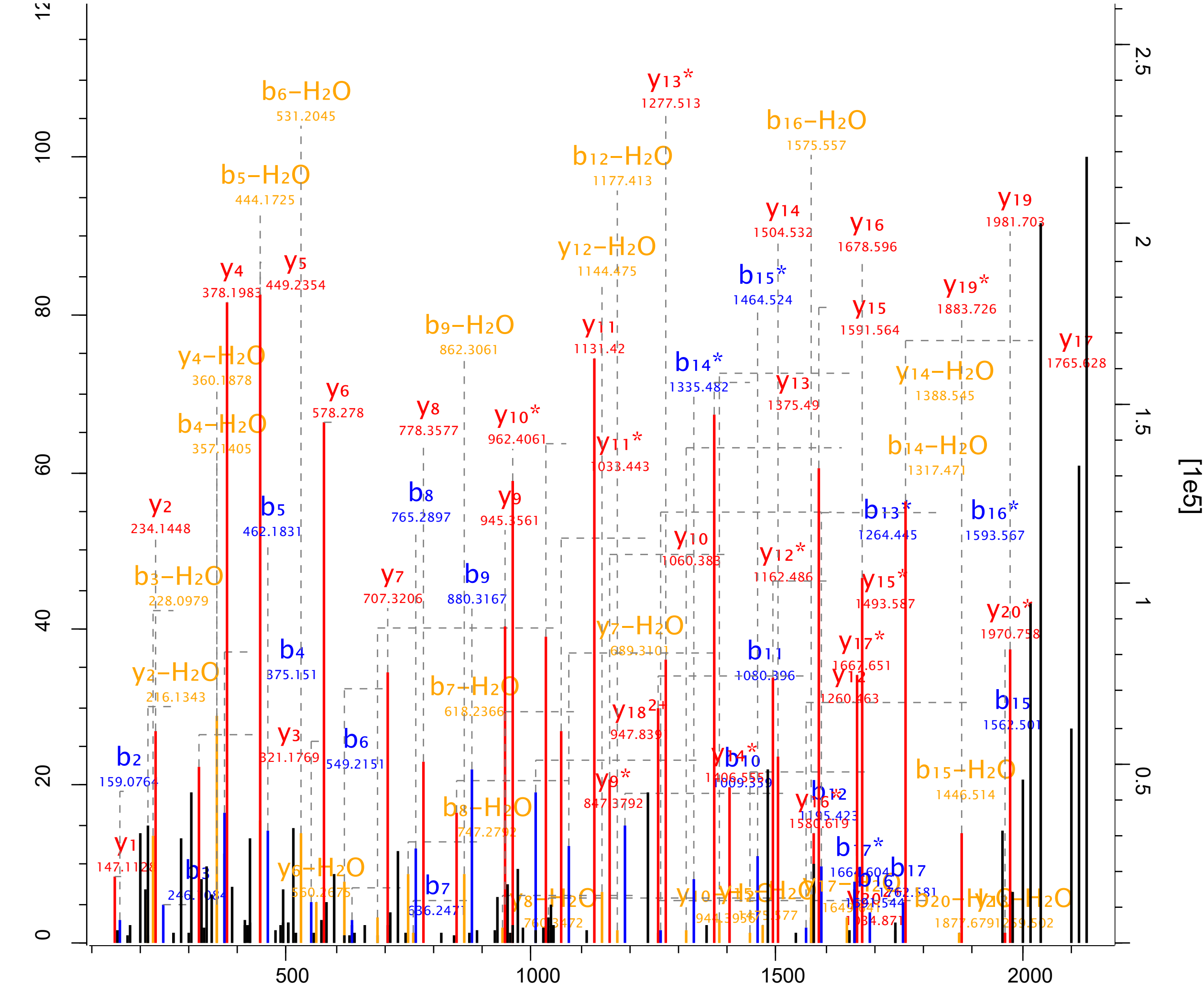


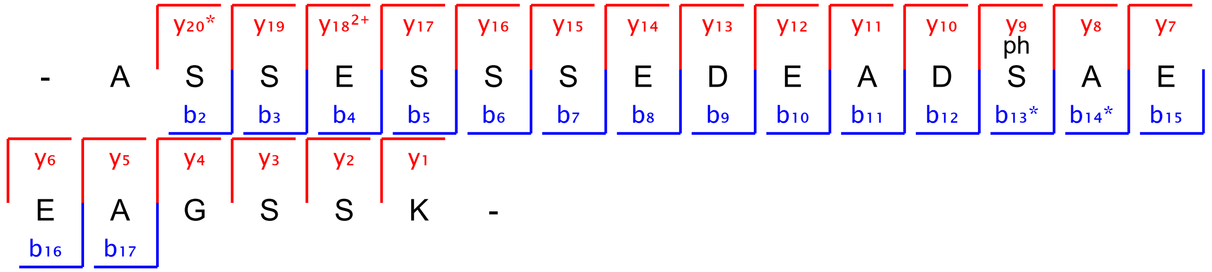

| Raw file | Scan | Method | Score | m/z |
| --- | --- | --- | --- | --- |
| A17_00_2 | 6069 | FTMS; HCD | 105.03 | 574.73 |

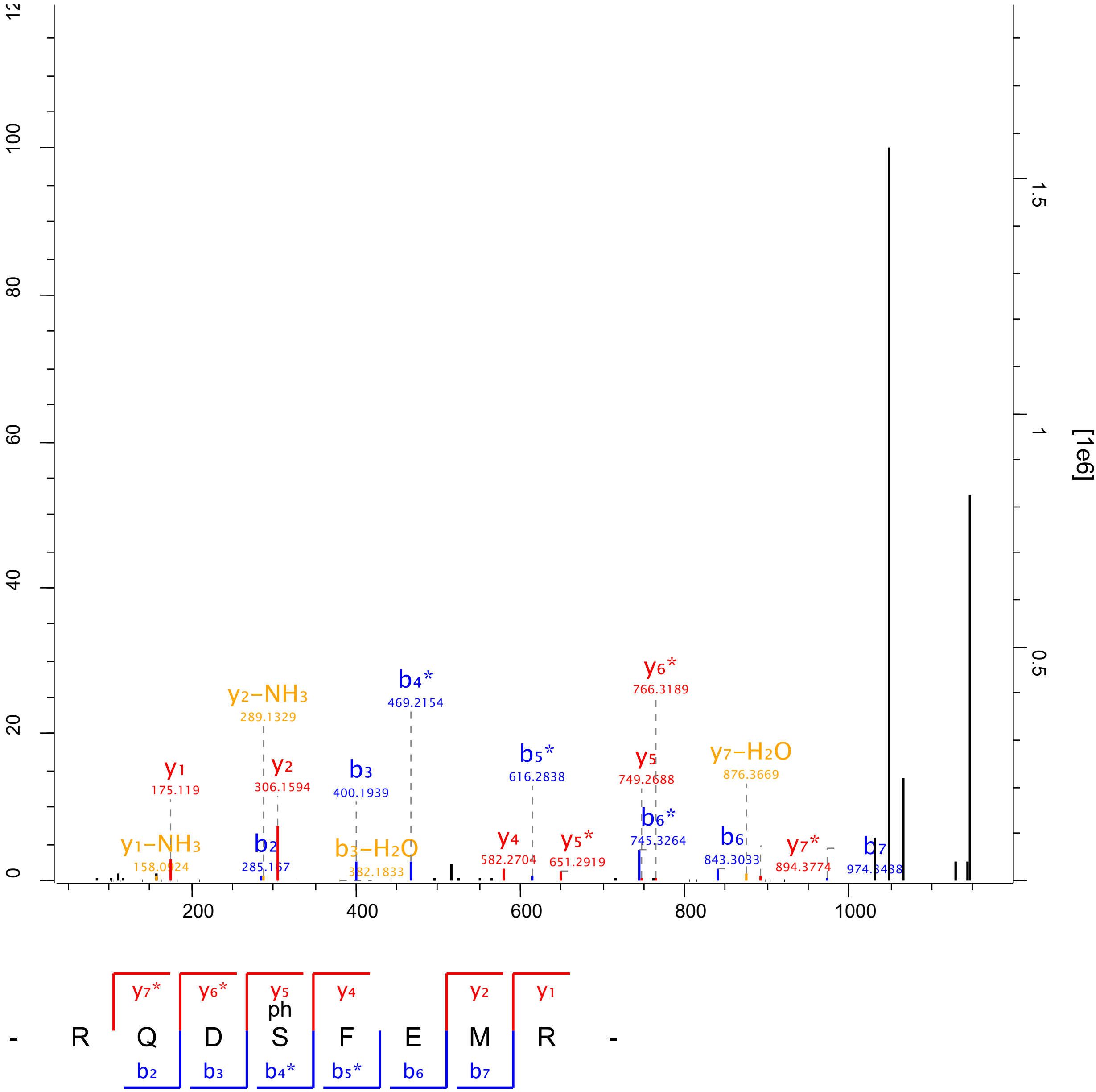

| Raw file | Scan | Method | Score | m/z |
| --- | --- | --- | --- | --- |
| A17_00_2 | 6095 | FTMS; HCD | 113.22 | 495.21 |

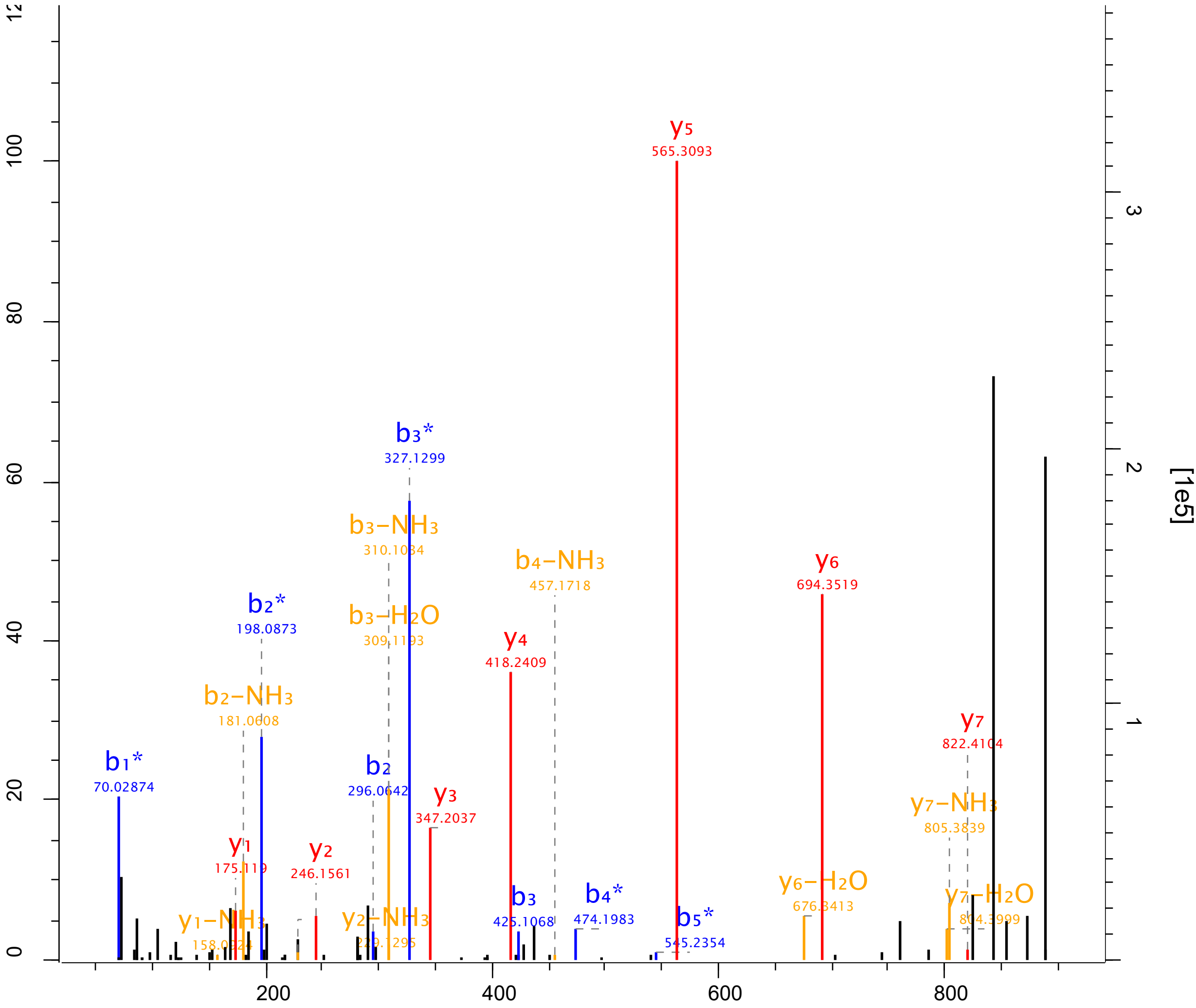


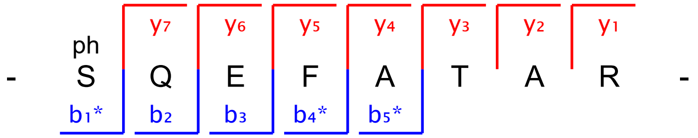

| Raw file | Scan | Method | Score | m/z |
|---|---|---|---|---|
| A17_00_2 | 6164 | FTMS; HCD | 41.09 | 532.57 |

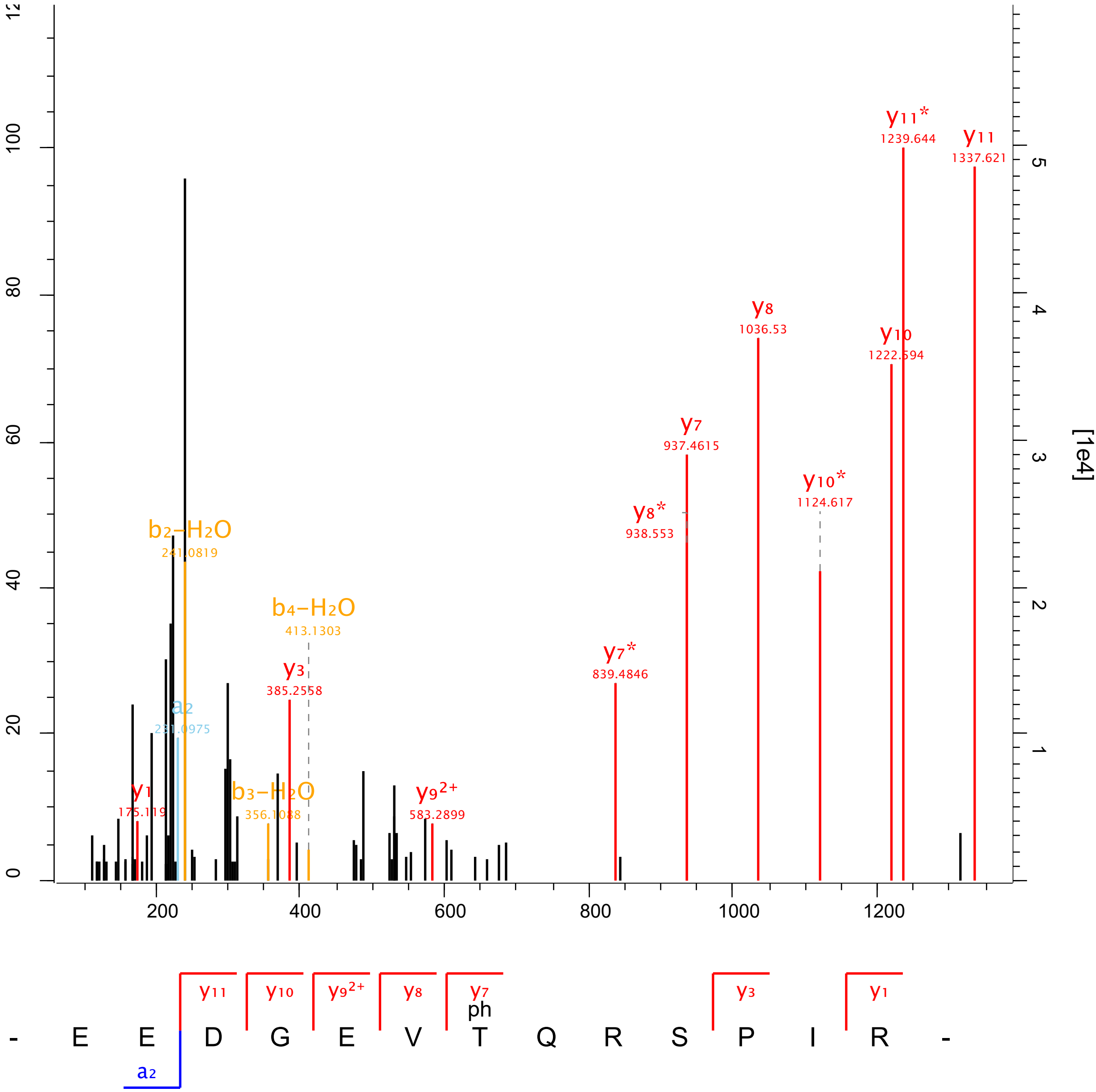

| Raw file | Scan | Method | Score | m/z |
|---|---|---|---|---|
| A17_00_2 | 6210 | FTMS; HCD | 63.68 | 462.2 |

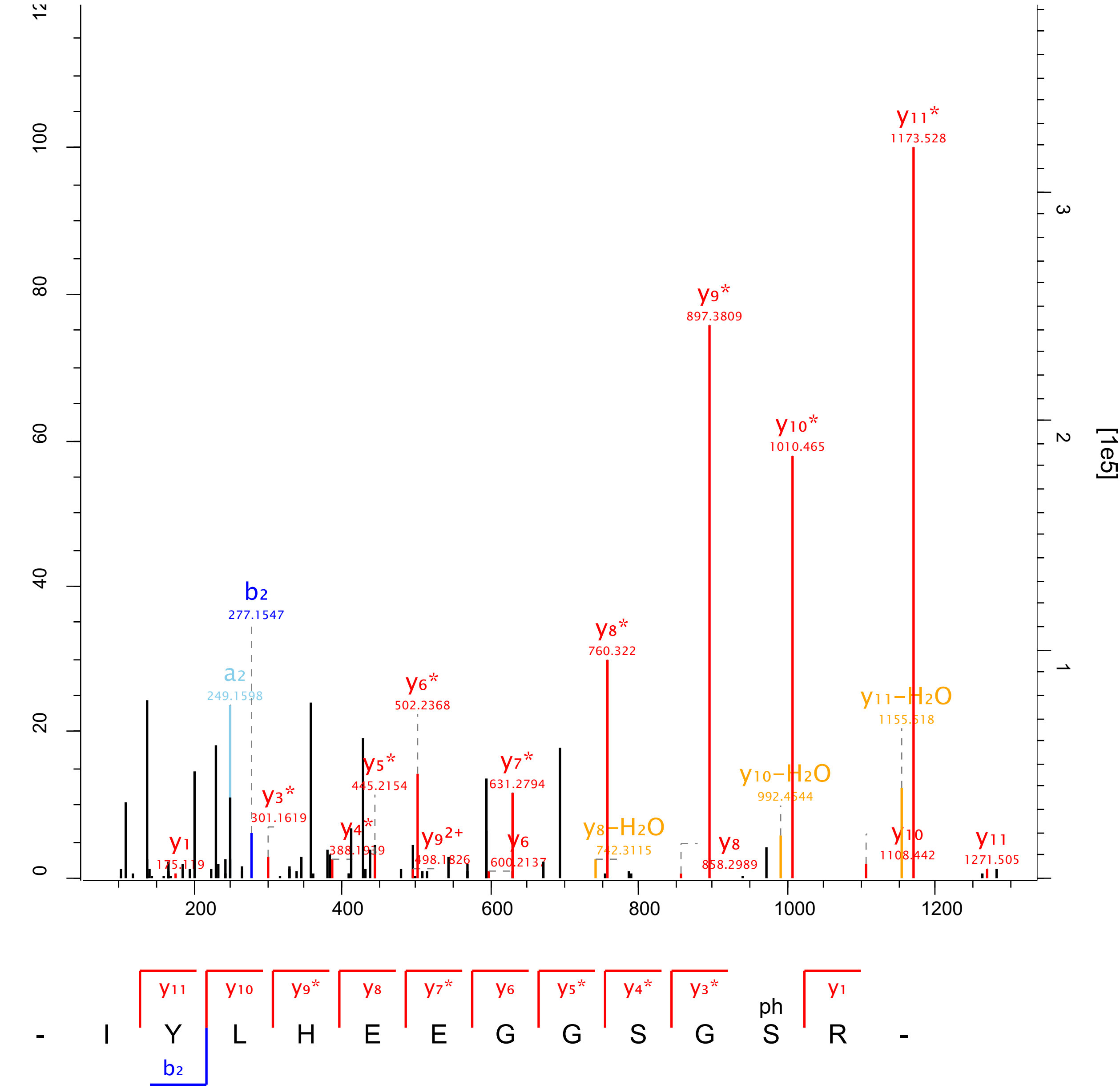

| Raw file | Scan | Method | Score | m/z |
| --- | --- | --- | --- | --- |
| A17_00_2 | 6260 | FTMS; HCD | 99.75 | 478.22 |

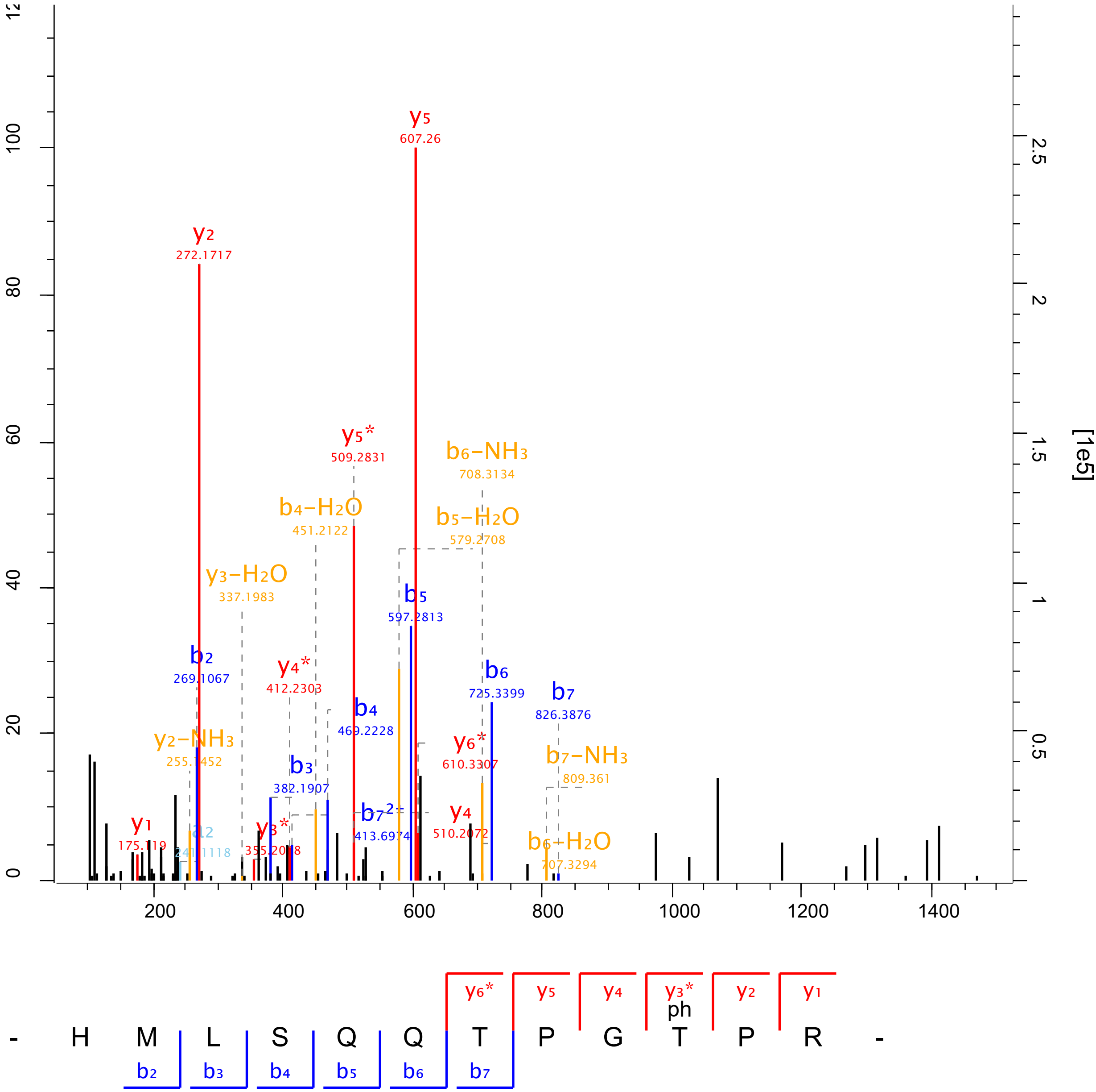

| Raw file | Scan | Method | Score | m/z |
|---|---|---|---|---|
| A17_00_2 | 6306 | FTMS; HCD | 92.19 | 546.24 |

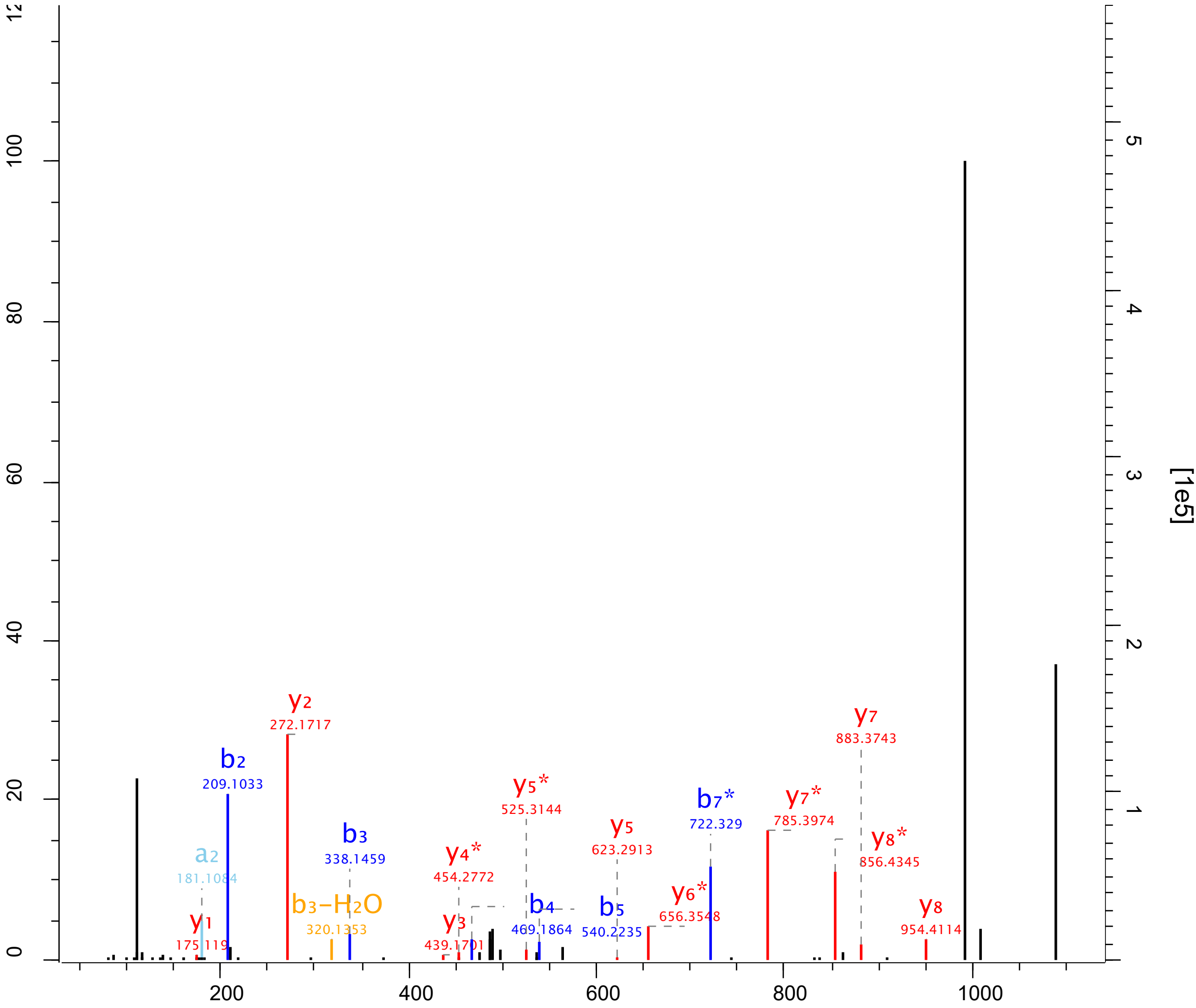


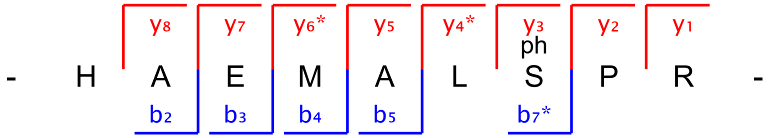

| Raw file | Scan | Method | Score | m/z |
|---|---|---|---|---|
| A17_00_2 | 6349 | FTMS; HCD | 56.12 | 847.35 |

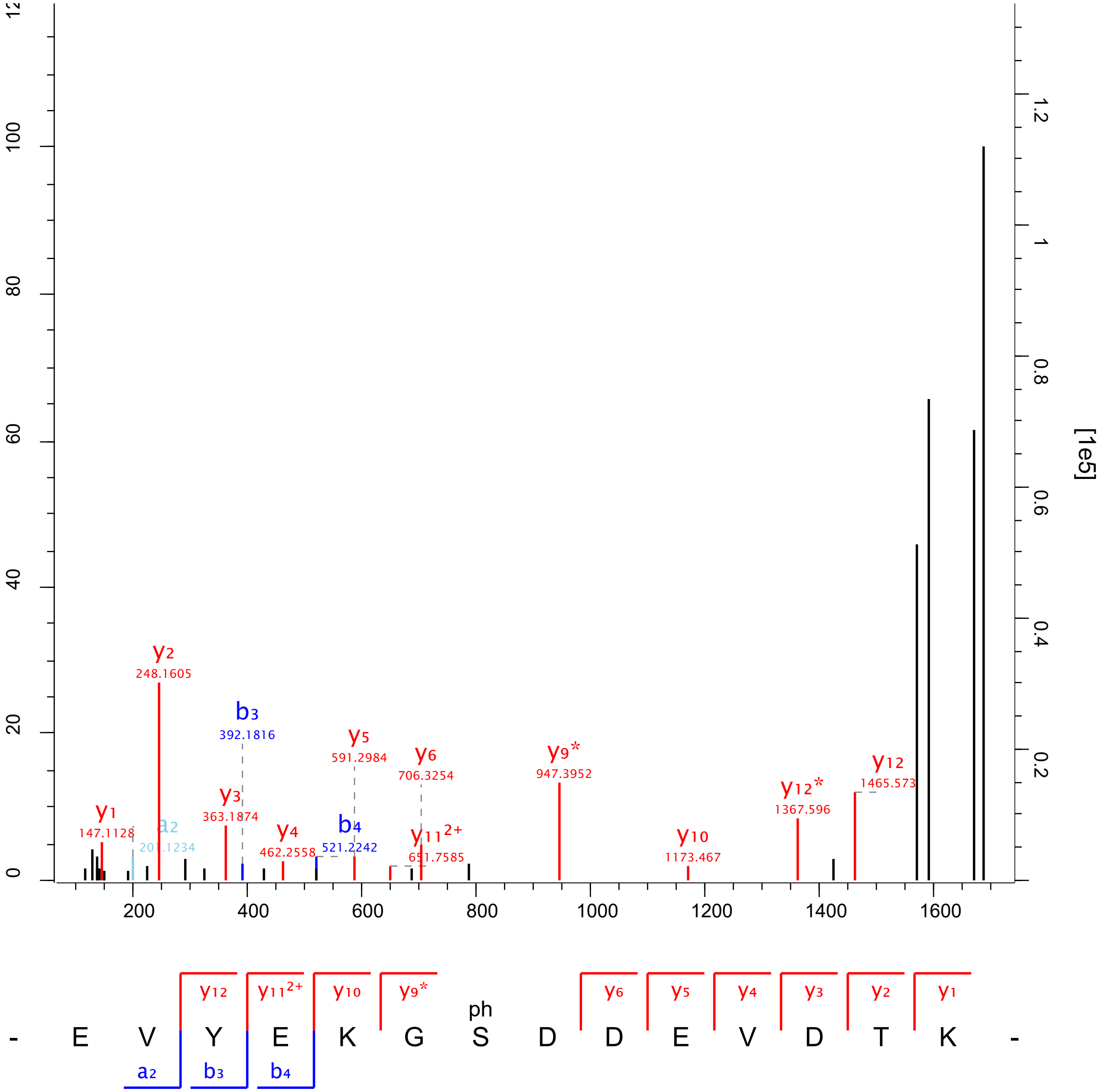

| Raw file | Scan | Method | Score | m/z |
| --- | --- | --- | --- | --- |
| A17_00_2 | 6362 | FTMS; HCD | 102.87 | 538.72 |

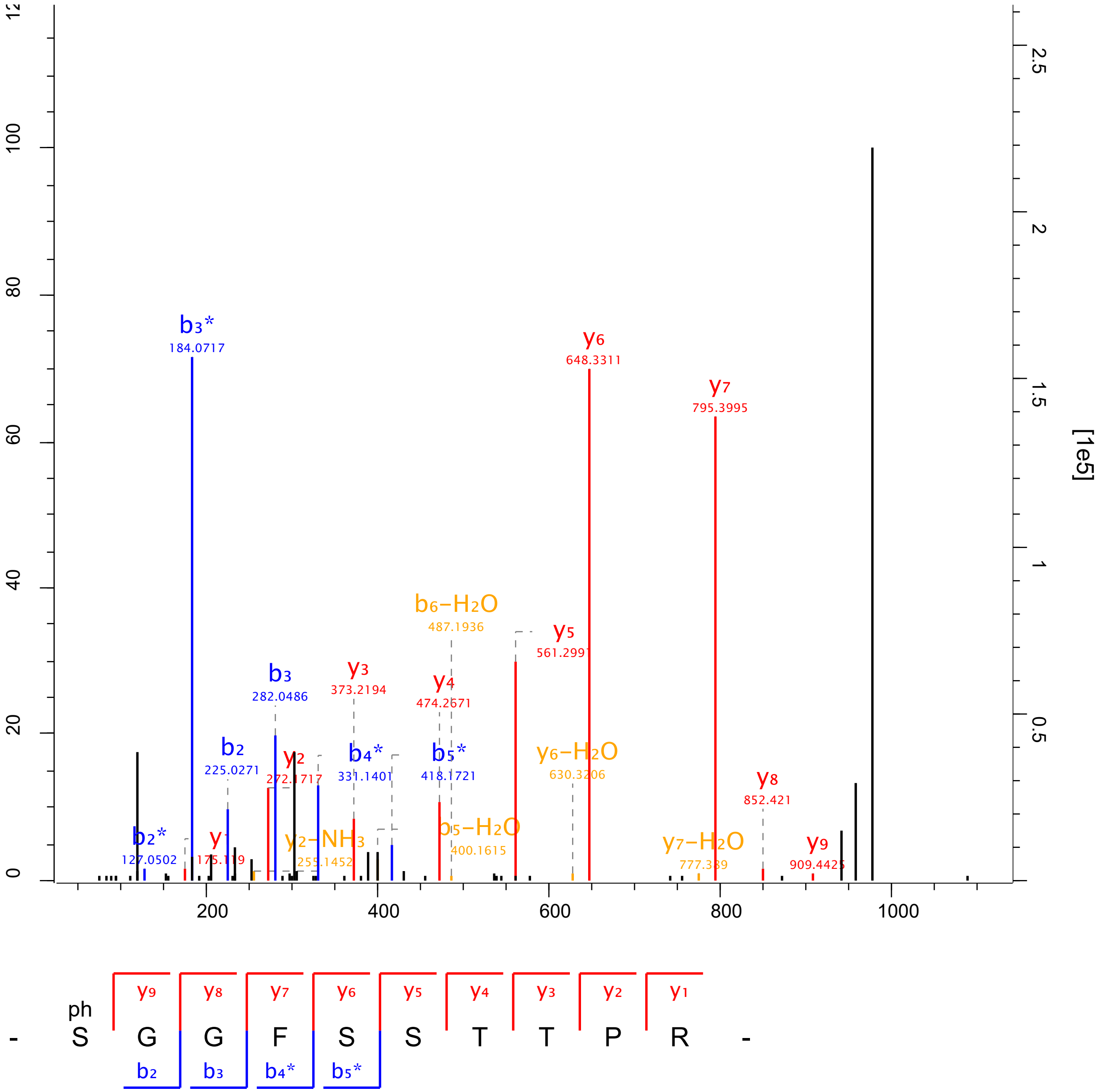

| Raw file | Scan | Method | Score | m/z |
| --- | --- | --- | --- | --- |
| A17_00_2 | 6586 | FTMS; HCD | 191.44 | 671.8 |

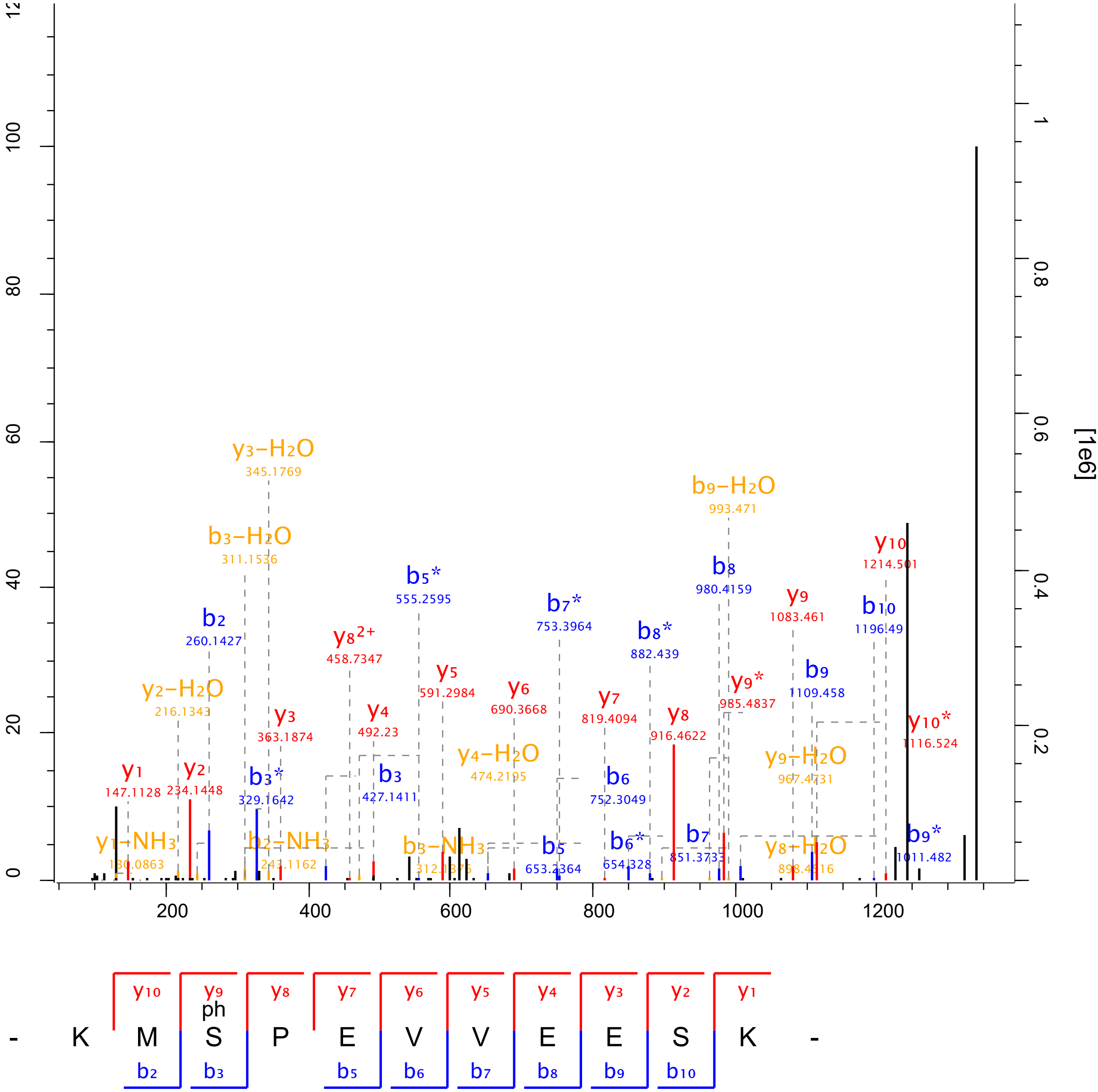

| Raw file | Scan | Method | Score | m/z |
|---|---|---|---|---|
| A17_00_2 | 6662 | FTMS; HCD | 48.62 | 638.63 |

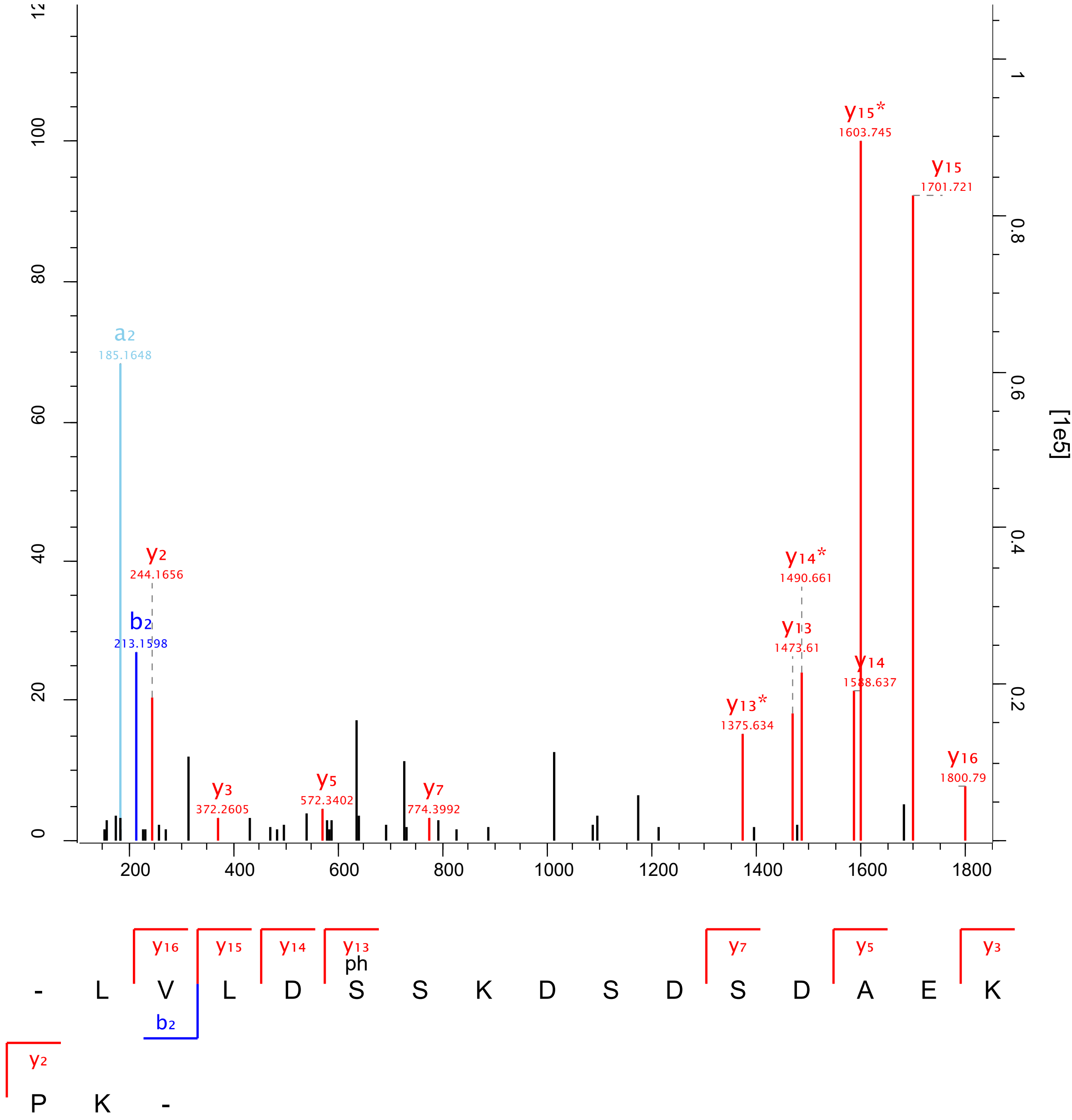

| Raw file | Scan | Method | Score | m/z |
|---|---|---|---|---|
| A17_00_2 | 7009 | FTMS; HCD | 52.77 | 613.94 |

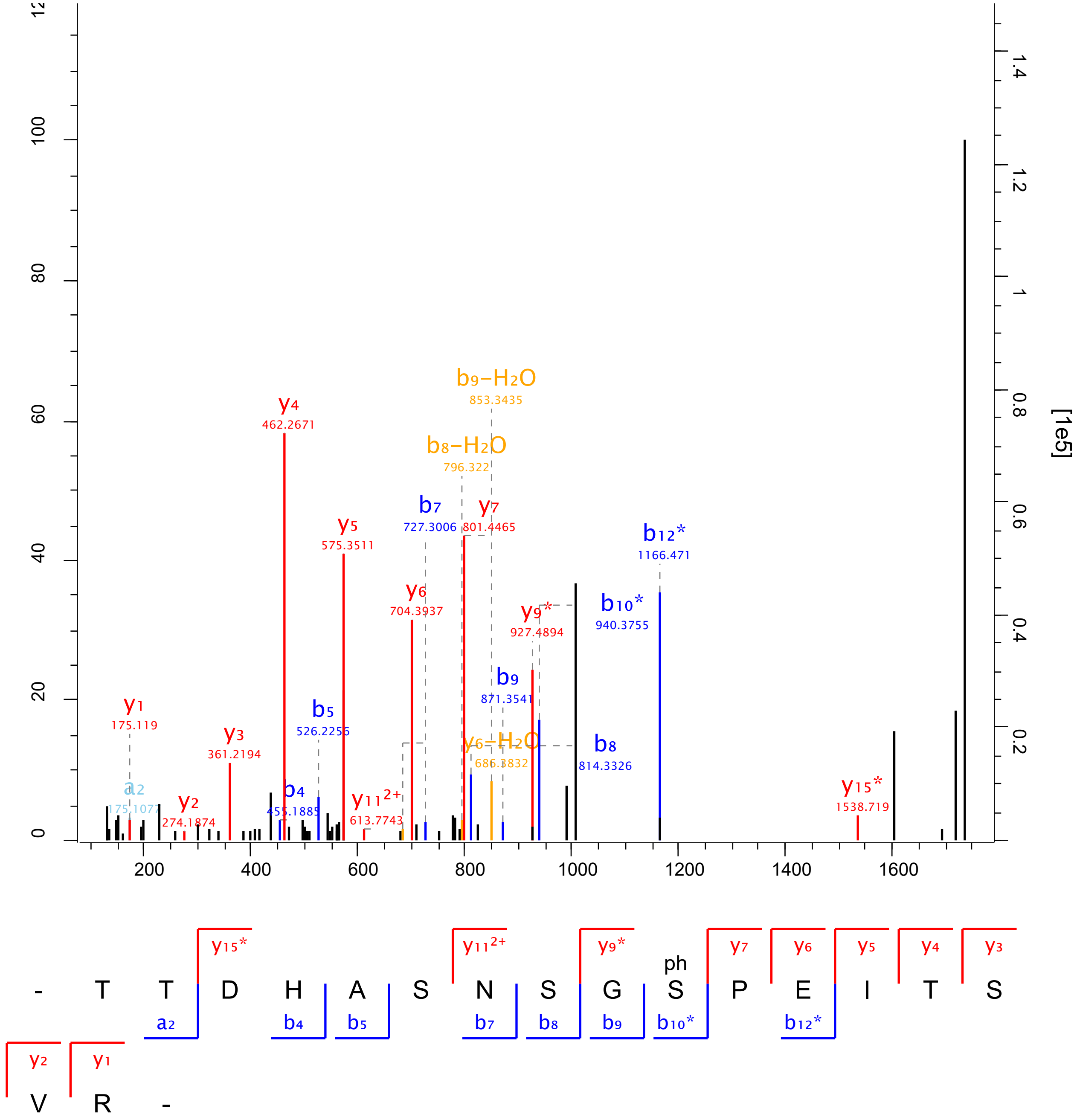

| Raw file | Scan | Method | Score | m/z |
| --- | --- | --- | --- | --- |
| A17_00_2 | 7097 | FTMS; HCD | 68.83 | 824 |

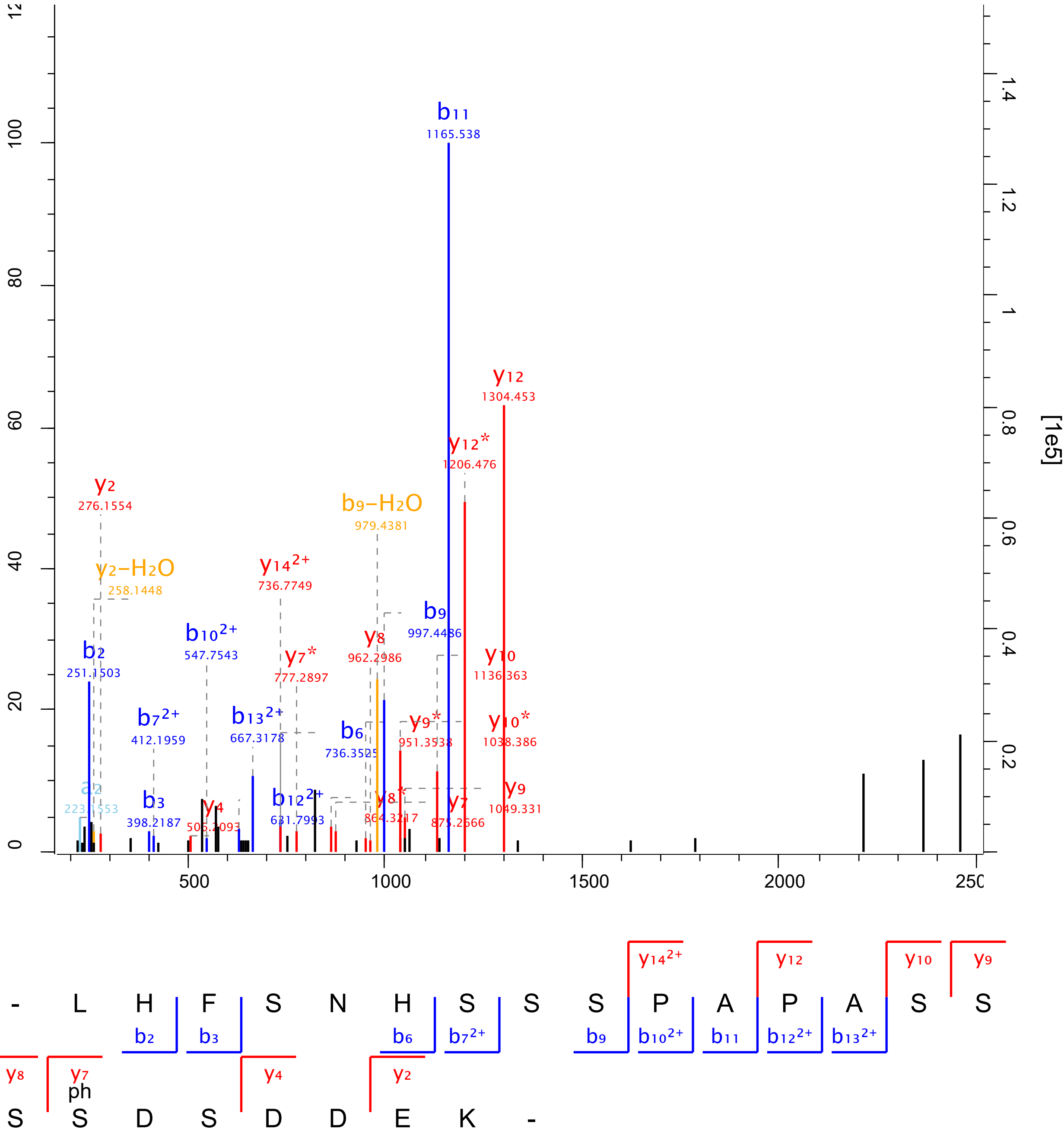

| Raw file | Scan | Method | Score | m/z |
| --- | --- | --- | --- | --- |
| A17_00_2 | 7129 | FTMS; HCD | 88.09 | 1011.84 |

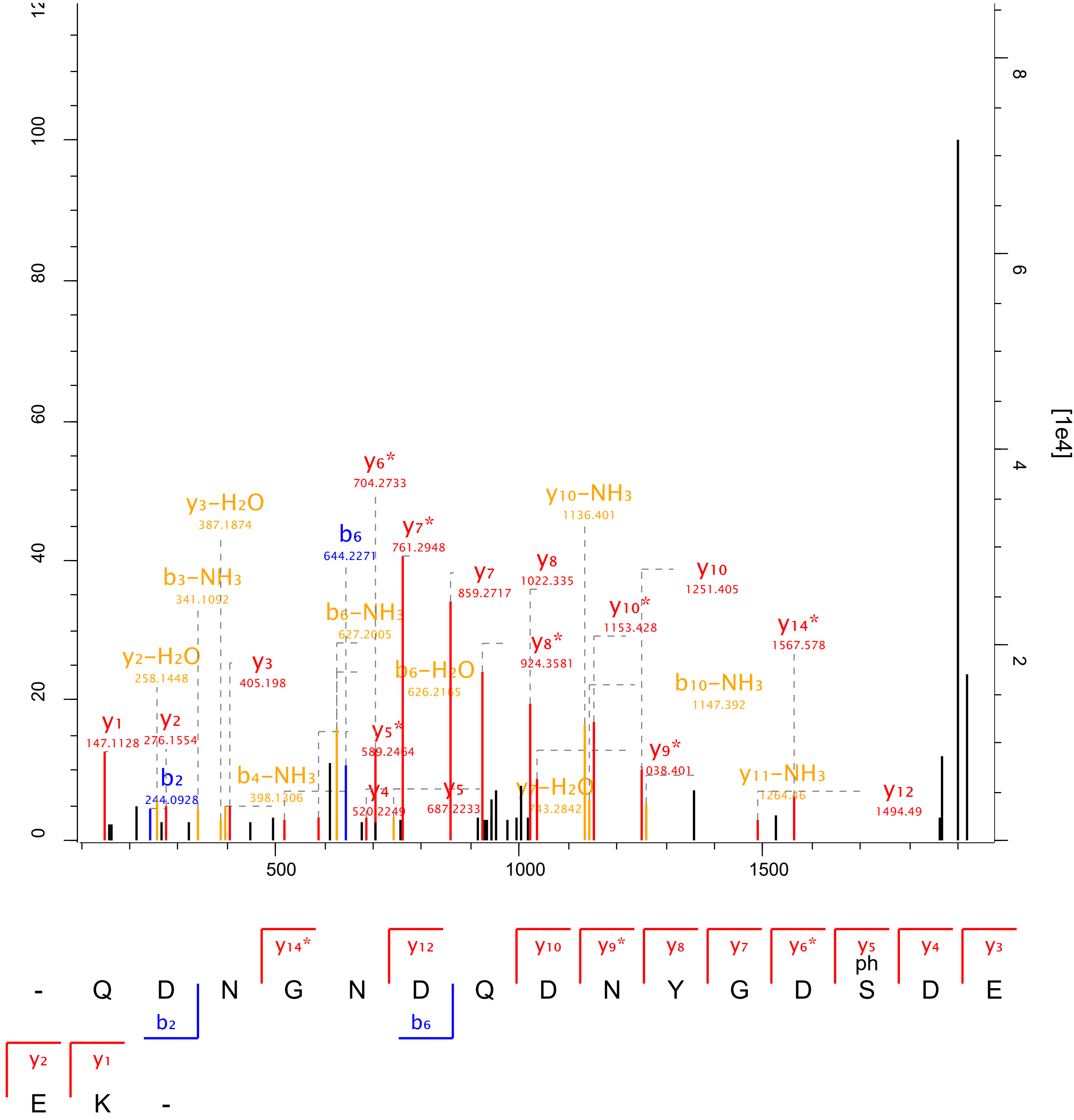

| Raw file | Scan | Method | Score | m/z |
|---|---|---|---|---|
| A17_00_2 | 7301 | FTMS; HCD | 52.5 | 672.28 |

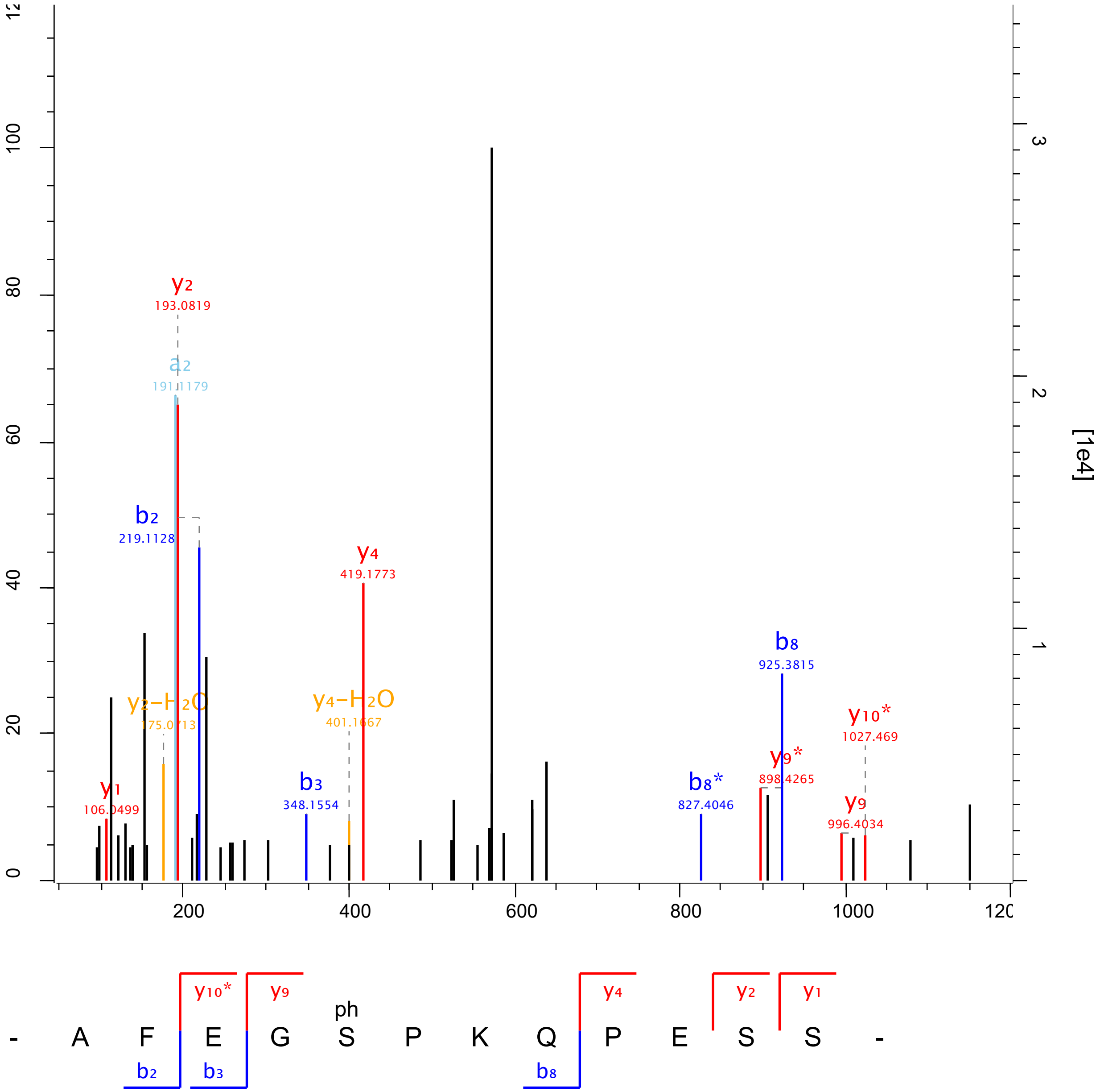

| Raw file | Scan | Method | Score | m/z |
|---|---|---|---|---|
| A17_00_2 | 7371 | FTMS; HCD | 67.14 | 485.87 |

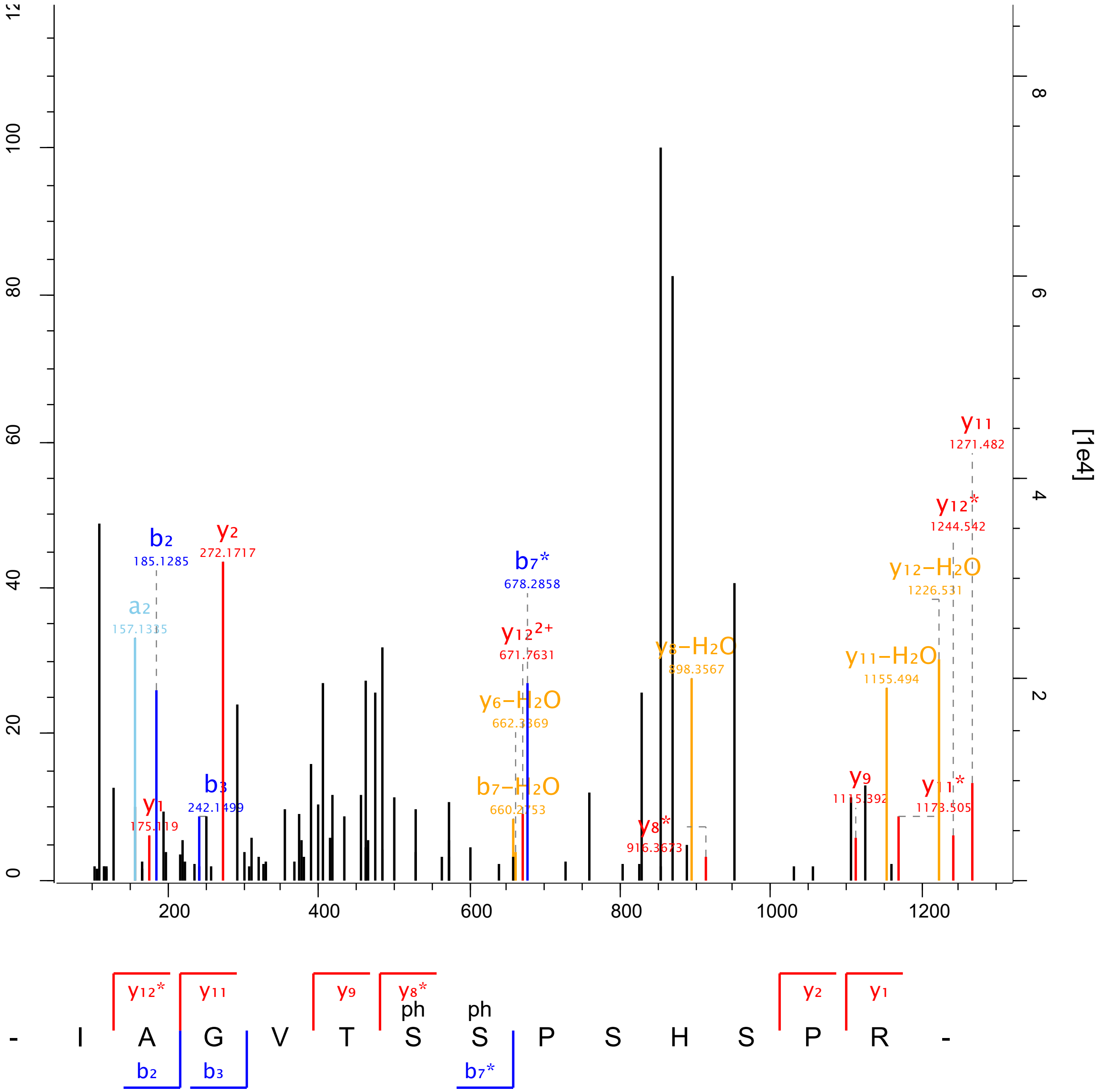

| Raw file | Scan | Method | Score | m/z |
|---|---|---|---|---|
| A17_00_2 | 7397 | FTMS; HCD | 59.23 | 634.28 |

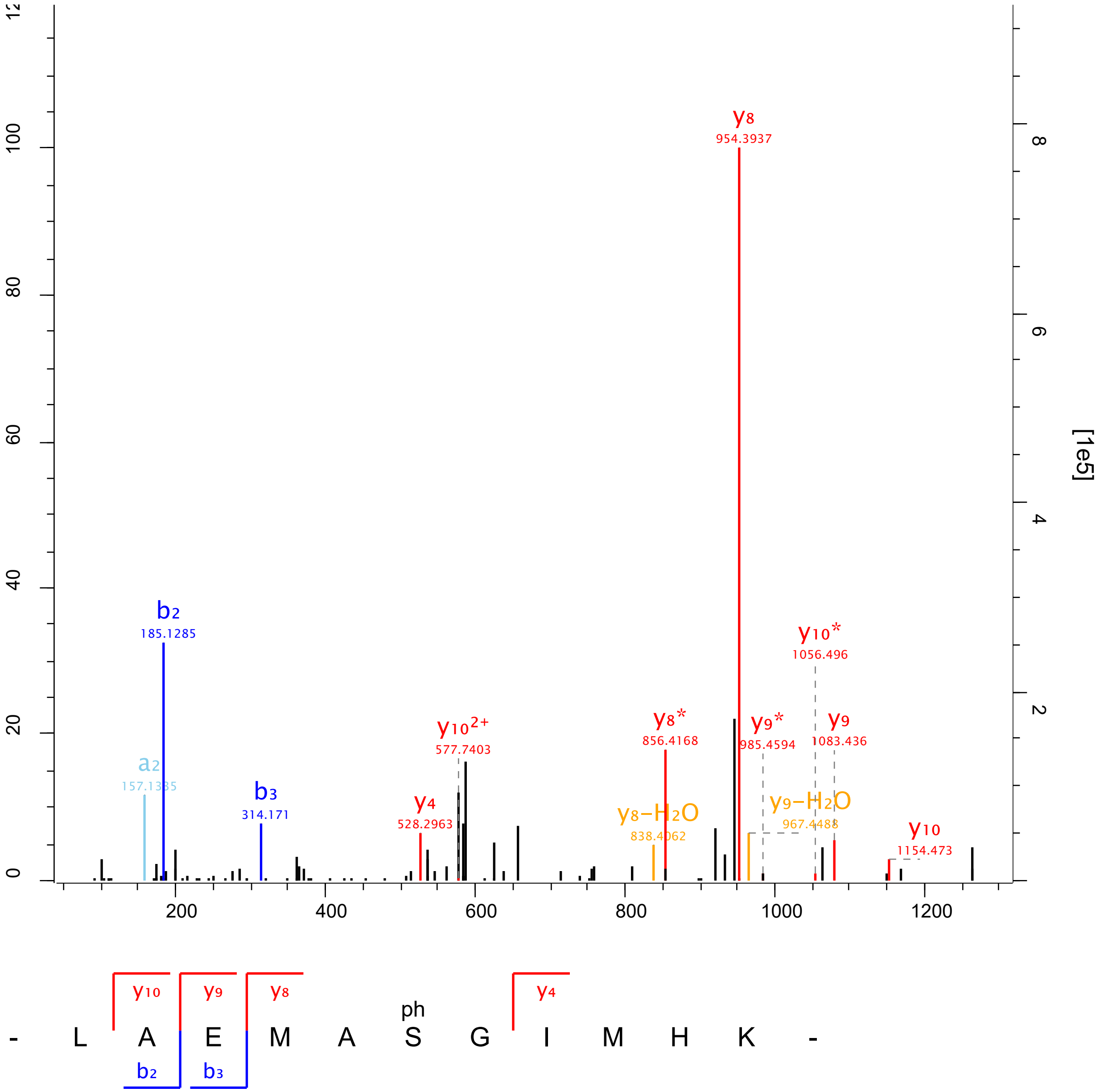

| Raw file | Scan | Method | Score | m/z |
|---|---|---|---|---|
| A17_00_2 | 7460 | FTMS; HCD | 107.09 | 622.75 |

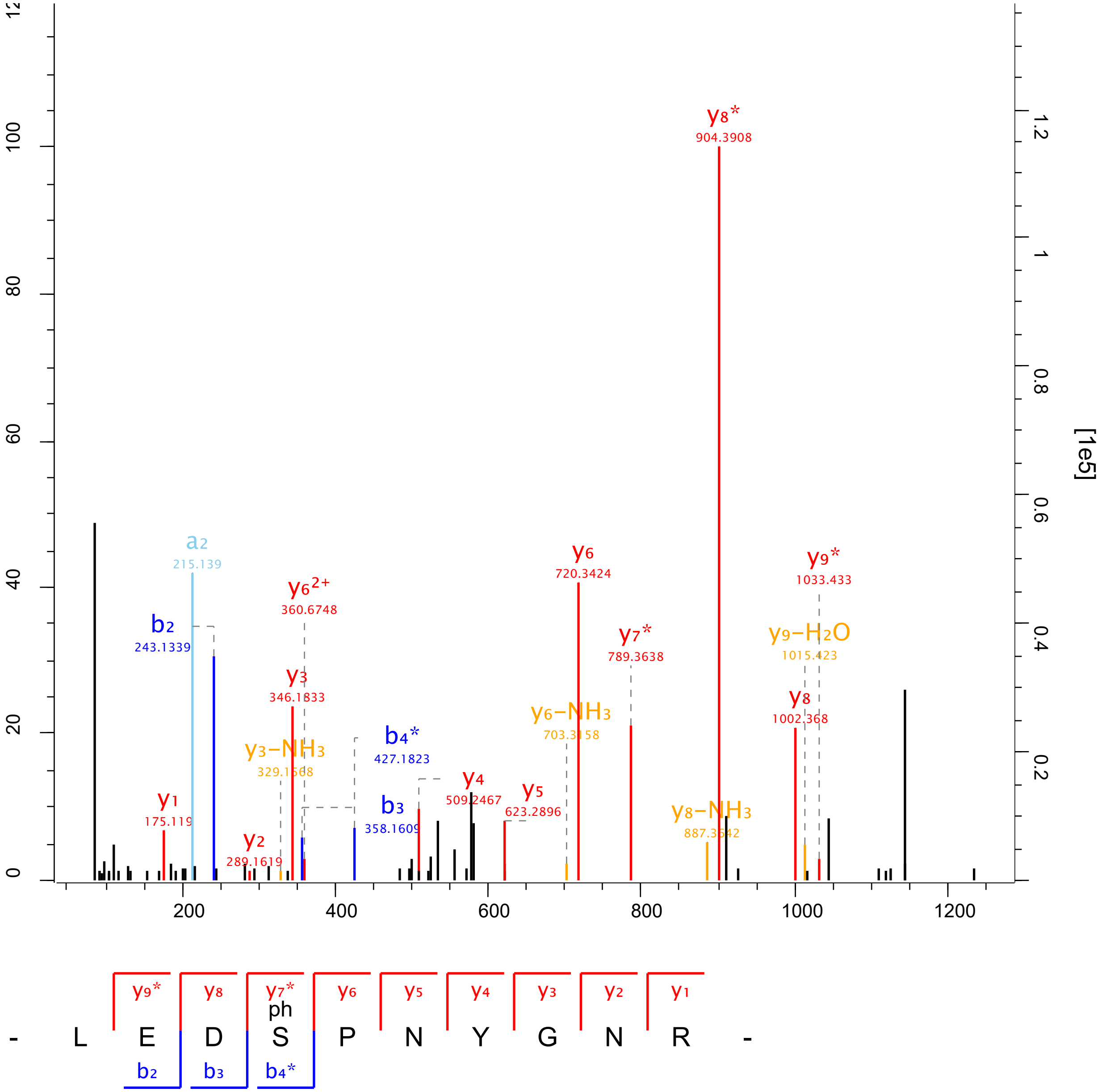

| Raw file | Scan | Method | Score | m/z |
|---|---|---|---|---|
| A17_00_2 | 7546 | FTMS; HCD | 114.96 | 541.89 |

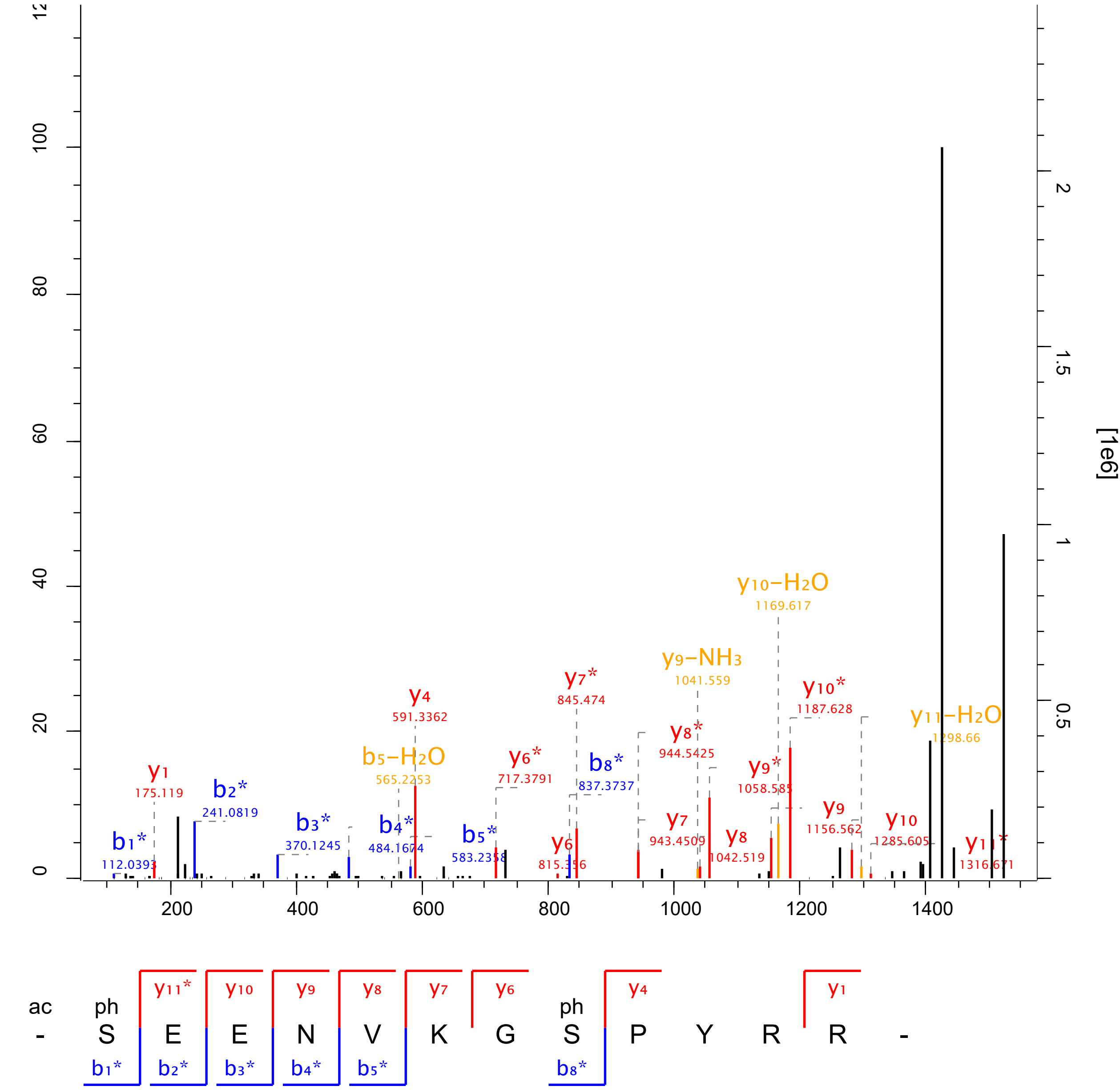

| Raw file | Scan | Method | Score | m/z |
|---|---|---|---|---|
| A17_00_2 | 7662 | FTMS; HCD | 55.72 | 651.81 |

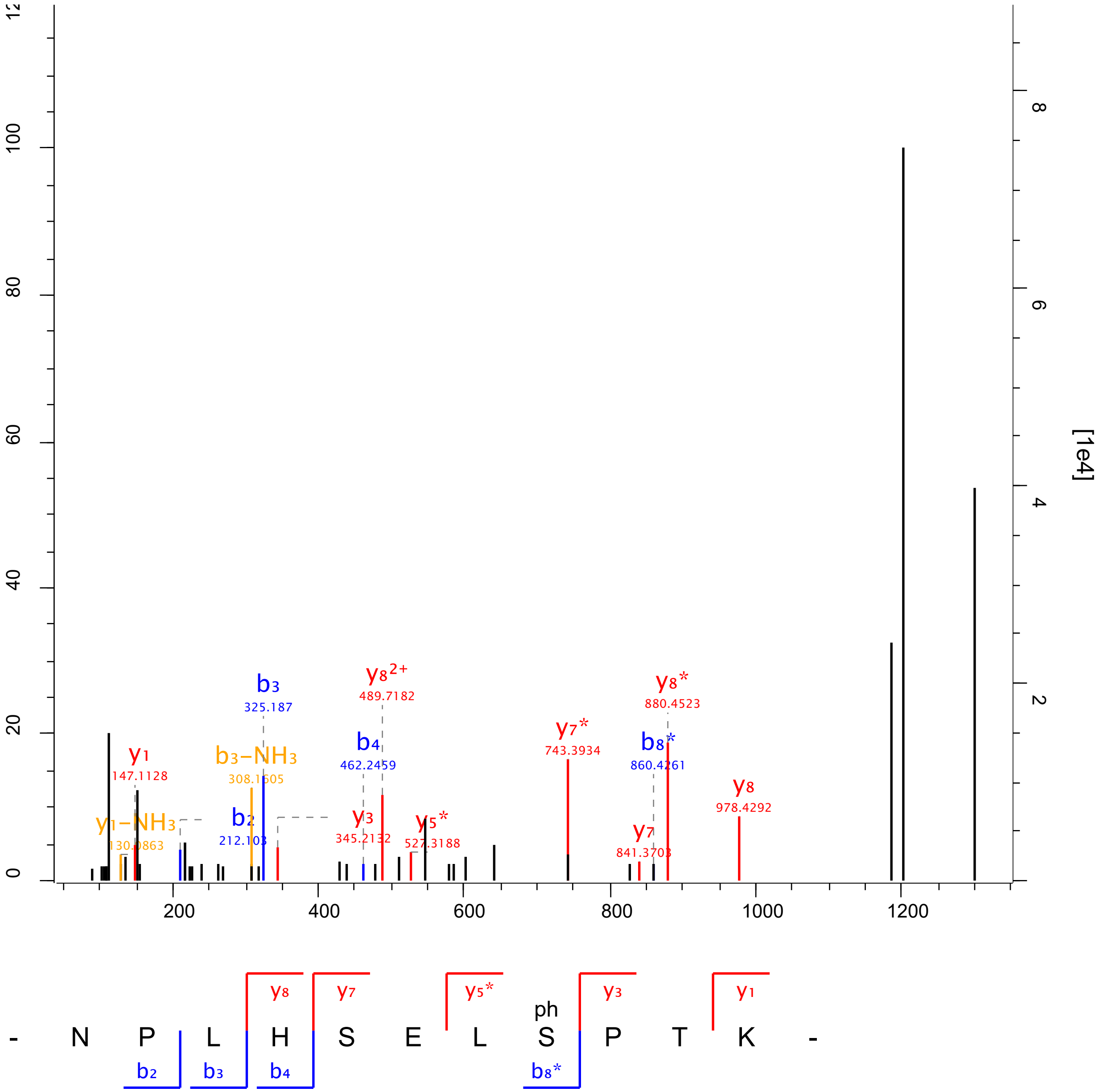

| Raw file | Scan | Method | Score | m/z |
| --- | --- | --- | --- | --- |
| A17_00_2 | 7690 | FTMS; HCD | 42.84 | 559.75 |

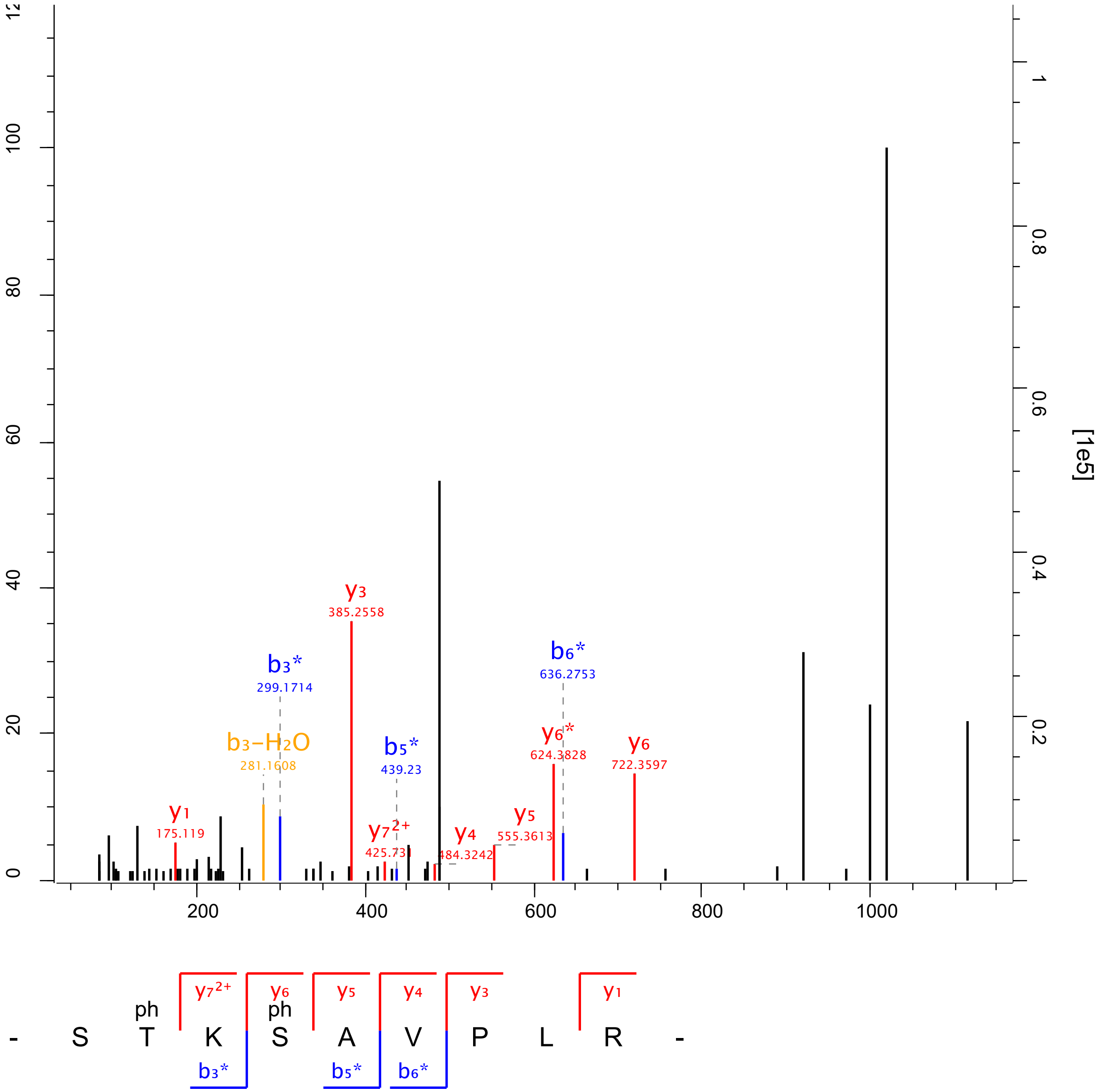

| Raw file | Scan | Method | Score | m/z |
| --- | --- | --- | --- | --- |
| A17_00_2 | 7754 | FTMS; HCD | 53.6 | 527.74 |

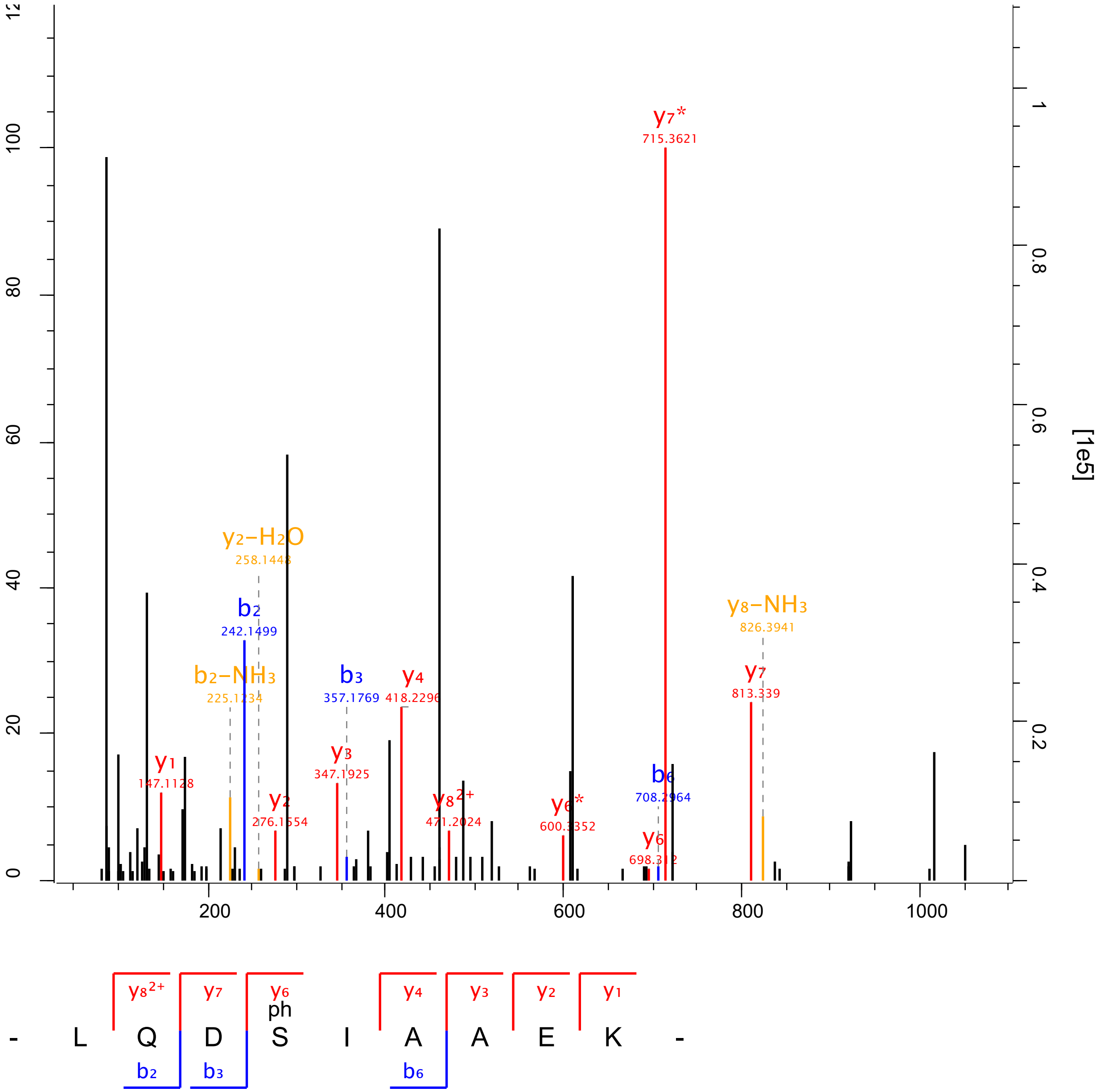

| Raw file | Scan | Method | Score | m/z |
| --- | --- | --- | --- | --- |
| A17_00_2 | 7824 | FTMS; HCD | 41.42 | 497.21 |

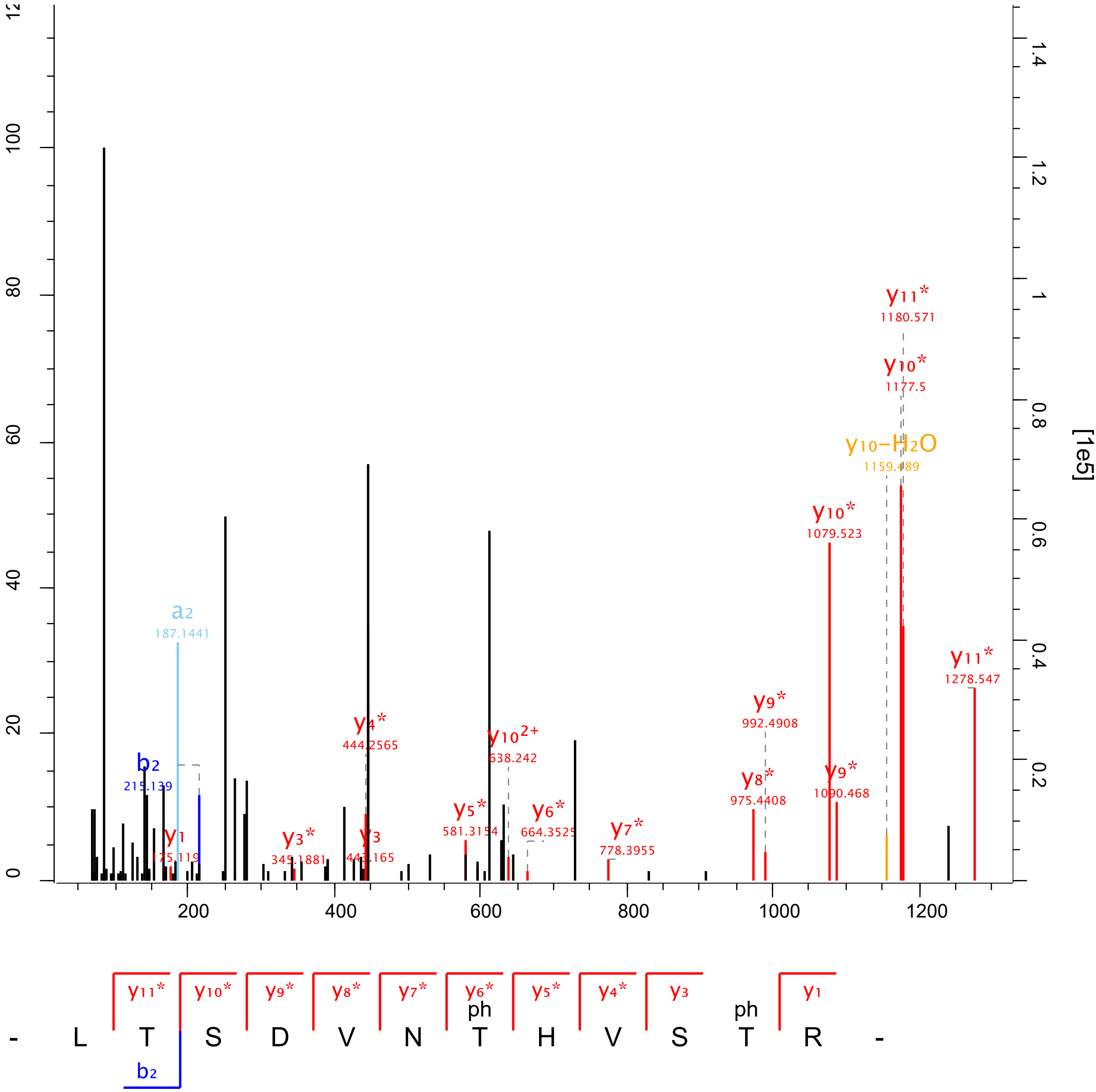

| Raw file | Scan | Method | Score | m/z |
|---|---|---|---|---|
| A17_00_2 | 7904 | FTMS; HCD | 57.17 | 540.22 |

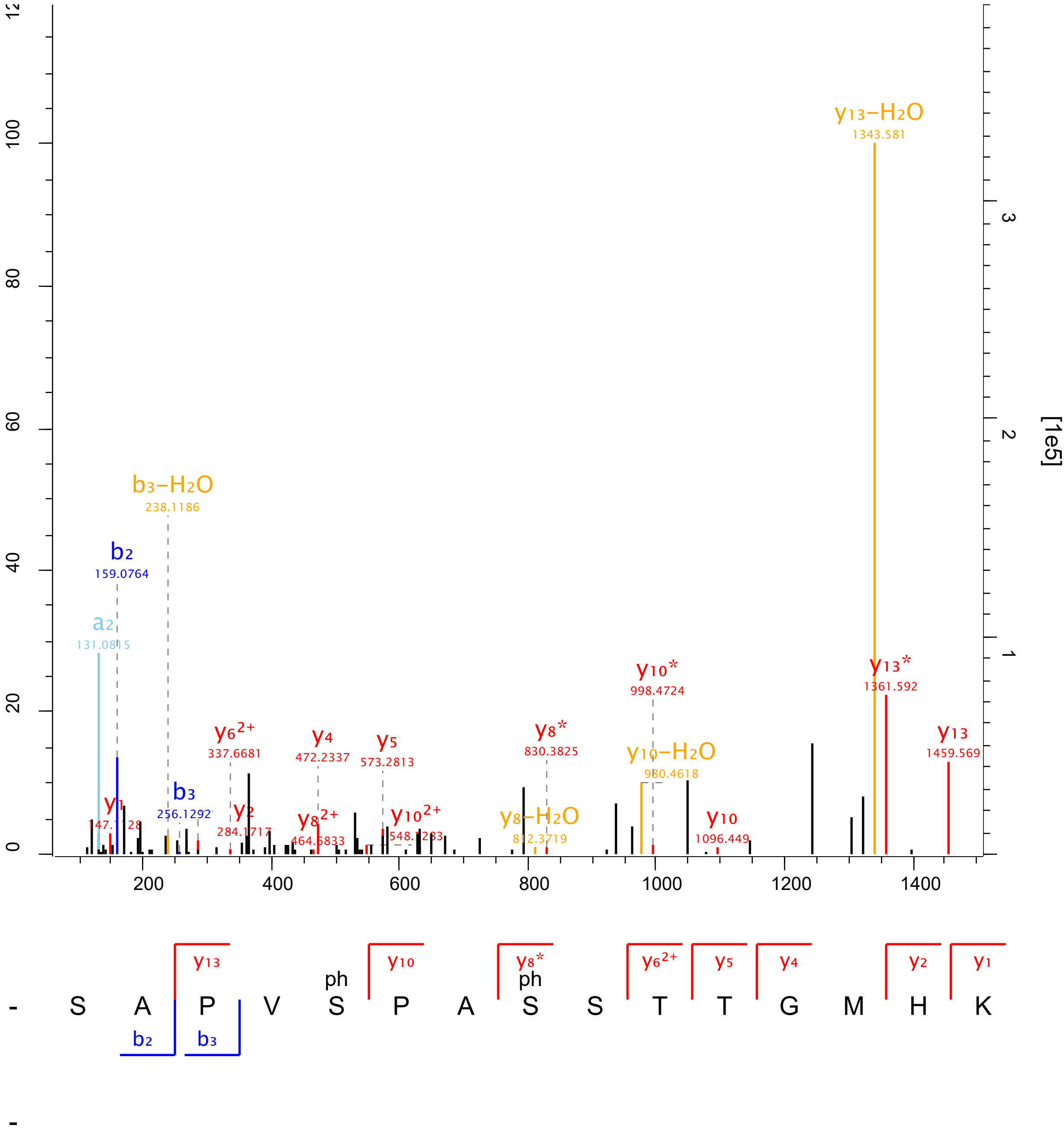

| Raw file | Scan | Method | Score | m/z |
|---|---|---|---|---|
| A17_00_2 | 8034 | FTMS; HCD | 81.66 | 448.22 |

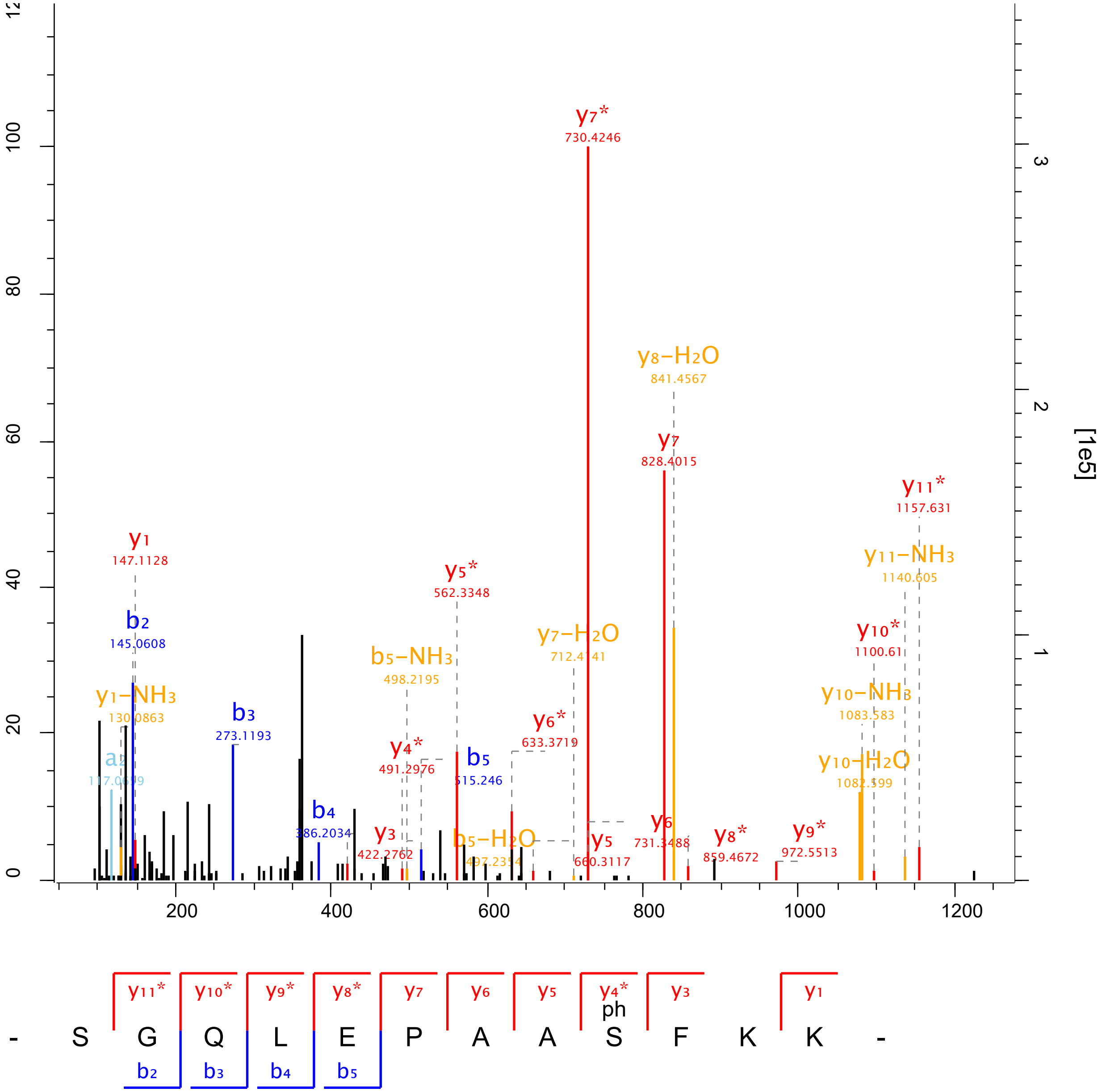

| Raw file | Scan | Method | Score | m/z |
| --- | --- | --- | --- | --- |
| A17_00_2 | 8165 | FTMS; HCD | 58.08 | 708.79 |

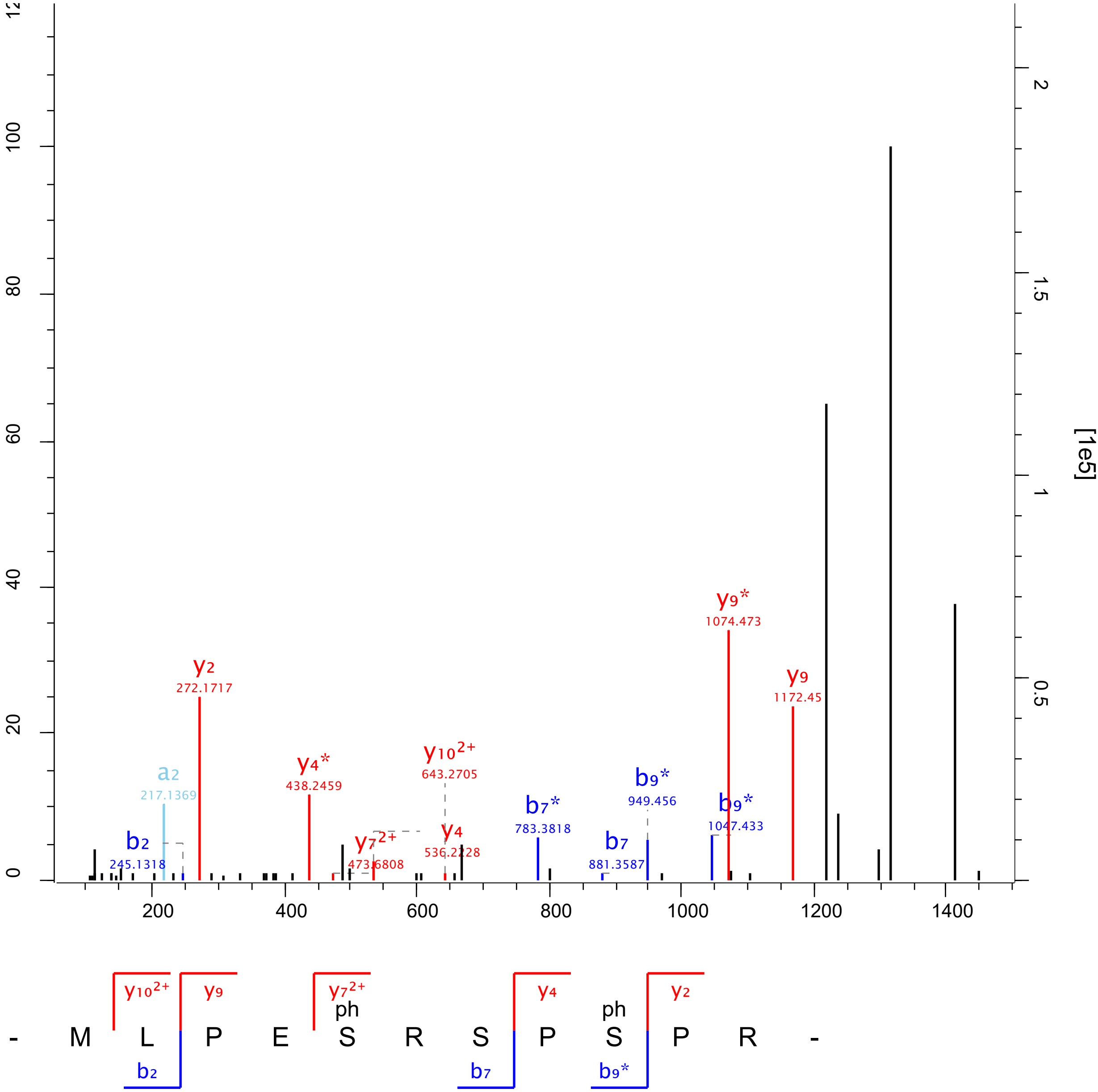

| Raw file | Scan | Method | Score | m/z |
|---|---|---|---|---|
| A17_00_2 | 8264 | FTMS; HCD | 135.18 | 567.6 |

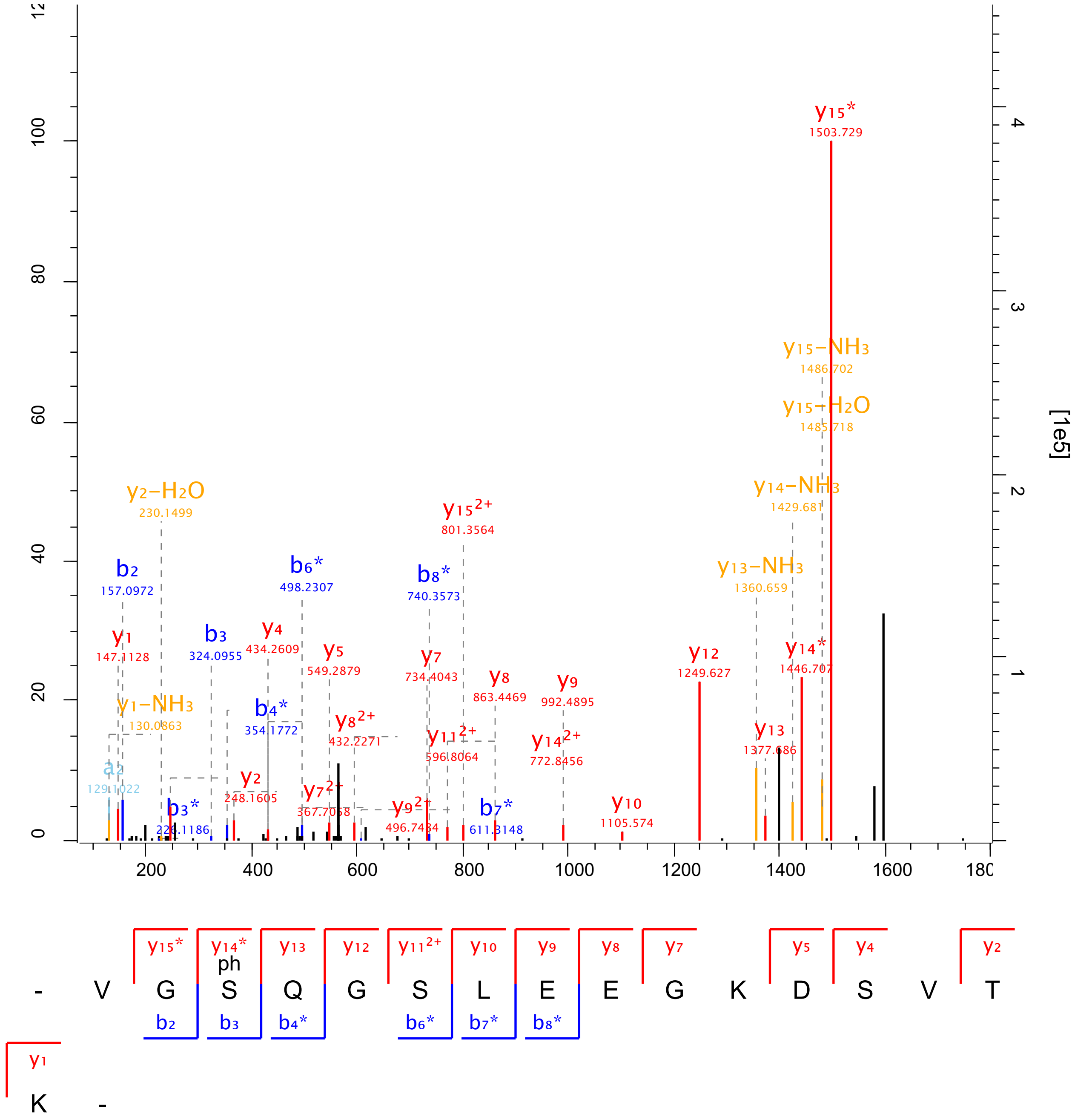

| Raw file | Scan | Method | Score | m/z |
|---|---|---|---|---|
| A17_00_2 | 8311 | FTMS; HCD | 45.99 | 633.29 |

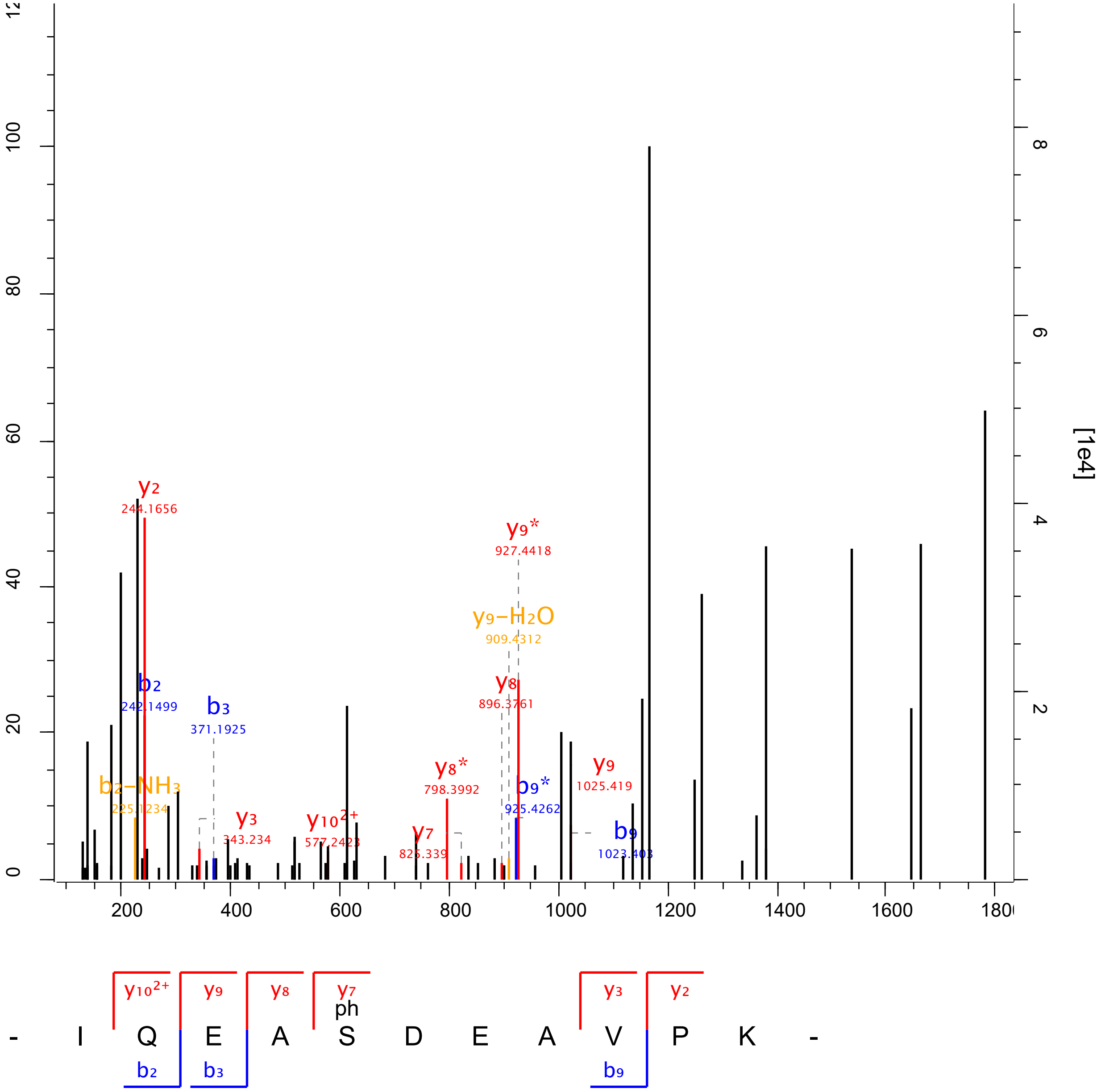

| Raw file | Scan | Method | Score | m/z |
|---|---|---|---|---|
| A17_00_2 | 8314 | FTMS; HCD | 82.37 | 516.91 |

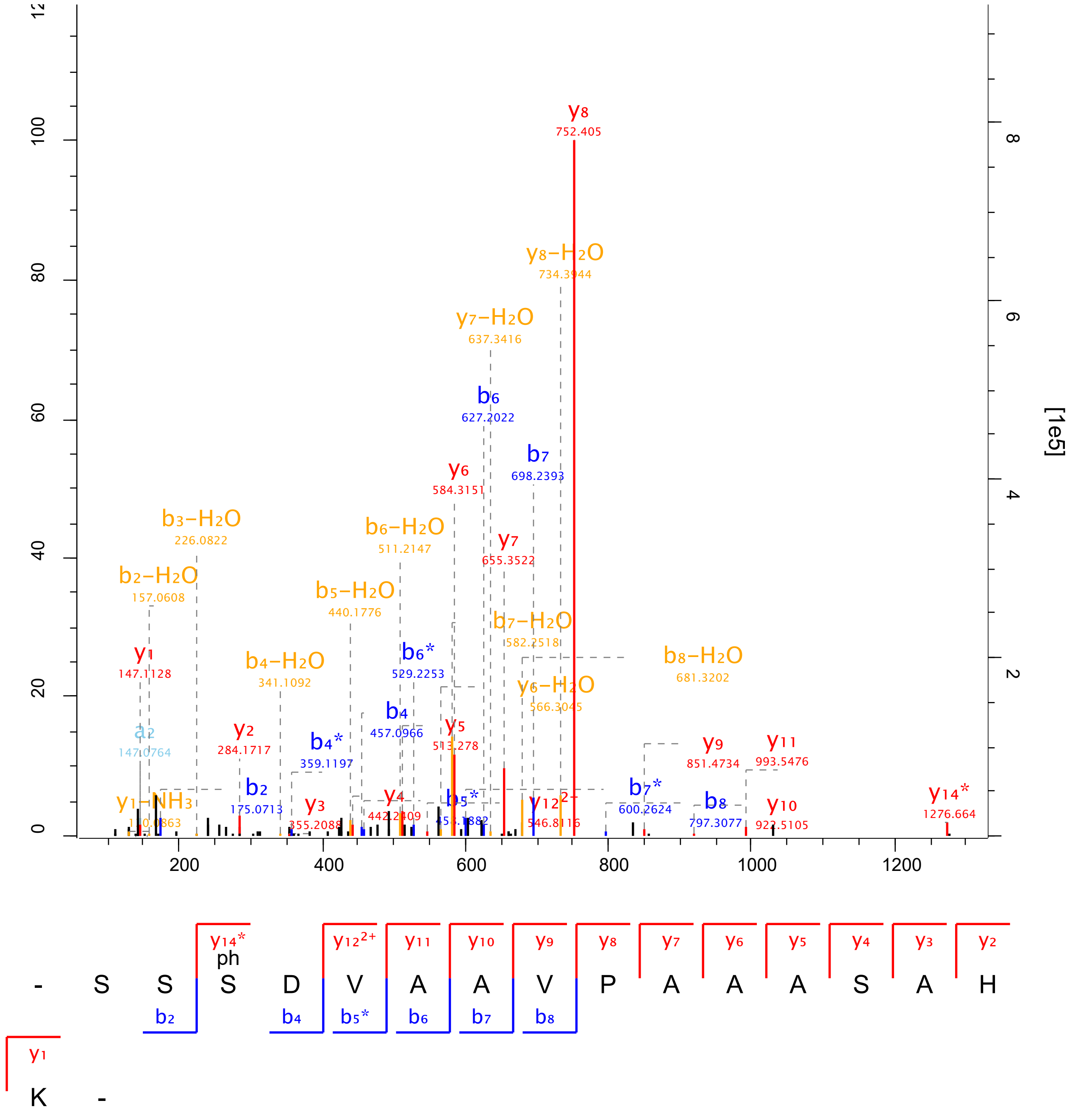

| Raw file | Scan | Method | Score | m/z |
|---|---|---|---|---|
| A17_00_2 | 8483 | FTMS; HCD | 66.99 | 614.72 |

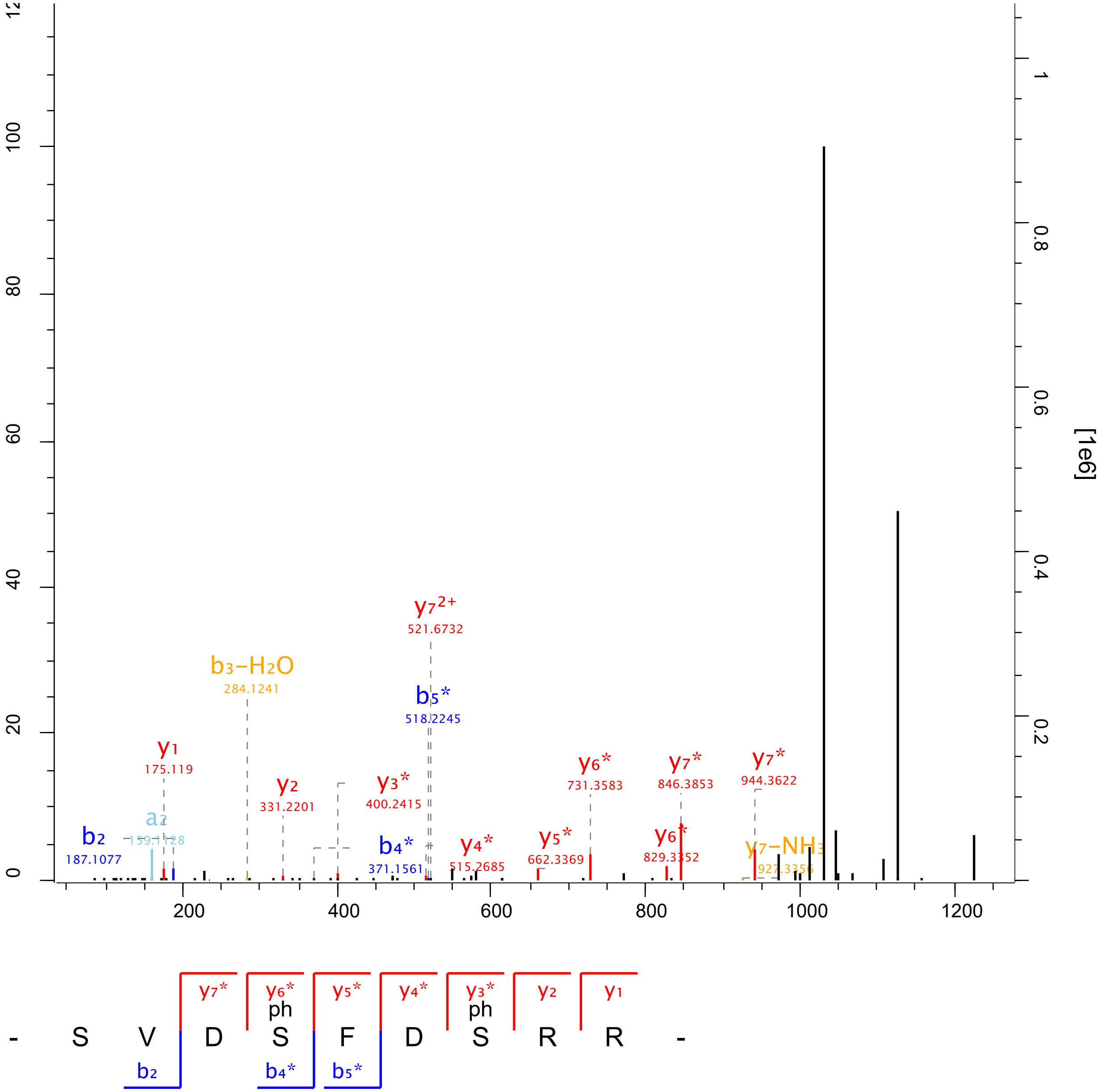

| Raw file | Scan | Method | Score | m/z |
| --- | --- | --- | --- | --- |
| A17_00_2 | 8507 | FTMS; HCD | 160.21 | 669.84 |

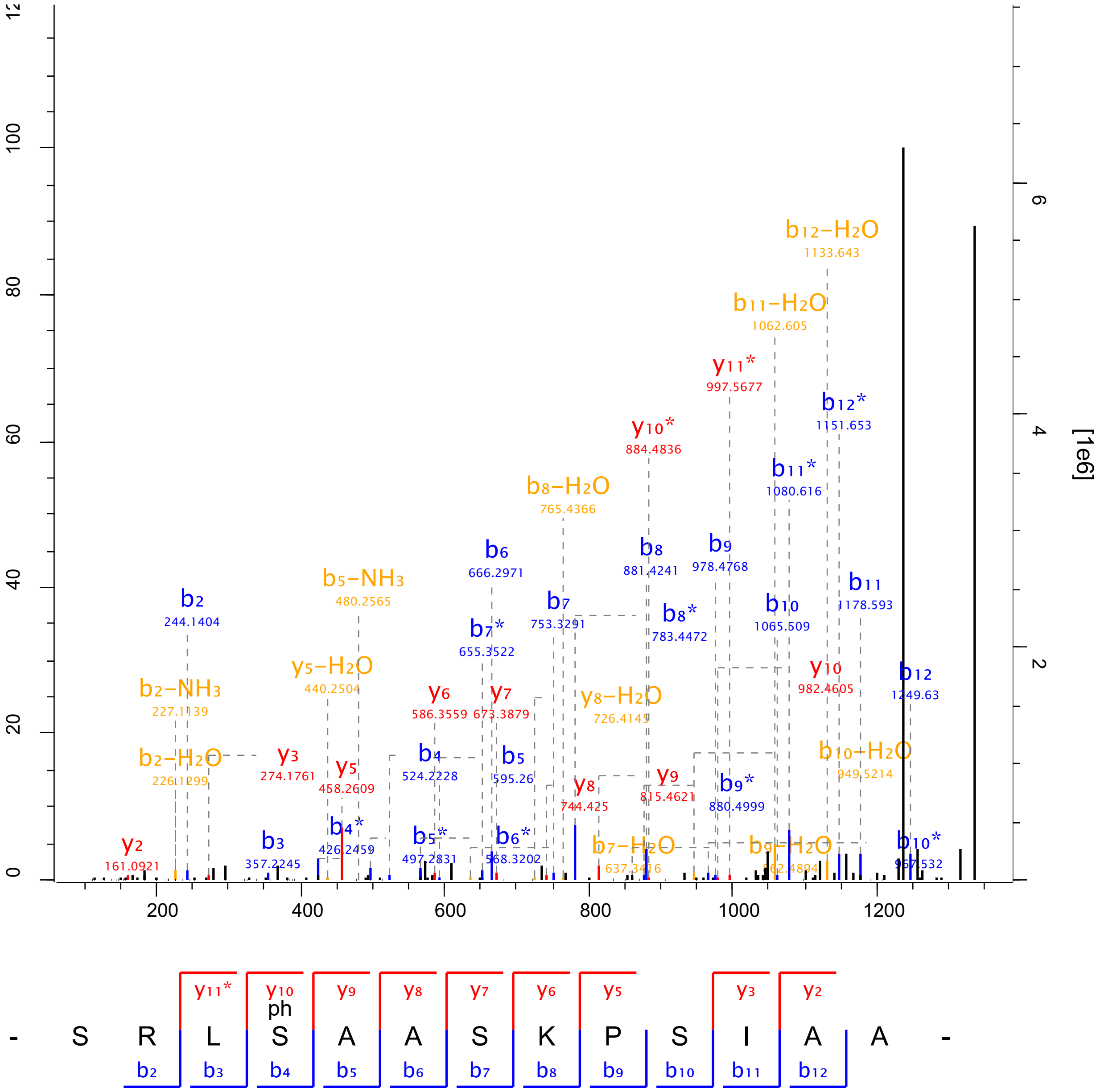

| Raw file | Scan | Method | Score | m/z |
|---|---|---|---|---|
| A17_00_2 | 8673 | FTMS; HCD | 152.7 | 478.7 |

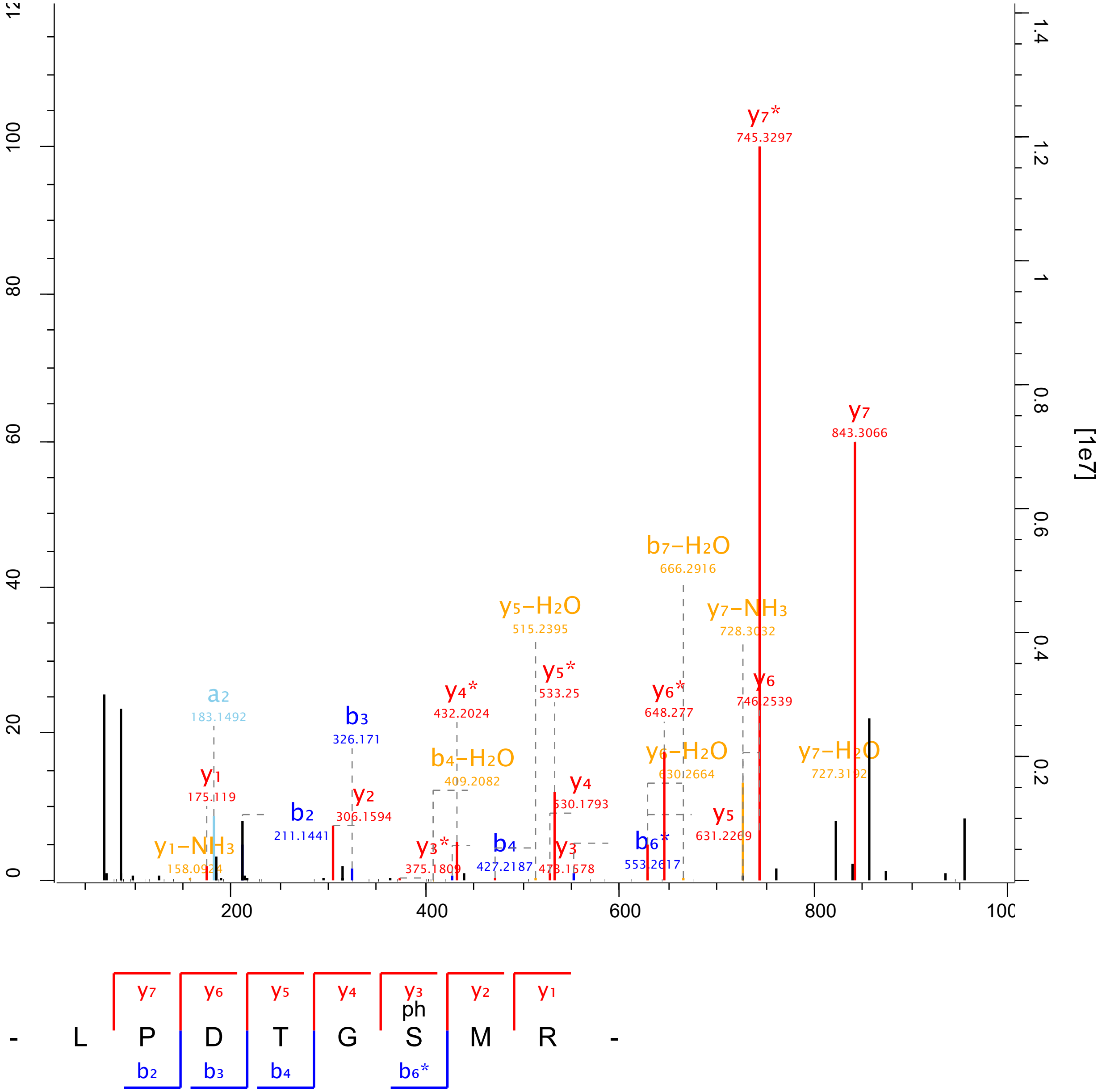

| Raw file | Scan | Method | Score | m/z |
| --- | --- | --- | --- | --- |
| A17_00_2 | 8809 | FTMS; HCD | 47.73 | 609.63 |

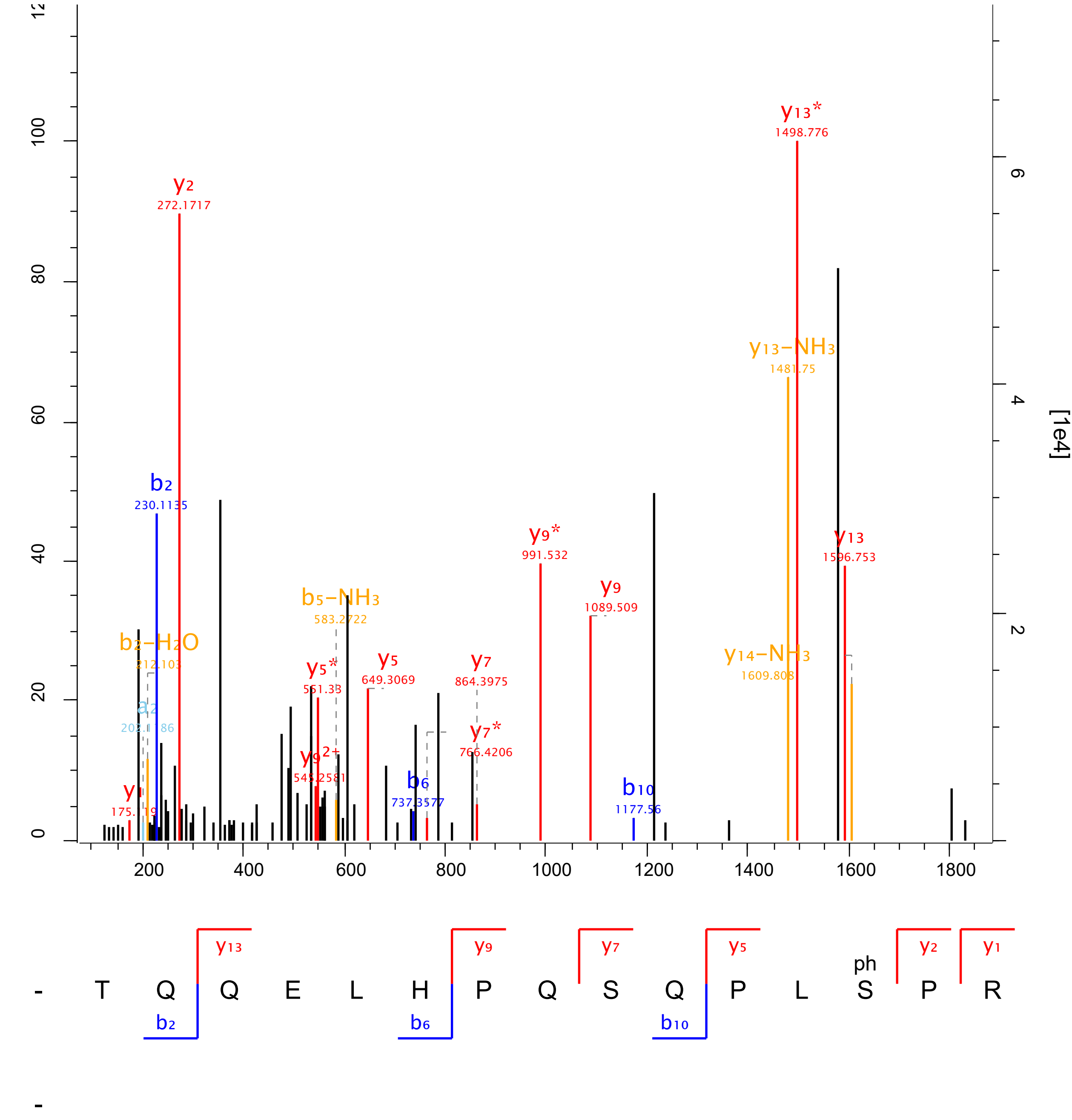

| Raw file | Scan | Method | Score | m/z |
|---|---|---|---|---|
| A17_00_2 | 8848 | FTMS; HCD | 63.82 | 425.2 |

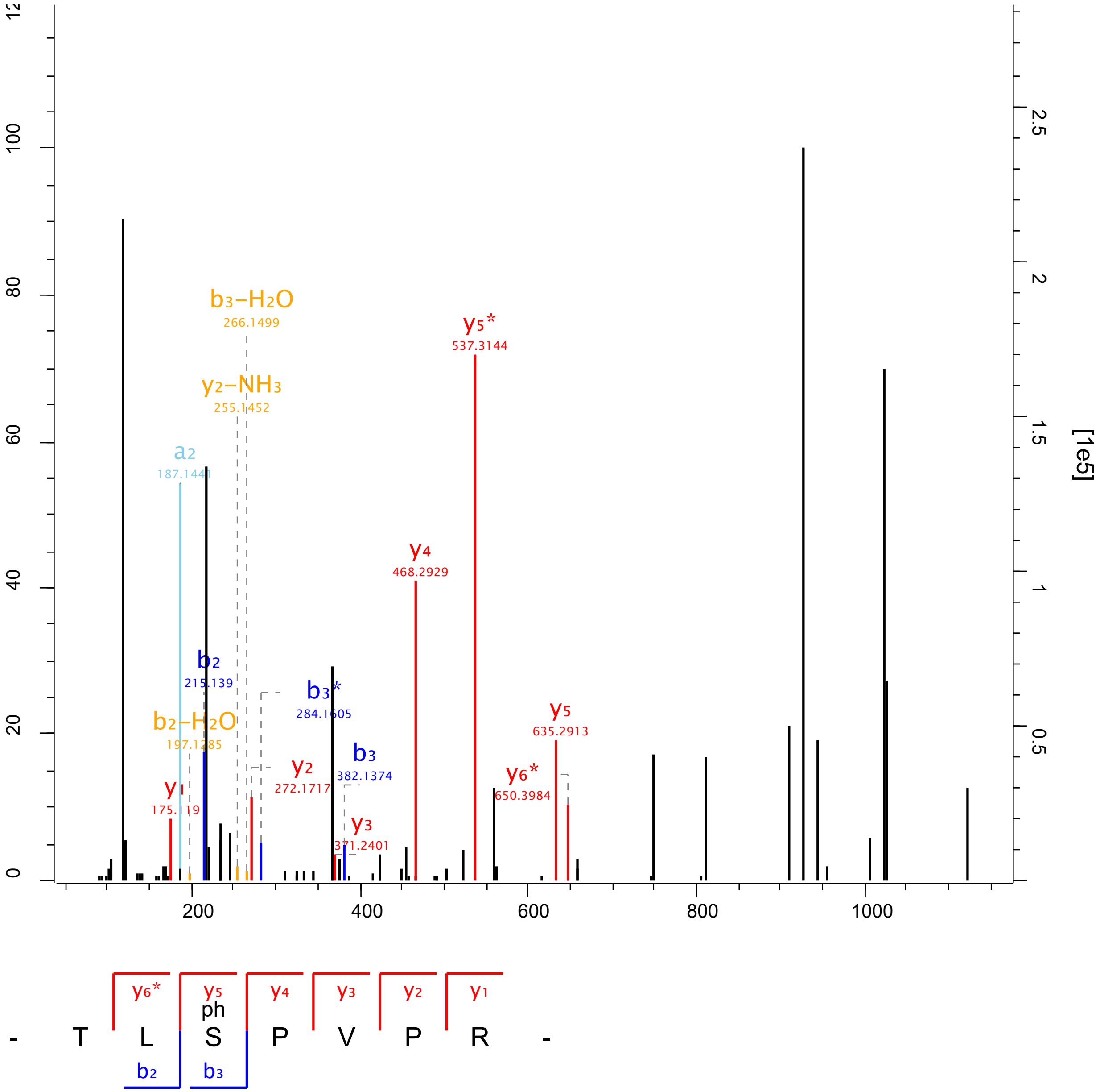

| Raw file | Scan | Method | Score | m/z |
|---|---|---|---|---|
| A17_00_2 | 8924 | FTMS; HCD | 105.96 | 578.58 |

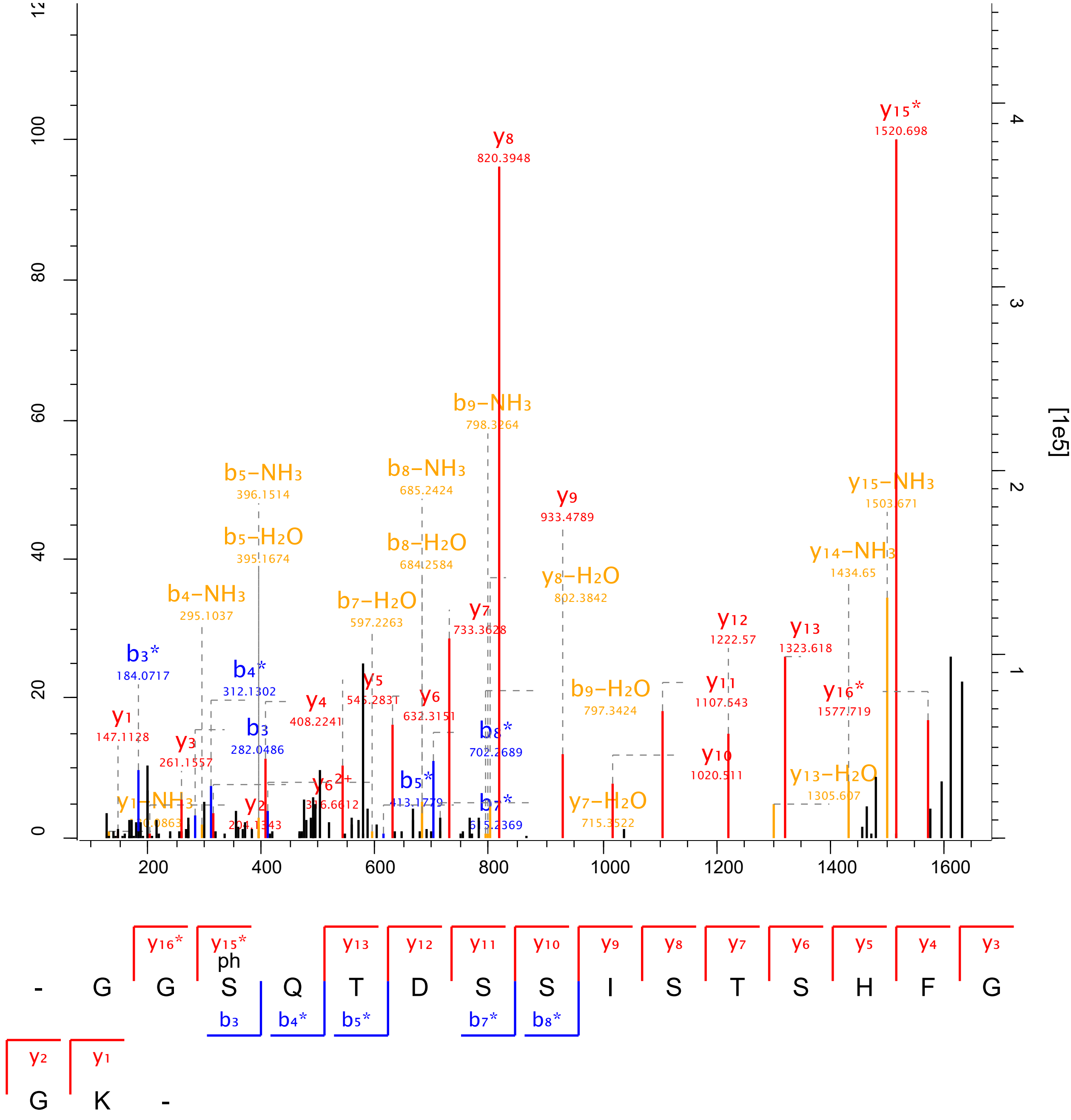

| Raw file | Scan | Method | Score | m/z |
|---|---|---|---|---|
| A17_00_2 | 9101 | FTMS; HCD | 86.8 | 557.24 |

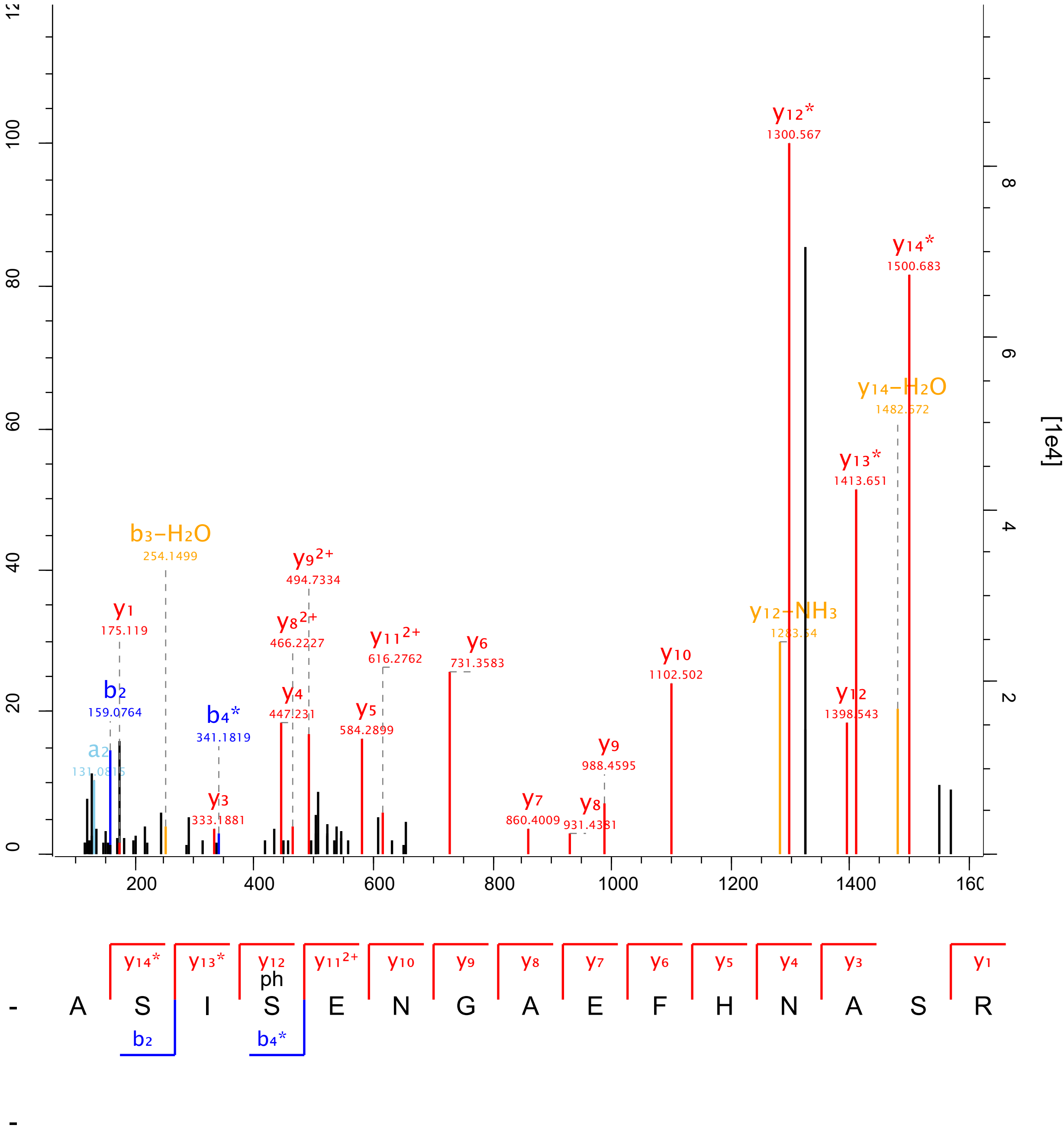

| Raw file | Scan | Method | Score | m/z |
| --- | --- | --- | --- | --- |
| A17_00_2 | 9154 | FTMS; HCD | 92.26 | 606.95 |

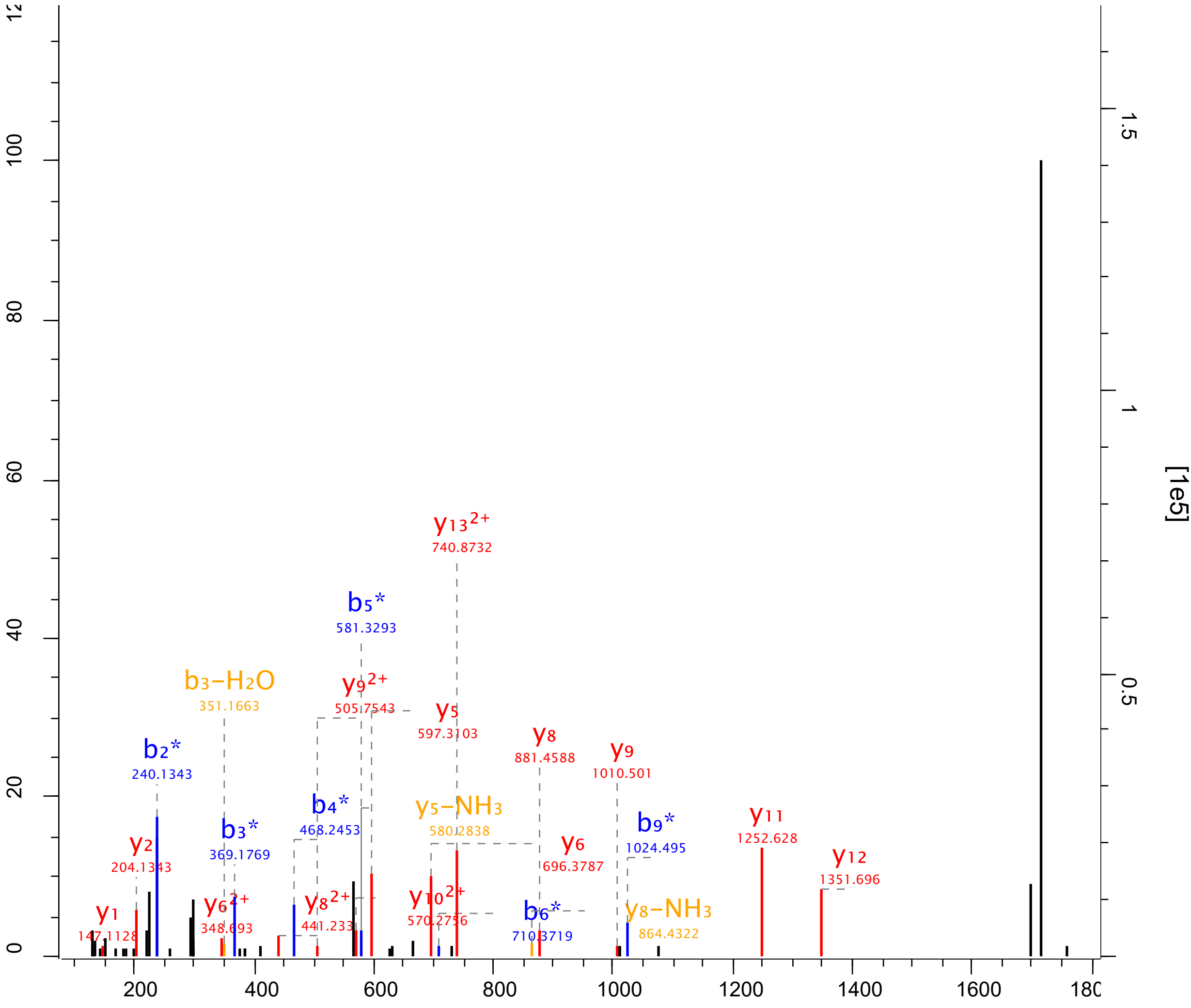


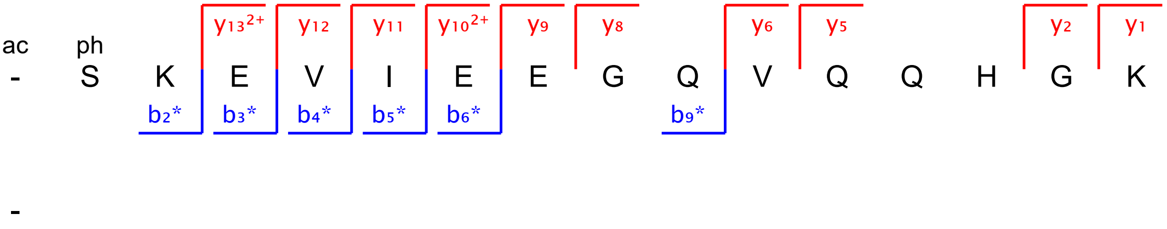

| Raw file | Scan | Method | Score | m/z |
|---|---|---|---|---|
| A17_00_2 | 9266 | FTMS; HCD | 48.47 | 545.9 |

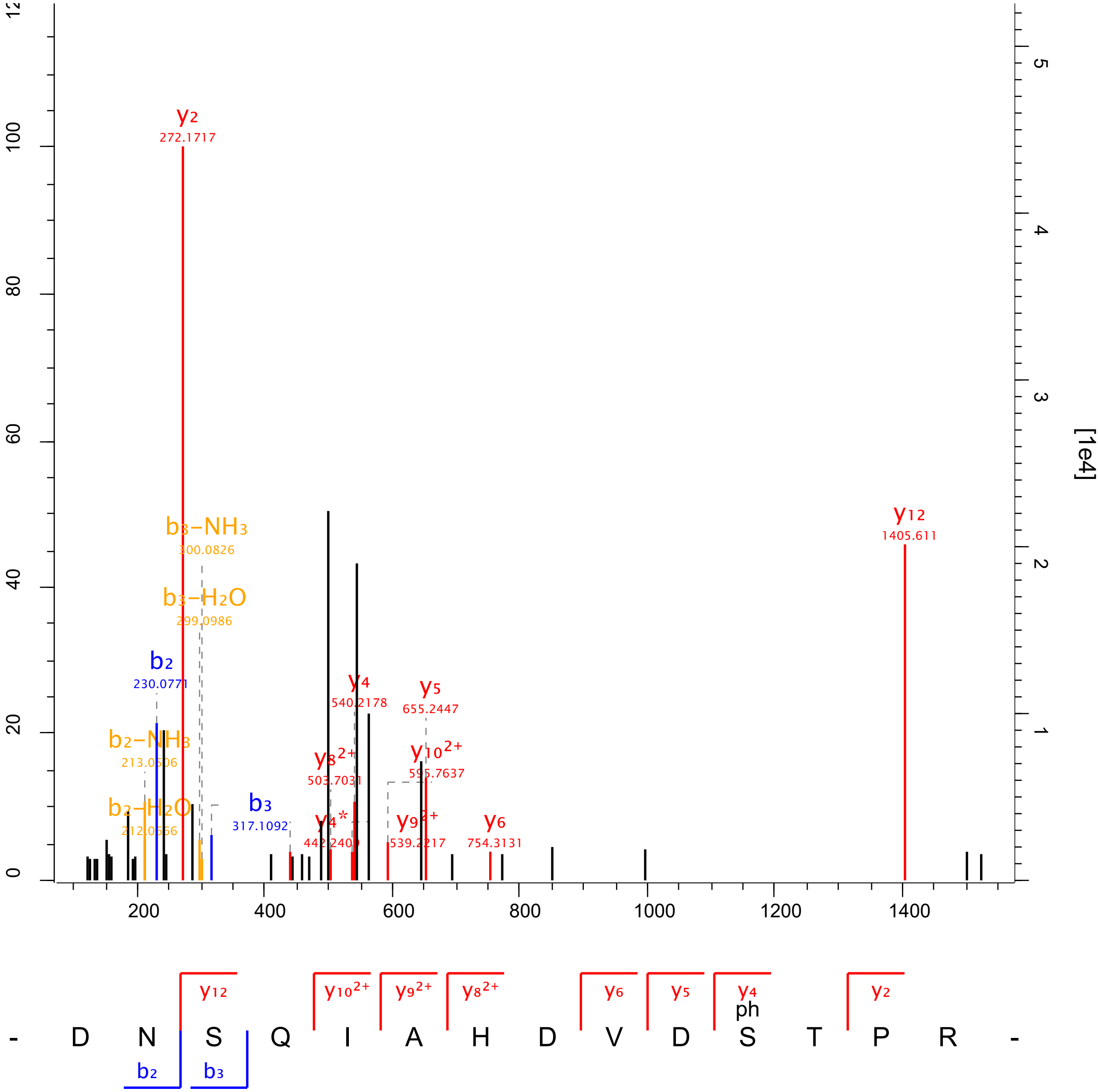

| Raw file | Scan | Method | Score | m/z |
| --- | --- | --- | --- | --- |
| A17_00_2 | 9410 | FTMS; HCD | 90.82 | 650.59 |

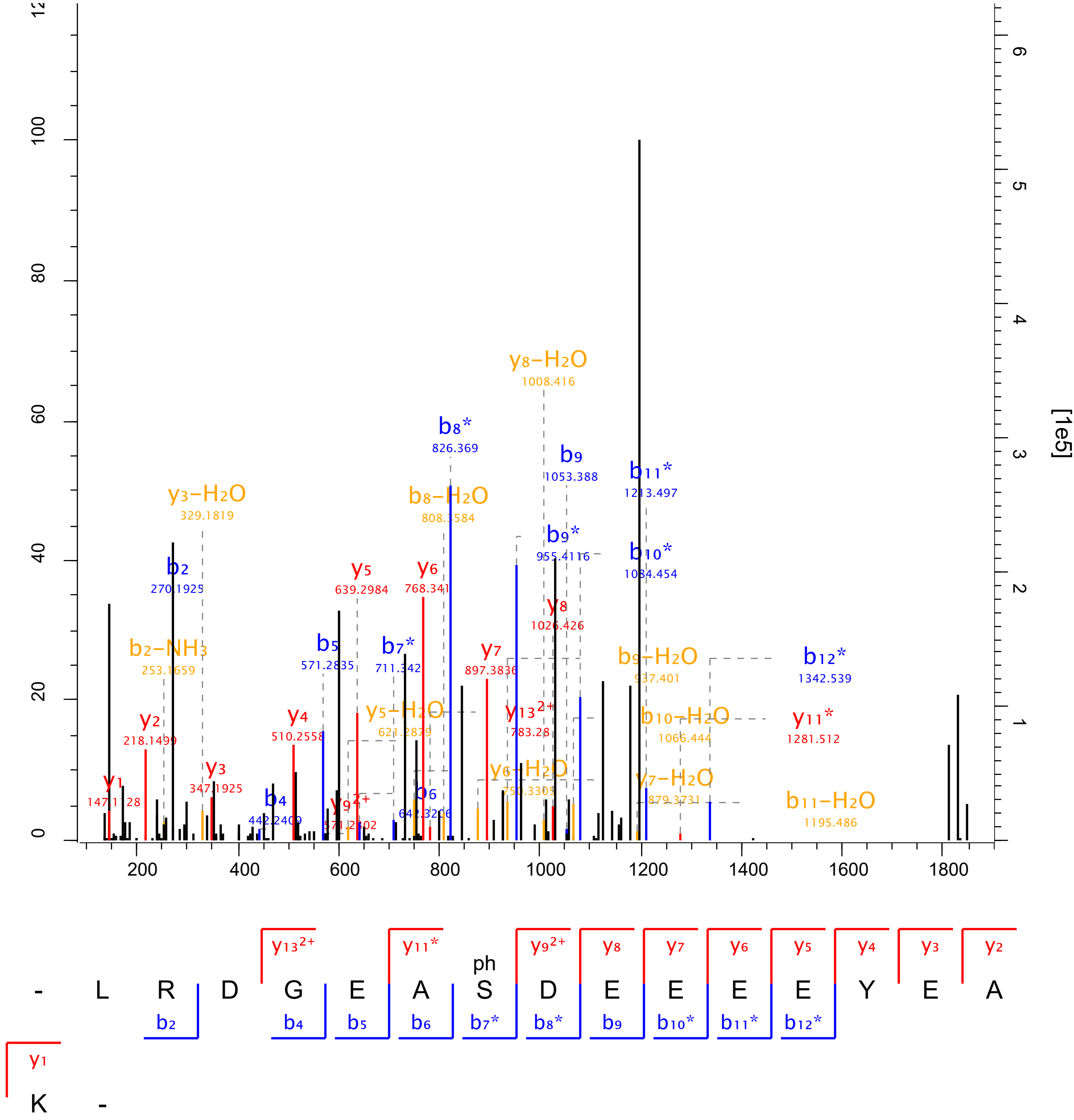

| Raw file | Scan | Method | Score | m/z |
|---|---|---|---|---|
| A17_00_2 | 9425 | FTMS; HCD | 113.36 | 982.4 |

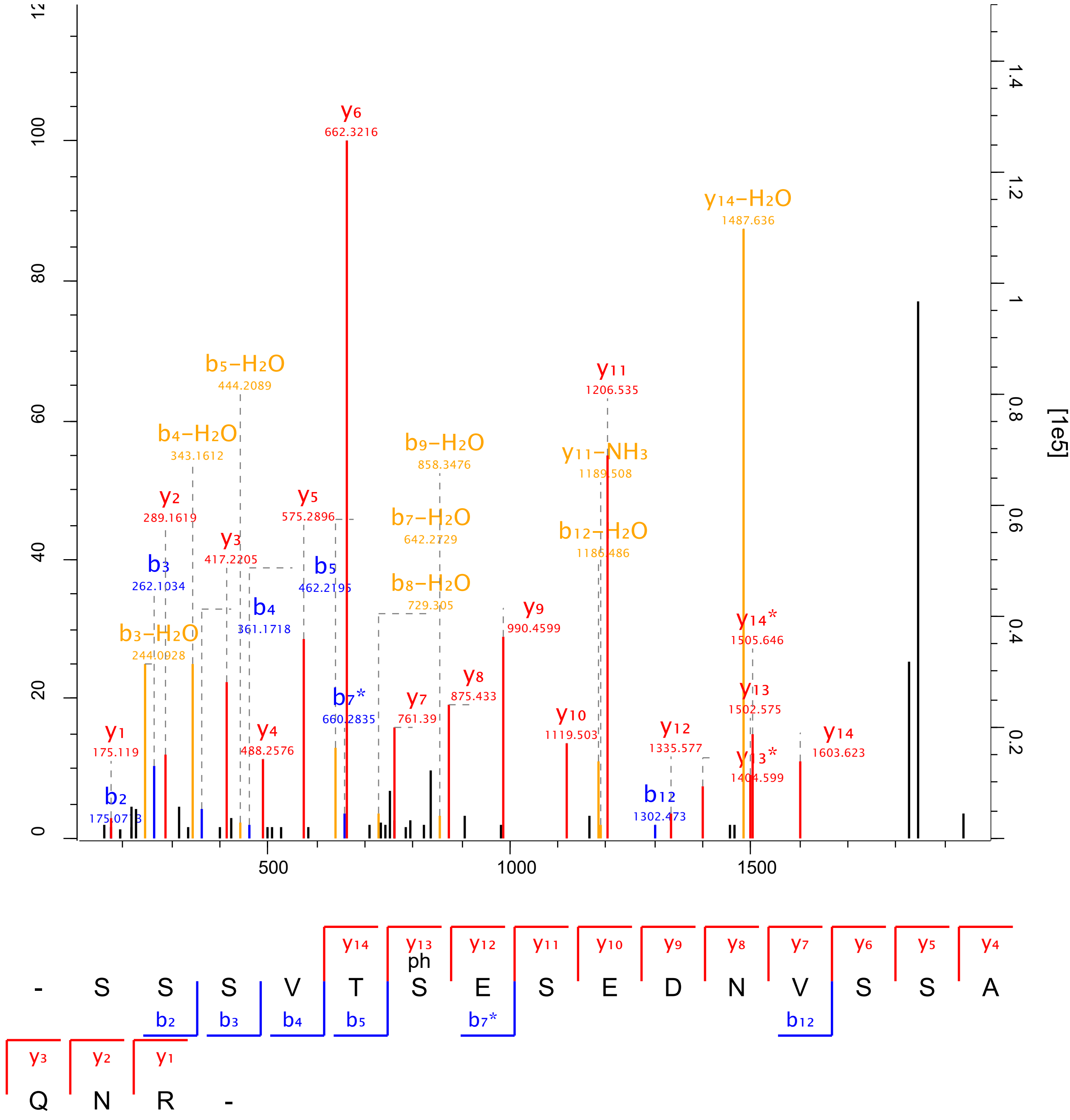

| Raw file | Scan | Method | Score | m/z |
|---|---|---|---|---|
| A17_00_2 | 9470 | FTMS; HCD | 190.53 | 566.25 |

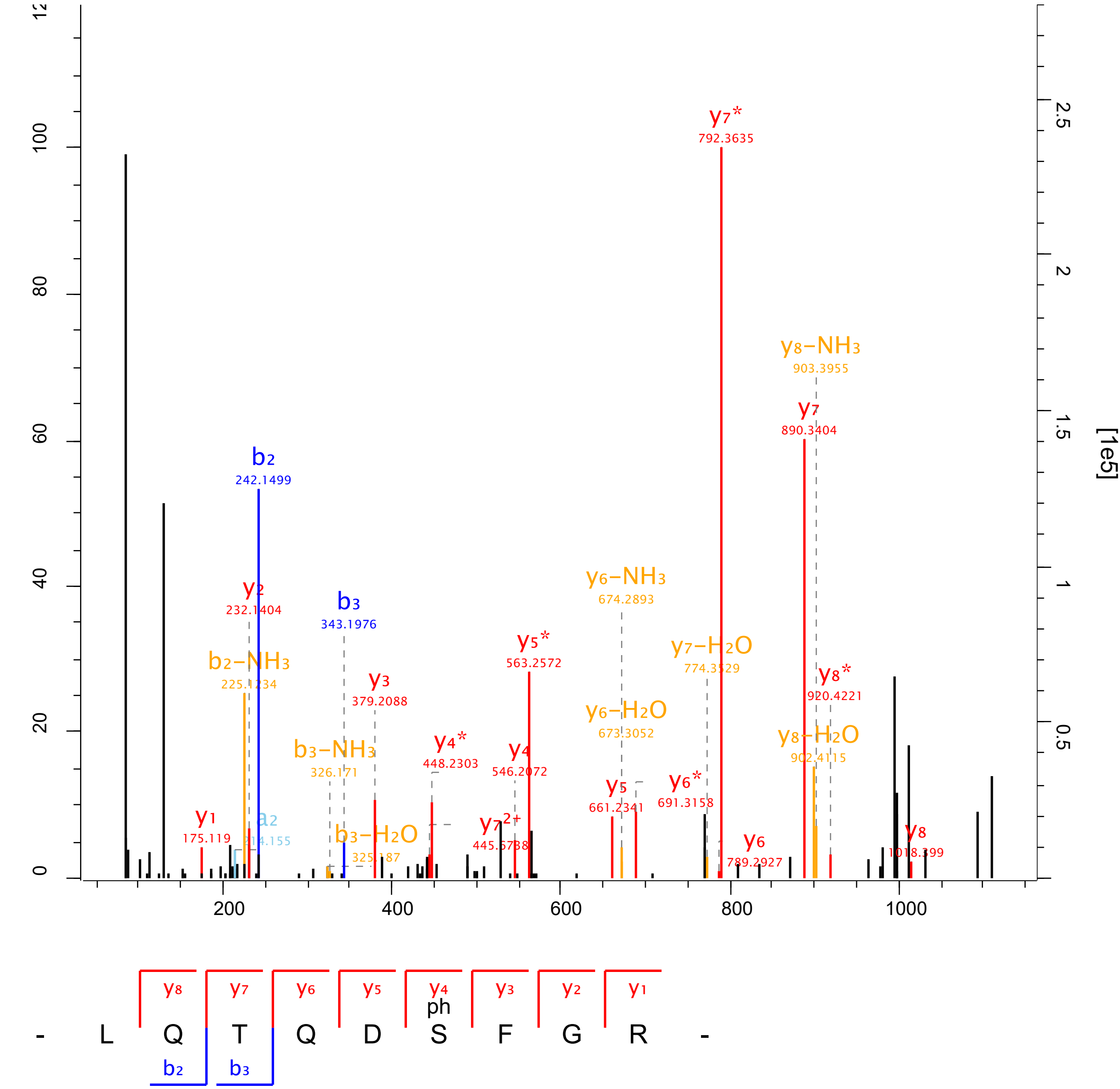

| Raw file | Scan | Method | Score | m/z |
|---|---|---|---|---|
| A17_00_2 | 9523 | FTMS; HCD | 120.06 | 524.69 |

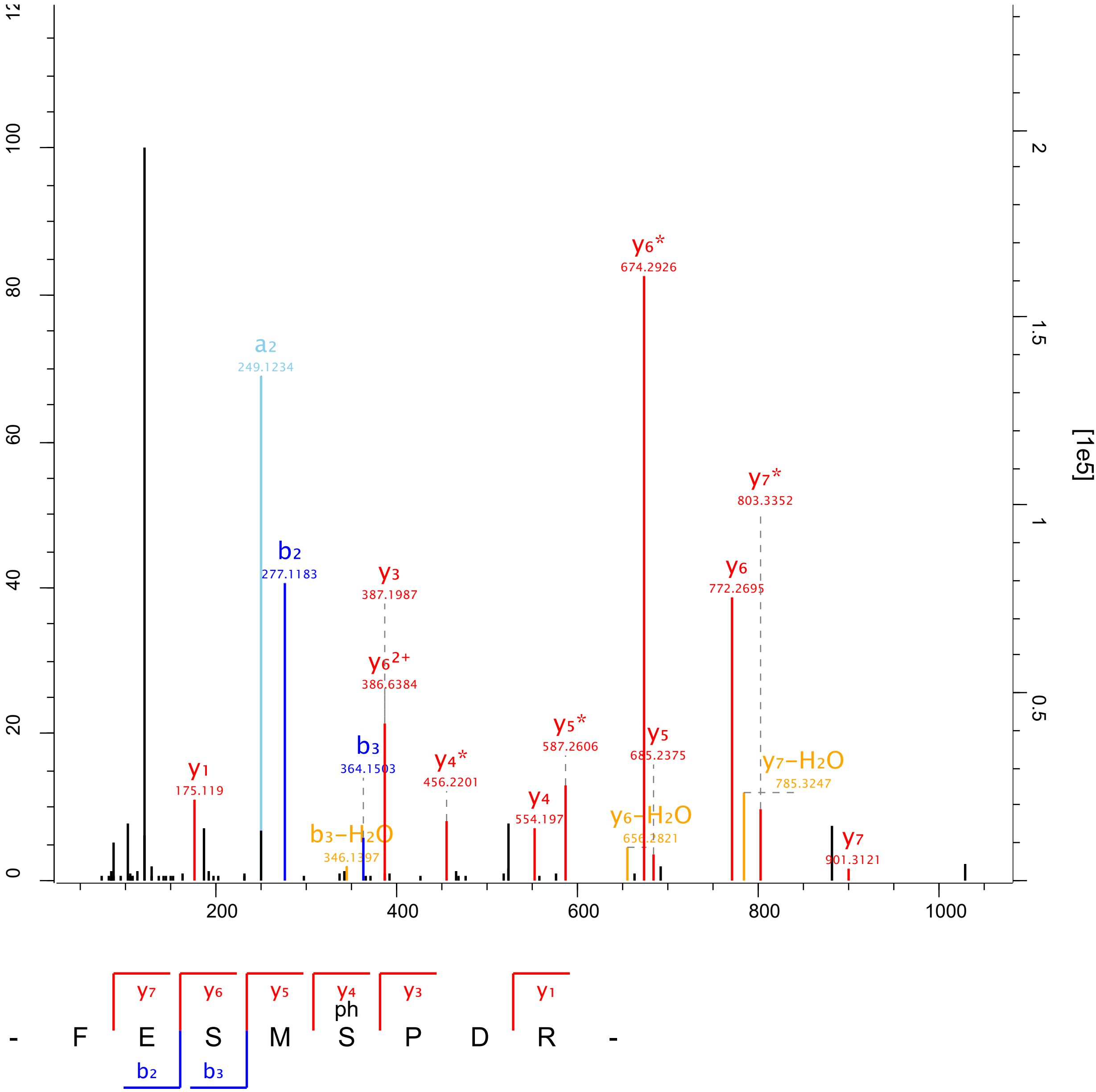

| Raw file | Scan | Method | Score | m/z |
|---|---|---|---|---|
| A17_00_2 | 9588 | FTMS; HCD | 119.46 | 717.76 |

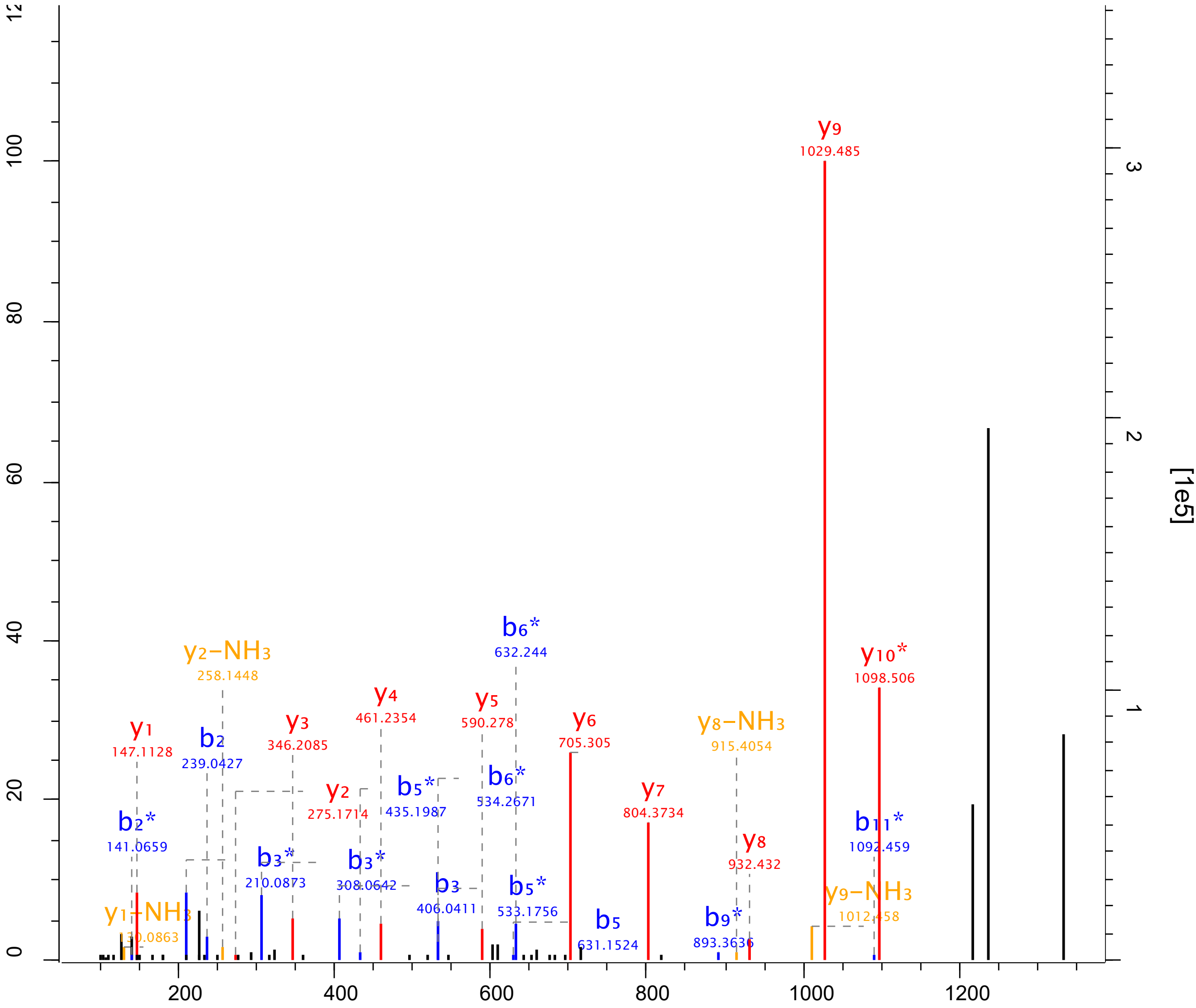


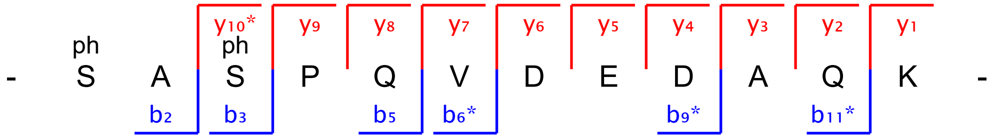

| Raw file | Scan | Method | Score | m/z |
|---|---|---|---|---|
| A17_00_2 | 9650 | FTMS; HCD | 143.04 | 650.93 |

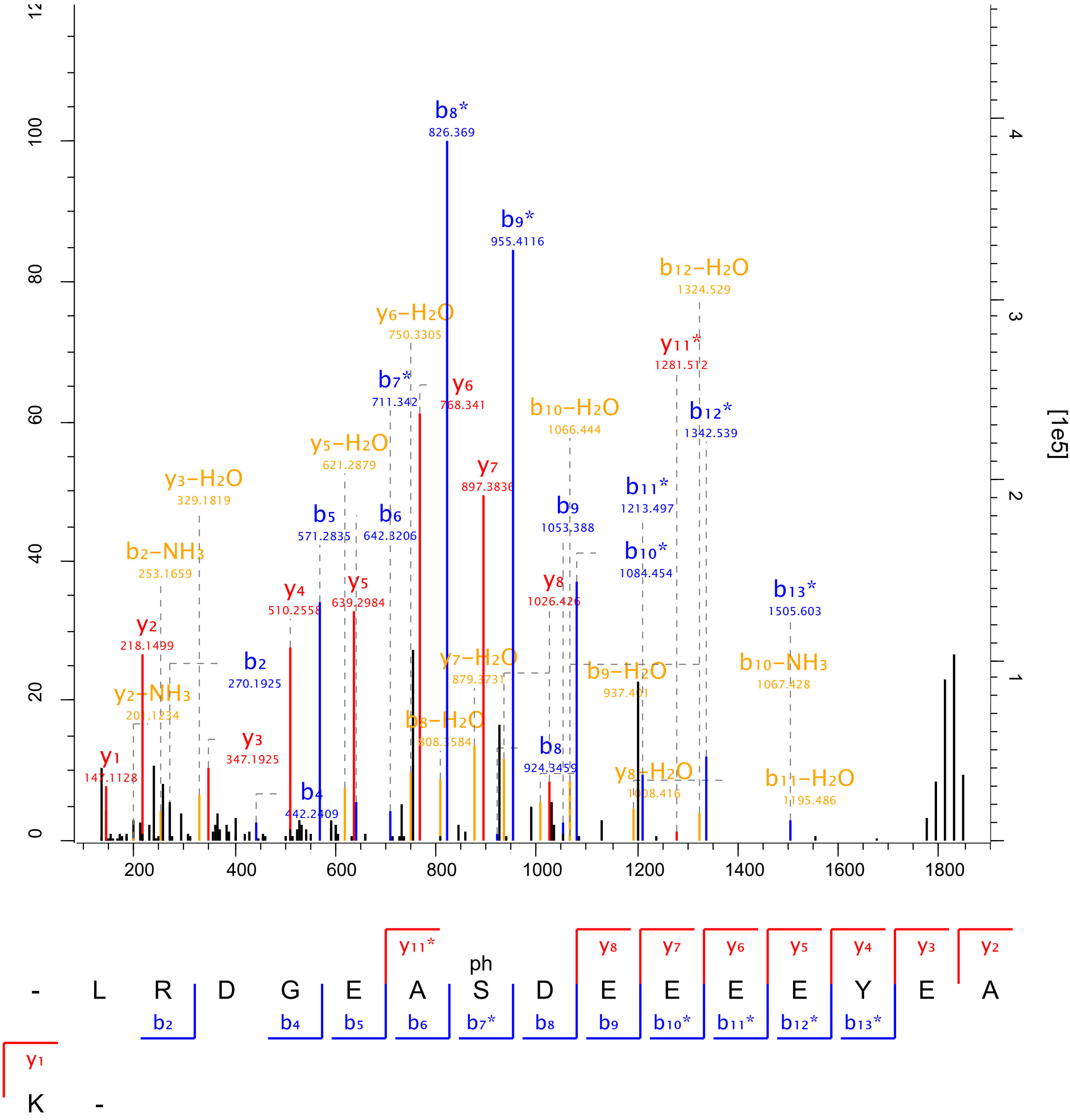

| Raw file | Scan | Method | Score | m/z |
| --- | --- | --- | --- | --- |
| A17_00_2 | 9672 | FTMS; HCD | 97.81 | 483.74 |

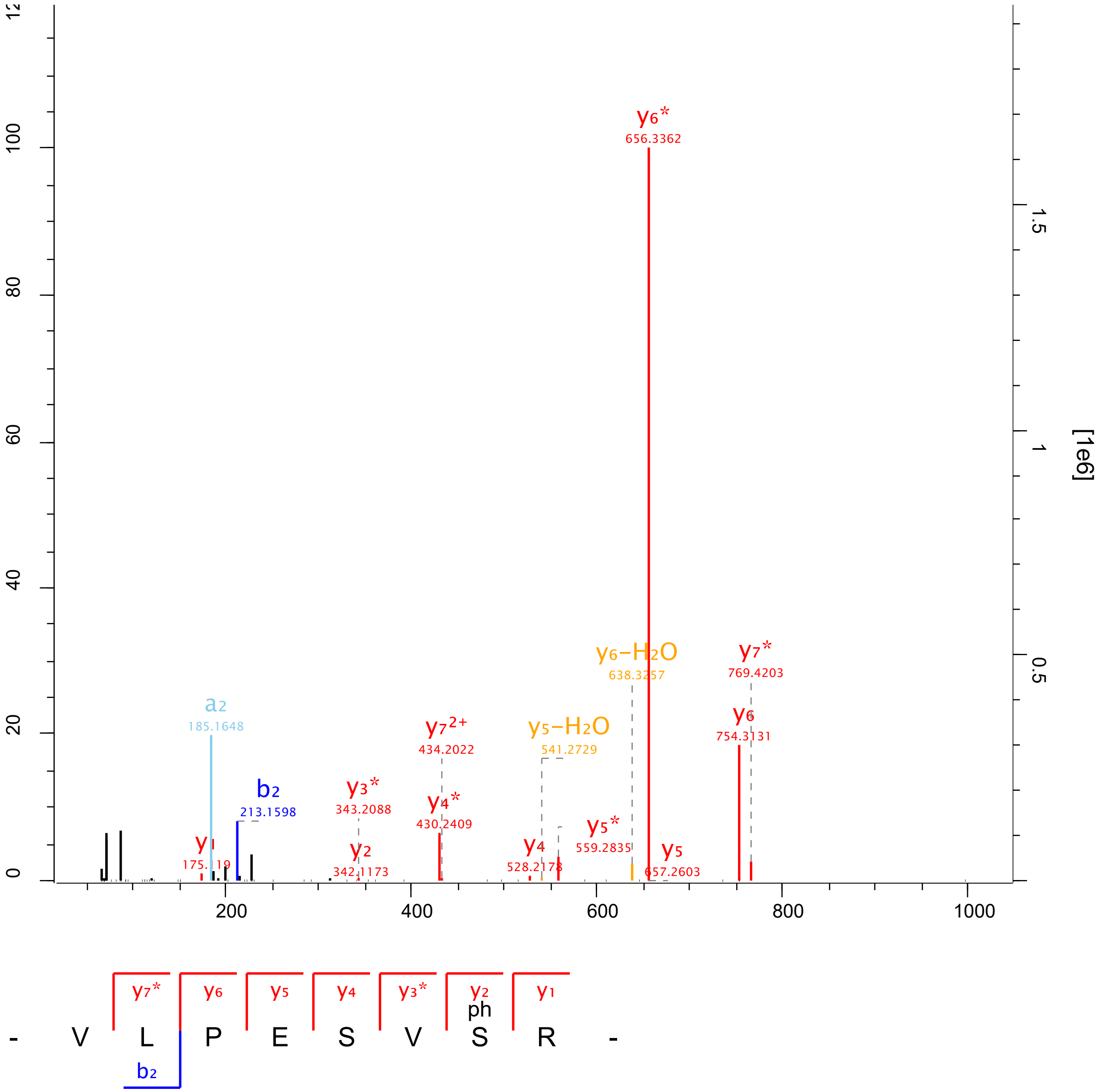

| Raw file | Scan | Method | Score | m/z |
|---|---|---|---|---|
| A17_00_2 | 9788 | FTMS; HCD | 176.25 | 695.27 |

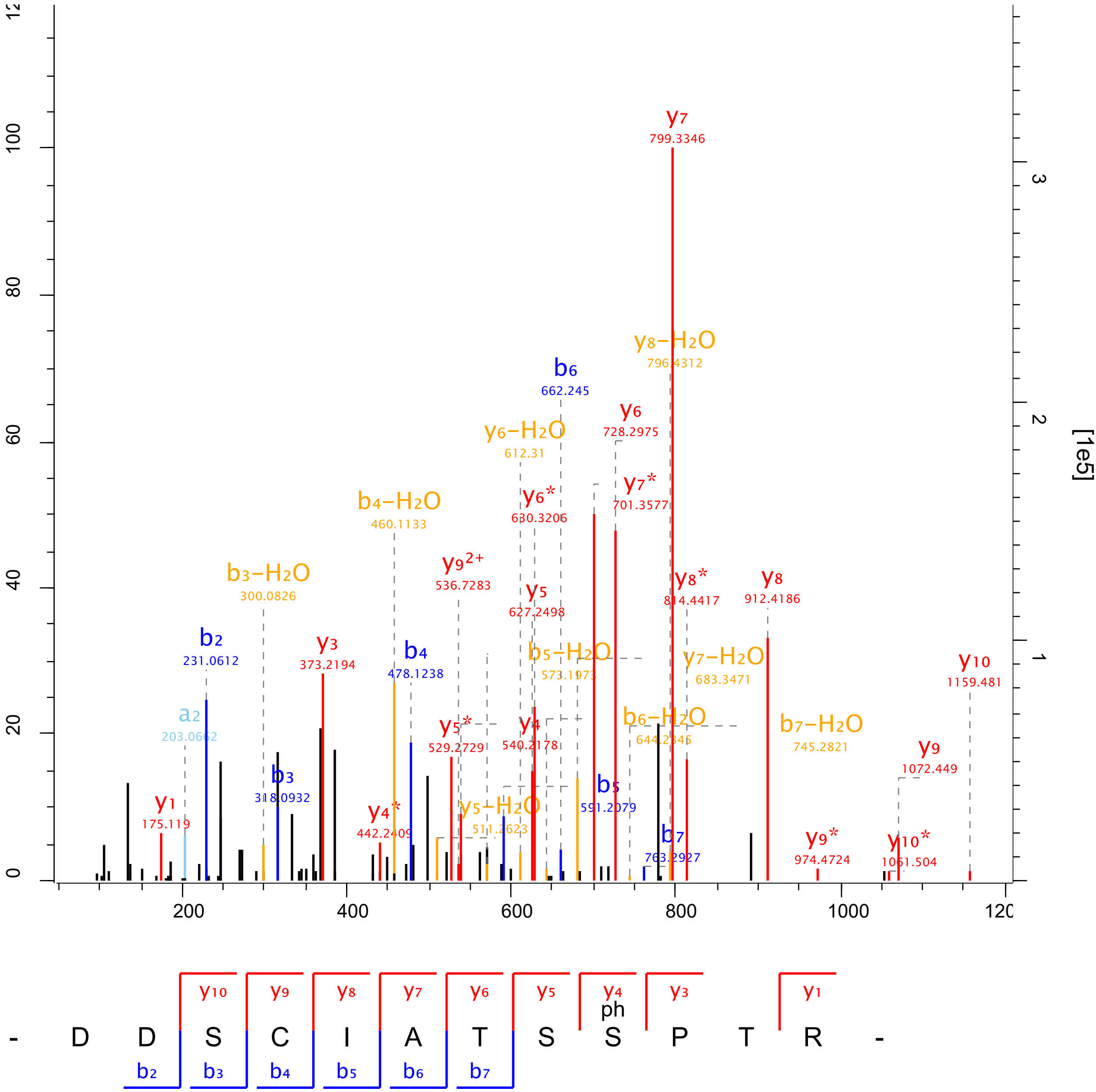

| Raw file | Scan | Method | Score | m/z |
|---|---|---|---|---|
| A17_00_3short | 16581 | FTMS; HCD | 134.68 | 541.71 |

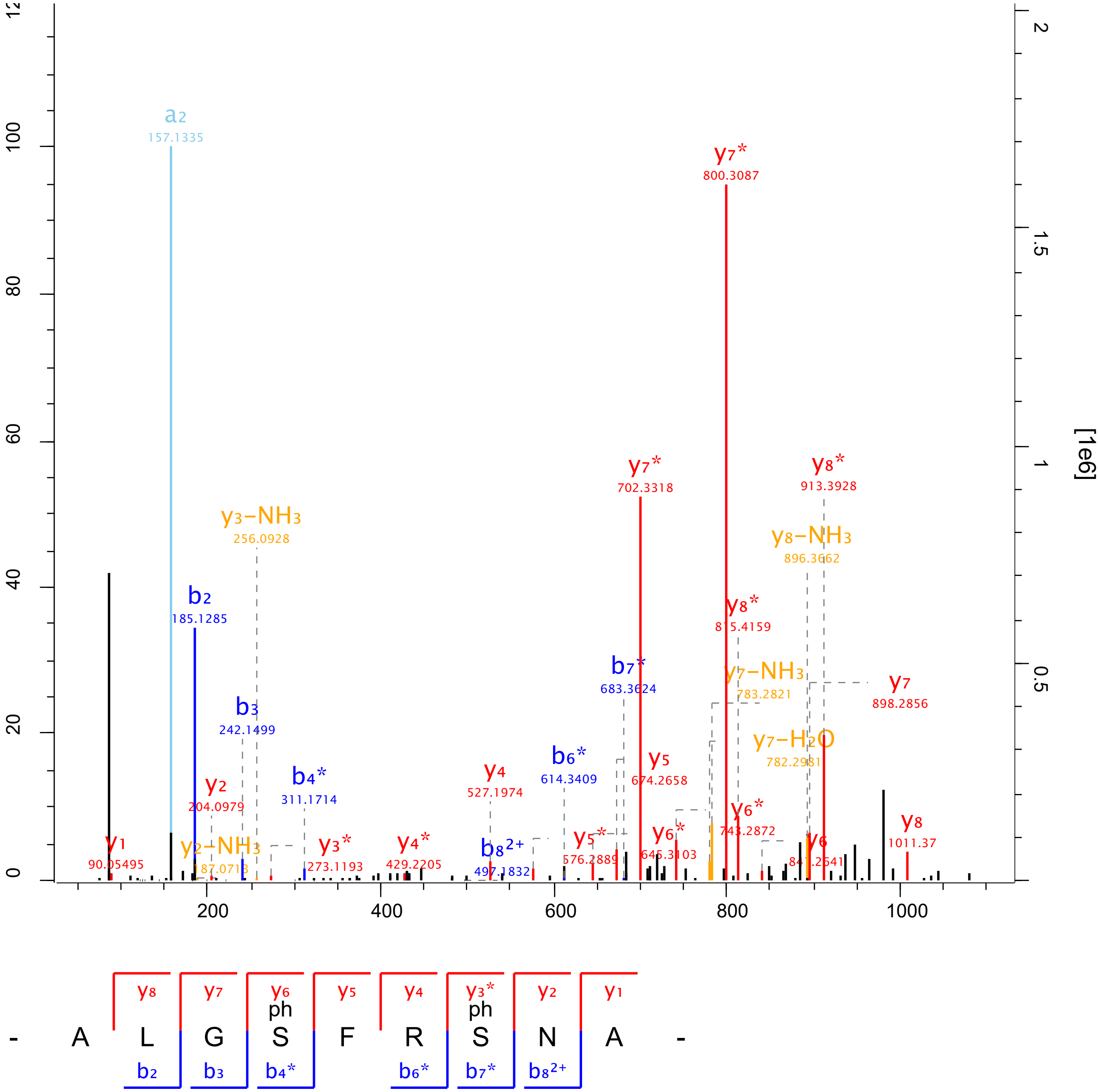

| Raw file | Scan | Method | Score | m/z |
| --- | --- | --- | --- | --- |
| A17_02_1 | 10357 | FTMS; HCD | 93.1 | 487.7 |

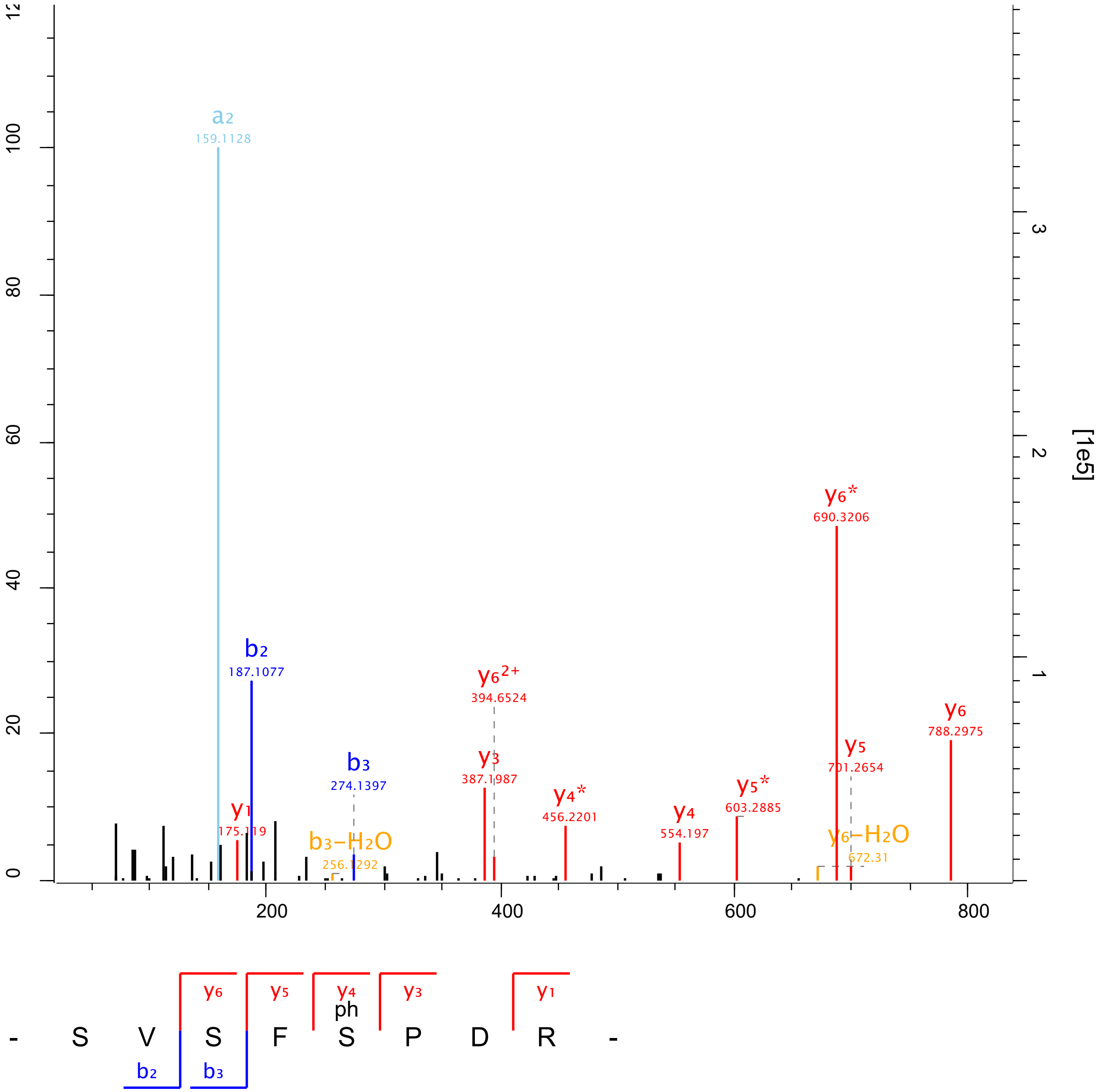

| Raw file | Scan | Method | Score | m/z |
|---|---|---|---|---|
| A17_02_1 | 10700 | FTMS; HCD | 187.47 | 770.33 |

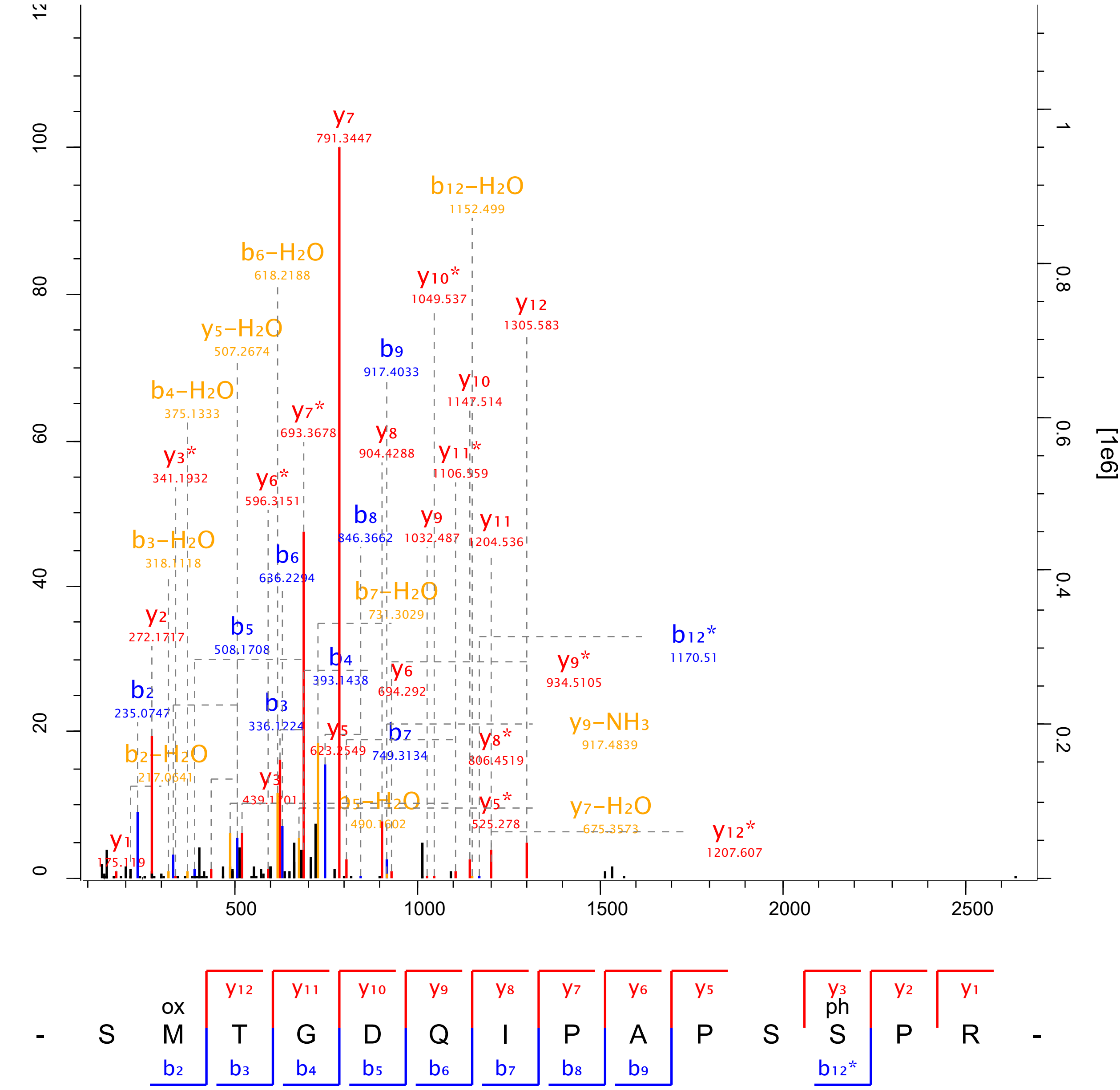

| Raw file | Scan | Method | Score | m/z |
|---|---|---|---|---|
| A17_02_1 | 10704 | FTMS; HCD | 139.58 | 697.28 |

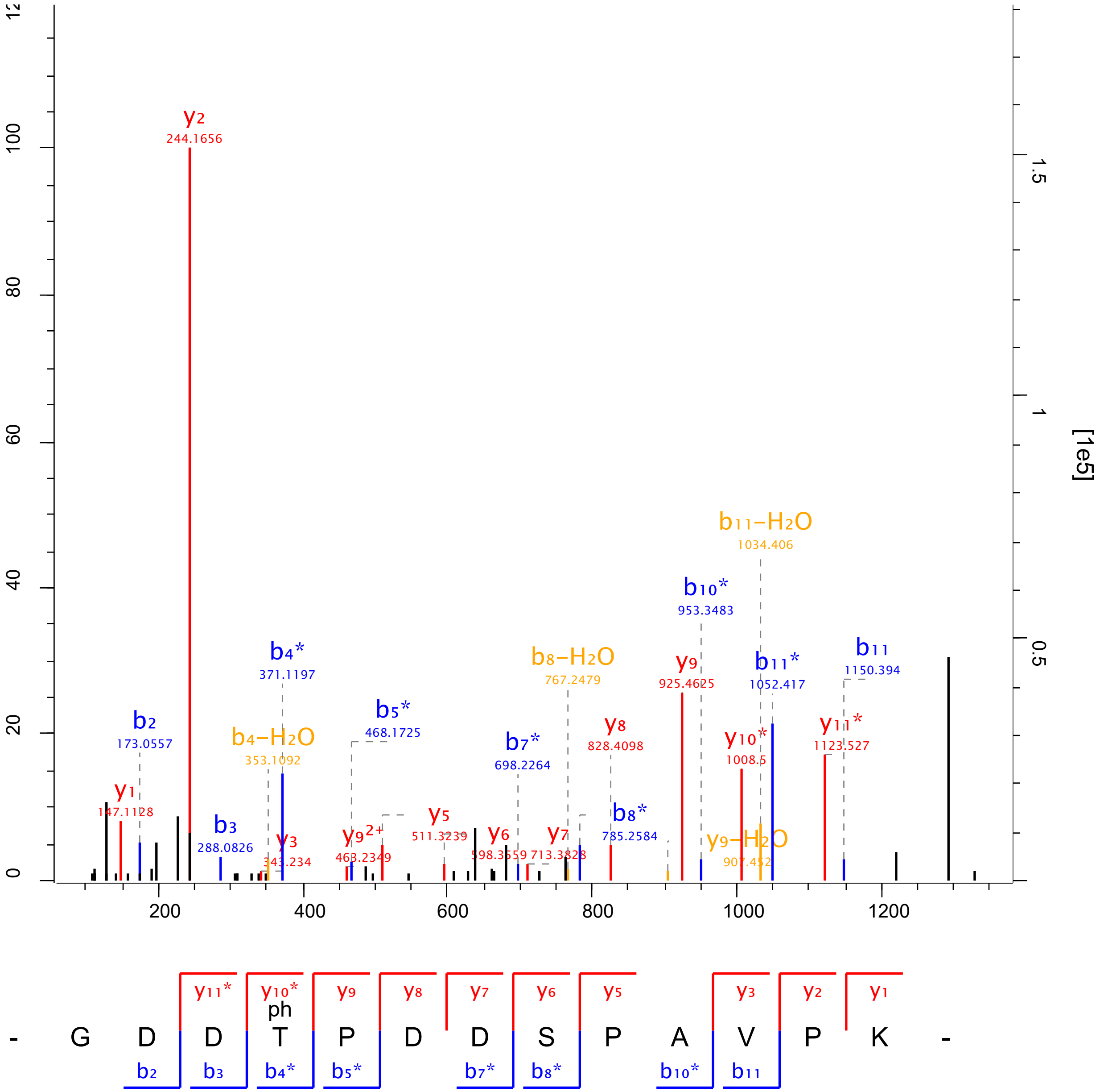

| Raw file | Scan | Method | Score | m/z |
|---|---|---|---|---|
| A17_02_1 | 10841 | FTMS; HCD | 44.93 | 755.85 |

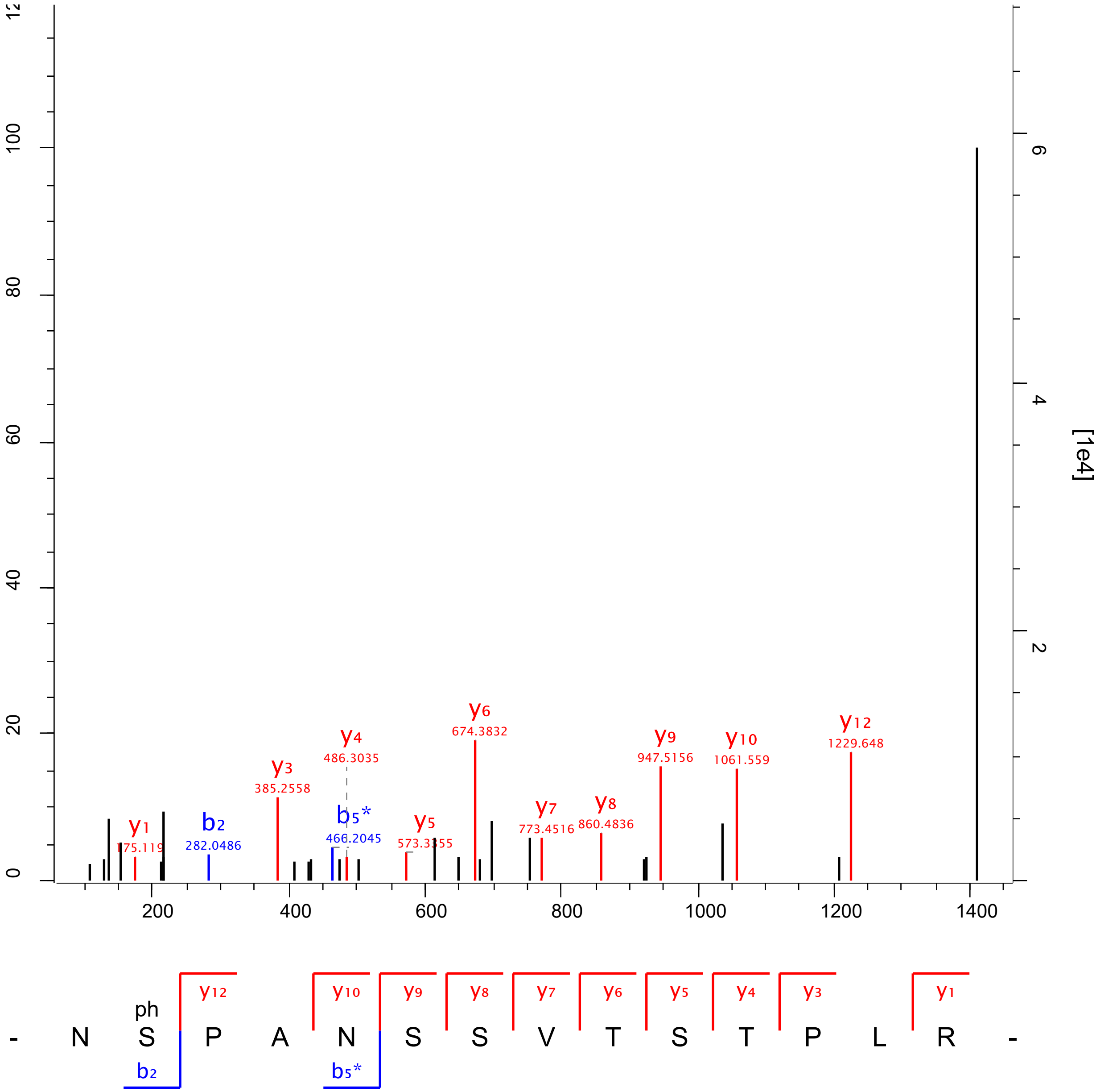

| Raw file | Scan | Method | Score | m/z |
|---|---|---|---|---|
| A17_02_1 | 11208 | FTMS; HCD | 78.32 | 586.75 |

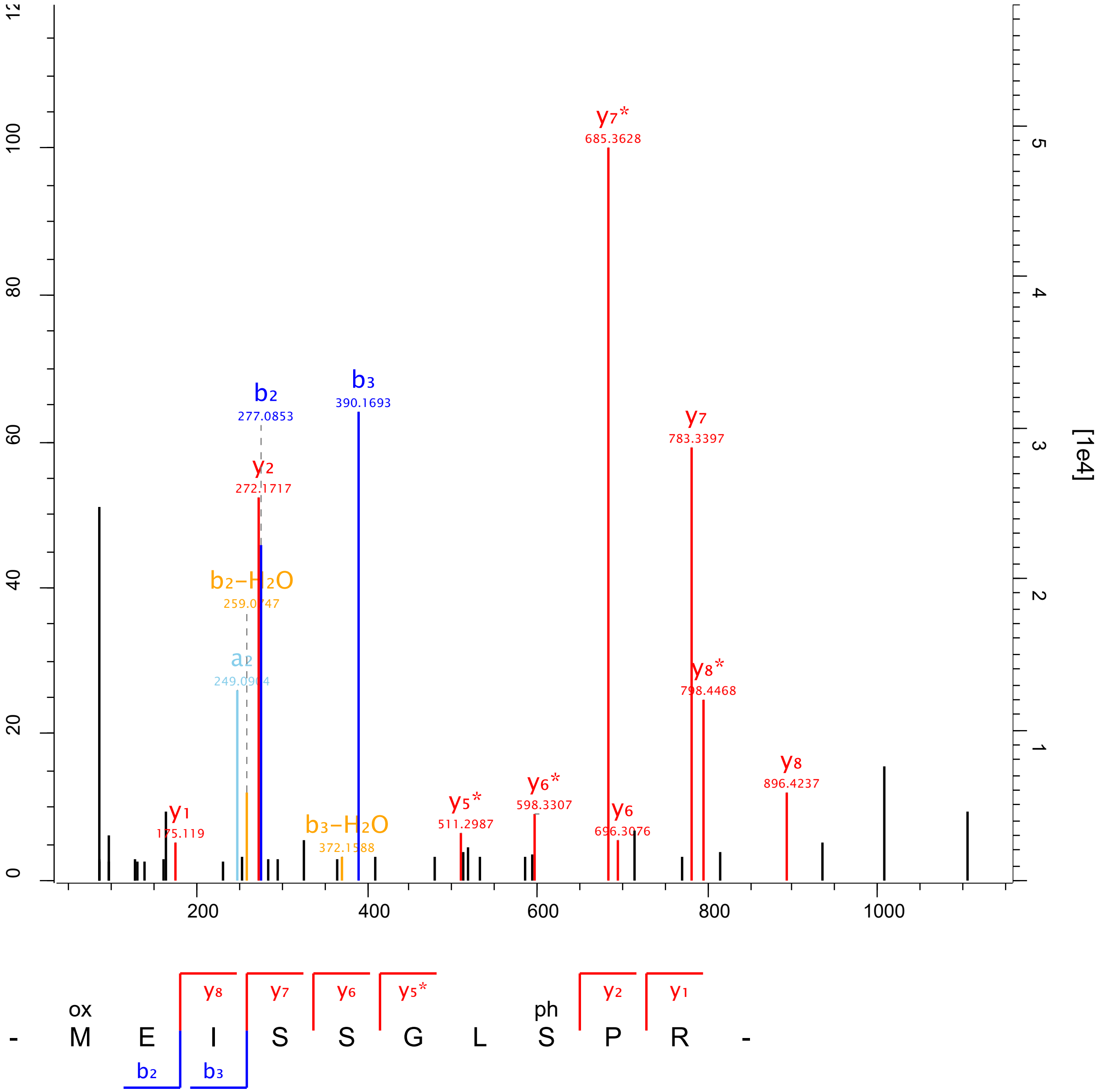

| Raw file | Scan | Method | Score | m/z |
| --- | --- | --- | --- | --- |
| A17_02_1 | 11507 | FTMS; HCD | 159.99 | 556.77 |

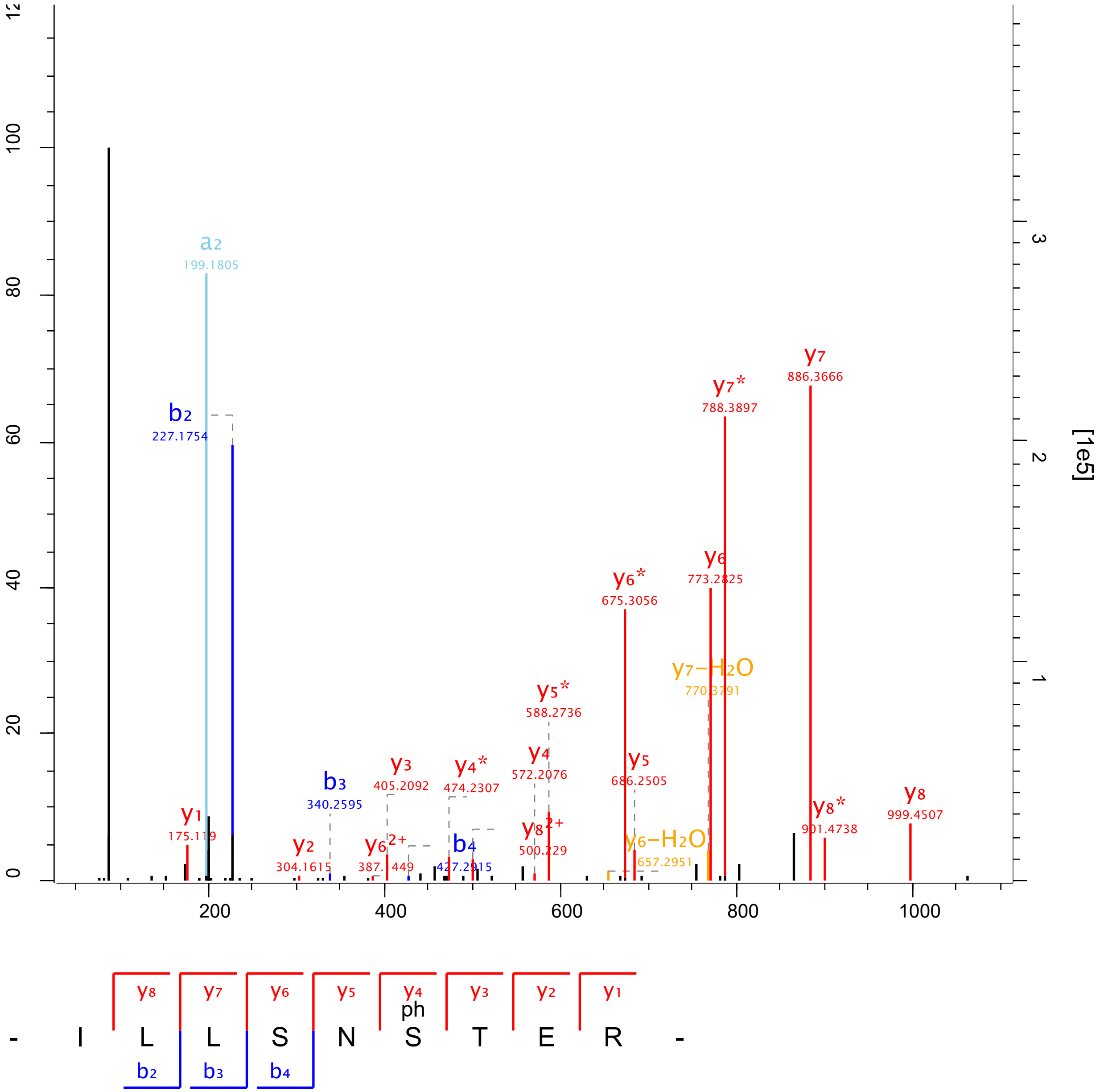

| Raw file | Scan | Method | Score | m/z |
| --- | --- | --- | --- | --- |
| A17_02_1 | 11653 | FTMS; HCD | 92.25 | 555.25 |

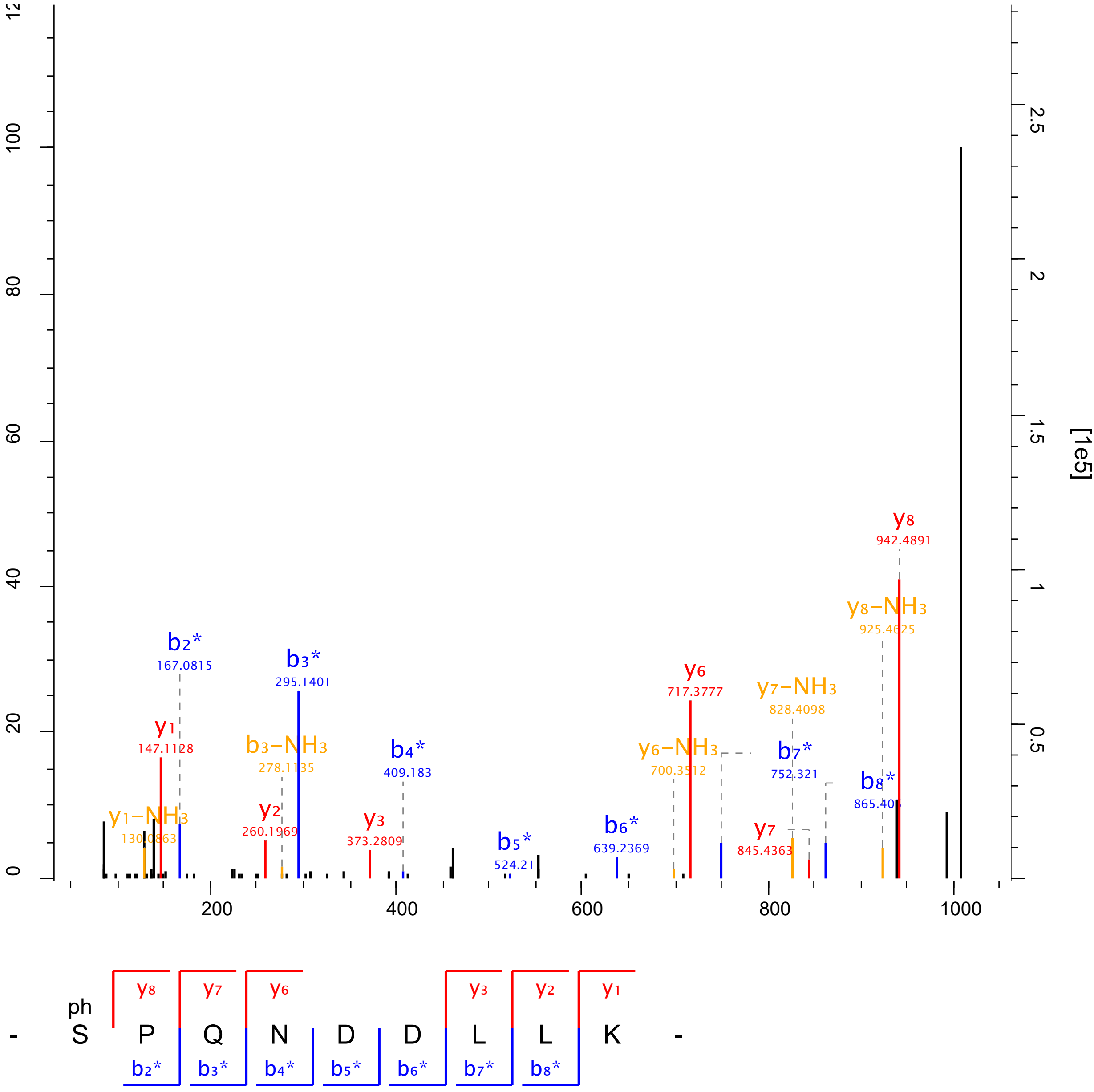

| Raw file | Scan | Method | Score | m/z |
|---|---|---|---|---|
| A17_02_1 | 11837 | FTMS; HCD | 95.48 | 435.7 |

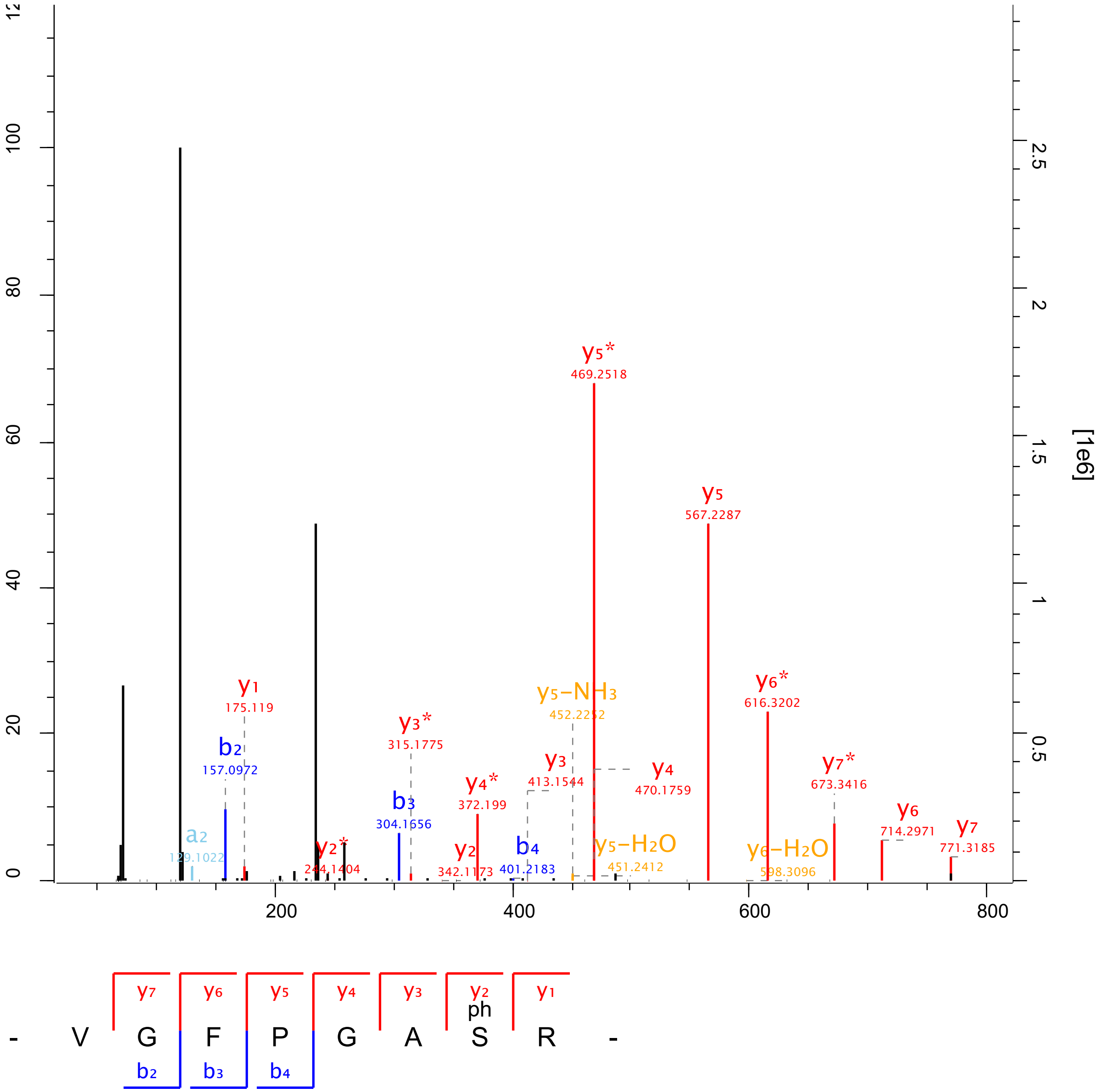

| Raw file | Scan | Method | Score | m/z |
|---|---|---|---|---|
| A17_02_1 | 12124 | FTMS; HCD | 62.2 | 773.33 |

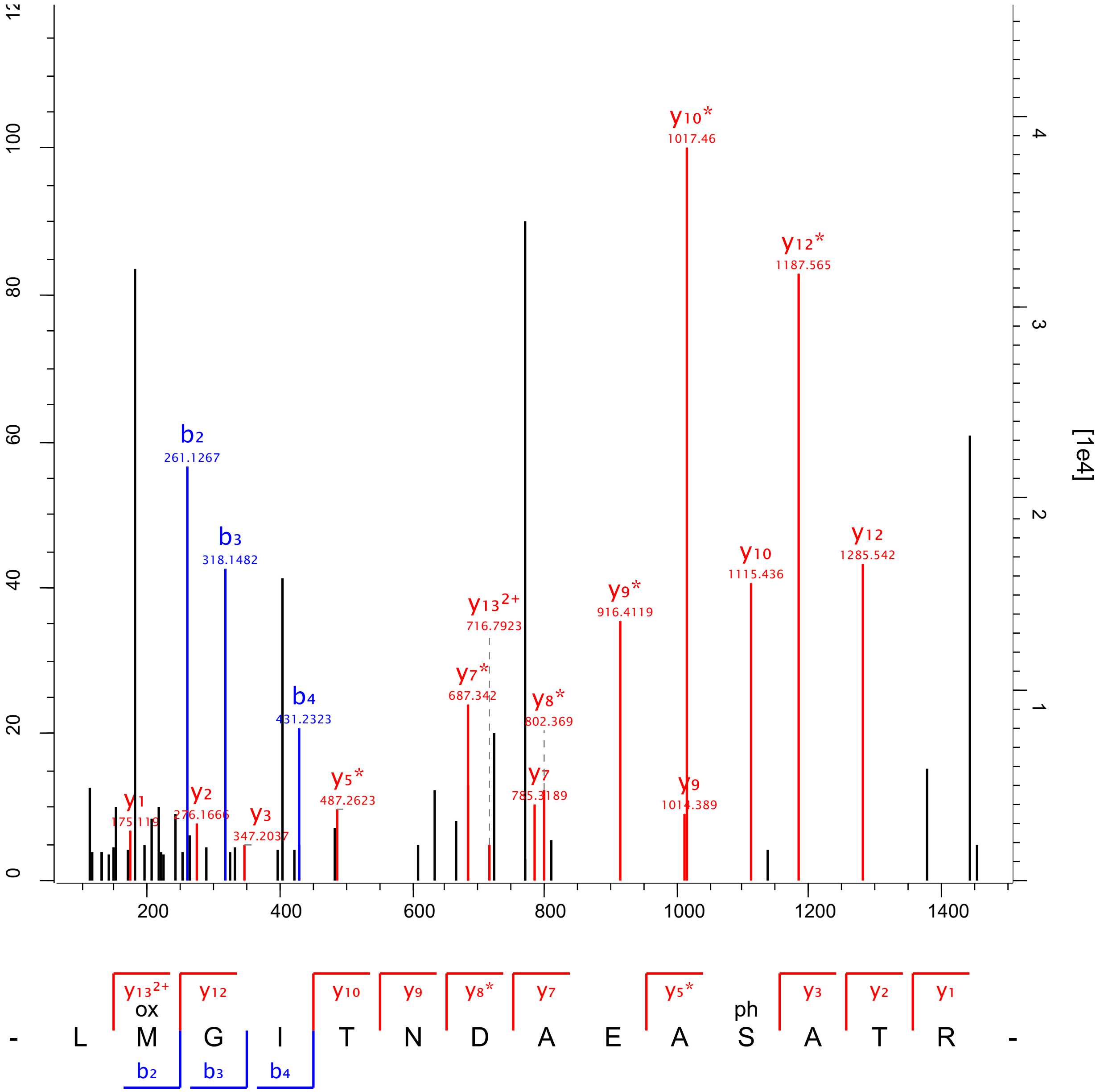

| Raw file | Scan | Method | Score | m/z |
| --- | --- | --- | --- | --- |
| A17_02_1 | 12197 | FTMS; HCD | 41.7 | 575.7 |

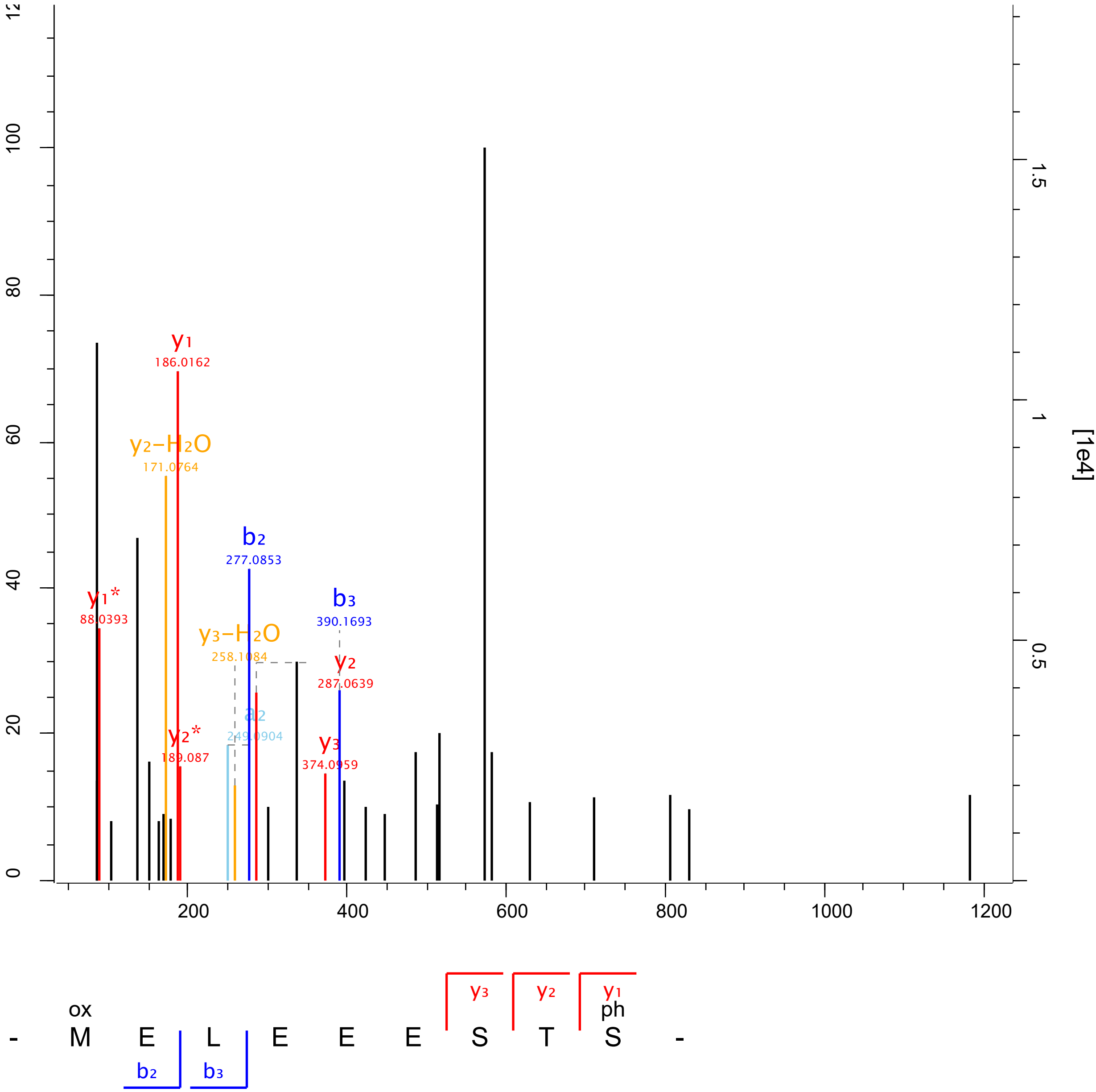

| Raw file | Scan | Method | Score | m/z |
|---|---|---|---|---|
| A17_02_1 | 12899 | FTMS; HCD | 184.27 | 547.24 |

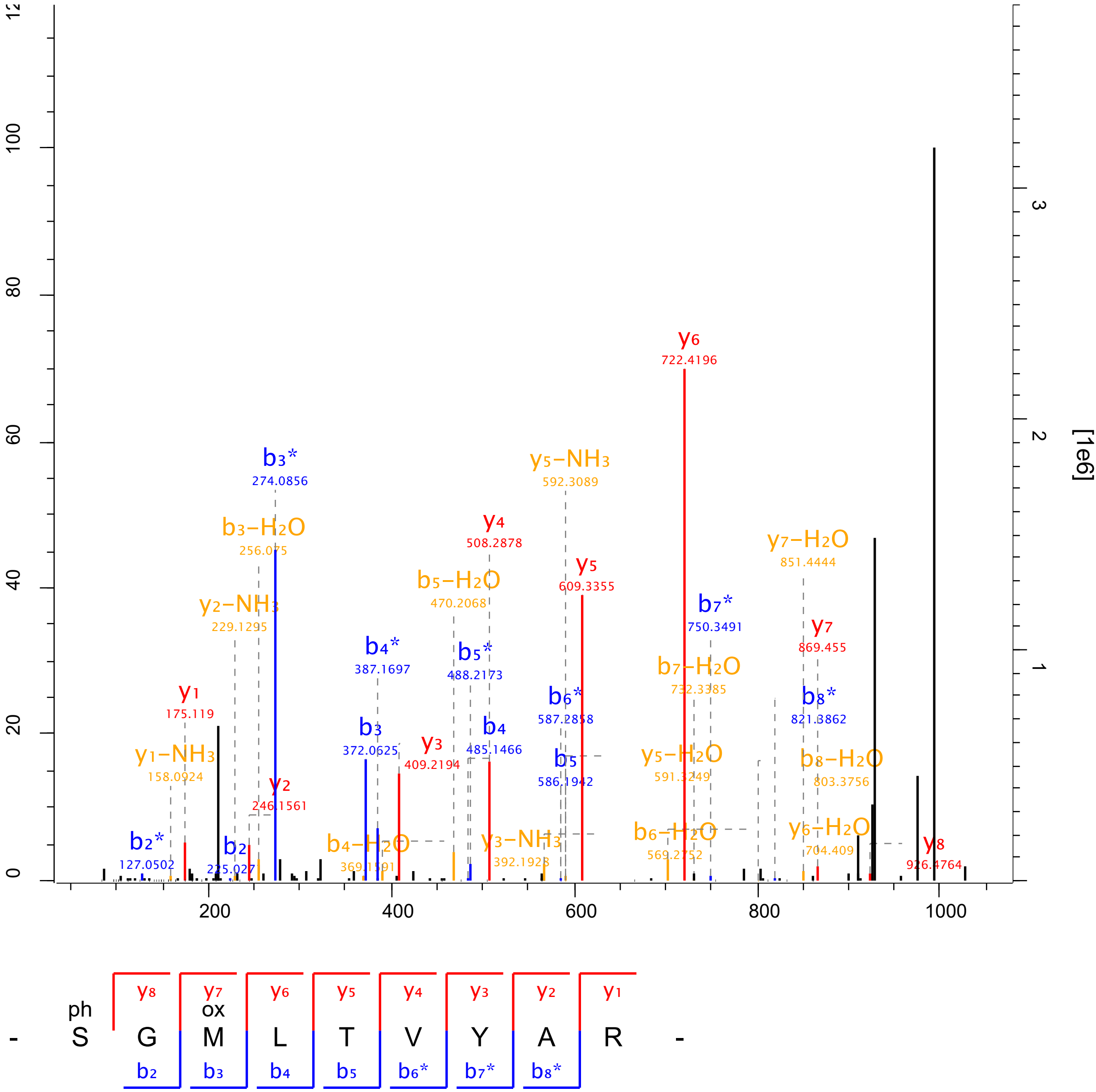

| Raw file | Scan | Method | Score | m/z |
|---|---|---|---|---|
| A17_02_1 | 13312 | FTMS; HCD | 327.55 | 682.77 |

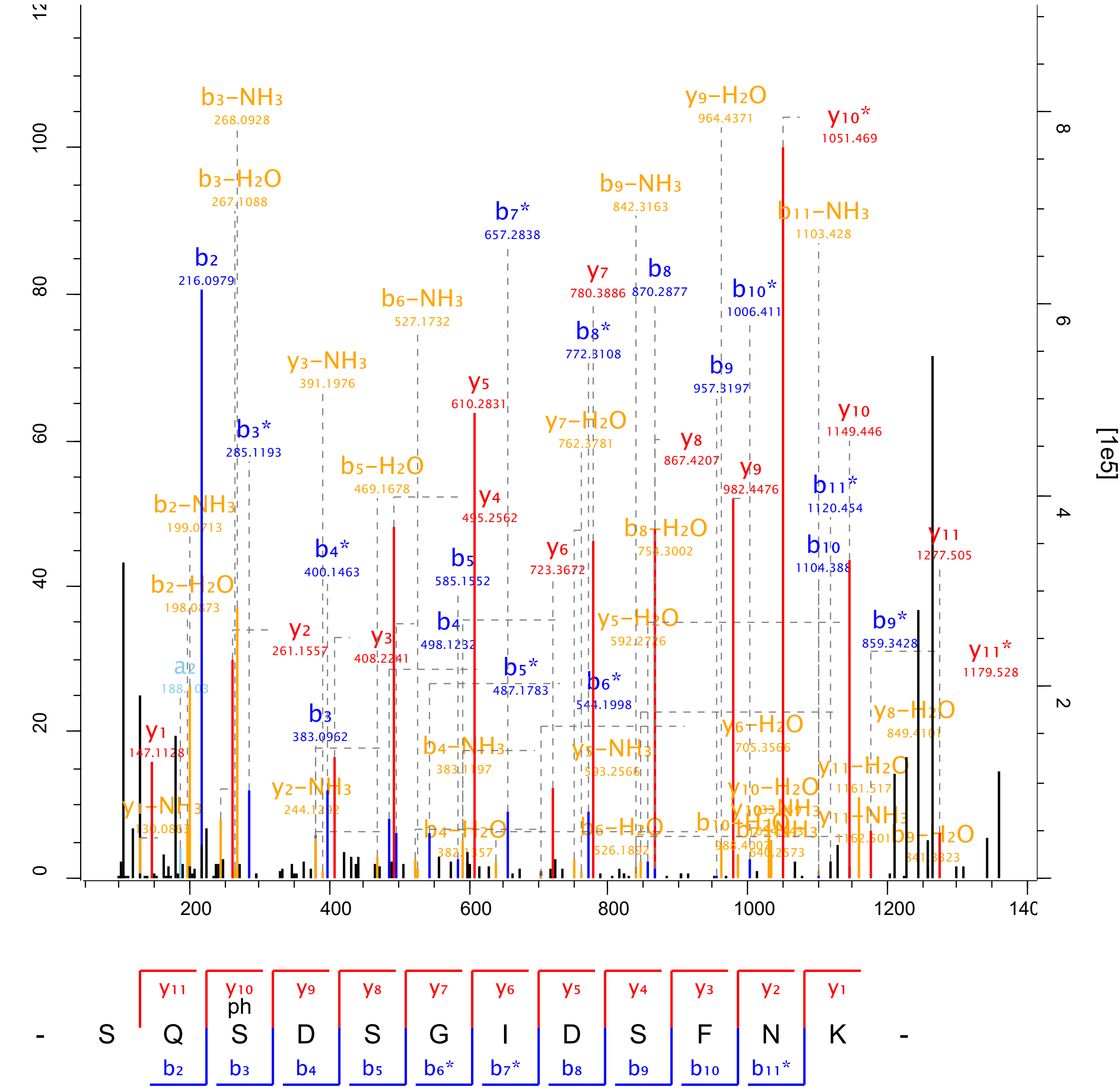

| Raw file | Scan | Method | Score | m/z |
|---|---|---|---|---|
| A17_02_1 | 13691 | FTMS; HCD | 62.47 | 515.22 |

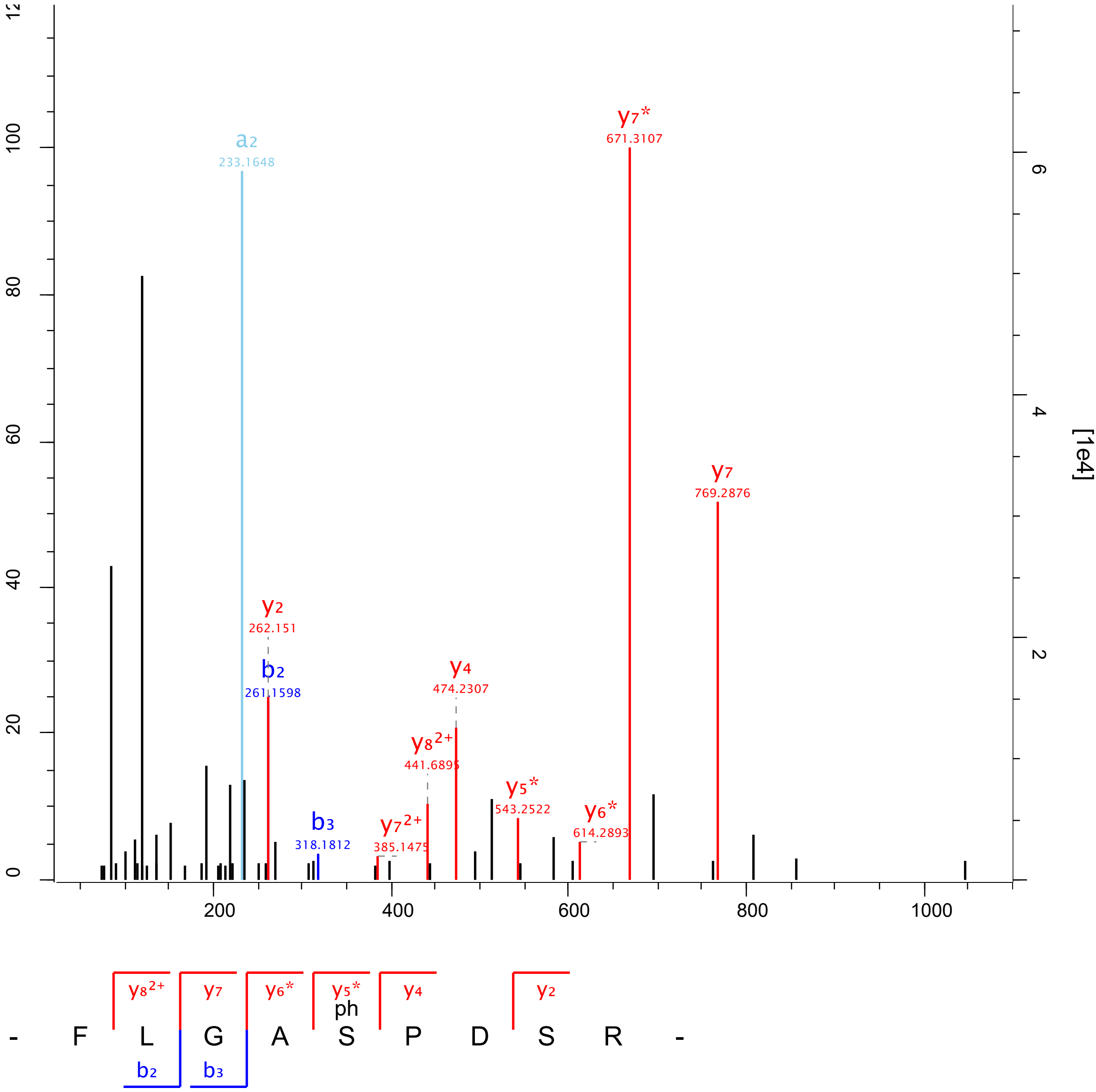

| Raw file | Scan | Method | Score | m/z |
|---|---|---|---|---|
| A17_02_1 | 13882 | FTMS; HCD | 269.53 | 977.91 |

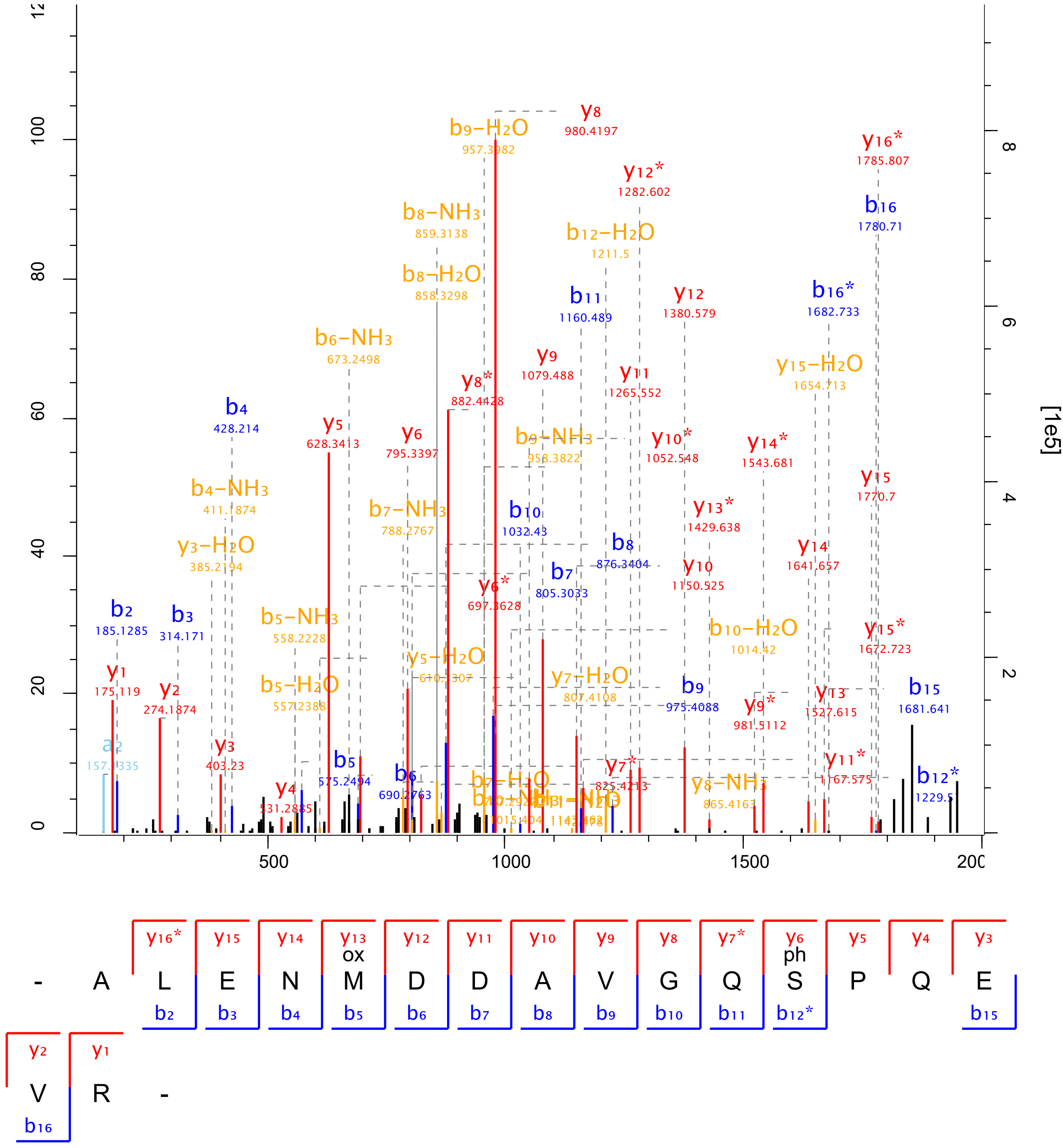

| Raw file | Scan | Method | Score | m/z |
|---|---|---|---|---|
| A17_02_1 | 14144 | FTMS; HCD | 228.76 | 780.32 |

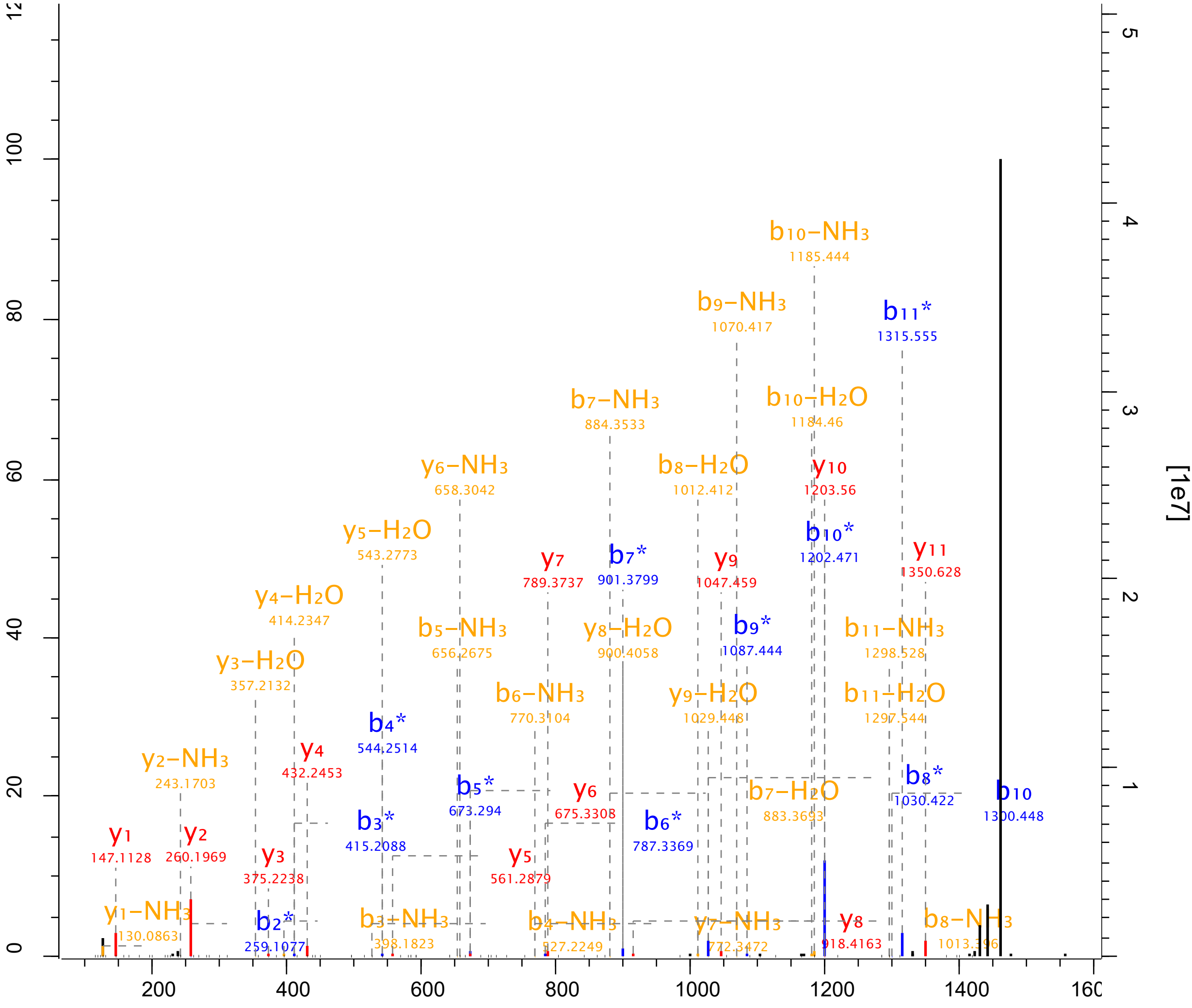


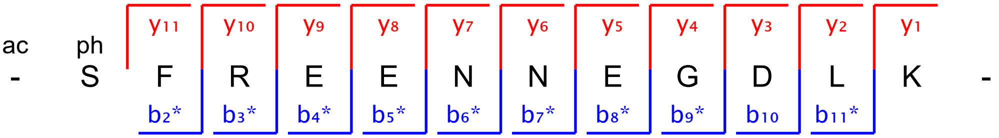

| Raw file | Scan | Method | Score | m/z |
|---|---|---|---|---|
| A17_02_1 | 15247 | FTMS; HCD | 109.83 | 680.31 |

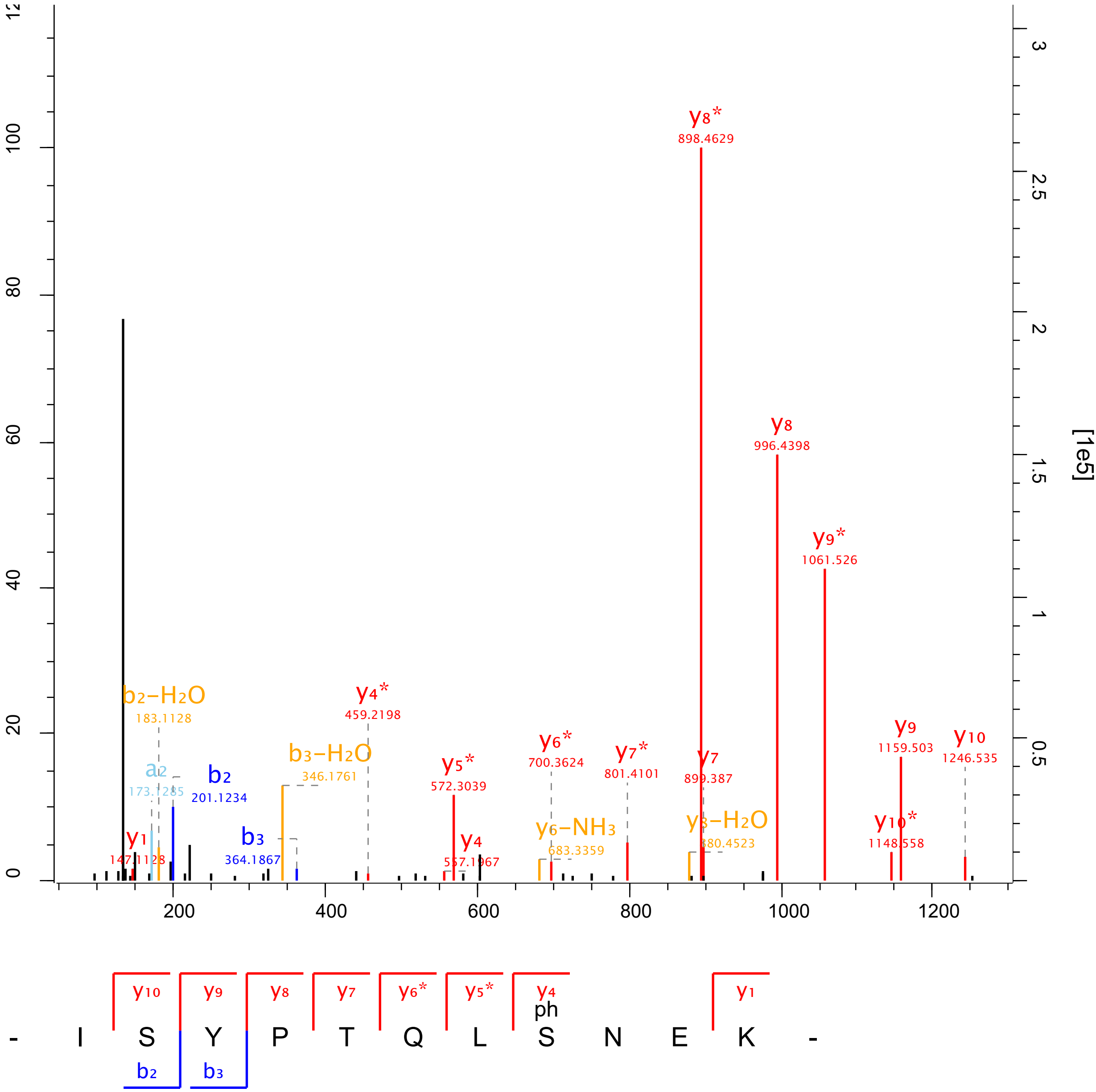

| Raw file | Scan | Method | Score | m/z |
| --- | --- | --- | --- | --- |
| A17_02_1 | 15665 | FTMS; HCD | 80.36 | 751.3 |

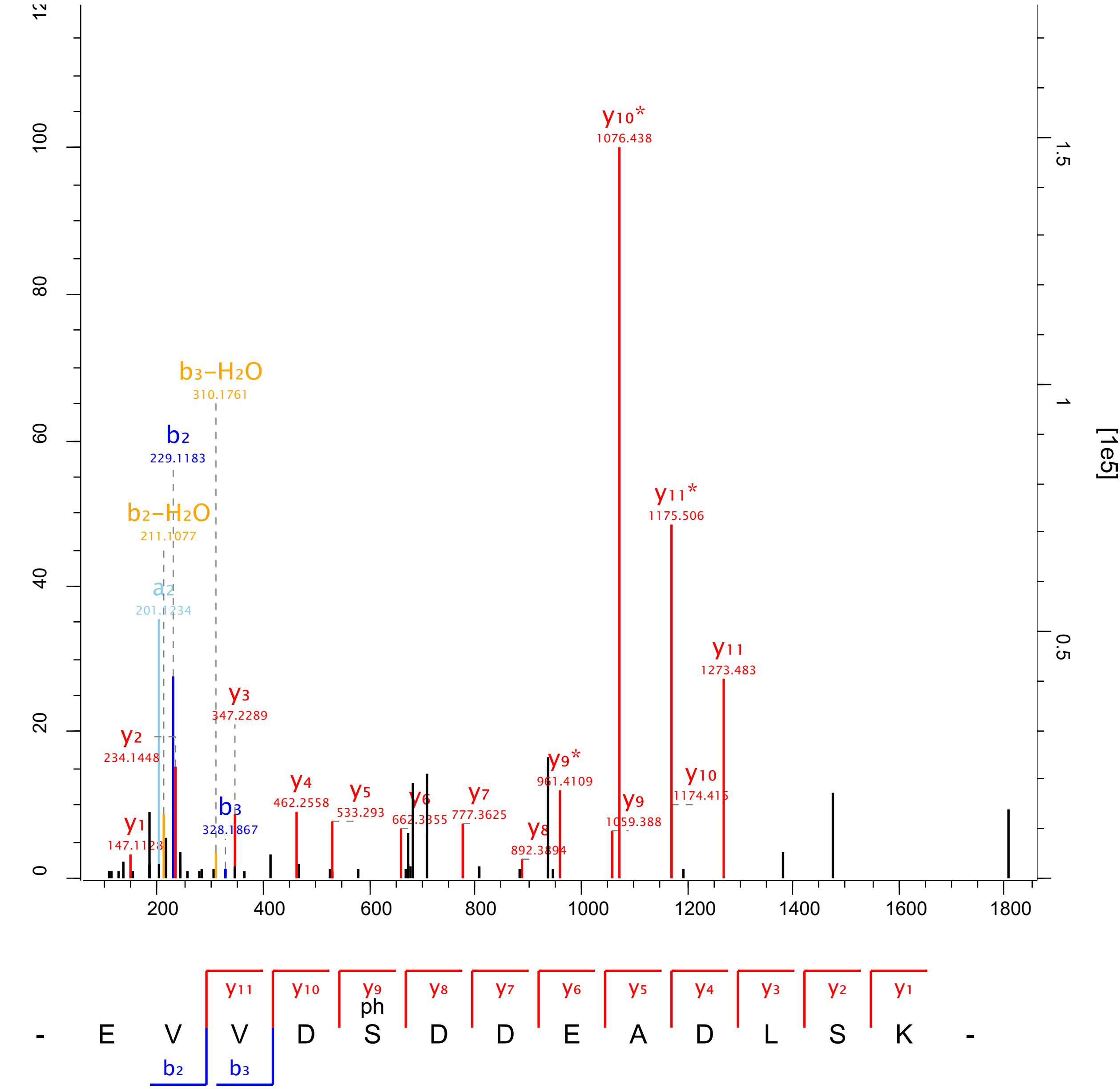

| Raw file | Scan | Method | Score | m/z |
| --- | --- | --- | --- | --- |
| A17_02_1 | 15858 | FTMS; HCD | 91.4 | 681.31 |

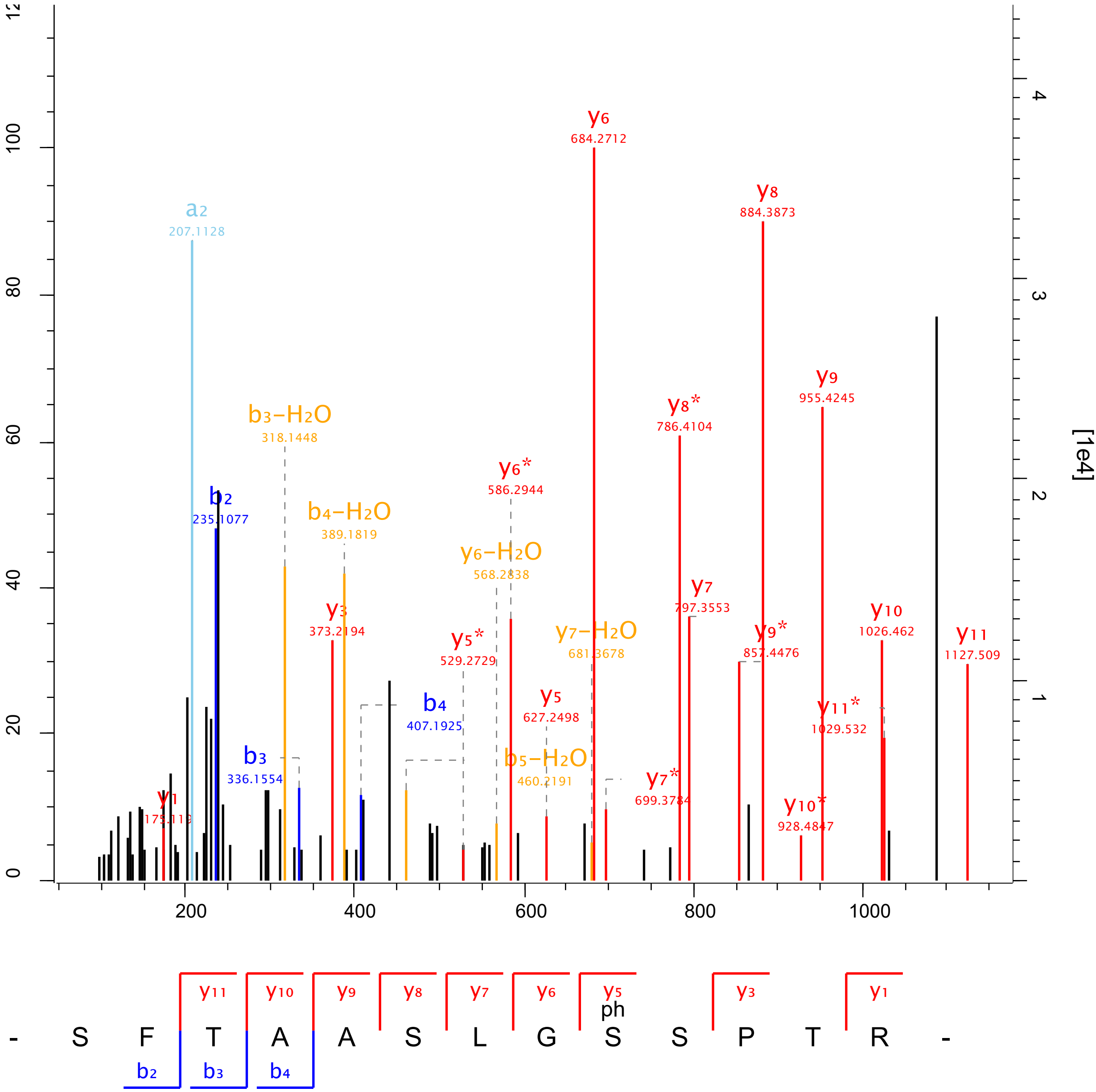

| Raw file | Scan | Method | Score | m/z |
|---|---|---|---|---|
| A17_02_1 | 1589 | FTMS; HCD | 59.2 | 572.72 |

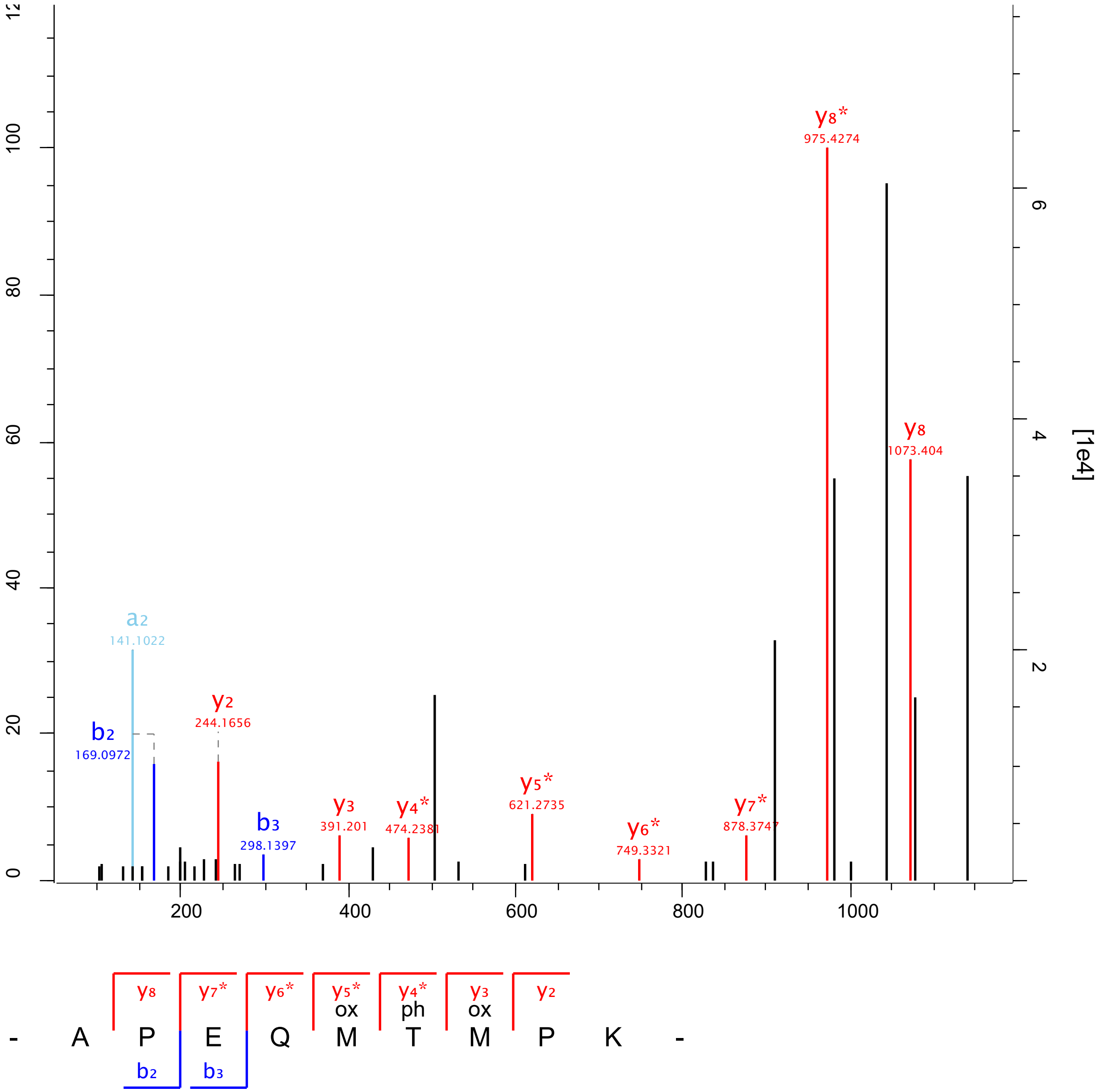

| Raw file | Scan | Method | Score | m/z |
| --- | --- | --- | --- | --- |
| A17_02_1 | 16126 | FTMS; HCD | 76.28 | 481.7 |

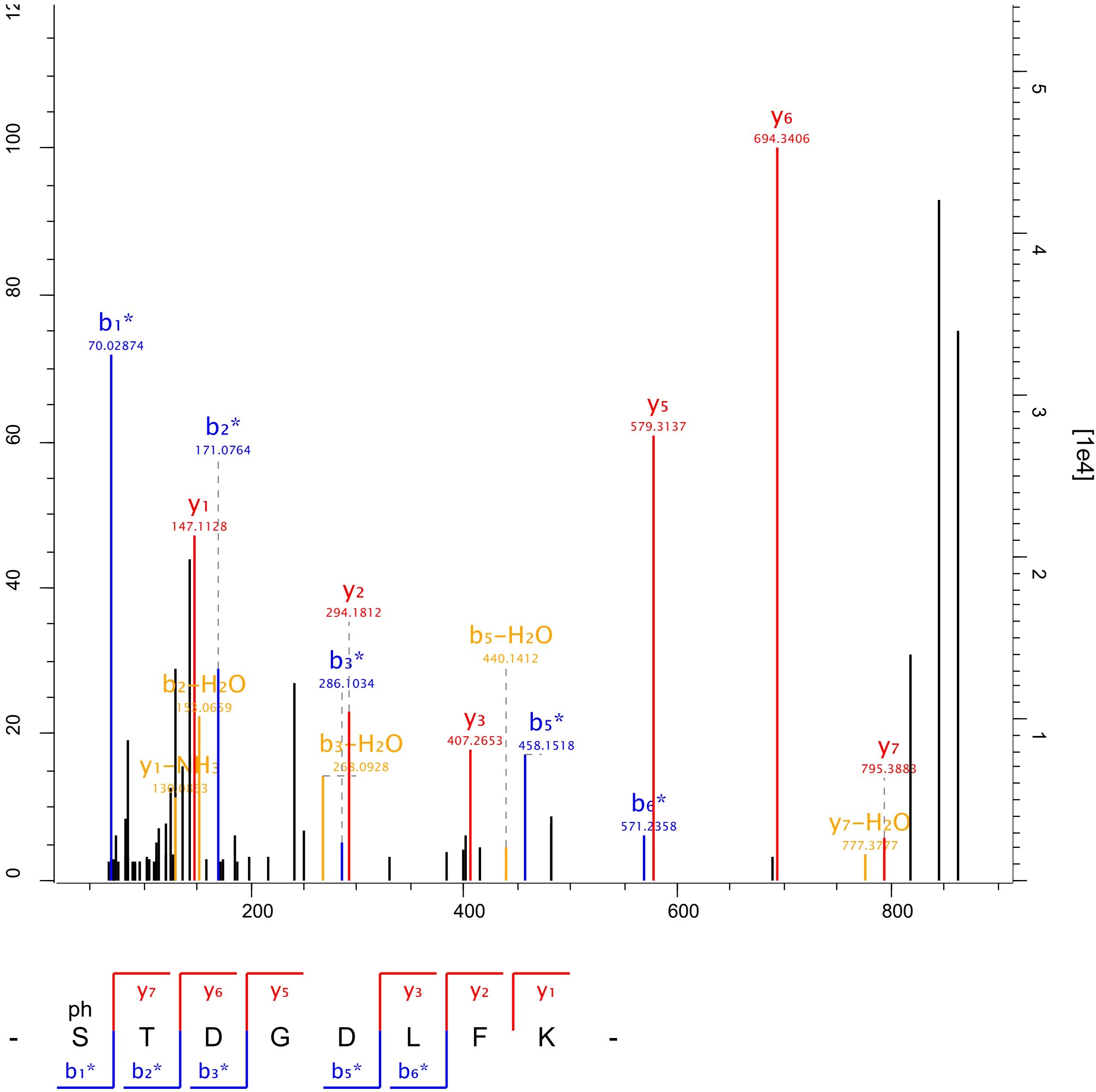

| Raw file | Scan | Method | Score | m/z |
|---|---|---|---|---|
| A17_02_1 | 16231 | FTMS; HCD | 115.91 | 690.3 |

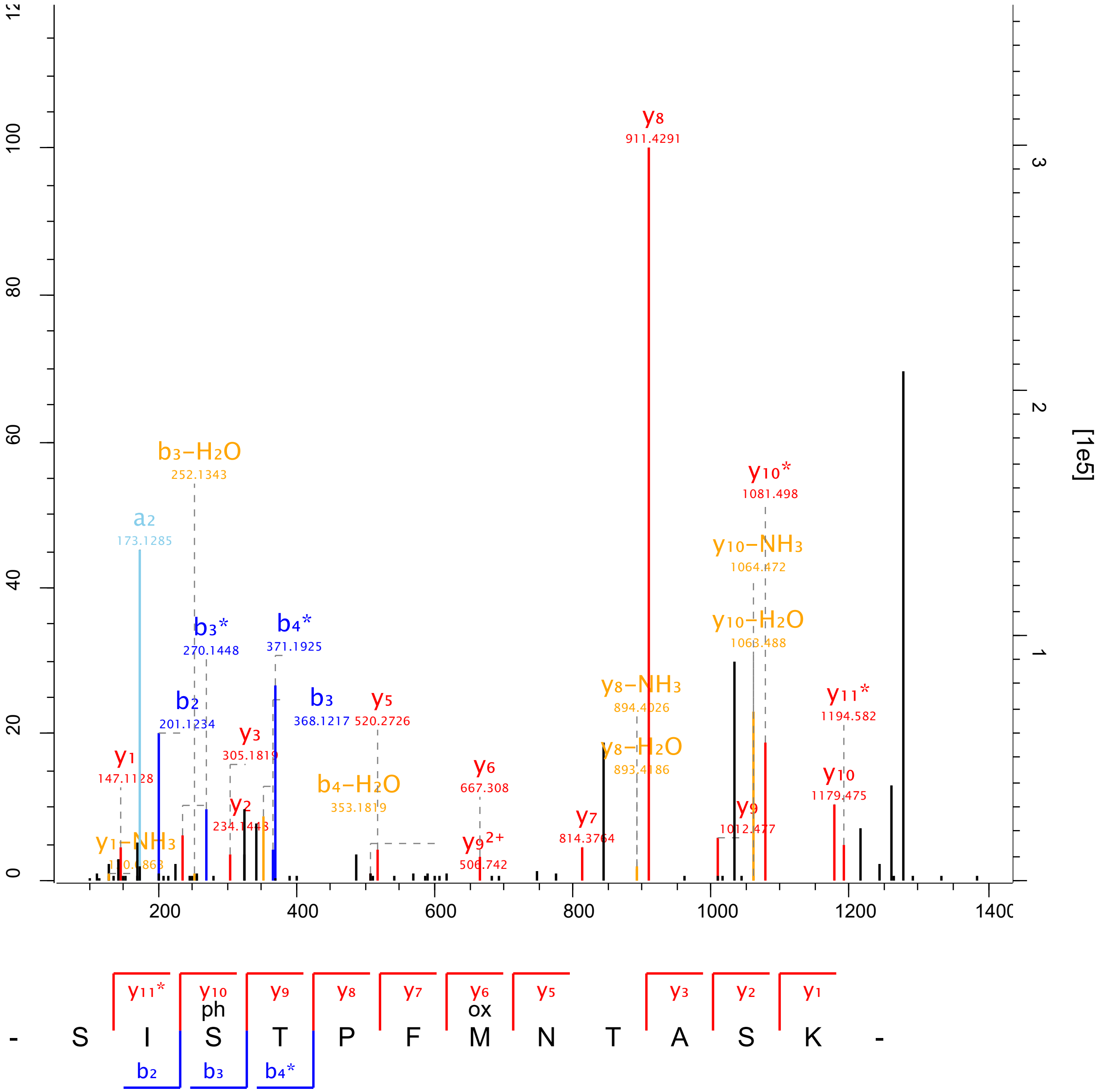

| Raw file | Scan | Method | Score | m/z |
| --- | --- | --- | --- | --- |
| A17_02_1 | 16325 | FTMS; HCD | 68.54 | 589.26 |

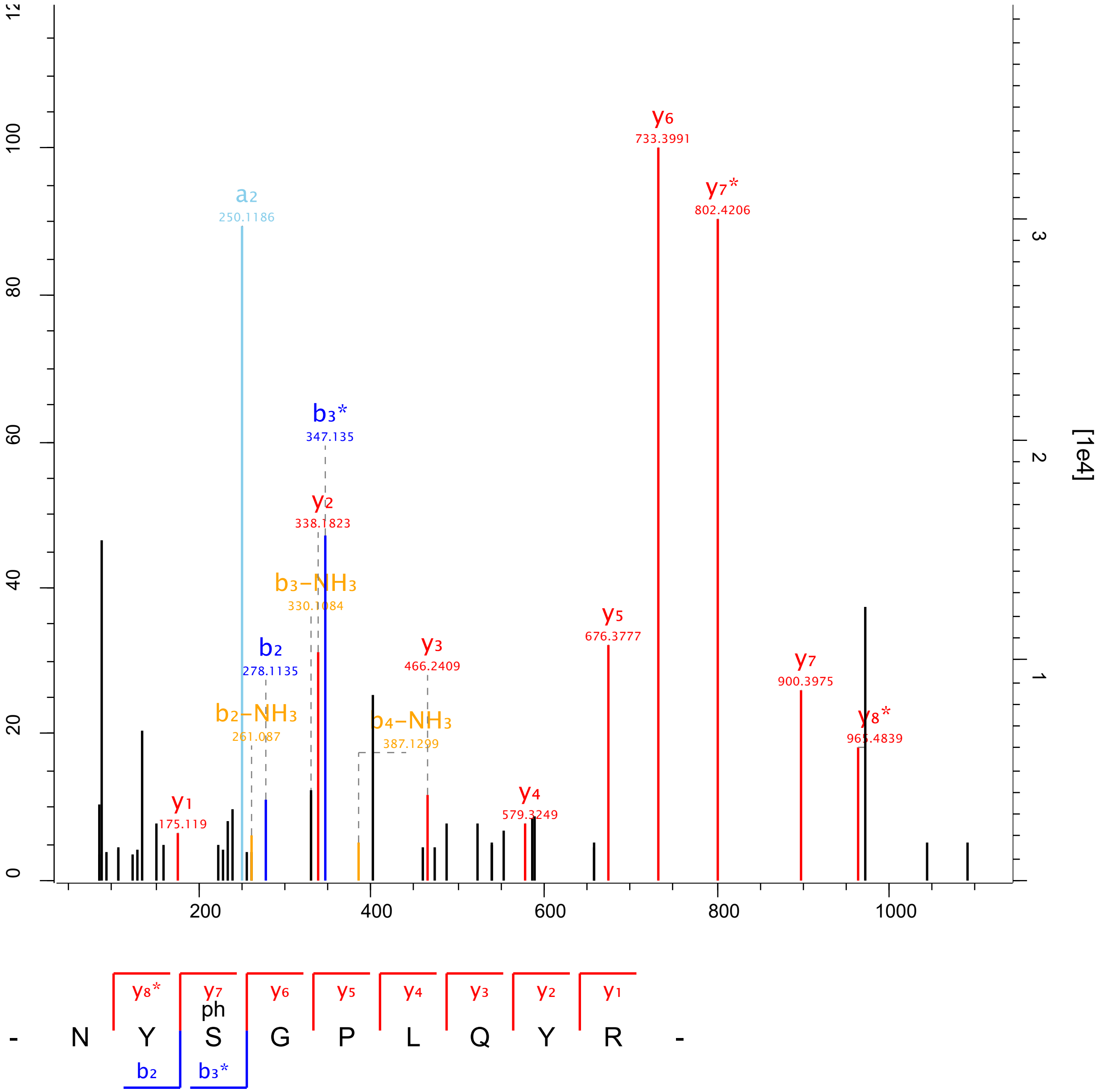

| Raw file | Scan | Method | Score | m/z |
|---|---|---|---|---|
| A17_02_1 | 16419 | FTMS; HCD | 70.89 | 557.22 |

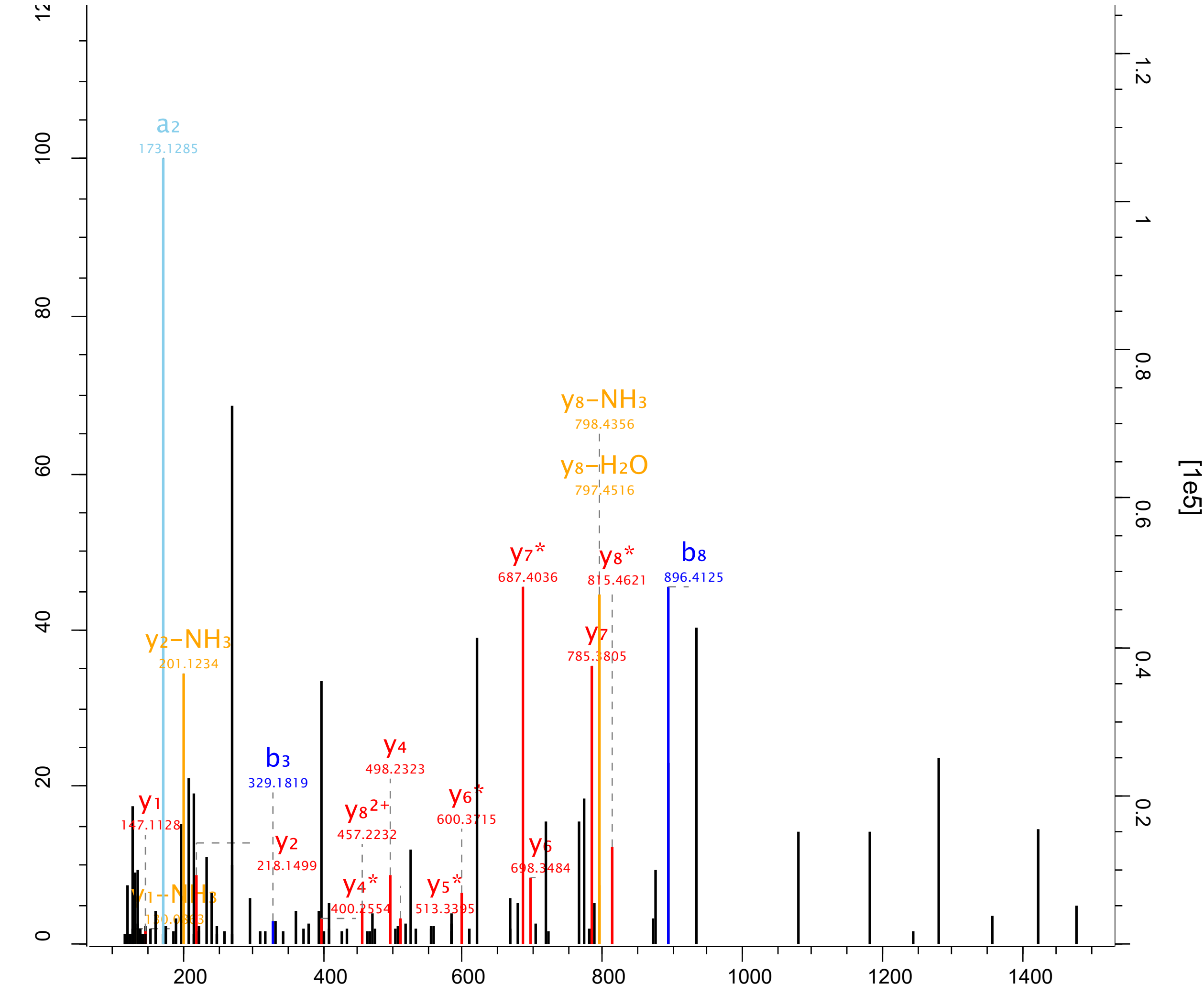


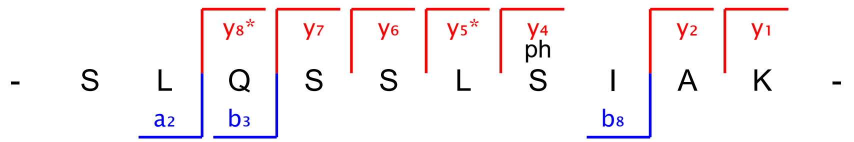

| Raw file | Scan | Method | Score | m/z |
|---|---|---|---|---|
| A17_02_1 | 16427 | FTMS; HCD | 63.57 | 527.27 |

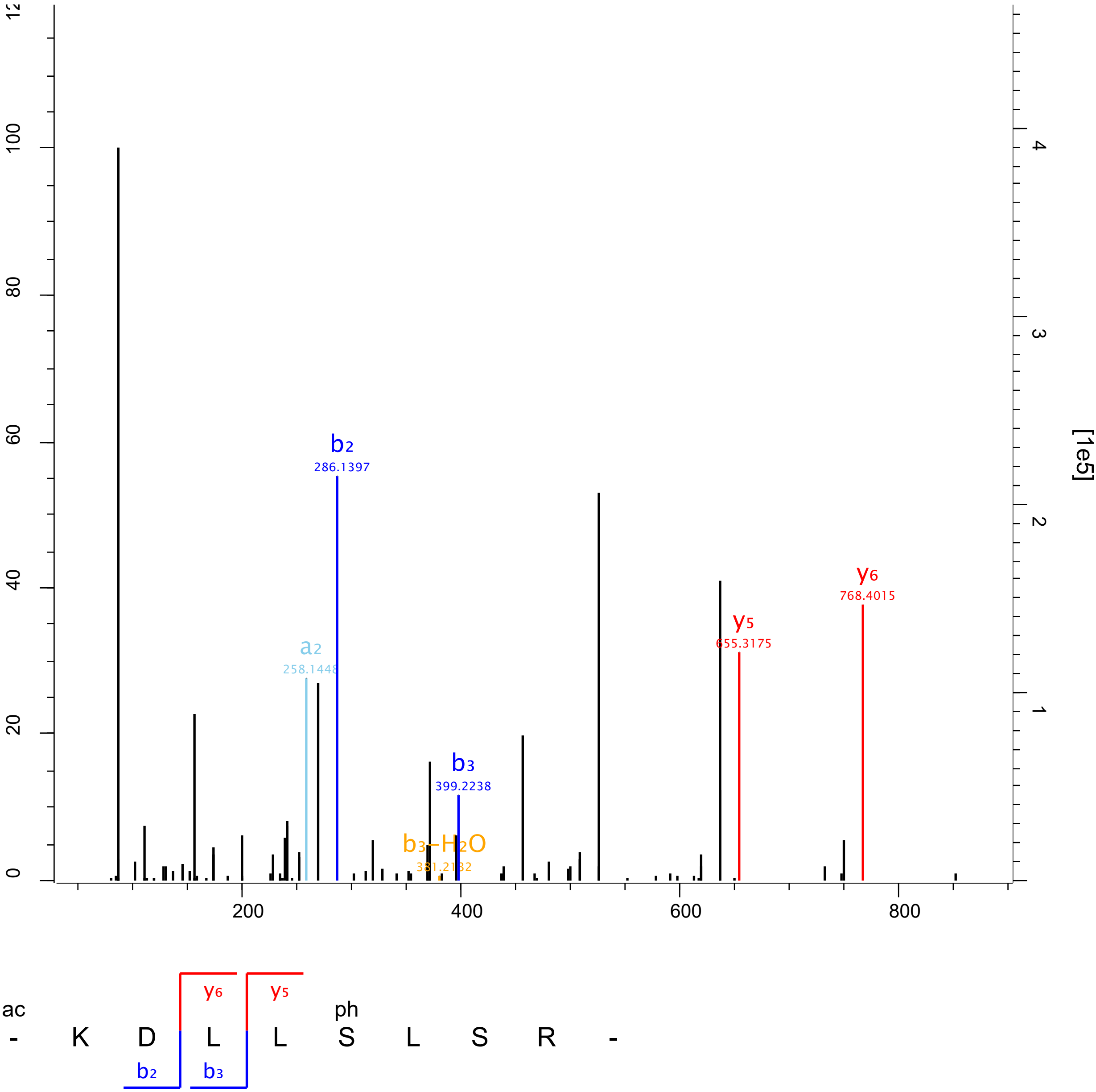

| Raw file | Scan | Method | Score | m/z |
| --- | --- | --- | --- | --- |
| A17_02_1 | 16806 | FTMS; HCD | 54.76 | 832.83 |

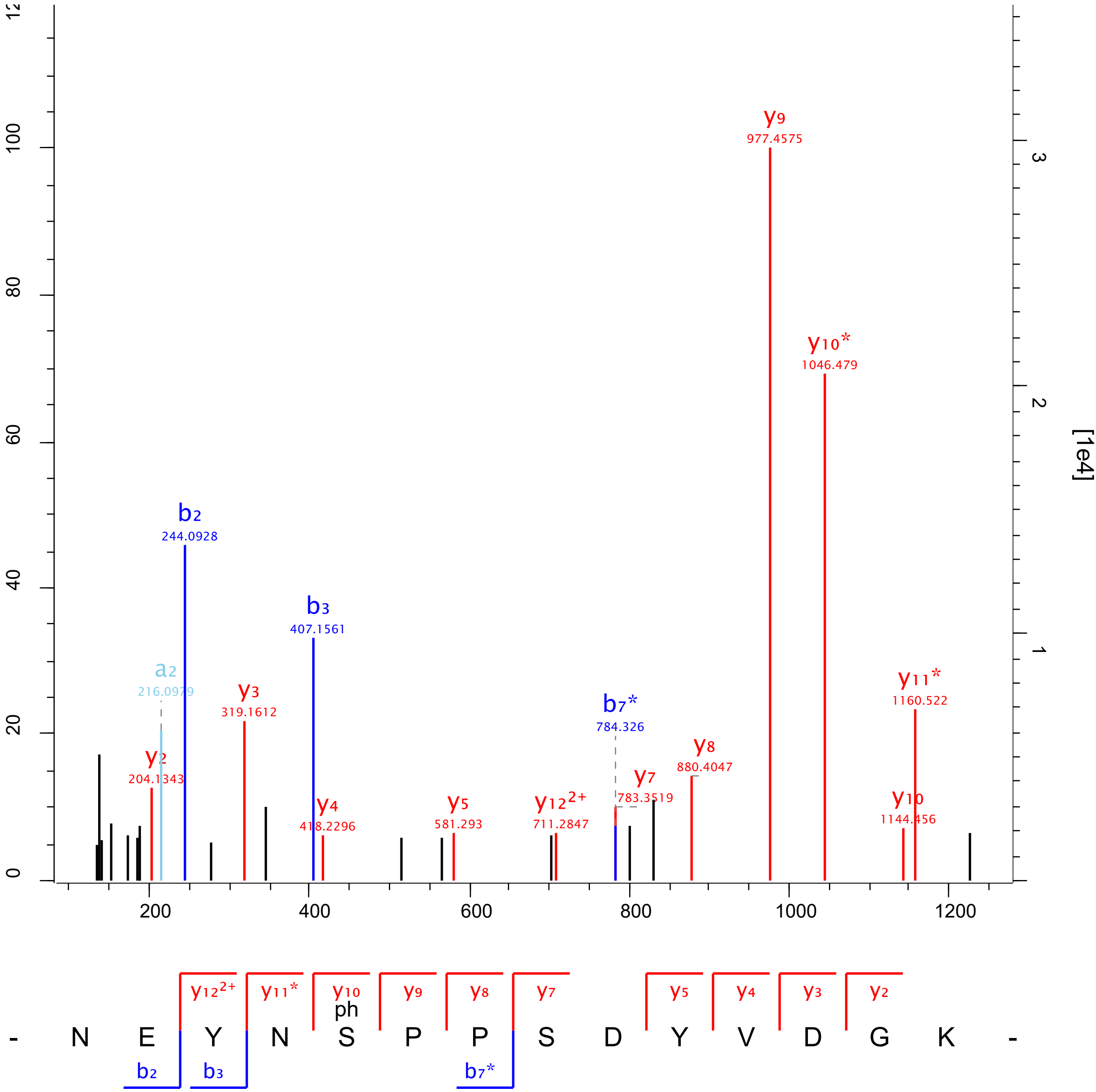

| Raw file | Scan | Method | Score | m/z |
|---|---|---|---|---|
| A17_02_1 | 17197 | FTMS; HCD | 122.49 | 728.84 |

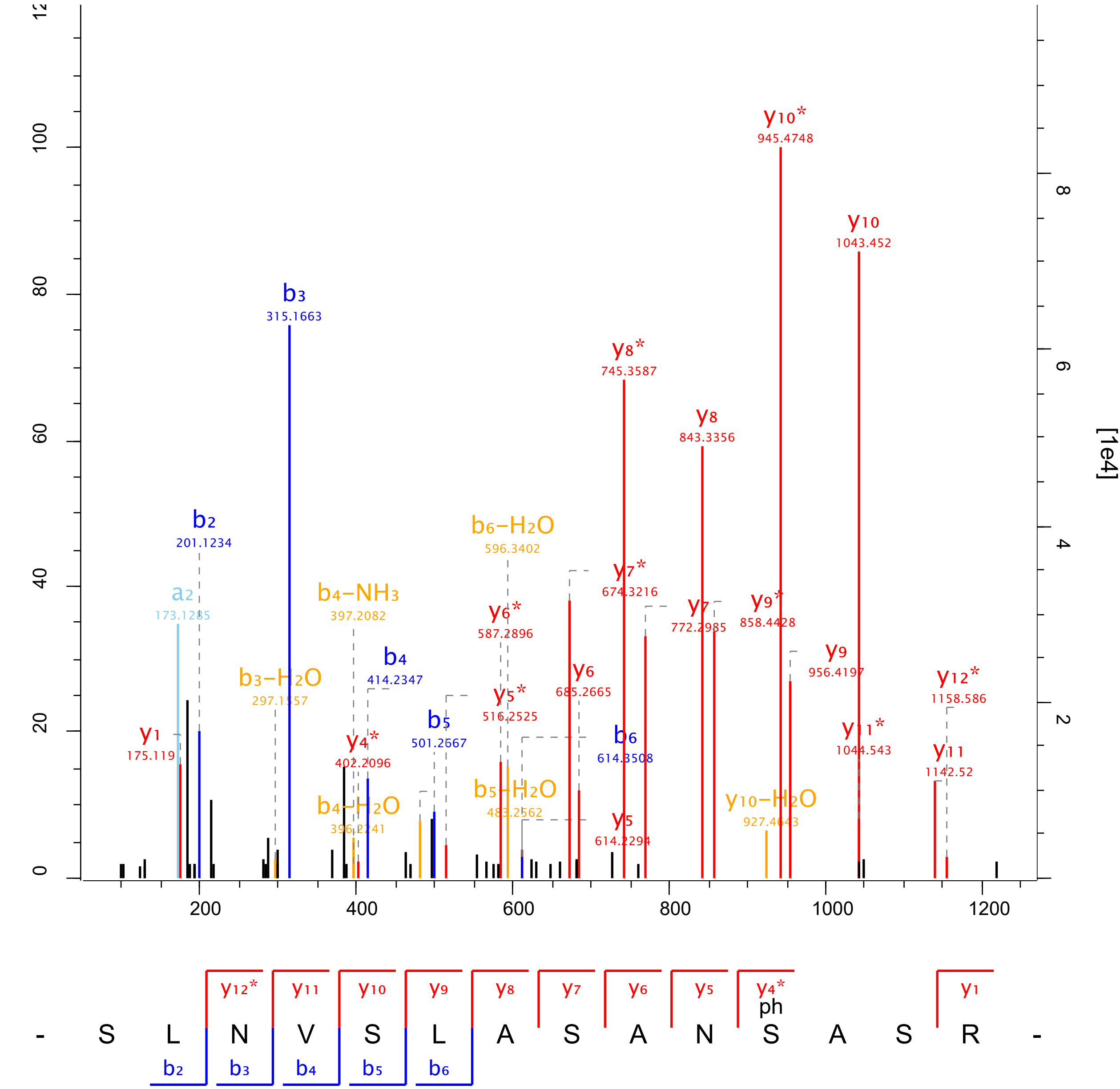

| Raw file | Scan | Method | Score | m/z |
| --- | --- | --- | --- | --- |
| A17_02_1 | 17369 | FTMS; HCD | 145.91 | 806.34 |

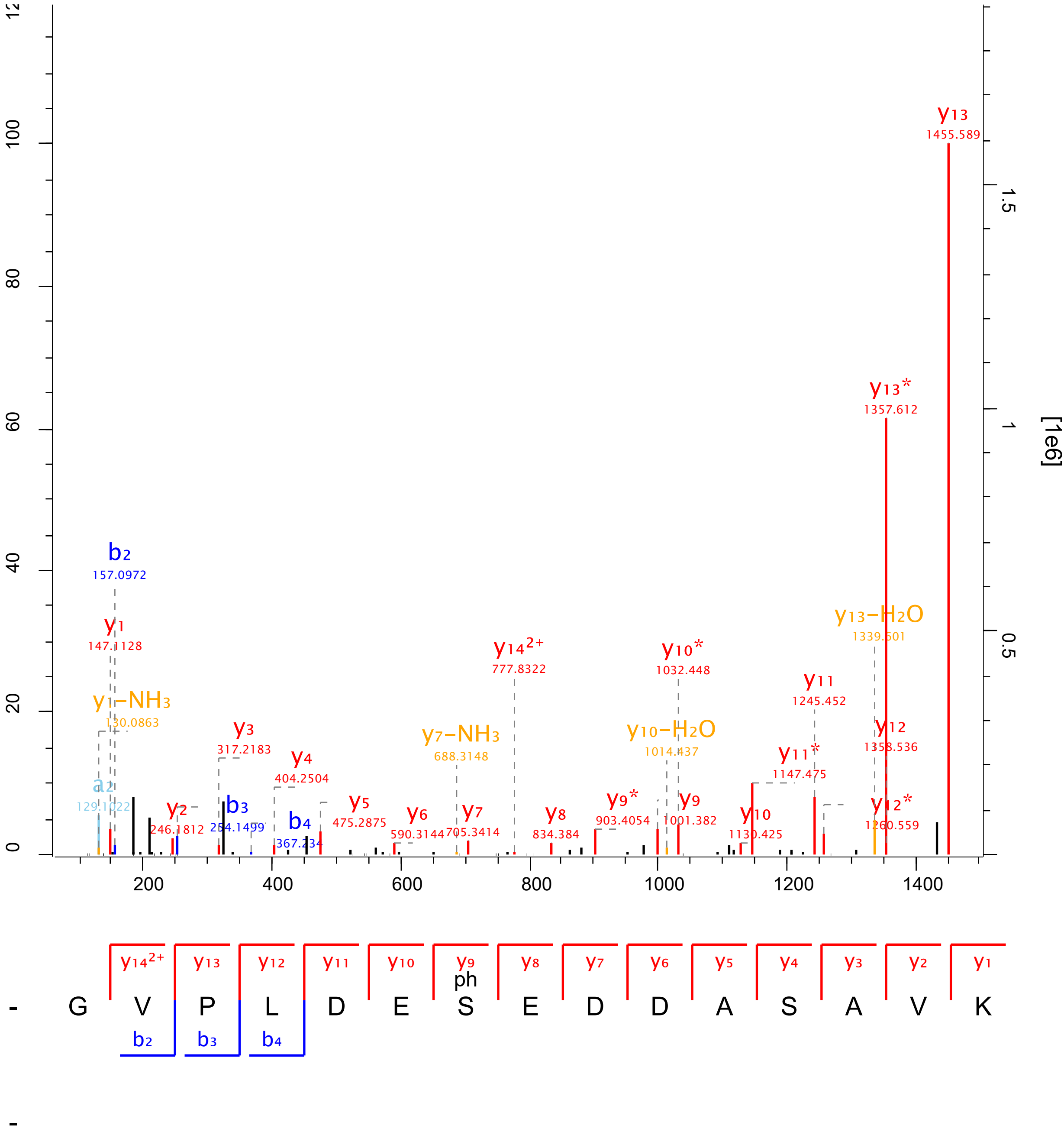

| Raw file | Scan | Method | Score | m/z |
|---|---|---|---|---|
| A17_02_1 | 17488 | FTMS; HCD | 81.63 | 744.77 |

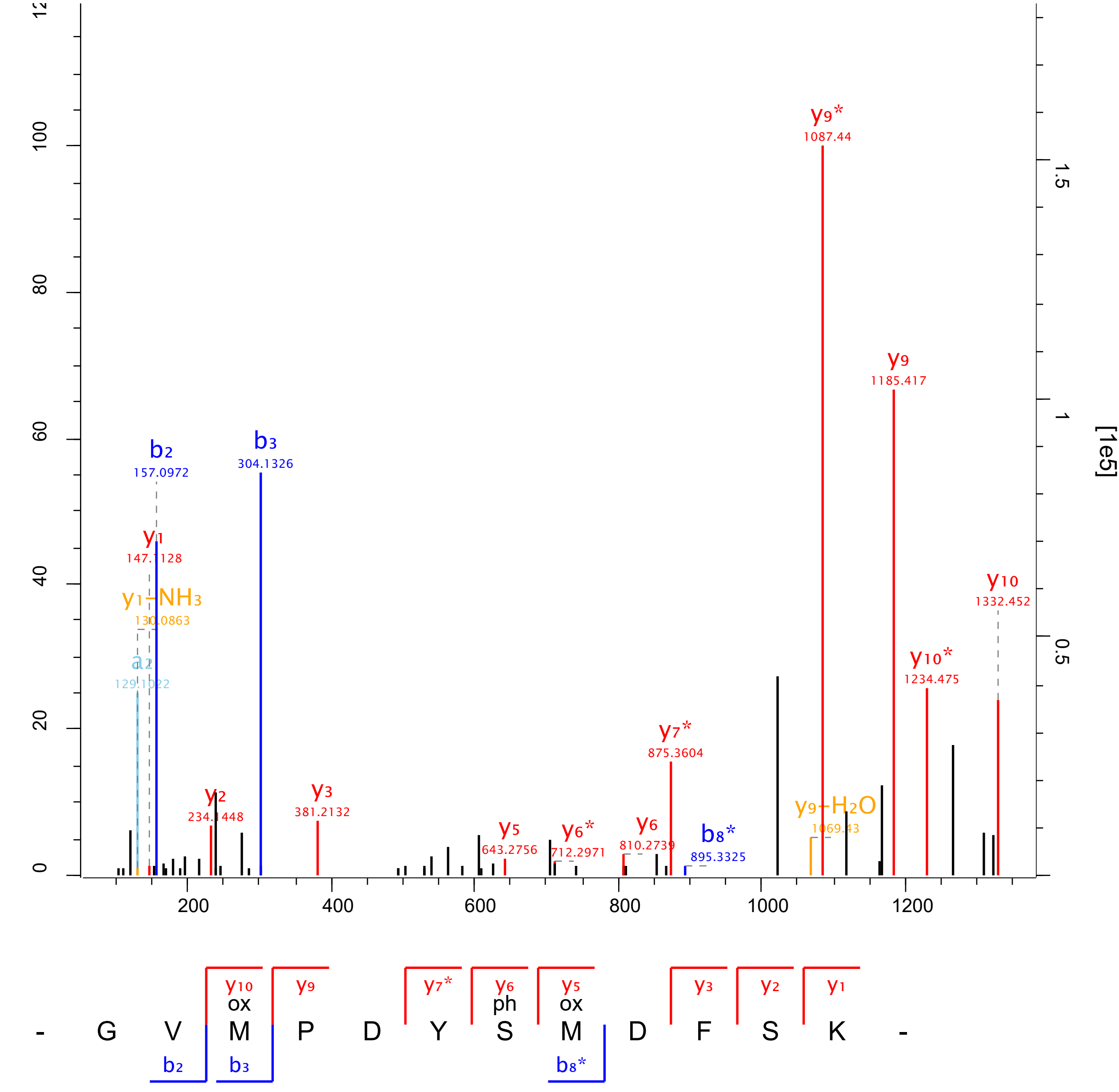

| Raw file | Scan | Method | Score | m/z |
|---|---|---|---|---|
| A17_02_1 | 17527 | FTMS; HCD | 190.37 | 635.28 |

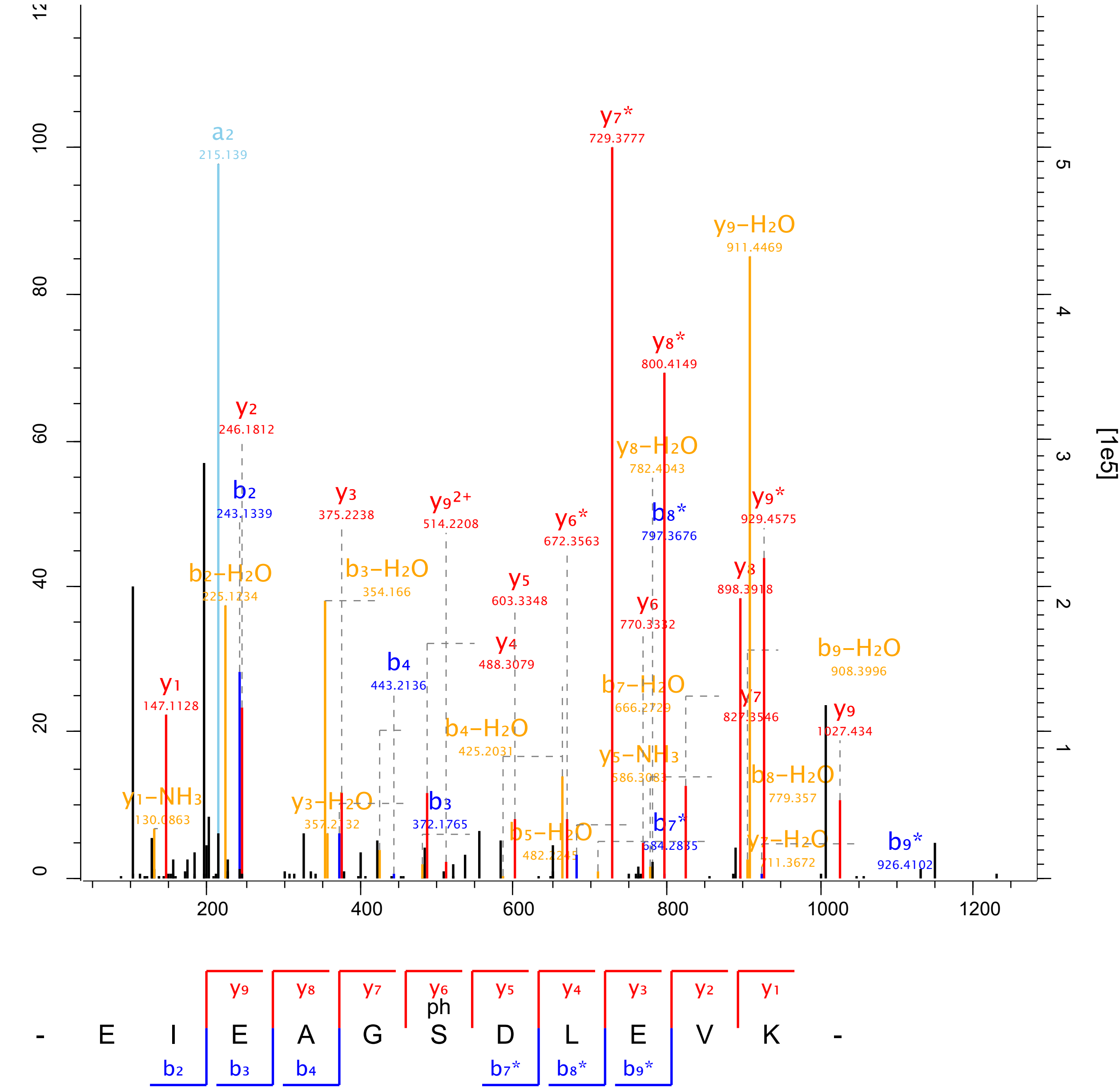

| Raw file | Scan | Method | Score | m/z |
|---|---|---|---|---|
| A17_02_1 | 17916 | FTMS; HCD | 193.42 | 1057.92 |

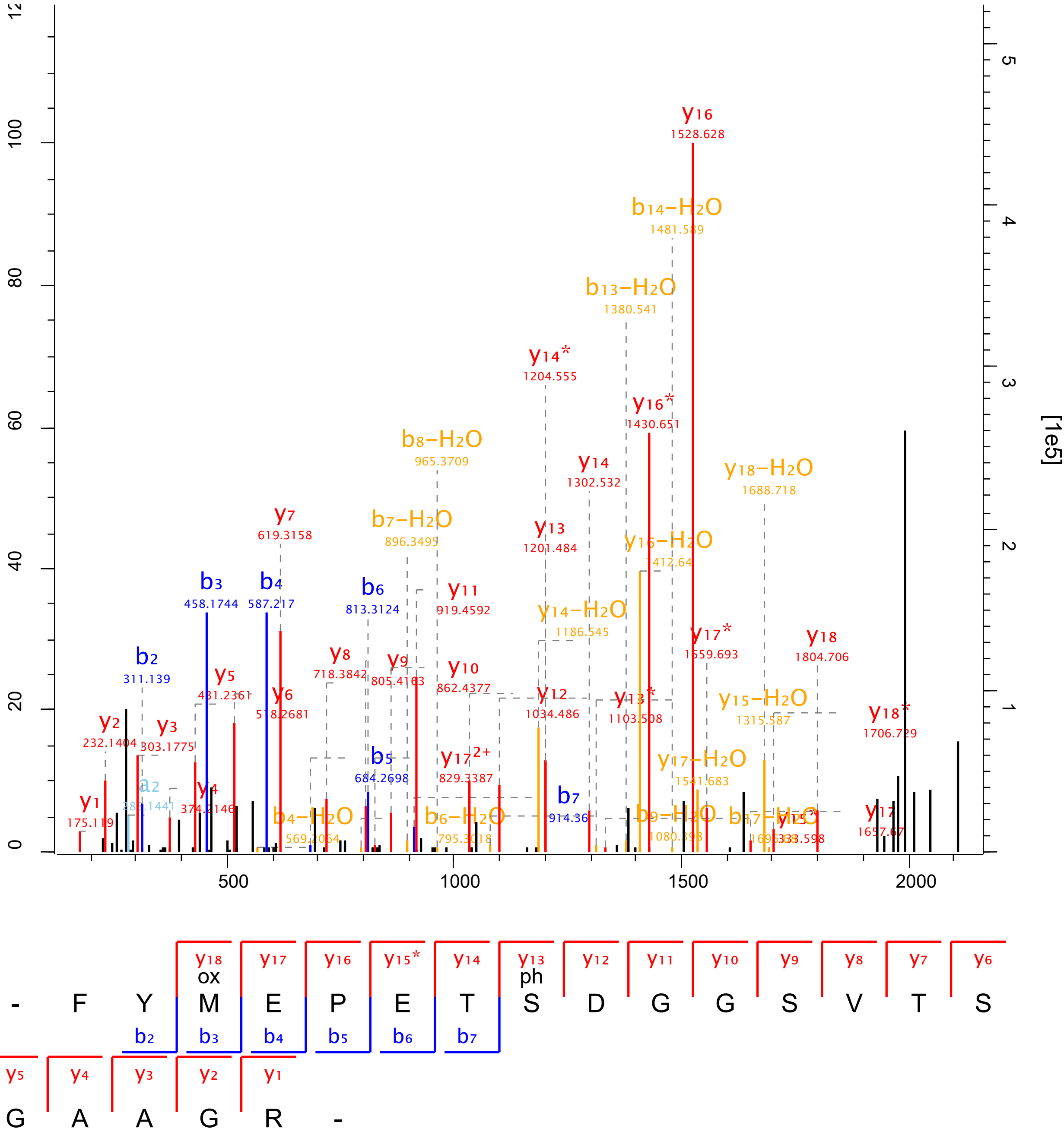

| Raw file | Scan | Method | Score | m/z |
|---|---|---|---|---|
| A17_02_1 | 18147 | FTMS; HCD | 49.45 | 617.74 |

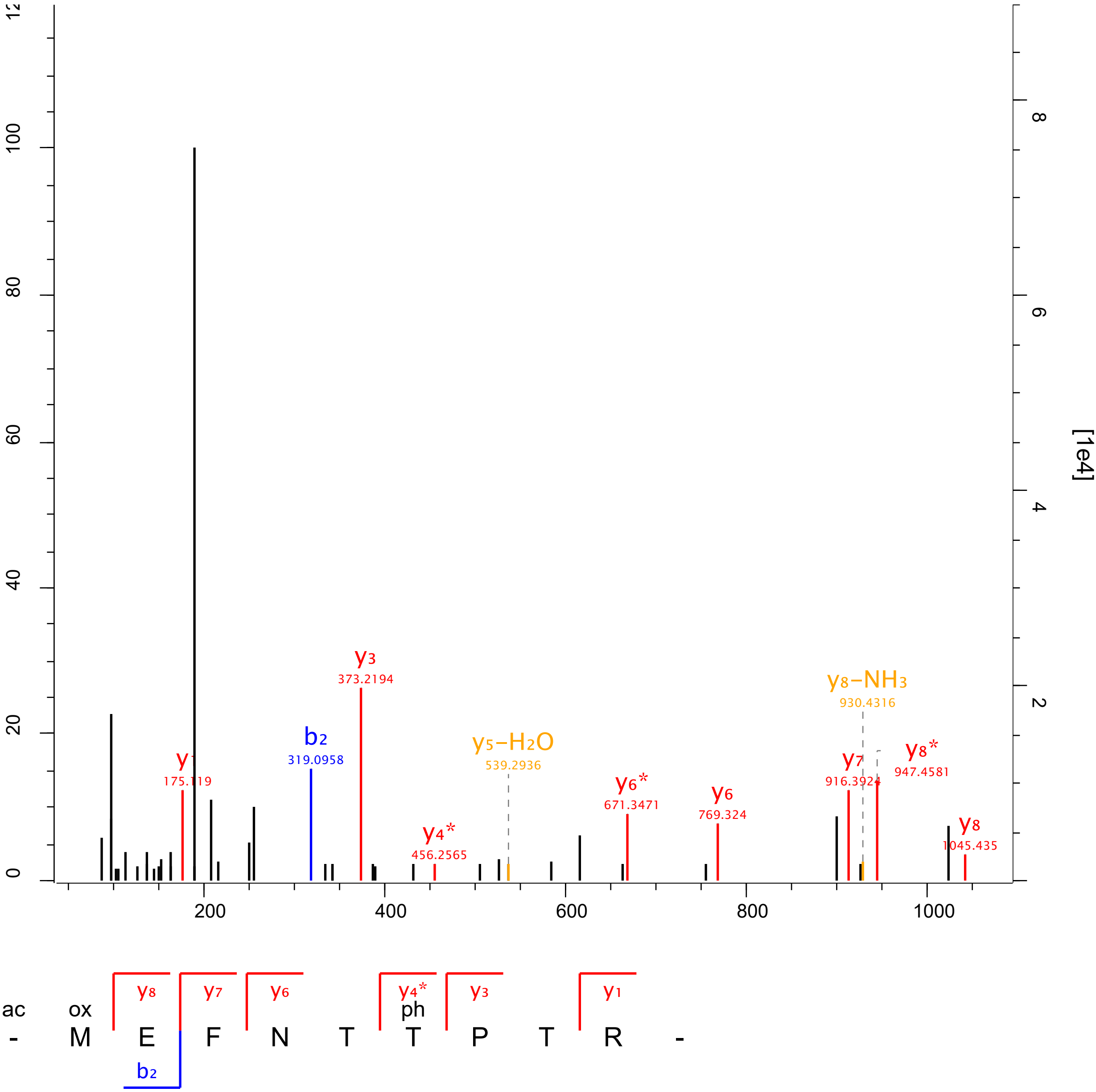

| Raw file | Scan | Method | Score | m/z |
|---|---|---|---|---|
| A17_02_1 | 18797 | FTMS; HCD | 42.21 | 599.27 |

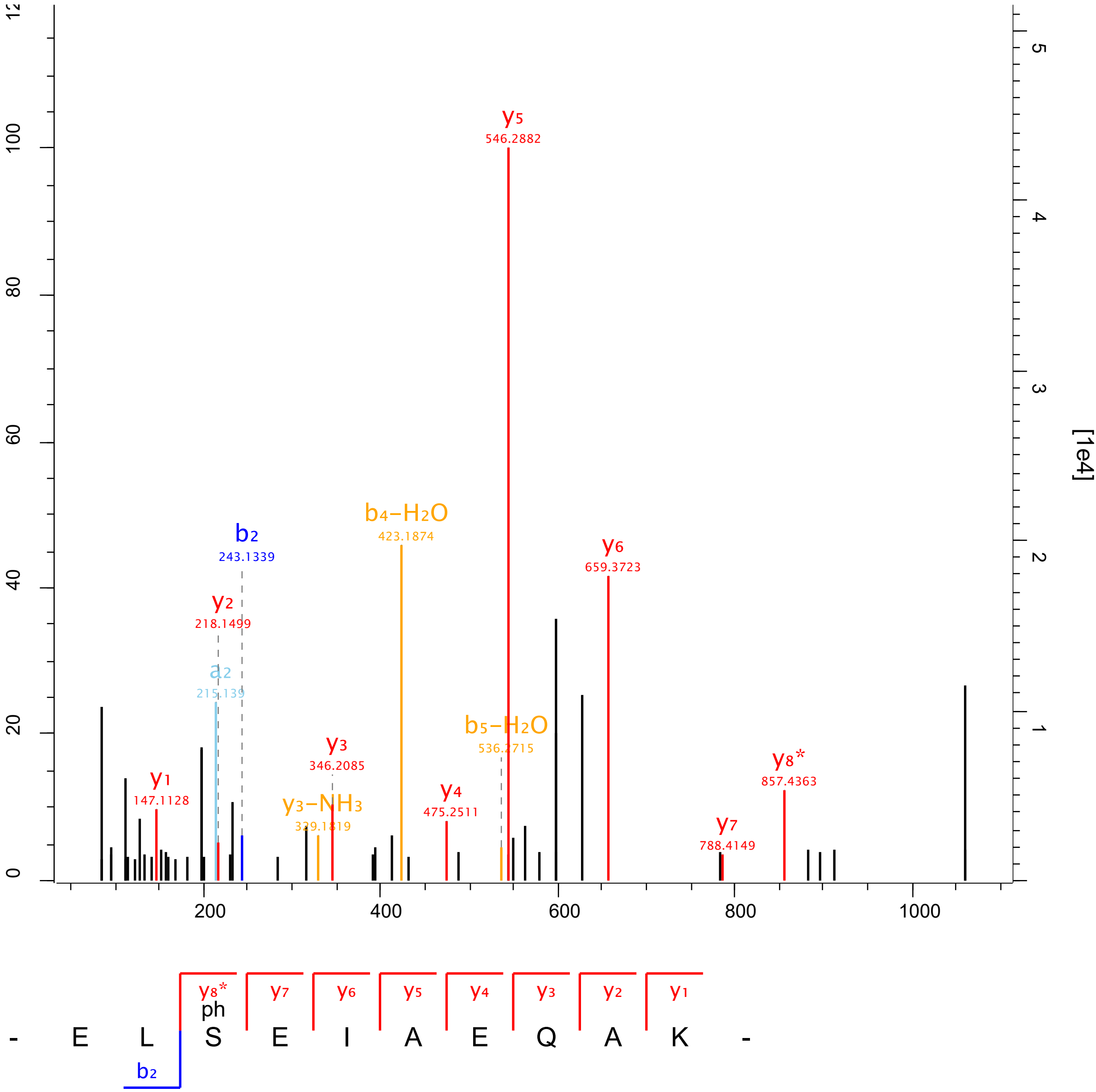

| Raw file | Scan | Method | Score | m/z |
| --- | --- | --- | --- | --- |
| A17_02_1 | 19097 | FTMS; HCD | 97.64 | 433.23 |

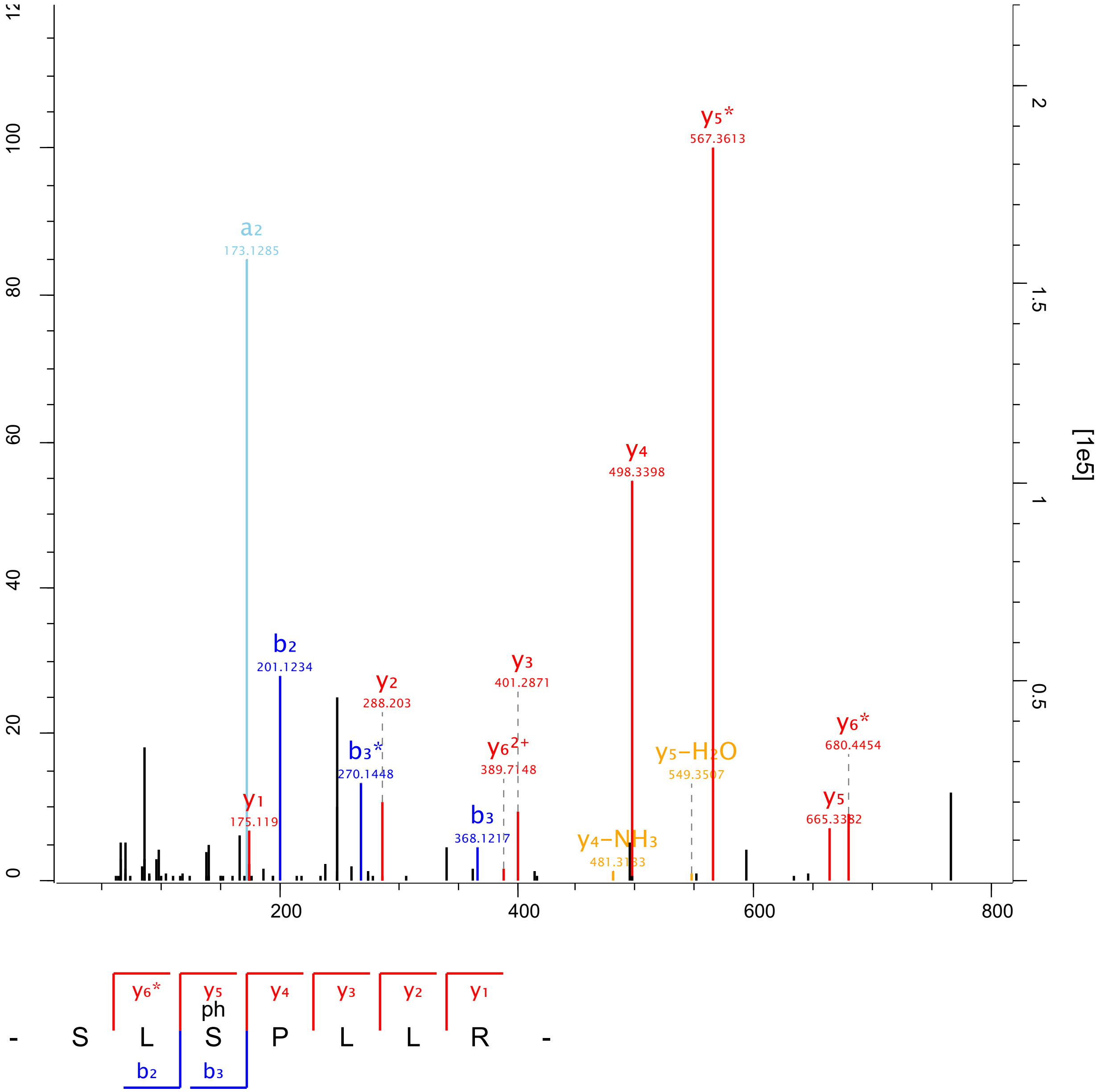

| Raw file | Scan | Method | Score | m/z |
|---|---|---|---|---|
| A17_02_1 | 19336 | FTMS; HCD | 212.43 | 696.31 |

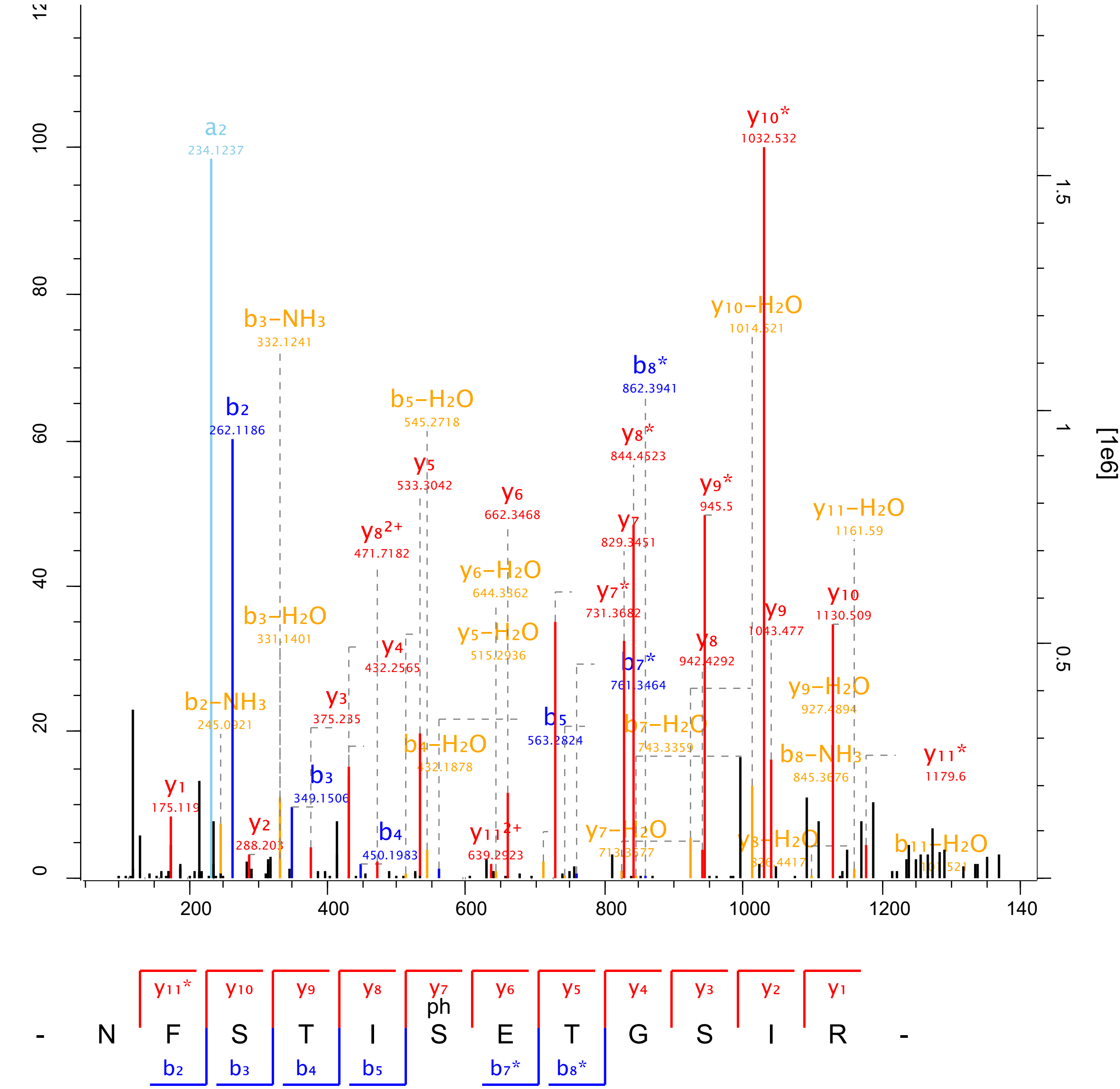

| Raw file | Scan | Method | Score | m/z |
| --- | --- | --- | --- | --- |
| A17_02_1 | 19397 | FTMS; HCD | 119.45 | 693.82 |

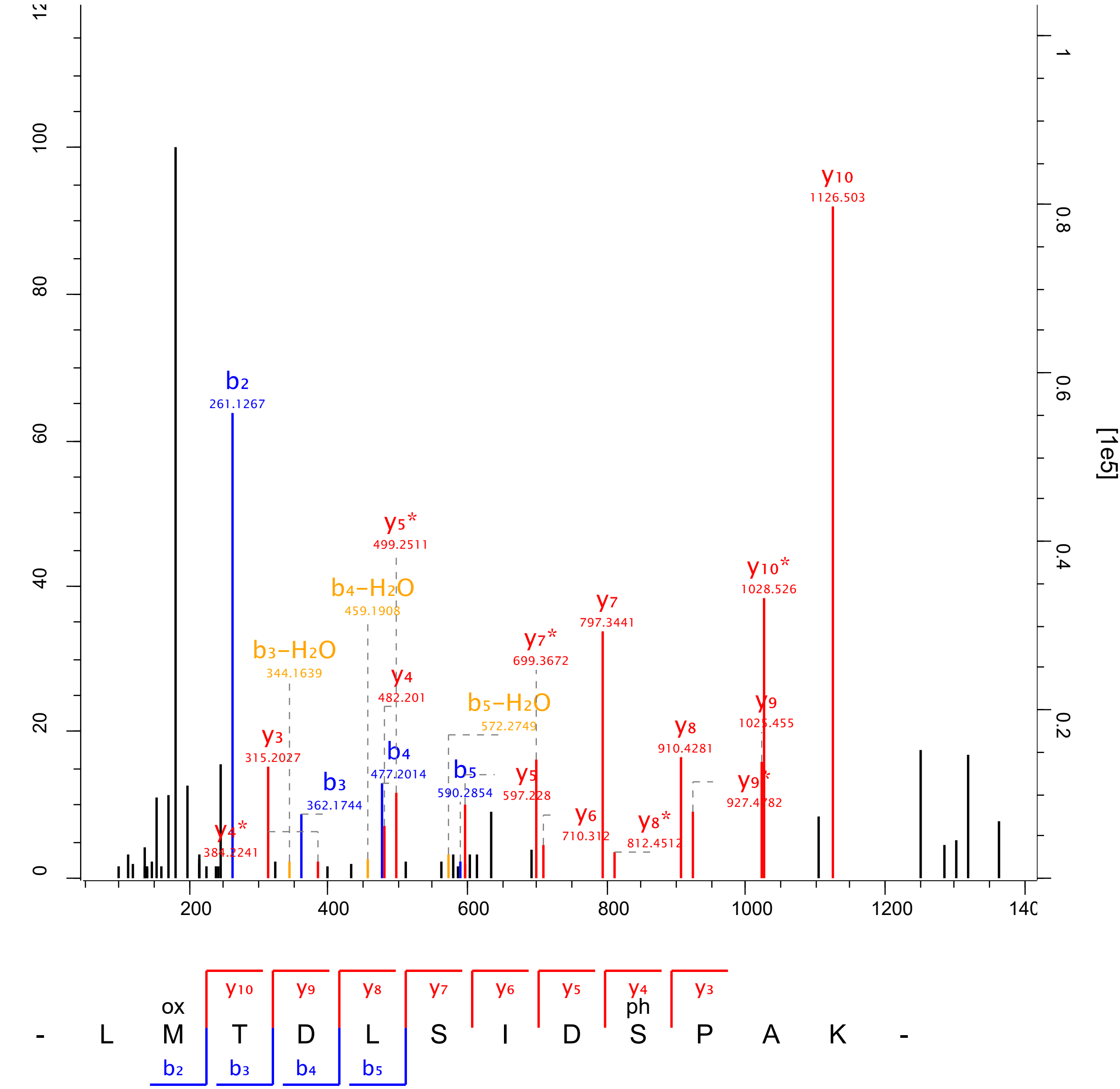

| Raw file | Scan | Method | Score | m/z |
|---|---|---|---|---|
| A17_02_1 | 19531 | FTMS; HCD | 52.58 | 654.25 |

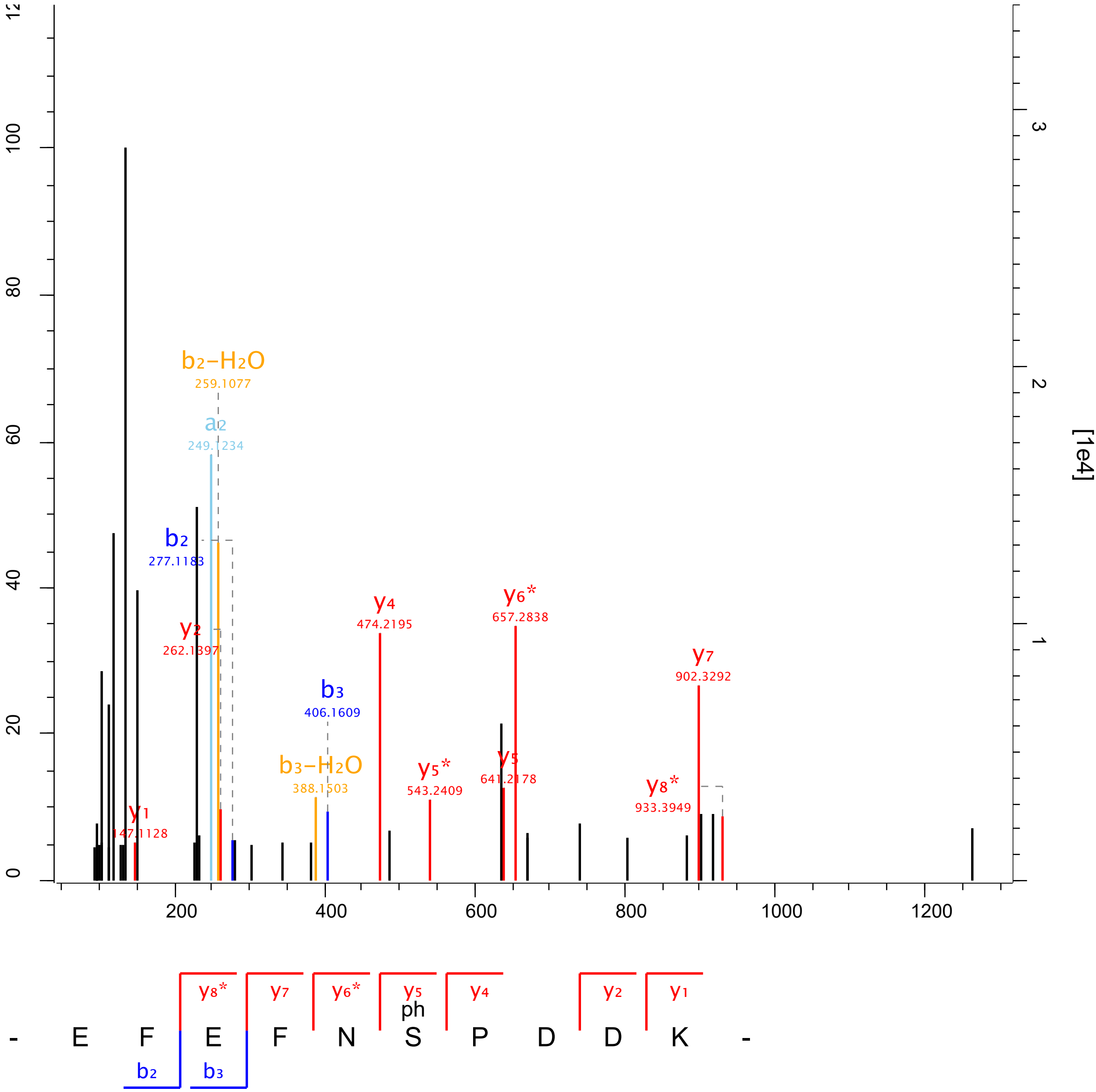

| Raw file | Scan | Method | Score | m/z |
| --- | --- | --- | --- | --- |
| A17_02_1 | 20033 | FTMS; HCD | 178.22 | 628.77 |

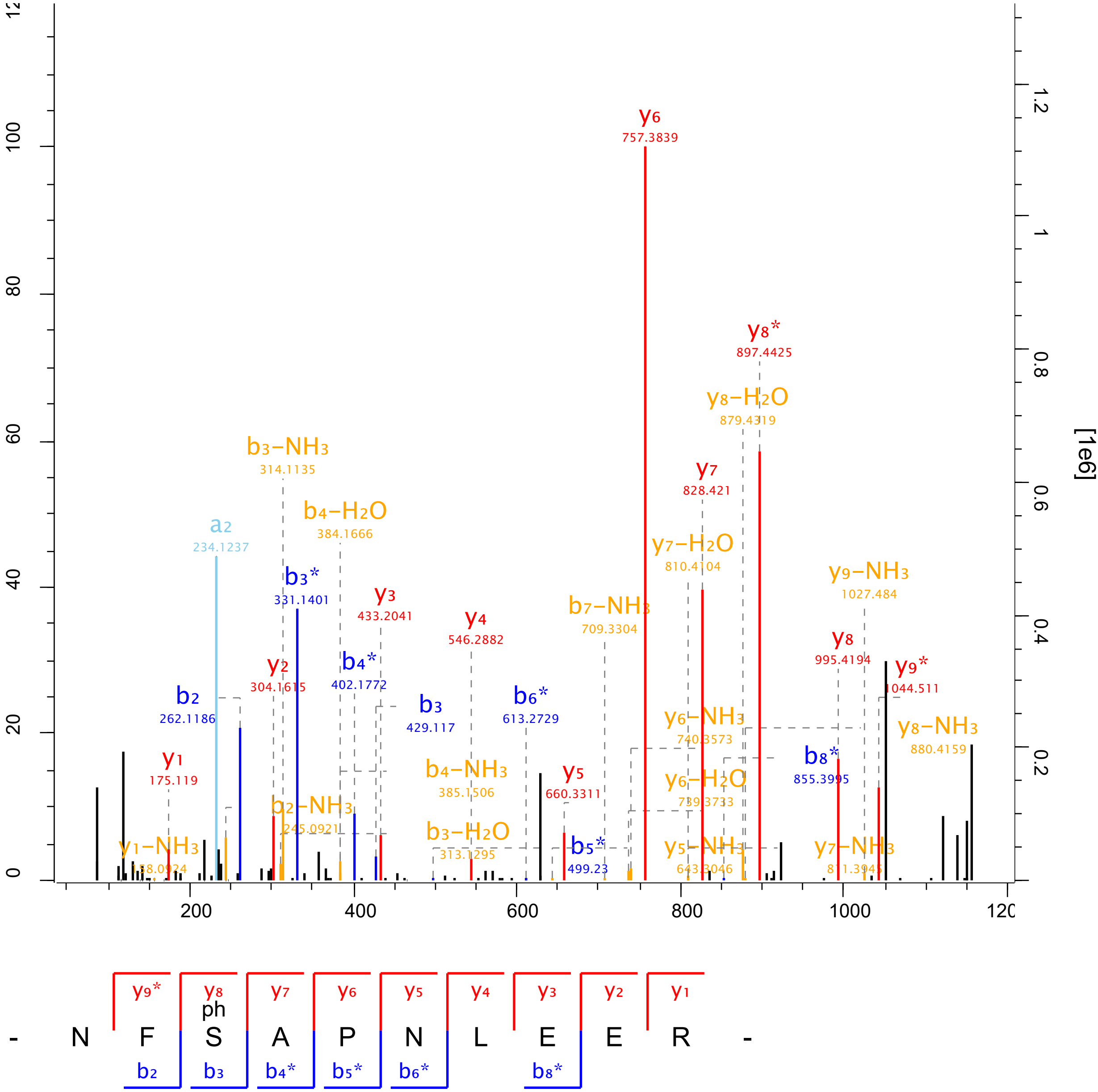

| Raw file | Scan | Method | Score | m/z |
| --- | --- | --- | --- | --- |
| A17_02_1 | 20354 | FTMS; HCD | 124.2 | 812.34 |

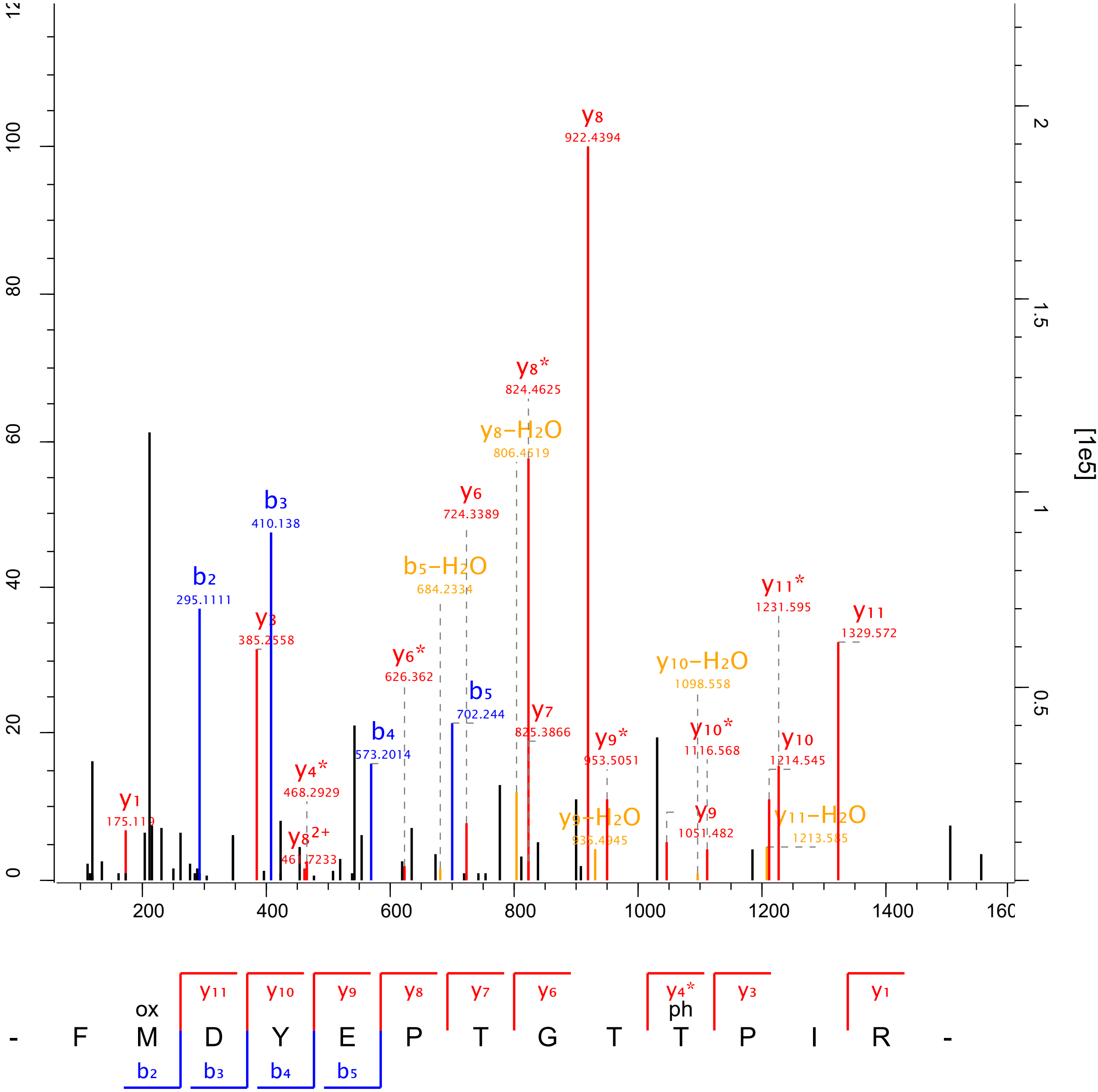

| Raw file | Scan | Method | Score | m/z |
|---|---|---|---|---|
| A17_02_1 | 2036 | FTMS; HCD | 71.88 | 572.22 |

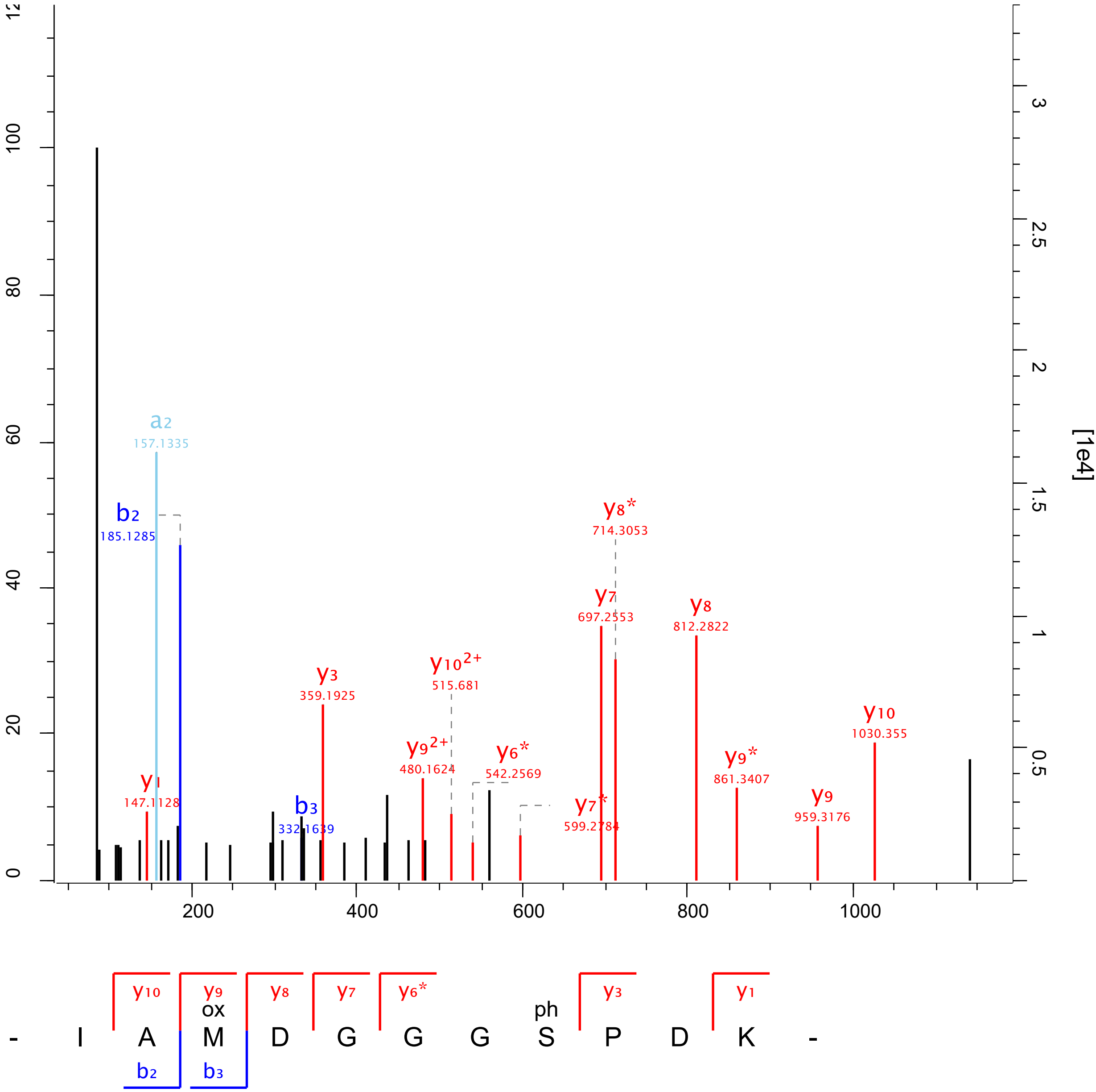

| Raw file | Scan | Method | Score | m/z |
|---|---|---|---|---|
| A17_02_1 | 20381 | FTMS; HCD | 43.21 | 715.28 |

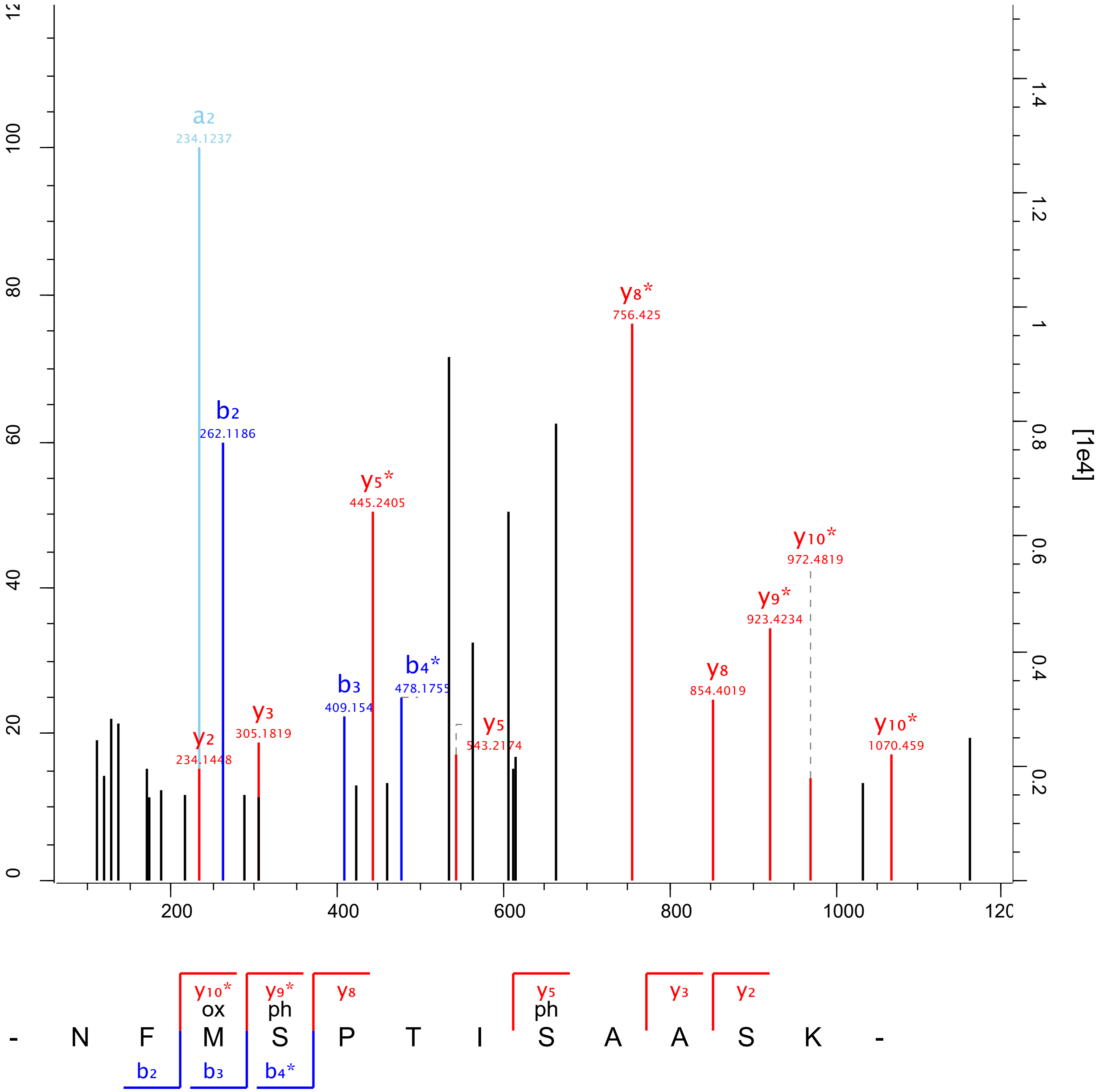

| Raw file | Scan | Method | Score | m/z |
| --- | --- | --- | --- | --- |
| A17_02_1 | 20973 | FTMS; HCD | 57.43 | 673.78 |

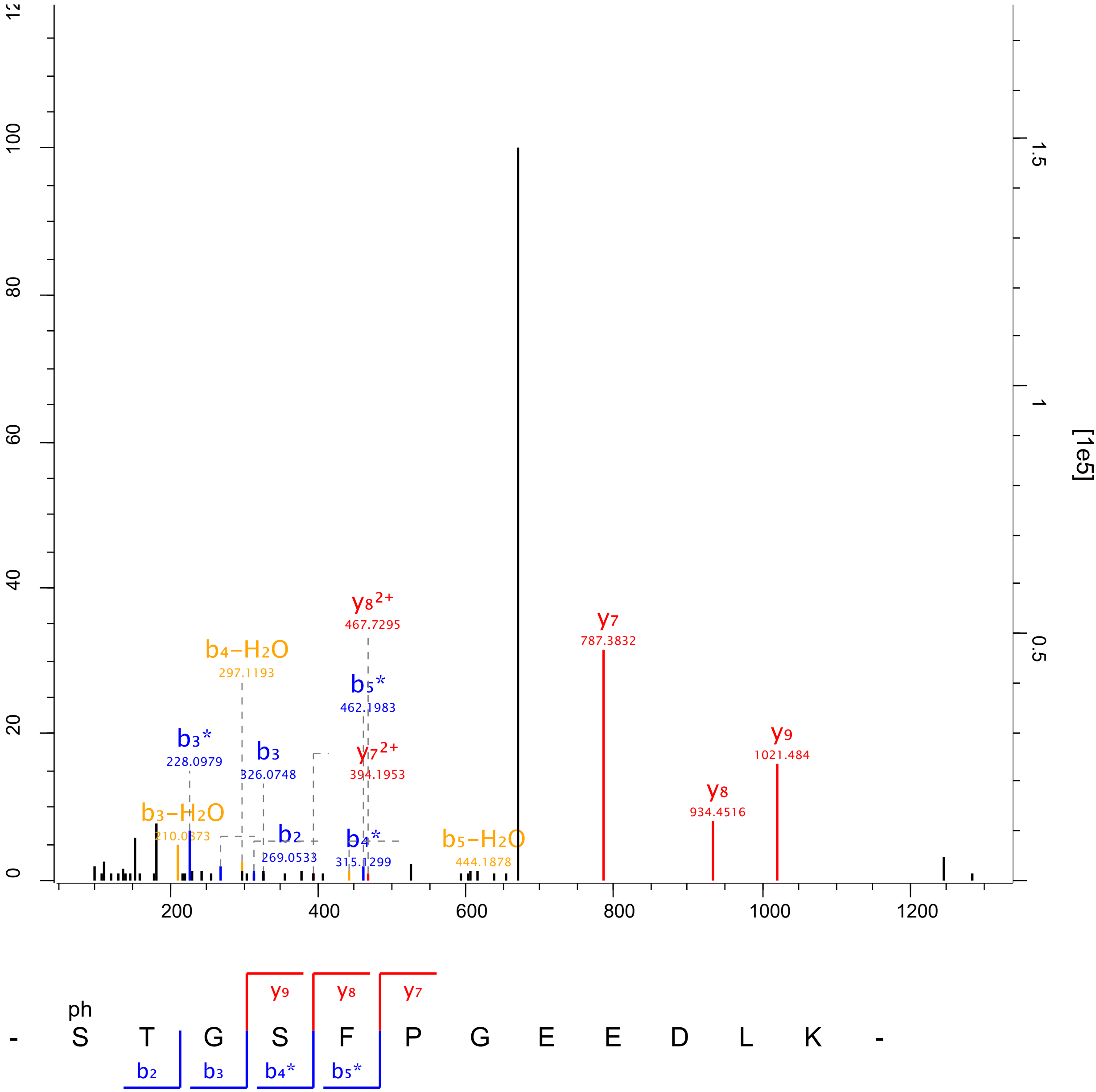

| Raw file | Scan | Method | Score | m/z |
|---|---|---|---|---|
| A17_02_1 | 21187 | FTMS; HCD | 65.43 | 719.34 |

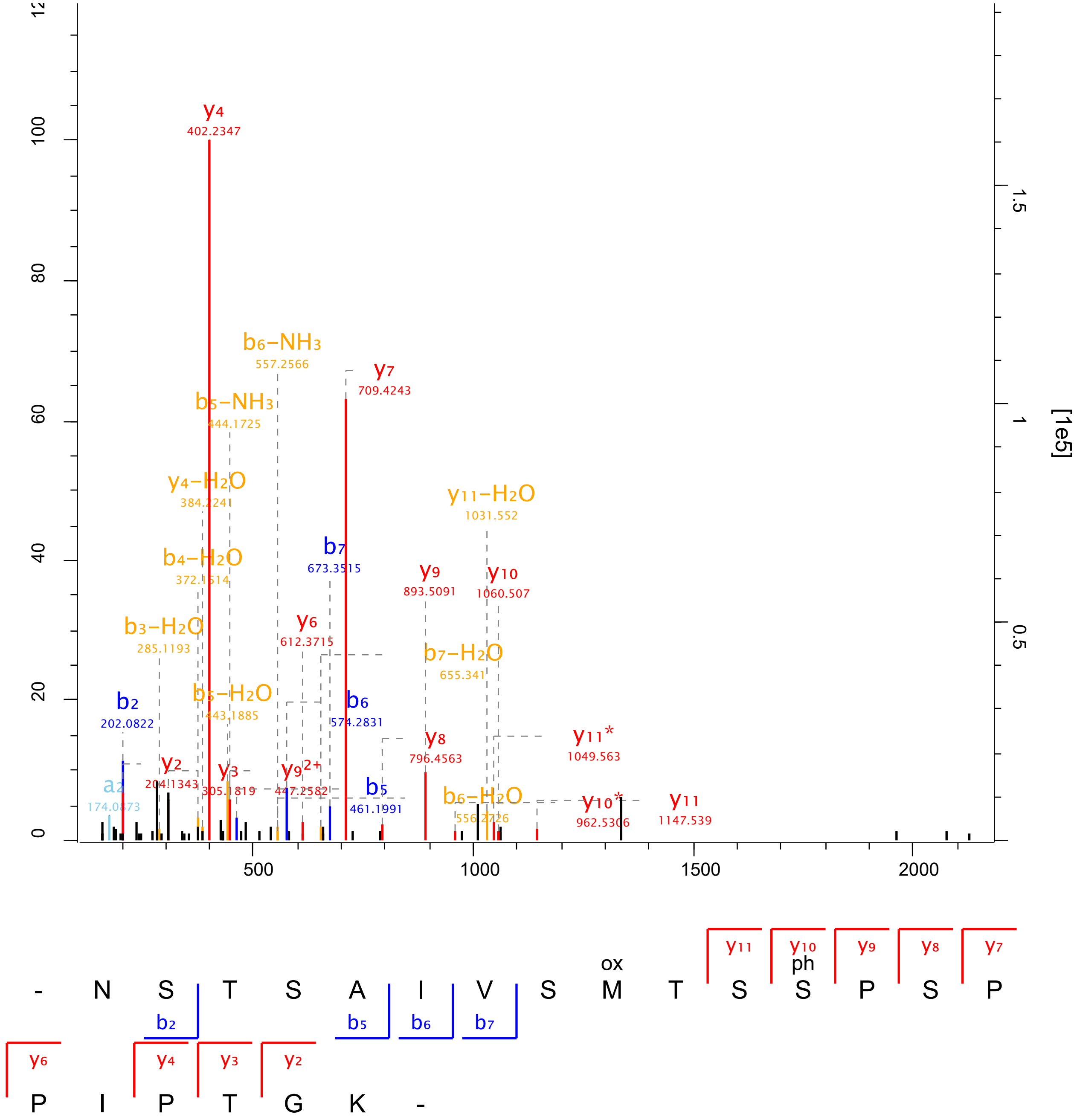

| Raw file | Scan | Method | Score | m/z |
|---|---|---|---|---|
| A17_02_1 | 21379 | FTMS; HCD | 136.66 | 697.86 |

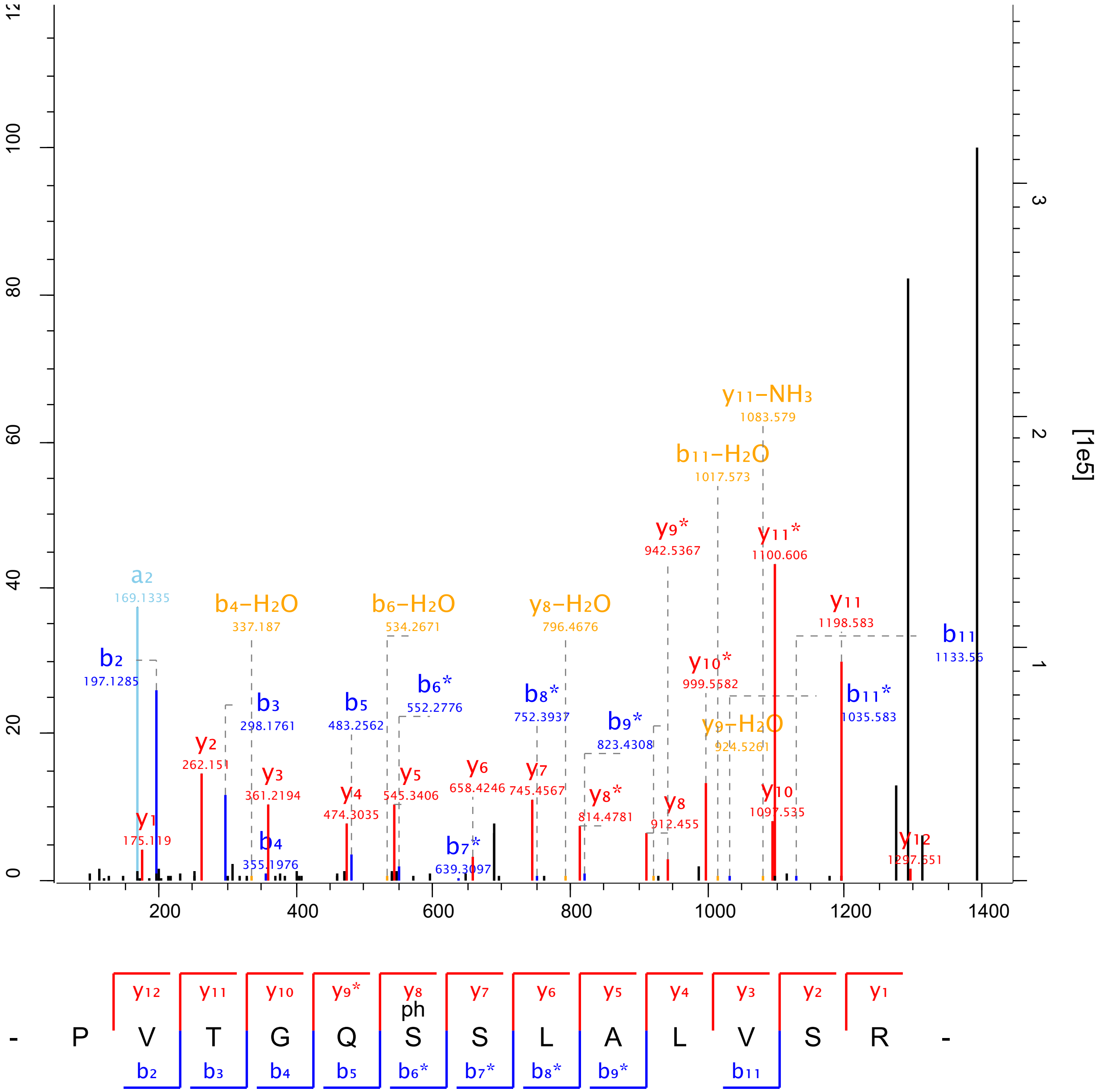

| Raw file | Scan | Method | Score | m/z |
|---|---|---|---|---|
| A17_02_1 | 2159 | FTMS; HCD | 60.4 | 425.16 |

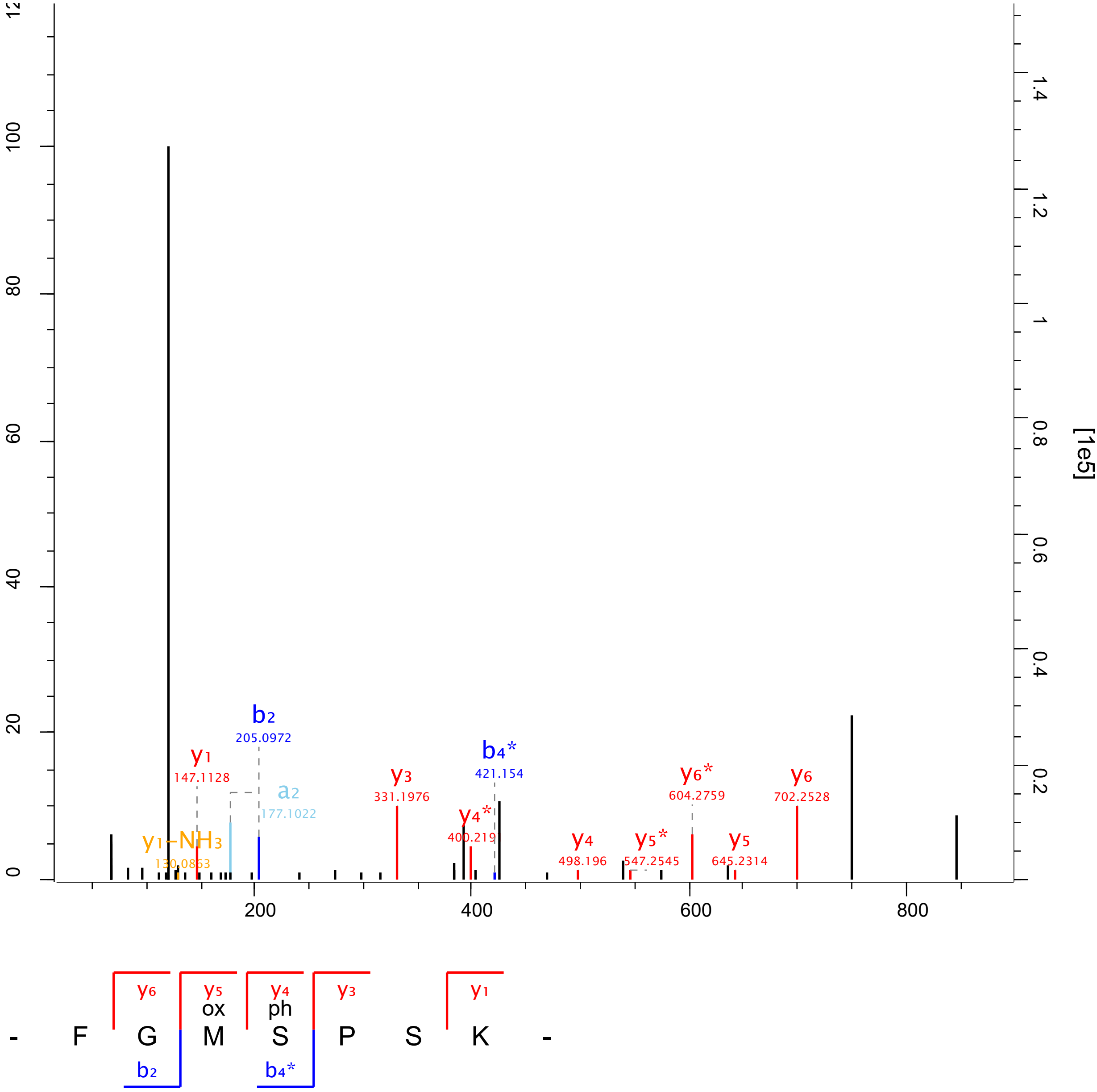

| Raw file | Scan | Method | Score | m/z |
|---|---|---|---|---|
| A17_02_1 | 22457 | FTMS; HCD | 121.01 | 762.36 |

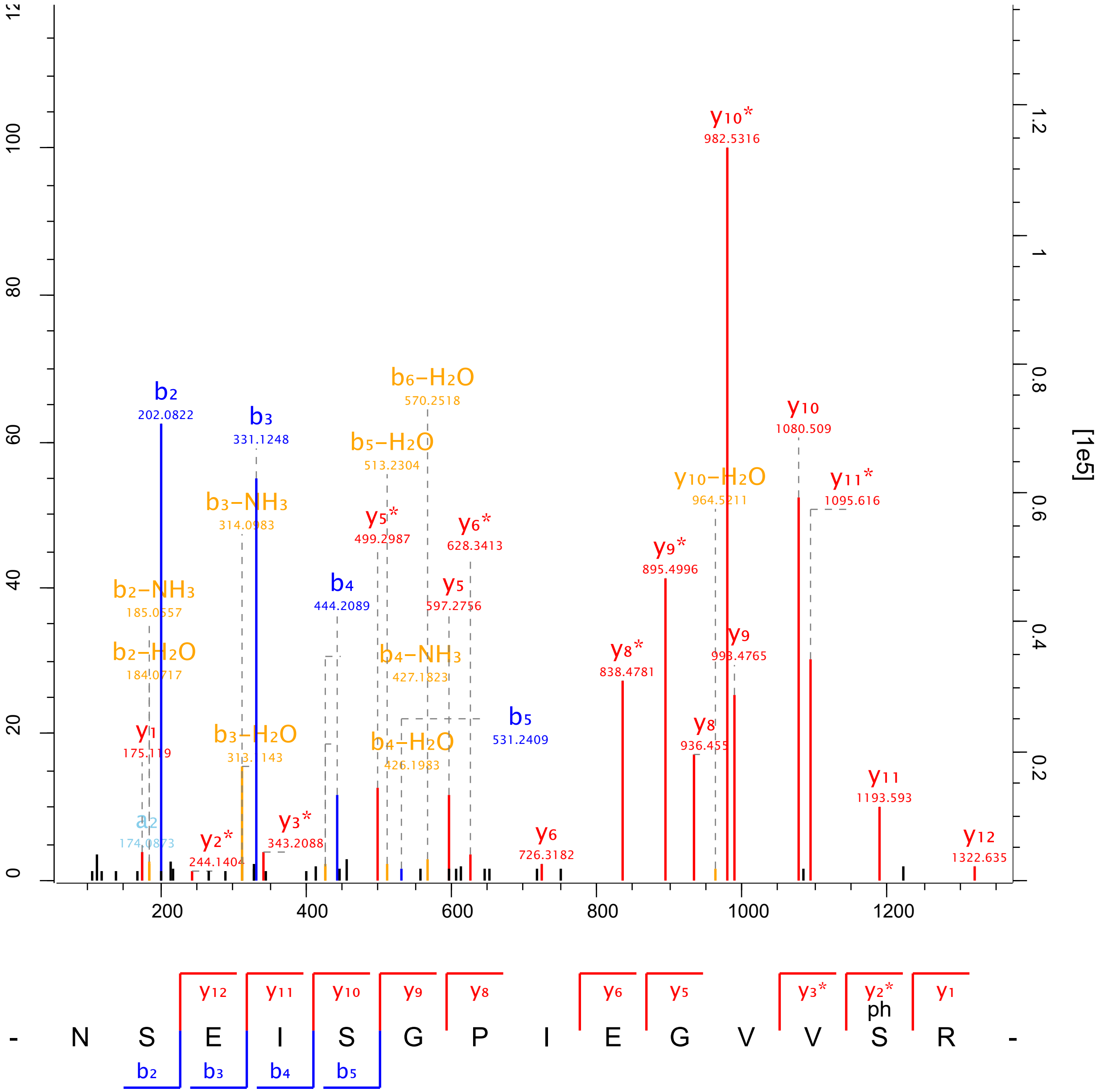

| Raw file | Scan | Method | Score | m/z |
| --- | --- | --- | --- | --- |
| A17_02_1 | 22913 | FTMS; HCD | 91.36 | 730.31 |

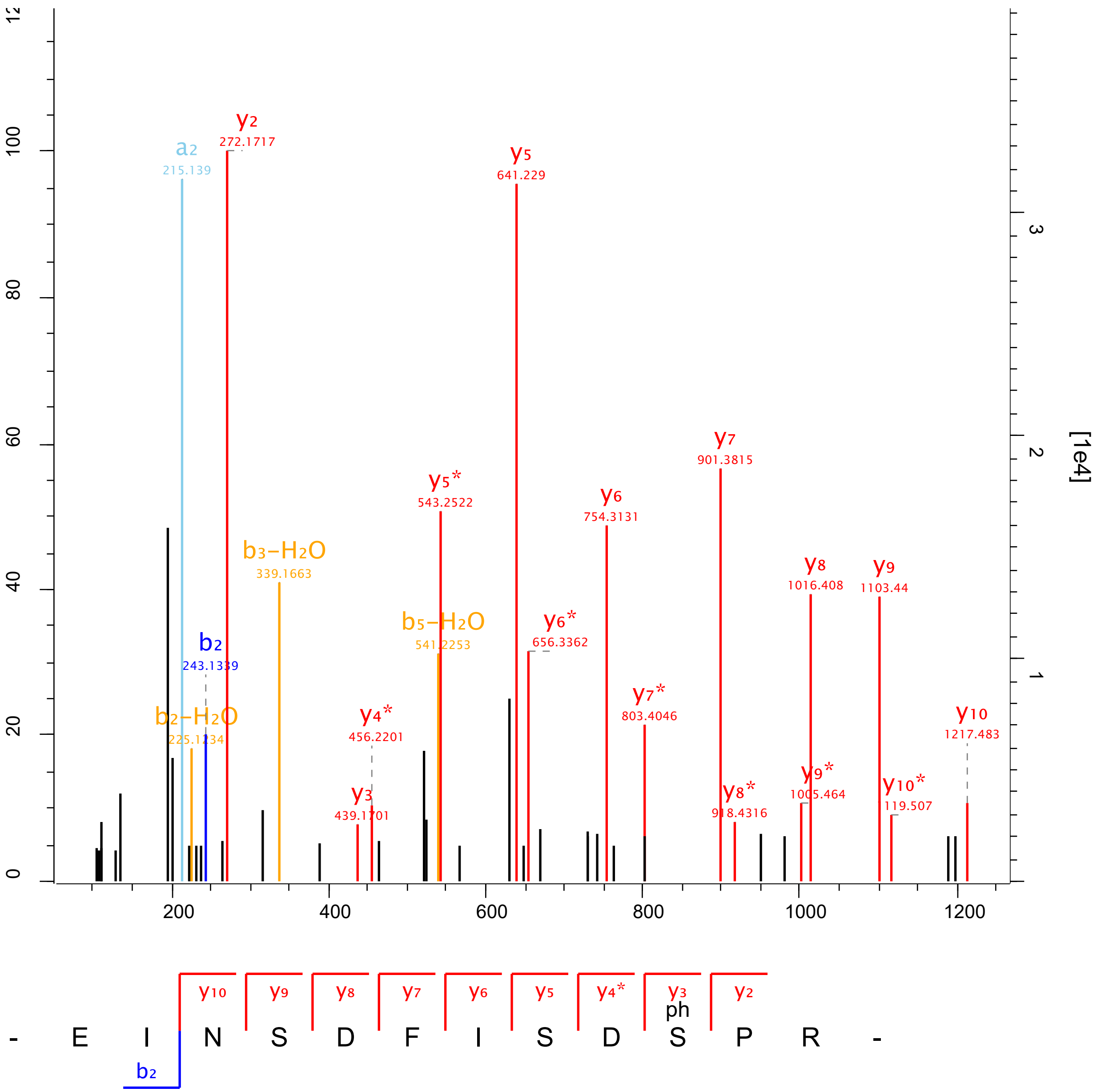

| Raw file | Scan | Method | Score | m/z |
|---|---|---|---|---|
| A17_02_1 | 23039 | FTMS; HCD | 50.83 | 747.87 |

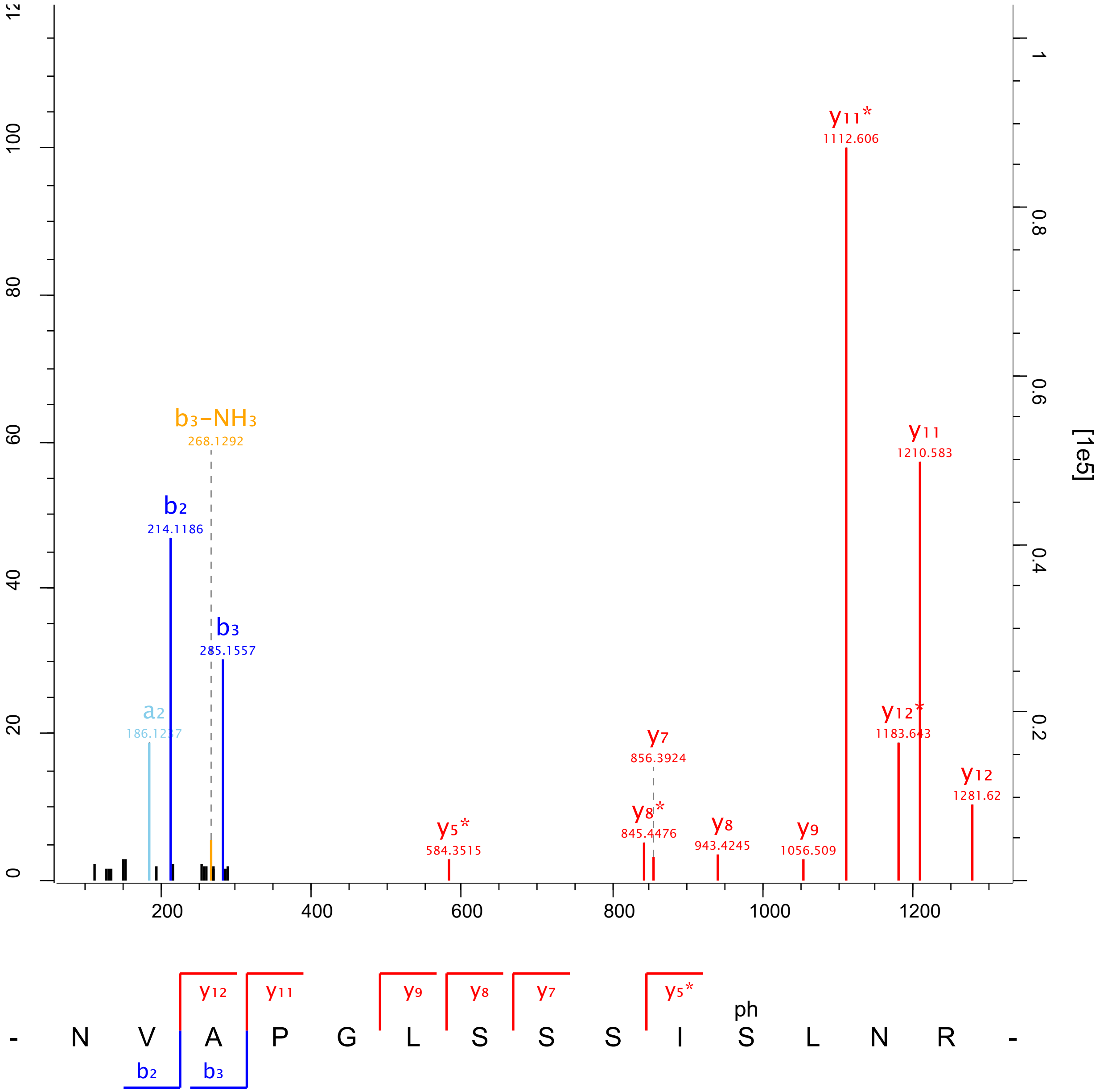

| Raw file | Scan | Method | Score | m/z |
|---|---|---|---|---|
| A17_02_1 | 23061 | FTMS; HCD | 141.34 | 656.76 |

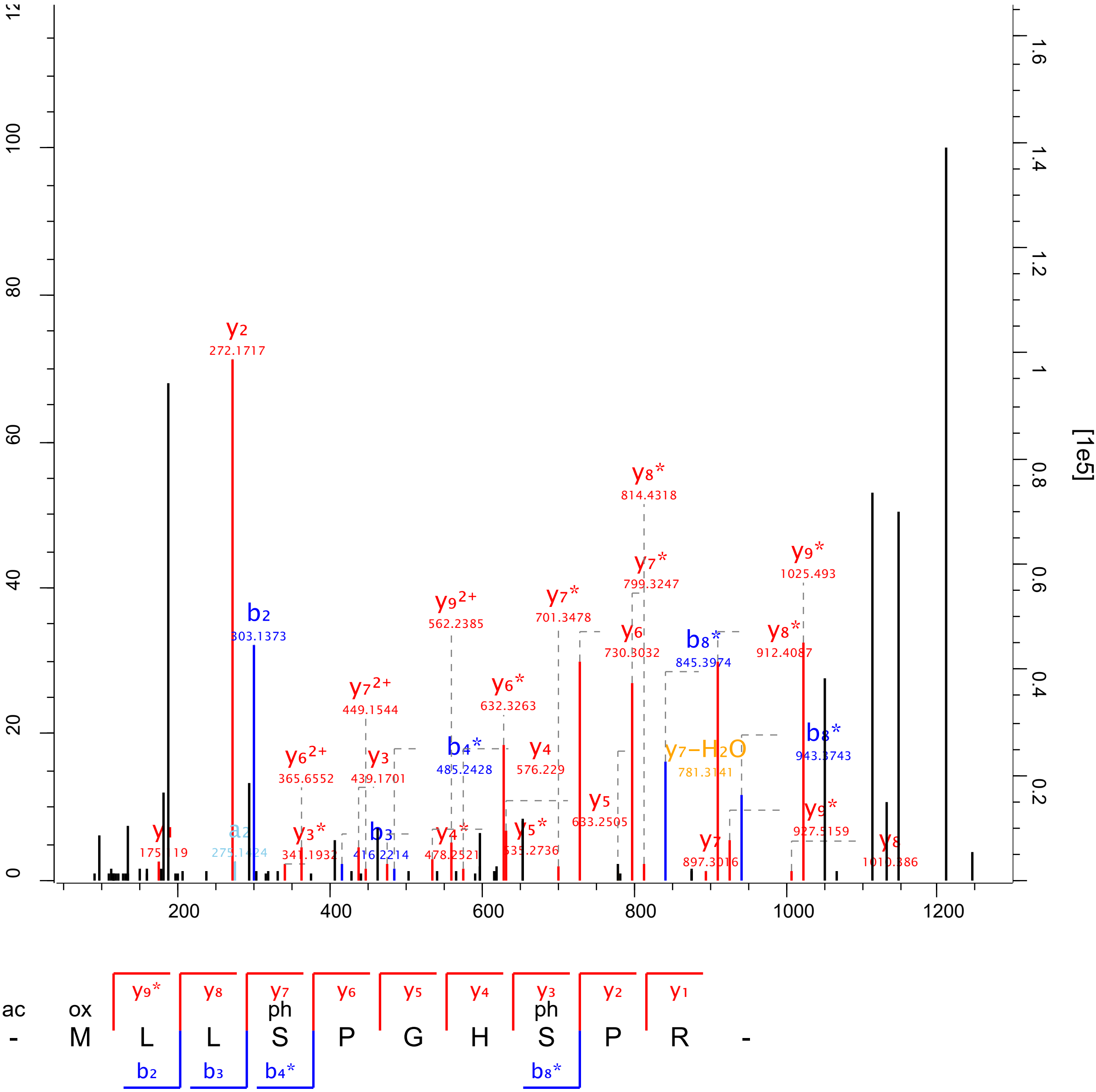

| Raw file | Scan | Method | Score | m/z |
|---|---|---|---|---|
| A17_02_1 | 23168 | FTMS; HCD | 155.2 | 825.85 |

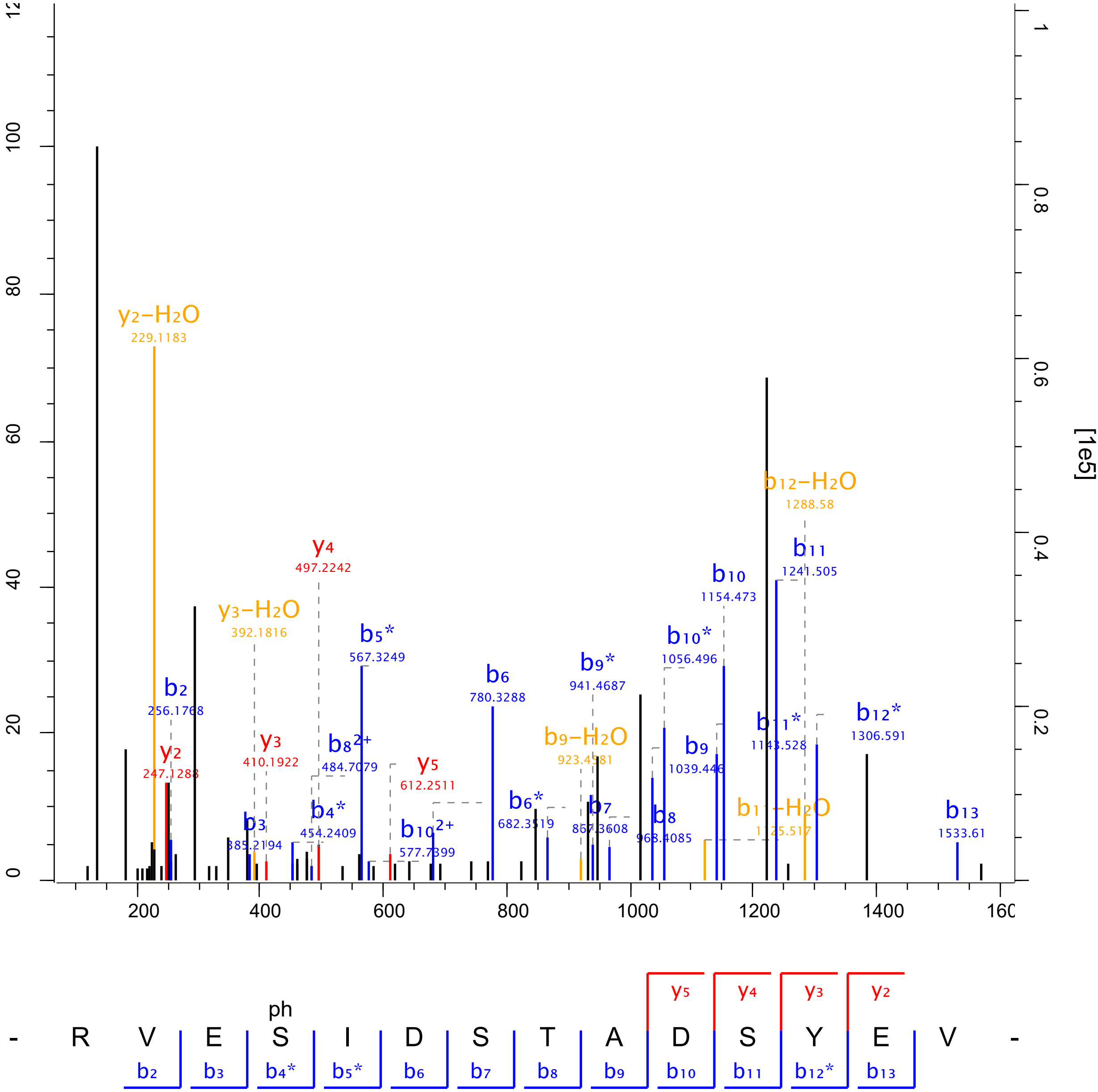

| Raw file | Scan | Method | Score | m/z |
| --- | --- | --- | --- | --- |
| A17_02_1 | 23586 | FTMS; HCD | 224.15 | 685.3 |

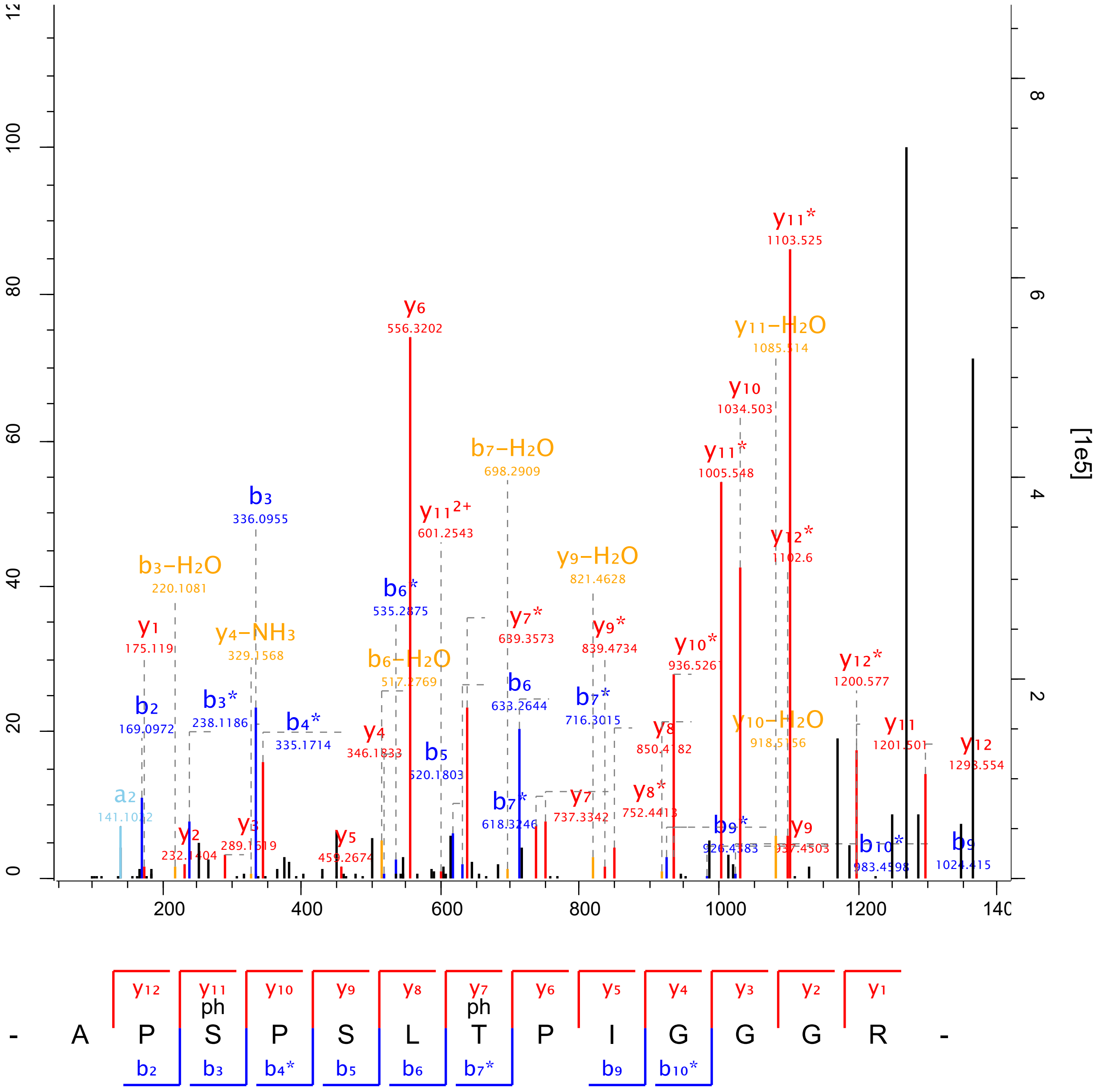

| Raw file | Scan | Method | Score | m/z |
|---|---|---|---|---|
| A17_02_1 | 23855 | FTMS; HCD | 57.43 | 687.31 |

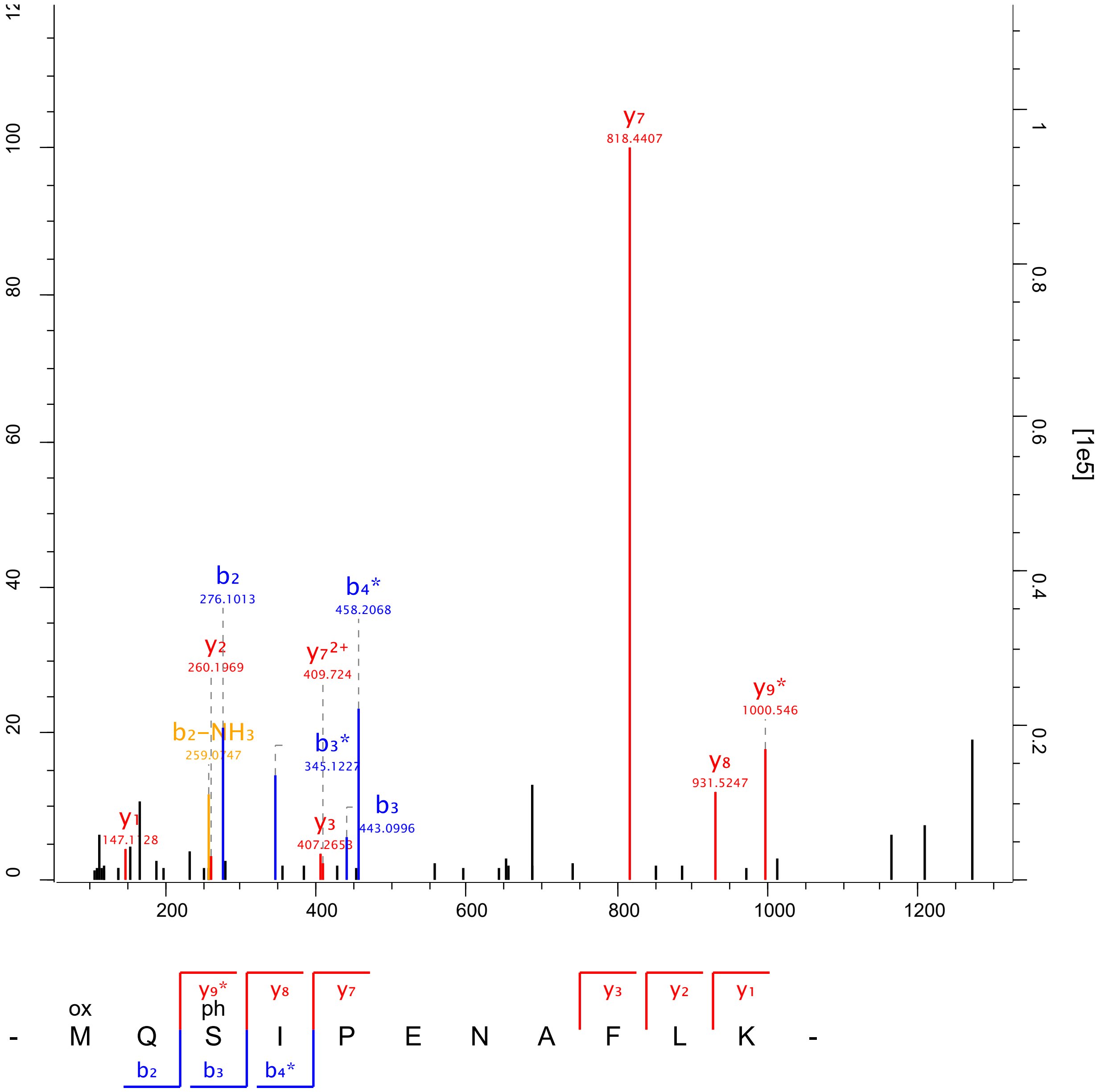

| Raw file | Scan | Method | Score | m/z |
| --- | --- | --- | --- | --- |
| A17_02_1 | 24312 | FTMS; HCD | 61.96 | 667.28 |

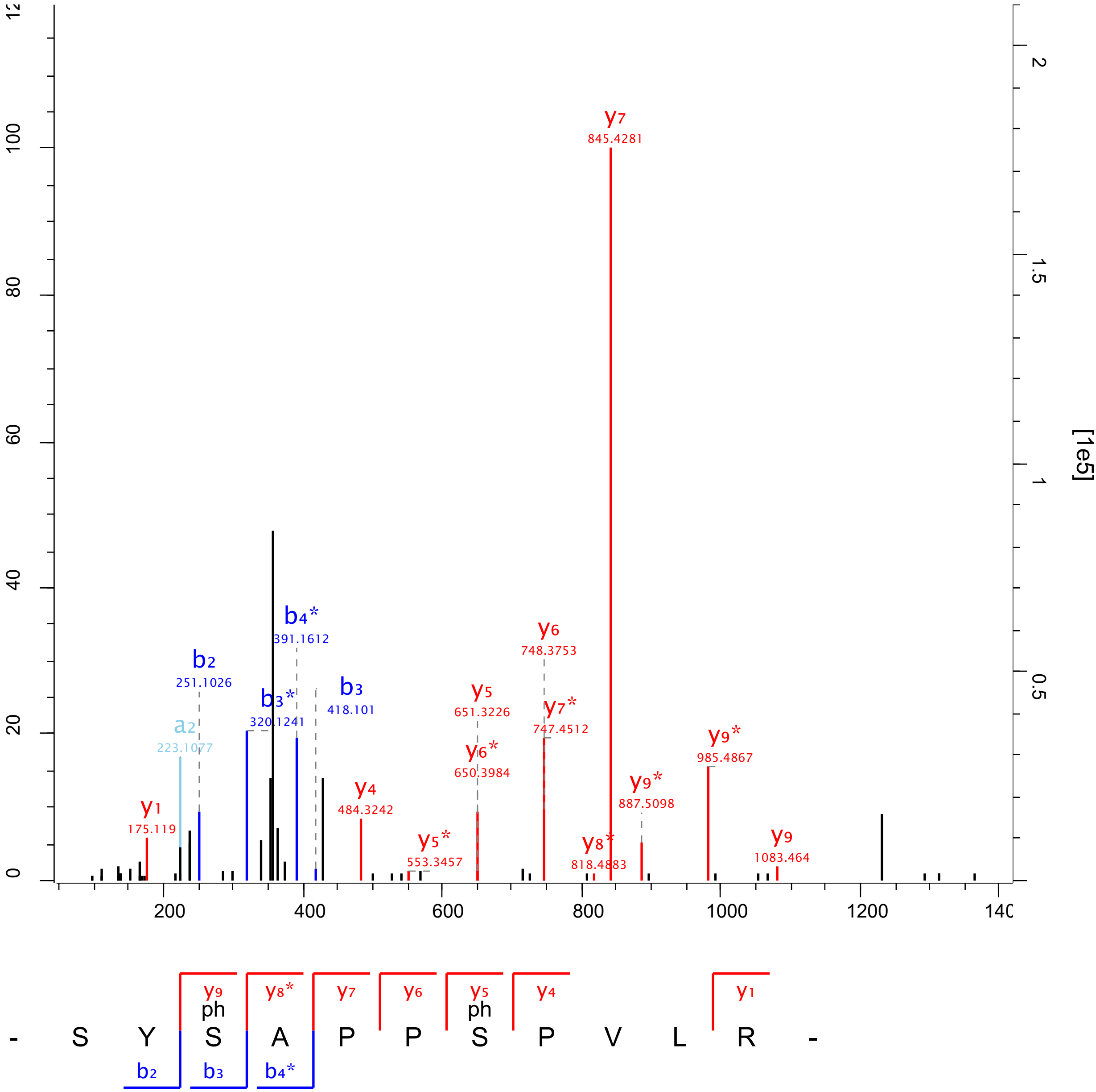

| Raw file | Scan | Method | Score | m/z |
|---|---|---|---|---|
| A17_02_1 | 24350 | FTMS; HCD | 132.64 | 792.35 |

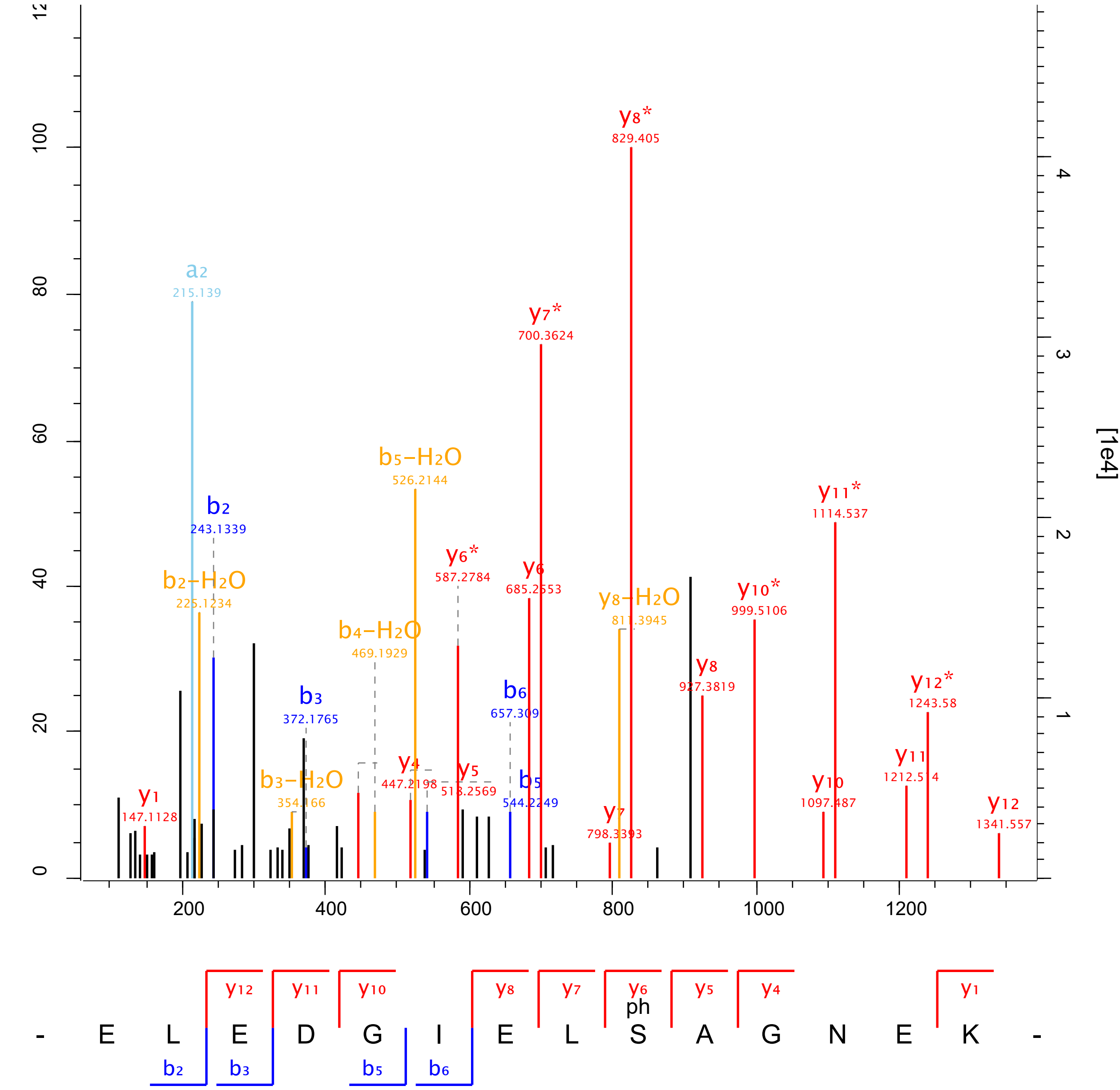

| Raw file | Scan | Method | Score | m/z |
| --- | --- | --- | --- | --- |
| A17_02_1 | 24633 | FTMS; HCD | 79.47 | 871.9 |

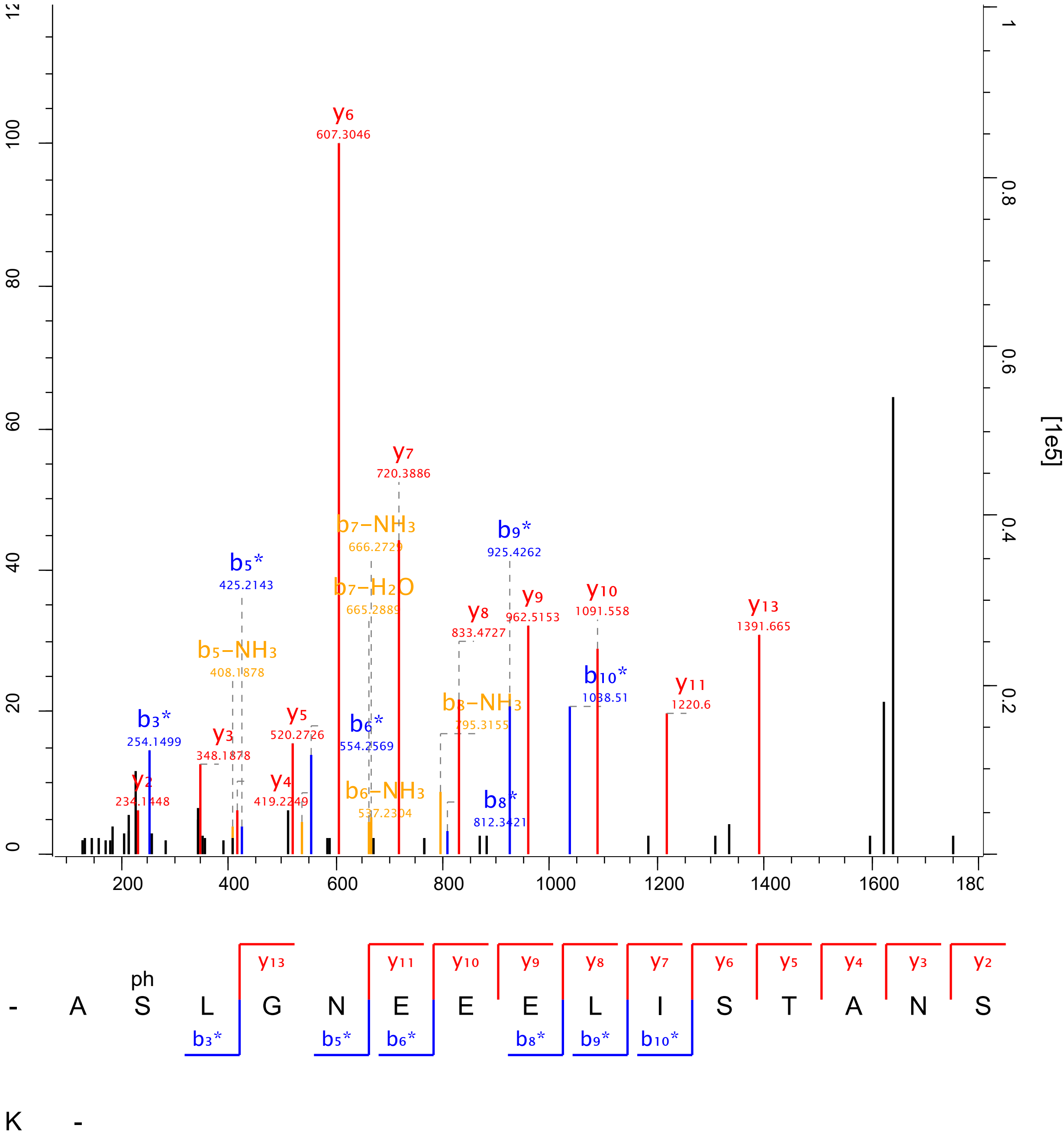

| Raw file | Scan | Method | Score | m/z |
| --- | --- | --- | --- | --- |
| A17_02_1 | 24793 | FTMS; HCD | 51.83 | 1060.94 |

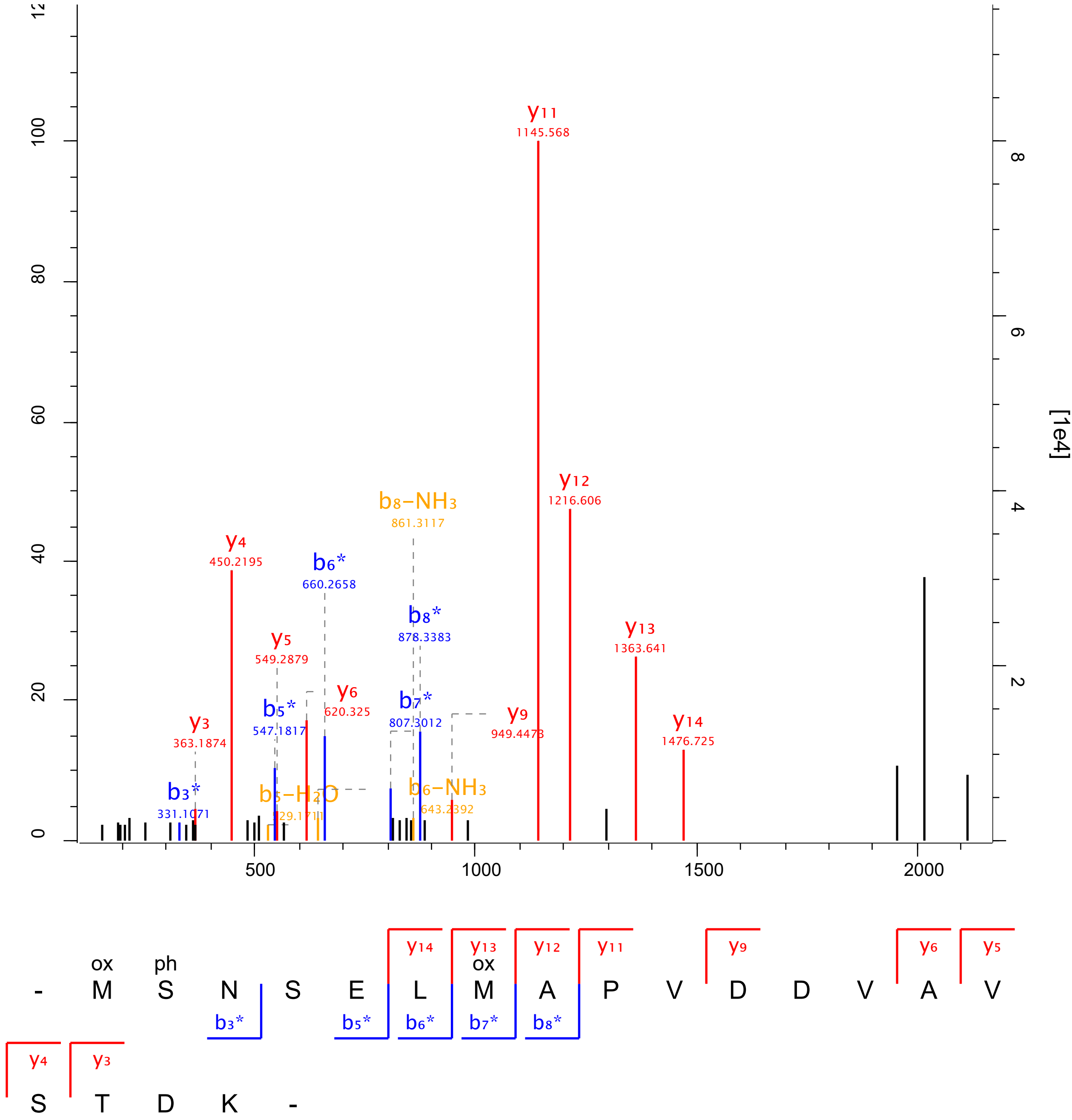

| Raw file | Scan | Method | Score | m/z |
| --- | --- | --- | --- | --- |
| A17_02_1 | 24826 | FTMS; HCD | 139.86 | 633.31 |

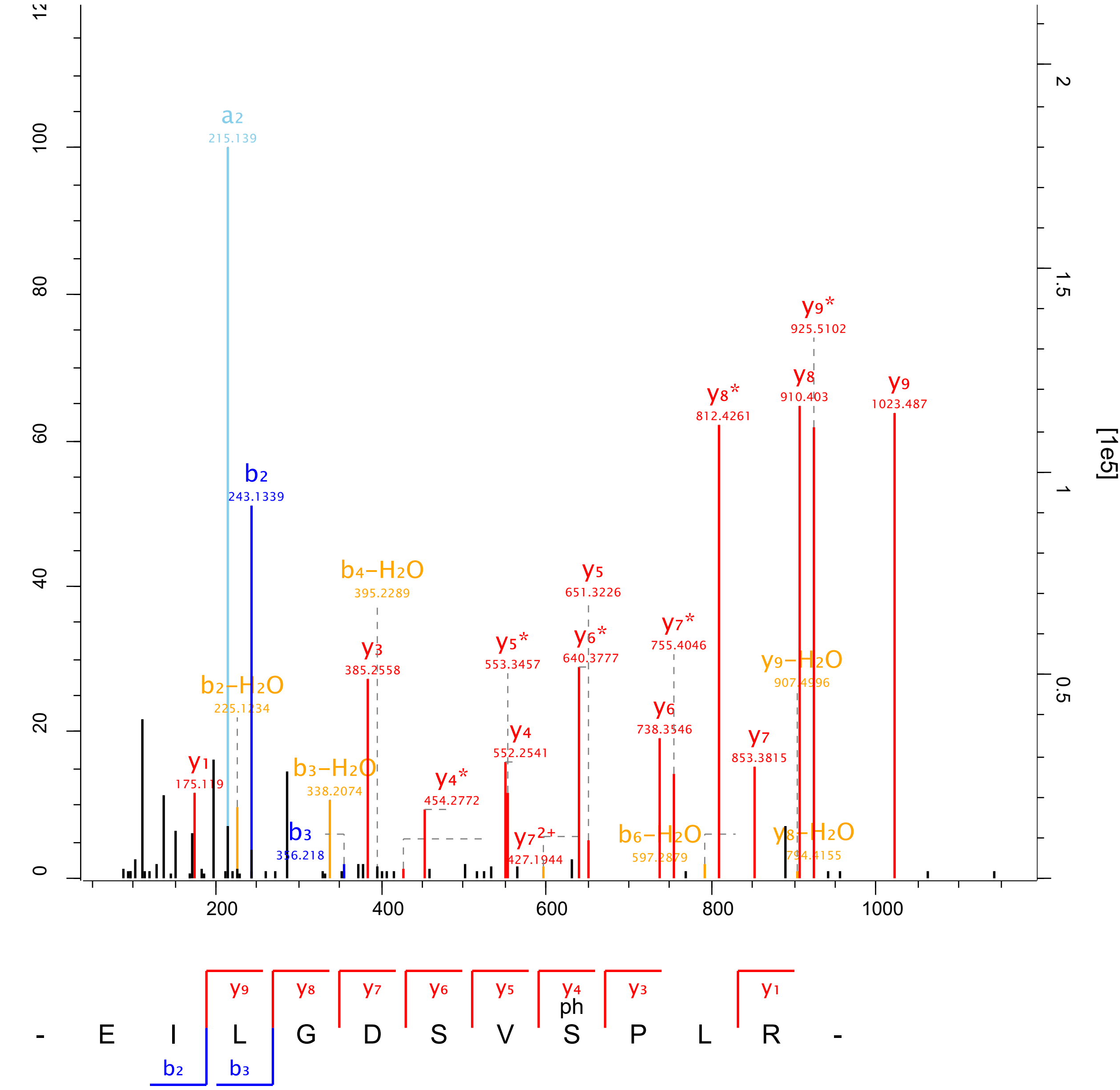

| Raw file | Scan | Method | Score | m/z |
|---|---|---|---|---|
| A17_02_1 | 25250 | FTMS; HCD | 252.52 | 852.82 |

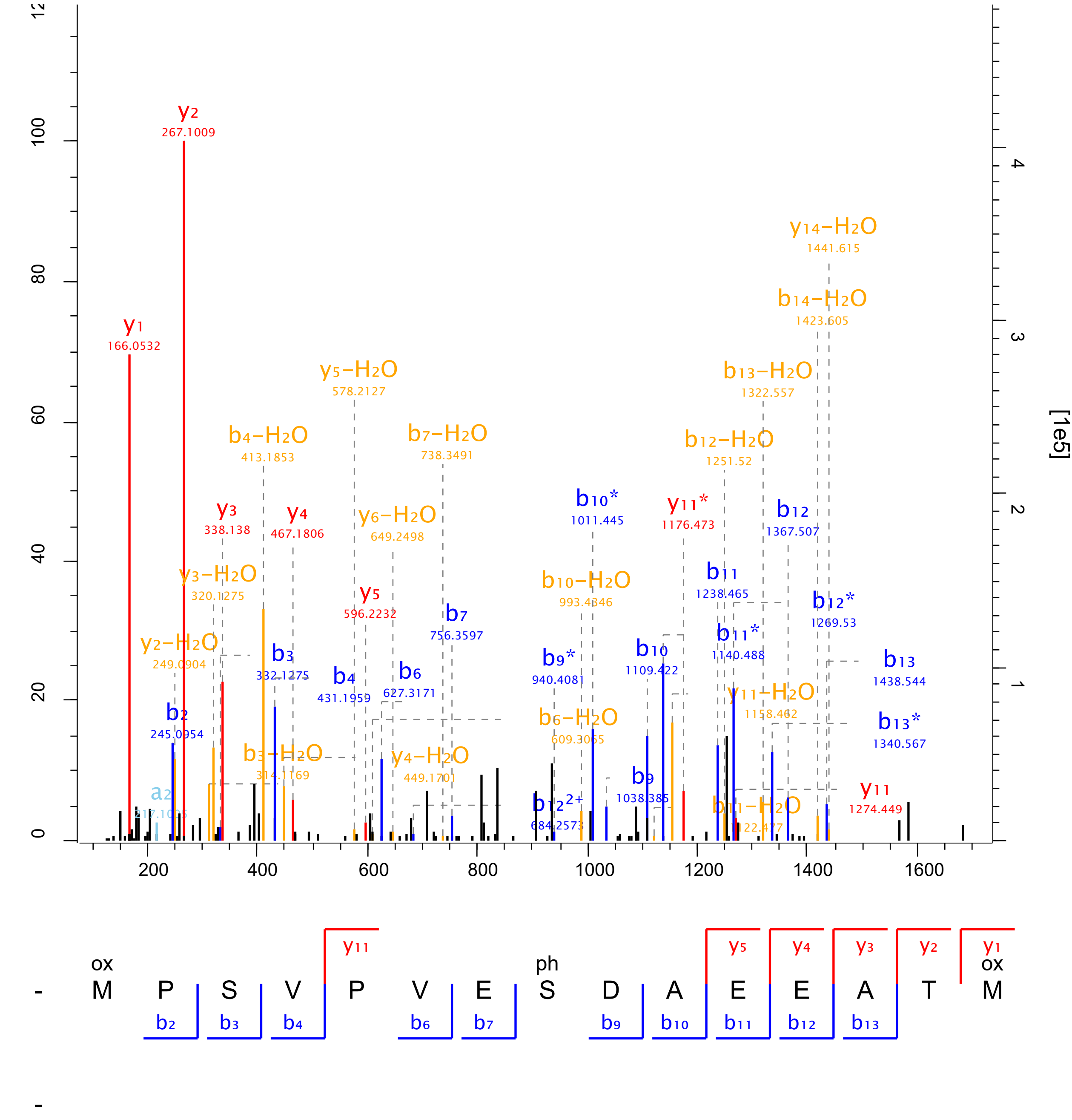

| Raw file | Scan | Method | Score | m/z |
| --- | --- | --- | --- | --- |
| A17_02_1 | 25301 | FTMS; HCD | 139.66 | 645.29 |

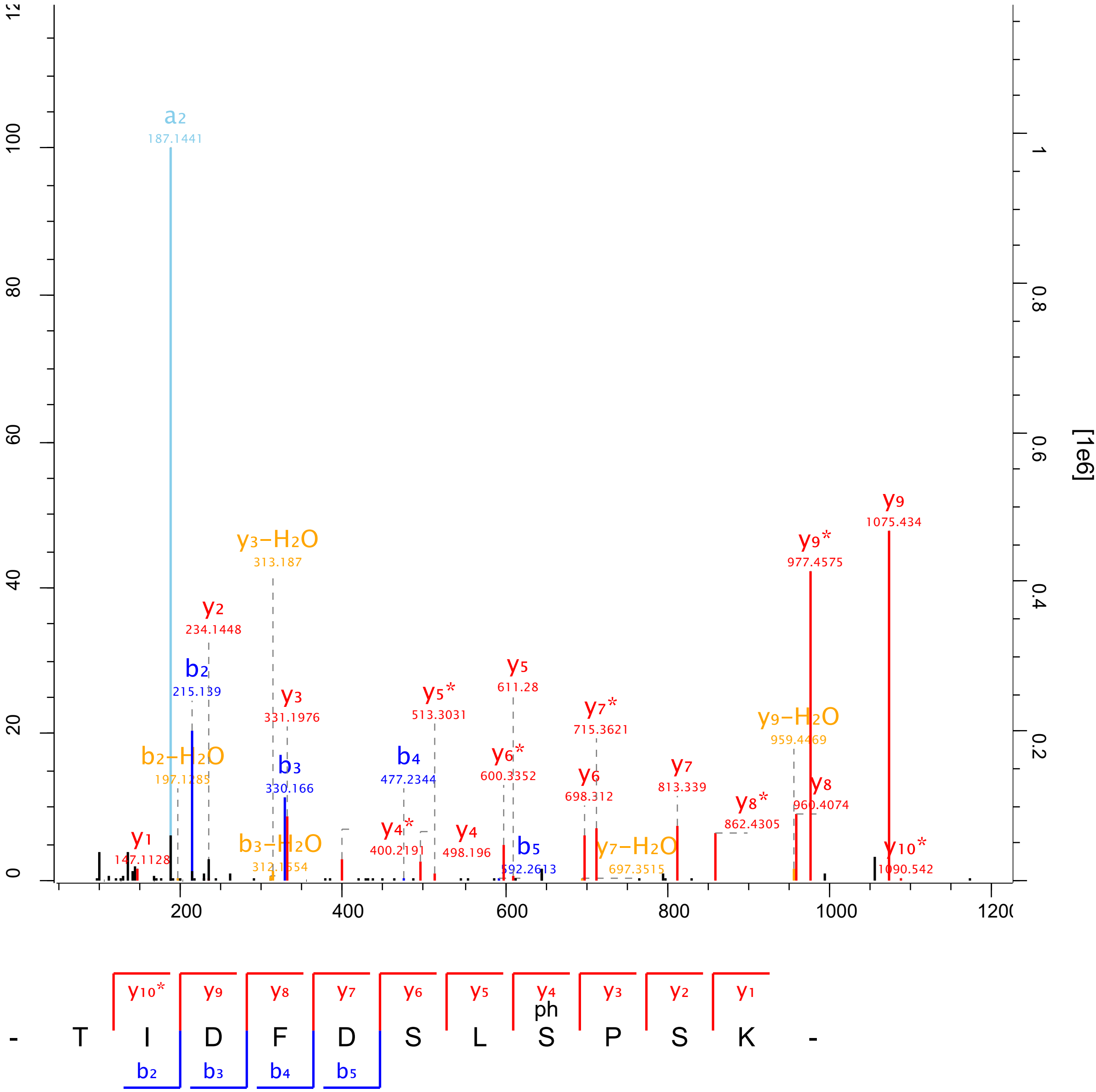

| Raw file | Scan | Method | Score | m/z |
| --- | --- | --- | --- | --- |
| A17_02_1 | 26373 | FTMS; HCD | 148.75 | 921.42 |

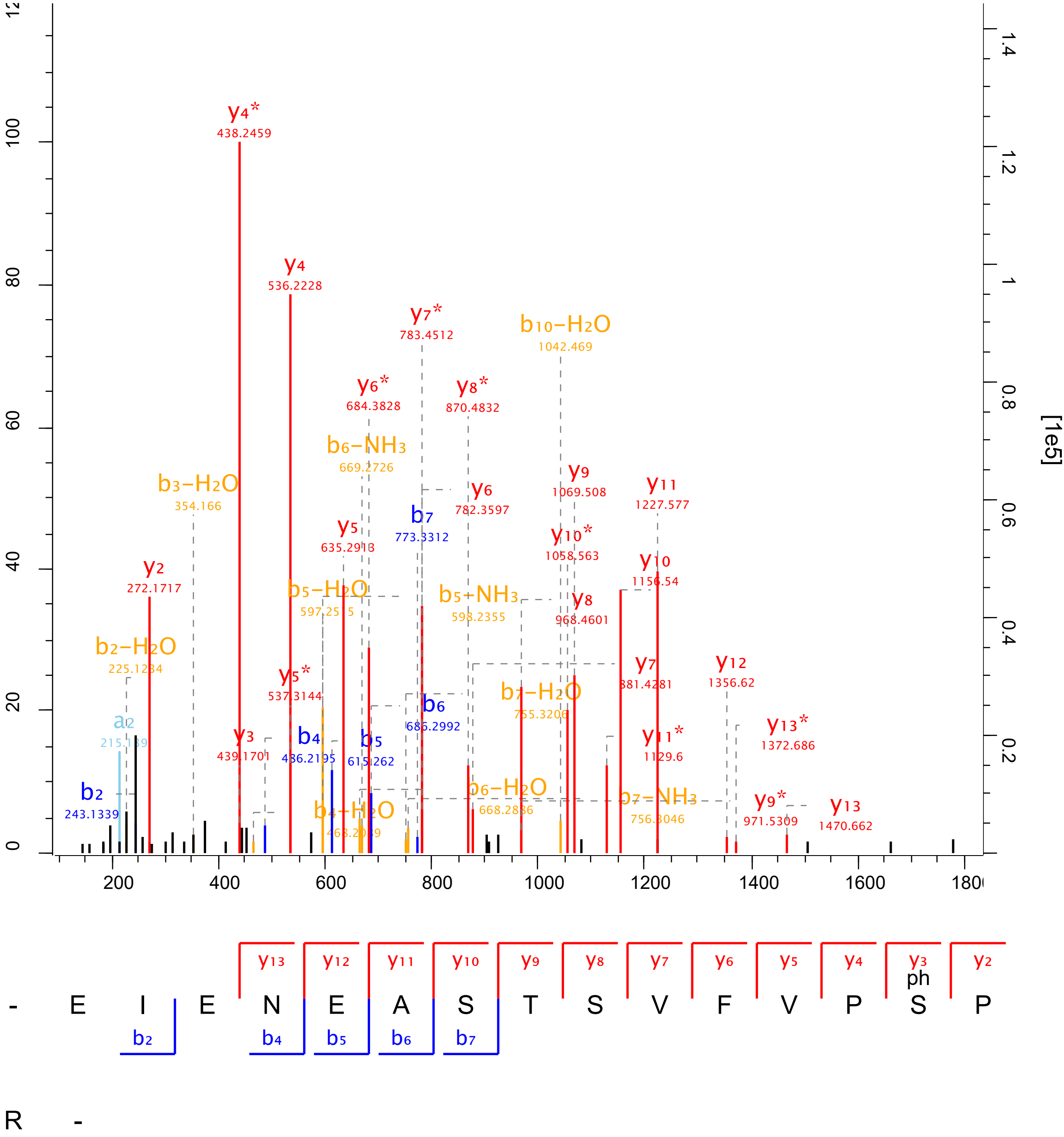

| Raw file | Scan | Method | Score | m/z |
|---|---|---|---|---|
| A17_02_1 | 26693 | FTMS; HCD | 169.67 | 760.87 |

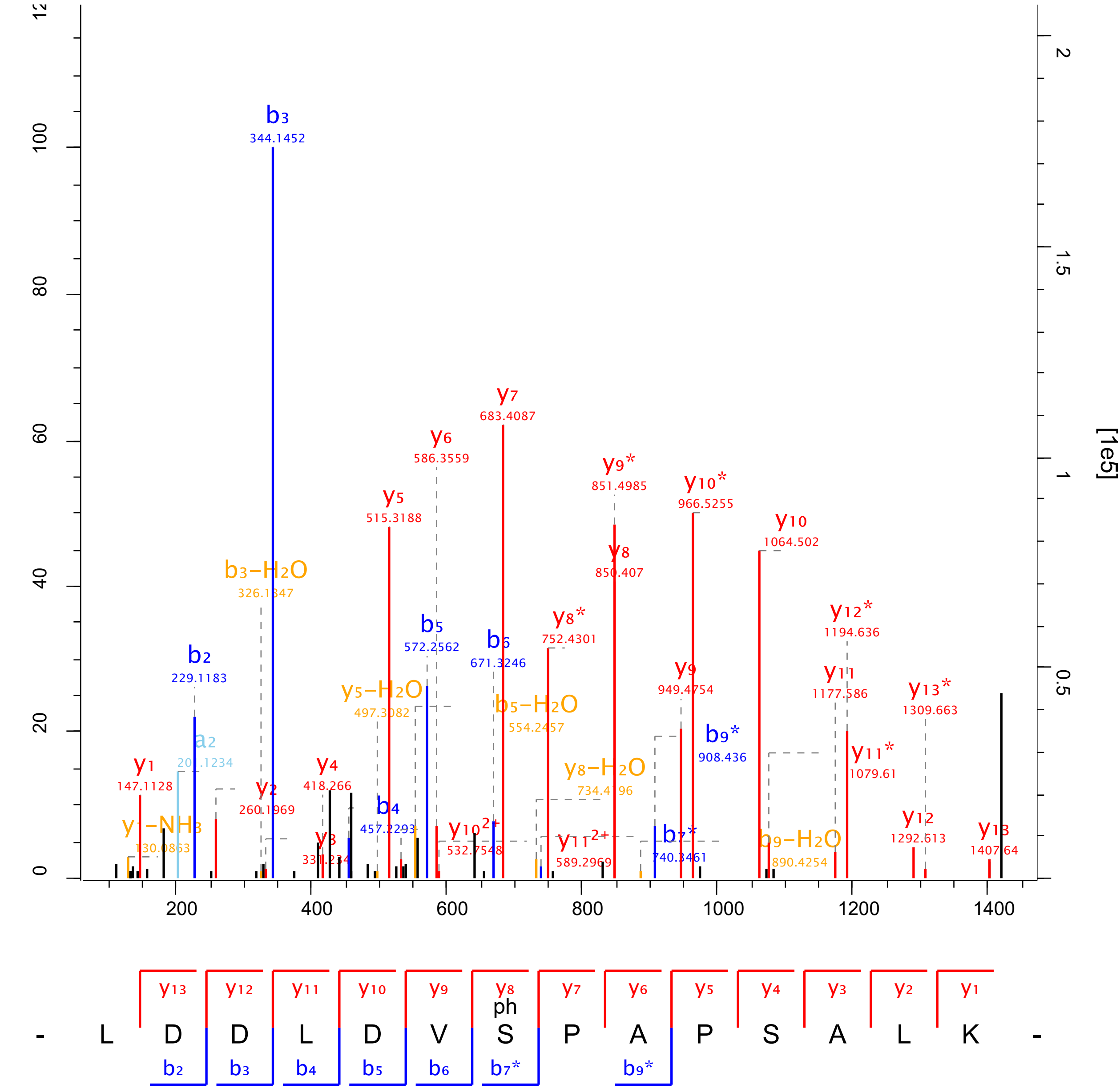

| Raw file | Scan | Method | Score | m/z |
| --- | --- | --- | --- | --- |
| A17_02_1 | 2738 | FTMS; HCD | 72.33 | 437.18 |

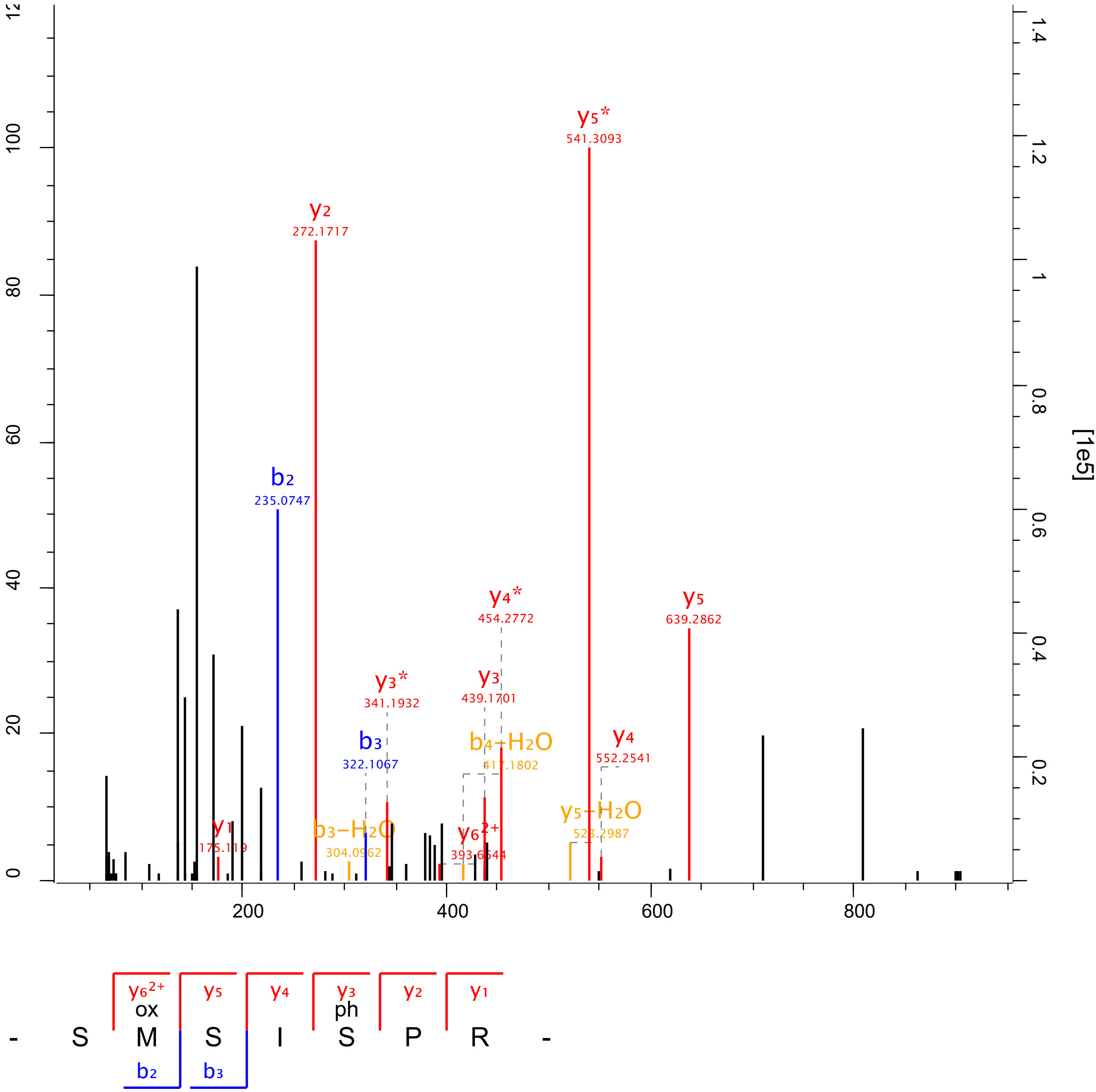

| Raw file | Scan | Method | Score | m/z |
| --- | --- | --- | --- | --- |
| A17_02_1 | 27674 | FTMS; HCD | 61.28 | 945.9 |

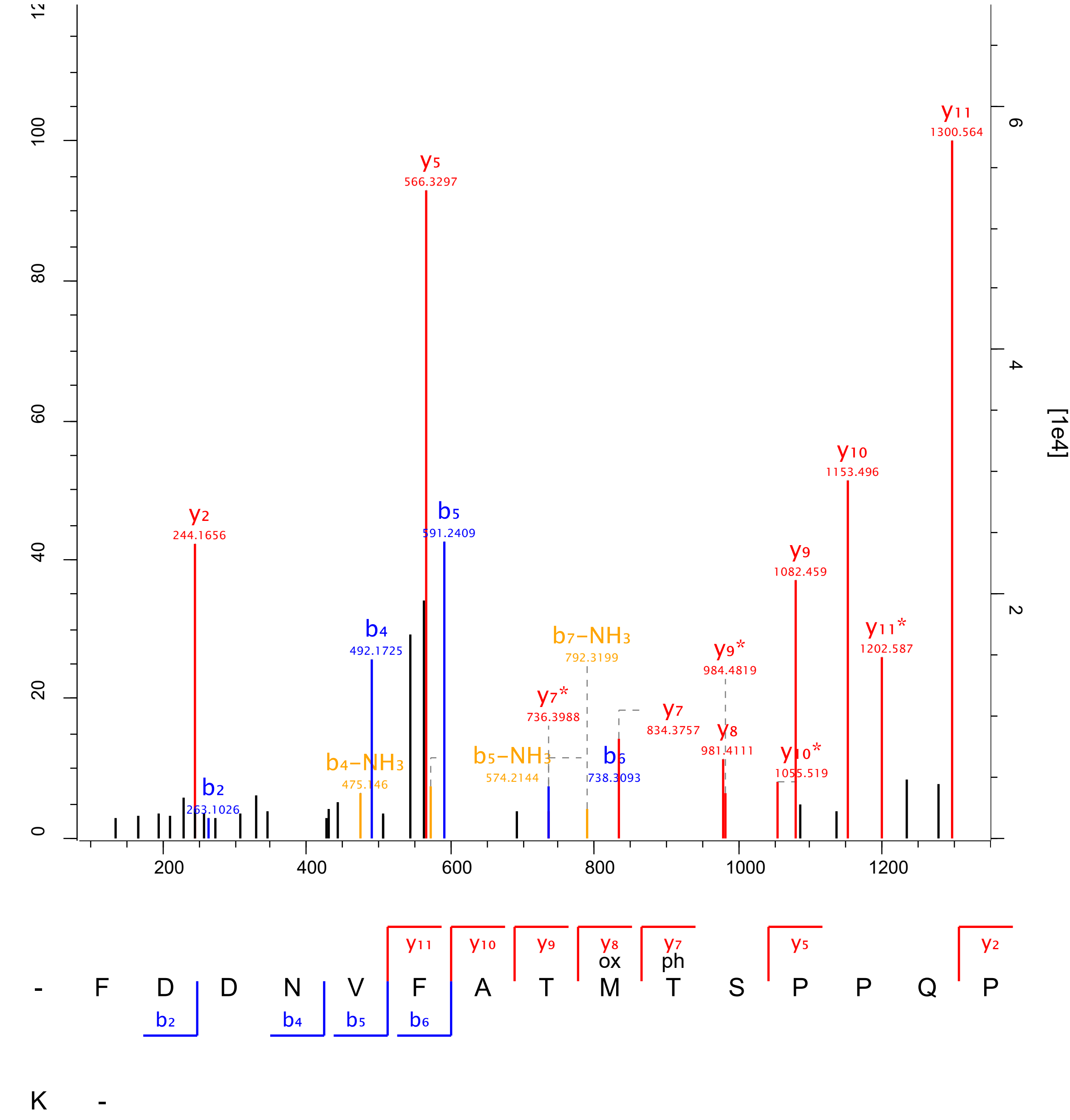

| Raw file | Scan | Method | Score | m/z |
| --- | --- | --- | --- | --- |
| A17_02_1 | 2818 | FTMS; HCD | 130.01 | 655.78 |

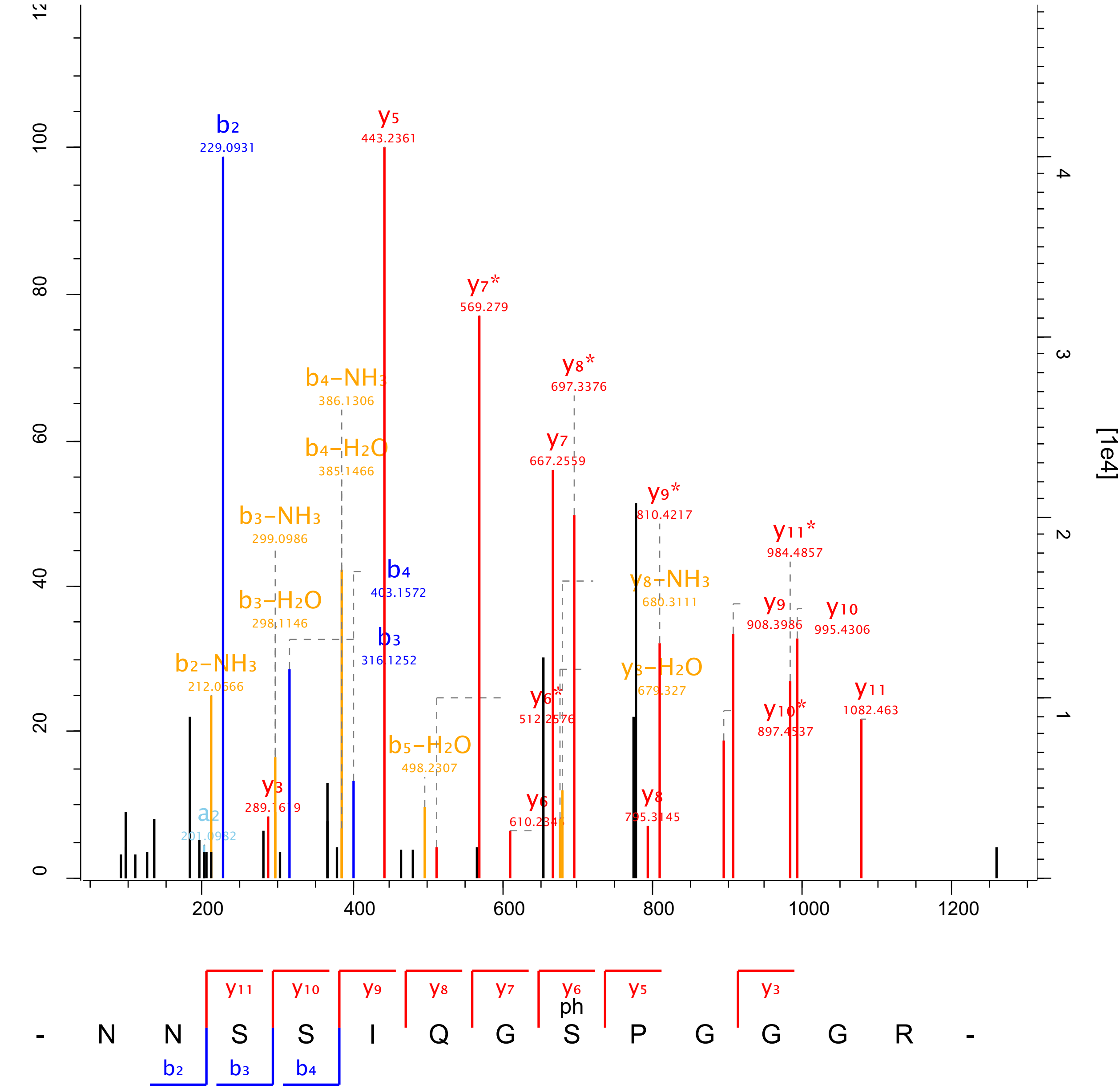

| Raw file | Scan | Method | Score | m/z |
|---|---|---|---|---|
| A17_02_1 | 28185 | FTMS; HCD | 44.93 | 729.34 |

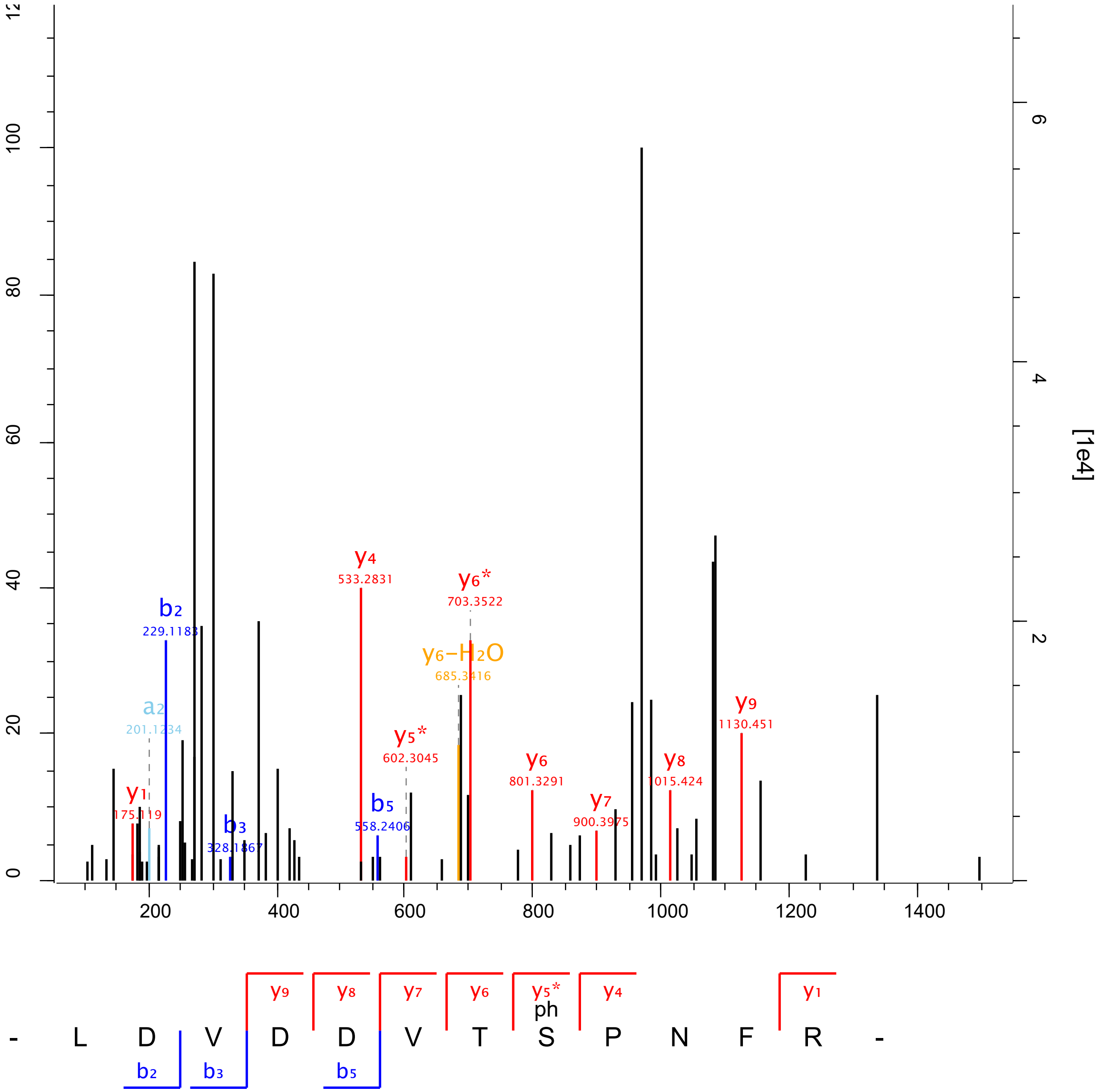

| Raw file | Scan | Method | Score | m/z |
|---|---|---|---|---|
| A17_02_1 | 28232 | FTMS; HCD | 70.94 | 515.26 |

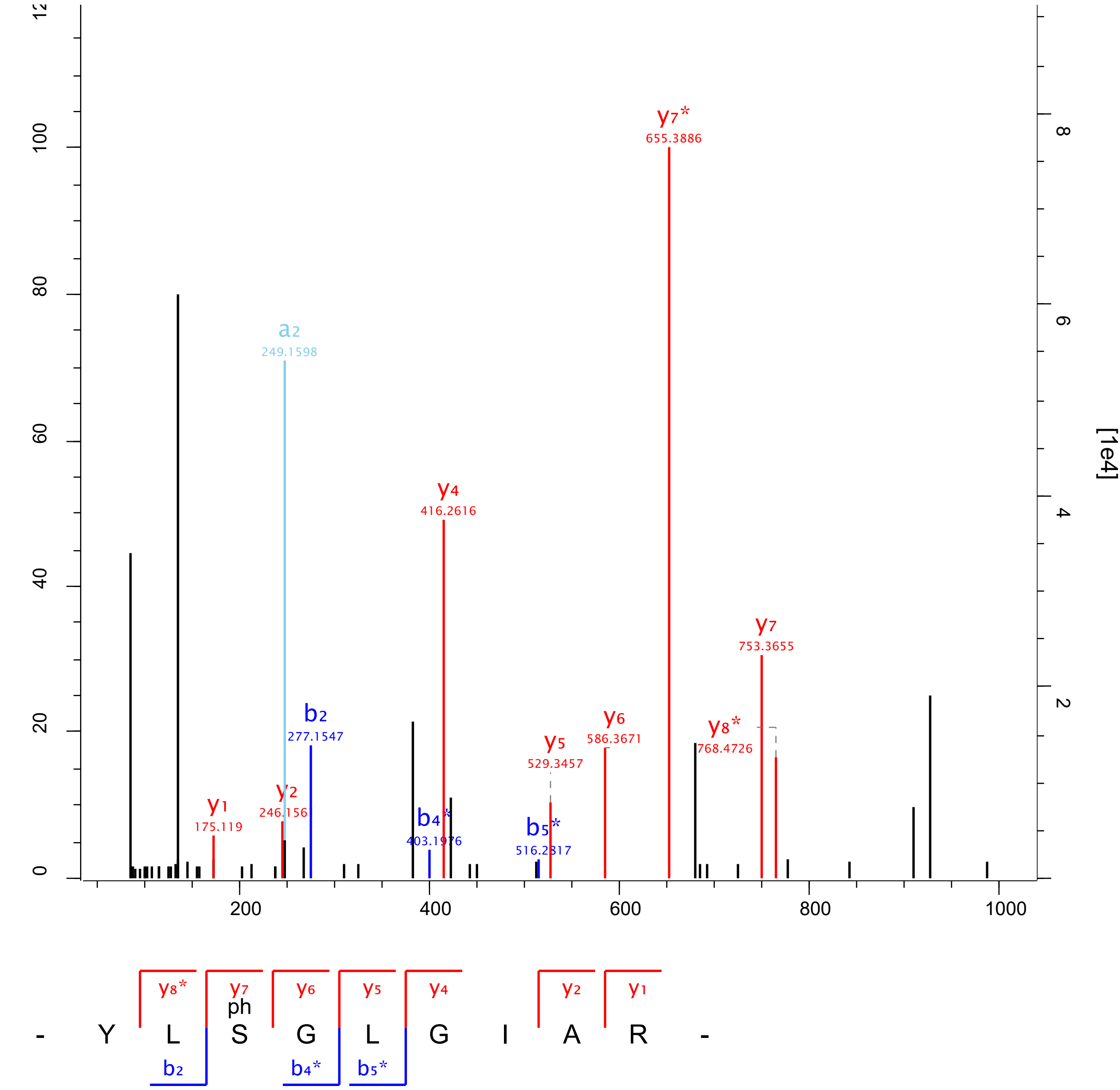

| Raw file | Scan | Method | Score | m/z |
|---|---|---|---|---|
| A17_02_1 | 2948 | FTMS; HCD | 53.24 | 645.27 |

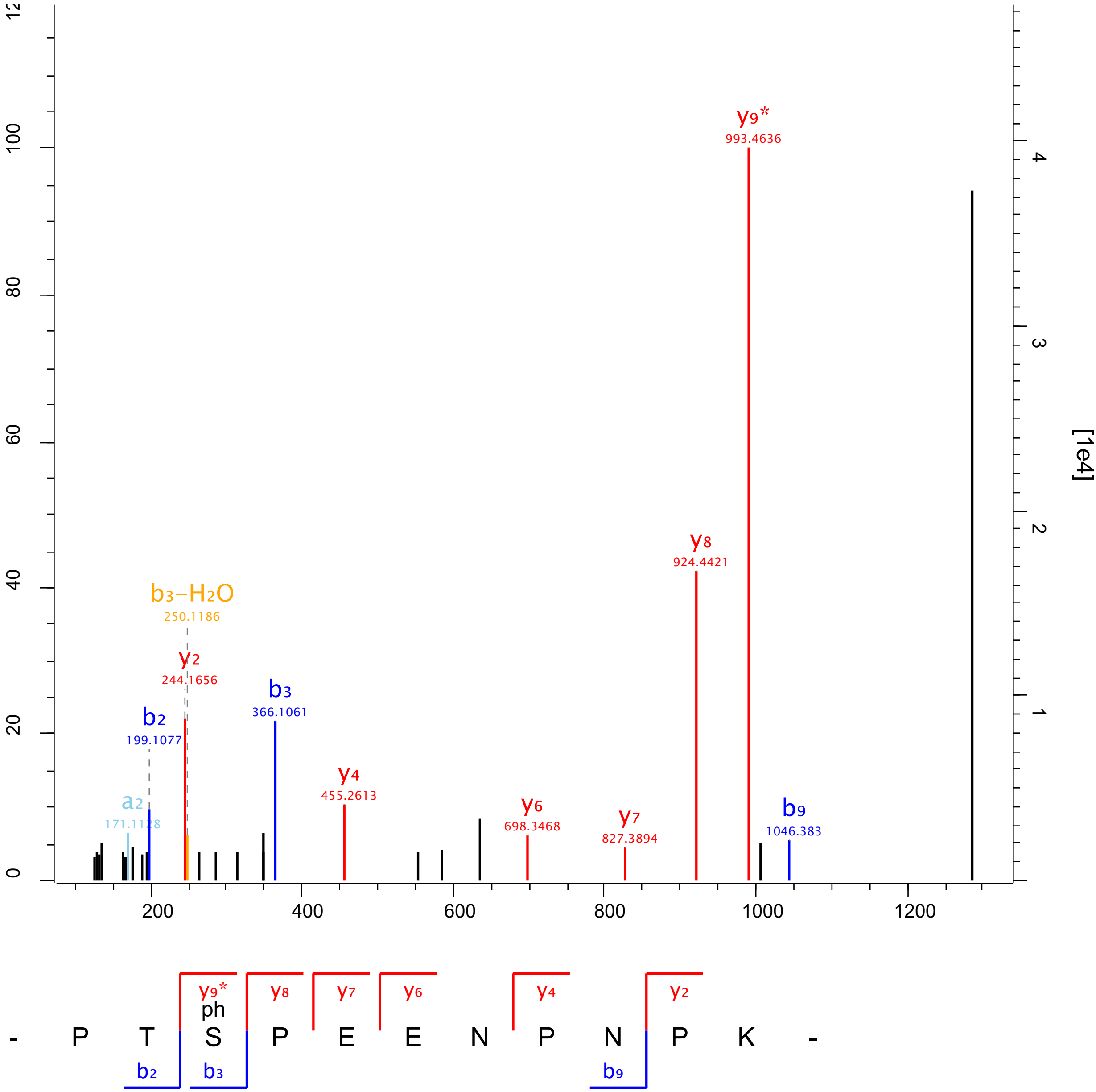

| Raw file | Scan | Method | Score | m/z |
|---|---|---|---|---|
| A17_02_1 | 29653 | FTMS; HCD | 165.45 | 793.85 |

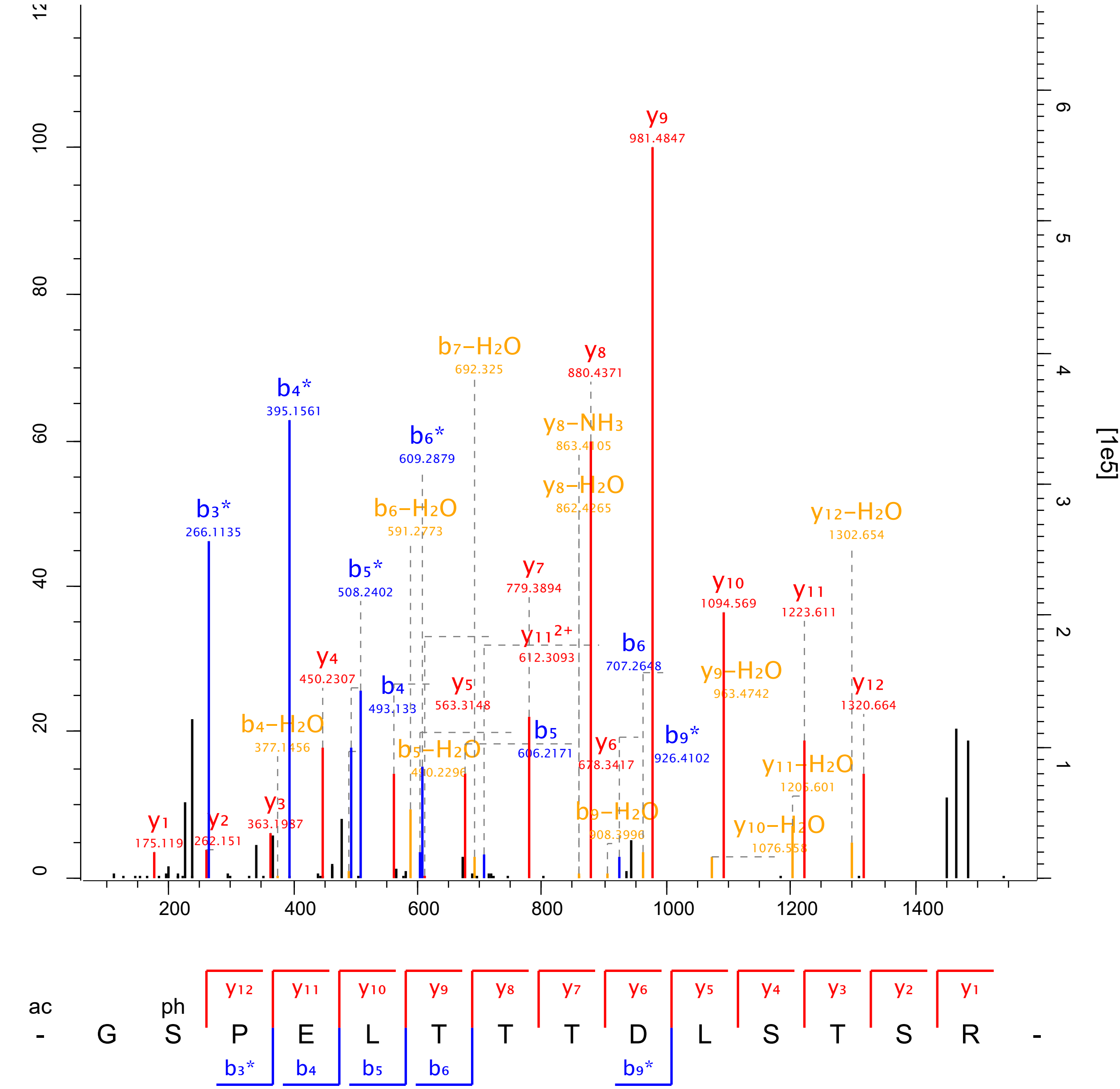

| Raw file | Scan | Method | Score | m/z |
|---|---|---|---|---|
| A17_02_1 | 29712 | FTMS; HCD | 50.9 | 685.32 |

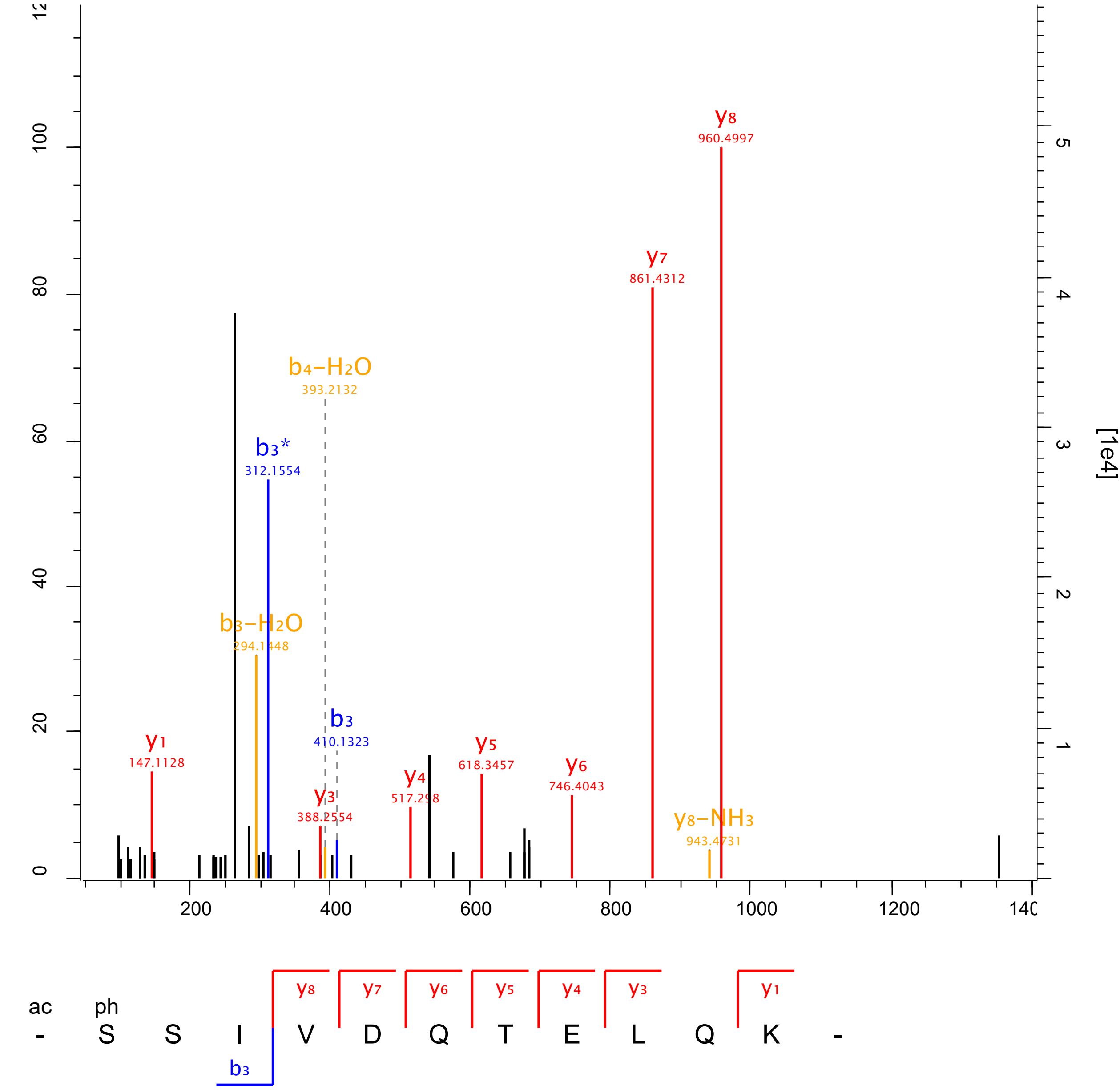

| Raw file | Scan | Method | Score | m/z |
|---|---|---|---|---|
| A17_02_1 | 30384 | FTMS; HCD | 52.72 | 869.37 |

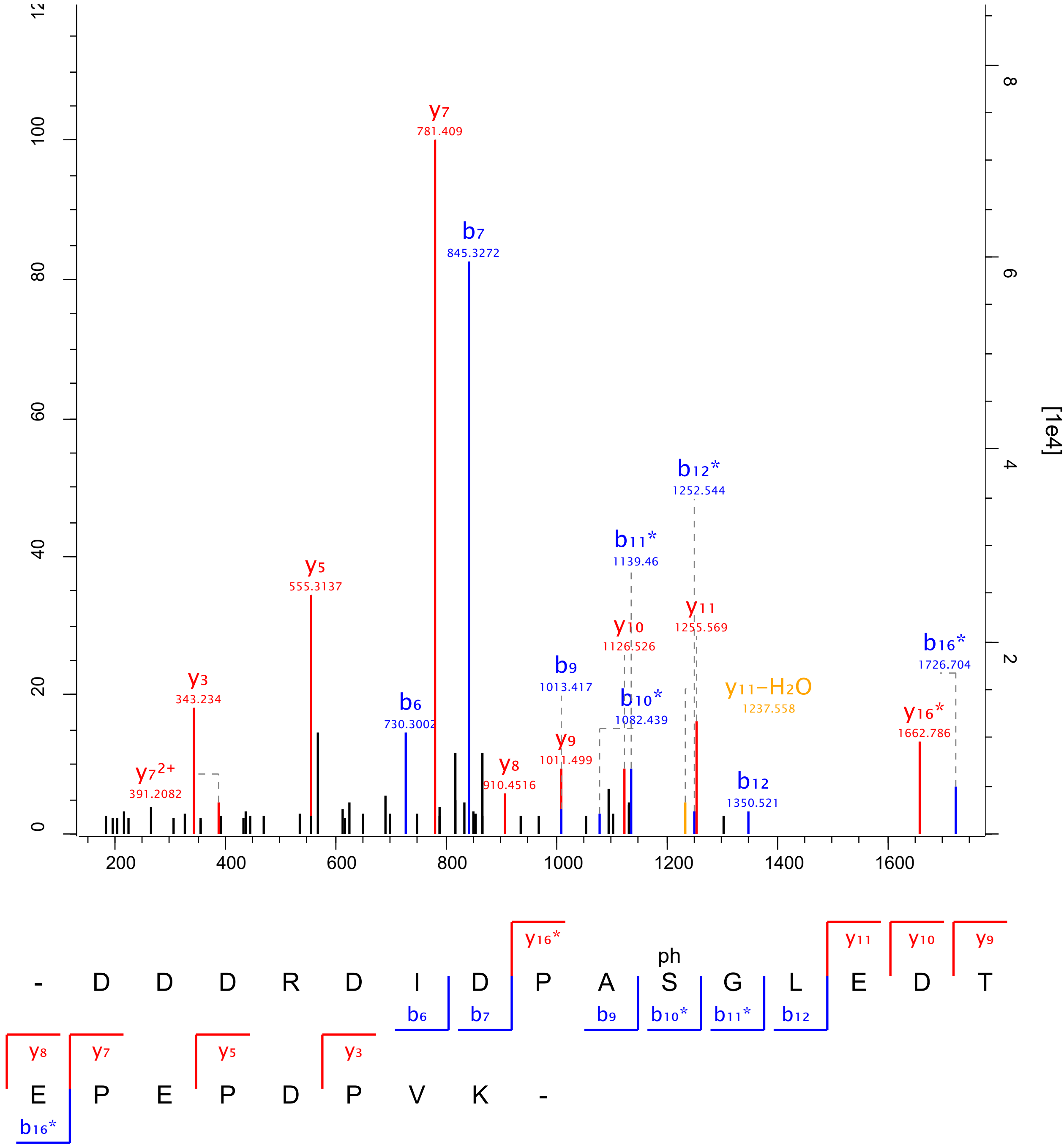

| Raw file | Scan | Method | Score | m/z |
|---|---|---|---|---|
| A17_02_1 | 30435 | FTMS; HCD | 190.59 | 779.37 |

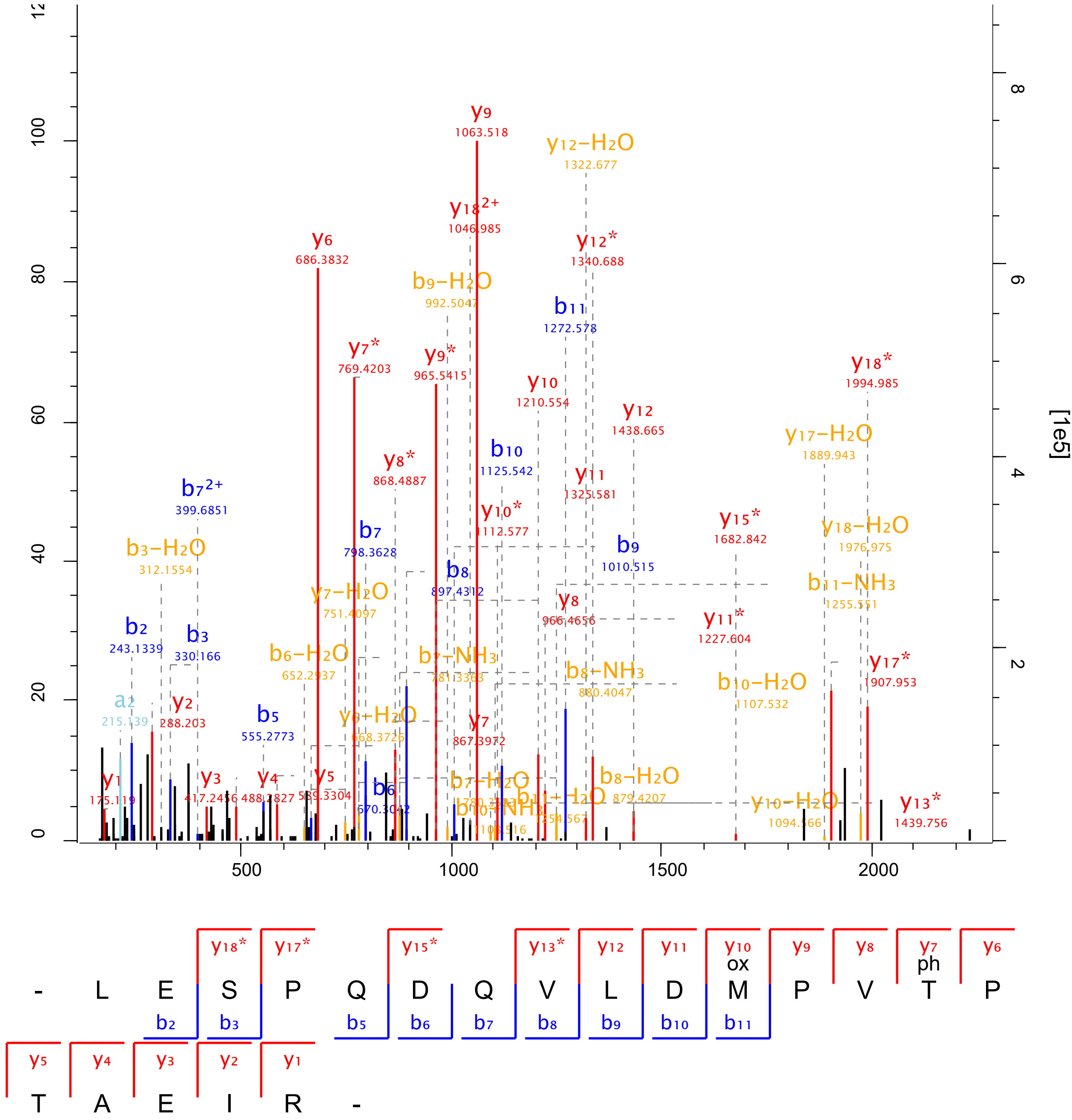

| Raw file | Scan | Method | Score | m/z |
|---|---|---|---|---|
| A17_02_1 | 30883 | FTMS; HCD | 118.31 | 832.89 |

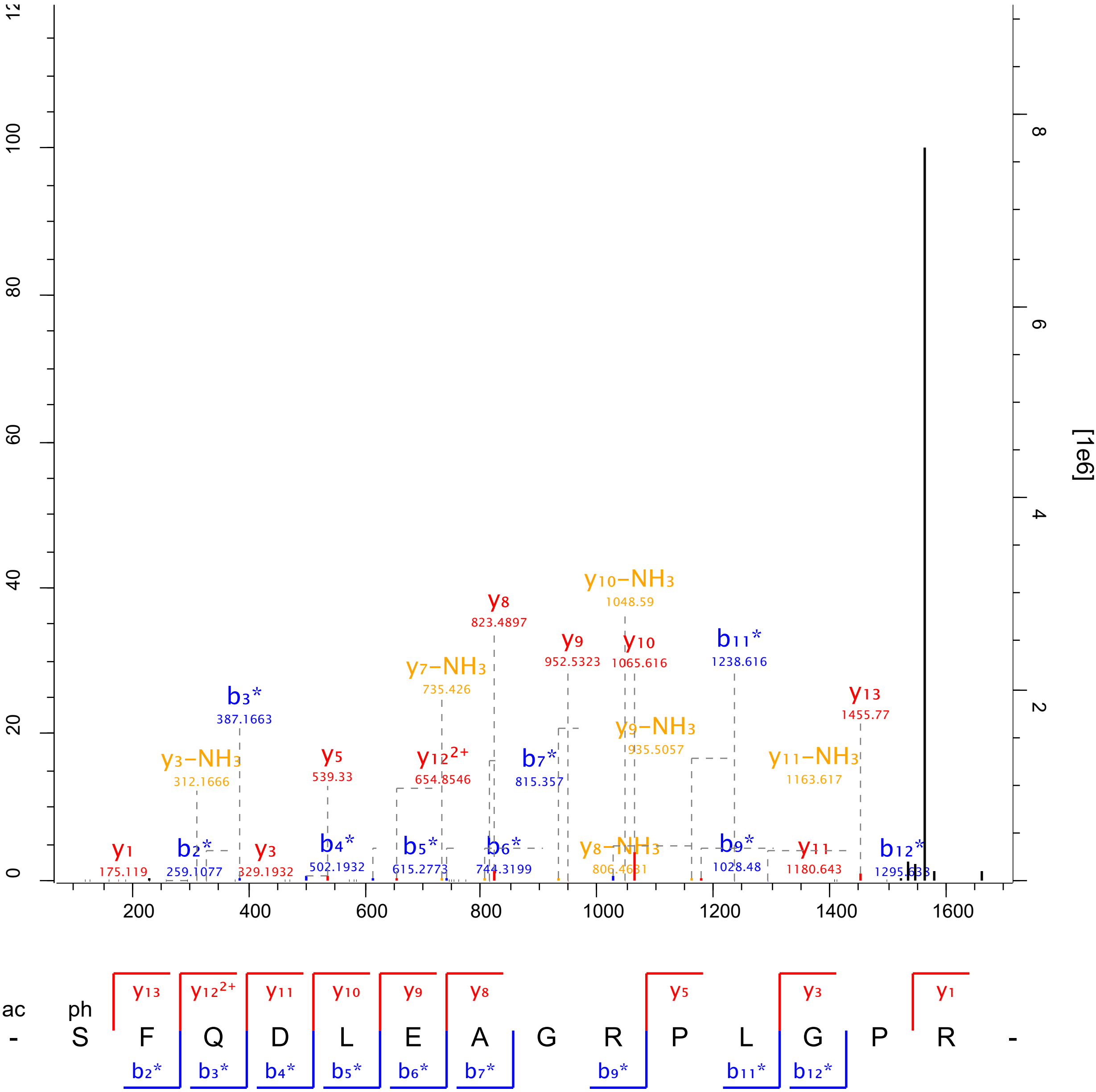

| Raw file | Scan | Method | Score | m/z |
|---|---|---|---|---|
| A17_02_1 | 31302 | FTMS; HCD | 73.67 | 1110.43 |

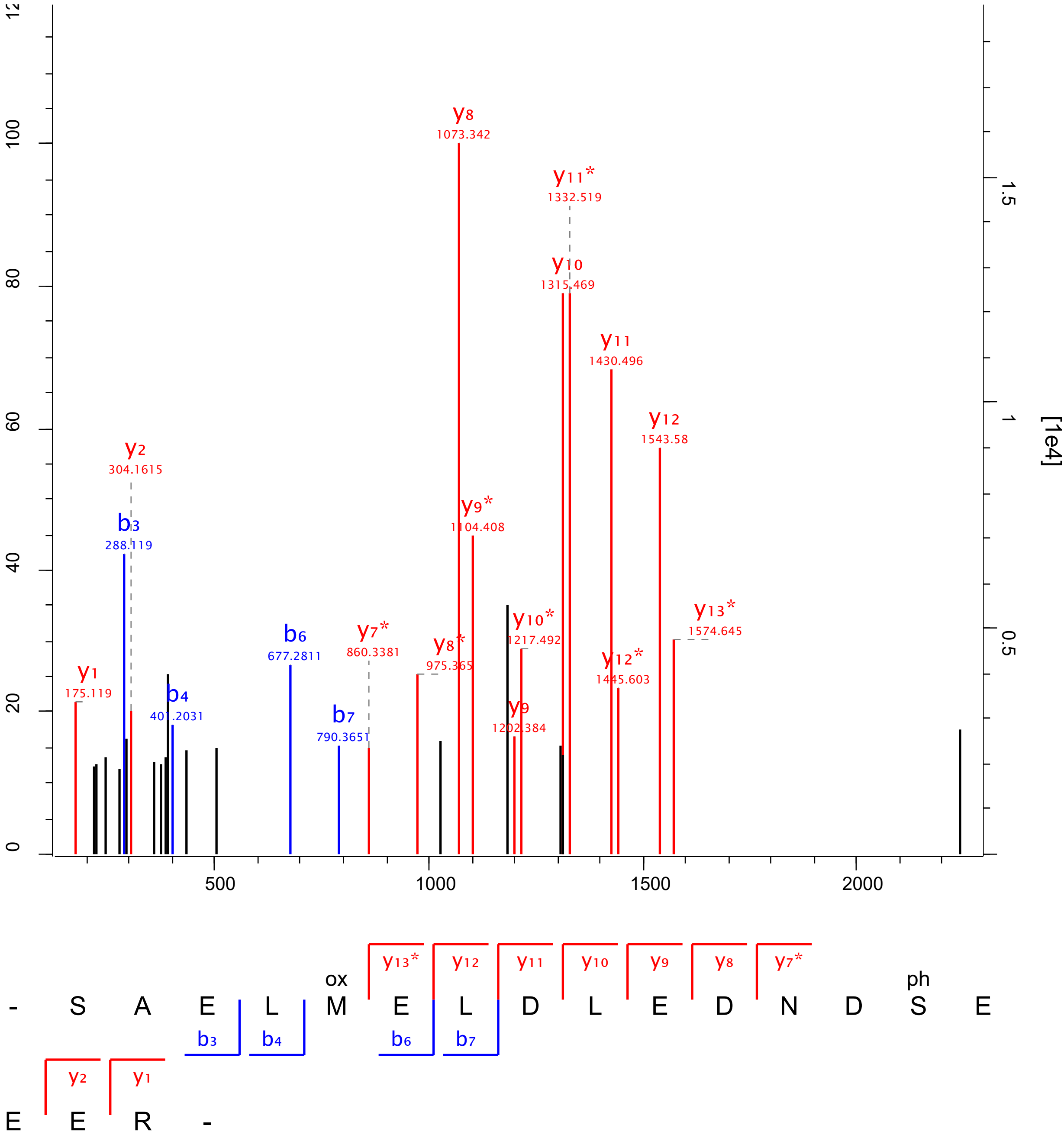

| Raw file | Scan | Method | Score | m/z |
|---|---|---|---|---|
| A17_02_1 | 3159 | FTMS; HCD | 46.32 | 630.75 |

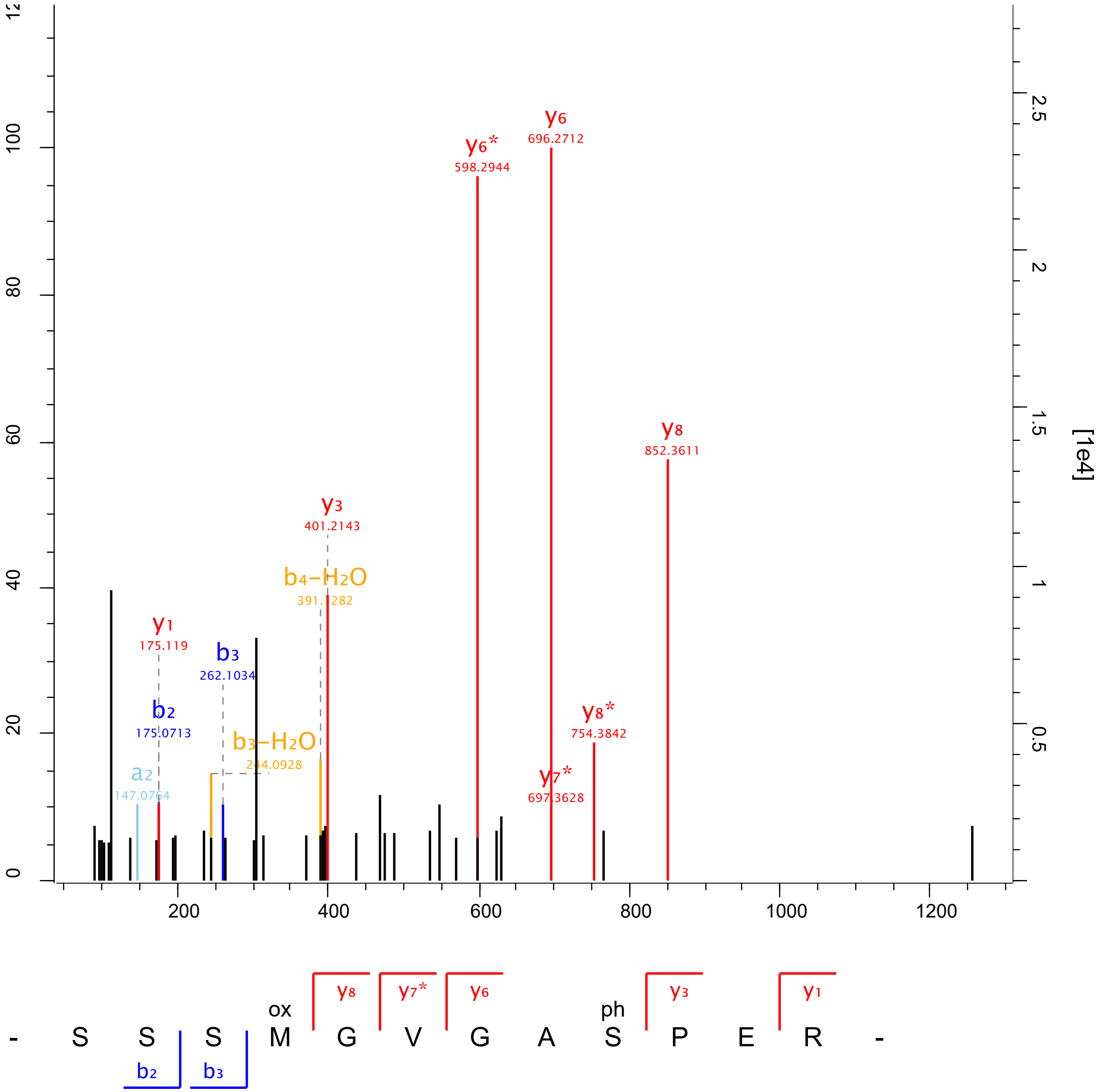

| Raw file | Scan | Method | Score | m/z |
|---|---|---|---|---|
| A17_02_1 | 32301 | FTMS; HCD | 55.31 | 869.72 |

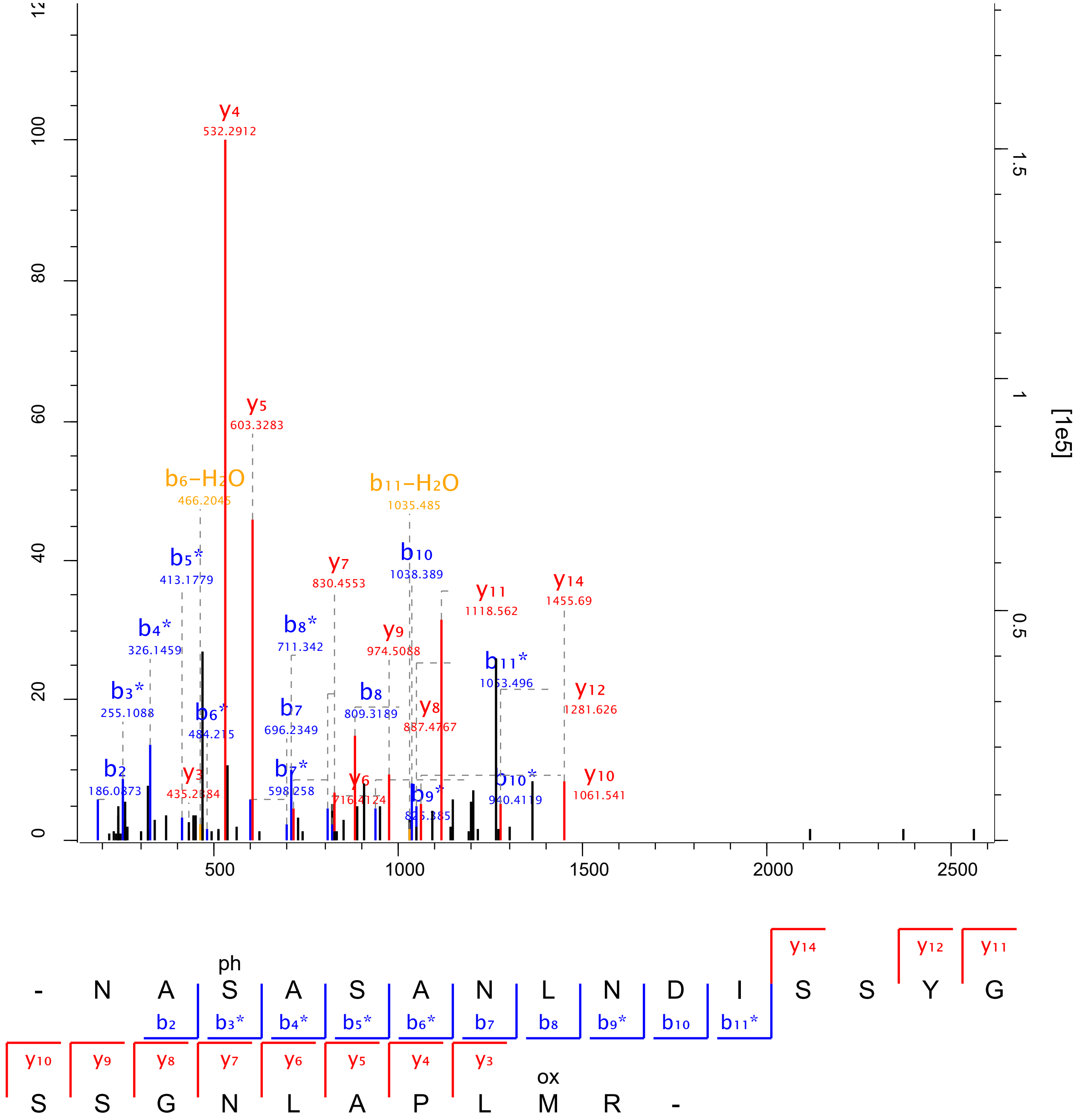

| Raw file | Scan | Method | Score | m/z |
| --- | --- | --- | --- | --- |
| A17_02_1 | 32504 | FTMS; HCD | 141.82 | 650.79 |

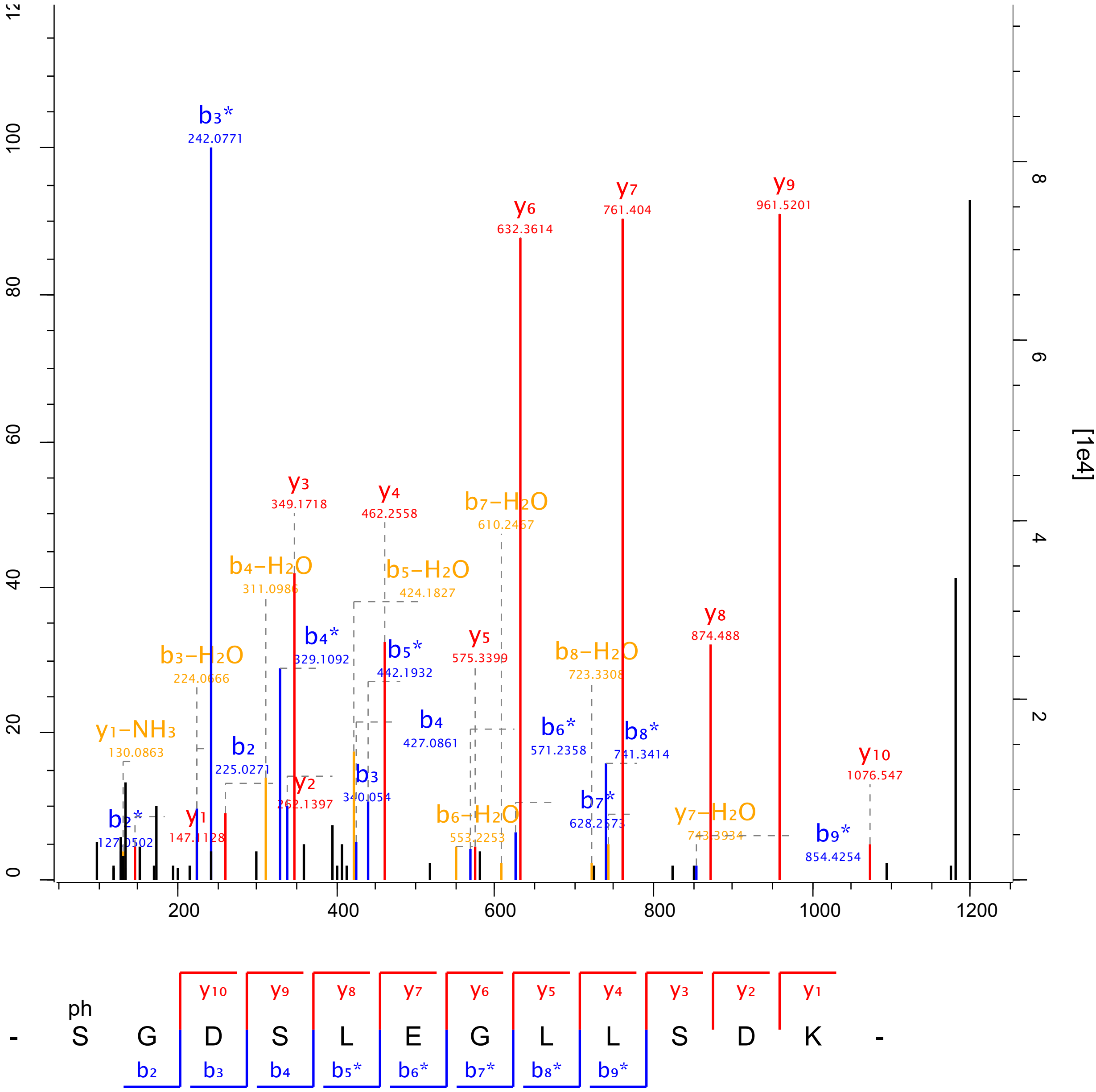

| Raw file | Scan | Method | Score | m/z |
|---|---|---|---|---|
| A17_02_1 | 32532 | FTMS; HCD | 311.1 | 843.87 |

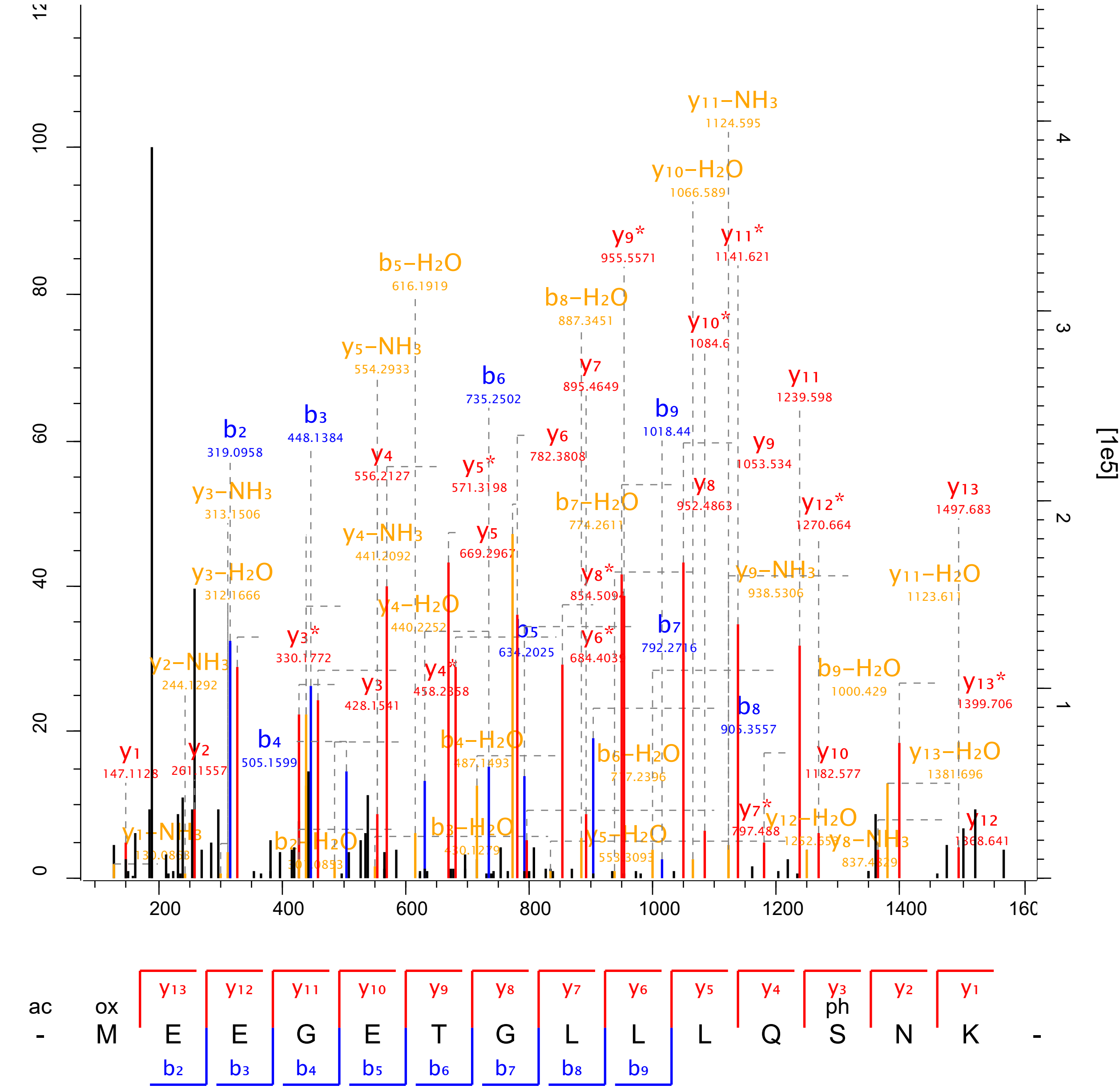

| Raw file | Scan | Method | Score | m/z |
|---|---|---|---|---|
| A17_02_1 | 32632 | FTMS; HCD | 53.56 | 696.82 |

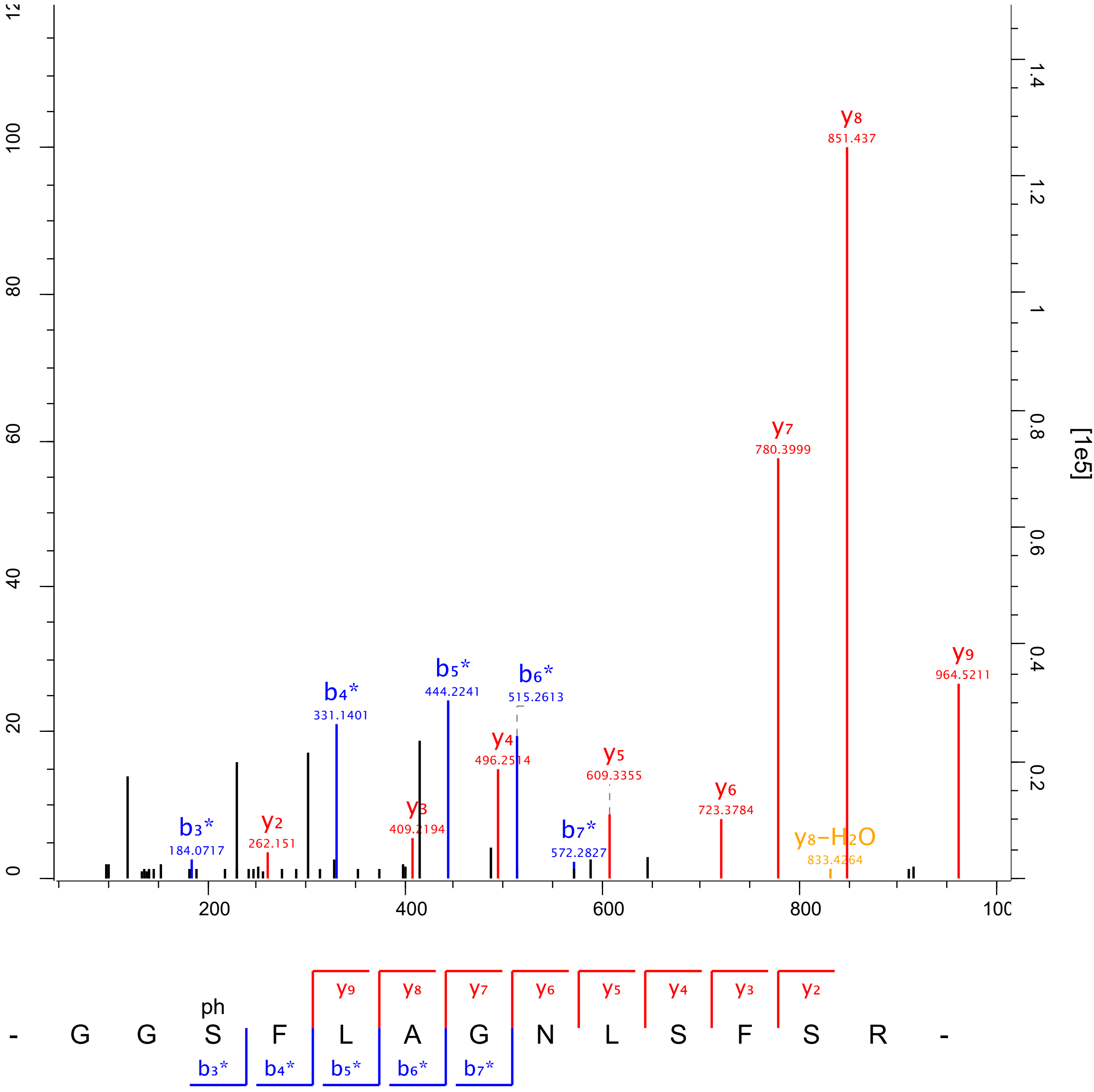

| Raw file | Scan | Method | Score | m/z |
|---|---|---|---|---|
| A17_02_1 | 32737 | FTMS; HCD | 115.8 | 744.89 |

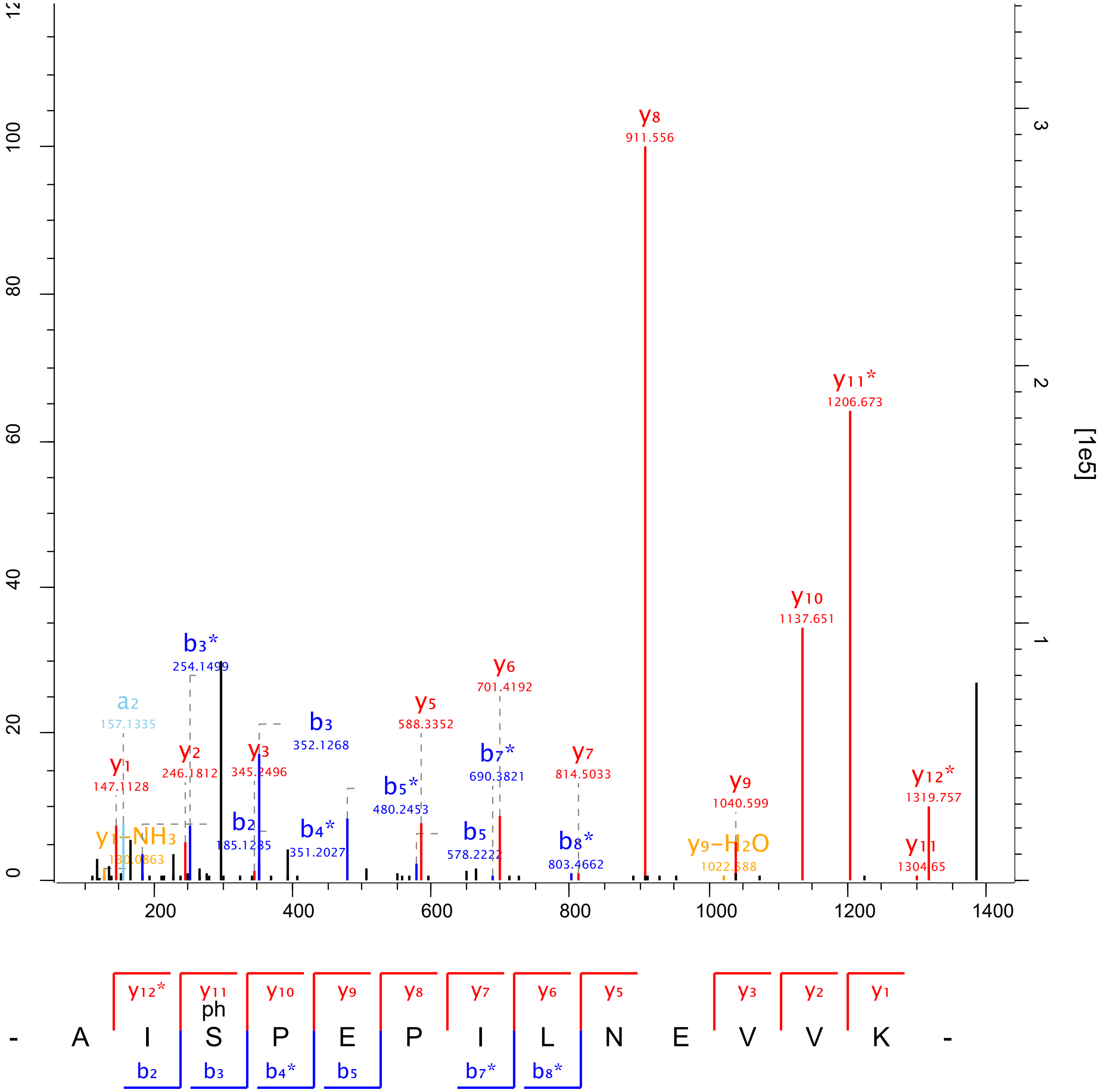

| Raw file | Scan | Method | Score | m/z |
| --- | --- | --- | --- | --- |
| A17_02_1 | 33133 | FTMS; HCD | 87.68 | 733.86 |

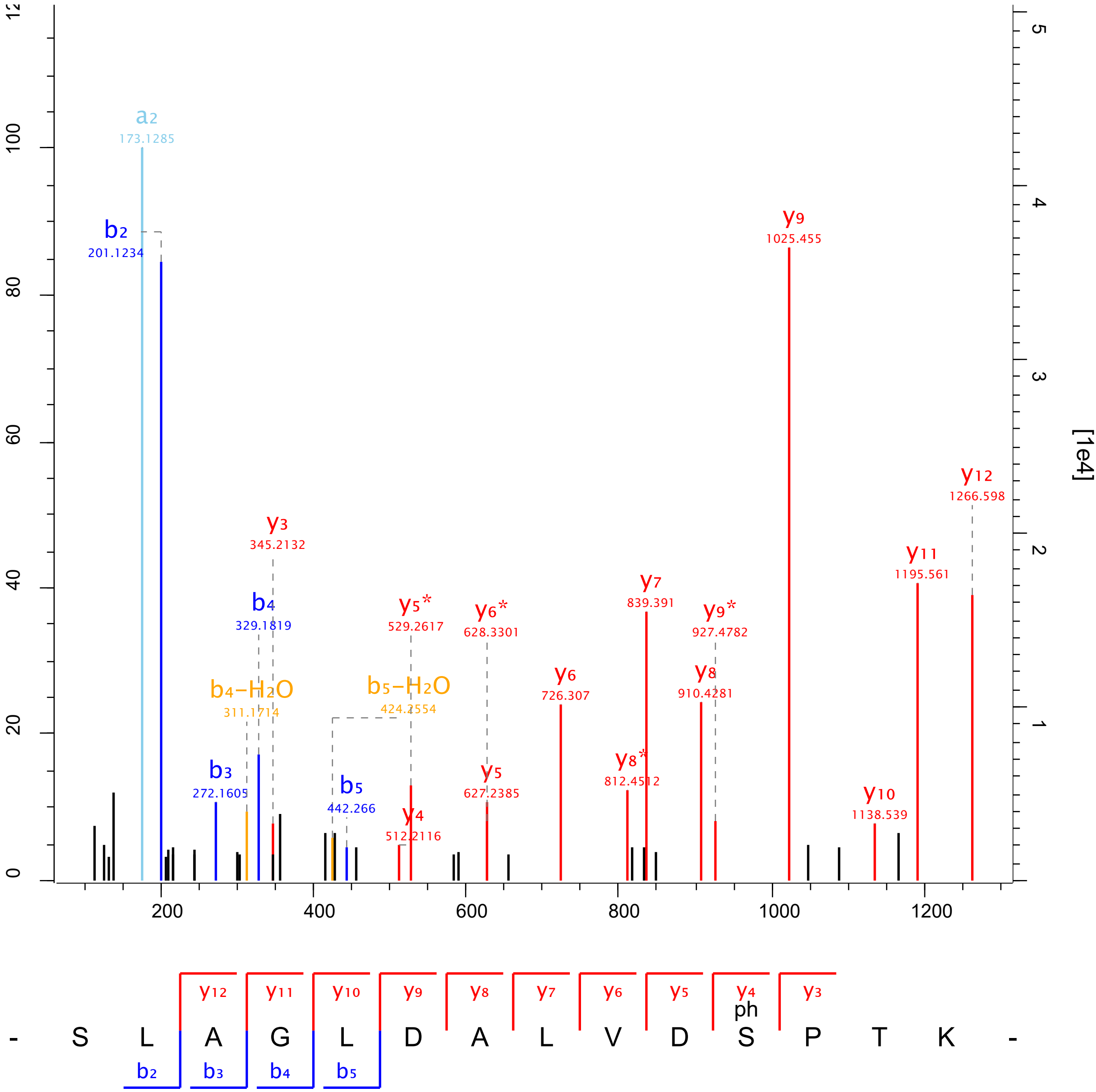

| Raw file | Scan | Method | Score | m/z |
|---|---|---|---|---|
| A17_02_1 | 33621 | FTMS; HCD | 133.76 | 812.36 |

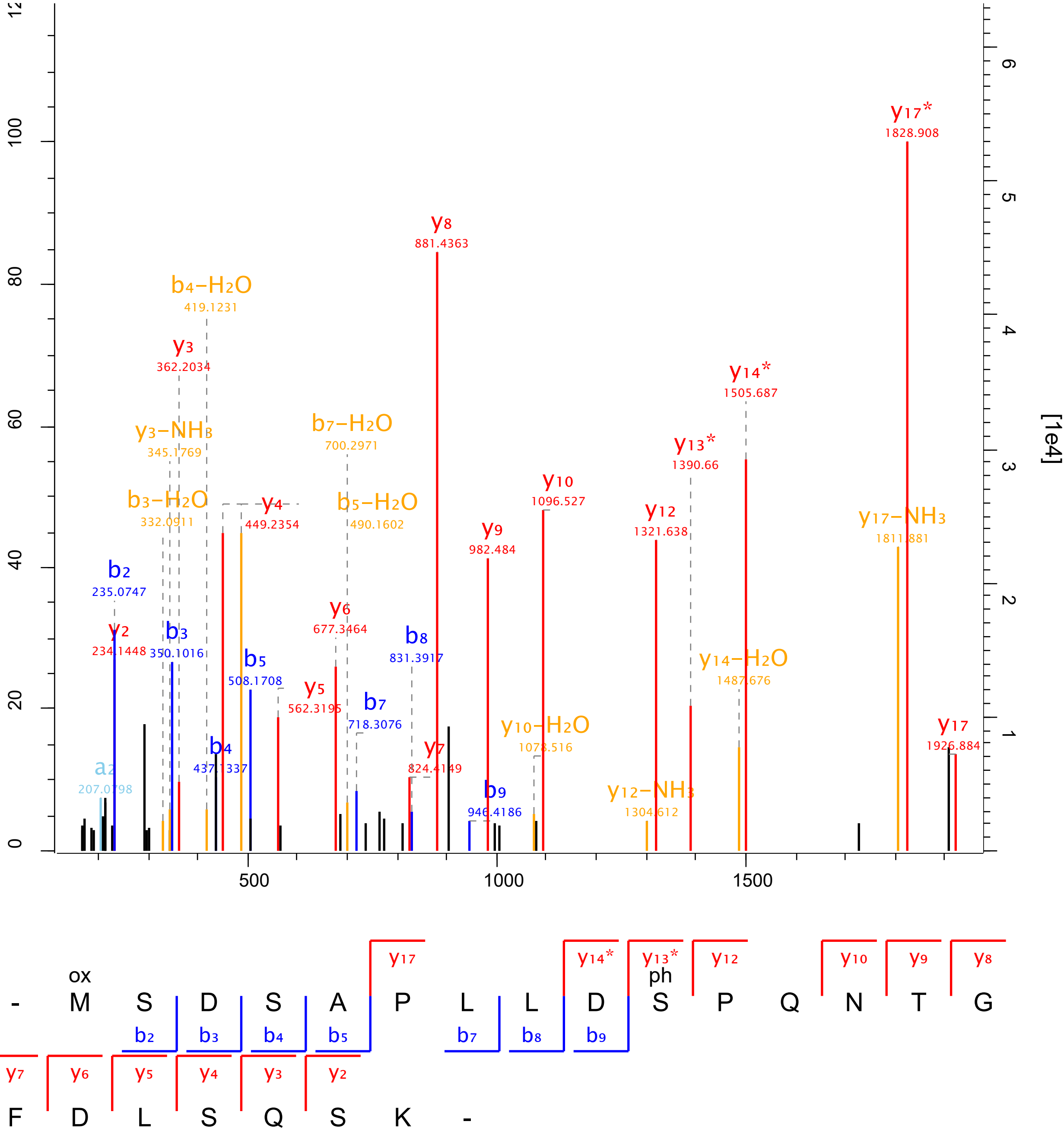

| Raw file | Scan | Method | Score | m/z |
|---|---|---|---|---|
| A17_02_1 | 33653 | FTMS; HCD | 68.45 | 1006.48 |

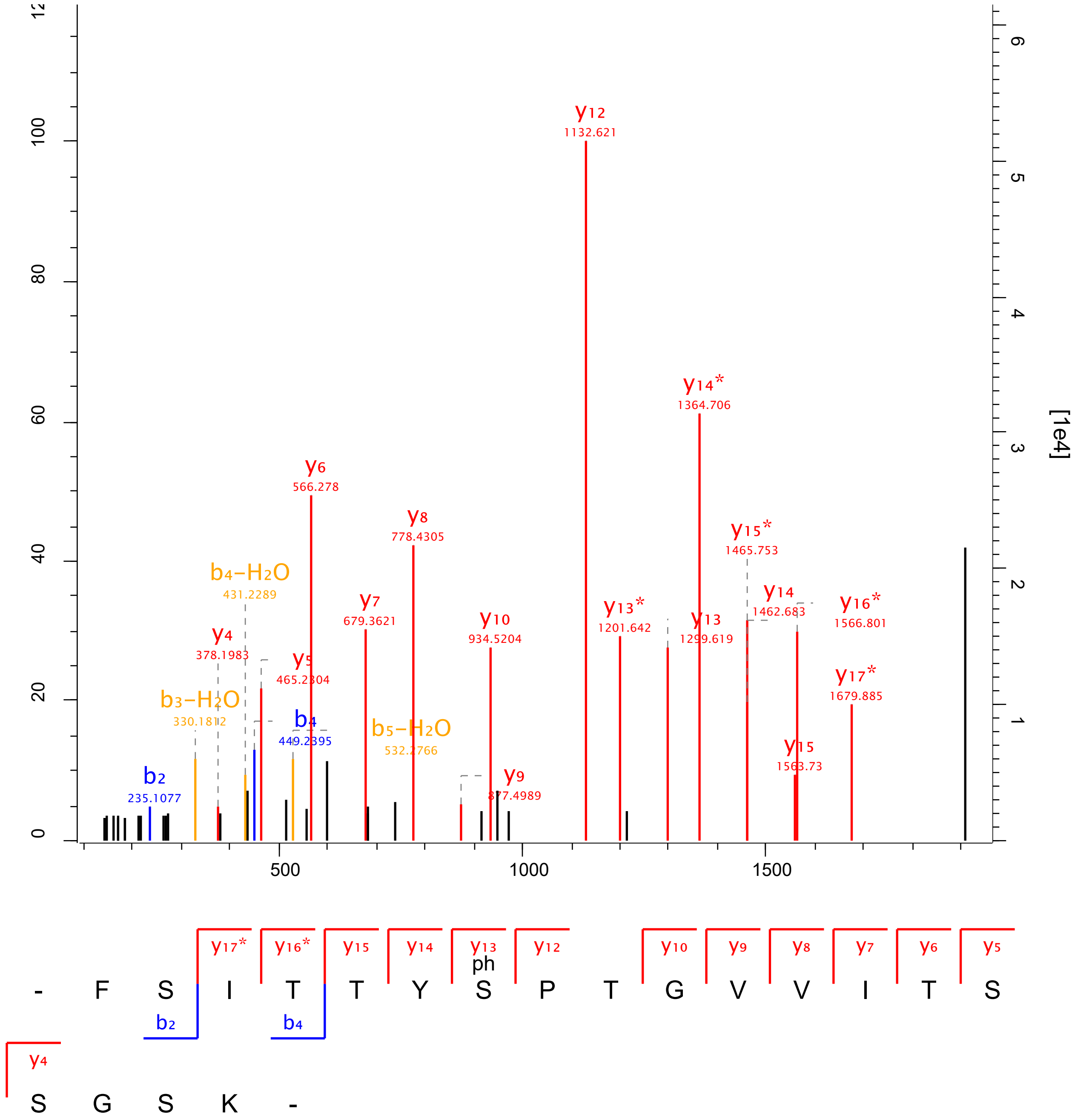

| Raw file | Scan | Method | Score | m/z |
|---|---|---|---|---|
| A17_02_1 | 33667 | FTMS; HCD | 183.31 | 742.83 |

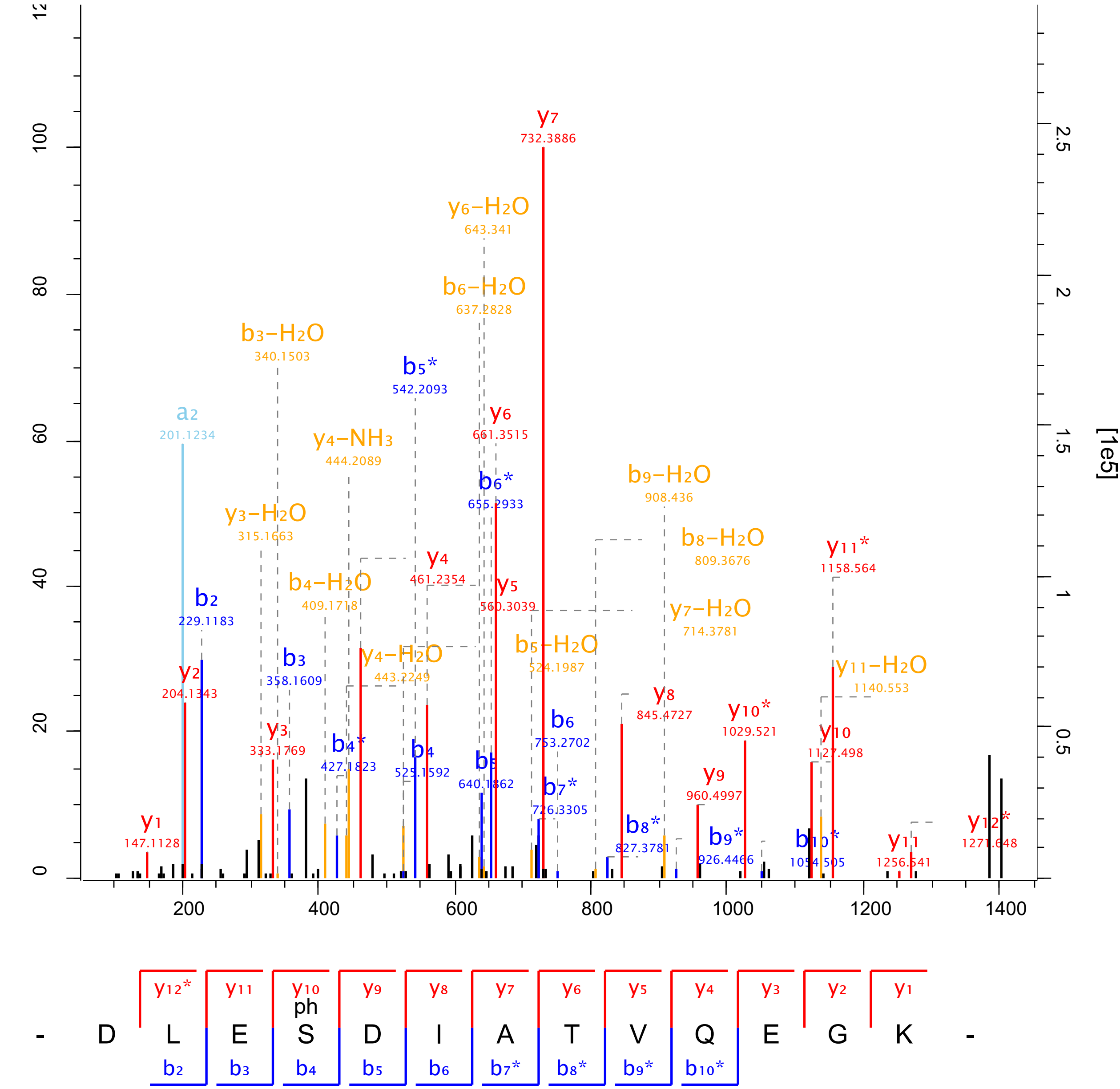

| Raw file | Scan | Method | Score | m/z |
|---|---|---|---|---|
| A17_02_1 | 33676 | FTMS; HCD | 111.64 | 659.82 |

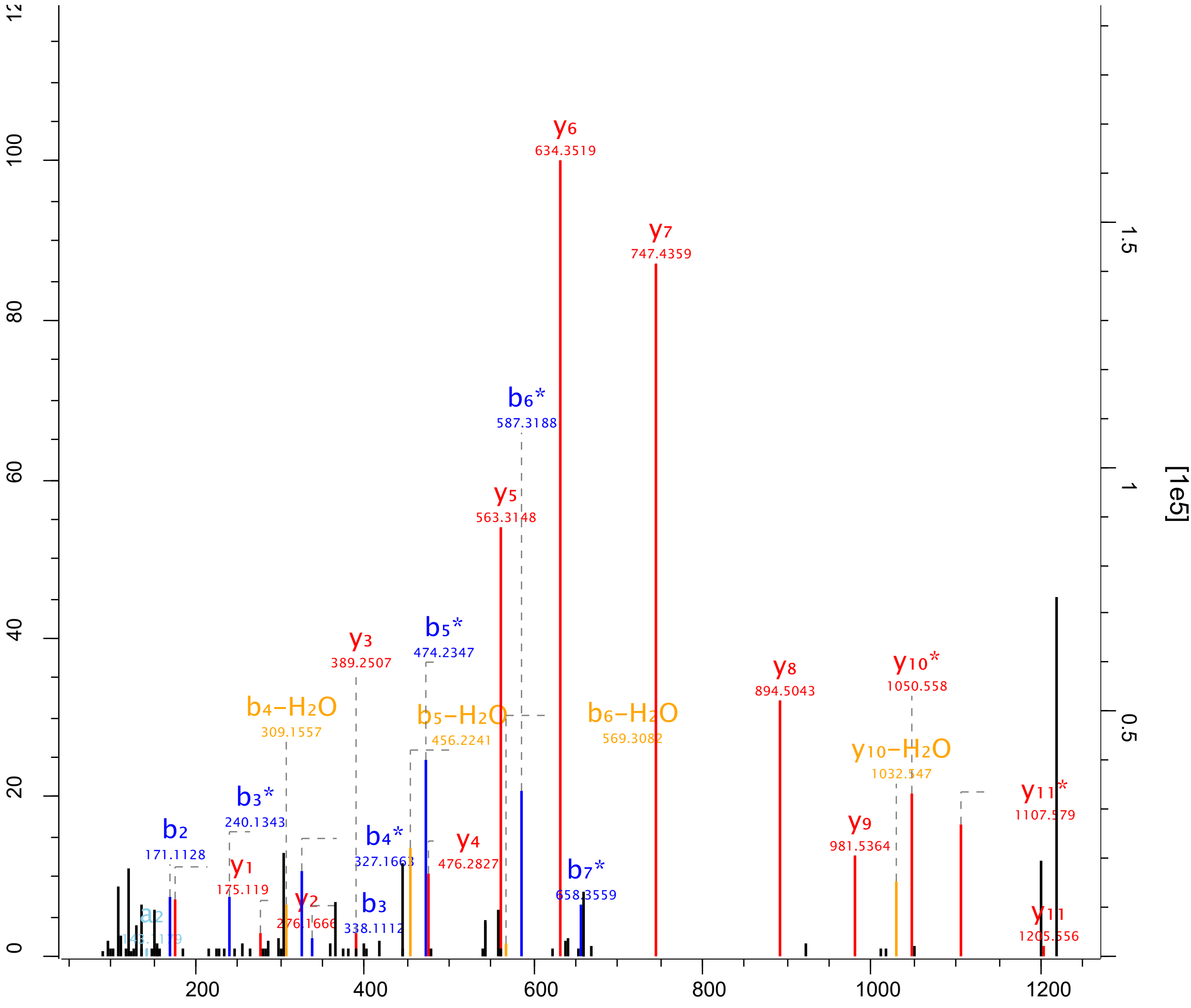


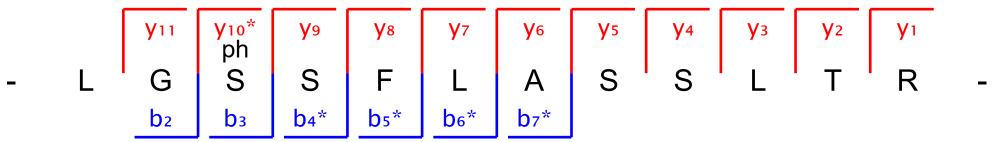

| Raw file | Scan | Method | Score | m/z |
| --- | --- | --- | --- | --- |
| A17_02_1 | 33959 | FTMS; HCD | 66.02 | 725.83 |

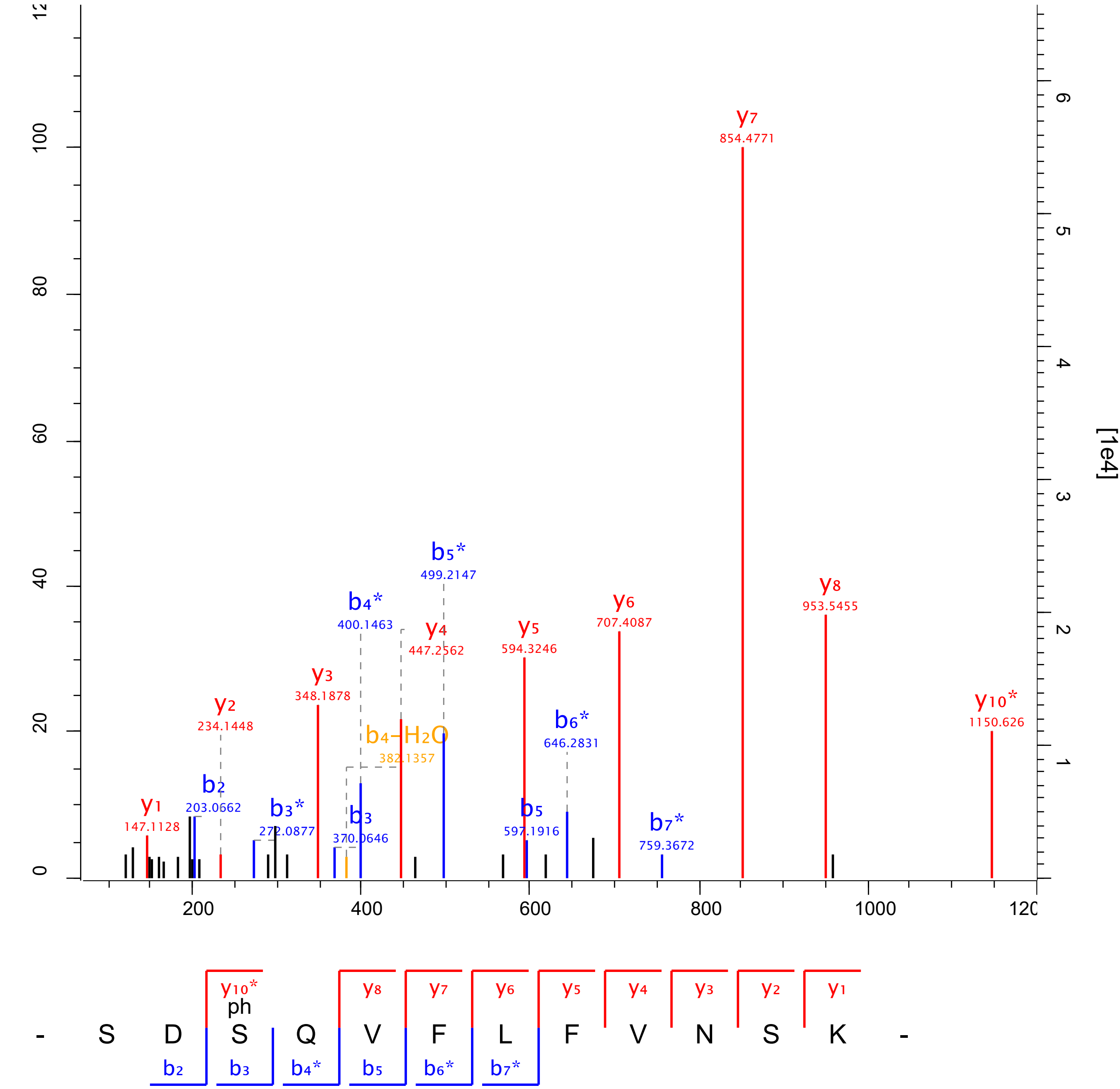

| Raw file | Scan | Method | Score | m/z |
|---|---|---|---|---|
| A17_02_1 | 34188 | FTMS; HCD | 65.24 | 927.73 |

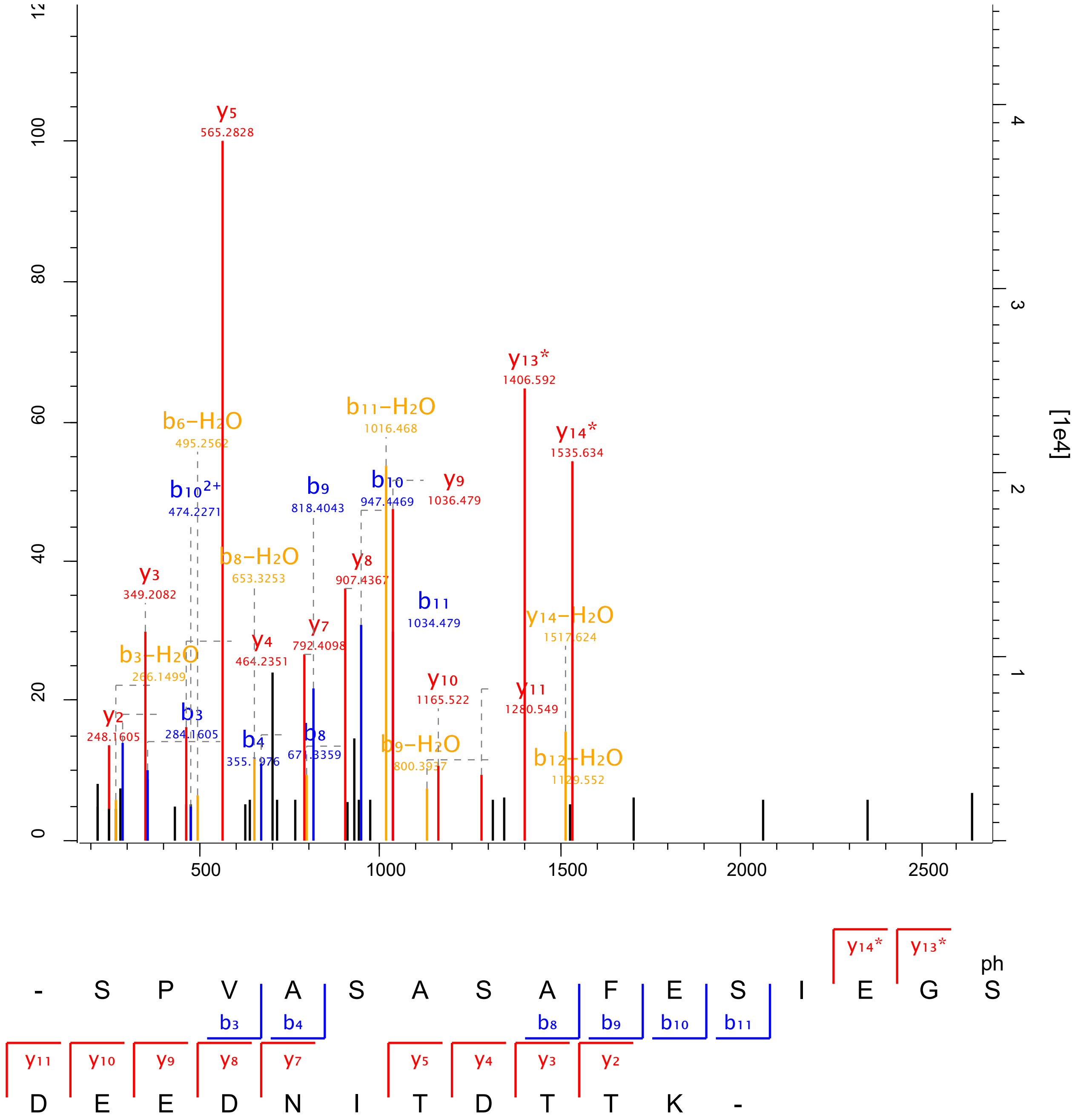

| Raw file | Scan | Method | Score | m/z |
|---|---|---|---|---|
| A17_02_1 | 36331 | FTMS; HCD | 124.51 | 1107.5 |

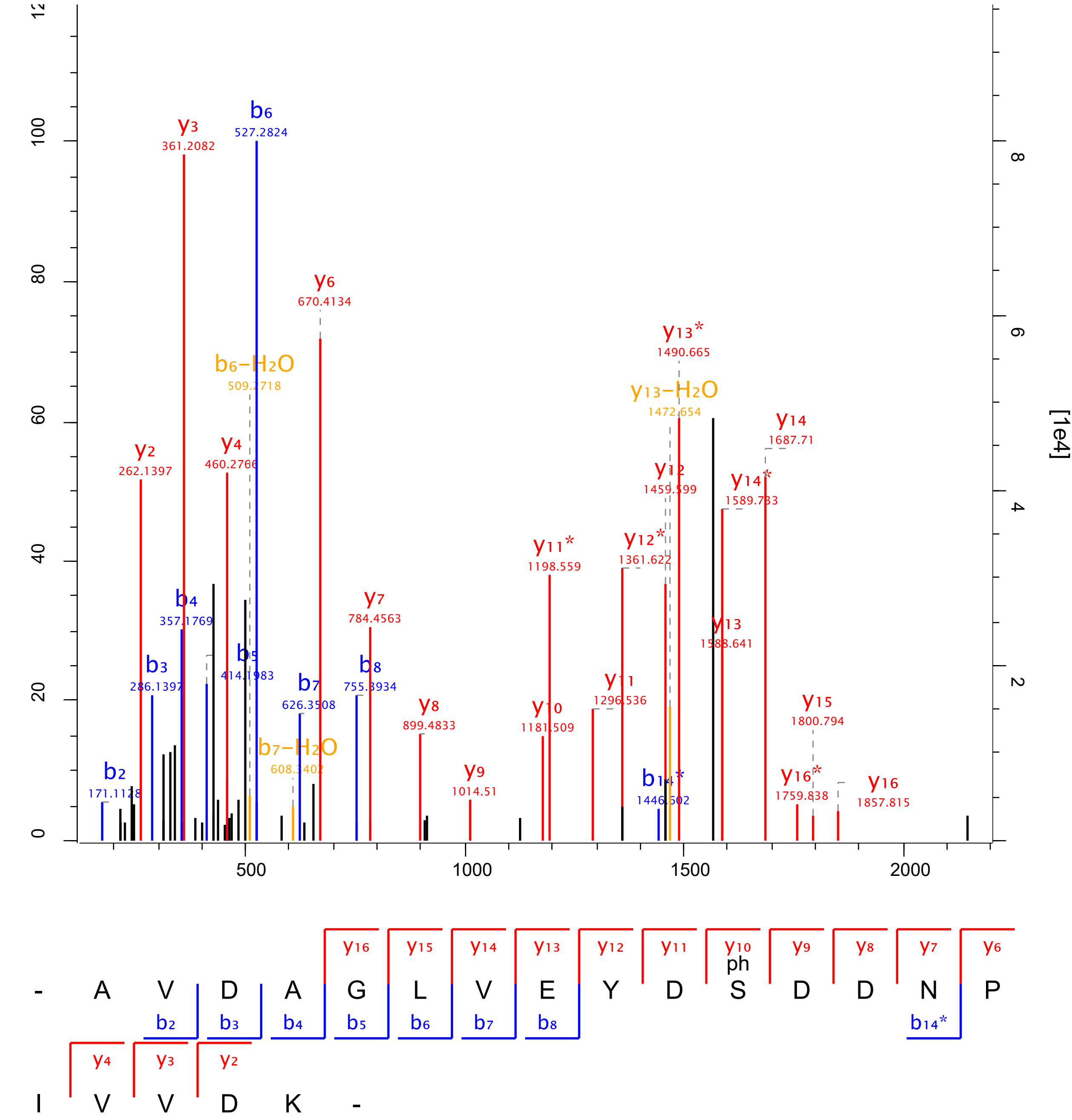

| Raw file | Scan | Method | Score | m/z |
|---|---|---|---|---|
| A17_02_1 | 3681 | FTMS; HCD | 103.08 | 485.71 |

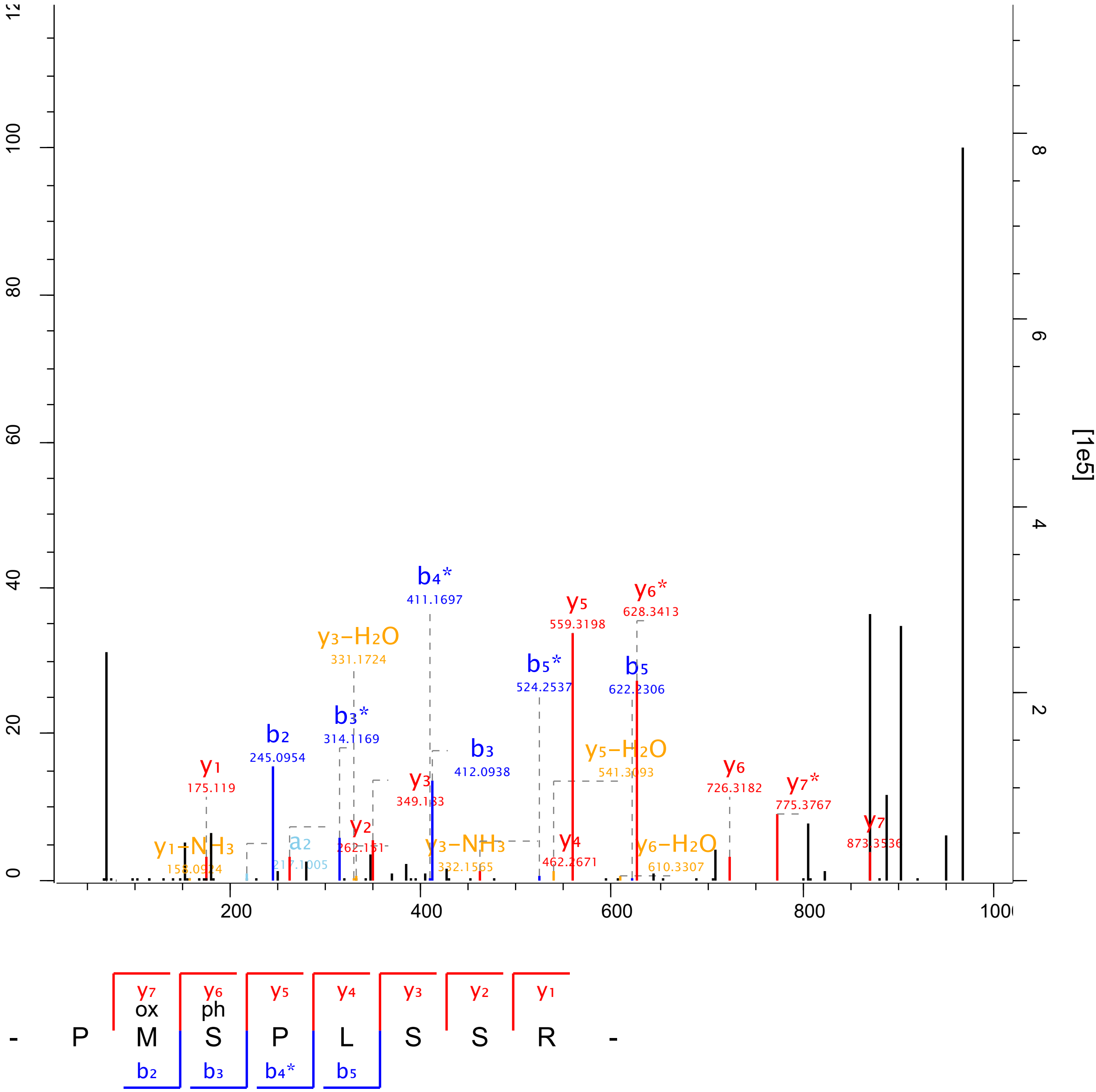

| Raw file | Scan | Method | Score | m/z |
|---|---|---|---|---|
| A17_02_1 | 37424 | FTMS; HCD | 43.31 | 806.02 |

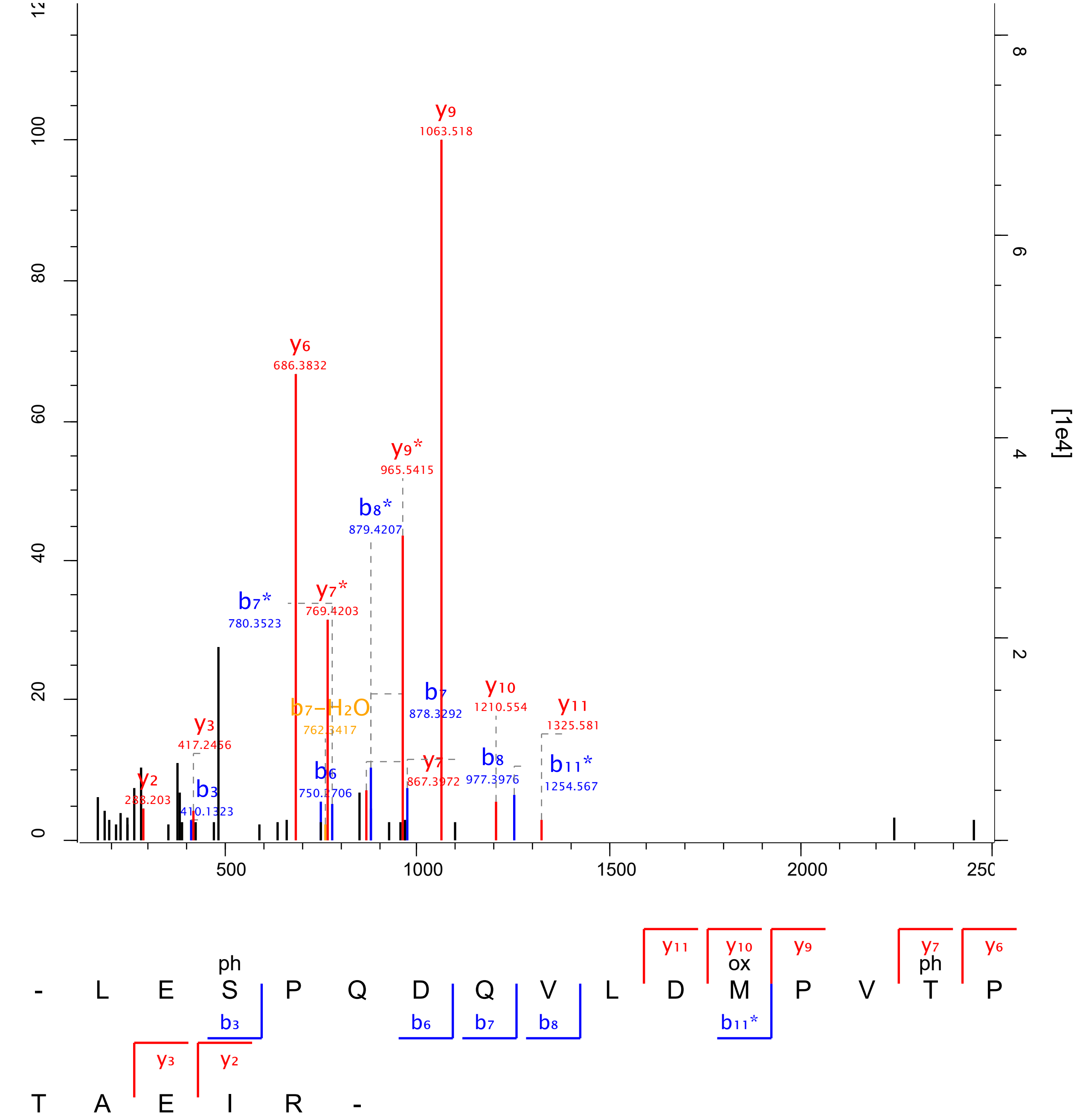

| Raw file | Scan | Method | Score | m/z |
|---|---|---|---|---|
| A17_02_1 | 37934 | FTMS; HCD | 139.11 | 934.94 |

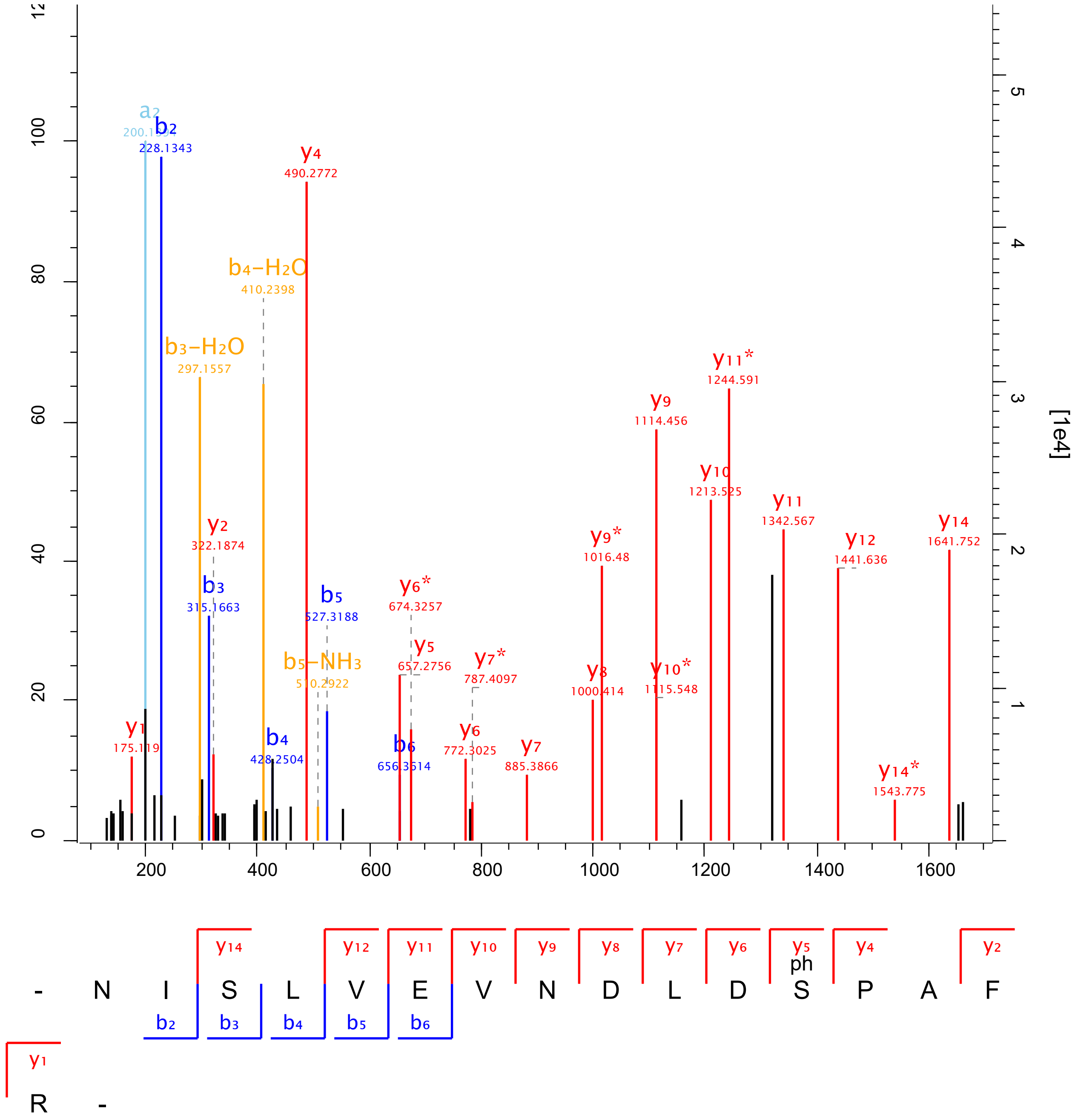

| Raw file | Scan | Method | Score | m/z |
| --- | --- | --- | --- | --- |
| A17_02_1 | 38328 | FTMS; HCD | 126.77 | 796.82 |

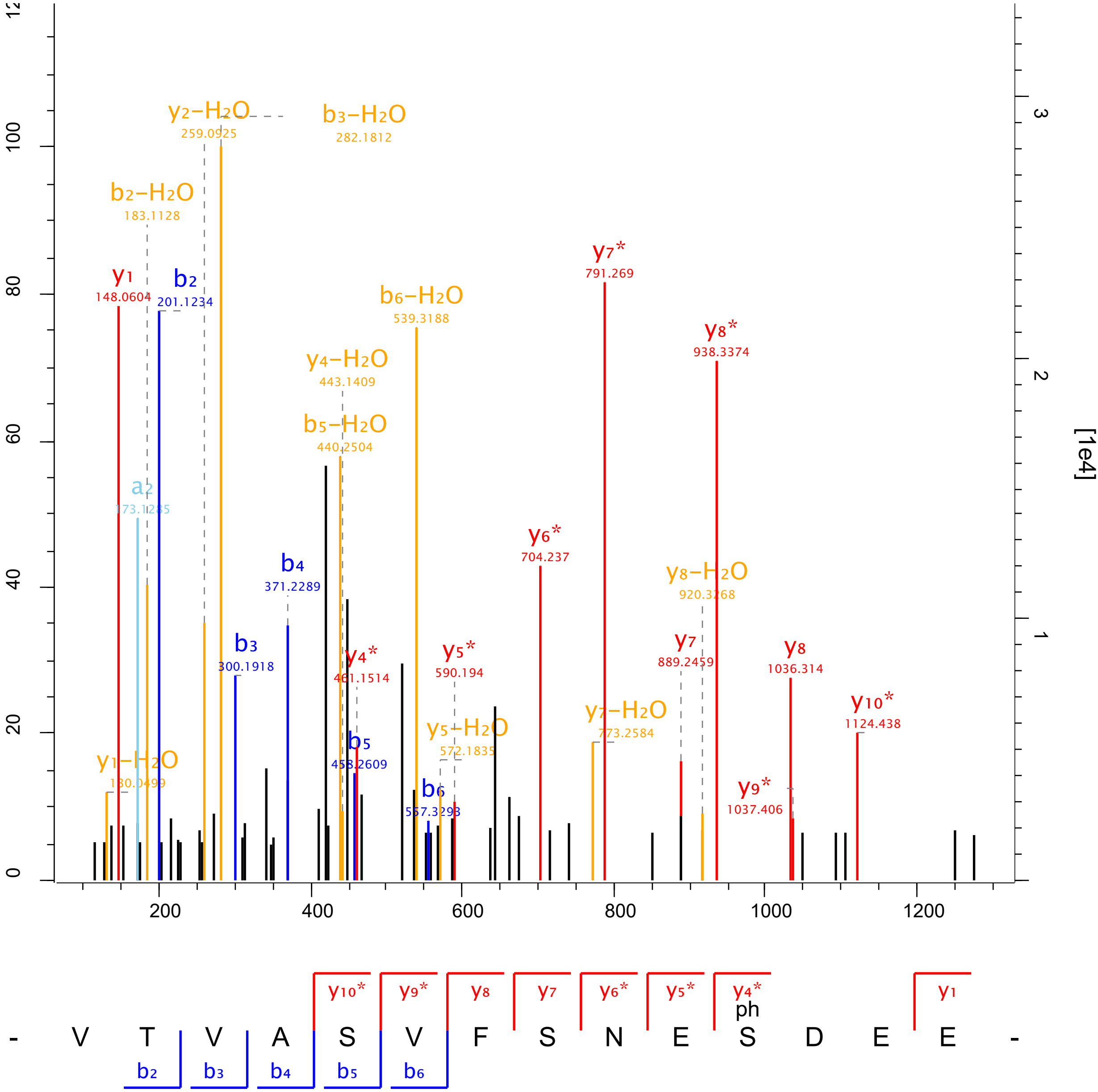

| Raw file | Scan | Method | Score | m/z |
| --- | --- | --- | --- | --- |
| A17_02_1 | 38972 | FTMS; HCD | 40.97 | 861.36 |

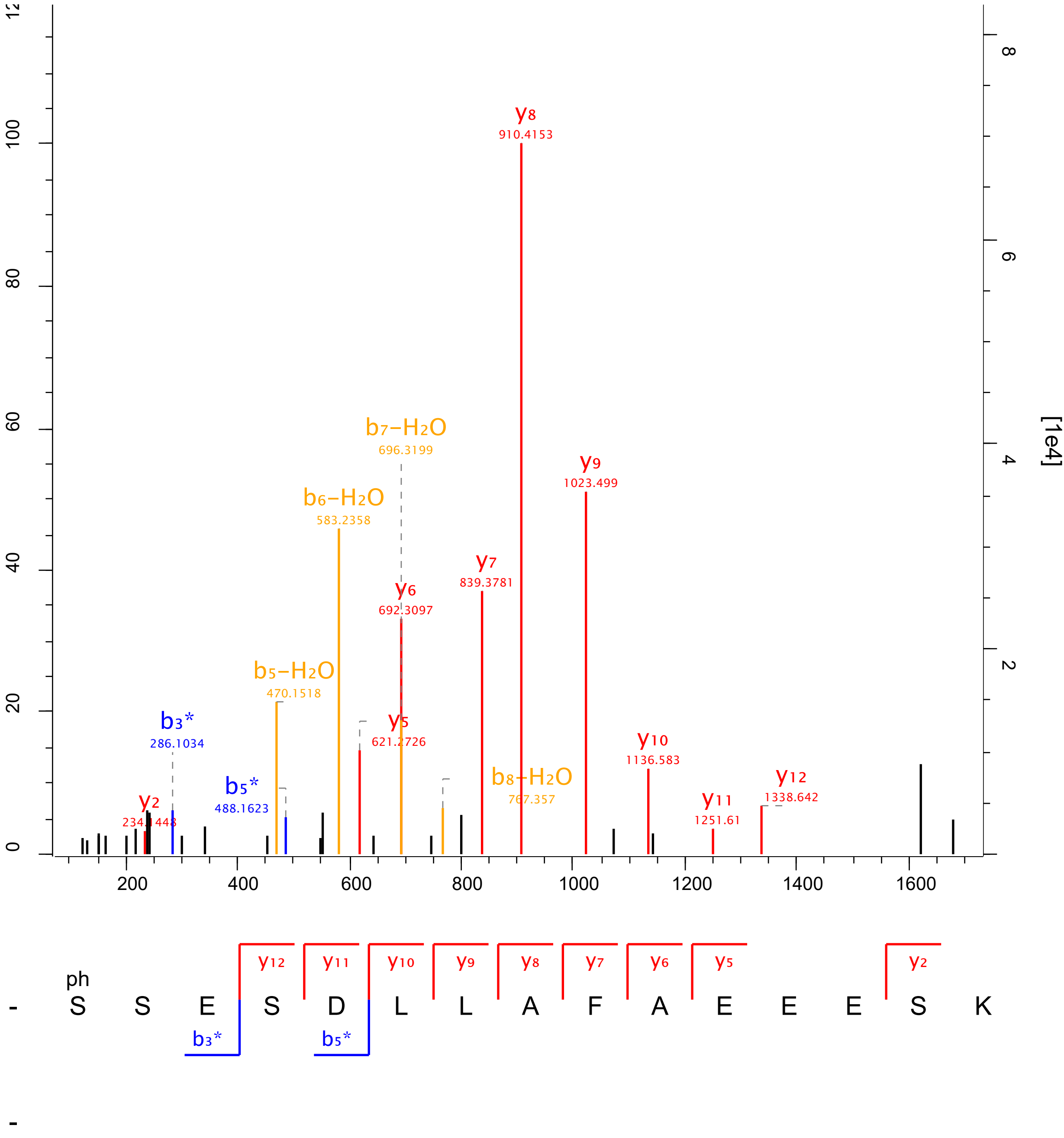

| Raw file | Scan | Method | Score | m/z |
|---|---|---|---|---|
| A17_02_1 | 39372 | FTMS; HCD | 44.25 | 701.39 |

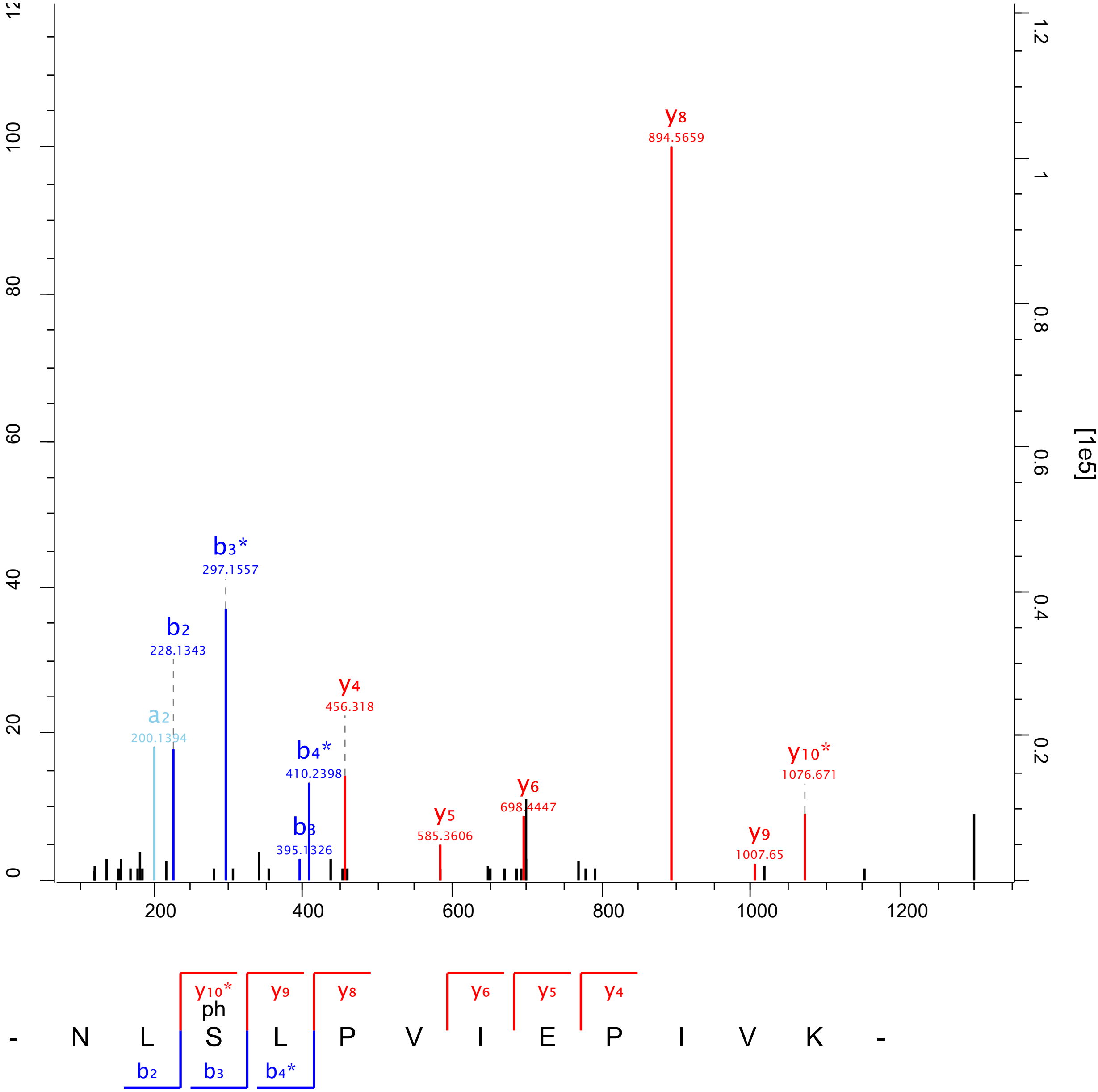

| Raw file | Scan | Method | Score | m/z |
|---|---|---|---|---|
| A17_02_1 | 39973 | FTMS; HCD | 126.51 | 721.31 |

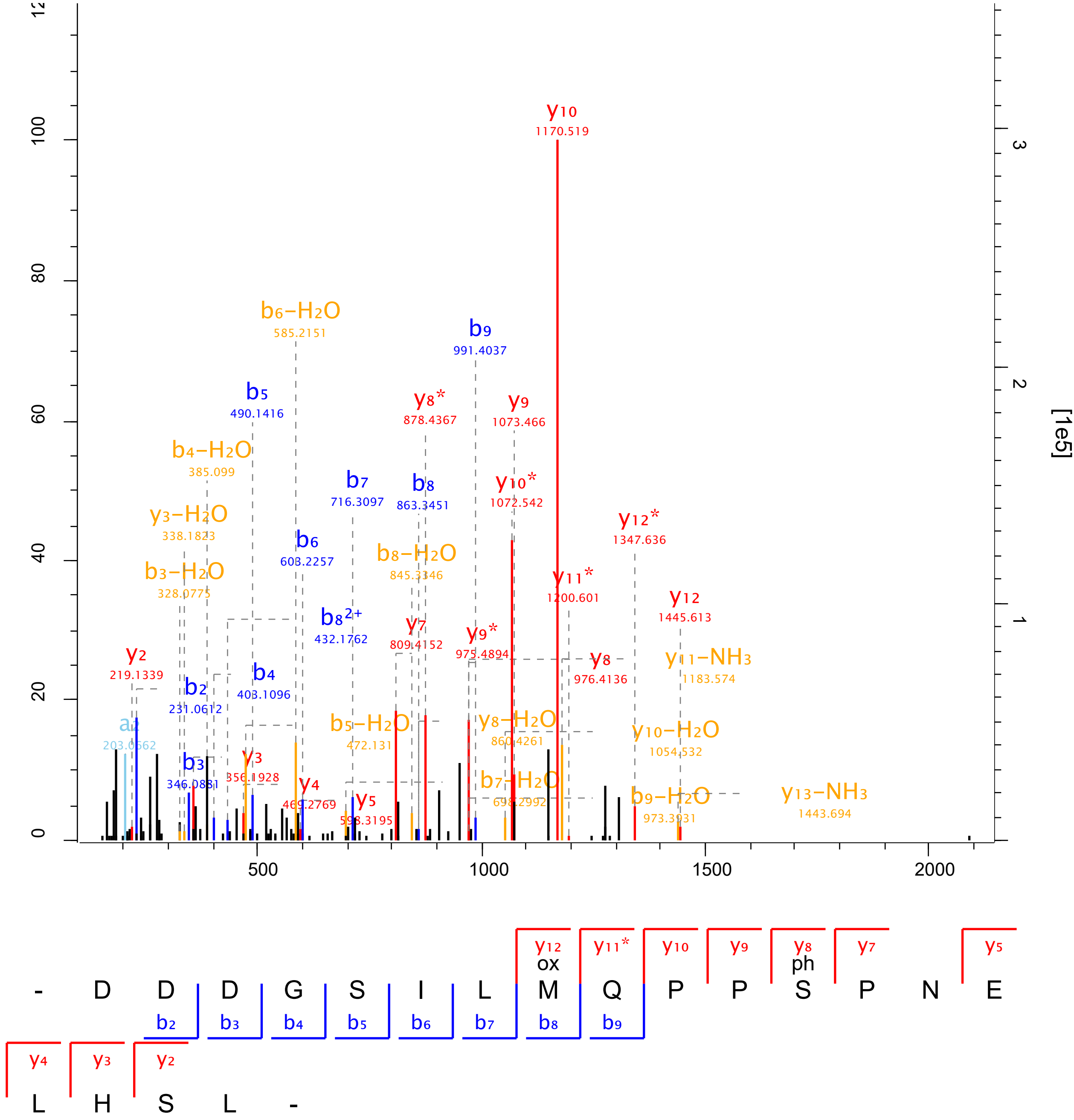

| Raw file | Scan | Method | Score | m/z |
|---|---|---|---|---|
| A17_02_1 | 40390 | FTMS; HCD | 170.89 | 913.44 |

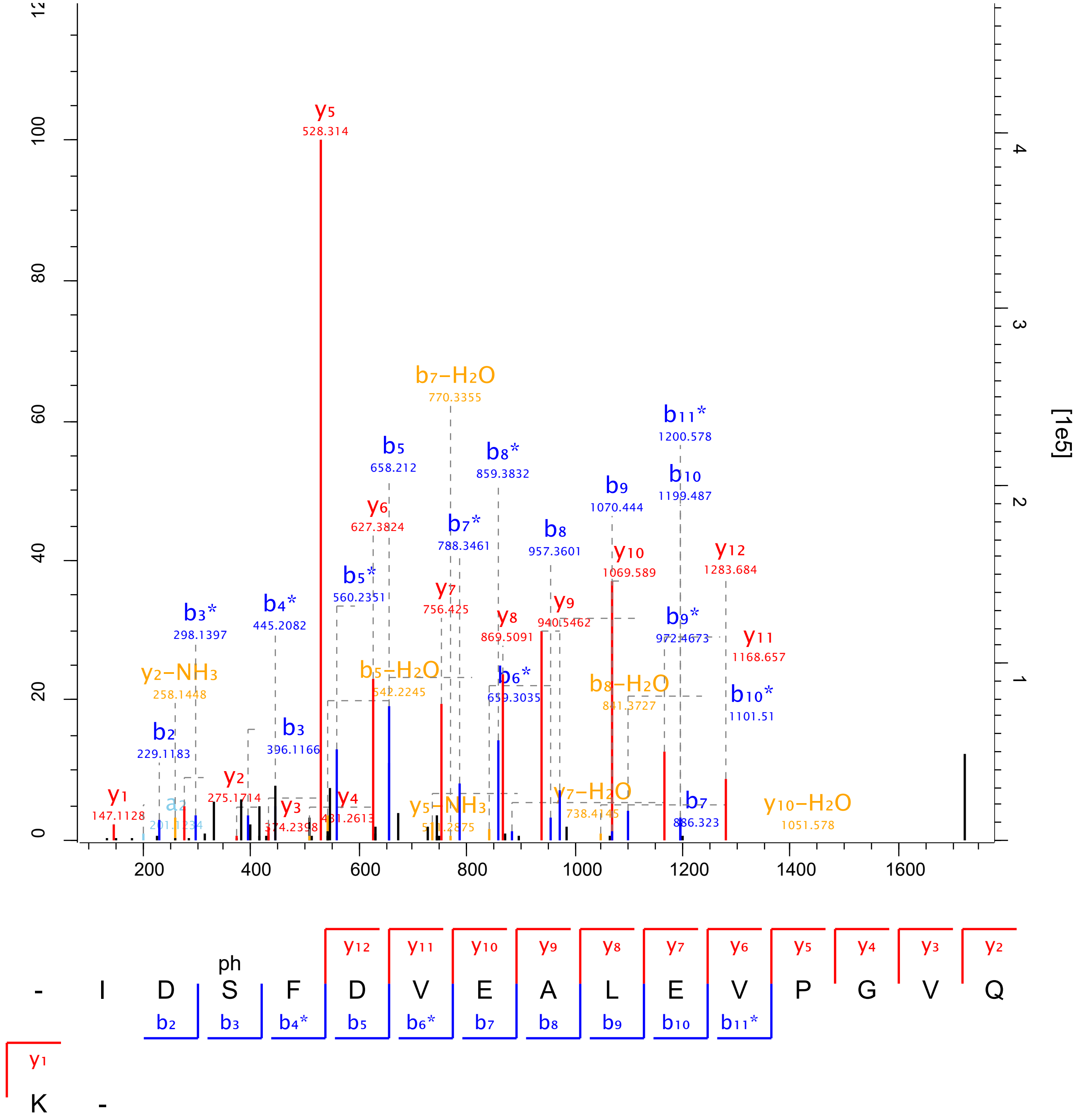

| Raw file | Scan | Method | Score | m/z |
| --- | --- | --- | --- | --- |
| A17_02_1 | 41104 | FTMS; HCD | 120.15 | 1025.99 |

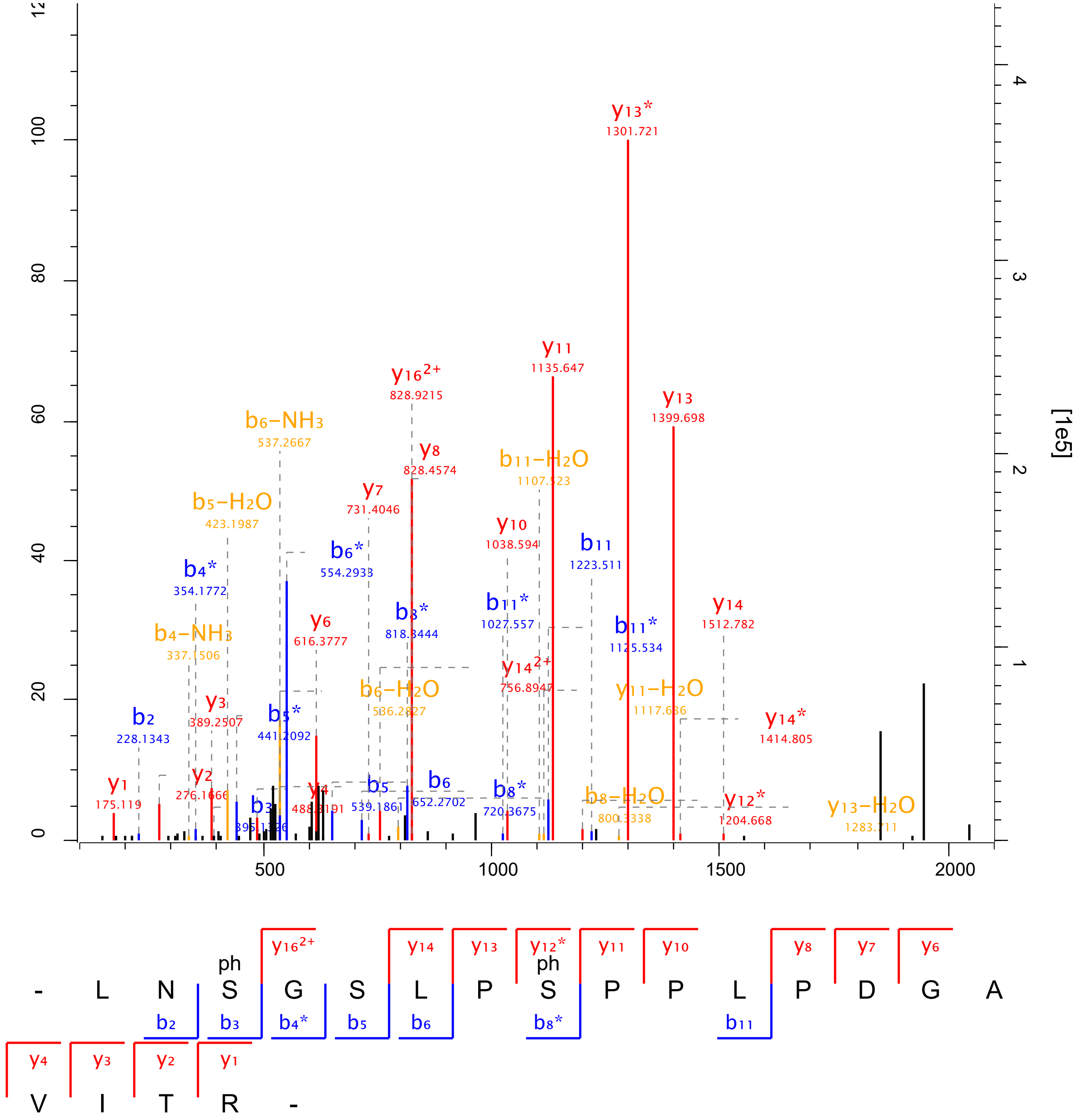

| Raw file | Scan | Method | Score | m/z |
|---|---|---|---|---|
| A17_02_1 | 4238 | FTMS; HCD | 156.72 | 645.27 |

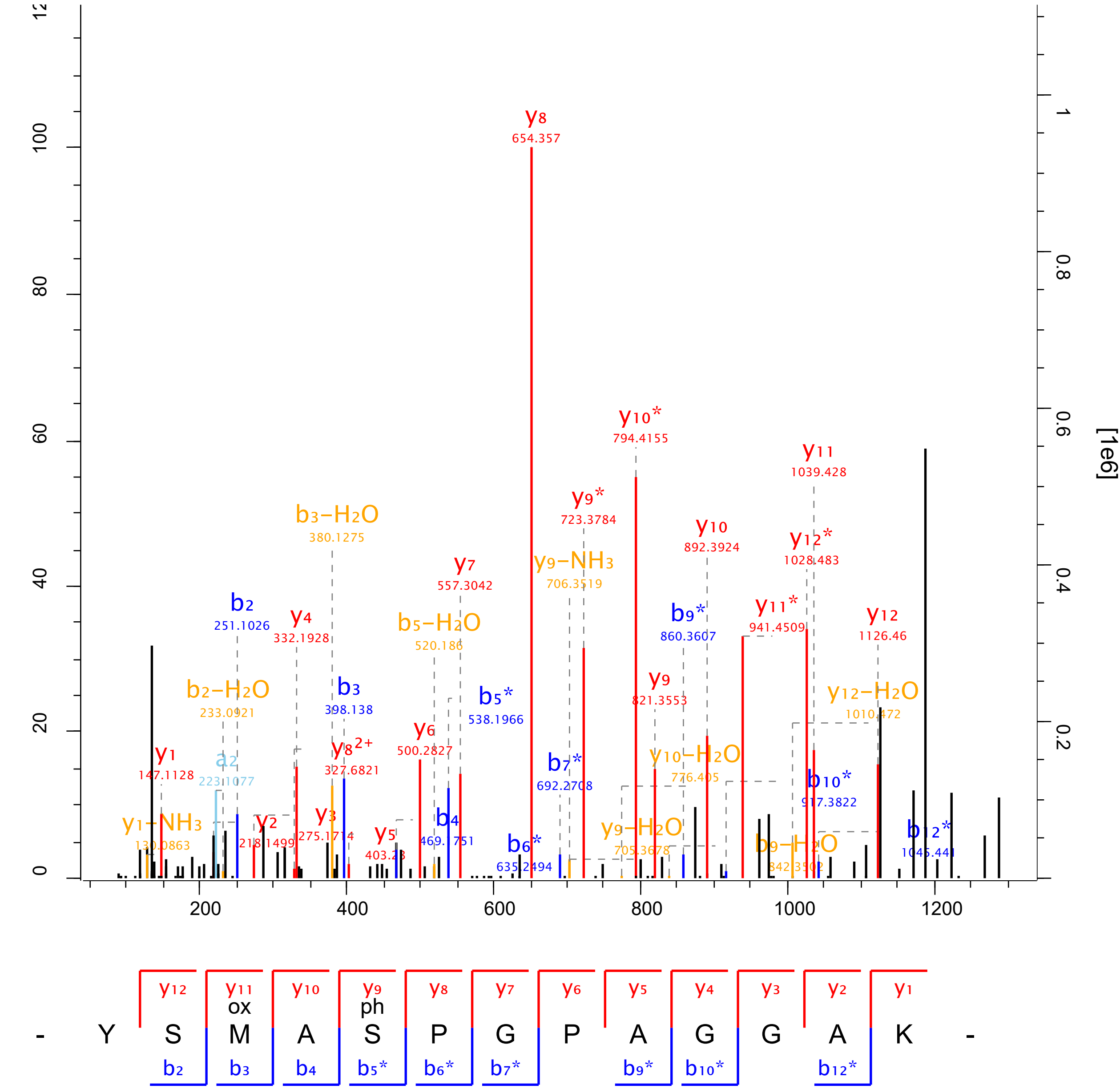

| Raw file | Scan | Method | Score | m/z |
|---|---|---|---|---|
| A17_02_1 | 42705 | FTMS; HCD | 305.19 | 773.86 |

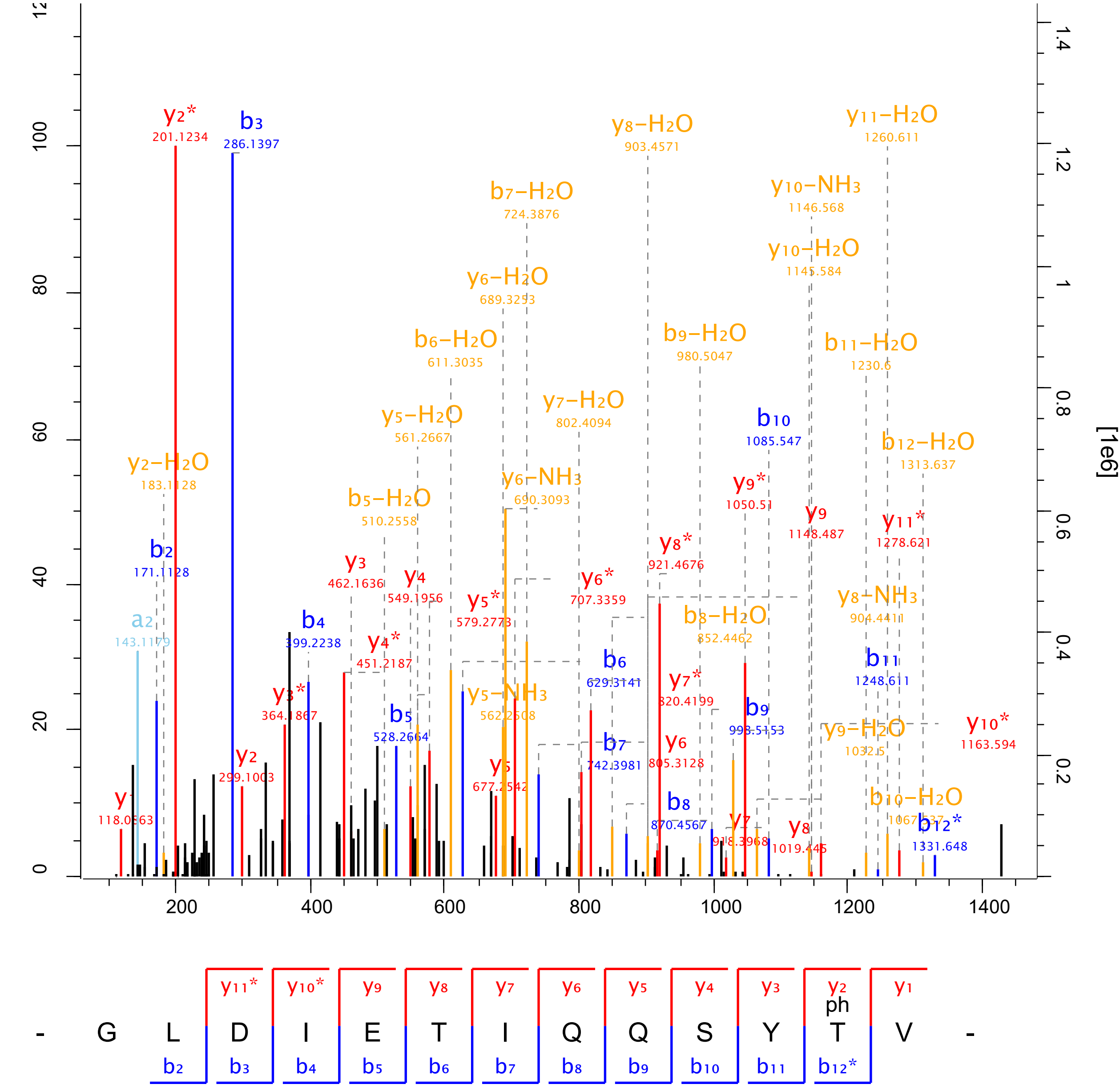

| Raw file | Scan | Method | Score | m/z |
| --- | --- | --- | --- | --- |
| A17_02_1 | 43257 | FTMS; HCD | 41.62 | 602.78 |

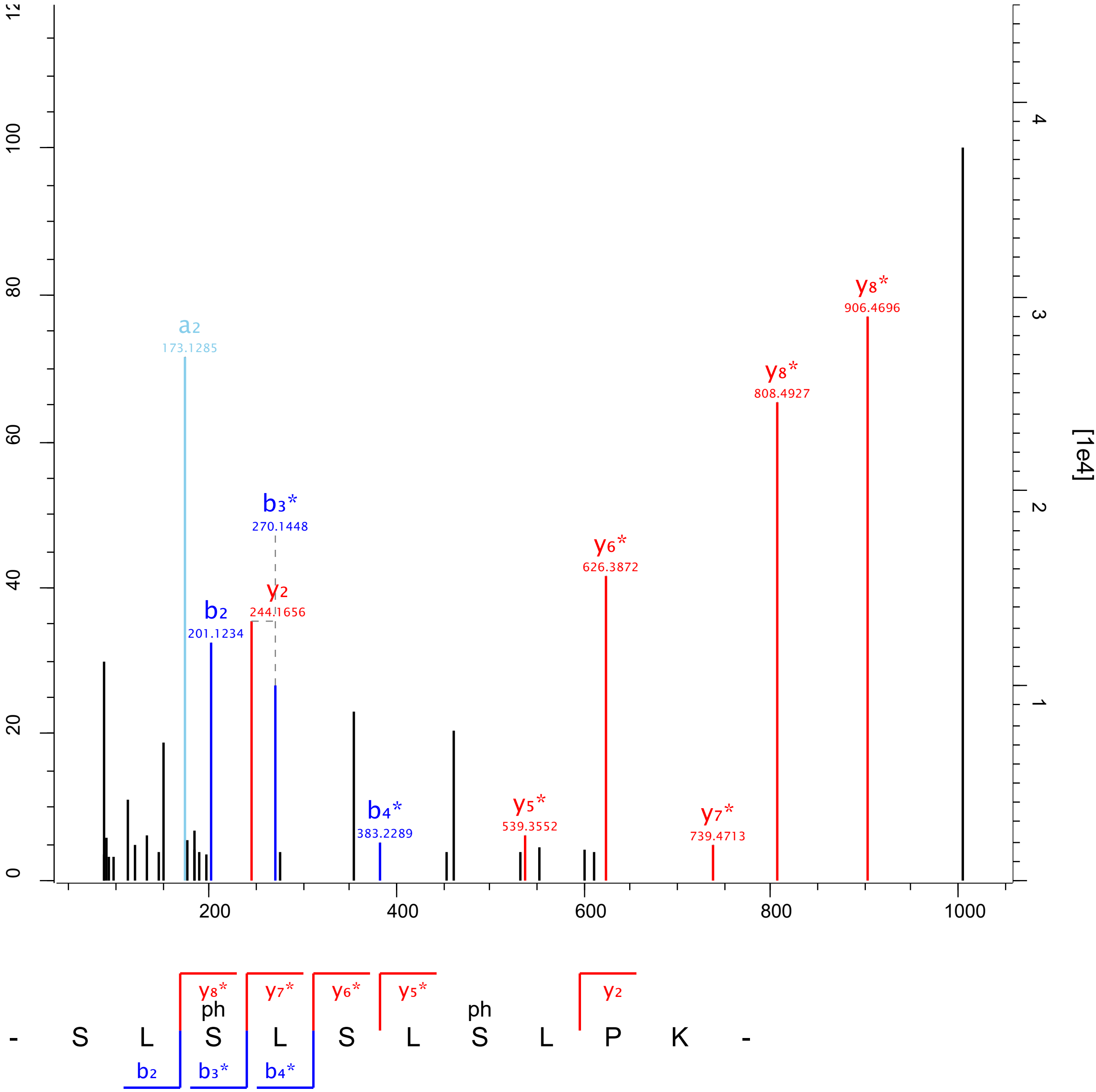

| Raw file | Scan | Method | Score | m/z |
|---|---|---|---|---|
| A17_02_1 | 43272 | FTMS; HCD | 118.68 | 941.08 |

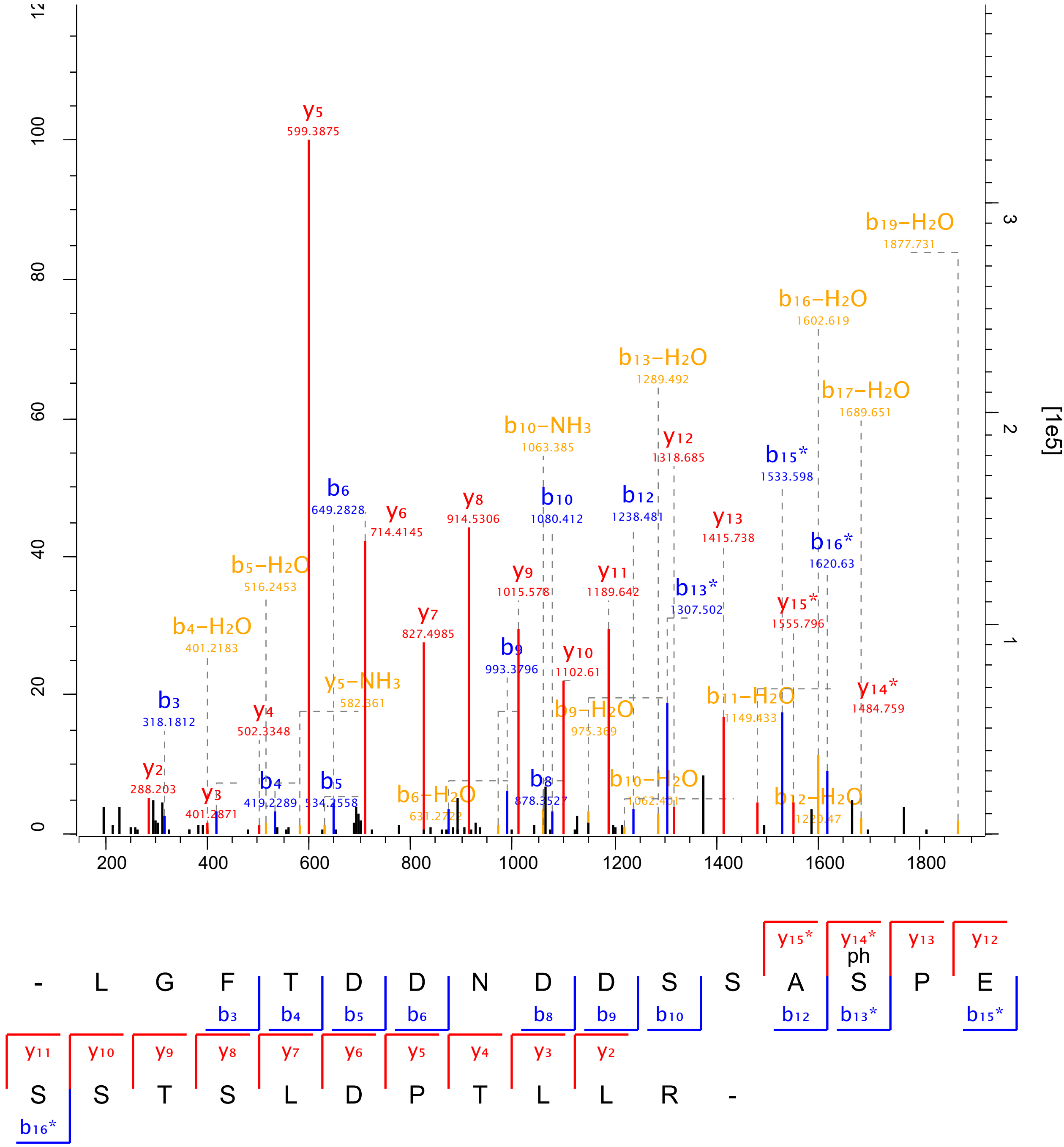

| Raw file | Scan | Method | Score | m/z |
| --- | --- | --- | --- | --- |
| A17_02_1 | 43274 | FTMS; HCD | 153.72 | 901.82 |

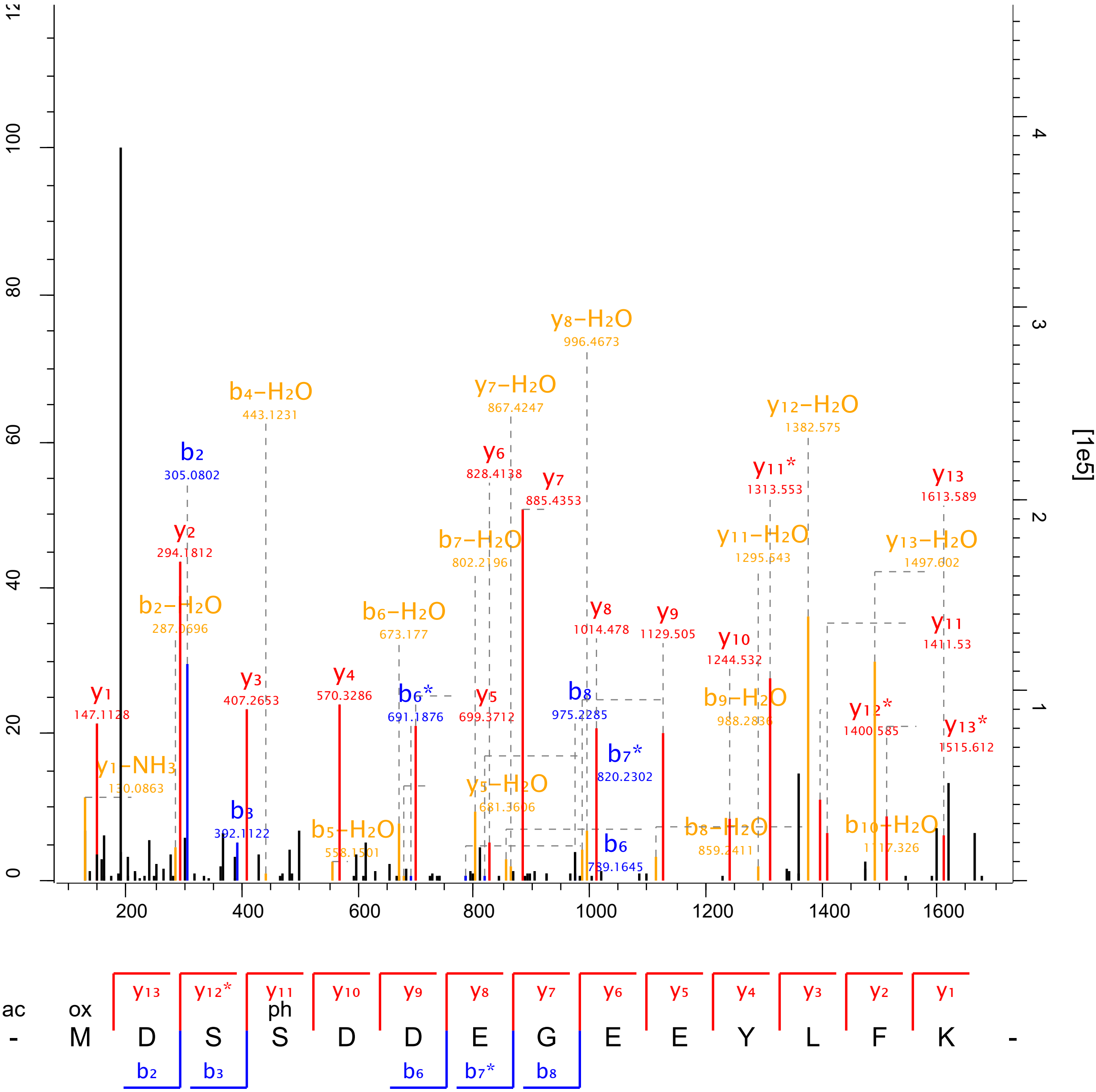

| Raw file | Scan | Method | Score | m/z |
|---|---|---|---|---|
| A17_02_1 | 43320 | FTMS; HCD | 40.85 | 848.09 |

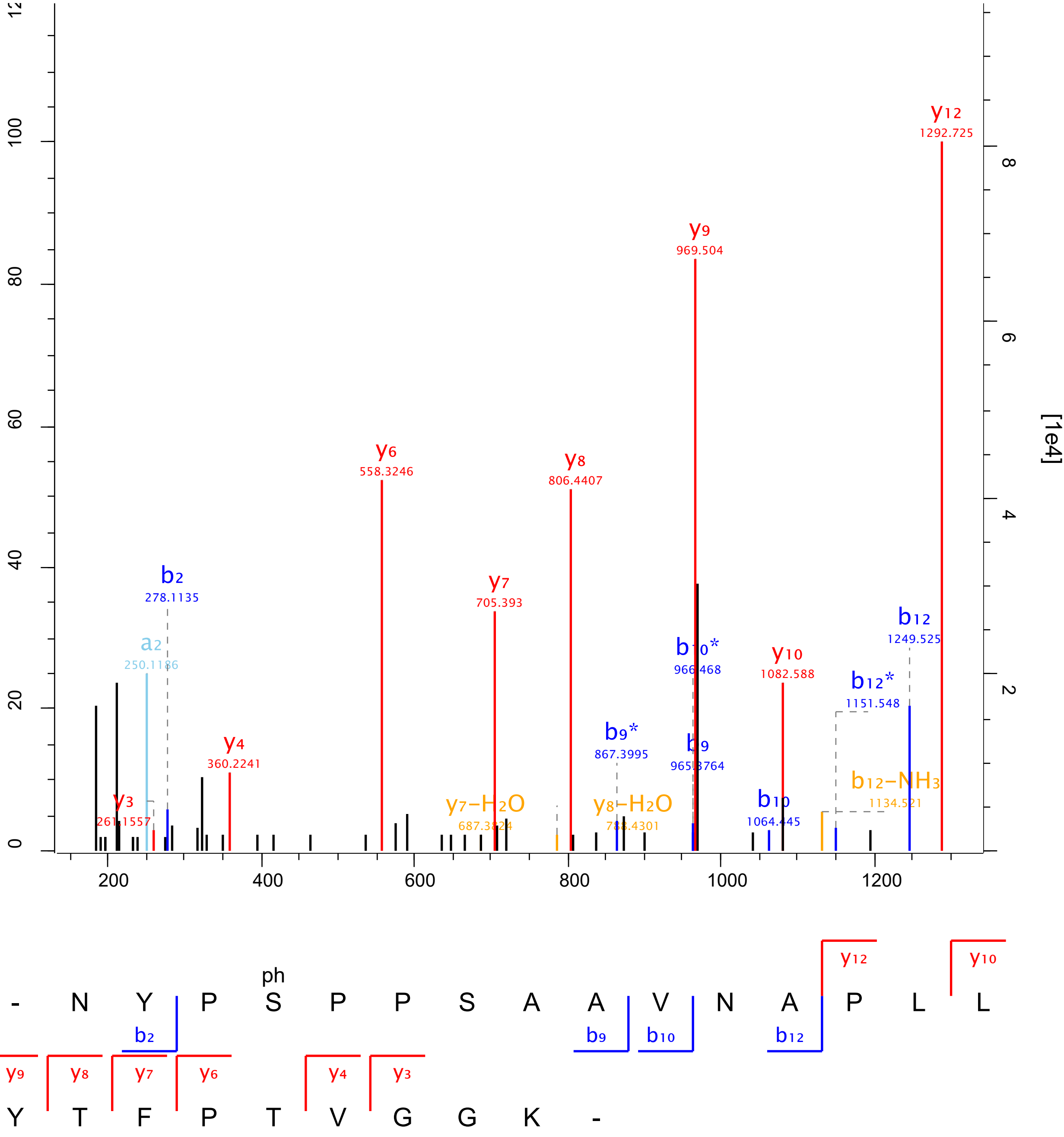

| Raw file | Scan | Method | Score | m/z |
|---|---|---|---|---|
| A17_02_1 | 44613 | FTMS; HCD | 82.64 | 783.7 |

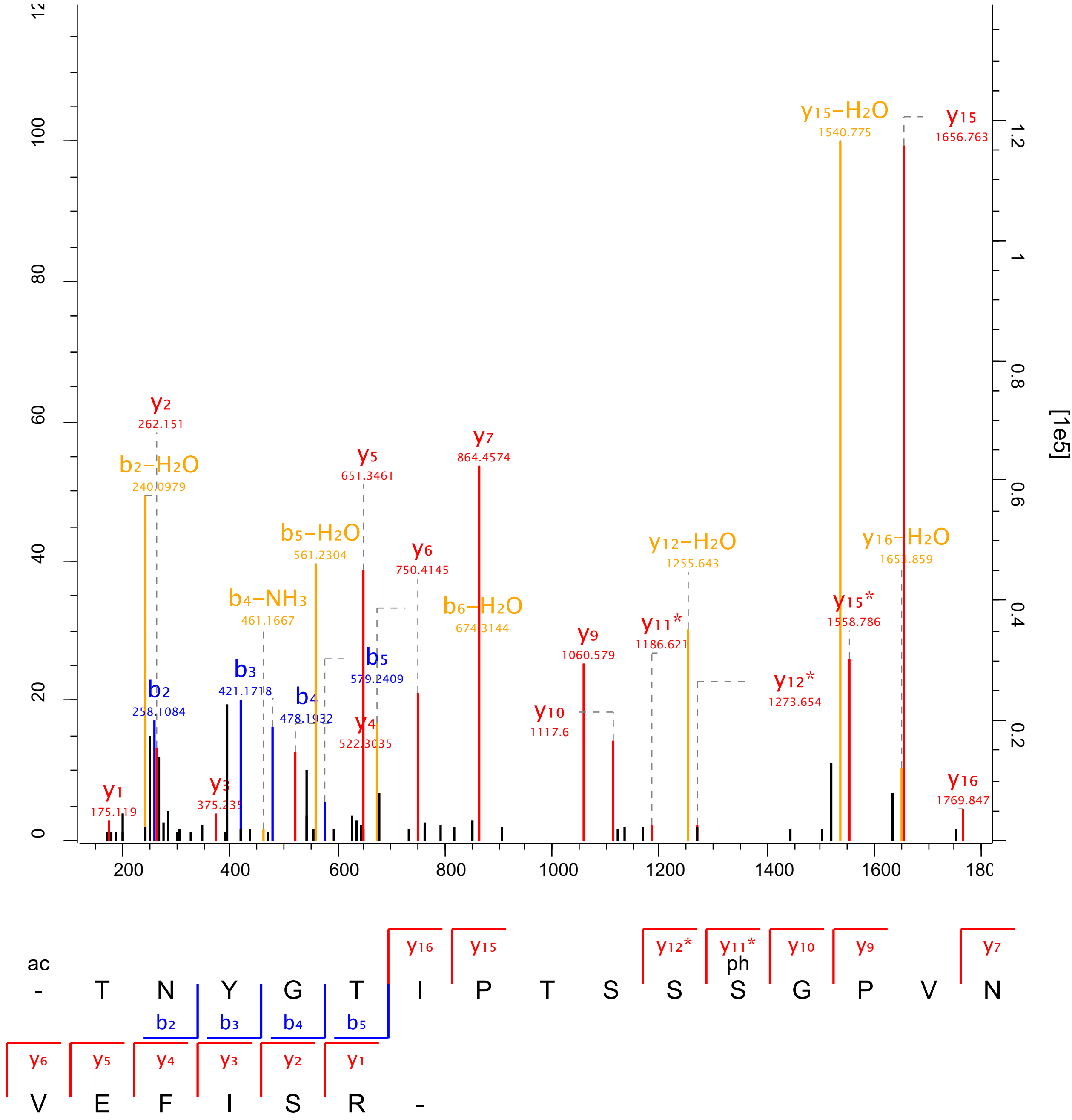

| Raw file | Scan | Method | Score | m/z |
|---|---|---|---|---|
| A17_02_1 | 45330 | FTMS; HCD | 51.03 | 733.35 |

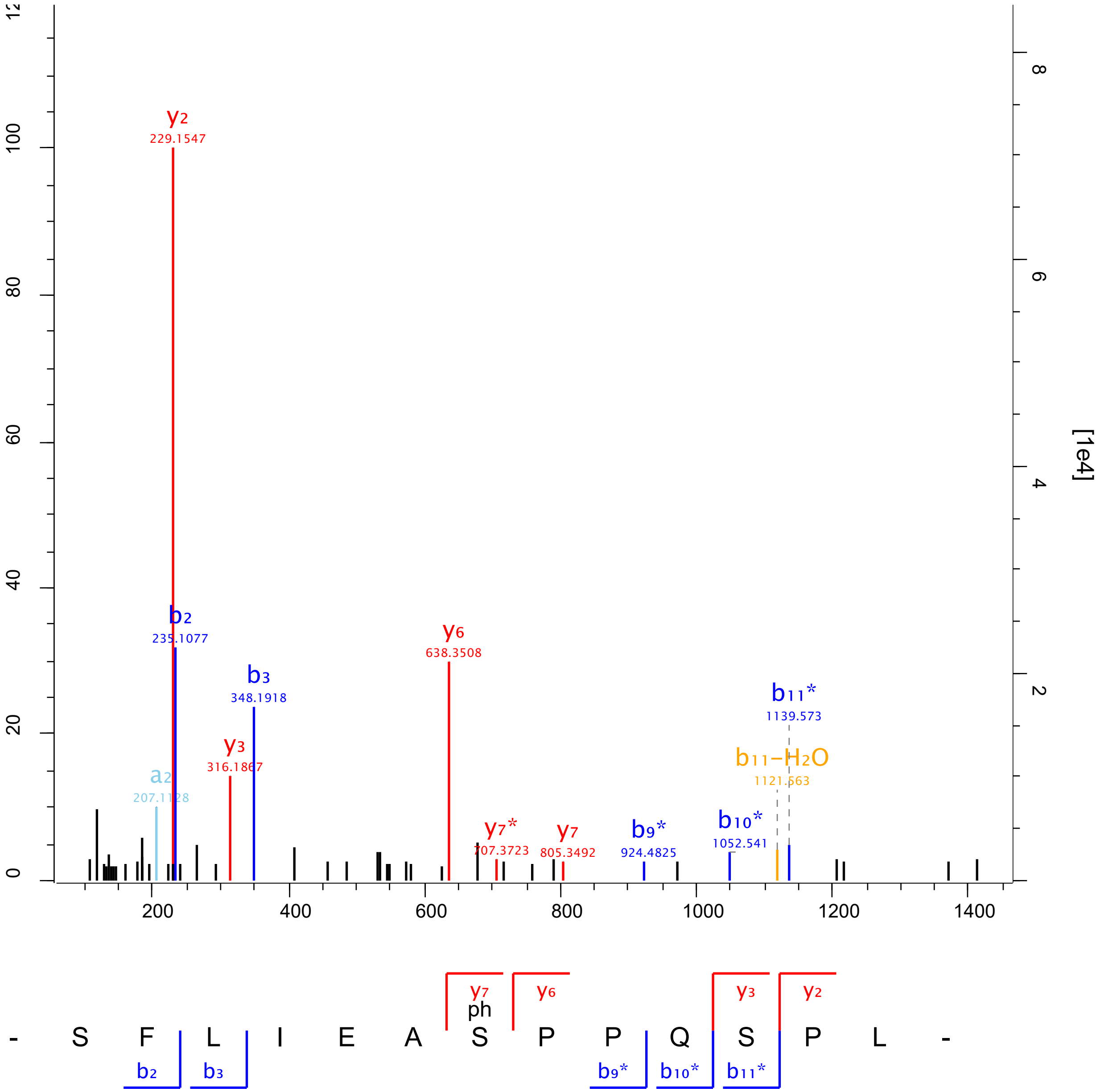

| Raw file | Scan | Method | Score | m/z |
|---|---|---|---|---|
| A17_02_1 | 45925 | FTMS; HCD | 184.82 | 846.36 |

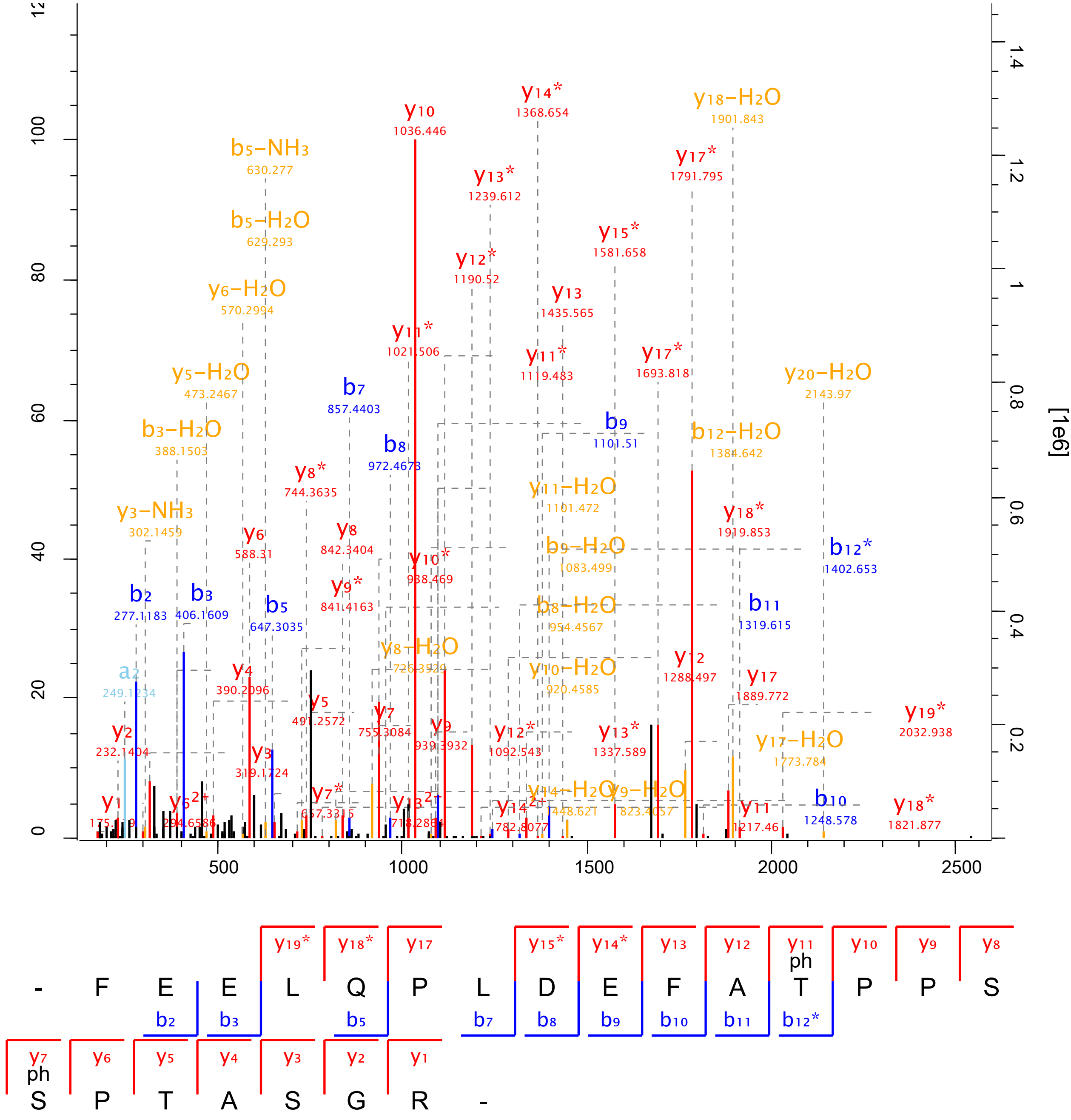

| Raw file | Scan | Method | Score | m/z |
|---|---|---|---|---|
| A17_02_1 | 46162 | FTMS; HCD | 50.96 | 910.43 |

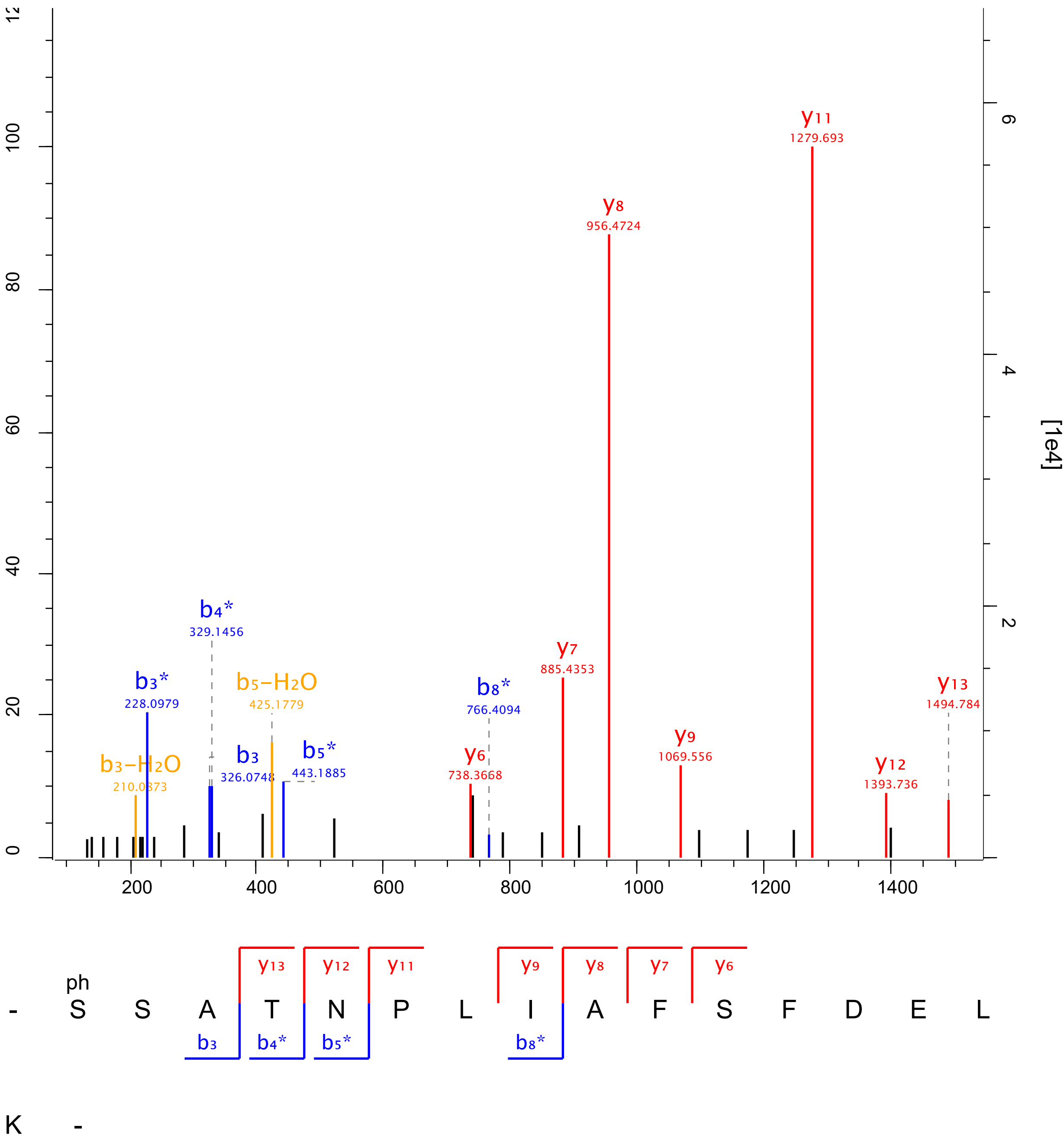

| Raw file | Scan | Method | Score | m/z |
| --- | --- | --- | --- | --- |
| A17_02_1 | 47279 | FTMS; HCD | 155.86 | 937.37 |

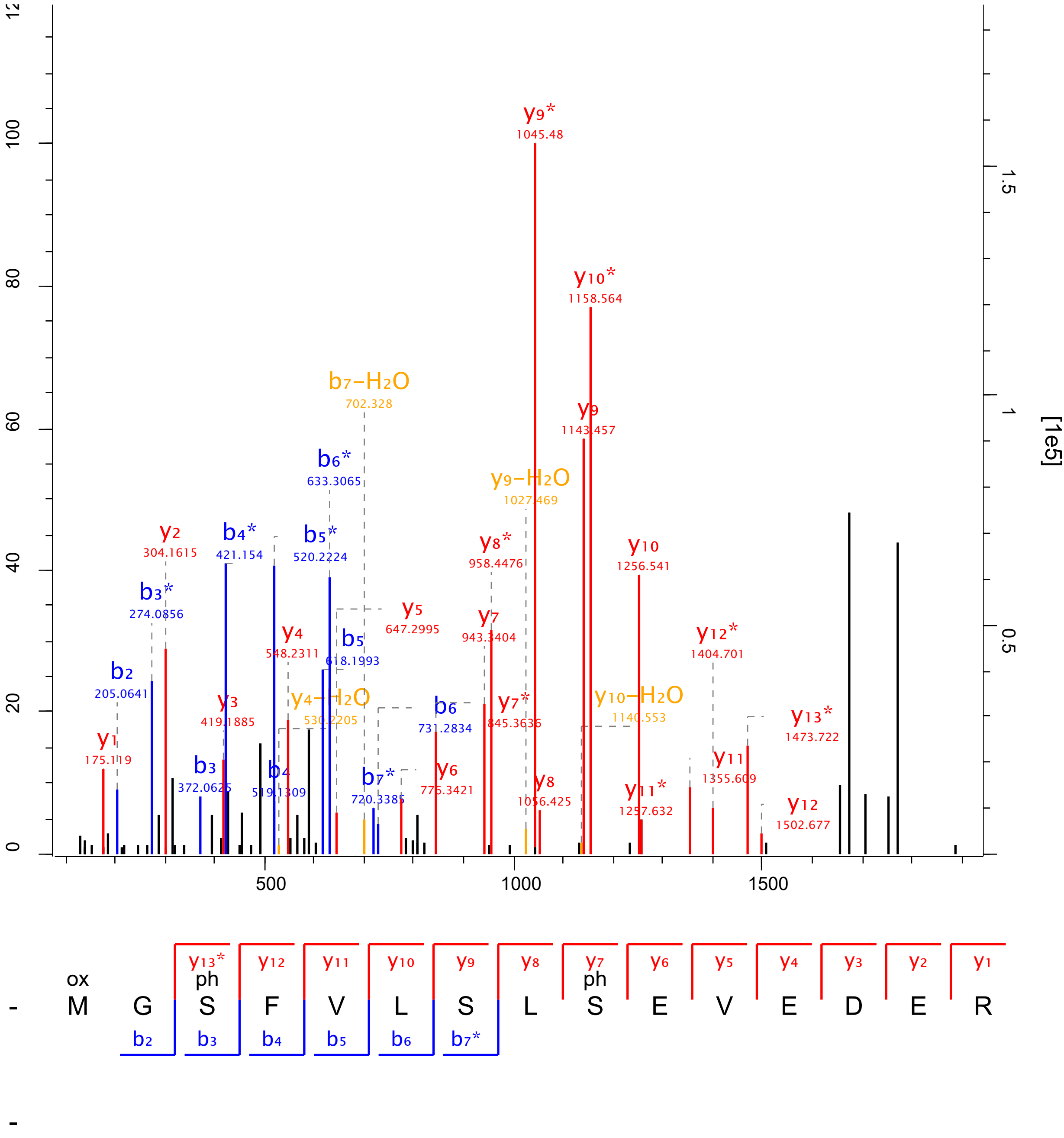

| Raw file | Scan | Method | Score | m/z |
|---|---|---|---|---|
| A17_02_1 | 47763 | FTMS; HCD | 78.32 | 963.41 |

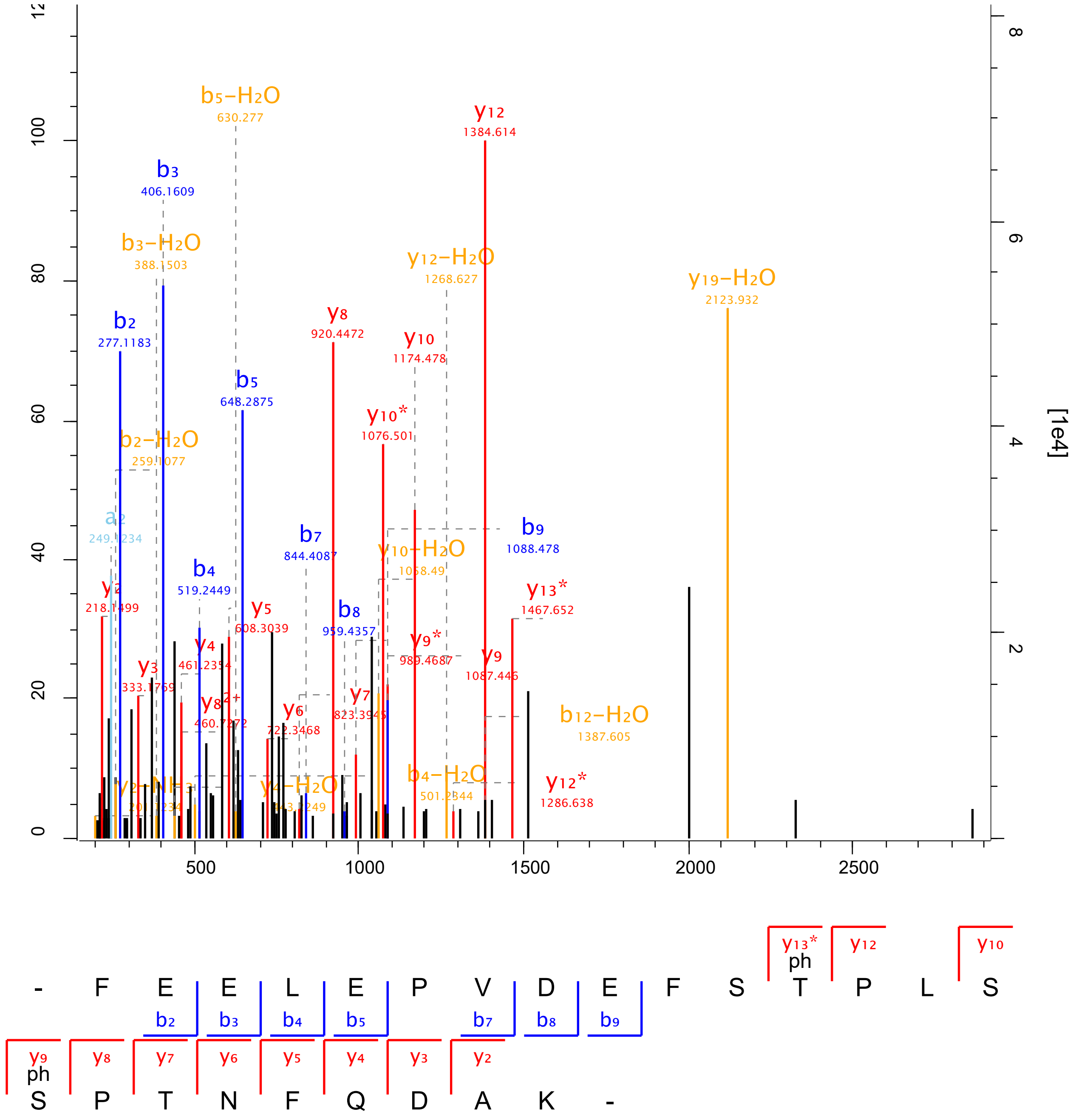

| Raw file | Scan | Method | Score | m/z |
|---|---|---|---|---|
| A17_02_1 | 48393 | FTMS; HCD | 45.56 | 1097.8 |

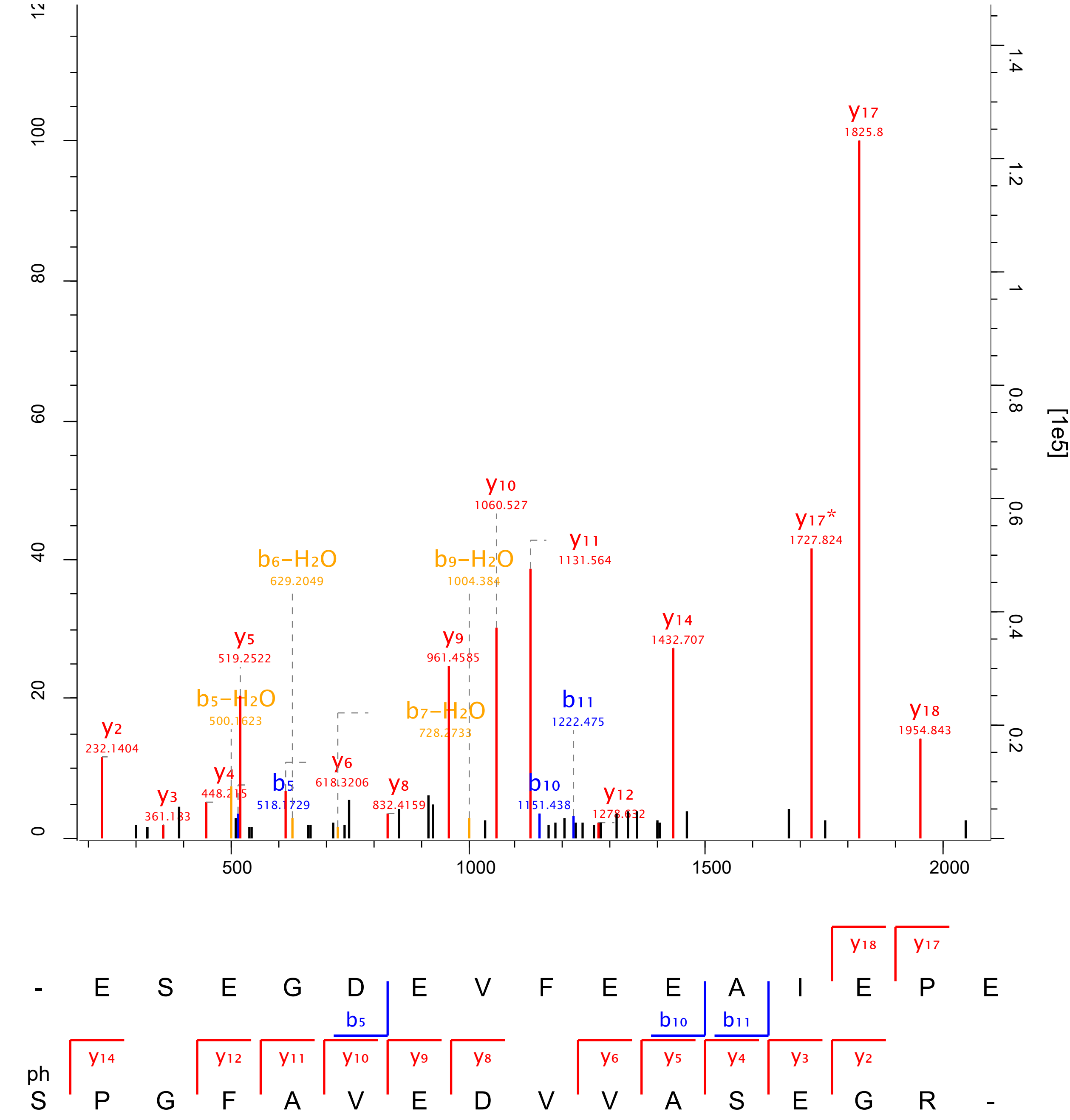

| Raw file | Scan | Method | Score | m/z |
|---|---|---|---|---|
| A17_02_1 | 48768 | FTMS; HCD | 136.48 | 982.11 |

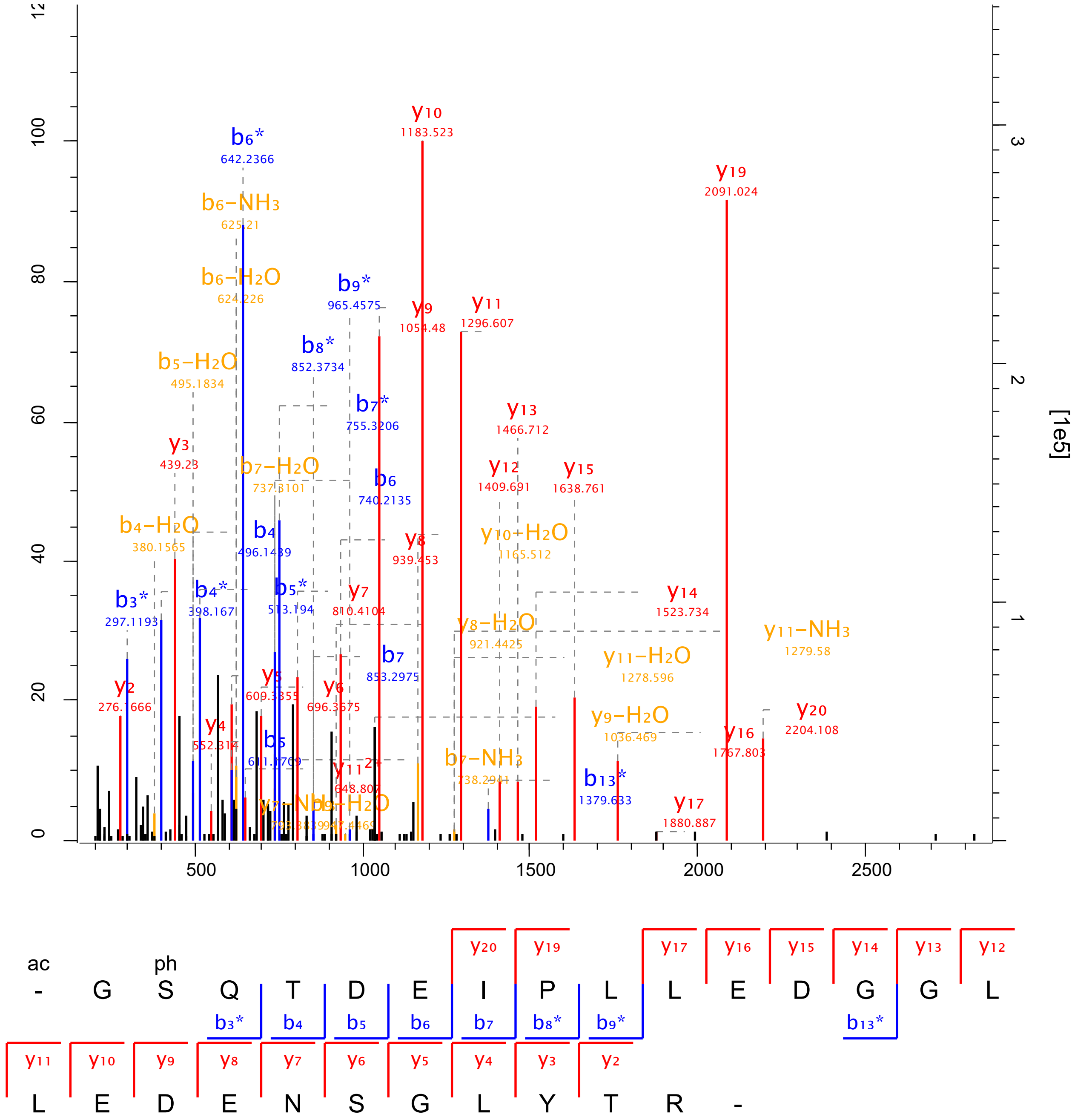

| Raw file | Scan | Method | Score | m/z |
|---|---|---|---|---|
| A17_02_1 | 4942 | FTMS; HCD | 115.71 | 450.2 |

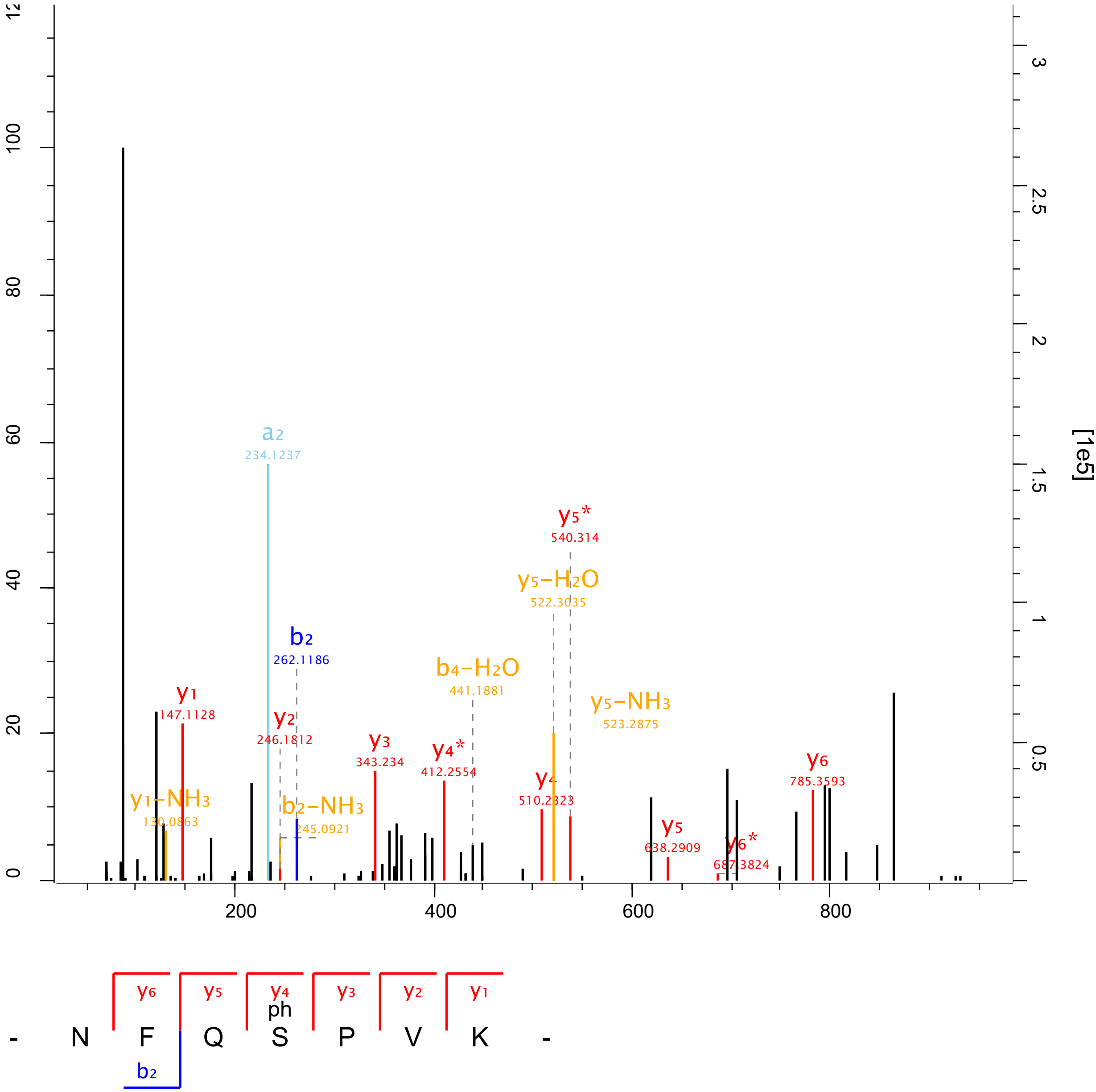

| Raw file | Scan | Method | Score | m/z |
| --- | --- | --- | --- | --- |
| A17_02_1 | 5045 | FTMS; HCD | 125.97 | 591.76 |

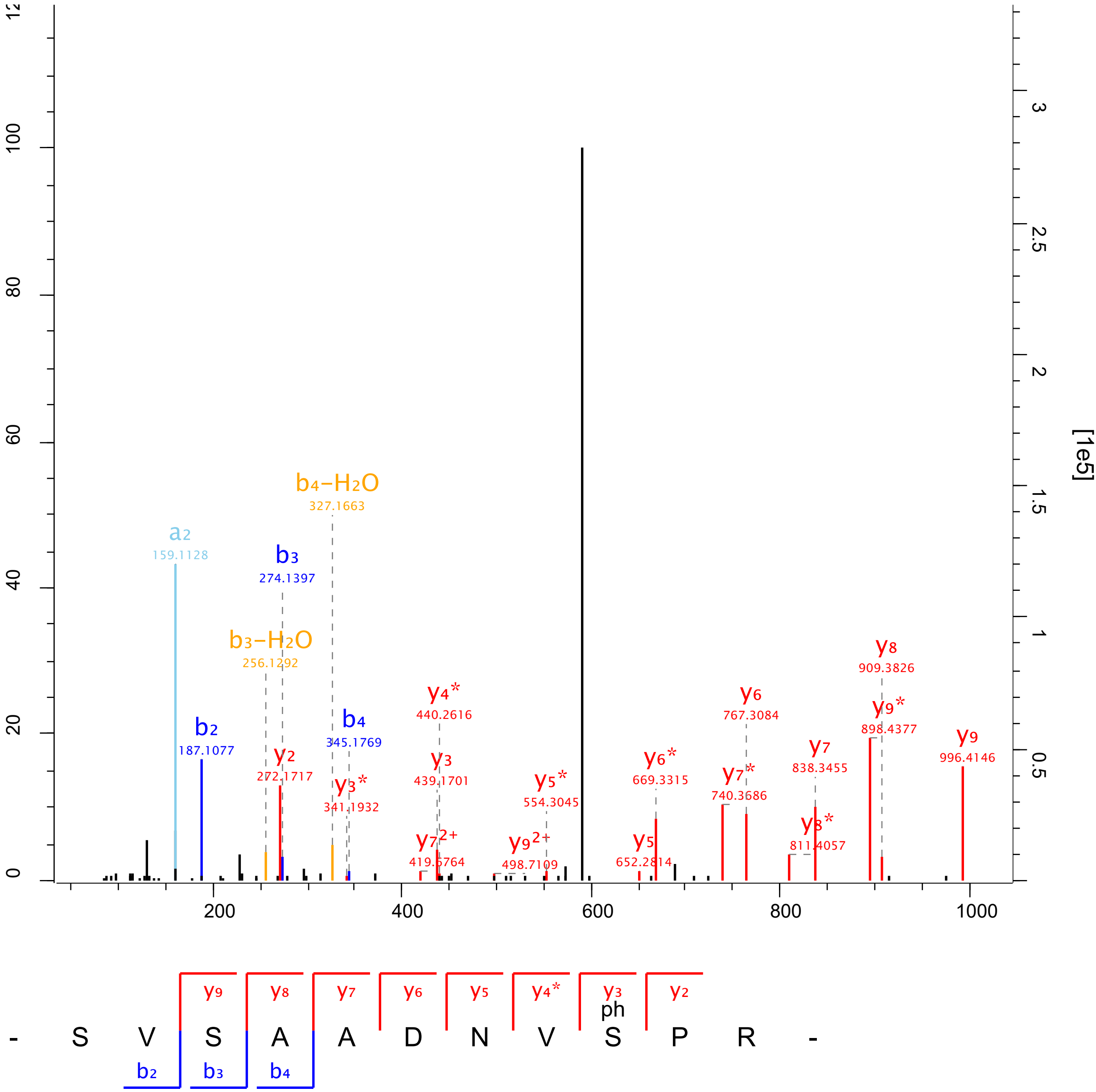

| Raw file | Scan | Method | Score | m/z |
| --- | --- | --- | --- | --- |
| A17_02_1 | 5080 | FTMS; HCD | 94.47 | 513.75 |

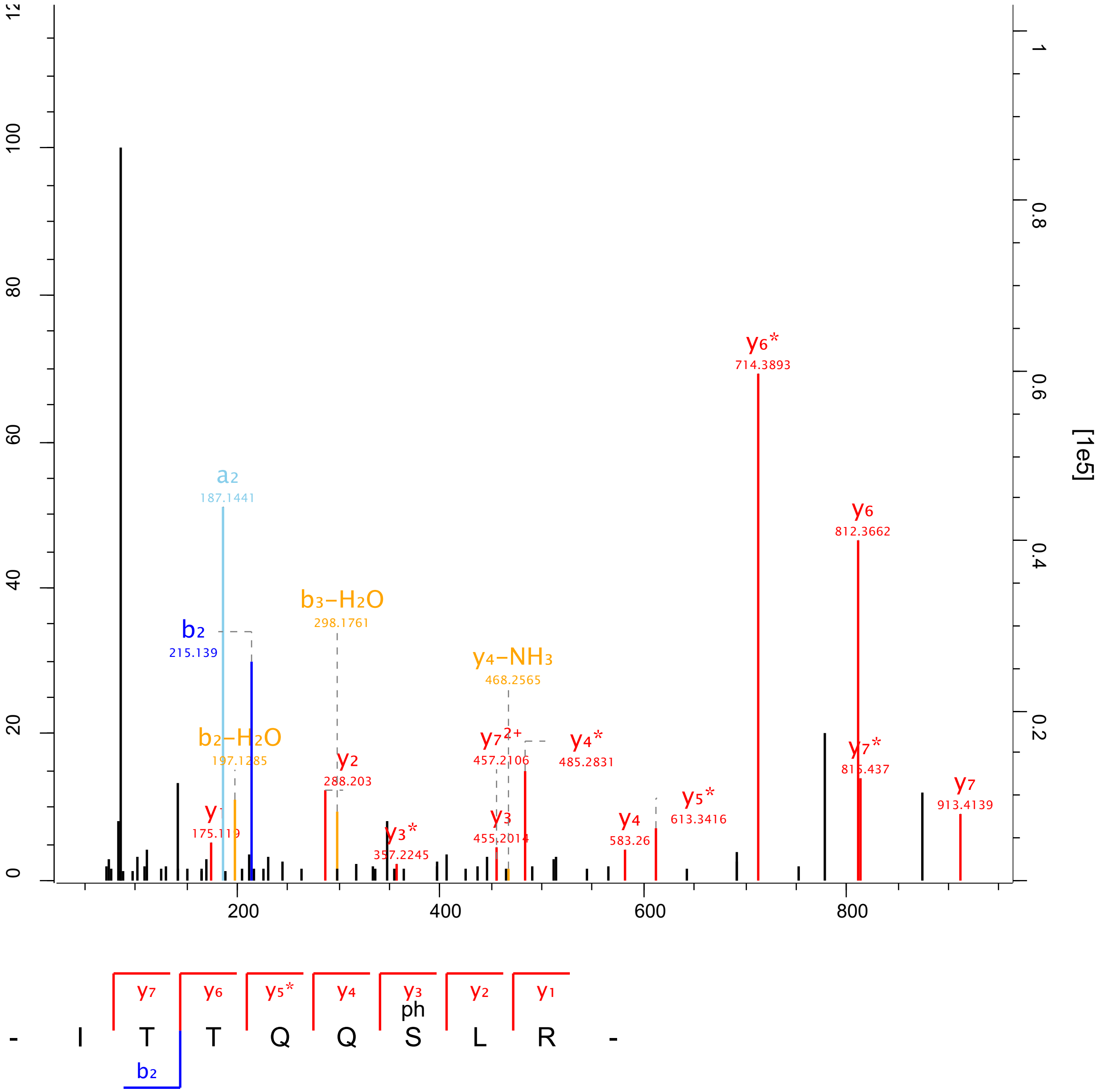

| Raw file | Scan | Method | Score | m/z |
| --- | --- | --- | --- | --- |
| A17_02_1 | 5152 | FTMS; HCD | 73.93 | 583.76 |

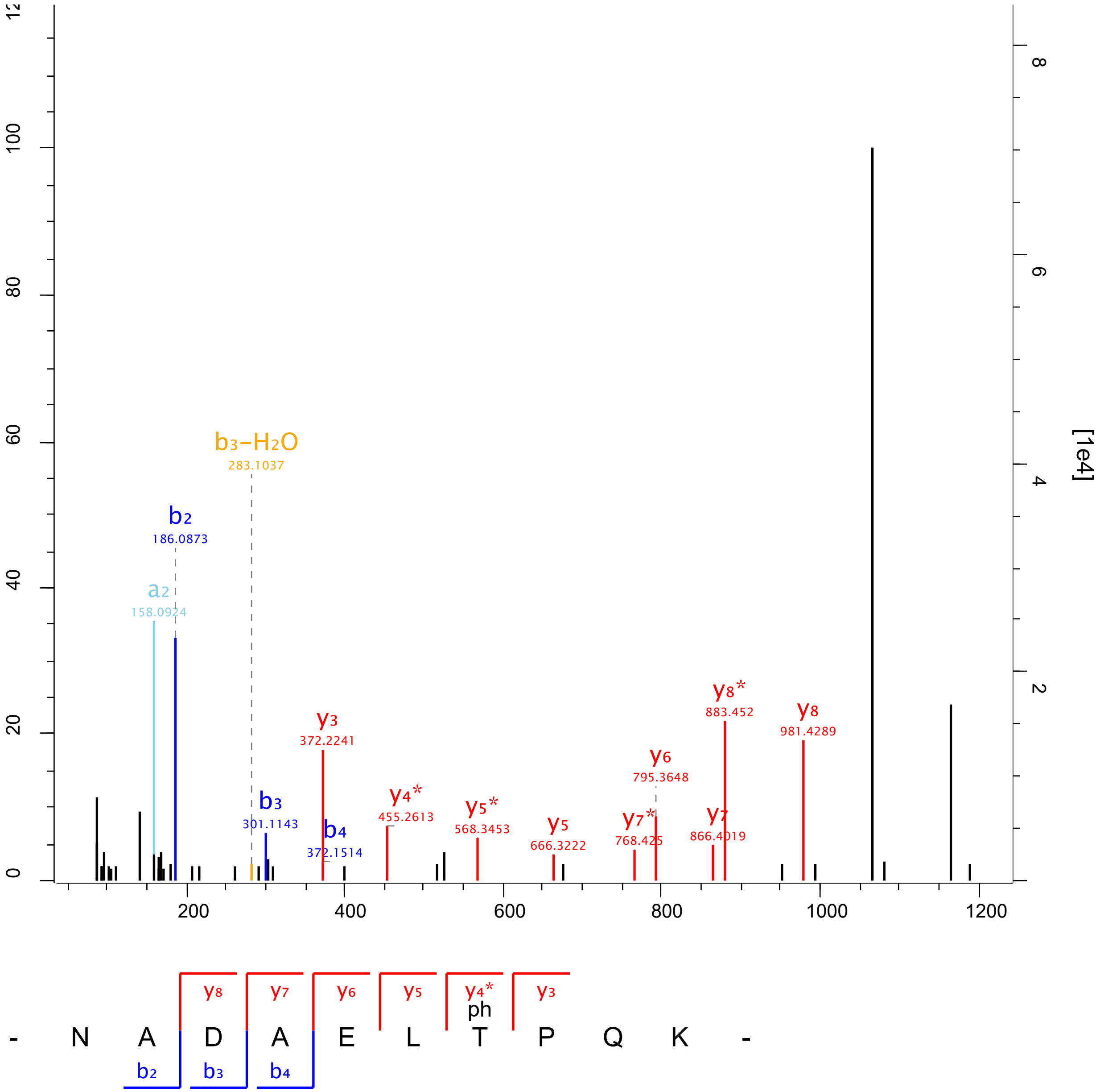

| Raw file | Scan | Method | Score | m/z |
| --- | --- | --- | --- | --- |
| A17_02_1 | 5193 | FTMS; HCD | 108.26 | 436.67 |

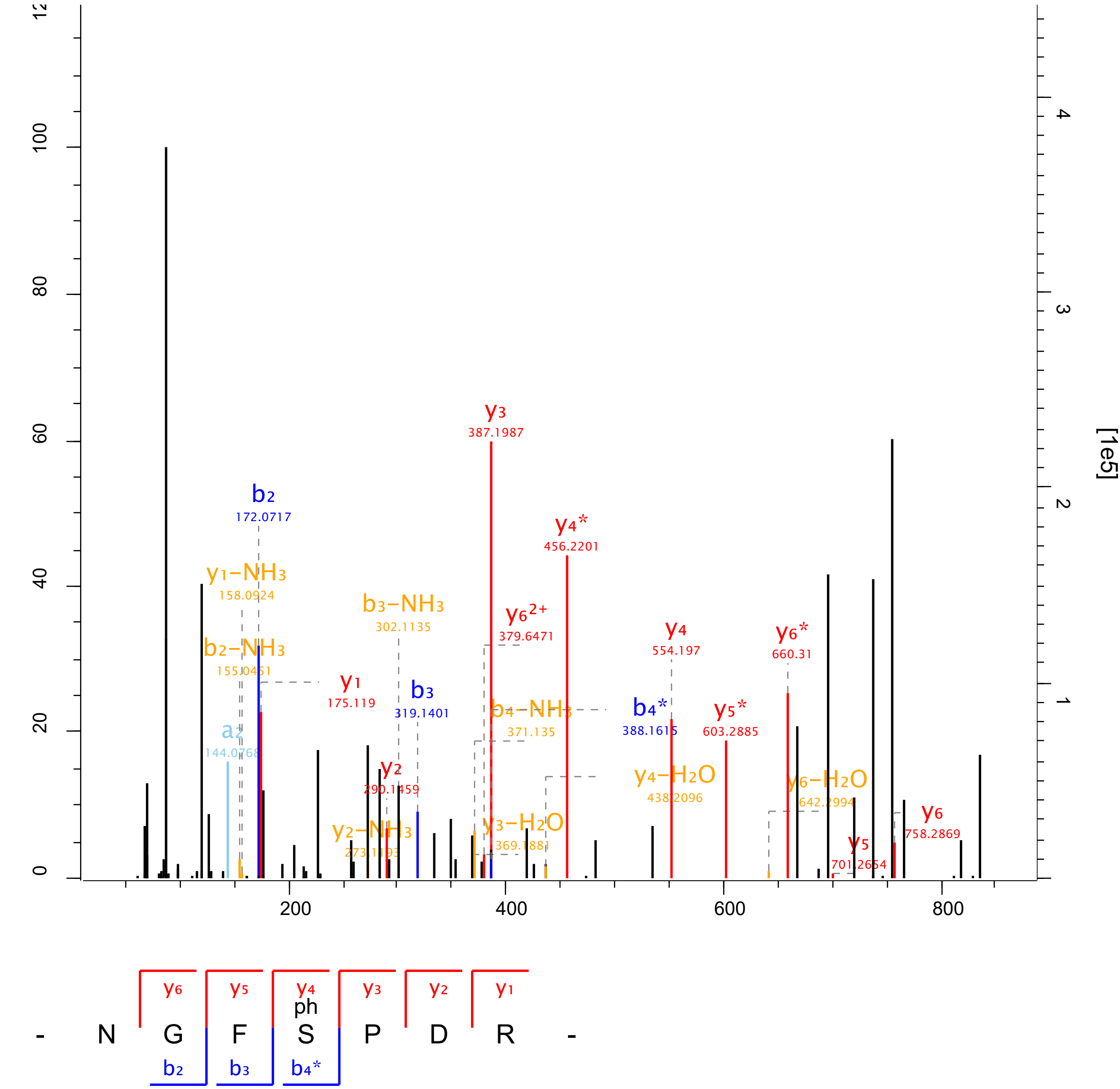

| Raw file | Scan | Method | Score | m/z |
|---|---|---|---|---|
| A17_02_1 | 5410 | FTMS; HCD | 93.24 | 593.75 |

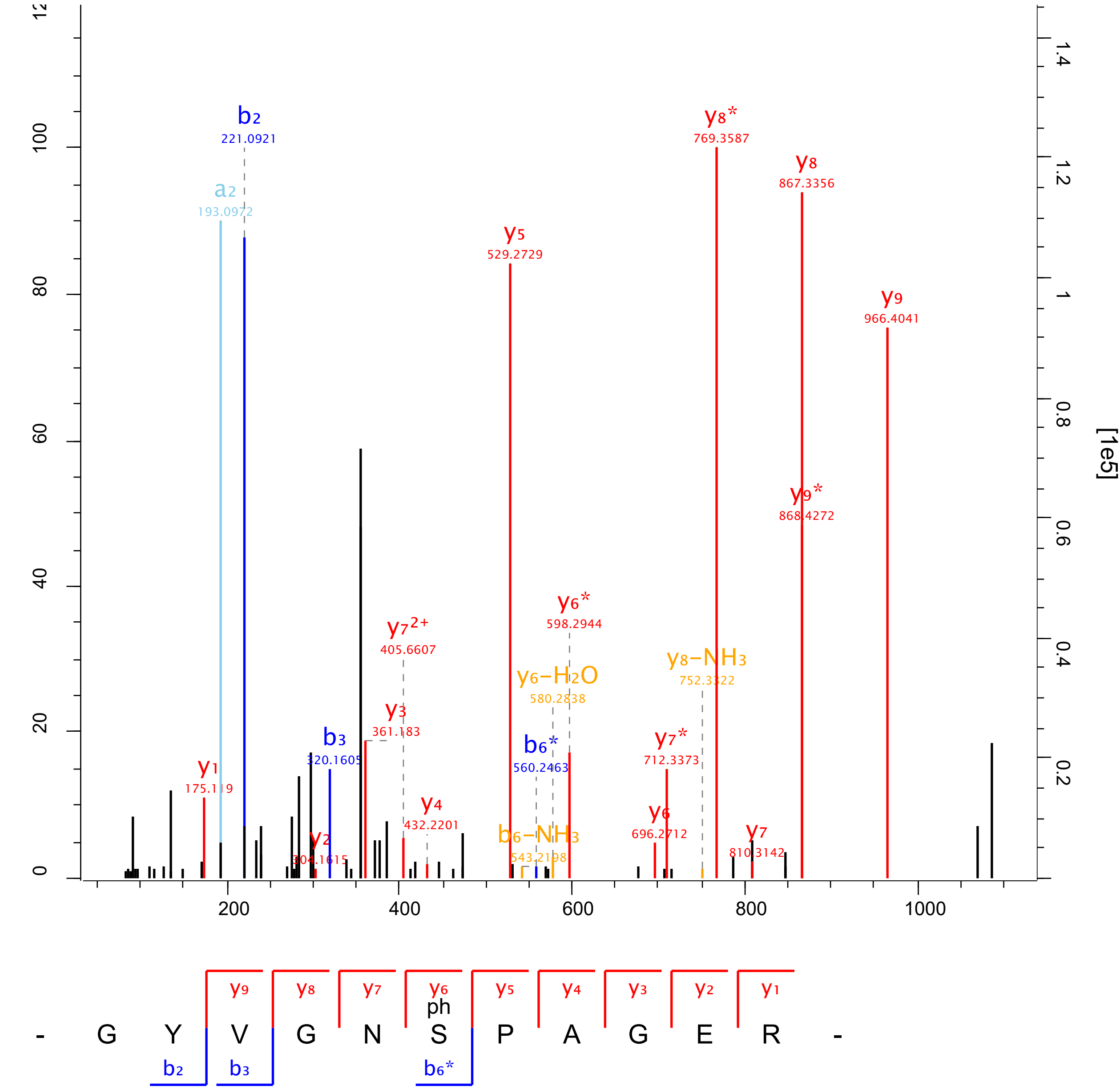

| Raw file | Scan | Method | Score | m/z |
|---|---|---|---|---|
| A17_02_1 | 5623 | FTMS; HCD | 177.83 | 593.76 |

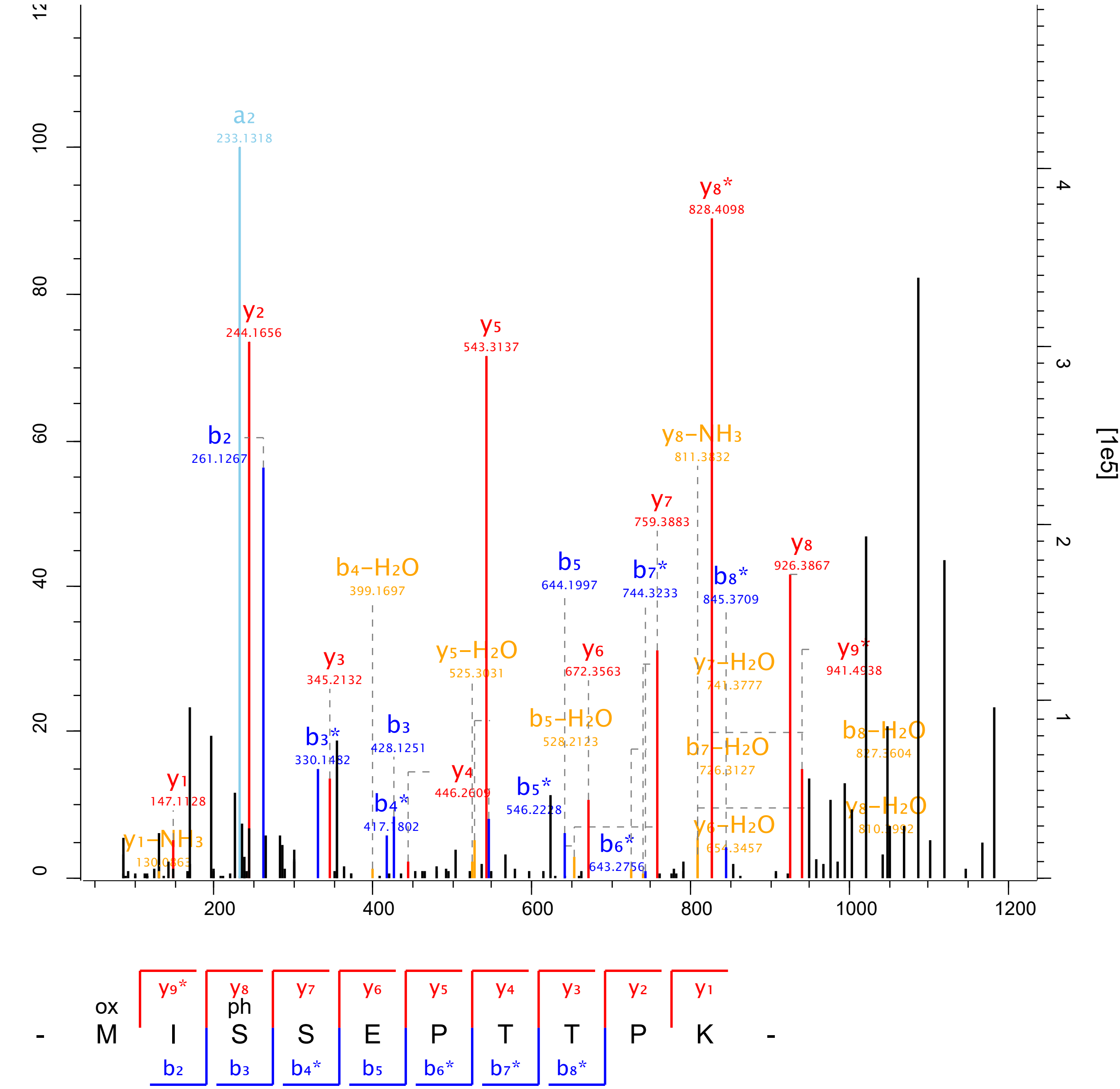

| Raw file | Scan | Method | Score | m/z |
|---|---|---|---|---|
| A17_02_1 | 5732 | FTMS; HCD | 177.1 | 539.75 |

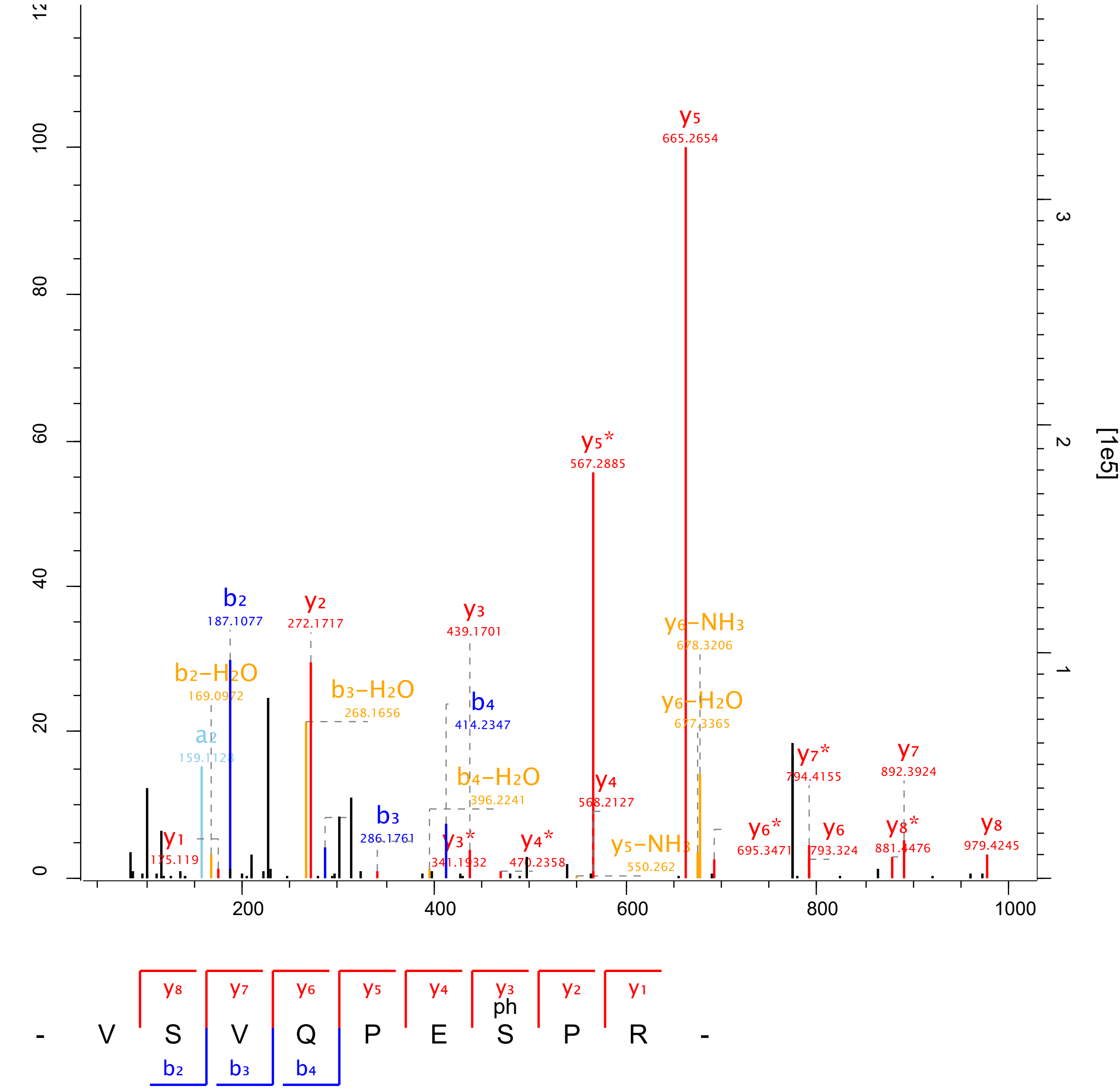

| Raw file | Scan | Method | Score | m/z |
|---|---|---|---|---|
| A17_02_1 | 5839 | FTMS; HCD | 143.99 | 785.31 |

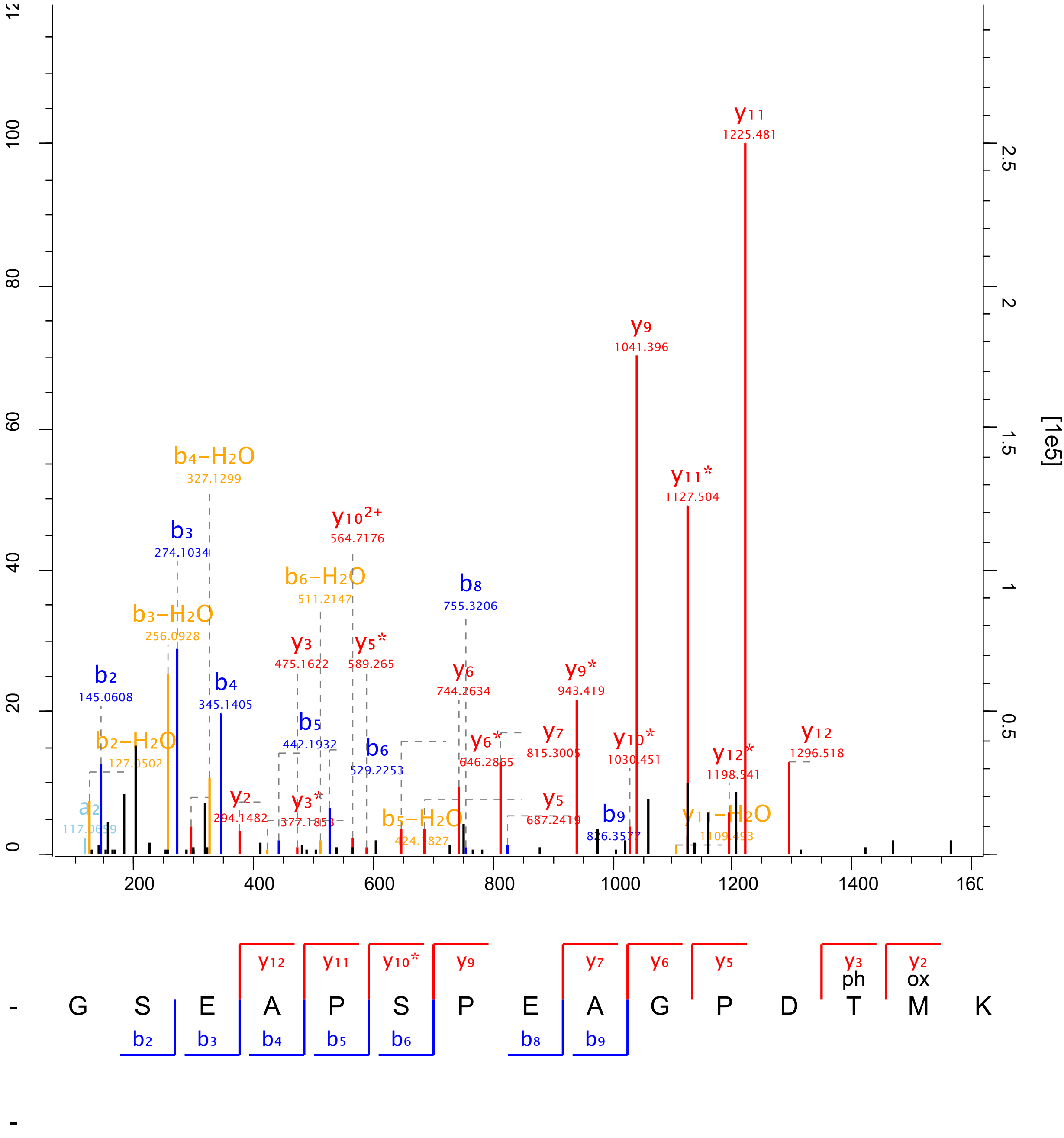

| Raw file | Scan | Method | Score | m/z |
| --- | --- | --- | --- | --- |
| A17_02_1 | 6173 | FTMS; HCD | 86.77 | 541.73 |

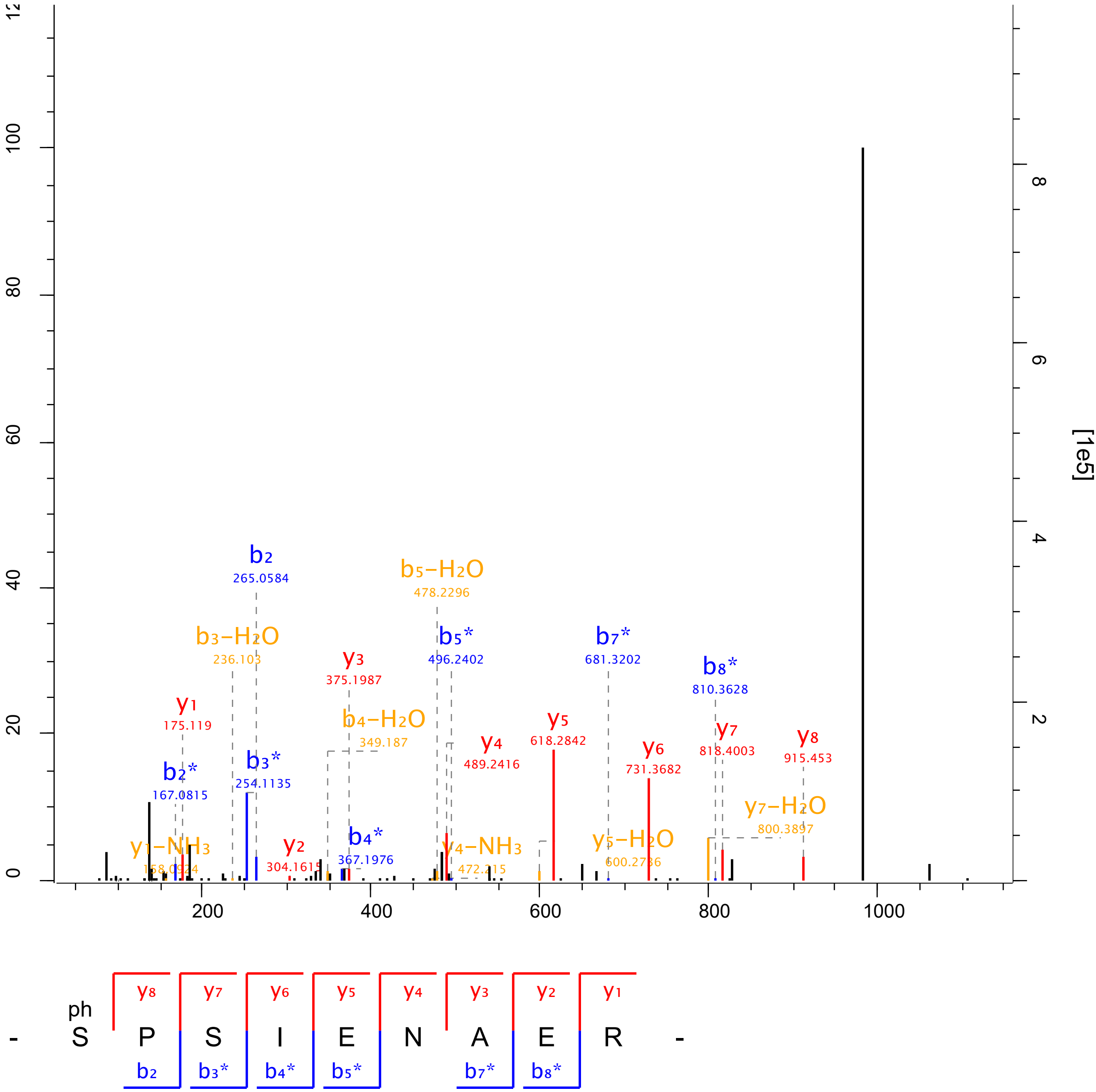

| Raw file | Scan | Method | Score | m/z |
|---|---|---|---|---|
| A17_02_1 | 6608 | FTMS; HCD | 42.42 | 550.74 |

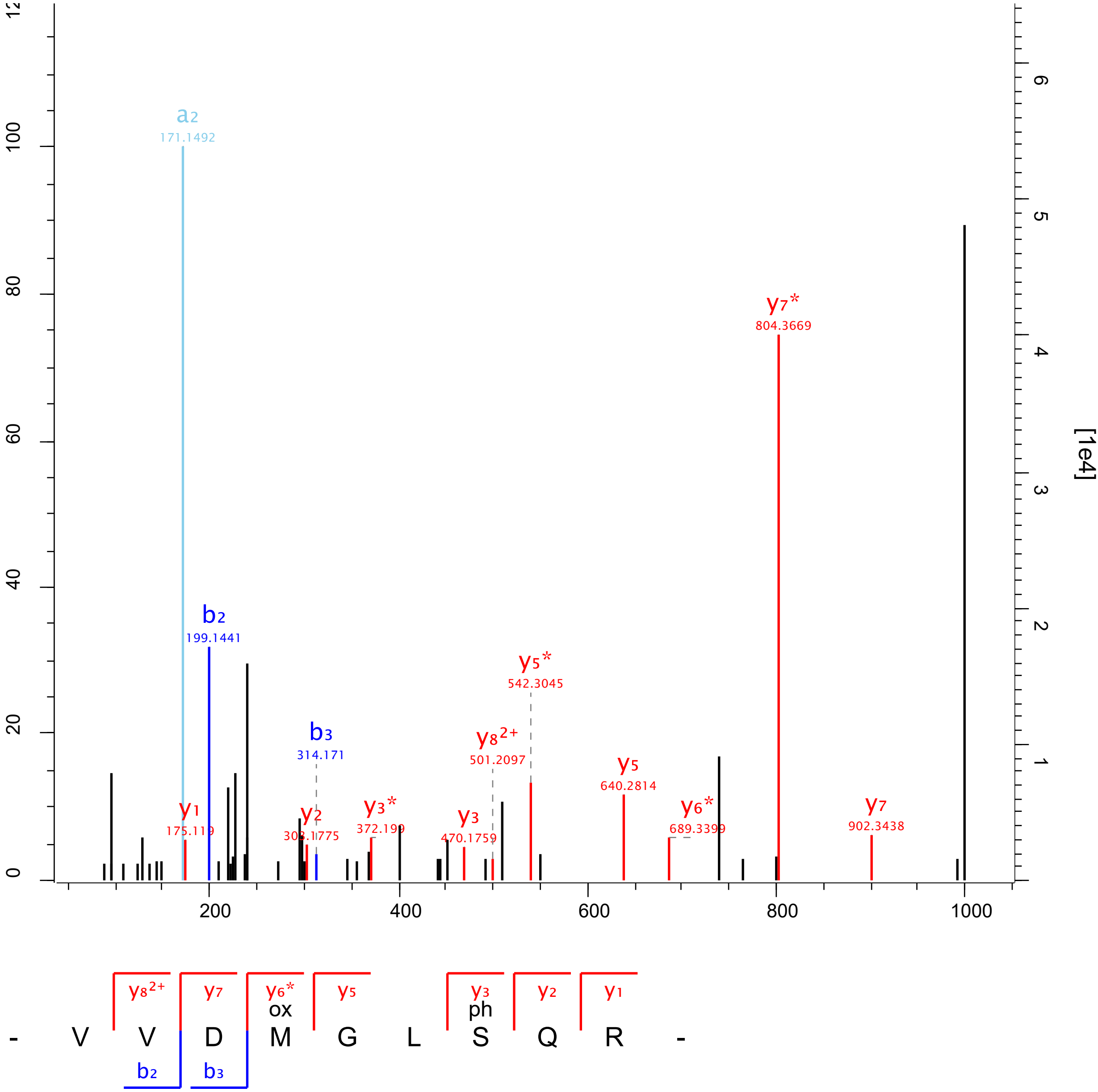

| Raw file | Scan | Method | Score | m/z |
|---|---|---|---|---|
| A17_02_1 | 7243 | FTMS; HCD | 261.58 | 826.33 |

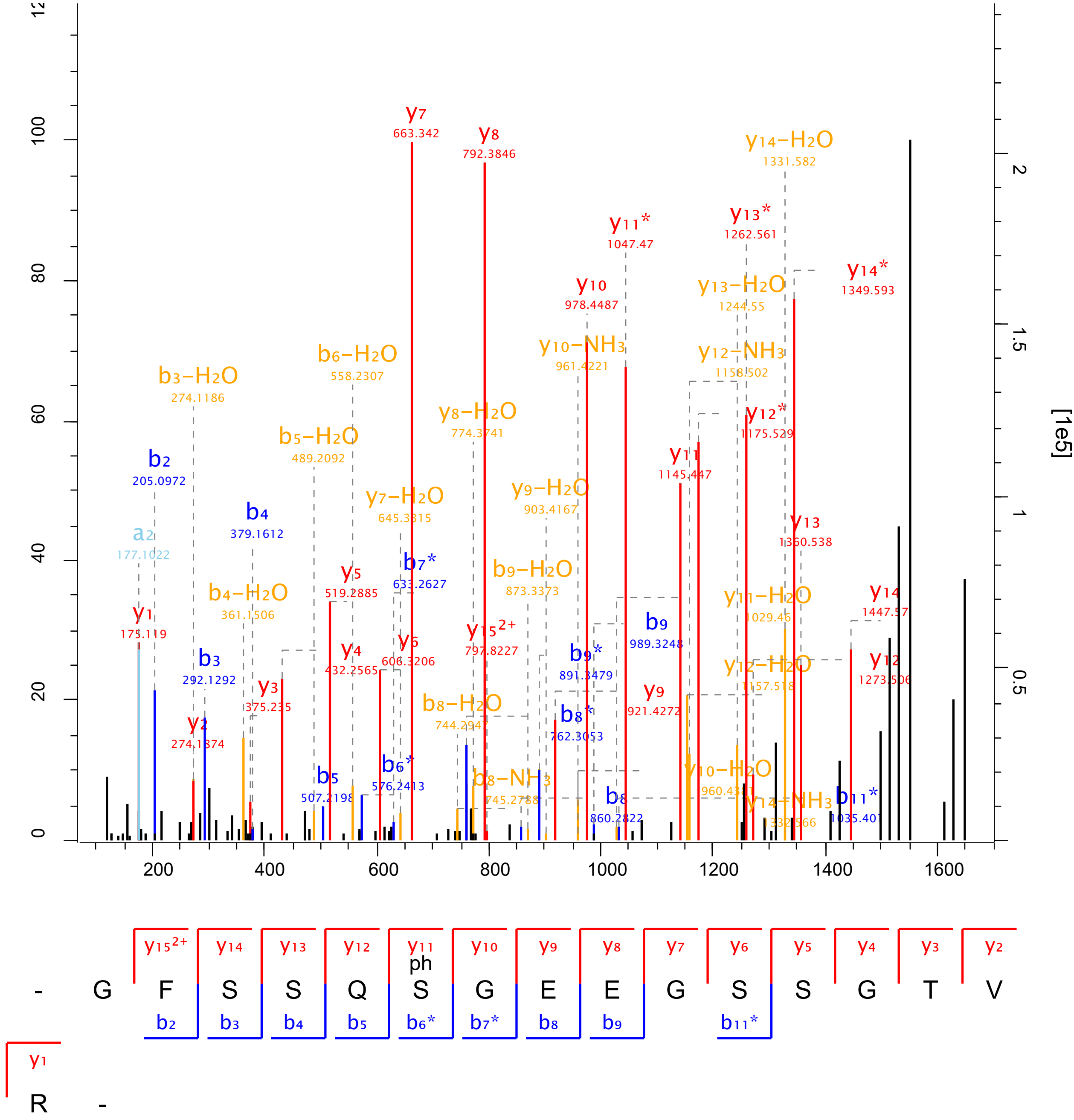

| Raw file | Scan | Method | Score | m/z |
|---|---|---|---|---|
| A17_02_1 | 7342 | FTMS; HCD | 111.5 | 515.22 |

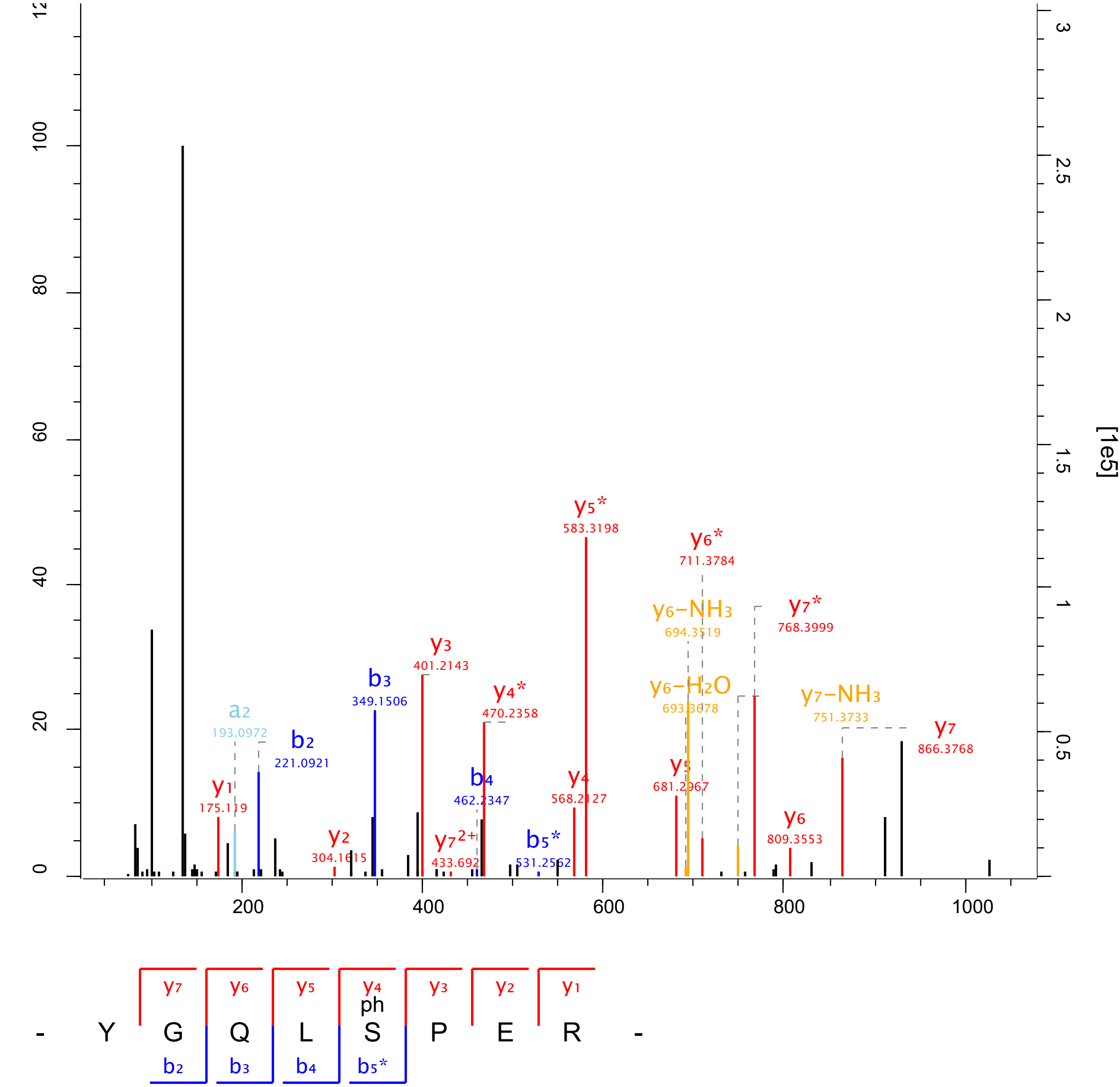

| Raw file | Scan | Method | Score | m/z |
|---|---|---|---|---|
| A17_02_1 | 7505 | FTMS; HCD | 144.18 | 636.74 |

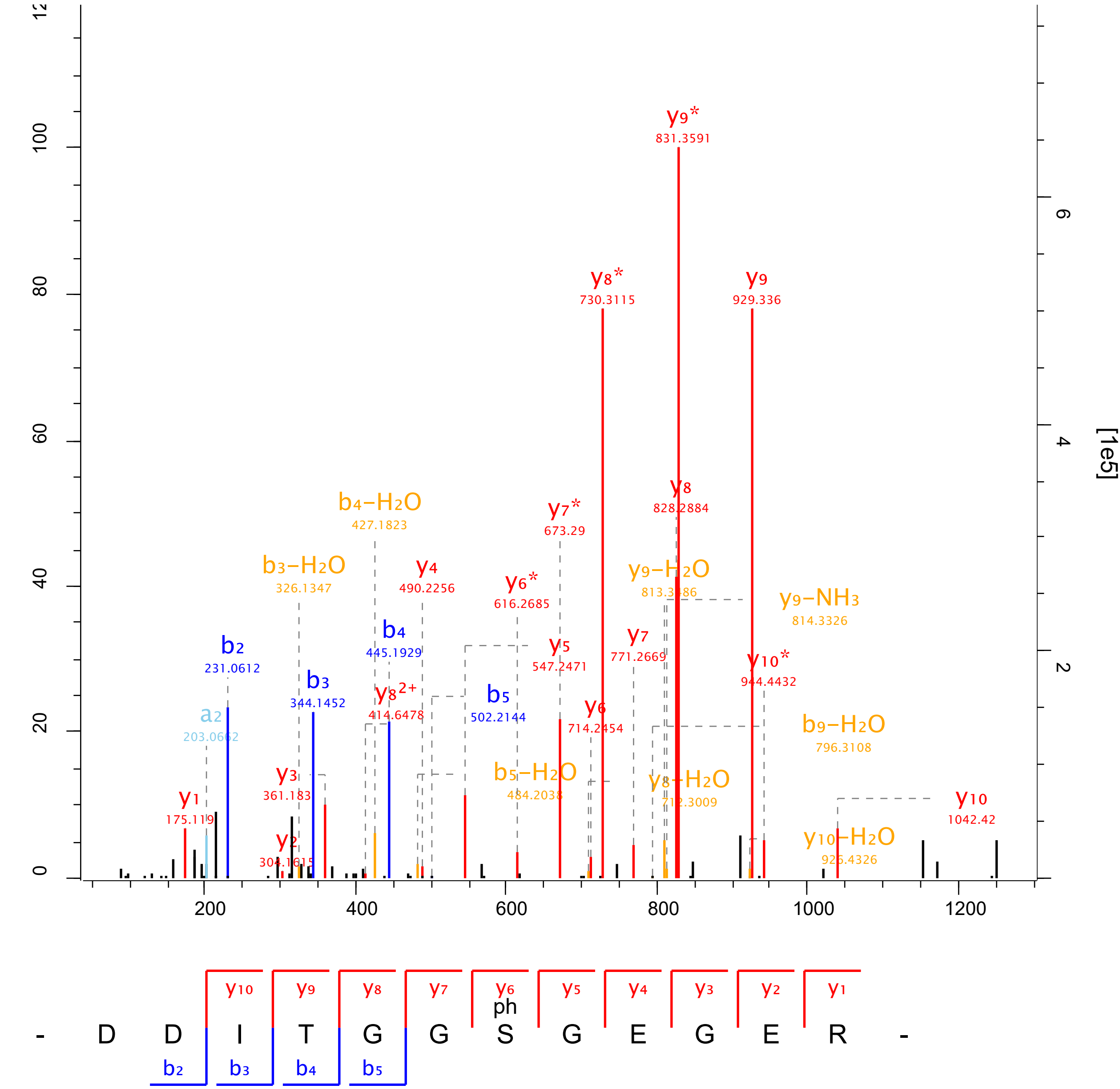

| Raw file | Scan | Method | Score | m/z |
|---|---|---|---|---|
| A17_02_1 | 7685 | FTMS; HCD | 94.77 | 529.22 |

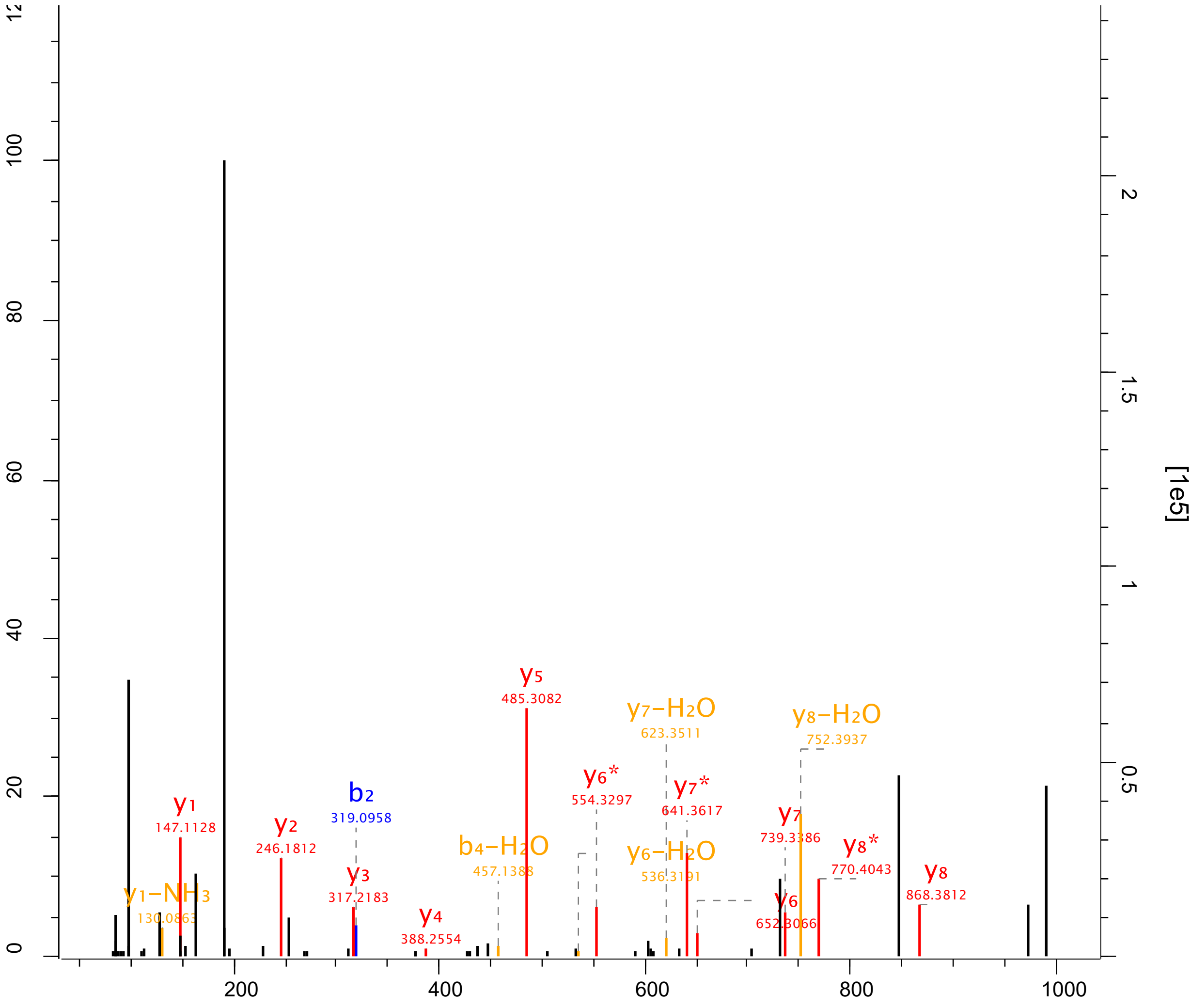


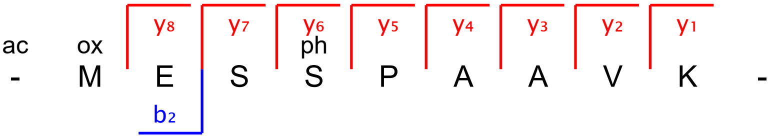

| Raw file | Scan | Method | Score | m/z |
|---|---|---|---|---|
| A17_02_1 | 8222 | FTMS; HCD | 146.88 | 782.82 |

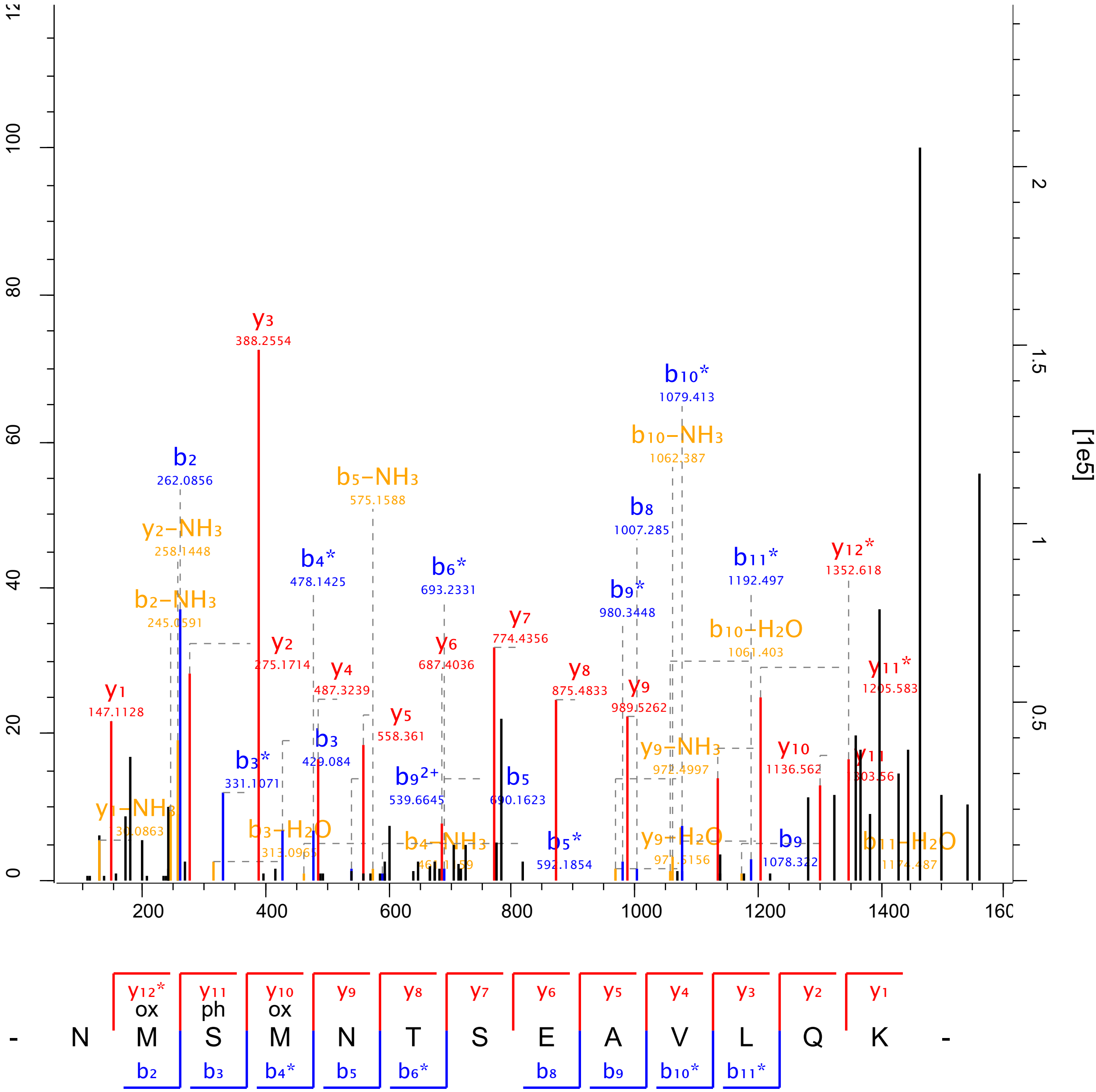

| Raw file | Scan | Method | Score | m/z |
| --- | --- | --- | --- | --- |
| A17_02_1 | 8281 | FTMS; HCD | 63.19 | 648.26 |

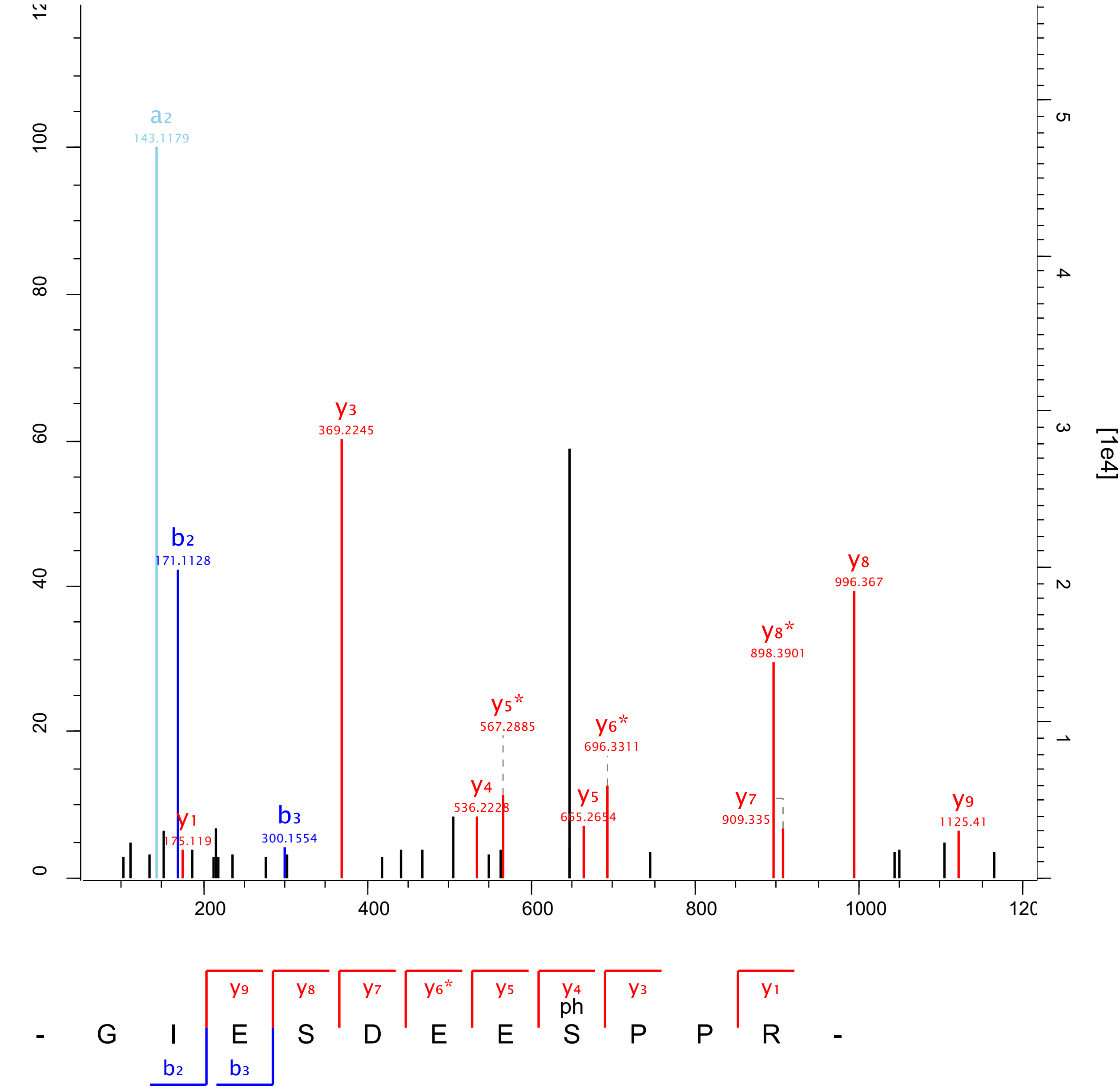

| Raw file | Scan | Method | Score | m/z |
| --- | --- | --- | --- | --- |
| A17_02_1 | 8413 | FTMS; HCD | 118.21 | 707.77 |

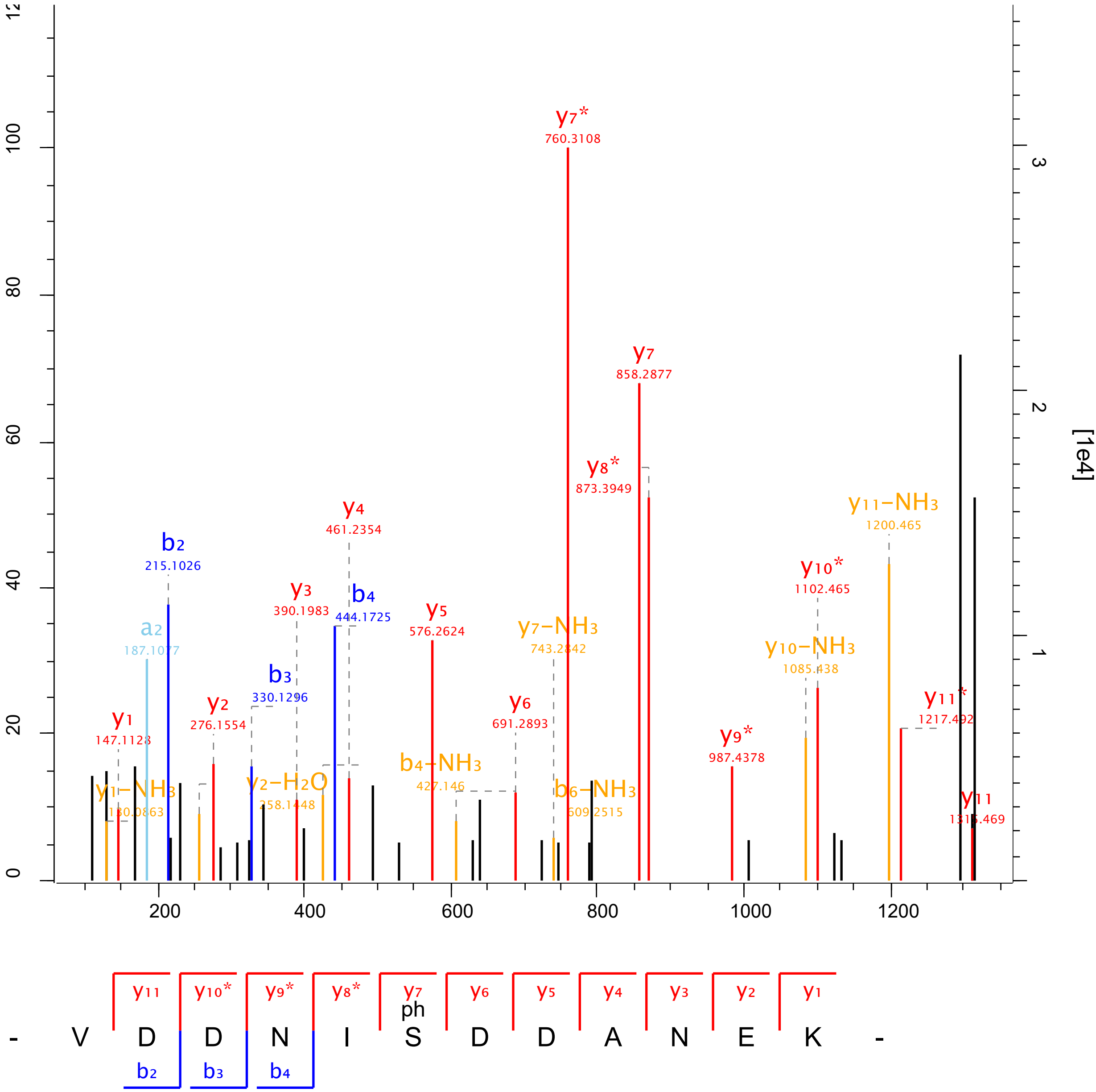

| Raw file | Scan | Method | Score | m/z |
| --- | --- | --- | --- | --- |
| A17_02_1 | 8427 | FTMS; HCD | 53.45 | 542.76 |

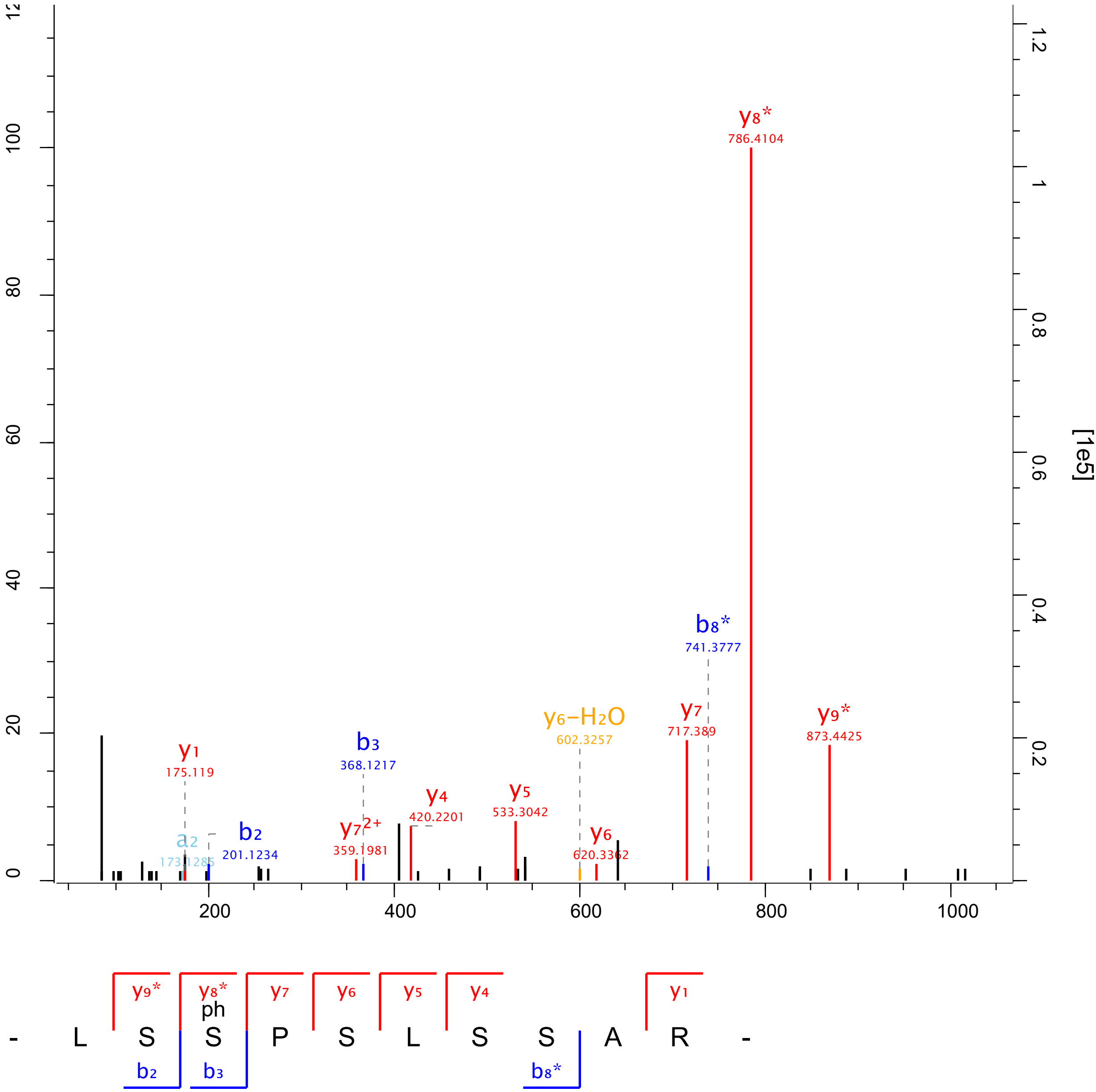

| Raw file | Scan | Method | Score | m/z |
| --- | --- | --- | --- | --- |
| A17_02_1 | 8791 | FTMS; HCD | 326.02 | 759.29 |

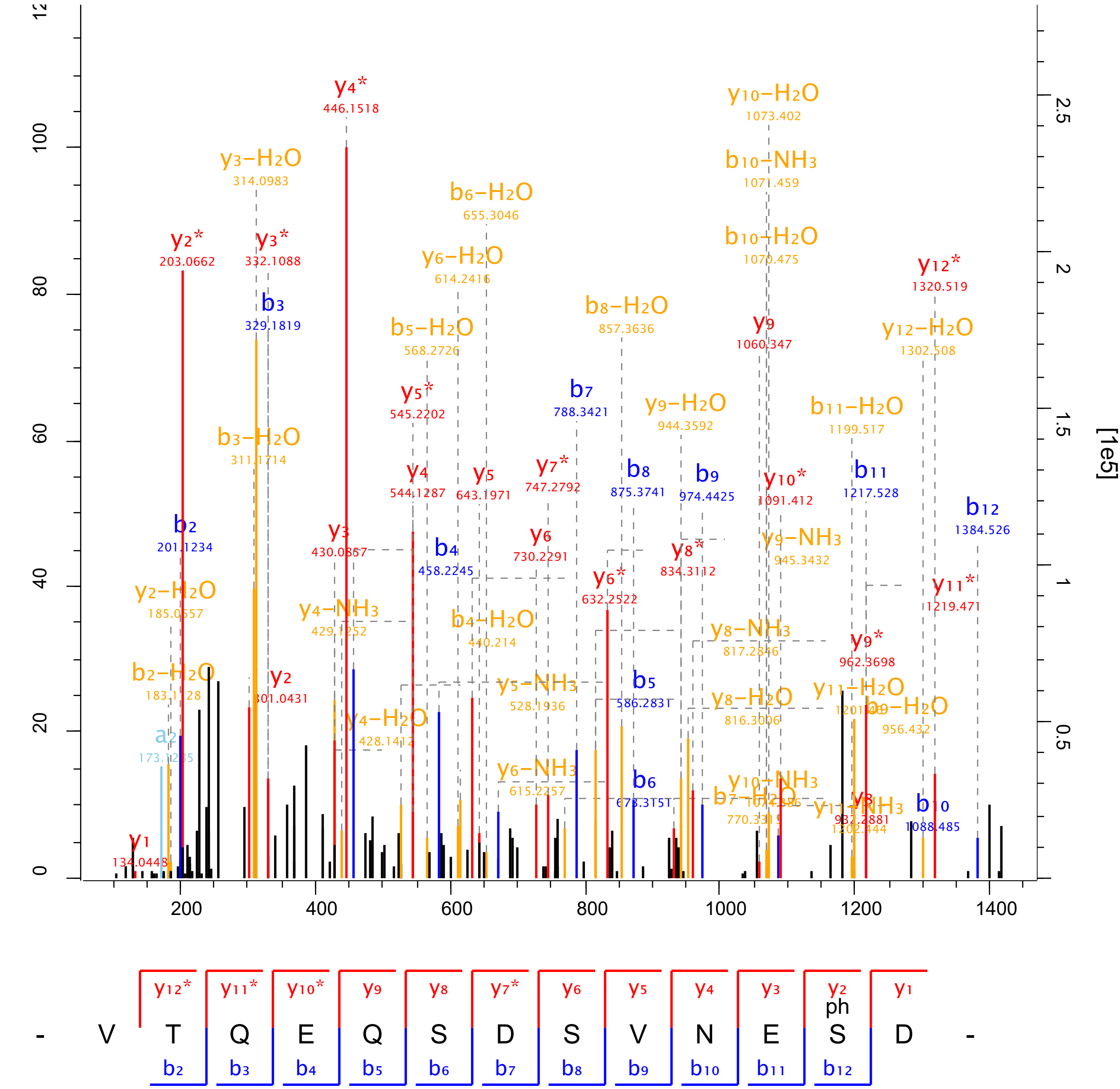

| Raw file | Scan | Method | Score | m/z |
|---|---|---|---|---|
| A17_02_1 | 8834 | FTMS; HCD | 264.97 | 812.85 |

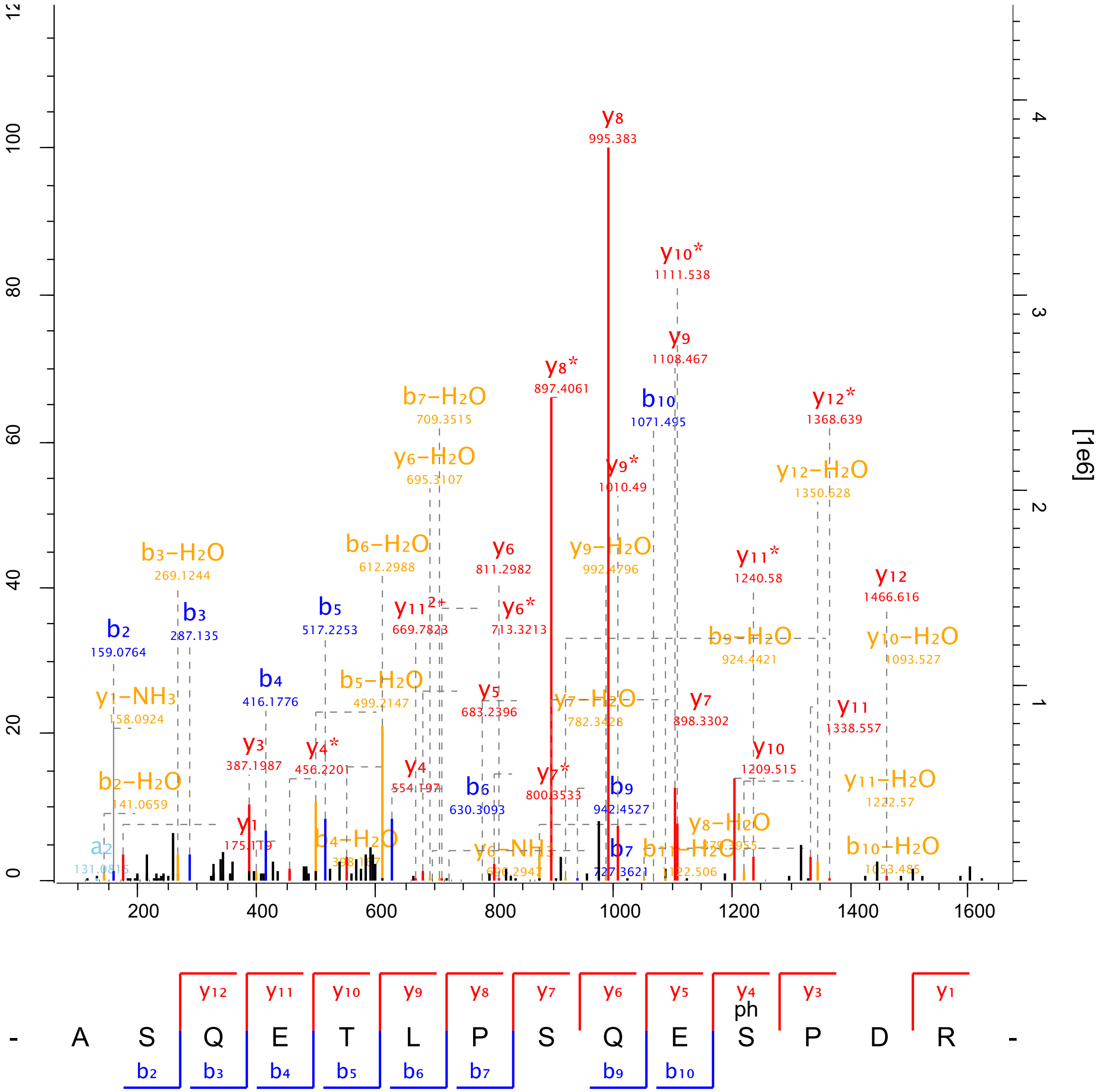

| Raw file | Scan | Method | Score | m/z |
|---|---|---|---|---|
| A17_02_1 | 9121 | FTMS; HCD | 156 | 592.27 |

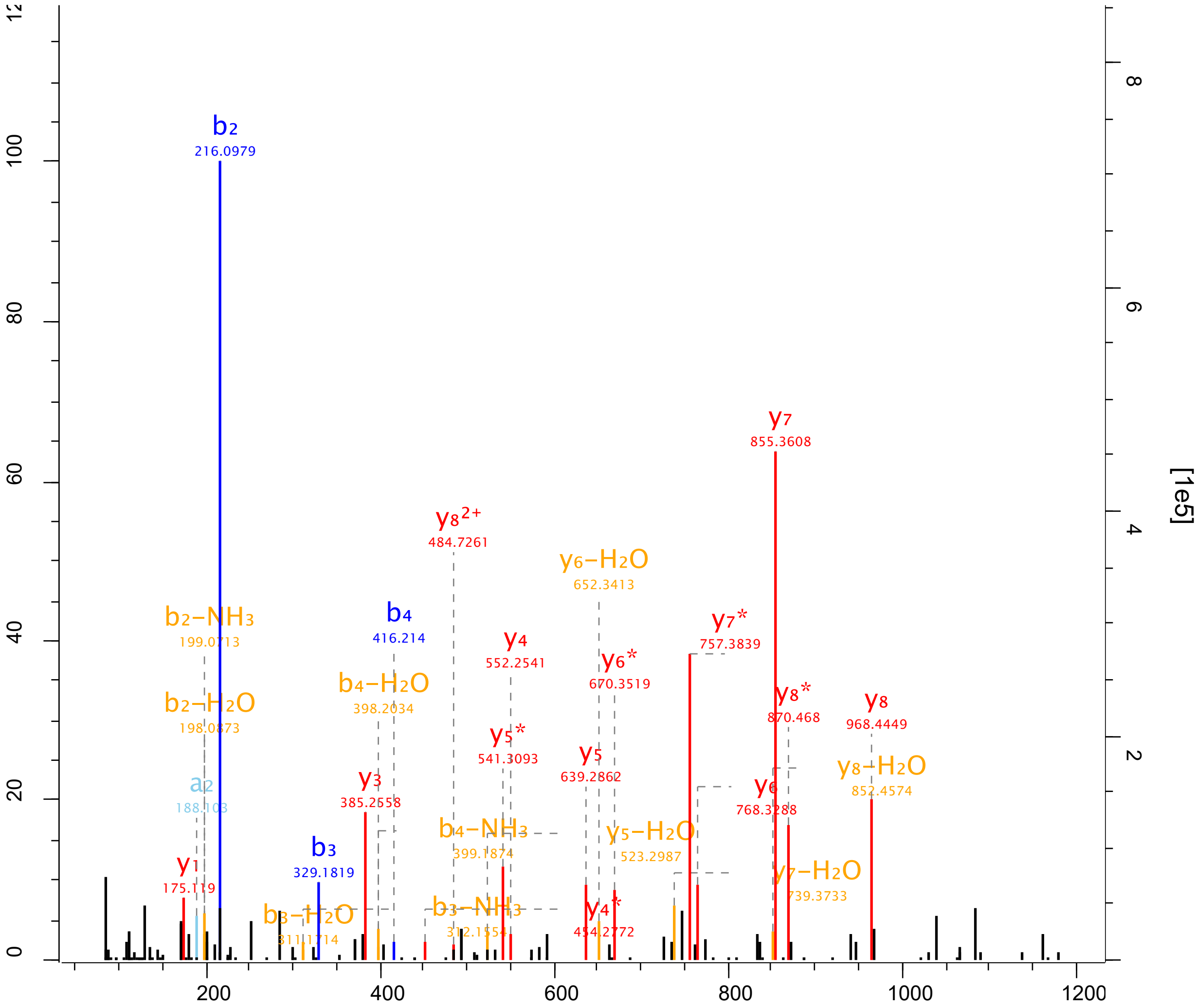


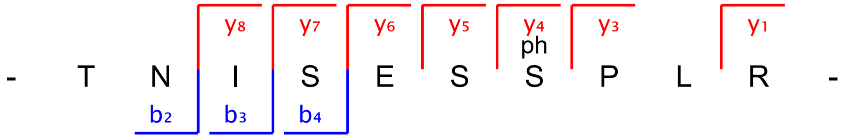

| Raw file | Scan | Method | Score | m/z |
|---|---|---|---|---|
| A17_02_1 | 9125 | FTMS; HCD | 51.5 | 767.84 |

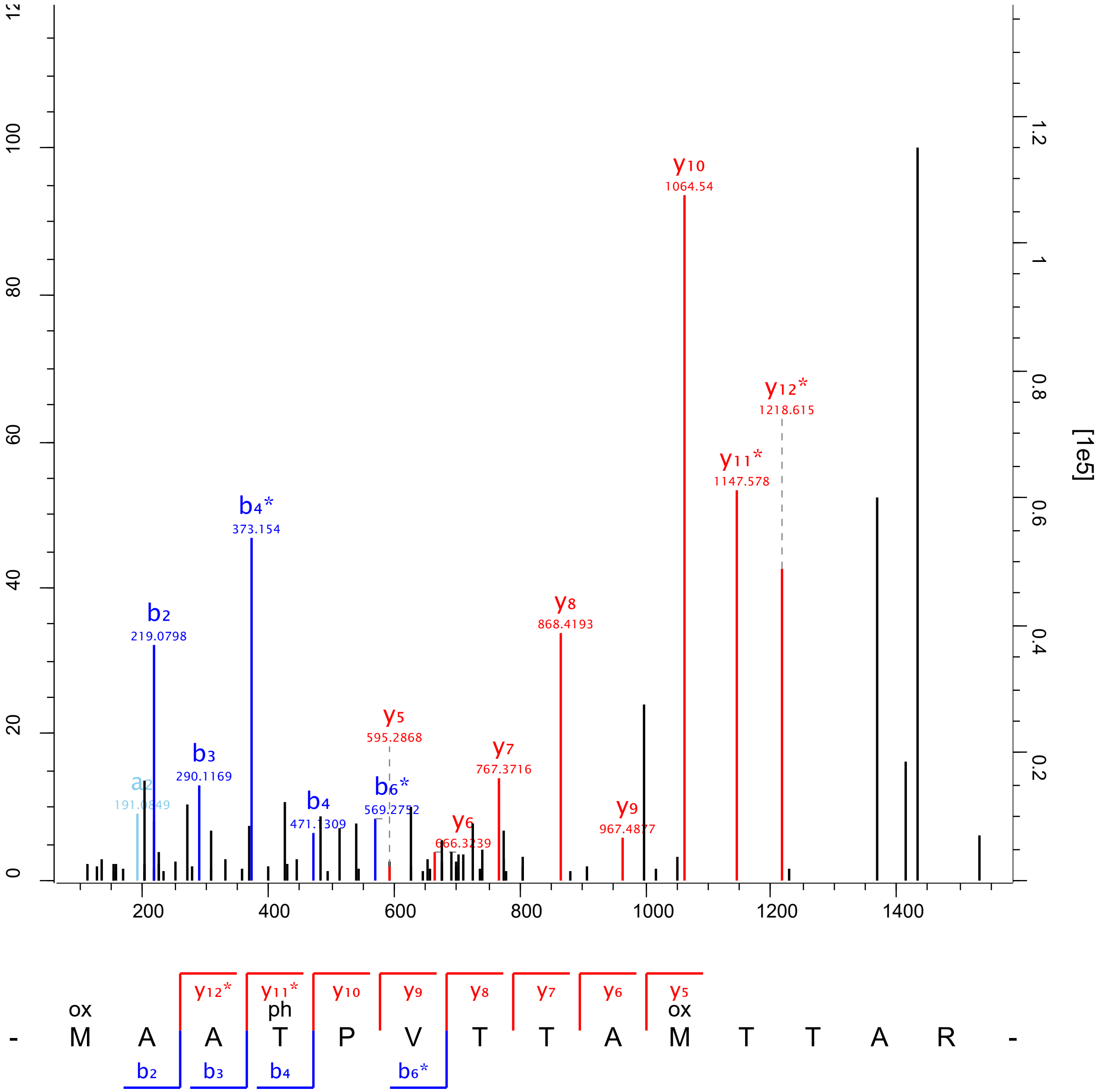

| Raw file | Scan | Method | Score | m/z |
|---|---|---|---|---|
| A17_02_1 | 9219 | FTMS; HCD | 104.81 | 611.74 |

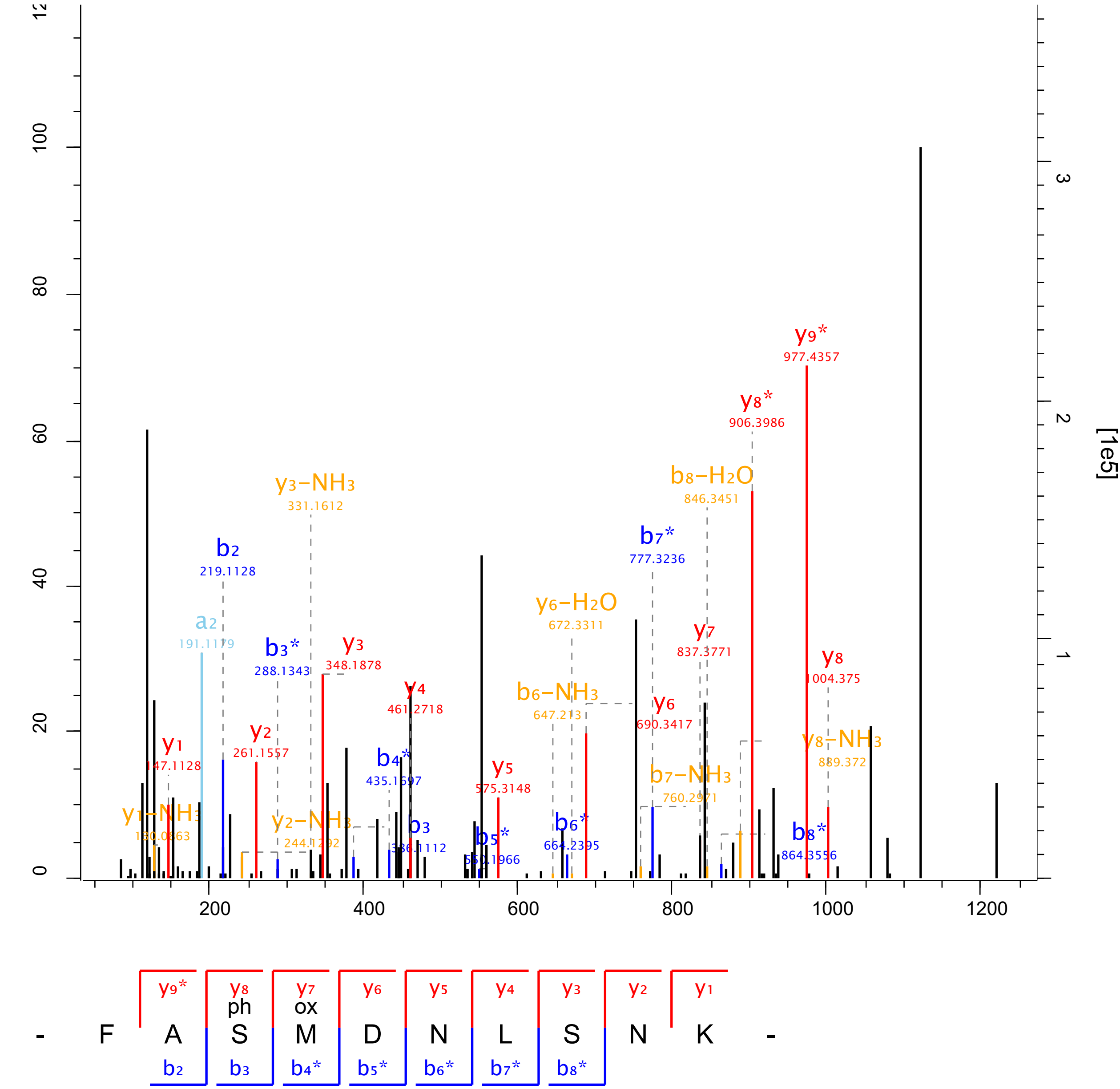

| Raw file | Scan | Method | Score | m/z |
|---|---|---|---|---|
| A17_02_1 | 9351 | FTMS; HCD | 62.29 | 627.78 |

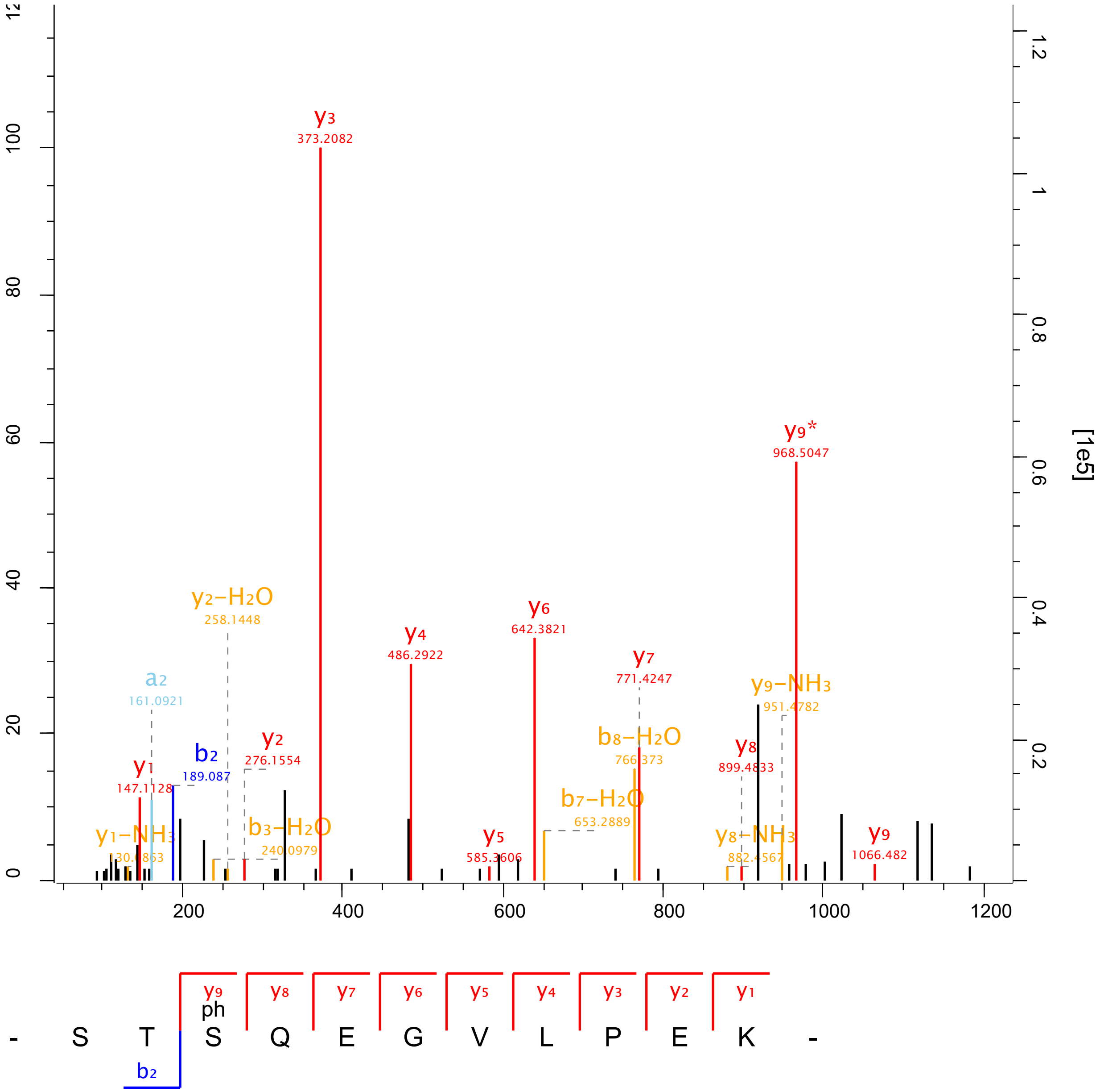

| Raw file | Scan | Method | Score | m/z |
|---|---|---|---|---|
| A17_02_1 | 9453 | FTMS; HCD | 53.76 | 497.76 |

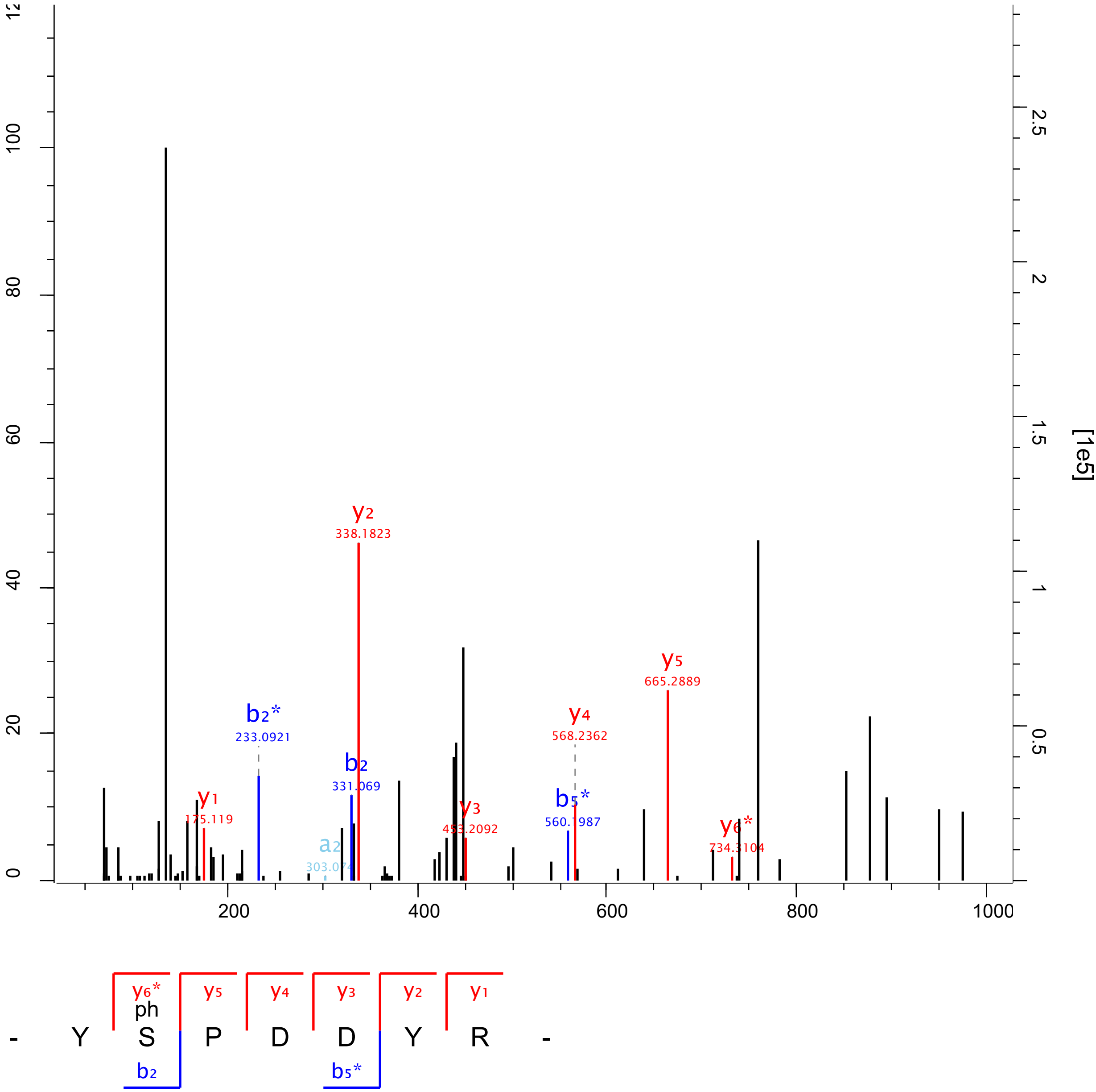

| Raw file | Scan | Method | Score | m/z |
|---|---|---|---|---|
| A17_02_1 | 9594 | FTMS; HCD | 130.56 | 819.86 |

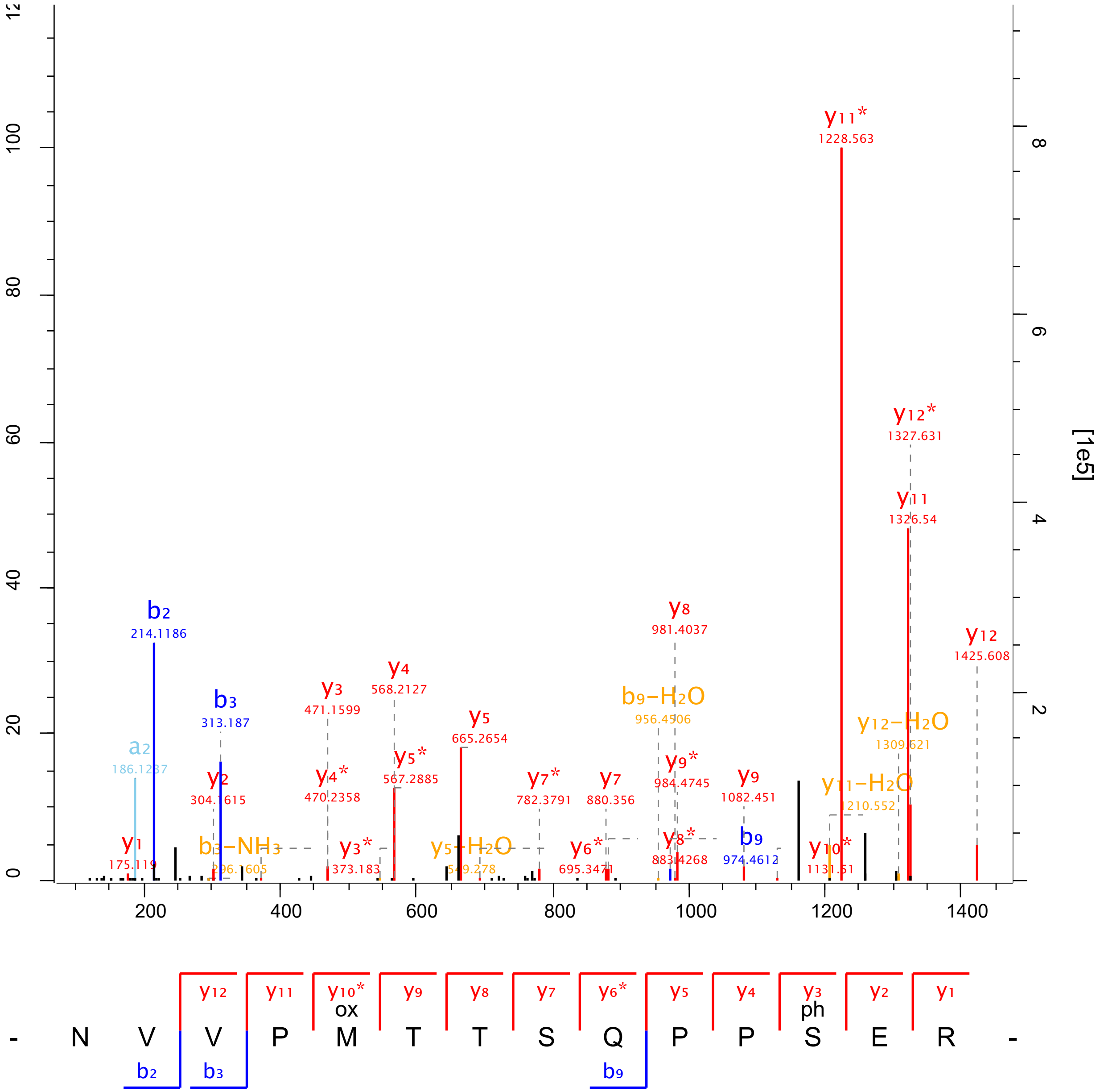

| Raw file | Scan | Method | Score | m/z |
|---|---|---|---|---|
| A17_02_1 | 9633 | FTMS; HCD | 103.13 | 544.25 |

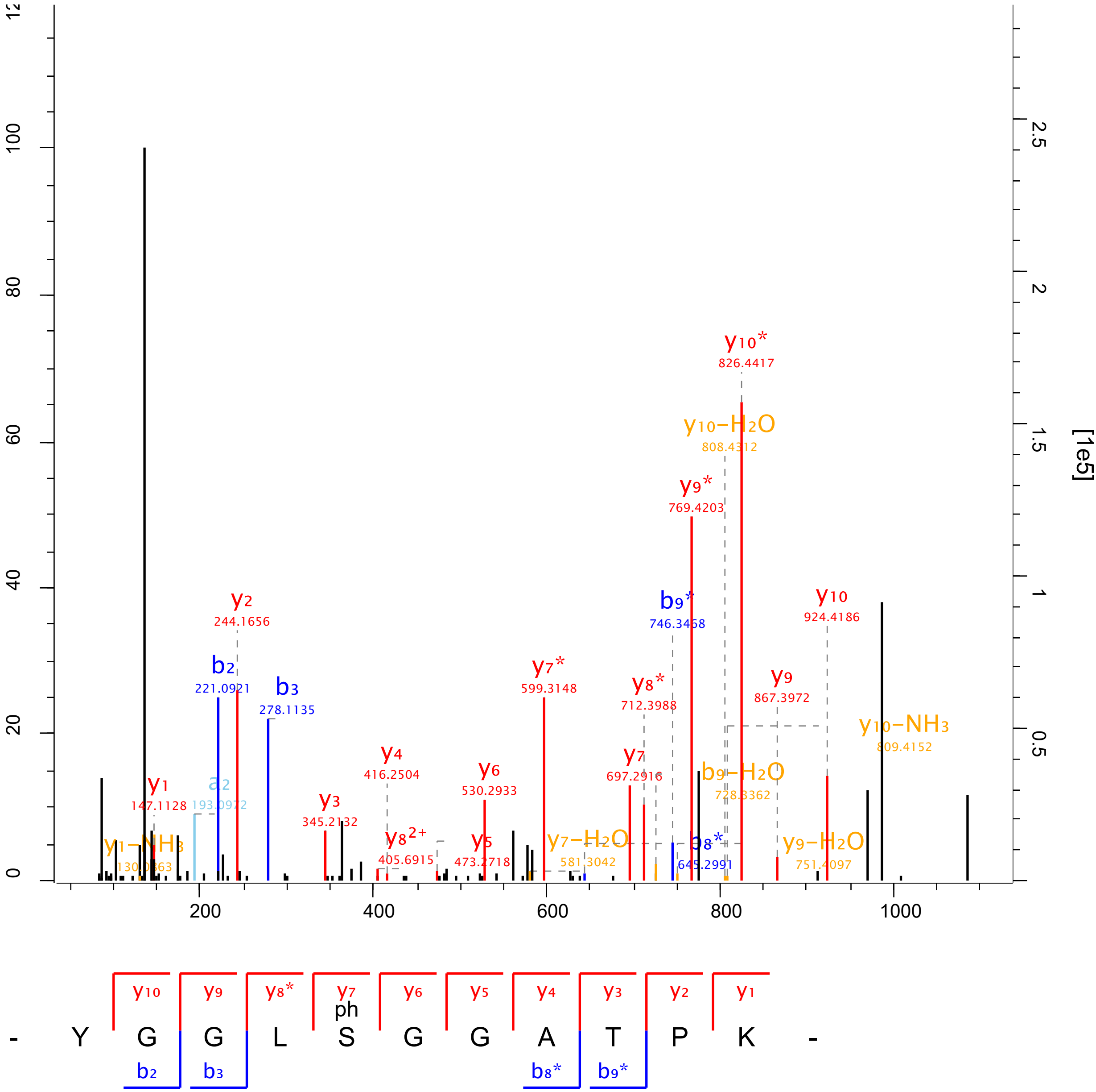

| Raw file | Scan | Method | Score | m/z |
| --- | --- | --- | --- | --- |
| A17_02_1 | 9661 | FTMS; HCD | 97.43 | 436.19 |

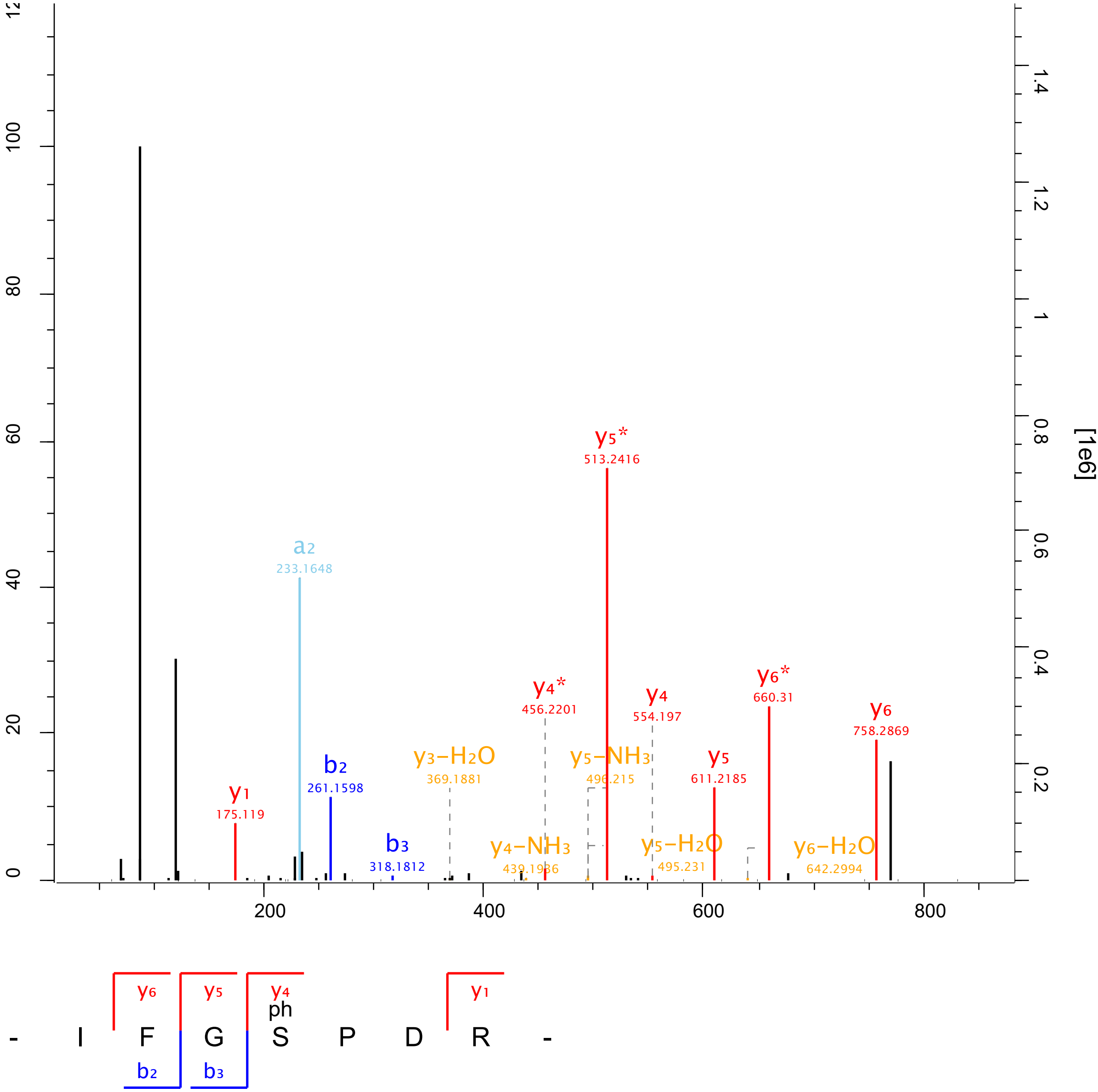

| Raw file | Scan | Method | Score | m/z |
|---|---|---|---|---|
| A17_05_1 | 27989 | FTMS; HCD | 152.86 | 516.71 |

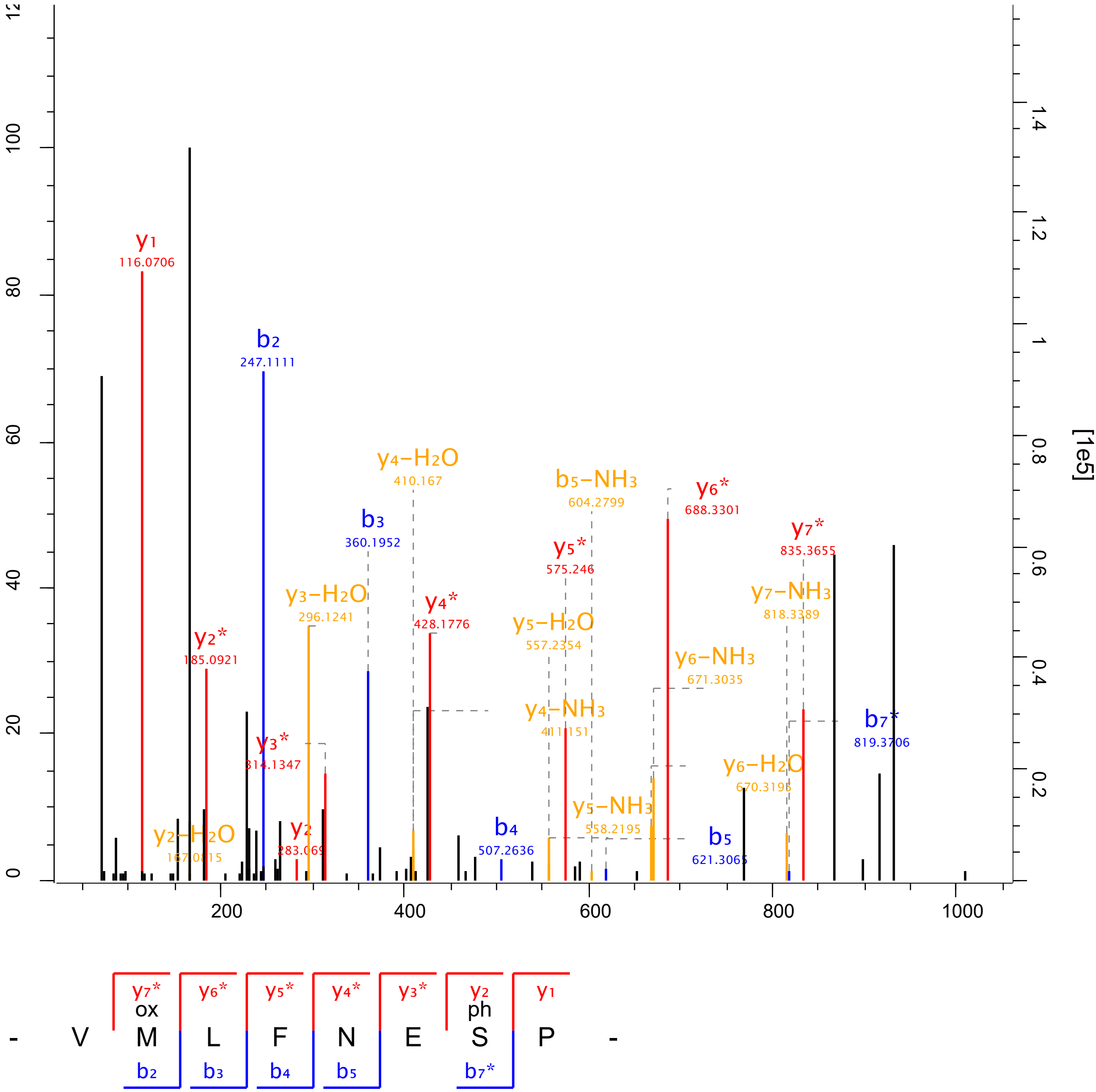

| Raw file | Scan | Method | Score | m/z |
|---|---|---|---|---|
| A17_05_1 | 5077 | FTMS; HCD | 102.95 | 427.7 |

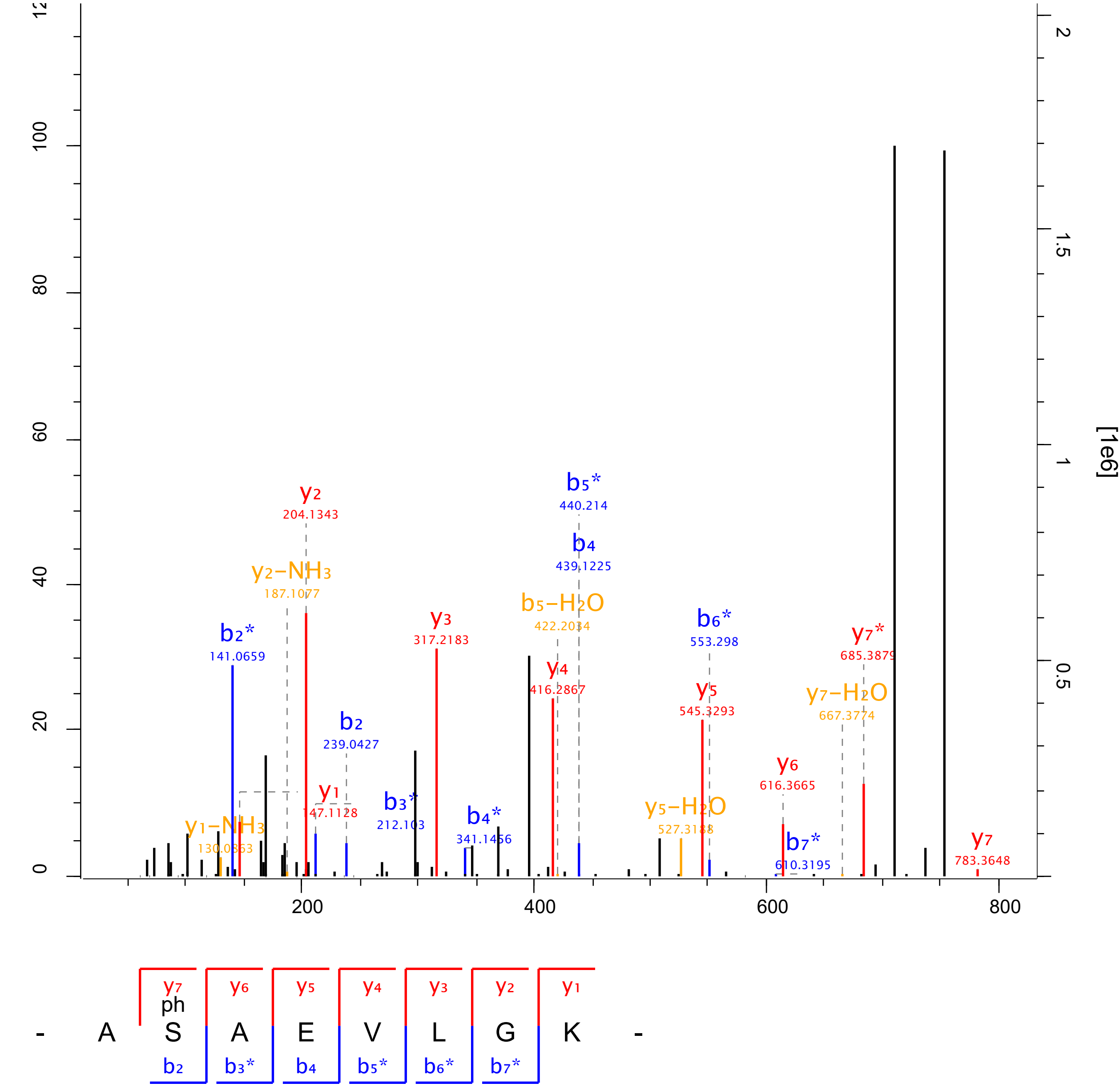

| Raw file | Scan | Method | Score | m/z |
|---|---|---|---|---|
| A17_05_2 | 18853 | FTMS; HCD | 55.35 | 646.3 |

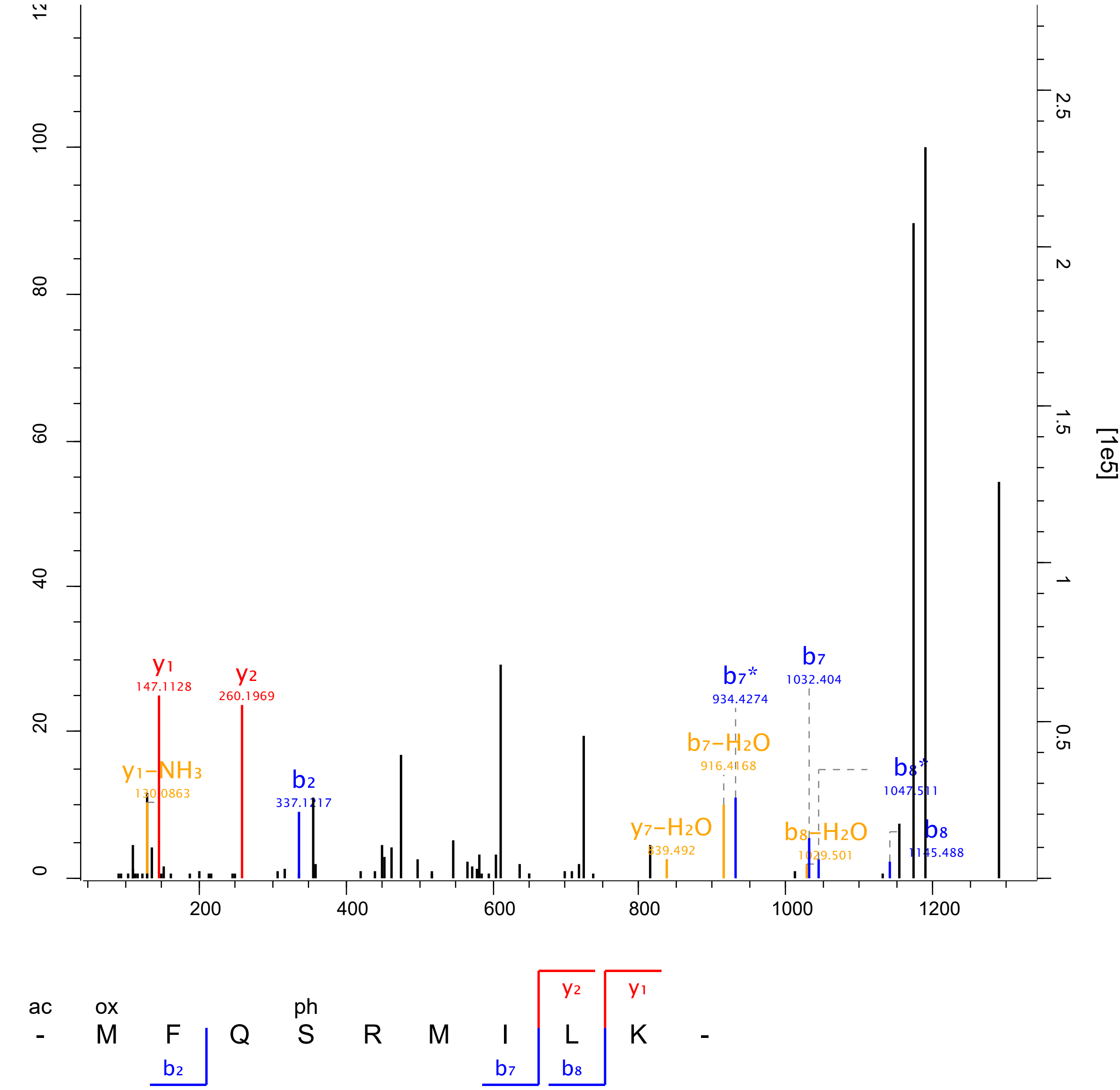

| Raw file | Scan | Method | Score | m/z |
|---|---|---|---|---|
| A17_05_2 | 5663 | FTMS; HCD | 107.06 | 551.73 |

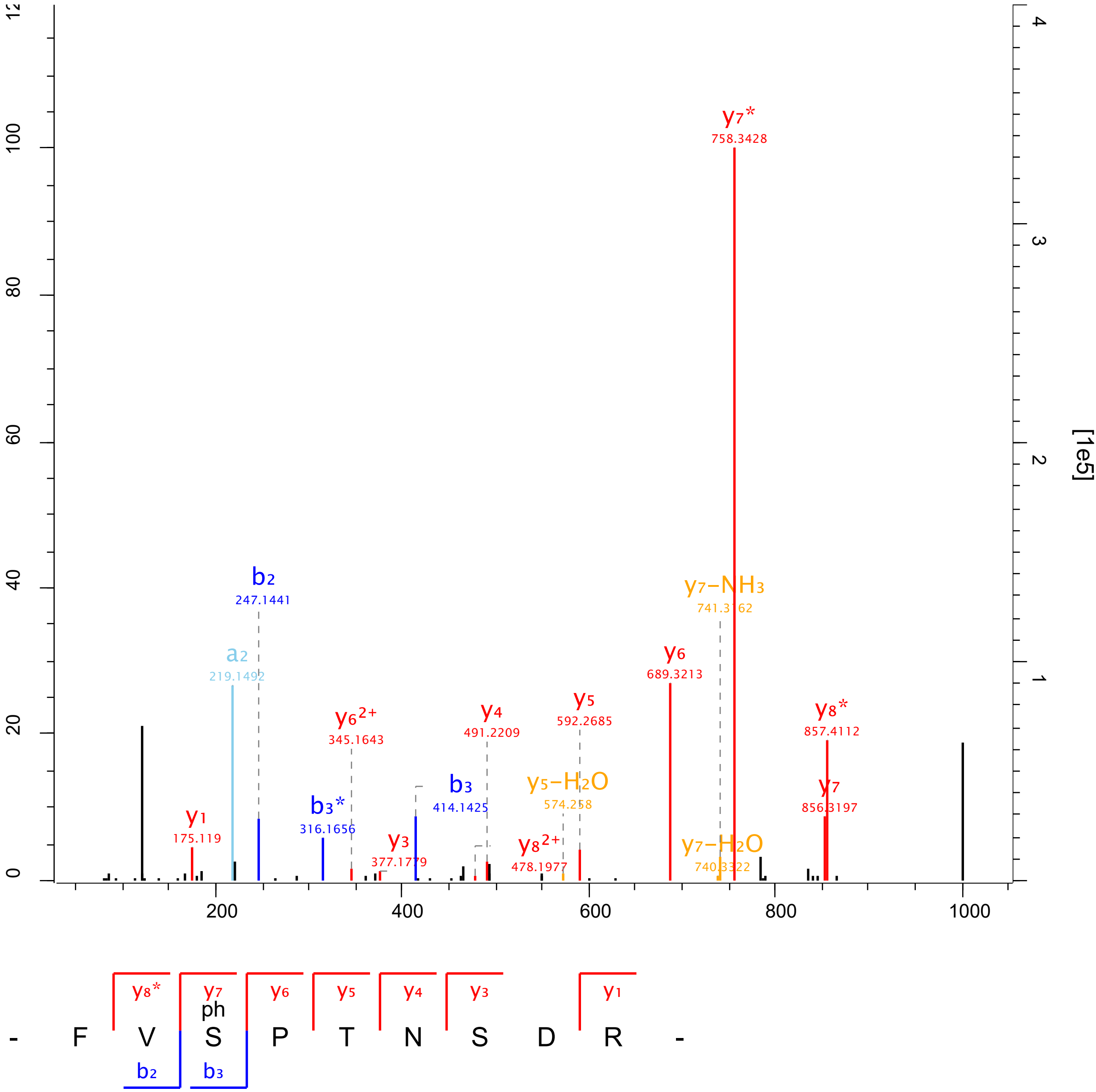

| Raw file | Scan | Method | Score | m/z |
|---|---|---|---|---|
| A17_15_1 | 10153 | FTMS; HCD | 92.26 | 743.83 |

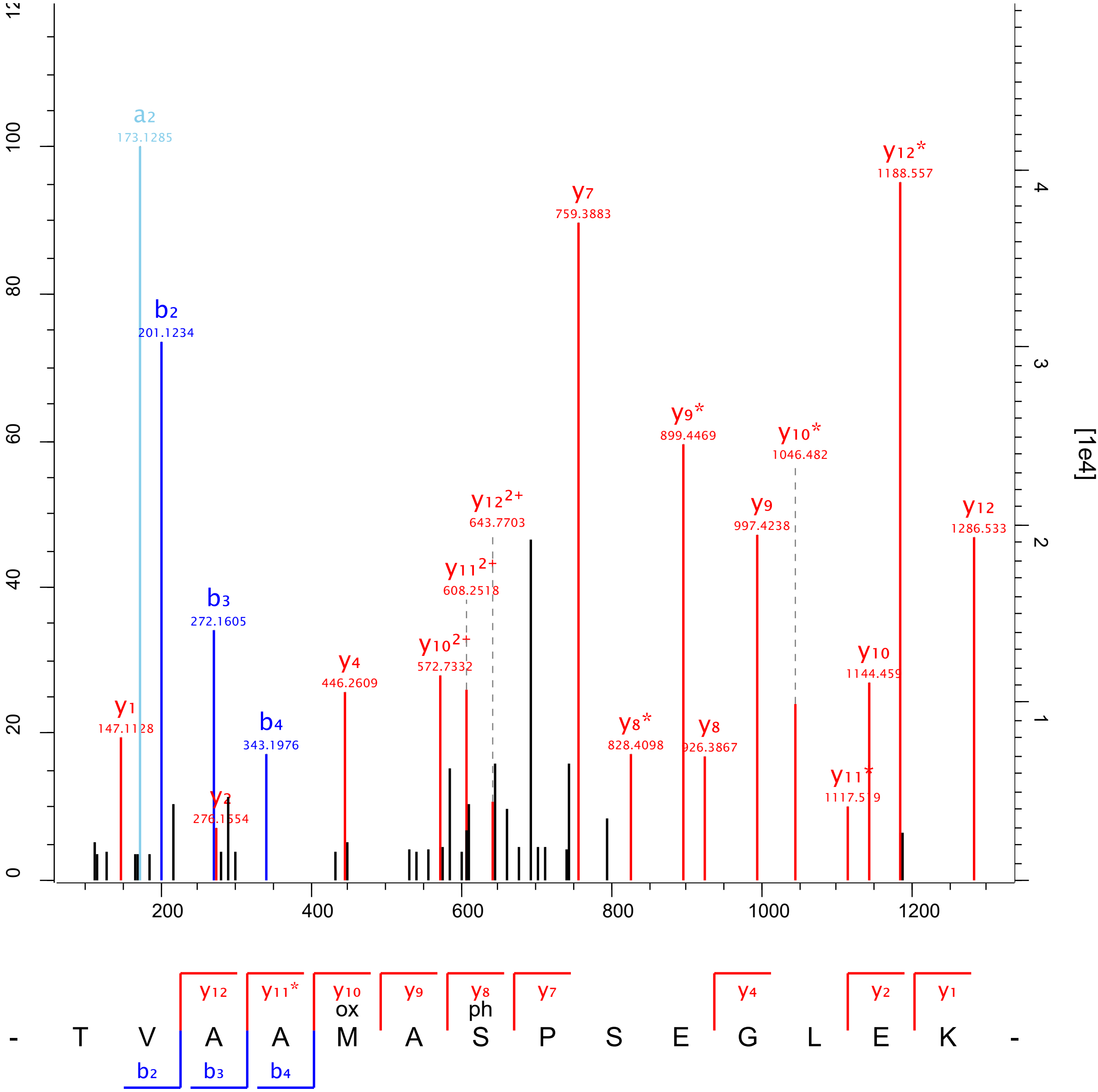

| Raw file | Scan | Method | Score | m/z |
|---|---|---|---|---|
| A17_15_1 | 10169 | FTMS; HCD | 56.57 | 677.77 |

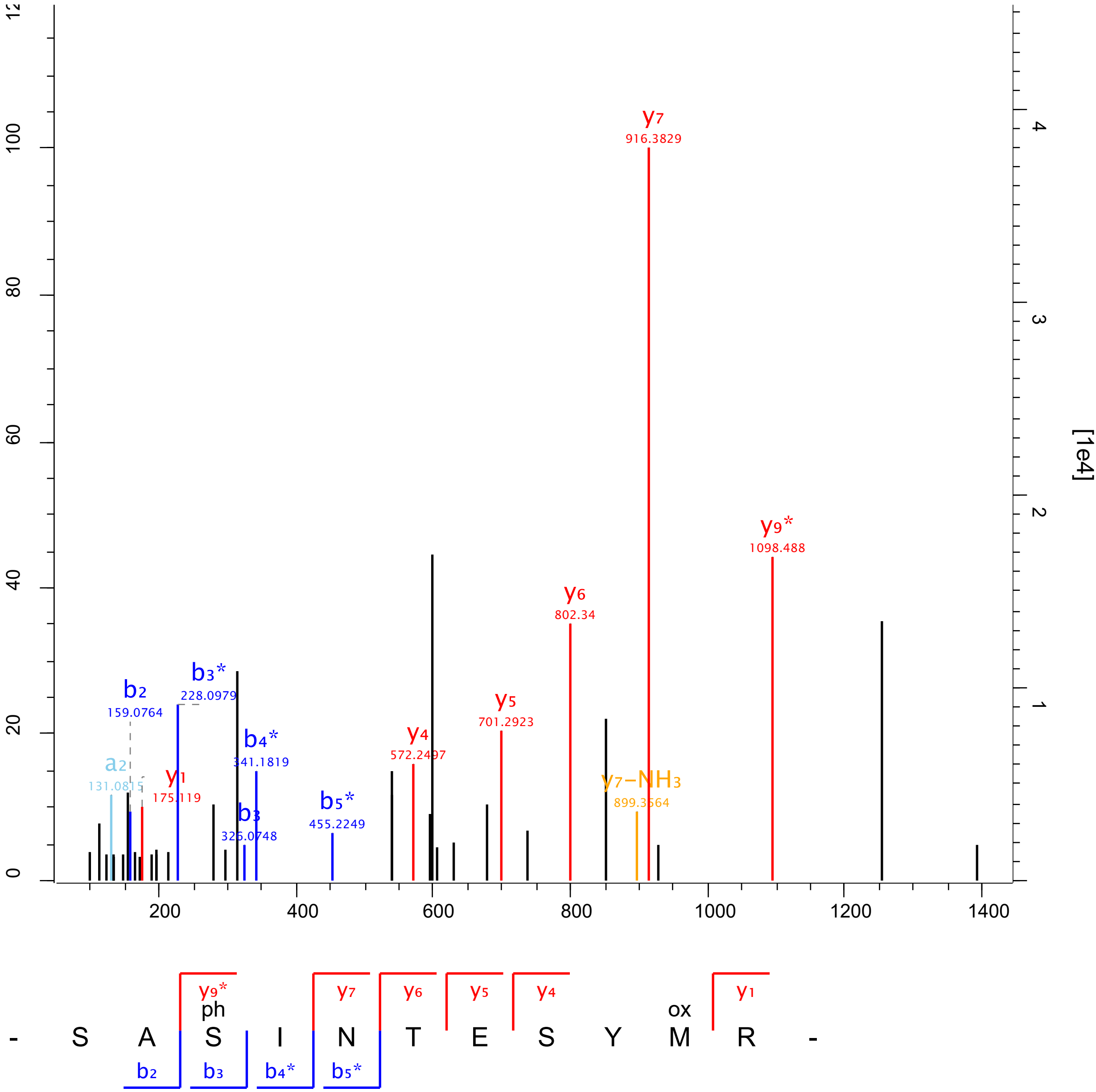

| Raw file | Scan | Method | Score | m/z |
| --- | --- | --- | --- | --- |
| A17_15_1 | 10345 | FTMS; HCD | 130.2 | 636.78 |

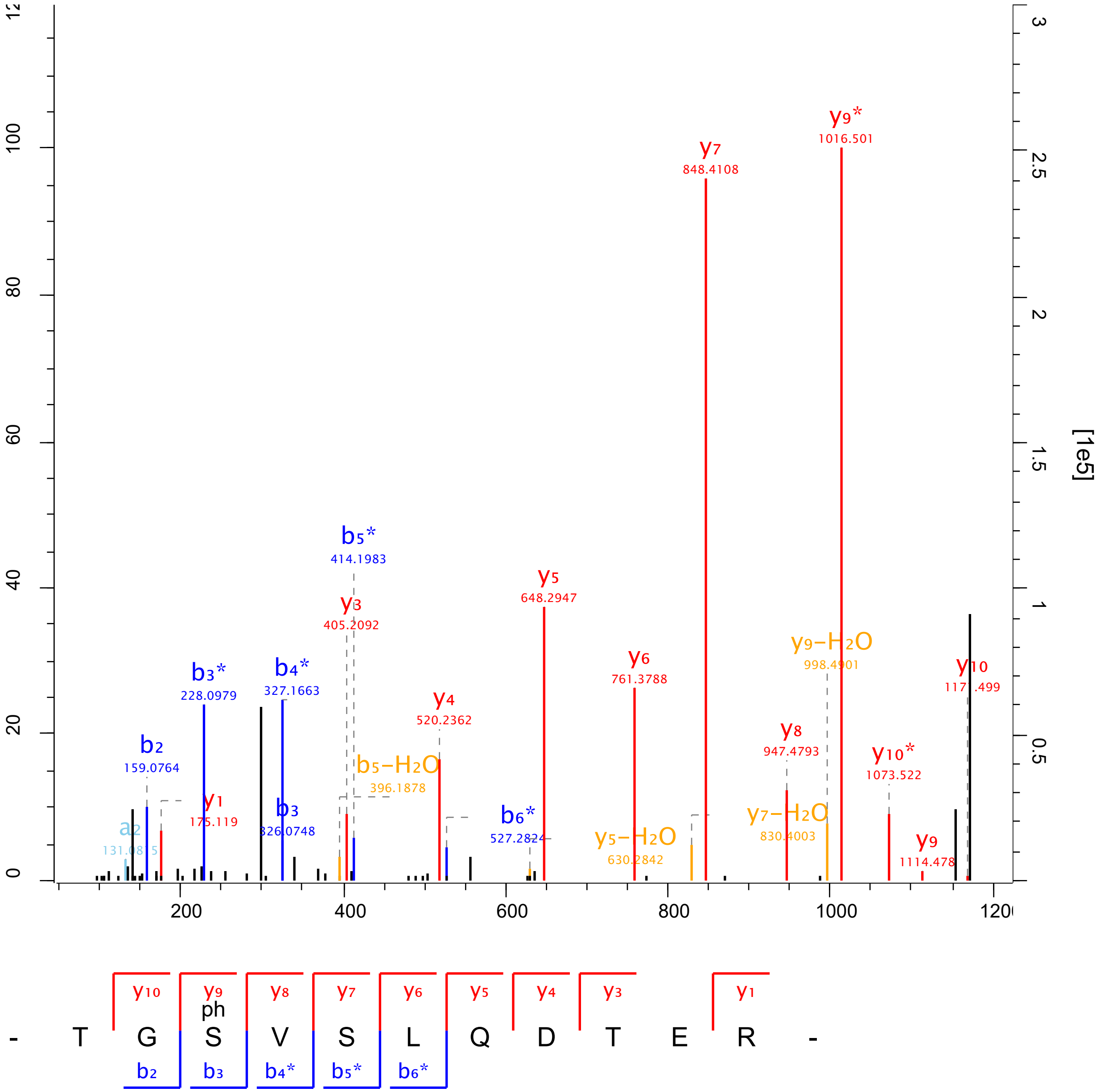

| Raw file | Scan | Method | Score | m/z |
| --- | --- | --- | --- | --- |
| A17_15_1 | 10346 | FTMS; HCD | 86.55 | 680.26 |

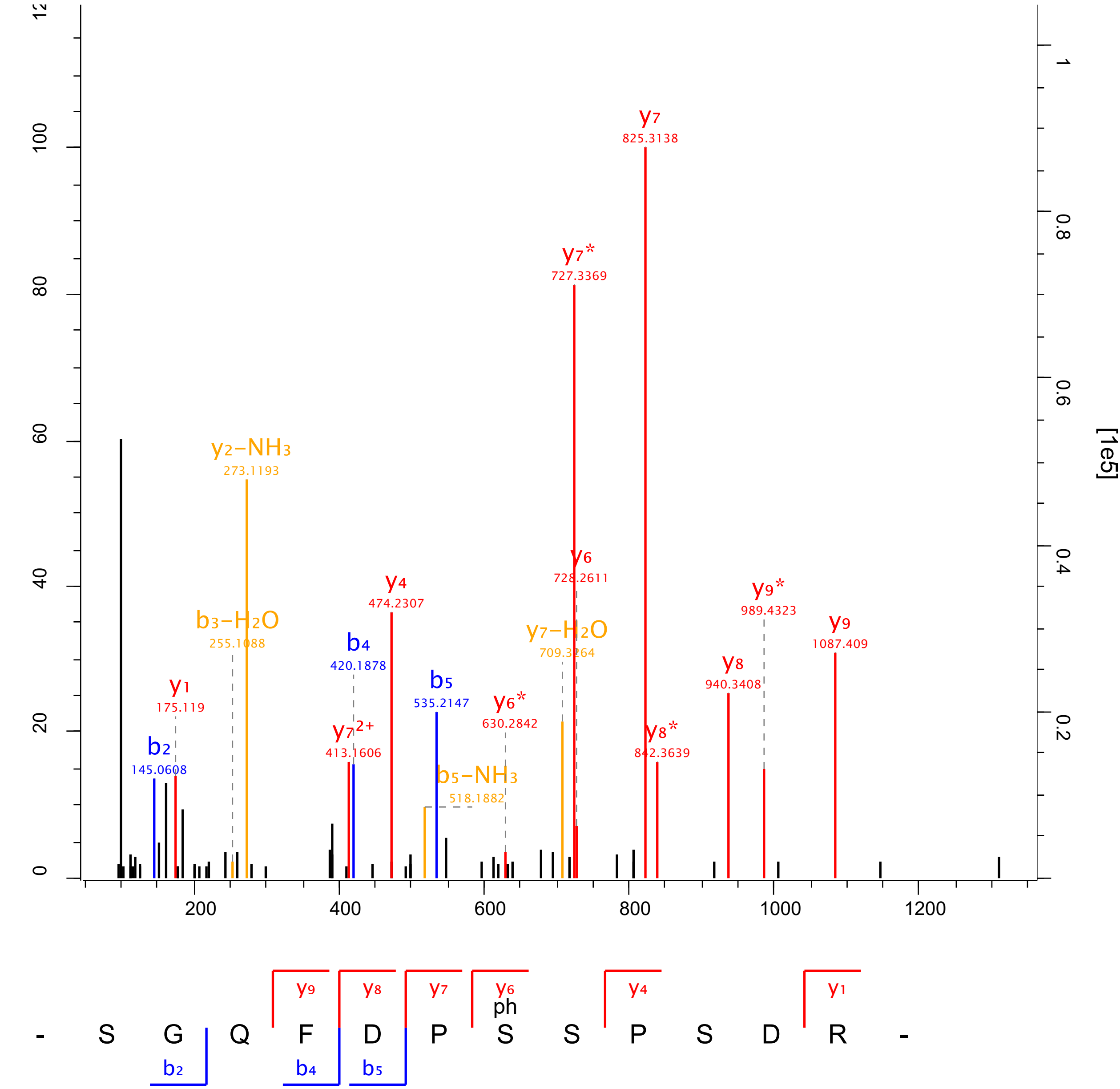

| Raw file | Scan | Method | Score | m/z |
|---|---|---|---|---|
| A17_15_1 | 10542 | FTMS; HCD | 198.49 | 877.38 |

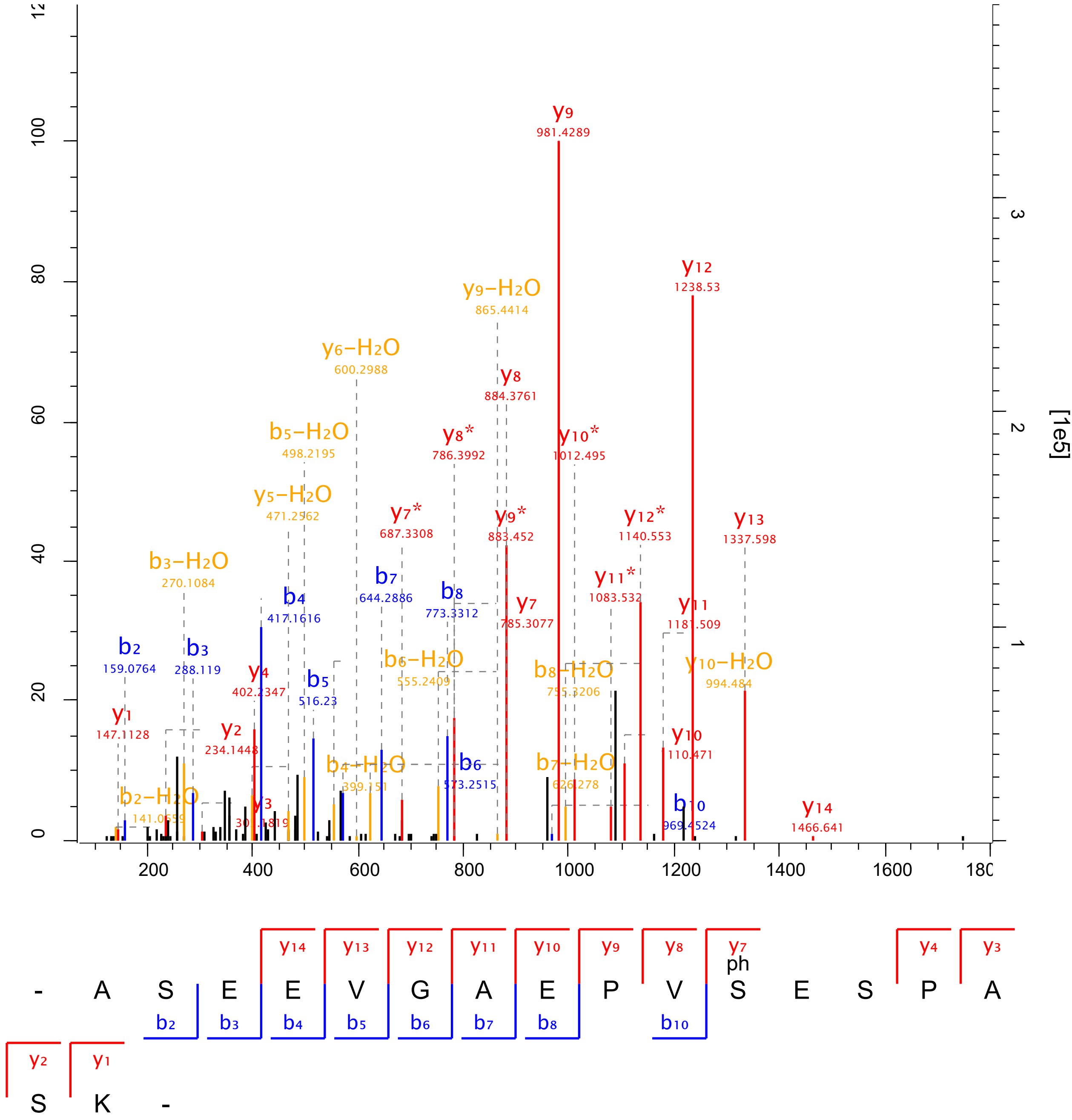

| Raw file | Scan | Method | Score | m/z |
|---|---|---|---|---|
| A17_15_1 | 10719 | FTMS; HCD | 234.04 | 866.8 |

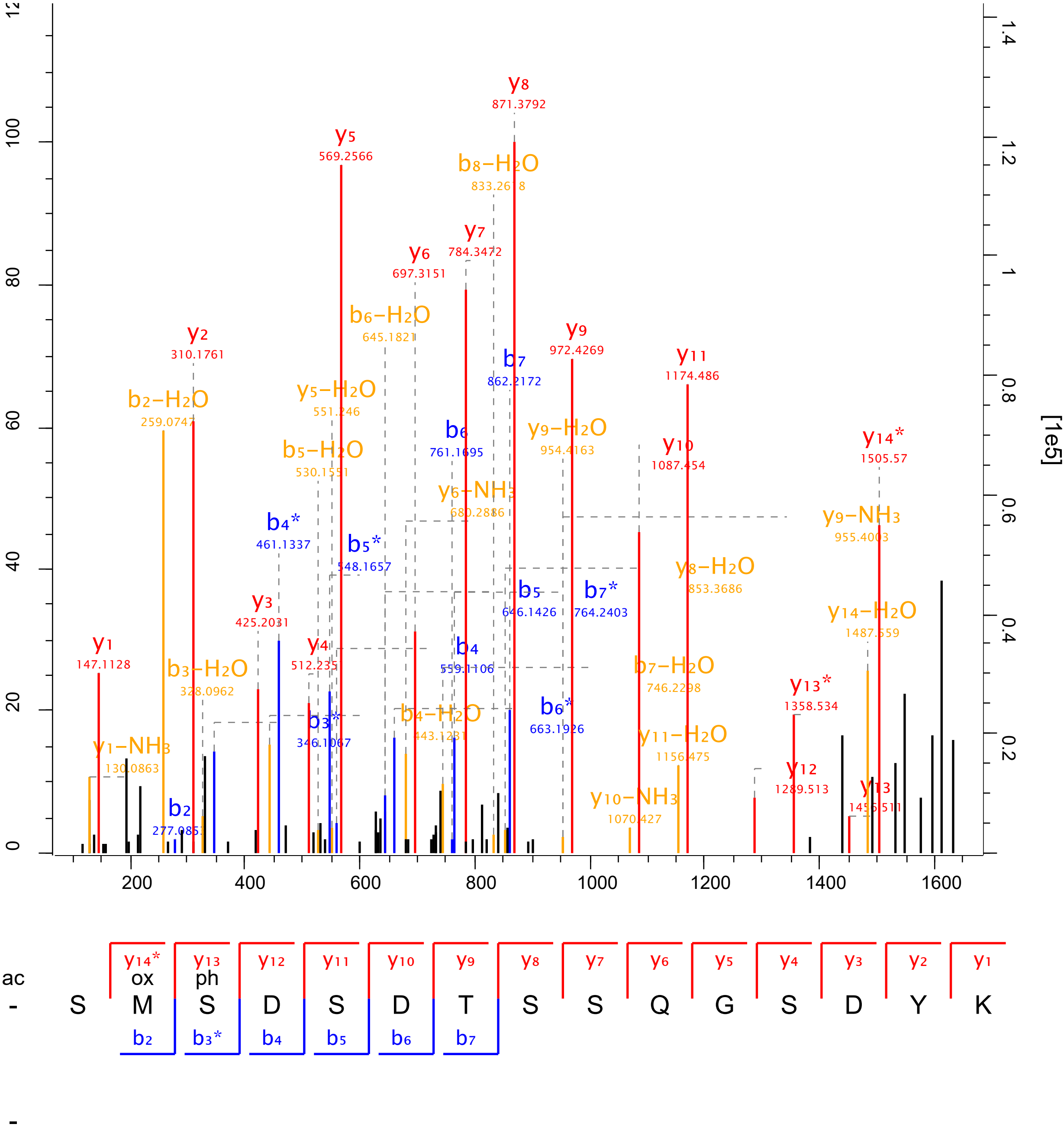

| Raw file | Scan | Method | Score | m/z |
| --- | --- | --- | --- | --- |
| A17_15_1 | 10949 | FTMS; HCD | 63.3 | 579.56 |

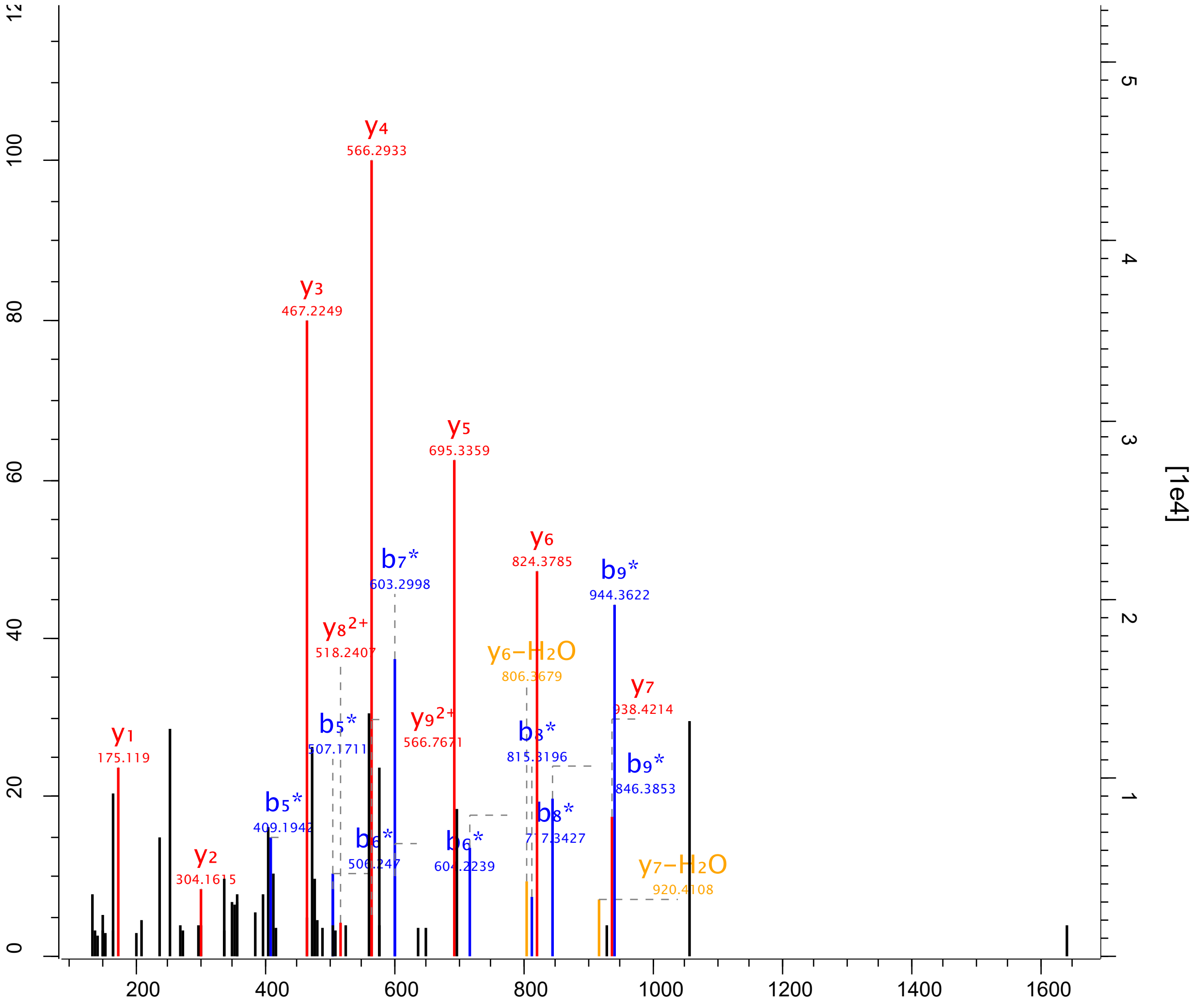


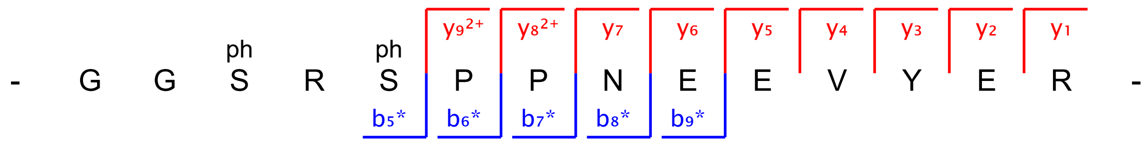

| Raw file | Scan | Method | Score | m/z |
|---|---|---|---|---|
| A17_15_1 | 11105 | FTMS; HCD | 78.71 | 825.29 |

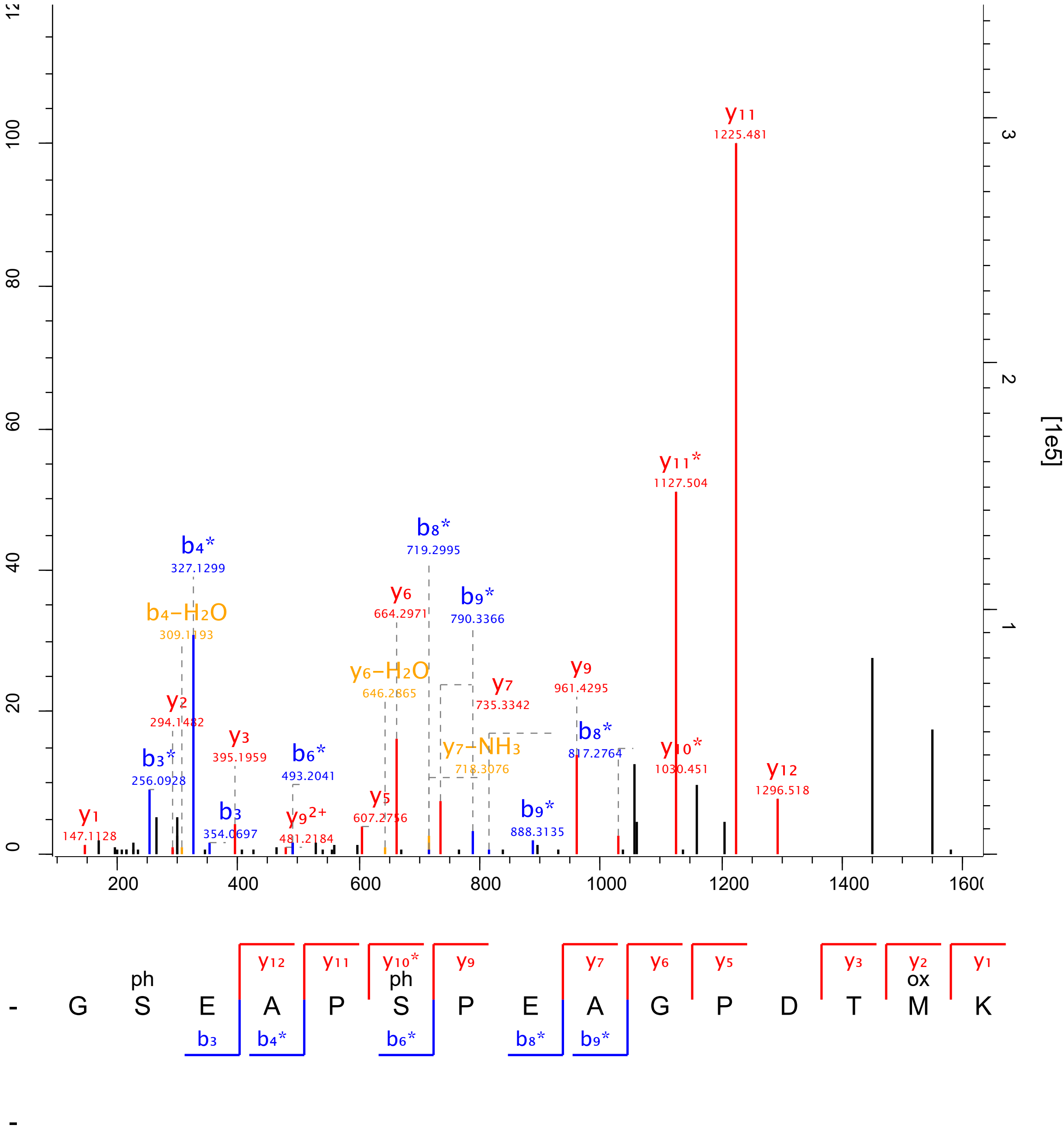

| Raw file | Scan | Method | Score | m/z |
|---|---|---|---|---|
| A17_15_1 | 11235 | FTMS; HCD | 50.35 | 538.21 |

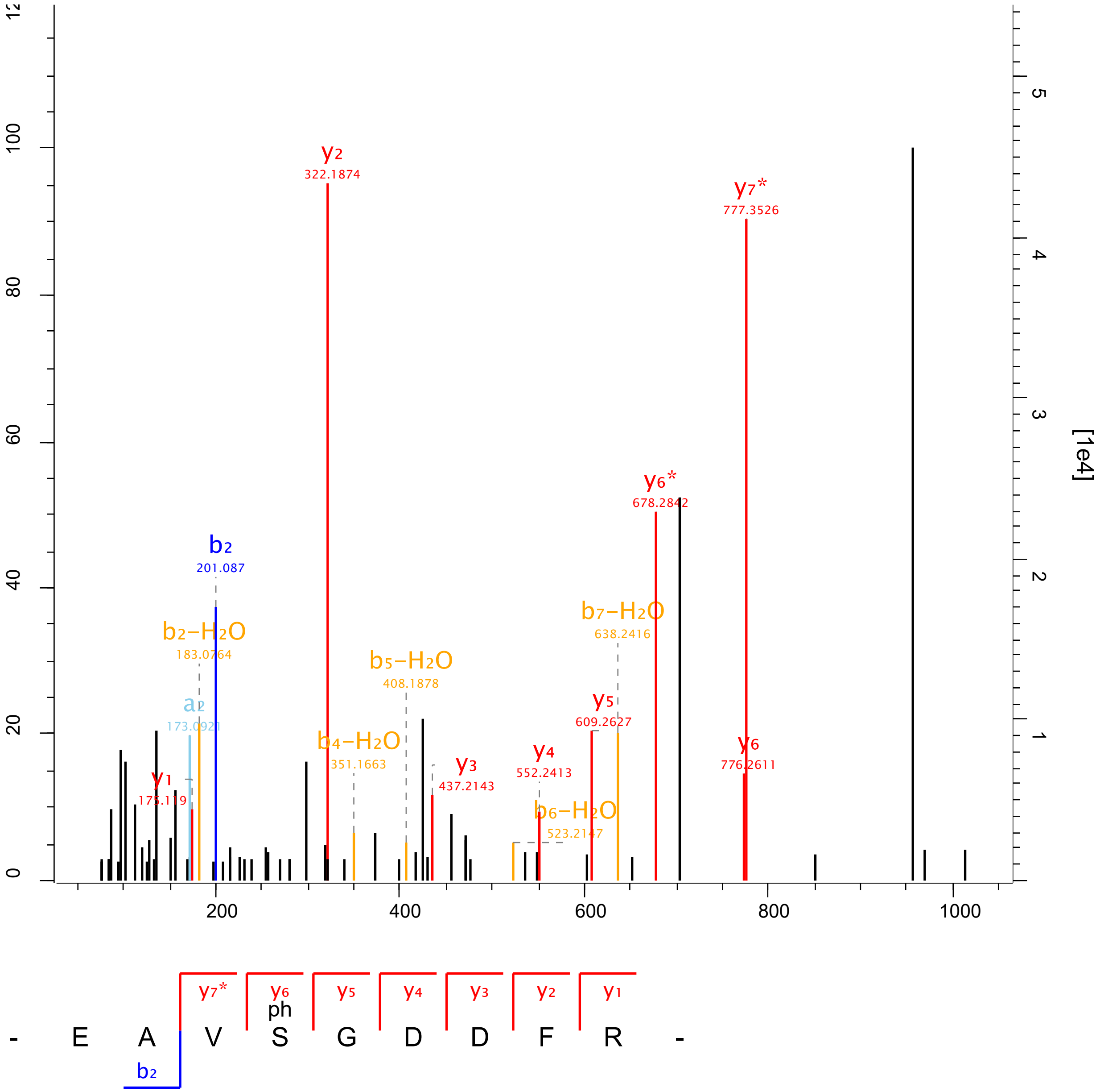

| Raw file | Scan | Method | Score | m/z |
|---|---|---|---|---|
| A17_15_1 | 11283 | FTMS; HCD | 96.07 | 604.28 |

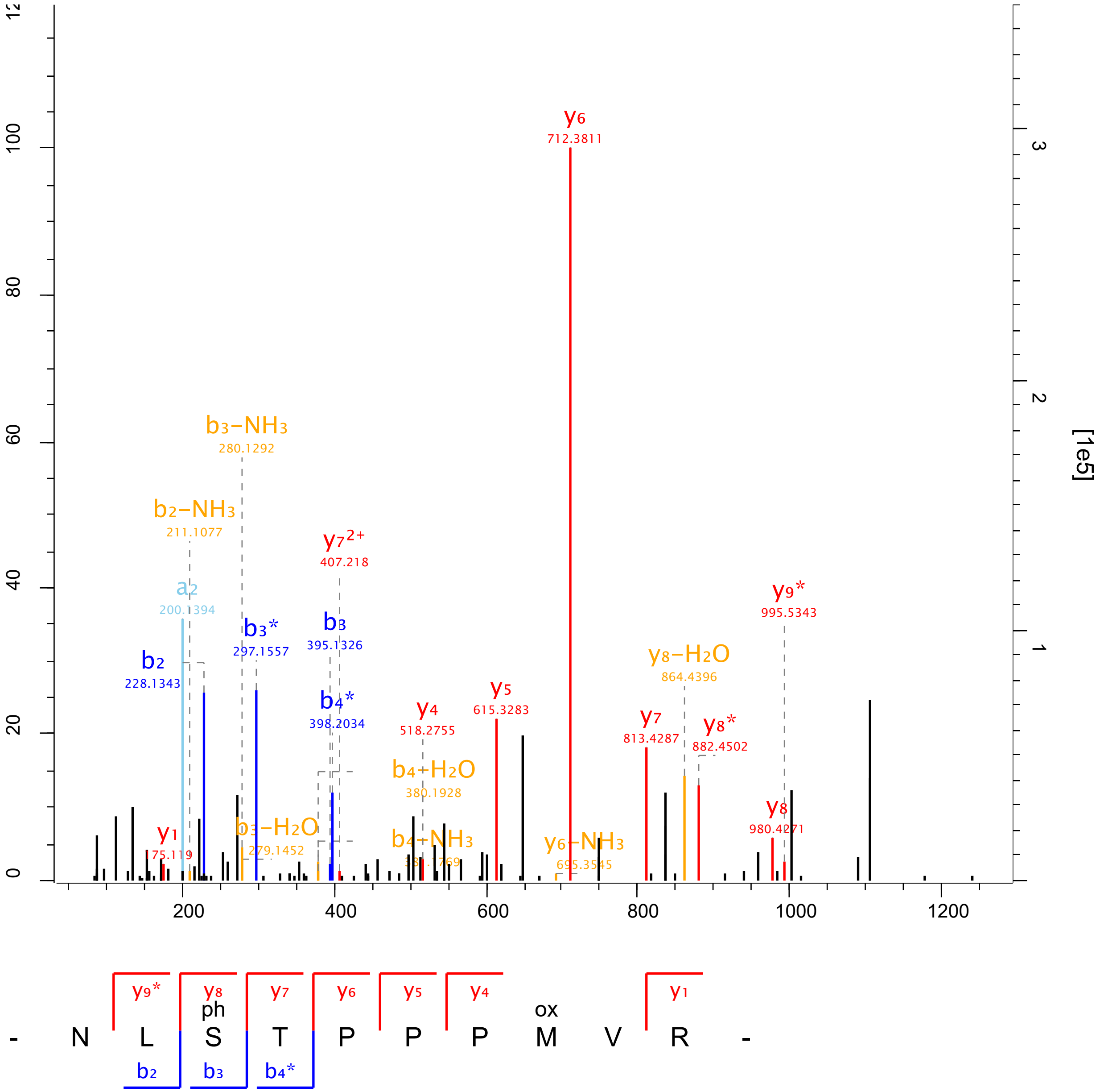

| Raw file | Scan | Method | Score | m/z |
| --- | --- | --- | --- | --- |
| A17_15_1 | 11292 | FTMS; HCD | 86.34 | 491.72 |

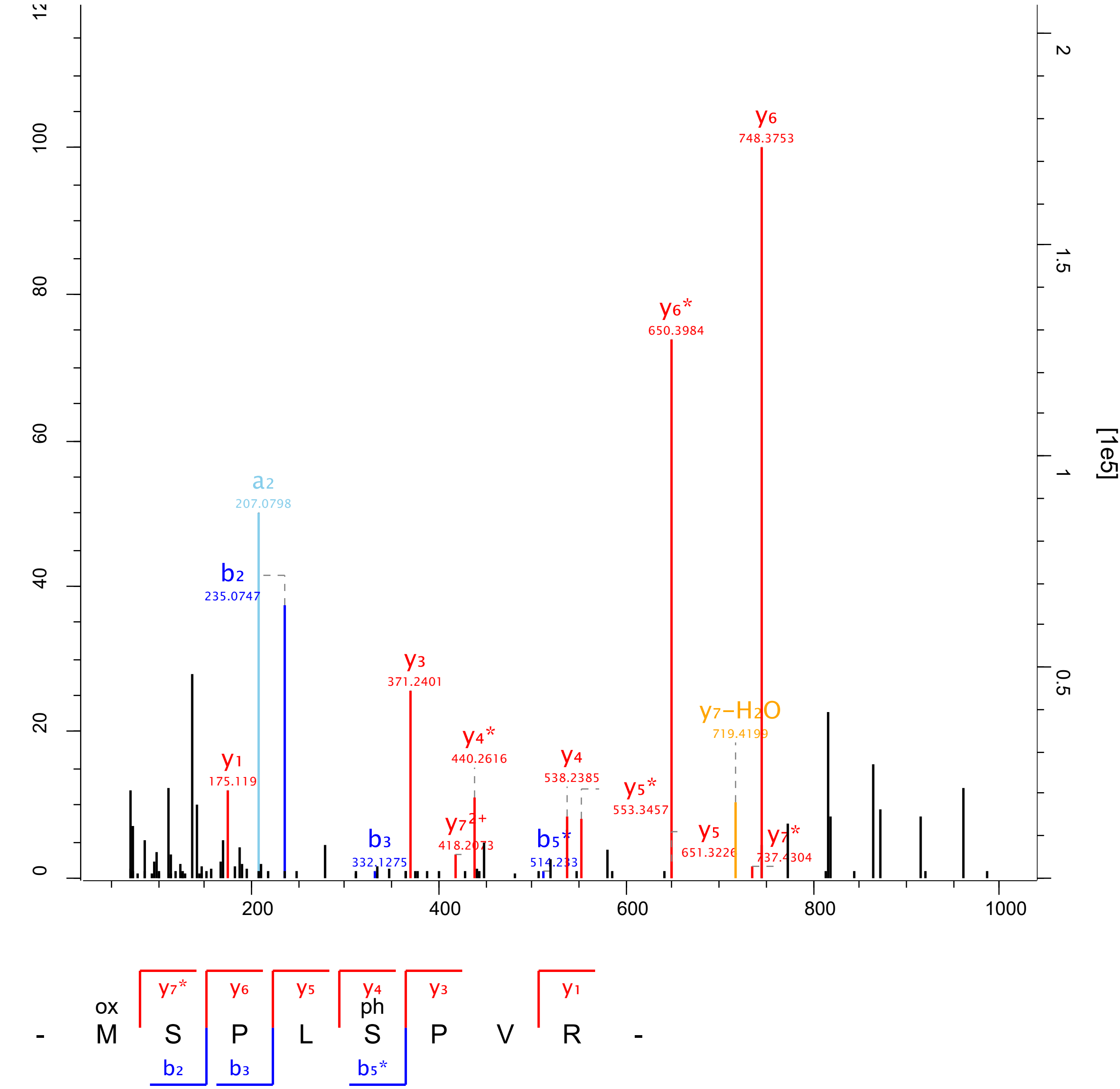

| Raw file | Scan | Method | Score | m/z |
|---|---|---|---|---|
| A17_15_1 | 11306 | FTMS; HCD | 266.39 | 795.34 |

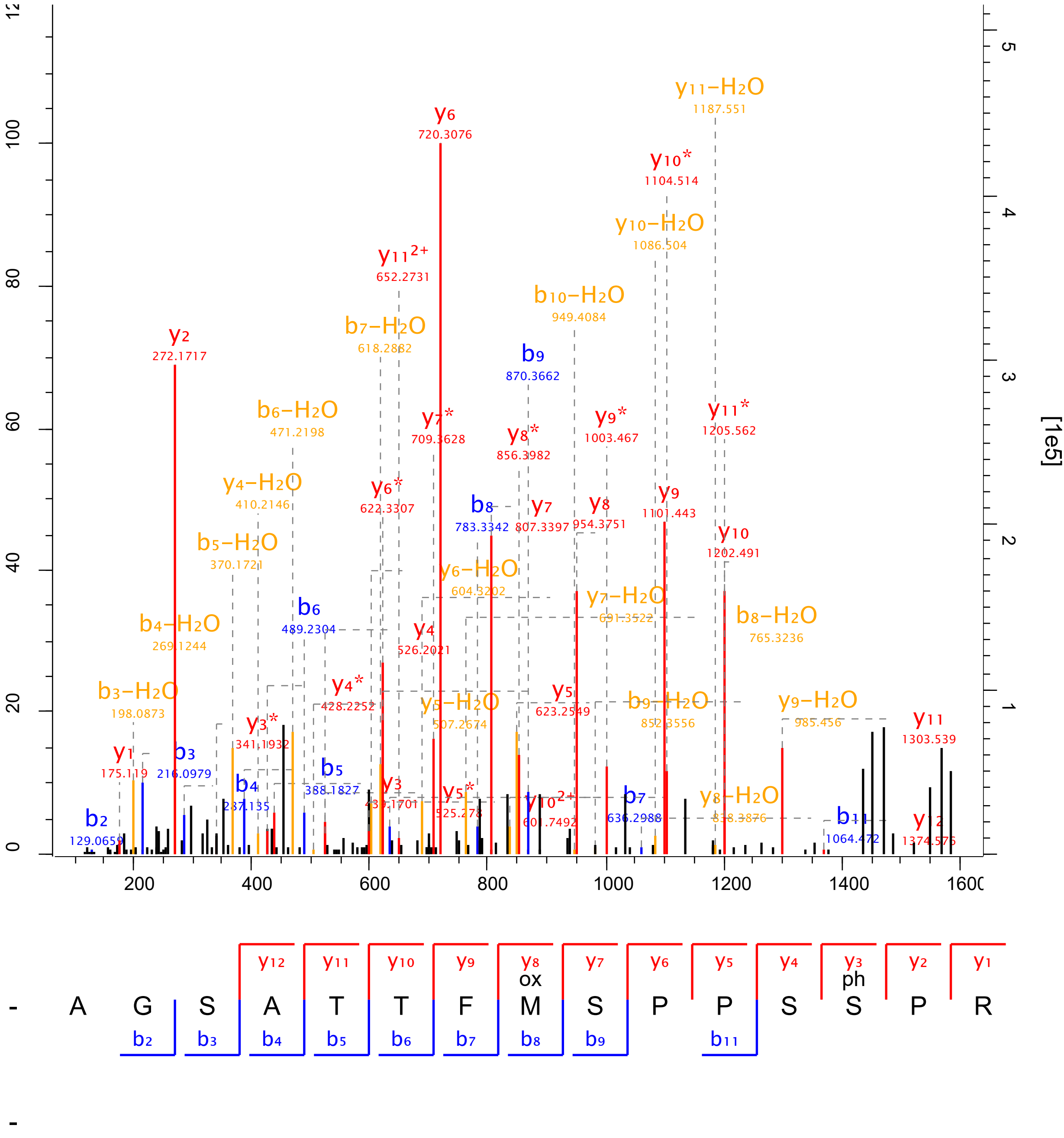

| Raw file | Scan | Method | Score | m/z |
|---|---|---|---|---|
| A17_15_1 | 11375 | FTMS; HCD | 62.81 | 797.85 |

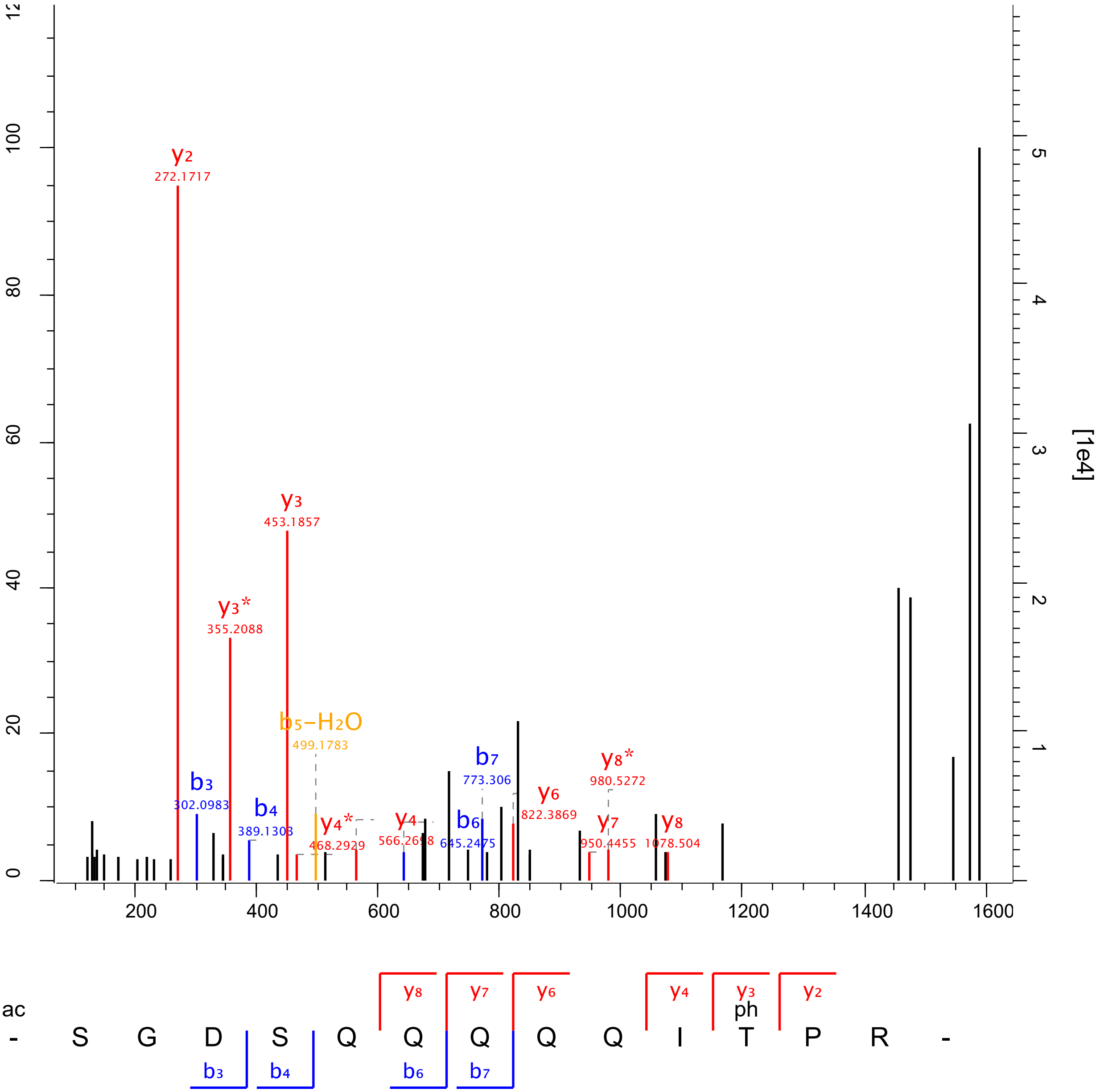

| Raw file | Scan | Method | Score | m/z |
|---|---|---|---|---|
| A17_15_1 | 11382 | FTMS; HCD | 86.77 | 615.26 |

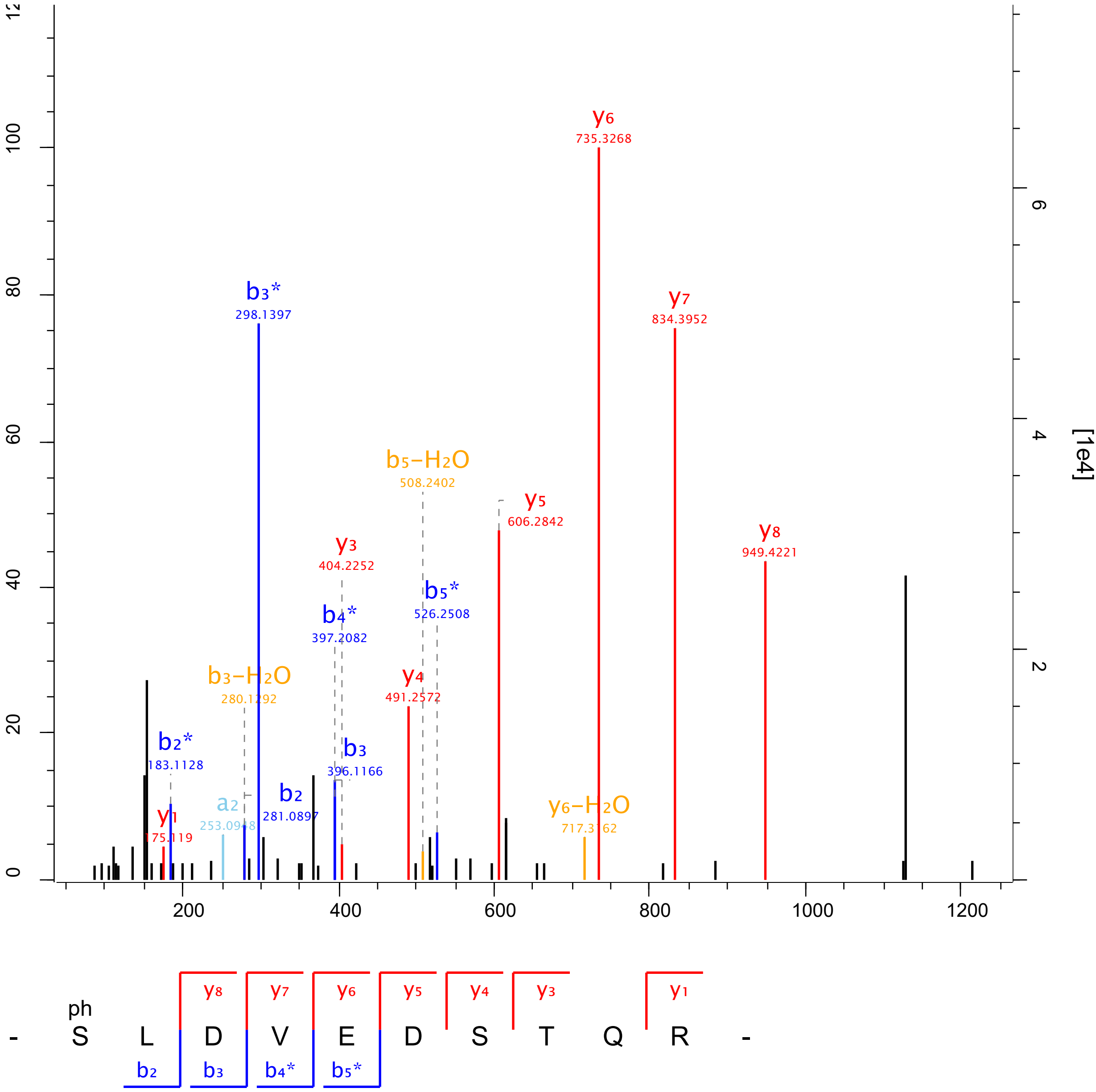

| Raw file | Scan | Method | Score | m/z |
| --- | --- | --- | --- | --- |
| A17_15_1 | 11572 | FTMS; HCD | 73.44 | 525.69 |

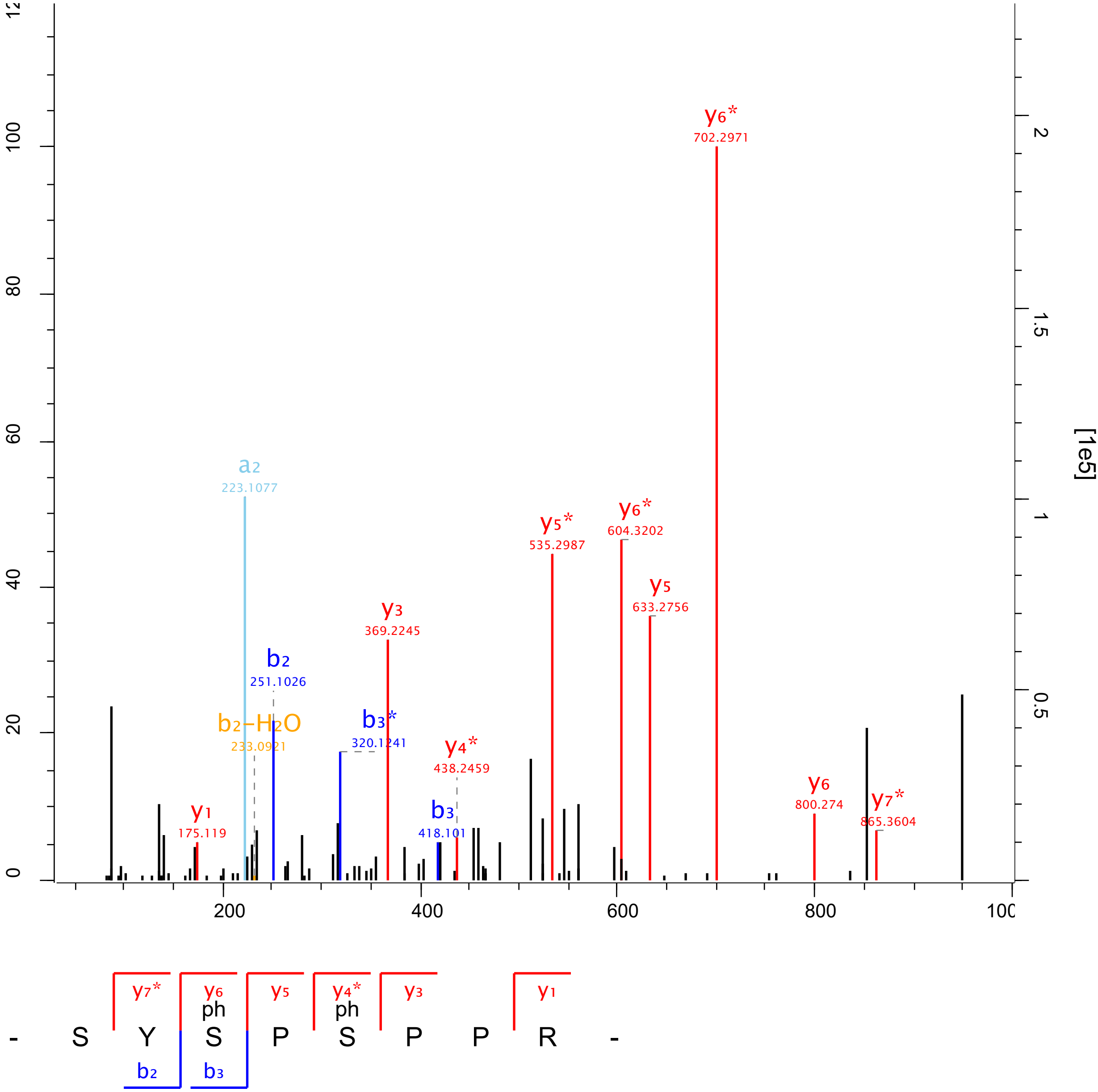

| Raw file | Scan | Method | Score | m/z |
| --- | --- | --- | --- | --- |
| A17_15_1 | 11597 | FTMS; HCD | 120.63 | 767.82 |

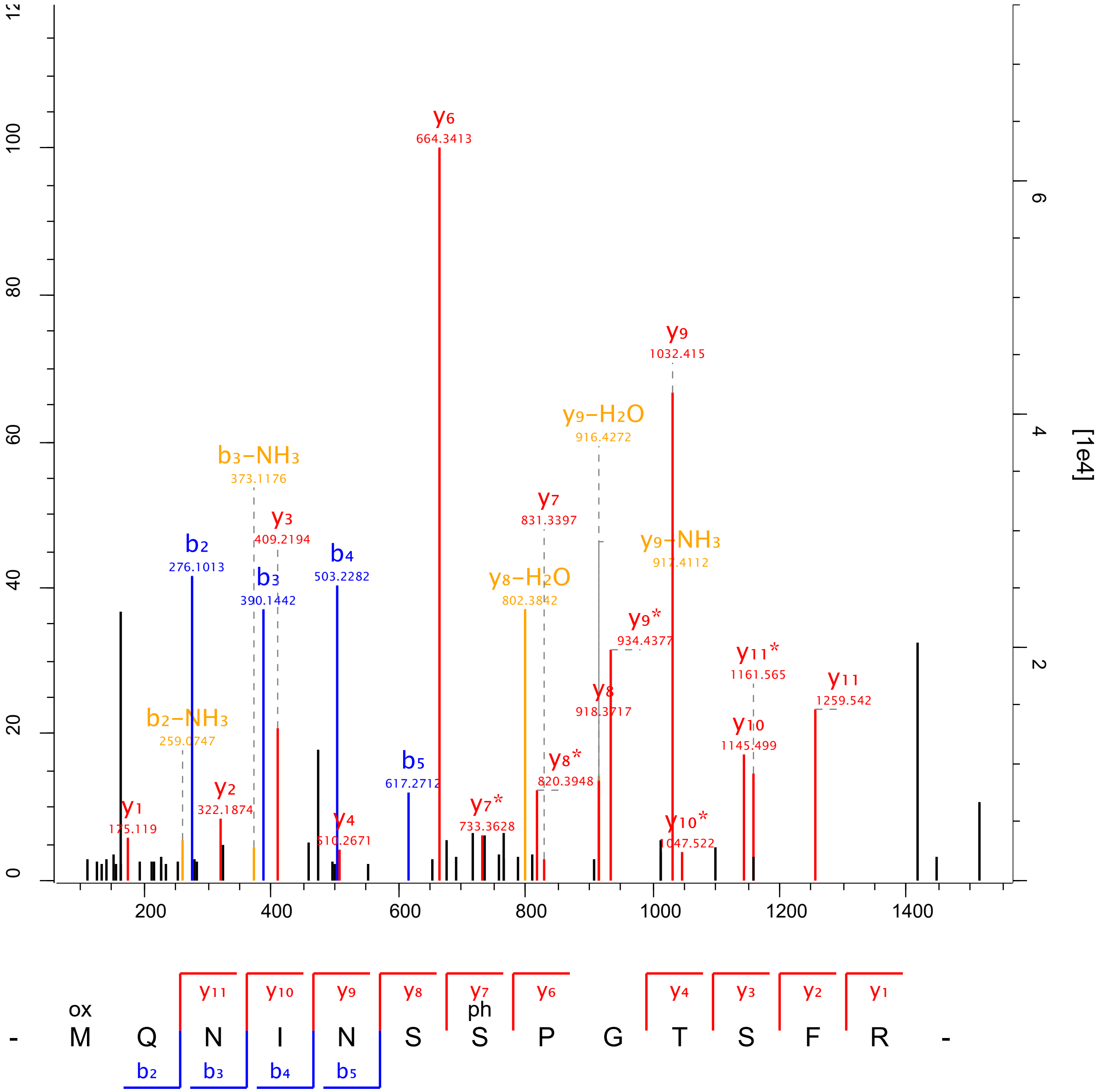

| Raw file | Scan | Method | Score | m/z |
|---|---|---|---|---|
| A17_15_1 | 11750 | FTMS; HCD | 53.56 | 800.84 |

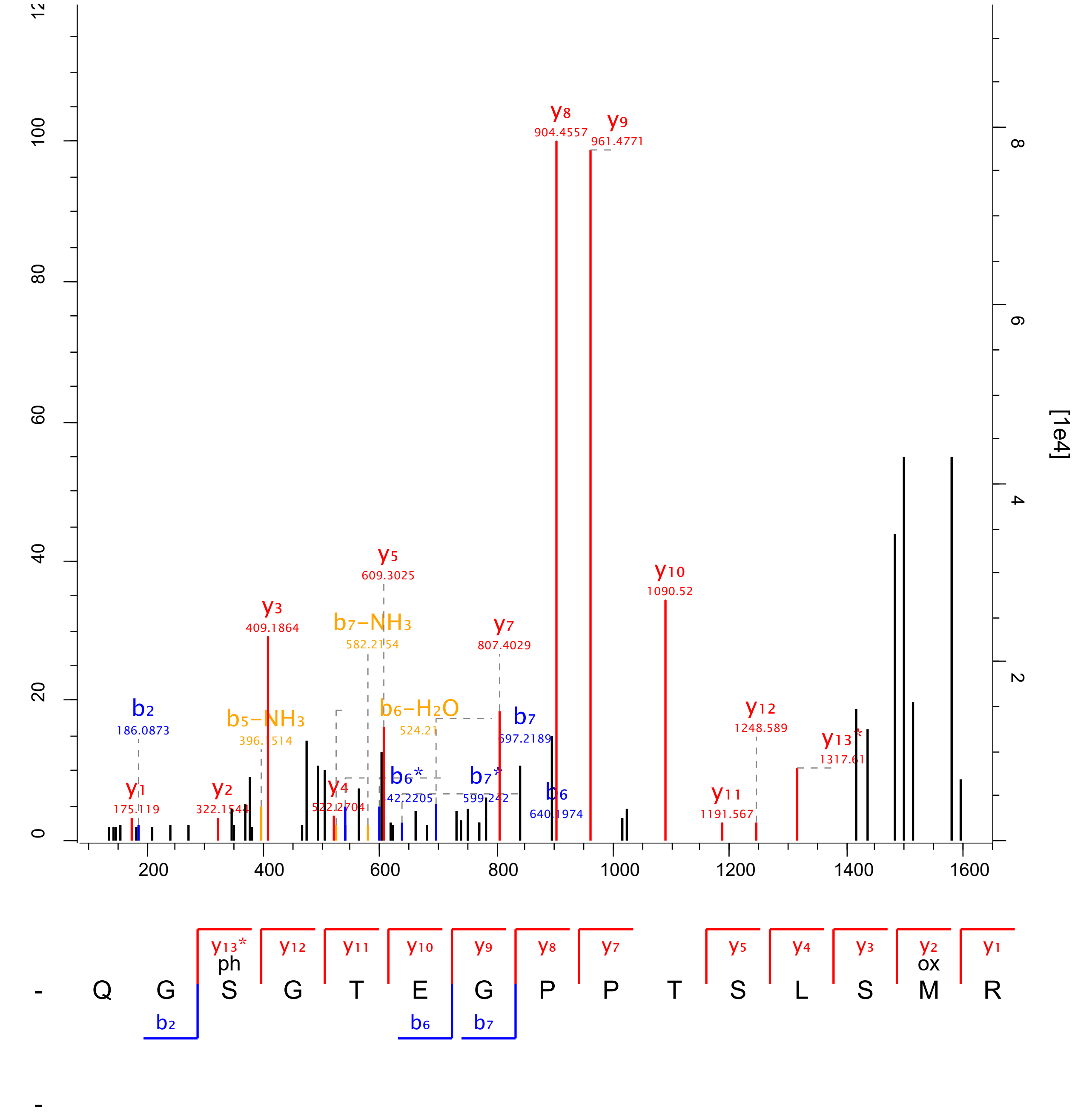

| Raw file | Scan | Method | Score | m/z |
|---|---|---|---|---|
| A17_15_1 | 11753 | FTMS; HCD | 46.07 | 622.77 |

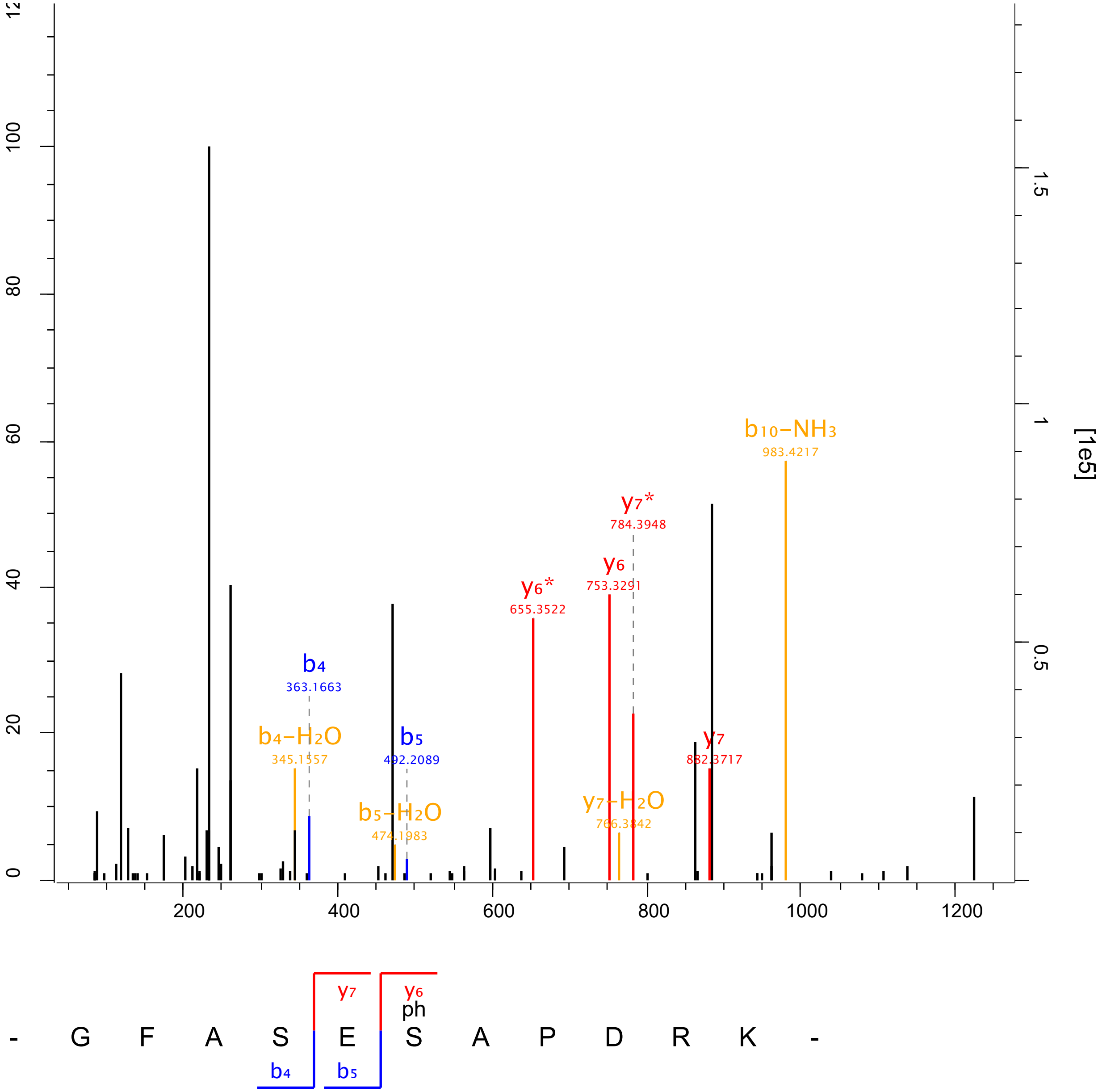

| Raw file | Scan | Method | Score | m/z |
| --- | --- | --- | --- | --- |
| A17_15_1 | 11808 | FTMS; HCD | 198.29 | 603.29 |

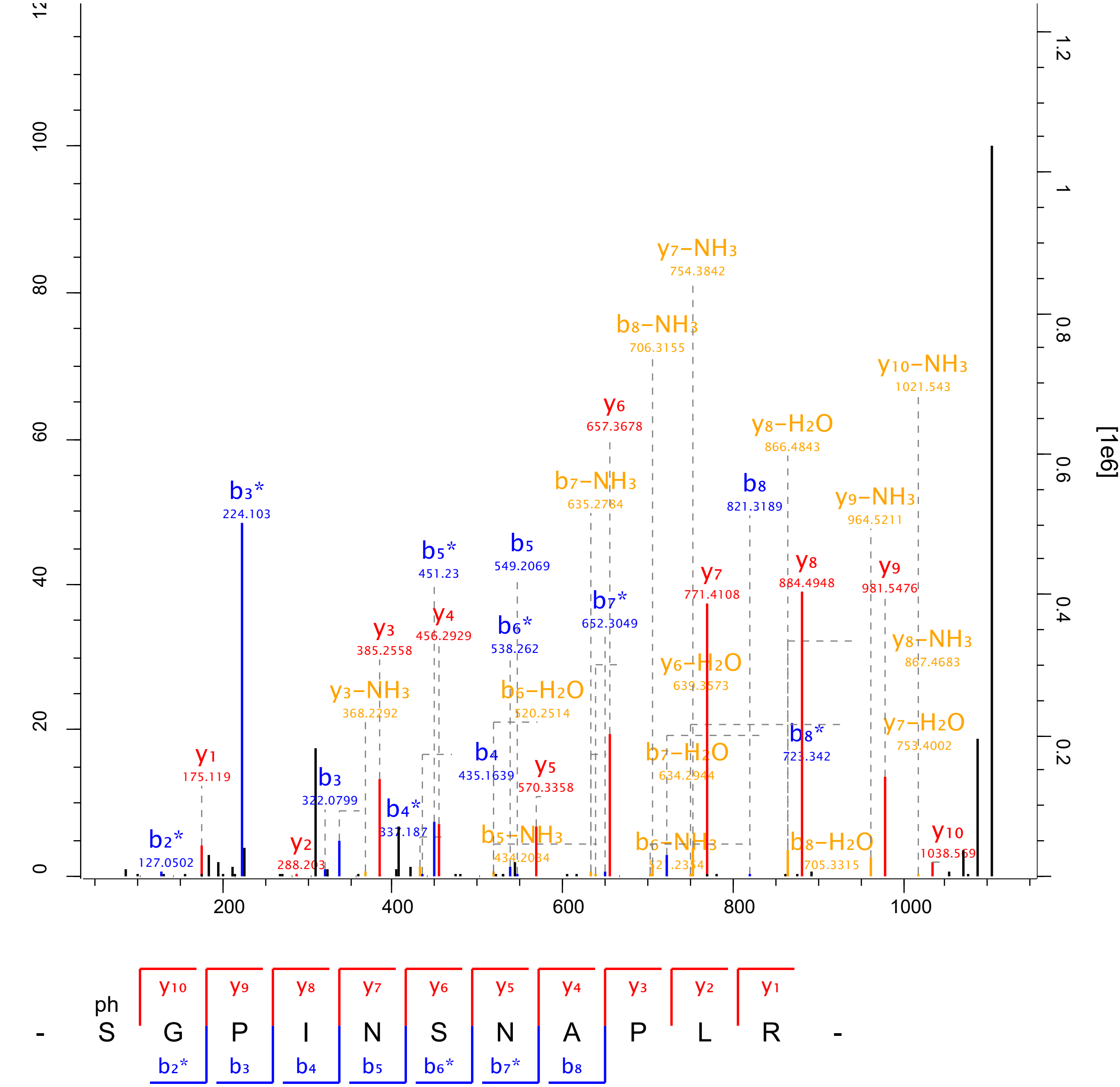

| Raw file | Scan | Method | Score | m/z |
|---|---|---|---|---|
| A17_15_1 | 11924 | FTMS; HCD | 56.72 | 718.79 |

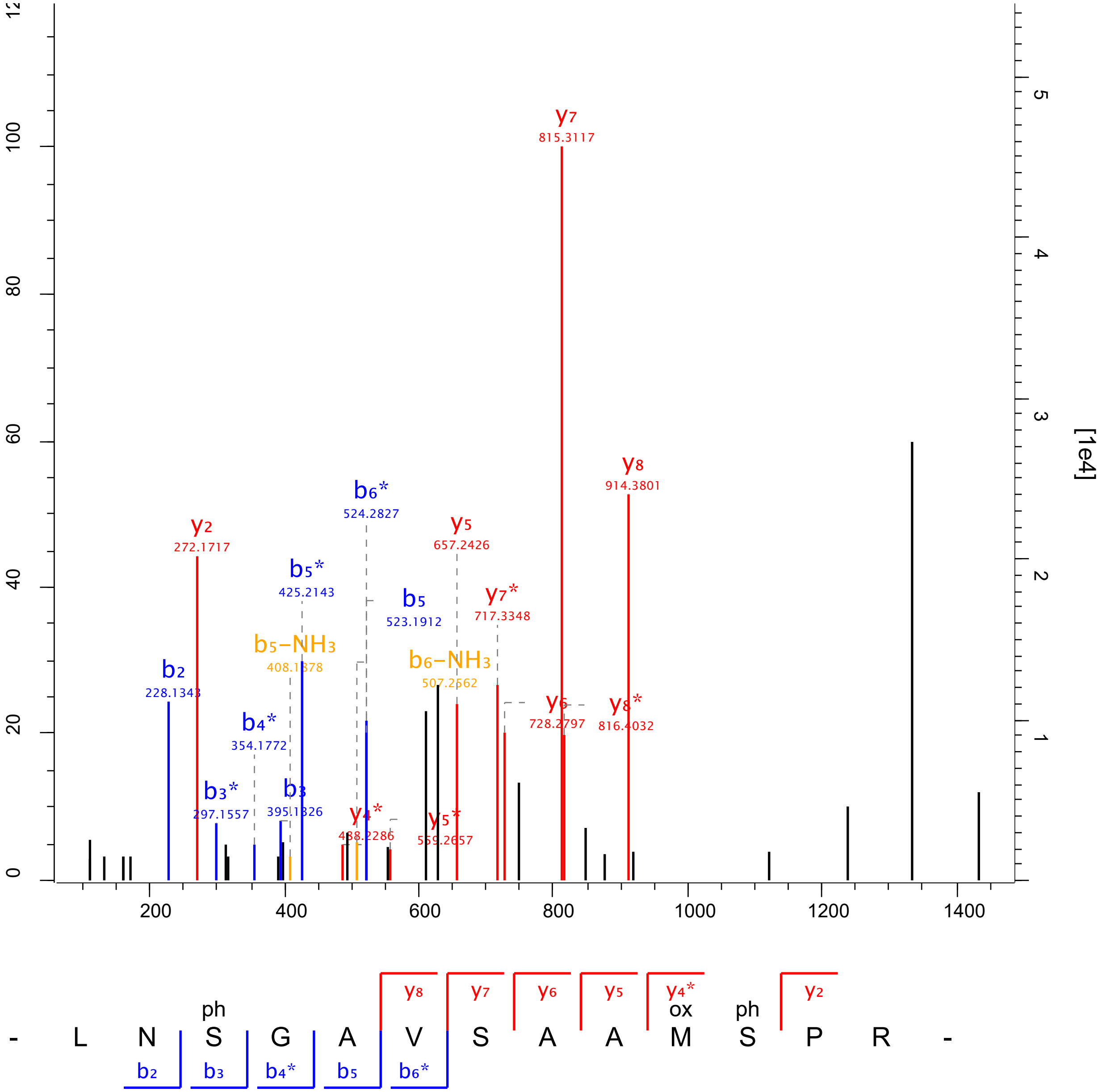

| Raw file | Scan | Method | Score | m/z |
|---|---|---|---|---|
| A17_15_1 | 11958 | FTMS; HCD | 96.38 | 823.85 |

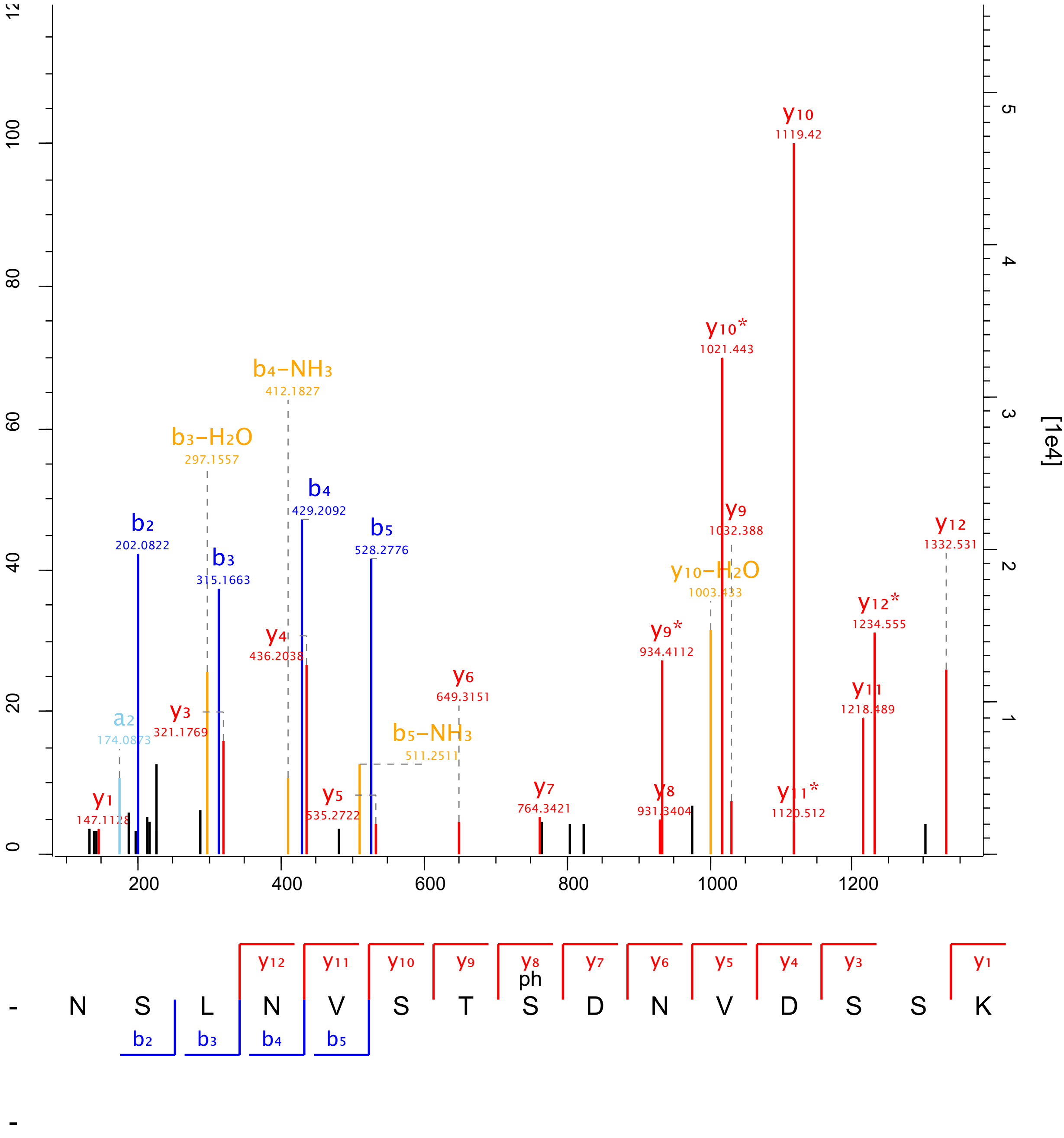

| Raw file | Scan | Method | Score | m/z |
| --- | --- | --- | --- | --- |
| A17_15_1 | 12057 | FTMS; HCD | 55.26 | 511.75 |

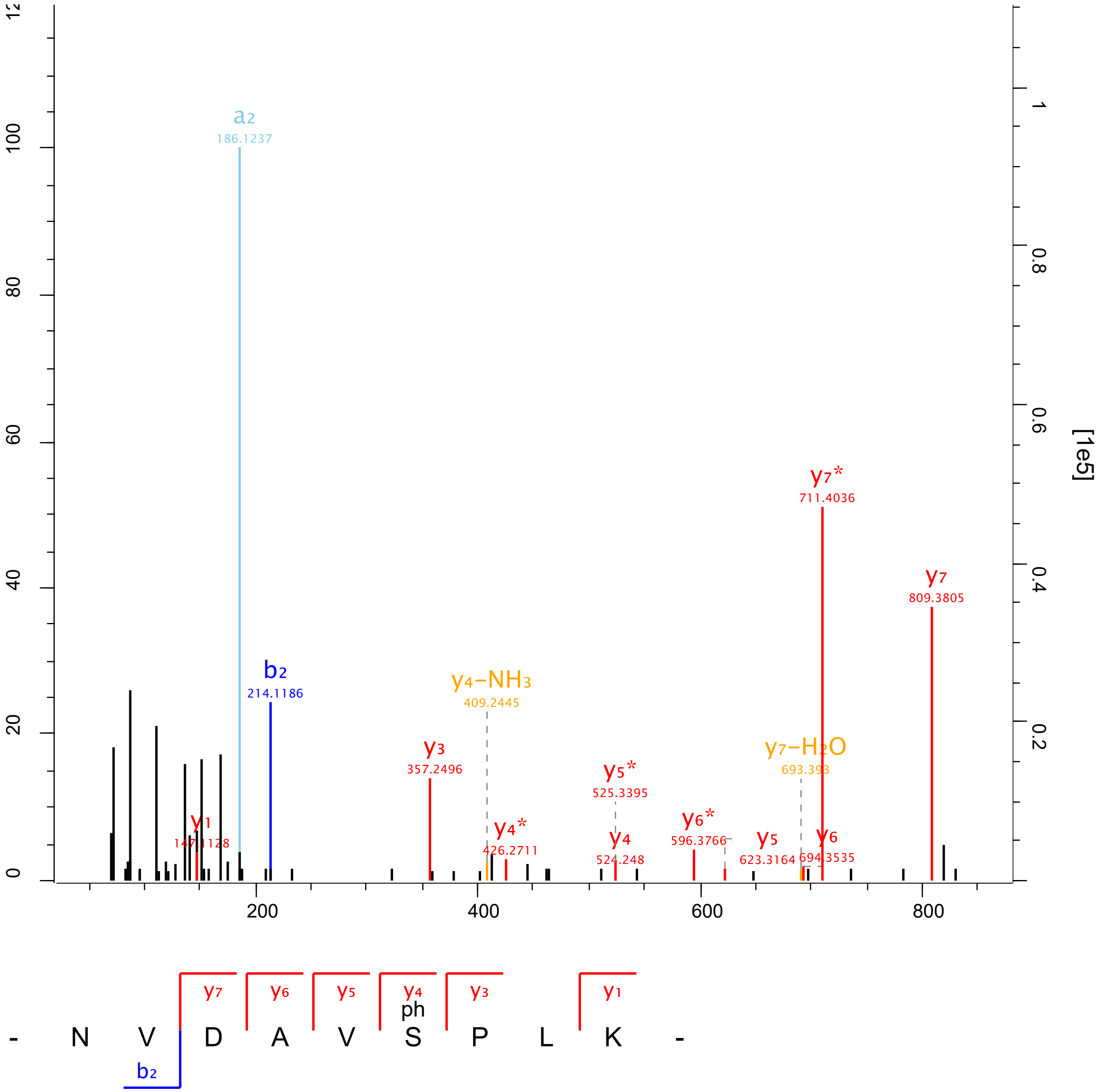

| Raw file | Scan | Method | Score | m/z |
| --- | --- | --- | --- | --- |
| A17_15_1 | 12133 | FTMS; HCD | 47.04 | 772.34 |

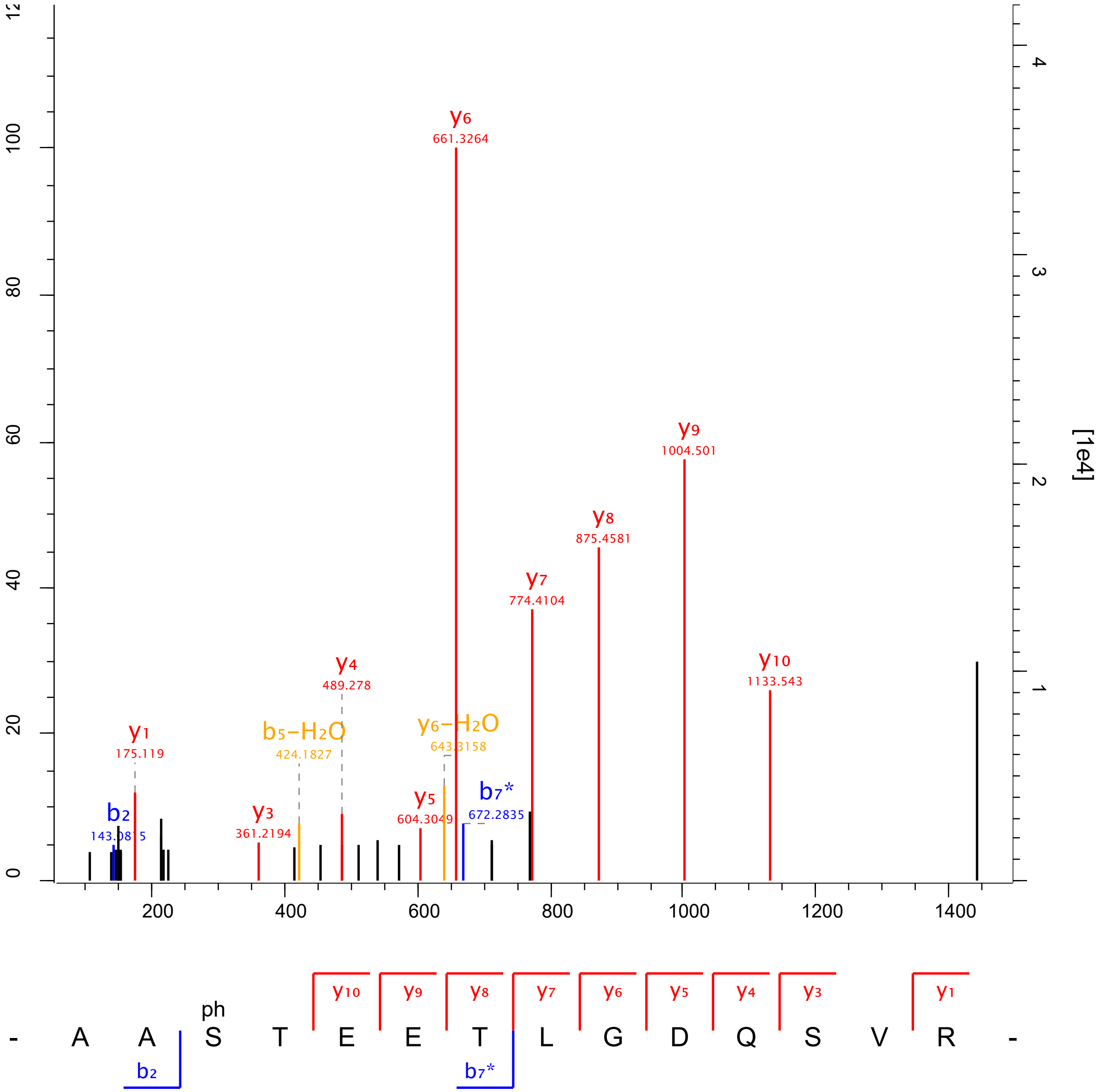

| Raw file | Scan | Method | Score | m/z |
|---|---|---|---|---|
| A17_15_1 | 12197 | FTMS; HCD | 114.99 | 759.79 |

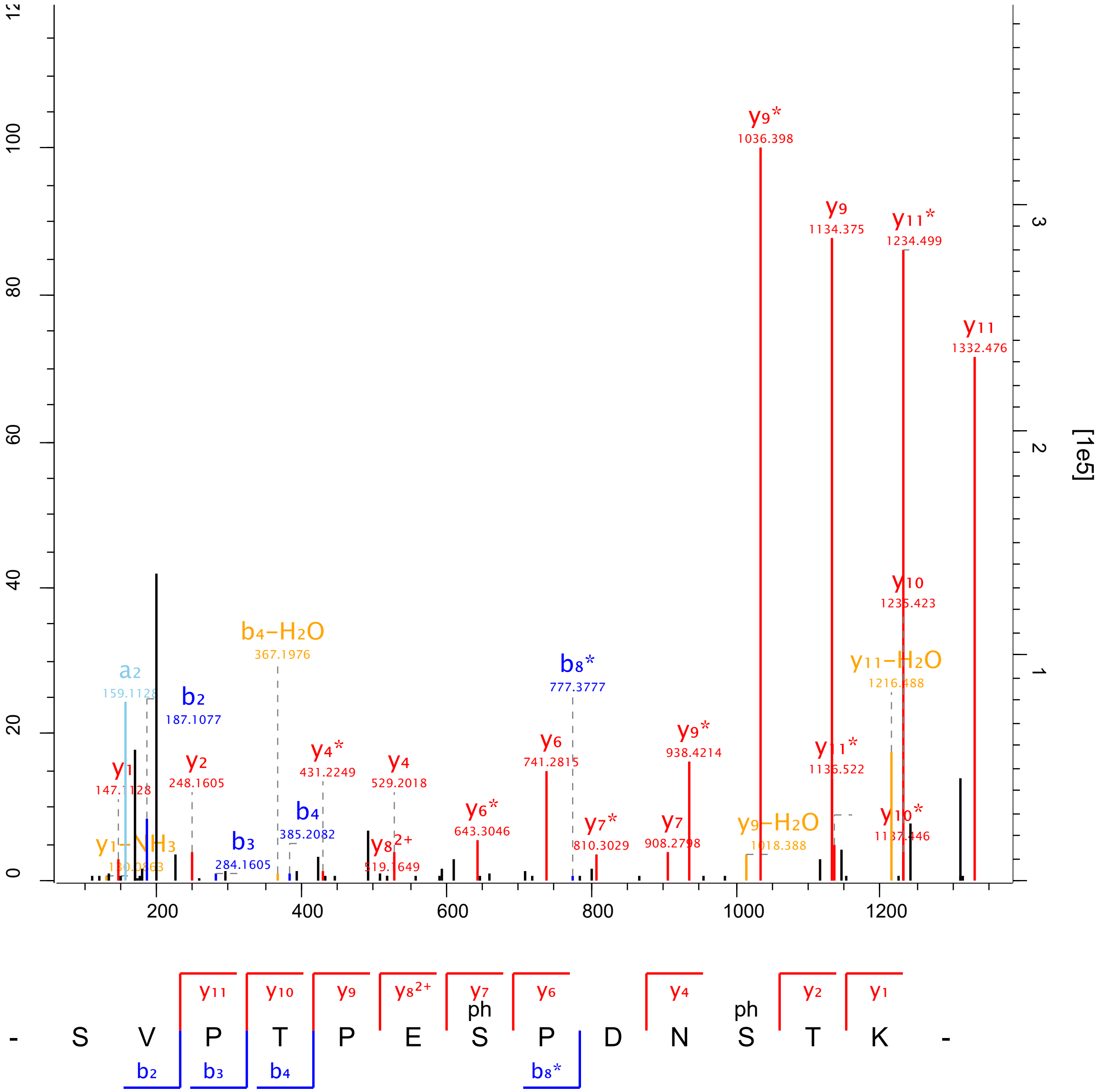

| Raw file | Scan | Method | Score | m/z |
|---|---|---|---|---|
| A17_15_1 | 12273 | FTMS; HCD | 126.26 | 855.89 |

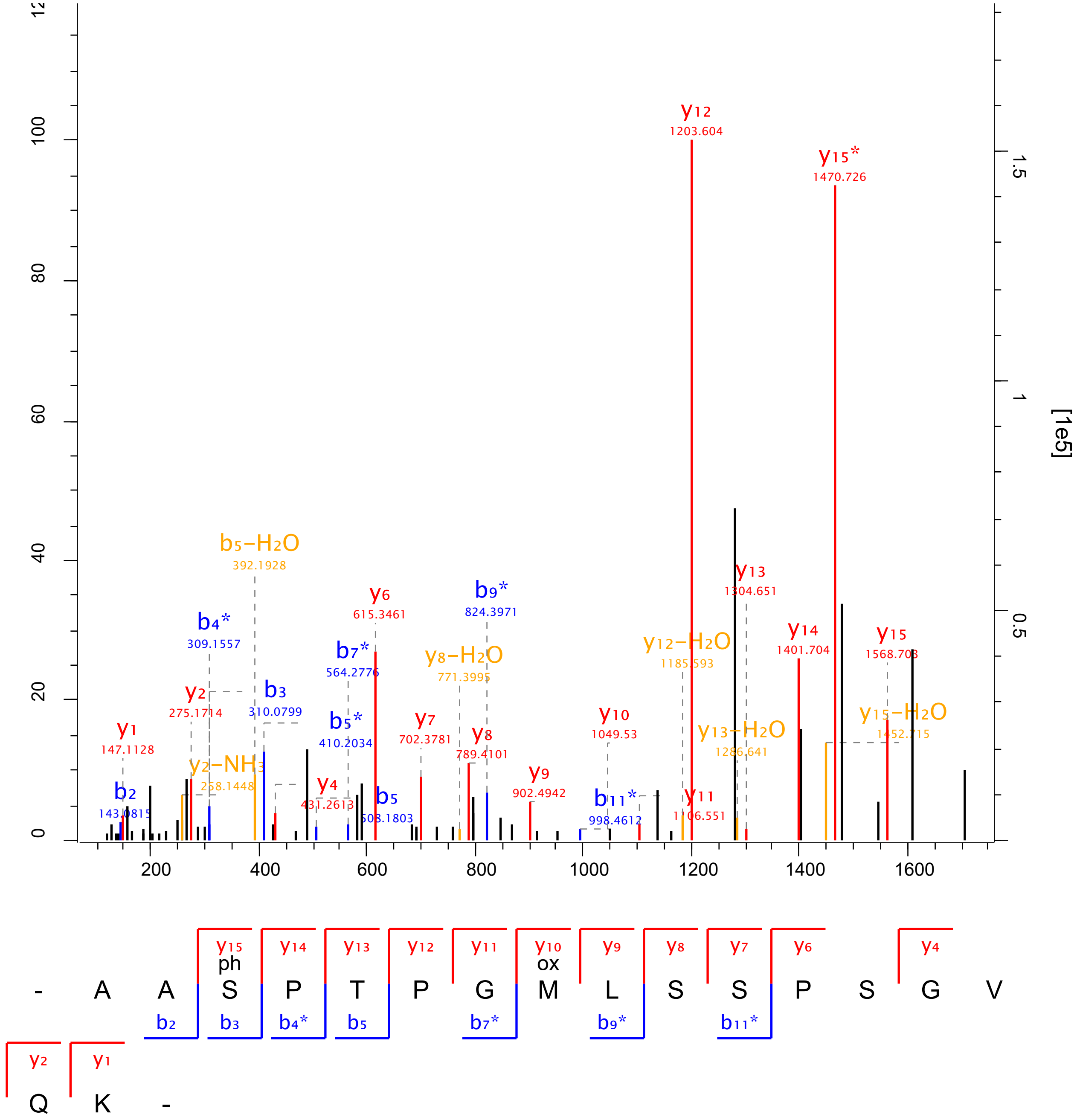

| Raw file | Scan | Method | Score | m/z |
|---|---|---|---|---|
| A17_15_1 | 12756 | FTMS; HCD | 62.2 | 860.84 |

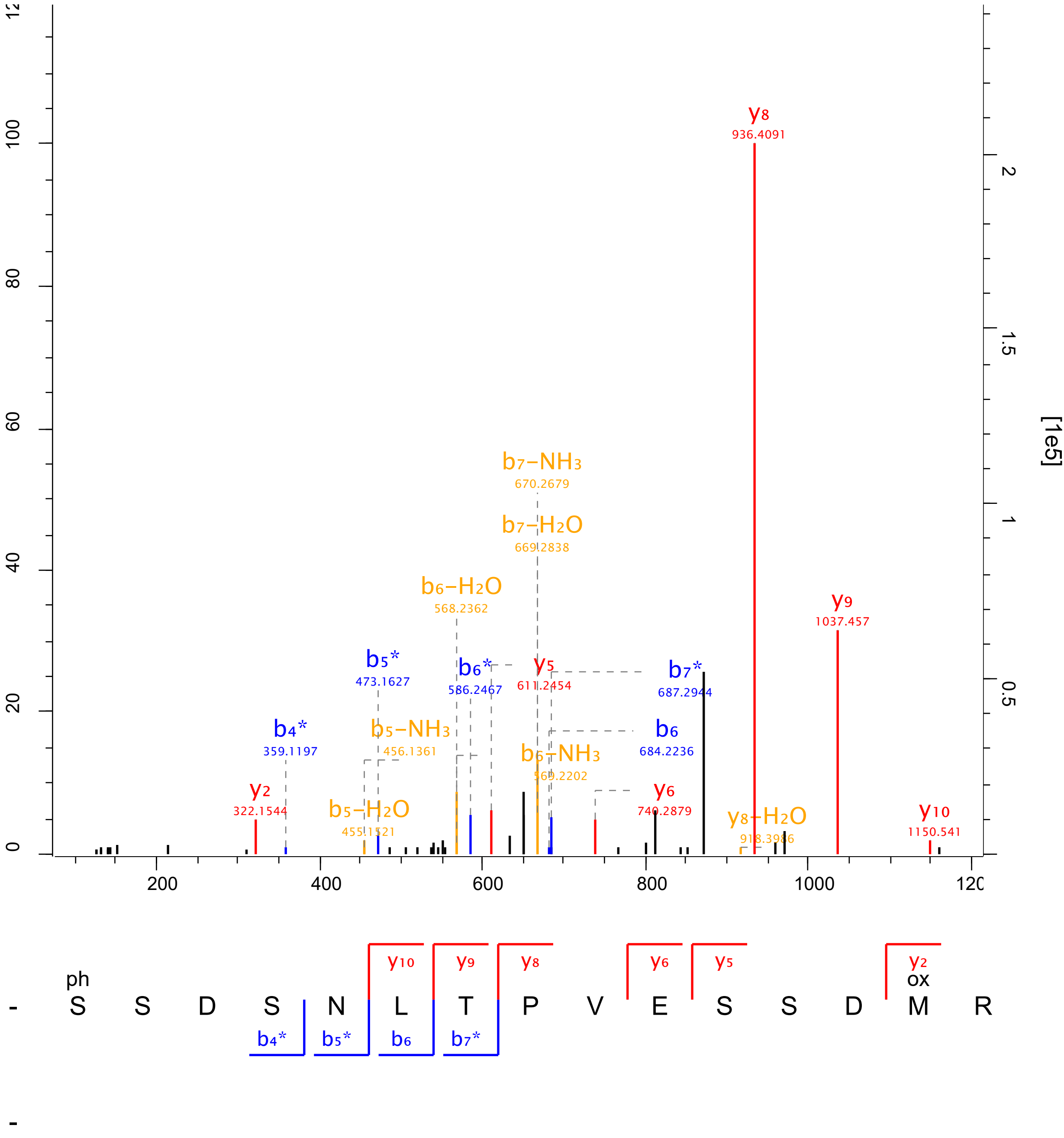

| Raw file | Scan | Method | Score | m/z |
| --- | --- | --- | --- | --- |
| A17_15_1 | 13009 | FTMS; HCD | 58.17 | 550.74 |

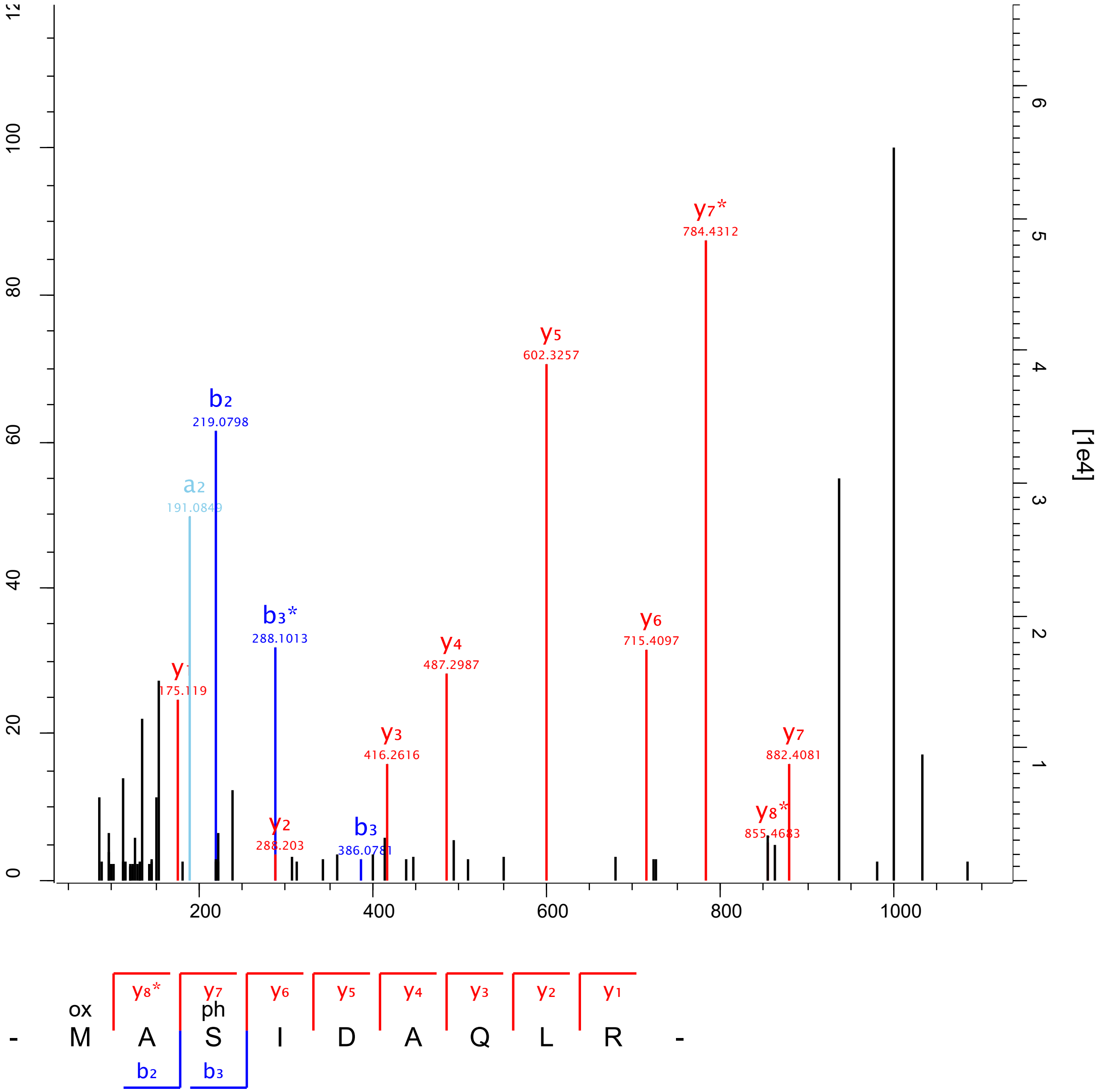

| Raw file | Scan | Method | Score | m/z |
| --- | --- | --- | --- | --- |
| A17_15_1 | 13182 | FTMS; HCD | 166.52 | 669.78 |

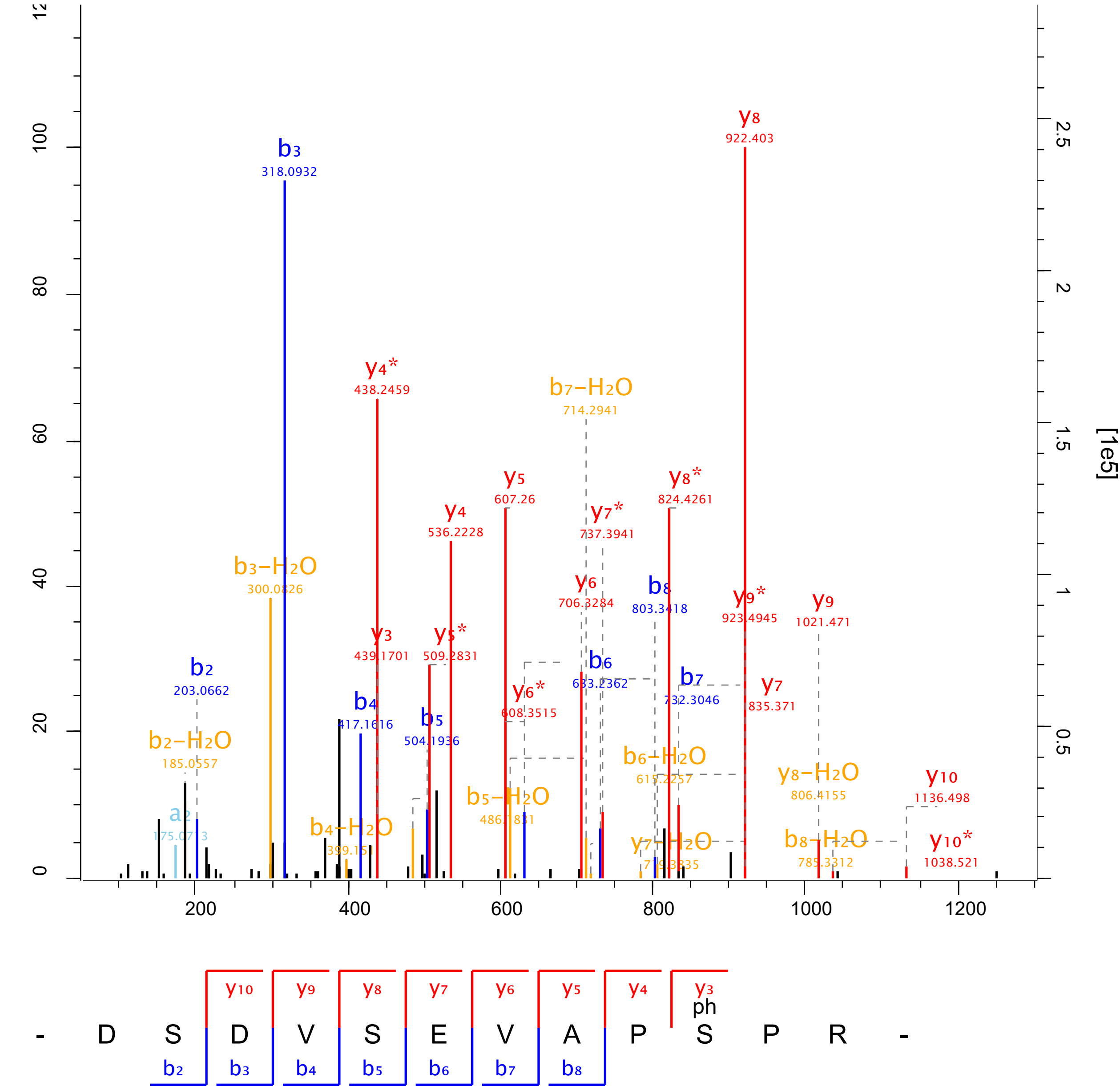

| Raw file | Scan | Method | Score | m/z |
| --- | --- | --- | --- | --- |
| A17_15_1 | 13274 | FTMS; HCD | 152.88 | 1088.92 |

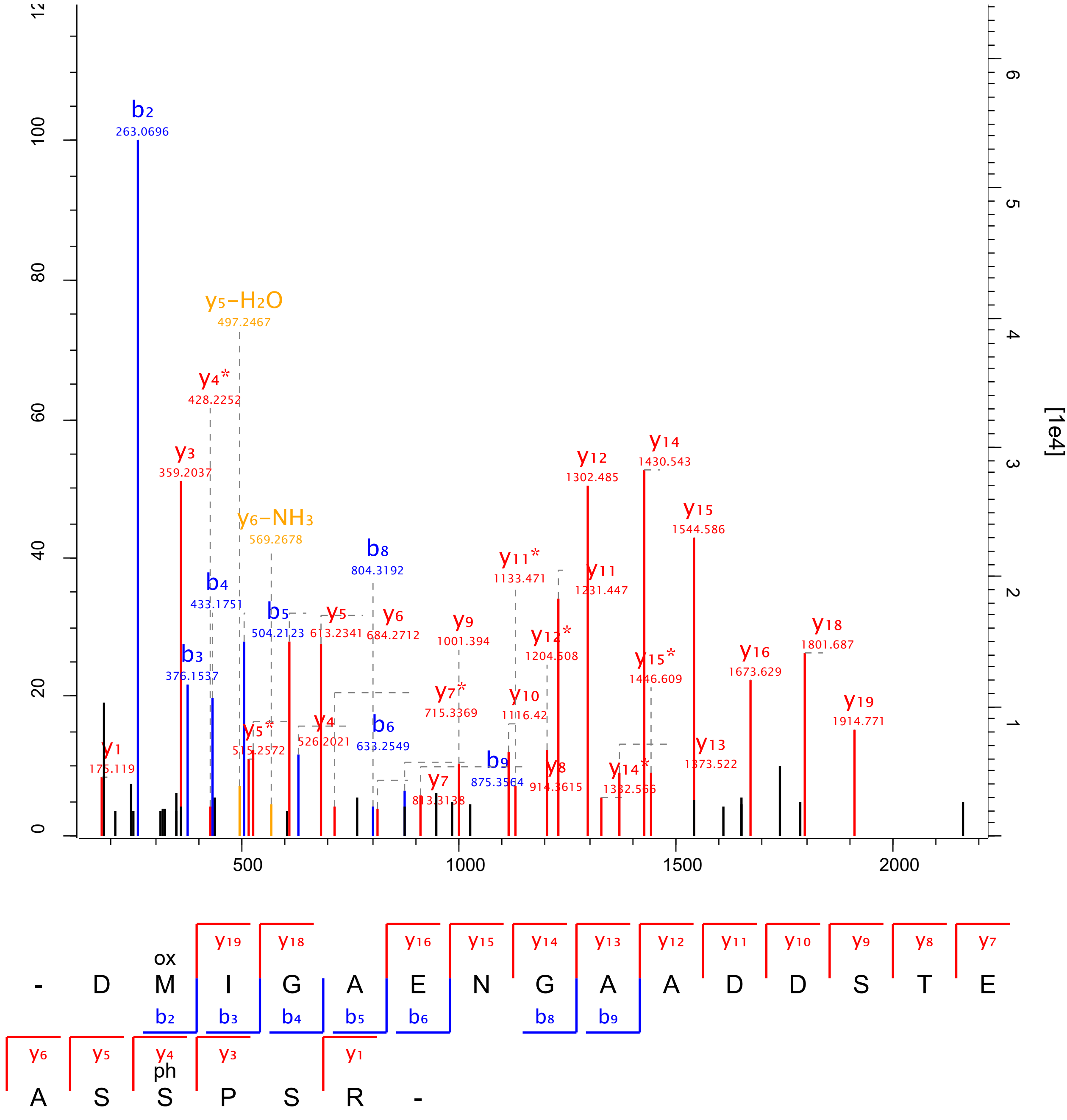

| Raw file | Scan | Method | Score | m/z |
|---|---|---|---|---|
| A17_15_1 | 13315 | FTMS; HCD | 130.29 | 699.28 |

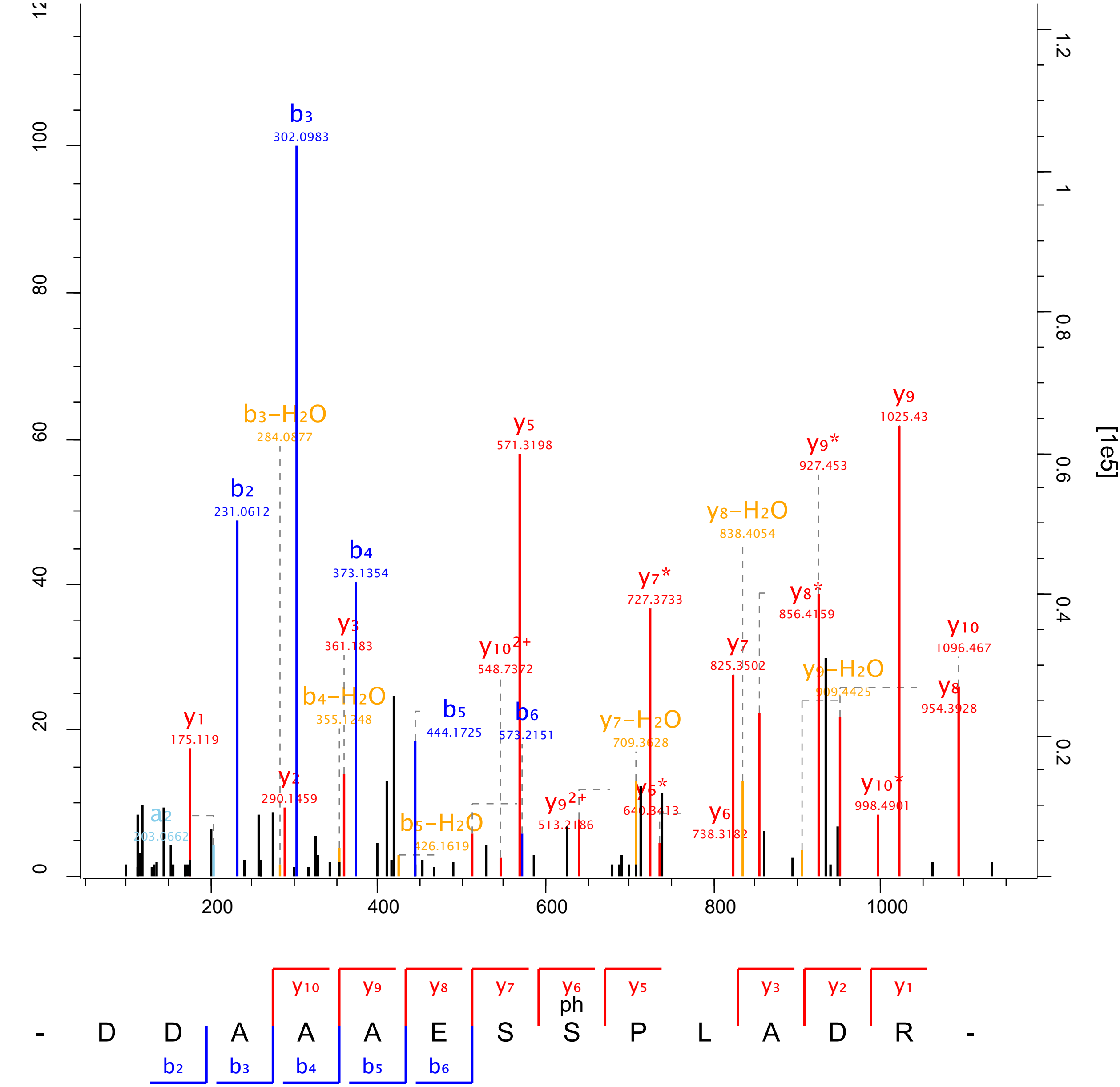

| Raw file | Scan | Method | Score | m/z |
| --- | --- | --- | --- | --- |
| A17_15_1 | 13852 | FTMS; HCD | 79.49 | 491.24 |

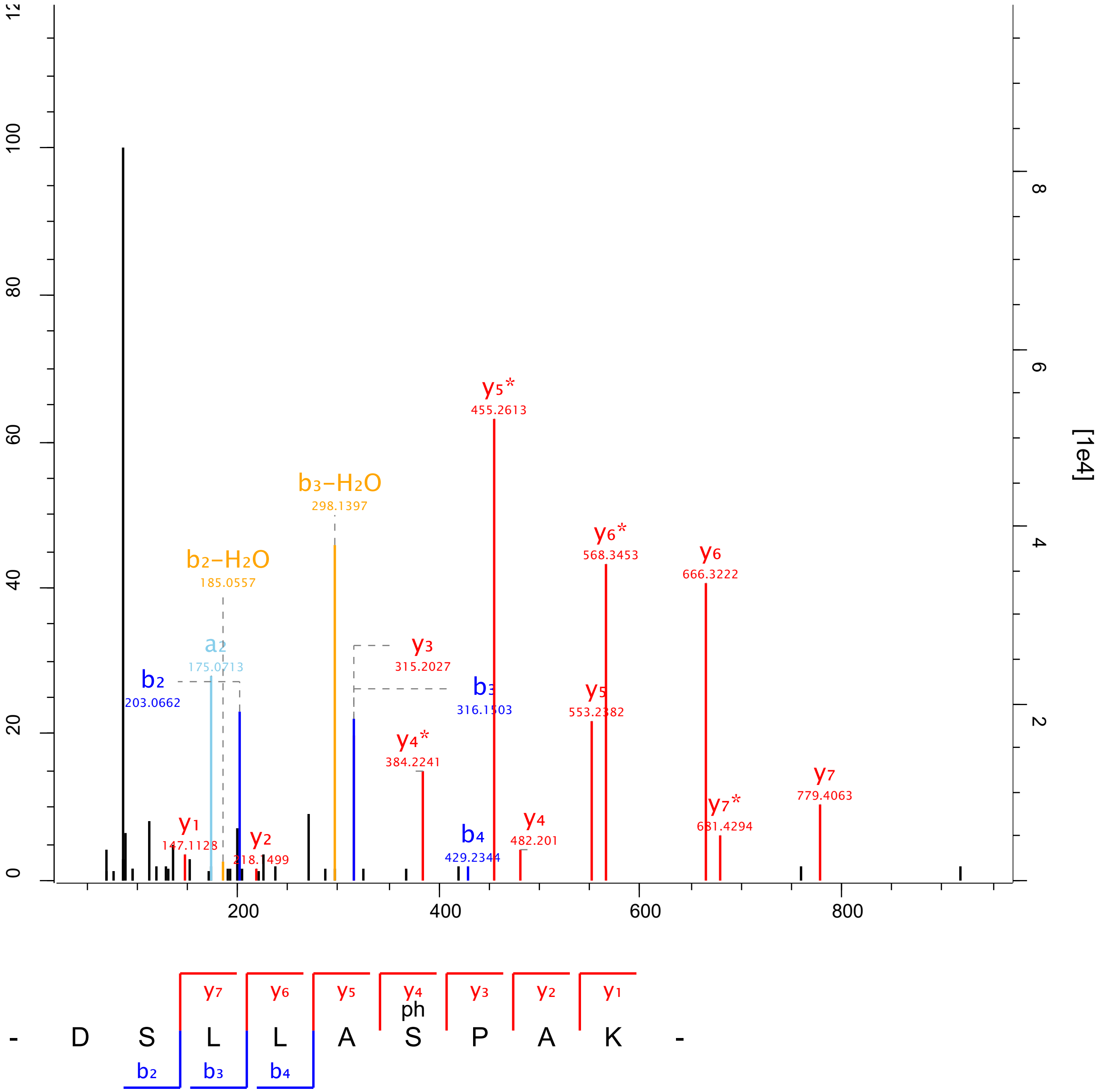

| Raw file | Scan | Method | Score | m/z |
|---|---|---|---|---|
| A17_15_1 | 14106 | FTMS; HCD | 77.96 | 852.83 |

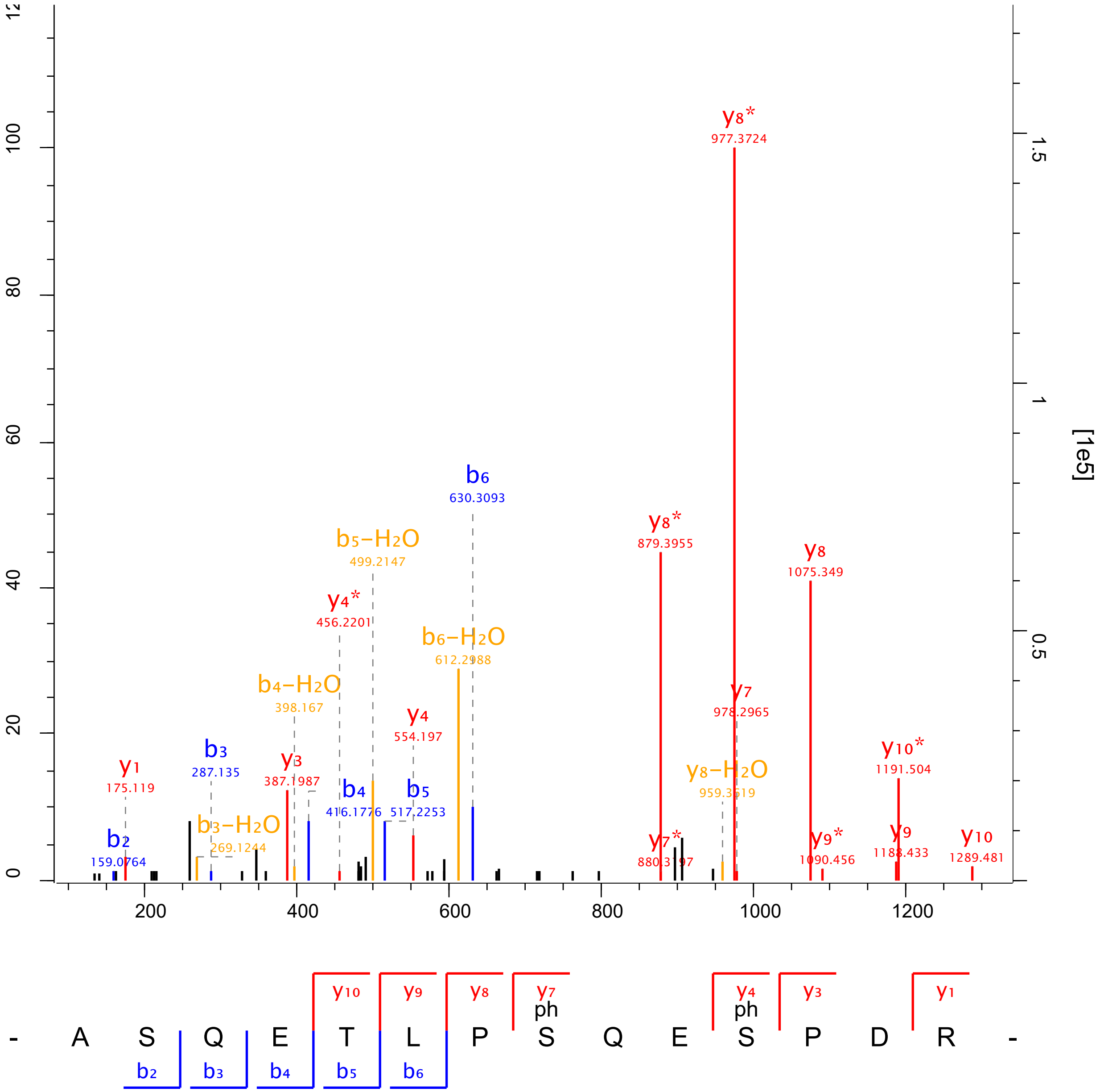

| Raw file | Scan | Method | Score | m/z |
| --- | --- | --- | --- | --- |
| A17_15_1 | 14203 | FTMS; HCD | 54.34 | 610.23 |

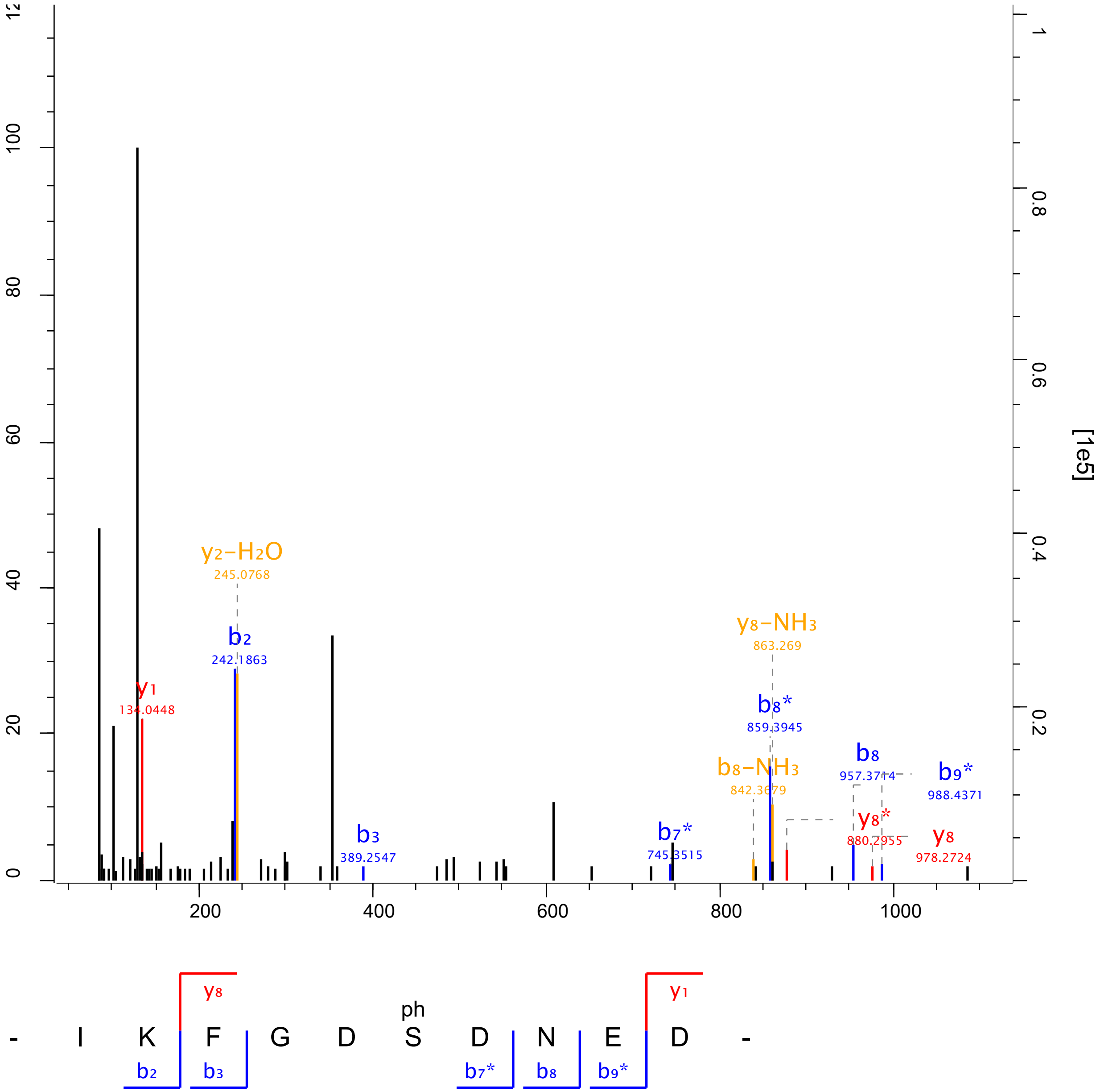

| Raw file | Scan | Method | Score | m/z |
| --- | --- | --- | --- | --- |
| A17_15_1 | 14294 | FTMS; HCD | 47.68 | 746.29 |

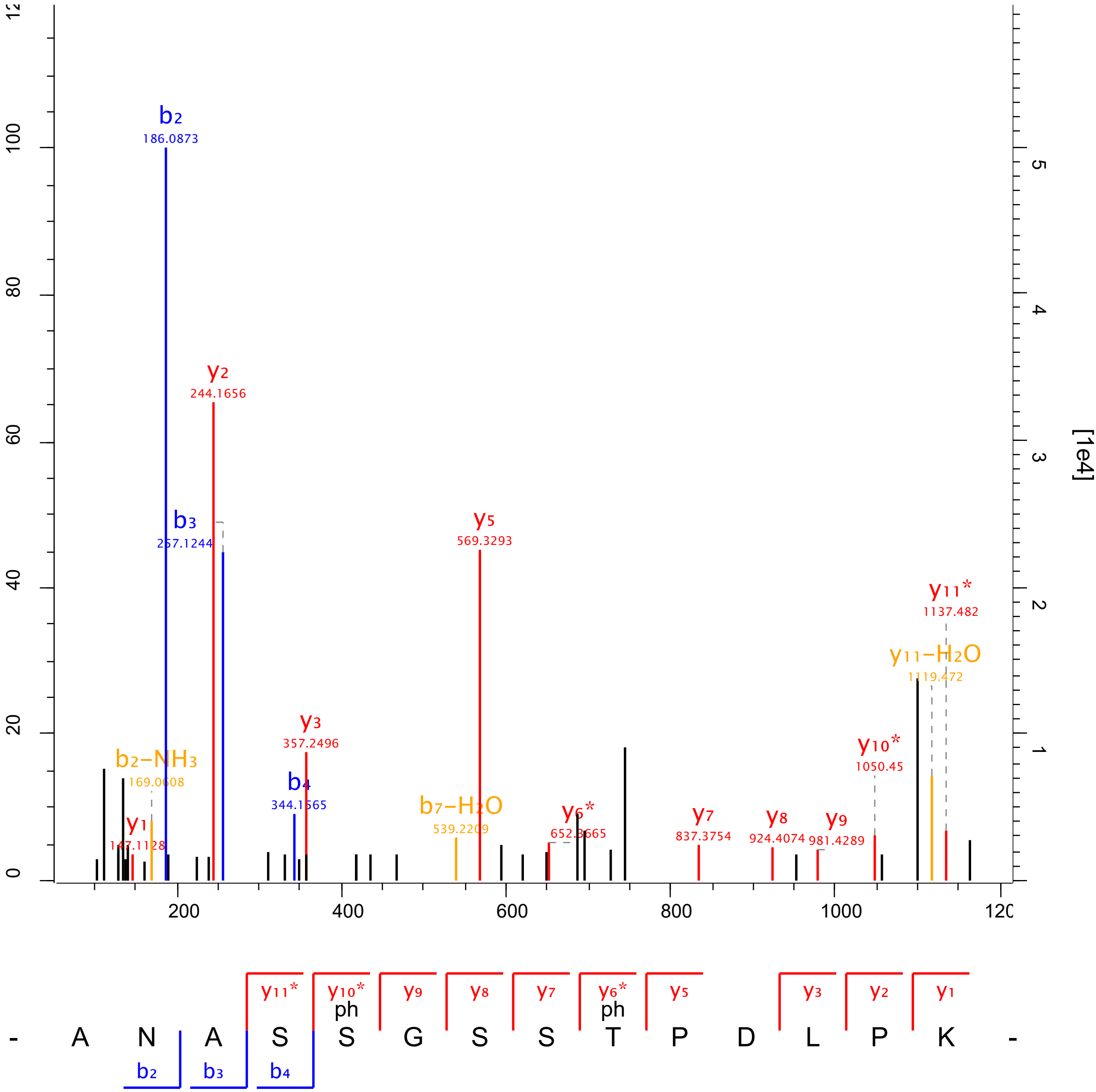

| Raw file | Scan | Method | Score | m/z |
|---|---|---|---|---|
| A17_15_1 | 14409 | FTMS; HCD | 199.19 | 760.82 |

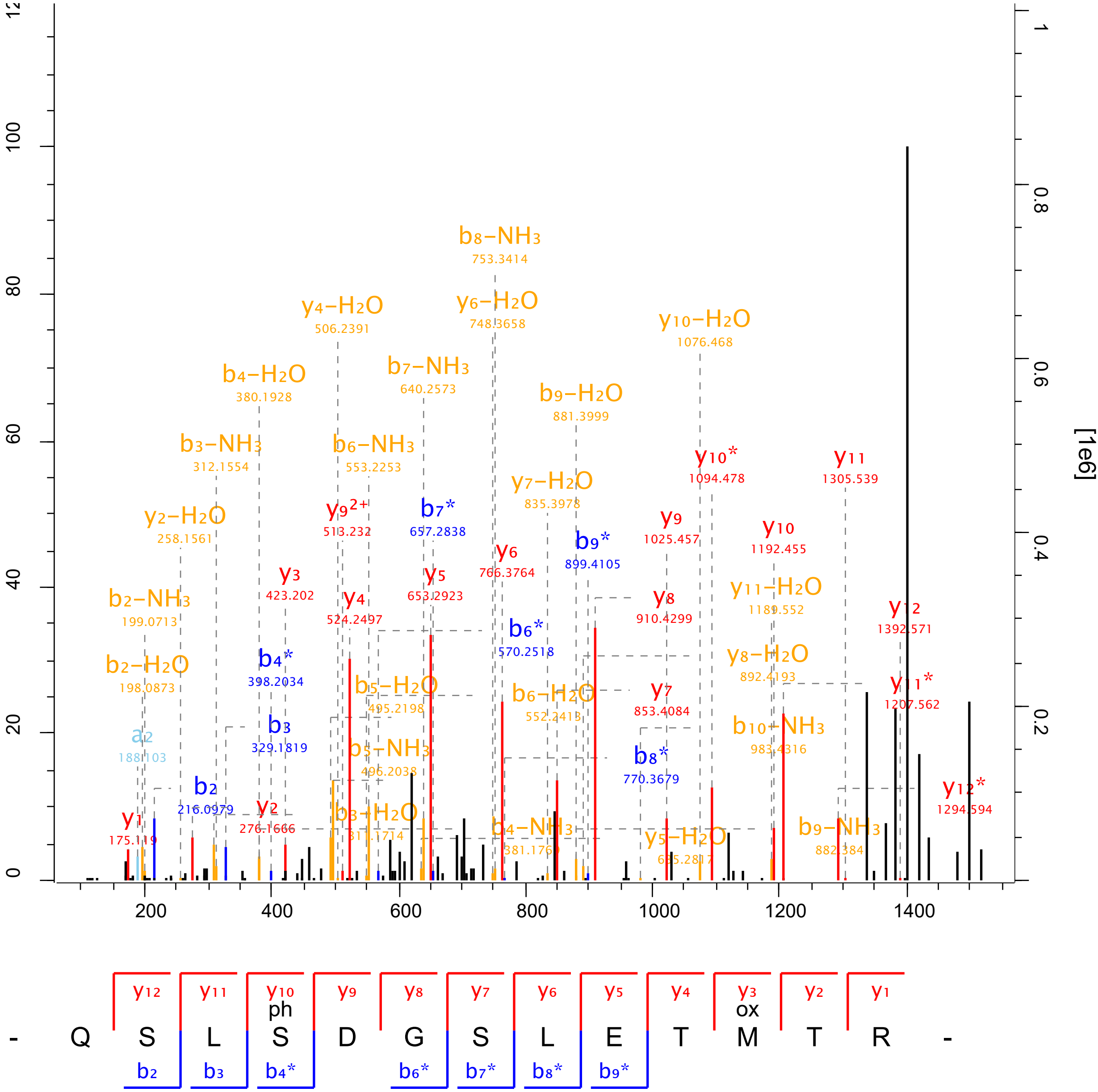

| Raw file | Scan | Method | Score | m/z |
| --- | --- | --- | --- | --- |
| A17_15_1 | 14828 | FTMS; HCD | 53.95 | 657.75 |

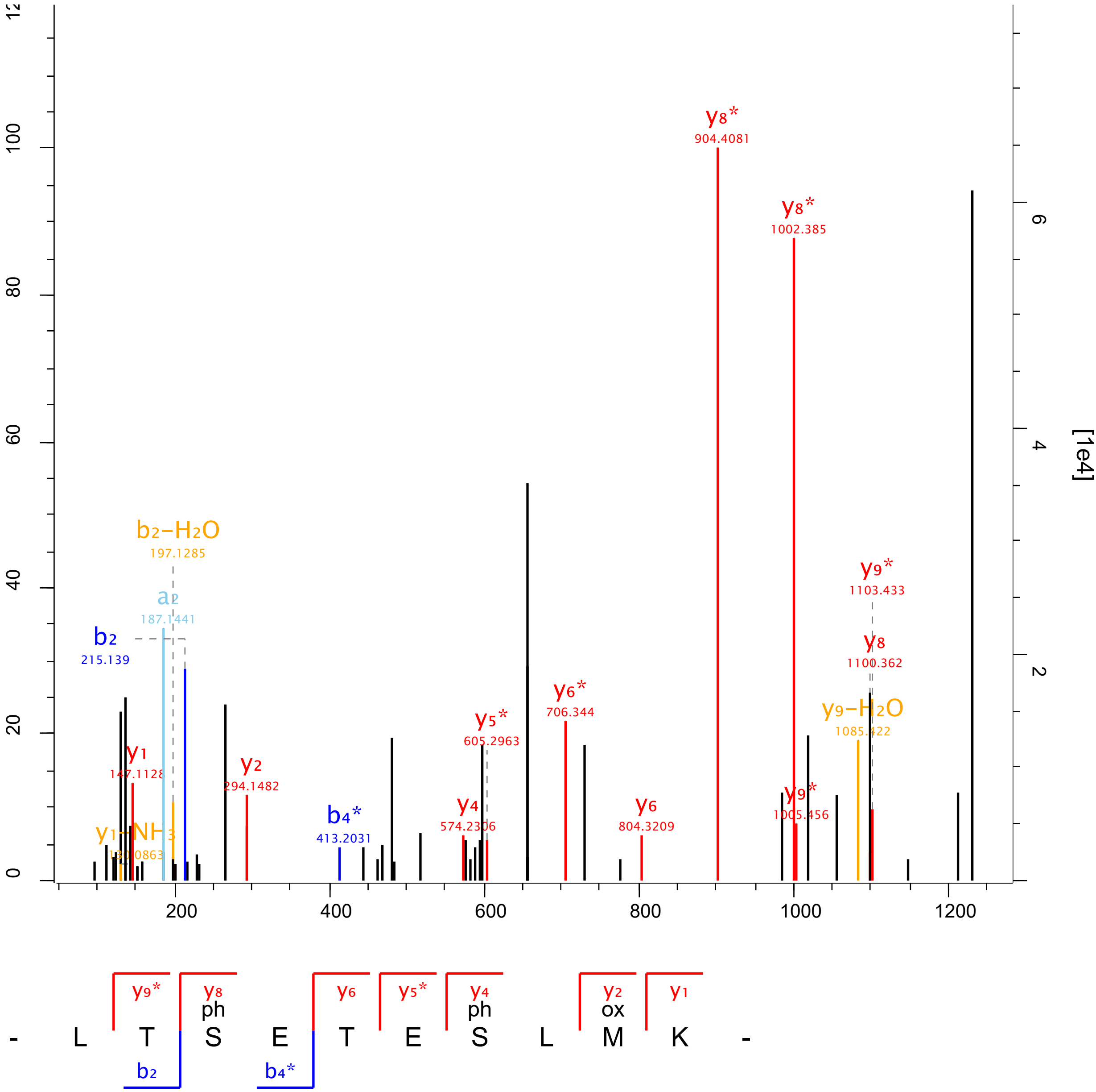

| Raw file | Scan | Method | Score | m/z |
| --- | --- | --- | --- | --- |
| A17_15_1 | 14879 | FTMS; HCD | 85.56 | 707.32 |

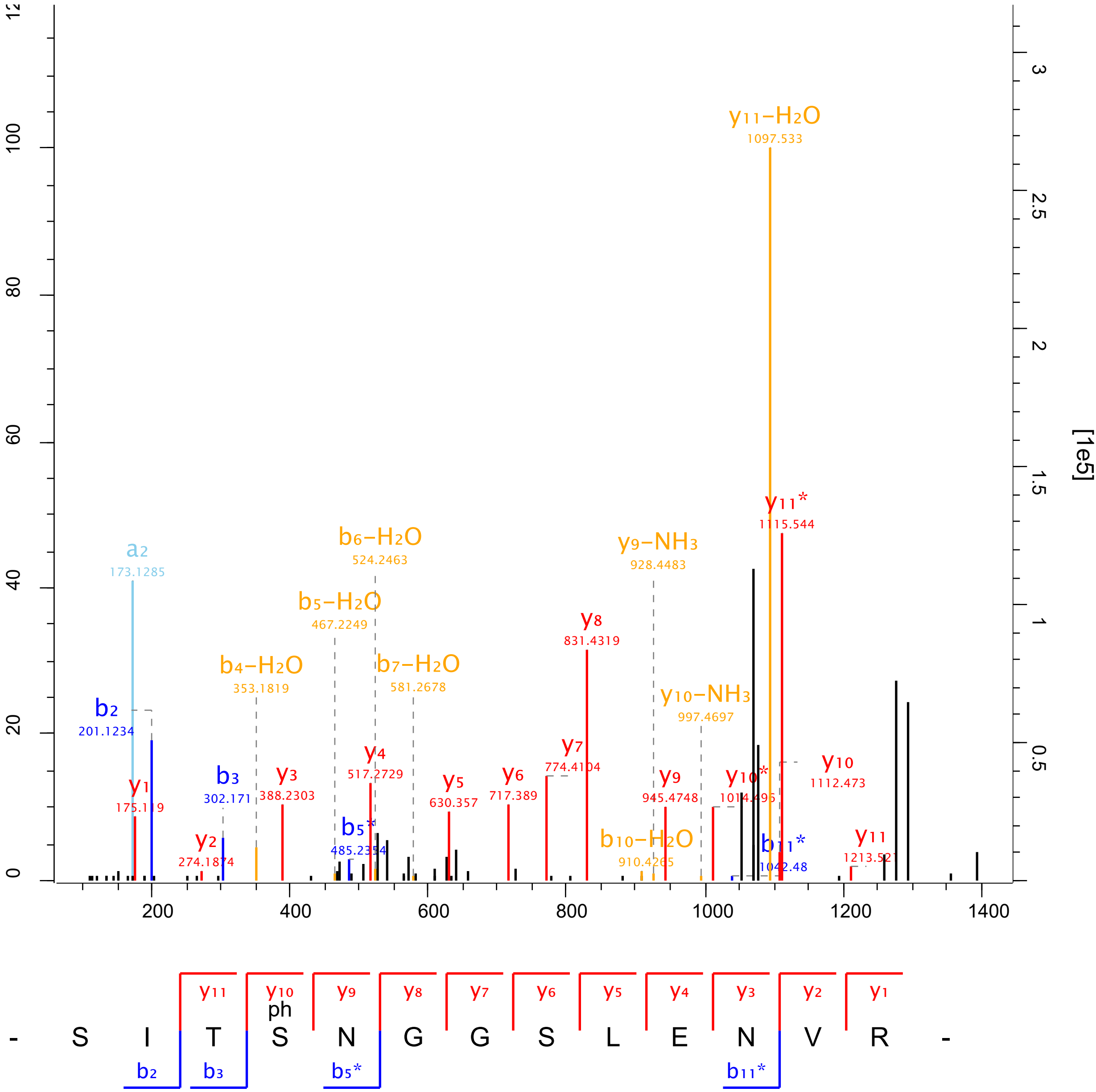

| Raw file | Scan | Method | Score | m/z |
| --- | --- | --- | --- | --- |
| A17_15_1 | 15149 | FTMS; HCD | 40.86 | 701.8 |

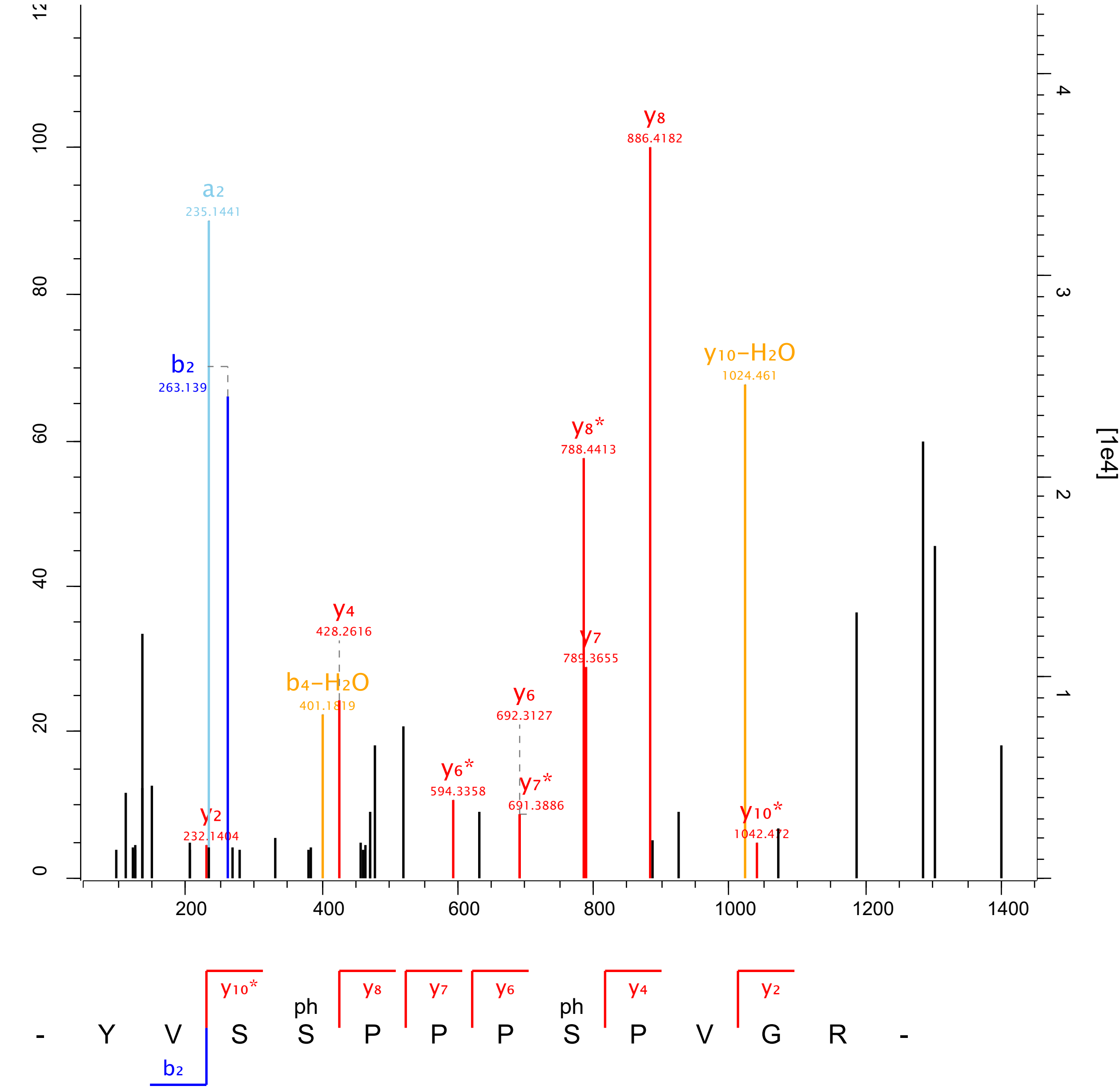

| Raw file | Scan | Method | Score | m/z |
| --- | --- | --- | --- | --- |
| A17_15_1 | 15504 | FTMS; HCD | 199.37 | 694.79 |

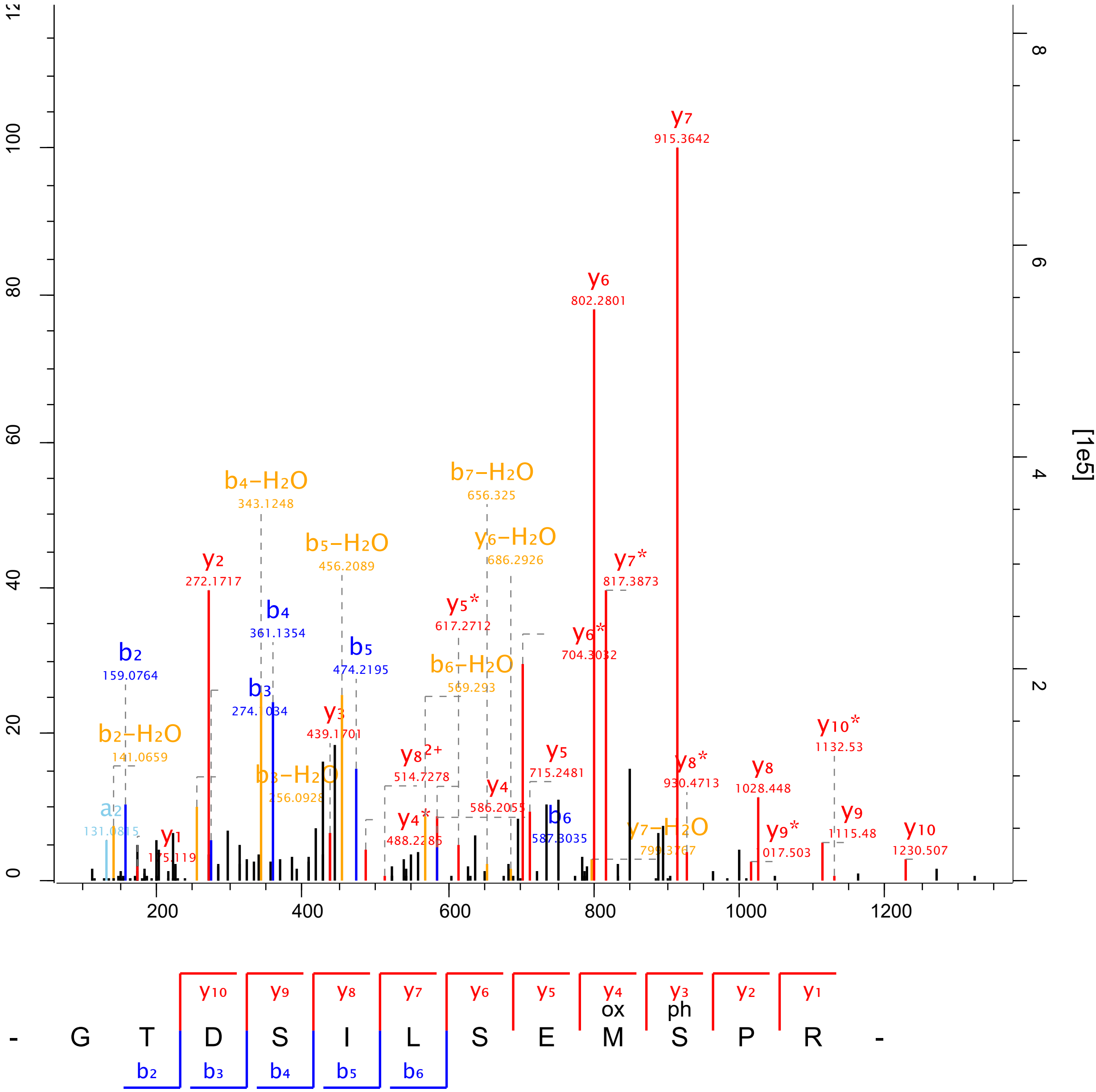

| Raw file | Scan | Method | Score | m/z |
|---|---|---|---|---|
| A17_15_1 | 1577 | FTMS; HCD | 108.47 | 537.71 |

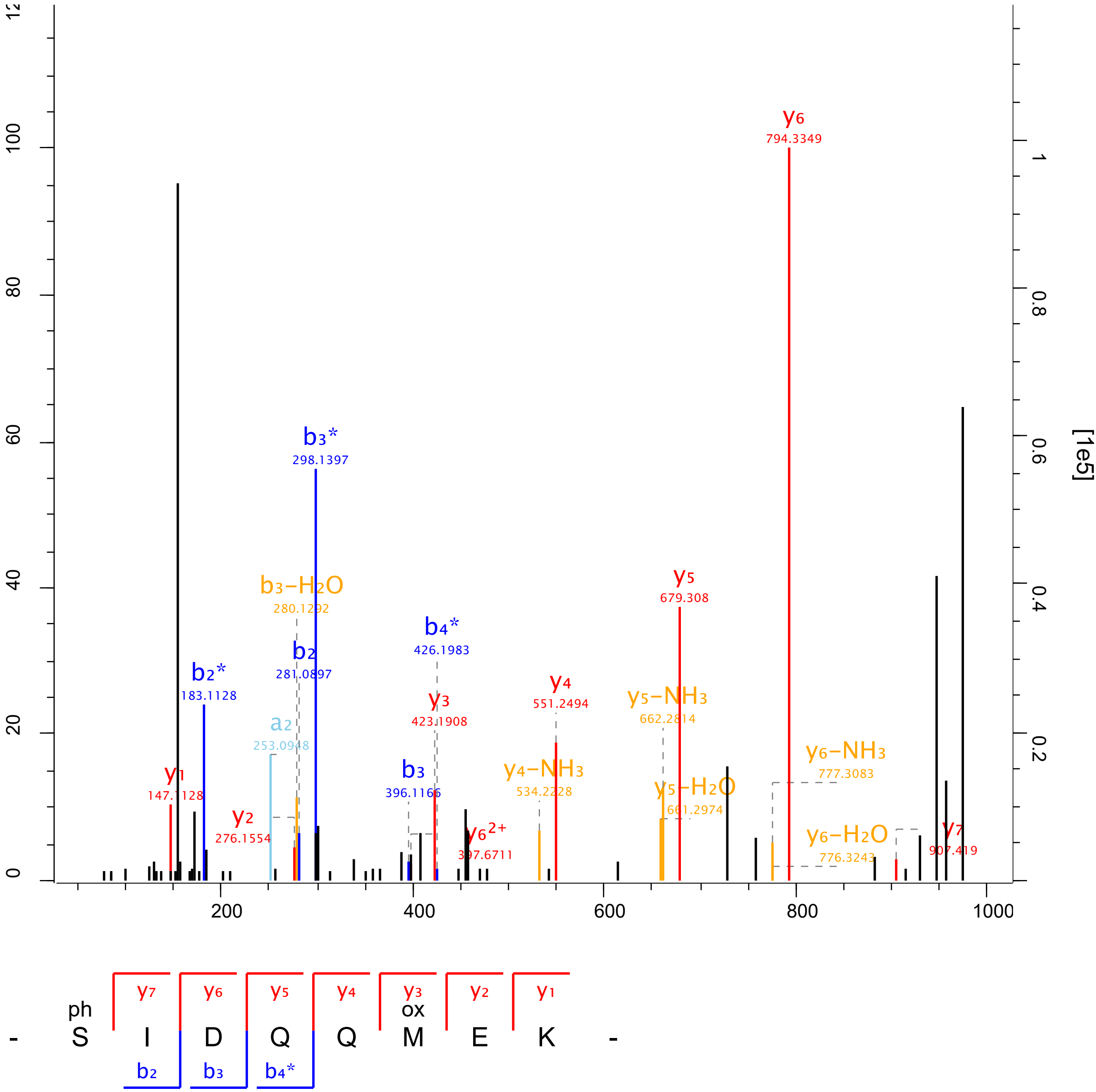

| Raw file | Scan | Method | Score | m/z |
|---|---|---|---|---|
| A17_15_1 | 16033 | FTMS; HCD | 157.2 | 551.24 |

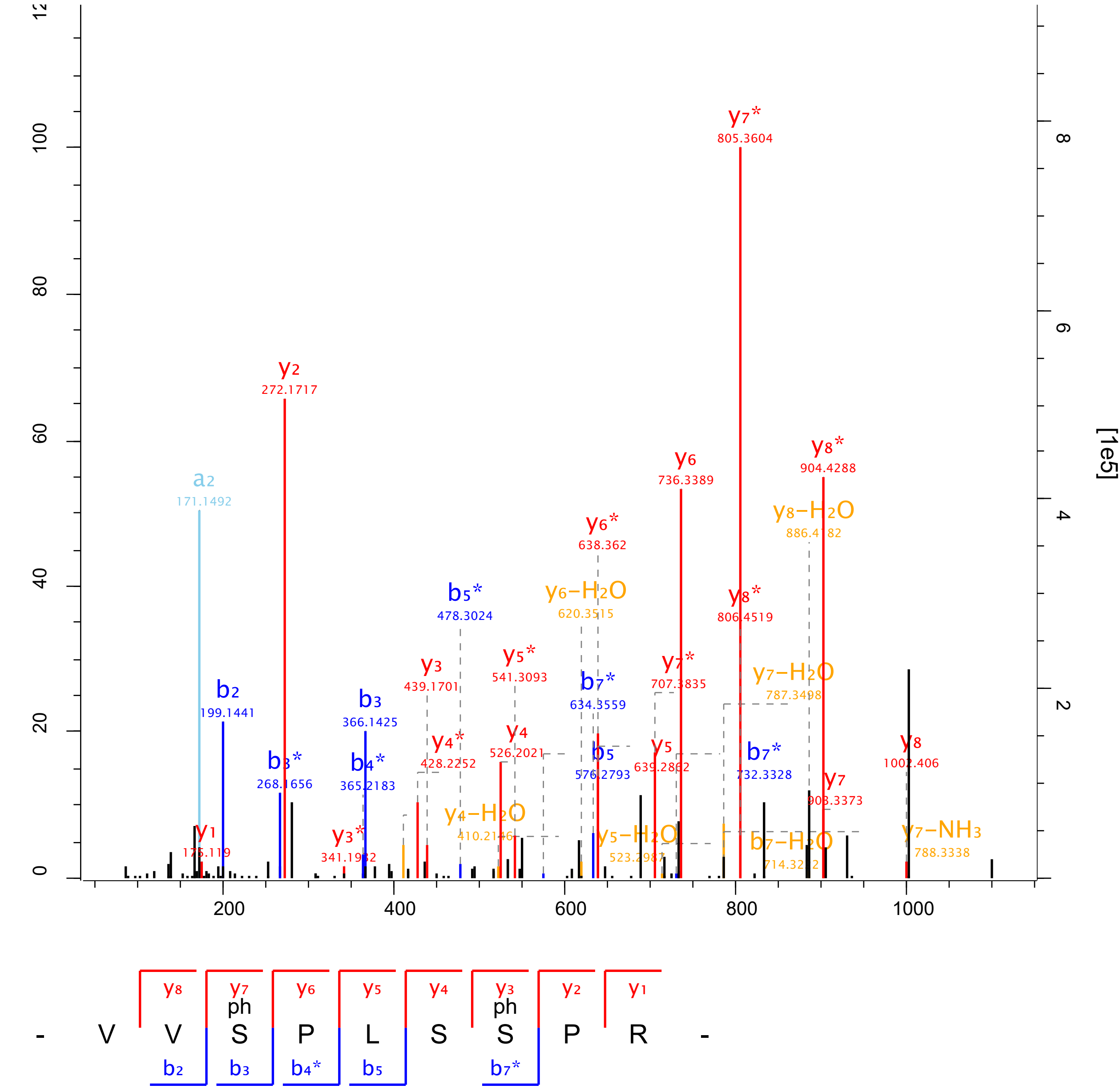

| Raw file | Scan | Method | Score | m/z |
| --- | --- | --- | --- | --- |
| A17_15_1 | 16145 | FTMS; HCD | 53.3 | 626.78 |

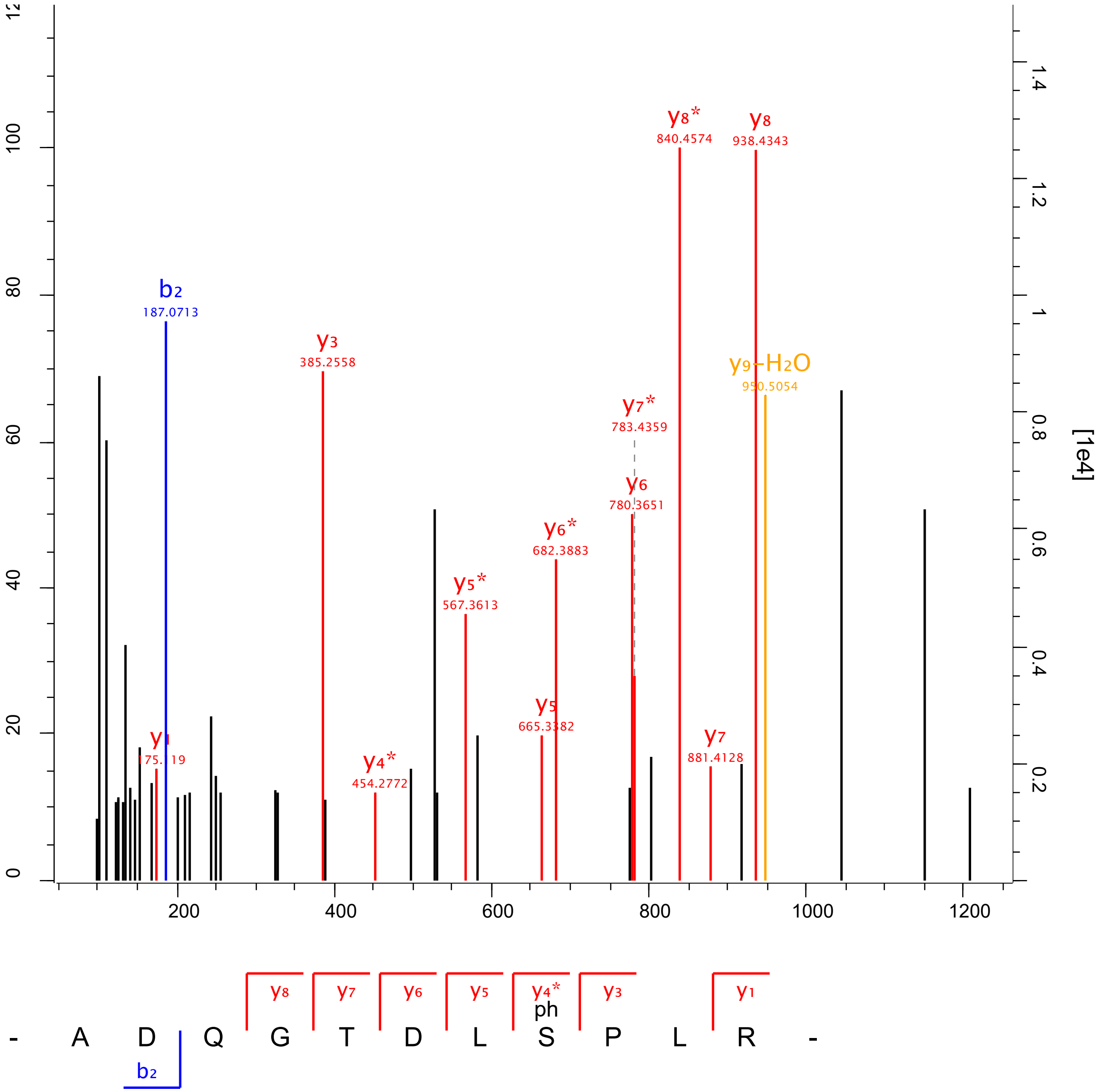

| Raw file | Scan | Method | Score | m/z |
|---|---|---|---|---|
| A17_15_1 | 16271 | FTMS; HCD | 71.18 | 500.25 |

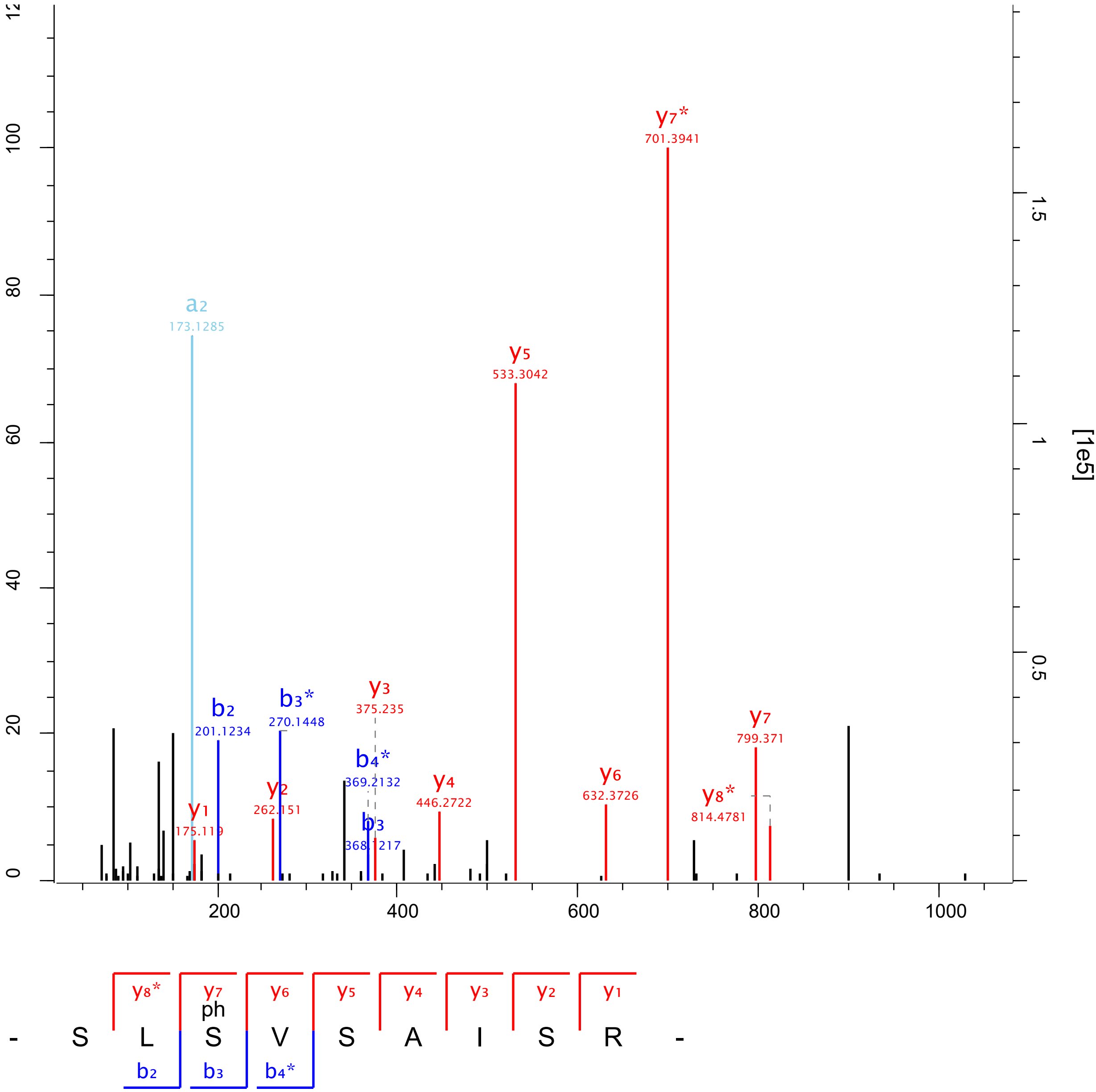

| Raw file | Scan | Method | Score | m/z |
|---|---|---|---|---|
| A17_15_1 | 16357 | FTMS; HCD | 102.95 | 507.72 |

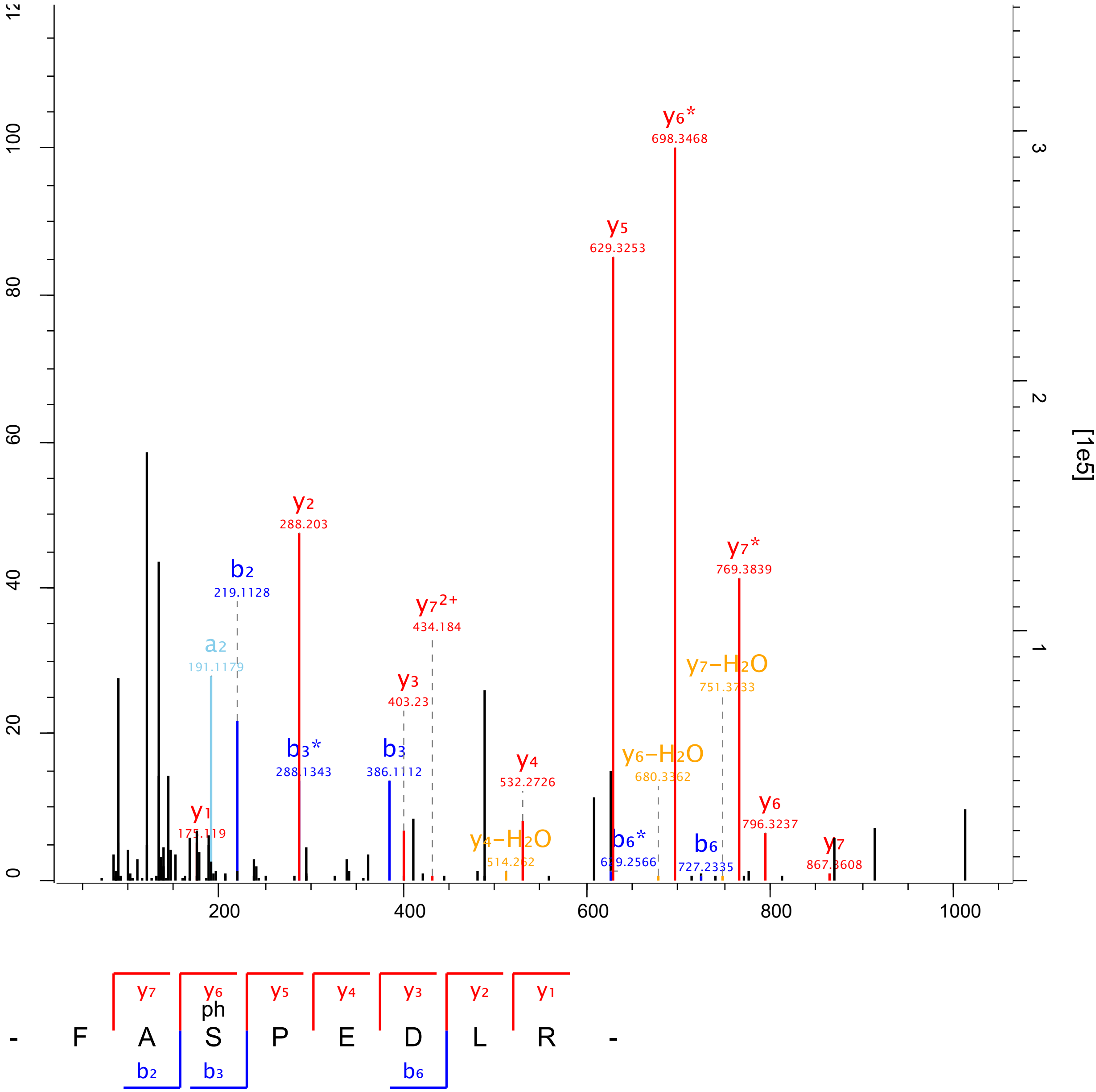

| Raw file | Scan | Method | Score | m/z |
|---|---|---|---|---|
| A17_15_1 | 1641 | FTMS; HCD | 96.67 | 531.53 |

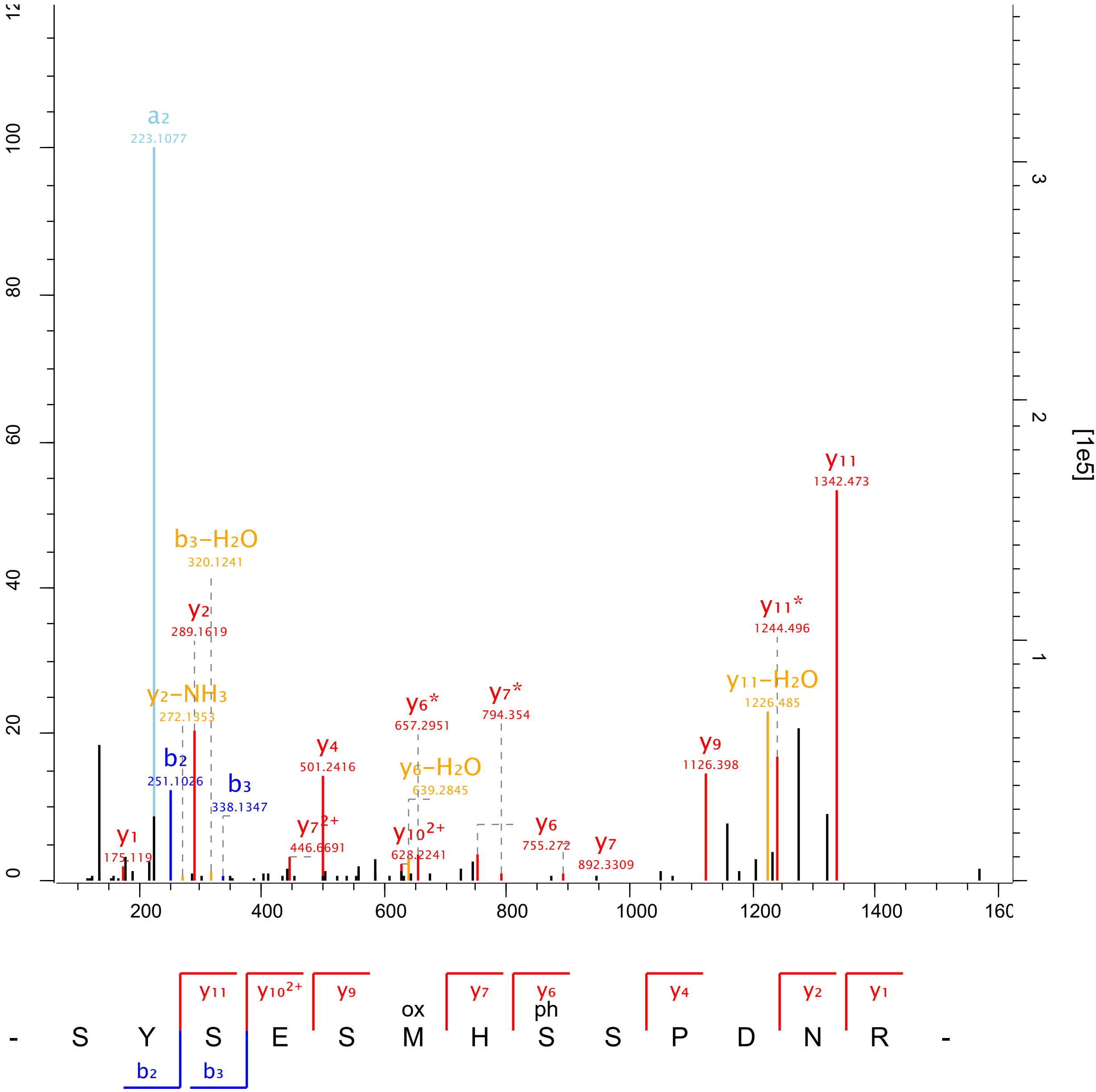

| Raw file | Scan | Method | Score | m/z |
|---|---|---|---|---|
| A17_15_1 | 16500 | FTMS; HCD | 152.34 | 688.24 |

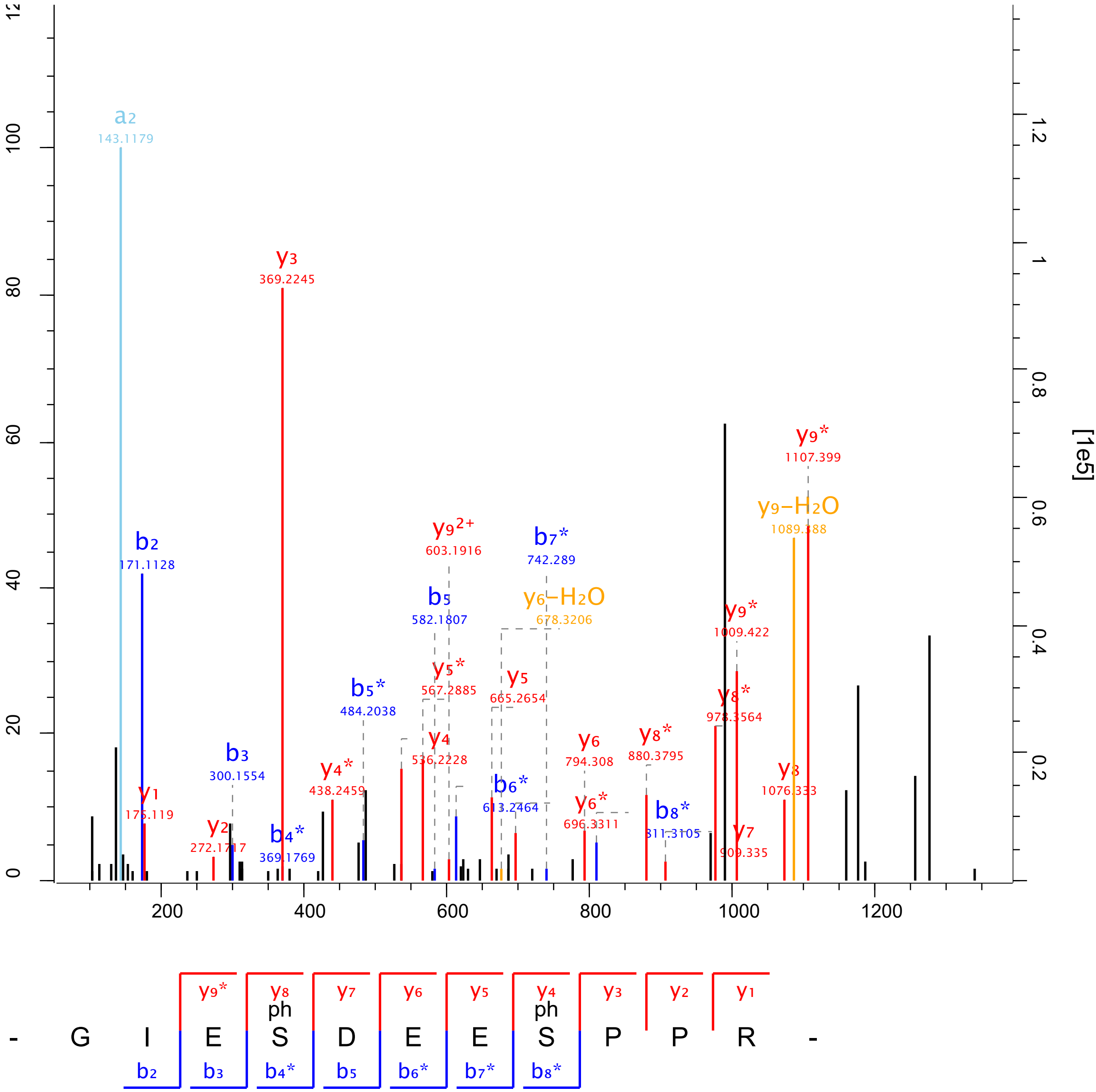

| Raw file | Scan | Method | Score | m/z |
|---|---|---|---|---|
| A17_15_1 | 16505 | FTMS; HCD | 113.65 | 792.02 |

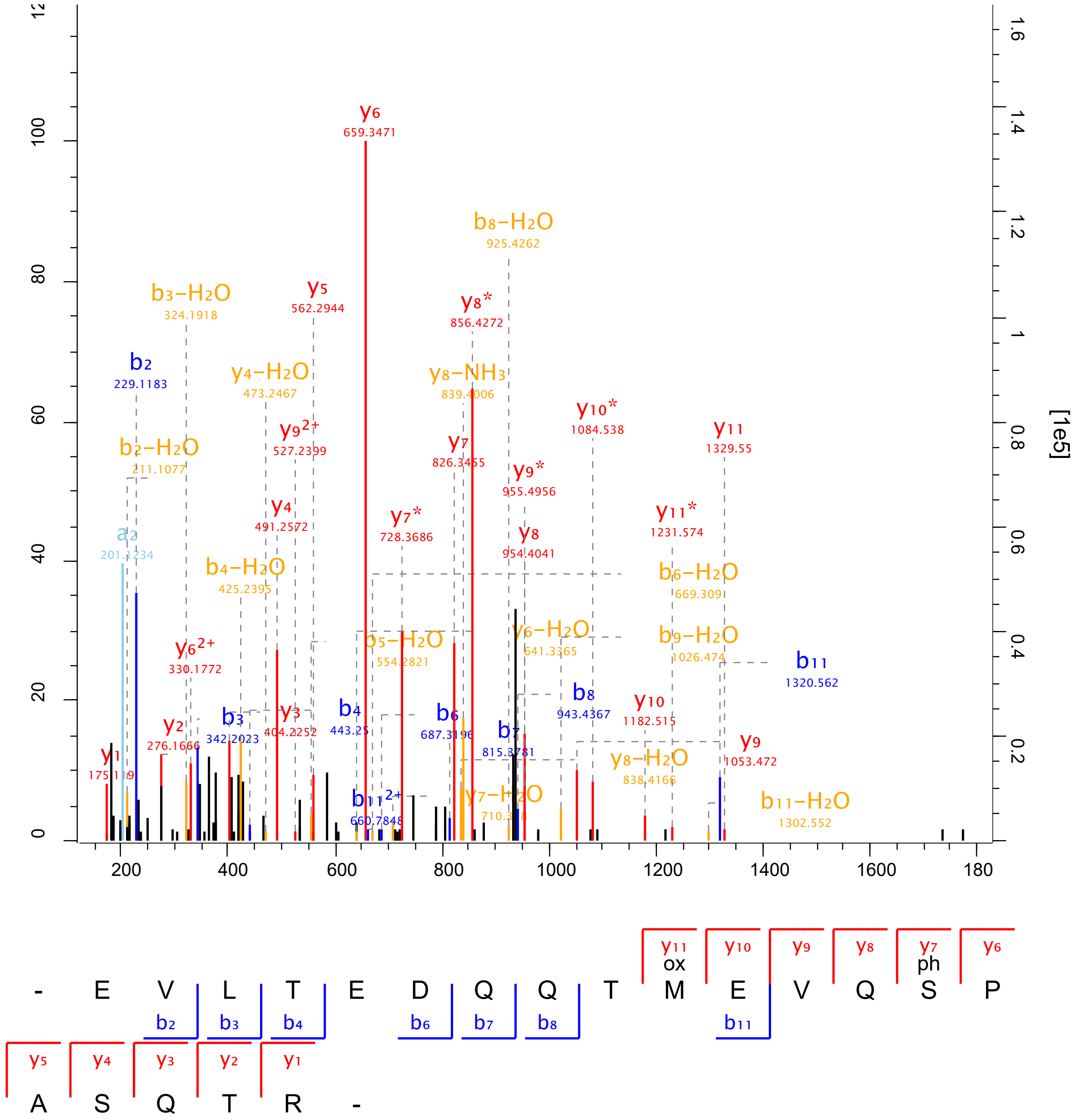

| Raw file | Scan | Method | Score | m/z |
|---|---|---|---|---|
| A17_15_1 | 16568 | FTMS; HCD | 88.08 | 665.78 |

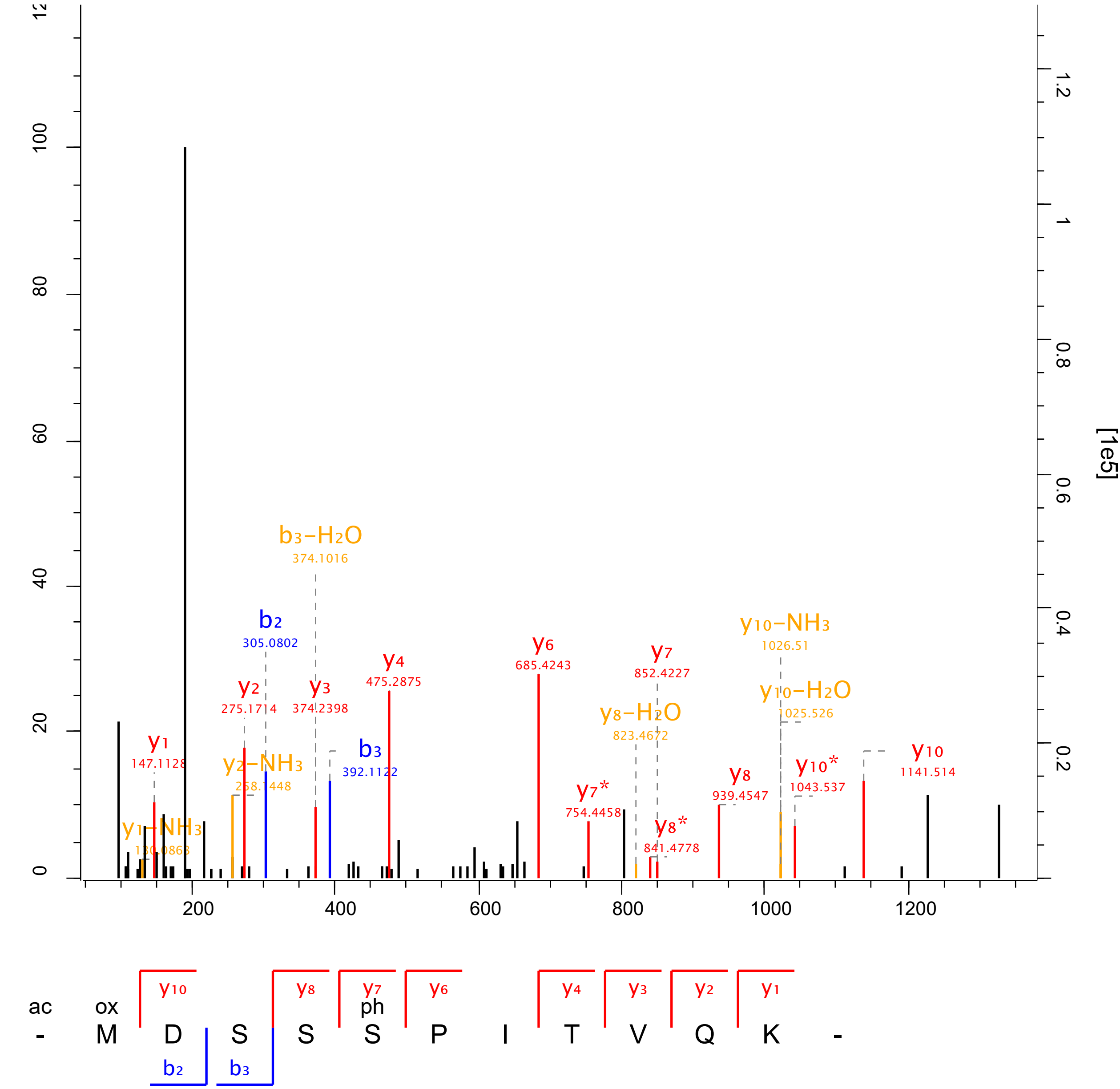

| Raw file | Scan | Method | Score | m/z |
|---|---|---|---|---|
| A17_15_1 | 16674 | FTMS; HCD | 55.26 | 530.25 |

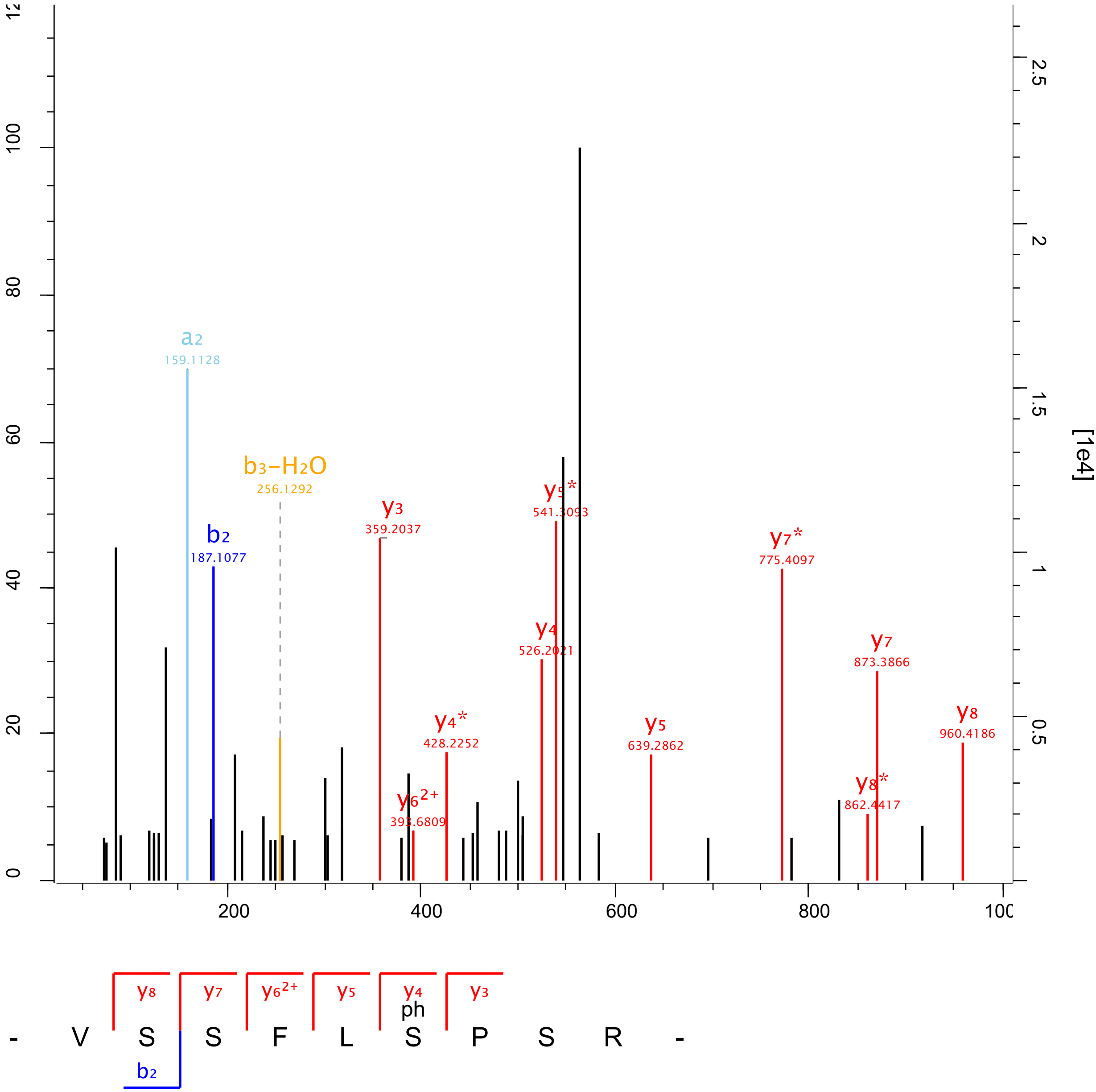

| Raw file | Scan | Method | Score | m/z |
|---|---|---|---|---|
| A17_15_1 | 16821 | FTMS; HCD | 96.38 | 918.86 |

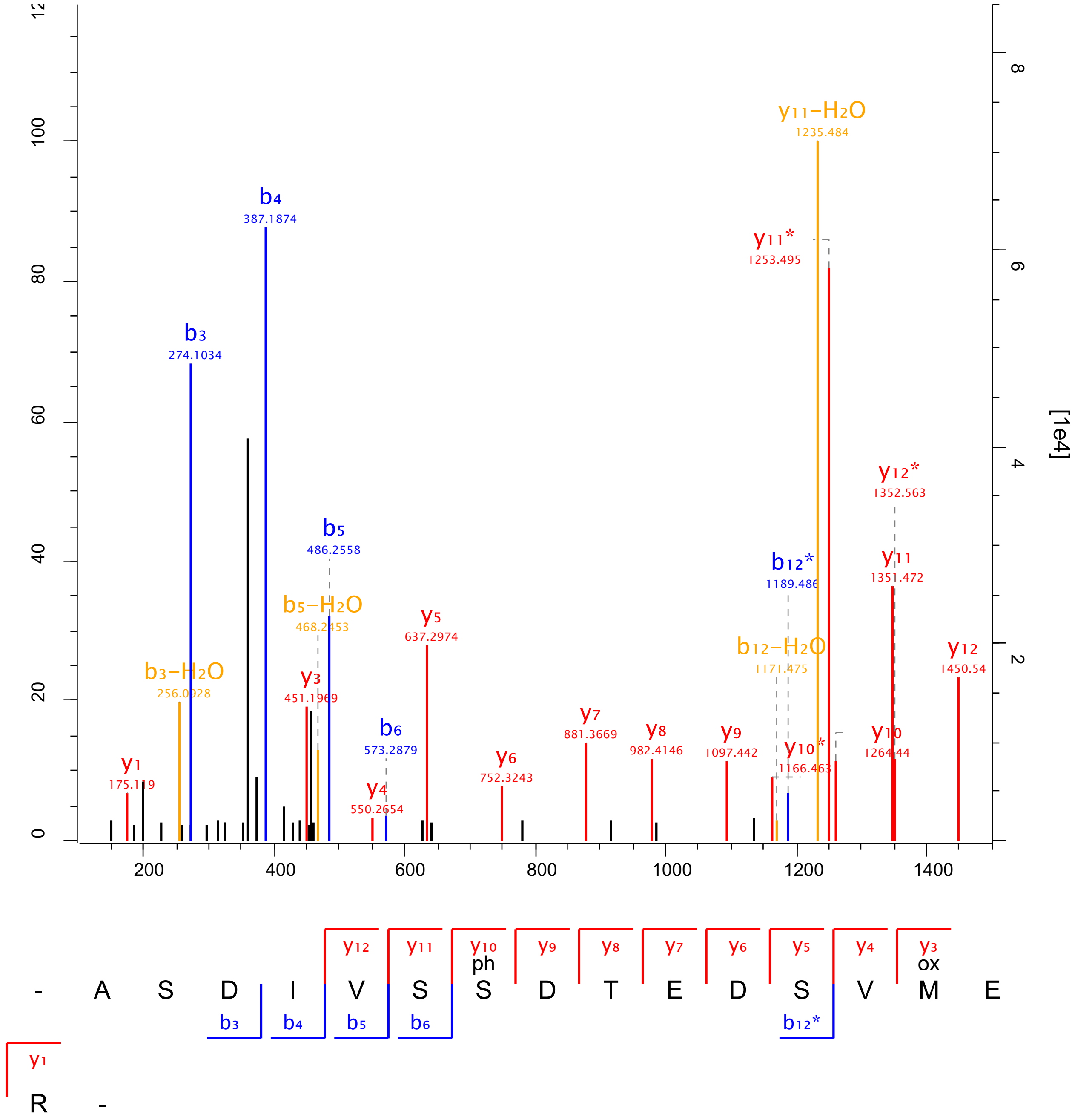

| Raw file | Scan | Method | Score | m/z |
|---|---|---|---|---|
| A17_15_1 | 17003 | FTMS; HCD | 167.89 | 946.37 |

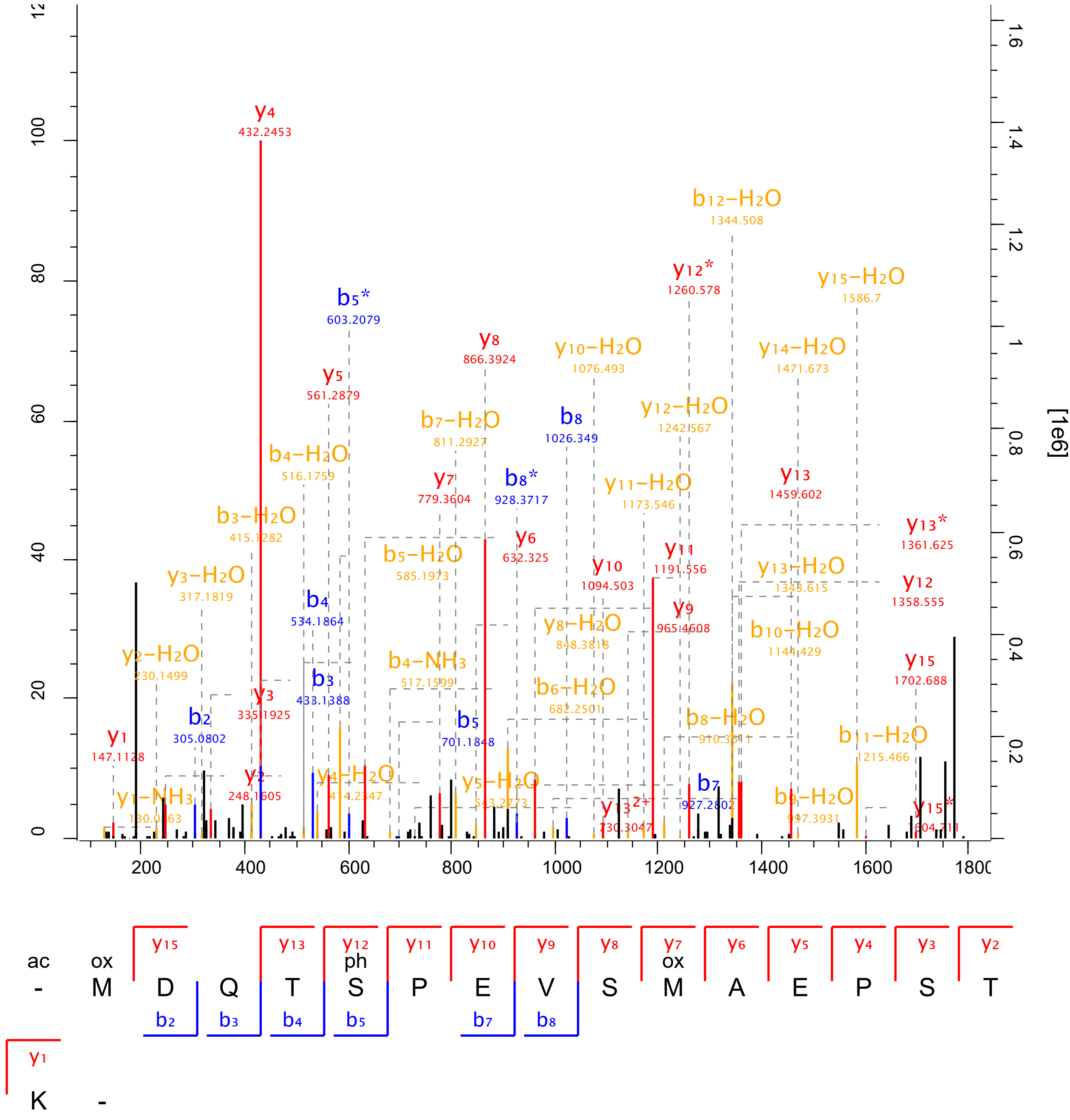

| Raw file | Scan | Method | Score | m/z |
|---|---|---|---|---|
| A17_15_1 | 17013 | FTMS; HCD | 66.66 | 533.22 |

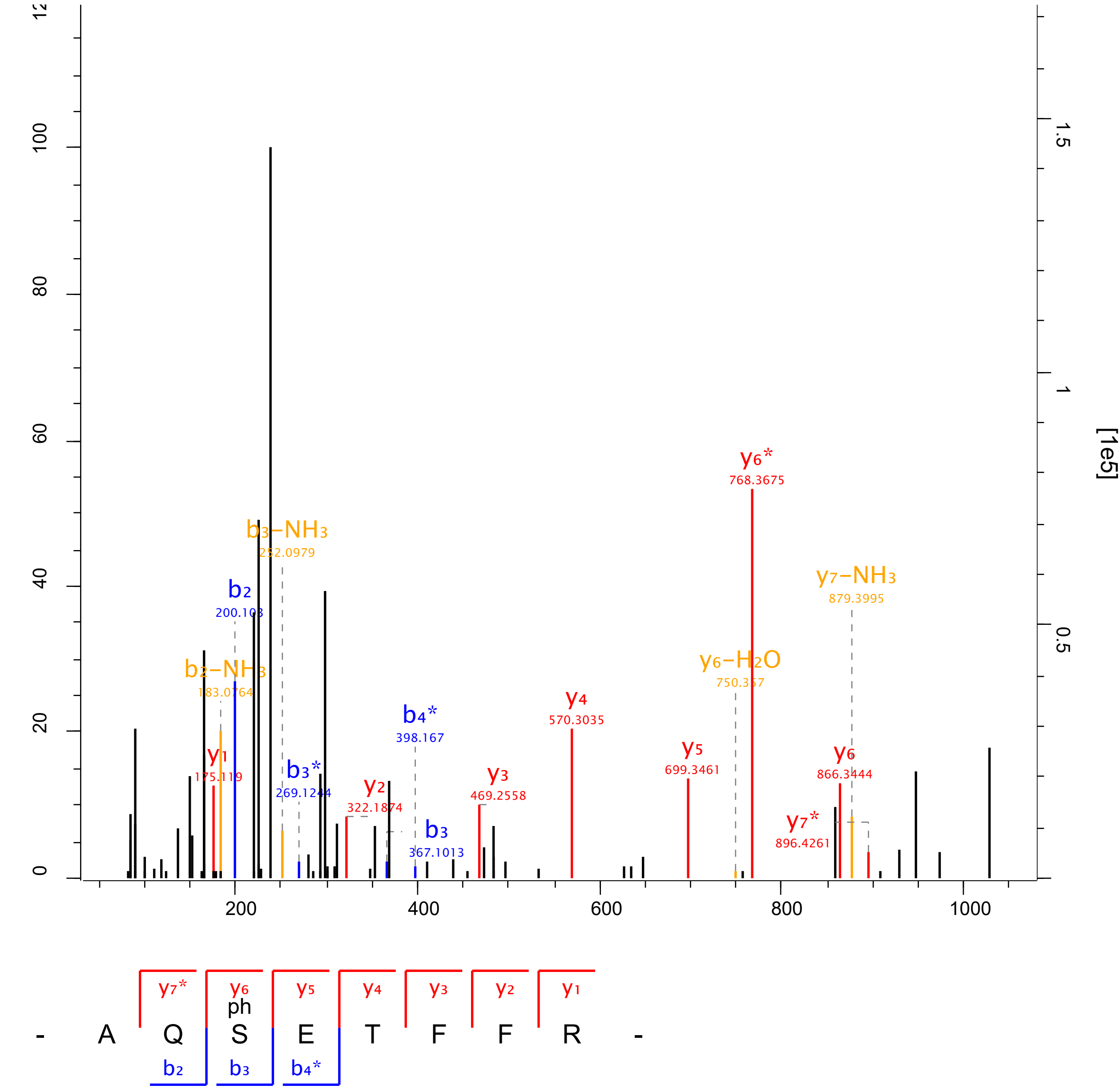

| Raw file | Scan | Method | Score | m/z |
|---|---|---|---|---|
| A17_15_1 | 17067 | FTMS; HCD | 222.46 | 895.88 |

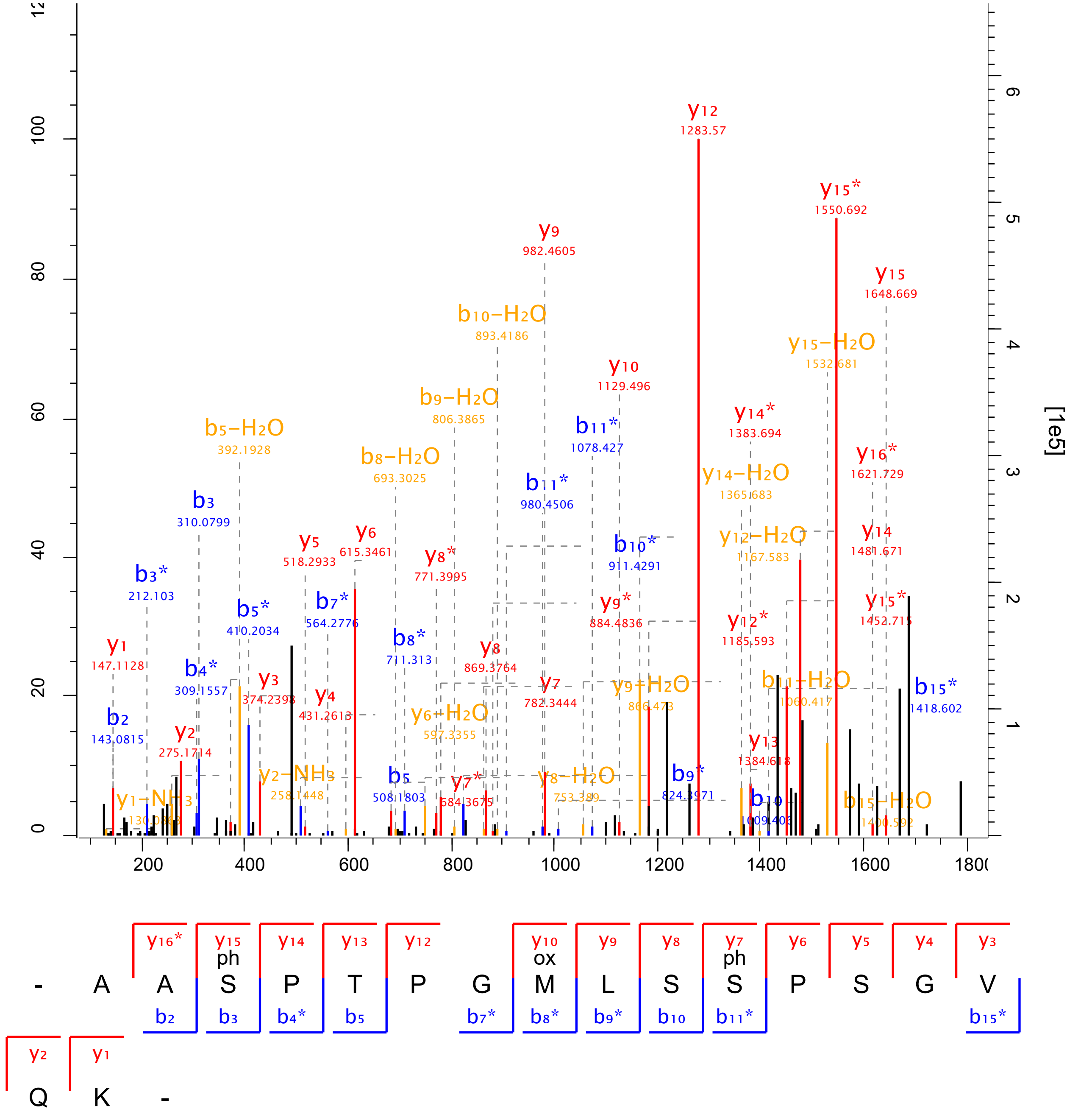

| Raw file | Scan | Method | Score | m/z |
|---|---|---|---|---|
| A17_15_1 | 17119 | FTMS; HCD | 135.62 | 669.95 |

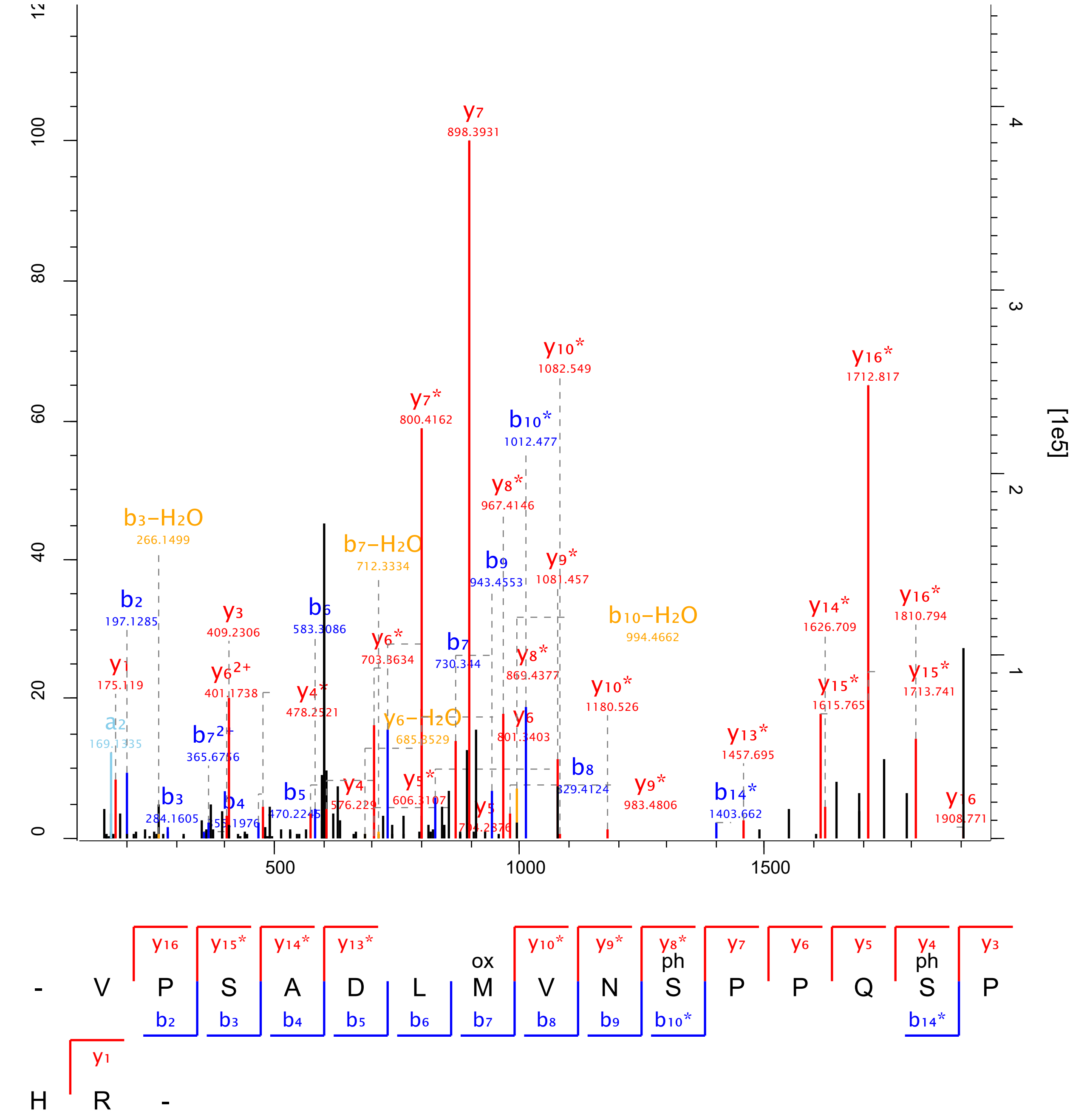

| Raw file | Scan | Method | Score | m/z |
| --- | --- | --- | --- | --- |
| A17_15_1 | 17435 | FTMS; HCD | 112.55 | 933.85 |

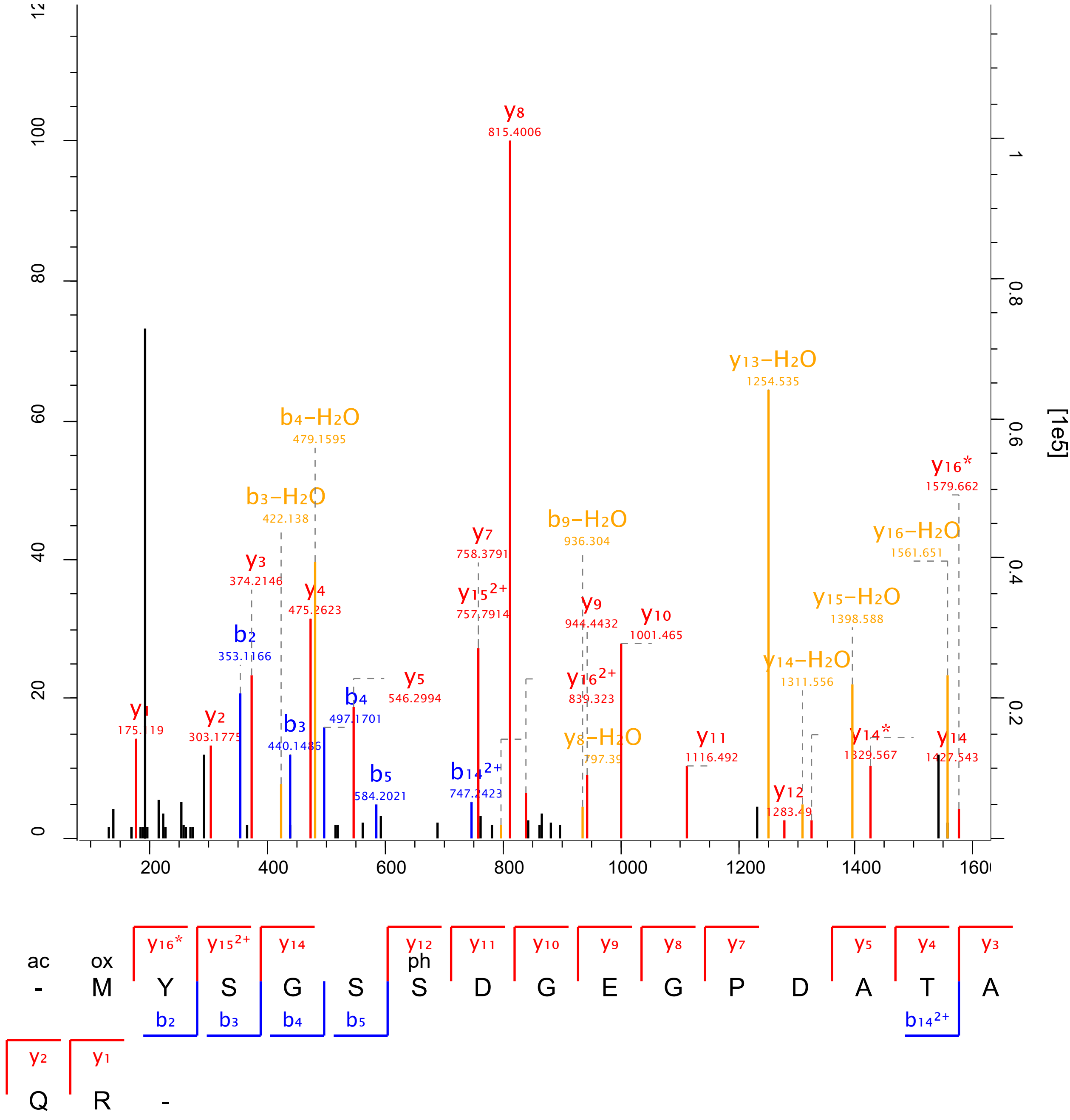

| Raw file | Scan | Method | Score | m/z |
| --- | --- | --- | --- | --- |
| A17_15_1 | 17740 | FTMS; HCD | 149.31 | 733.79 |

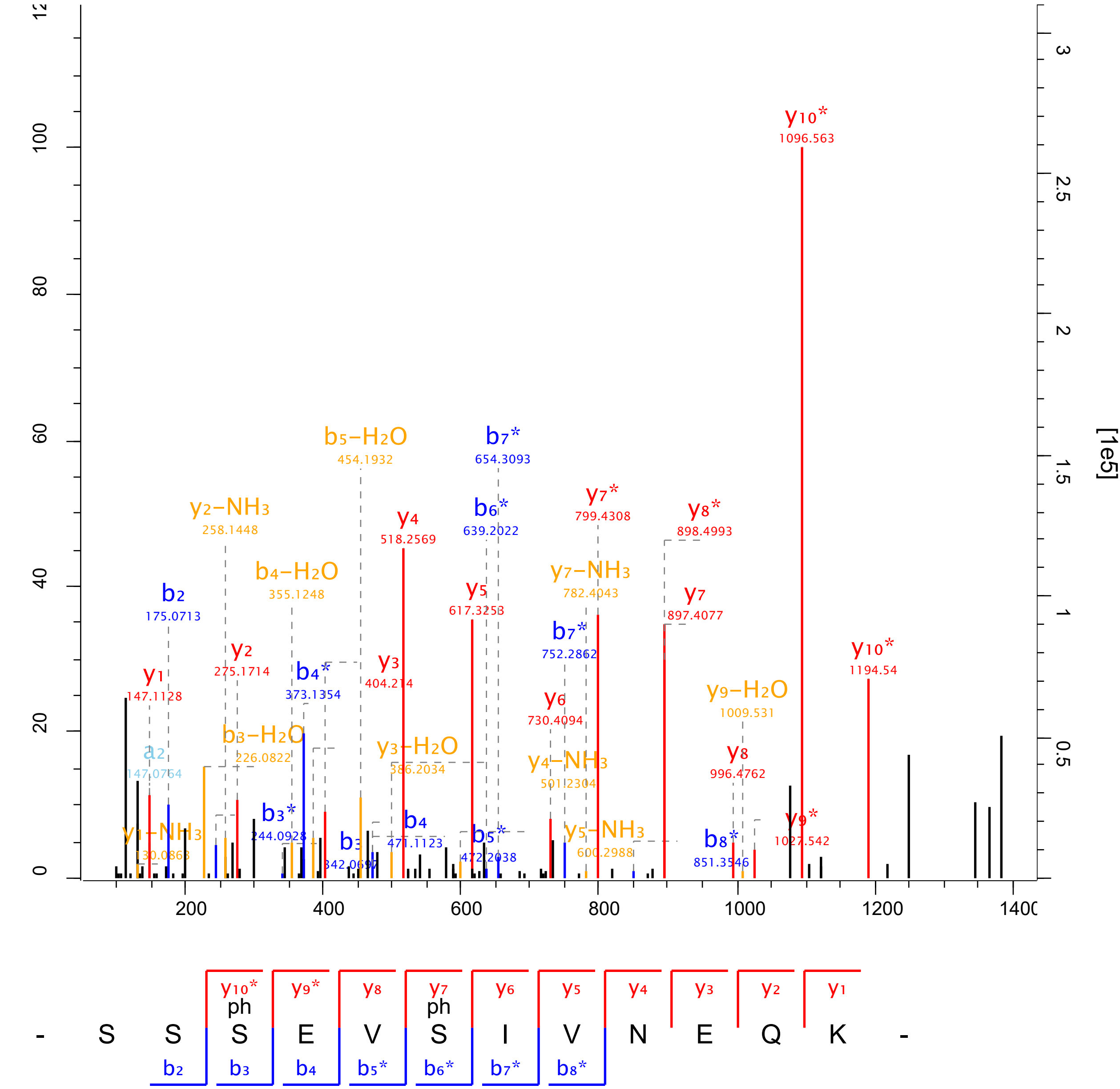

| Raw file | Scan | Method | Score | m/z |
| --- | --- | --- | --- | --- |
| A17_15_1 | 17814 | FTMS; HCD | 51.88 | 664.97 |

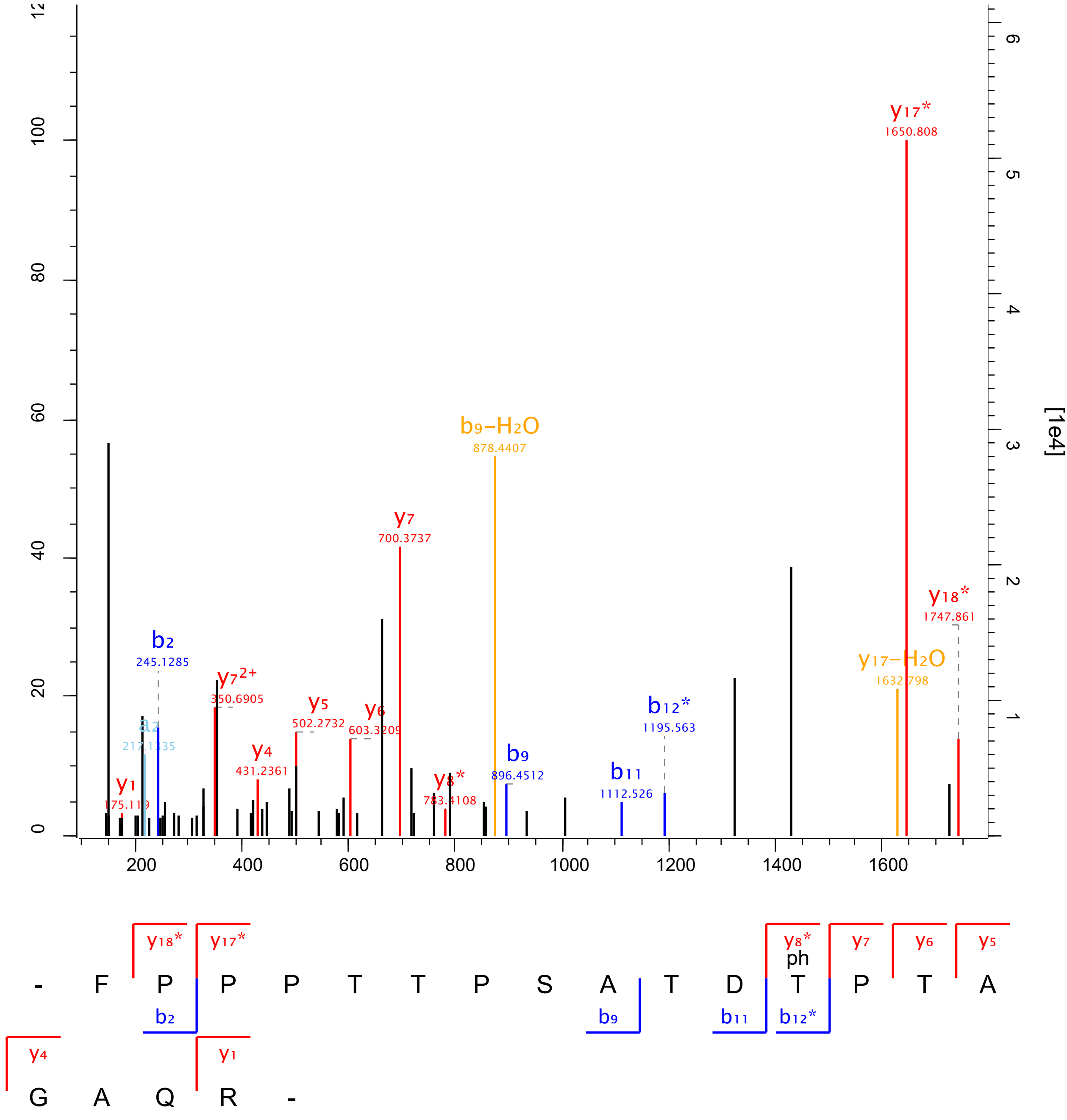

| Raw file | Scan | Method | Score | m/z |
|---|---|---|---|---|
| A17_15_1 | 18044 | FTMS; HCD | 183.23 | 588.26 |

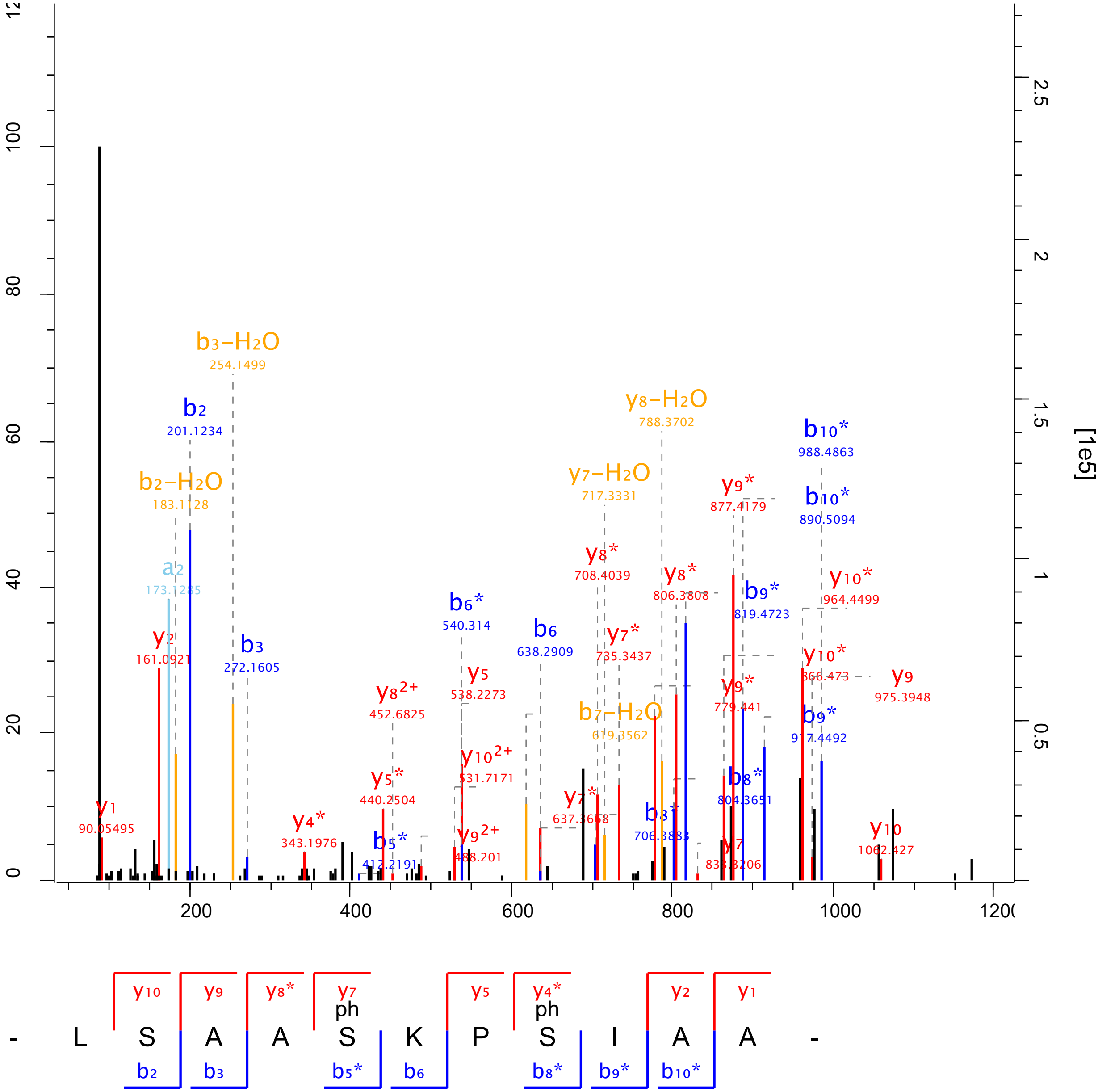

| Raw file | Scan | Method | Score | m/z |
|---|---|---|---|---|
| A17_15_1 | 18238 | FTMS; HCD | 112.34 | 671.3 |

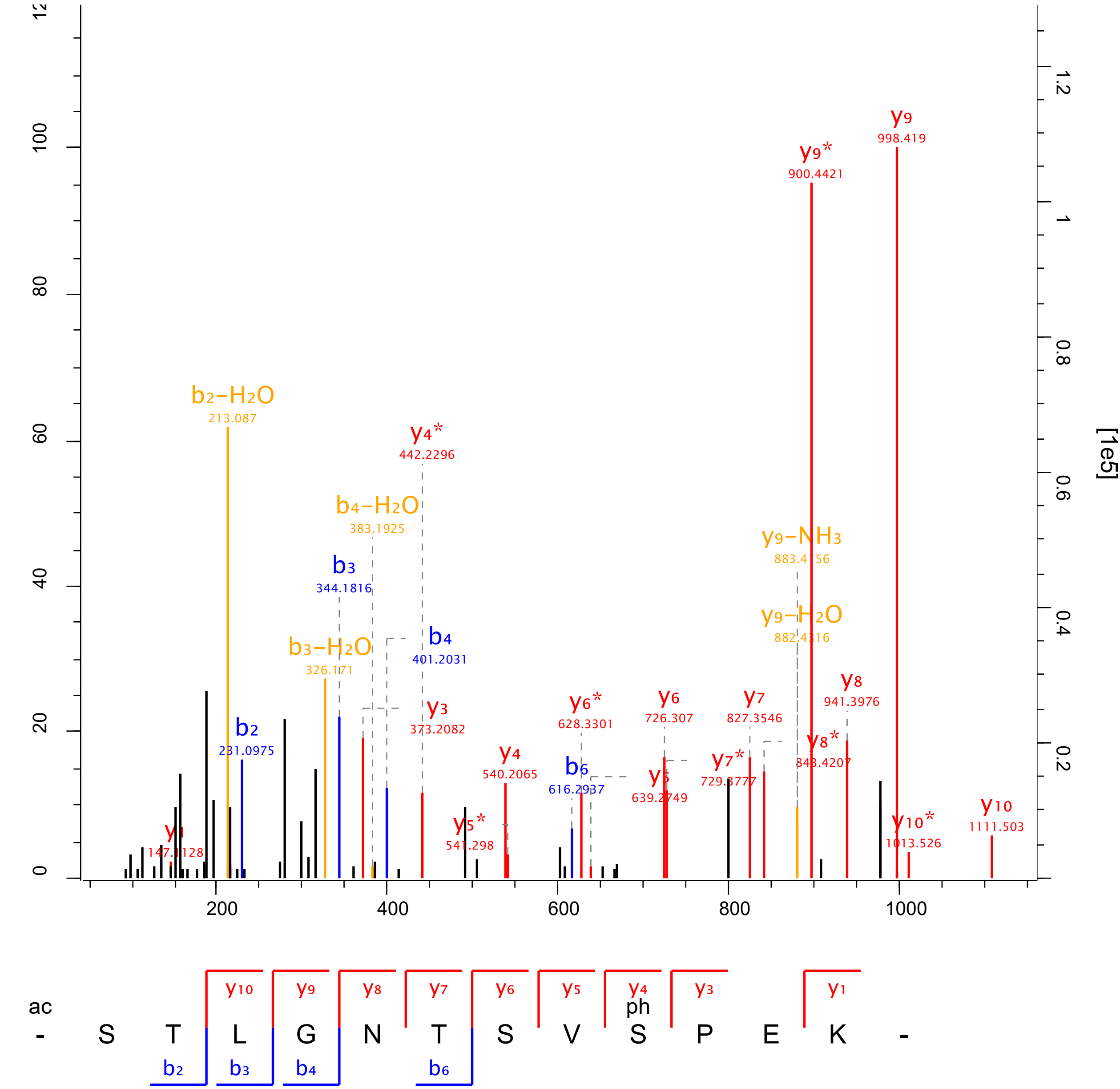

| Raw file | Scan | Method | Score | m/z |
|---|---|---|---|---|
| A17_15_1 | 18552 | FTMS; HCD | 84.33 | 1109.91 |

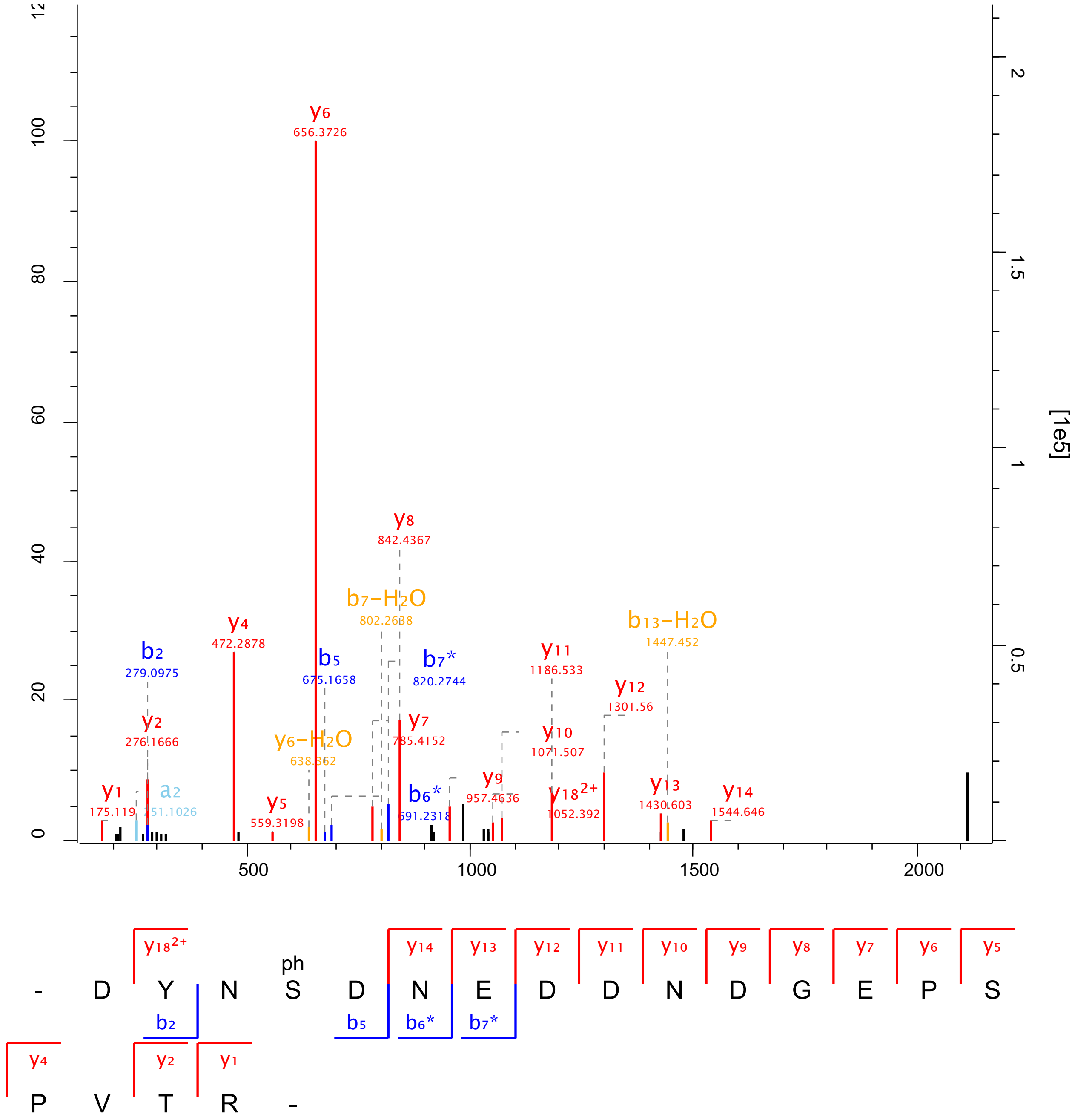

| Raw file | Scan | Method | Score | m/z |
| --- | --- | --- | --- | --- |
| A17_15_1 | 18601 | FTMS; HCD | 69.98 | 839.87 |

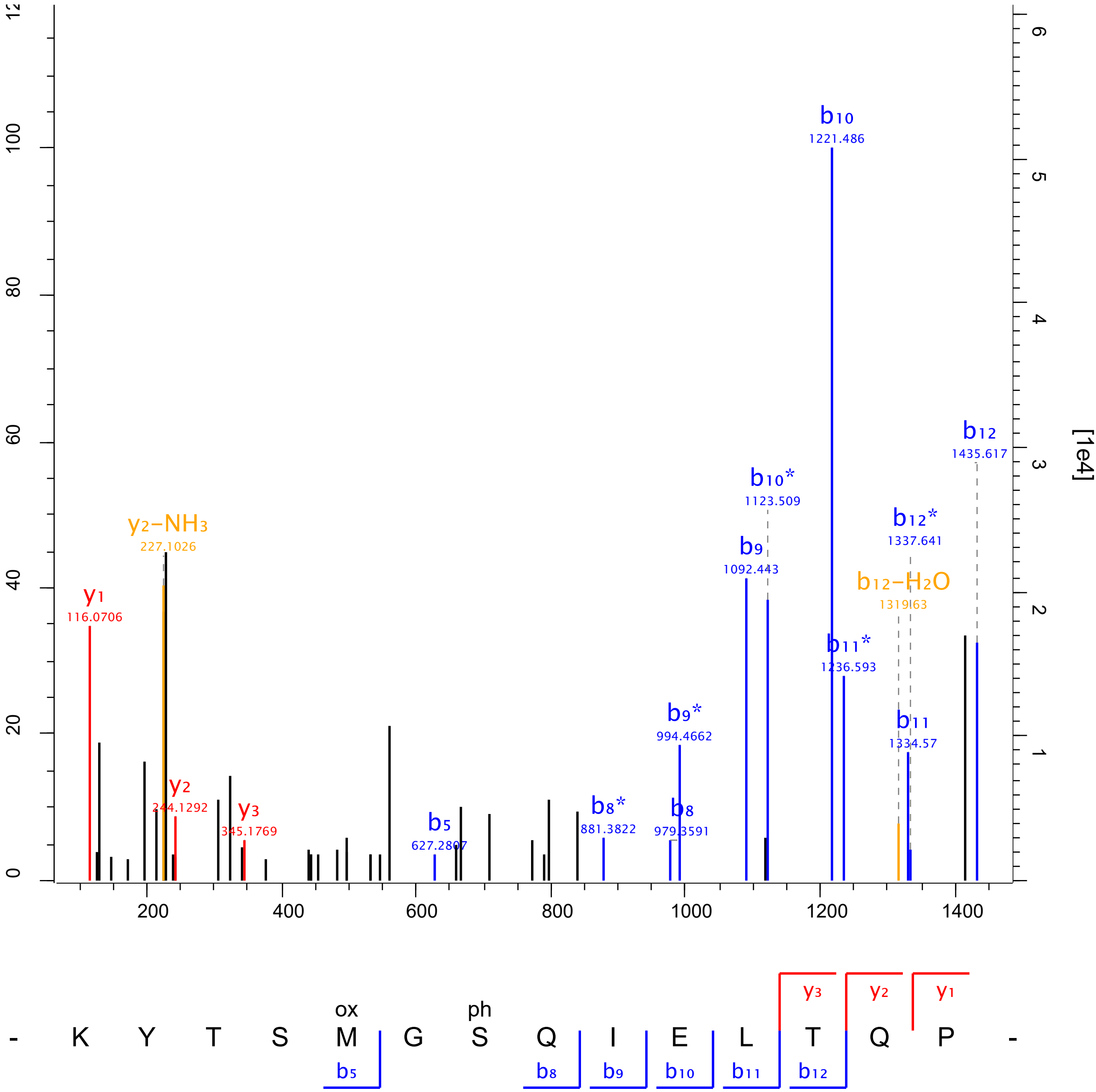

| Raw file | Scan | Method | Score | m/z |
| --- | --- | --- | --- | --- |
| A17_15_1 | 18665 | FTMS; HCD | 63.87 | 937.89 |

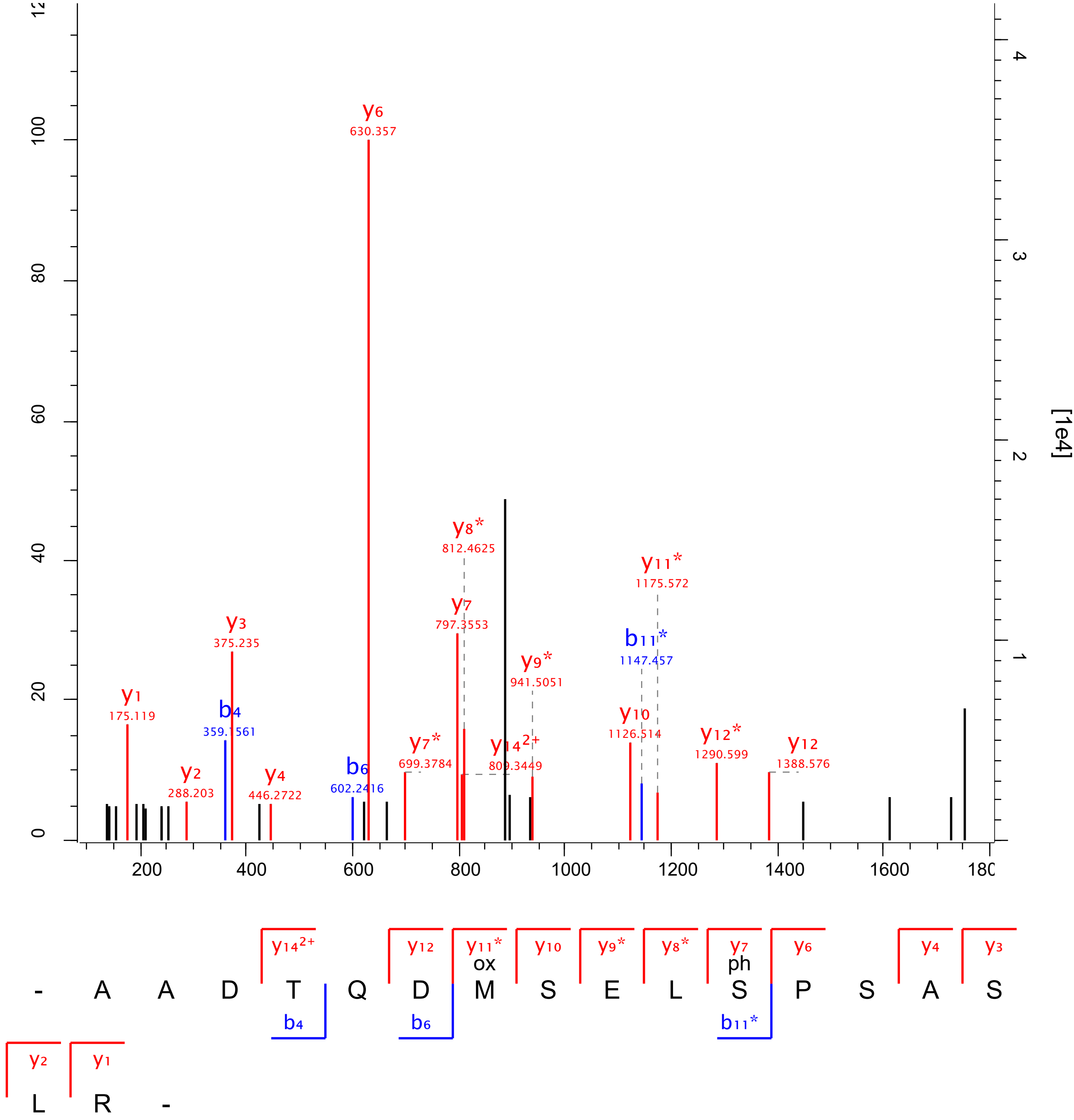

| Raw file | Scan | Method | Score | m/z |
|---|---|---|---|---|
| A17_15_1 | 19187 | FTMS; HCD | 93.35 | 580.74 |

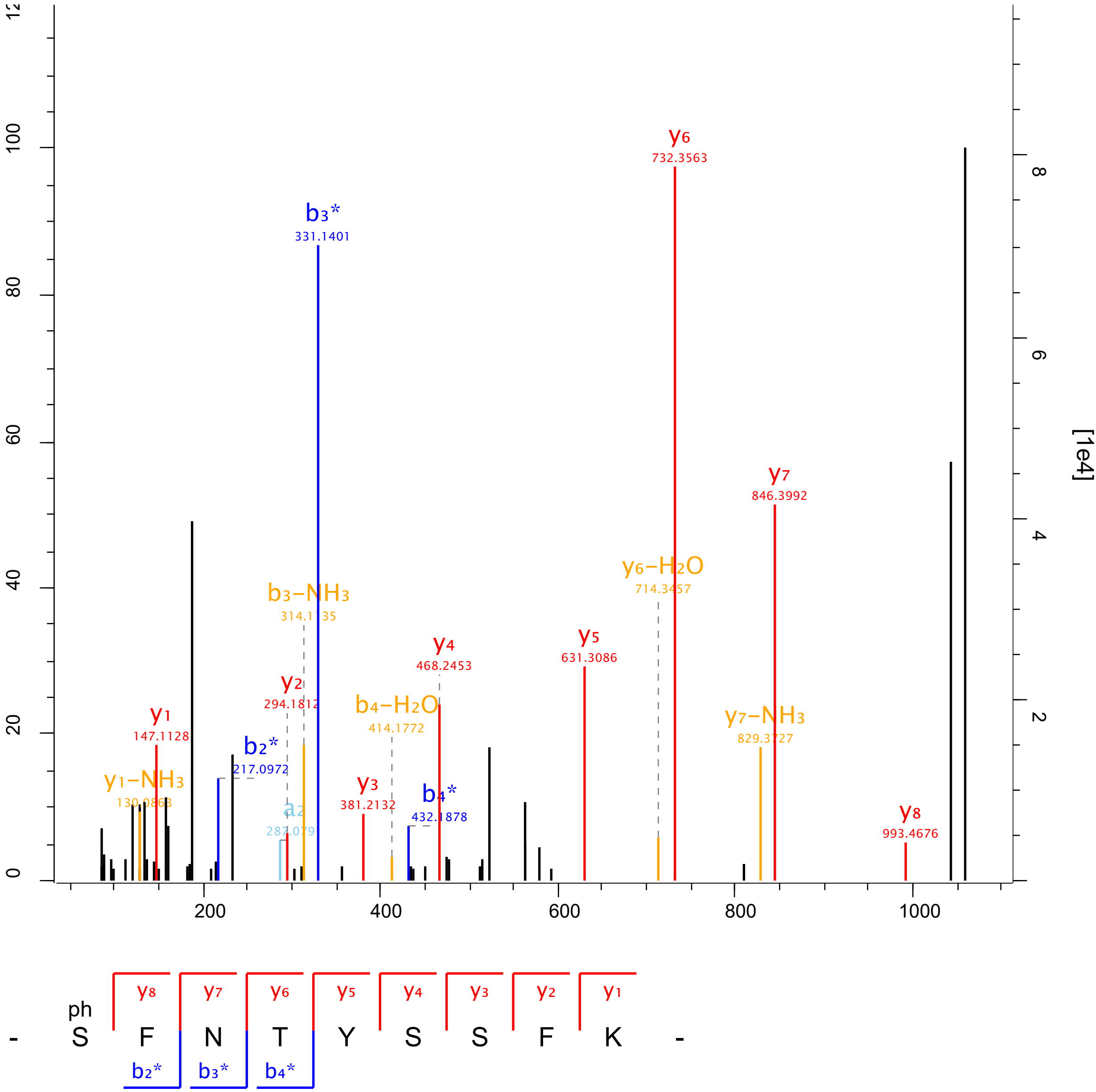

| Raw file | Scan | Method | Score | m/z |
|---|---|---|---|---|
| A17_15_1 | 19189 | FTMS; HCD | 55.26 | 572.7 |

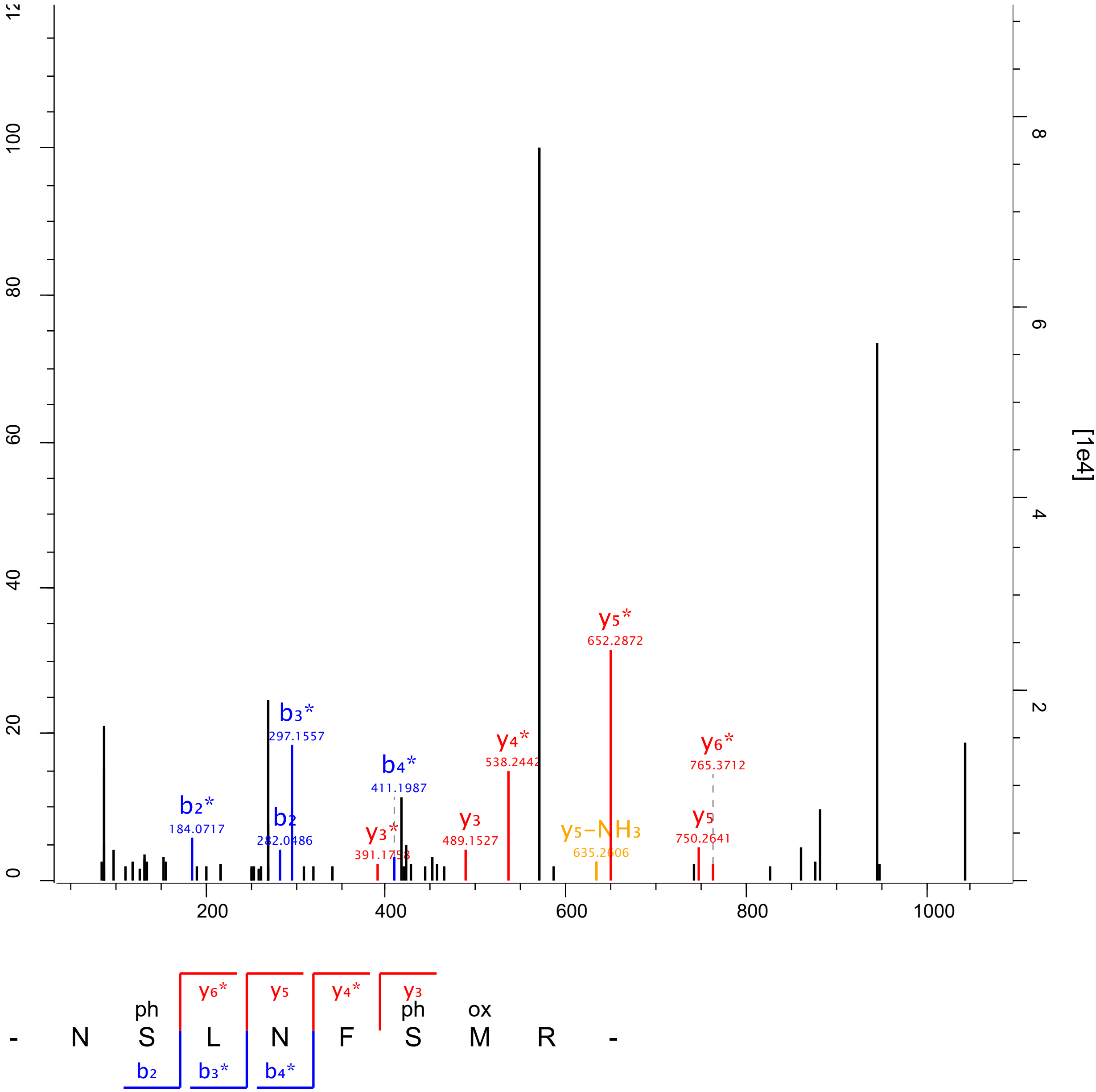

| Raw file | Scan | Method | Score | m/z |
|---|---|---|---|---|
| A17_15_1 | 19418 | FTMS; HCD | 188.96 | 787.81 |

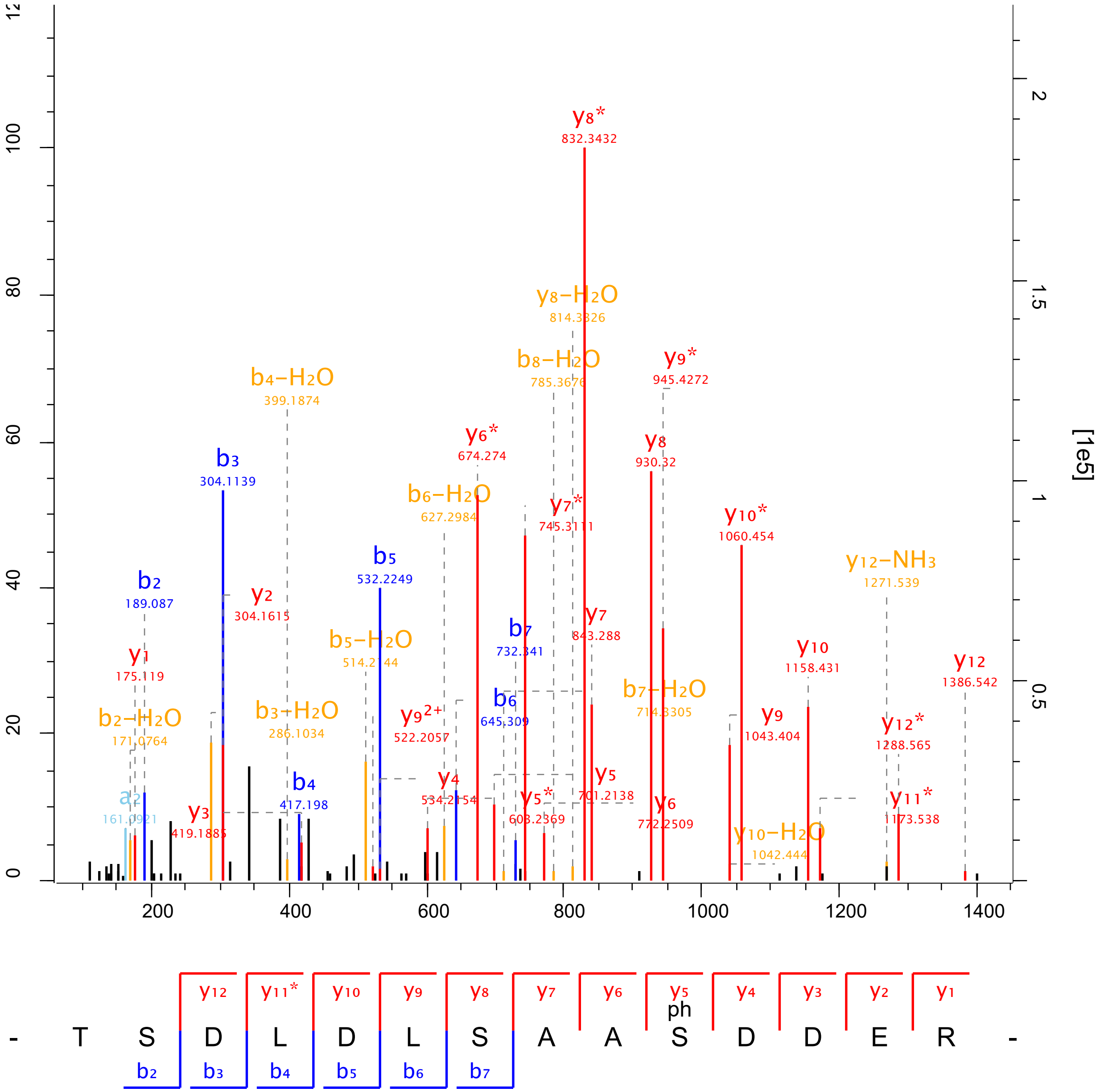

| Raw file | Scan | Method | Score | m/z |
|---|---|---|---|---|
| A17_15_1 | 19755 | FTMS; HCD | 53.3 | 889.89 |

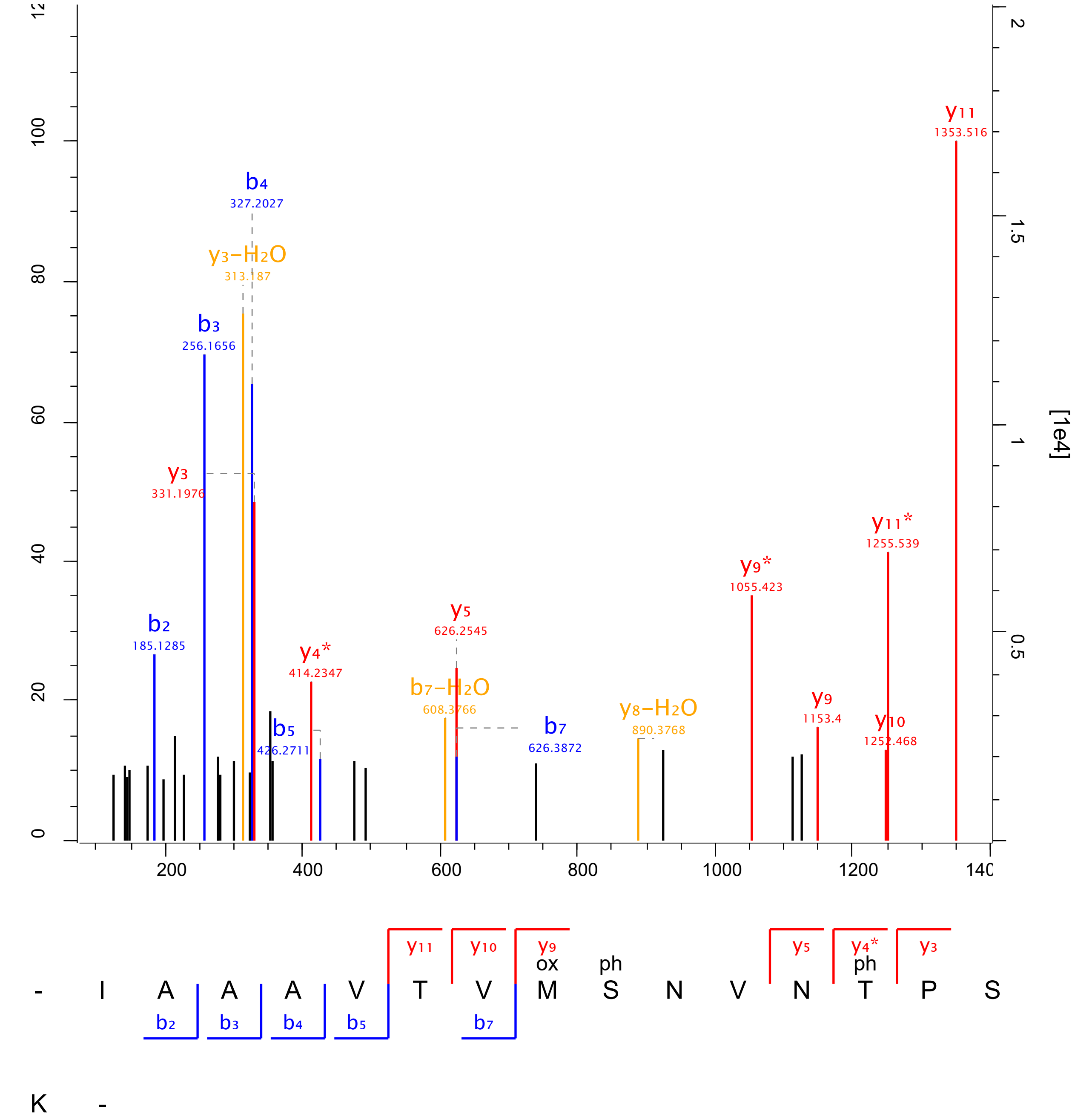

| Raw file | Scan | Method | Score | m/z |
|---|---|---|---|---|
| A17_15_1 | 19893 | FTMS; HCD | 46.89 | 724.3 |

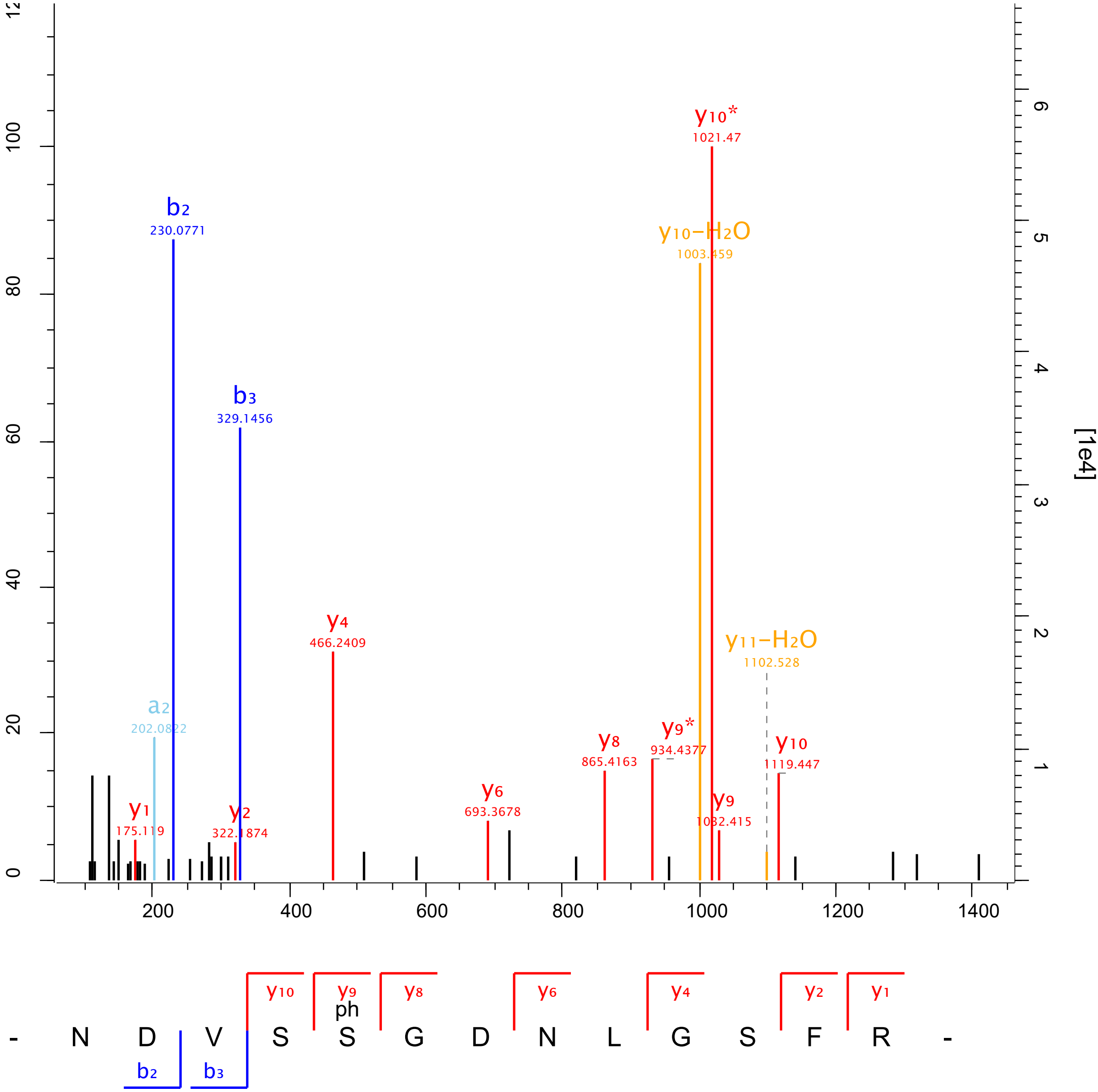

| Raw file | Scan | Method | Score | m/z |
|---|---|---|---|---|
| A17_15_1 | 20009 | FTMS; HCD | 68.17 | 843.31 |

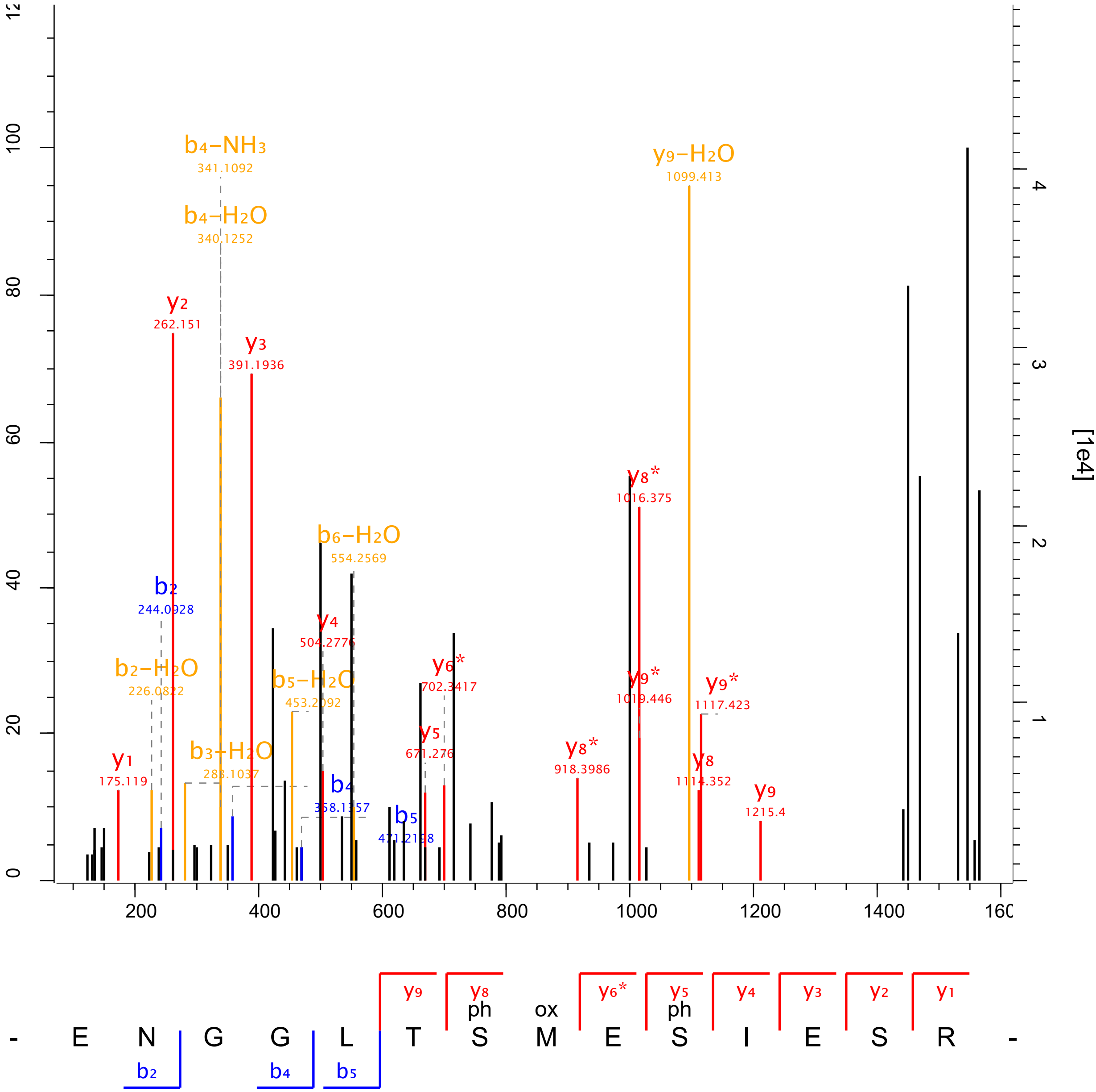

| Raw file | Scan | Method | Score | m/z |
|---|---|---|---|---|
| A17_15_1 | 20087 | FTMS; HCD | 140.48 | 939.44 |

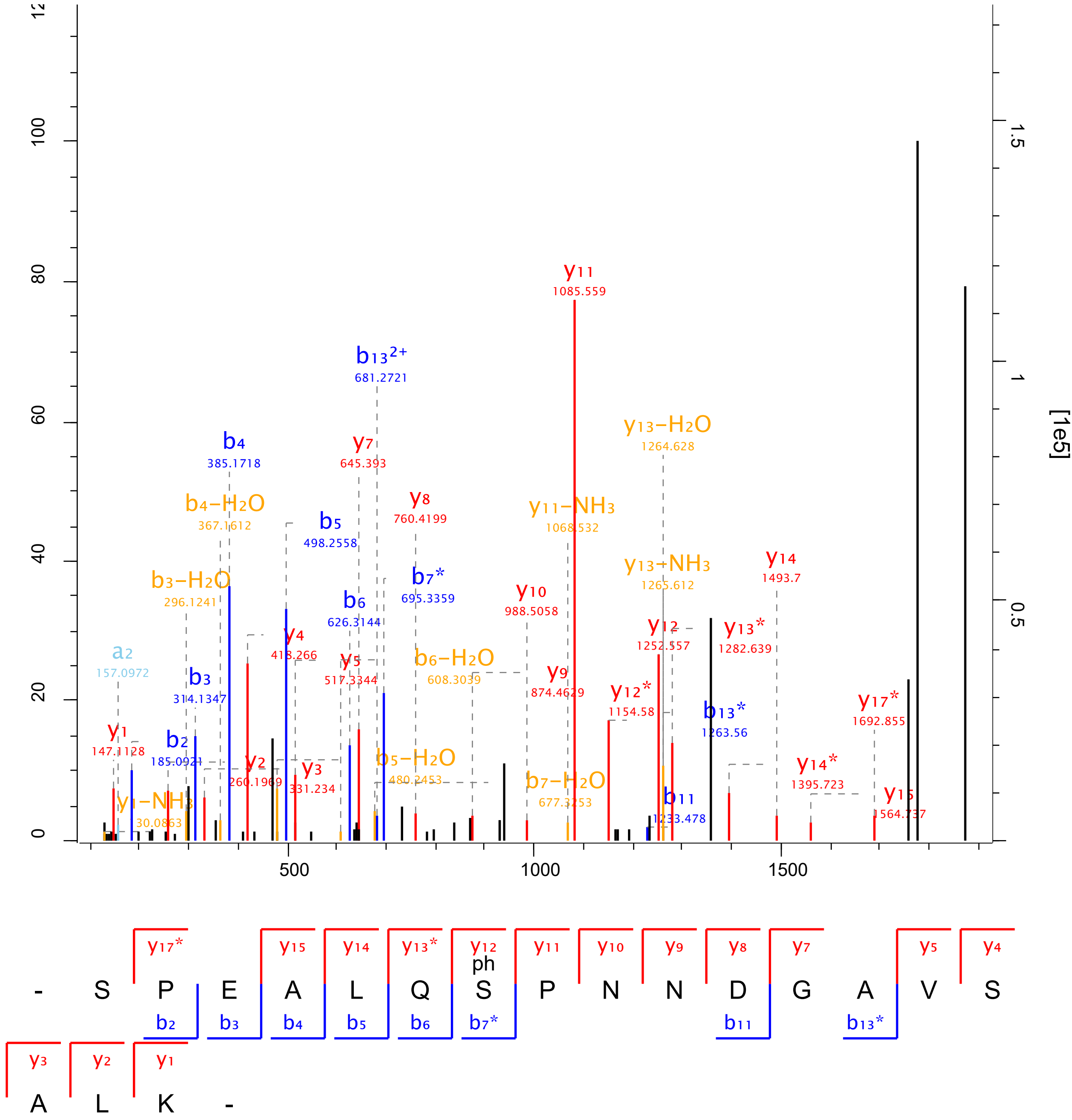

| Raw file | Scan | Method | Score | m/z |
|---|---|---|---|---|
| A17_15_1 | 20315 | FTMS; HCD | 71.59 | 770.32 |

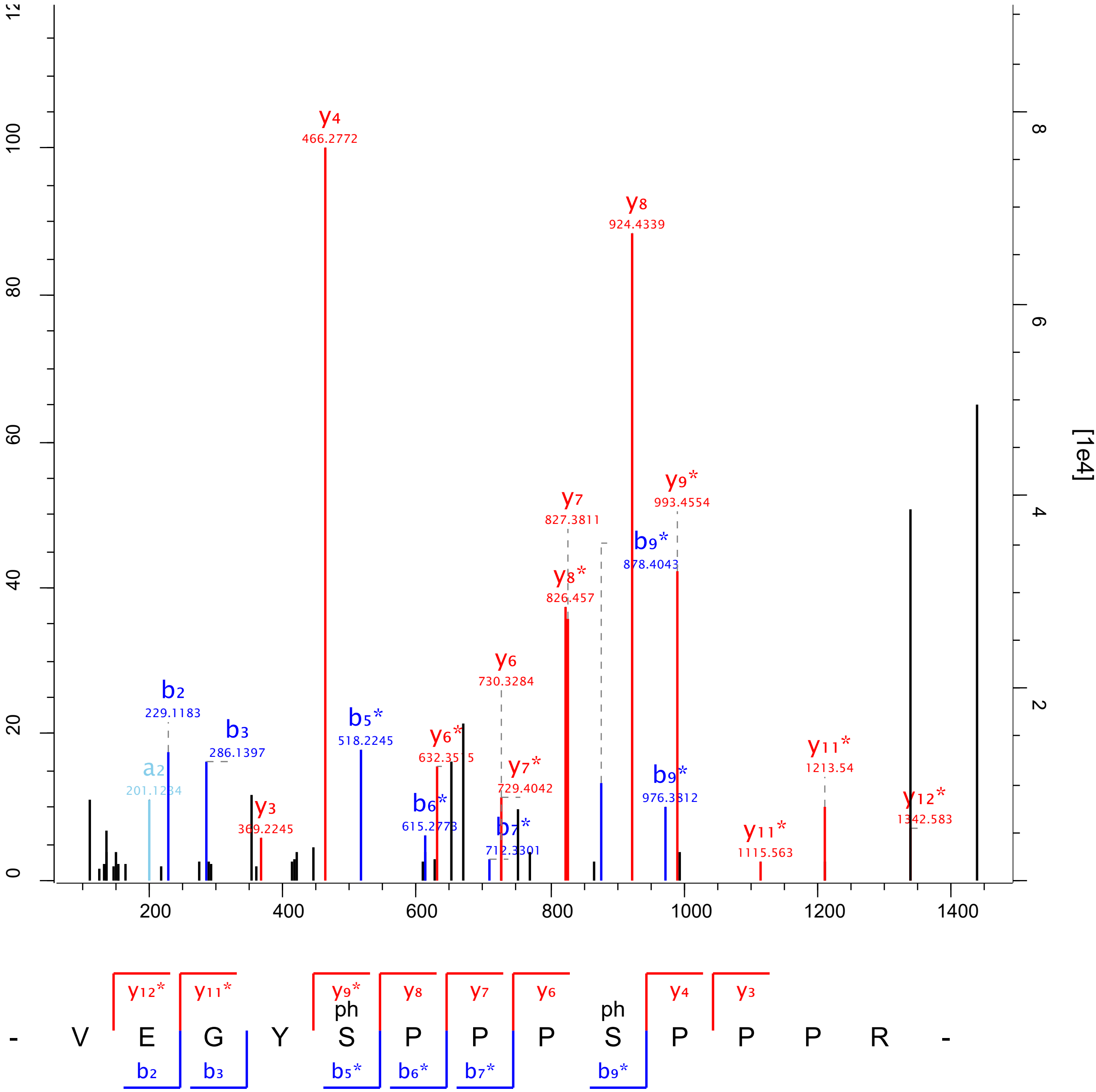

| Raw file | Scan | Method | Score | m/z |
|---|---|---|---|---|
| A17_15_1 | 20326 | FTMS; HCD | 114.55 | 746.31 |

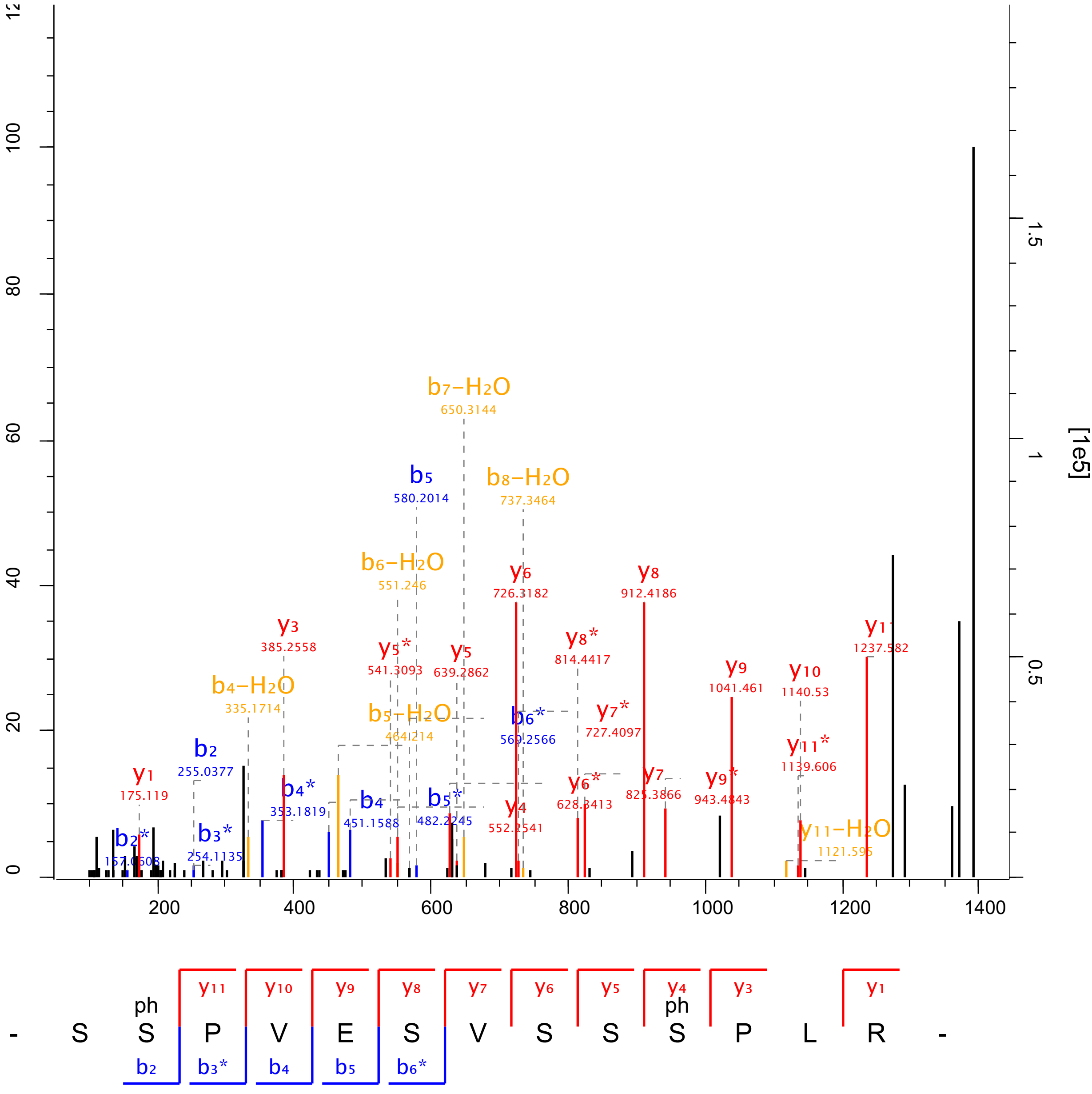

| Raw file | Scan | Method | Score | m/z |
|---|---|---|---|---|
| A17_15_1 | 20663 | FTMS; HCD | 112.55 | 787.89 |

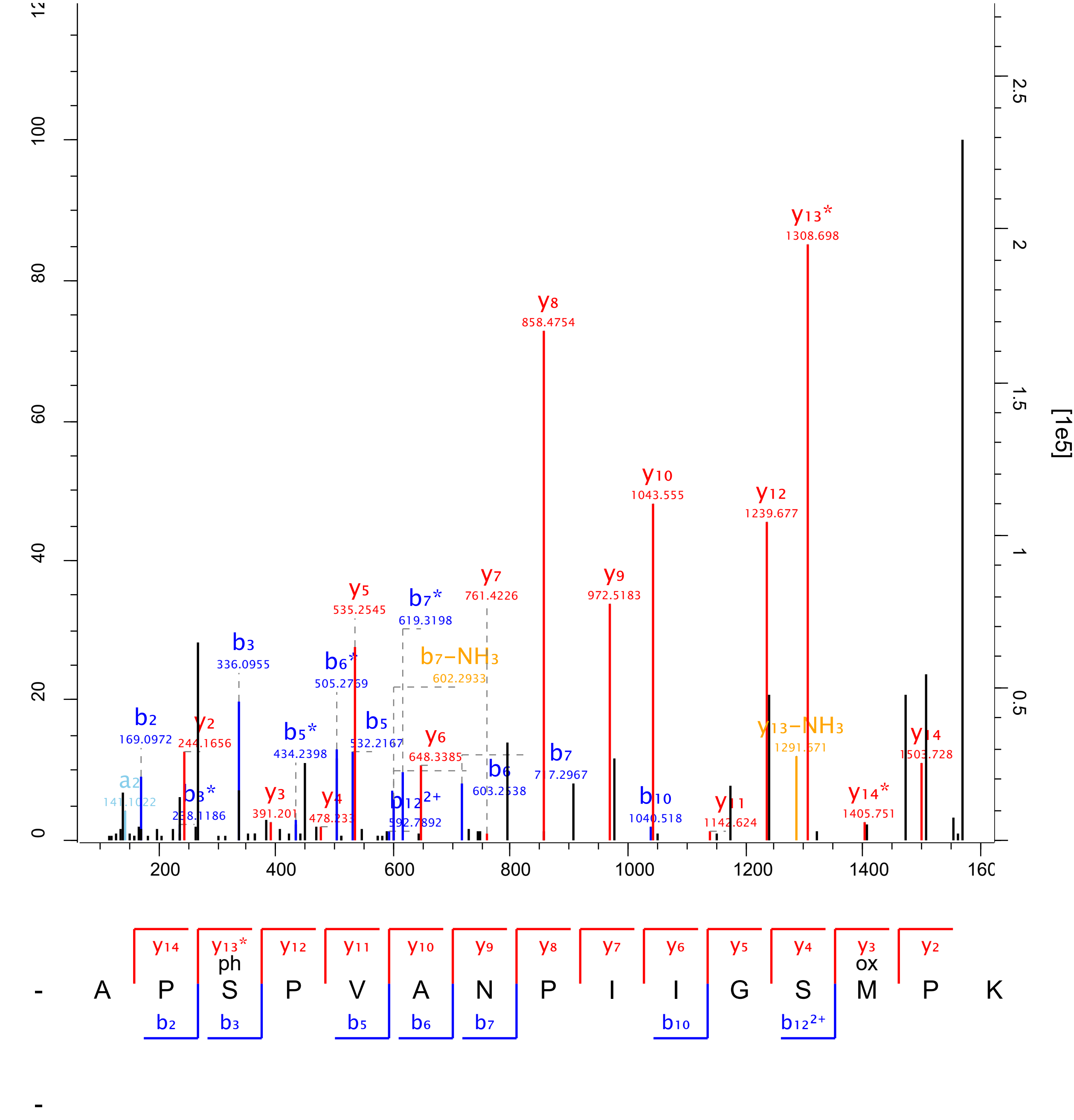

| Raw file | Scan | Method | Score | m/z |
|---|---|---|---|---|
| A17_15_1 | 21296 | FTMS; HCD | 72.62 | 674.25 |

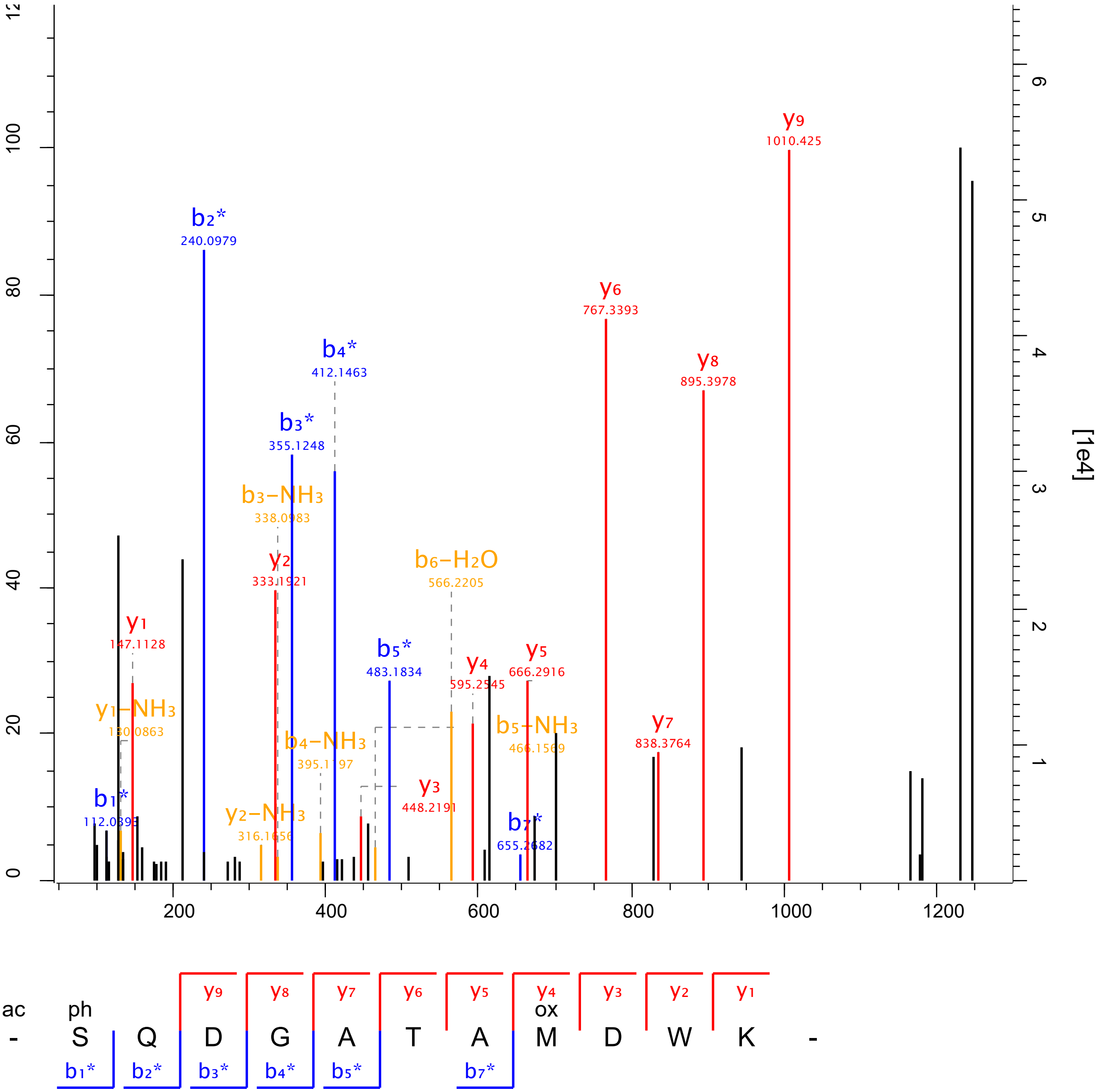

| Raw file | Scan | Method | Score | m/z |
|---|---|---|---|---|
| A17_15_1 | 21330 | FTMS; HCD | 64.59 | 697.31 |

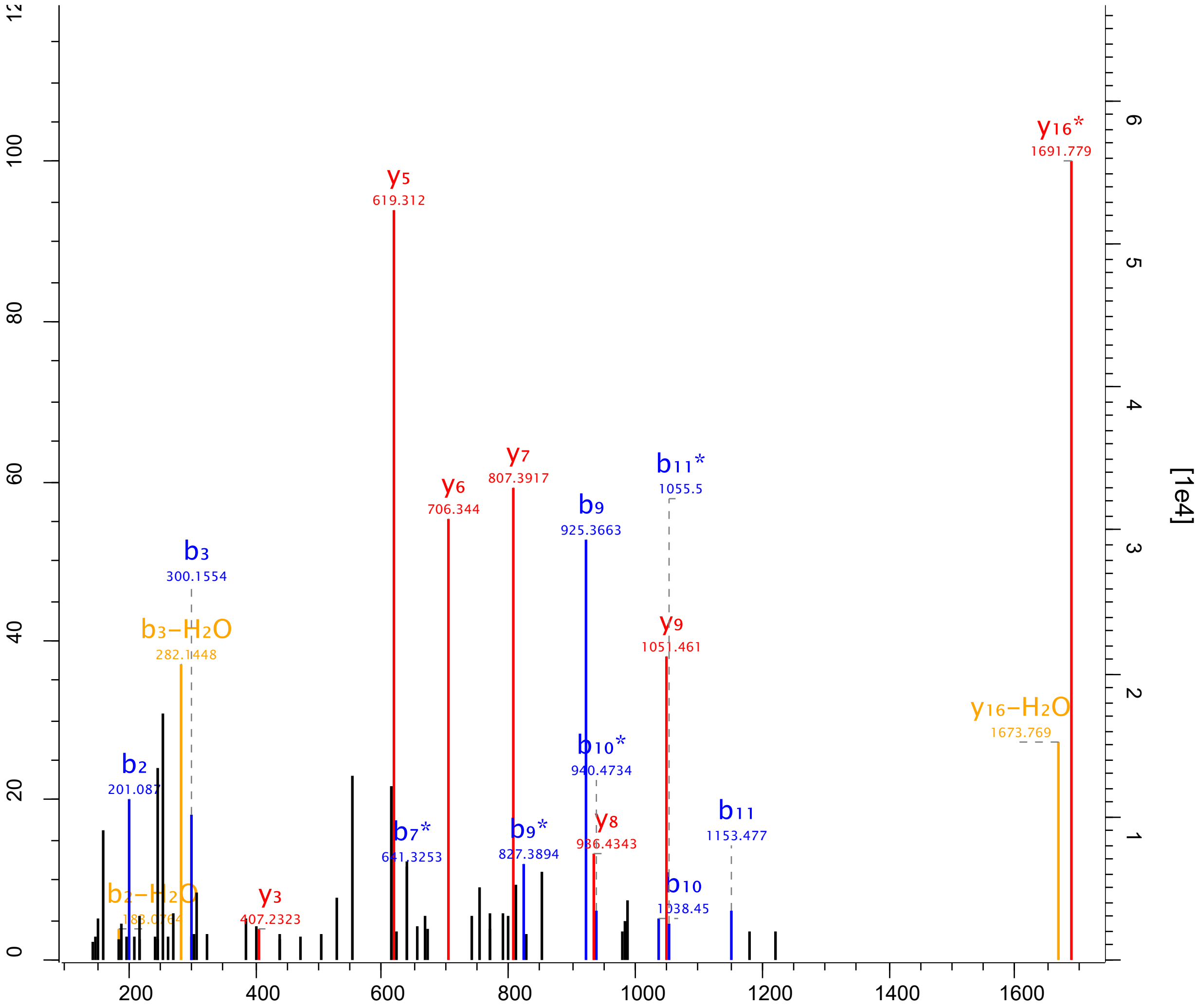


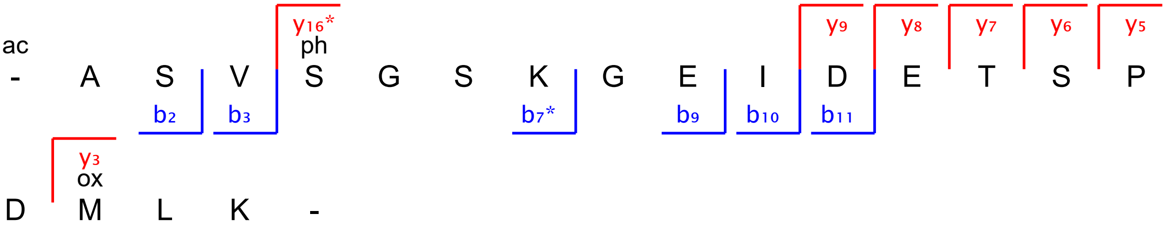

| Raw file | Scan | Method | Score | m/z |
|---|---|---|---|---|
| A17_15_1 | 21369 | FTMS; HCD | 159.98 | 937.41 |

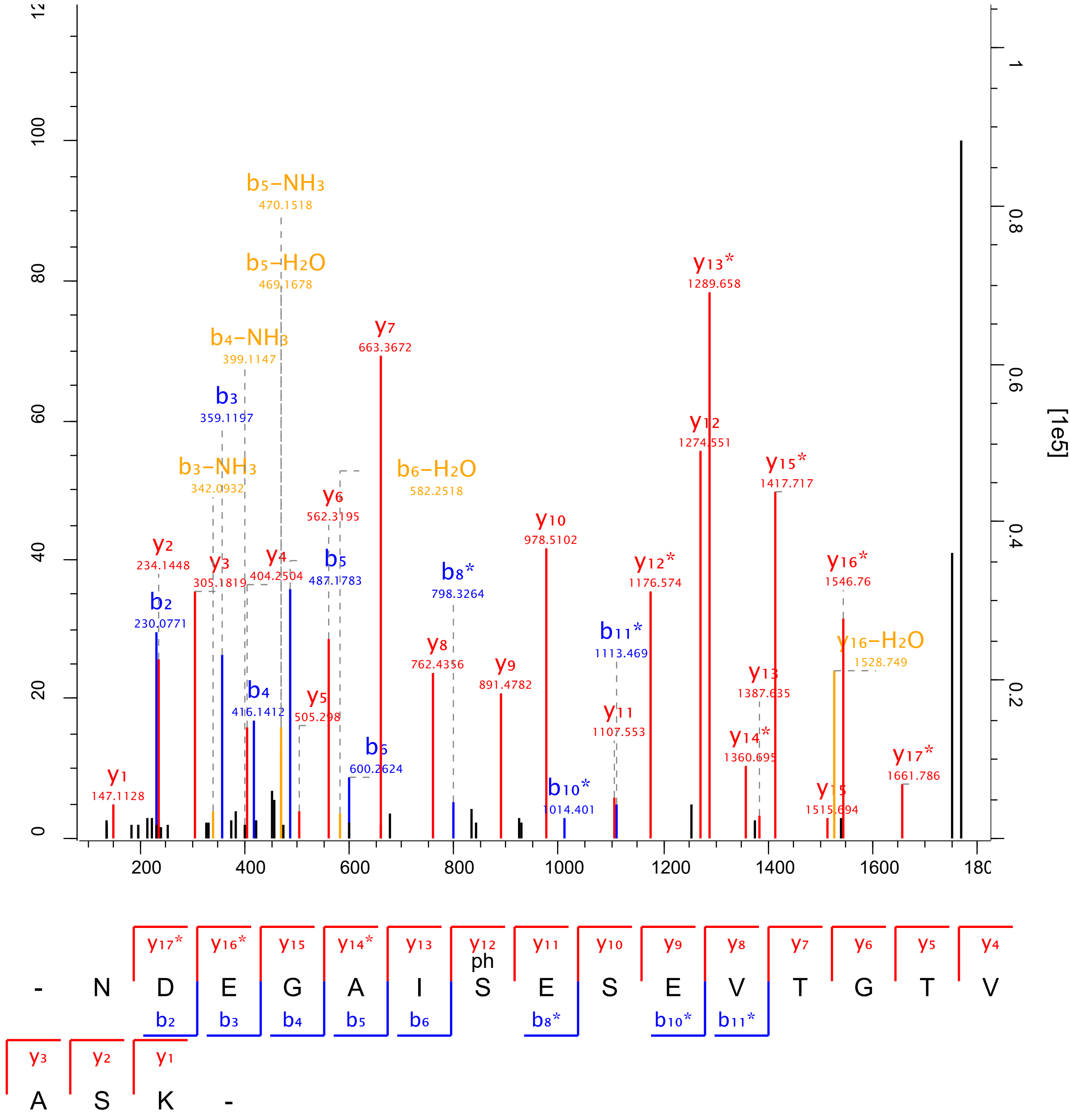

| Raw file | Scan | Method | Score | m/z |
|---|---|---|---|---|
| A17_15_1 | 21757 | FTMS; HCD | 41.7 | 739.85 |

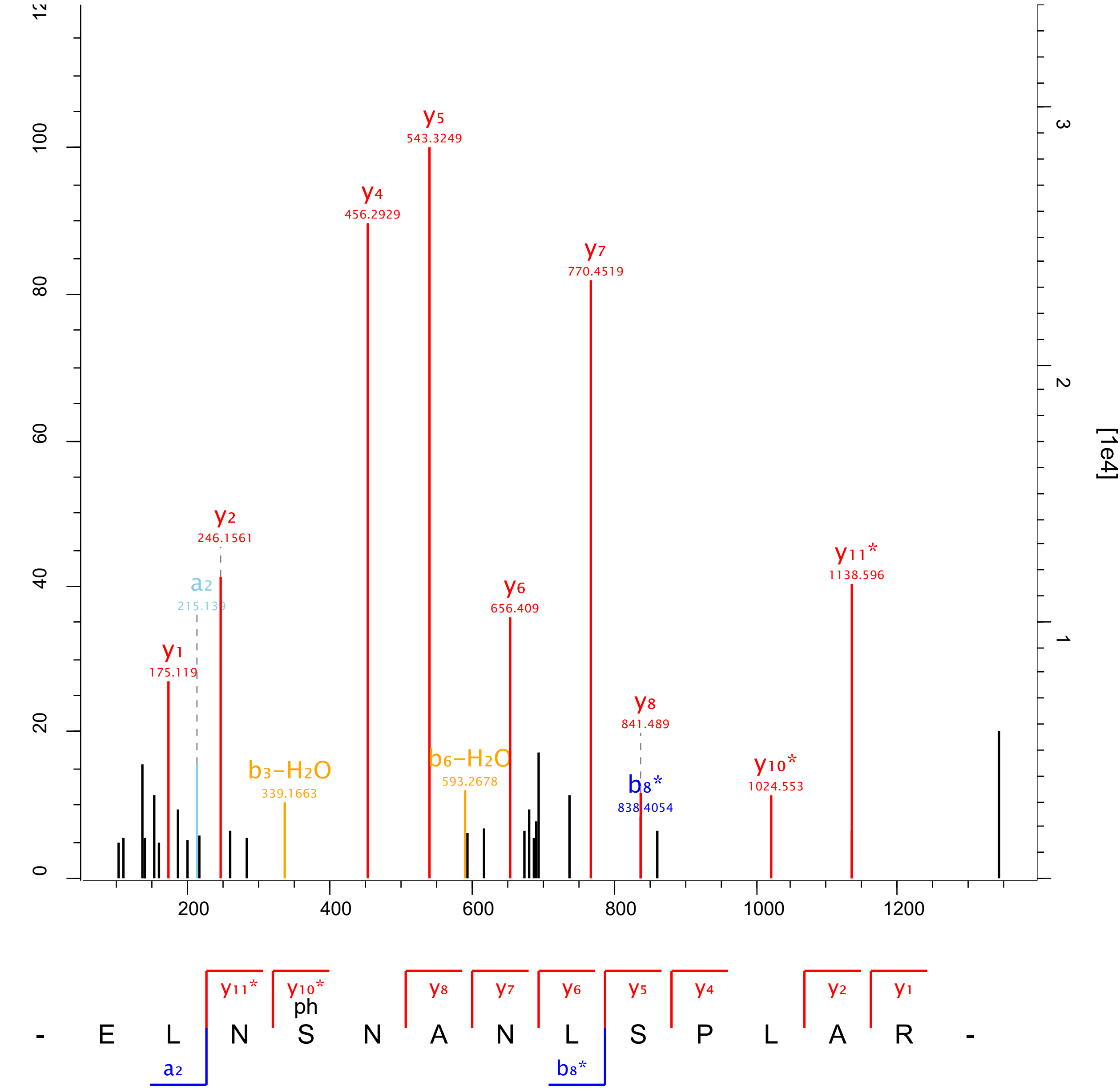

| Raw file | Scan | Method | Score | m/z |
|---|---|---|---|---|
| A17_15_1 | 22163 | FTMS; HCD | 103.44 | 553.29 |

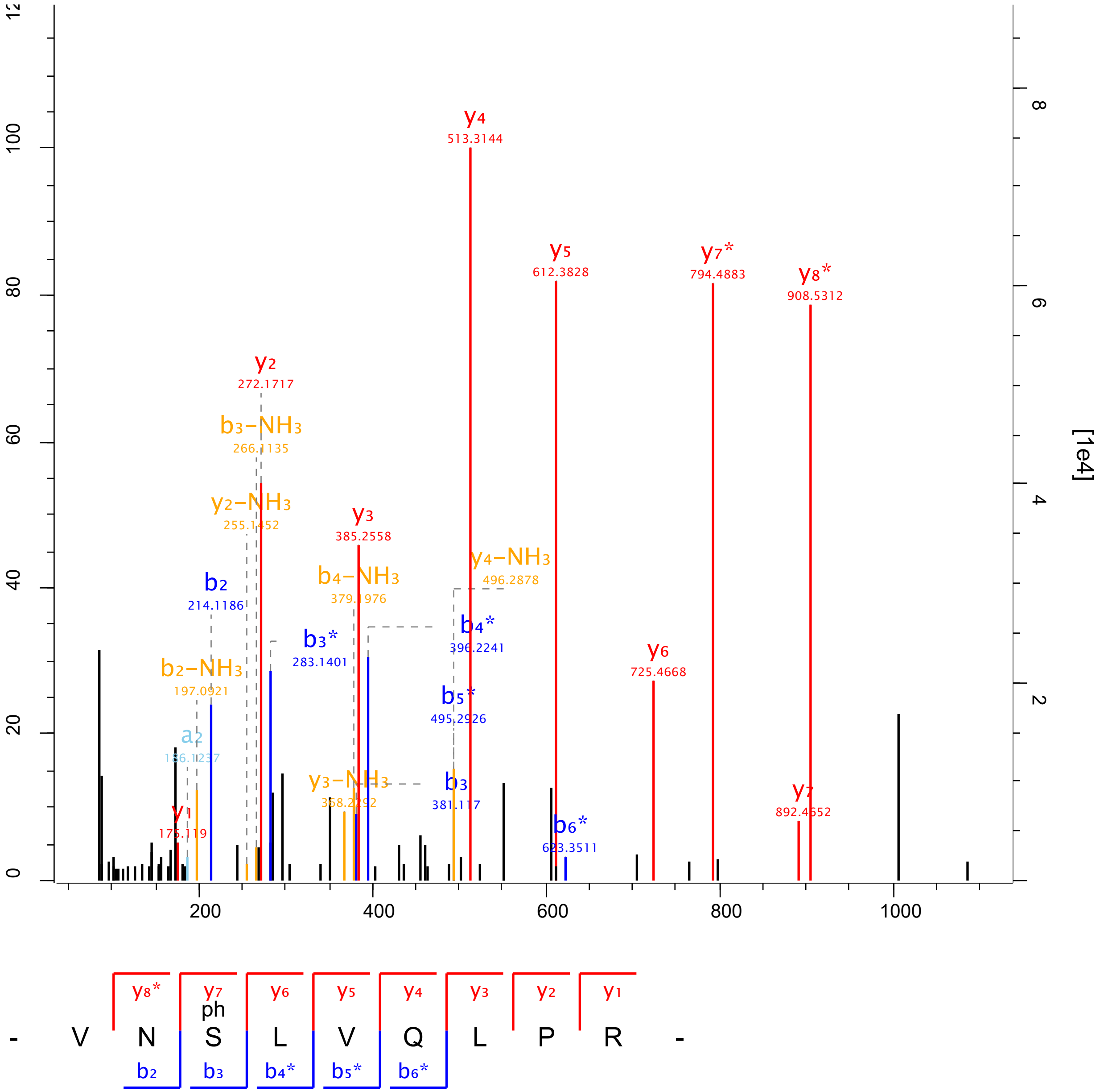

| Raw file | Scan | Method | Score | m/z |
|---|---|---|---|---|
| A17_15_1 | 22186 | FTMS; HCD | 70.2 | 682.78 |

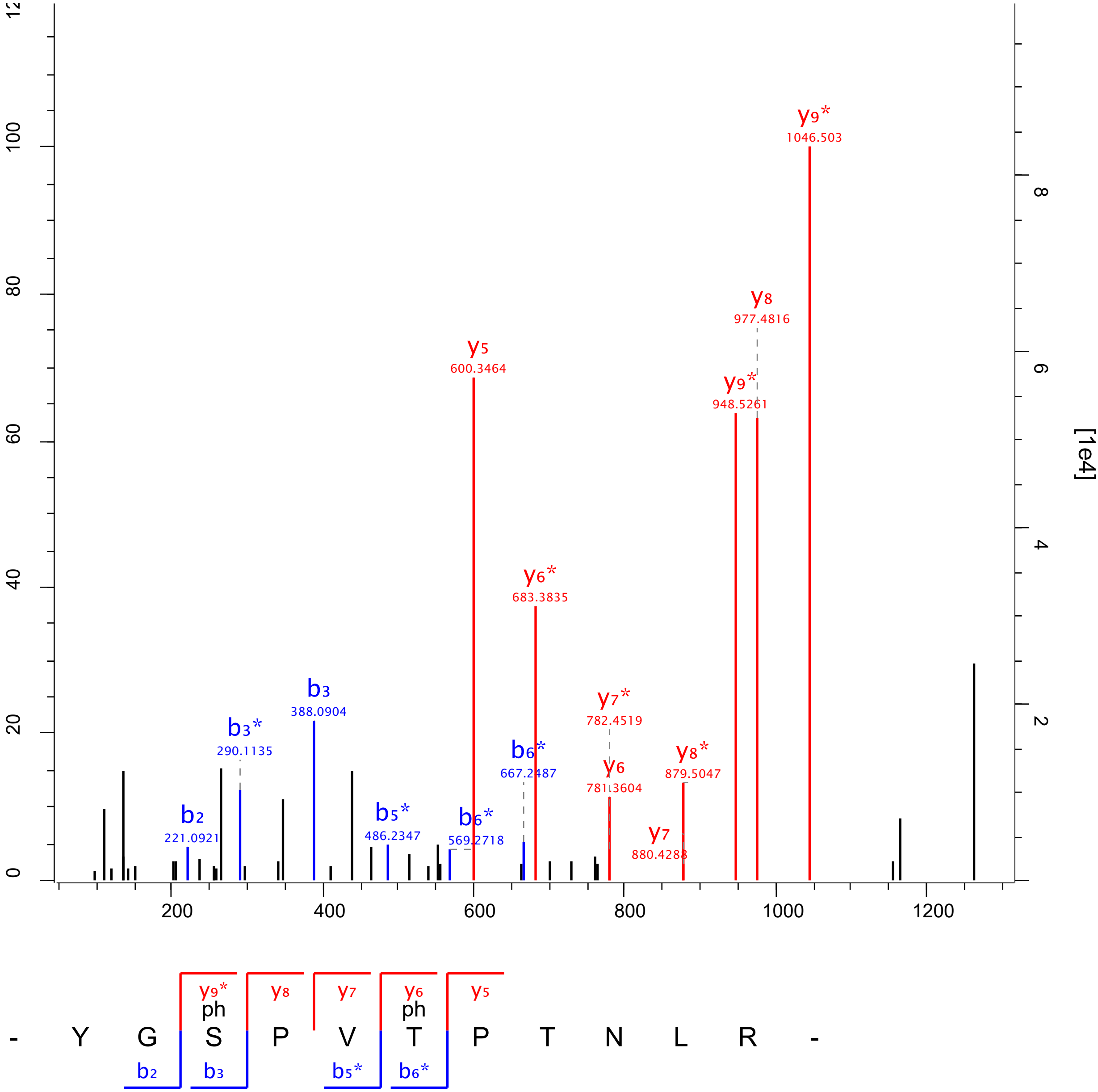

| Raw file | Scan | Method | Score | m/z |
|---|---|---|---|---|
| A17_15_1 | 22337 | FTMS; HCD | 57.17 | 700.81 |

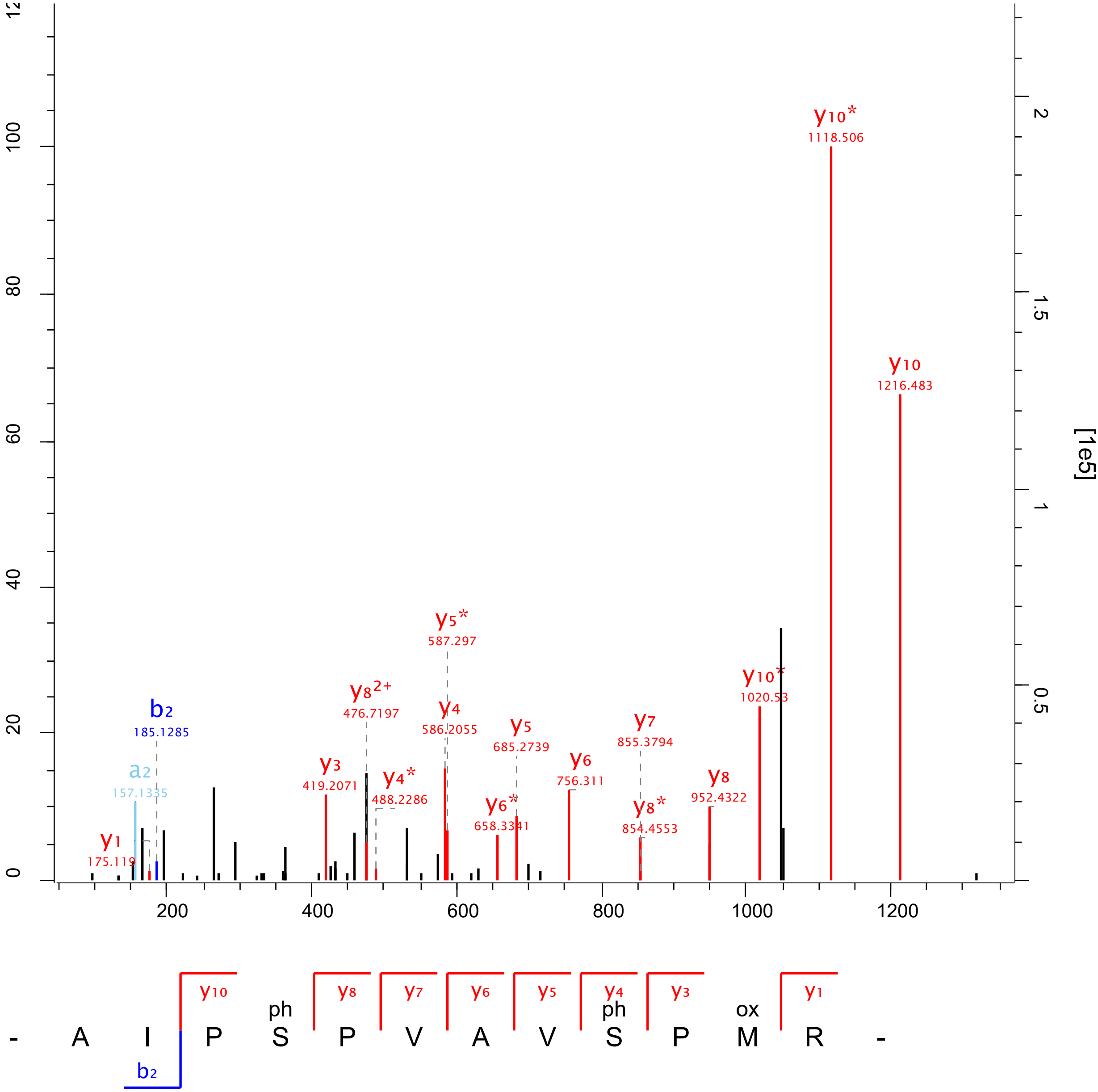

| Raw file | Scan | Method | Score | m/z |
|---|---|---|---|---|
| A17_15_1 | 22462 | FTMS; HCD | 60.76 | 747.31 |

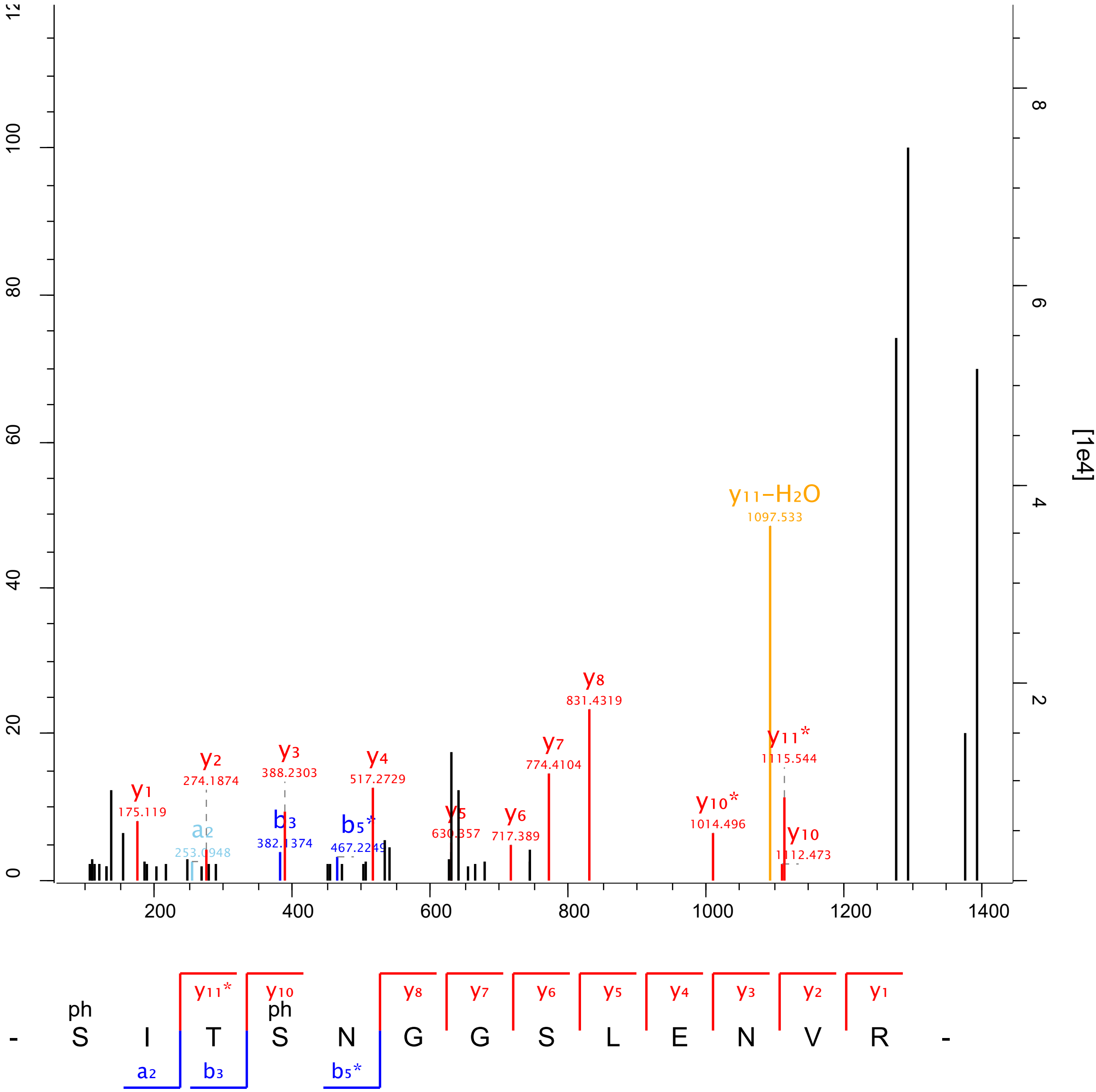

| Raw file | Scan | Method | Score | m/z |
|---|---|---|---|---|
| A17_15_1 | 22471 | FTMS; HCD | 83.76 | 517.91 |

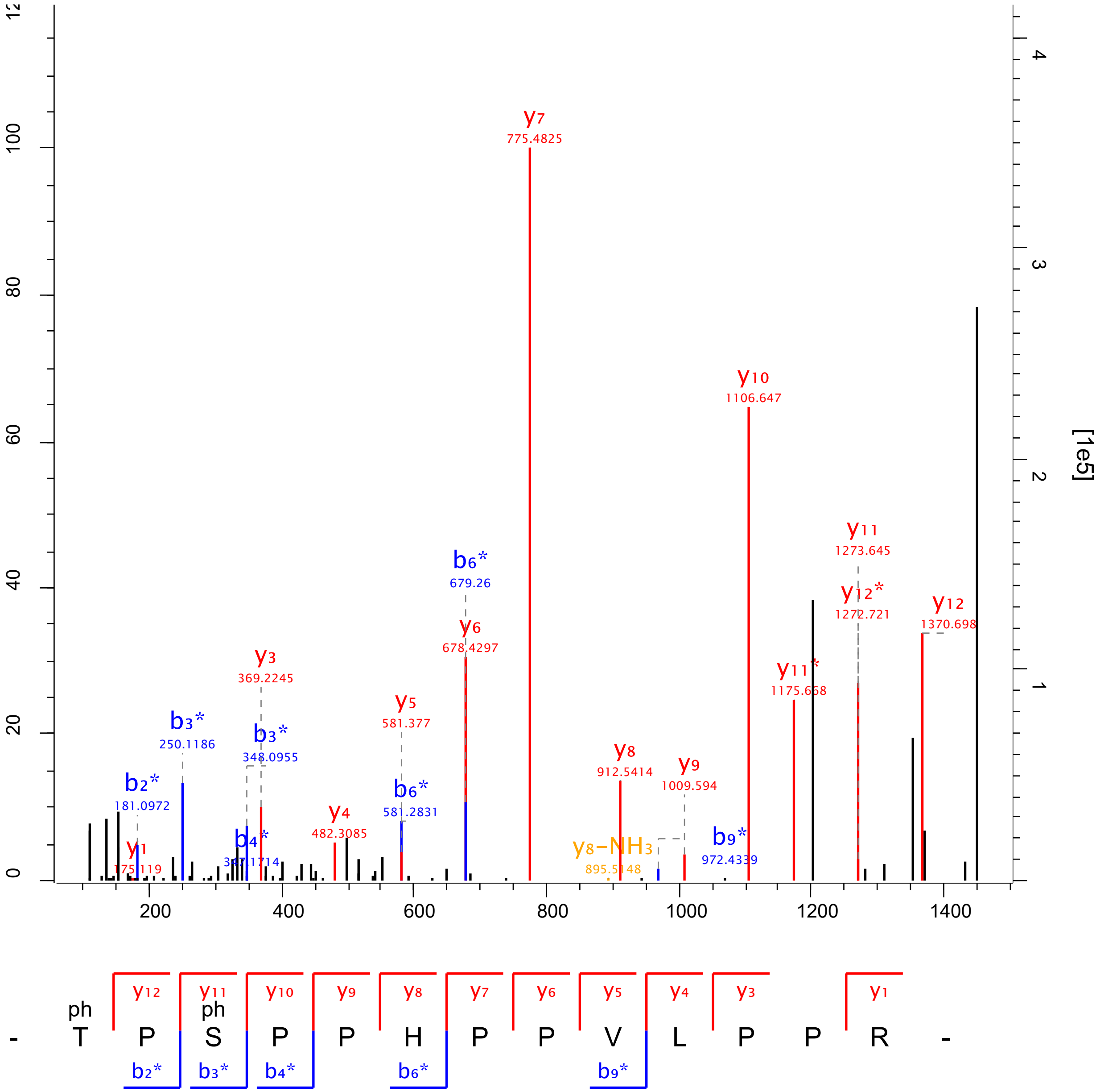

| Raw file | Scan | Method | Score | m/z |
|---|---|---|---|---|
| A17_15_1 | 22637 | FTMS; HCD | 45 | 680.75 |

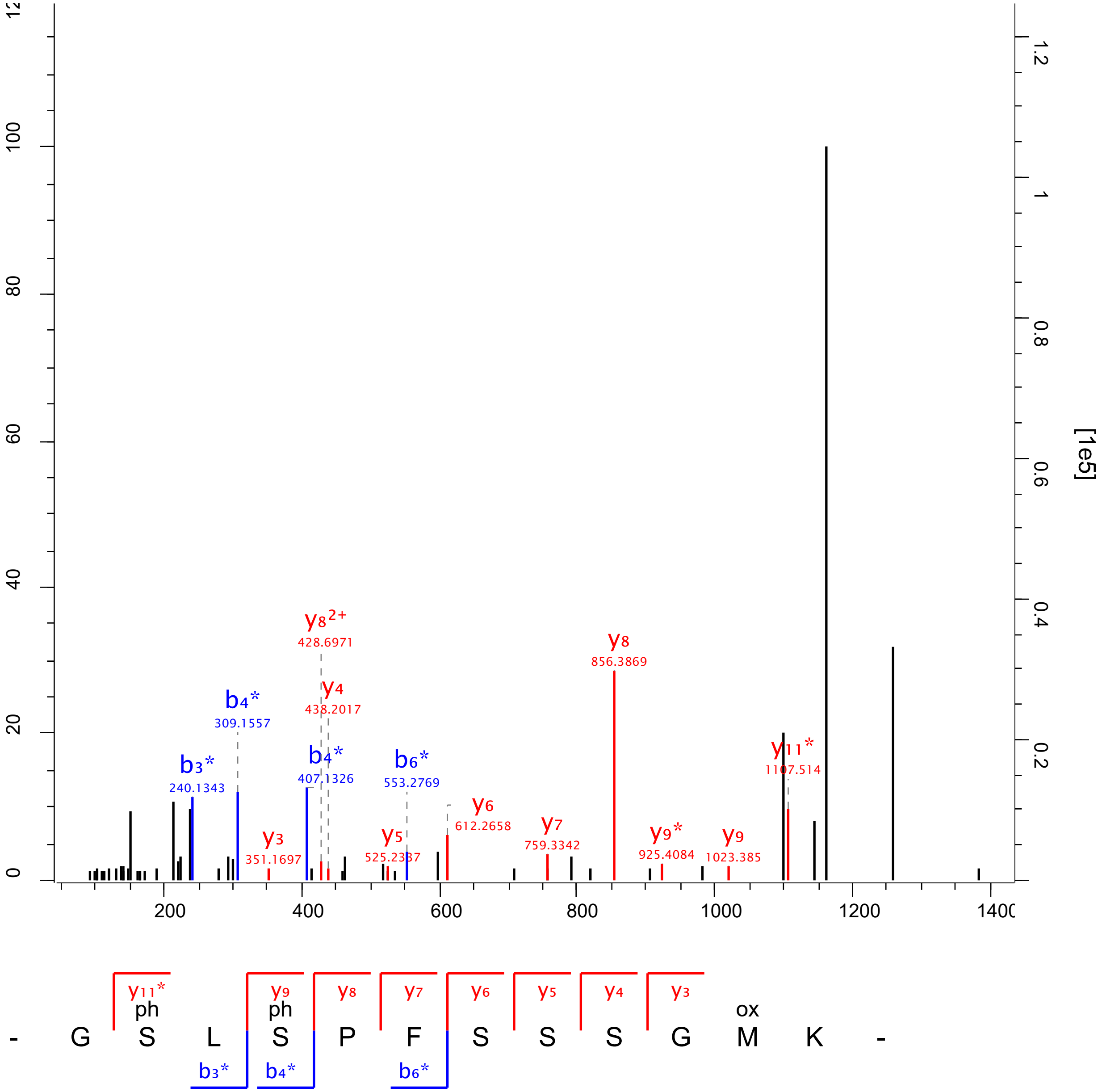

| Raw file | Scan | Method | Score | m/z |
| --- | --- | --- | --- | --- |
| A17_15_1 | 22675 | FTMS; HCD | 87.43 | 768.83 |

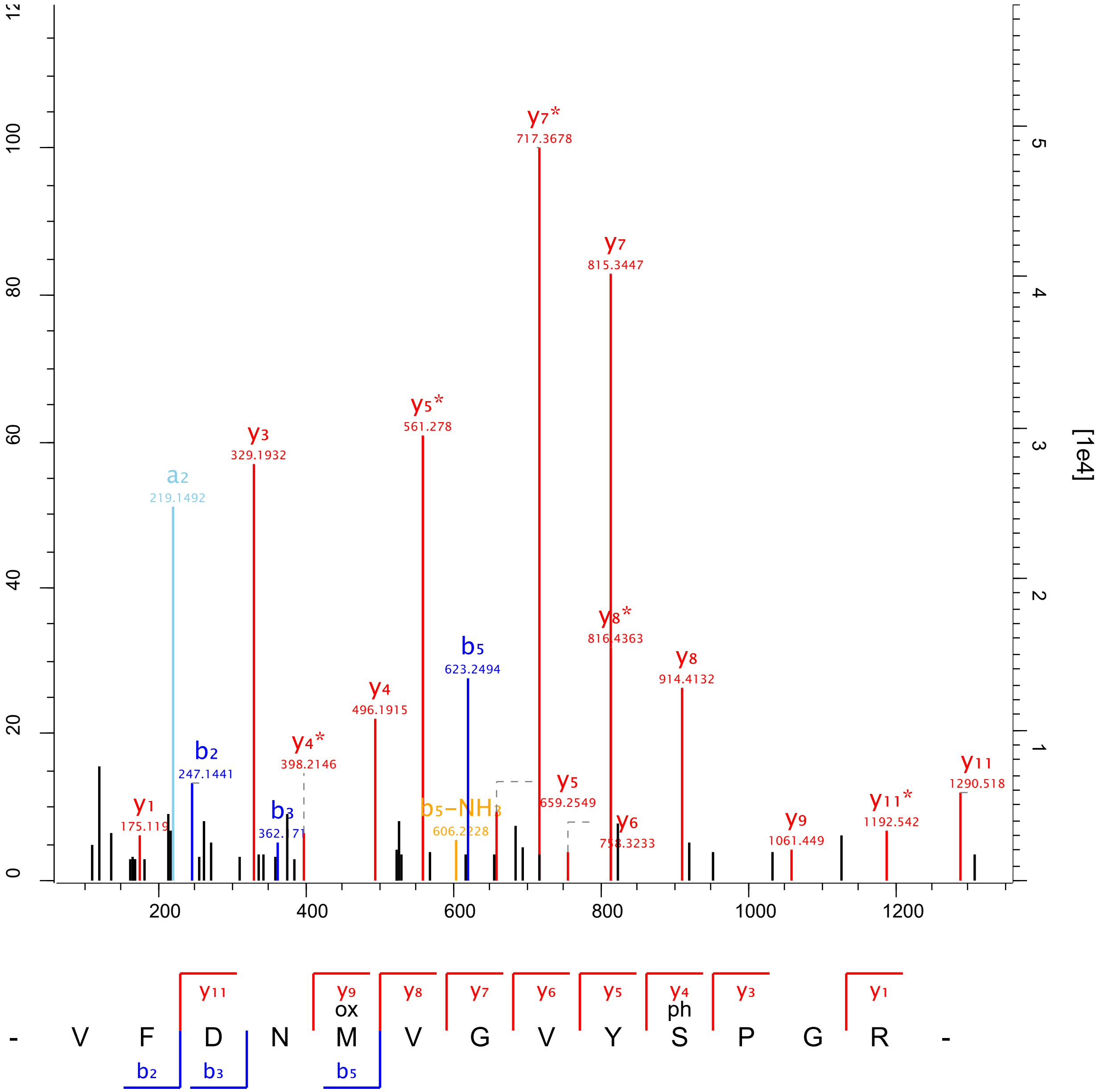

| Raw file | Scan | Method | Score | m/z |
|---|---|---|---|---|
| A17_15_1 | 22943 | FTMS; HCD | 49.19 | 676.98 |

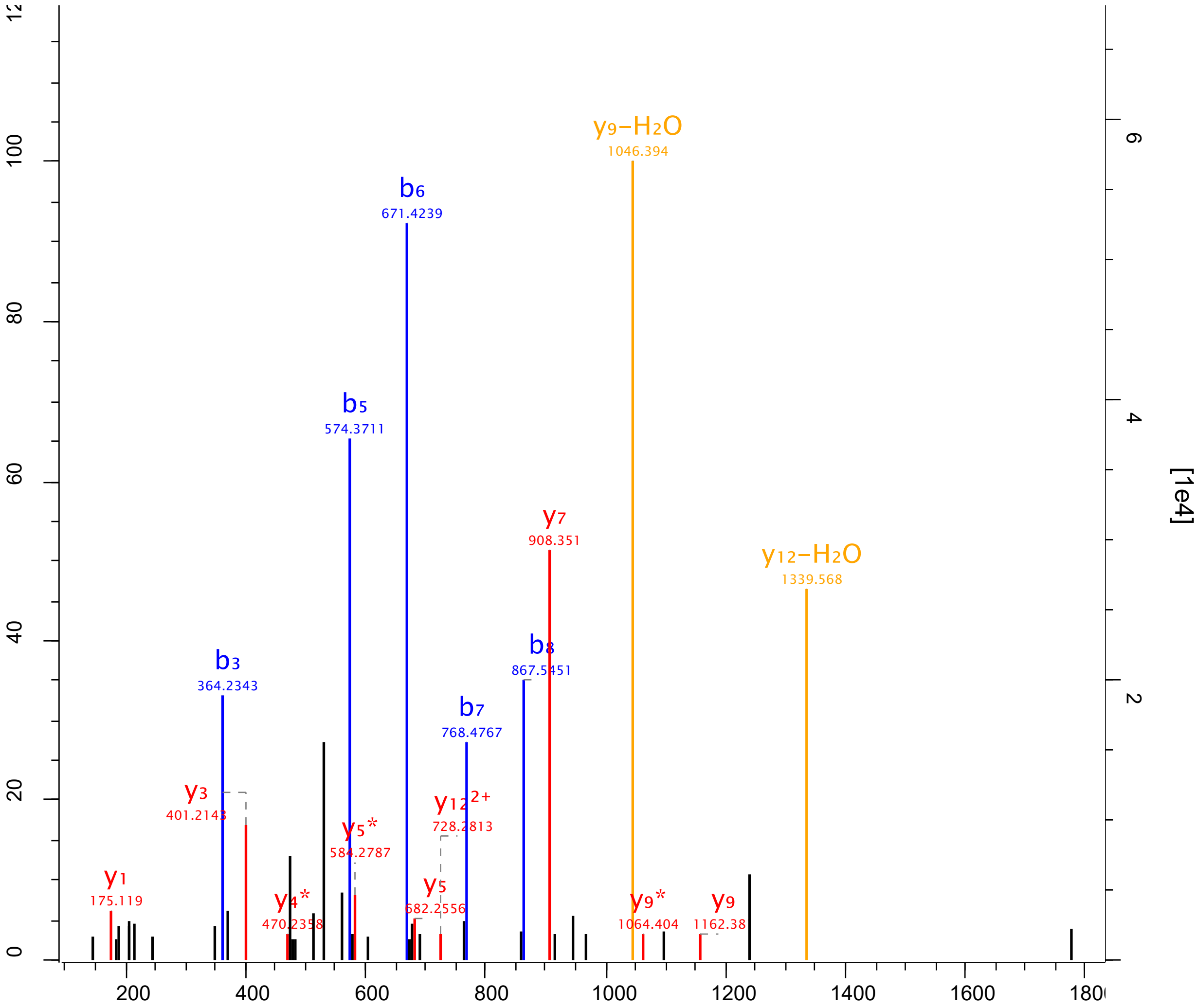


- L I H P I P P V S S P E N S P
E R -

(ph on S, ph on S; fragment ions: b3, b5, b6, b7, b8; y12 2+, y9, y7, y5, y4*, y3, y1)

| Raw file | Scan | Method | Score | m/z |
| --- | --- | --- | --- | --- |
| A17_15_1 | 23043 | FTMS; HCD | 200.58 | 799.85 |

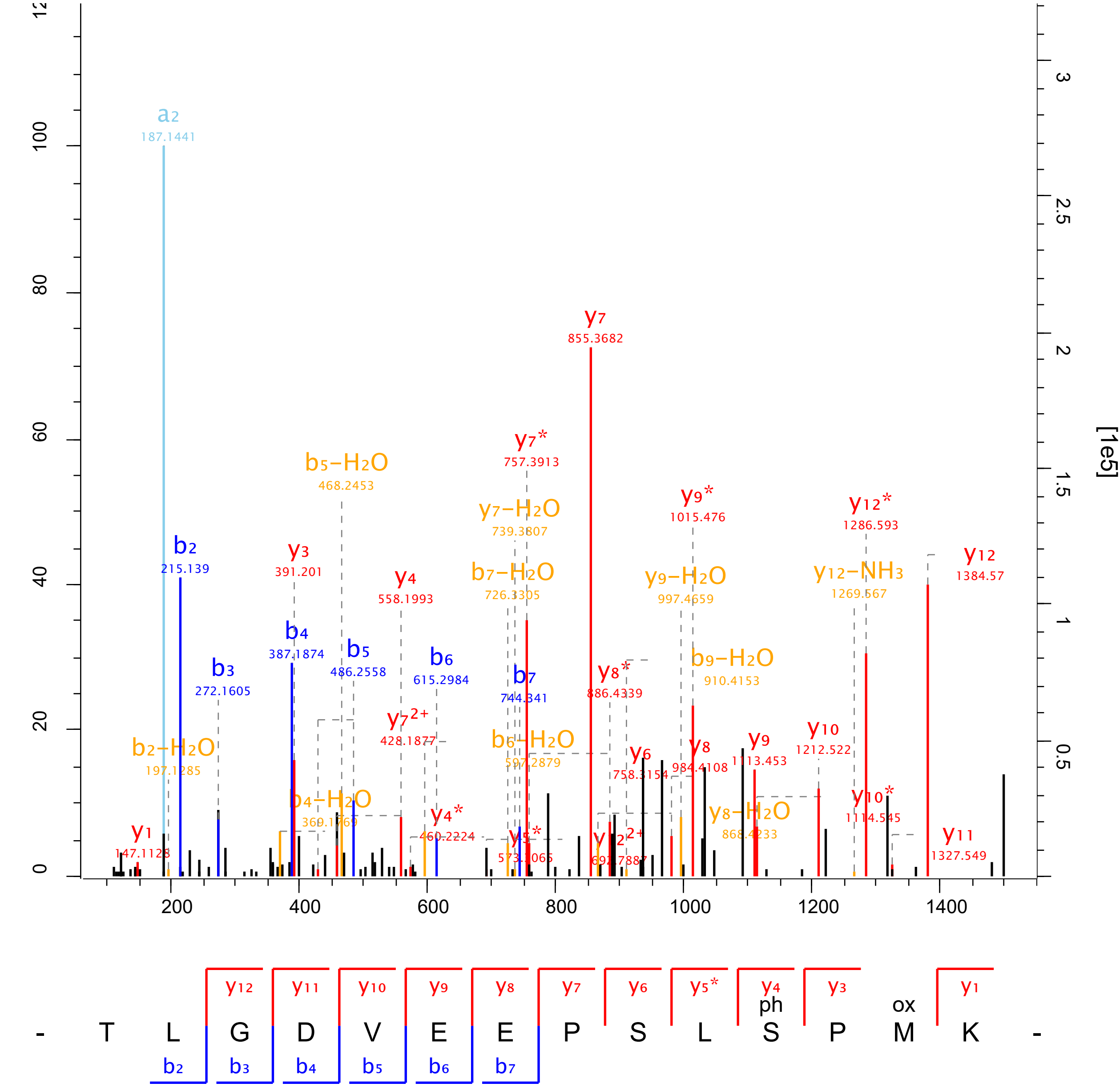

| Raw file | Scan | Method | Score | m/z |
| --- | --- | --- | --- | --- |
| A17_15_1 | 23058 | FTMS; HCD | 89.17 | 520.72 |

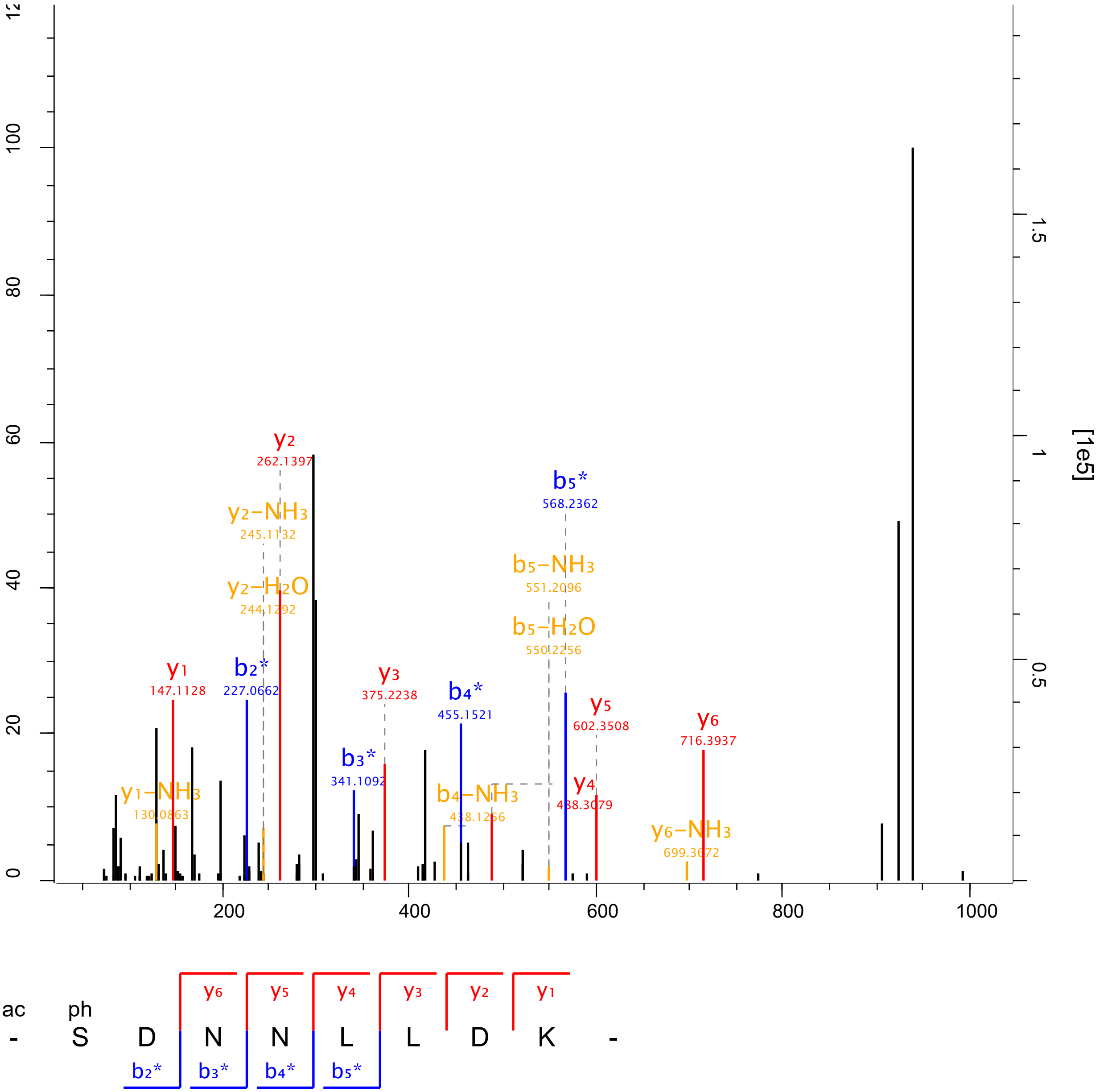

| Raw file | Scan | Method | Score | m/z |
|---|---|---|---|---|
| A17_15_1 | 23255 | FTMS; HCD | 333.68 | 920.4 |

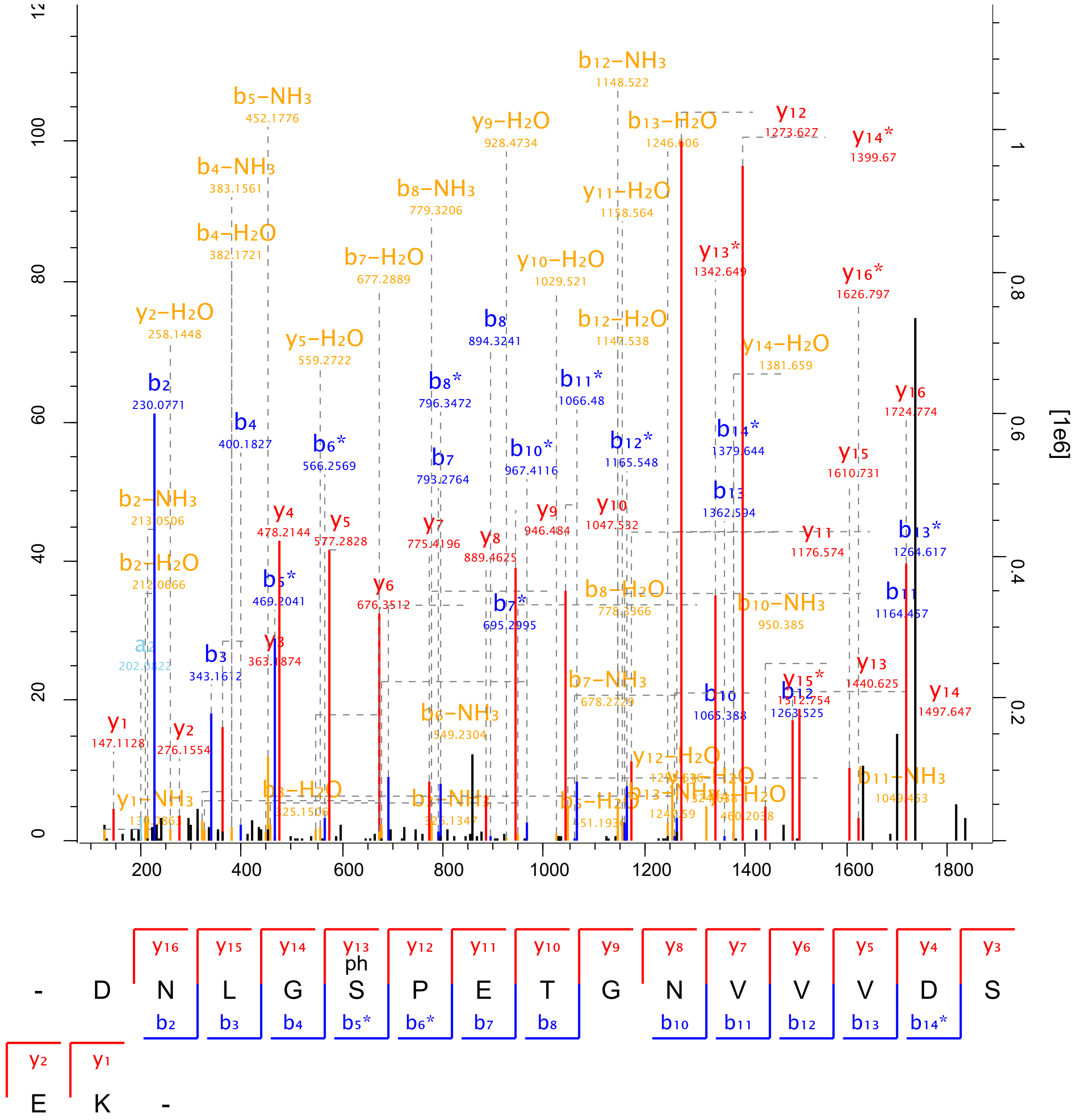

| Raw file | Scan | Method | Score | m/z |
| --- | --- | --- | --- | --- |
| A17_15_1 | 23312 | FTMS; HCD | 55.13 | 774.34 |

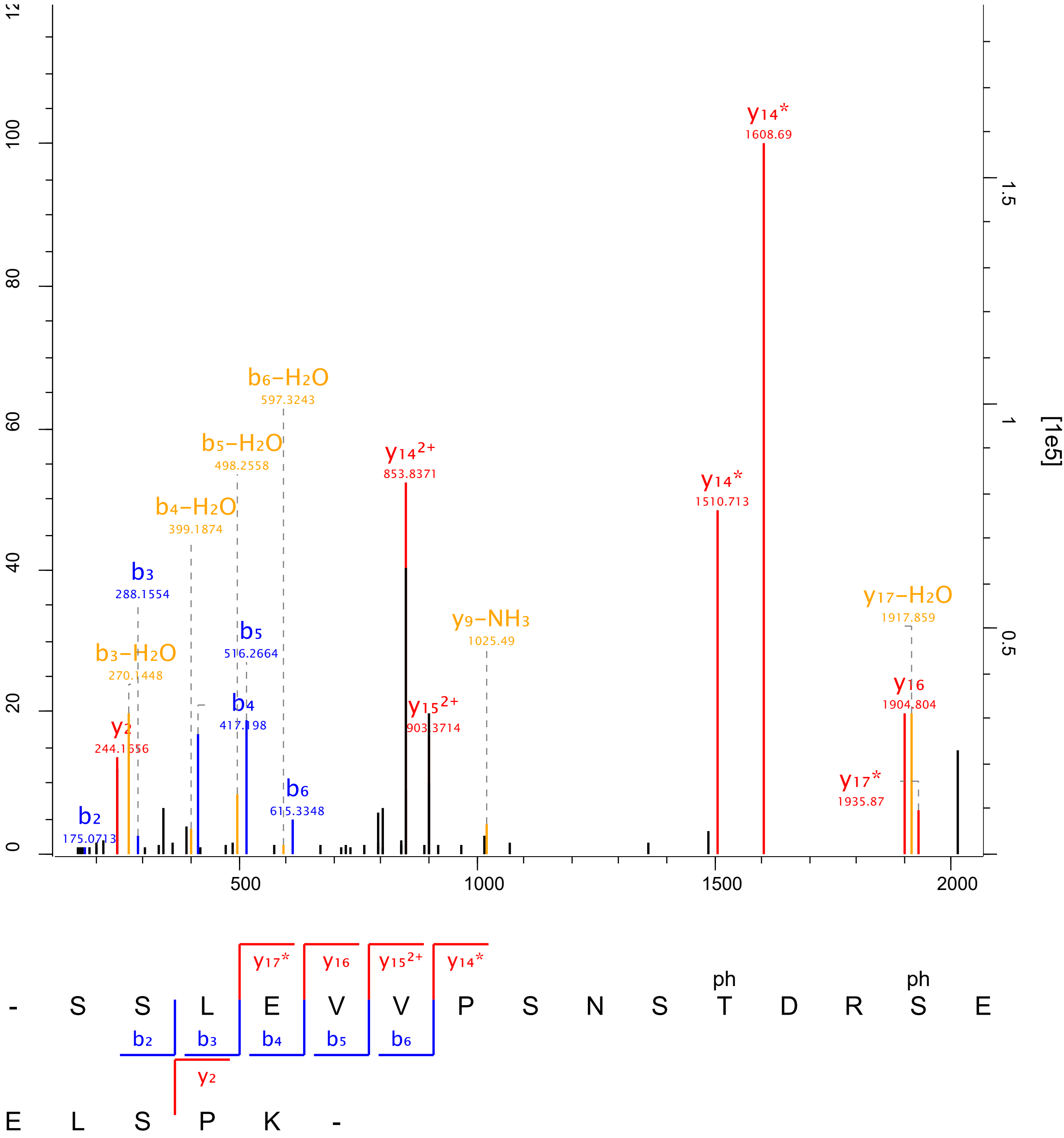

| Raw file | Scan | Method | Score | m/z |
|---|---|---|---|---|
| A17_15_1 | 23473 | FTMS; HCD | 102.89 | 824.37 |

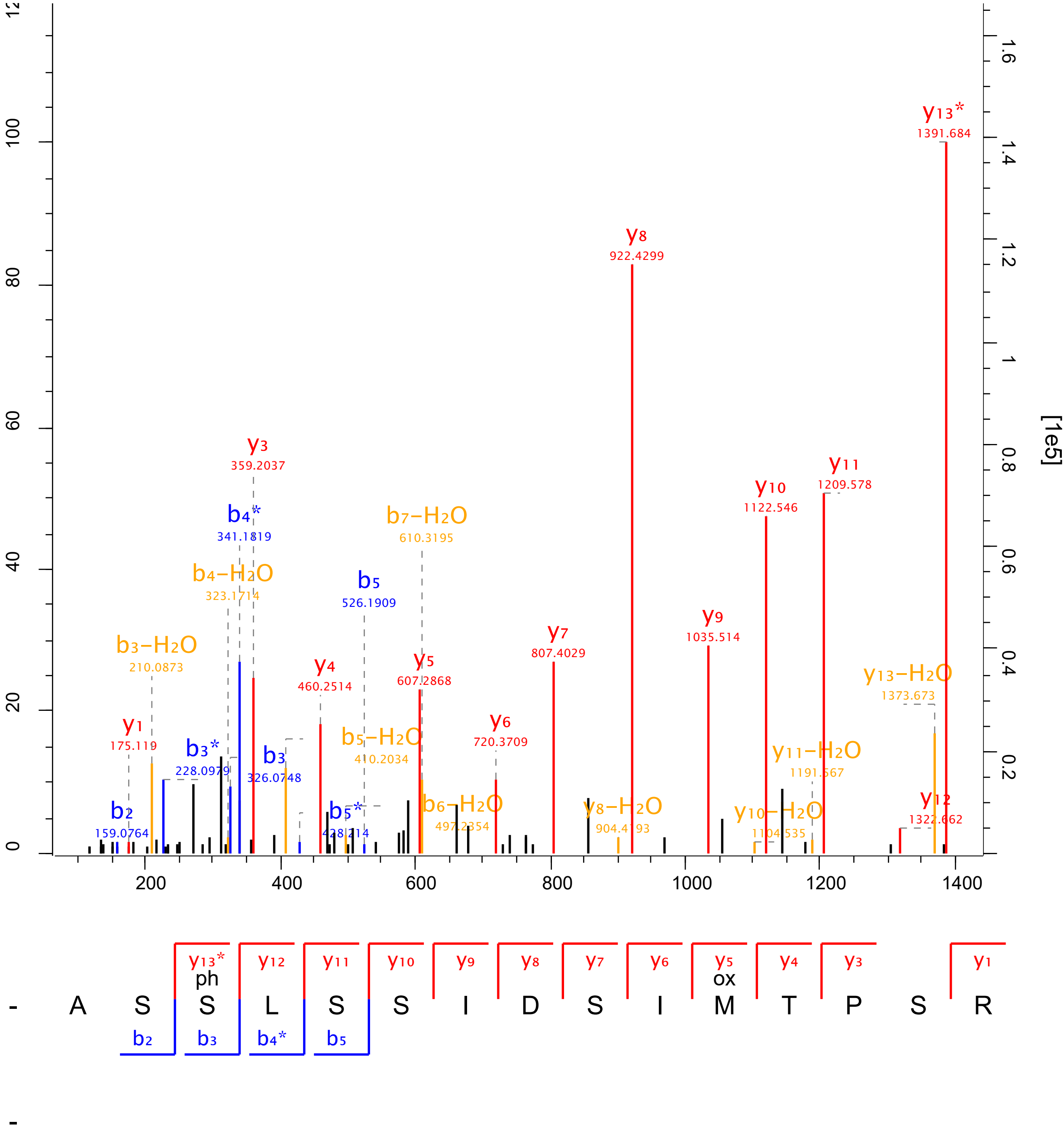

| Raw file | Scan | Method | Score | m/z |
|---|---|---|---|---|
| A17_15_1 | 23514 | FTMS; HCD | 63.62 | 693.8 |

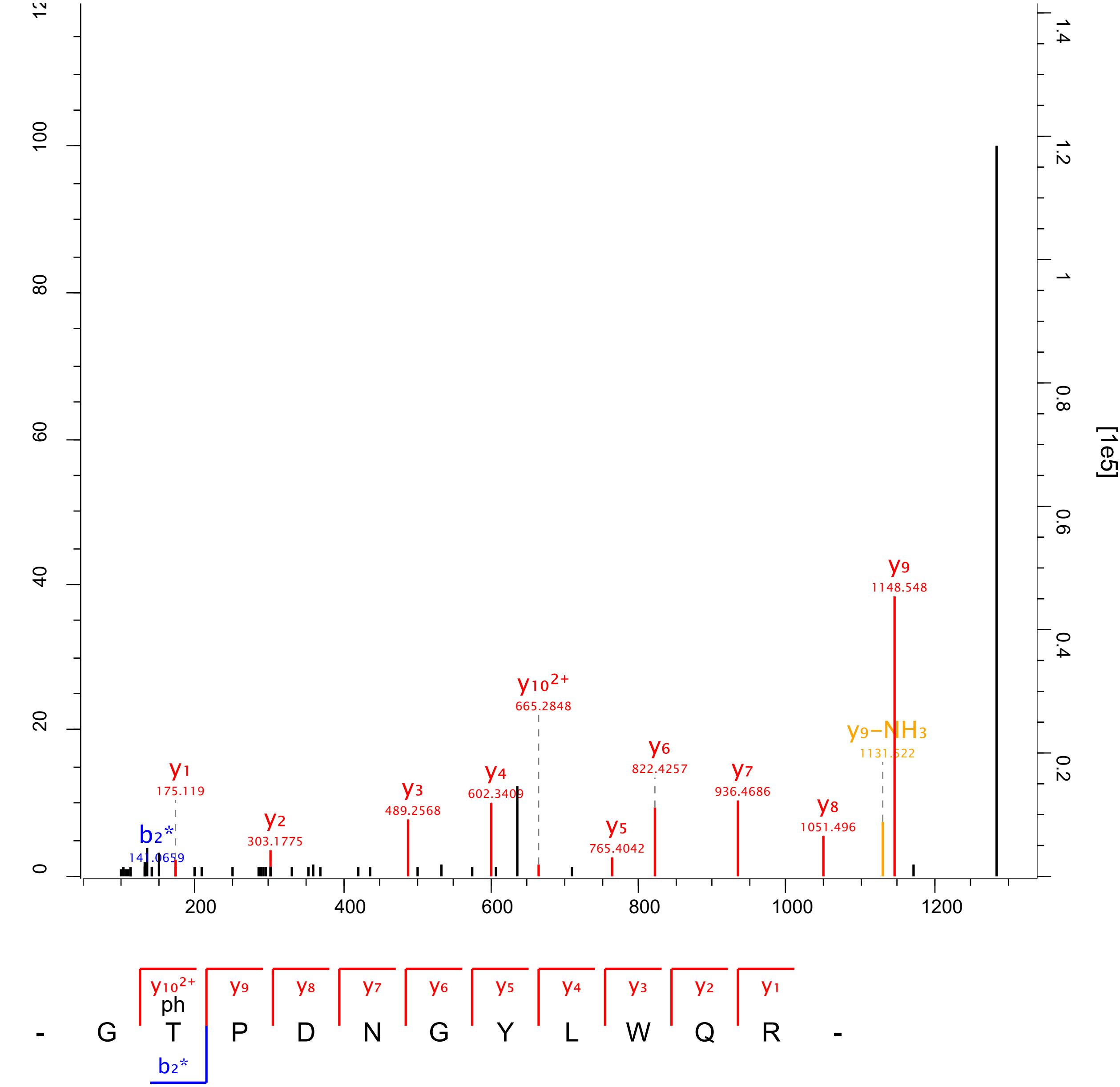

| Raw file | Scan | Method | Score | m/z |
|---|---|---|---|---|
| A17_15_1 | 24204 | FTMS; HCD | 104.76 | 763.82 |

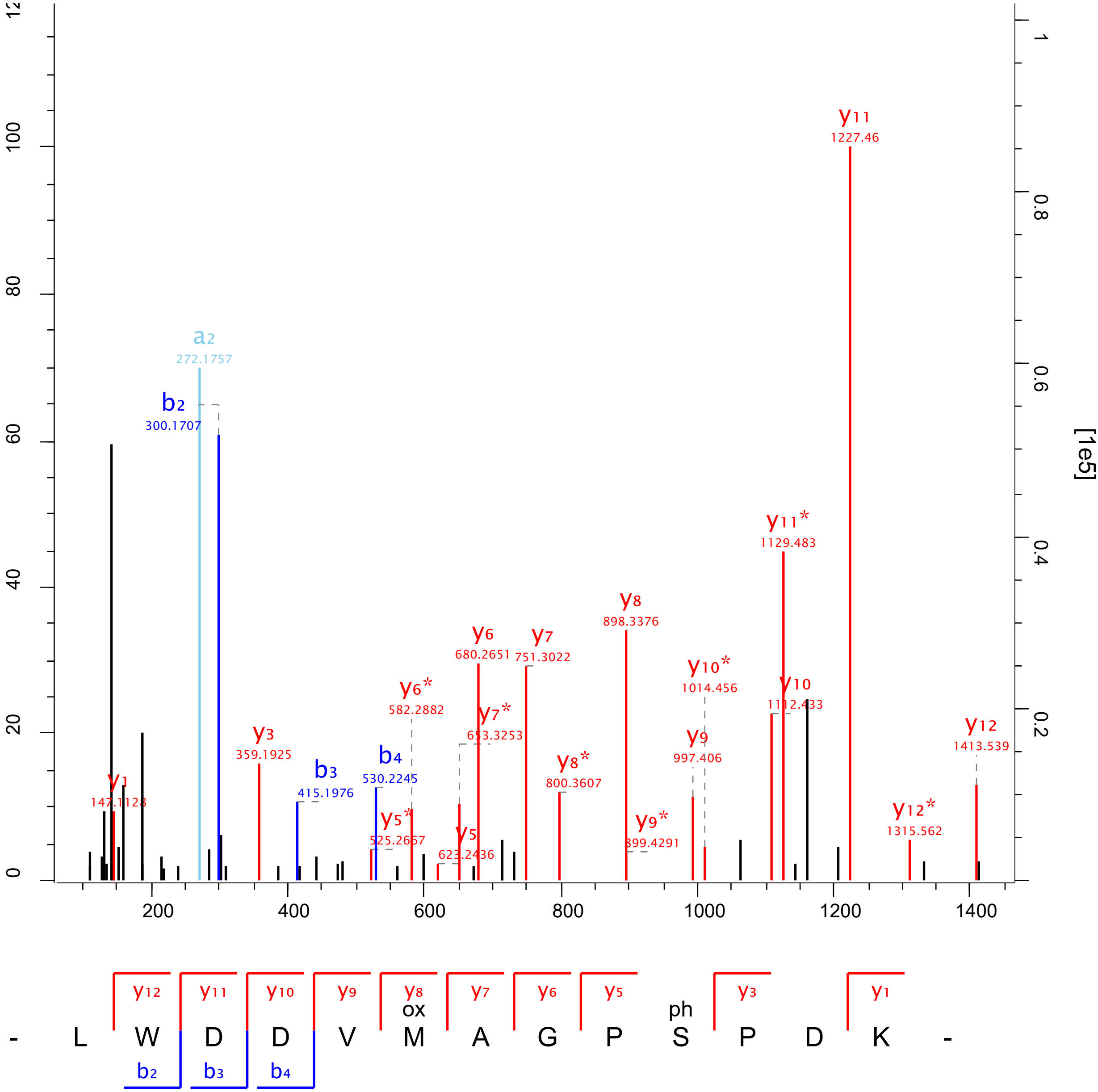

| Raw file | Scan | Method | Score | m/z |
|---|---|---|---|---|
| A17_15_1 | 24395 | FTMS; HCD | 80.96 | 675.29 |

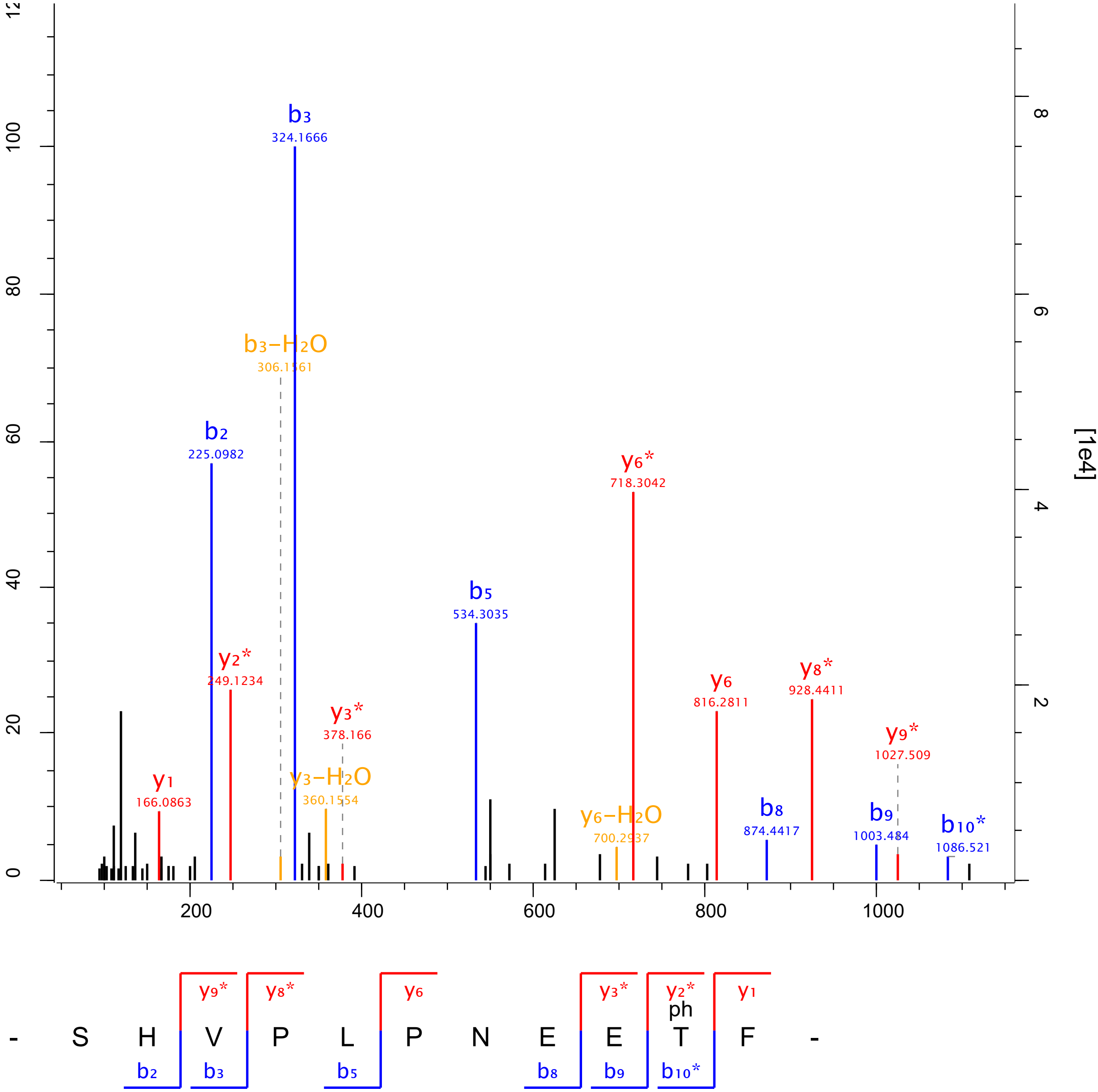

| Raw file | Scan | Method | Score | m/z |
|---|---|---|---|---|
| A17_15_1 | 24571 | FTMS; HCD | 92.38 | 710.79 |

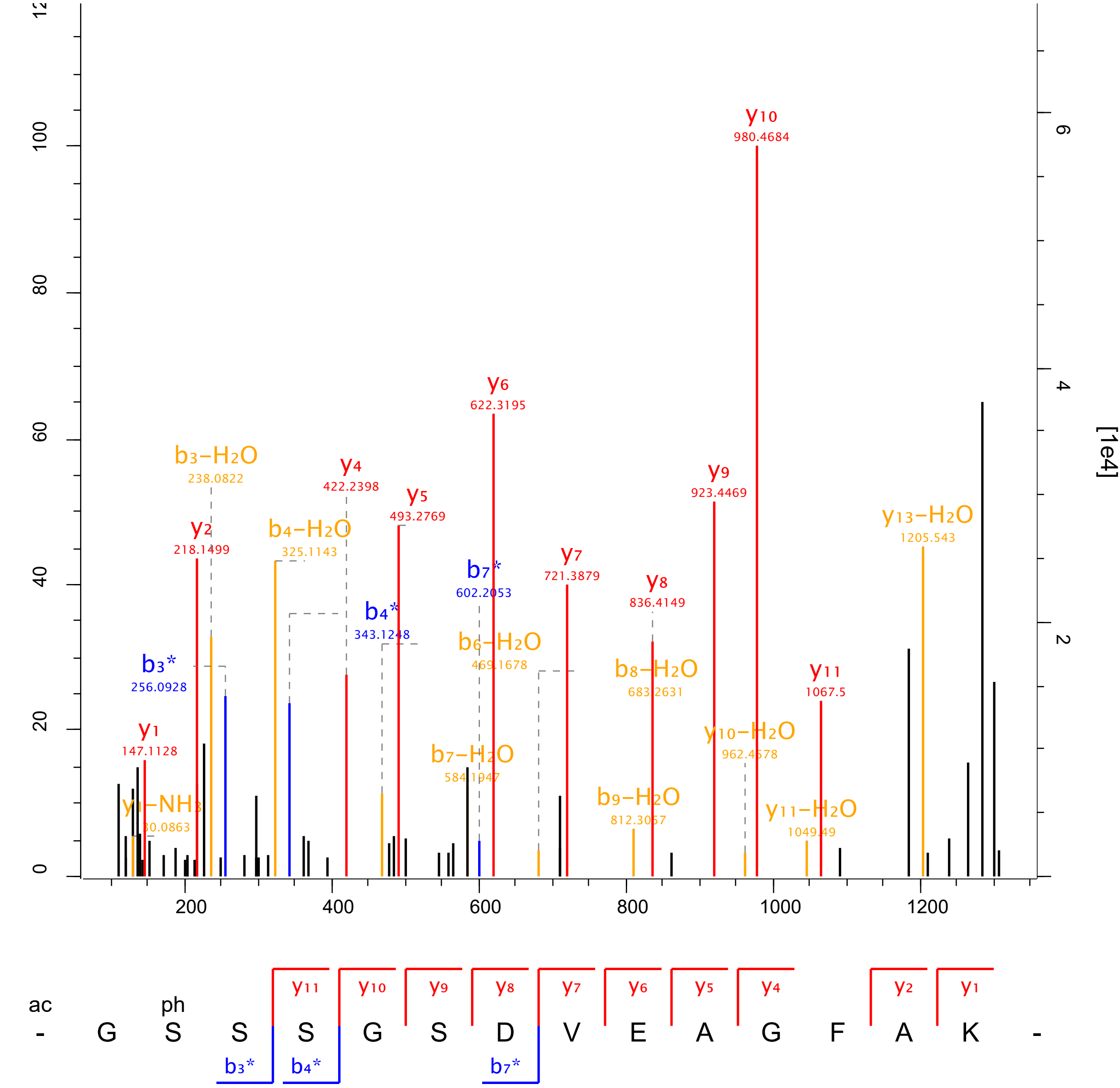

| Raw file | Scan | Method | Score | m/z |
|---|---|---|---|---|
| A17_15_1 | 2470 | FTMS; HCD | 98.9 | 554.24 |

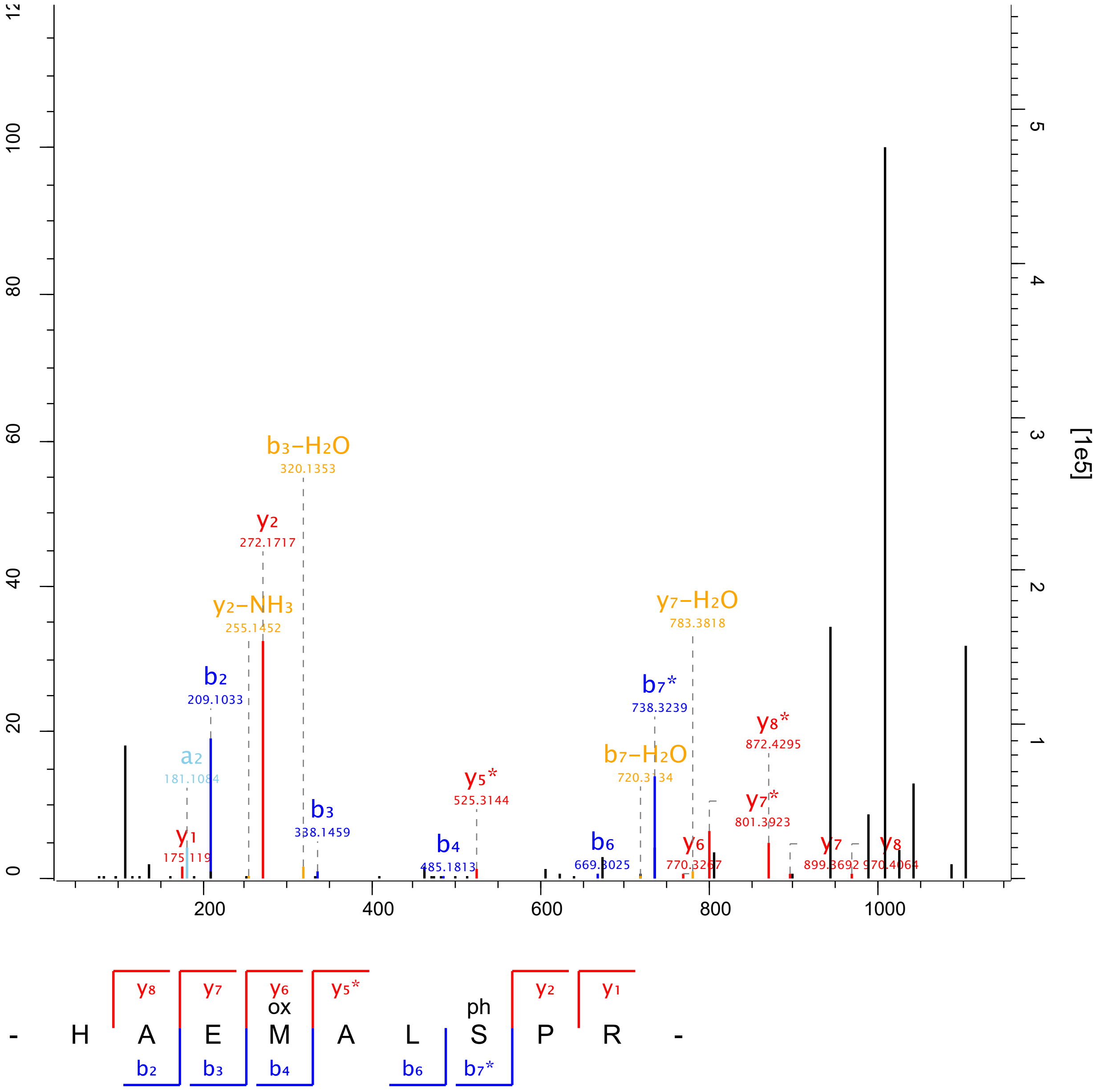

| Raw file | Scan | Method | Score | m/z |
|---|---|---|---|---|
| A17_15_1 | 24709 | FTMS; HCD | 147.06 | 817.4 |

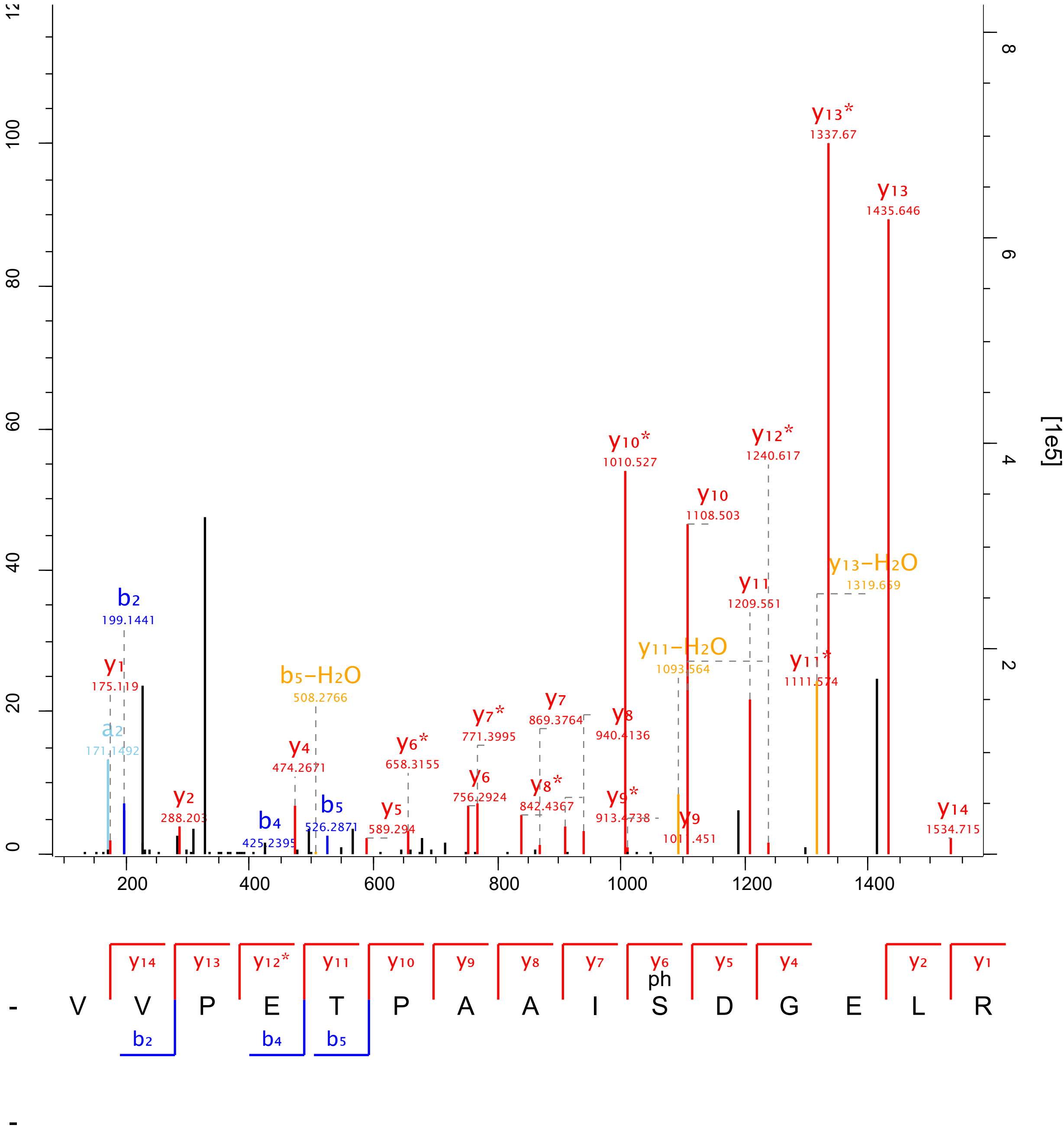

| Raw file | Scan | Method | Score | m/z |
|---|---|---|---|---|
| A17_15_1 | 24743 | FTMS; HCD | 44.85 | 867.02 |

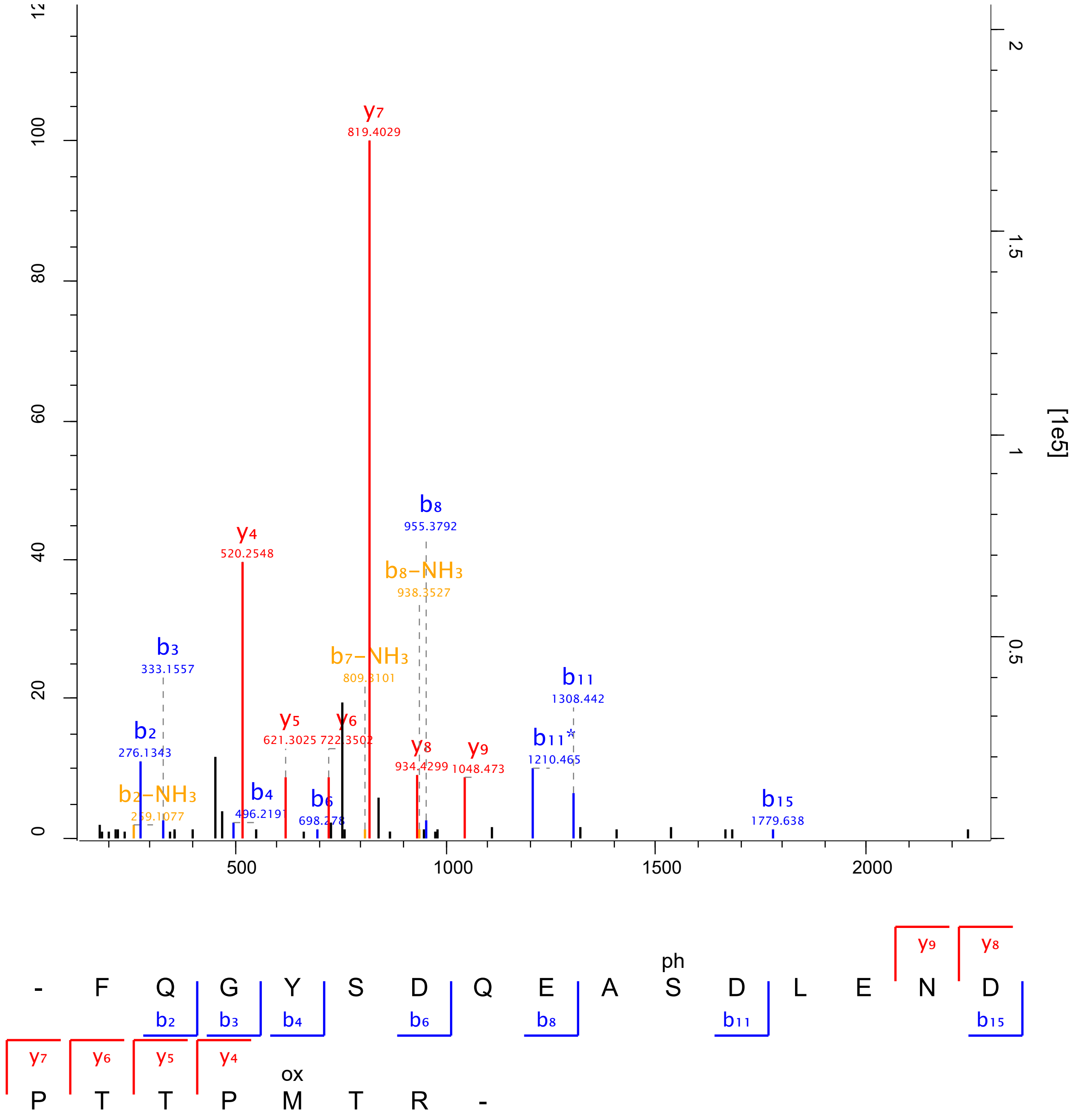

| Raw file | Scan | Method | Score | m/z |
| --- | --- | --- | --- | --- |
| A17_15_1 | 24906 | FTMS; HCD | 63.57 | 497.72 |

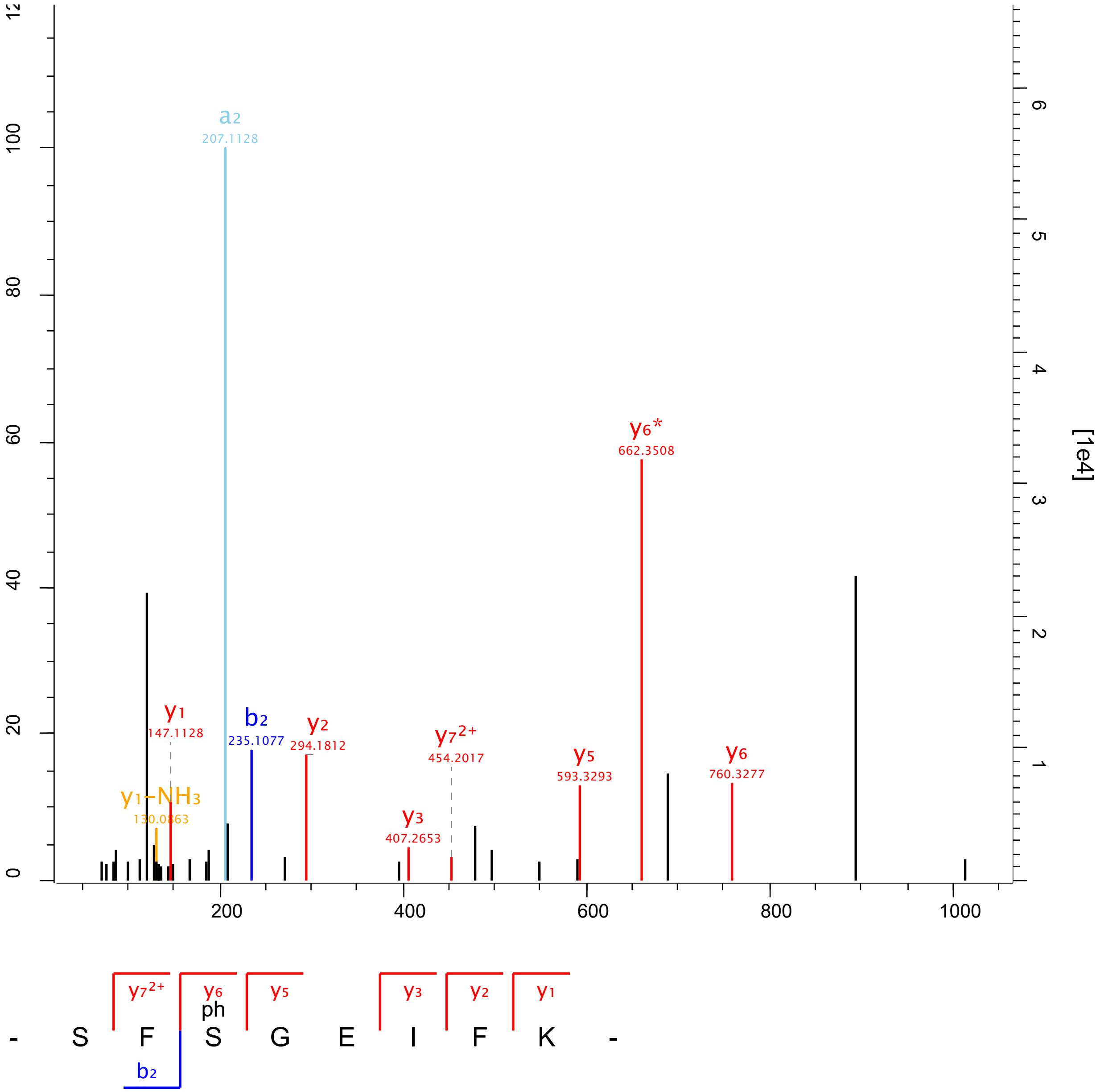

| Raw file | Scan | Method | Score | m/z |
|---|---|---|---|---|
| A17_15_1 | 25172 | FTMS; HCD | 143.57 | 916.86 |

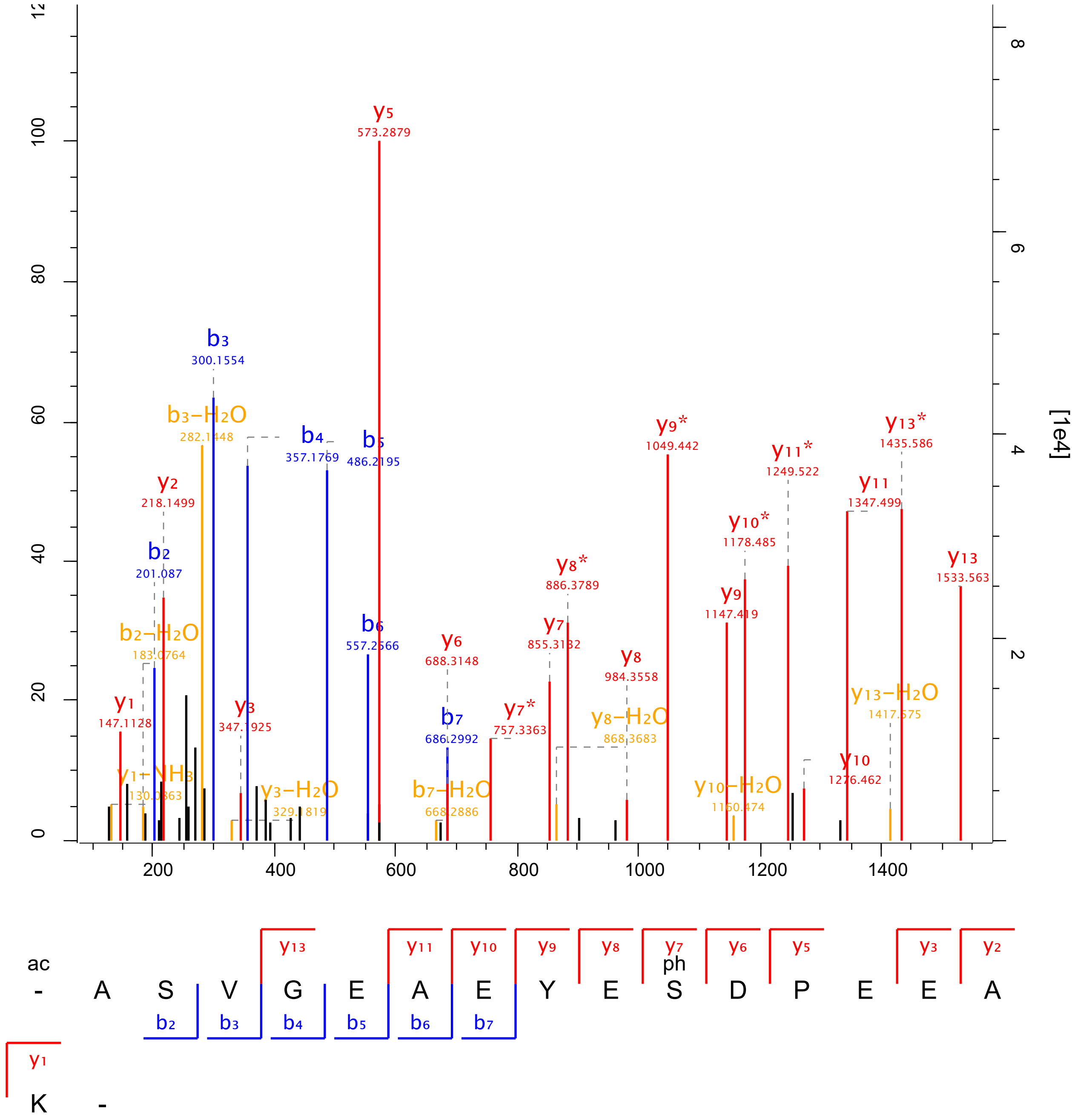

| Raw file | Scan | Method | Score | m/z |
|---|---|---|---|---|
| A17_15_1 | 25460 | FTMS; HCD | 40.94 | 606.77 |

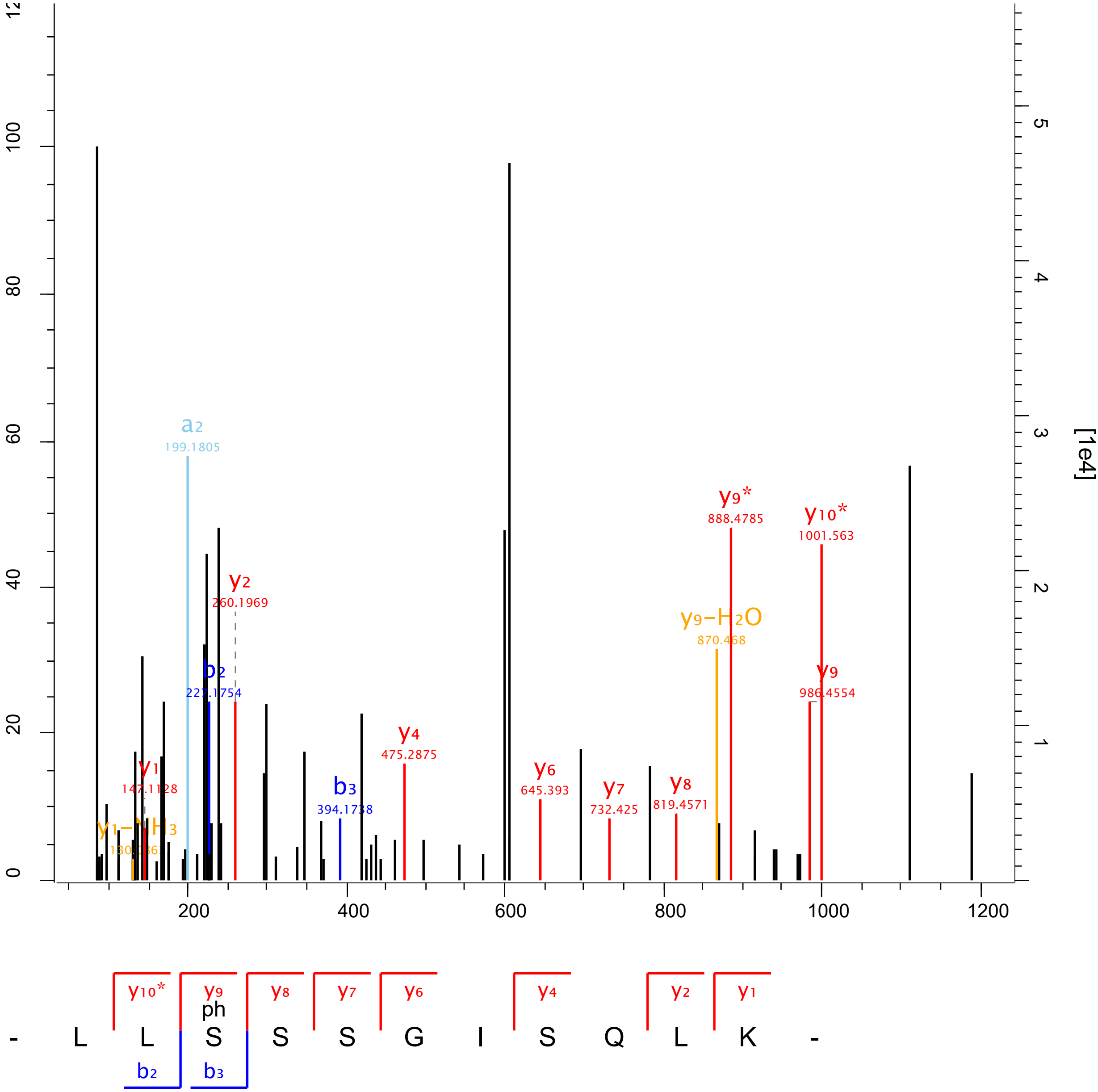

| Raw file | Scan | Method | Score | m/z |
|---|---|---|---|---|
| A17_15_1 | 25567 | FTMS; HCD | 47.19 | 485.71 |

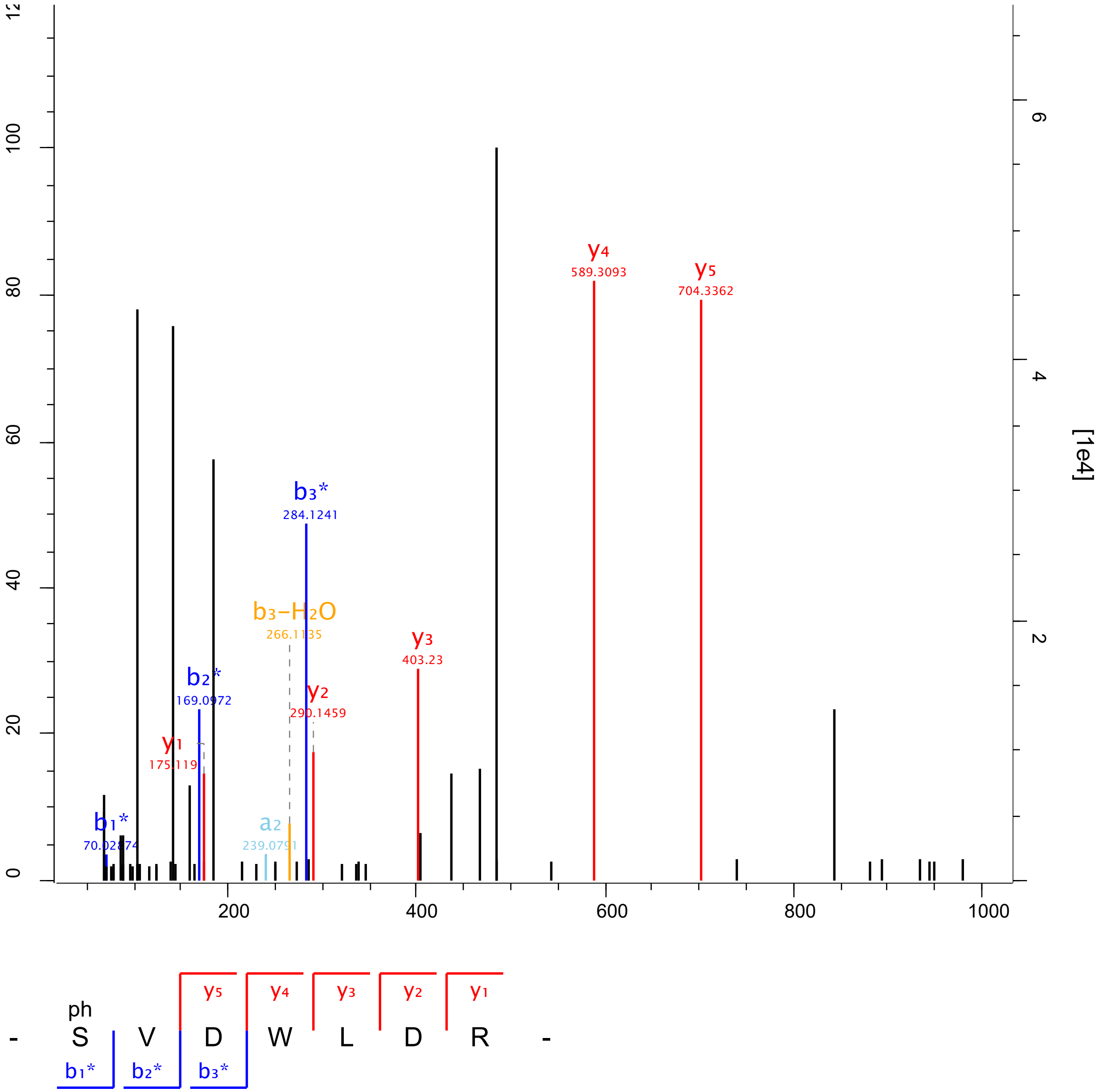

| Raw file | Scan | Method | Score | m/z |
|---|---|---|---|---|
| A17_15_1 | 26301 | FTMS; HCD | 76.07 | 620.72 |

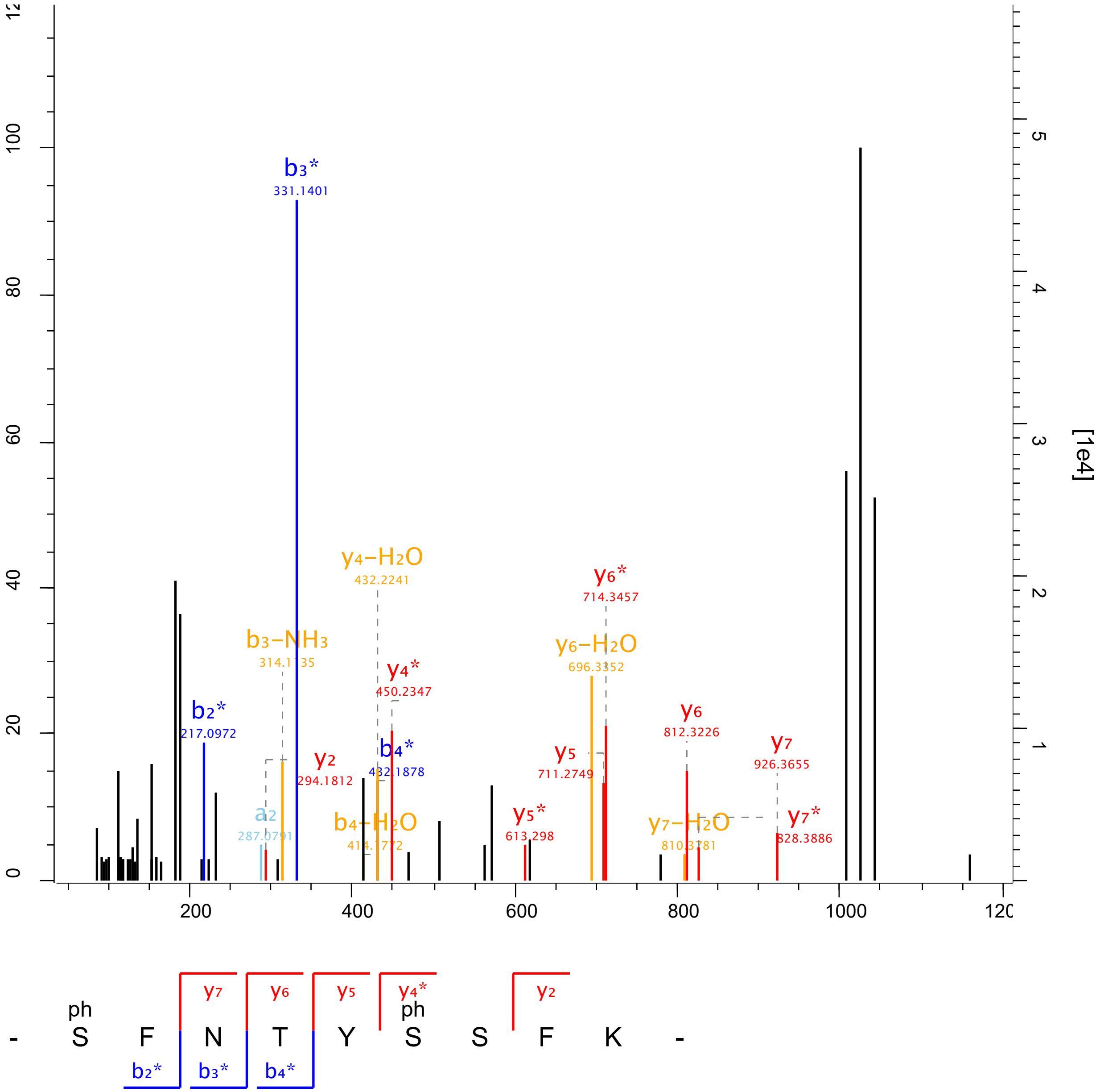

| Raw file | Scan | Method | Score | m/z |
| --- | --- | --- | --- | --- |
| A17_15_1 | 26310 | FTMS; HCD | 59.43 | 897.38 |

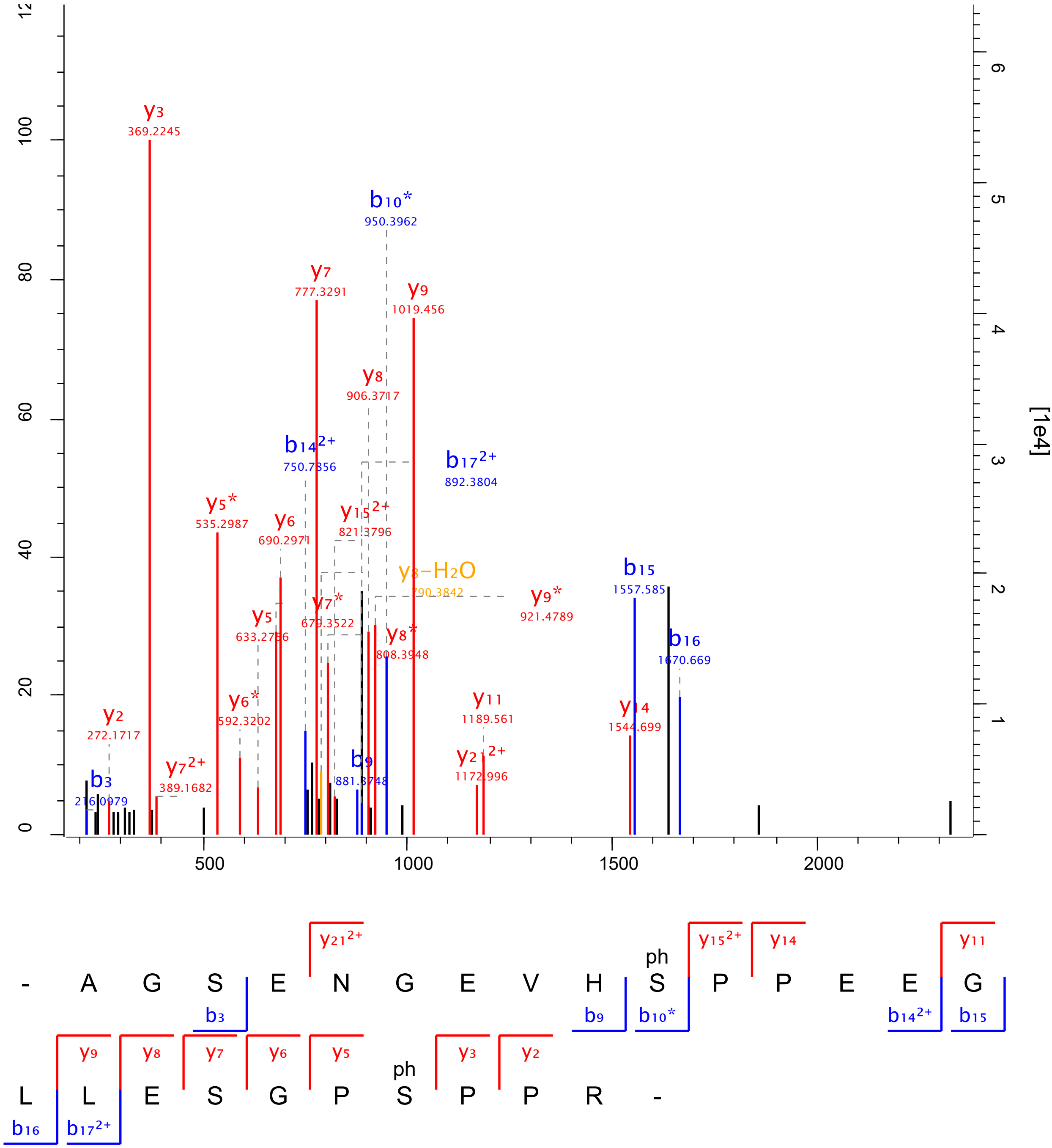

| Raw file | Scan | Method | Score | m/z |
|---|---|---|---|---|
| A17_15_1 | 26372 | FTMS; HCD | 199.38 | 688.27 |

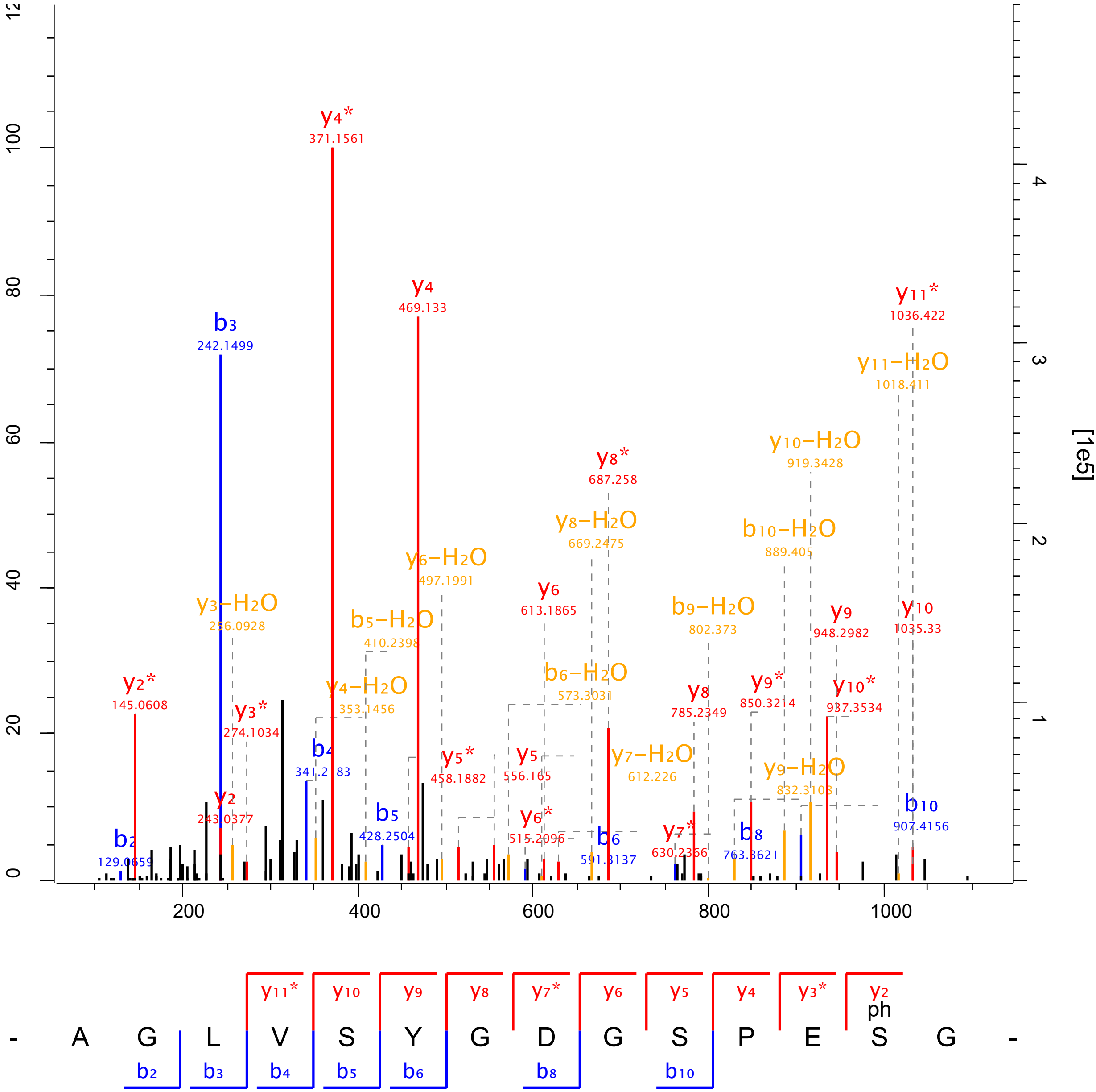

| Raw file | Scan | Method | Score | m/z |
|---|---|---|---|---|
| A17_15_1 | 26898 | FTMS; HCD | 46.49 | 944.4 |

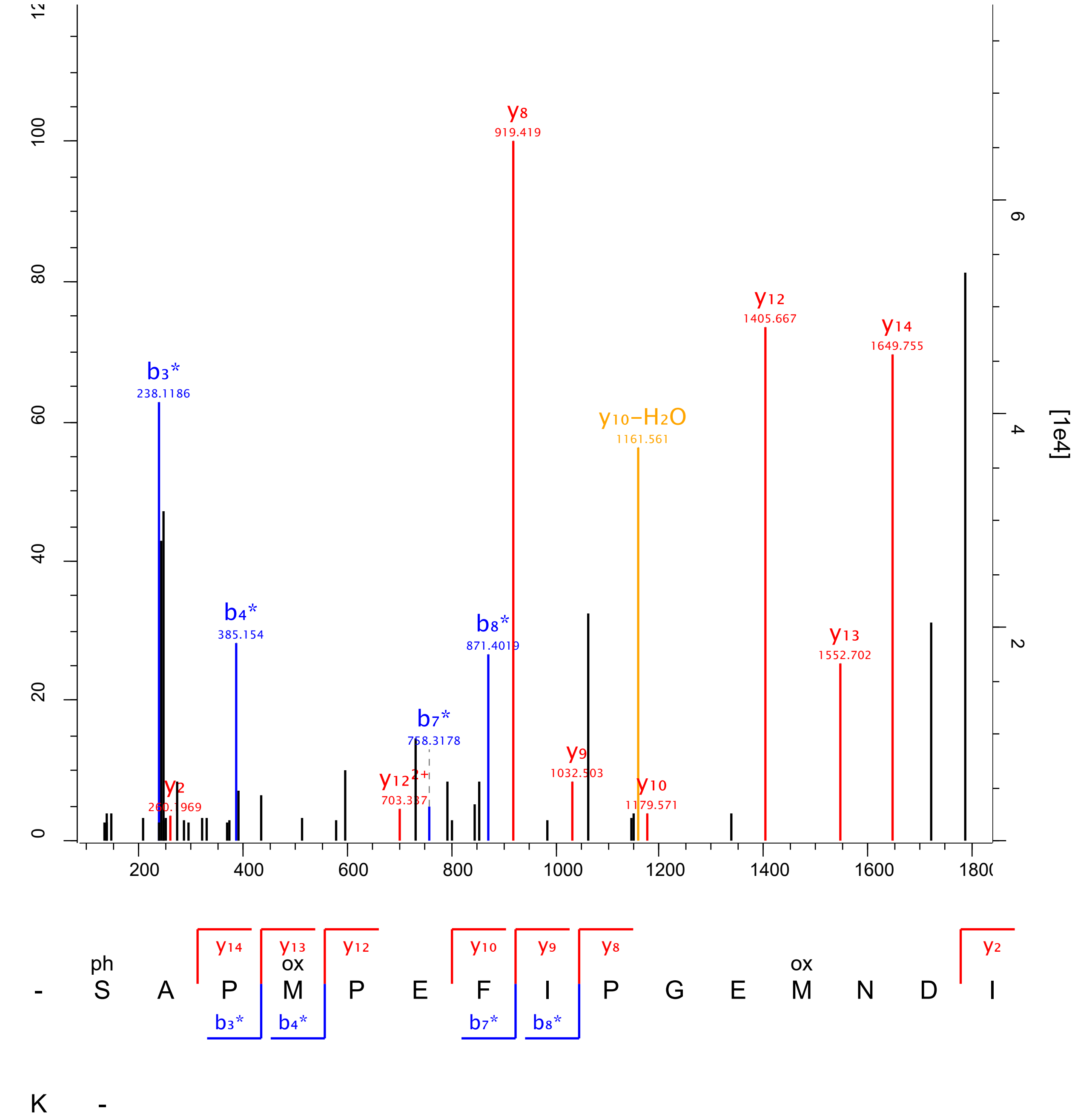

| Raw file | Scan | Method | Score | m/z |
|---|---|---|---|---|
| A17_15_1 | 26992 | FTMS; HCD | 47.33 | 814.7 |

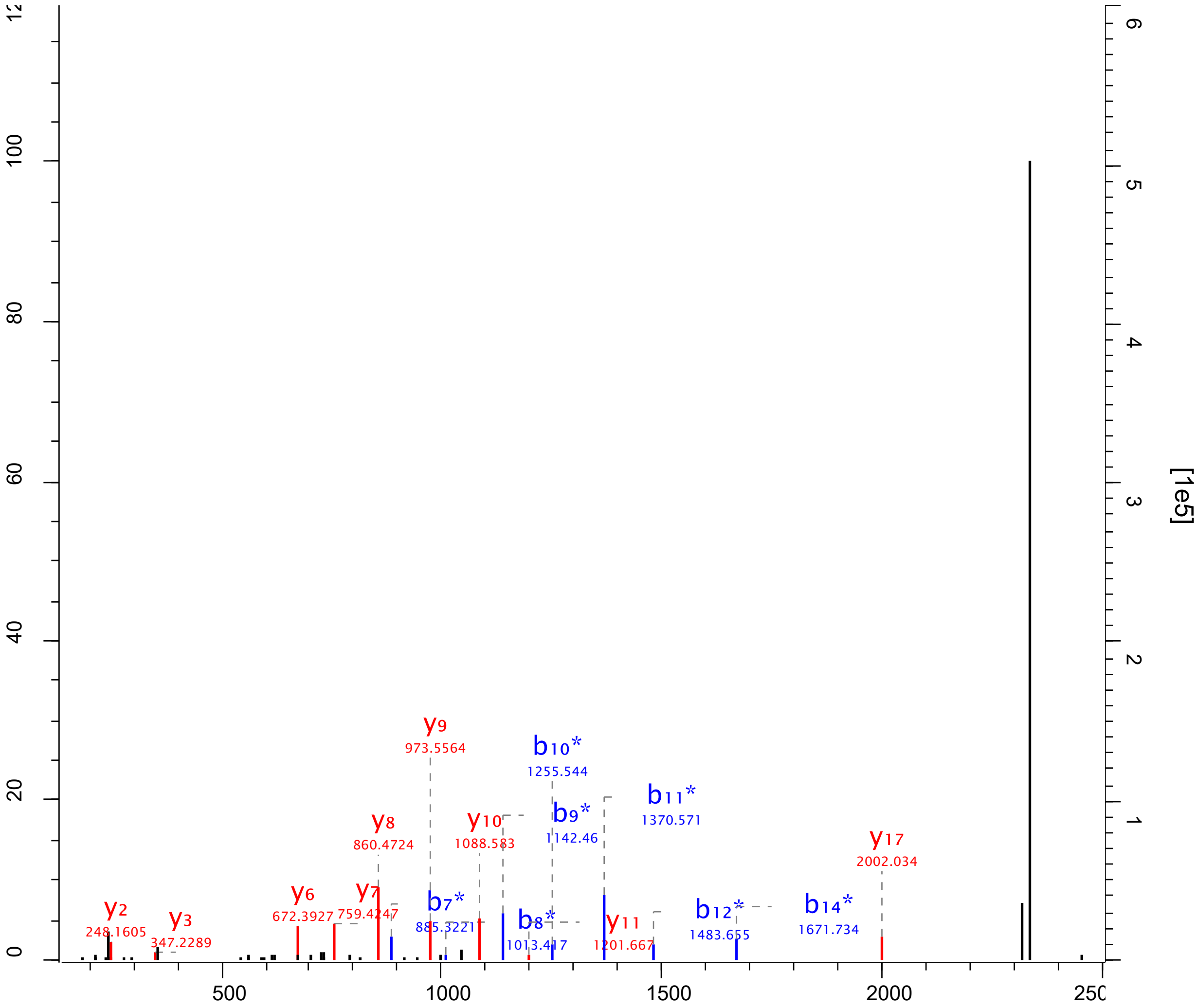


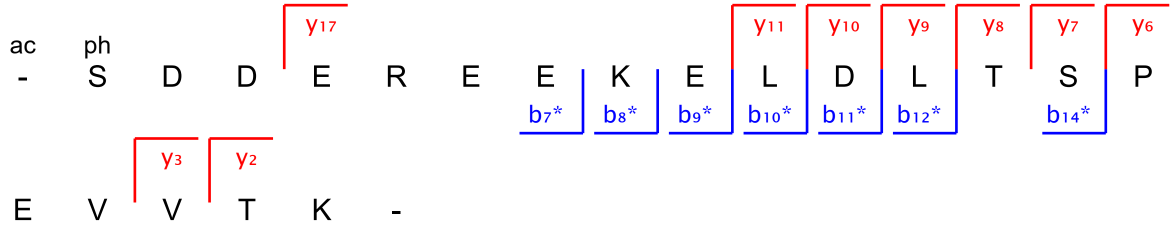

| Raw file | Scan | Method | Score | m/z |
|---|---|---|---|---|
| A17_15_1 | 27235 | FTMS; HCD | 52.58 | 628.3 |

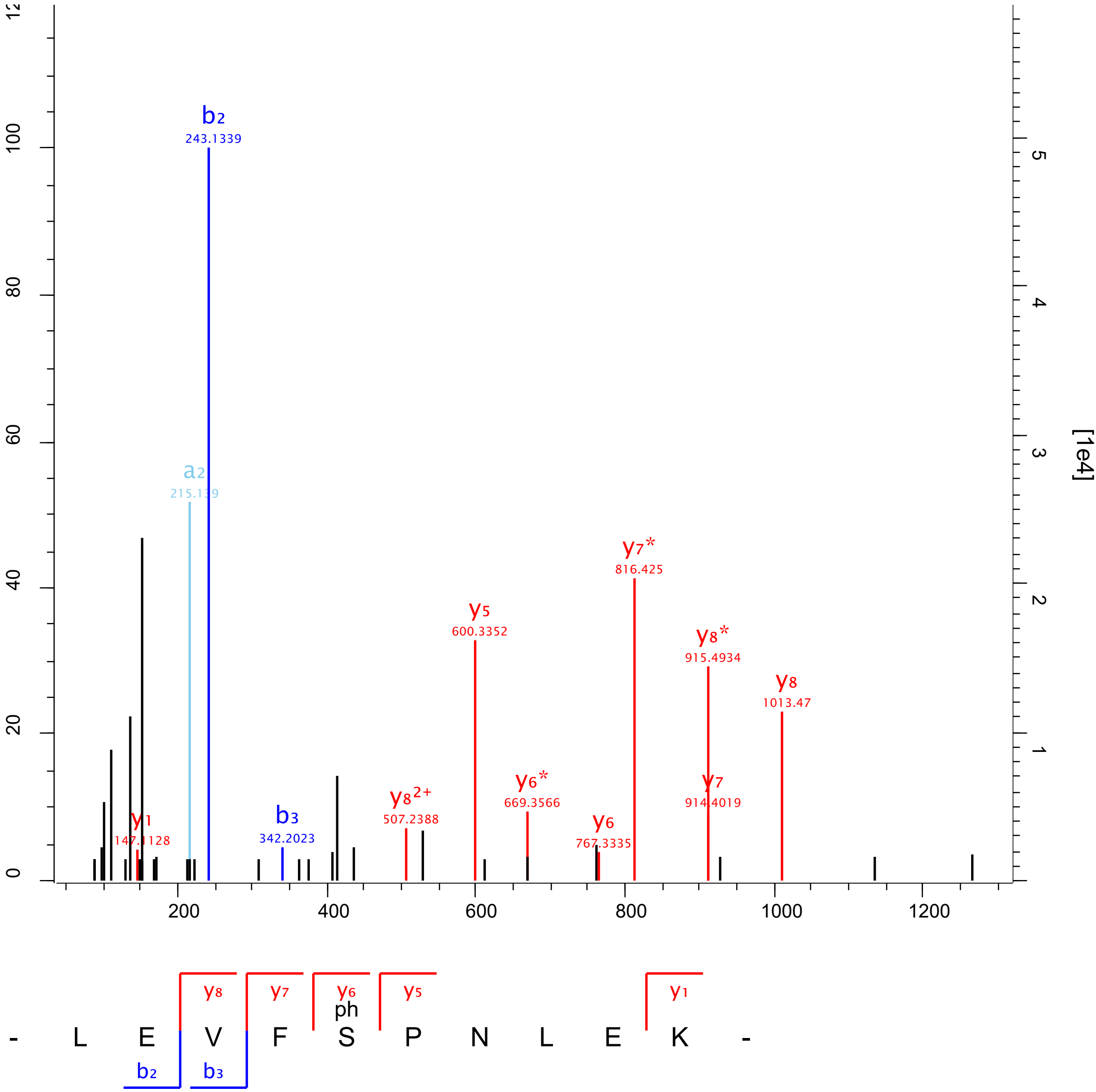

| Raw file | Scan | Method | Score | m/z |
|---|---|---|---|---|
| A17_15_1 | 27295 | FTMS; HCD | 84.51 | 701.82 |

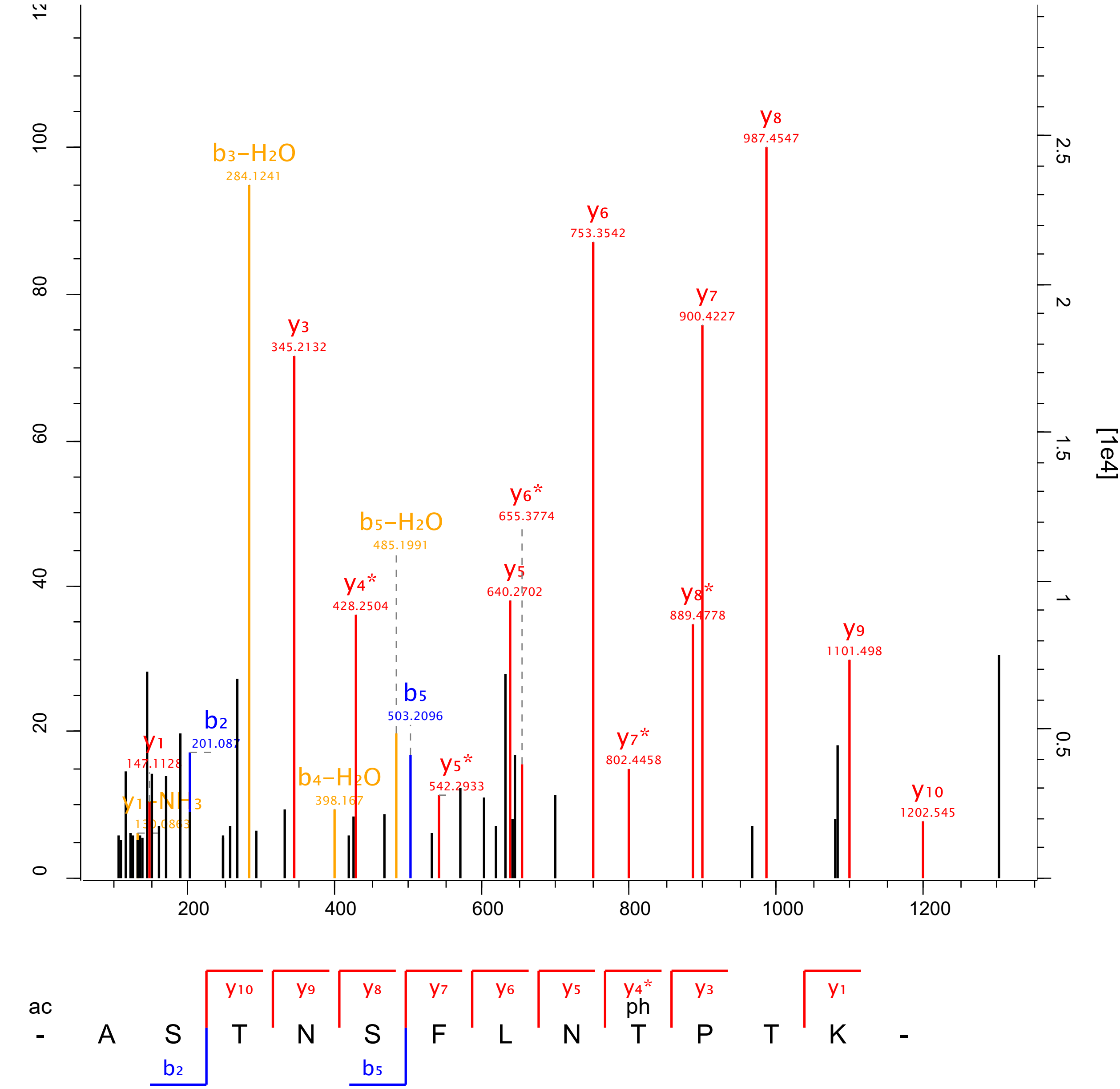

| Raw file | Scan | Method | Score | m/z |
|---|---|---|---|---|
| A17_15_1 | 27424 | FTMS; HCD | 112.44 | 632.79 |

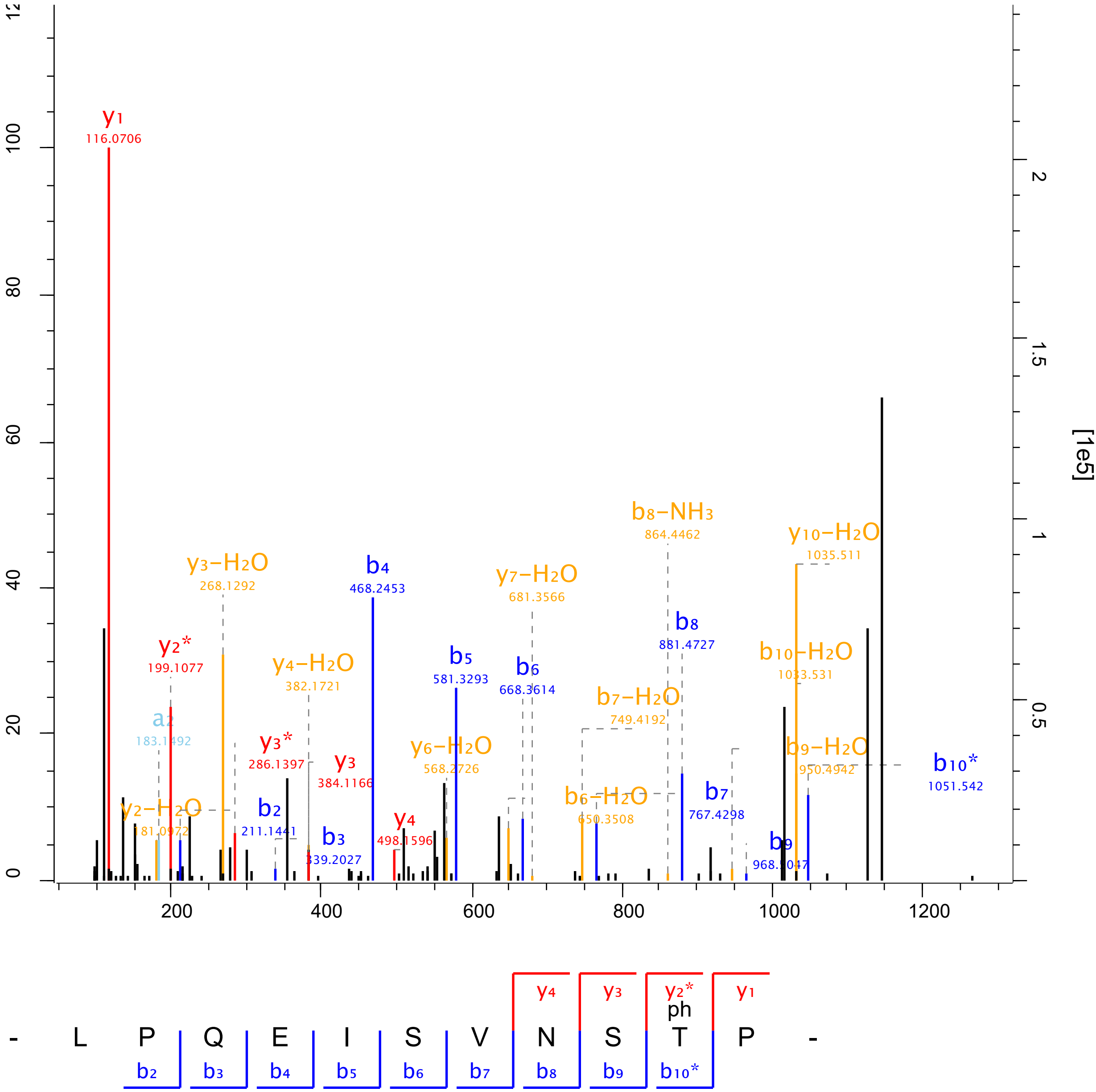

| Raw file | Scan | Method | Score | m/z |
| --- | --- | --- | --- | --- |
| A17_15_1 | 27493 | FTMS; HCD | 117.48 | 933.44 |

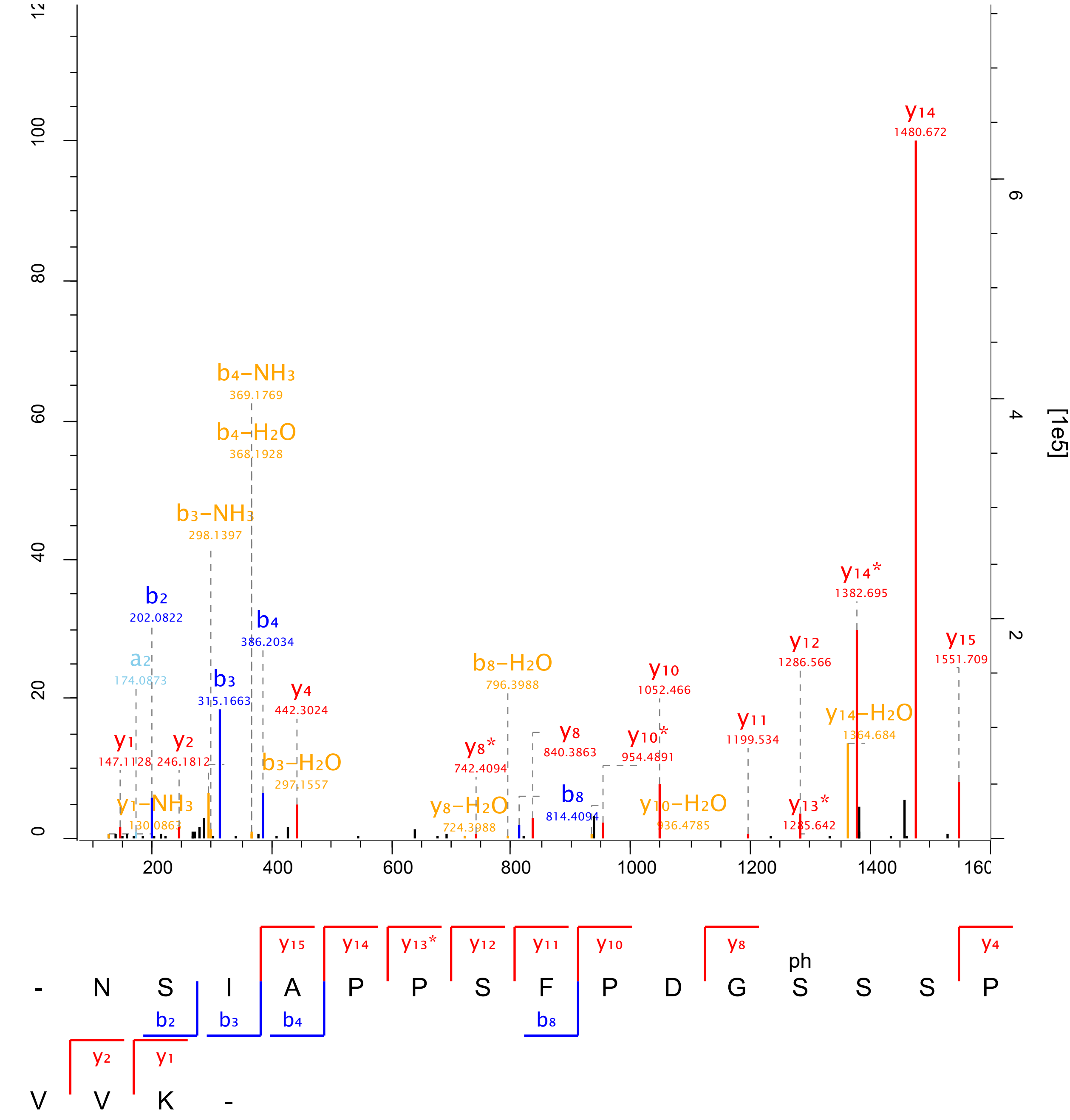

| Raw file | Scan | Method | Score | m/z |
| --- | --- | --- | --- | --- |
| A17_15_1 | 28229 | FTMS; HCD | 41.62 | 701.33 |

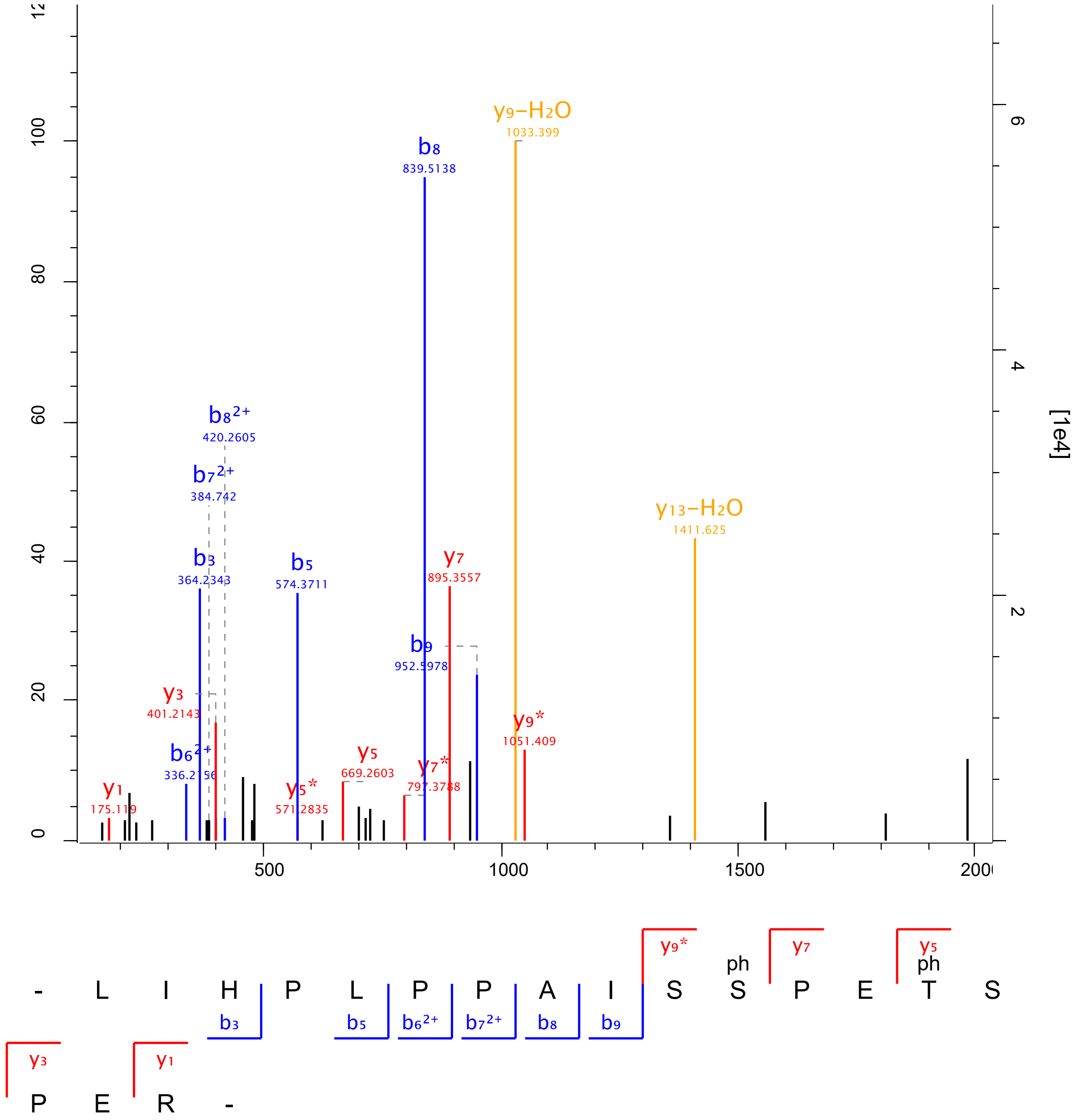

| Raw file | Scan | Method | Score | m/z |
|---|---|---|---|---|
| A17_15_1 | 28380 | FTMS; HCD | 137.73 | 721.32 |

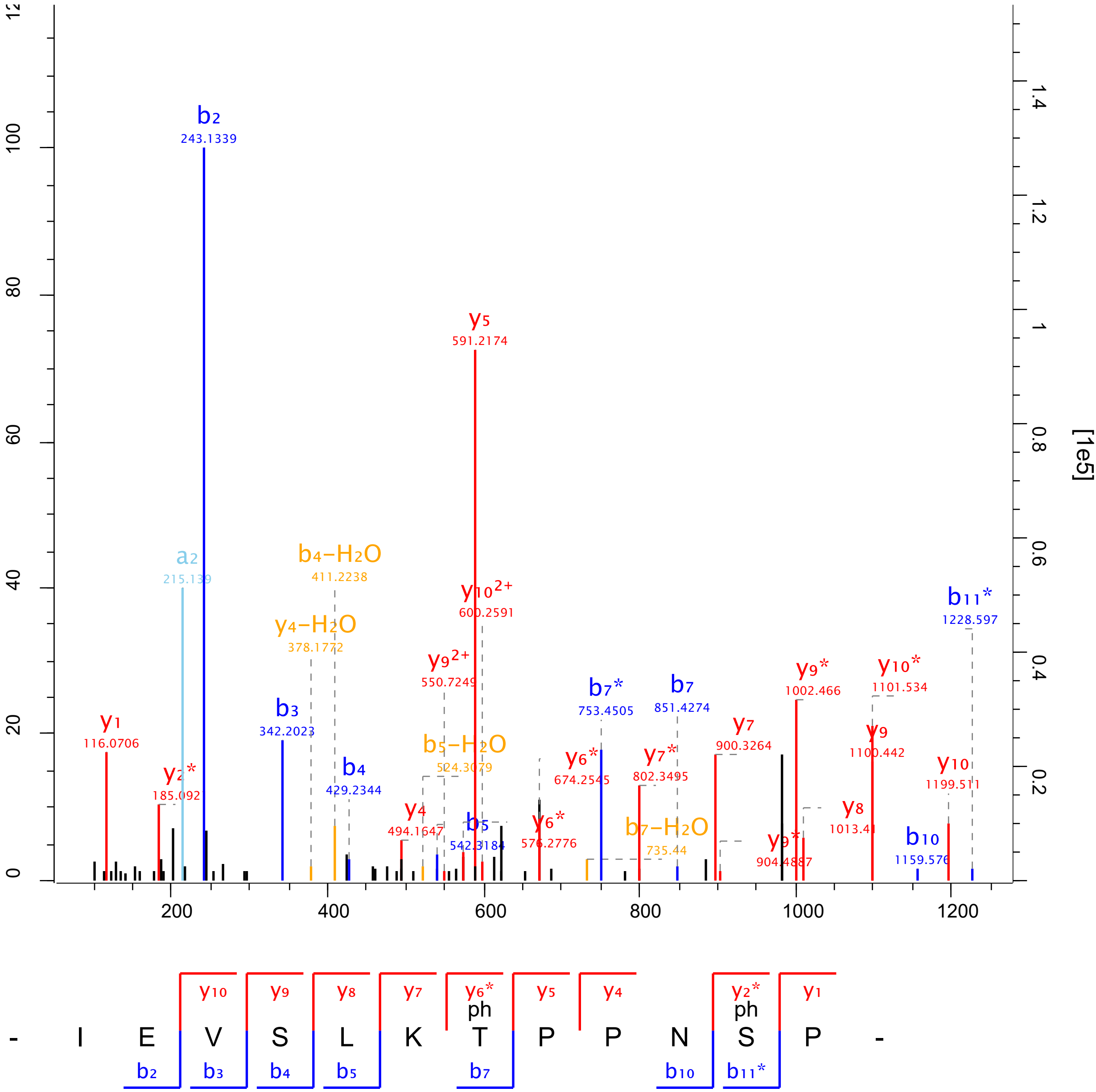

| Raw file | Scan | Method | Score | m/z |
|---|---|---|---|---|
| A17_15_1 | 29102 | FTMS; HCD | 40.62 | 835.33 |

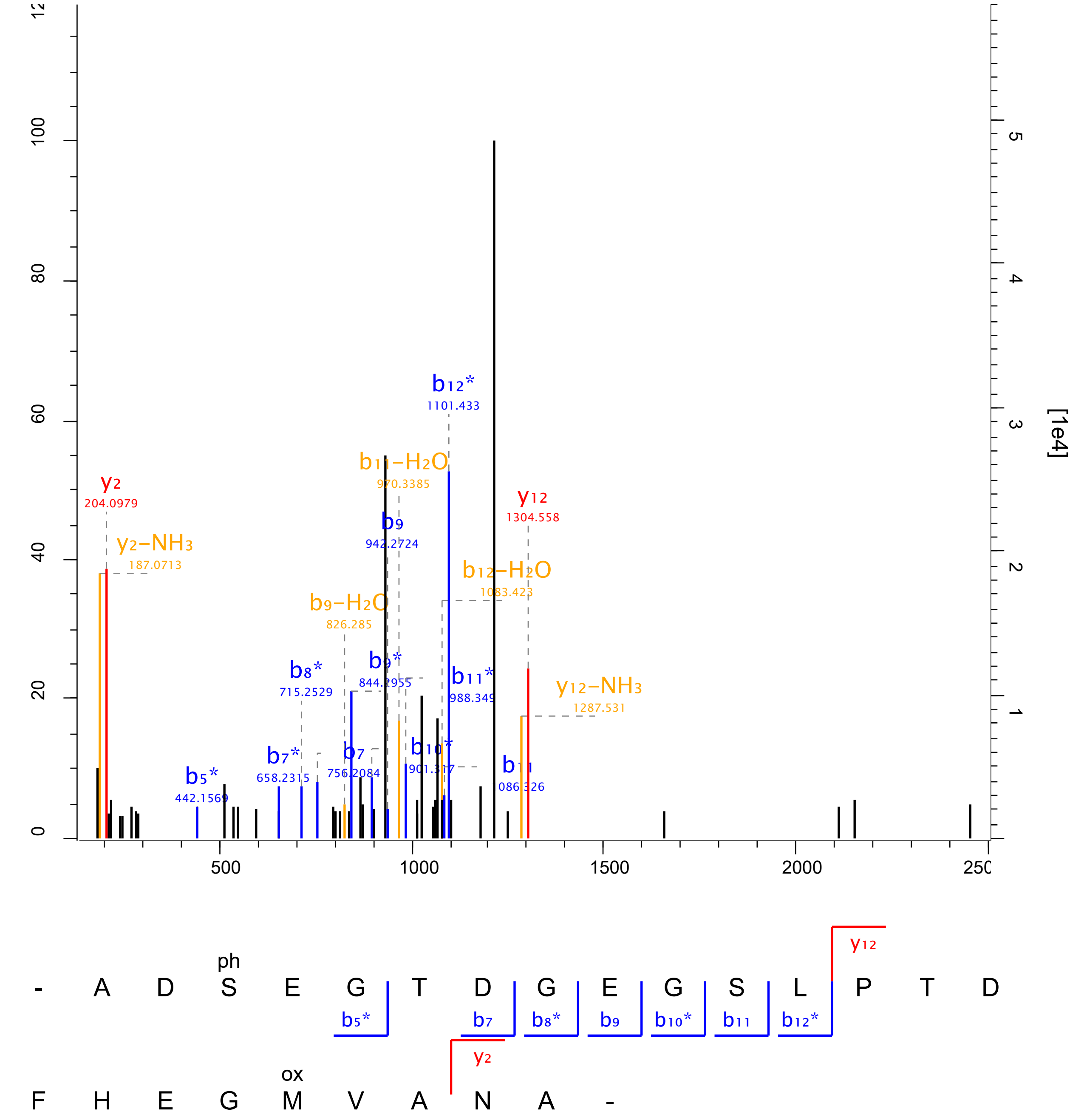

| Raw file | Scan | Method | Score | m/z |
|---|---|---|---|---|
| A17_15_1 | 29271 | FTMS; HCD | 161.04 | 846.4 |

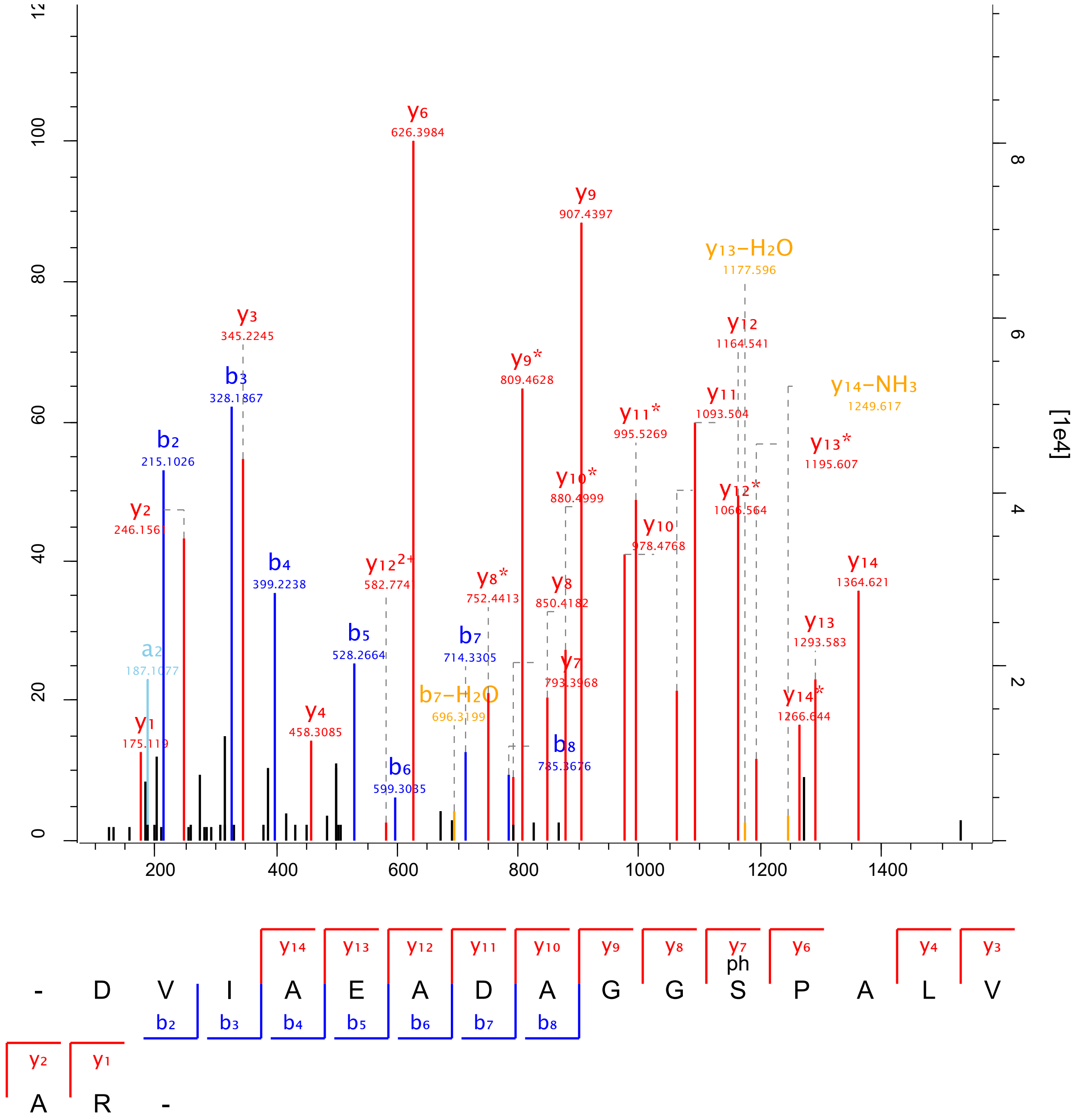

| Raw file | Scan | Method | Score | m/z |
|---|---|---|---|---|
| A17_15_1 | 29834 | FTMS; HCD | 71.85 | 823.37 |

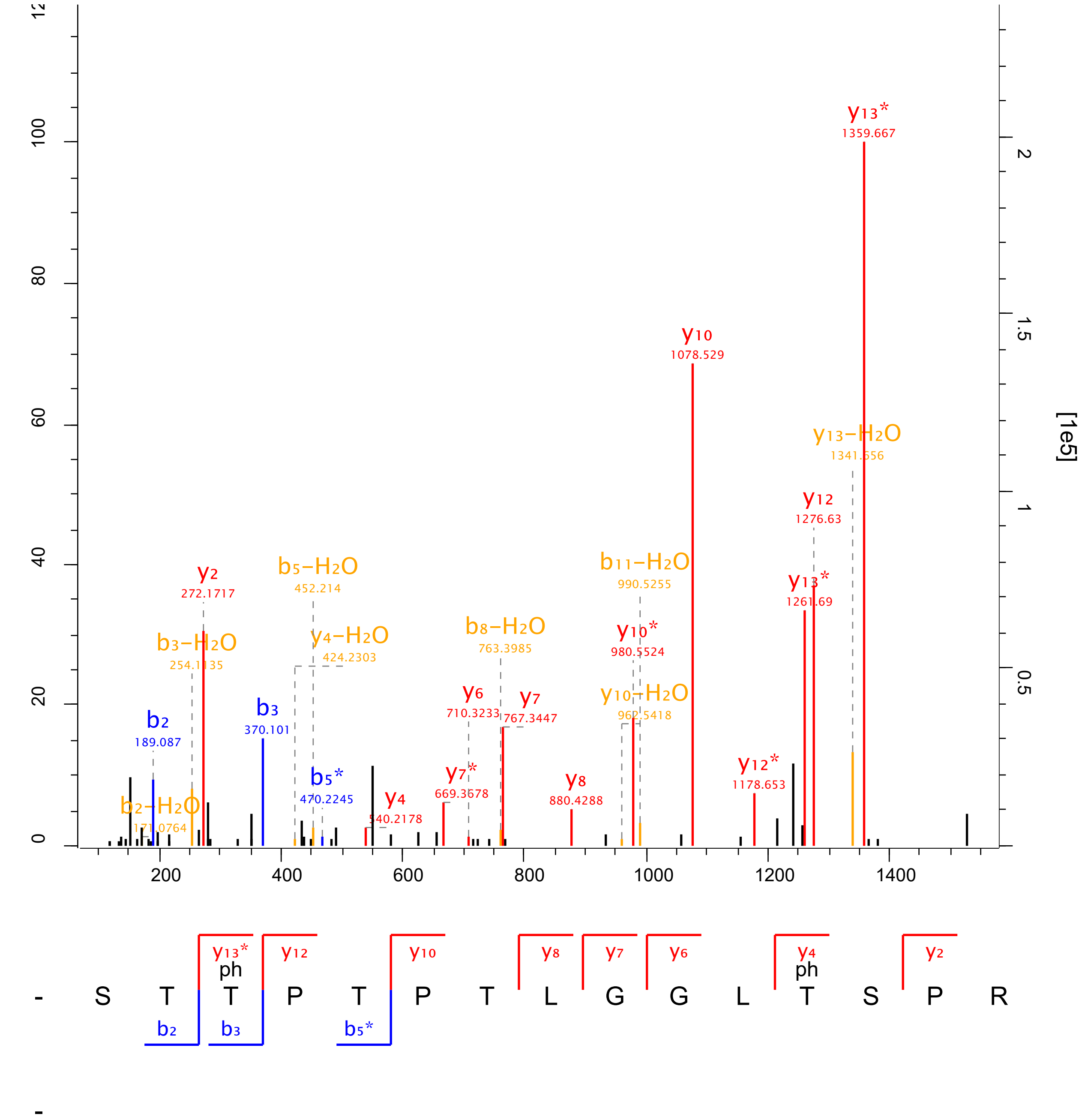

| Raw file | Scan | Method | Score | m/z |
|---|---|---|---|---|
| A17_15_1 | 30538 | FTMS; HCD | 48.9 | 748.27 |

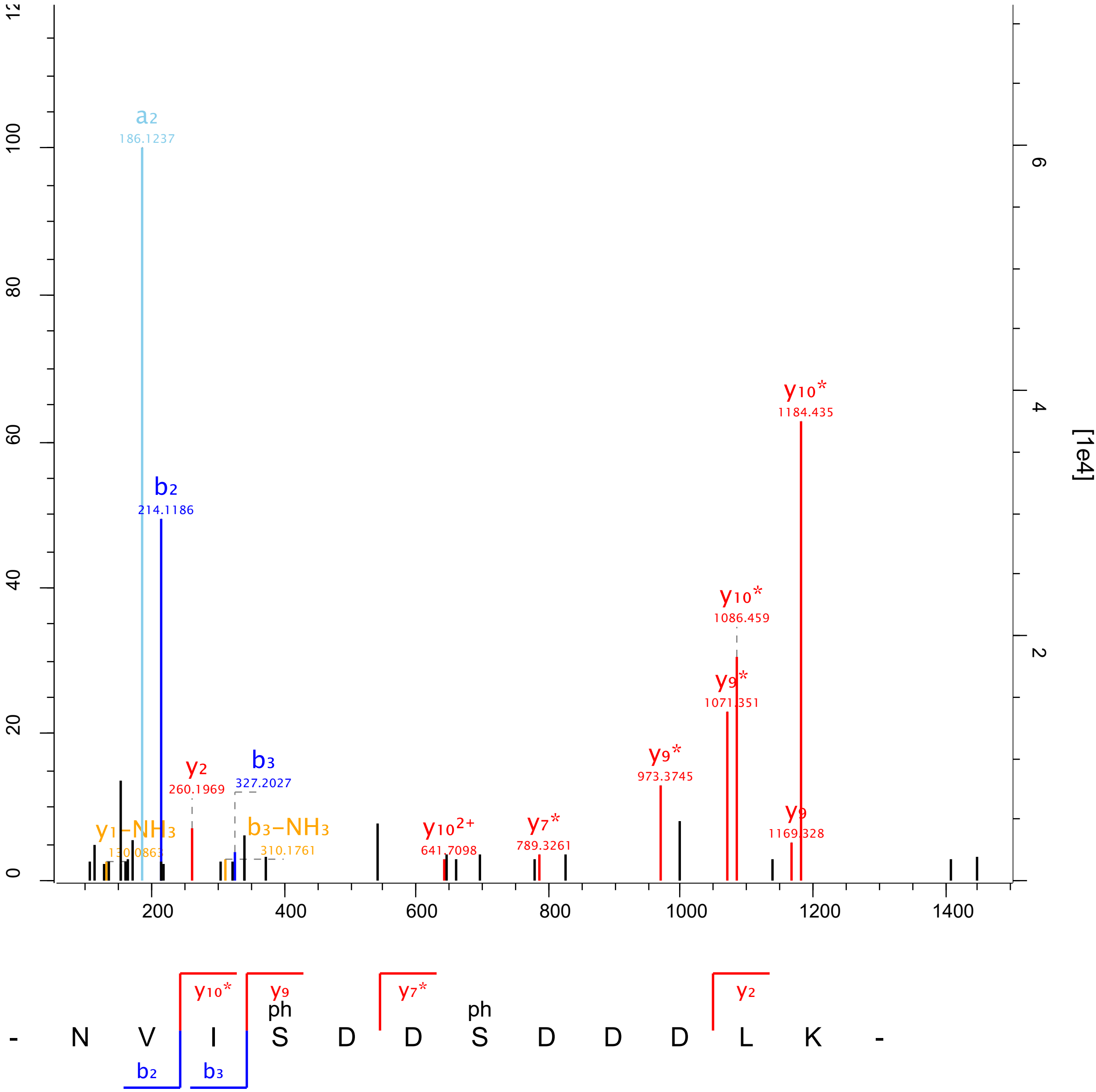

| Raw file | Scan | Method | Score | m/z |
| --- | --- | --- | --- | --- |
| A17_15_1 | 3072 | FTMS; HCD | 149.93 | 560.72 |

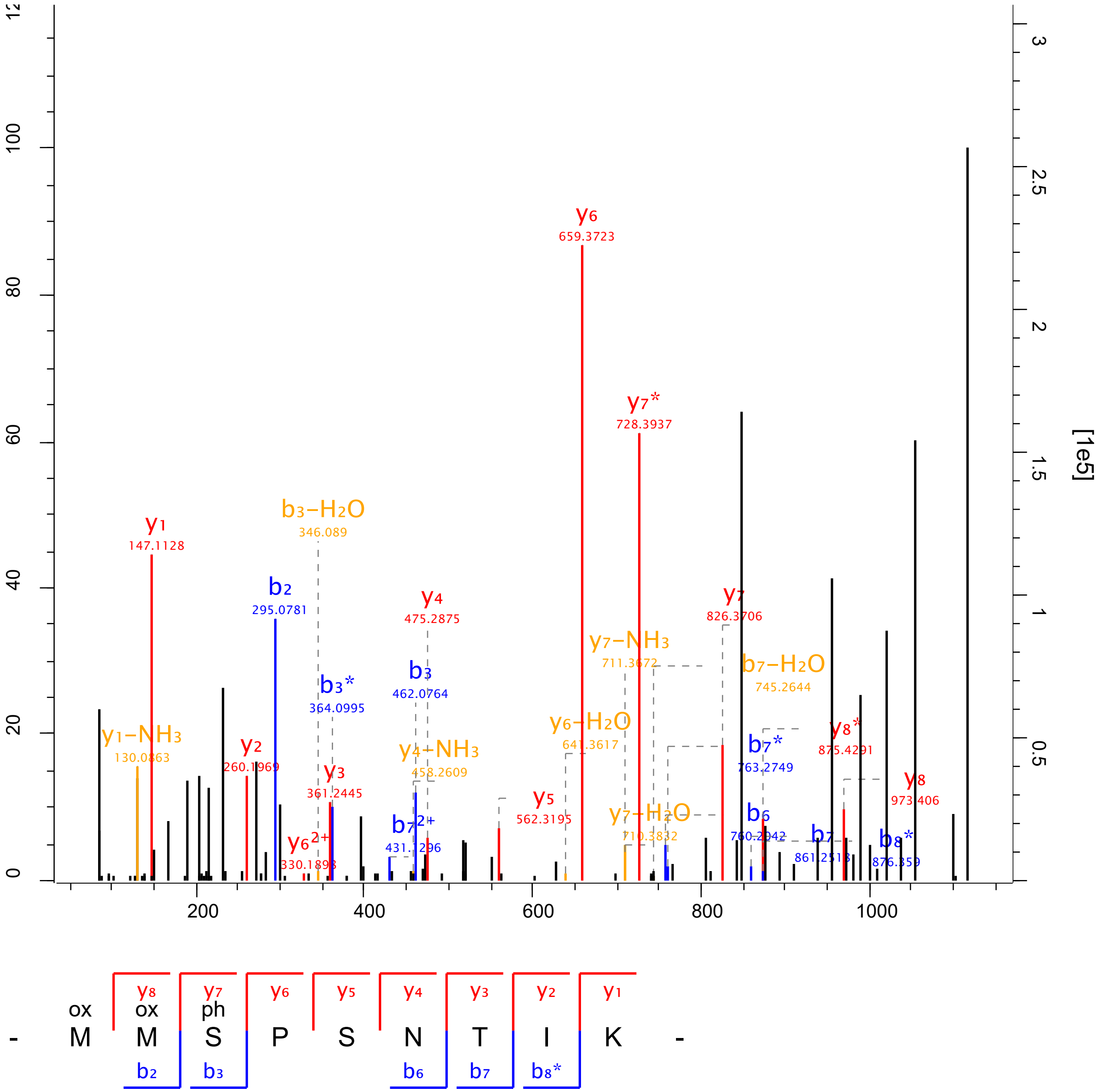

| Raw file | Scan | Method | Score | m/z |
|---|---|---|---|---|
| A17_15_1 | 30759 | FTMS; HCD | 231.17 | 774.35 |

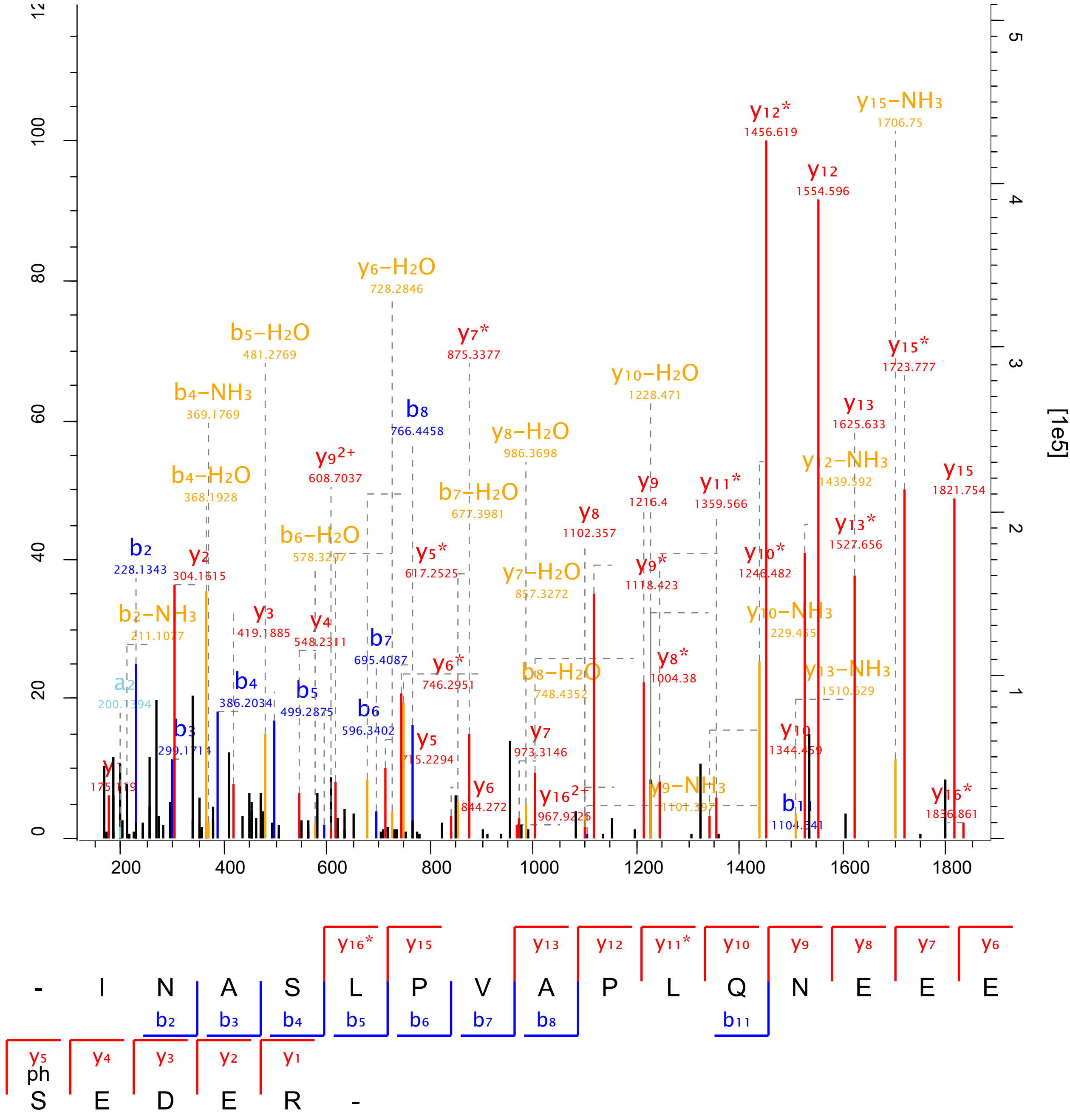

| Raw file | Scan | Method | Score | m/z |
|---|---|---|---|---|
| A17_15_1 | 30788 | FTMS; HCD | 55.44 | 855.88 |

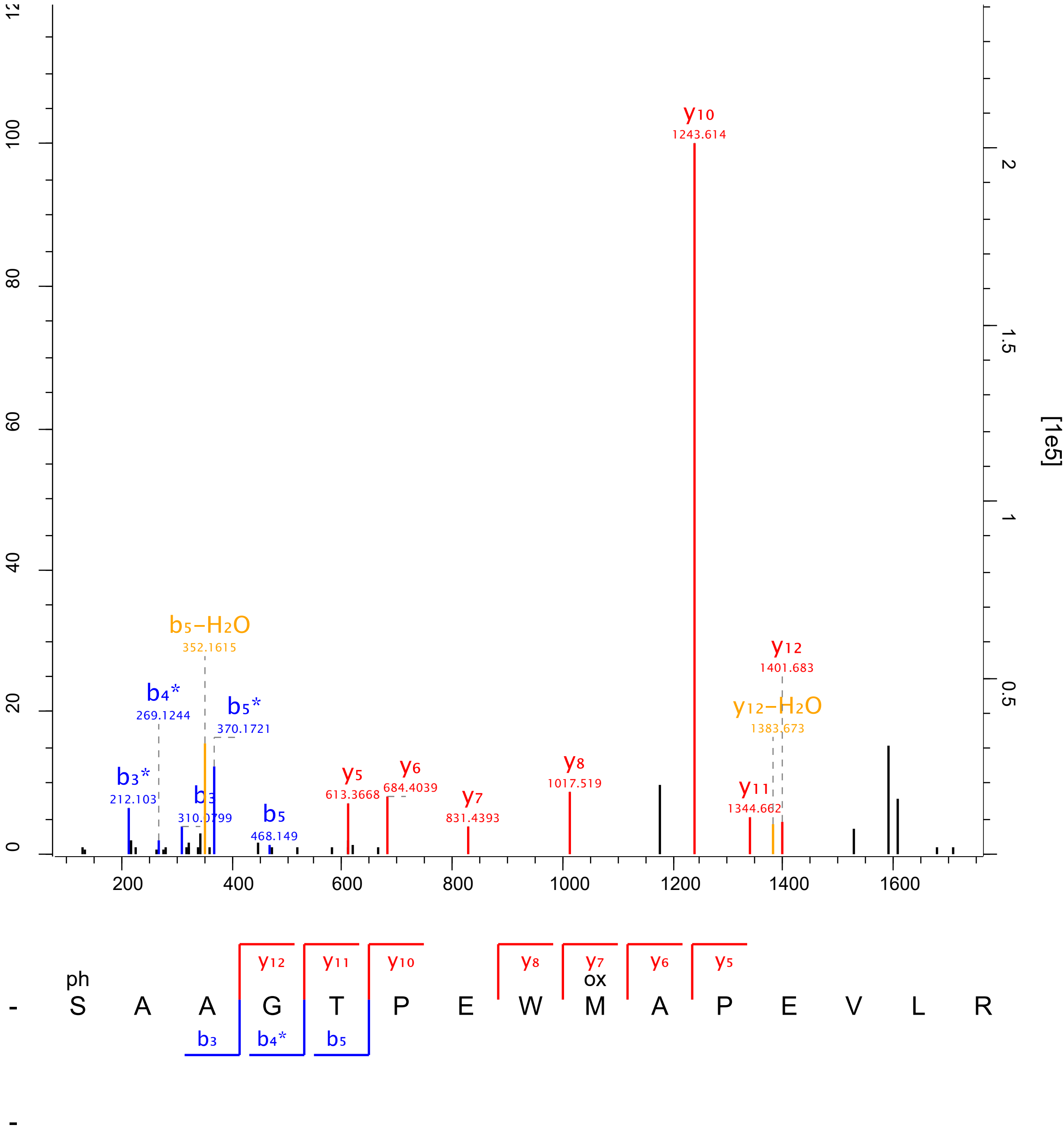

| Raw file | Scan | Method | Score | m/z |
| --- | --- | --- | --- | --- |
| A17_15_1 | 3082 | FTMS; HCD | 130.21 | 736.32 |

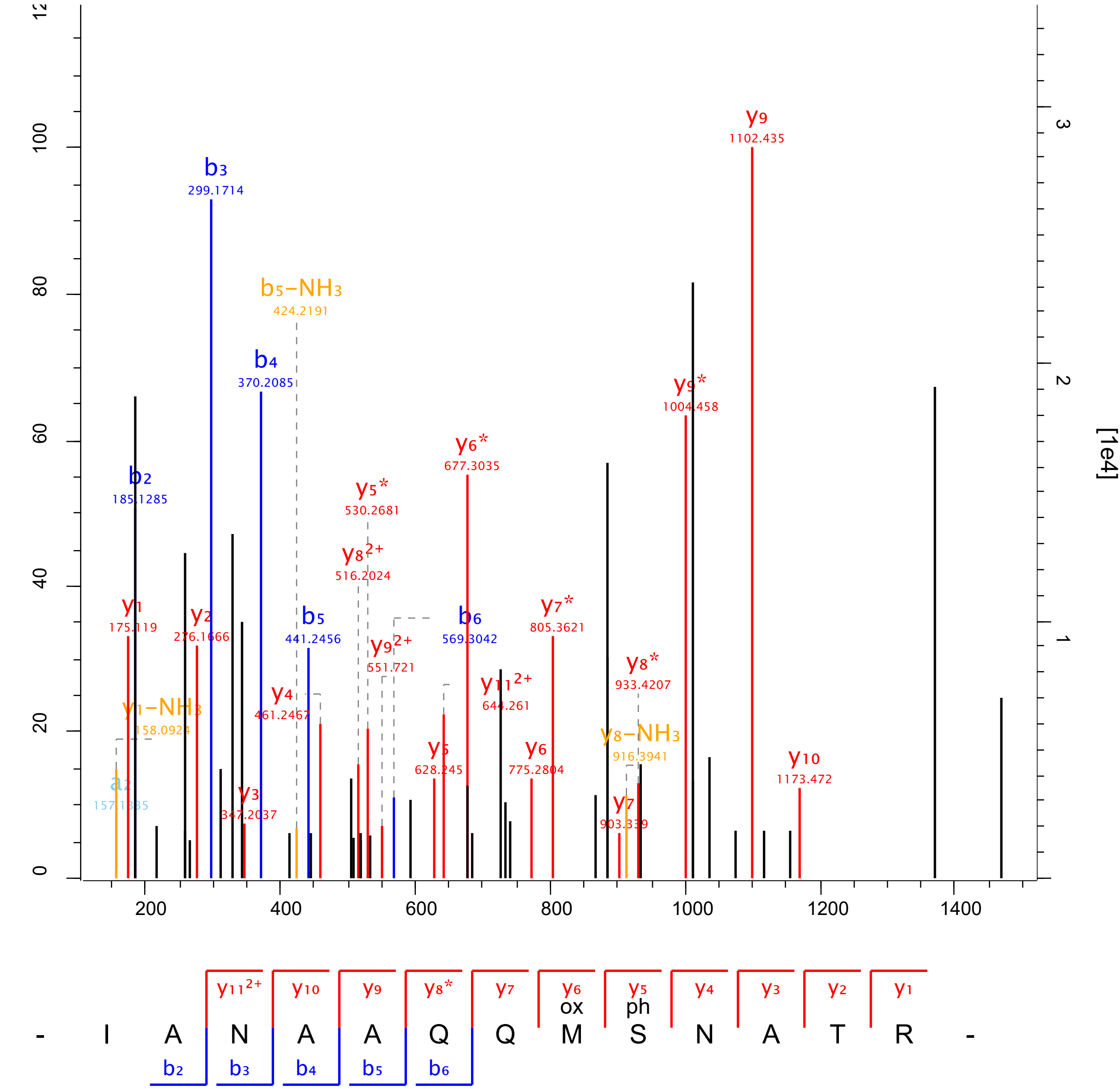

| Raw file | Scan | Method | Score | m/z |
|---|---|---|---|---|
| A17_15_1 | 31190 | FTMS; HCD | 148.6 | 828.03 |

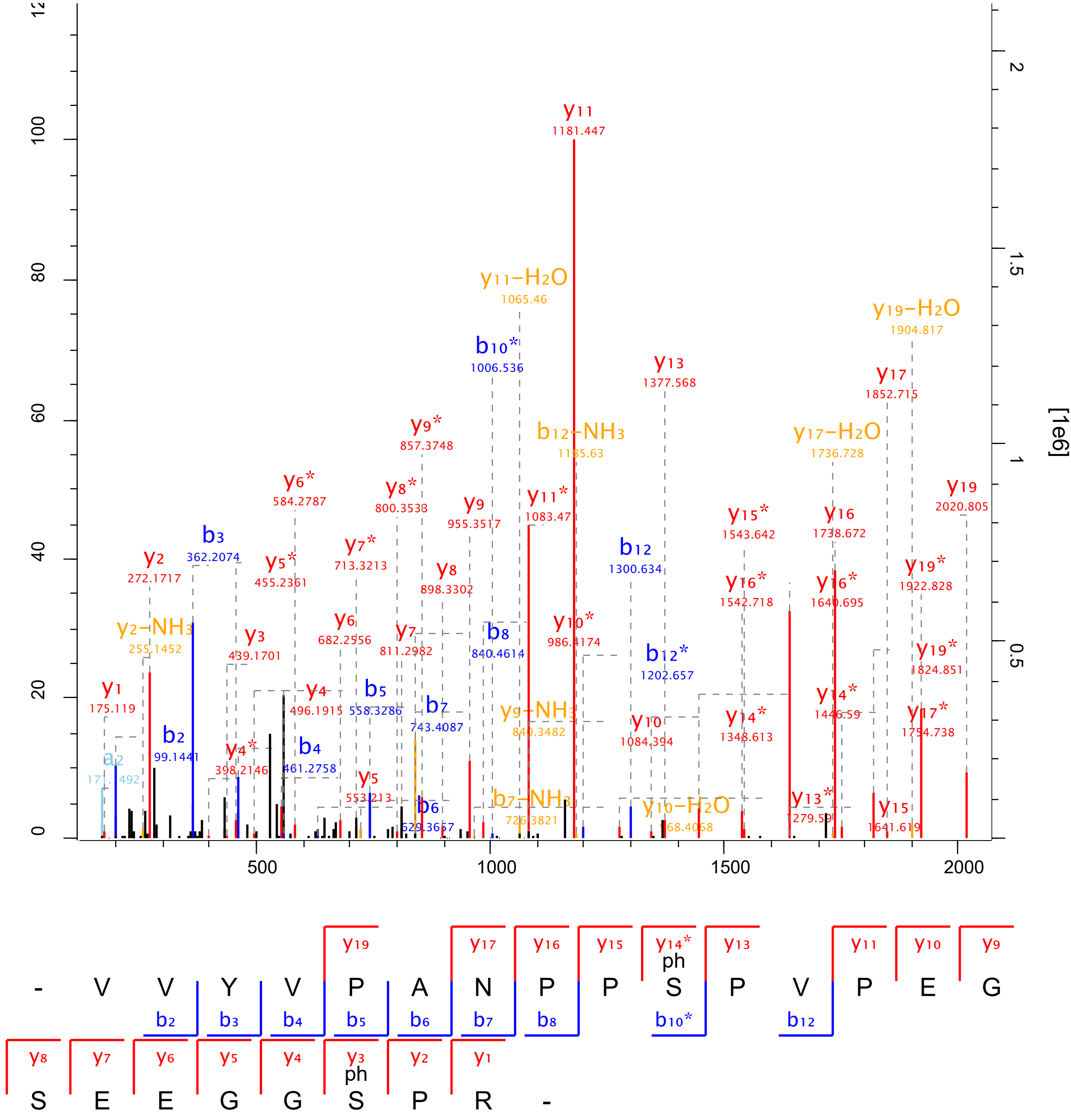

| Raw file | Scan | Method | Score | m/z |
|---|---|---|---|---|
| A17_15_1 | 3146 | FTMS; HCD | 49.54 | 887.81 |

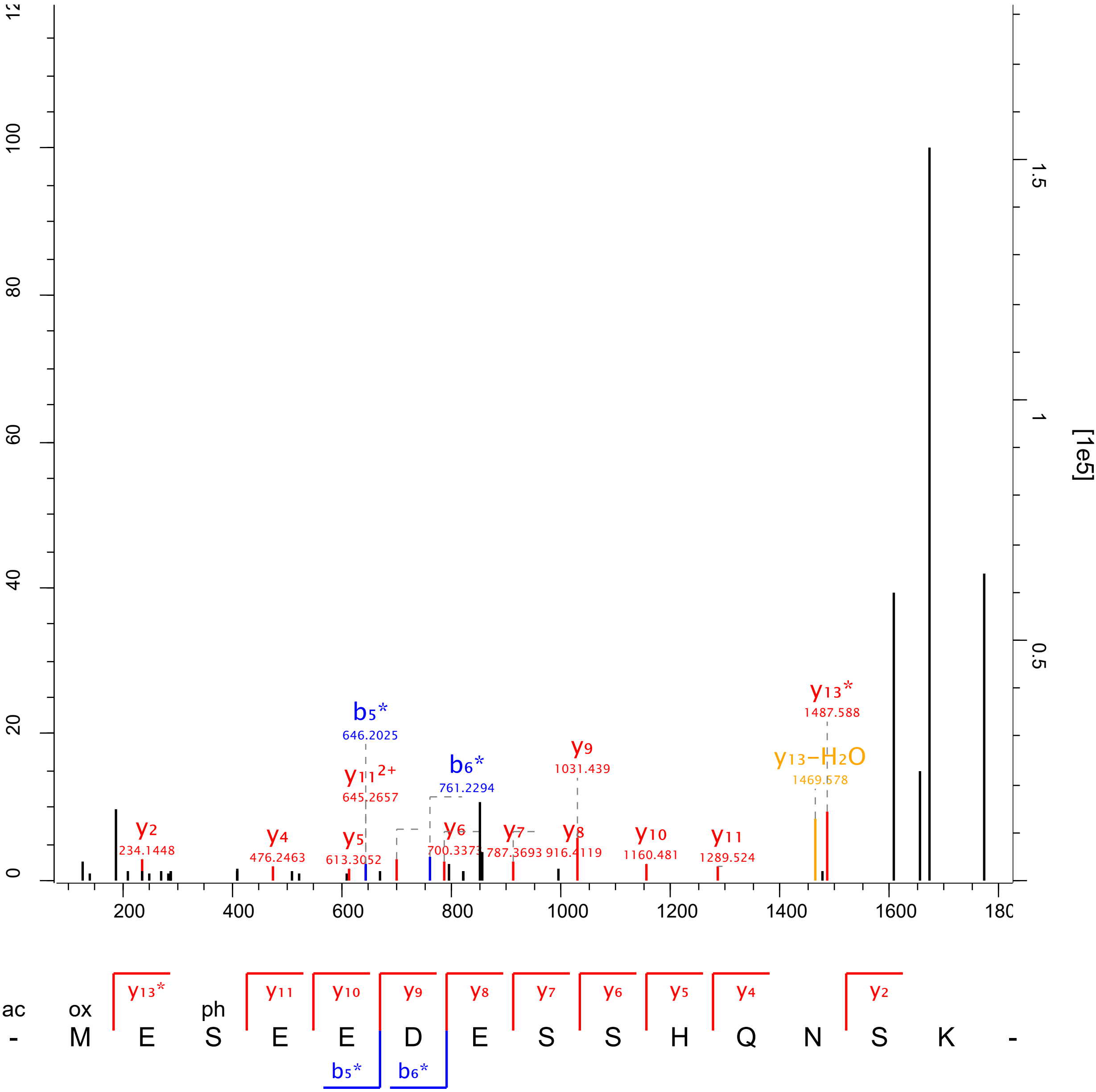

| Raw file | Scan | Method | Score | m/z |
| --- | --- | --- | --- | --- |
| A17_15_1 | 32344 | FTMS; HCD | 69.85 | 1107.46 |

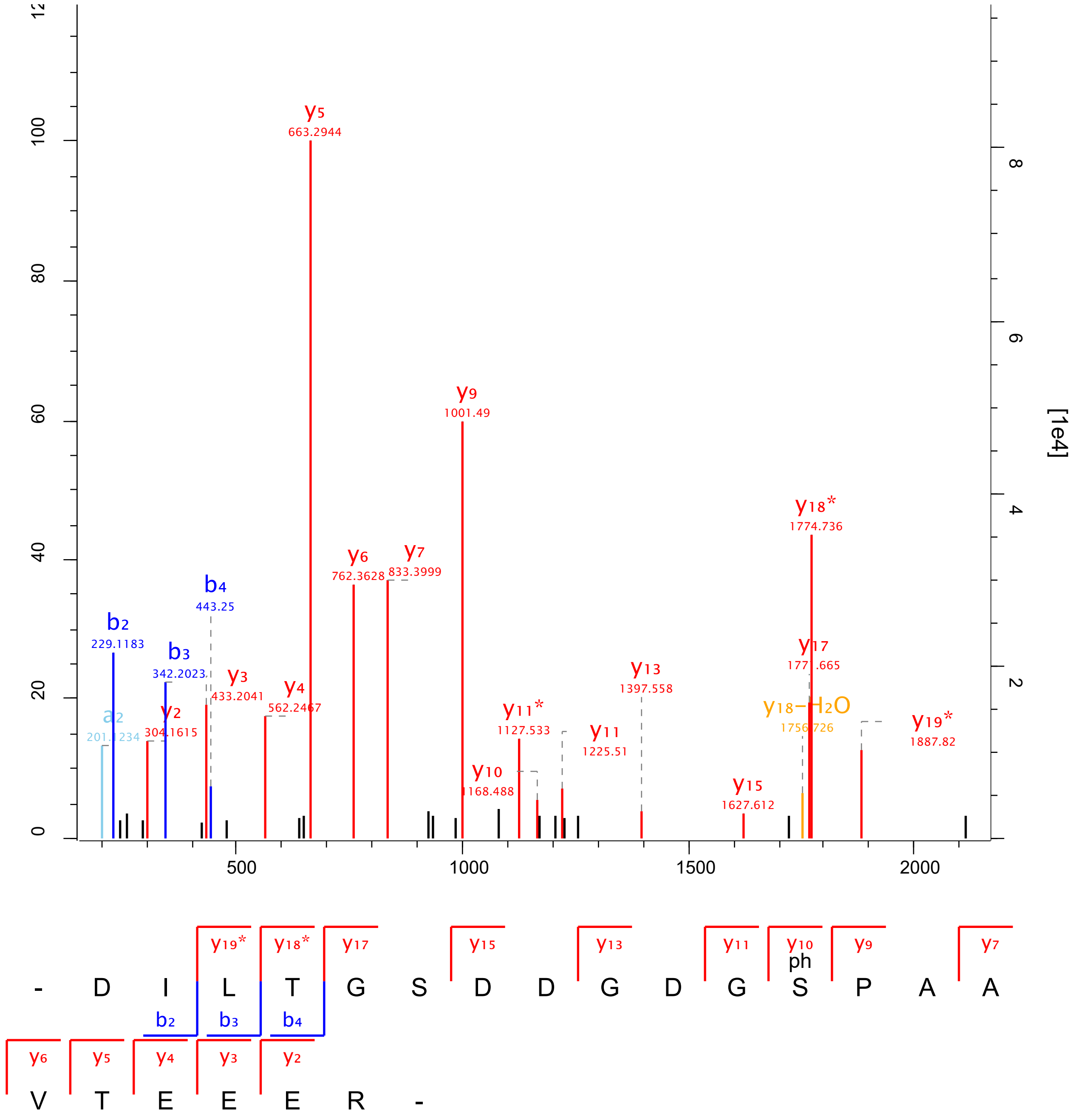

| Raw file | Scan | Method | Score | m/z |
|---|---|---|---|---|
| A17_15_1 | 32643 | FTMS; HCD | 76.89 | 1136.47 |

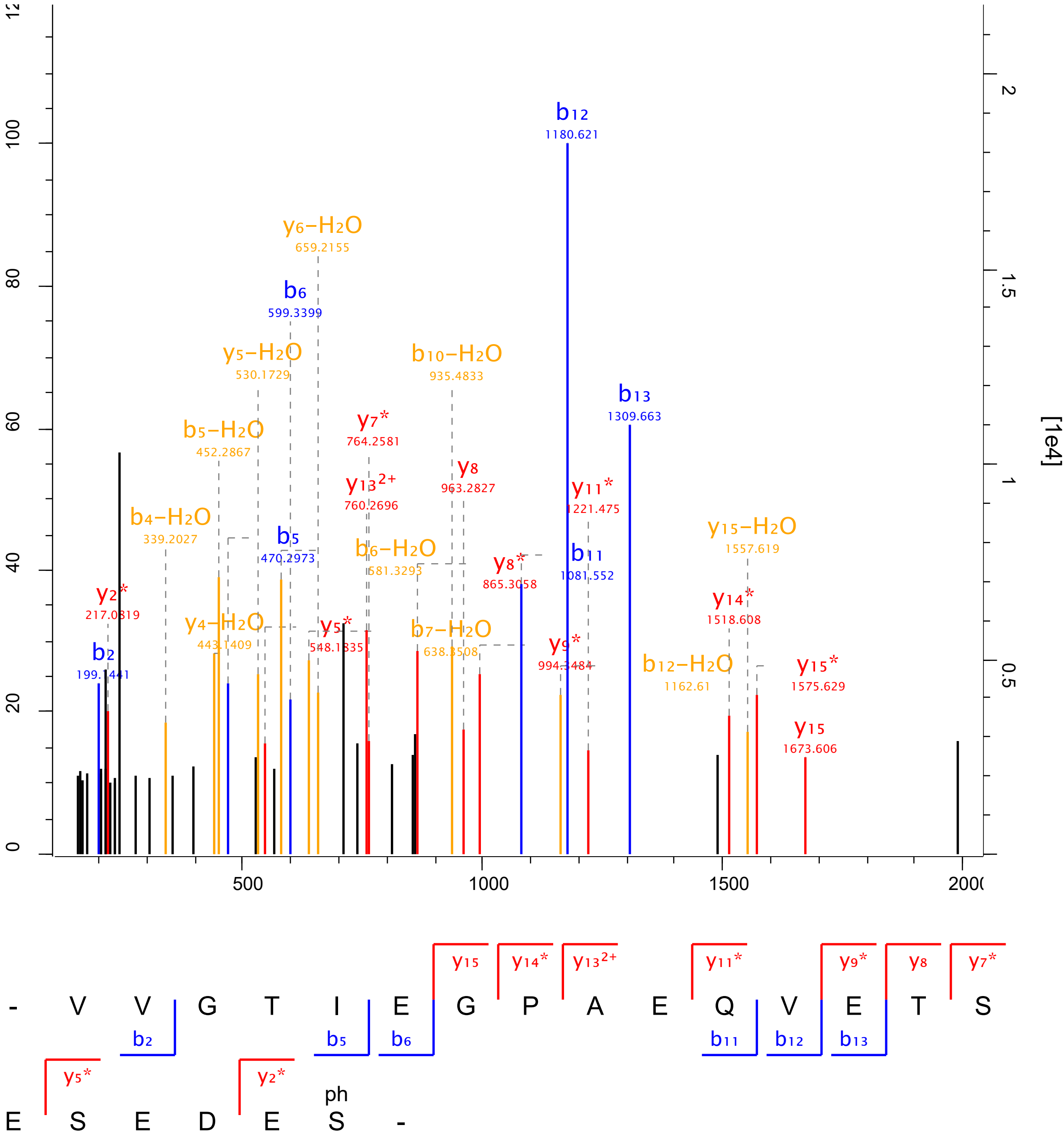

| Raw file | Scan | Method | Score | m/z |
| --- | --- | --- | --- | --- |
| A17_15_1 | 32760 | FTMS; HCD | 118.71 | 469.22 |

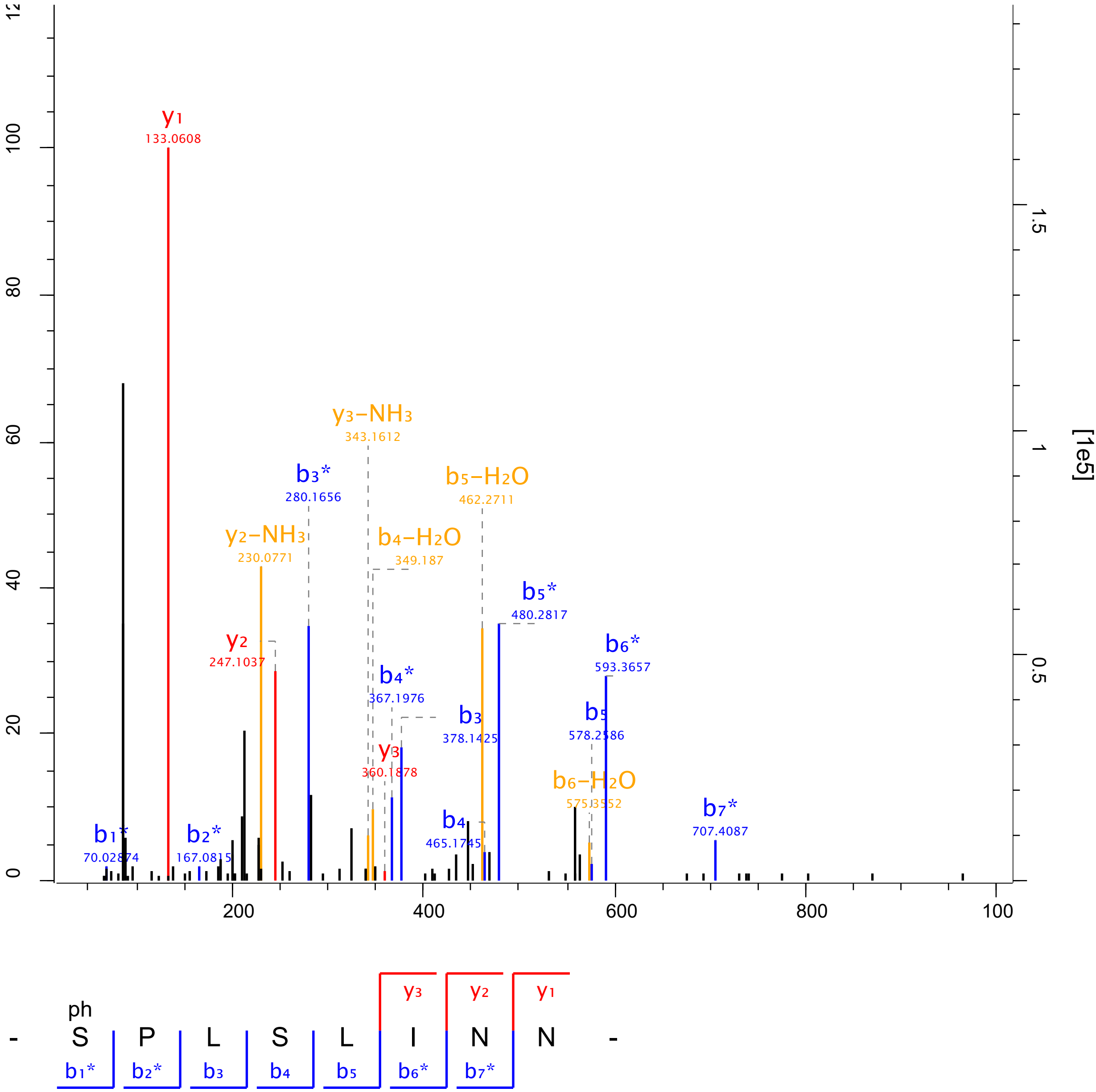

| Raw file | Scan | Method | Score | m/z |
| --- | --- | --- | --- | --- |
| A17_15_1 | 32992 | FTMS; HCD | 145.69 | 683.27 |

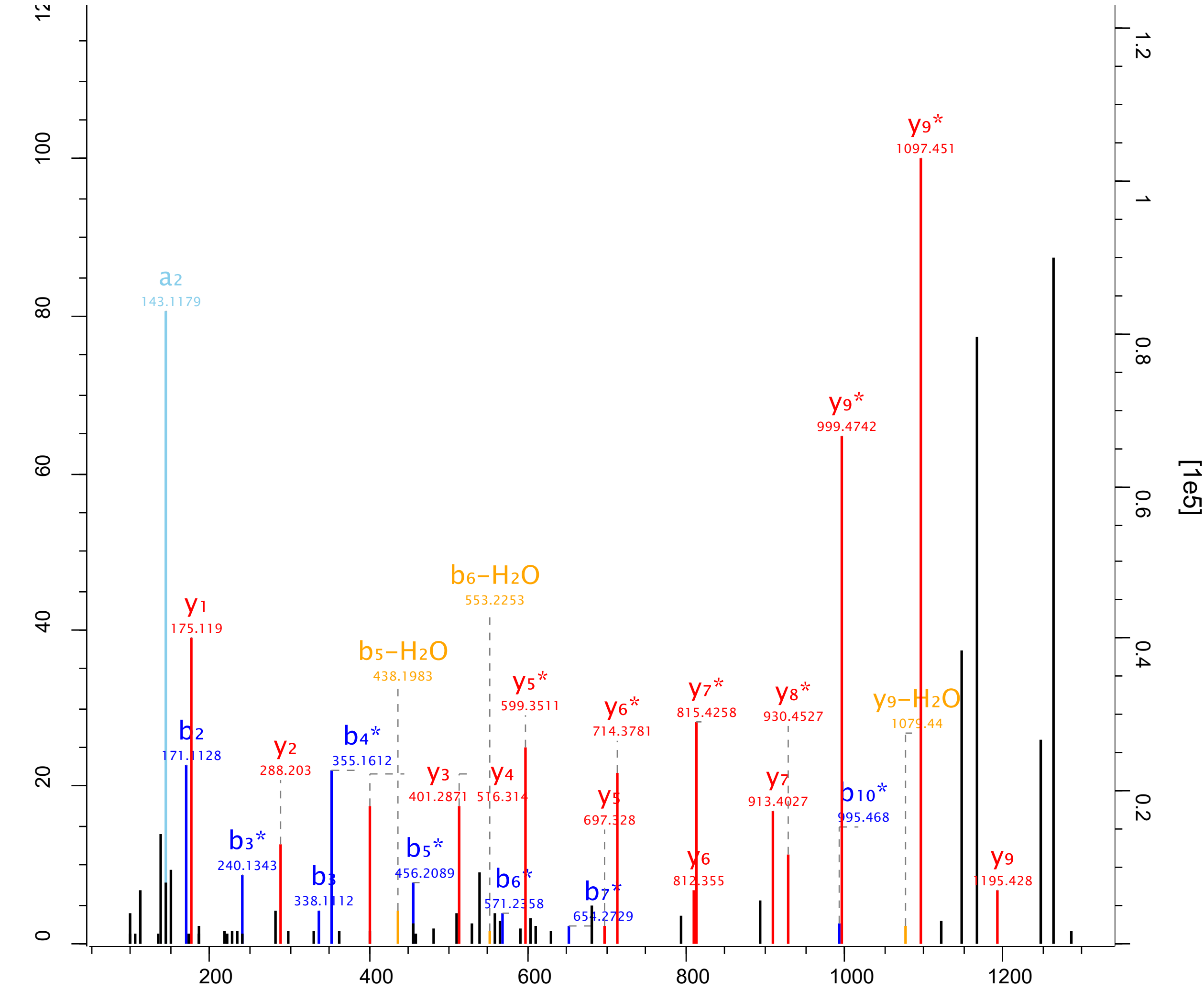


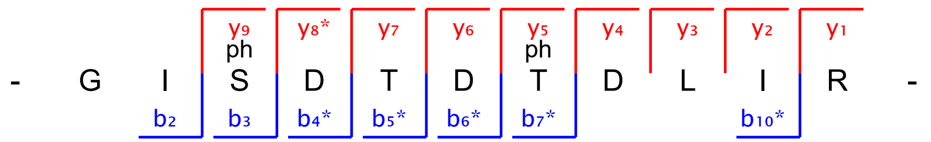

| Raw file | Scan | Method | Score | m/z |
|---|---|---|---|---|
| A17_15_1 | 33778 | FTMS; HCD | 70.84 | 974.45 |

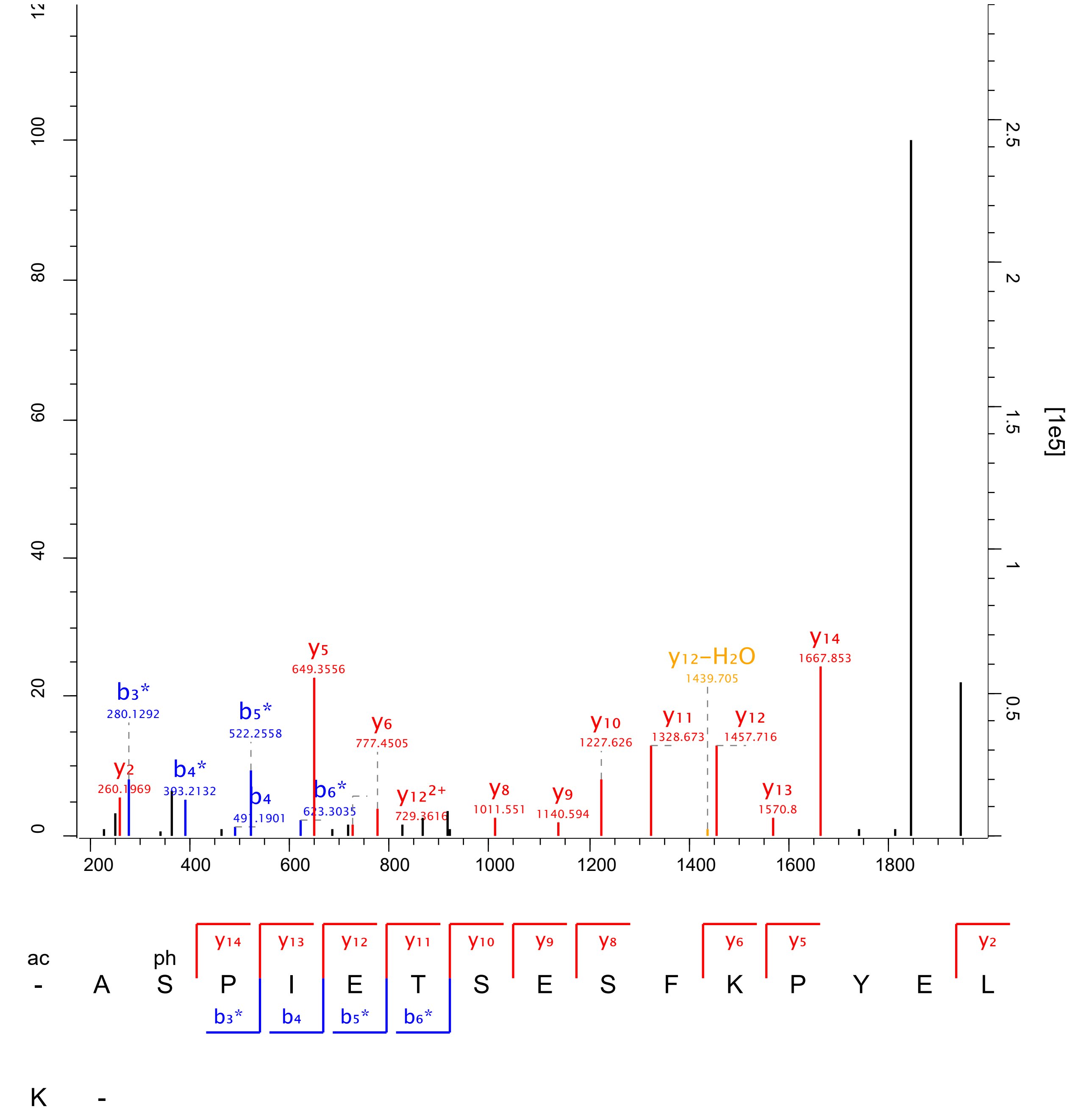

| Raw file | Scan | Method | Score | m/z |
| --- | --- | --- | --- | --- |
| A17_15_1 | 33787 | FTMS; HCD | 162.71 | 1102.93 |

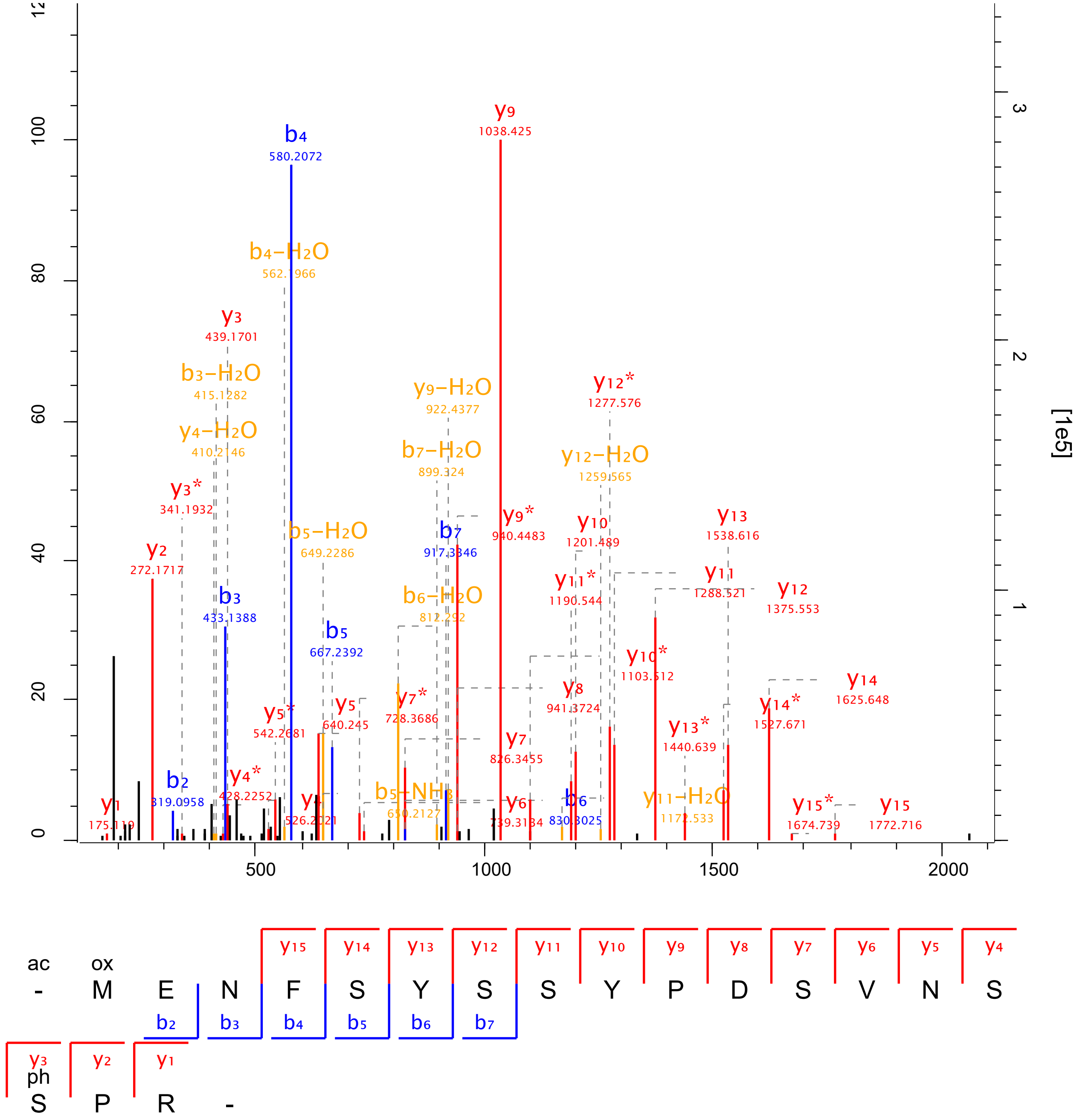

| Raw file | Scan | Method | Score | m/z |
| --- | --- | --- | --- | --- |
| A17_15_1 | 33837 | FTMS; HCD | 136.64 | 1052.46 |

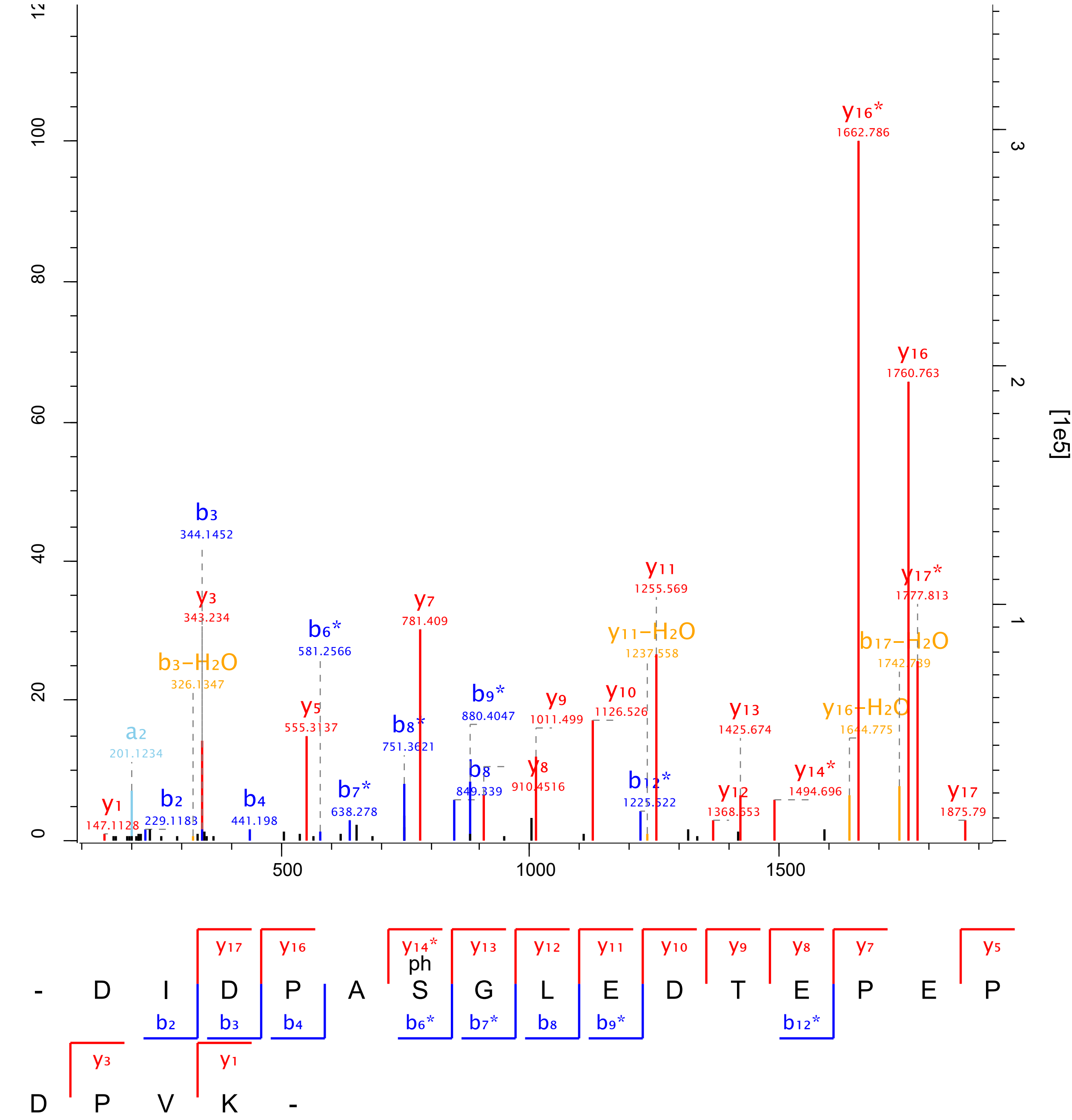

| Raw file | Scan | Method | Score | m/z |
| --- | --- | --- | --- | --- |
| A17_15_1 | 34473 | FTMS; HCD | 153.6 | 924.03 |

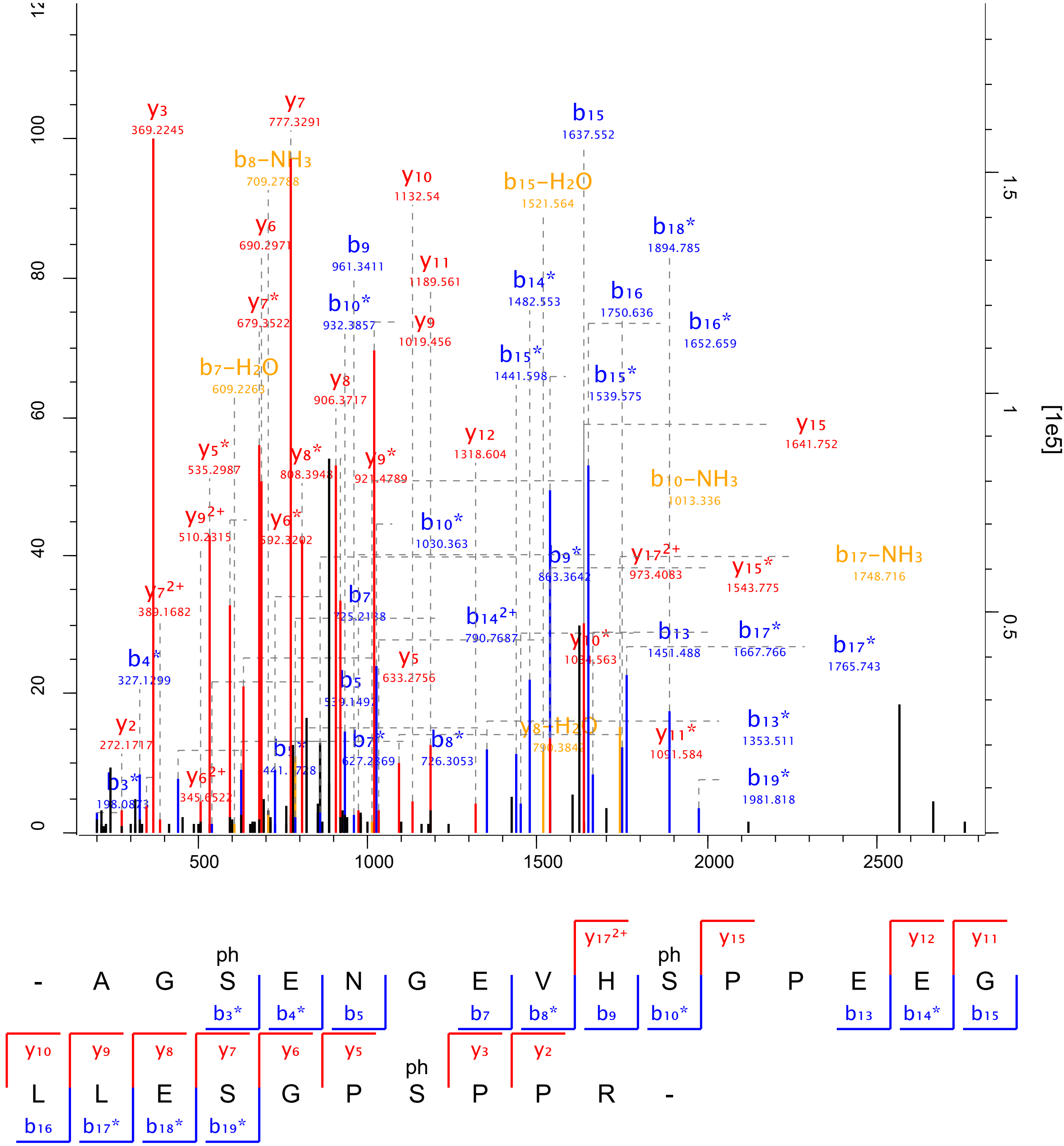

| Raw file | Scan | Method | Score | m/z |
|---|---|---|---|---|
| A17_15_1 | 35020 | FTMS; HCD | 83.56 | 894.75 |

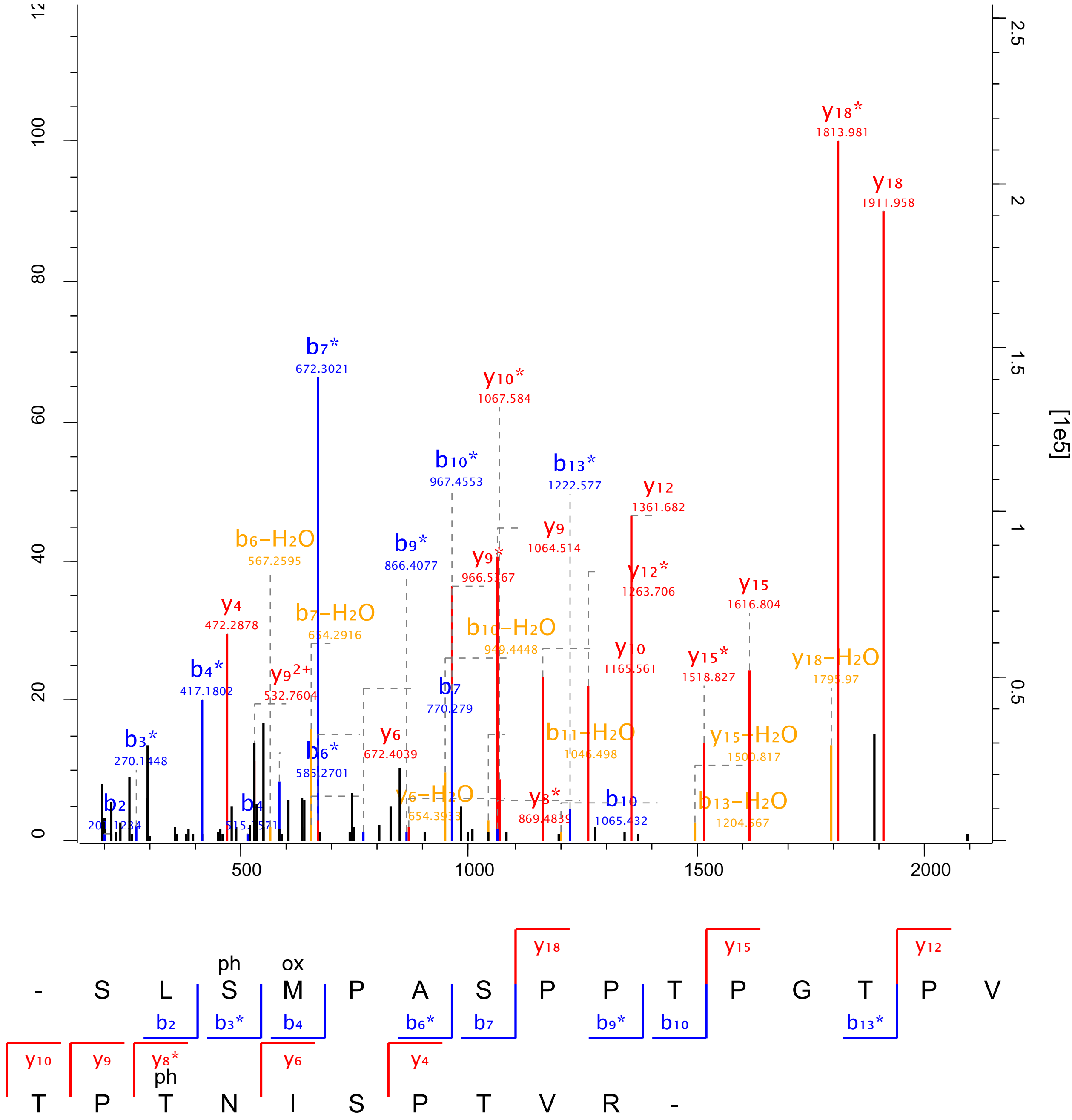

| Raw file | Scan | Method | Score | m/z |
|---|---|---|---|---|
| A17_15_1 | 35967 | FTMS; HCD | 85.09 | 927.86 |

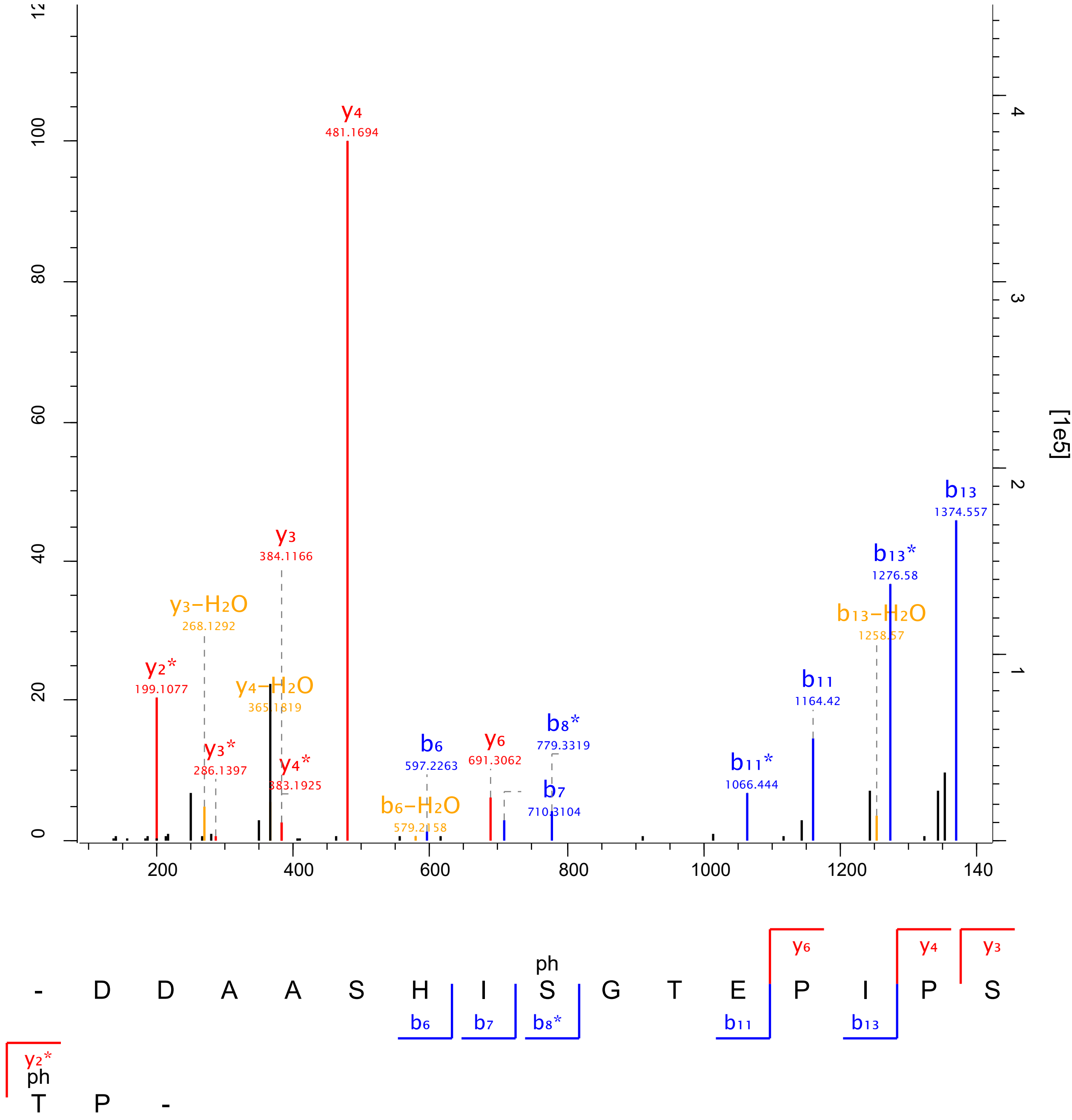

| Raw file | Scan | Method | Score | m/z |
|---|---|---|---|---|
| A17_15_1 | 37291 | FTMS; HCD | 69.34 | 620.3 |

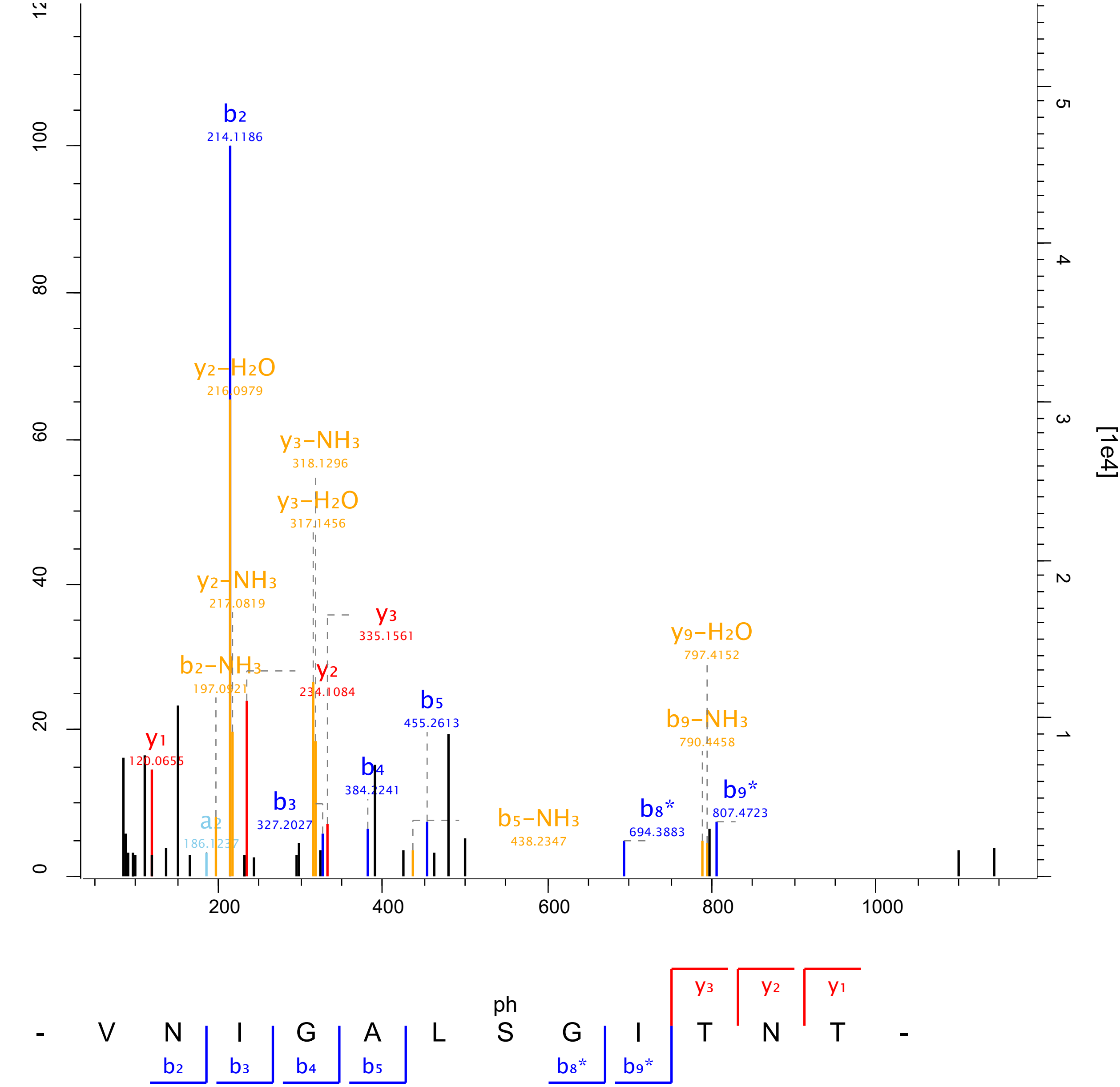

| Raw file | Scan | Method | Score | m/z |
| --- | --- | --- | --- | --- |
| A17_15_1 | 3928 | FTMS; HCD | 211.09 | 758.81 |

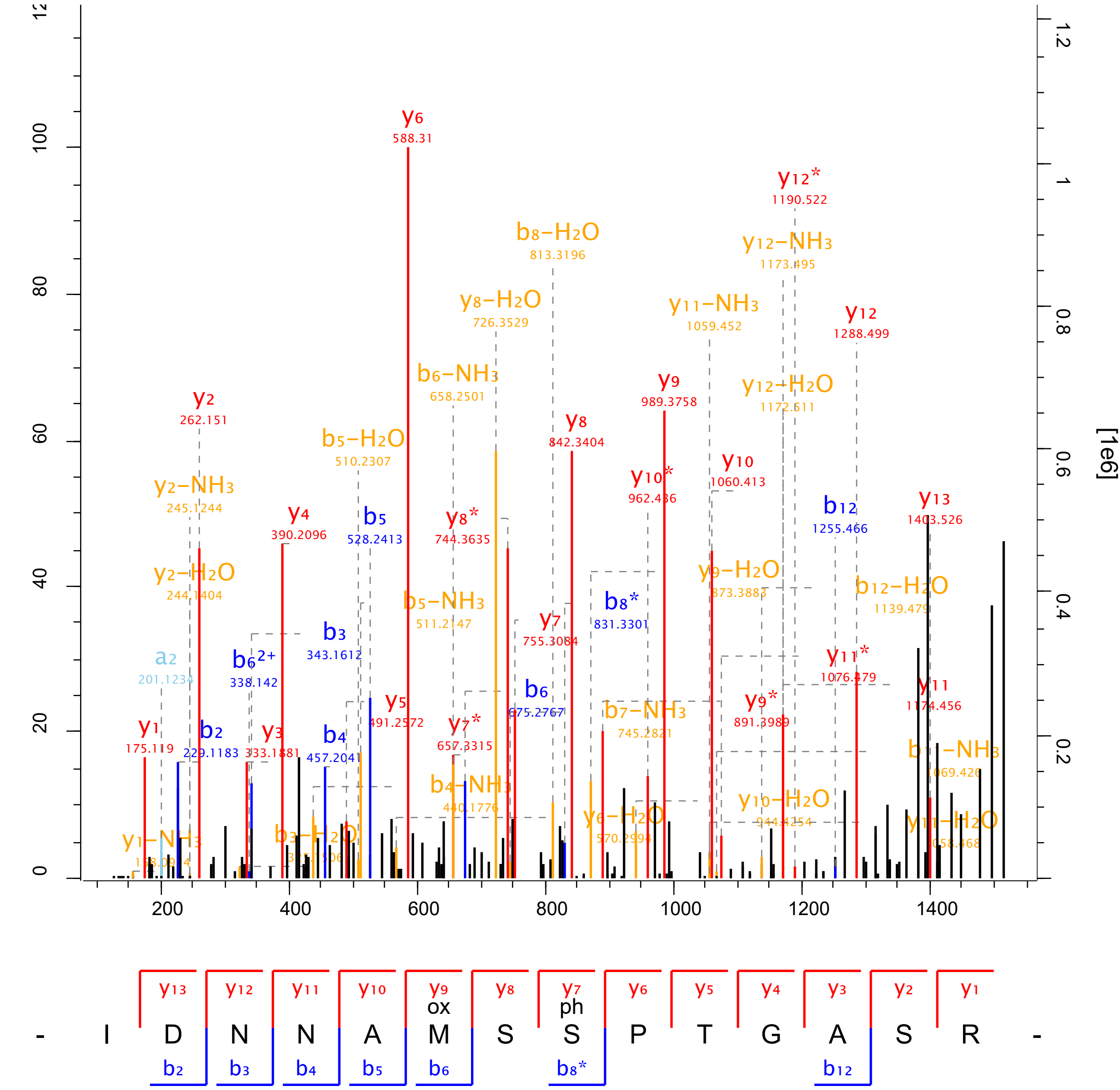

| Raw file | Scan | Method | Score | m/z |
|---|---|---|---|---|
| A17_15_1 | 4255 | FTMS; HCD | 43.03 | 621.76 |

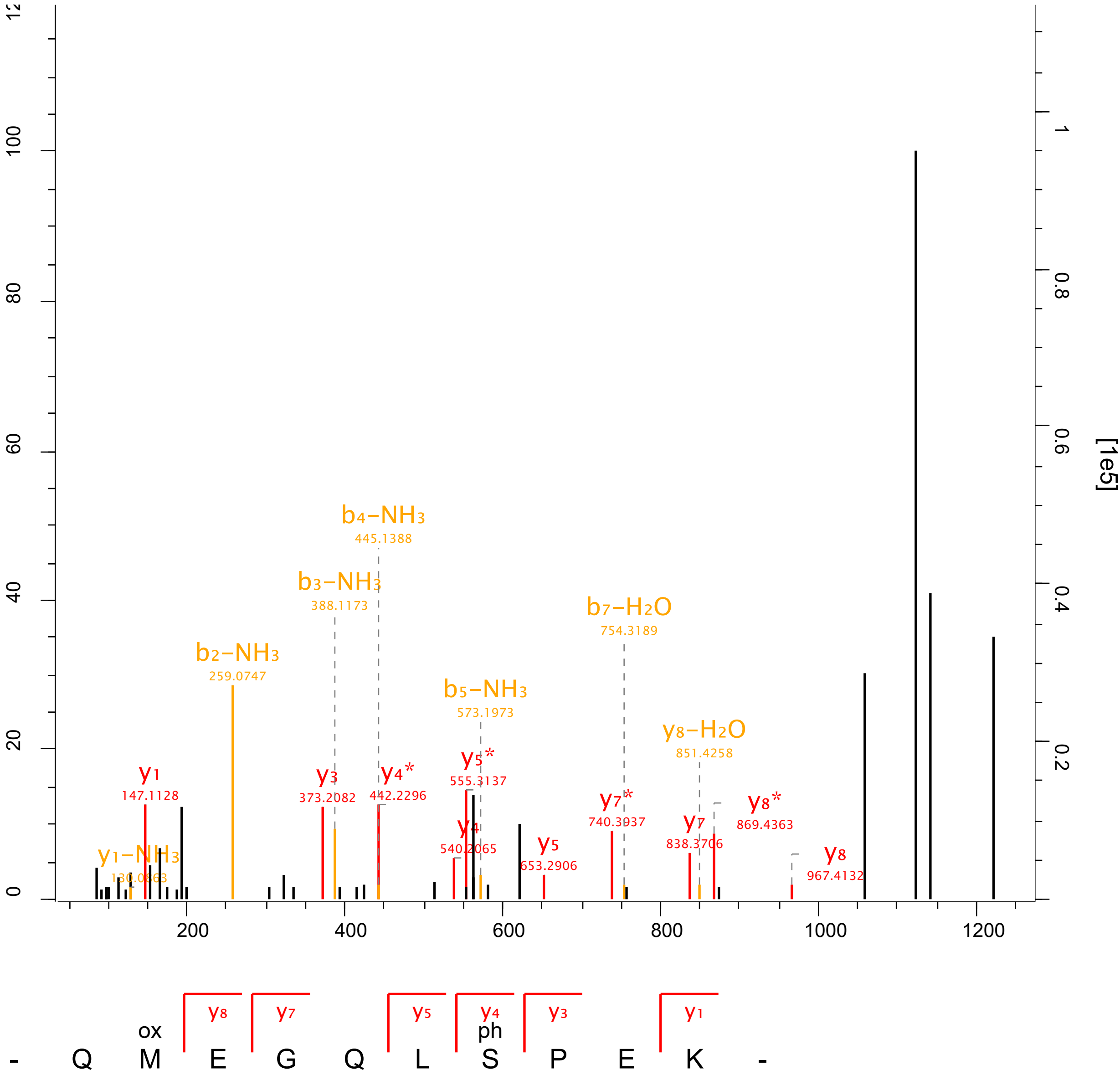

| Raw file | Scan | Method | Score | m/z |
|---|---|---|---|---|
| A17_15_1 | 4344 | FTMS; HCD | 78.51 | 594.22 |

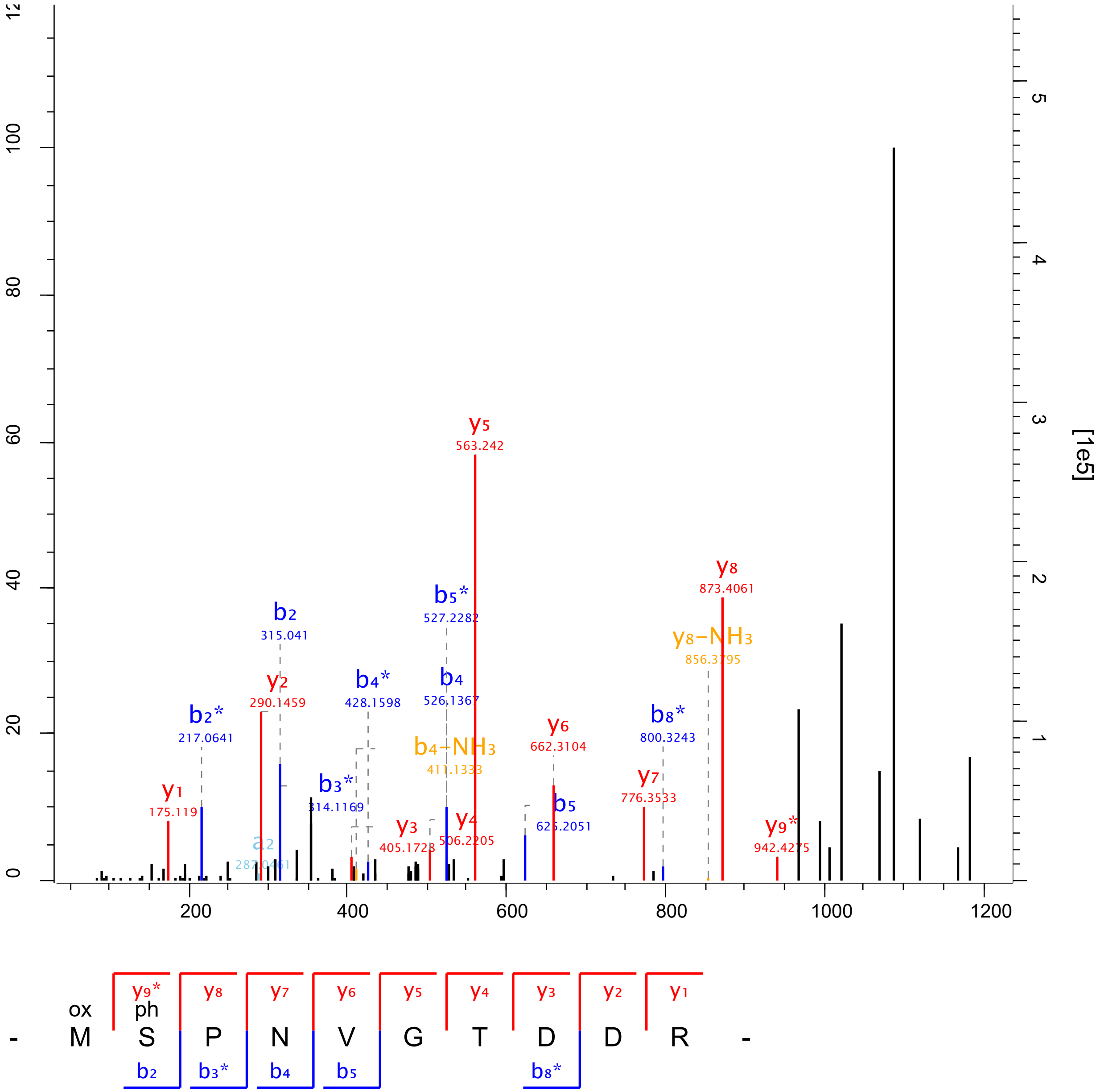

| Raw file | Scan | Method | Score | m/z |
|---|---|---|---|---|
| A17_15_1 | 44264 | FTMS; HCD | 58.98 | 838.32 |

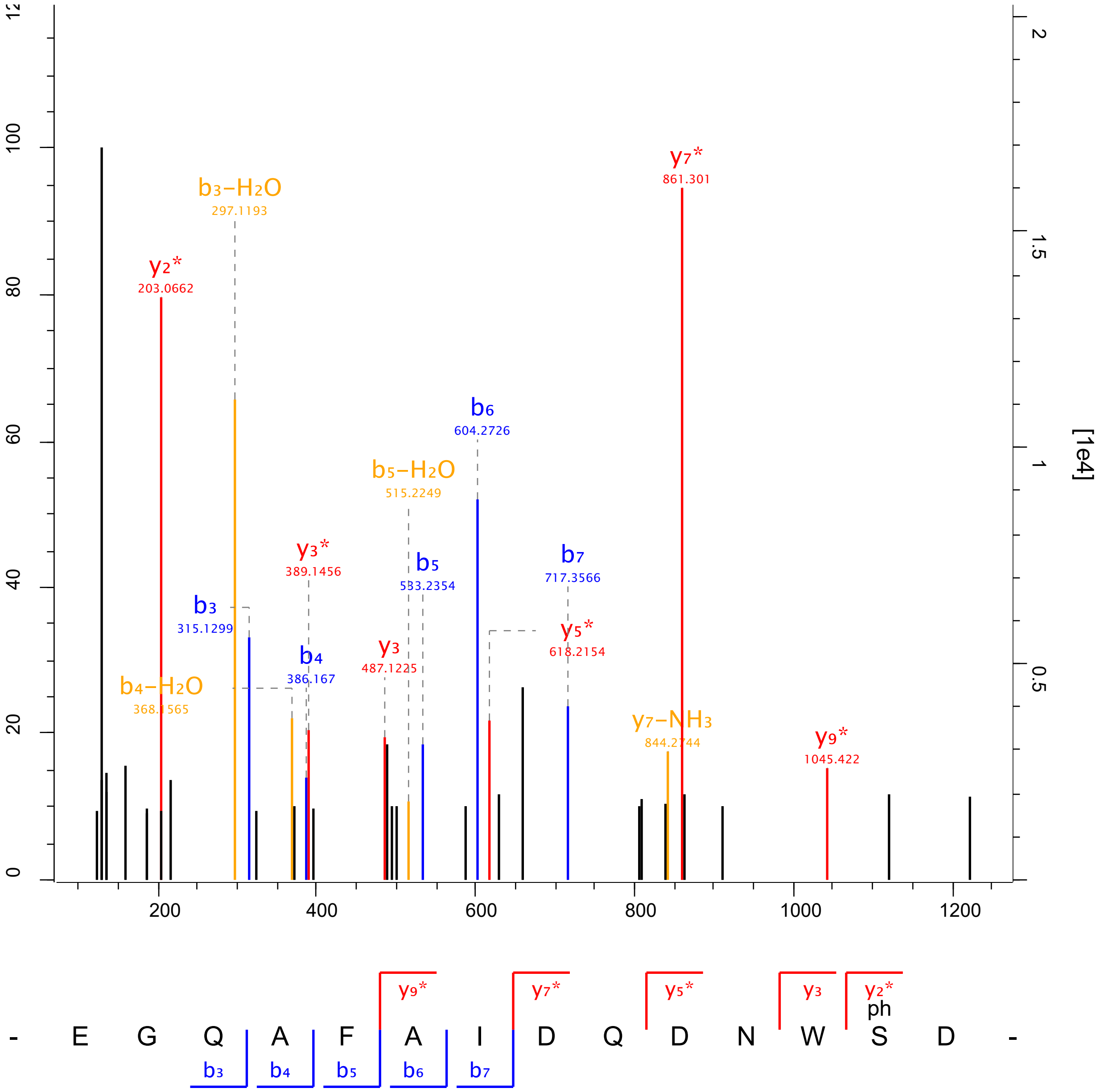

| Raw file | Scan | Method | Score | m/z |
| --- | --- | --- | --- | --- |
| A17_15_1 | 4544 | FTMS; HCD | 53.24 | 483.71 |

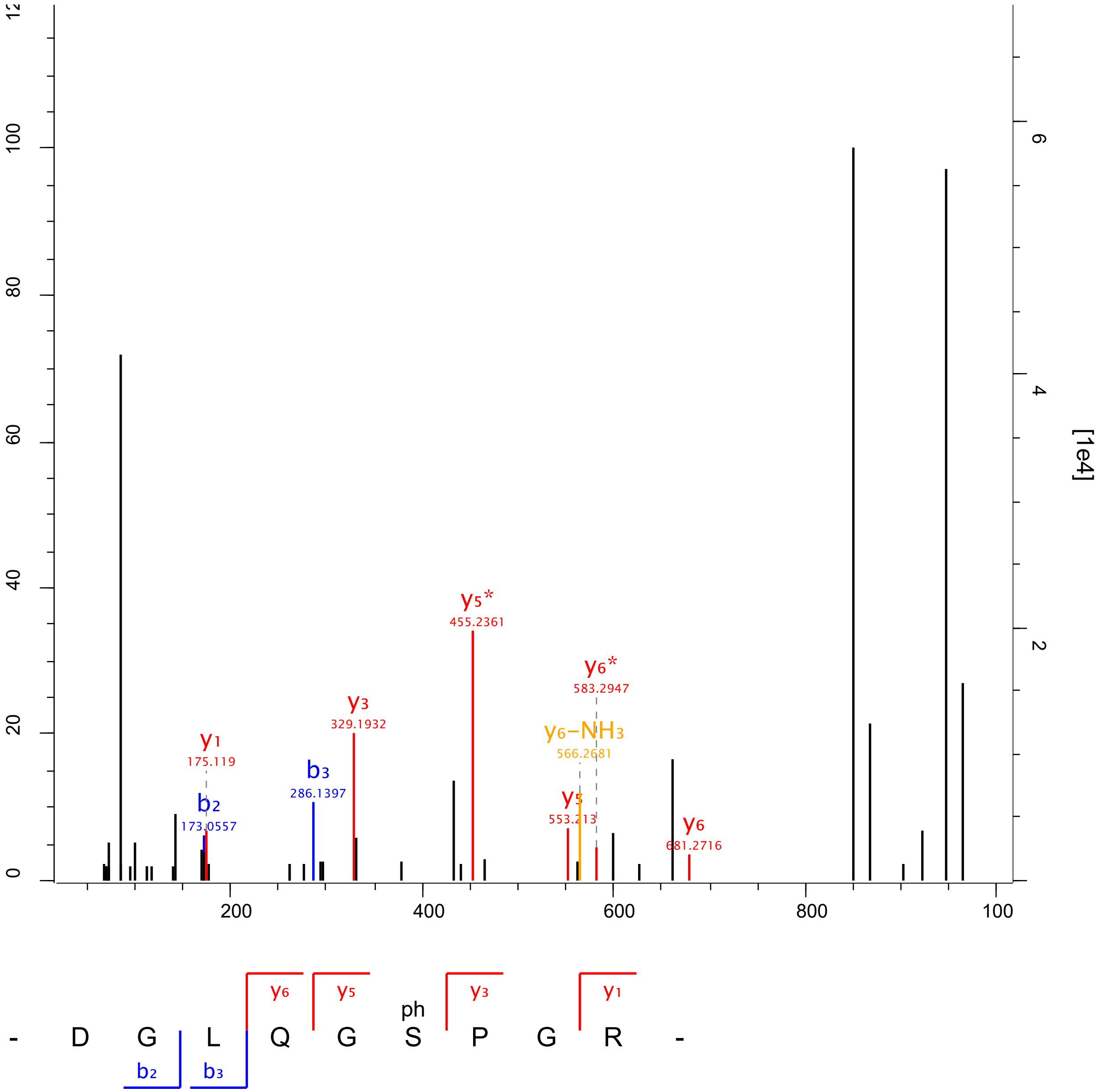

| Raw file | Scan | Method | Score | m/z |
|---|---|---|---|---|
| A17_15_1 | 45557 | FTMS; HCD | 79.84 | 974.32 |

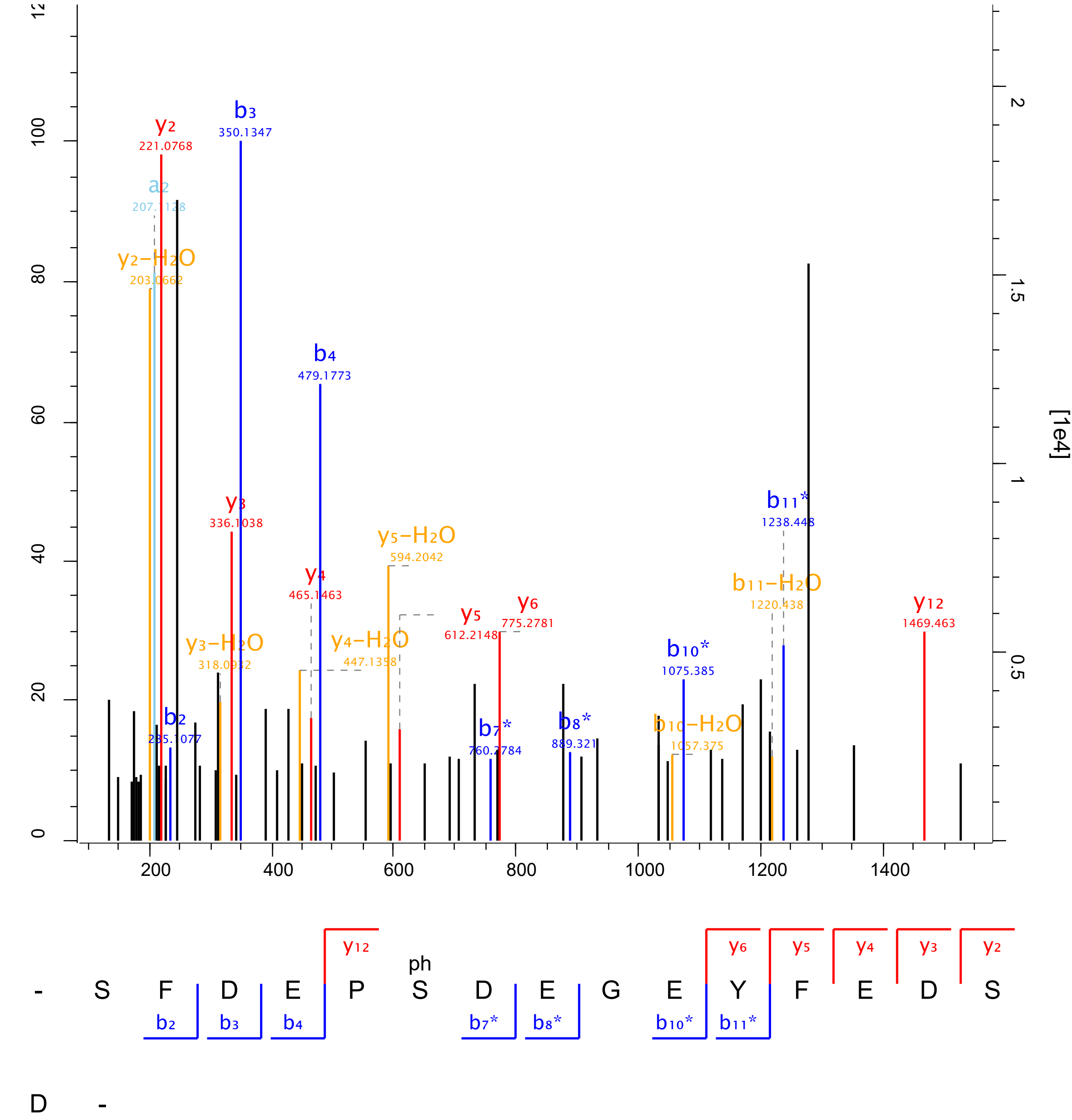

| Raw file | Scan | Method | Score | m/z |
|---|---|---|---|---|
| A17_15_1 | 45697 | FTMS; HCD | 94.92 | 728.26 |

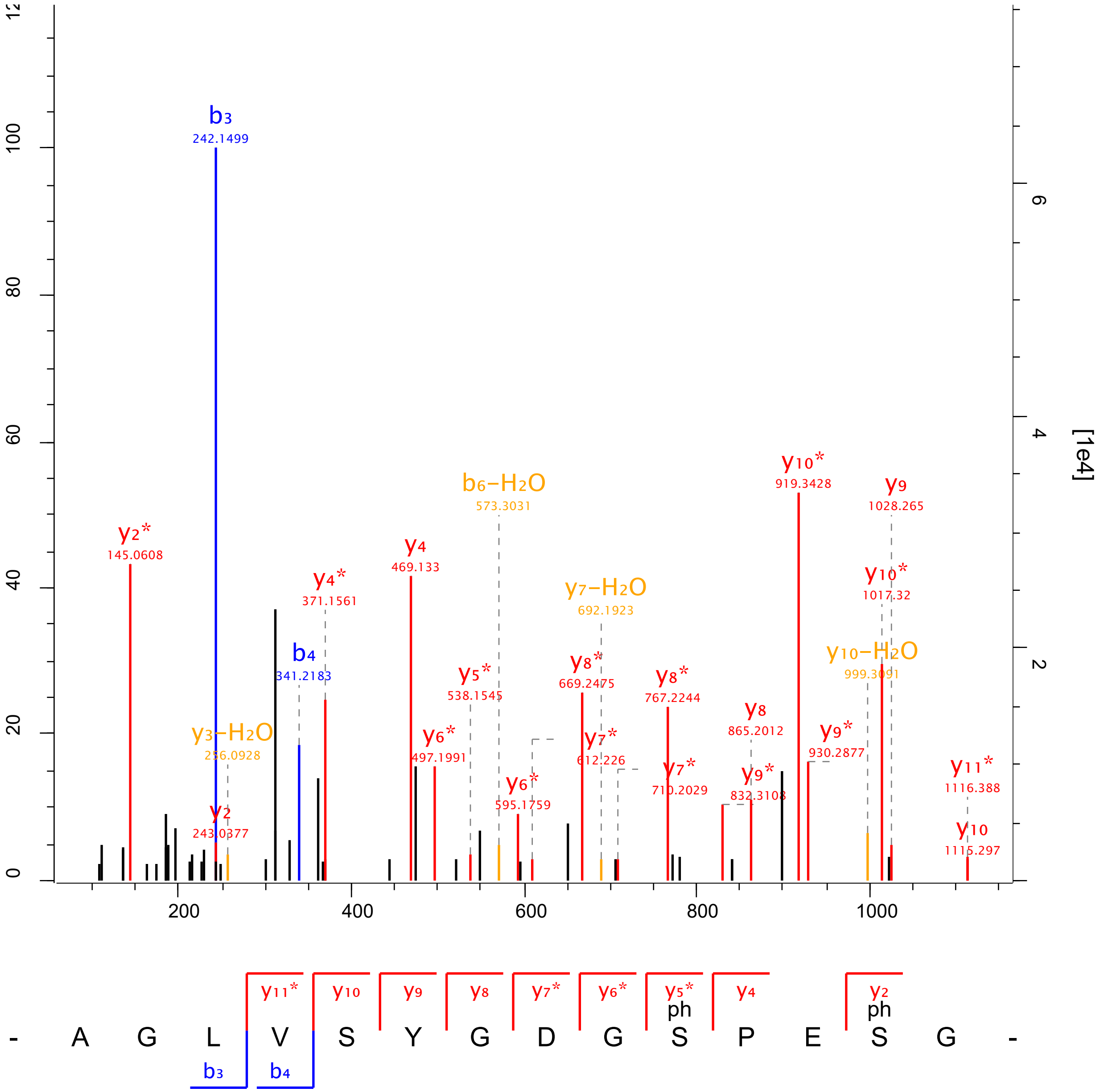

| Raw file | Scan | Method | Score | m/z |
|---|---|---|---|---|
| A17_15_1 | 4606 | FTMS; HCD | 62.34 | 499.23 |

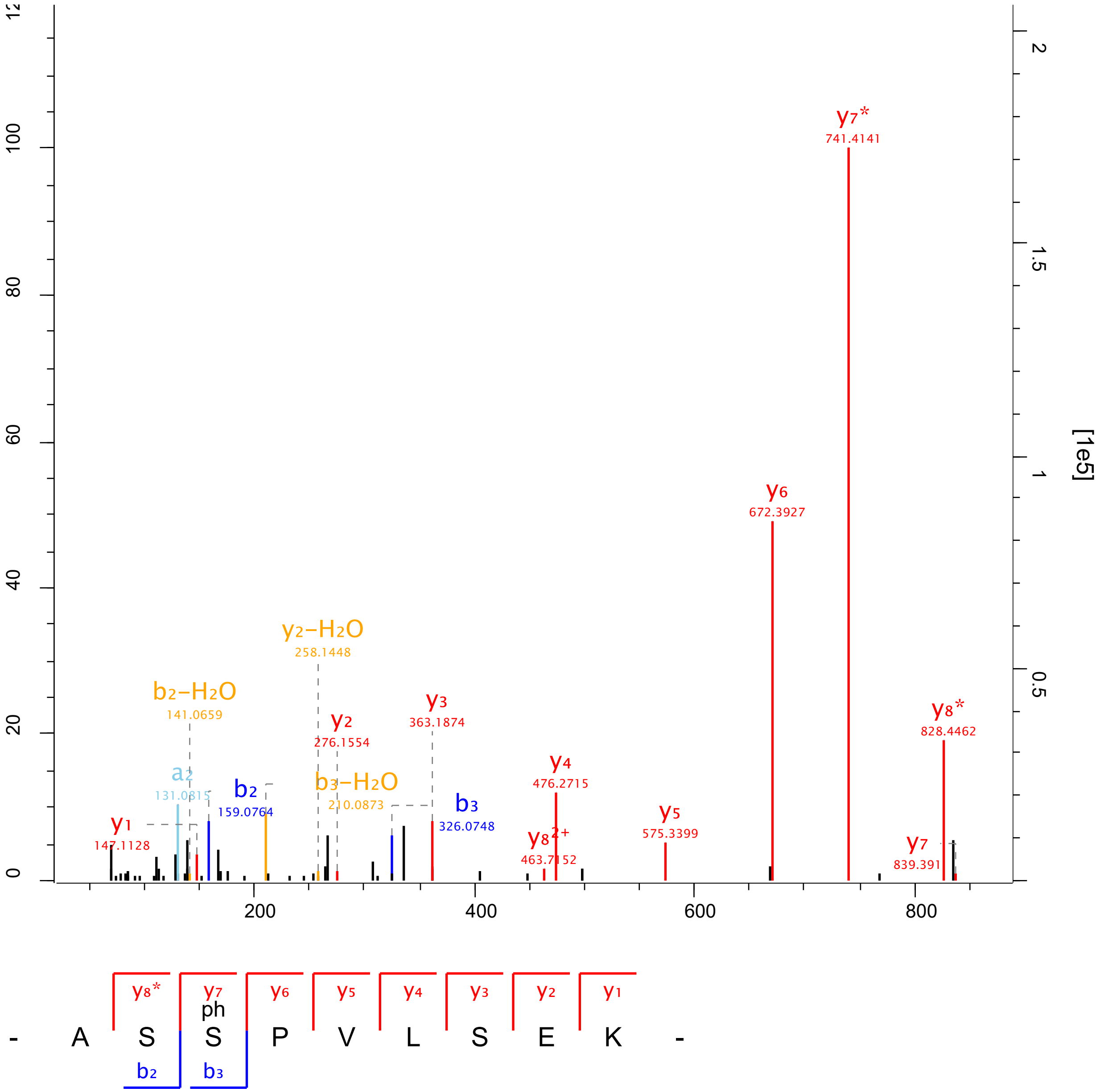

| Raw file | Scan | Method | Score | m/z |
|---|---|---|---|---|
| A17_15_1 | 4905 | FTMS; HCD | 45 | 488.52 |

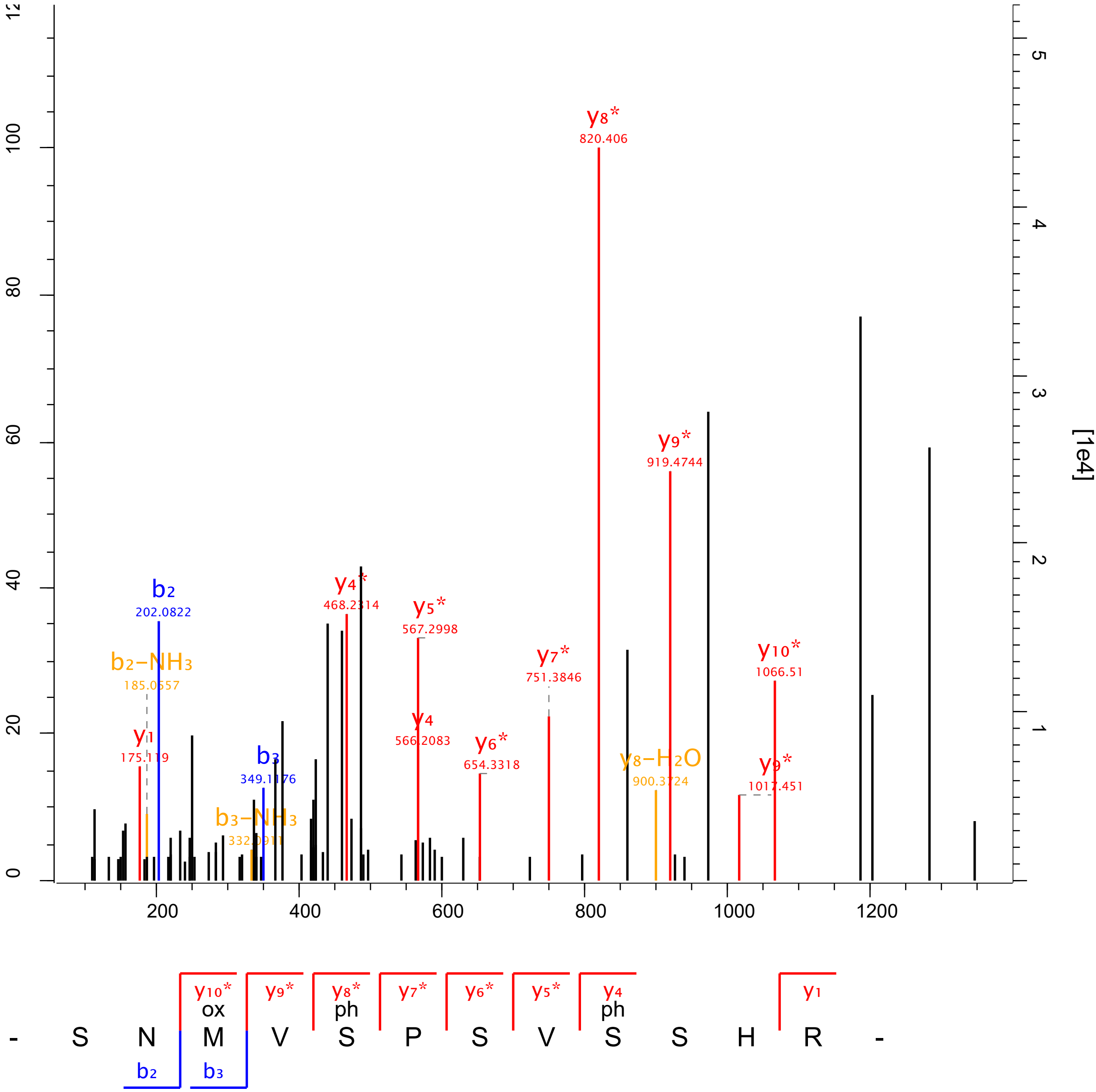

| Raw file | Scan | Method | Score | m/z |
|---|---|---|---|---|
| A17_15_1 | 4933 | FTMS; HCD | 121.18 | 550.74 |

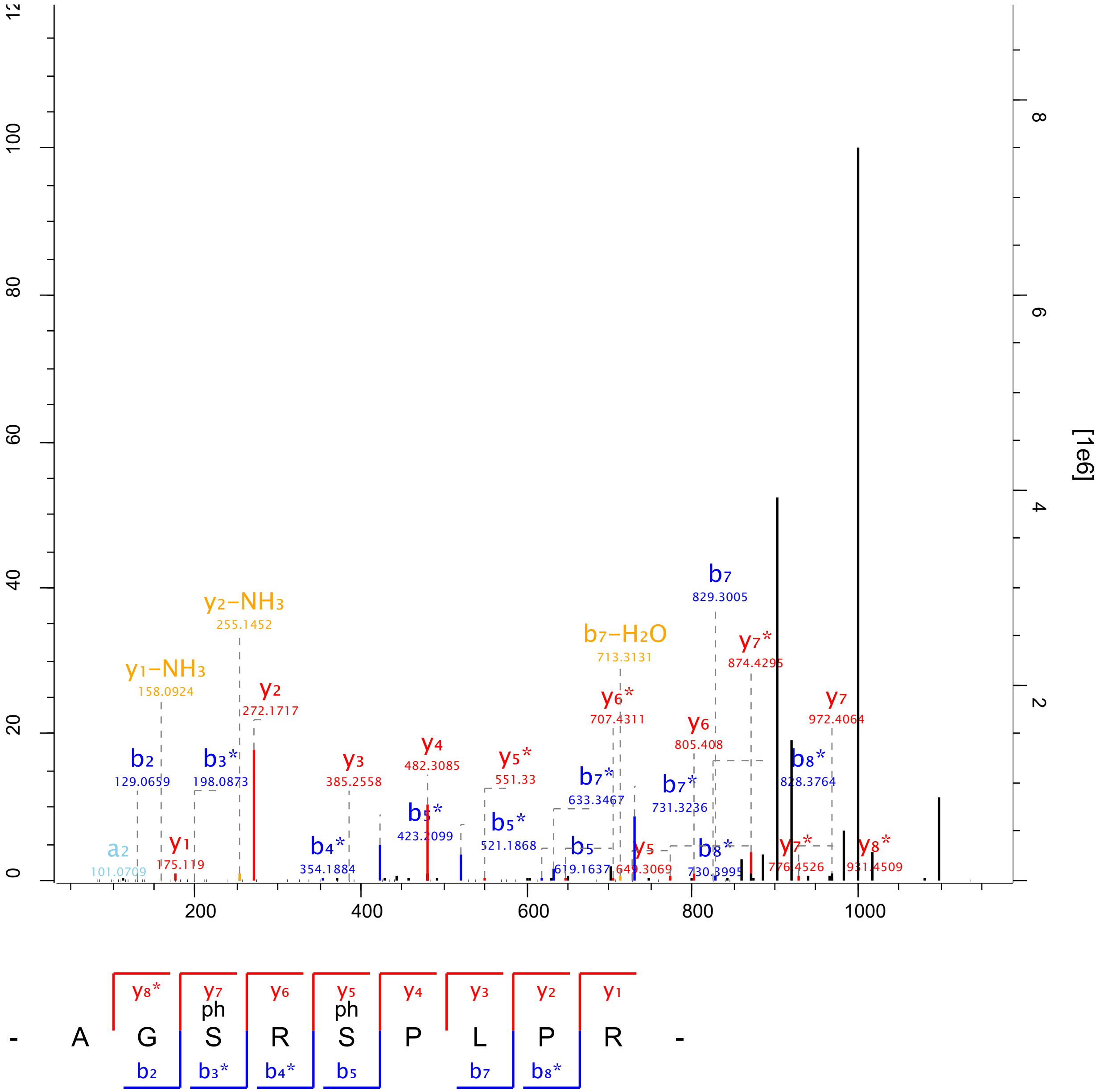

| Raw file | Scan | Method | Score | m/z |
|---|---|---|---|---|
| A17_15_1 | 5003 | FTMS; HCD | 49.3 | 481.21 |

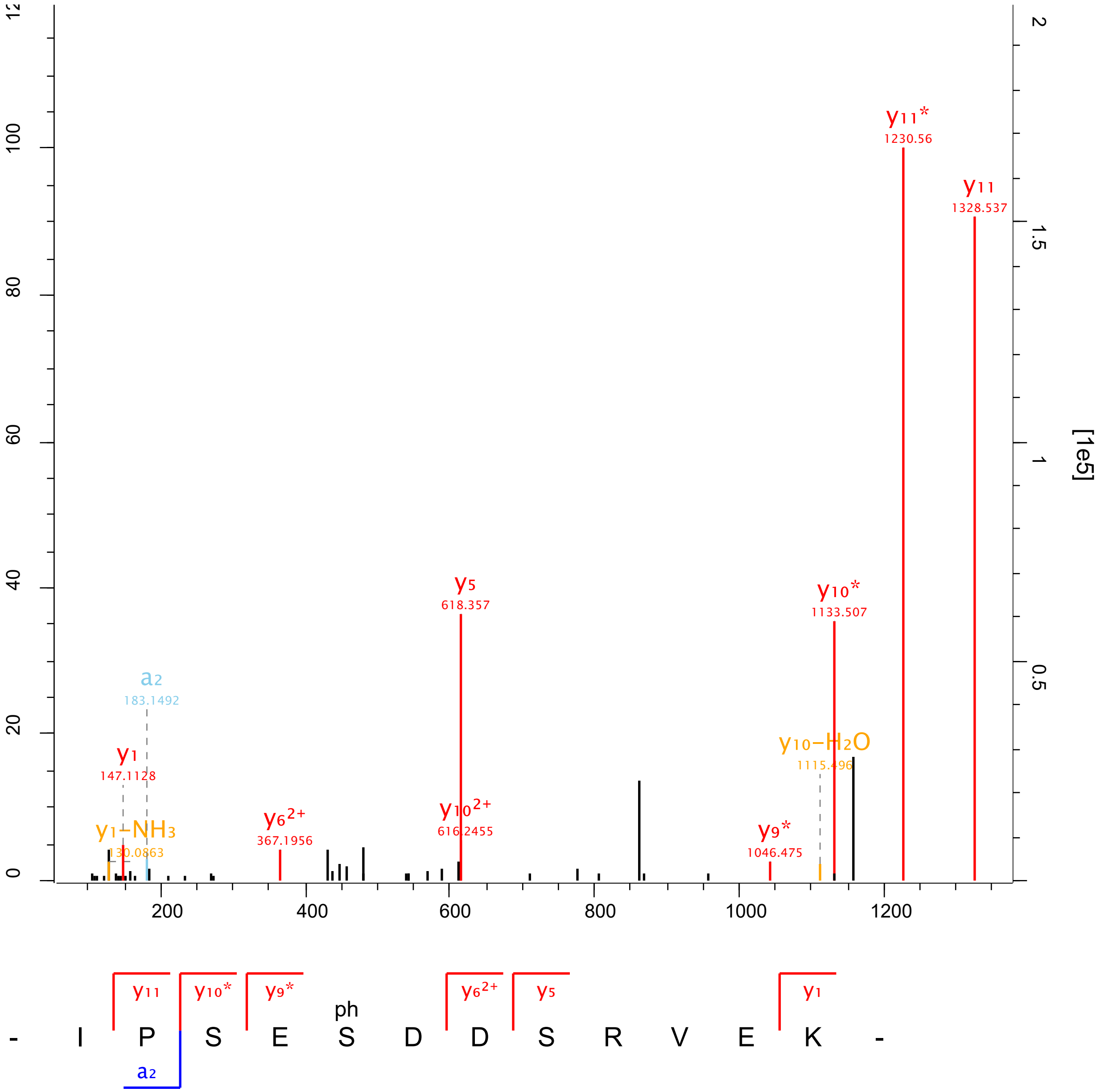

| Raw file | Scan | Method | Score | m/z |
| --- | --- | --- | --- | --- |
| A17_15_1 | 5090 | FTMS; HCD | 89.8 | 484.71 |

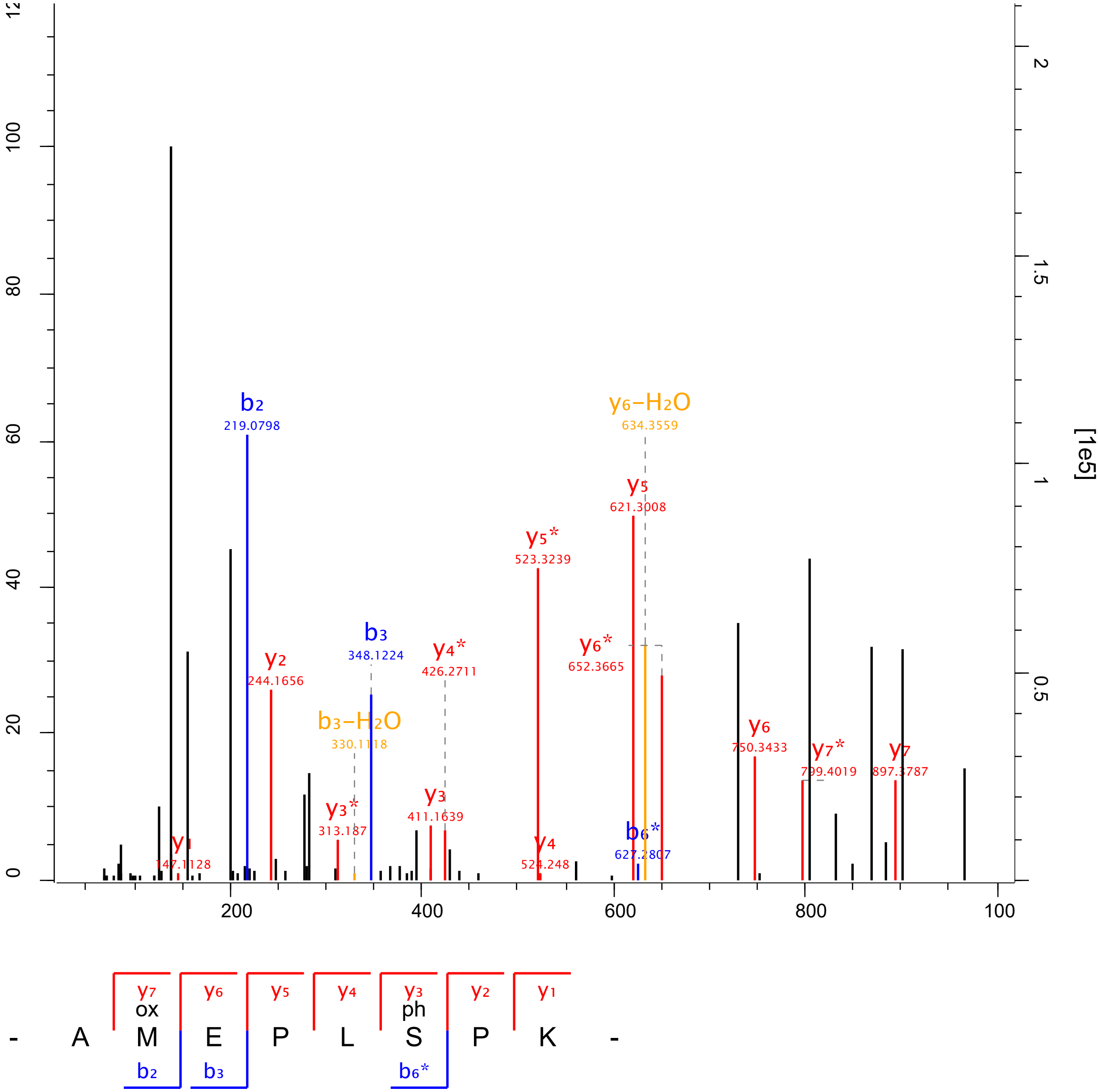

| Raw file | Scan | Method | Score | m/z |
|---|---|---|---|---|
| A17_15_1 | 5418 | FTMS; HCD | 67.14 | 632.25 |

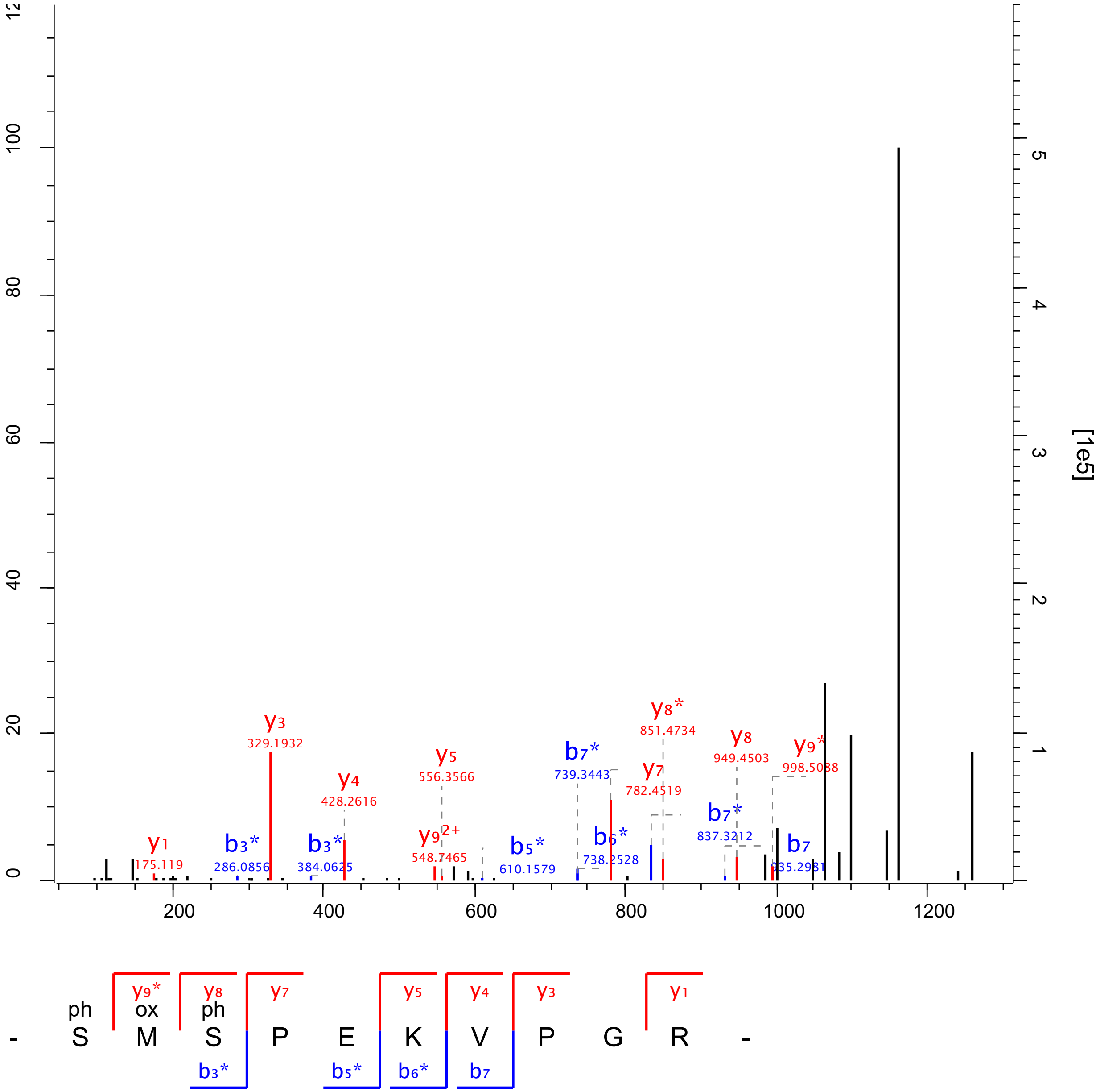

| Raw file | Scan | Method | Score | m/z |
| --- | --- | --- | --- | --- |
| A17_15_1 | 5438 | FTMS; HCD | 42.64 | 941.34 |

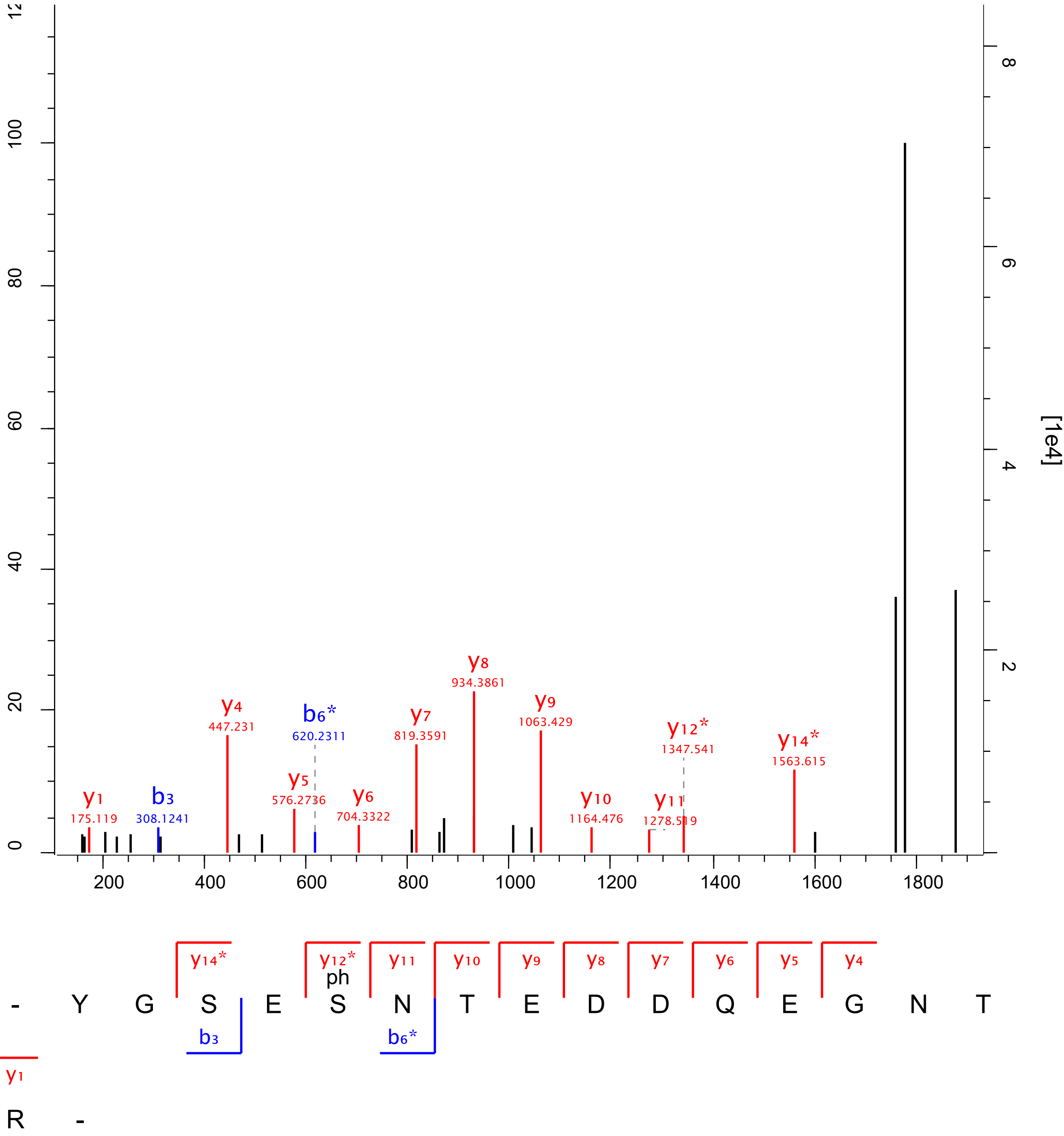

| Raw file | Scan | Method | Score | m/z |
|---|---|---|---|---|
| A17_15_1 | 5634 | FTMS; HCD | 92.11 | 522.72 |

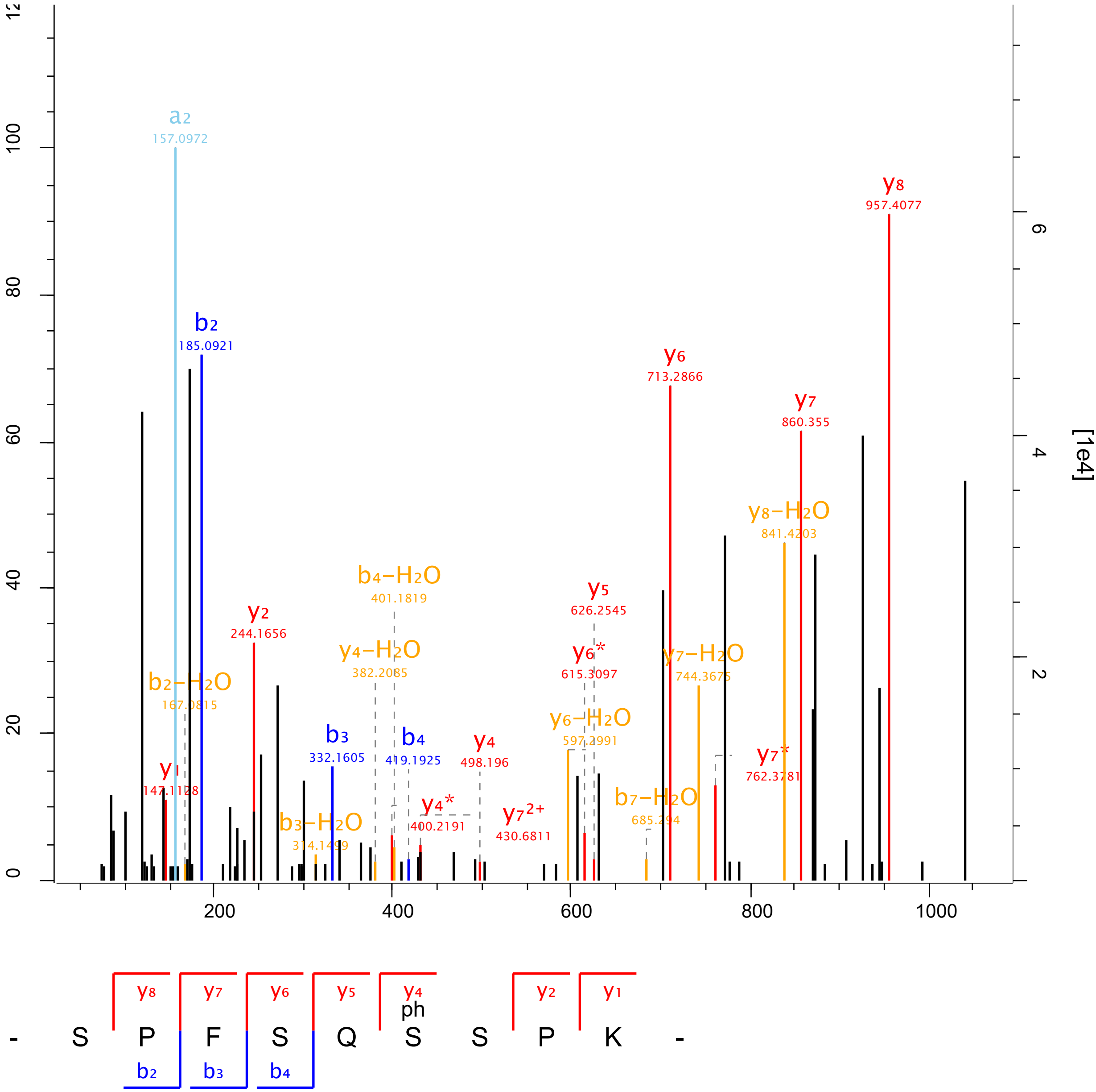

| Raw file | Scan | Method | Score | m/z |
| --- | --- | --- | --- | --- |
| A17_15_1 | 6016 | FTMS; HCD | 89.8 | 477.16 |

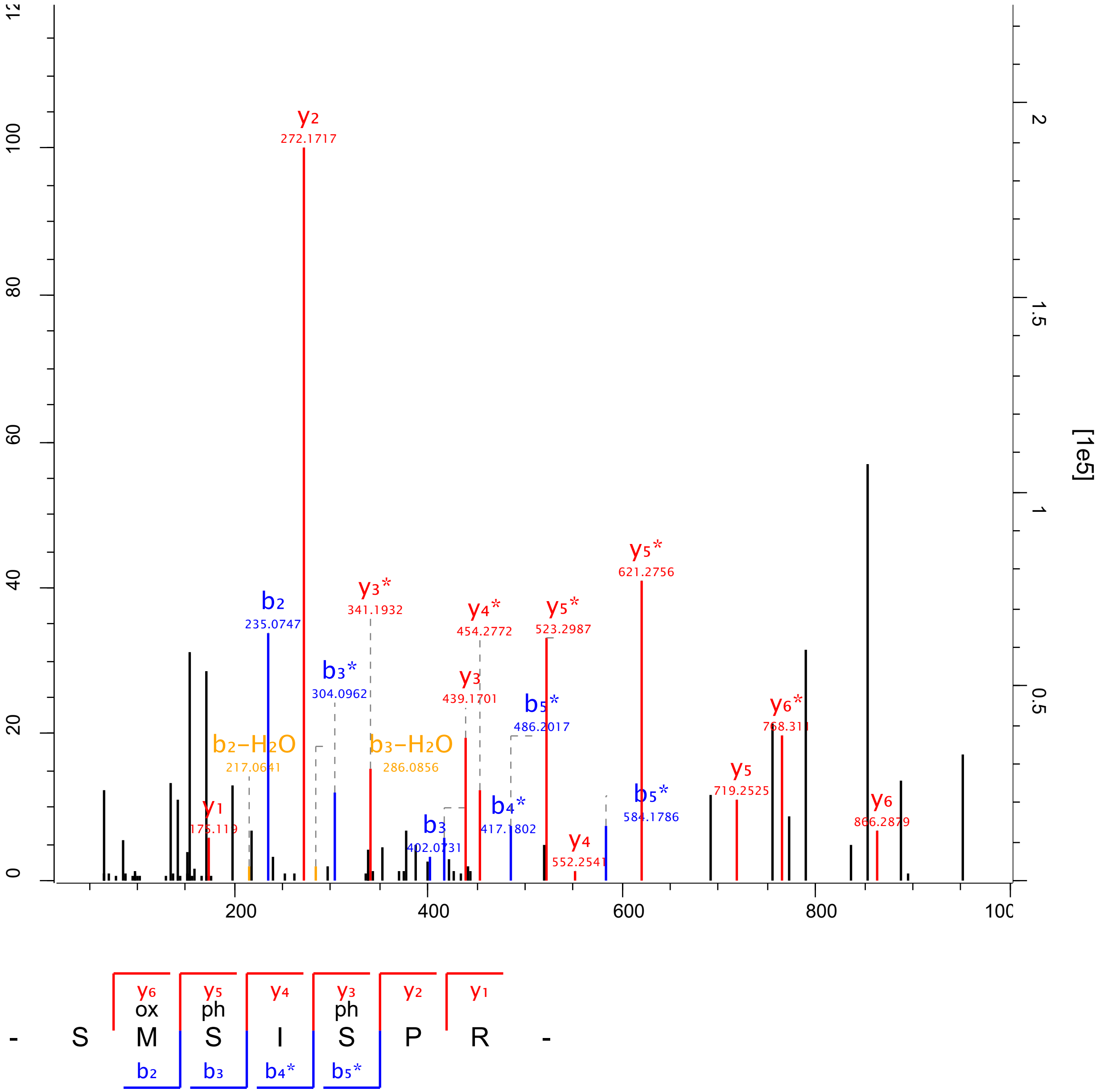

| Raw file | Scan | Method | Score | m/z |
|---|---|---|---|---|
| A17_15_1 | 6325 | FTMS; HCD | 53.24 | 540.22 |

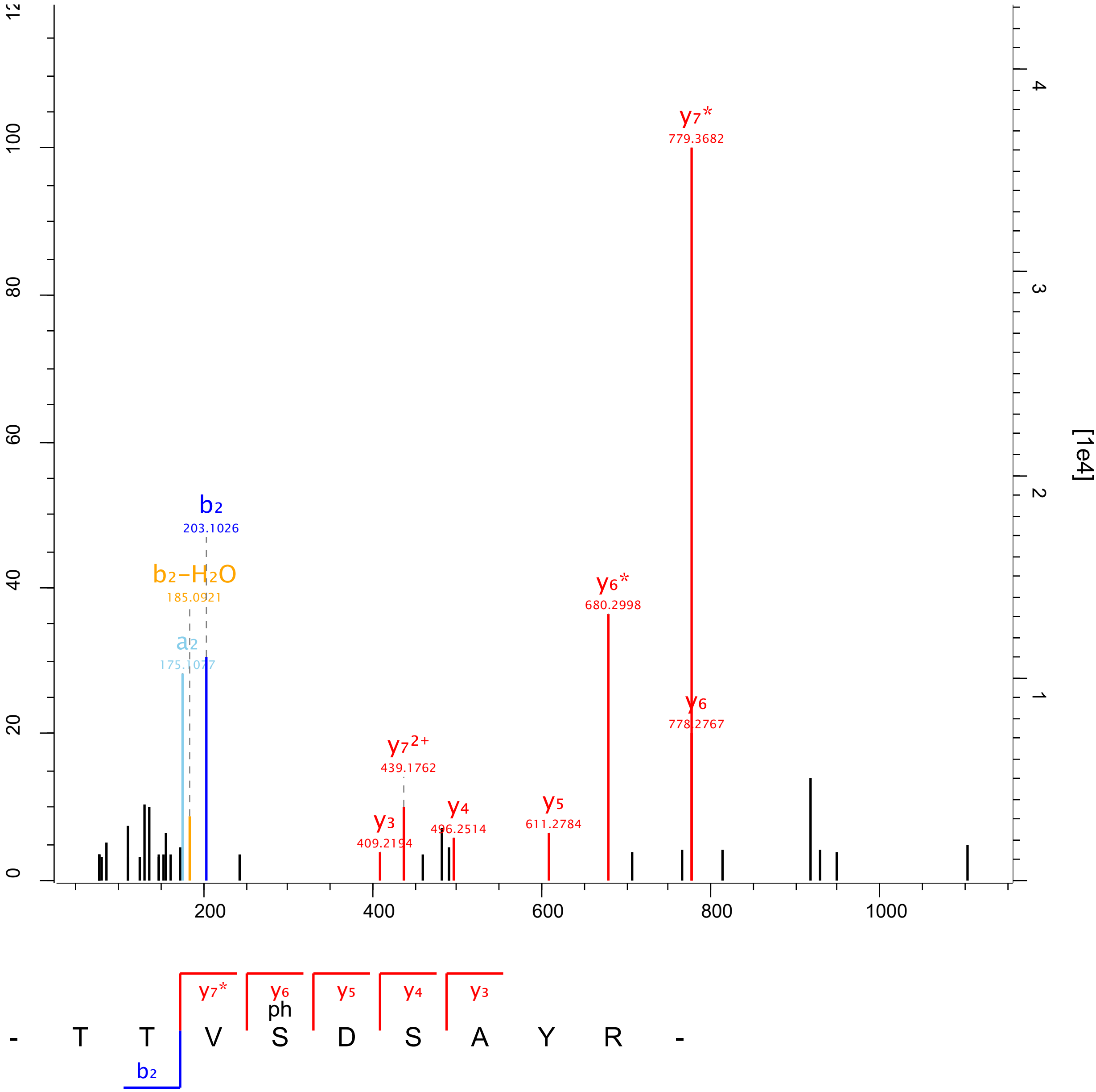

| Raw file | Scan | Method | Score | m/z |
| --- | --- | --- | --- | --- |
| A17_15_1 | 6563 | FTMS; HCD | 104.66 | 583.59 |

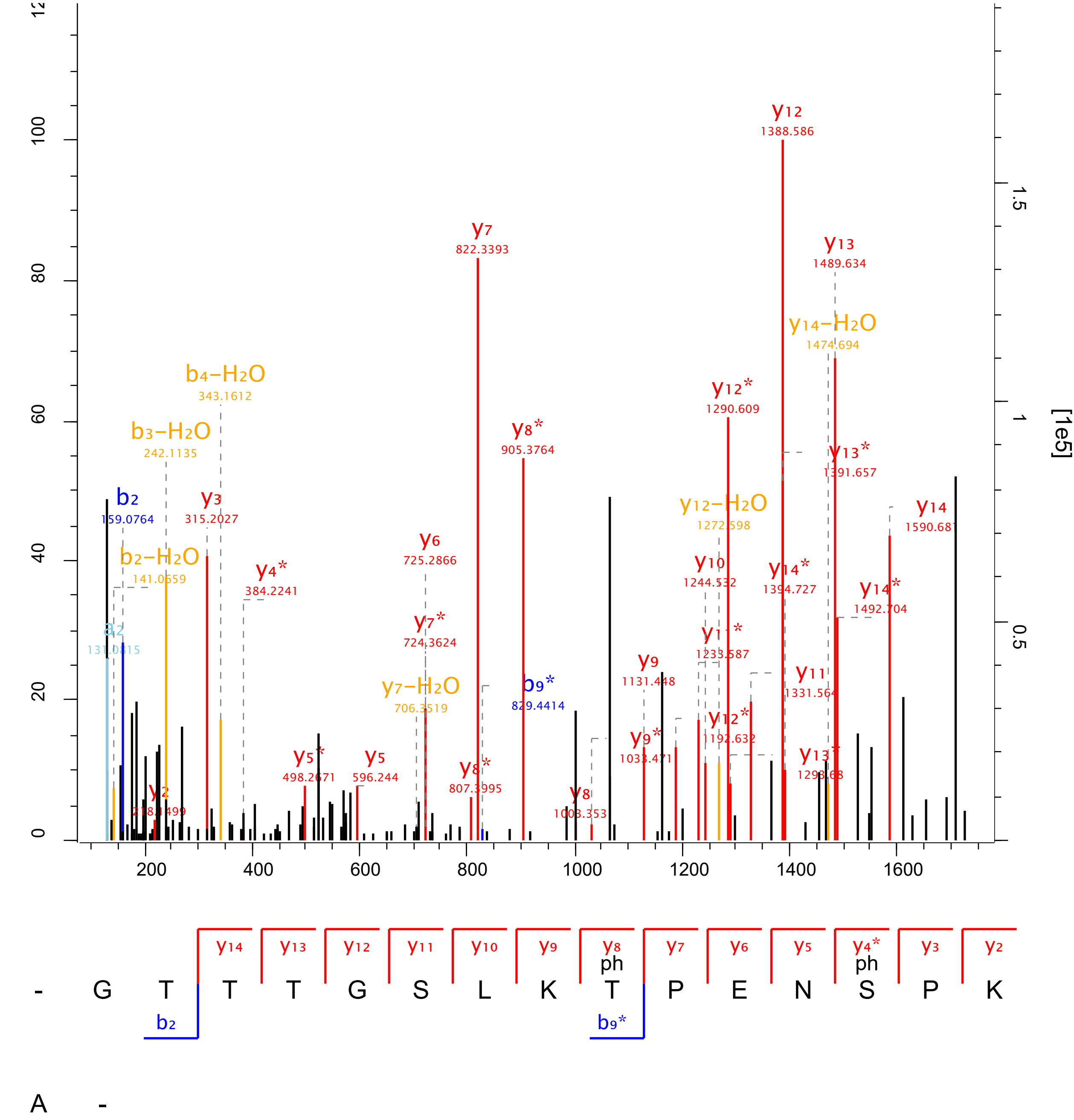

| Raw file | Scan | Method | Score | m/z |
|---|---|---|---|---|
| A17_15_1 | 6631 | FTMS; HCD | 161.75 | 705.78 |

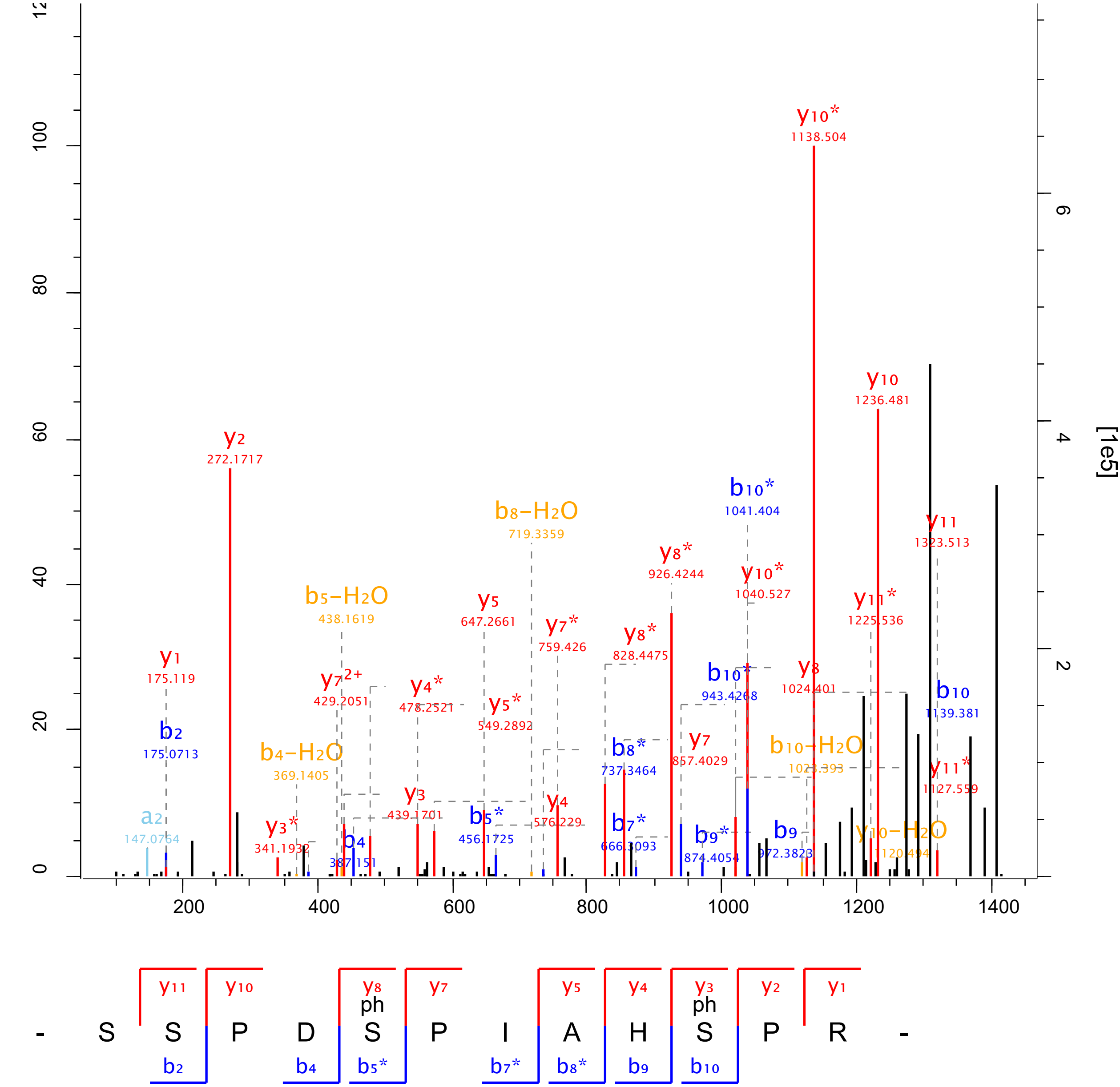

| Raw file | Scan | Method | Score | m/z |
|---|---|---|---|---|
| A17_15_1 | 6773 | FTMS; HCD | 139.86 | 716.26 |

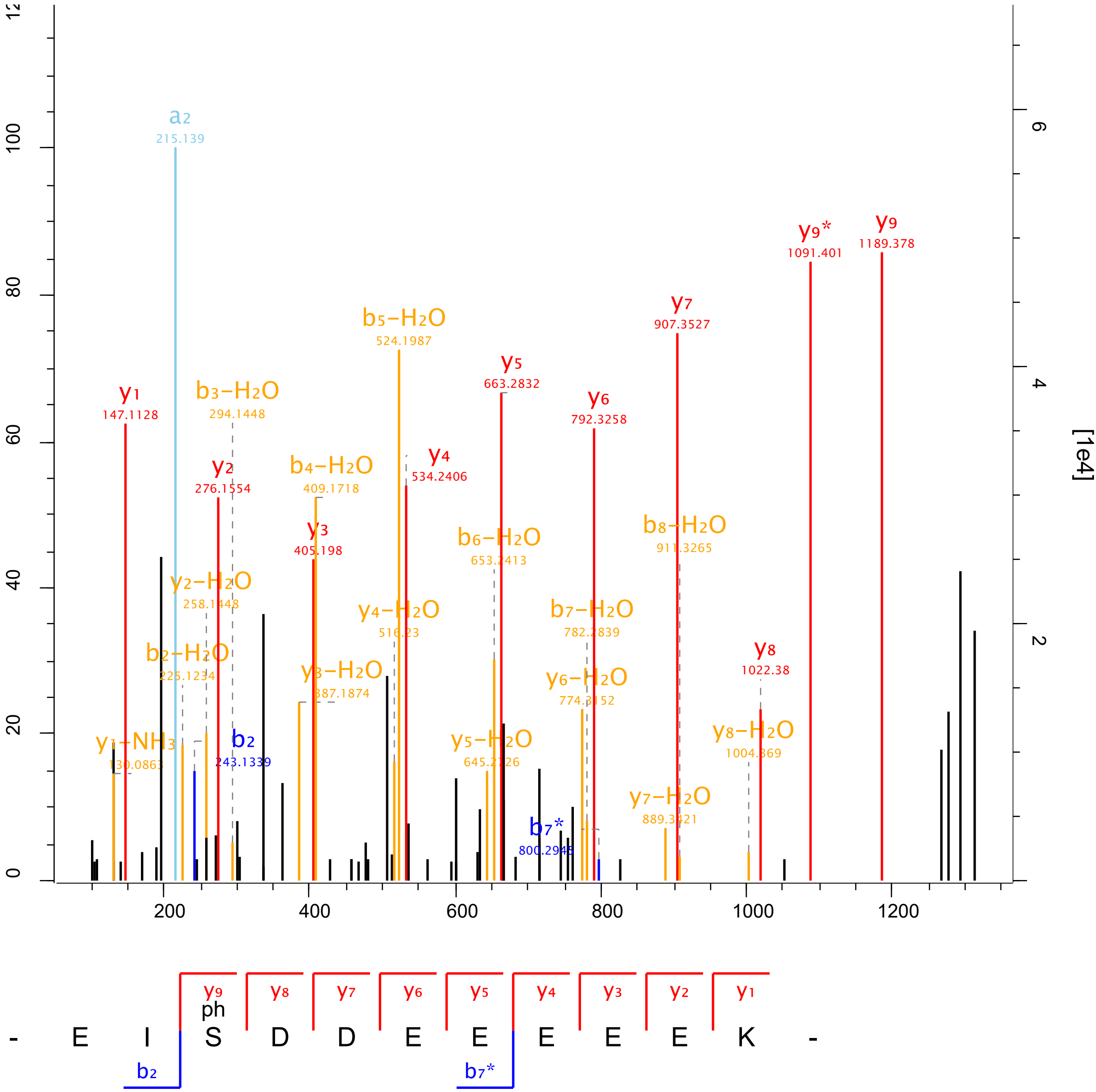

| Raw file | Scan | Method | Score | m/z |
|---|---|---|---|---|
| A17_15_1 | 6827 | FTMS; HCD | 138.22 | 493.22 |

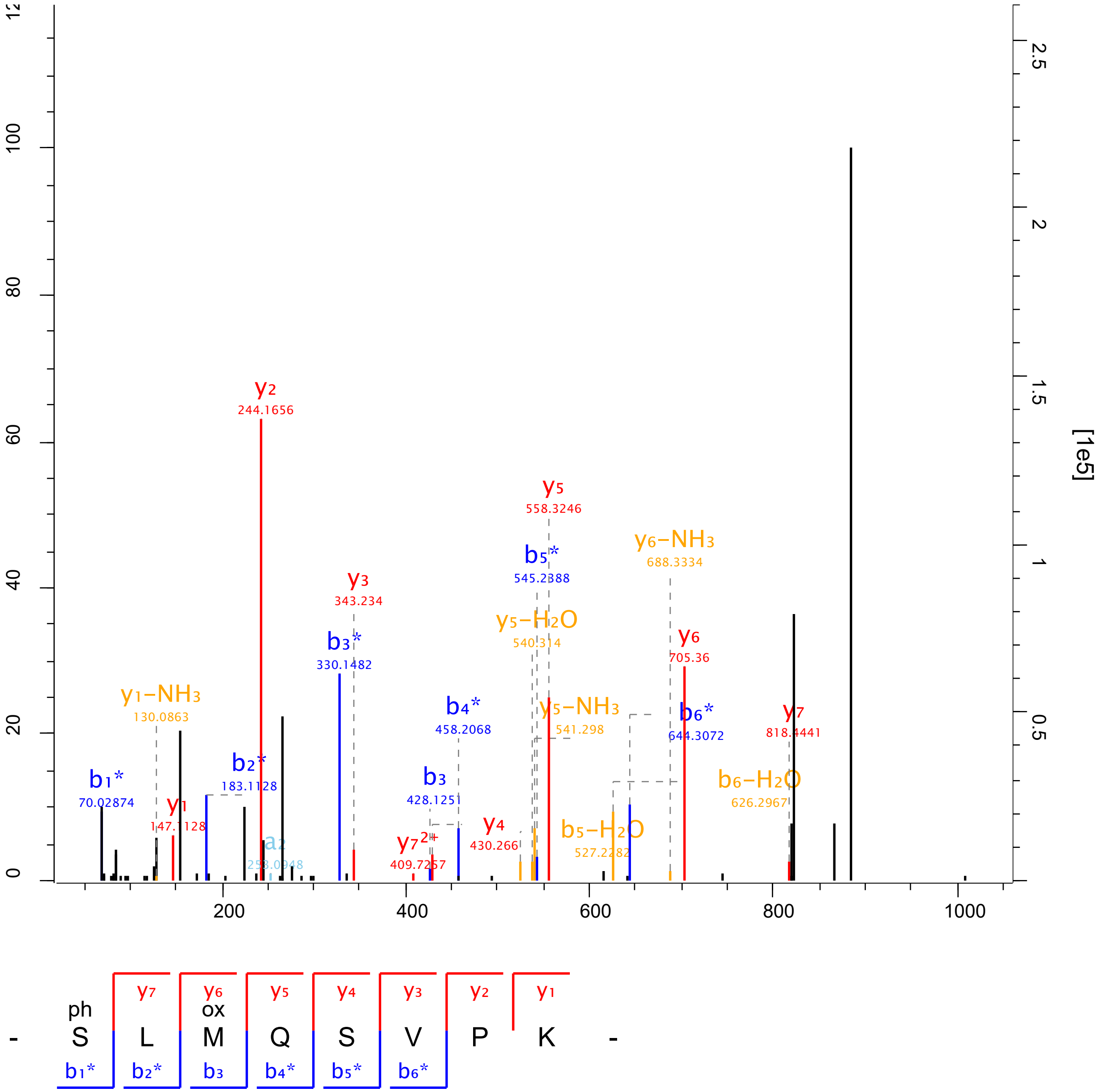

| Raw file | Scan | Method | Score | m/z |
| --- | --- | --- | --- | --- |
| A17_15_1 | 7045 | FTMS; HCD | 123.1 | 576.73 |

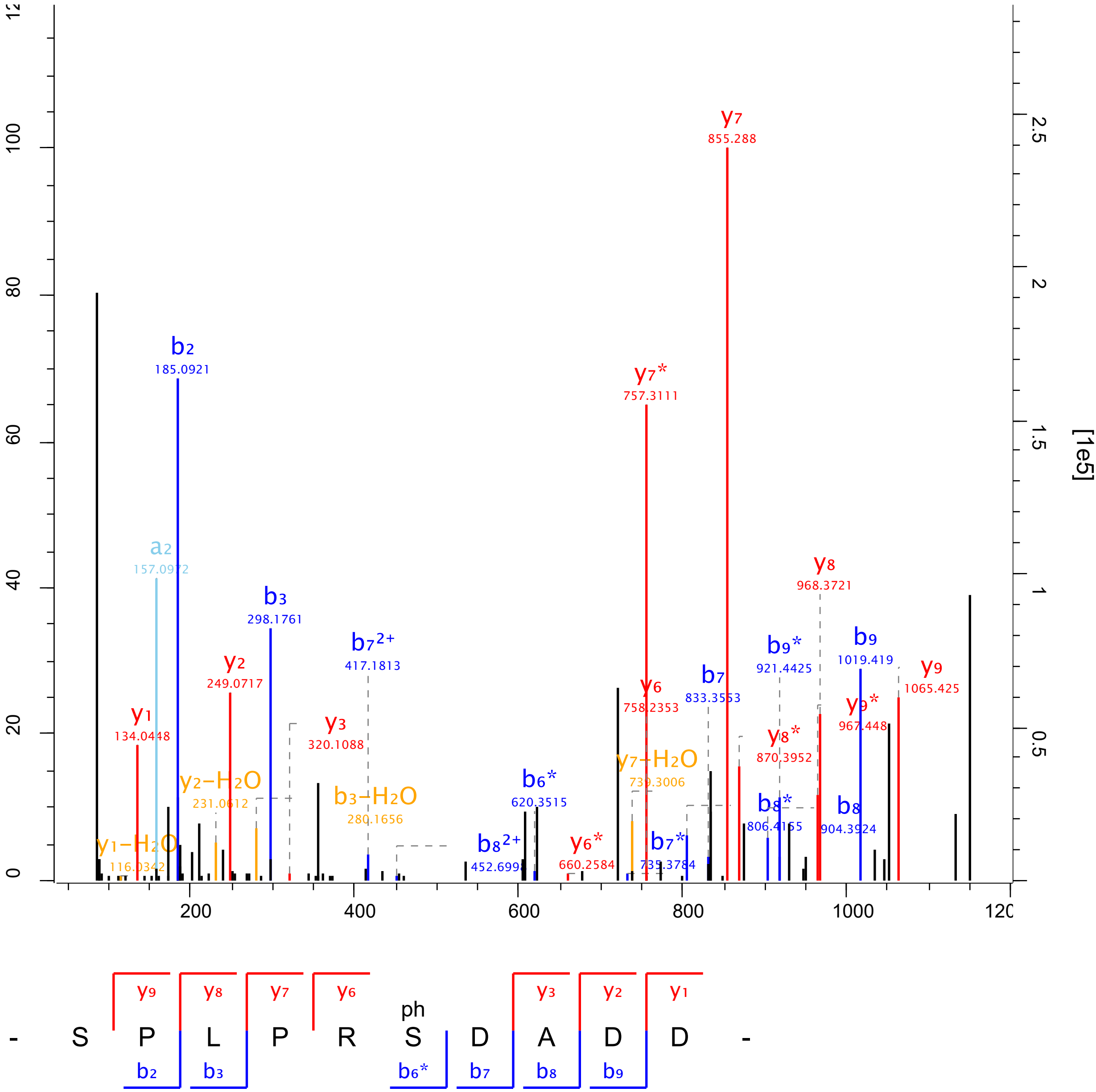

| Raw file | Scan | Method | Score | m/z |
|---|---|---|---|---|
| A17_15_1 | 7099 | FTMS; HCD | 57.05 | 535.21 |

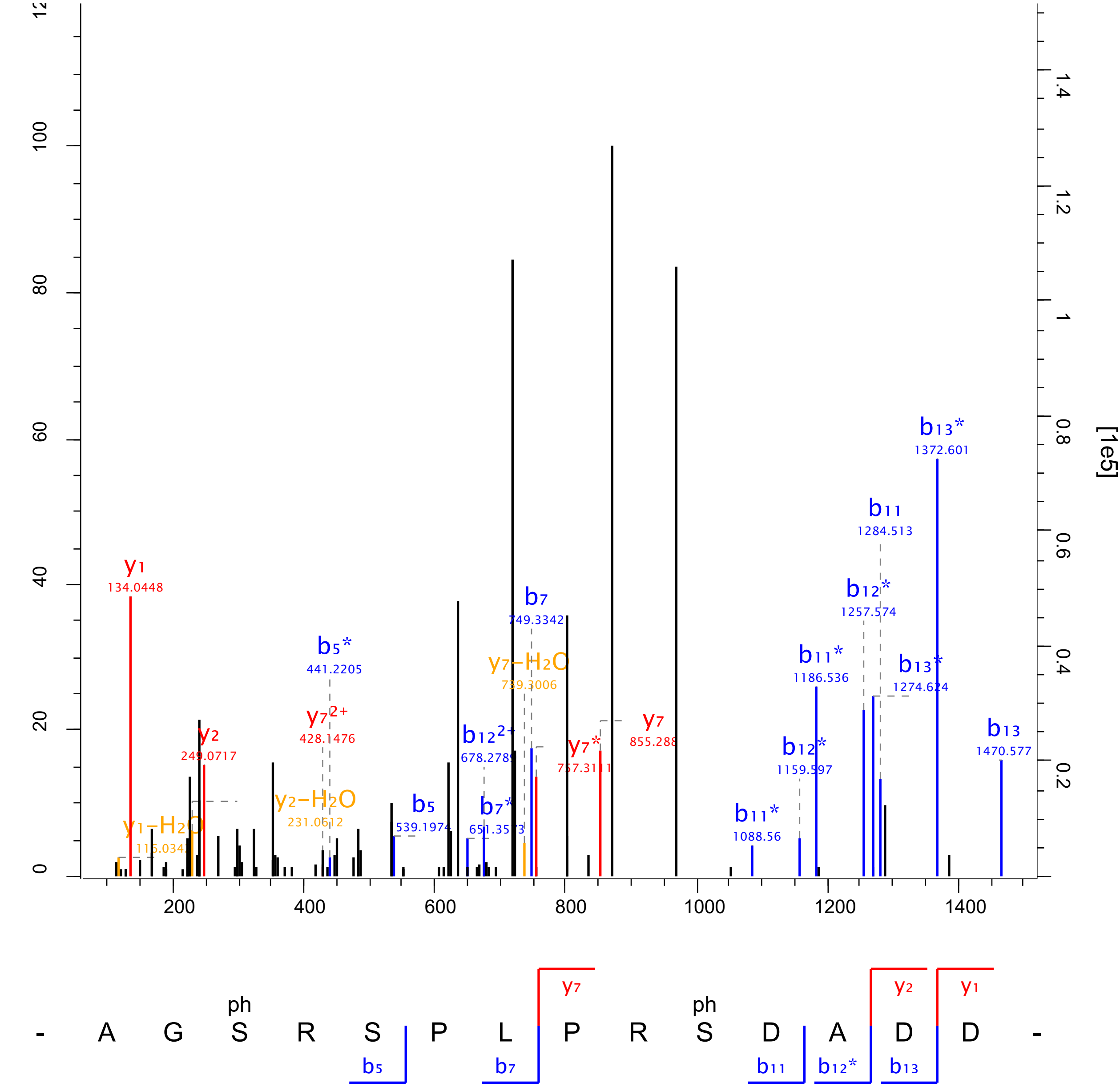

| Raw file | Scan | Method | Score | m/z |
| --- | --- | --- | --- | --- |
| A17_15_1 | 7180 | FTMS; HCD | 102.52 | 612.31 |

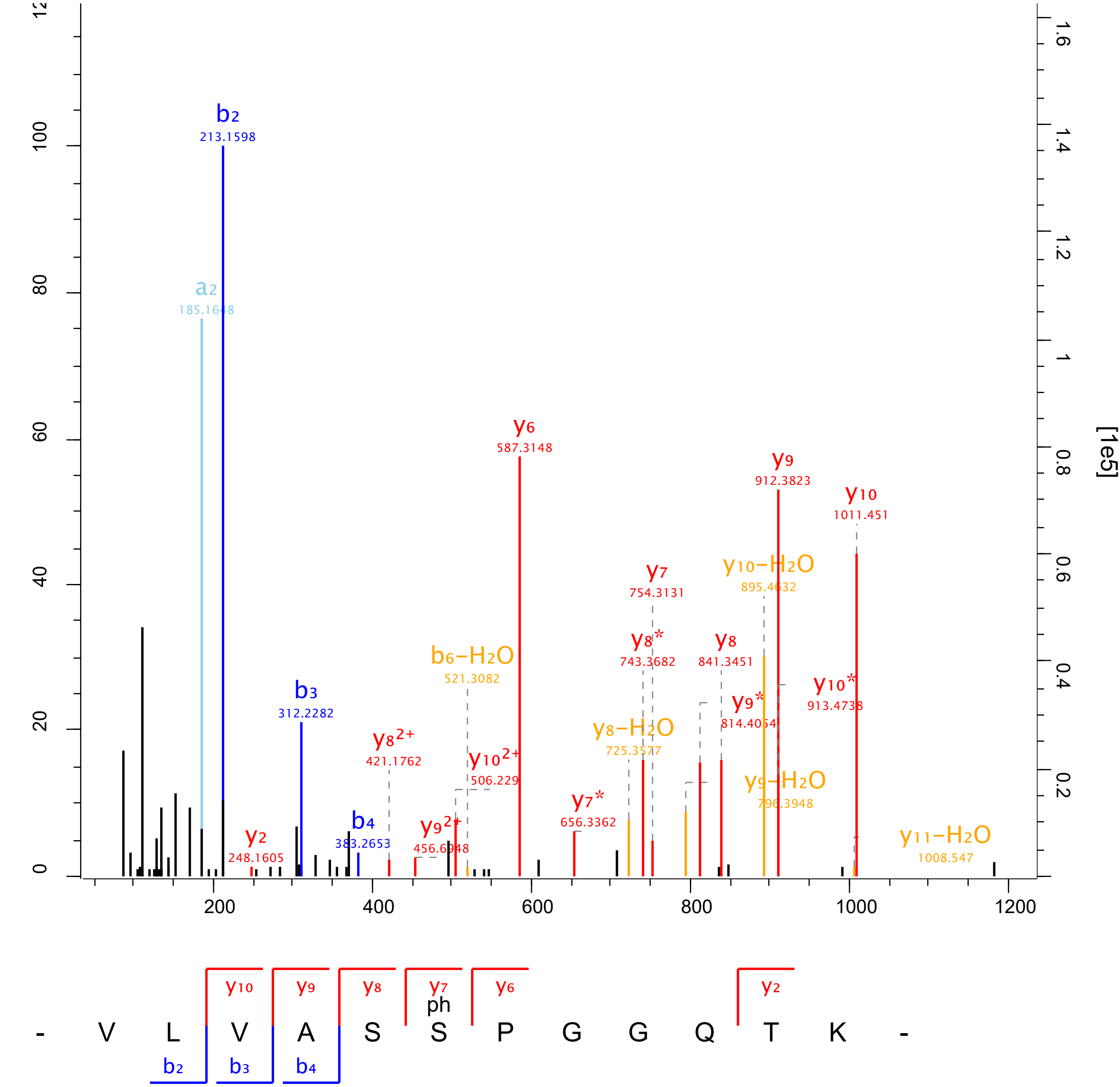

| Raw file | Scan | Method | Score | m/z |
|---|---|---|---|---|
| A17_15_1 | 7225 | FTMS; HCD | 64 | 761.33 |

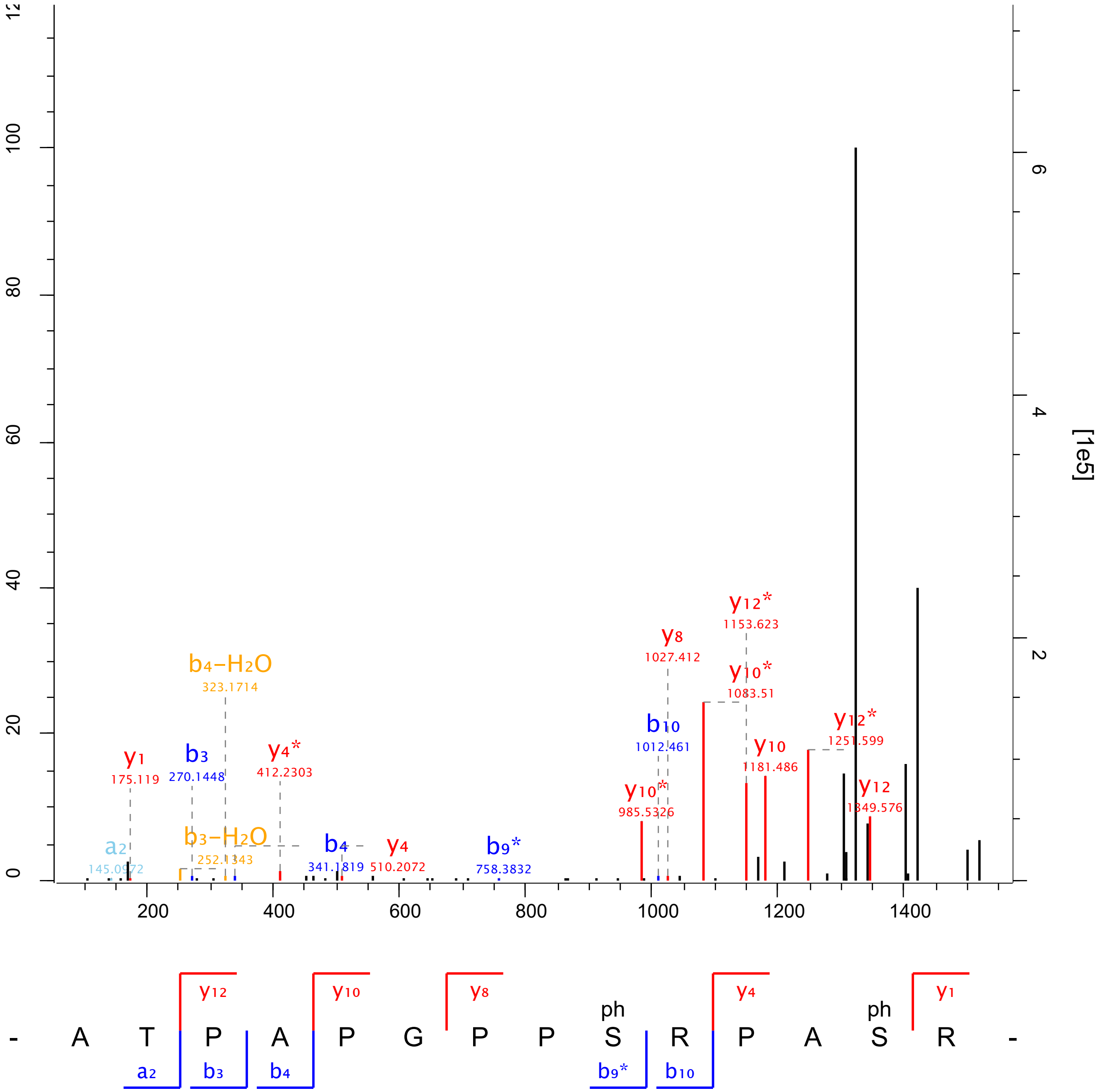

| Raw file | Scan | Method | Score | m/z |
|---|---|---|---|---|
| A17_15_1 | 7285 | FTMS; HCD | 46.32 | 753.83 |

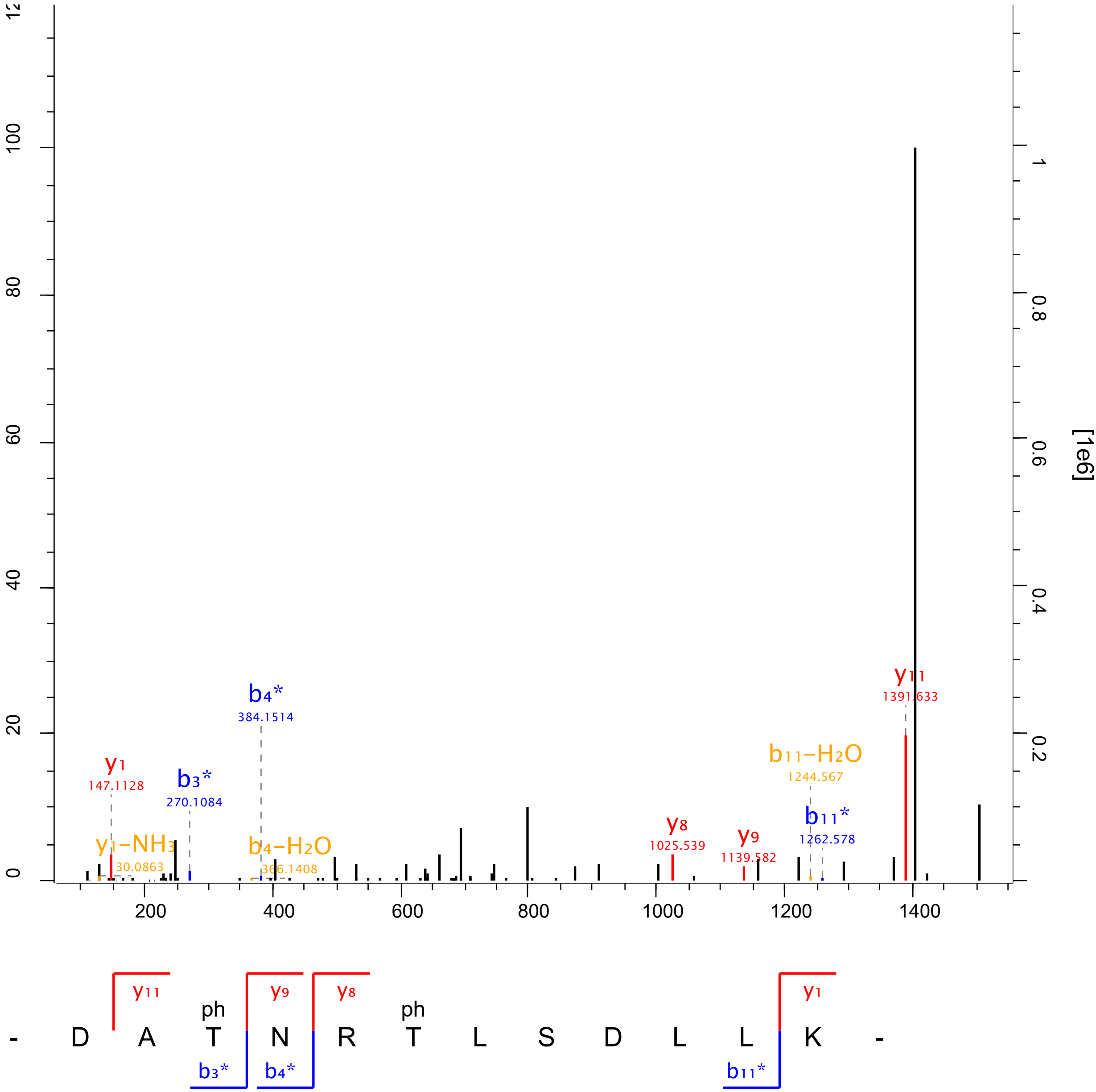

| Raw file | Scan | Method | Score | m/z |
|---|---|---|---|---|
| A17_15_1 | 7325 | FTMS; HCD | 102.64 | 652.27 |

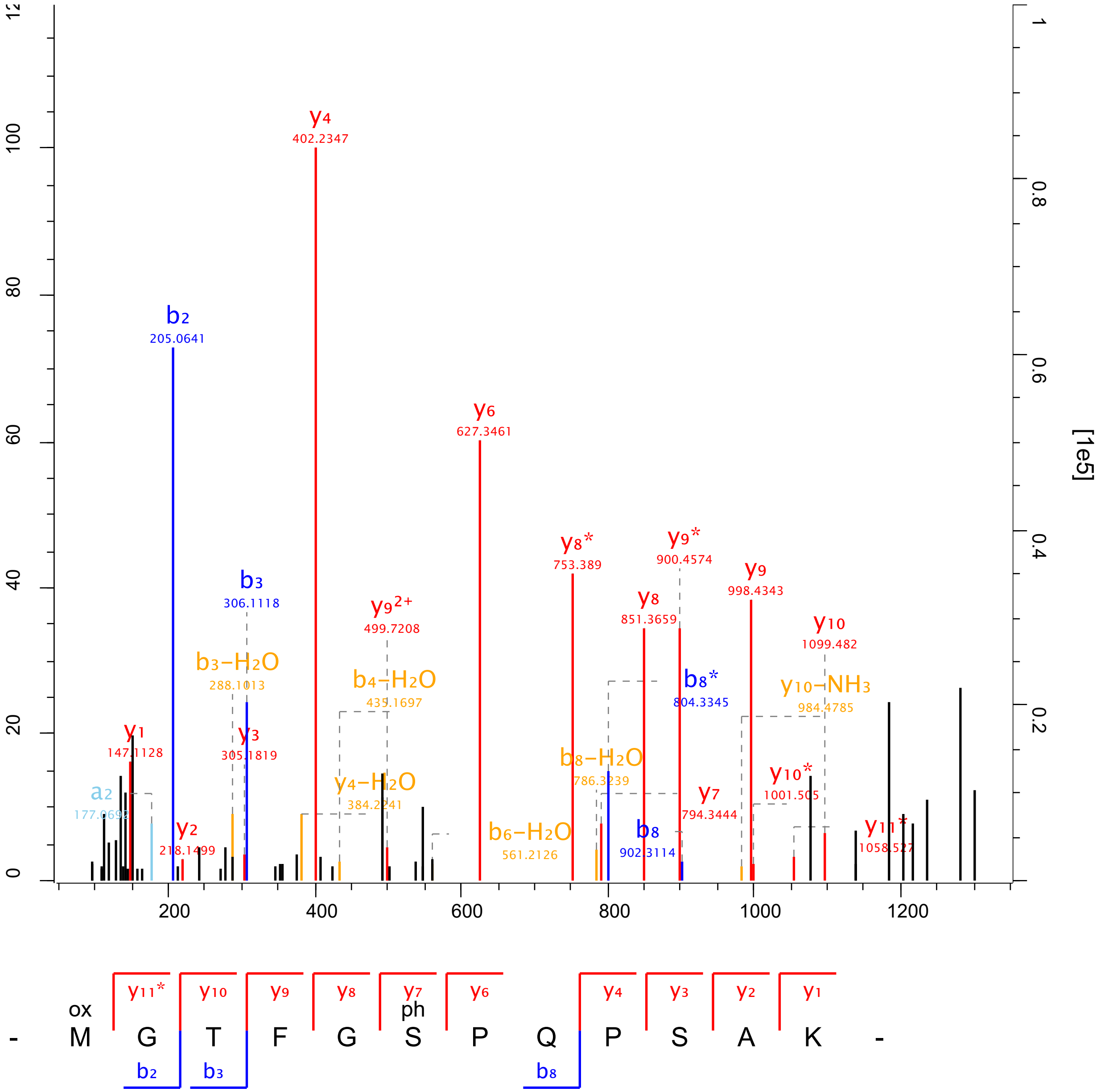

| Raw file | Scan | Method | Score | m/z |
| --- | --- | --- | --- | --- |
| A17_15_1 | 7401 | FTMS; HCD | 122.78 | 628.28 |

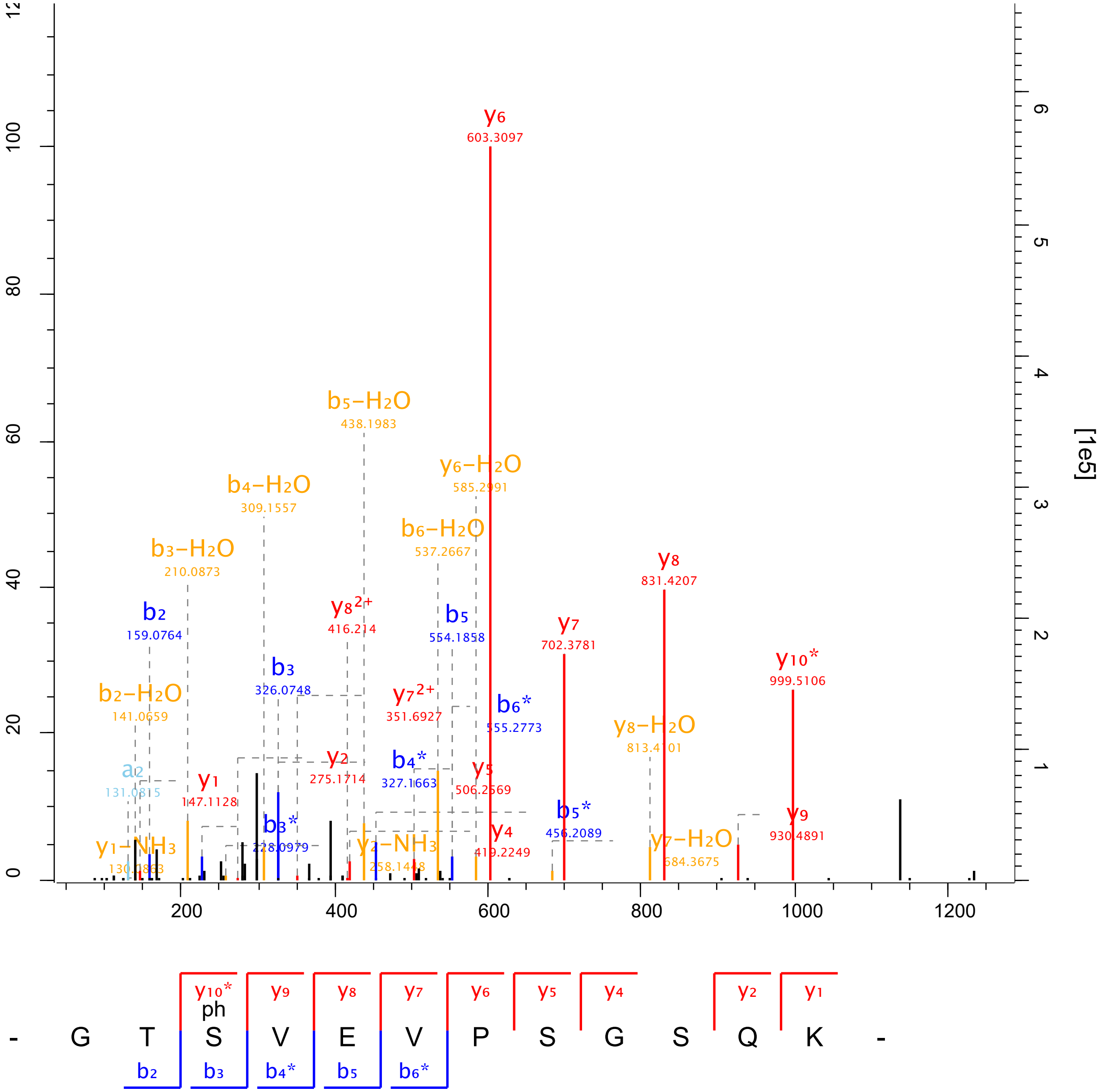

| Raw file | Scan | Method | Score | m/z |
|---|---|---|---|---|
| A17_15_1 | 7461 | FTMS; HCD | 73.45 | 553.9 |

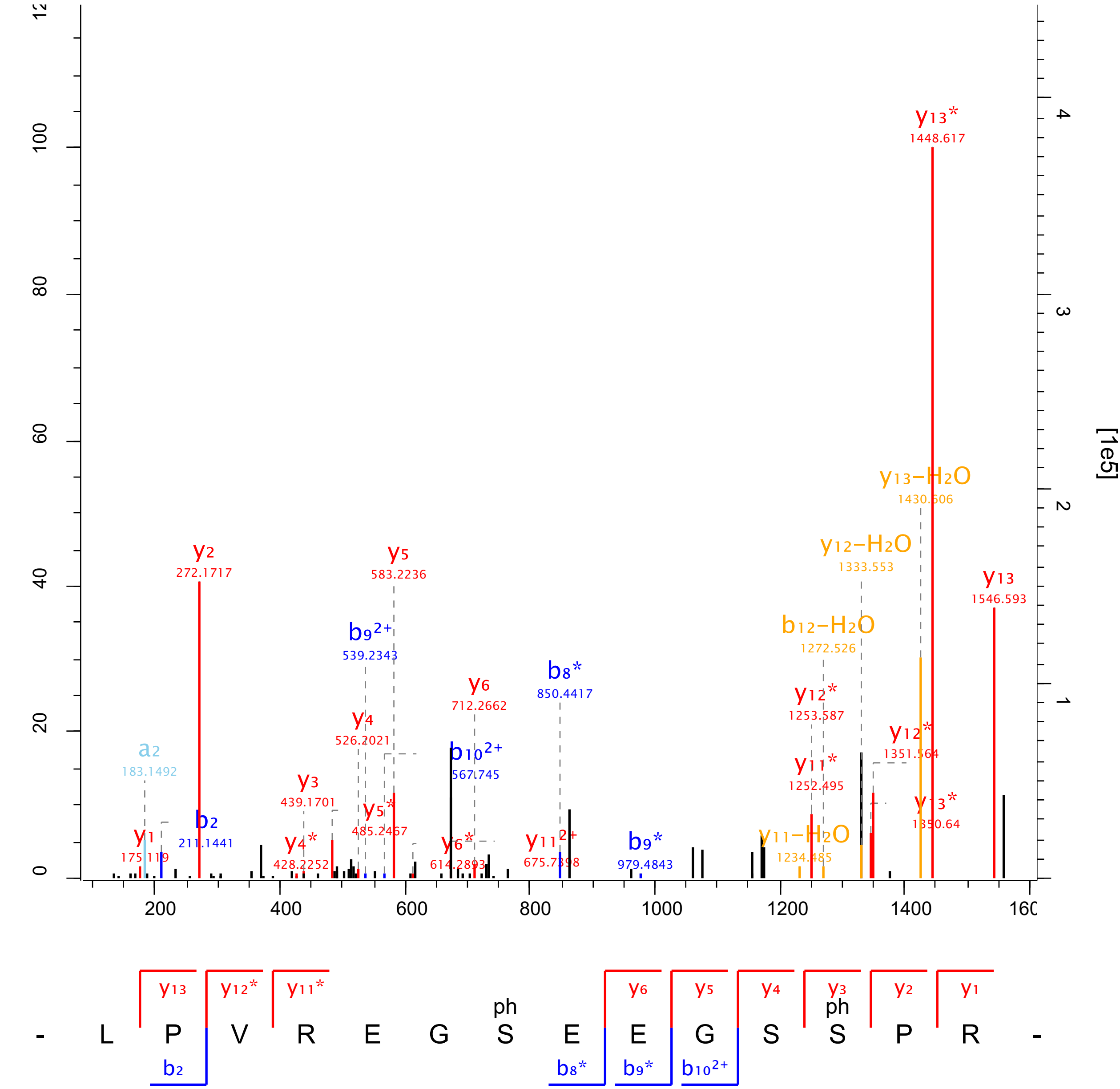

| Raw file | Scan | Method | Score | m/z |
|---|---|---|---|---|
| A17_15_1 | 7669 | FTMS; HCD | 64.24 | 683.8 |

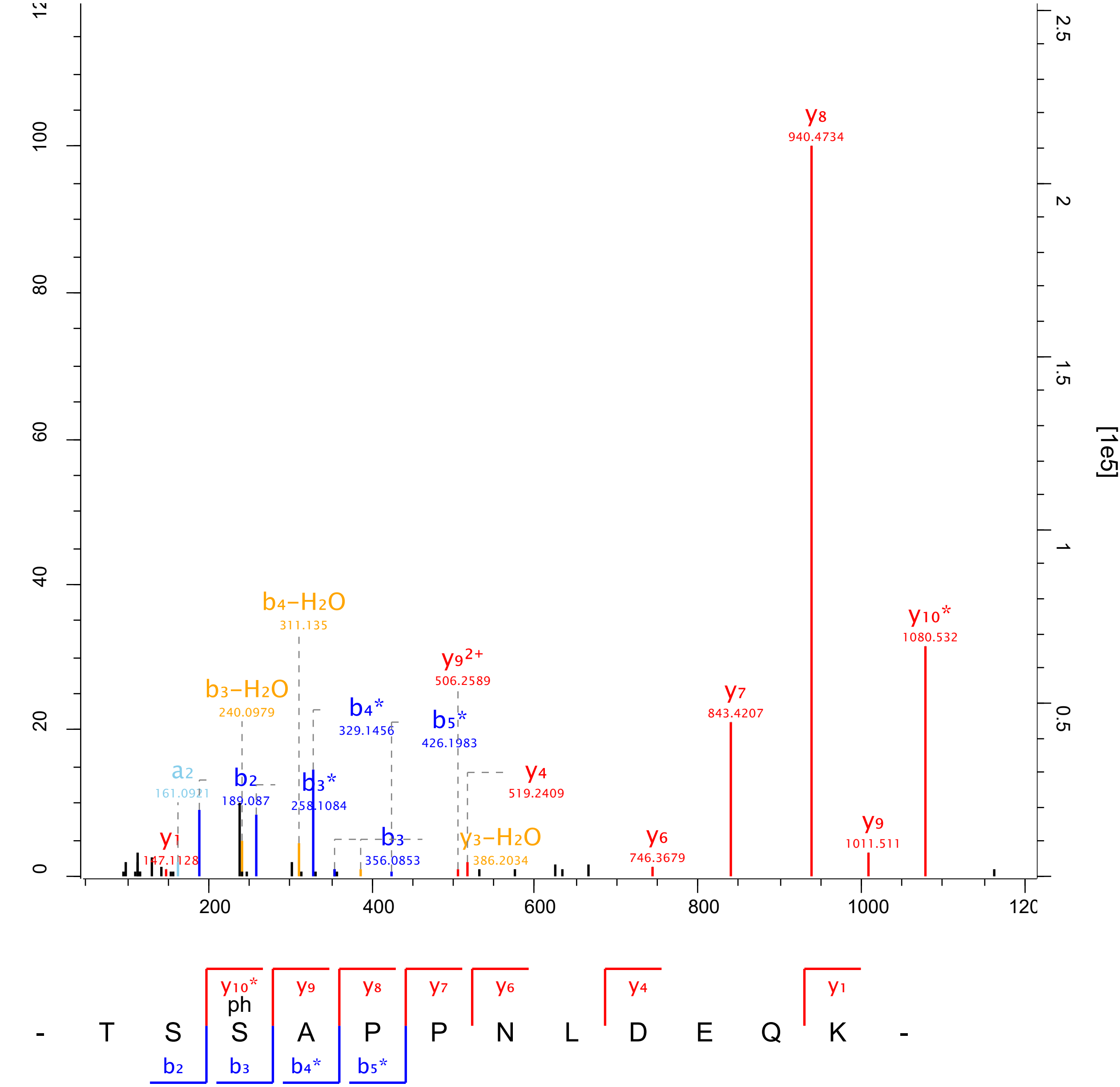

| Raw file | Scan | Method | Score | m/z |
|---|---|---|---|---|
| A17_15_1 | 7703 | FTMS; HCD | 168.47 | 719.81 |

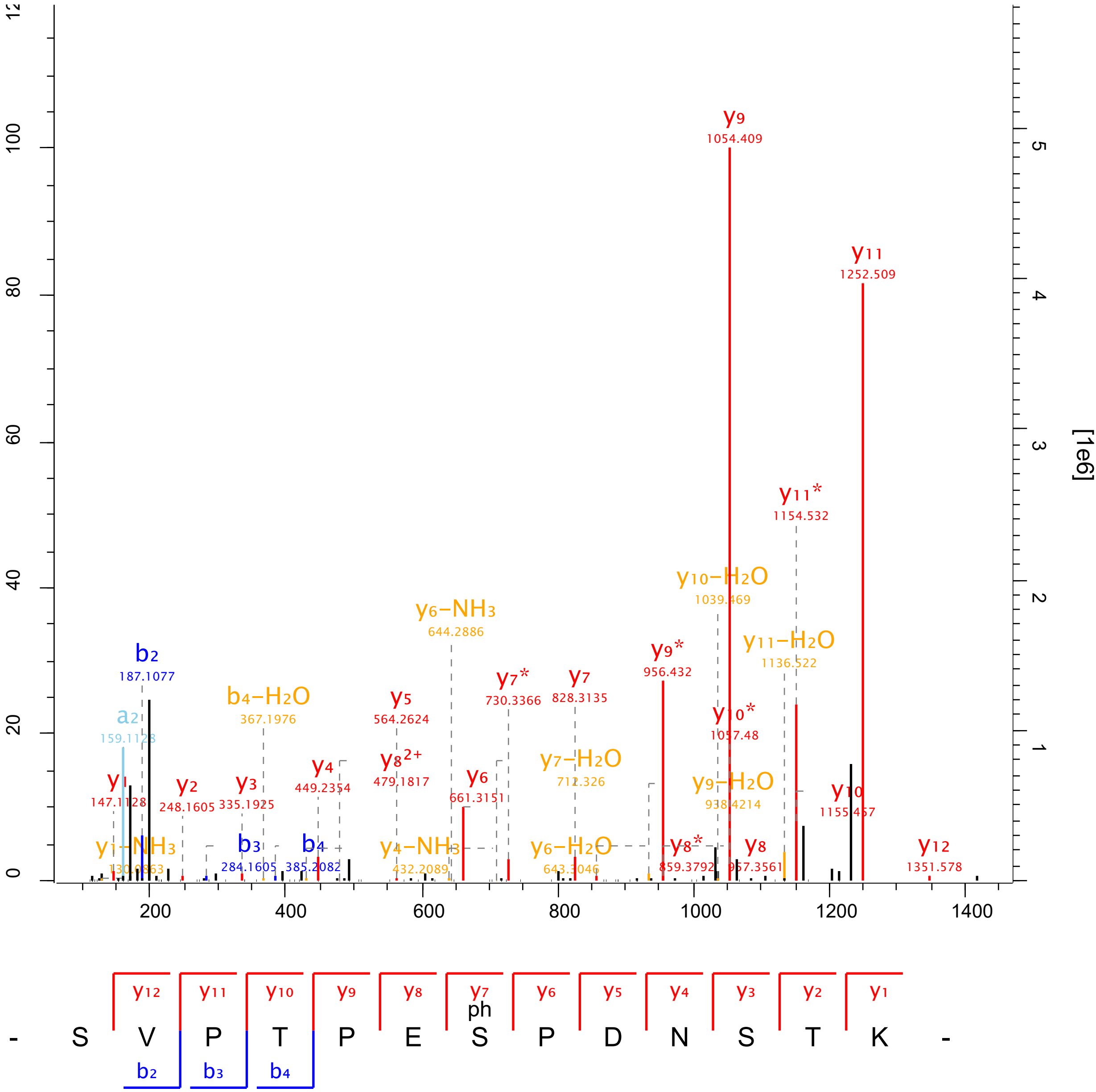

| Raw file | Scan | Method | Score | m/z |
|---|---|---|---|---|
| A17_15_1 | 7832 | FTMS; HCD | 111.94 | 576.23 |

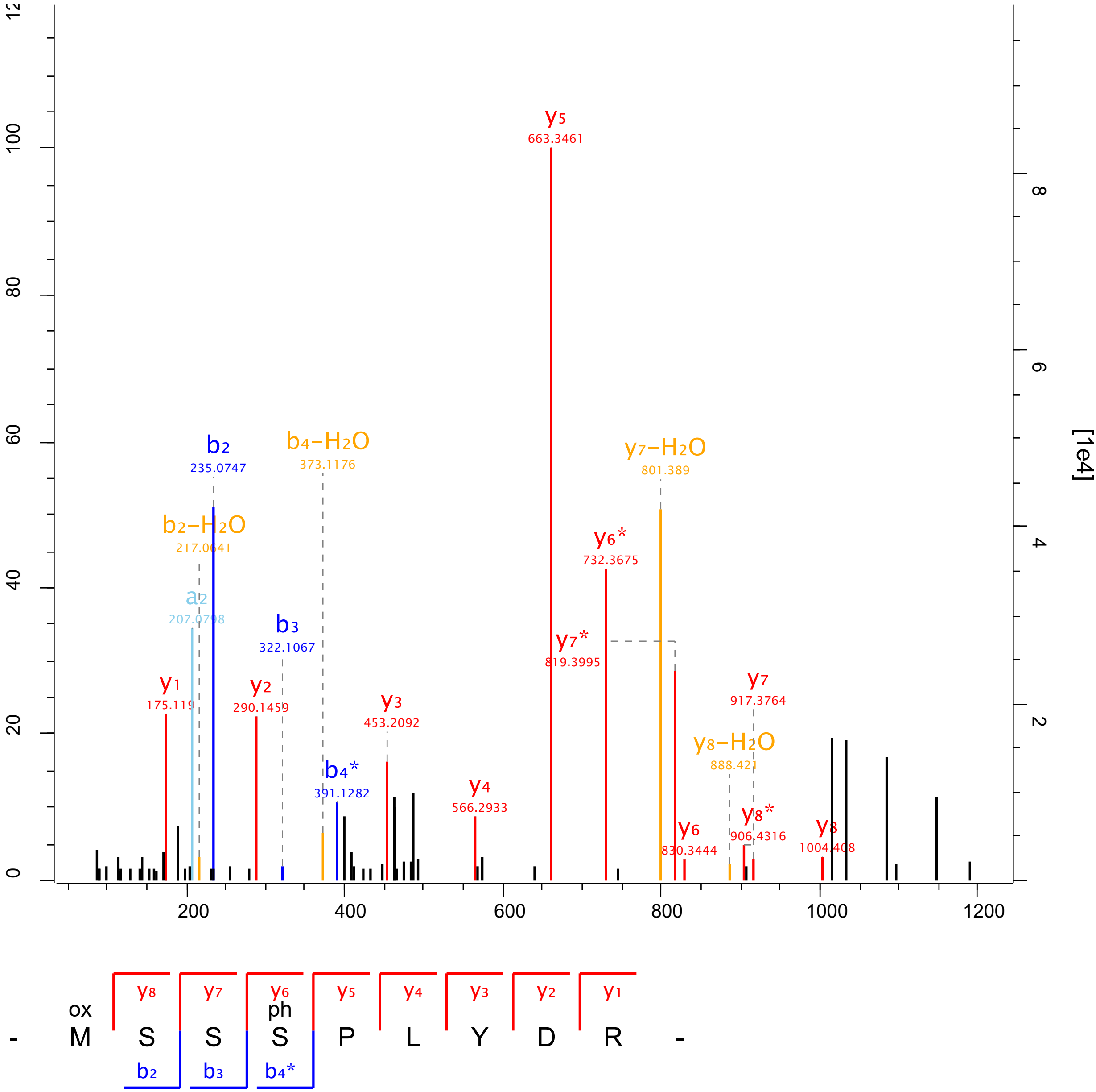

| Raw file | Scan | Method | Score | m/z |
| --- | --- | --- | --- | --- |
| A17_15_1 | 7998 | FTMS; HCD | 83.05 | 686.24 |

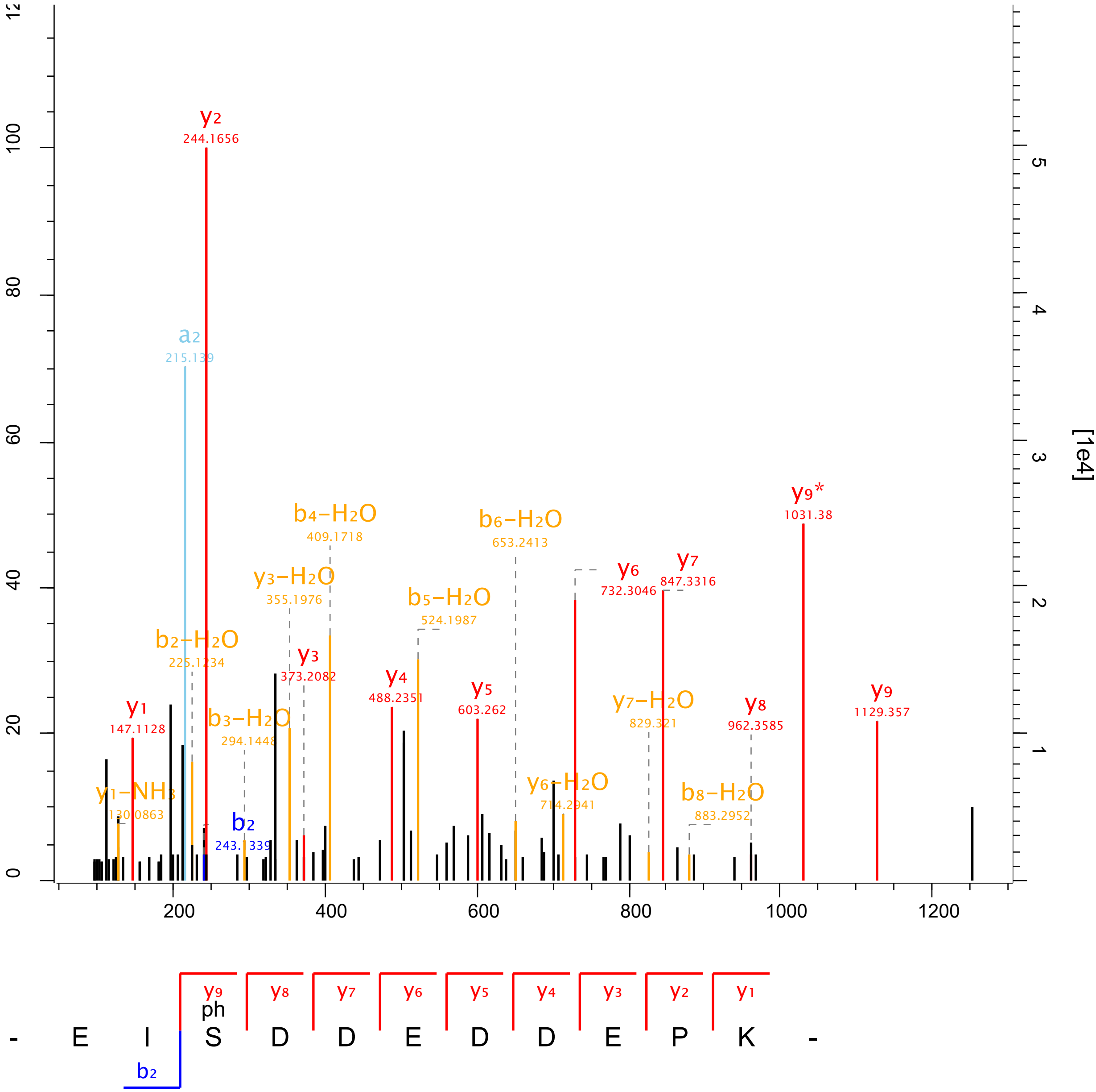

| Raw file | Scan | Method | Score | m/z |
|---|---|---|---|---|
| A17_15_1 | 8027 | FTMS; HCD | 122.44 | 798.8 |

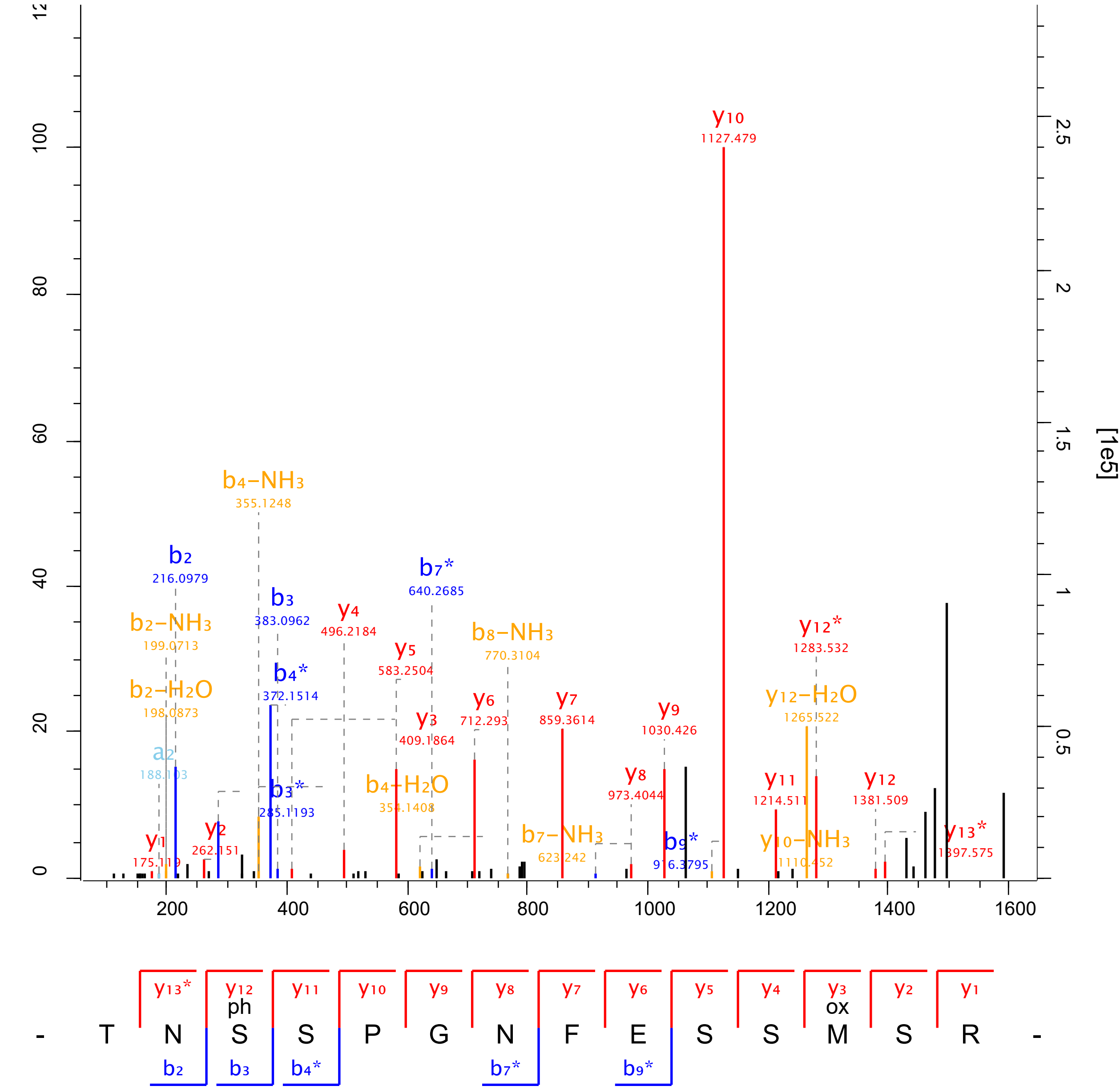

| Raw file | Scan | Method | Score | m/z |
|---|---|---|---|---|
| A17_15_1 | 8112 | FTMS; HCD | 57.53 | 665.29 |

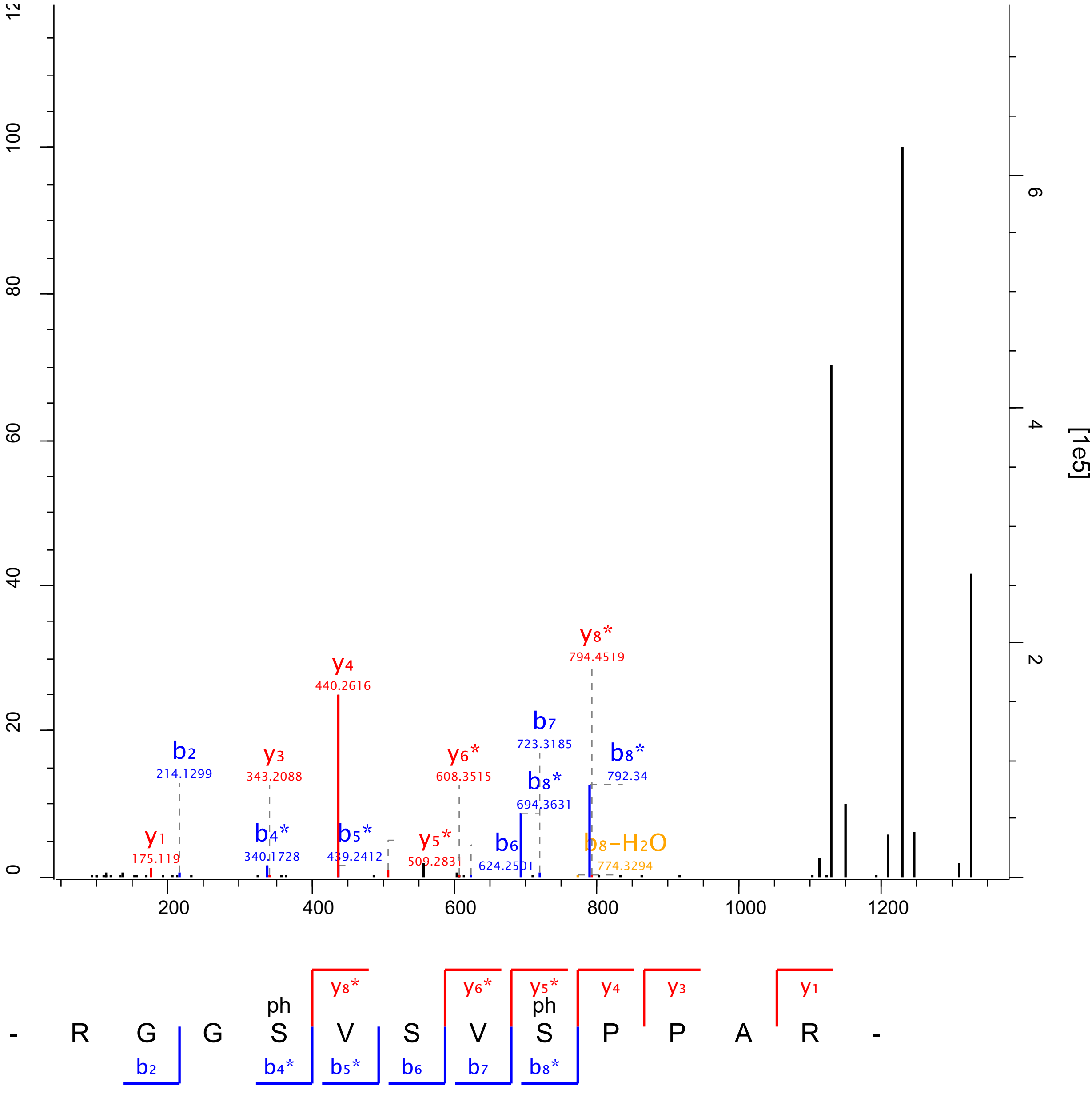

| Raw file | Scan | Method | Score | m/z |
| --- | --- | --- | --- | --- |
| A17_15_1 | 8287 | FTMS; HCD | 62.09 | 607.27 |

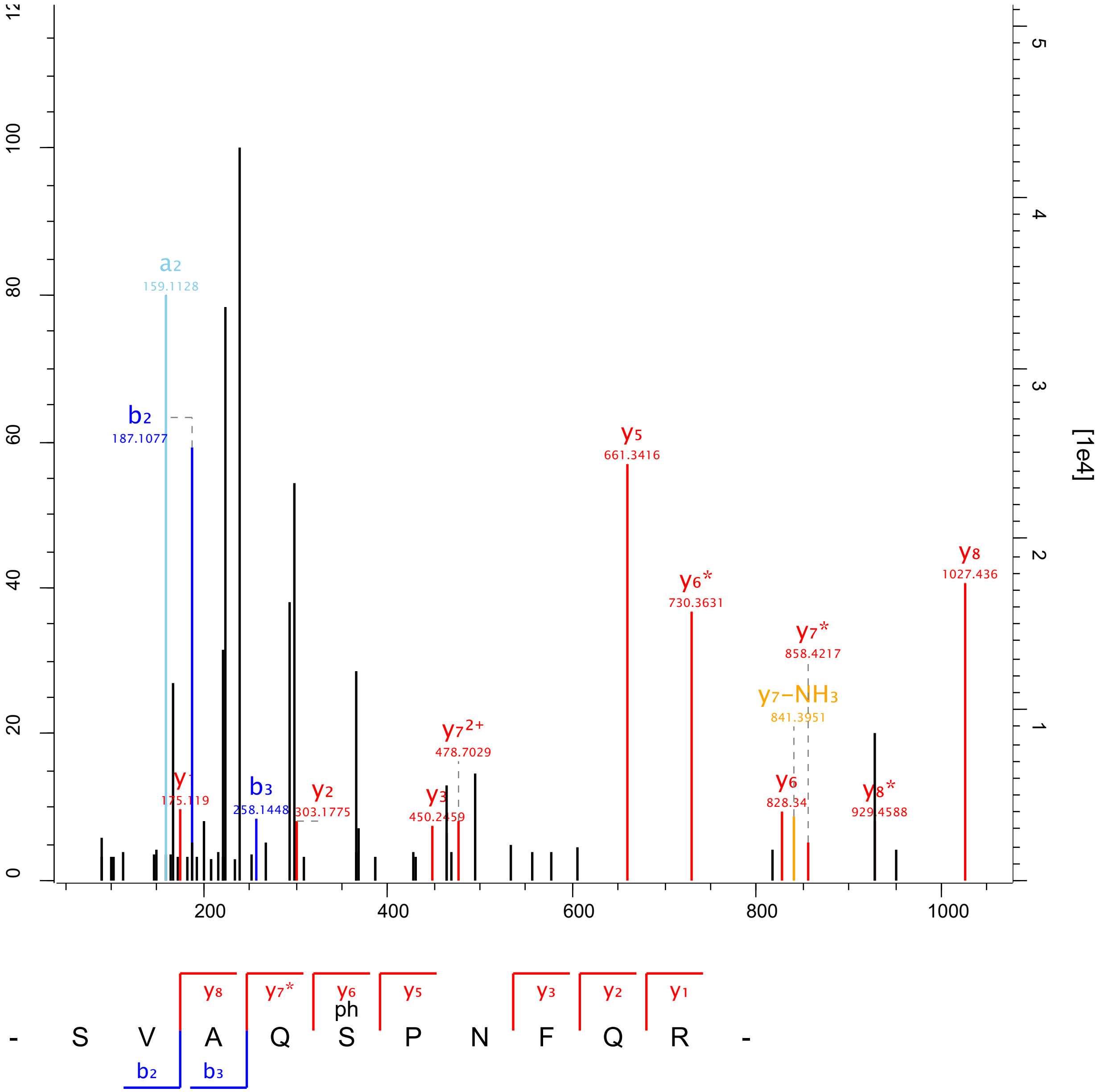

| Raw file | Scan | Method | Score | m/z |
|---|---|---|---|---|
| A17_15_1 | 8364 | FTMS; HCD | 199.33 | 847.79 |

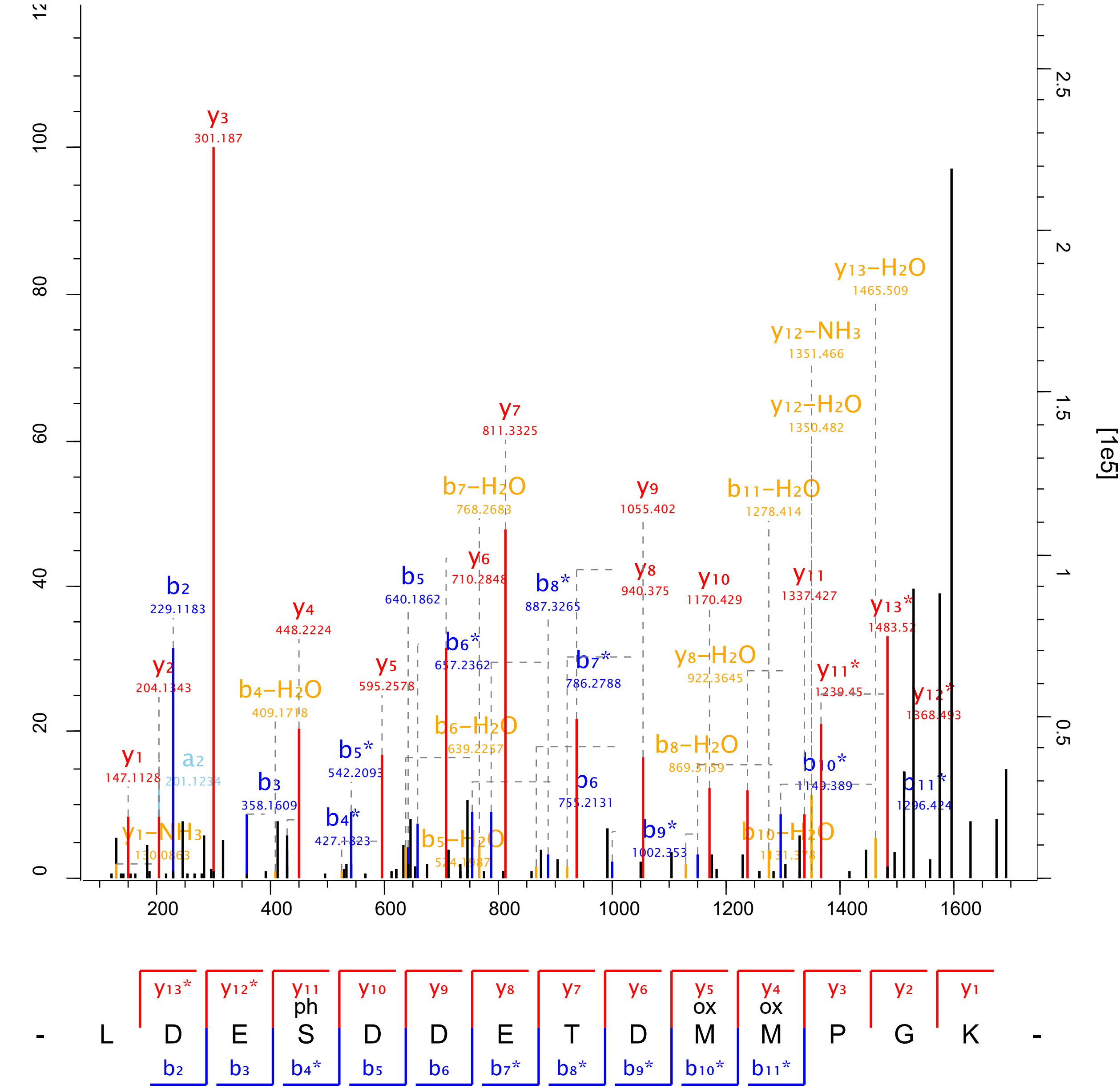

| Raw file | Scan | Method | Score | m/z |
|---|---|---|---|---|
| A17_15_1 | 8401 | FTMS; HCD | 42.07 | 750.32 |

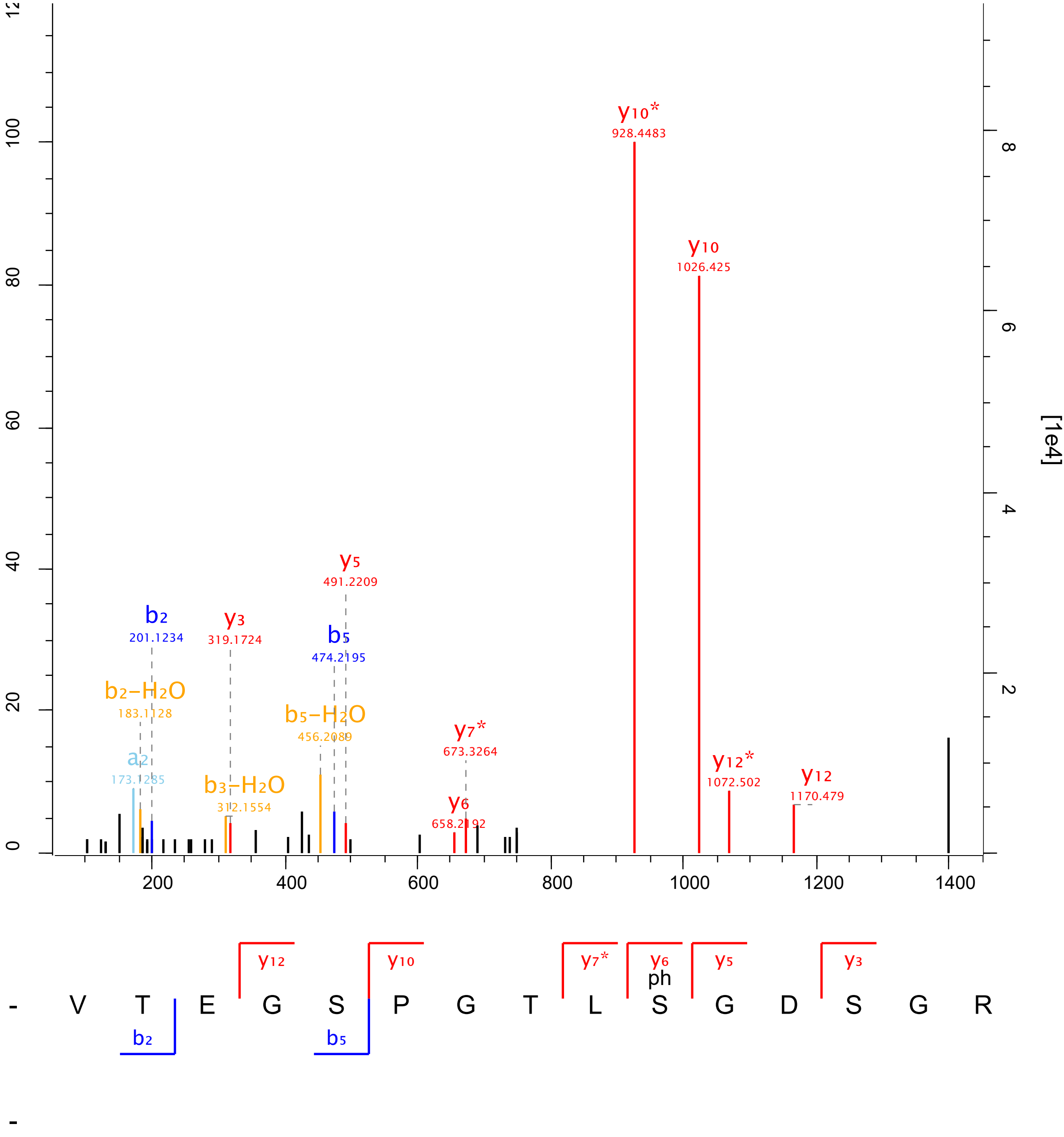

| Raw file | Scan | Method | Score | m/z |
|---|---|---|---|---|
| A17_15_1 | 8438 | FTMS; HCD | 97.06 | 711.62 |

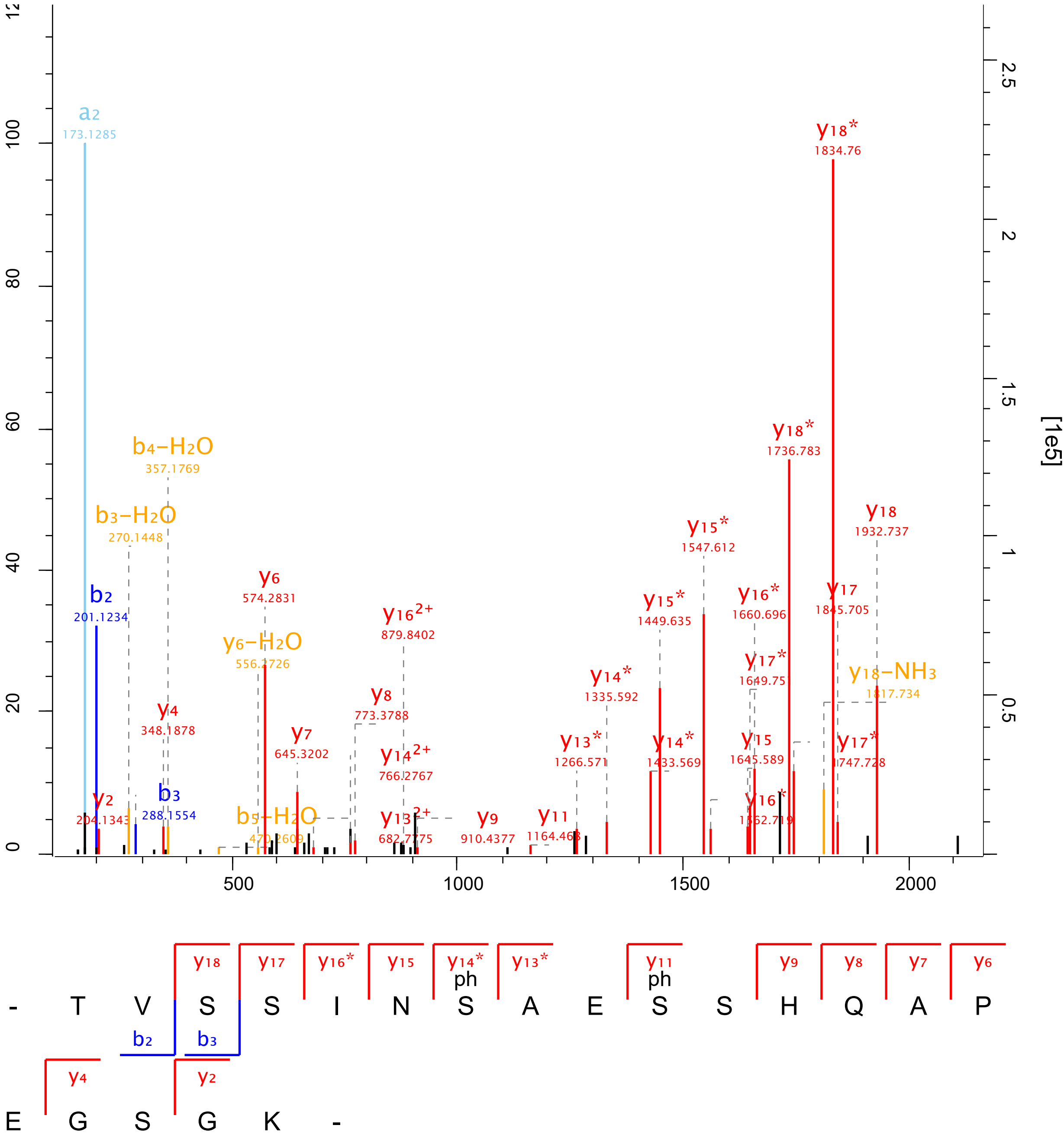

| Raw file | Scan | Method | Score | m/z |
|---|---|---|---|---|
| A17_15_1 | 8470 | FTMS; HCD | 117.86 | 588.74 |

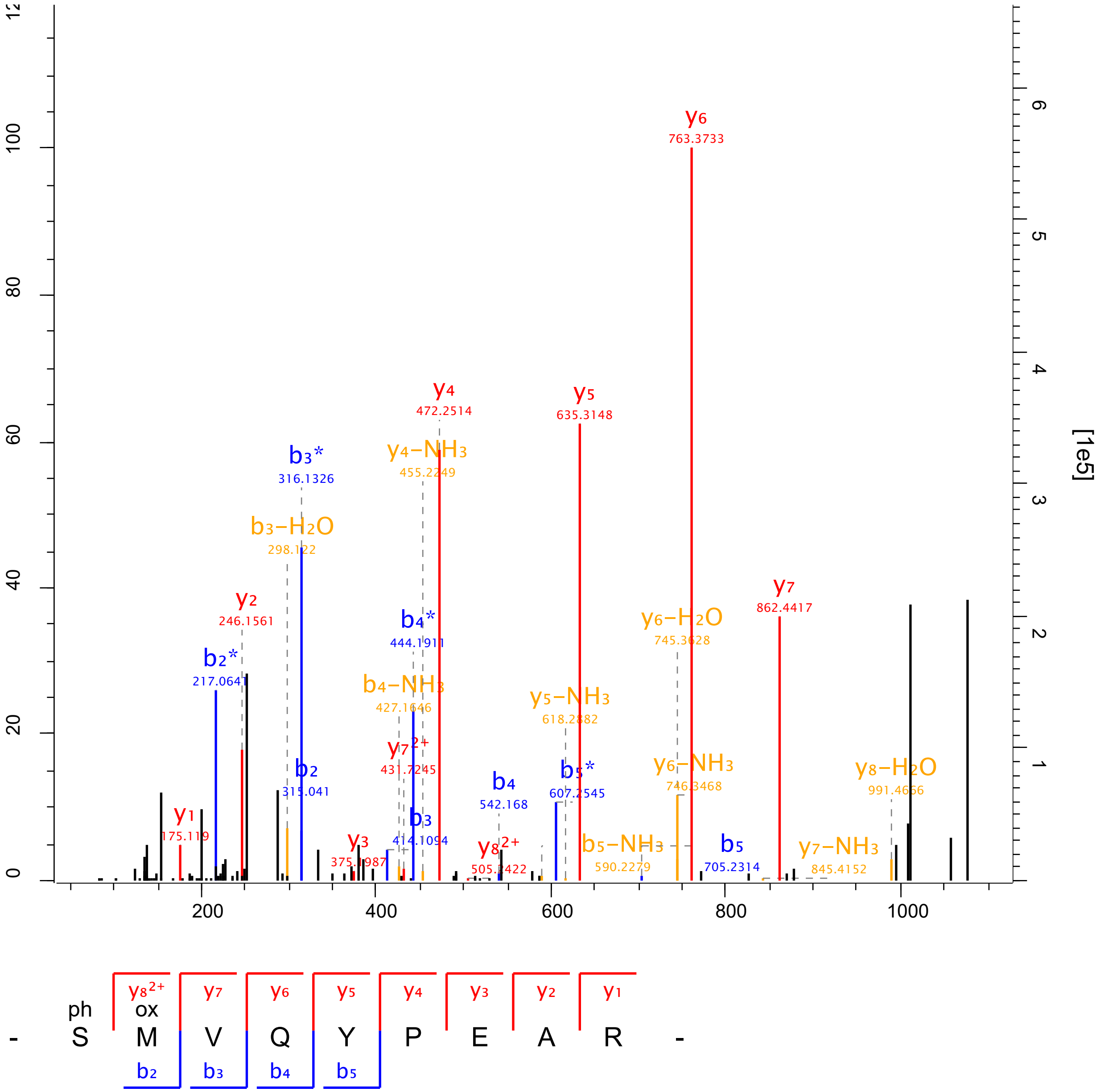

| Raw file | Scan | Method | Score | m/z |
| --- | --- | --- | --- | --- |
| A17_15_1 | 8595 | FTMS; HCD | 43.79 | 753.3 |

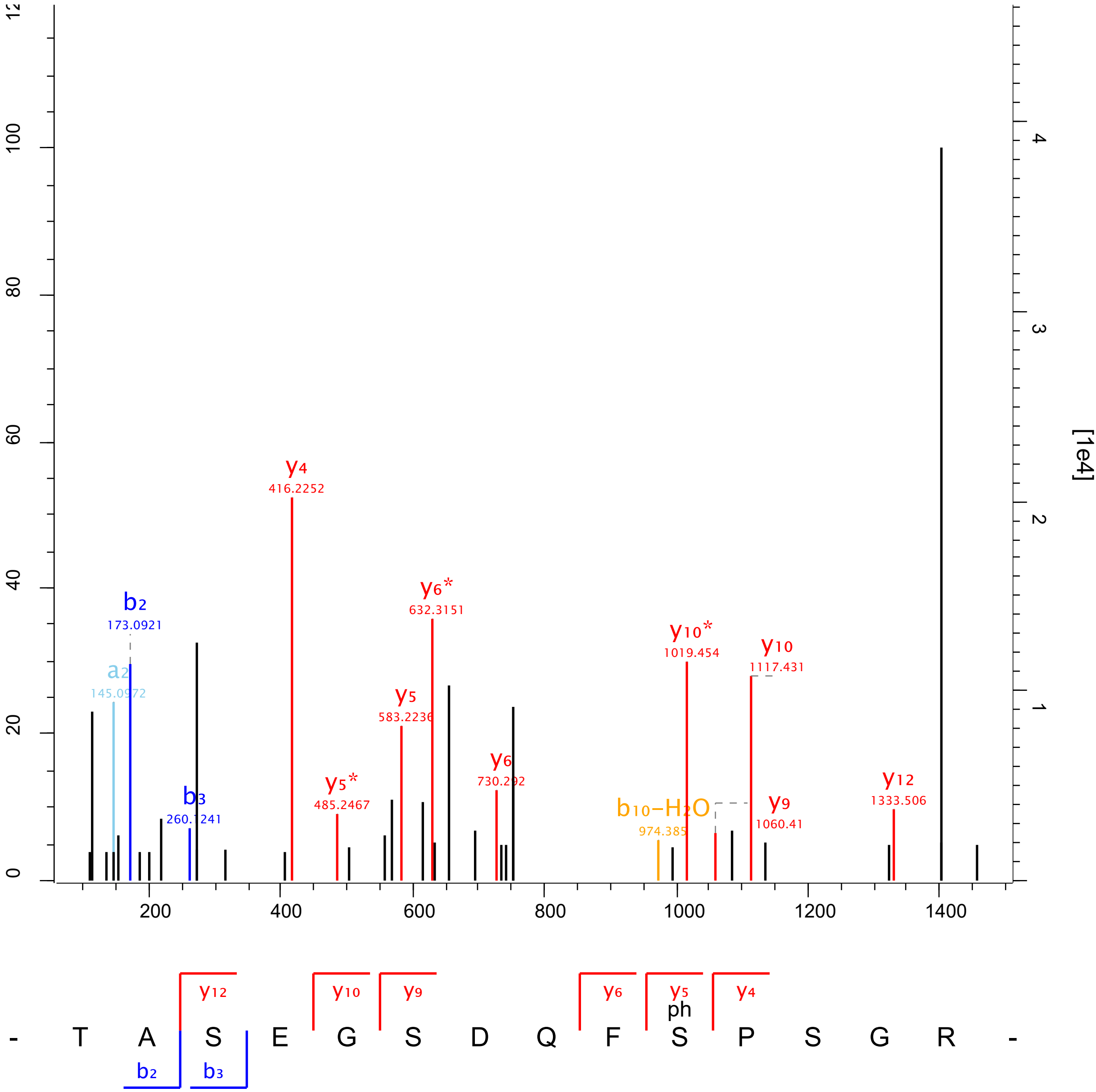

| Raw file | Scan | Method | Score | m/z |
| --- | --- | --- | --- | --- |
| A17_15_1 | 8626 | FTMS; HCD | 58.71 | 525.88 |

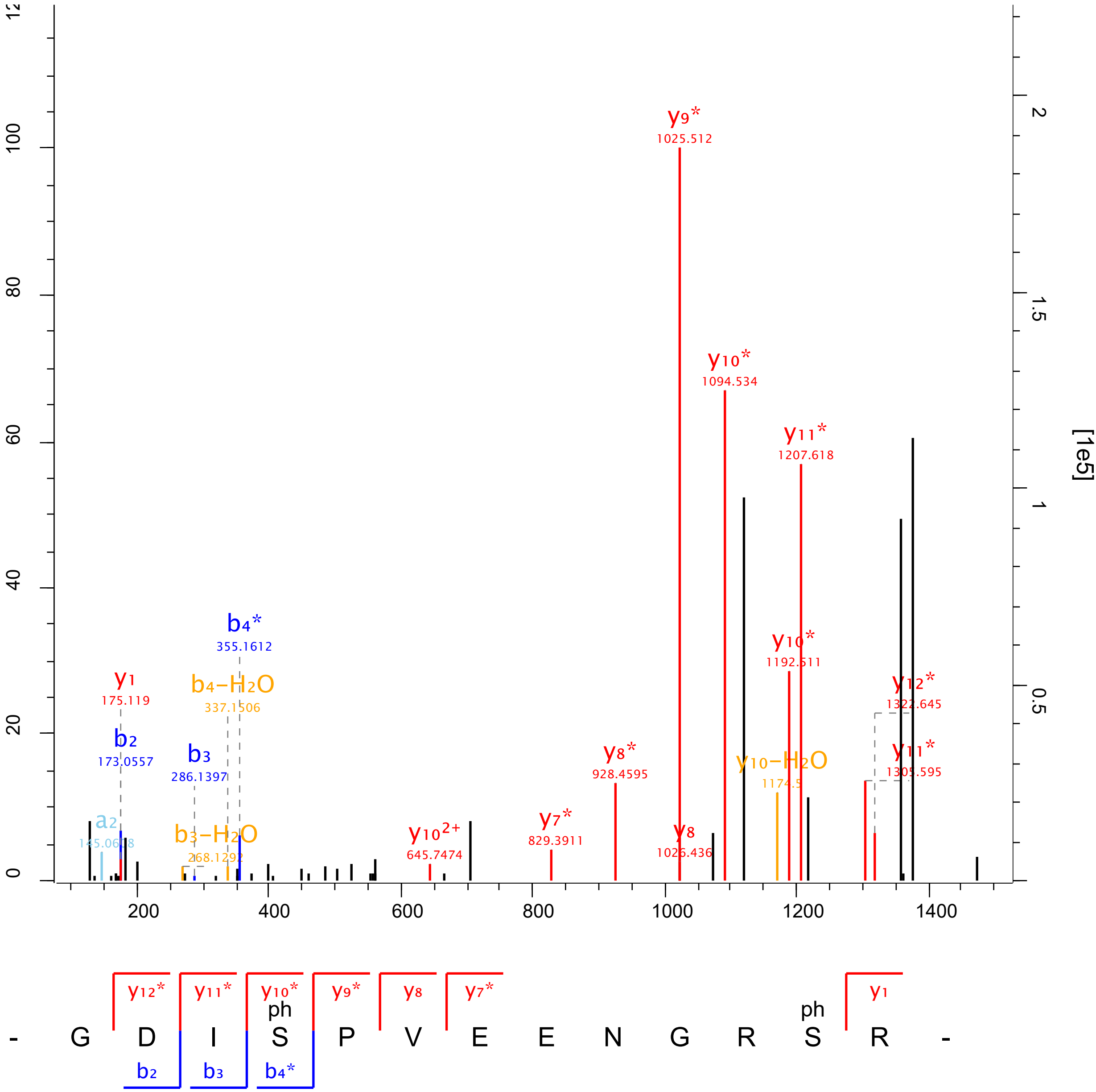

| Raw file | Scan | Method | Score | m/z |
| --- | --- | --- | --- | --- |
| A17_15_1 | 8672 | FTMS; HCD | 79.66 | 534.22 |

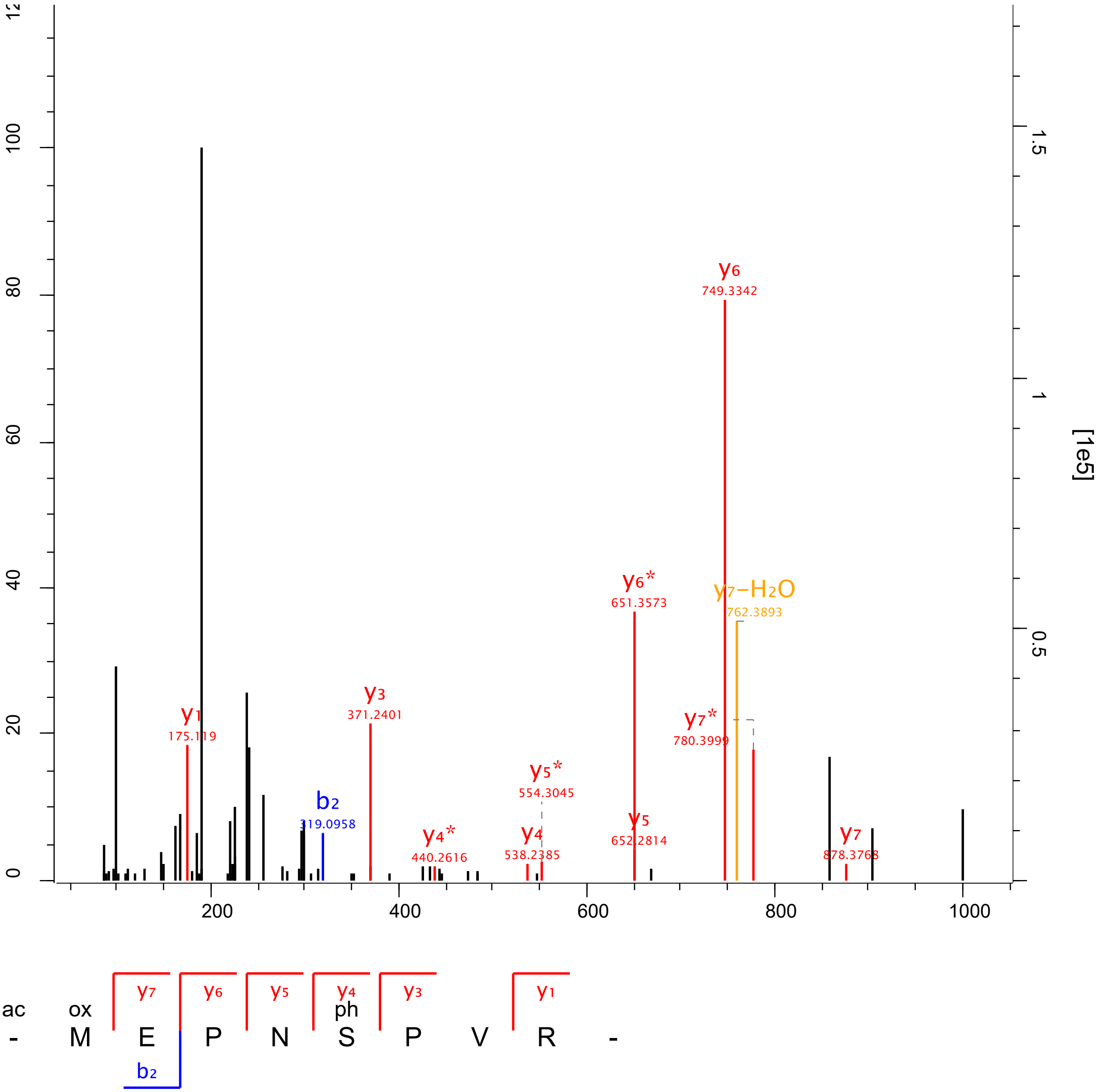

| Raw file | Scan | Method | Score | m/z |
|---|---|---|---|---|
| A17_15_1 | 8725 | FTMS; HCD | 163.99 | 684.79 |

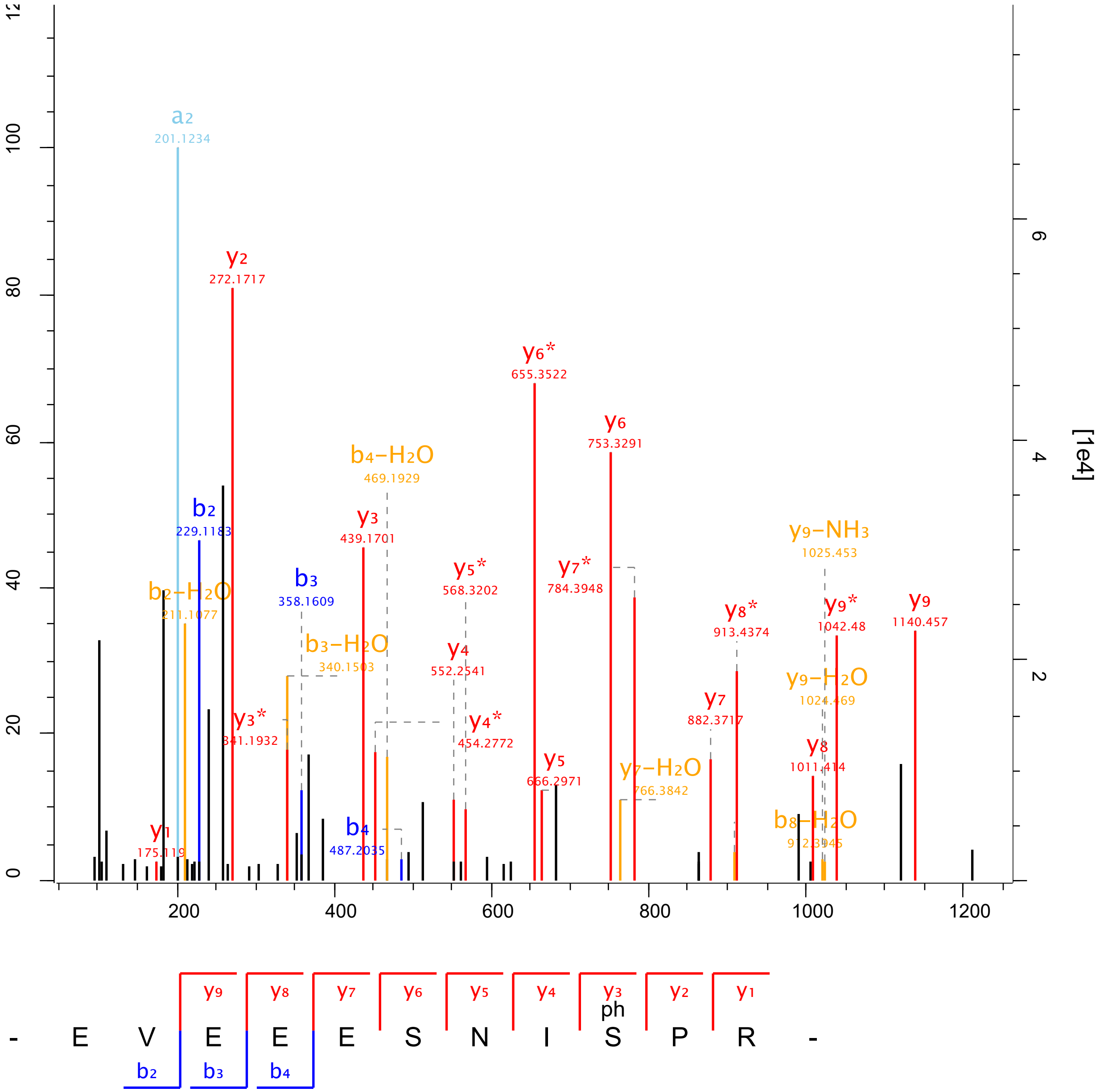

| Raw file | Scan | Method | Score | m/z |
| --- | --- | --- | --- | --- |
| A17_15_1 | 8839 | FTMS; HCD | 78.34 | 515.72 |

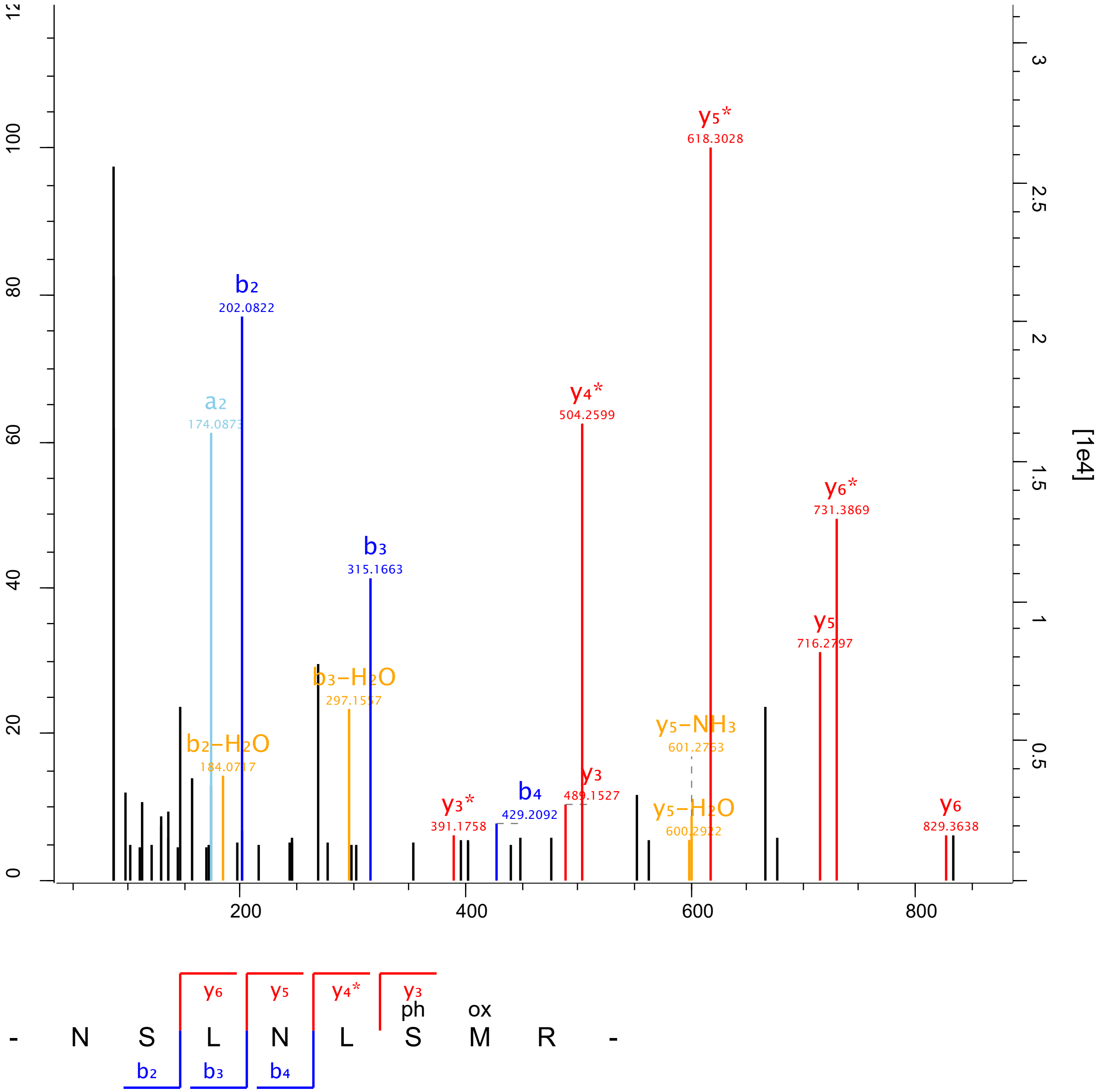

| Raw file | Scan | Method | Score | m/z |
|---|---|---|---|---|
| A17_15_1 | 9051 | FTMS; HCD | 57.37 | 867.85 |

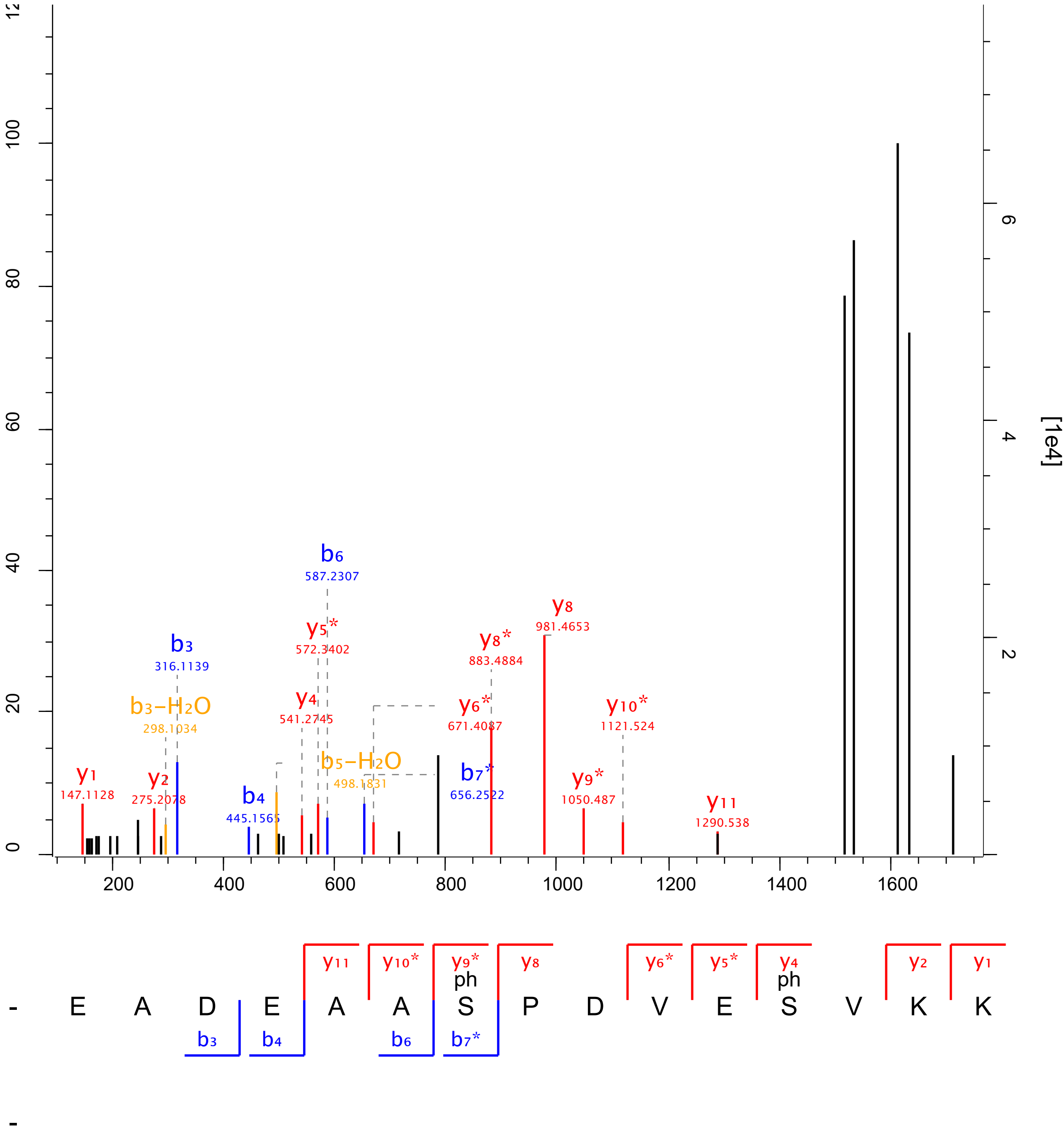

| Raw file | Scan | Method | Score | m/z |
| --- | --- | --- | --- | --- |
| A17_15_1 | 9079 | FTMS; HCD | 94.45 | 561.27 |

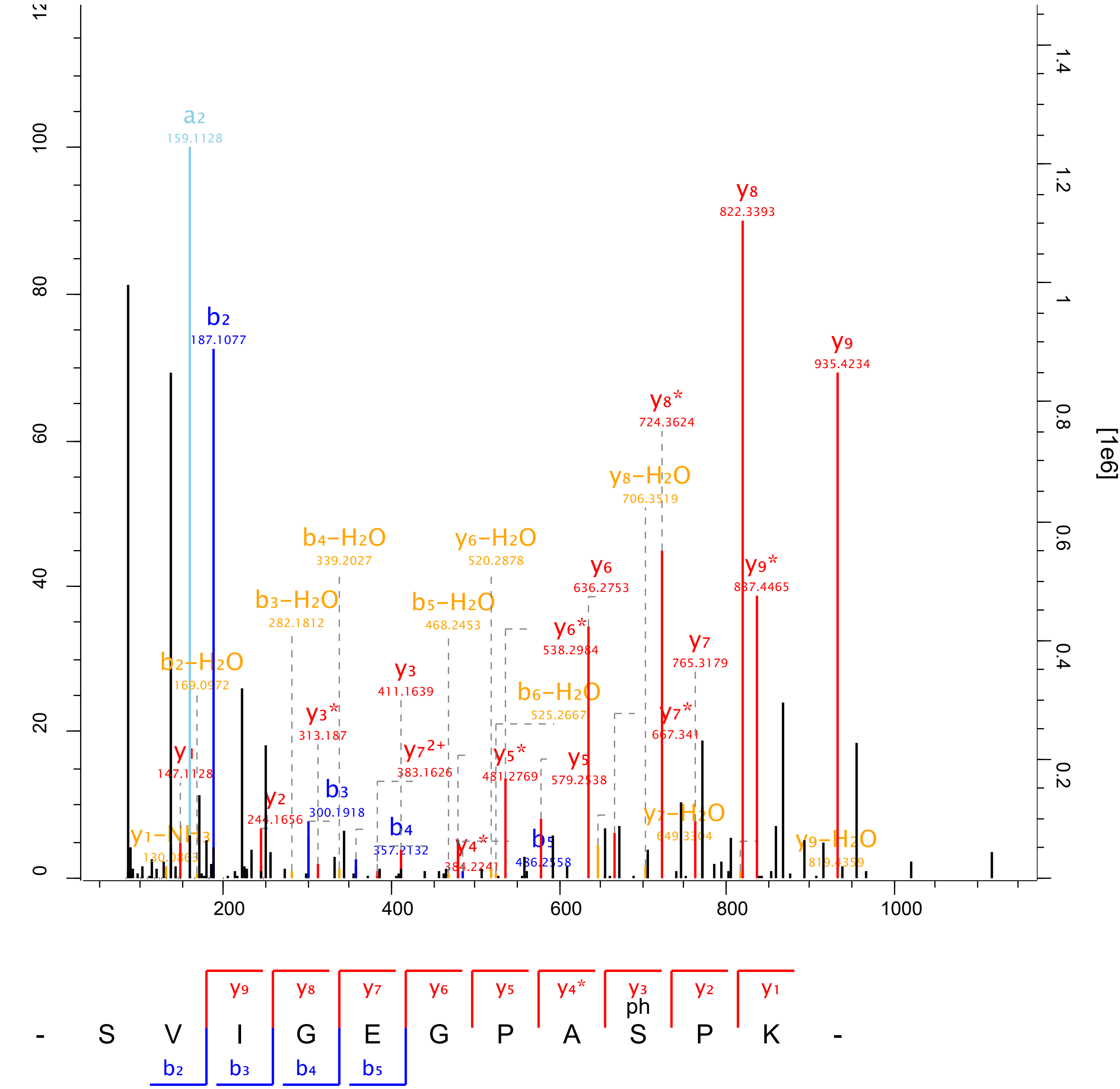

| Raw file | Scan | Method | Score | m/z |
|---|---|---|---|---|
| A17_15_1 | 9397 | FTMS; HCD | 58.98 | 763.33 |

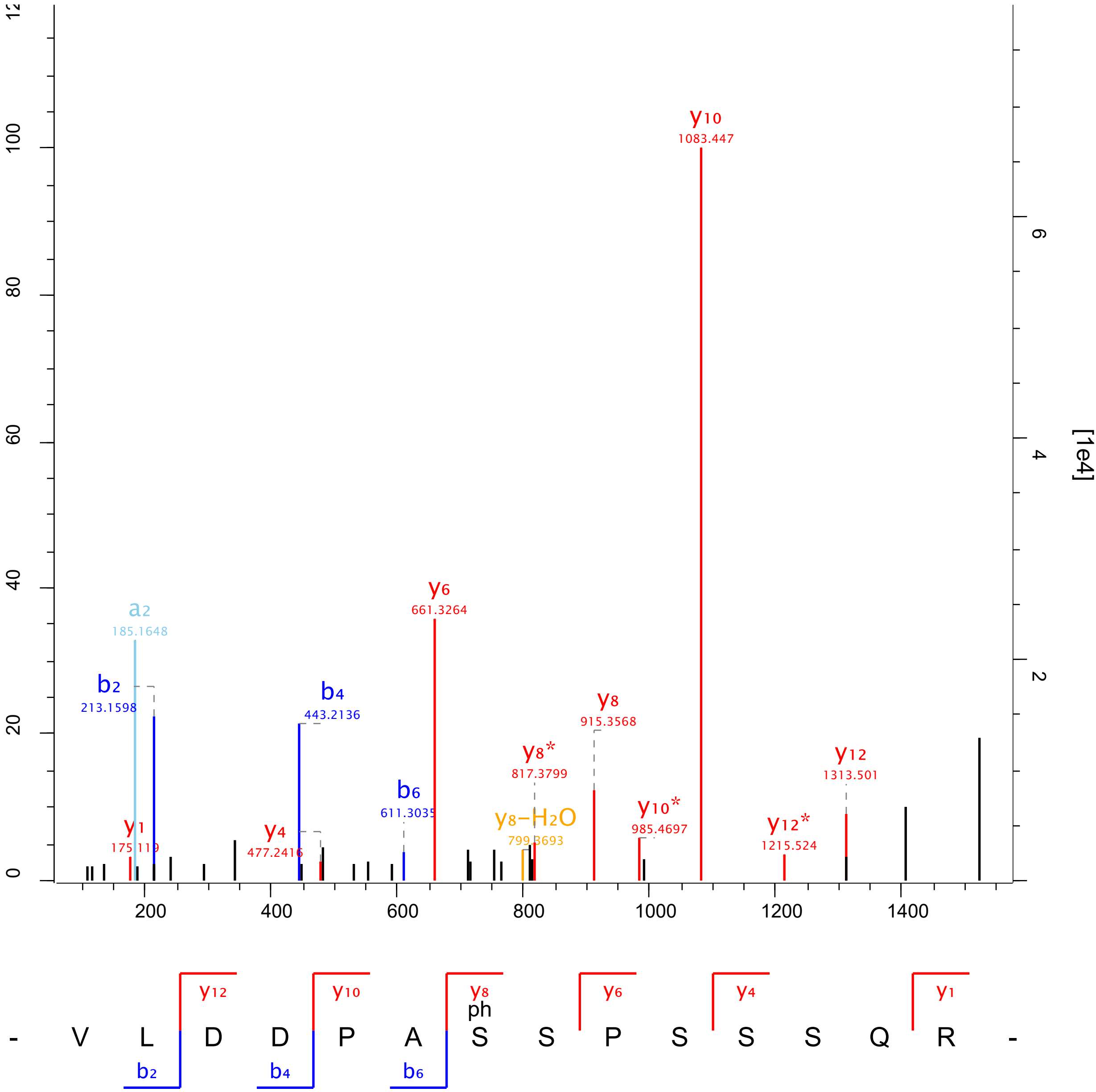

| Raw file | Scan | Method | Score | m/z |
|---|---|---|---|---|
| A17_15_1 | 9619 | FTMS; HCD | 108.96 | 794.31 |

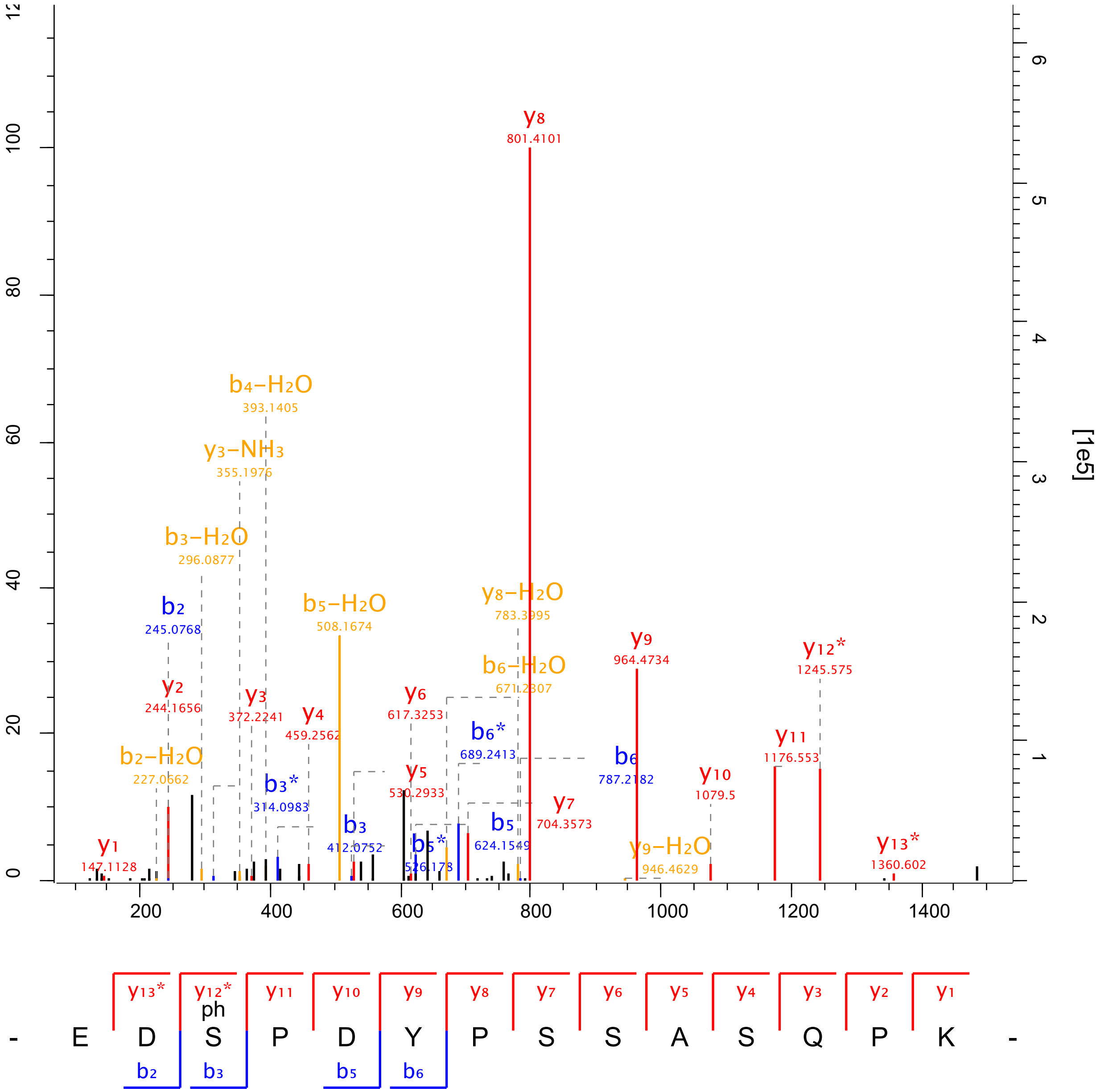

| Raw file | Scan | Method | Score | m/z |
|---|---|---|---|---|
| A17_15_1 | 9790 | FTMS; HCD | 87.88 | 642.94 |

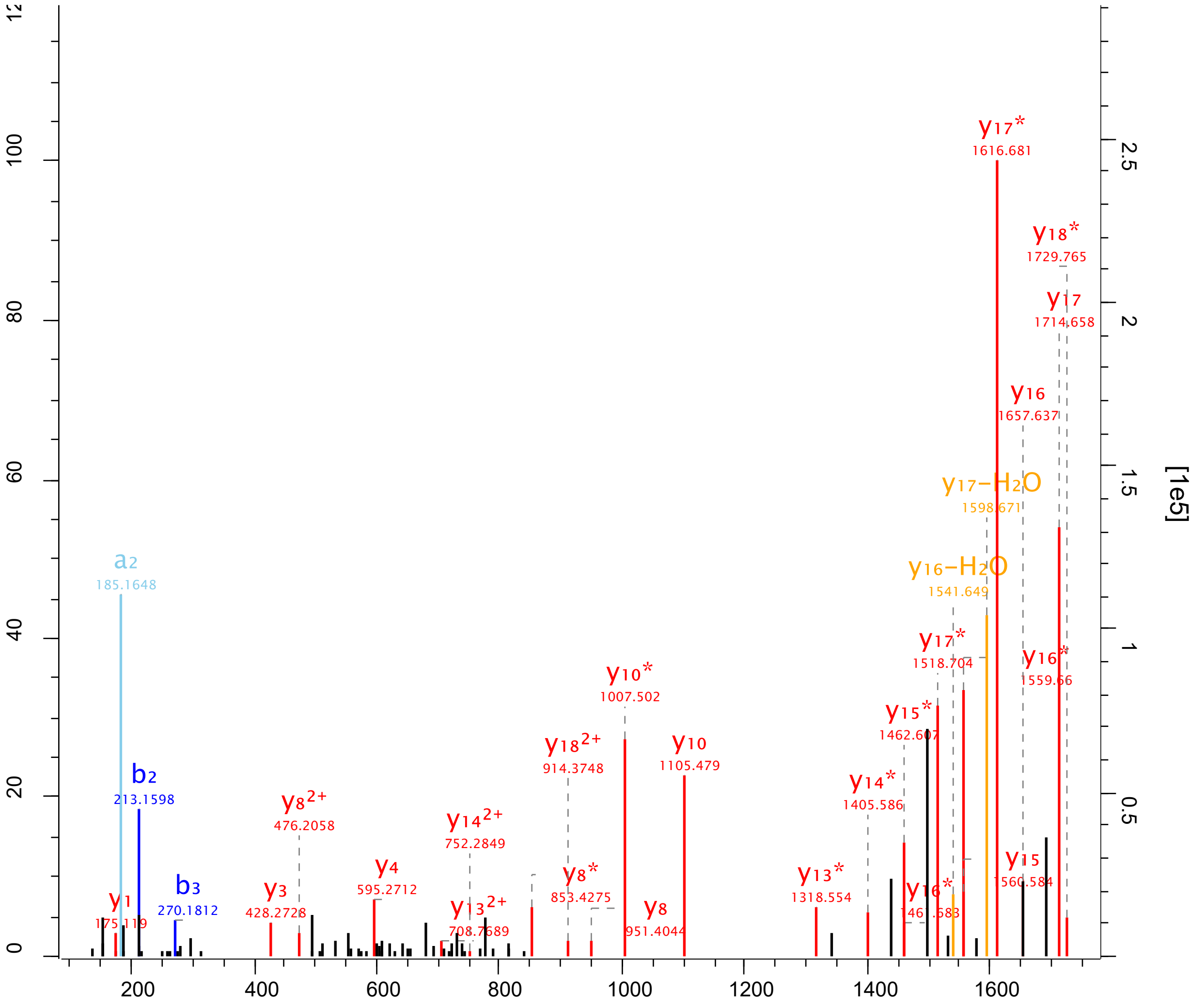


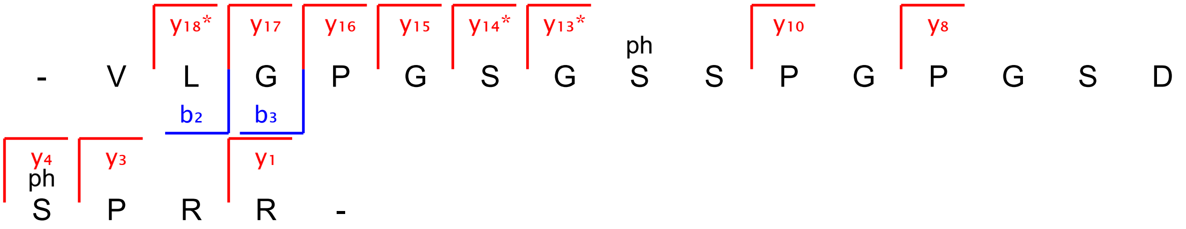

| Raw file | Scan | Method | Score | m/z |
|---|---|---|---|---|
| A17_15_1 | 9977 | FTMS; HCD | 74.99 | 487.21 |

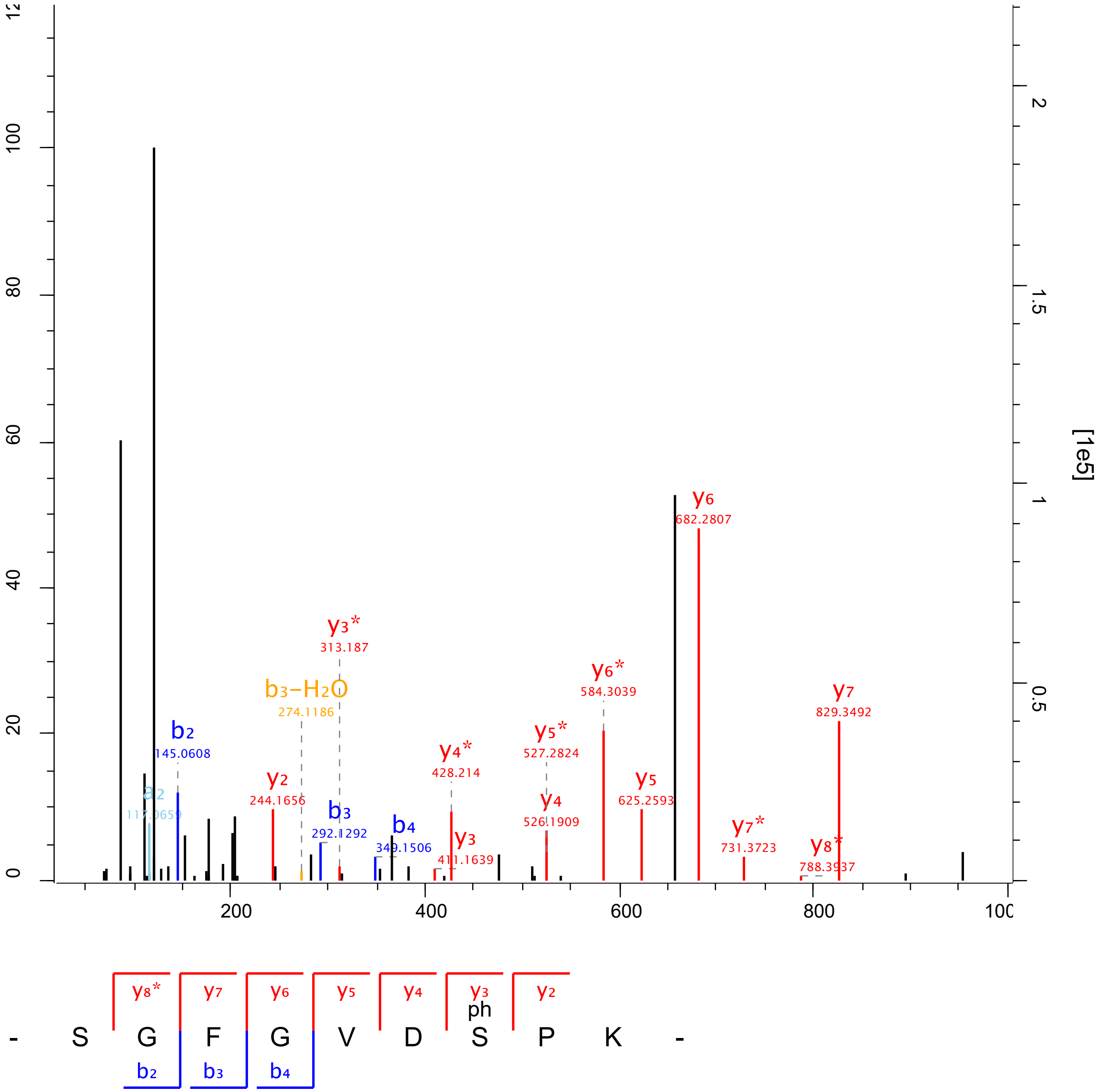

| Raw file | Scan | Method | Score | m/z |
|---|---|---|---|---|
| A17_15_3 | 6306 | FTMS; HCD | 192.78 | 625.29 |

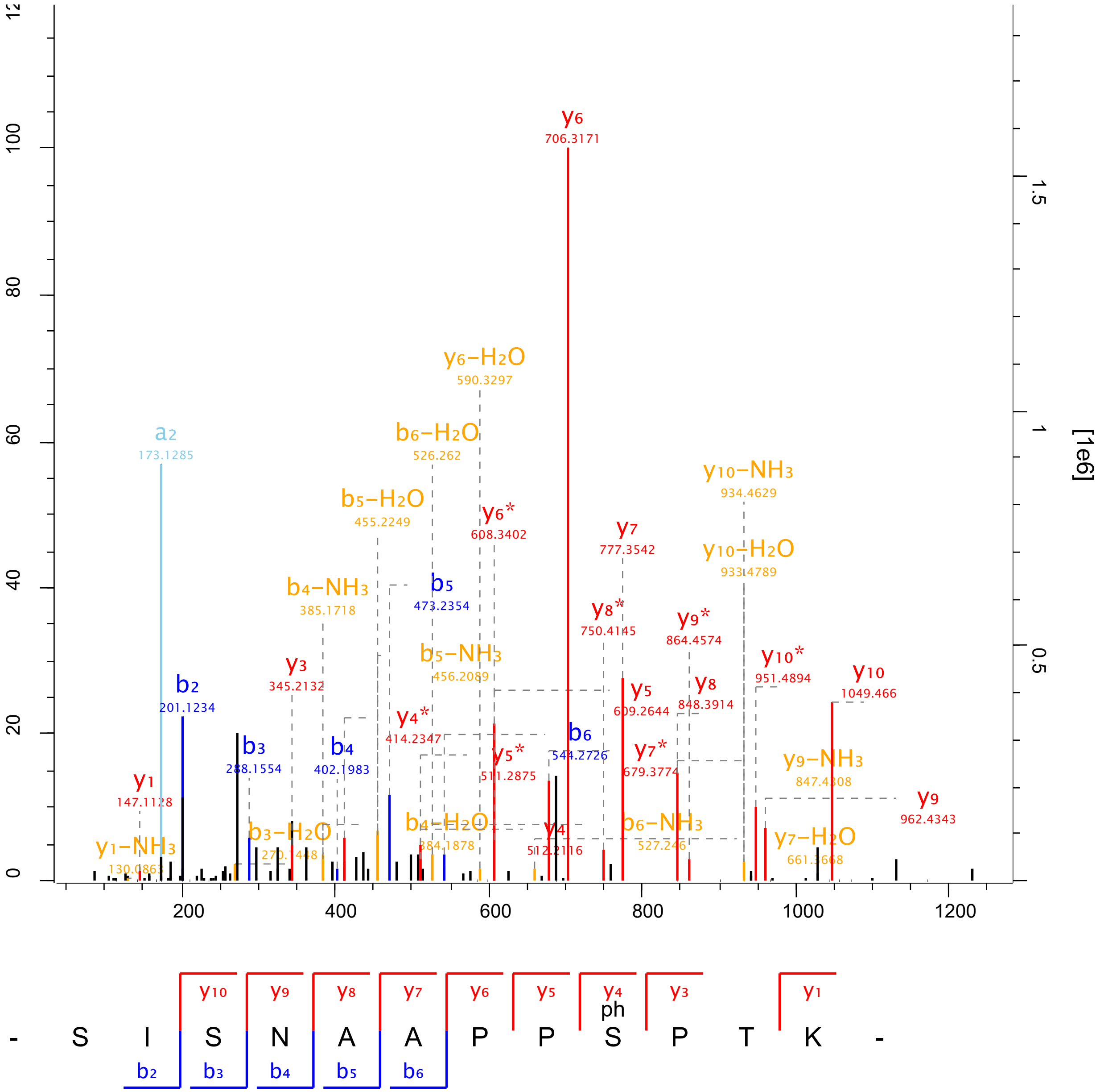

| Raw file | Scan | Method | Score | m/z |
| --- | --- | --- | --- | --- |
| A17_30_1 | 15446 | FTMS; HCD | 109.65 | 859.85 |

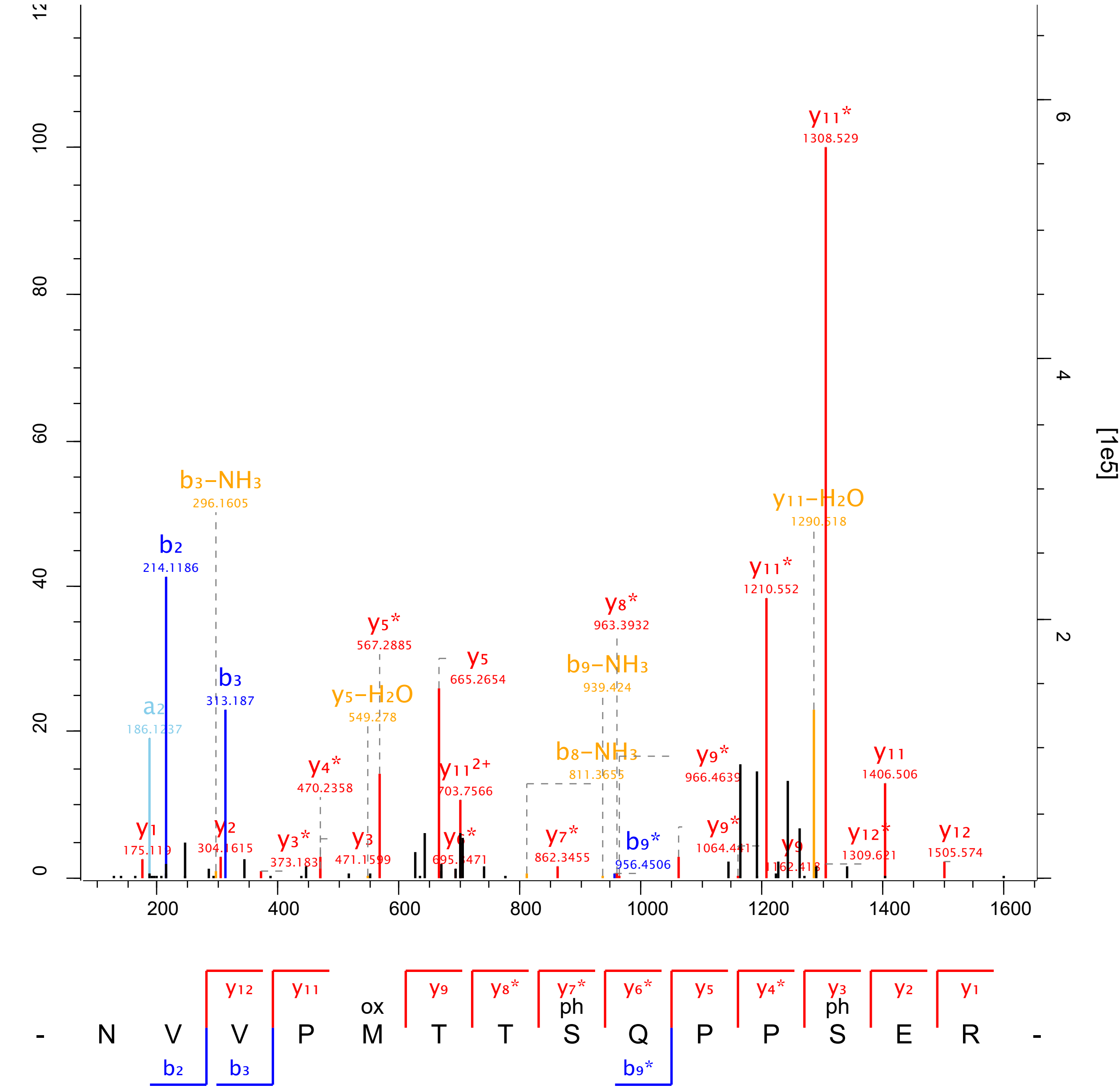

| Raw file | Scan | Method | Score | m/z |
|---|---|---|---|---|
| A17_30_1 | 20082 | FTMS; HCD | 134.14 | 744.83 |

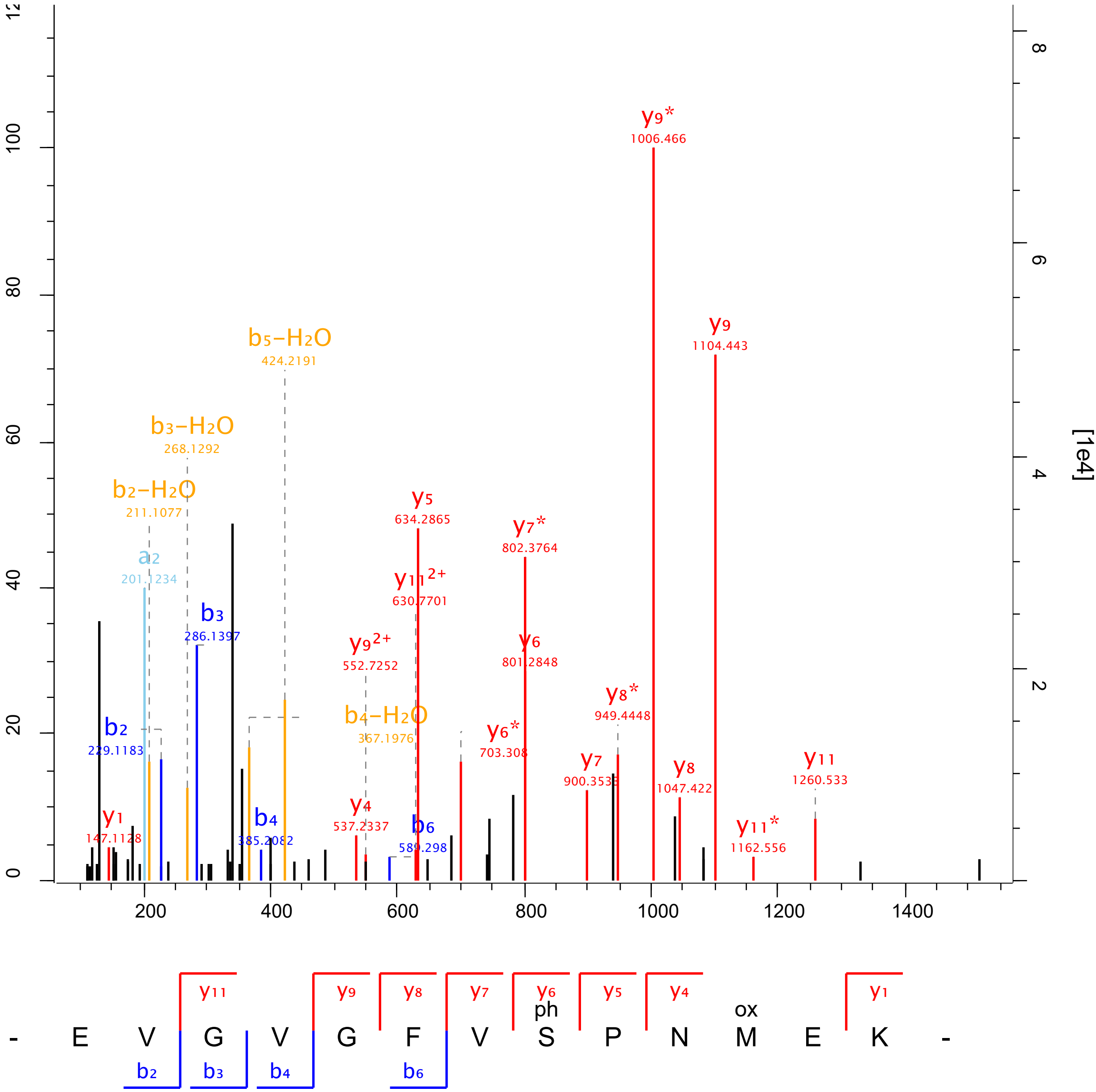

| Raw file | Scan | Method | Score | m/z |
|---|---|---|---|---|
| A17_30_1 | 20314 | FTMS; HCD | 40.94 | 663.79 |

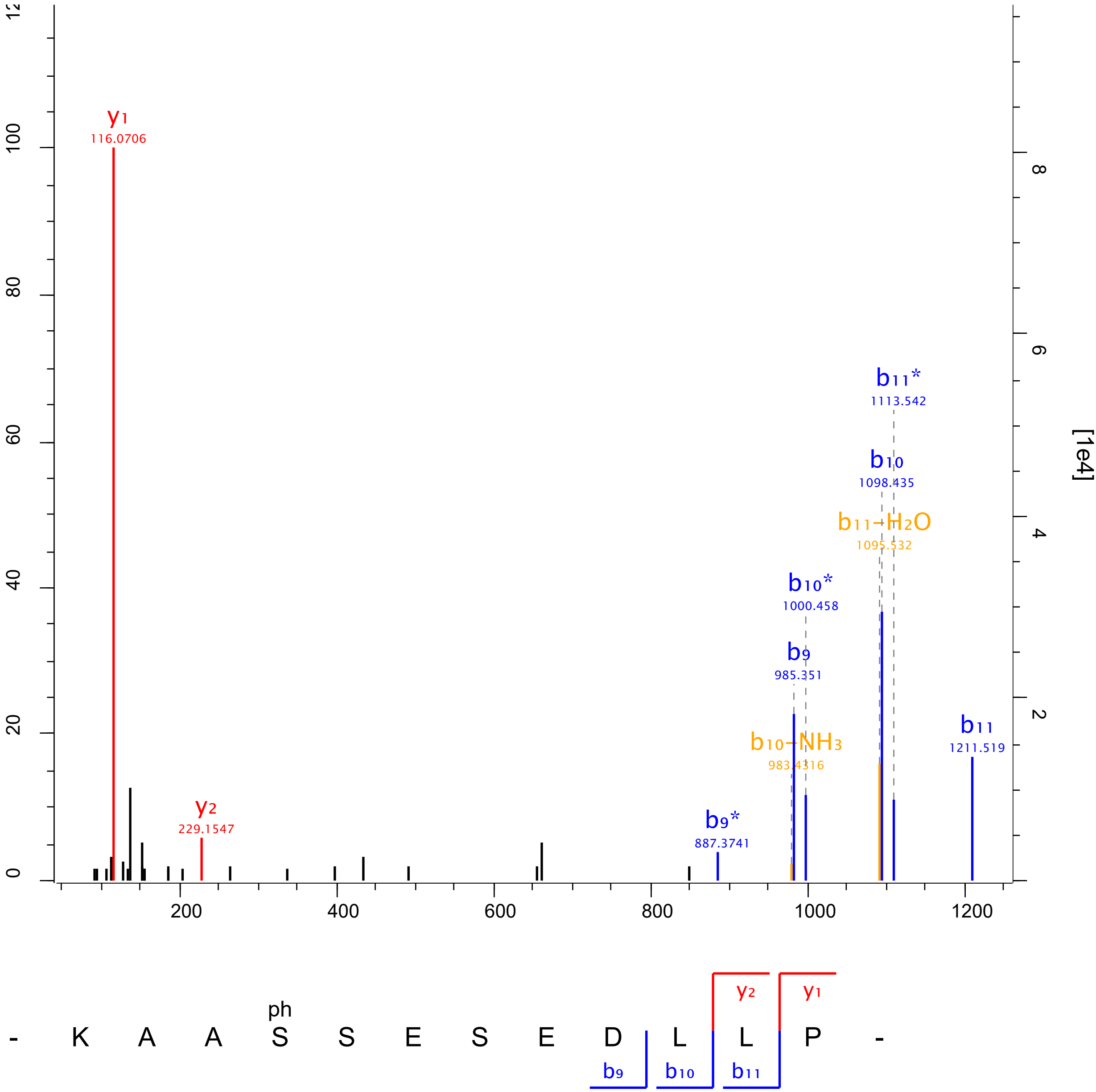

| Raw file | Scan | Method | Score | m/z |
|---|---|---|---|---|
| A17_30_1 | 21537 | FTMS; HCD | 154.69 | 966.39 |

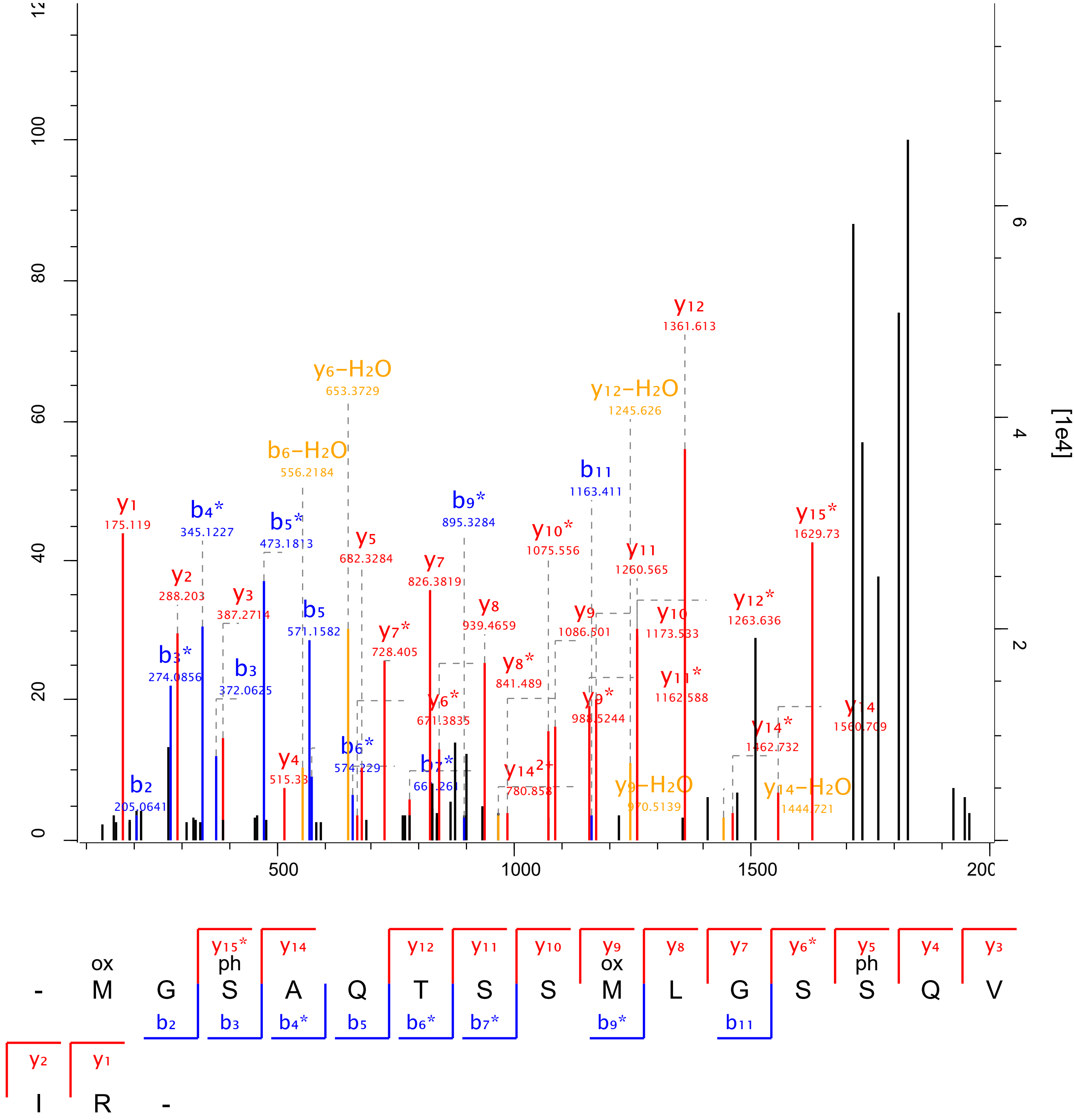

| Raw file | Scan | Method | Score | m/z |
|---|---|---|---|---|
| A17_30_1 | 22464 | FTMS; HCD | 240.63 | 788.88 |

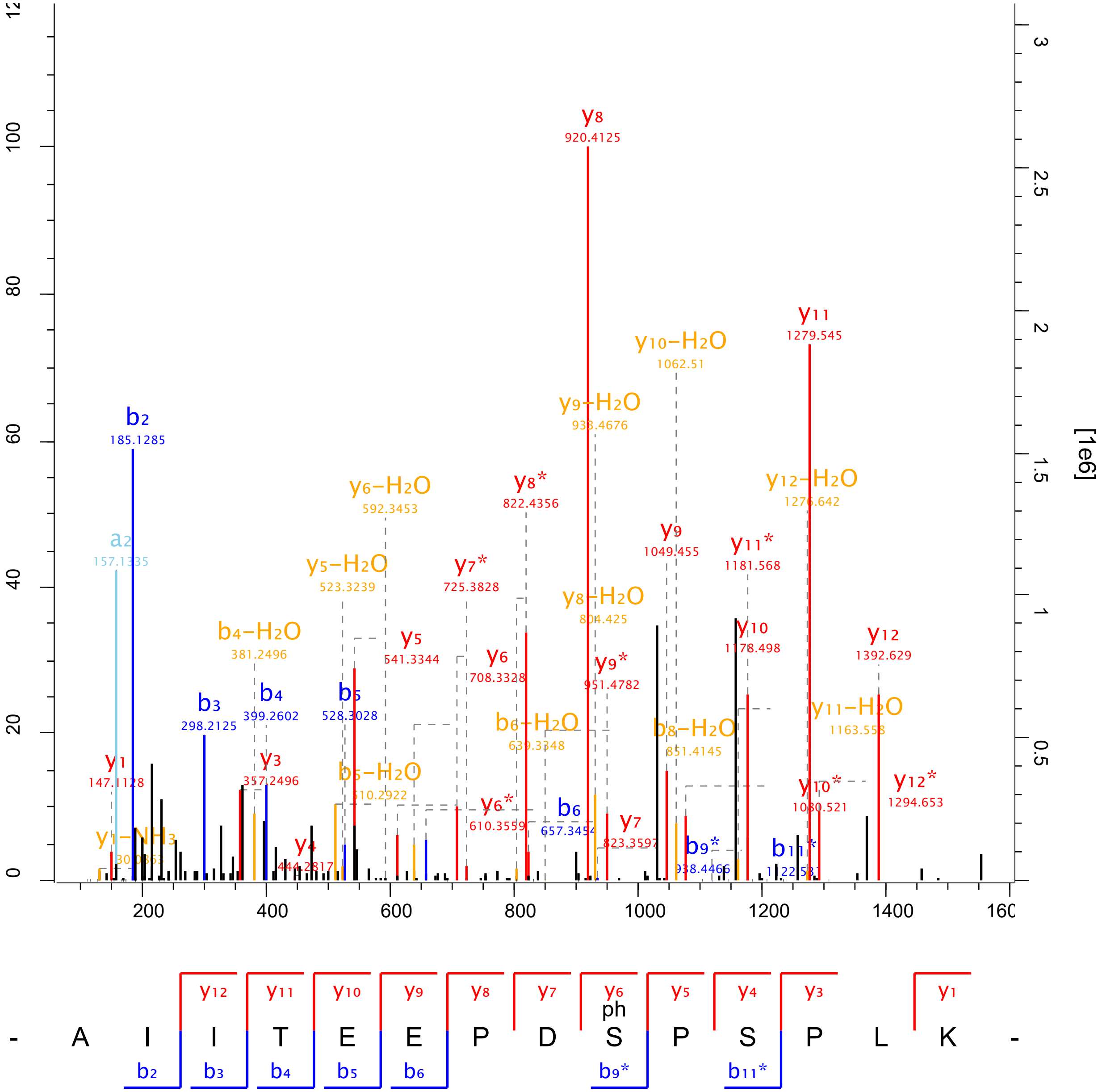

| Raw file | Scan | Method | Score | m/z |
|---|---|---|---|---|
| A17_30_1 | 2360 | FTMS; HCD | 52.58 | 472.74 |

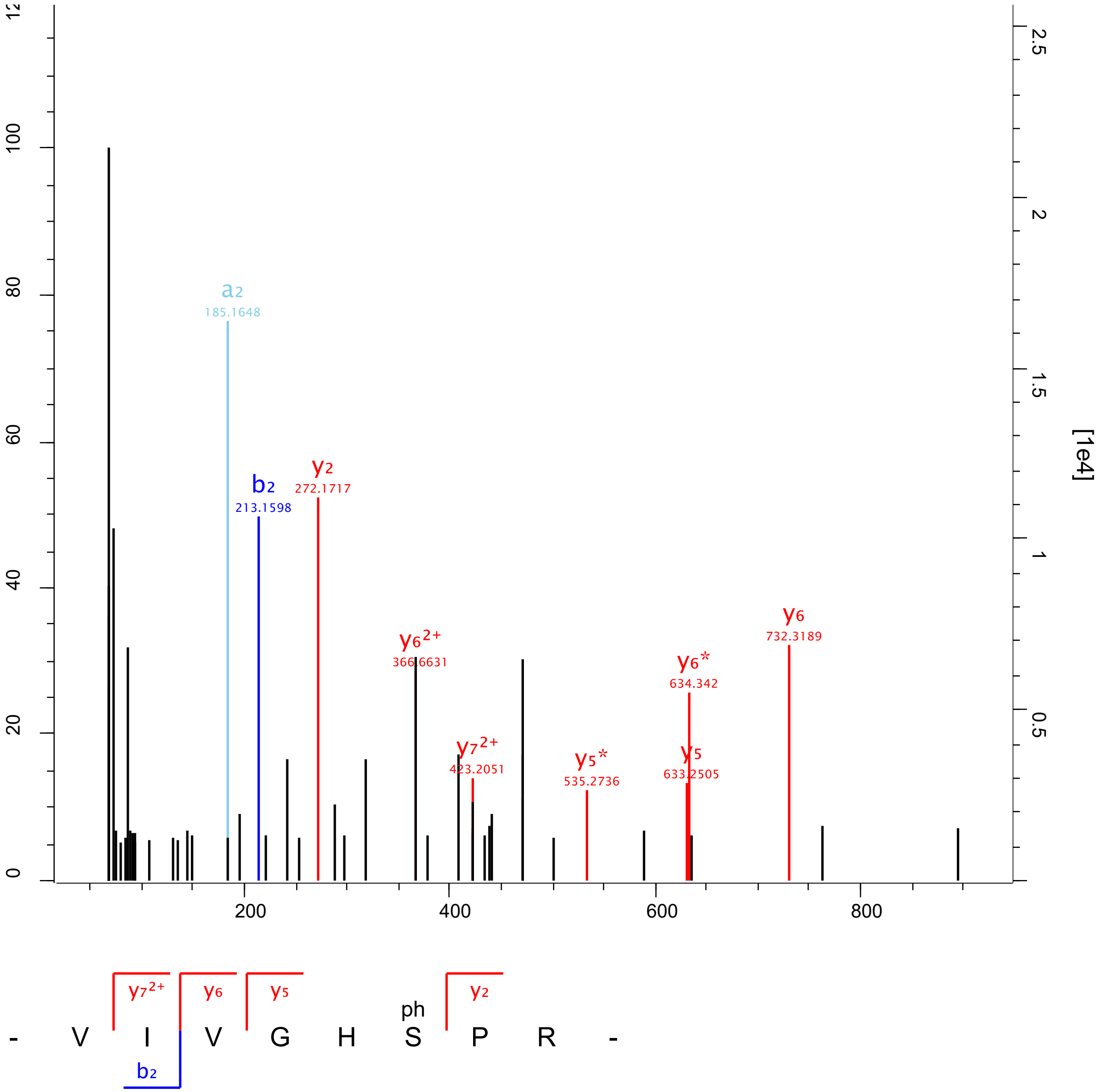

| Raw file | Scan | Method | Score | m/z |
|---|---|---|---|---|
| A17_30_1 | 25373 | FTMS; HCD | 242.8 | 809.35 |

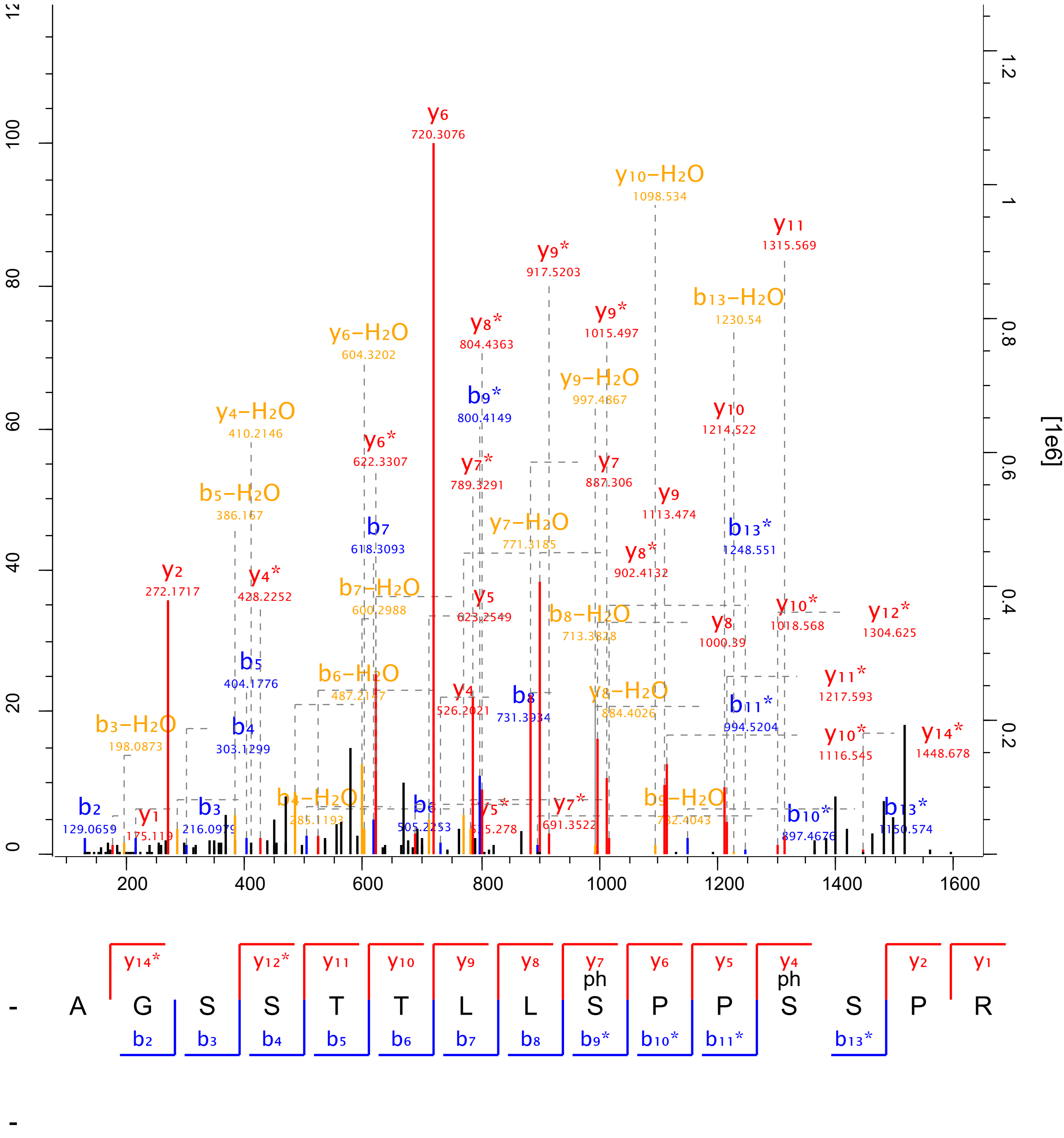

| Raw file | Scan | Method | Score | m/z |
|---|---|---|---|---|
| A17_30_1 | 26746 | FTMS; HCD | 44.25 | 694.84 |

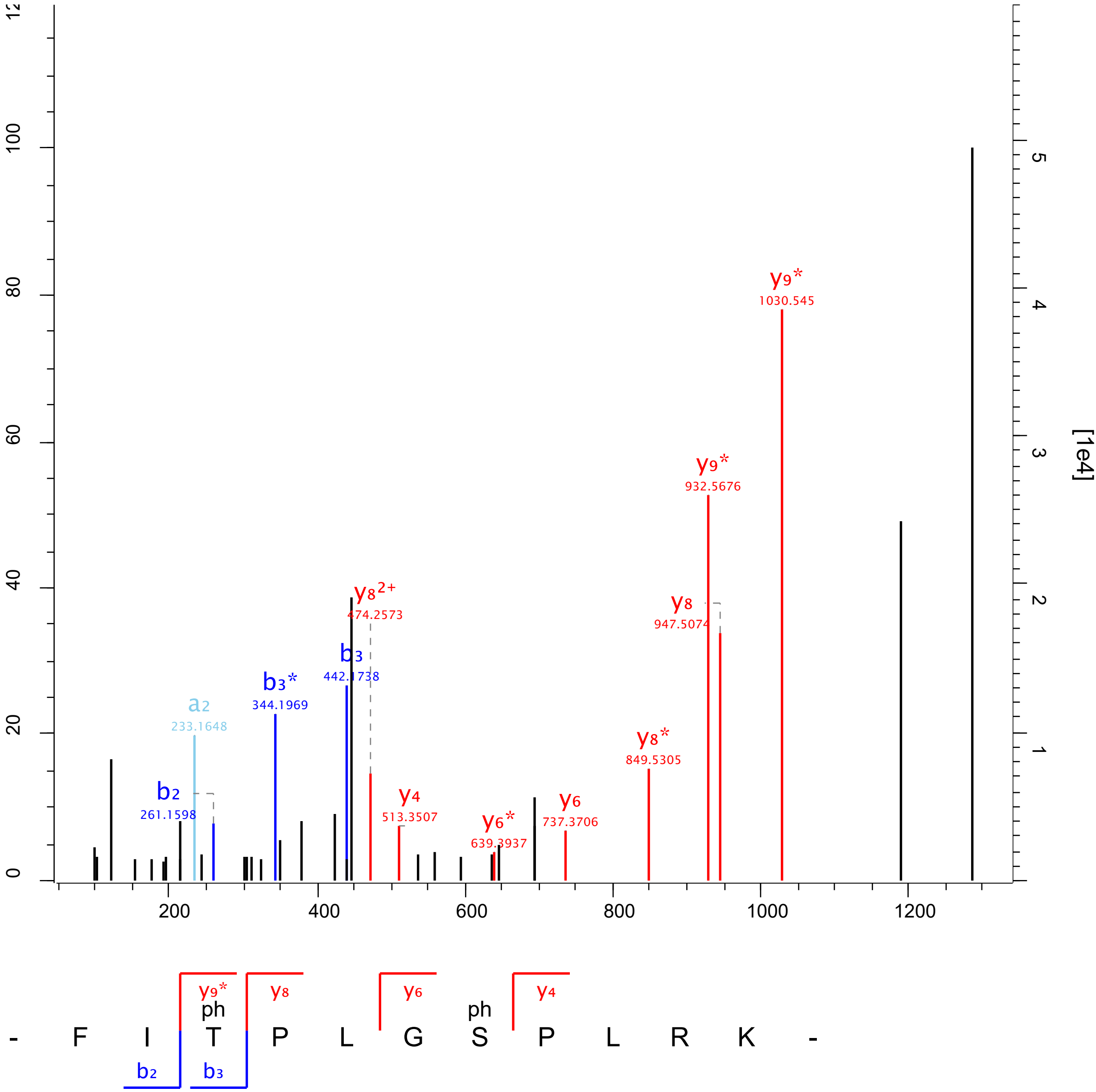

| Raw file | Scan | Method | Score | m/z |
|---|---|---|---|---|
| A17_30_1 | 27233 | FTMS; HCD | 86.77 | 893.73 |

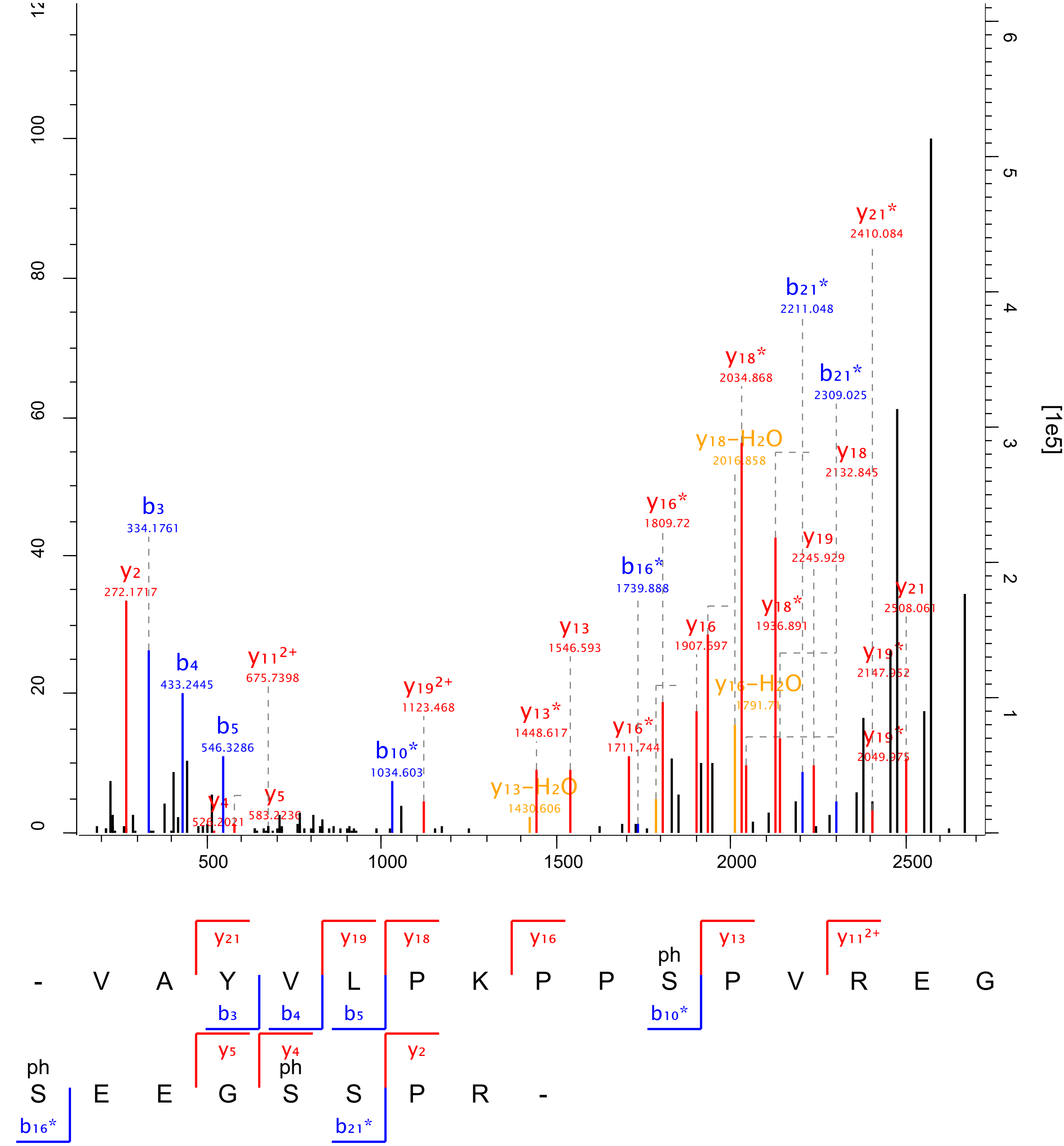

| Raw file | Scan | Method | Score | m/z |
|---|---|---|---|---|
| A17_30_1 | 27389 | FTMS; HCD | 155.9 | 1053.77 |

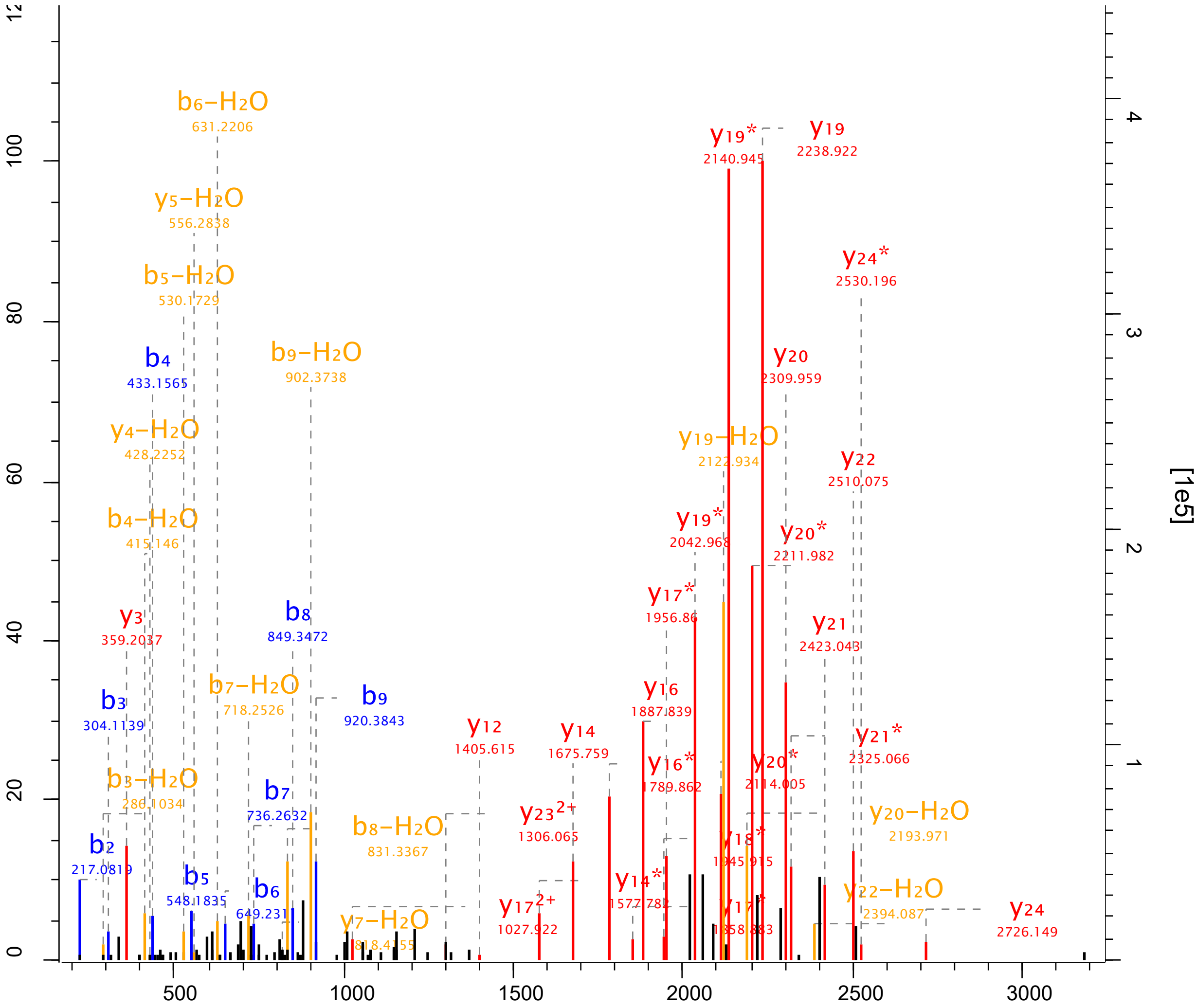


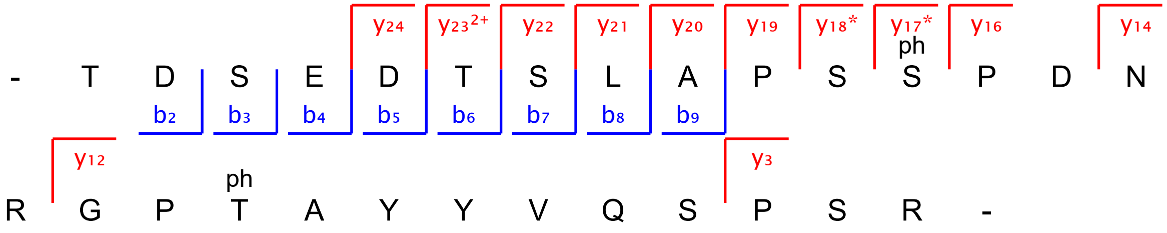

| Raw file | Scan | Method | Score | m/z |
| --- | --- | --- | --- | --- |
| A17_30_1 | 27827 | FTMS; HCD | 147.7 | 692.34 |

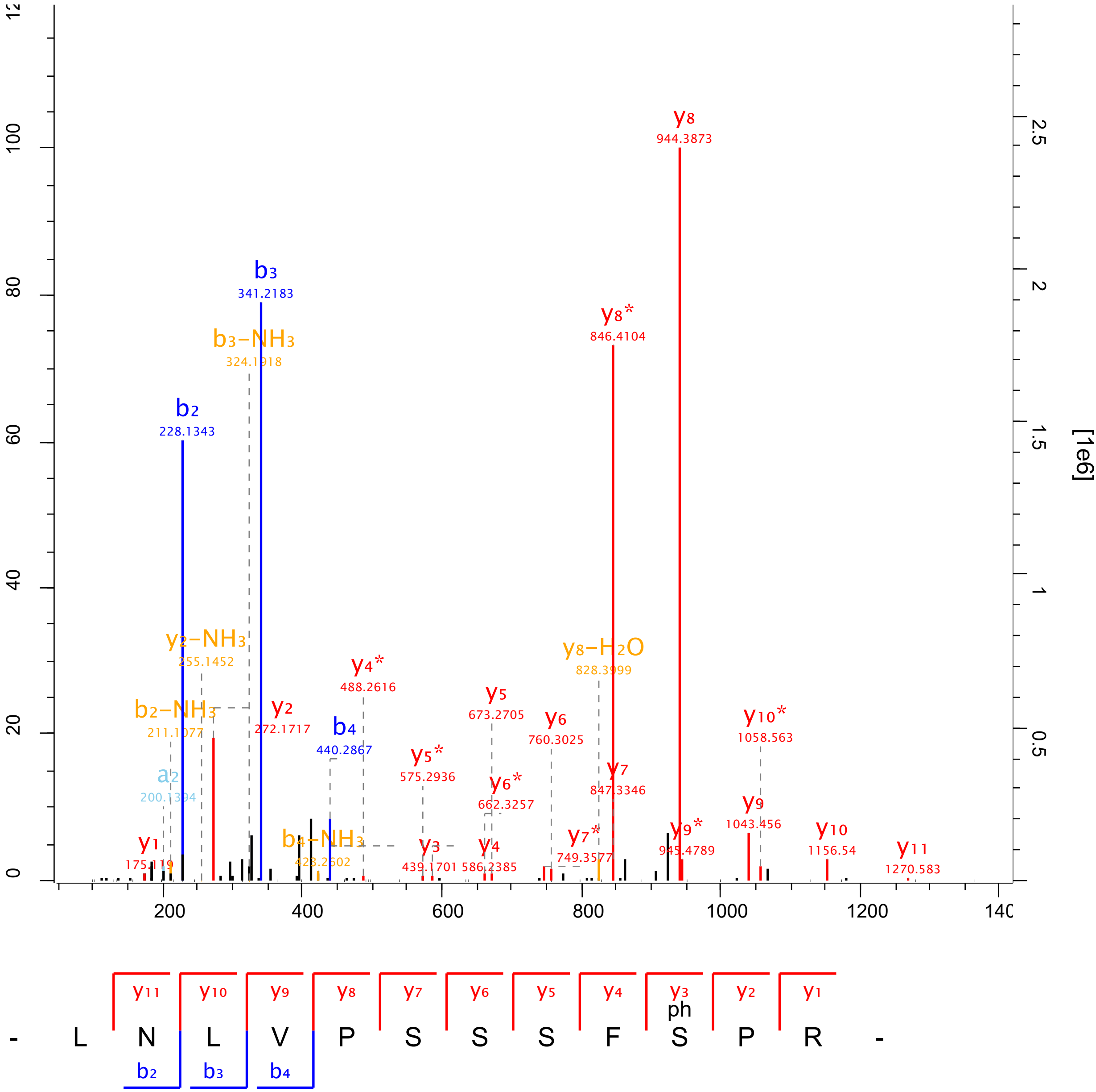

| Raw file | Scan | Method | Score | m/z |
|---|---|---|---|---|
| A17_30_1 | 3433 | FTMS; HCD | 97.73 | 567.74 |

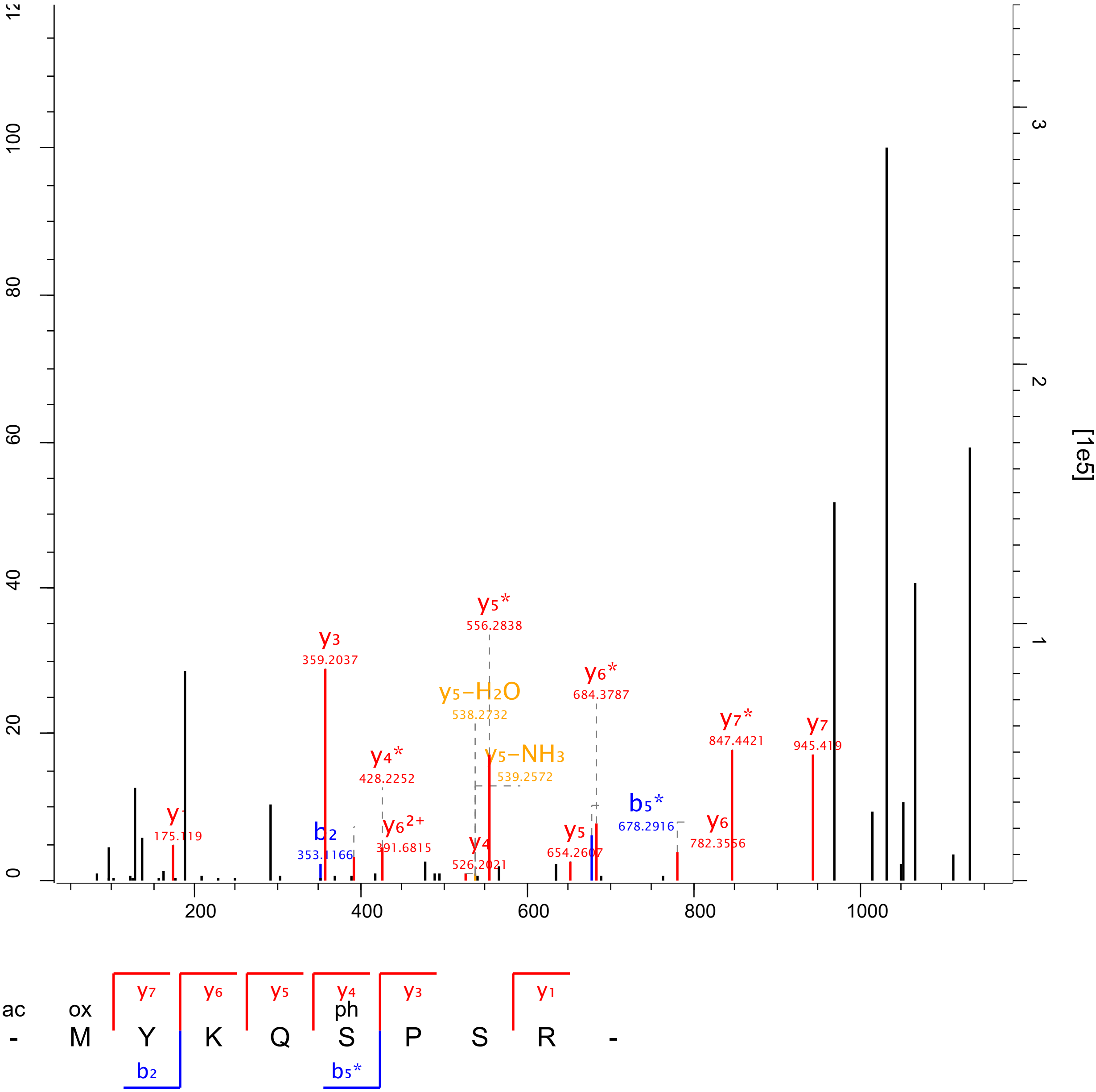

| Raw file | Scan | Method | Score | m/z |
|---|---|---|---|---|
| A17_30_1 | 4053 | FTMS; HCD | 40.94 | 616.26 |

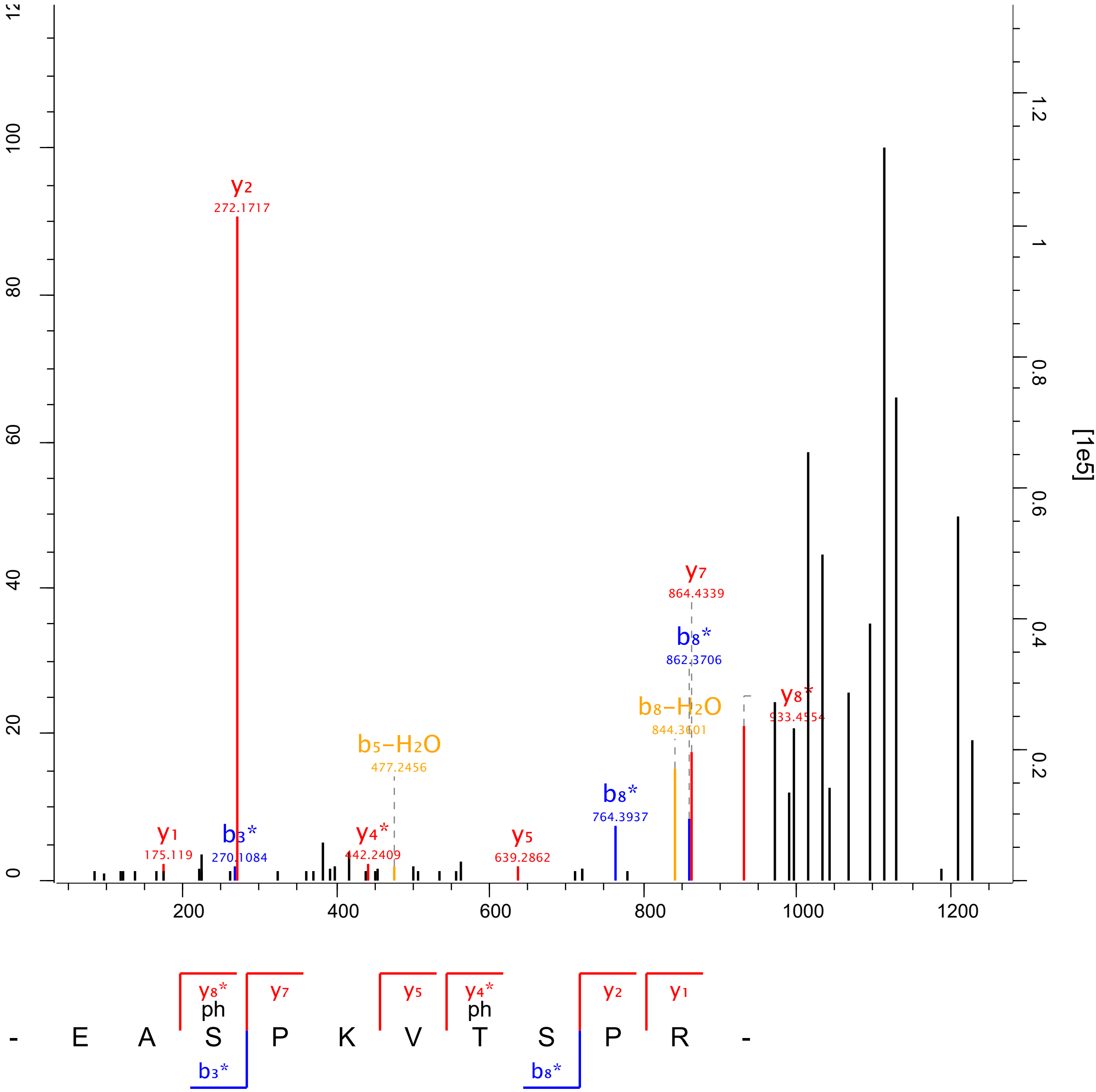

| Raw file | Scan | Method | Score | m/z |
| --- | --- | --- | --- | --- |
| A17_30_1 | 5074 | FTMS; HCD | 55.35 | 654.75 |

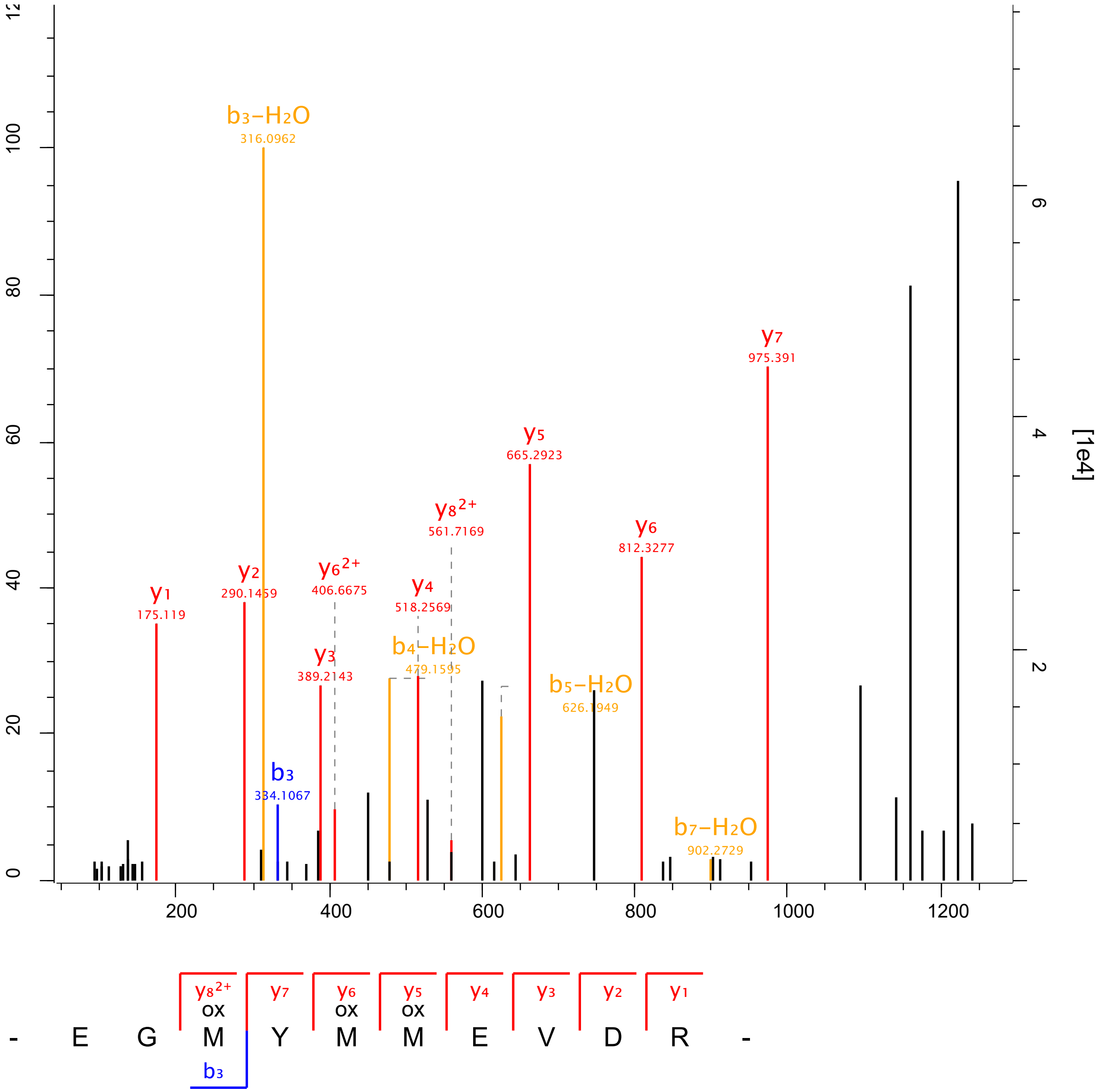

| Raw file | Scan | Method | Score | m/z |
| --- | --- | --- | --- | --- |
| A17_30_1 | 5871 | FTMS; HCD | 73.5 | 737.76 |

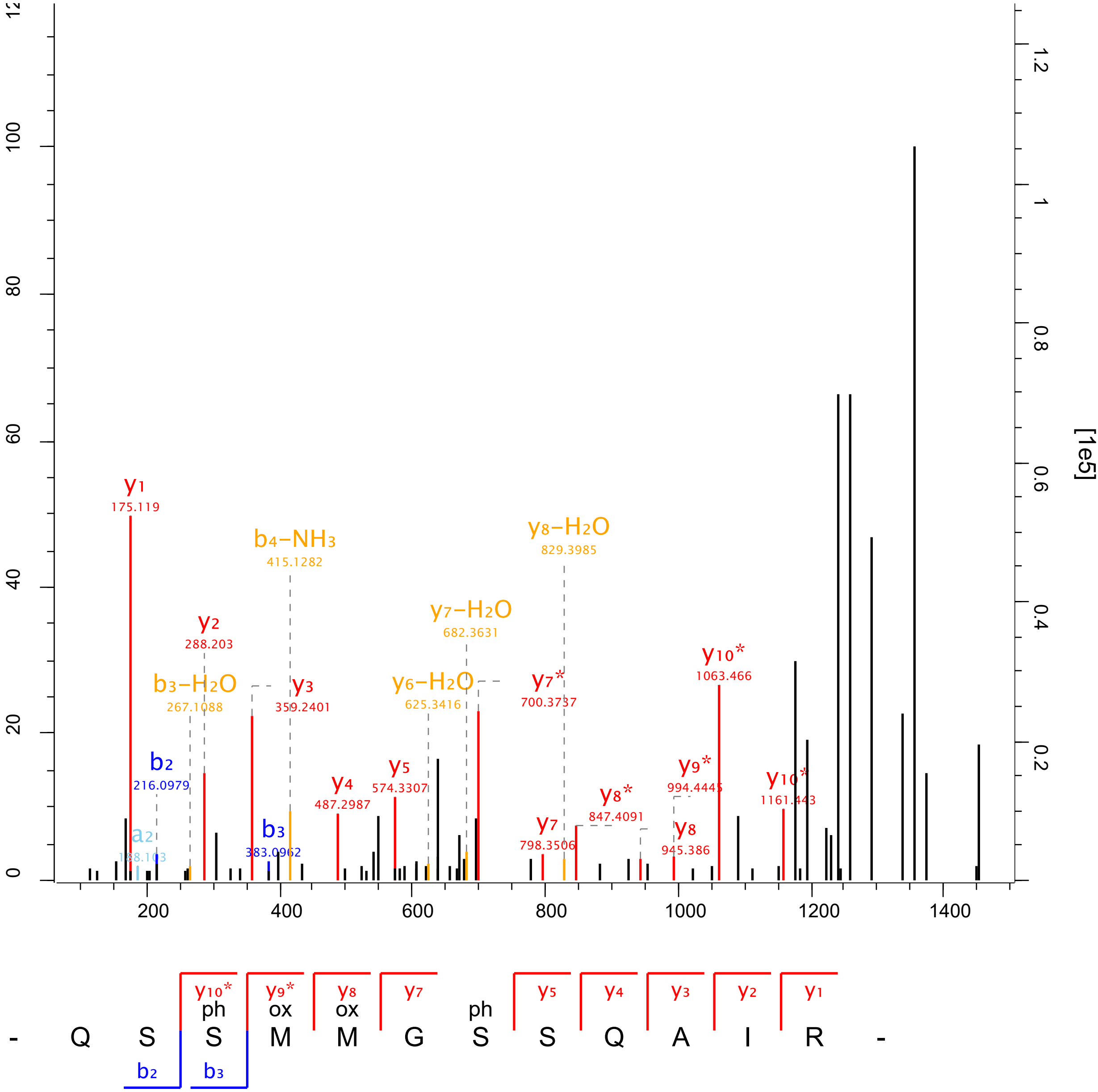

| Raw file | Scan | Method | Score | m/z |
|---|---|---|---|---|
| A17_30_1 | 7251 | FTMS; HCD | 119.39 | 606.72 |

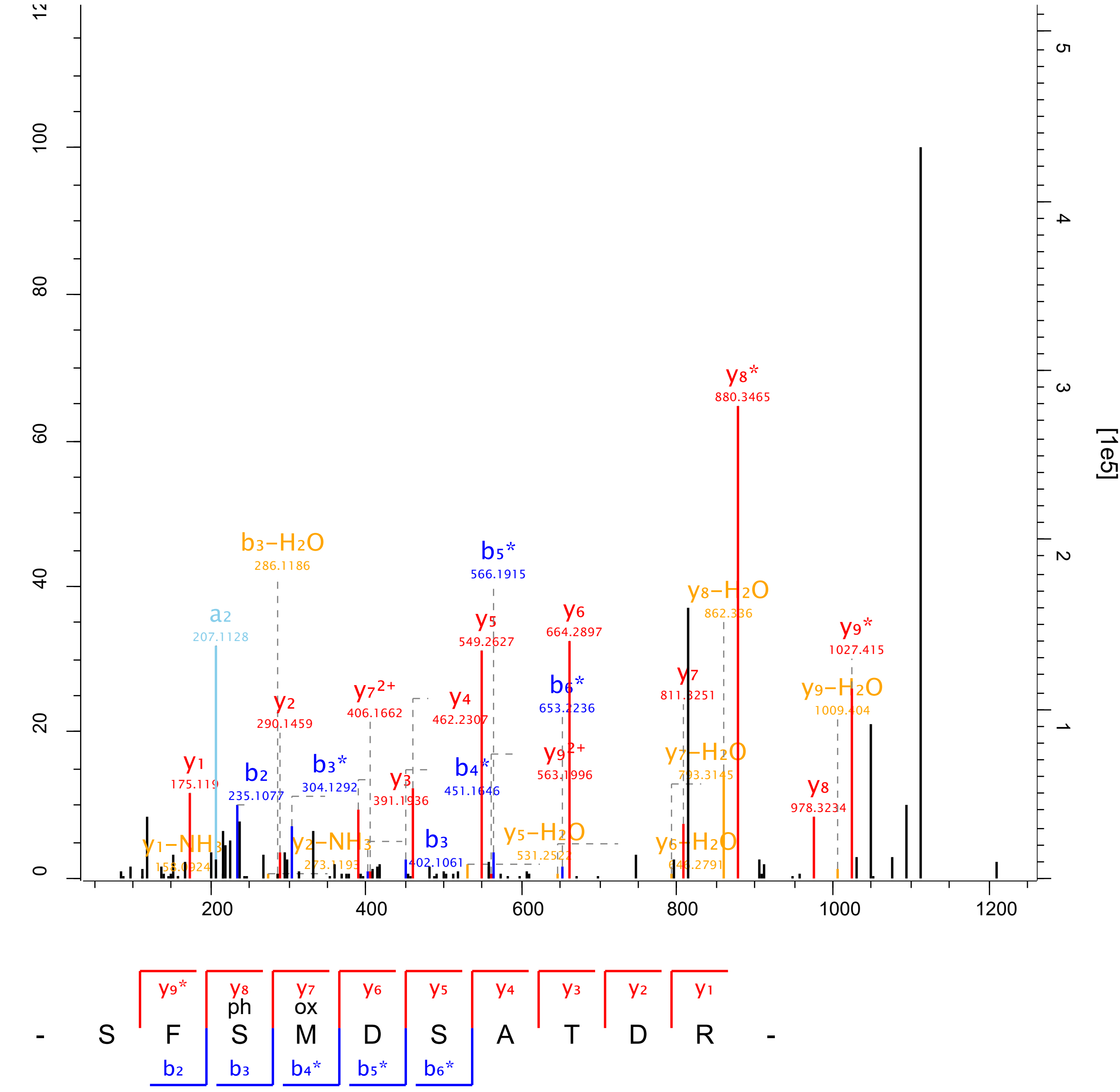

| Raw file | Scan | Method | Score | m/z |
|---|---|---|---|---|
| A17_30_1 | 7789 | FTMS; HCD | 114.89 | 456.16 |

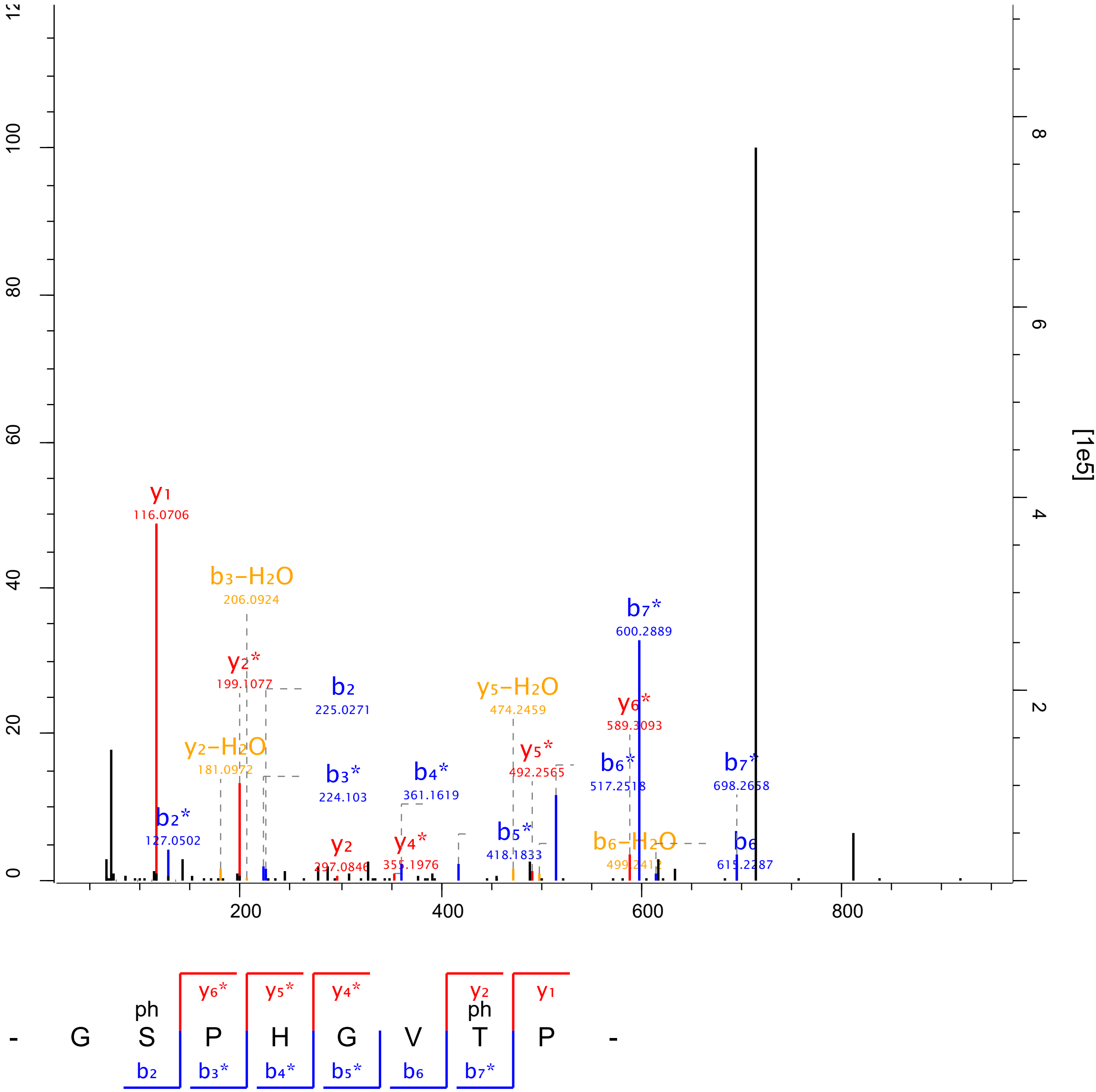

| Raw file | Scan | Method | Score | m/z |
|---|---|---|---|---|
| A17_30_1 | 8299 | FTMS; HCD | 127.07 | 722.3 |

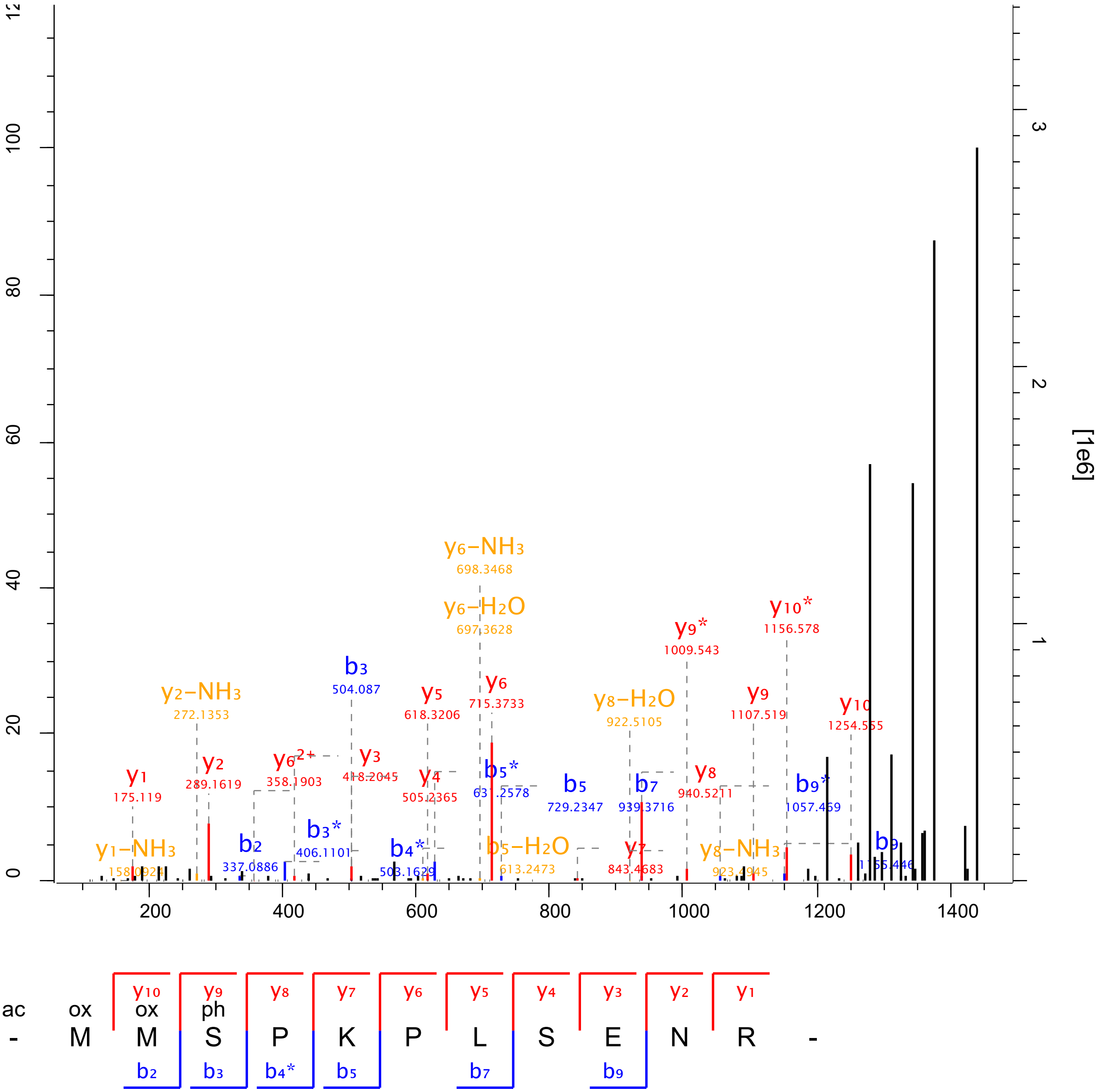

| Raw file | Scan | Method | Score | m/z |
| --- | --- | --- | --- | --- |
| A17_30_2 | 10014 | FTMS; HCD | 79.61 | 621.27 |

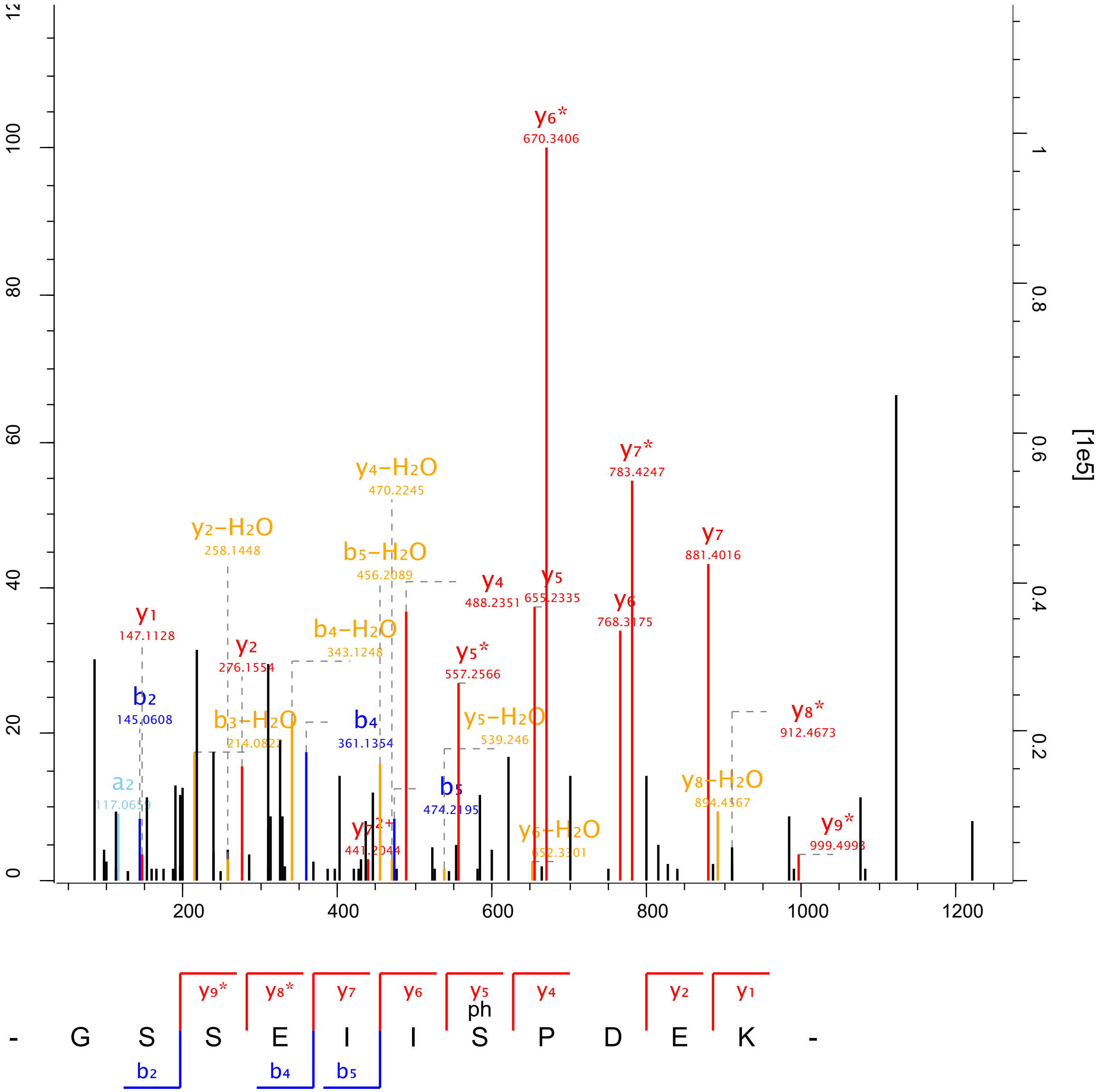

| Raw file | Scan | Method | Score | m/z |
|---|---|---|---|---|
| A17_30_2 | 10873 | FTMS; HCD | 239.58 | 756.34 |

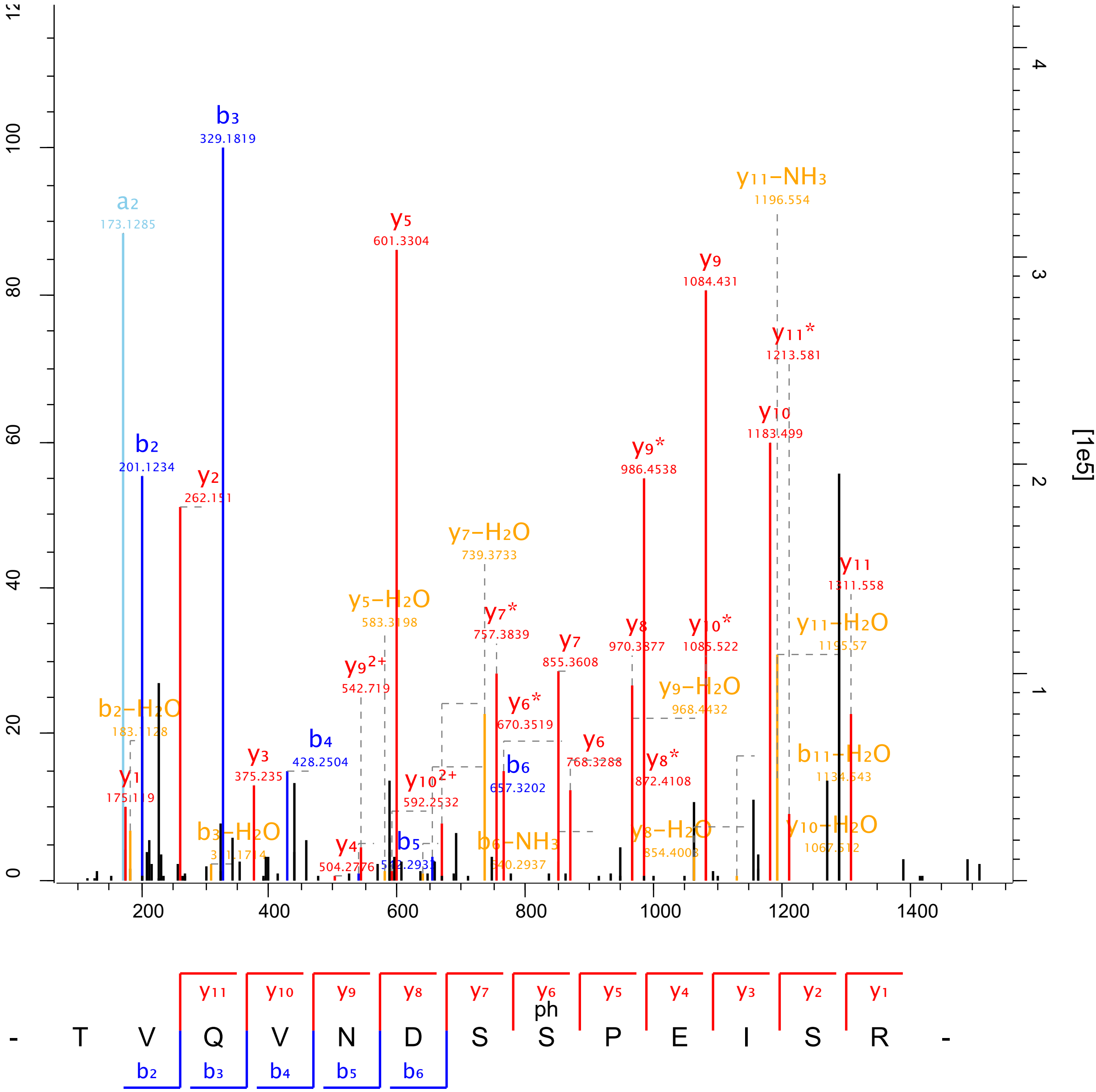

| Raw file | Scan | Method | Score | m/z |
|---|---|---|---|---|
| A17_30_2 | 11543 | FTMS; HCD | 189.17 | 514.22 |

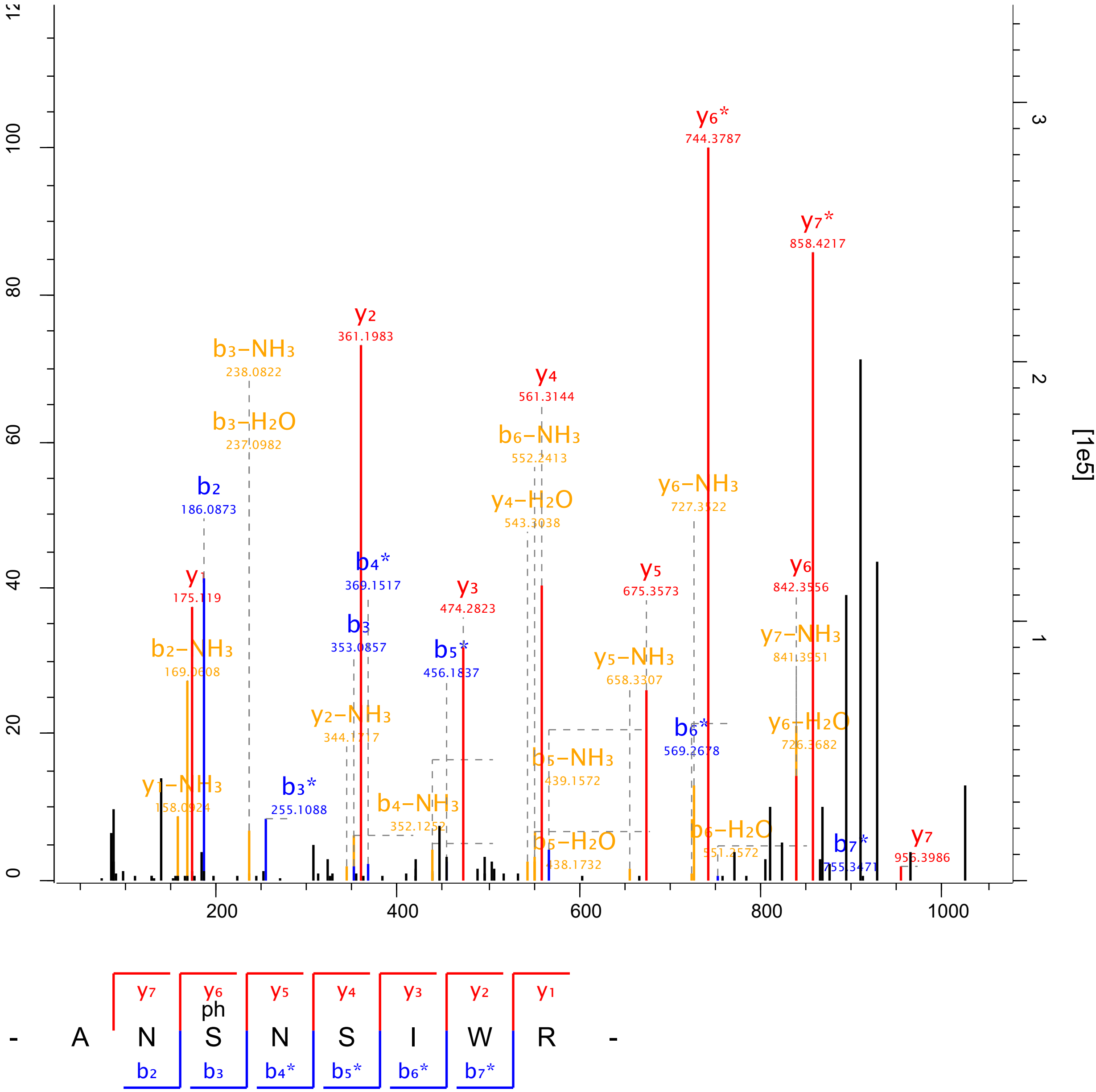

| Raw file | Scan | Method | Score | m/z |
|---|---|---|---|---|
| A17_30_2 | 11578 | FTMS; HCD | 137.69 | 614.29 |

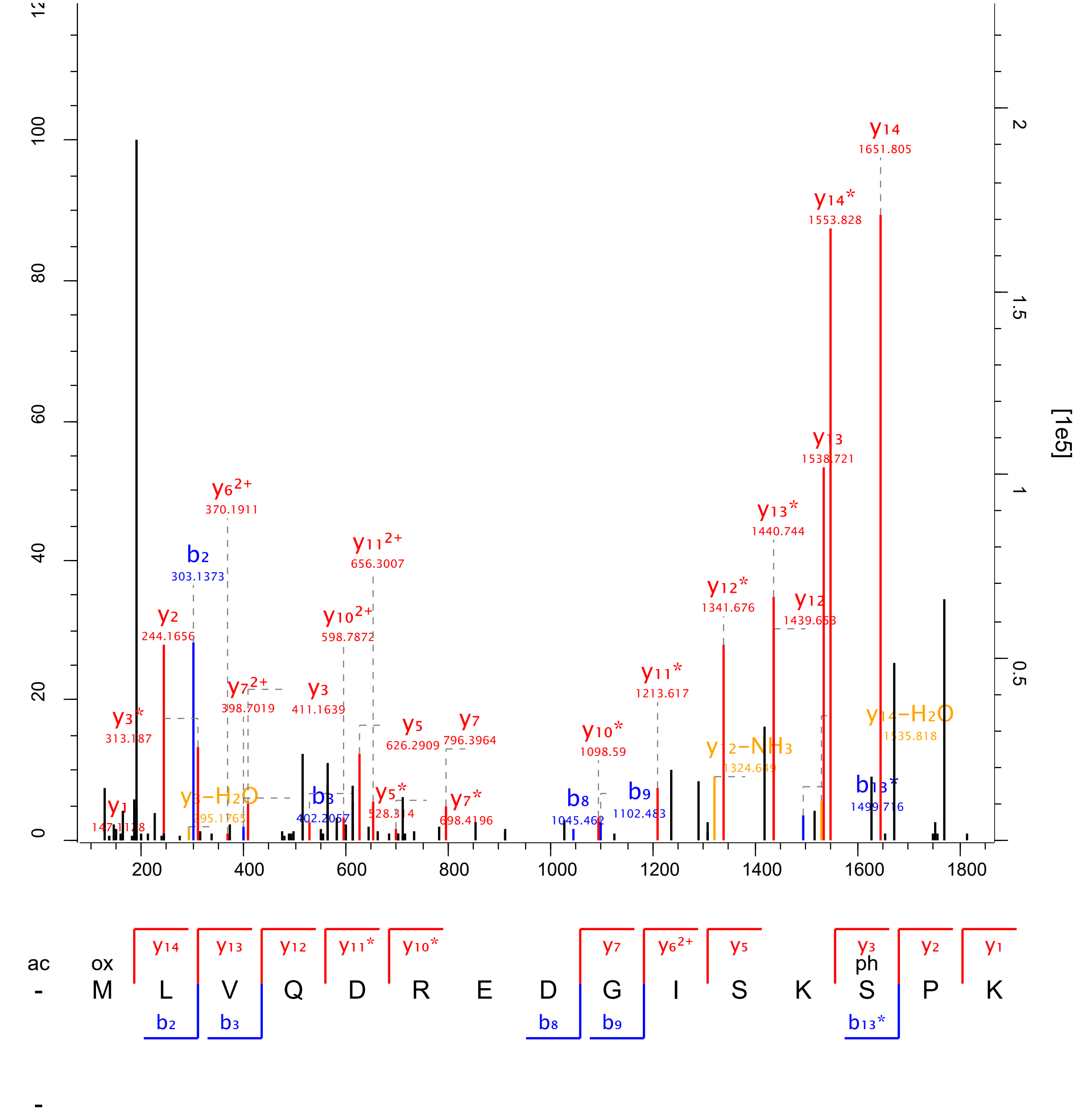

| Raw file | Scan | Method | Score | m/z |
| --- | --- | --- | --- | --- |
| A17_30_2 | 11809 | FTMS; HCD | 144.88 | 590.21 |

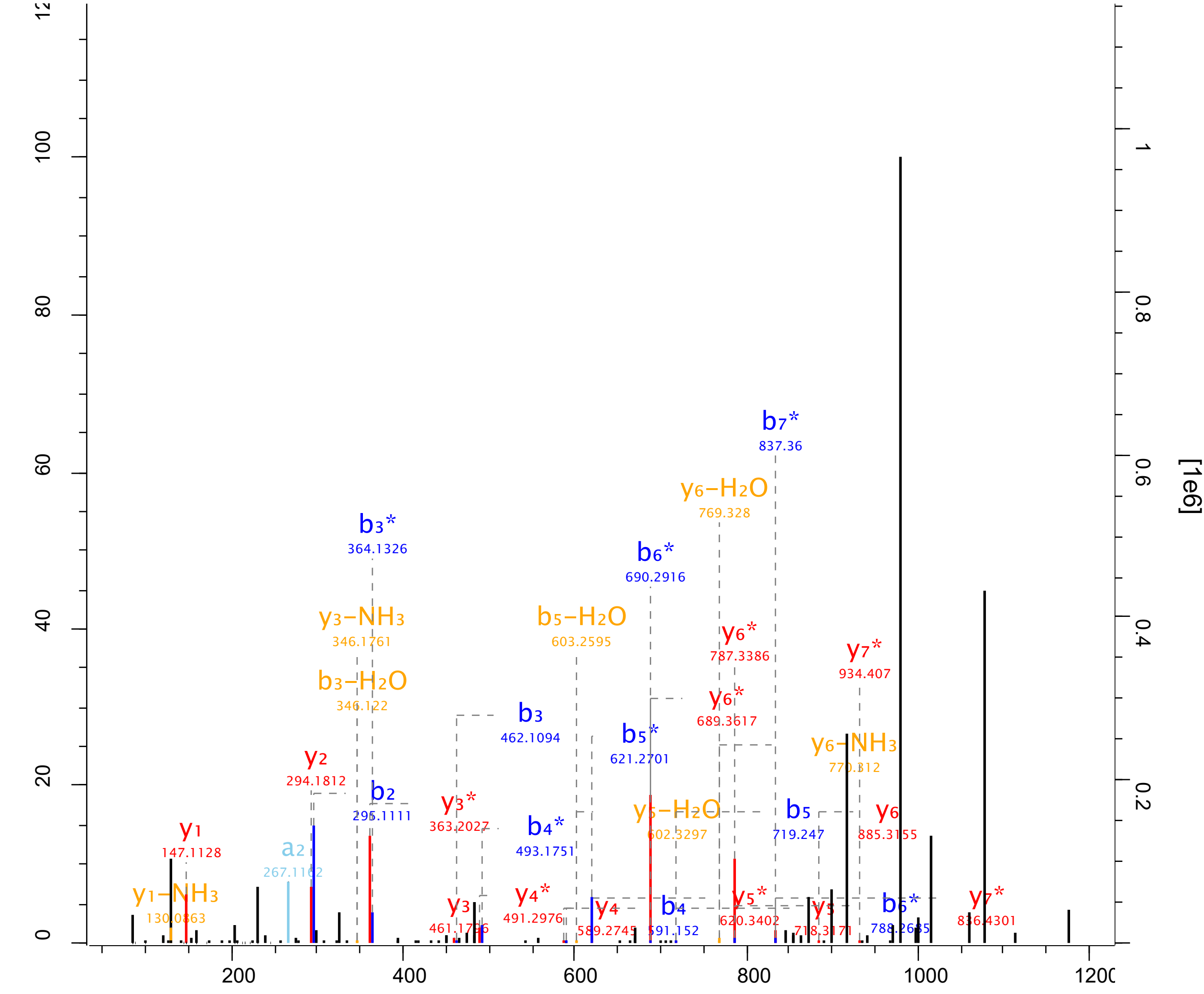


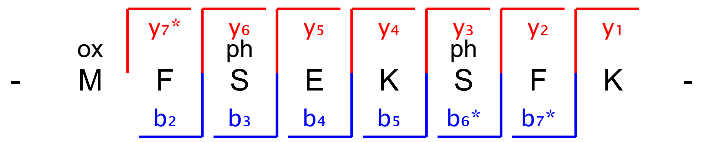

| Raw file | Scan | Method | Score | m/z |
| --- | --- | --- | --- | --- |
| A17_30_2 | 12601 | FTMS; HCD | 87.65 | 714.31 |

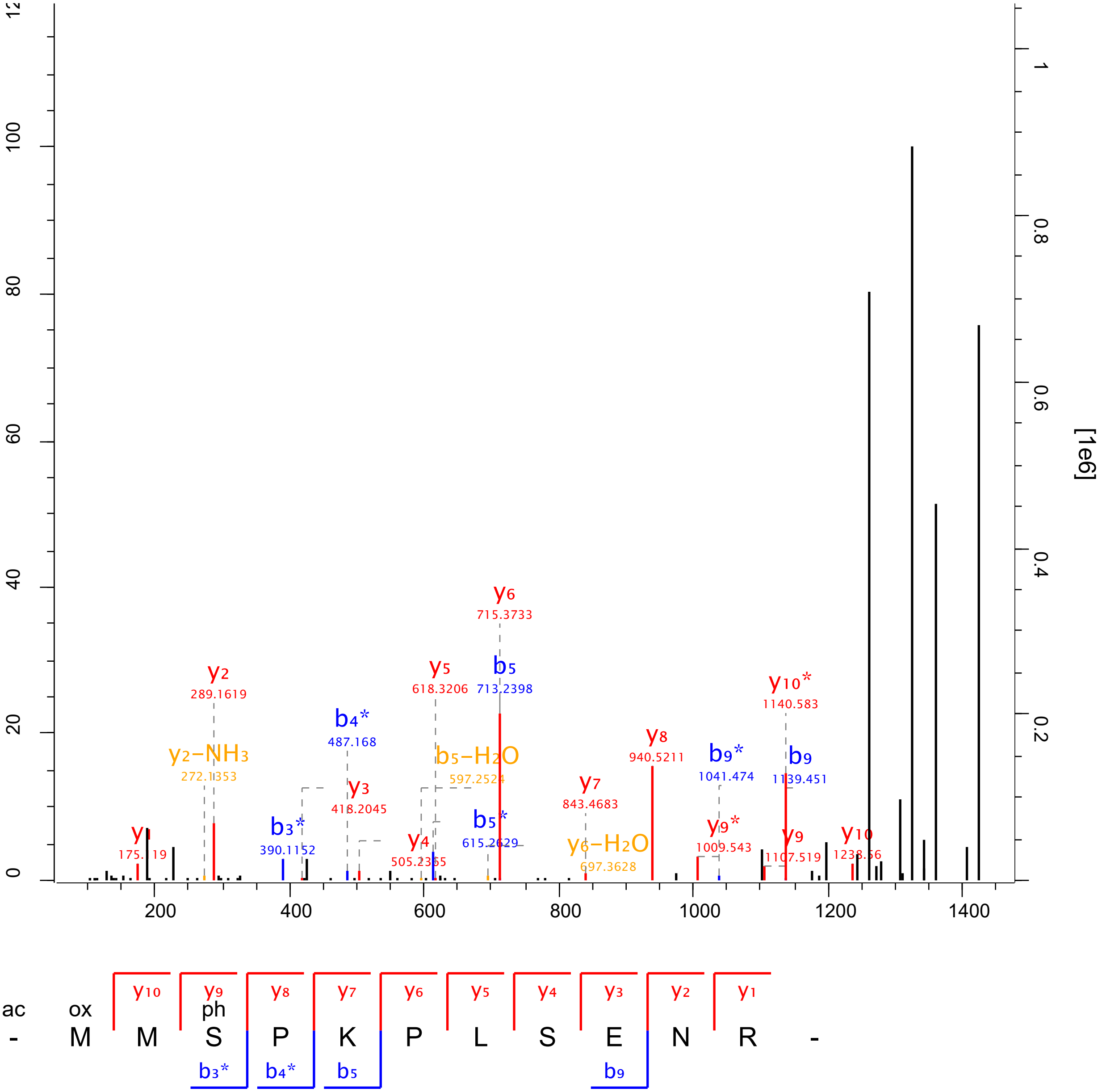

| Raw file | Scan | Method | Score | m/z |
|---|---|---|---|---|
| A17_30_2 | 13066 | FTMS; HCD | 99.06 | 656.27 |

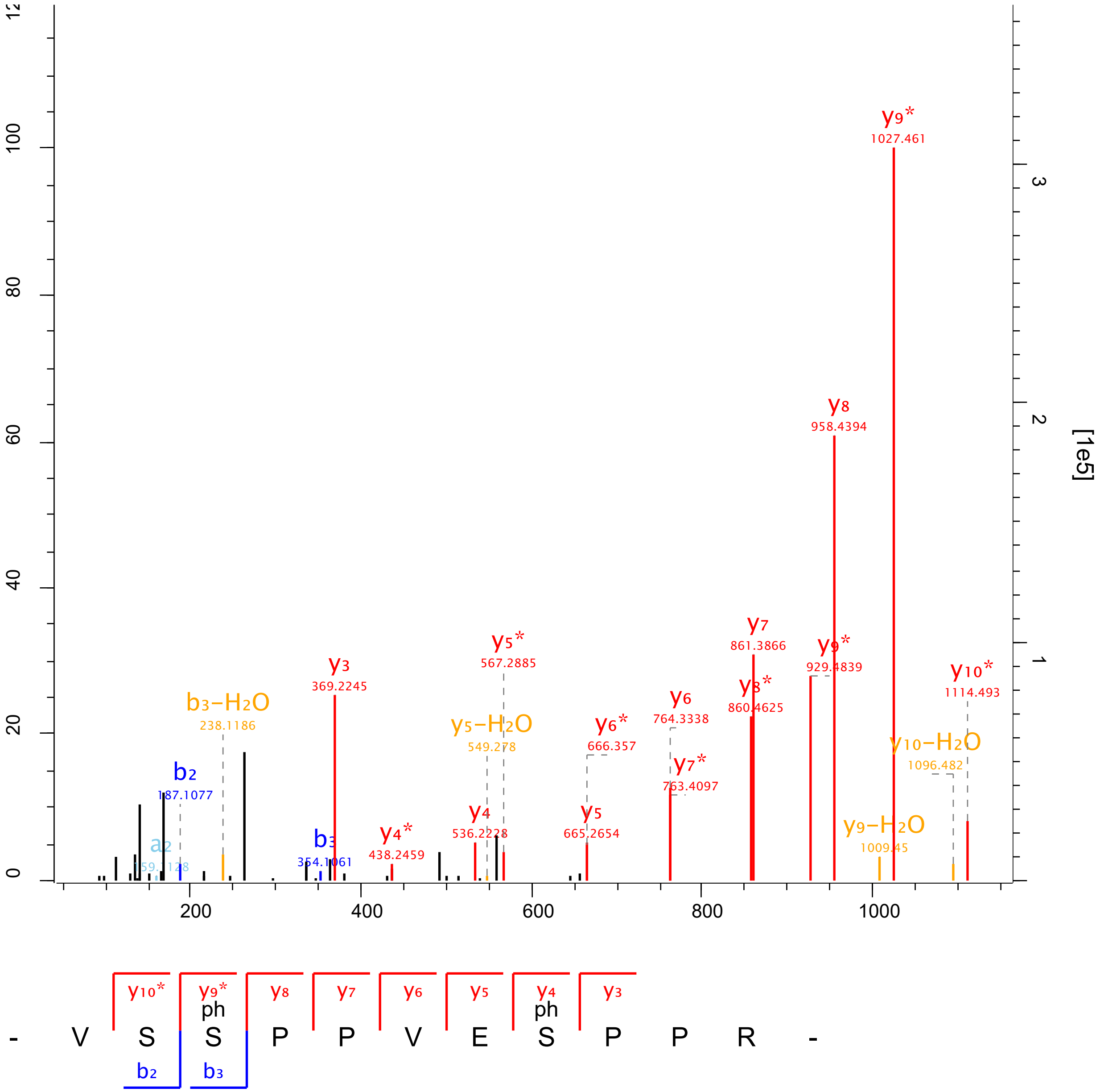

| Raw file | Scan | Method | Score | m/z |
| --- | --- | --- | --- | --- |
| A17_30_2 | 13068 | FTMS; HCD | 98.58 | 610.27 |

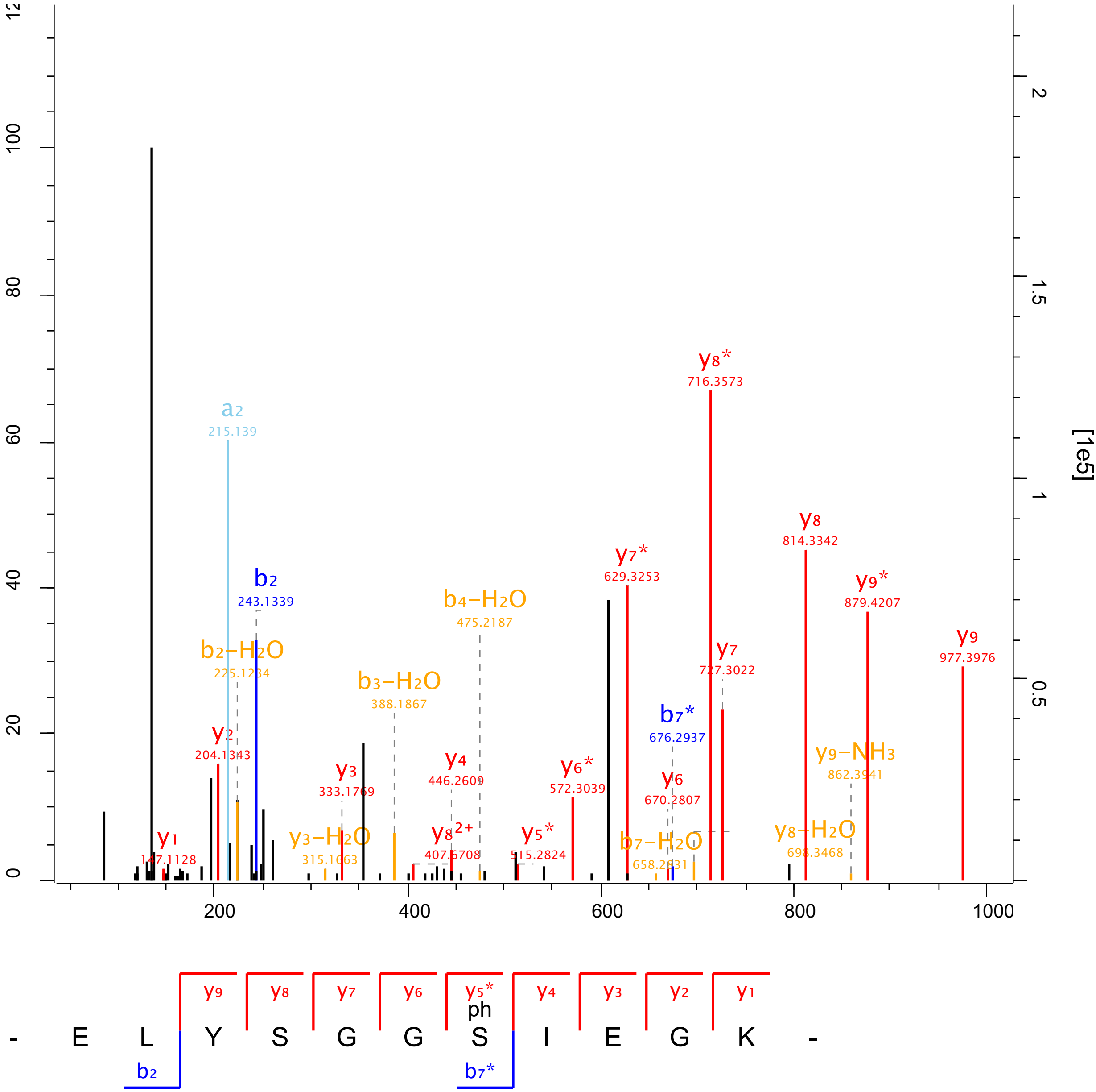

| Raw file | Scan | Method | Score | m/z |
| --- | --- | --- | --- | --- |
| A17_30_2 | 13789 | FTMS; HCD | 102.87 | 587.76 |

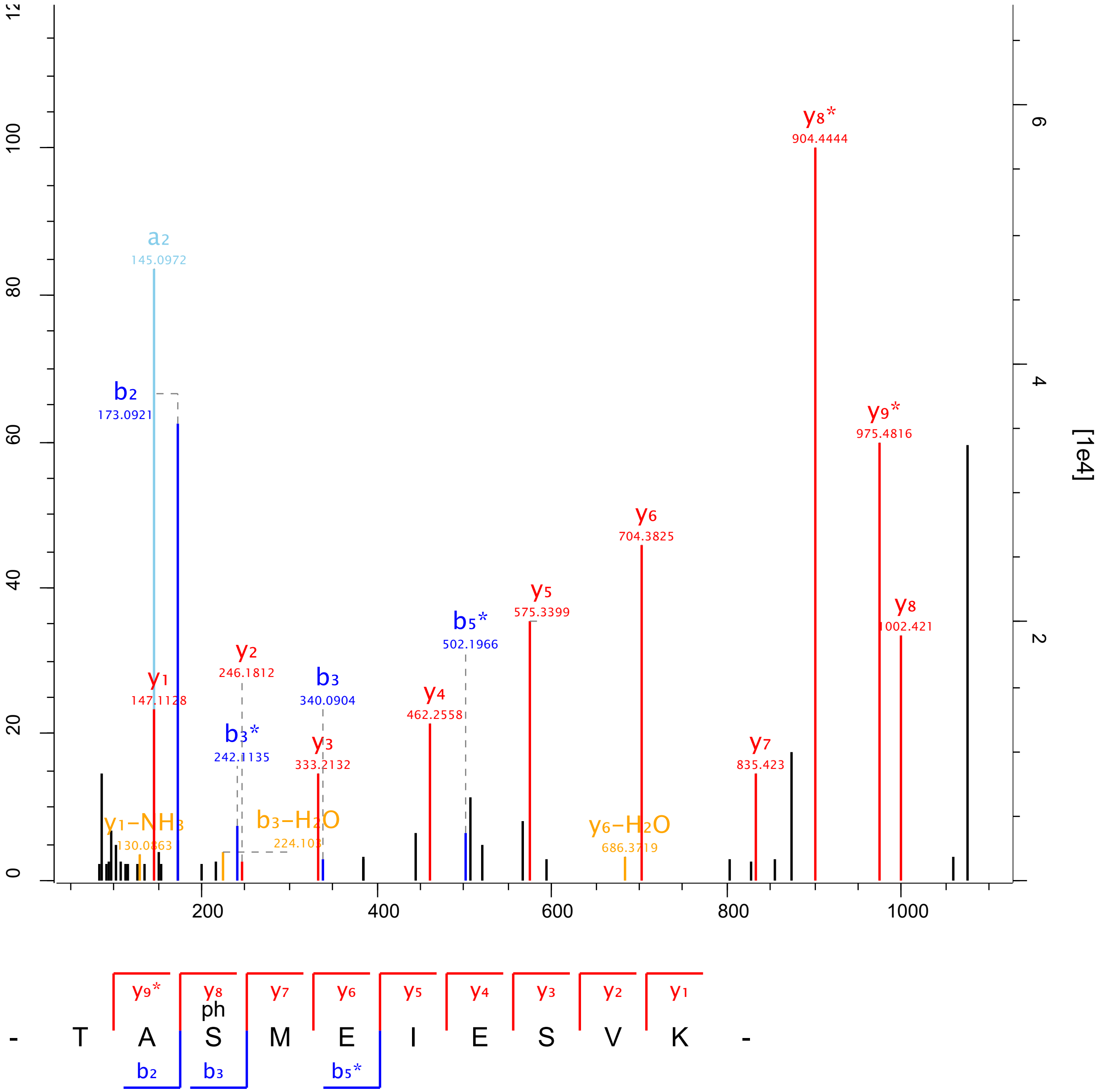

| Raw file | Scan | Method | Score | m/z |
|---|---|---|---|---|
| A17_30_2 | 13811 | FTMS; HCD | 125.73 | 811.87 |

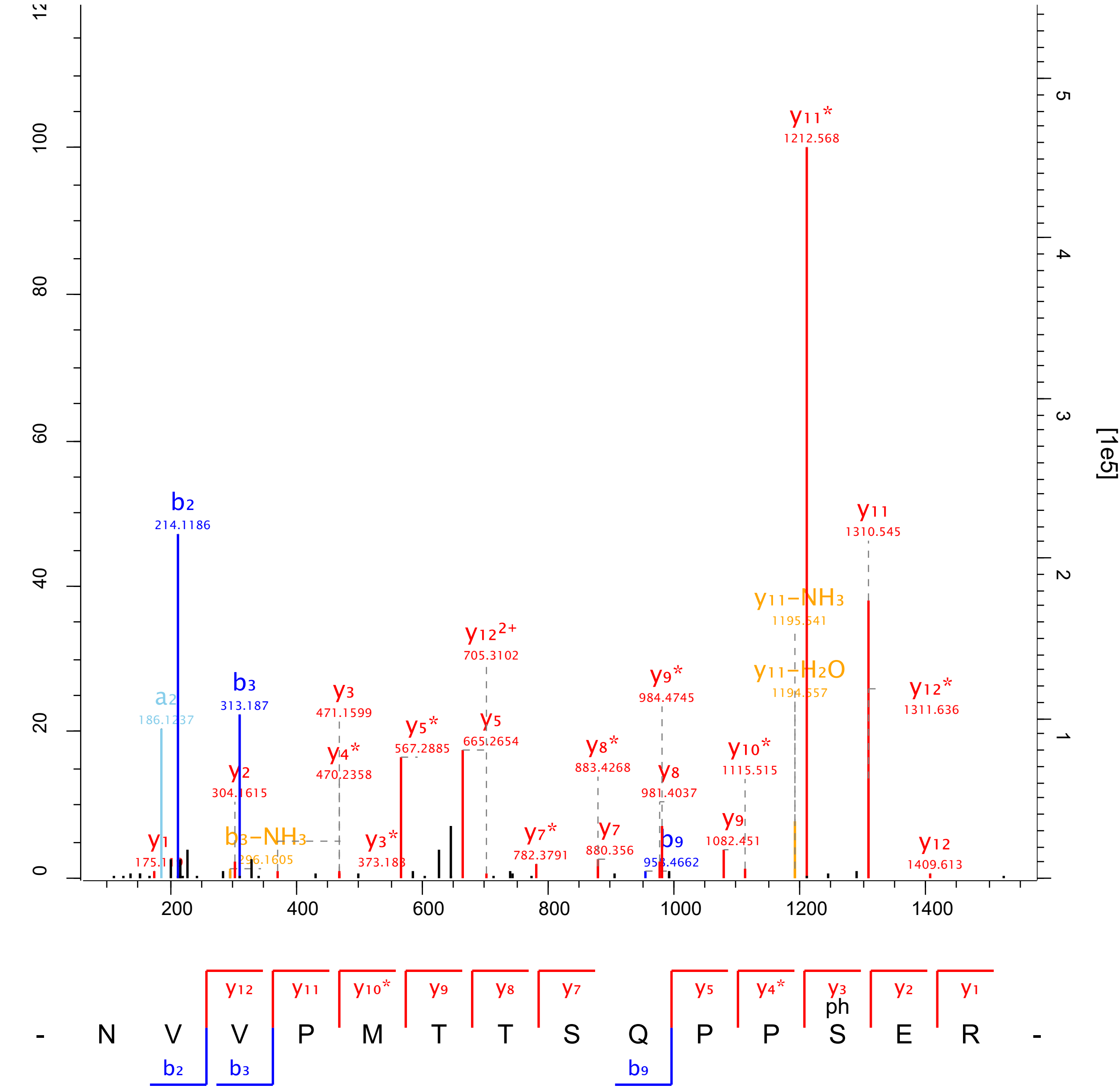

| Raw file | Scan | Method | Score | m/z |
|---|---|---|---|---|
| A17_30_2 | 14734 | FTMS; HCD | 91.22 | 500.75 |

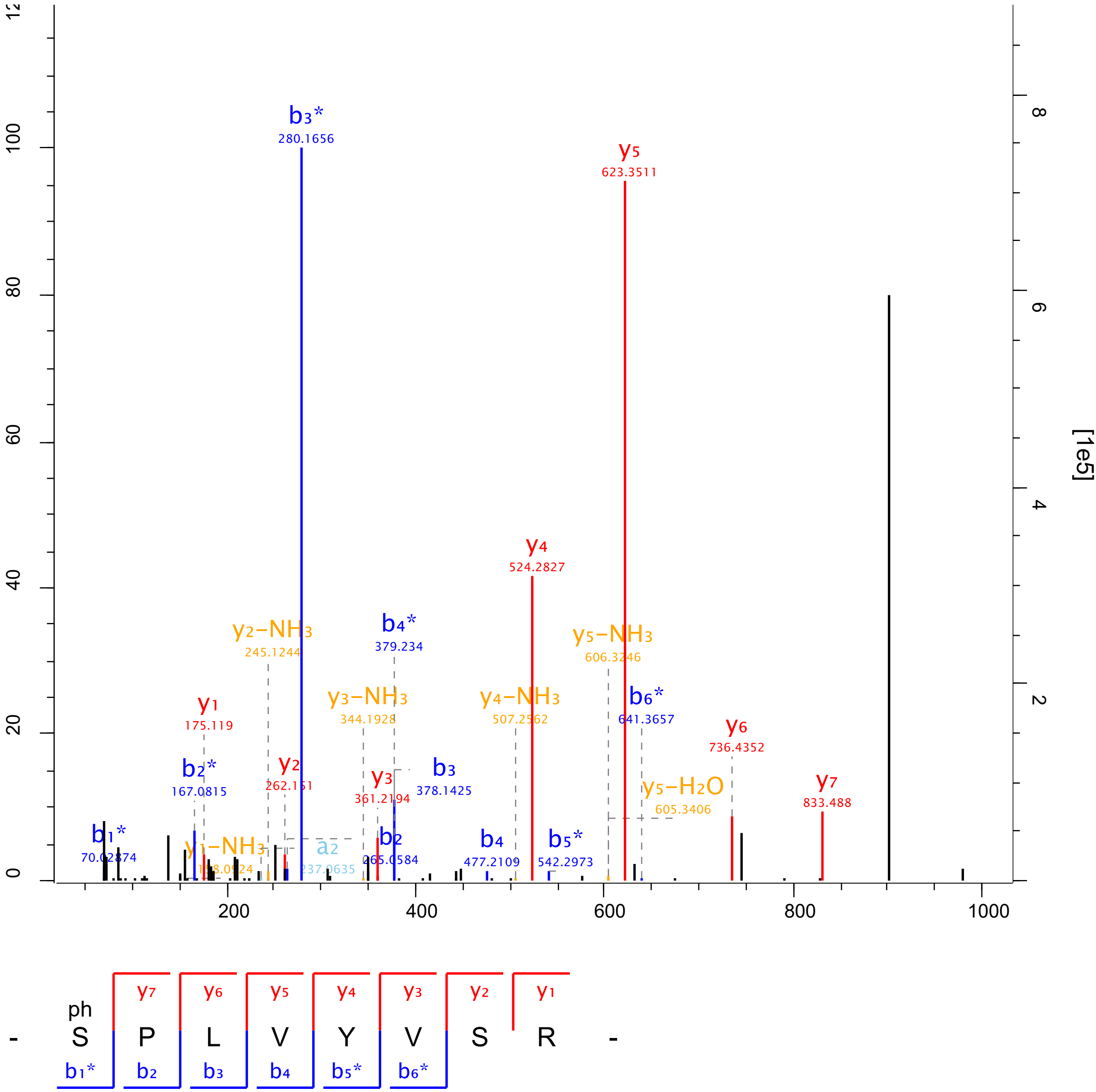

| Raw file | Scan | Method | Score | m/z |
|---|---|---|---|---|
| A17_30_2 | 14900 | FTMS; HCD | 117.09 | 480.2 |

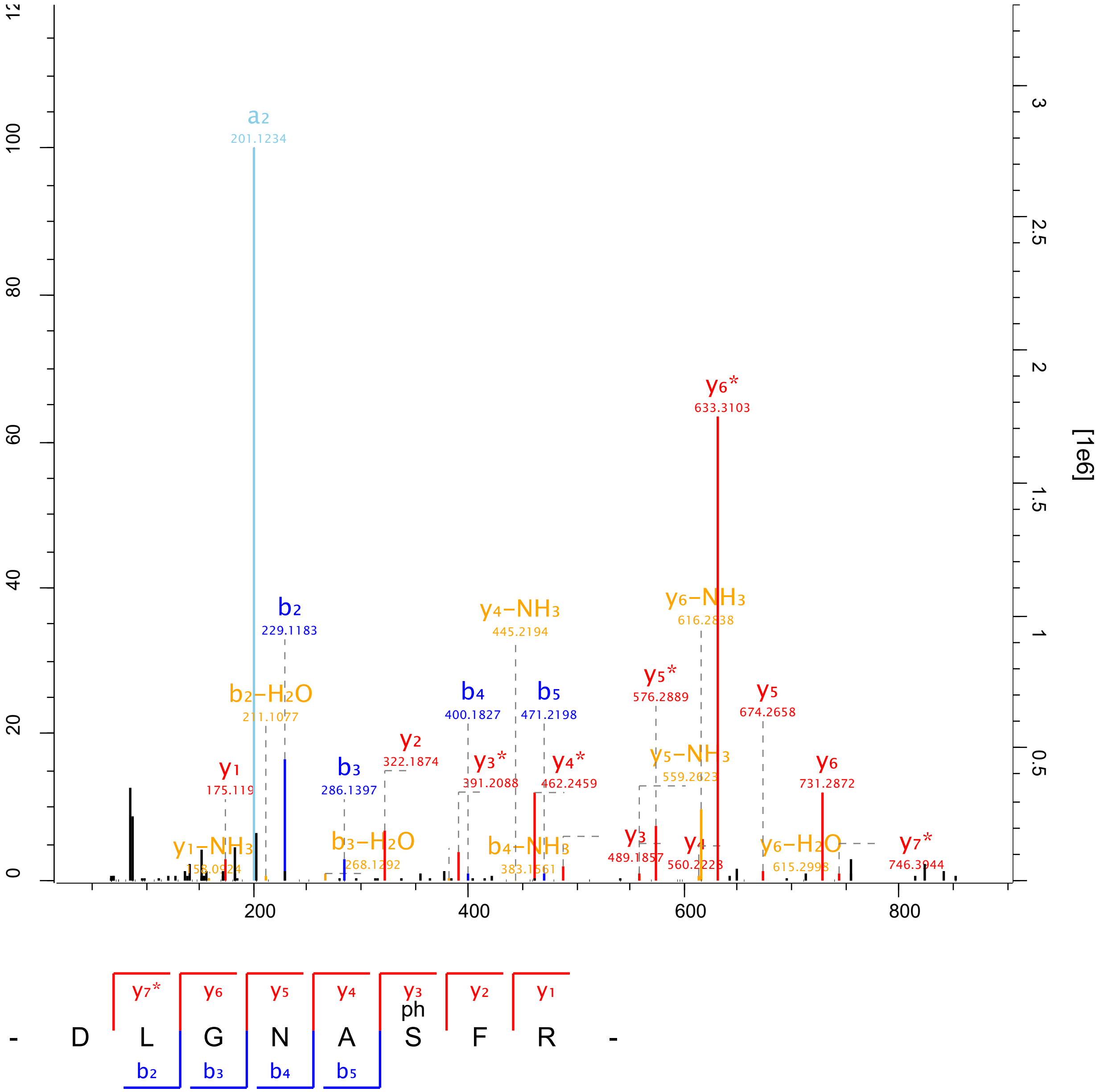

| Raw file | Scan | Method | Score | m/z |
| --- | --- | --- | --- | --- |
| A17_30_2 | 15502 | FTMS; HCD | 158.89 | 618.72 |

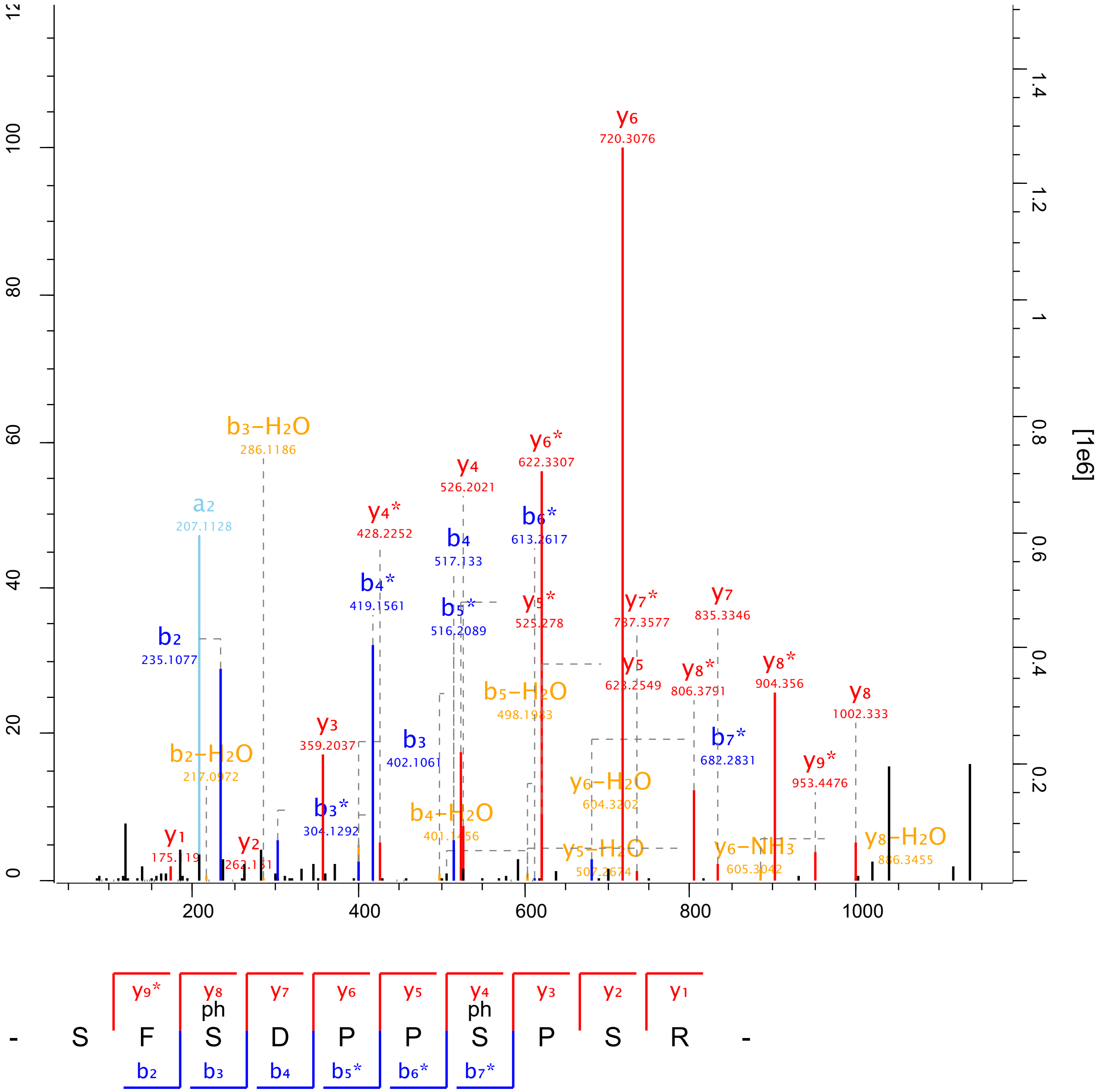

| Raw file | Scan | Method | Score | m/z |
|---|---|---|---|---|
| A17_30_2 | 18103 | FTMS; HCD | 106.93 | 723.27 |

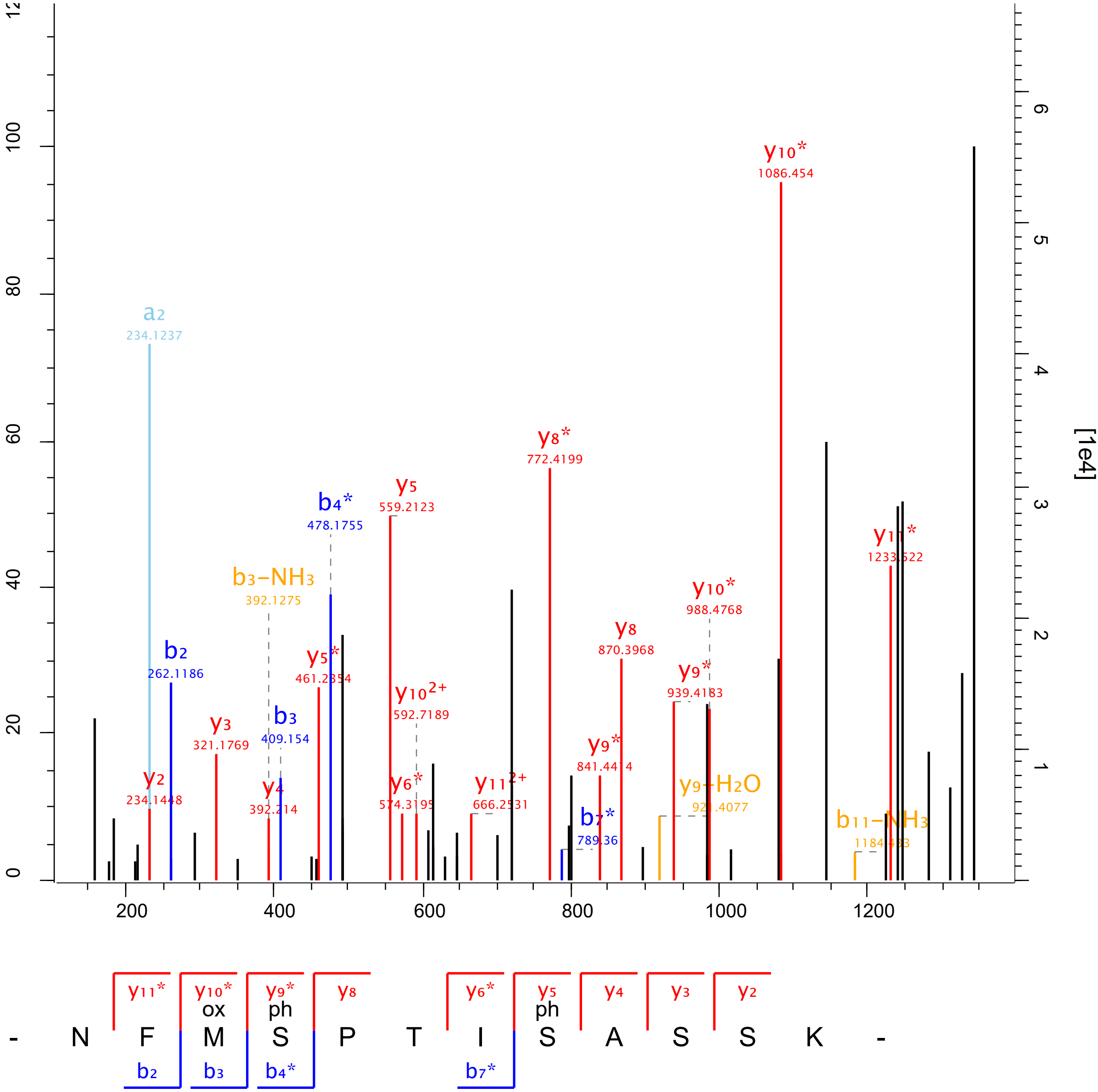

| Raw file | Scan | Method | Score | m/z |
| --- | --- | --- | --- | --- |
| A17_30_2 | 19406 | FTMS; HCD | 43.9 | 620.8 |

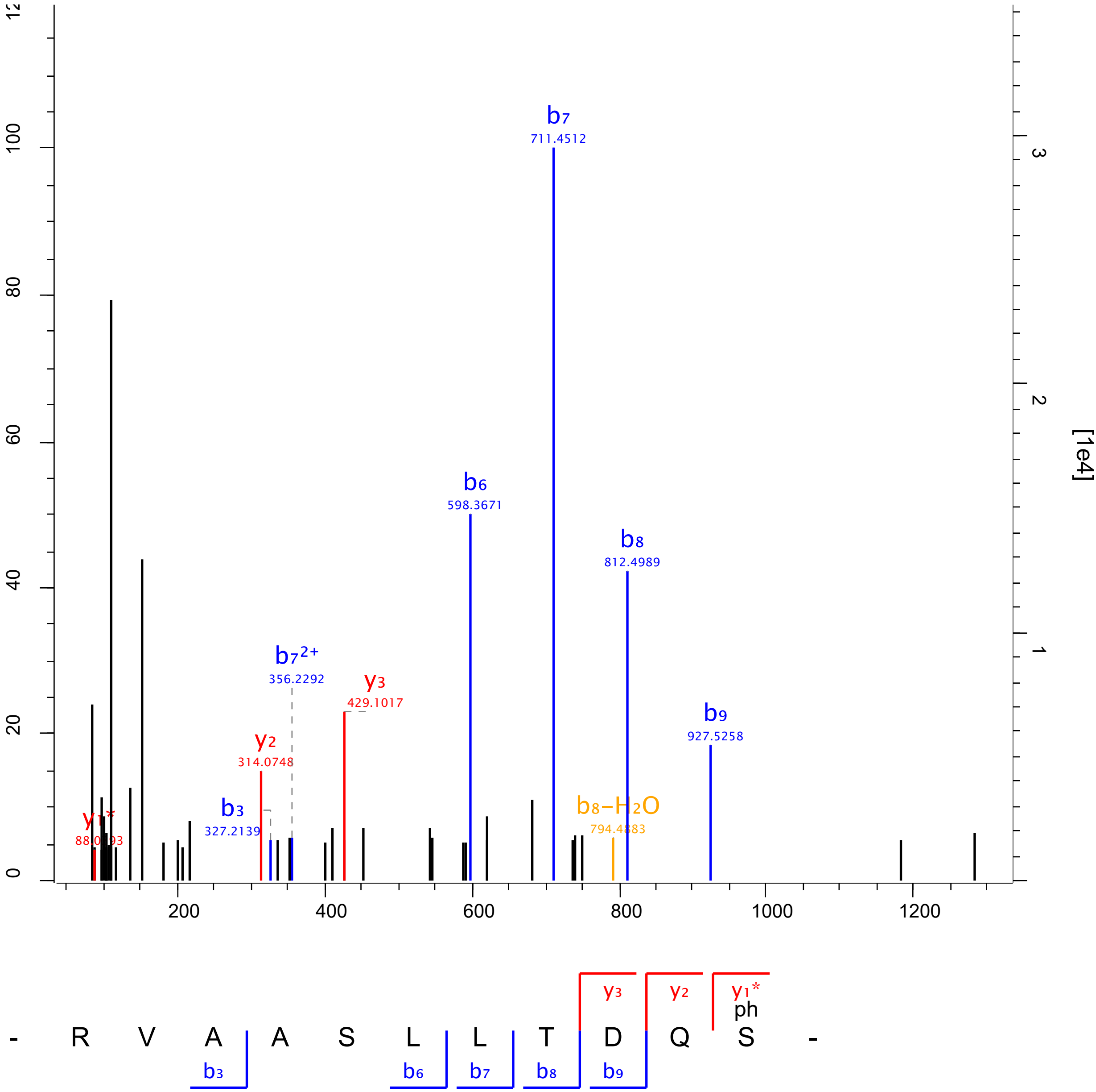

| Raw file | Scan | Method | Score | m/z |
|---|---|---|---|---|
| A17_30_2 | 20789 | FTMS; HCD | 97.24 | 748.85 |

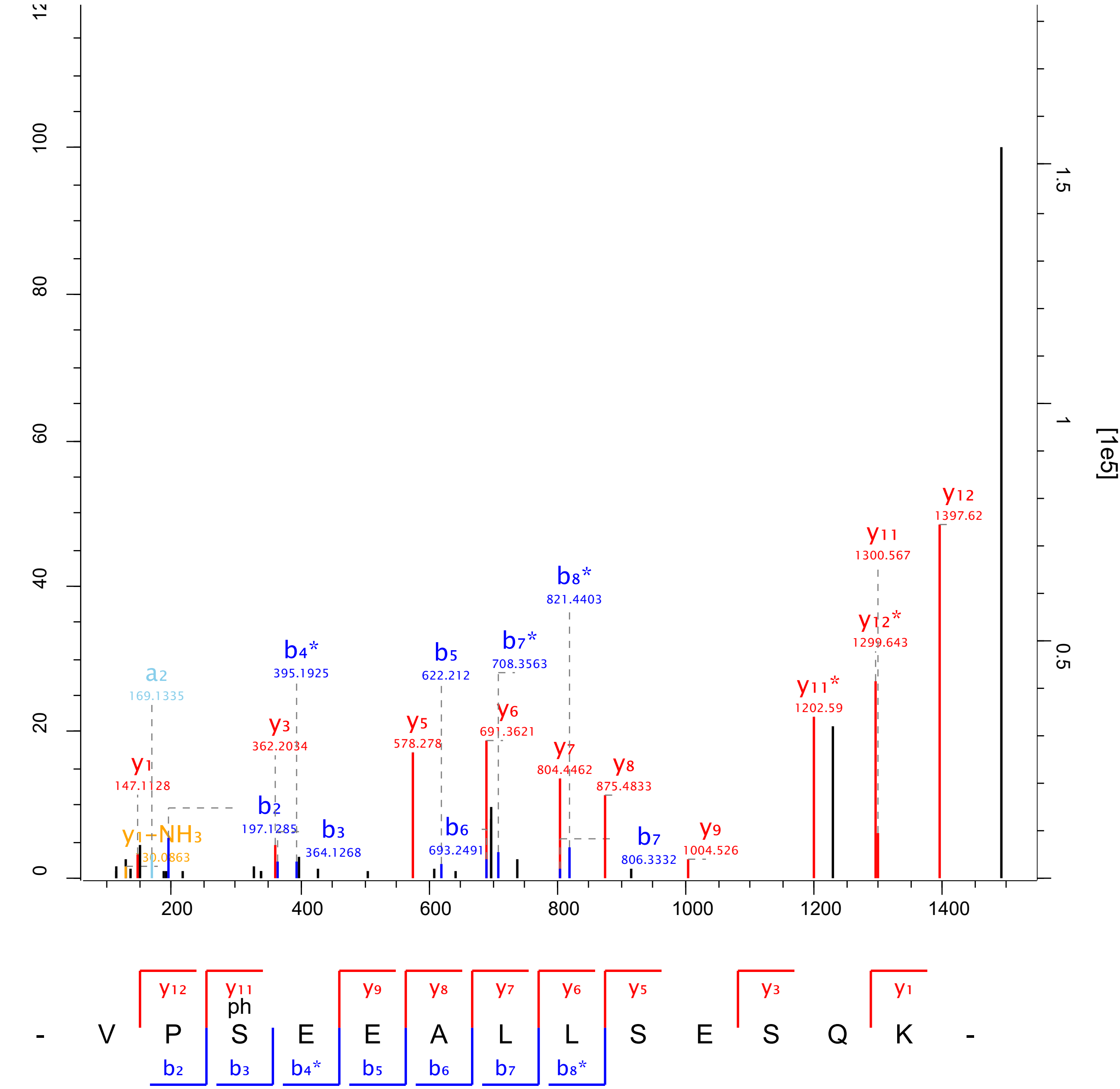

| Raw file | Scan | Method | Score | m/z |
| --- | --- | --- | --- | --- |
| A17_30_2 | 21707 | FTMS; HCD | 48.28 | 525.73 |

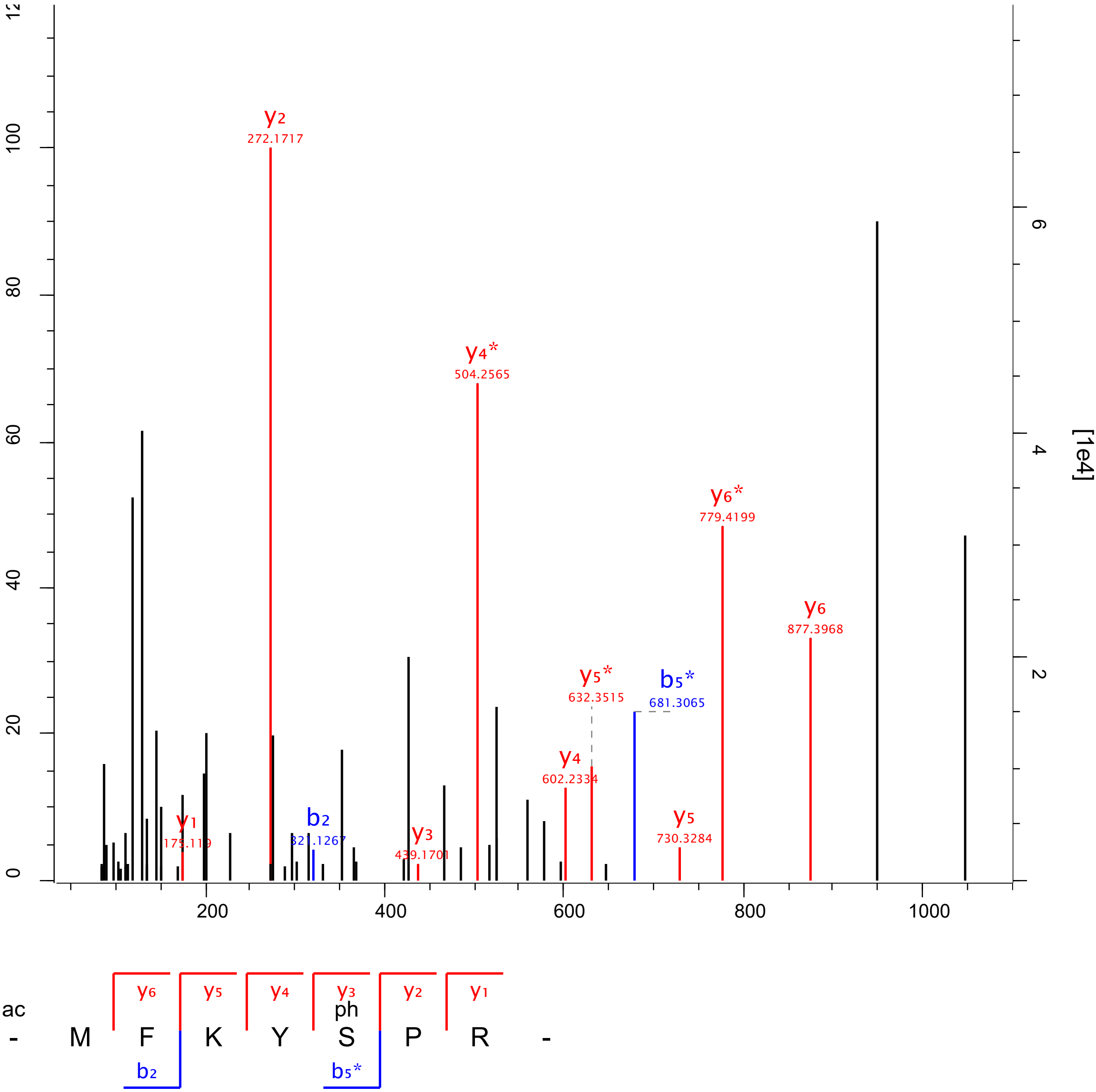

| Raw file | Scan | Method | Score | m/z |
|---|---|---|---|---|
| A17_30_2 | 24372 | FTMS; HCD | 91.24 | 1006.44 |

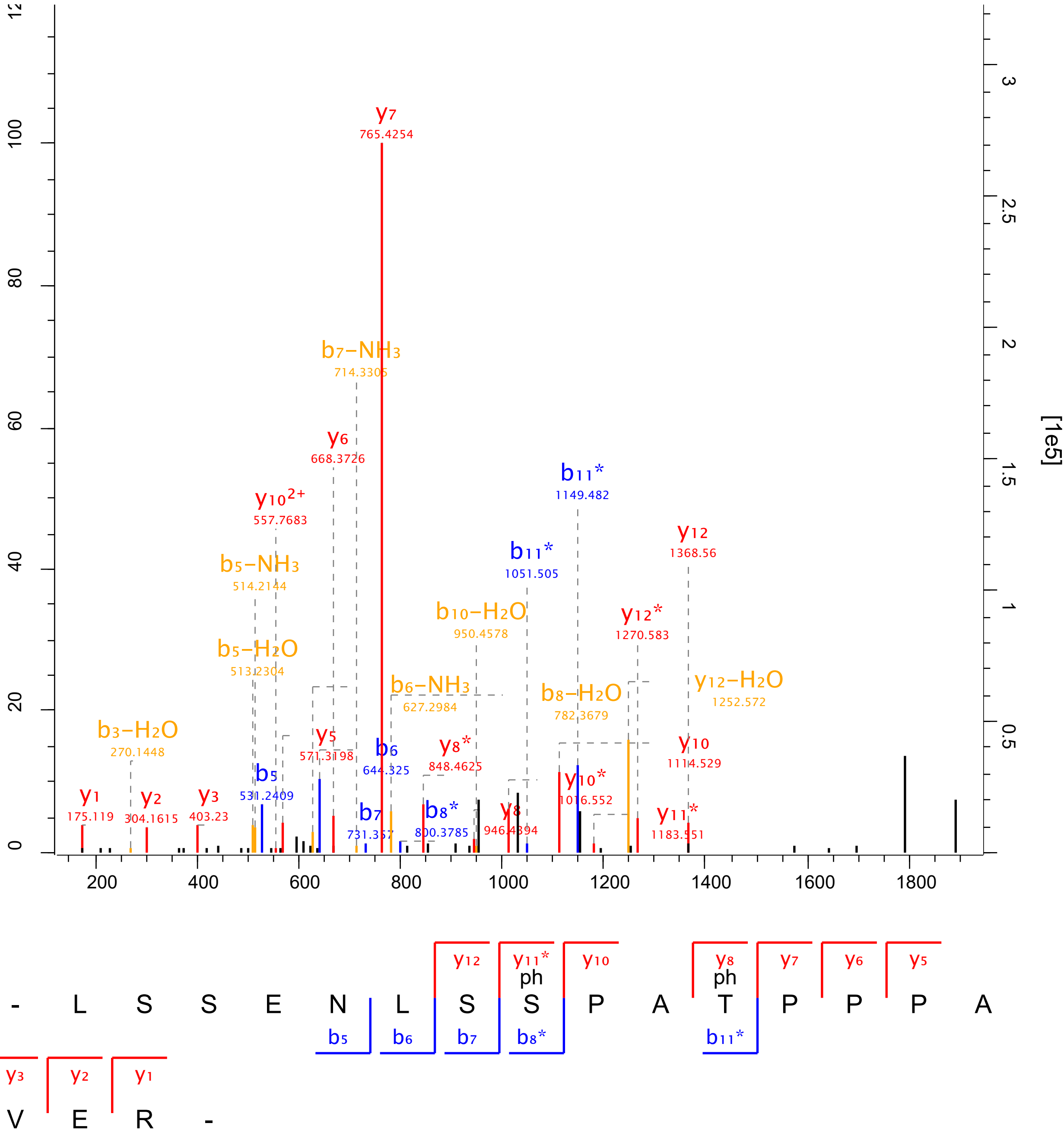

| Raw file | Scan | Method | Score | m/z |
|---|---|---|---|---|
| A17_30_2 | 25589 | FTMS; HCD | 110.18 | 499.7 |

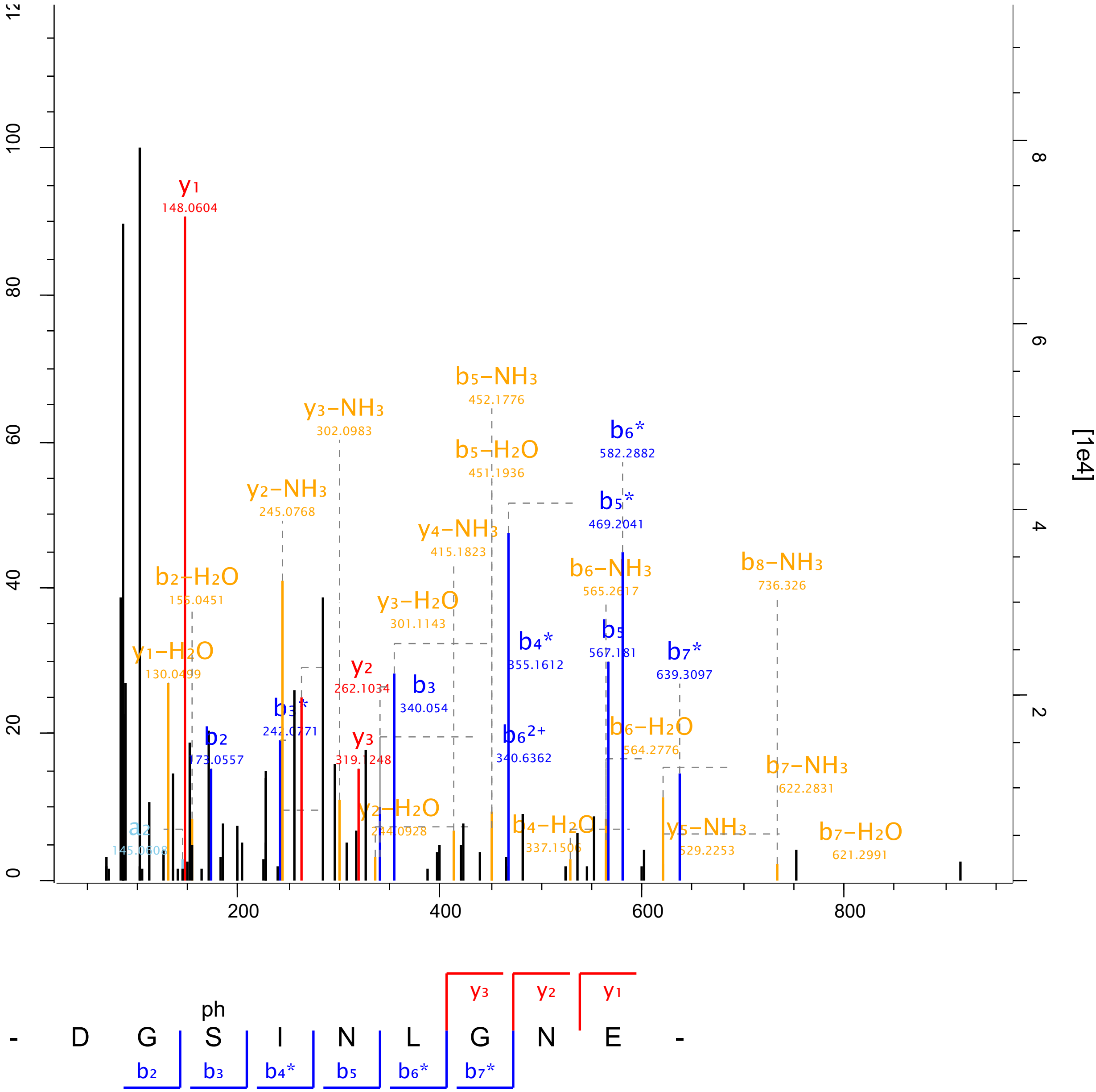

| Raw file | Scan | Method | Score | m/z |
| --- | --- | --- | --- | --- |
| A17_30_2 | 28689 | FTMS; HCD | 48.05 | 798.32 |

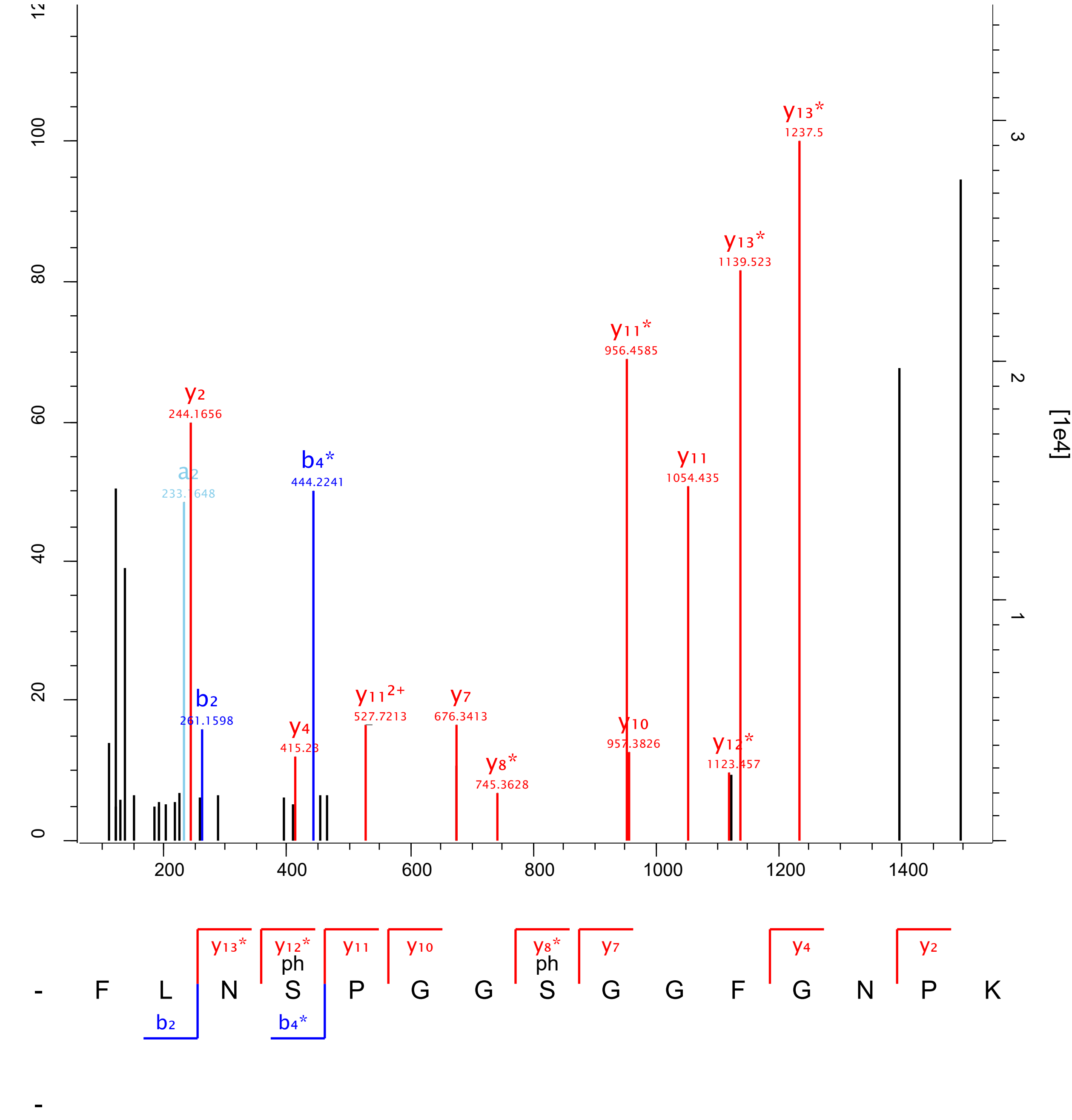

| Raw file | Scan | Method | Score | m/z |
| --- | --- | --- | --- | --- |
| A17_30_2 | 2928 | FTMS; HCD | 40.35 | 506.18 |

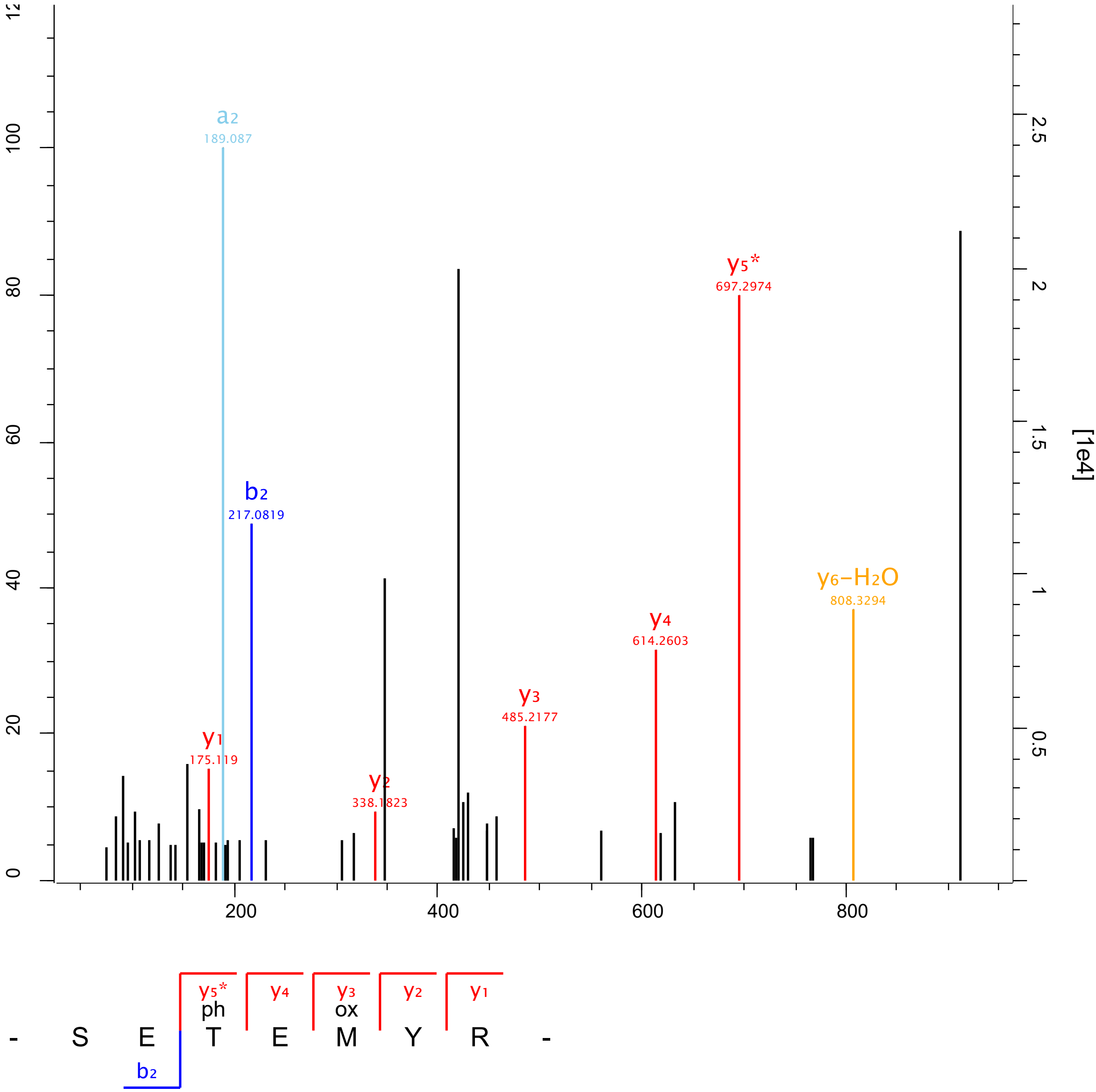

| Raw file | Scan | Method | Score | m/z |
|---|---|---|---|---|
| A17_30_2 | 2940 | FTMS; HCD | 82.64 | 416.17 |

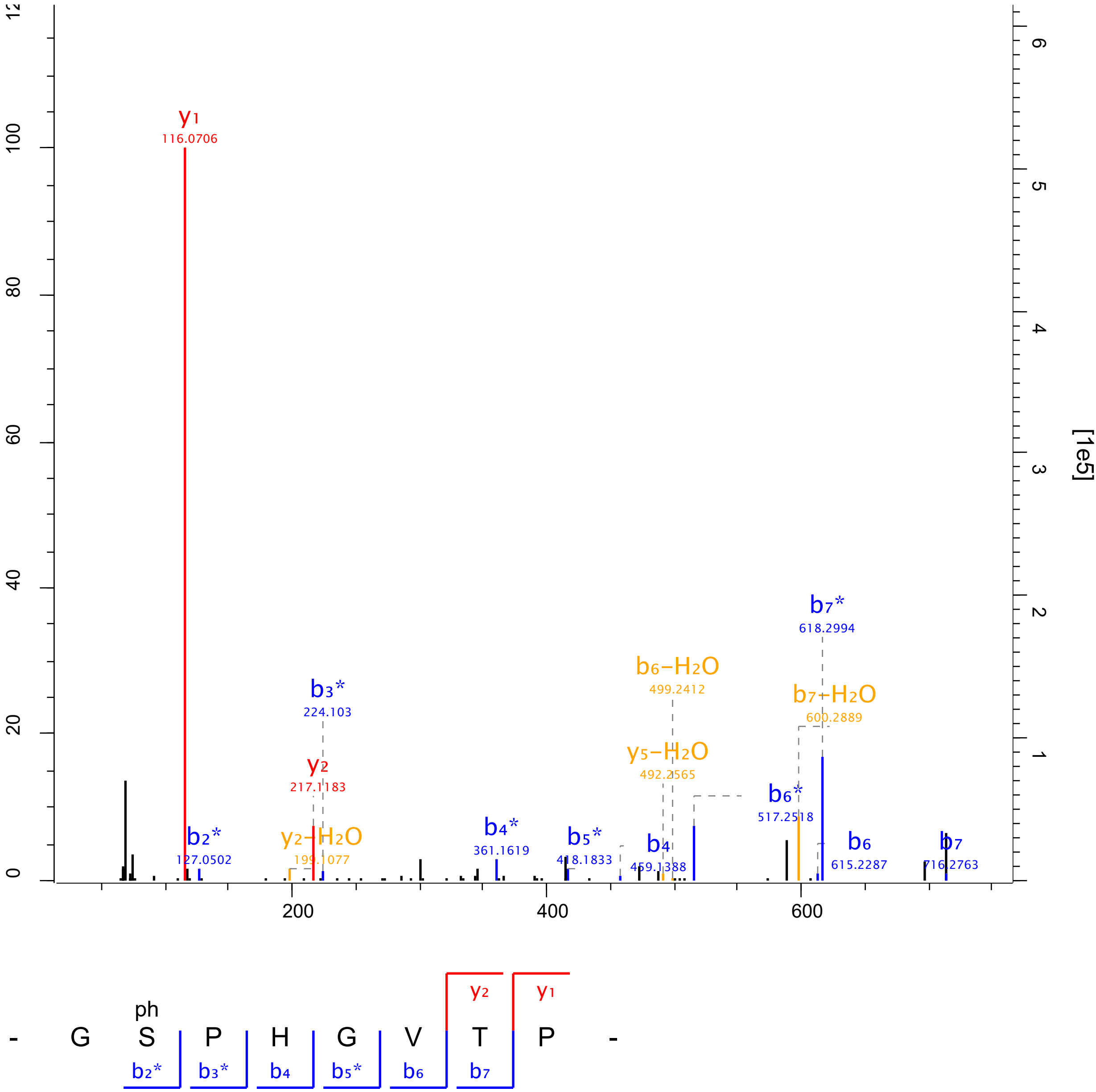

| Raw file | Scan | Method | Score | m/z |
|---|---|---|---|---|
| A17_30_2 | 3369 | FTMS; HCD | 94.09 | 483.55 |

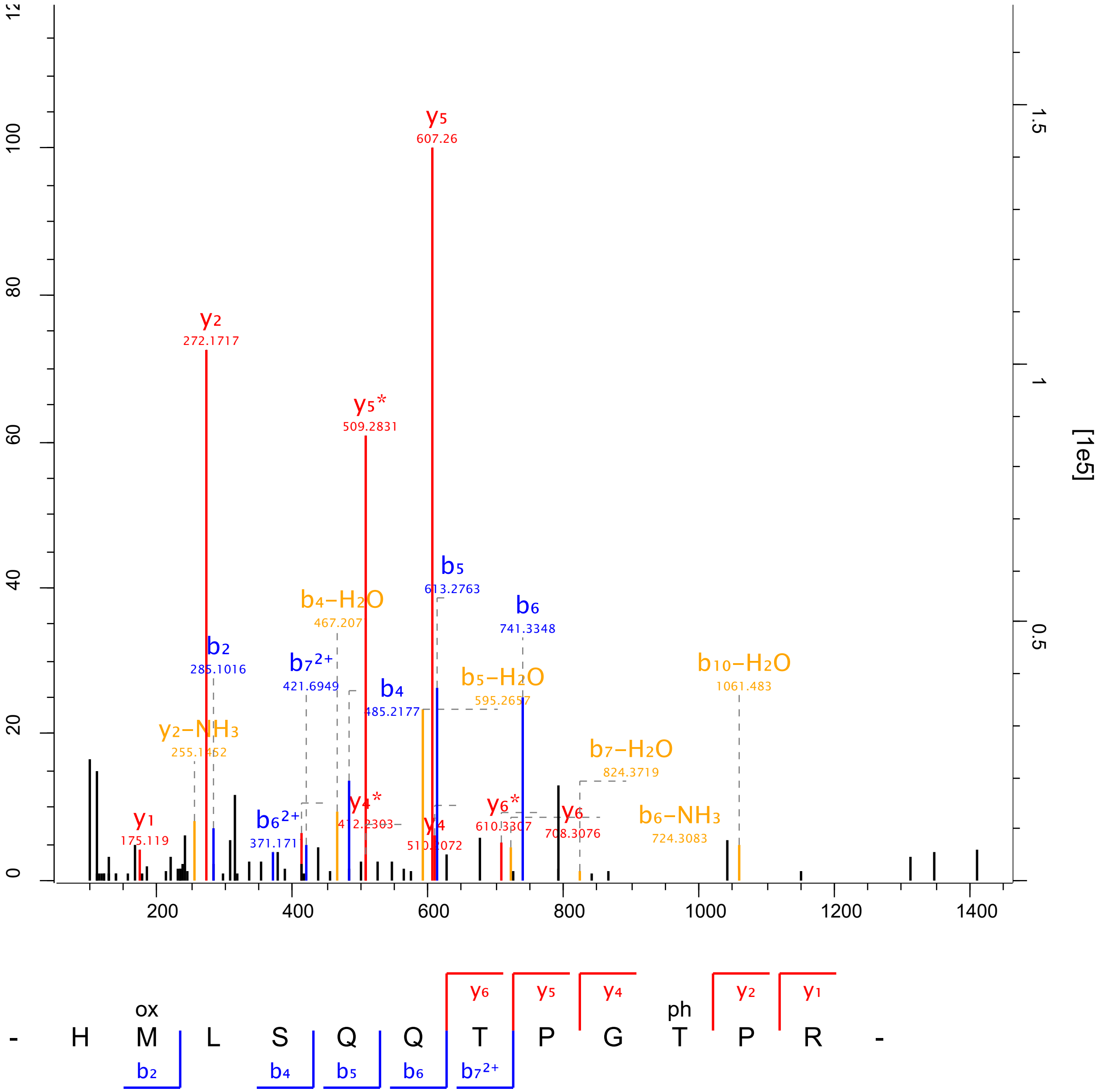

| Raw file | Scan | Method | Score | m/z |
|---|---|---|---|---|
| A17_30_2 | 38122 | FTMS; HCD | 233.02 | 699.77 |

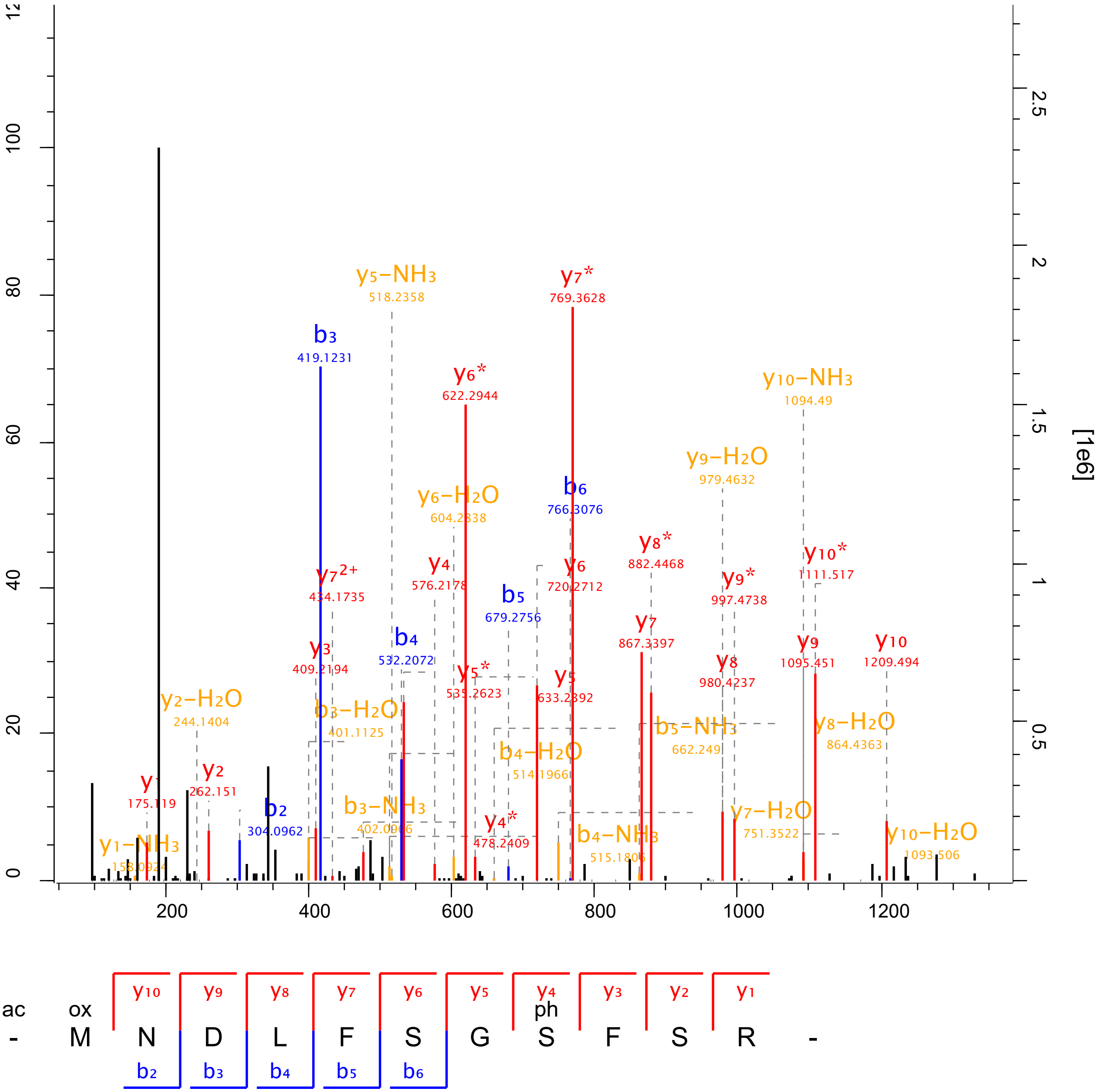

| Raw file | Scan | Method | Score | m/z |
| --- | --- | --- | --- | --- |
| A17_30_2 | 41838 | FTMS; HCD | 48.79 | 507.22 |

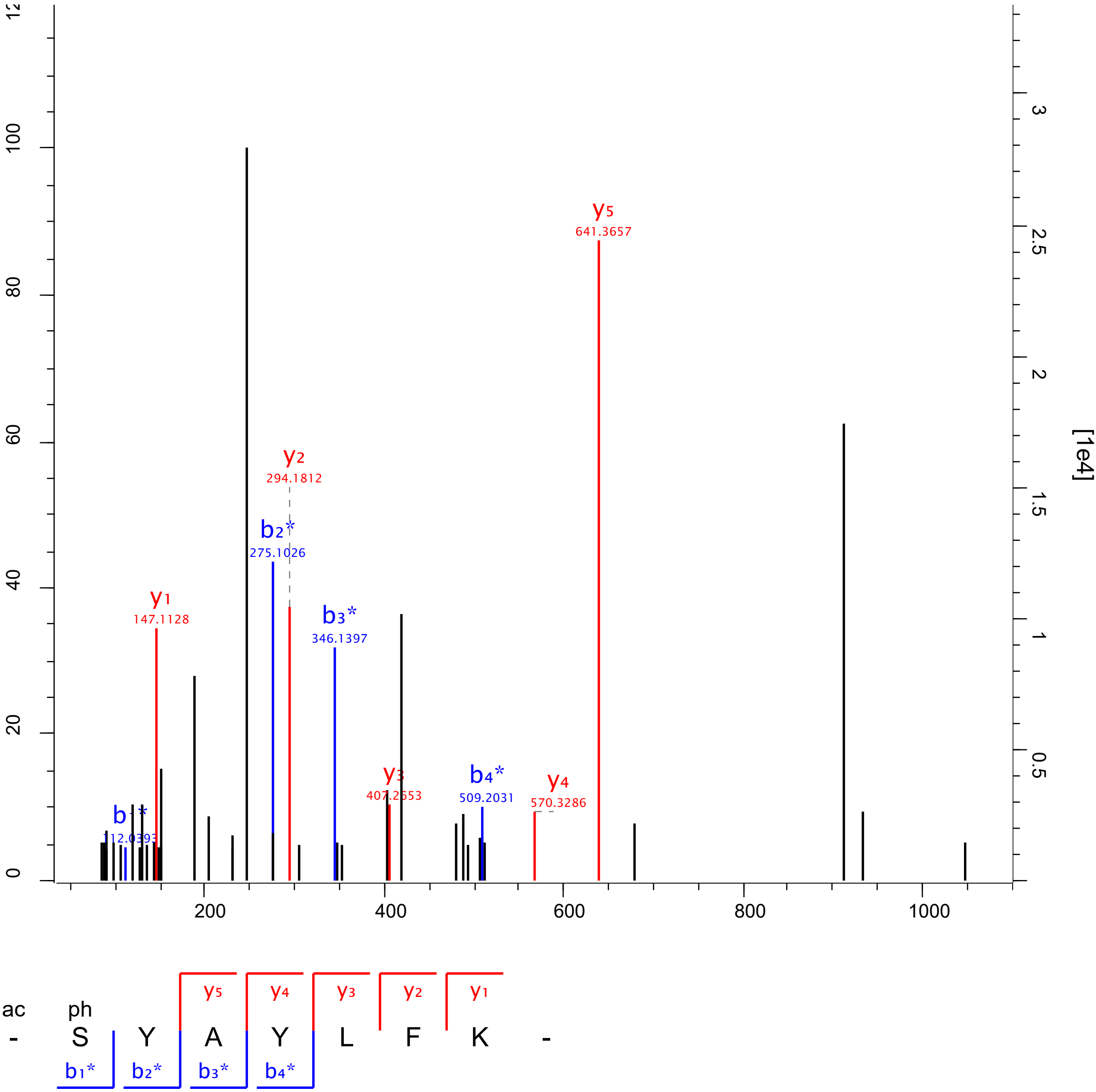

| Raw file | Scan | Method | Score | m/z |
|---|---|---|---|---|
| A17_30_2 | 4352 | FTMS; HCD | 55.75 | 628.28 |

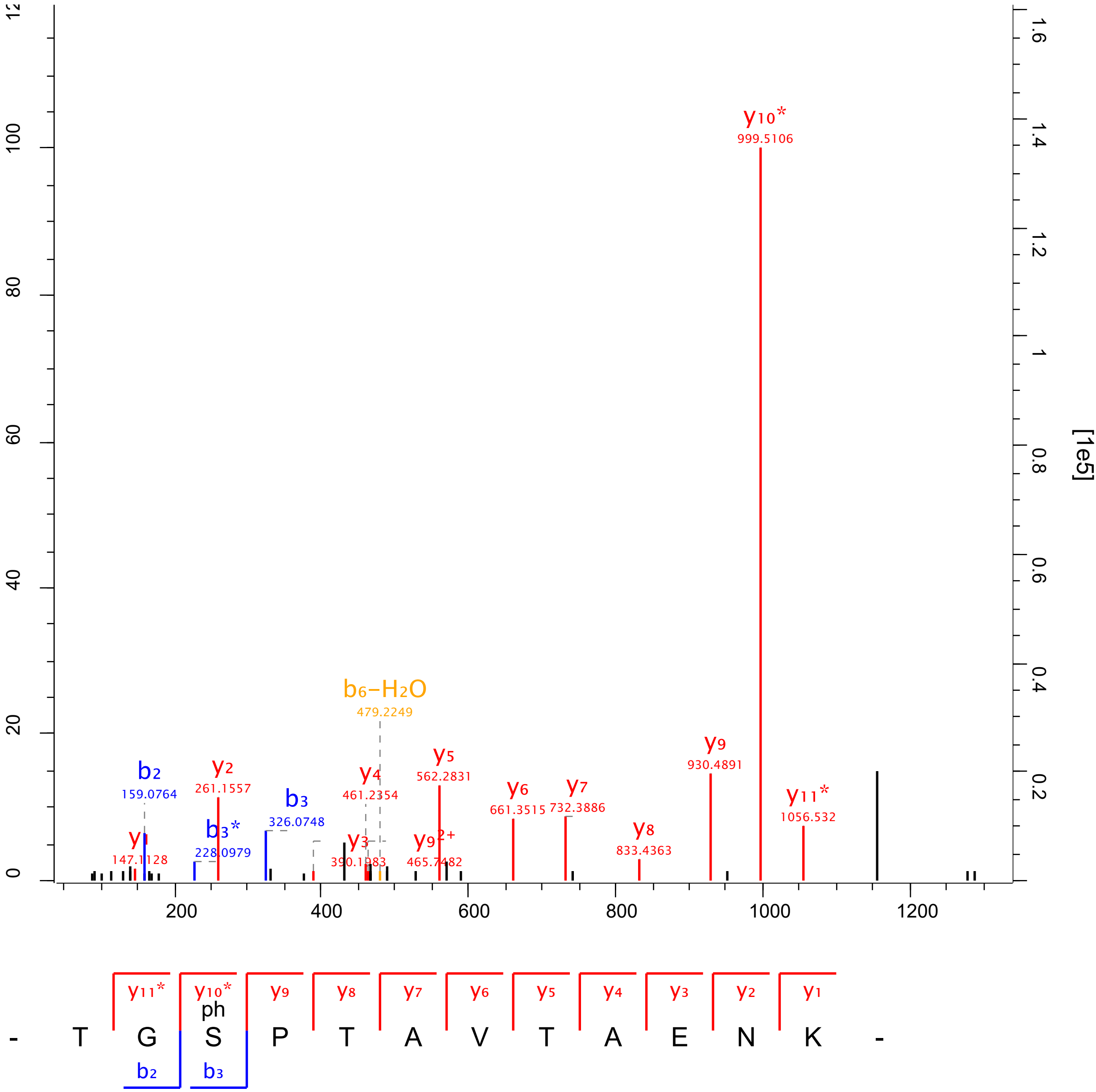

| Raw file | Scan | Method | Score | m/z |
| --- | --- | --- | --- | --- |
| A17_30_2 | 5983 | FTMS; HCD | 117.93 | 608.24 |

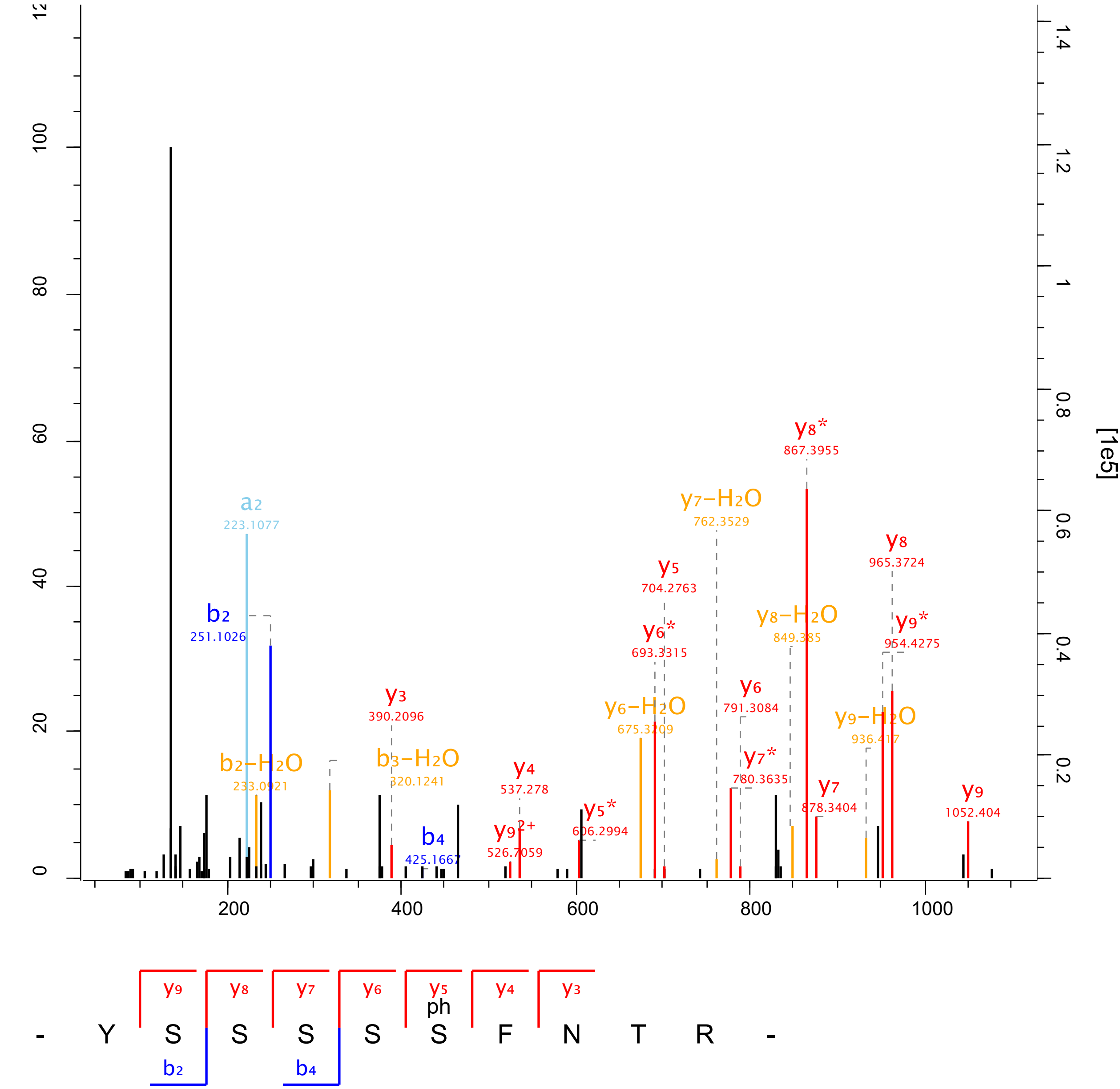

| Raw file | Scan | Method | Score | m/z |
|---|---|---|---|---|
| A17_30_2 | 6189 | FTMS; HCD | 66 | 538.24 |

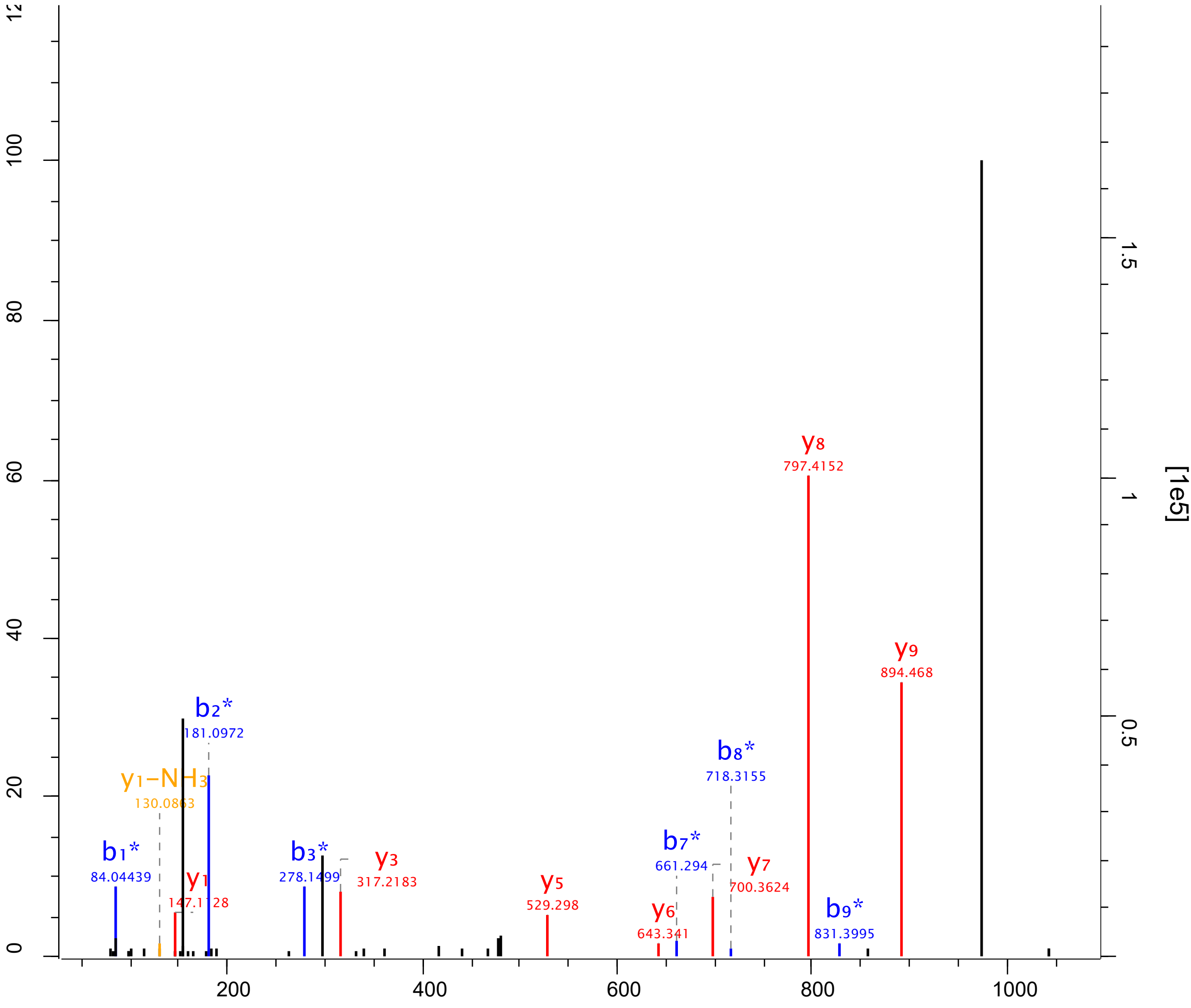


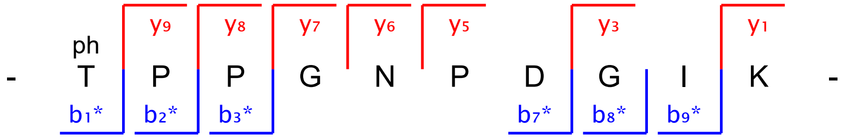

| Raw file | Scan | Method | Score | m/z |
|---|---|---|---|---|
| A17_30_2 | 6800 | FTMS; HCD | 74.79 | 649.78 |

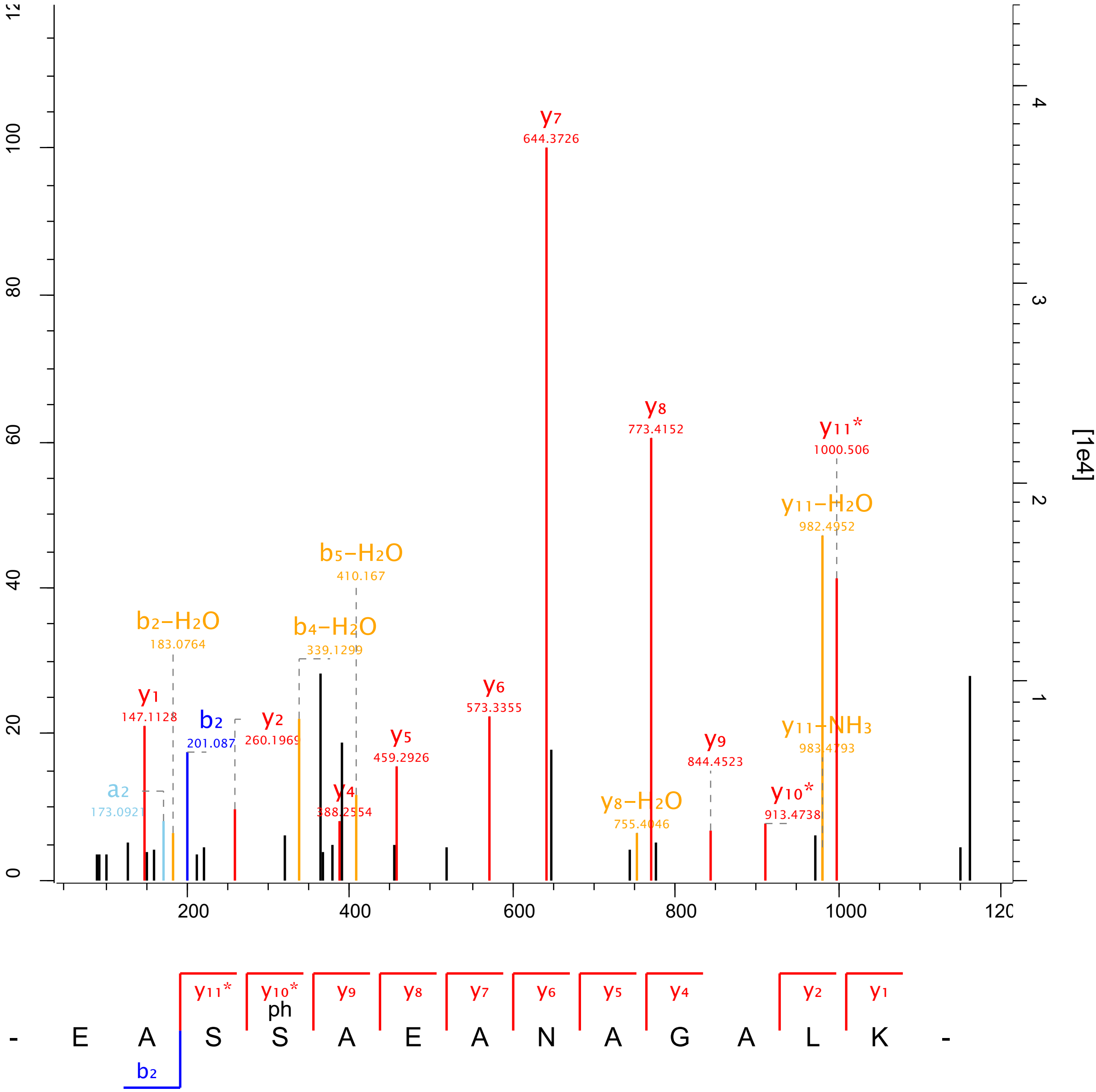

| Raw file | Scan | Method | Score | m/z |
|---|---|---|---|---|
| A17_30_2 | 7031 | FTMS; HCD | 42.86 | 433.19 |

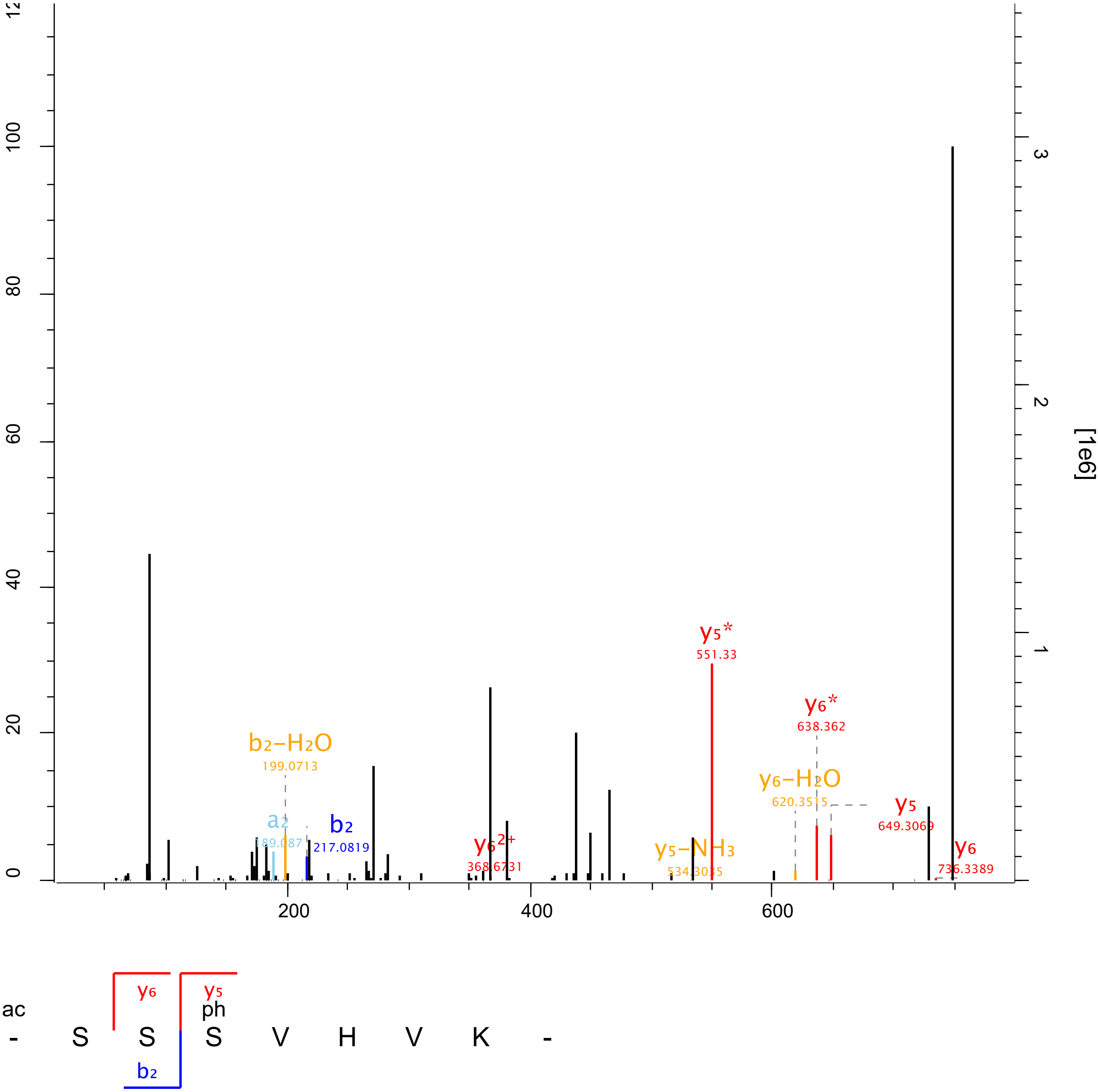

| Raw file | Scan | Method | Score | m/z |
| --- | --- | --- | --- | --- |
| A17_30_2 | 7864 | FTMS; HCD | 51.73 | 649.31 |

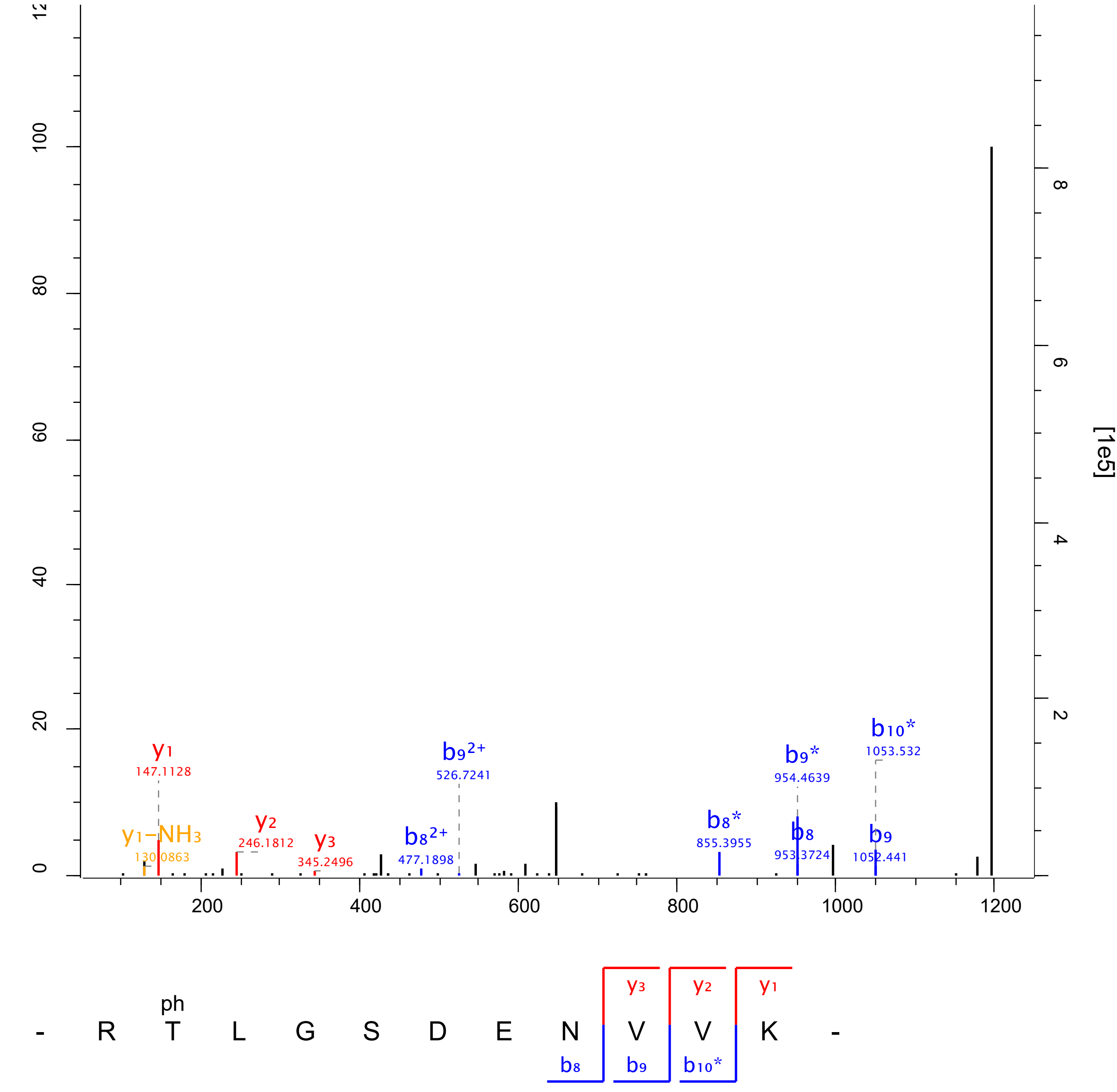

| Raw file | Scan | Method | Score | m/z |
| --- | --- | --- | --- | --- |
| A17_30_2 | 8586 | FTMS; HCD | 149.93 | 508.22 |

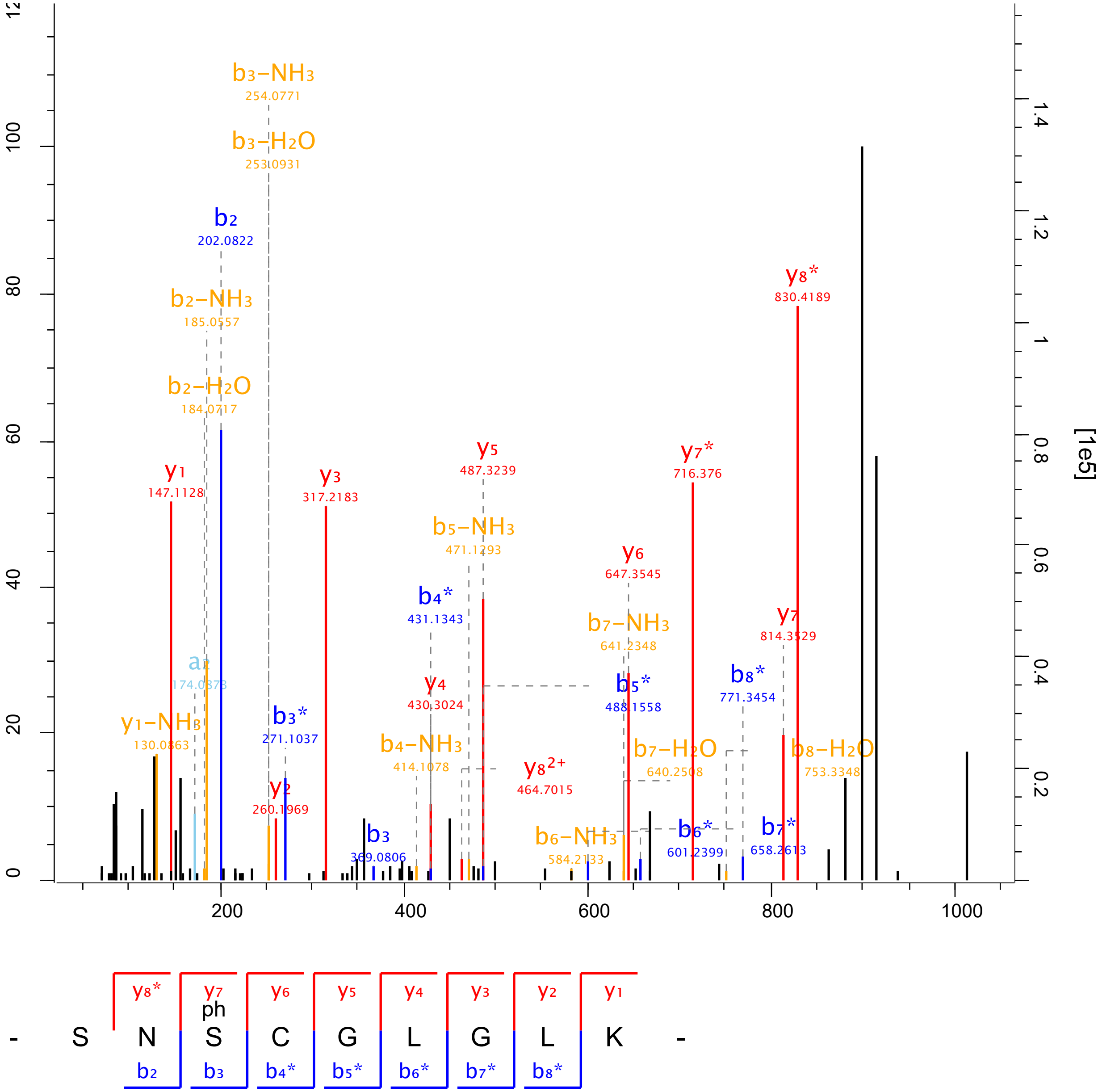

| Raw file | Scan | Method | Score | m/z |
|---|---|---|---|---|
| A17_30_2 | 9142 | FTMS; HCD | 159.58 | 591.28 |

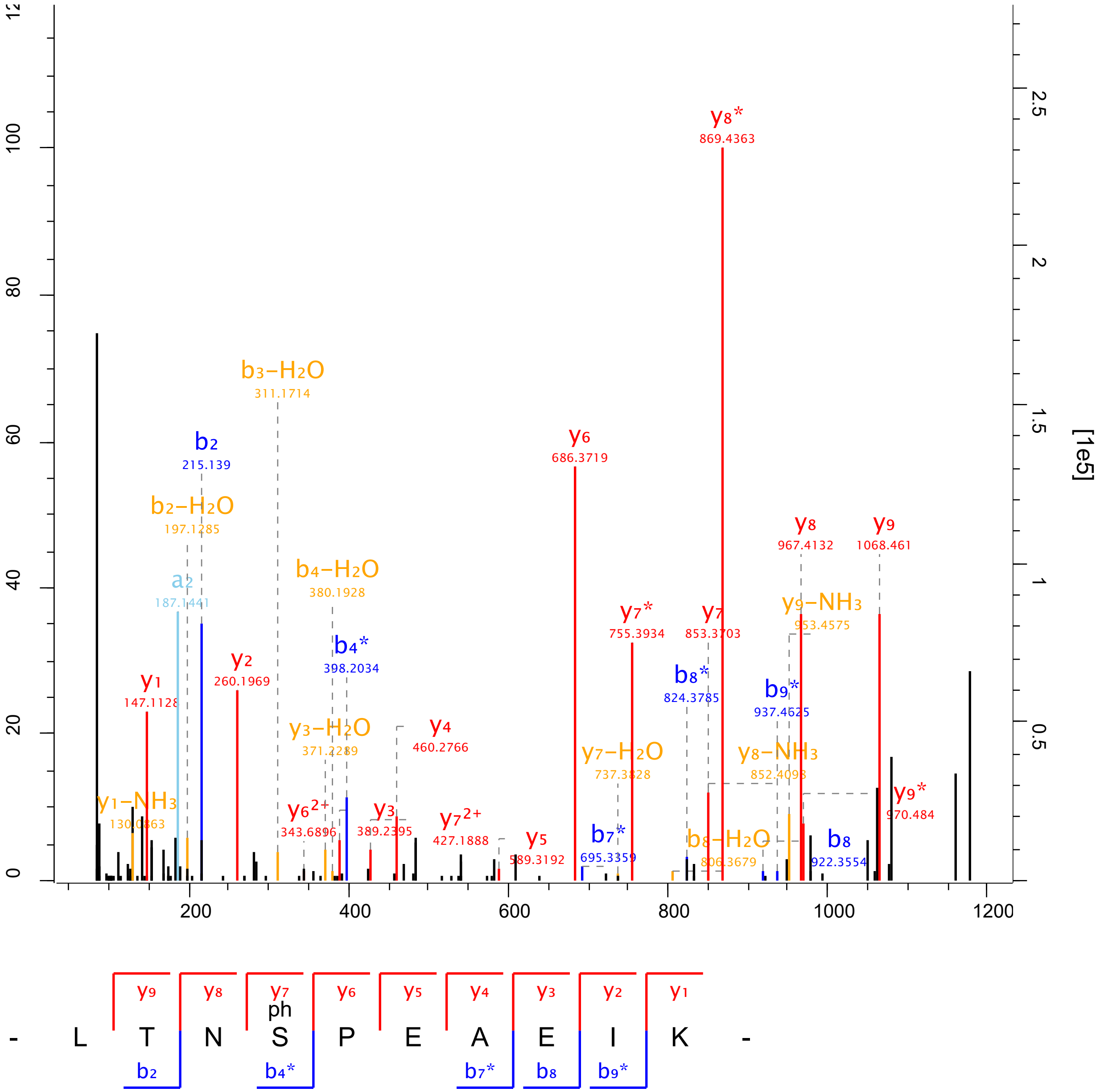

| Raw file | Scan | Method | Score | m/z |
| --- | --- | --- | --- | --- |
| A17_30_2 | 9630 | FTMS; HCD | 103.55 | 454.22 |

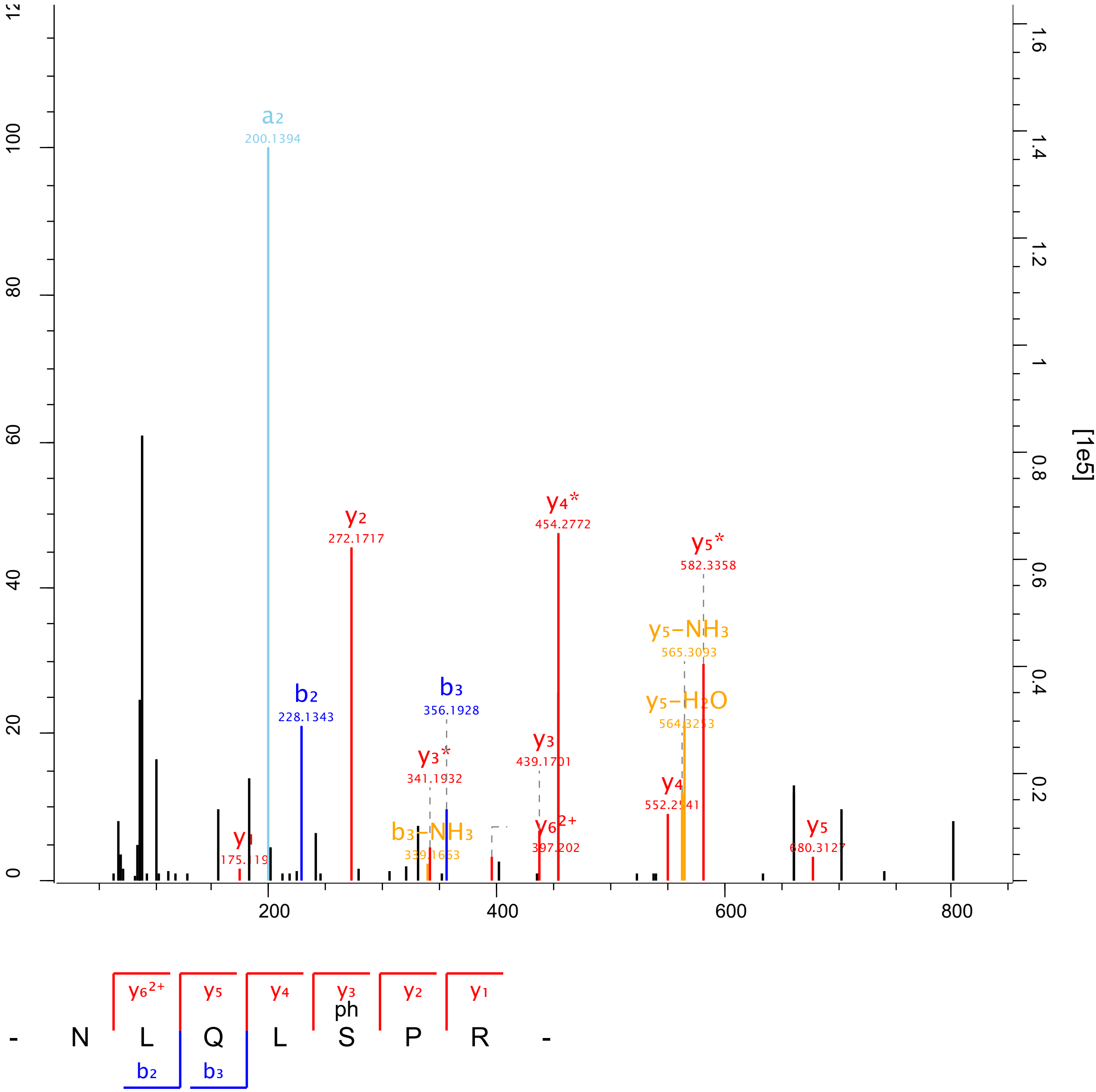

| Raw file | Scan | Method | Score | m/z |
| --- | --- | --- | --- | --- |
| A17_30_2 | 9969 | FTMS; HCD | 57.27 | 437.68 |

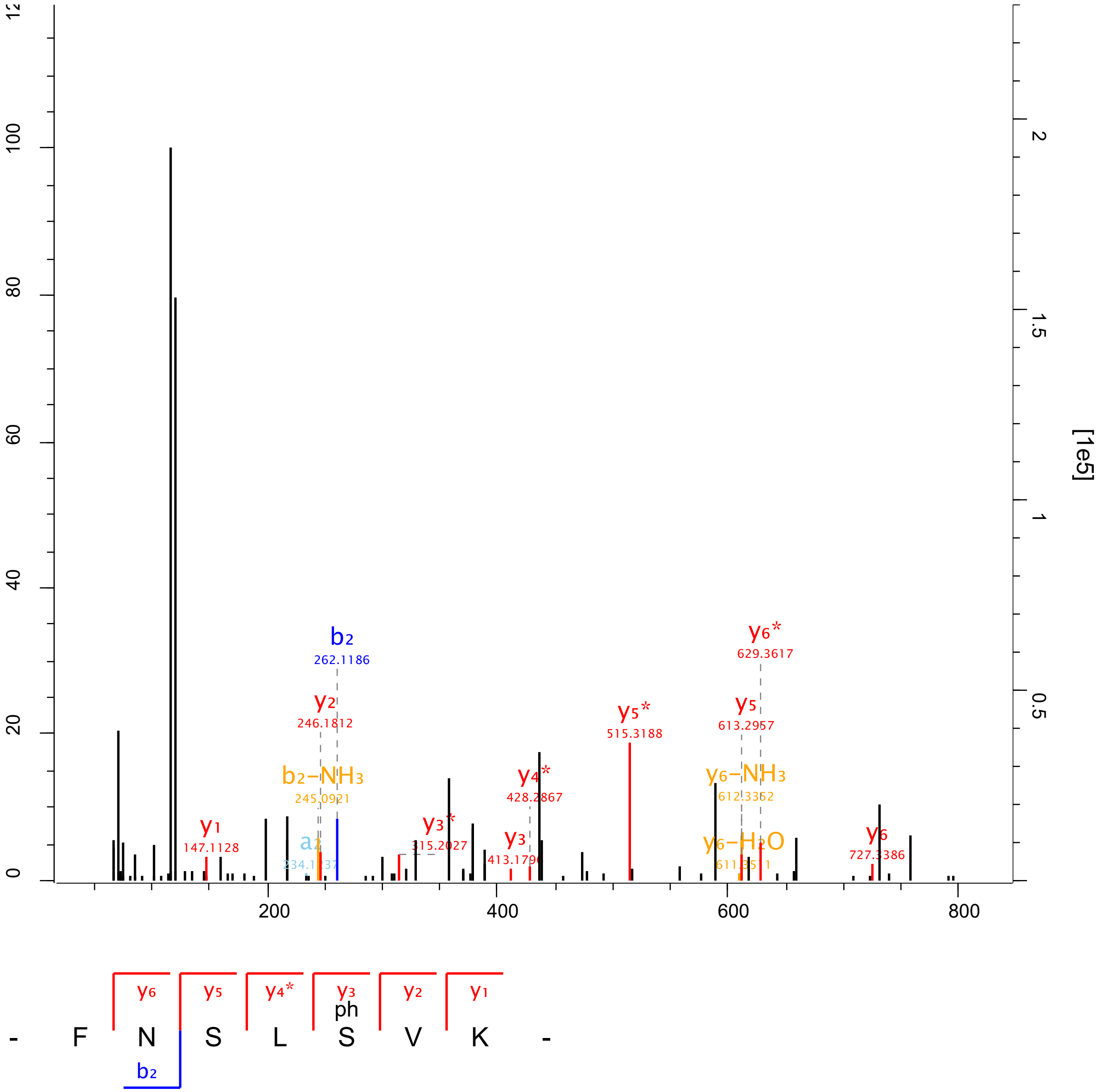

| Raw file | Scan | Method | Score | m/z |
|---|---|---|---|---|
| sys_00_1 | 12829 | FTMS; HCD | 62.94 | 769.86 |

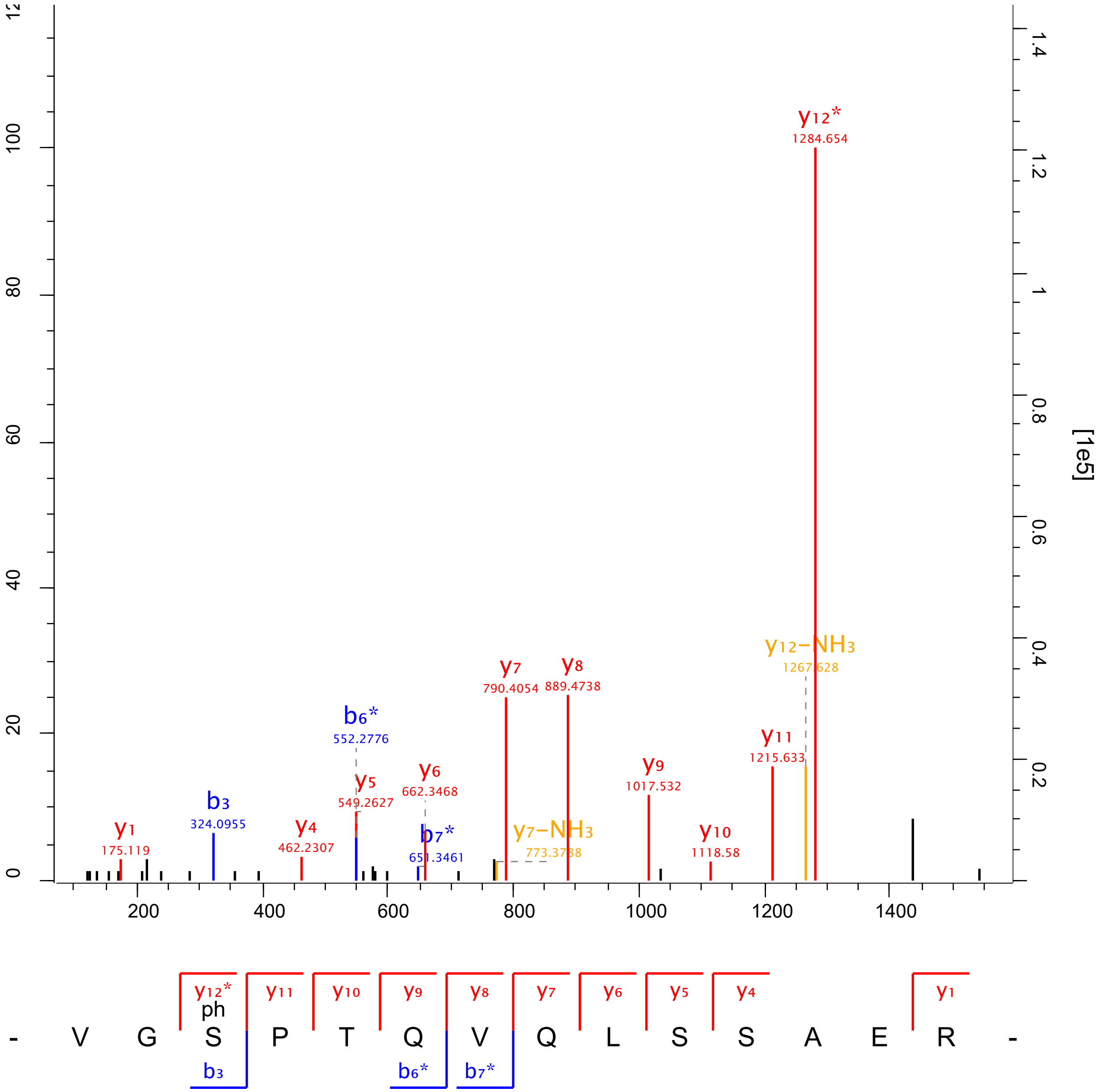

| Raw file | Scan | Method | Score | m/z |
| --- | --- | --- | --- | --- |
| sys_00_1 | 1551 | FTMS; HCD | 62.46 | 480.19 |

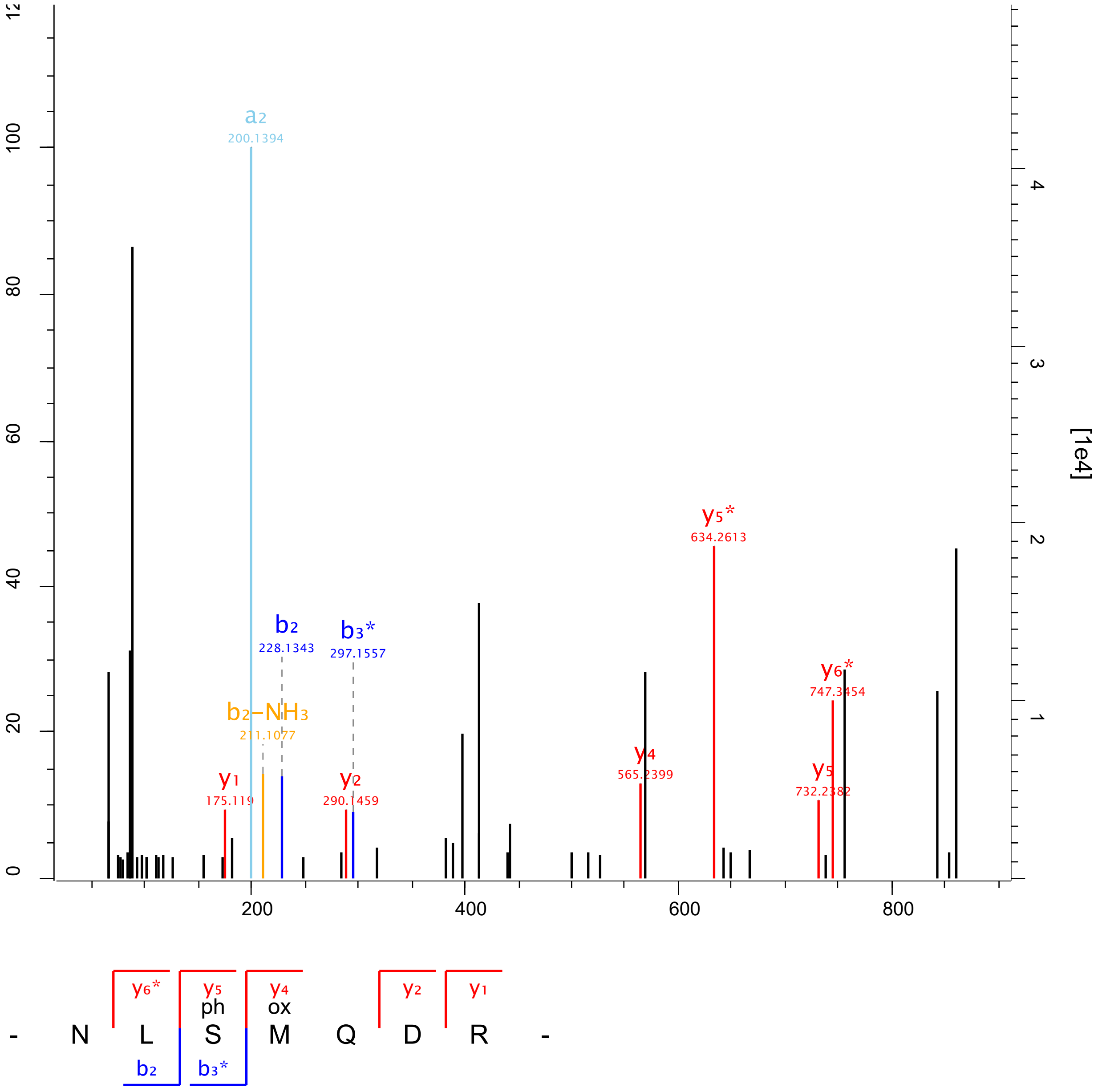

| Raw file | Scan | Method | Score | m/z |
| --- | --- | --- | --- | --- |
| sys_00_1 | 1755 | FTMS; HCD | 50.08 | 721.8 |

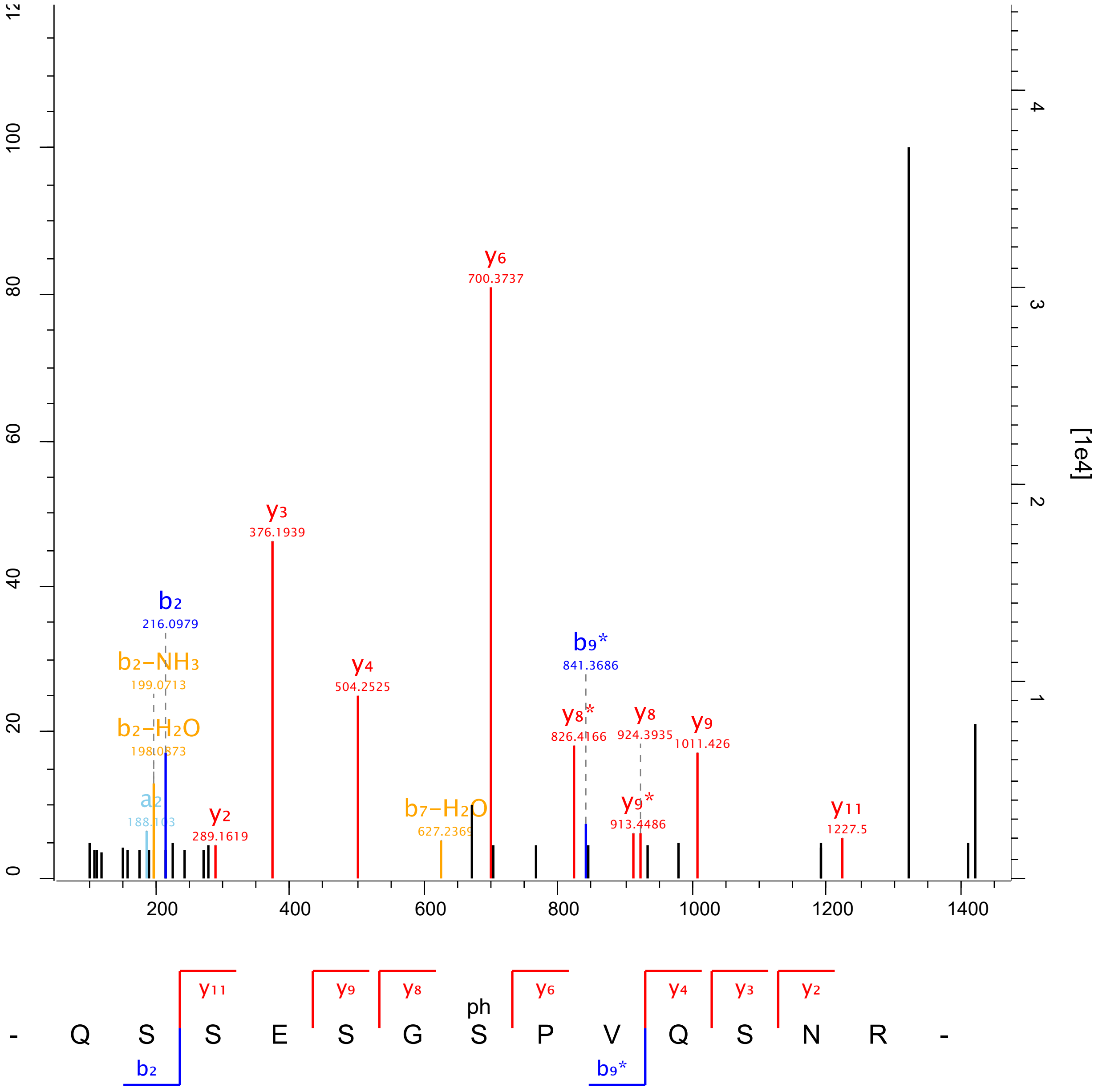

| Raw file | Scan | Method | Score | m/z |
| --- | --- | --- | --- | --- |
| sys_00_1 | 2700 | FTMS; HCD | 51.28 | 506.73 |

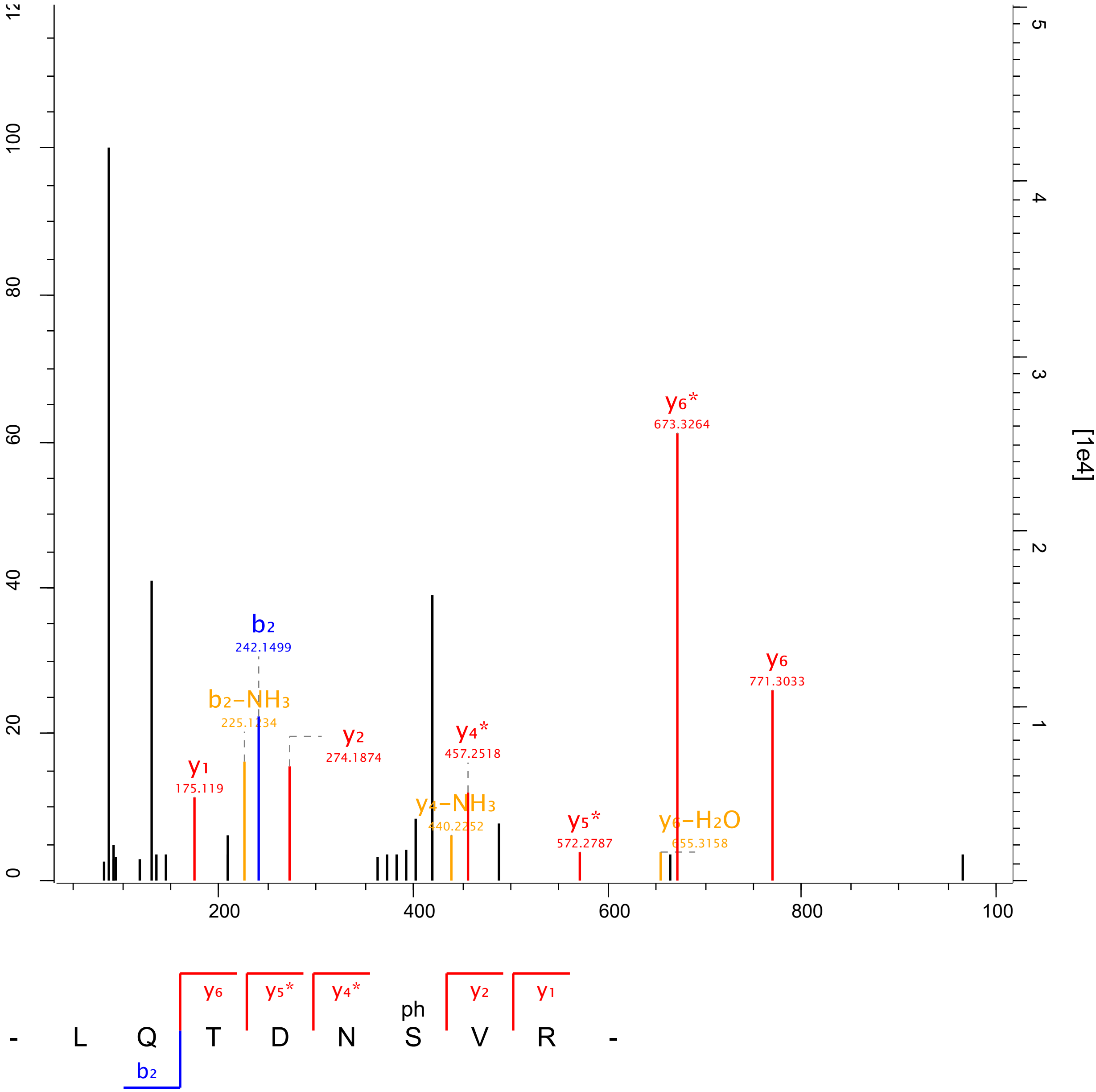

| Raw file | Scan | Method | Score | m/z |
|---|---|---|---|---|
| sys_00_1 | 2897 | FTMS; HCD | 46.8 | 680.74 |

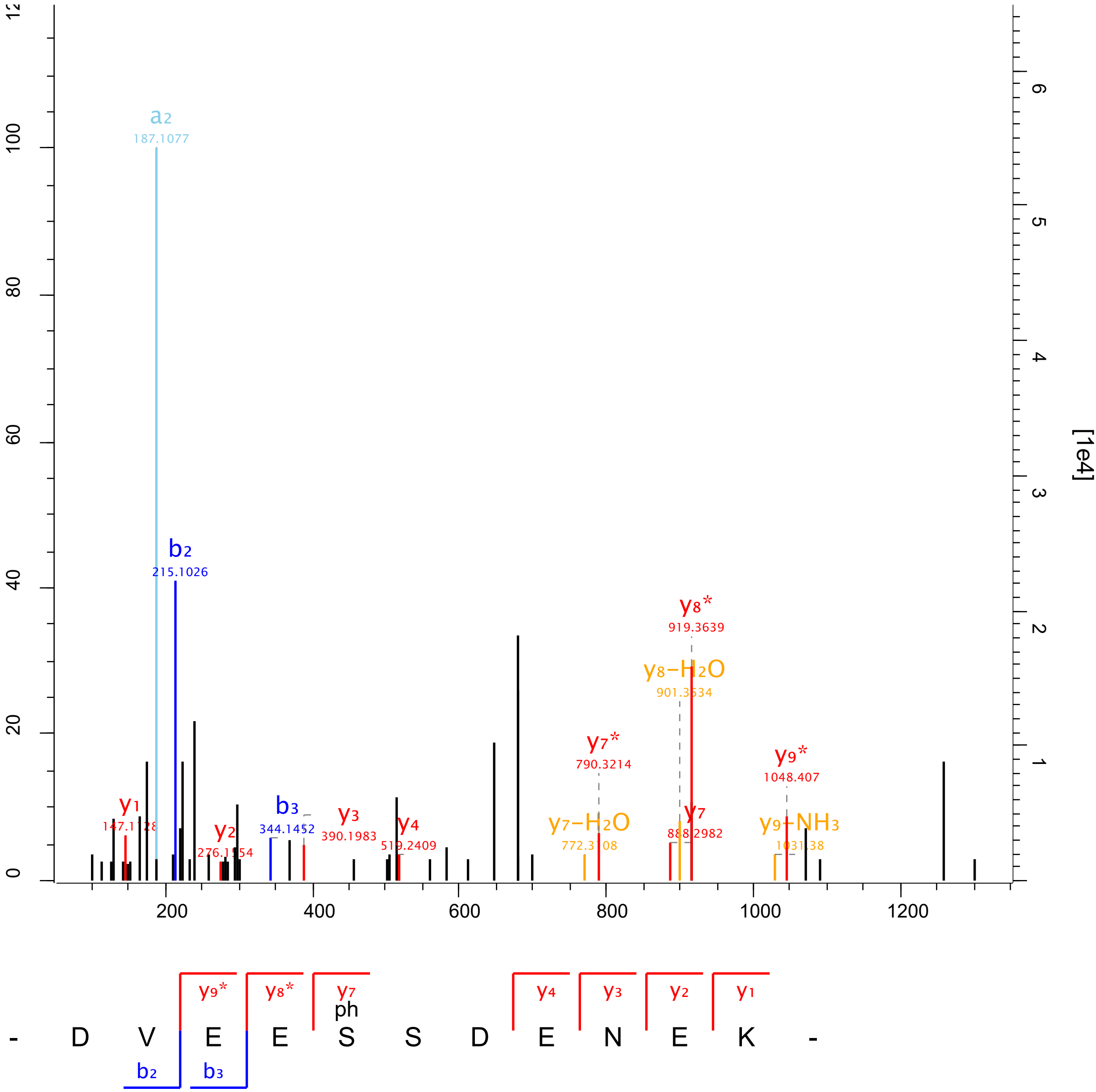

| Raw file | Scan | Method | Score | m/z |
|---|---|---|---|---|
| sys_00_1 | 3740 | FTMS; HCD | 139.31 | 486.7 |

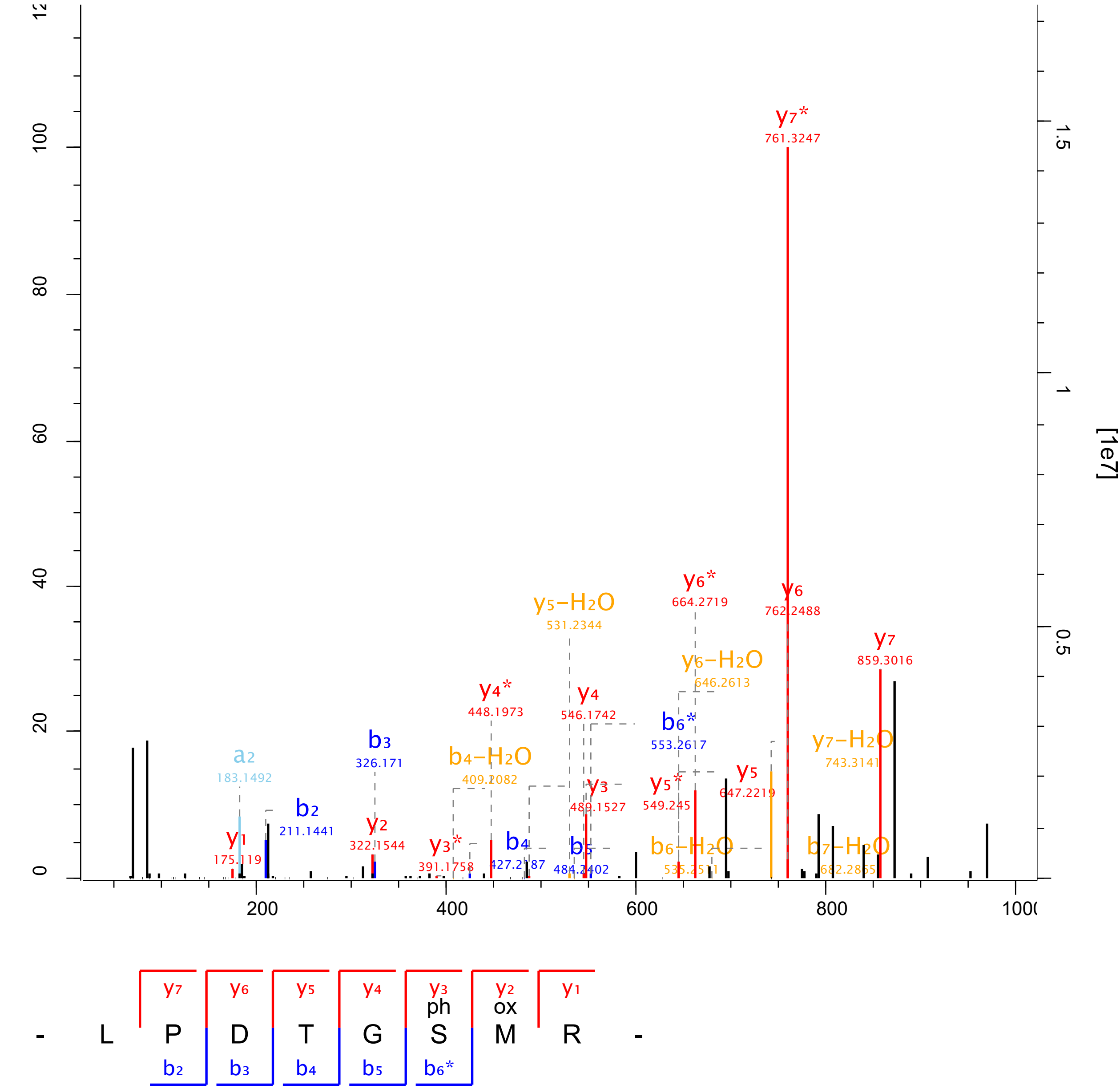

| Raw file | Scan | Method | Score | m/z |
|---|---|---|---|---|
| sys_00_1 | 3784 | FTMS; HCD | 54.09 | 616.7 |

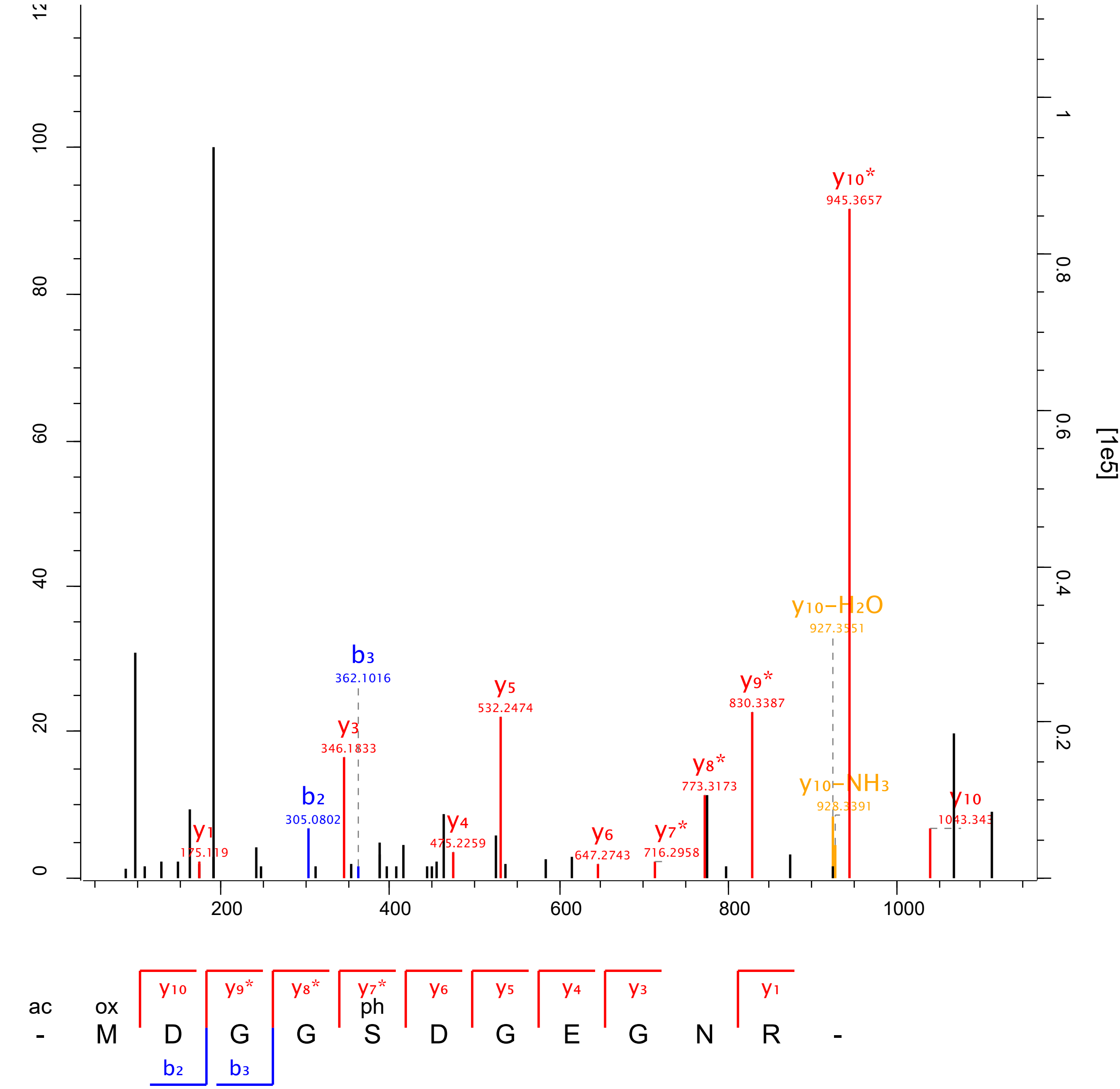

| Raw file | Scan | Method | Score | m/z |
|---|---|---|---|---|
| sys_00_1 | 4842 | FTMS; HCD | 68.66 | 421.7 |

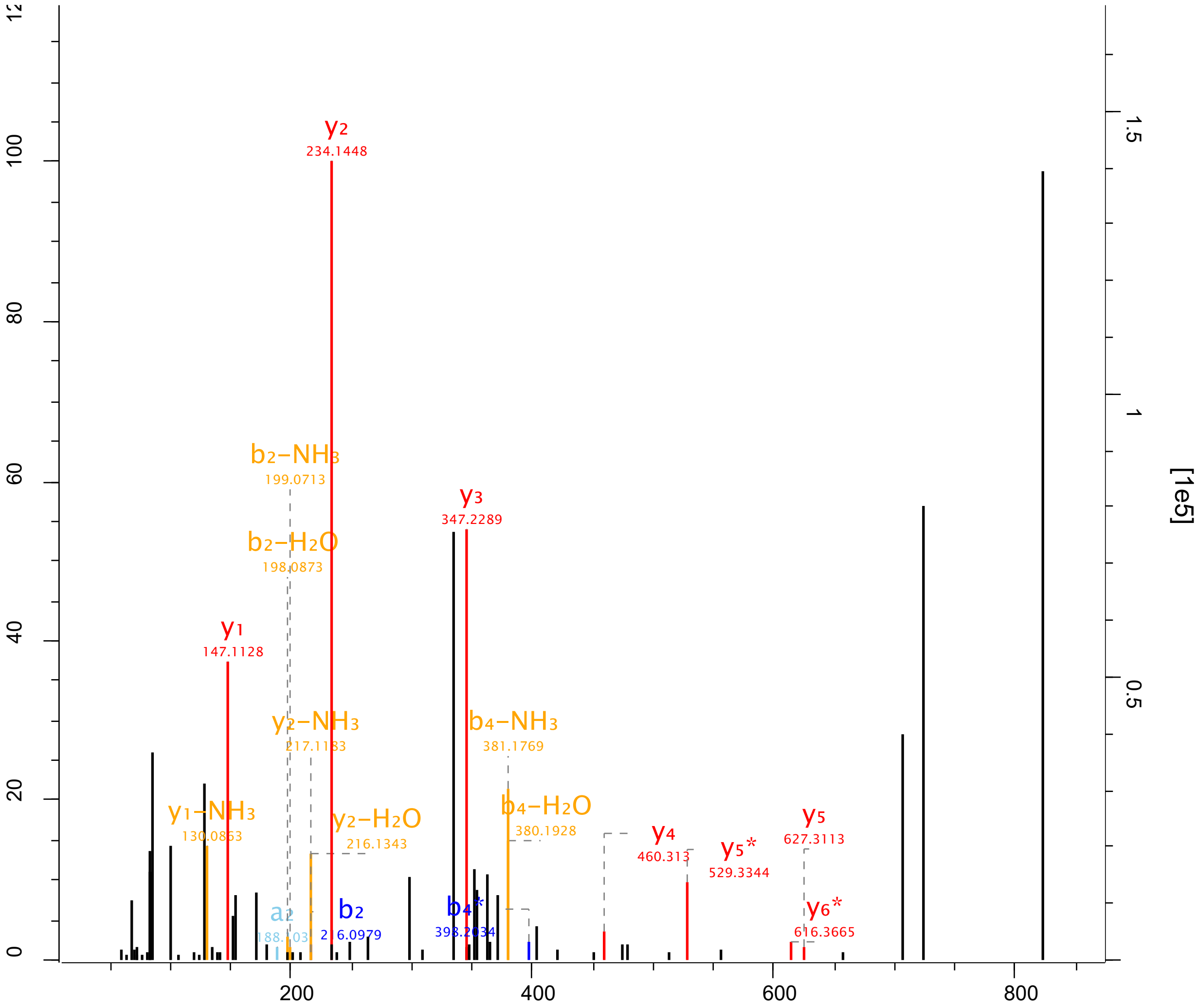


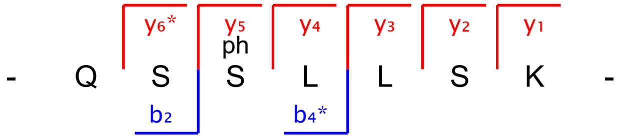

| Raw file | Scan | Method | Score | m/z |
| --- | --- | --- | --- | --- |
| sys_00_1 | 5069 | FTMS; HCD | 51.29 | 522.73 |

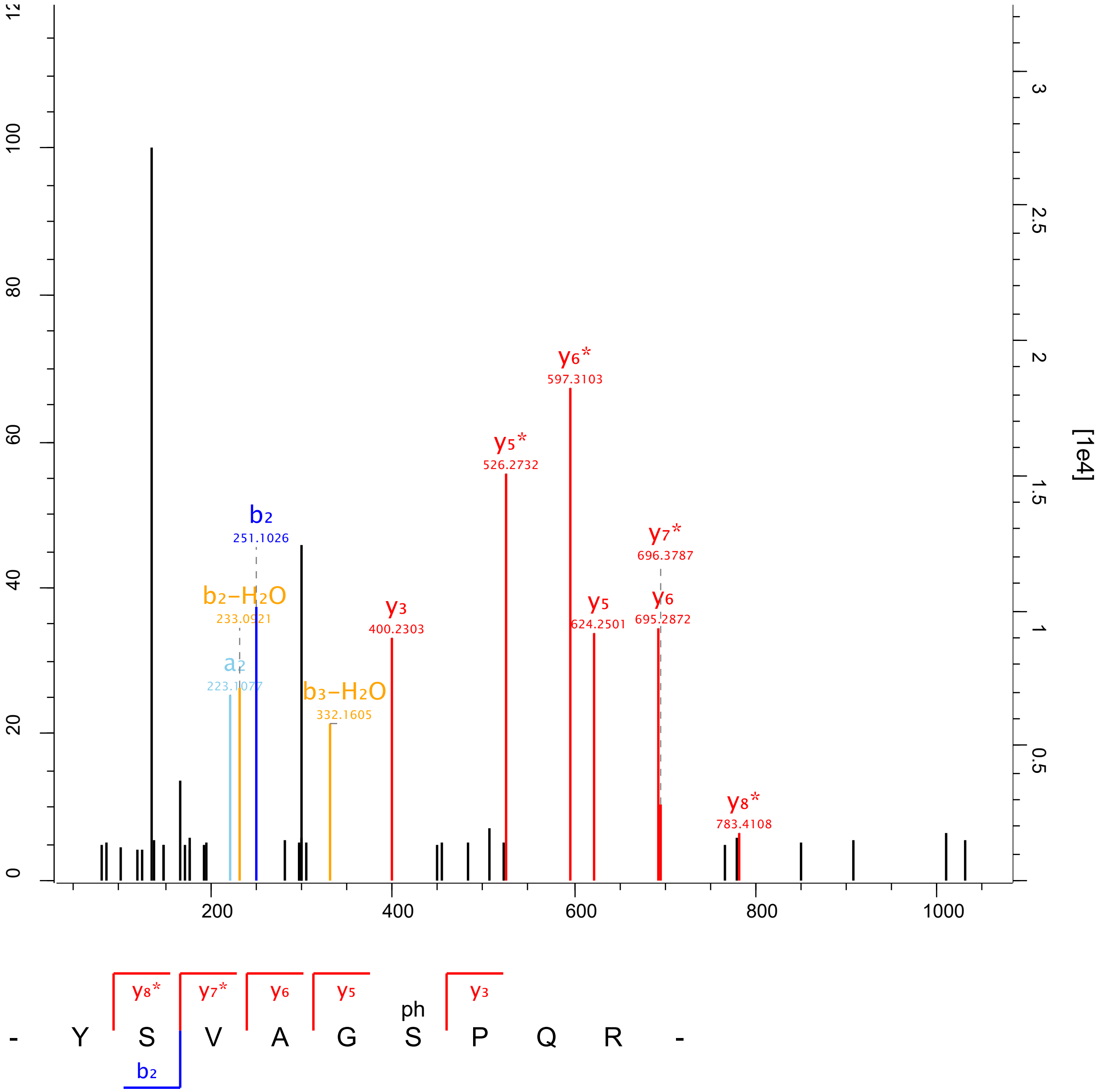

| Raw file | Scan | Method | Score | m/z |
|---|---|---|---|---|
| sys_00_1 | 6749 | FTMS; HCD | 114.86 | 498.23 |

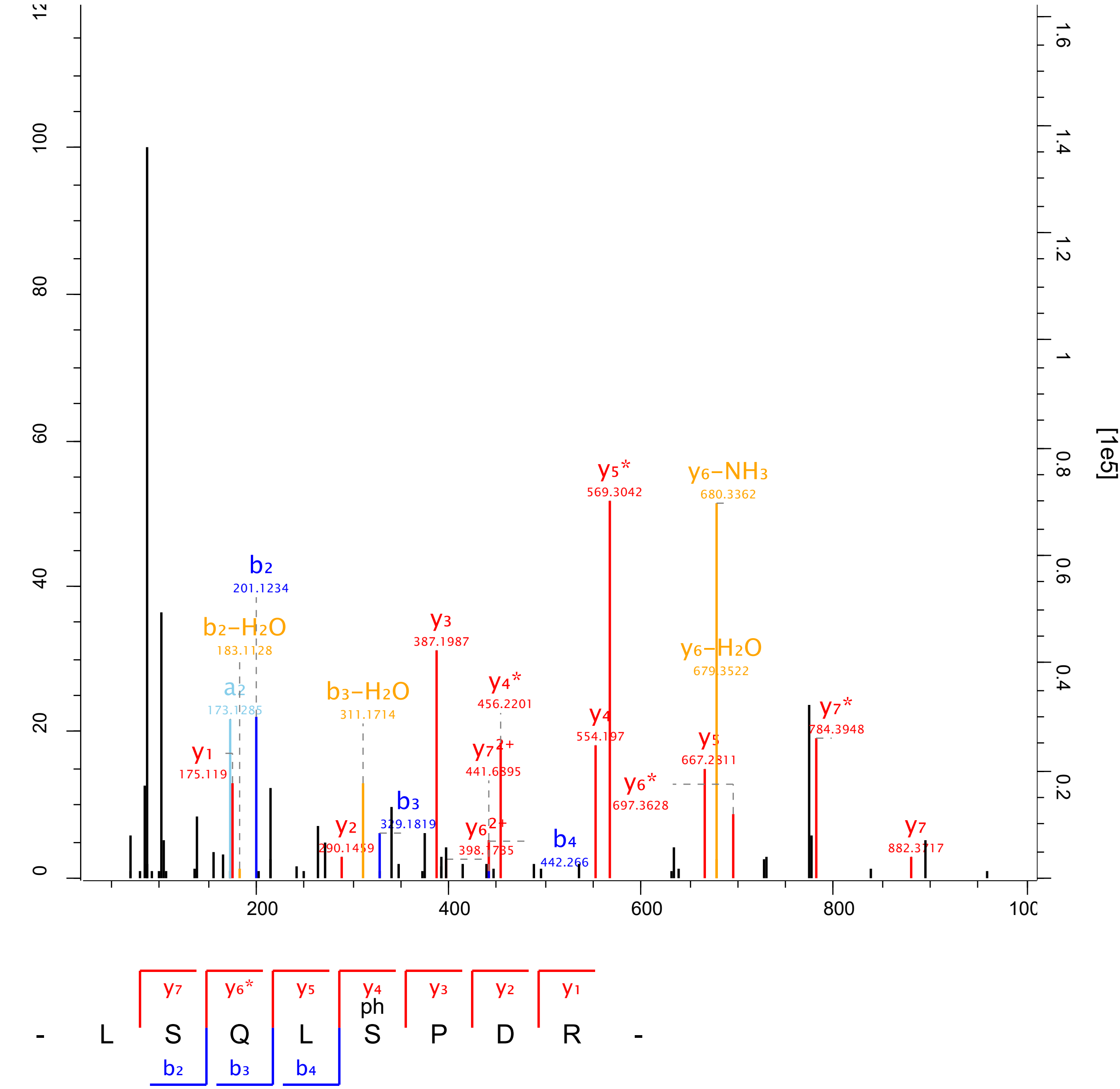

| Raw file | Scan | Method | Score | m/z |
| --- | --- | --- | --- | --- |
| sys_00_1 | 7359 | FTMS; HCD | 115.57 | 430.7 |

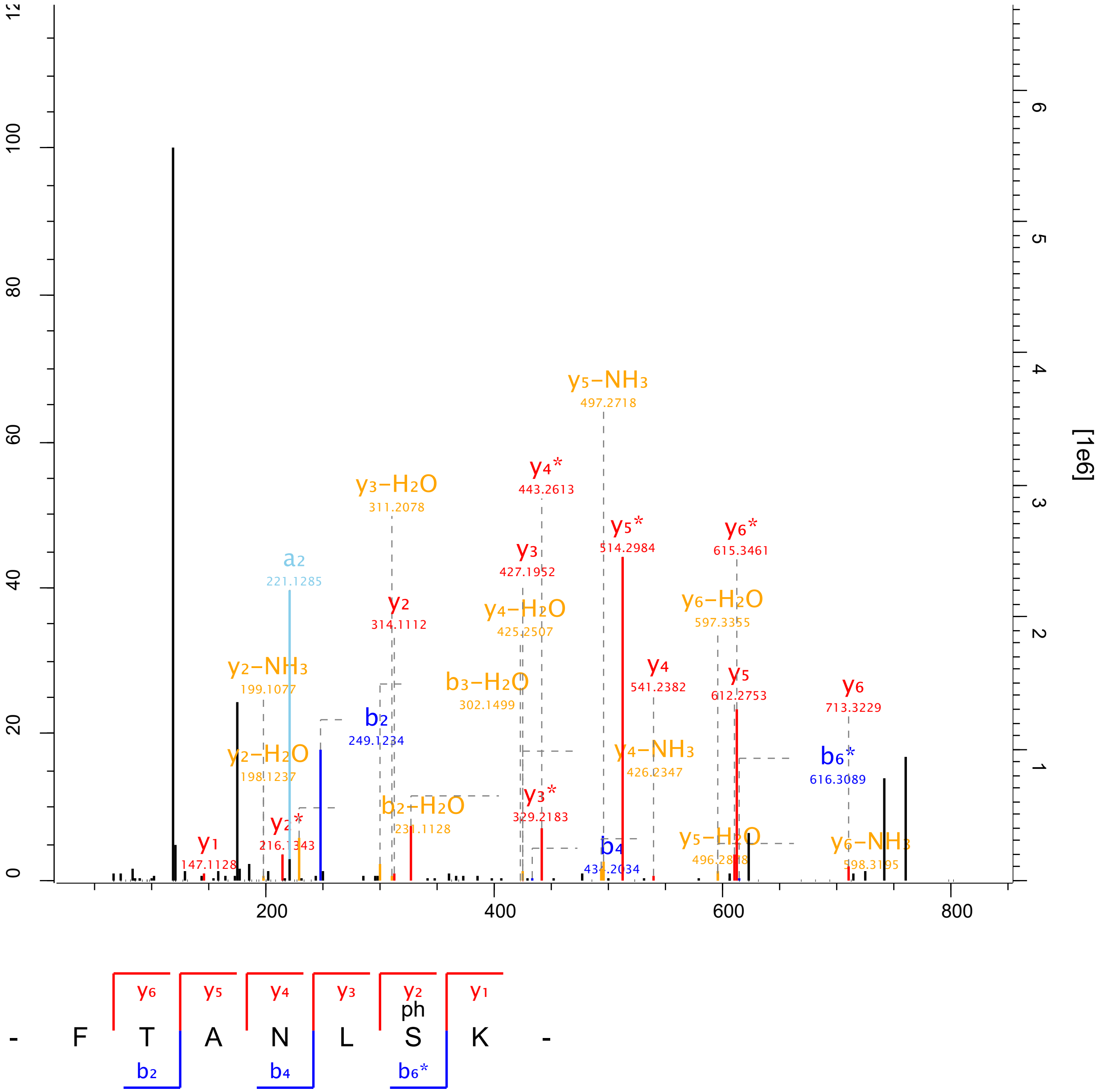

| Raw file | Scan | Method | Score | m/z |
|---|---|---|---|---|
| sys_00_1 | 7613 | FTMS; HCD | 81.63 | 650.75 |

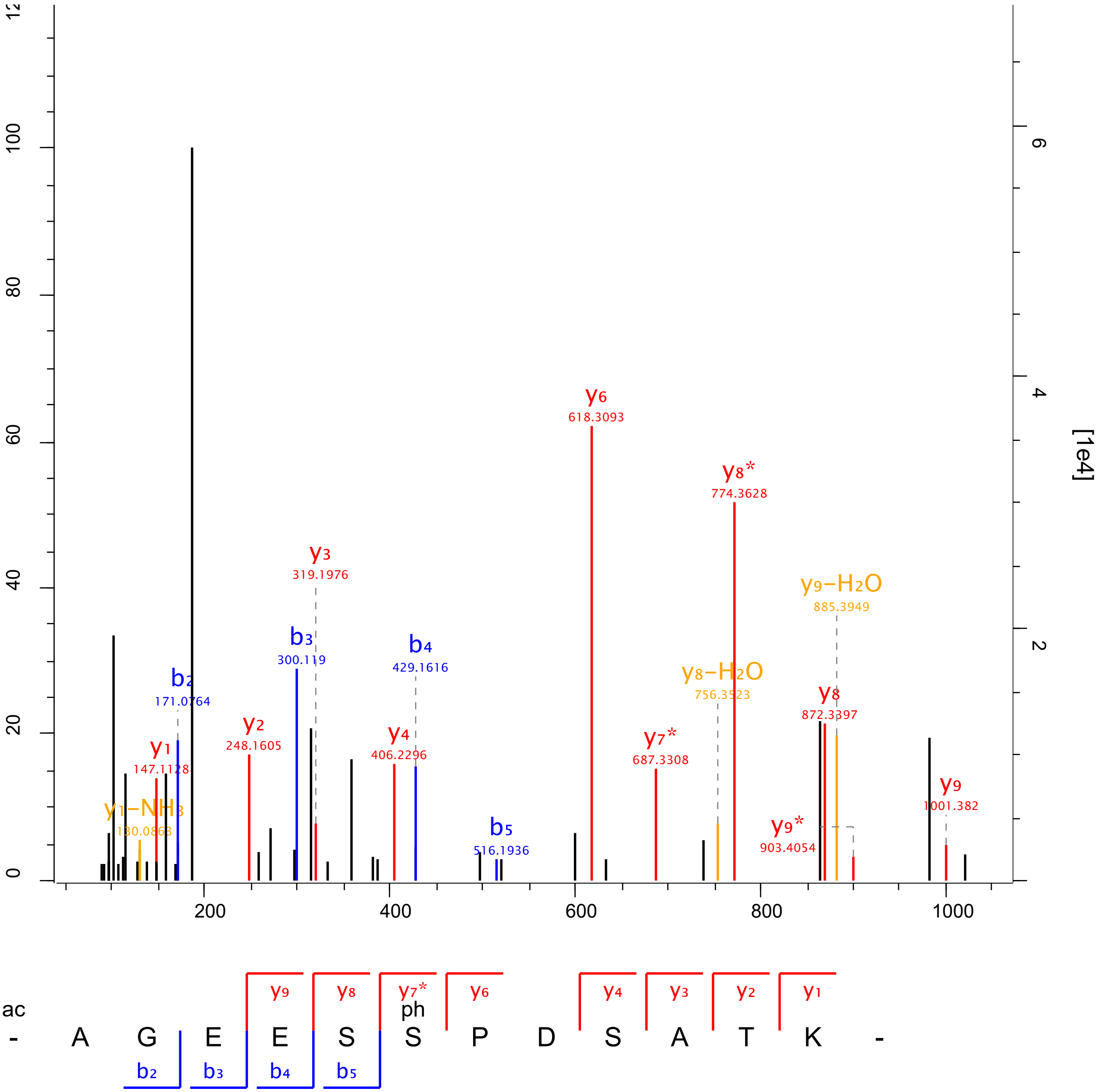

| Raw file | Scan | Method | Score | m/z |
|---|---|---|---|---|
| sys_00_1 | 8284 | FTMS; HCD | 97.69 | 601.75 |

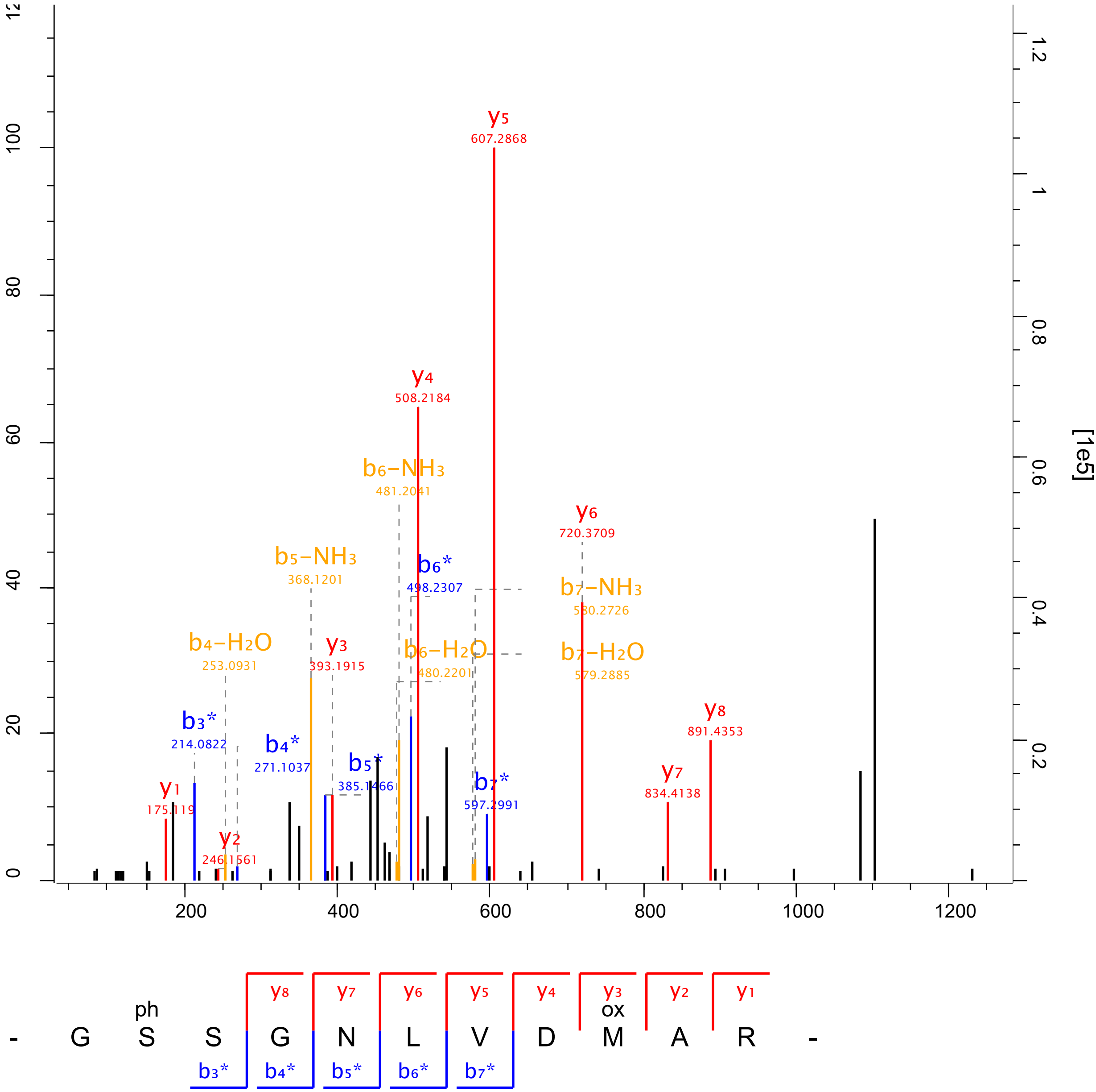

| Raw file | Scan | Method | Score | m/z |
|---|---|---|---|---|
| sys_00_1 | 8470 | FTMS; HCD | 83.18 | 628.76 |

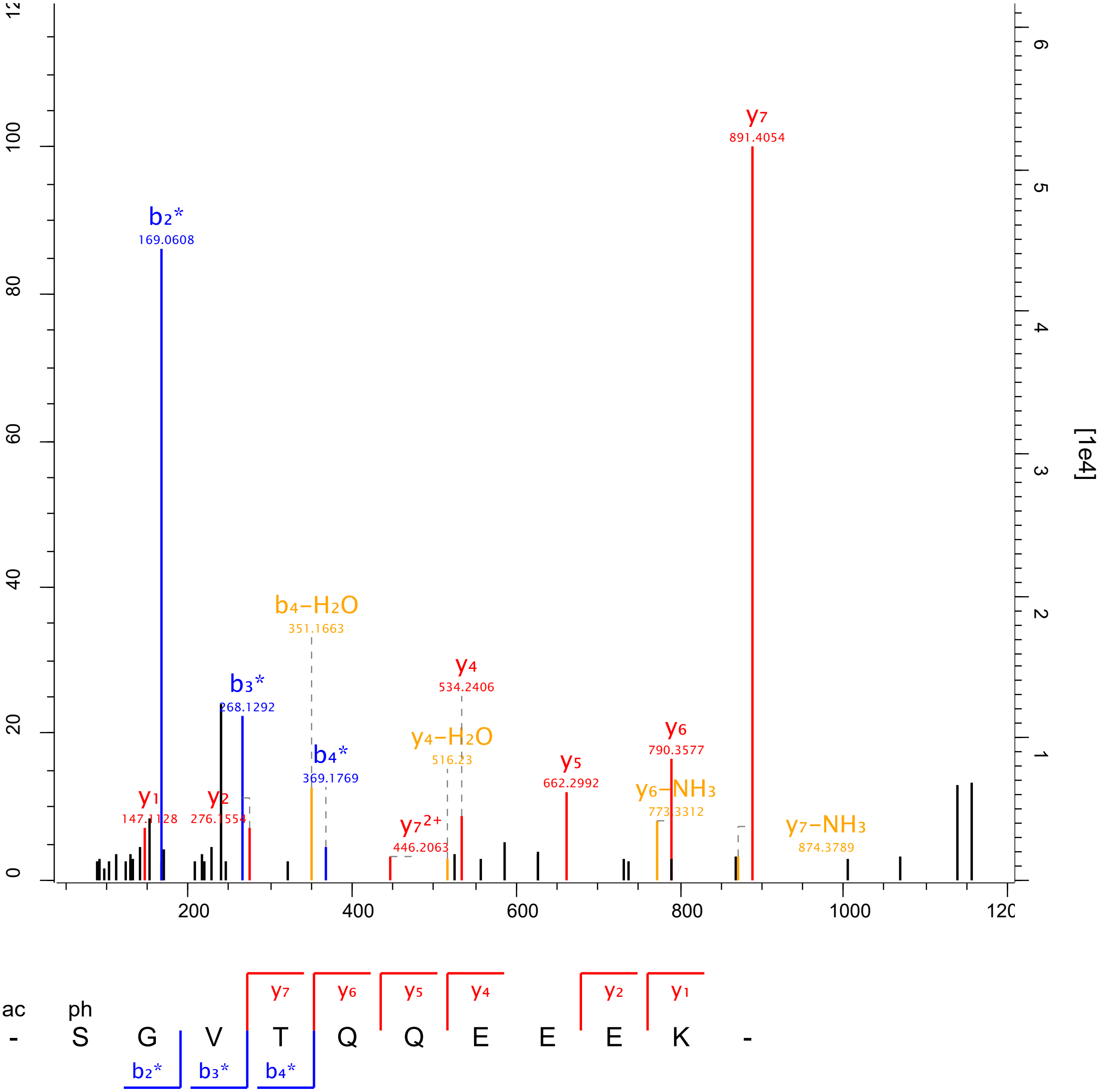

| Raw file | Scan | Method | Score | m/z |
| --- | --- | --- | --- | --- |
| sys_00_2 | 10058 | FTMS; HCD | 67.11 | 483.18 |

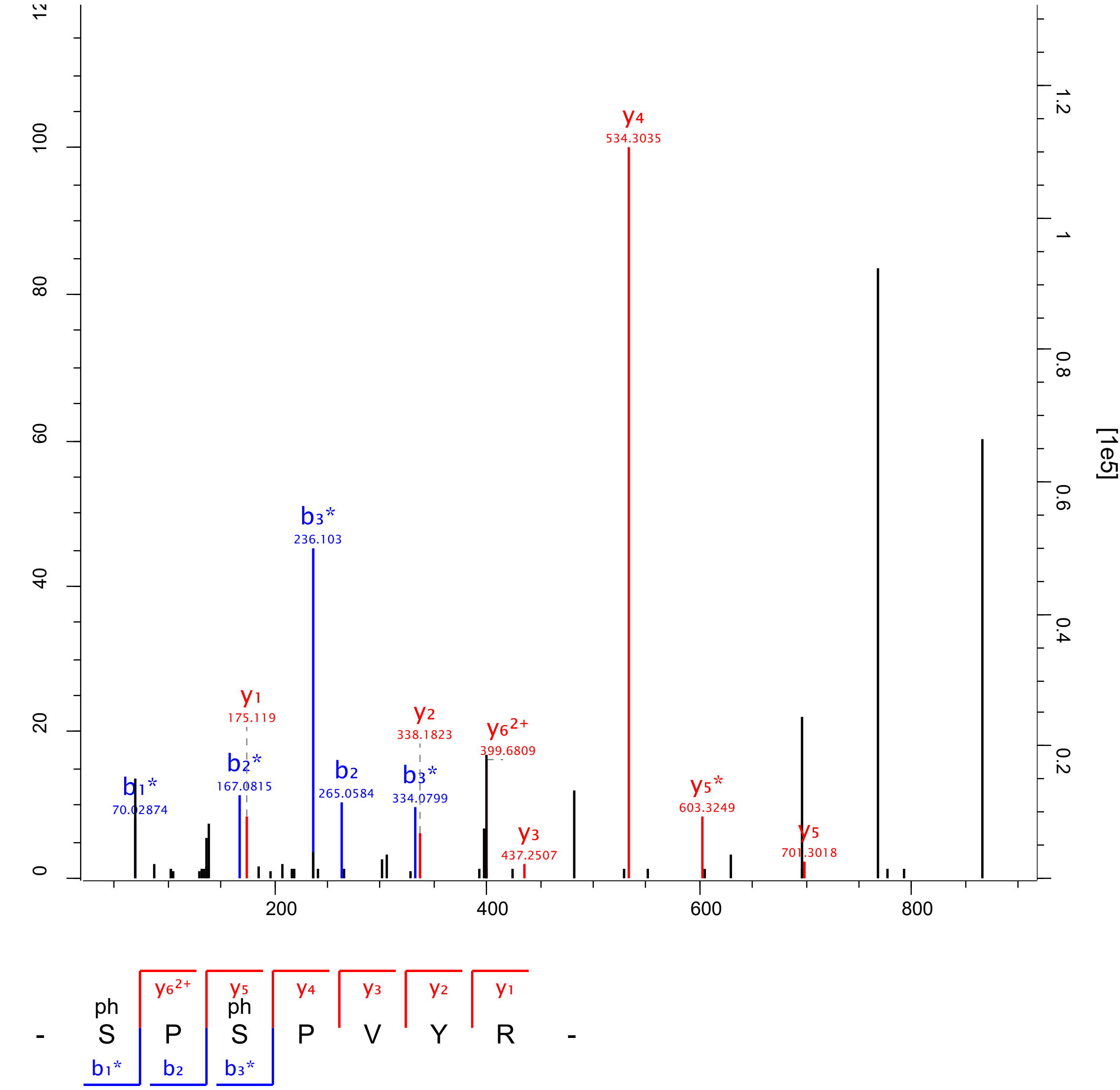

| Raw file | Scan | Method | Score | m/z |
|---|---|---|---|---|
| sys_00_2 | 10457 | FTMS; HCD | 44.25 | 644.27 |

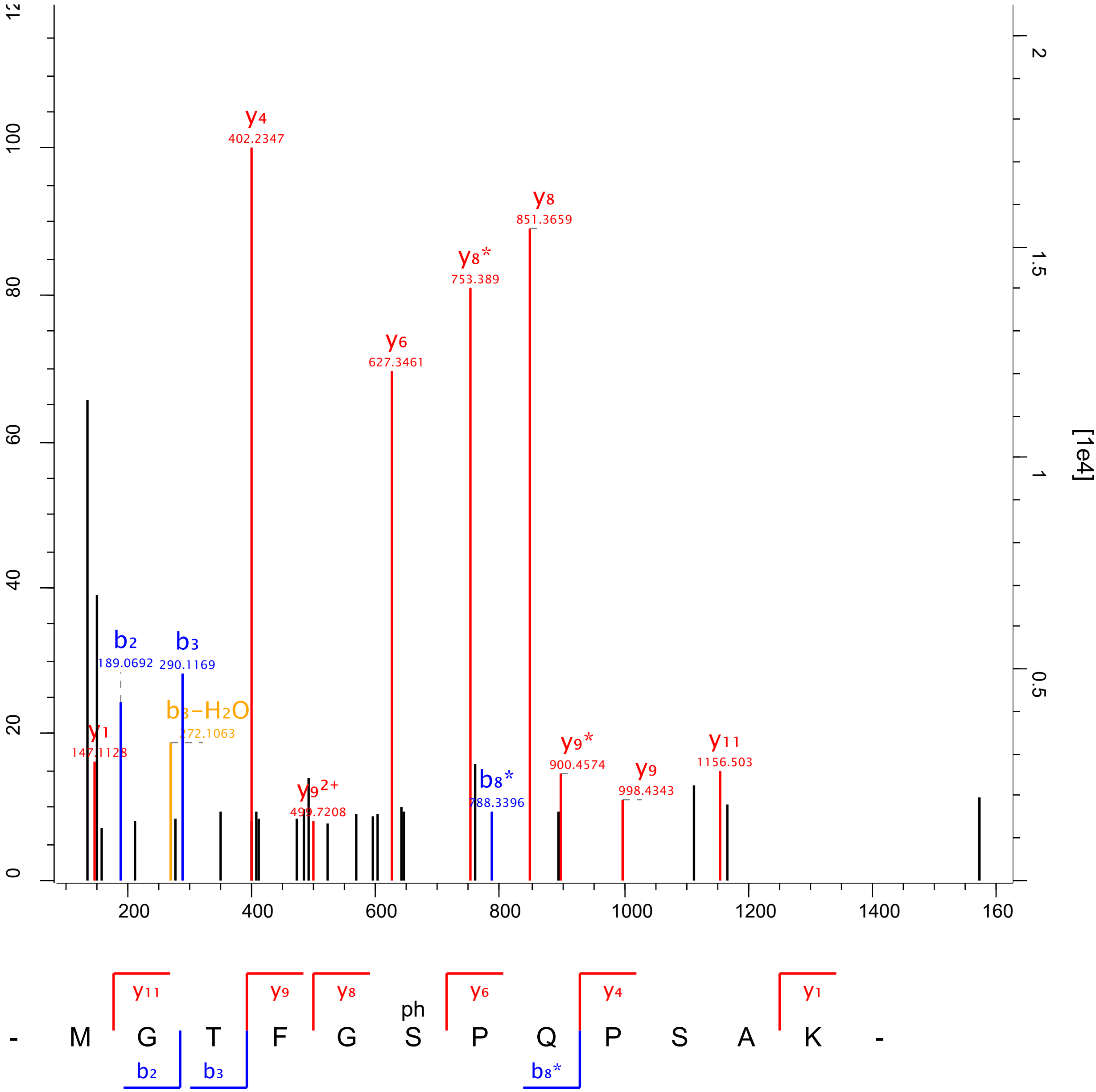

| Raw file | Scan | Method | Score | m/z |
|---|---|---|---|---|
| sys_00_2 | 10468 | FTMS; HCD | 54.16 | 490.71 |

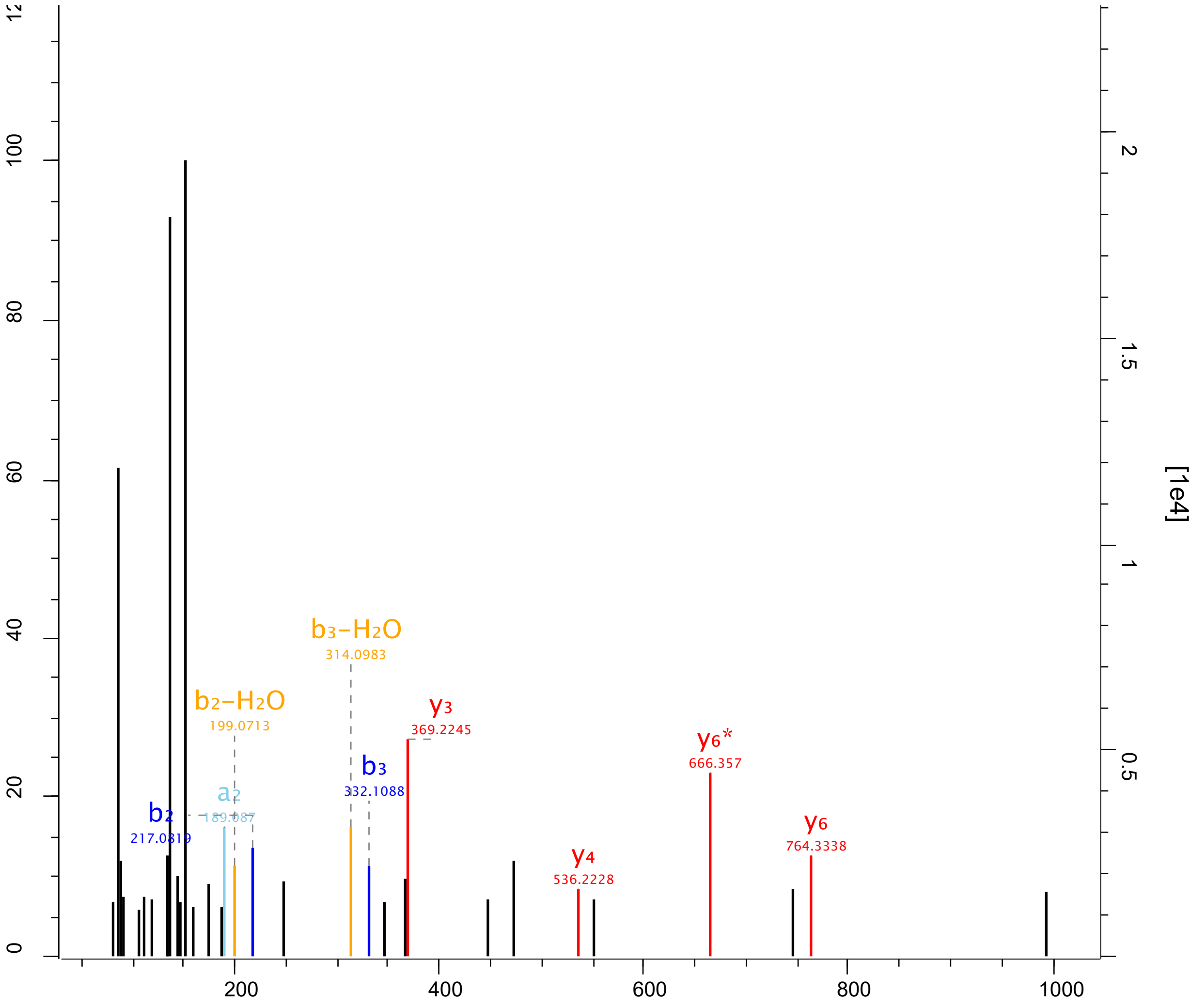


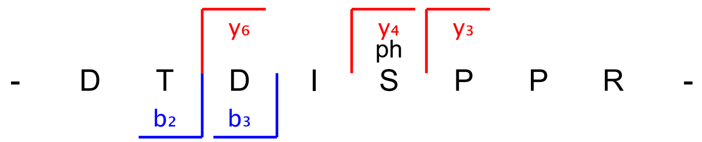

| Raw file | Scan | Method | Score | m/z |
|---|---|---|---|---|
| sys_00_2 | 10876 | FTMS; HCD | 147.62 | 456.73 |

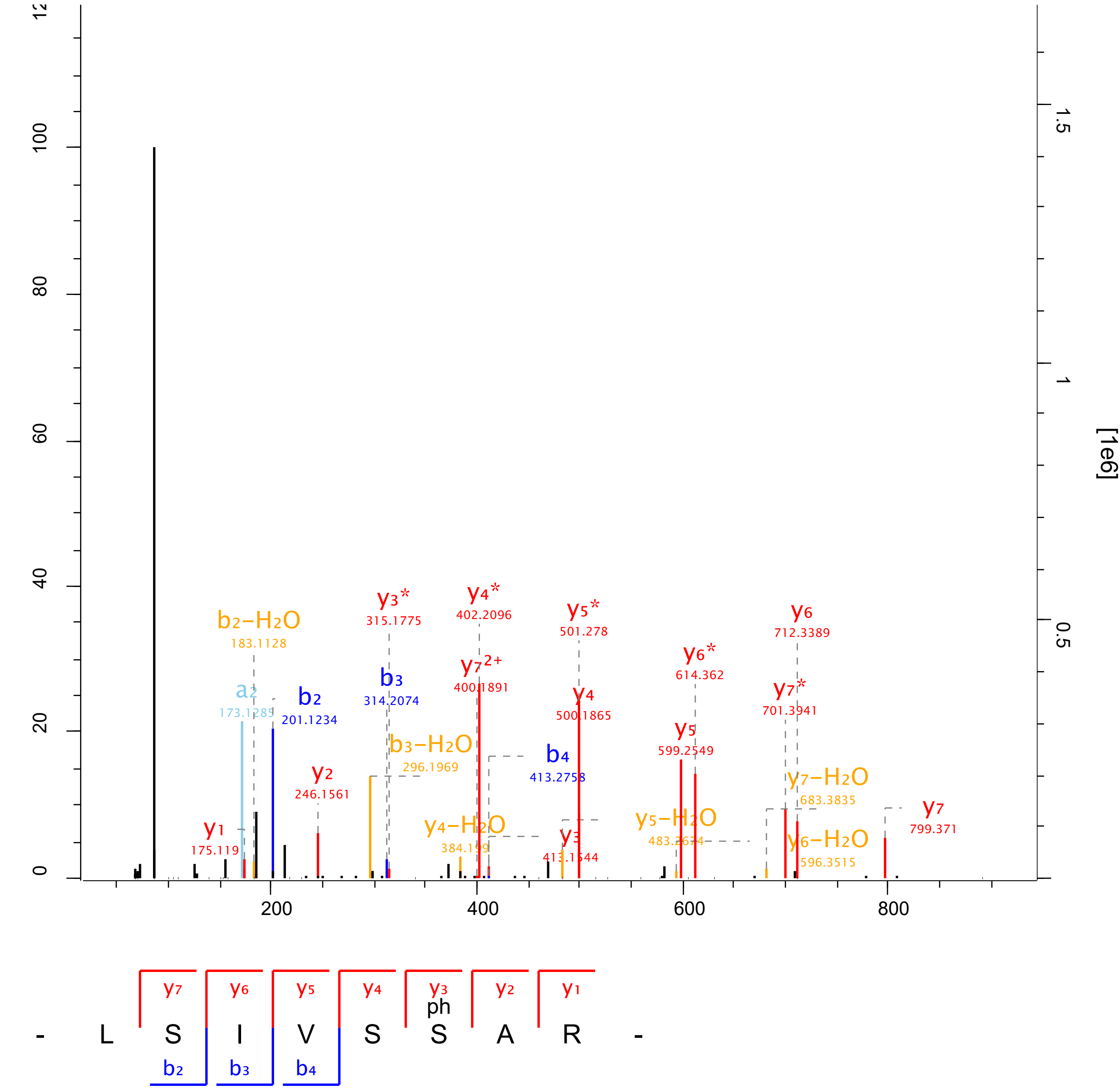

| Raw file | Scan | Method | Score | m/z |
|---|---|---|---|---|
| sys_00_2 | 11570 | FTMS; HCD | 109.04 | 657.3 |

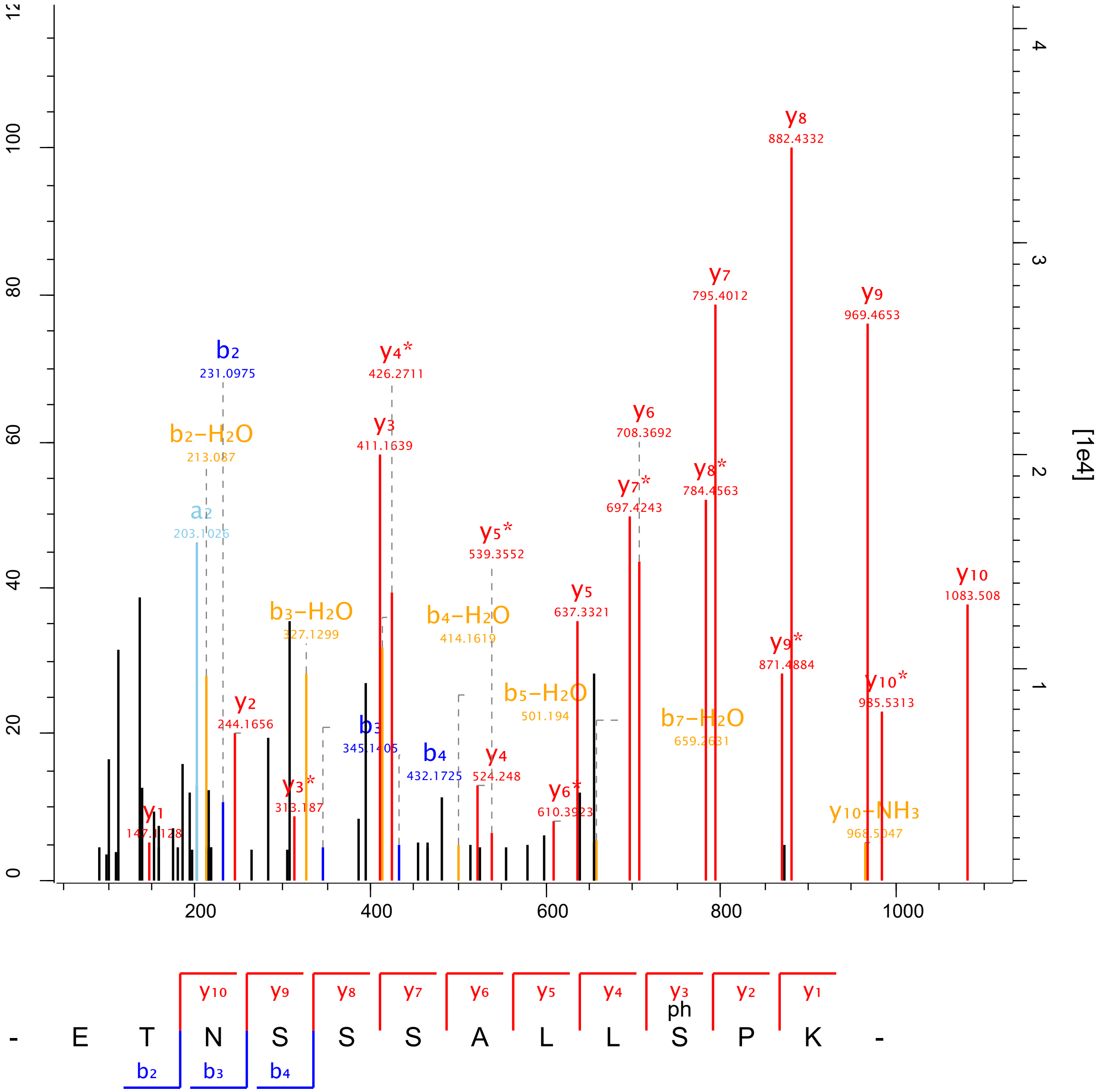

| Raw file | Scan | Method | Score | m/z |
| --- | --- | --- | --- | --- |
| sys_00_2 | 13502 | FTMS; HCD | 40.43 | 582.23 |

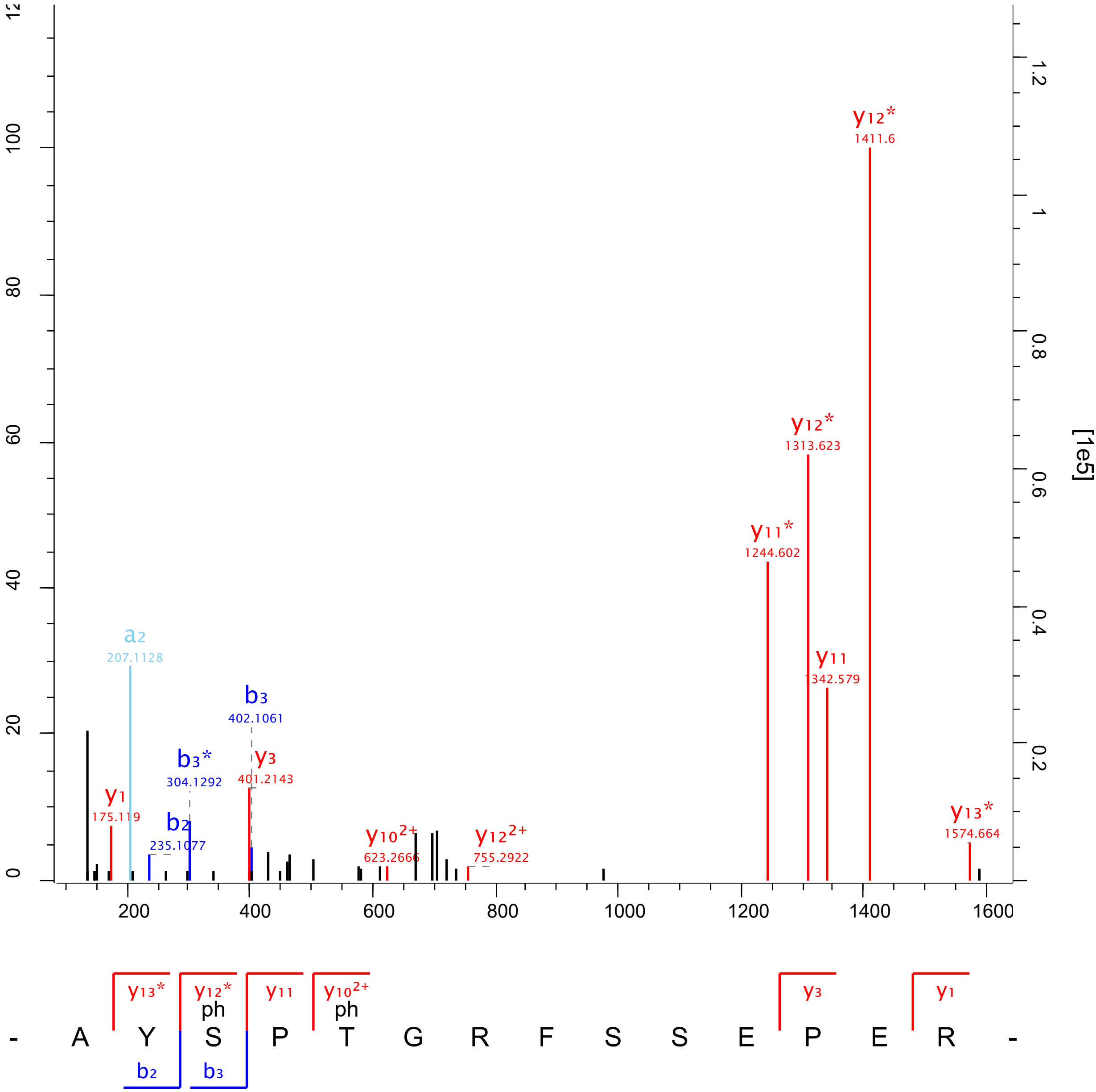

| Raw file | Scan | Method | Score | m/z |
| --- | --- | --- | --- | --- |
| sys_00_2 | 14148 | FTMS; HCD | 69.72 | 514.24 |

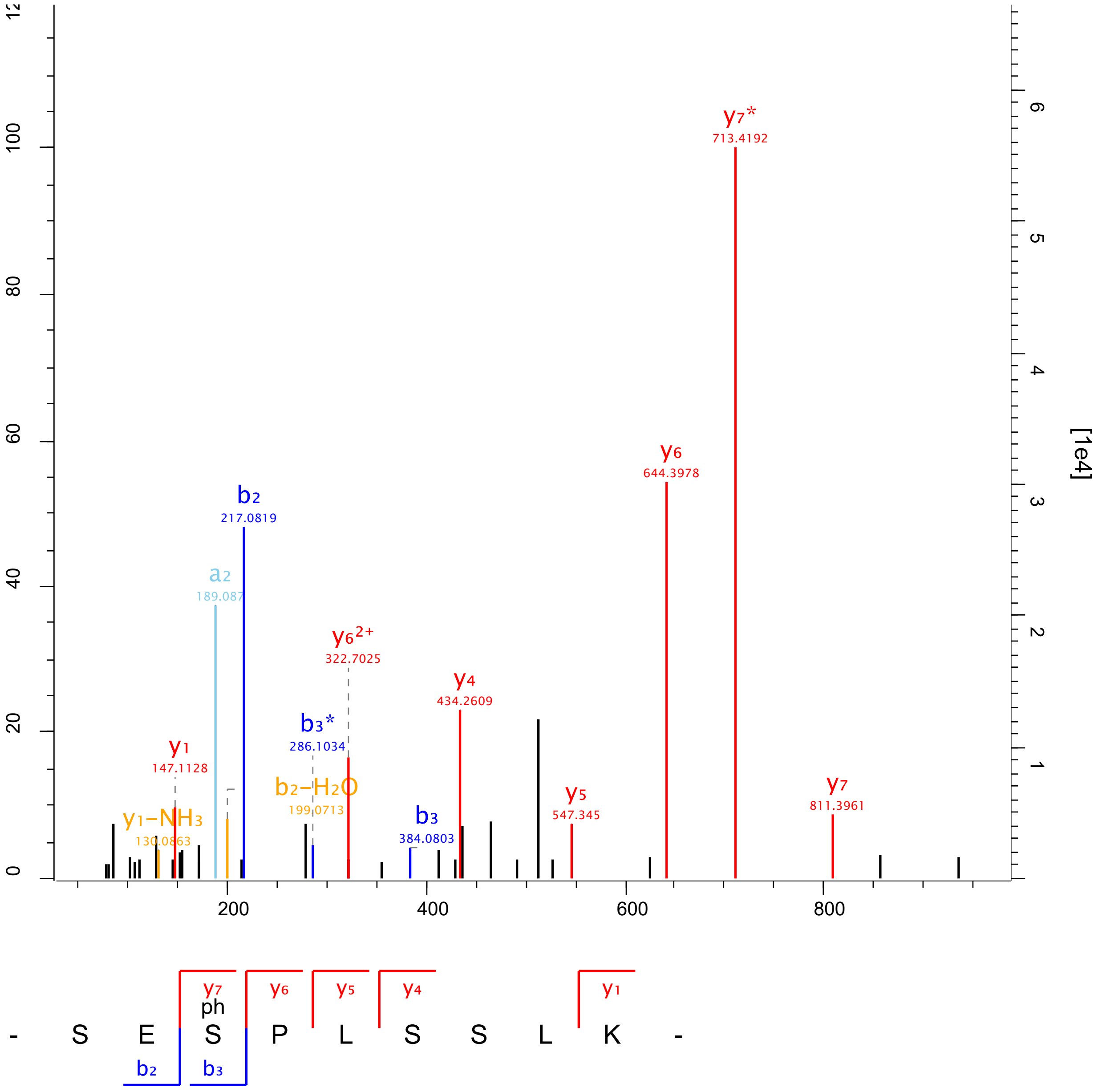

| Raw file | Scan | Method | Score | m/z |
|---|---|---|---|---|
| sys_00_2 | 14643 | FTMS; HCD | 54.28 | 759.83 |

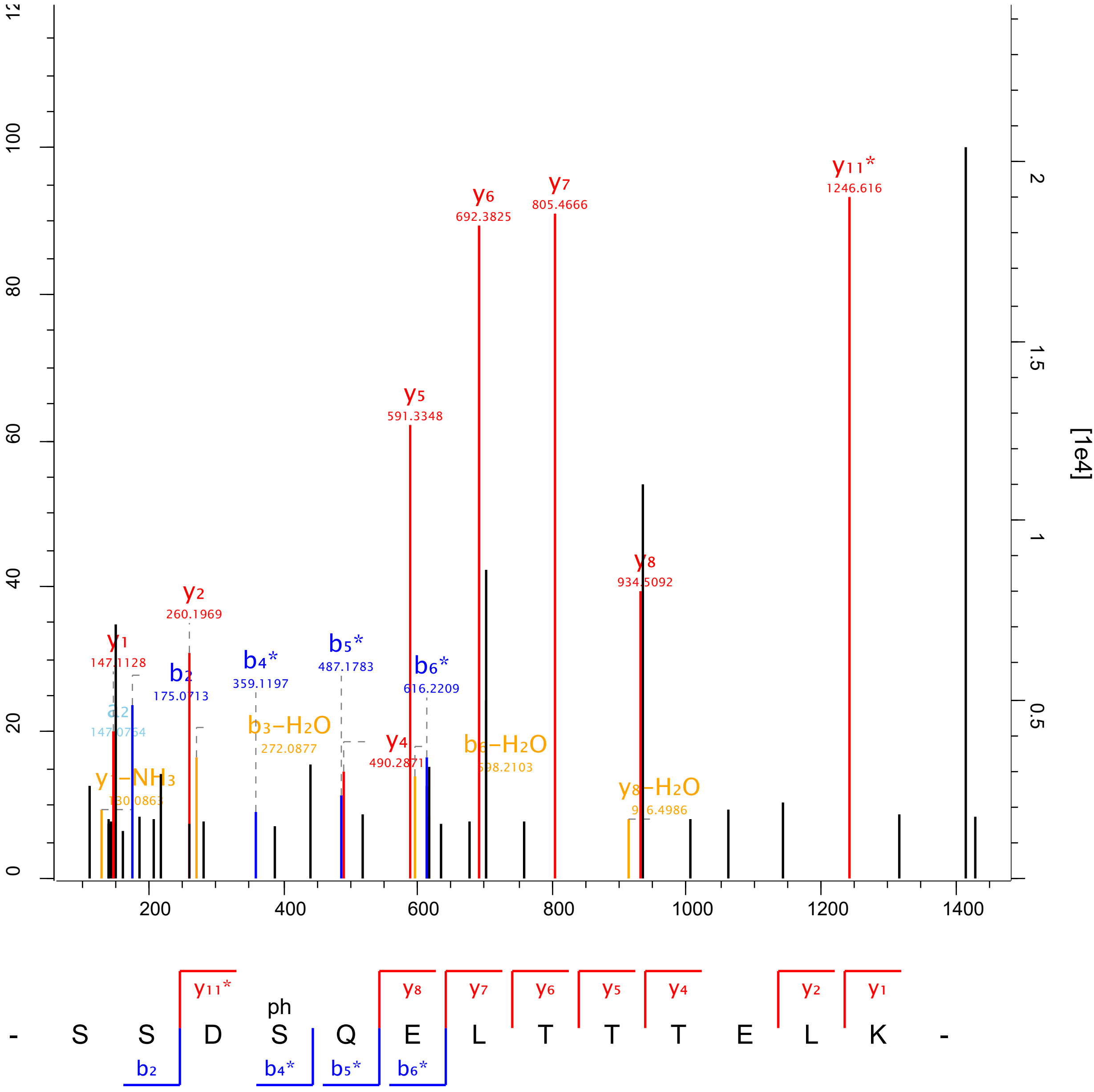

| Raw file | Scan | Method | Score | m/z |
|---|---|---|---|---|
| sys_00_2 | 14960 | FTMS; HCD | 305.22 | 756.29 |

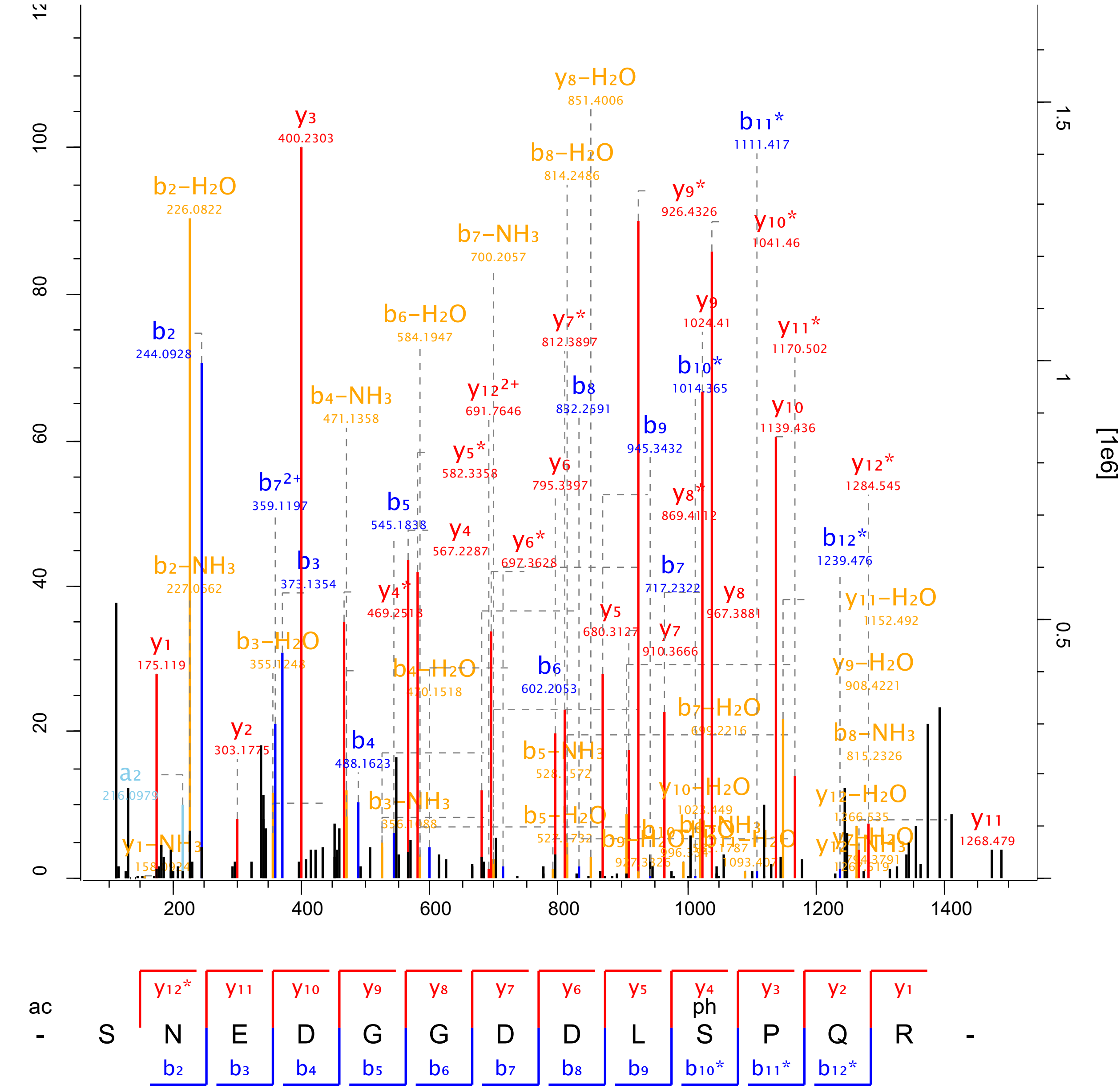

| Raw file | Scan | Method | Score | m/z |
| --- | --- | --- | --- | --- |
| sys_00_2 | 1497 | FTMS; HCD | 43.77 | 685.93 |

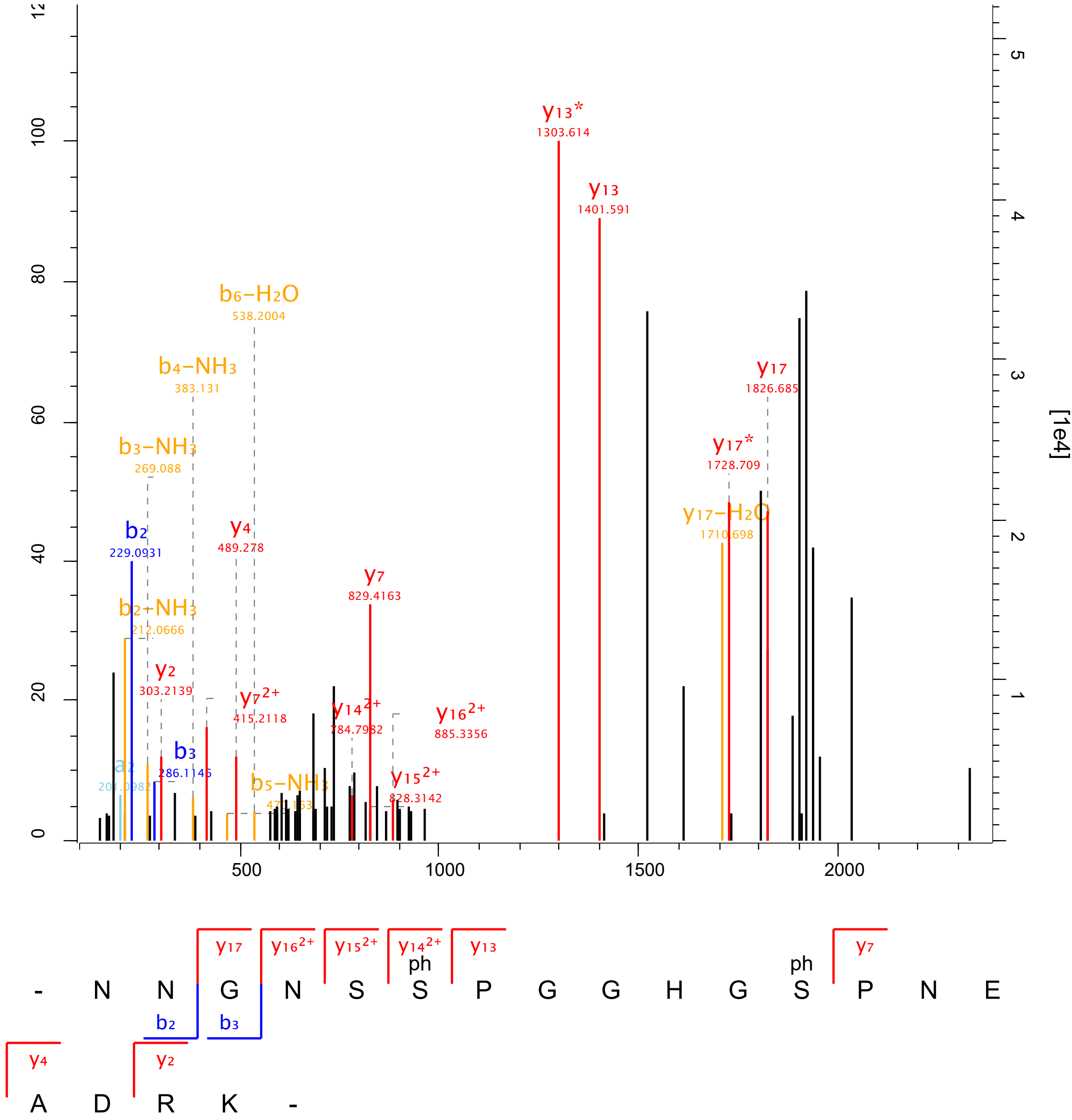

| Raw file | Scan | Method | Score | m/z |
|---|---|---|---|---|
| sys_00_2 | 15999 | FTMS; HCD | 85.55 | 603.75 |

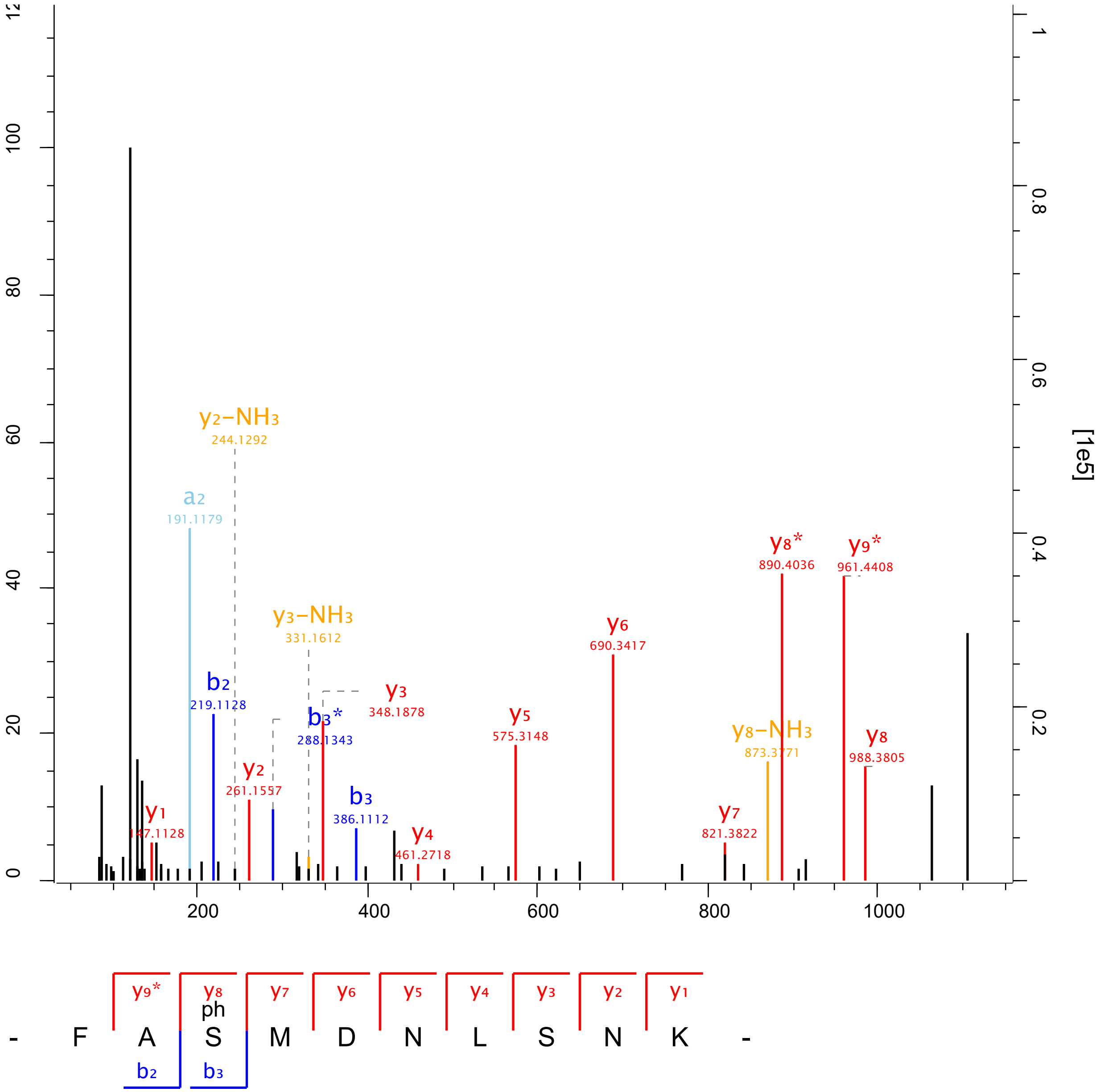

| Raw file | Scan | Method | Score | m/z |
|---|---|---|---|---|
| sys_00_2 | 16007 | FTMS; HCD | 117.86 | 553.24 |

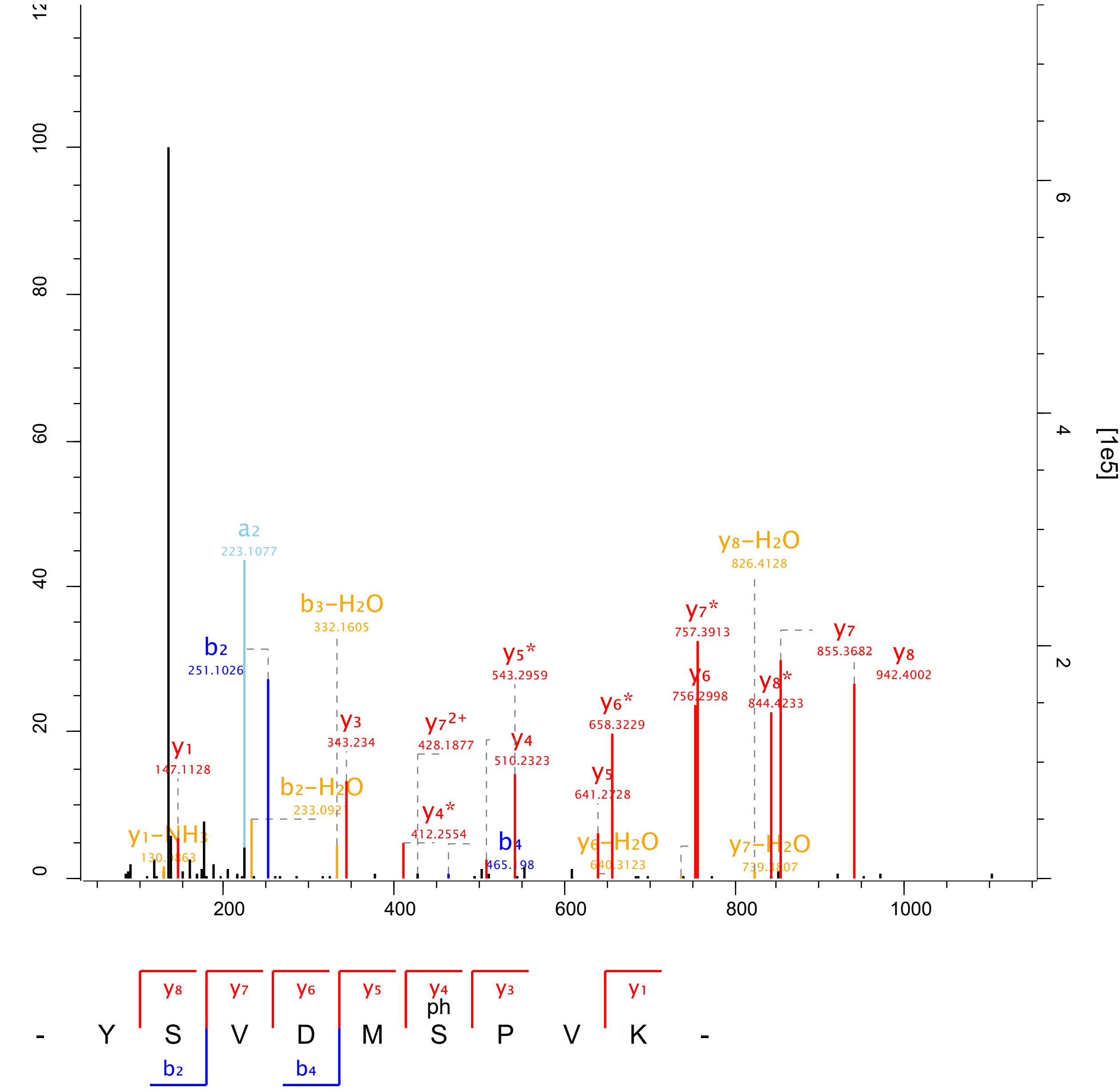

| Raw file | Scan | Method | Score | m/z |
|---|---|---|---|---|
| sys_00_2 | 17622 | FTMS; HCD | 177.42 | 649.25 |

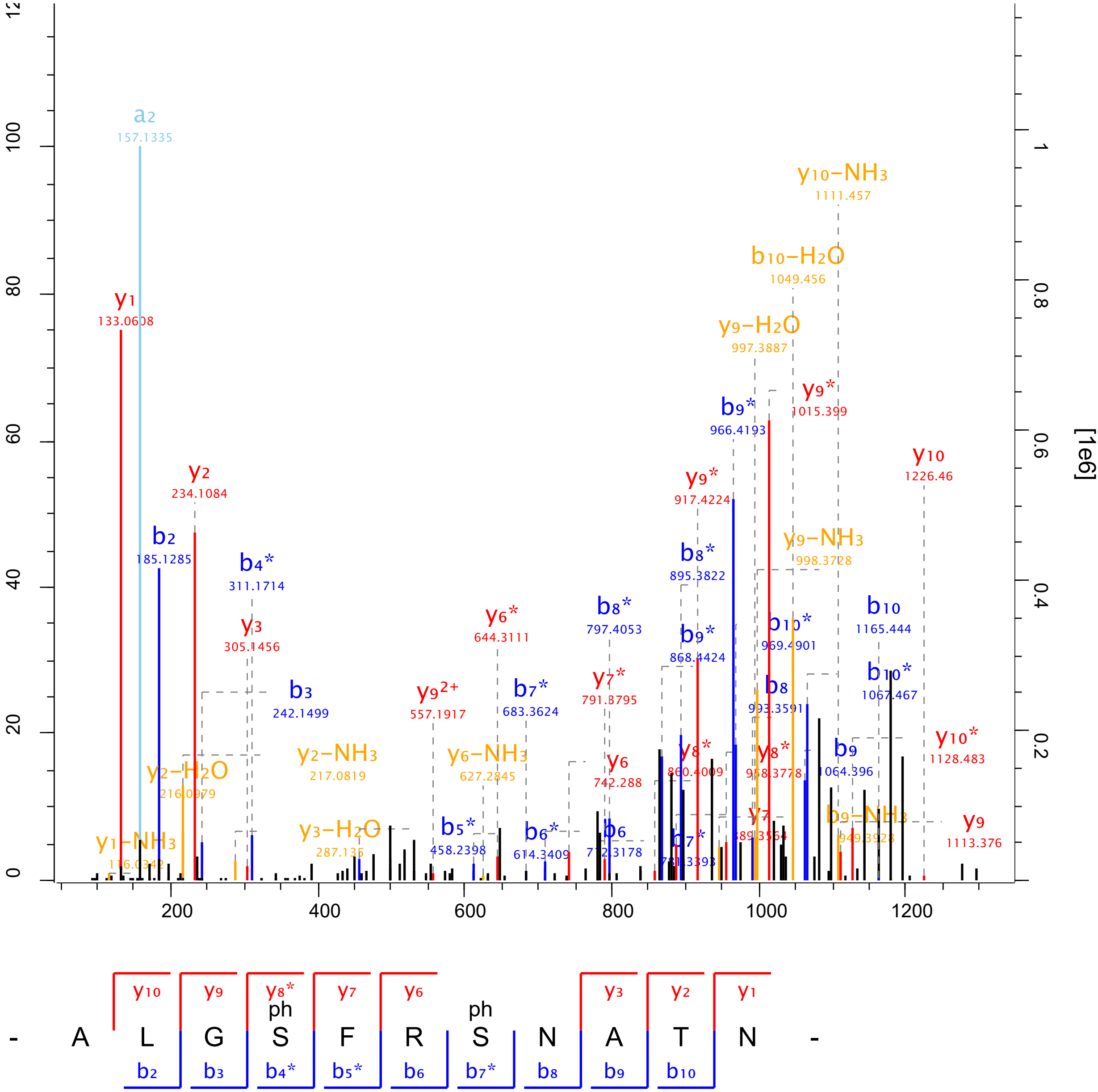

| Raw file | Scan | Method | Score | m/z |
| --- | --- | --- | --- | --- |
| sys_00_2 | 19780 | FTMS; HCD | 46.89 | 751.84 |

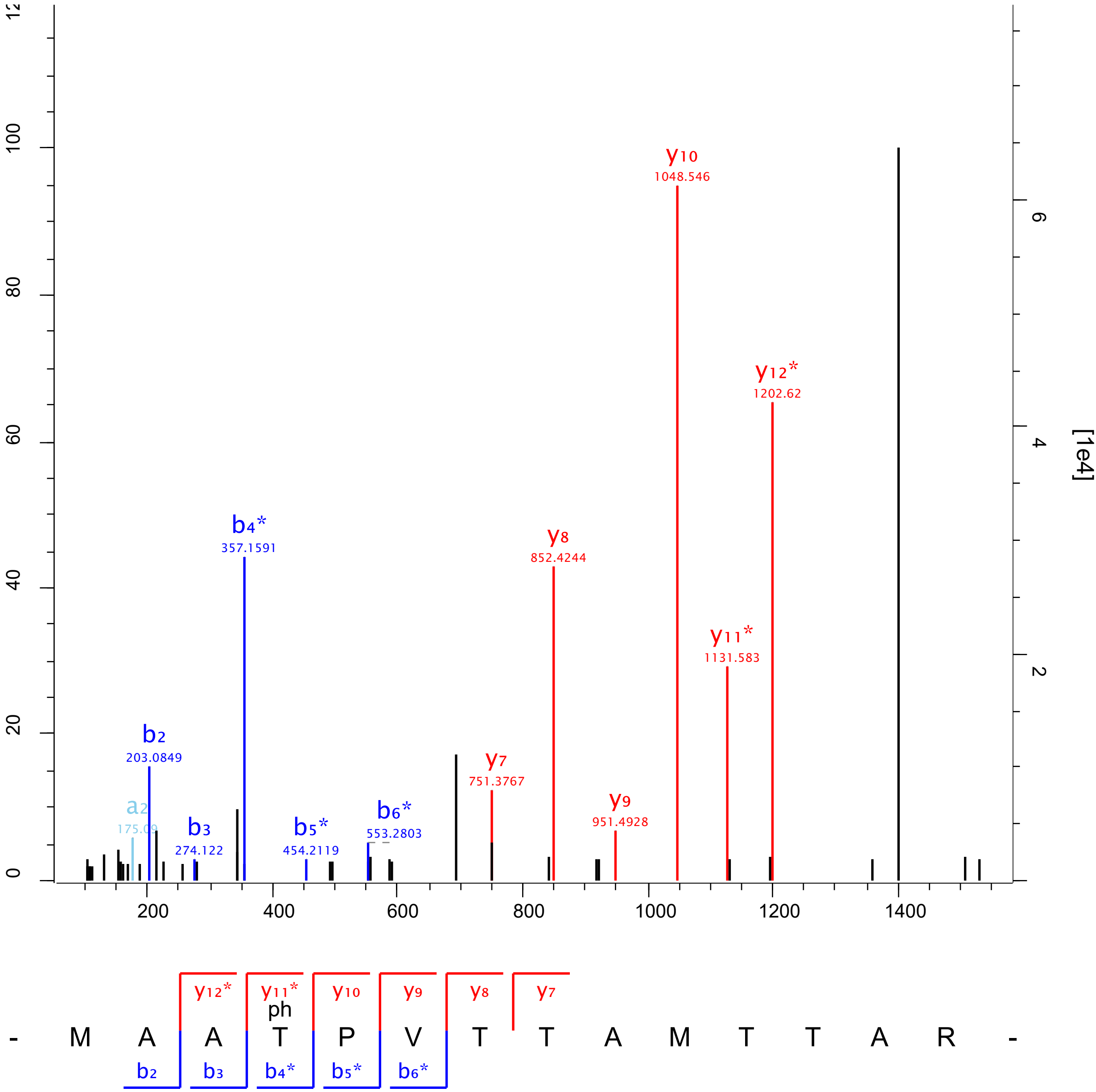

| Raw file | Scan | Method | Score | m/z |
| --- | --- | --- | --- | --- |
| sys_00_2 | 20133 | FTMS; HCD | 210.64 | 969.92 |

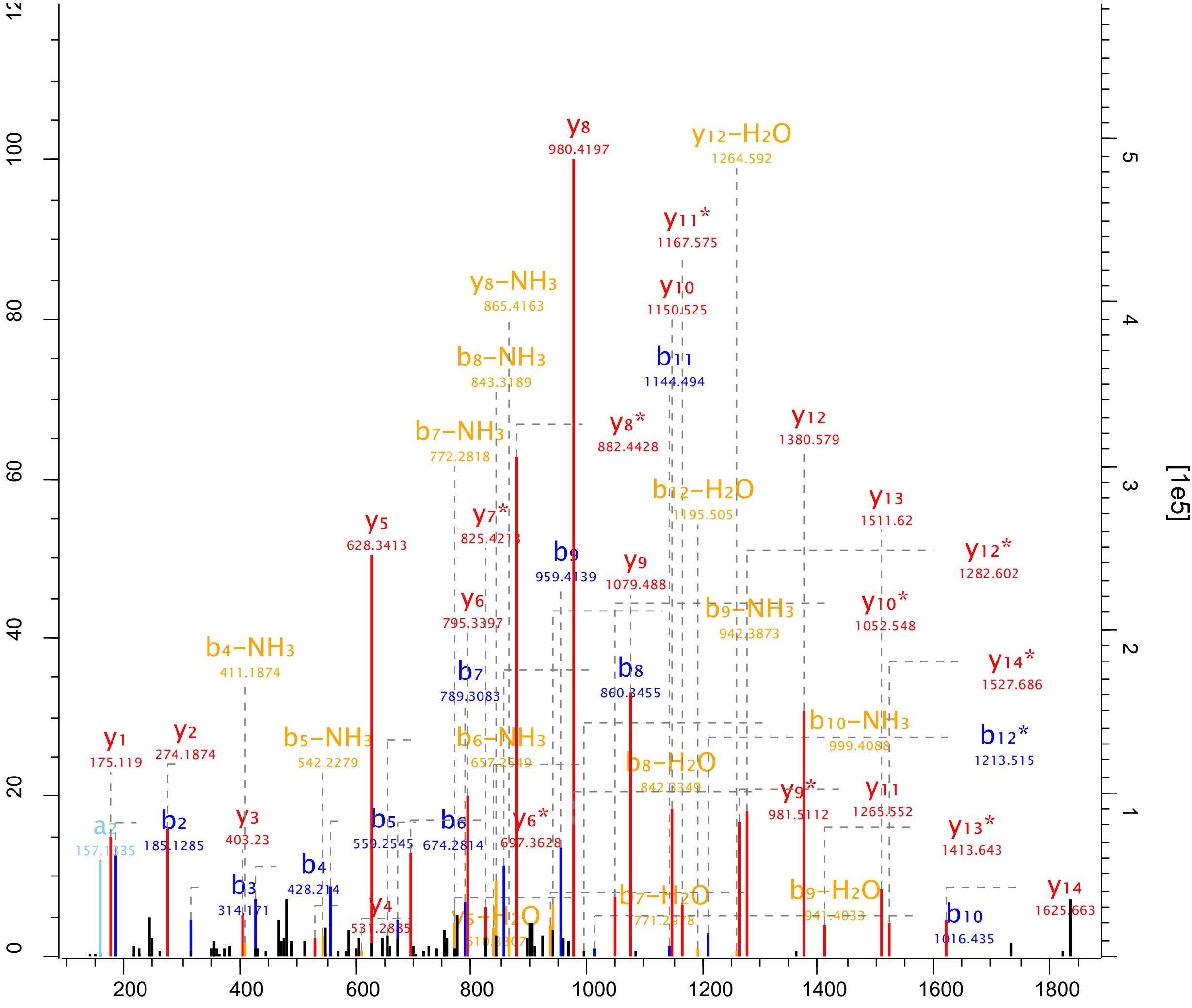


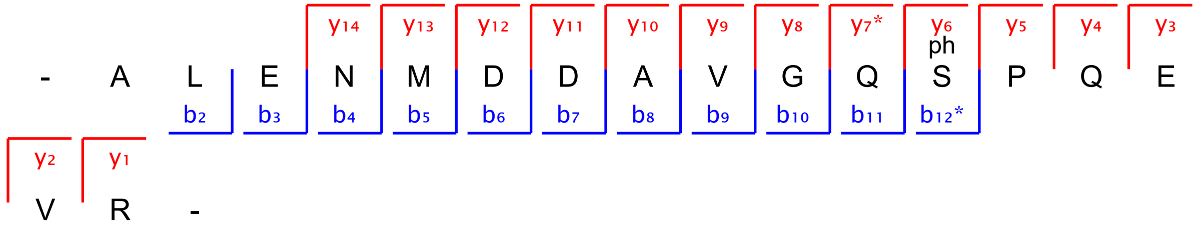

| Raw file | Scan | Method | Score | m/z |
|---|---|---|---|---|
| sys_00_2 | 21664 | FTMS; HCD | 201.29 | 887.88 |

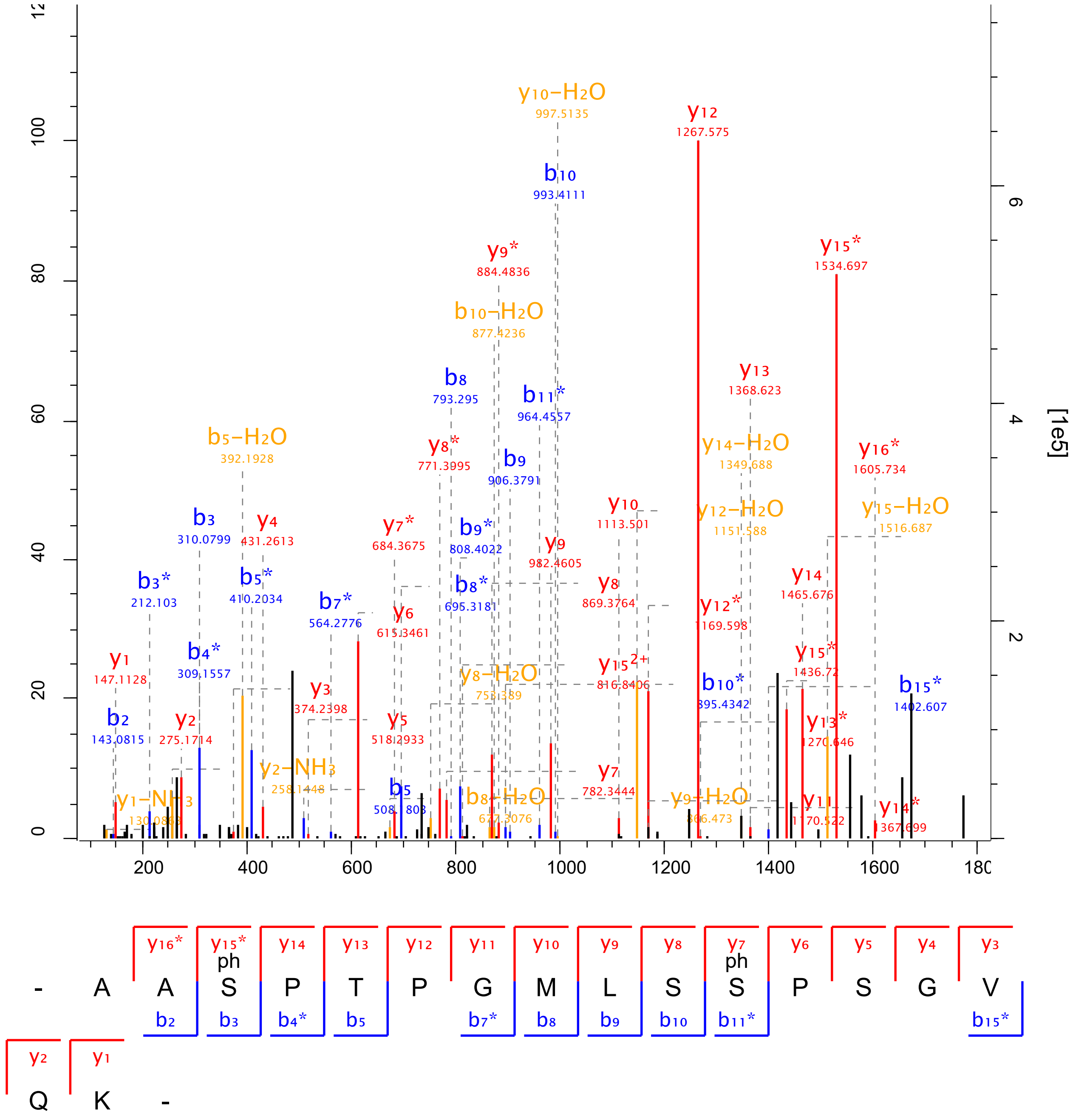

| Raw file | Scan | Method | Score | m/z |
|---|---|---|---|---|
| sys_00_2 | 22688 | FTMS; HCD | 241.79 | 718.86 |

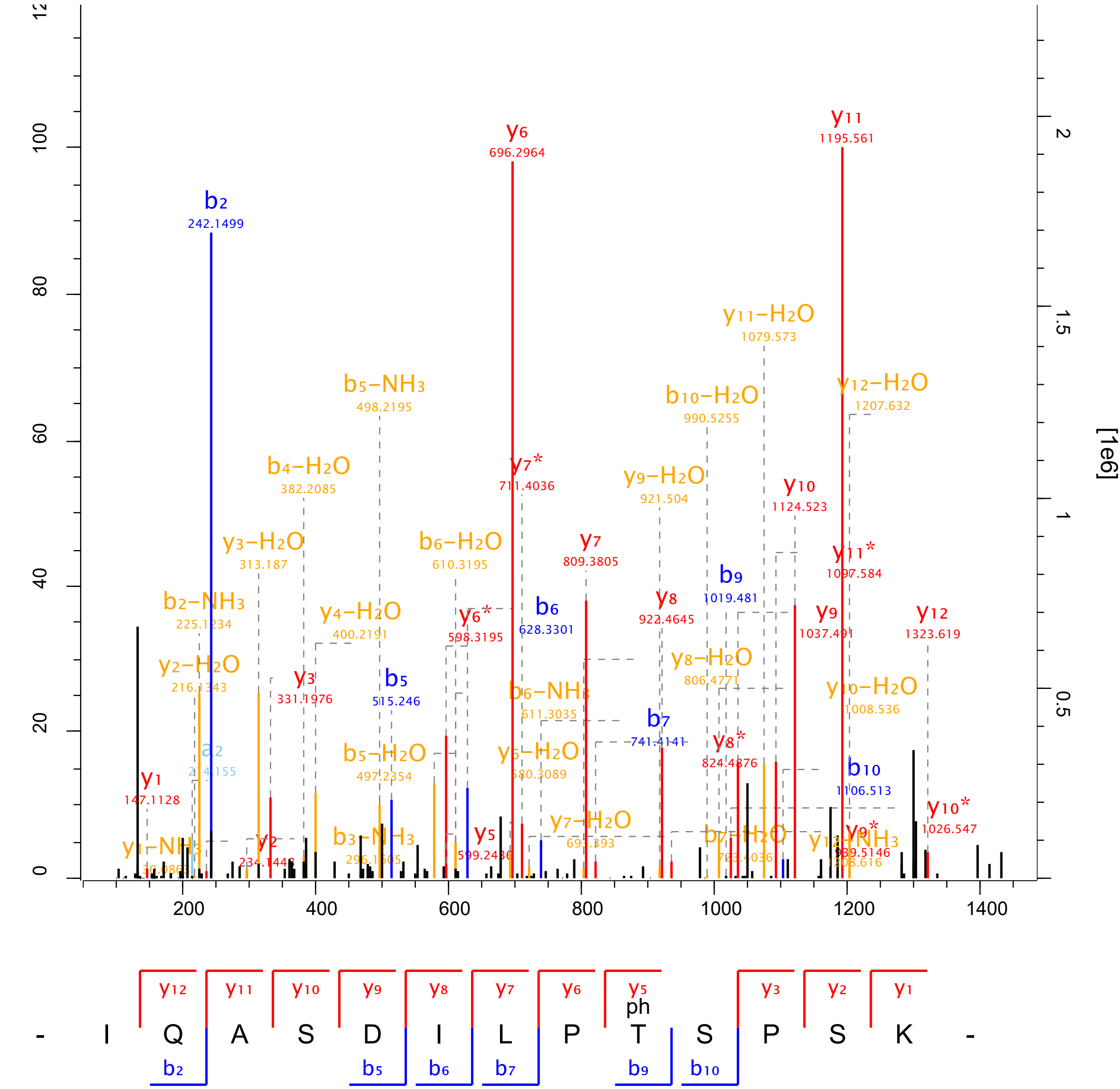

| Raw file | Scan | Method | Score | m/z |
|---|---|---|---|---|
| sys_00_2 | 23125 | FTMS; HCD | 134.68 | 643.29 |

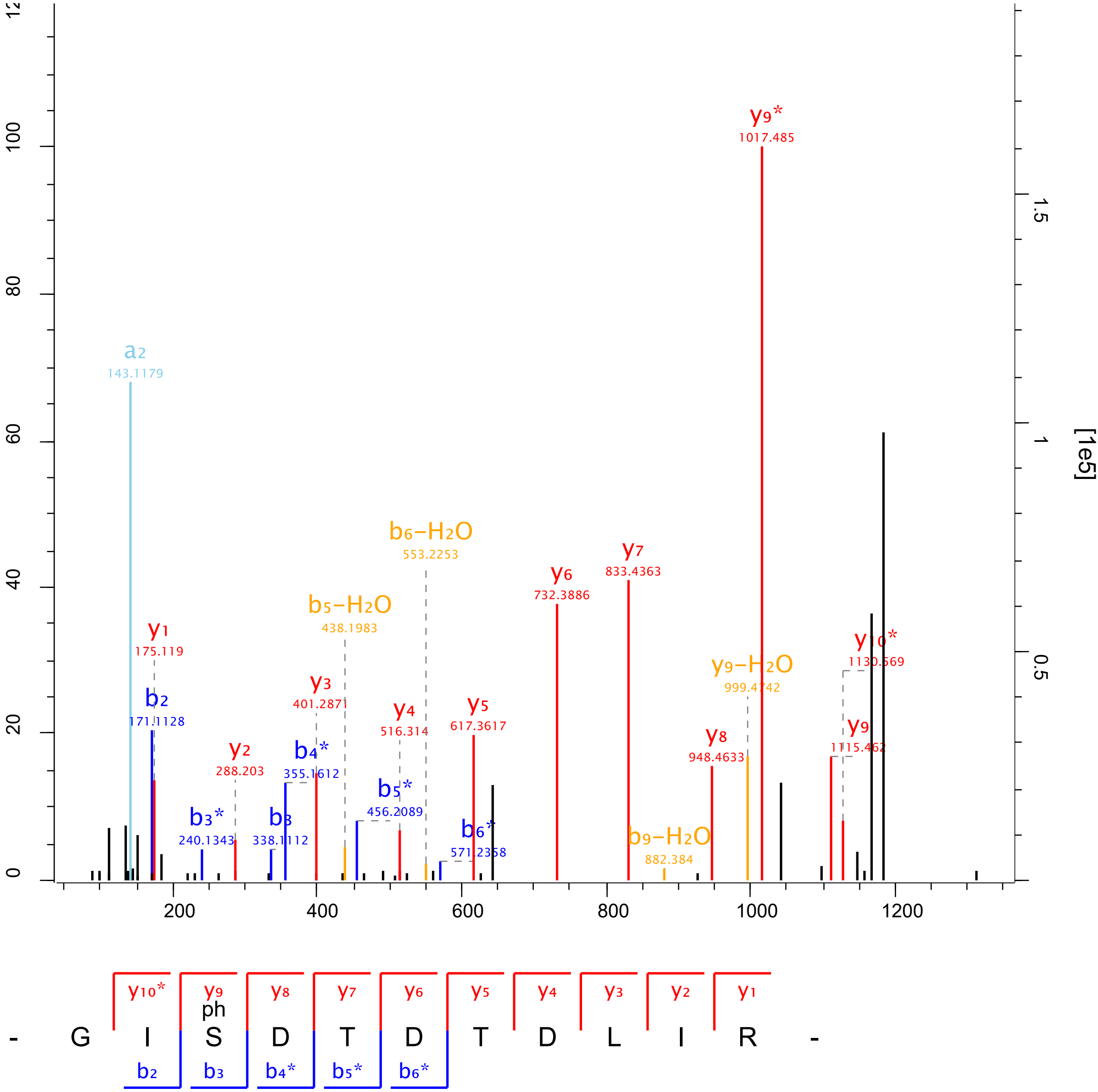

| Raw file | Scan | Method | Score | m/z |
|---|---|---|---|---|
| sys_00_2 | 25449 | FTMS; HCD | 167.63 | 886.36 |

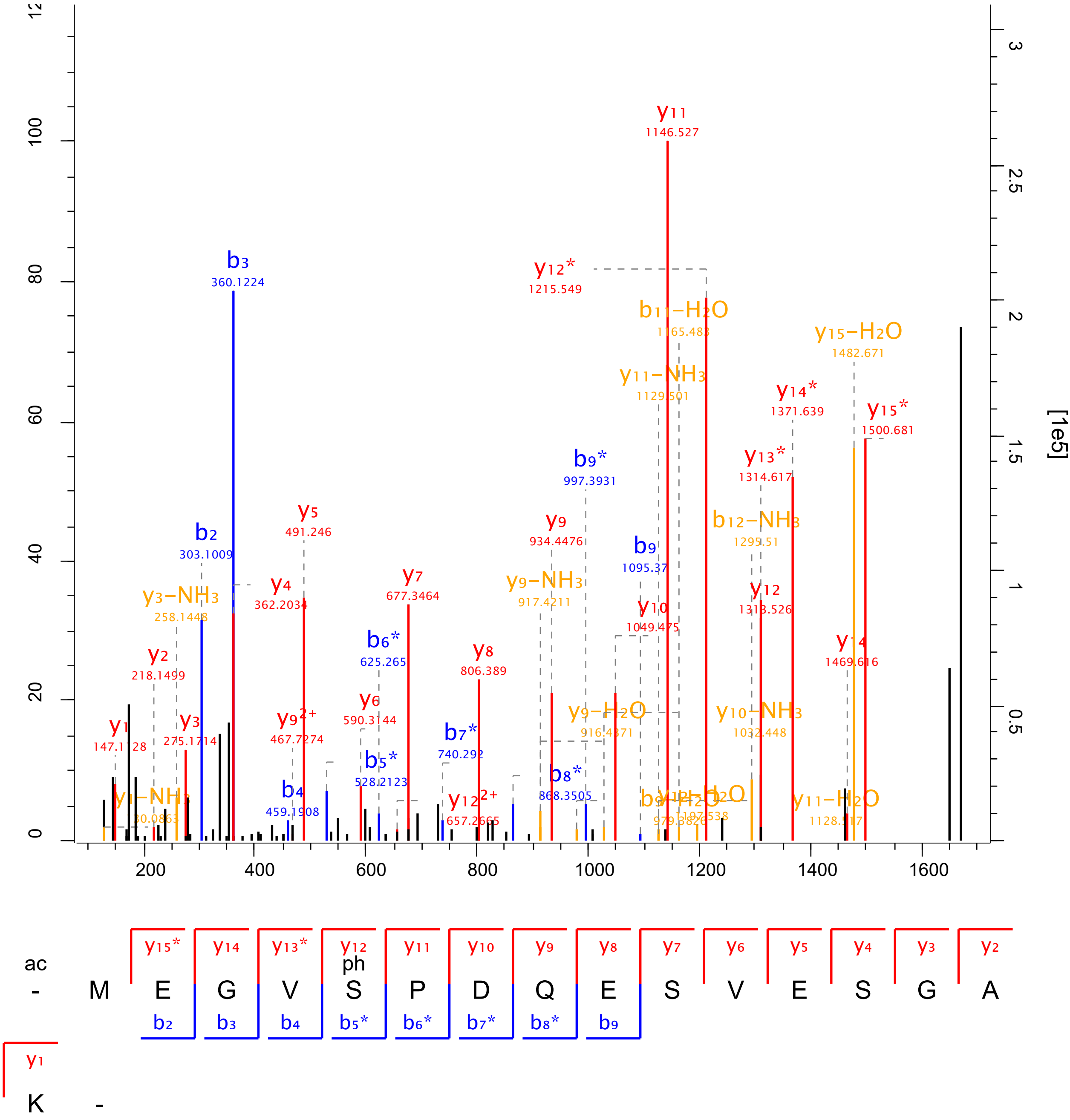

| Raw file | Scan | Method | Score | m/z |
|---|---|---|---|---|
| sys_00_2 | 28719 | FTMS; HCD | 135.21 | 930.38 |

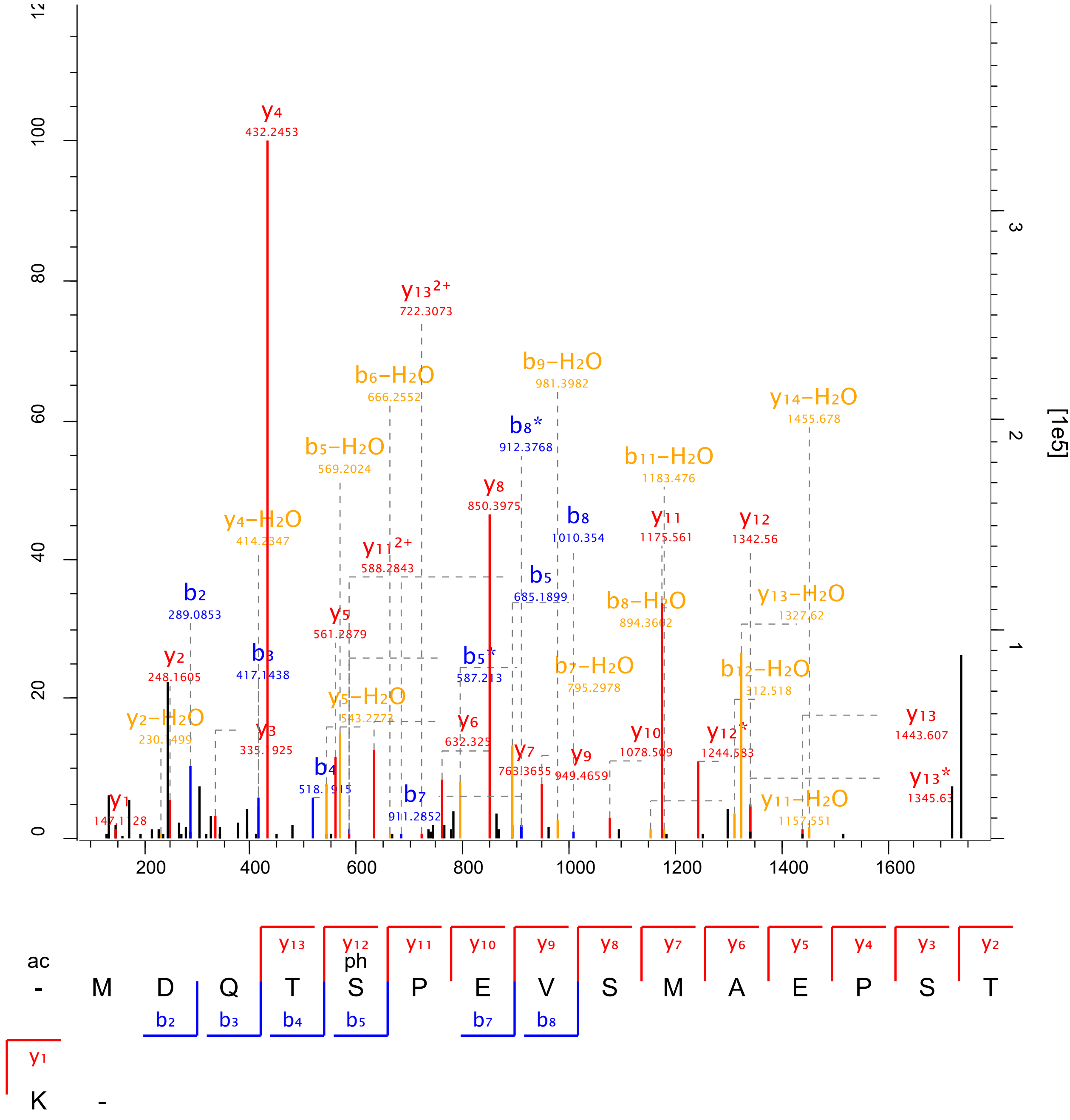

| Raw file | Scan | Method | Score | m/z |
| --- | --- | --- | --- | --- |
| sys_00_2 | 30261 | FTMS; HCD | 131.79 | 800.34 |

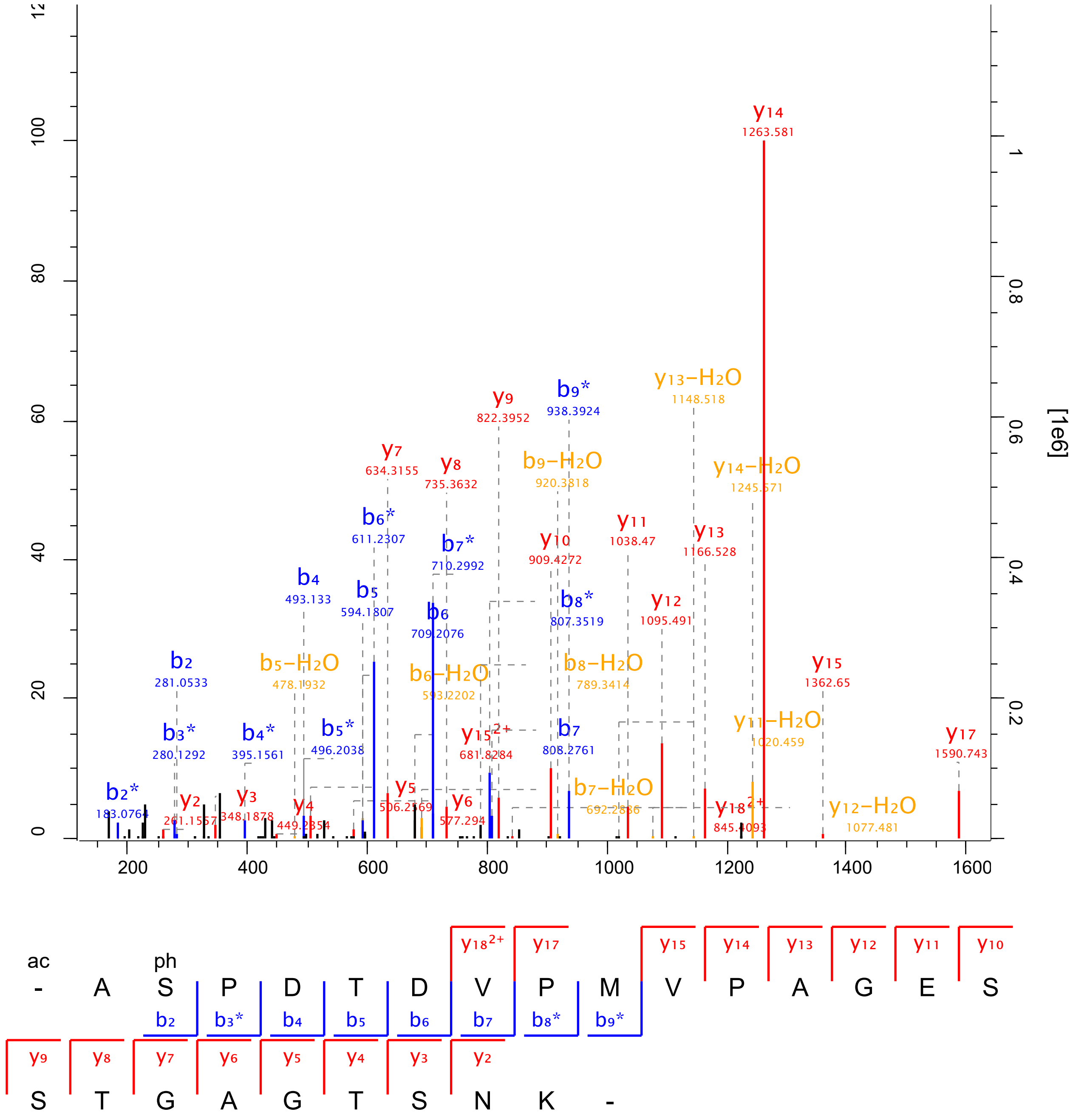

| Raw file | Scan | Method | Score | m/z |
|---|---|---|---|---|
| sys_00_2 | 3484 | FTMS; HCD | 121.25 | 980.36 |

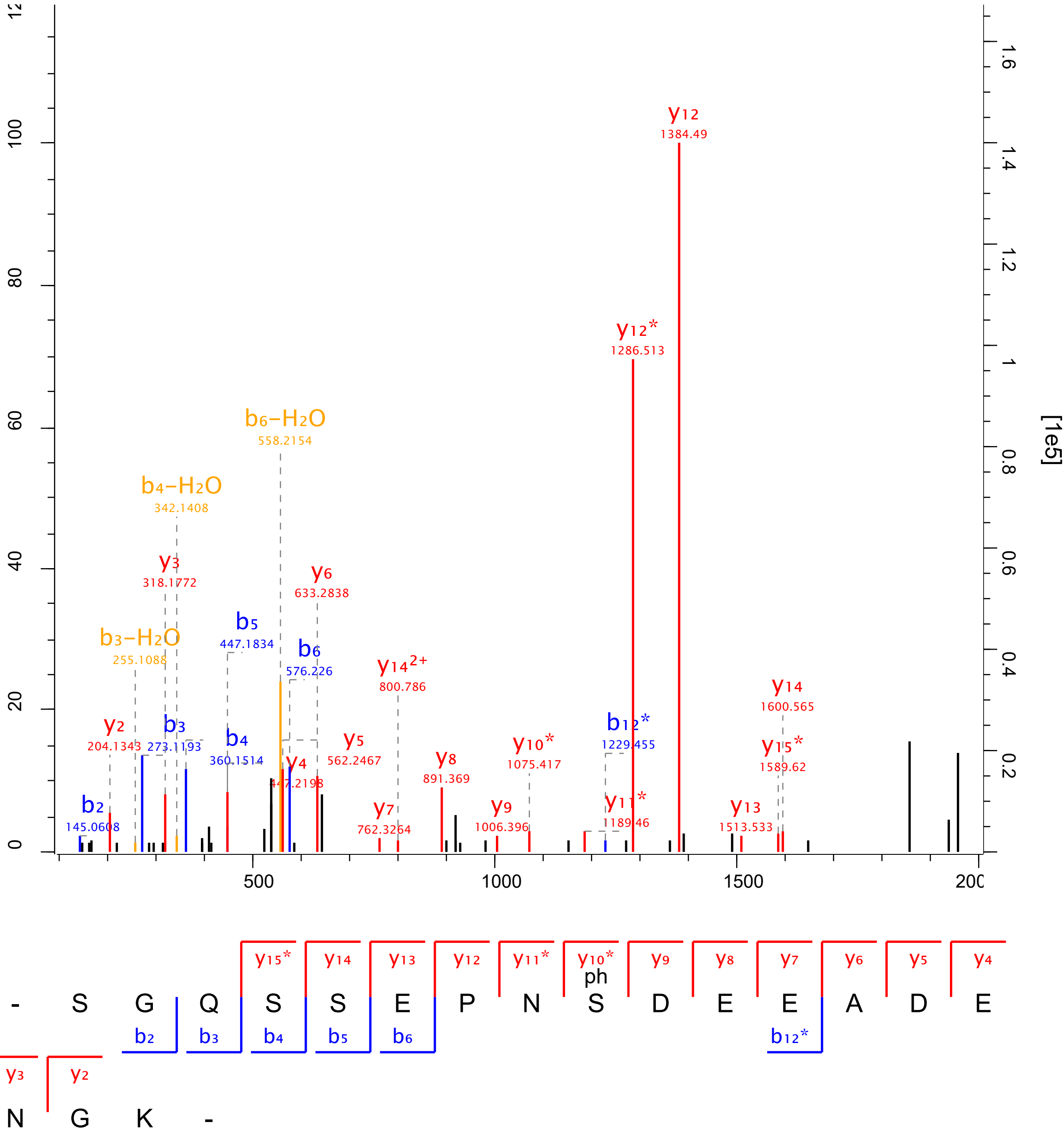

| Raw file | Scan | Method | Score | m/z |
|---|---|---|---|---|
| sys_00_2 | 36121 | FTMS; HCD | 76.07 | 666.27 |

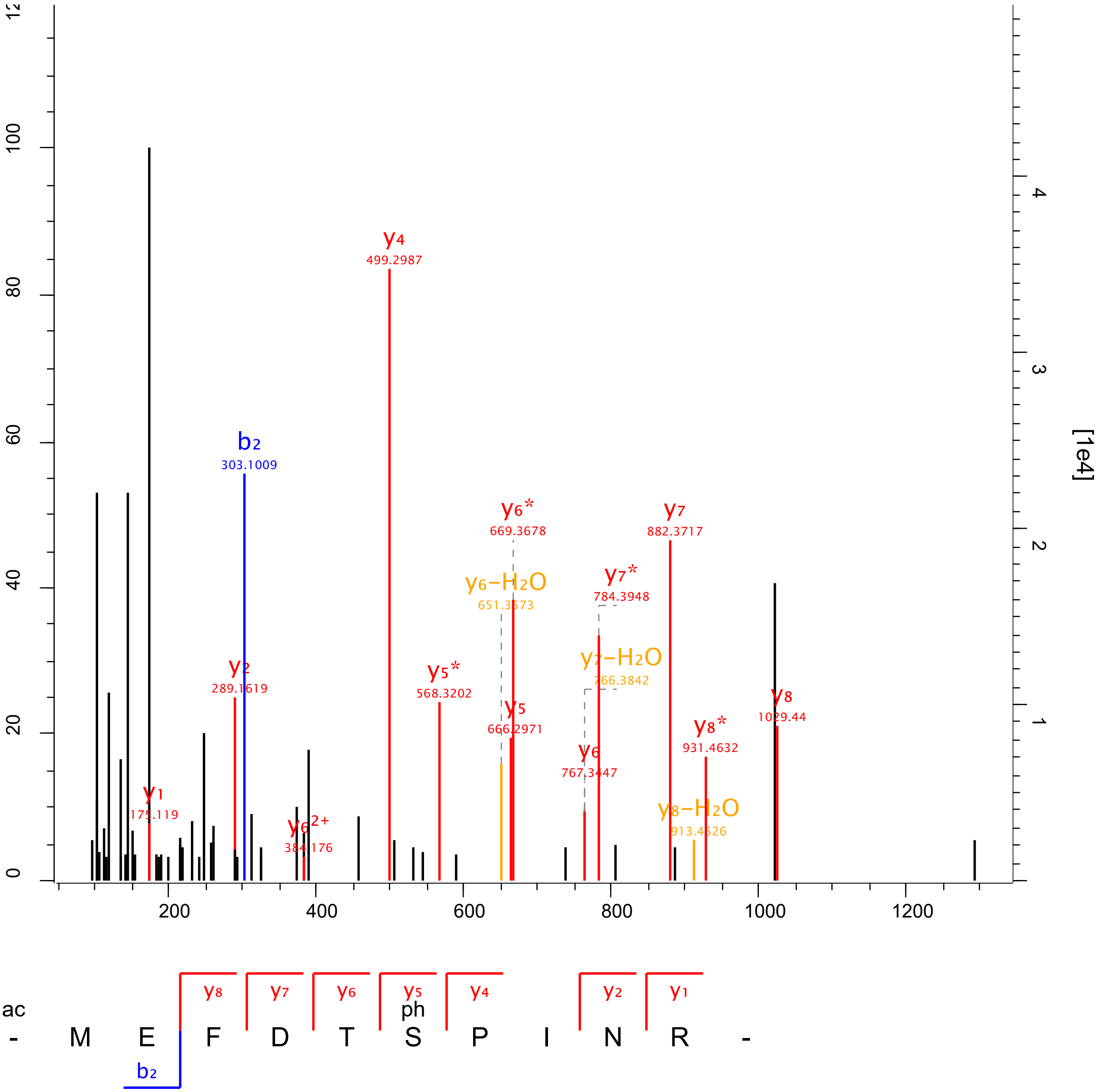

| Raw file | Scan | Method | Score | m/z |
| --- | --- | --- | --- | --- |
| sys_00_2 | 3993 | FTMS; HCD | 50.11 | 456.22 |

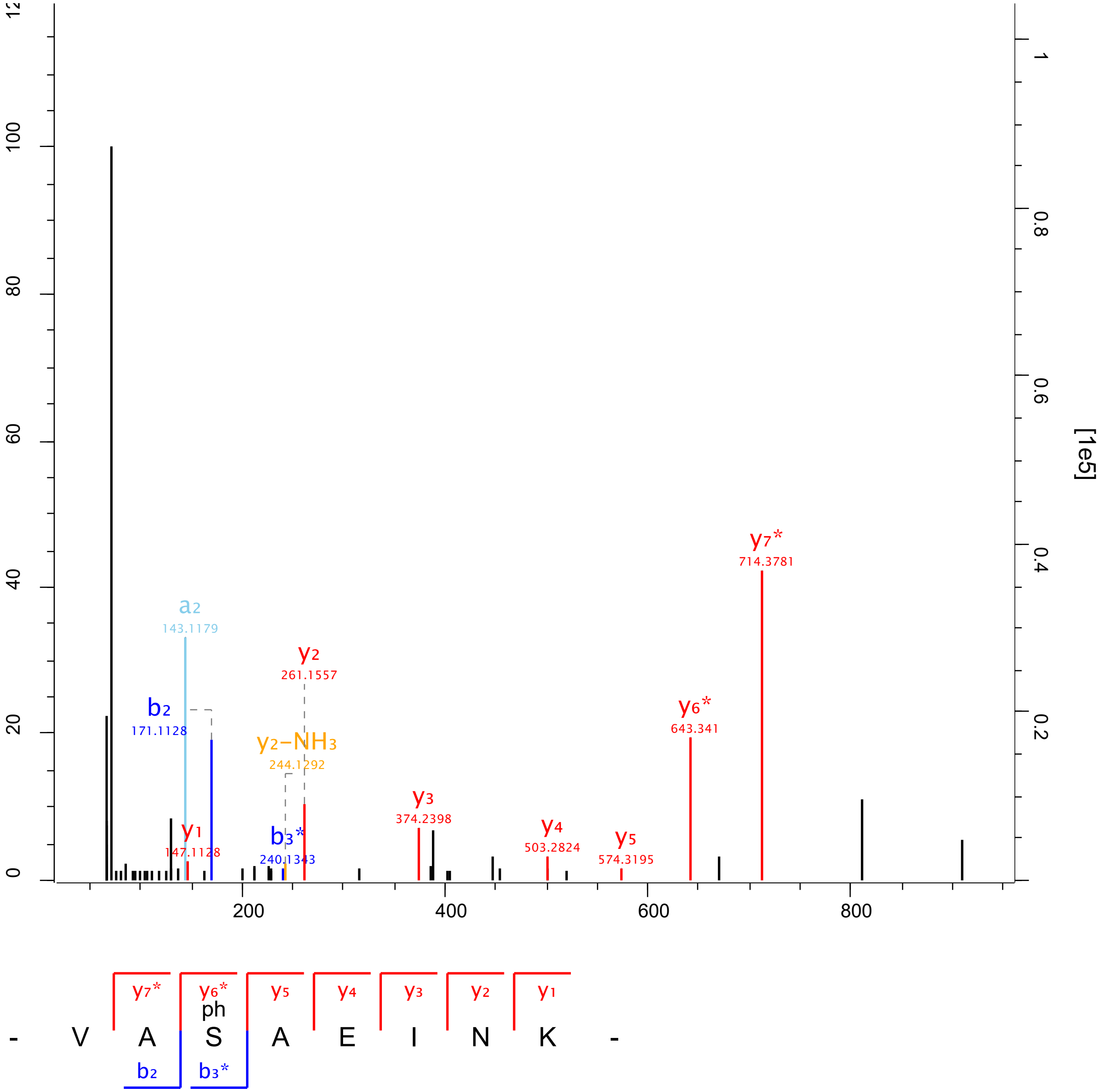

| Raw file | Scan | Method | Score | m/z |
|---|---|---|---|---|
| sys_00_2 | 4203 | FTMS; HCD | 110.97 | 677.78 |

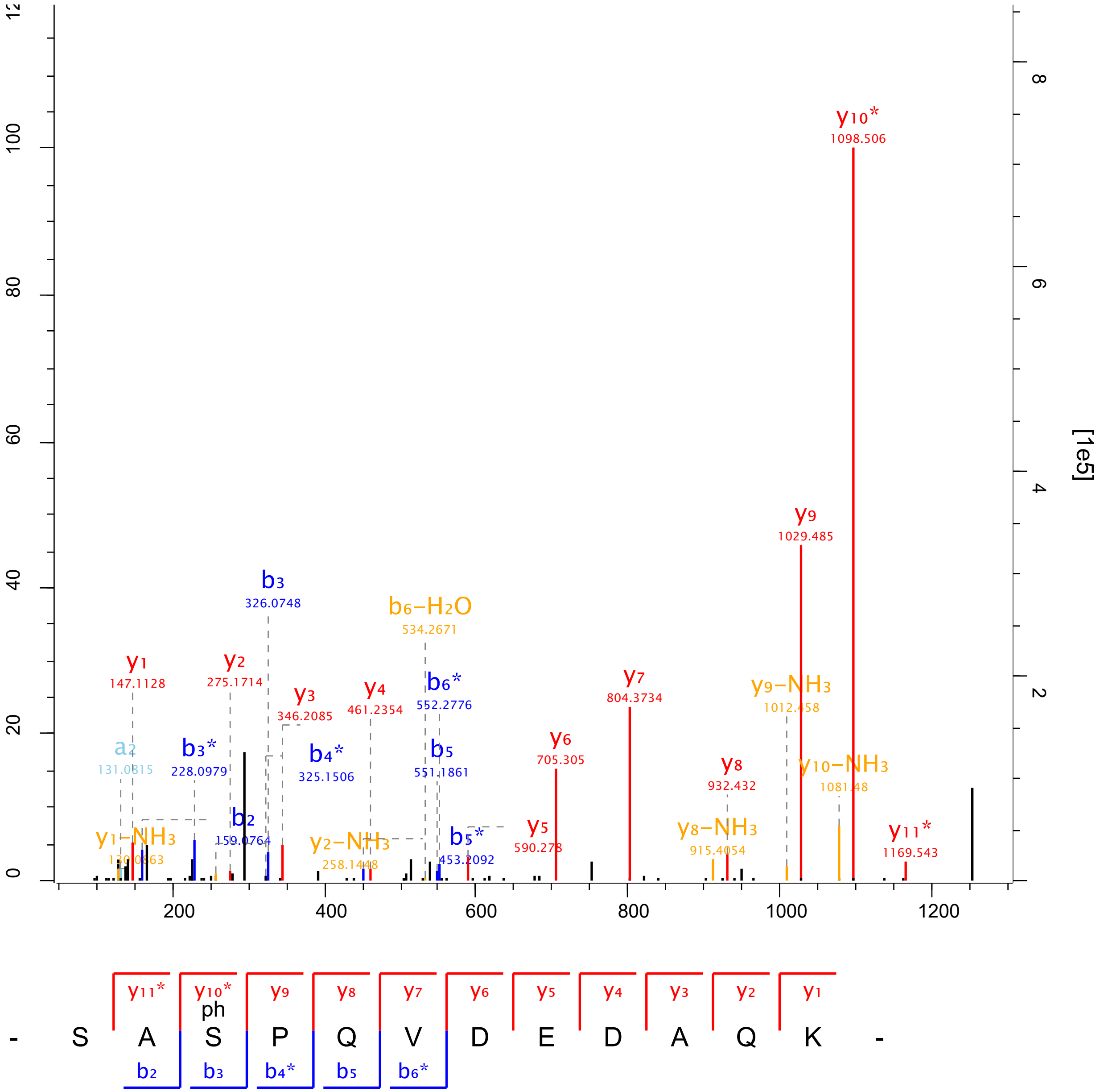

| Raw file | Scan | Method | Score | m/z |
|---|---|---|---|---|
| sys_00_2 | 4253 | FTMS; HCD | 62.46 | 413.17 |

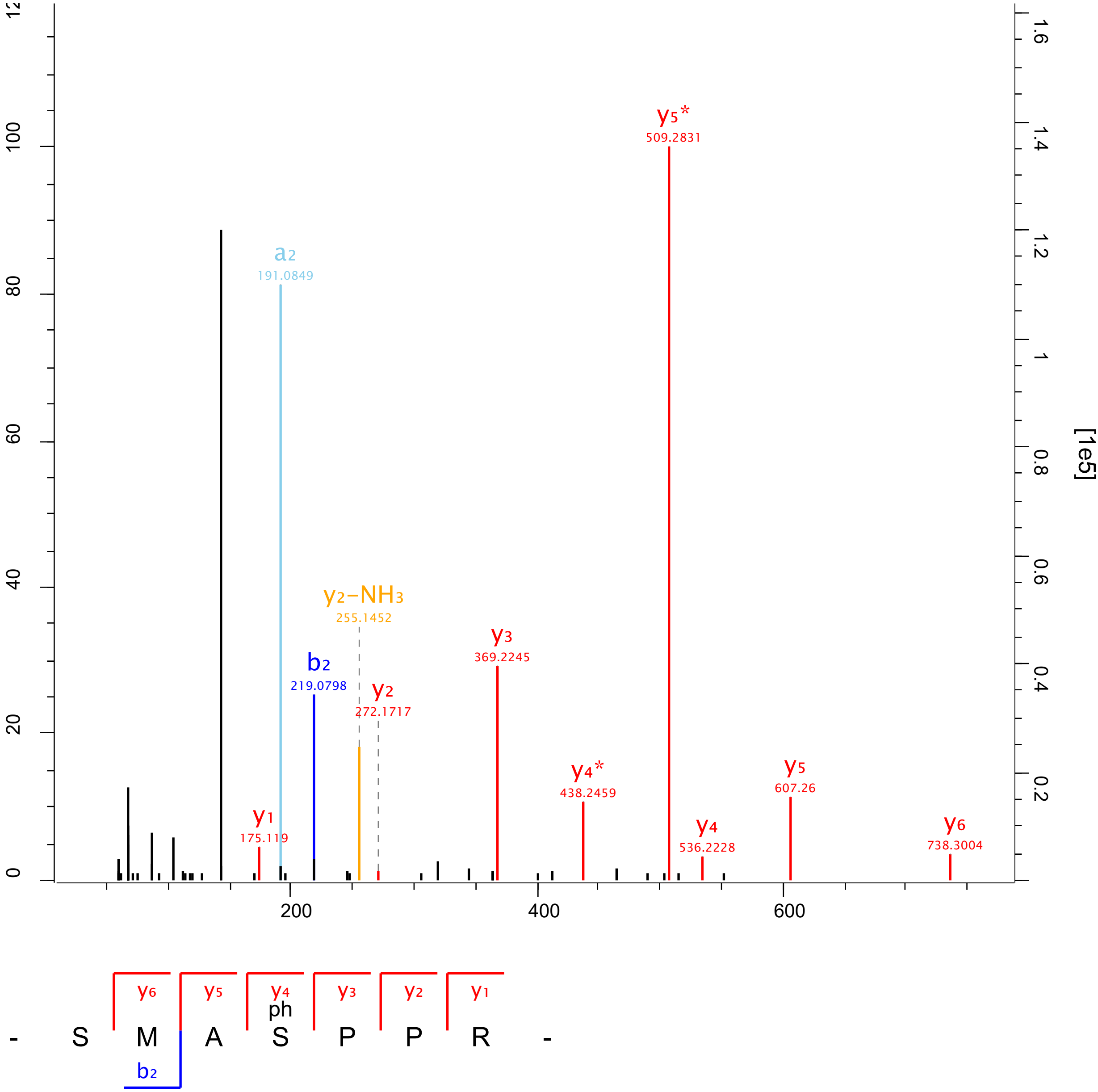

| Raw file | Scan | Method | Score | m/z |
|---|---|---|---|---|
| sys_00_2 | 4275 | FTMS; HCD | 113.22 | 492.72 |

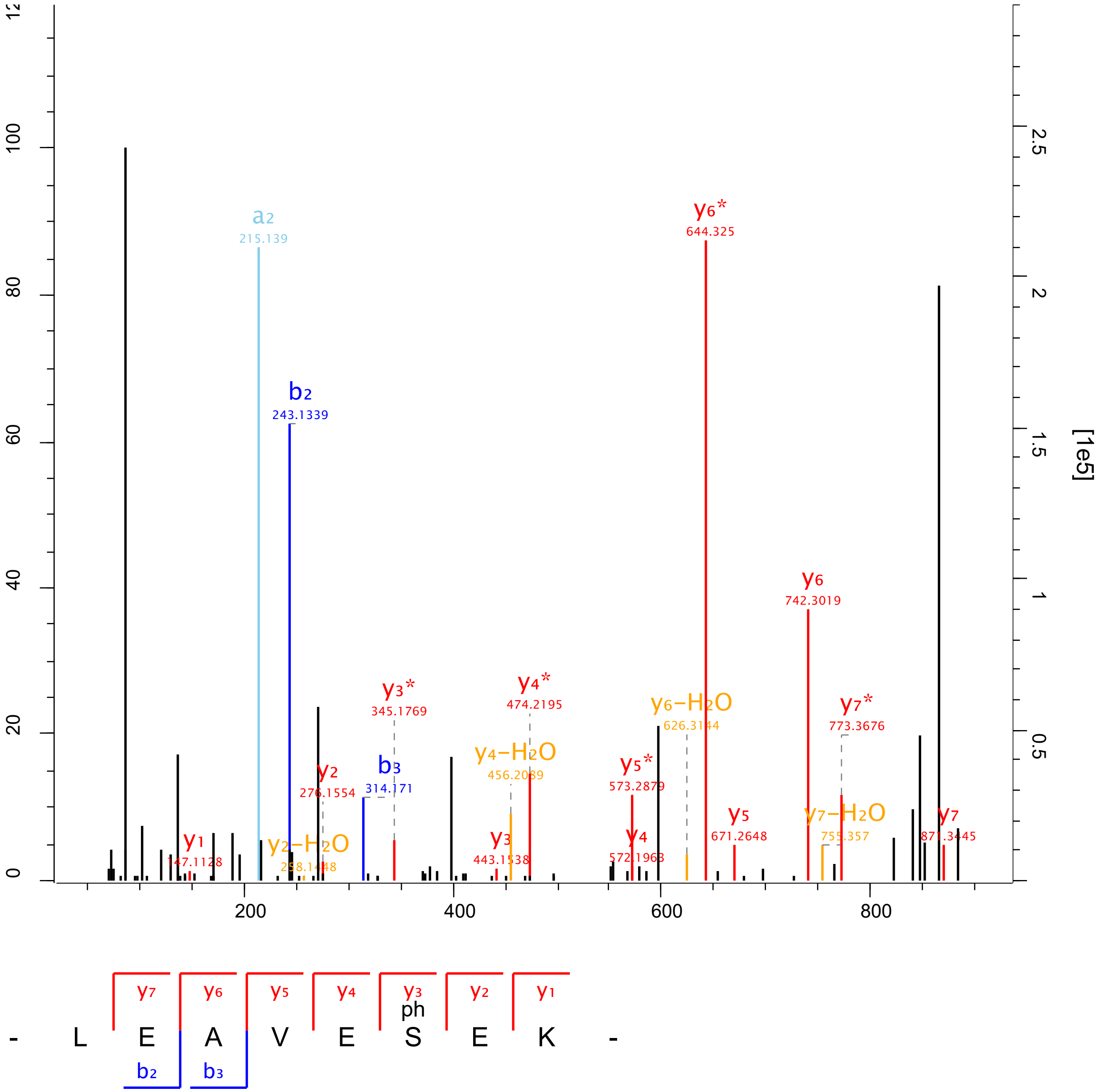

| Raw file | Scan | Method | Score | m/z |
|---|---|---|---|---|
| sys_00_2 | 4437 | FTMS; HCD | 42.71 | 980.37 |

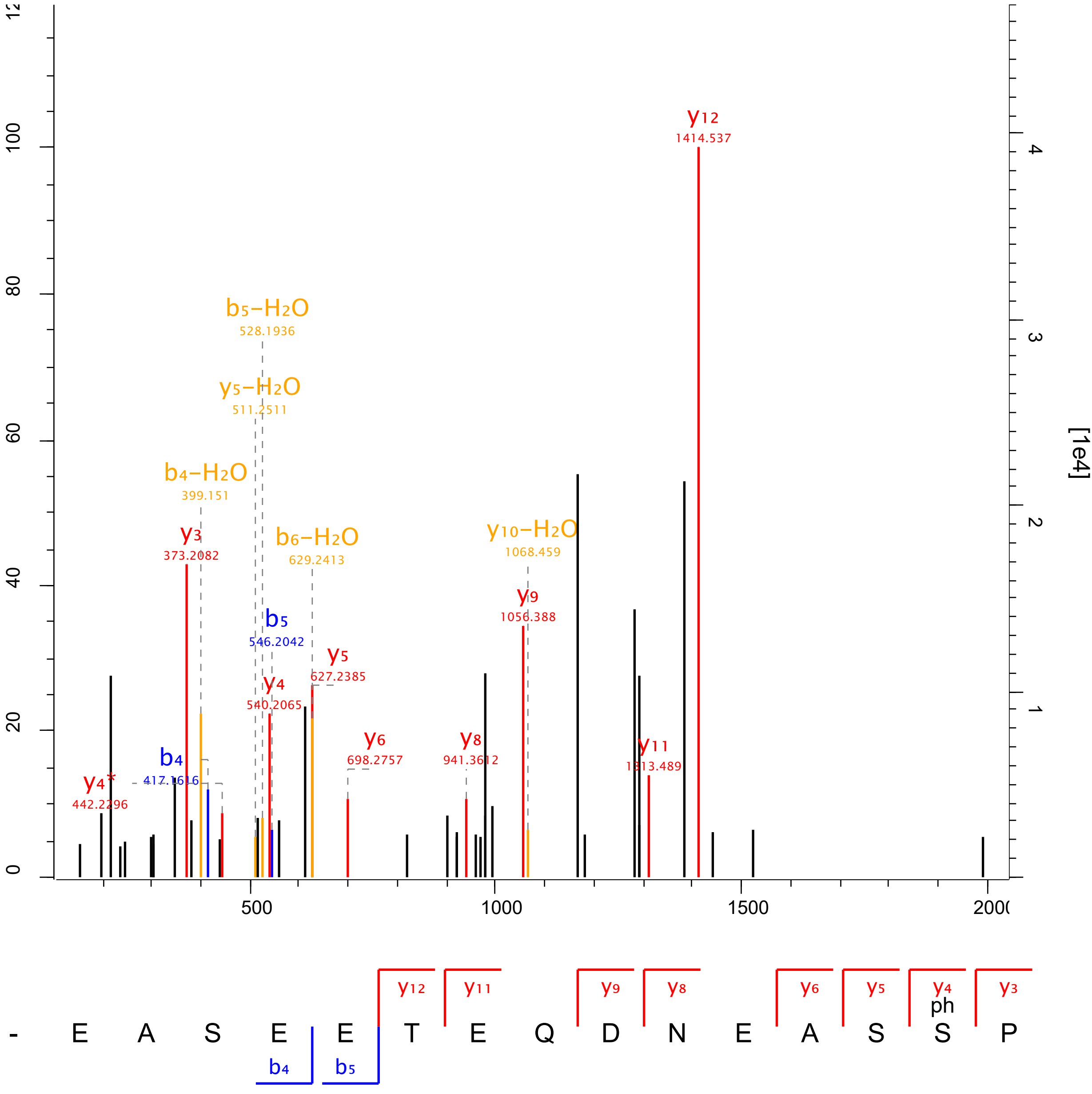


- E A S E E T E Q D N E A S S P

E K -

| Raw file | Scan | Method | Score | m/z |
|---|---|---|---|---|
| sys_00_2 | 44682 | FTMS; HCD | 72.36 | 861.88 |

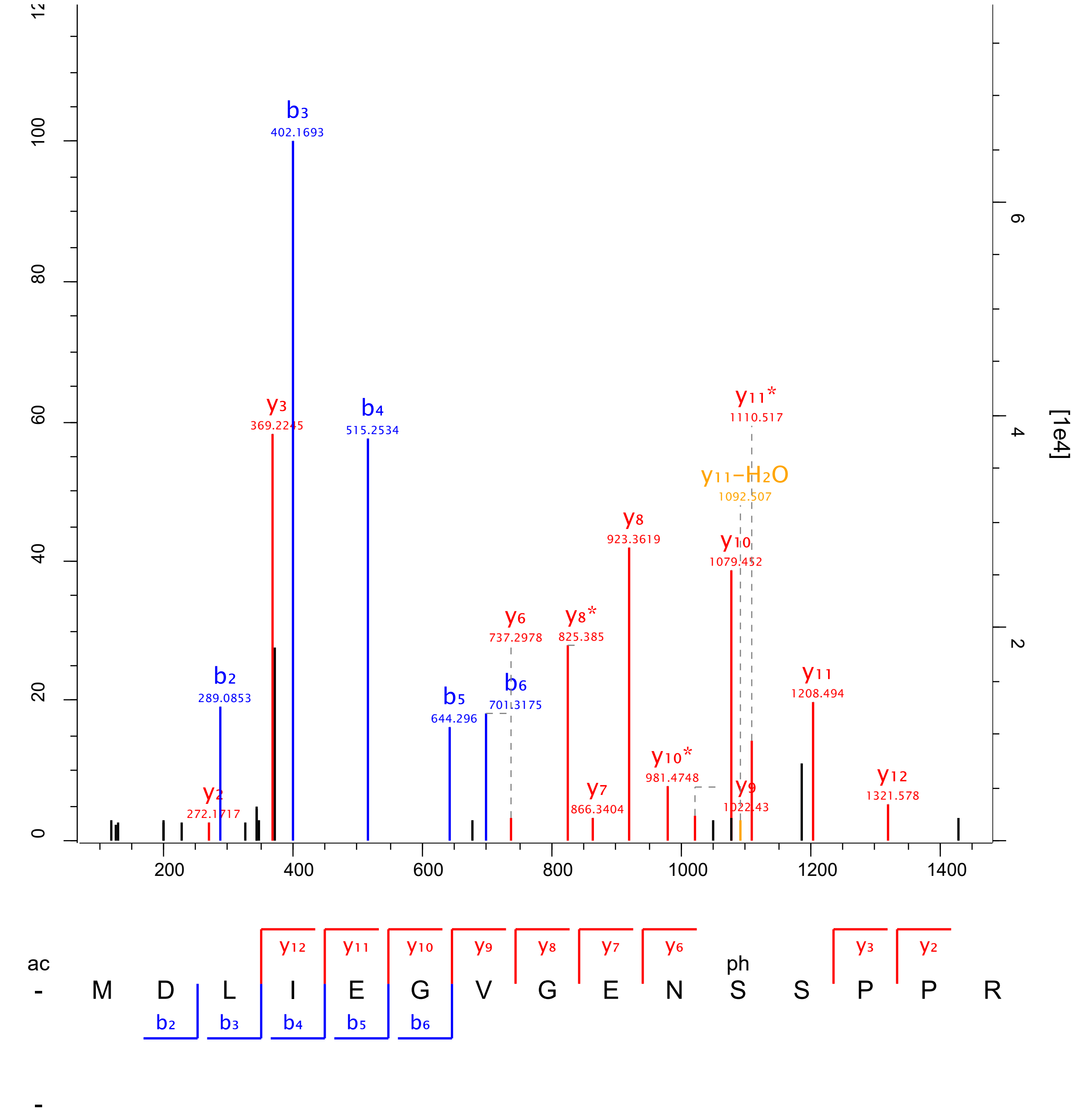

| Raw file | Scan | Method | Score | m/z |
| --- | --- | --- | --- | --- |
| sys_00_2 | 4939 | FTMS; HCD | 72.7 | 541.25 |

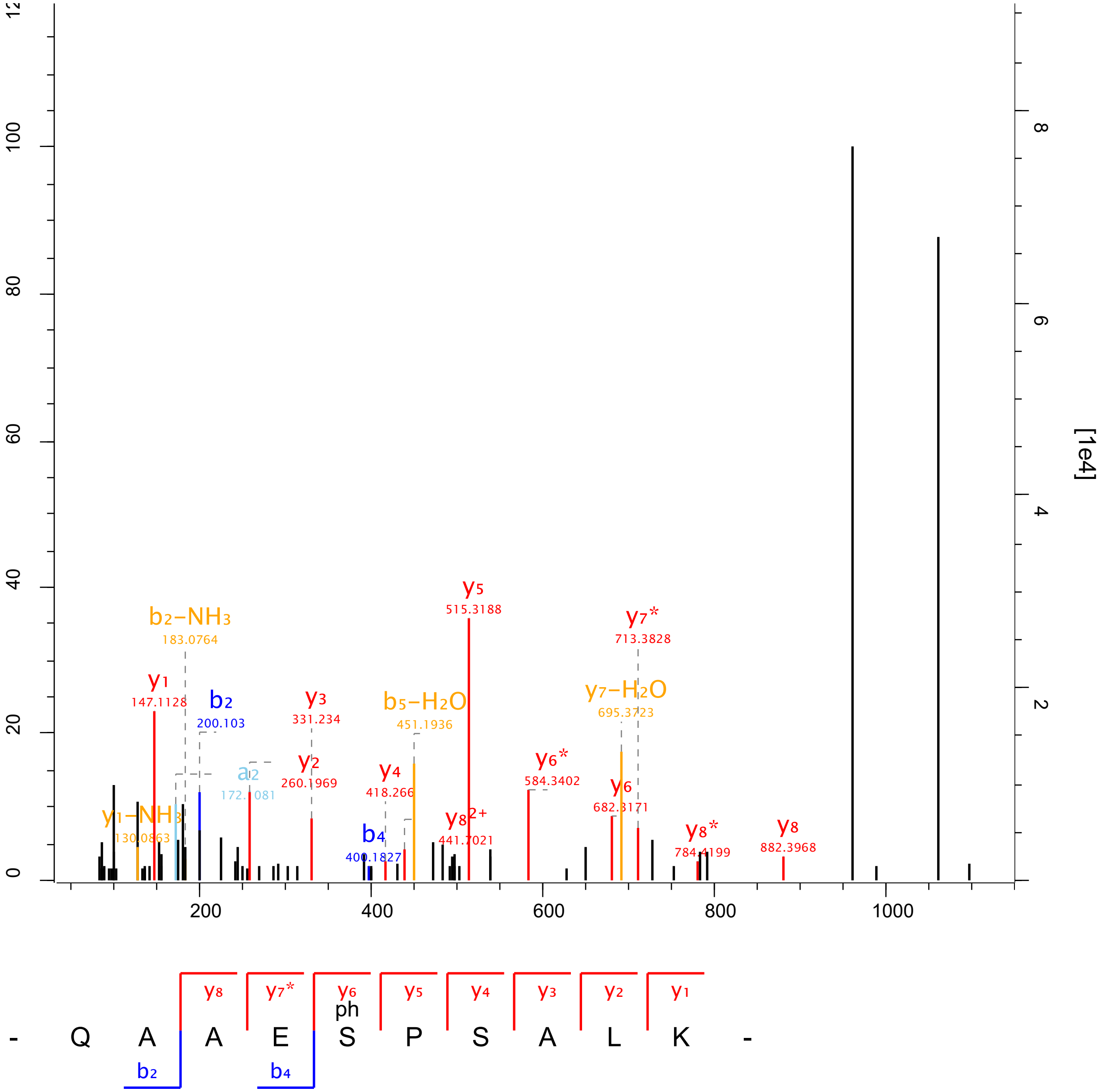

| Raw file | Scan | Method | Score | m/z |
|---|---|---|---|---|
| sys_00_2 | 5489 | FTMS; HCD | 77.66 | 529.72 |

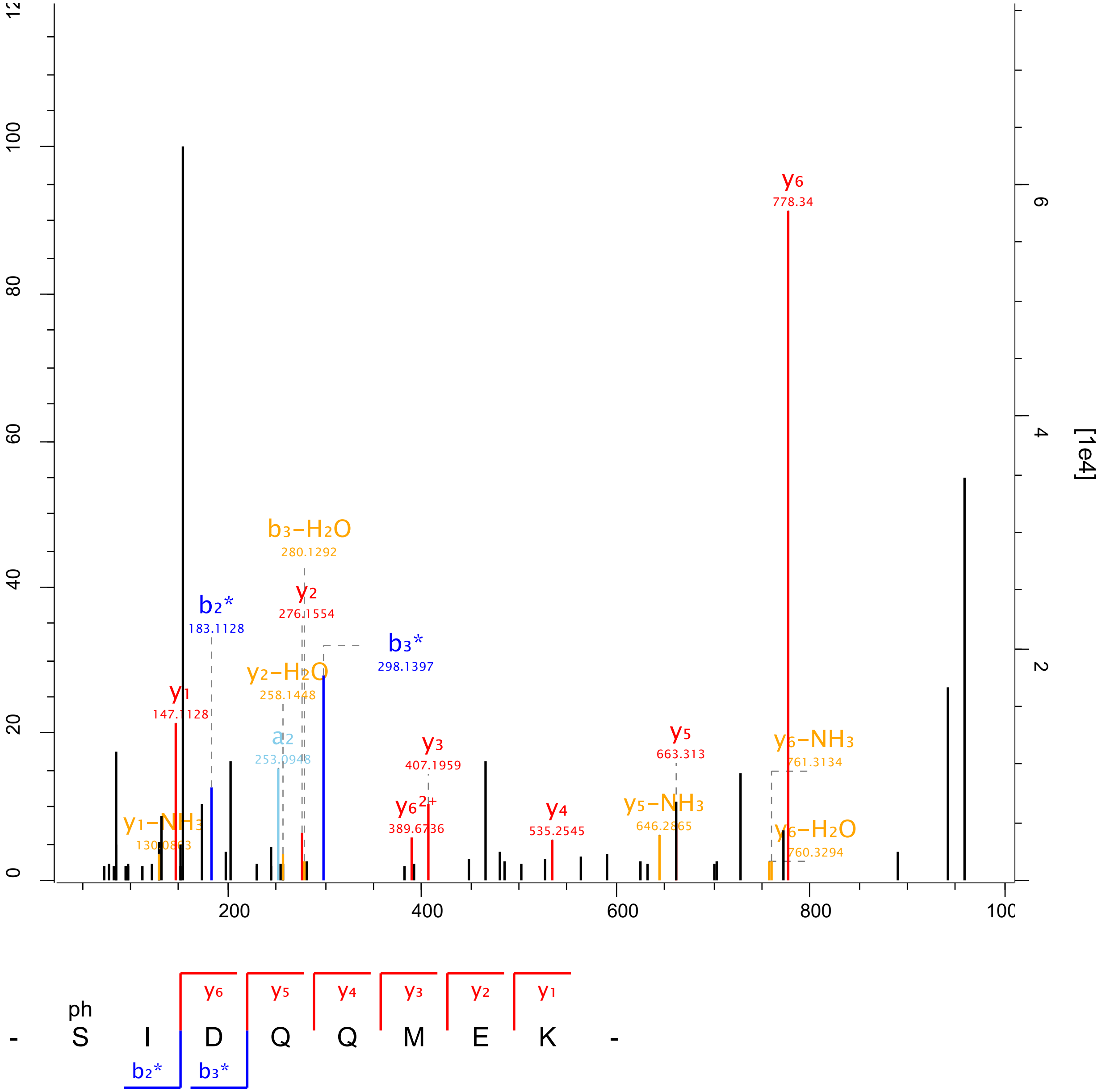

| Raw file | Scan | Method | Score | m/z |
| --- | --- | --- | --- | --- |
| sys_00_2 | 5669 | FTMS; HCD | 55.31 | 556.24 |

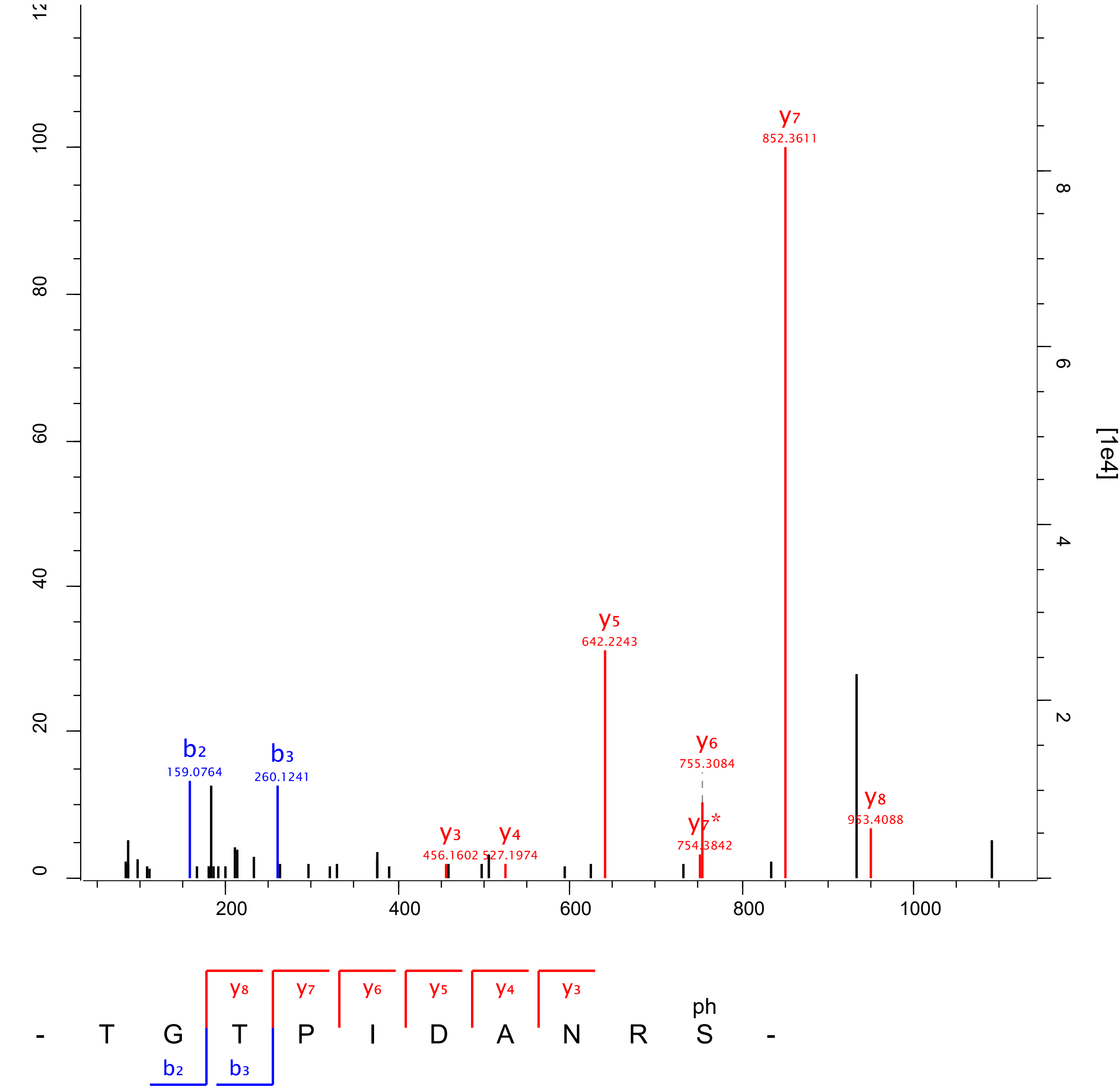

| Raw file | Scan | Method | Score | m/z |
|---|---|---|---|---|
| sys_00_2 | 5851 | FTMS; HCD | 99.97 | 469.72 |

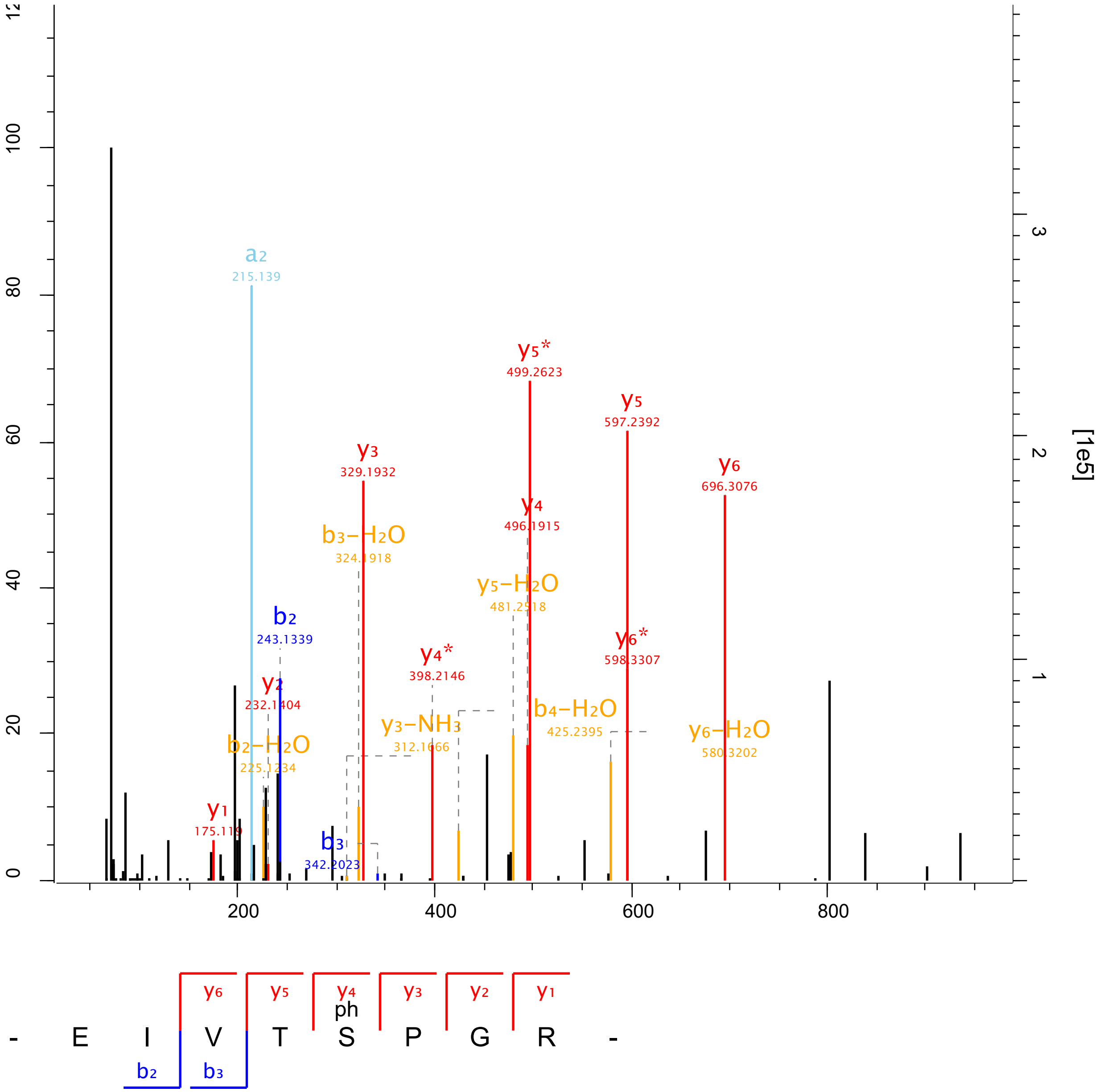

| Raw file | Scan | Method | Score | m/z |
|---|---|---|---|---|
| sys_00_2 | 6408 | FTMS; HCD | 91.09 | 419.2 |

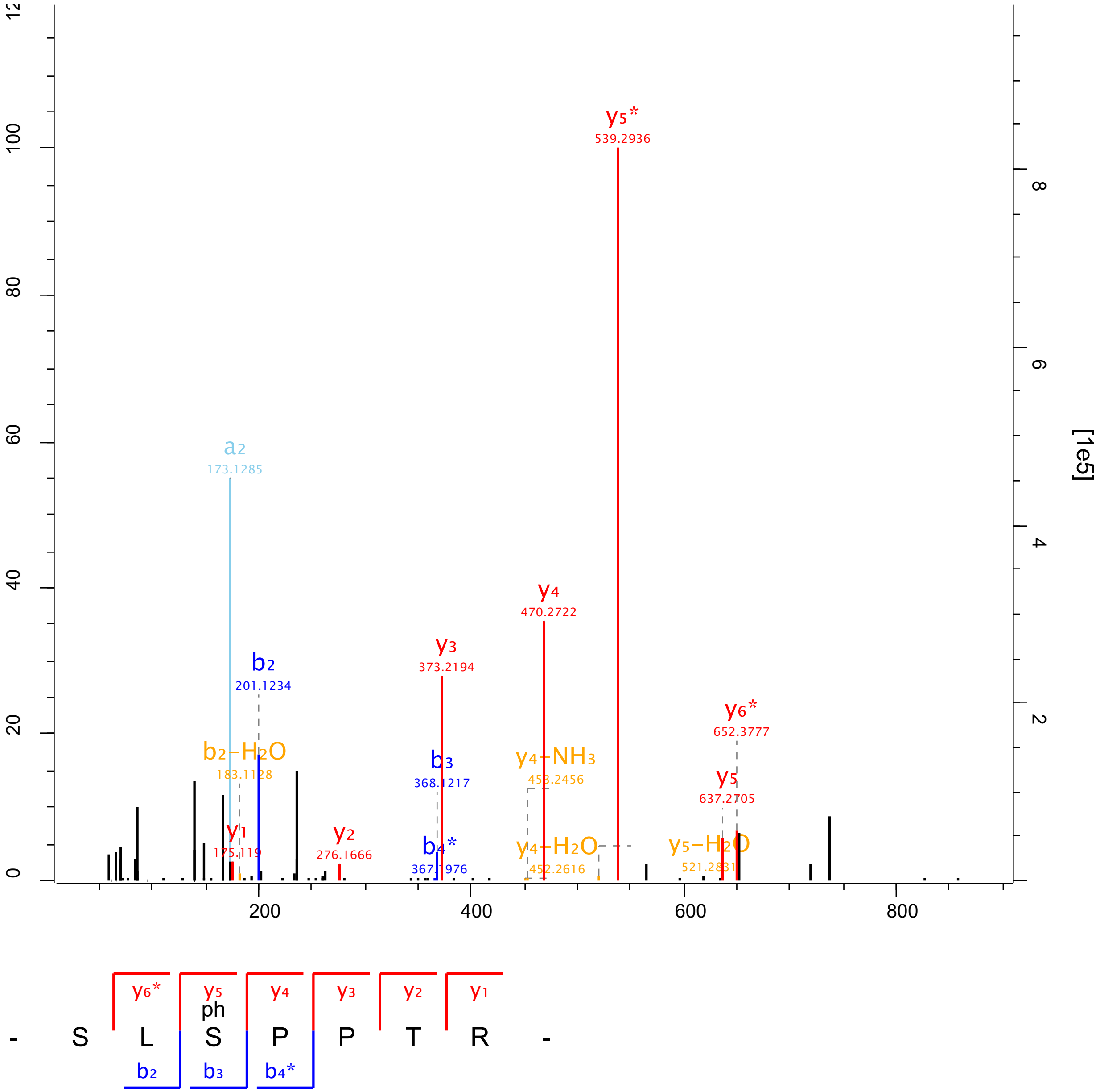

| Raw file | Scan | Method | Score | m/z |
| --- | --- | --- | --- | --- |
| sys_00_2 | 6656 | FTMS; HCD | 42.84 | 513.23 |

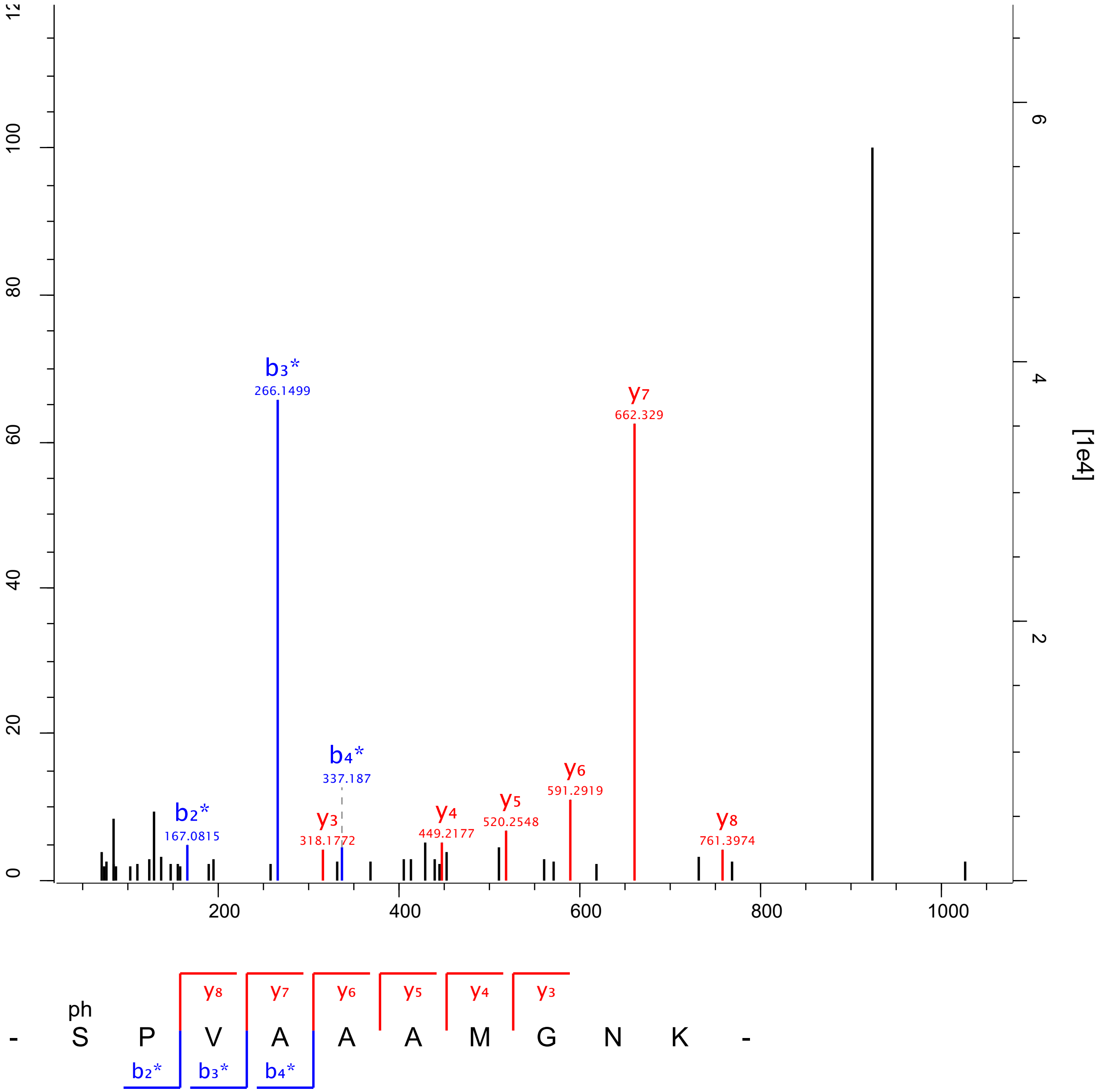

| Raw file | Scan | Method | Score | m/z |
| --- | --- | --- | --- | --- |
| sys_00_2 | 6933 | FTMS; HCD | 78.1 | 429.18 |

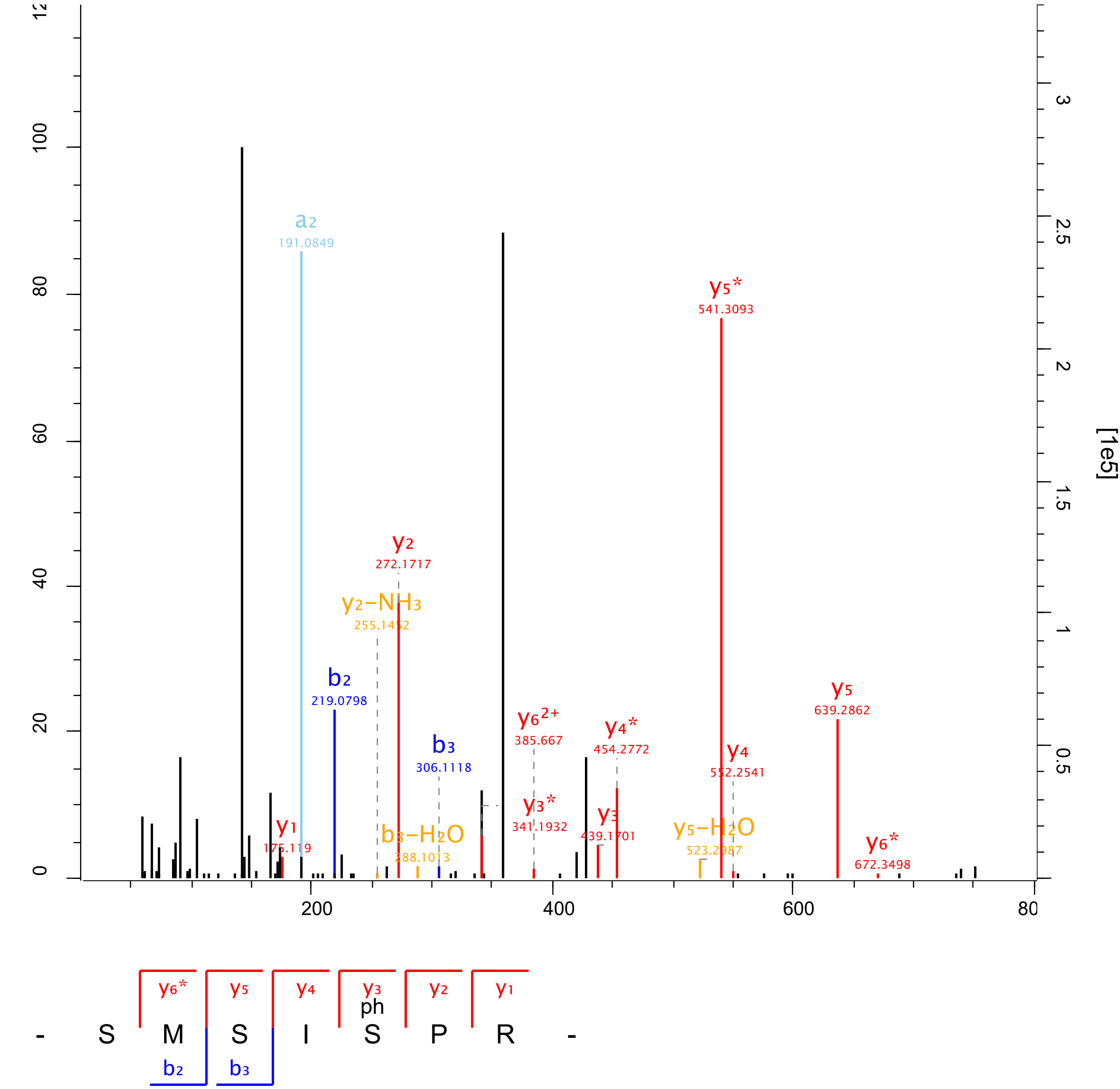

| Raw file | Scan | Method | Score | m/z |
| --- | --- | --- | --- | --- |
| sys_00_2 | 7343 | FTMS; HCD | 91.31 | 586.23 |

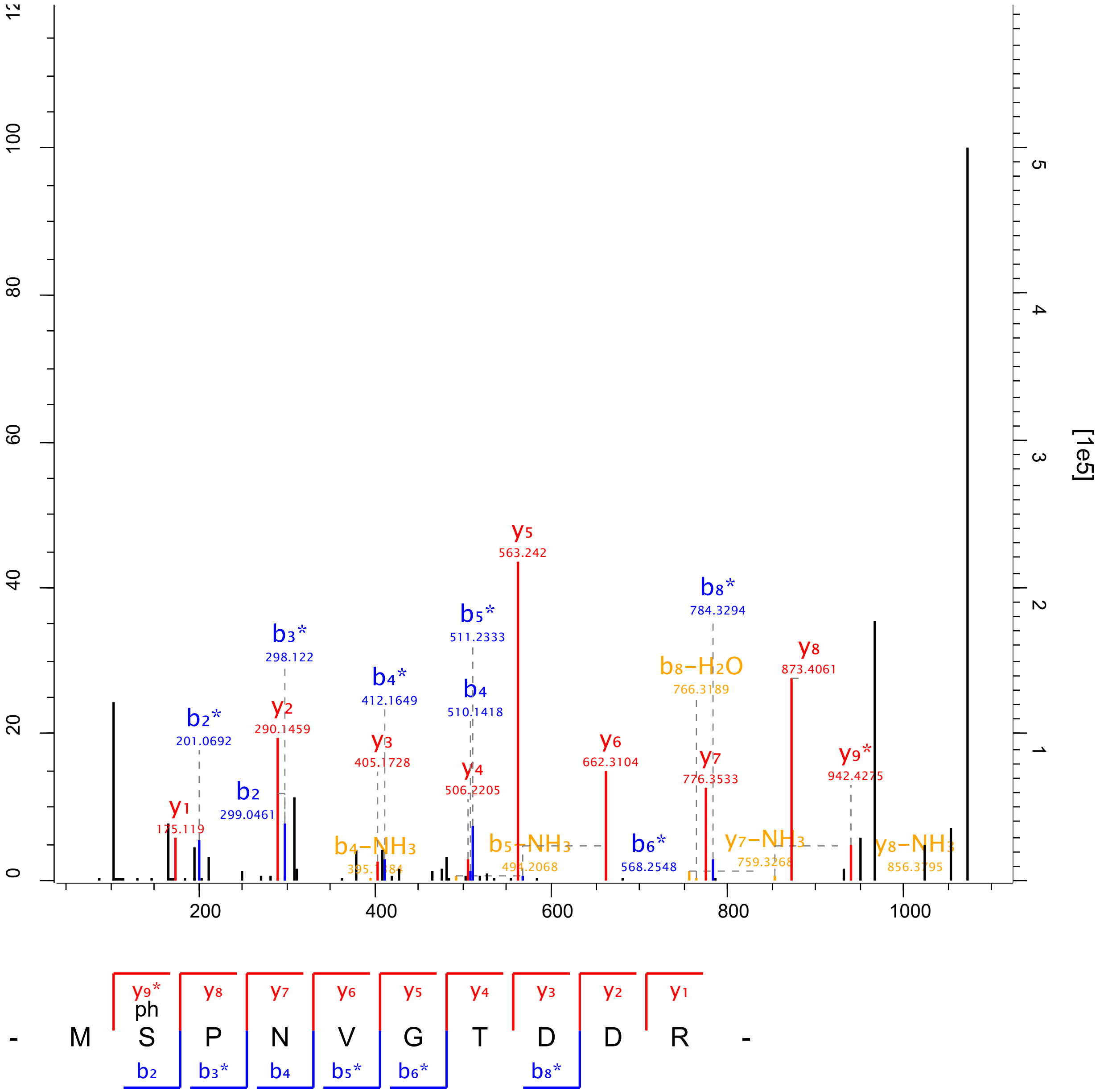

| Raw file | Scan | Method | Score | m/z |
|---|---|---|---|---|
| sys_00_2 | 7606 | FTMS; HCD | 62.86 | 728.32 |

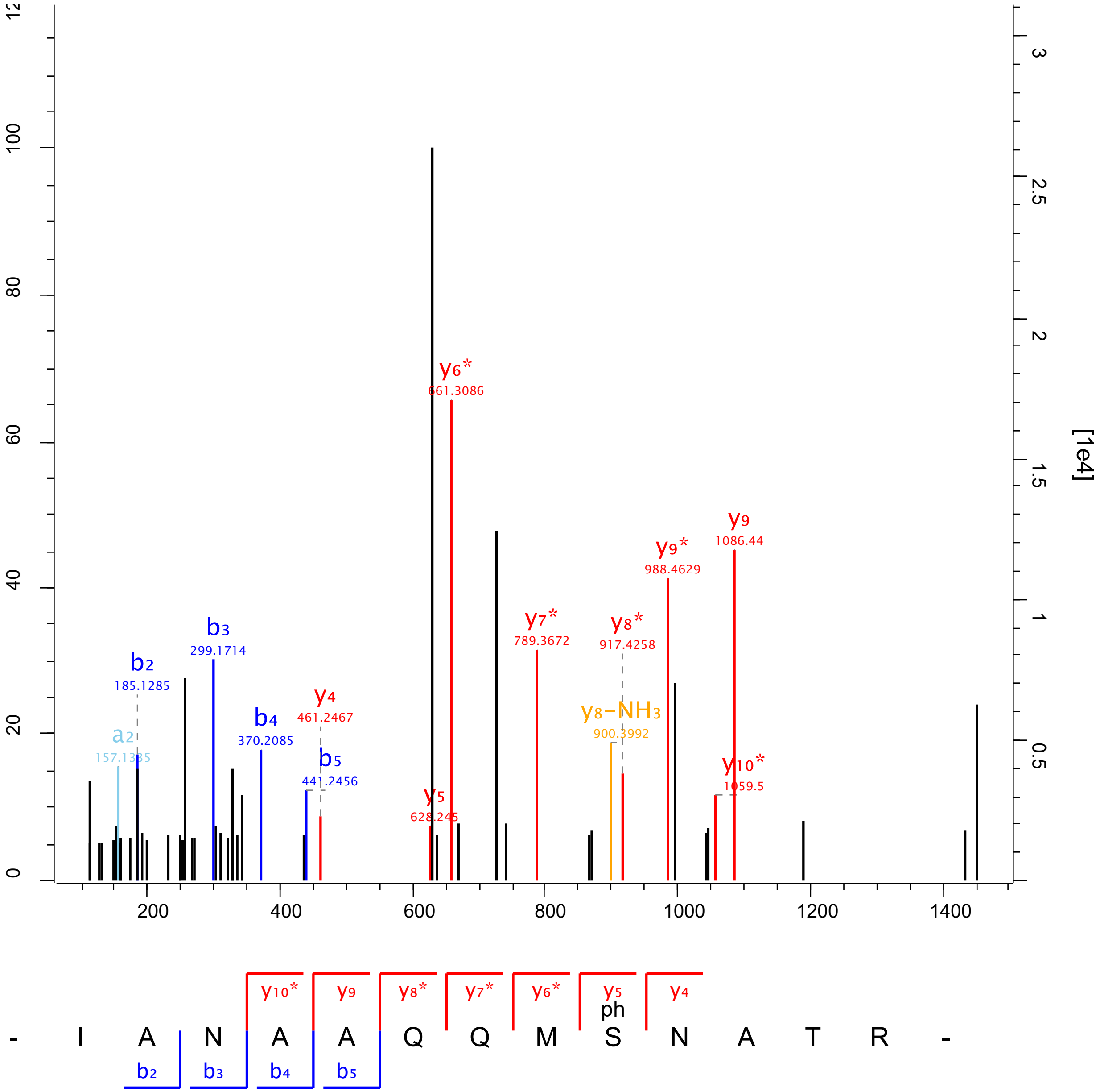

| Raw file | Scan | Method | Score | m/z |
|---|---|---|---|---|
| sys_00_2 | 8276 | FTMS; HCD | 126.09 | 626.76 |

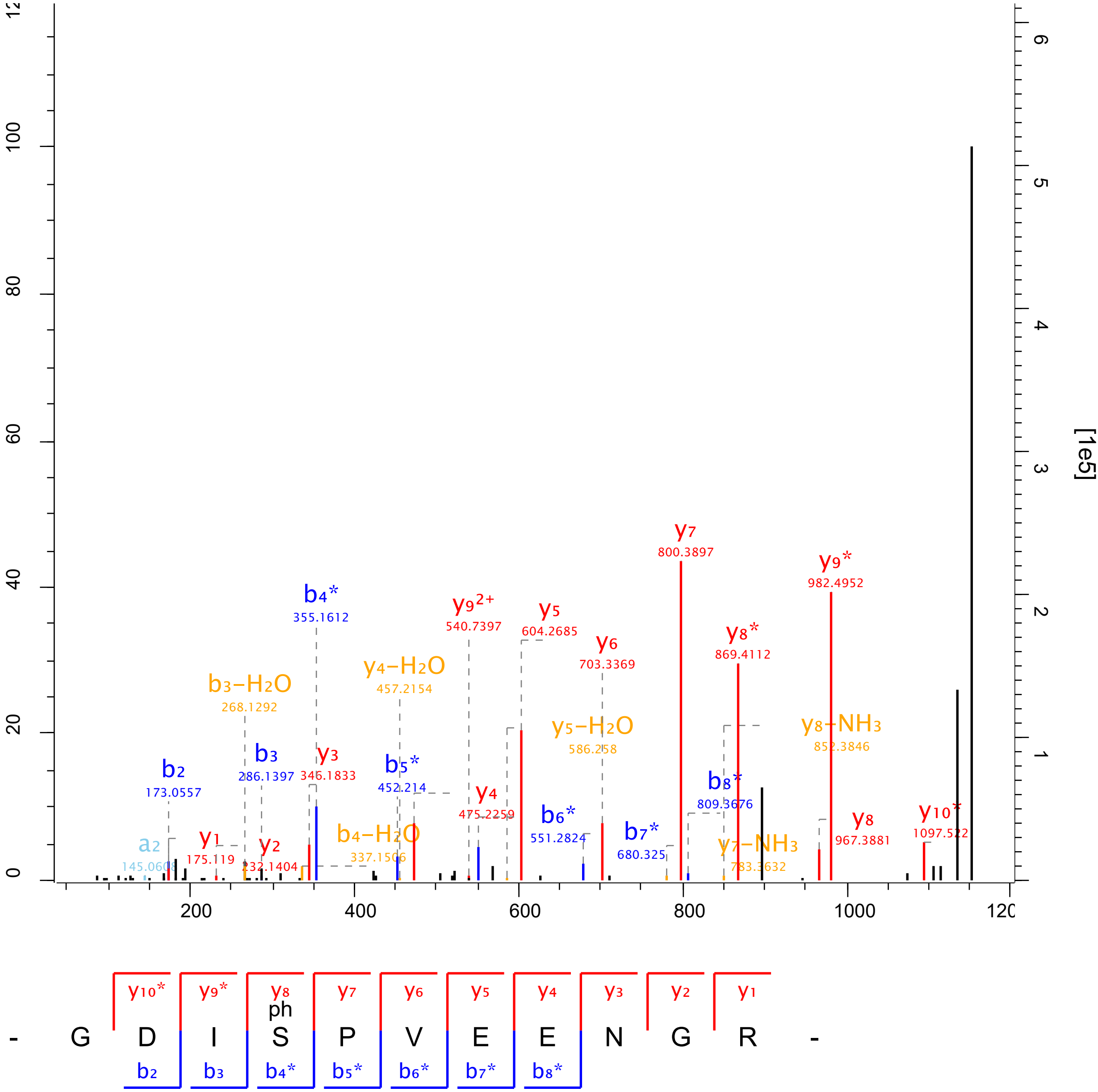

| Raw file | Scan | Method | Score | m/z |
|---|---|---|---|---|
| sys_00_2 | 8553 | FTMS; HCD | 139.7 | 561.27 |

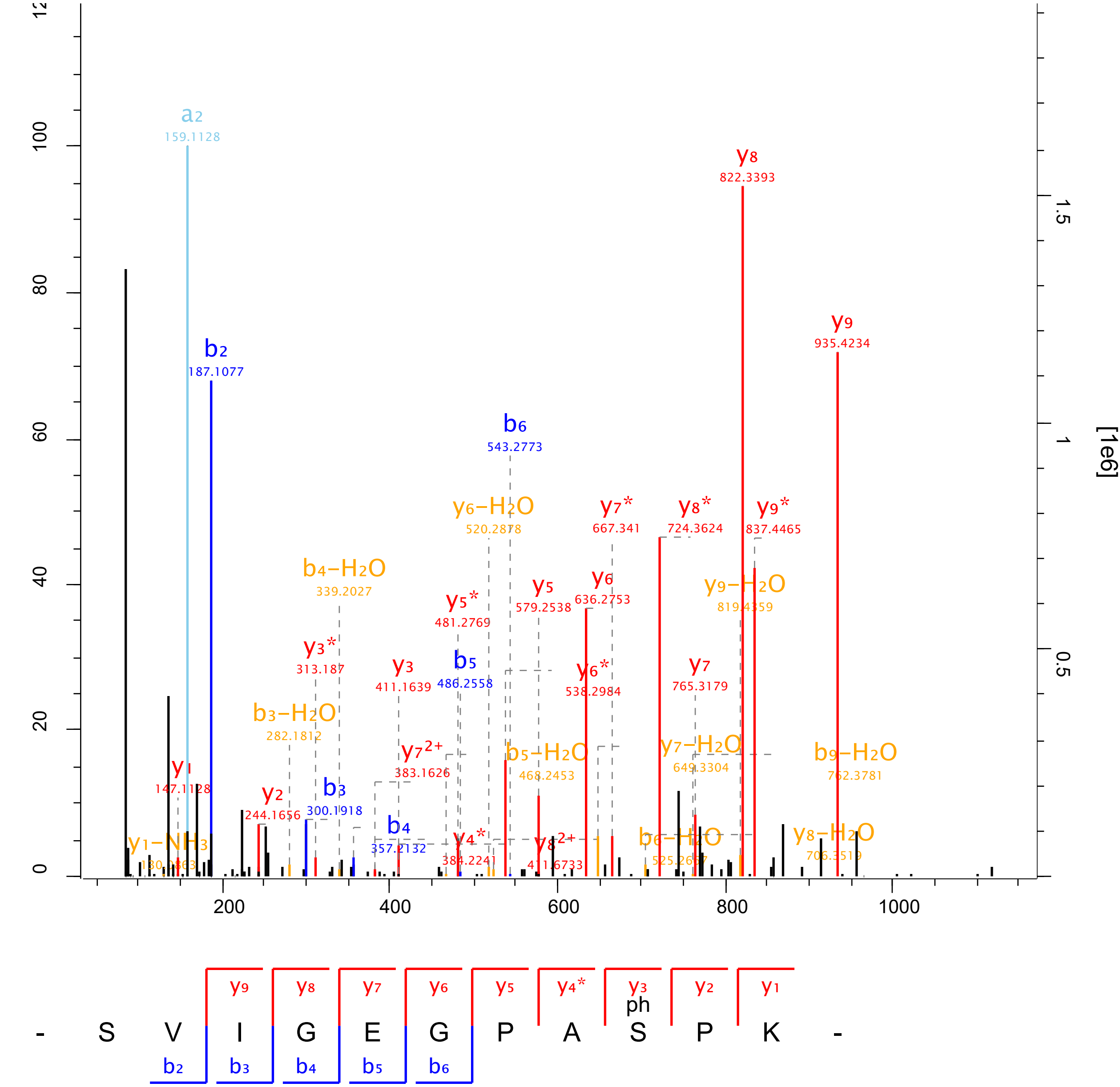

| Raw file | Scan | Method | Score | m/z |
|---|---|---|---|---|
| sys_00_2 | 8753 | FTMS; HCD | 148.18 | 562.76 |

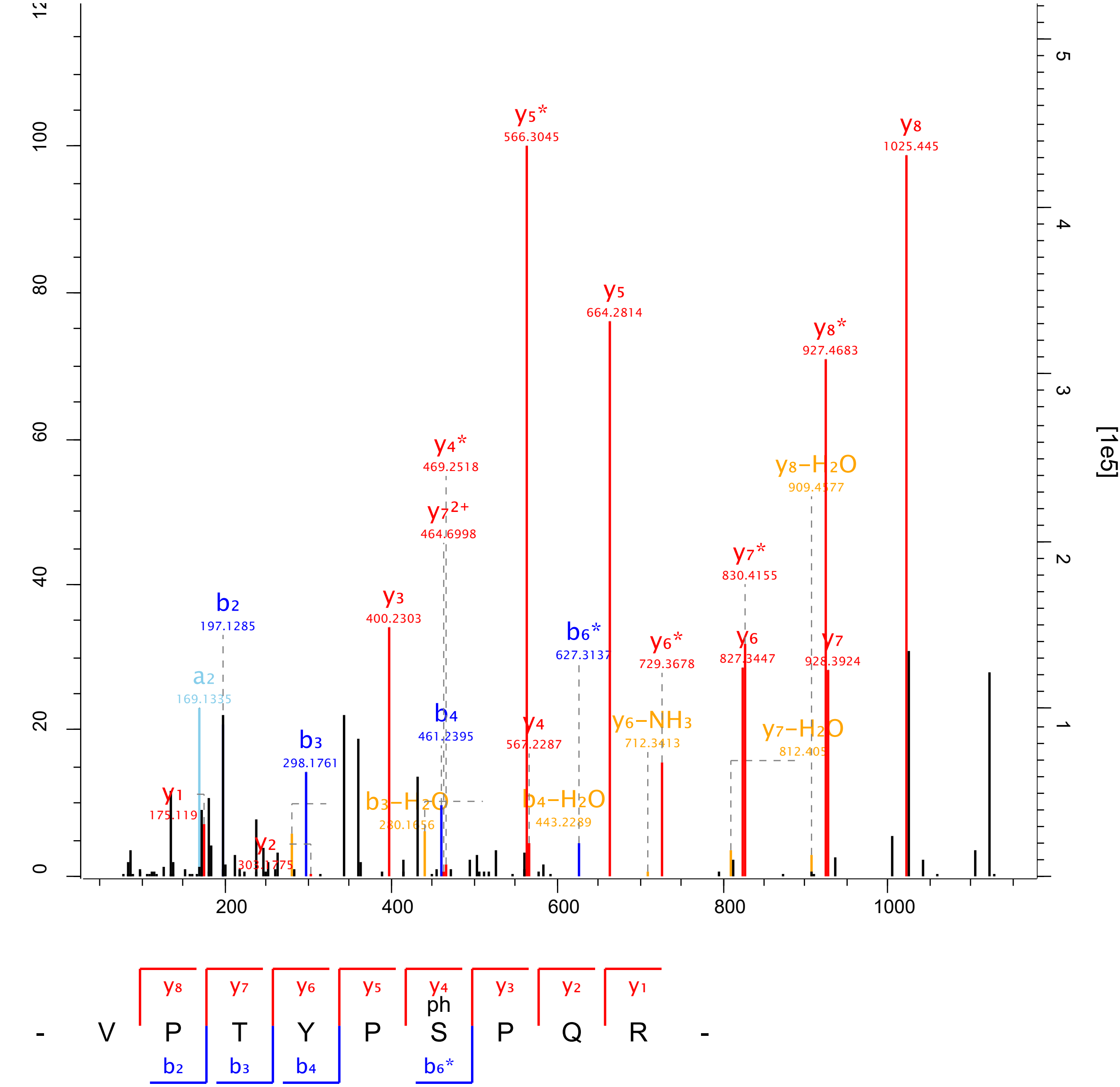

| Raw file | Scan | Method | Score | m/z |
| --- | --- | --- | --- | --- |
| sys_00_2 | 8885 | FTMS; HCD | 40.94 | 651.26 |

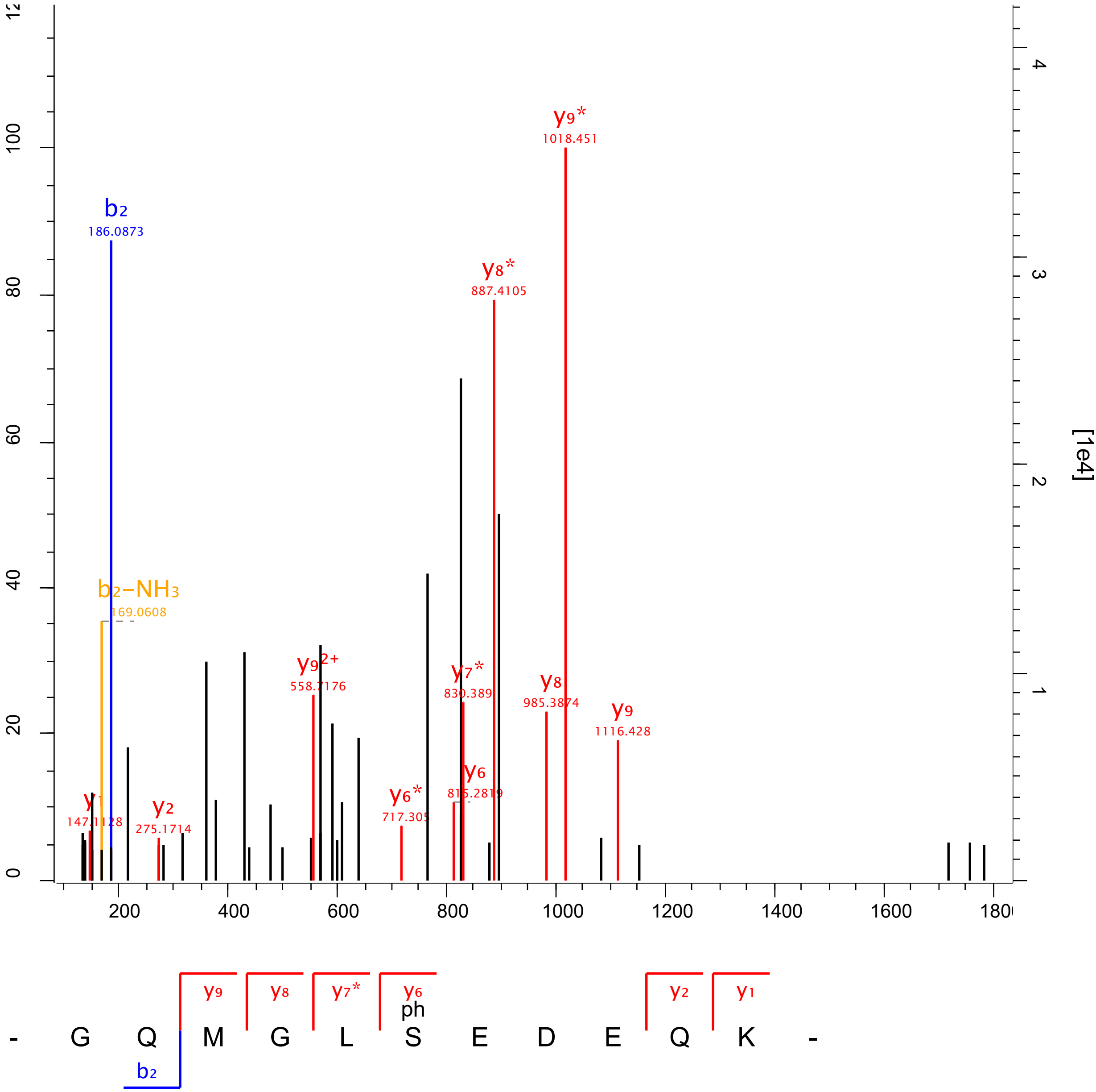

| Raw file | Scan | Method | Score | m/z |
| --- | --- | --- | --- | --- |
| sys_00_2 | 9070 | FTMS; HCD | 179.67 | 570.77 |

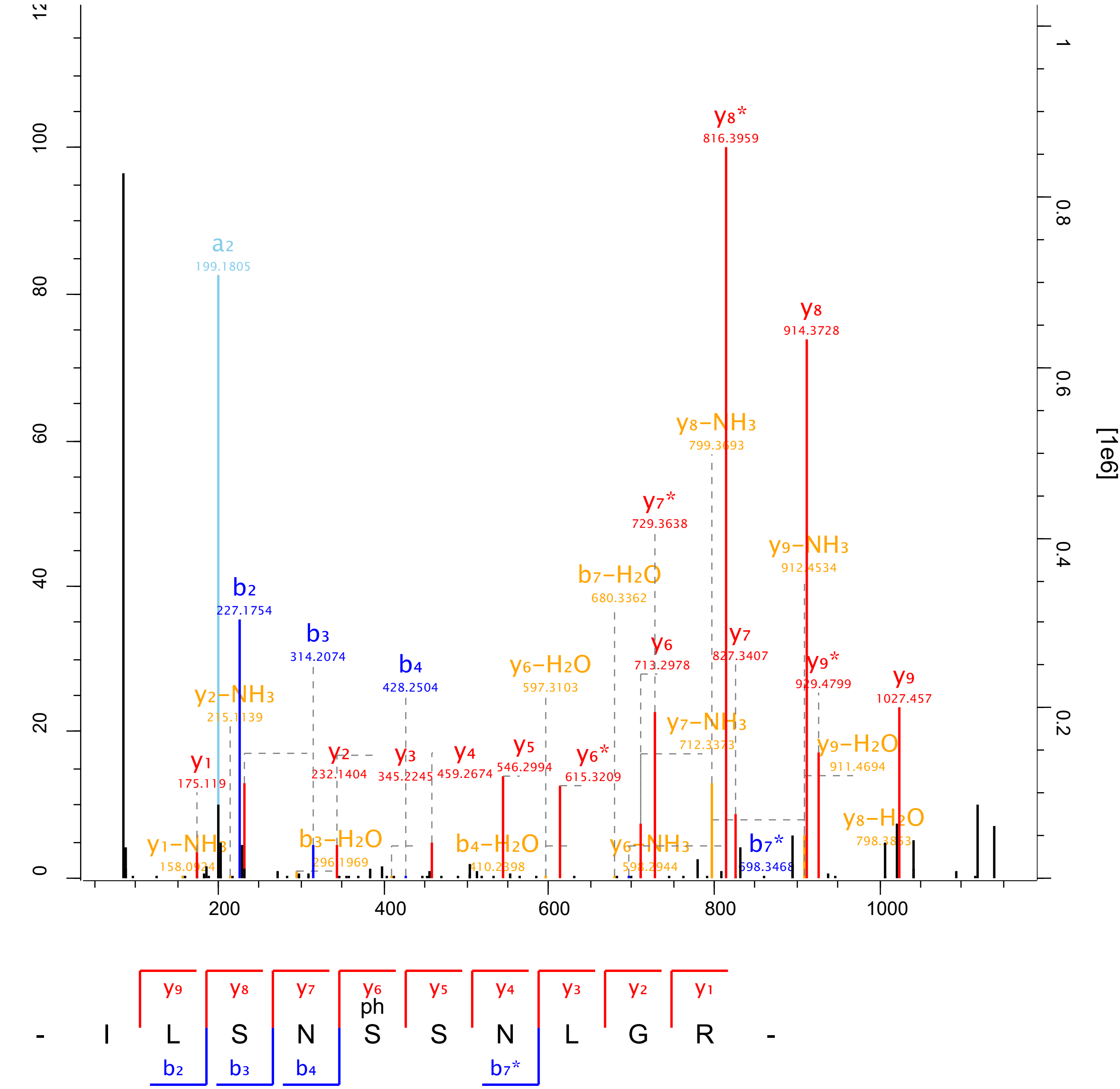

| Raw file | Scan | Method | Score | m/z |
|---|---|---|---|---|
| sys_00_2 | 9125 | FTMS; HCD | 72.61 | 591.28 |

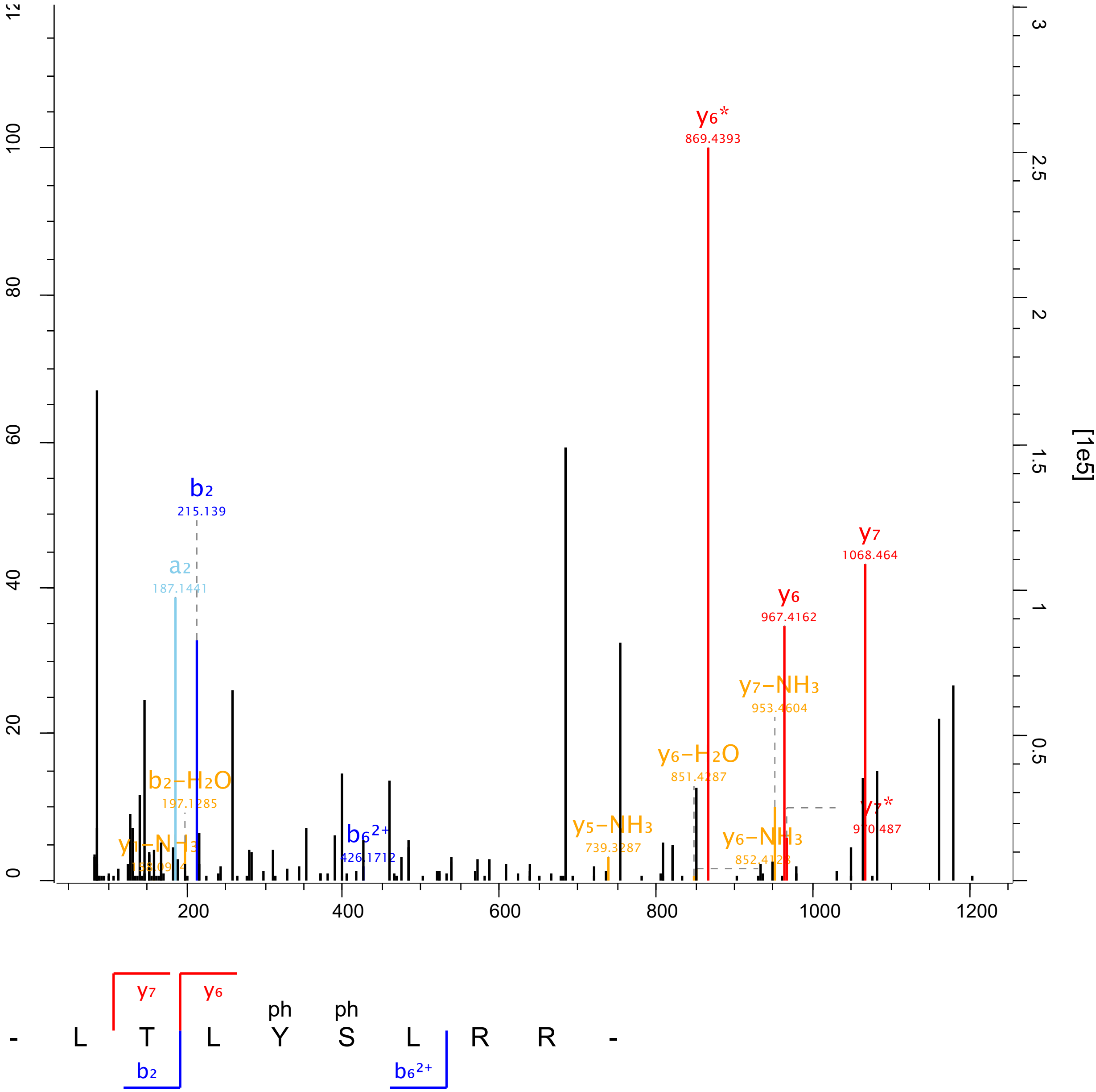

| Raw file | Scan | Method | Score | m/z |
|---|---|---|---|---|
| sys_00_2 | 9319 | FTMS; HCD | 118.28 | 476.71 |

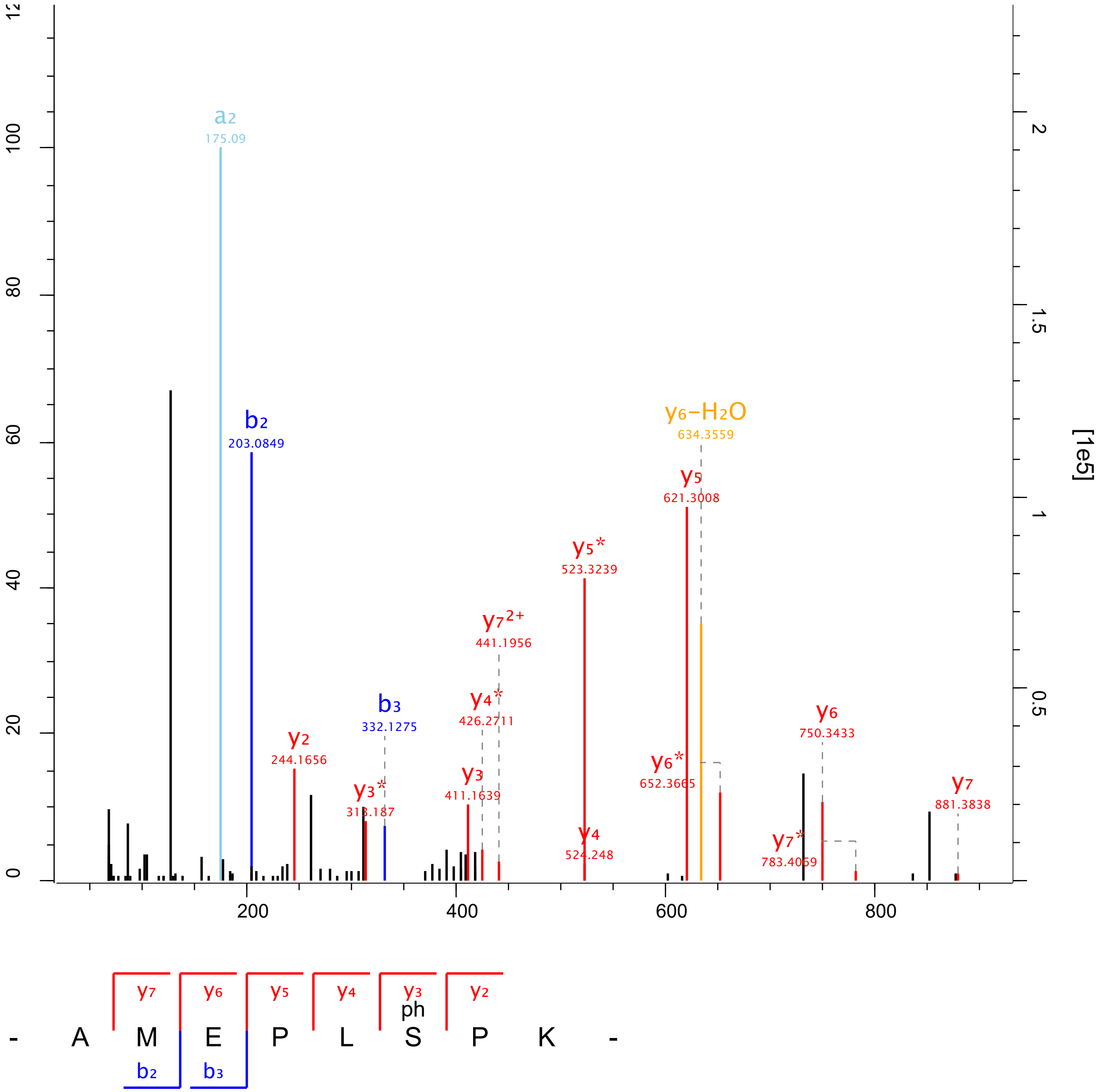

| Raw file | Scan | Method | Score | m/z |
| --- | --- | --- | --- | --- |
| sys_00_2 | 9677 | FTMS; HCD | 137.58 | 609.27 |

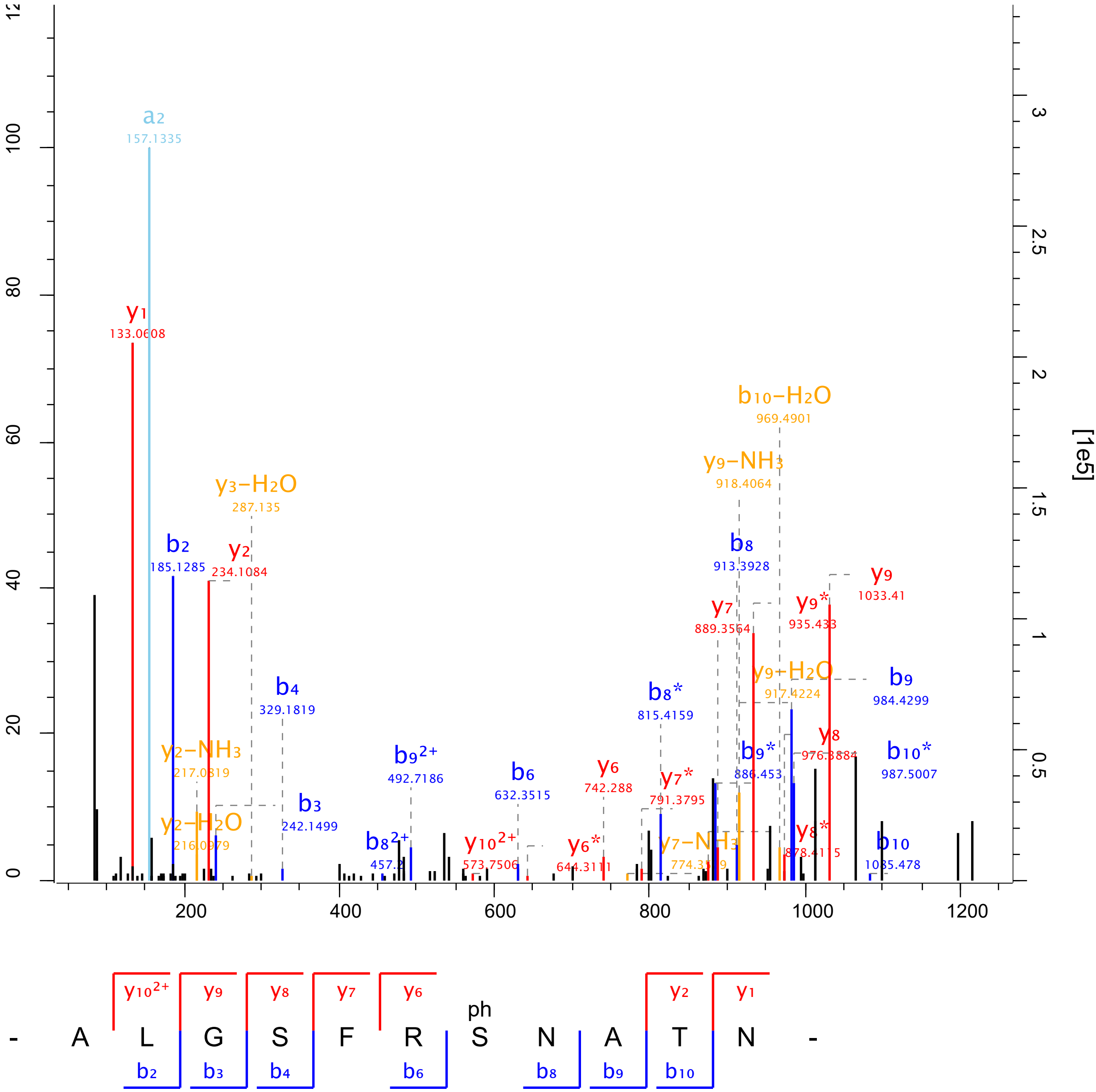

| Raw file | Scan | Method | Score | m/z |
|---|---|---|---|---|
| sys_00_2 | 9738 | FTMS; HCD | 94.77 | 524.22 |

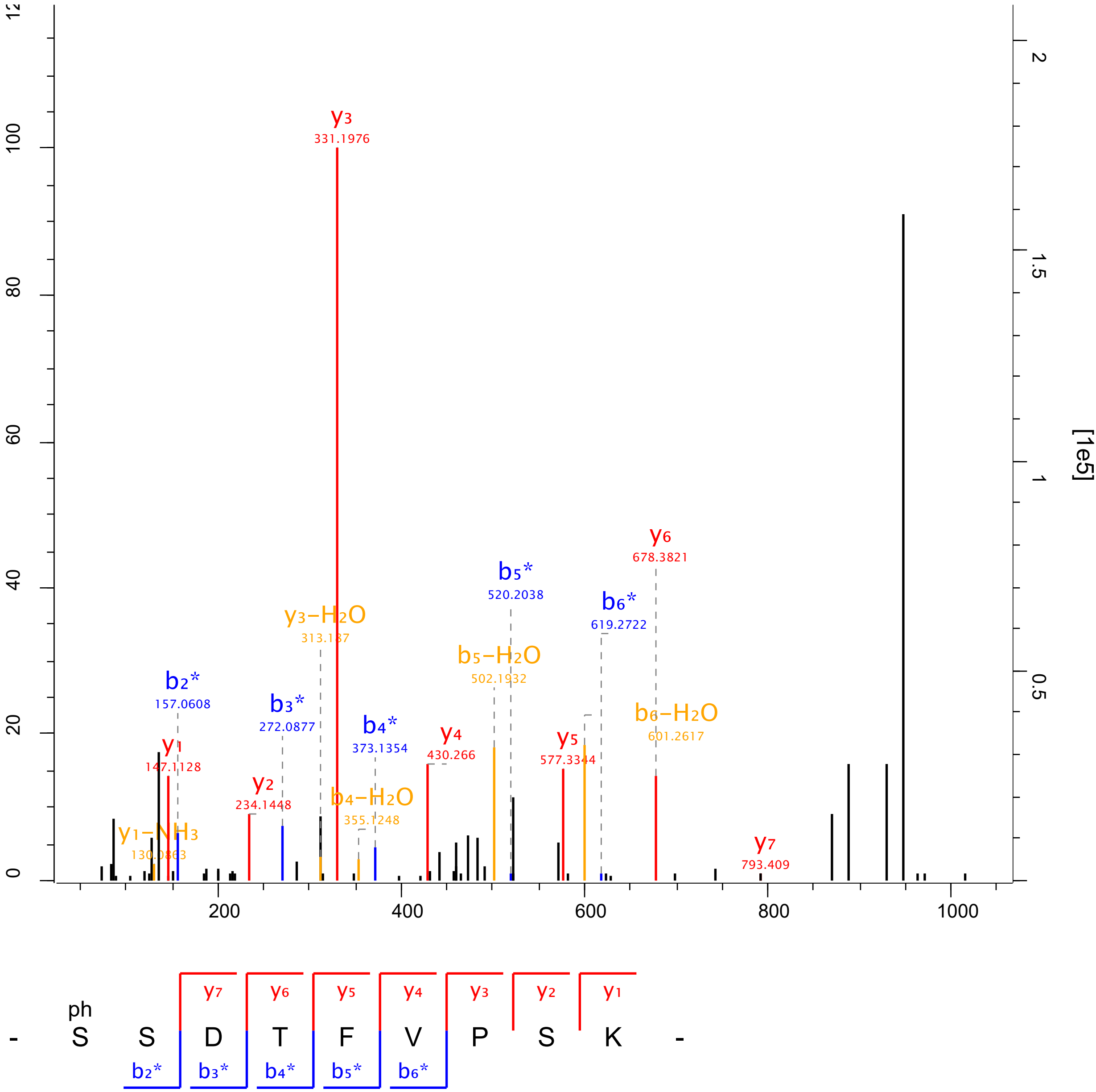

| Raw file | Scan | Method | Score | m/z |
|---|---|---|---|---|
| sys_00_2 | 9959 | FTMS; HCD | 79.2 | 544.73 |

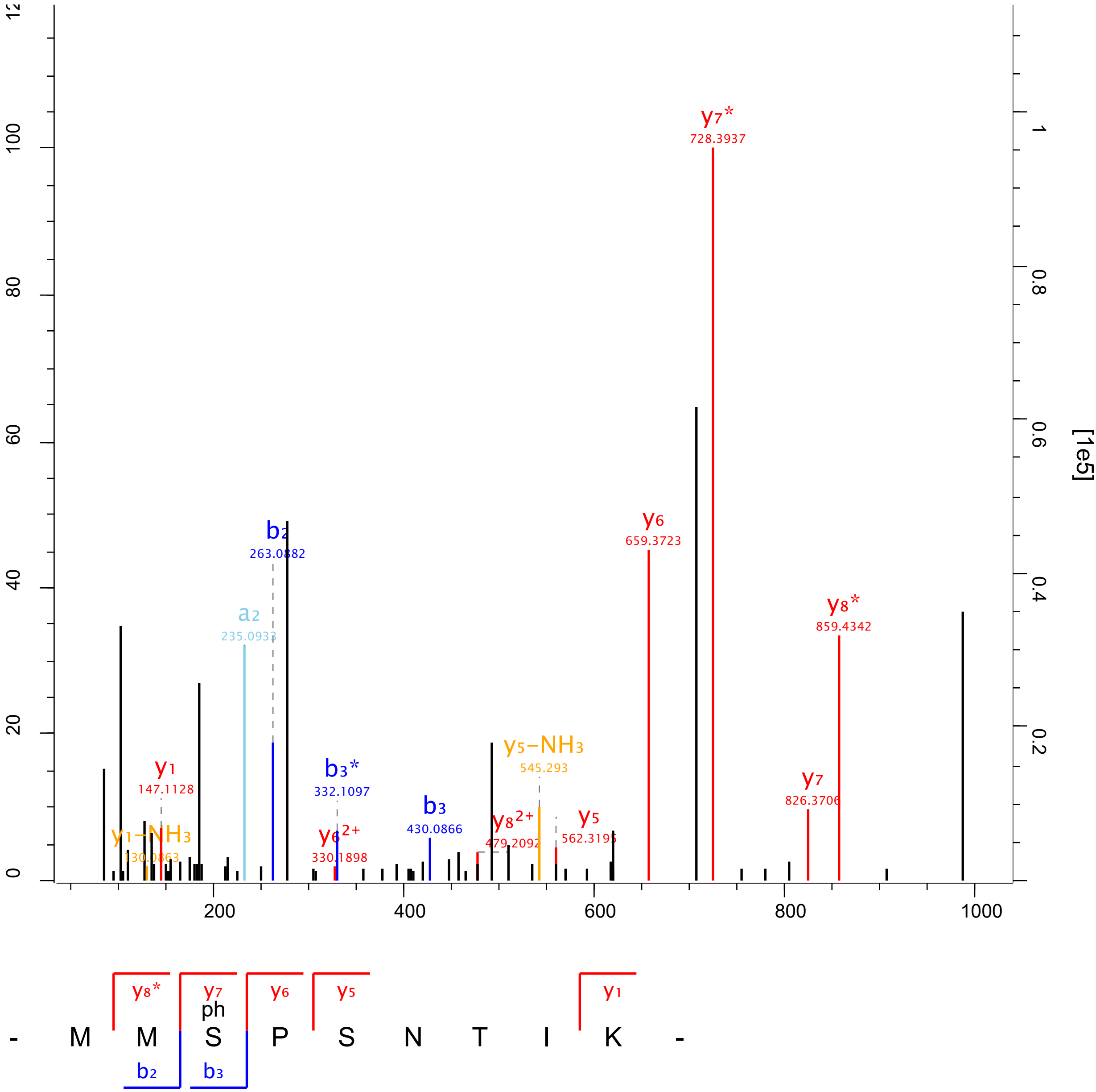

| Raw file | Scan | Method | Score | m/z |
| --- | --- | --- | --- | --- |
| sys_00_3short | 10195 | FTMS; HCD | 66.27 | 511.26 |

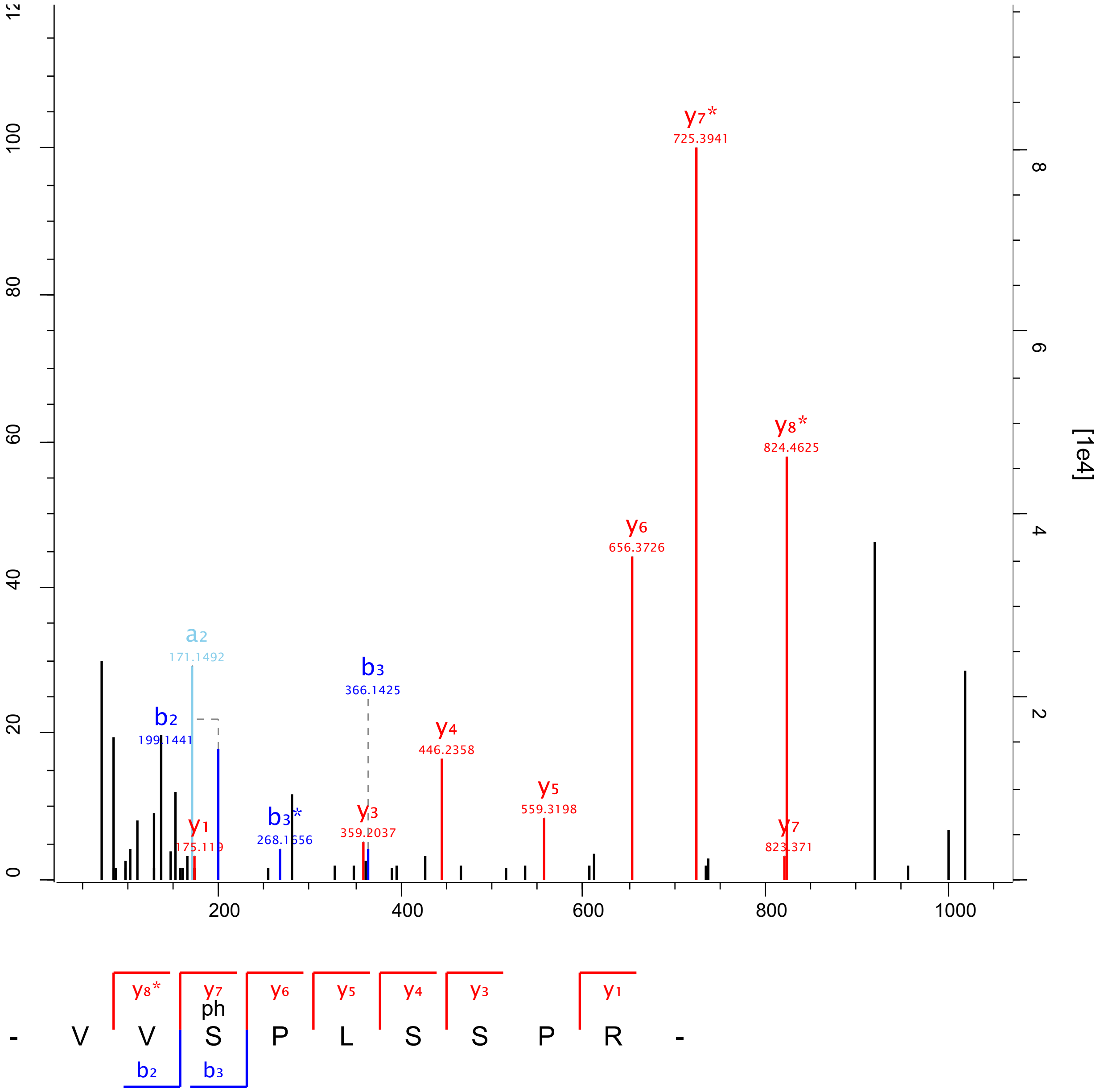

| Raw file | Scan | Method | Score | m/z |
|---|---|---|---|---|
| sys_00_3short | 10211 | FTMS; HCD | 84.05 | 495.75 |

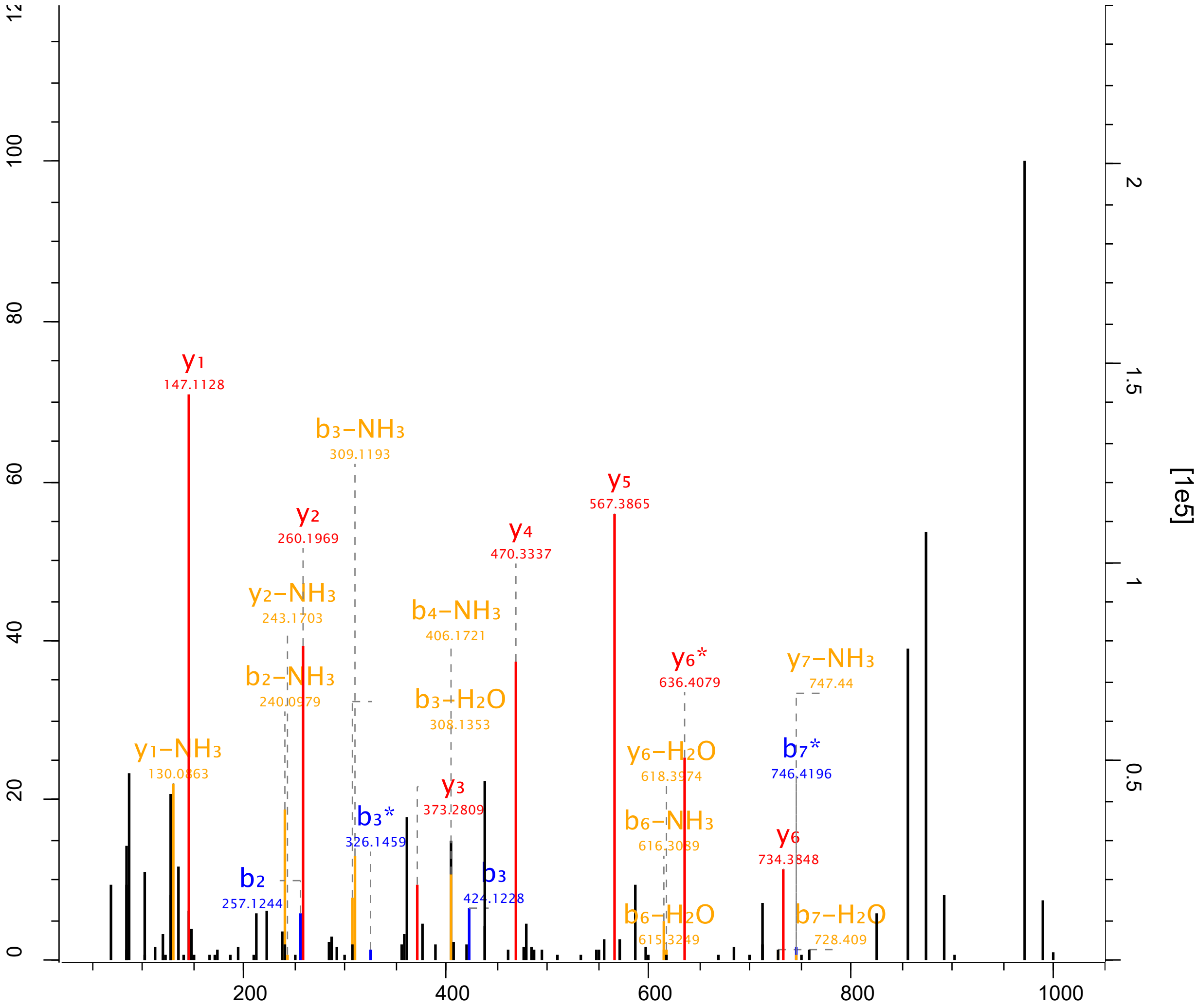


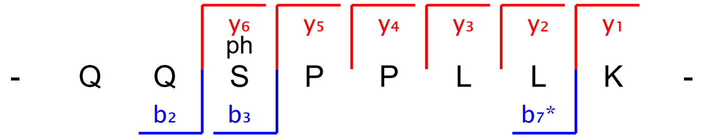

| Raw file | Scan | Method | Score | m/z |
|---|---|---|---|---|
| sys_00_3short | 10304 | FTMS; HCD | 55.01 | 714.8 |

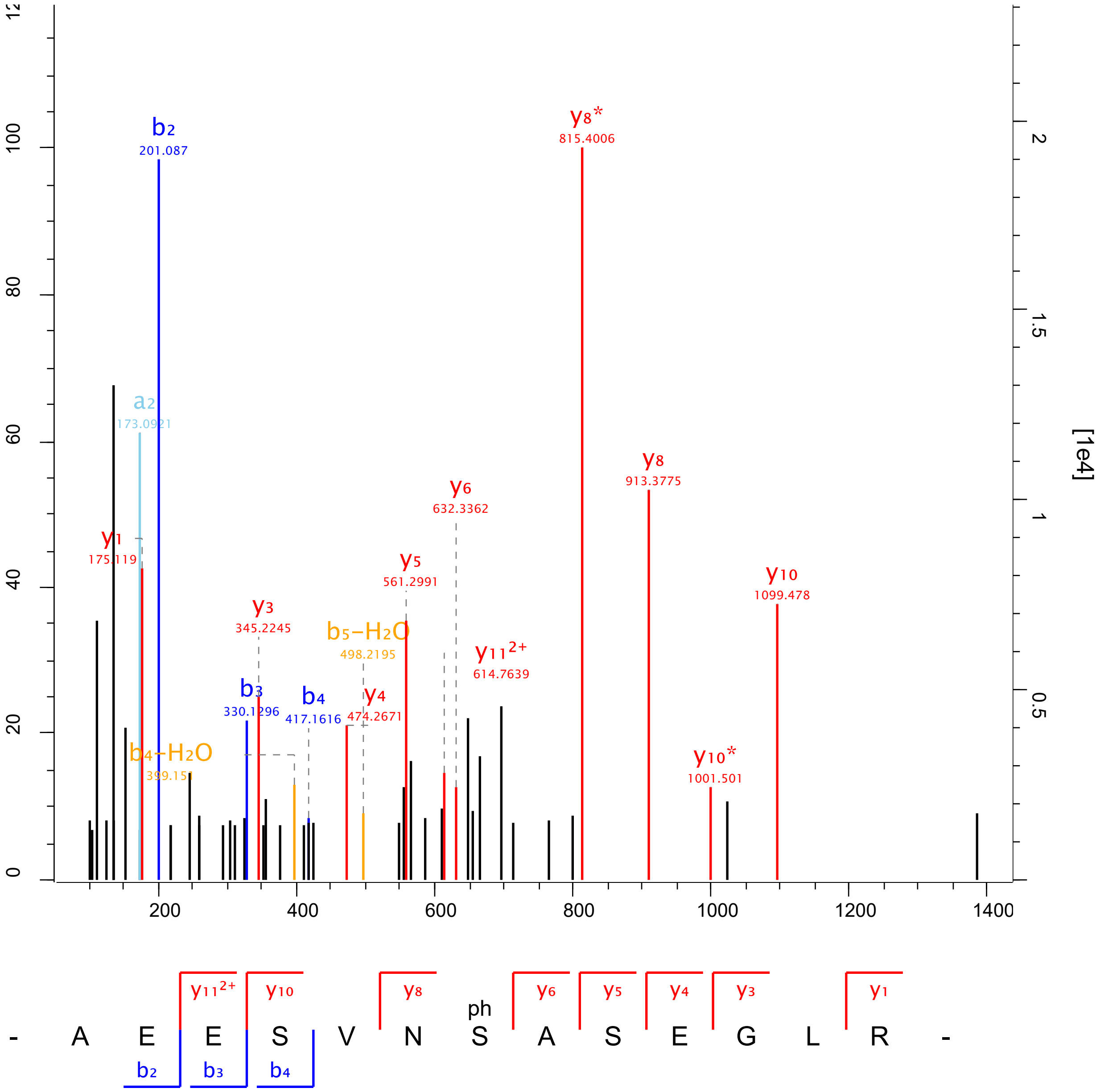

| Raw file | Scan | Method | Score | m/z |
|---|---|---|---|---|
| sys_00_3short | 10520 | FTMS; HCD | 62.47 | 478.71 |

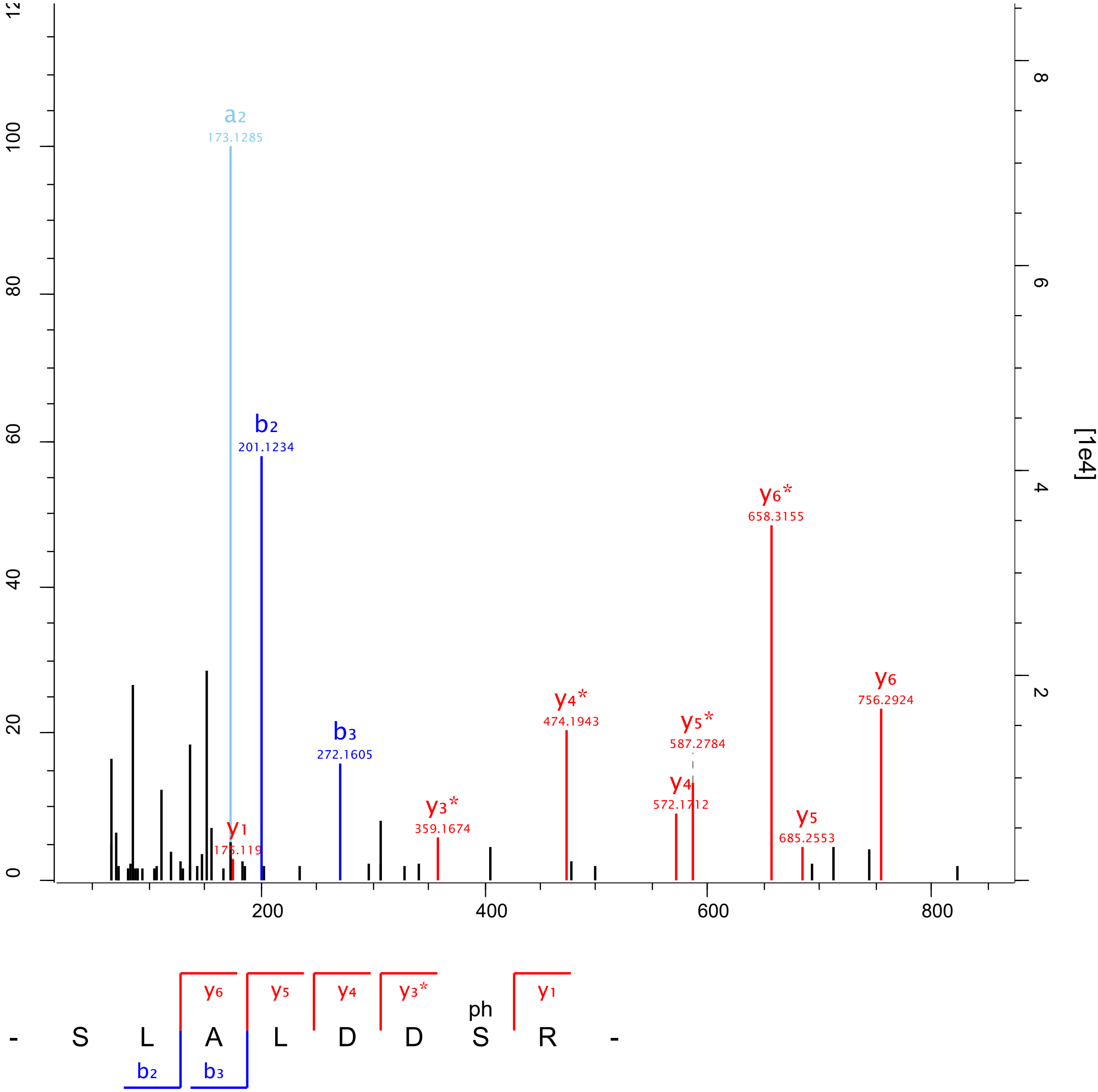

| Raw file | Scan | Method | Score | m/z |
|---|---|---|---|---|
| sys_00_3short | 10603 | FTMS; HCD | 53.26 | 712.33 |

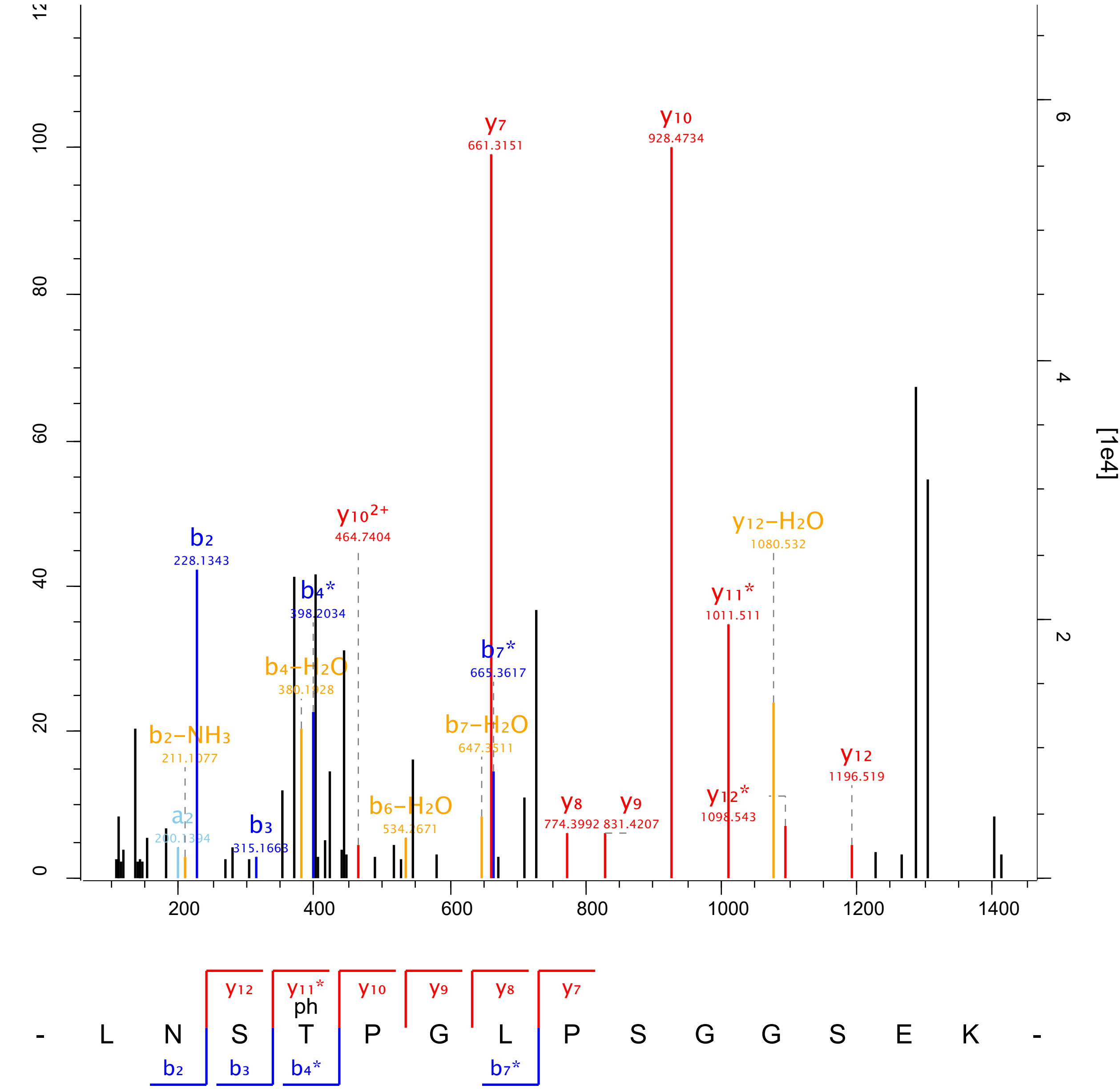

| Raw file | Scan | Method | Score | m/z |
| --- | --- | --- | --- | --- |
| sys_00_3short | 10989 | FTMS; HCD | 58.08 | 698.82 |

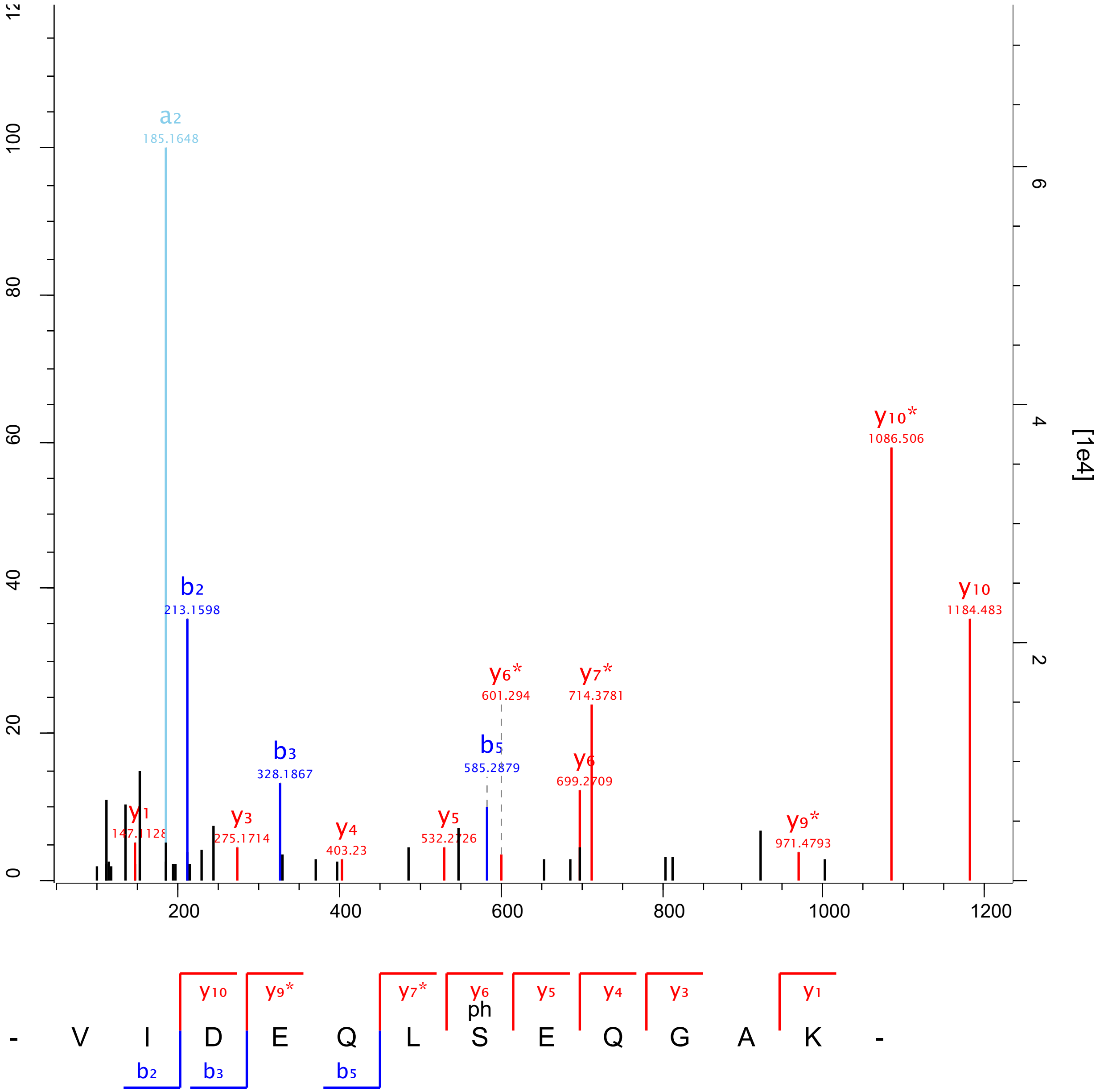

| Raw file | Scan | Method | Score | m/z |
|---|---|---|---|---|
| sys_00_3short | 11016 | FTMS; HCD | 147.7 | 662.31 |

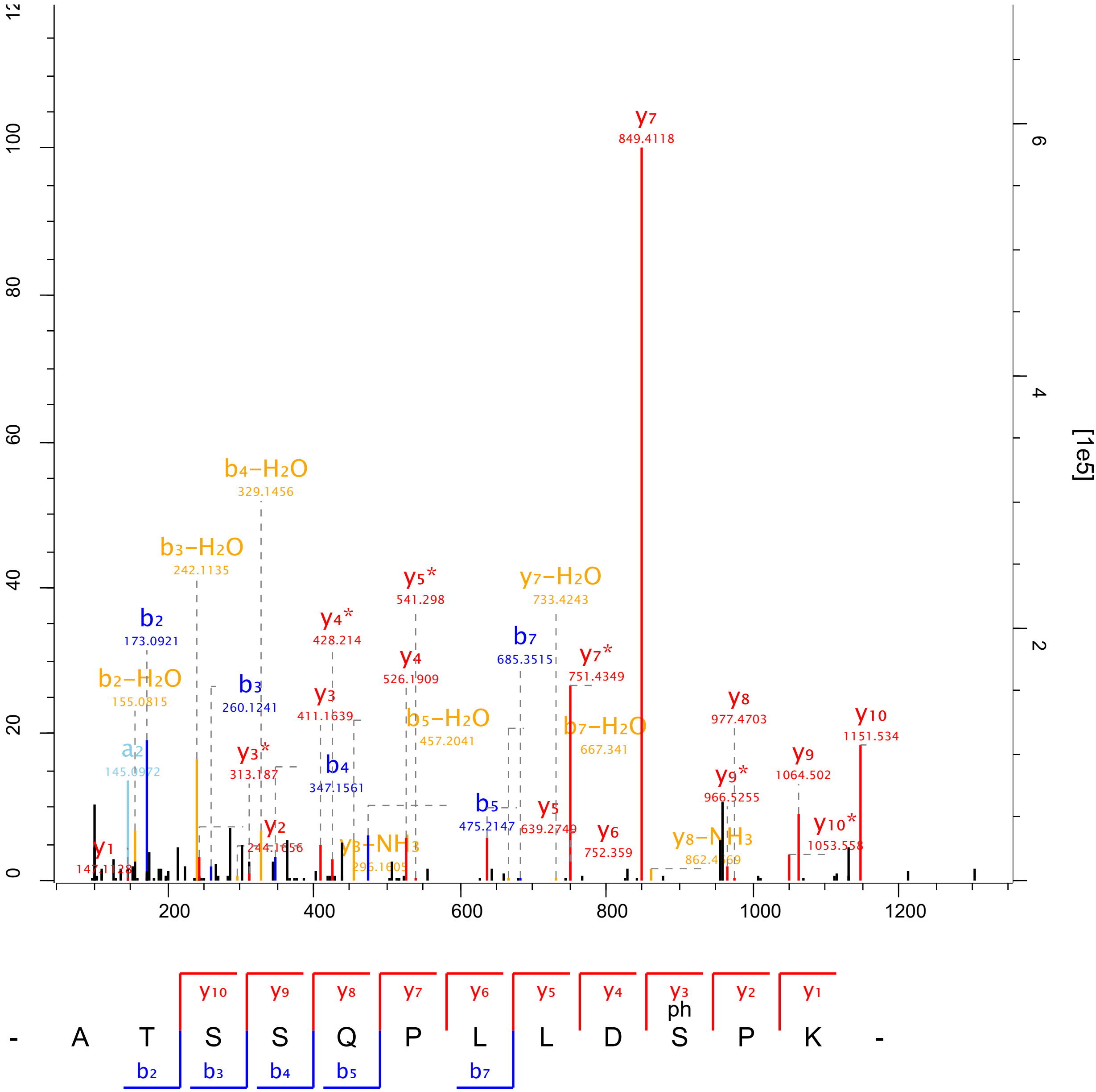

| Raw file | Scan | Method | Score | m/z |
|---|---|---|---|---|
| sys_00_3short | 11136 | FTMS; HCD | 42.34 | 433.7 |

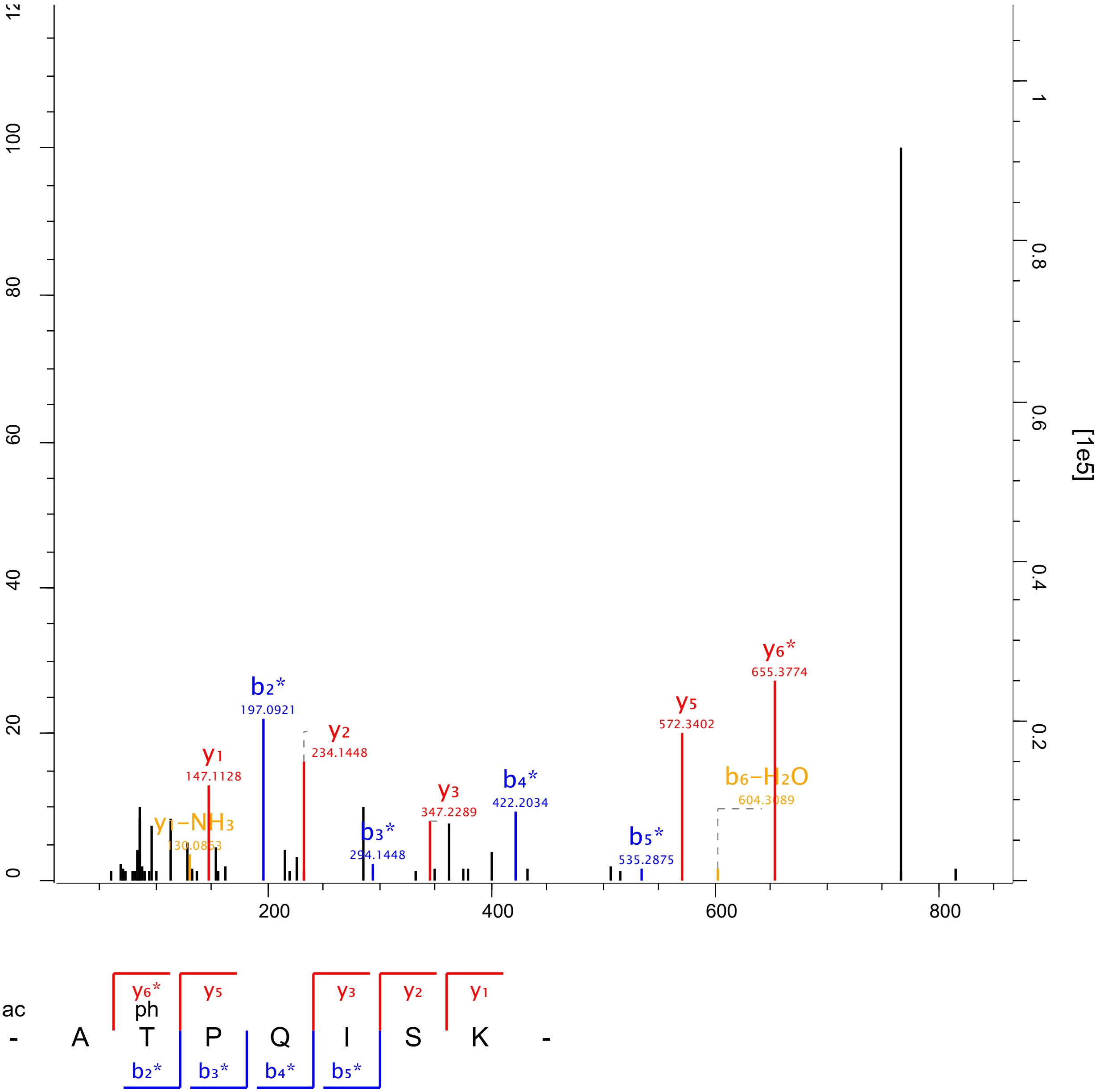

| Raw file | Scan | Method | Score | m/z |
|---|---|---|---|---|
| sys_00_3short | 11182 | FTMS; HCD | 95.42 | 510.72 |

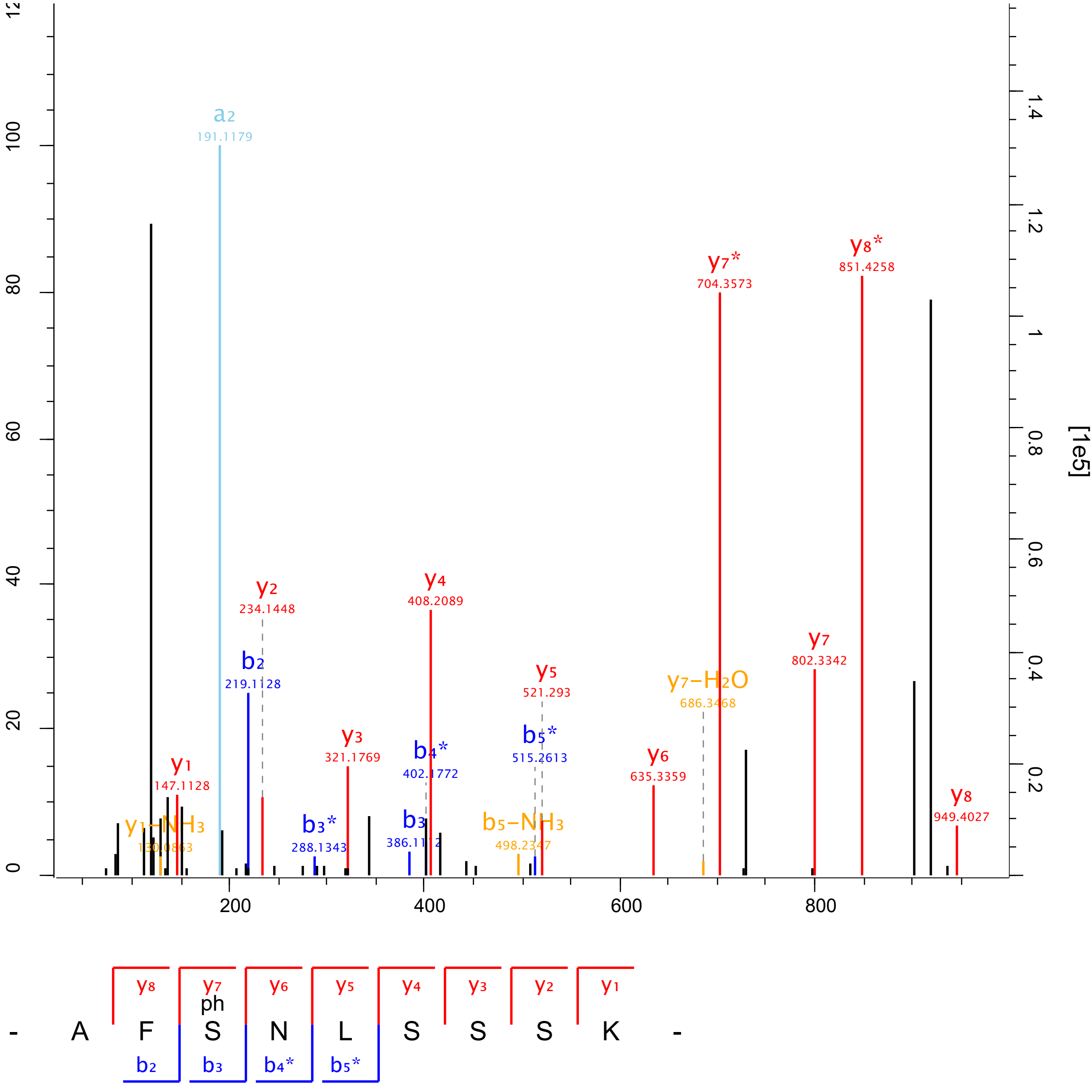

| Raw file | Scan | Method | Score | m/z |
| --- | --- | --- | --- | --- |
| sys_00_3short | 11403 | FTMS; HCD | 125.97 | 452.71 |

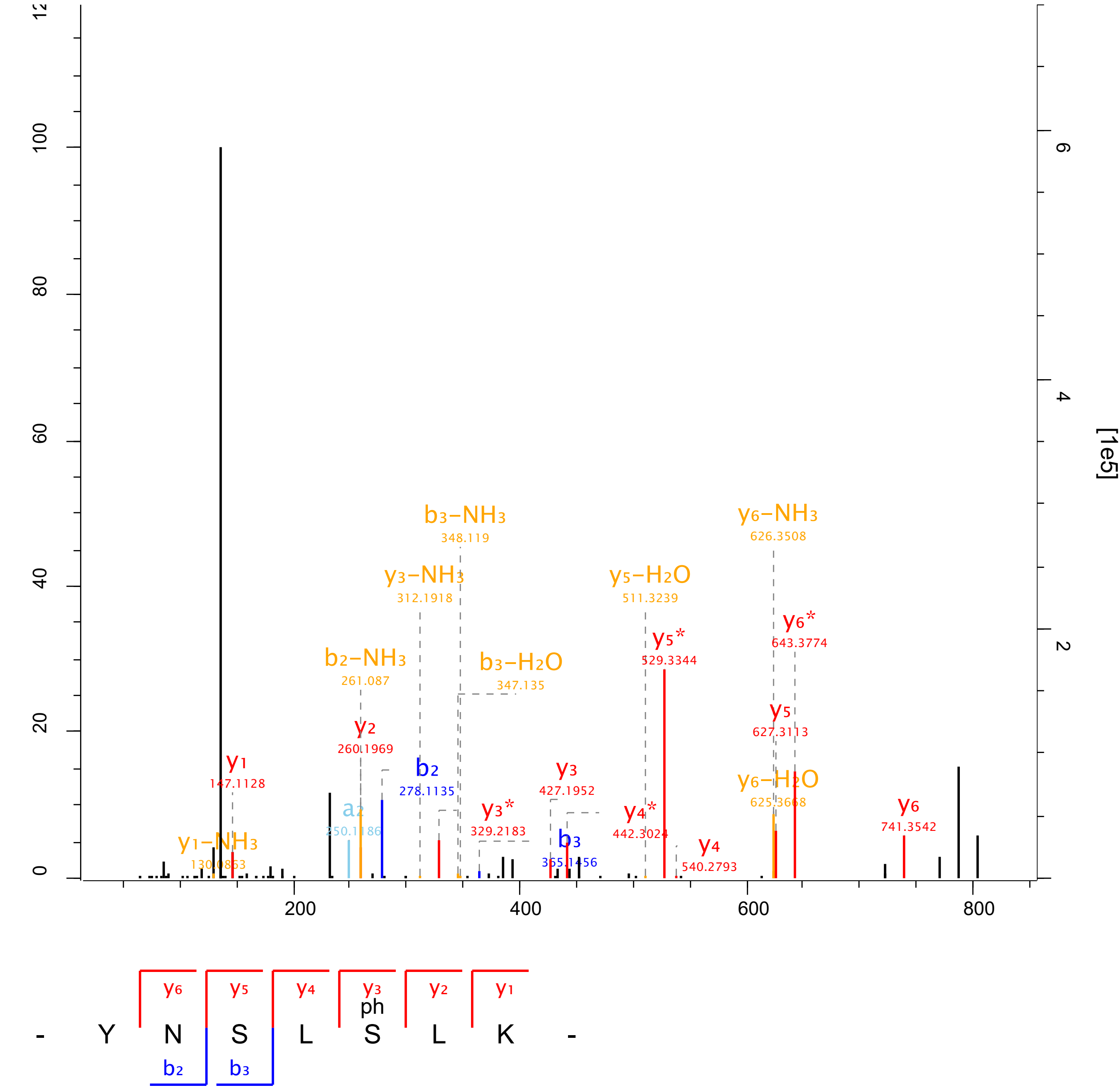

| Raw file | Scan | Method | Score | m/z |
|---|---|---|---|---|
| sys_00_3short | 11629 | FTMS; HCD | 52.58 | 630.73 |

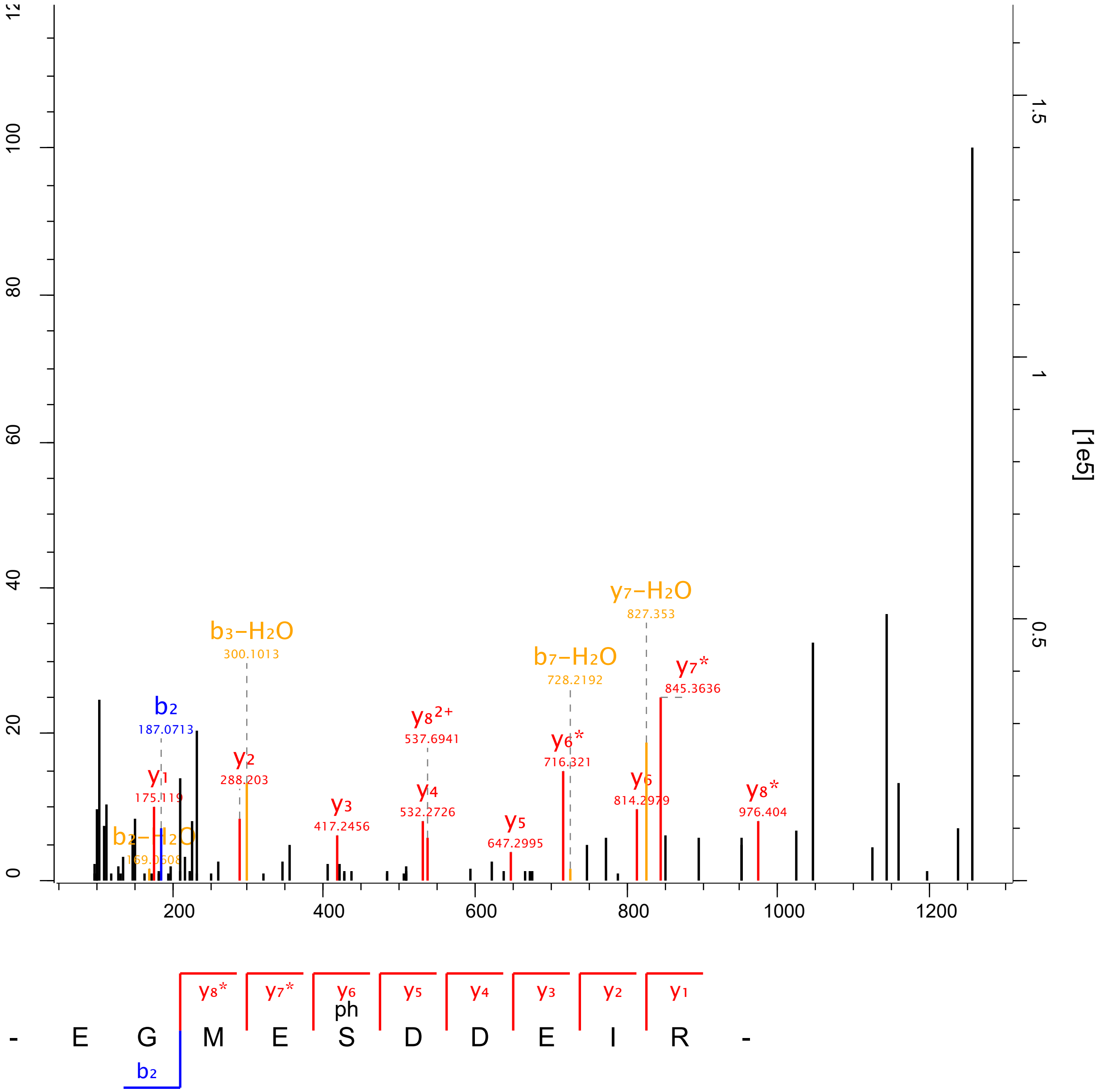

| Raw file | Scan | Method | Score | m/z |
|---|---|---|---|---|
| sys_00_3short | 12013 | FTMS; HCD | 116.24 | 568.23 |

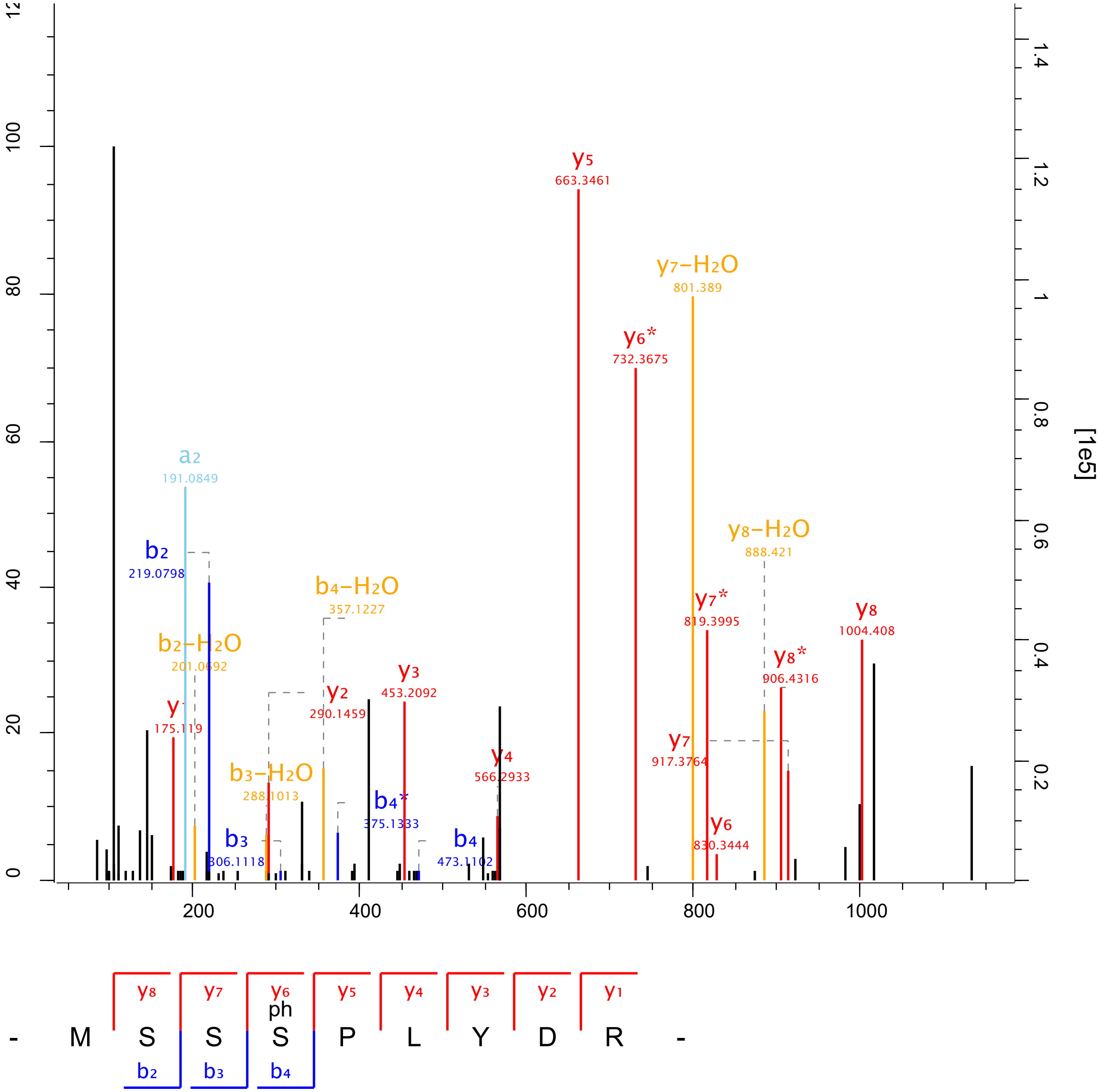

| Raw file | Scan | Method | Score | m/z |
|---|---|---|---|---|
| sys_00_3short | 12110 | FTMS; HCD | 136.39 | 686.31 |

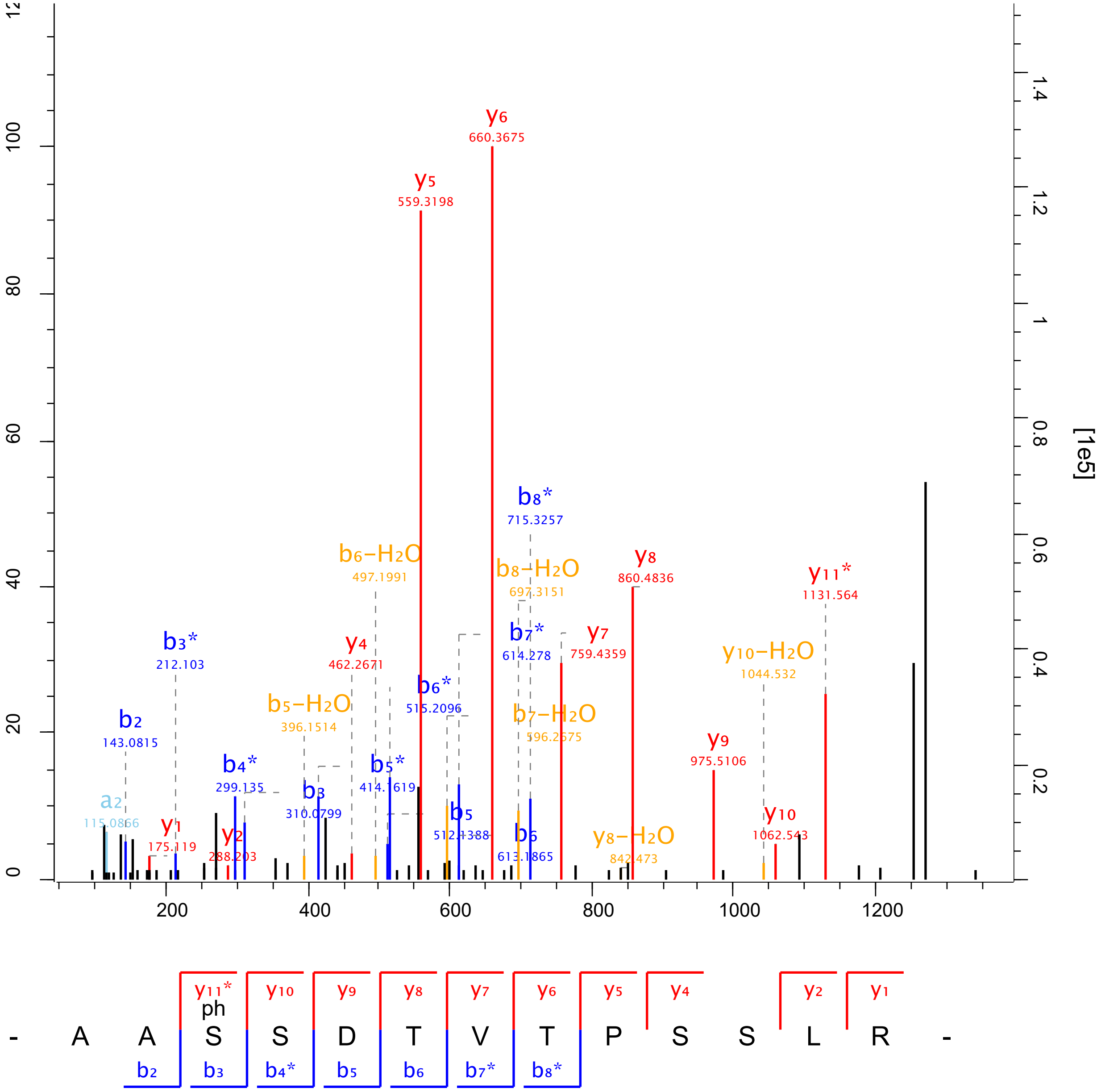

| Raw file | Scan | Method | Score | m/z |
| --- | --- | --- | --- | --- |
| sys_00_3short | 12113 | FTMS; HCD | 146.16 | 515.24 |

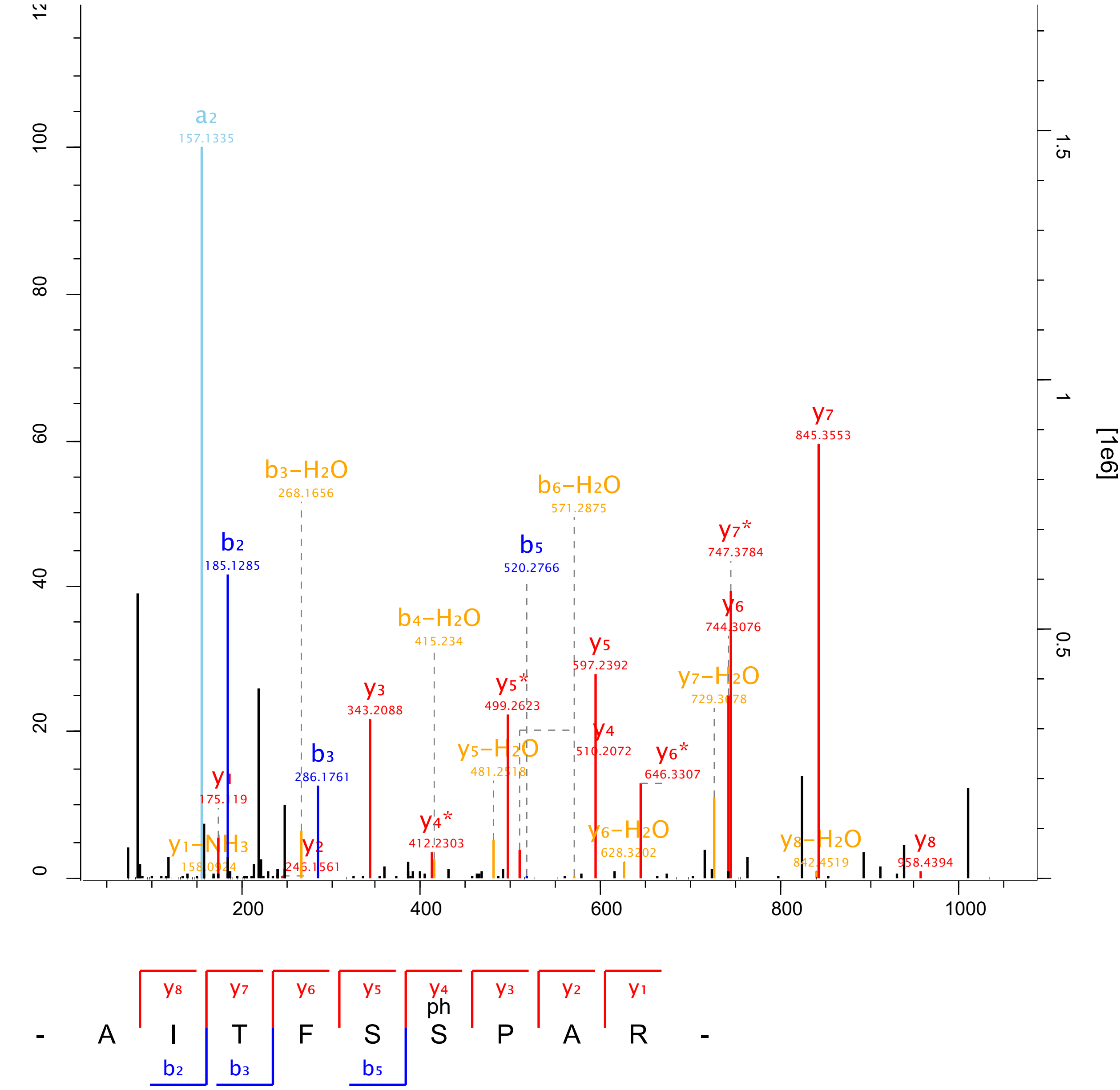

| Raw file | Scan | Method | Score | m/z |
|---|---|---|---|---|
| sys_00_3short | 12650 | FTMS; HCD | 148.62 | 445.19 |

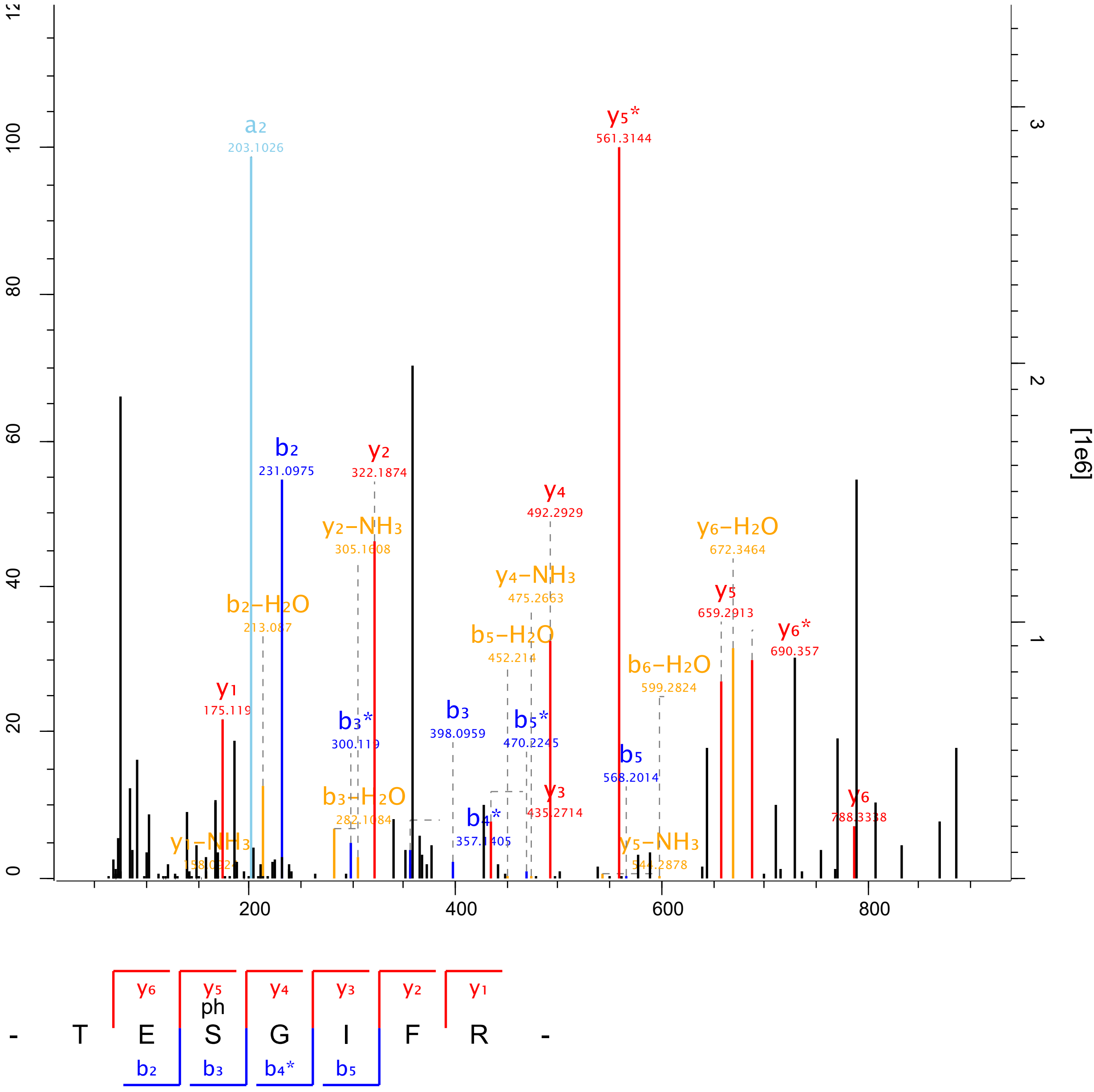

| Raw file | Scan | Method | Score | m/z |
|---|---|---|---|---|
| sys_00_3short | 12797 | FTMS; HCD | 178.51 | 598.28 |

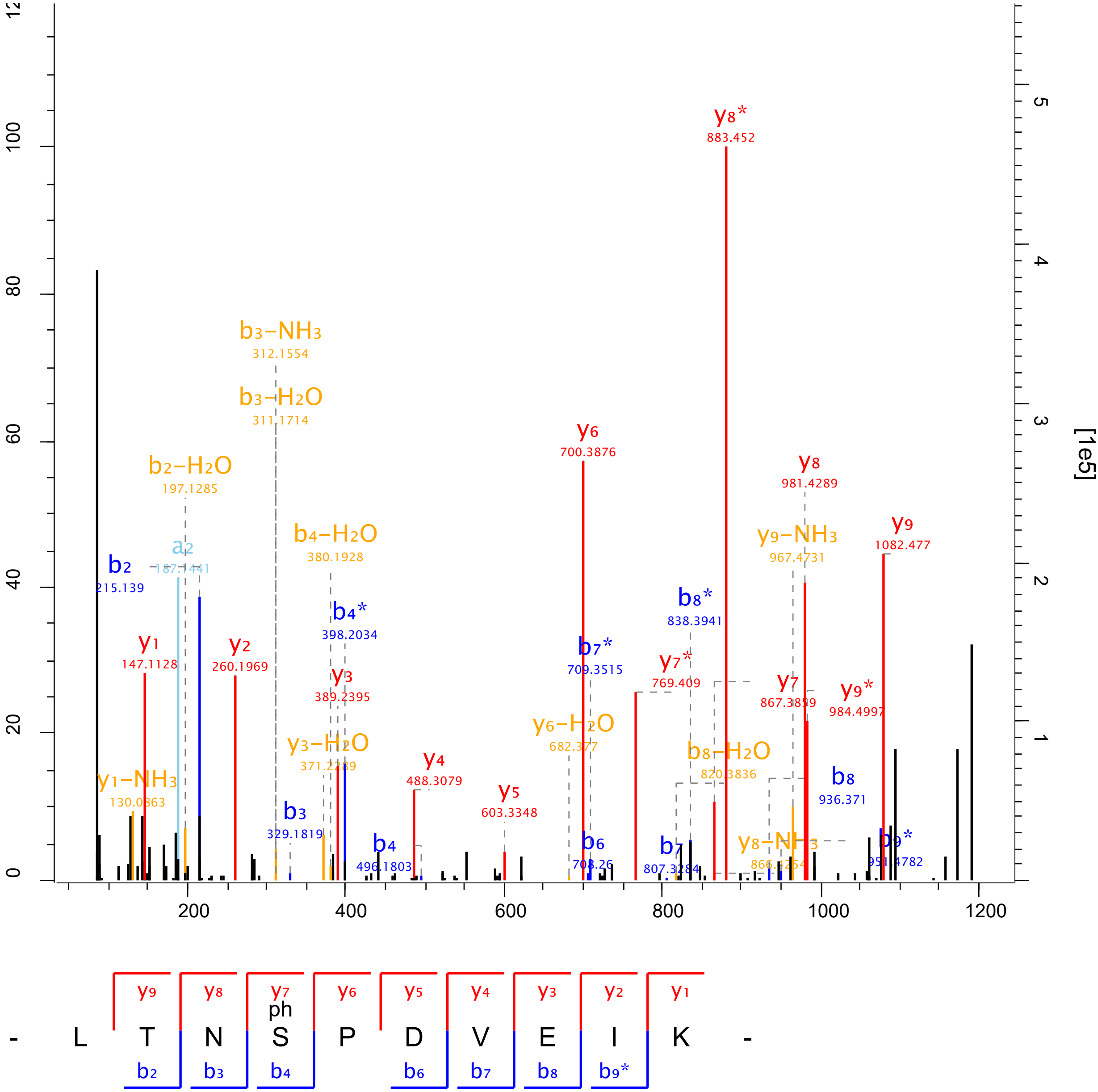

| Raw file | Scan | Method | Score | m/z |
| --- | --- | --- | --- | --- |
| sys_00_3short | 12943 | FTMS; HCD | 123.75 | 562.75 |

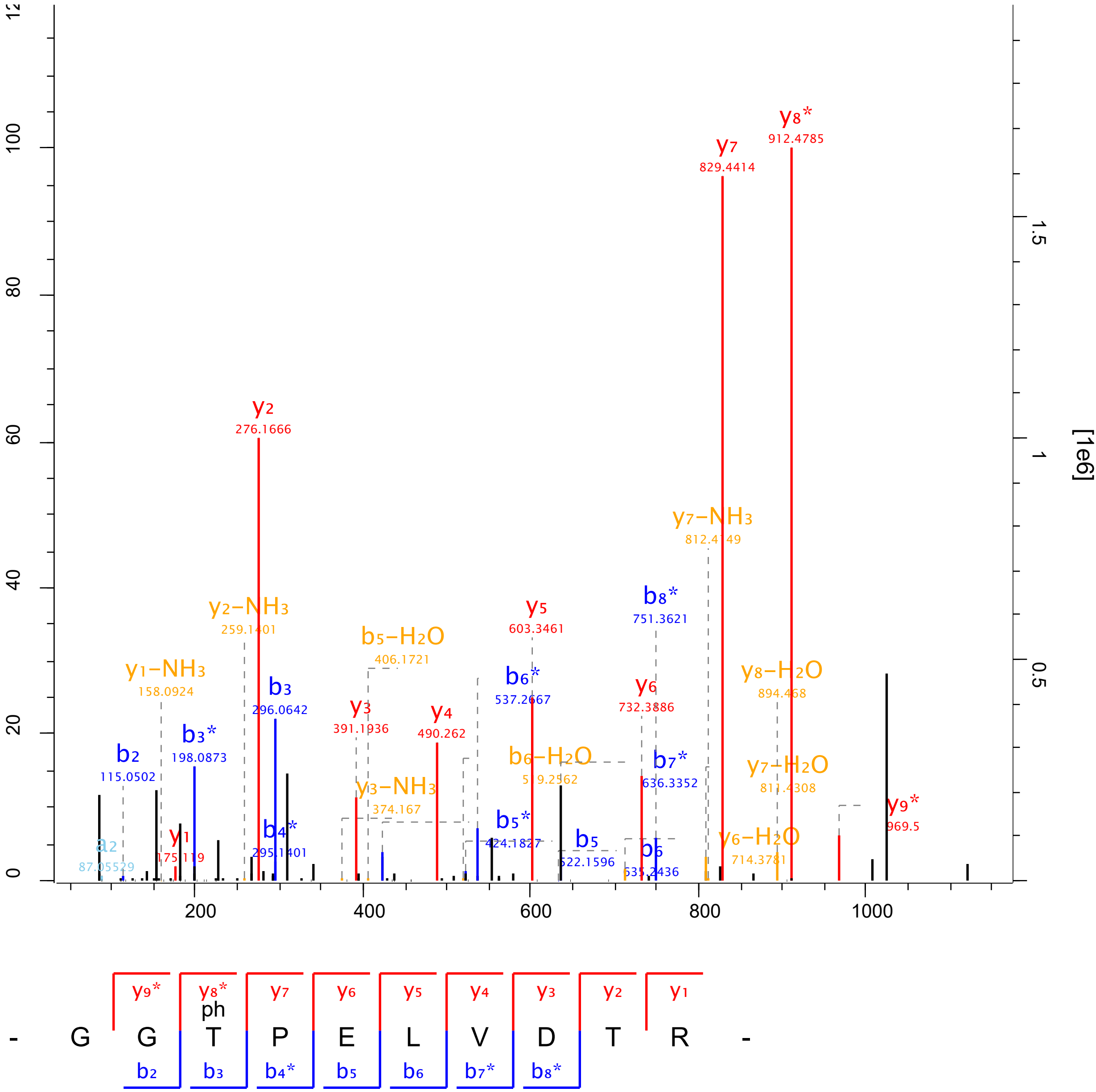

| Raw file | Scan | Method | Score | m/z |
| --- | --- | --- | --- | --- |
| sys_00_3short | 12999 | FTMS; HCD | 111.63 | 524.23 |

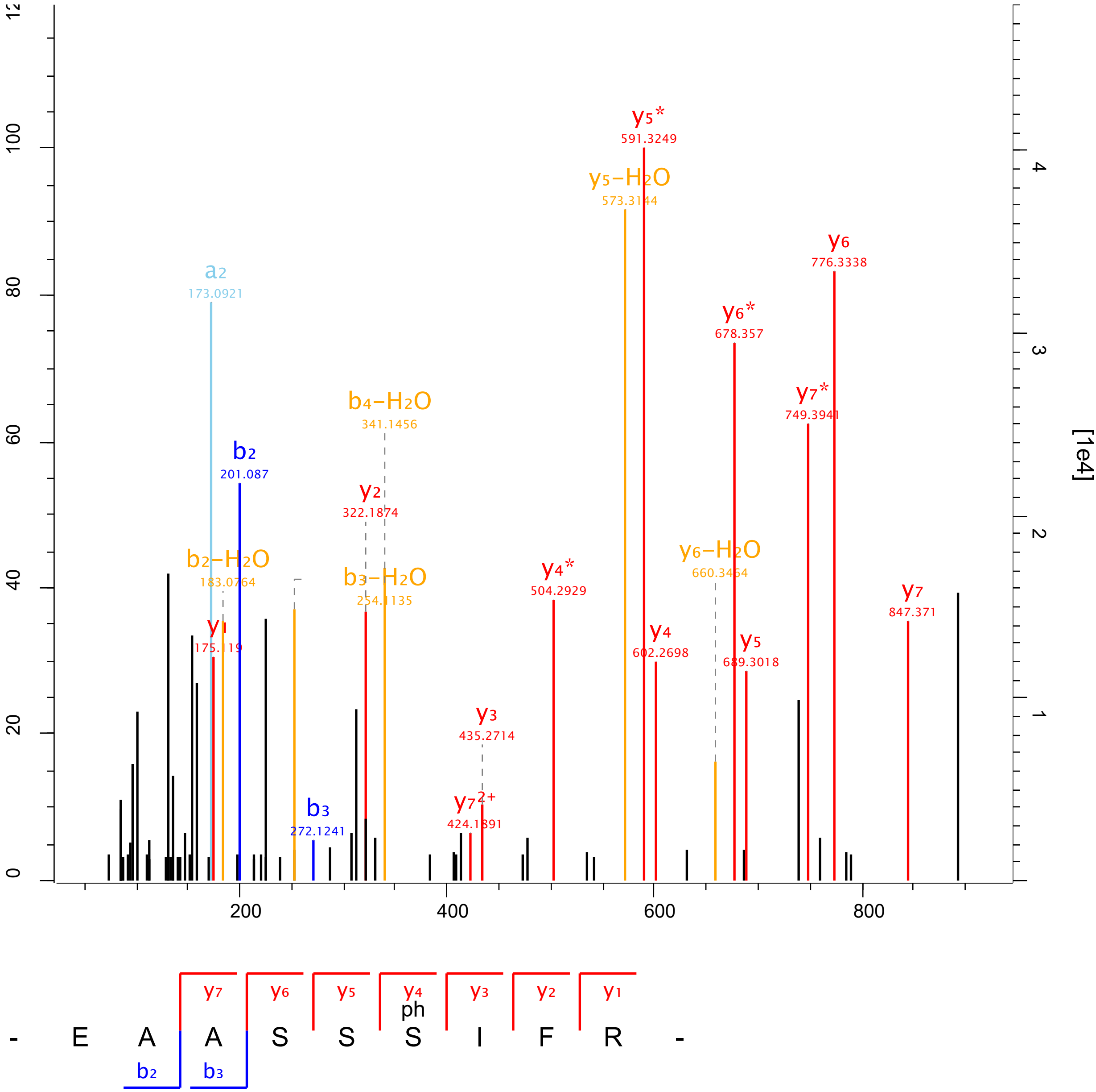

| Raw file | Scan | Method | Score | m/z |
|---|---|---|---|---|
| sys_00_3short | 13034 | FTMS; HCD | 51.73 | 560.23 |

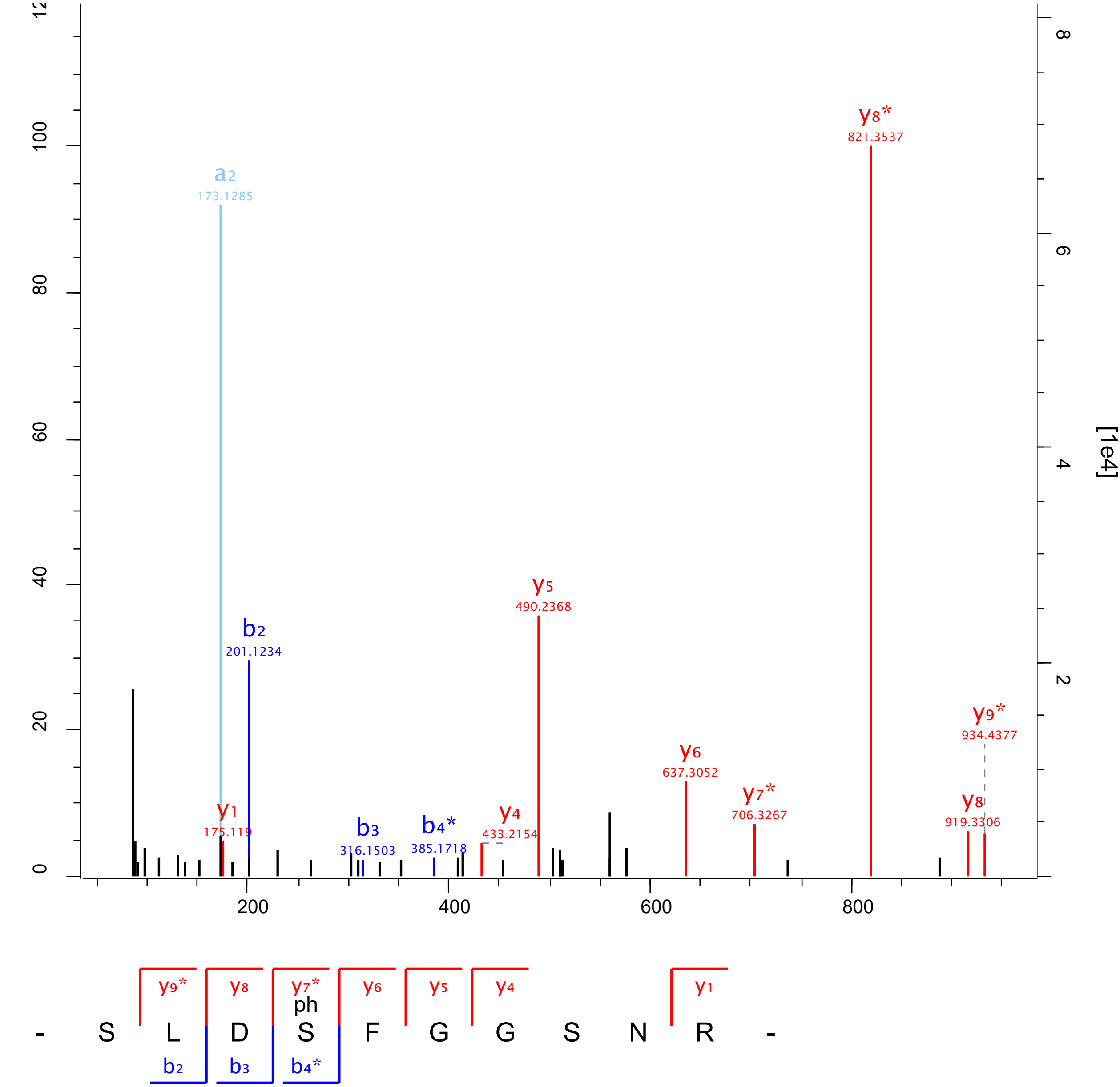

| Raw file | Scan | Method | Score | m/z |
| --- | --- | --- | --- | --- |
| sys_00_3short | 13244 | FTMS; HCD | 55.26 | 542.25 |

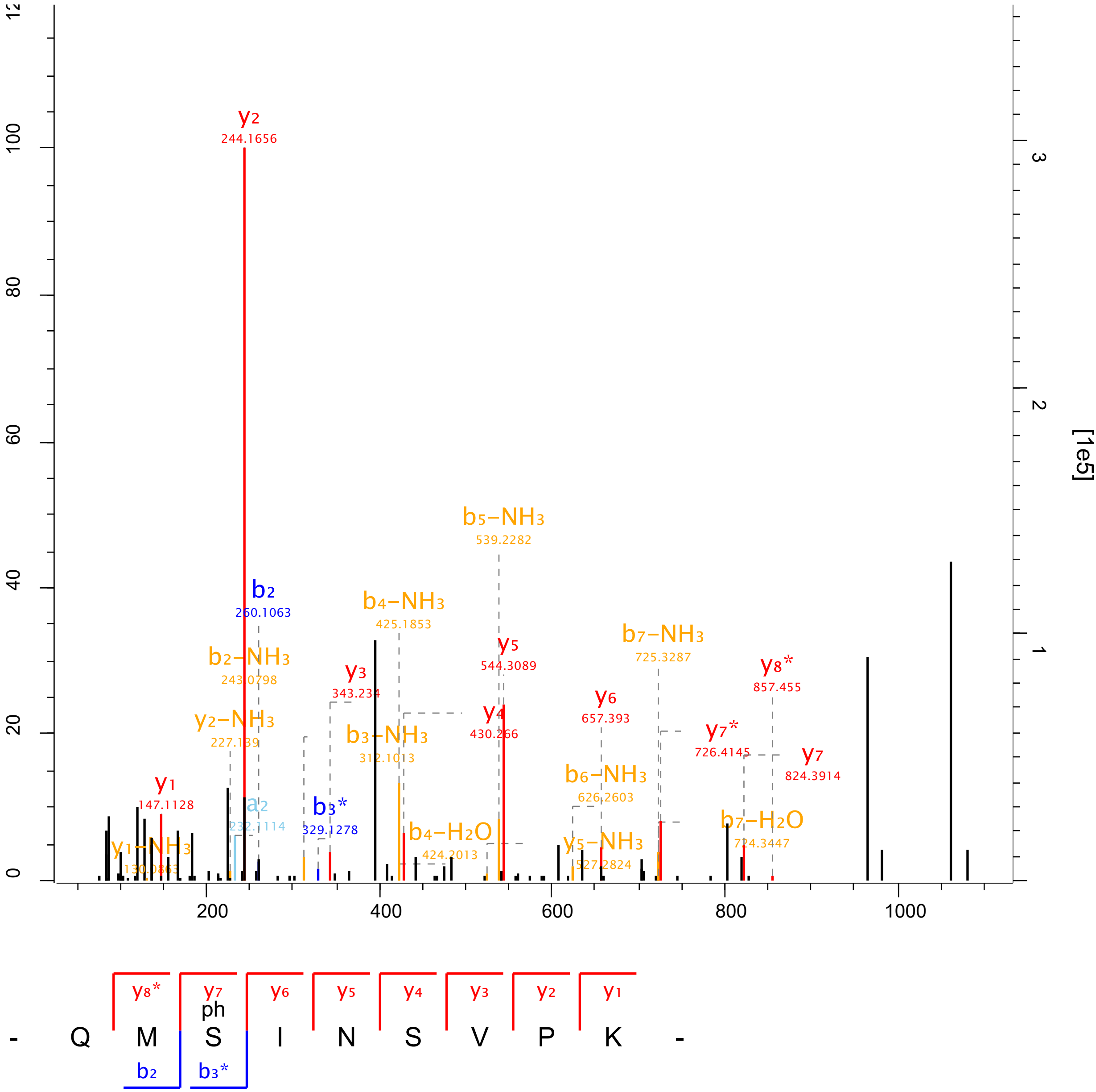

| Raw file | Scan | Method | Score | m/z |
| --- | --- | --- | --- | --- |
| sys_00_3short | 13707 | FTMS; HCD | 61.24 | 564.27 |

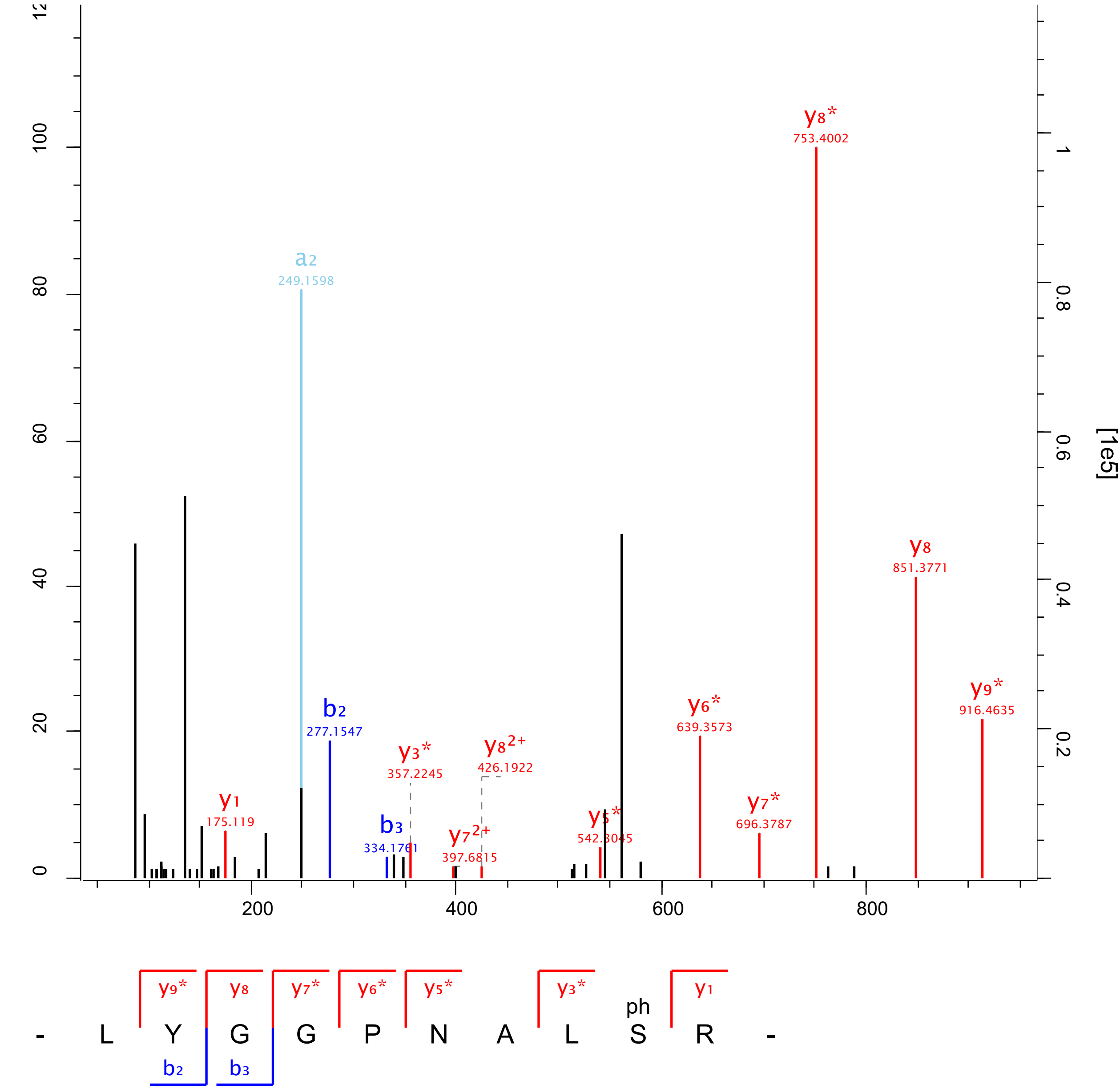

| Raw file | Scan | Method | Score | m/z |
| --- | --- | --- | --- | --- |
| sys_00_3short | 13717 | FTMS; HCD | 148.78 | 550.76 |

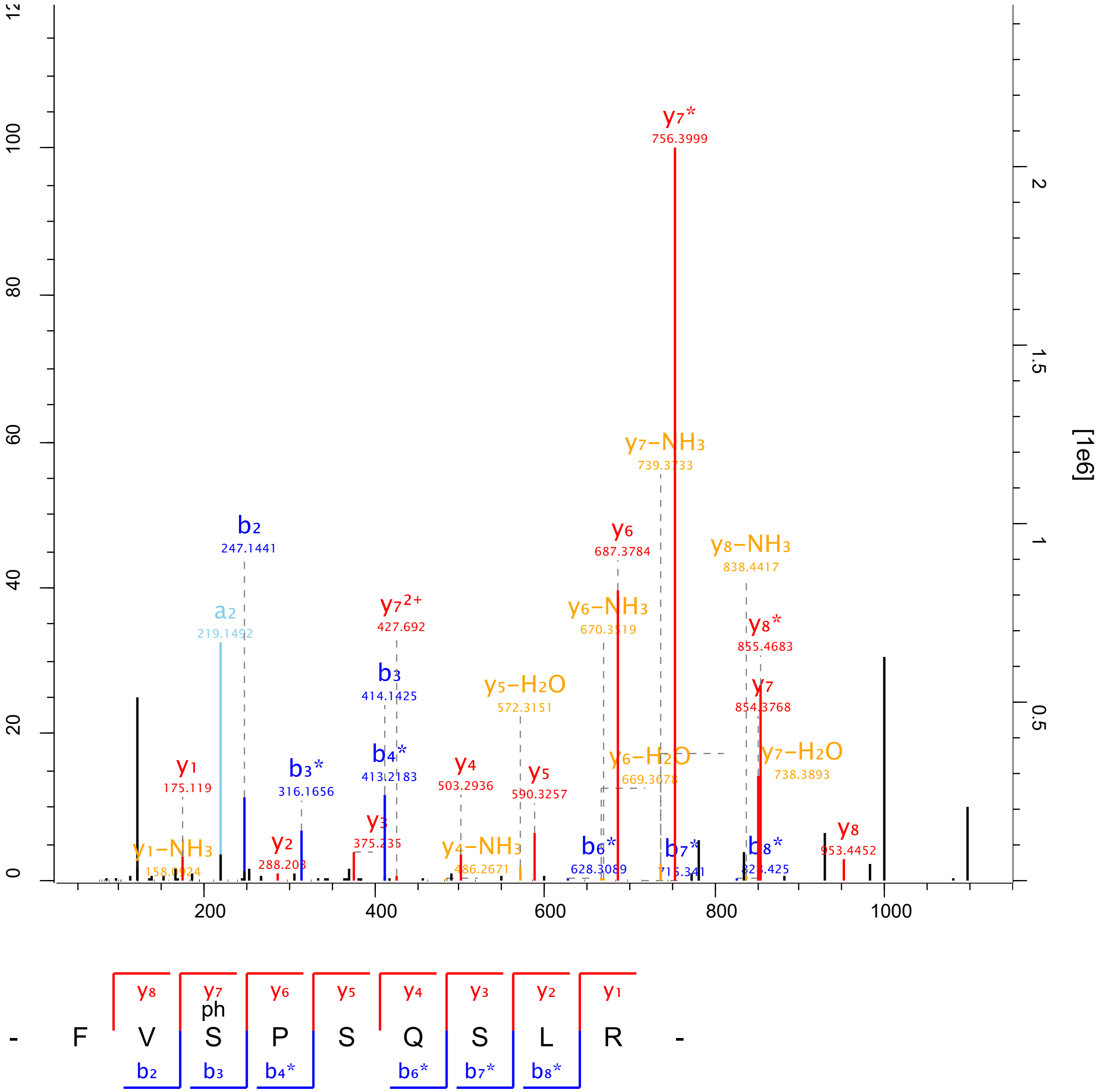

| Raw file | Scan | Method | Score | m/z |
|---|---|---|---|---|
| sys_00_3short | 13838 | FTMS; HCD | 67.52 | 541.75 |

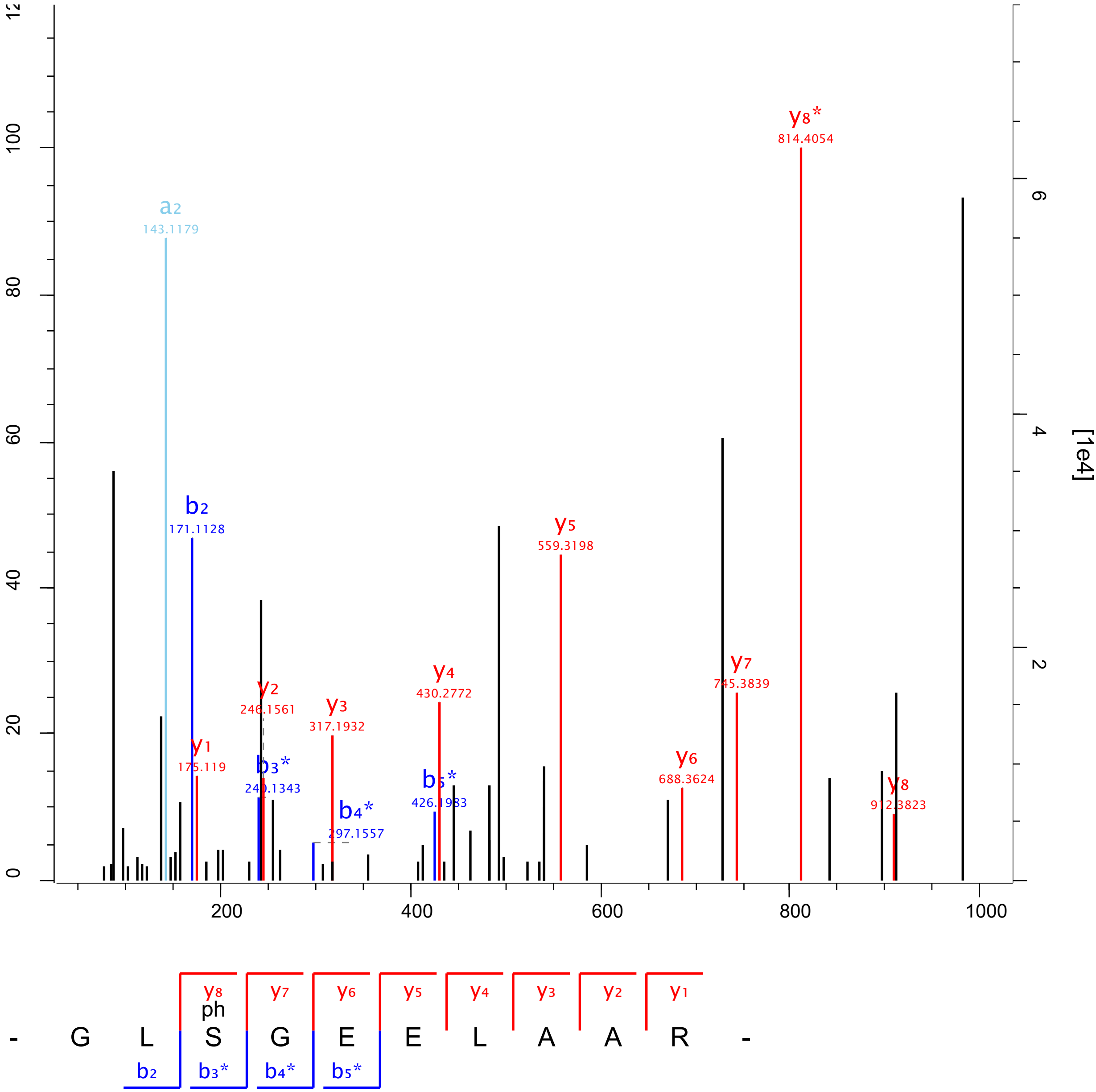

| Raw file | Scan | Method | Score | m/z |
|---|---|---|---|---|
| sys_00_3short | 13862 | FTMS; HCD | 84.51 | 758.31 |

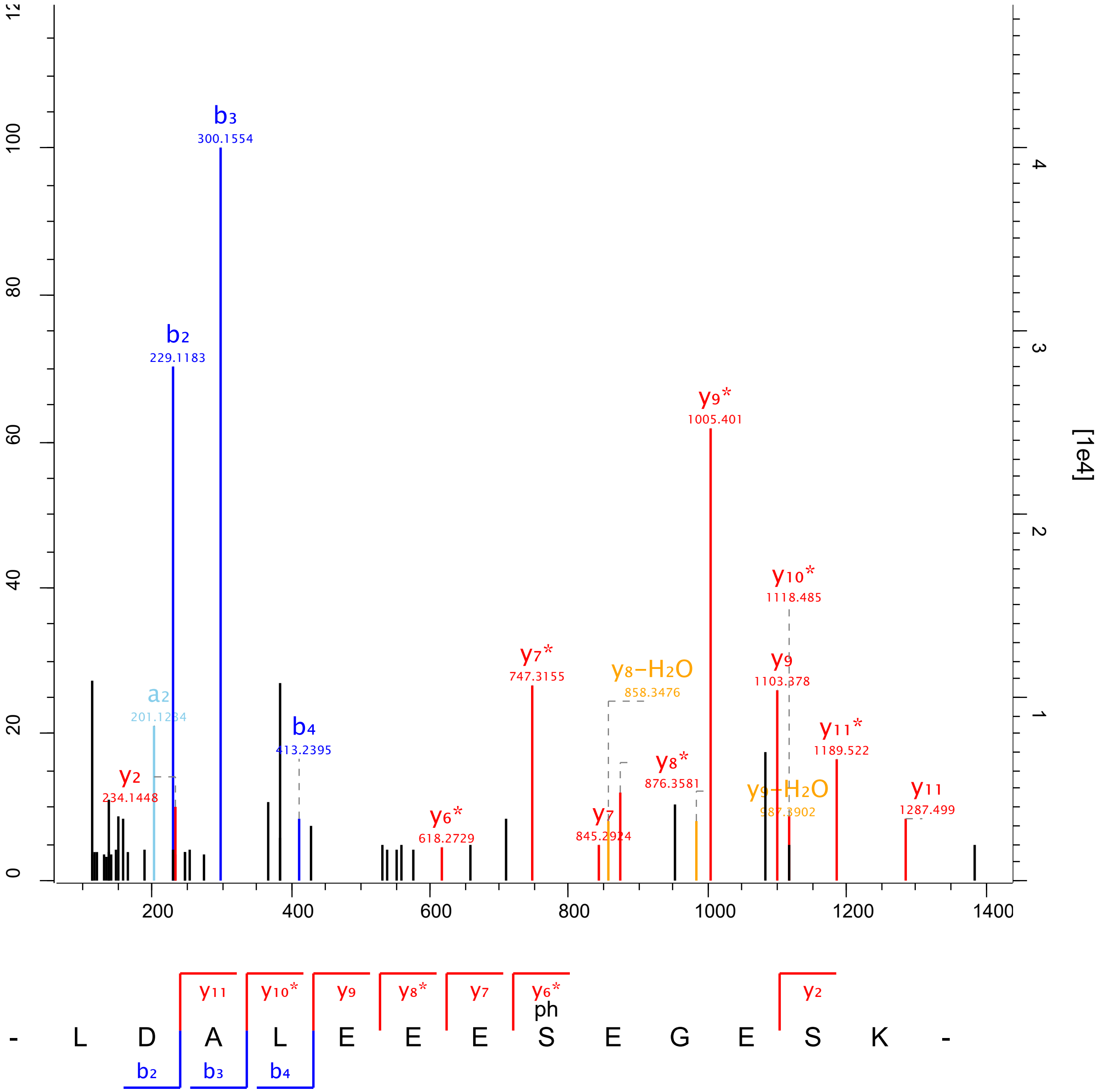

| Raw file | Scan | Method | Score | m/z |
| --- | --- | --- | --- | --- |
| sys_00_3short | 14037 | FTMS; HCD | 89.51 | 561.79 |

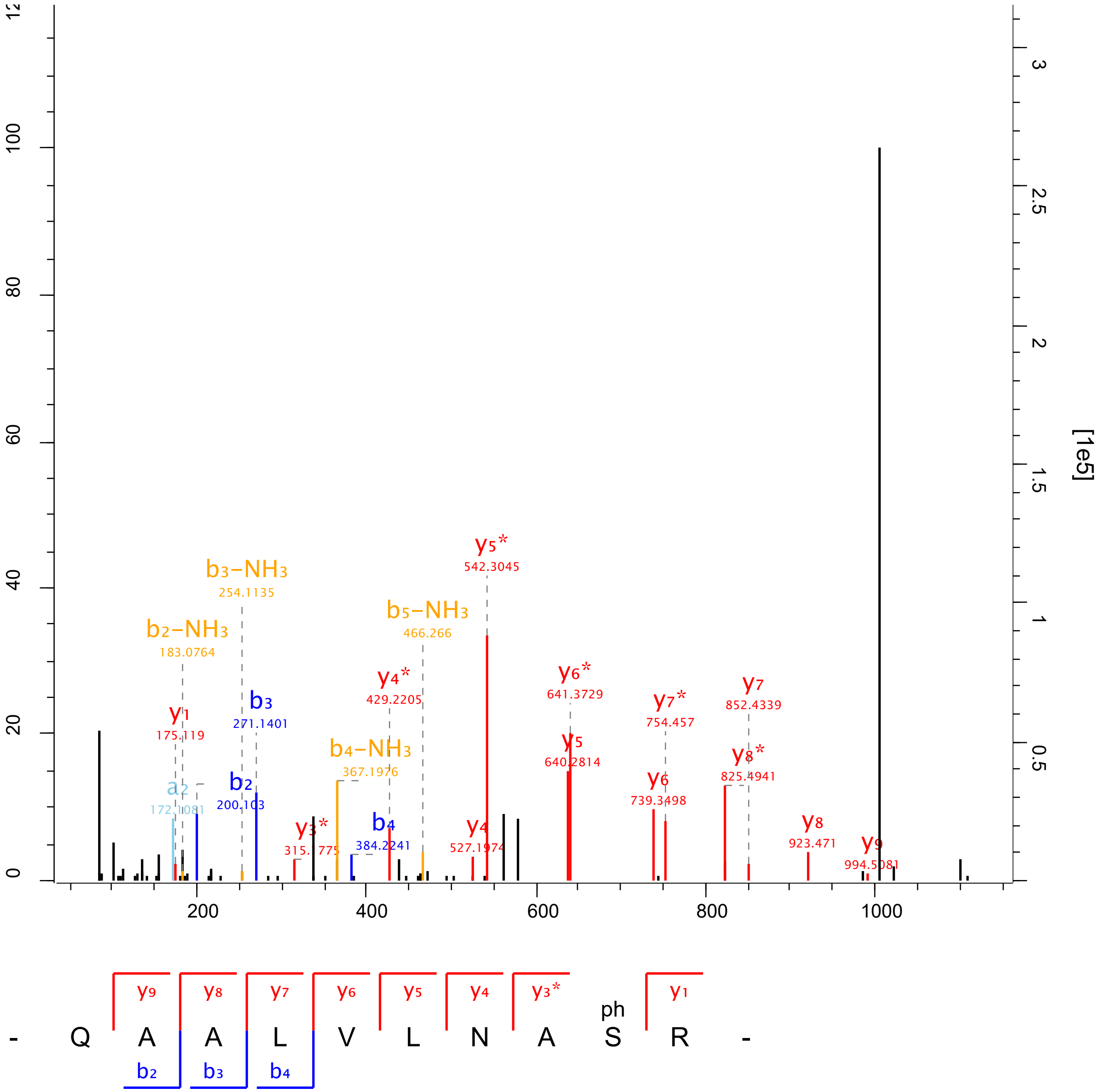

| Raw file | Scan | Method | Score | m/z |
| --- | --- | --- | --- | --- |
| sys_00_3short | 14152 | FTMS; HCD | 82.35 | 695.3 |

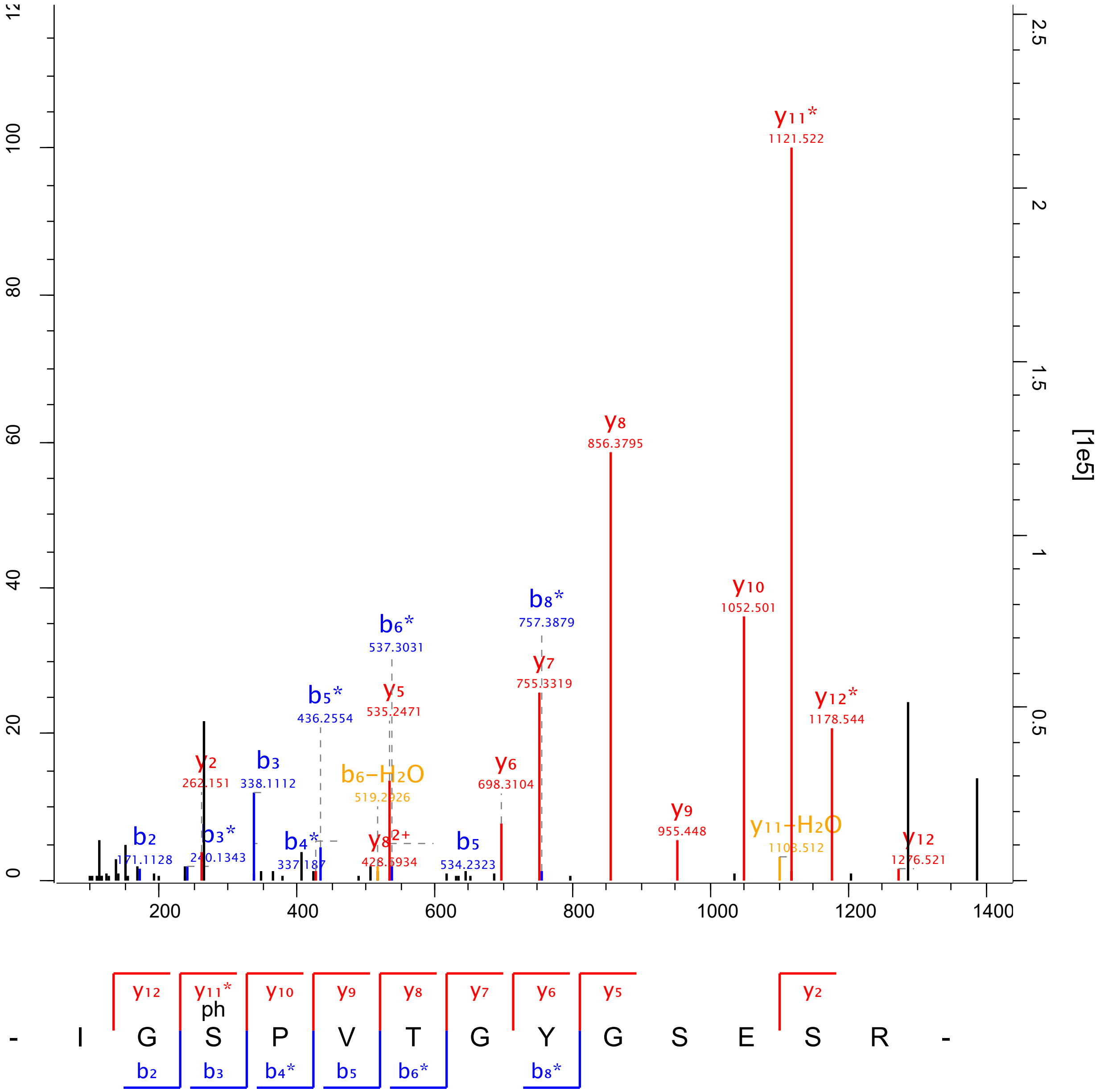

| Raw file | Scan | Method | Score | m/z |
| --- | --- | --- | --- | --- |
| sys_00_3short | 14176 | FTMS; HCD | 200.58 | 769.79 |

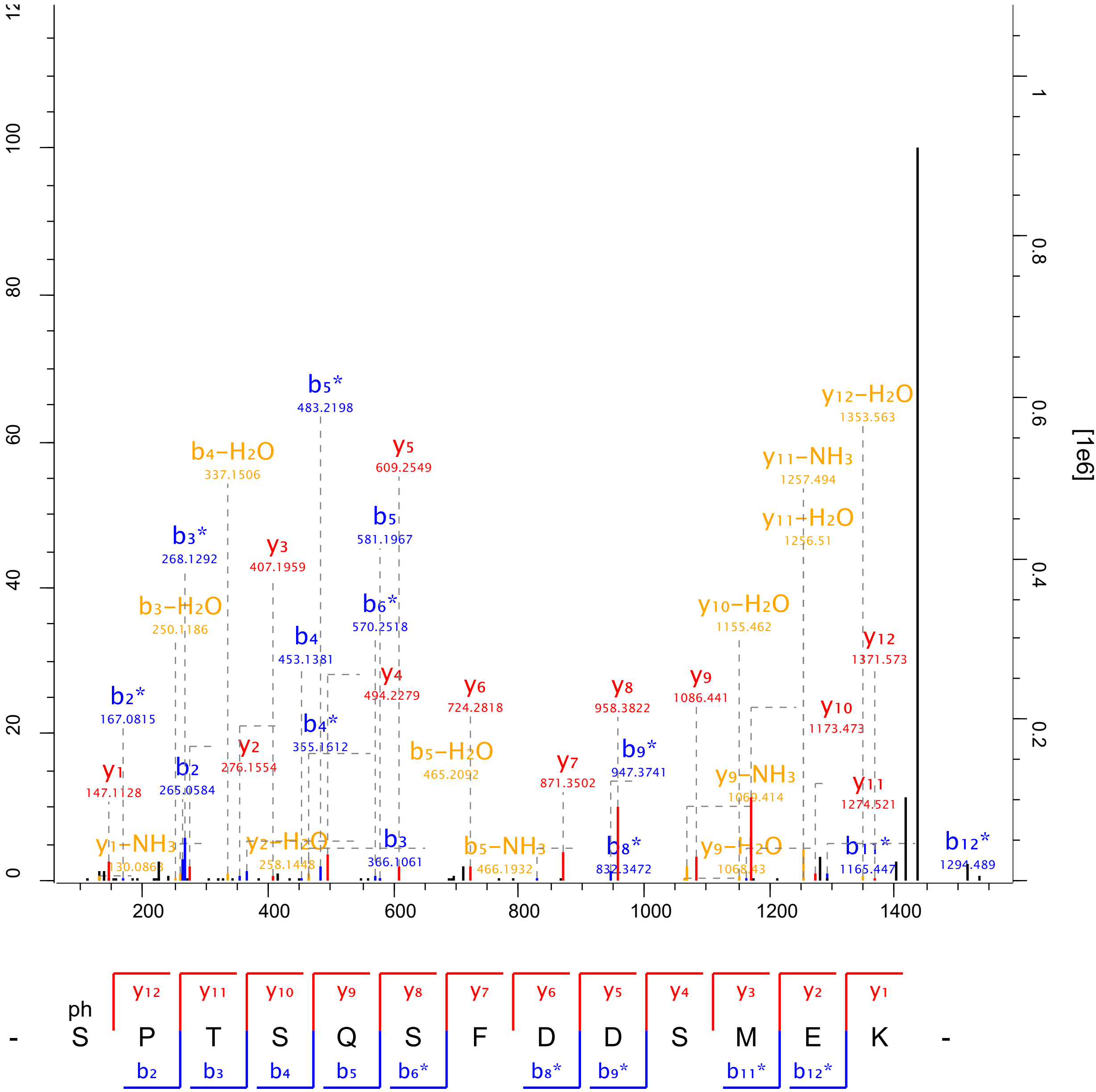

| Raw file | Scan | Method | Score | m/z |
| --- | --- | --- | --- | --- |
| sys_00_3short | 14218 | FTMS; HCD | 108.72 | 528.73 |

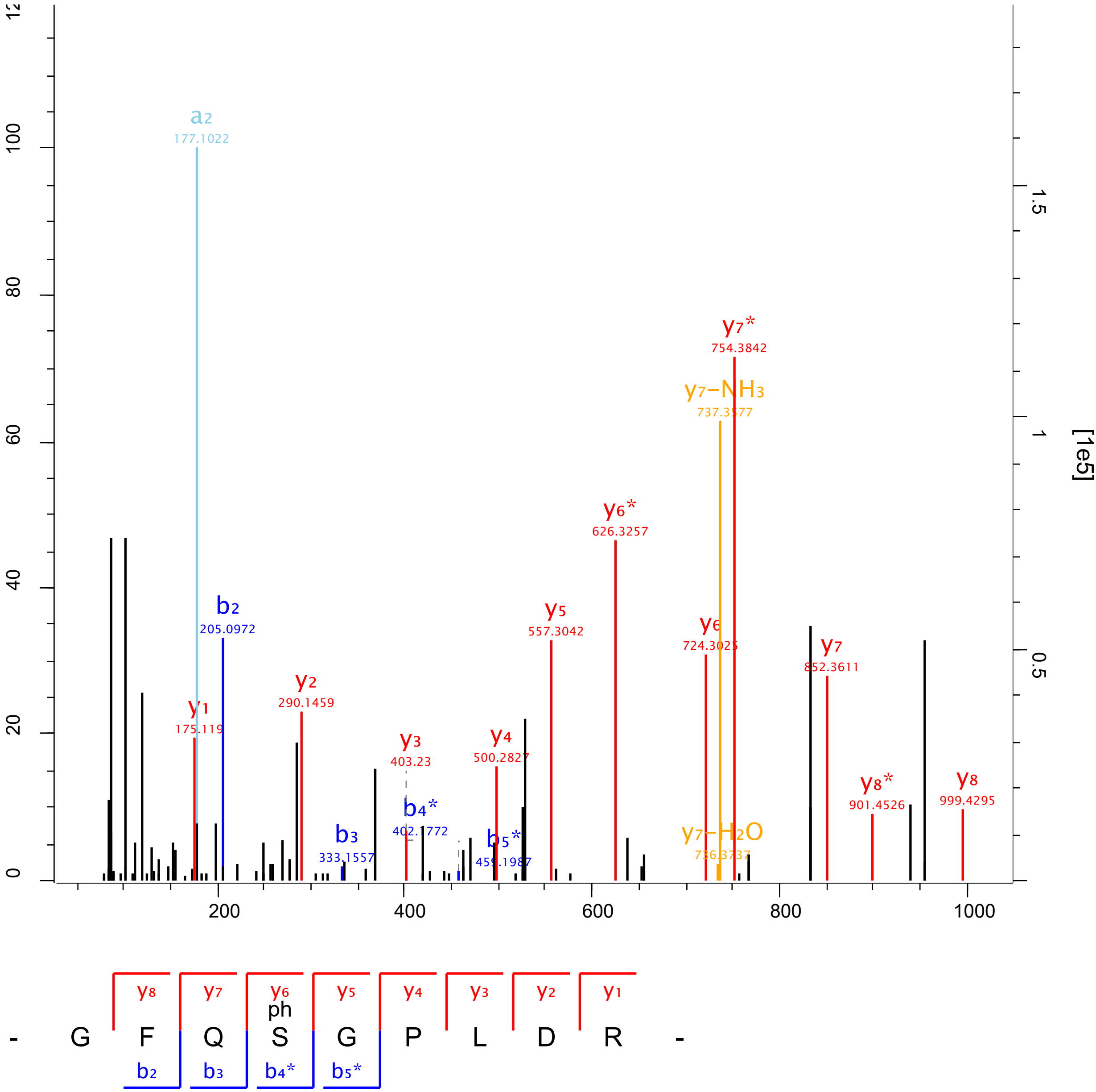

| Raw file | Scan | Method | Score | m/z |
| --- | --- | --- | --- | --- |
| sys_00_3short | 14426 | FTMS; HCD | 98.05 | 512.73 |

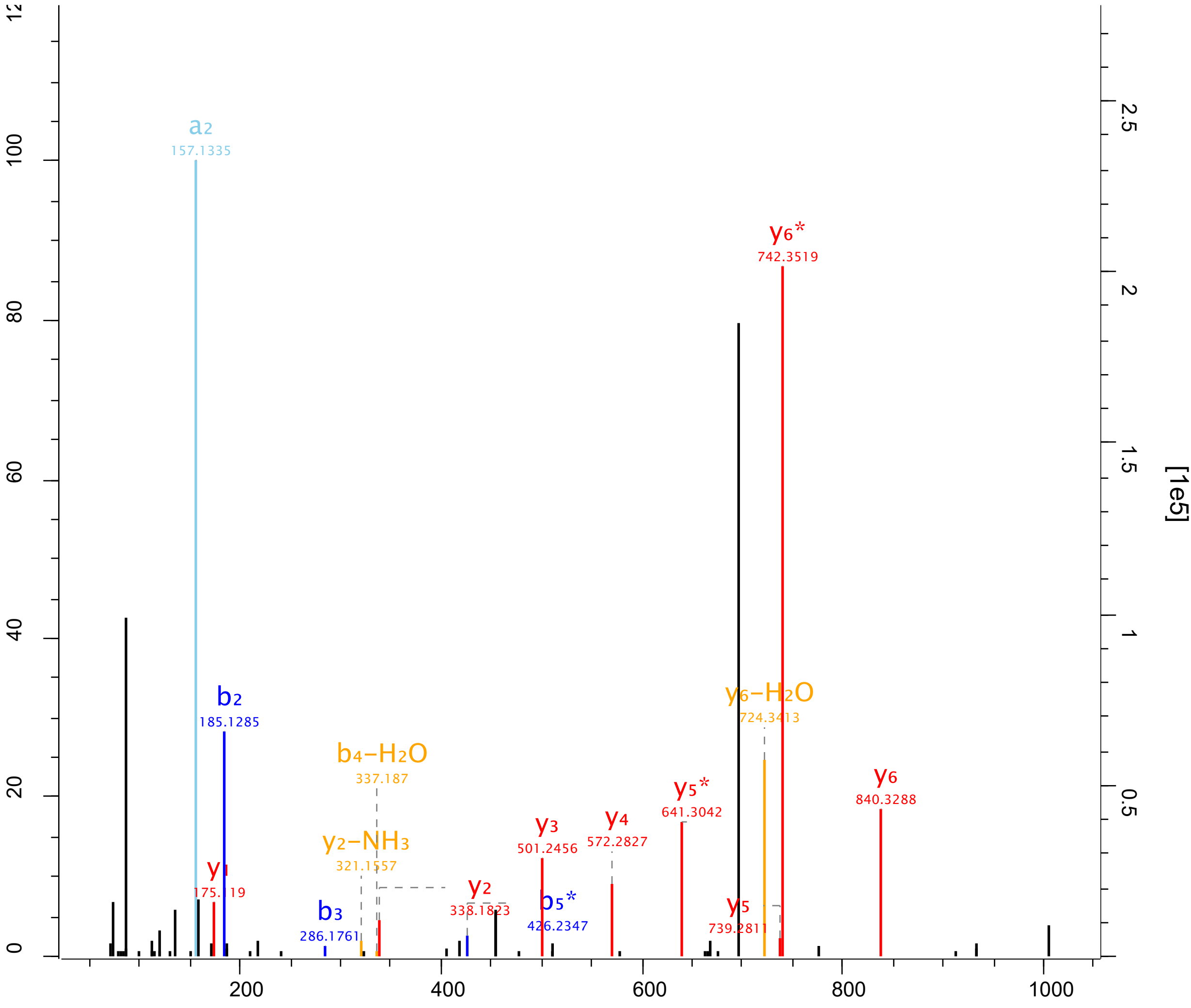


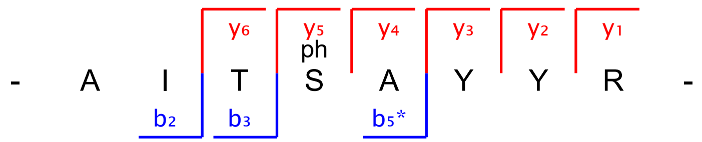

| Raw file | Scan | Method | Score | m/z |
| --- | --- | --- | --- | --- |
| sys_00_3short | 14444 | FTMS; HCD | 117.18 | 802.32 |

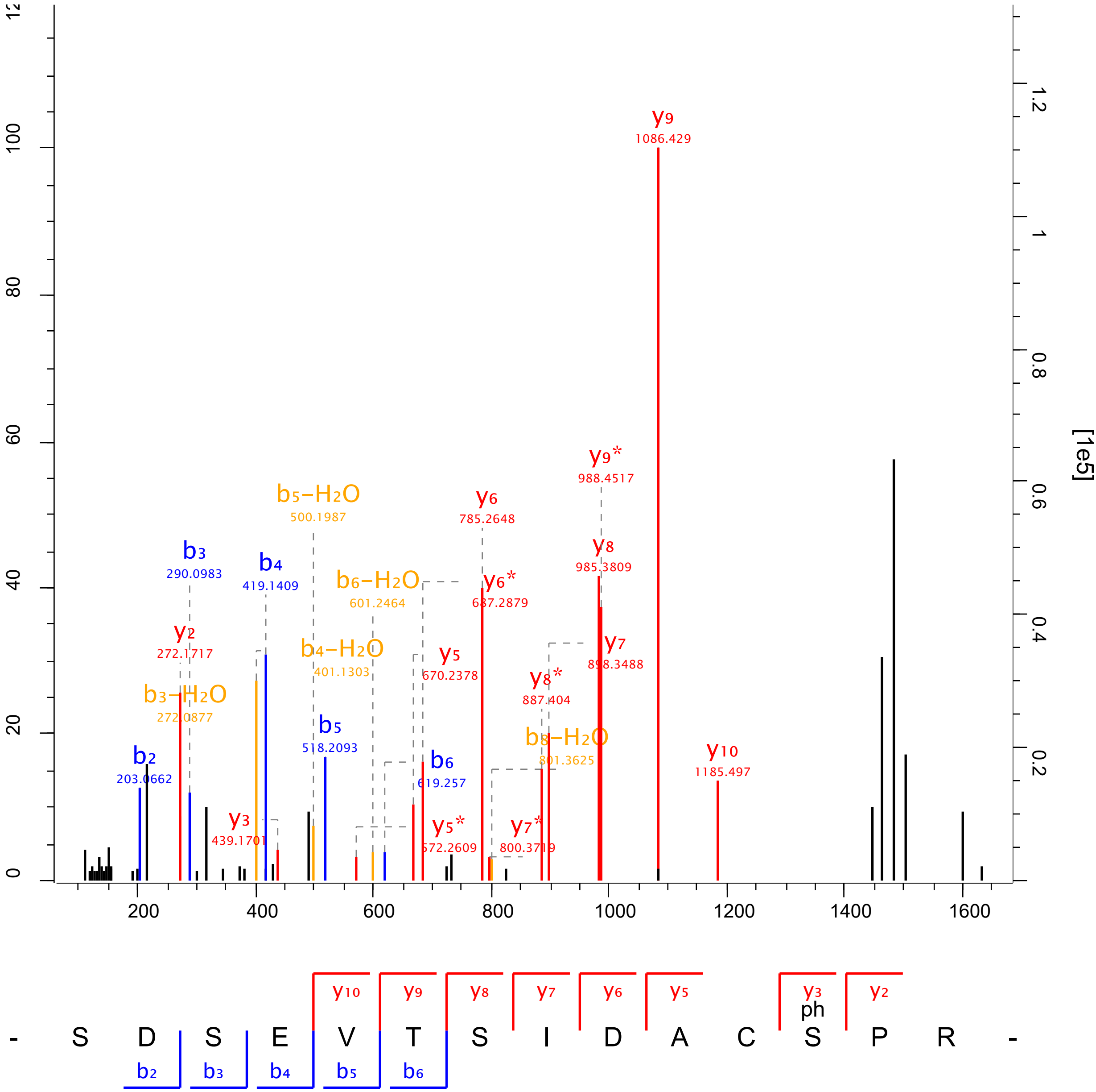

| Raw file | Scan | Method | Score | m/z |
|---|---|---|---|---|
| sys_00_3short | 14450 | FTMS; HCD | 82.07 | 563.76 |

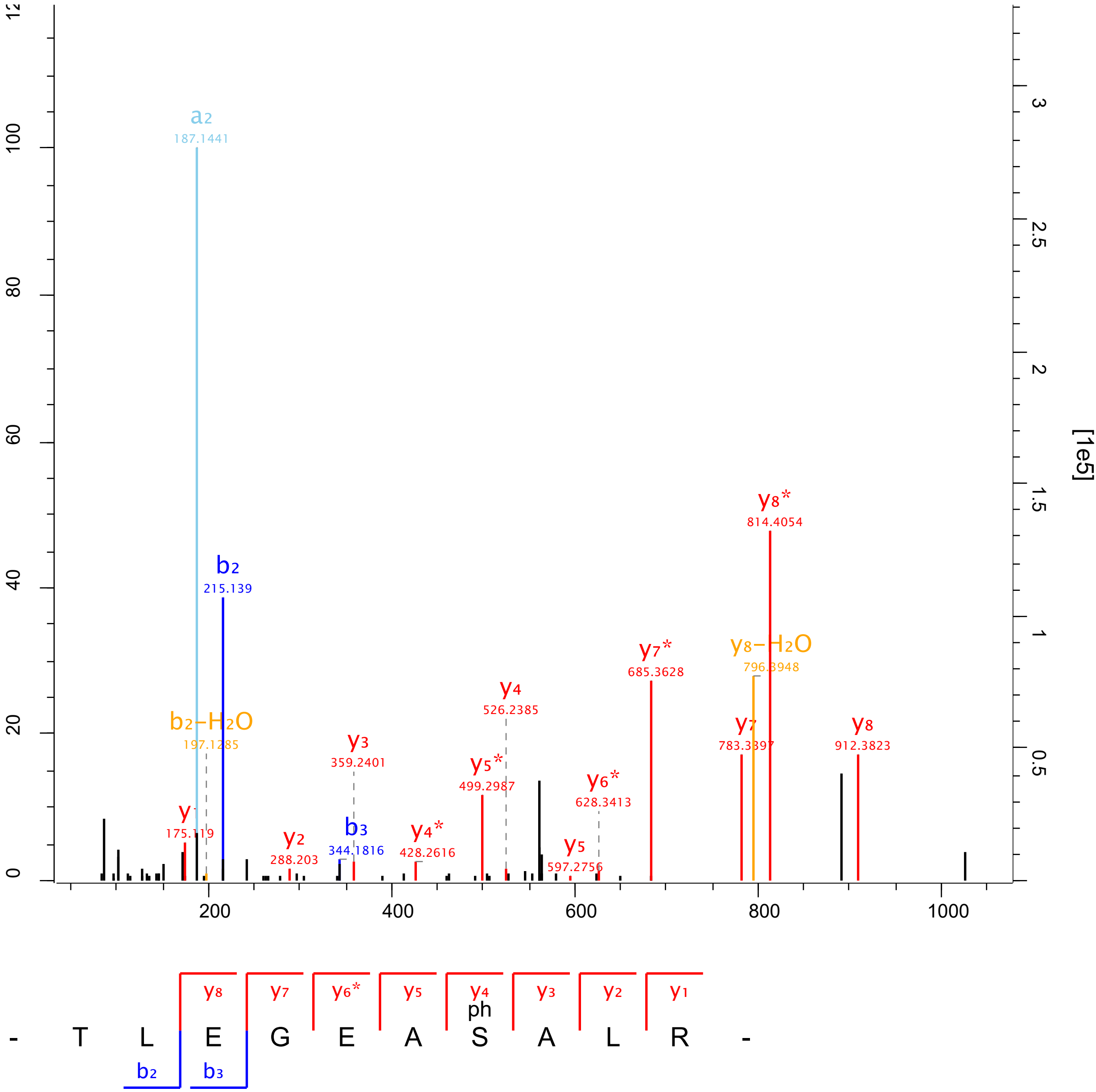

| Raw file | Scan | Method | Score | m/z |
| --- | --- | --- | --- | --- |
| sys_00_3short | 14488 | FTMS; HCD | 45.89 | 506.21 |

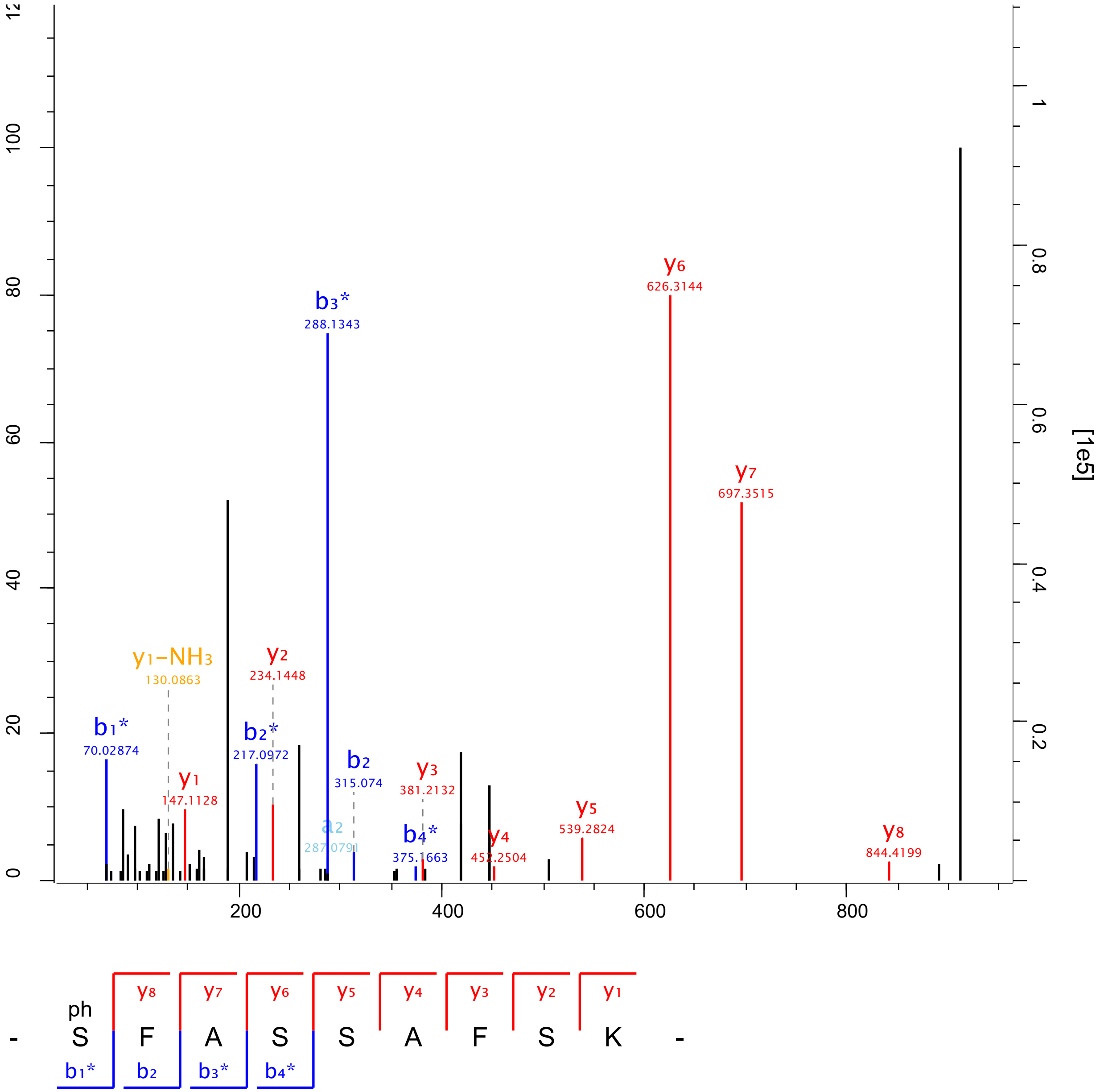

| Raw file | Scan | Method | Score | m/z |
|---|---|---|---|---|
| sys_00_3short | 14628 | FTMS; HCD | 185.96 | 511.23 |

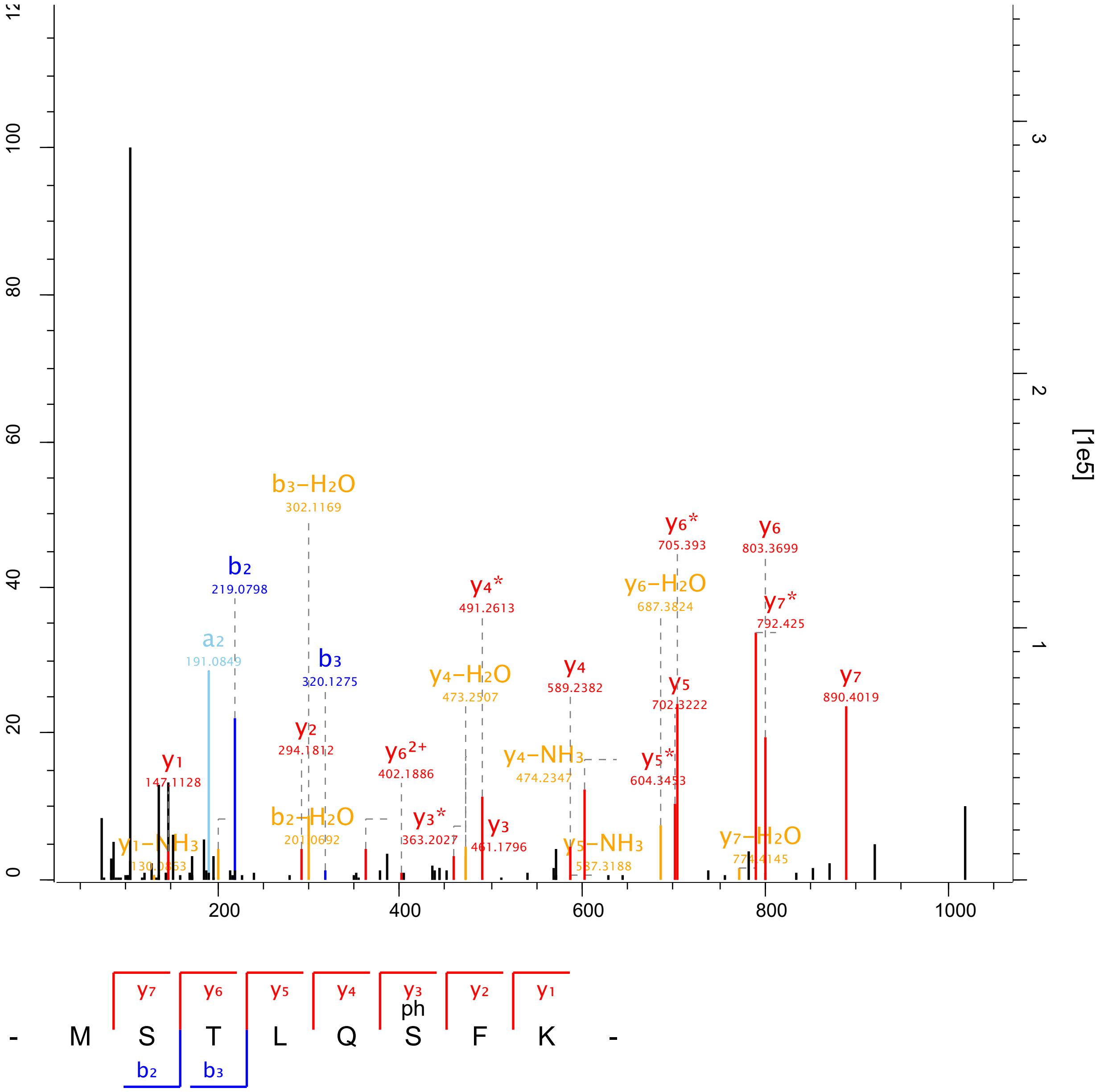

| Raw file | Scan | Method | Score | m/z |
| --- | --- | --- | --- | --- |
| sys_00_3short | 14744 | FTMS; HCD | 96.46 | 562.26 |

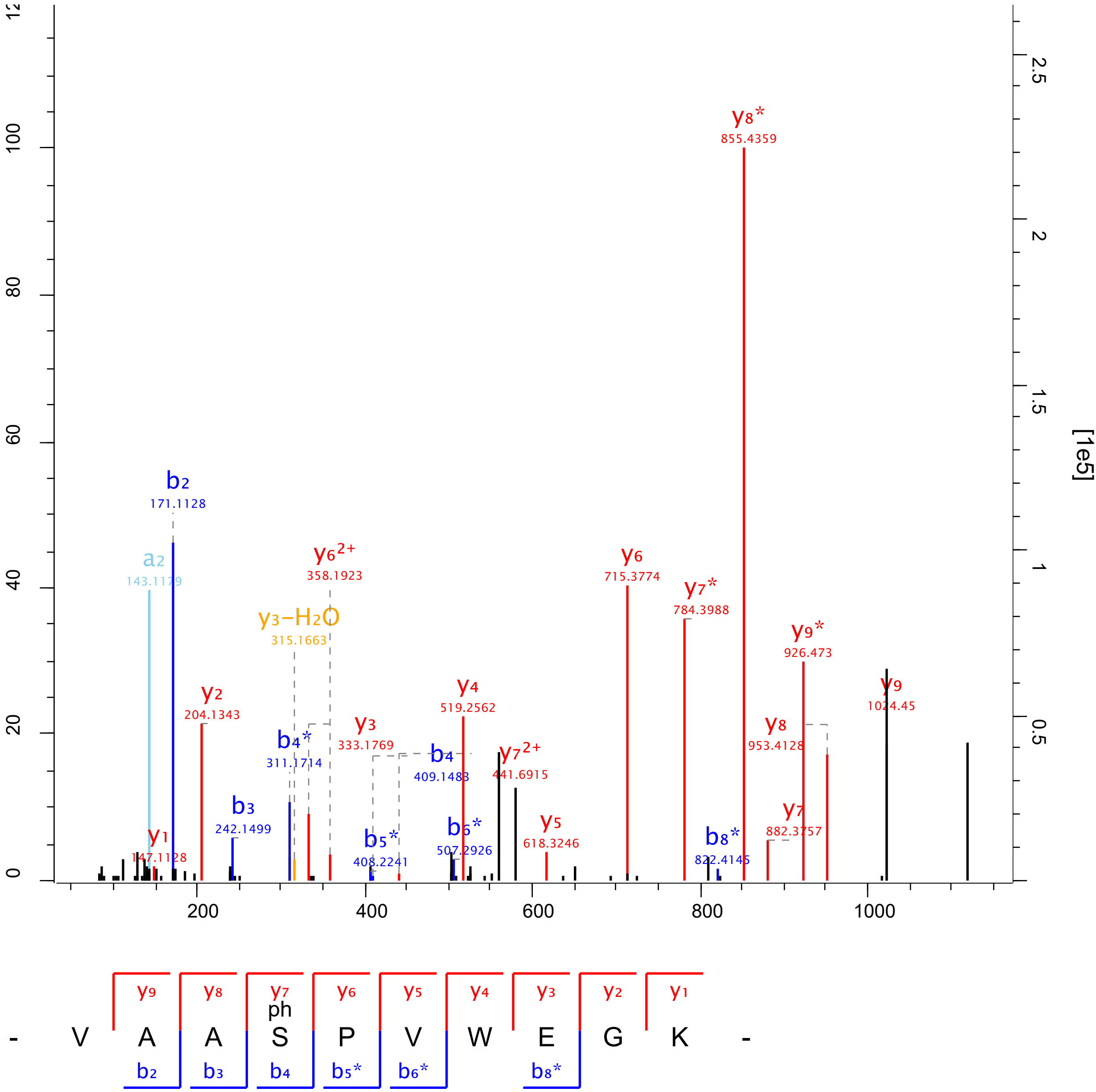

| Raw file | Scan | Method | Score | m/z |
|---|---|---|---|---|
| sys_00_3short | 15024 | FTMS; HCD | 117.93 | 568.75 |

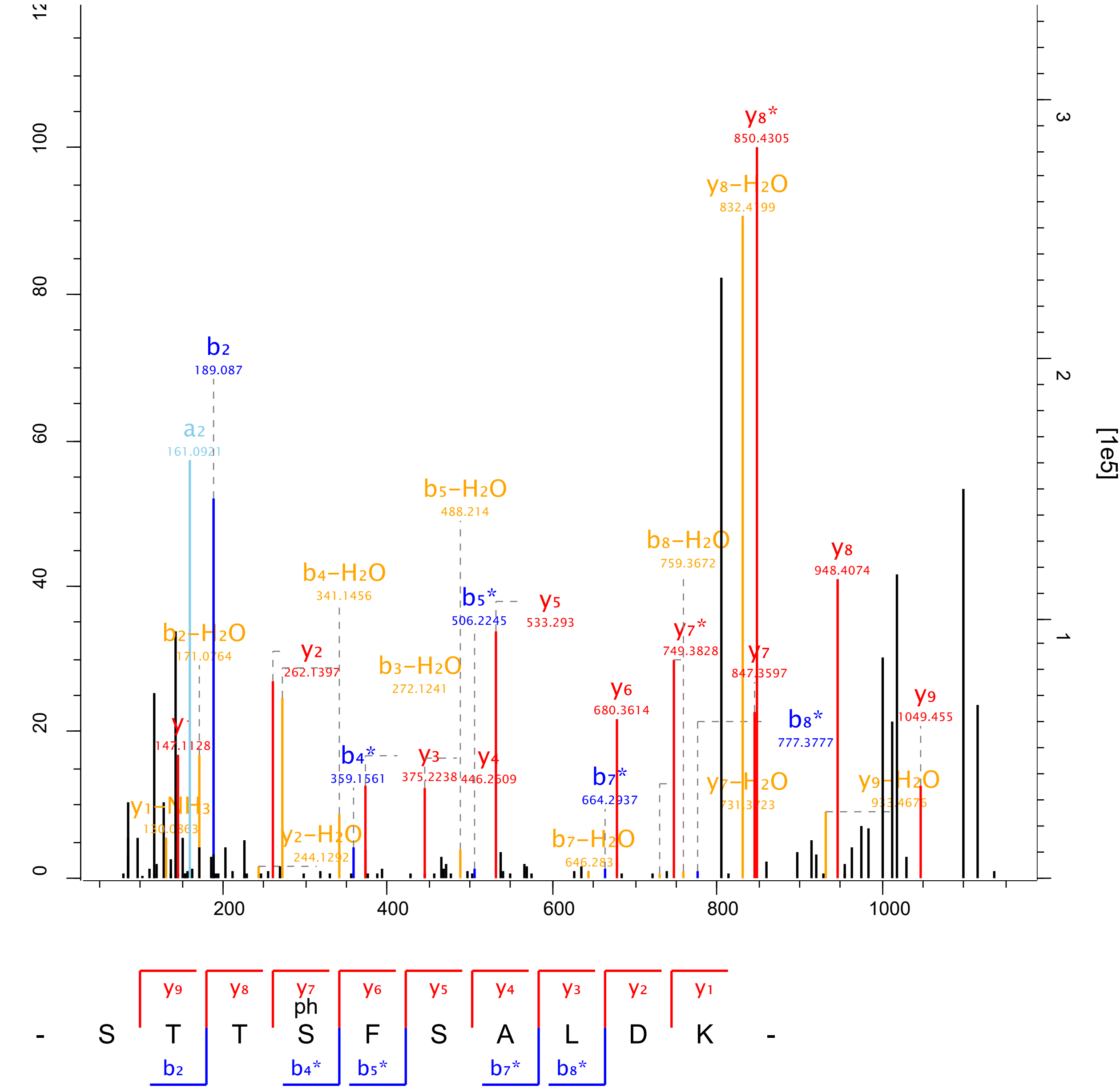

| Raw file | Scan | Method | Score | m/z |
| --- | --- | --- | --- | --- |
| sys_00_3short | 15426 | FTMS; HCD | 44.85 | 482.73 |

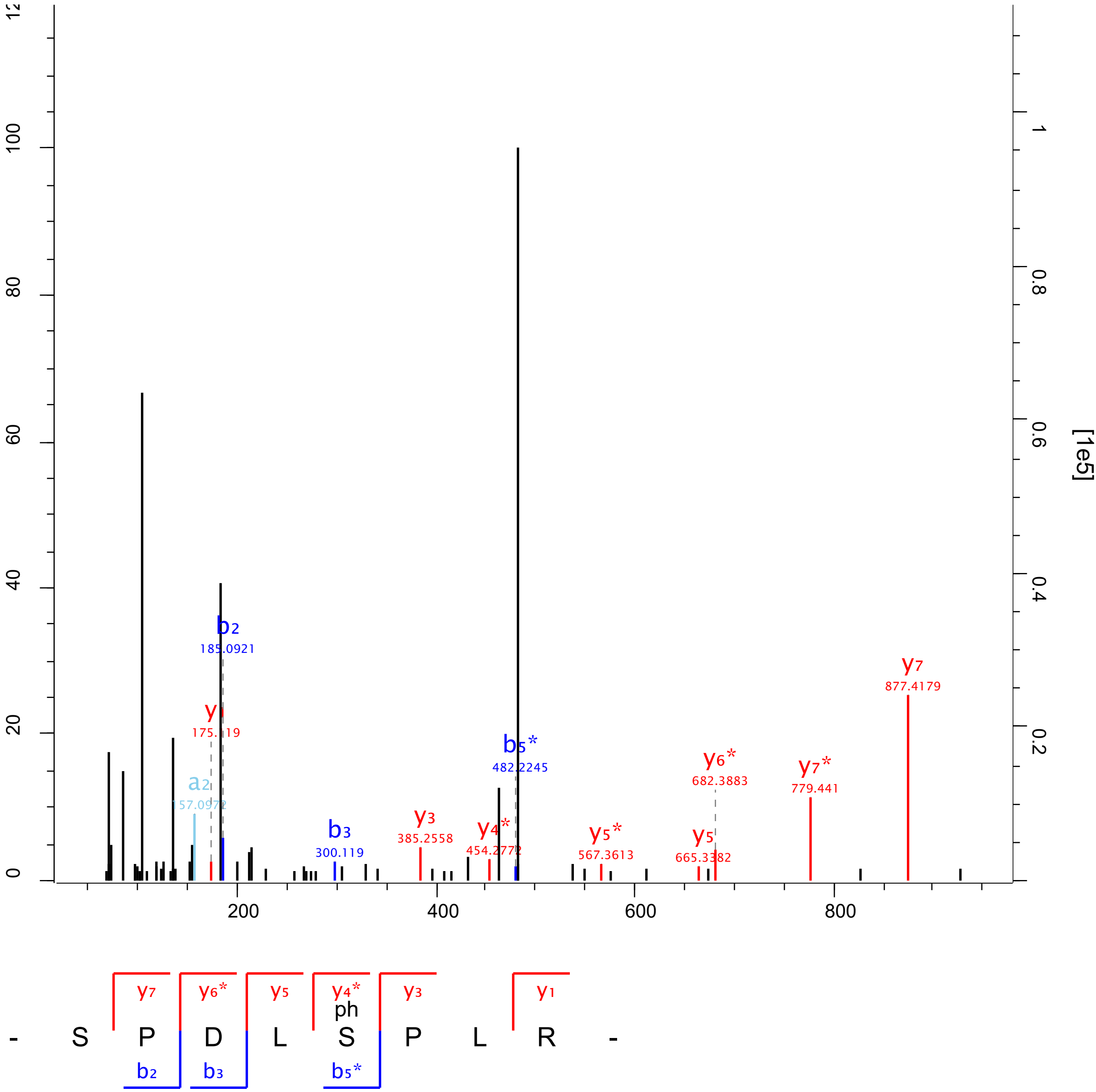

| Raw file | Scan | Method | Score | m/z |
| --- | --- | --- | --- | --- |
| sys_00_3short | 15684 | FTMS; HCD | 85.81 | 697.78 |

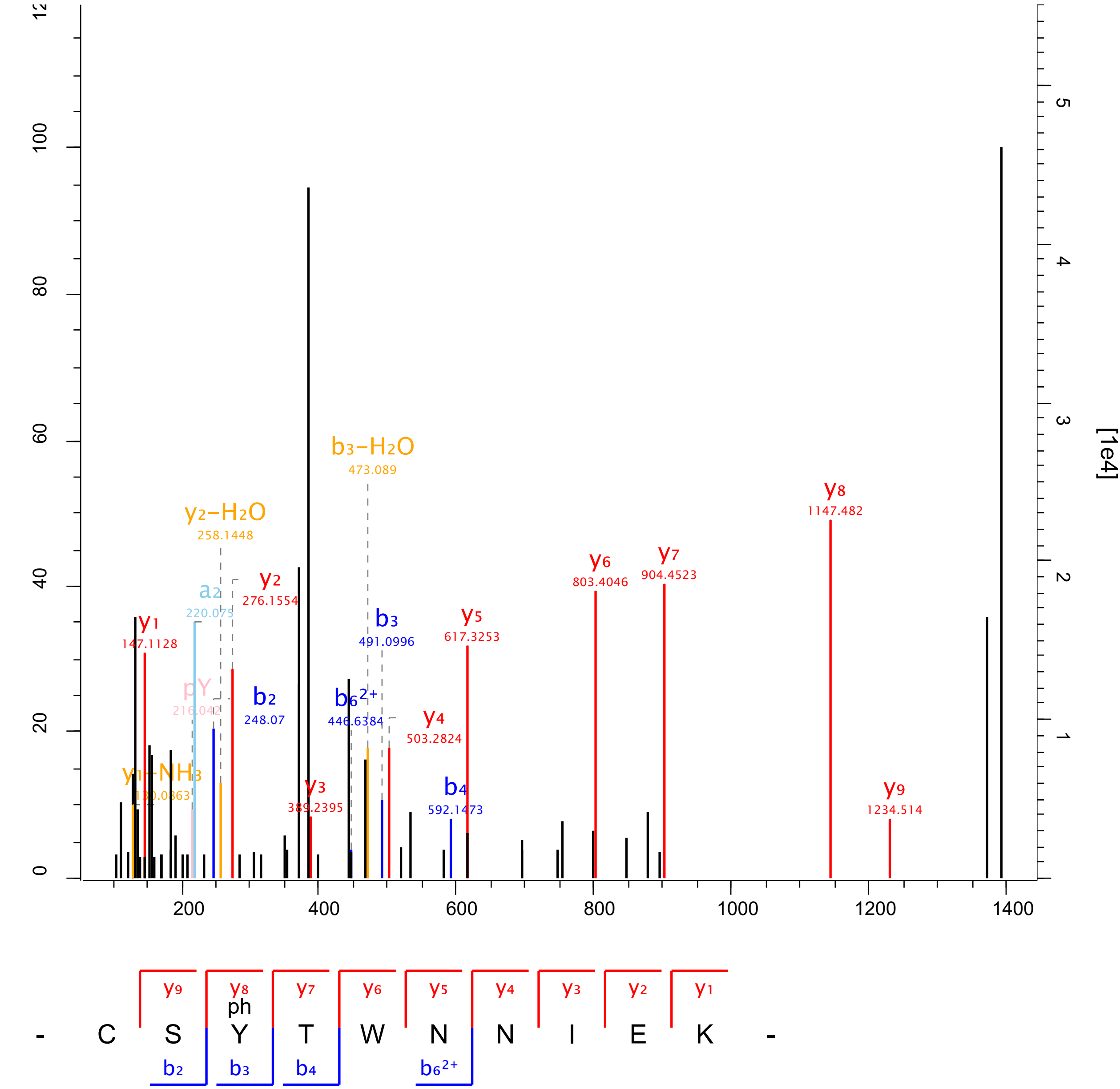

| Raw file | Scan | Method | Score | m/z |
| --- | --- | --- | --- | --- |
| sys_00_3short | 15821 | FTMS; HCD | 105.03 | 613.29 |

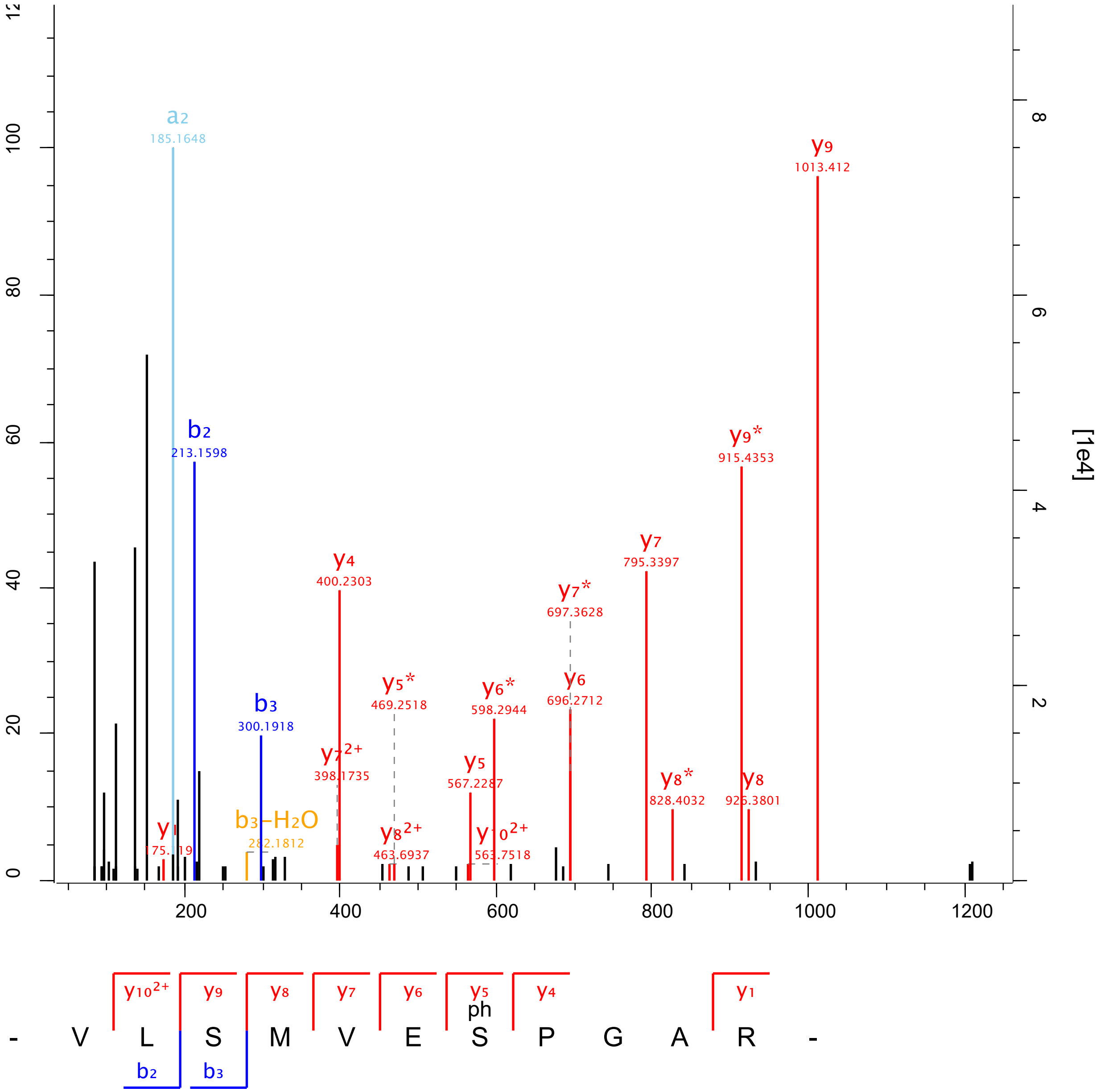

| Raw file | Scan | Method | Score | m/z |
|---|---|---|---|---|
| sys_00_3short | 15846 | FTMS; HCD | 163.99 | 700.81 |

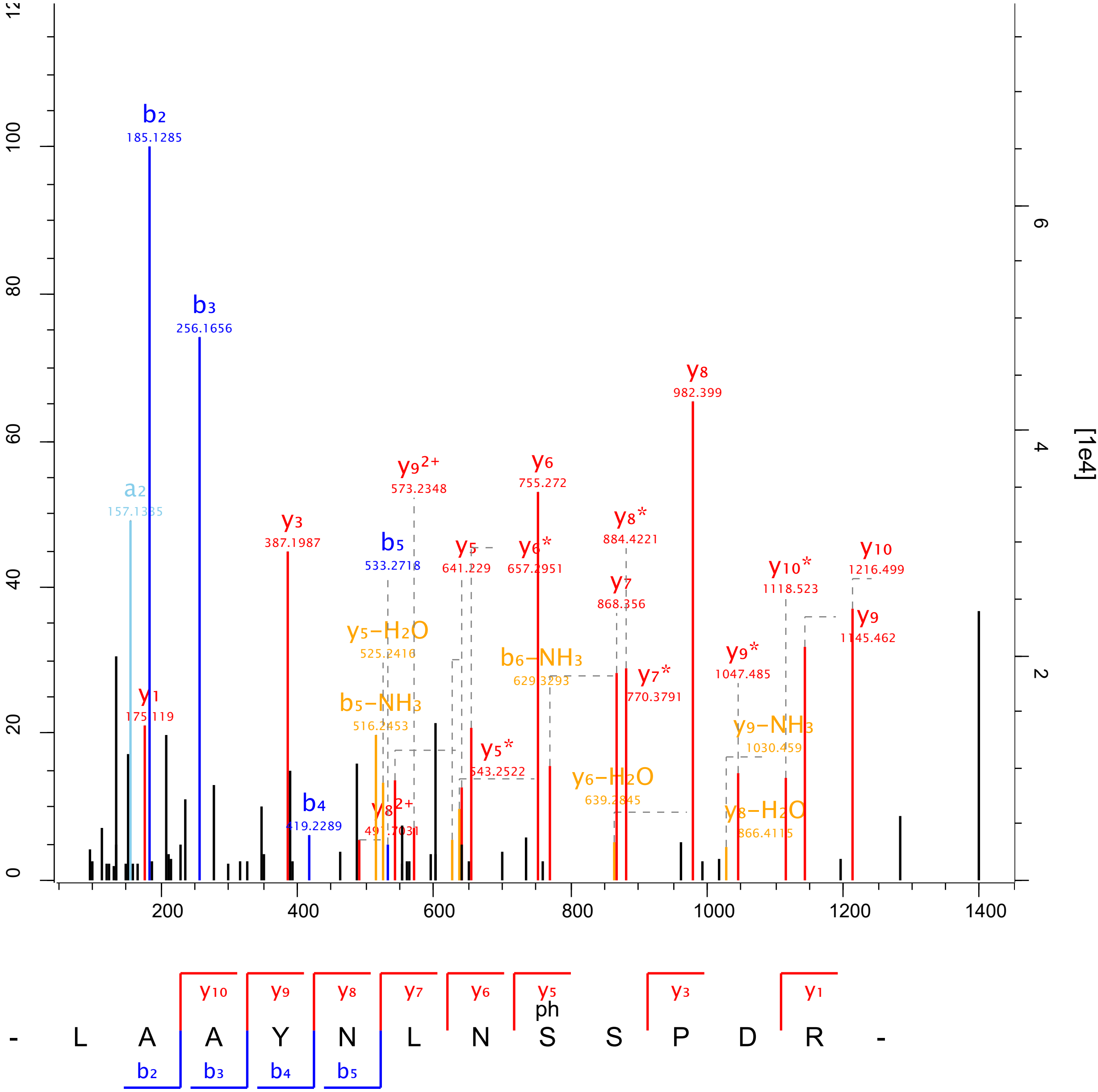

| Raw file | Scan | Method | Score | m/z |
| --- | --- | --- | --- | --- |
| sys_00_3short | 15878 | FTMS; HCD | 46.84 | 513.25 |

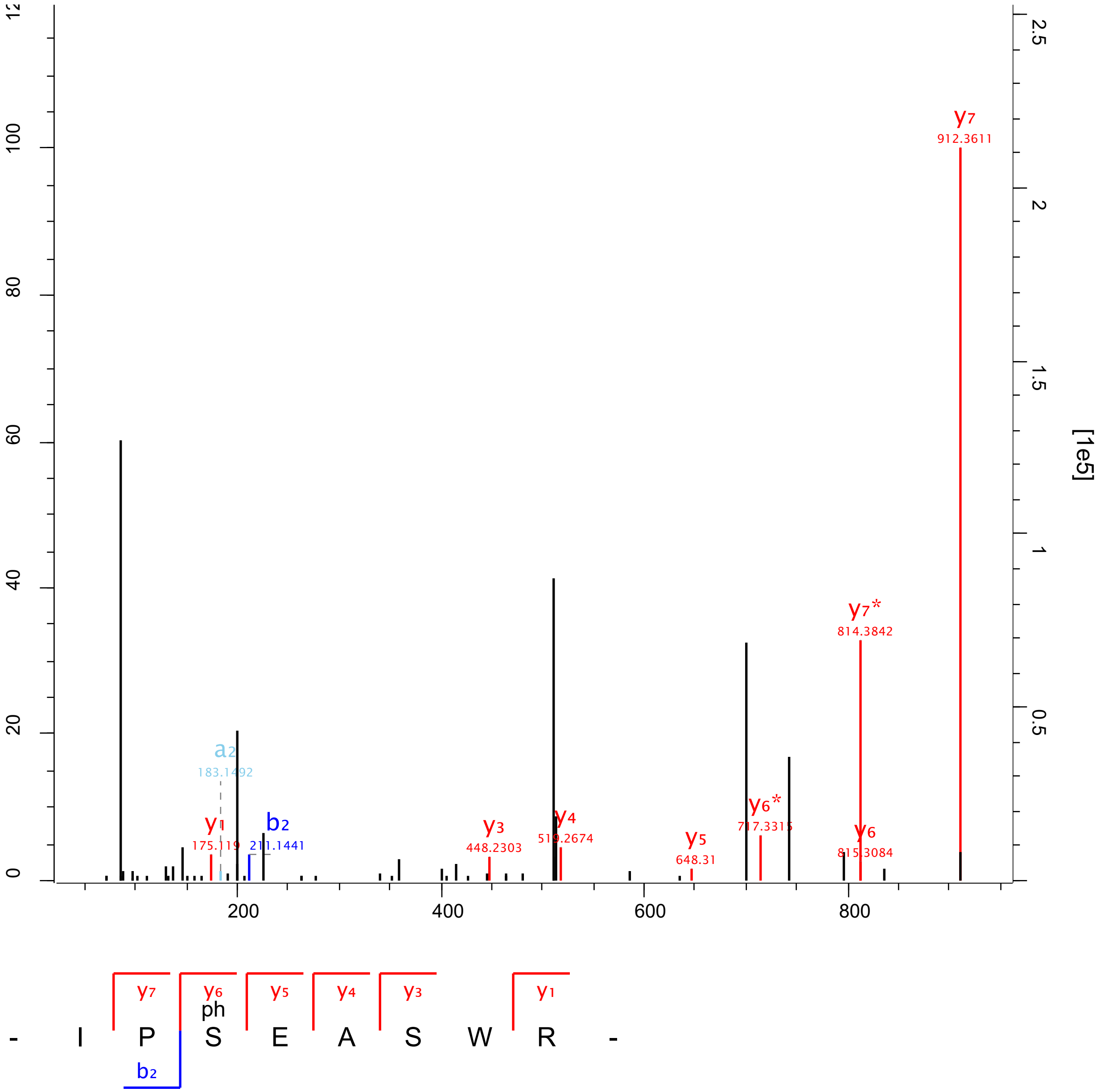

| Raw file | Scan | Method | Score | m/z |
| --- | --- | --- | --- | --- |
| sys_00_3short | 15891 | FTMS; HCD | 69.72 | 483.73 |

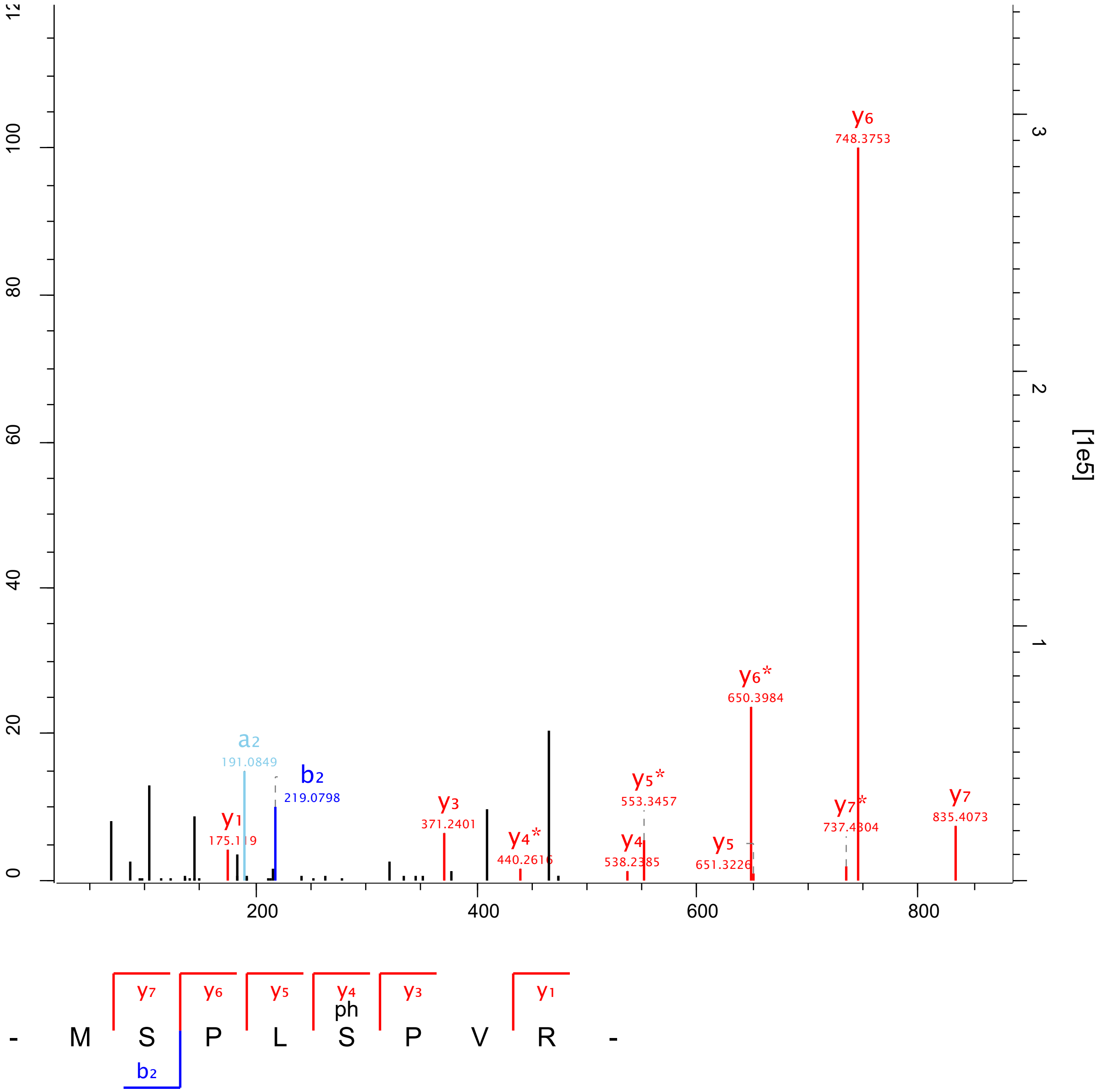

| Raw file | Scan | Method | Score | m/z |
|---|---|---|---|---|
| sys_00_3short | 15895 | FTMS; HCD | 117.7 | 620.77 |

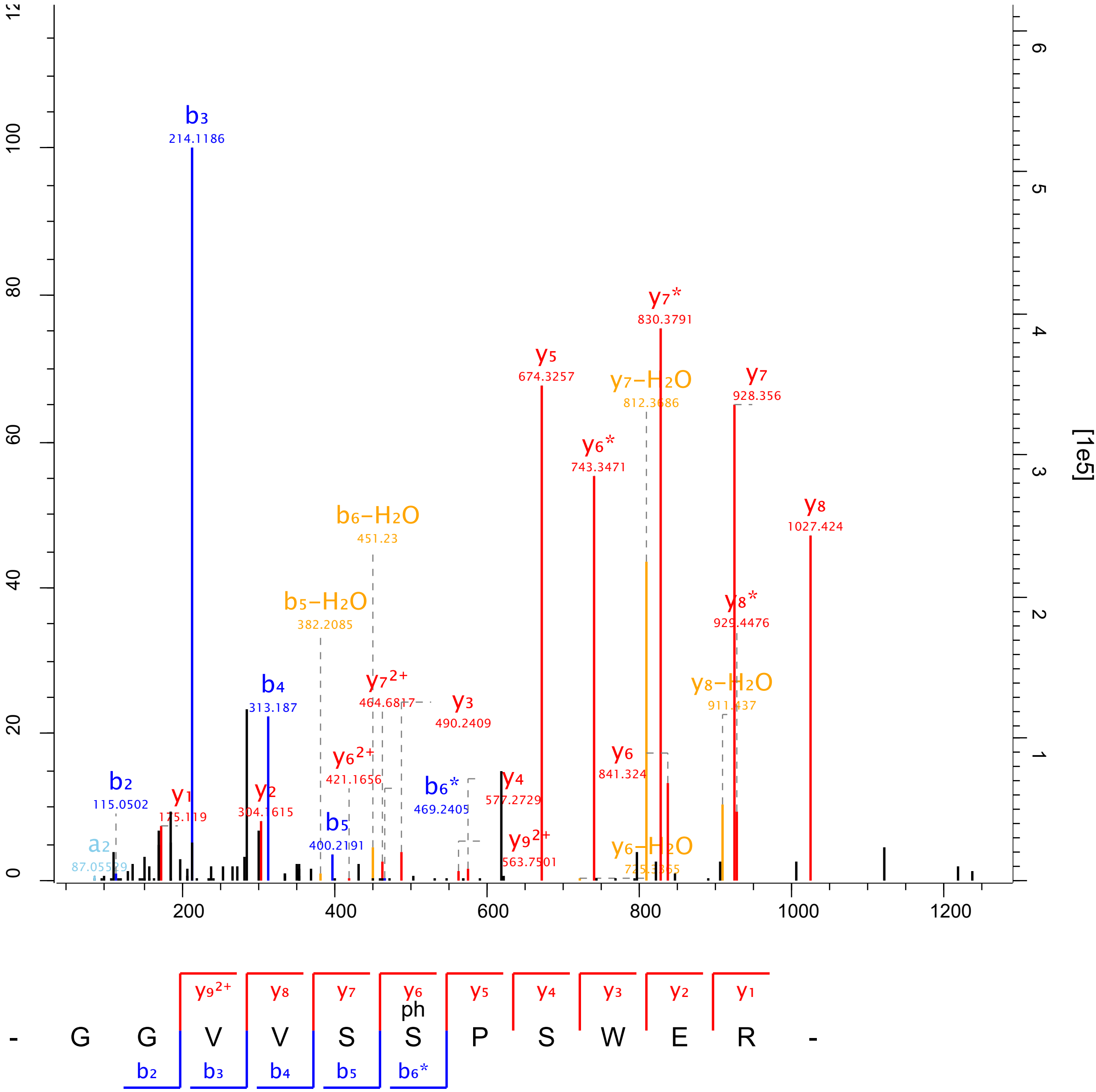

| Raw file | Scan | Method | Score | m/z |
| --- | --- | --- | --- | --- |
| sys_00_3short | 16065 | FTMS; HCD | 128.59 | 761.84 |

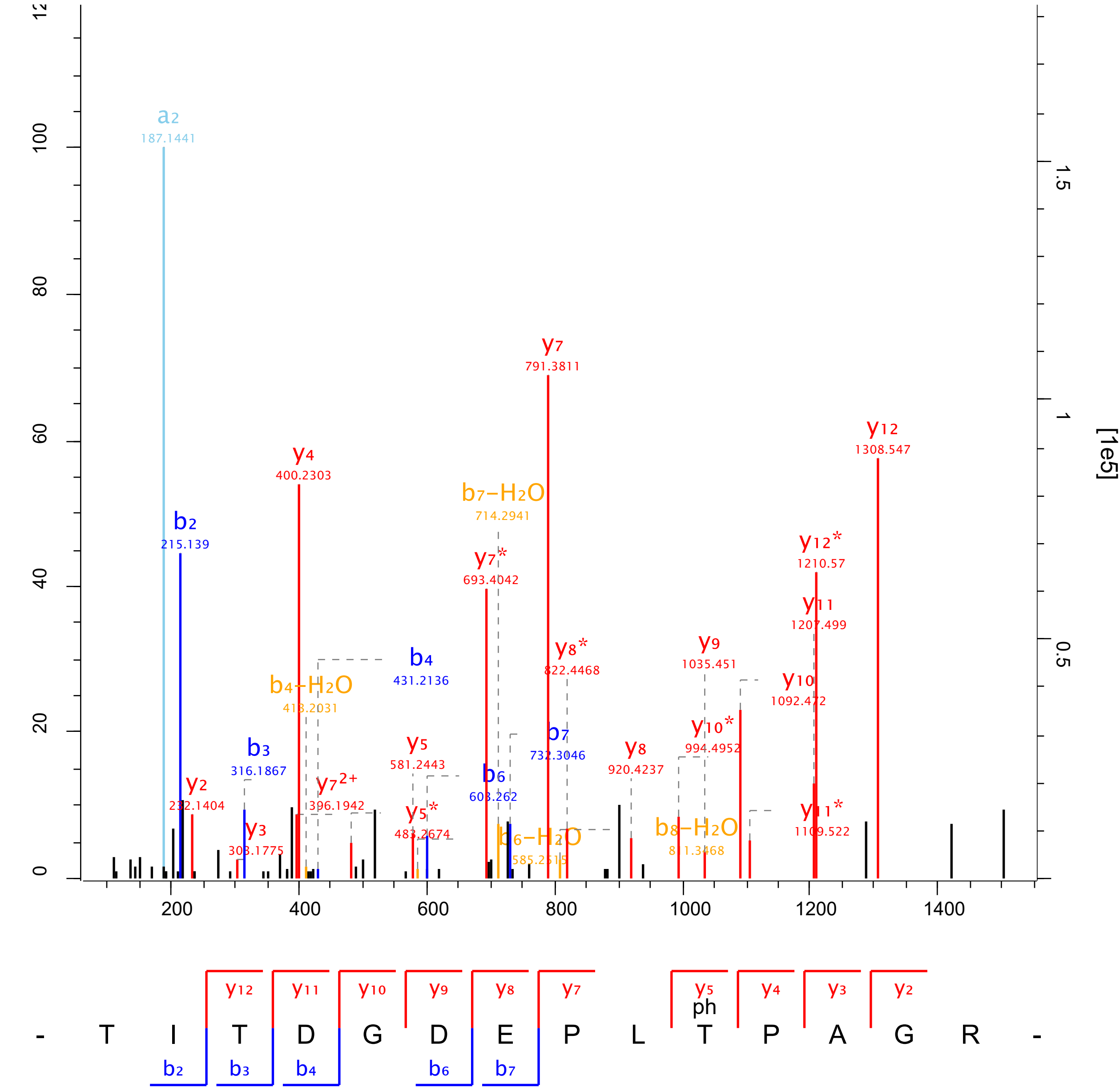

| Raw file | Scan | Method | Score | m/z |
|---|---|---|---|---|
| sys_00_3short | 16229 | FTMS; HCD | 47.5 | 934.43 |

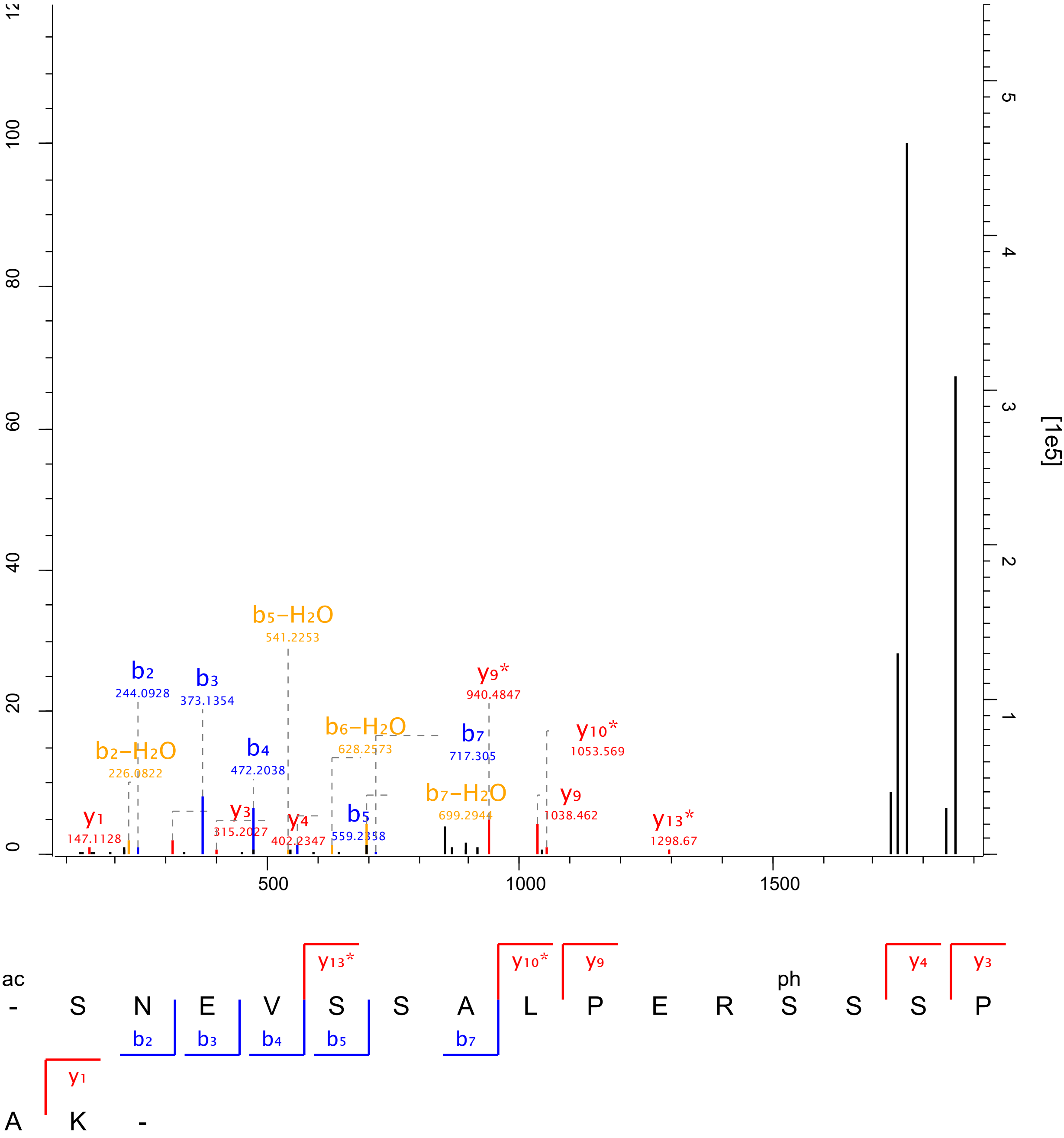

| Raw file | Scan | Method | Score | m/z |
|---|---|---|---|---|
| sys_00_3short | 16276 | FTMS; HCD | 71.88 | 692.28 |

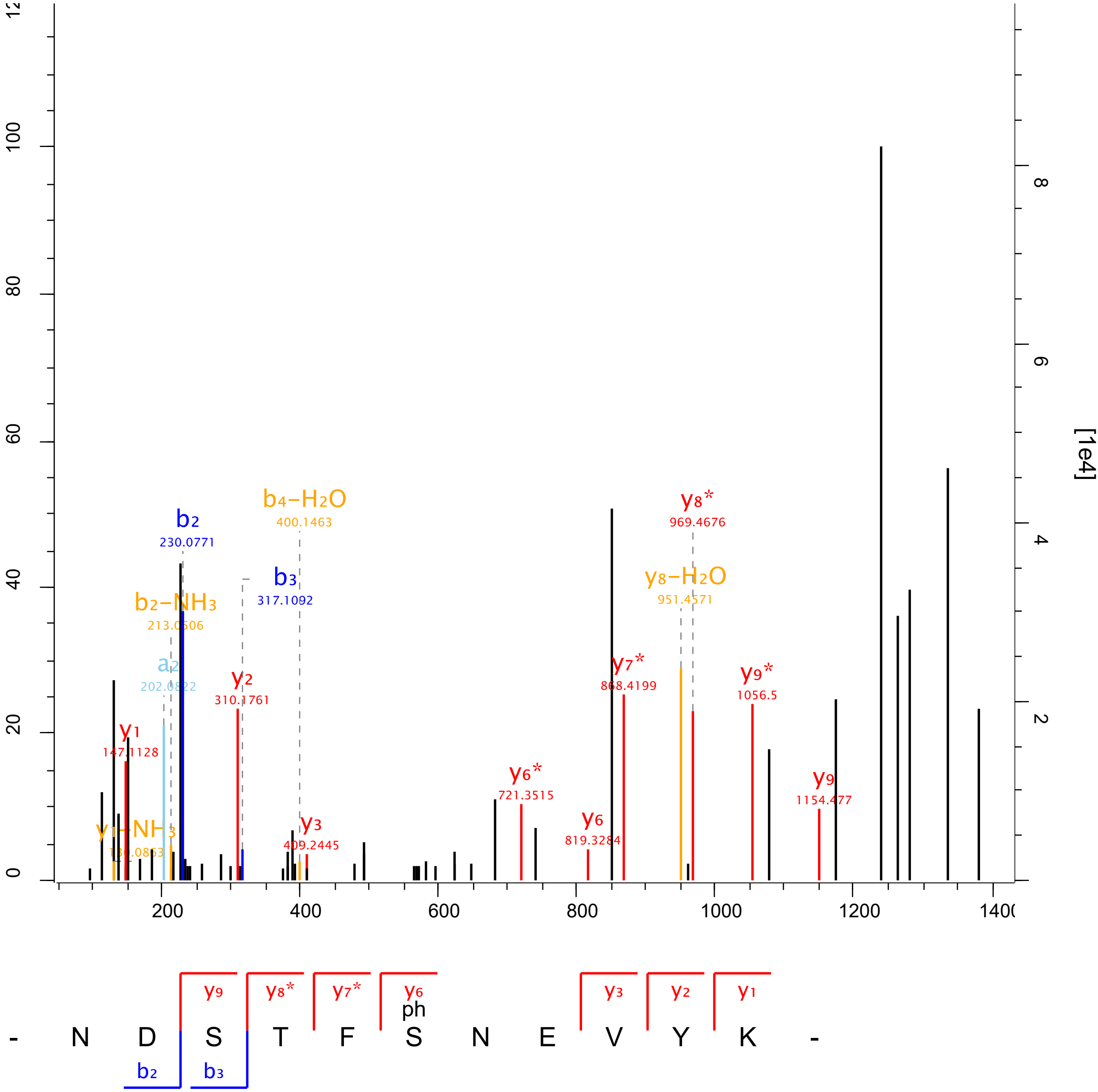

| Raw file | Scan | Method | Score | m/z |
|---|---|---|---|---|
| sys_00_3short | 16361 | FTMS; HCD | 47.6 | 452.18 |

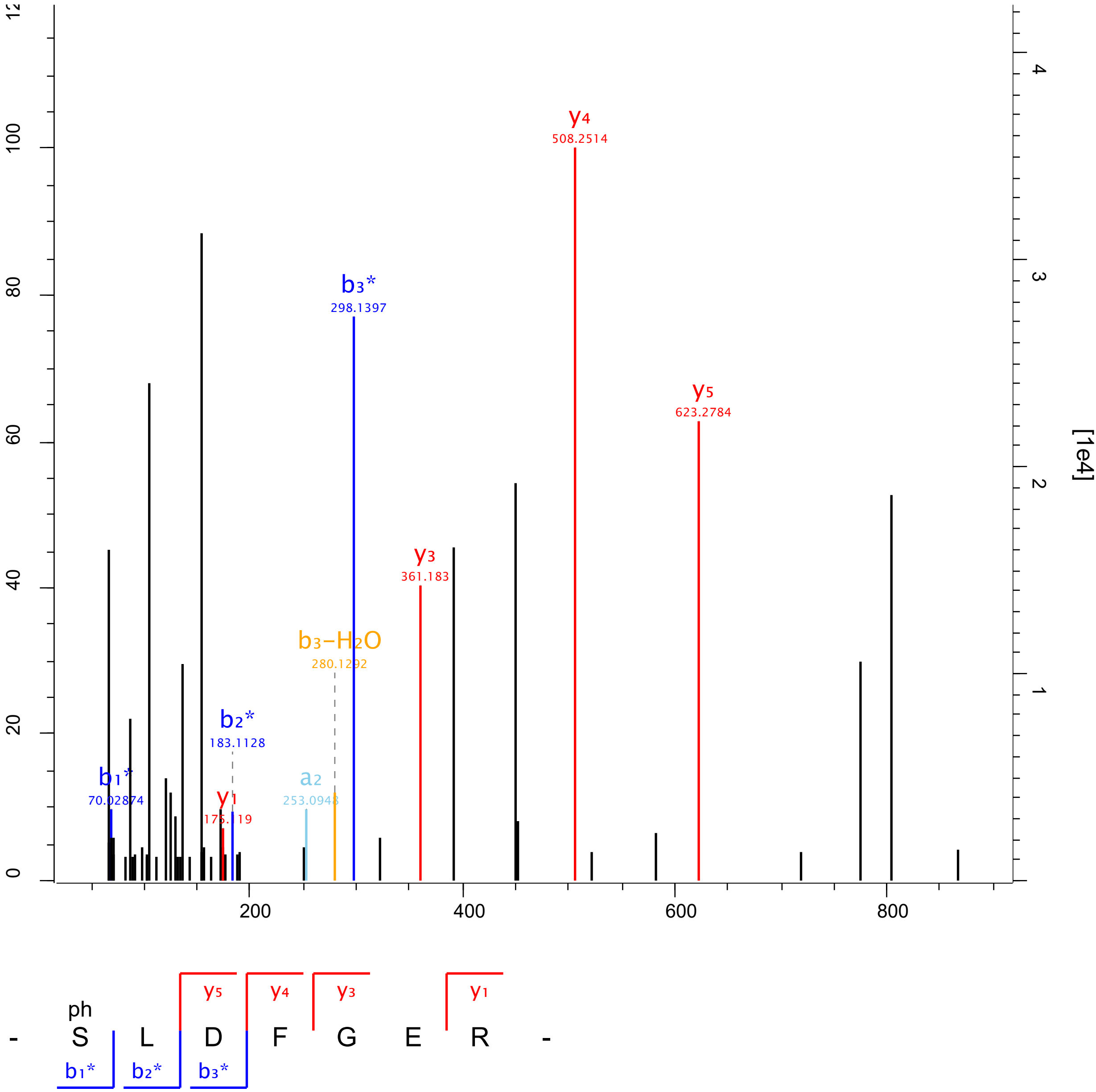

| Raw file | Scan | Method | Score | m/z |
|---|---|---|---|---|
| sys_00_3short | 16439 | FTMS; HCD | 52.58 | 602.74 |

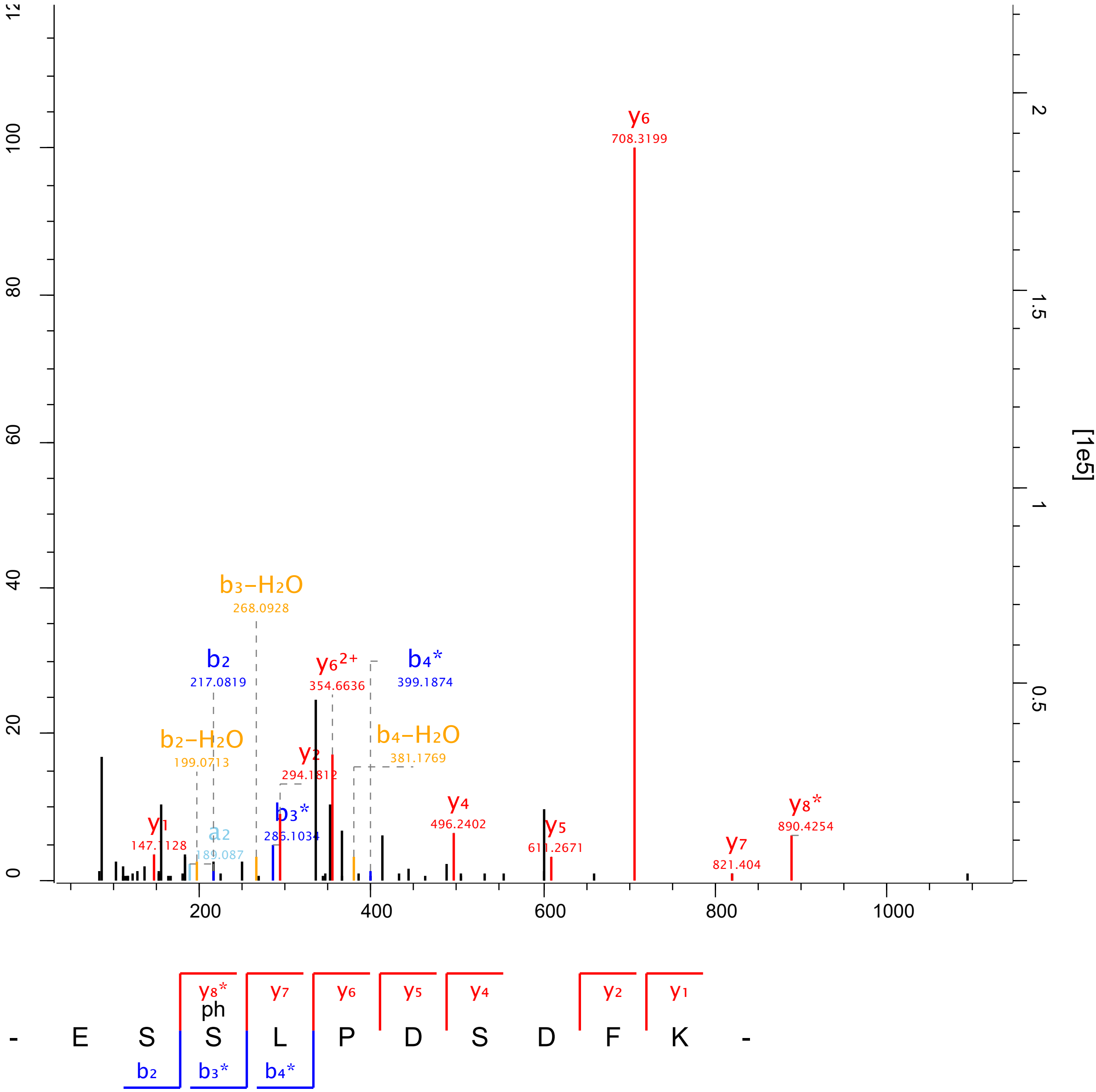

| Raw file | Scan | Method | Score | m/z |
| --- | --- | --- | --- | --- |
| sys_00_3short | 16535 | FTMS; HCD | 218.83 | 750.33 |

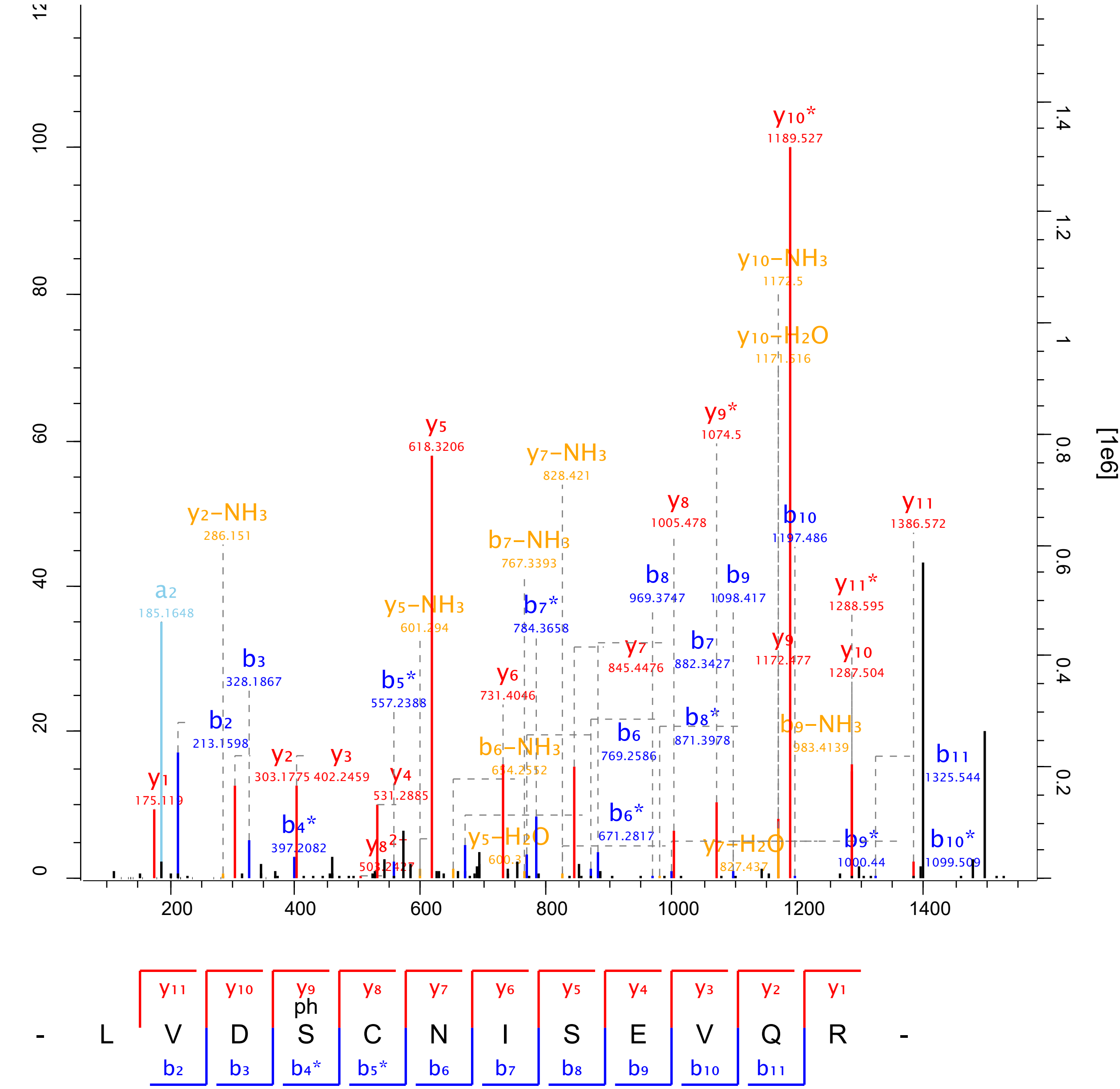

| Raw file | Scan | Method | Score | m/z |
|---|---|---|---|---|
| sys_00_3short | 16599 | FTMS; HCD | 70.2 | 702.83 |

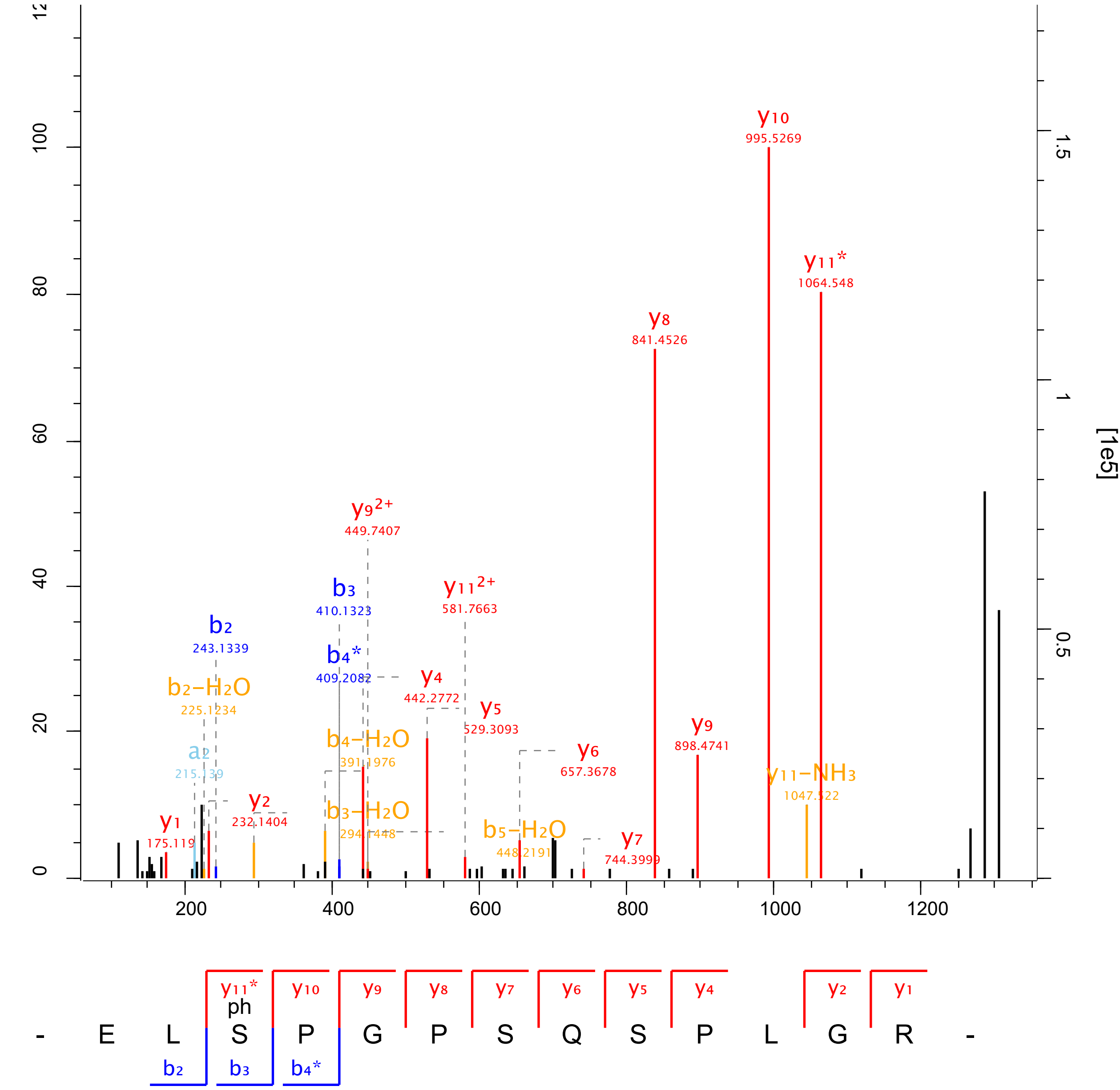

| Raw file | Scan | Method | Score | m/z |
| --- | --- | --- | --- | --- |
| sys_00_3short | 16873 | FTMS; HCD | 75.55 | 748.36 |

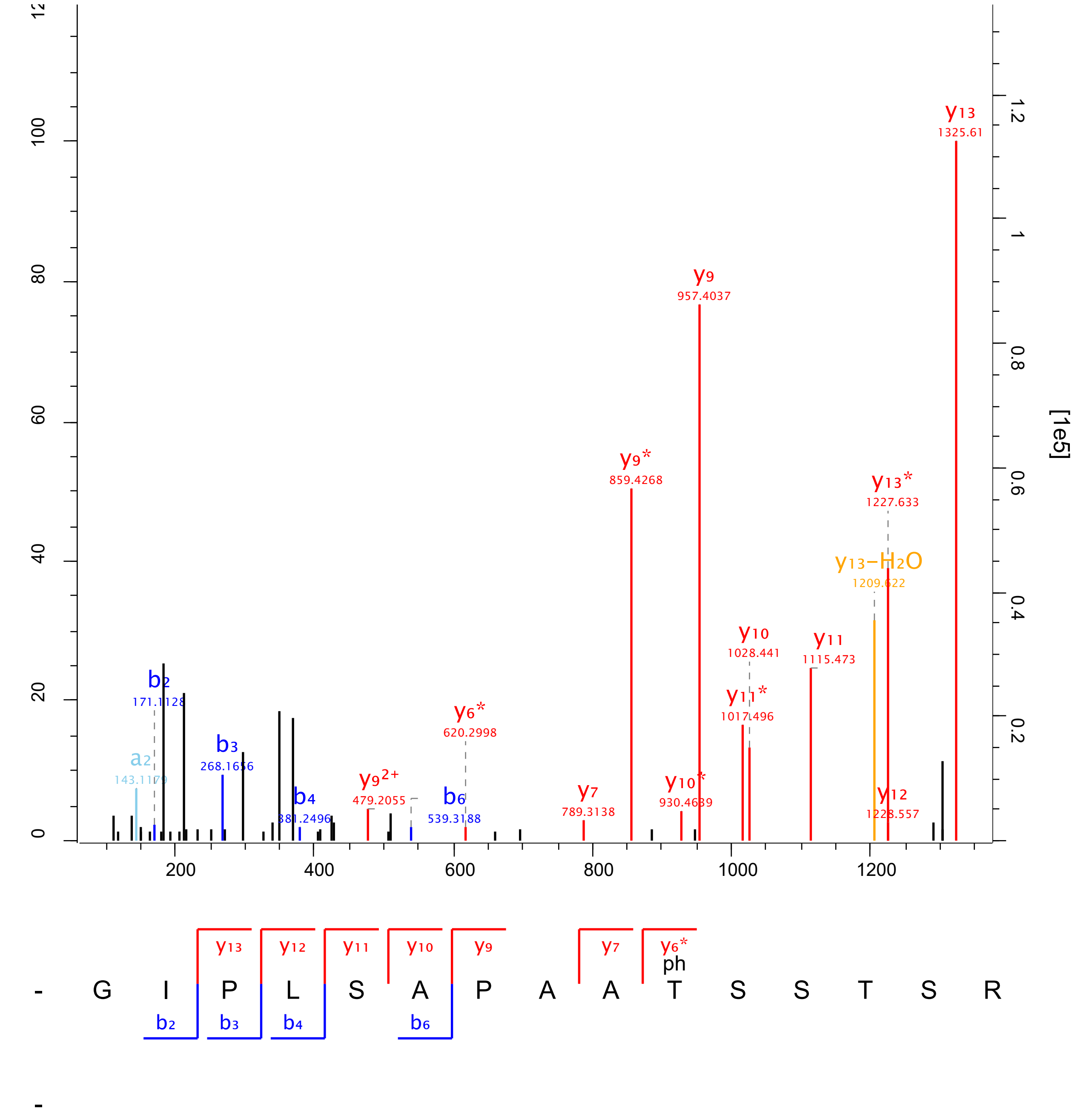

| Raw file | Scan | Method | Score | m/z |
| --- | --- | --- | --- | --- |
| sys_00_3short | 16971 | FTMS; HCD | 79.49 | 740.33 |

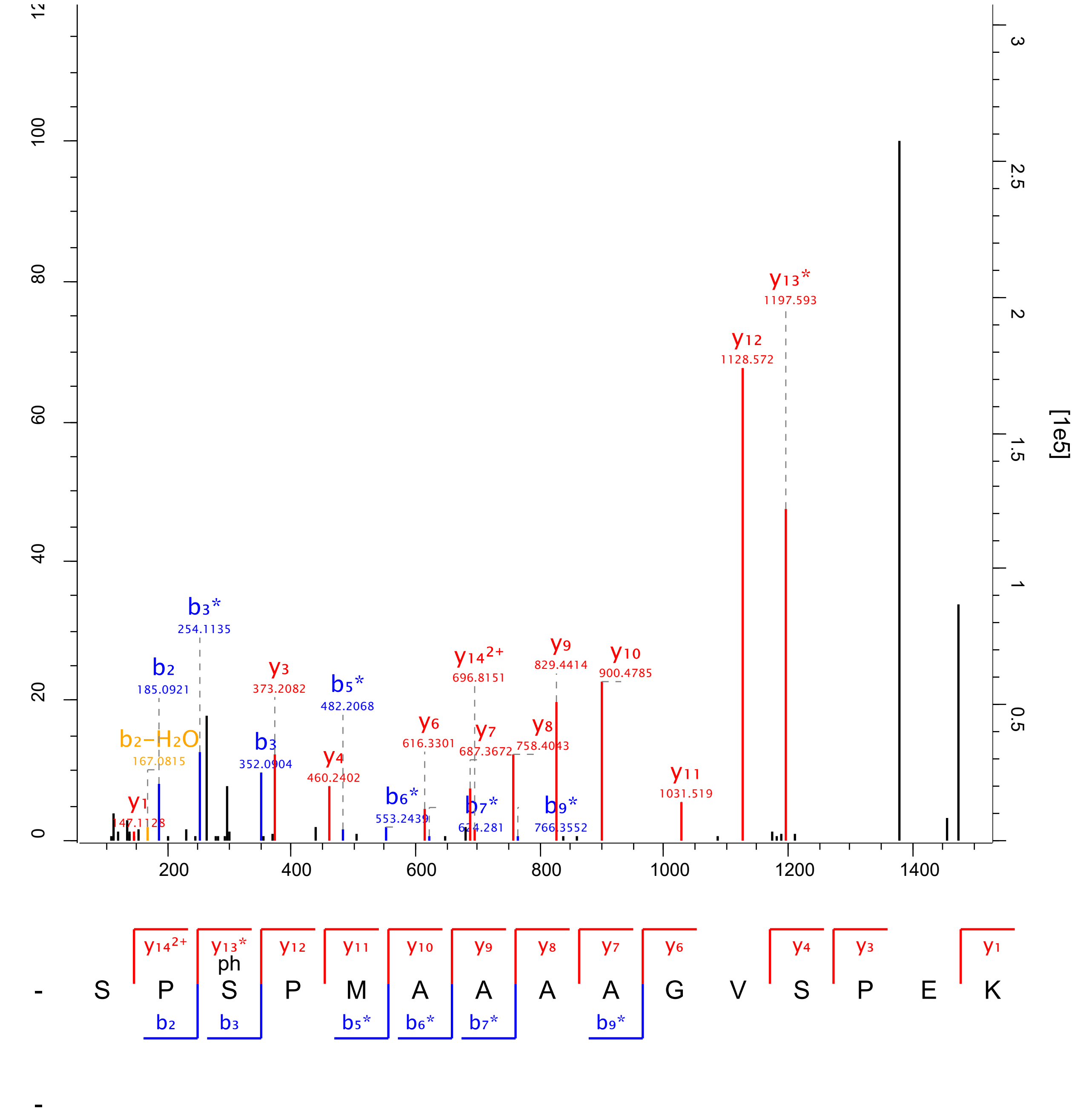

| Raw file | Scan | Method | Score | m/z |
| --- | --- | --- | --- | --- |
| sys_00_3short | 17196 | FTMS; HCD | 85.84 | 606.77 |

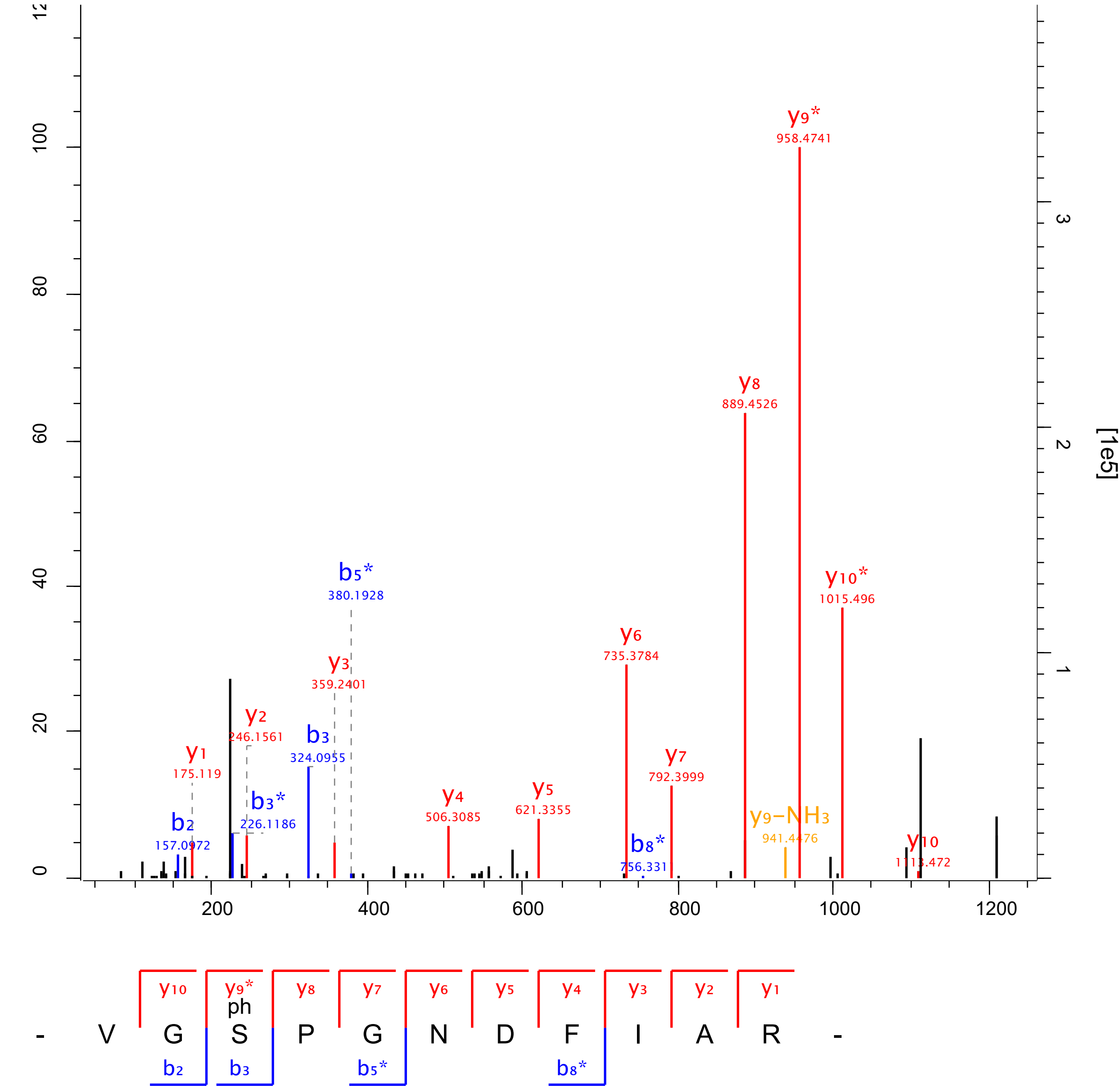

| Raw file | Scan | Method | Score | m/z |
|---|---|---|---|---|
| sys_00_3short | 17246 | FTMS; HCD | 40.95 | 887.41 |

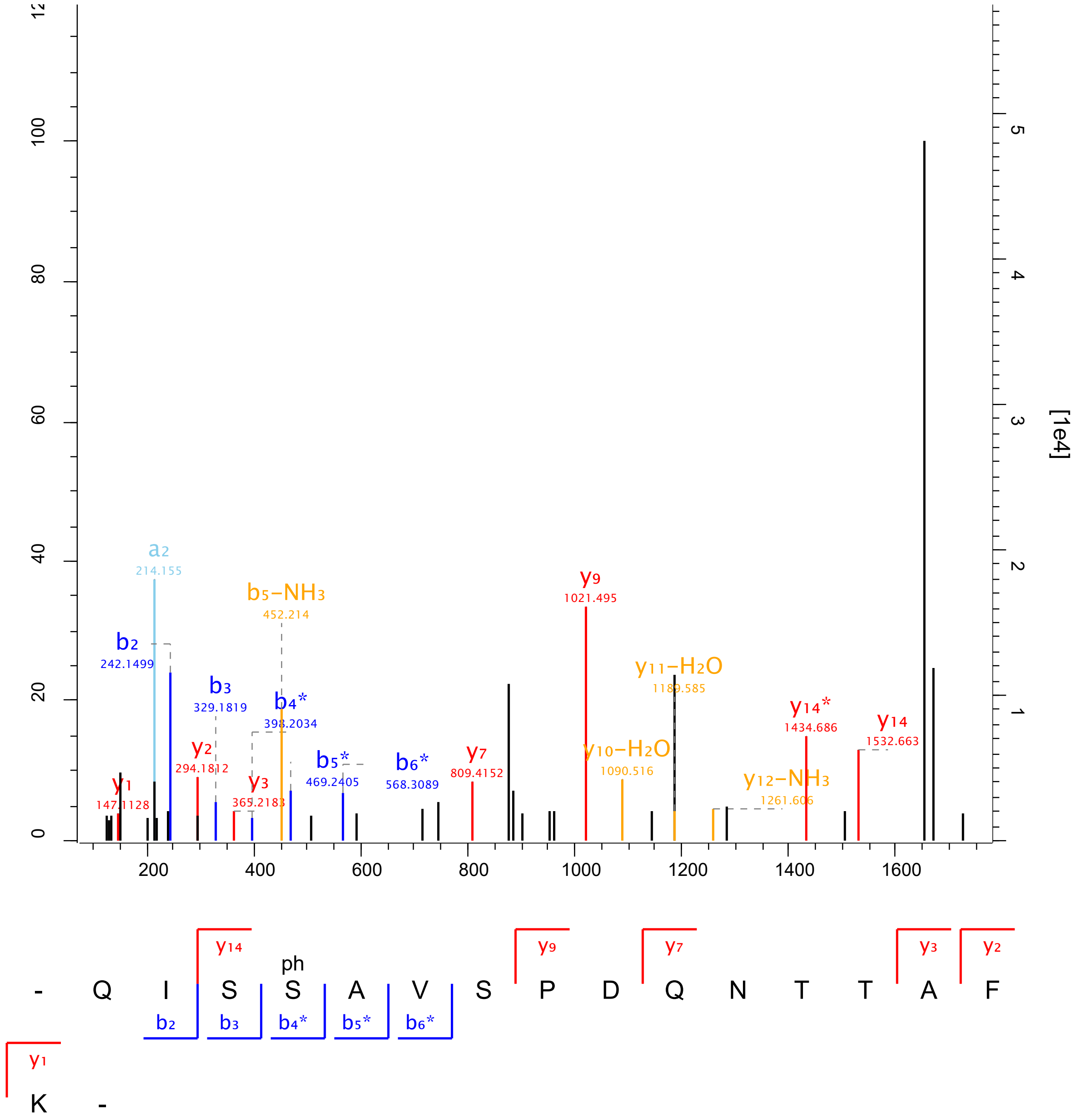

| Raw file | Scan | Method | Score | m/z |
|---|---|---|---|---|
| sys_00_3short | 17323 | FTMS; HCD | 51.45 | 573.74 |

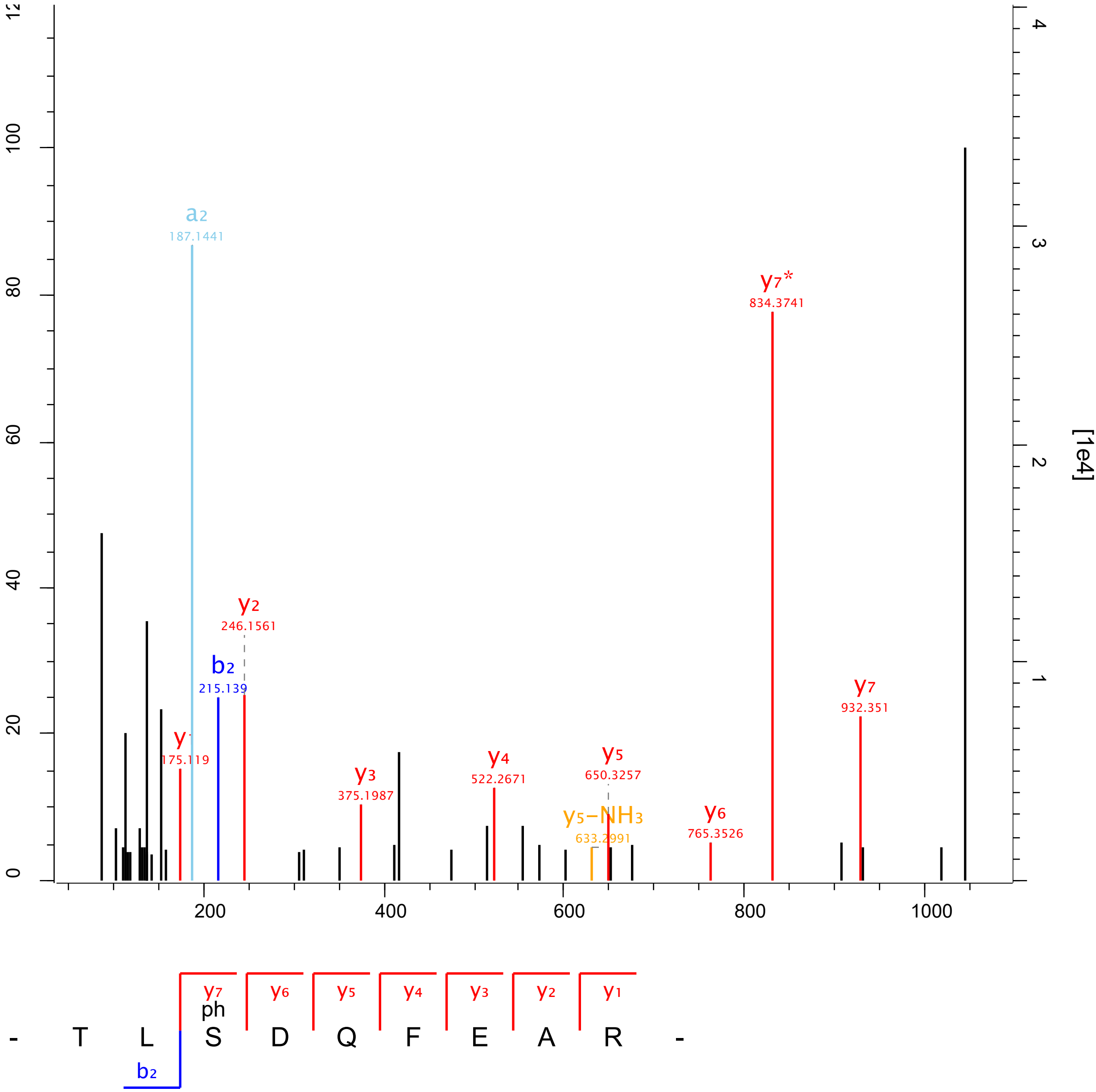

| Raw file | Scan | Method | Score | m/z |
| --- | --- | --- | --- | --- |
| sys_00_3short | 17561 | FTMS; HCD | 98.59 | 578.76 |

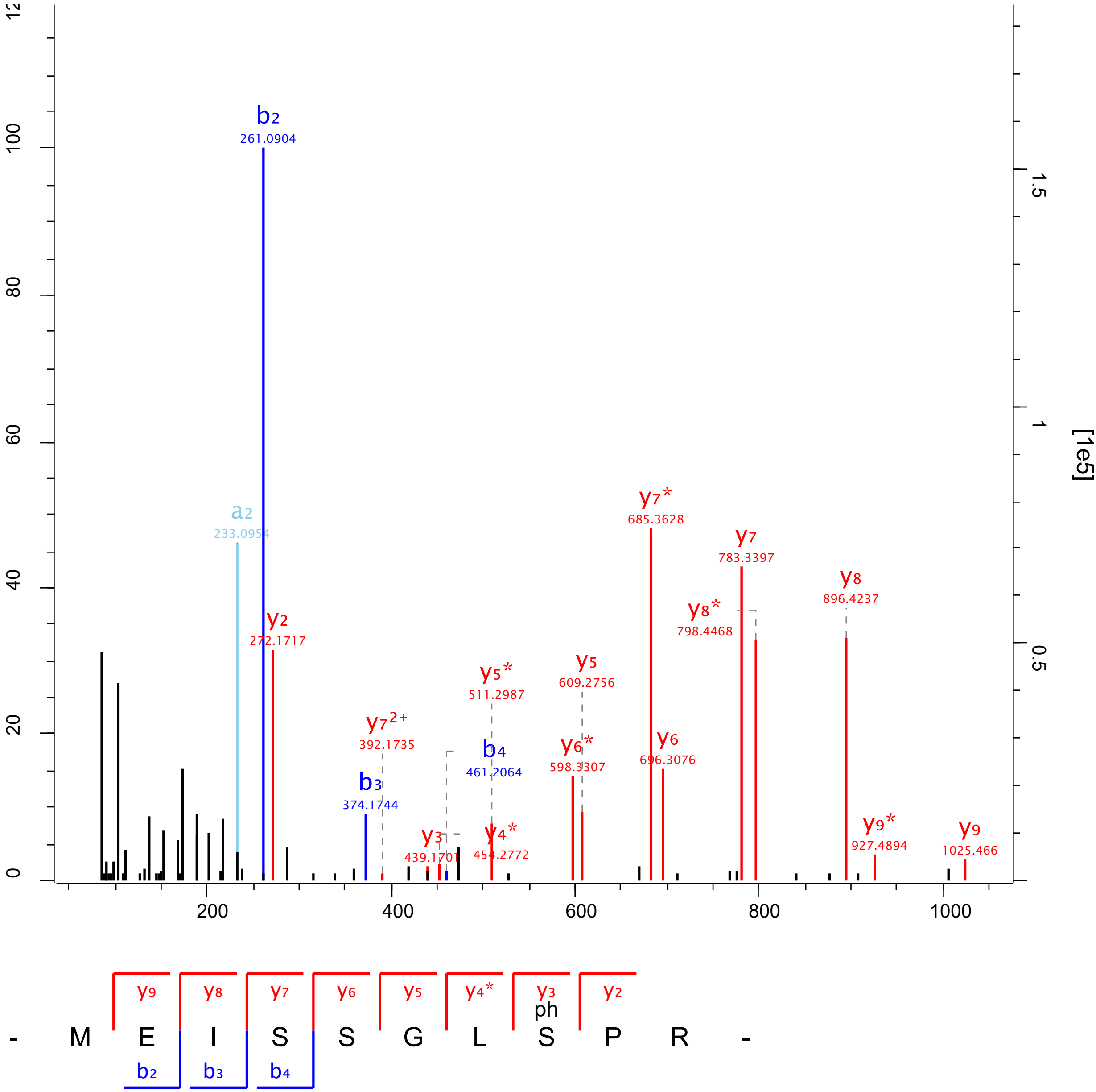

| Raw file | Scan | Method | Score | m/z |
|---|---|---|---|---|
| sys_00_3short | 17610 | FTMS; HCD | 55.72 | 651.77 |

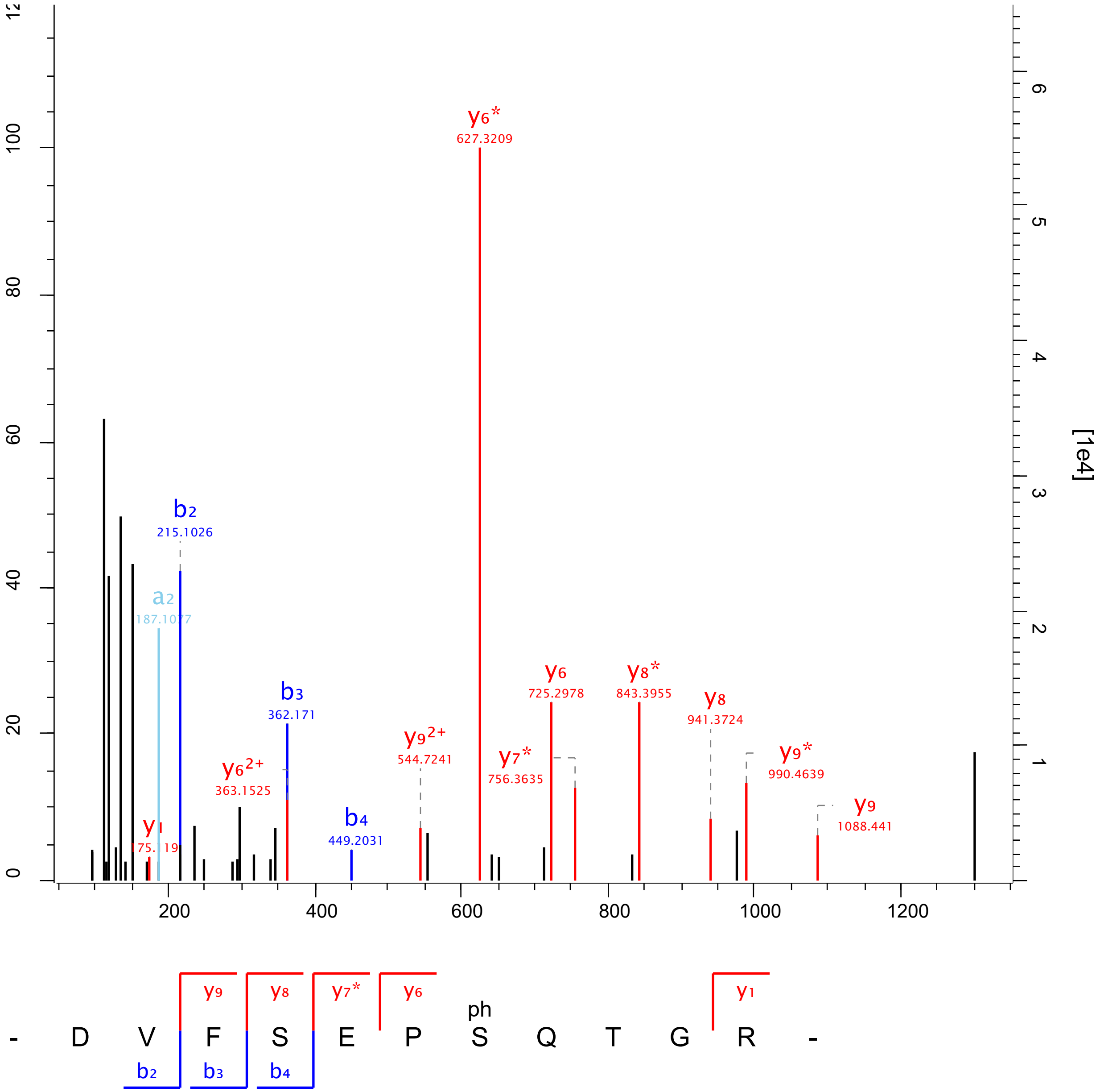

| Raw file | Scan | Method | Score | m/z |
|---|---|---|---|---|
| sys_00_3short | 17639 | FTMS; HCD | 157.98 | 838.85 |

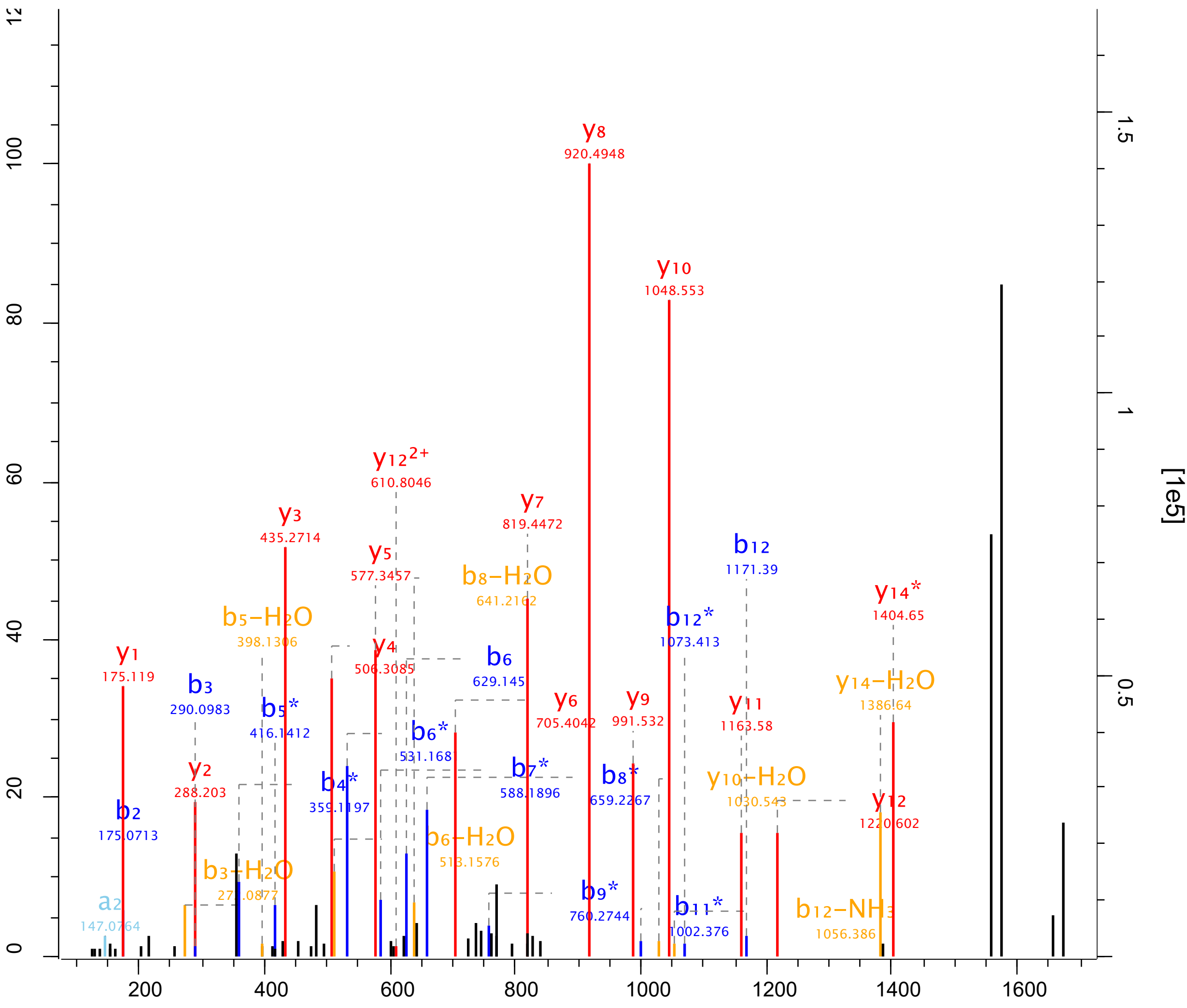


| | | | | | | | | | | | | | | | |
|---|---|---|---|---|---|---|---|---|---|---|---|---|---|---|---|
| | | | y14* | | y12 | y11 | y10 | y9 | y8 | y7 | y6 | y5 | y4 | y3 | y2 |
| | | | | ph | | | | | | | | | | | |
| - | S | S | D | S | G | D | G | A | T | N | Q | A | A | F | I |
| | | b2 | b3 | b4* | b5* | b6 | b7* | b8* | b9* | | b11* | b12 | | | |
| y1 | | | | | | | | | | | | | | | |
| R | - | | | | | | | | | | | | | | |

| Raw file | Scan | Method | Score | m/z |
|---|---|---|---|---|
| sys_00_3short | 17705 | FTMS; HCD | 70.03 | 515.22 |

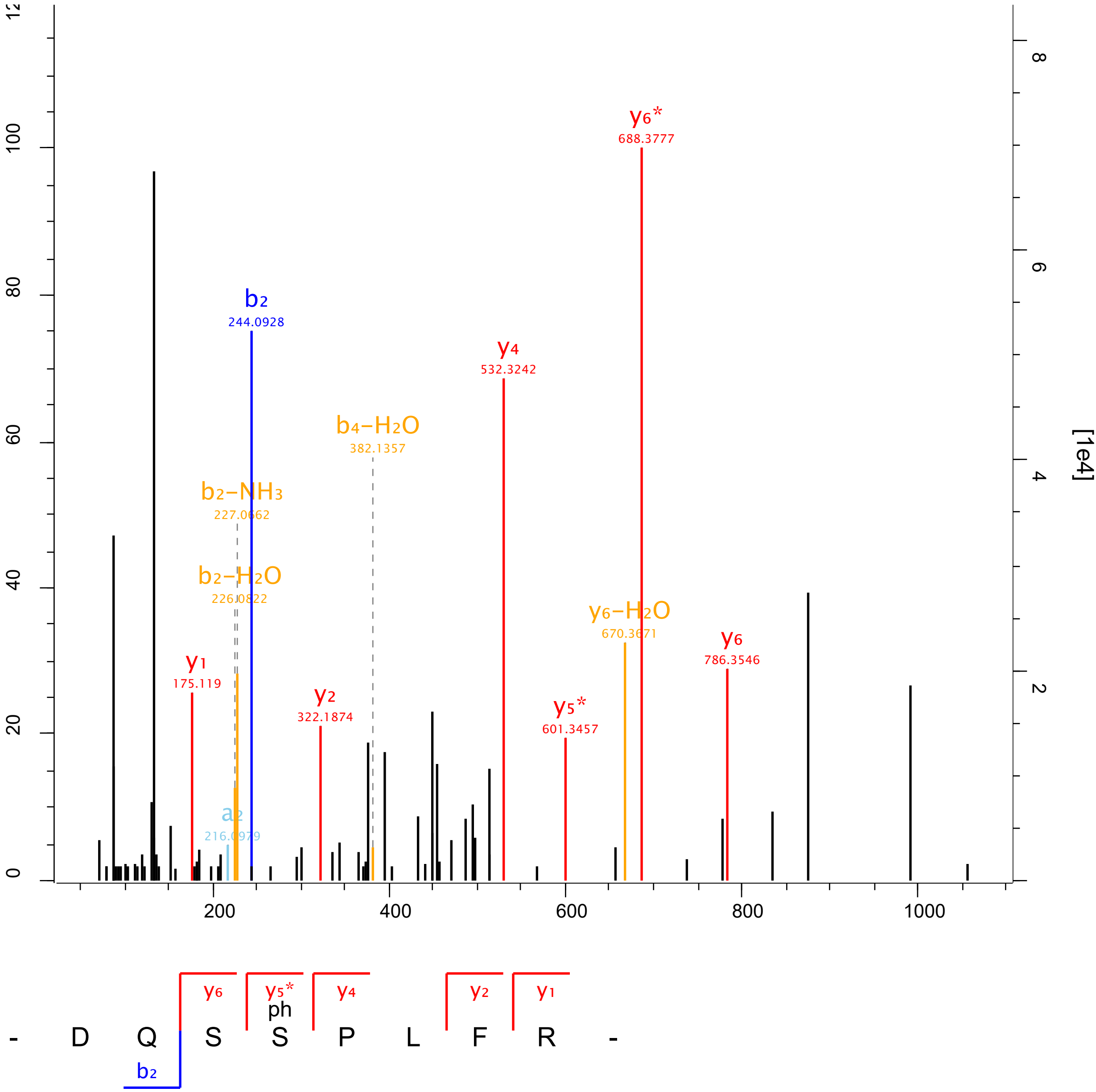

| Raw file | Scan | Method | Score | m/z |
| --- | --- | --- | --- | --- |
| sys_00_3short | 17742 | FTMS; HCD | 143.25 | 572.76 |

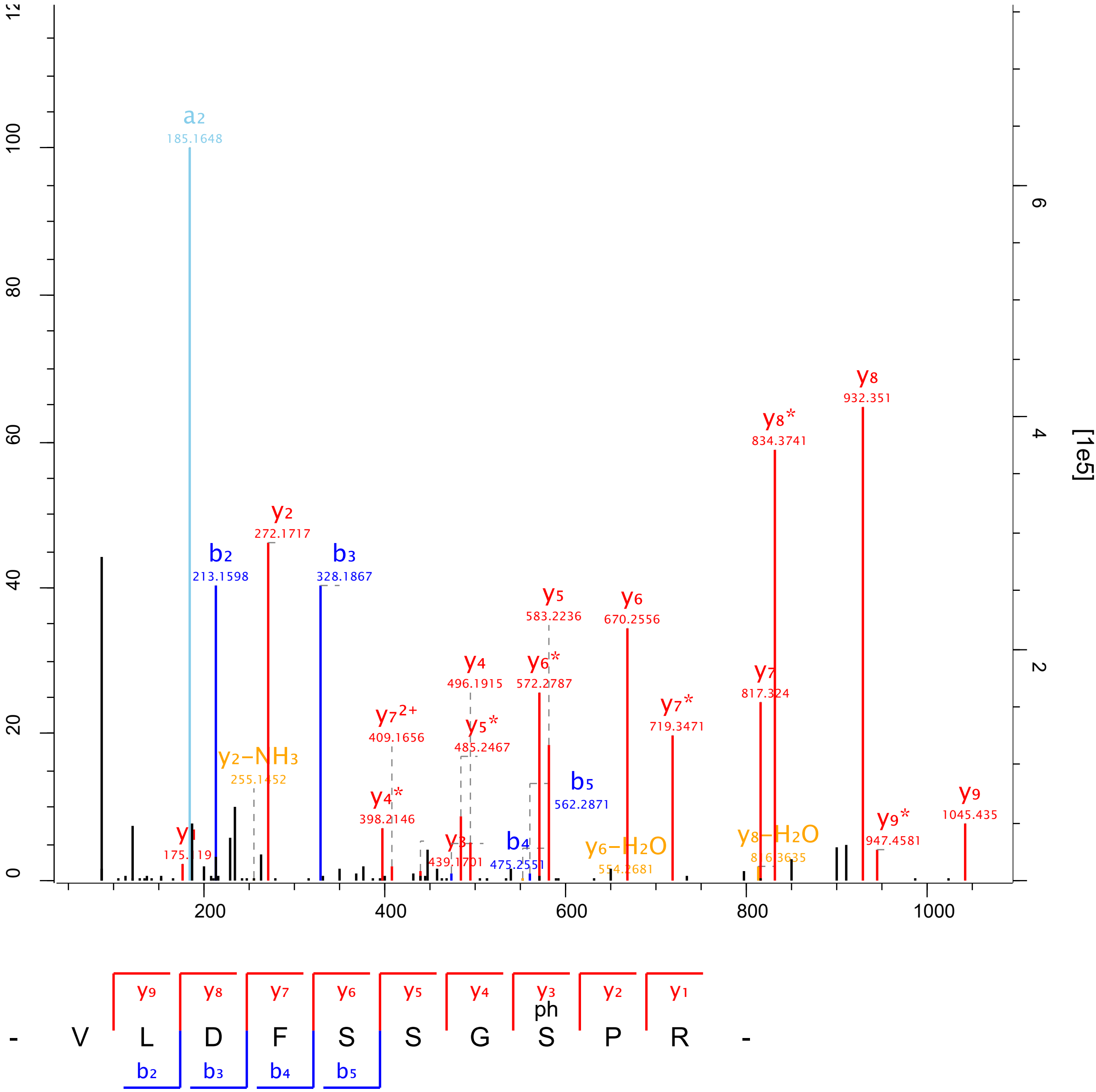

| Raw file | Scan | Method | Score | m/z |
| --- | --- | --- | --- | --- |
| sys_00_3short | 17750 | FTMS; HCD | 73.93 | 648.3 |

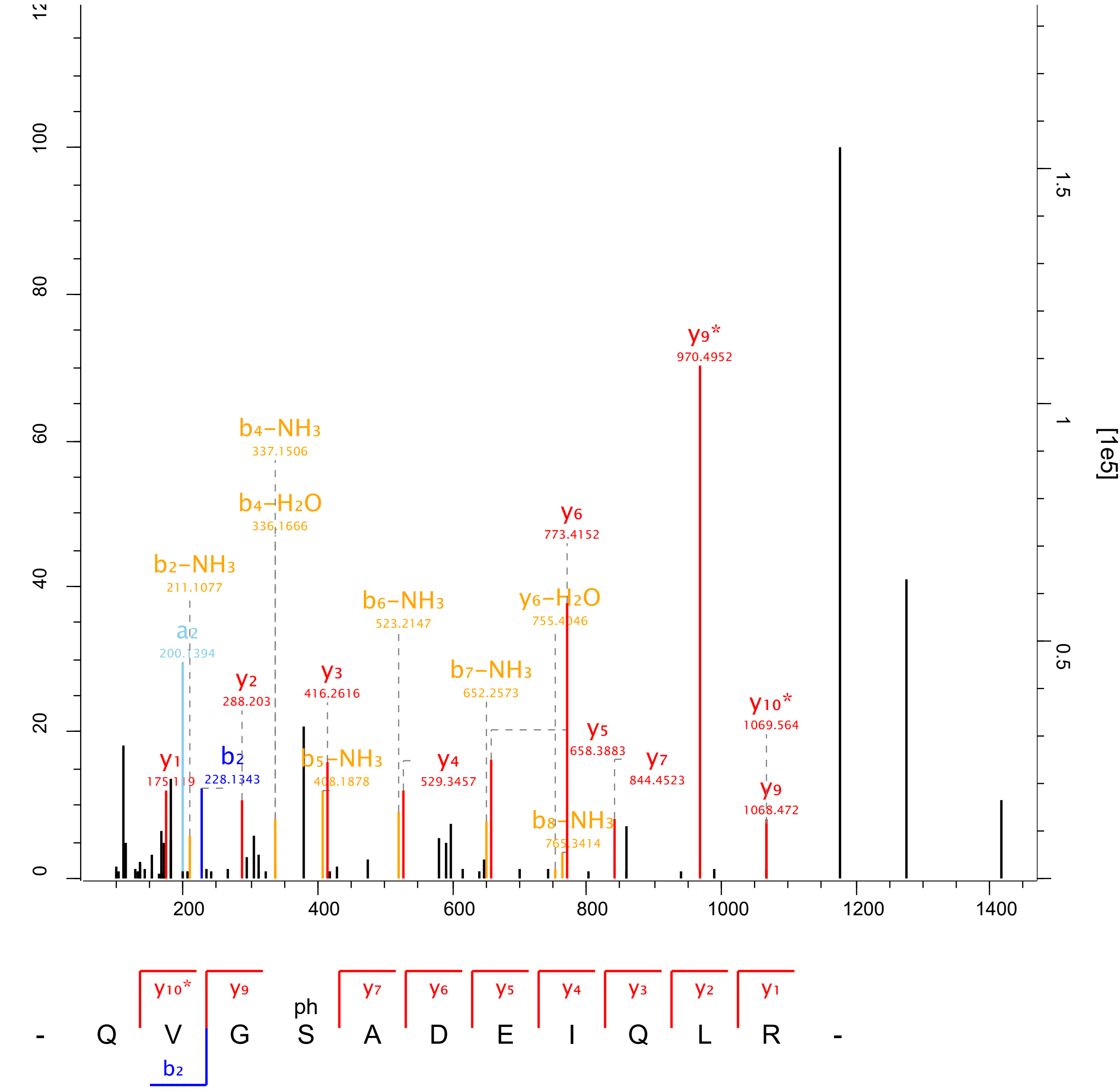

| Raw file | Scan | Method | Score | m/z |
| --- | --- | --- | --- | --- |
| sys_00_3short | 17870 | FTMS; HCD | 84.36 | 525.78 |

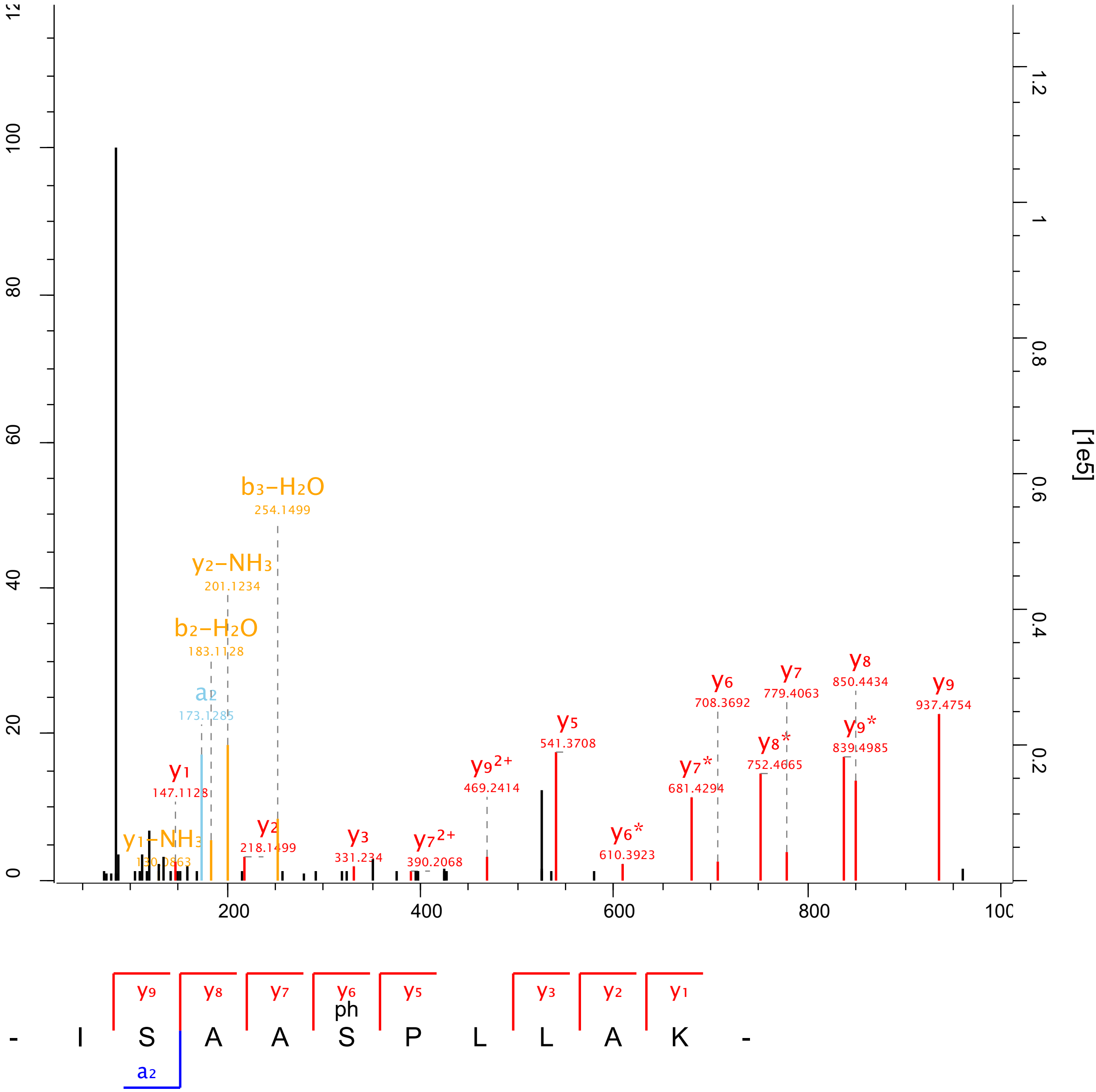

| Raw file | Scan | Method | Score | m/z |
|---|---|---|---|---|
| sys_00_3short | 17947 | FTMS; HCD | 130.15 | 607.74 |

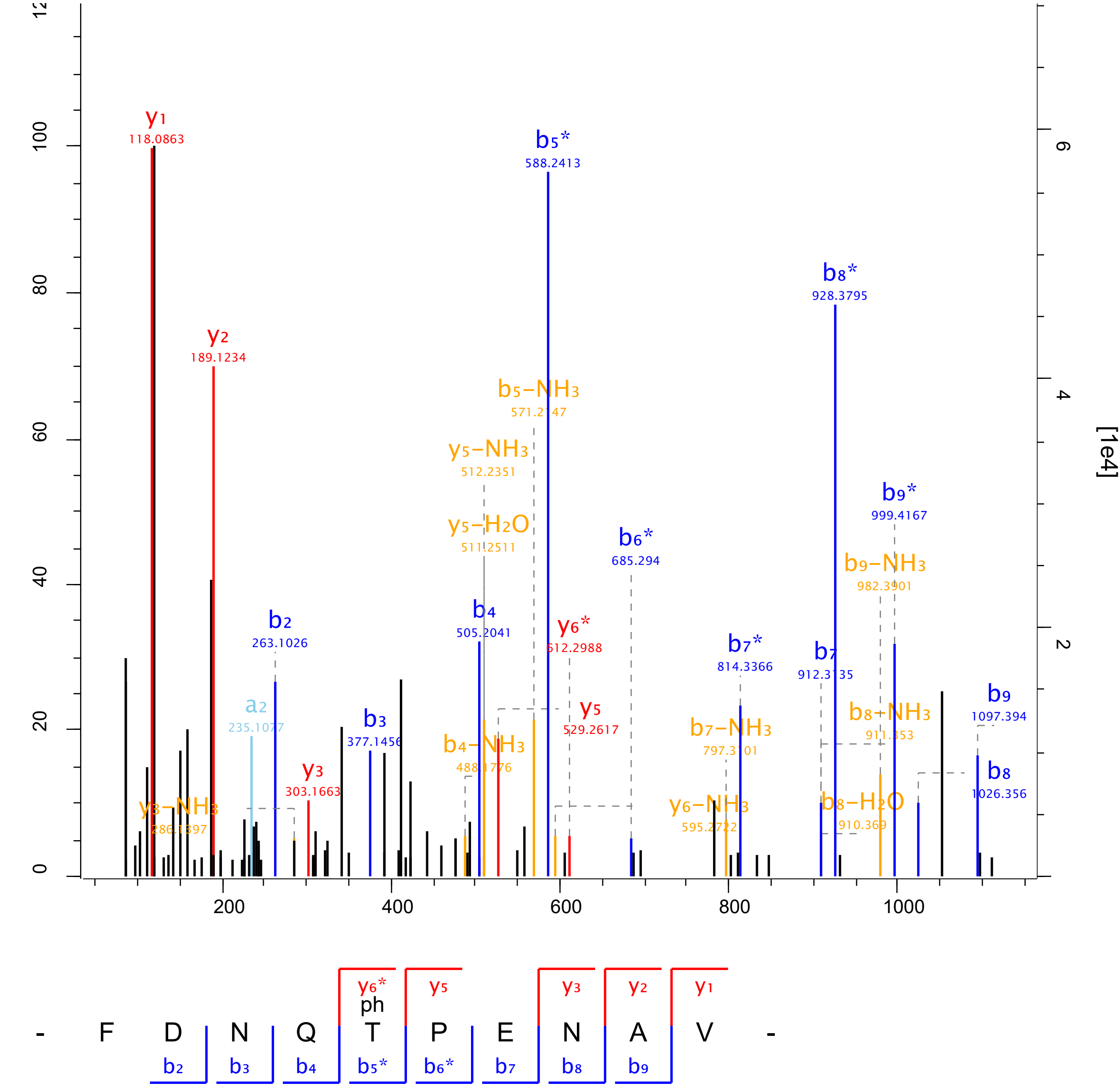

| Raw file | Scan | Method | Score | m/z |
| --- | --- | --- | --- | --- |
| sys_00_3short | 18233 | FTMS; HCD | 60.16 | 581.3 |

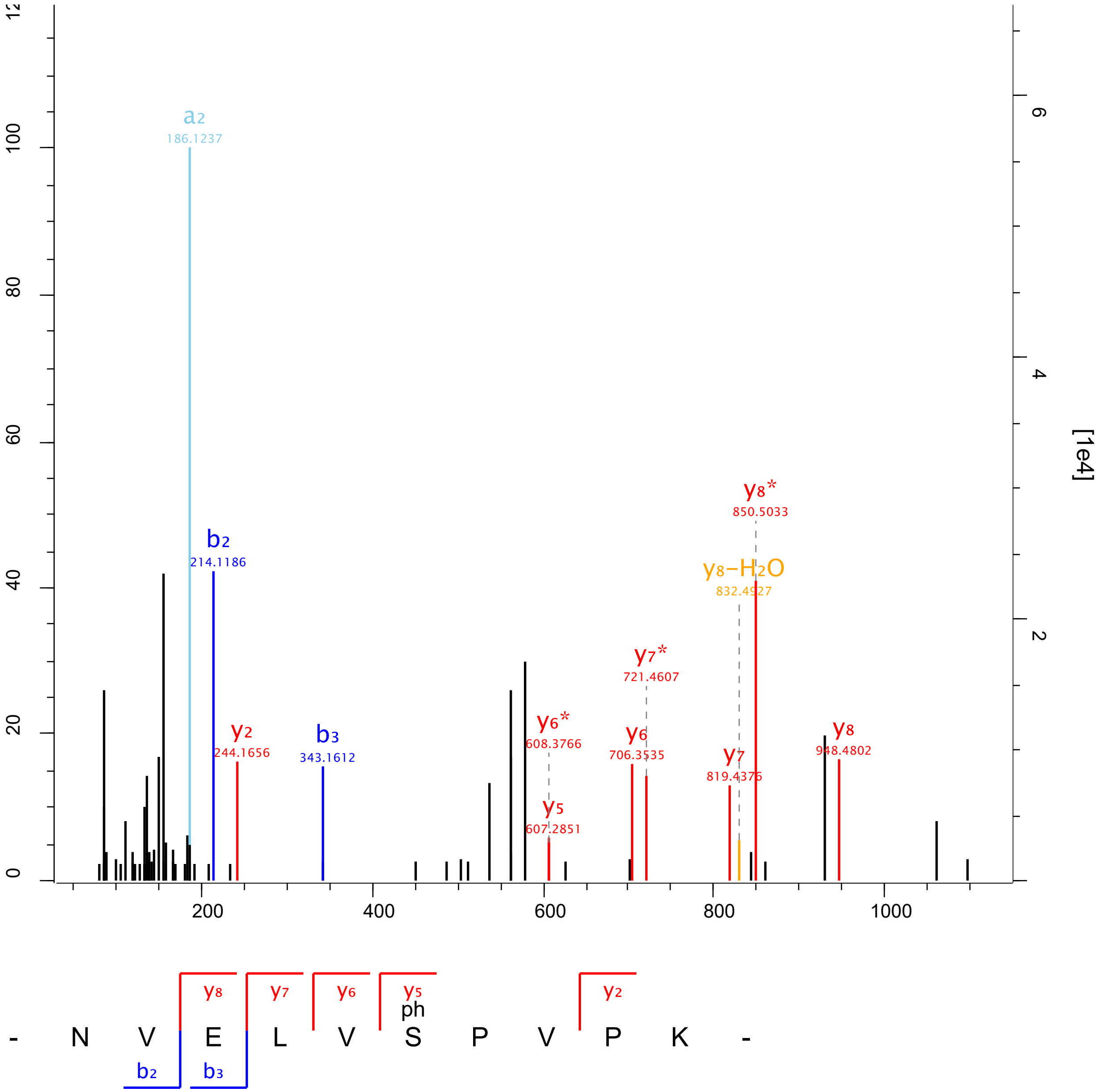

| Raw file | Scan | Method | Score | m/z |
|---|---|---|---|---|
| sys_00_3short | 18297 | FTMS; HCD | 150.11 | 657.79 |

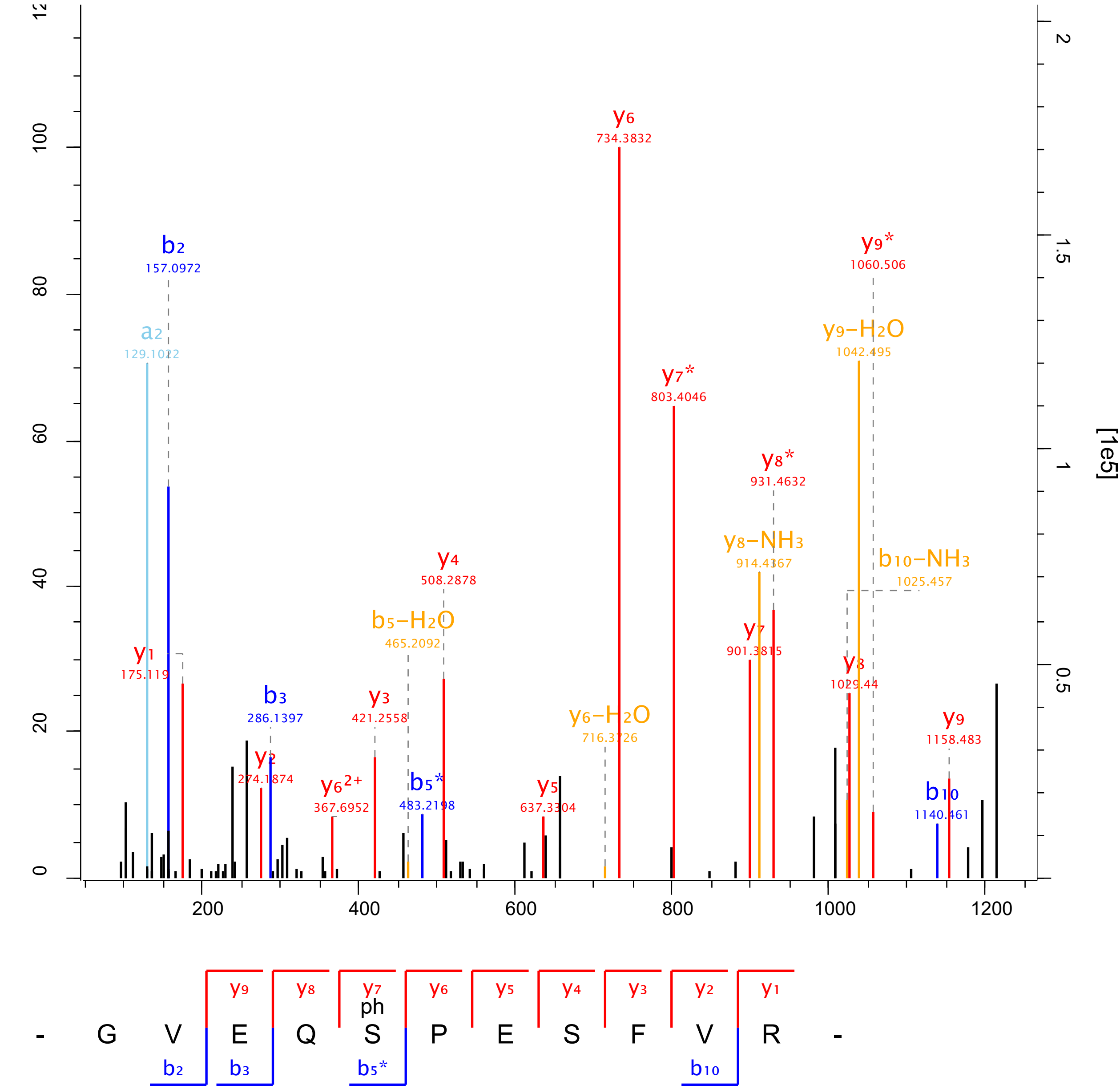

| Raw file | Scan | Method | Score | m/z |
|---|---|---|---|---|
| sys_00_3short | 18580 | FTMS; HCD | 49.33 | 562.76 |

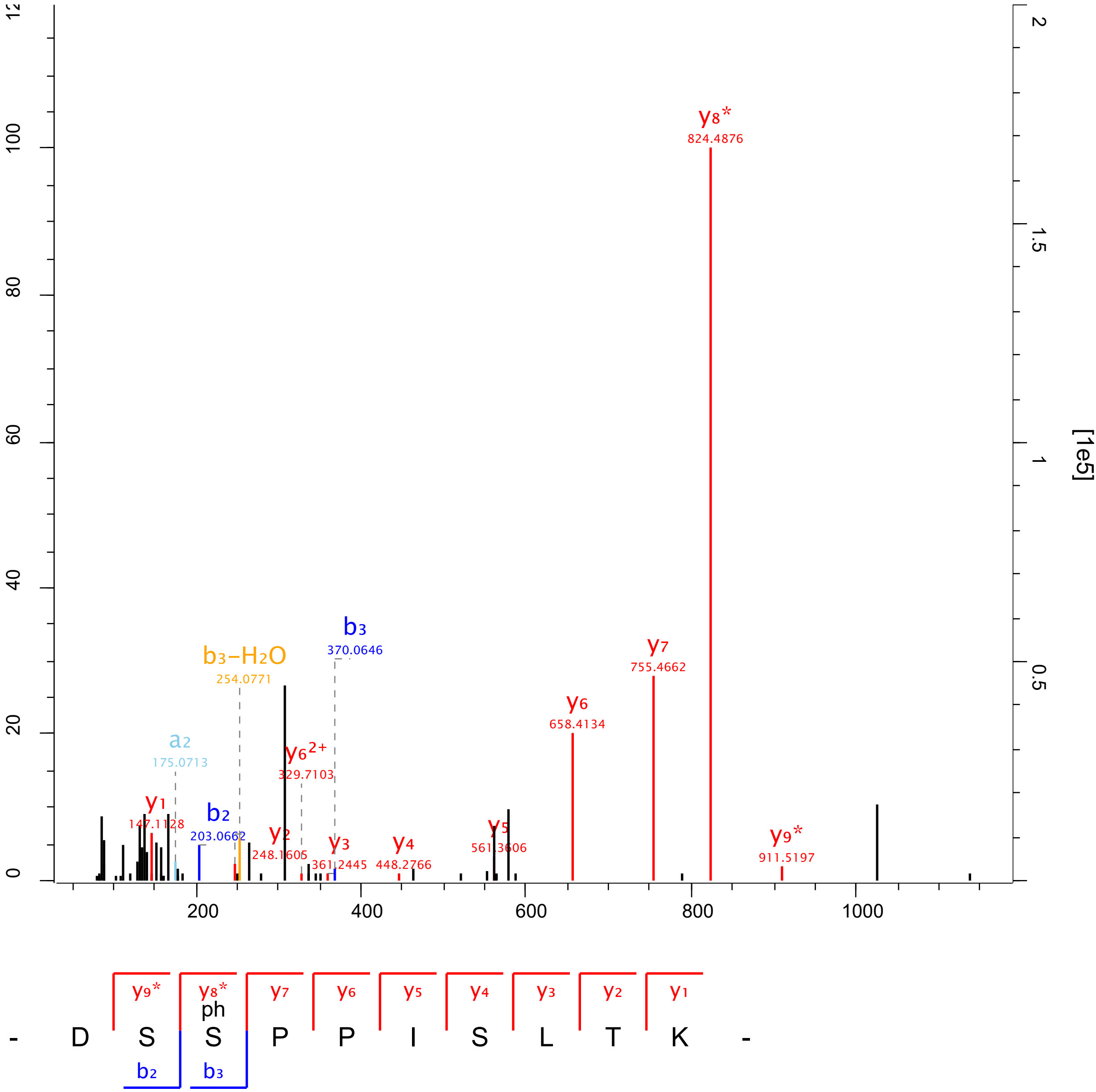

| Raw file | Scan | Method | Score | m/z |
|---|---|---|---|---|
| sys_00_3short | 18685 | FTMS; HCD | 98.57 | 775.36 |

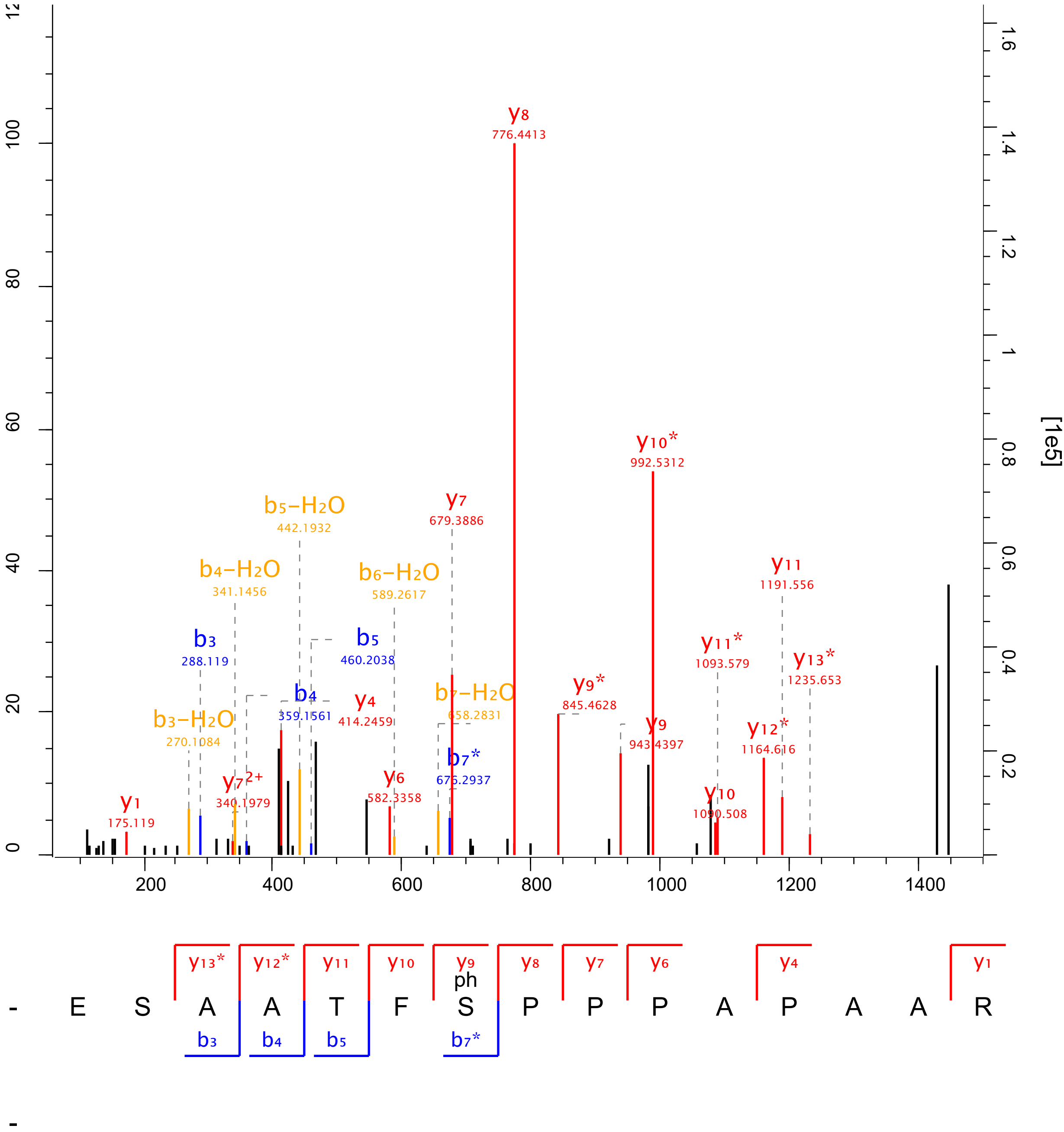

| Raw file | Scan | Method | Score | m/z |
|---|---|---|---|---|
| sys_00_3short | 18691 | FTMS; HCD | 59.71 | 602.77 |

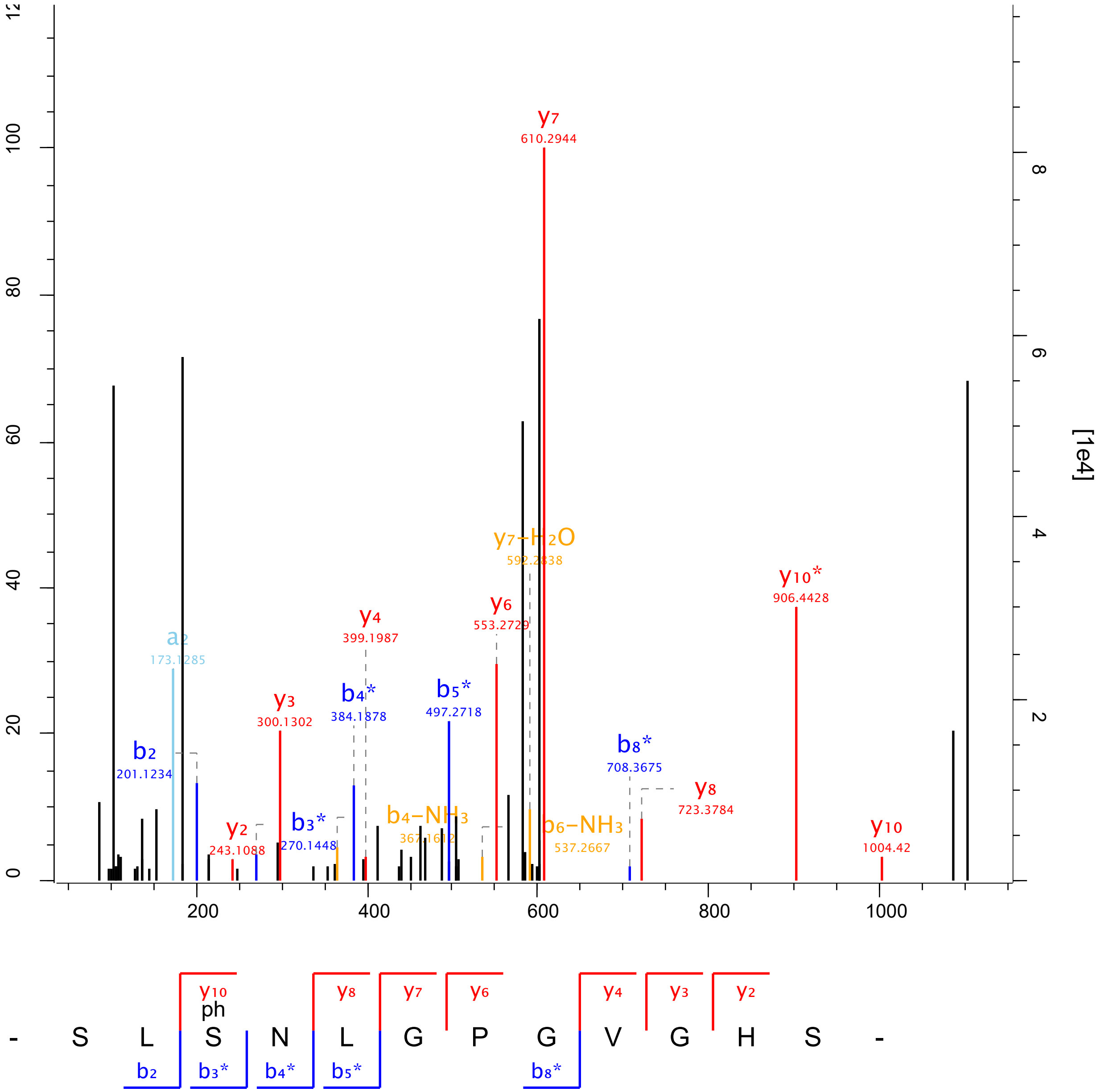

| Raw file | Scan | Method | Score | m/z |
| --- | --- | --- | --- | --- |
| sys_00_3short | 1871 | FTMS; HCD | 70.81 | 562.75 |

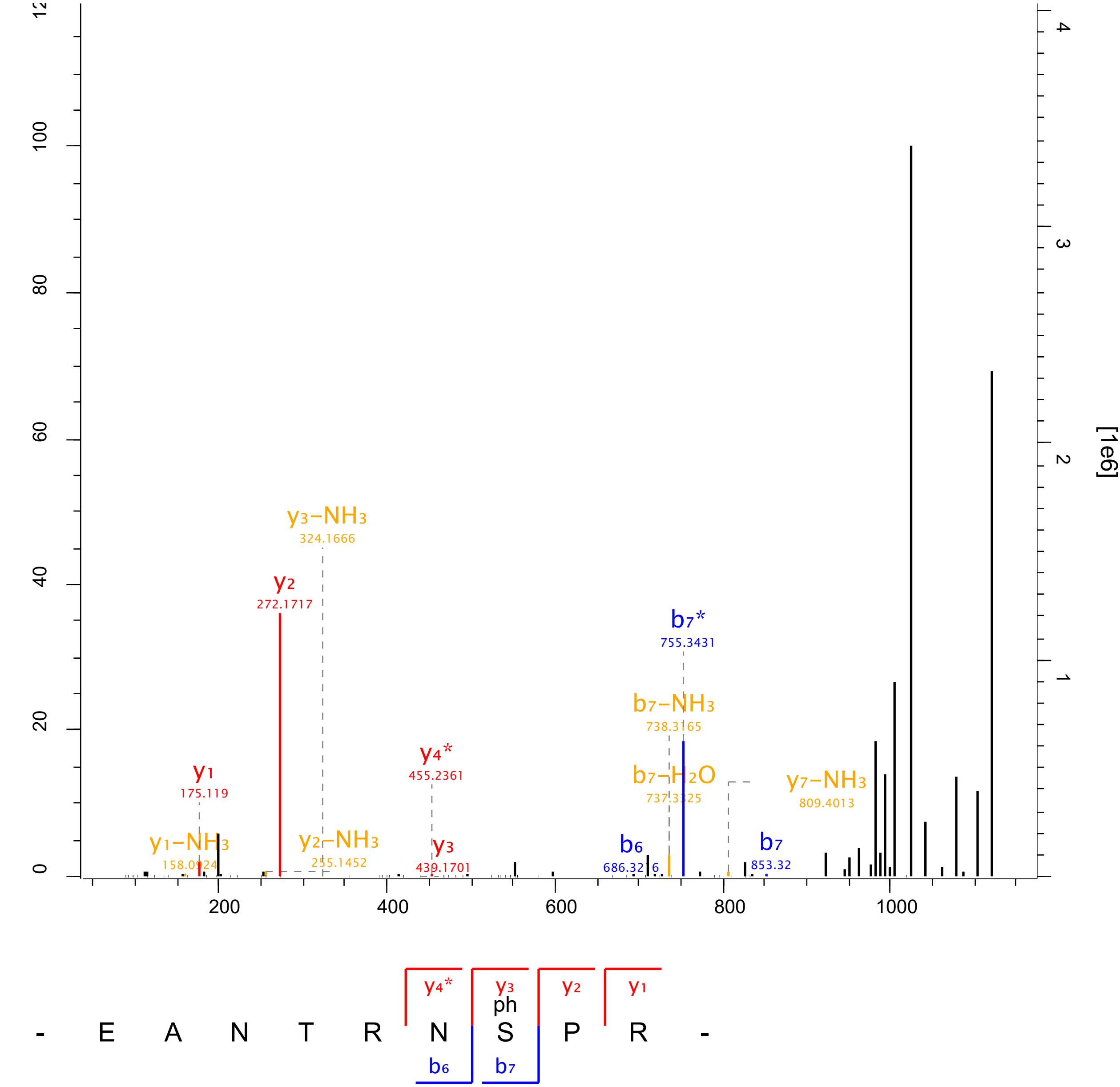

| Raw file | Scan | Method | Score | m/z |
| --- | --- | --- | --- | --- |
| sys_00_3short | 19002 | FTMS; HCD | 62.09 | 772.87 |

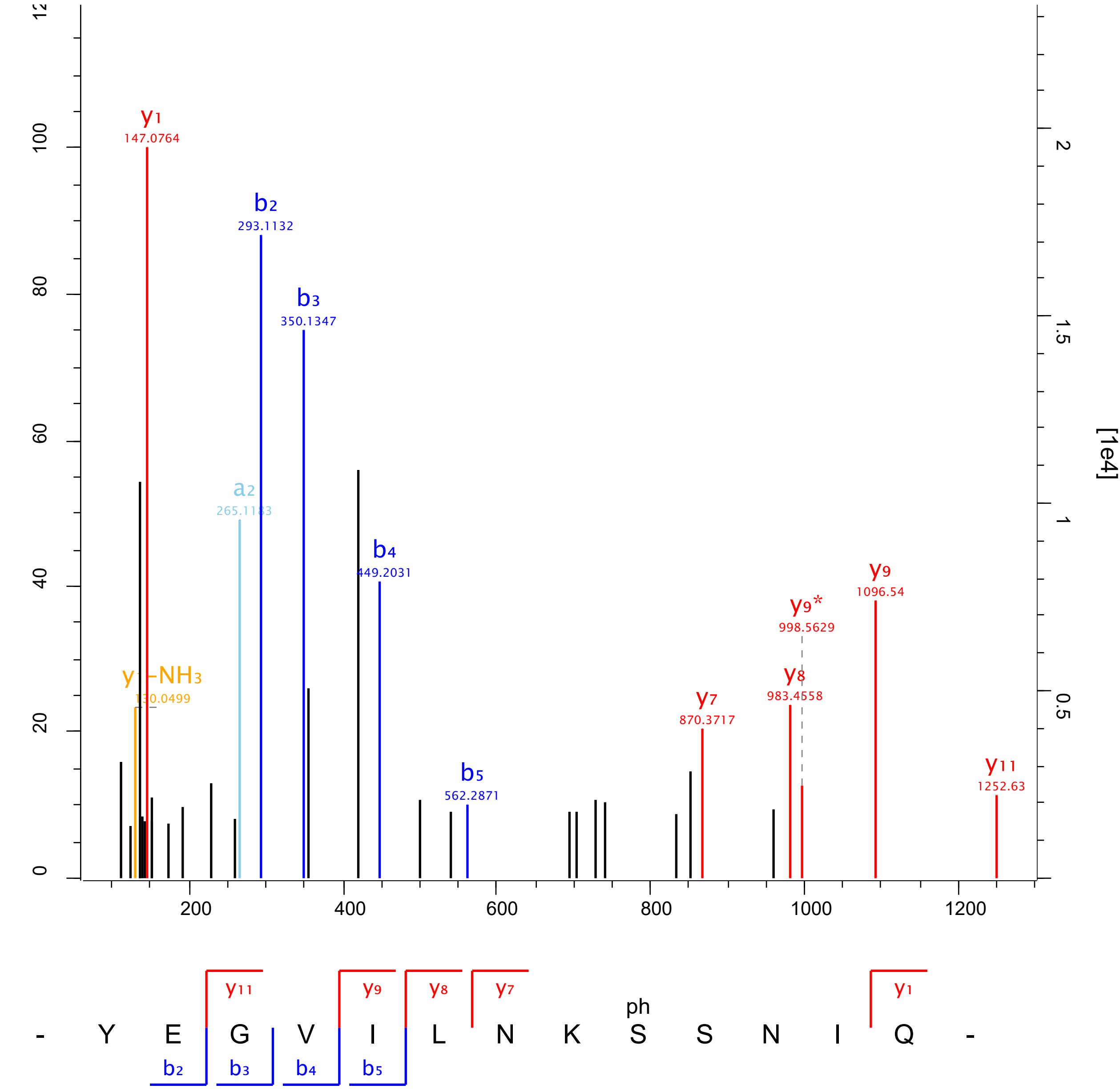

| Raw file | Scan | Method | Score | m/z |
|---|---|---|---|---|
| sys_00_3short | 19224 | FTMS; HCD | 54.26 | 516.75 |

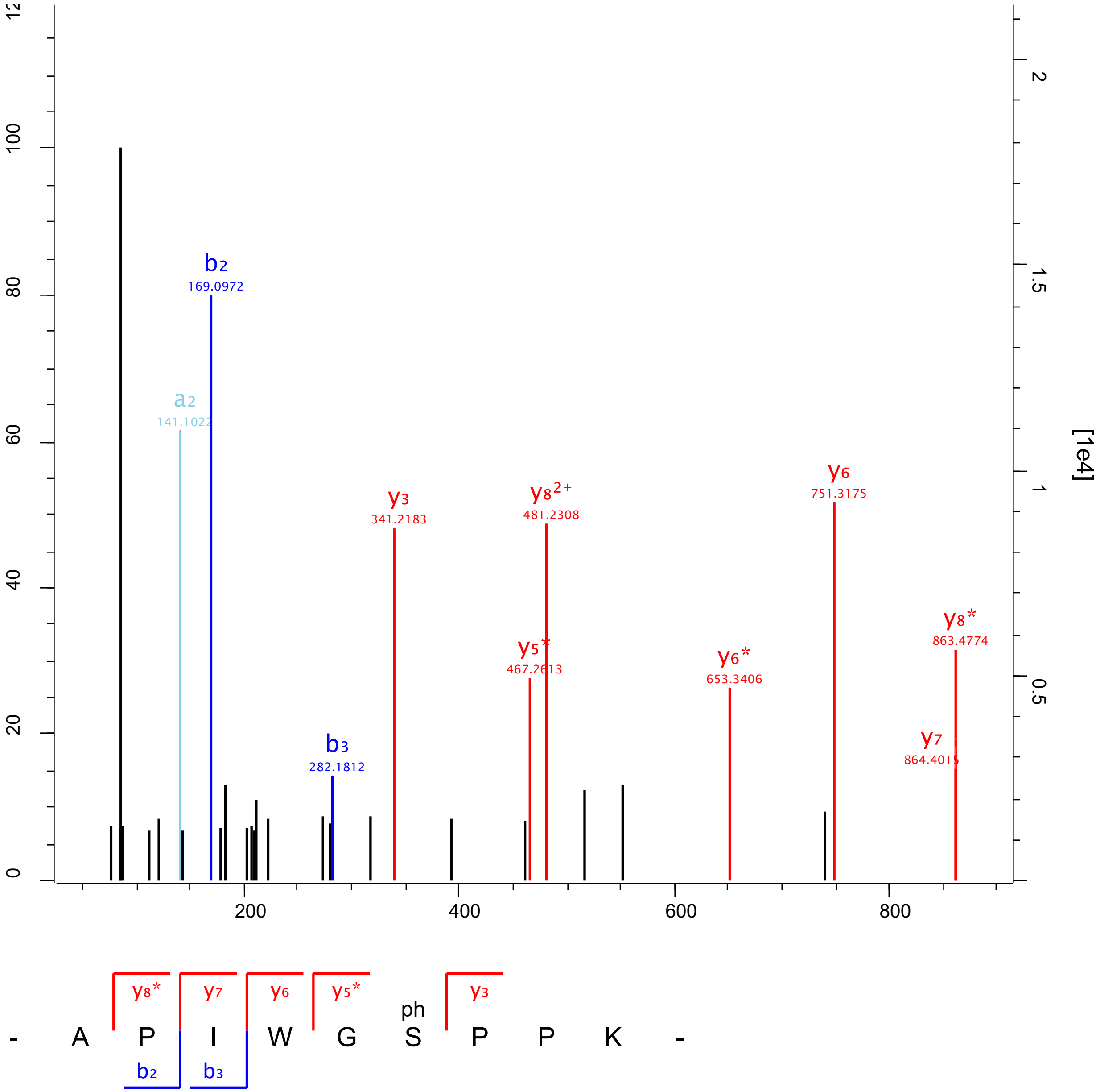

| Raw file | Scan | Method | Score | m/z |
|---|---|---|---|---|
| sys_00_3short | 19356 | FTMS; HCD | 149.42 | 805.36 |

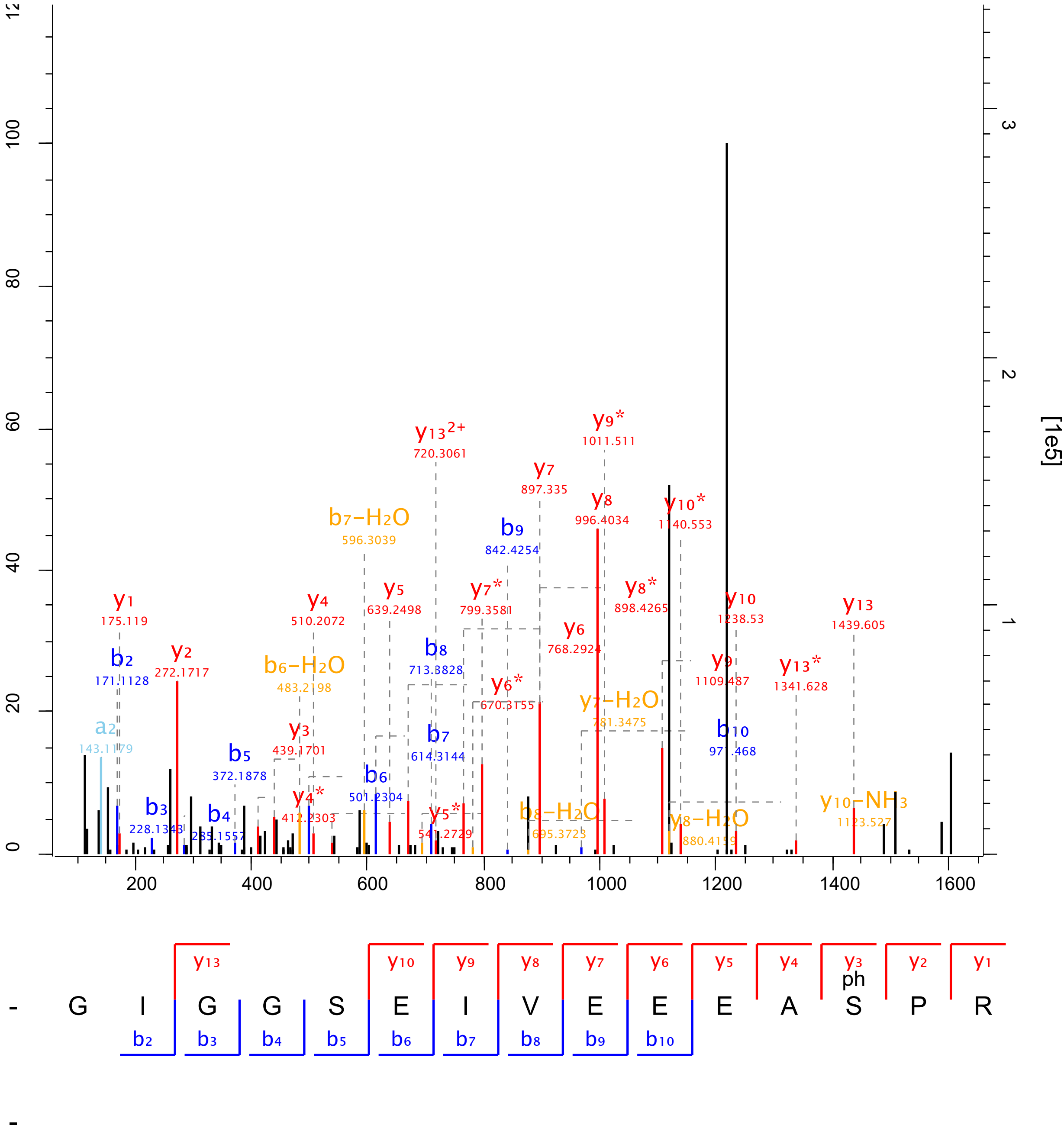

| Raw file | Scan | Method | Score | m/z |
| --- | --- | --- | --- | --- |
| sys_00_3short | 19433 | FTMS; HCD | 75.06 | 966.46 |

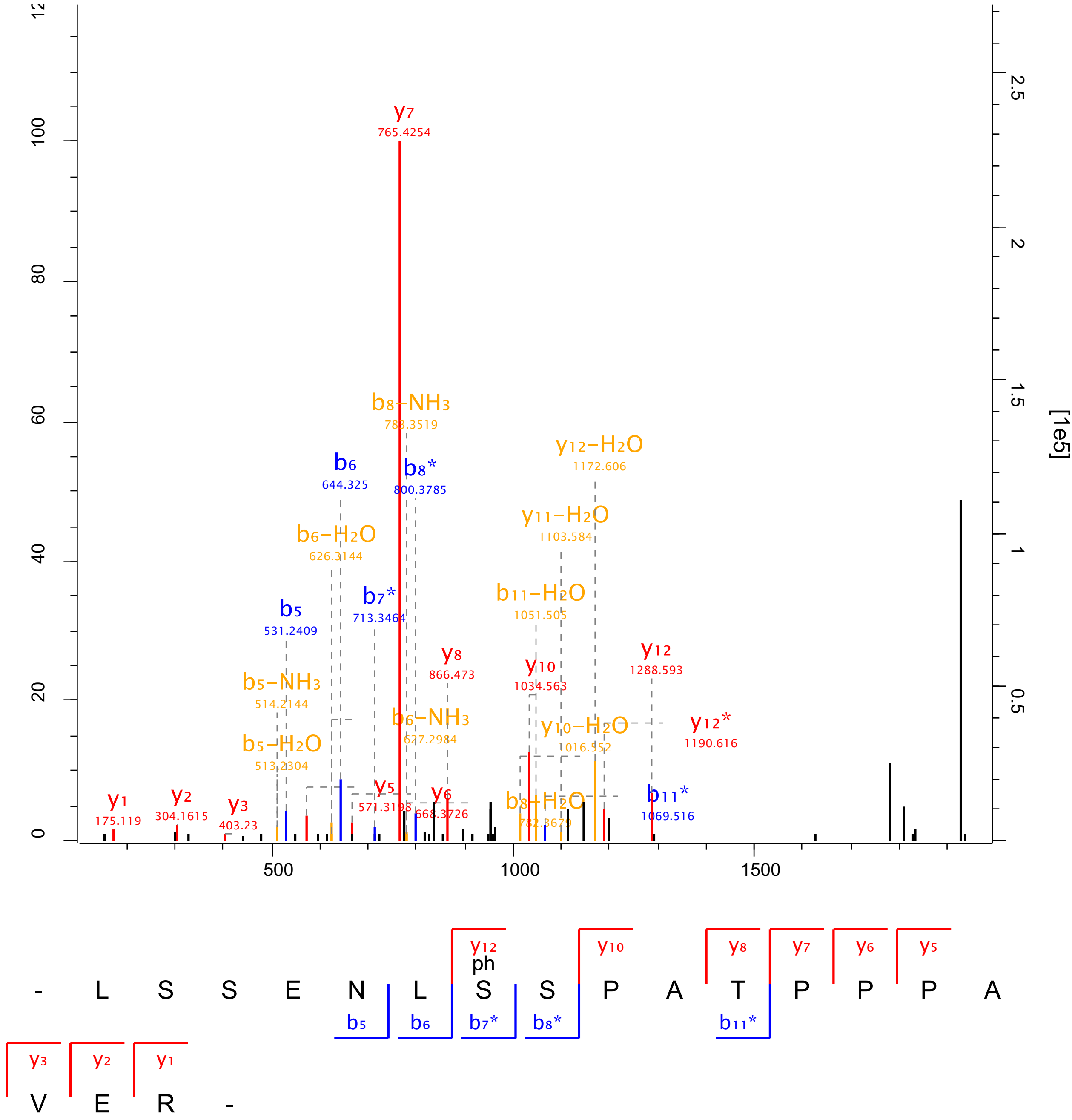

| Raw file | Scan | Method | Score | m/z |
|---|---|---|---|---|
| sys_00_3short | 19525 | FTMS; HCD | 74.84 | 439.72 |

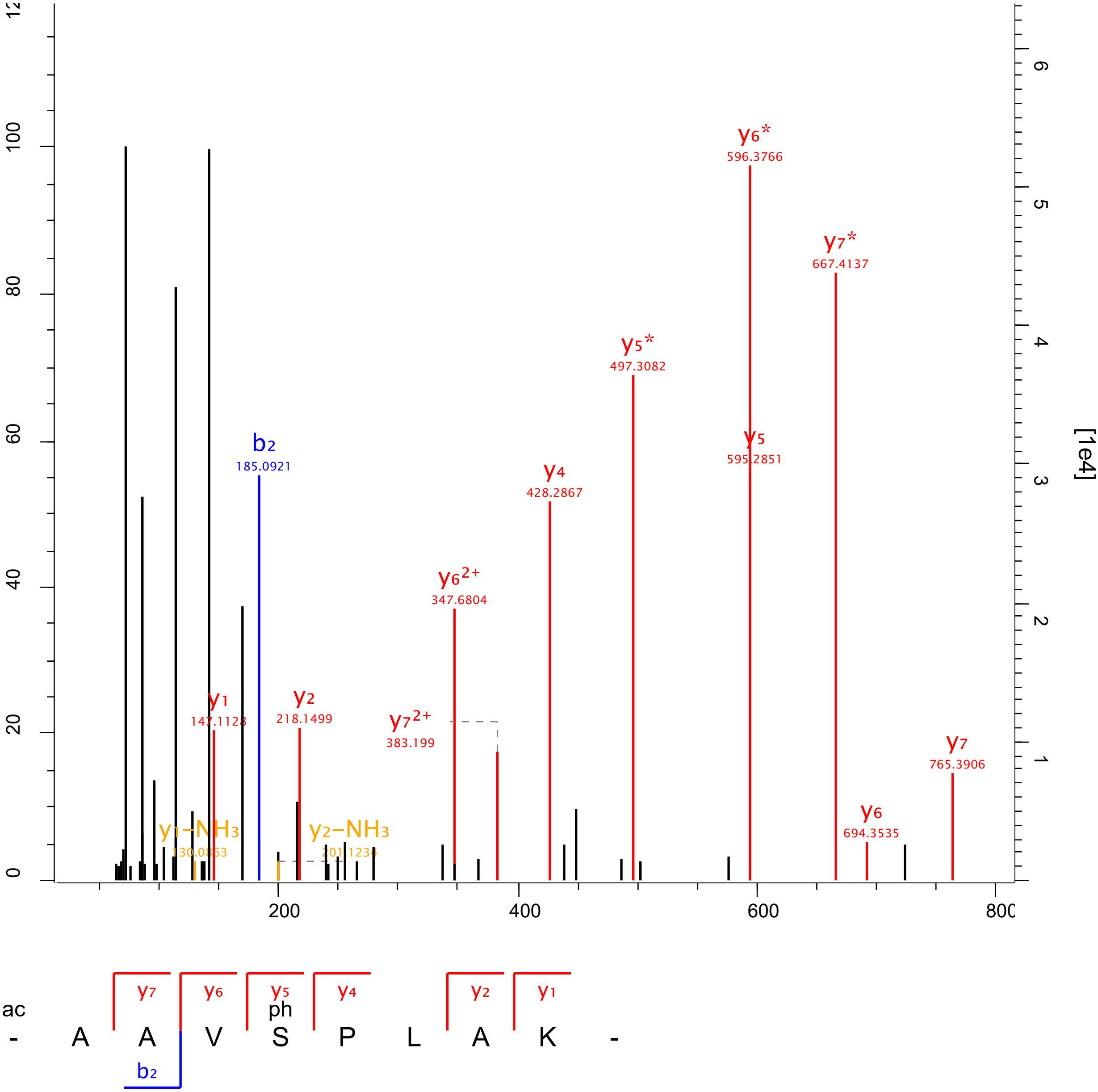

| Raw file | Scan | Method | Score | m/z |
|---|---|---|---|---|
| sys_00_3short | 19584 | FTMS; HCD | 82.64 | 470.23 |

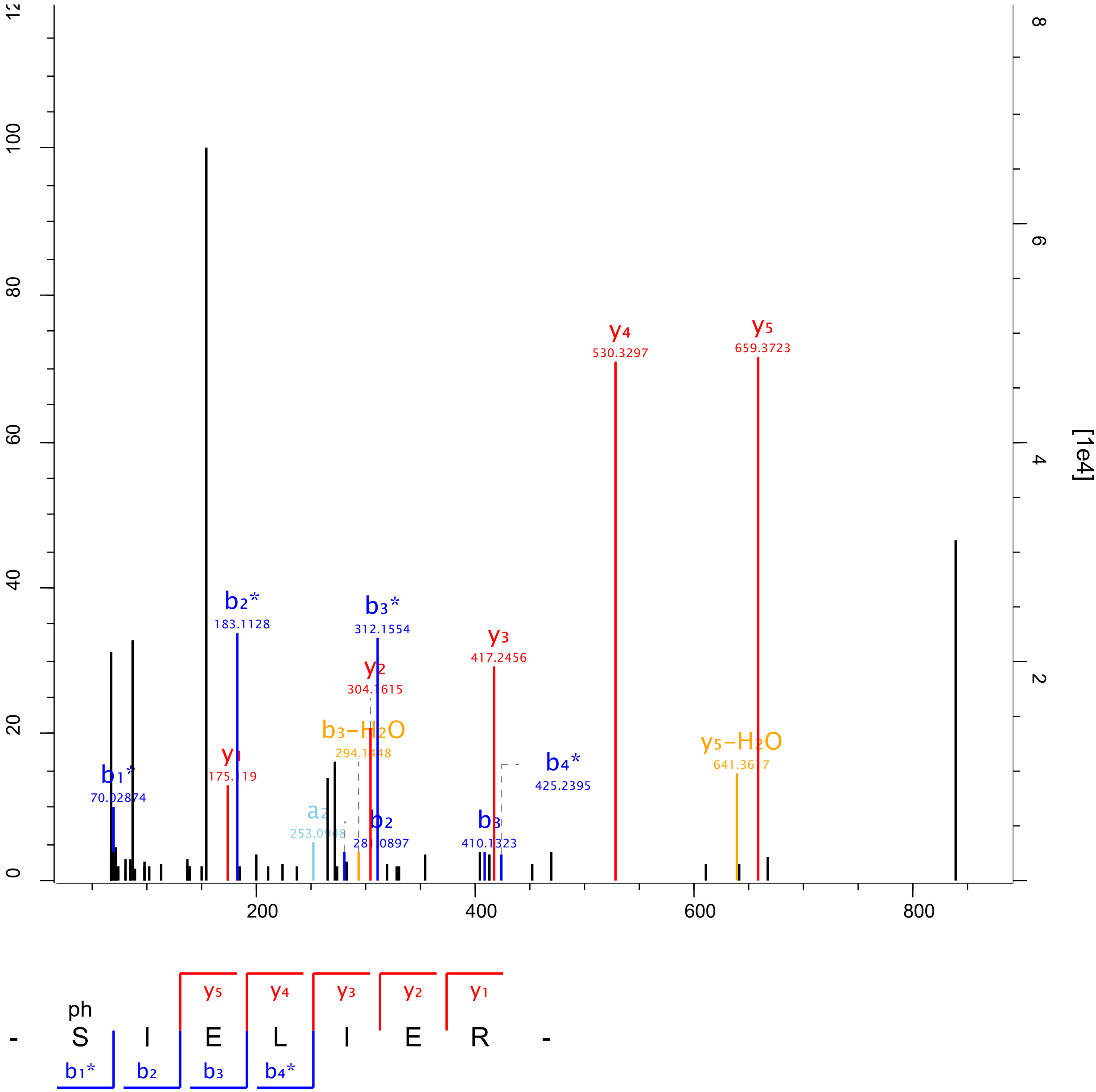

| Raw file | Scan | Method | Score | m/z |
| --- | --- | --- | --- | --- |
| sys_00_3short | 19632 | FTMS; HCD | 125.98 | 734.28 |

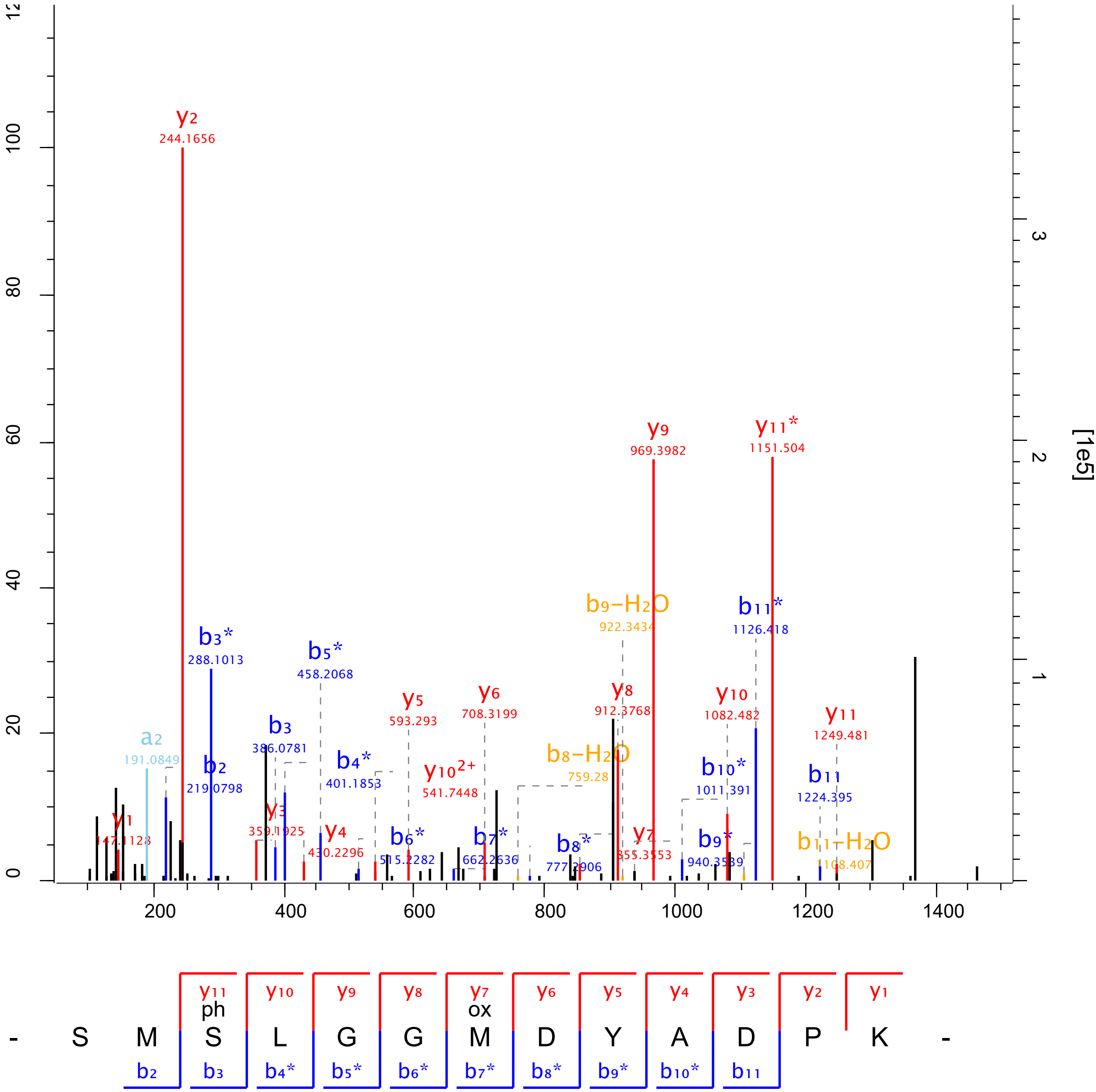

| Raw file | Scan | Method | Score | m/z |
|---|---|---|---|---|
| sys_00_3short | 19679 | FTMS; HCD | 99.41 | 758.34 |

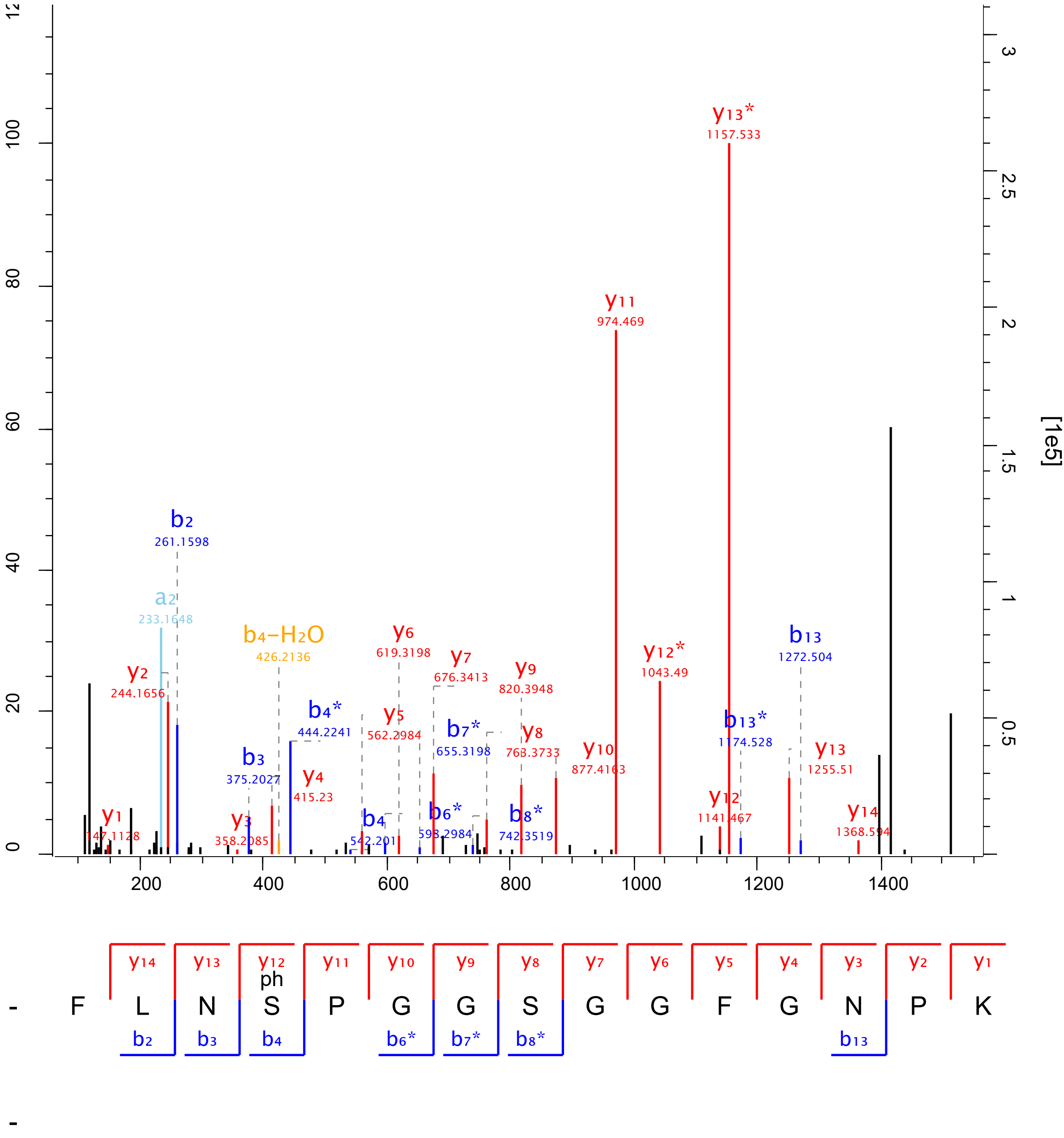

| Raw file | Scan | Method | Score | m/z |
|---|---|---|---|---|
| sys_00_3short | 19768 | FTMS; HCD | 79.84 | 729.32 |

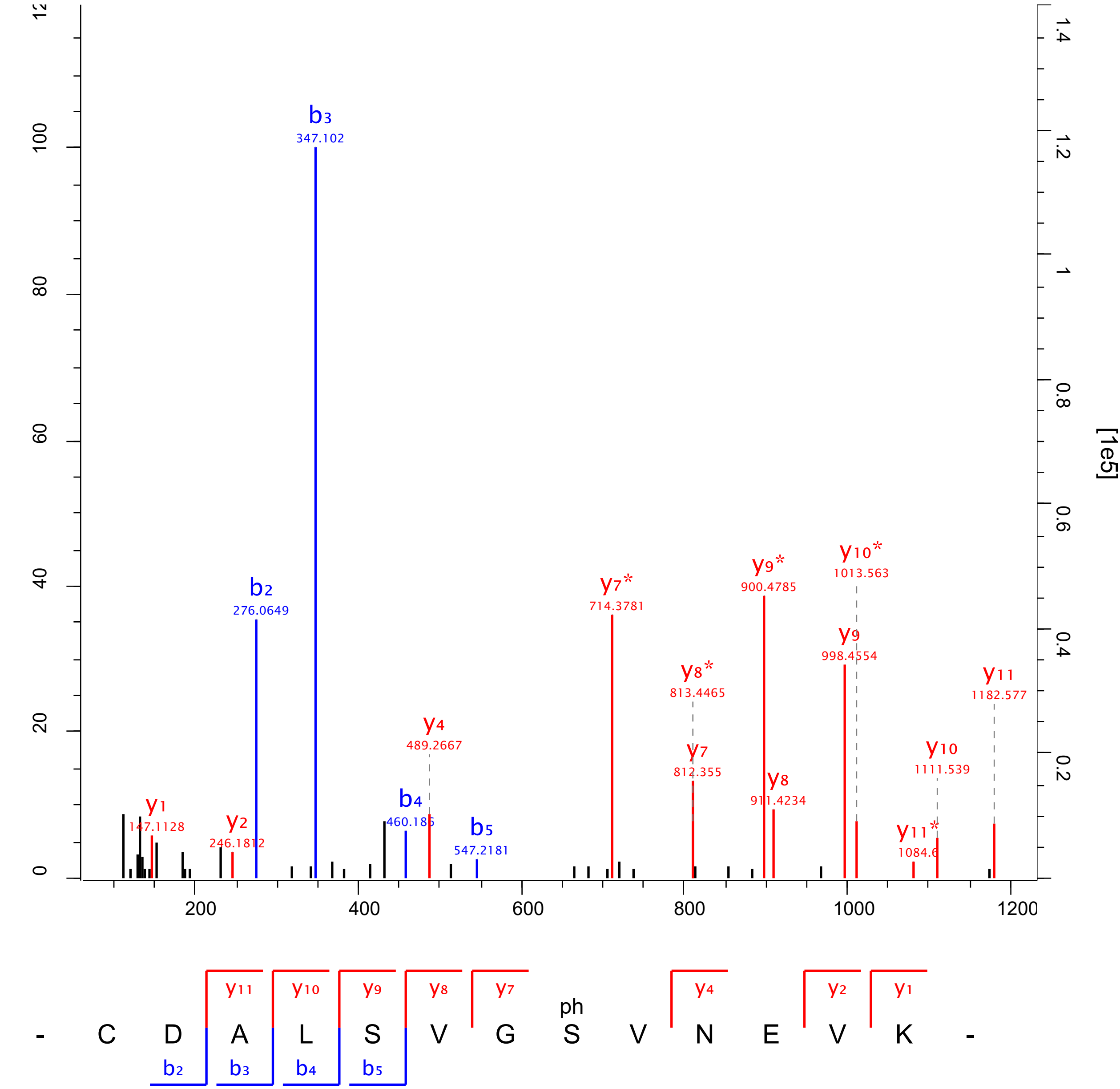

| Raw file | Scan | Method | Score | m/z |
|---|---|---|---|---|
| sys_00_3short | 19769 | FTMS; HCD | 90.61 | 533.77 |

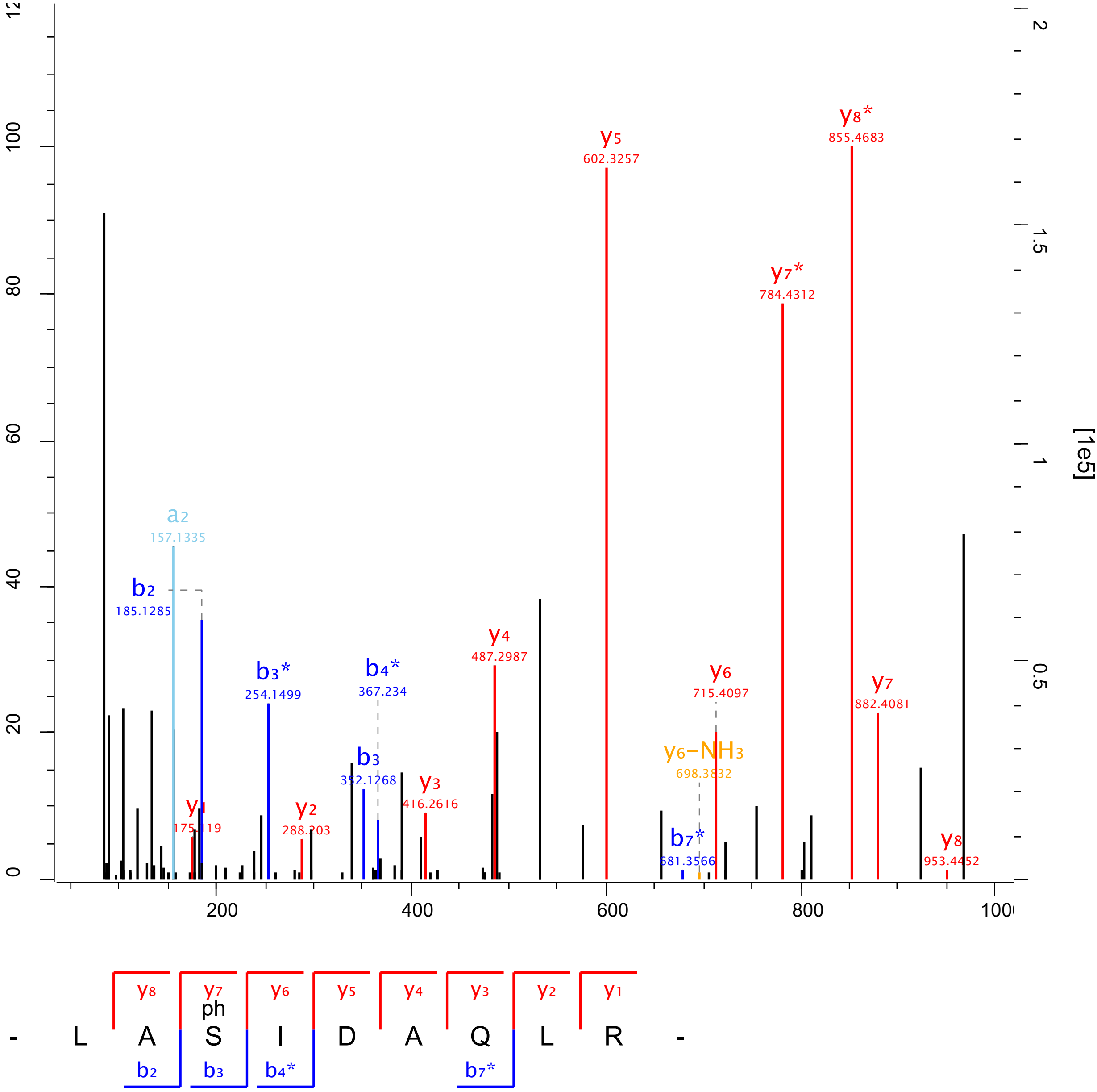

| Raw file | Scan | Method | Score | m/z |
|---|---|---|---|---|
| sys_00_3short | 19783 | FTMS; HCD | 85.52 | 698.79 |

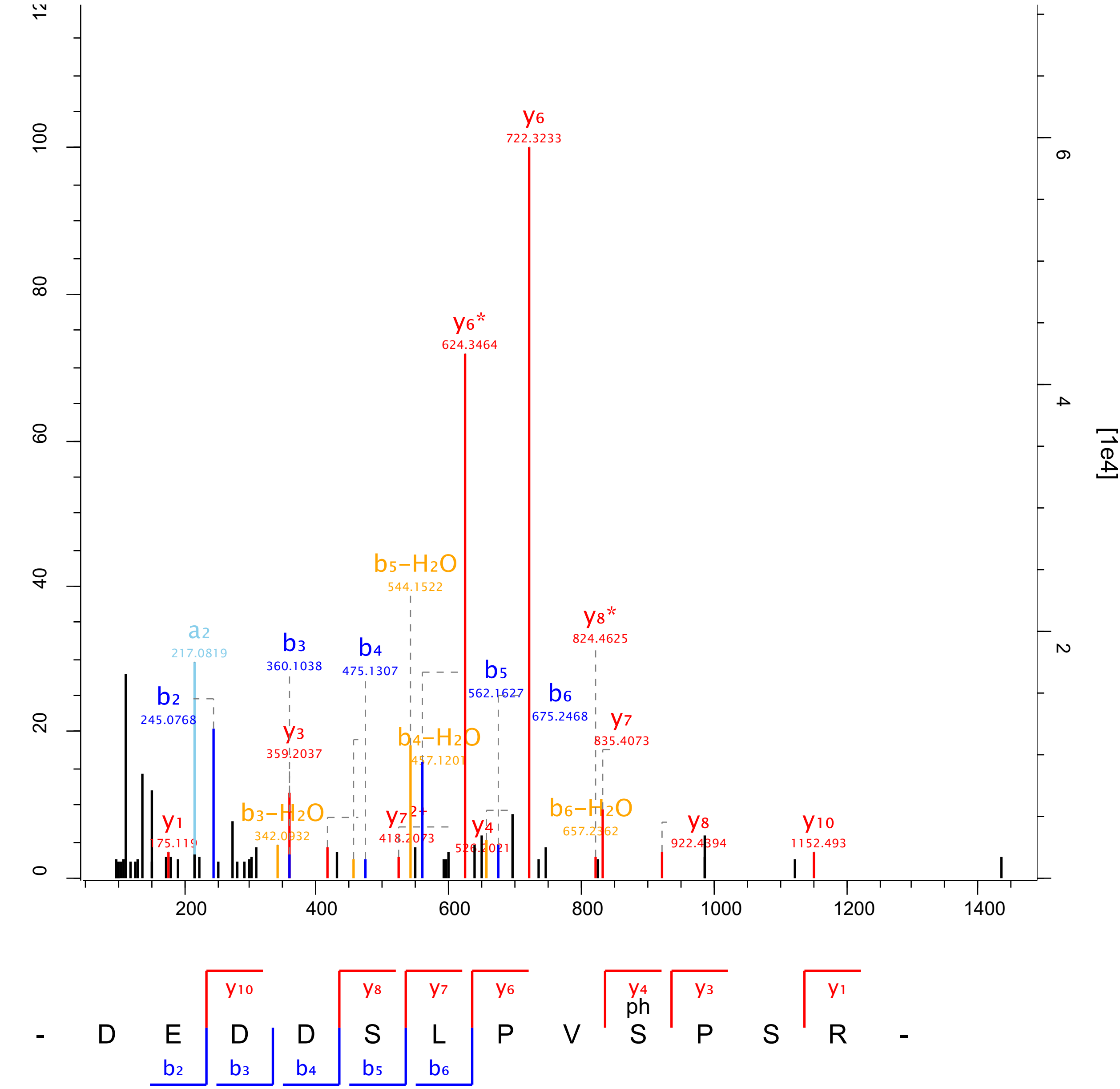

| Raw file | Scan | Method | Score | m/z |
| --- | --- | --- | --- | --- |
| sys_00_3short | 19824 | FTMS; HCD | 93.24 | 706.31 |

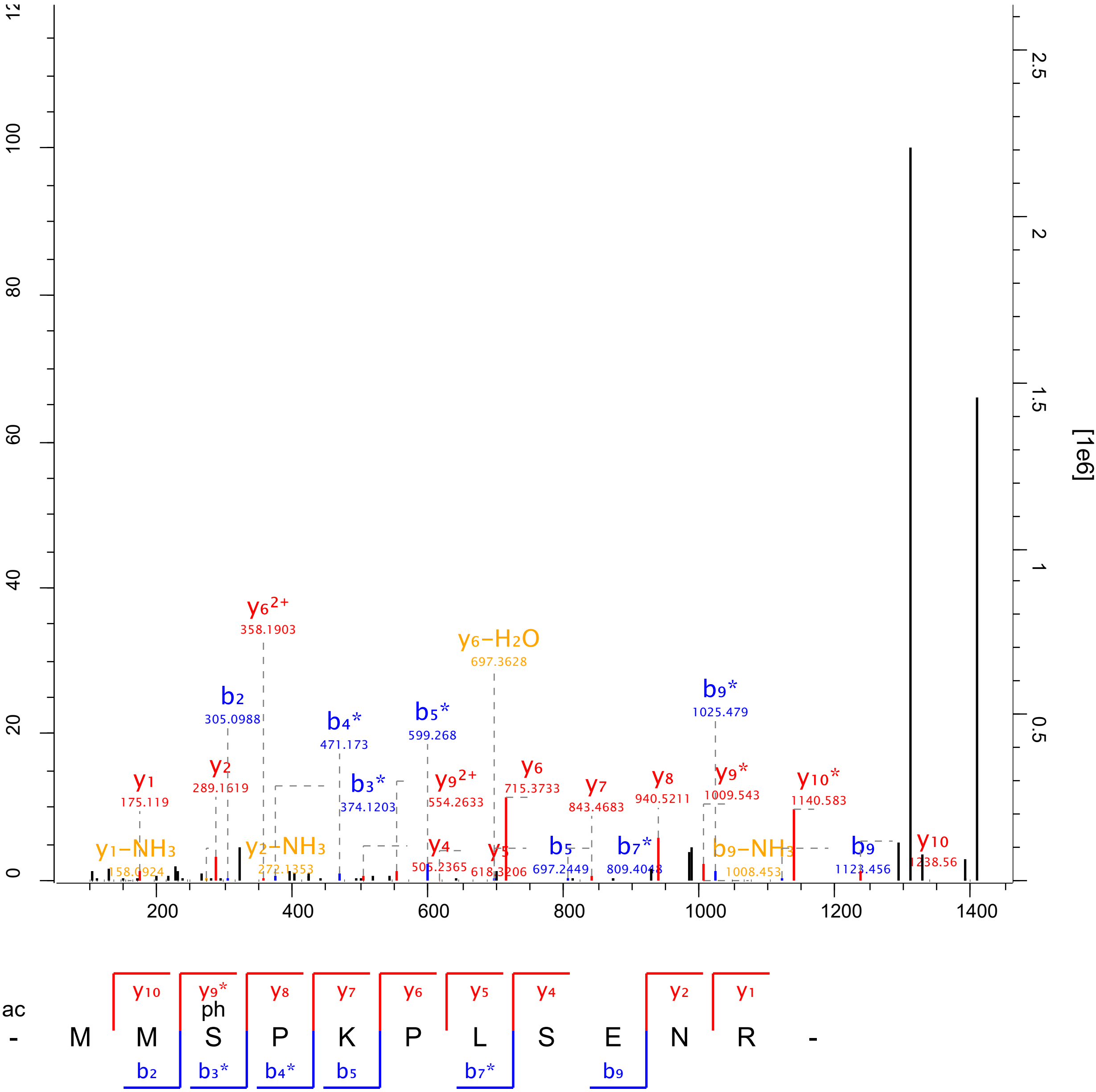

| Raw file | Scan | Method | Score | m/z |
| --- | --- | --- | --- | --- |
| sys_00_3short | 19831 | FTMS; HCD | 97.16 | 475.76 |

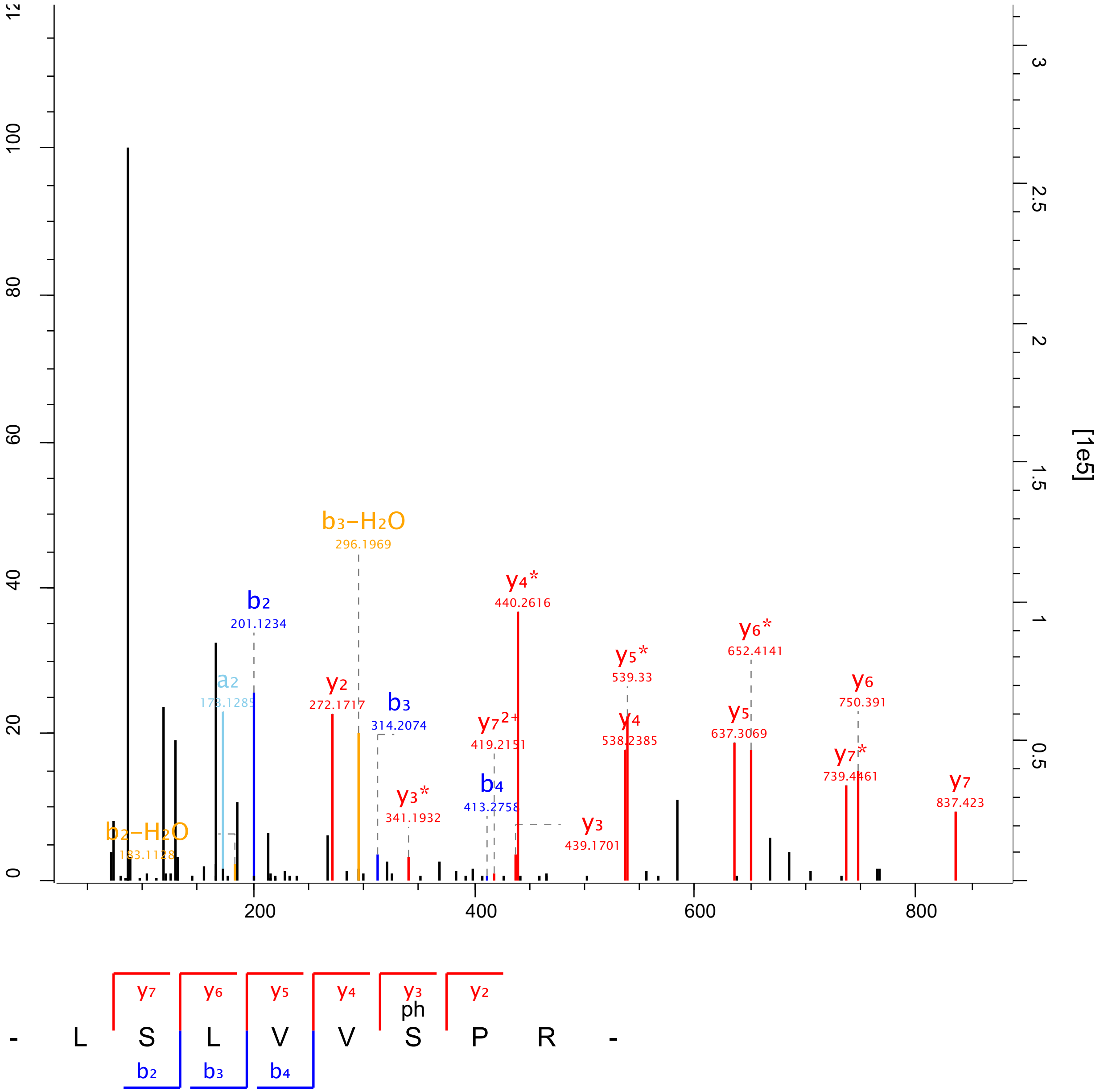

| Raw file | Scan | Method | Score | m/z |
|---|---|---|---|---|
| sys_00_3short | 20043 | FTMS; HCD | 82.01 | 853.86 |

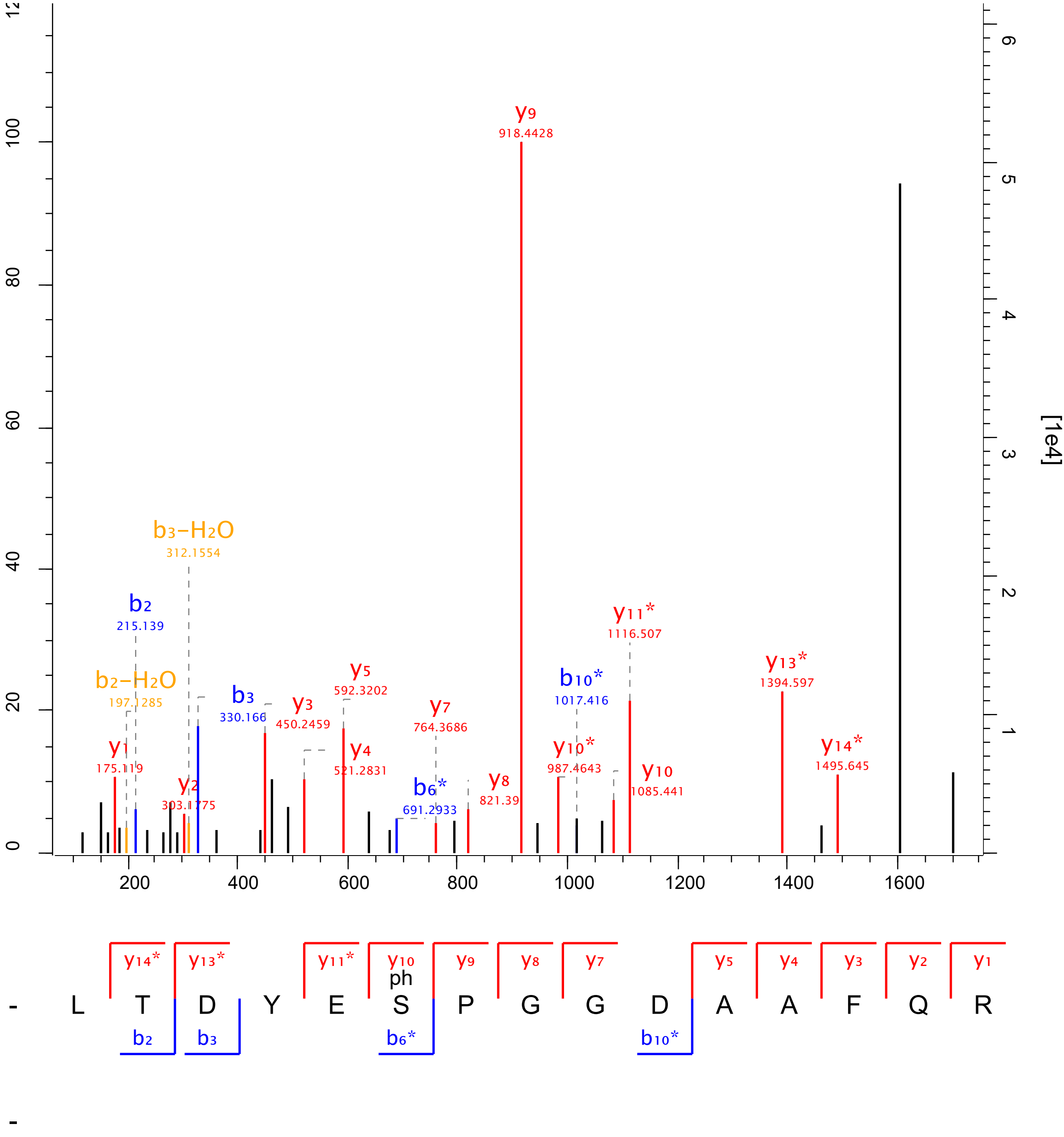

| Raw file | Scan | Method | Score | m/z |
|---|---|---|---|---|
| sys_00_3short | 20186 | FTMS; HCD | 178.71 | 807.36 |

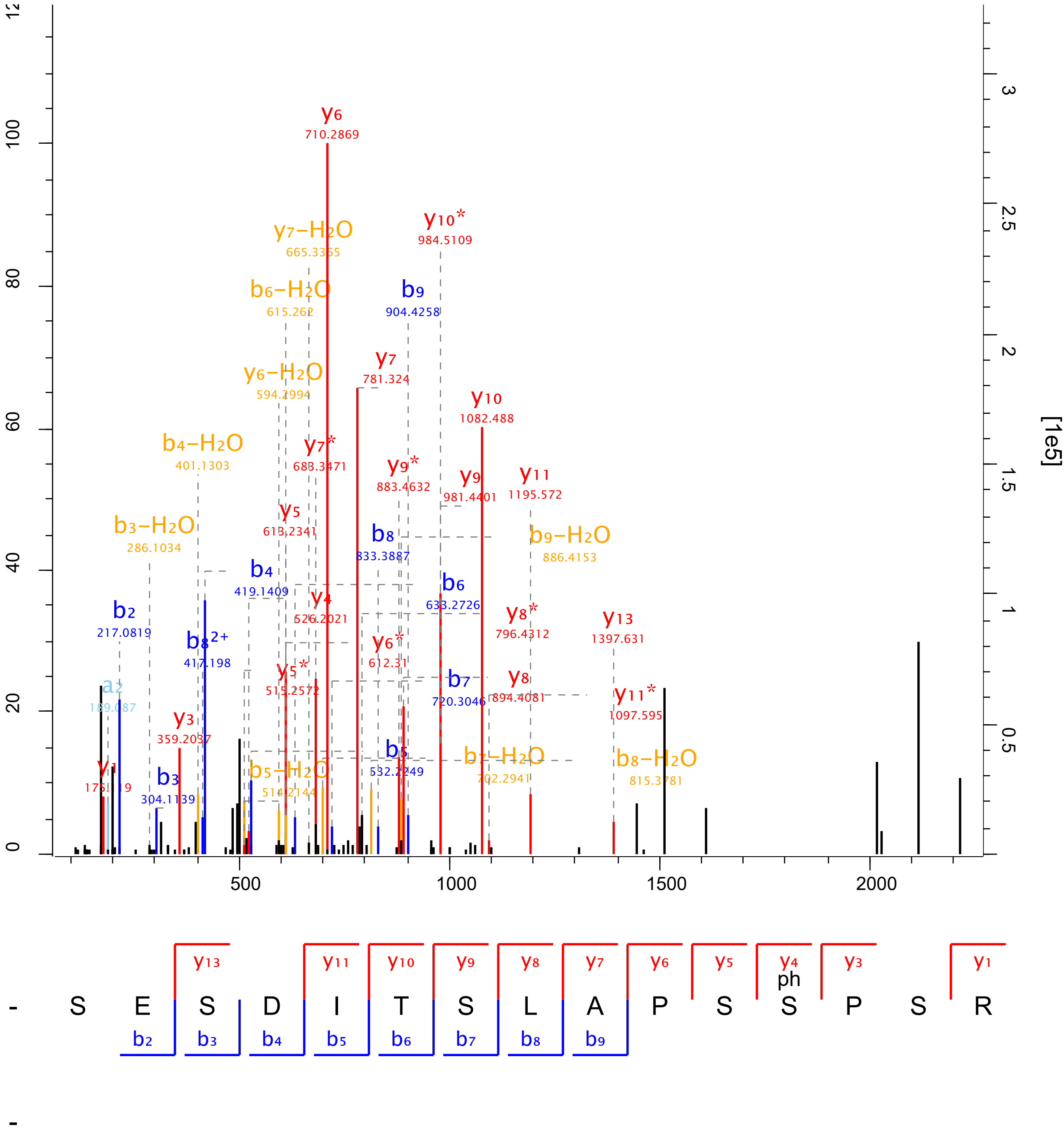

| Raw file | Scan | Method | Score | m/z |
|---|---|---|---|---|
| sys_00_3short | 20462 | FTMS; HCD | 72.9 | 646.24 |

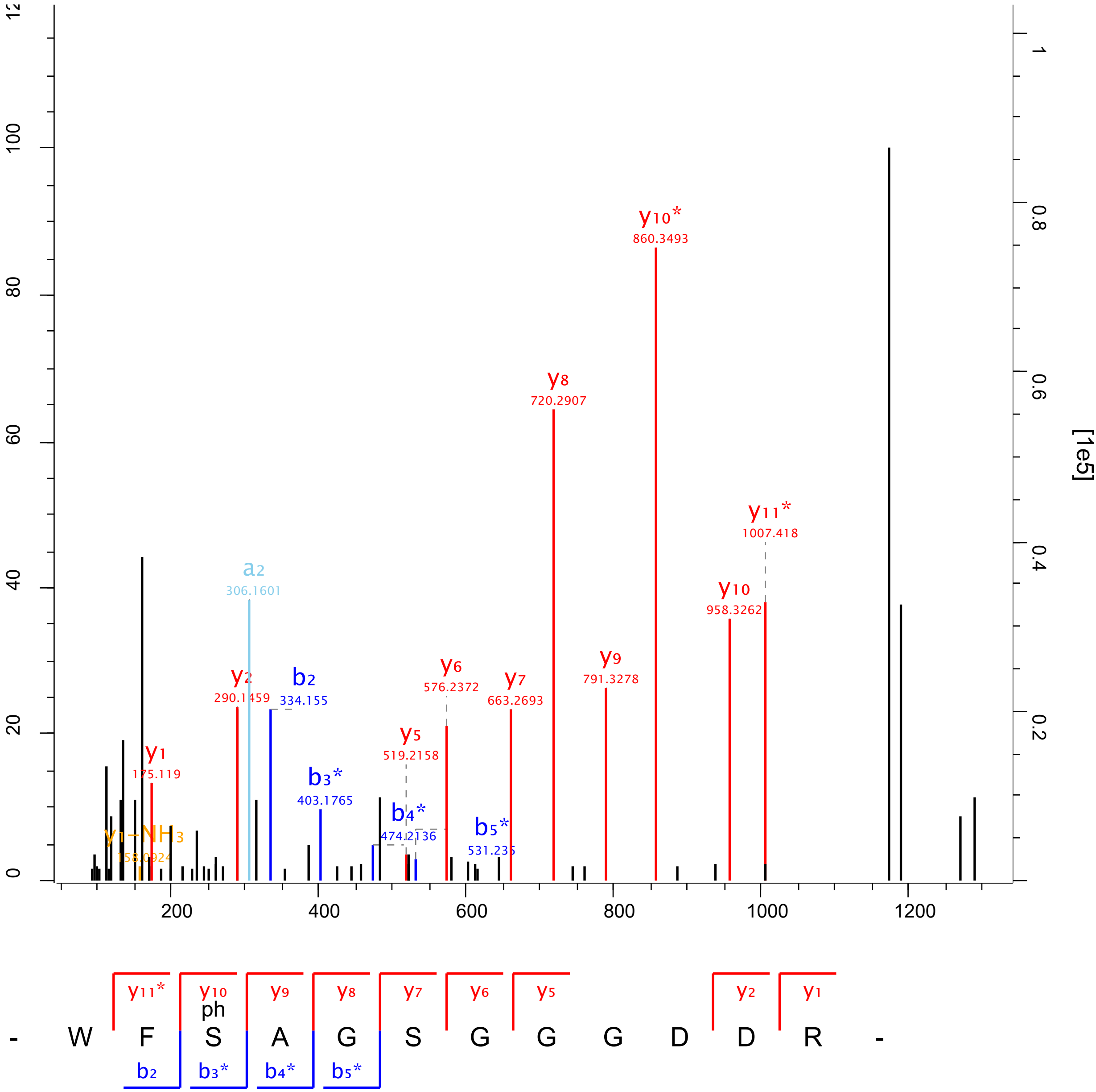

| Raw file | Scan | Method | Score | m/z |
|---|---|---|---|---|
| sys_00_3short | 20627 | FTMS; HCD | 66.27 | 529.24 |

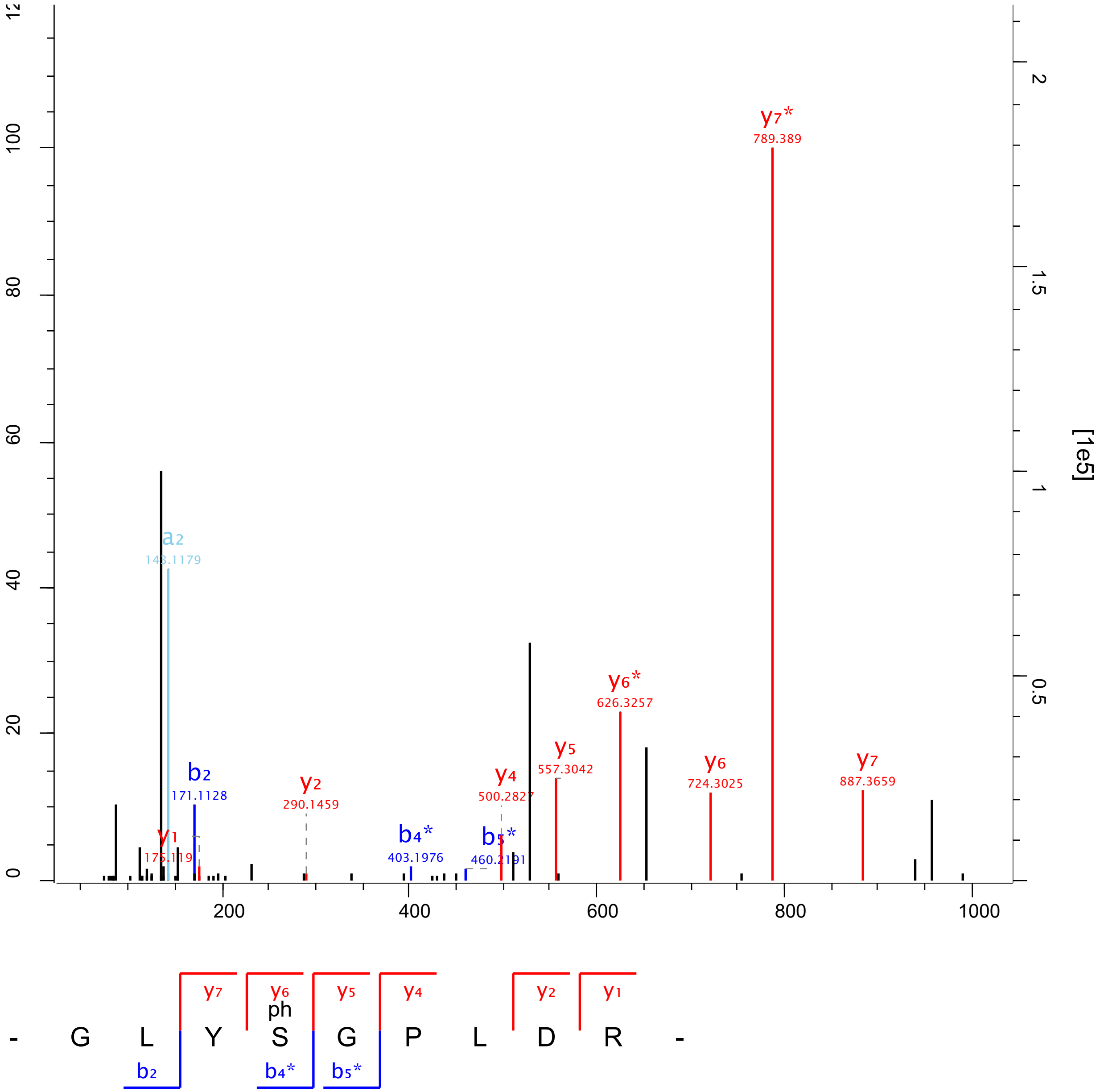

| Raw file | Scan | Method | Score | m/z |
|---|---|---|---|---|
| sys_00_3short | 20689 | FTMS; HCD | 70.47 | 675.29 |

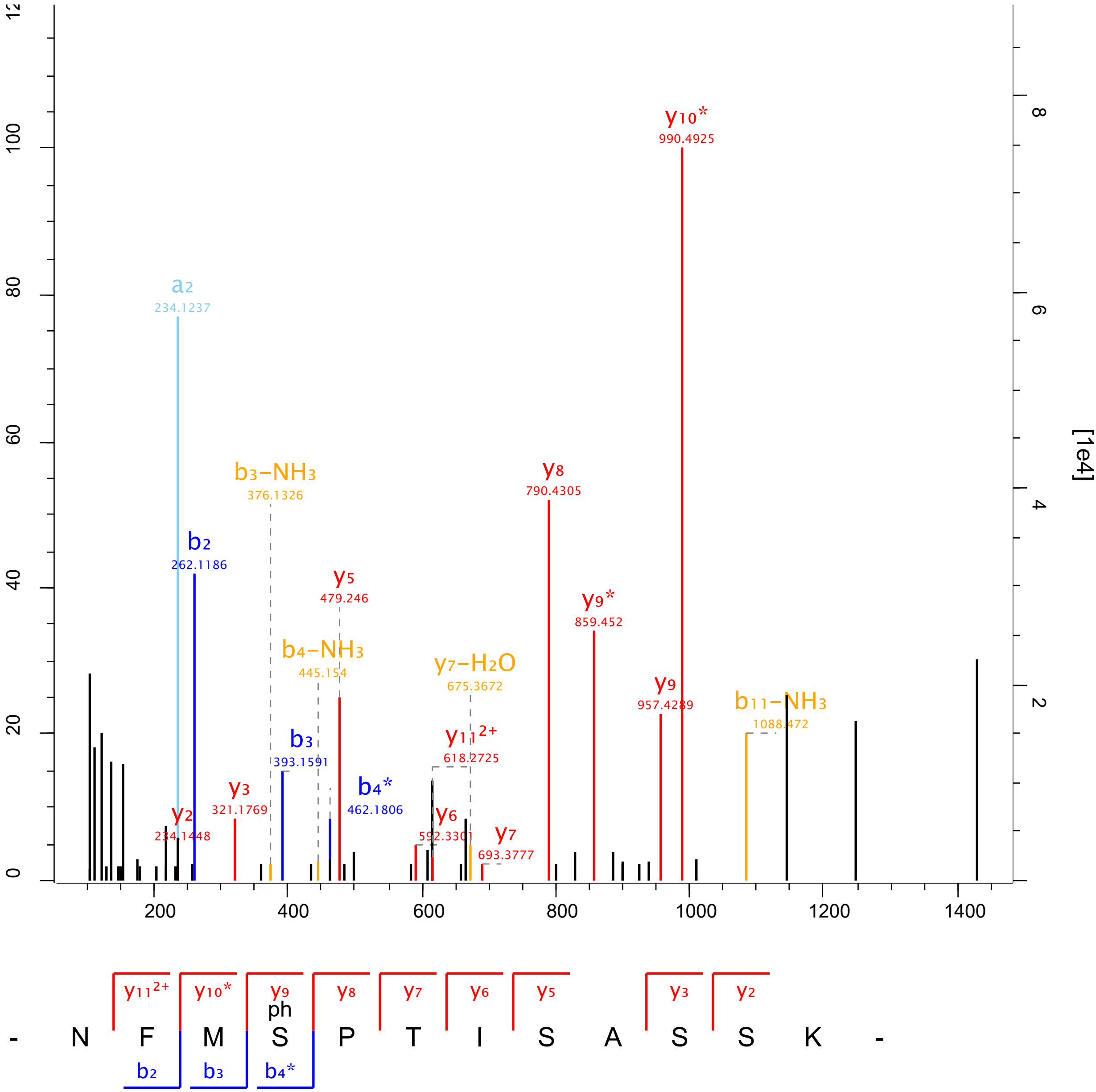

| Raw file | Scan | Method | Score | m/z |
|---|---|---|---|---|
| sys_00_3short | 20716 | FTMS; HCD | 131.26 | 709.3 |

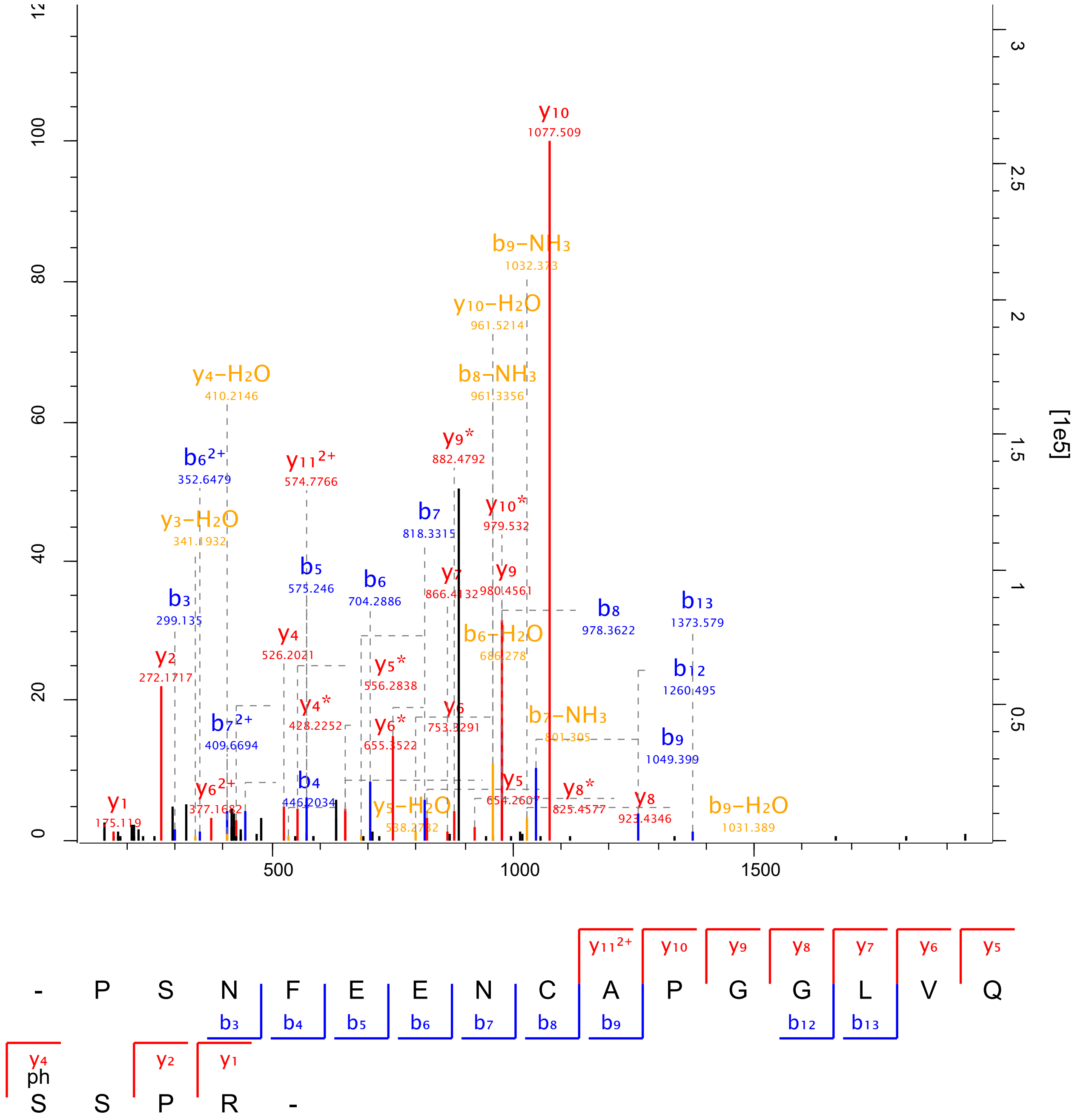

| Raw file | Scan | Method | Score | m/z |
|---|---|---|---|---|
| sys_00_3short | 20790 | FTMS; HCD | 80.24 | 572.76 |

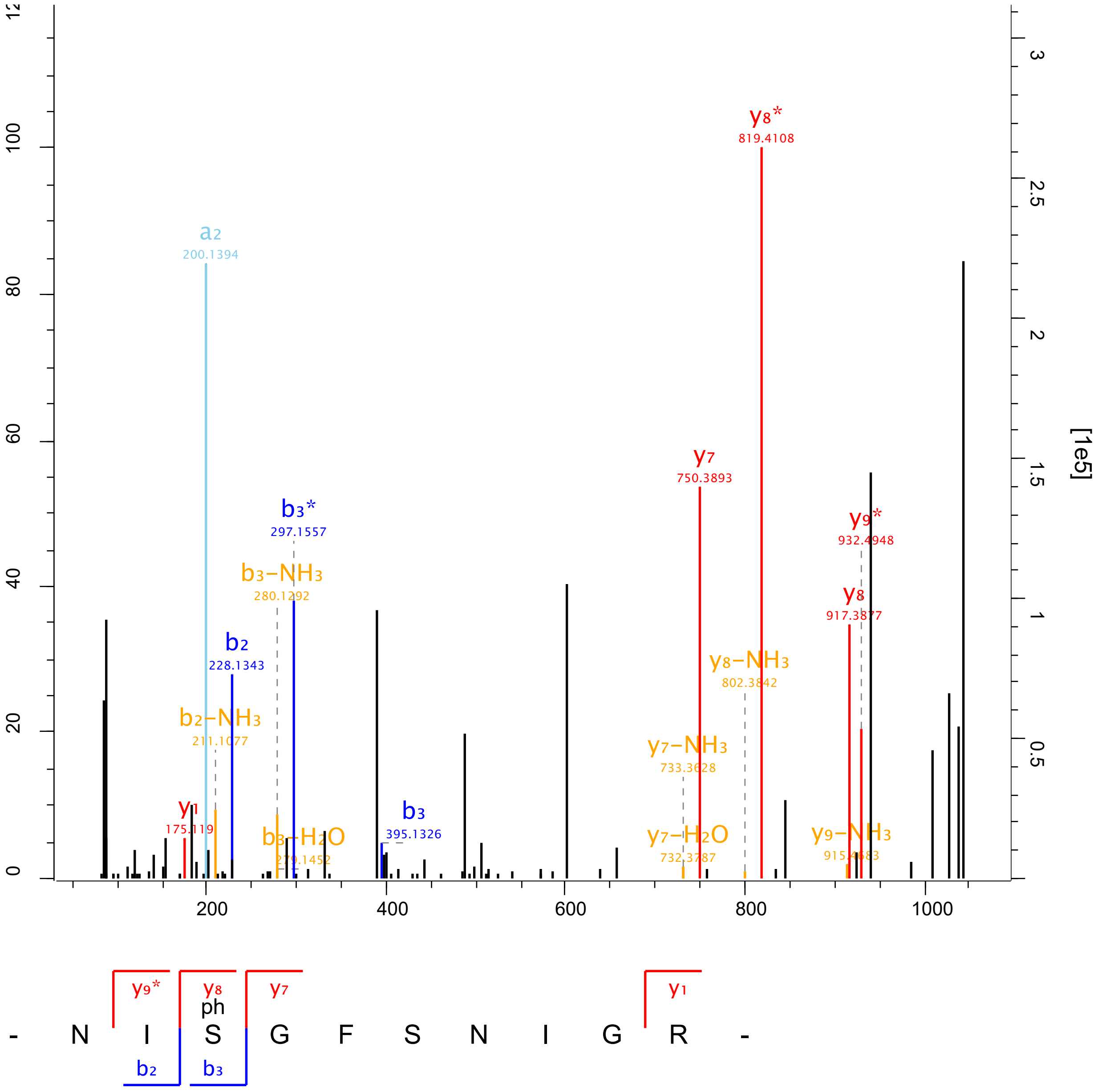

| Raw file | Scan | Method | Score | m/z |
|---|---|---|---|---|
| sys_00_3short | 20825 | FTMS; HCD | 123.14 | 754.8 |

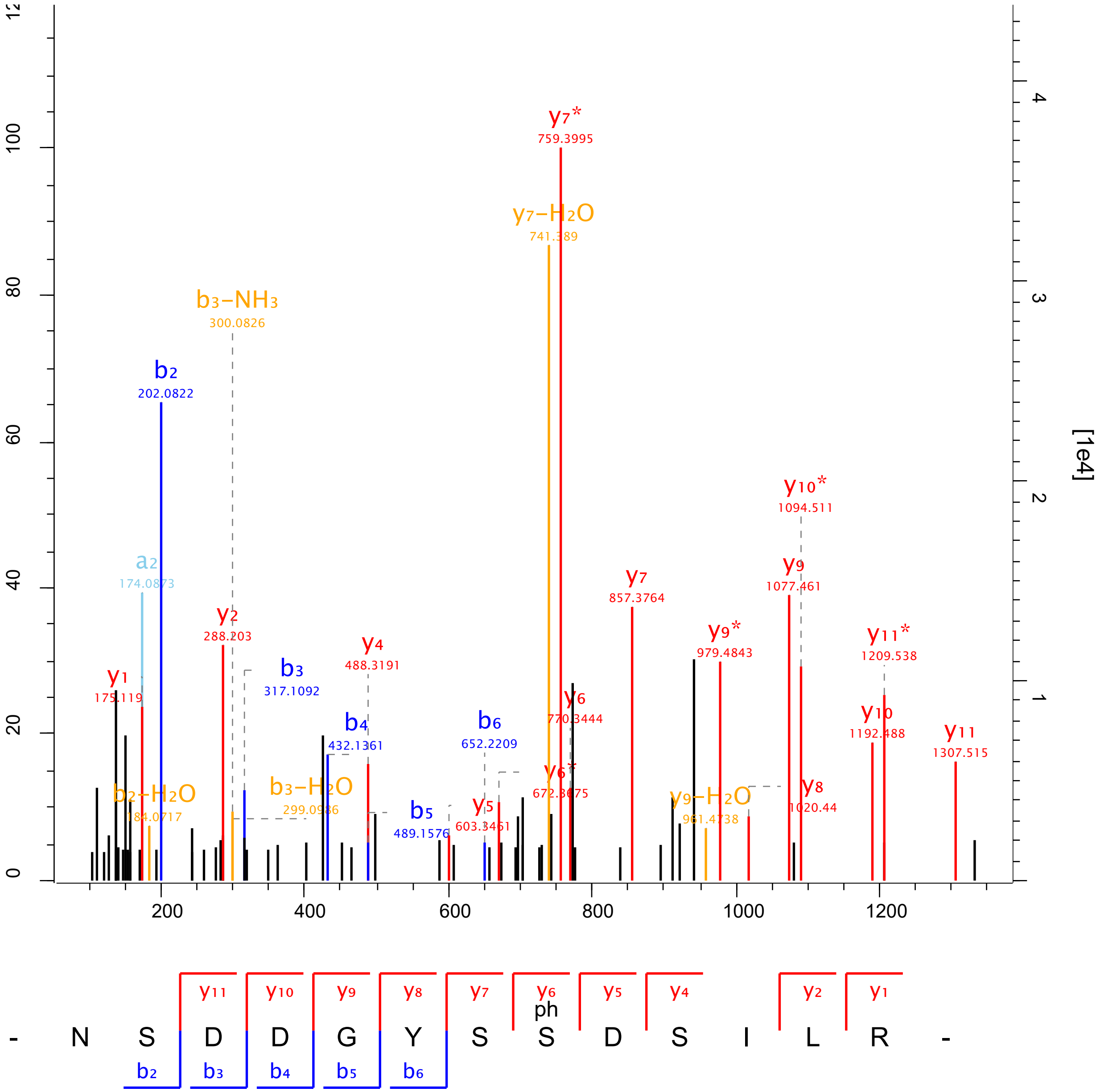

| Raw file | Scan | Method | Score | m/z |
|---|---|---|---|---|
| sys_00_3short | 21140 | FTMS; HCD | 71.15 | 543.27 |

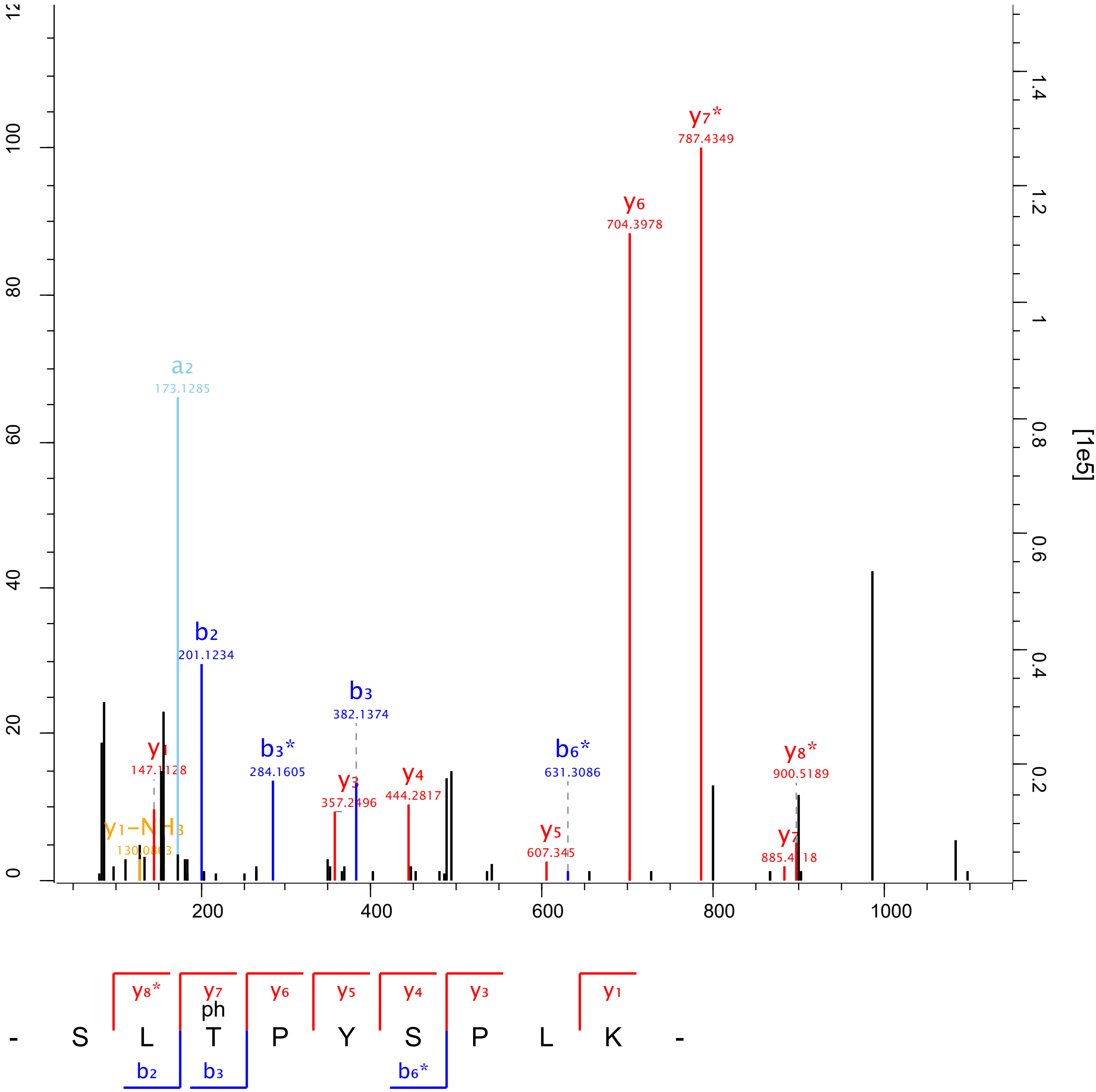

| Raw file | Scan | Method | Score | m/z |
|---|---|---|---|---|
| sys_00_3short | 21501 | FTMS; HCD | 109.72 | 869.86 |

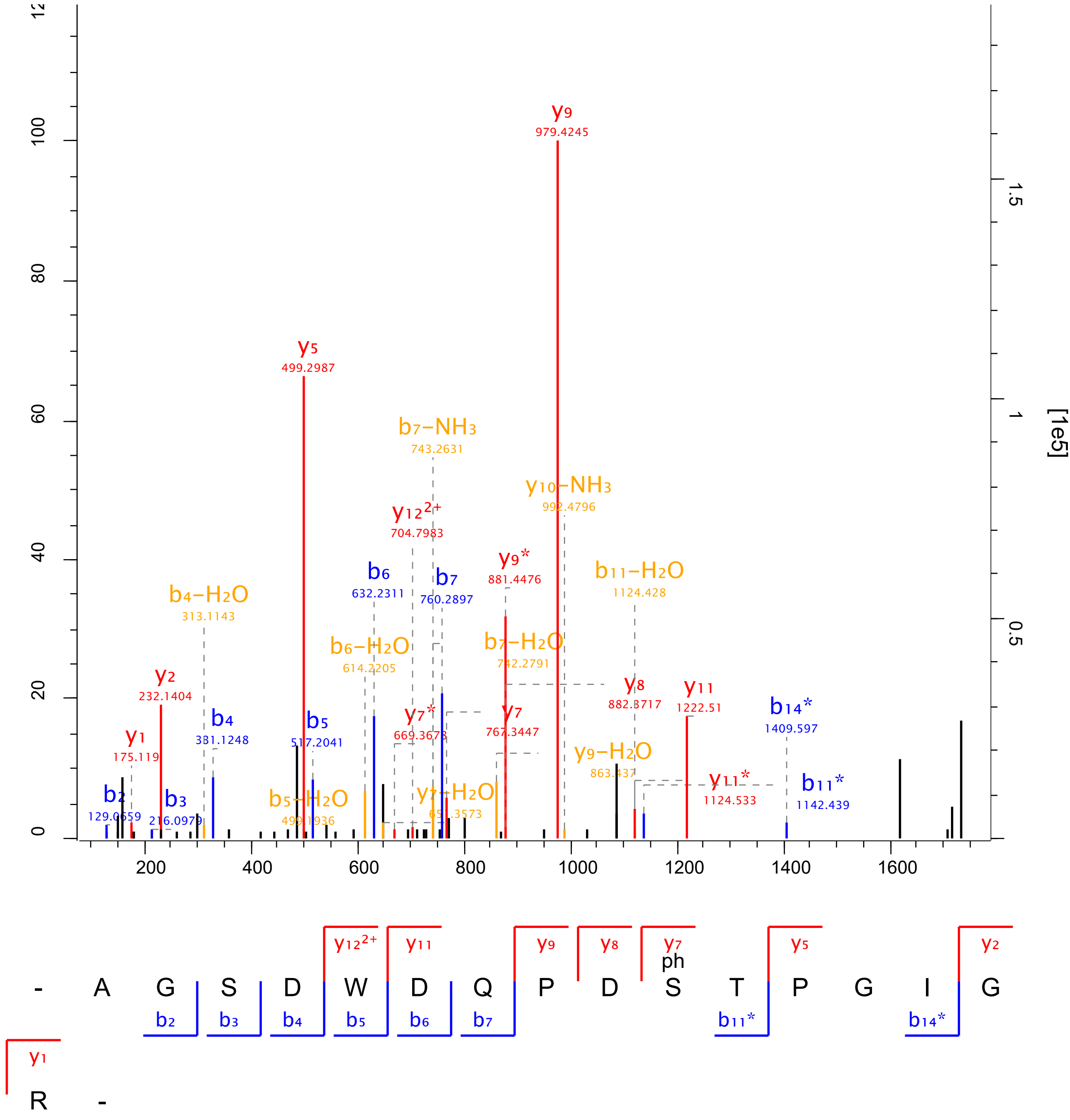

| Raw file | Scan | Method | Score | m/z |
|---|---|---|---|---|
| sys_00_3short | 21749 | FTMS; HCD | 49.59 | 732.32 |

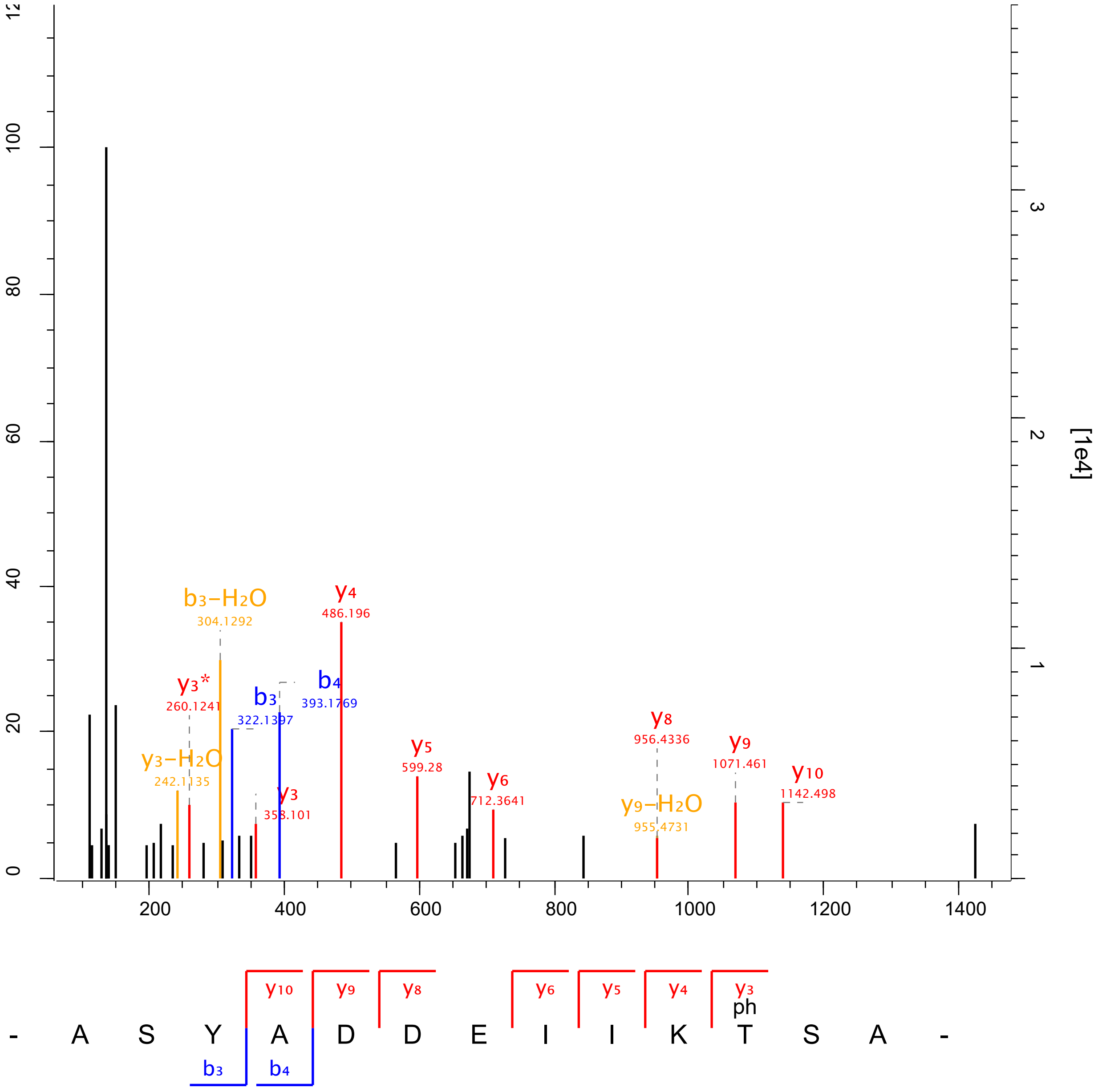

| Raw file | Scan | Method | Score | m/z |
|---|---|---|---|---|
| sys_00_3short | 21801 | FTMS; HCD | 73.44 | 481.26 |

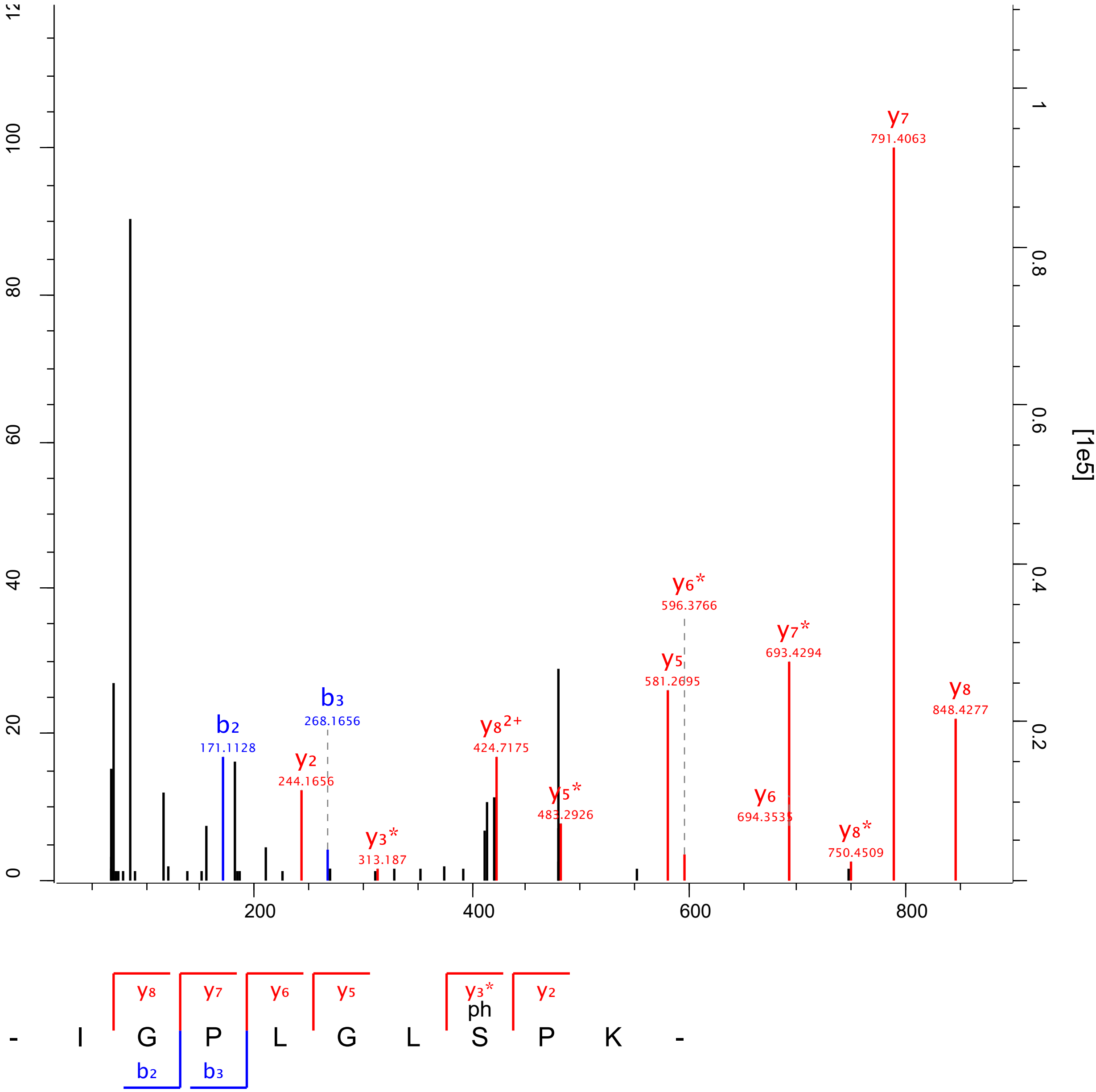

| Raw file | Scan | Method | Score | m/z |
|---|---|---|---|---|
| sys_00_3short | 21929 | FTMS; HCD | 203.2 | 691.79 |

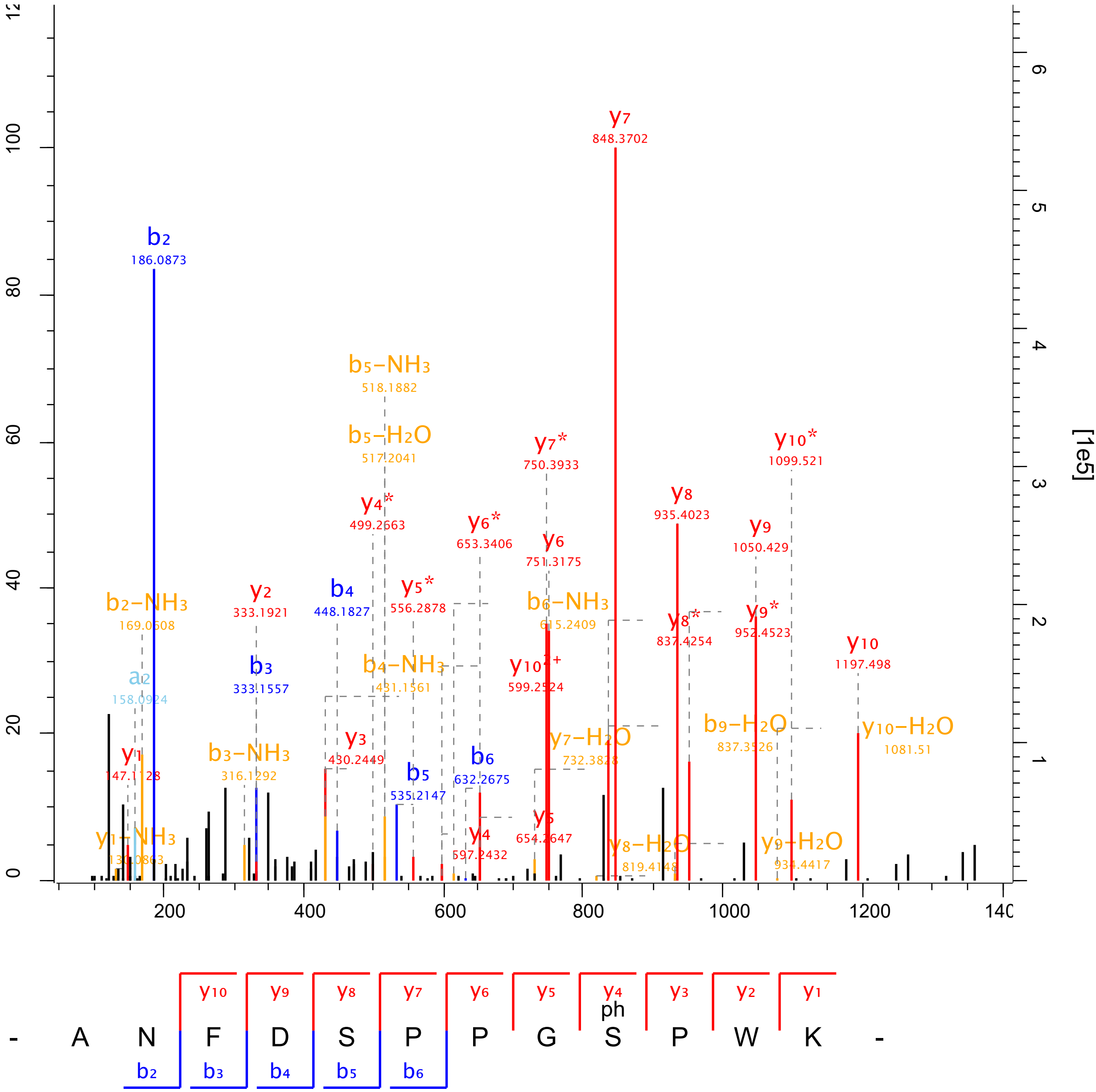

| Raw file | Scan | Method | Score | m/z |
|---|---|---|---|---|
| sys_00_3short | 21997 | FTMS; HCD | 42.31 | 667.3 |

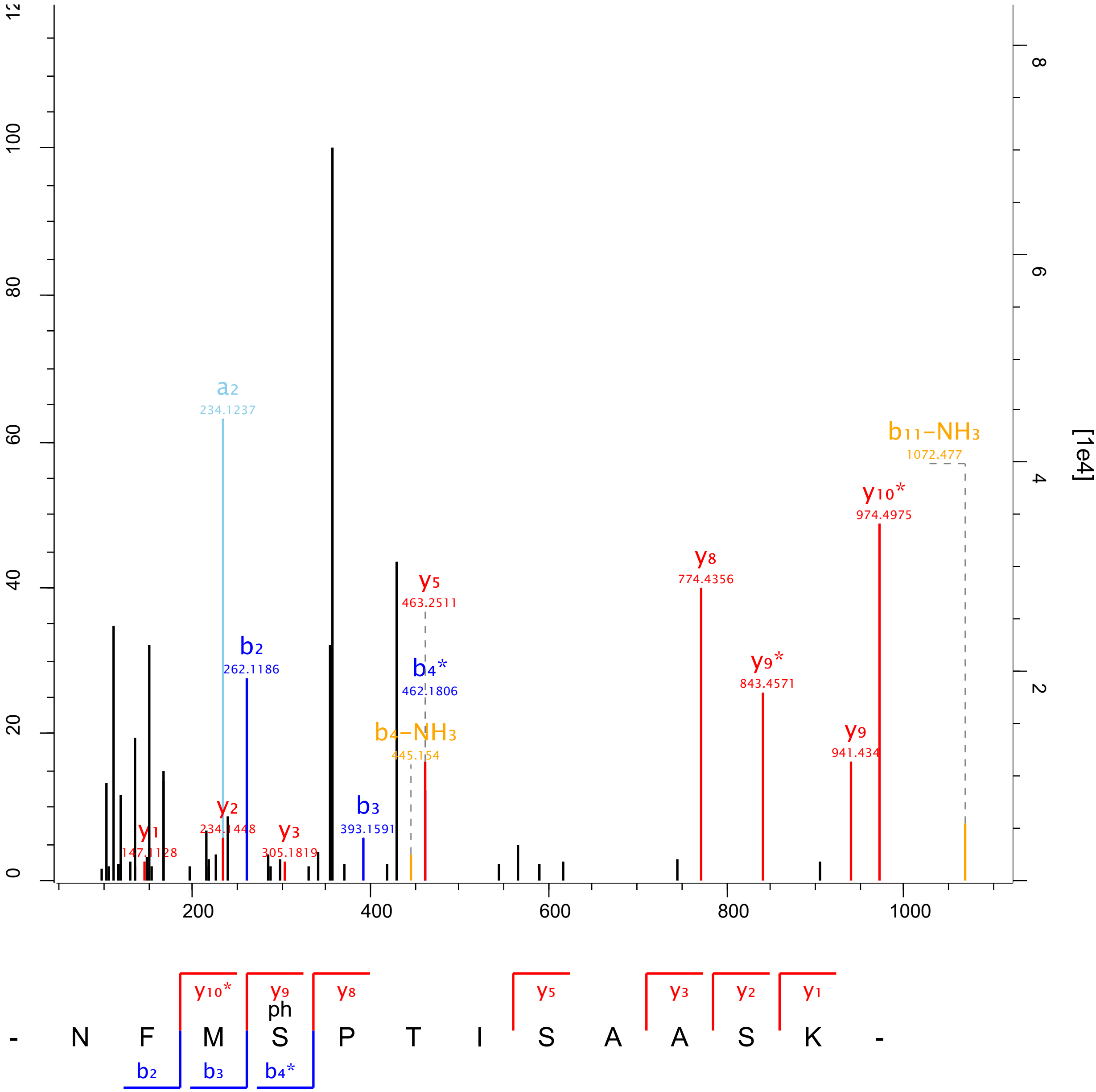

| Raw file | Scan | Method | Score | m/z |
|---|---|---|---|---|
| sys_00_3short | 22149 | FTMS; HCD | 63.69 | 732.82 |

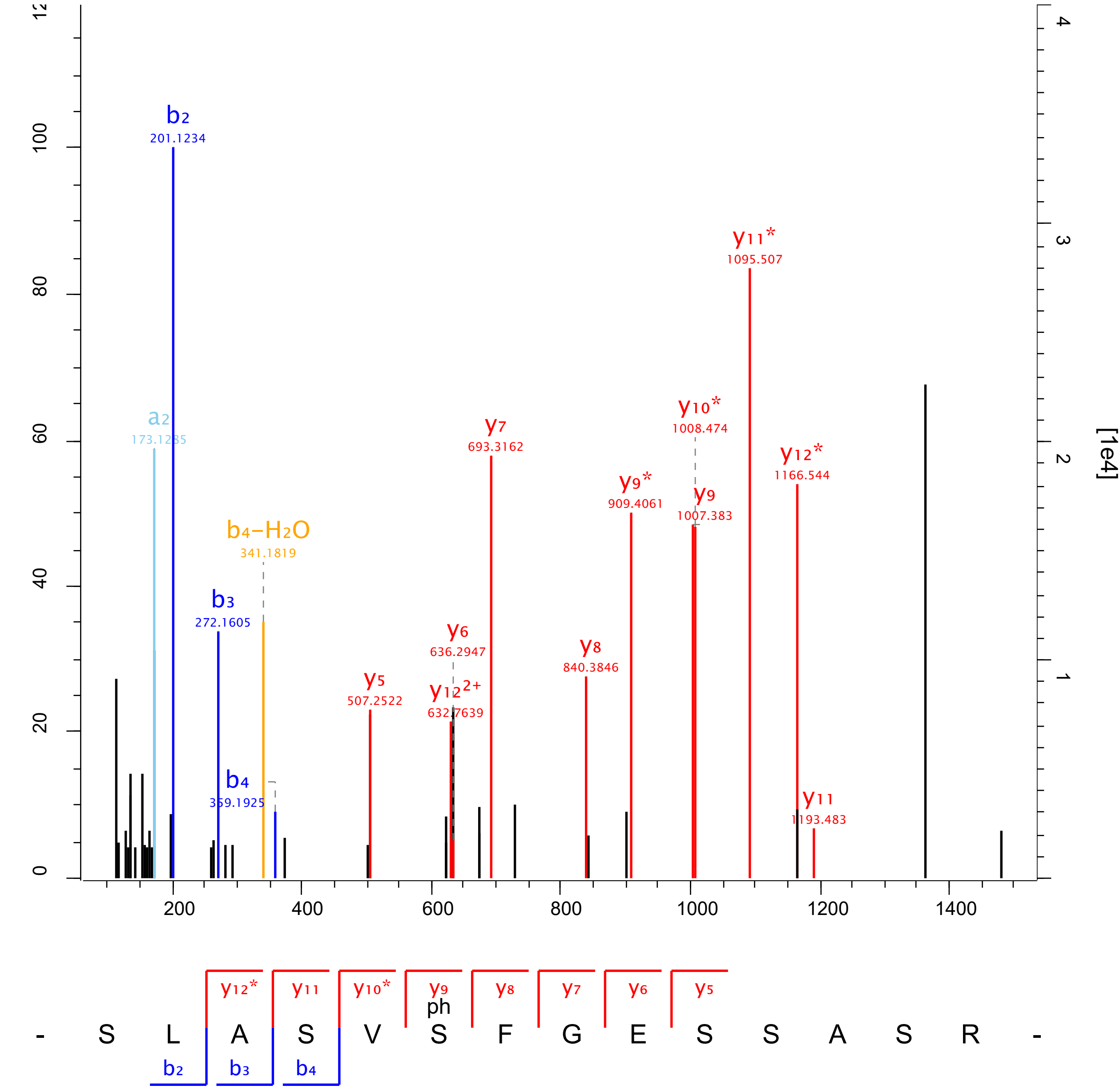

| Raw file | Scan | Method | Score | m/z |
|---|---|---|---|---|
| sys_00_3short | 22190 | FTMS; HCD | 83.75 | 767.3 |

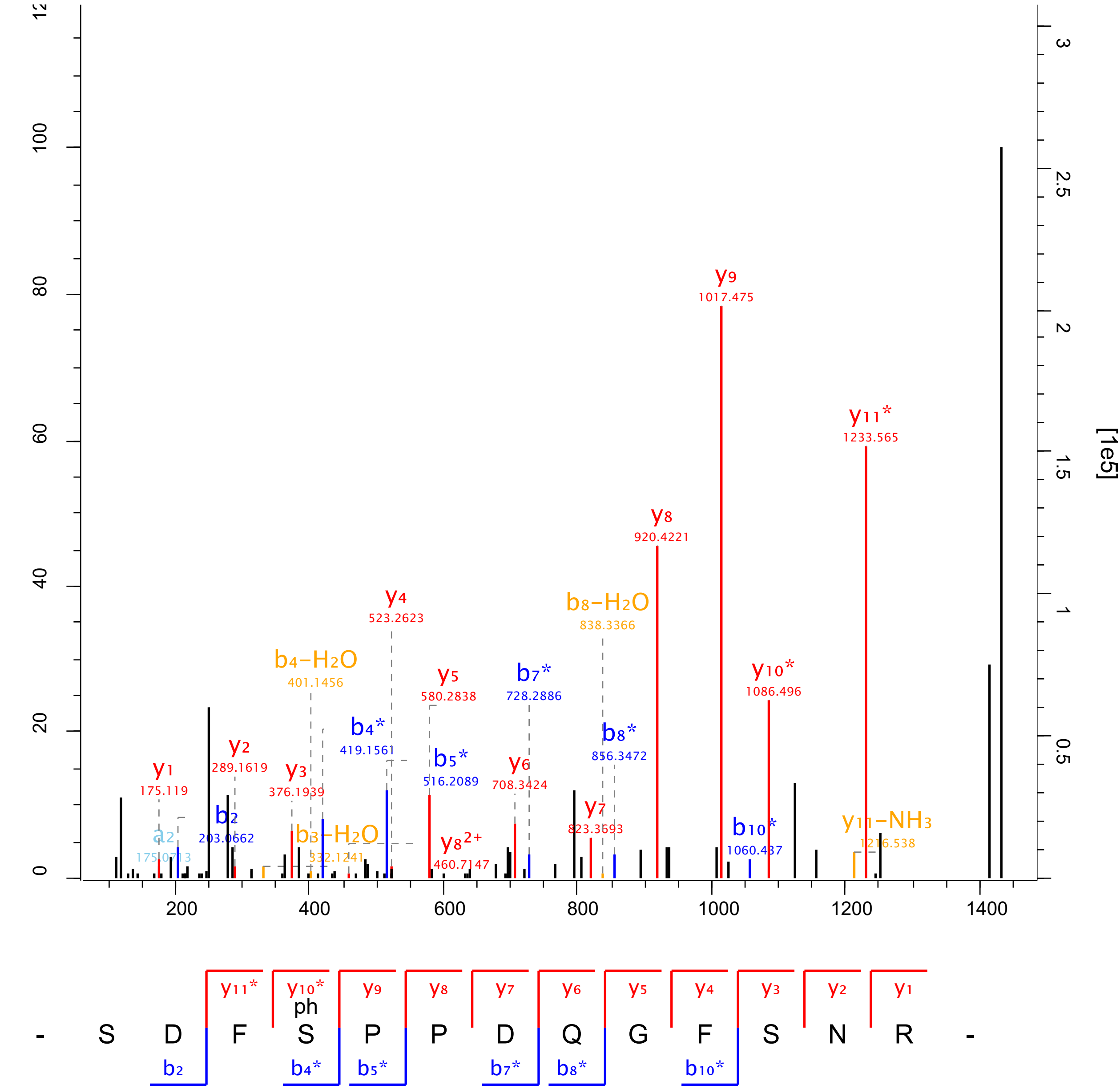

| Raw file | Scan | Method | Score | m/z |
|---|---|---|---|---|
| sys_00_3short | 22242 | FTMS; HCD | 117.37 | 819.87 |

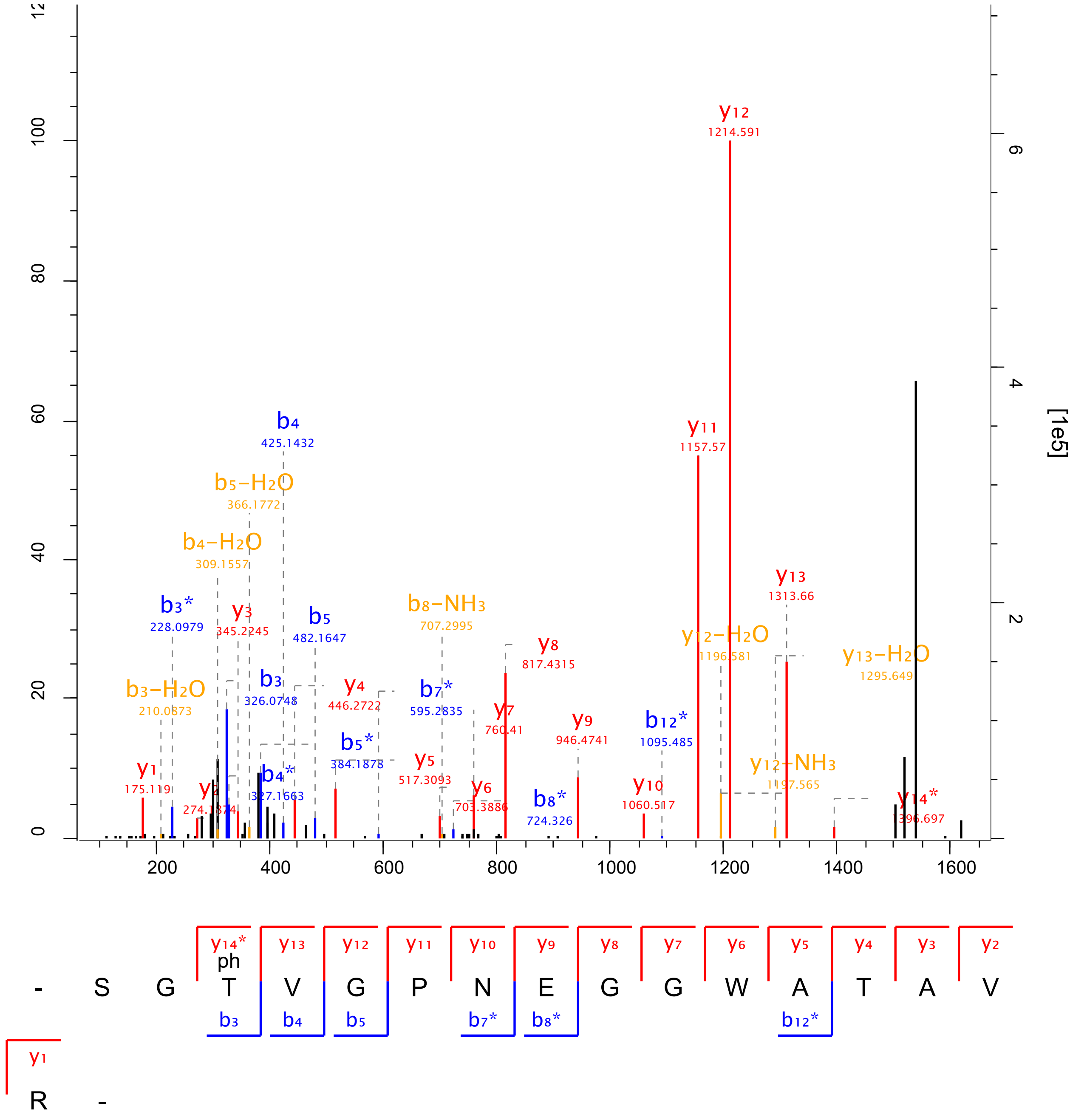

| Raw file | Scan | Method | Score | m/z |
|---|---|---|---|---|
| sys_00_3short | 22337 | FTMS; HCD | 40.28 | 799.87 |

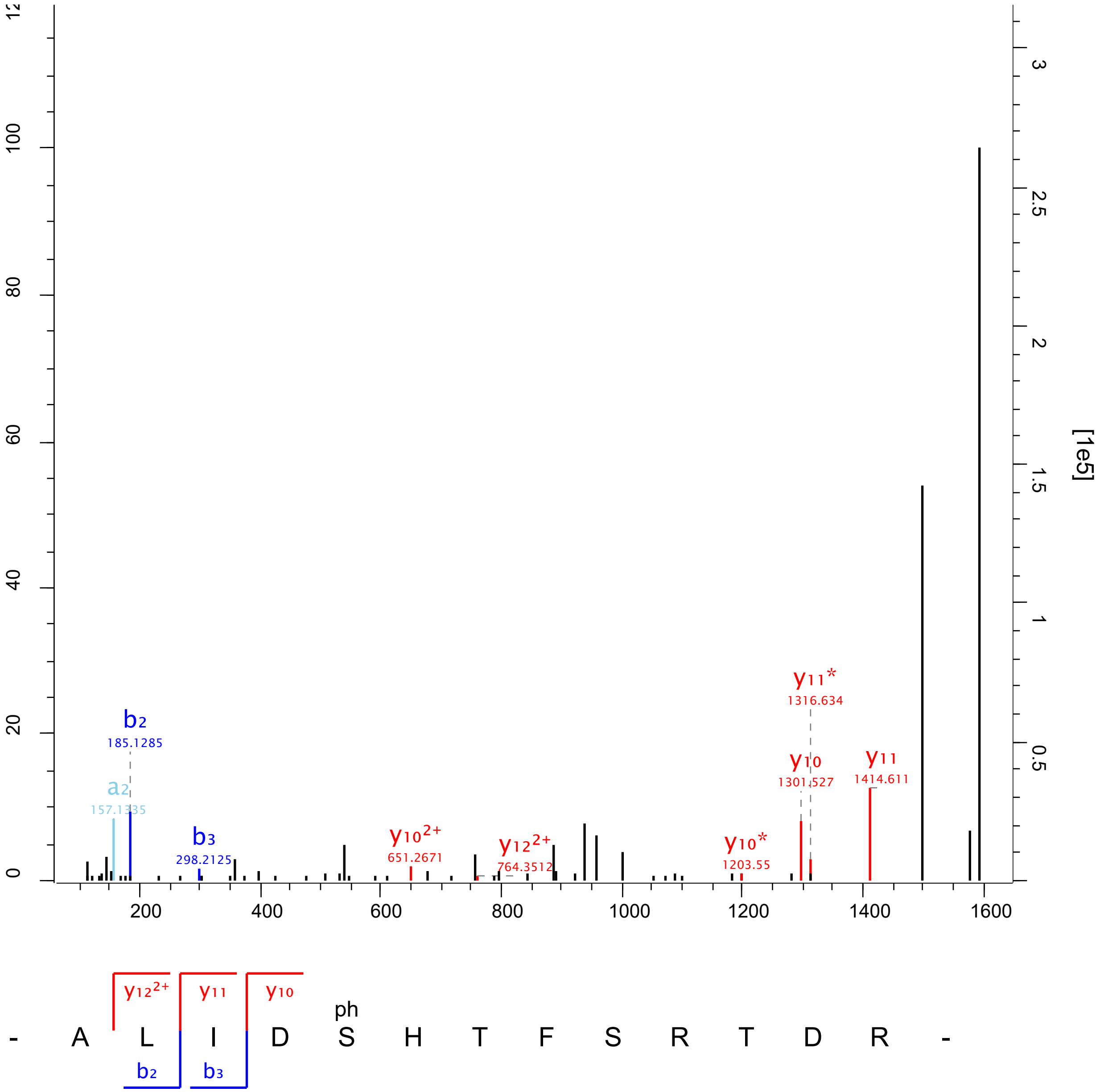

| Raw file | Scan | Method | Score | m/z |
| --- | --- | --- | --- | --- |
| sys_00_3short | 22454 | FTMS; HCD | 225.29 | 782.89 |

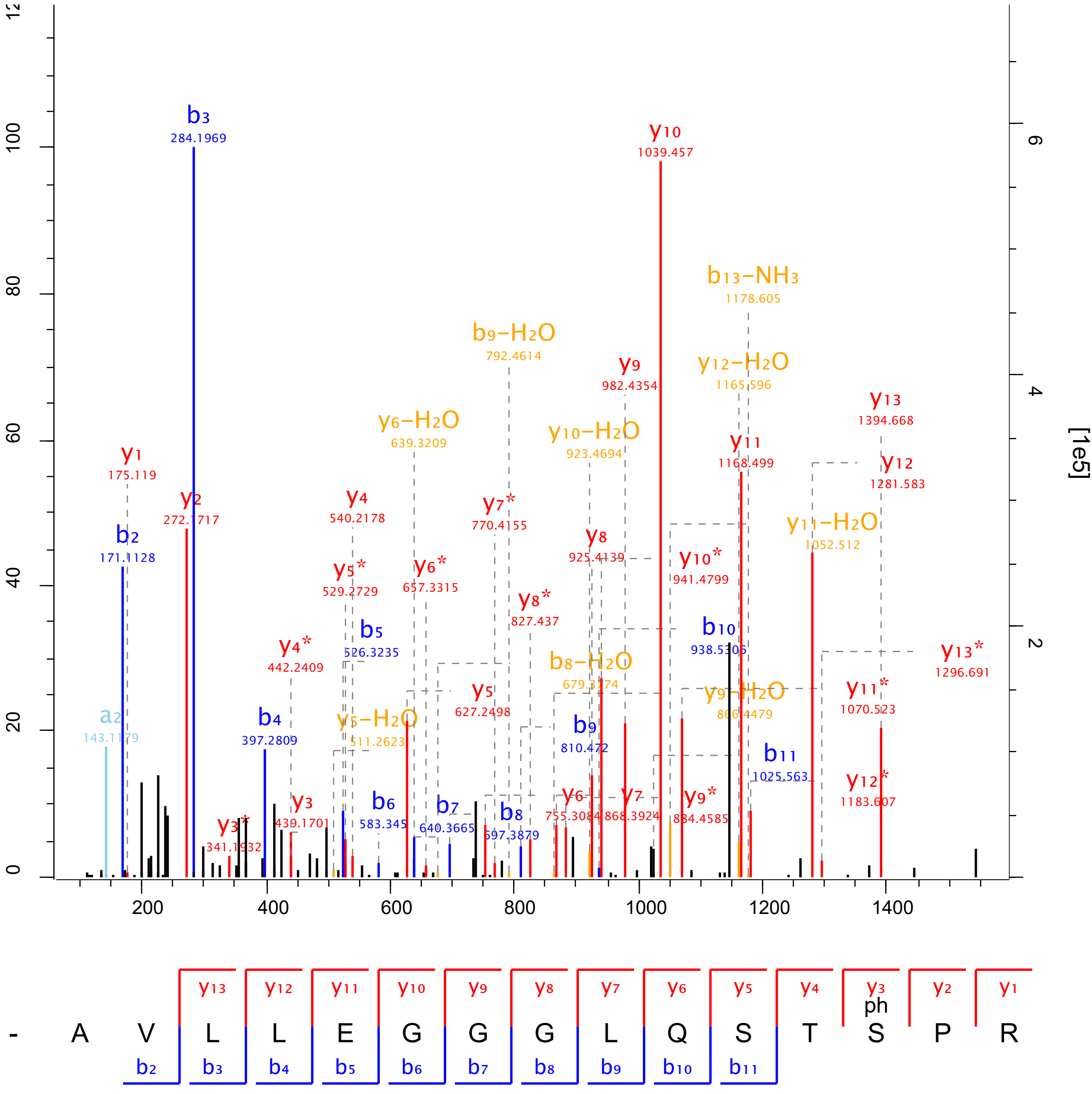

| Raw file | Scan | Method | Score | m/z |
| --- | --- | --- | --- | --- |
| sys_00_3short | 22513 | FTMS; HCD | 65.42 | 795.33 |

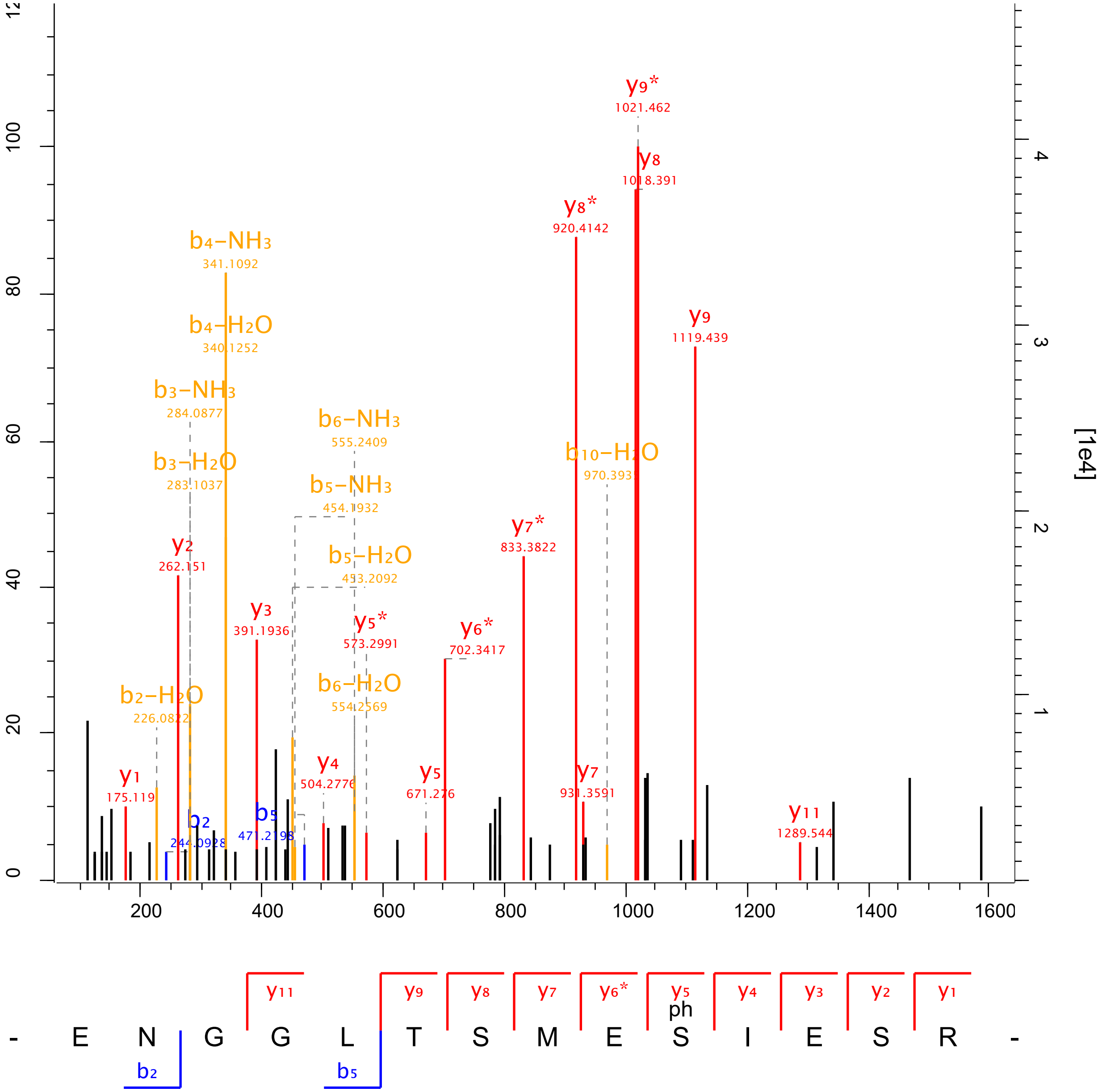

| Raw file | Scan | Method | Score | m/z |
|---|---|---|---|---|
| sys_00_3short | 22647 | FTMS; HCD | 159.18 | 715.3 |

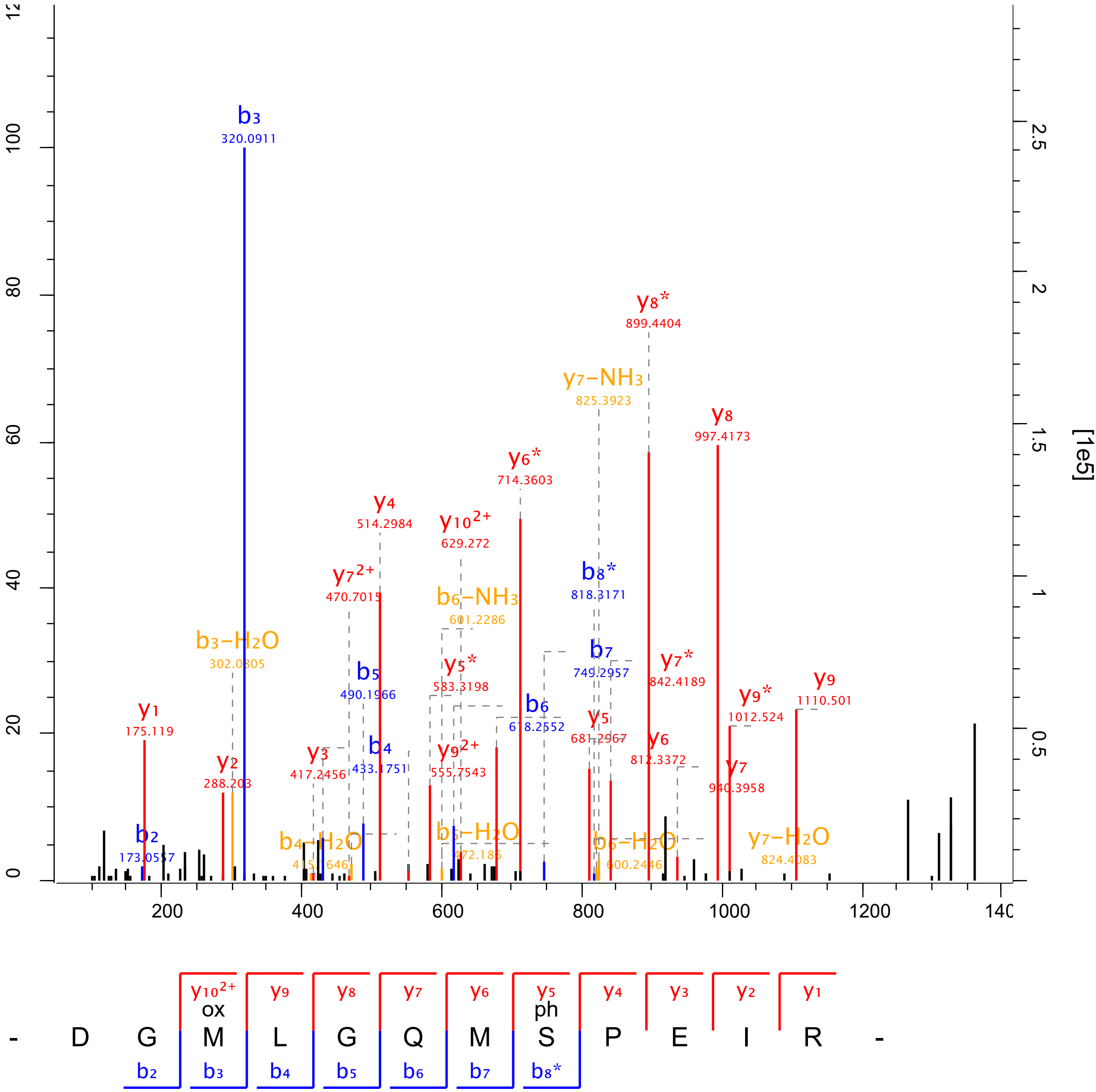

| Raw file | Scan | Method | Score | m/z |
|---|---|---|---|---|
| sys_00_3short | 22870 | FTMS; HCD | 159.64 | 713.83 |

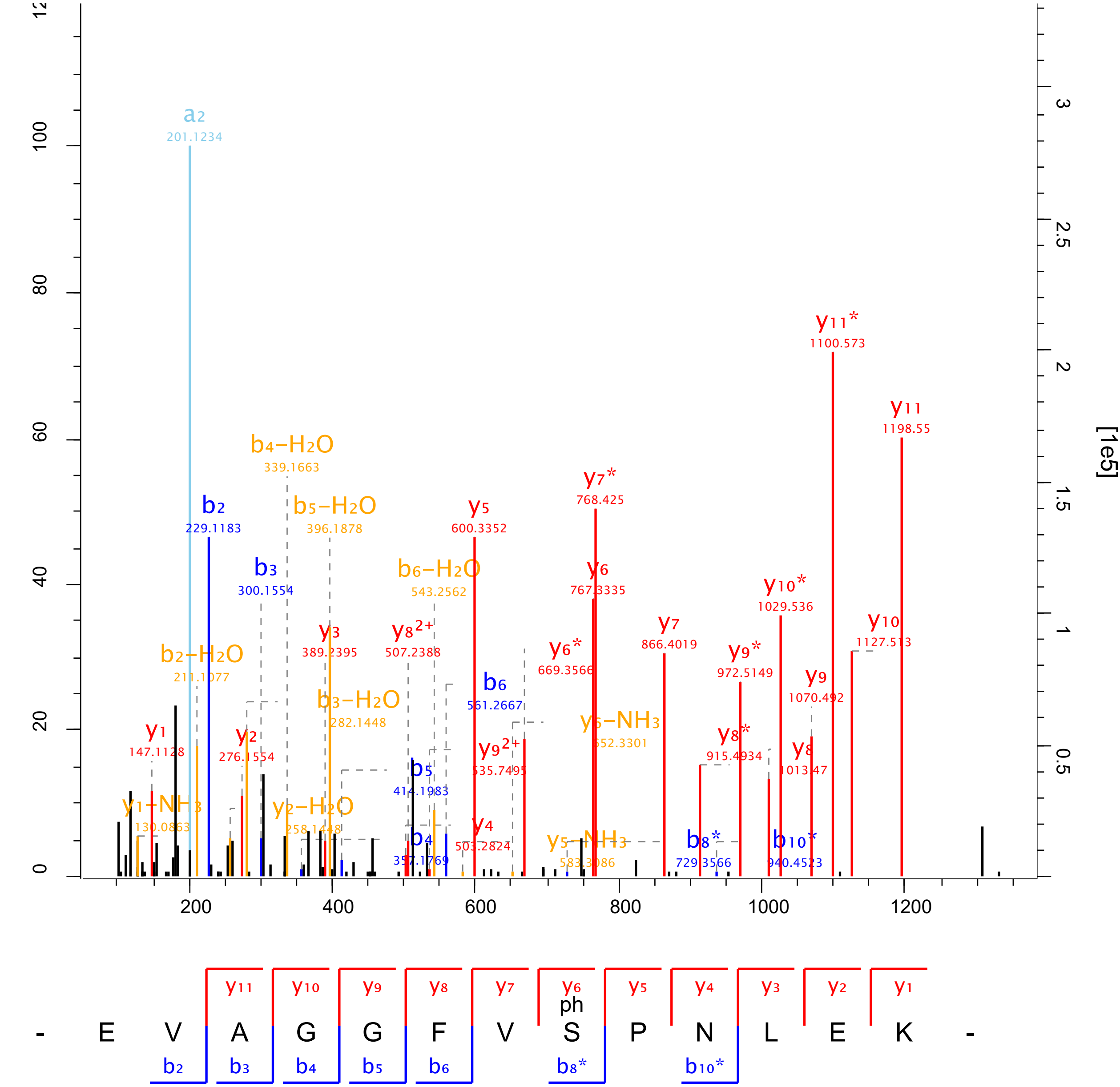

| Raw file | Scan | Method | Score | m/z |
| --- | --- | --- | --- | --- |
| sys_00_3short | 23033 | FTMS; HCD | 119.18 | 938.37 |

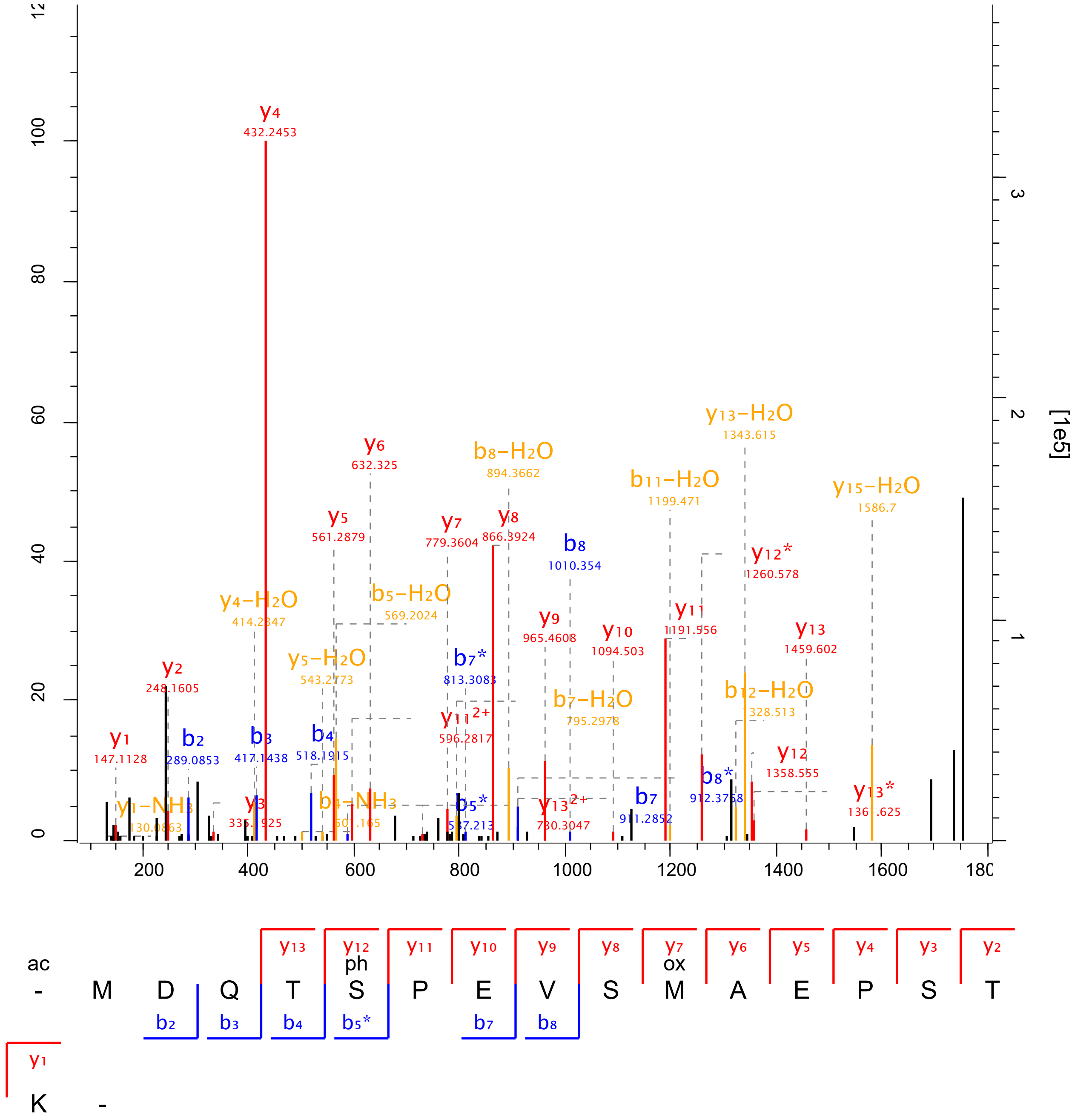

| Raw file | Scan | Method | Score | m/z |
|---|---|---|---|---|
| sys_00_3short | 23098 | FTMS; HCD | 43.76 | 700.81 |

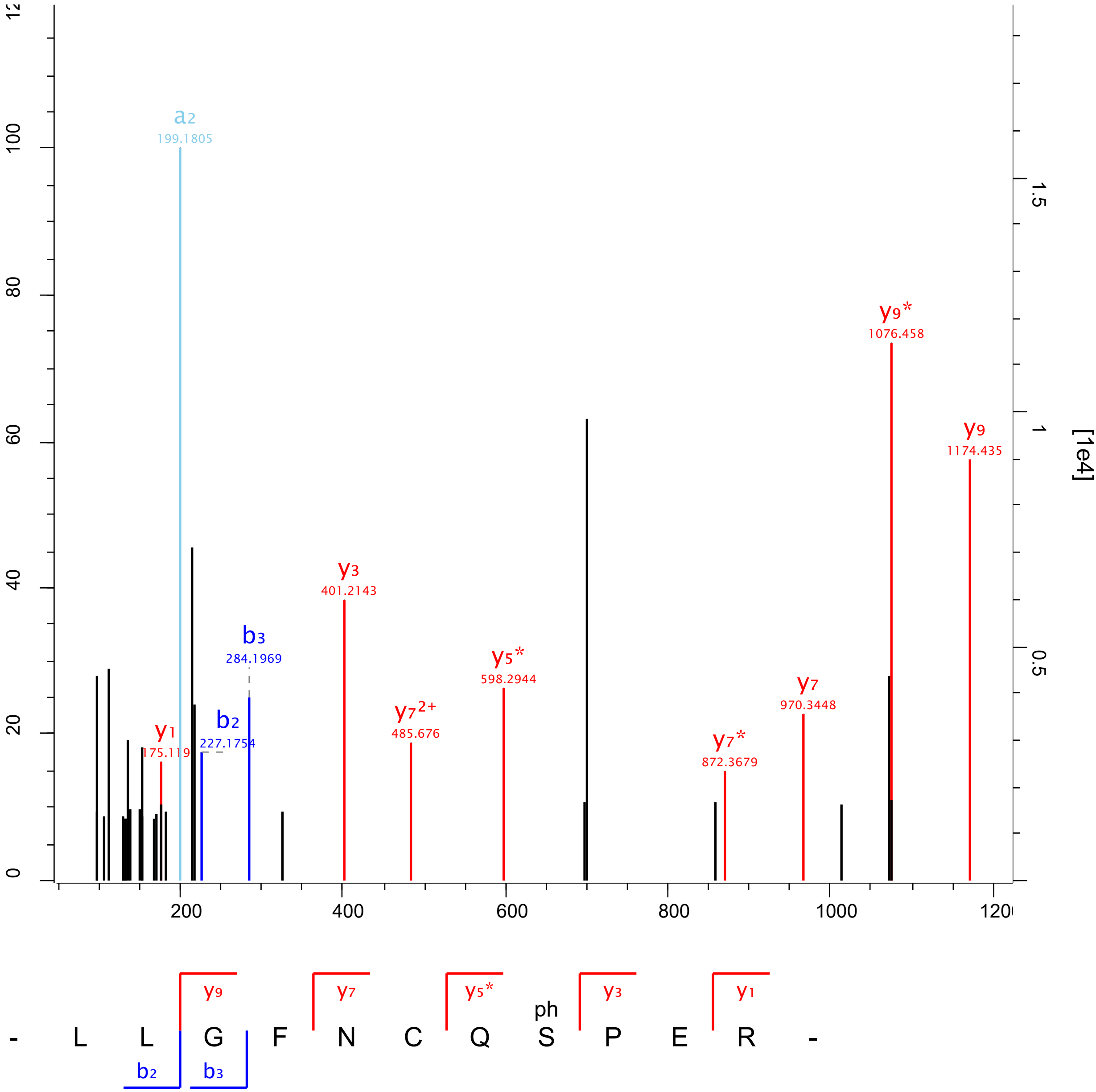

| Raw file | Scan | Method | Score | m/z |
|---|---|---|---|---|
| sys_00_3short | 23160 | FTMS; HCD | 69.92 | 598.3 |

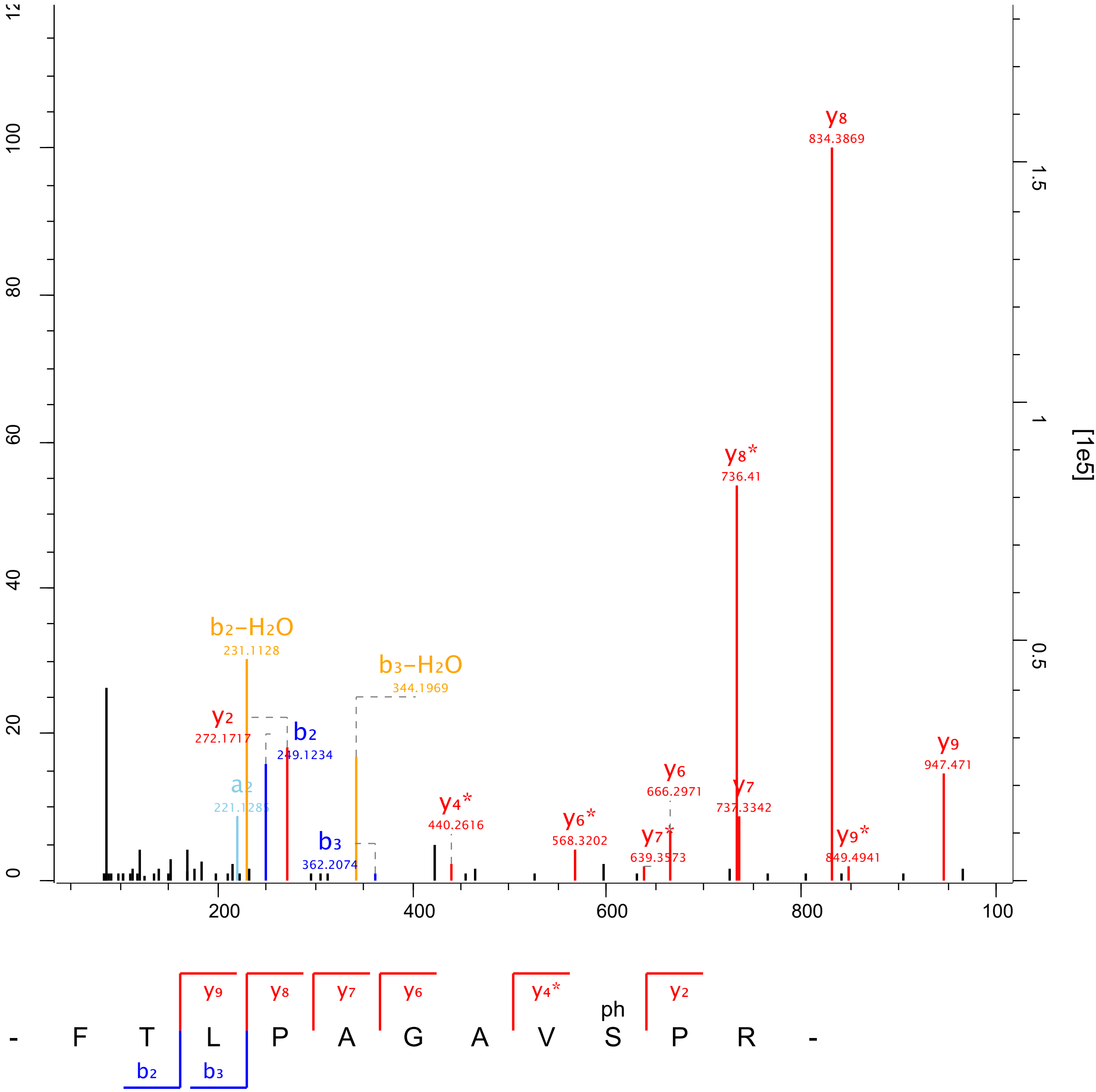

| Raw file | Scan | Method | Score | m/z |
|---|---|---|---|---|
| sys_00_3short | 24112 | FTMS; HCD | 147.12 | 794.37 |

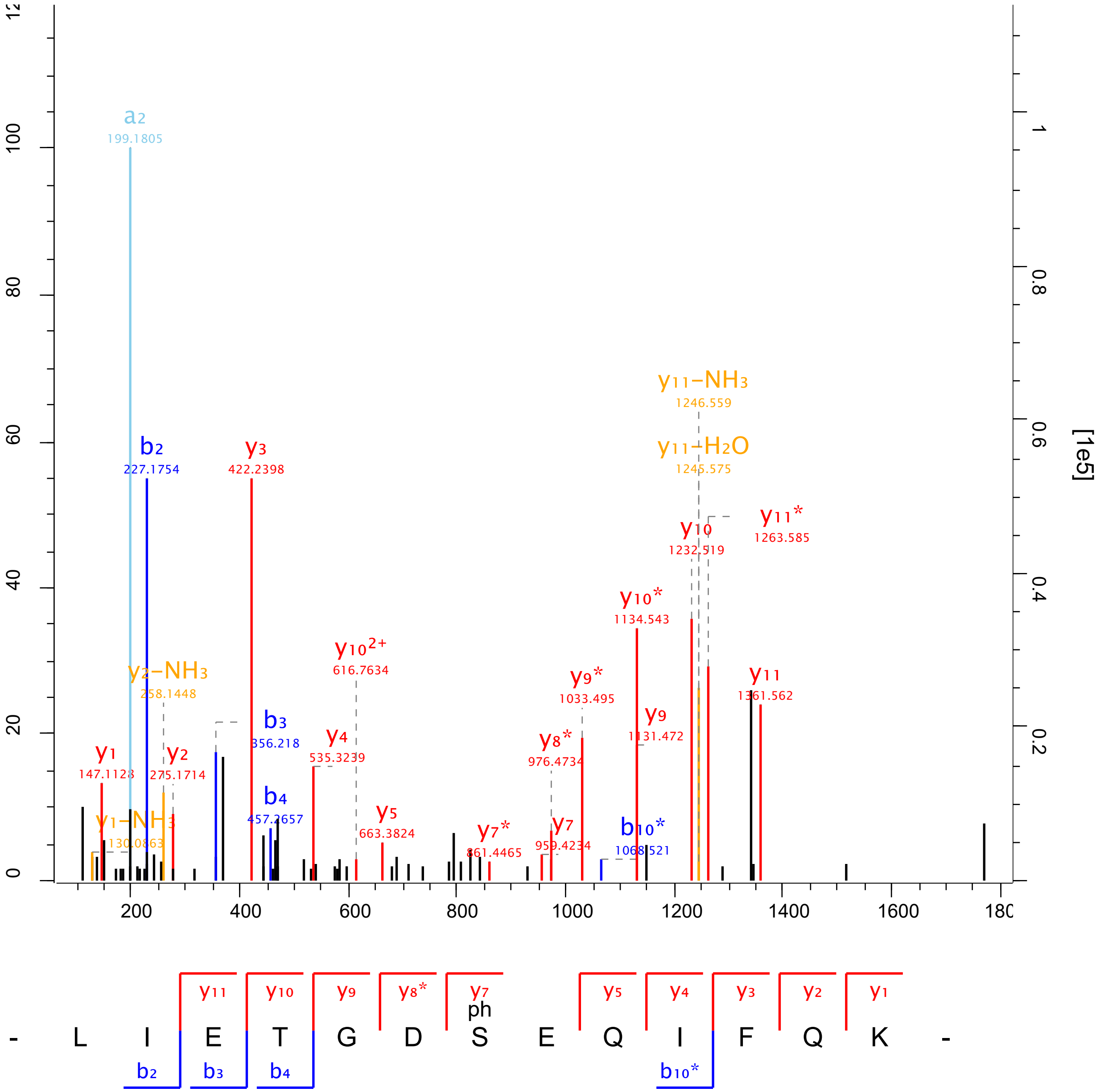

| Raw file | Scan | Method | Score | m/z |
|---|---|---|---|---|
| sys_00_3short | 24350 | FTMS; HCD | 142.45 | 1049.93 |

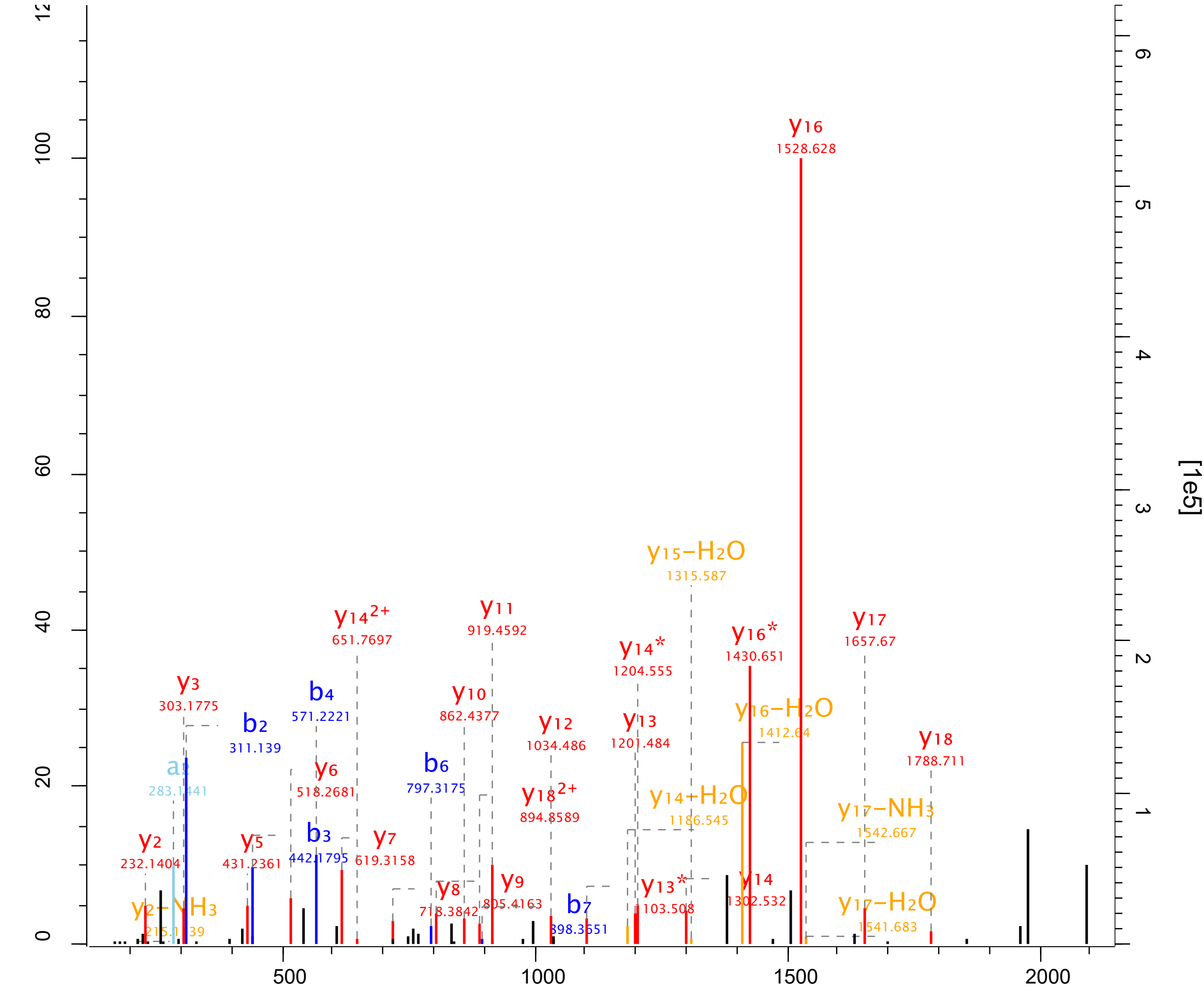


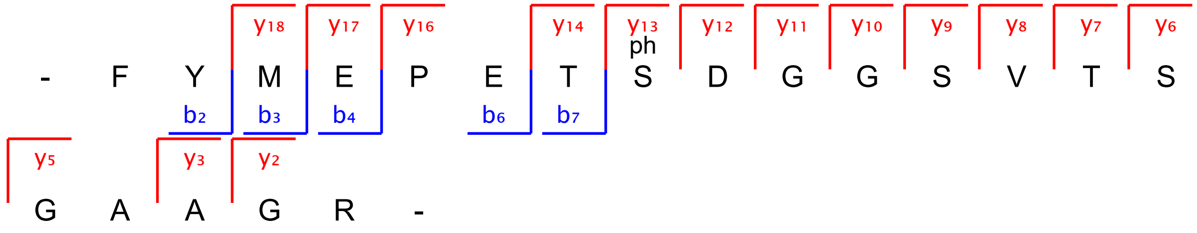

| Raw file | Scan | Method | Score | m/z |
|---|---|---|---|---|
| sys_00_3short | 24723 | FTMS; HCD | 180.27 | 706.34 |

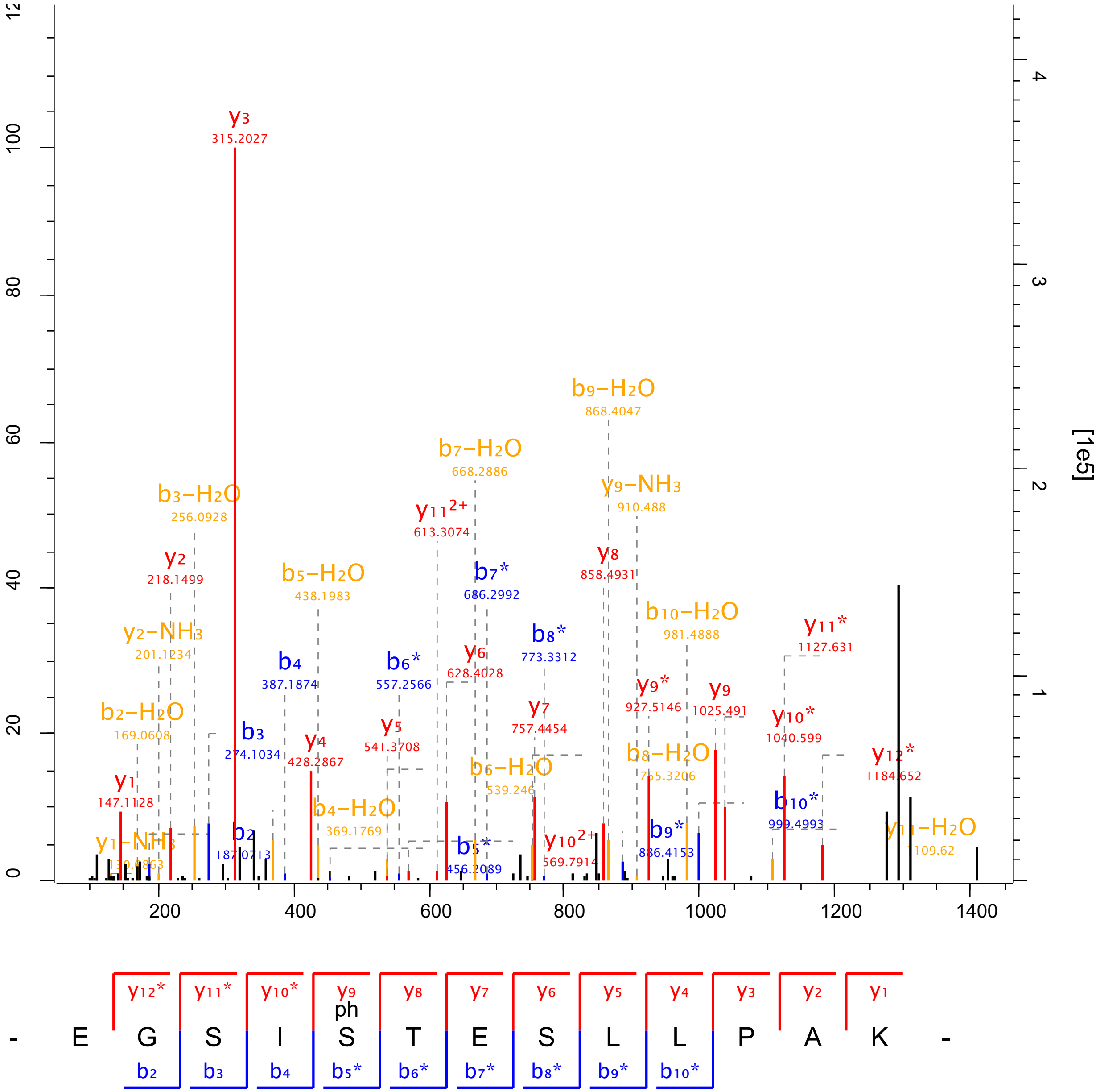

| Raw file | Scan | Method | Score | m/z |
|---|---|---|---|---|
| sys_00_3short | 24865 | FTMS; HCD | 151.44 | 736.83 |

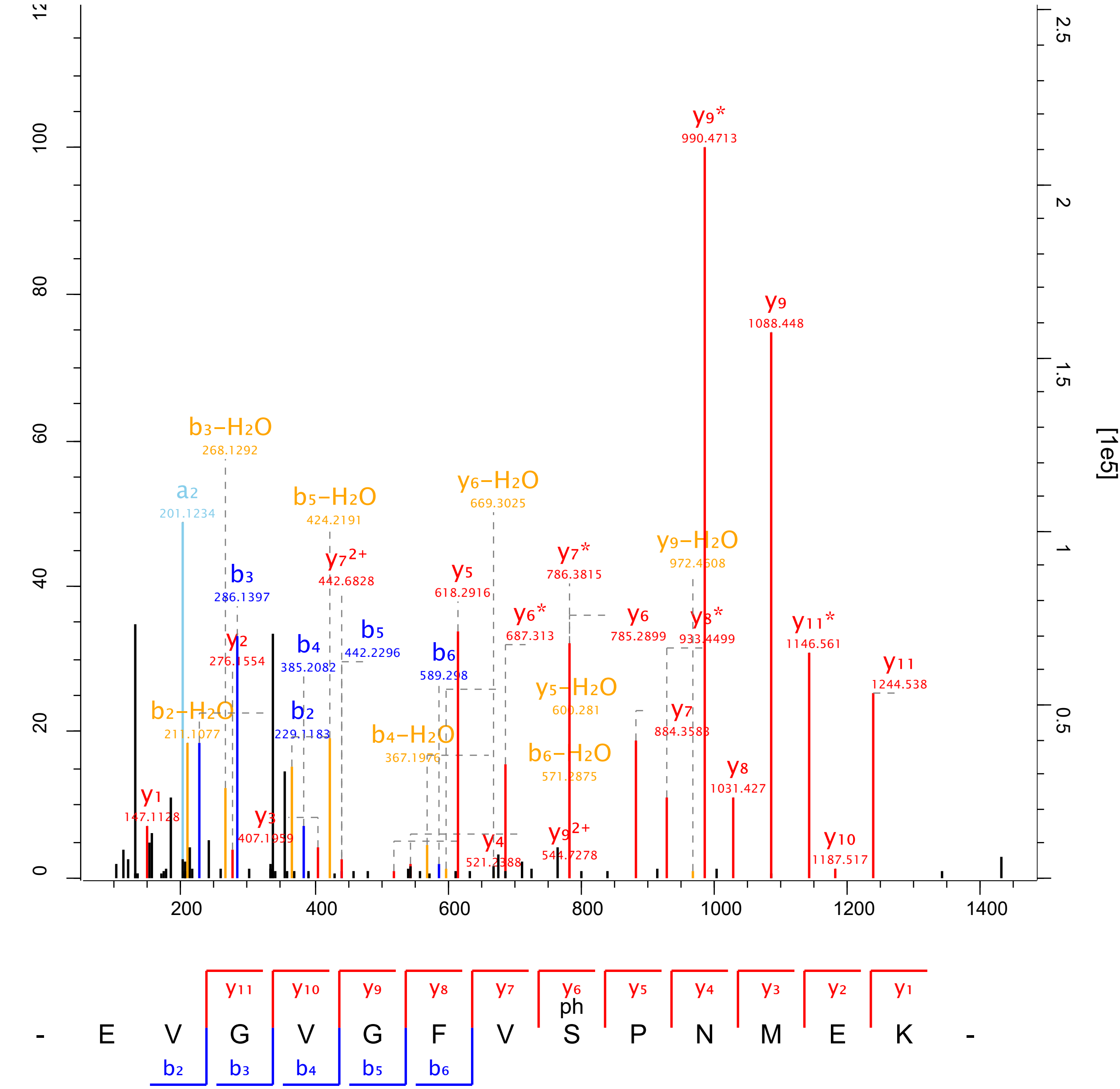

| Raw file | Scan | Method | Score | m/z |
| --- | --- | --- | --- | --- |
| sys_00_3short | 24868 | FTMS; HCD | 116.86 | 683.32 |

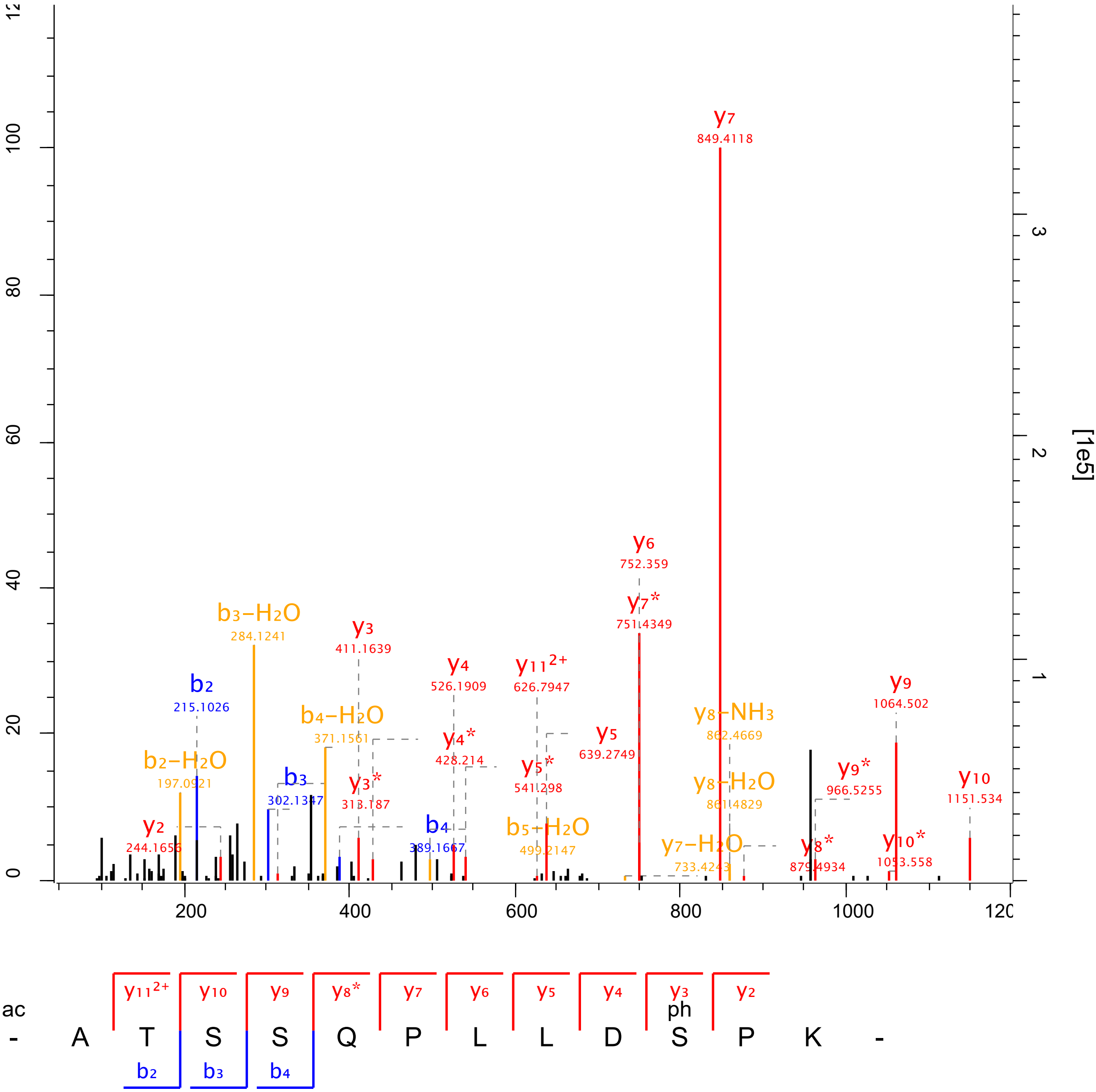

| Raw file | Scan | Method | Score | m/z |
| --- | --- | --- | --- | --- |
| sys_00_3short | 24908 | FTMS; HCD | 58.98 | 831.88 |

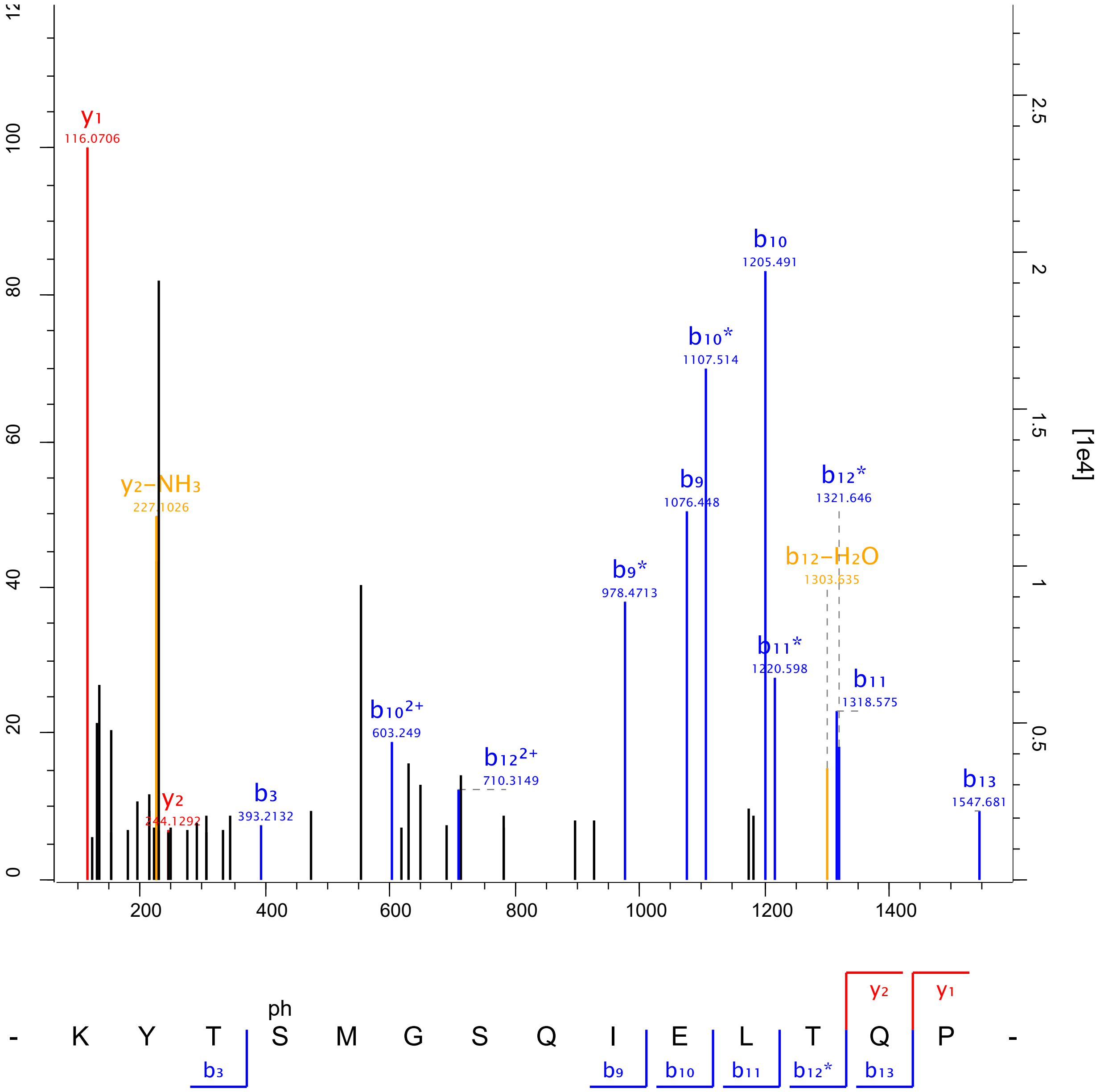

| Raw file | Scan | Method | Score | m/z |
|---|---|---|---|---|
| sys_00_3short | 25096 | FTMS; HCD | 160.78 | 710.33 |

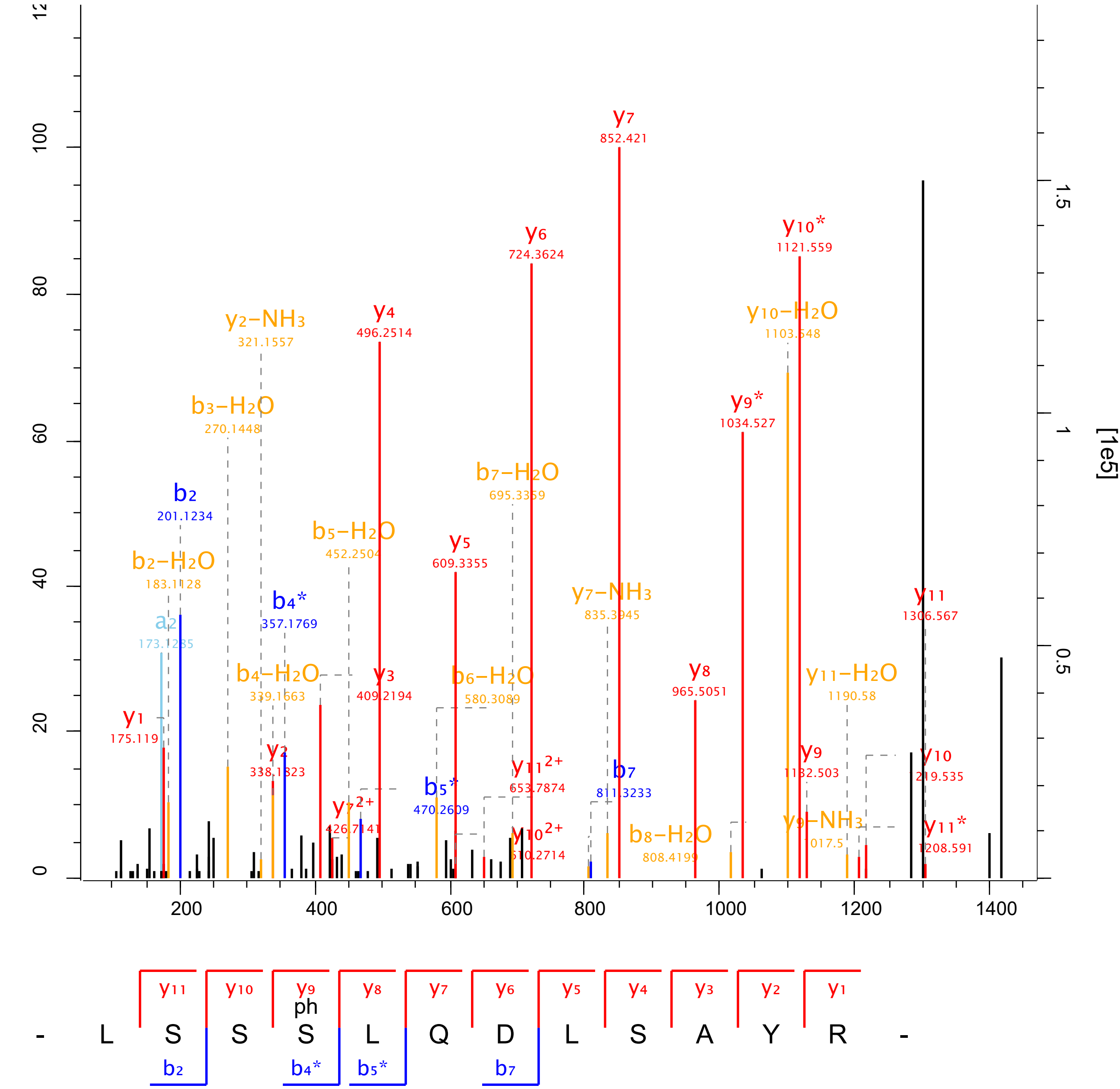

| Raw file | Scan | Method | Score | m/z |
|---|---|---|---|---|
| sys_00_3short | 25587 | FTMS; HCD | 218.11 | 799.9 |

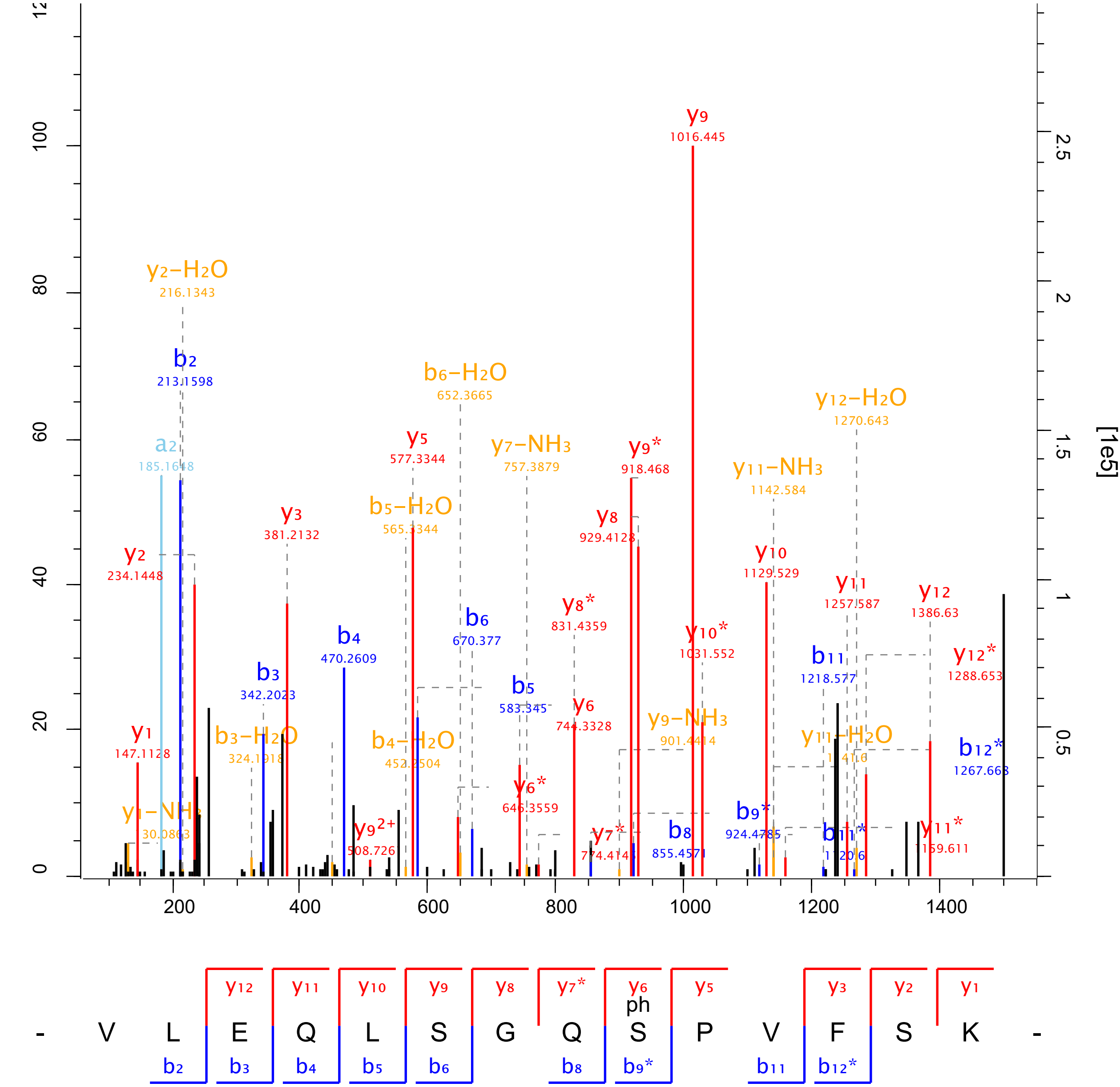

| Raw file | Scan | Method | Score | m/z |
| --- | --- | --- | --- | --- |
| sys_00_3short | 25716 | FTMS; HCD | 124.25 | 795.35 |

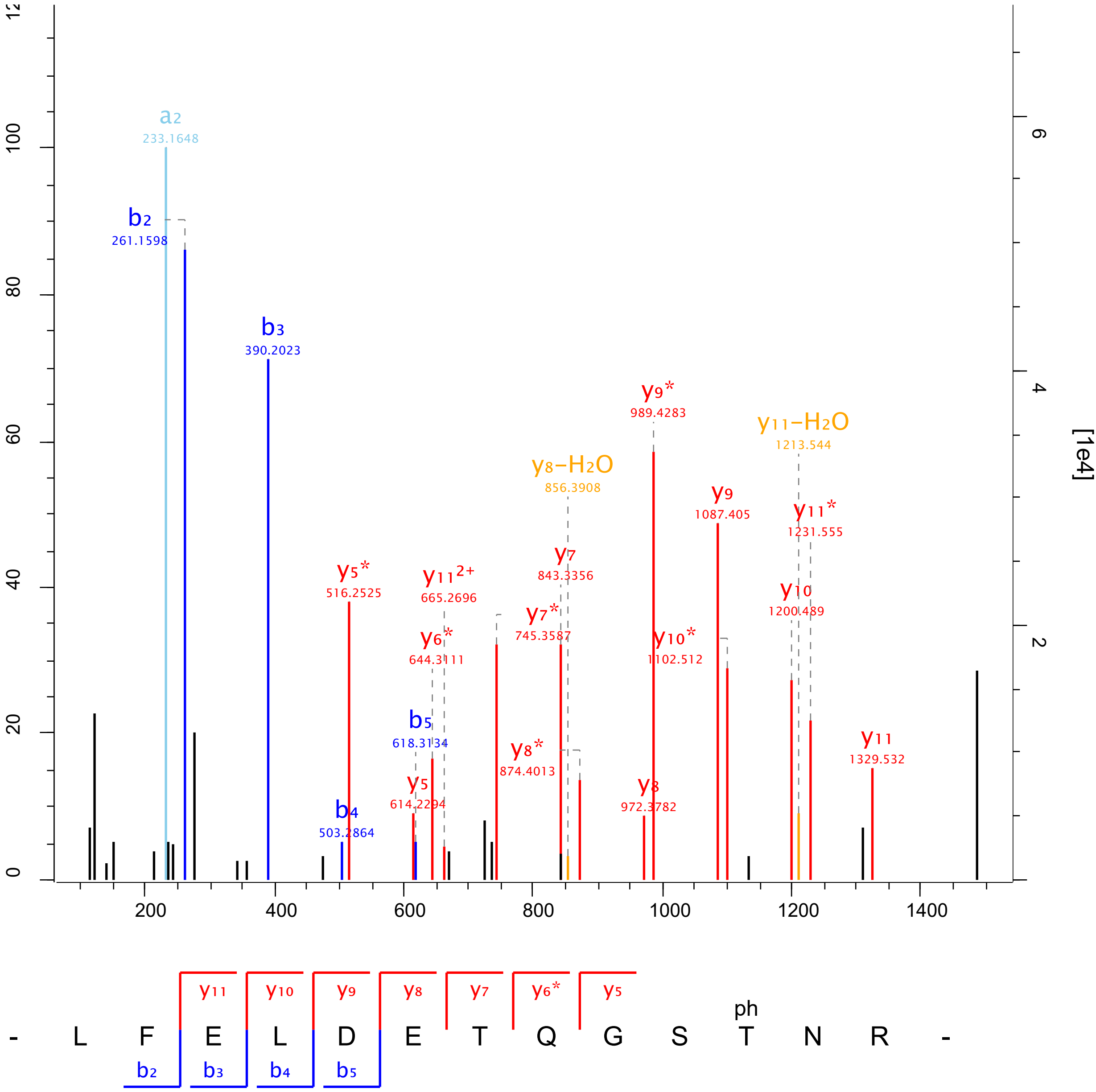

| Raw file | Scan | Method | Score | m/z |
|---|---|---|---|---|
| sys_00_3short | 25762 | FTMS; HCD | 71.88 | 588.81 |

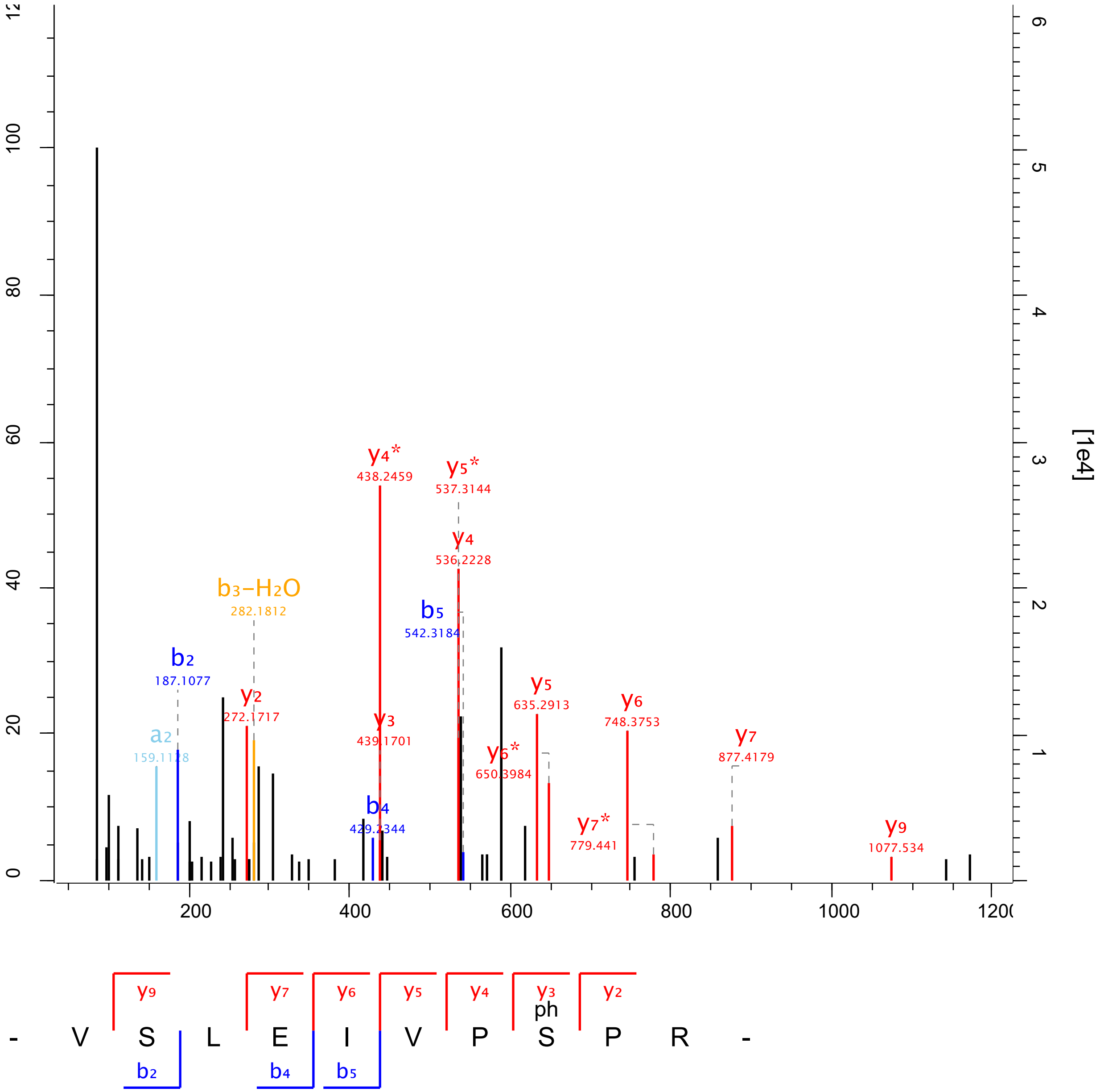

| Raw file | Scan | Method | Score | m/z |
| --- | --- | --- | --- | --- |
| sys_00_3short | 25778 | FTMS; HCD | 161.21 | 598.3 |

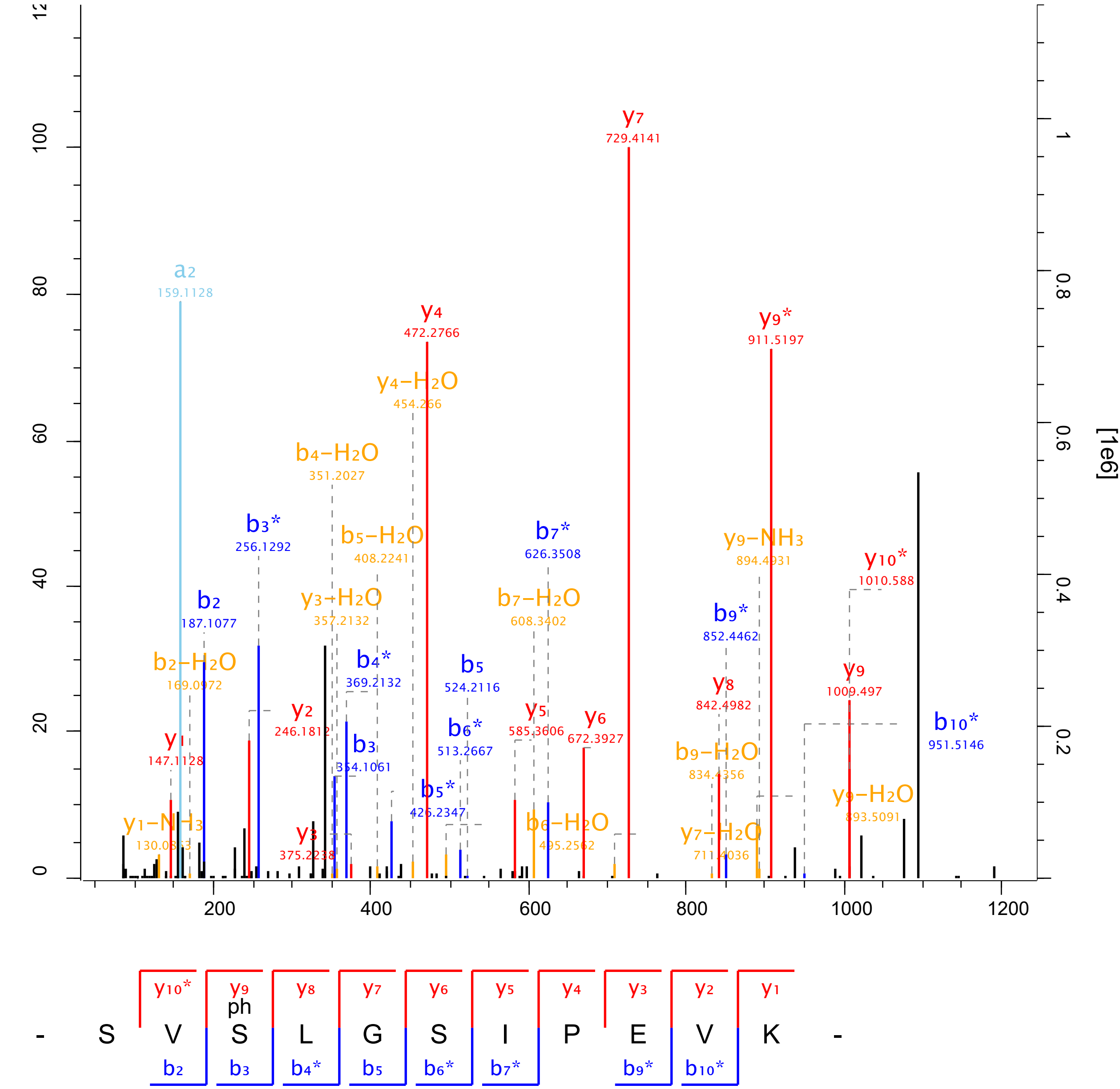

| Raw file | Scan | Method | Score | m/z |
| --- | --- | --- | --- | --- |
| sys_00_3short | 25897 | FTMS; HCD | 73.69 | 729.69 |

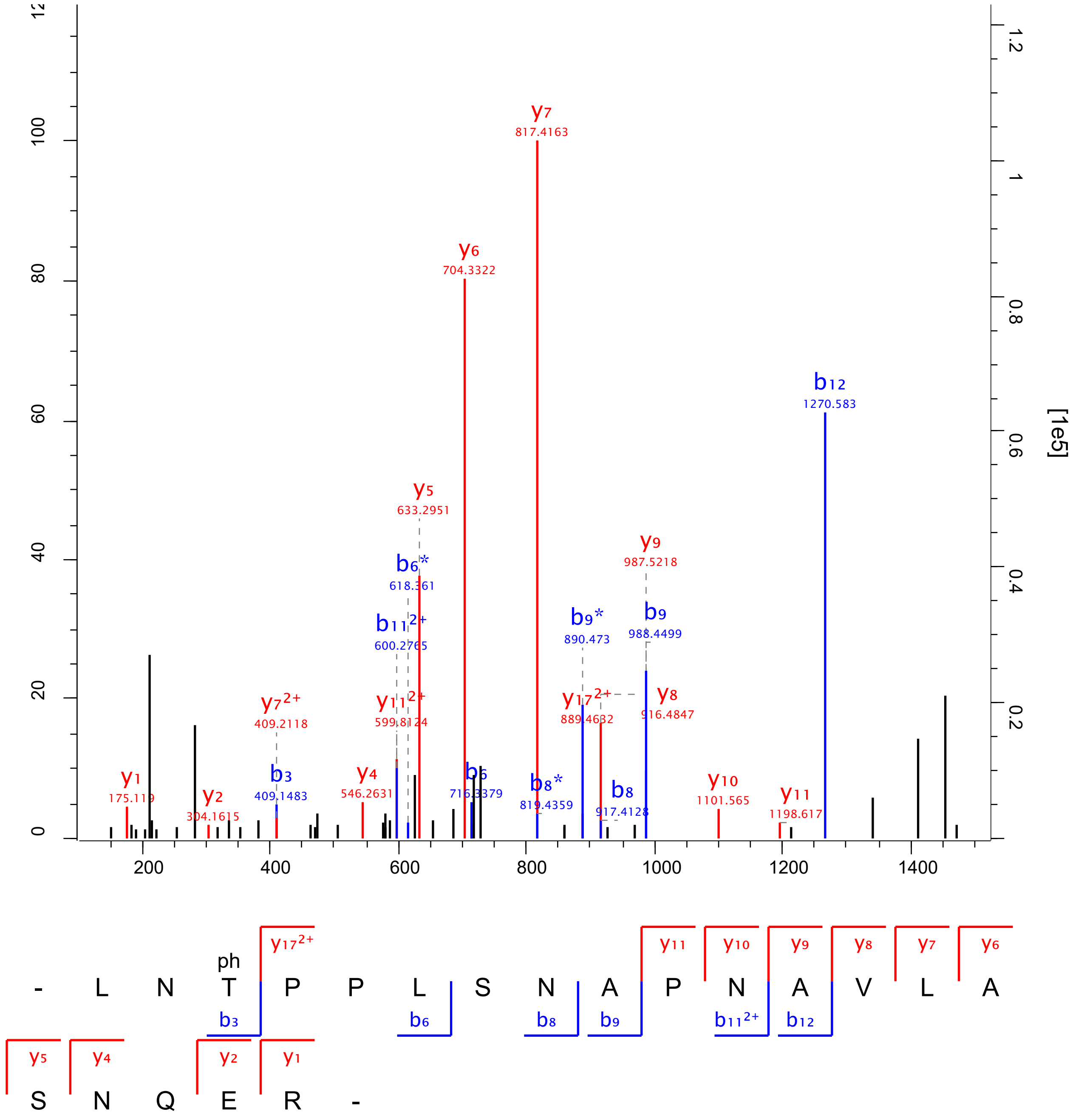

| Raw file | Scan | Method | Score | m/z |
|---|---|---|---|---|
| sys_00_3short | 25963 | FTMS; HCD | 109.44 | 639.75 |

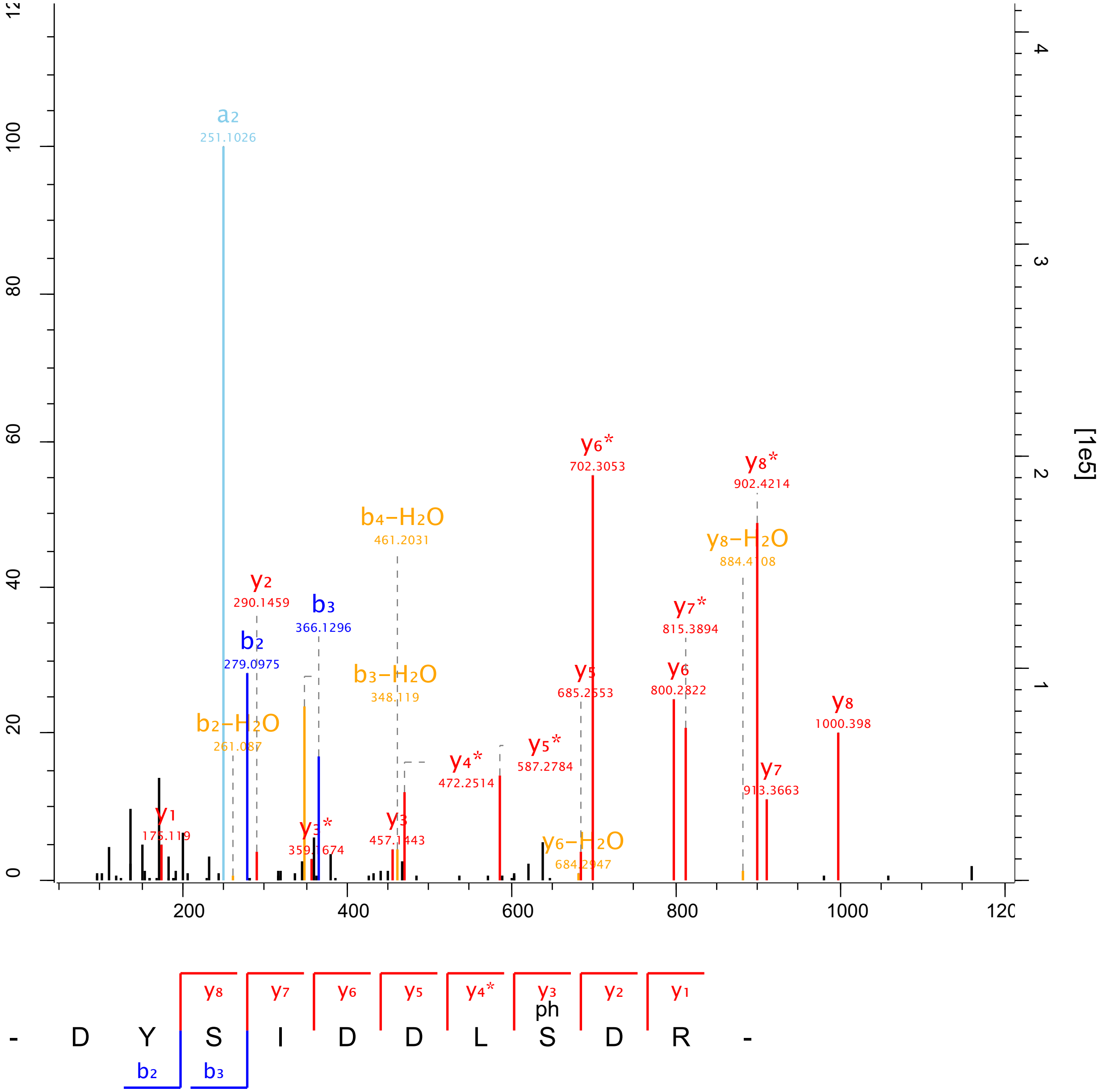

| Raw file | Scan | Method | Score | m/z |
| --- | --- | --- | --- | --- |
| sys_00_3short | 25996 | FTMS; HCD | 58.32 | 601.28 |

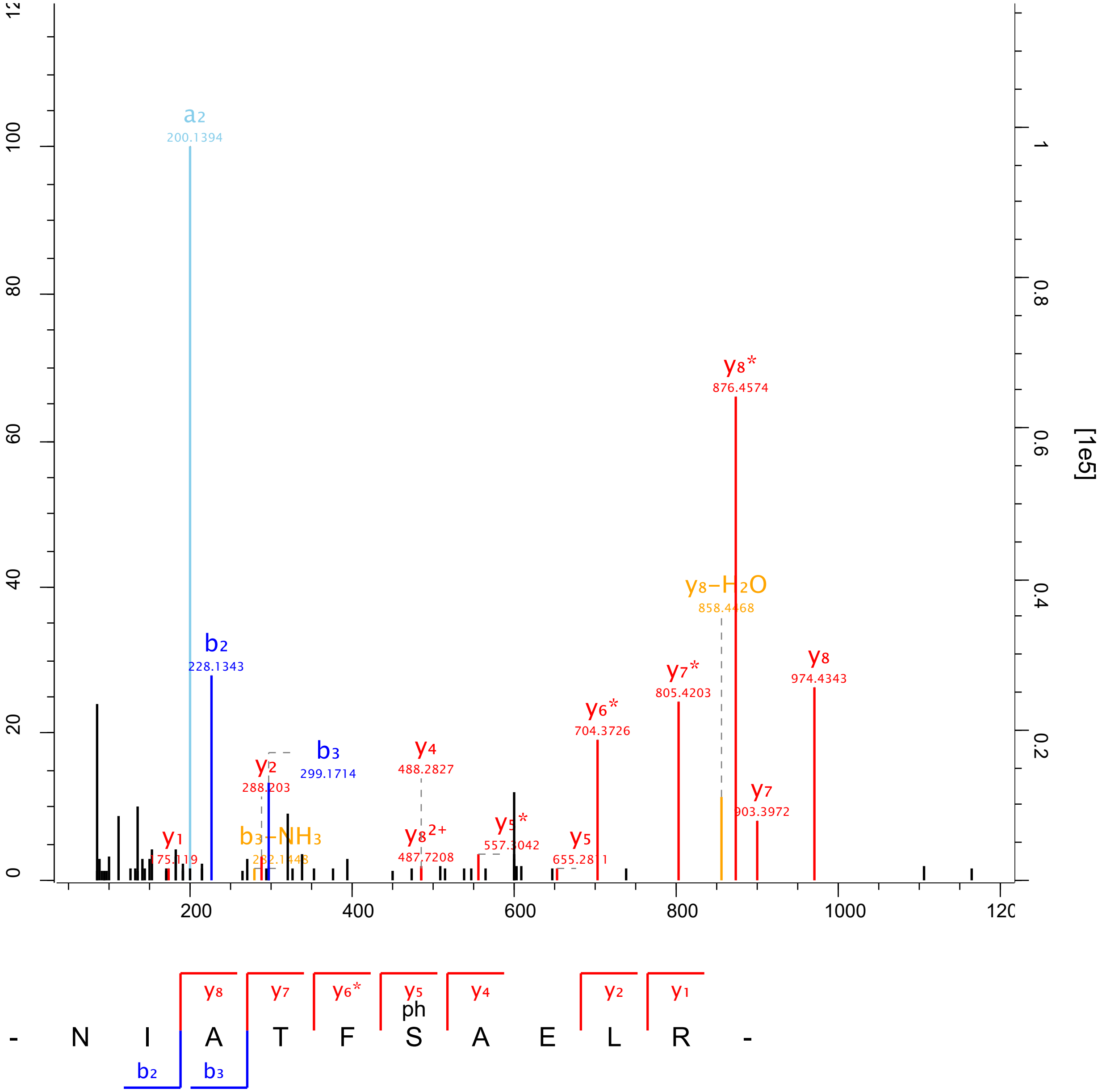

| Raw file | Scan | Method | Score | m/z |
|---|---|---|---|---|
| sys_00_3short | 26045 | FTMS; HCD | 45.22 | 744.82 |

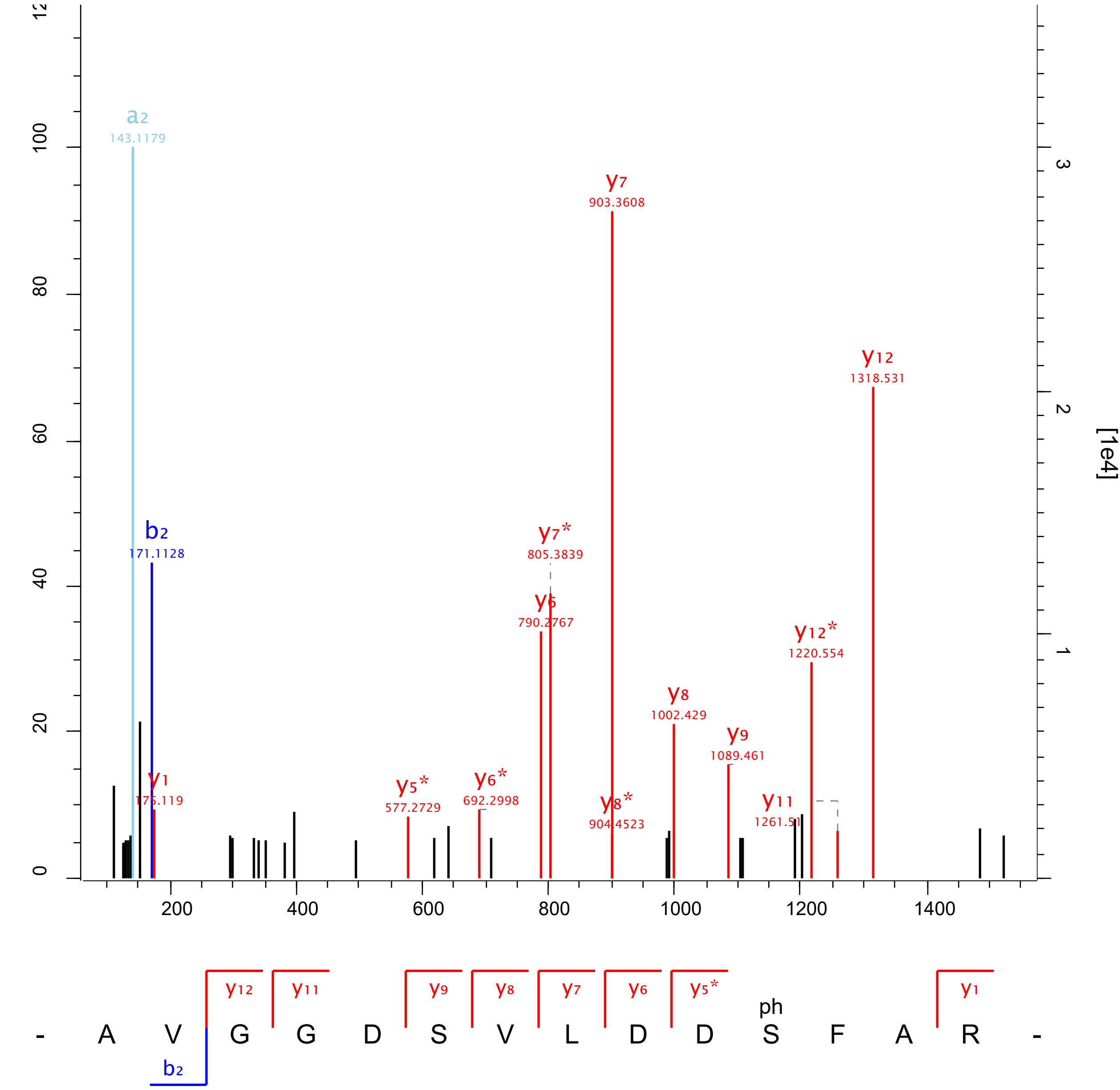

| Raw file | Scan | Method | Score | m/z |
|---|---|---|---|---|
| sys_00_3short | 26120 | FTMS; HCD | 69.33 | 776.84 |

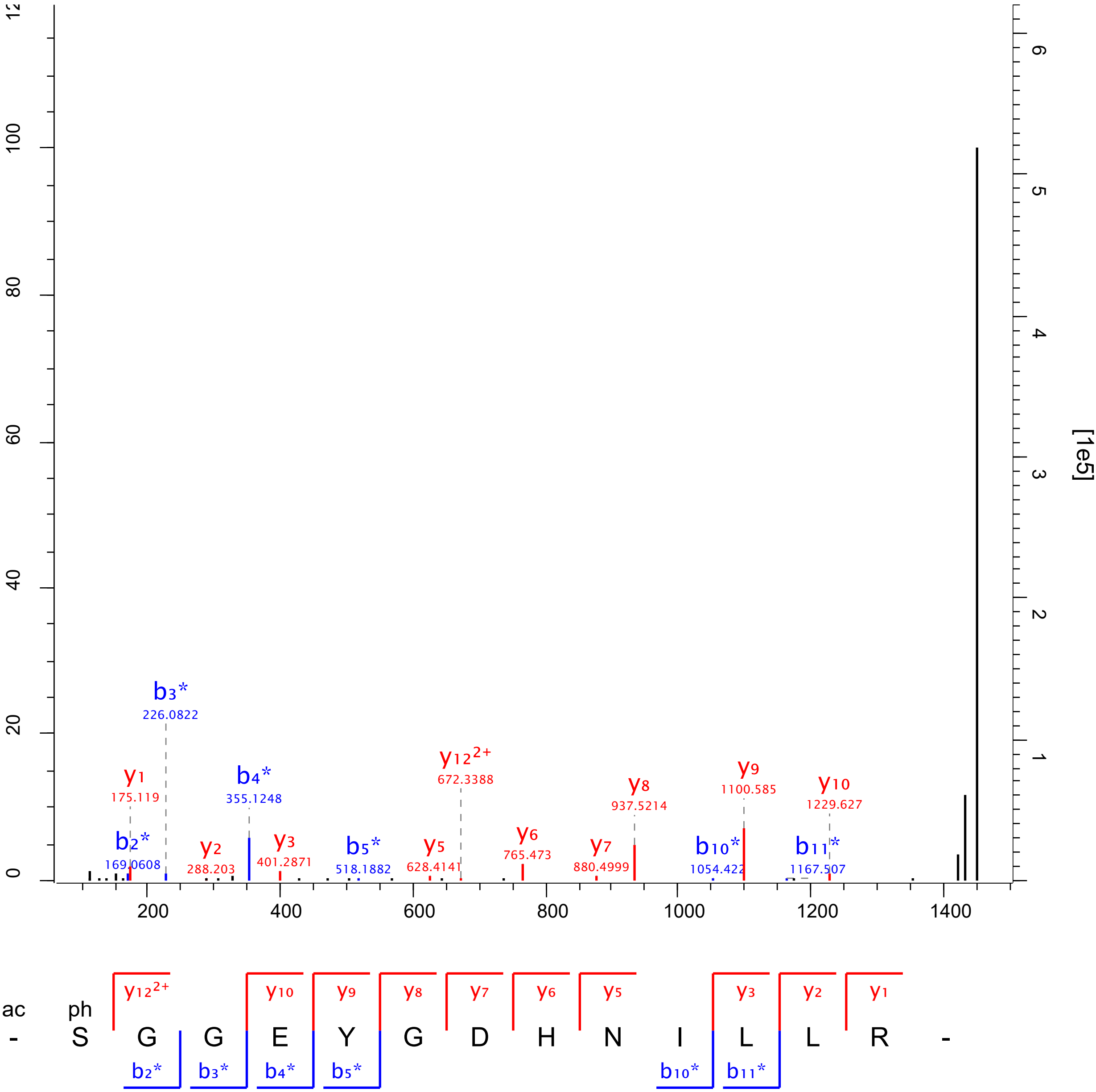

| Raw file | Scan | Method | Score | m/z |
|---|---|---|---|---|
| sys_00_3short | 26443 | FTMS; HCD | 110.12 | 726.28 |

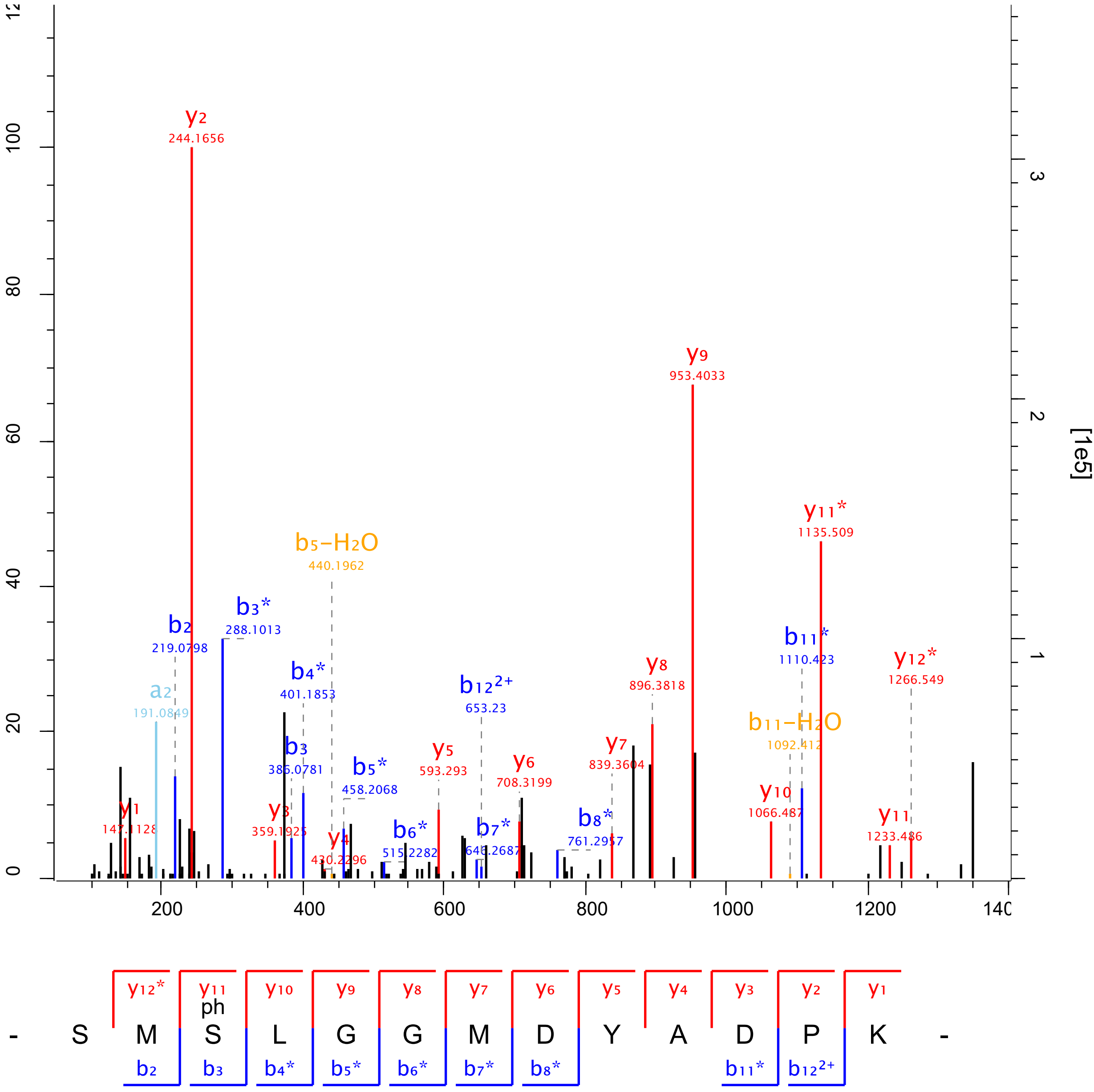

| Raw file | Scan | Method | Score | m/z |
| --- | --- | --- | --- | --- |
| sys_00_3short | 26476 | FTMS; HCD | 108.63 | 895.82 |

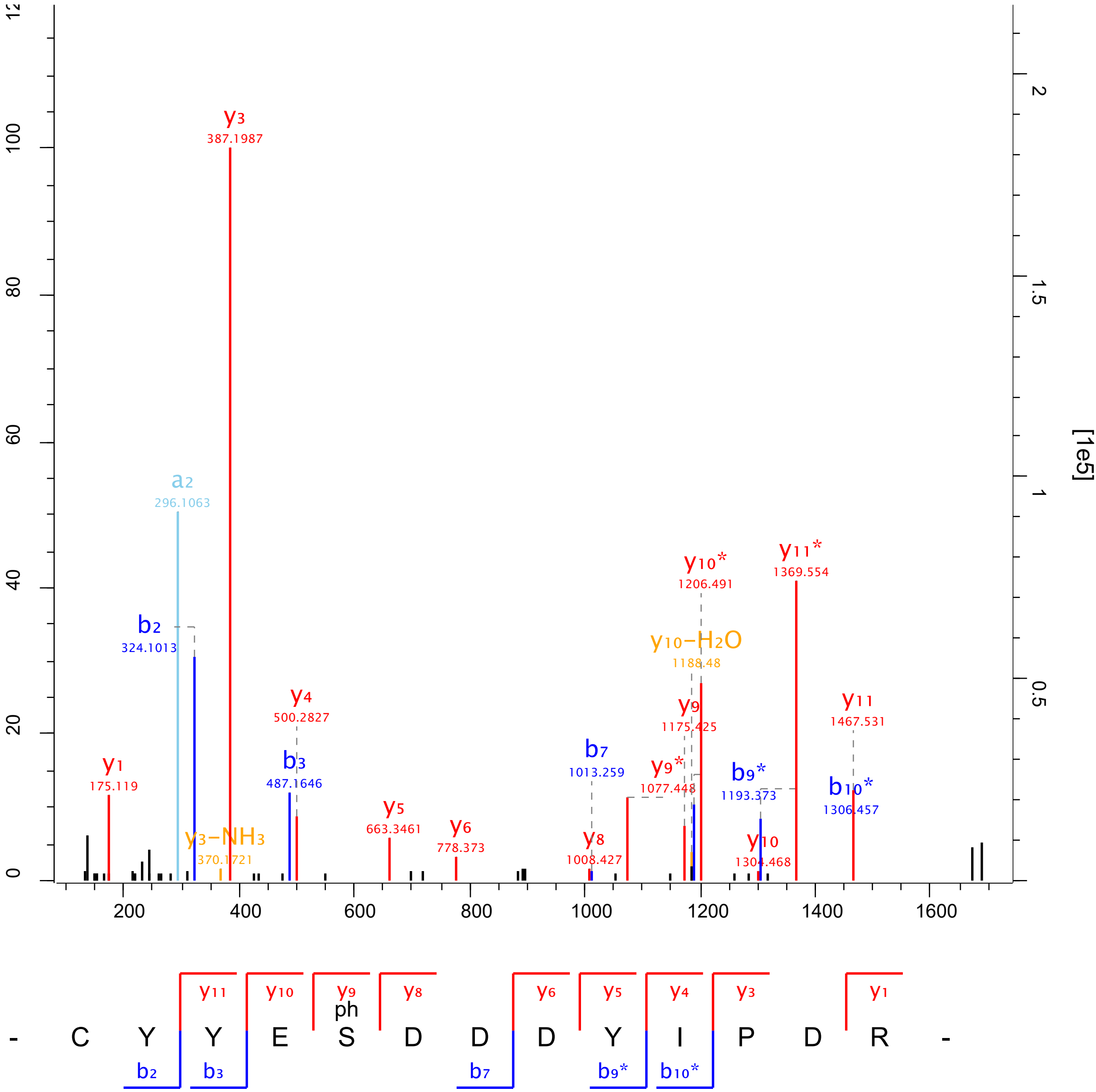

| Raw file | Scan | Method | Score | m/z |
|---|---|---|---|---|
| sys_00_3short | 26954 | FTMS; HCD | 168.84 | 833.41 |

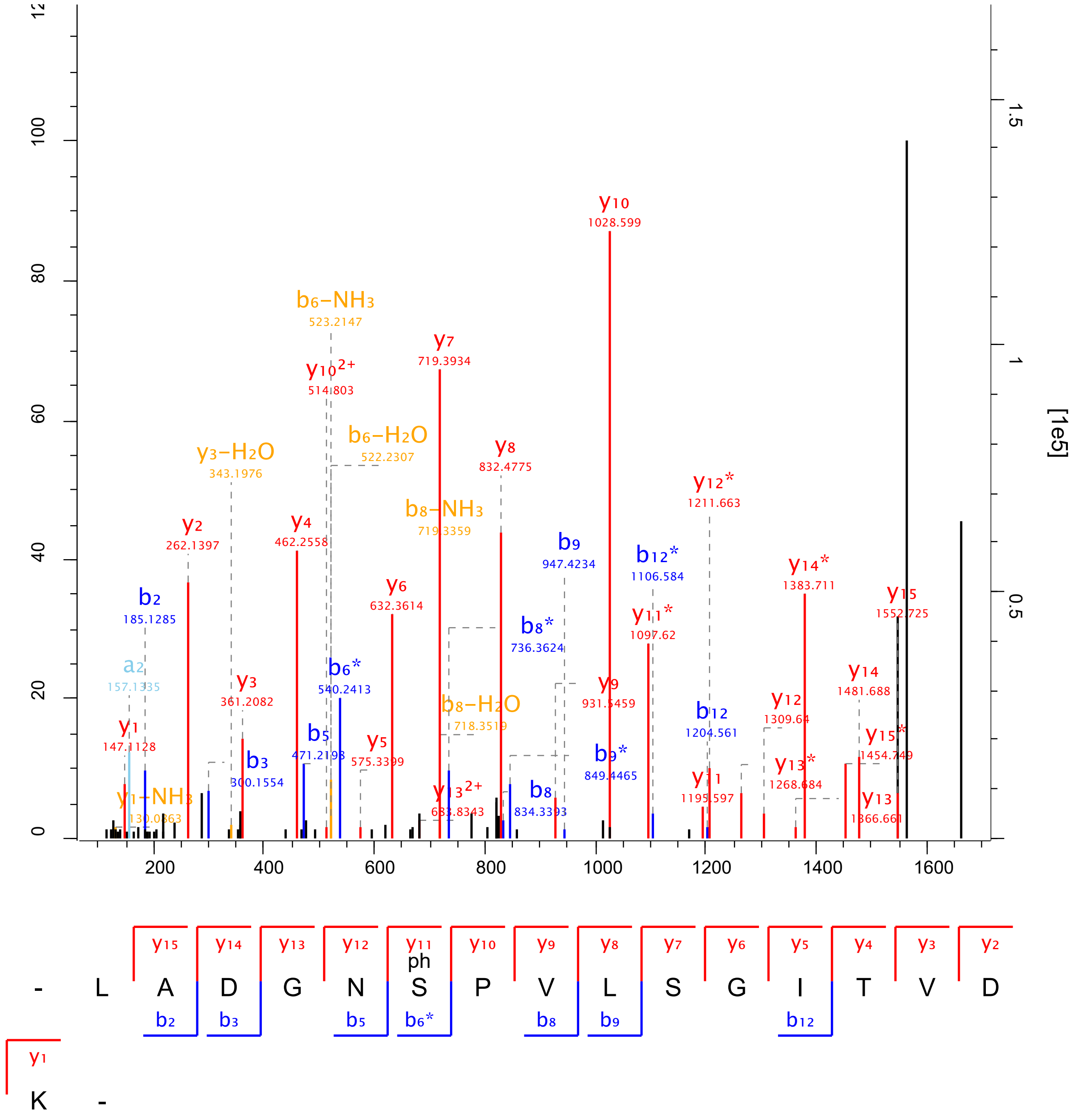

| Raw file | Scan | Method | Score | m/z |
|---|---|---|---|---|
| sys_00_3short | 27519 | FTMS; HCD | 64.52 | 621.29 |

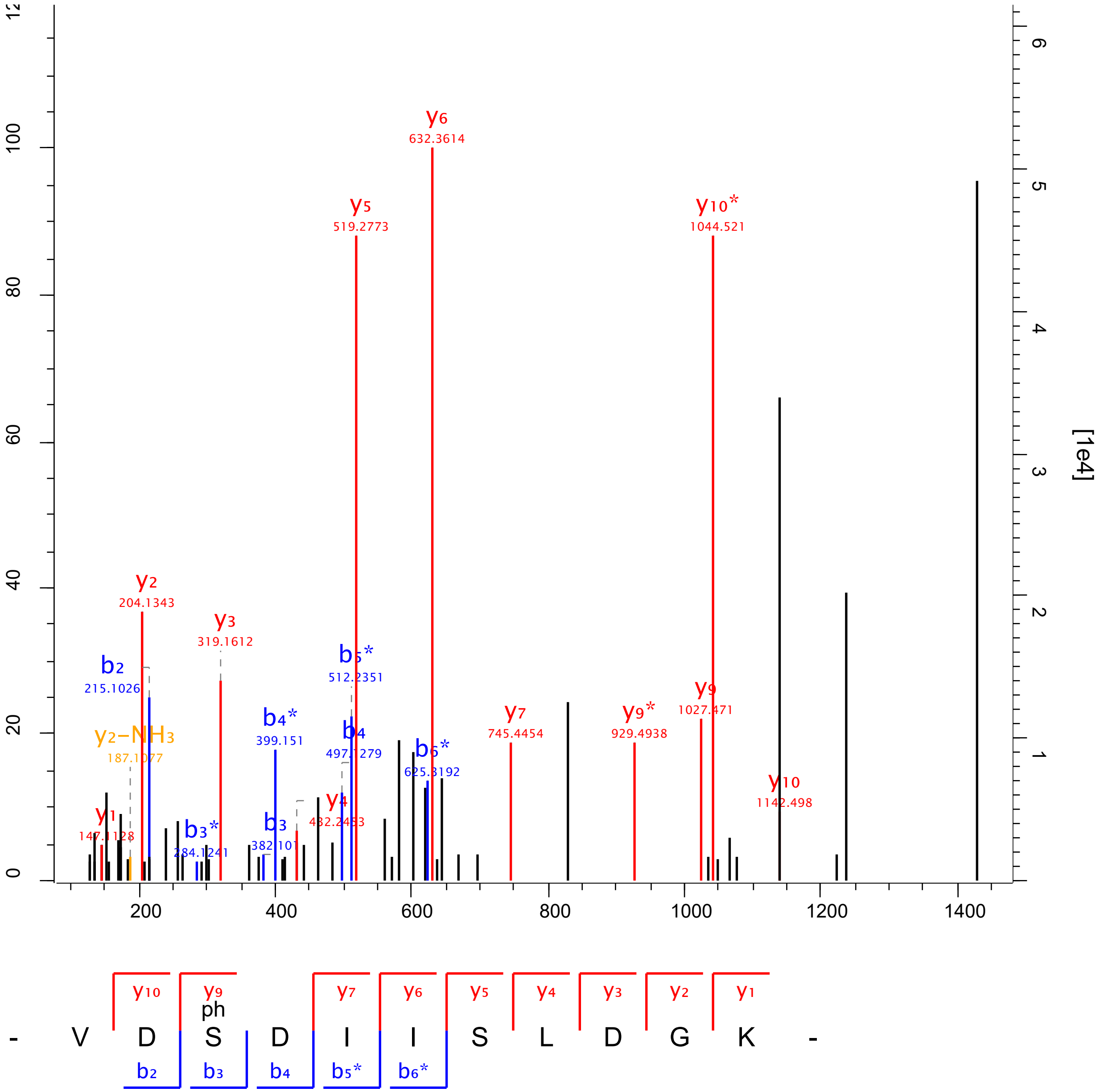

| Raw file | Scan | Method | Score | m/z |
|---|---|---|---|---|
| sys_00_3short | 27536 | FTMS; HCD | 62.09 | 618.78 |

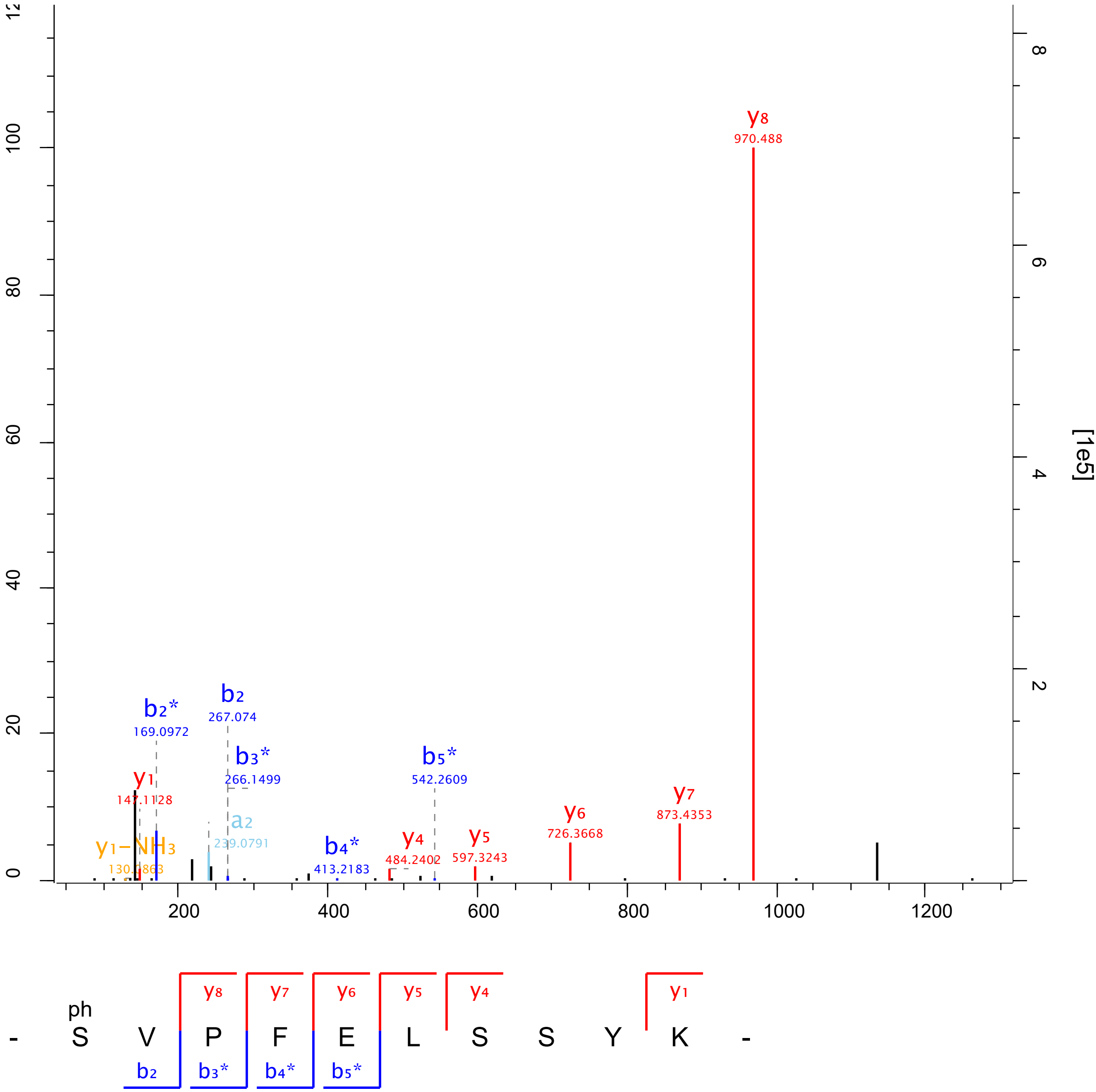

| Raw file | Scan | Method | Score | m/z |
|---|---|---|---|---|
| sys_00_3short | 2774 | FTMS; HCD | 143.7 | 398.2 |

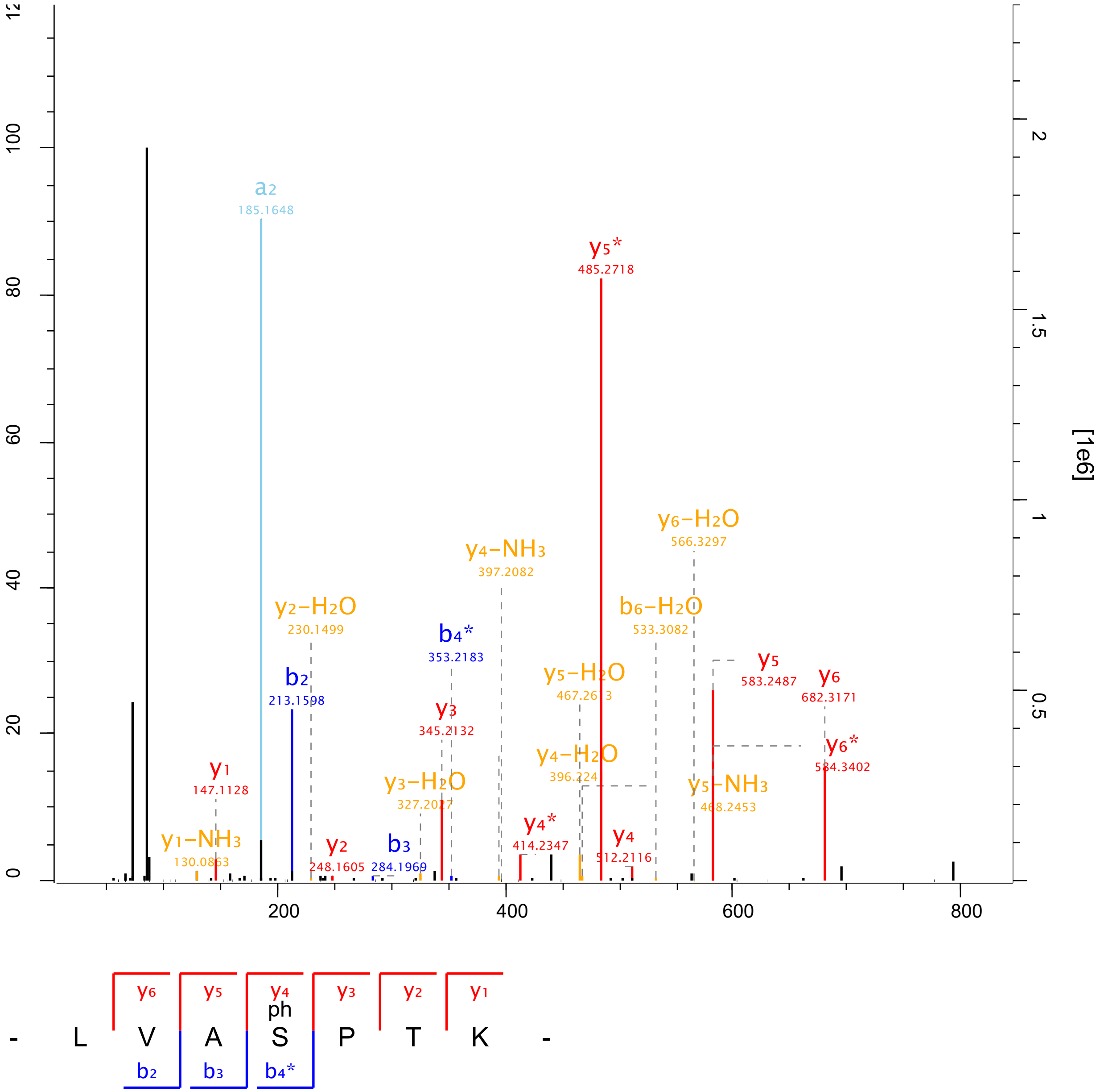

| Raw file | Scan | Method | Score | m/z |
|---|---|---|---|---|
| sys_00_3short | 27750 | FTMS; HCD | 49.12 | 739.68 |

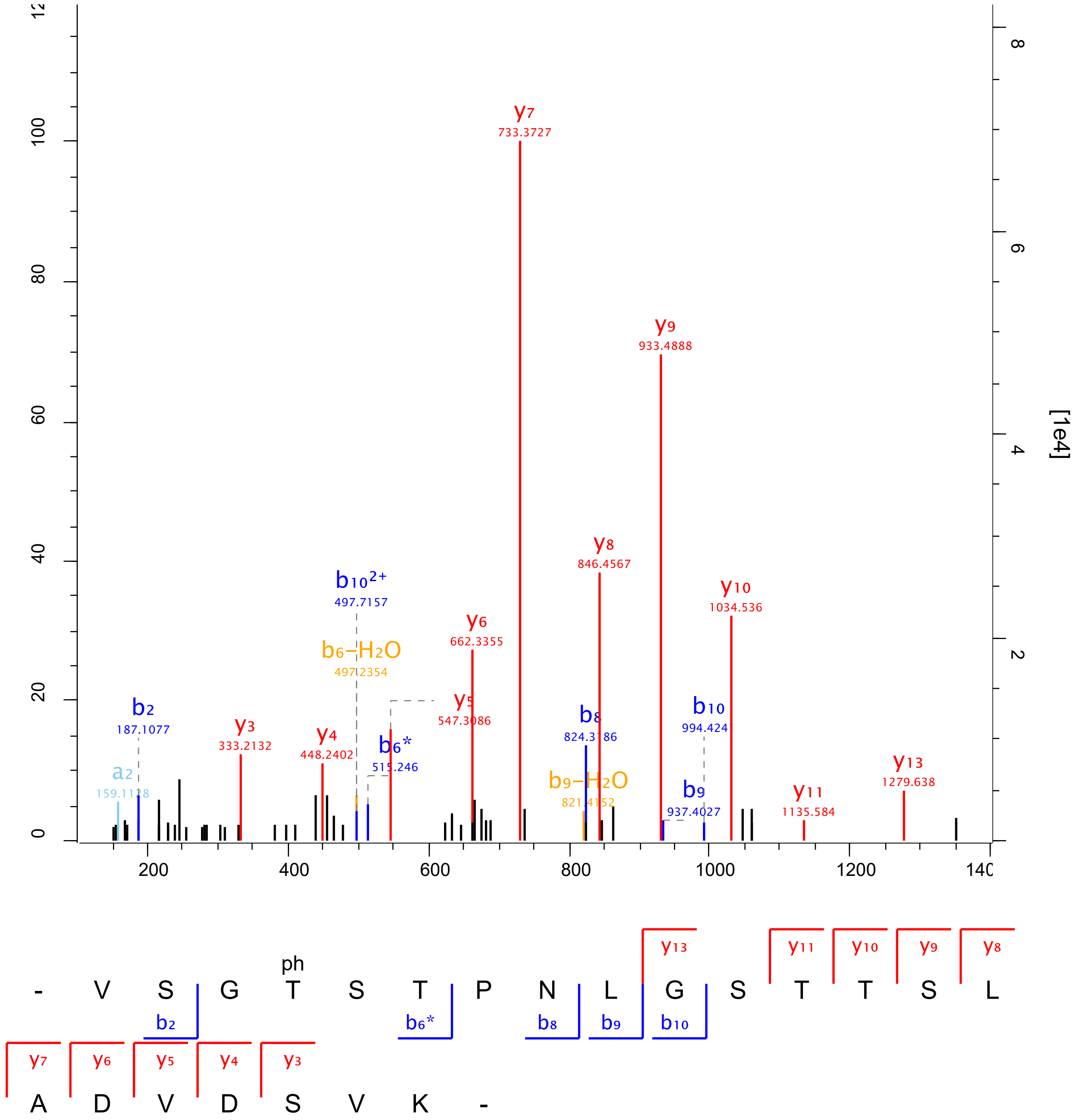

| Raw file | Scan | Method | Score | m/z |
| --- | --- | --- | --- | --- |
| sys_00_3short | 28318 | FTMS; HCD | 72.2 | 527.25 |

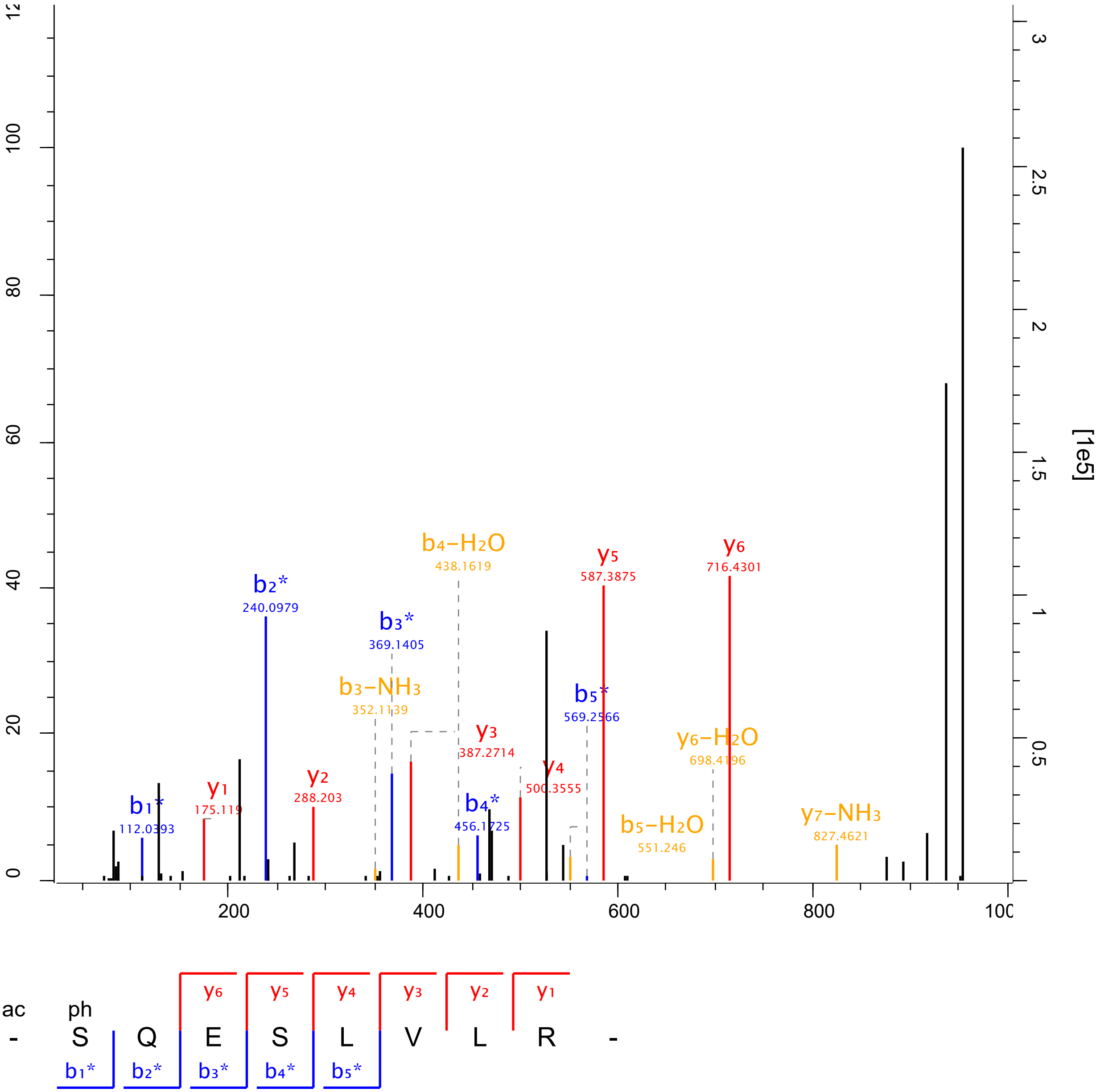

| Raw file | Scan | Method | Score | m/z |
|---|---|---|---|---|
| sys_00_3short | 28403 | FTMS; HCD | 45.58 | 1151.47 |

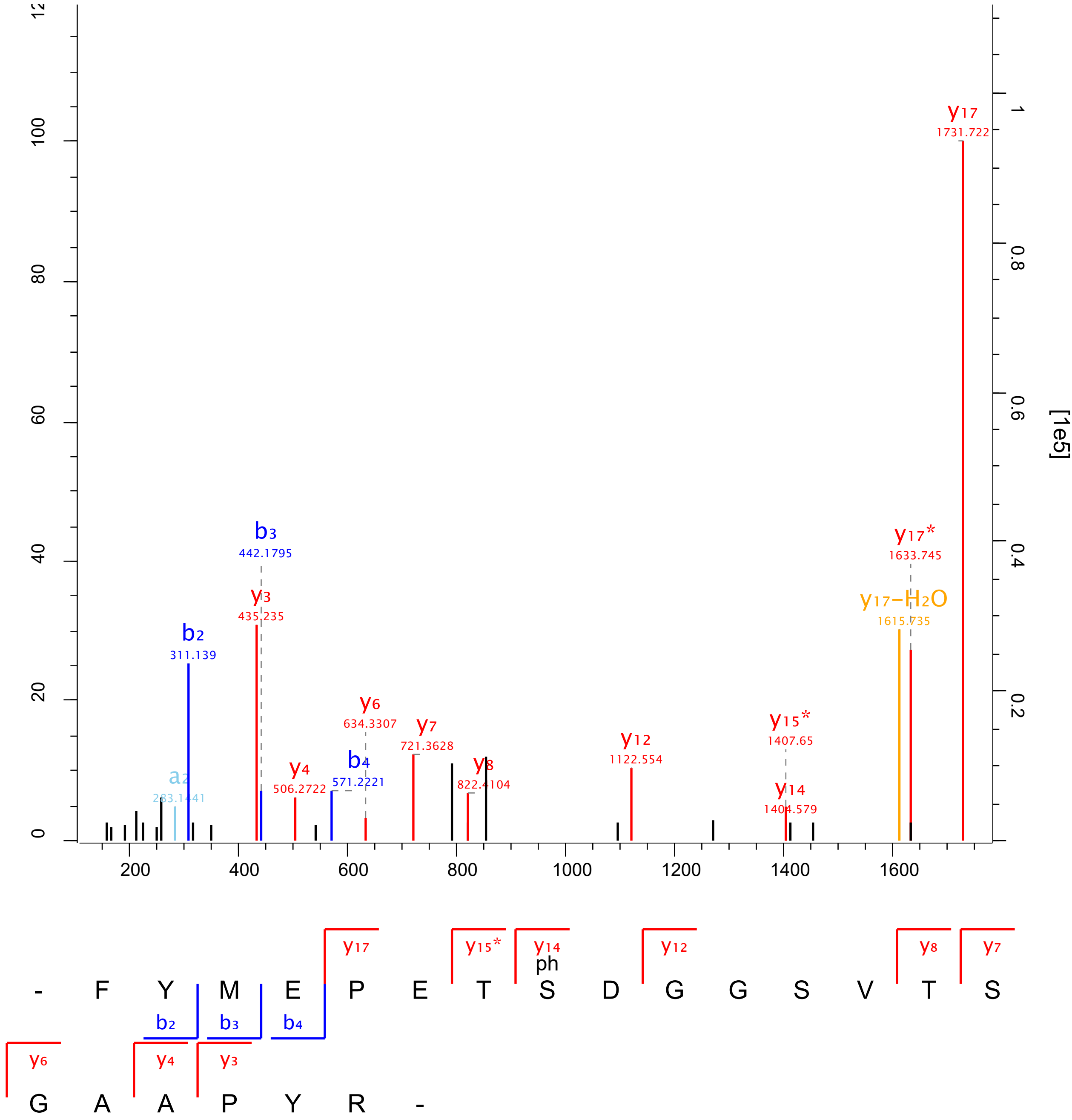

| Raw file | Scan | Method | Score | m/z |
|---|---|---|---|---|
| sys_00_3short | 28802 | FTMS; HCD | 136.6 | 649.8 |

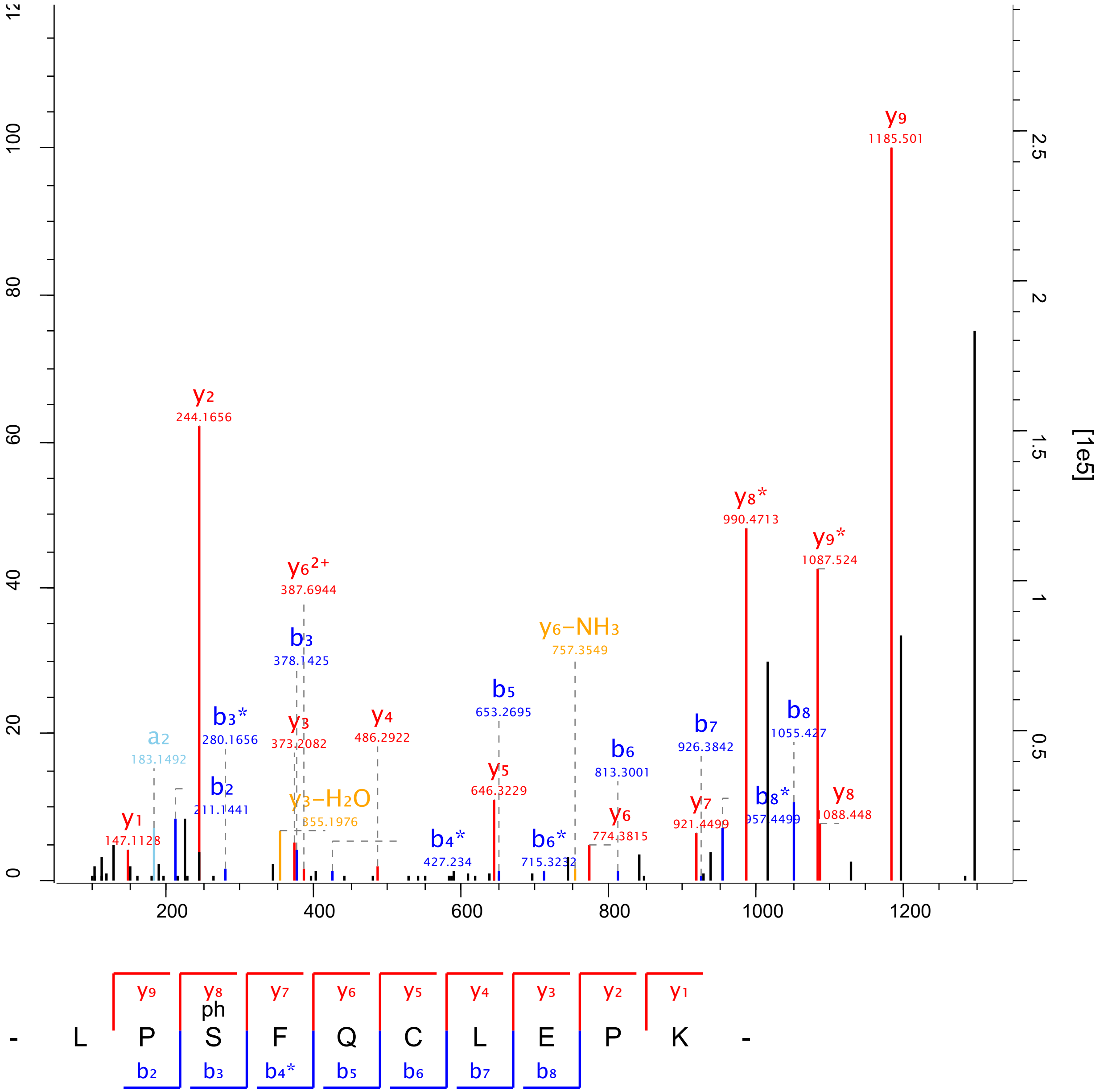

| Raw file | Scan | Method | Score | m/z |
|---|---|---|---|---|
| sys_00_3short | 29095 | FTMS; HCD | 186.94 | 736.3 |

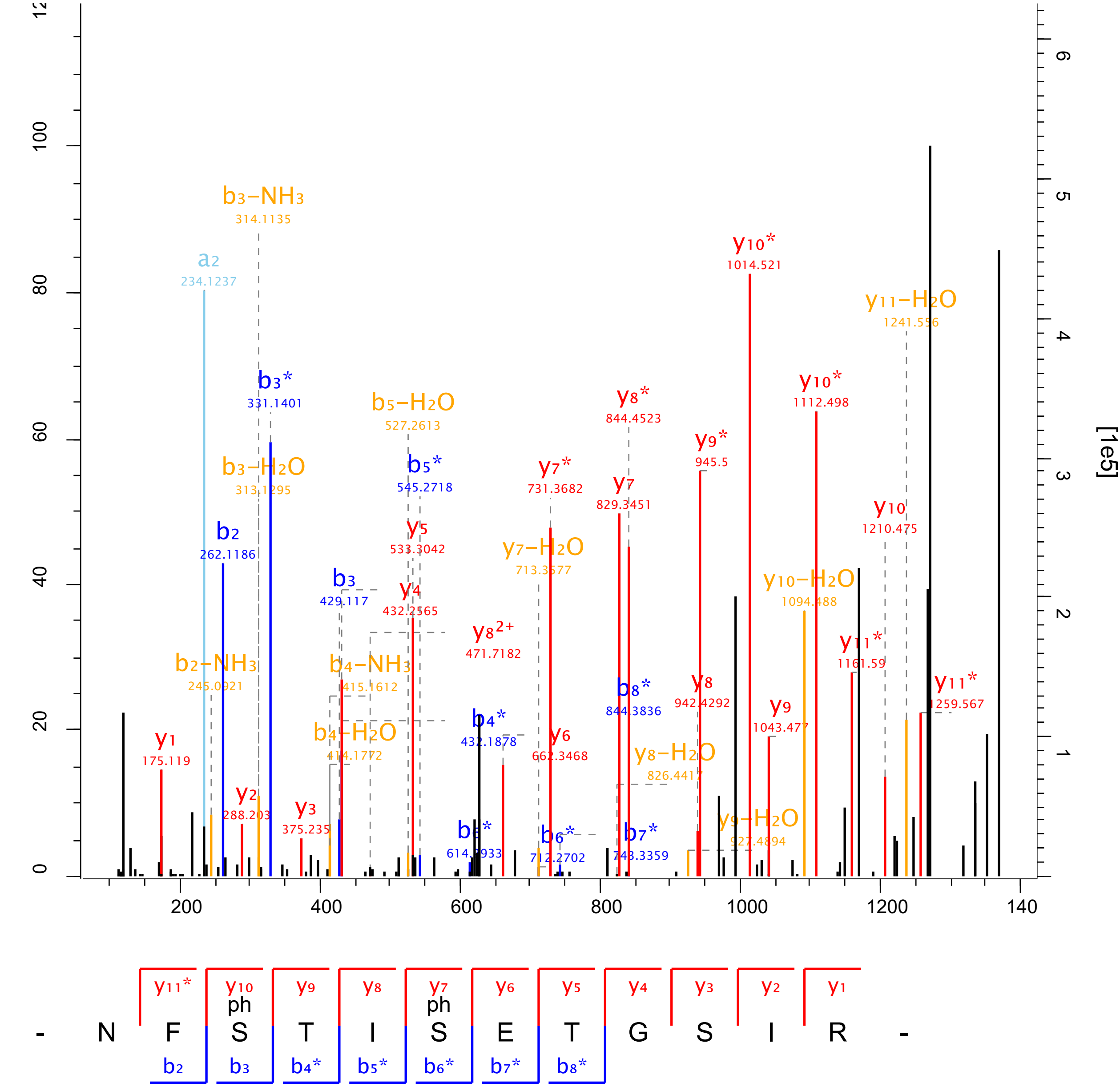

| Raw file | Scan | Method | Score | m/z |
|---|---|---|---|---|
| sys_00_3short | 29176 | FTMS; HCD | 143.28 | 750.87 |

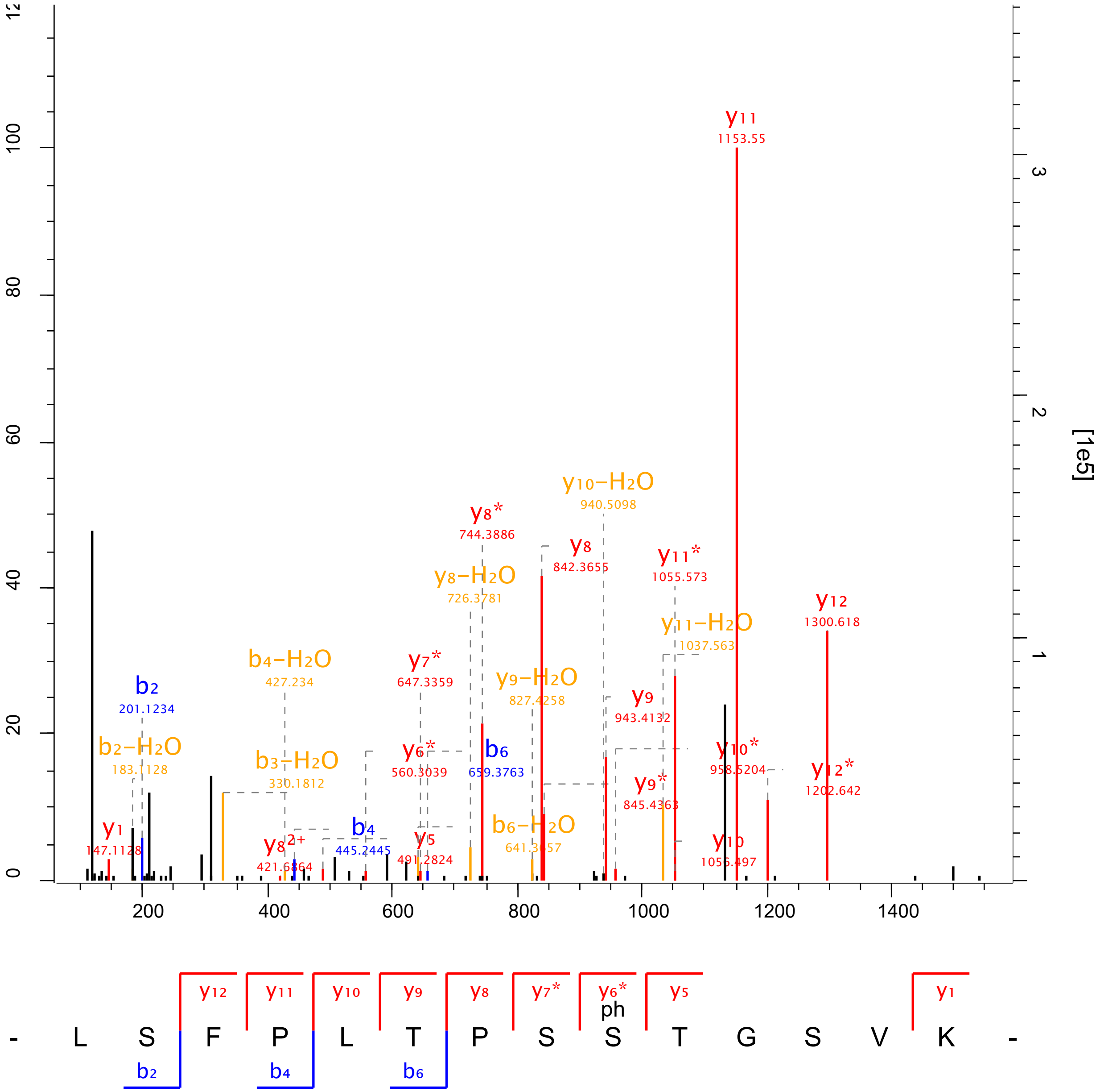

| Raw file | Scan | Method | Score | m/z |
|---|---|---|---|---|
| sys_00_3short | 2951 | FTMS; HCD | 119.39 | 522.22 |

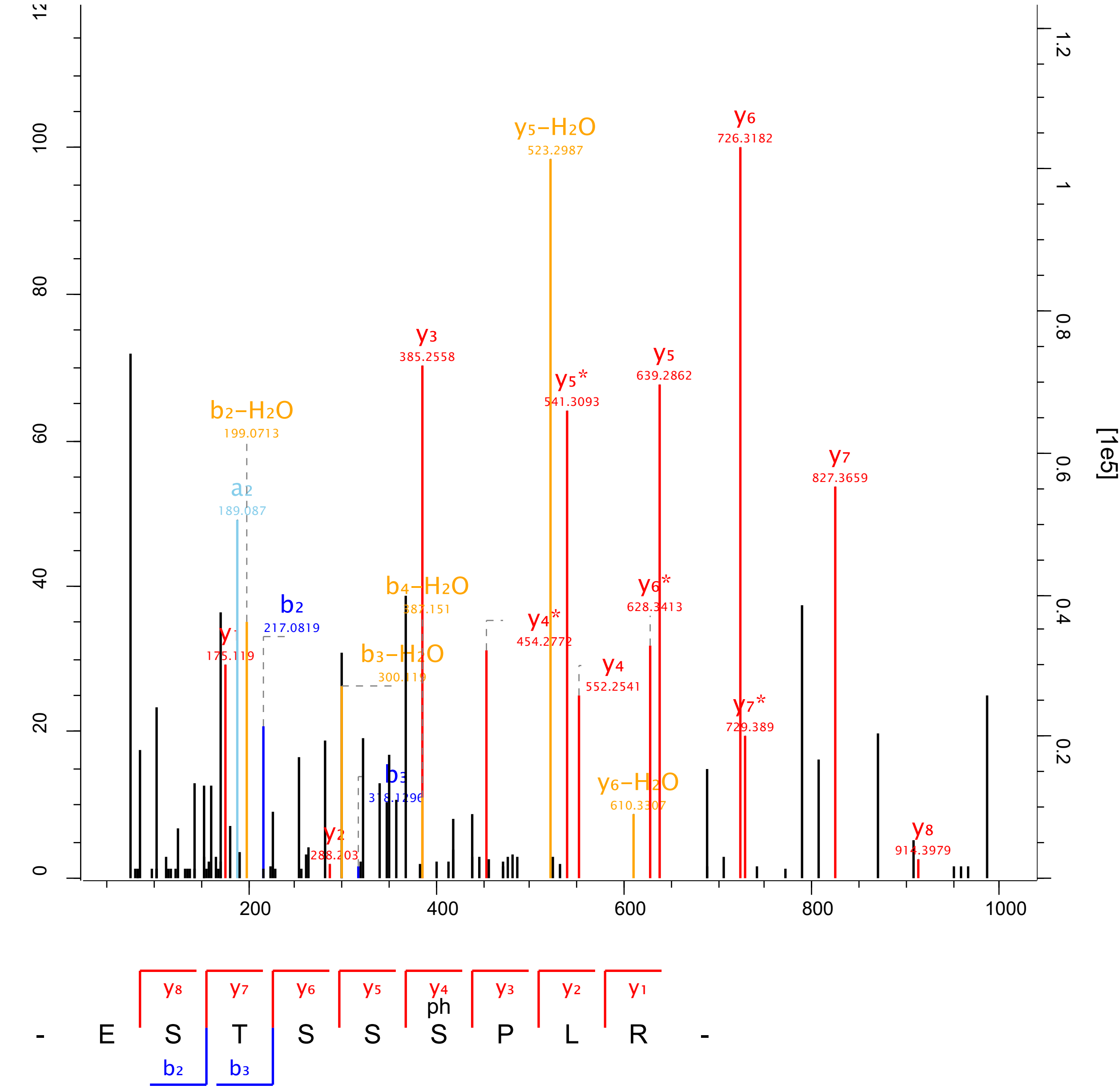

| Raw file | Scan | Method | Score | m/z |
| --- | --- | --- | --- | --- |
| sys_00_3short | 29759 | FTMS; HCD | 68.48 | 692.33 |

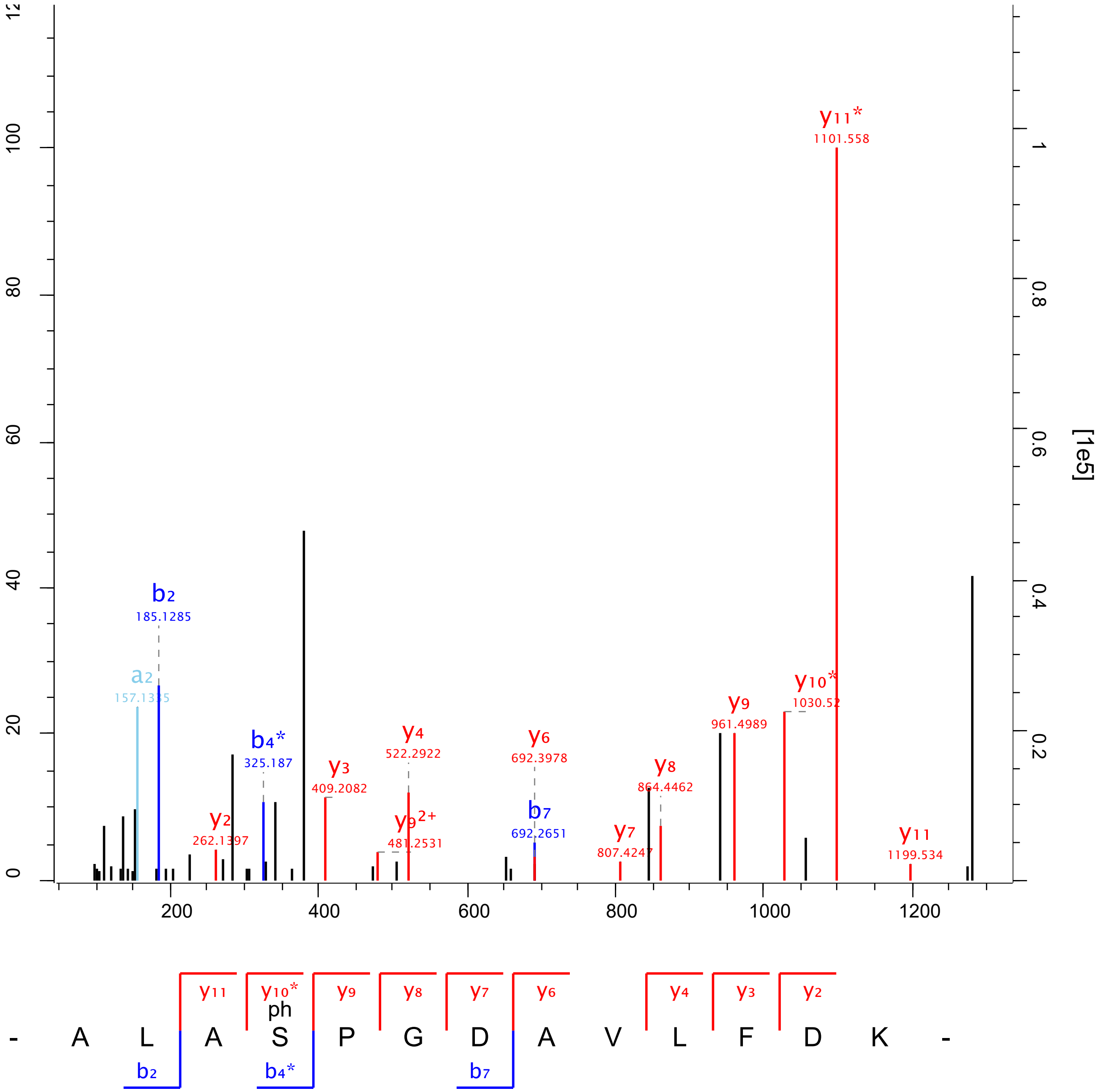

| Raw file | Scan | Method | Score | m/z |
|---|---|---|---|---|
| sys_00_3short | 29829 | FTMS; HCD | 62.2 | 760.83 |

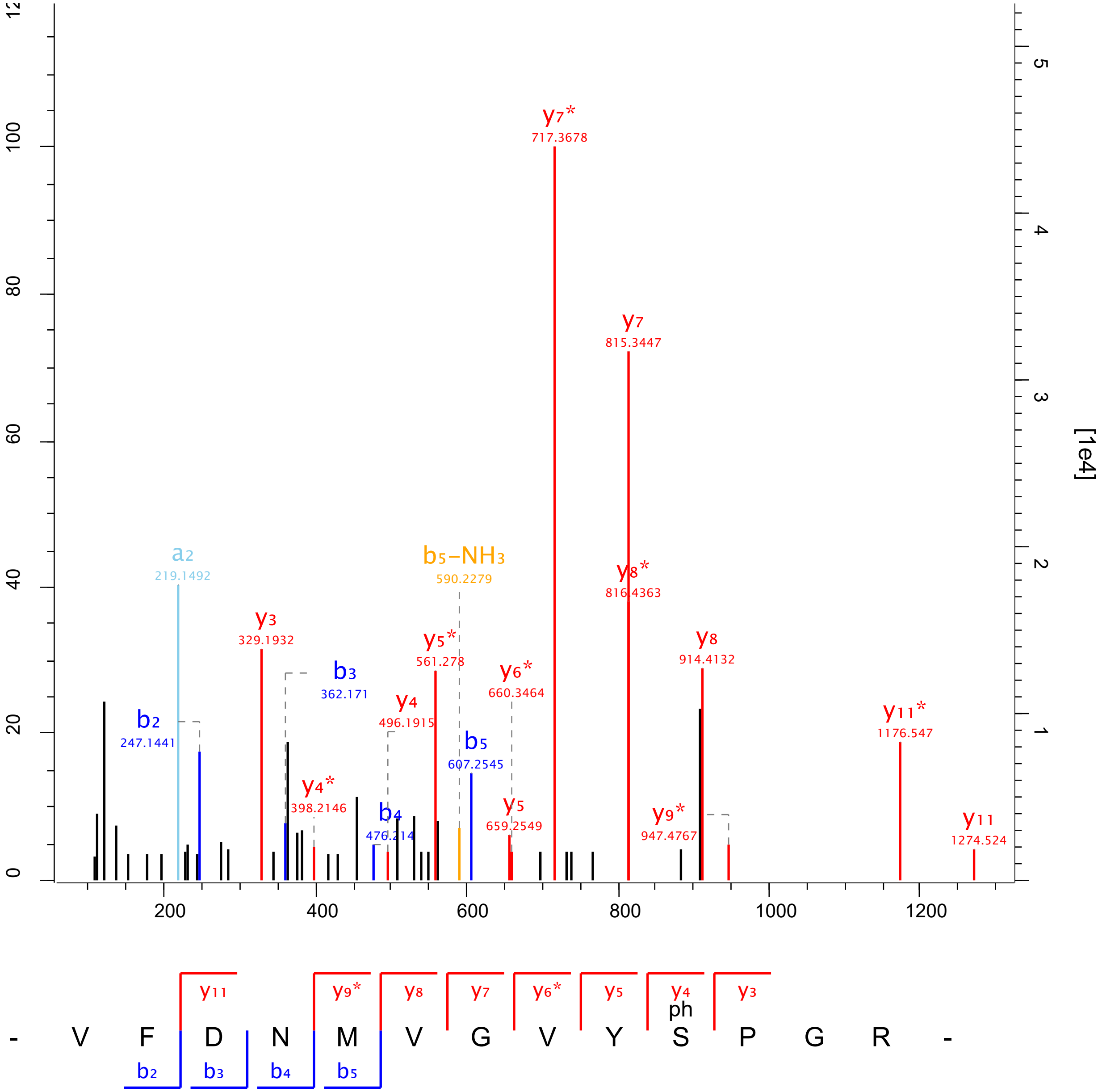

| Raw file | Scan | Method | Score | m/z |
|---|---|---|---|---|
| sys_00_3short | 29941 | FTMS; HCD | 113.38 | 573.73 |

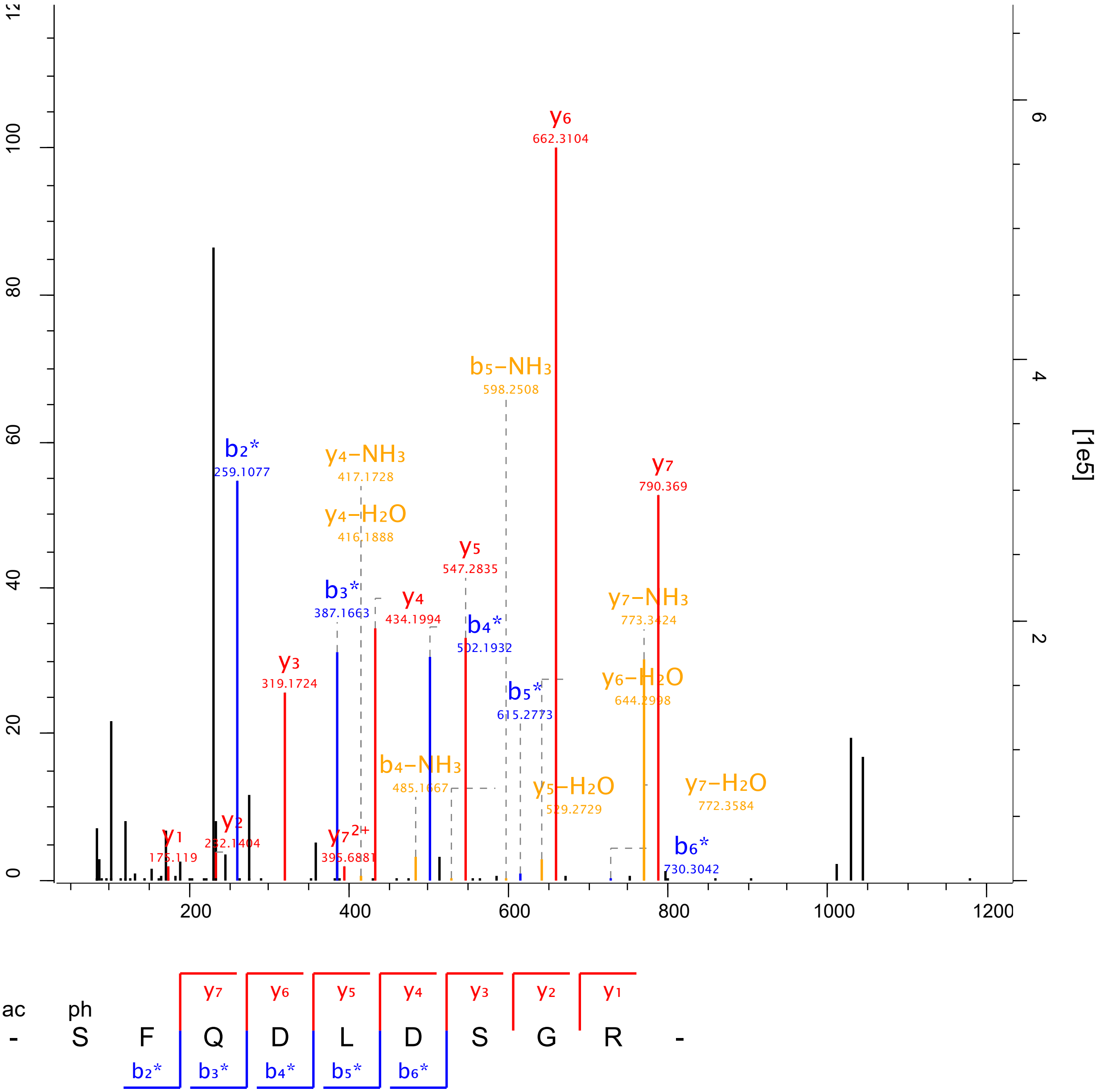

| Raw file | Scan | Method | Score | m/z |
| --- | --- | --- | --- | --- |
| sys_00_3short | 30029 | FTMS; HCD | 80.31 | 590.81 |

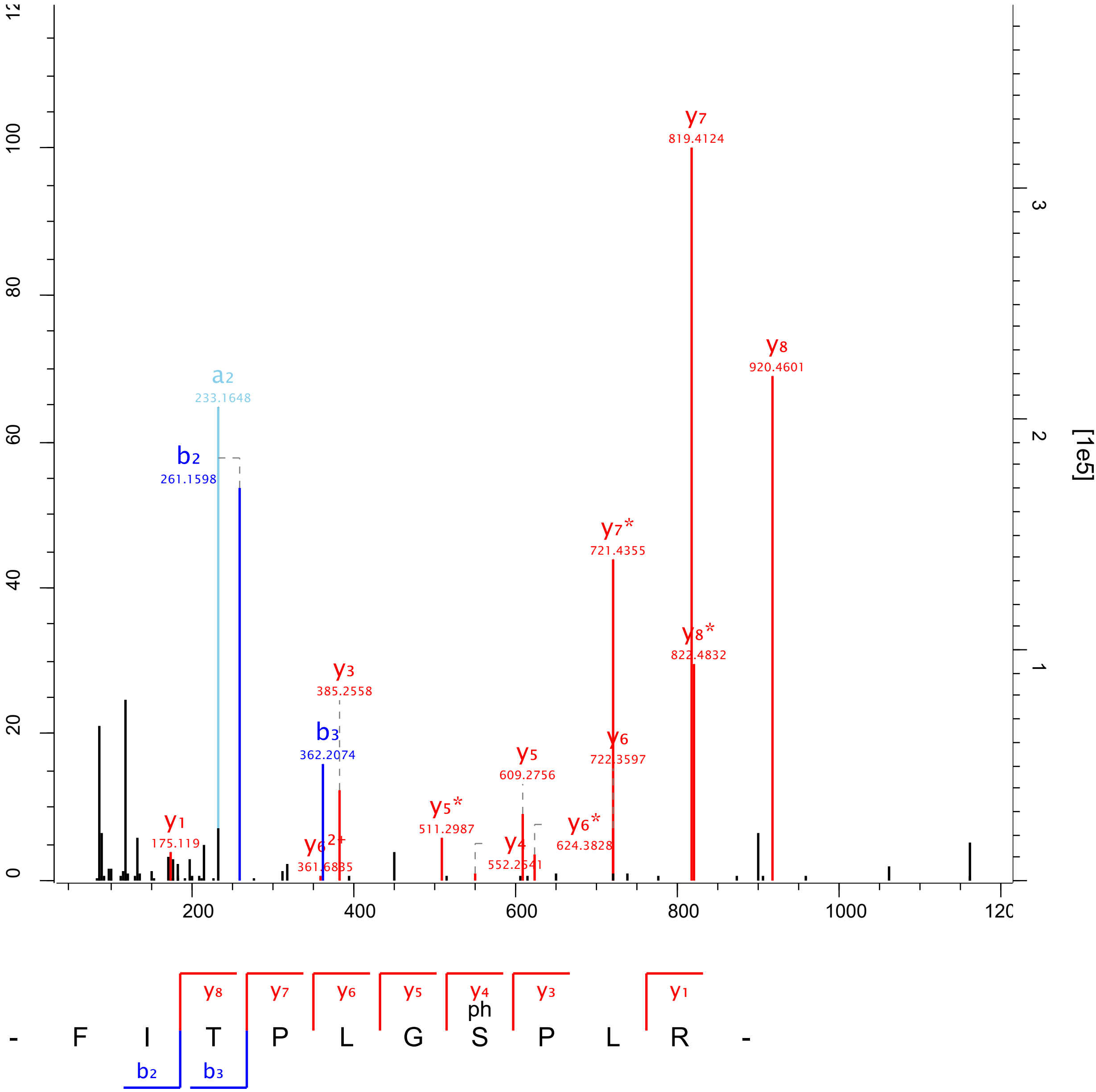

| Raw file | Scan | Method | Score | m/z |
|---|---|---|---|---|
| sys_00_3short | 30170 | FTMS; HCD | 141.55 | 707.3 |

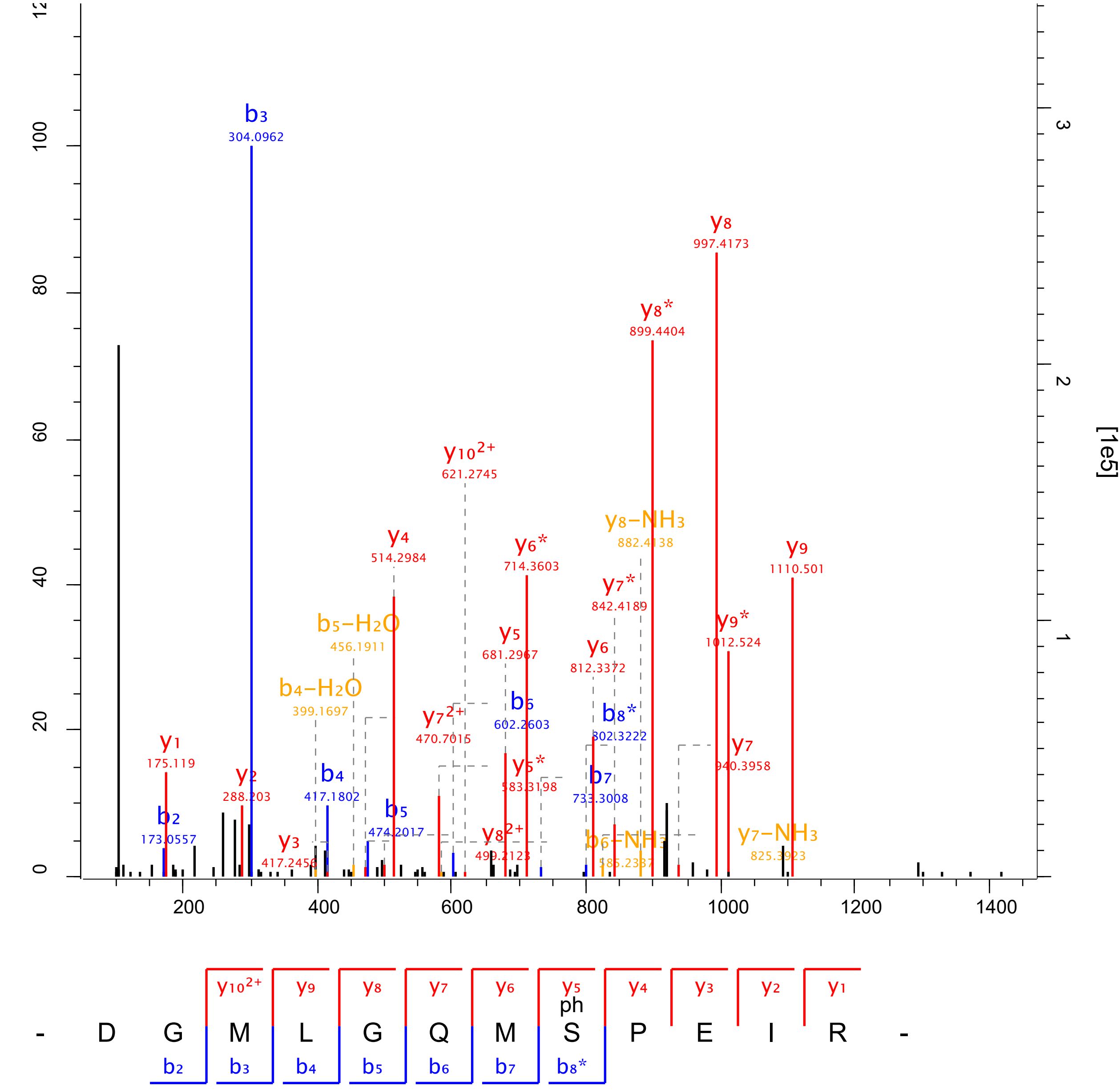

| Raw file | Scan | Method | Score | m/z |
|---|---|---|---|---|
| sys_00_3short | 30458 | FTMS; HCD | 48.05 | 842.89 |

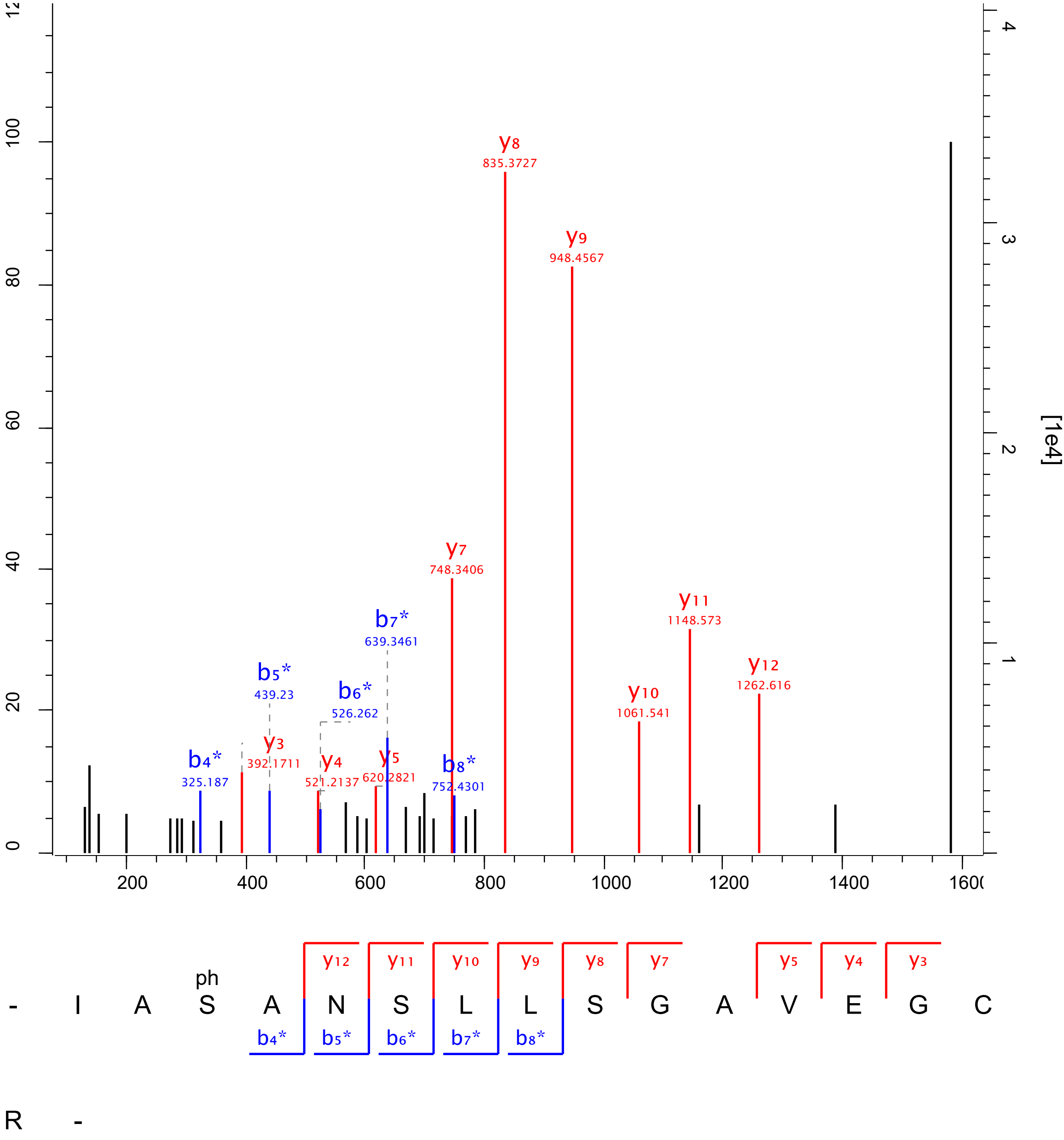

| Raw file | Scan | Method | Score | m/z |
|---|---|---|---|---|
| sys_00_3short | 30506 | FTMS; HCD | 96.14 | 865.36 |

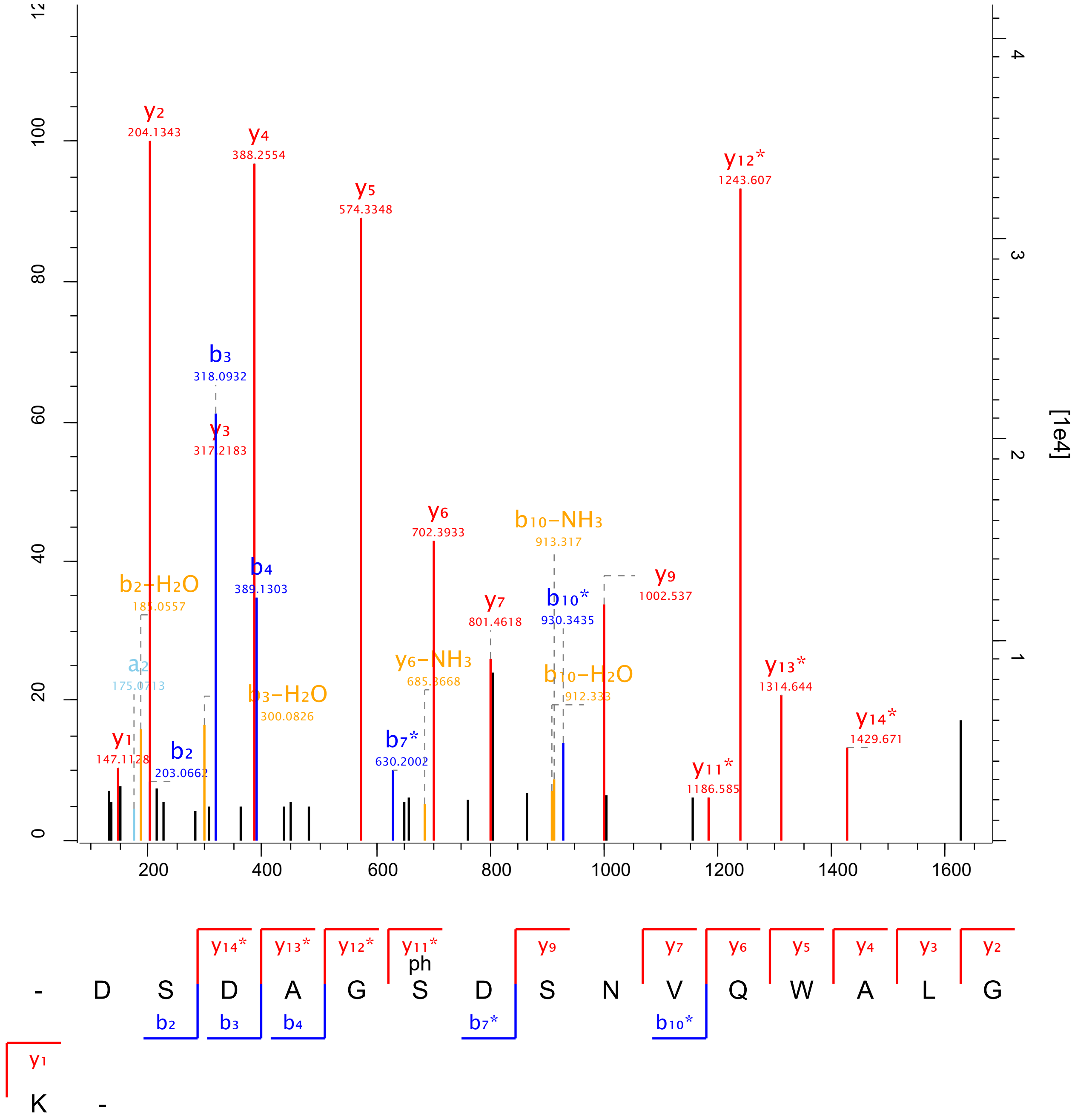

| Raw file | Scan | Method | Score | m/z |
|---|---|---|---|---|
| sys_00_3short | 30548 | FTMS; HCD | 106.57 | 769.9 |

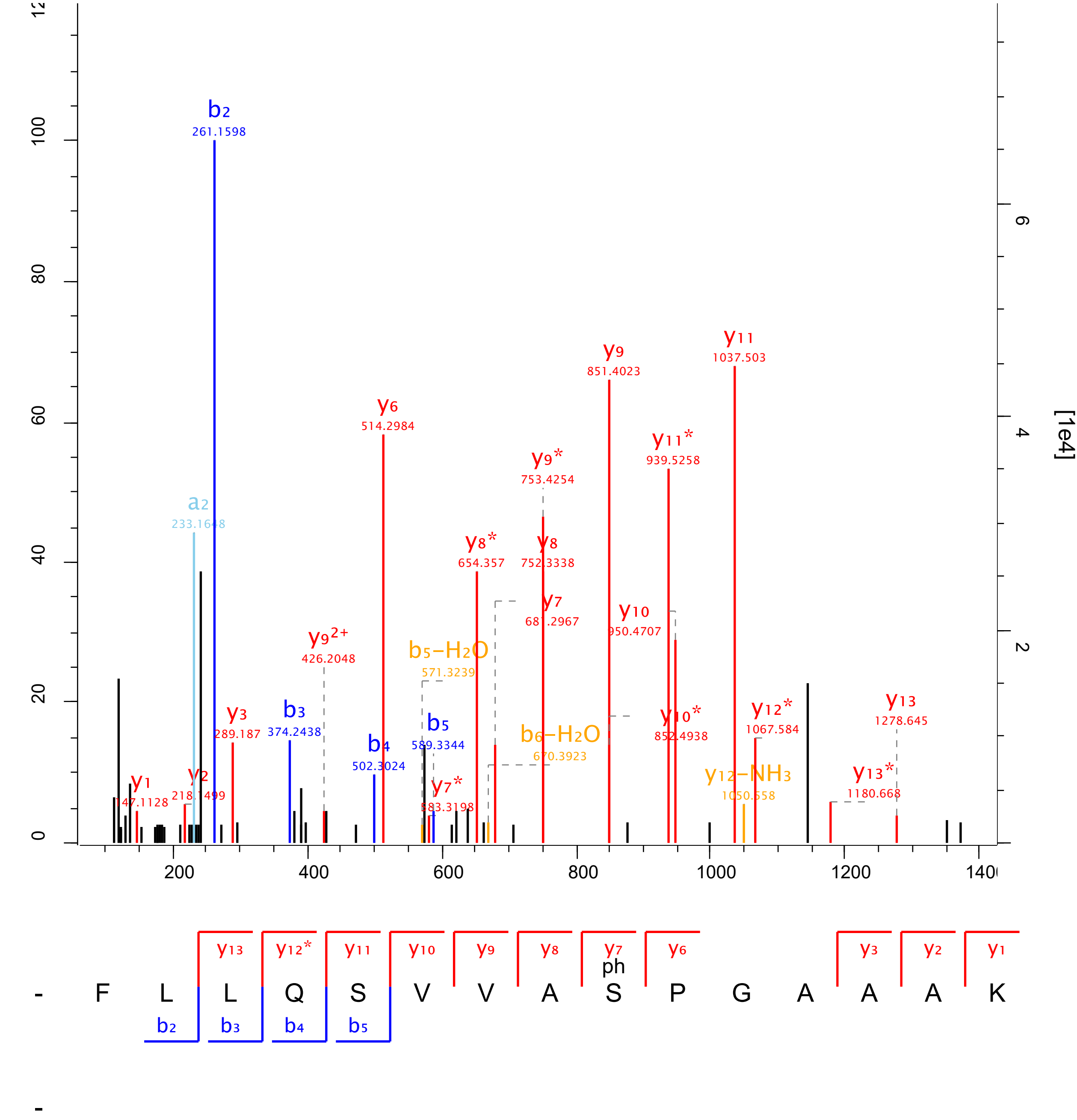

| Raw file | Scan | Method | Score | m/z |
|---|---|---|---|---|
| sys_00_3short | 30791 | FTMS; HCD | 222.81 | 637.79 |

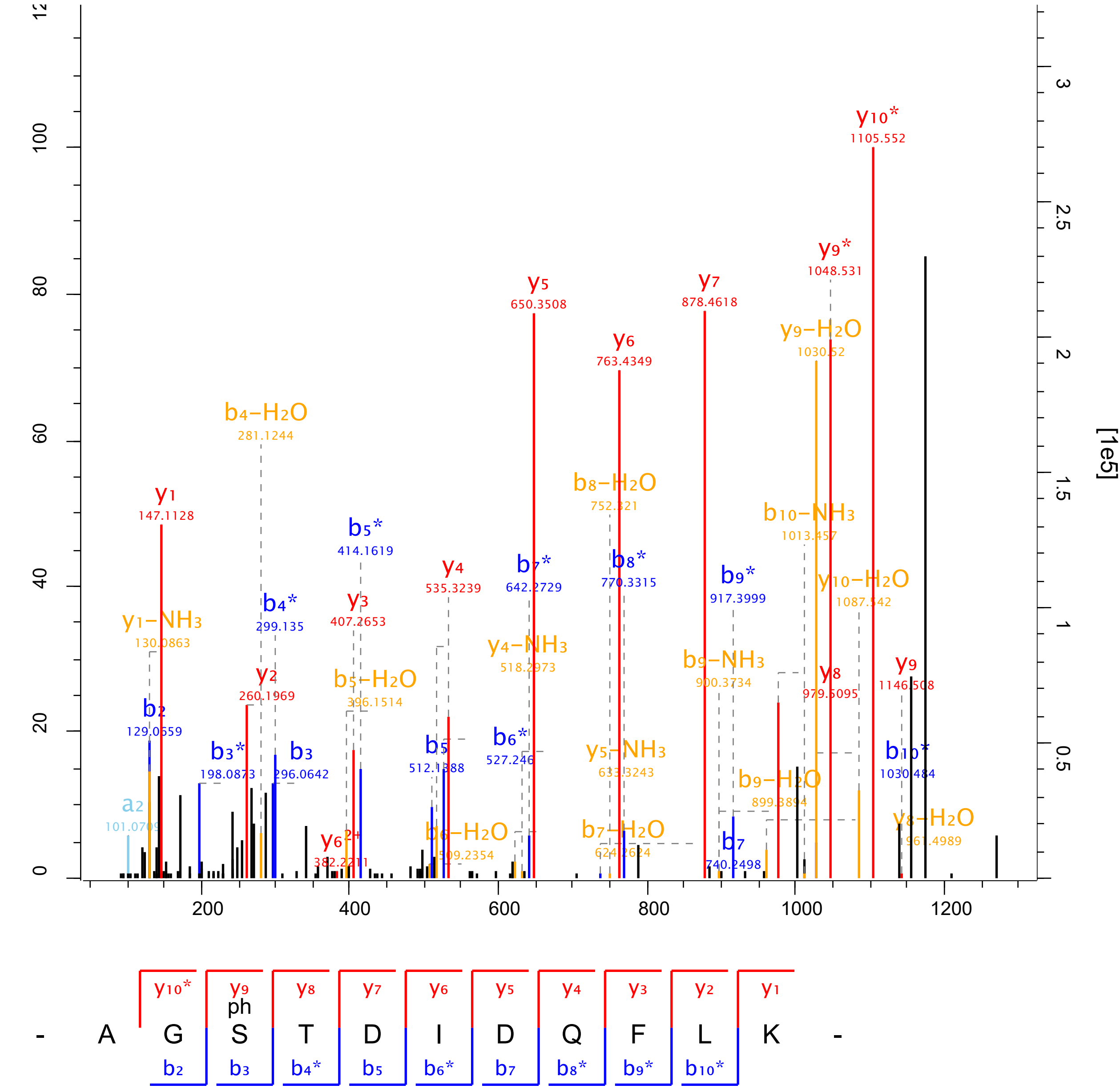

| Raw file | Scan | Method | Score | m/z |
|---|---|---|---|---|
| sys_00_3short | 30987 | FTMS; HCD | 126.71 | 721.33 |

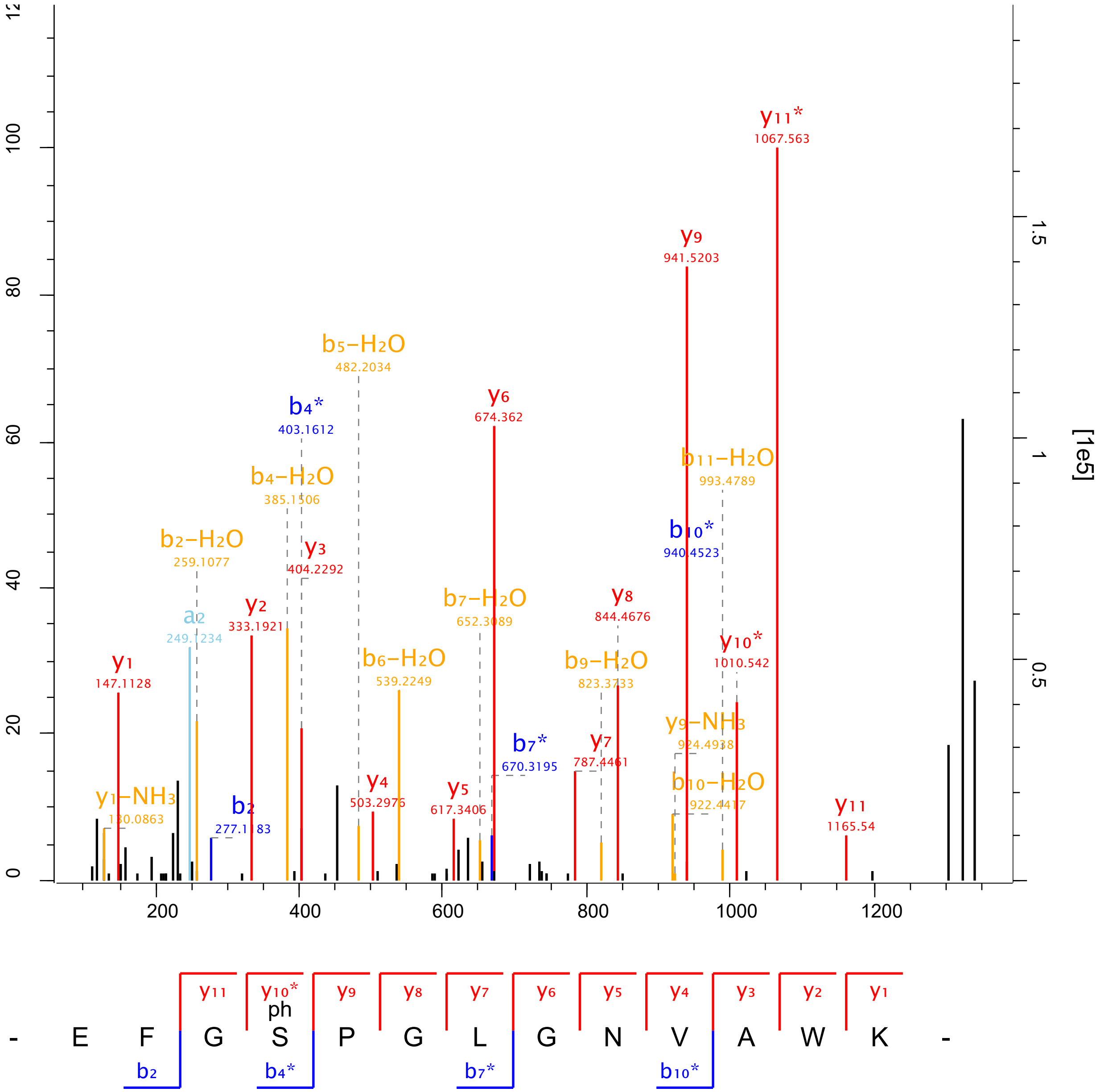

| Raw file | Scan | Method | Score | m/z |
|---|---|---|---|---|
| sys_00_3short | 31060 | FTMS; HCD | 80.75 | 626.81 |

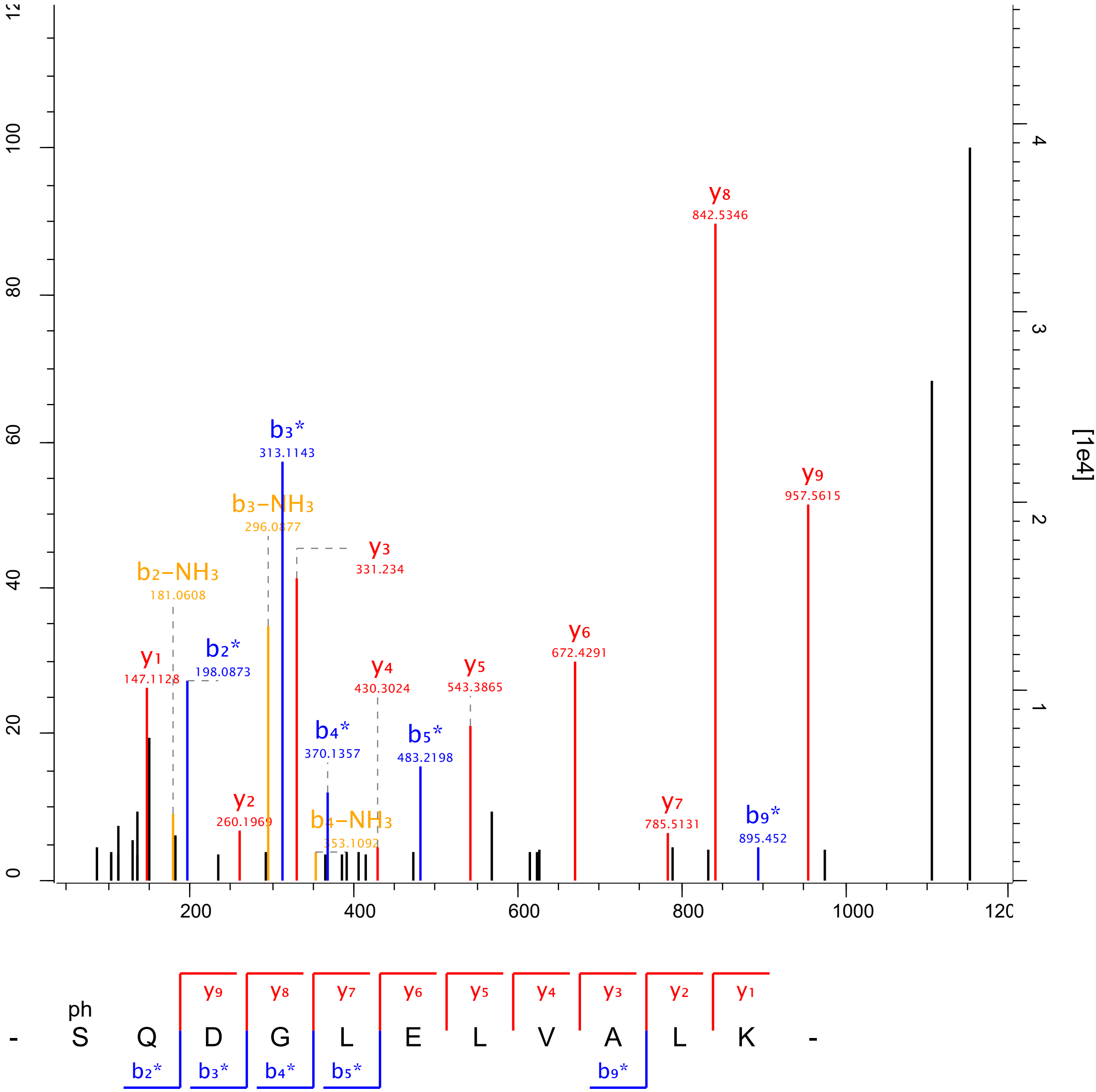

| Raw file | Scan | Method | Score | m/z |
|---|---|---|---|---|
| sys_00_3short | 31127 | FTMS; HCD | 206.62 | 837.33 |

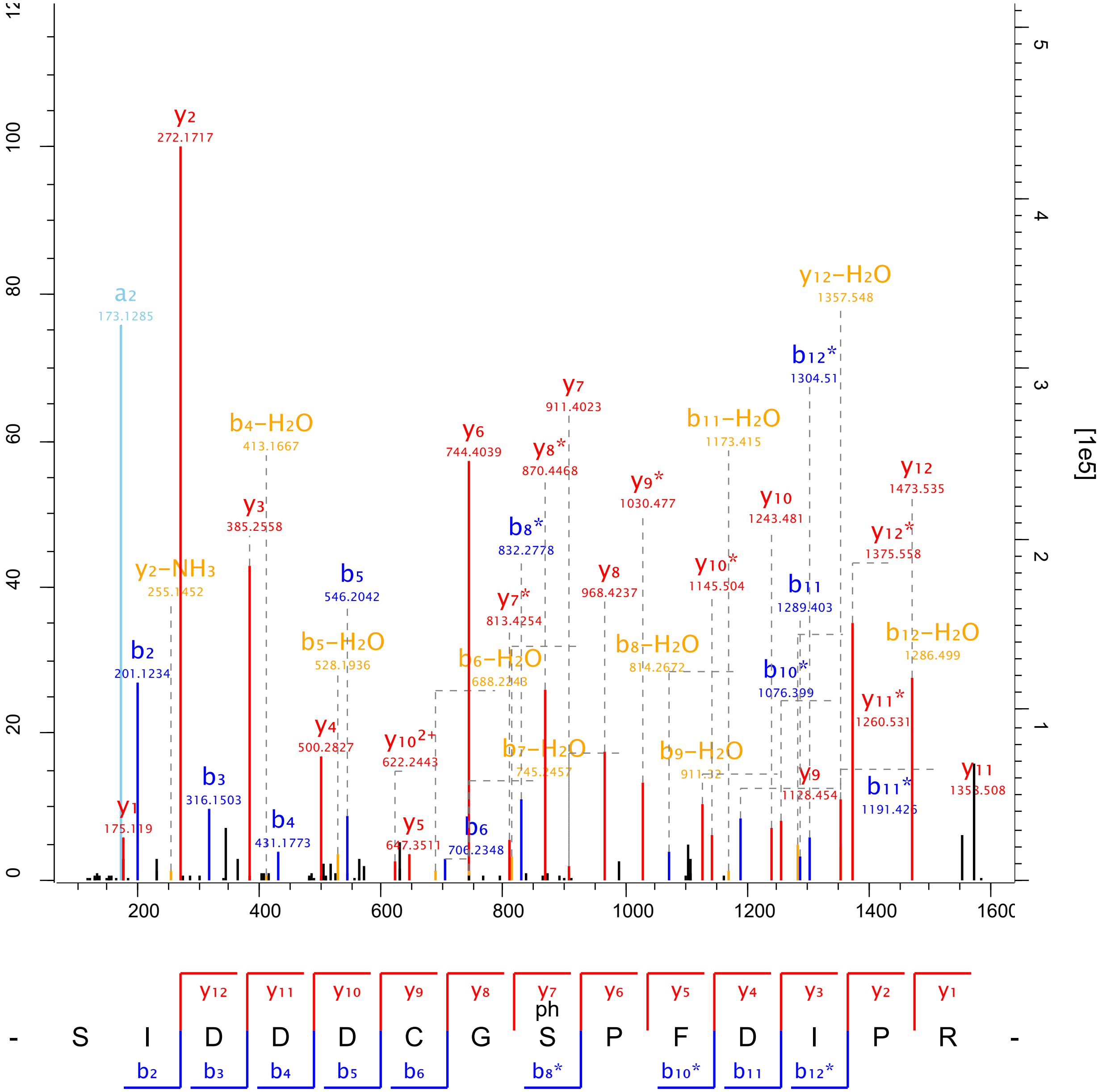

| Raw file | Scan | Method | Score | m/z |
|---|---|---|---|---|
| sys_00_3short | 31164 | FTMS; HCD | 272.72 | 844.83 |

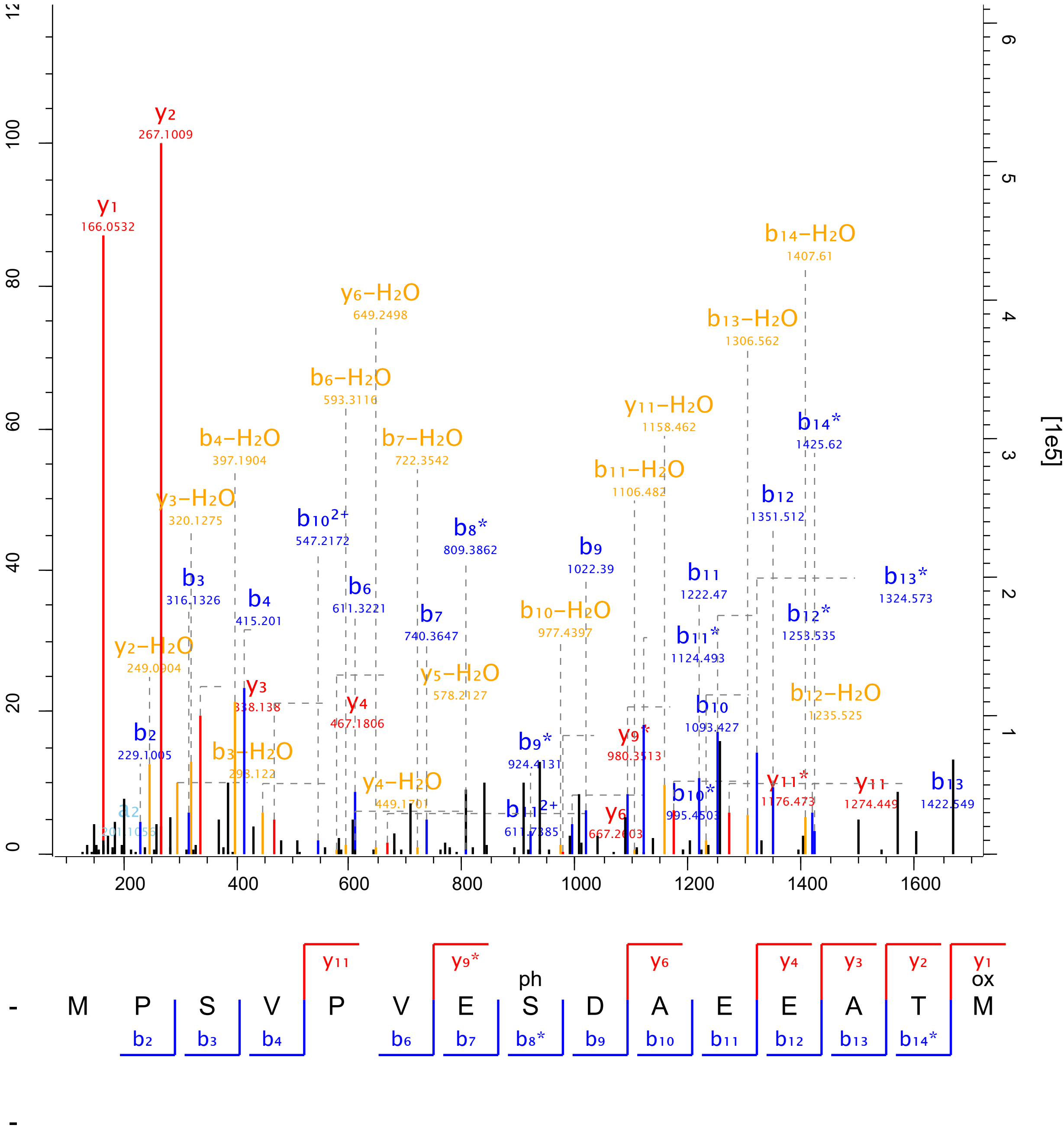

| Raw file | Scan | Method | Score | m/z |
|---|---|---|---|---|
| sys_00_3short | 31198 | FTMS; HCD | 67.65 | 816.37 |

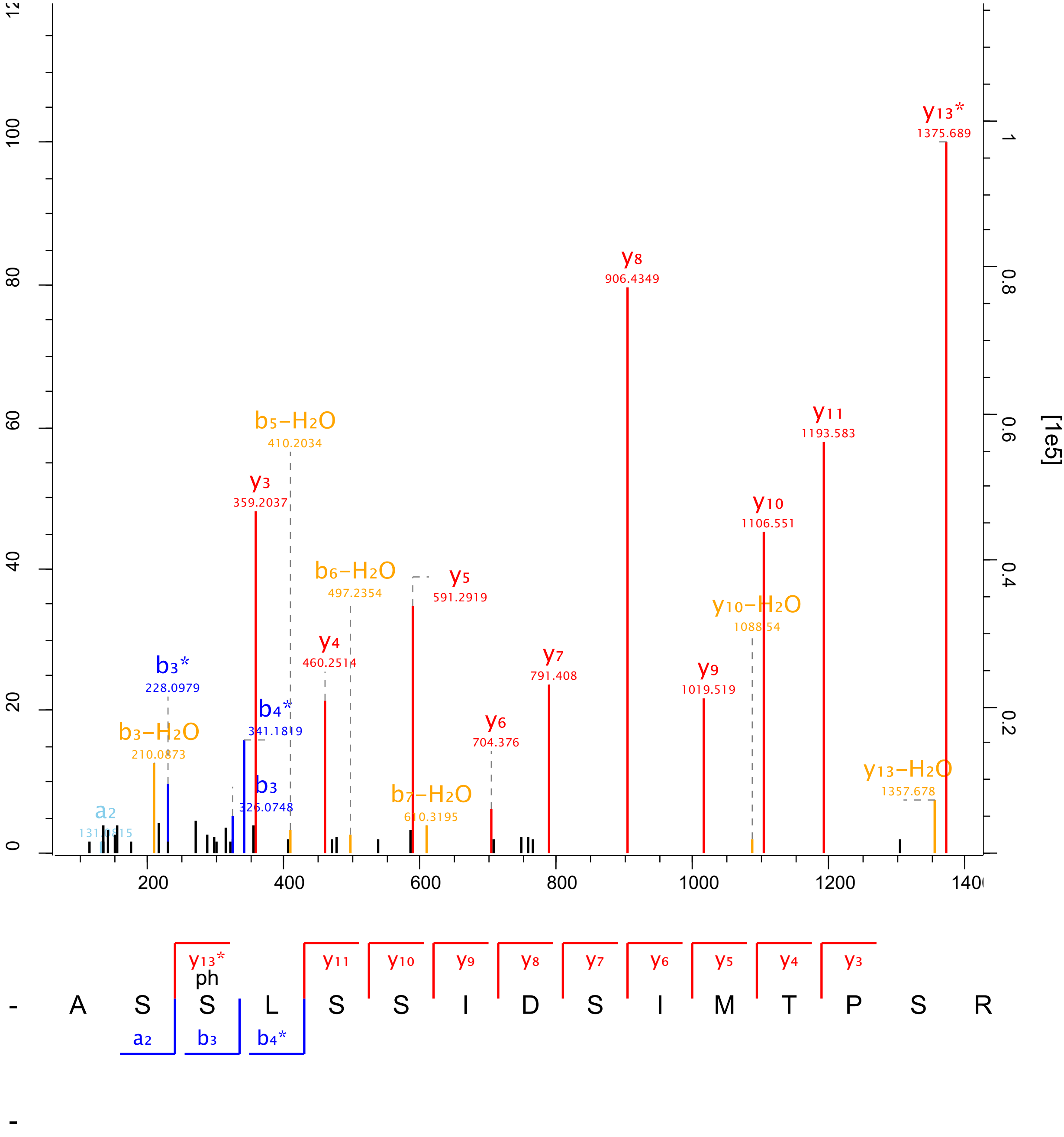

| Raw file | Scan | Method | Score | m/z |
|---|---|---|---|---|
| sys_00_3short | 31306 | FTMS; HCD | 54.34 | 617.77 |

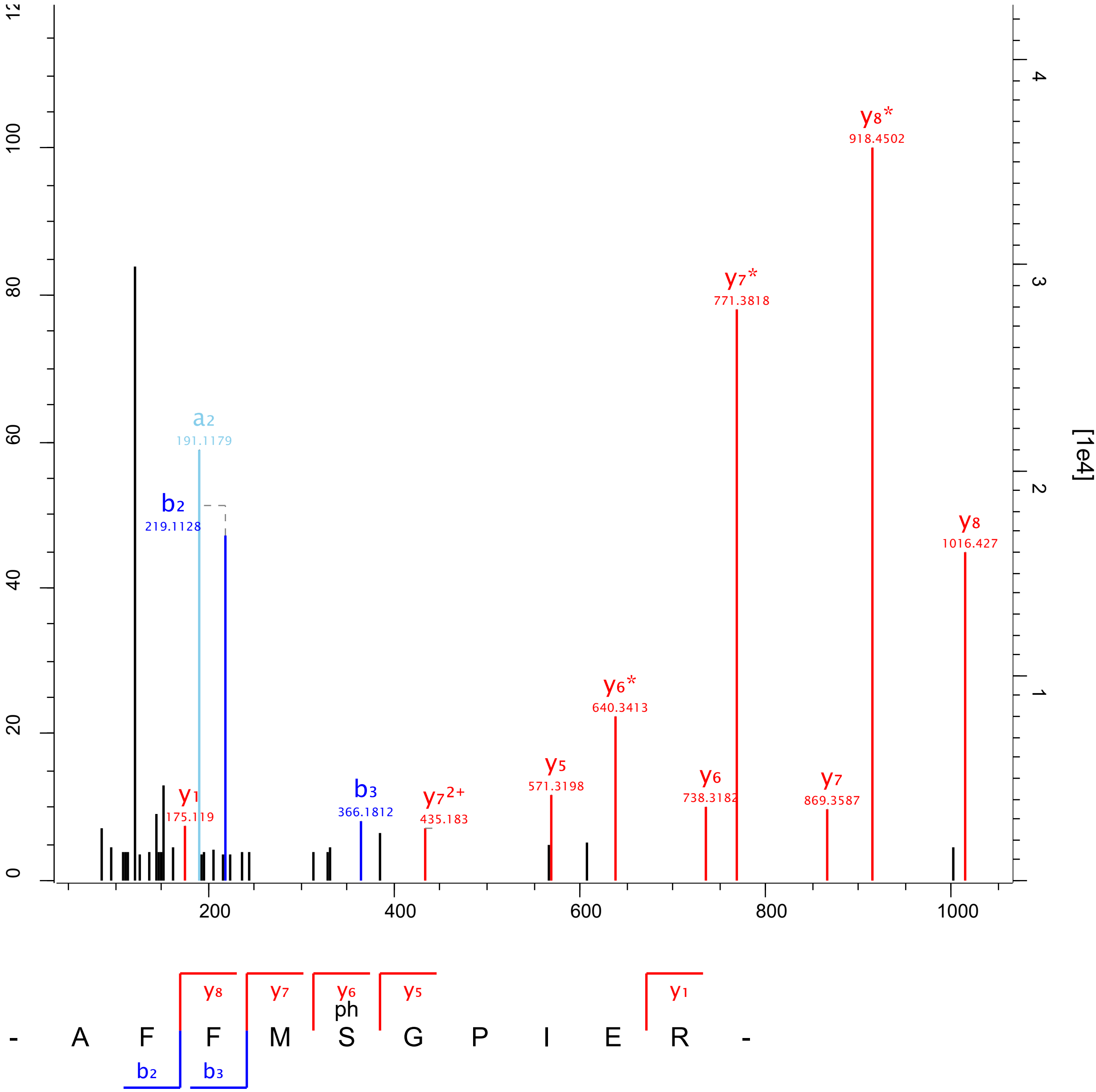

| Raw file | Scan | Method | Score | m/z |
| --- | --- | --- | --- | --- |
| sys_00_3short | 31579 | FTMS; HCD | 82.01 | 787.35 |

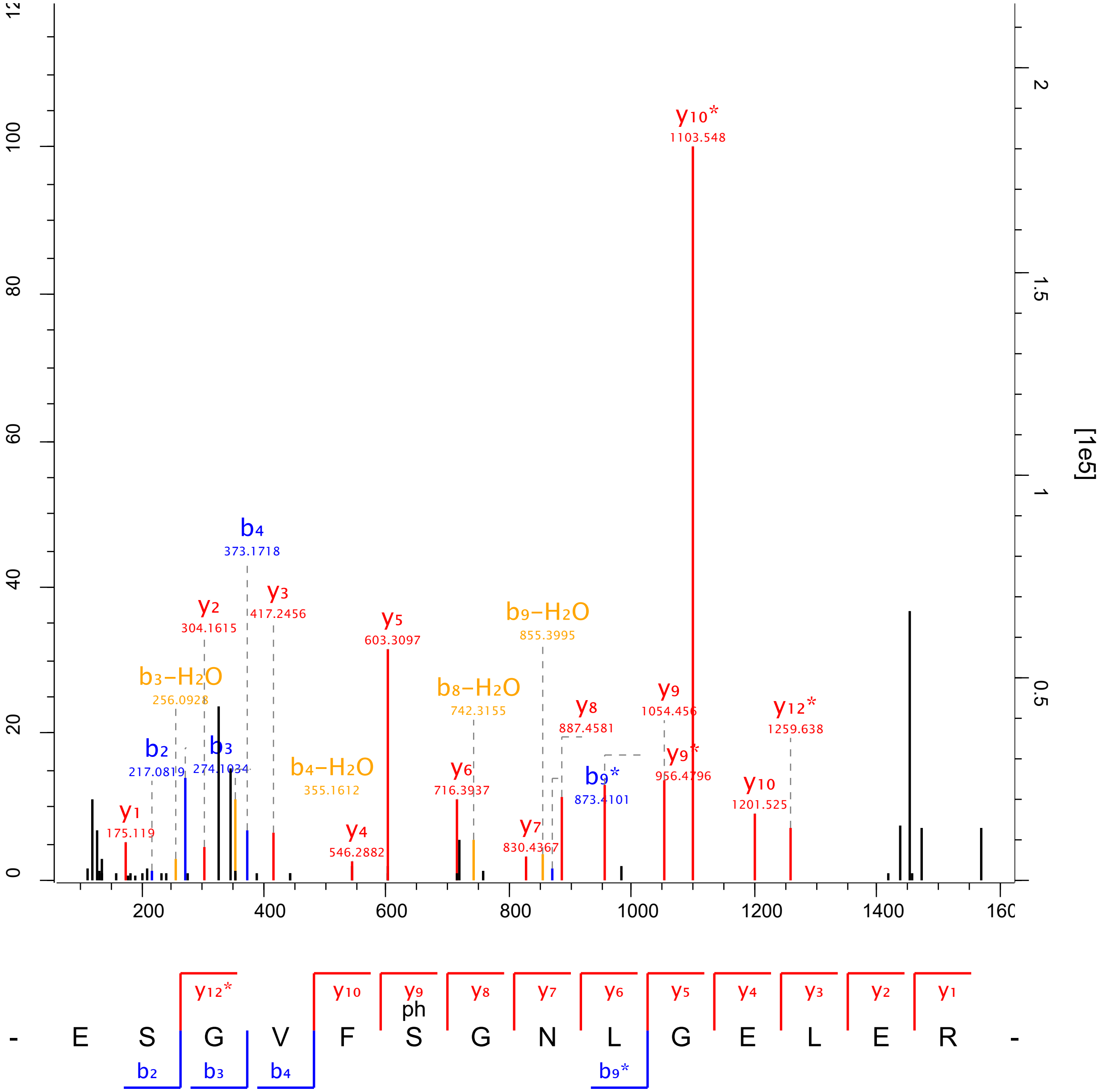

| Raw file | Scan | Method | Score | m/z |
| --- | --- | --- | --- | --- |
| sys_00_3short | 31724 | FTMS; HCD | 78.19 | 700.32 |

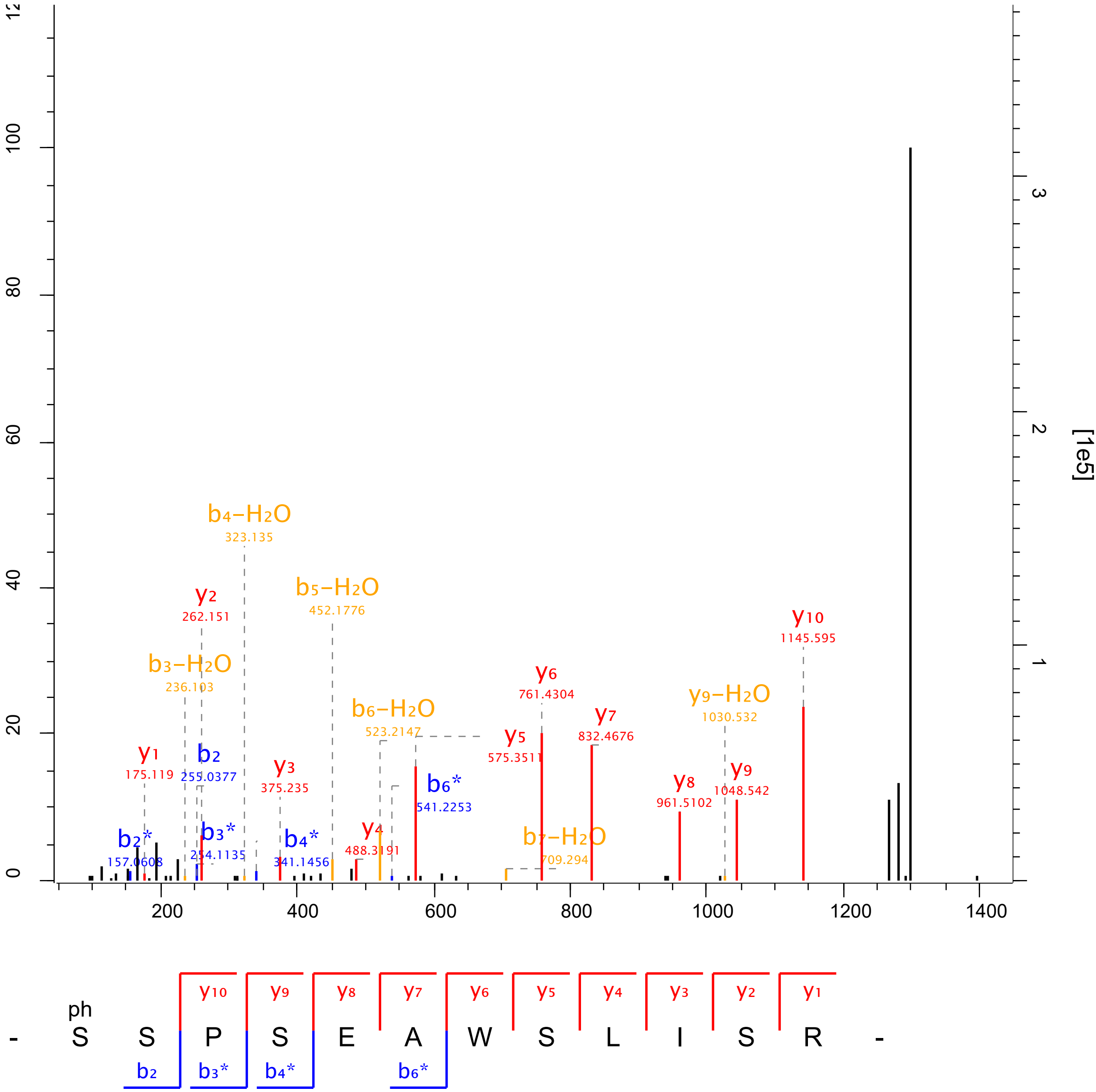

| Raw file | Scan | Method | Score | m/z |
|---|---|---|---|---|
| sys_00_3short | 31753 | FTMS; HCD | 332.56 | 868.69 |

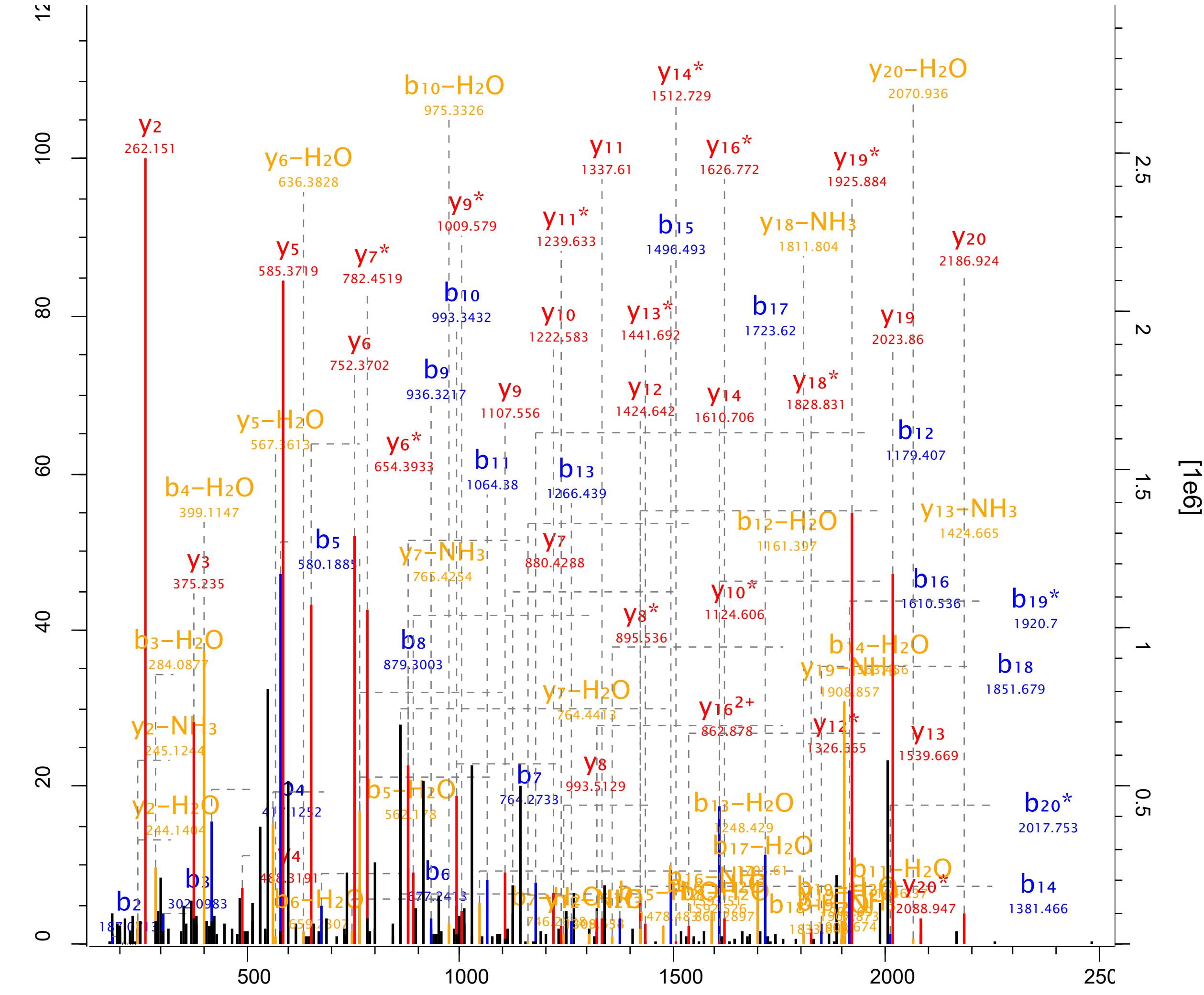


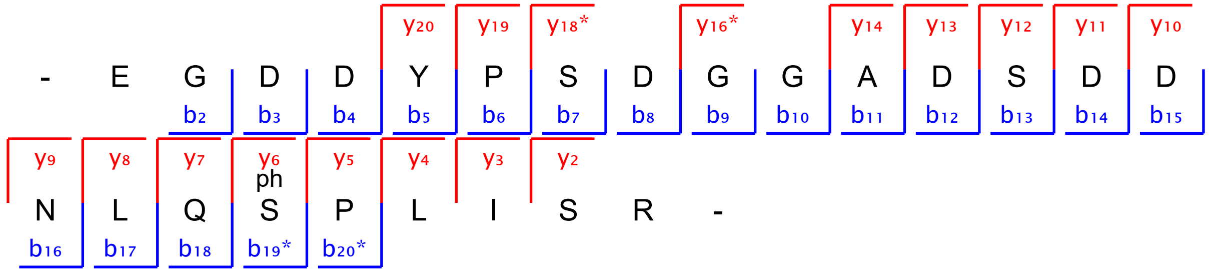

| Raw file | Scan | Method | Score | m/z |
|---|---|---|---|---|
| sys_00_3short | 32041 | FTMS; HCD | 65.18 | 599.75 |

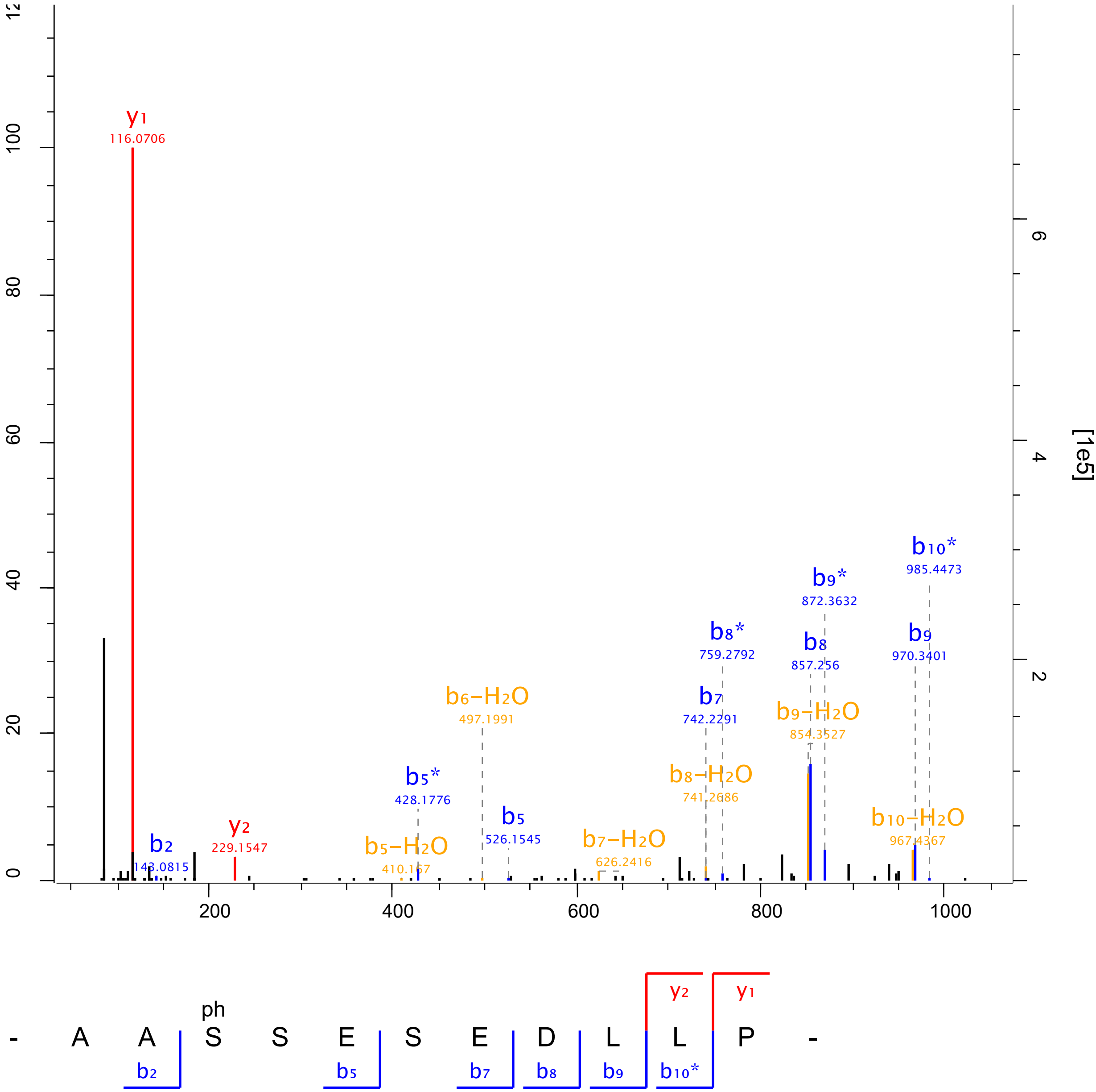

| Raw file | Scan | Method | Score | m/z |
| --- | --- | --- | --- | --- |
| sys_00_3short | 3291 | FTMS; HCD | 72.29 | 577.23 |

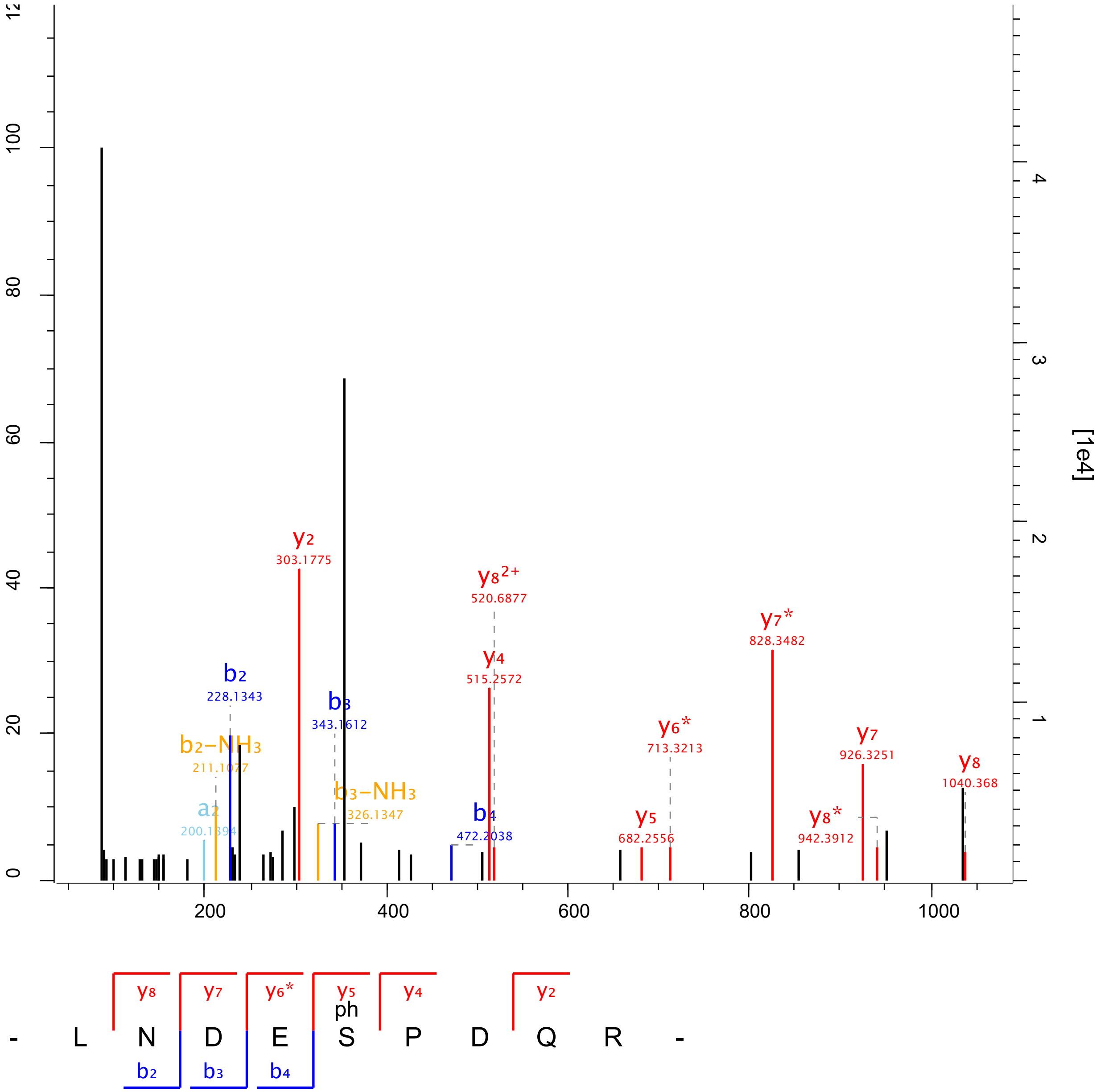

| Raw file | Scan | Method | Score | m/z |
|---|---|---|---|---|
| sys_00_3short | 33357 | FTMS; HCD | 53.24 | 651.29 |

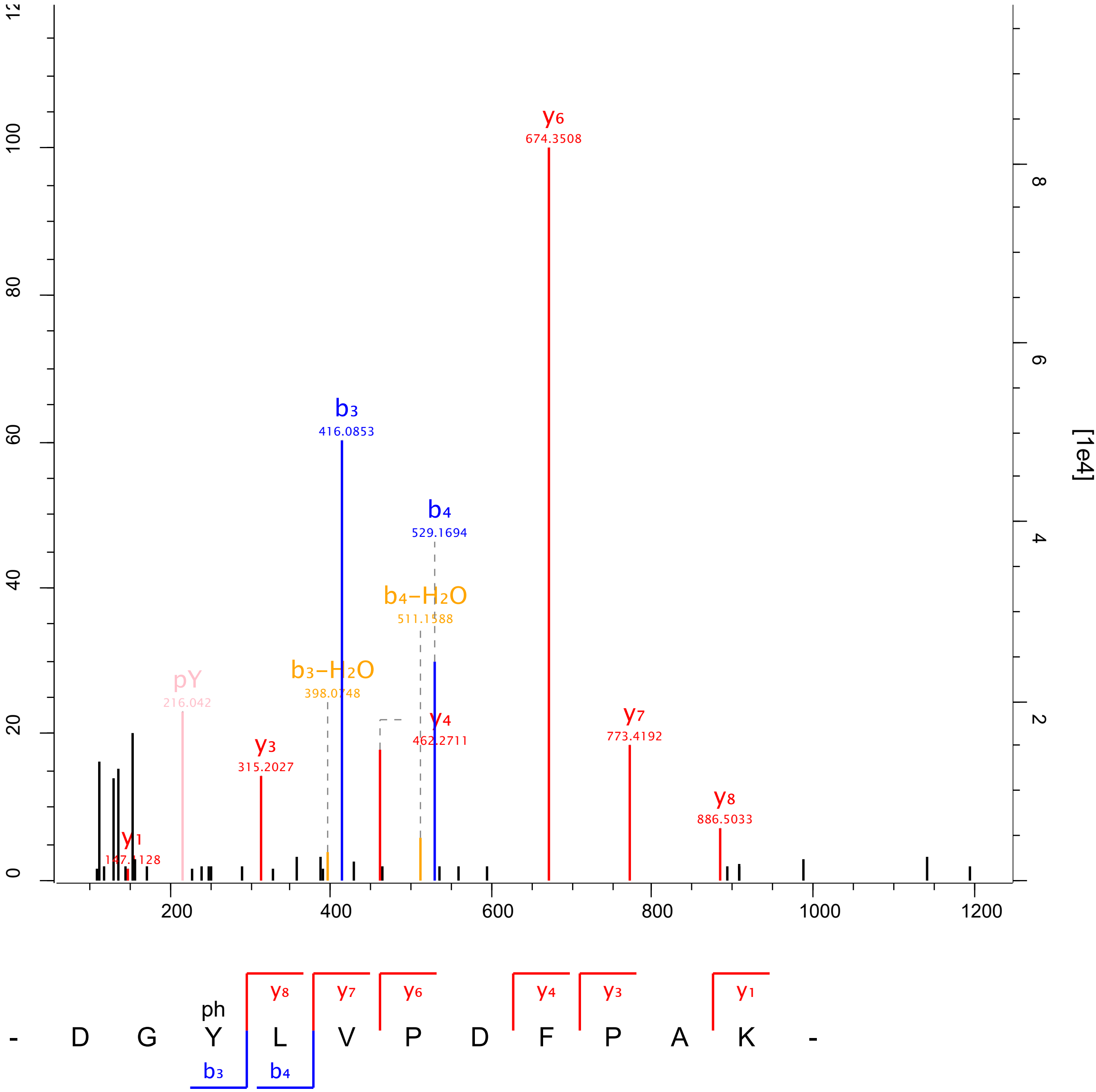

| Raw file | Scan | Method | Score | m/z |
|---|---|---|---|---|
| sys_00_3short | 33404 | FTMS; HCD | 130.89 | 997.38 |

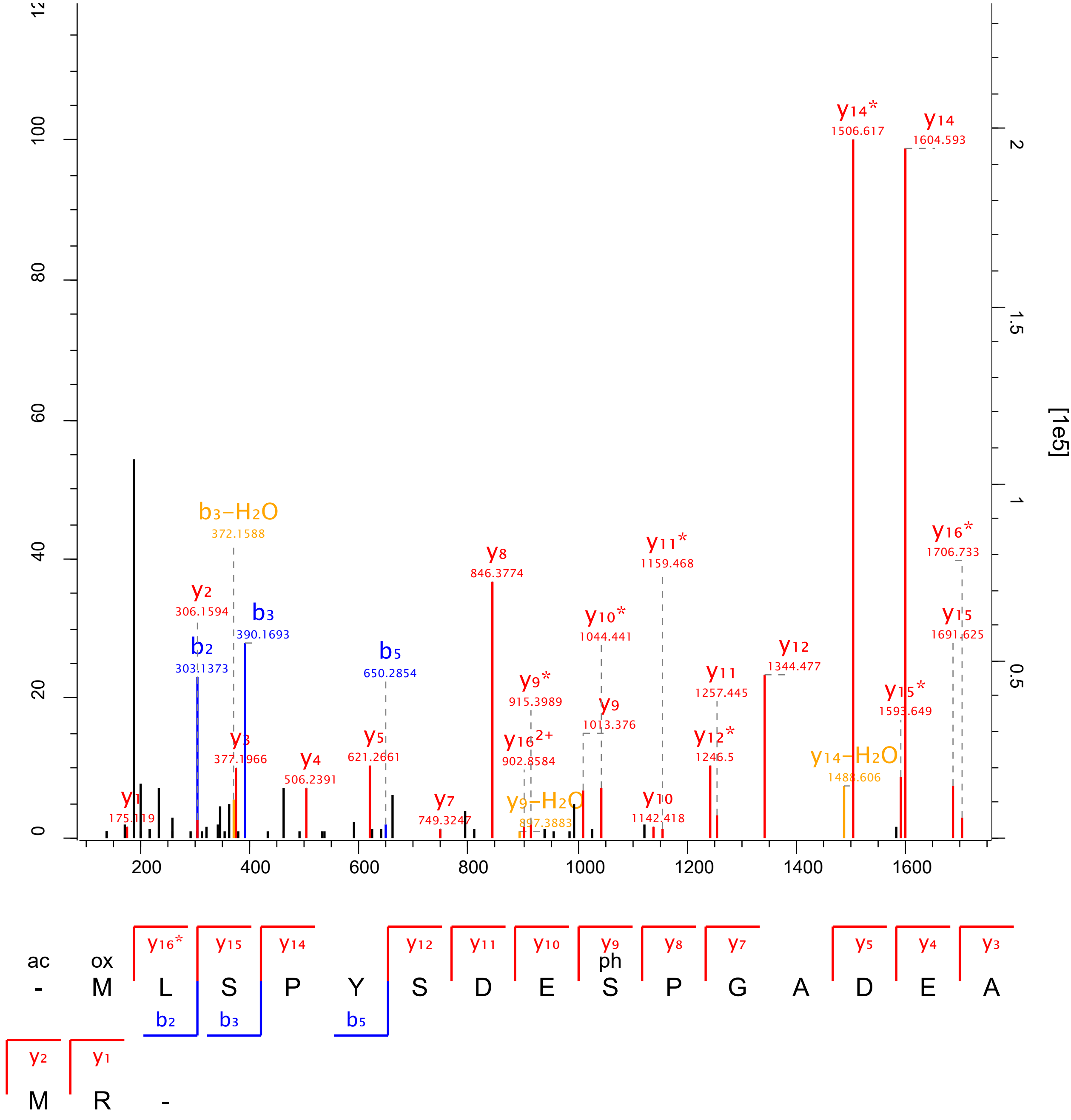

| Raw file | Scan | Method | Score | m/z |
| --- | --- | --- | --- | --- |
| sys_00_3short | 33606 | FTMS; HCD | 42.78 | 607.28 |

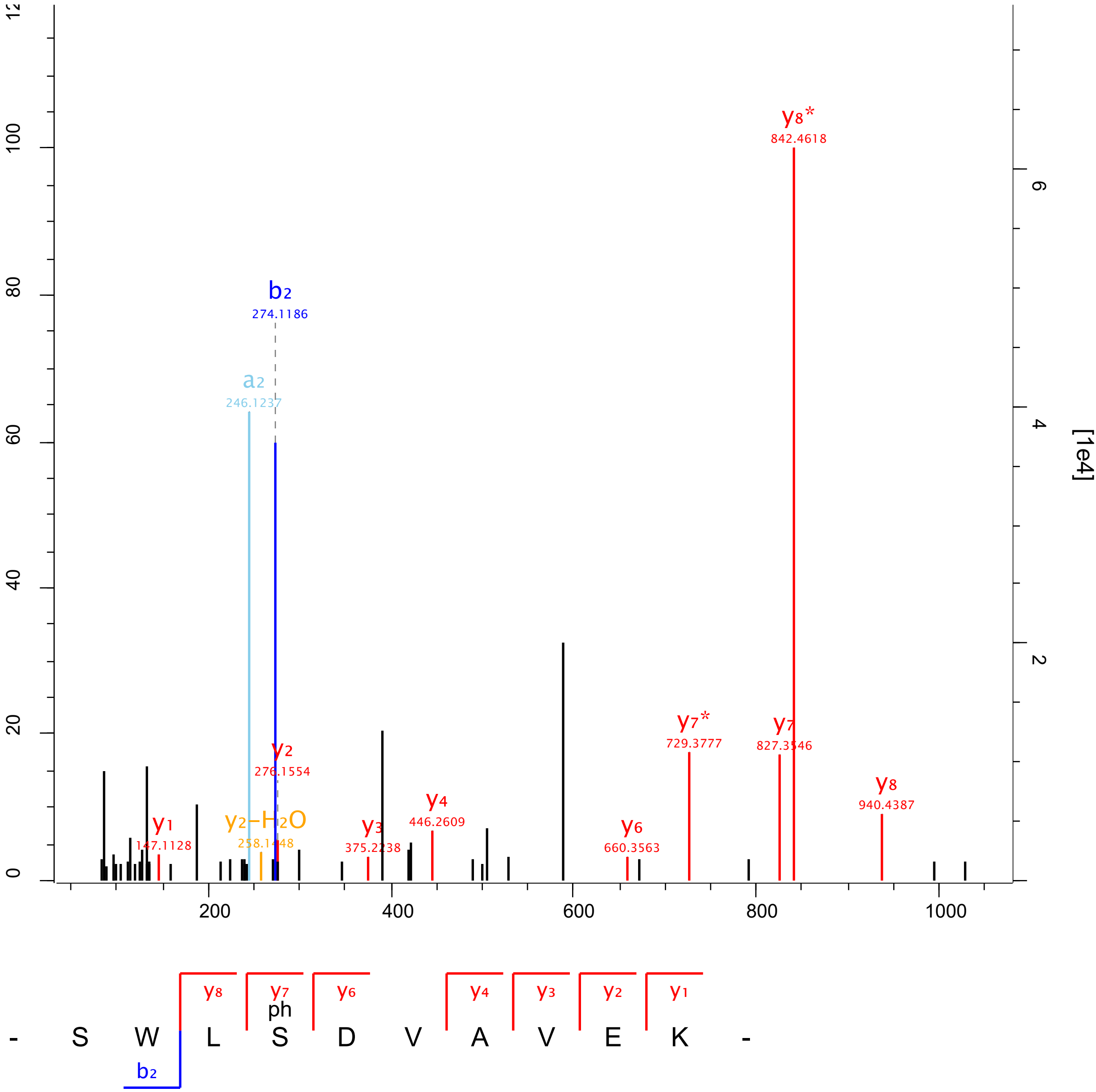

| Raw file | Scan | Method | Score | m/z |
|---|---|---|---|---|
| sys_00_3short | 33950 | FTMS; HCD | 57.29 | 796.36 |

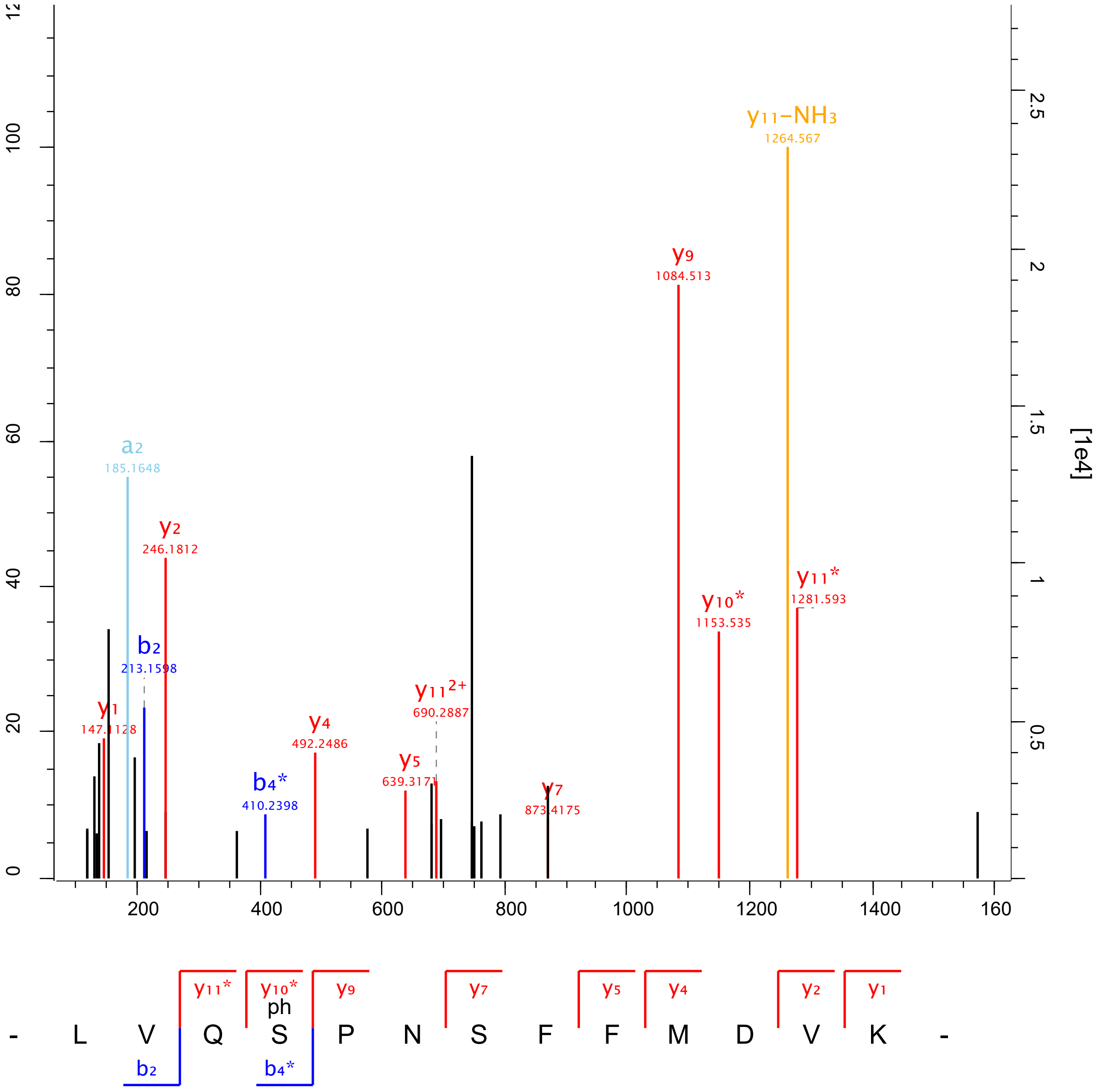

| Raw file | Scan | Method | Score | m/z |
|---|---|---|---|---|
| sys_00_3short | 34236 | FTMS; HCD | 135.91 | 599.76 |

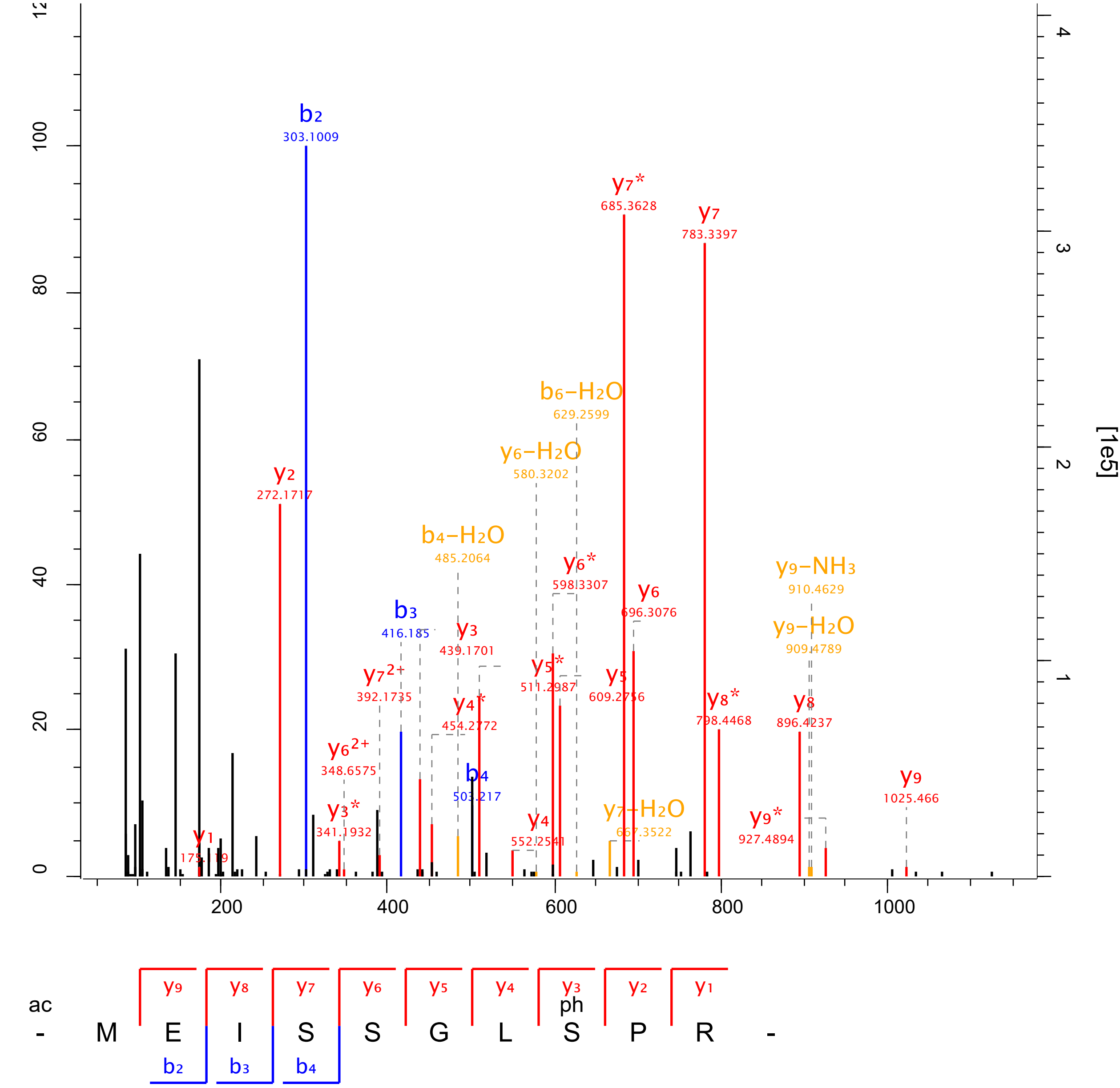

| Raw file | Scan | Method | Score | m/z |
|---|---|---|---|---|
| sys_00_3short | 3469 | FTMS; HCD | 83.2 | 526.74 |

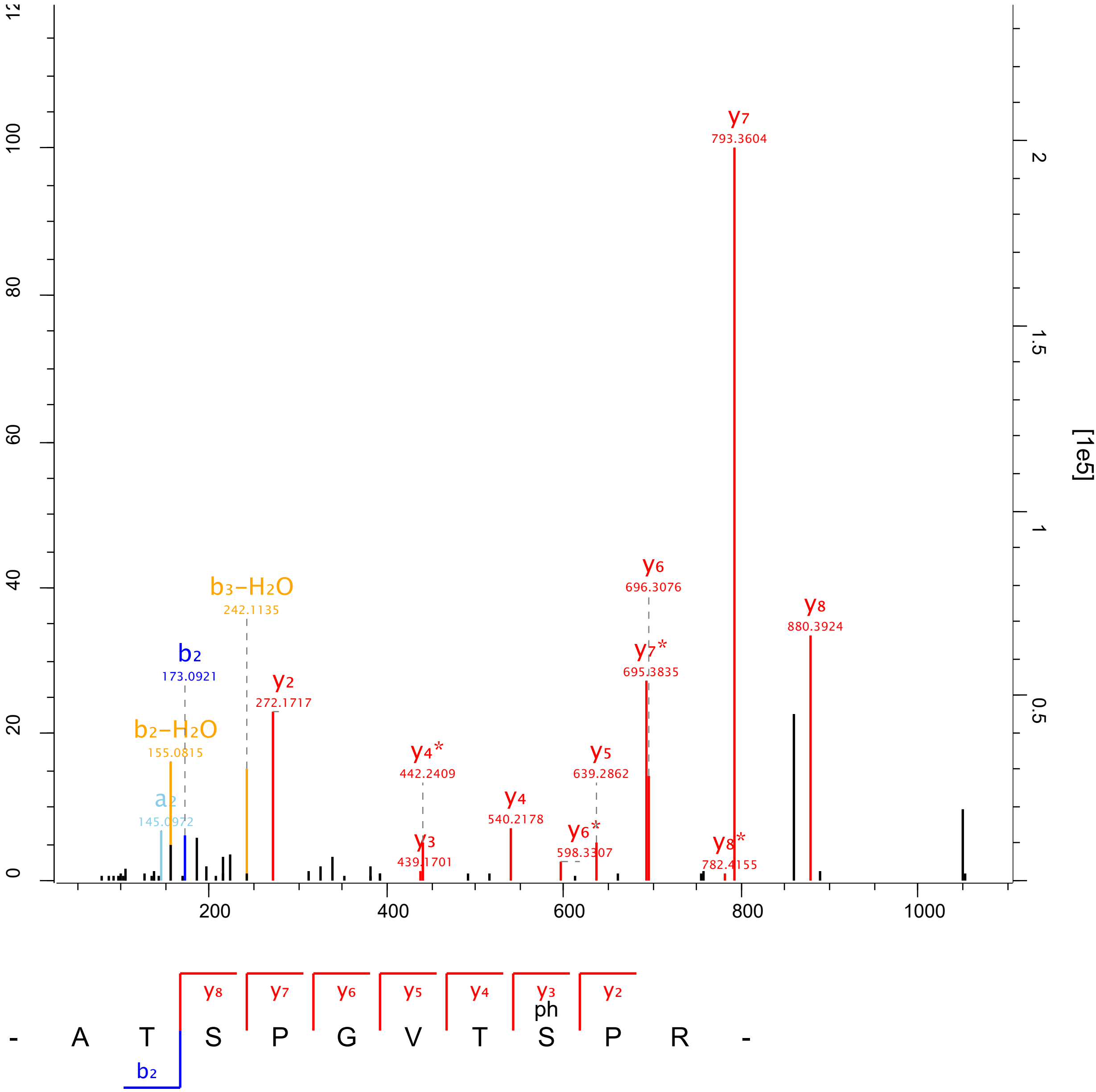

| Raw file | Scan | Method | Score | m/z |
|---|---|---|---|---|
| sys_00_3short | 34878 | FTMS; HCD | 219.95 | 836.83 |

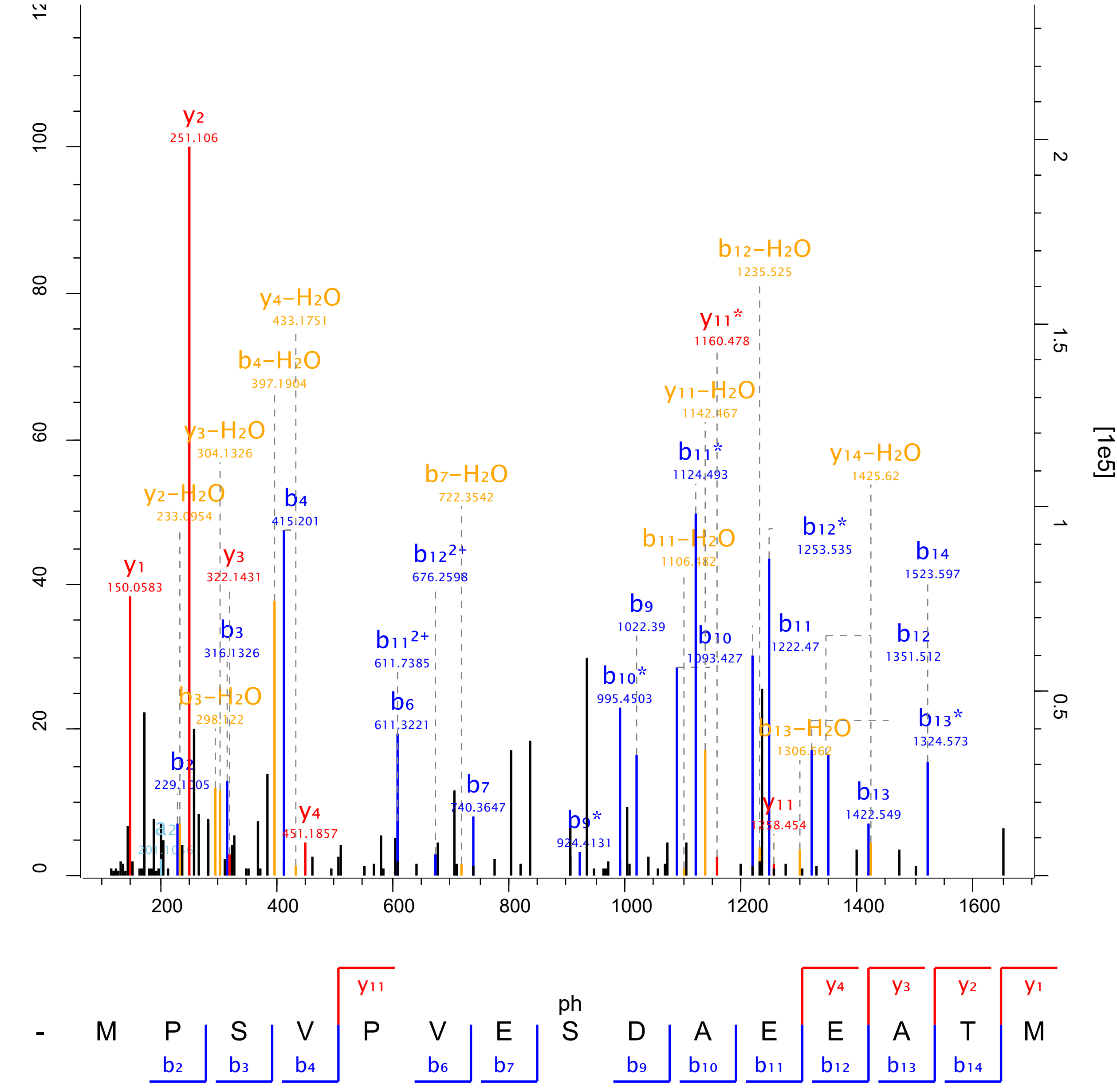

| Raw file | Scan | Method | Score | m/z |
|---|---|---|---|---|
| sys_00_3short | 3573 | FTMS; HCD | 71.61 | 482.19 |

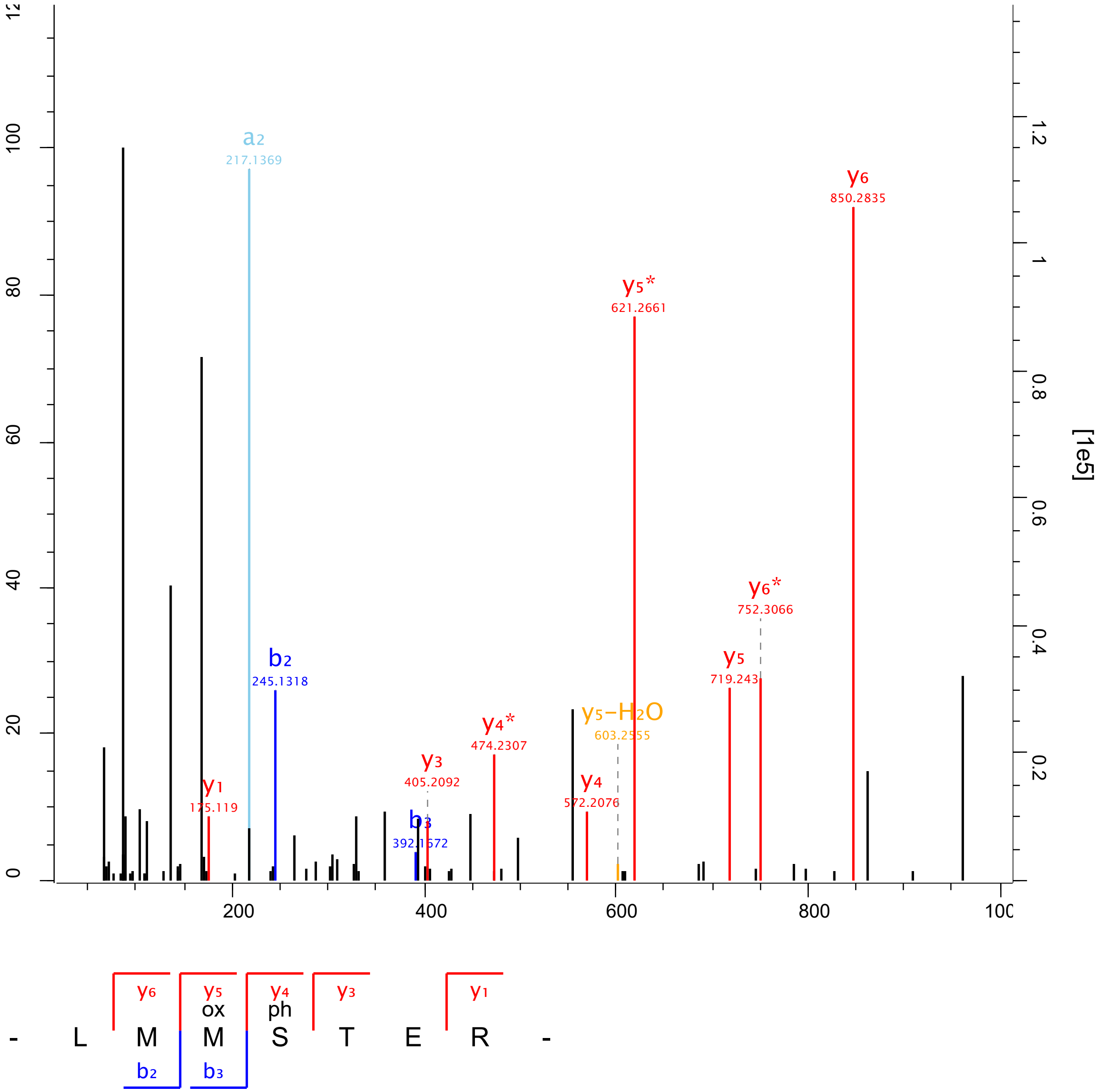

| Raw file | Scan | Method | Score | m/z |
|---|---|---|---|---|
| sys_00_3short | 3584 | FTMS; HCD | 68.52 | 457.21 |

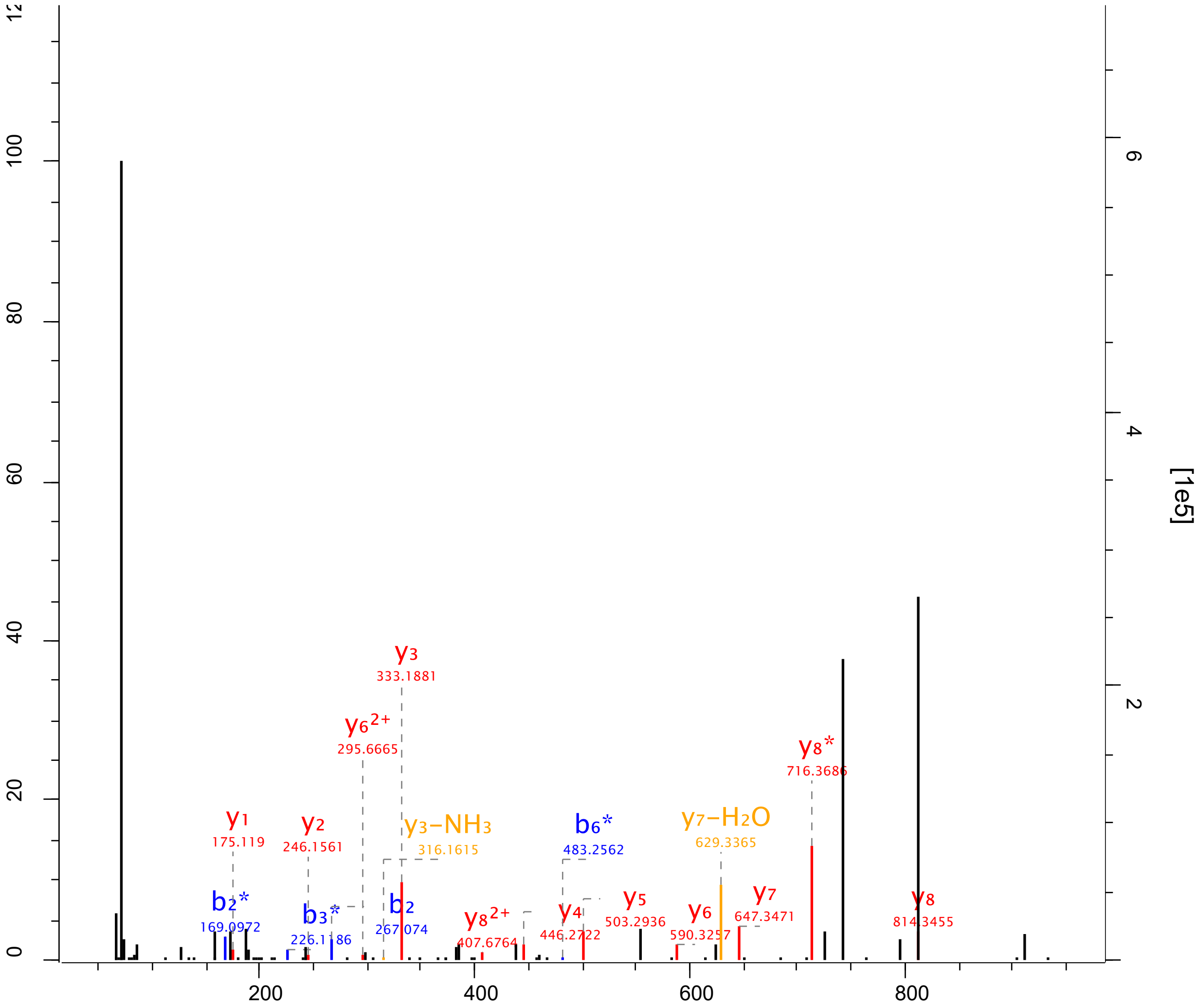


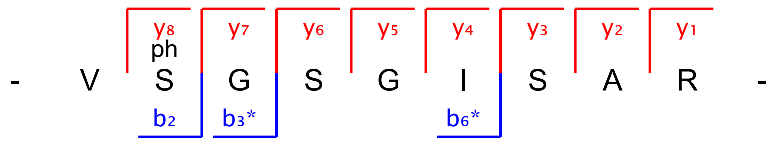

| Raw file | Scan | Method | Score | m/z |
|---|---|---|---|---|
| sys_00_3short | 3758 | FTMS; HCD | 119.51 | 636.76 |

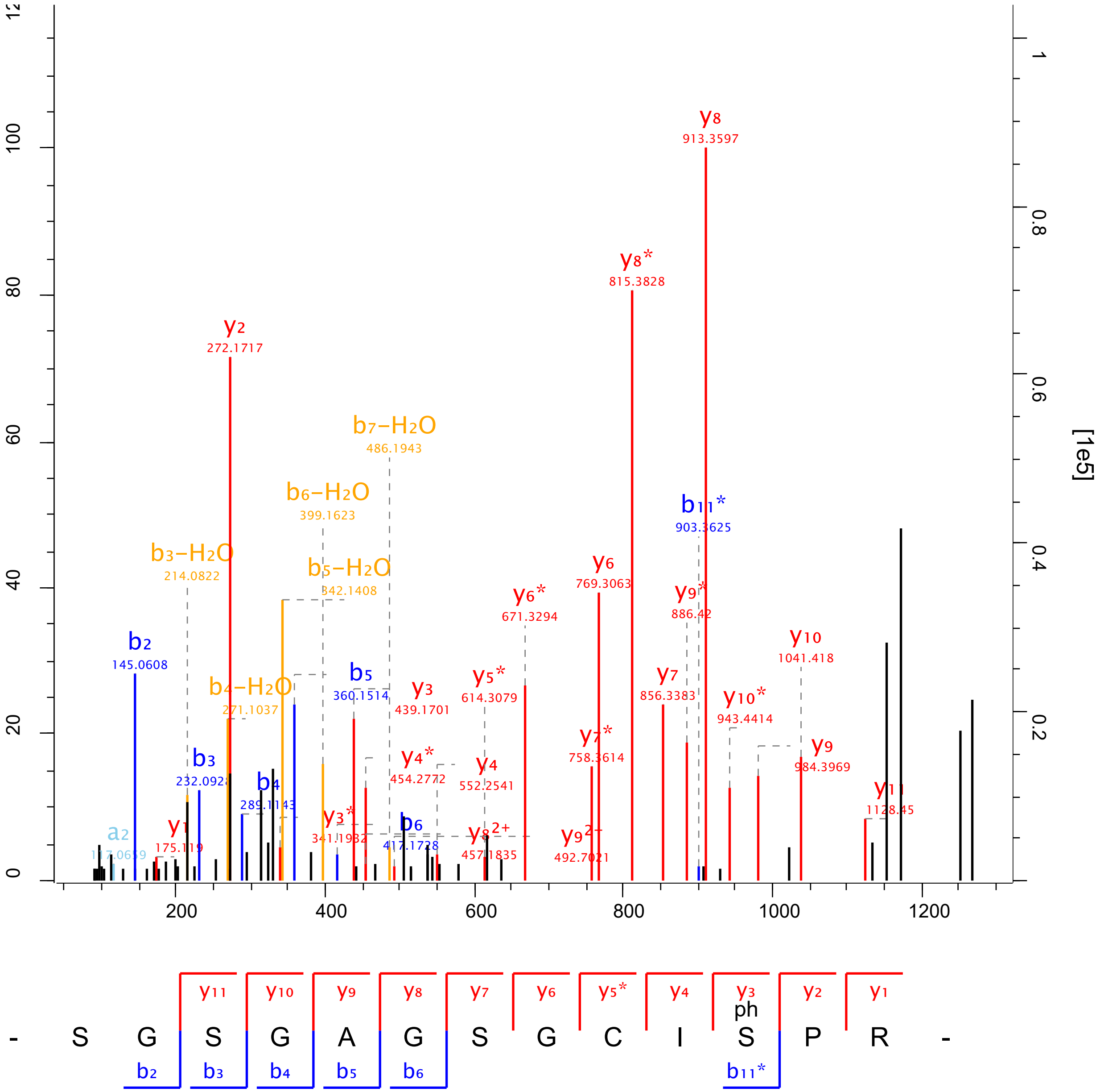

| Raw file | Scan | Method | Score | m/z |
|---|---|---|---|---|
| sys_00_3short | 4082 | FTMS; HCD | 107.06 | 536.25 |

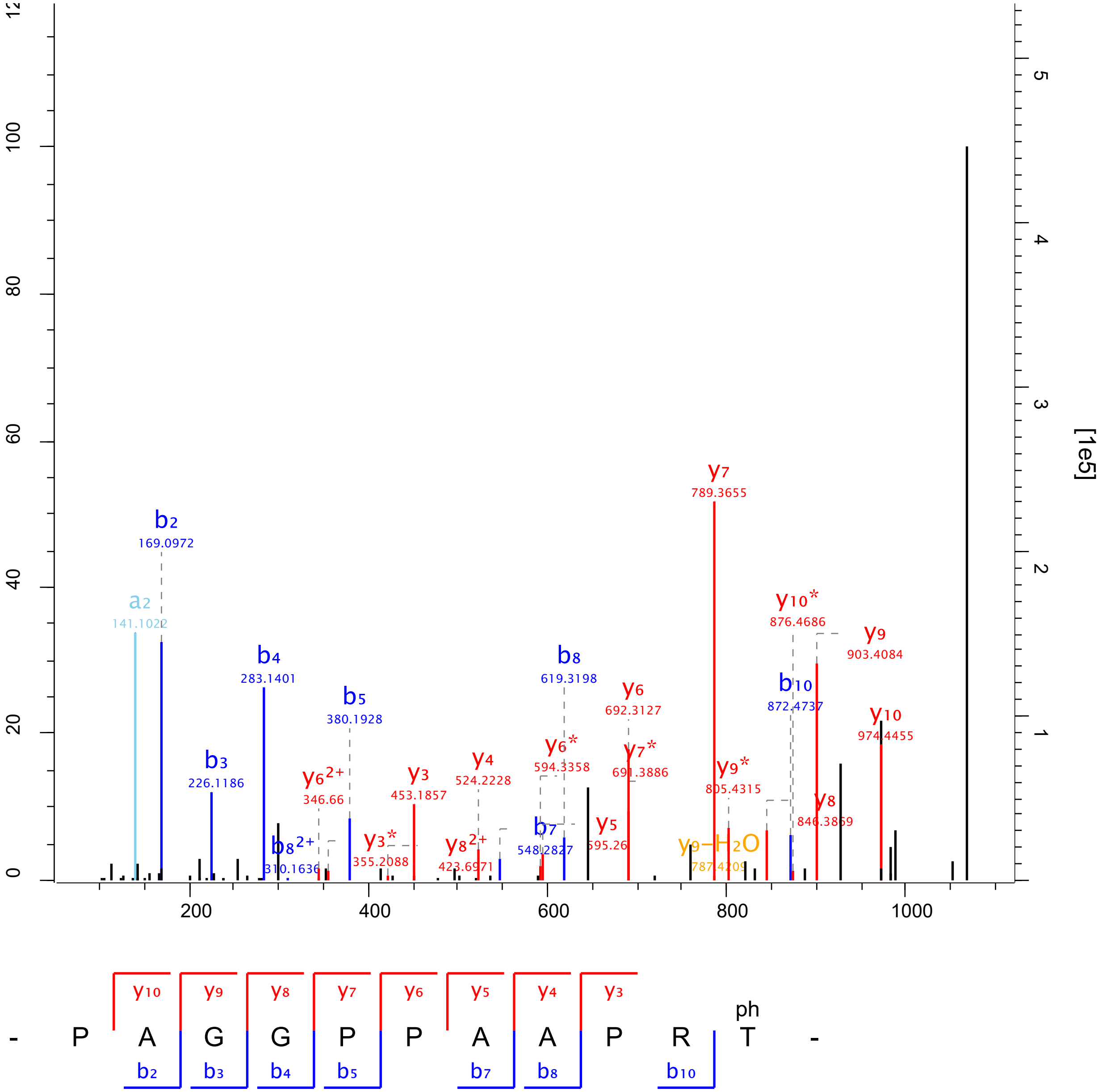

| Raw file | Scan | Method | Score | m/z |
|---|---|---|---|---|
| sys_00_3short | 4262 | FTMS; HCD | 48.57 | 742.78 |

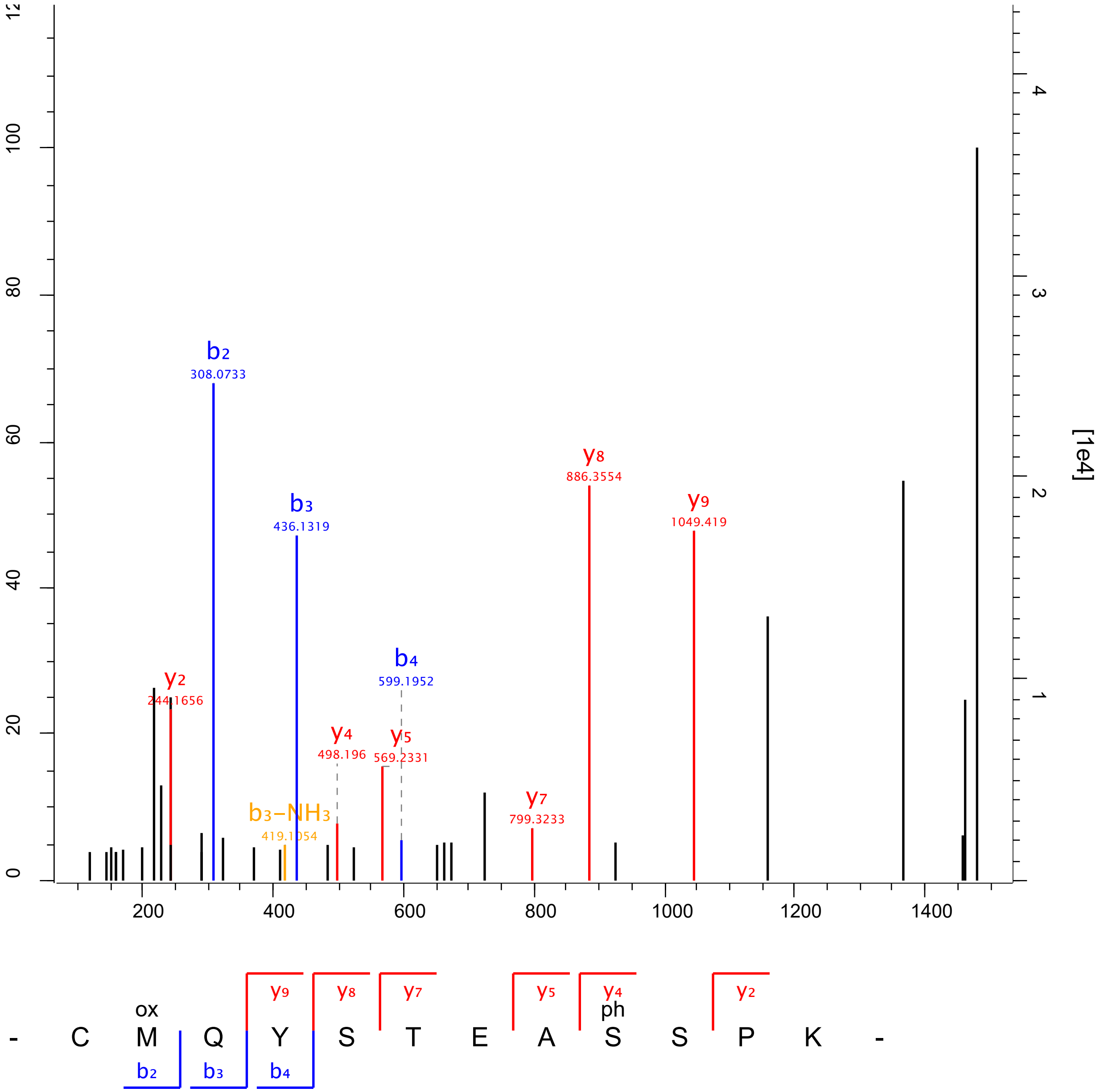

| Raw file | Scan | Method | Score | m/z |
|---|---|---|---|---|
| sys_00_3short | 4338 | FTMS; HCD | 60.16 | 614.75 |

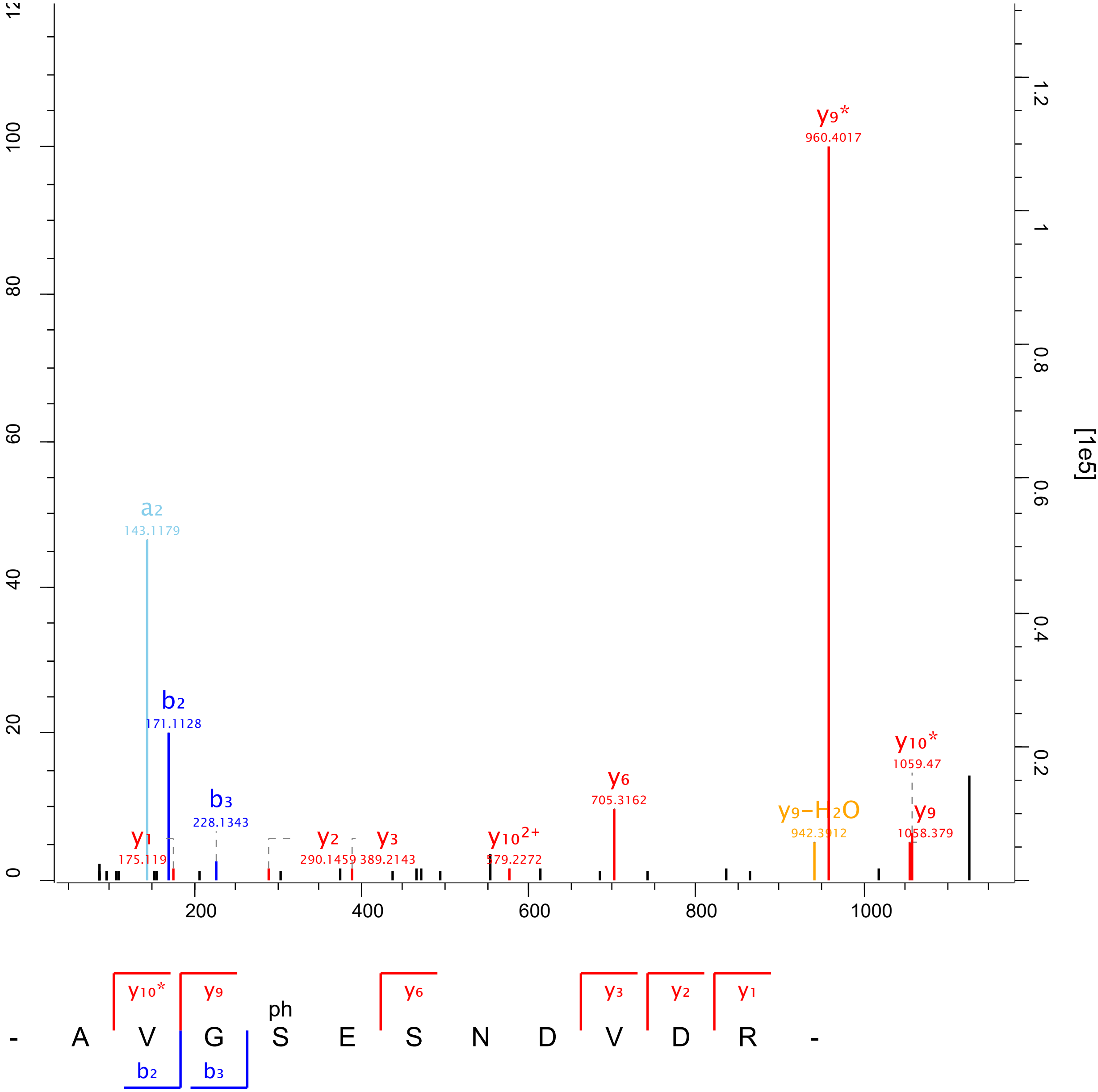

| Raw file | Scan | Method | Score | m/z |
|---|---|---|---|---|
| sys_00_3short | 4837 | FTMS; HCD | 234.28 | 665.26 |

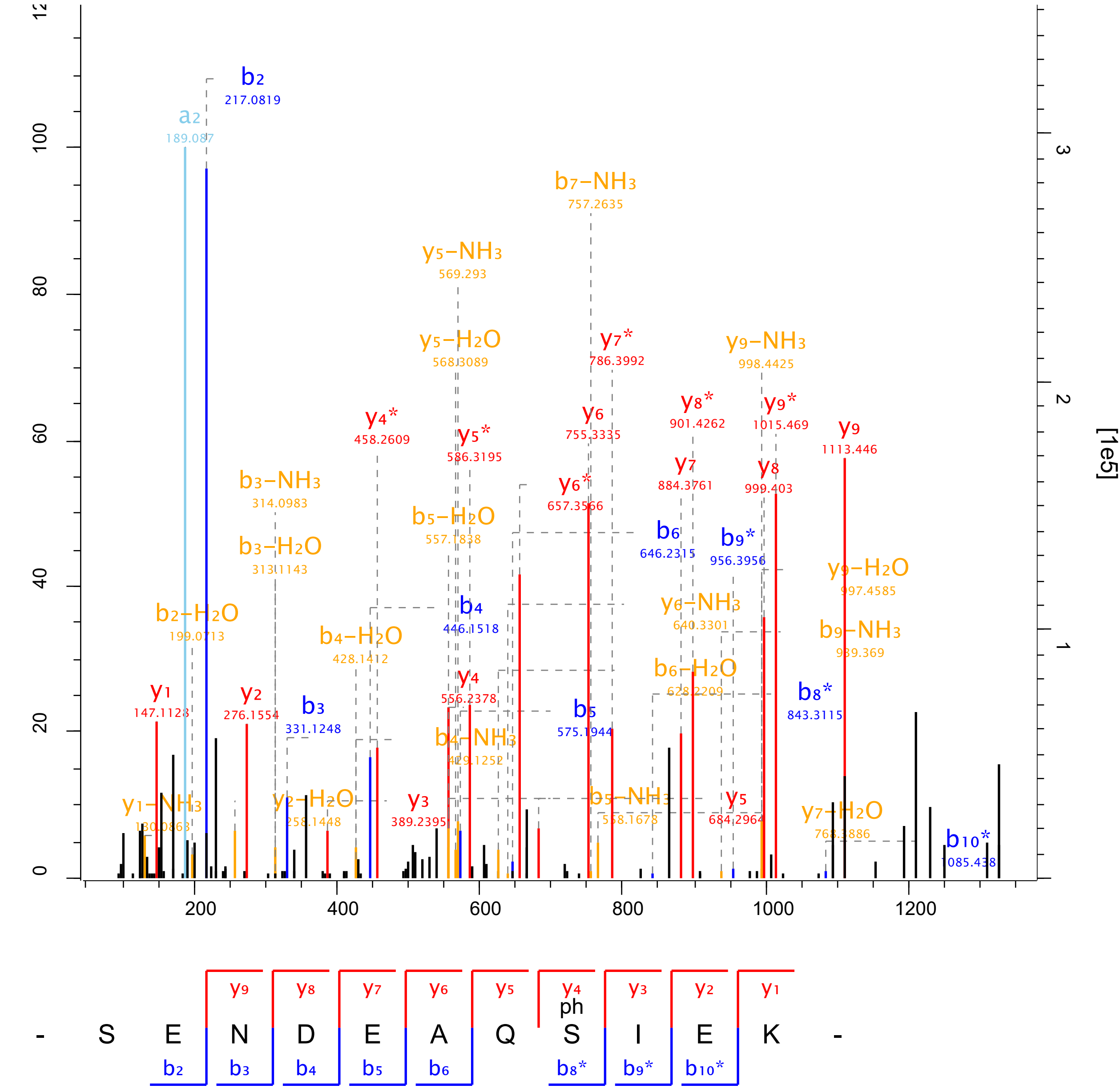

| Raw file | Scan | Method | Score | m/z |
|---|---|---|---|---|
| sys_00_3short | 4863 | FTMS; HCD | 83.86 | 492.23 |

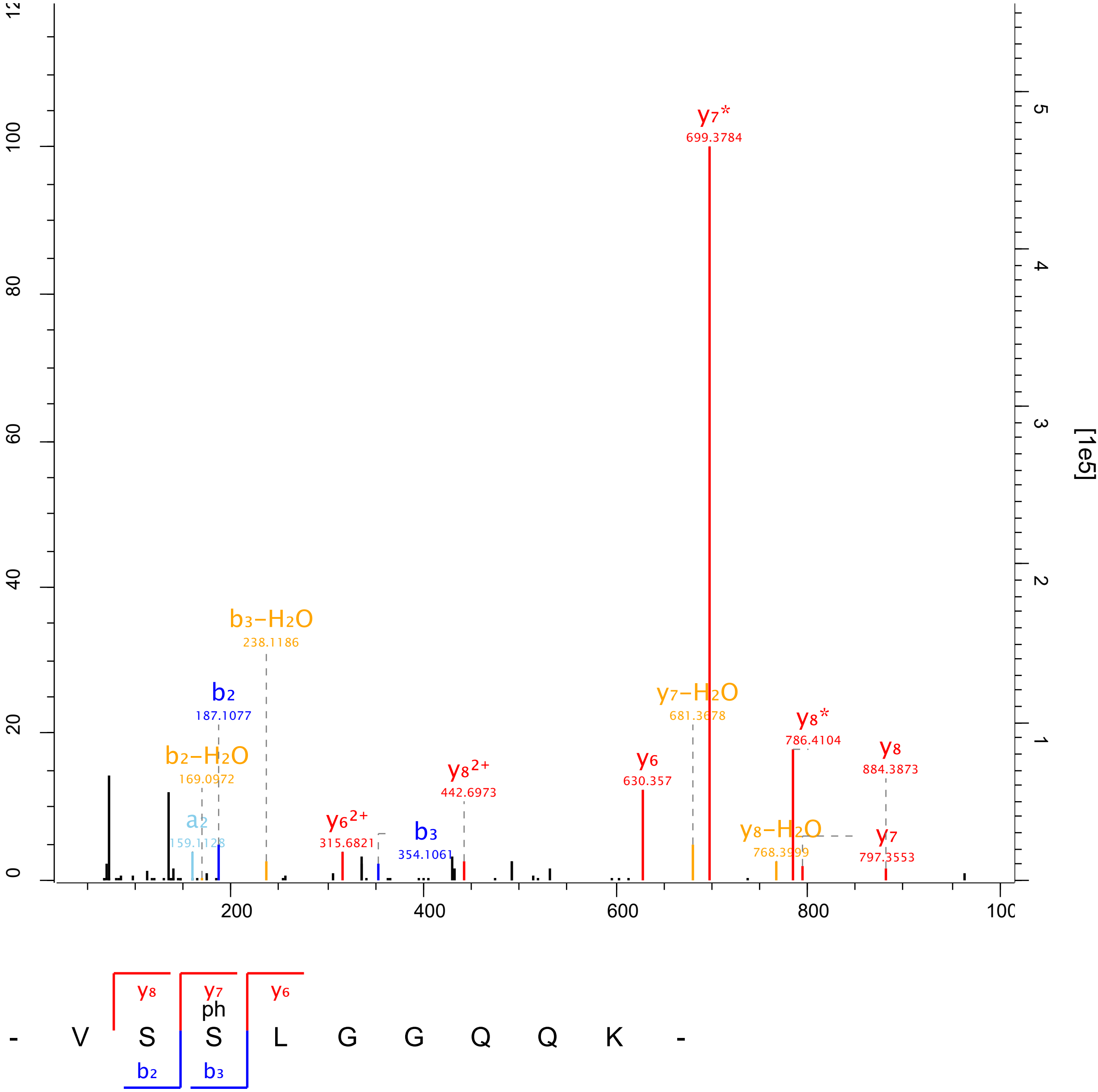

| Raw file | Scan | Method | Score | m/z |
| --- | --- | --- | --- | --- |
| sys_00_3short | 5304 | FTMS; HCD | 187.51 | 529.23 |

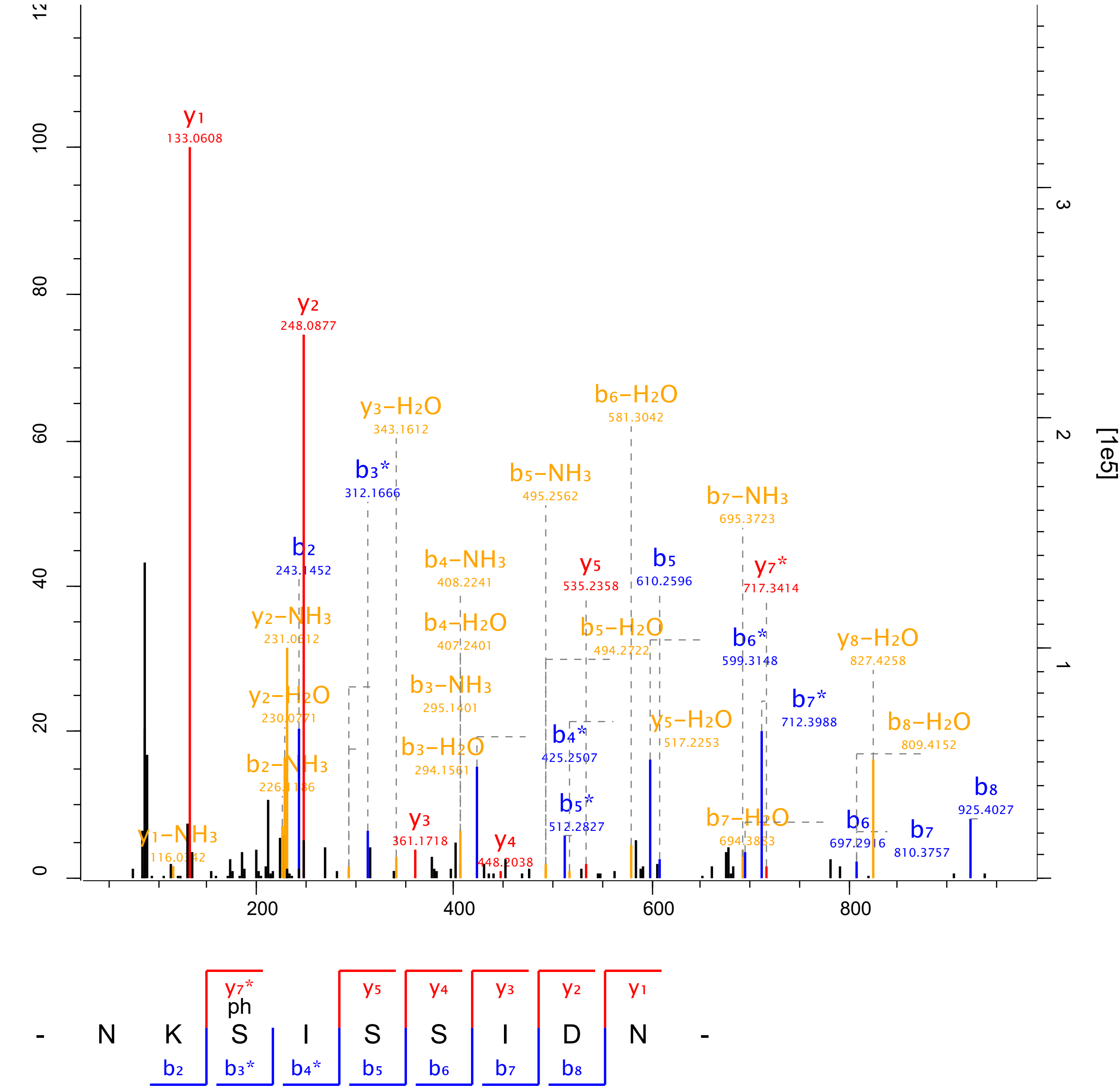

| Raw file | Scan | Method | Score | m/z |
|---|---|---|---|---|
| sys_00_3short | 5515 | FTMS; HCD | 113.74 | 437.2 |

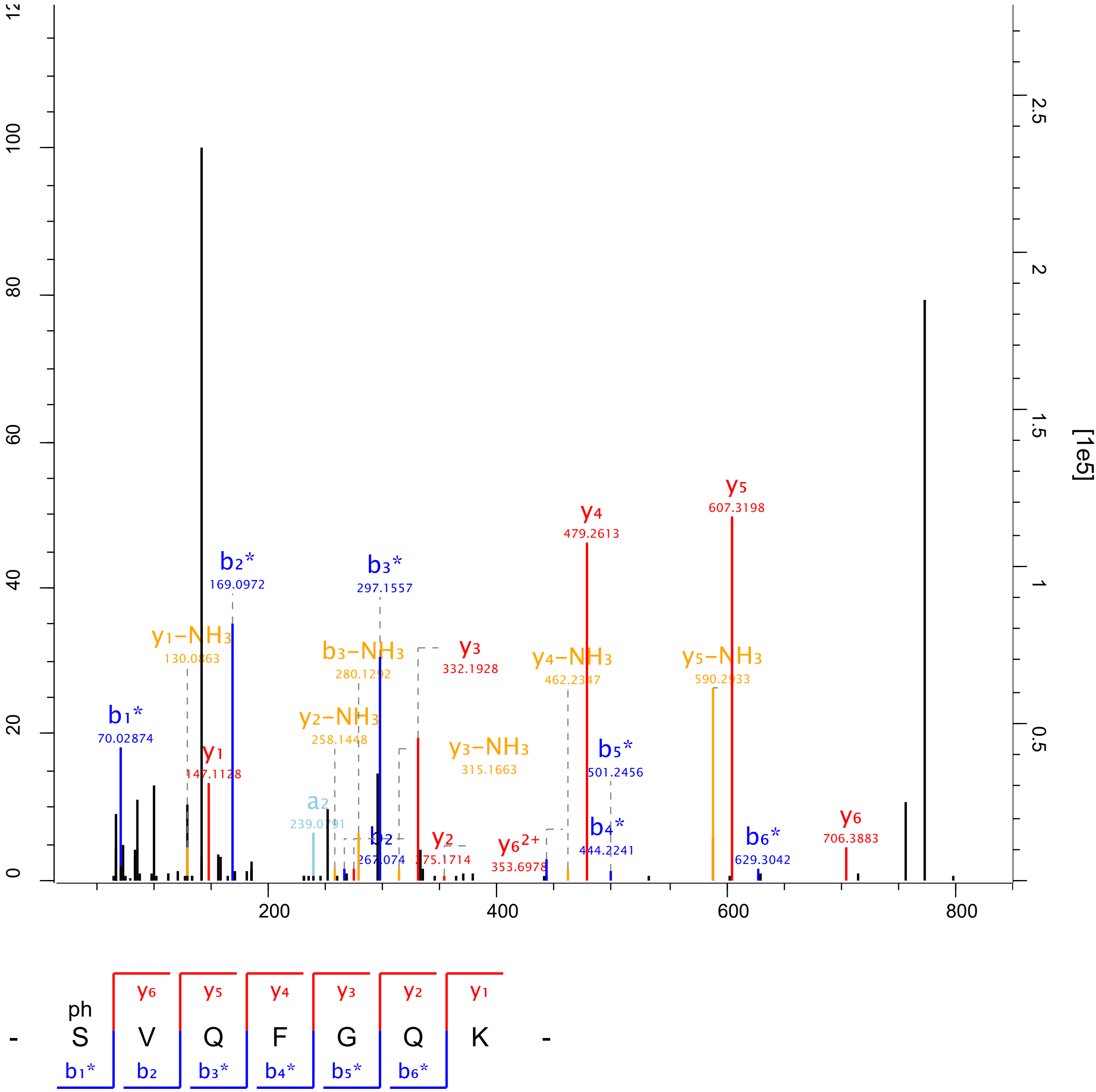

| Raw file | Scan | Method | Score | m/z |
|---|---|---|---|---|
| sys_00_3short | 5564 | FTMS; HCD | 119.25 | 620.22 |

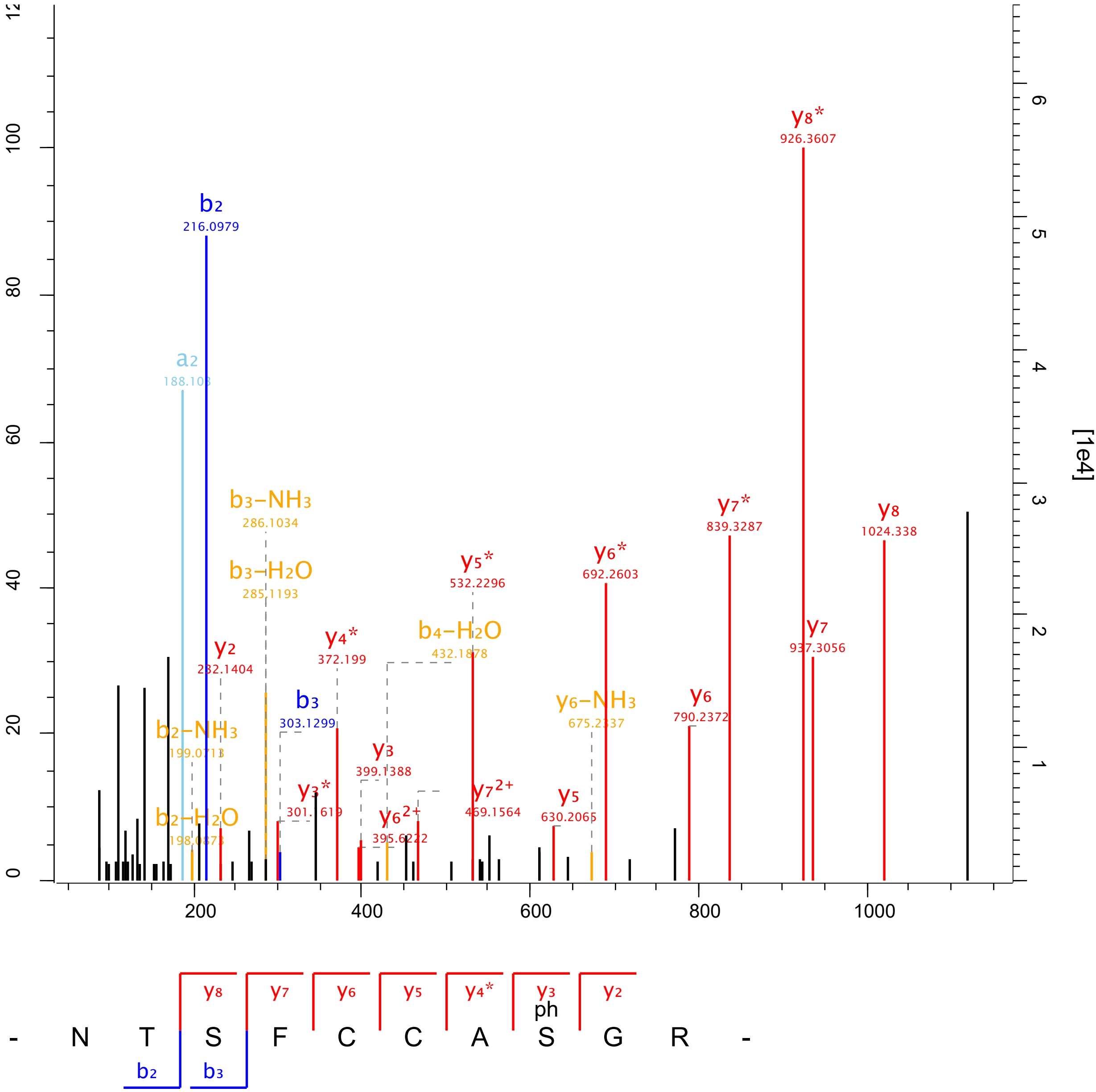

| Raw file | Scan | Method | Score | m/z |
|---|---|---|---|---|
| sys_00_3short | 5781 | FTMS; HCD | 71.88 | 584.28 |

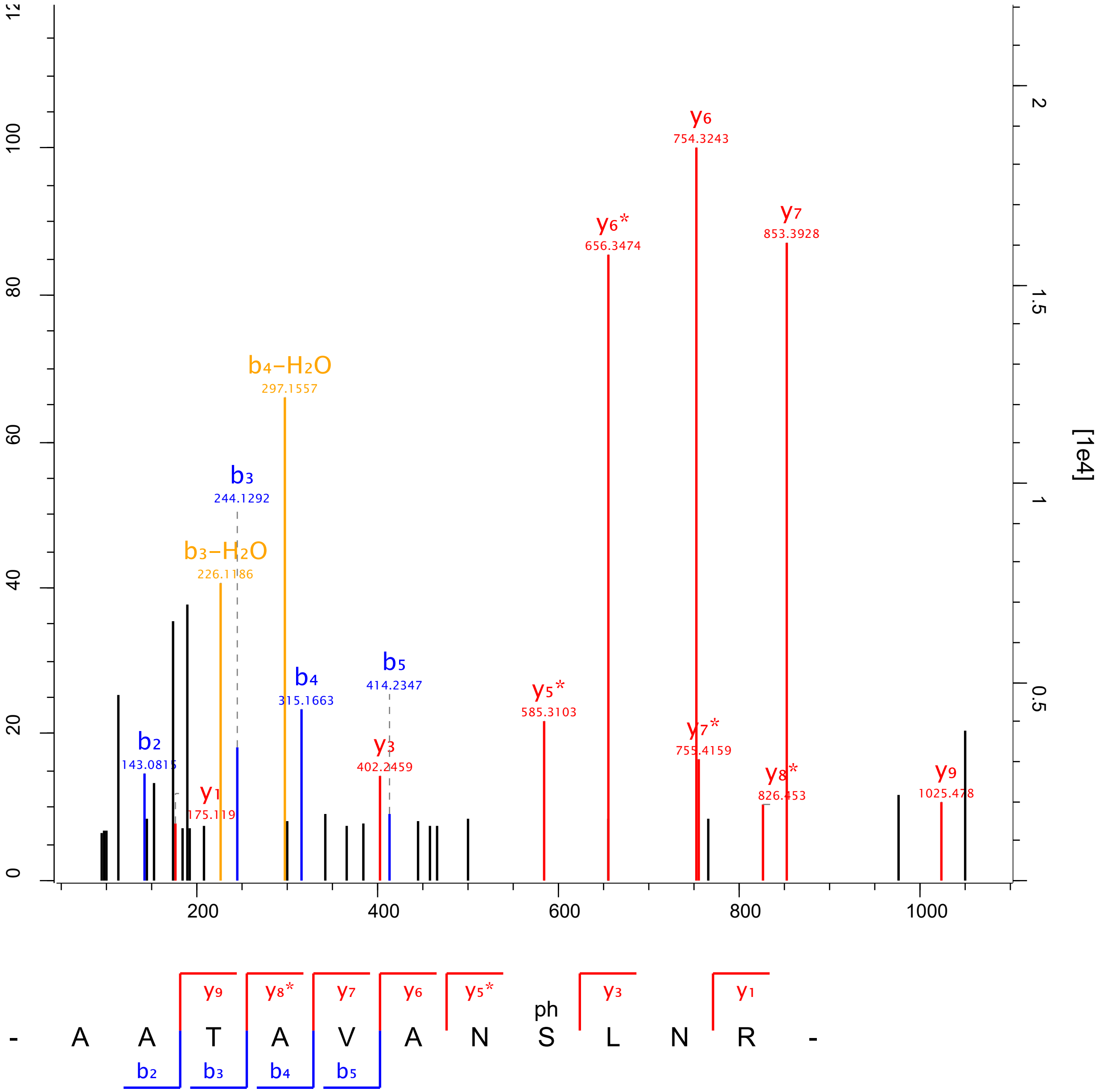

| Raw file | Scan | Method | Score | m/z |
| --- | --- | --- | --- | --- |
| sys_00_3short | 5903 | FTMS; HCD | 64.45 | 805.75 |

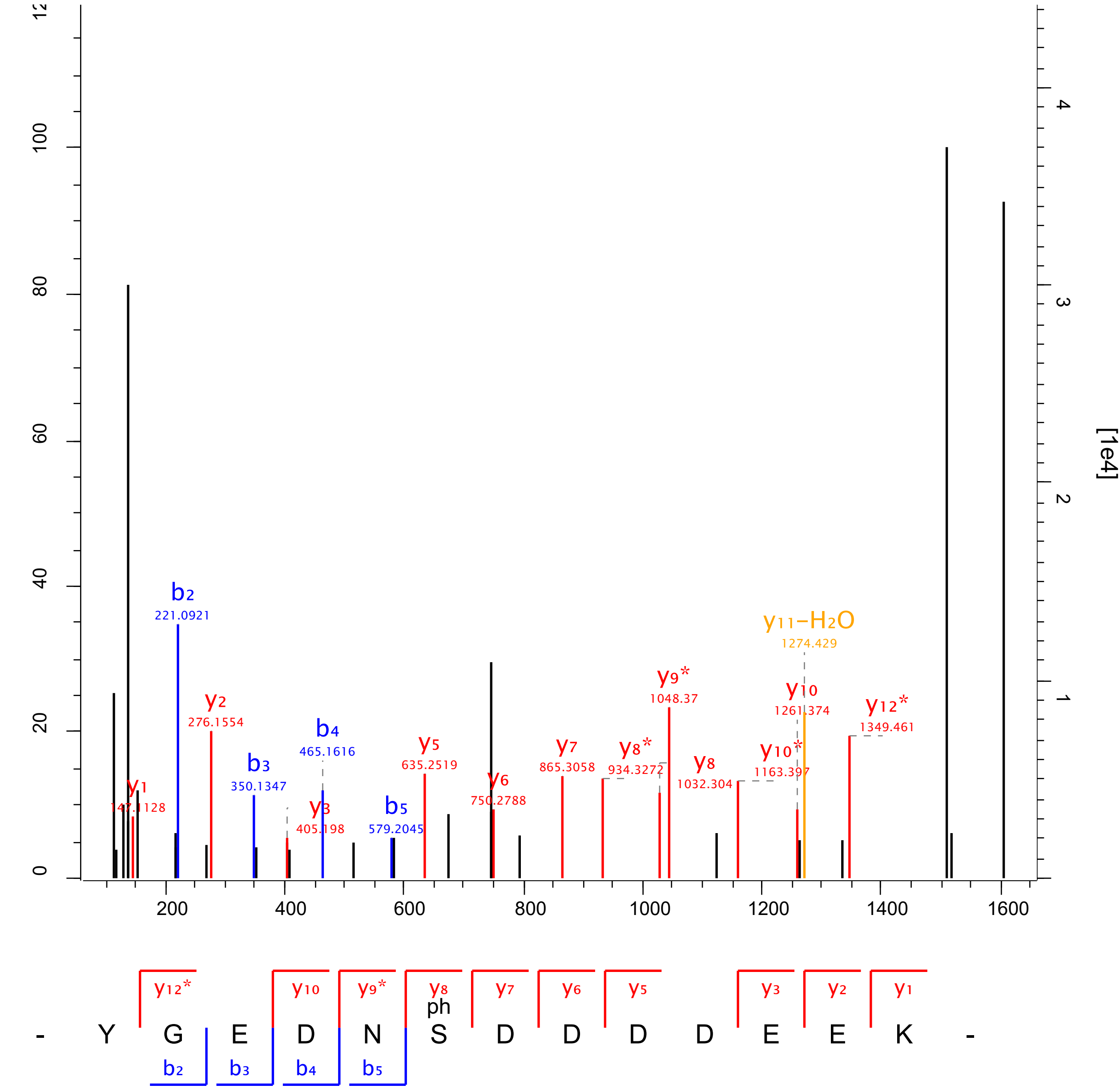

| Raw file | Scan | Method | Score | m/z |
|---|---|---|---|---|
| sys_00_3short | 6034 | FTMS; HCD | 130.05 | 851.82 |

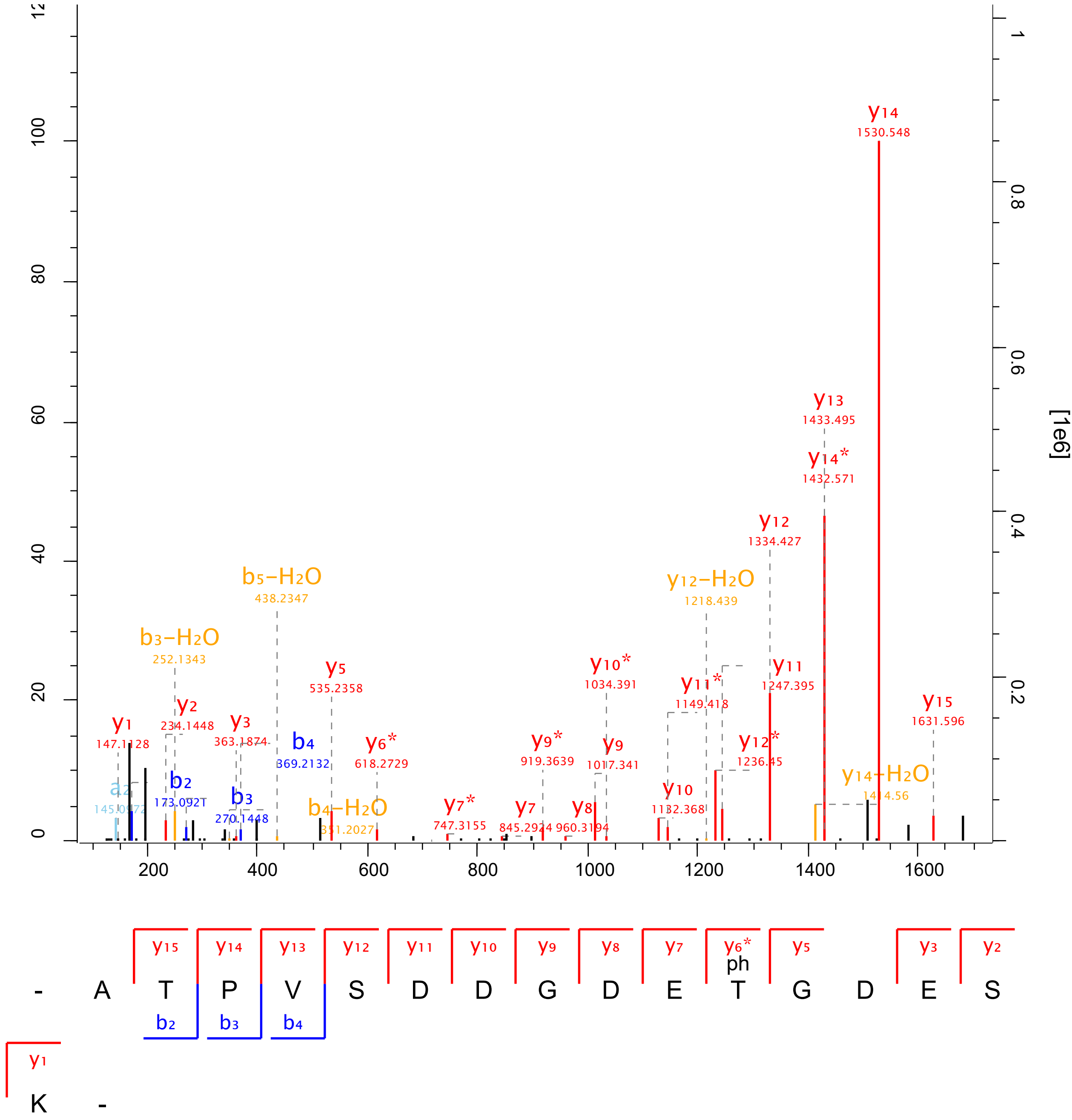

| Raw file | Scan | Method | Score | m/z |
| --- | --- | --- | --- | --- |
| sys_00_3short | 6557 | FTMS; HCD | 66.71 | 619.75 |

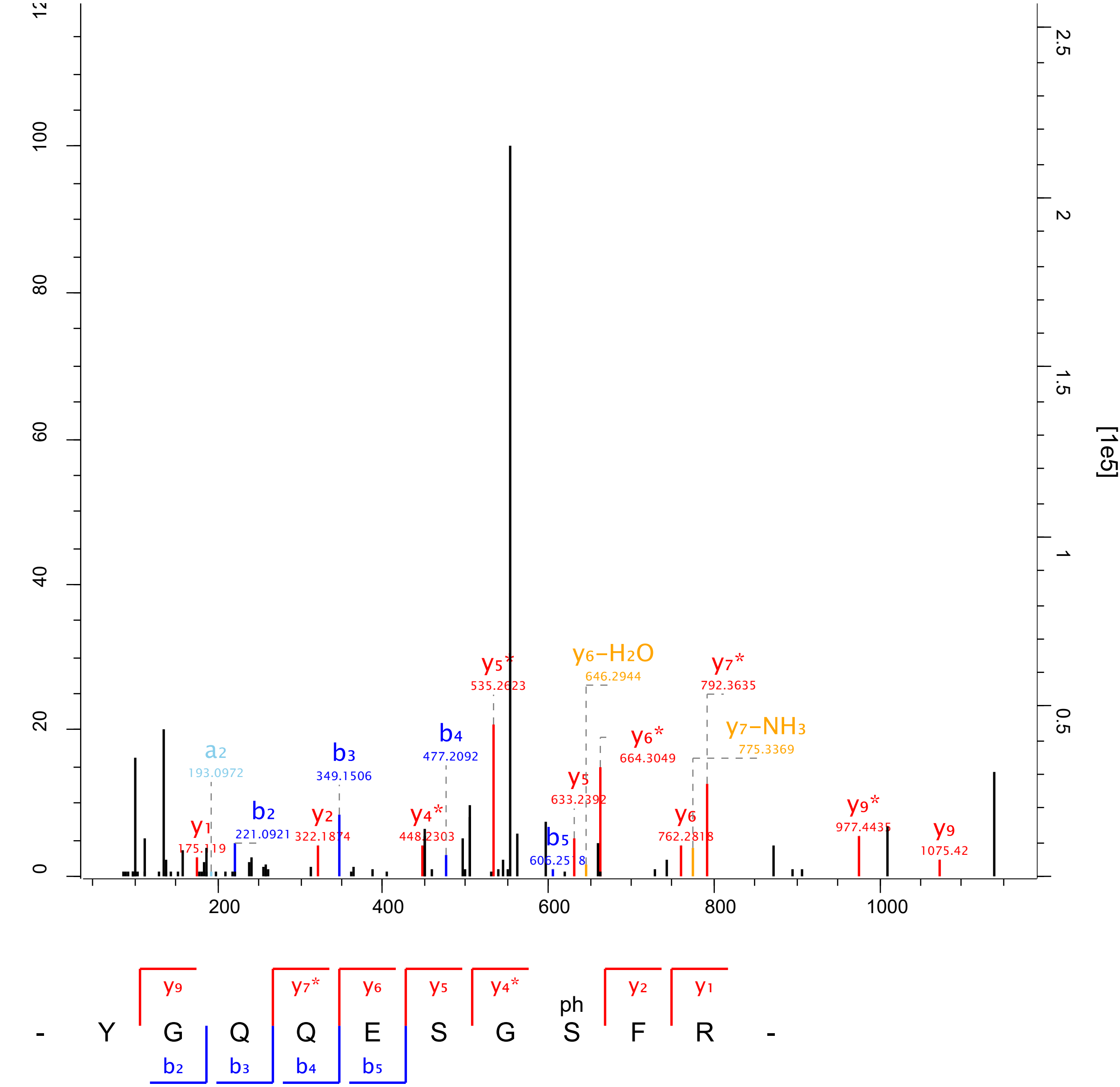

| Raw file | Scan | Method | Score | m/z |
| --- | --- | --- | --- | --- |
| sys_00_3short | 6563 | FTMS; HCD | 57.17 | 473.71 |

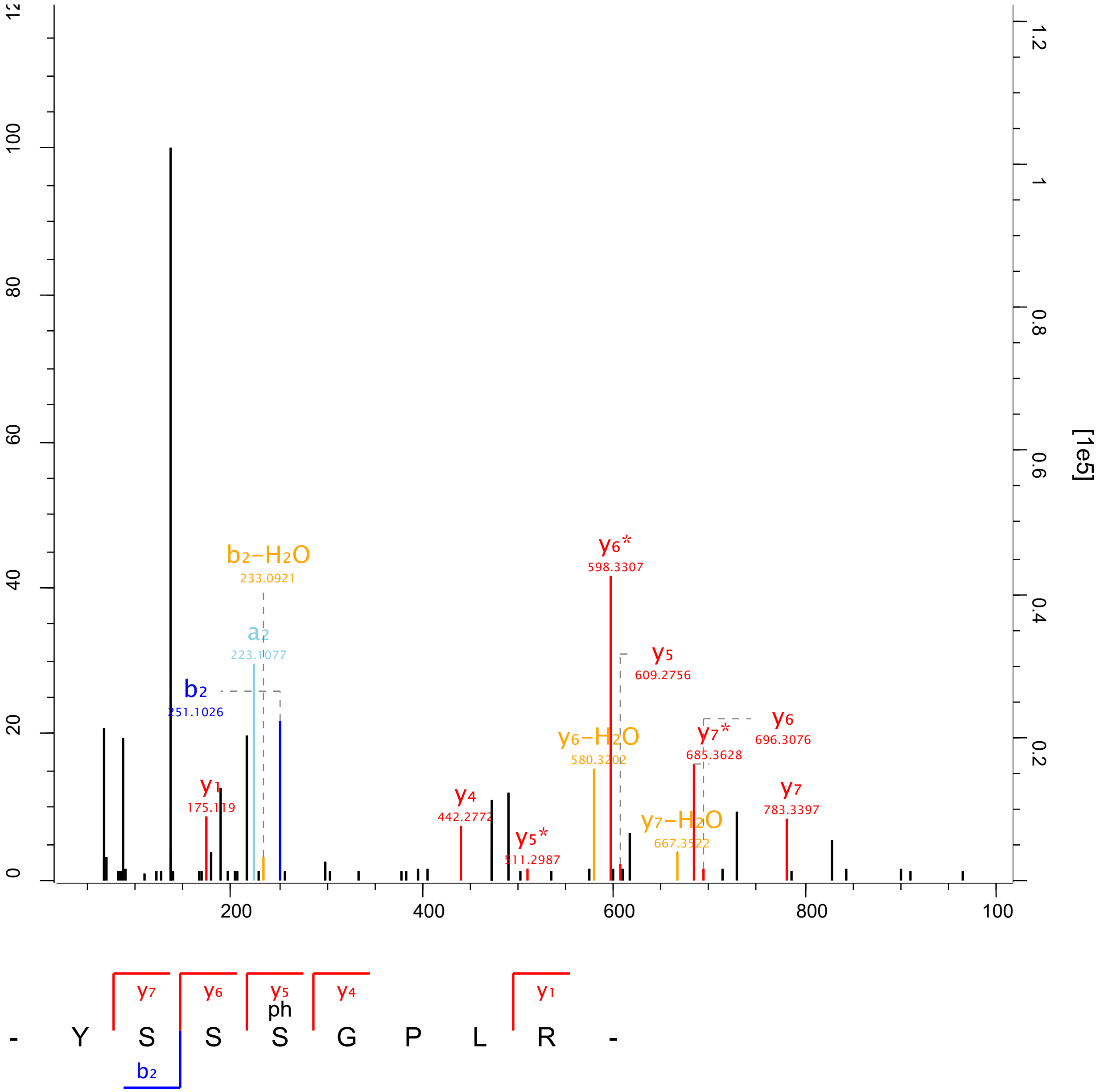

| Raw file | Scan | Method | Score | m/z |
|---|---|---|---|---|
| sys_00_3short | 6651 | FTMS; HCD | 98.17 | 540.76 |

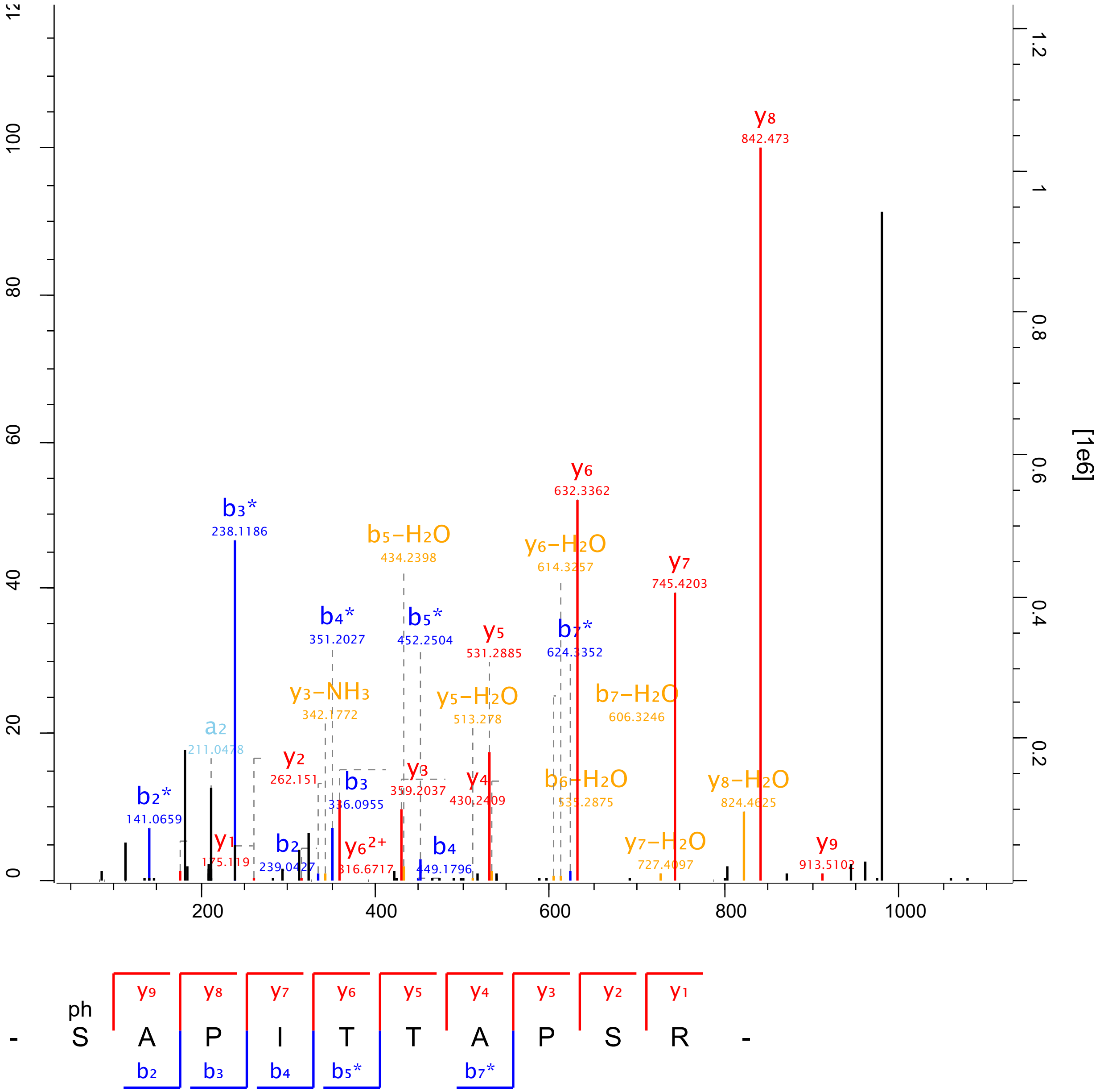

| Raw file | Scan | Method | Score | m/z |
| --- | --- | --- | --- | --- |
| sys_00_3short | 6827 | FTMS; HCD | 113.99 | 510.75 |

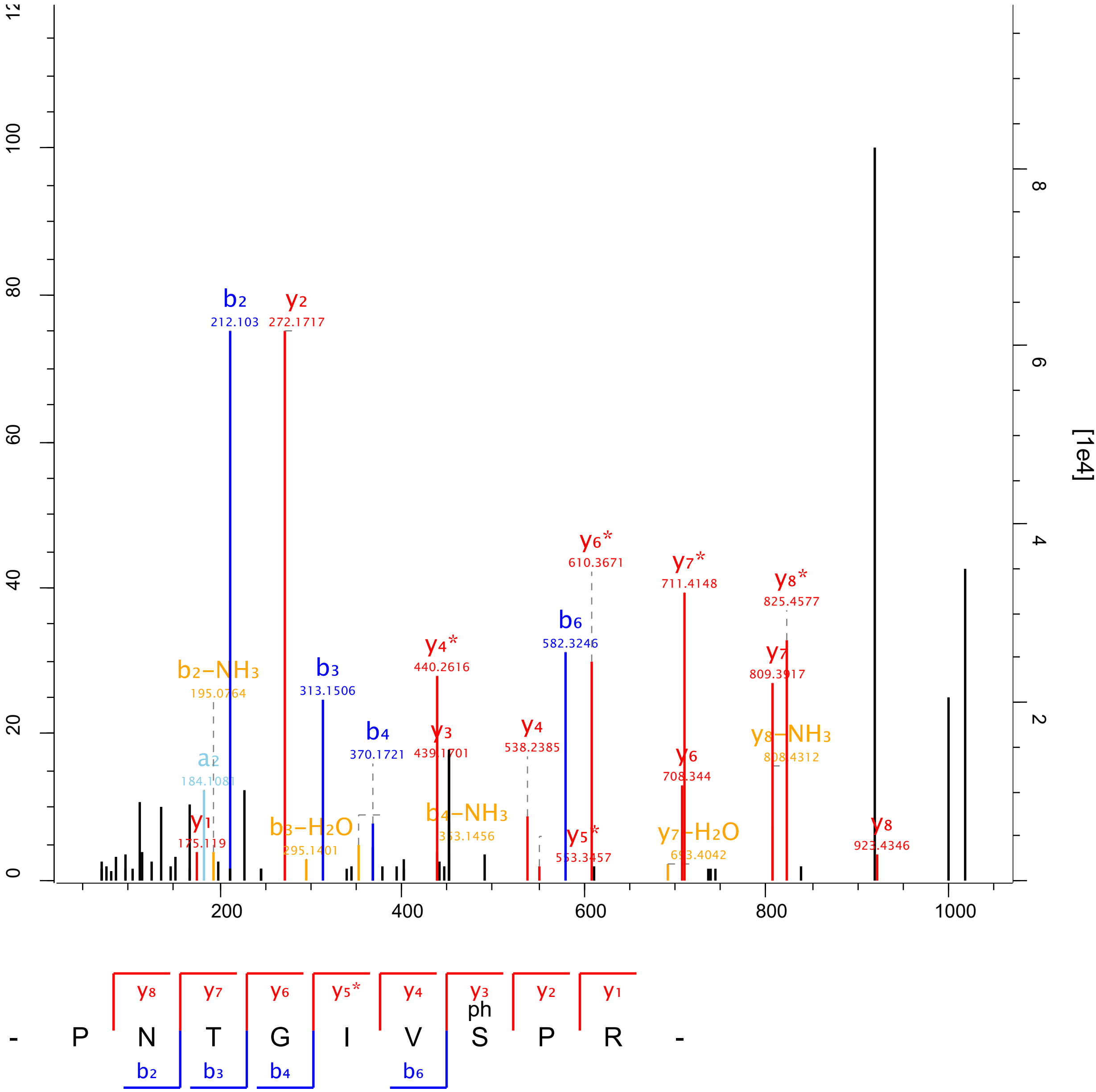

| Raw file | Scan | Method | Score | m/z |
|---|---|---|---|---|
| sys_00_3short | 6940 | FTMS; HCD | 146.59 | 634.28 |

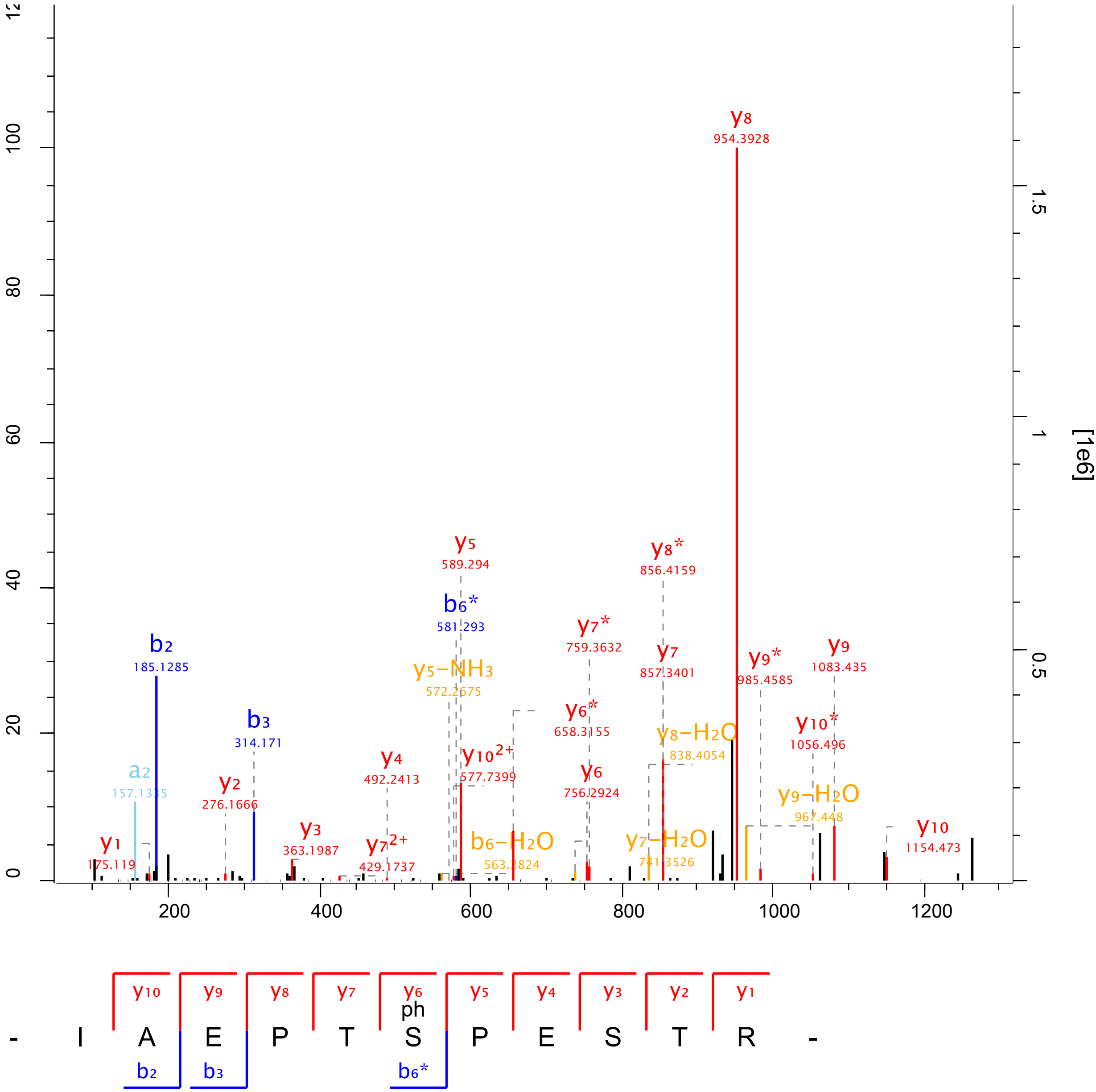

| Raw file | Scan | Method | Score | m/z |
|---|---|---|---|---|
| sys_00_3short | 7401 | FTMS; HCD | 54.9 | 737.29 |

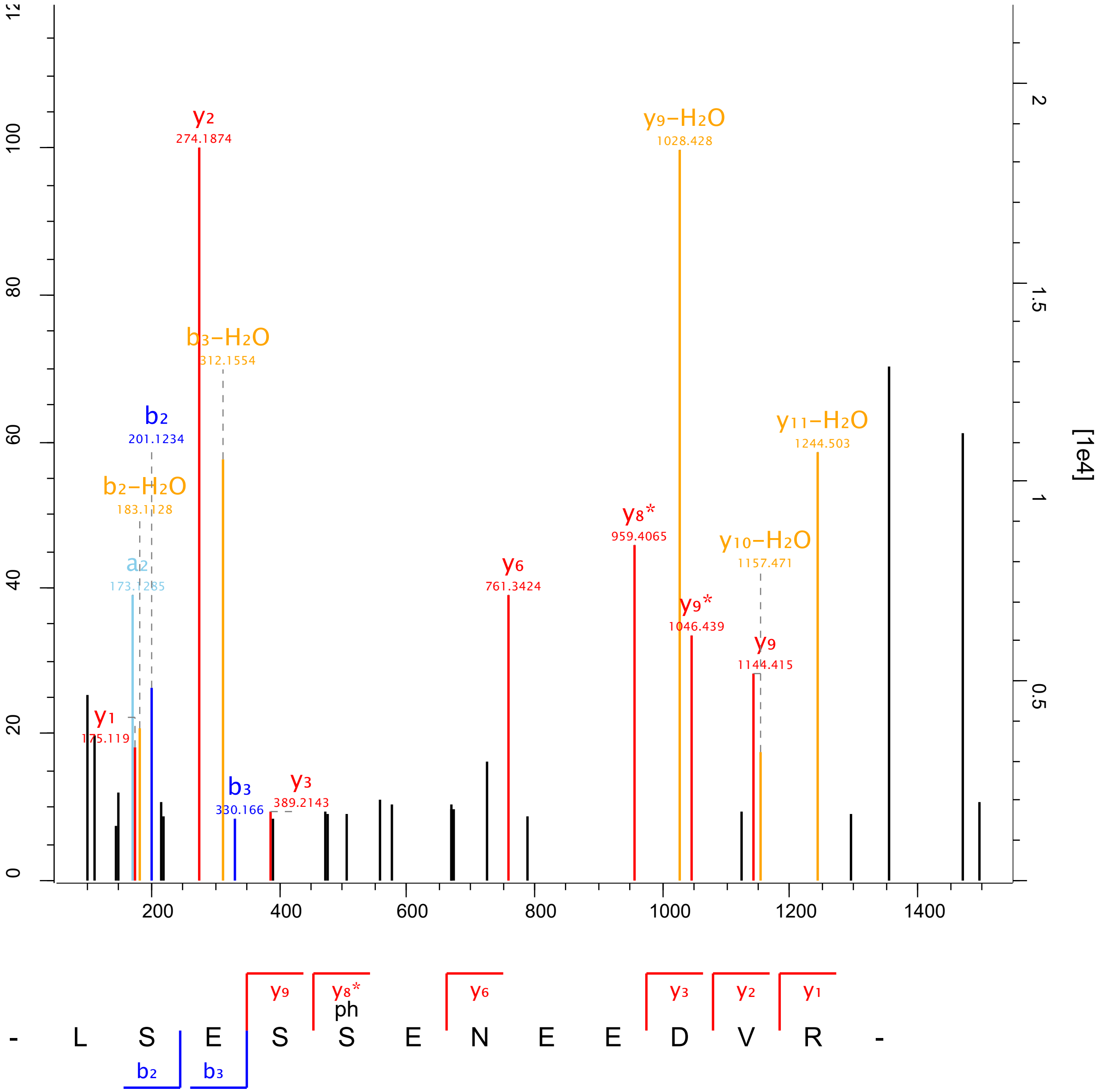

| Raw file | Scan | Method | Score | m/z |
|---|---|---|---|---|
| sys_00_3short | 7417 | FTMS; HCD | 252.66 | 694.3 |

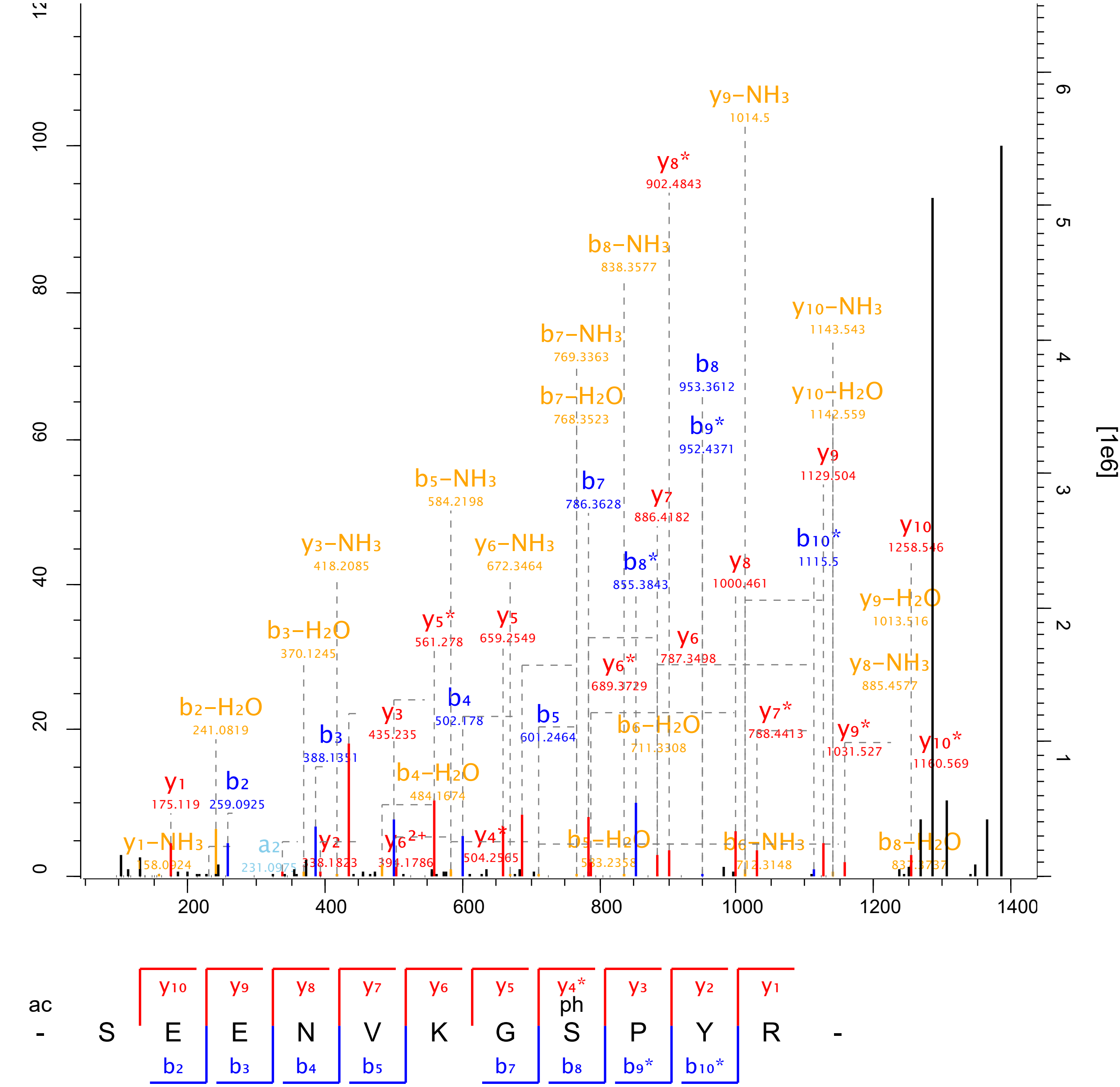

| Raw file | Scan | Method | Score | m/z |
| --- | --- | --- | --- | --- |
| sys_00_3short | 7649 | FTMS; HCD | 248.75 | 680.29 |

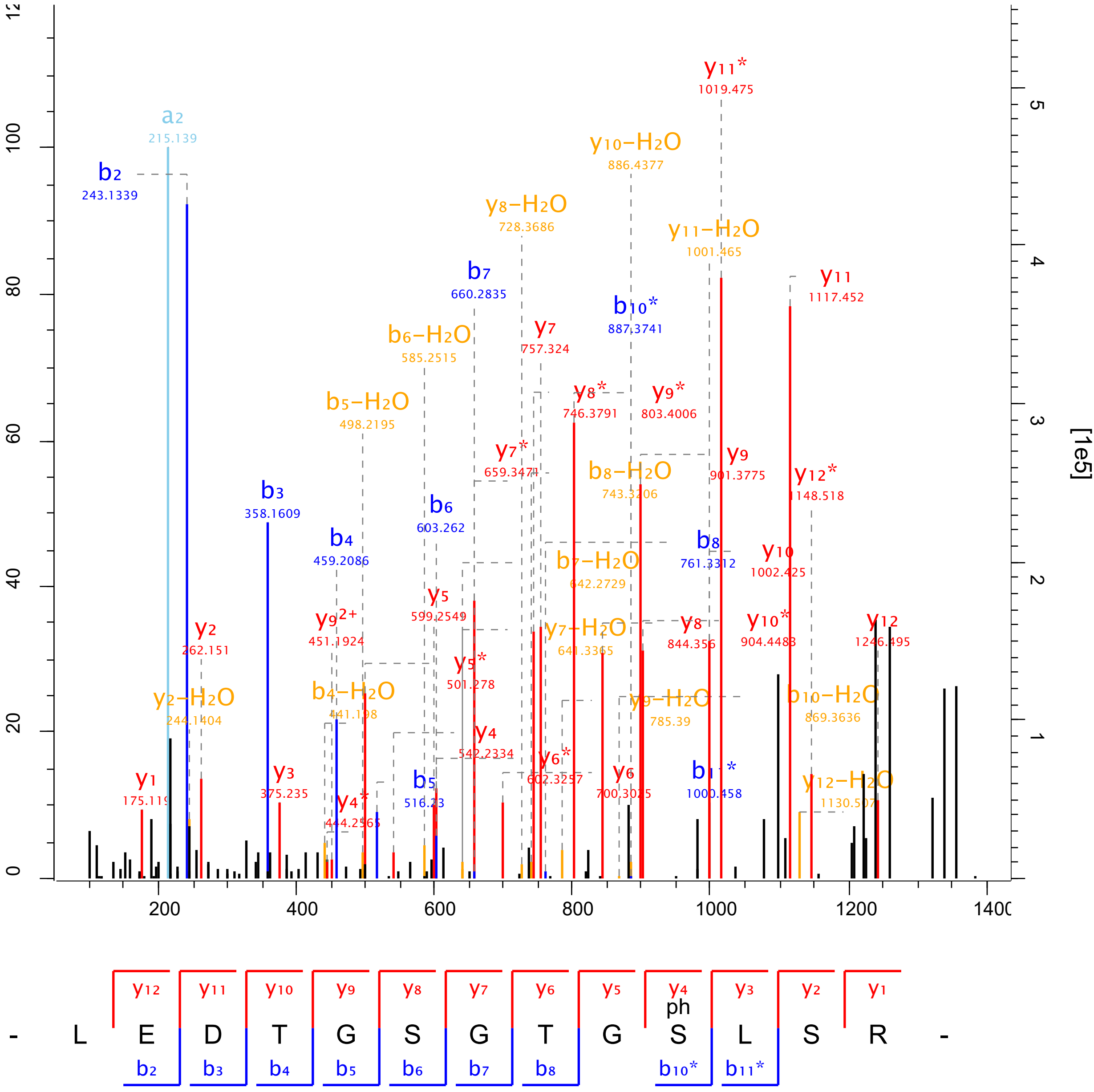

| Raw file | Scan | Method | Score | m/z |
|---|---|---|---|---|
| sys_00_3short | 7859 | FTMS; HCD | 152.04 | 474.2 |

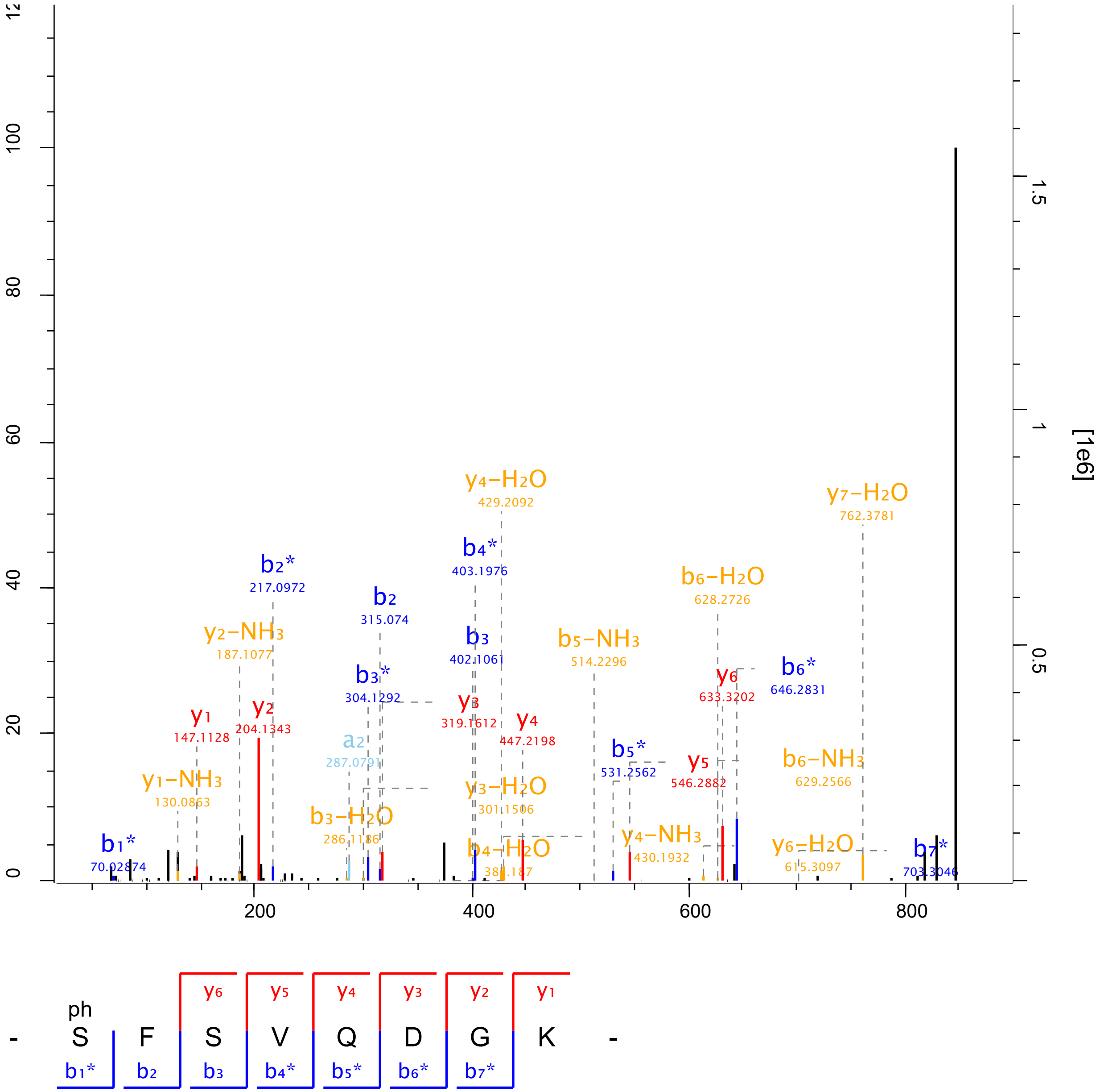

| Raw file | Scan | Method | Score | m/z |
| --- | --- | --- | --- | --- |
| sys_00_3short | 7980 | FTMS; HCD | 99.14 | 410.21 |

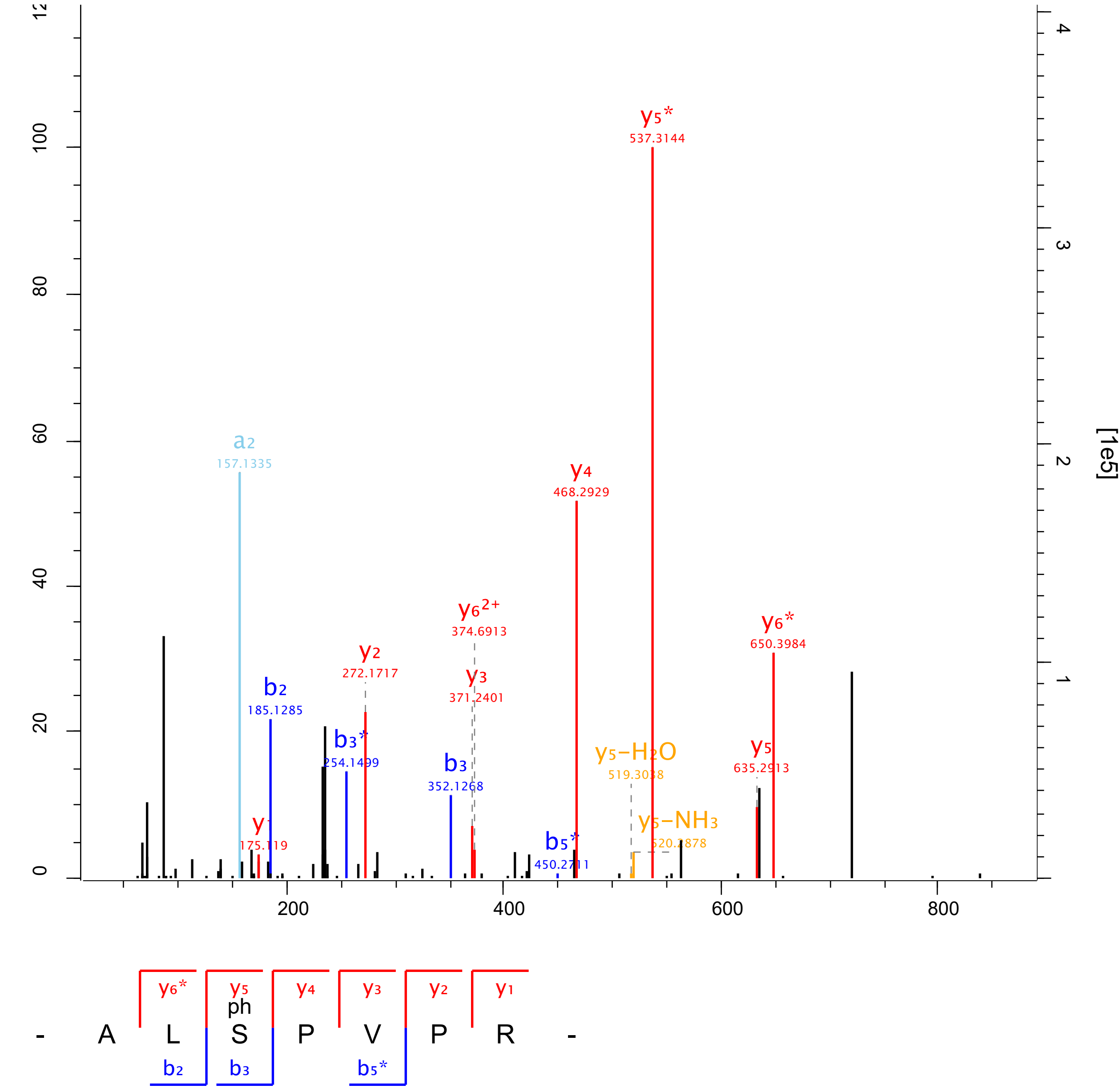

| Raw file | Scan | Method | Score | m/z |
|---|---|---|---|---|
| sys_00_3short | 8115 | FTMS; HCD | 79.66 | 477.71 |

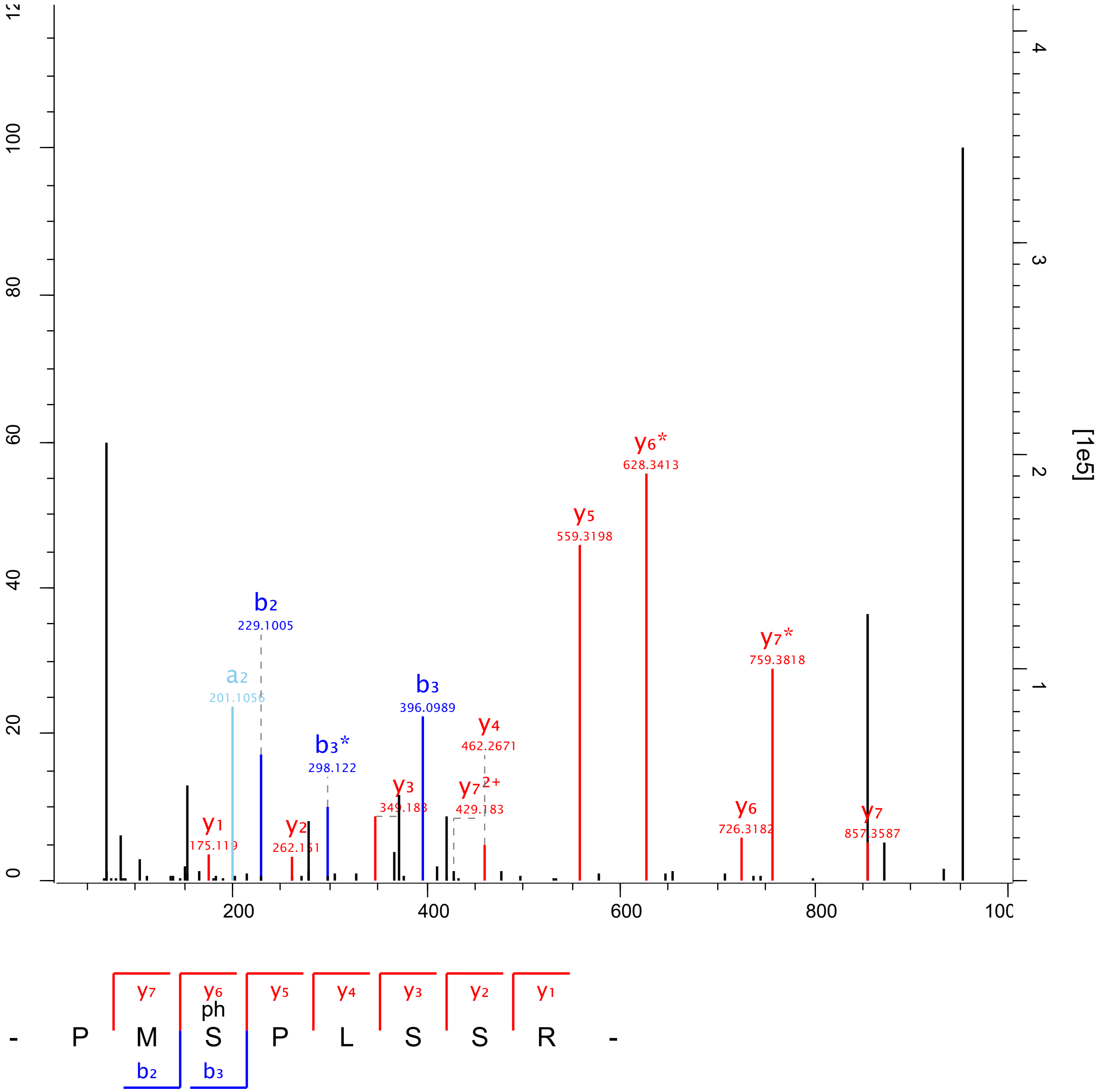

| Raw file | Scan | Method | Score | m/z |
| --- | --- | --- | --- | --- |
| sys_00_3short | 8275 | FTMS; HCD | 79.09 | 616.79 |

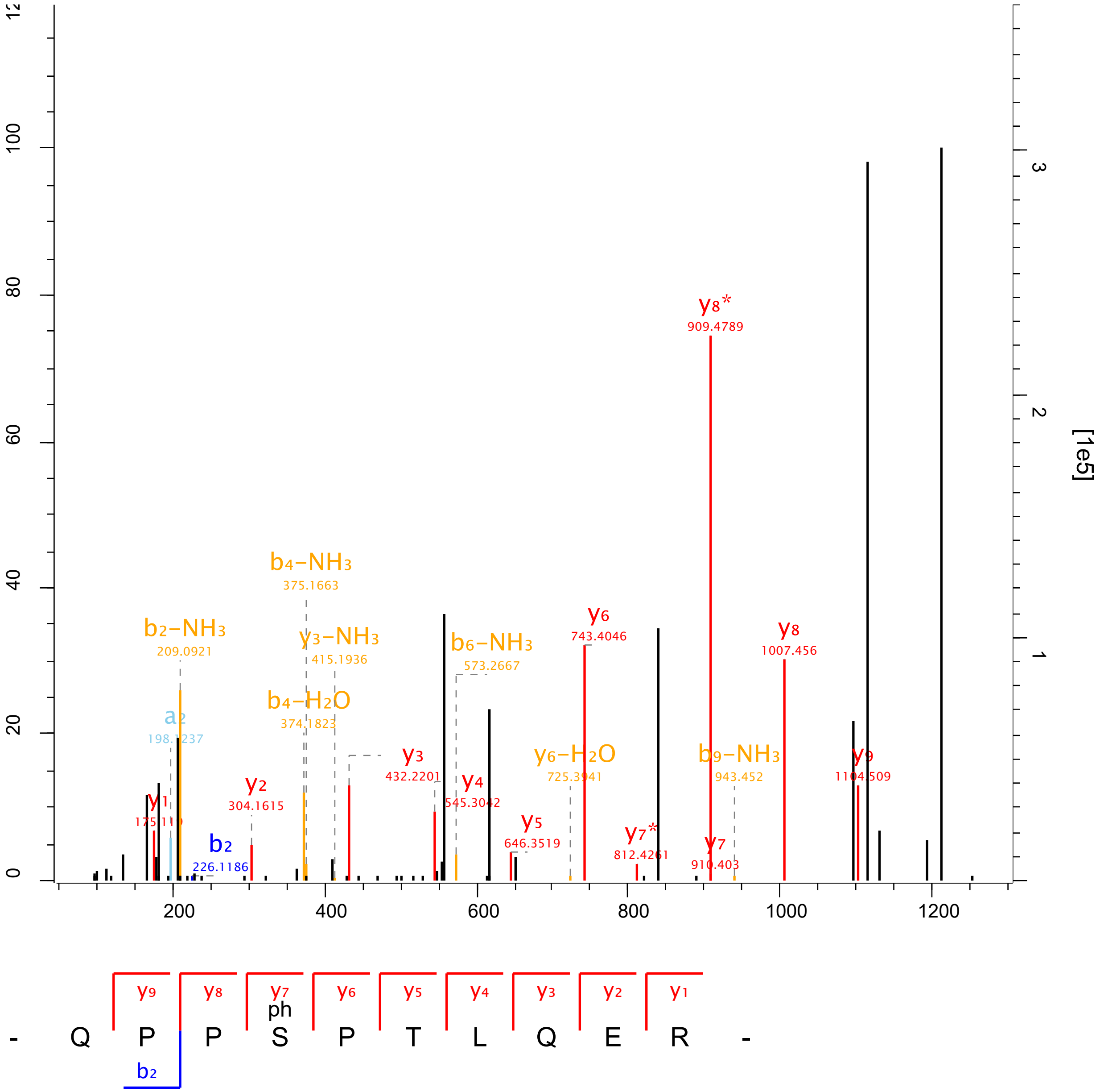

| Raw file | Scan | Method | Score | m/z |
|---|---|---|---|---|
| sys_00_3short | 8459 | FTMS; HCD | 147.33 | 521.73 |

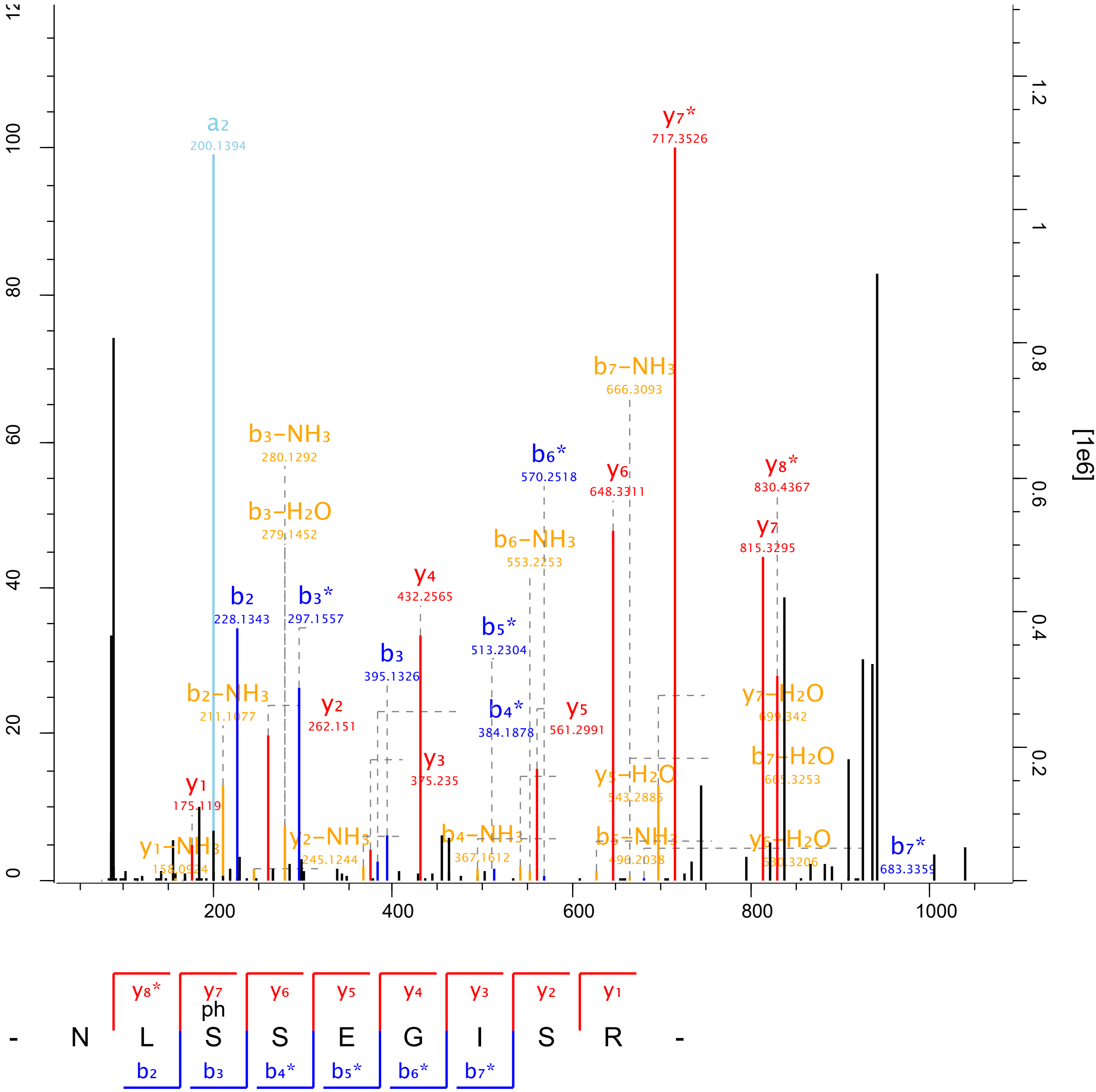

| Raw file | Scan | Method | Score | m/z |
| --- | --- | --- | --- | --- |
| sys_00_3short | 8496 | FTMS; HCD | 100.69 | 469.72 |

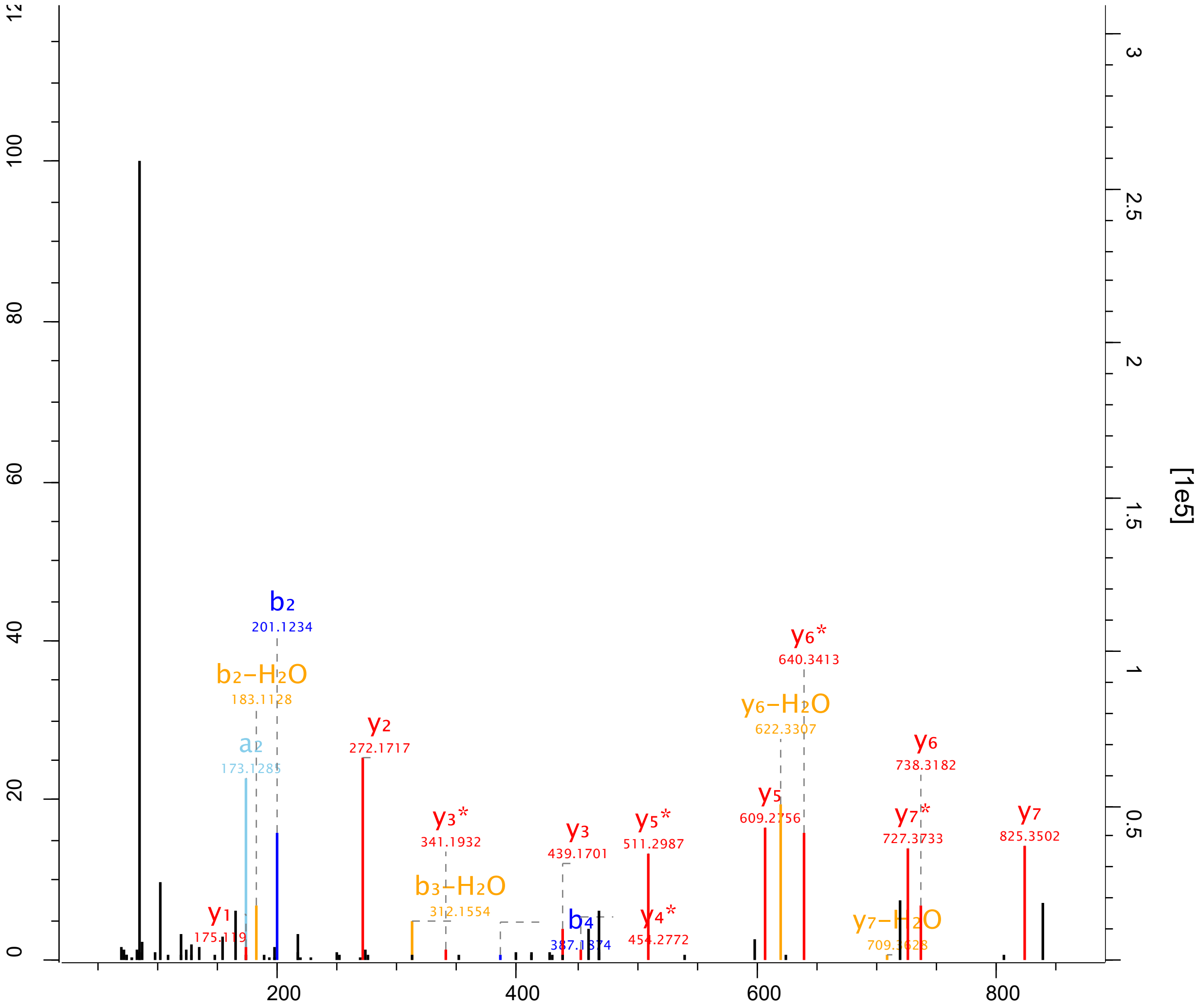


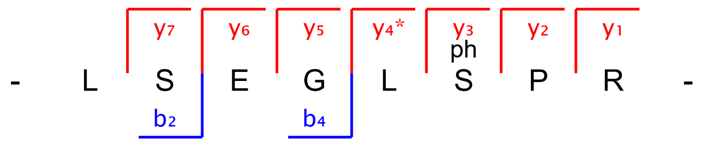

| Raw file | Scan | Method | Score | m/z |
|---|---|---|---|---|
| sys_00_3short | 8704 | FTMS; HCD | 46.13 | 536.29 |

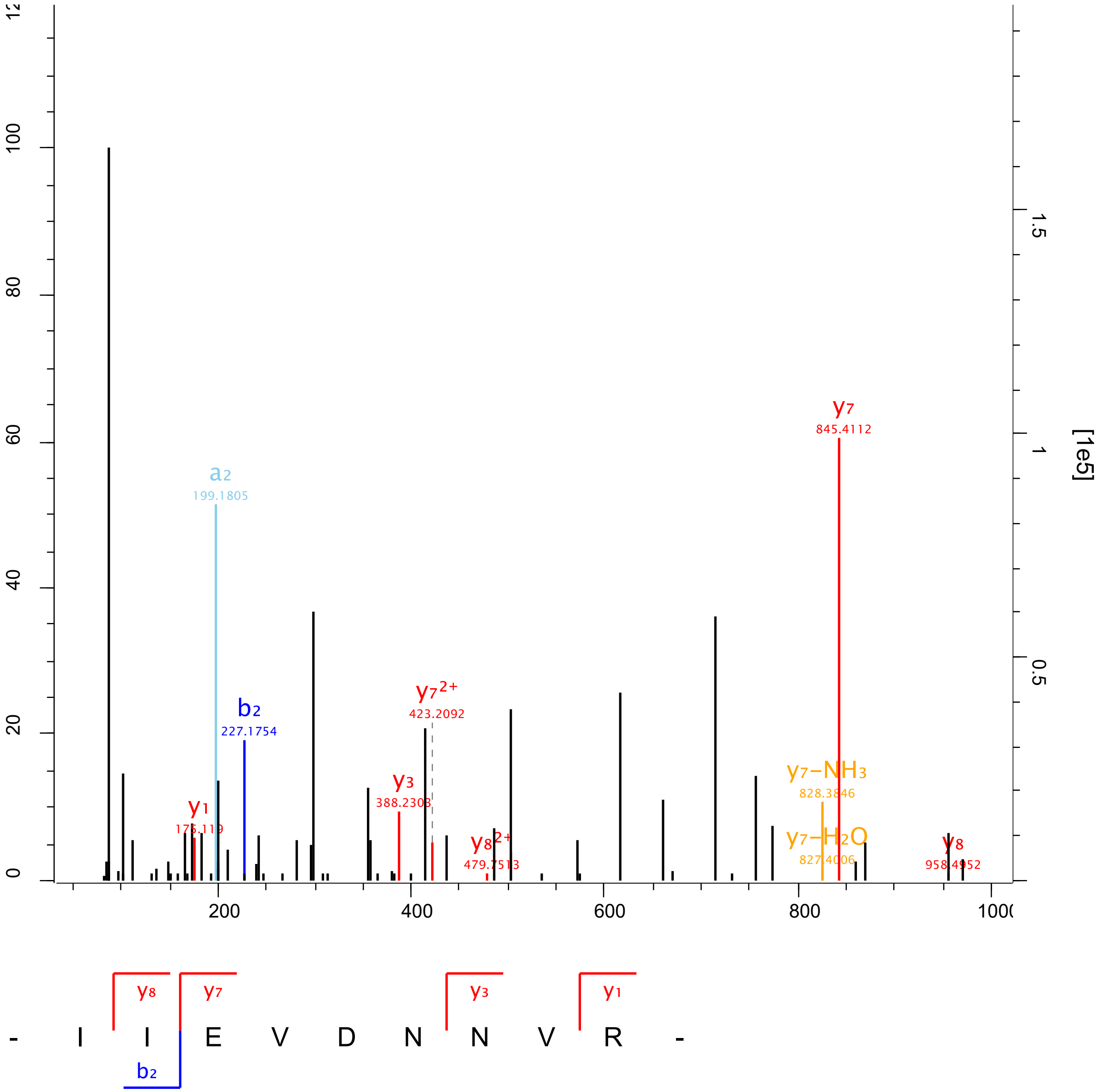

| Raw file | Scan | Method | Score | m/z |
| --- | --- | --- | --- | --- |
| sys_00_3short | 8721 | FTMS; HCD | 116.55 | 496.69 |

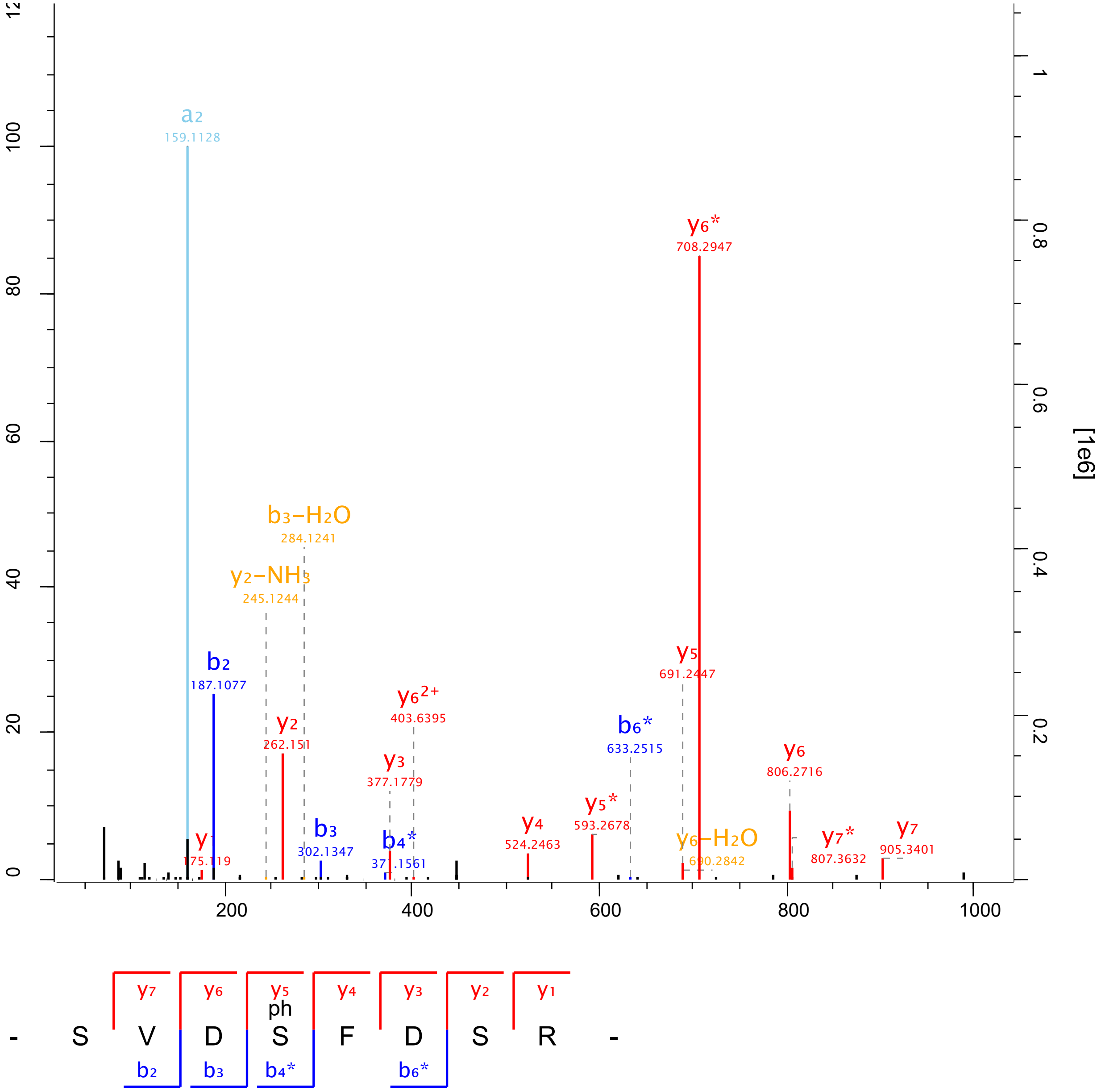

| Raw file | Scan | Method | Score | m/z |
|---|---|---|---|---|
| sys_00_3short | 8767 | FTMS; HCD | 74.99 | 604.24 |

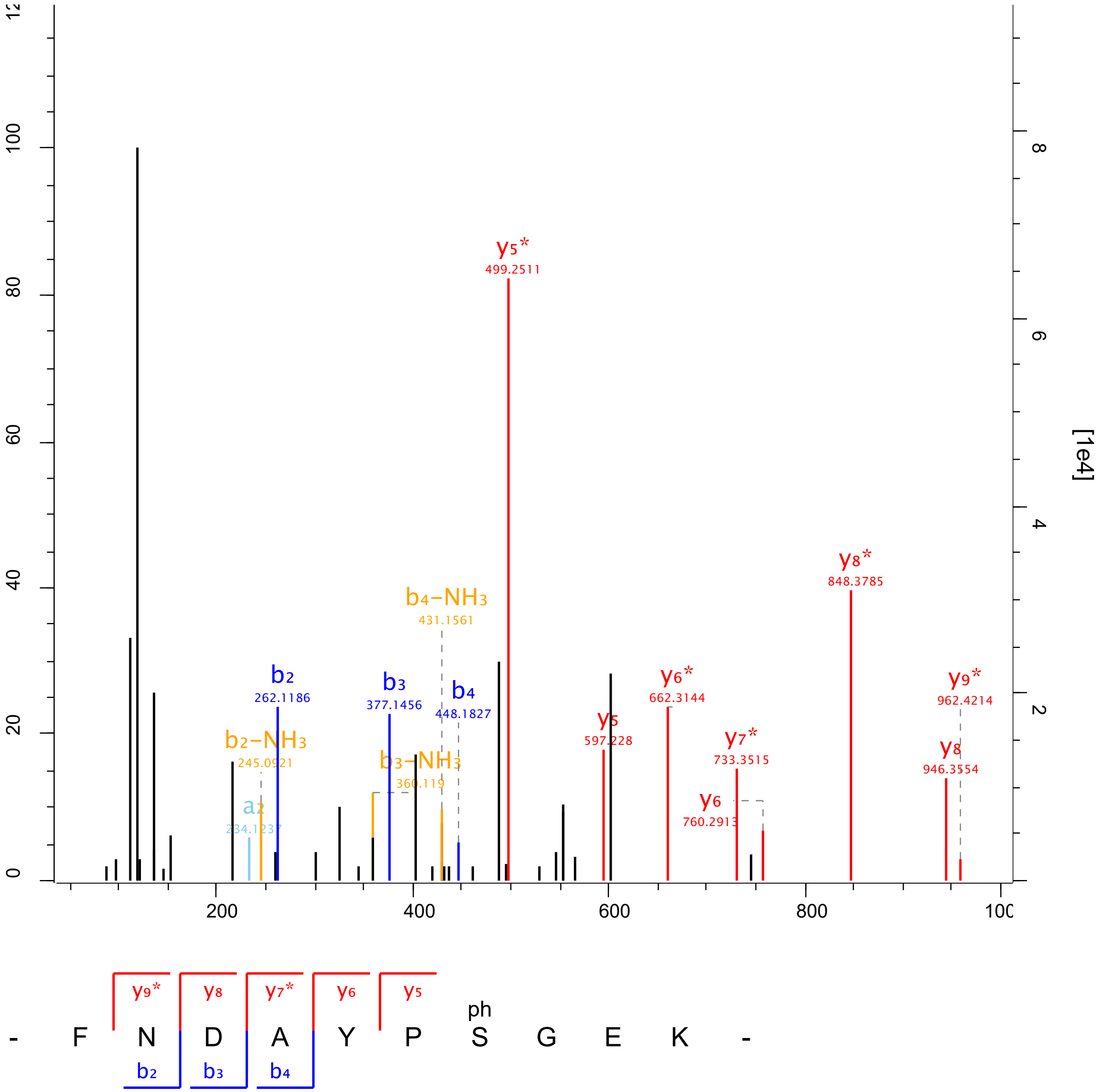

| Raw file | Scan | Method | Score | m/z |
|---|---|---|---|---|
| sys_00_3short | 8811 | FTMS; HCD | 68.66 | 474.19 |

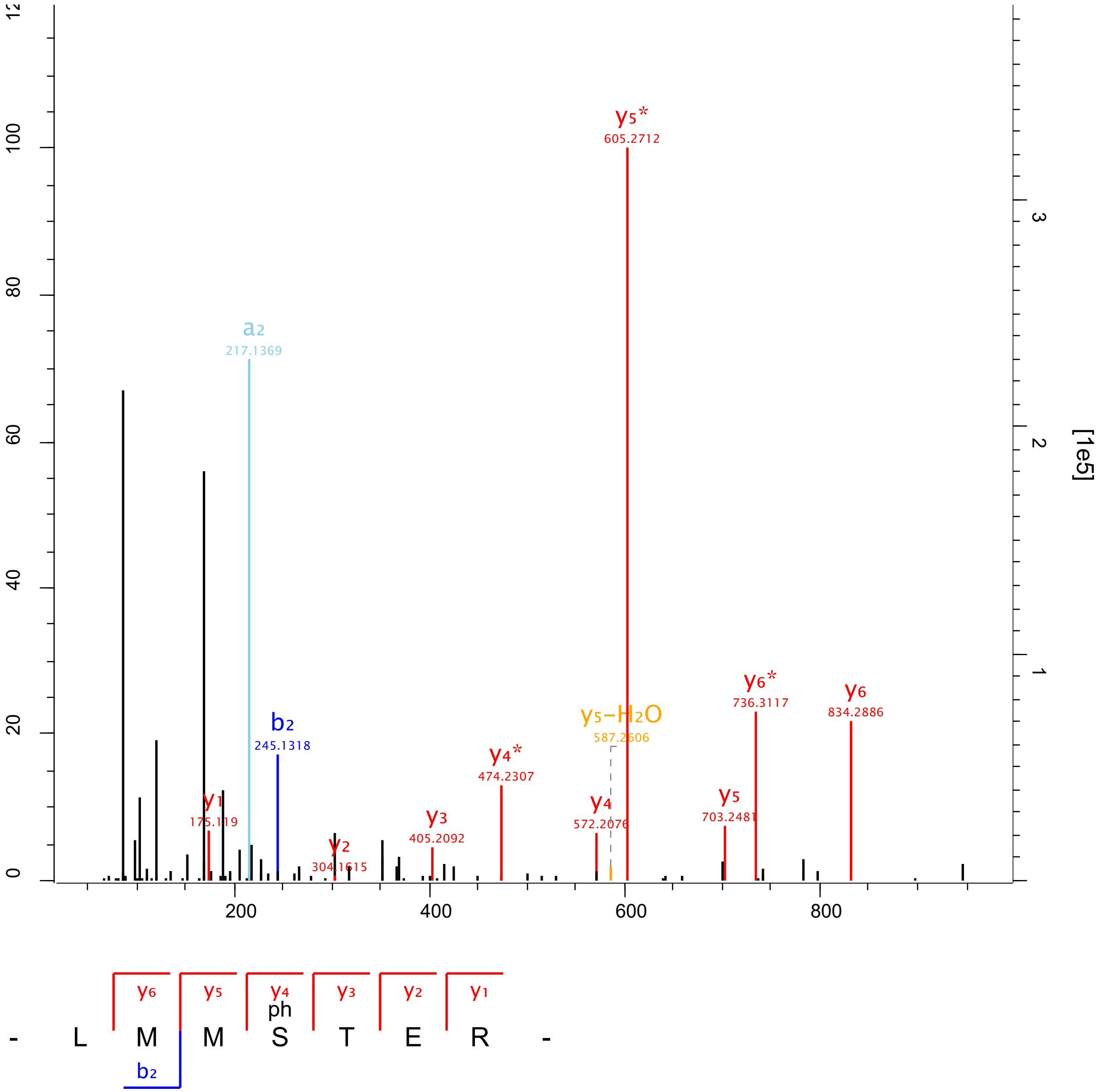

| Raw file | Scan | Method | Score | m/z |
| --- | --- | --- | --- | --- |
| sys_00_3short | 8857 | FTMS; HCD | 154.1 | 578.74 |

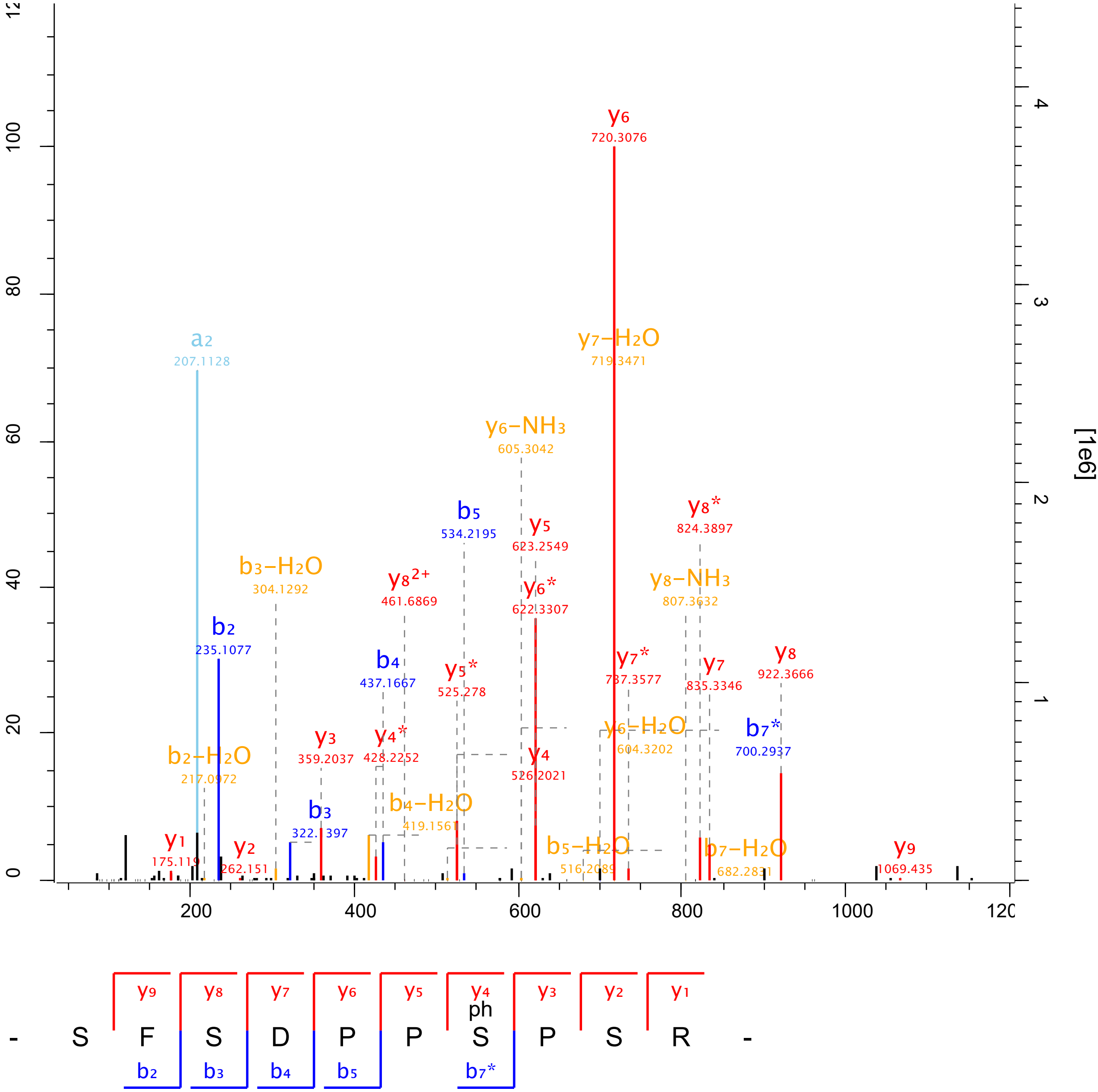

| Raw file | Scan | Method | Score | m/z |
|---|---|---|---|---|
| sys_00_3short | 8892 | FTMS; HCD | 99.5 | 627.24 |

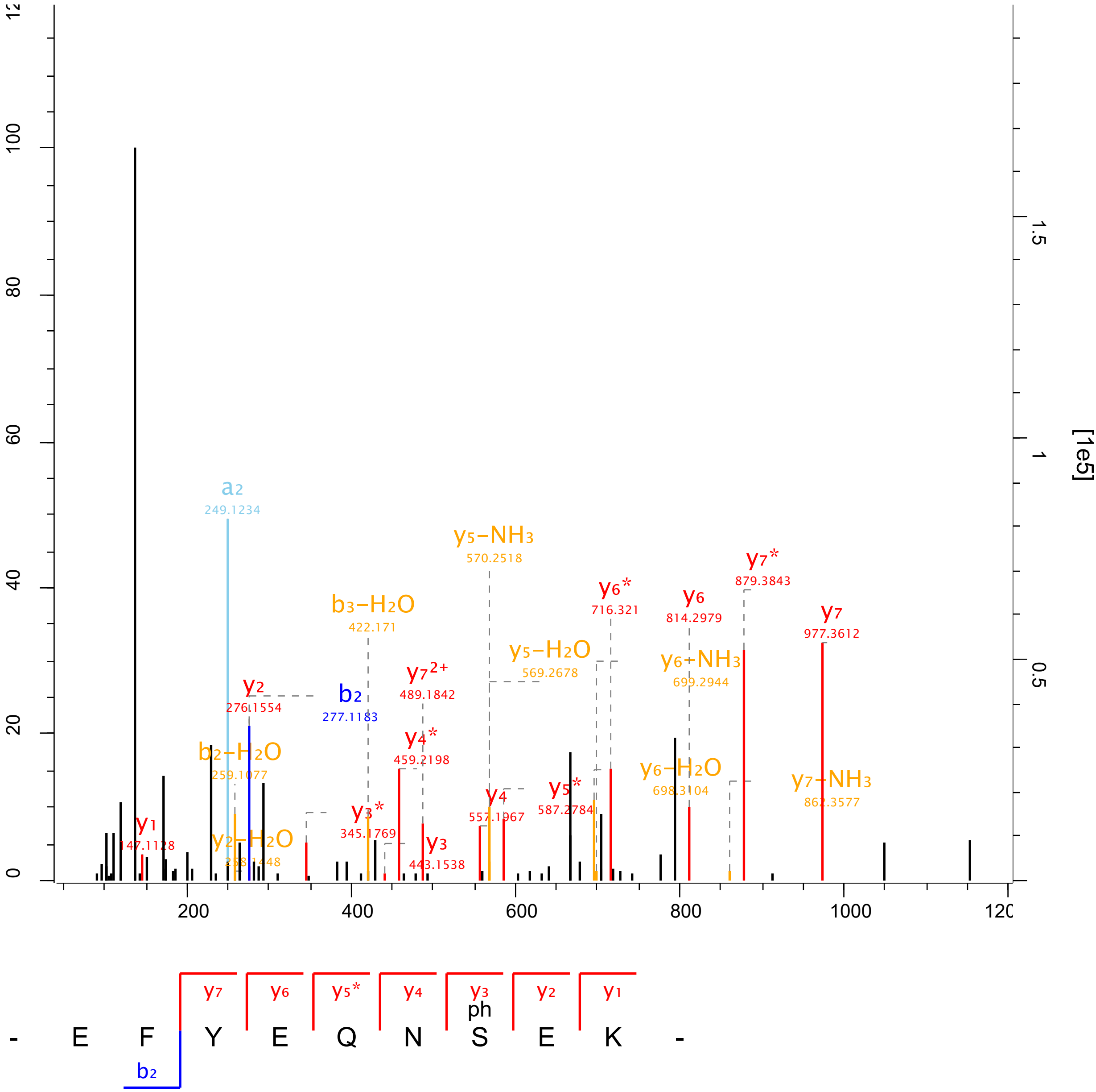

| Raw file | Scan | Method | Score | m/z |
|---|---|---|---|---|
| sys_00_3short | 8986 | FTMS; HCD | 175.08 | 637.78 |

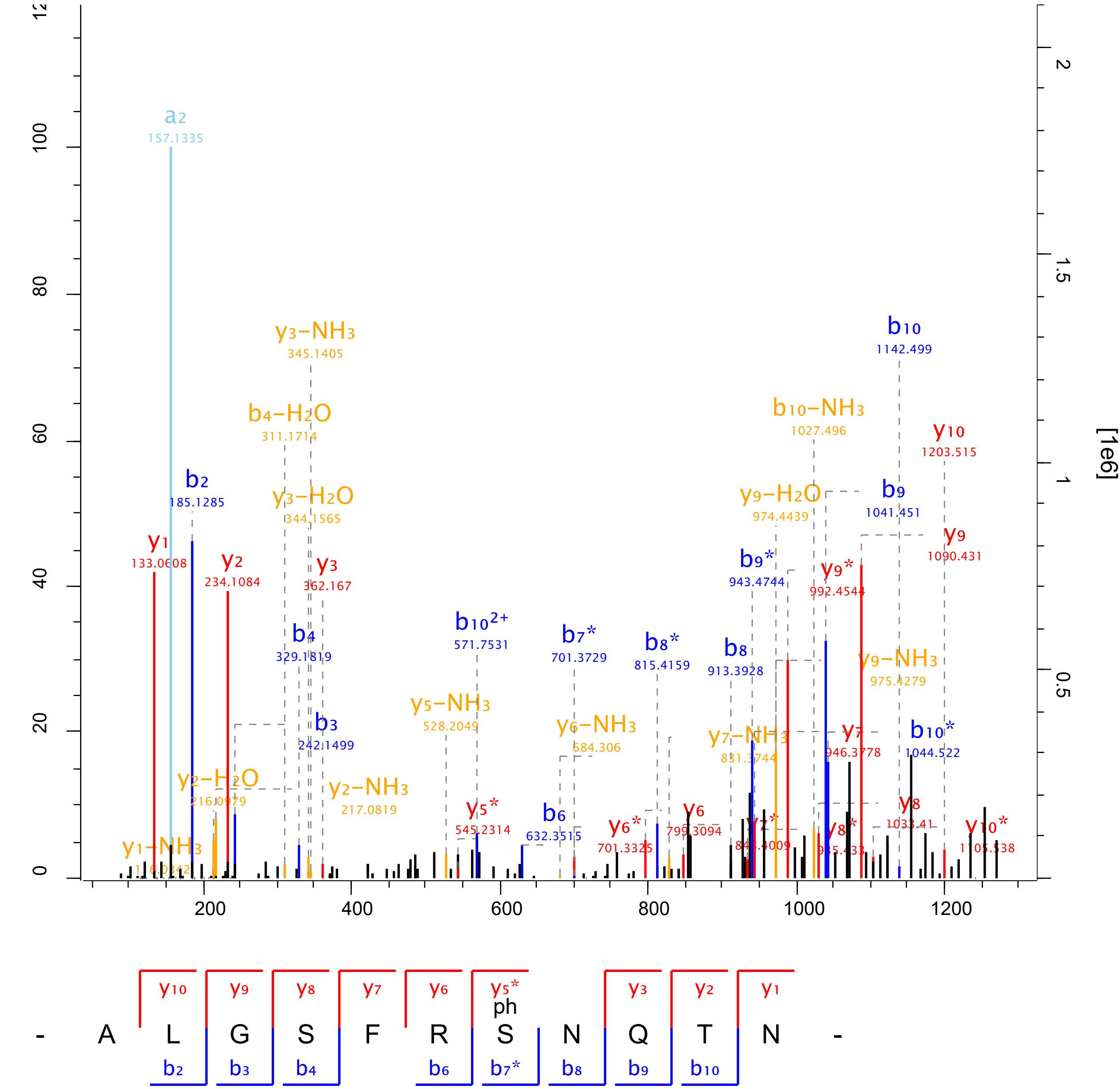

| Raw file | Scan | Method | Score | m/z |
|---|---|---|---|---|
| sys_00_3short | 9228 | FTMS; HCD | 105.58 | 620.3 |

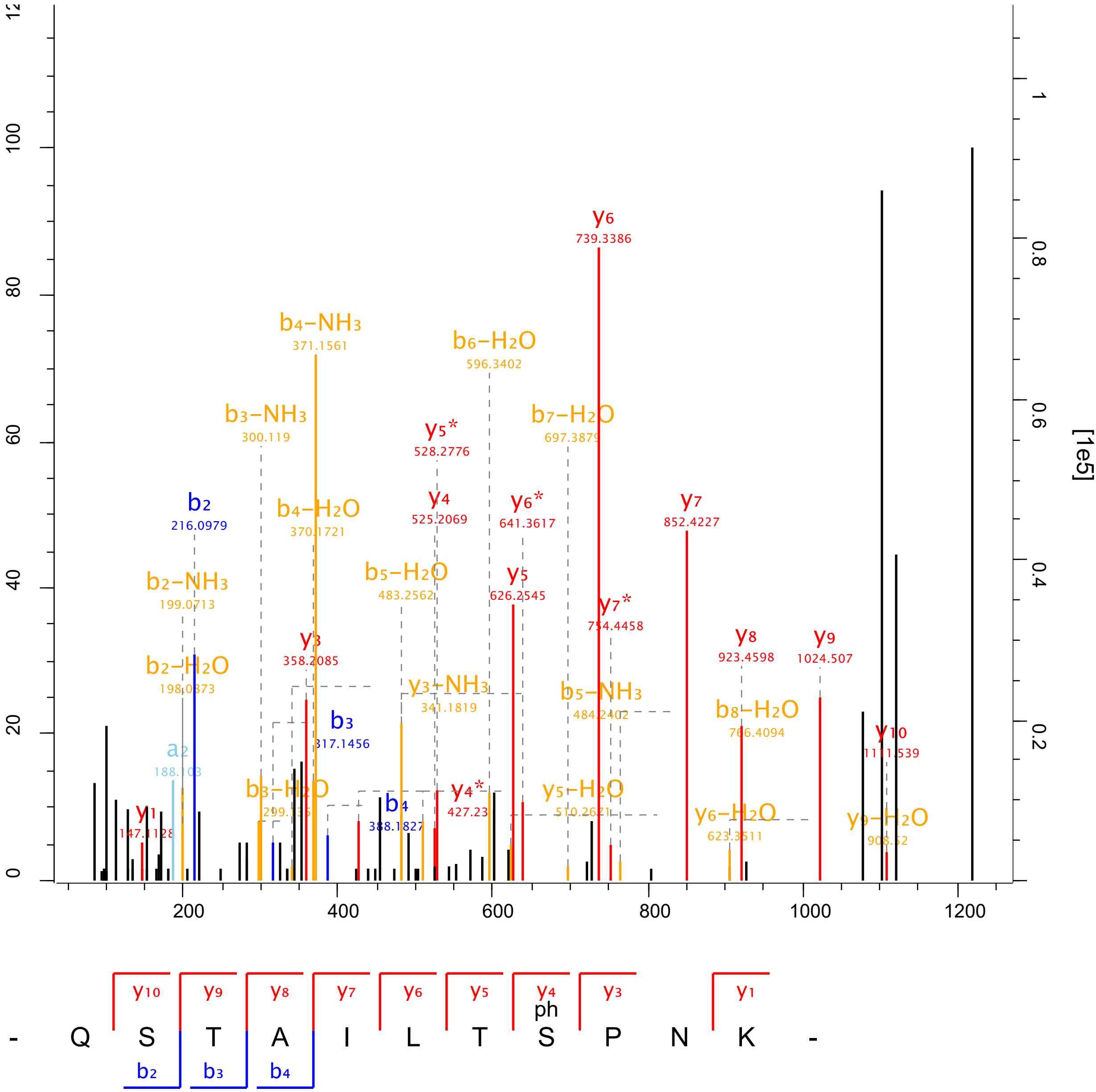

| Raw file | Scan | Method | Score | m/z |
|---|---|---|---|---|
| sys_00_3short | 9266 | FTMS; HCD | 80.23 | 398.19 |

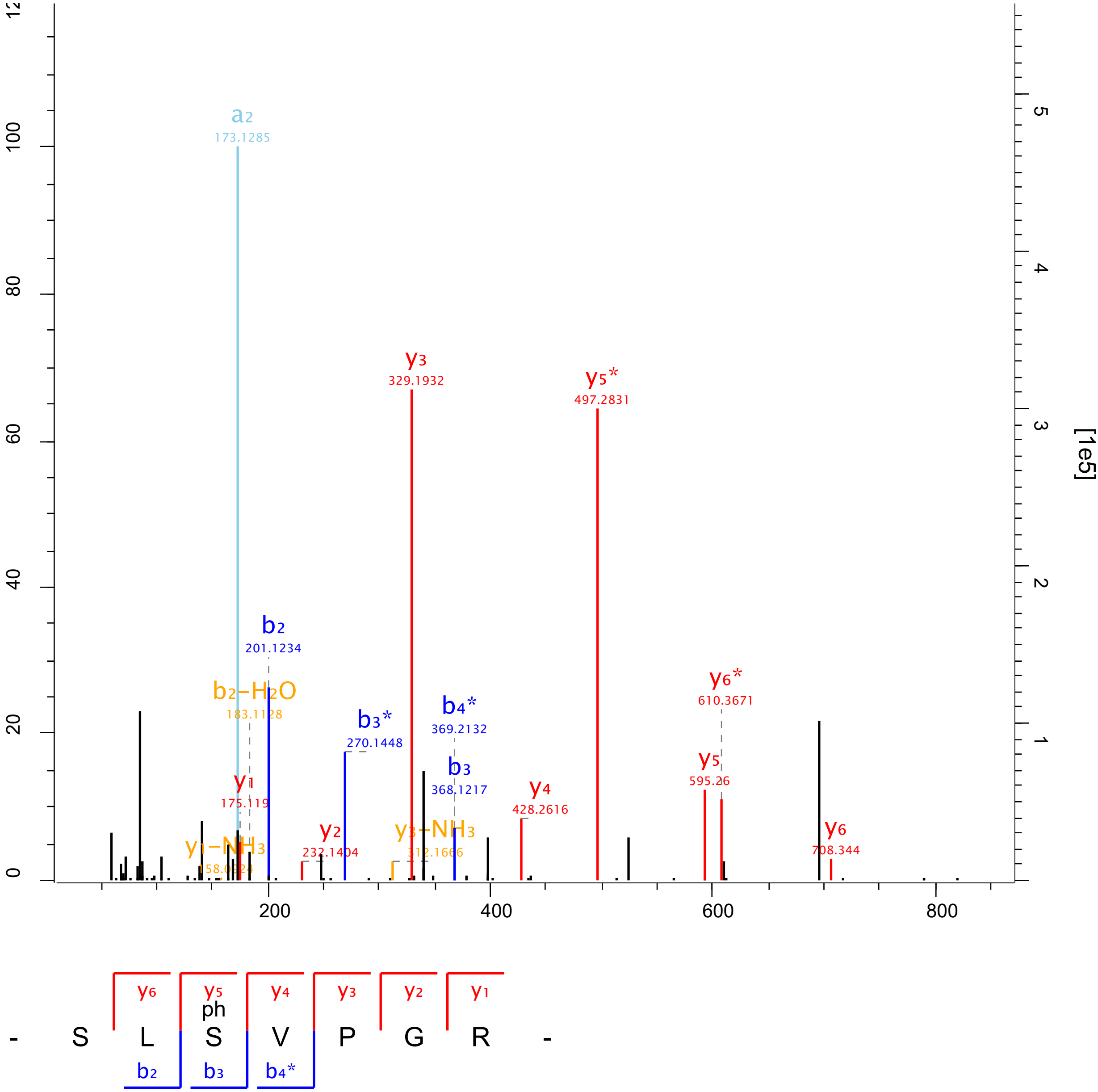

| Raw file | Scan | Method | Score | m/z |
|---|---|---|---|---|
| sys_00_3short | 9342 | FTMS; HCD | 98.9 | 538.76 |

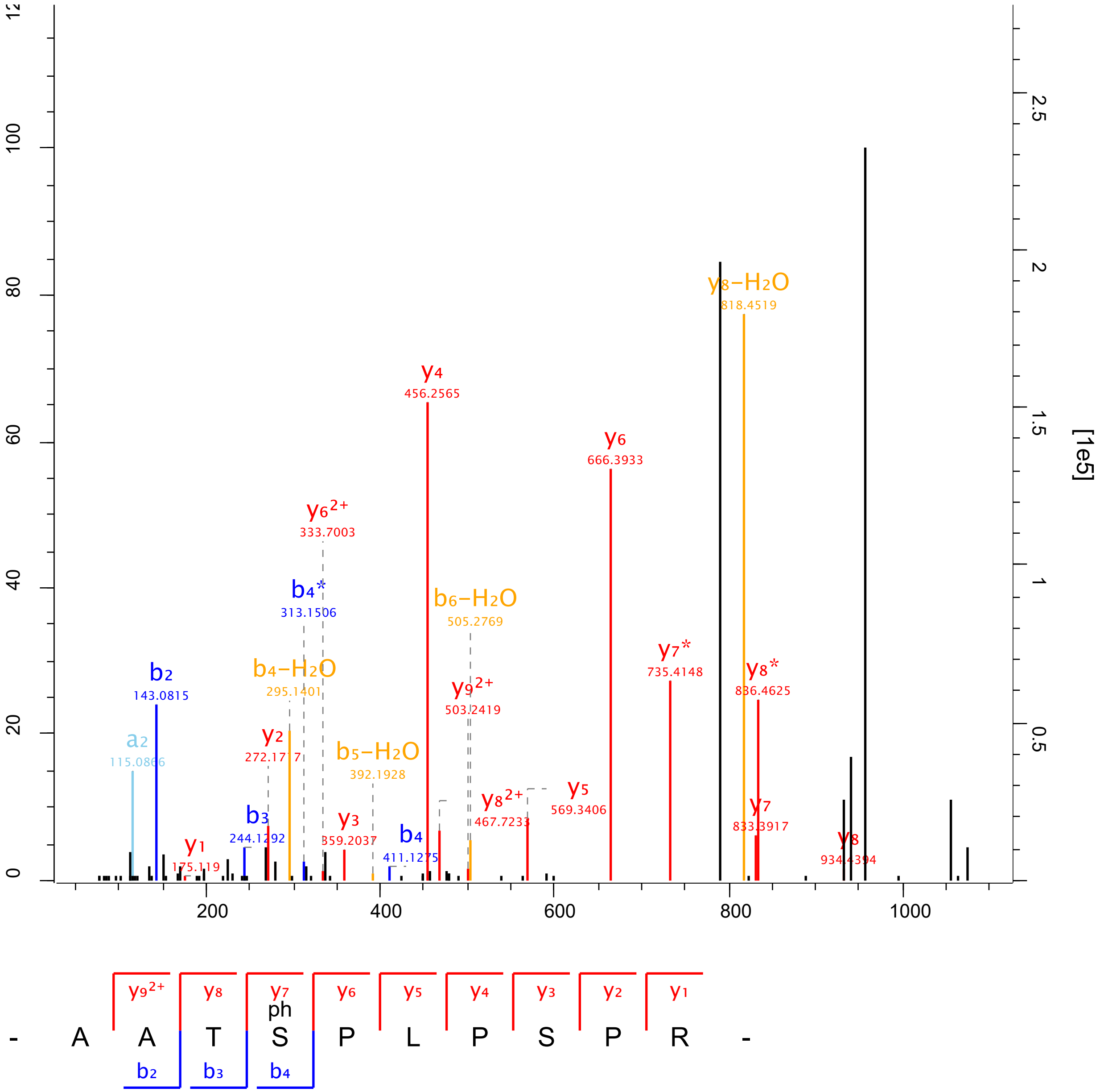

| Raw file | Scan | Method | Score | m/z |
|---|---|---|---|---|
| sys_00_3short | 9602 | FTMS; HCD | 41.4 | 406.7 |

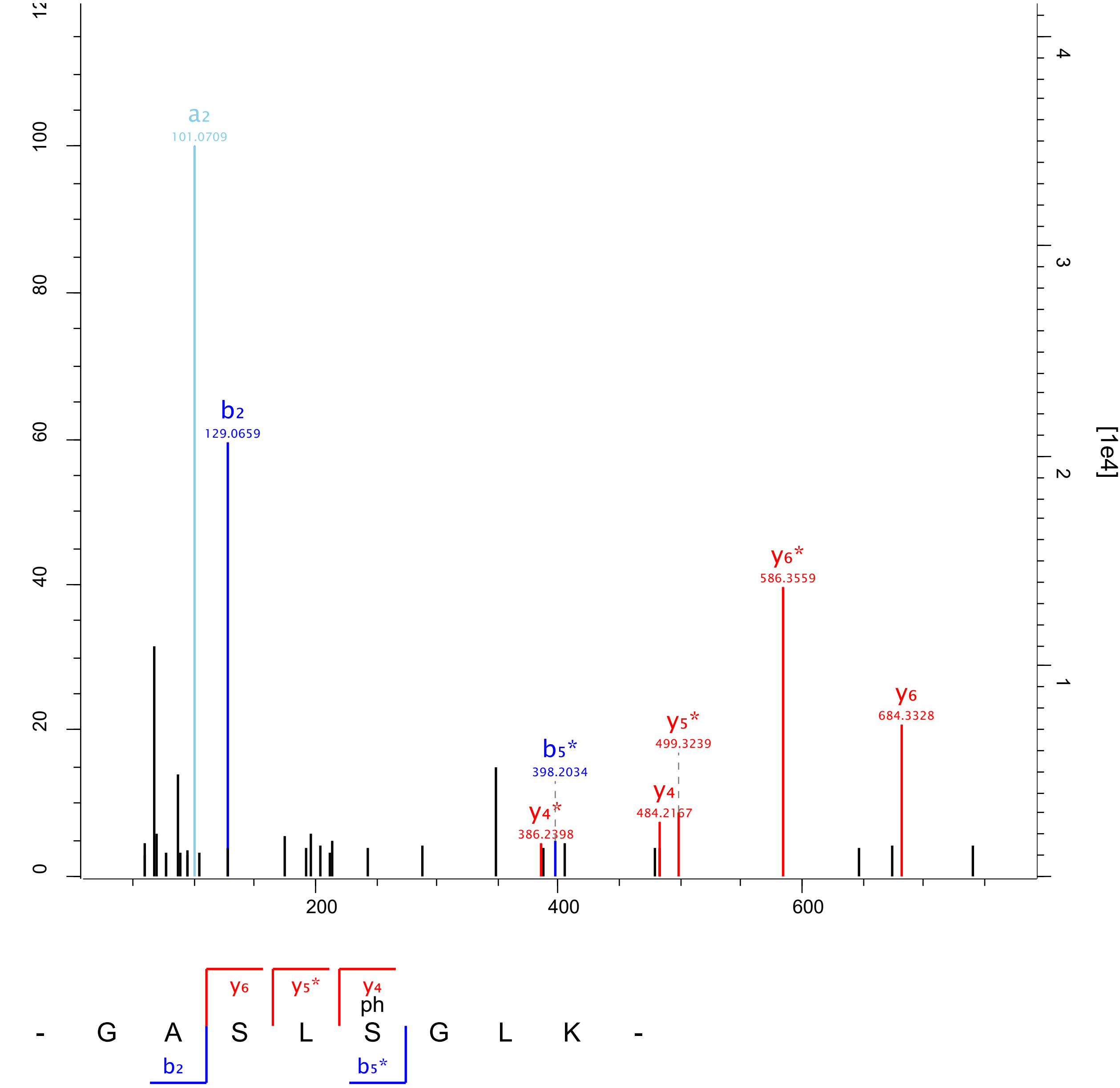

| Raw file | Scan | Method | Score | m/z |
|---|---|---|---|---|
| sys_00_3short | 9657 | FTMS; HCD | 119.04 | 676.8 |

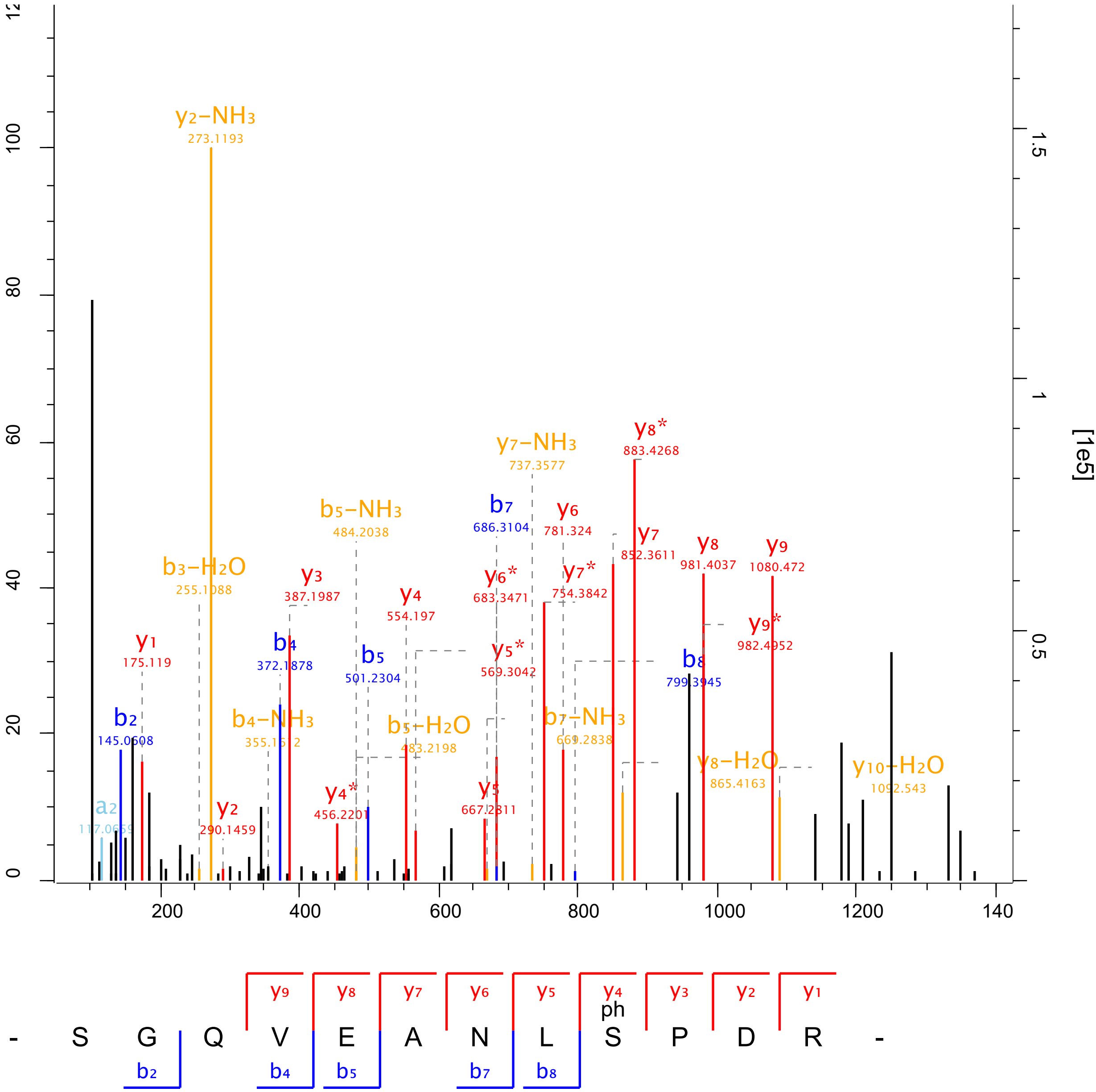

| Raw file | Scan | Method | Score | m/z |
| --- | --- | --- | --- | --- |
| sys_00_3short | 9951 | FTMS; HCD | 142.12 | 577.77 |

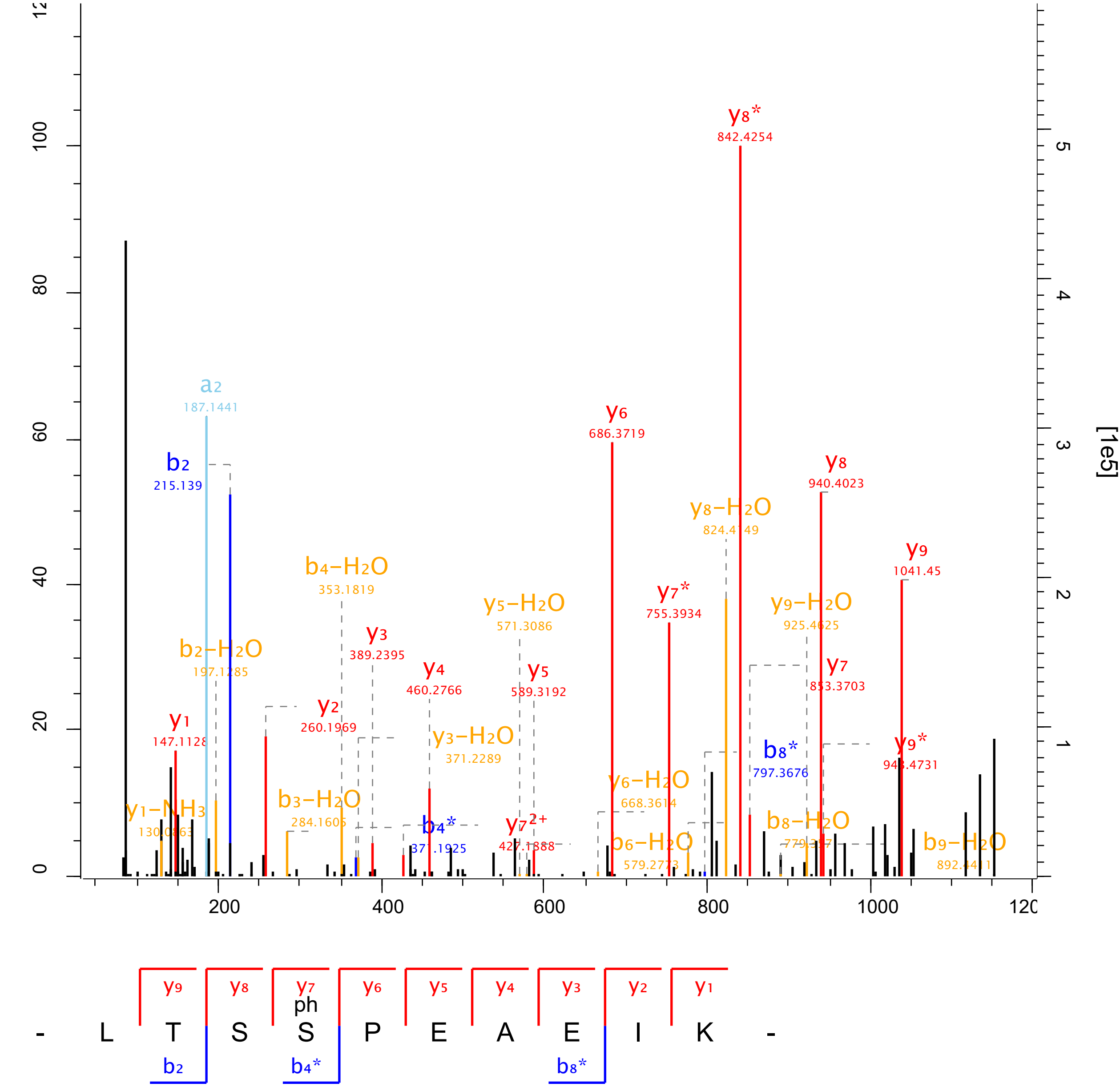

| Raw file | Scan | Method | Score | m/z |
|---|---|---|---|---|
| sys_02_2 | 10092 | FTMS; HCD | 50.9 | 502.24 |

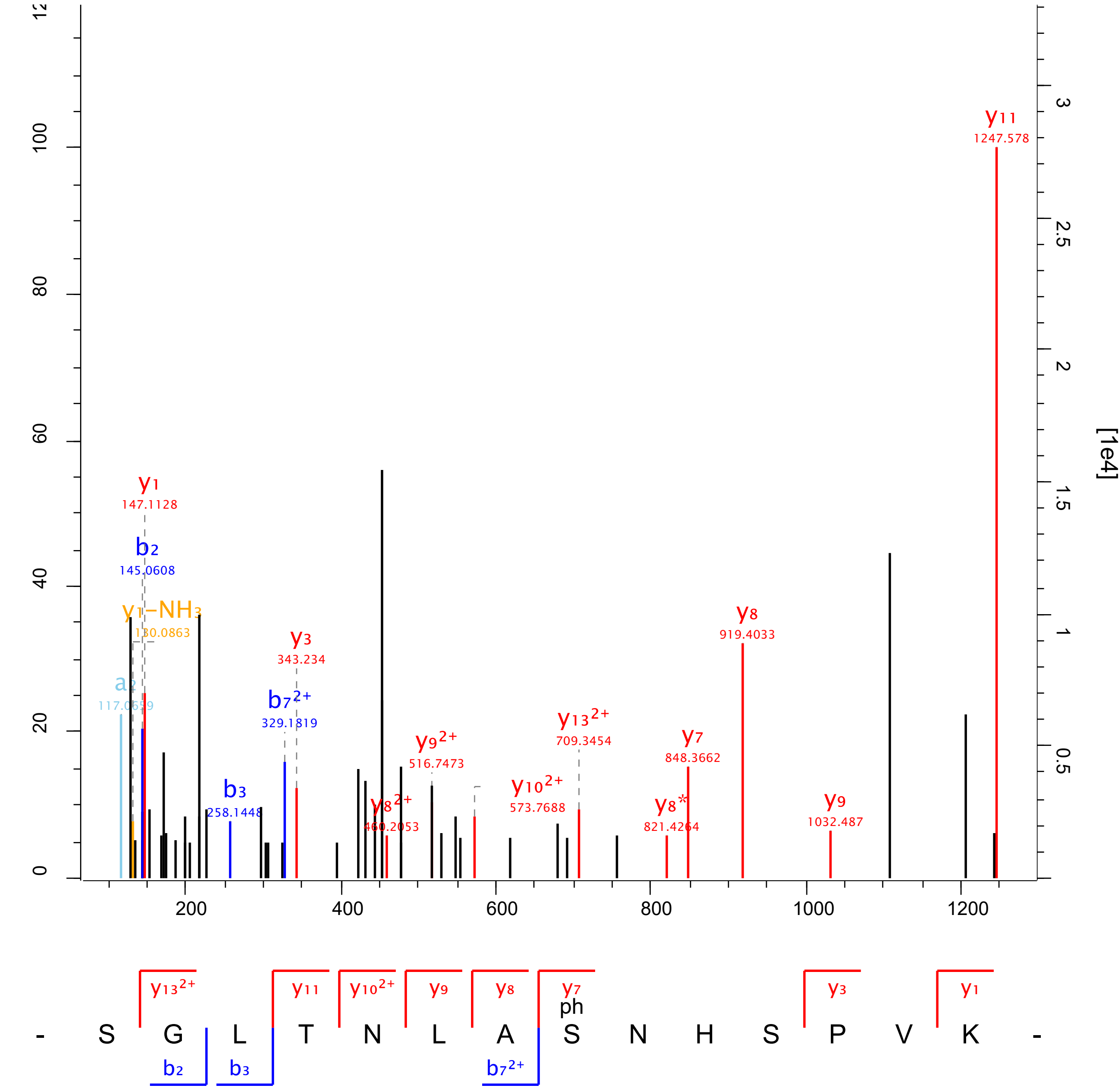

| Raw file | Scan | Method | Score | m/z |
| --- | --- | --- | --- | --- |
| sys_02_2 | 10197 | FTMS; HCD | 42.32 | 616.29 |

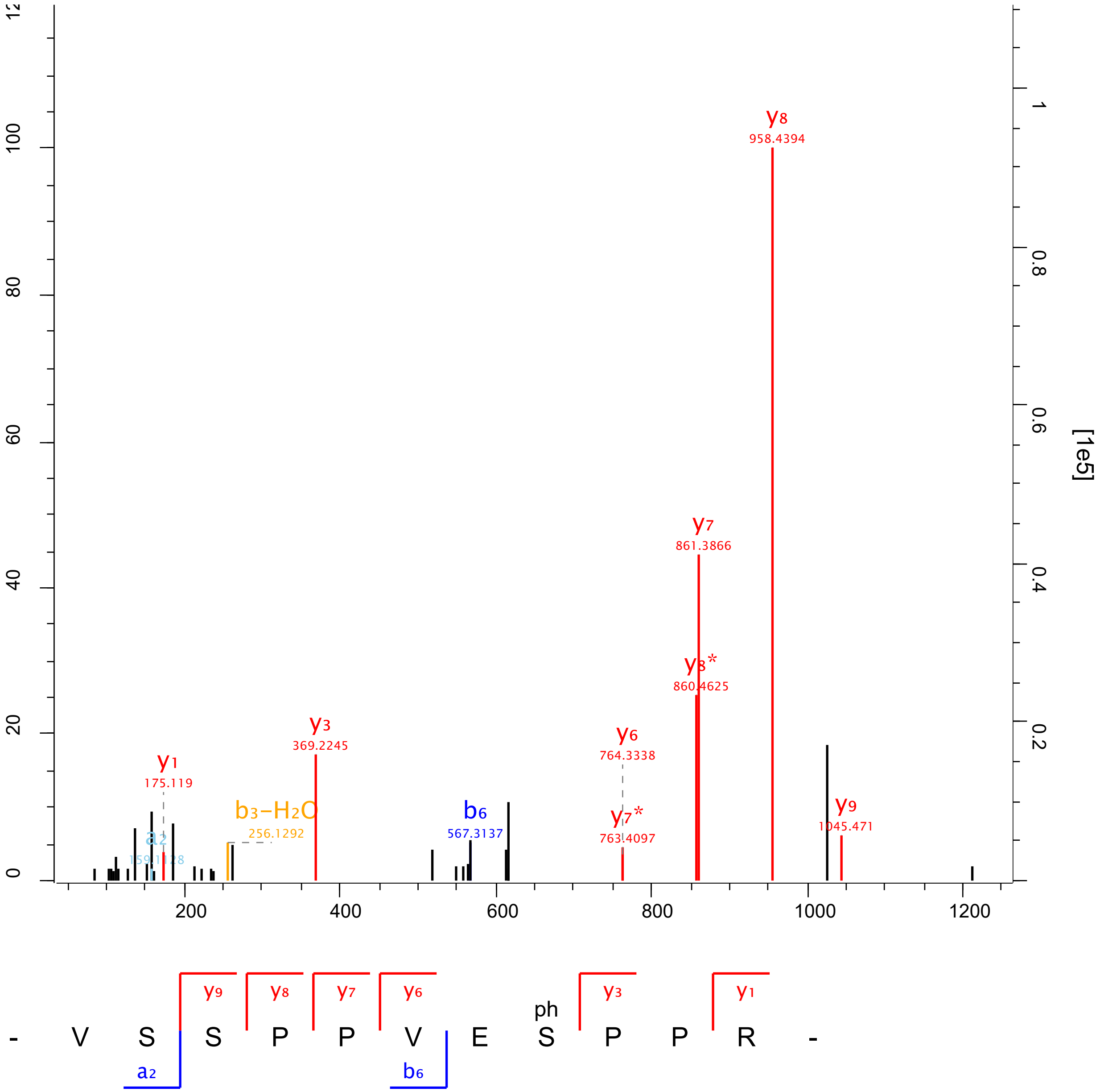

| Raw file | Scan | Method | Score | m/z |
|---|---|---|---|---|
| sys_02_2 | 10215 | FTMS; HCD | 94.12 | 560.78 |

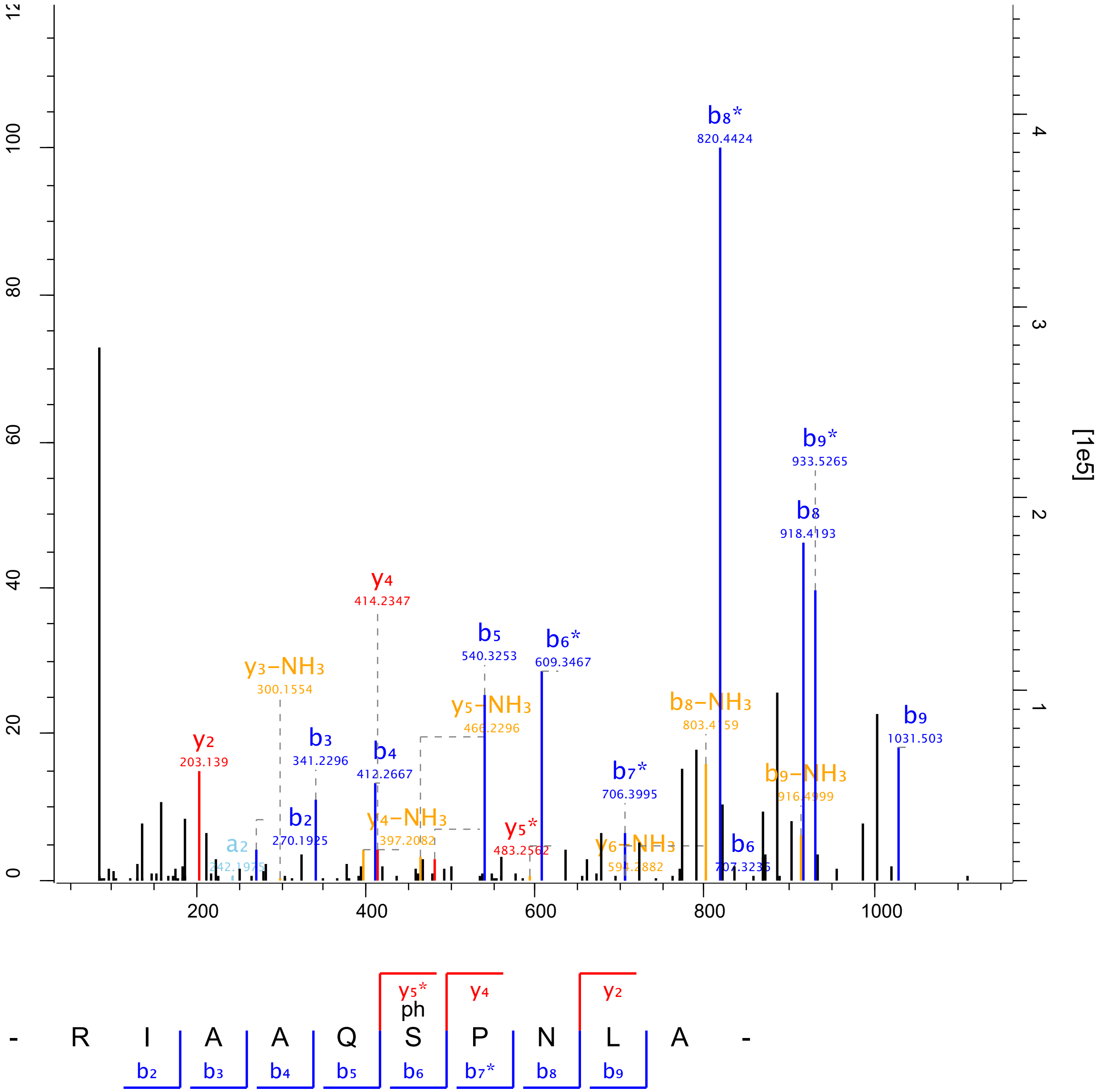

| Raw file | Scan | Method | Score | m/z |
| --- | --- | --- | --- | --- |
| sys_02_2 | 10248 | FTMS; HCD | 130.56 | 870.38 |

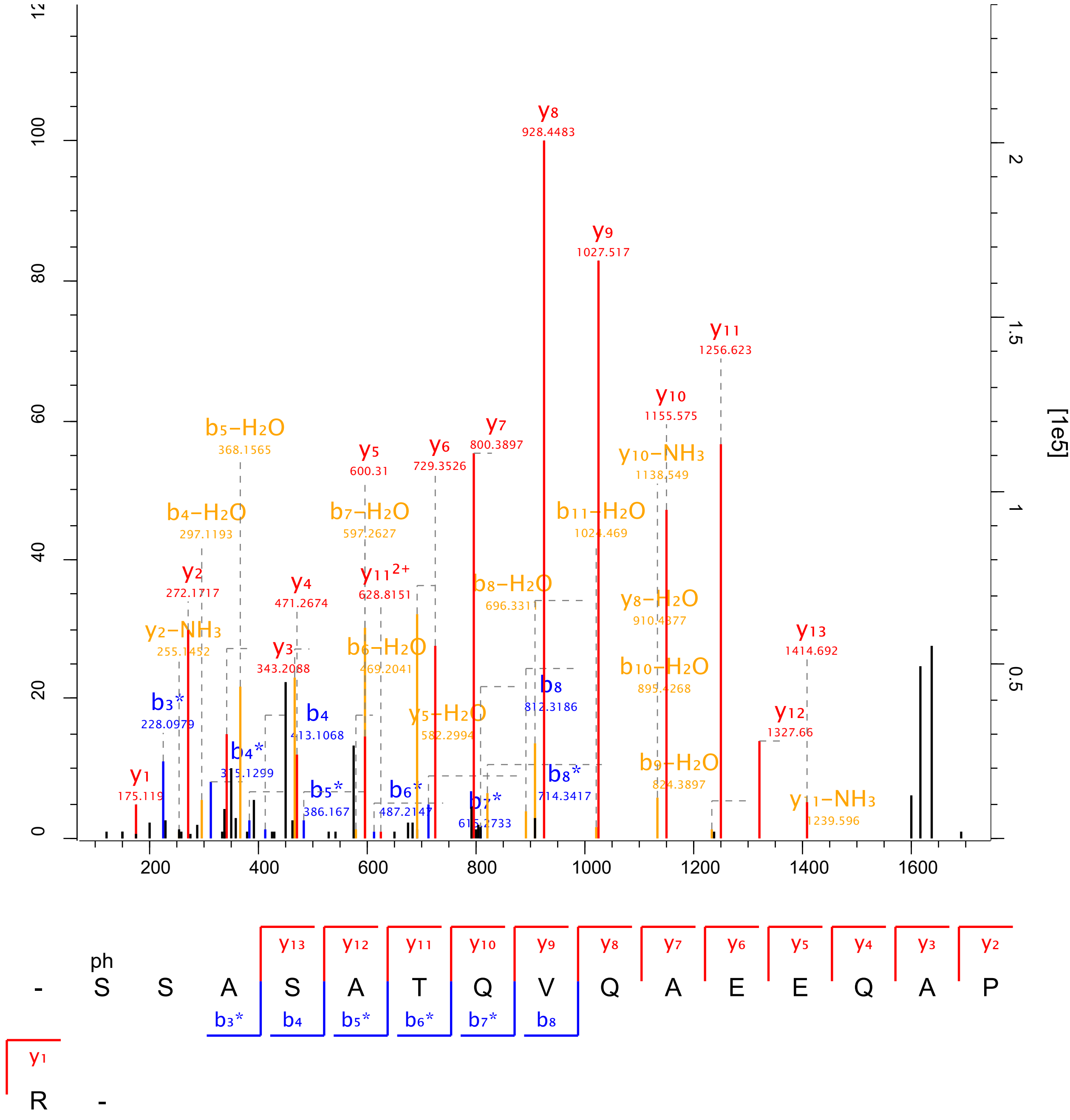

| Raw file | Scan | Method | Score | m/z |
|---|---|---|---|---|
| sys_02_2 | 10362 | FTMS; HCD | 47.77 | 393.86 |

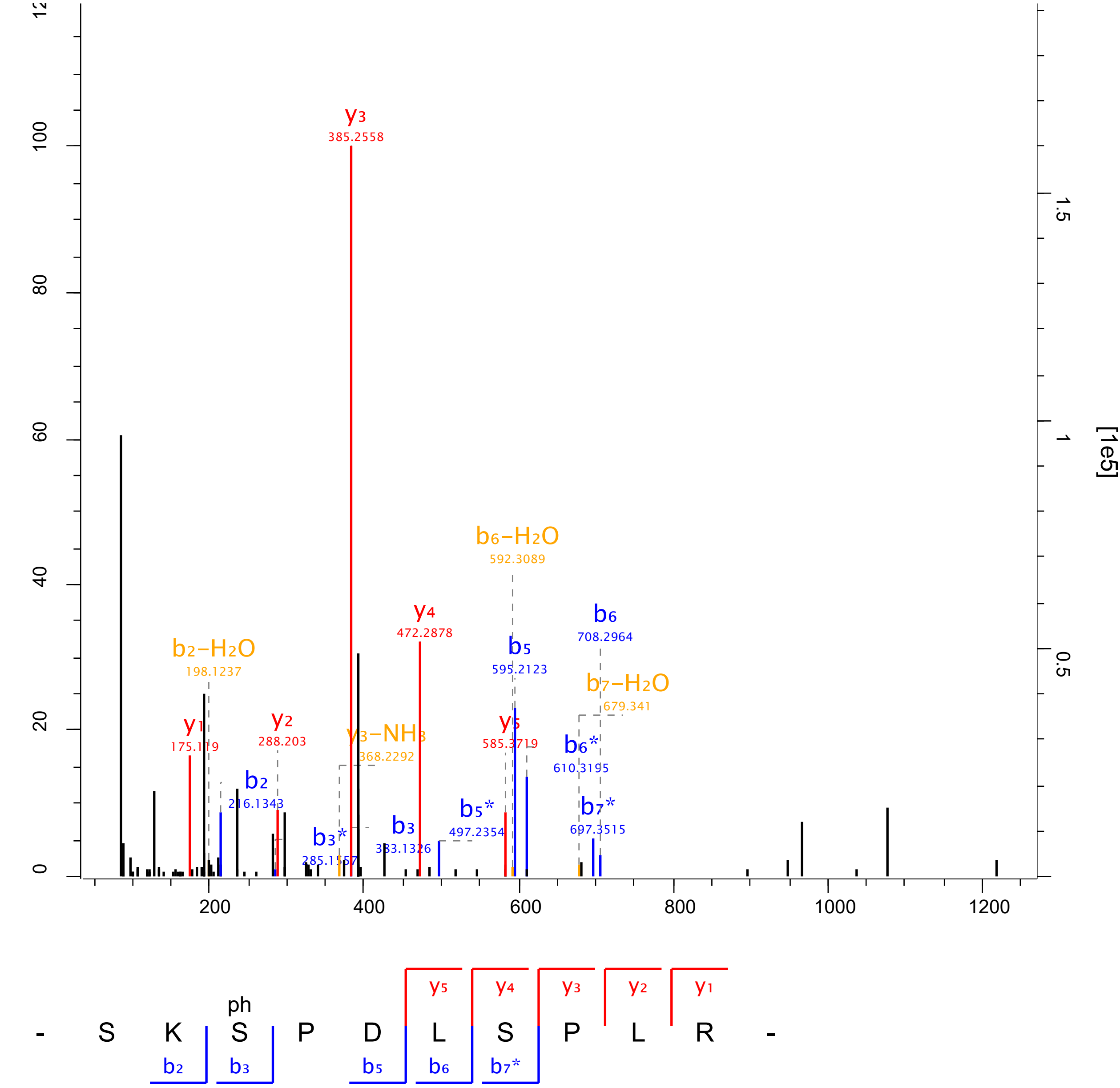

| Raw file | Scan | Method | Score | m/z |
|---|---|---|---|---|
| sys_02_2 | 10407 | FTMS; HCD | 248.95 | 706.31 |

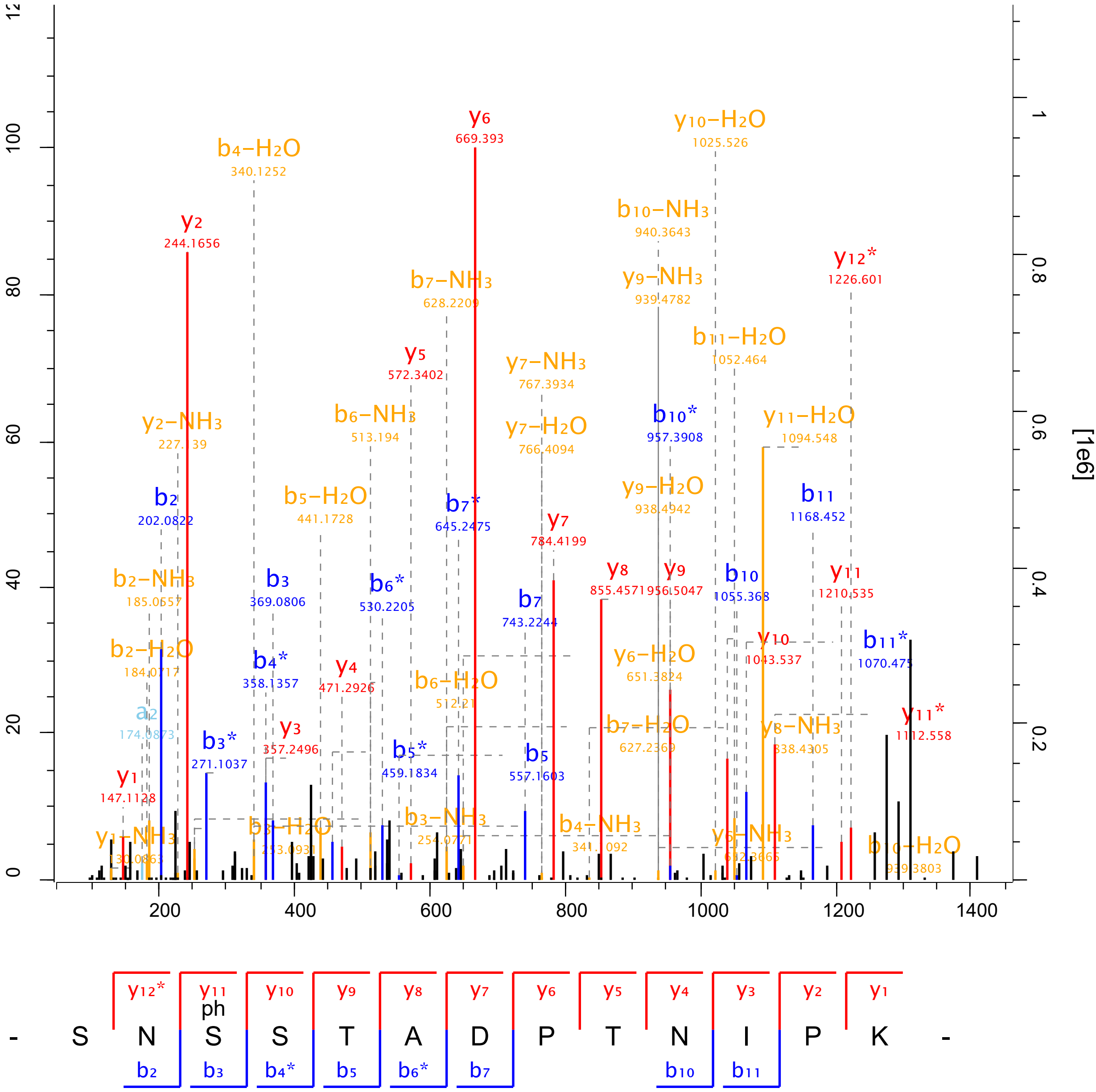

| Raw file | Scan | Method | Score | m/z |
| --- | --- | --- | --- | --- |
| sys_02_2 | 10429 | FTMS; HCD | 142.08 | 517.72 |

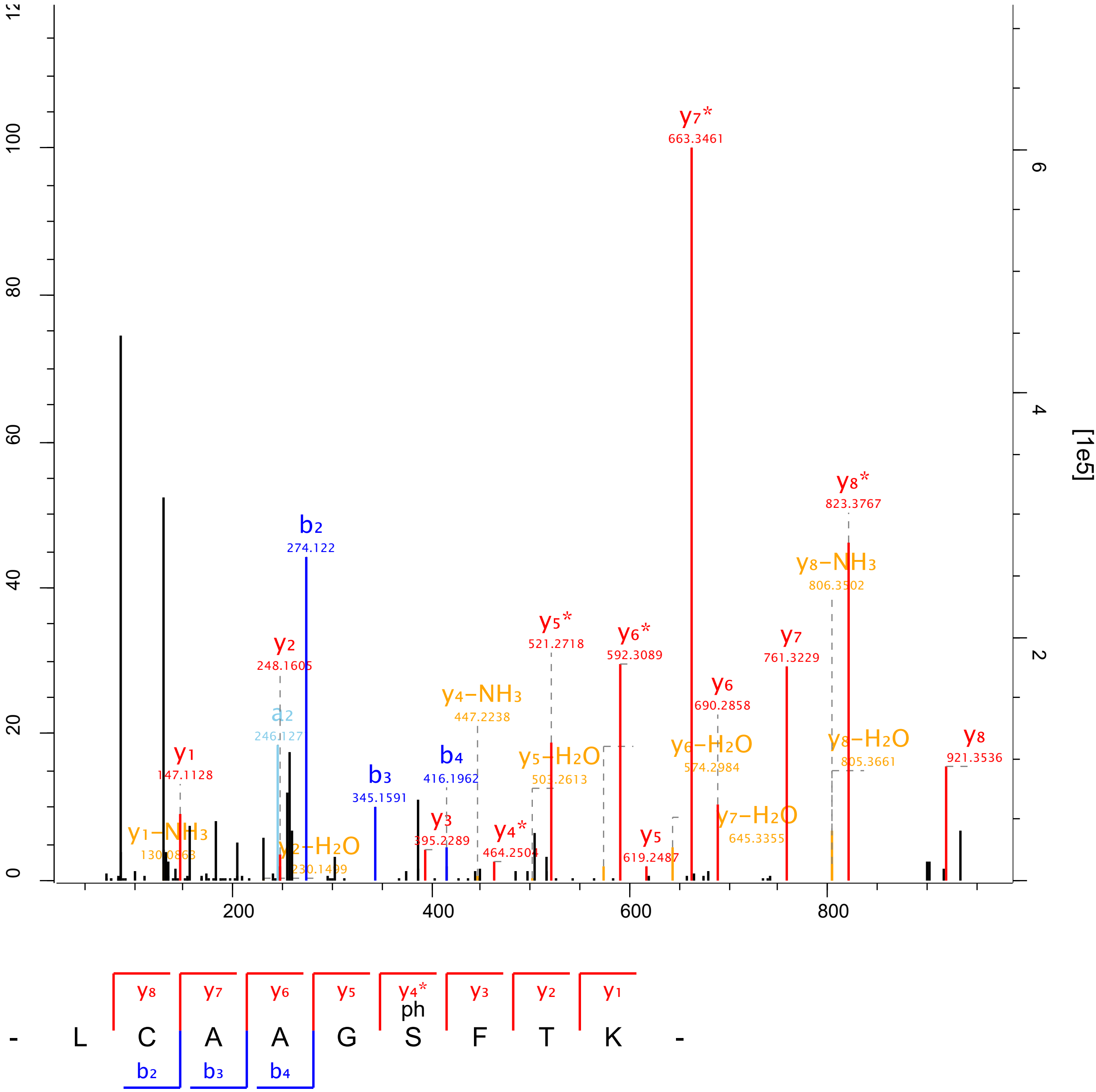

| Raw file | Scan | Method | Score | m/z |
|---|---|---|---|---|
| sys_02_2 | 10449 | FTMS; HCD | 70.26 | 463.22 |

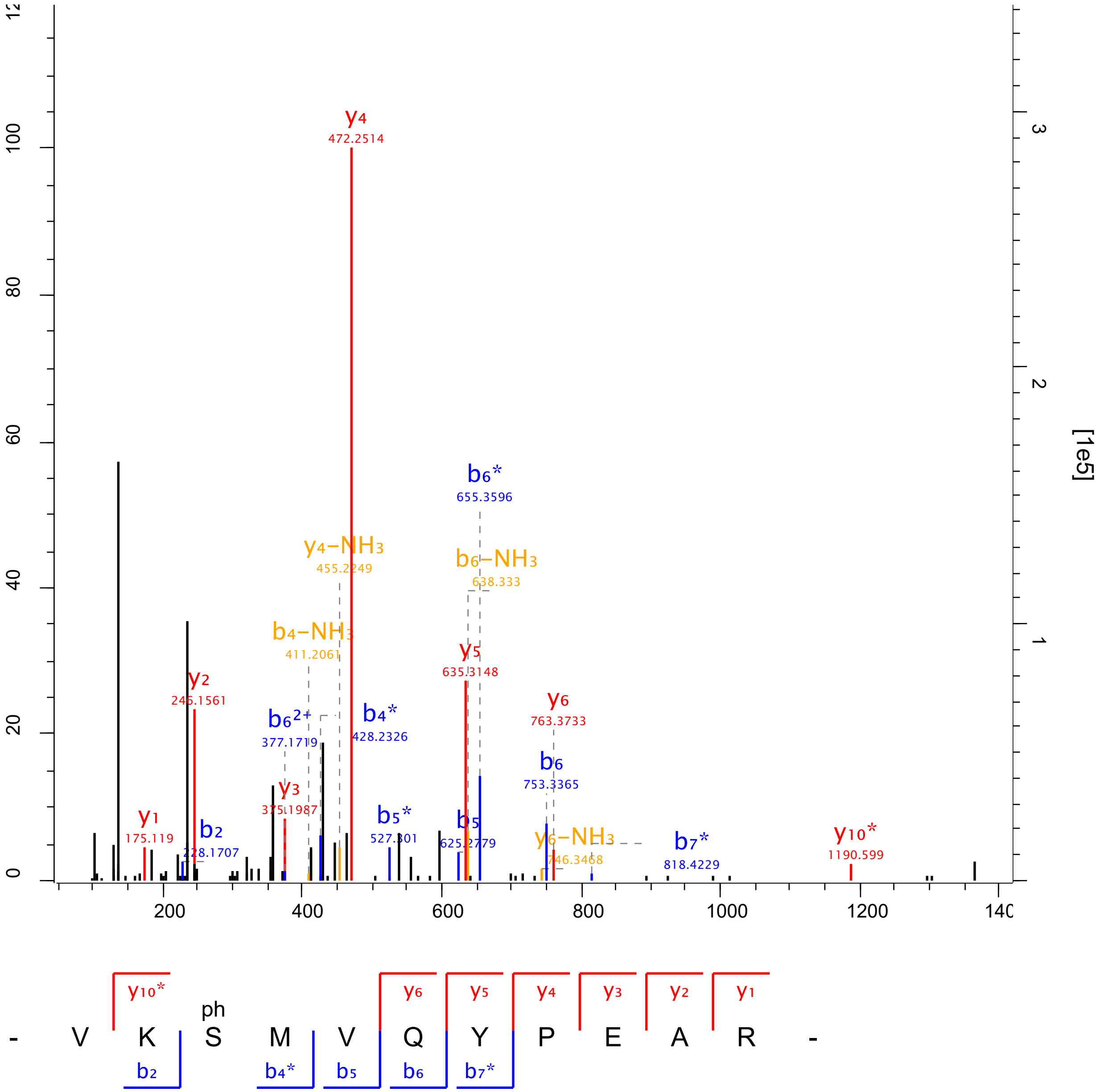

| Raw file | Scan | Method | Score | m/z |
| --- | --- | --- | --- | --- |
| sys_02_2 | 10482 | FTMS; HCD | 190.24 | 724.28 |

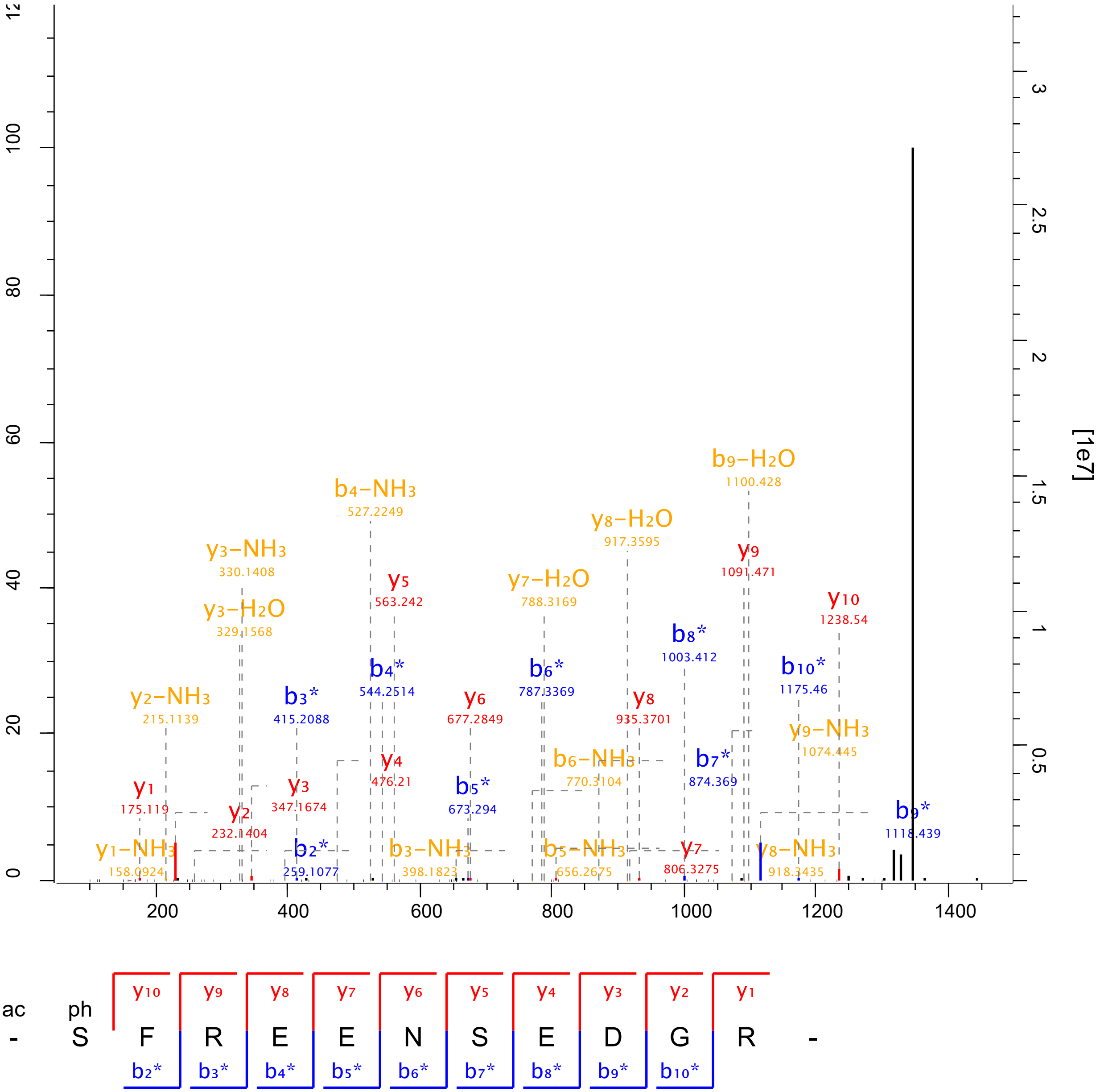

| Raw file | Scan | Method | Score | m/z |
| --- | --- | --- | --- | --- |
| sys_02_2 | 10839 | FTMS; HCD | 190.35 | 675.94 |

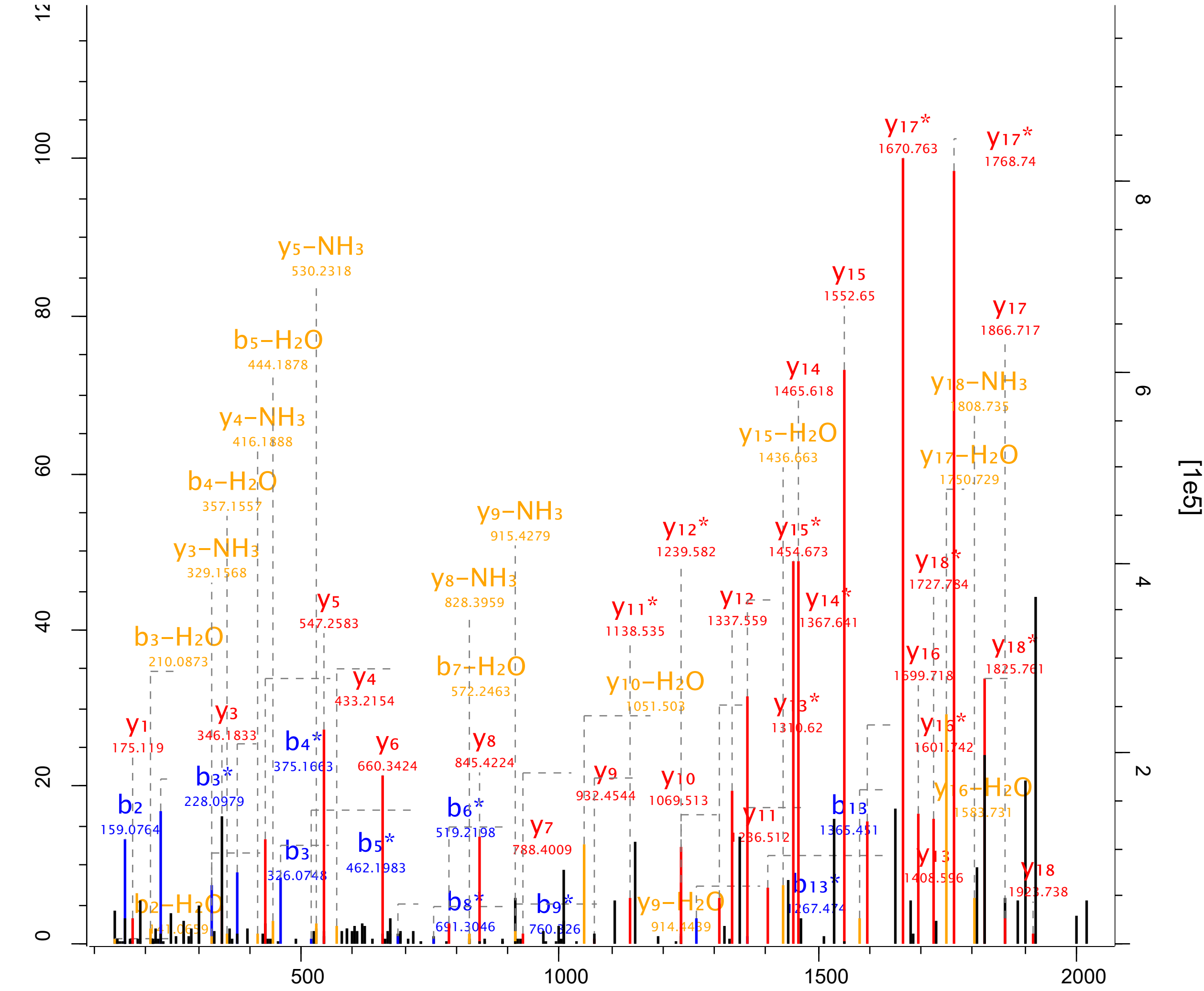


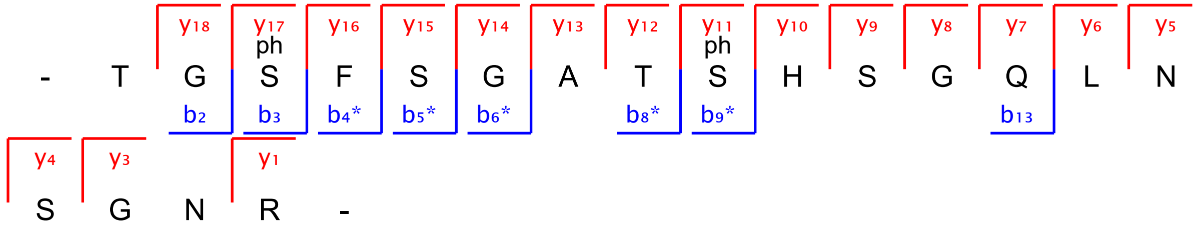

| Raw file | Scan | Method | Score | m/z |
|---|---|---|---|---|
| sys_02_2 | 10970 | FTMS; HCD | 230.41 | 440.72 |

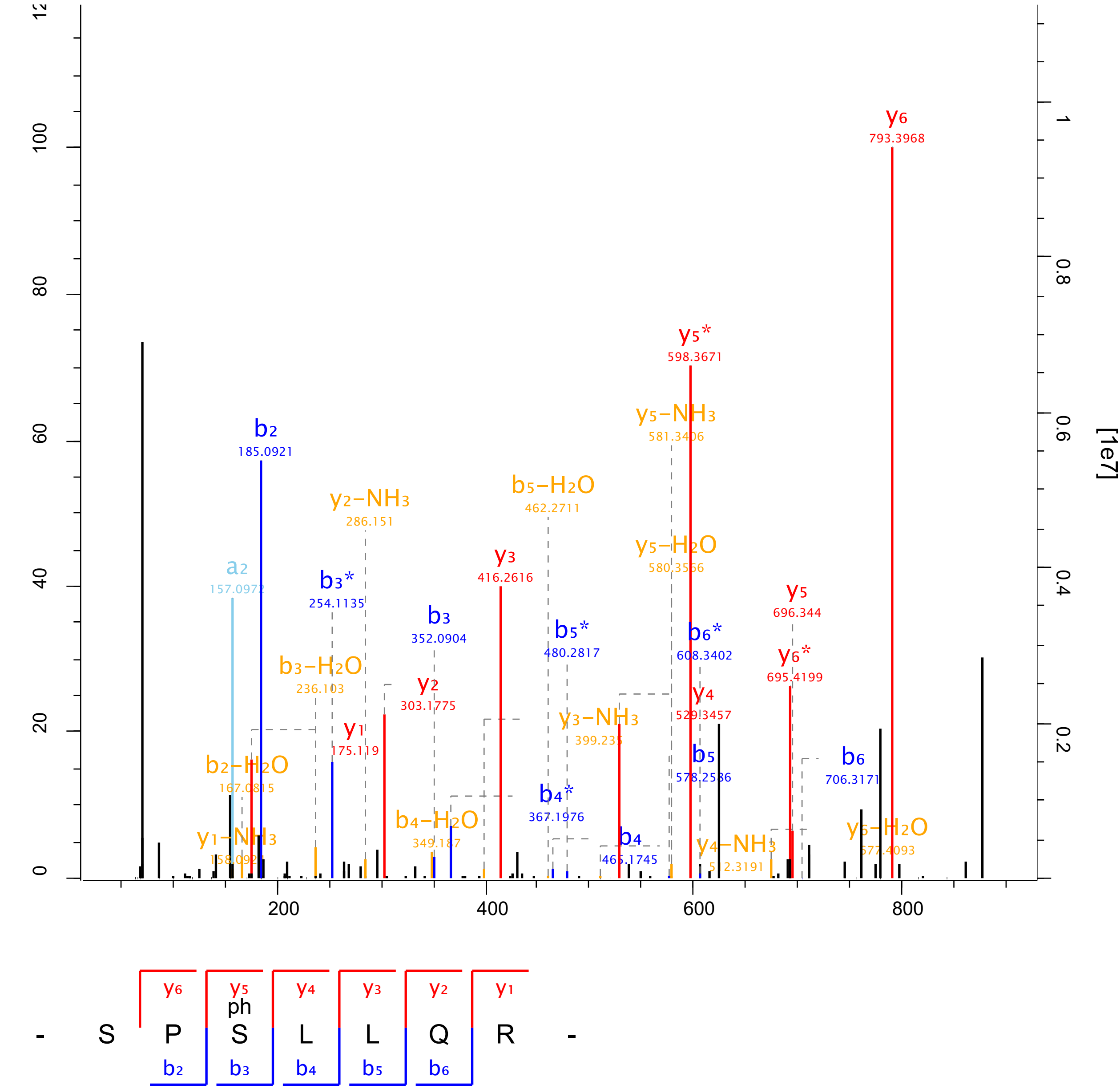

| Raw file | Scan | Method | Score | m/z |
|---|---|---|---|---|
| sys_02_2 | 10983 | FTMS; HCD | 47.62 | 576.75 |

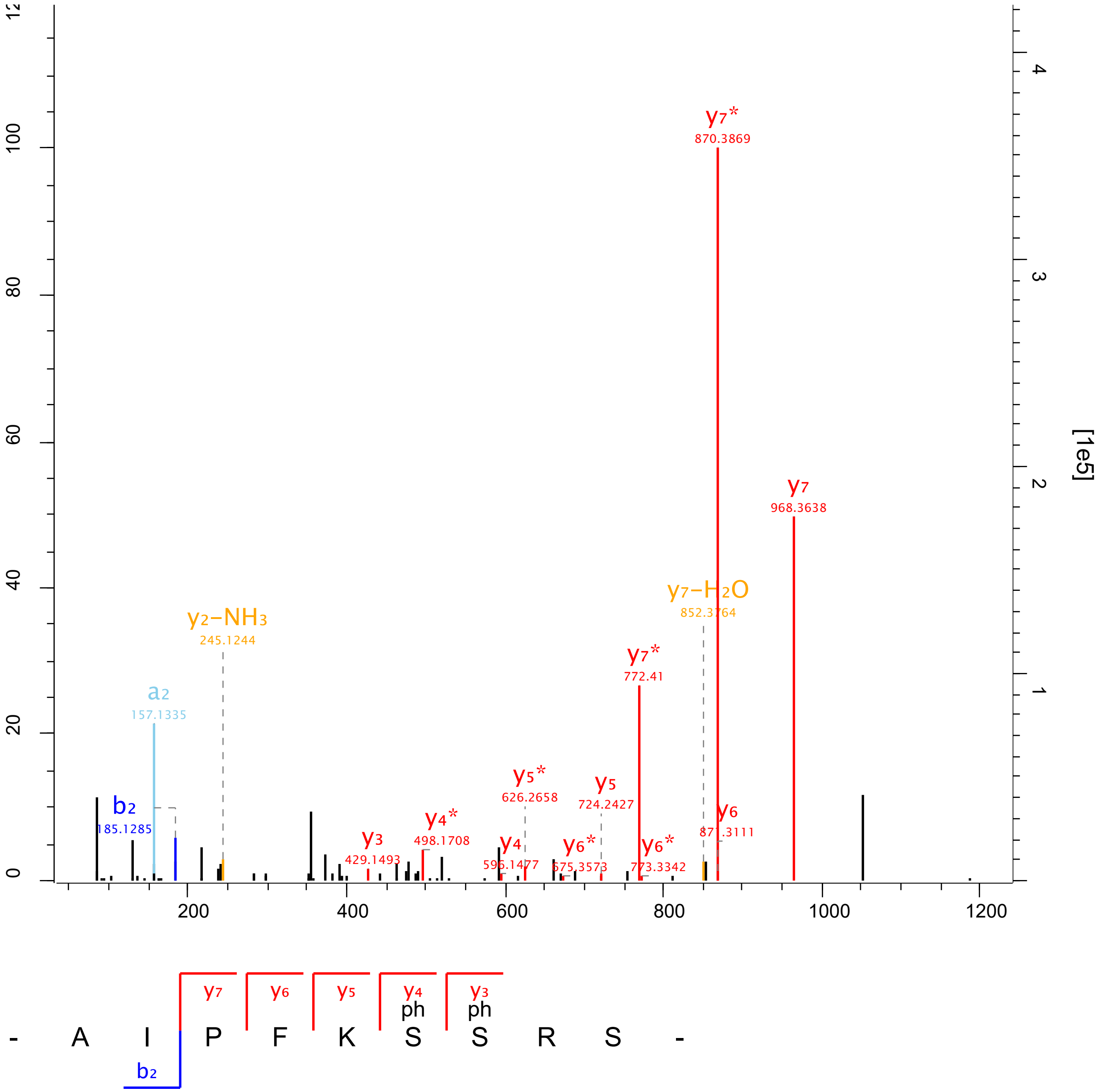

| Raw file | Scan | Method | Score | m/z |
| --- | --- | --- | --- | --- |
| sys_02_2 | 11014 | FTMS; HCD | 84.61 | 412.21 |

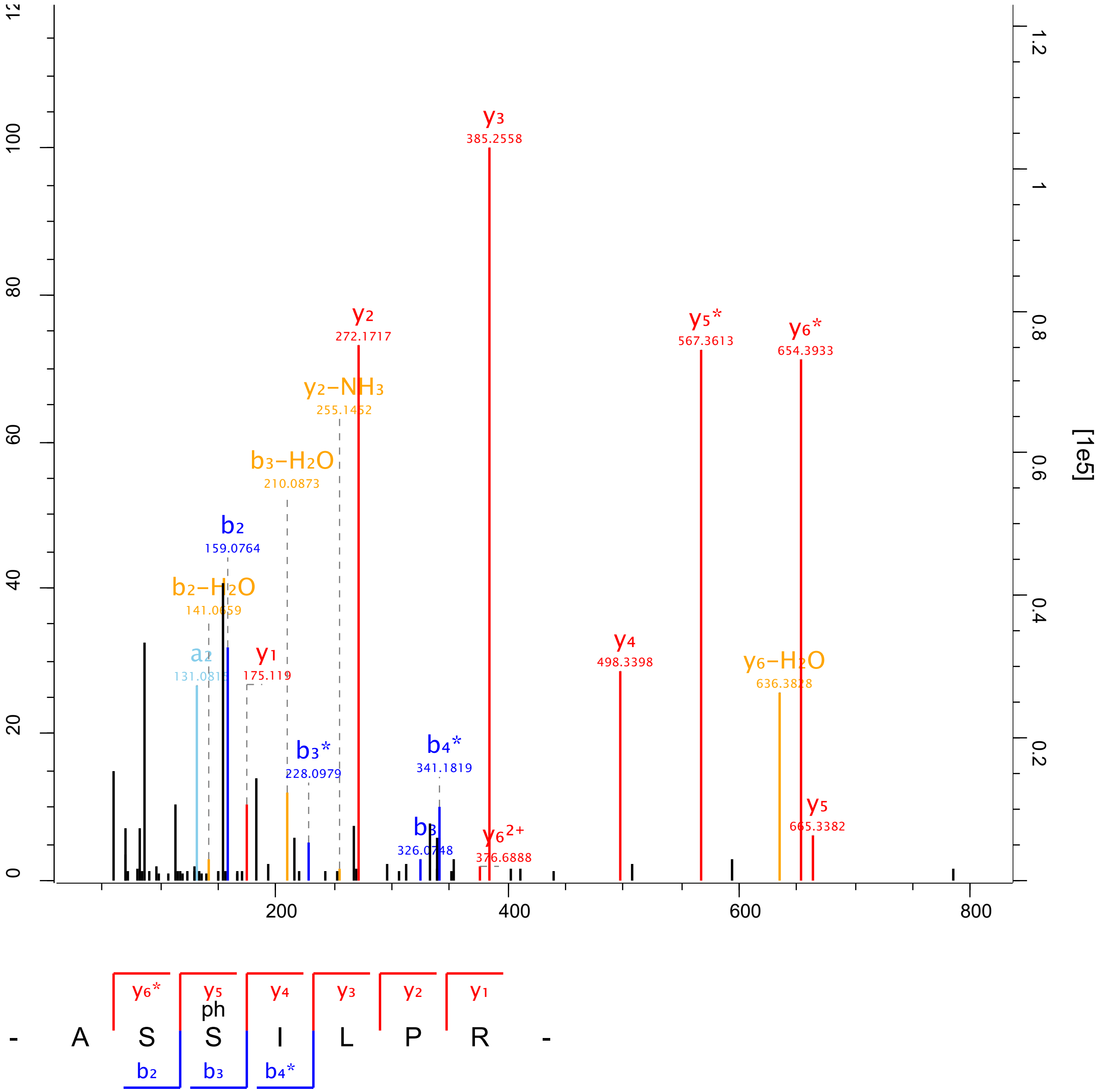

| Raw file | Scan | Method | Score | m/z |
|---|---|---|---|---|
| sys_02_2 | 11367 | FTMS; HCD | 196.37 | 678.3 |

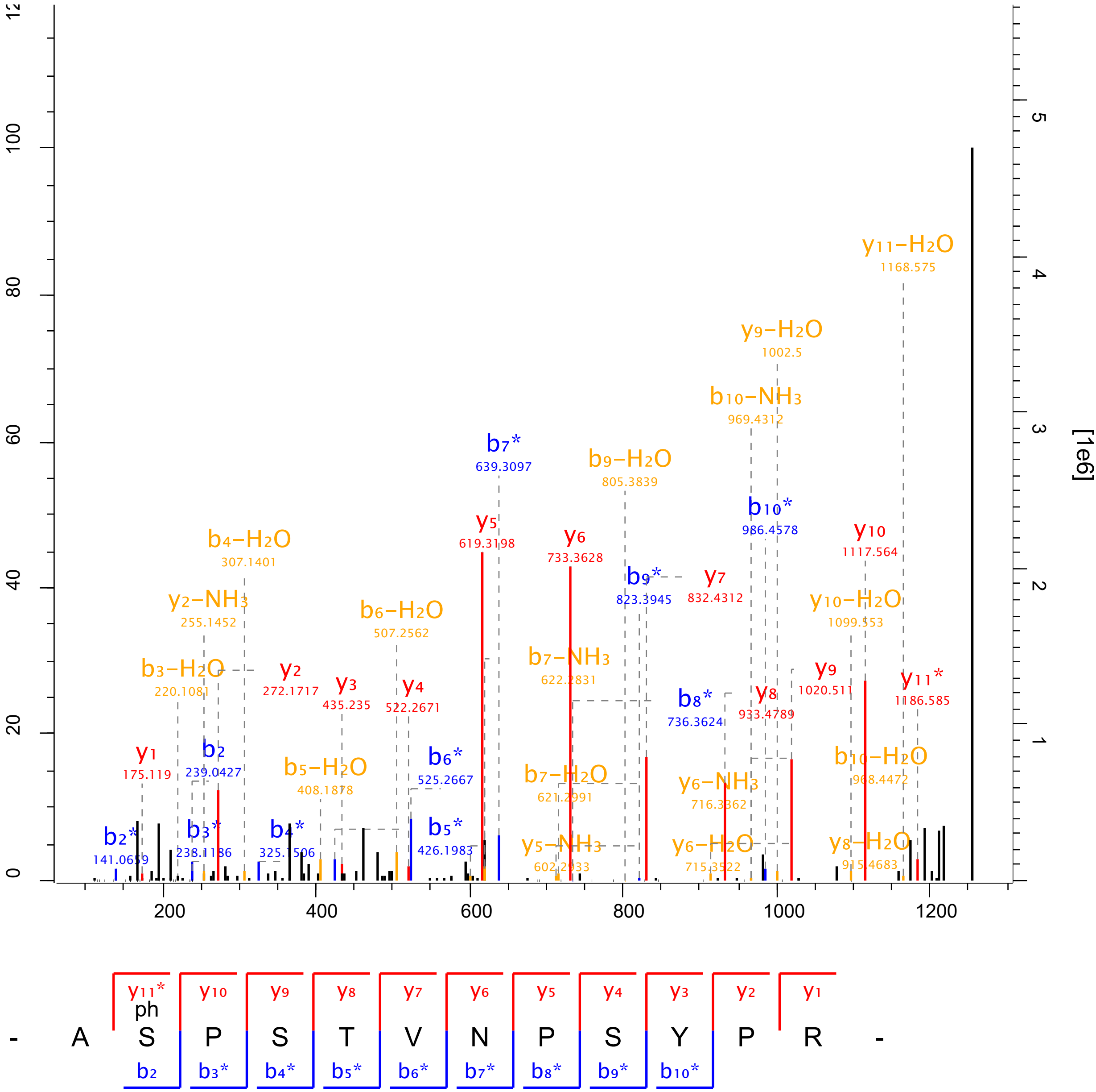

| Raw file | Scan | Method | Score | m/z |
|---|---|---|---|---|
| sys_02_2 | 11426 | FTMS; HCD | 46.89 | 496.25 |

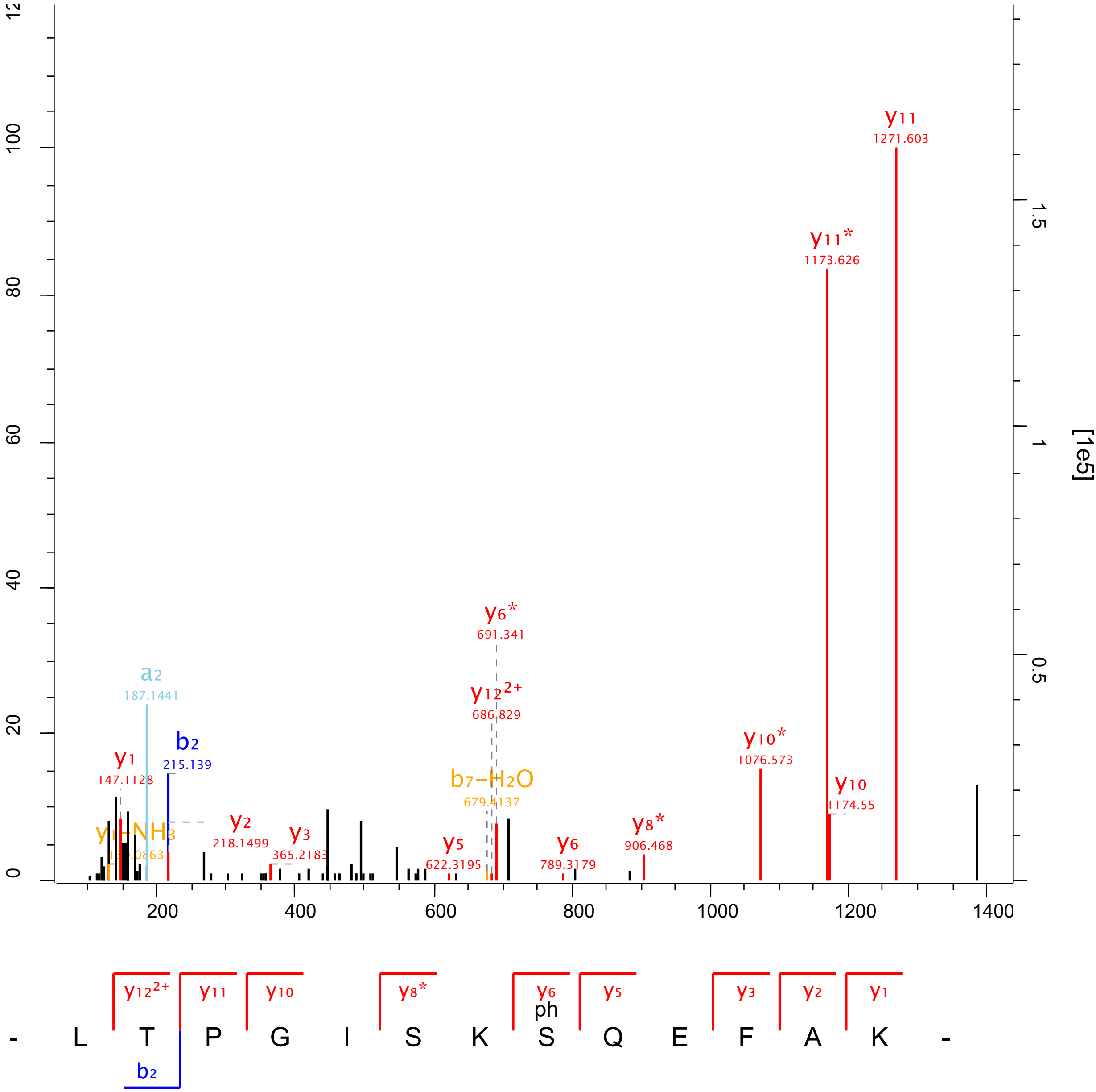

| Raw file | Scan | Method | Score | m/z |
|---|---|---|---|---|
| sys_02_2 | 11489 | FTMS; HCD | 99.34 | 627.6 |

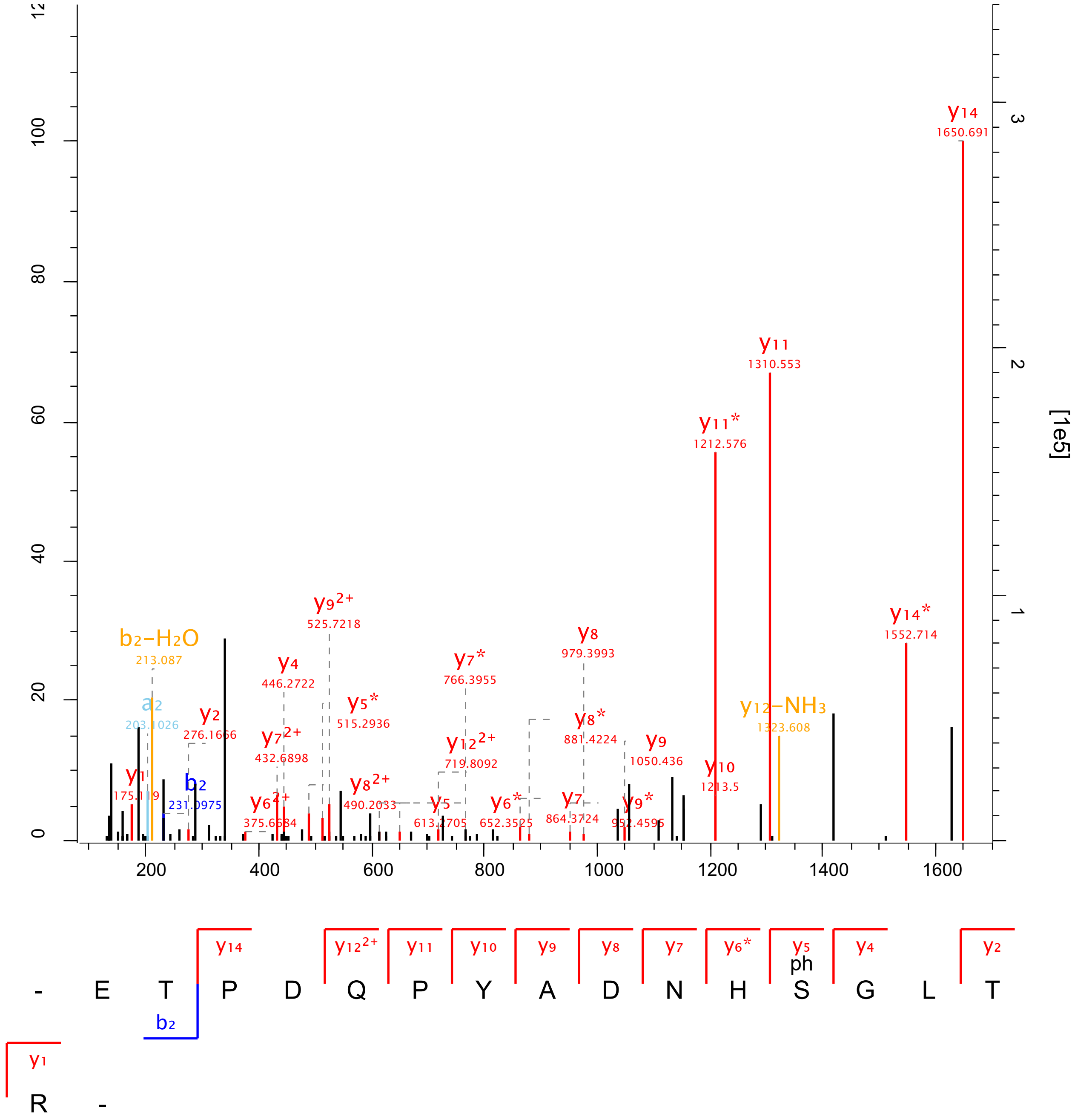

| Raw file | Scan | Method | Score | m/z |
|---|---|---|---|---|
| sys_02_2 | 11522 | FTMS; HCD | 51.52 | 540.94 |

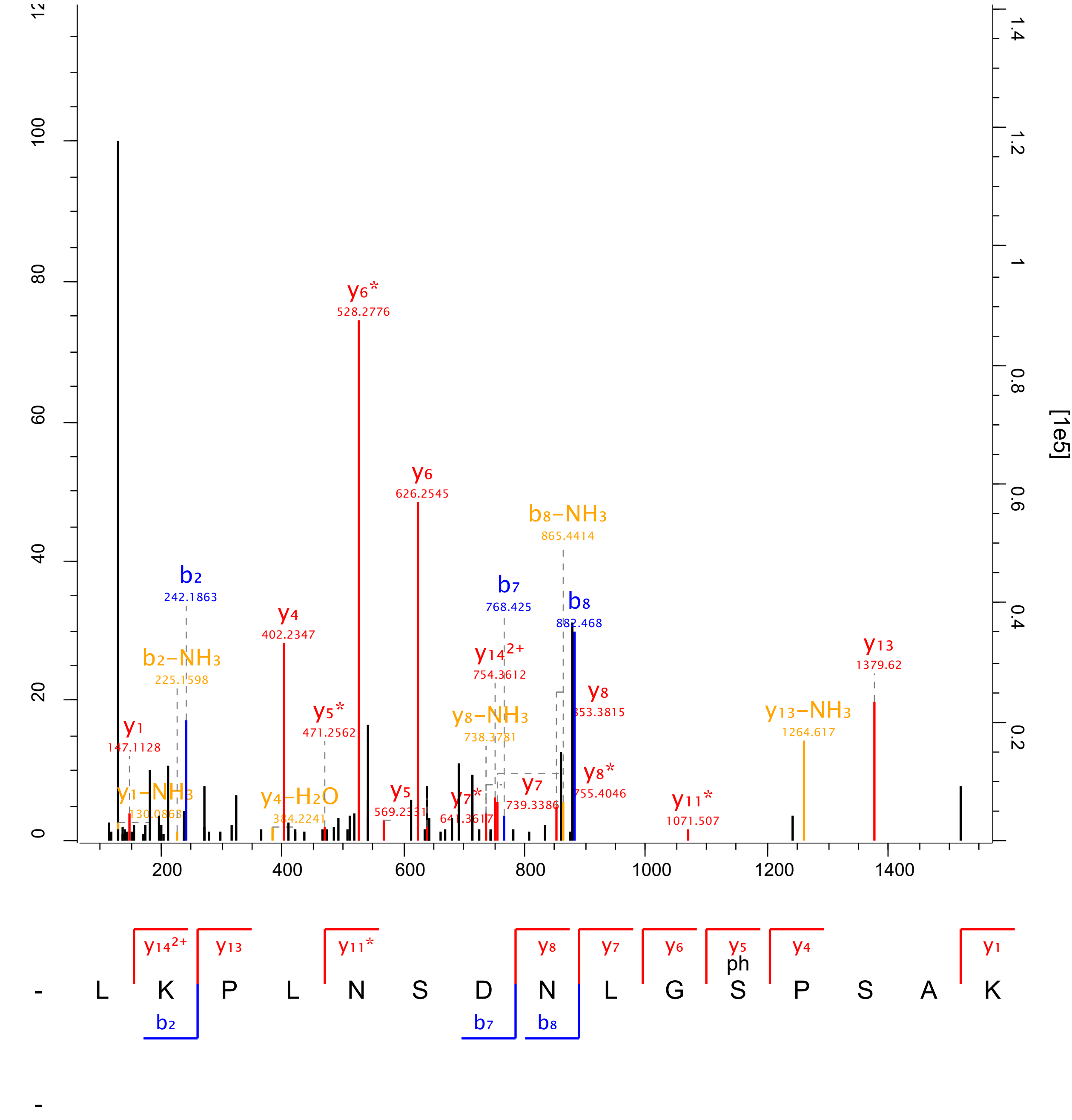

| Raw file | Scan | Method | Score | m/z |
|---|---|---|---|---|
| sys_02_2 | 11567 | FTMS; HCD | 72.89 | 625.3 |

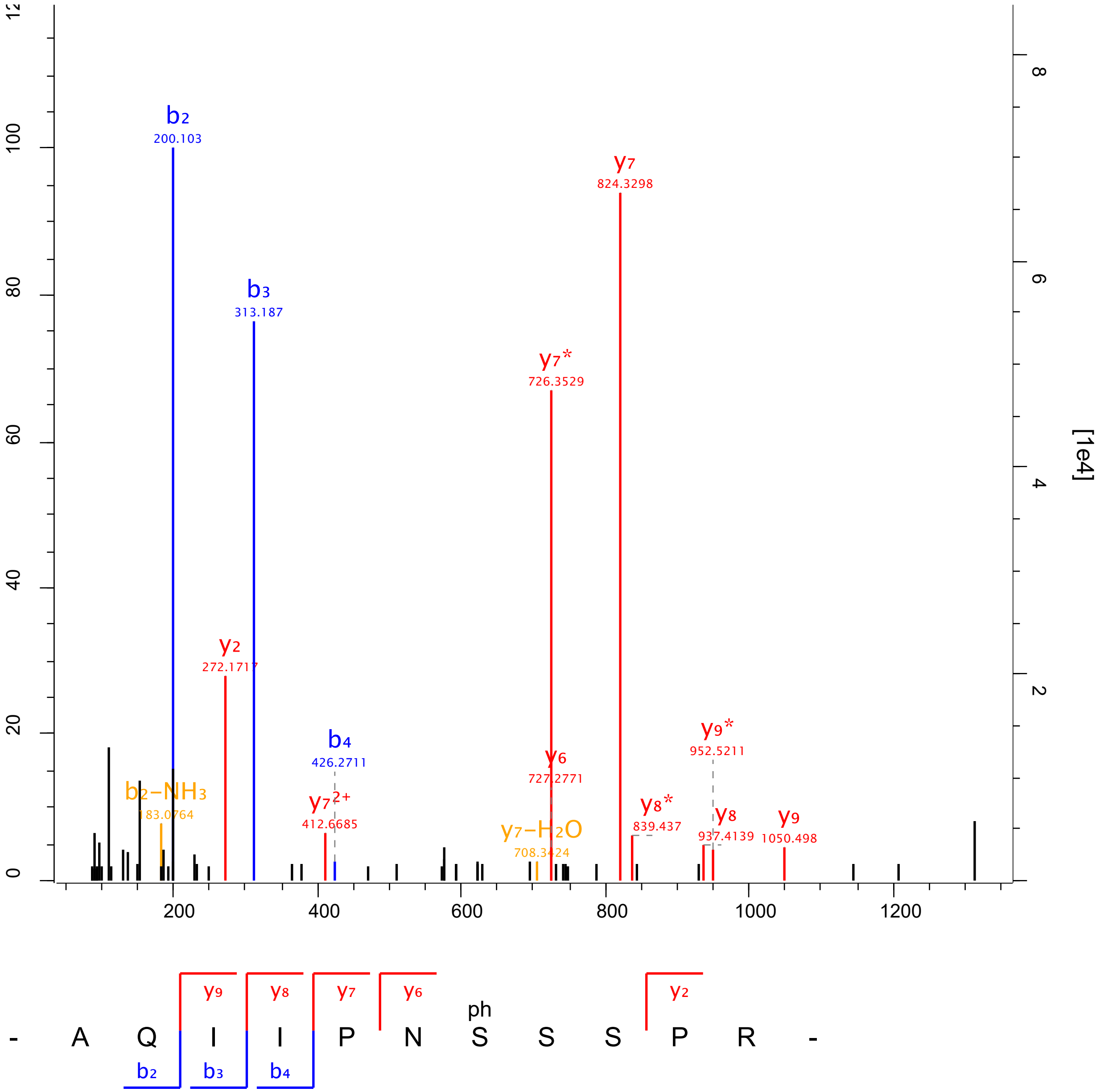

| Raw file | Scan | Method | Score | m/z |
| --- | --- | --- | --- | --- |
| sys_02_2 | 11568 | FTMS; HCD | 52.81 | 614.96 |

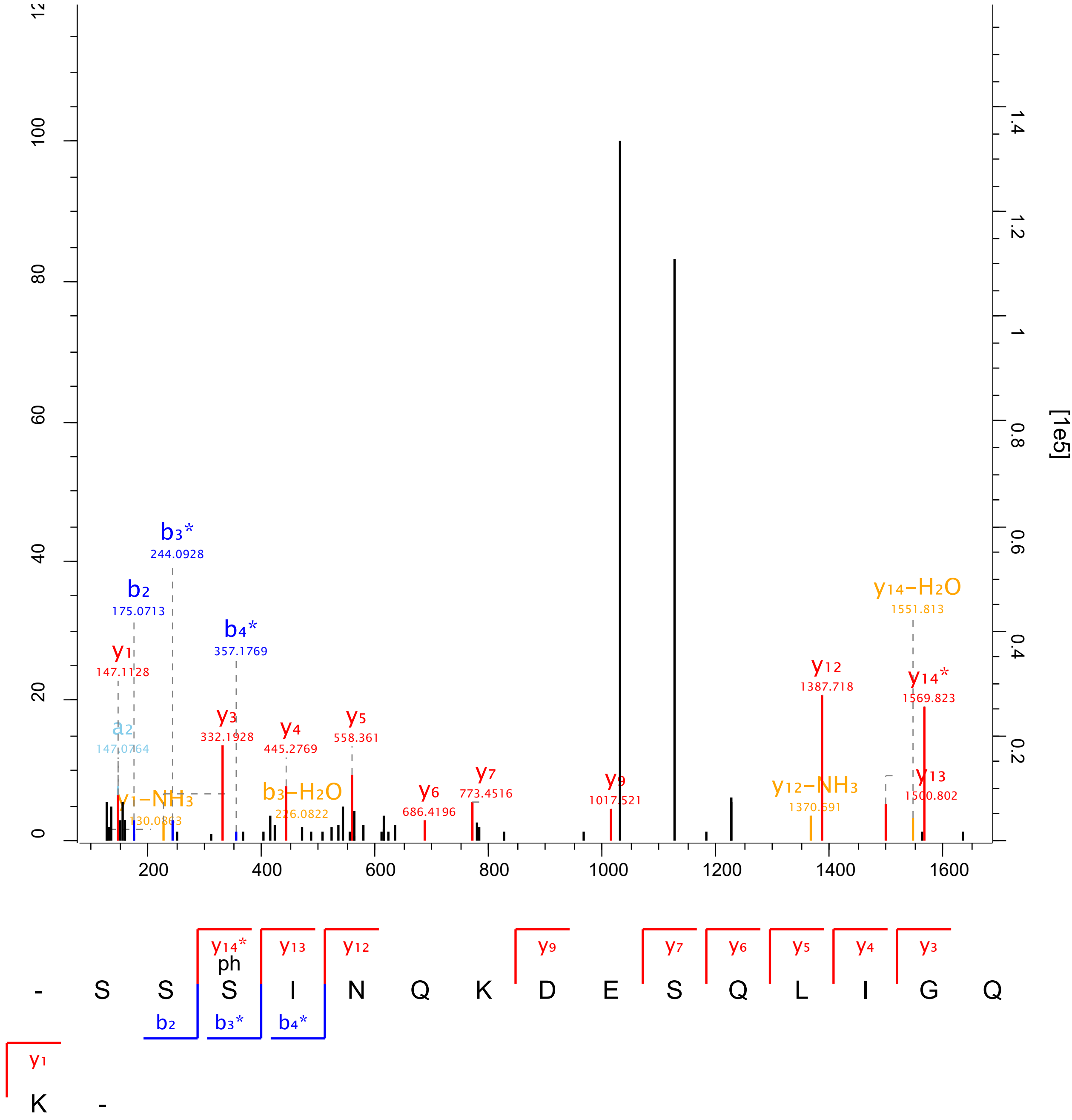

| Raw file | Scan | Method | Score | m/z |
|---|---|---|---|---|
| sys_02_2 | 11613 | FTMS; HCD | 44.25 | 835.35 |

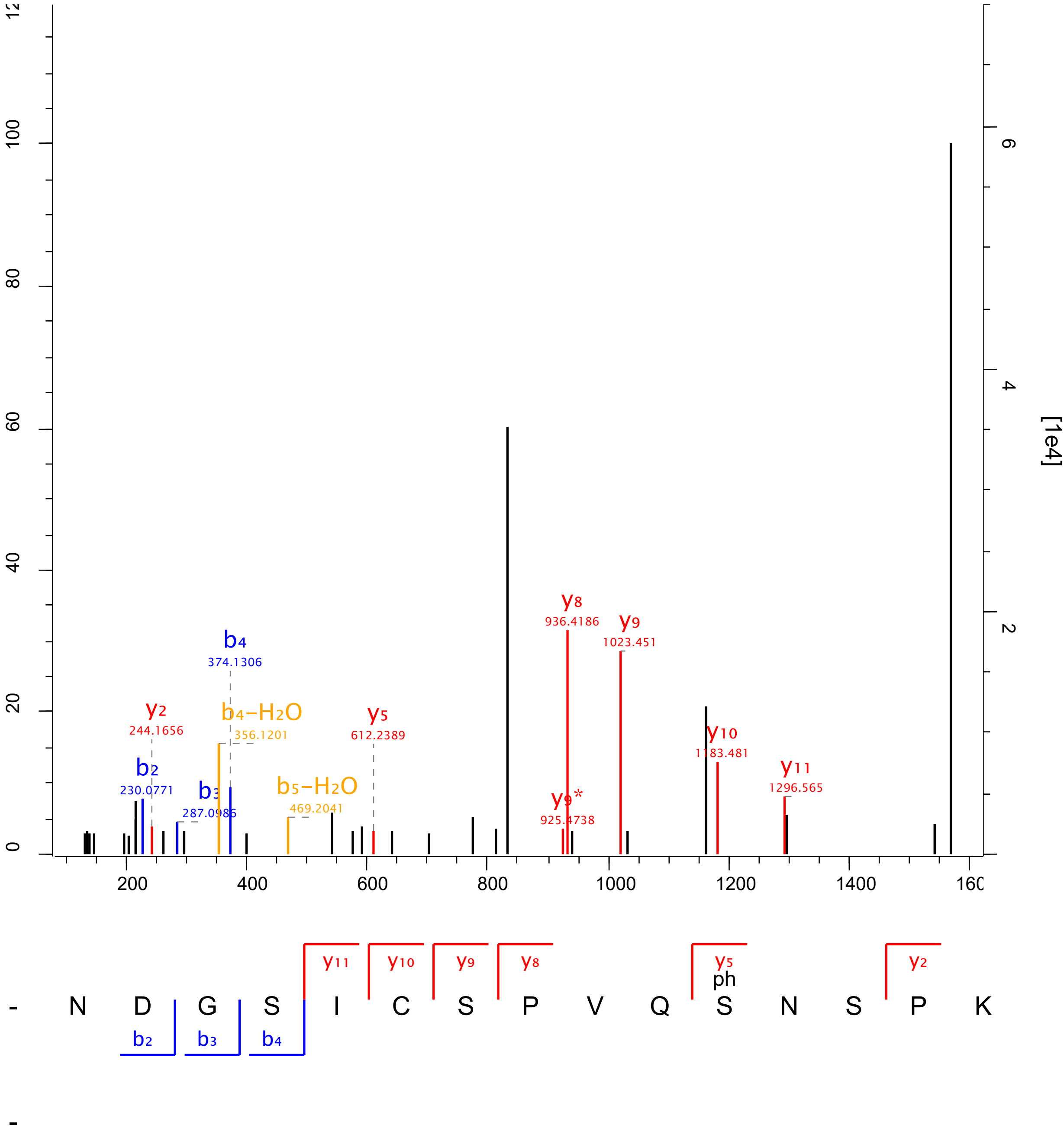

| Raw file | Scan | Method | Score | m/z |
|---|---|---|---|---|
| sys_02_2 | 11616 | FTMS; HCD | 77.42 | 665.79 |

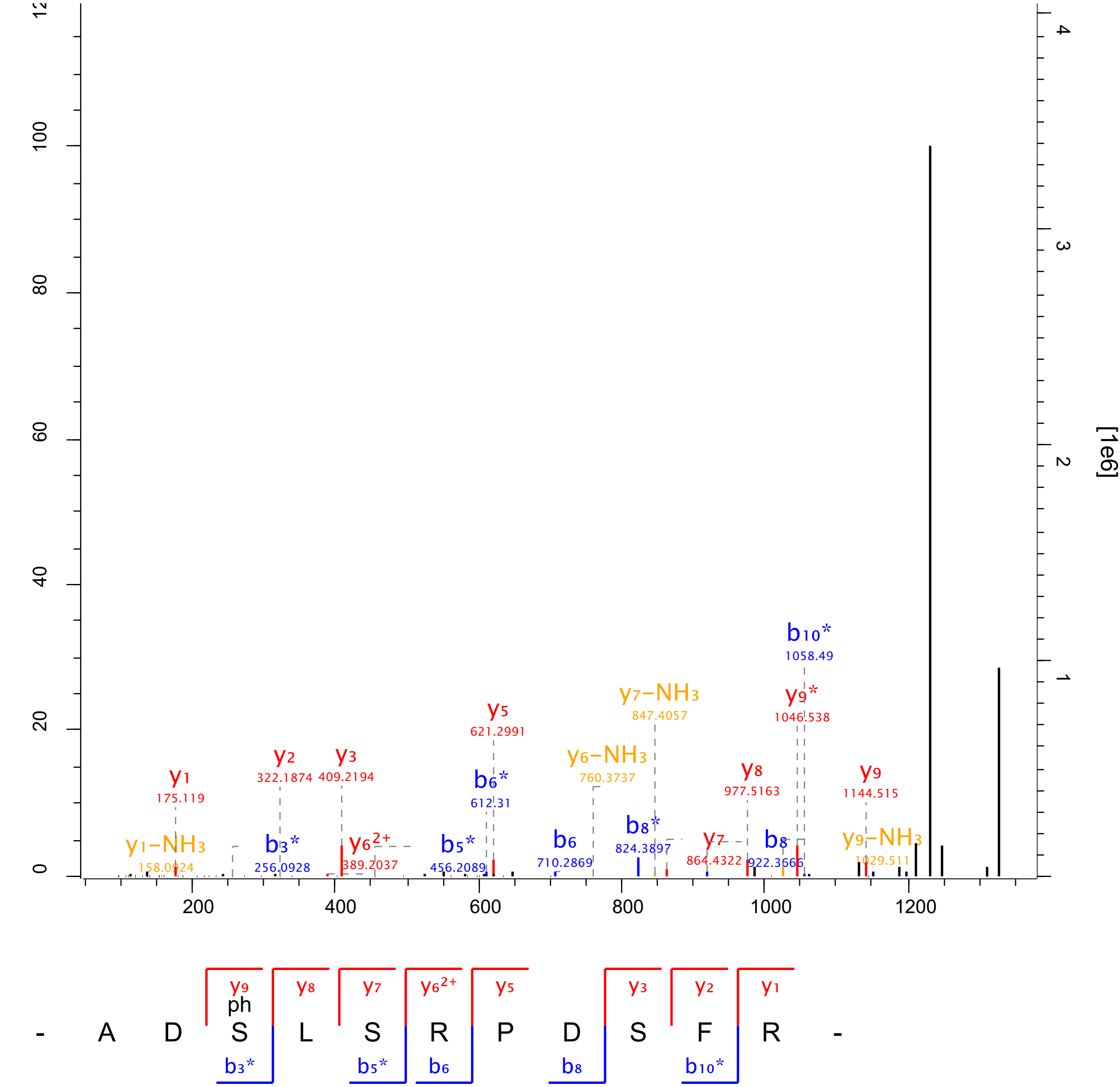

| Raw file | Scan | Method | Score | m/z |
|---|---|---|---|---|
| sys_02_2 | 11700 | FTMS; HCD | 56.96 | 536.26 |

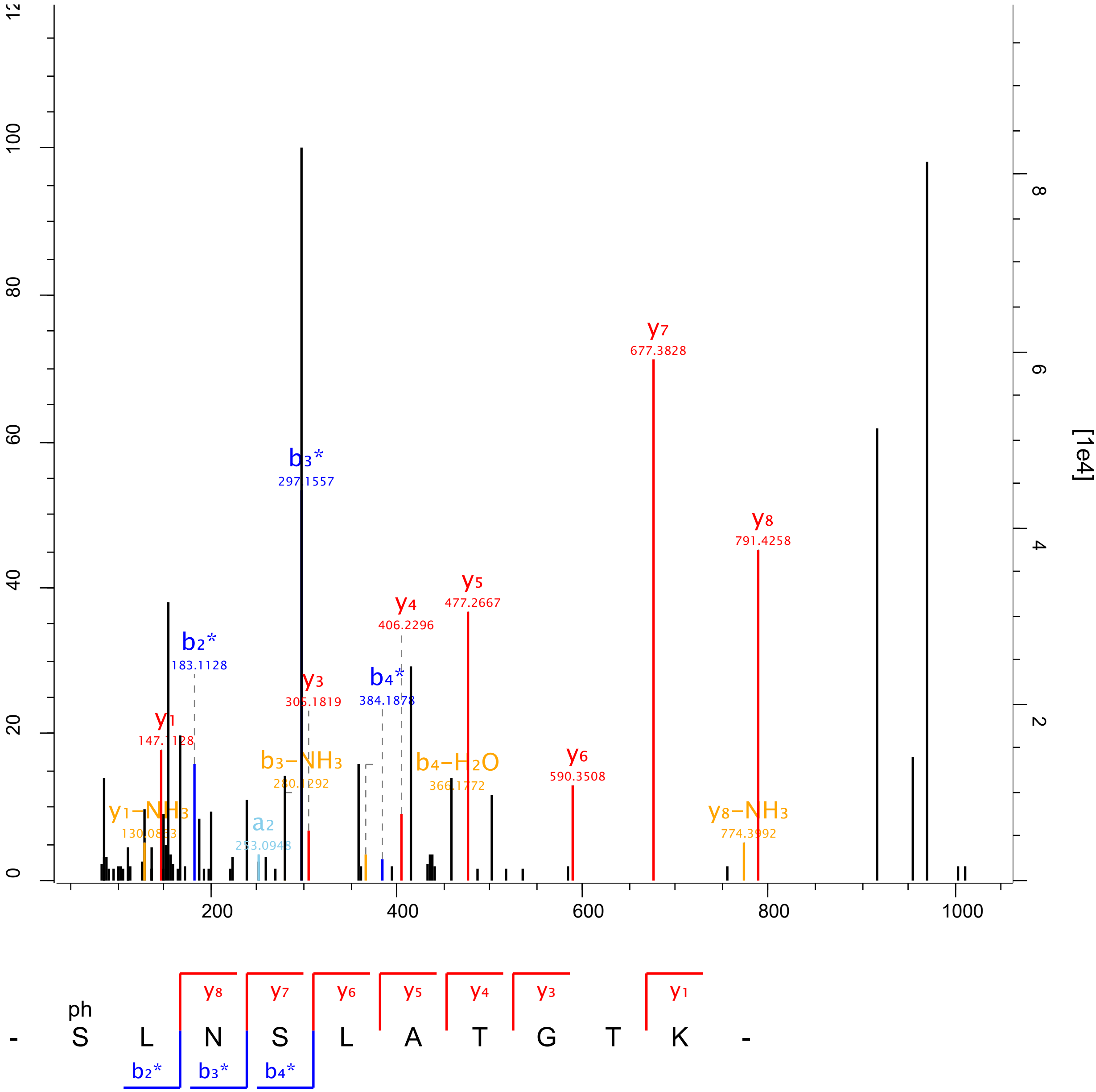

| Raw file | Scan | Method | Score | m/z |
|---|---|---|---|---|
| sys_02_2 | 11732 | FTMS; HCD | 42.31 | 546.25 |

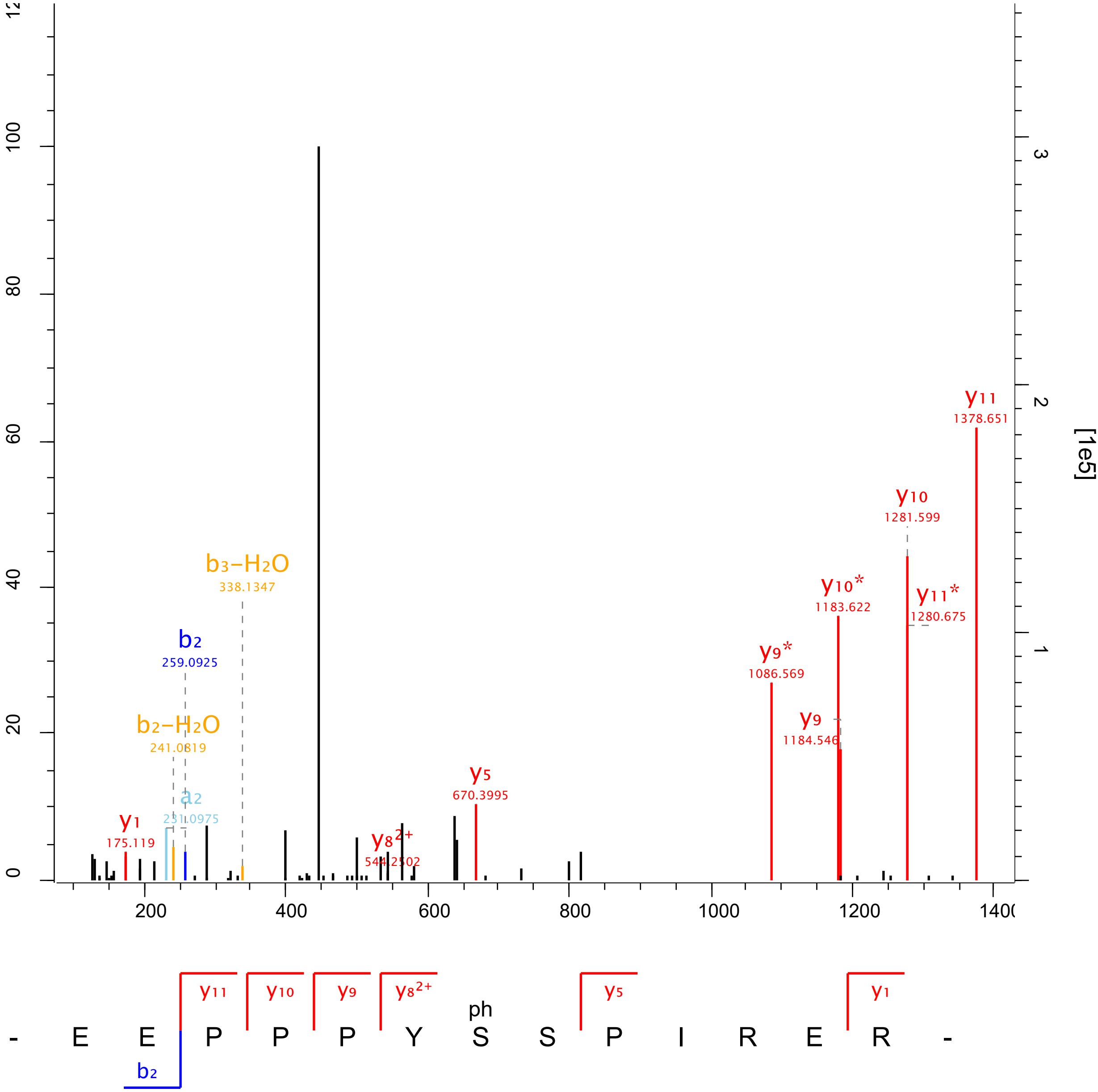

| Raw file | Scan | Method | Score | m/z |
| --- | --- | --- | --- | --- |
| sys_02_2 | 11778 | FTMS; HCD | 138.91 | 672.29 |

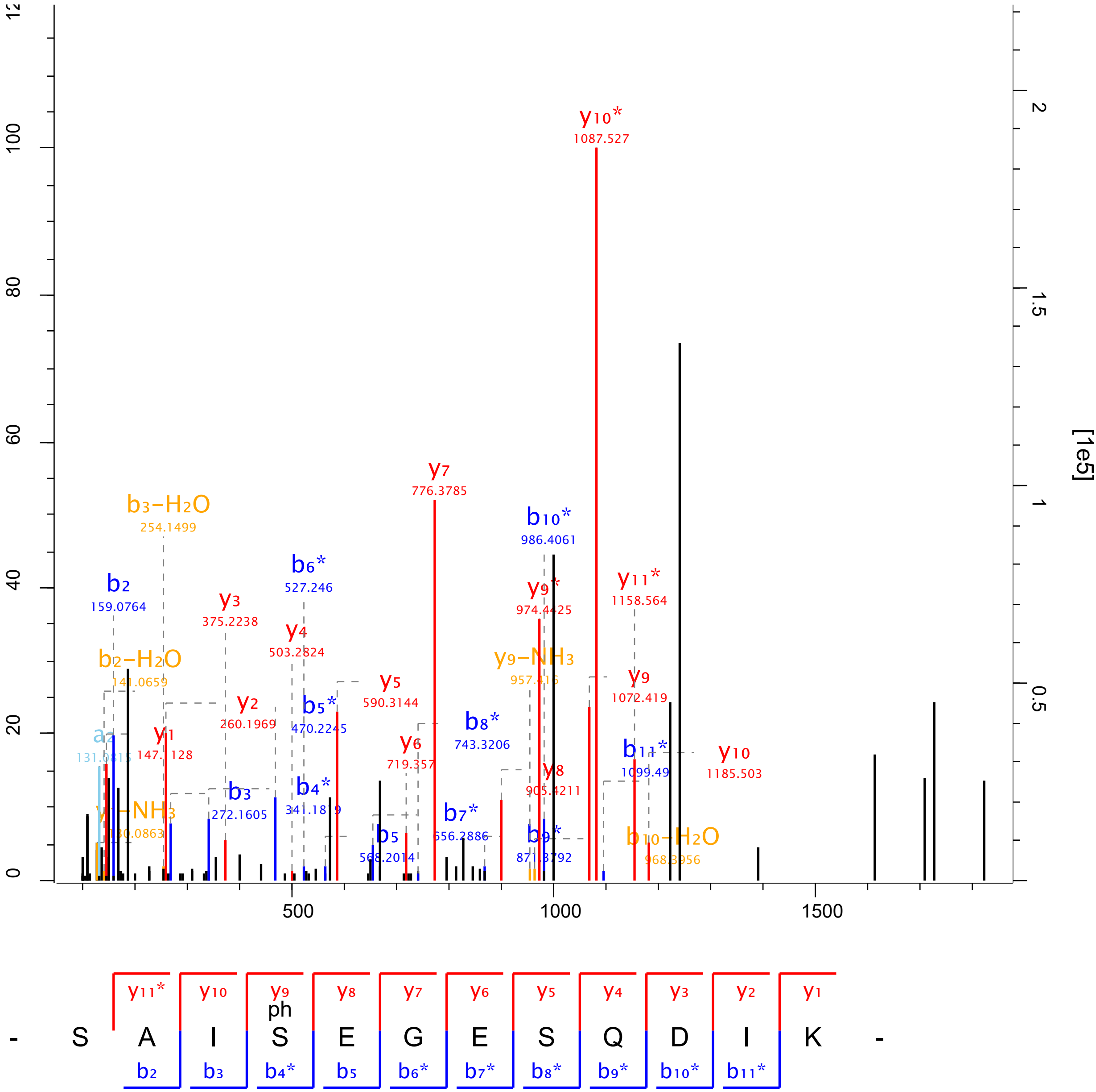

| Raw file | Scan | Method | Score | m/z |
|---|---|---|---|---|
| sys_02_2 | 11962 | FTMS; HCD | 56.72 | 535.25 |

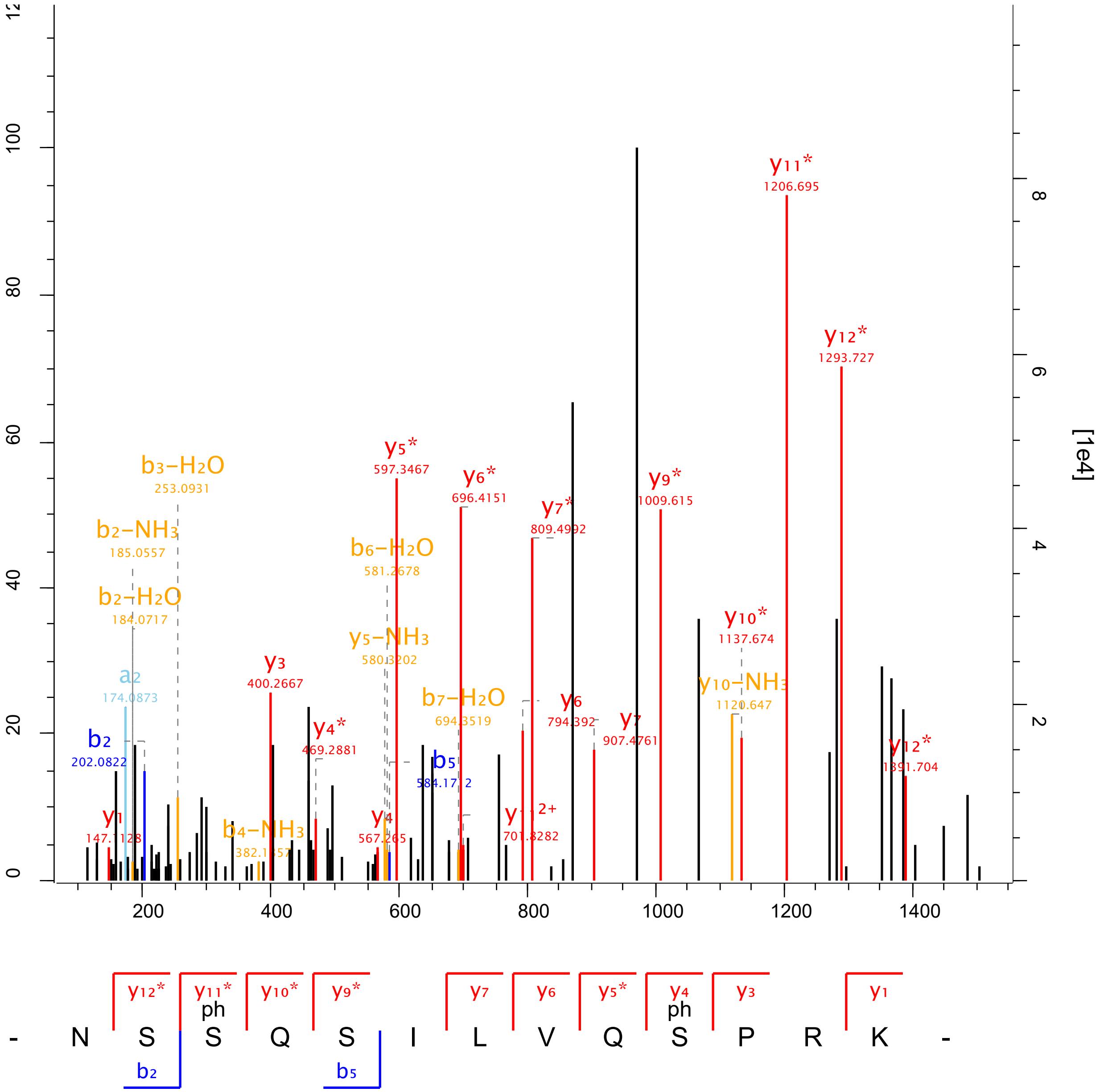

| Raw file | Scan | Method | Score | m/z |
|---|---|---|---|---|
| sys_02_2 | 12020 | FTMS; HCD | 108.55 | 690.98 |

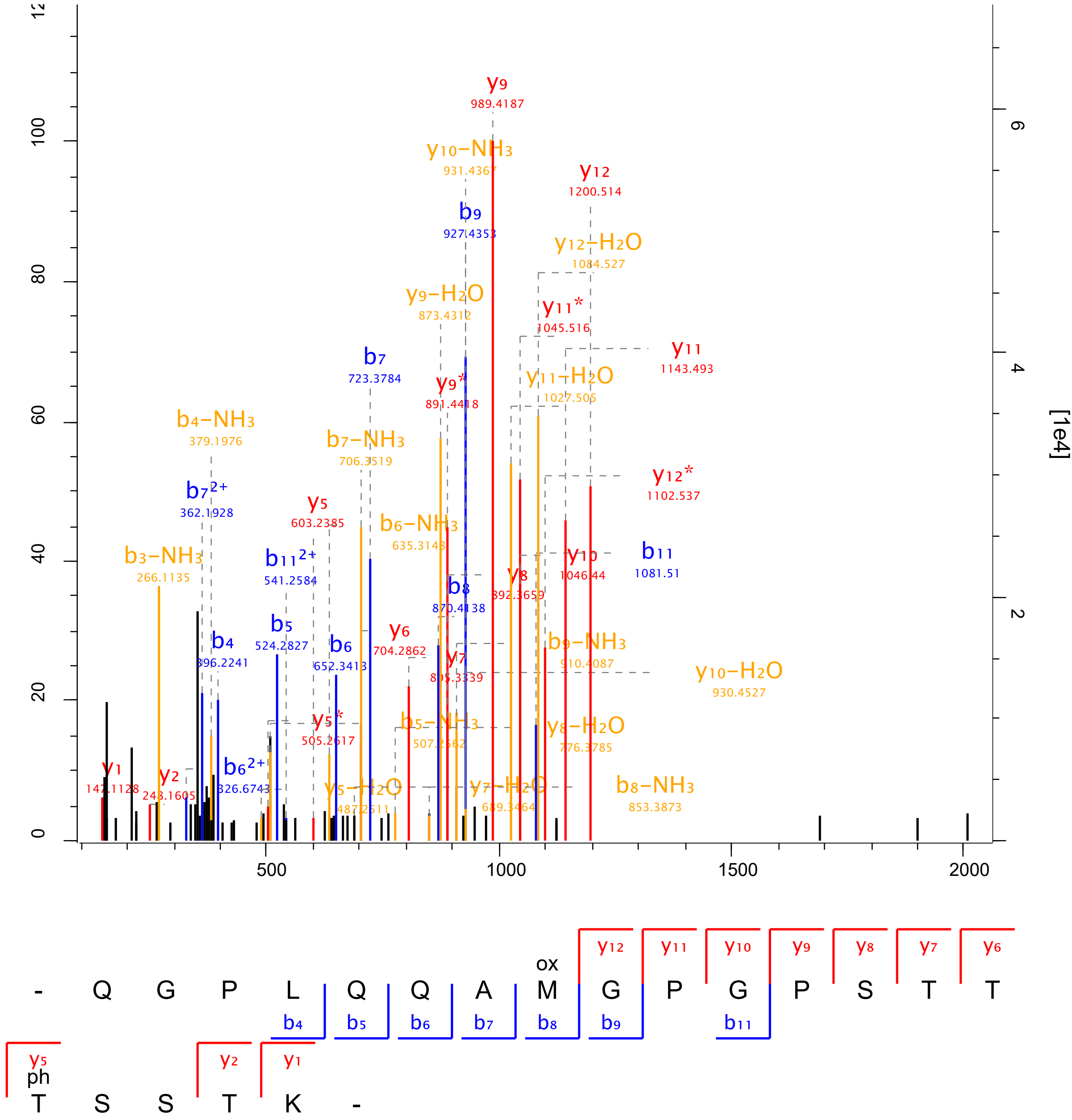

| Raw file | Scan | Method | Score | m/z |
| --- | --- | --- | --- | --- |
| sys_02_2 | 12390 | FTMS; HCD | 42.63 | 512.72 |

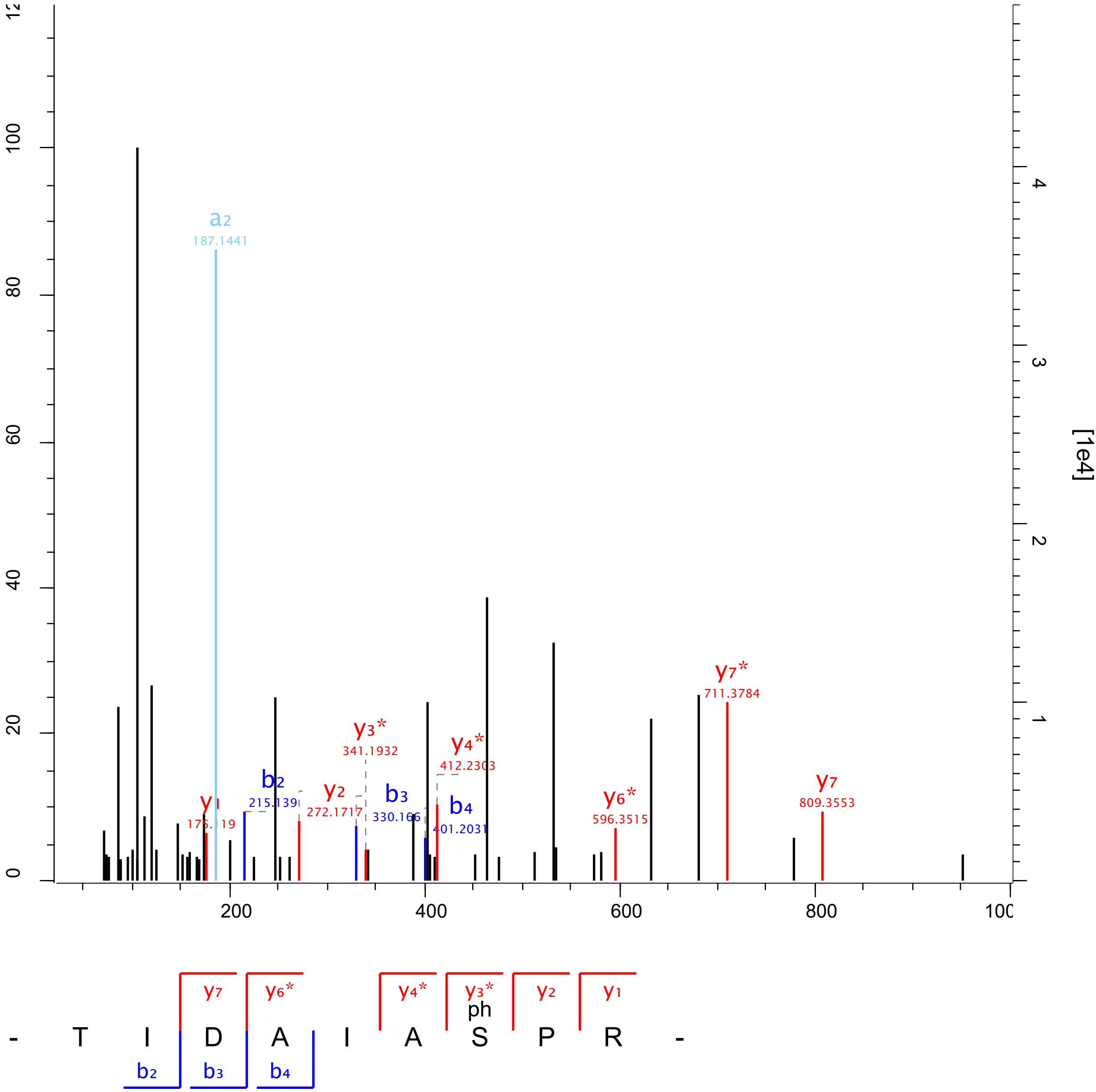

| Raw file | Scan | Method | Score | m/z |
|---|---|---|---|---|
| sys_02_2 | 12400 | FTMS; HCD | 72.47 | 466.56 |

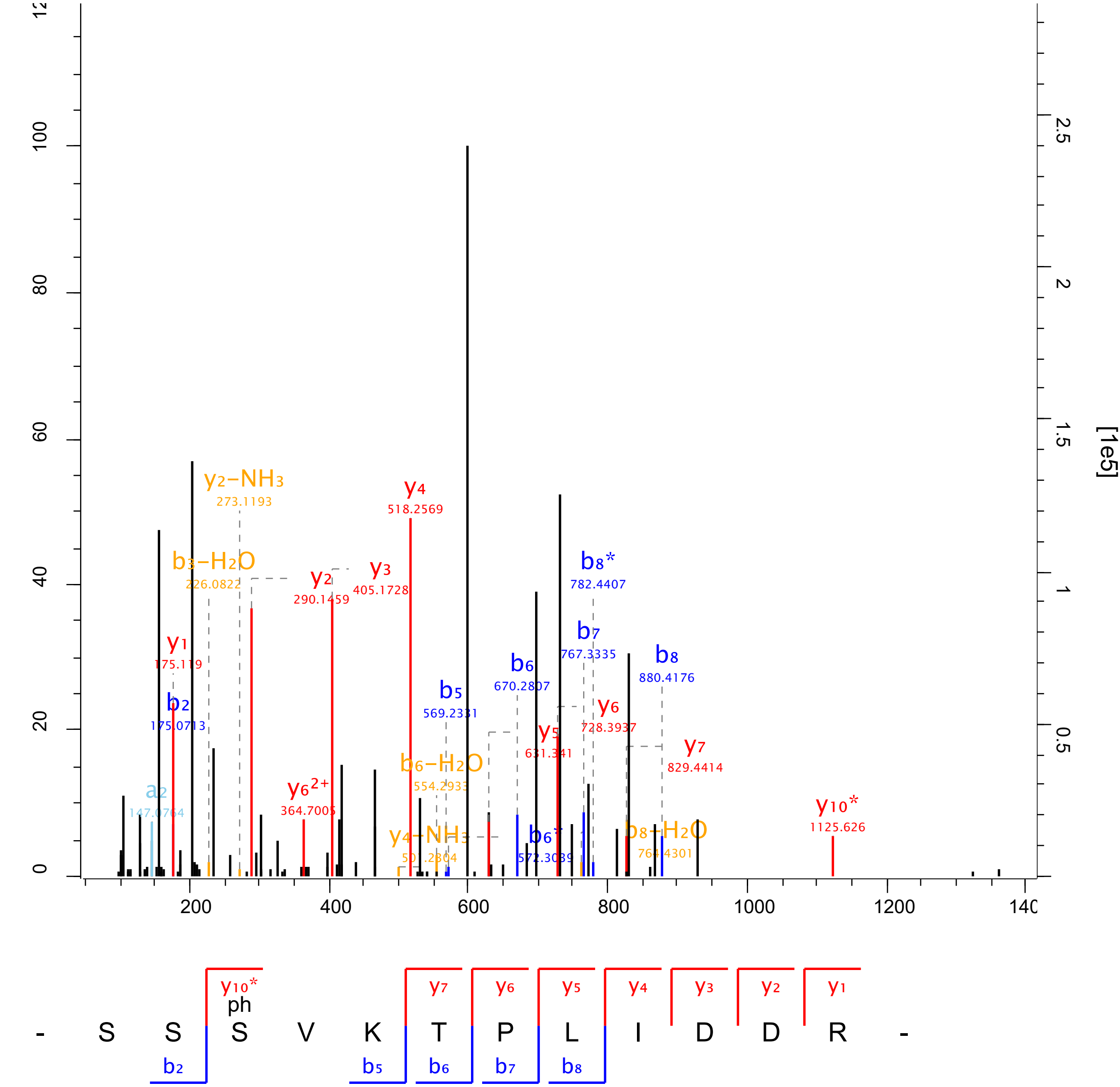

| Raw file | Scan | Method | Score | m/z |
|---|---|---|---|---|
| sys_02_2 | 12582 | FTMS; HCD | 73.5 | 424.2 |

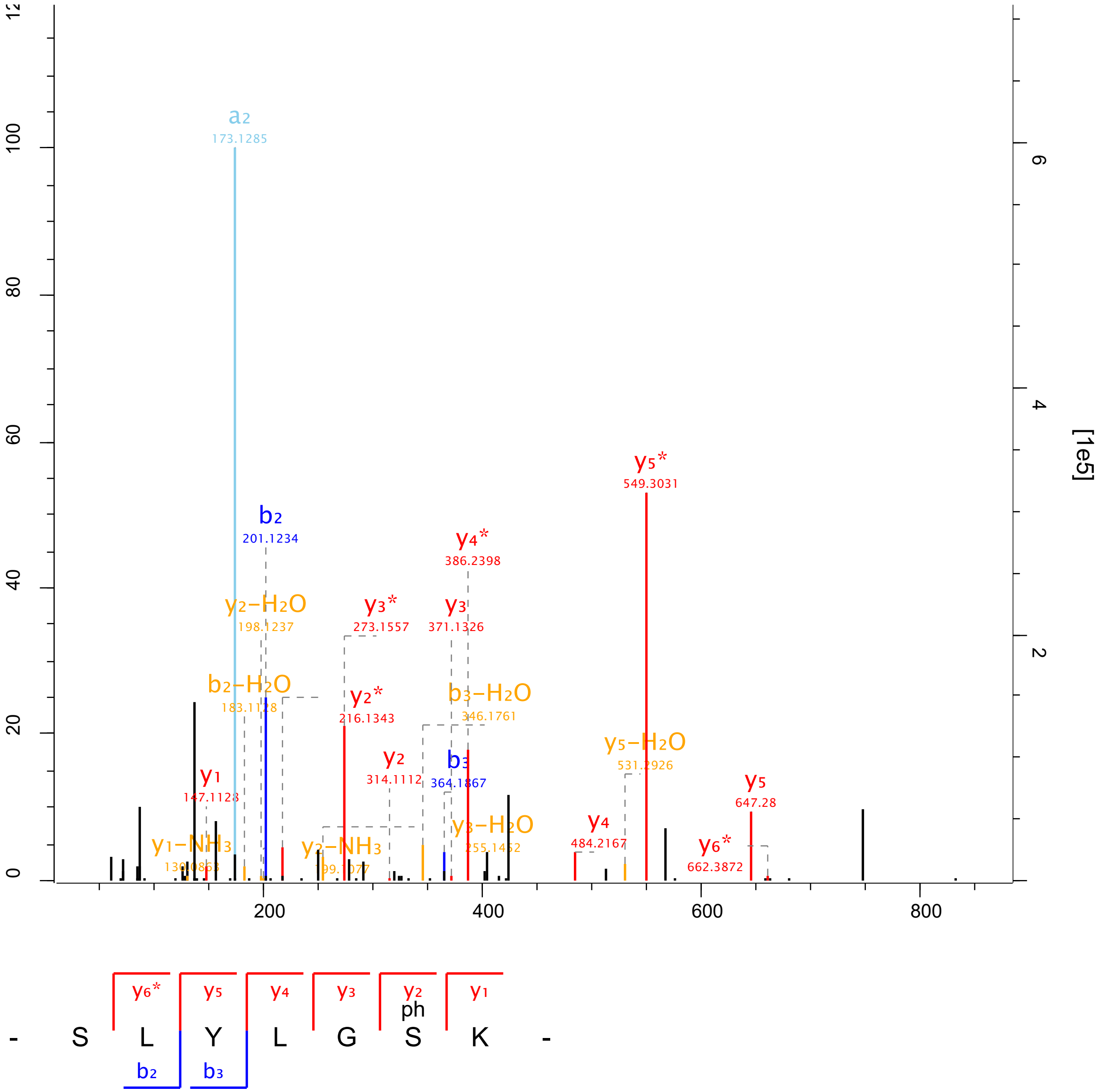

| Raw file | Scan | Method | Score | m/z |
| --- | --- | --- | --- | --- |
| sys_02_2 | 12986 | FTMS; HCD | 149.21 | 713.78 |

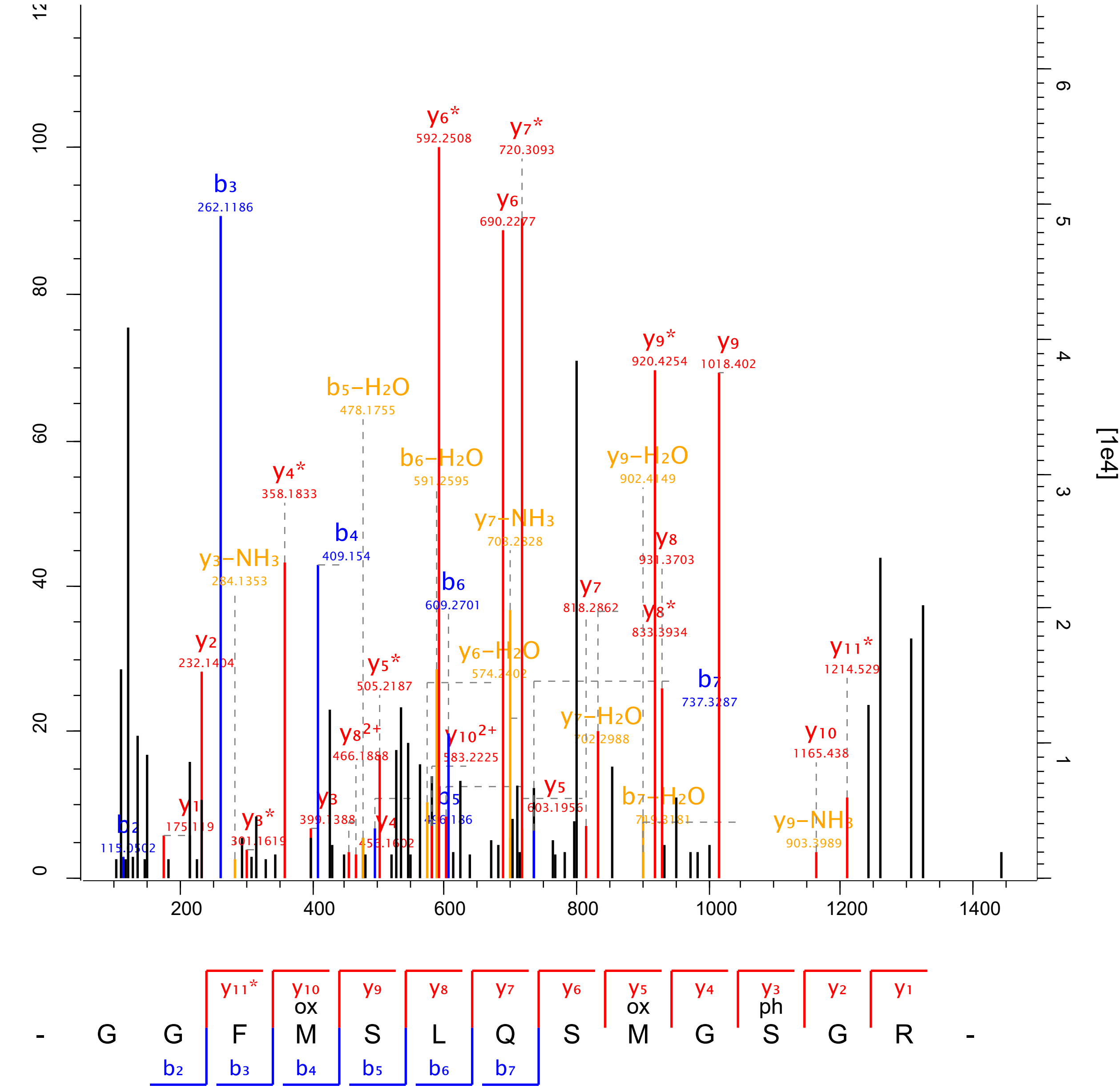

| Raw file | Scan | Method | Score | m/z |
|---|---|---|---|---|
| sys_02_2 | 13005 | FTMS; HCD | 47.62 | 563.76 |

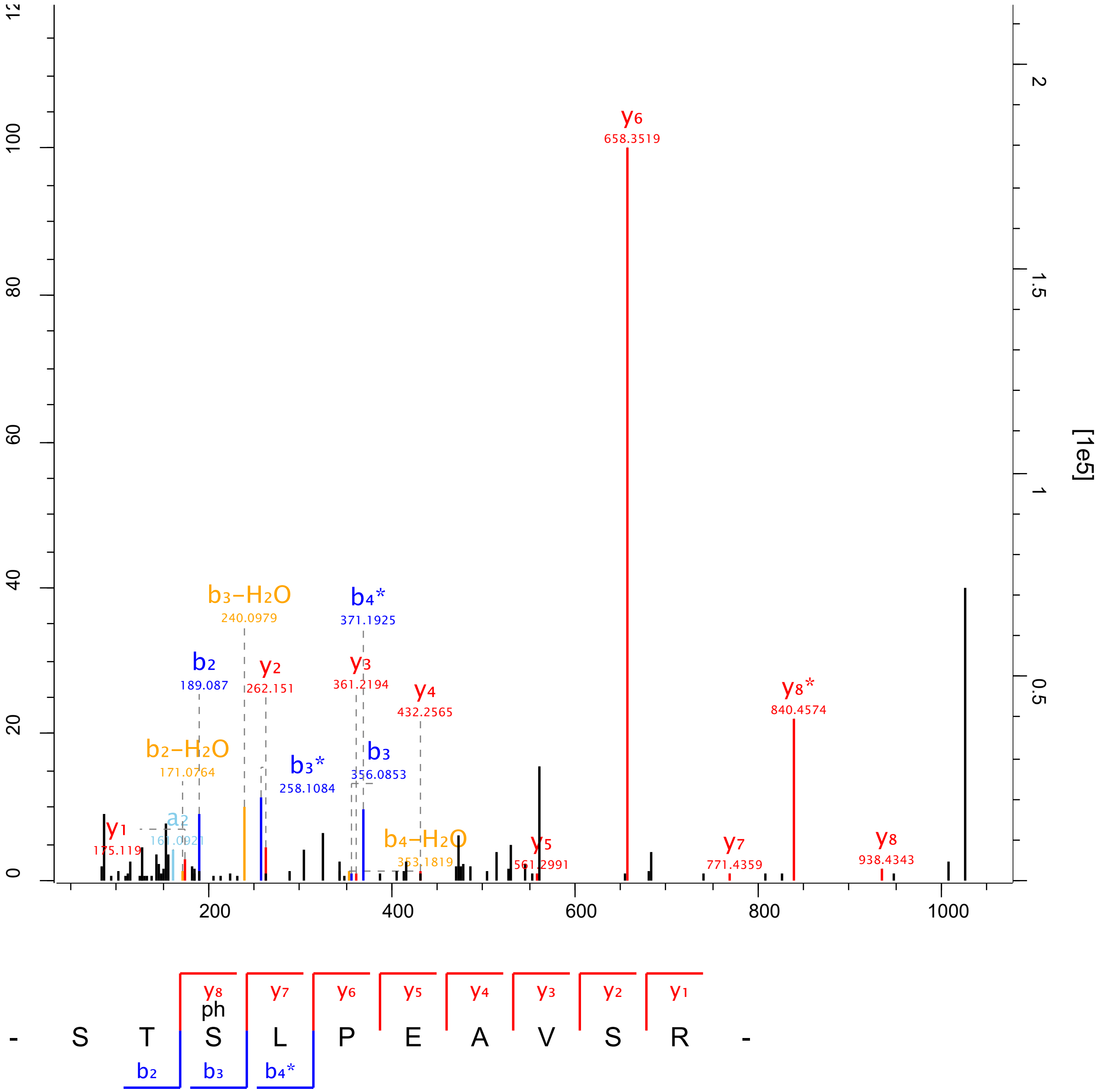

| Raw file | Scan | Method | Score | m/z |
| --- | --- | --- | --- | --- |
| sys_02_2 | 13044 | FTMS; HCD | 155.8 | 816.84 |

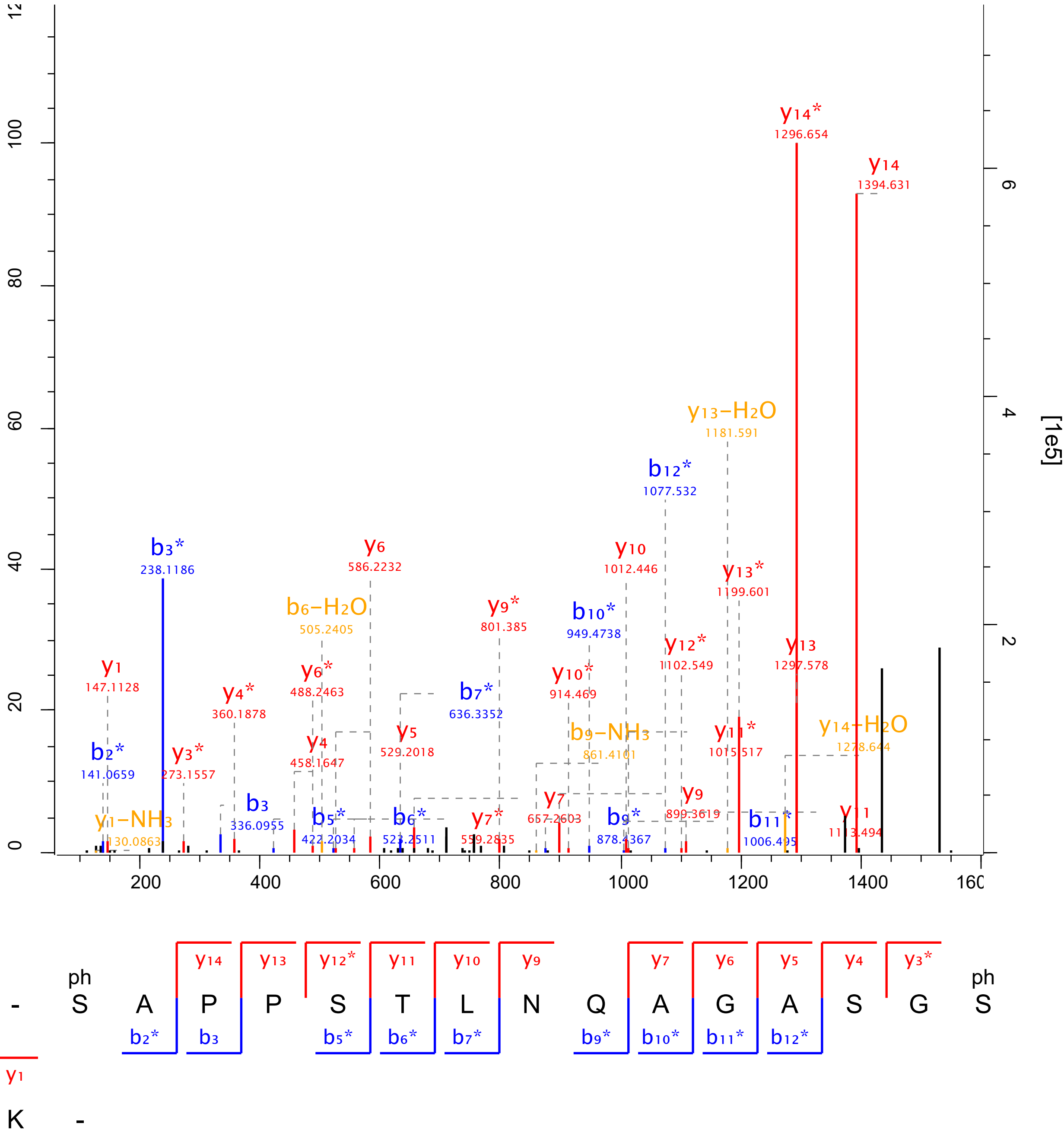

| Raw file | Scan | Method | Score | m/z |
|---|---|---|---|---|
| sys_02_2 | 13135 | FTMS; HCD | 42.07 | 737.3 |

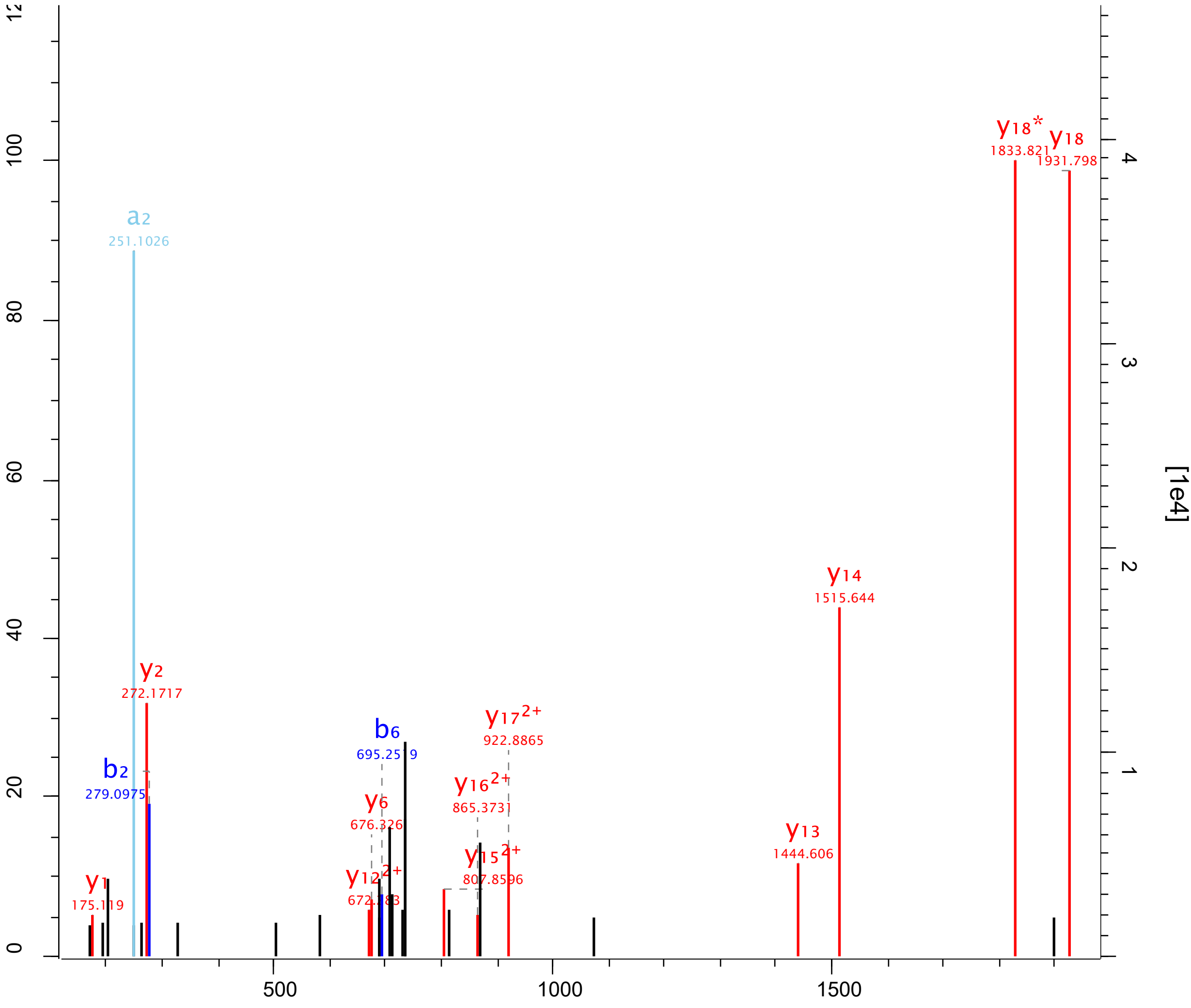


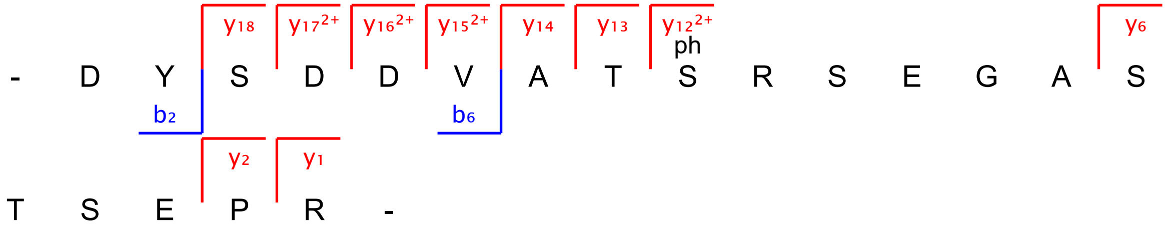

| Raw file | Scan | Method | Score | m/z |
|---|---|---|---|---|
| sys_02_2 | 13167 | FTMS; HCD | 139.49 | 564.27 |

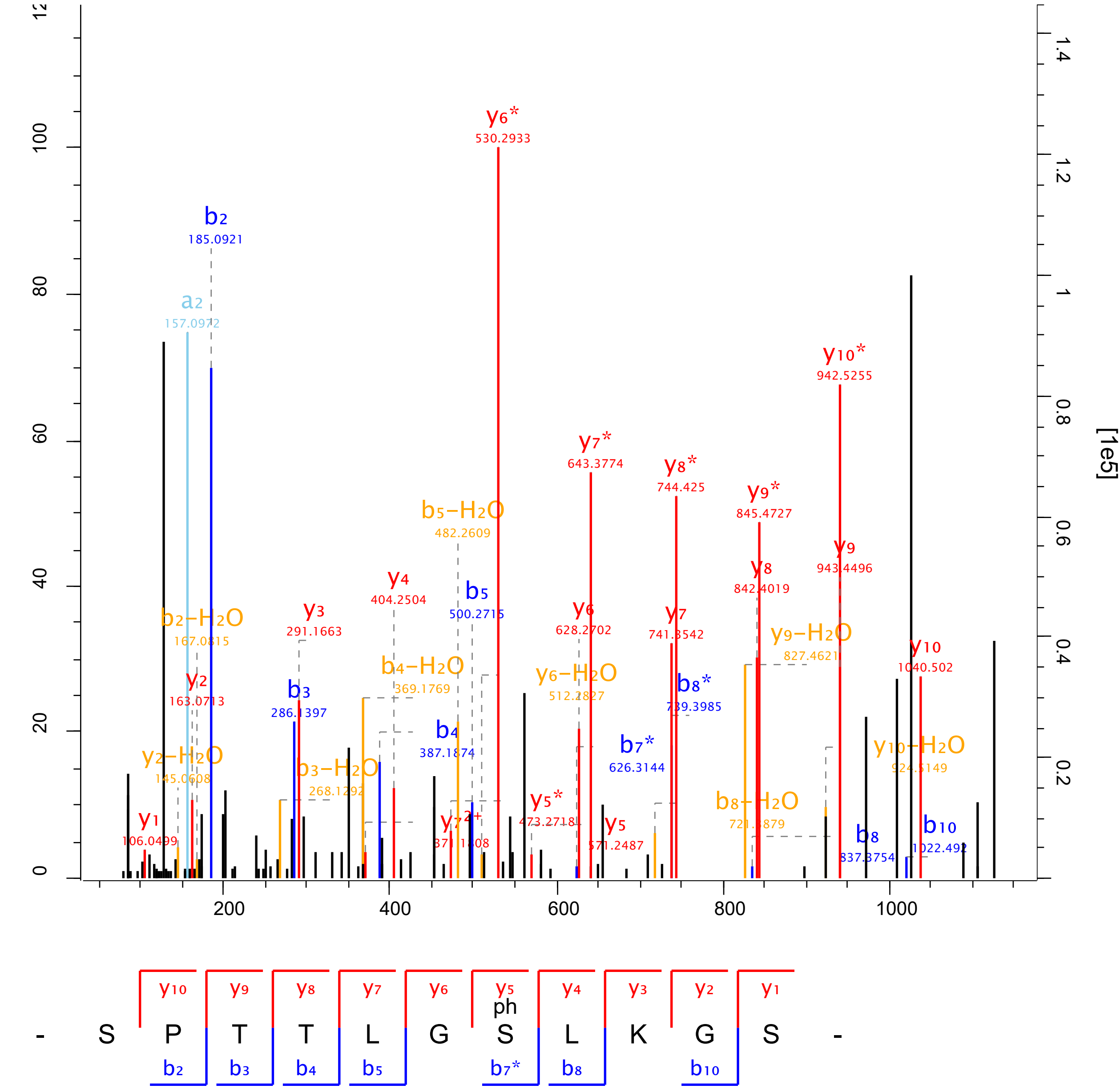

| Raw file | Scan | Method | Score | m/z |
|---|---|---|---|---|
| sys_02_2 | 13302 | FTMS; HCD | 94.96 | 538.94 |

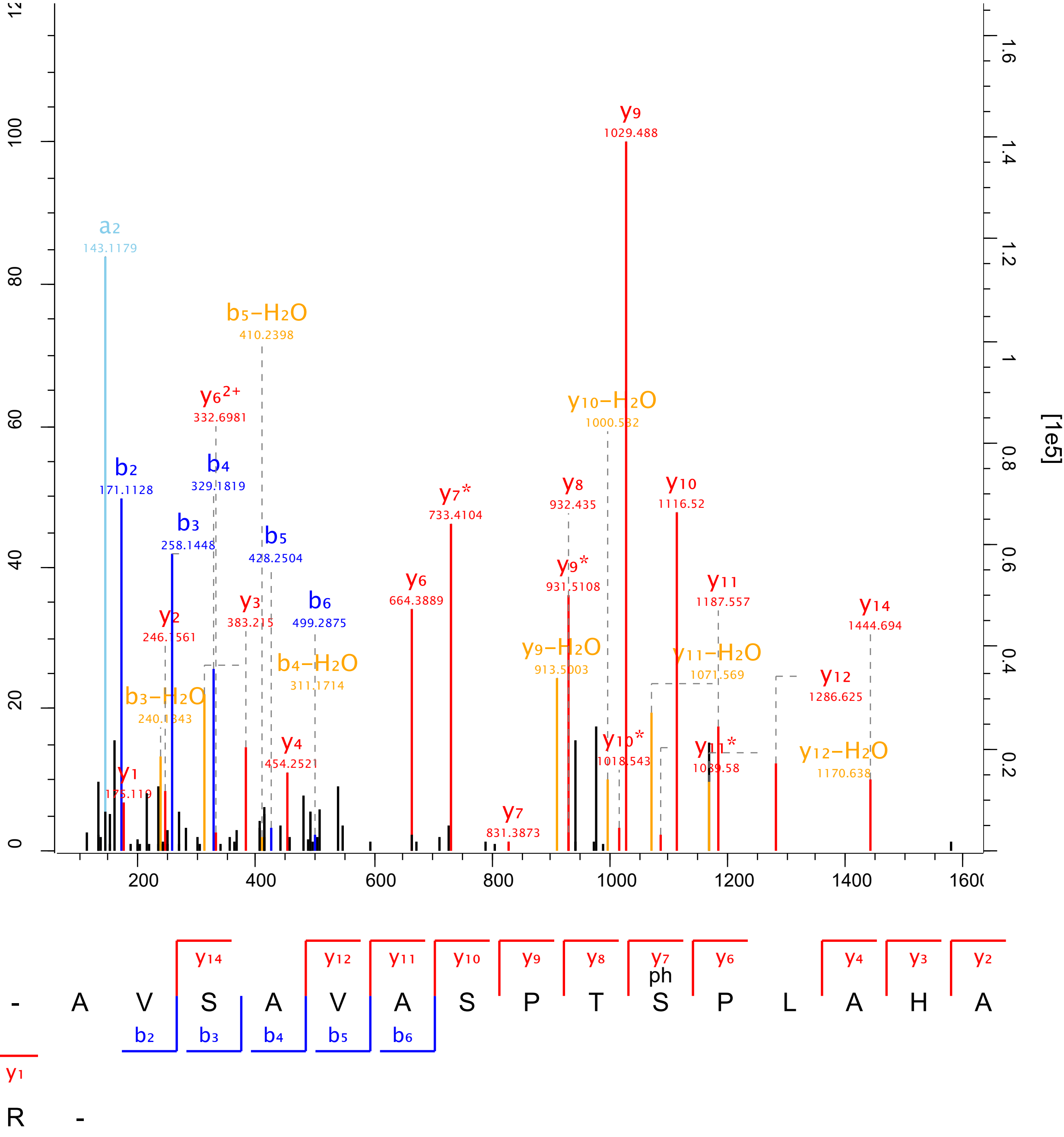

| Raw file | Scan | Method | Score | m/z |
|---|---|---|---|---|
| sys_02_2 | 13492 | FTMS; HCD | 120.56 | 563.93 |

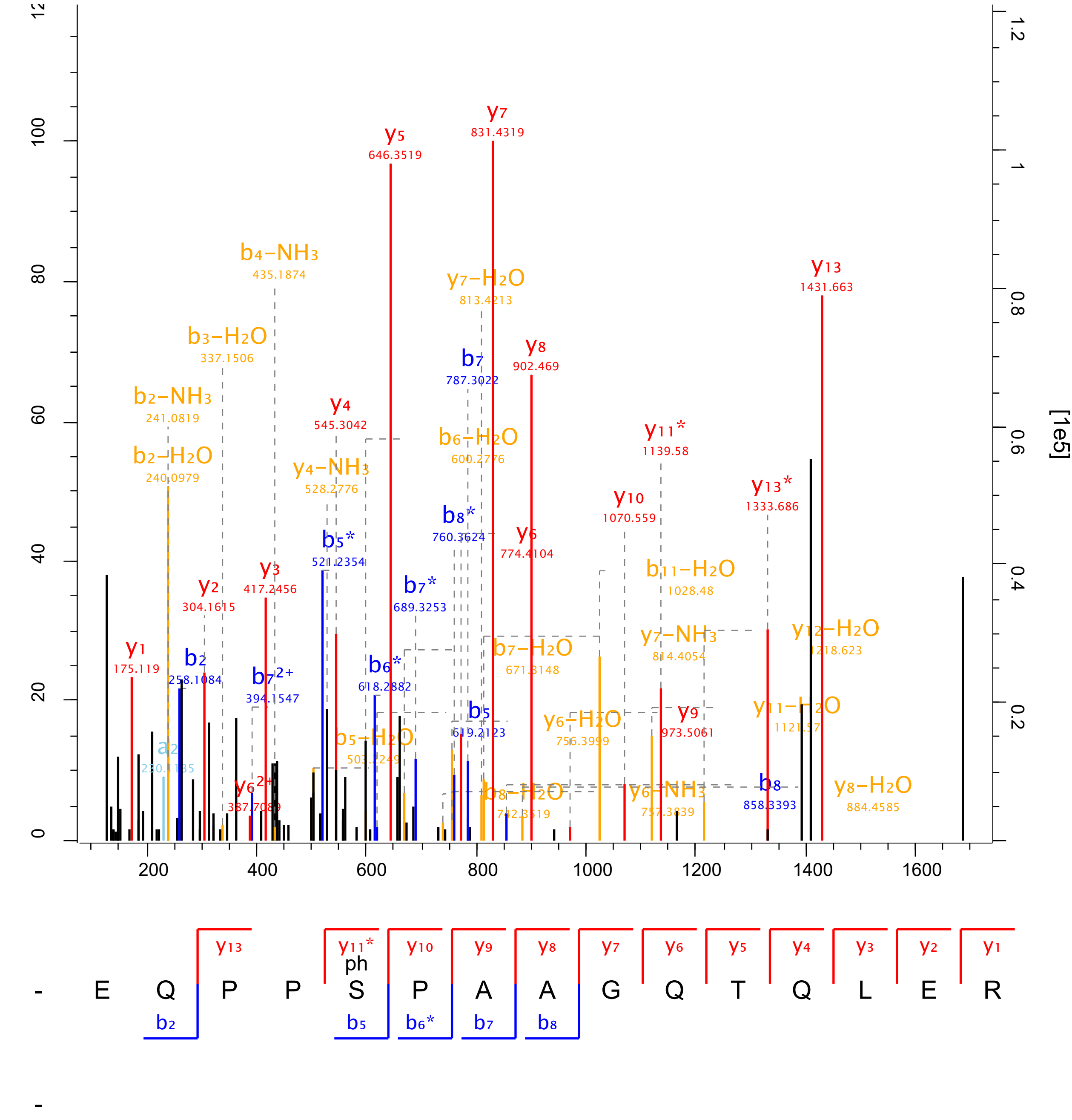

| Raw file | Scan | Method | Score | m/z |
|---|---|---|---|---|
| sys_02_2 | 13547 | FTMS; HCD | 113.47 | 774.83 |

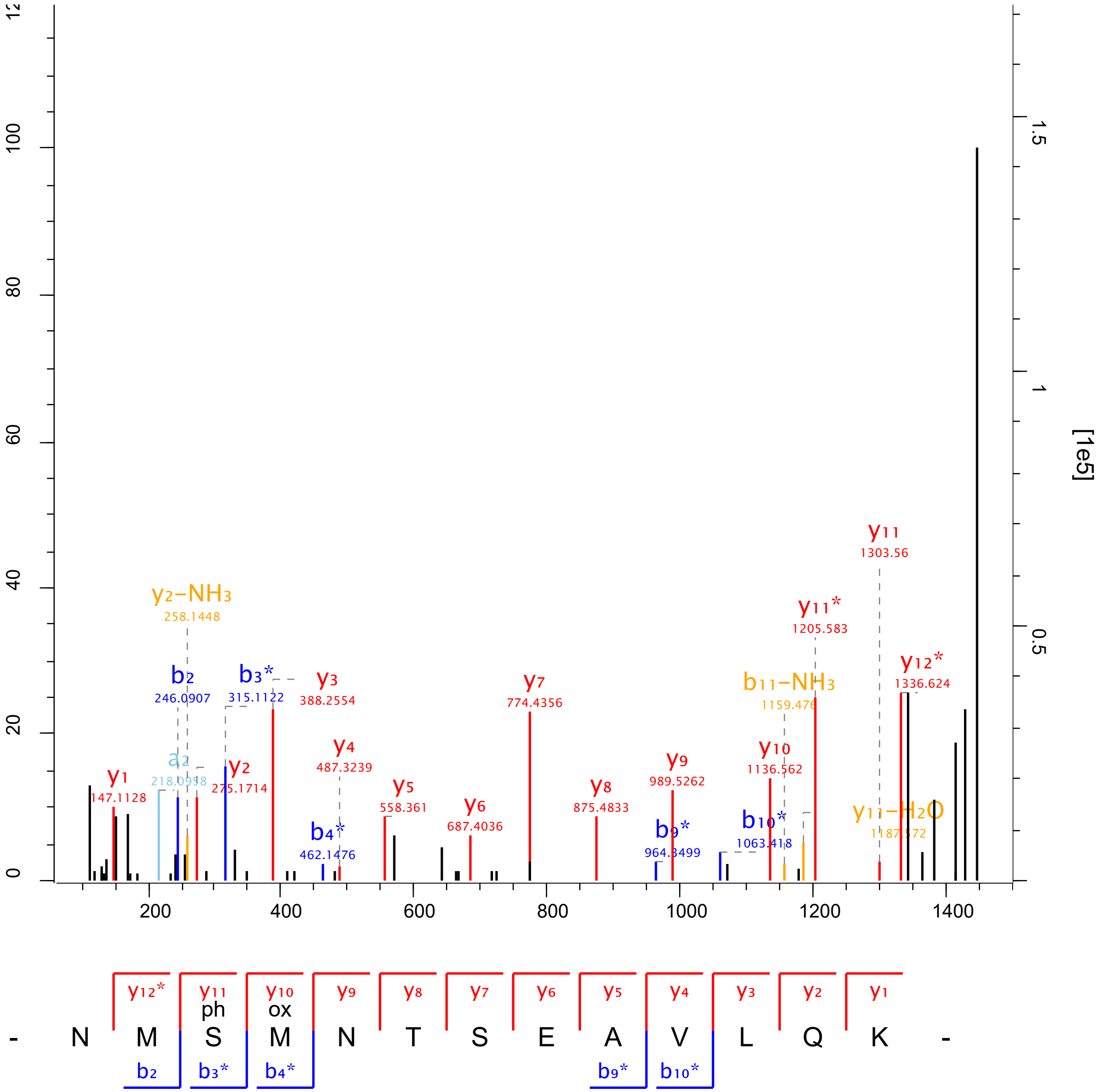

| Raw file | Scan | Method | Score | m/z |
| --- | --- | --- | --- | --- |
| sys_02_2 | 13627 | FTMS; HCD | 75.82 | 573.78 |

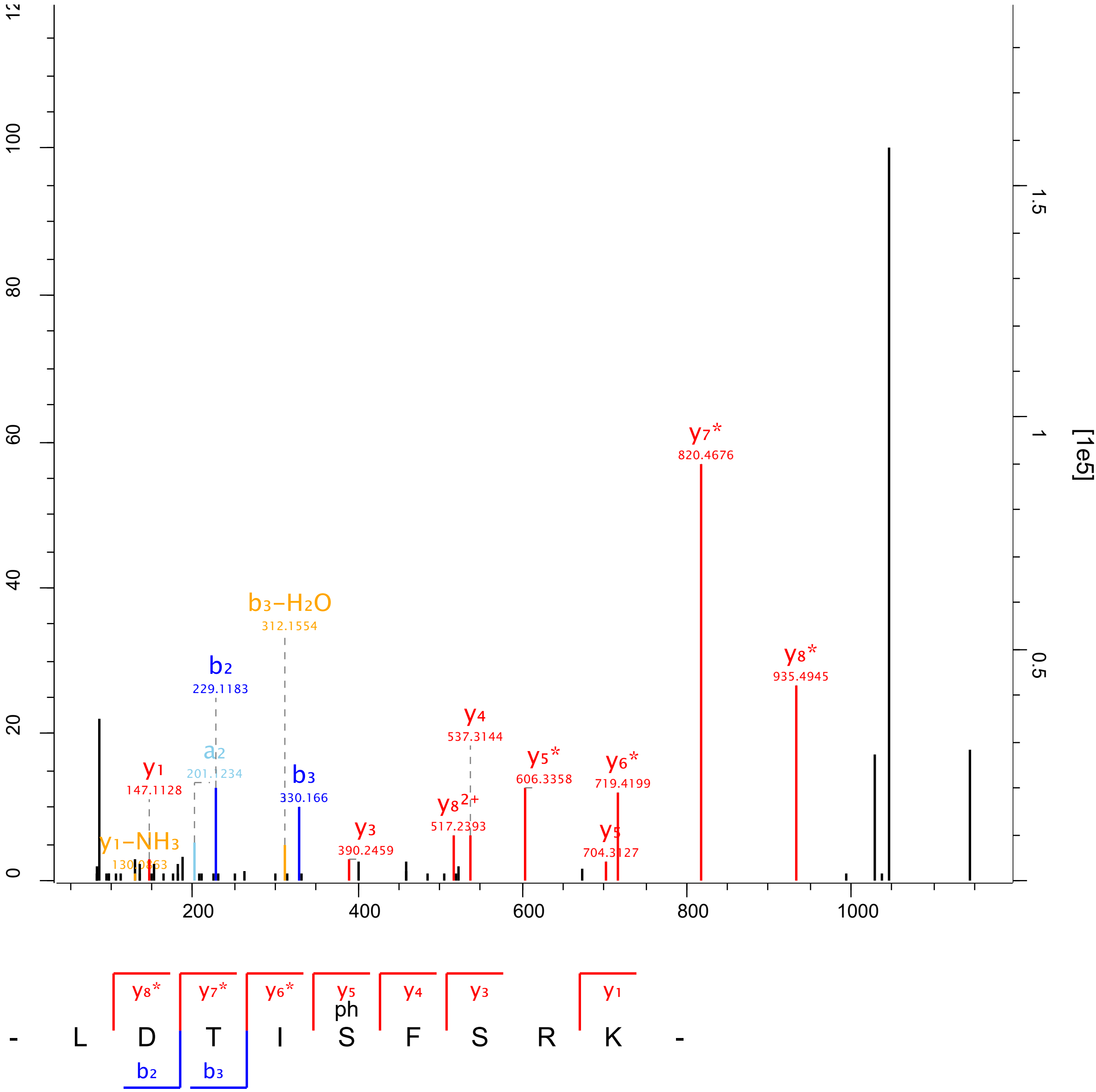

| Raw file | Scan | Method | Score | m/z |
|---|---|---|---|---|
| sys_02_2 | 13694 | FTMS; HCD | 289.87 | 698.33 |

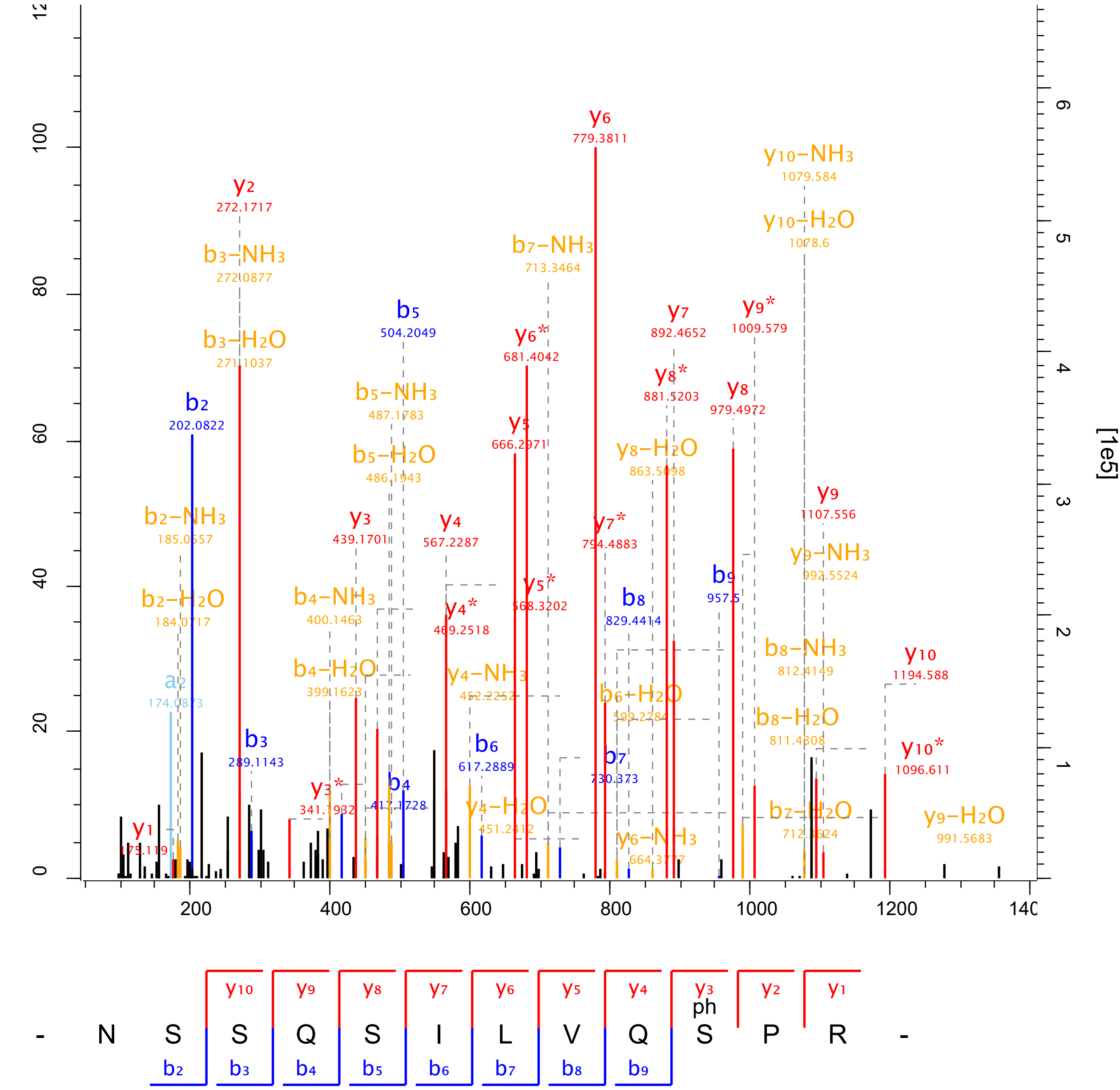

| Raw file | Scan | Method | Score | m/z |
| --- | --- | --- | --- | --- |
| sys_02_2 | 13814 | FTMS; HCD | 129.89 | 498.75 |

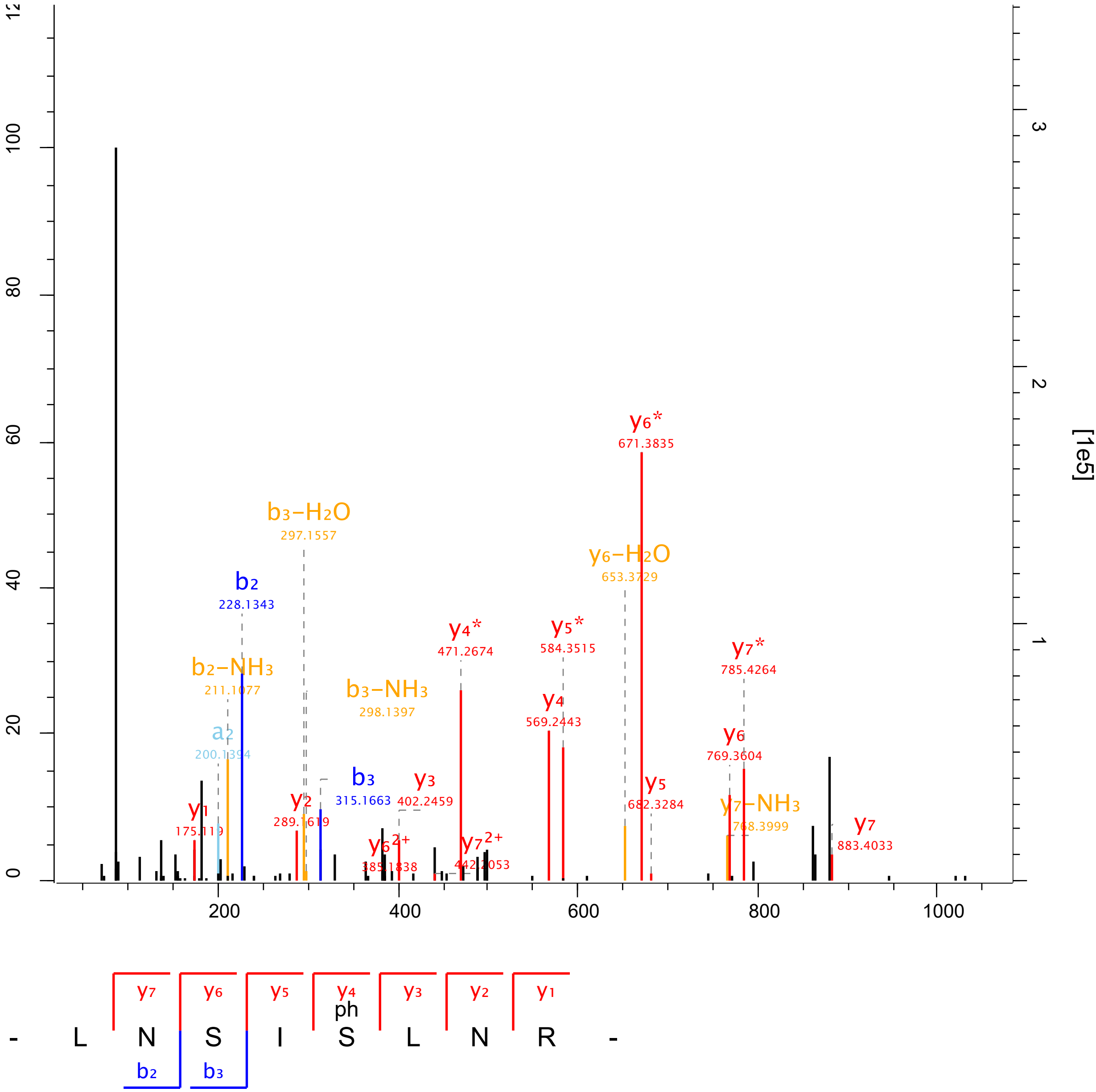

| Raw file | Scan | Method | Score | m/z |
| --- | --- | --- | --- | --- |
| sys_02_2 | 13826 | FTMS; HCD | 204.32 | 549.76 |

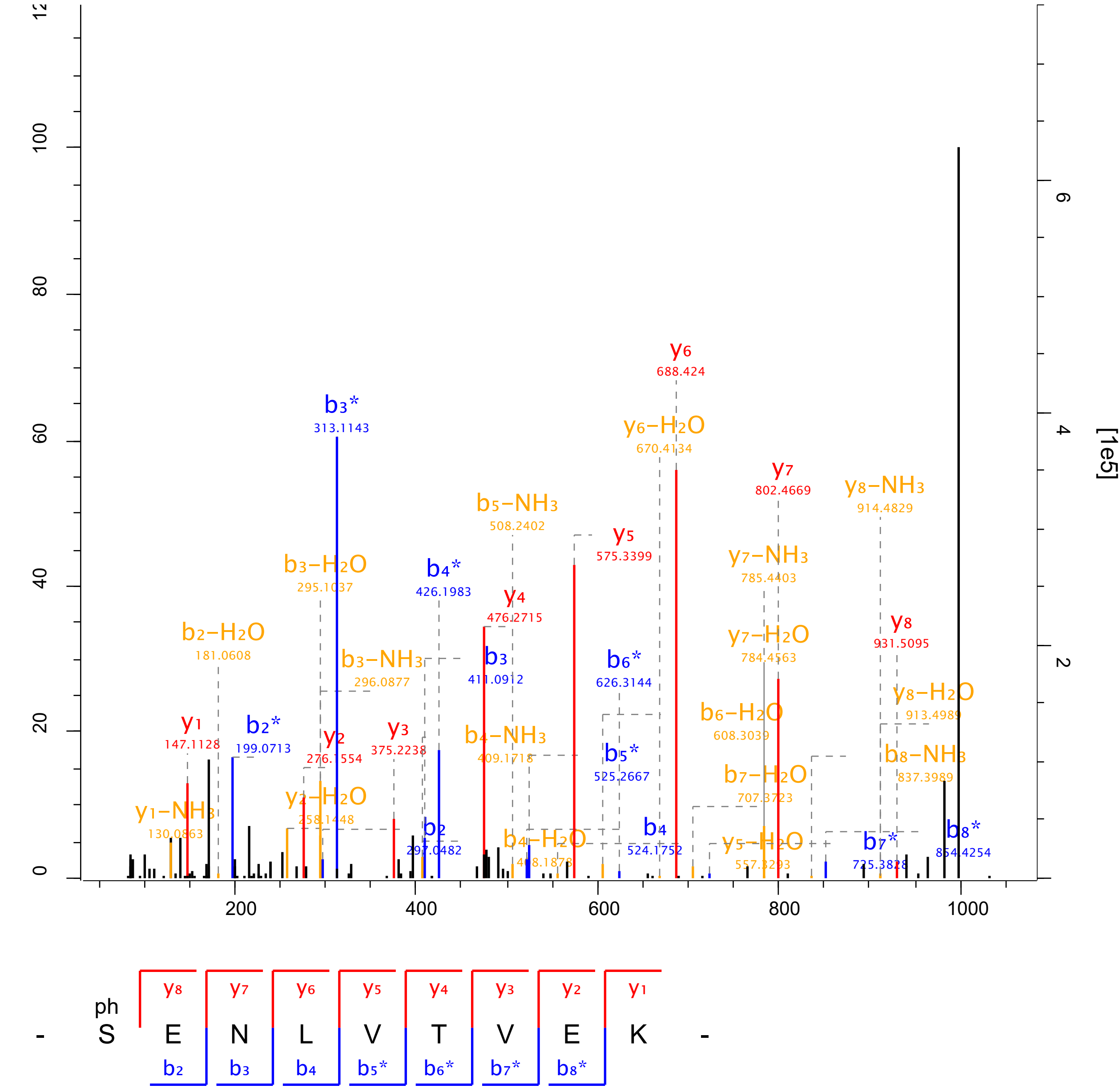

| Raw file | Scan | Method | Score | m/z |
|---|---|---|---|---|
| sys_02_2 | 13833 | FTMS; HCD | 148.15 | 745.33 |

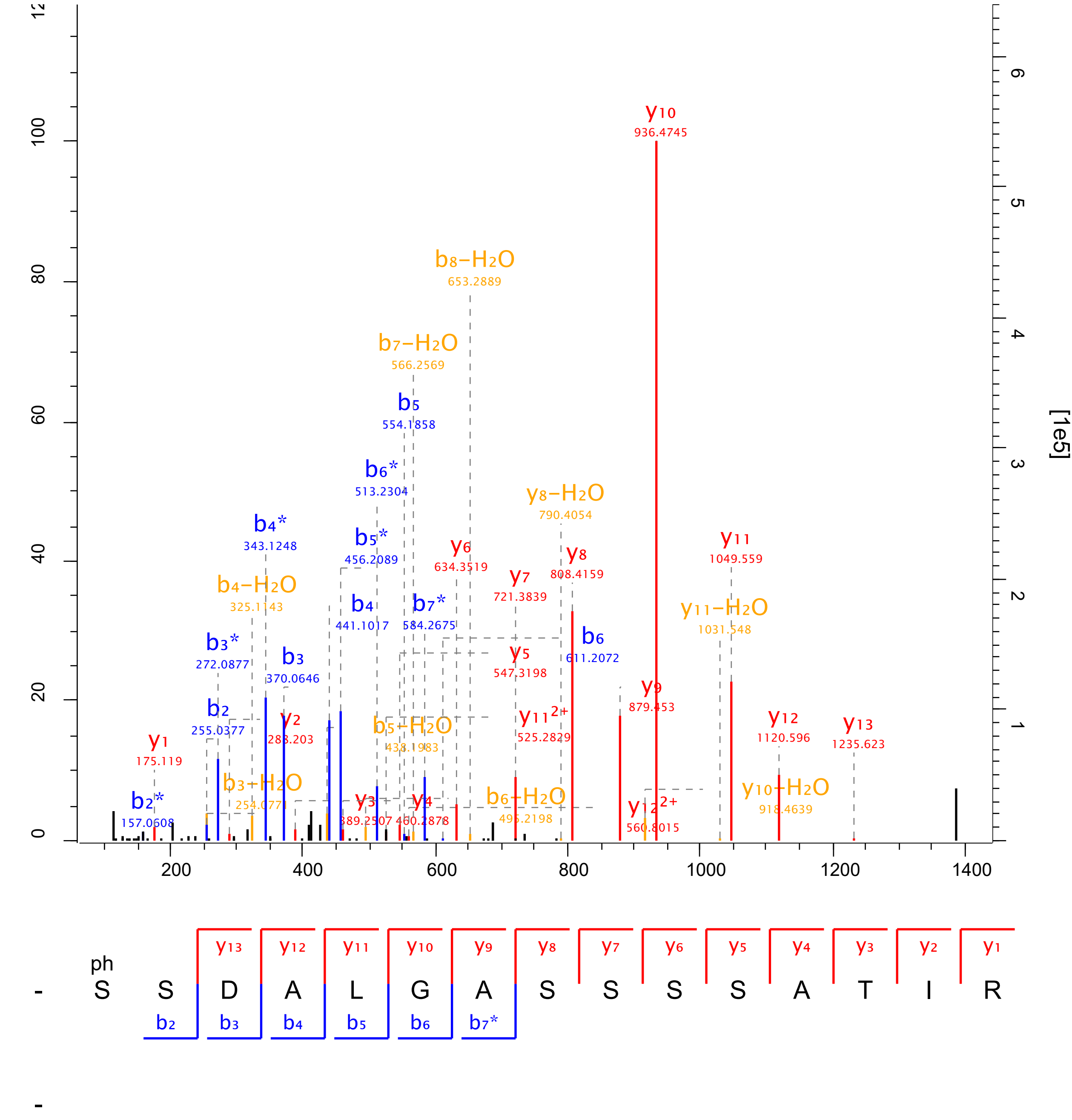

| Raw file | Scan | Method | Score | m/z |
|---|---|---|---|---|
| sys_02_2 | 14029 | FTMS; HCD | 50.19 | 468.25 |

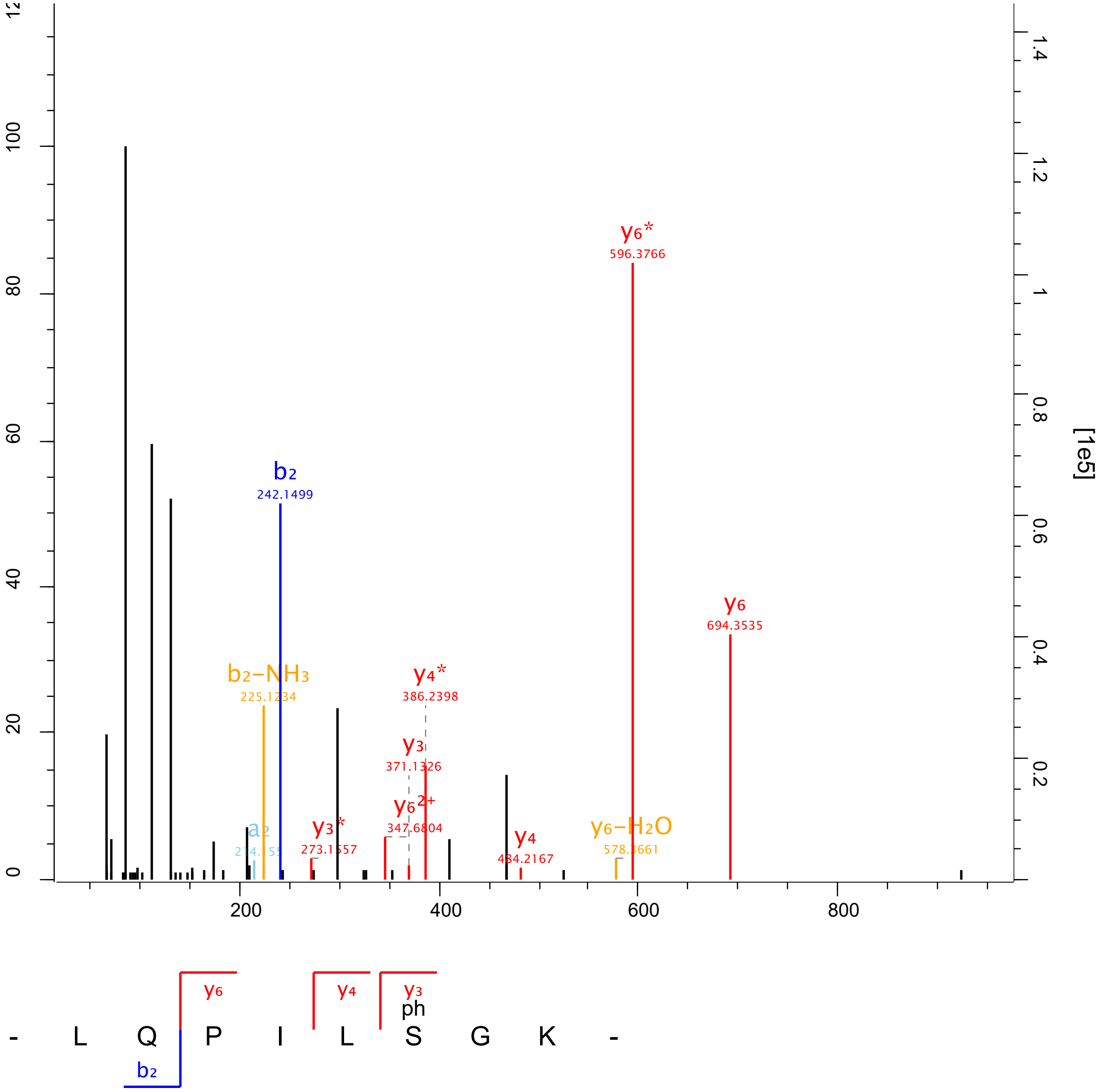

| Raw file | Scan | Method | Score | m/z |
|---|---|---|---|---|
| sys_02_2 | 14242 | FTMS; HCD | 136.68 | 554.22 |

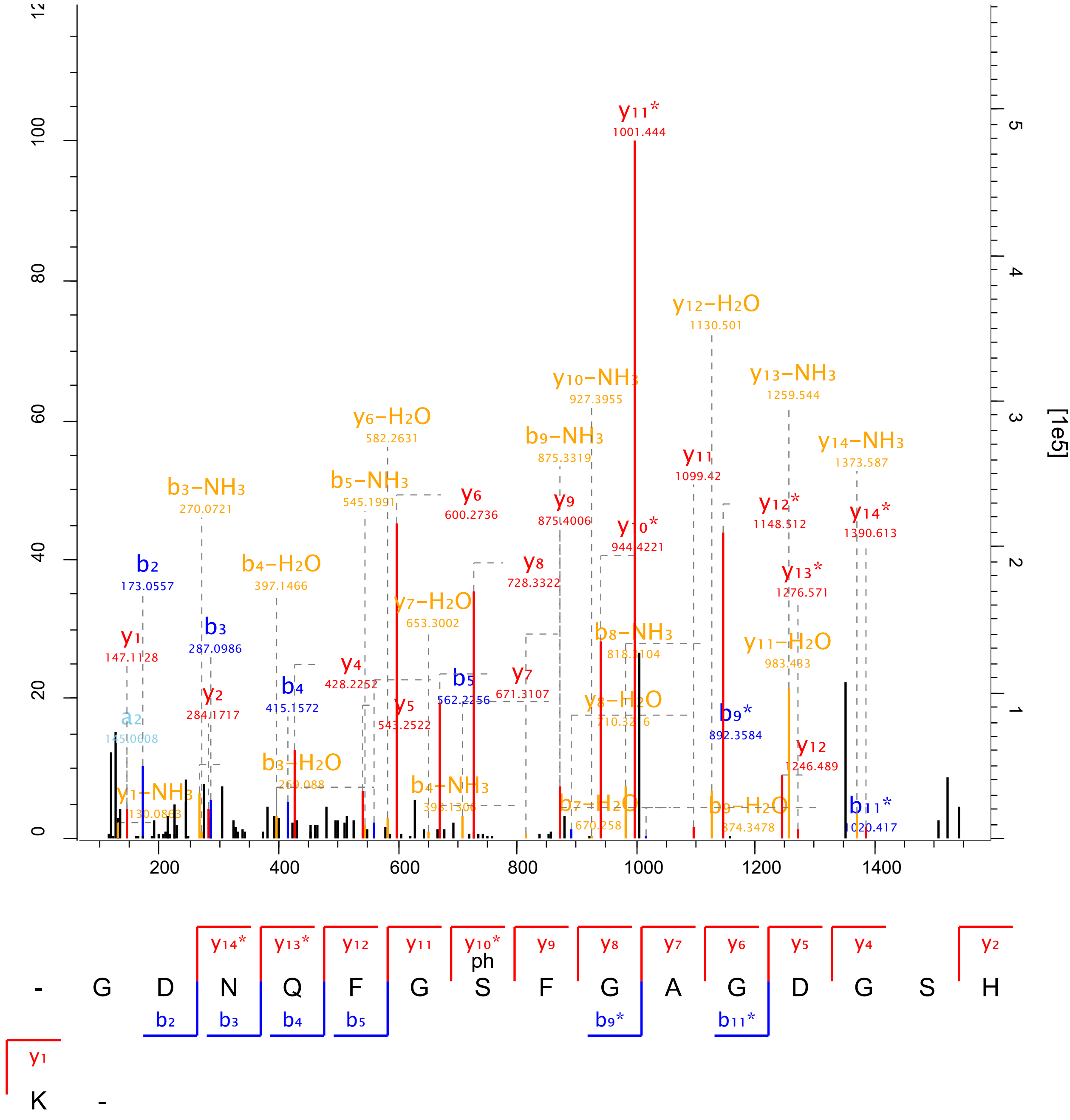

| Raw file | Scan | Method | Score | m/z |
|---|---|---|---|---|
| sys_02_2 | 14254 | FTMS; HCD | 111.79 | 580.74 |

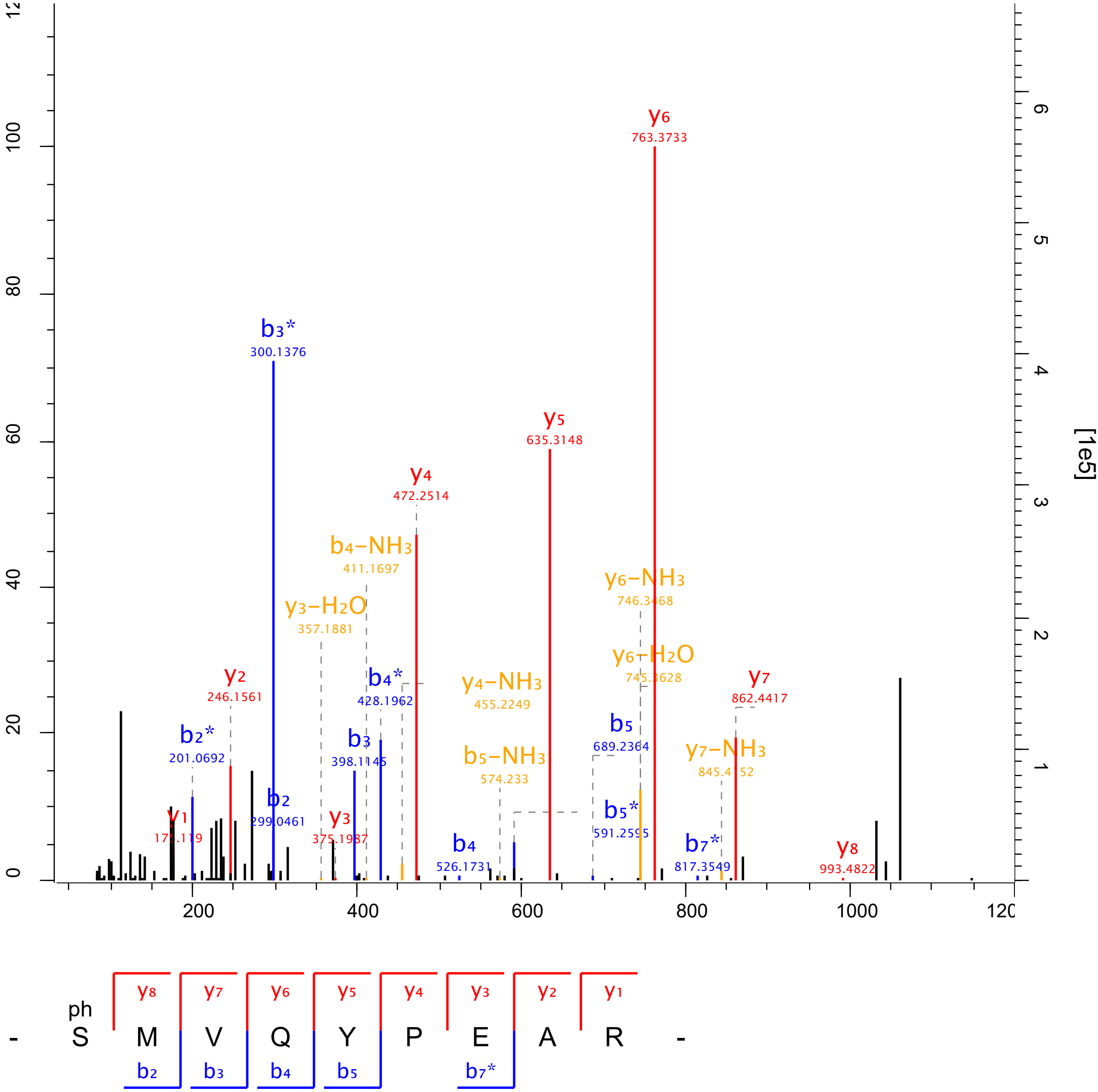

| Raw file | Scan | Method | Score | m/z |
|---|---|---|---|---|
| sys_02_2 | 14356 | FTMS; HCD | 69.35 | 562.75 |

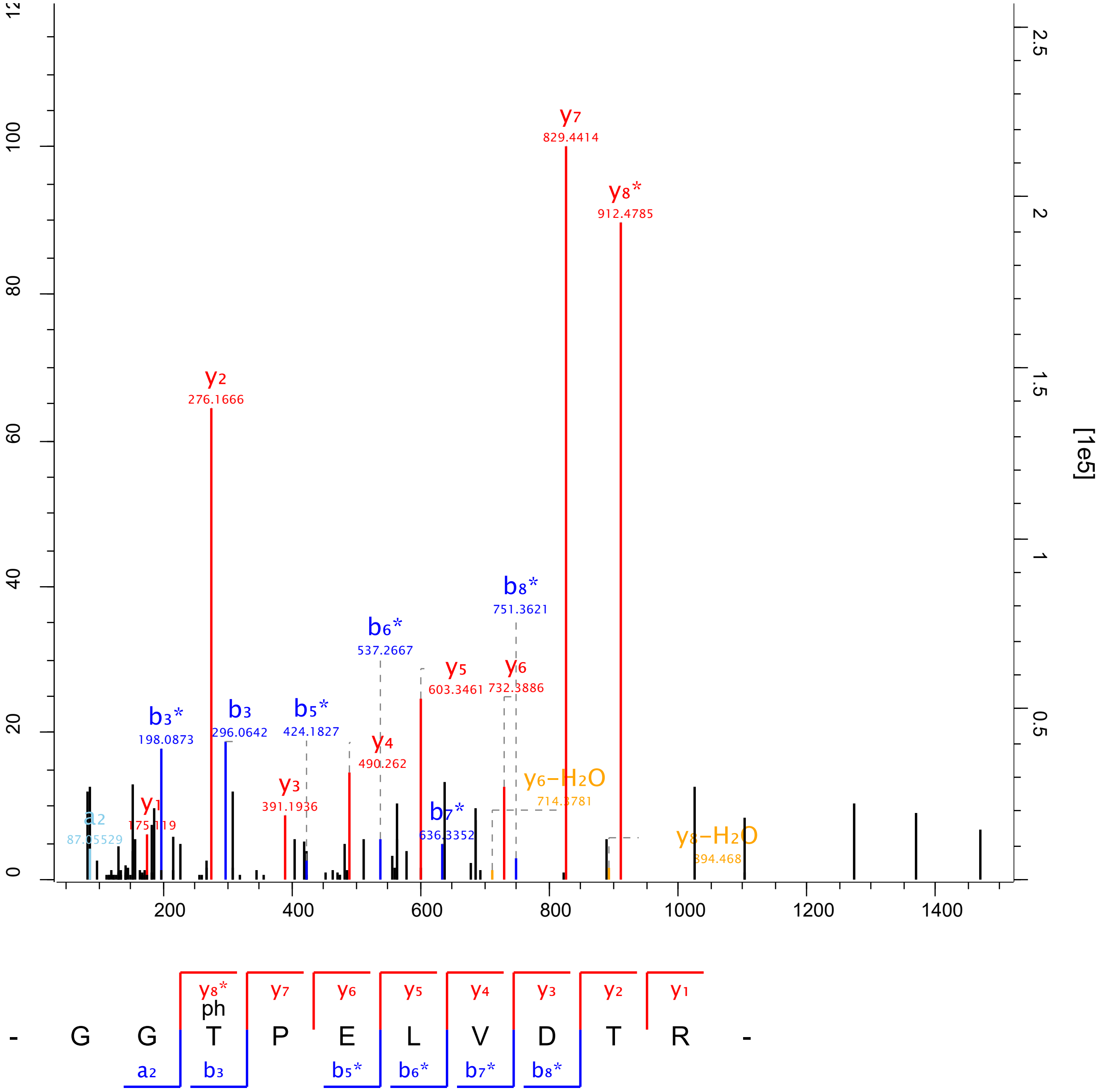

| Raw file | Scan | Method | Score | m/z |
|---|---|---|---|---|
| sys_02_2 | 14417 | FTMS; HCD | 111.22 | 760.8 |

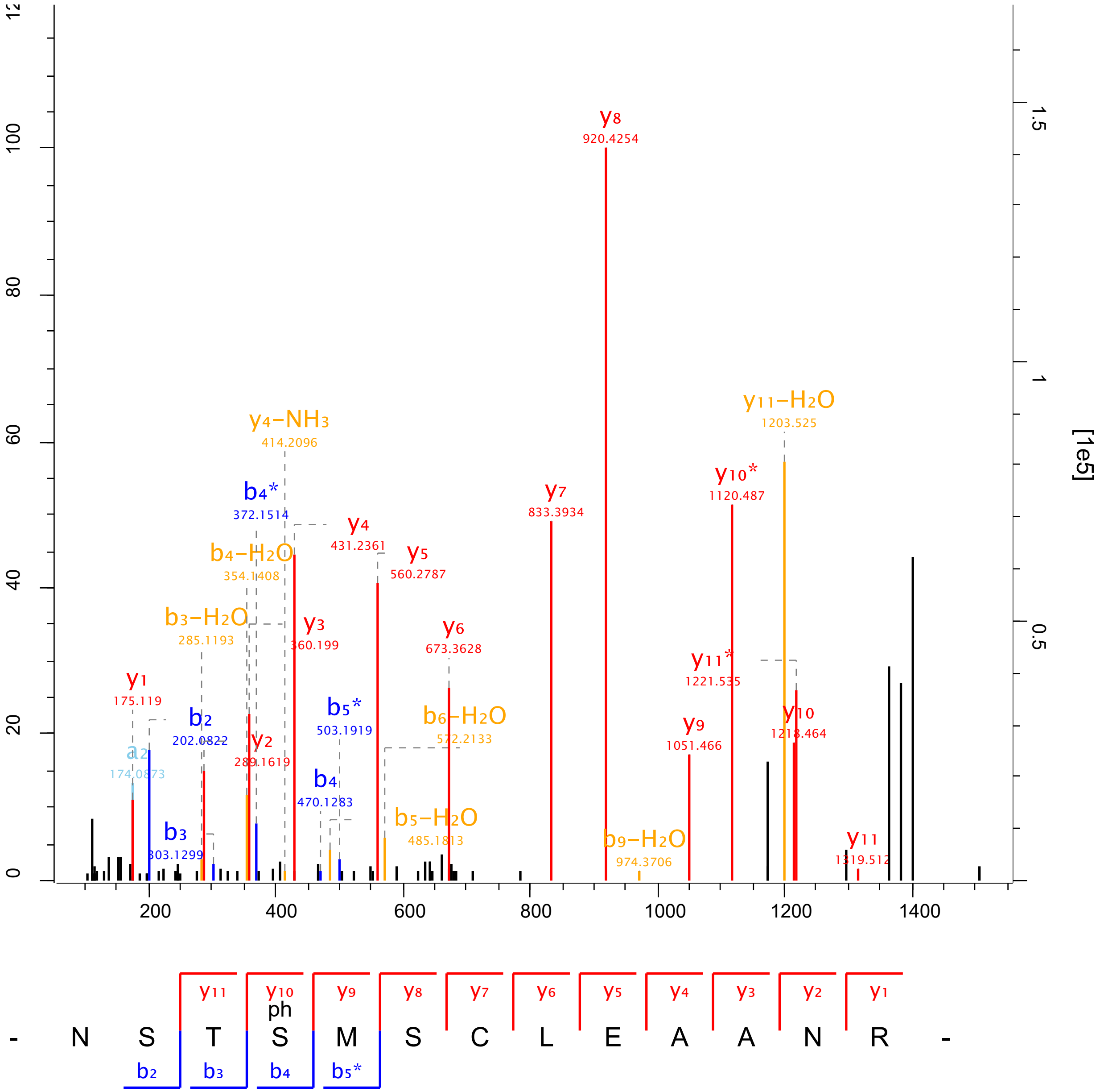

| Raw file | Scan | Method | Score | m/z |
|---|---|---|---|---|
| sys_02_2 | 1461 | FTMS; HCD | 48.53 | 440.86 |

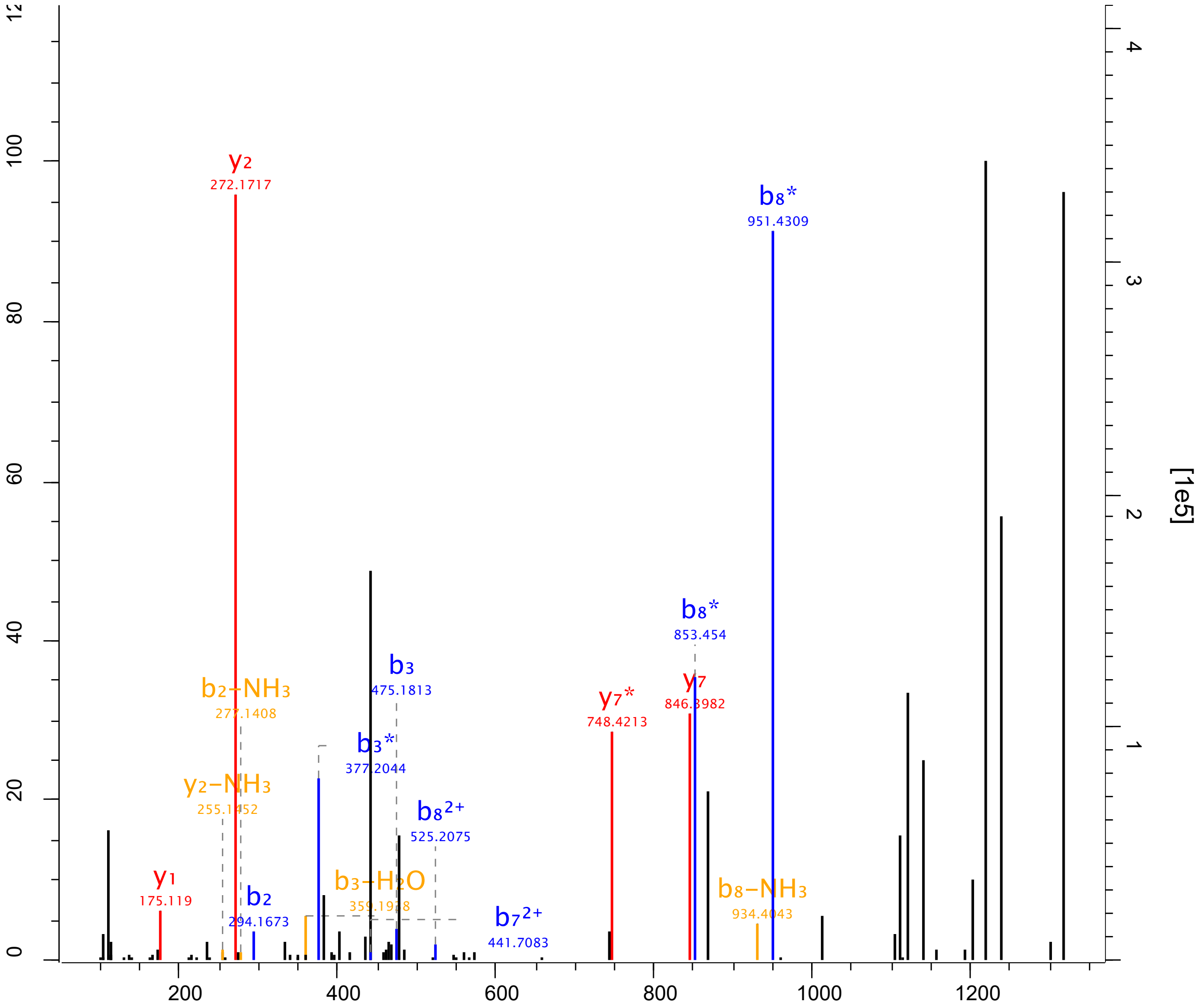


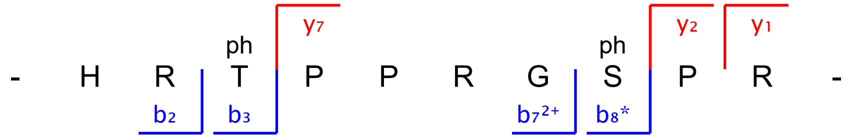

| Raw file | Scan | Method | Score | m/z |
|---|---|---|---|---|
| sys_02_2 | 14752 | FTMS; HCD | 58.32 | 601.26 |

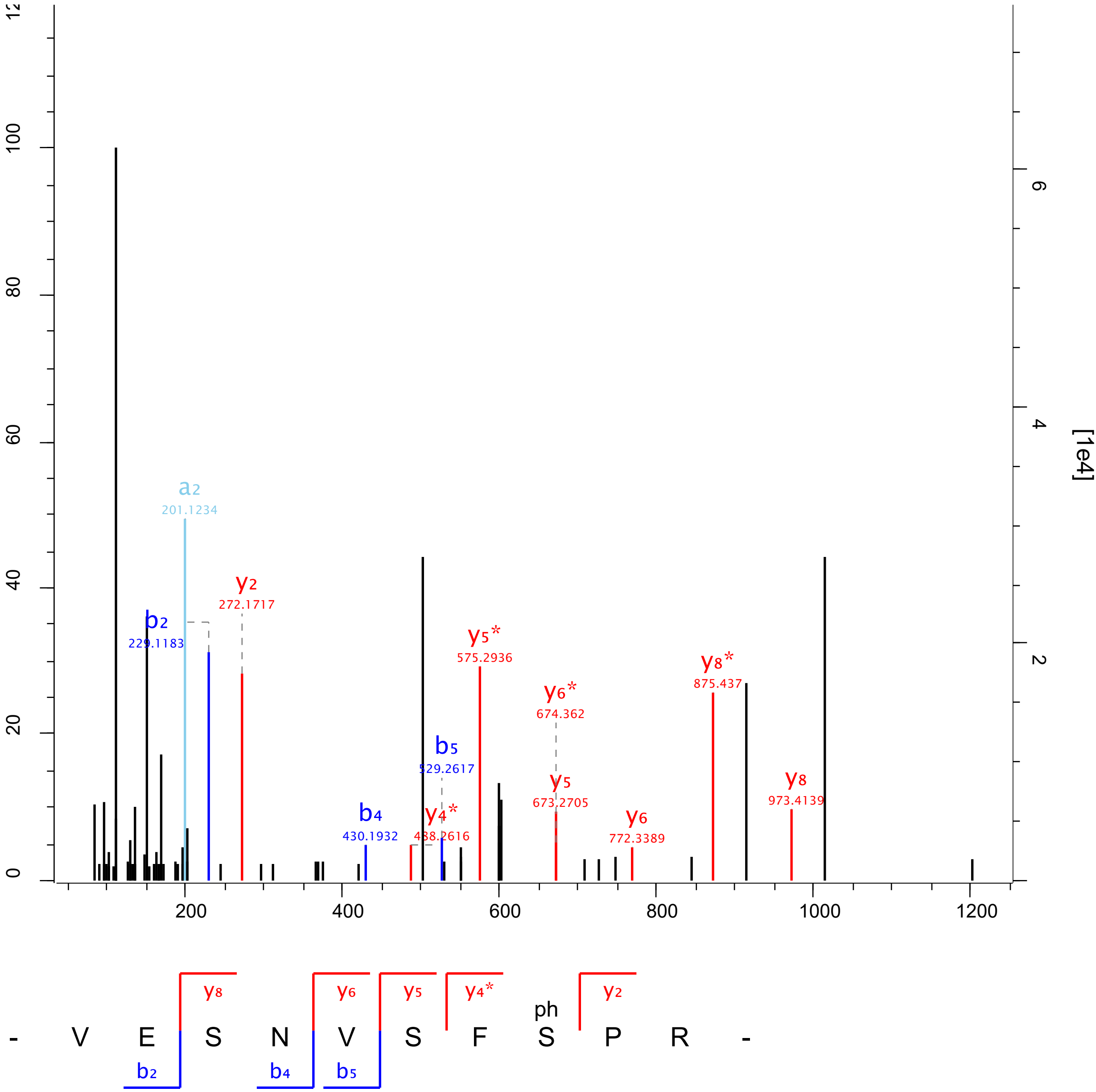

| Raw file | Scan | Method | Score | m/z |
|---|---|---|---|---|
| sys_02_2 | 1498 | FTMS; HCD | 43.03 | 427.16 |

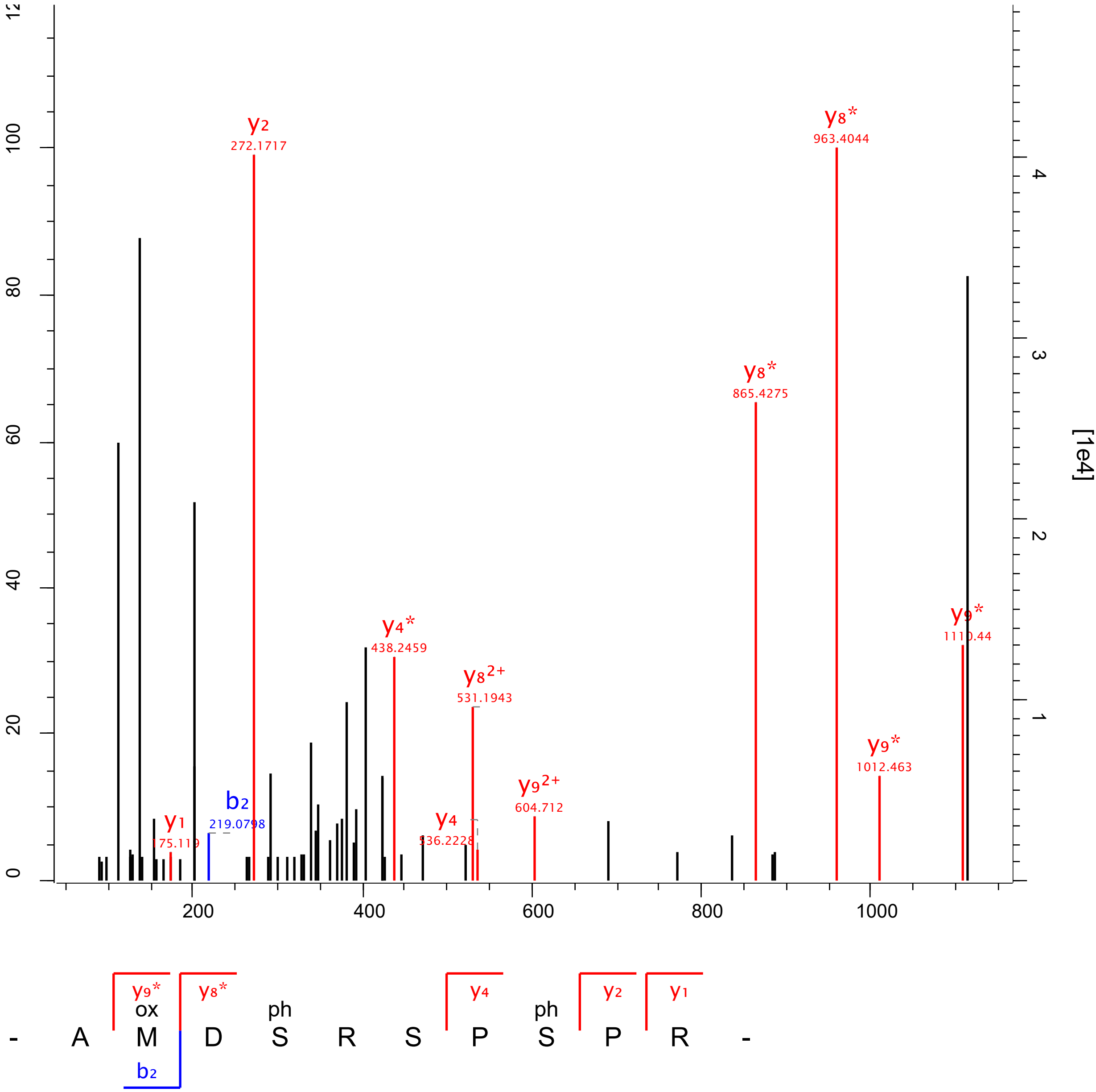

| Raw file | Scan | Method | Score | m/z |
| --- | --- | --- | --- | --- |
| sys_02_2 | 15114 | FTMS; HCD | 74.12 | 685.67 |

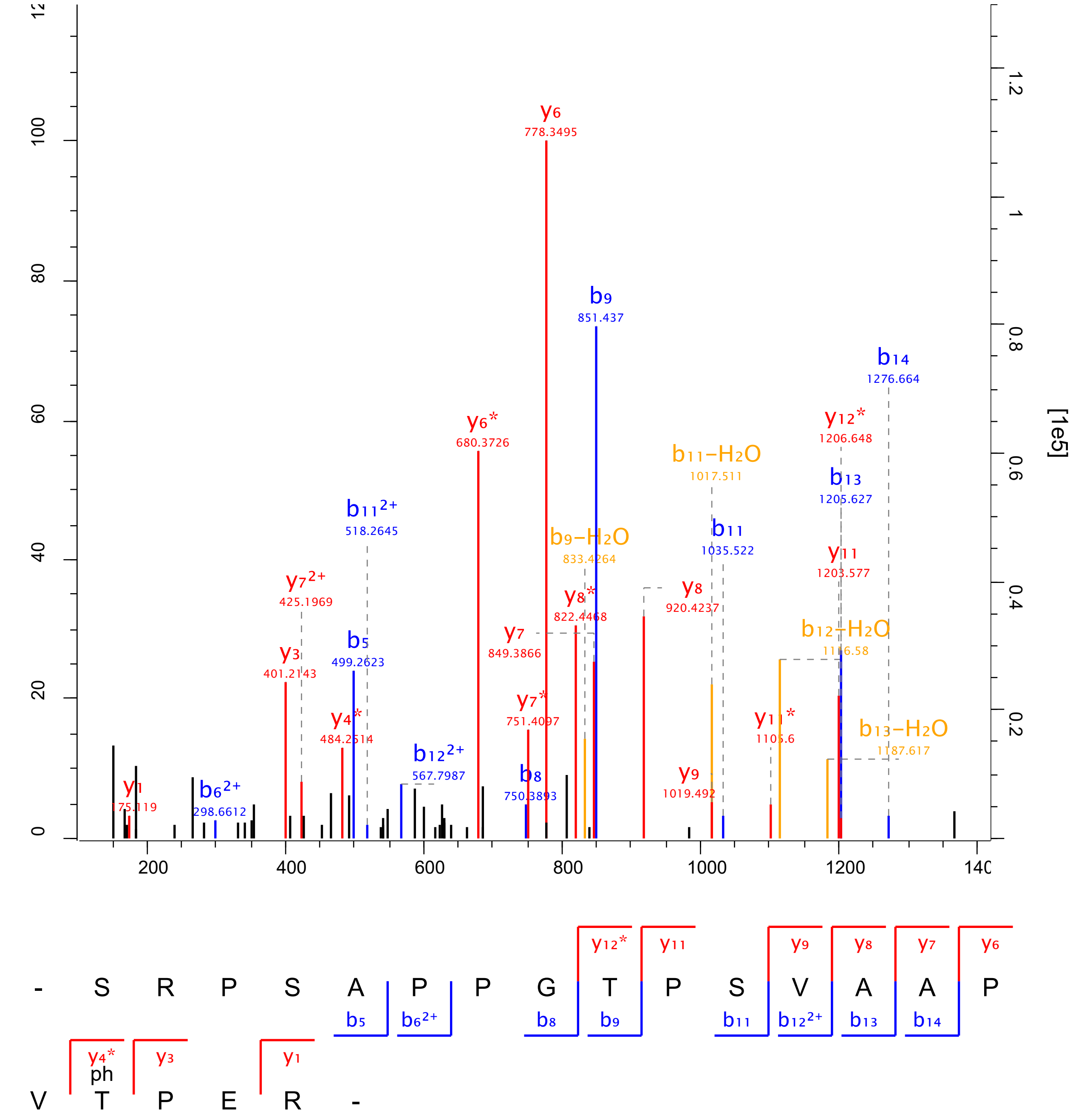

| Raw file | Scan | Method | Score | m/z |
|---|---|---|---|---|
| sys_02_2 | 15423 | FTMS; HCD | 160.27 | 584.95 |

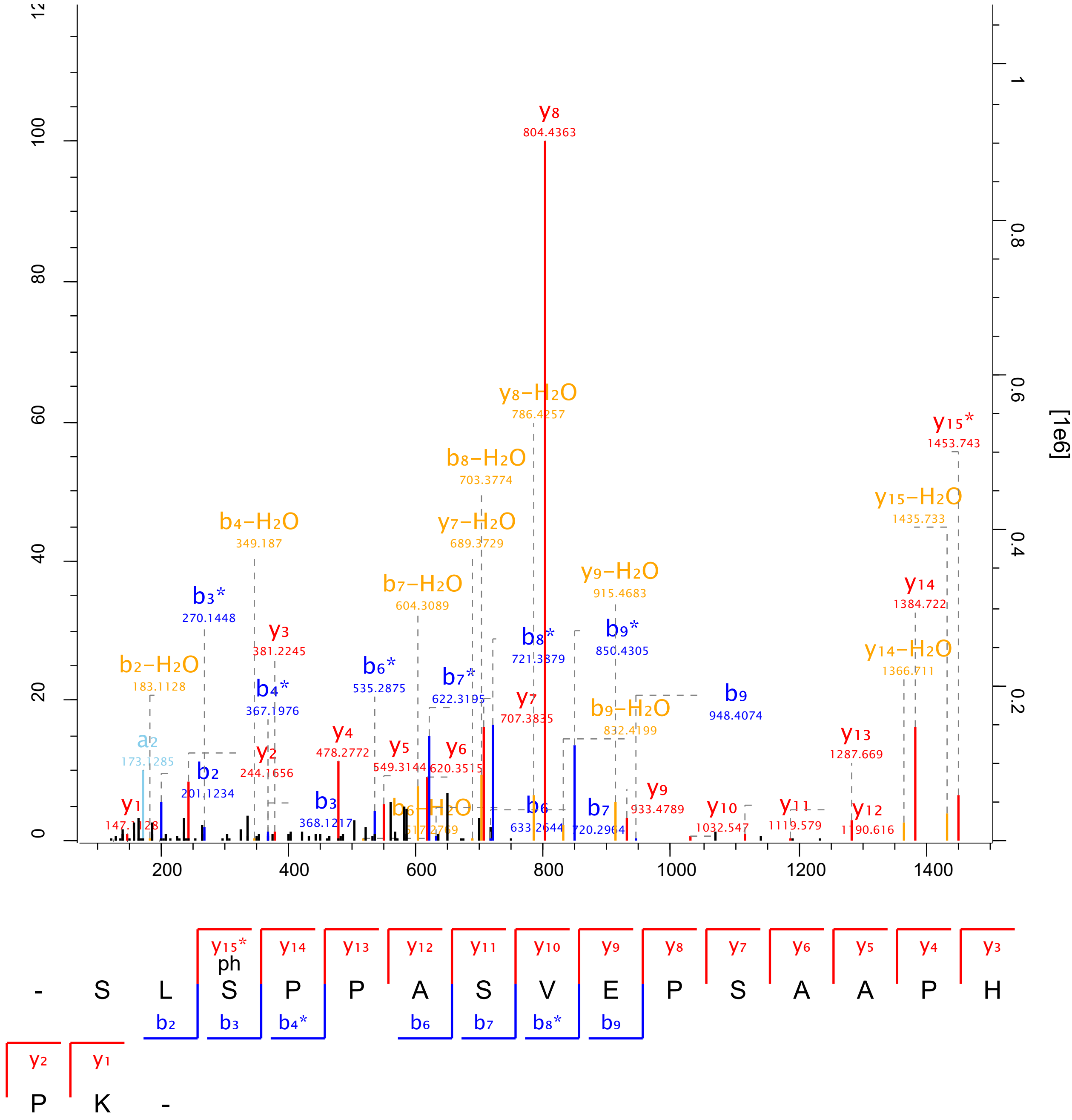

| Raw file | Scan | Method | Score | m/z |
|---|---|---|---|---|
| sys_02_2 | 1550 | FTMS; HCD | 68.41 | 462.23 |

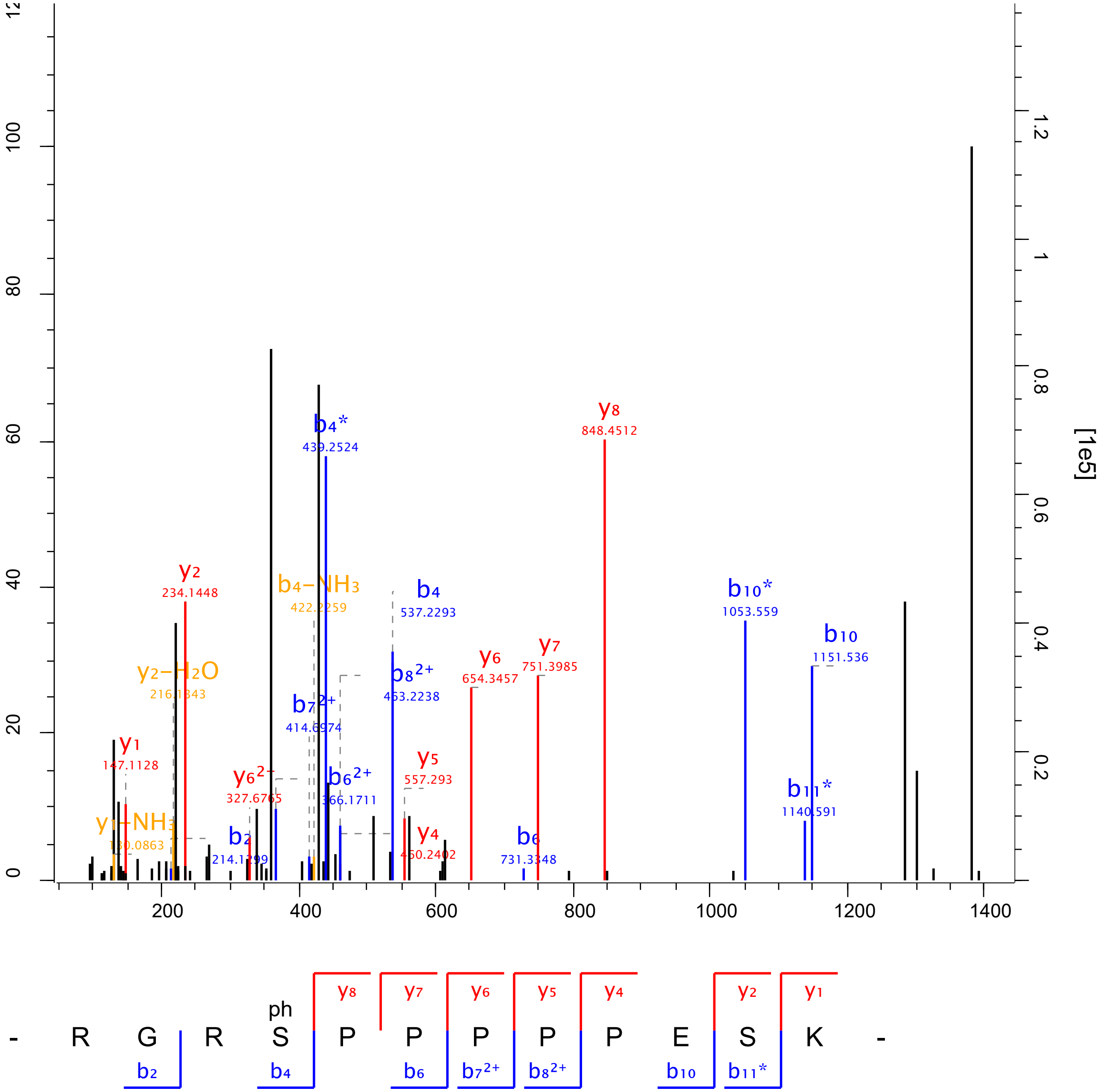

| Raw file | Scan | Method | Score | m/z |
| --- | --- | --- | --- | --- |
| sys_02_2 | 15620 | FTMS; HCD | 69.48 | 530.24 |

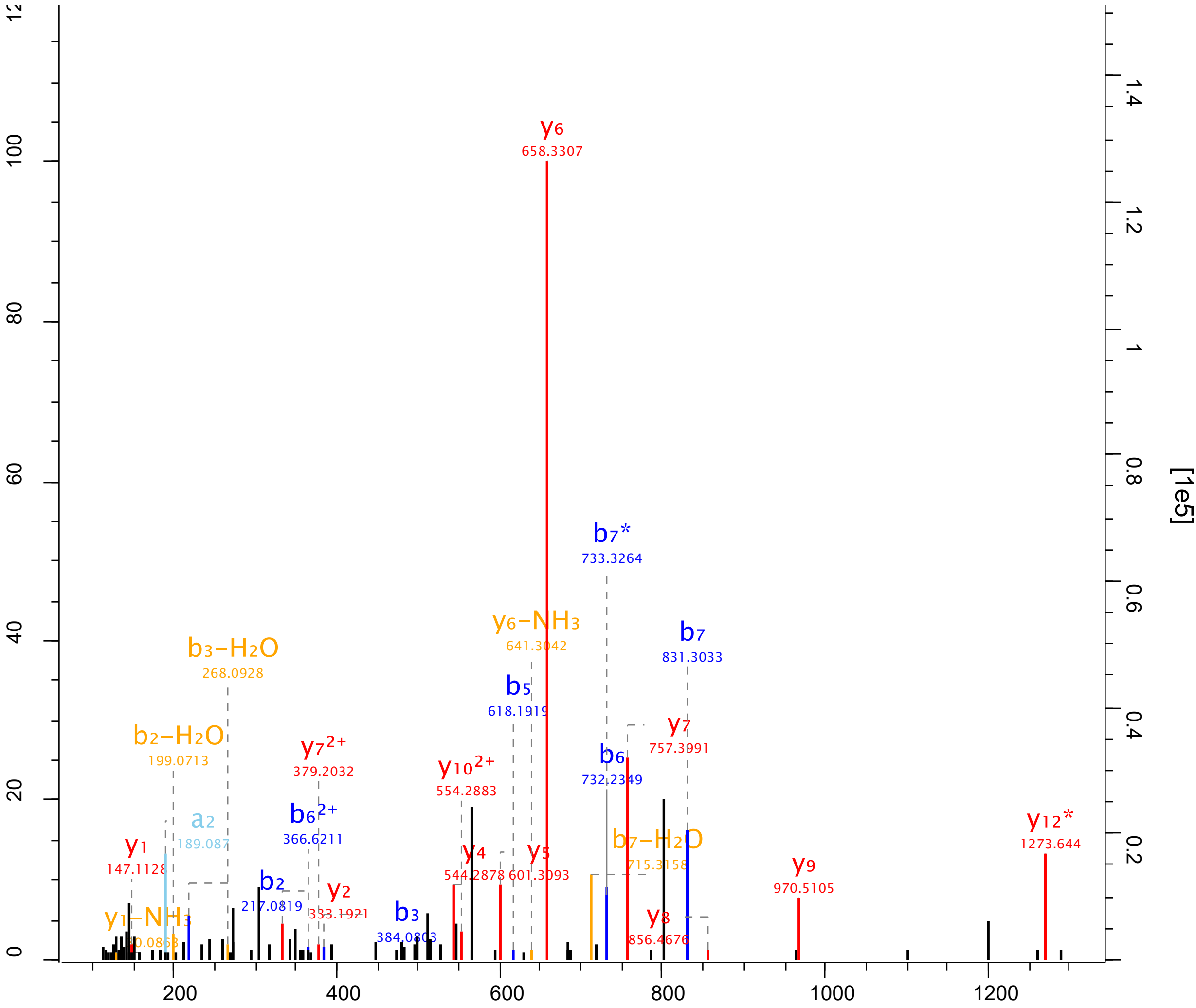


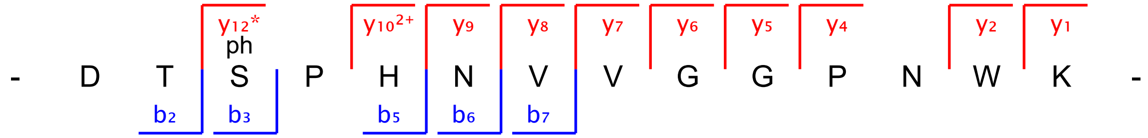

| Raw file | Scan | Method | Score | m/z |
|---|---|---|---|---|
| sys_02_2 | 15637 | FTMS; HCD | 80.86 | 790.81 |

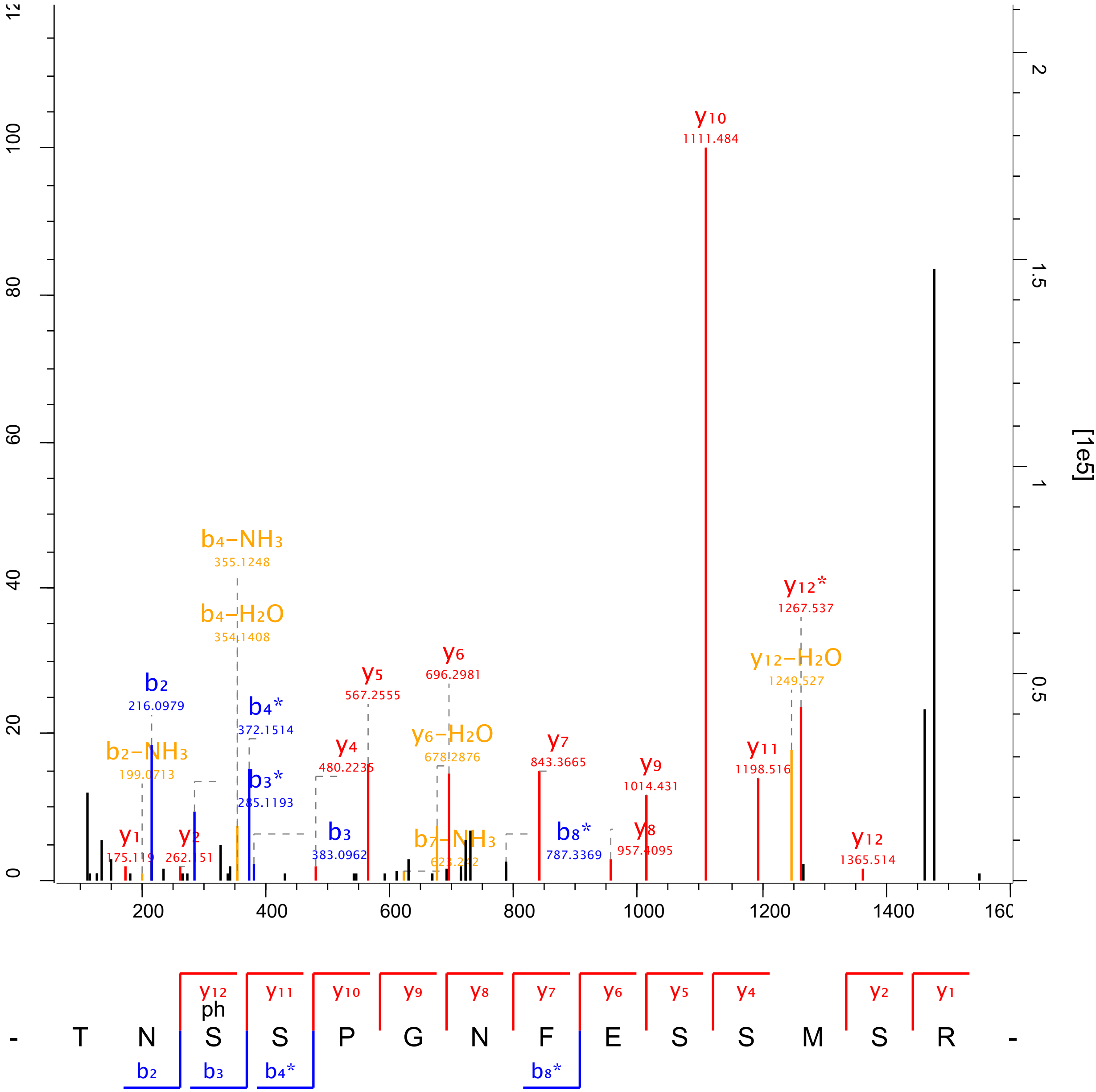

| Raw file | Scan | Method | Score | m/z |
|---|---|---|---|---|
| sys_02_2 | 15819 | FTMS; HCD | 46.41 | 533.2 |

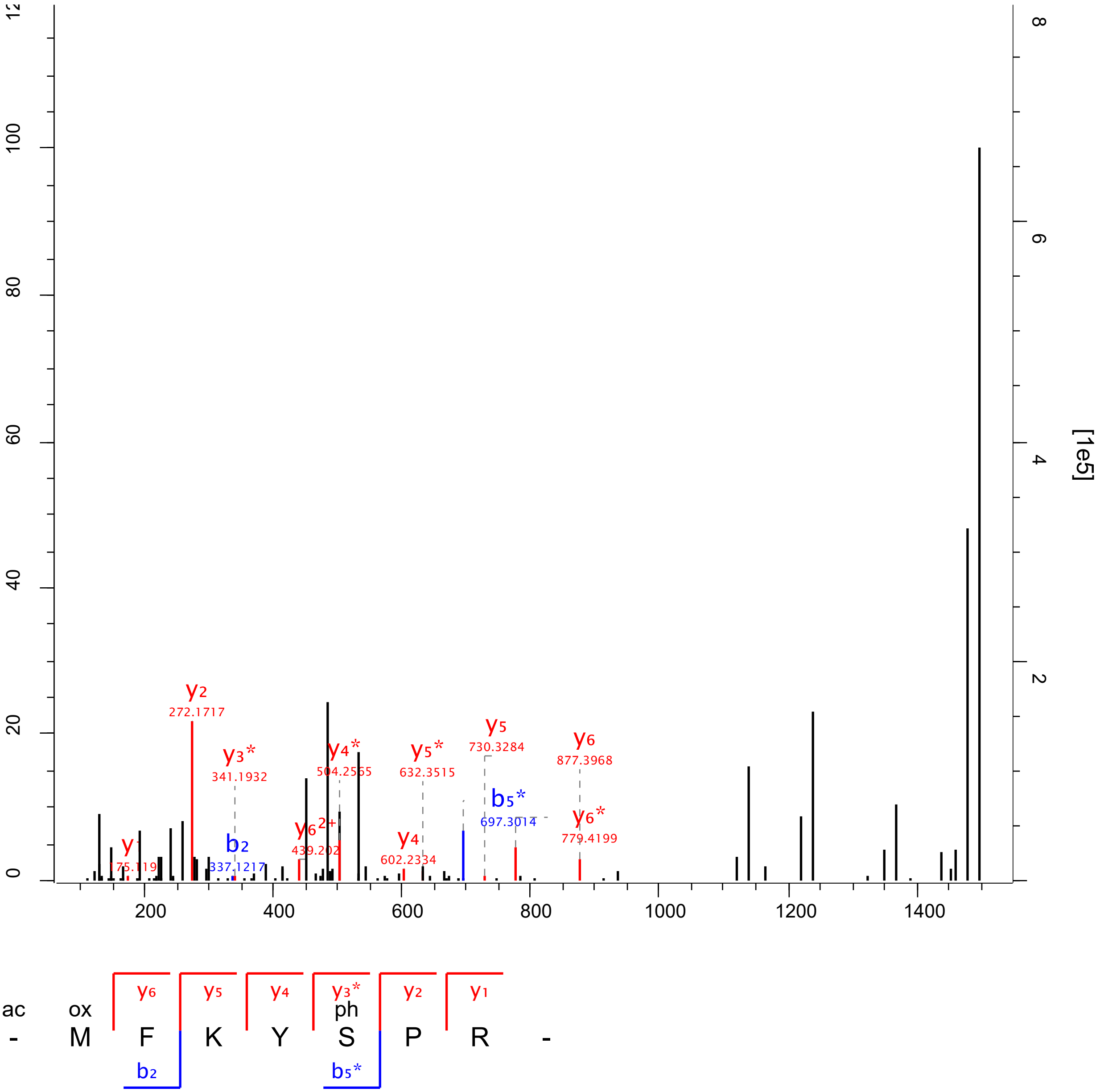

| Raw file | Scan | Method | Score | m/z |
| --- | --- | --- | --- | --- |
| sys_02_2 | 15980 | FTMS; HCD | 66.02 | 675.28 |

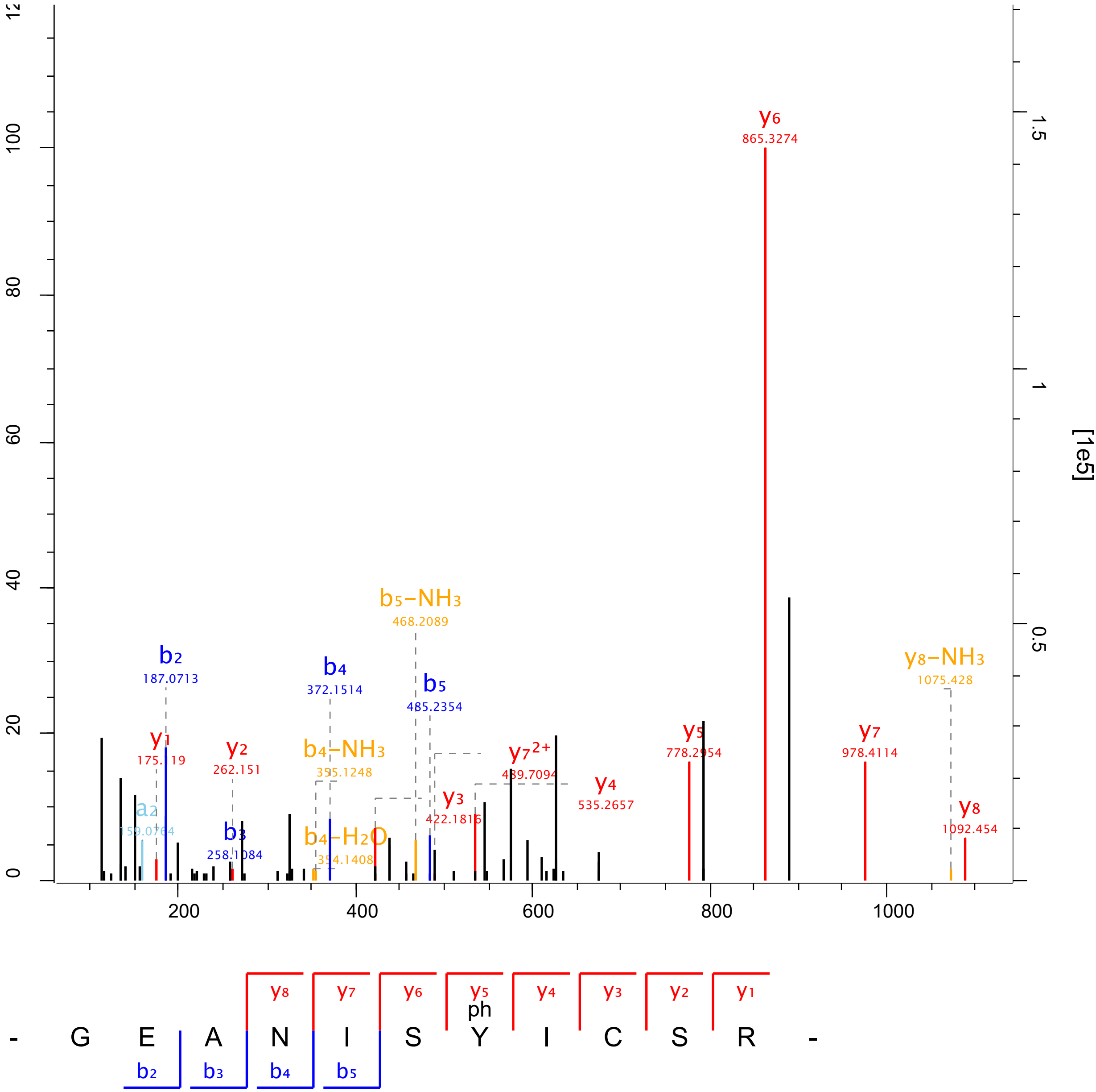

| Raw file | Scan | Method | Score | m/z |
|---|---|---|---|---|
| sys_02_2 | 16047 | FTMS; HCD | 135.23 | 620.79 |

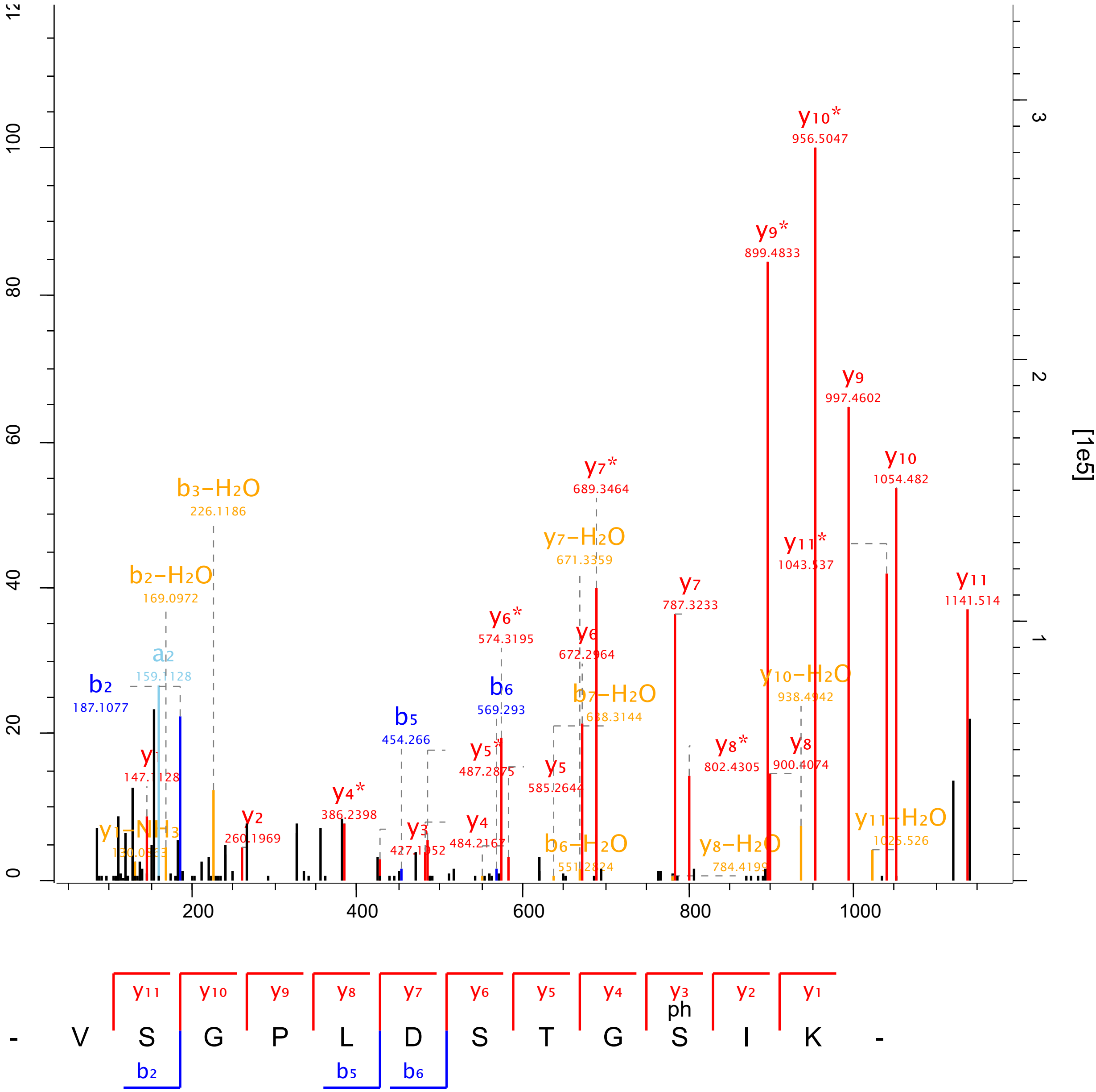

| Raw file | Scan | Method | Score | m/z |
|---|---|---|---|---|
| sys_02_2 | 16060 | FTMS; HCD | 98.06 | 507.59 |

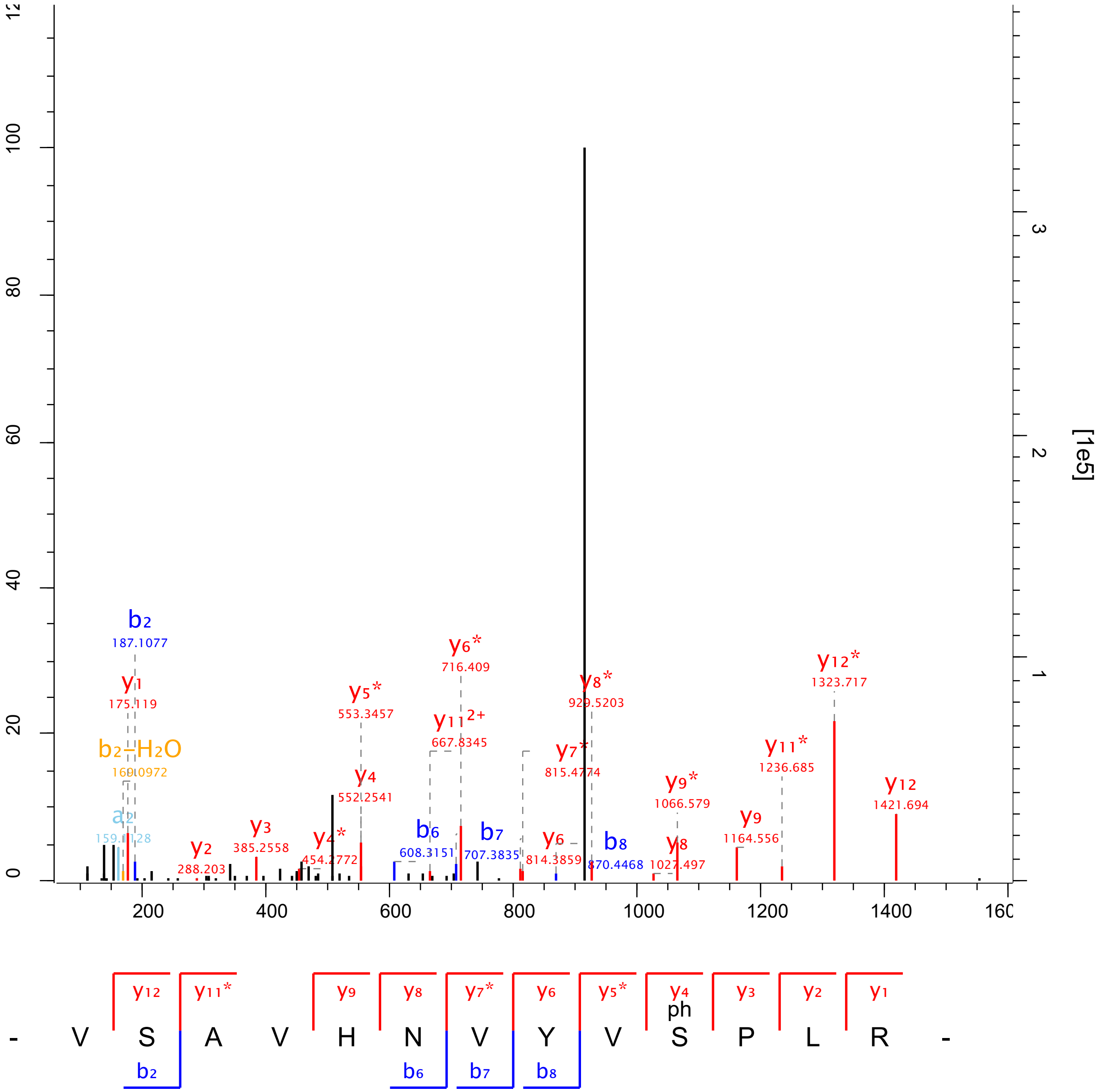

| Raw file | Scan | Method | Score | m/z |
|---|---|---|---|---|
| sys_02_2 | 16108 | FTMS; HCD | 257.58 | 557.27 |

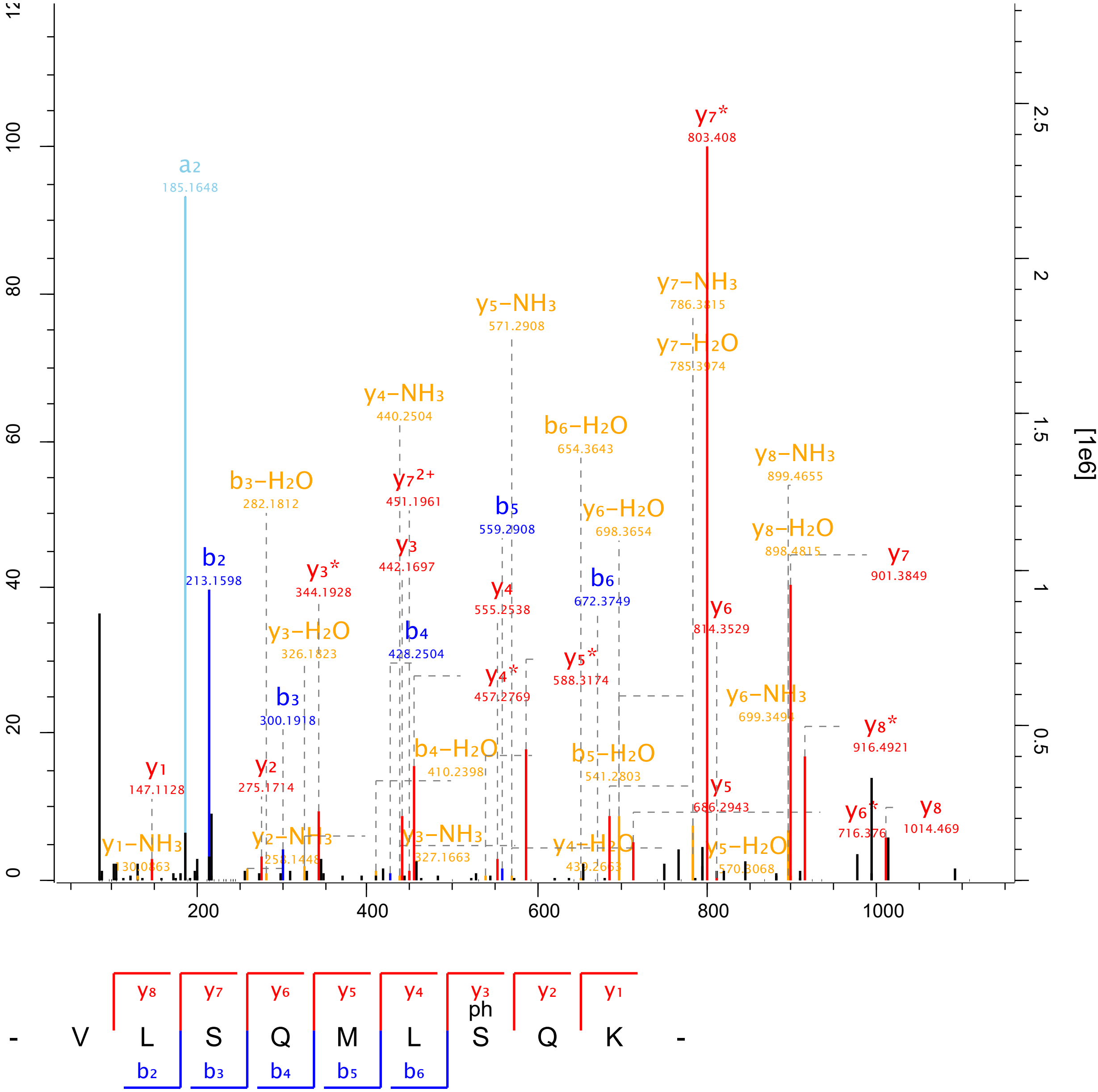

| Raw file | Scan | Method | Score | m/z |
|---|---|---|---|---|
| sys_02_2 | 16141 | FTMS; HCD | 98.94 | 540.76 |

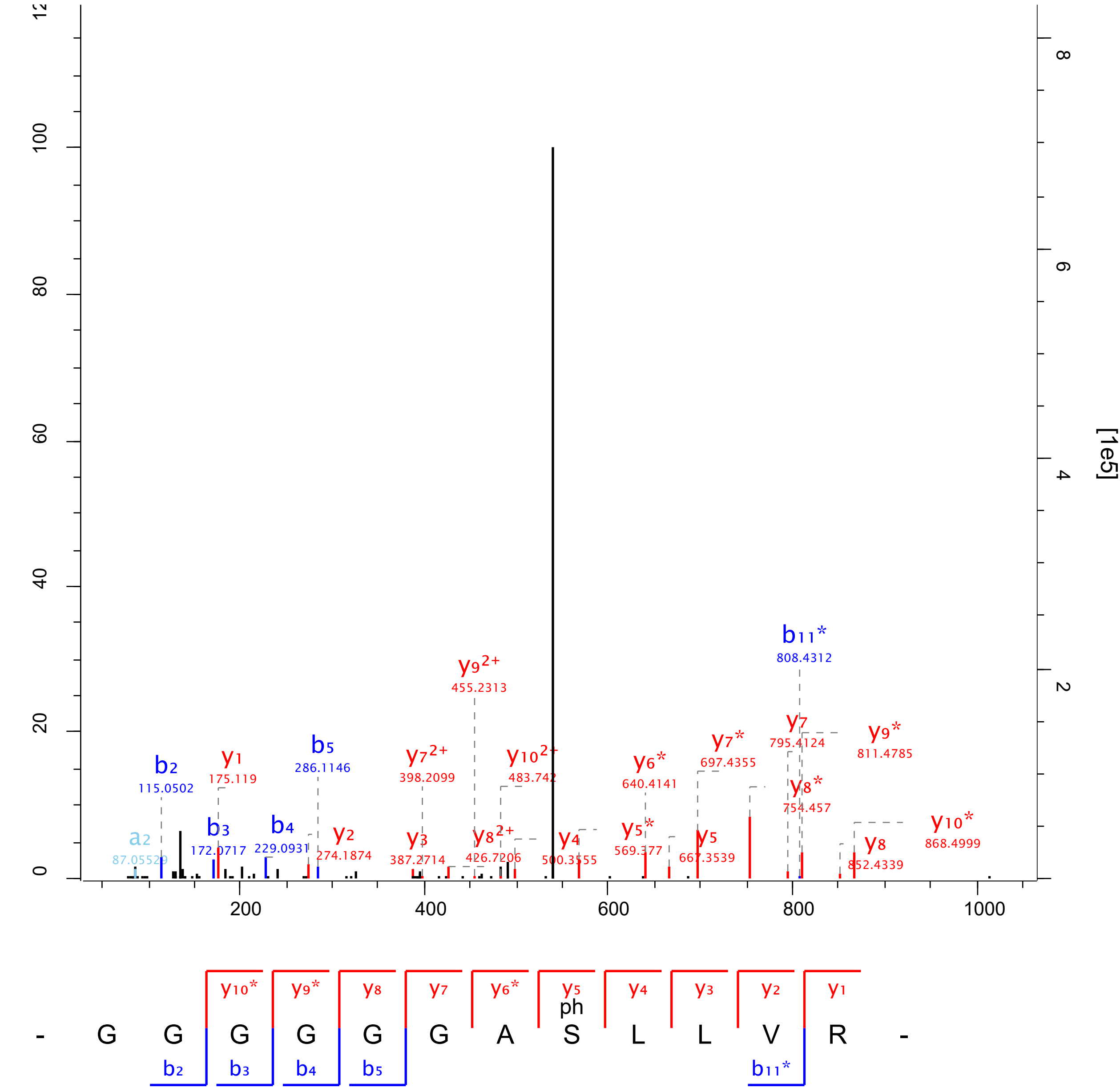

| Raw file | Scan | Method | Score | m/z |
| --- | --- | --- | --- | --- |
| sys_02_2 | 16209 | FTMS; HCD | 57.11 | 416.55 |

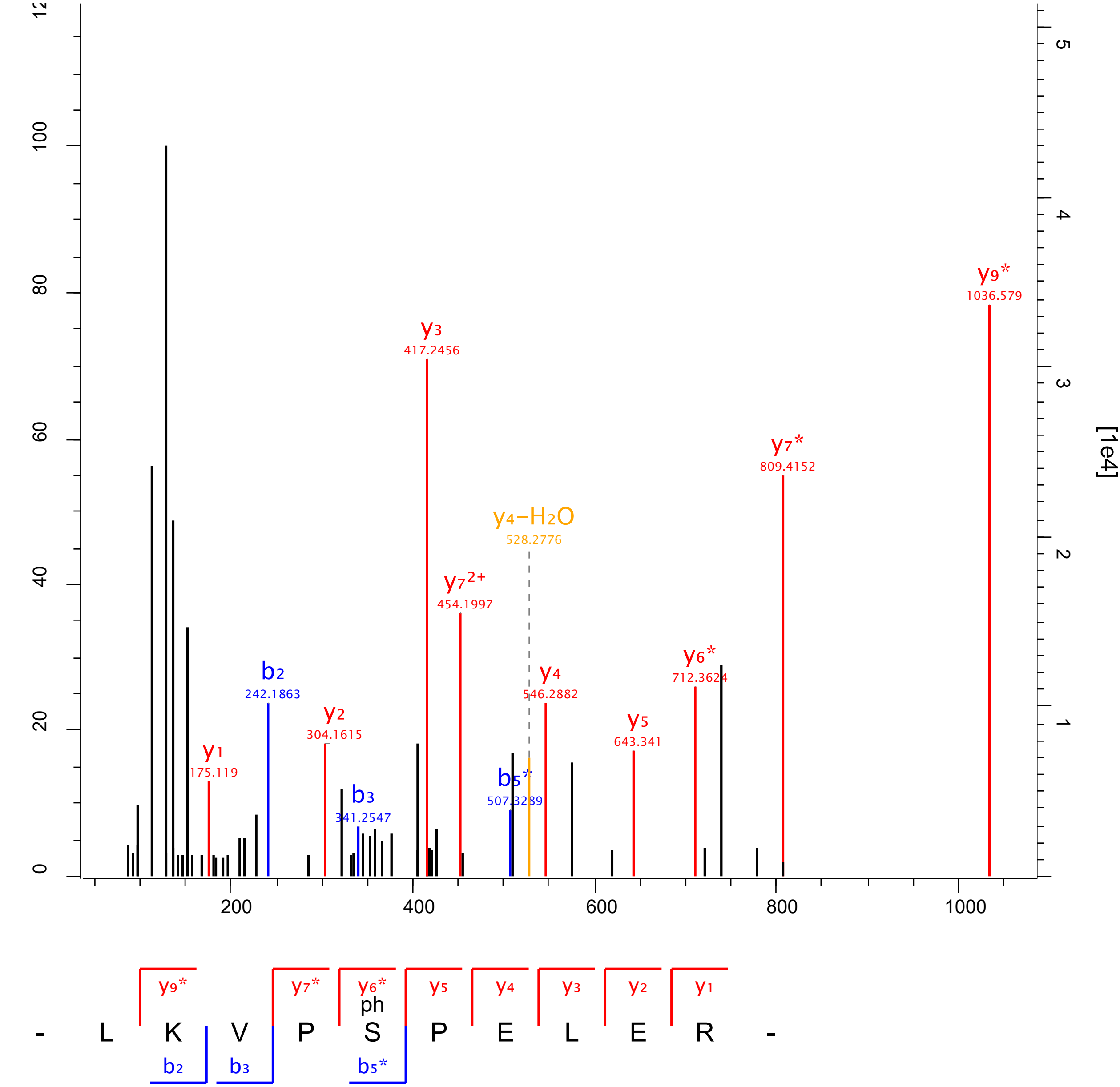

| Raw file | Scan | Method | Score | m/z |
| --- | --- | --- | --- | --- |
| sys_02_2 | 1630 | FTMS; HCD | 41.68 | 450.22 |

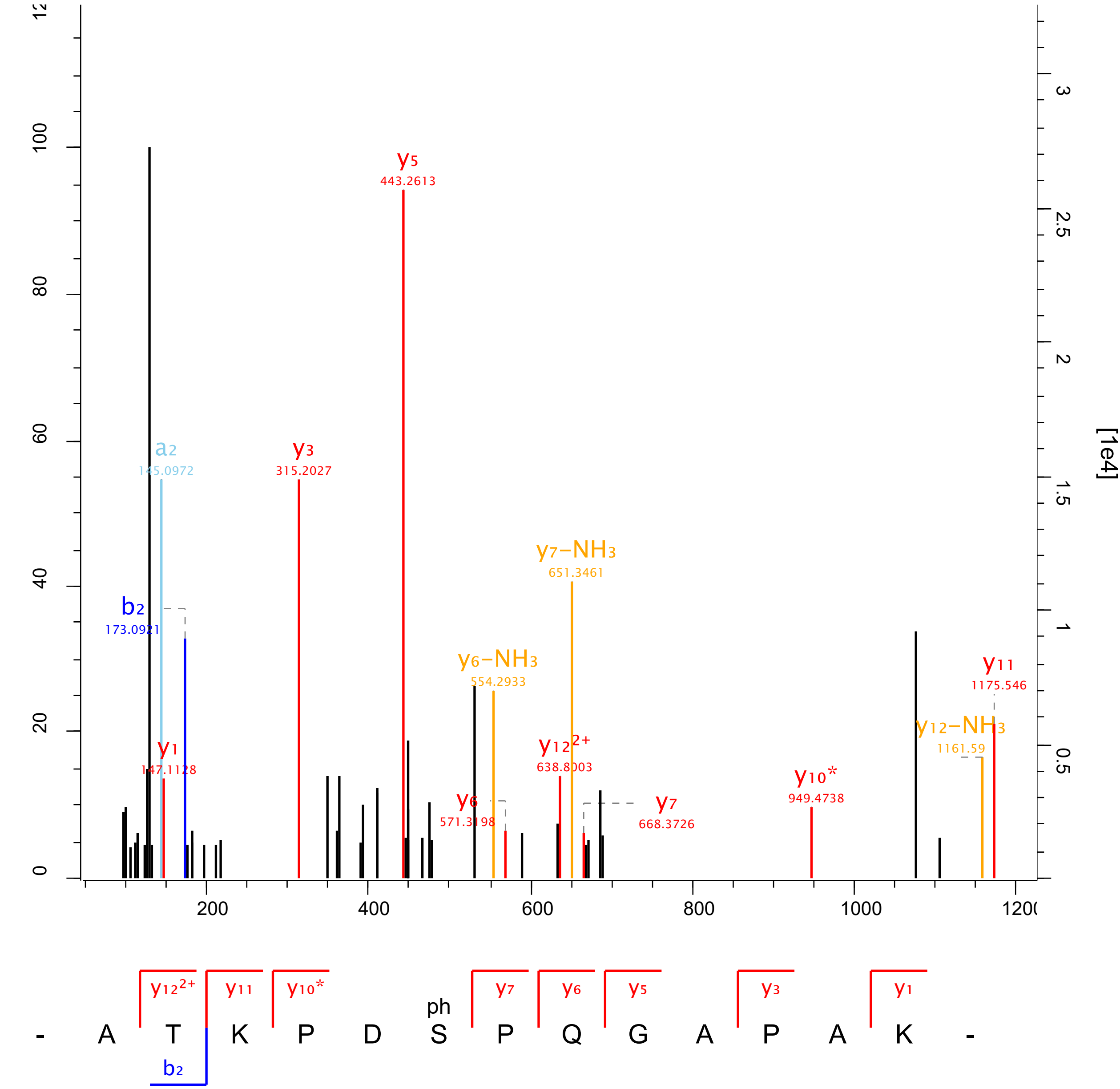

| Raw file | Scan | Method | Score | m/z |
| --- | --- | --- | --- | --- |
| sys_02_2 | 16381 | FTMS; HCD | 43.3 | 421.88 |

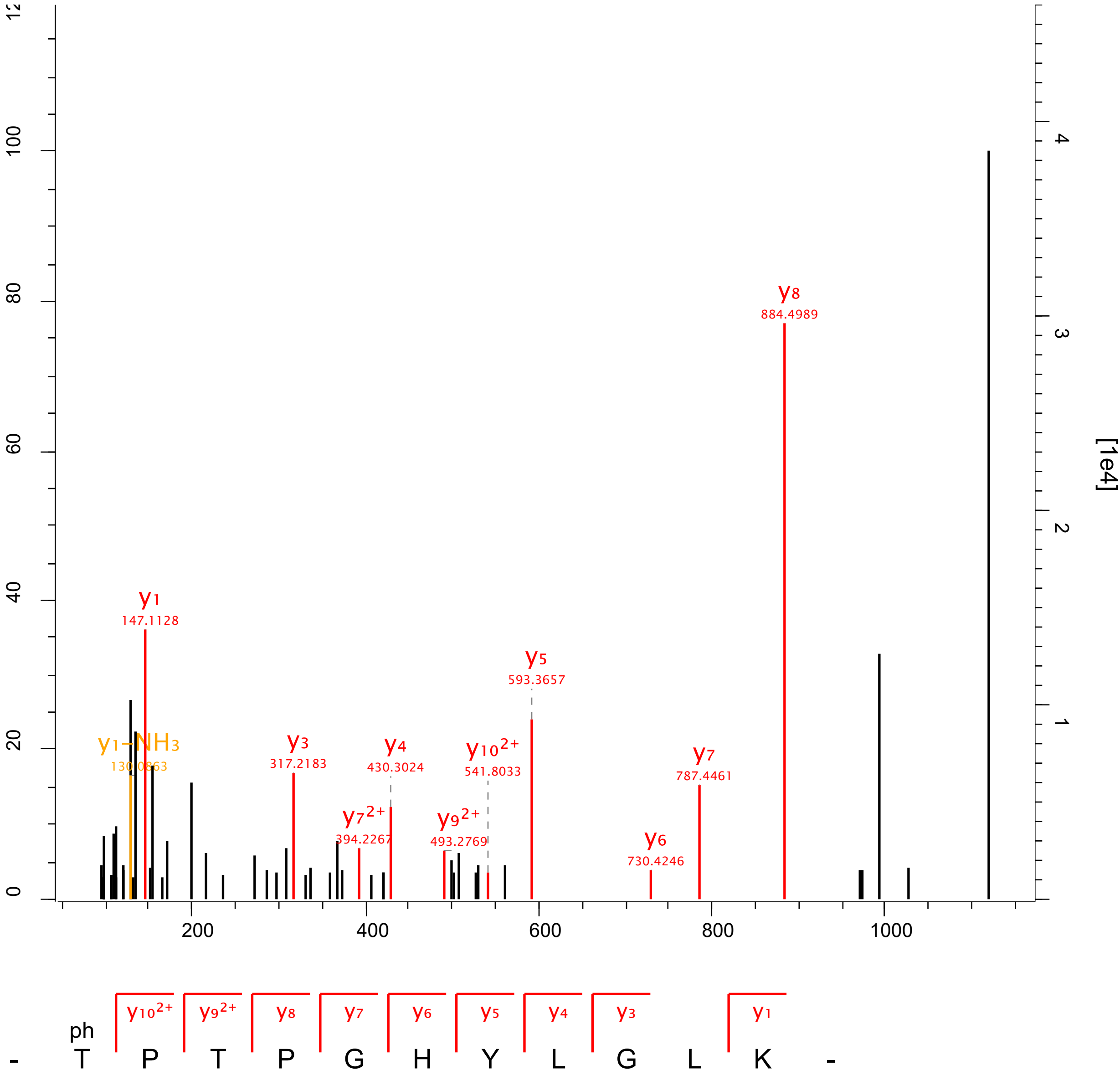

| Raw file | Scan | Method | Score | m/z |
|---|---|---|---|---|
| sys_02_2 | 16438 | FTMS; HCD | 81.66 | 727.82 |

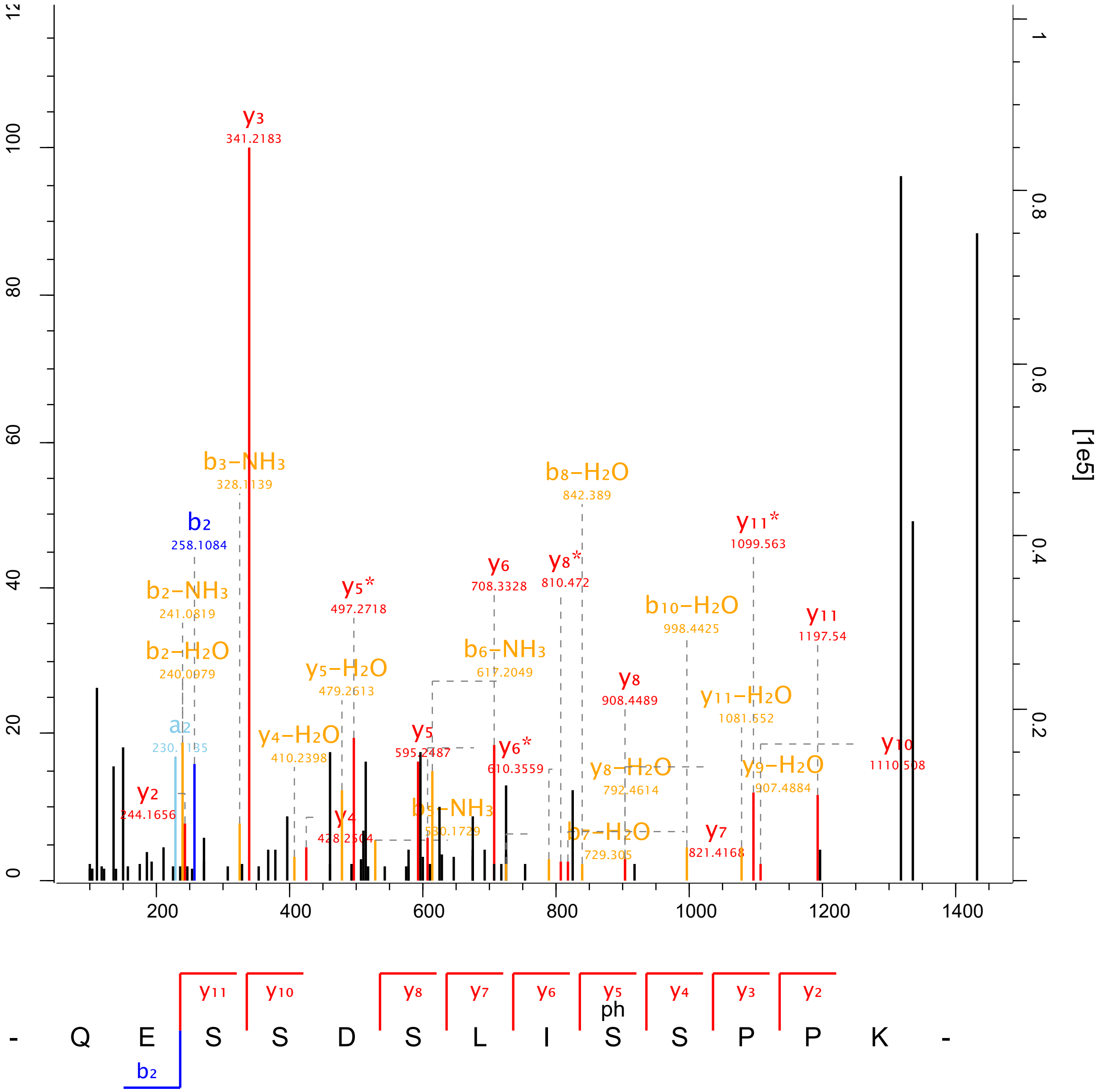

| Raw file | Scan | Method | Score | m/z |
|---|---|---|---|---|
| sys_02_2 | 16870 | FTMS; HCD | 102.65 | 672.79 |

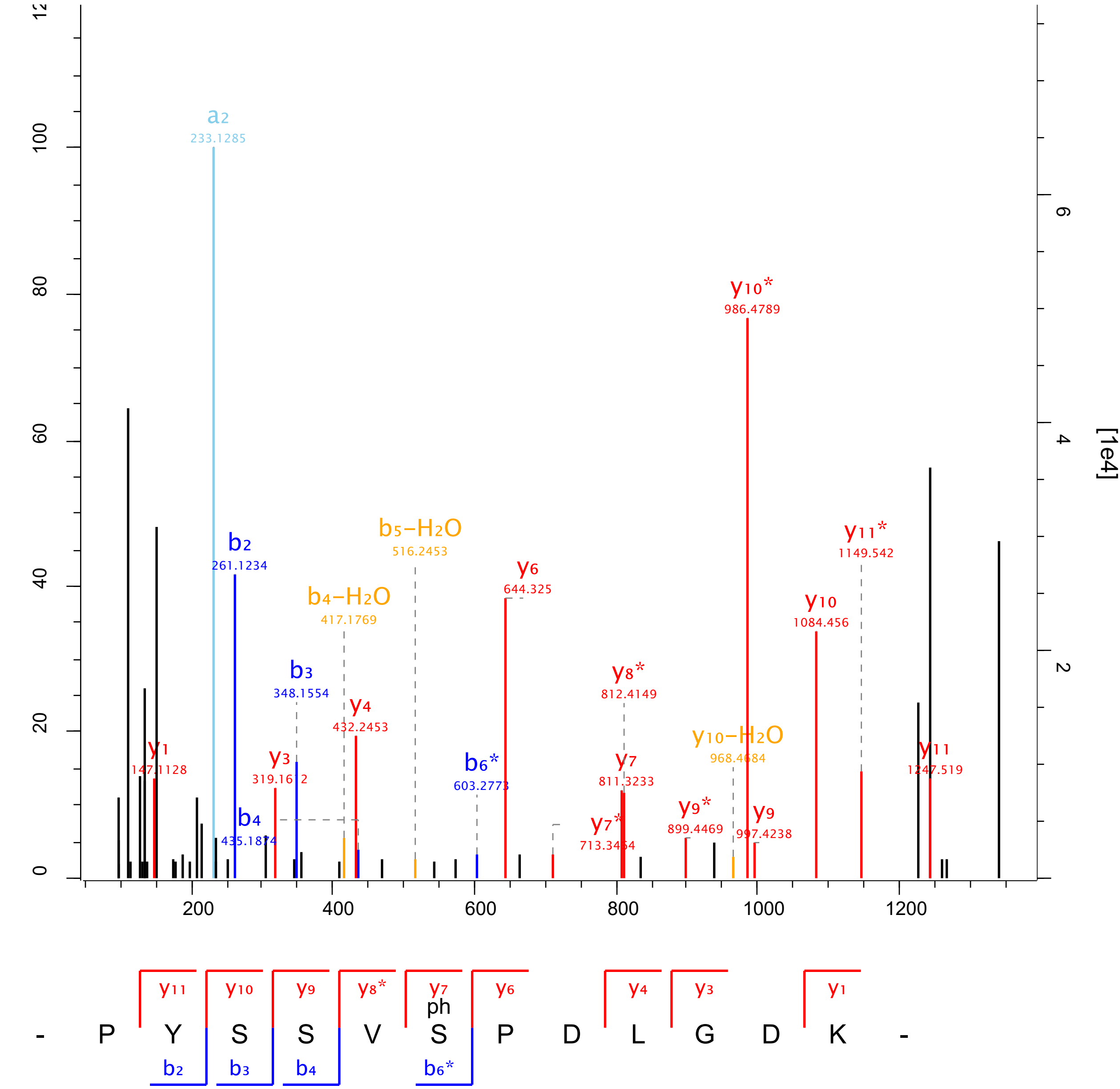

| Raw file | Scan | Method | Score | m/z |
| --- | --- | --- | --- | --- |
| sys_02_2 | 16985 | FTMS; HCD | 60.8 | 689.31 |

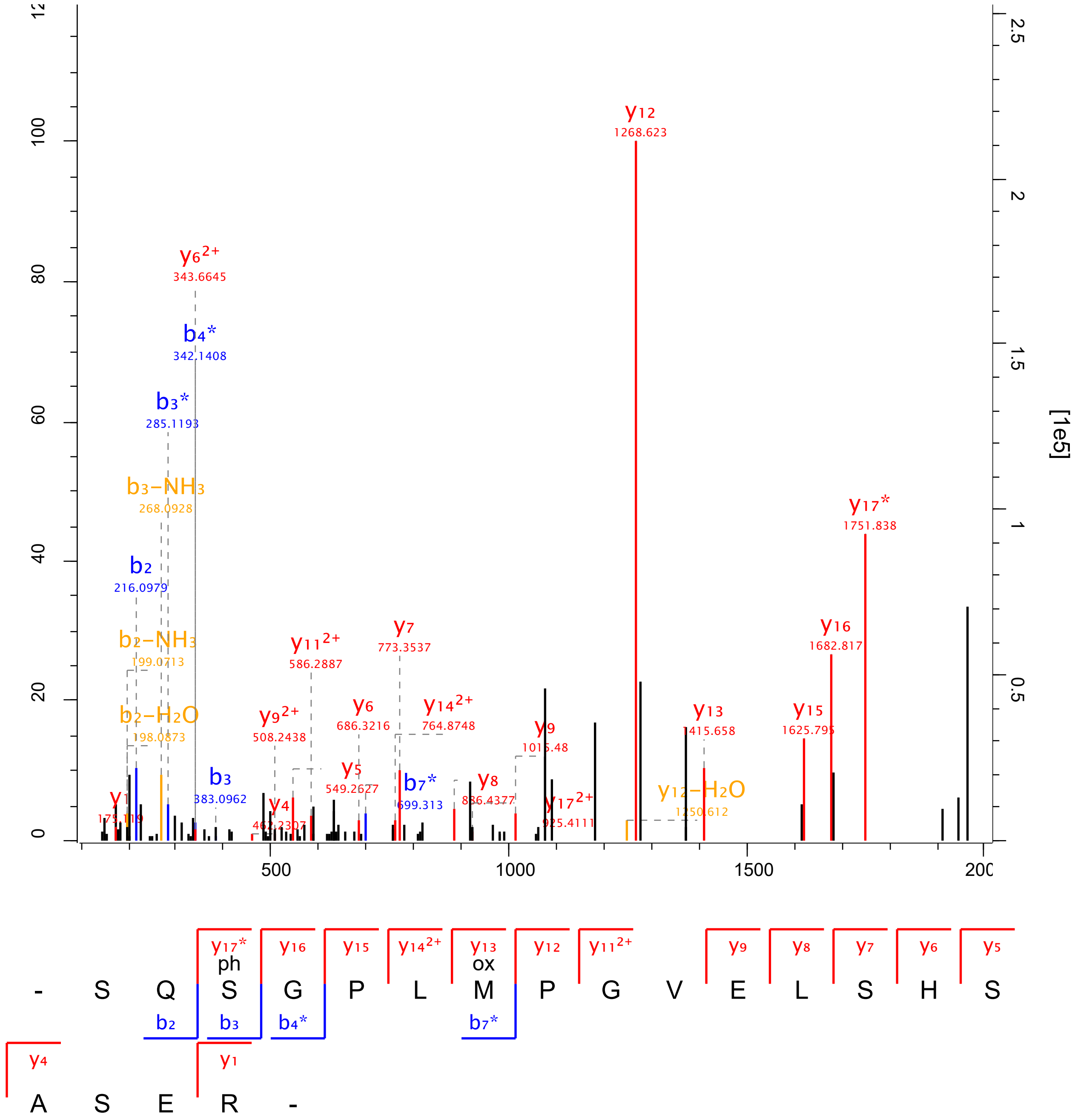

| Raw file | Scan | Method | Score | m/z |
|---|---|---|---|---|
| sys_02_2 | 17105 | FTMS; HCD | 80.36 | 712 |

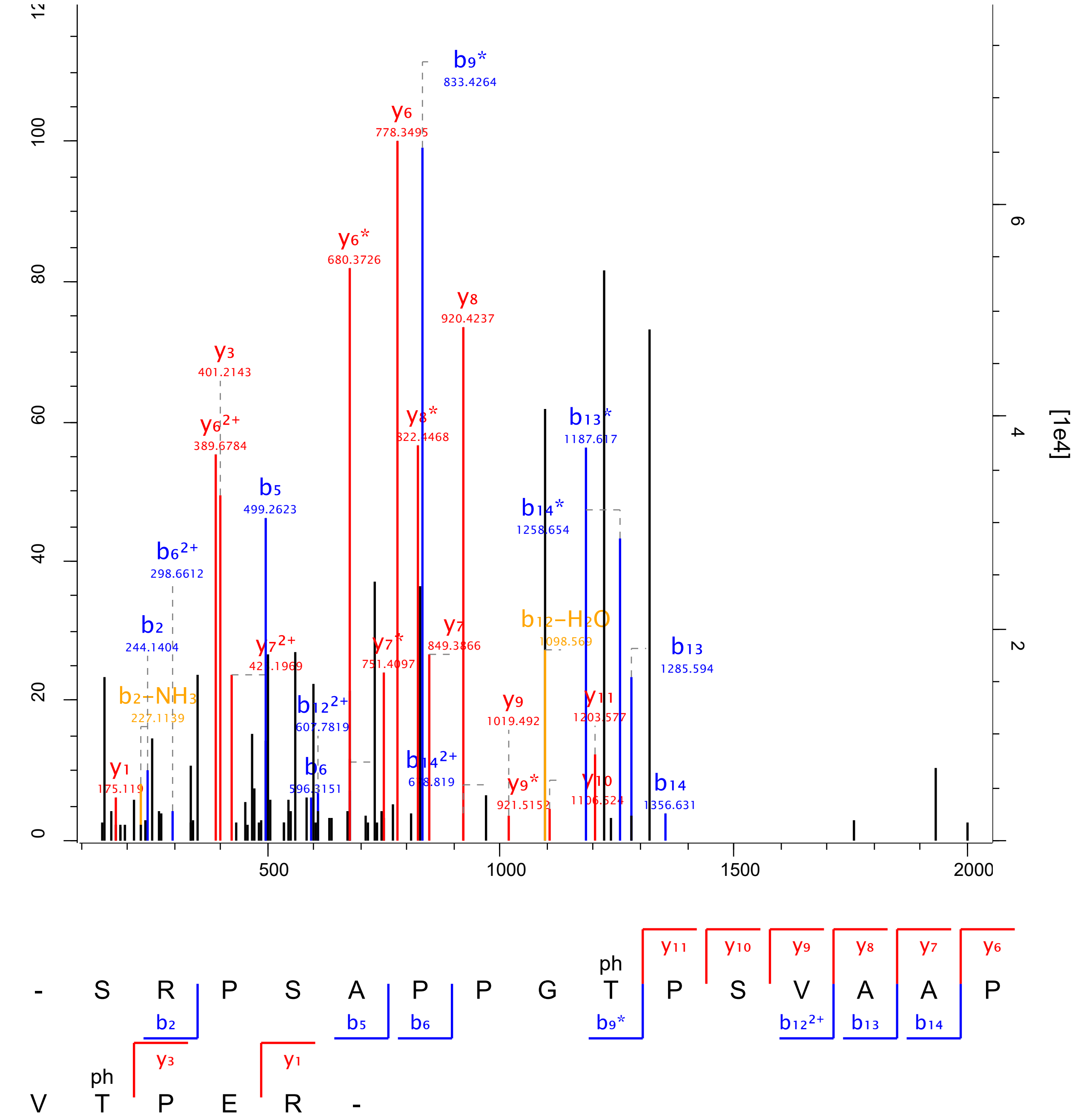

| Raw file | Scan | Method | Score | m/z |
|---|---|---|---|---|
| sys_02_2 | 17108 | FTMS; HCD | 144.2 | 574.26 |

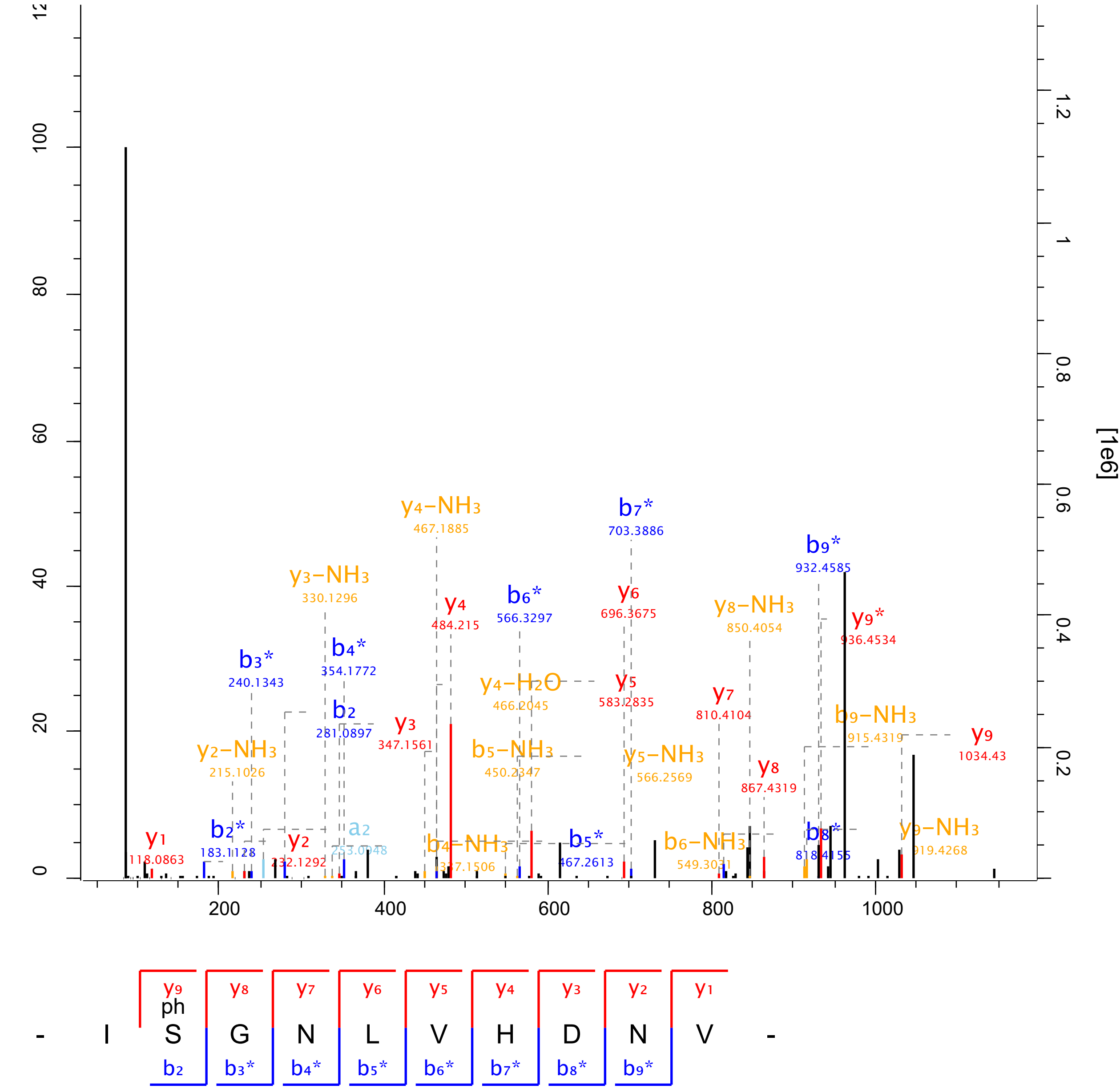

| Raw file | Scan | Method | Score | m/z |
|---|---|---|---|---|
| sys_02_2 | 17225 | FTMS; HCD | 73.5 | 635.26 |

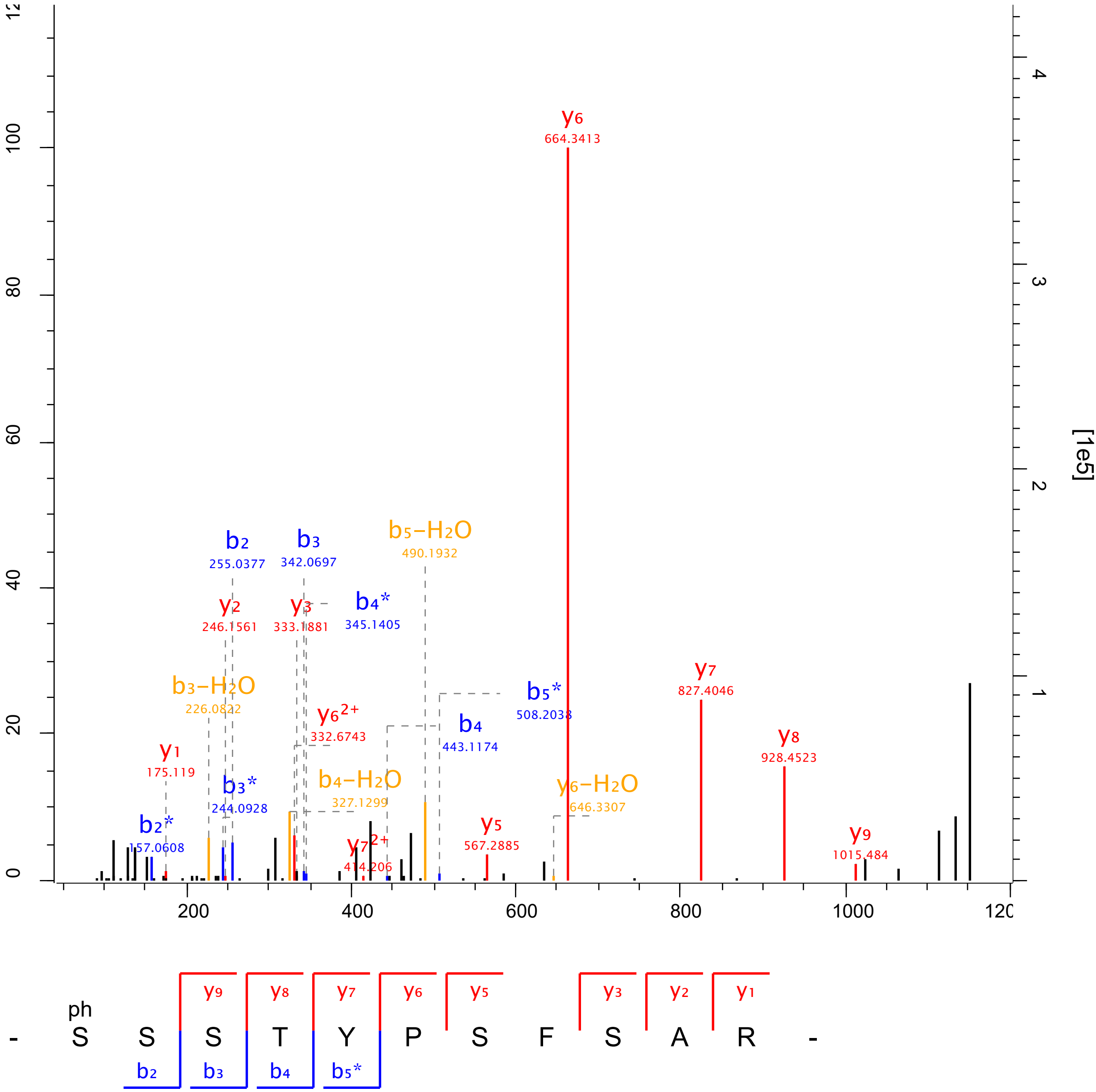

| Raw file | Scan | Method | Score | m/z |
|---|---|---|---|---|
| sys_02_2 | 17233 | FTMS; HCD | 137.64 | 913.9 |

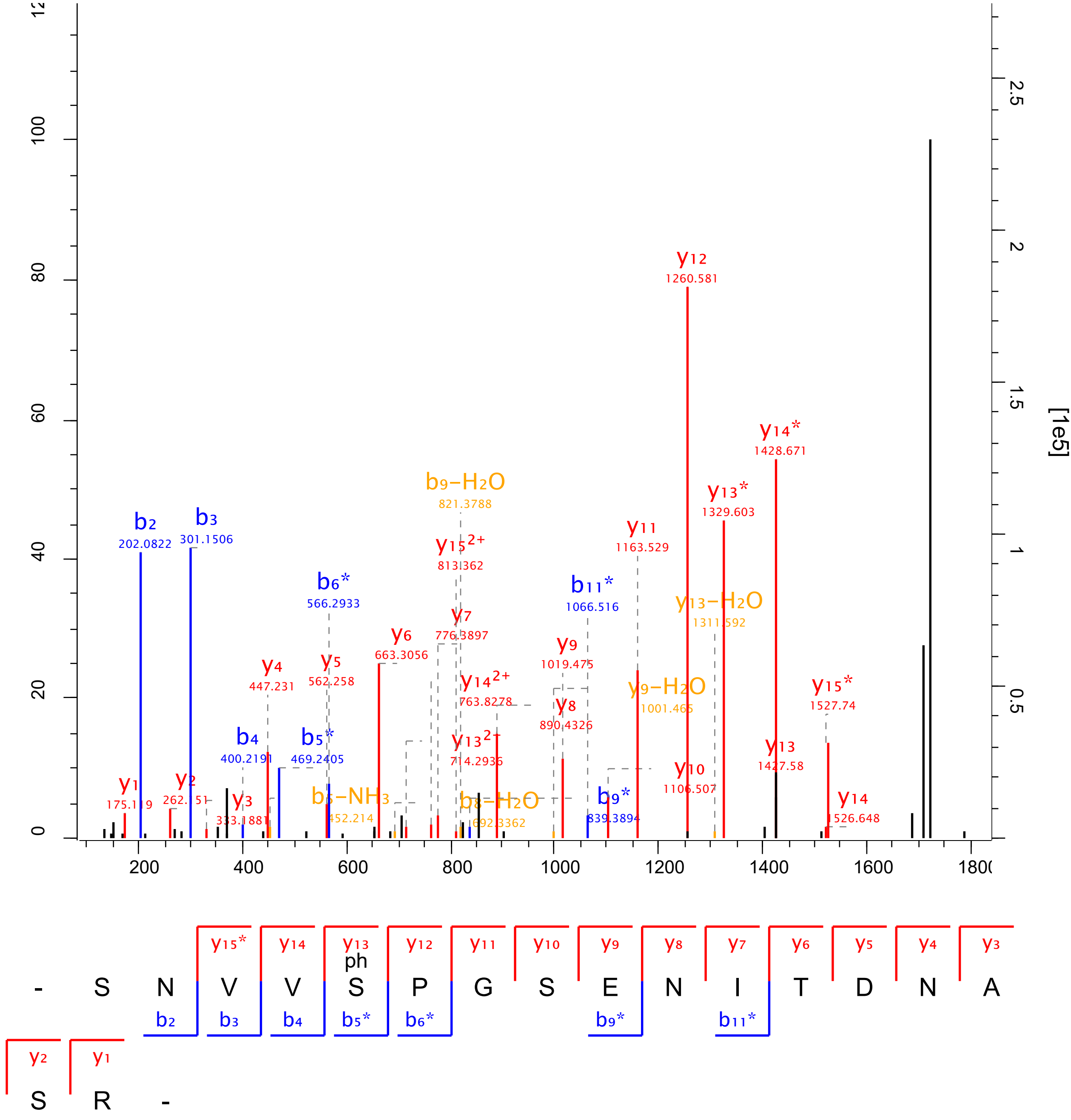

| Raw file | Scan | Method | Score | m/z |
|---|---|---|---|---|
| sys_02_2 | 17287 | FTMS; HCD | 260.97 | 865.85 |

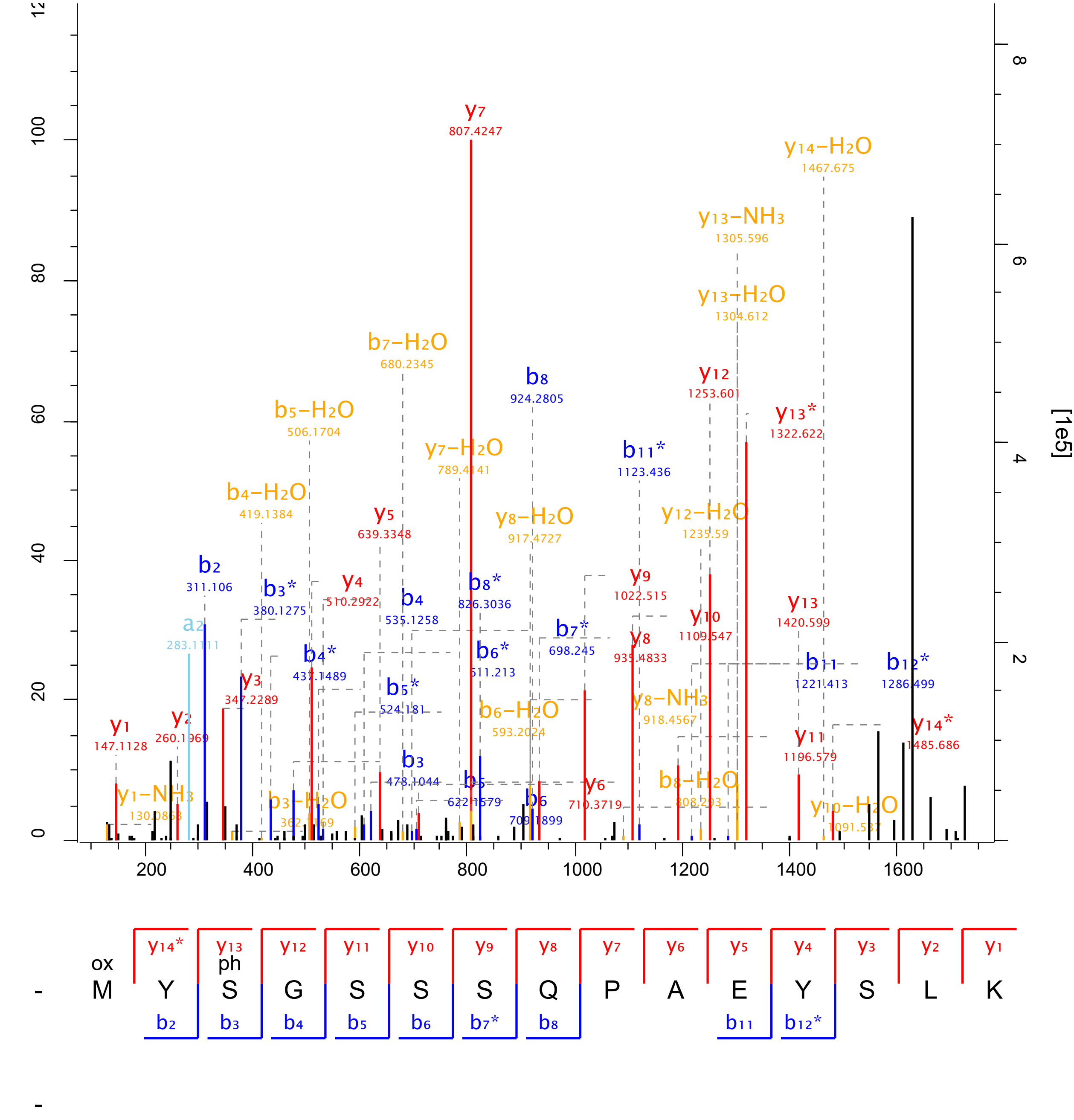

| Raw file | Scan | Method | Score | m/z |
| --- | --- | --- | --- | --- |
| sys_02_2 | 17409 | FTMS; HCD | 42.52 | 631.83 |

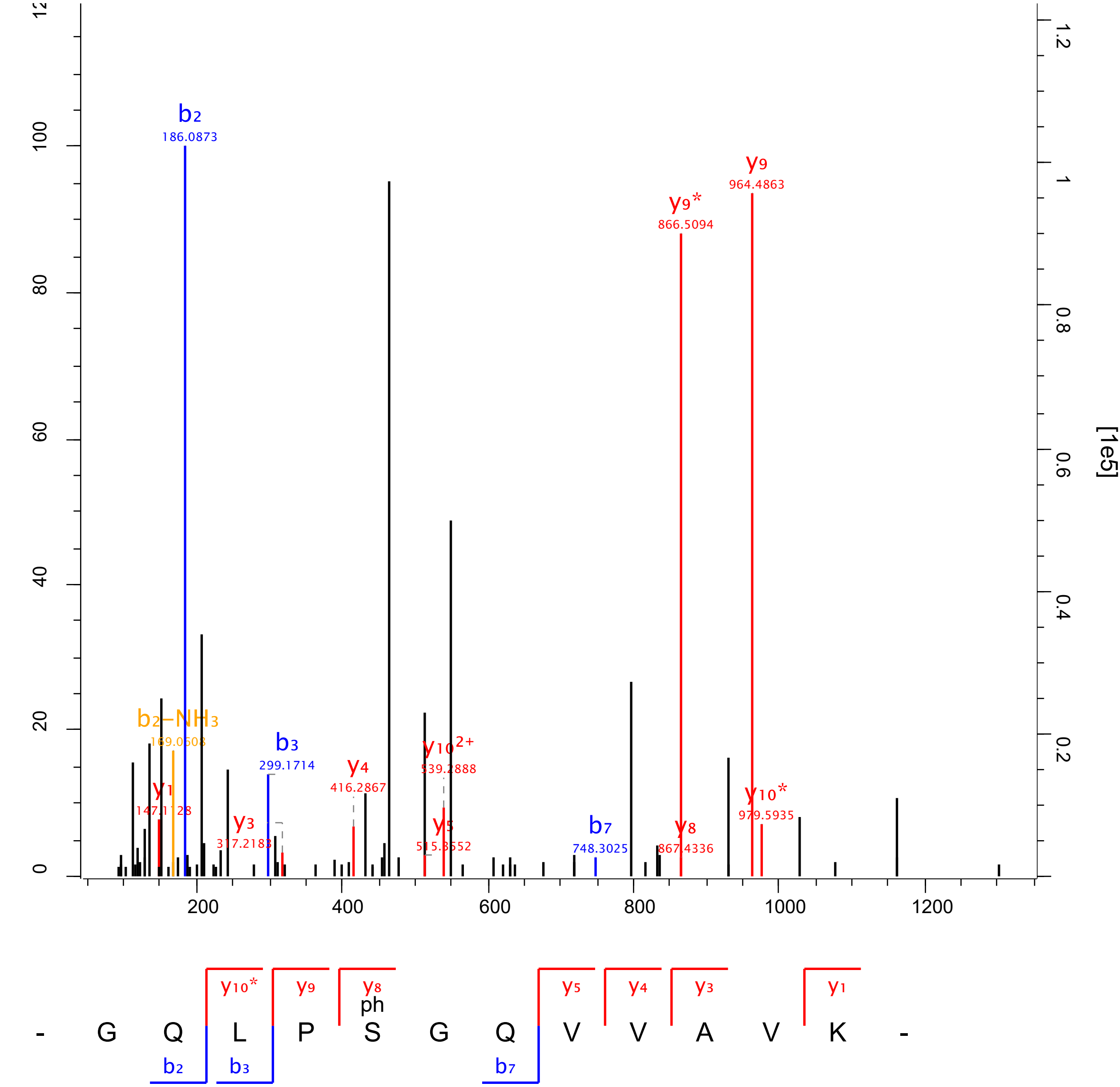

| Raw file | Scan | Method | Score | m/z |
|---|---|---|---|---|
| sys_02_2 | 17513 | FTMS; HCD | 171.08 | 607.78 |

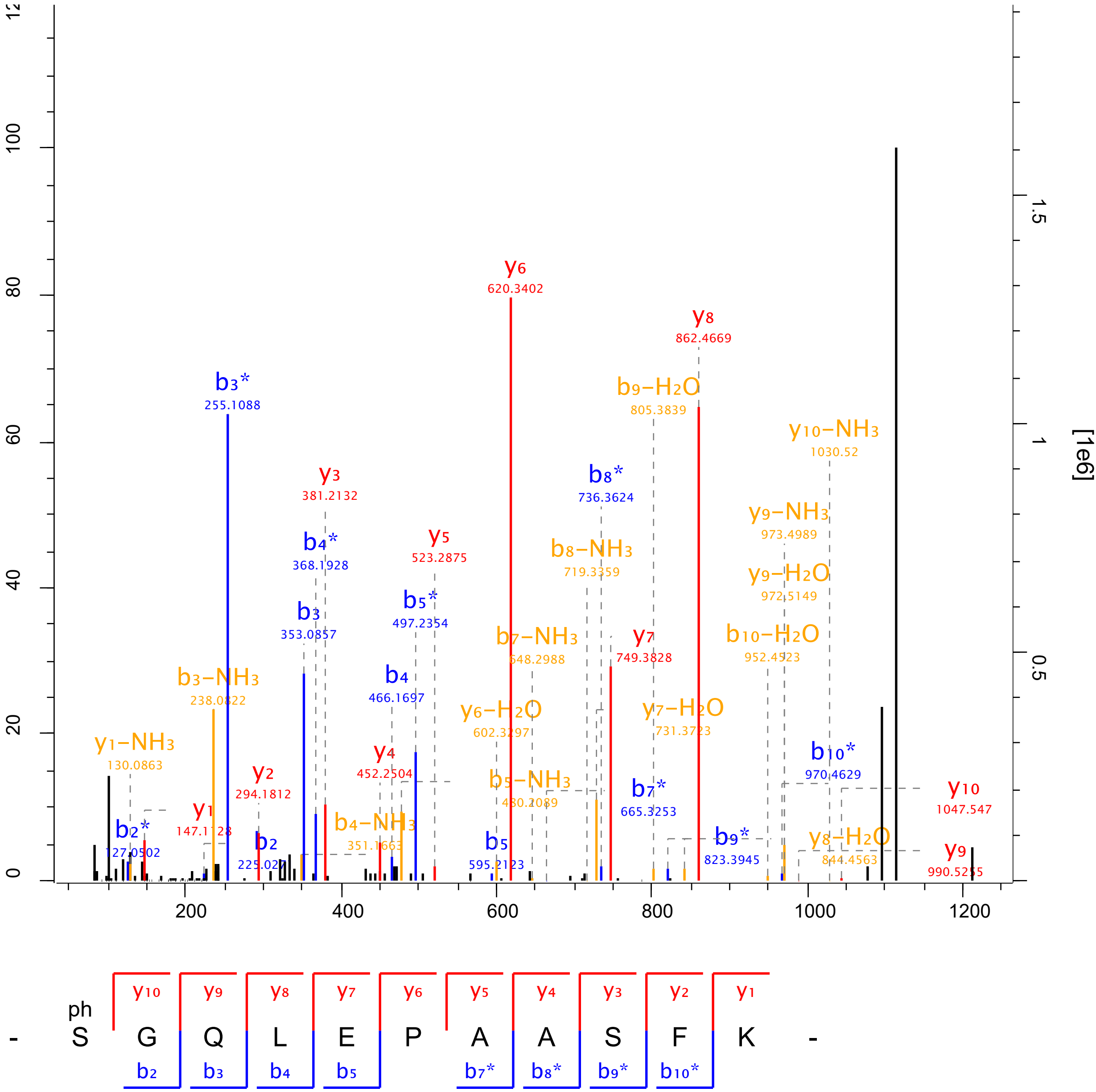

| Raw file | Scan | Method | Score | m/z |
|---|---|---|---|---|
| sys_02_2 | 17756 | FTMS; HCD | 318.08 | 923.42 |

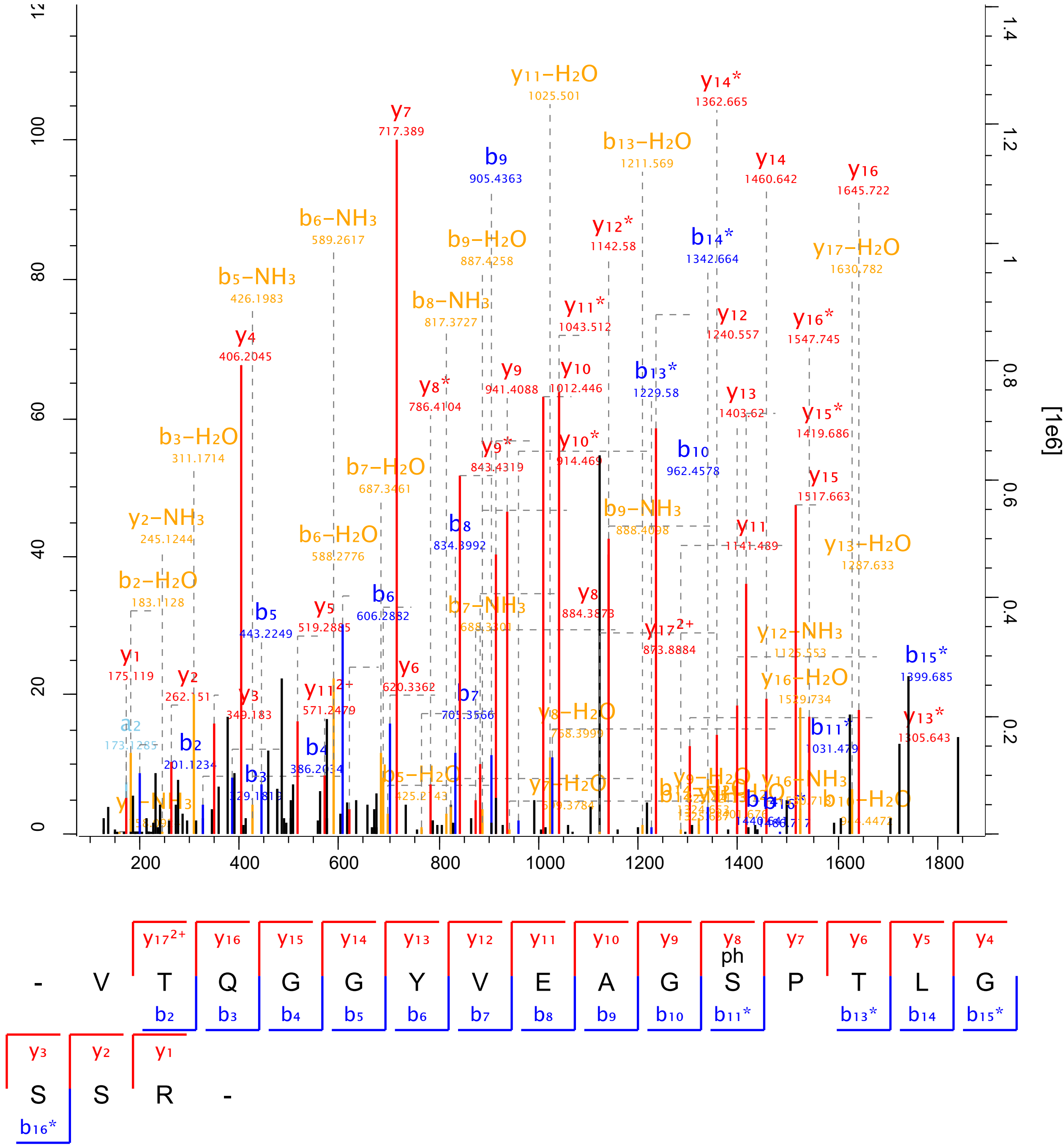

| Raw file | Scan | Method | Score | m/z |
|---|---|---|---|---|
| sys_02_2 | 18161 | FTMS; HCD | 45.35 | 799.38 |

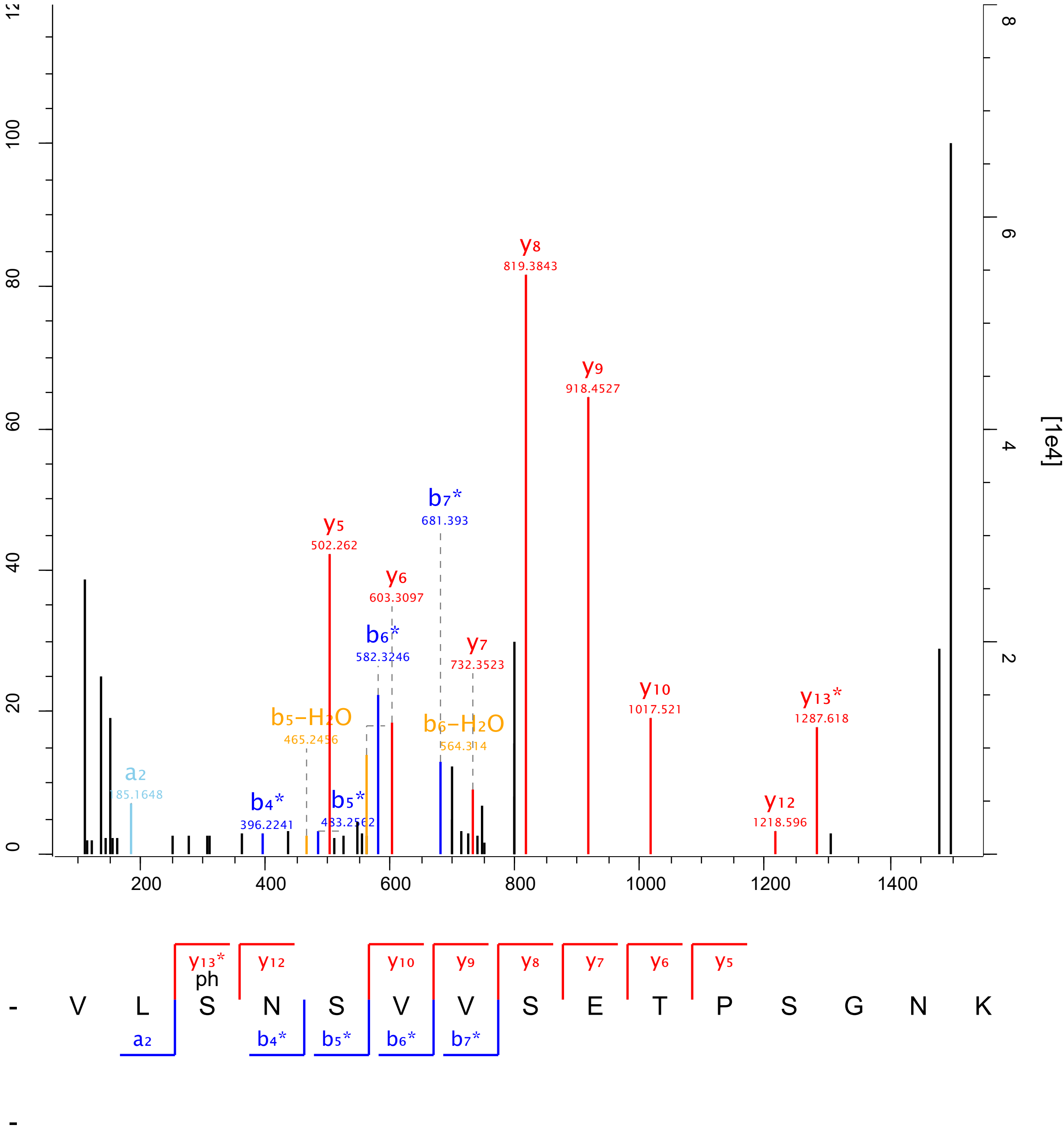

| Raw file | Scan | Method | Score | m/z |
|---|---|---|---|---|
| sys_02_2 | 18166 | FTMS; HCD | 97.9 | 528.6 |

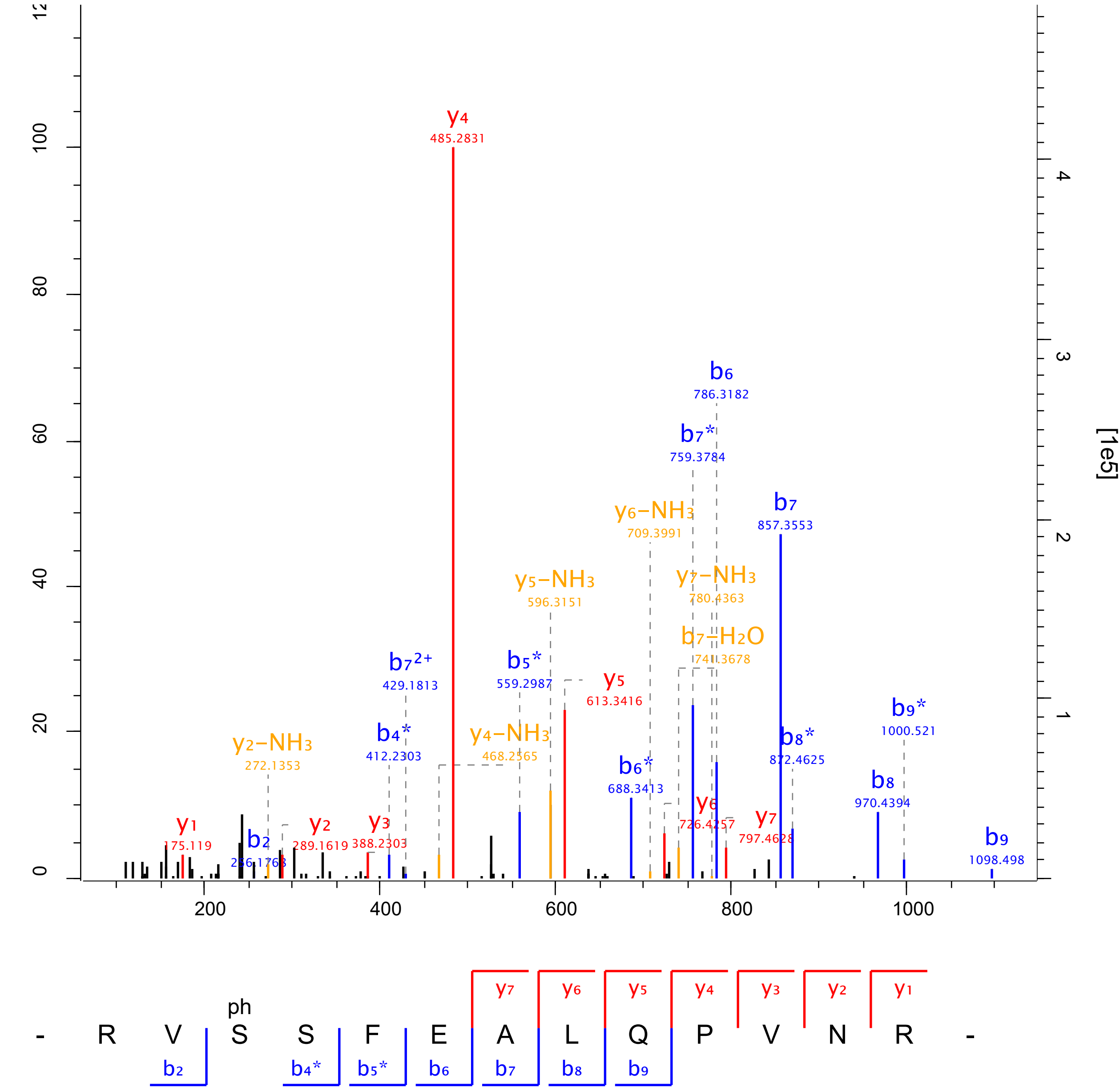

| Raw file | Scan | Method | Score | m/z |
| --- | --- | --- | --- | --- |
| sys_02_2 | 18195 | FTMS; HCD | 218.98 | 927.35 |

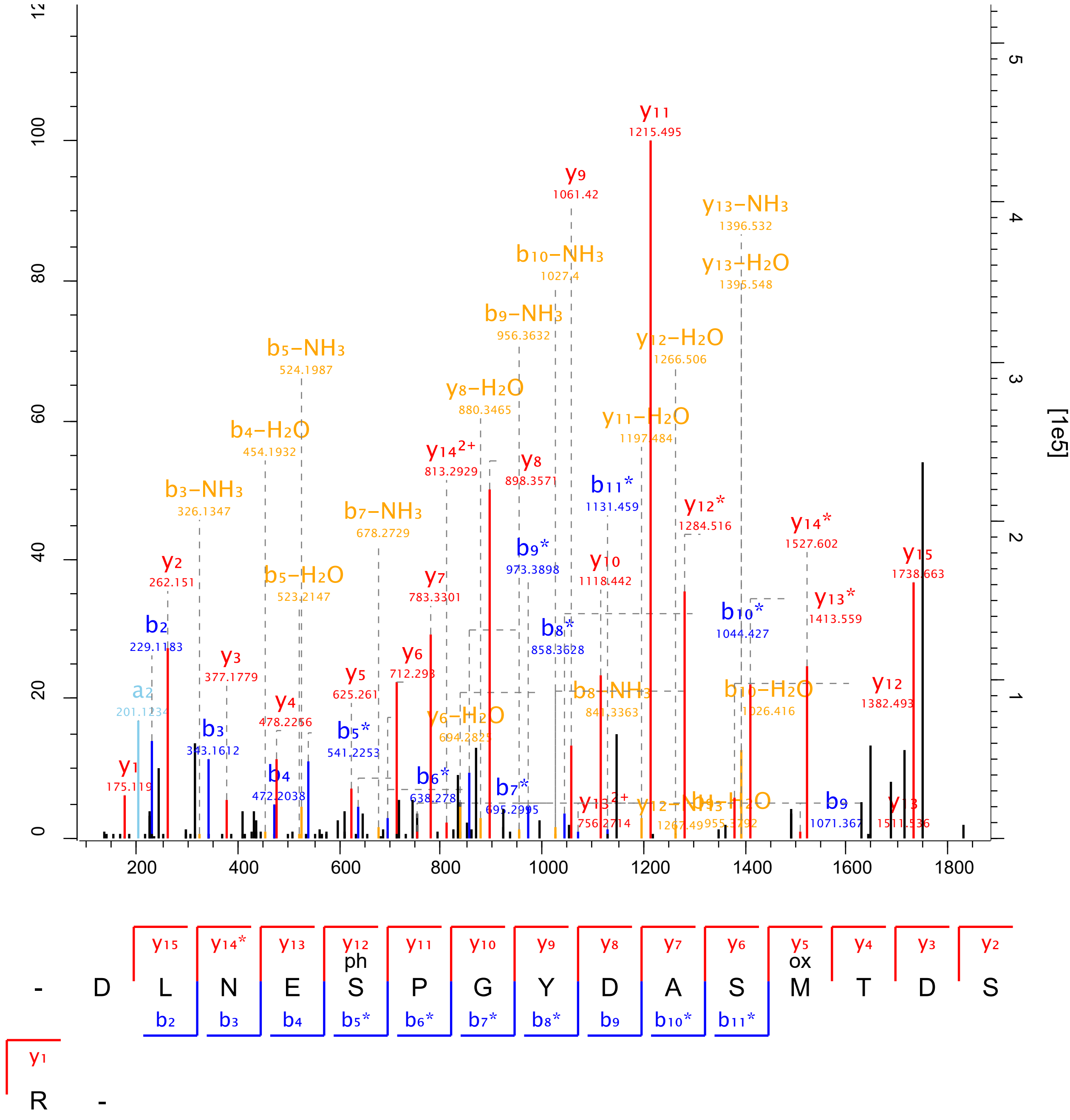

| Raw file | Scan | Method | Score | m/z |
|---|---|---|---|---|
| sys_02_2 | 18388 | FTMS; HCD | 60.52 | 685.32 |

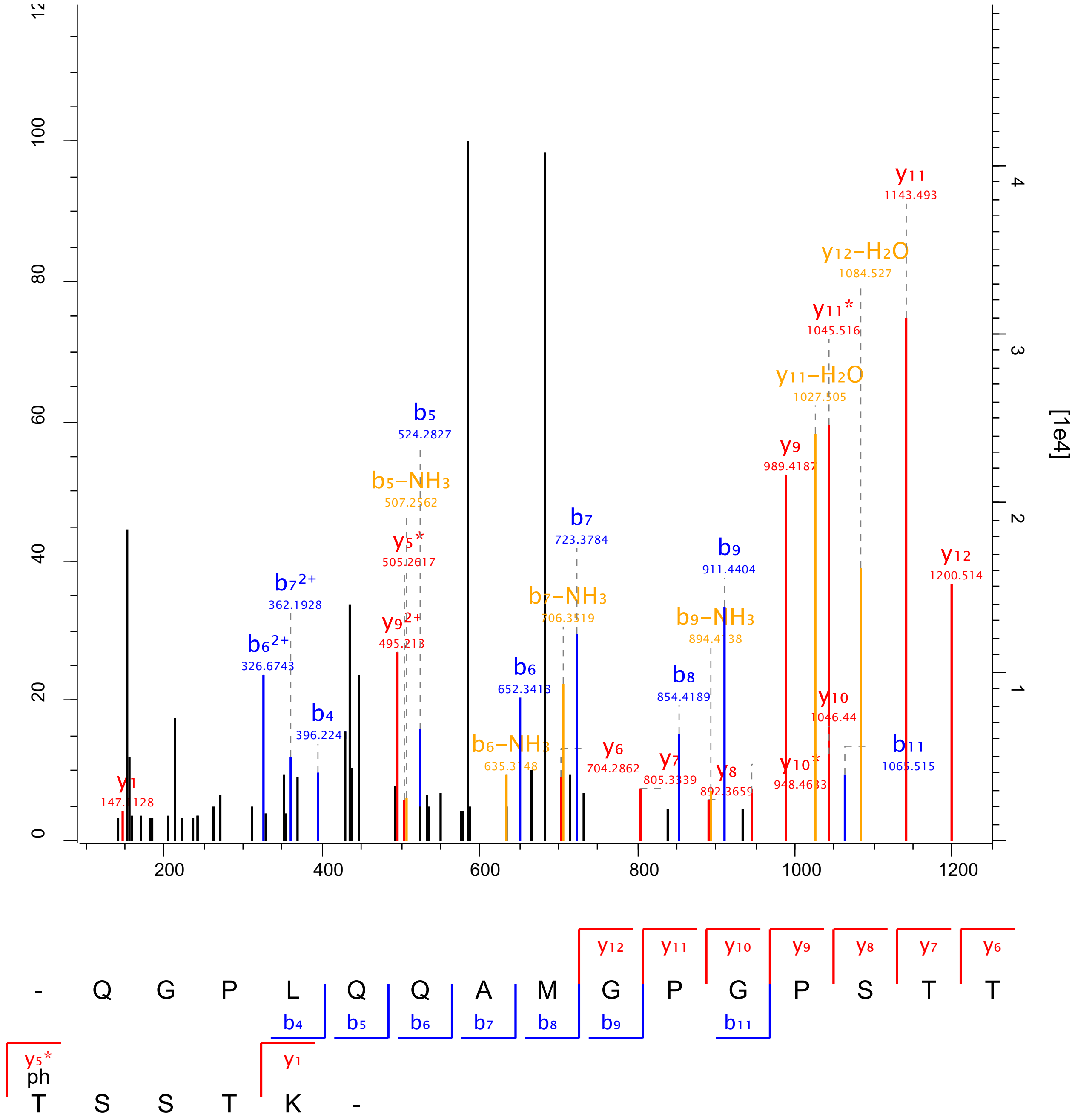

| Raw file | Scan | Method | Score | m/z |
| --- | --- | --- | --- | --- |
| sys_02_2 | 18424 | FTMS; HCD | 124.08 | 585.6 |

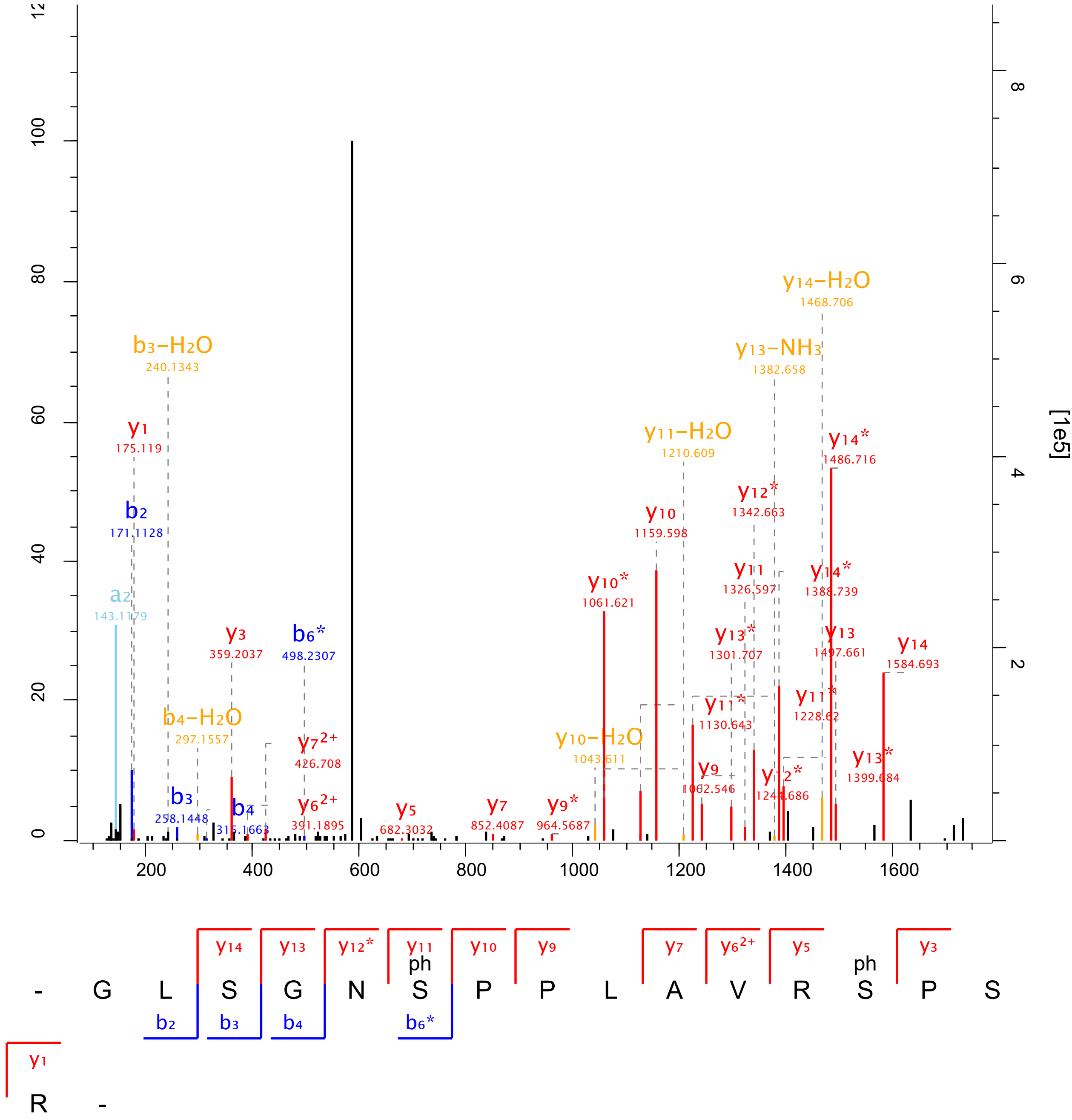

| Raw file | Scan | Method | Score | m/z |
| --- | --- | --- | --- | --- |
| sys_02_2 | 18544 | FTMS; HCD | 140.53 | 605.25 |

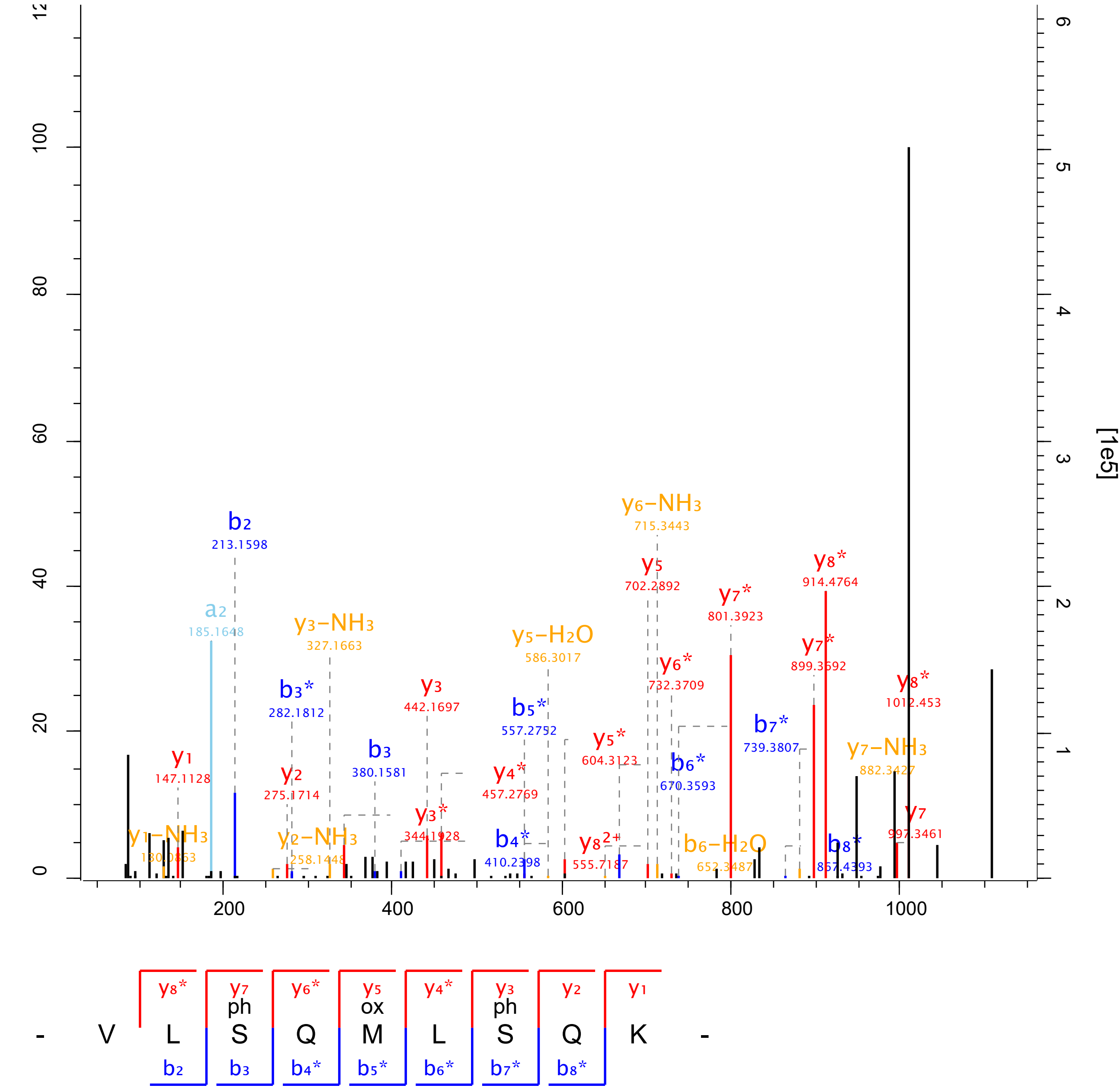

| Raw file | Scan | Method | Score | m/z |
|---|---|---|---|---|
| sys_02_2 | 18579 | FTMS; HCD | 56.29 | 593.75 |

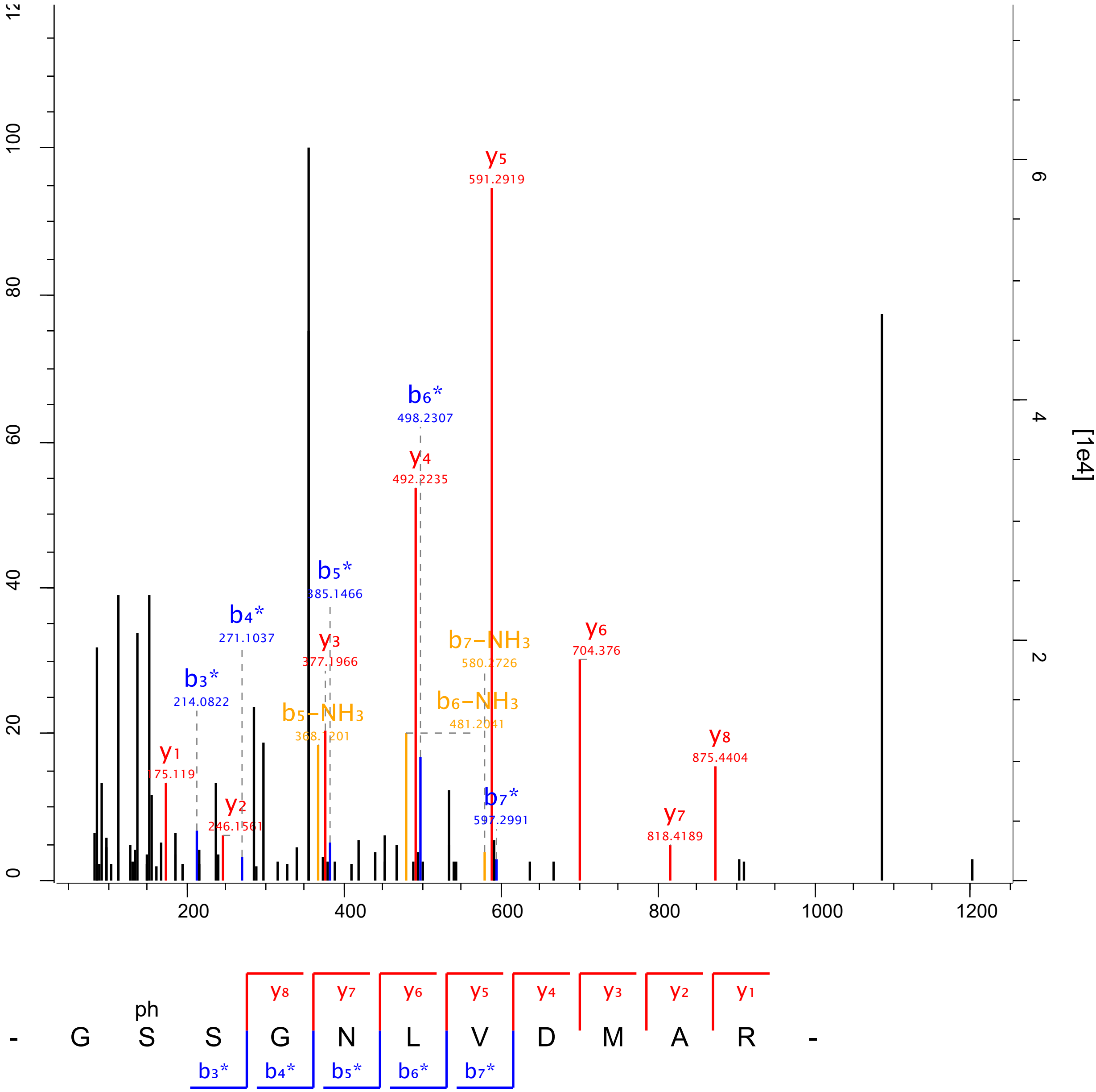

| Raw file | Scan | Method | Score | m/z |
|---|---|---|---|---|
| sys_02_2 | 18807 | FTMS; HCD | 128.54 | 567.76 |

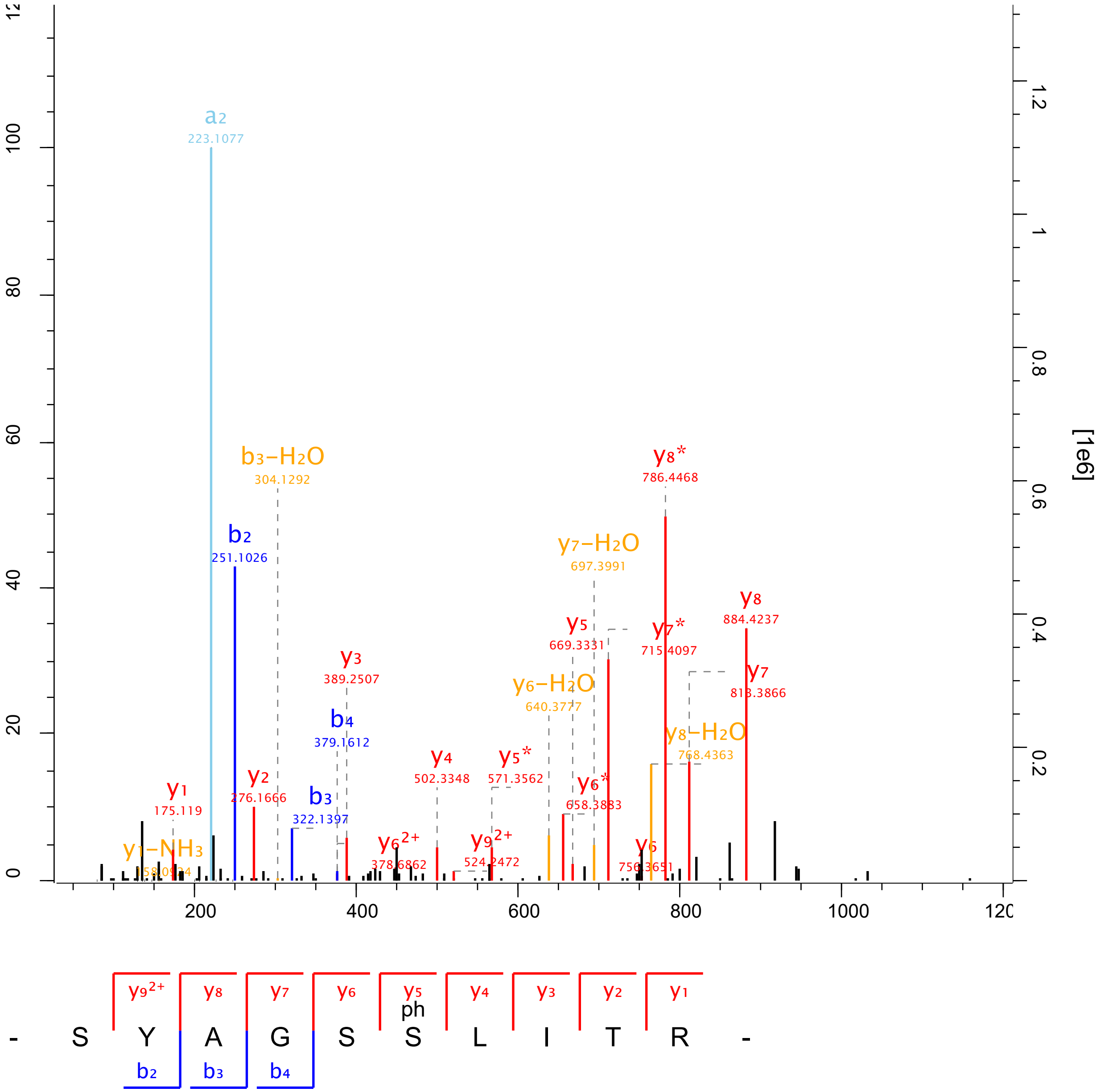

| Raw file | Scan | Method | Score | m/z |
|---|---|---|---|---|
| sys_02_2 | 1884 | FTMS; HCD | 42.96 | 615.27 |

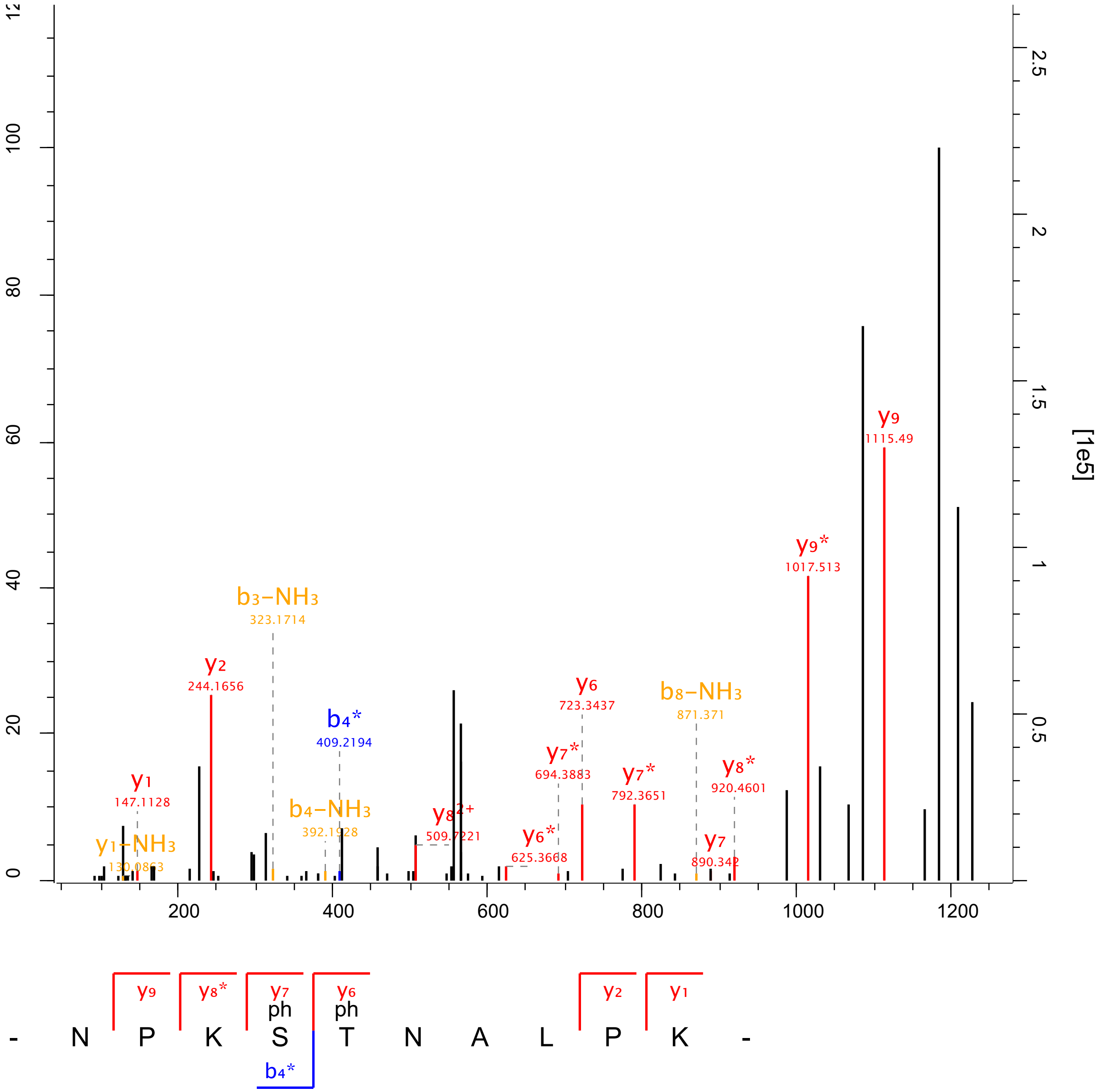

| Raw file | Scan | Method | Score | m/z |
|---|---|---|---|---|
| sys_02_2 | 1898 | FTMS; HCD | 125.47 | 427.53 |

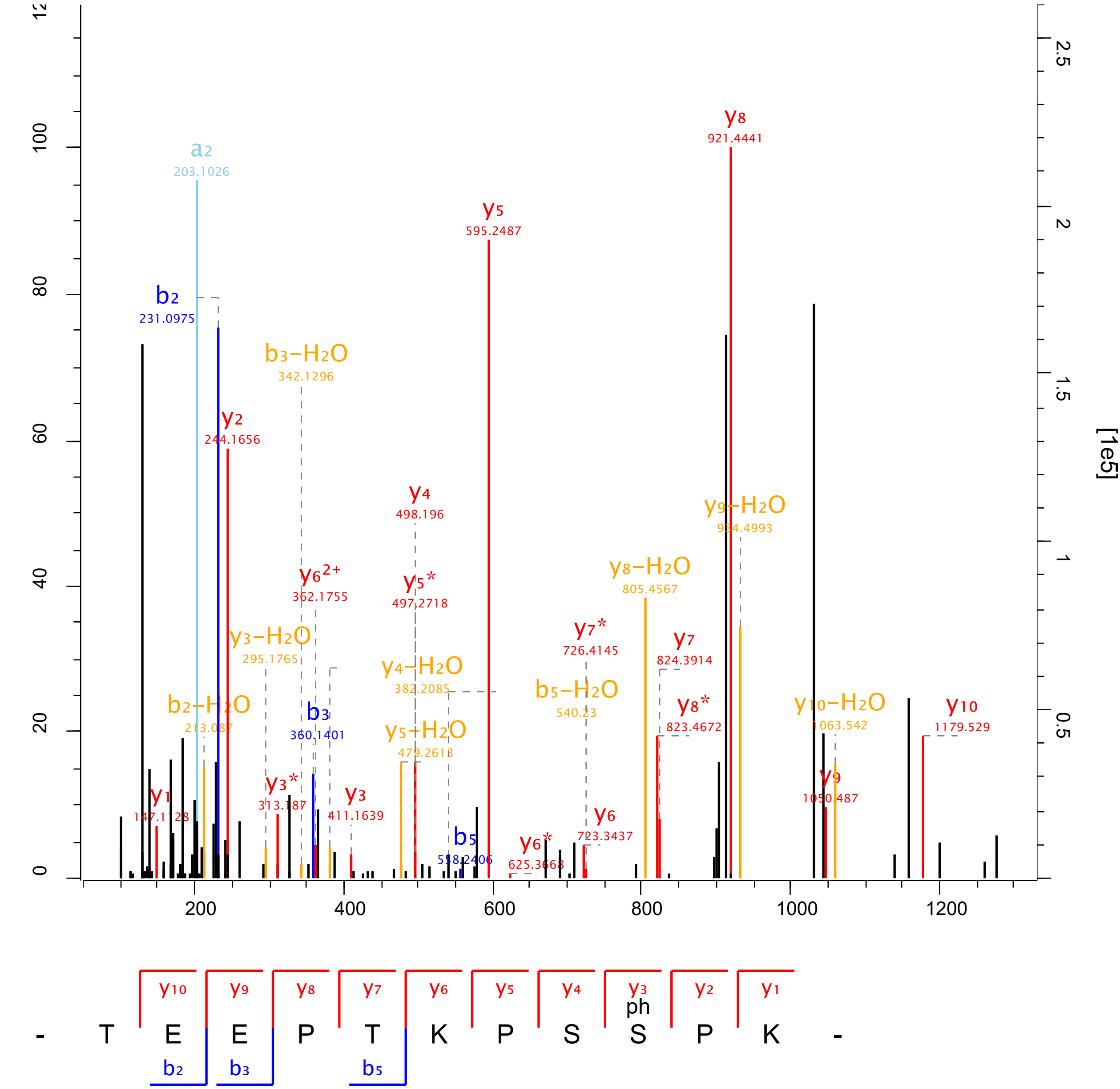

| Raw file | Scan | Method | Score | m/z |
|---|---|---|---|---|
| sys_02_2 | 19480 | FTMS; HCD | 155.8 | 574.77 |

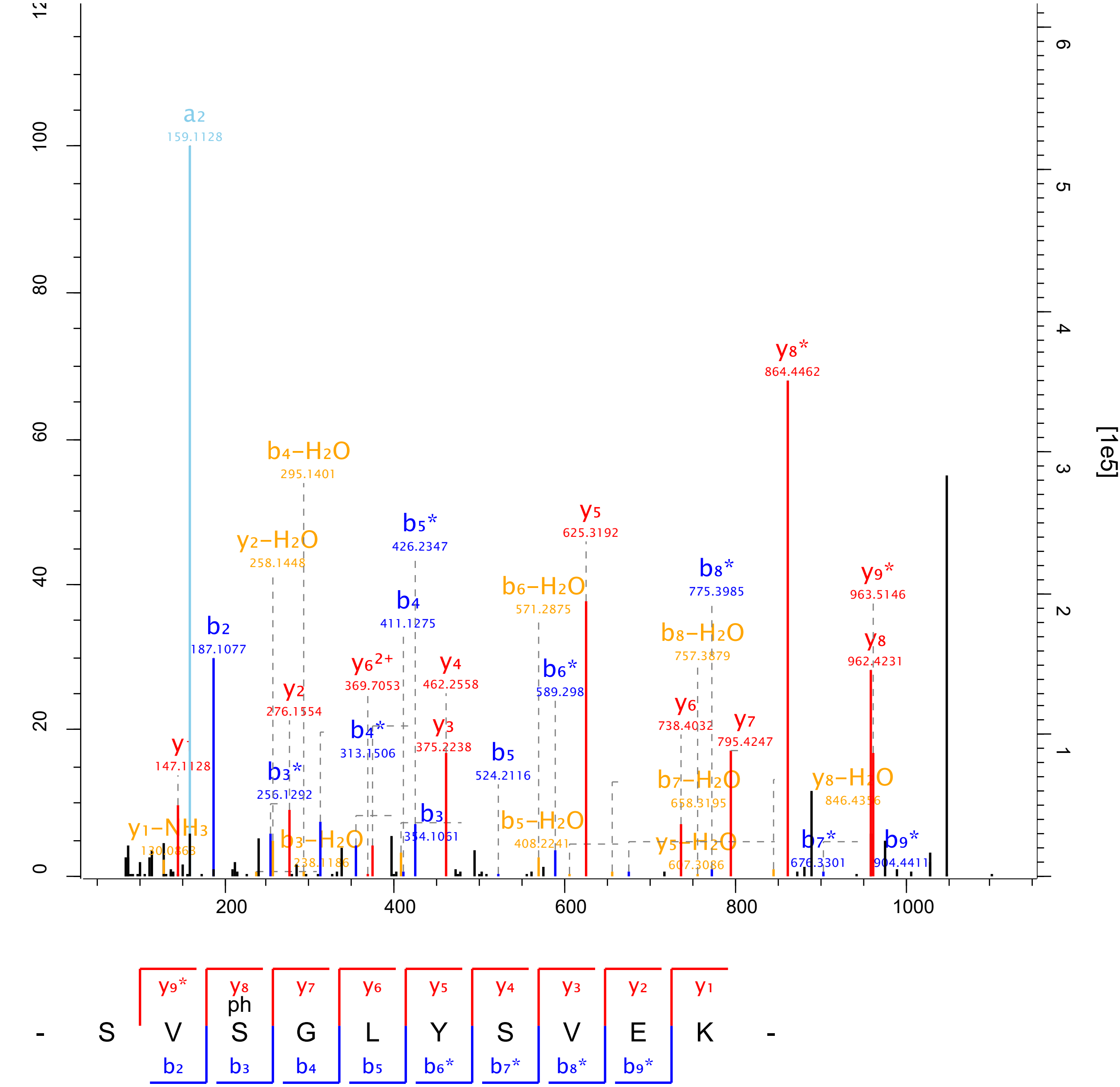

| Raw file | Scan | Method | Score | m/z |
| --- | --- | --- | --- | --- |
| sys_02_2 | 19756 | FTMS; HCD | 47.51 | 698.79 |

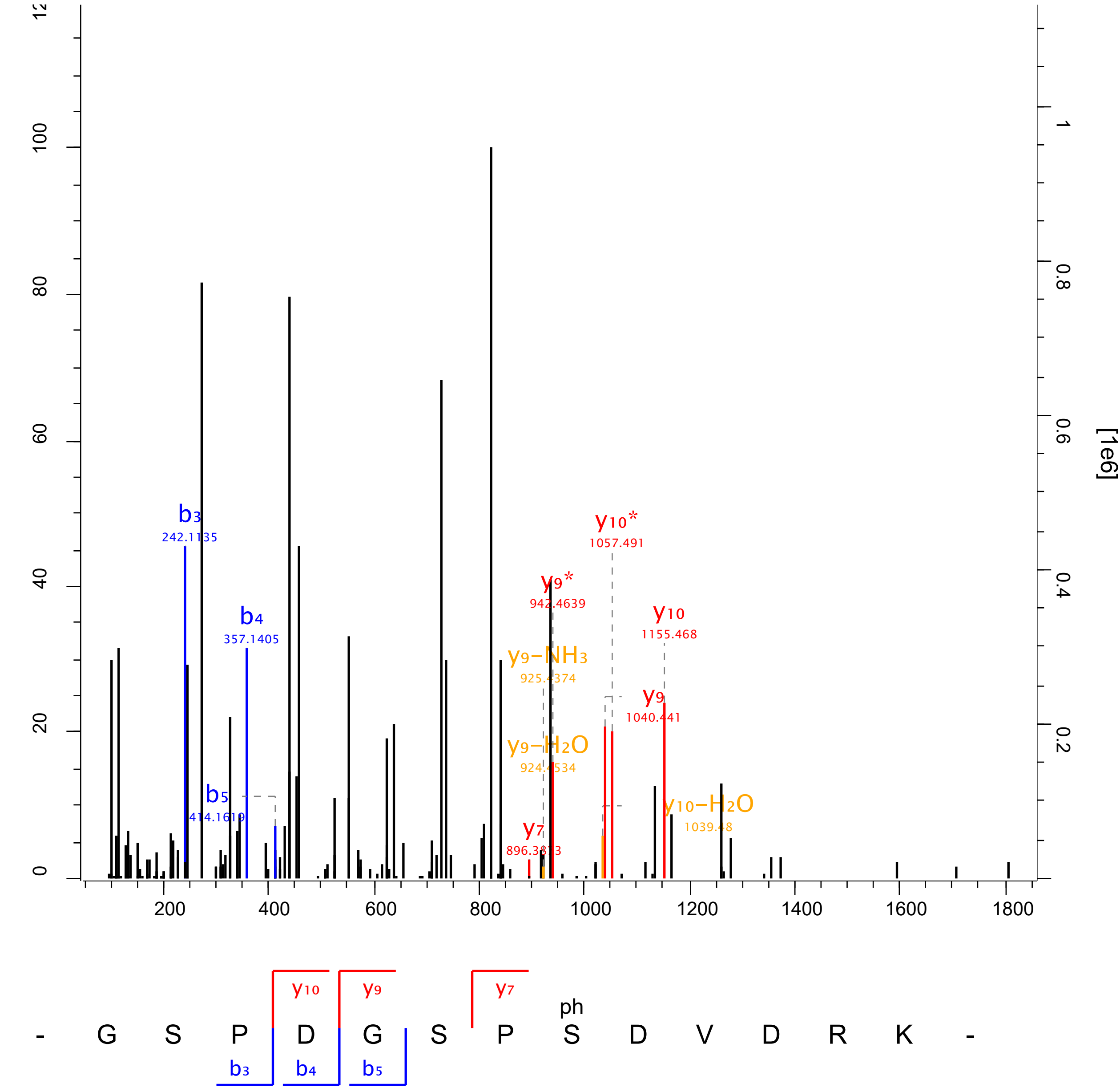

| Raw file | Scan | Method | Score | m/z |
|---|---|---|---|---|
| sys_02_2 | 19846 | FTMS; HCD | 62.34 | 534.75 |

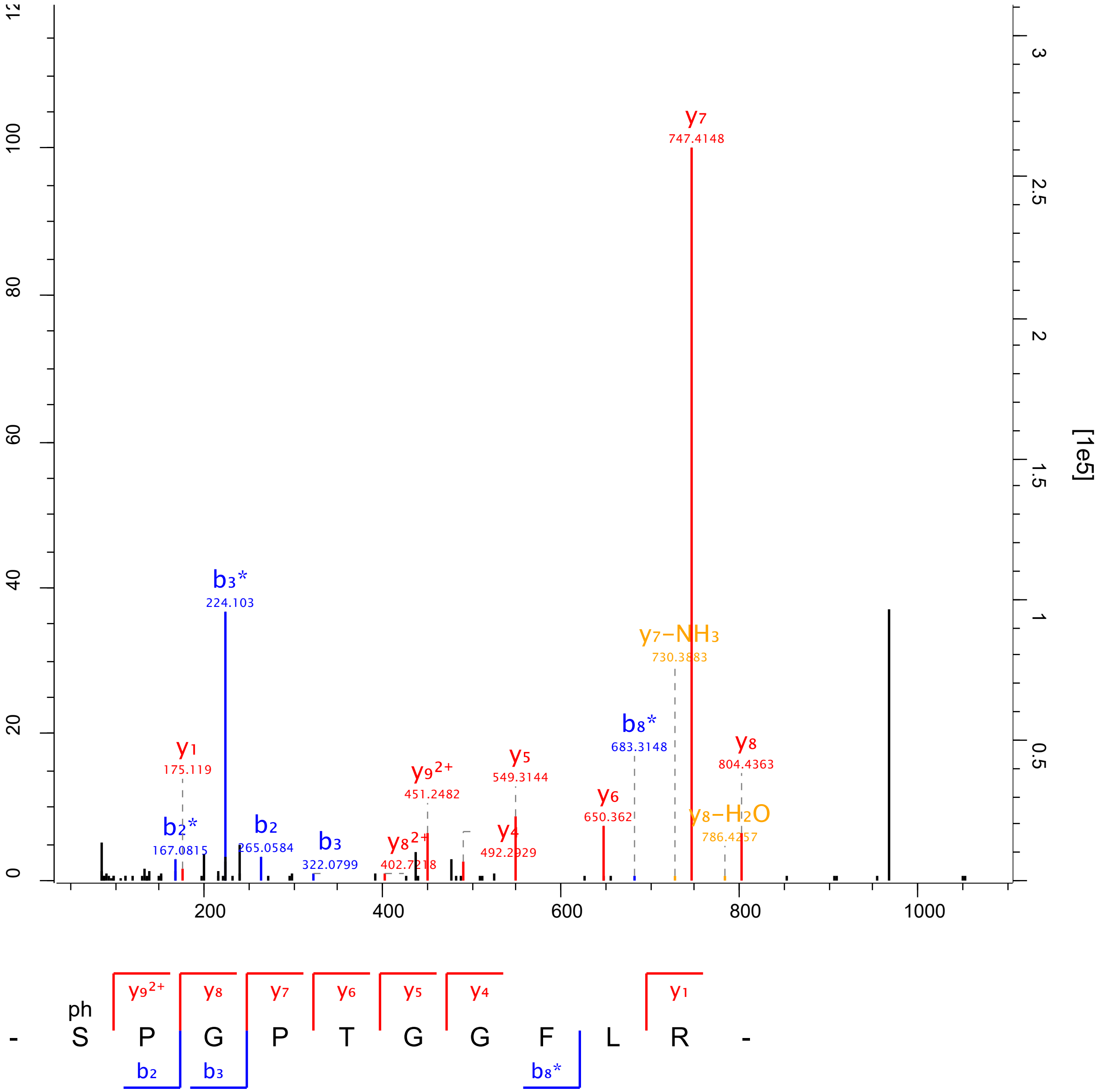

| Raw file | Scan | Method | Score | m/z |
|---|---|---|---|---|
| sys_02_2 | 20188 | FTMS; HCD | 72.72 | 946.42 |

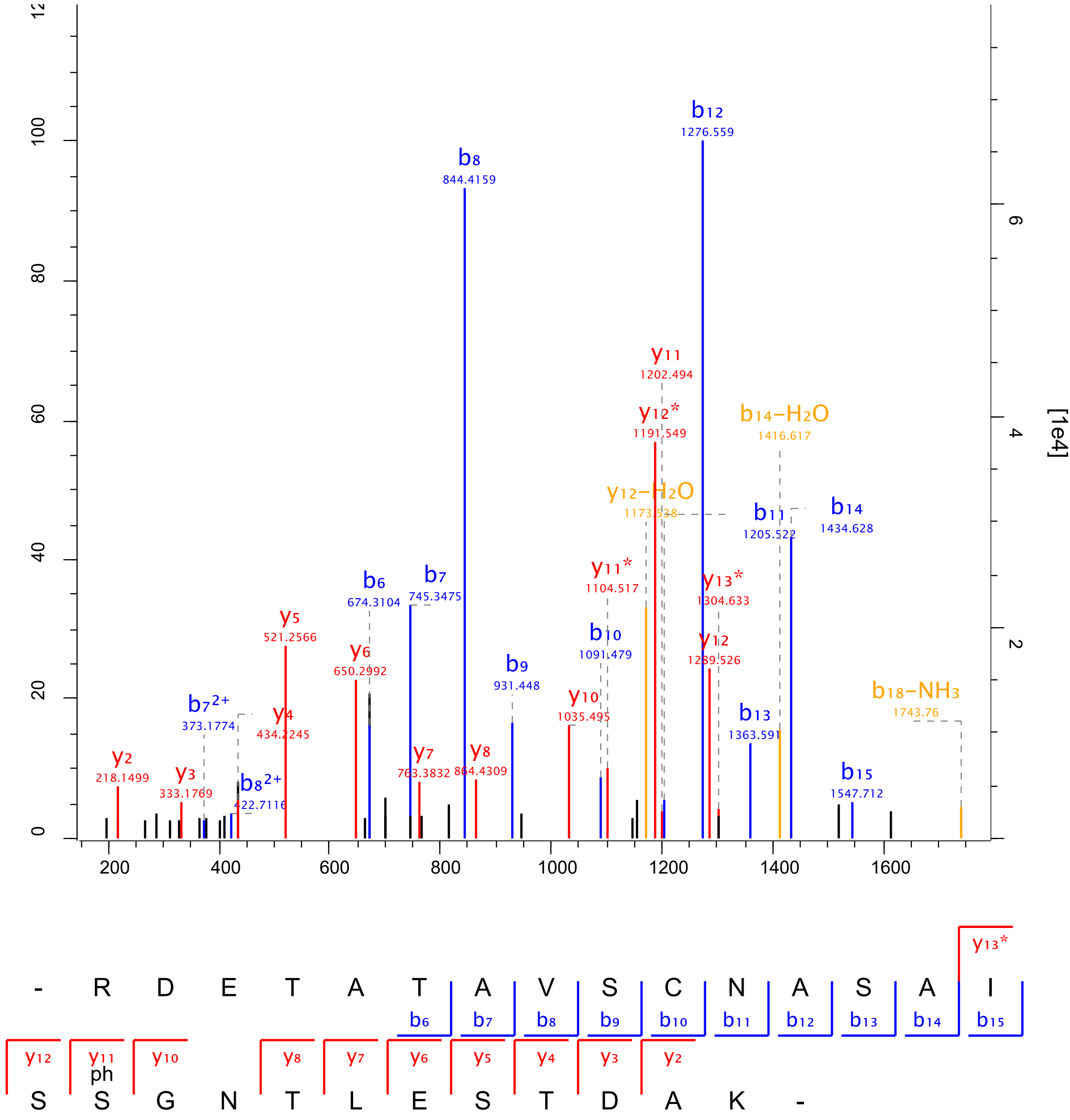

| Raw file | Scan | Method | Score | m/z |
|---|---|---|---|---|
| sys_02_2 | 20583 | FTMS; HCD | 141.31 | 862.86 |

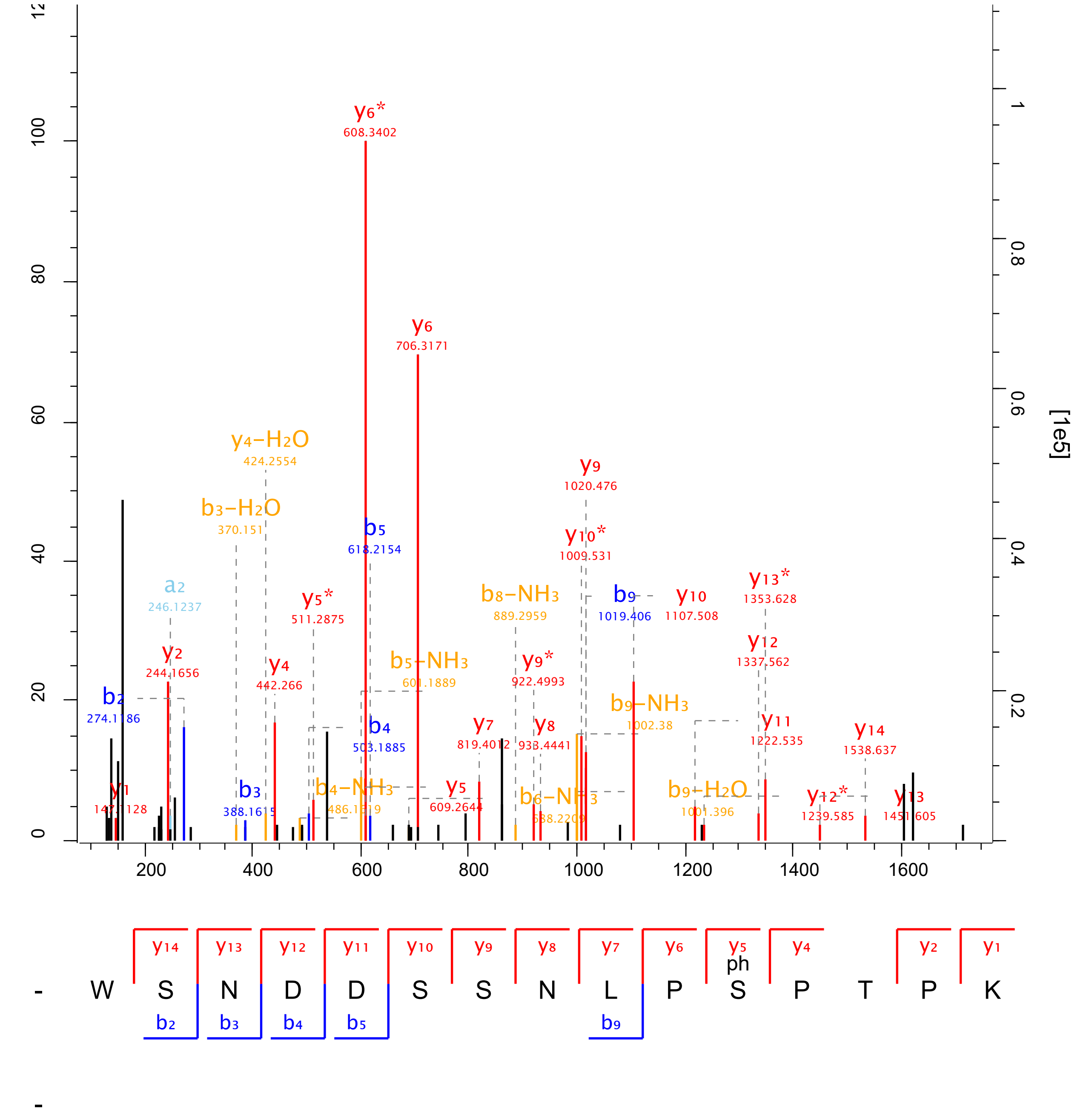

| Raw file | Scan | Method | Score | m/z |
|---|---|---|---|---|
| sys_02_2 | 2095 | FTMS; HCD | 45.28 | 597.78 |

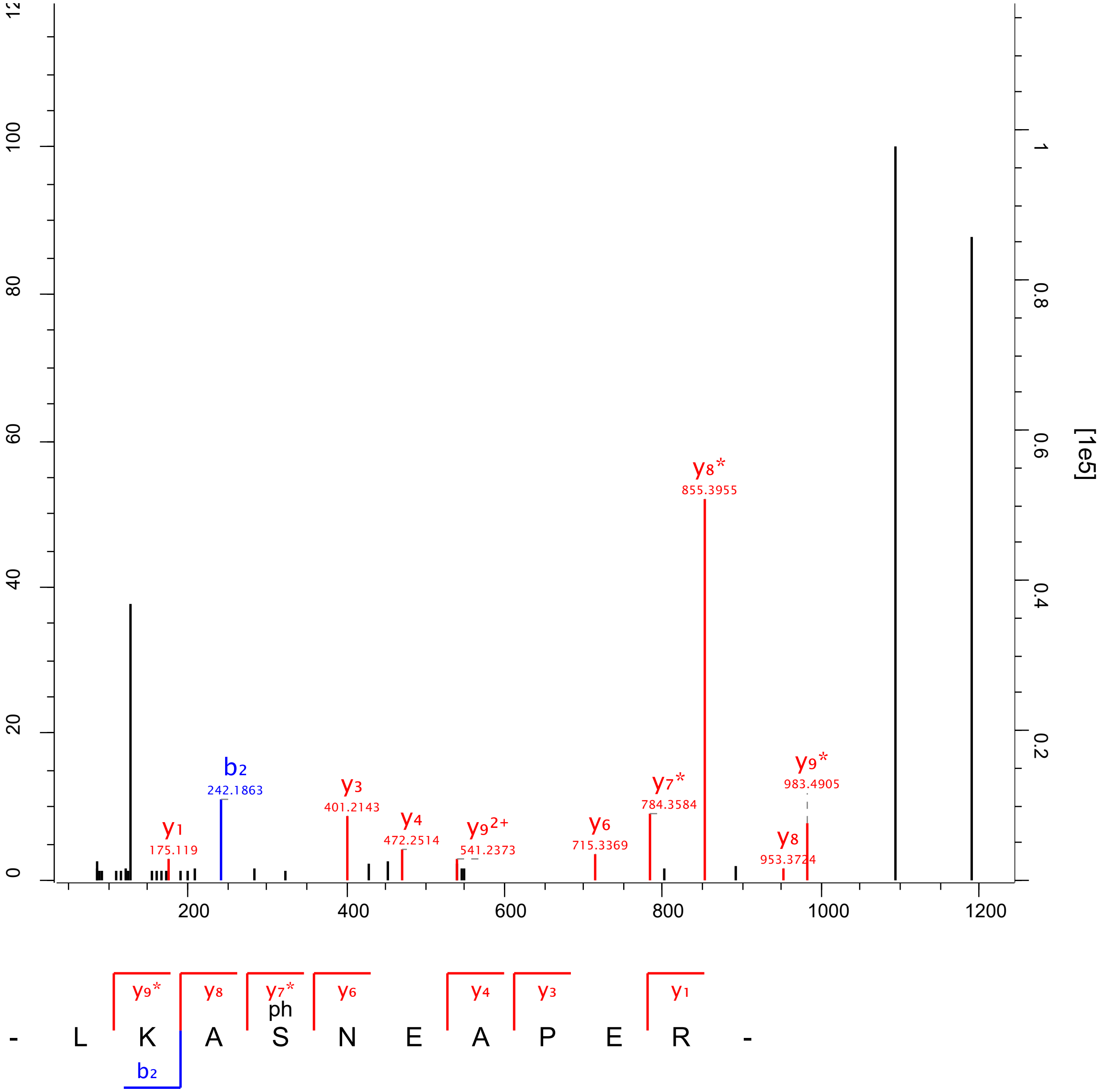

| Raw file | Scan | Method | Score | m/z |
|---|---|---|---|---|
| sys_02_2 | 21265 | FTMS; HCD | 44.93 | 637.62 |

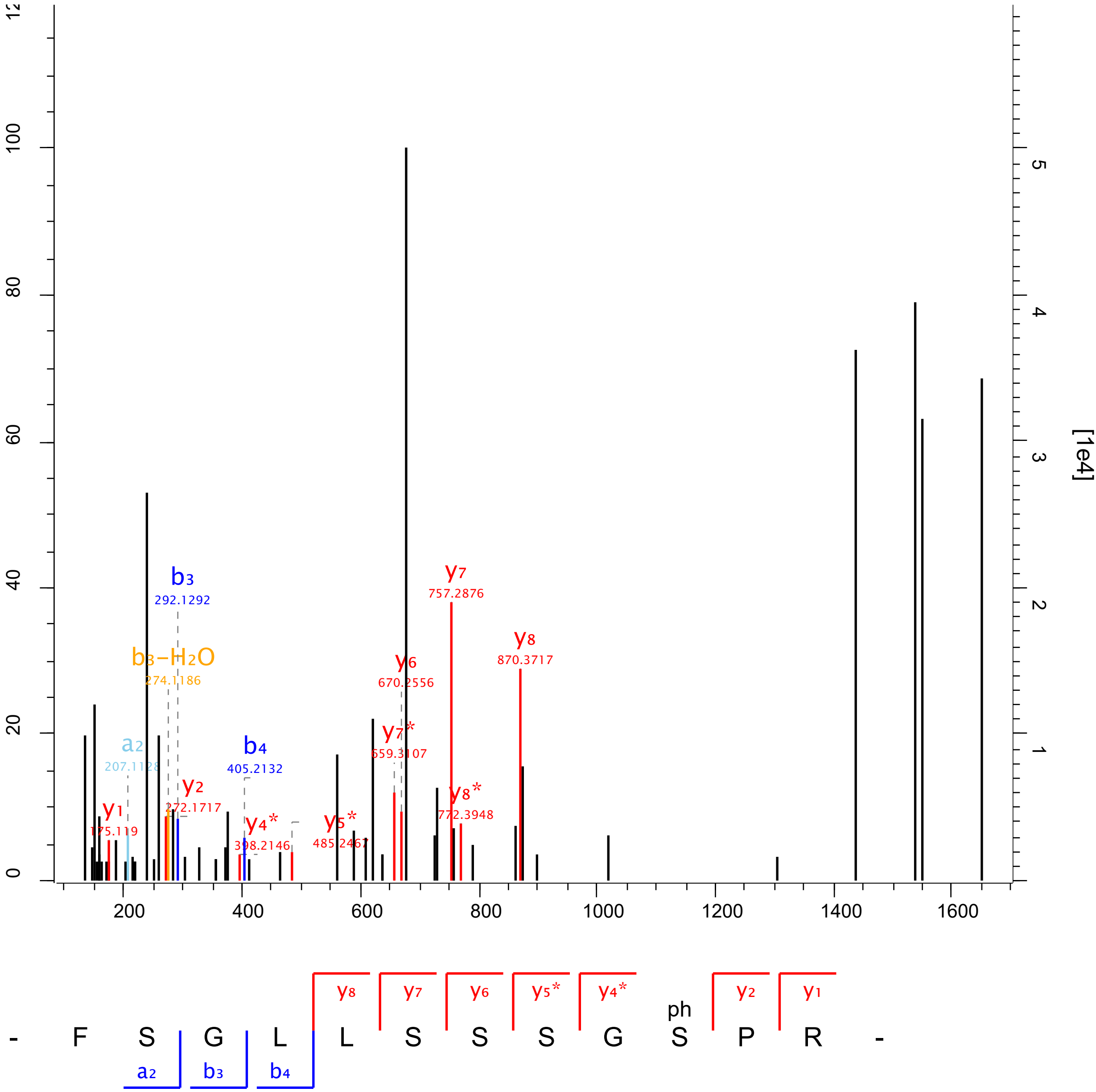

| Raw file | Scan | Method | Score | m/z |
|---|---|---|---|---|
| sys_02_2 | 2137 | FTMS; HCD | 128.21 | 535.27 |

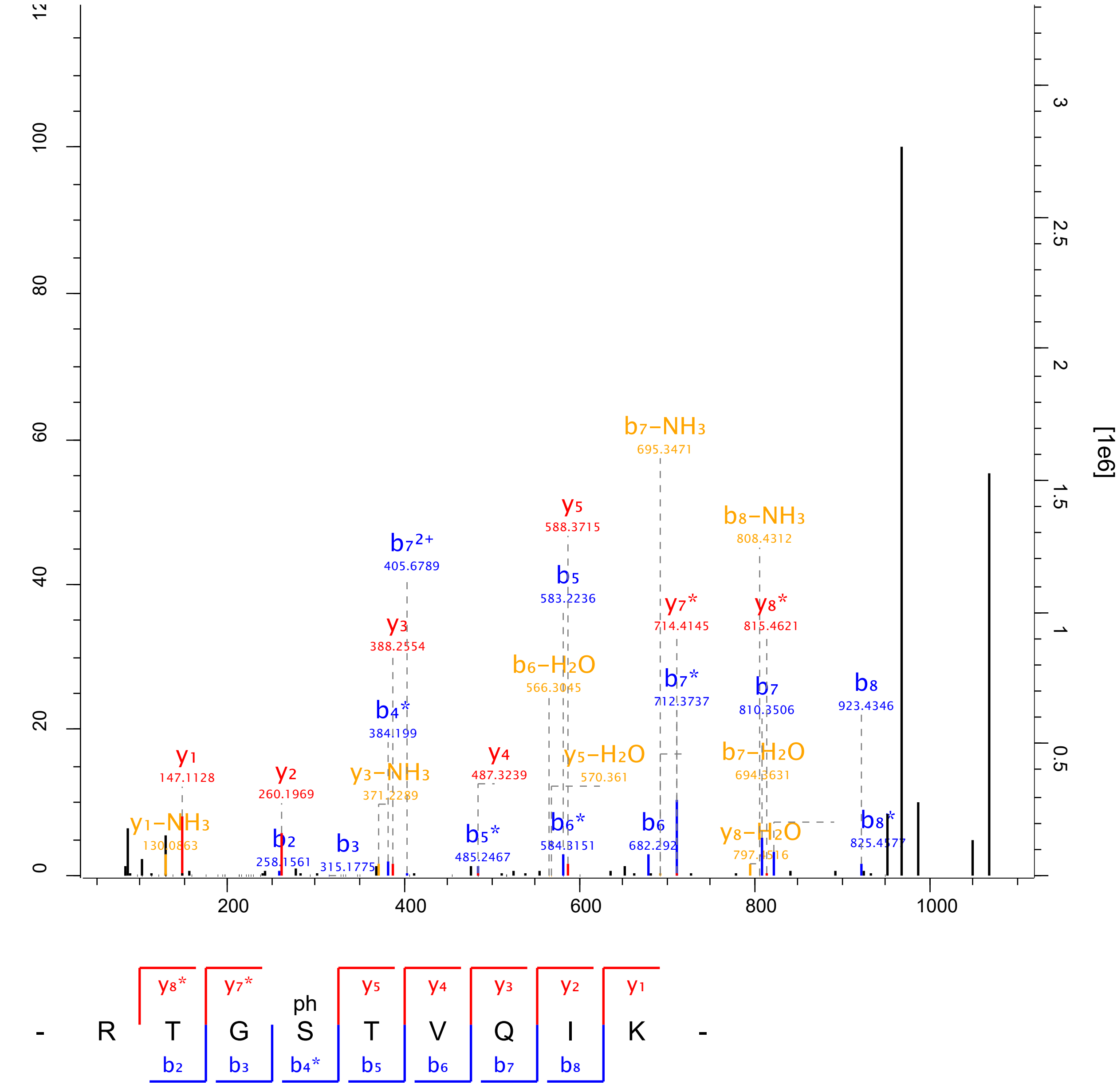

| Raw file | Scan | Method | Score | m/z |
| --- | --- | --- | --- | --- |
| sys_02_2 | 2159 | FTMS; HCD | 40.5 | 340.84 |

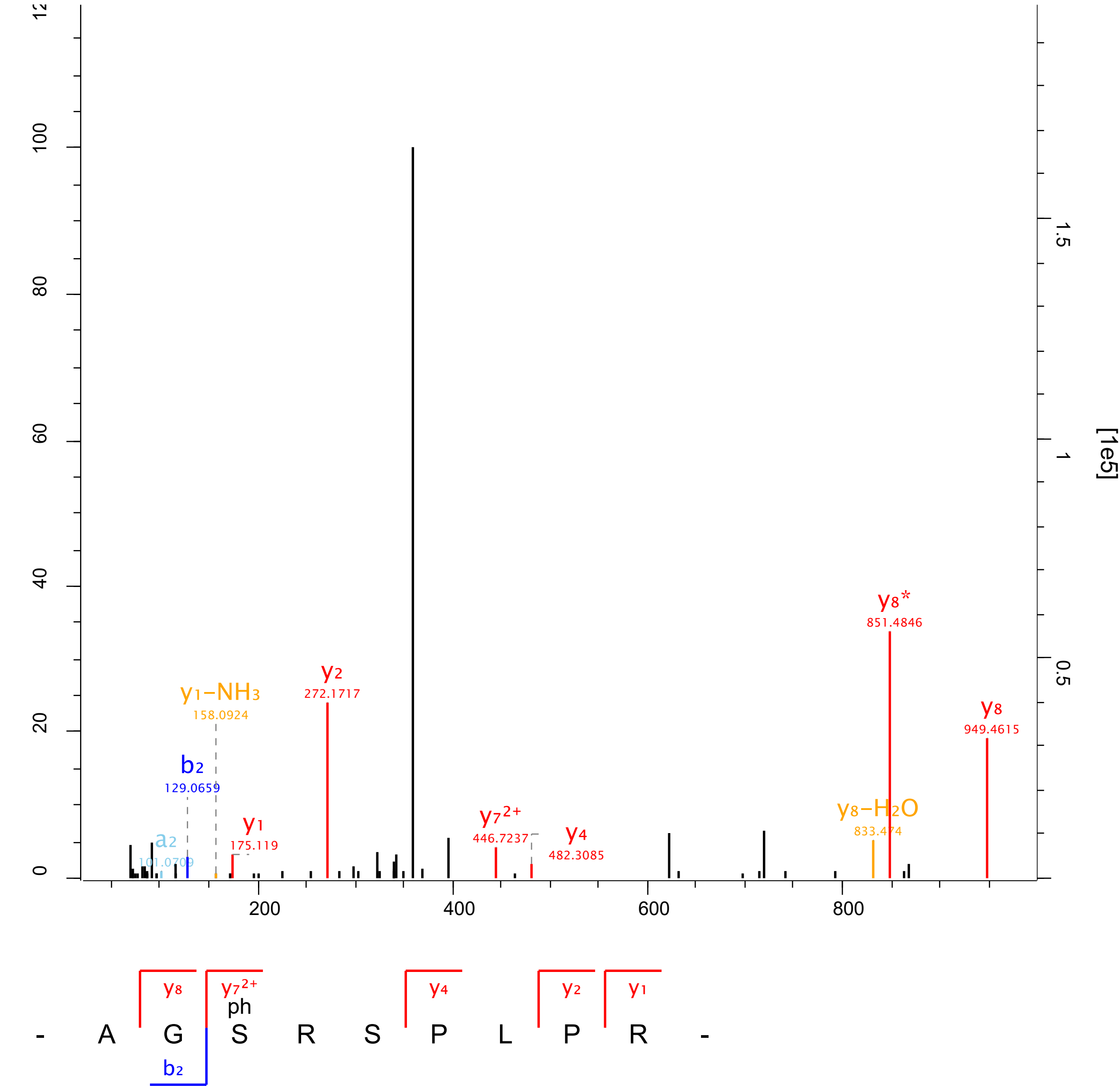

| Raw file | Scan | Method | Score | m/z |
| --- | --- | --- | --- | --- |
| sys_02_2 | 21604 | FTMS; HCD | 108.56 | 551.75 |

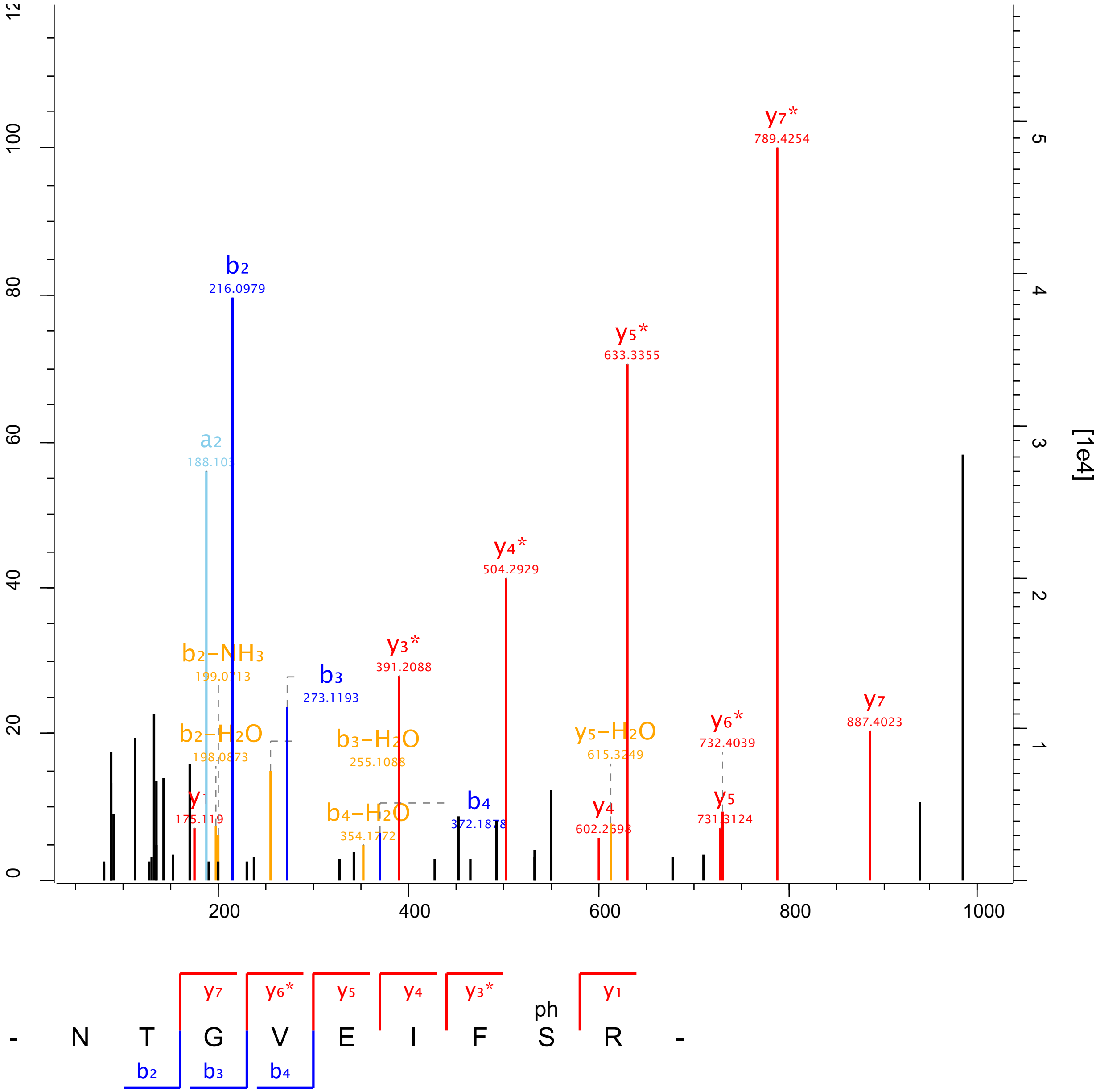

| Raw file | Scan | Method | Score | m/z |
|---|---|---|---|---|
| sys_02_2 | 21758 | FTMS; HCD | 54.34 | 538.76 |

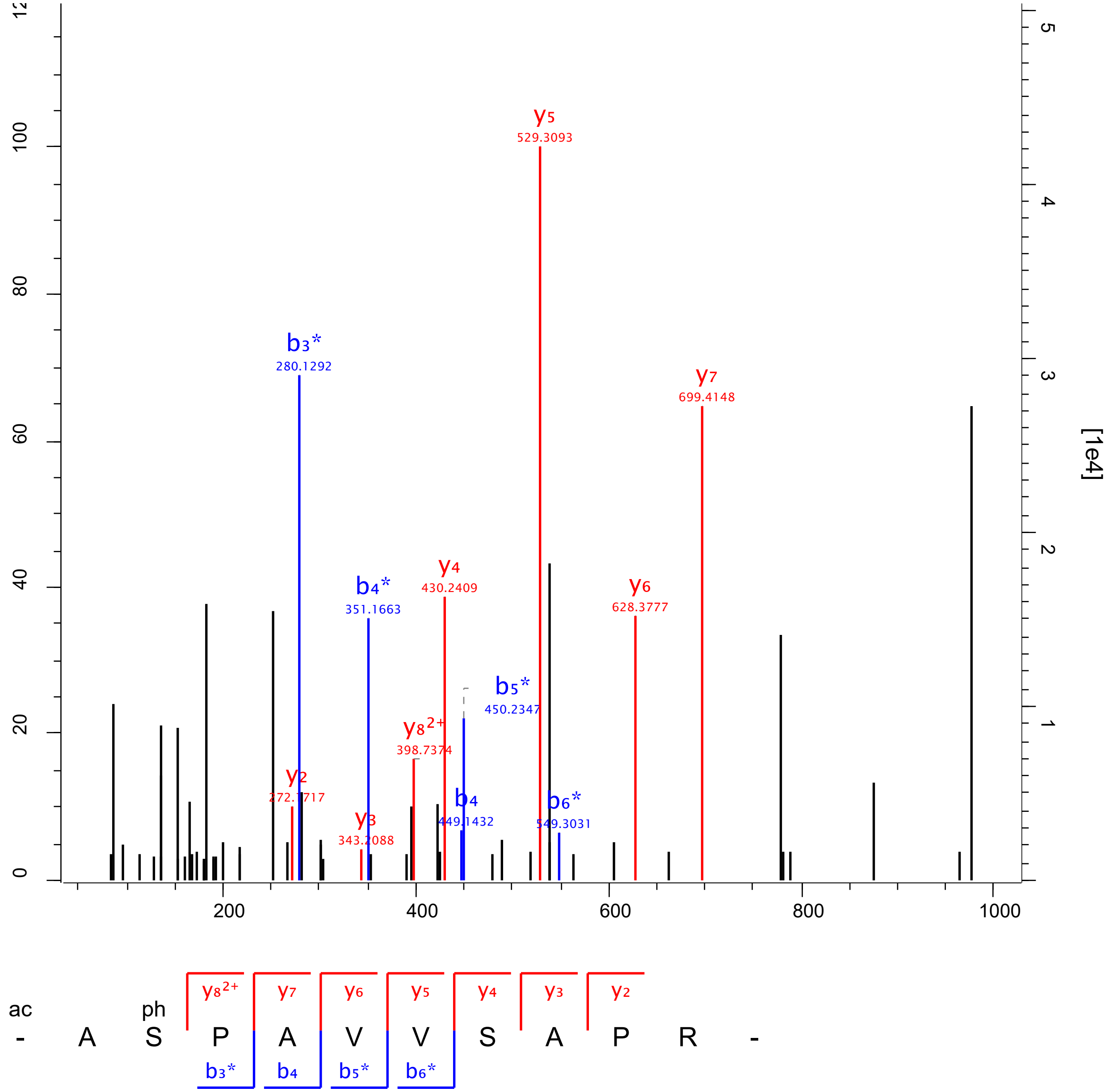

| Raw file | Scan | Method | Score | m/z |
|---|---|---|---|---|
| sys_02_2 | 21784 | FTMS; HCD | 54.22 | 850.4 |

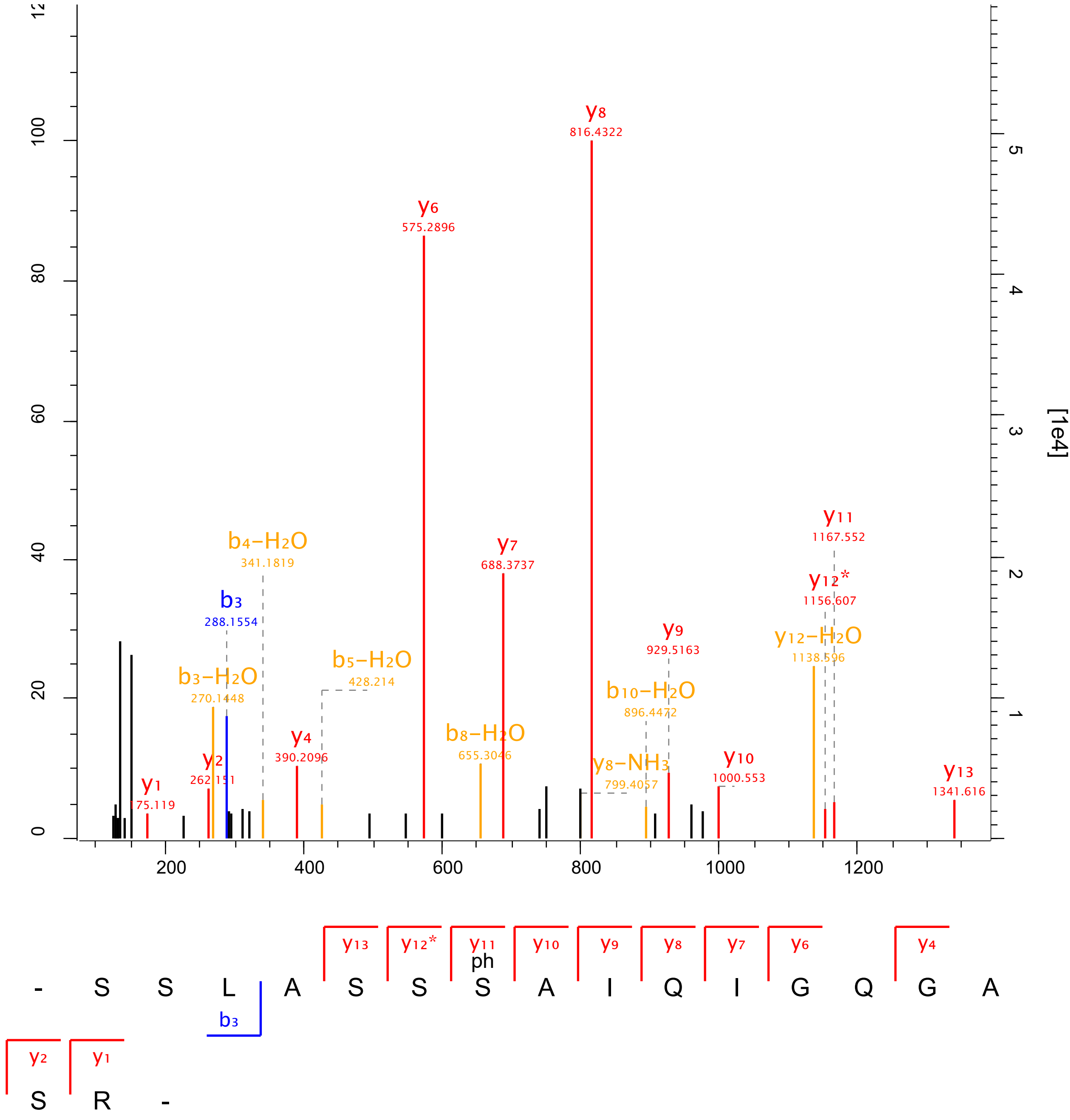

| Raw file | Scan | Method | Score | m/z |
|---|---|---|---|---|
| sys_02_2 | 21948 | FTMS; HCD | 82.88 | 587.63 |

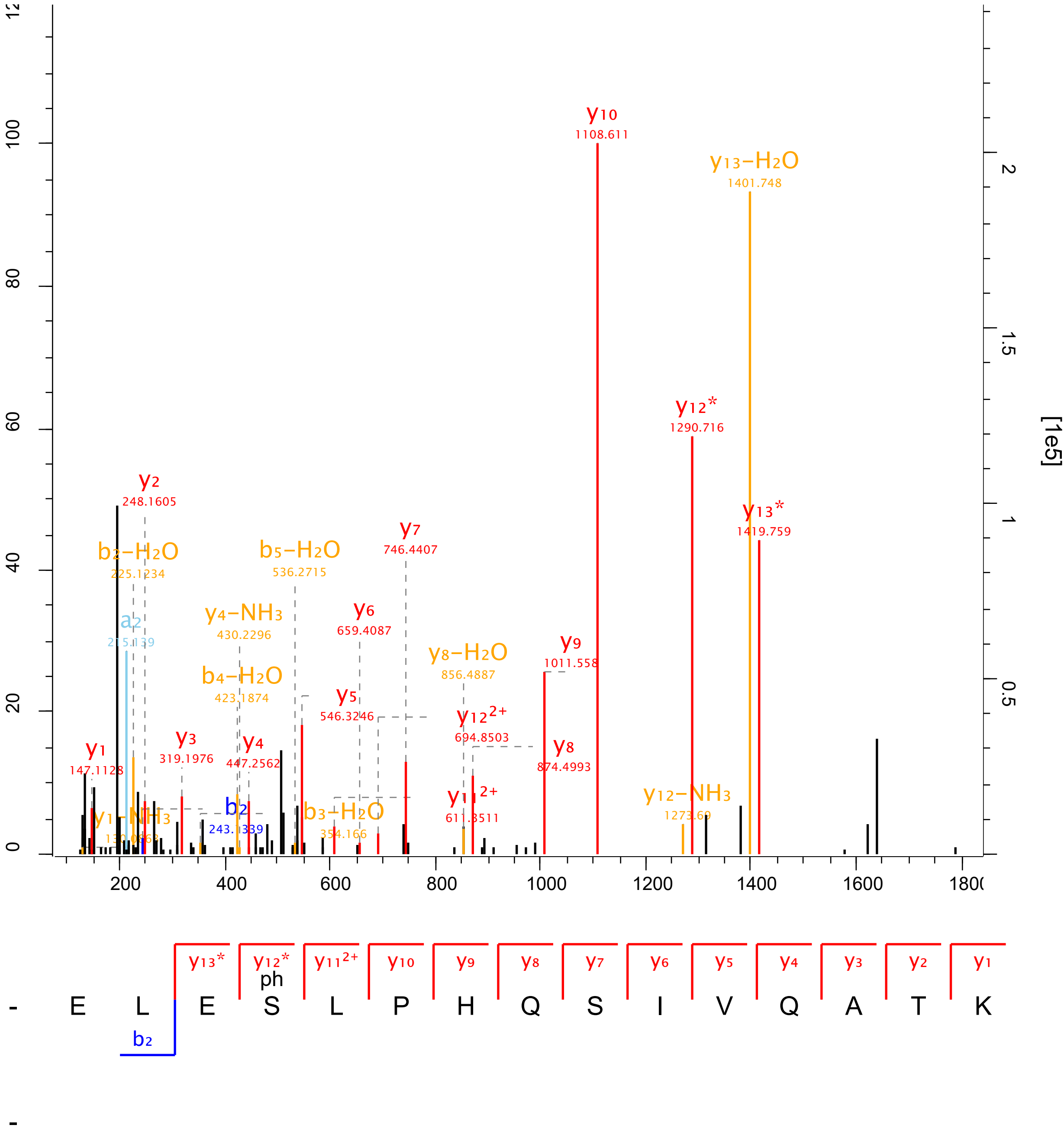

| Raw file | Scan | Method | Score | m/z |
| --- | --- | --- | --- | --- |
| sys_02_2 | 22037 | FTMS; HCD | 48.62 | 720.79 |

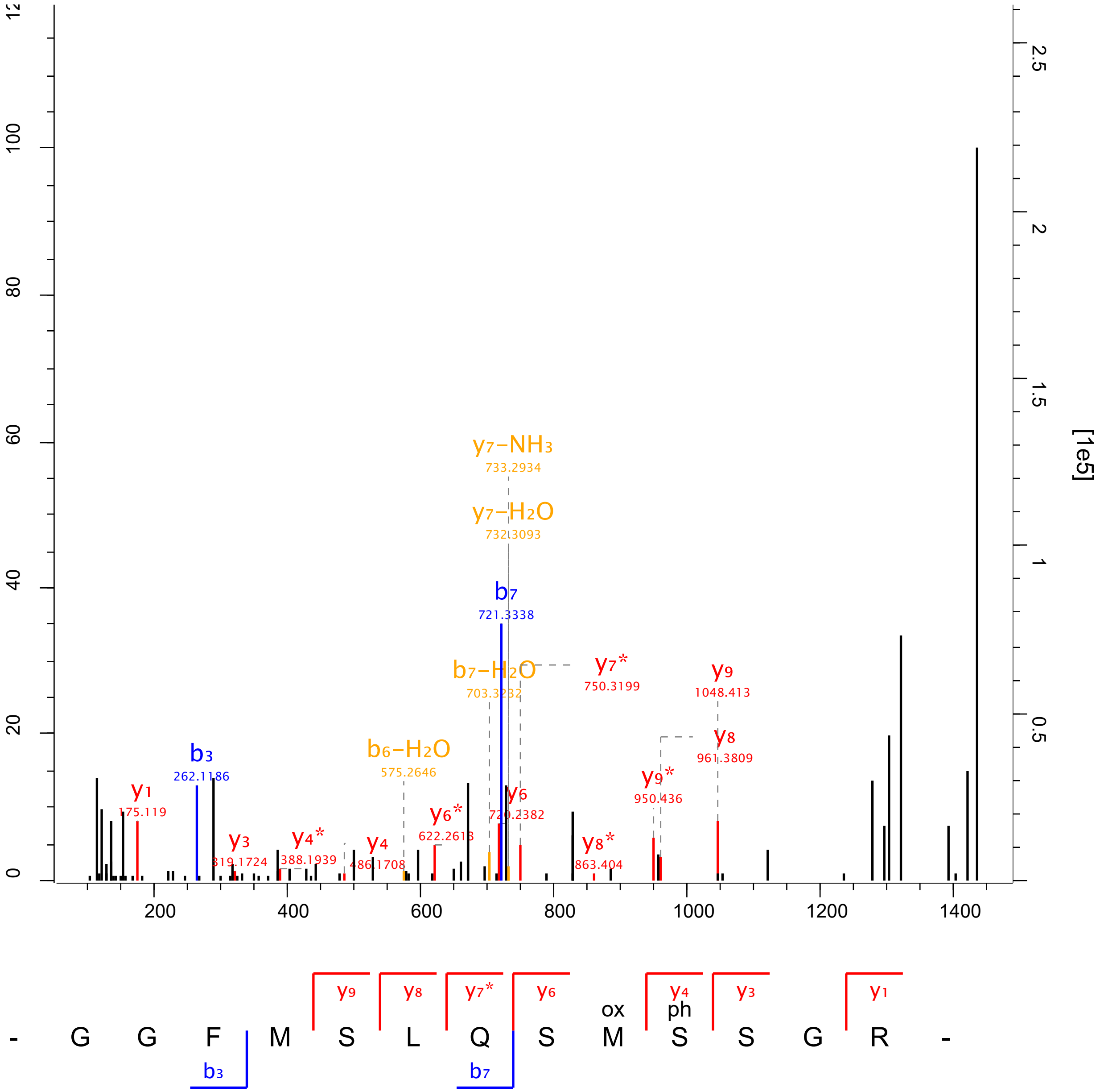

| Raw file | Scan | Method | Score | m/z |
|---|---|---|---|---|
| sys_02_2 | 2210 | FTMS; HCD | 112.7 | 543.72 |

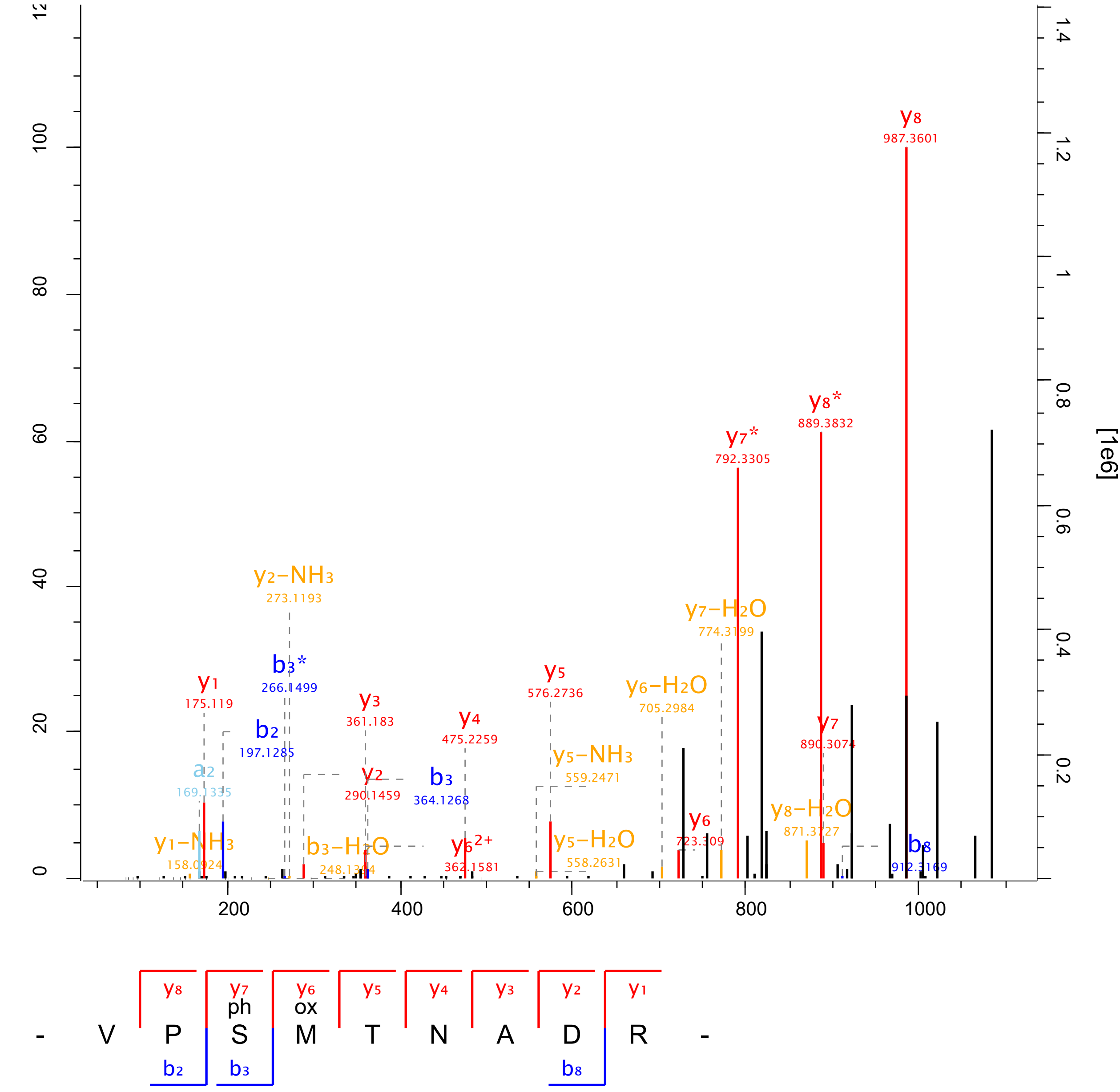

| Raw file | Scan | Method | Score | m/z |
|---|---|---|---|---|
| sys_02_2 | 22151 | FTMS; HCD | 42.71 | 705.78 |

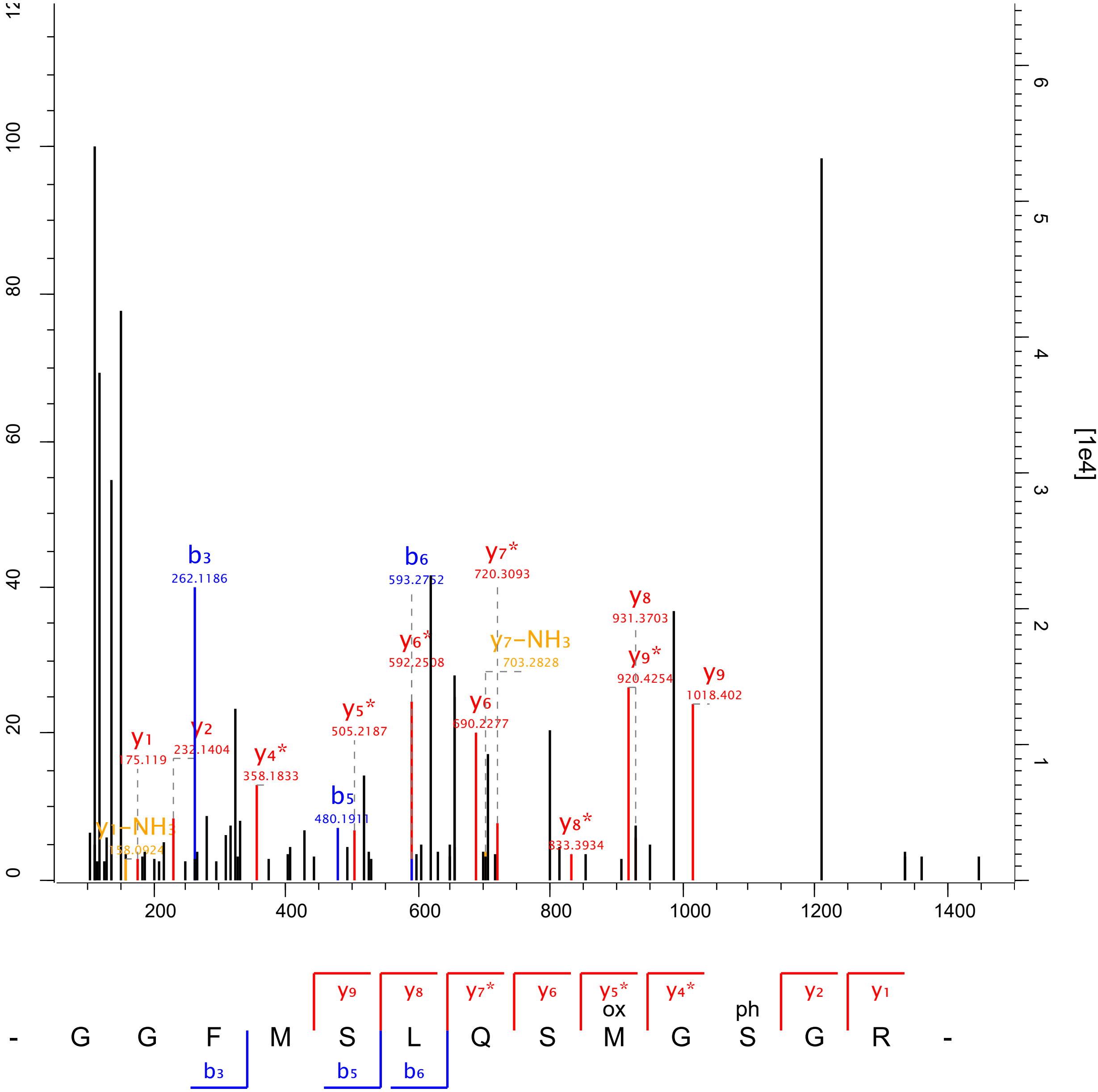

| Raw file | Scan | Method | Score | m/z |
|---|---|---|---|---|
| sys_02_2 | 22821 | FTMS; HCD | 49.66 | 683.98 |

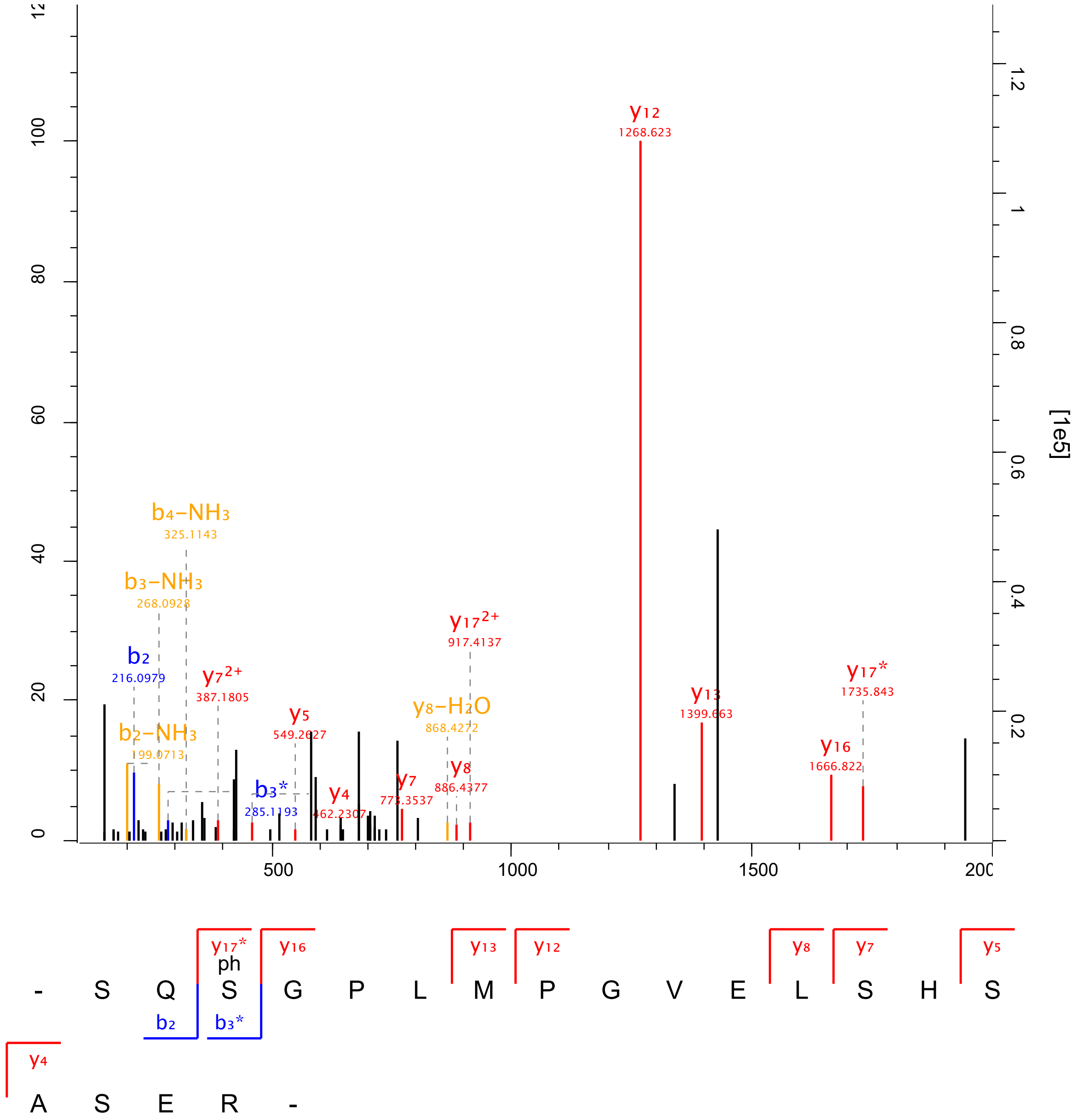

| Raw file | Scan | Method | Score | m/z |
| --- | --- | --- | --- | --- |
| sys_02_2 | 22888 | FTMS; HCD | 81.48 | 664.95 |

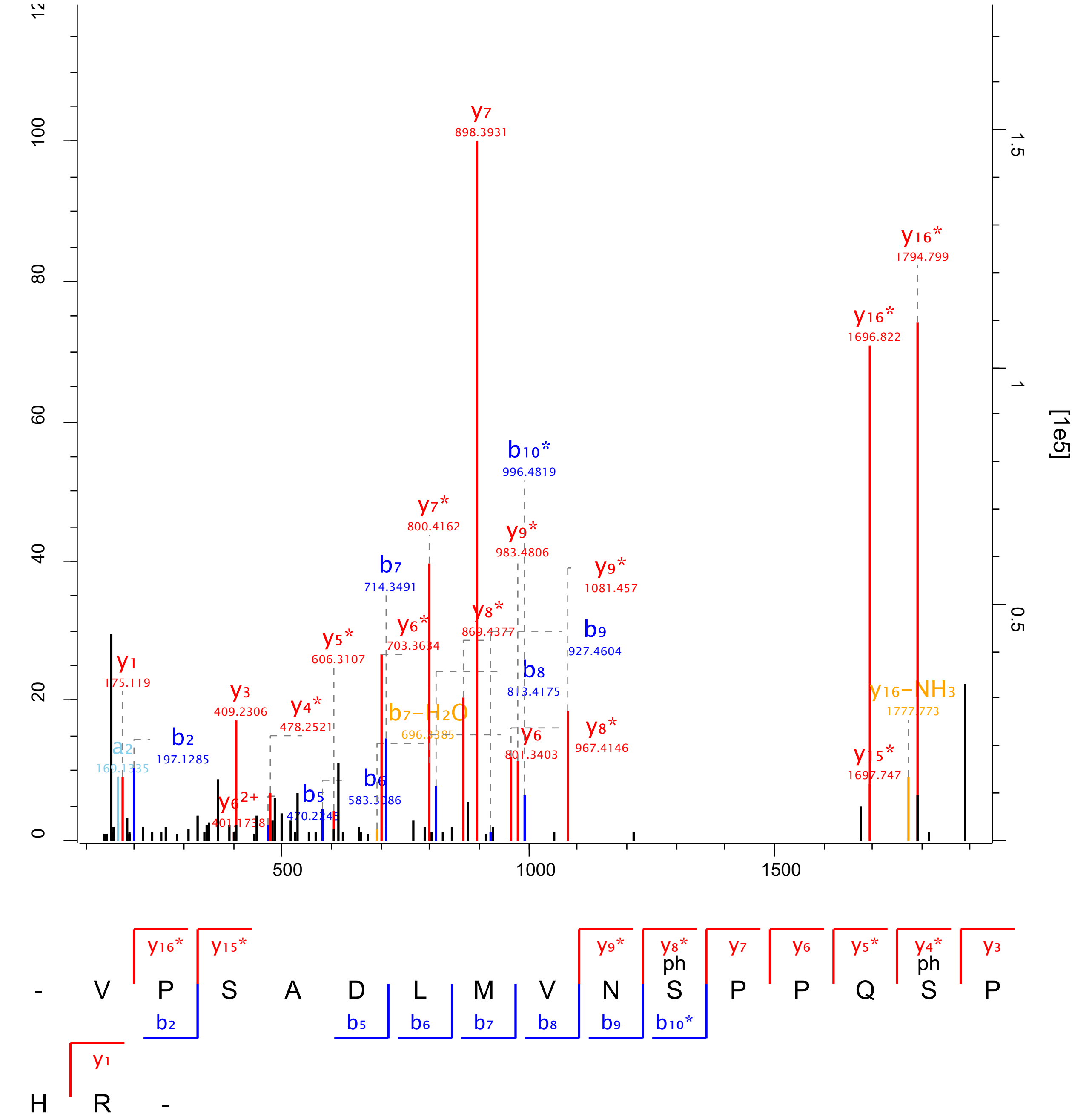

| Raw file | Scan | Method | Score | m/z |
|---|---|---|---|---|
| sys_02_2 | 22939 | FTMS; HCD | 56.08 | 653.98 |

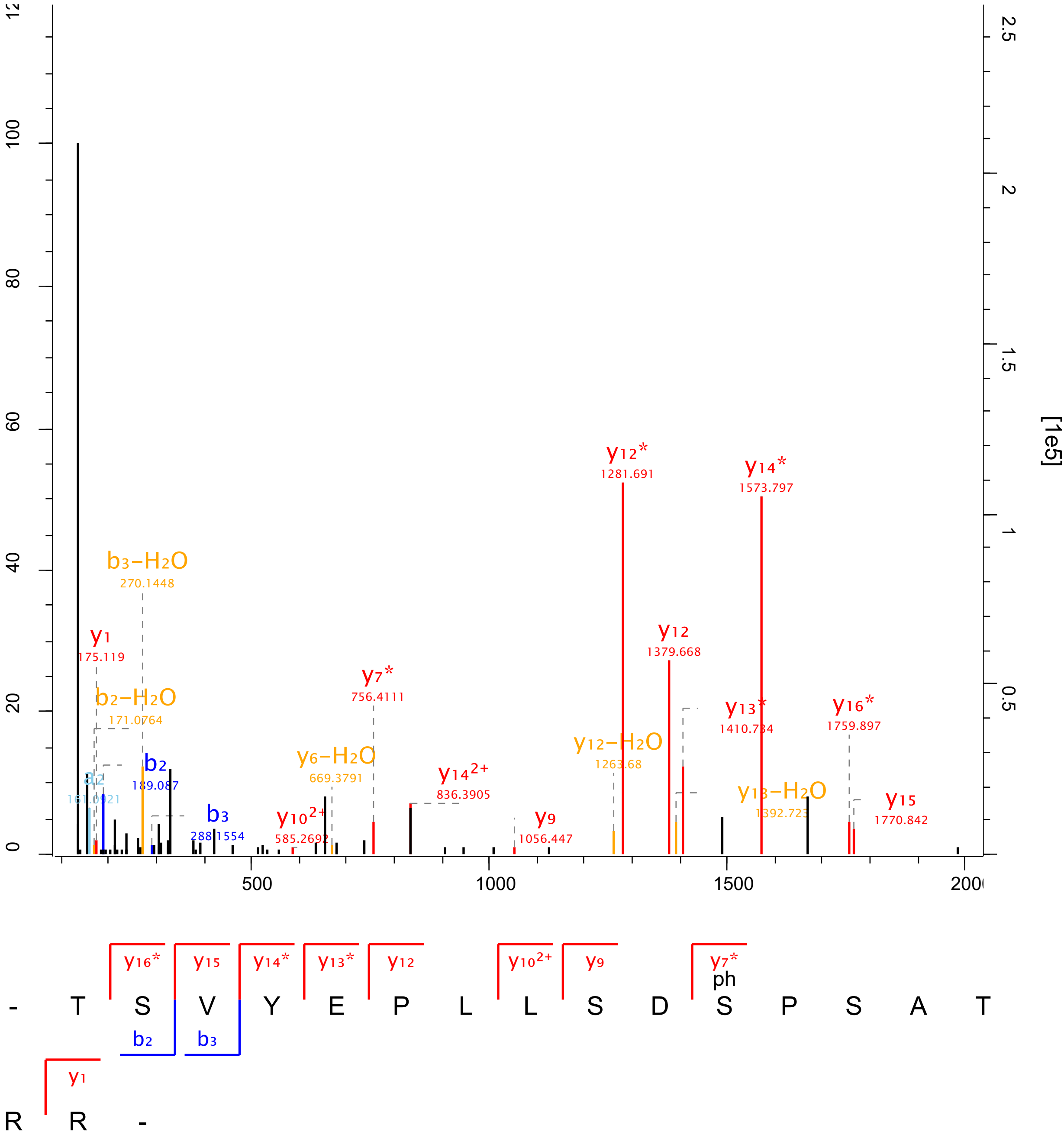

| Raw file | Scan | Method | Score | m/z |
|---|---|---|---|---|
| sys_02_2 | 23023 | FTMS; HCD | 138.75 | 762.36 |

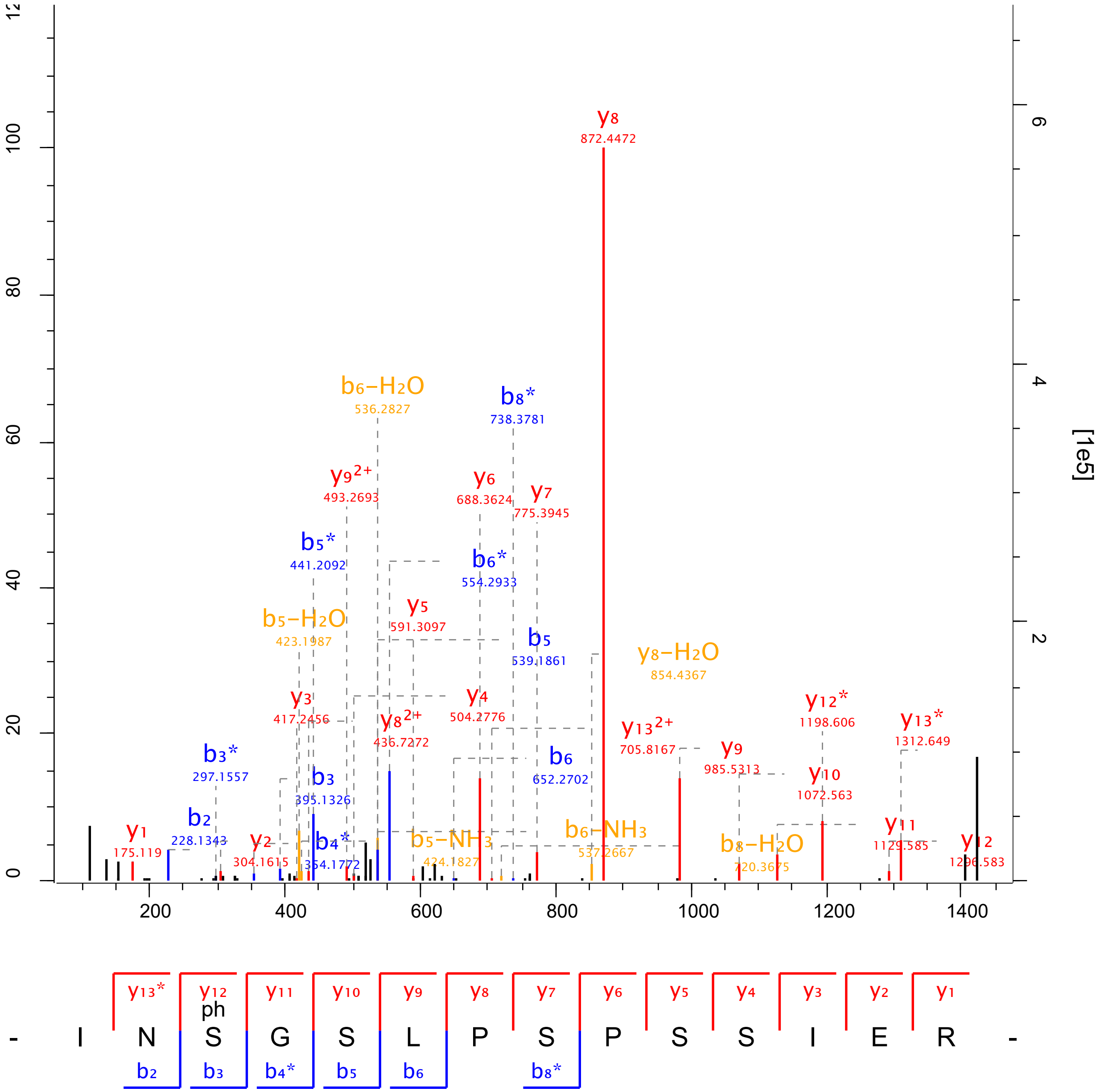

| Raw file | Scan | Method | Score | m/z |
|---|---|---|---|---|
| sys_02_2 | 23101 | FTMS; HCD | 75.56 | 786.84 |

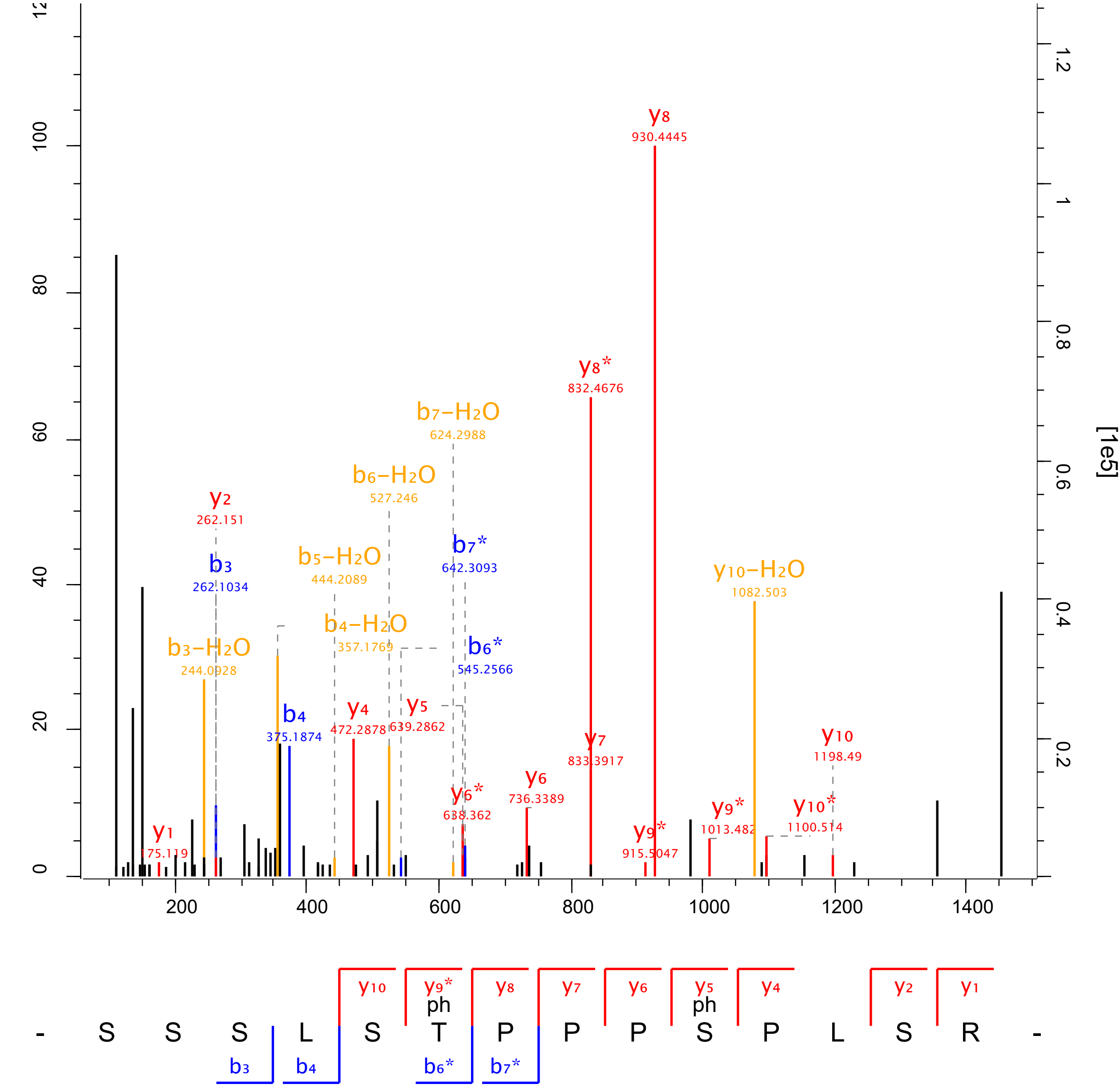

| Raw file | Scan | Method | Score | m/z |
|---|---|---|---|---|
| sys_02_2 | 23556 | FTMS; HCD | 124.32 | 933.06 |

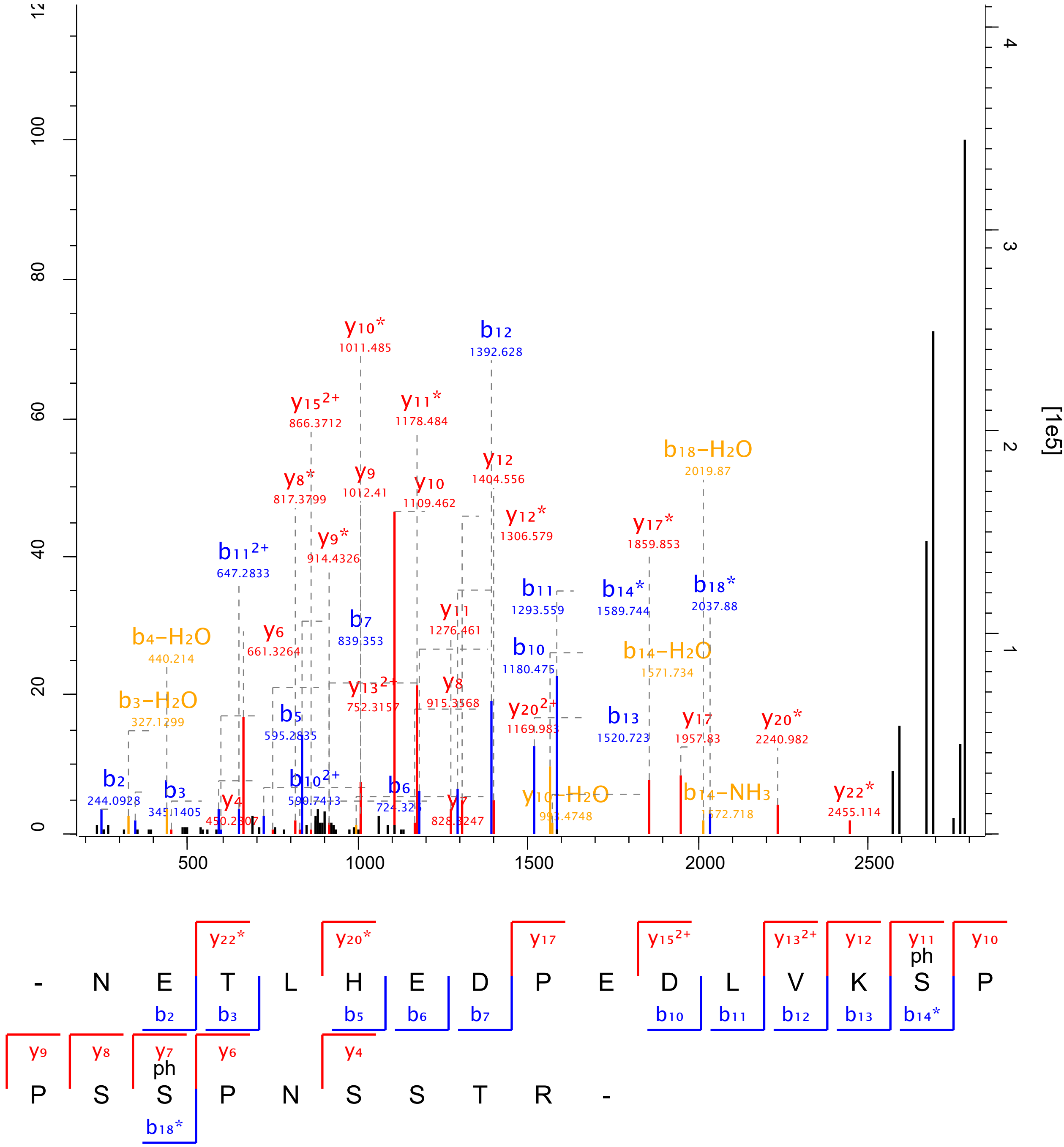

| Raw file | Scan | Method | Score | m/z |
|---|---|---|---|---|
| sys_02_2 | 23588 | FTMS; HCD | 117.7 | 636.31 |

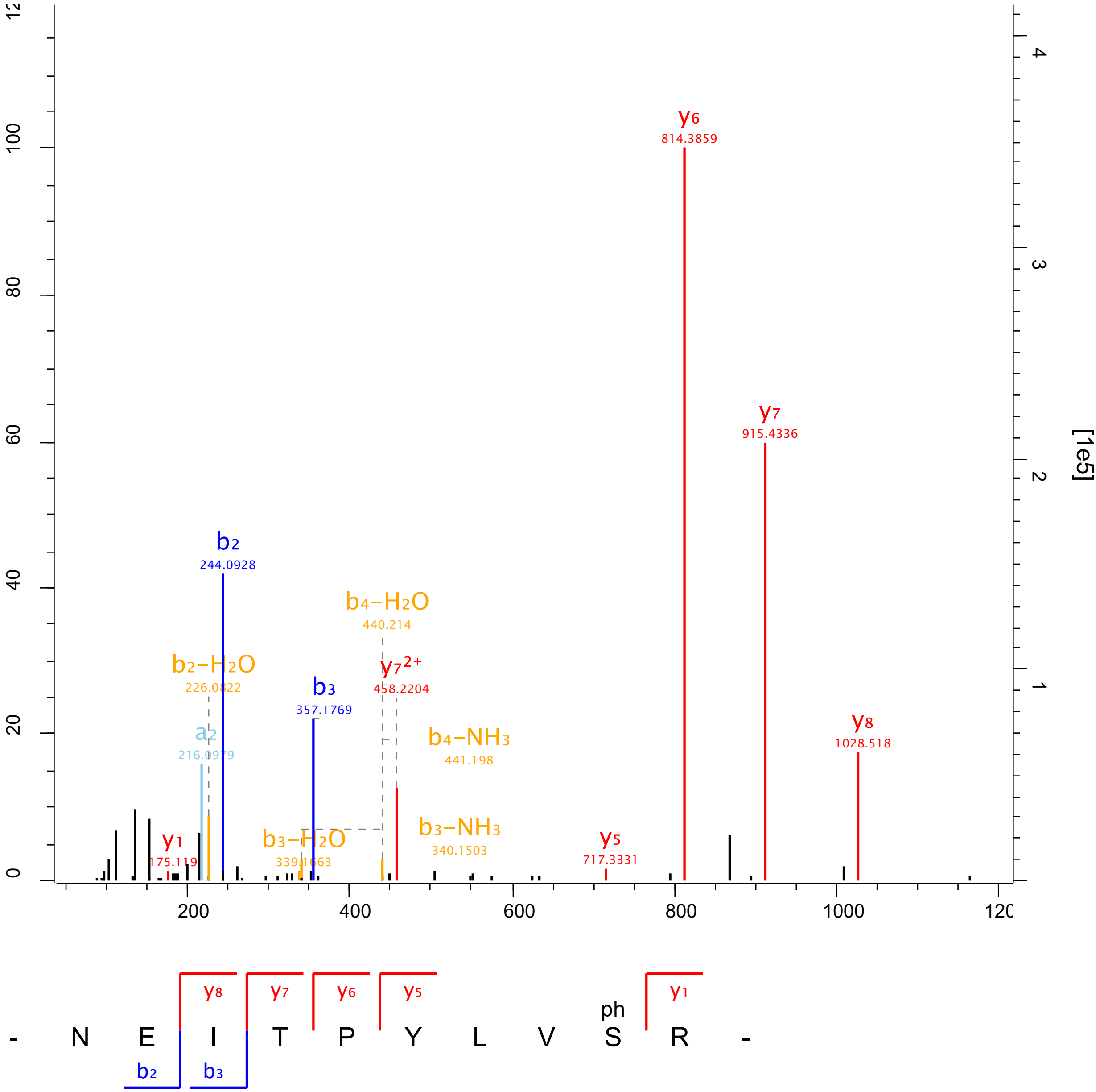

| Raw file | Scan | Method | Score | m/z |
|---|---|---|---|---|
| sys_02_2 | 23839 | FTMS; HCD | 89.66 | 939.41 |

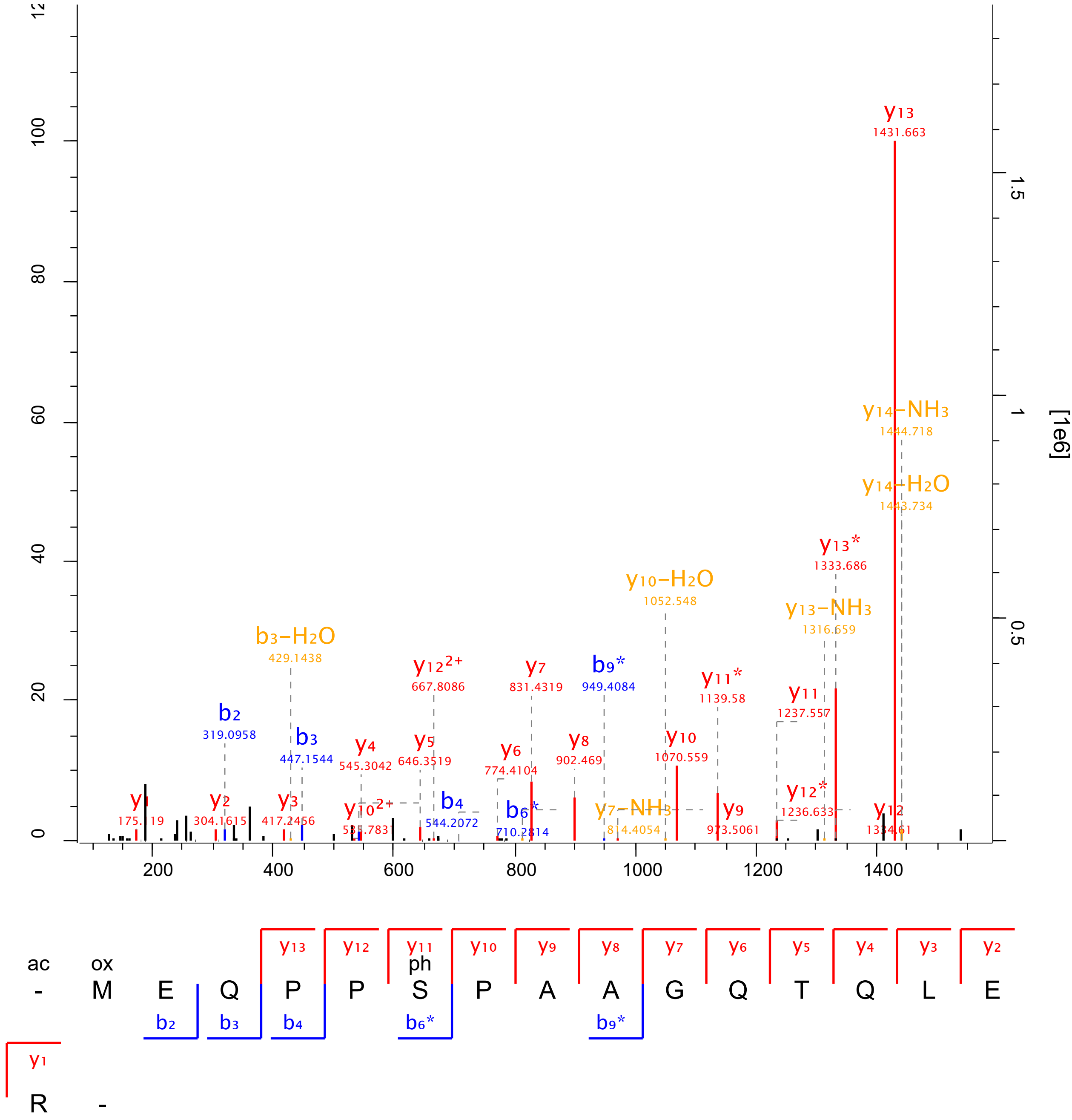

| Raw file | Scan | Method | Score | m/z |
|---|---|---|---|---|
| sys_02_2 | 23919 | FTMS; HCD | 158.25 | 537.77 |

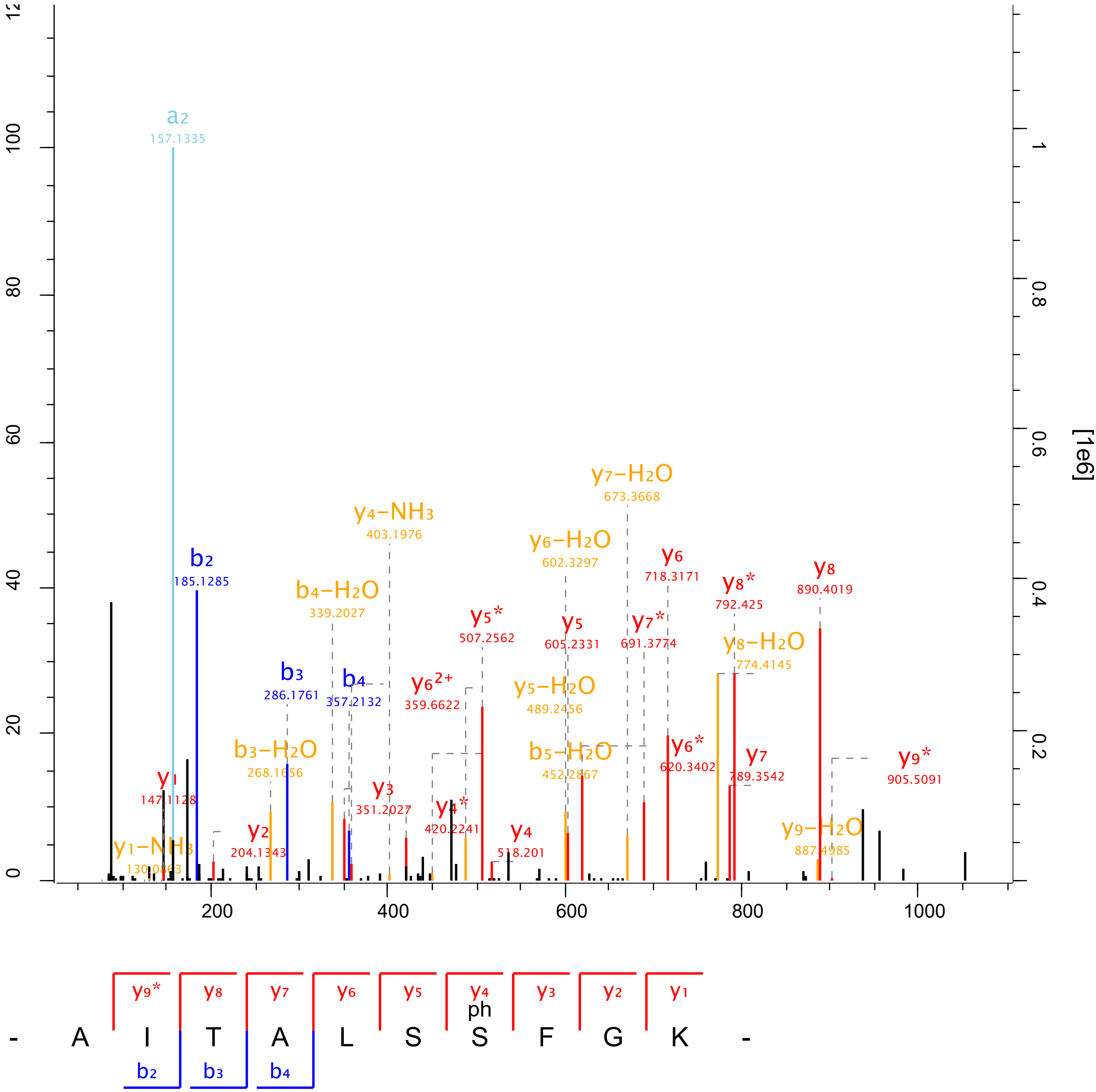

| Raw file | Scan | Method | Score | m/z |
|---|---|---|---|---|
| sys_02_2 | 24054 | FTMS; HCD | 229.51 | 920.39 |

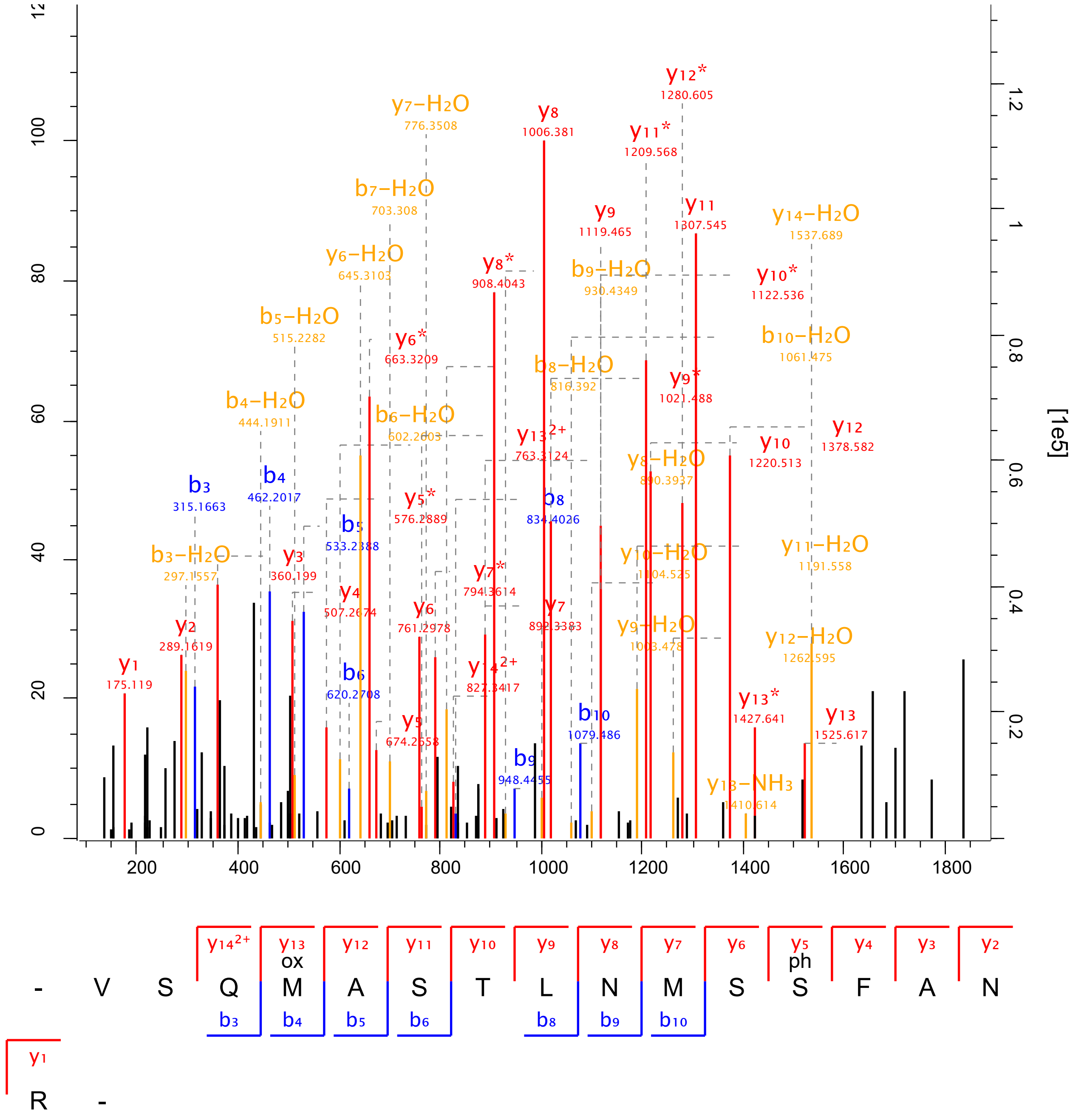

| Raw file | Scan | Method | Score | m/z |
| --- | --- | --- | --- | --- |
| sys_02_2 | 24114 | FTMS; HCD | 44.34 | 790.72 |

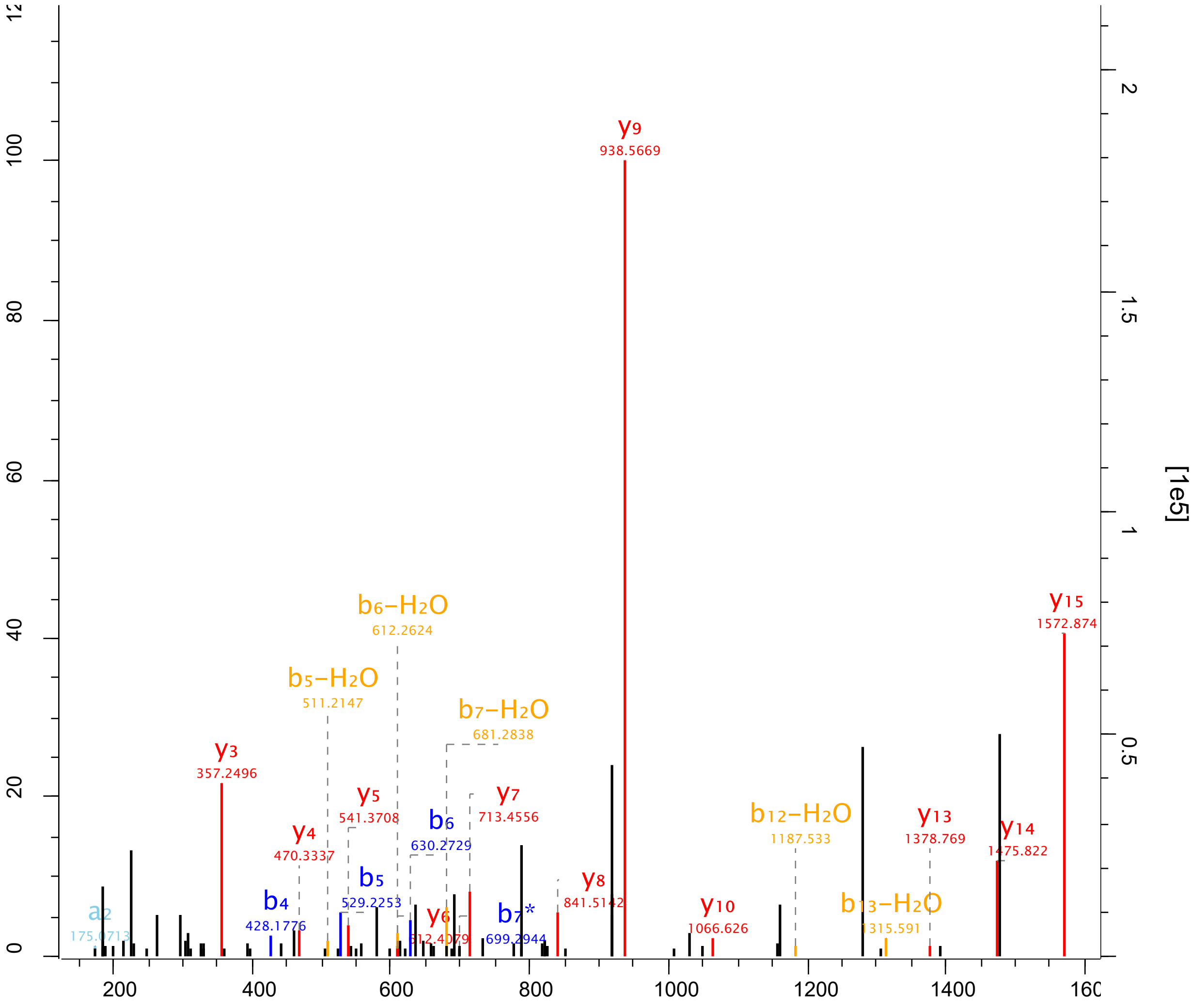


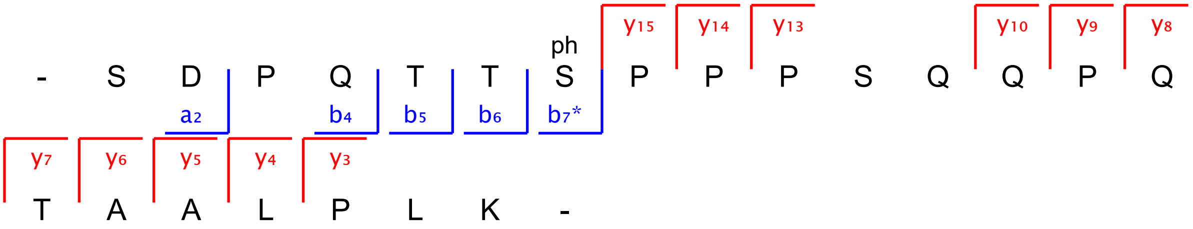

| Raw file | Scan | Method | Score | m/z |
|---|---|---|---|---|
| sys_02_2 | 24136 | FTMS; HCD | 67.4 | 880.9 |

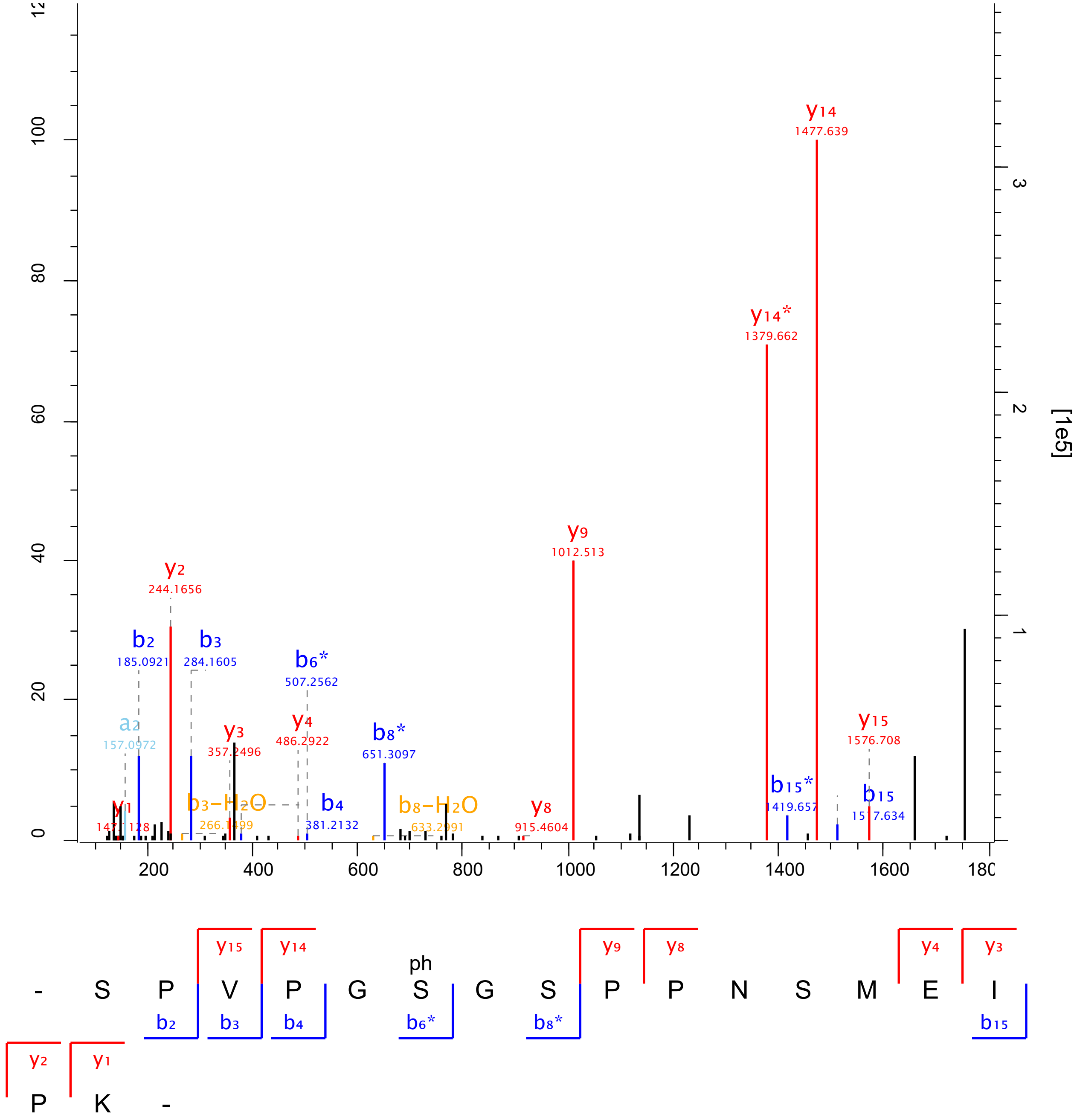

| Raw file | Scan | Method | Score | m/z |
|---|---|---|---|---|
| sys_02_2 | 24208 | FTMS; HCD | 45.28 | 572.76 |

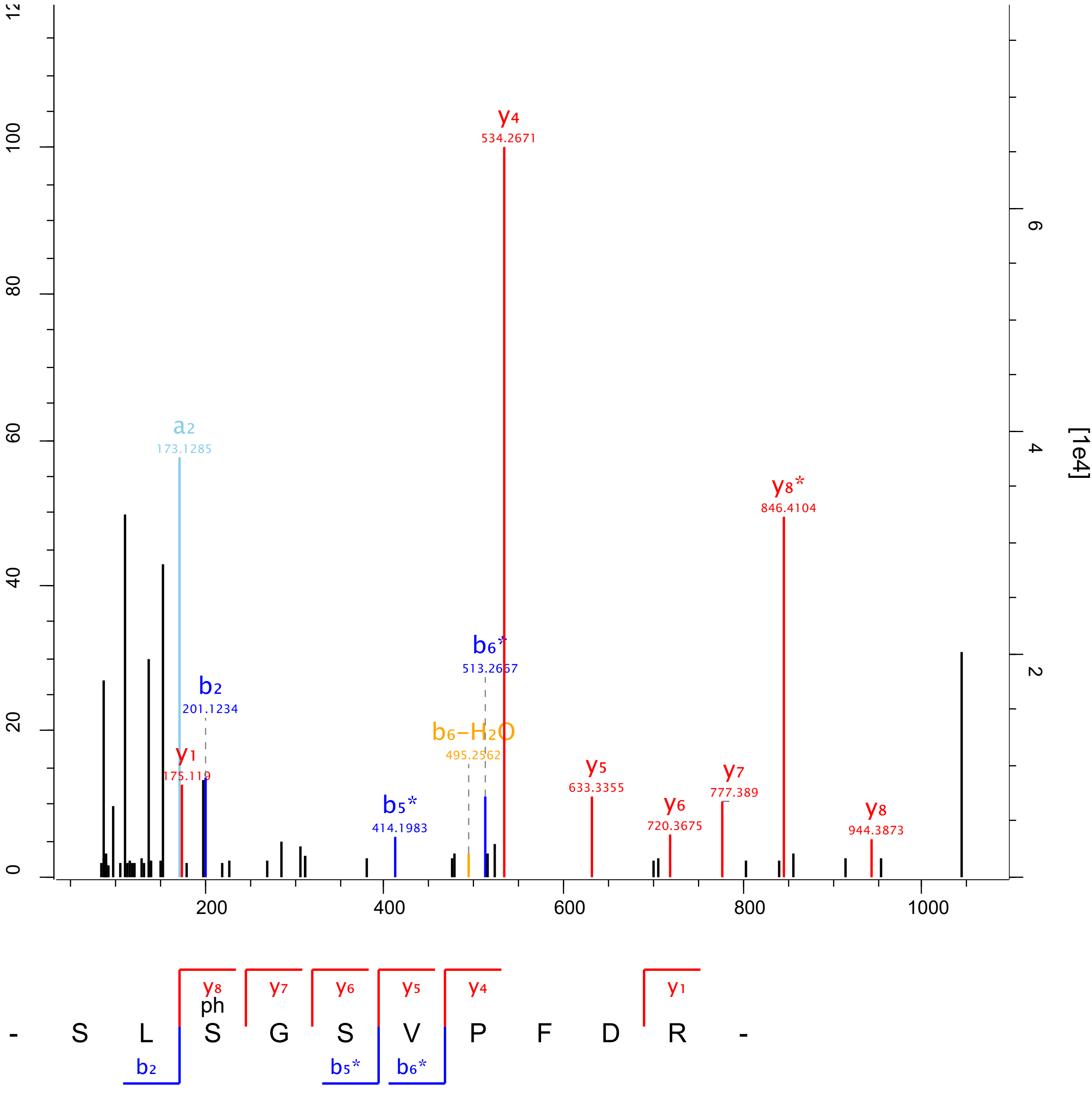

| Raw file | Scan | Method | Score | m/z |
|---|---|---|---|---|
| sys_02_2 | 2445 | FTMS; HCD | 42.34 | 525.68 |

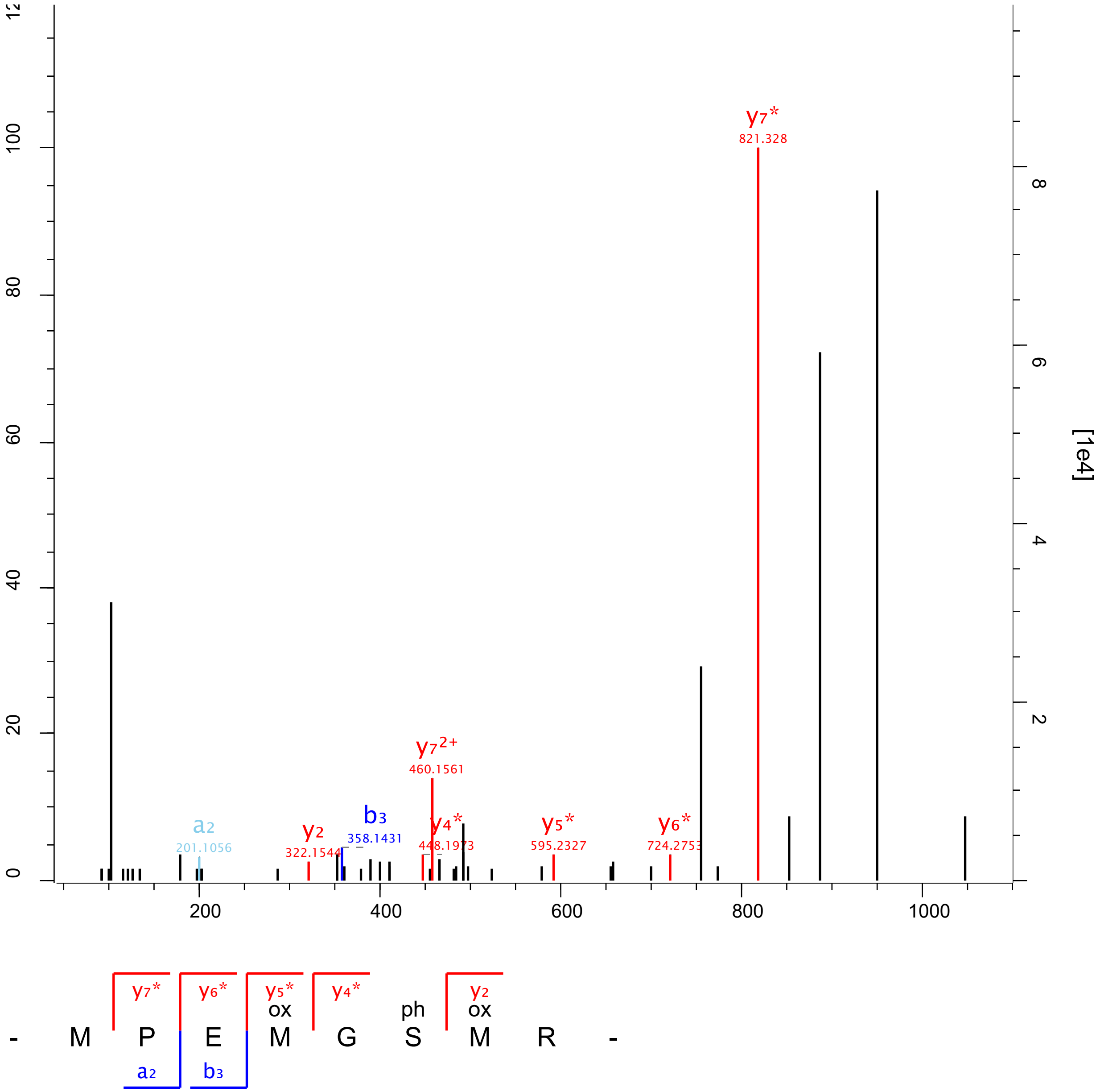

| Raw file | Scan | Method | Score | m/z |
|---|---|---|---|---|
| sys_02_2 | 24655 | FTMS; HCD | 115.42 | 499.26 |

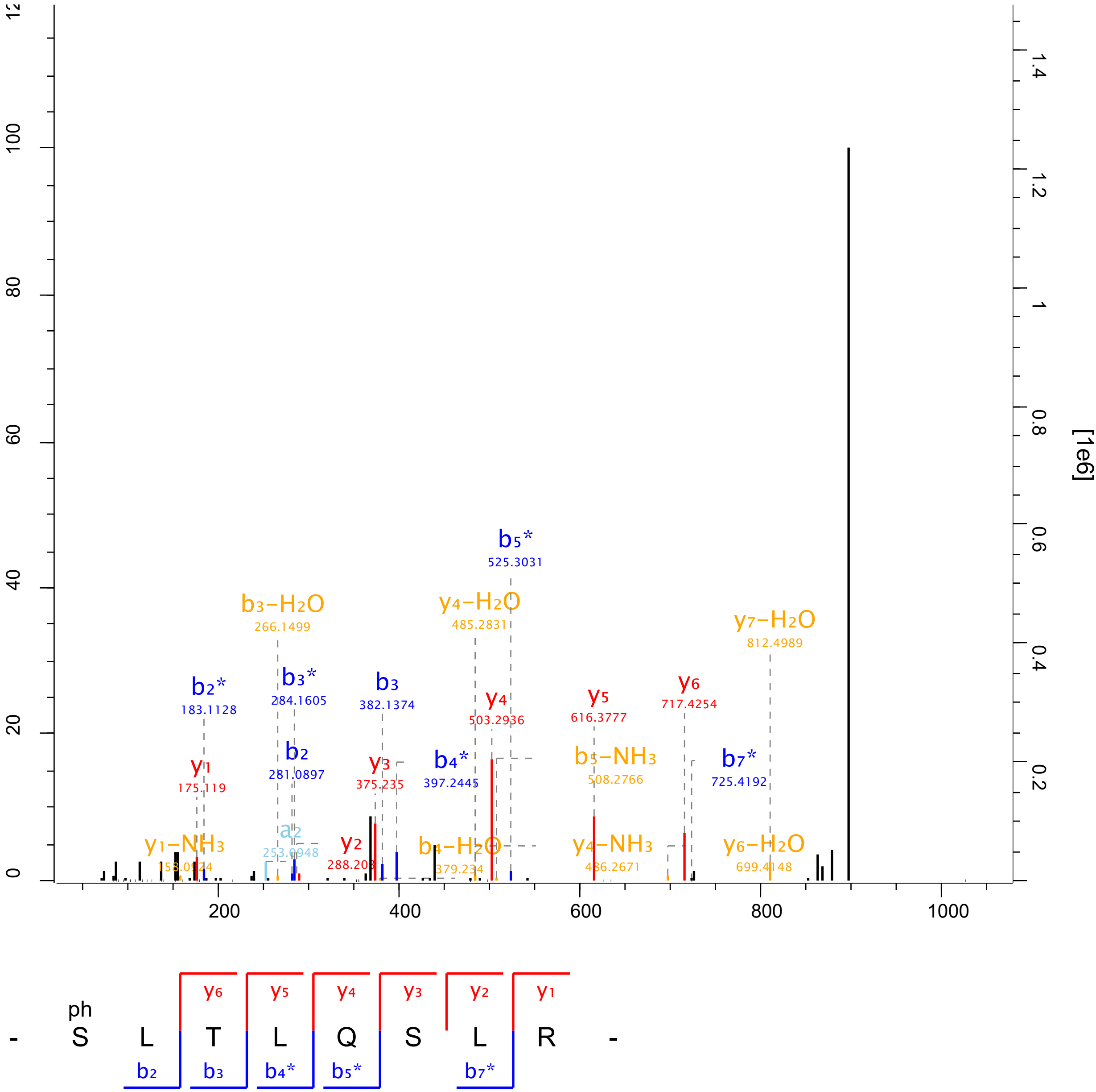

| Raw file | Scan | Method | Score | m/z |
|---|---|---|---|---|
| sys_02_2 | 25351 | FTMS; HCD | 127.2 | 1027.12 |

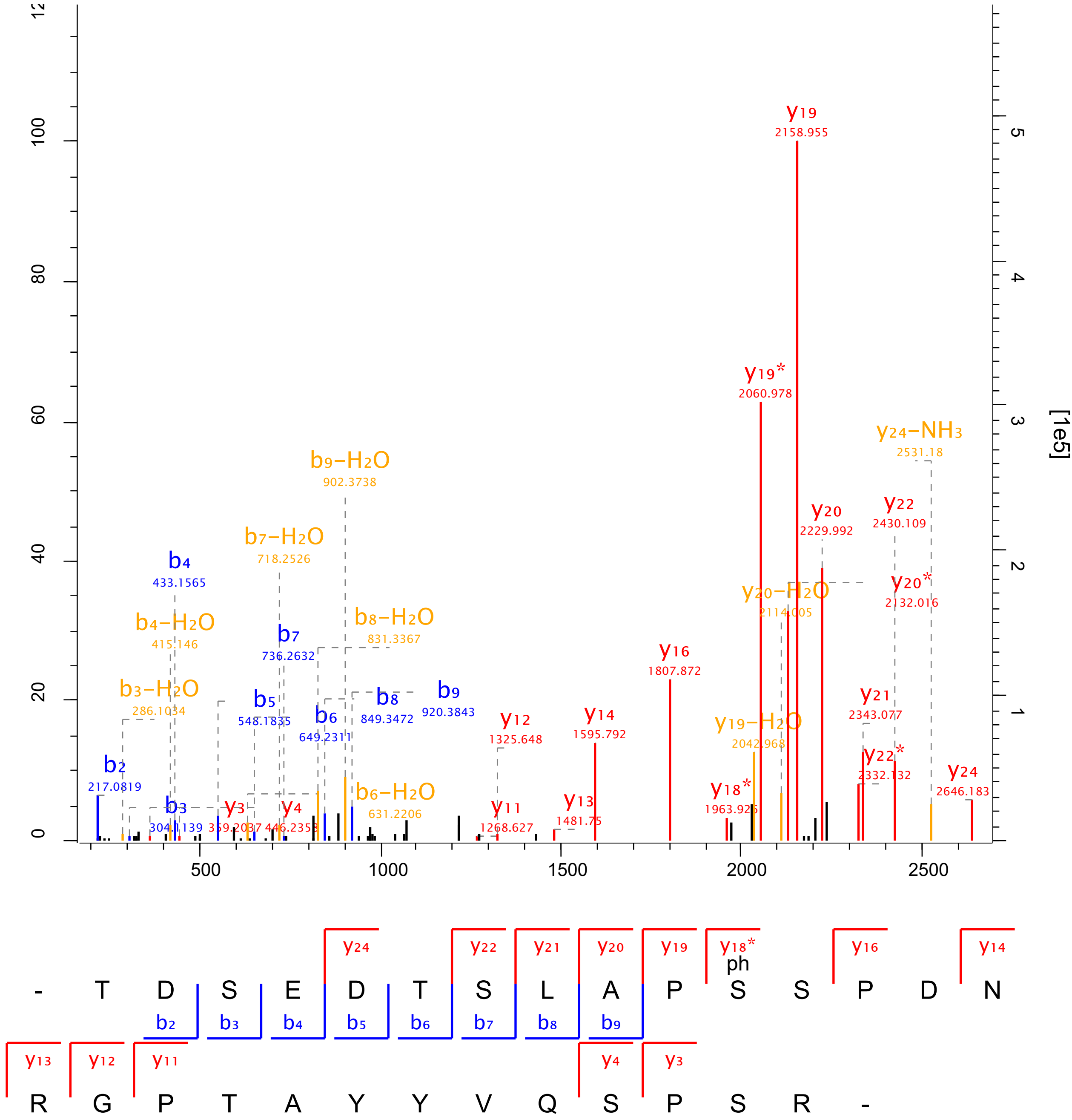

| Raw file | Scan | Method | Score | m/z |
|---|---|---|---|---|
| sys_02_2 | 25751 | FTMS; HCD | 199.09 | 782.87 |

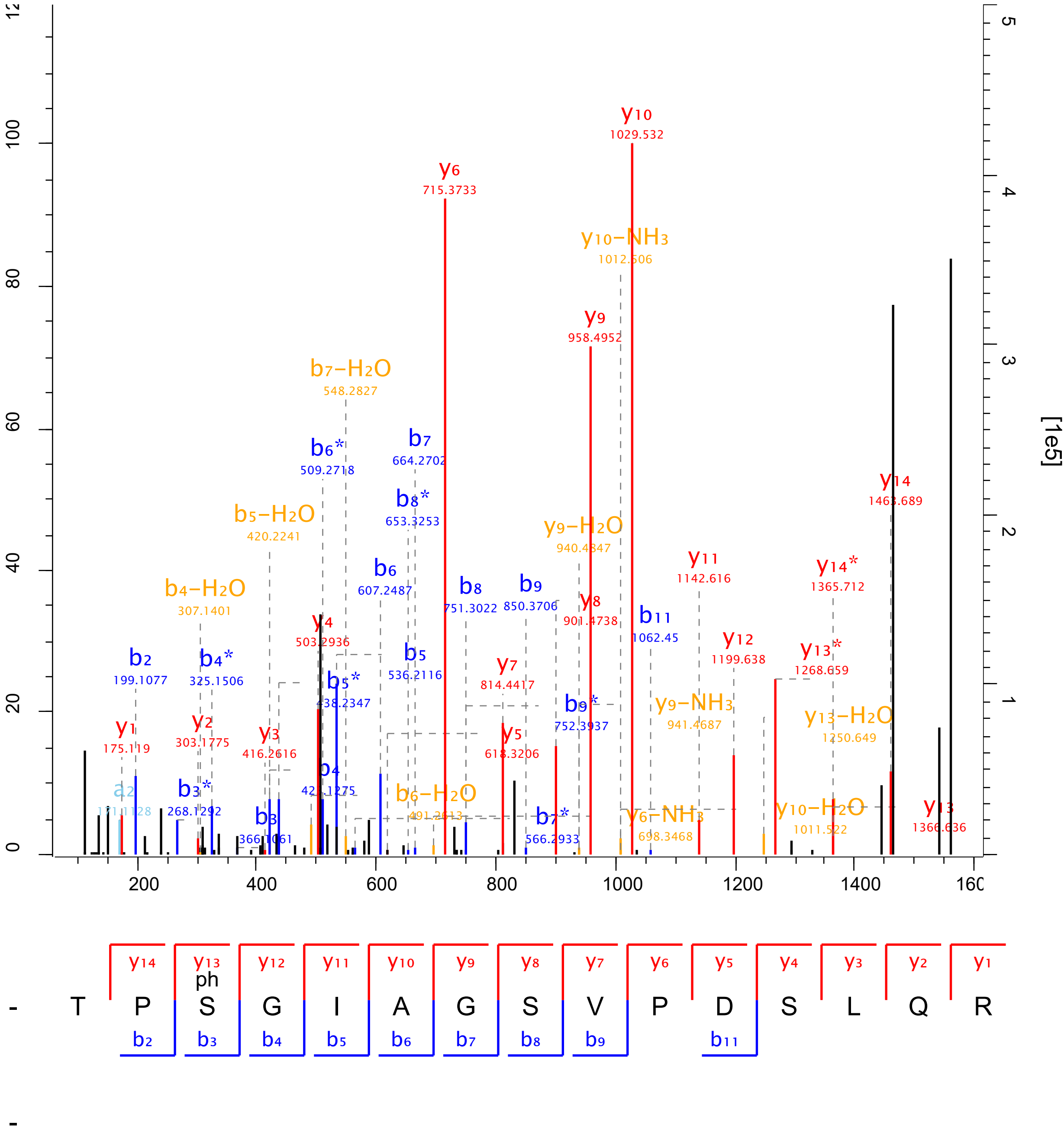

| Raw file | Scan | Method | Score | m/z |
|---|---|---|---|---|
| sys_02_2 | 25862 | FTMS; HCD | 219.23 | 682.35 |

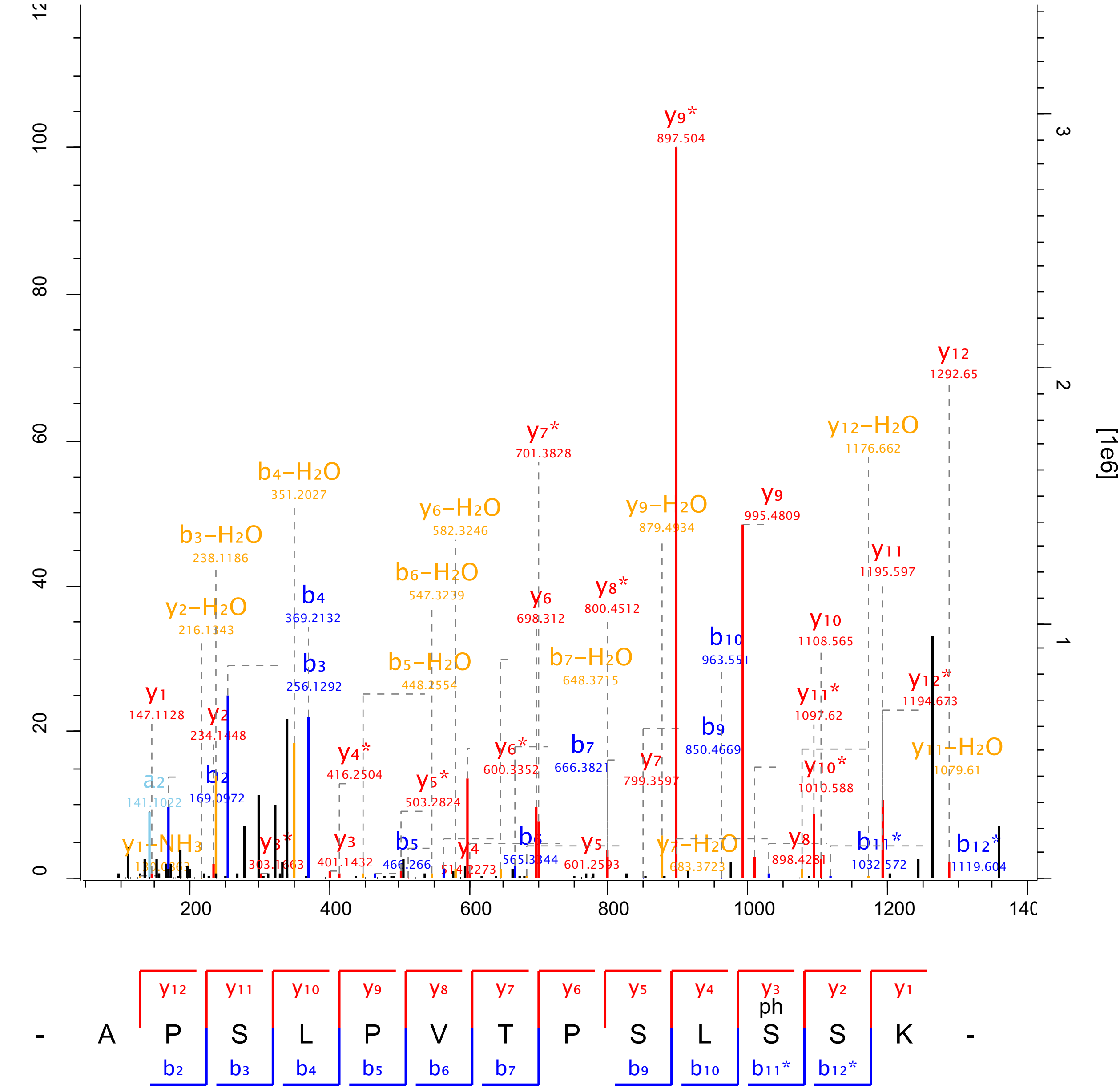

| Raw file | Scan | Method | Score | m/z |
| --- | --- | --- | --- | --- |
| sys_02_2 | 2598 | FTMS; HCD | 119.54 | 625.8 |

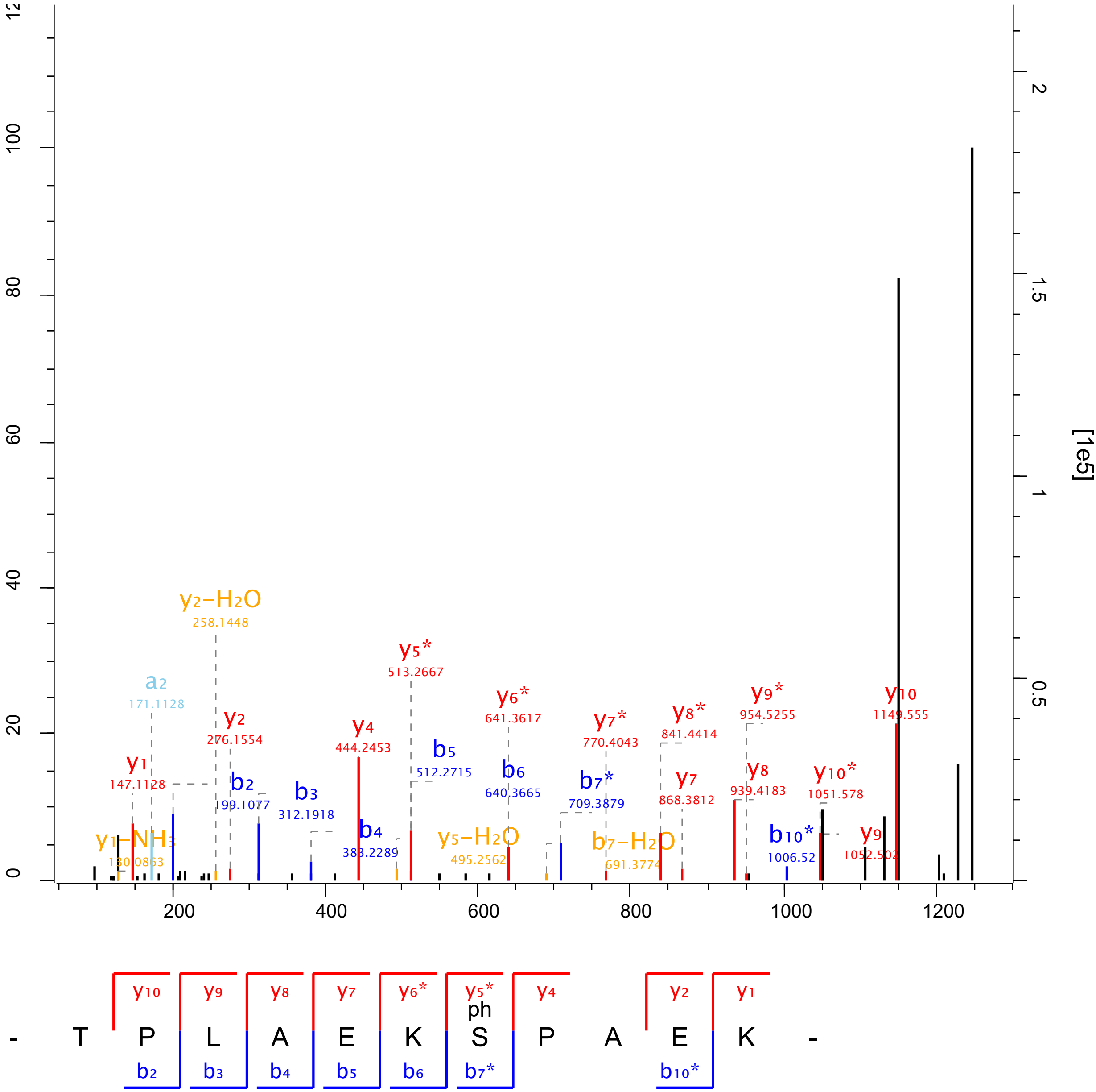

| Raw file | Scan | Method | Score | m/z |
|---|---|---|---|---|
| sys_02_2 | 26009 | FTMS; HCD | 55.26 | 550.77 |

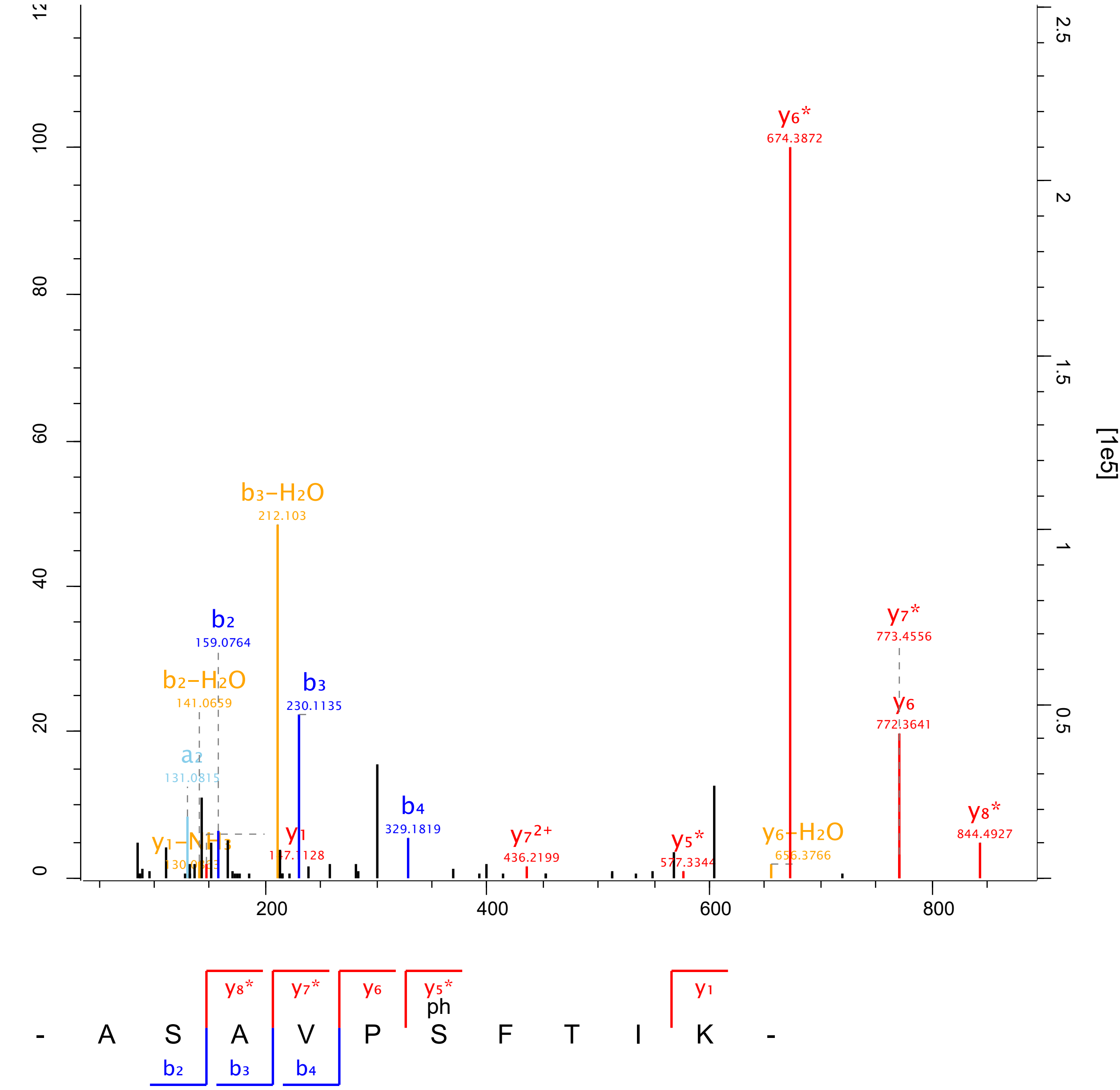

| Raw file | Scan | Method | Score | m/z |
| --- | --- | --- | --- | --- |
| sys_02_2 | 2624 | FTMS; HCD | 43.9 | 399.17 |

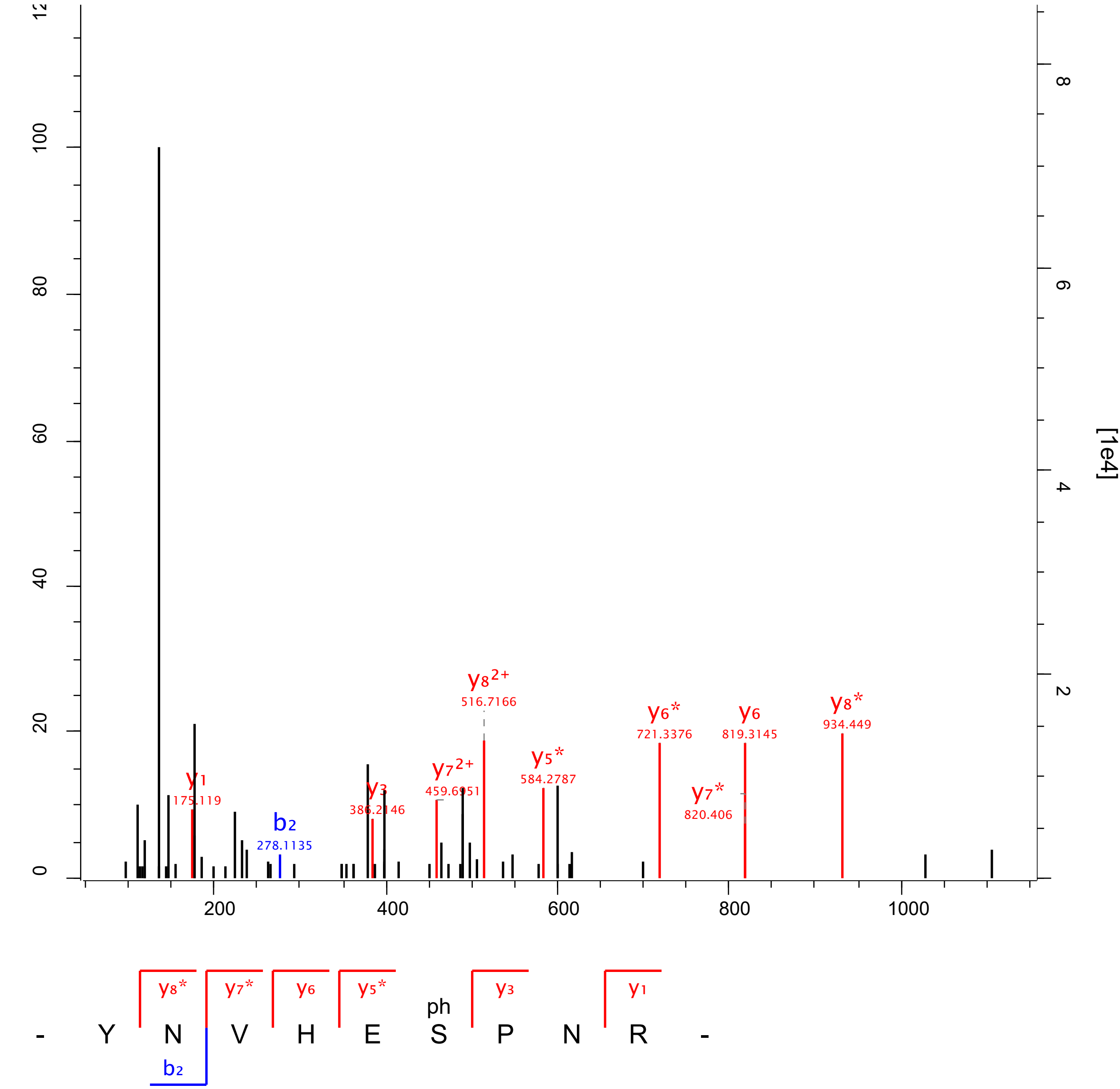

| Raw file | Scan | Method | Score | m/z |
|---|---|---|---|---|
| sys_02_2 | 26446 | FTMS; HCD | 188.15 | 590.75 |

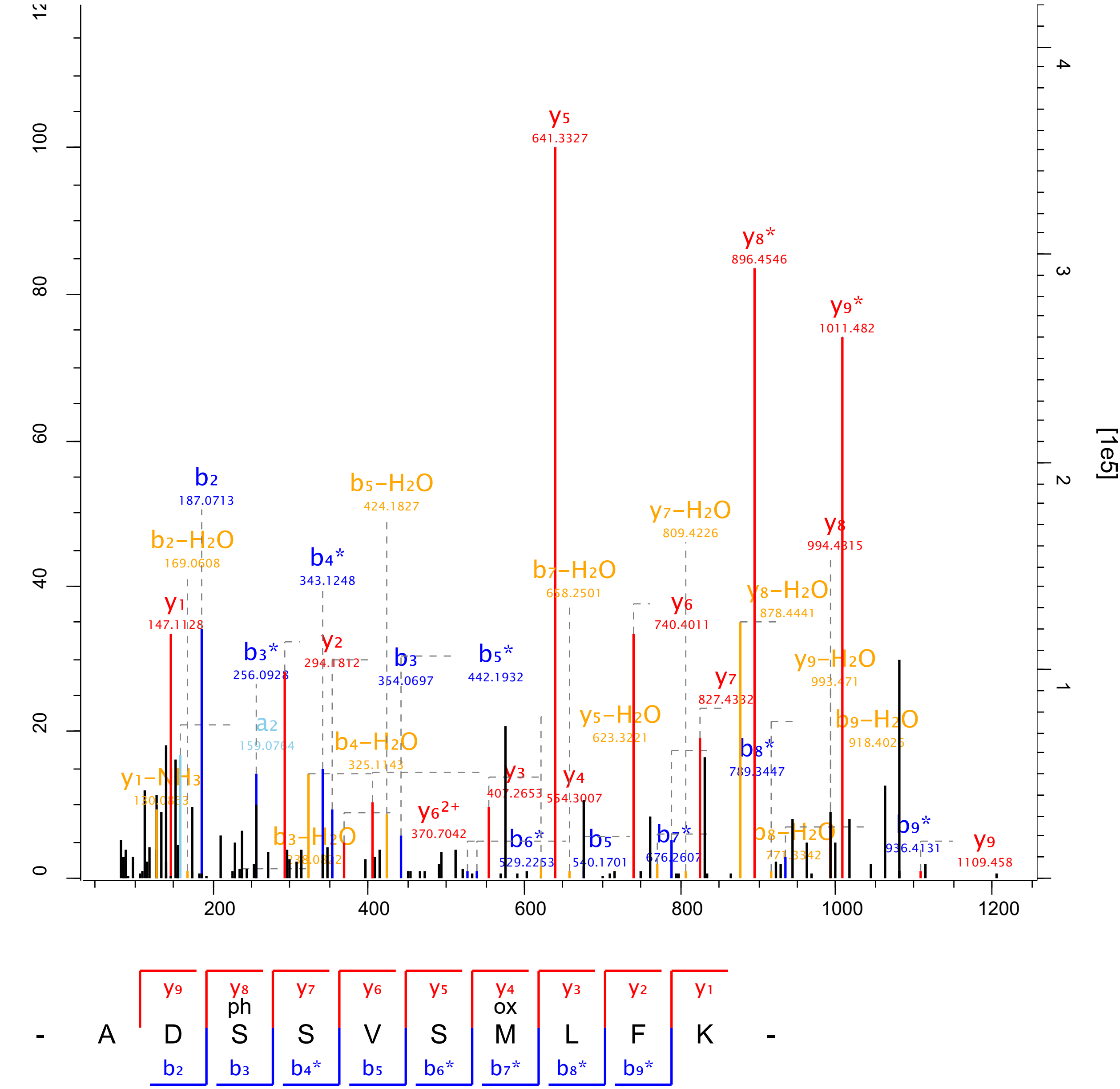

| Raw file | Scan | Method | Score | m/z |
|---|---|---|---|---|
| sys_02_2 | 2645 | FTMS; HCD | 66.27 | 395.86 |

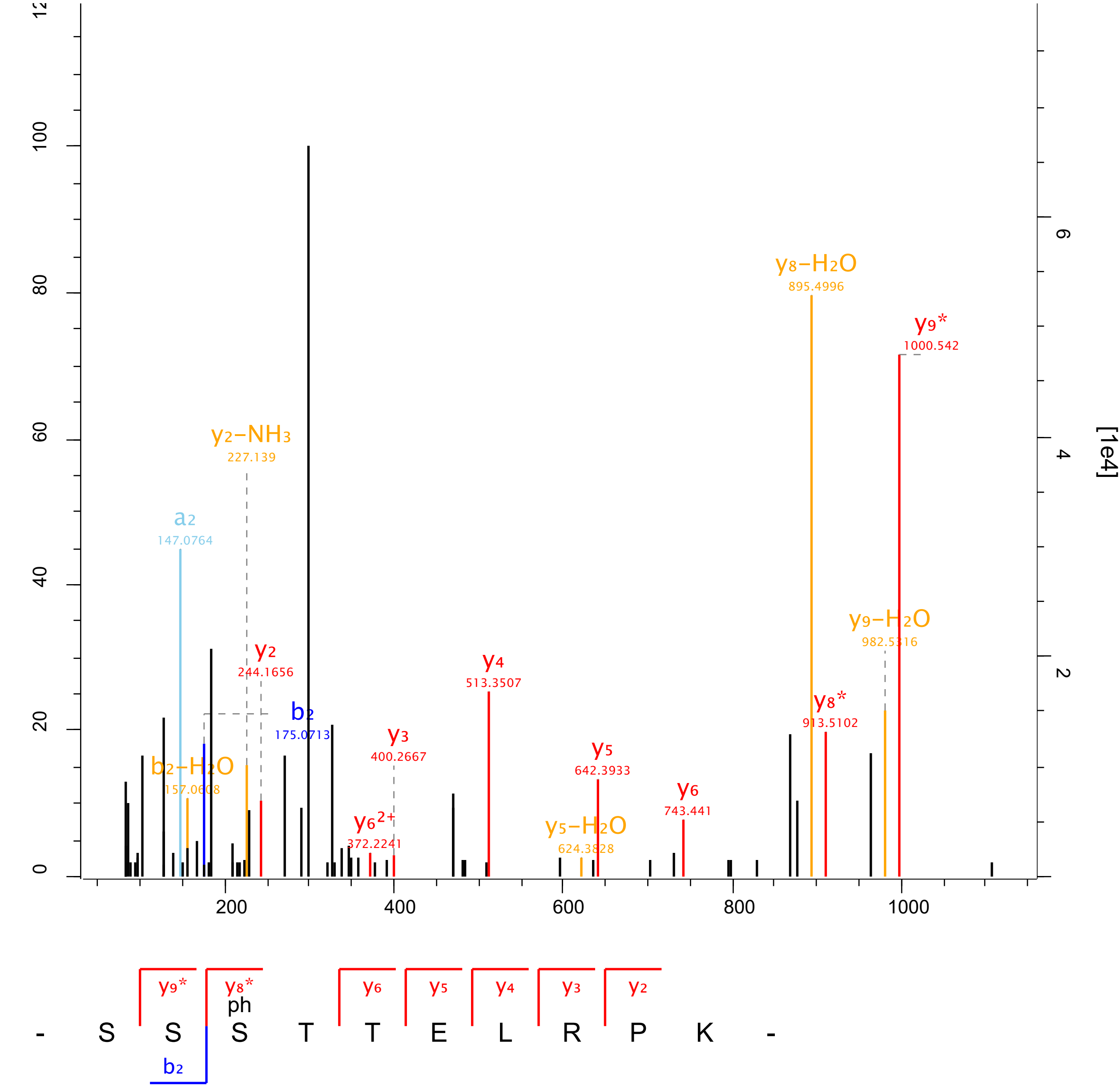

| Raw file | Scan | Method | Score | m/z |
|---|---|---|---|---|
| sys_02_2 | 26563 | FTMS; HCD | 95.12 | 722.96 |

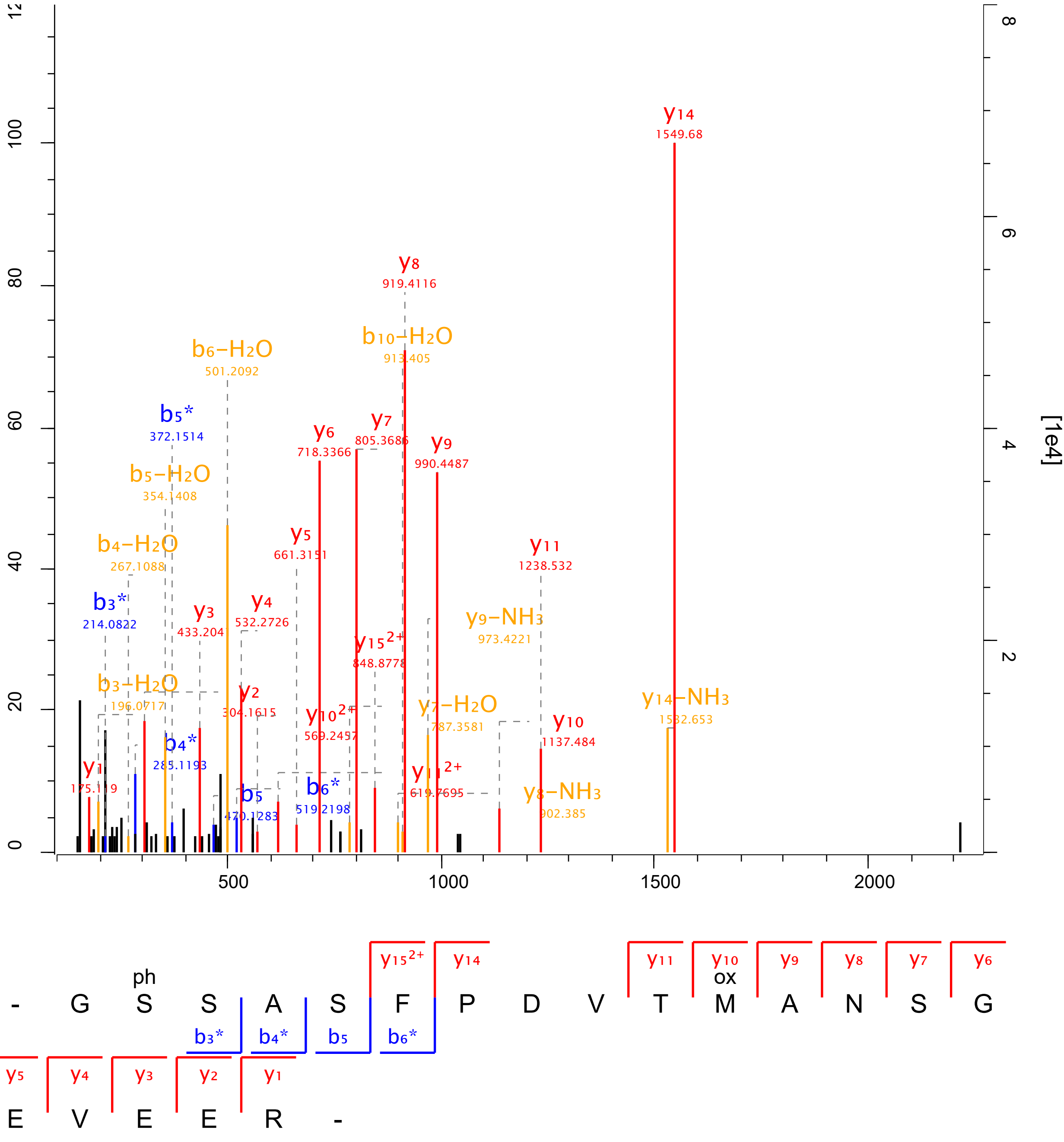

| Raw file | Scan | Method | Score | m/z |
|---|---|---|---|---|
| sys_02_2 | 27519 | FTMS; HCD | 42.07 | 811.39 |

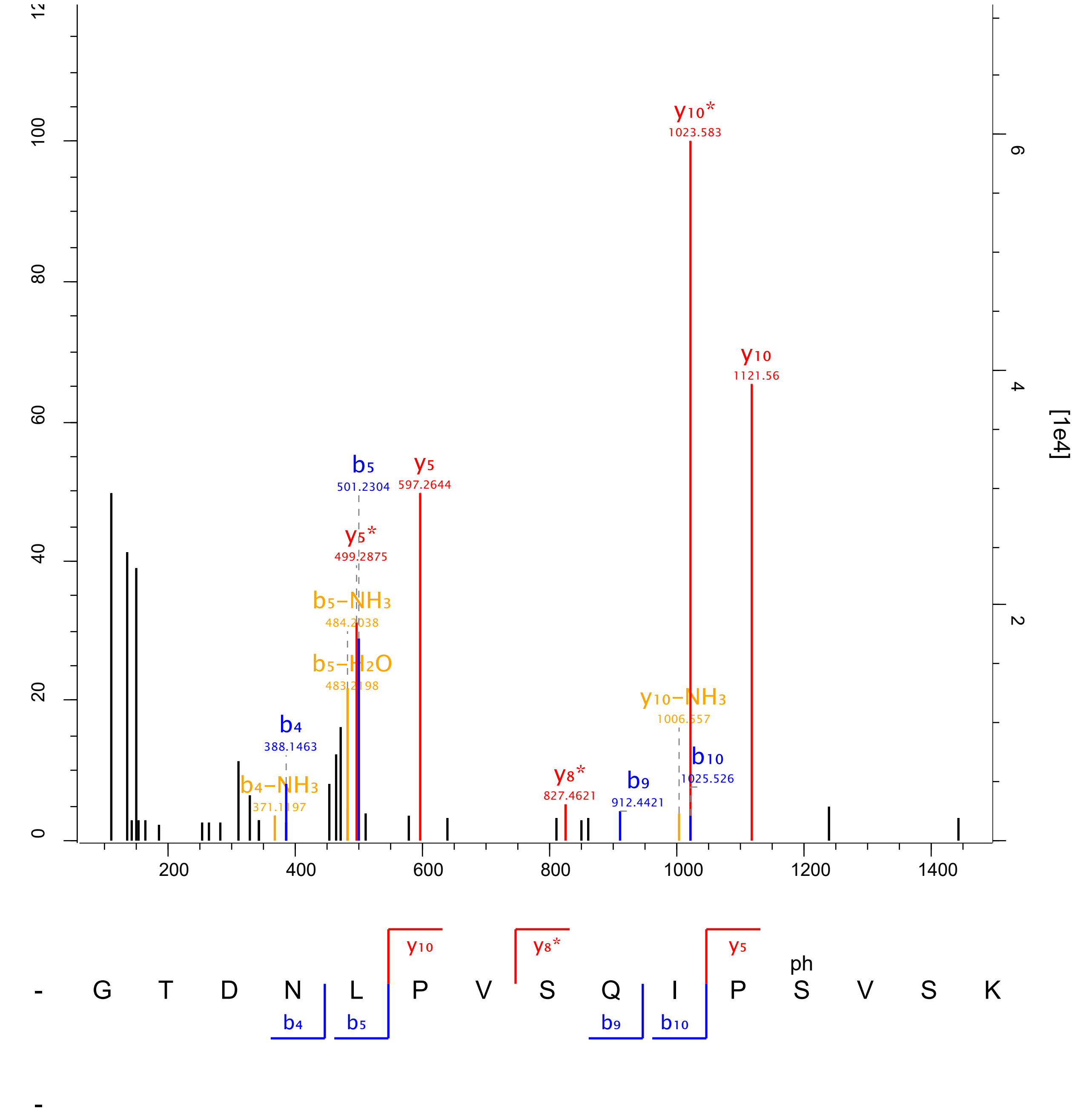

| Raw file | Scan | Method | Score | m/z |
|---|---|---|---|---|
| sys_02_2 | 27621 | FTMS; HCD | 61.13 | 741.32 |

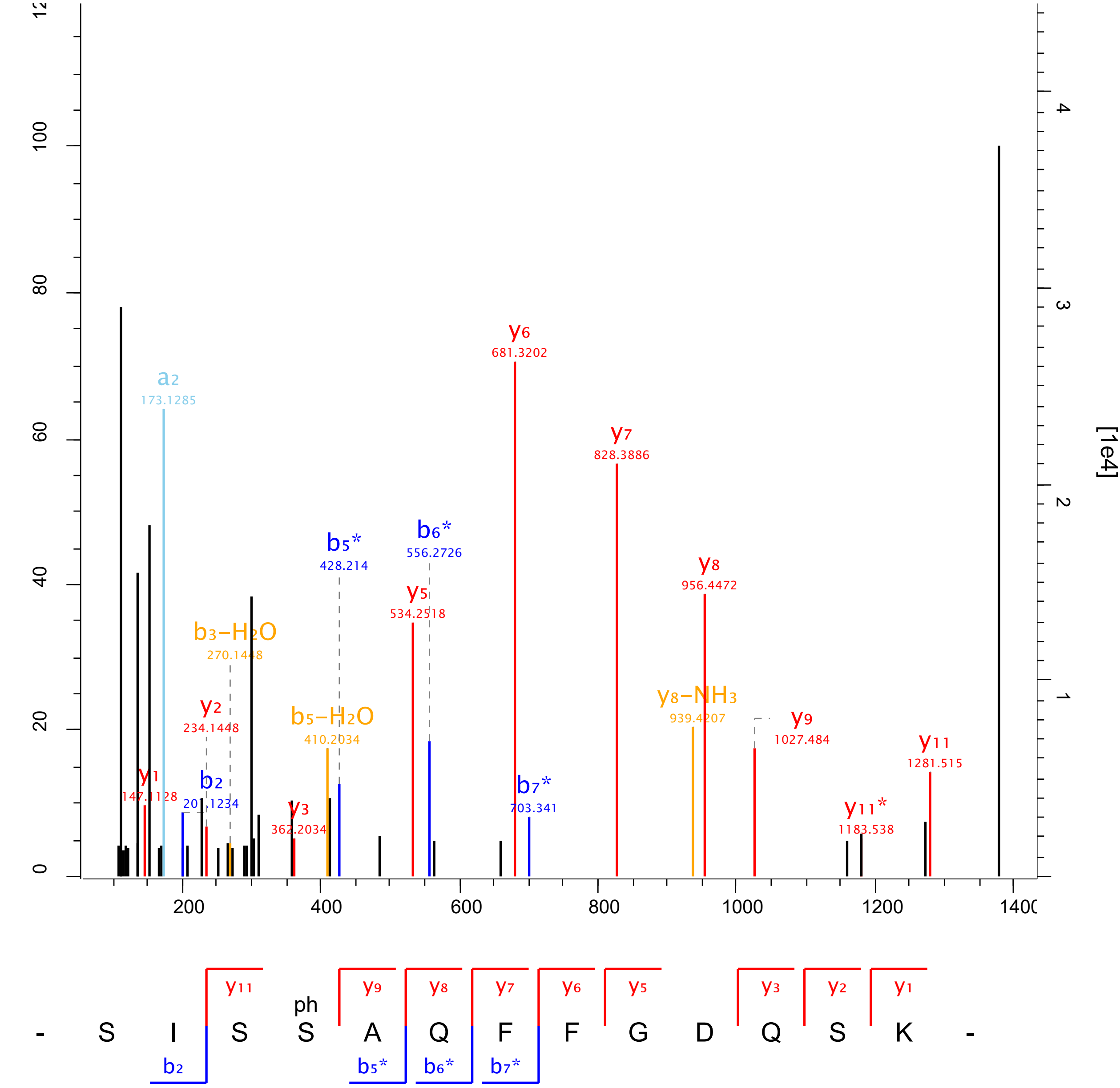

| Raw file | Scan | Method | Score | m/z |
|---|---|---|---|---|
| sys_02_2 | 2792 | FTMS; HCD | 75.97 | 517.87 |

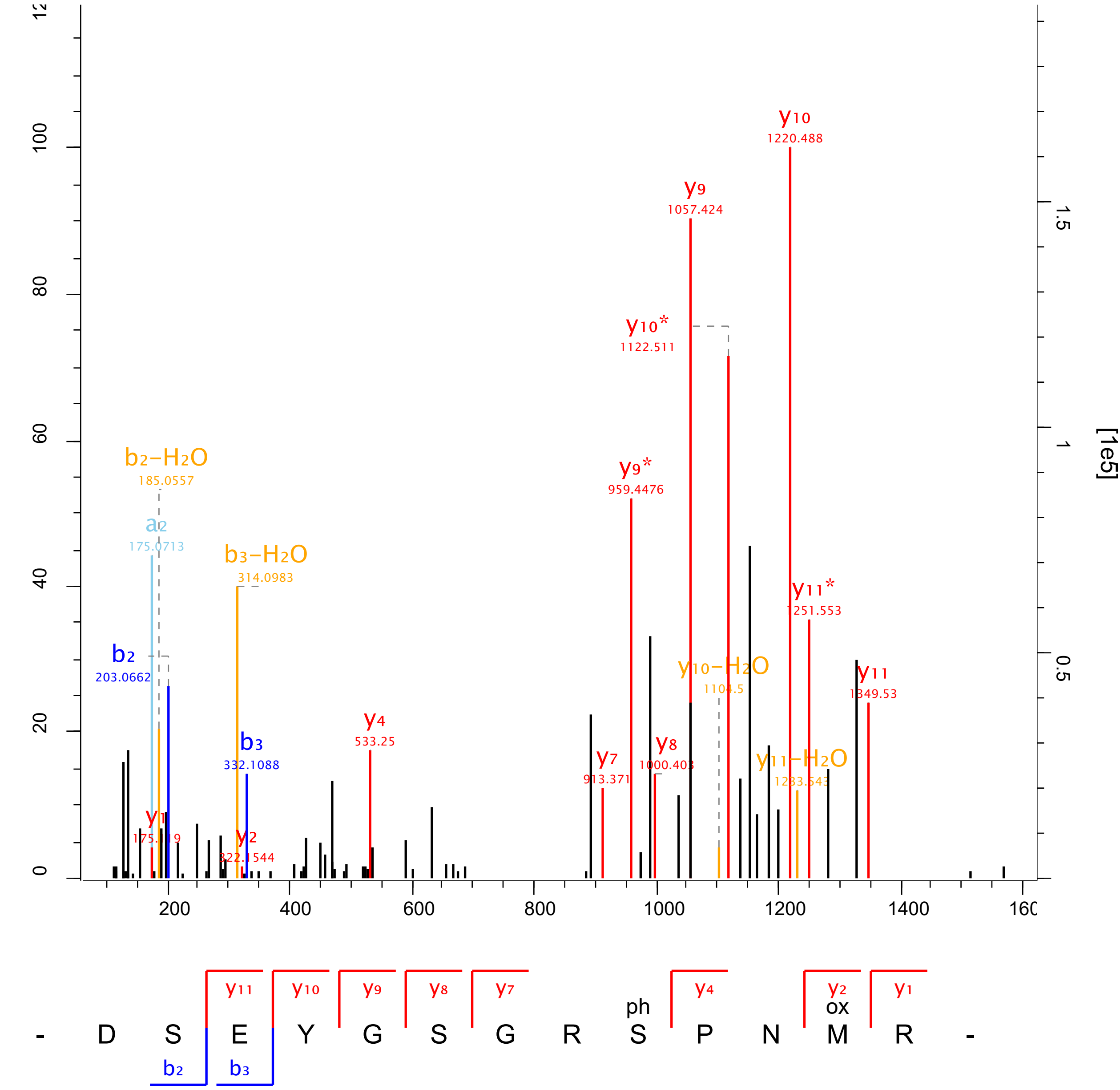

| Raw file | Scan | Method | Score | m/z |
|---|---|---|---|---|
| sys_02_2 | 2798 | FTMS; HCD | 70.44 | 588.79 |

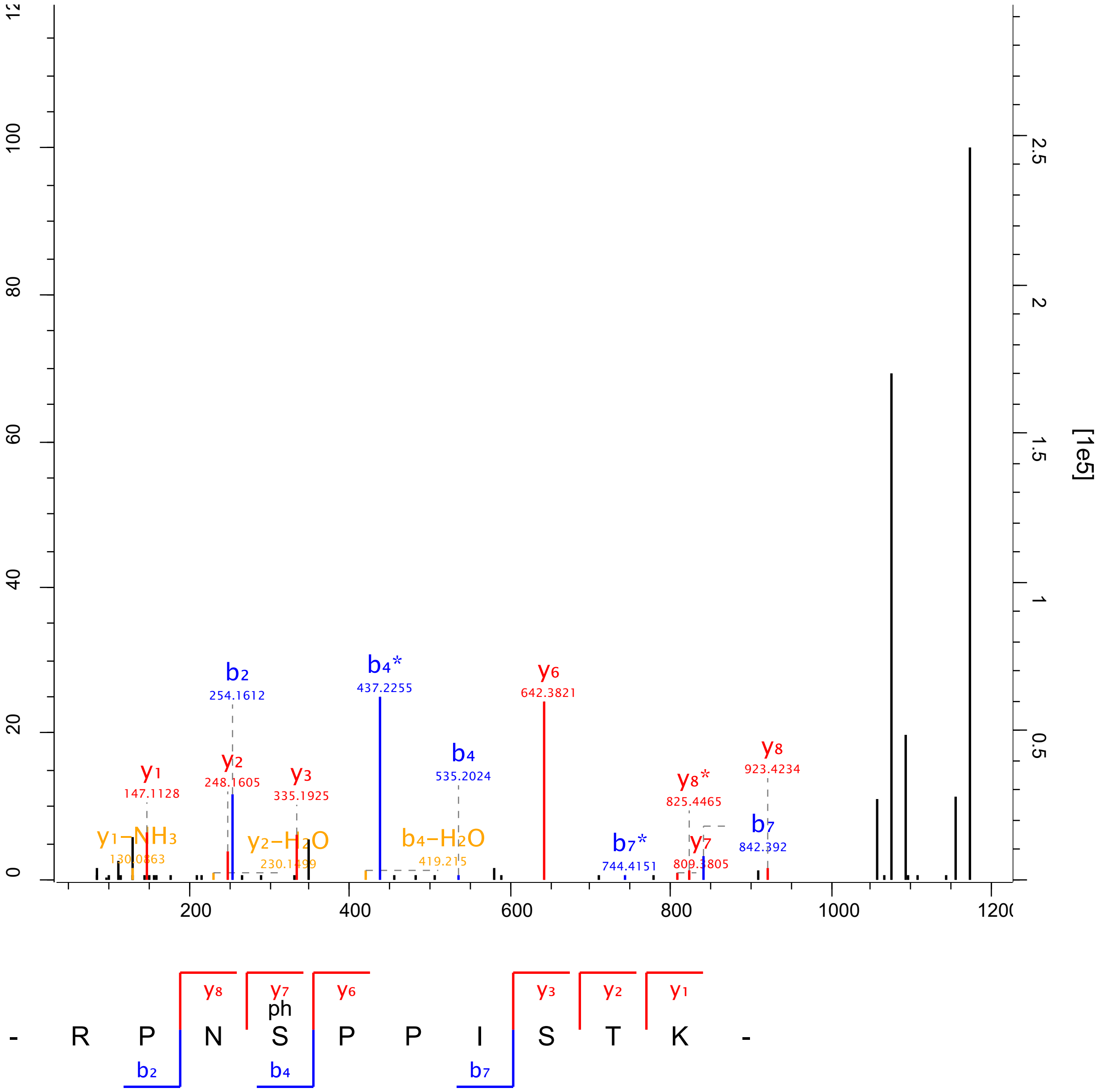

| Raw file | Scan | Method | Score | m/z |
|---|---|---|---|---|
| sys_02_2 | 2803 | FTMS; HCD | 116.51 | 670.3 |

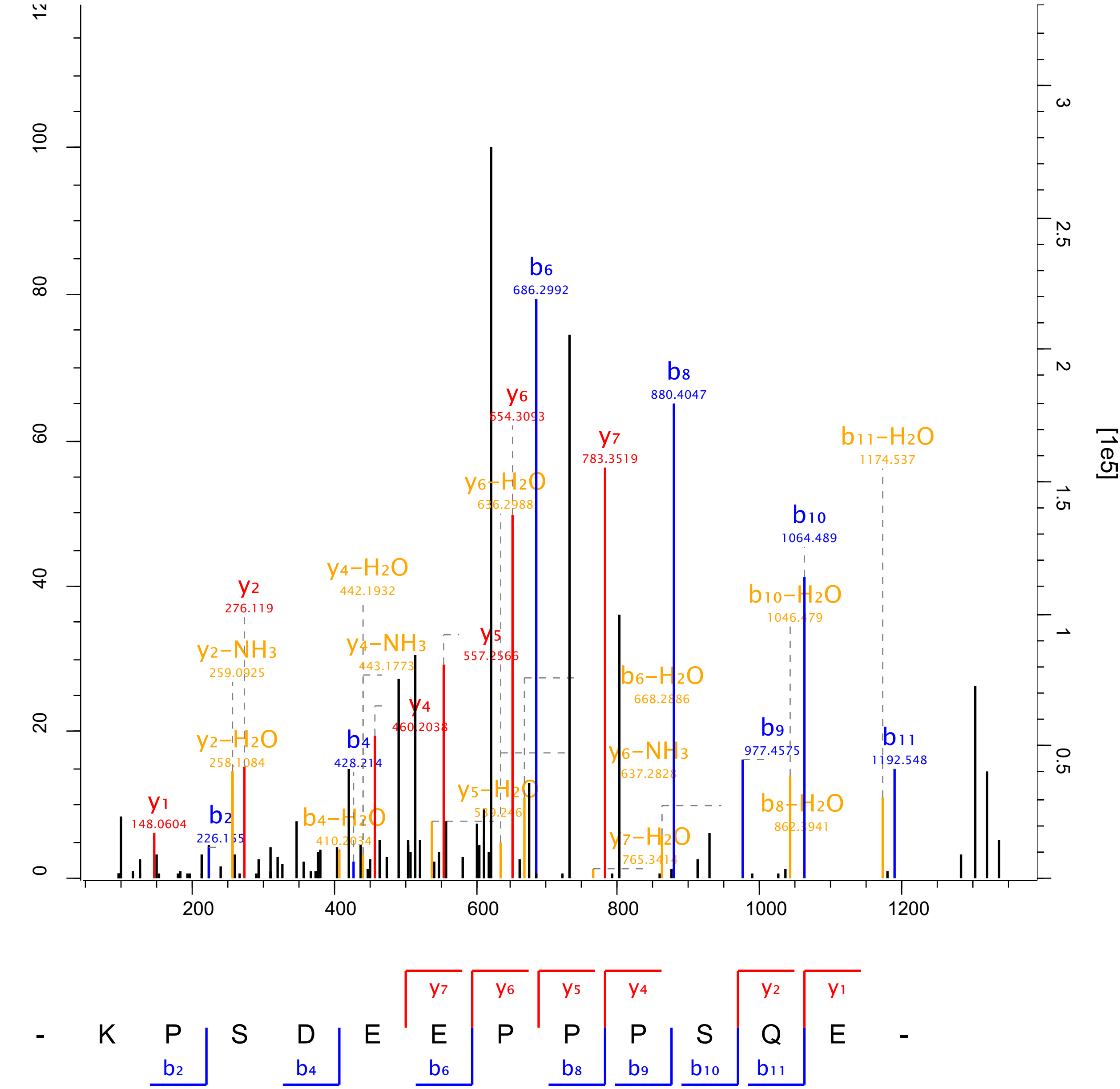

| Raw file | Scan | Method | Score | m/z |
|---|---|---|---|---|
| sys_02_2 | 2844 | FTMS; HCD | 43.03 | 439.53 |

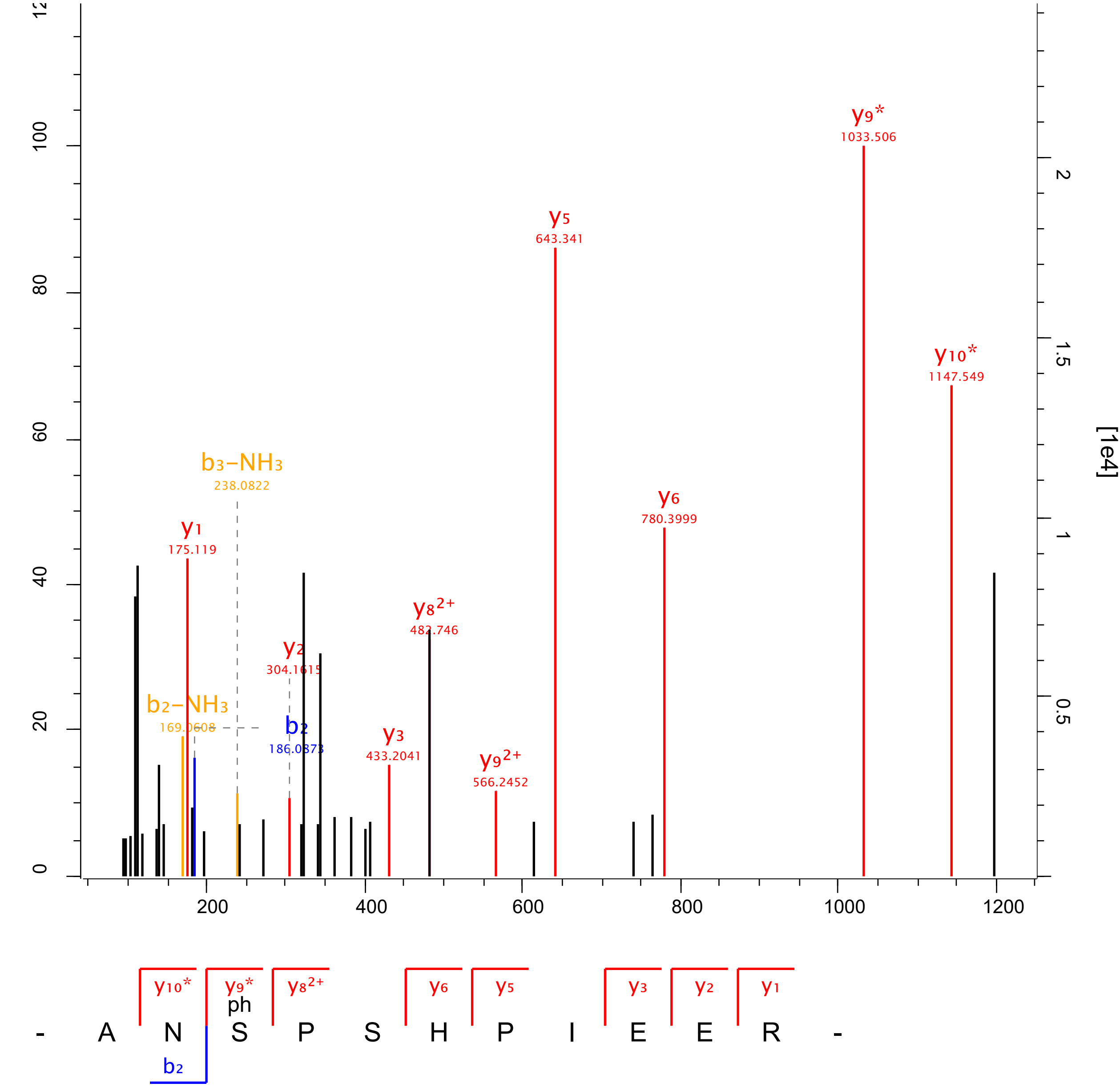

| Raw file | Scan | Method | Score | m/z |
|---|---|---|---|---|
| sys_02_2 | 28733 | FTMS; HCD | 46.35 | 525.78 |

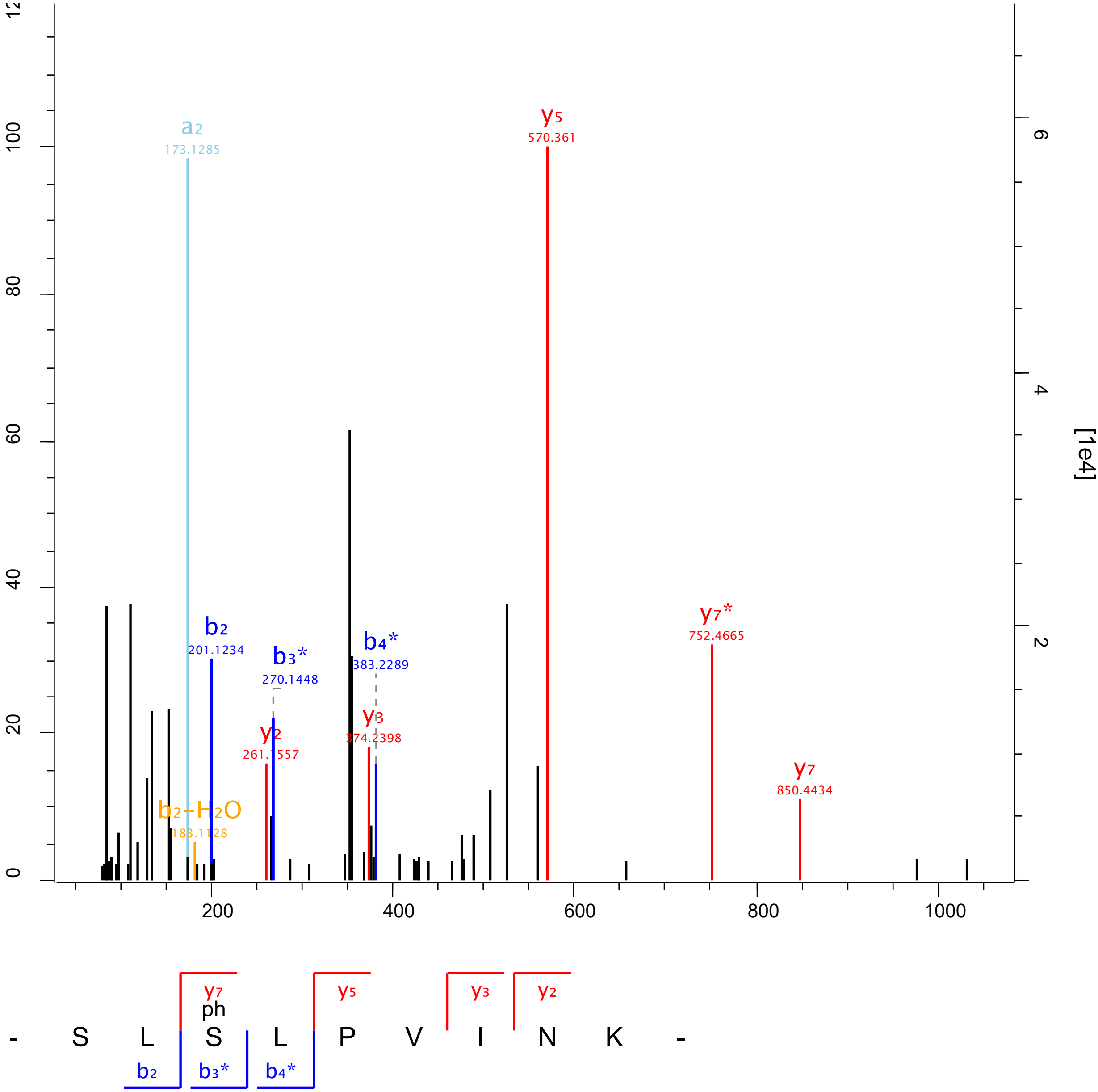

| Raw file | Scan | Method | Score | m/z |
|---|---|---|---|---|
| sys_02_2 | 2886 | FTMS; HCD | 62.47 | 474.24 |

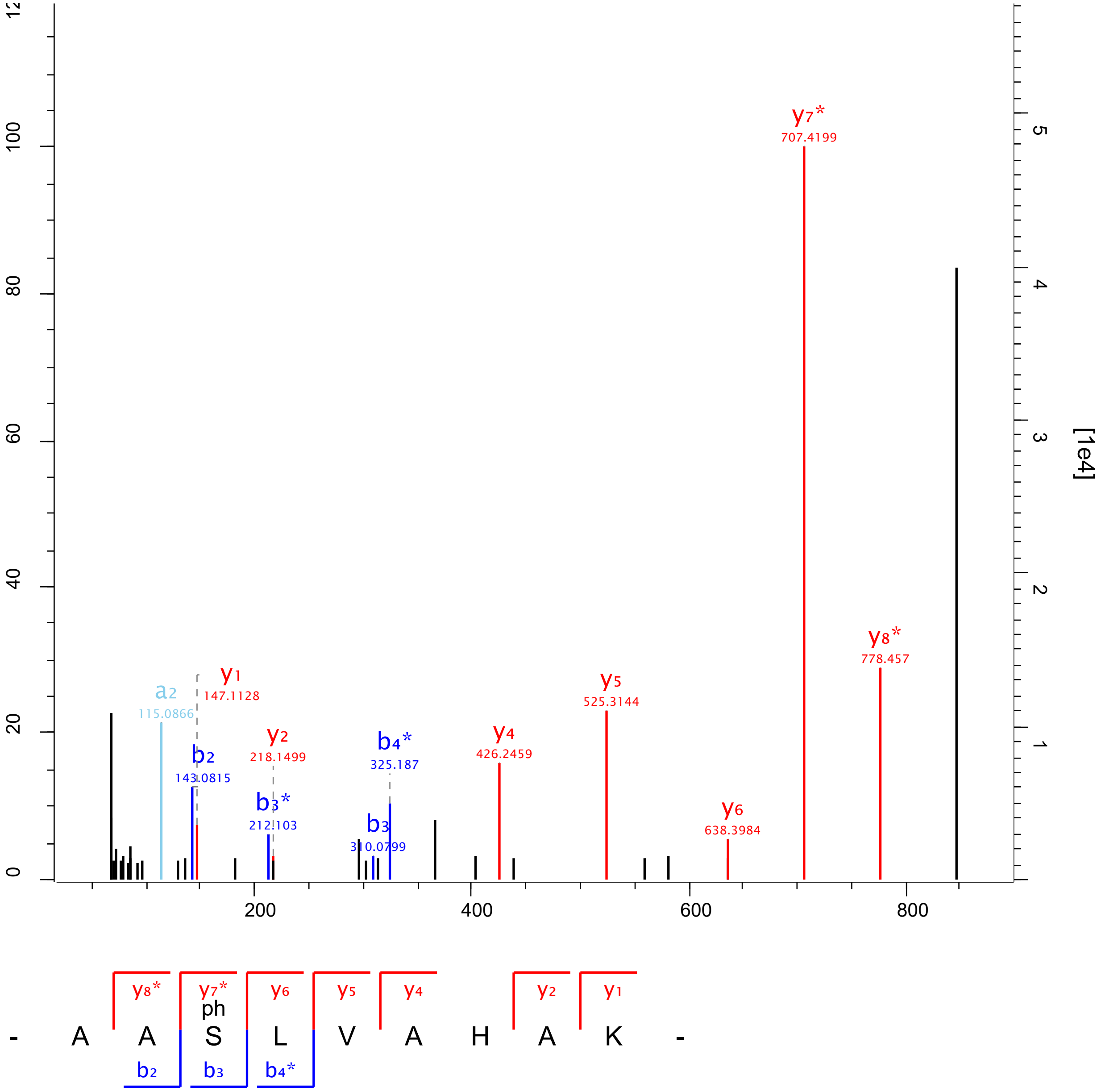

| Raw file | Scan | Method | Score | m/z |
|---|---|---|---|---|
| sys_02_2 | 2949 | FTMS; HCD | 148.62 | 596.26 |

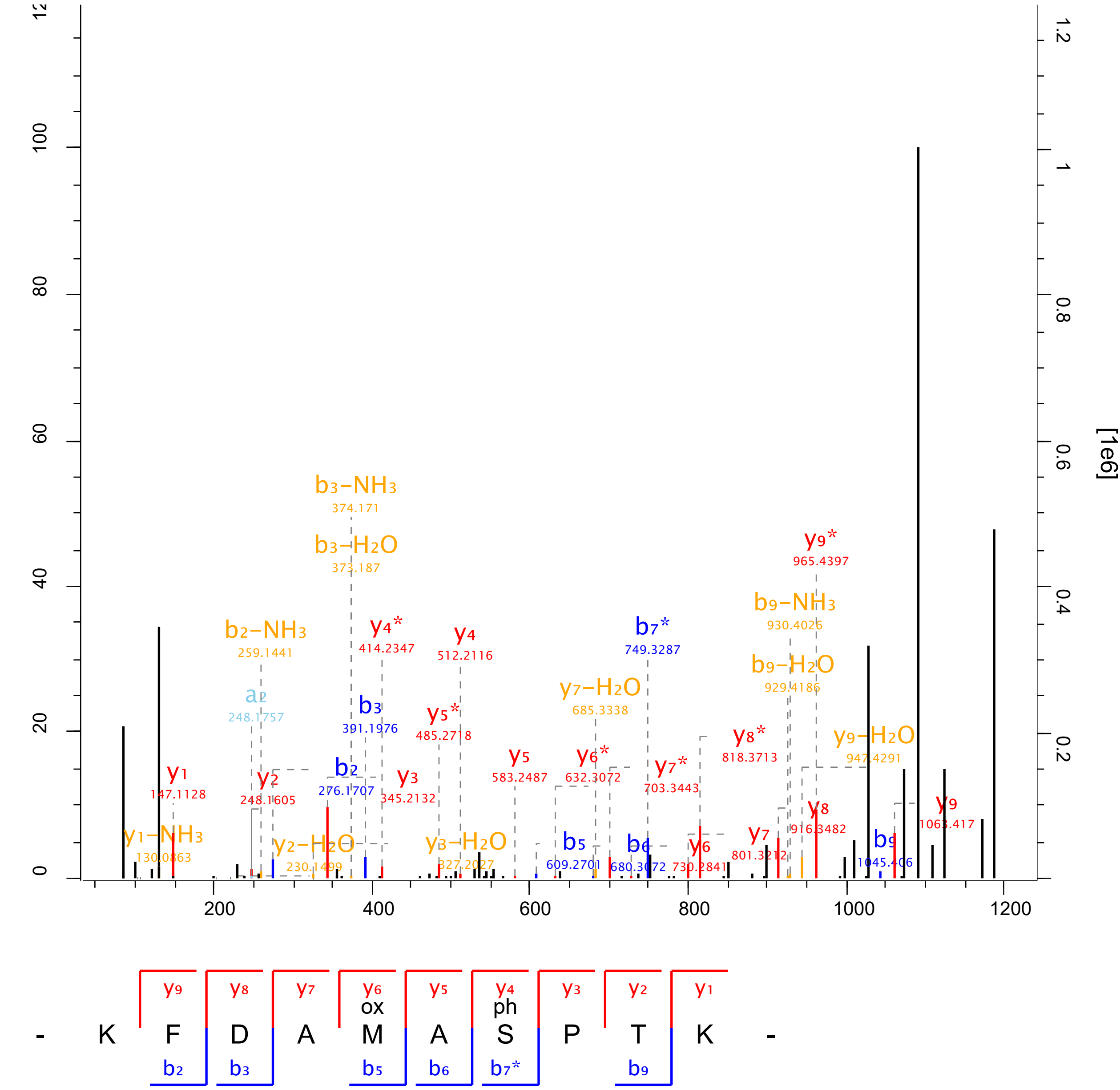

| Raw file | Scan | Method | Score | m/z |
| --- | --- | --- | --- | --- |
| sys_02_2 | 3026 | FTMS; HCD | 95.42 | 566.74 |

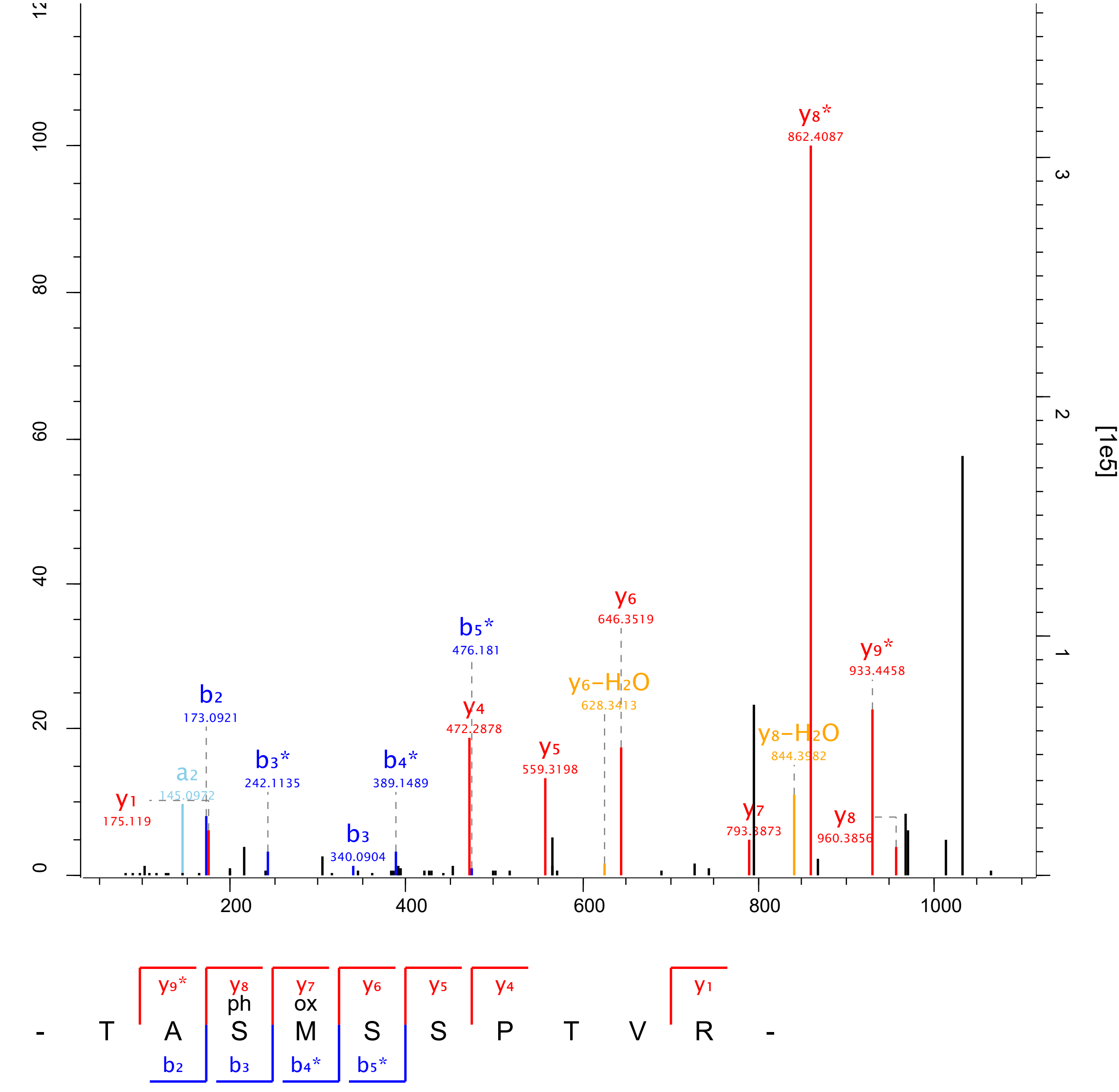

| Raw file | Scan | Method | Score | m/z |
|---|---|---|---|---|
| sys_02_2 | 3061 | FTMS; HCD | 97.45 | 631.58 |

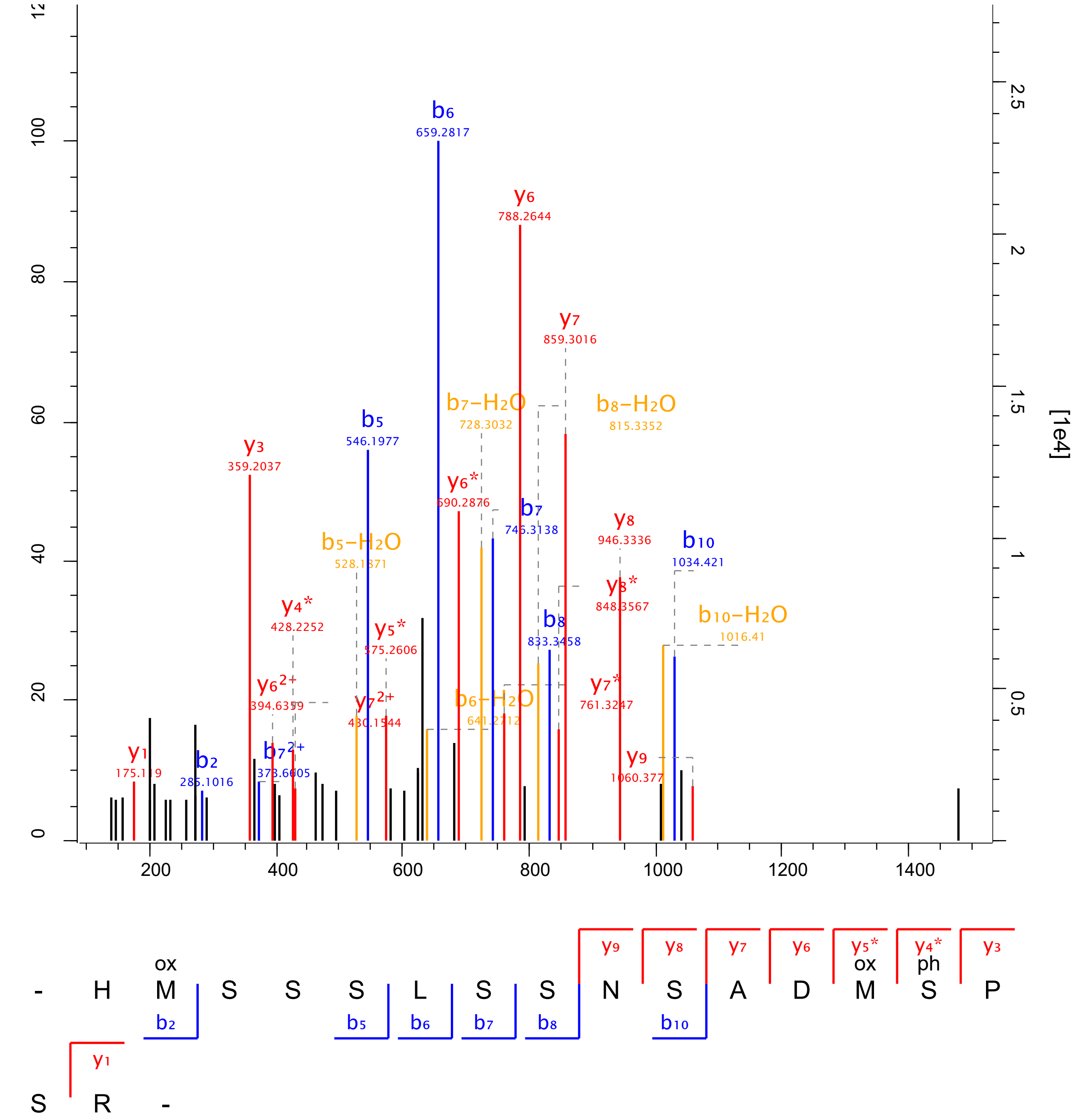

| Raw file | Scan | Method | Score | m/z |
|---|---|---|---|---|
| sys_02_2 | 3115 | FTMS; HCD | 58.17 | 571.74 |

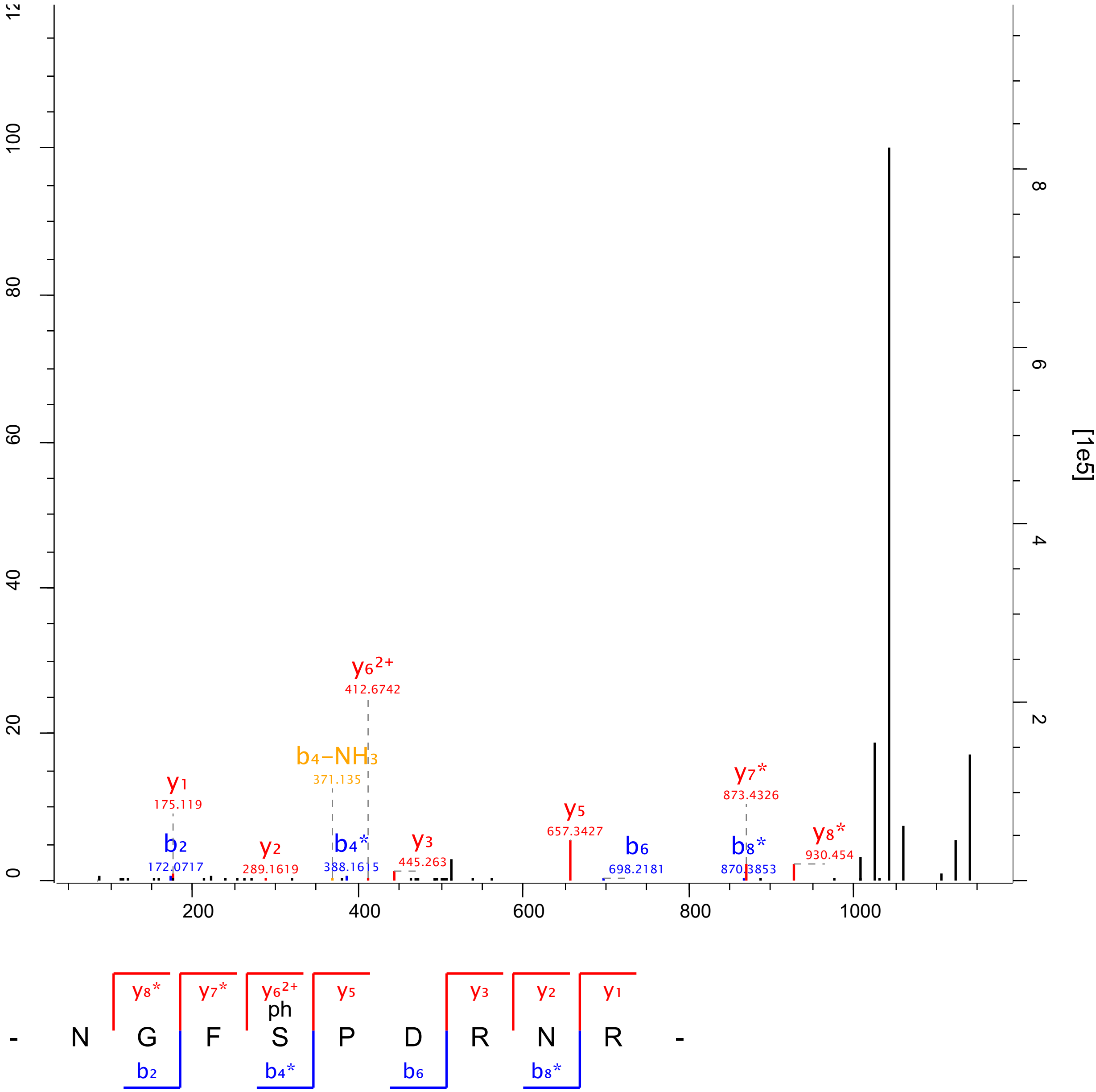

| Raw file | Scan | Method | Score | m/z |
|---|---|---|---|---|
| sys_02_2 | 3227 | FTMS; HCD | 50.39 | 533.21 |

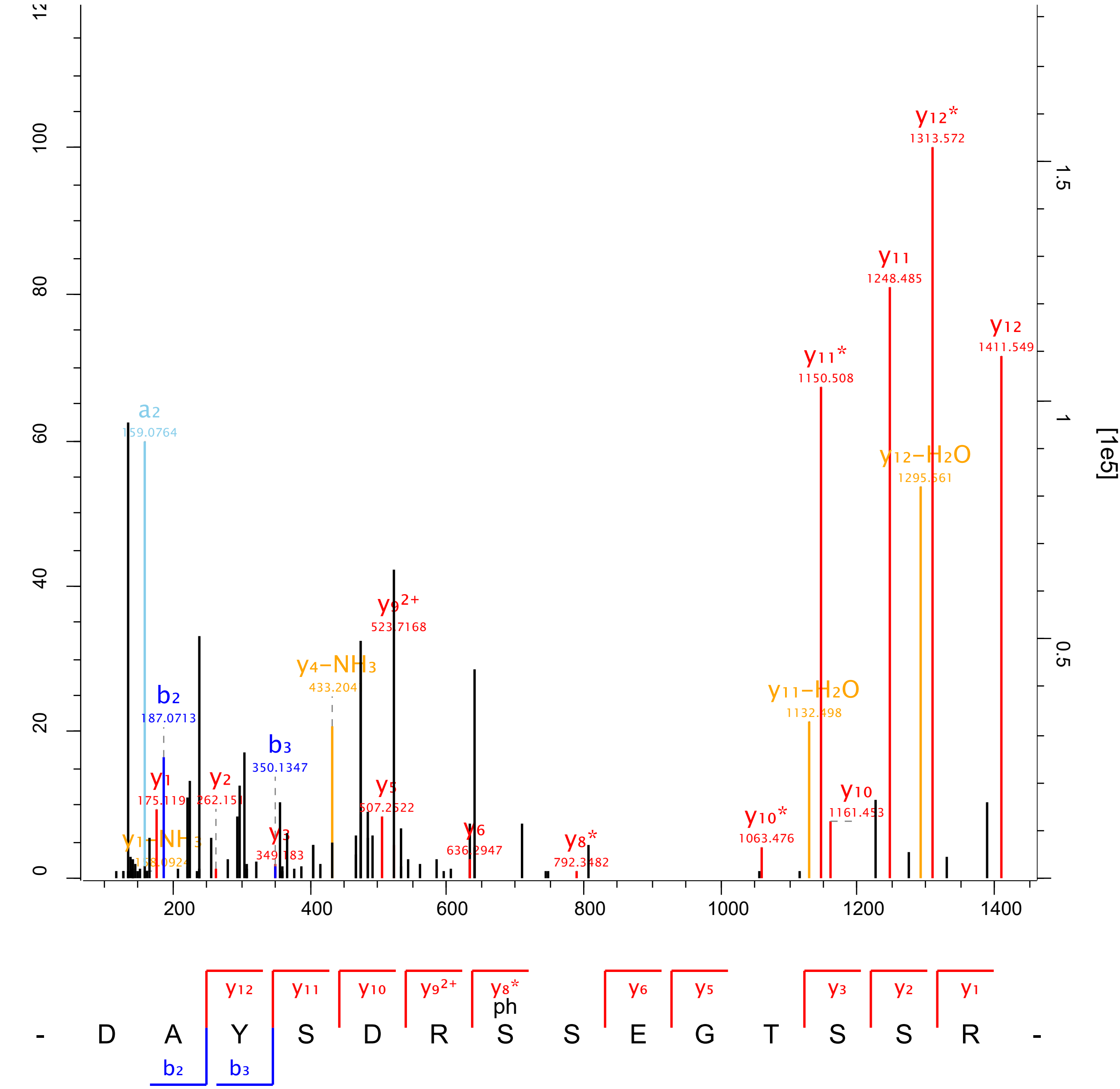

| Raw file | Scan | Method | Score | m/z |
| --- | --- | --- | --- | --- |
| sys_02_2 | 3258 | FTMS; HCD | 42.34 | 563.24 |

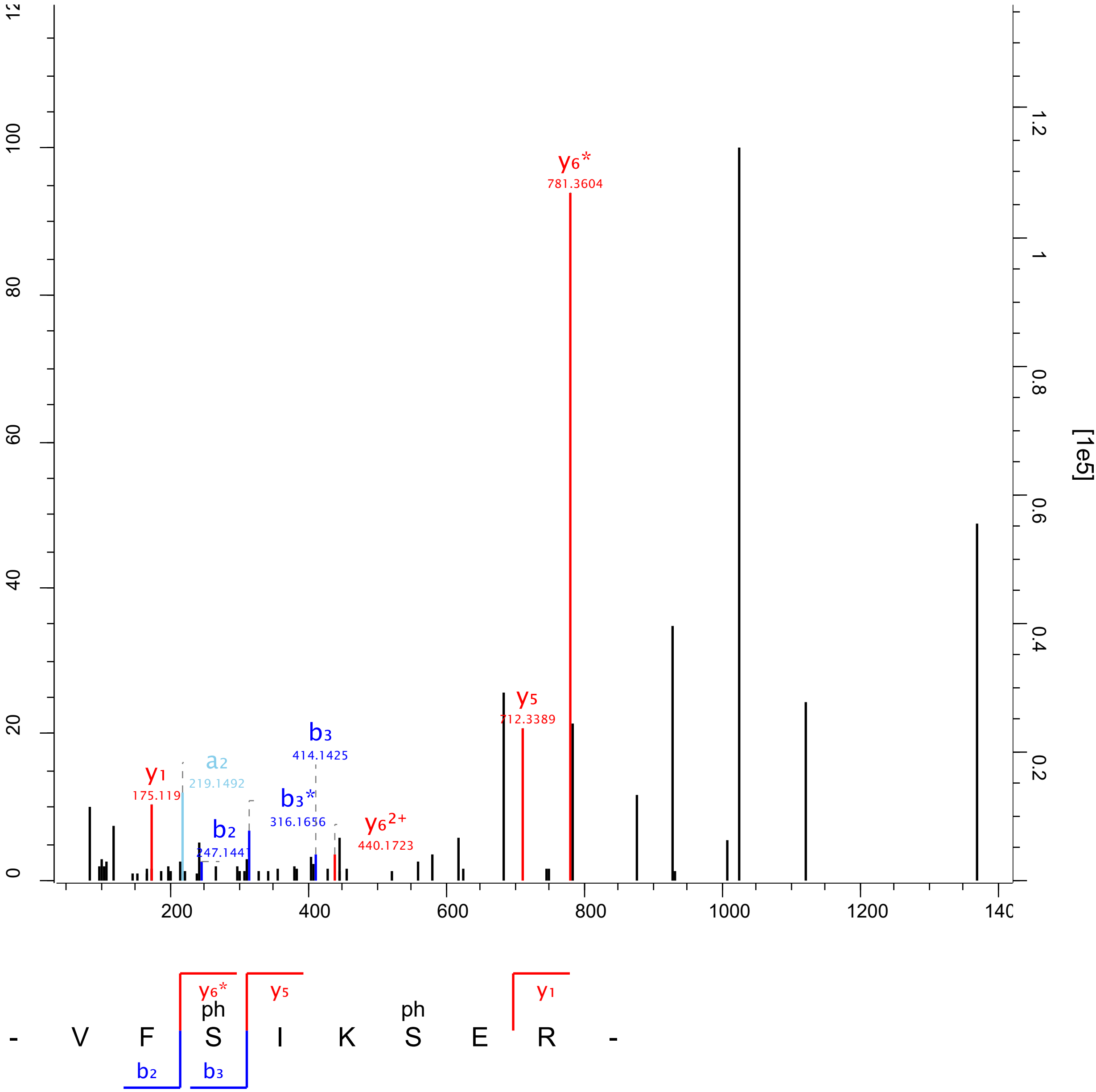

| Raw file | Scan | Method | Score | m/z |
|---|---|---|---|---|
| sys_02_2 | 32914 | FTMS; HCD | 230.82 | 850.36 |

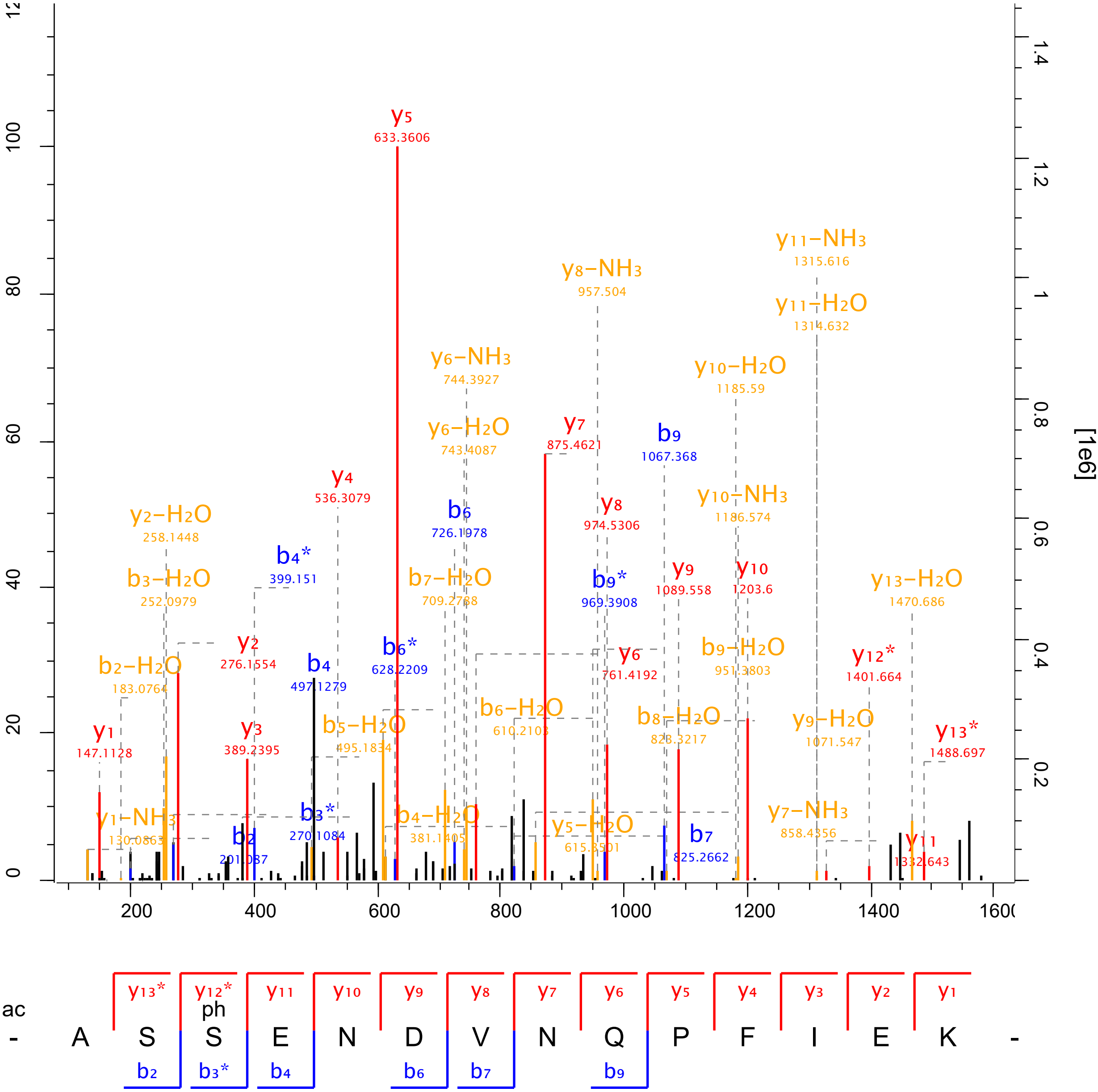

| Raw file | Scan | Method | Score | m/z |
| --- | --- | --- | --- | --- |
| sys_02_2 | 3307 | FTMS; HCD | 107.99 | 554.25 |

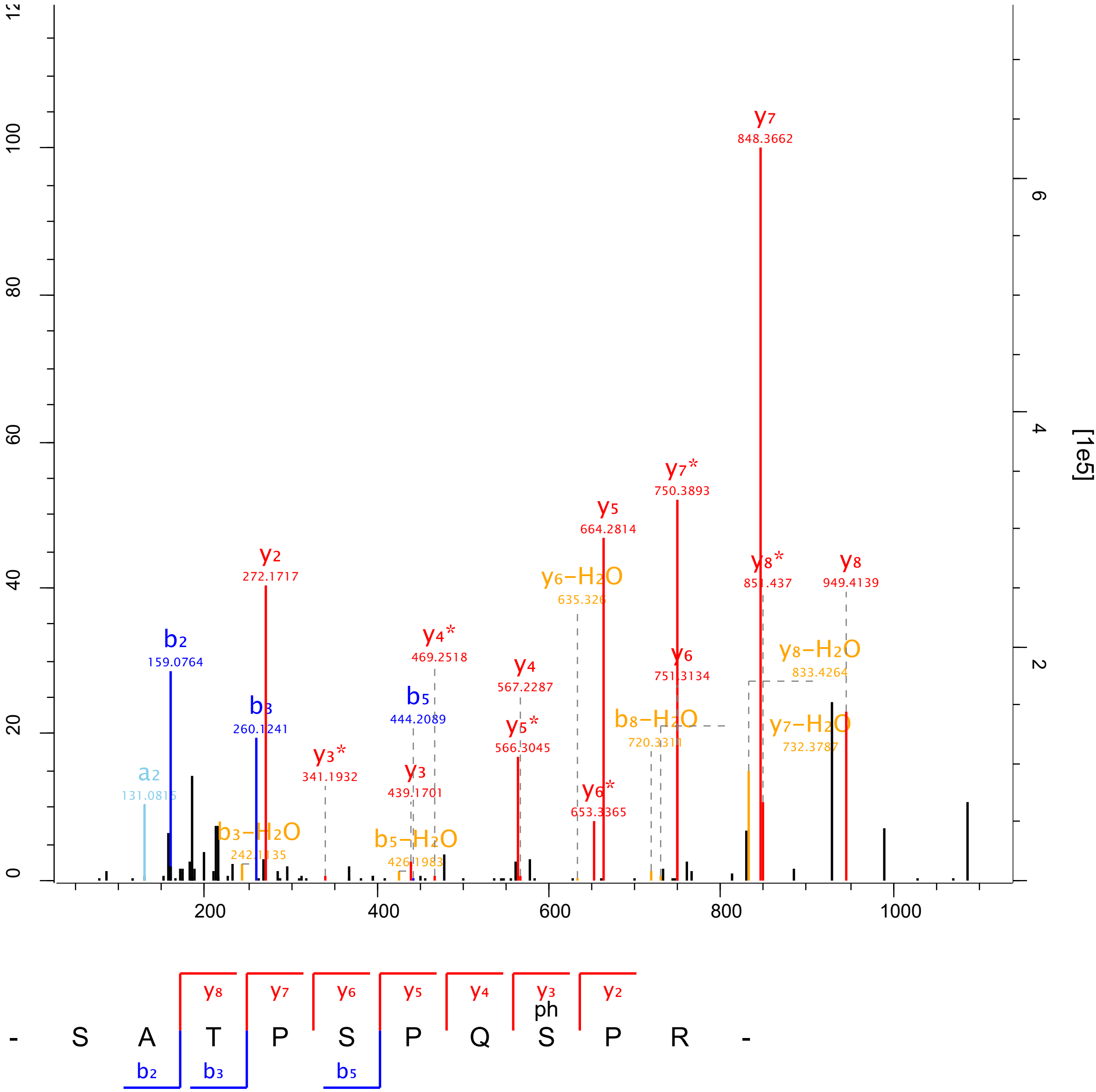

| Raw file | Scan | Method | Score | m/z |
|---|---|---|---|---|
| sys_02_2 | 3320 | FTMS; HCD | 60.49 | 431.87 |

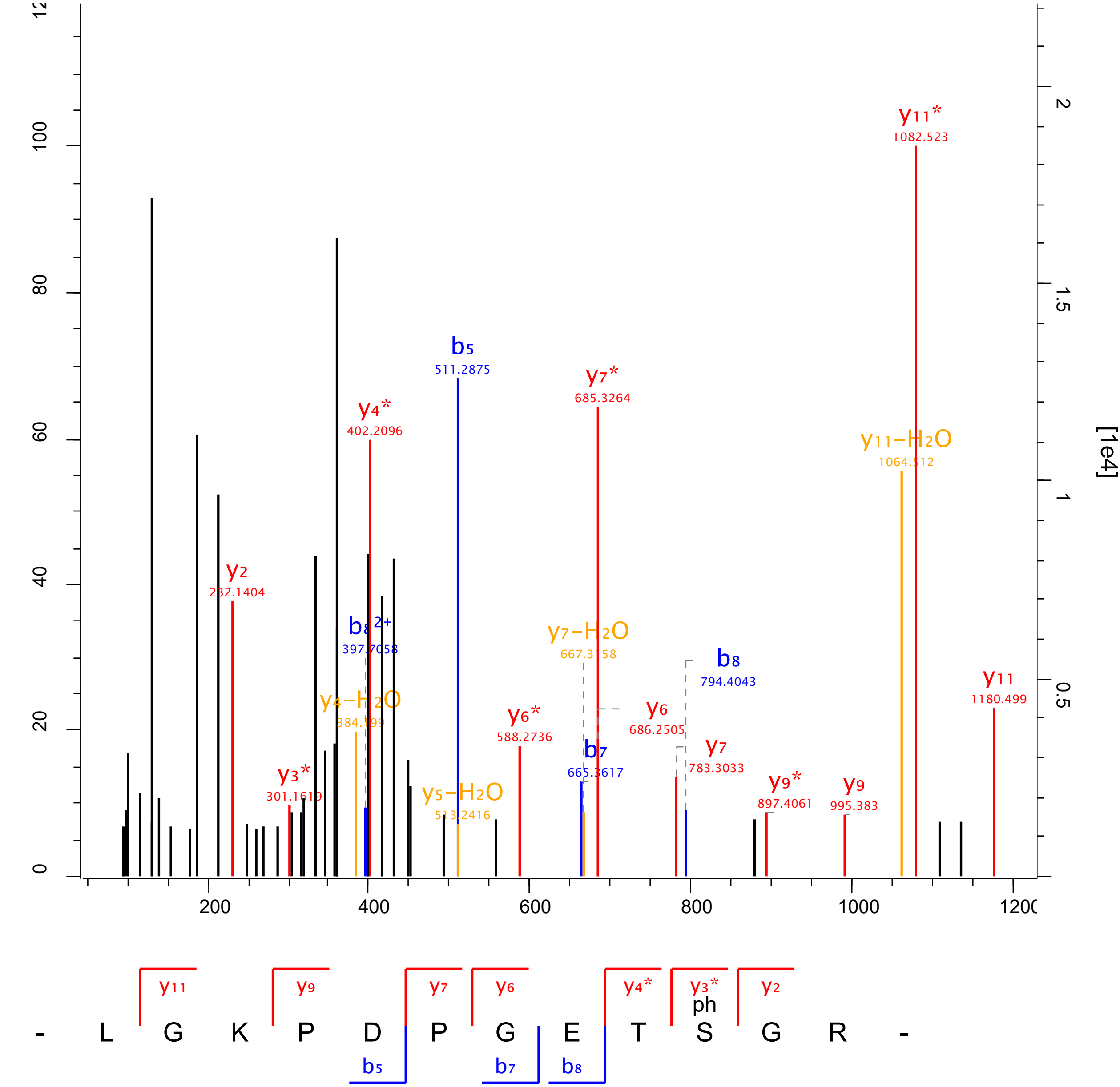

| Raw file | Scan | Method | Score | m/z |
|---|---|---|---|---|
| sys_02_2 | 3426 | FTMS; HCD | 240.42 | 587.58 |

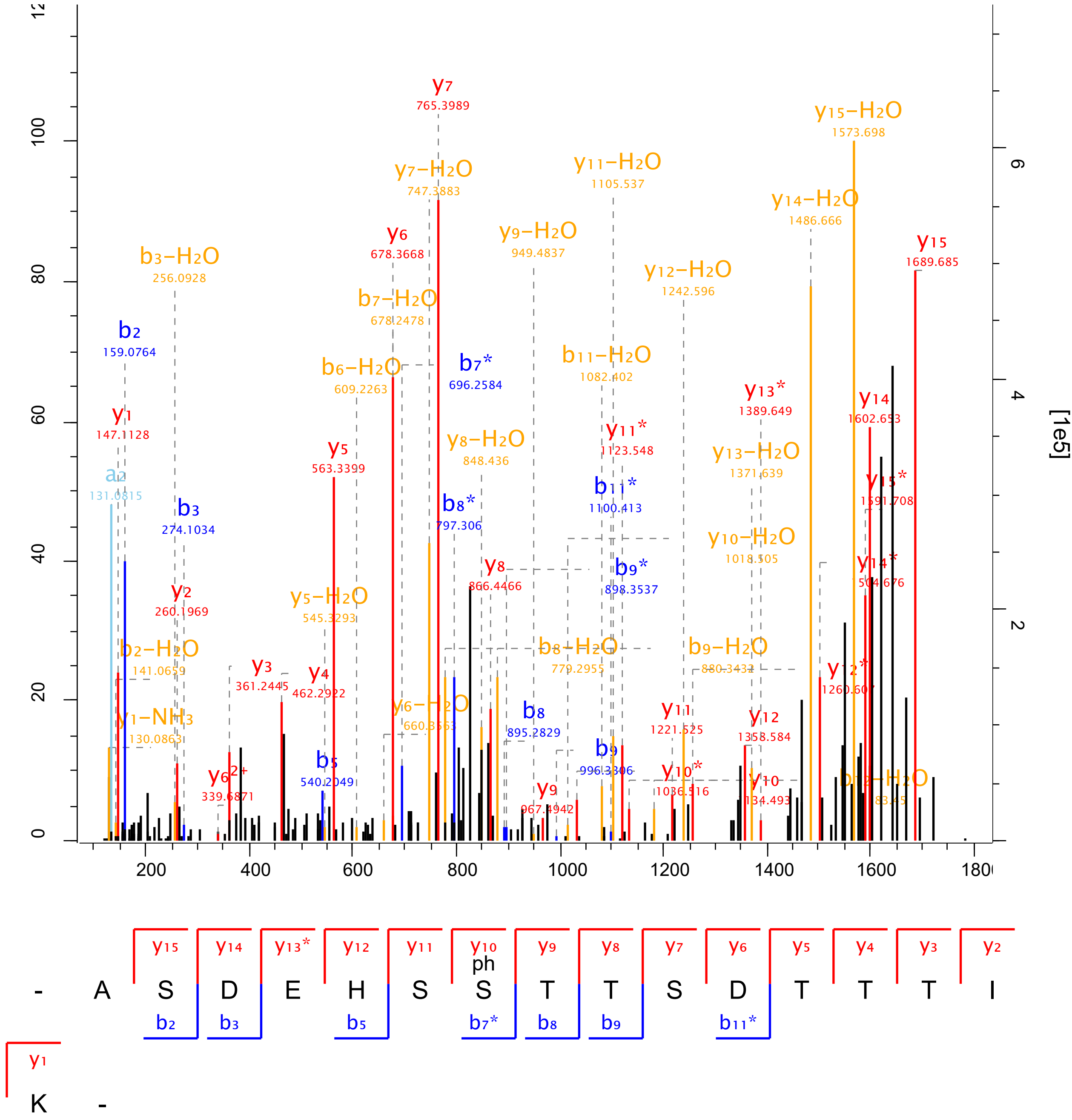

| Raw file | Scan | Method | Score | m/z |
| --- | --- | --- | --- | --- |
| sys_02_2 | 3437 | FTMS; HCD | 88.07 | 851.83 |

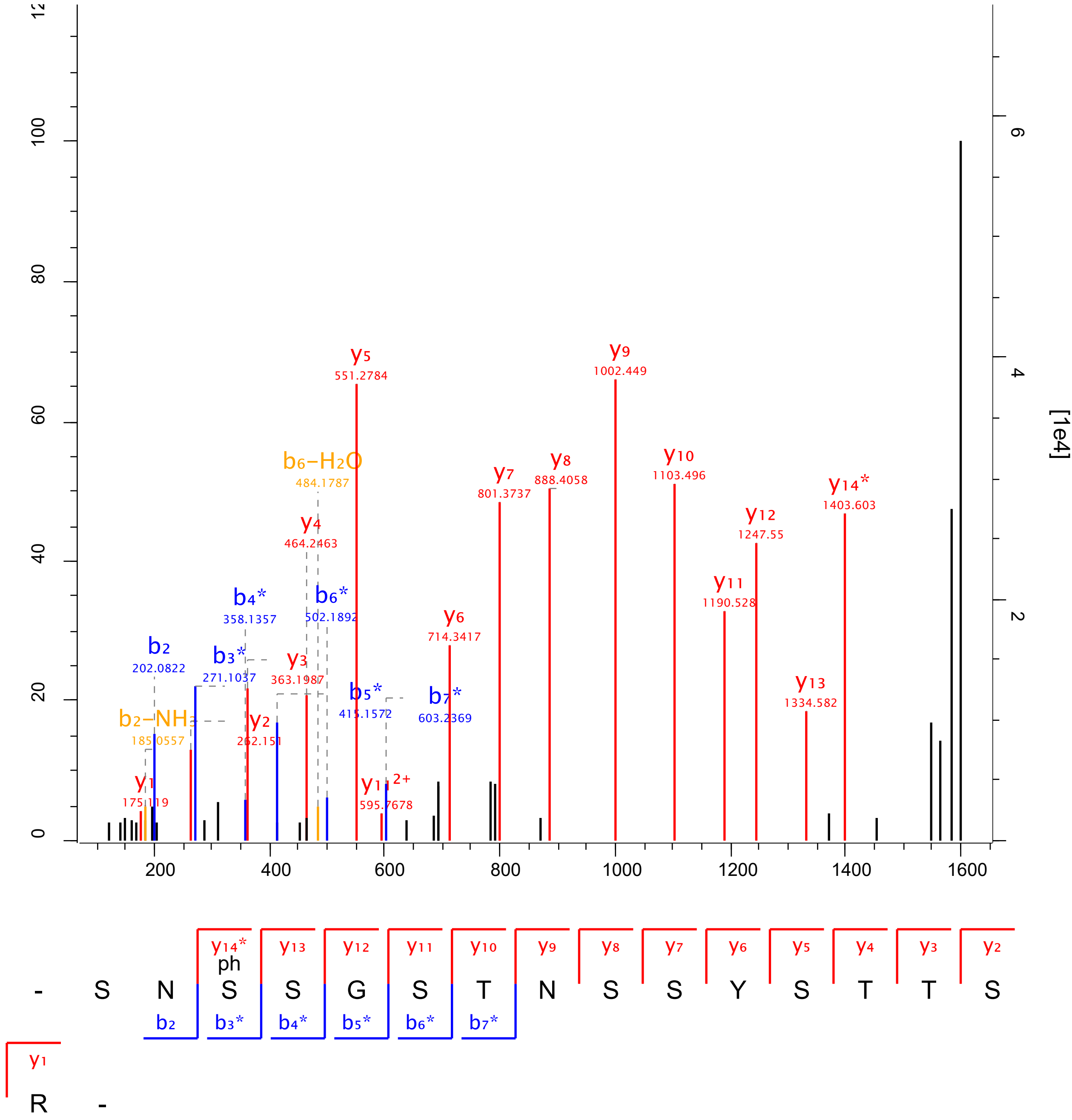

| Raw file | Scan | Method | Score | m/z |
|---|---|---|---|---|
| sys_02_2 | 3460 | FTMS; HCD | 168 | 611.77 |

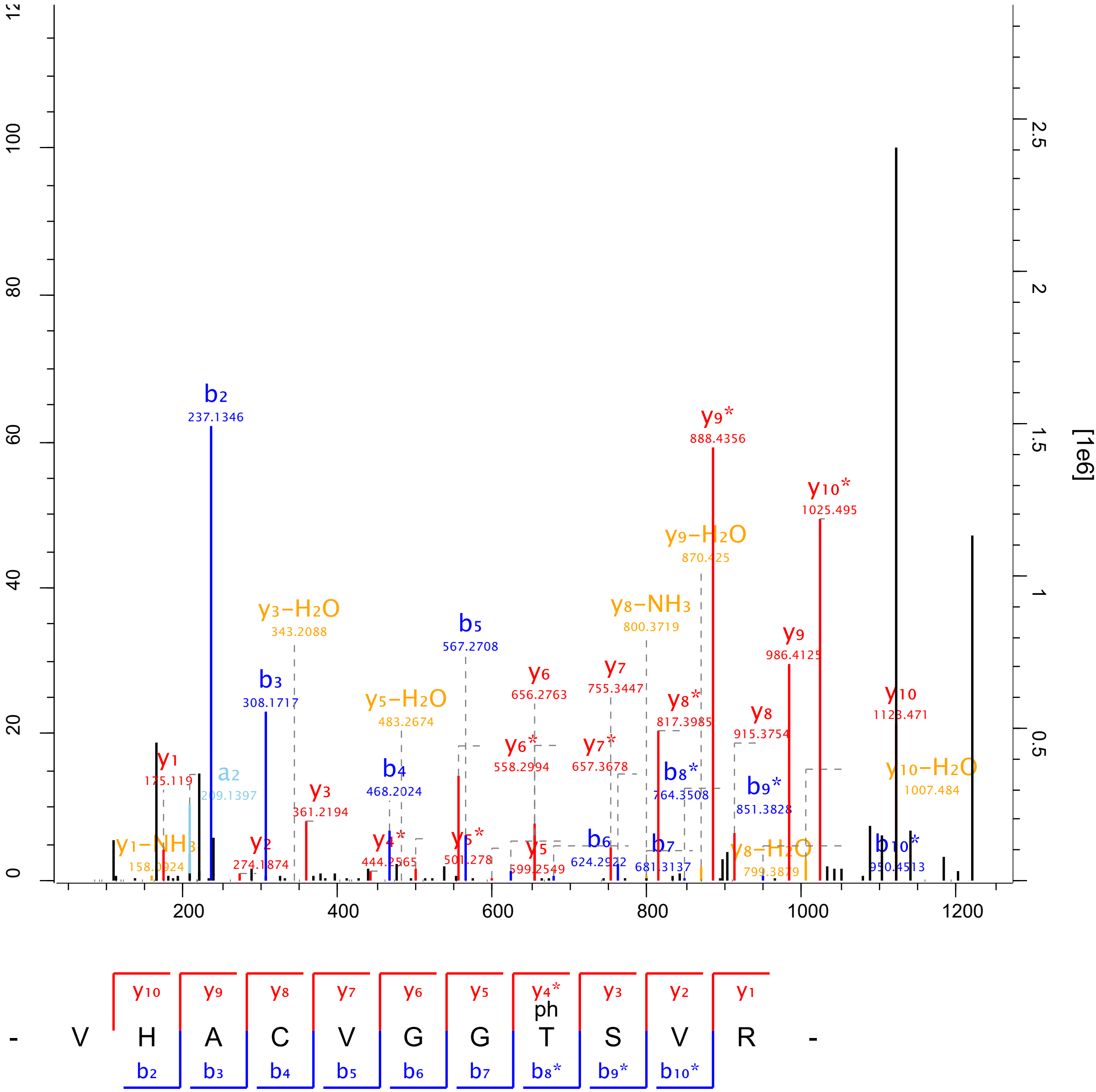

| Raw file | Scan | Method | Score | m/z |
|---|---|---|---|---|
| sys_02_2 | 3518 | FTMS; HCD | 80.24 | 649.61 |

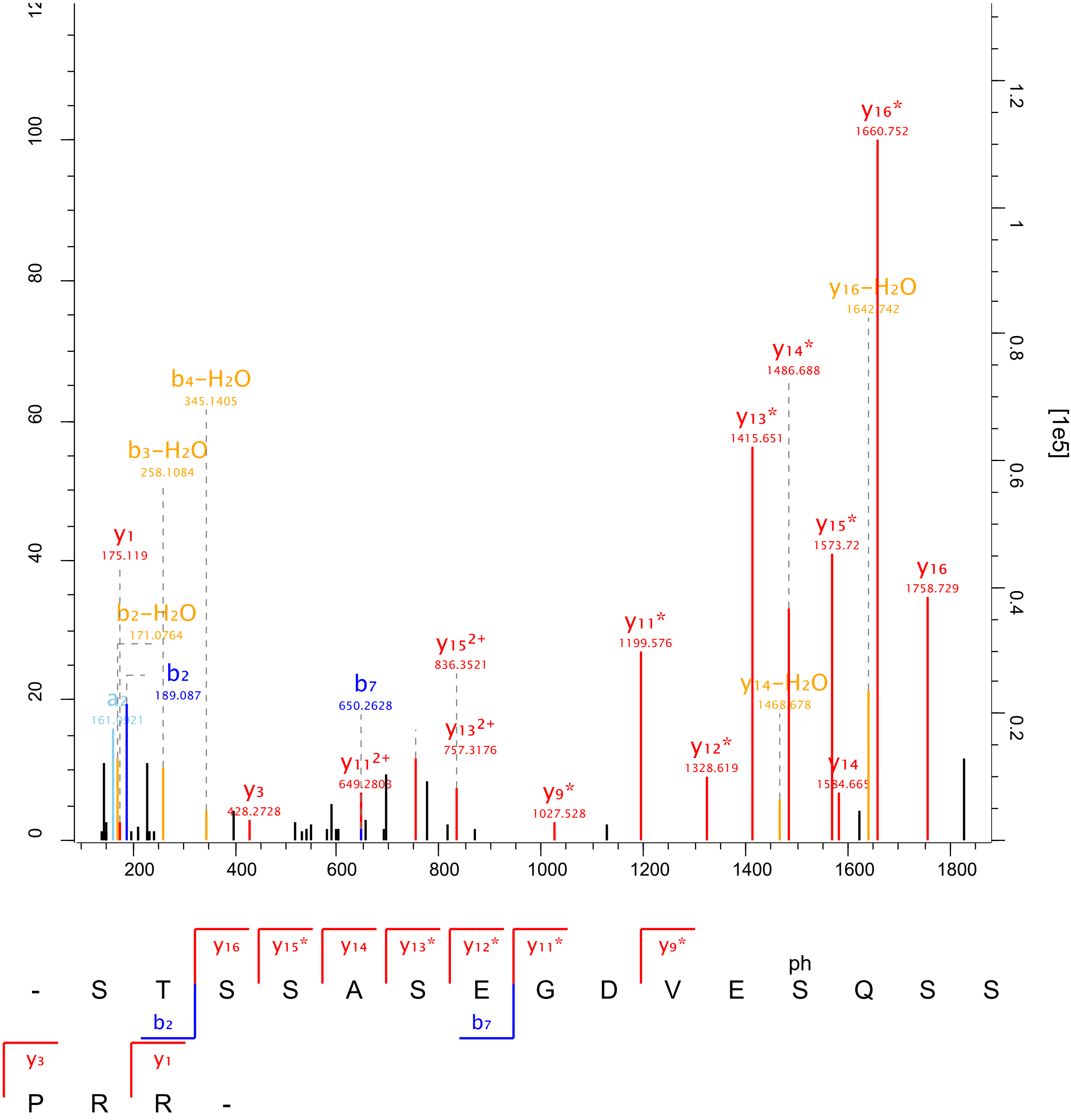

| Raw file | Scan | Method | Score | m/z |
| --- | --- | --- | --- | --- |
| sys_02_2 | 3556 | FTMS; HCD | 89.47 | 605.59 |

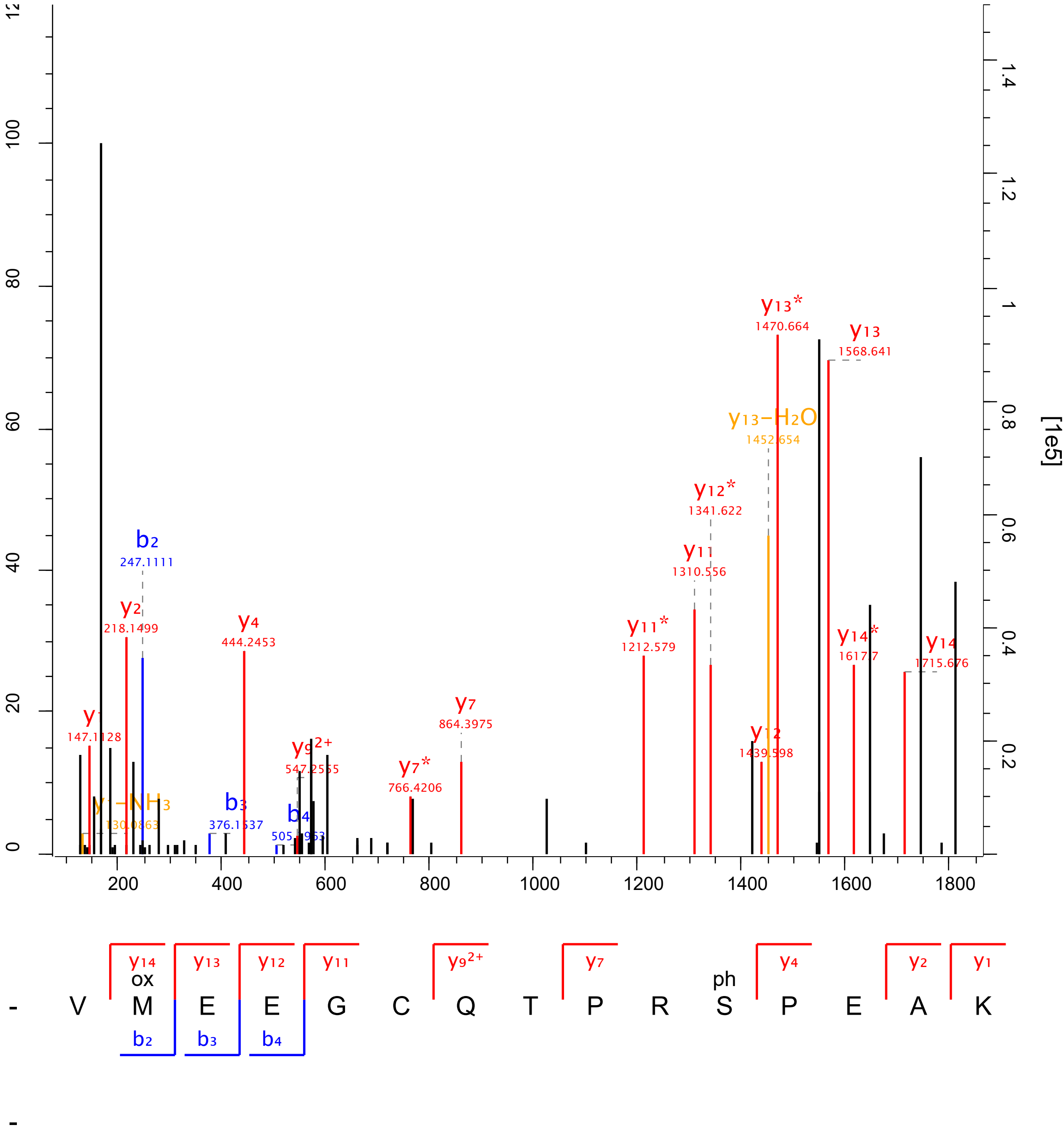

| Raw file | Scan | Method | Score | m/z |
|---|---|---|---|---|
| sys_02_2 | 3754 | FTMS; HCD | 62.86 | 526.2 |

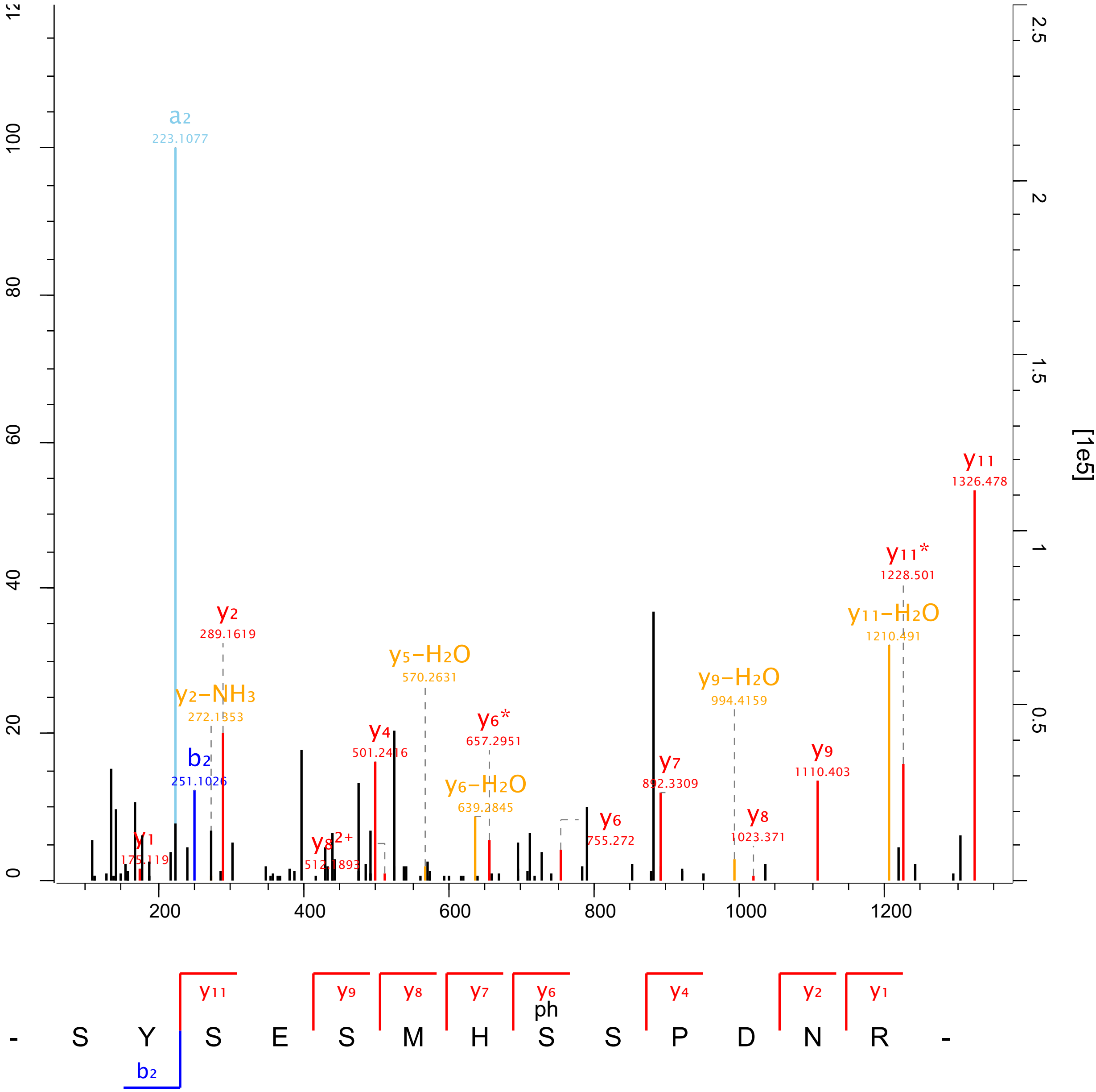

| Raw file | Scan | Method | Score | m/z |
|---|---|---|---|---|
| sys_02_2 | 3791 | FTMS; HCD | 168.43 | 586.27 |

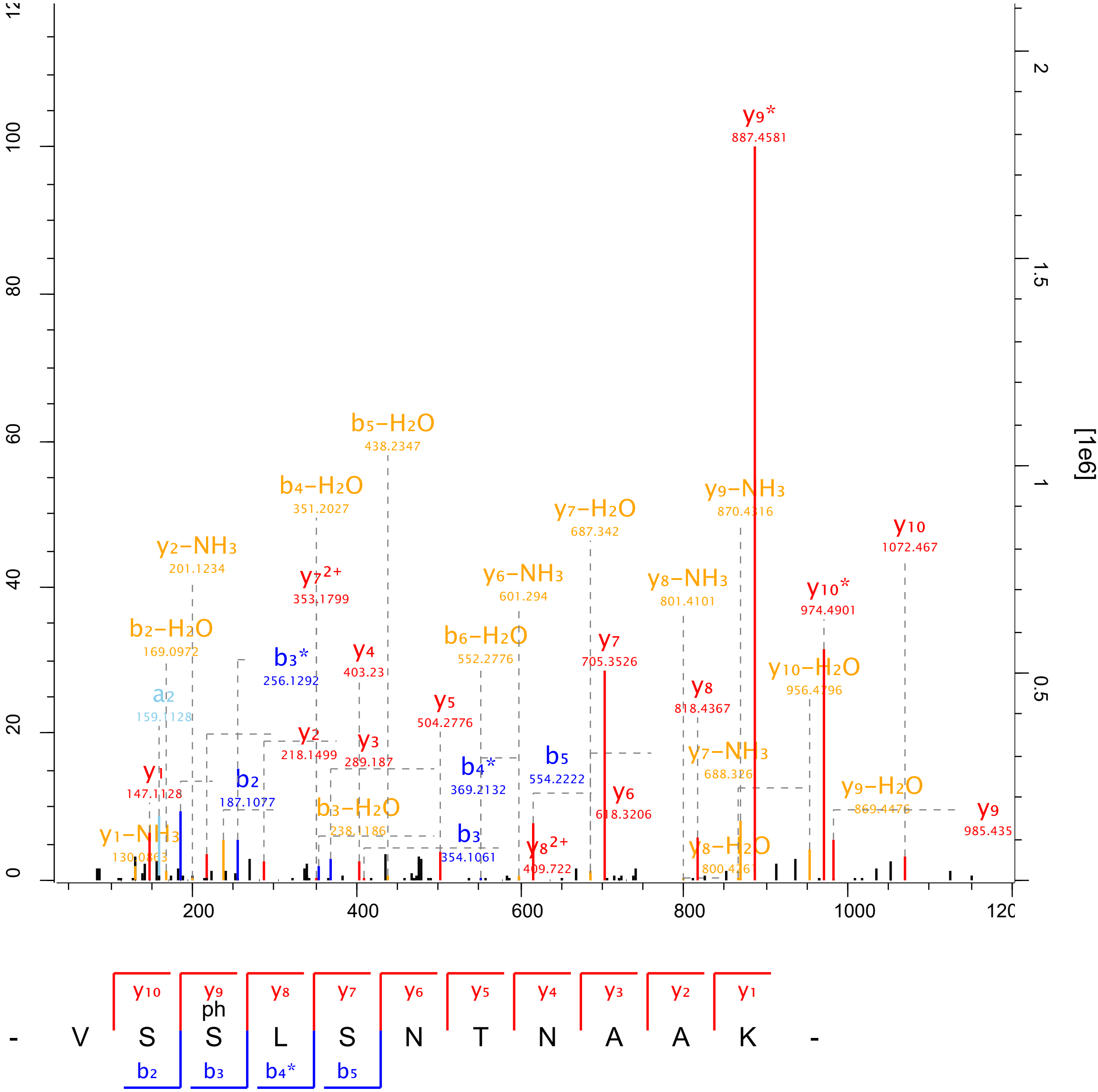

| Raw file | Scan | Method | Score | m/z |
| --- | --- | --- | --- | --- |
| sys_02_2 | 3795 | FTMS; HCD | 58.89 | 465.89 |

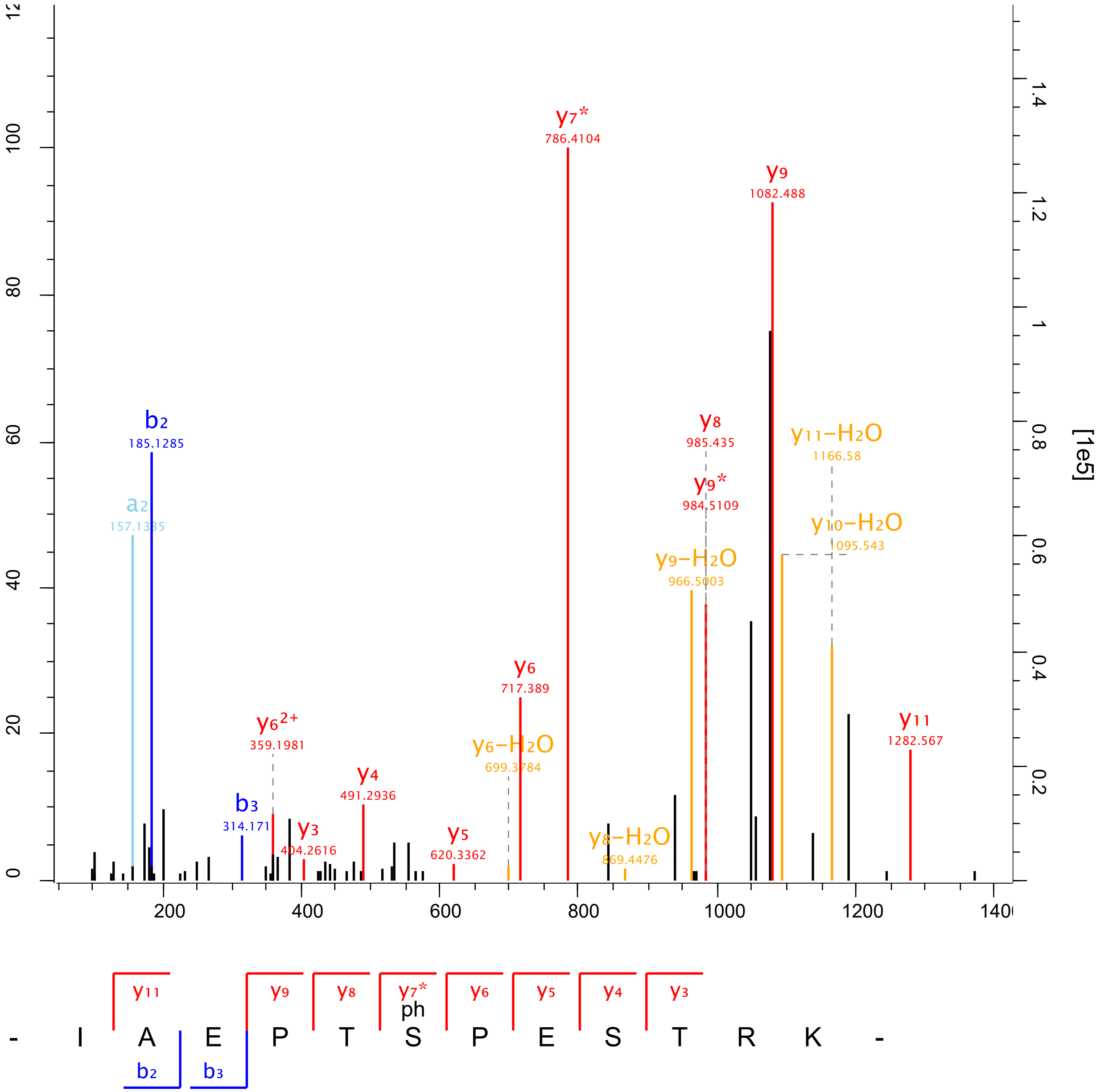

| Raw file | Scan | Method | Score | m/z |
|---|---|---|---|---|
| sys_02_2 | 38641 | FTMS; HCD | 206.49 | 1030.96 |

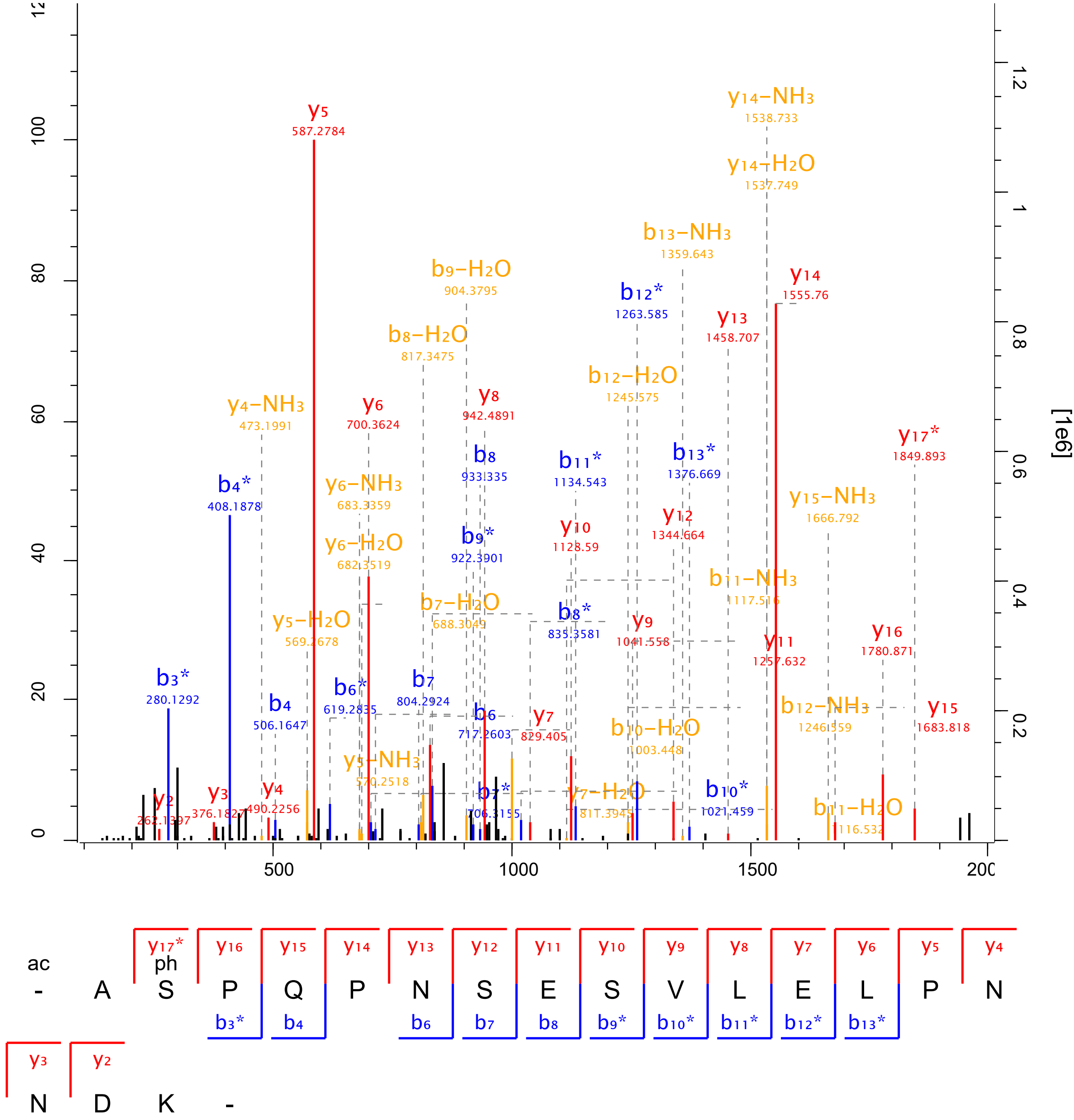

| Raw file | Scan | Method | Score | m/z |
|---|---|---|---|---|
| sys_02_2 | 3950 | FTMS; HCD | 40.48 | 615.27 |

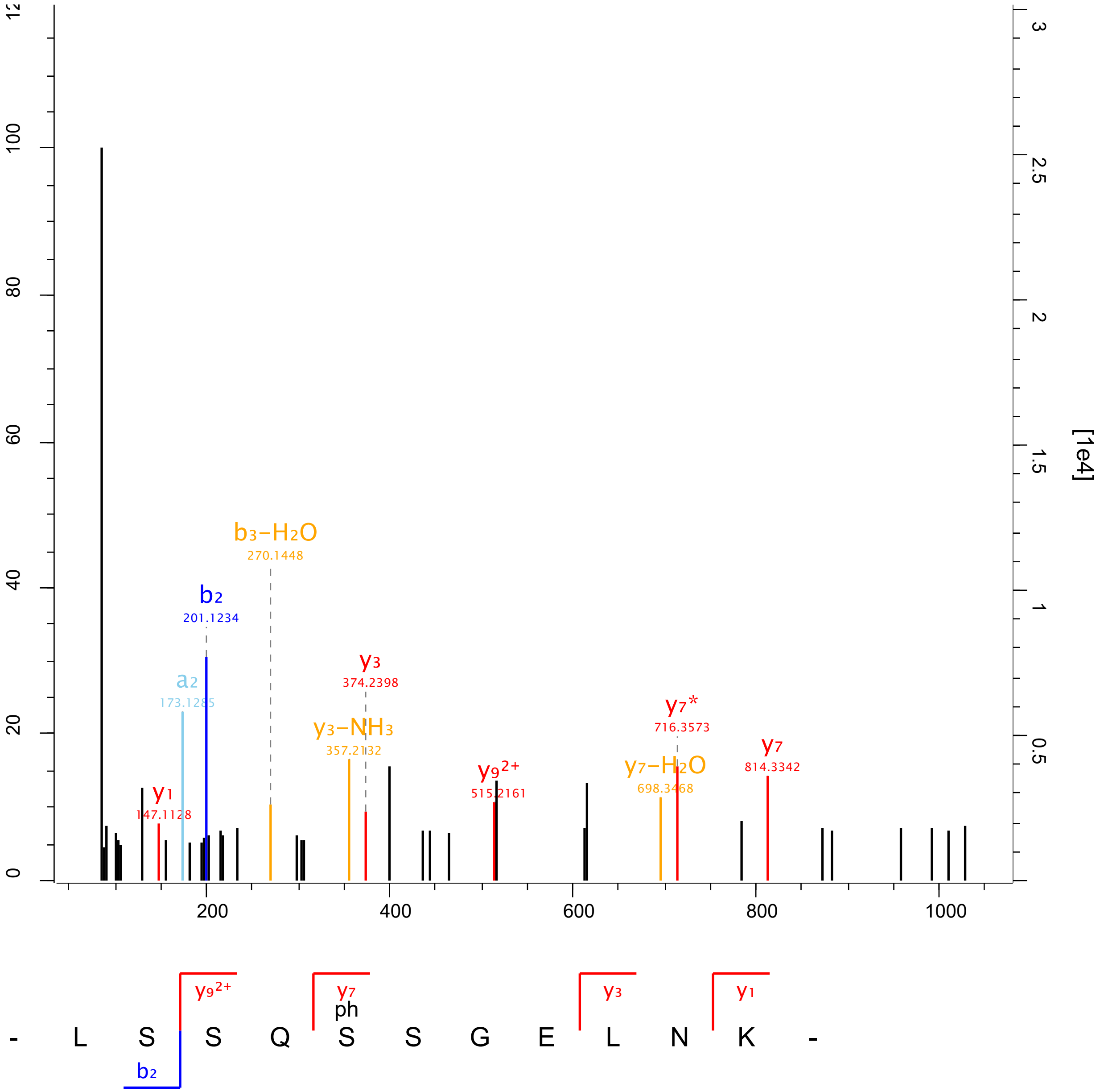

| Raw file | Scan | Method | Score | m/z |
|---|---|---|---|---|
| sys_02_2 | 3953 | FTMS; HCD | 124.6 | 402.19 |

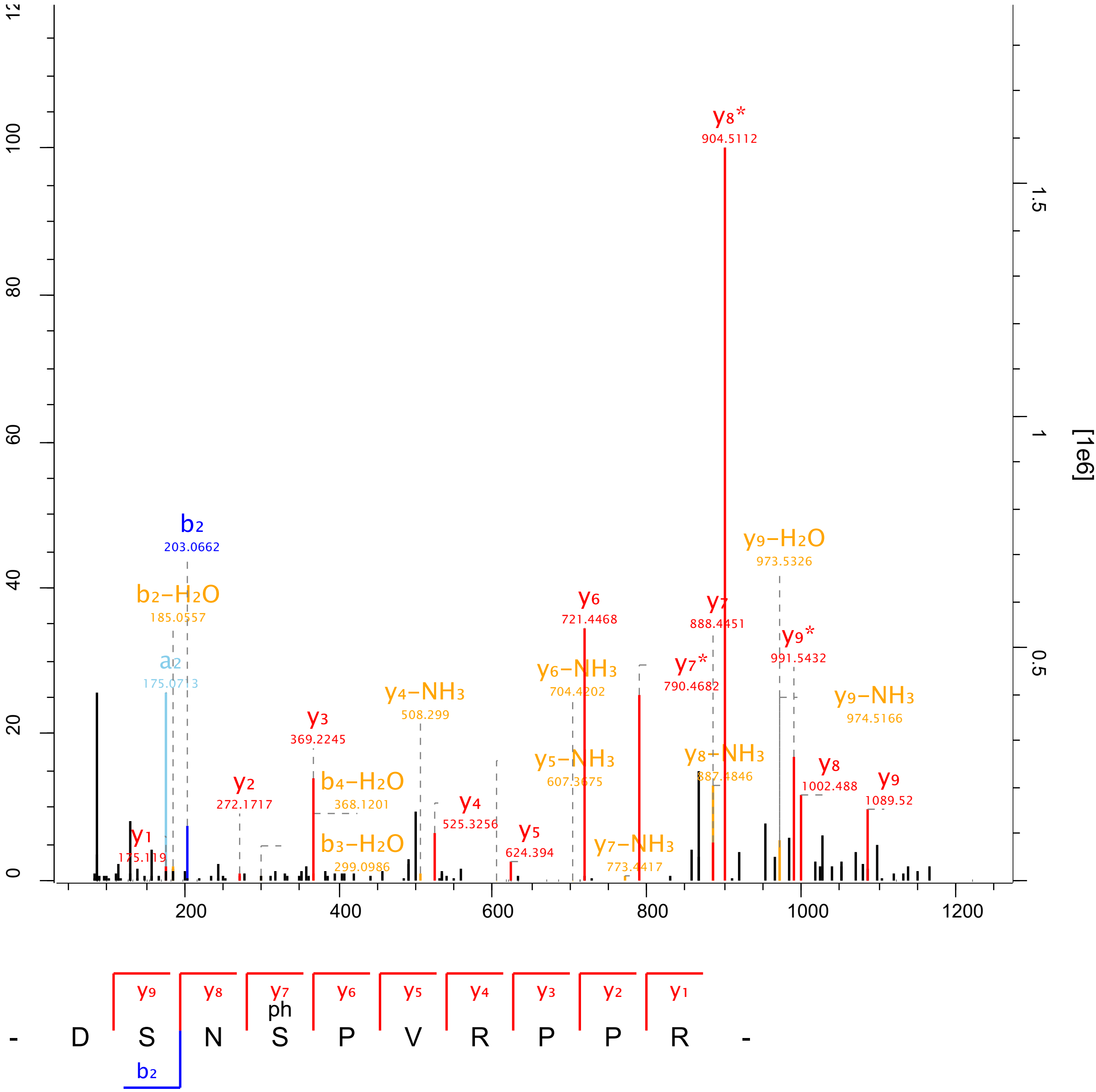

| Raw file | Scan | Method | Score | m/z |
|---|---|---|---|---|
| sys_02_2 | 4013 | FTMS; HCD | 70.2 | 738.79 |

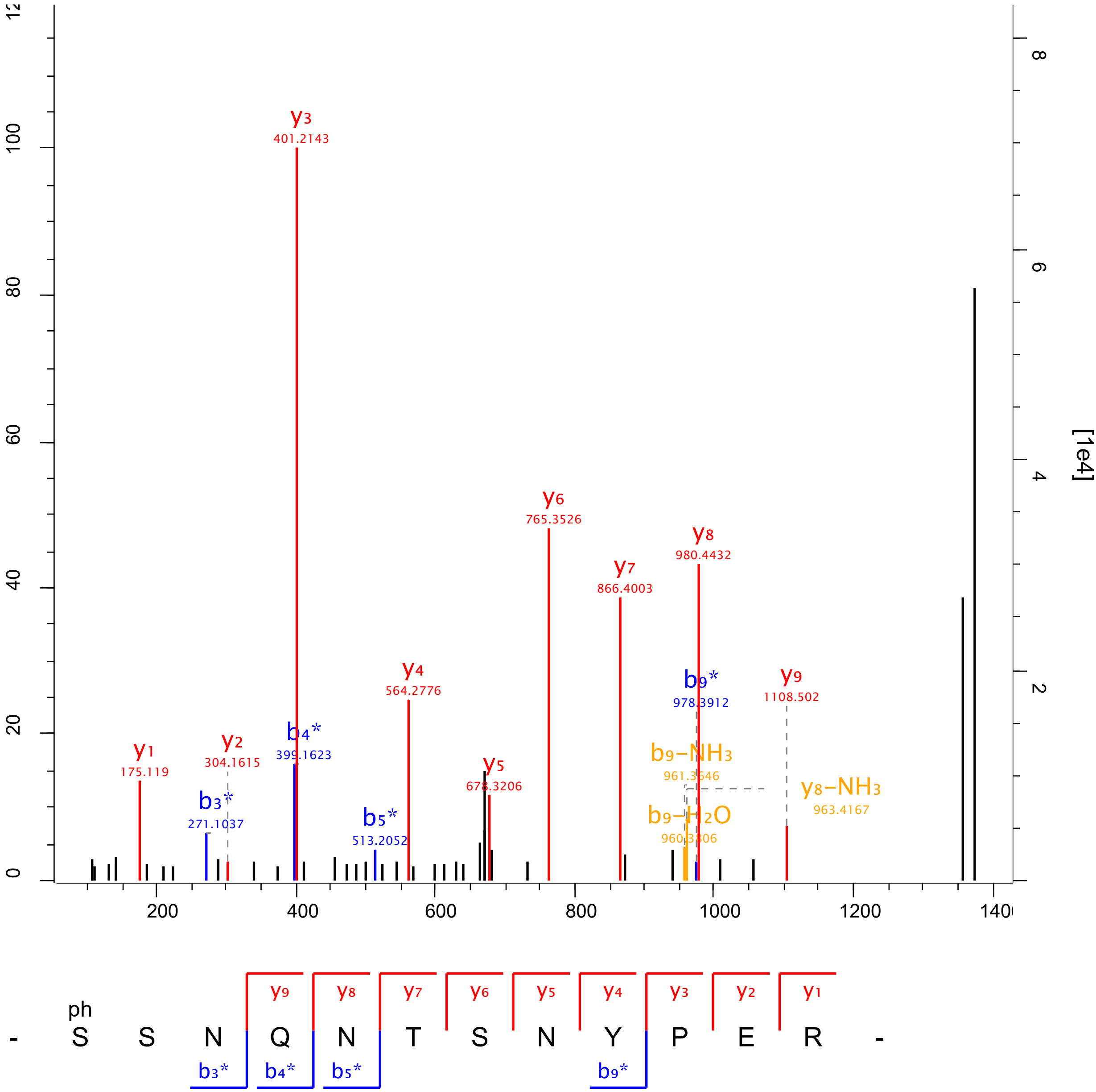

| Raw file | Scan | Method | Score | m/z |
|---|---|---|---|---|
| sys_02_2 | 4025 | FTMS; HCD | 149.83 | 546.78 |

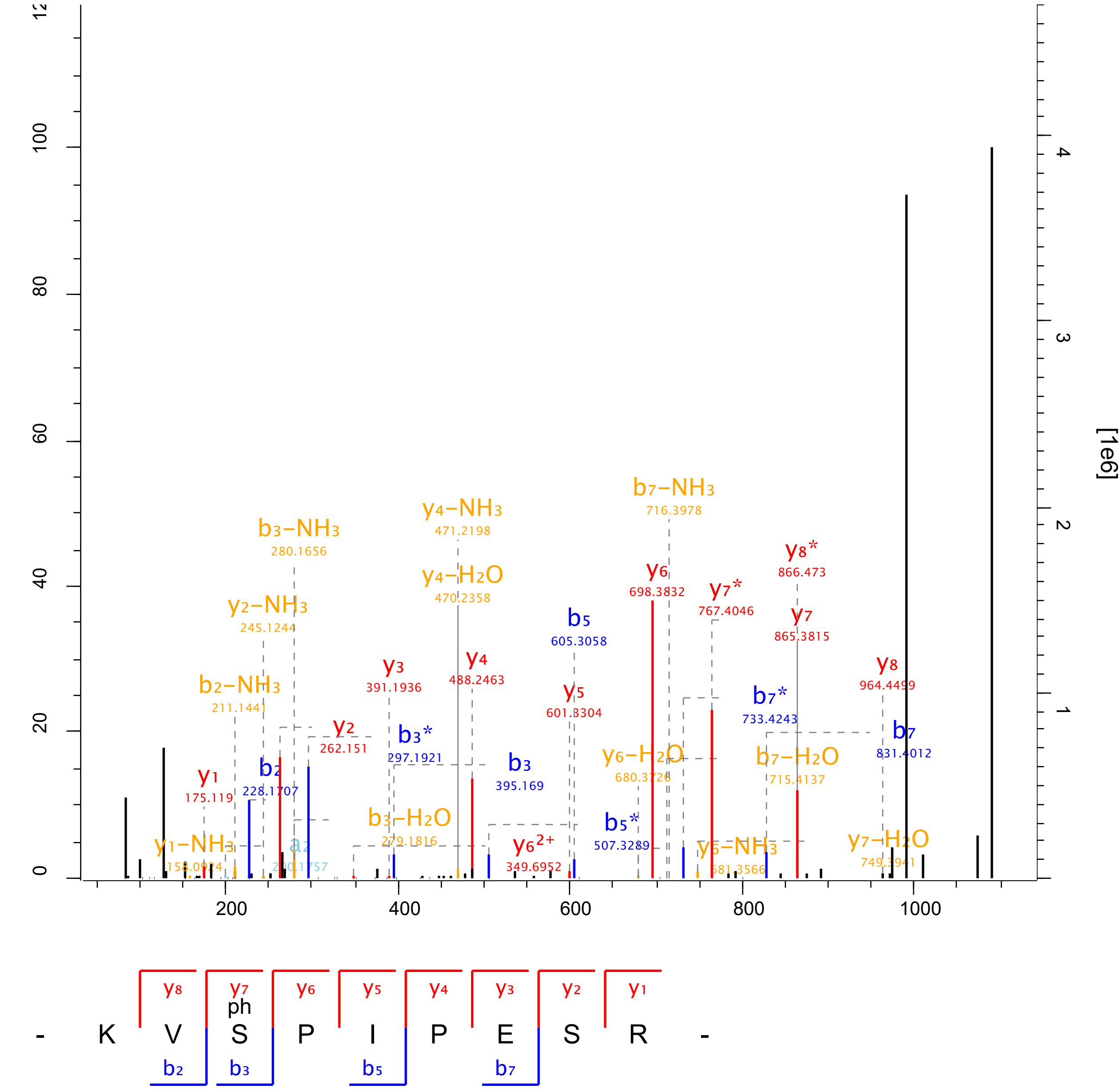

| Raw file | Scan | Method | Score | m/z |
|---|---|---|---|---|
| sys_02_2 | 4034 | FTMS; HCD | 90.56 | 464.88 |

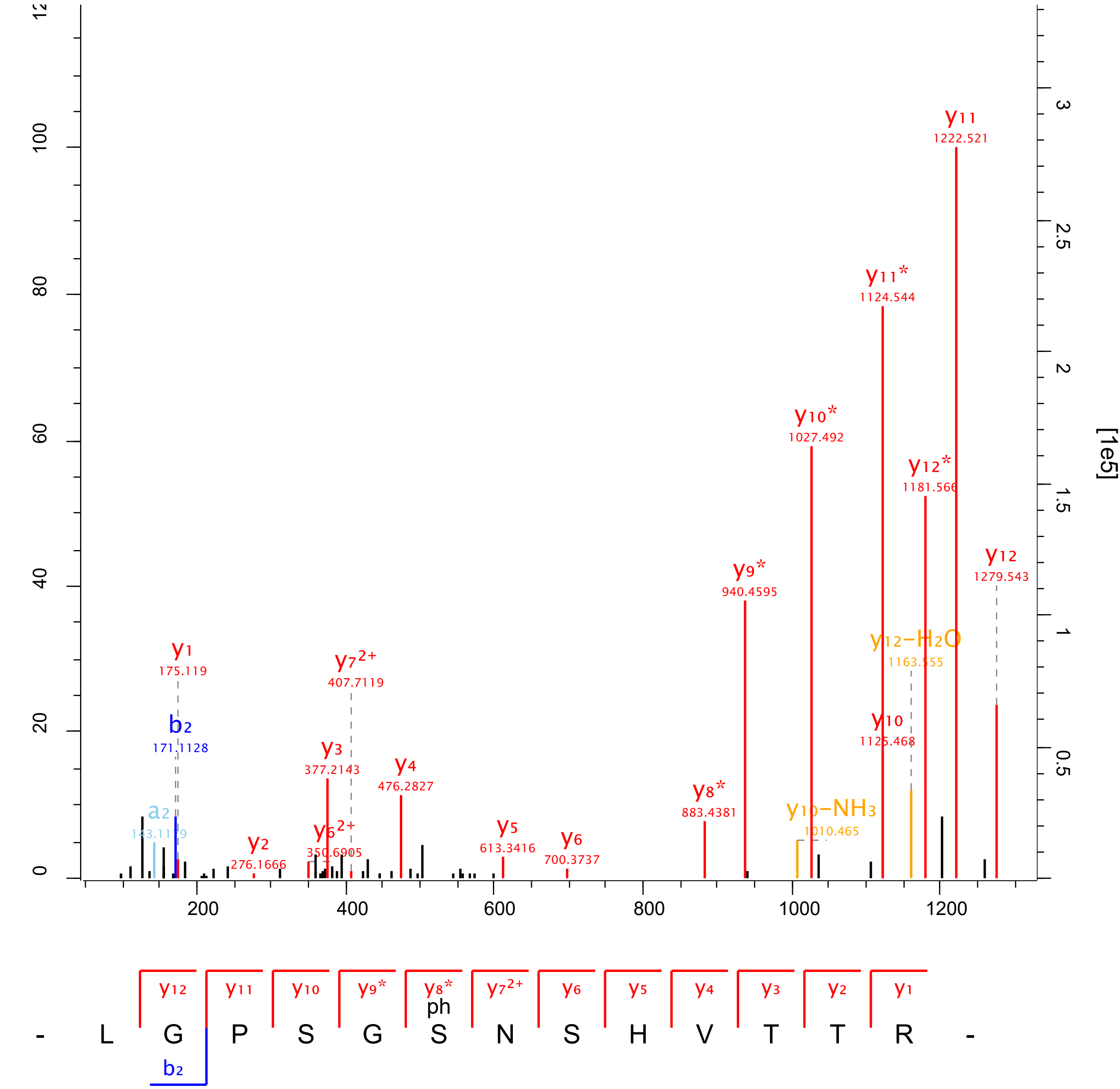

| Raw file | Scan | Method | Score | m/z |
|---|---|---|---|---|
| sys_02_2 | 4061 | FTMS; HCD | 68 | 569.26 |

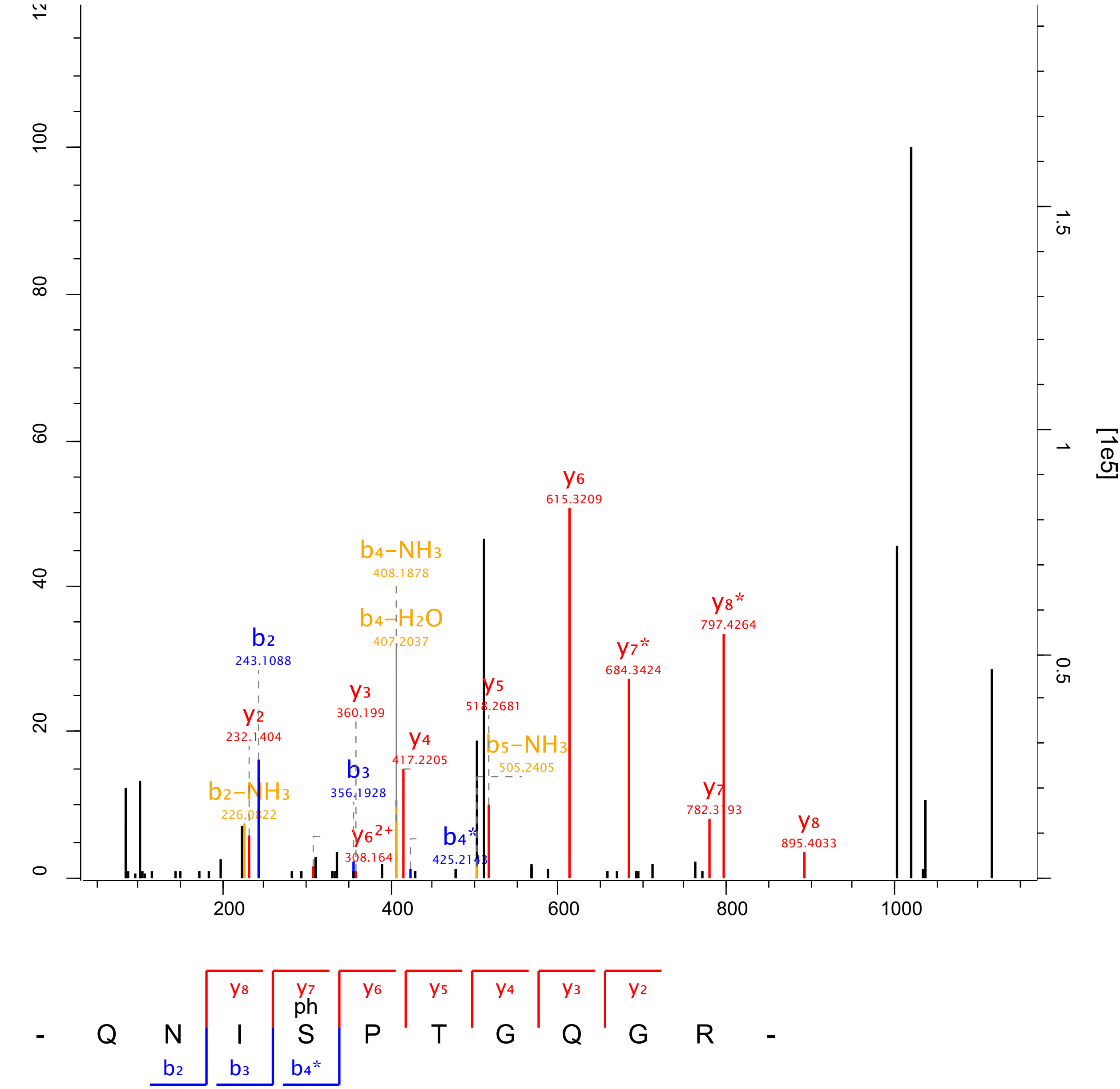

| Raw file | Scan | Method | Score | m/z |
|---|---|---|---|---|
| sys_02_2 | 4063 | FTMS; HCD | 49.59 | 654.29 |

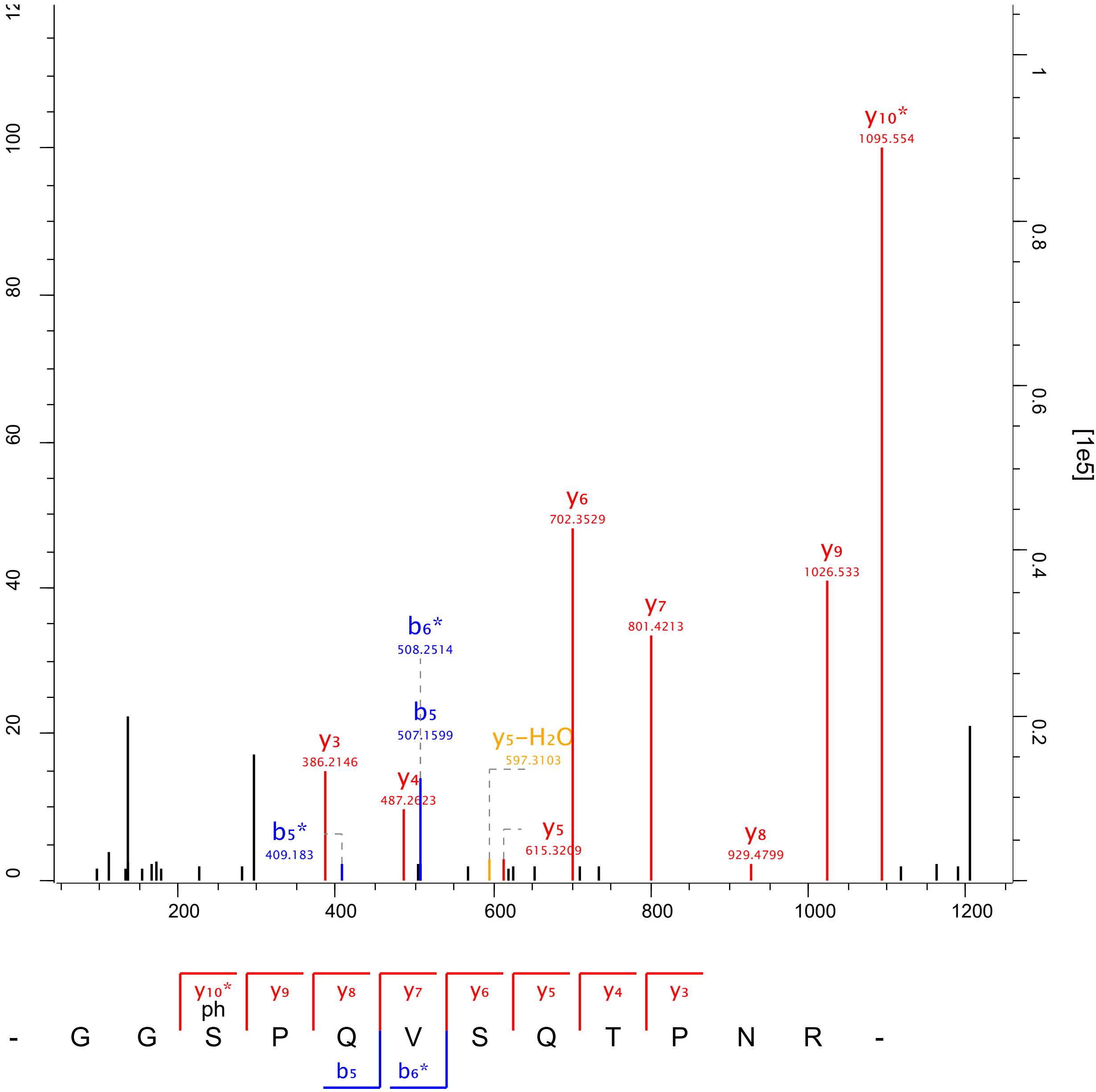

| Raw file | Scan | Method | Score | m/z |
|---|---|---|---|---|
| sys_02_2 | 4082 | FTMS; HCD | 52.86 | 381.49 |

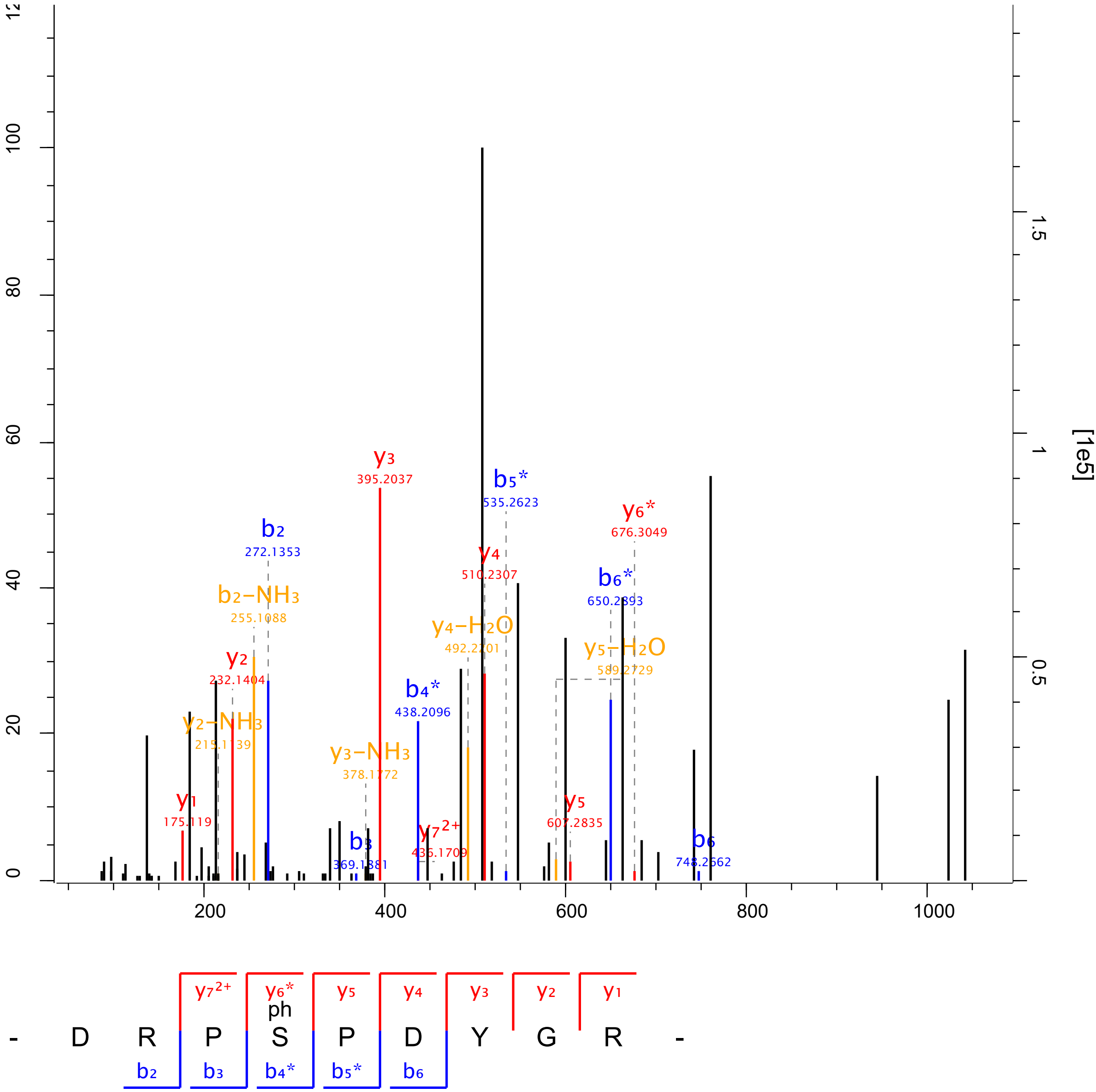

| Raw file | Scan | Method | Score | m/z |
|---|---|---|---|---|
| sys_02_2 | 4184 | FTMS; HCD | 133.3 | 693.64 |

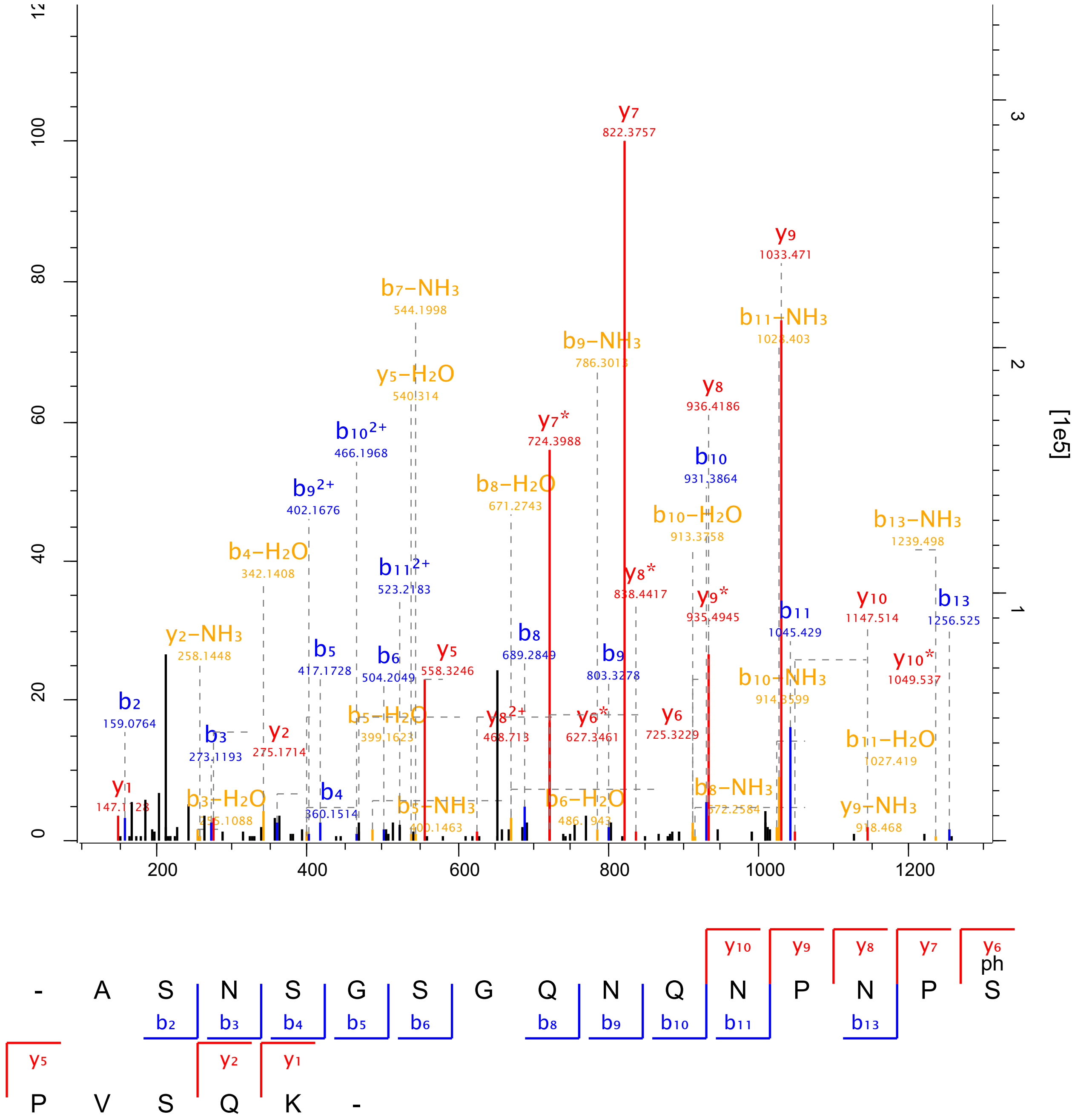

| Raw file | Scan | Method | Score | m/z |
|---|---|---|---|---|
| sys_02_2 | 4203 | FTMS; HCD | 81.57 | 566.24 |

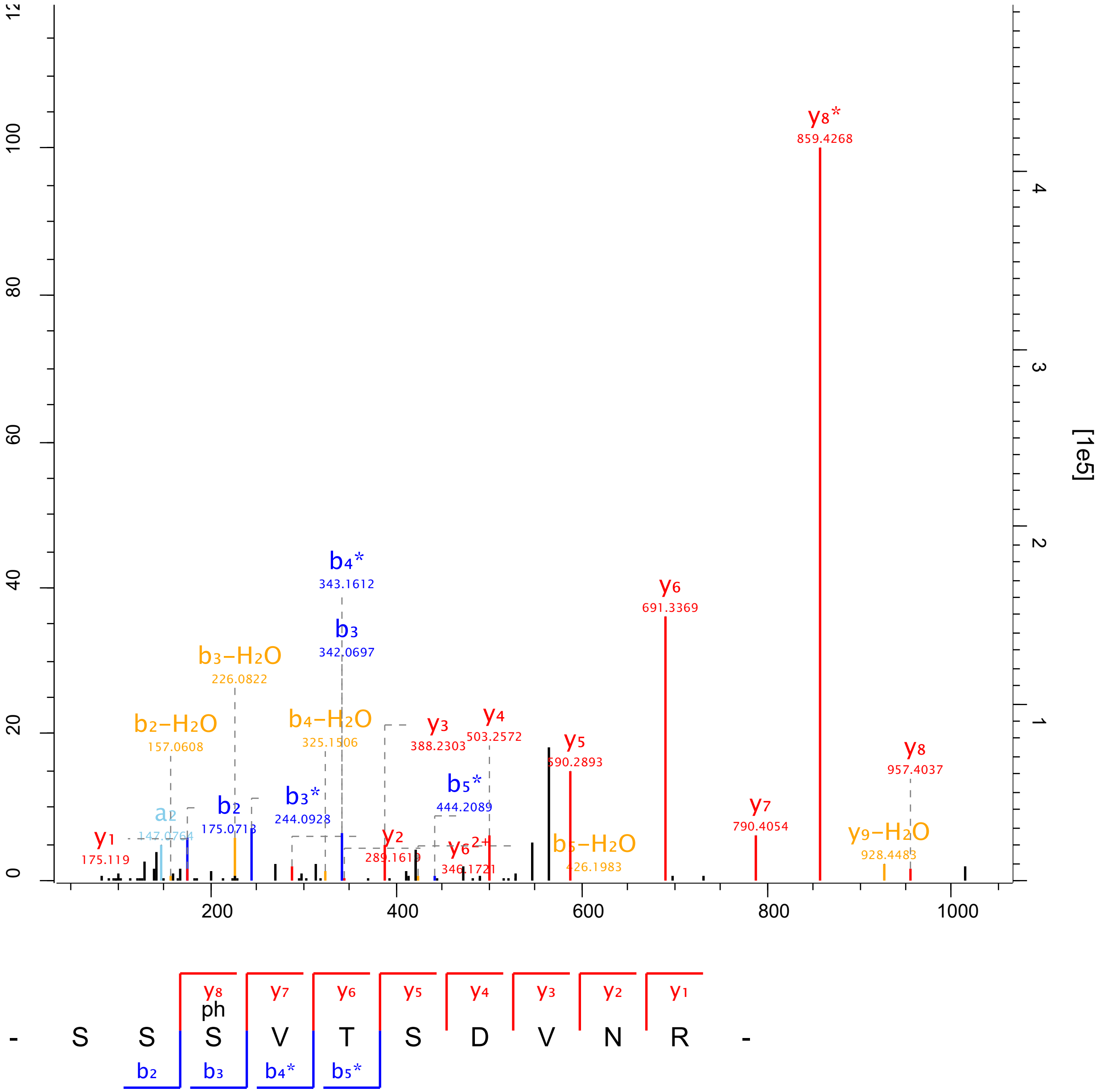

| Raw file | Scan | Method | Score | m/z |
| --- | --- | --- | --- | --- |
| sys_02_2 | 4414 | FTMS; HCD | 112.02 | 476.23 |

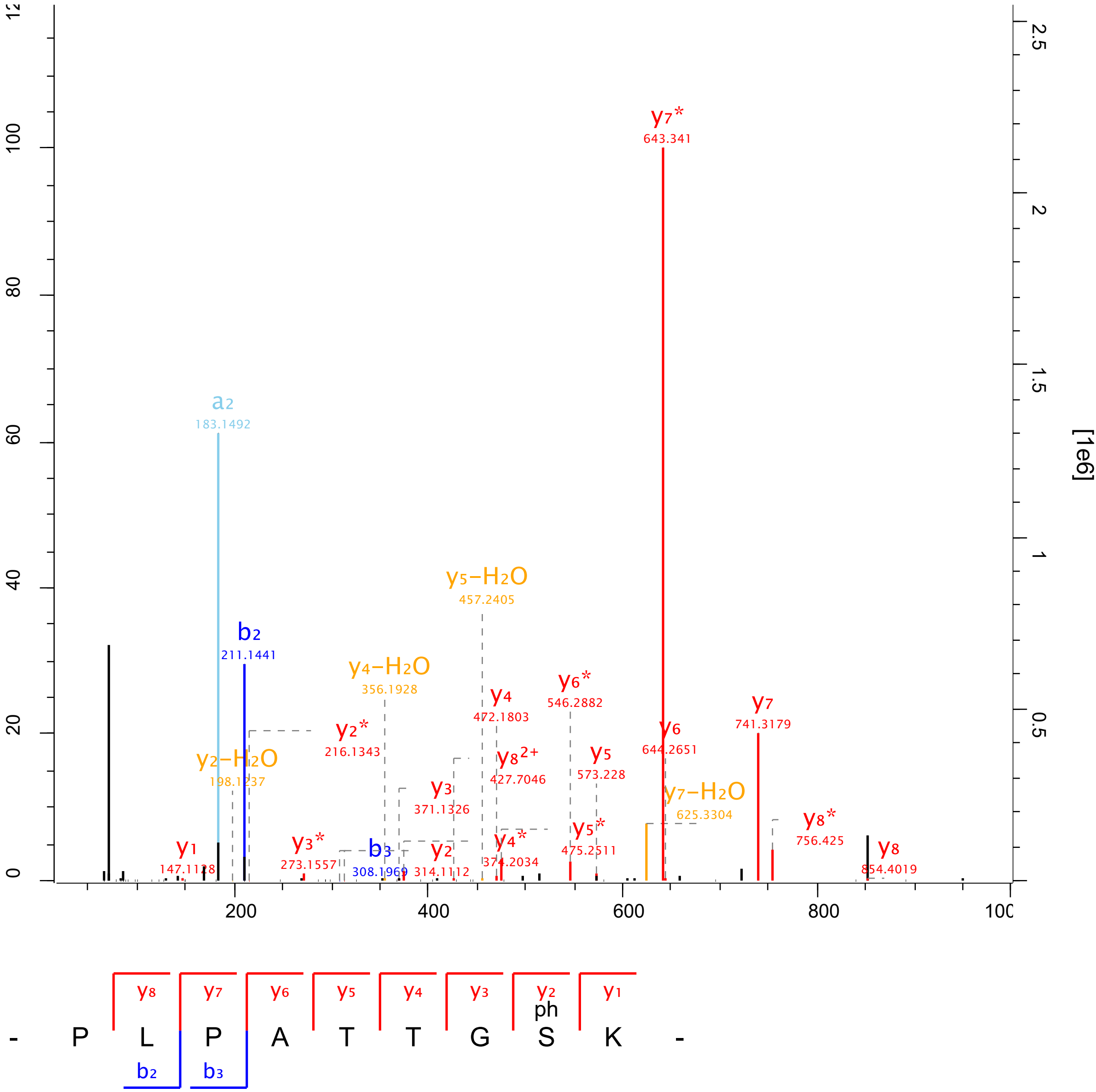

| Raw file | Scan | Method | Score | m/z |
| --- | --- | --- | --- | --- |
| sys_02_2 | 4429 | FTMS; HCD | 101.3 | 531.74 |

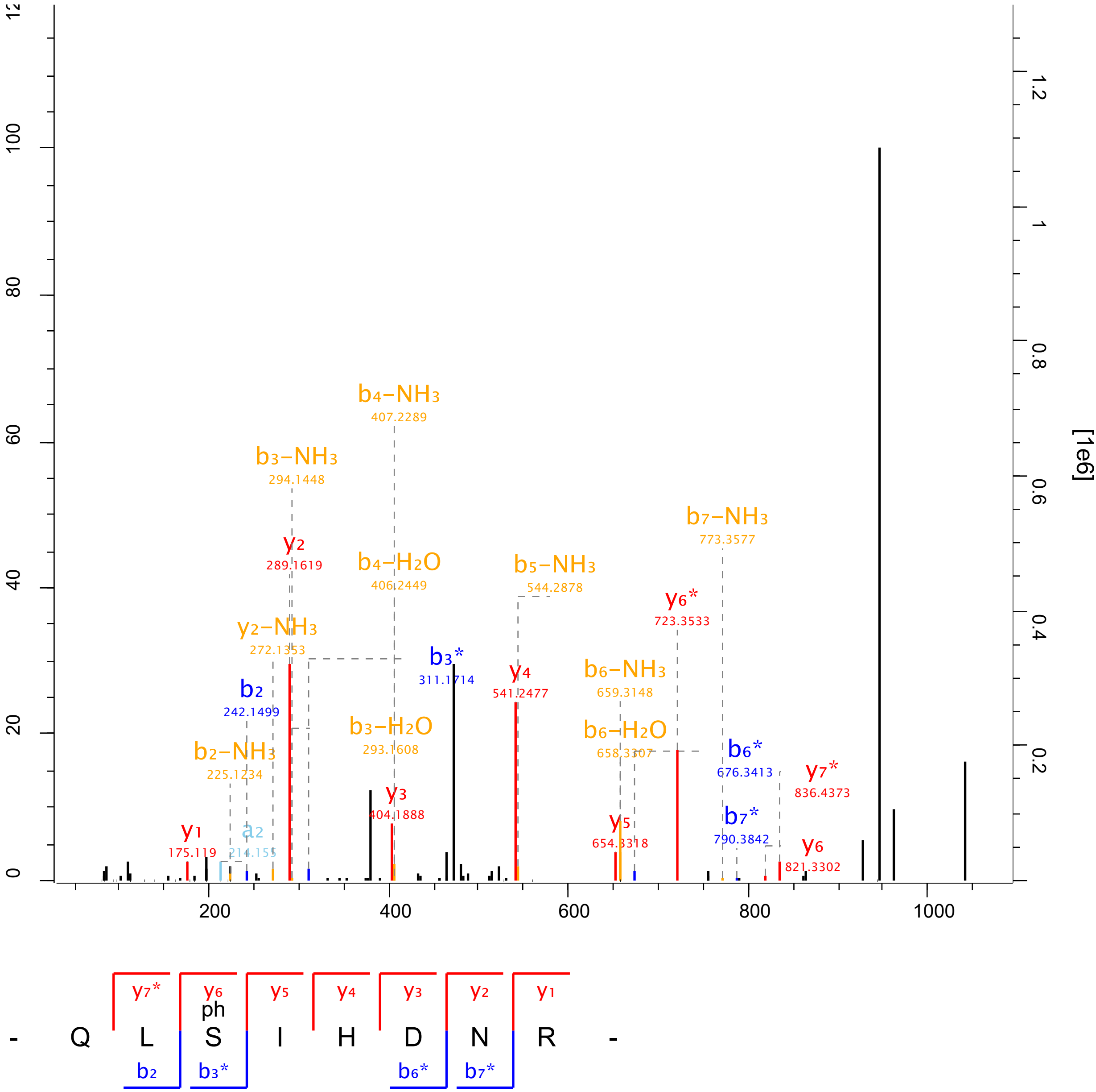

| Raw file | Scan | Method | Score | m/z |
|---|---|---|---|---|
| sys_02_2 | 4609 | FTMS; HCD | 67.22 | 702.32 |

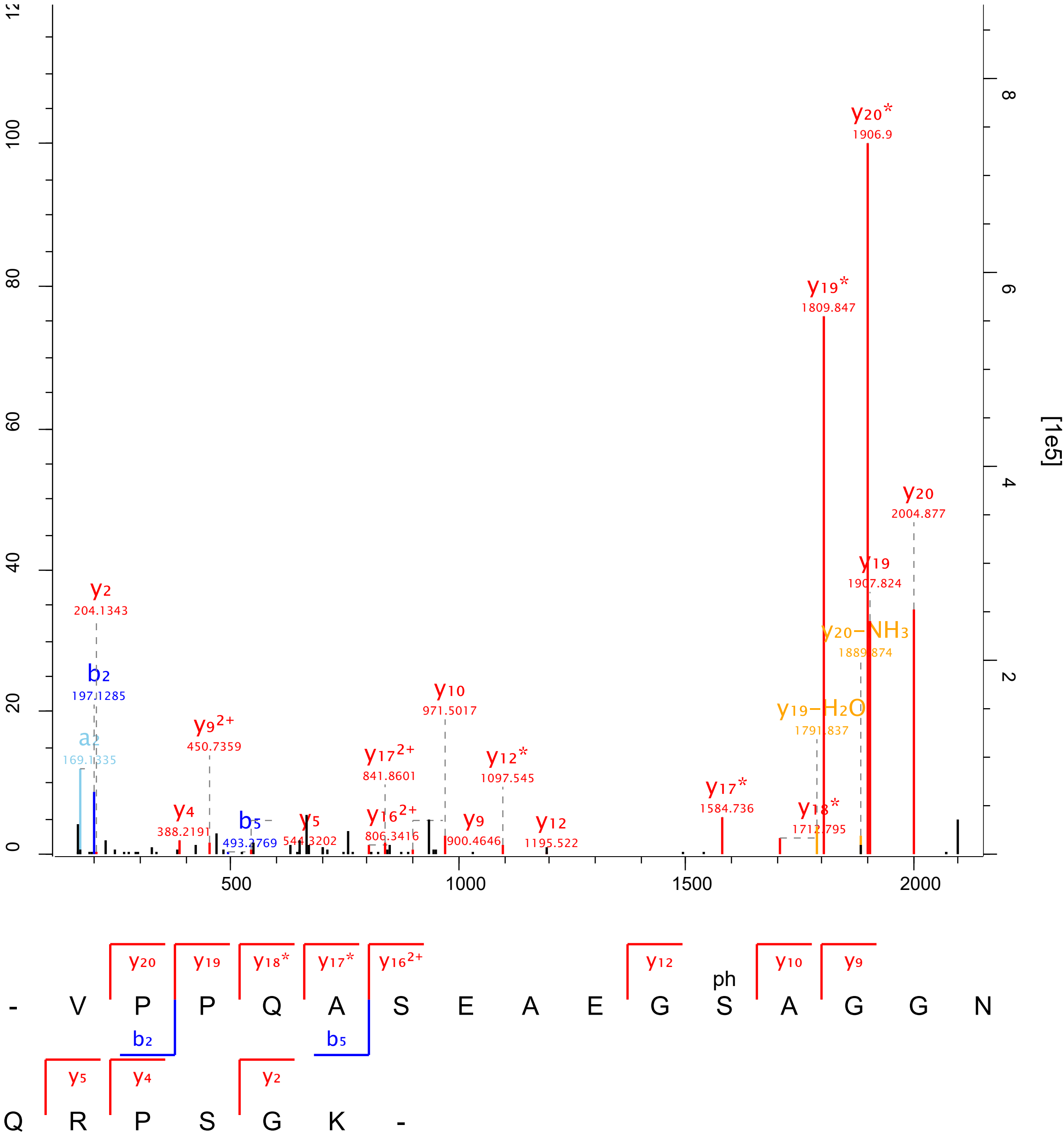

| Raw file | Scan | Method | Score | m/z |
| --- | --- | --- | --- | --- |
| sys_02_2 | 4627 | FTMS; HCD | 41.54 | 469.55 |

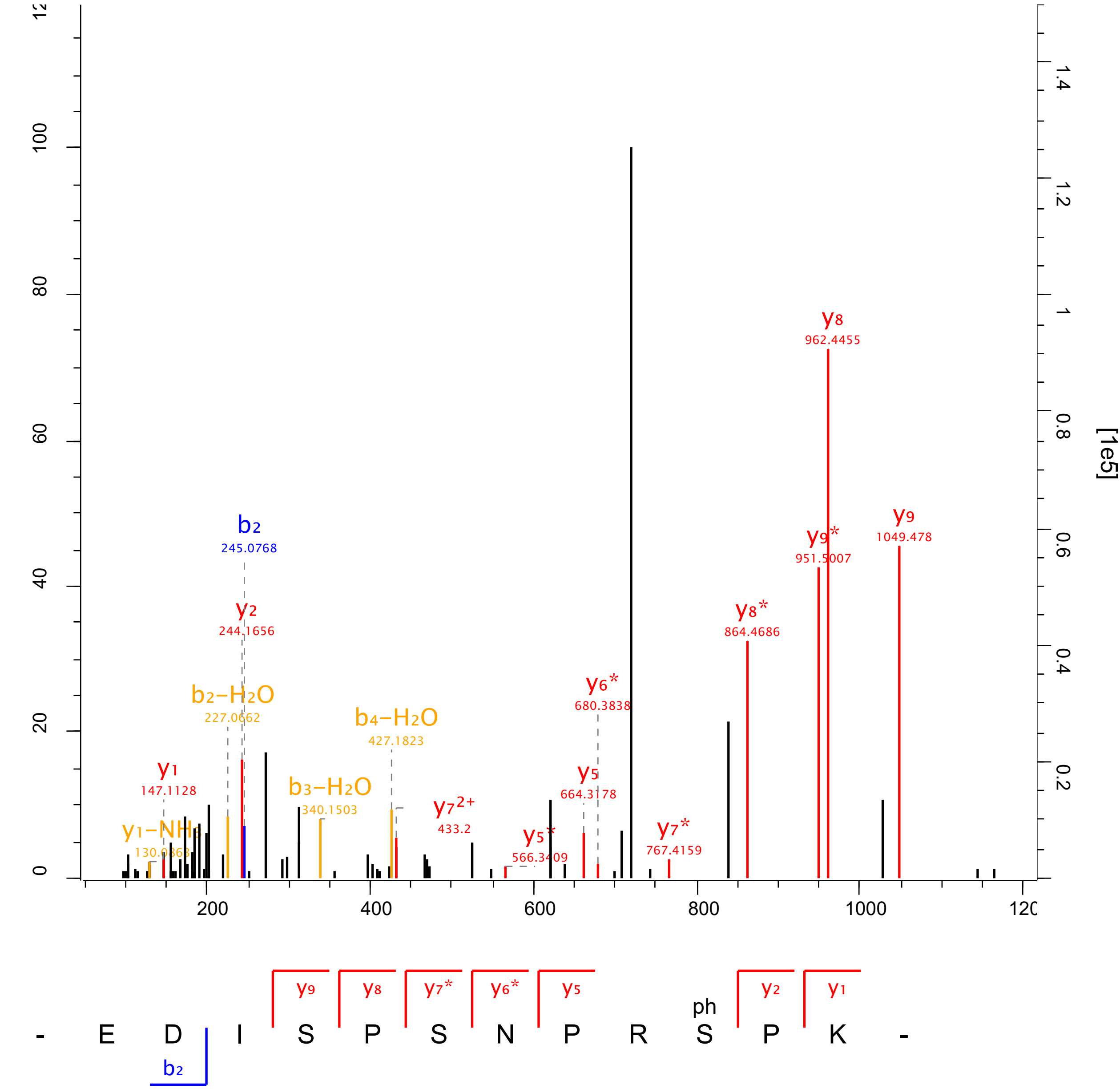

| Raw file | Scan | Method | Score | m/z |
|---|---|---|---|---|
| sys_02_2 | 4675 | FTMS; HCD | 76.07 | 605.78 |

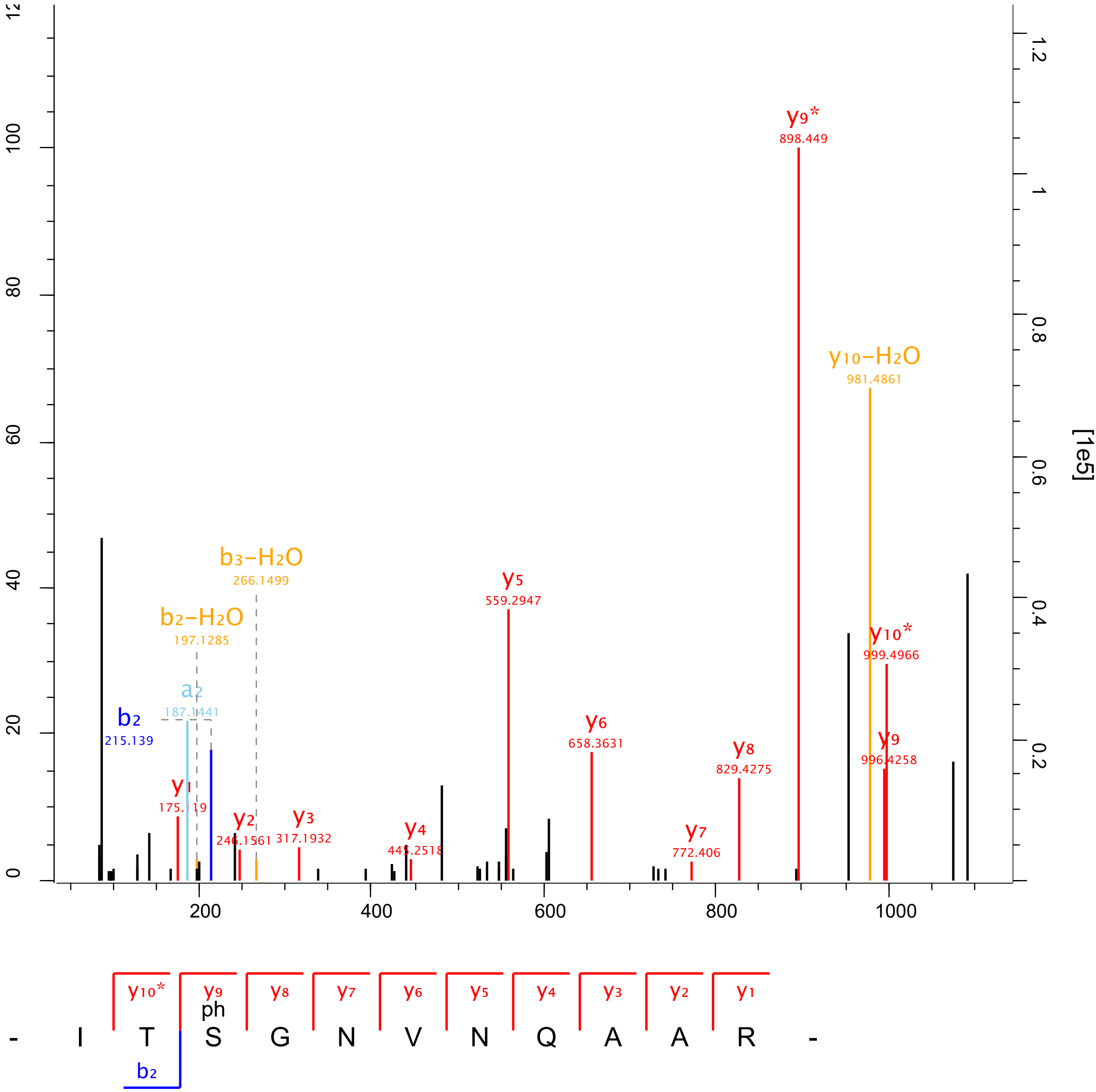

| Raw file | Scan | Method | Score | m/z |
|---|---|---|---|---|
| sys_02_2 | 4786 | FTMS; HCD | 41.09 | 469.23 |

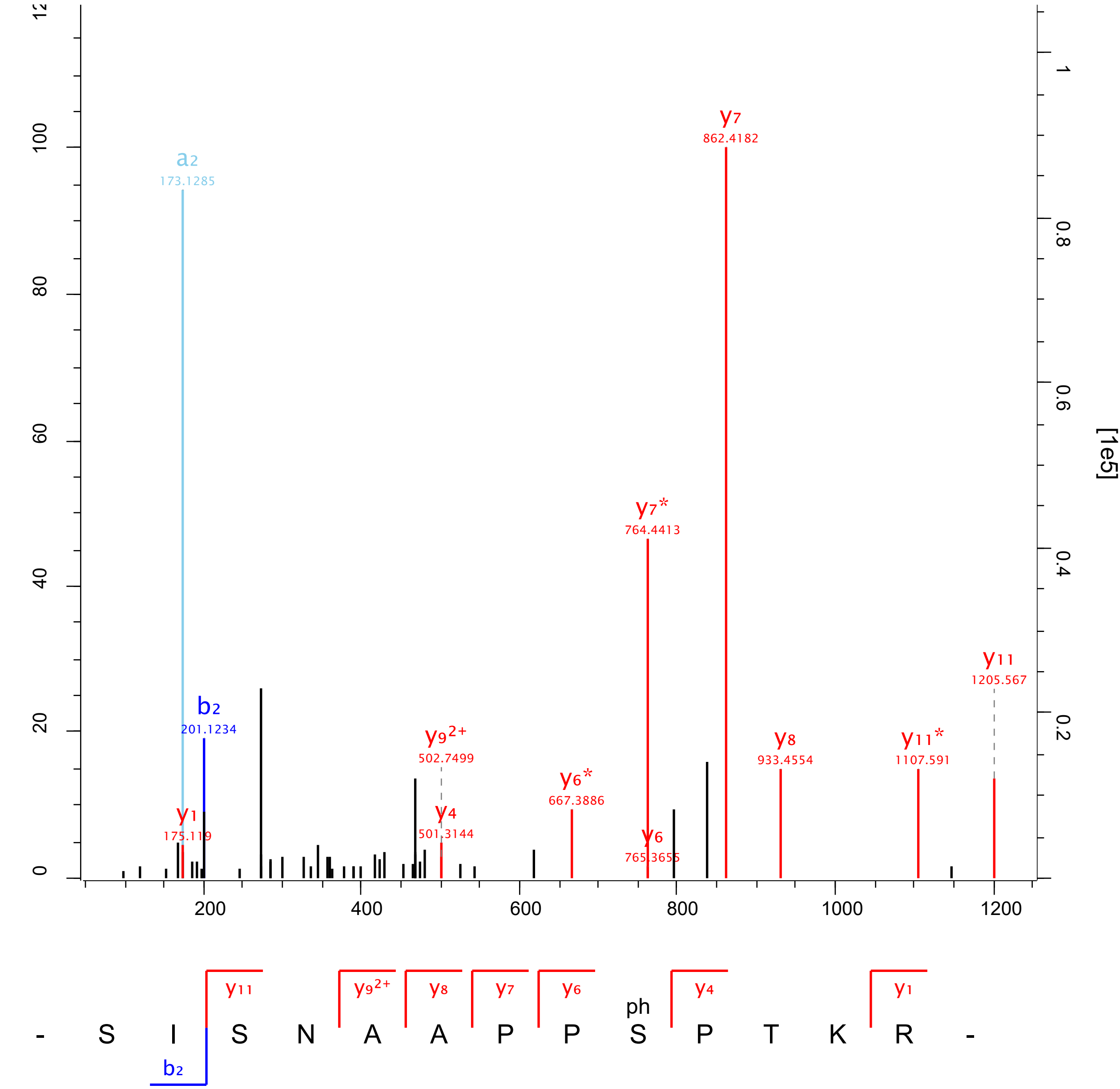

| Raw file | Scan | Method | Score | m/z |
|---|---|---|---|---|
| sys_02_2 | 4860 | FTMS; HCD | 50.09 | 617.79 |

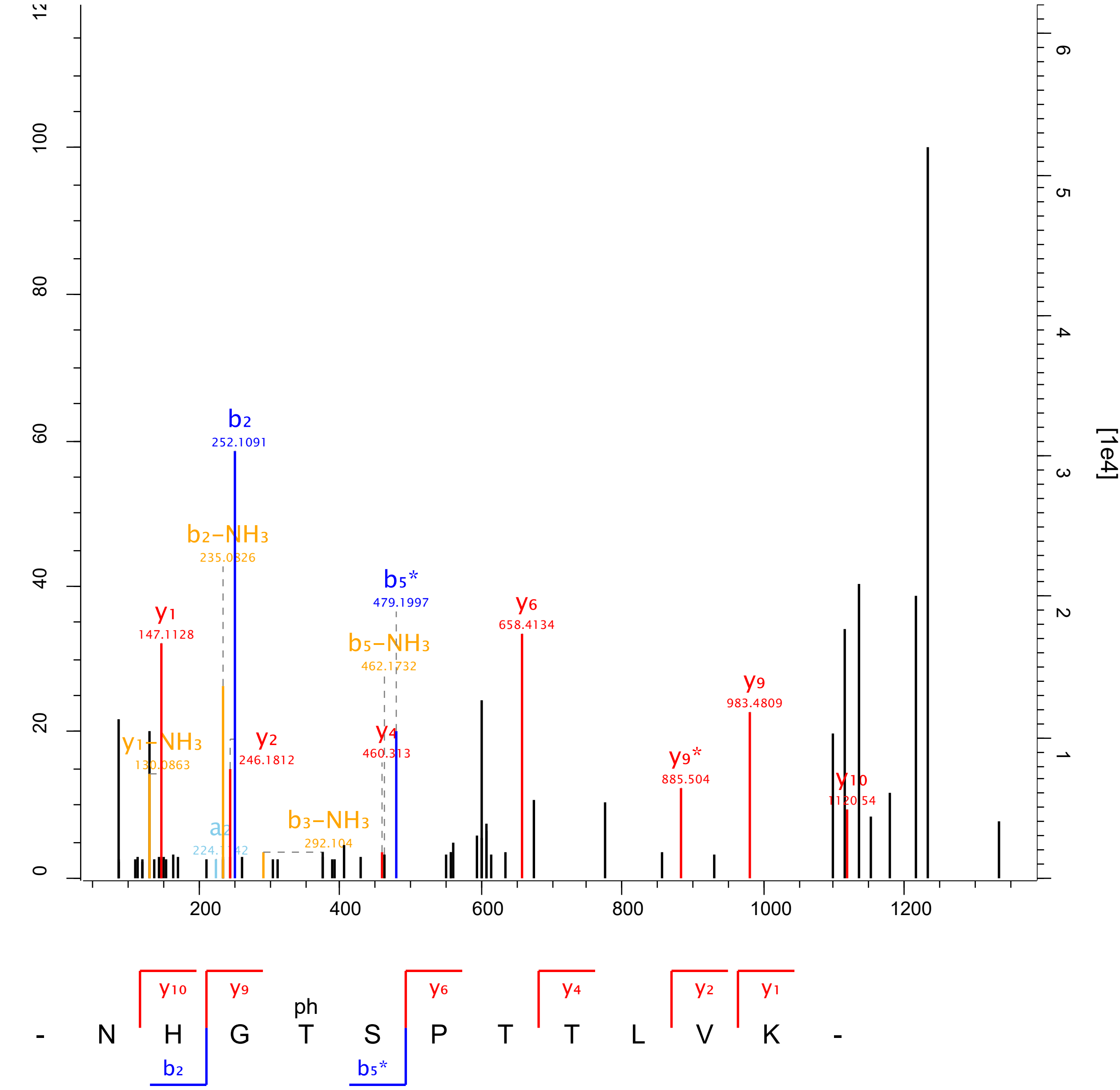

| Raw file | Scan | Method | Score | m/z |
|---|---|---|---|---|
| sys_02_2 | 4927 | FTMS; HCD | 64.64 | 471.21 |

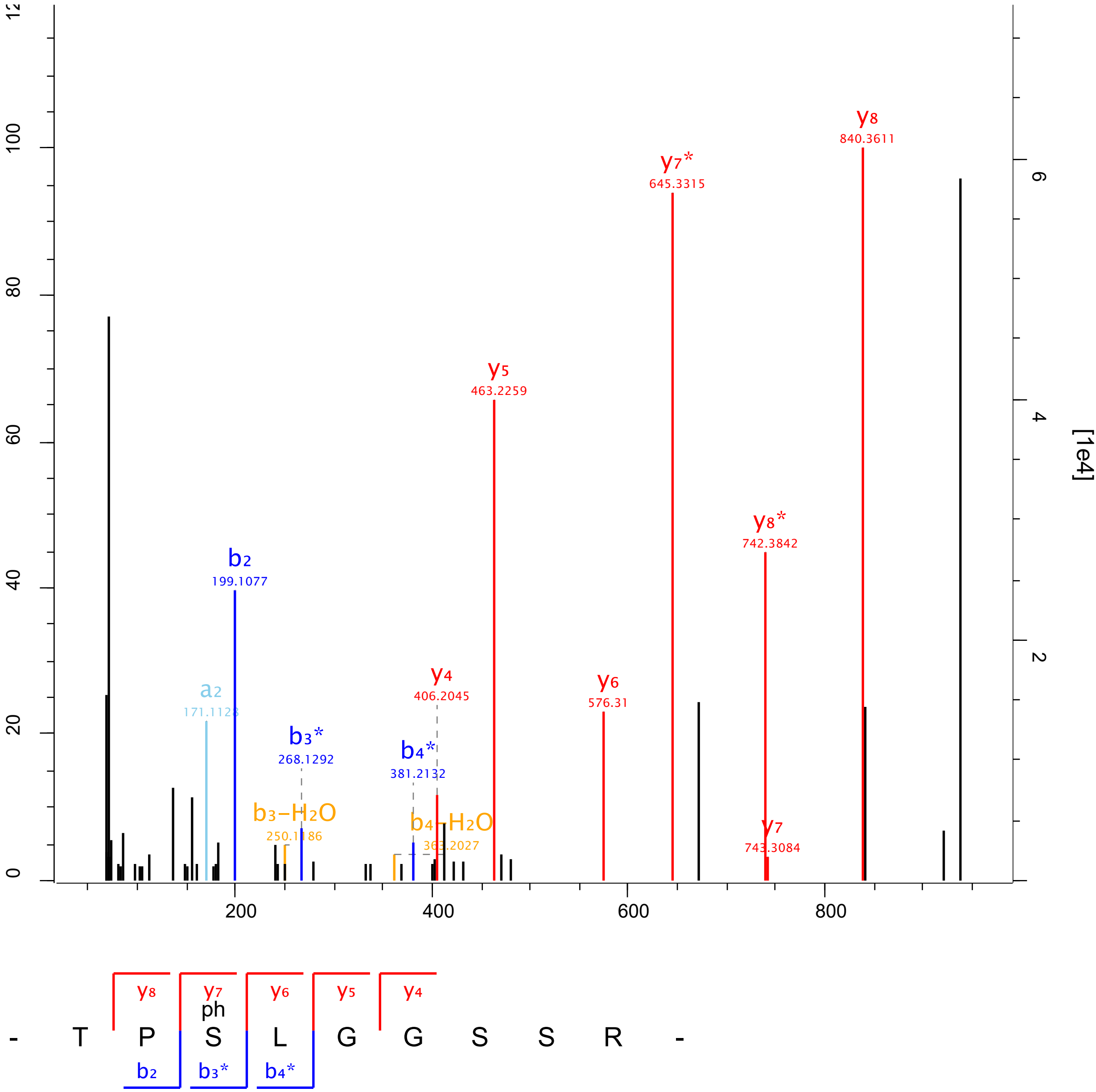

| Raw file | Scan | Method | Score | m/z |
|---|---|---|---|---|
| sys_02_2 | 4994 | FTMS; HCD | 95.85 | 627.28 |

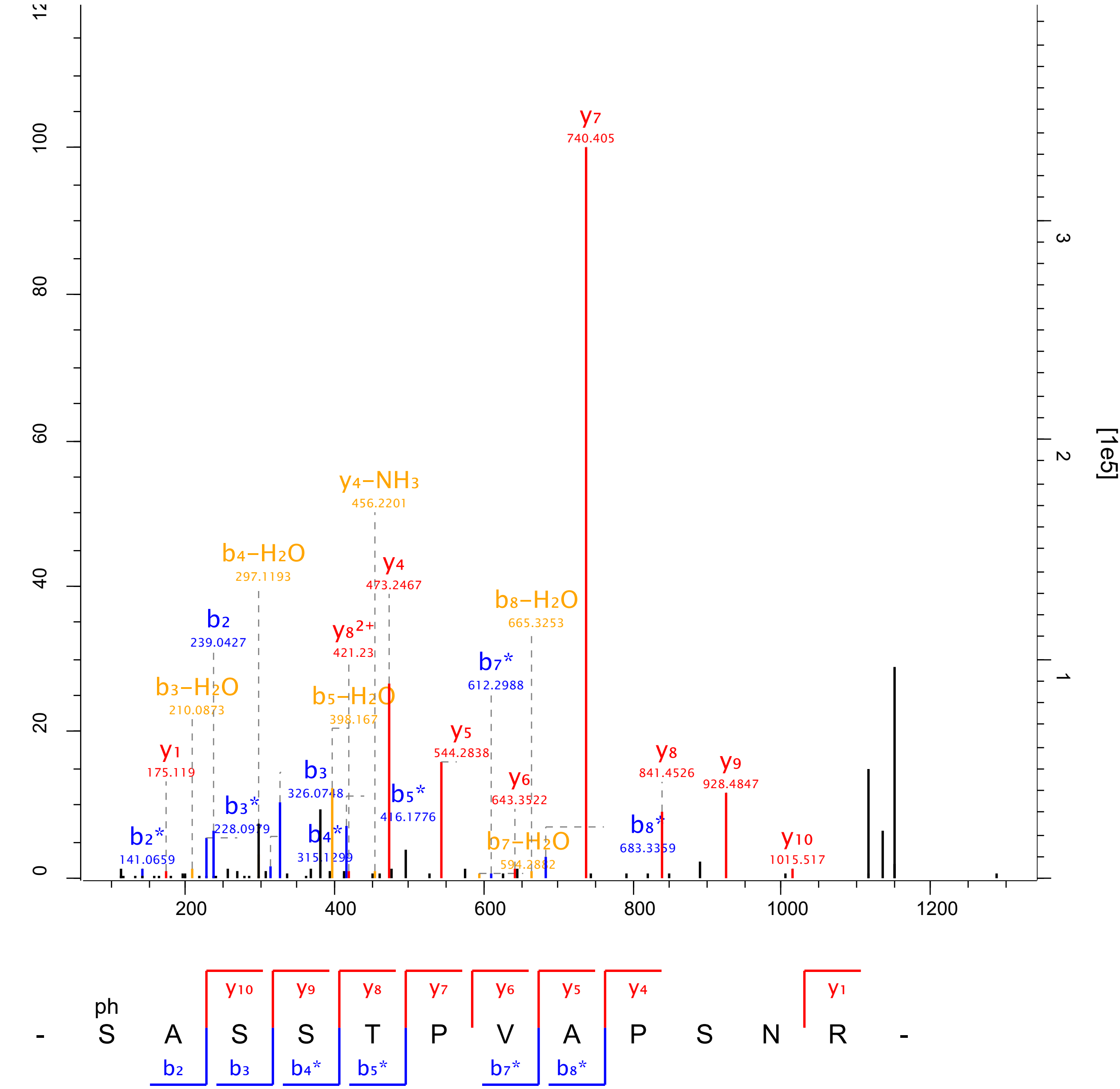

| Raw file | Scan | Method | Score | m/z |
|---|---|---|---|---|
| sys_02_2 | 5046 | FTMS; HCD | 155.49 | 879.86 |

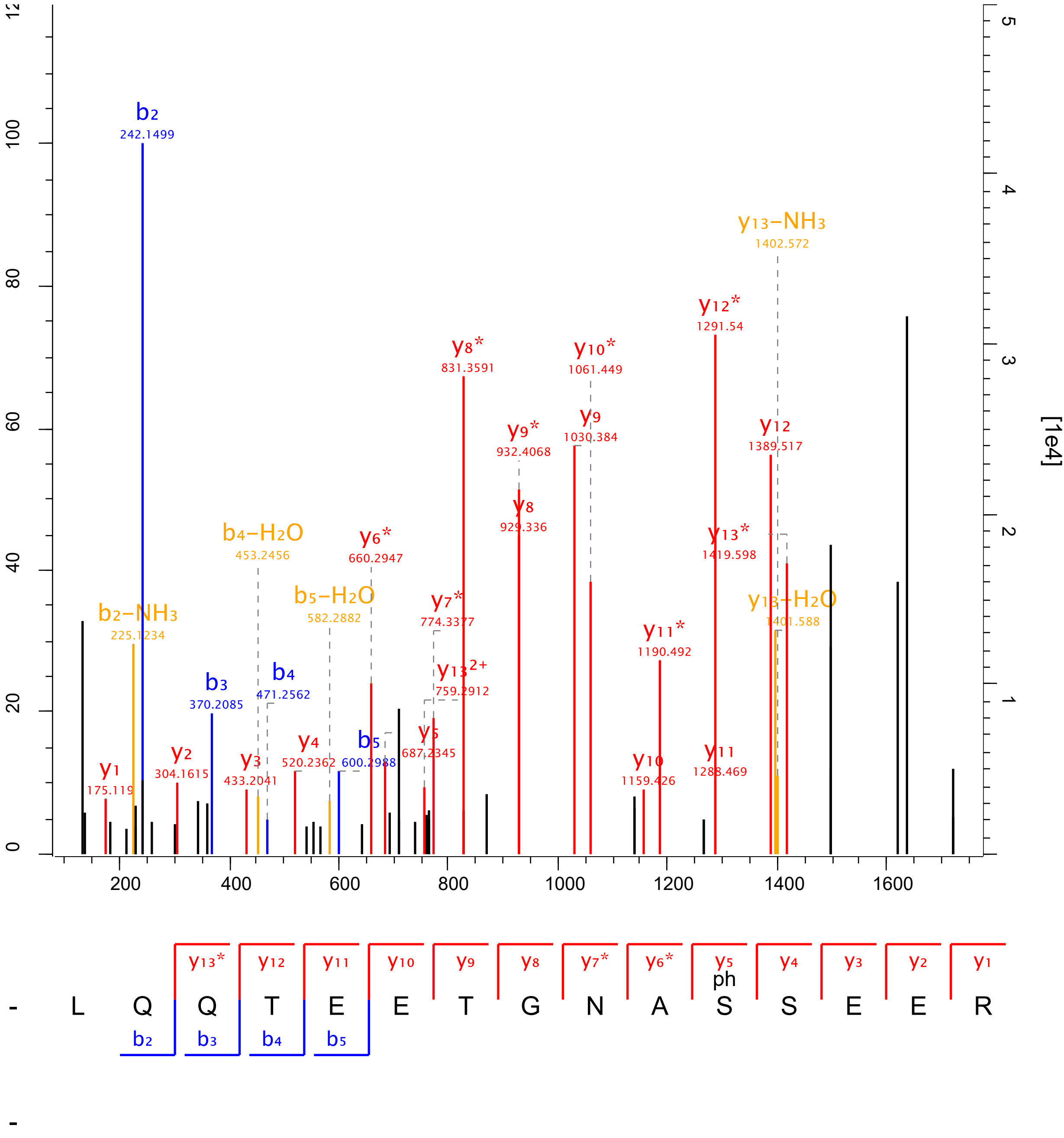

| Raw file | Scan | Method | Score | m/z |
|---|---|---|---|---|
| sys_02_2 | 5120 | FTMS; HCD | 43.21 | 600.59 |

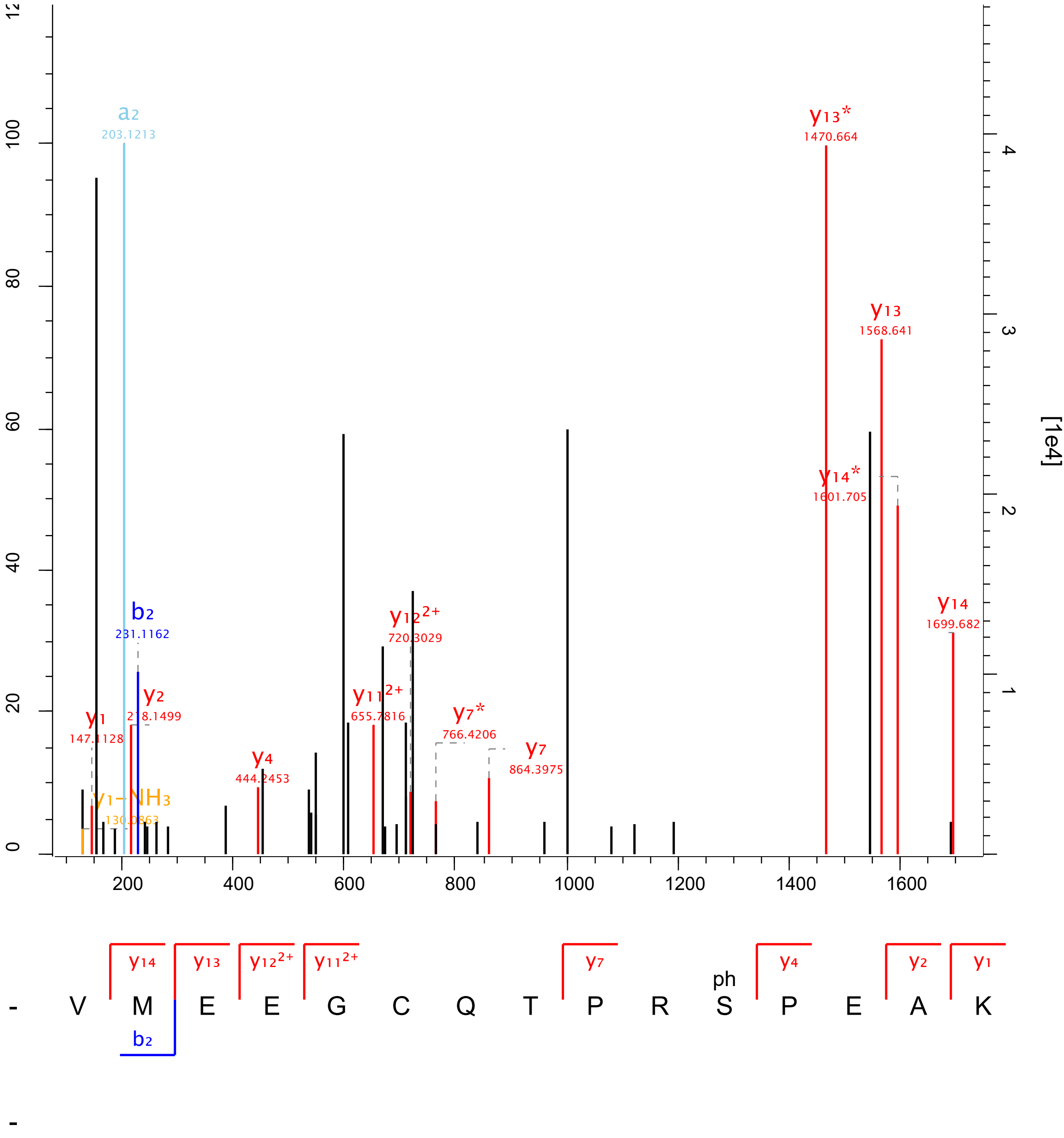

| Raw file | Scan | Method | Score | m/z |
|---|---|---|---|---|
| sys_02_2 | 5166 | FTMS; HCD | 77.64 | 569.25 |

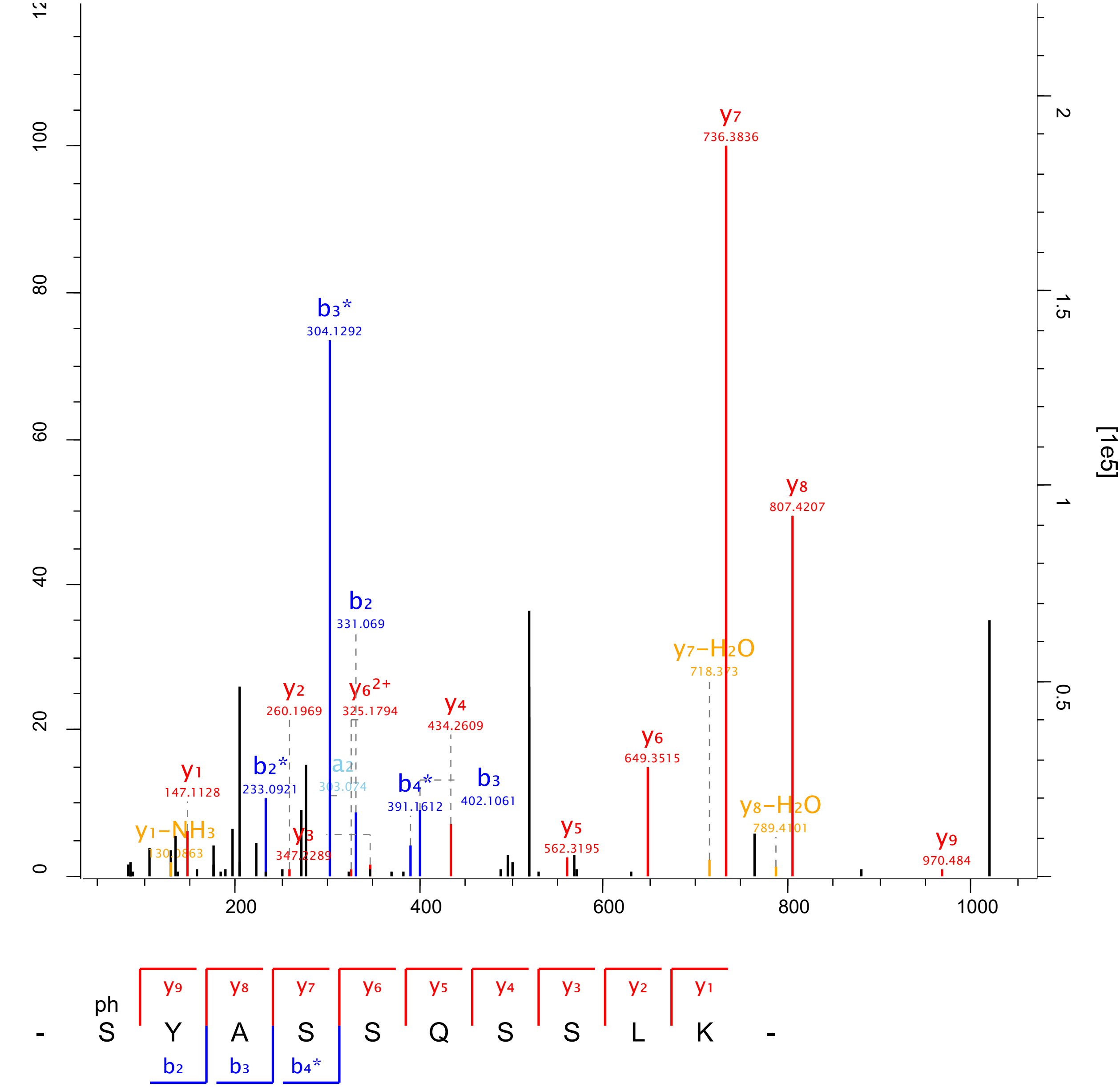

| Raw file | Scan | Method | Score | m/z |
|---|---|---|---|---|
| sys_02_2 | 5203 | FTMS; HCD | 71.98 | 659.29 |

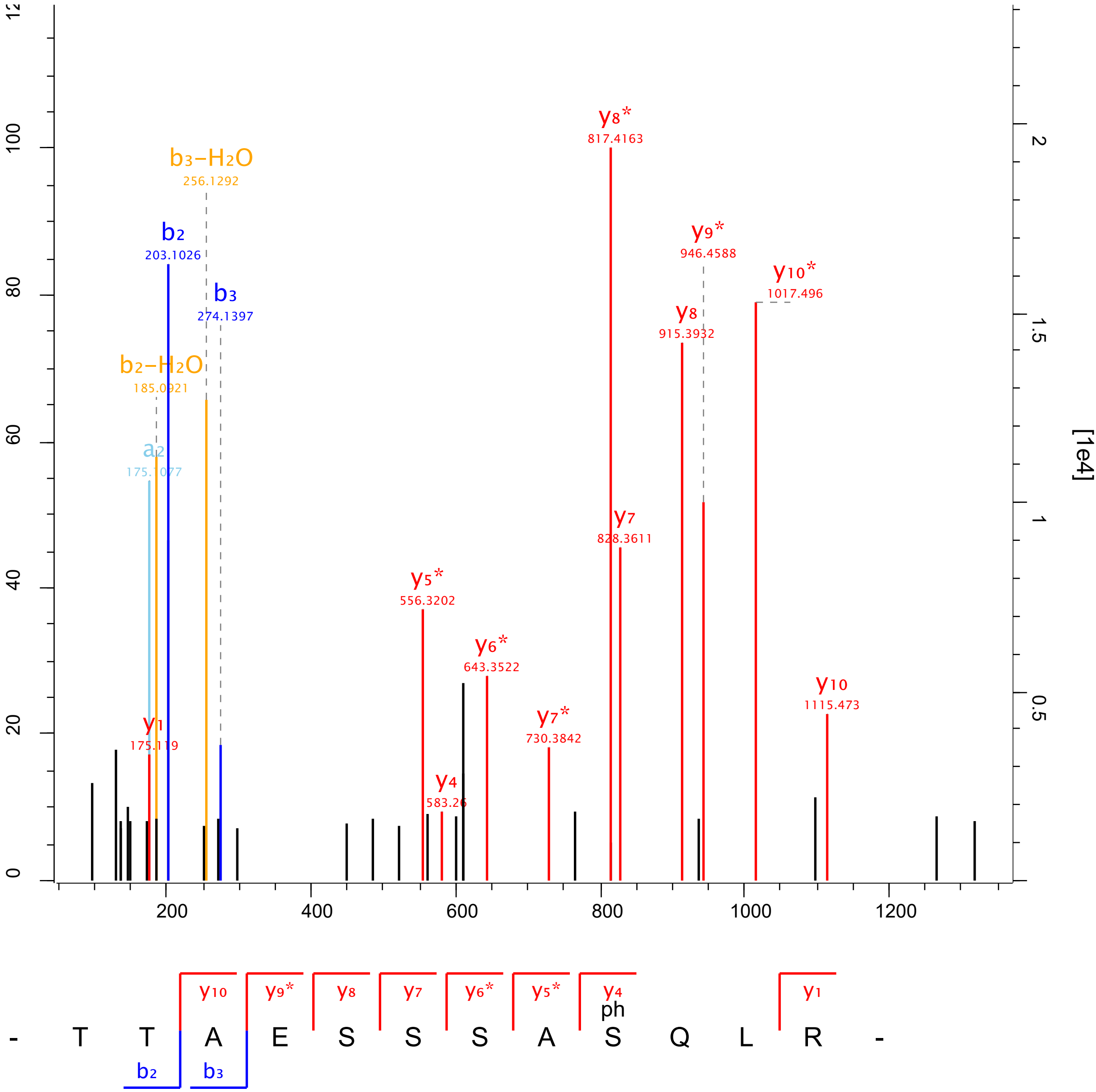

| Raw file | Scan | Method | Score | m/z |
|---|---|---|---|---|
| sys_02_2 | 5210 | FTMS; HCD | 40.28 | 628.79 |

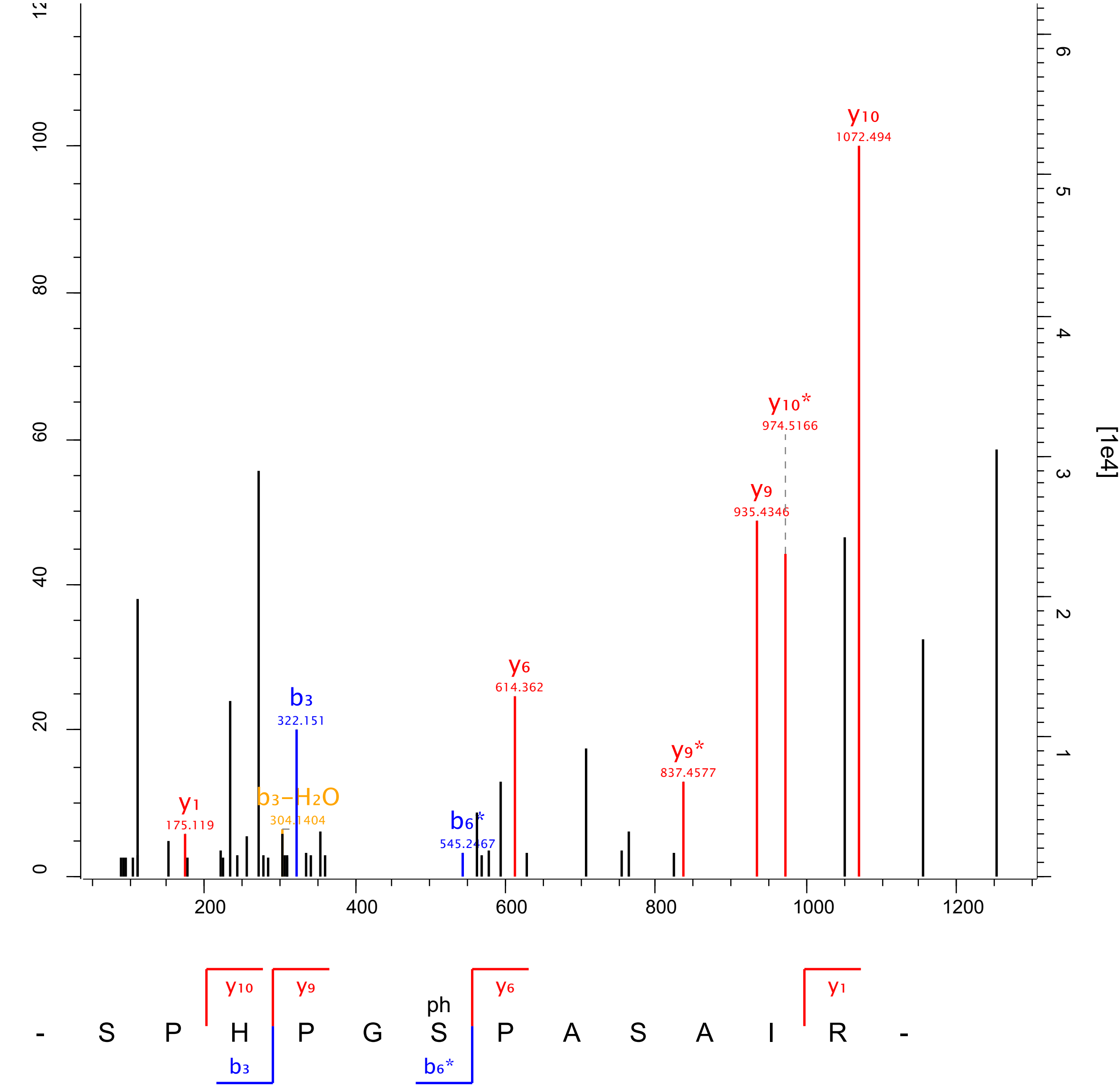

| Raw file | Scan | Method | Score | m/z |
|---|---|---|---|---|
| sys_02_2 | 5409 | FTMS; HCD | 143.57 | 834.34 |

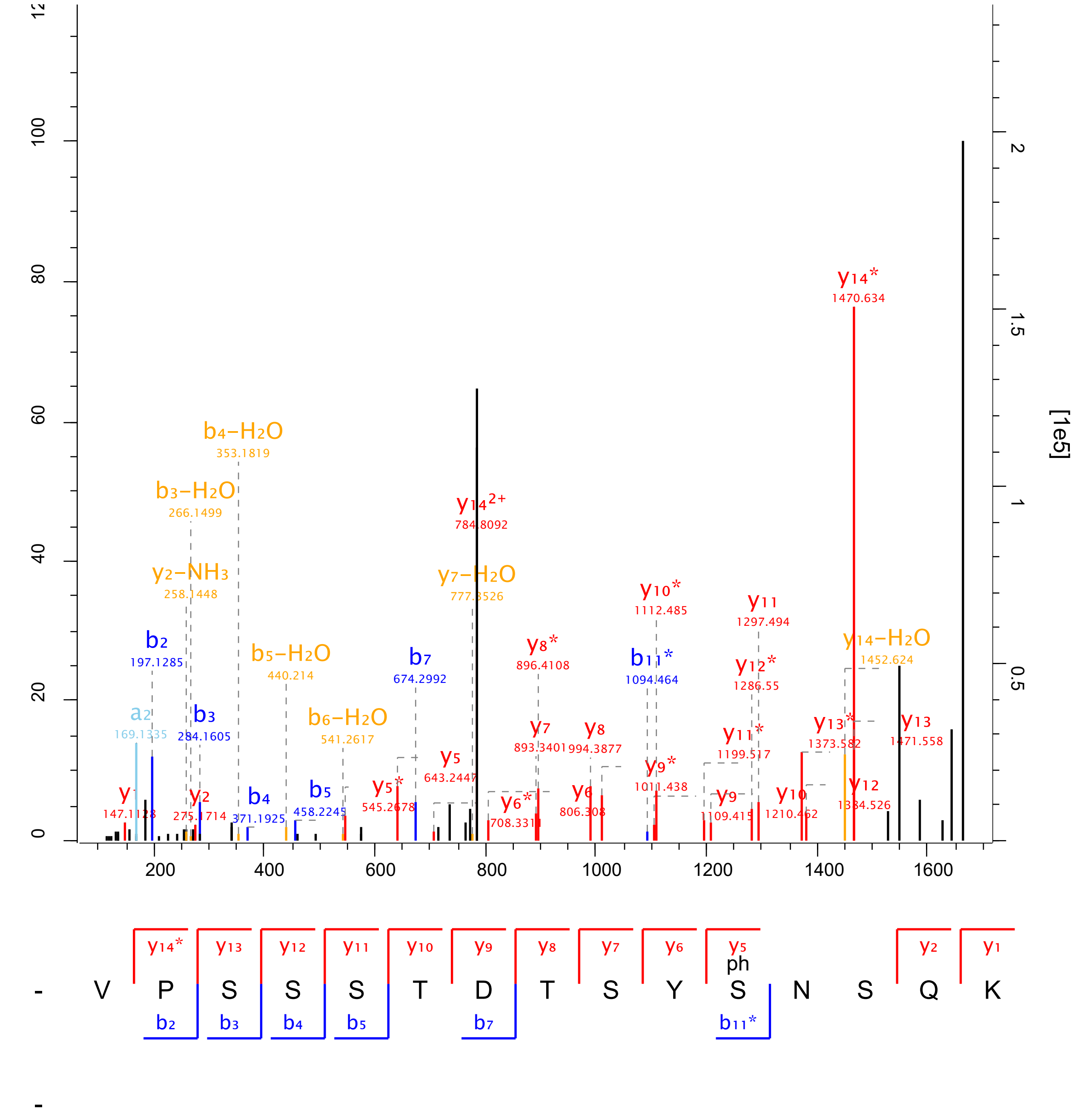

| Raw file | Scan | Method | Score | m/z |
|---|---|---|---|---|
| sys_02_2 | 5443 | FTMS; HCD | 52.25 | 588.22 |

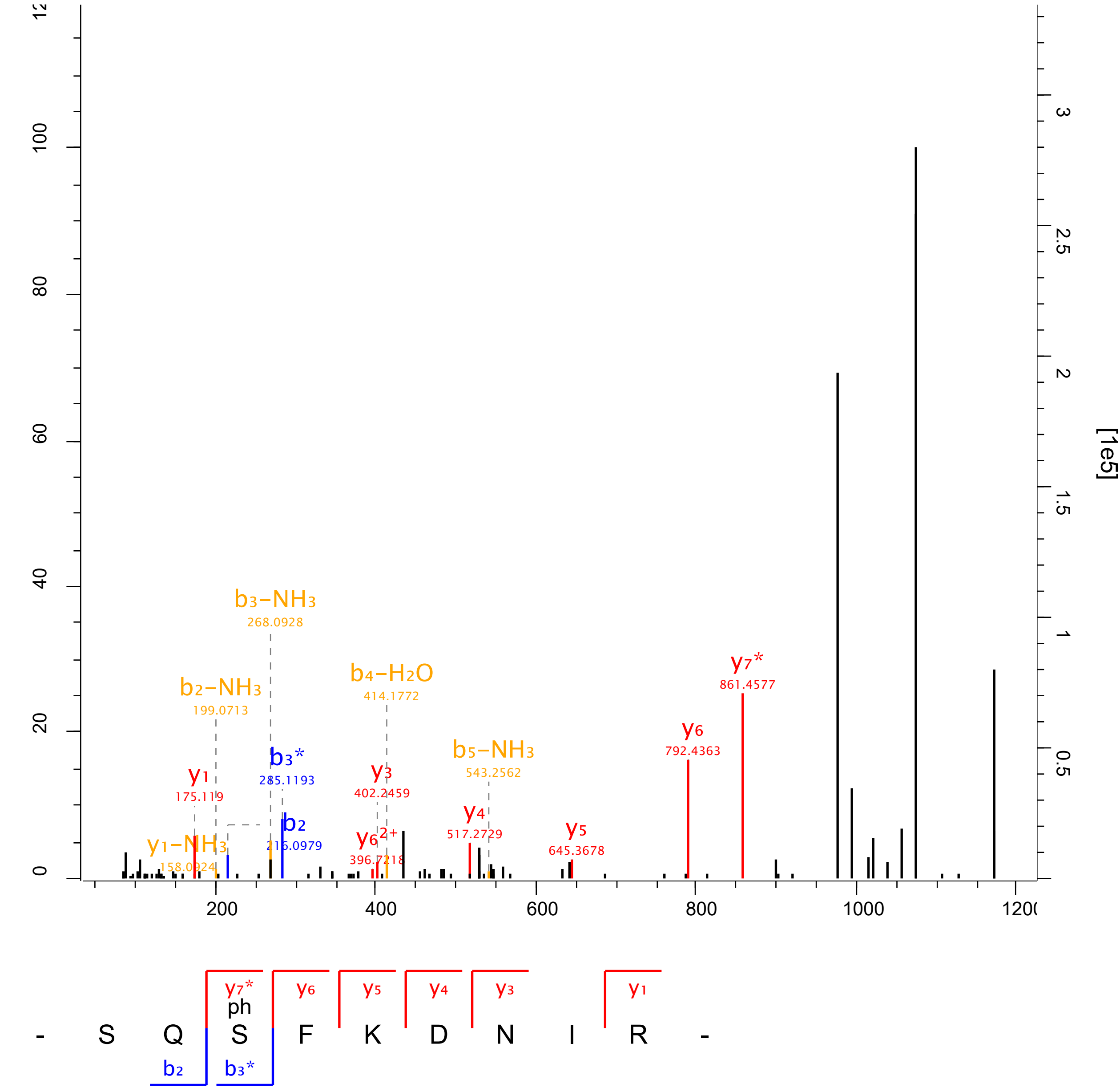

| Raw file | Scan | Method | Score | m/z |
|---|---|---|---|---|
| sys_02_2 | 5469 | FTMS; HCD | 211.9 | 713.33 |

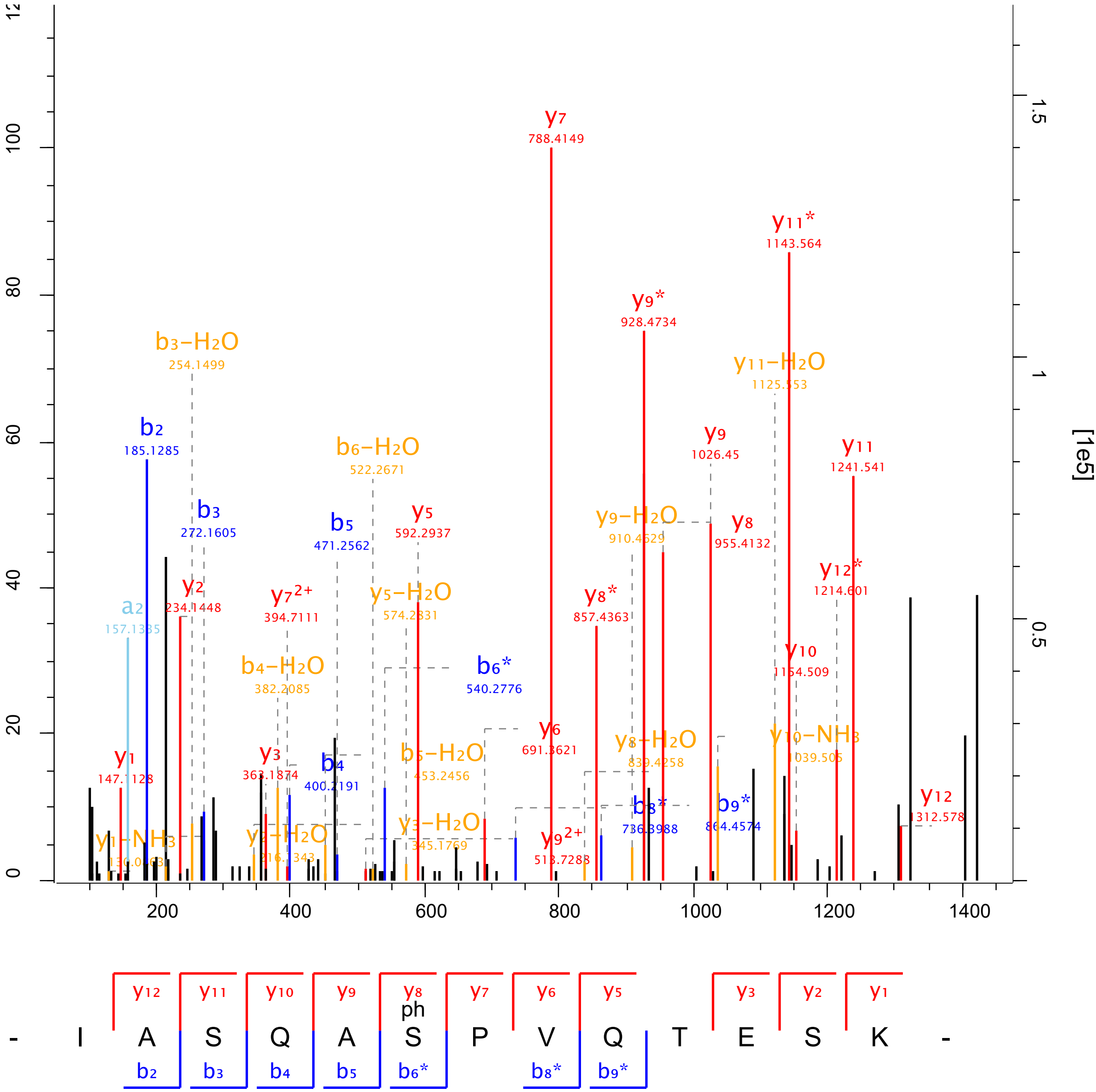

| Raw file | Scan | Method | Score | m/z |
| --- | --- | --- | --- | --- |
| sys_02_2 | 5475 | FTMS; HCD | 58.32 | 400.86 |

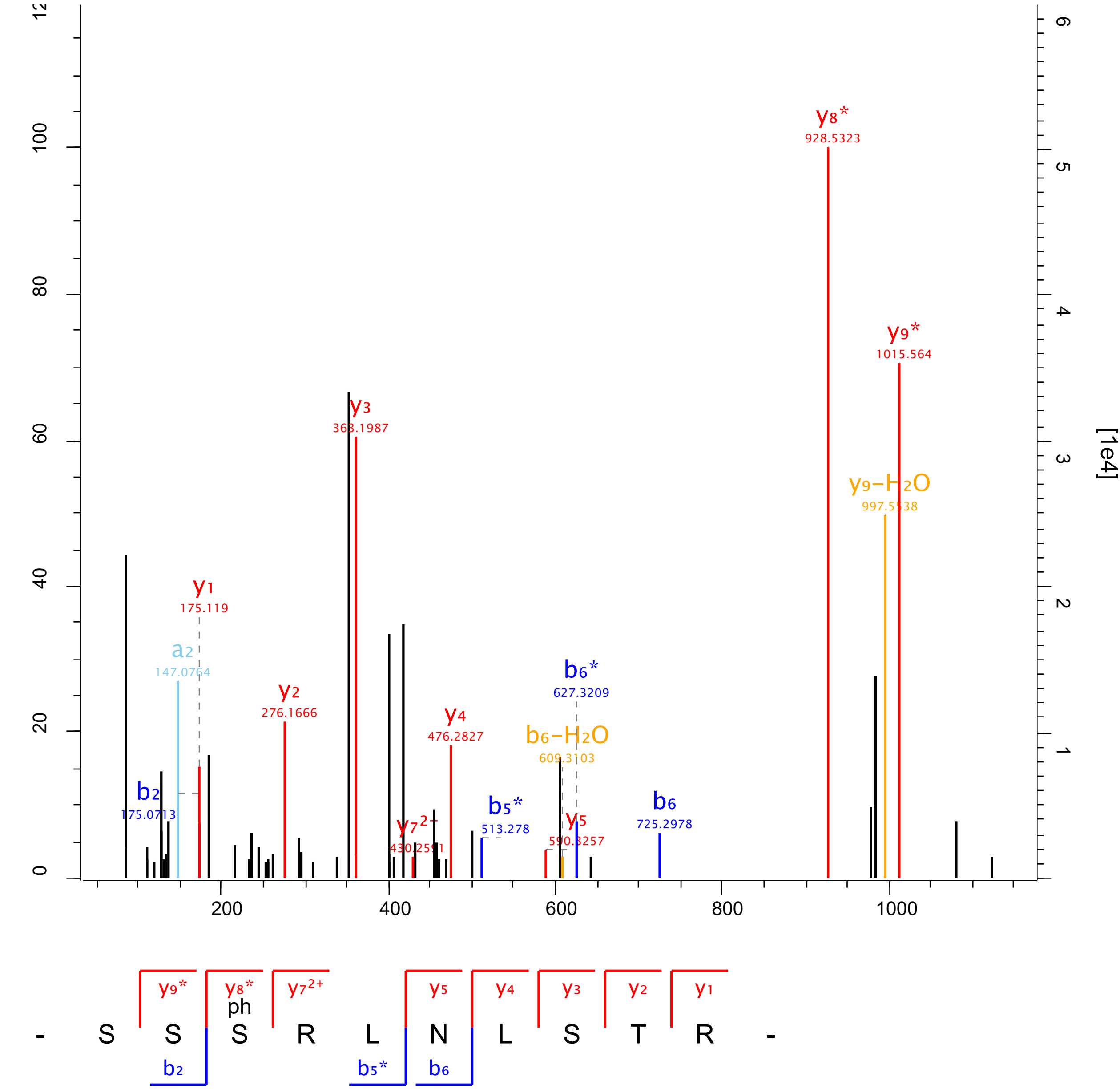

| Raw file | Scan | Method | Score | m/z |
|---|---|---|---|---|
| sys_02_2 | 5551 | FTMS; HCD | 206.96 | 695.65 |

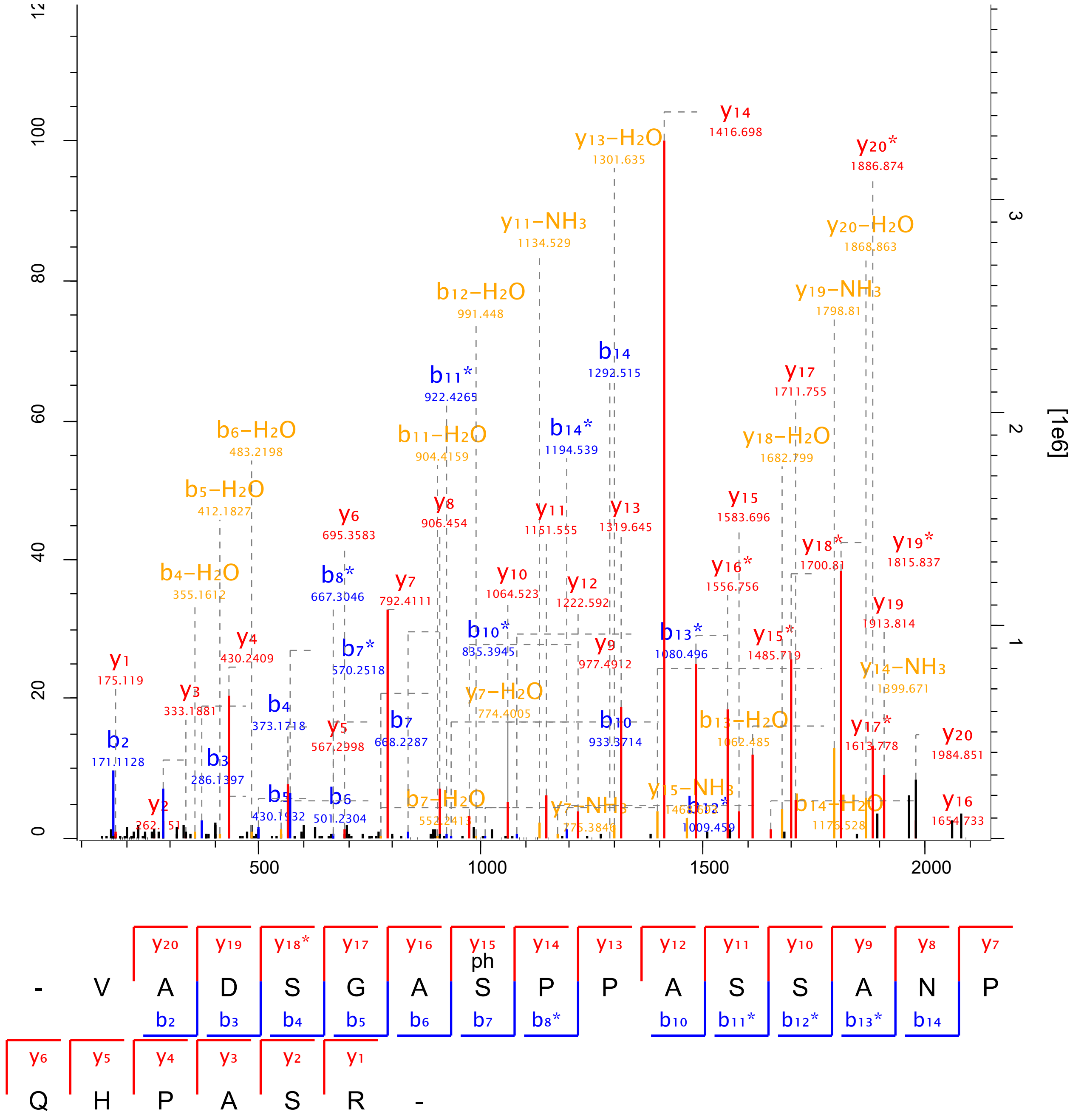

| Raw file | Scan | Method | Score | m/z |
| --- | --- | --- | --- | --- |
| sys_02_2 | 5622 | FTMS; HCD | 102.65 | 672.78 |

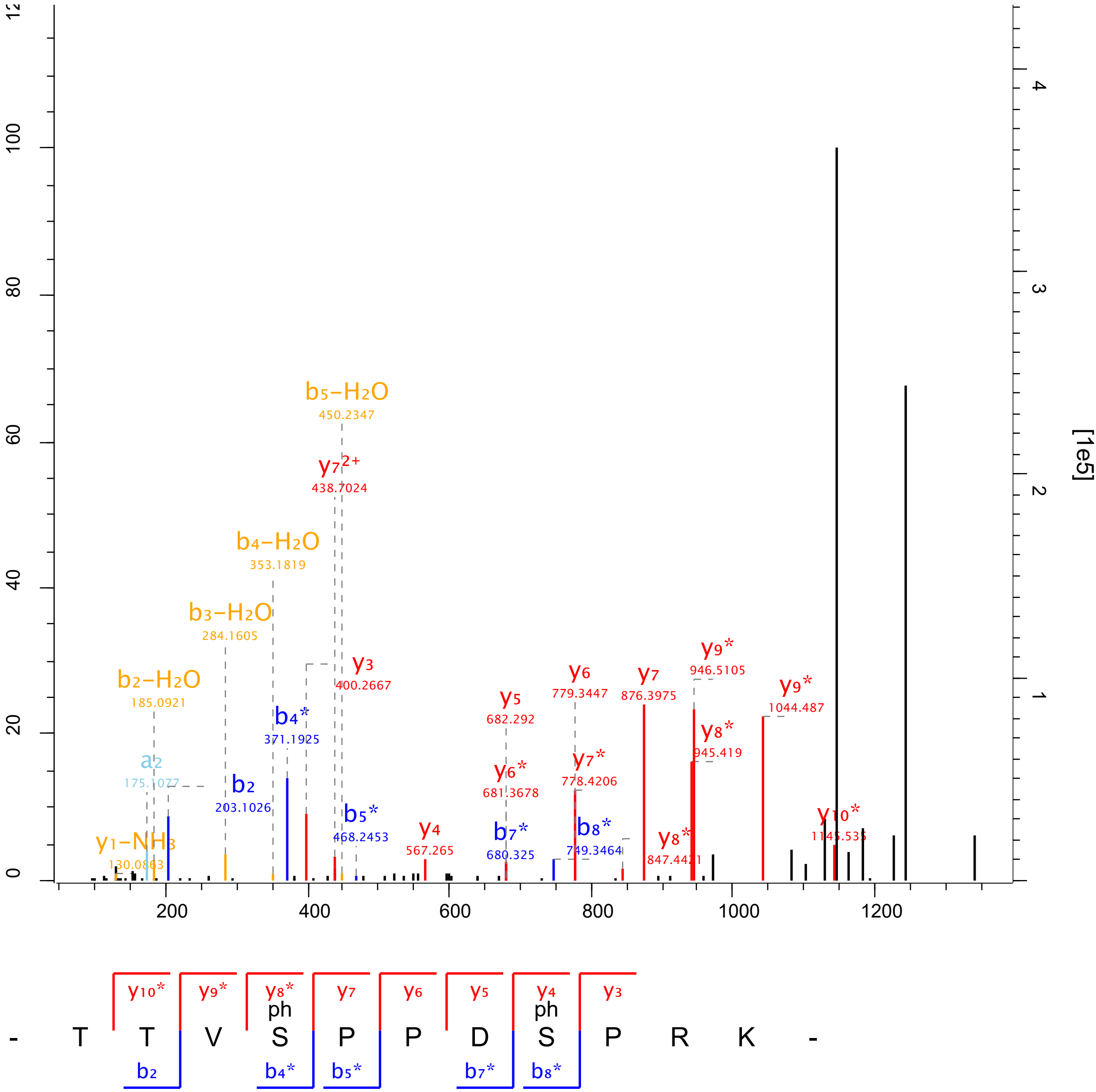

| Raw file | Scan | Method | Score | m/z |
| --- | --- | --- | --- | --- |
| sys_02_2 | 5766 | FTMS; HCD | 149.86 | 523.89 |

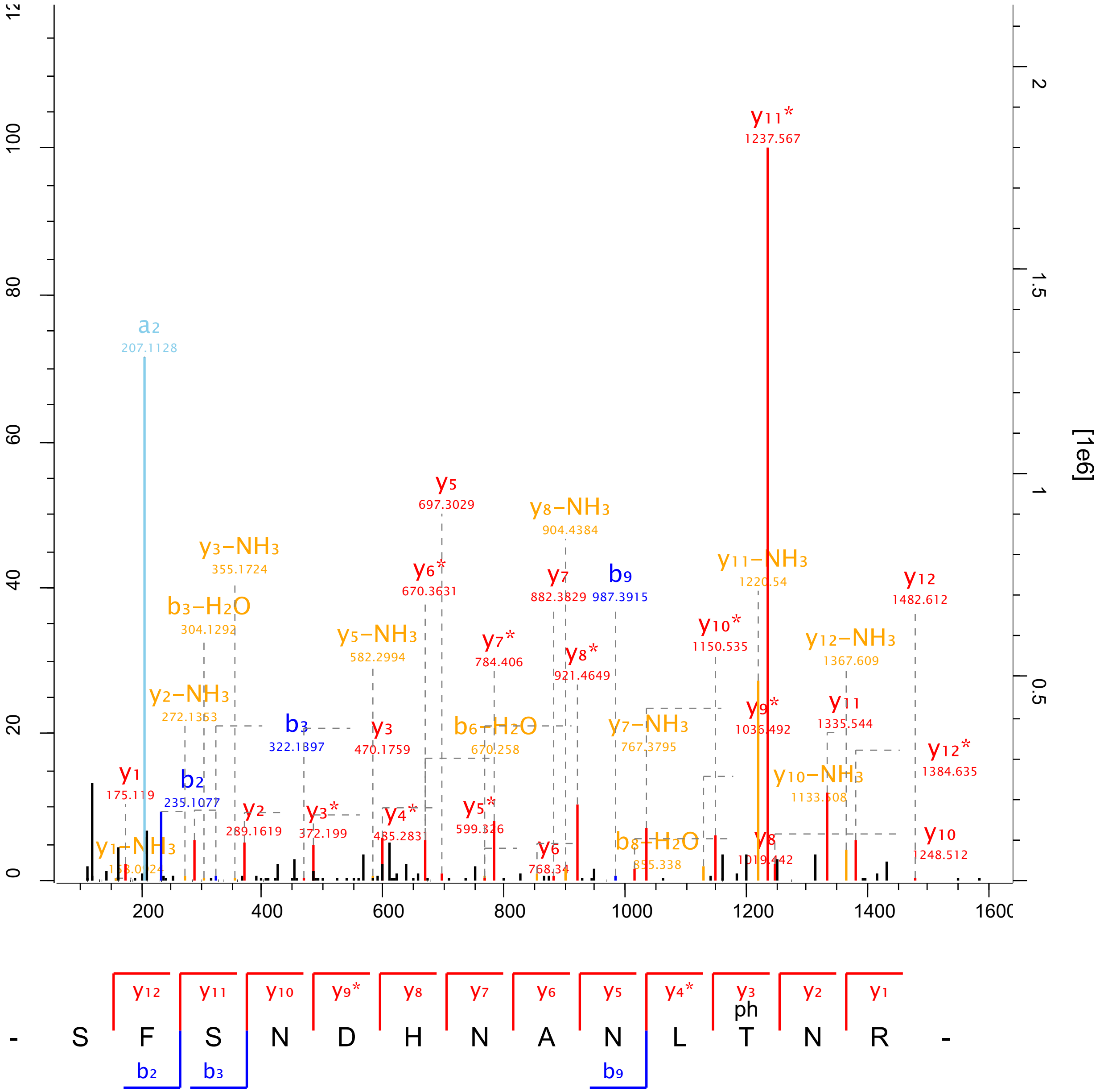

| Raw file | Scan | Method | Score | m/z |
|---|---|---|---|---|
| sys_02_2 | 5815 | FTMS; HCD | 74.96 | 457.22 |

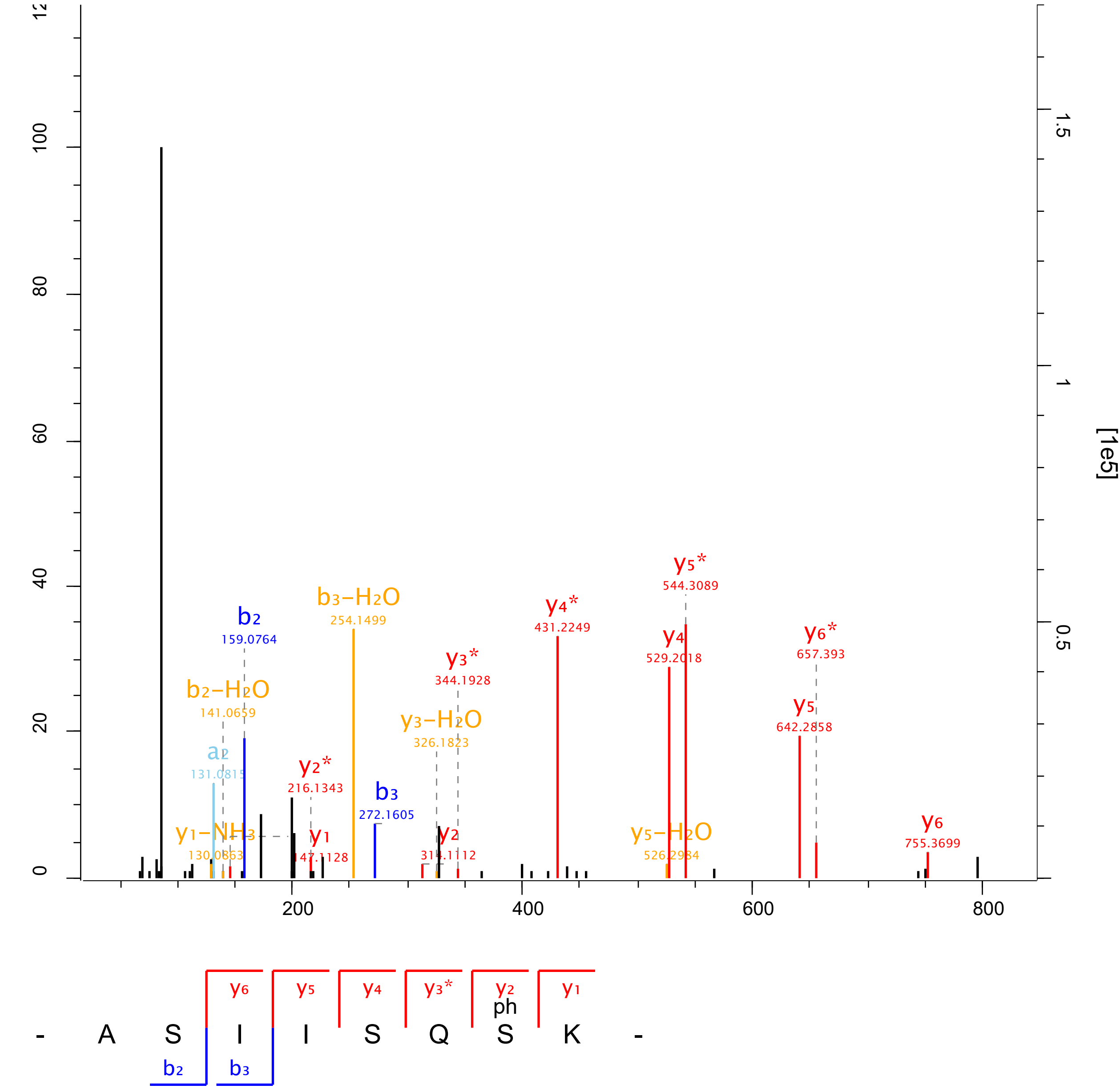

| Raw file | Scan | Method | Score | m/z |
|---|---|---|---|---|
| sys_02_2 | 5846 | FTMS; HCD | 66 | 506.74 |

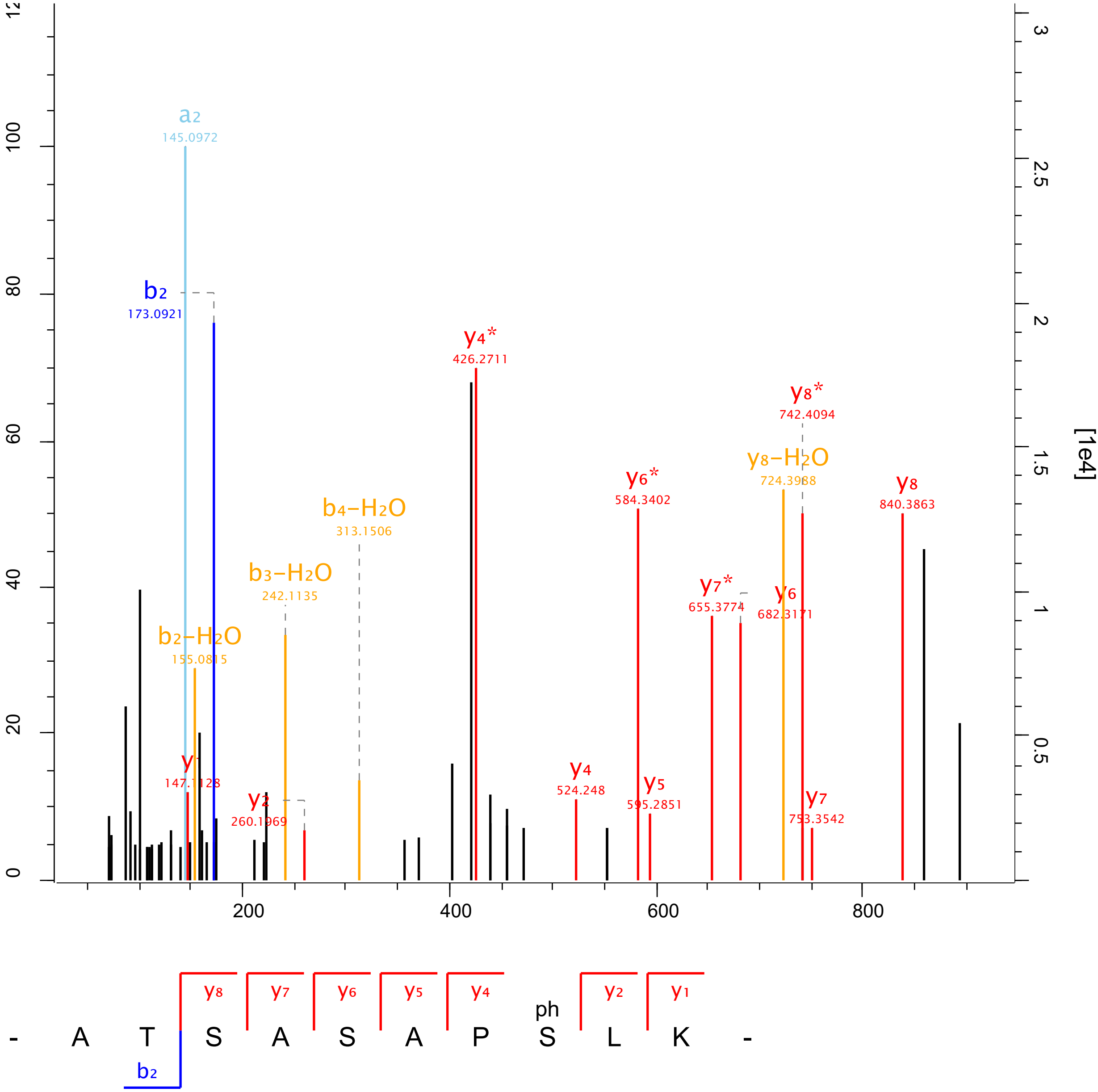

| Raw file | Scan | Method | Score | m/z |
| --- | --- | --- | --- | --- |
| sys_02_2 | 5899 | FTMS; HCD | 54.39 | 468.55 |

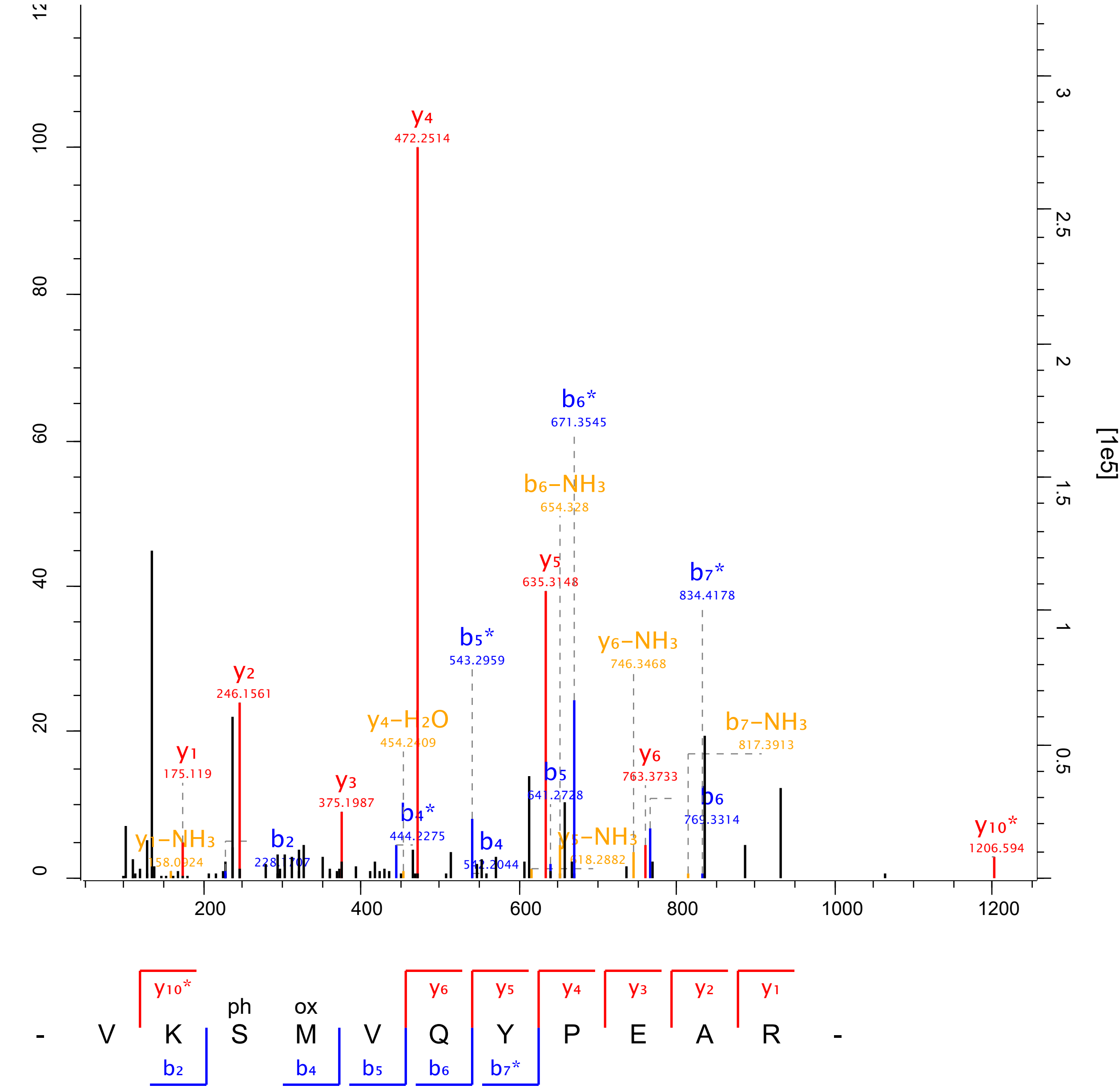

| Raw file | Scan | Method | Score | m/z |
|---|---|---|---|---|
| sys_02_2 | 5935 | FTMS; HCD | 68.41 | 626.24 |

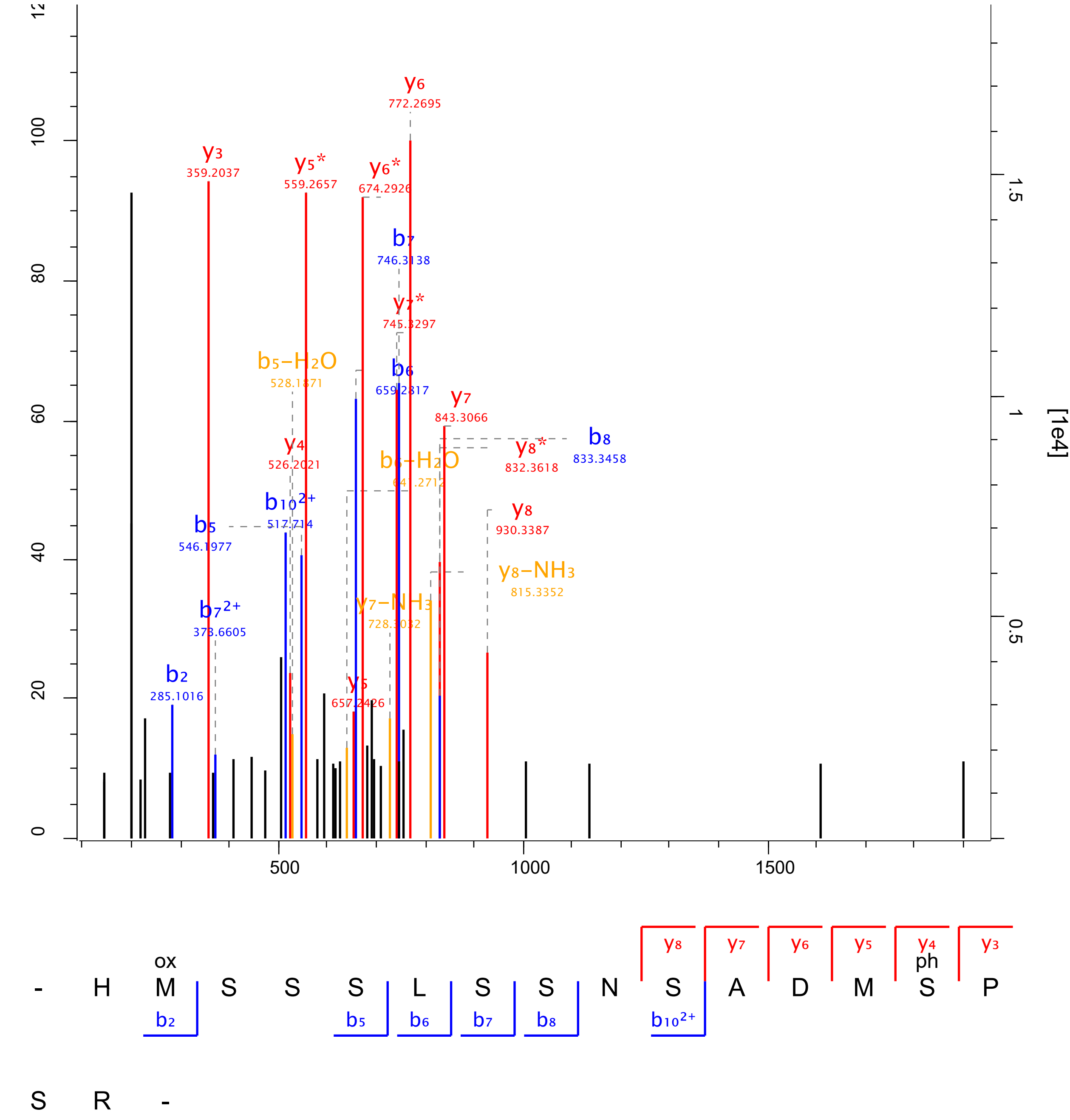

| Raw file | Scan | Method | Score | m/z |
|---|---|---|---|---|
| sys_02_2 | 5954 | FTMS; HCD | 209.73 | 656.26 |

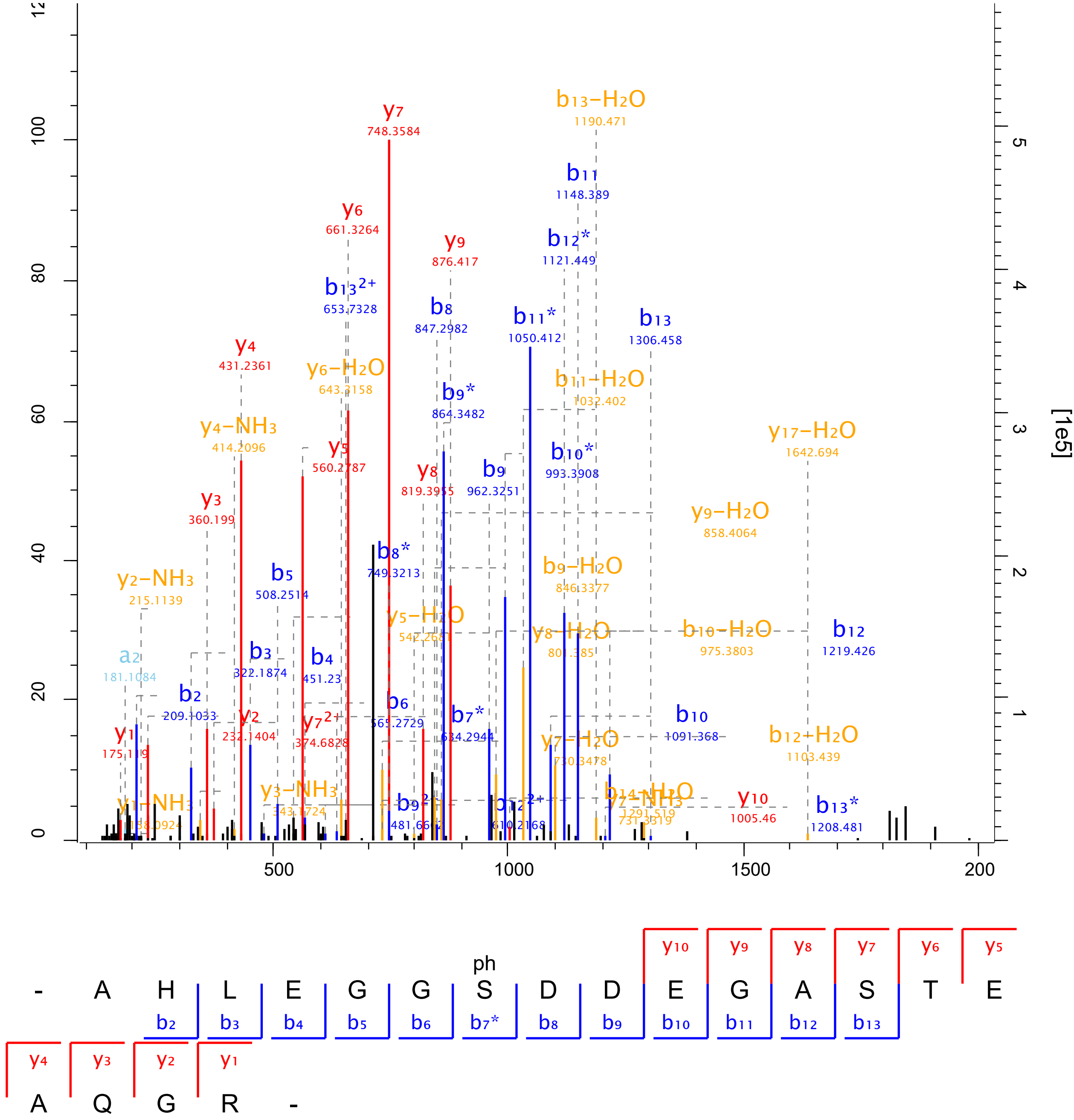

| Raw file | Scan | Method | Score | m/z |
|---|---|---|---|---|
| sys_02_2 | 5982 | FTMS; HCD | 76.07 | 434.87 |

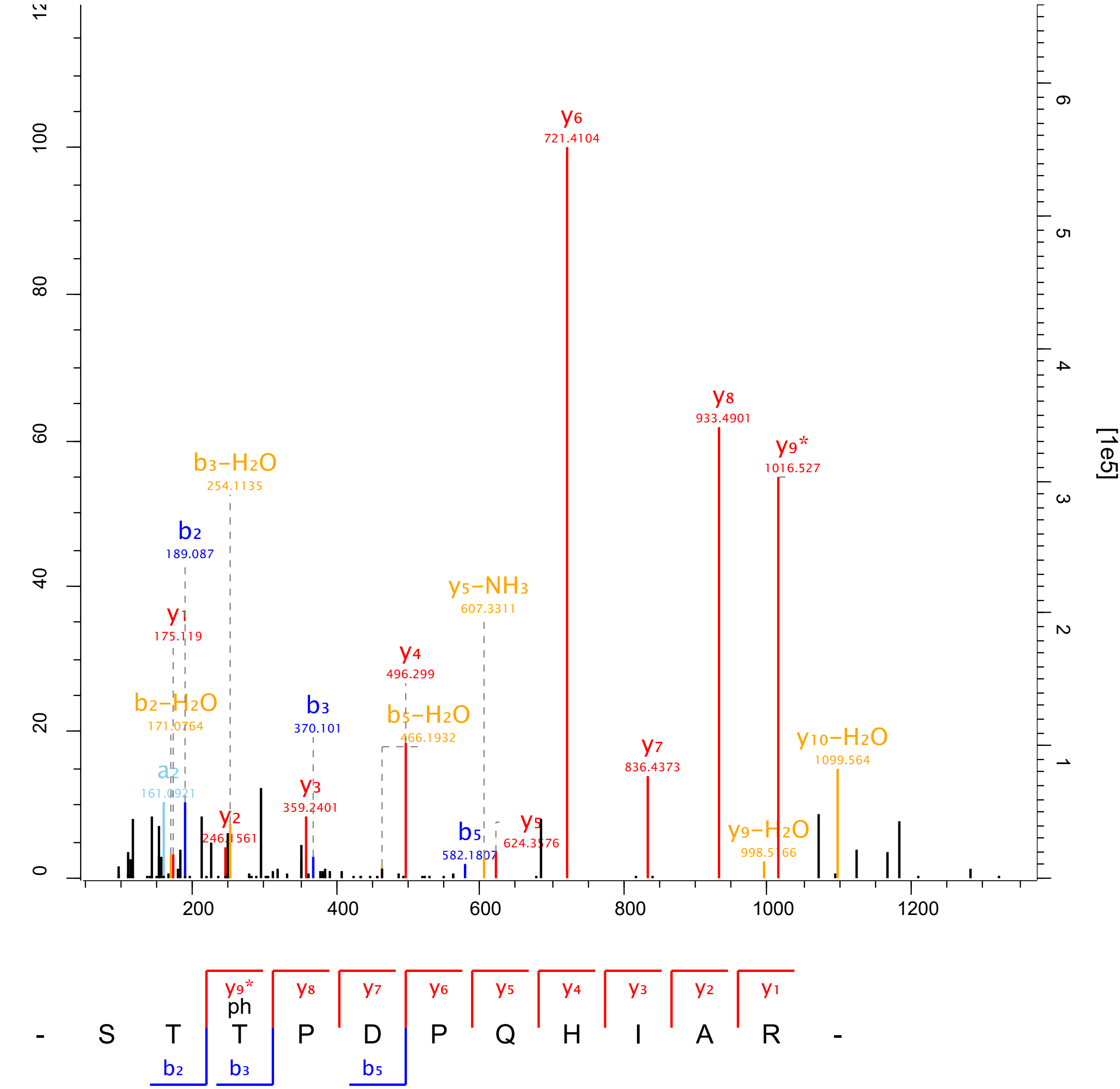

| Raw file | Scan | Method | Score | m/z |
|---|---|---|---|---|
| sys_02_2 | 6039 | FTMS; HCD | 44.83 | 614.23 |

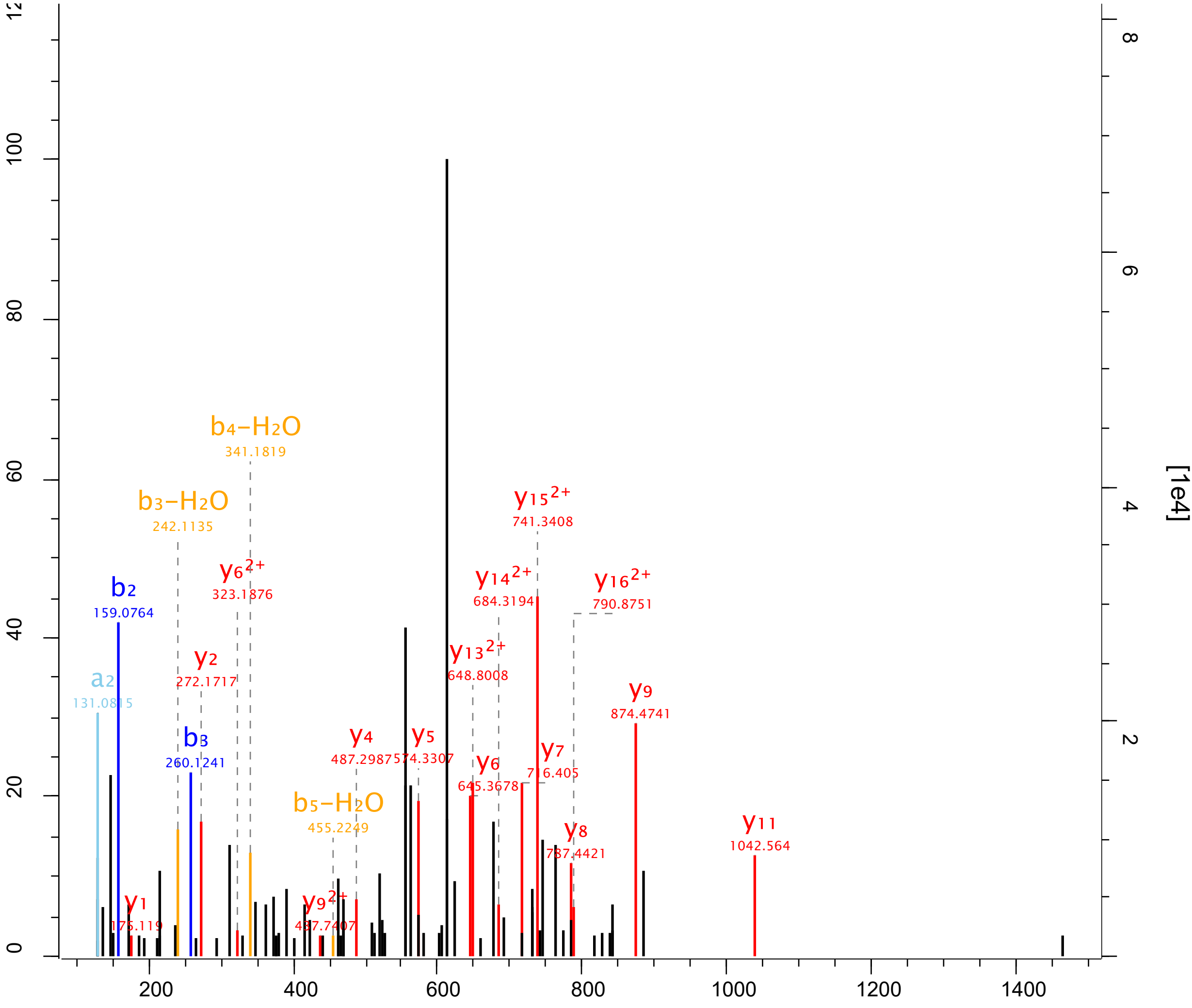


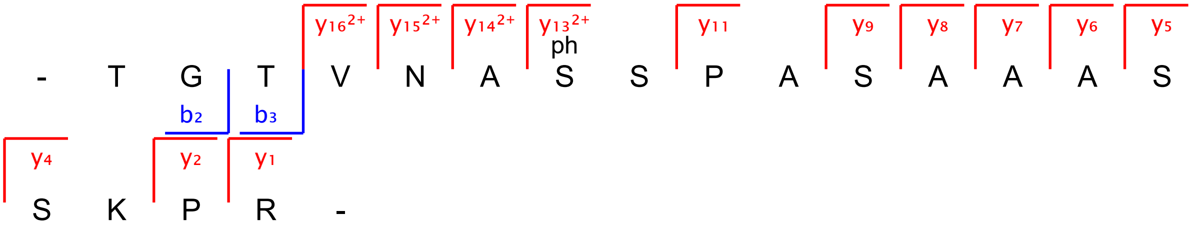

| Raw file | Scan | Method | Score | m/z |
| --- | --- | --- | --- | --- |
| sys_02_2 | 6086 | FTMS; HCD | 211.12 | 551.56 |

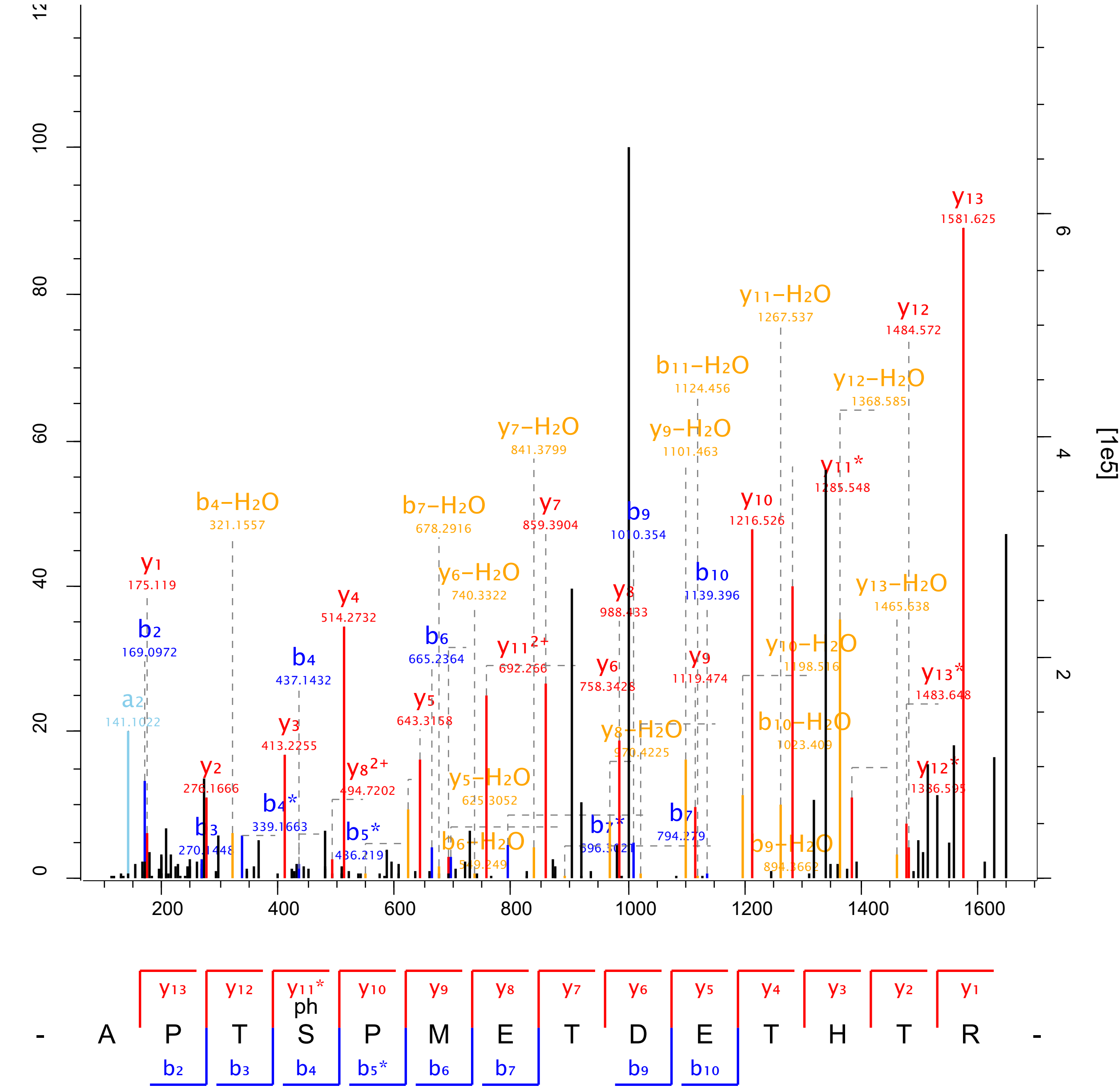

| Raw file | Scan | Method | Score | m/z |
|---|---|---|---|---|
| sys_02_2 | 6103 | FTMS; HCD | 84.51 | 452.2 |

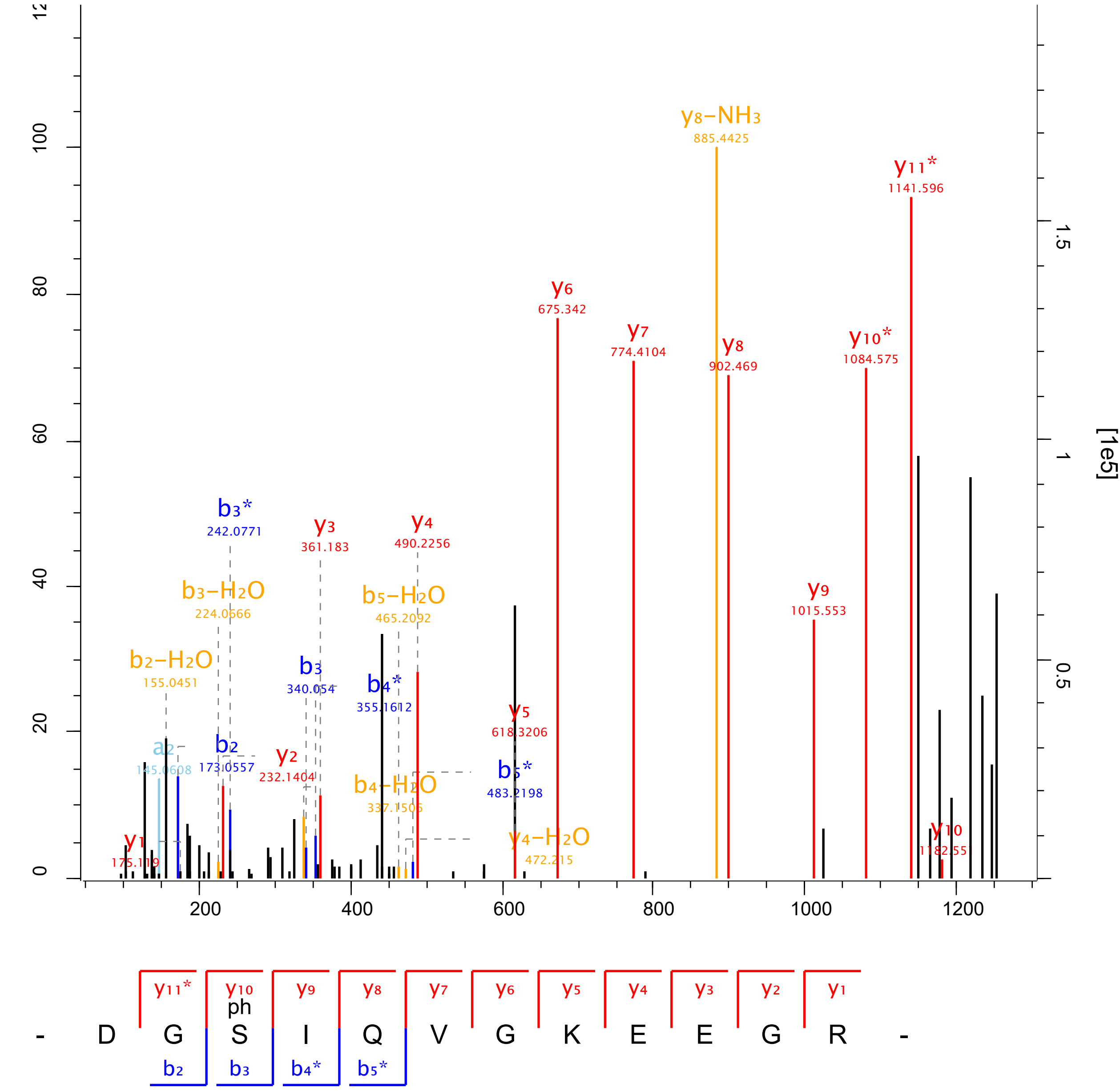

| Raw file | Scan | Method | Score | m/z |
|---|---|---|---|---|
| sys_02_2 | 6193 | FTMS; HCD | 159.75 | 632.63 |

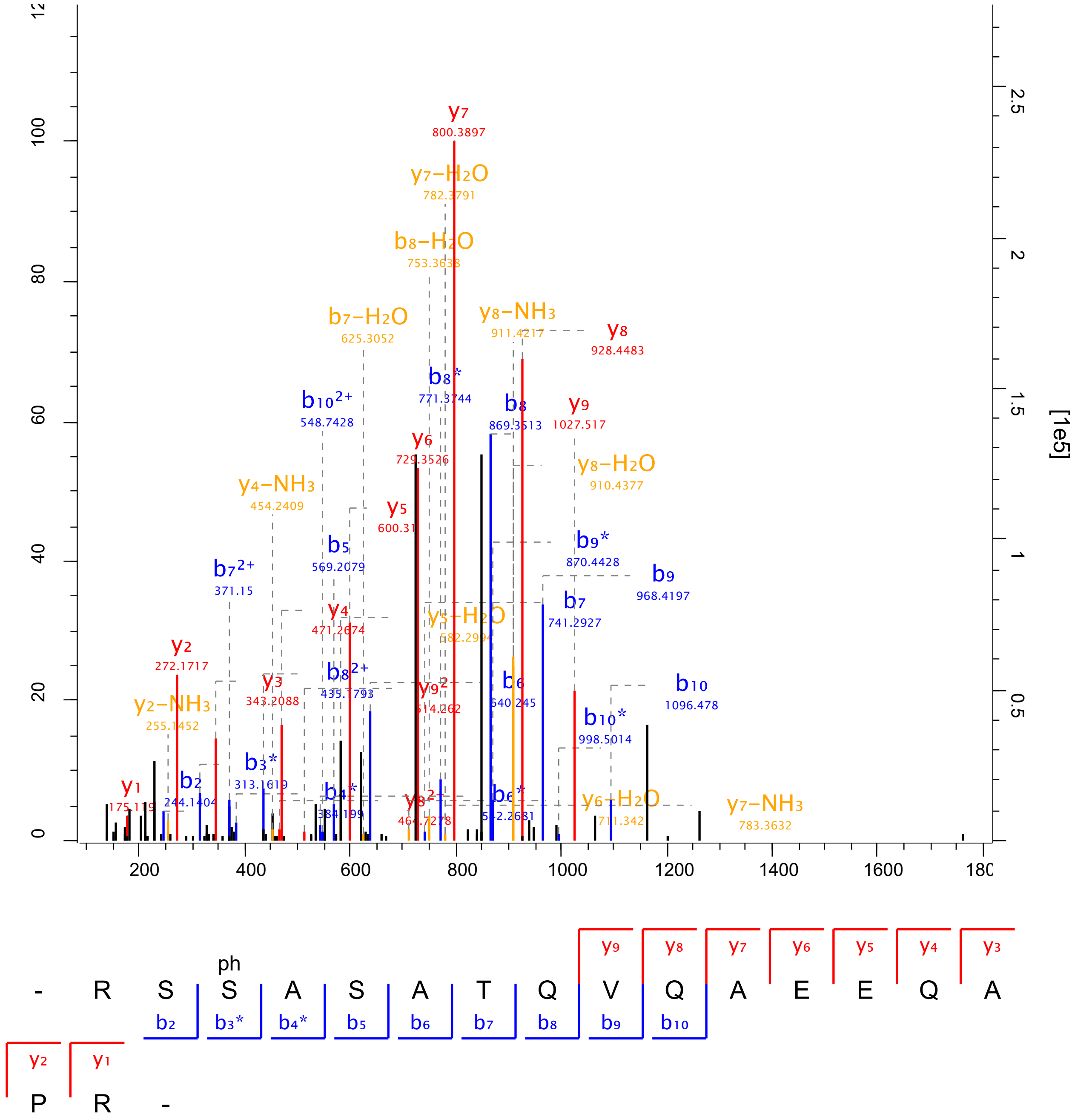

| Raw file | Scan | Method | Score | m/z |
|---|---|---|---|---|
| sys_02_2 | 6200 | FTMS; HCD | 115.29 | 462.73 |

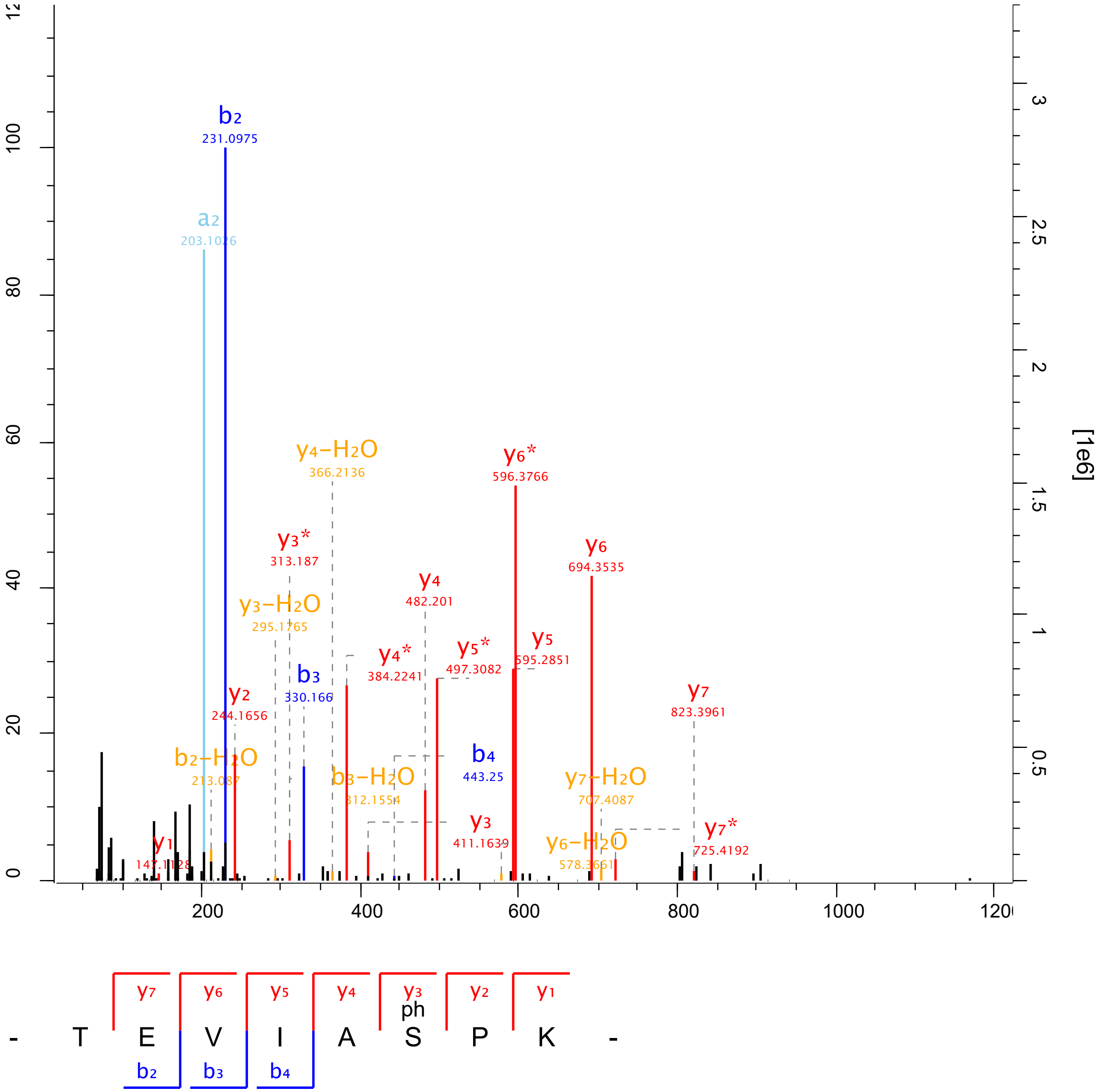

| Raw file | Scan | Method | Score | m/z |
|---|---|---|---|---|
| sys_02_2 | 6222 | FTMS; HCD | 132.79 | 764.82 |

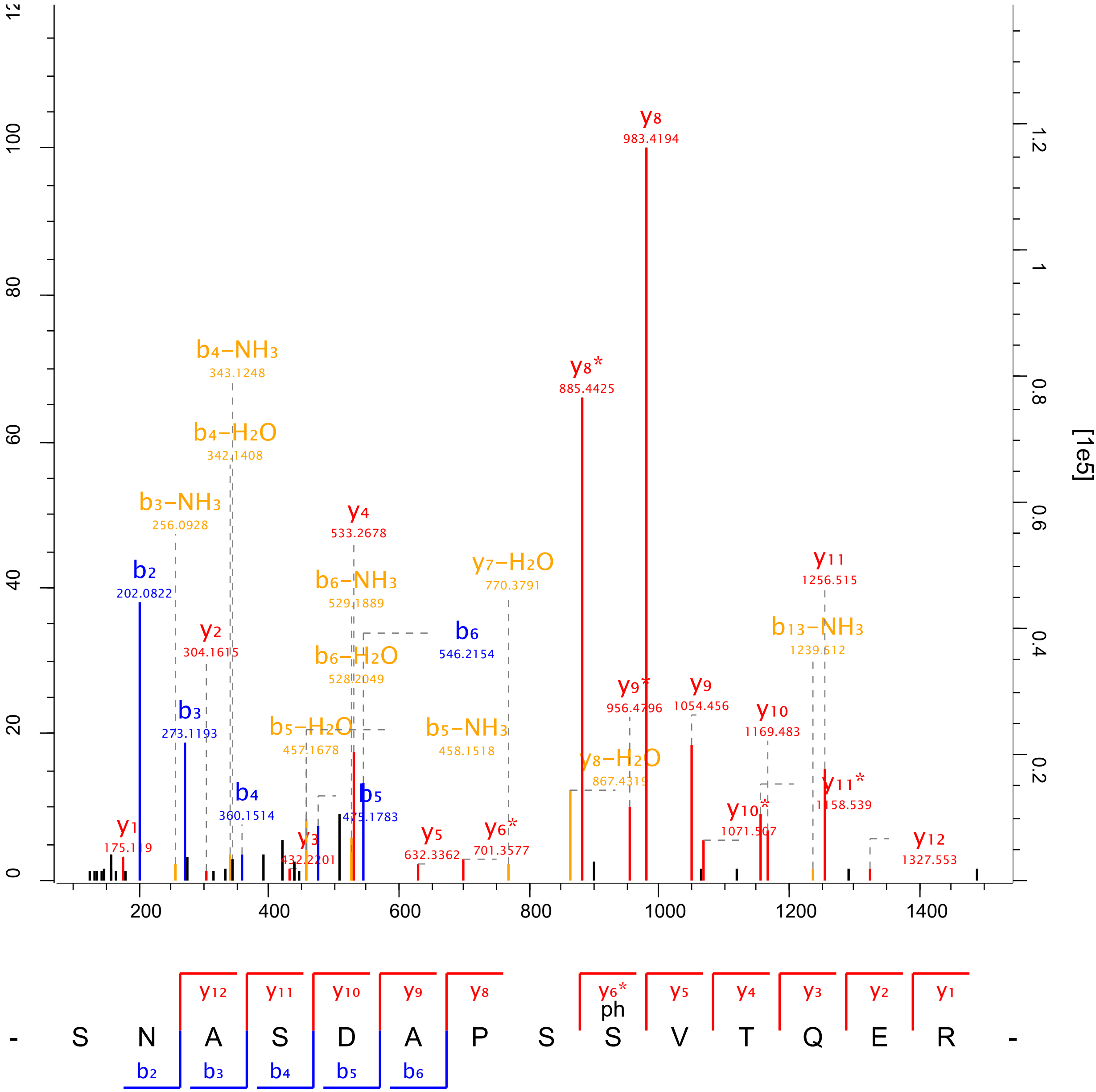

| Raw file | Scan | Method | Score | m/z |
| --- | --- | --- | --- | --- |
| sys_02_2 | 6226 | FTMS; HCD | 107.78 | 651.95 |

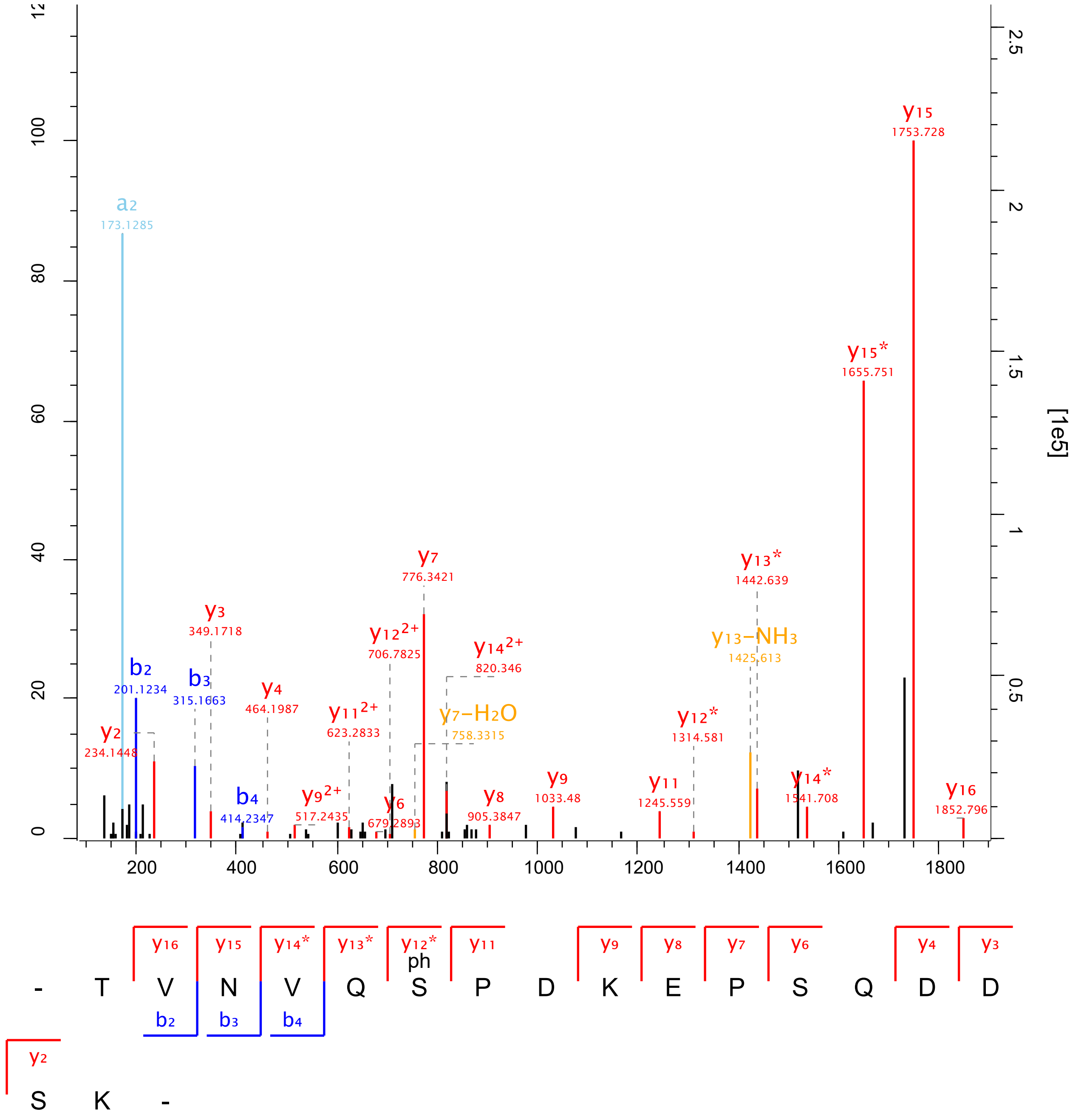

| Raw file | Scan | Method | Score | m/z |
|---|---|---|---|---|
| sys_02_2 | 6237 | FTMS; HCD | 90.7 | 602.77 |

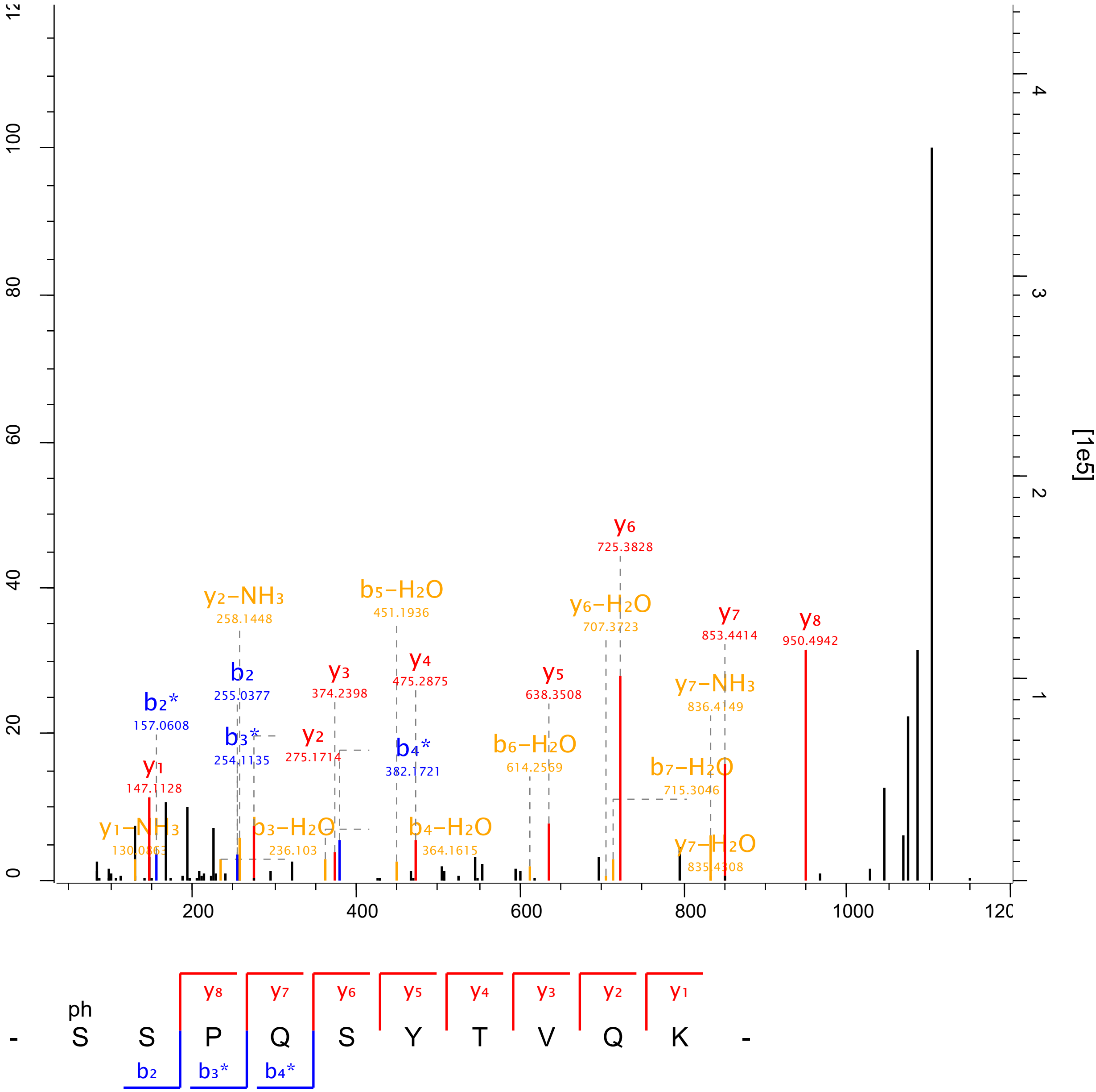

| Raw file | Scan | Method | Score | m/z |
|---|---|---|---|---|
| sys_02_2 | 6418 | FTMS; HCD | 42.03 | 535.23 |

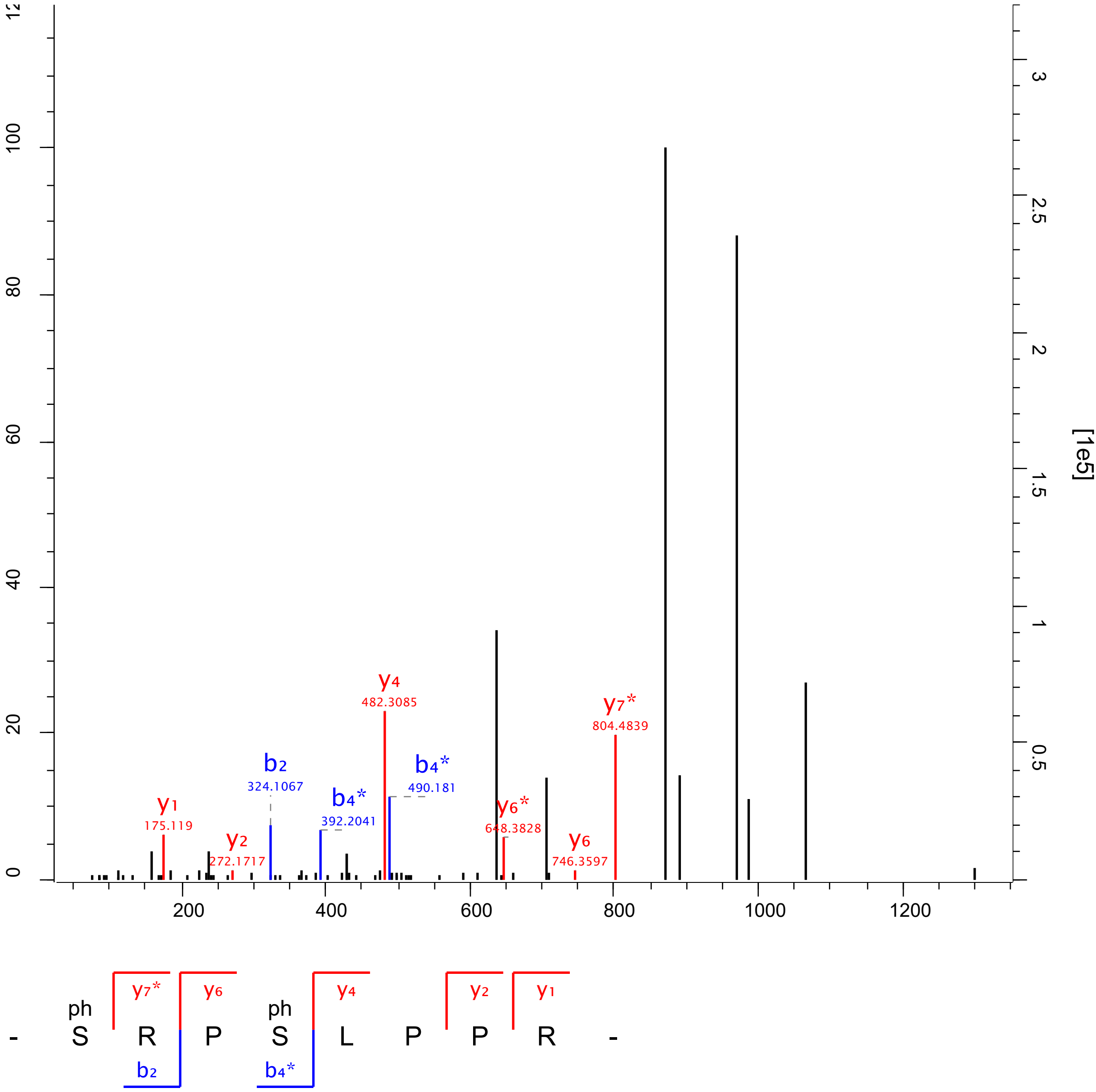

| Raw file | Scan | Method | Score | m/z |
| --- | --- | --- | --- | --- |
| sys_02_2 | 6447 | FTMS; HCD | 78.19 | 659.77 |

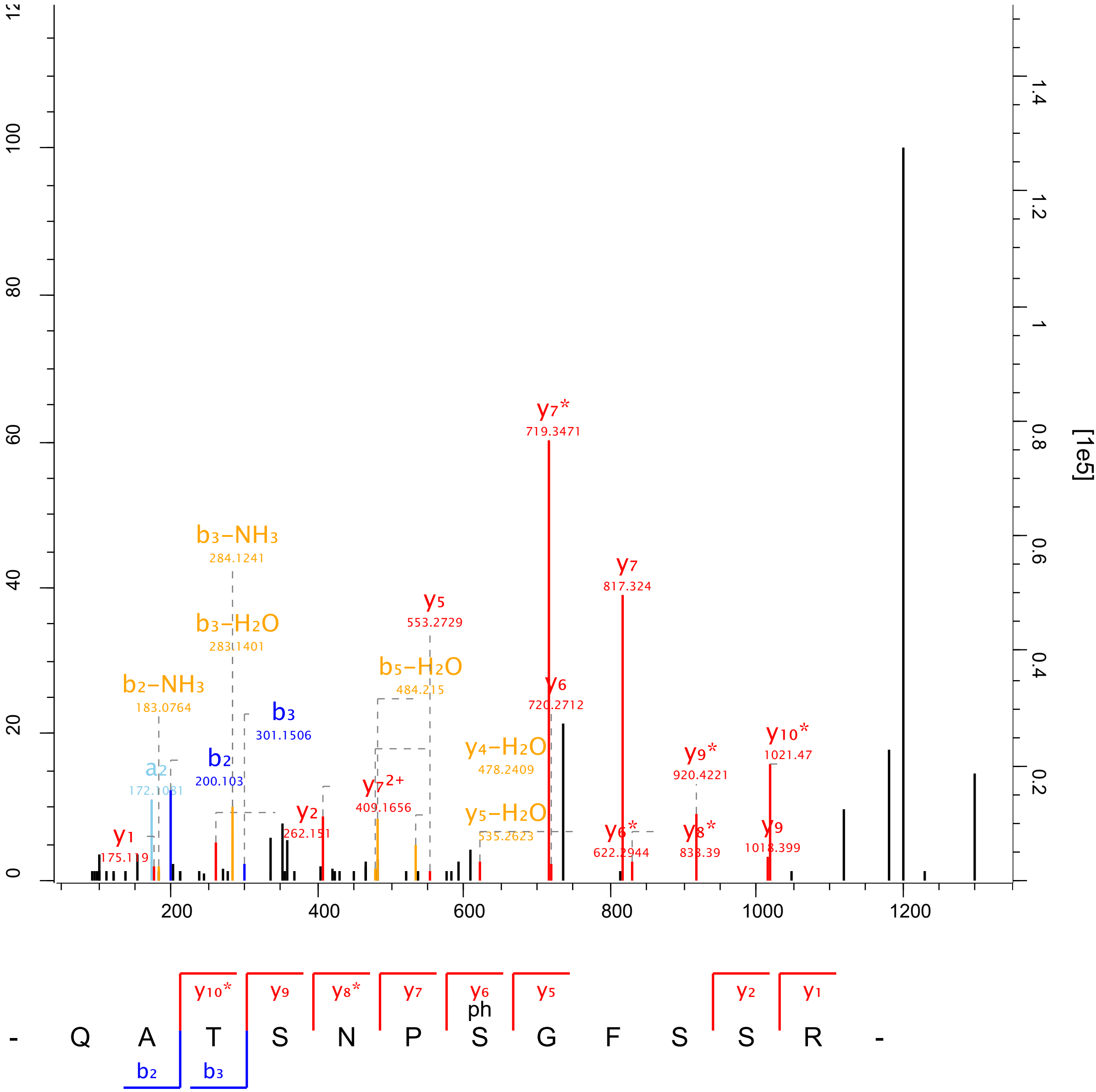

| Raw file | Scan | Method | Score | m/z |
|---|---|---|---|---|
| sys_02_2 | 6495 | FTMS; HCD | 98.42 | 588.26 |

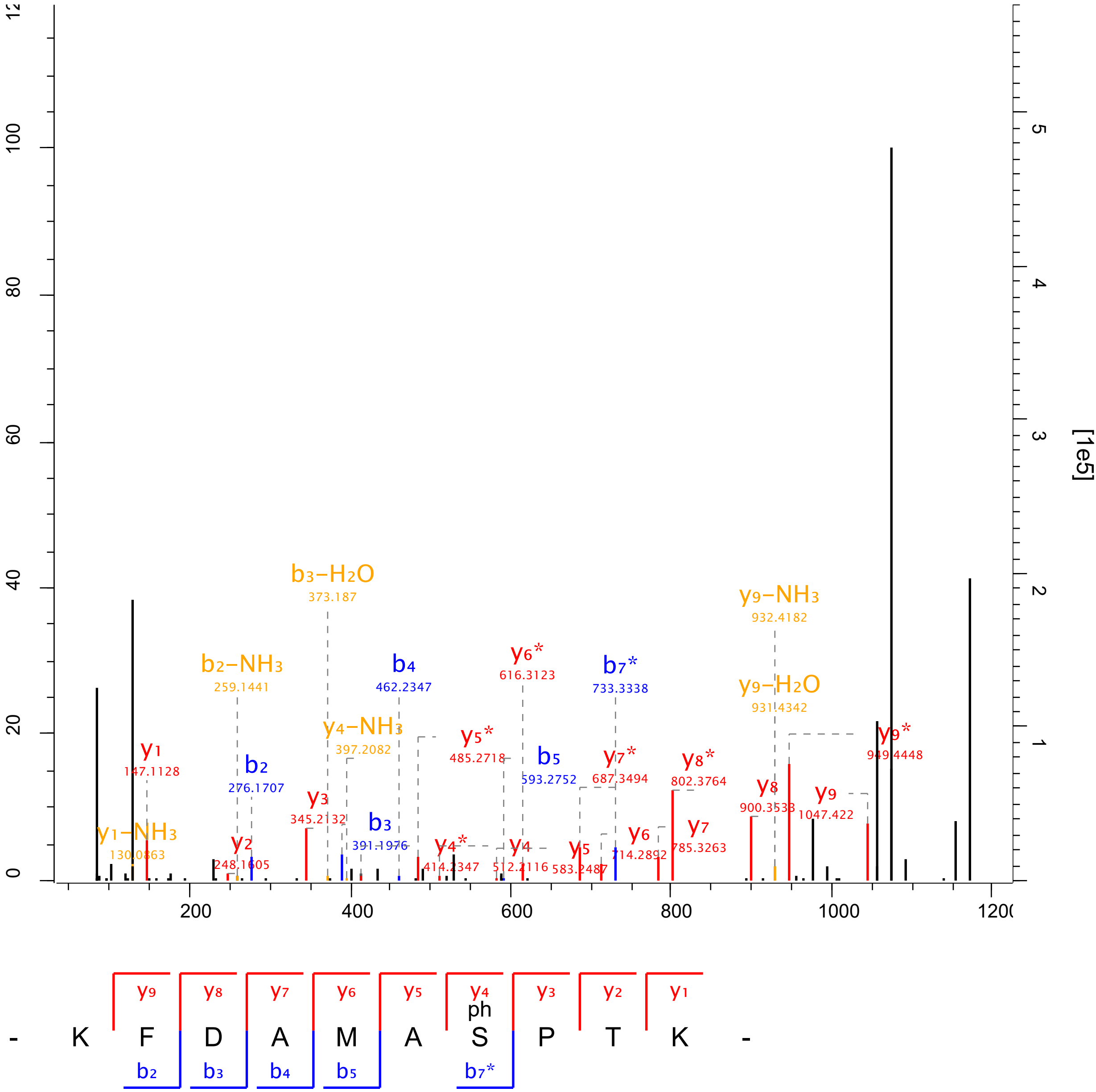

| Raw file | Scan | Method | Score | m/z |
|---|---|---|---|---|
| sys_02_2 | 6522 | FTMS; HCD | 62.59 | 628.29 |

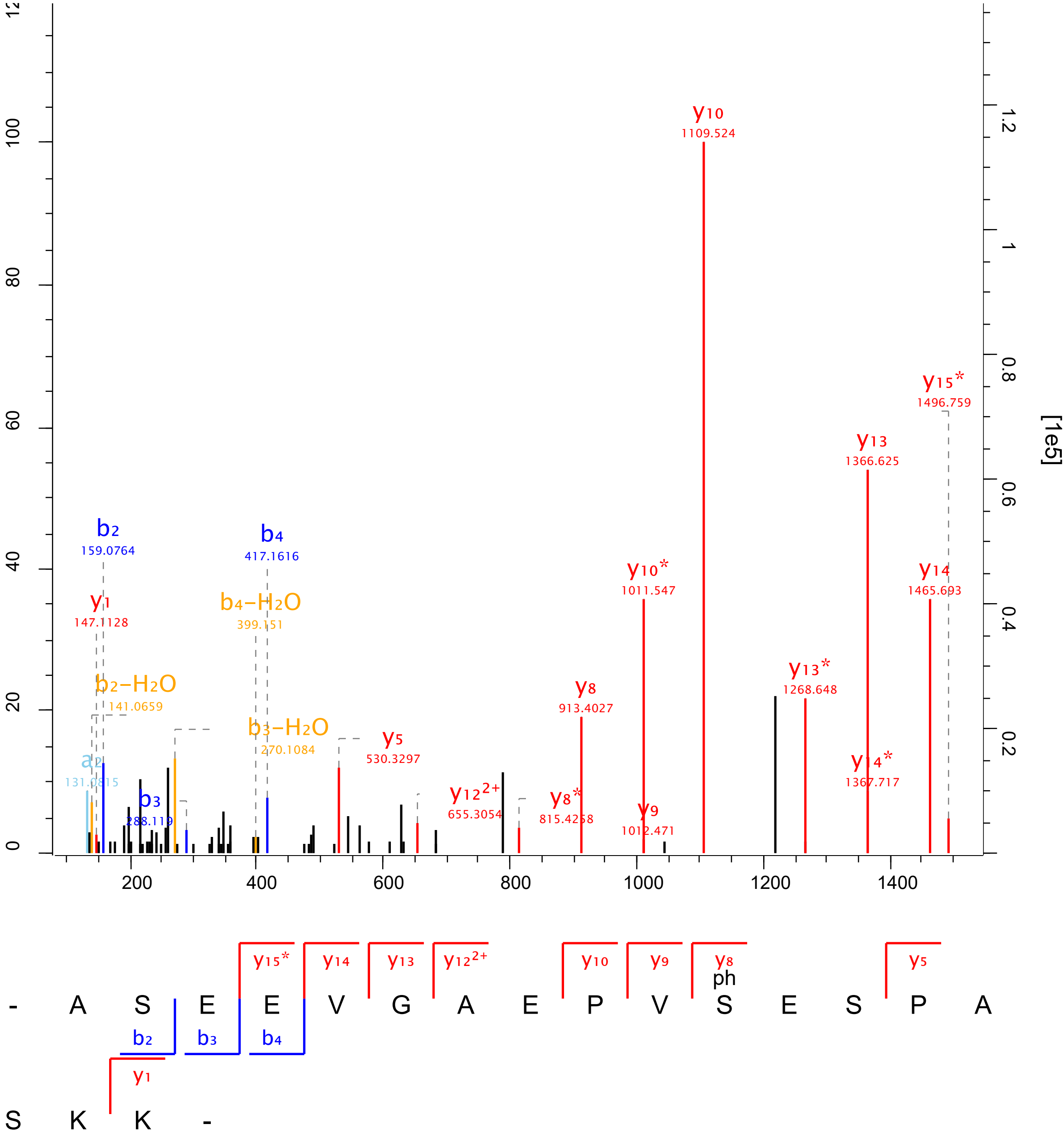

| Raw file | Scan | Method | Score | m/z |
| --- | --- | --- | --- | --- |
| sys_02_2 | 6699 | FTMS; HCD | 102.45 | 653.95 |

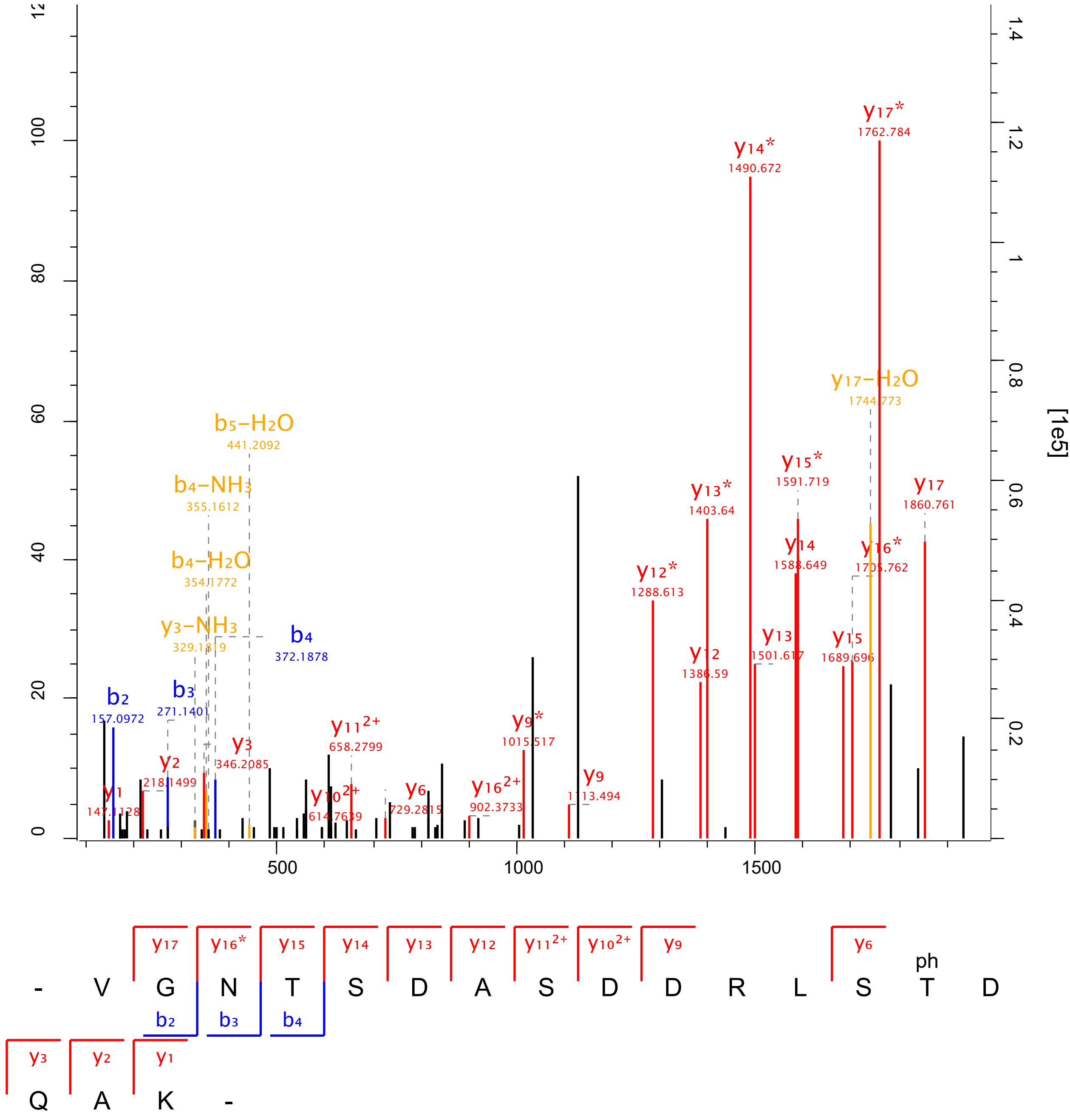

| Raw file | Scan | Method | Score | m/z |
|---|---|---|---|---|
| sys_02_2 | 6730 | FTMS; HCD | 47.71 | 609.78 |

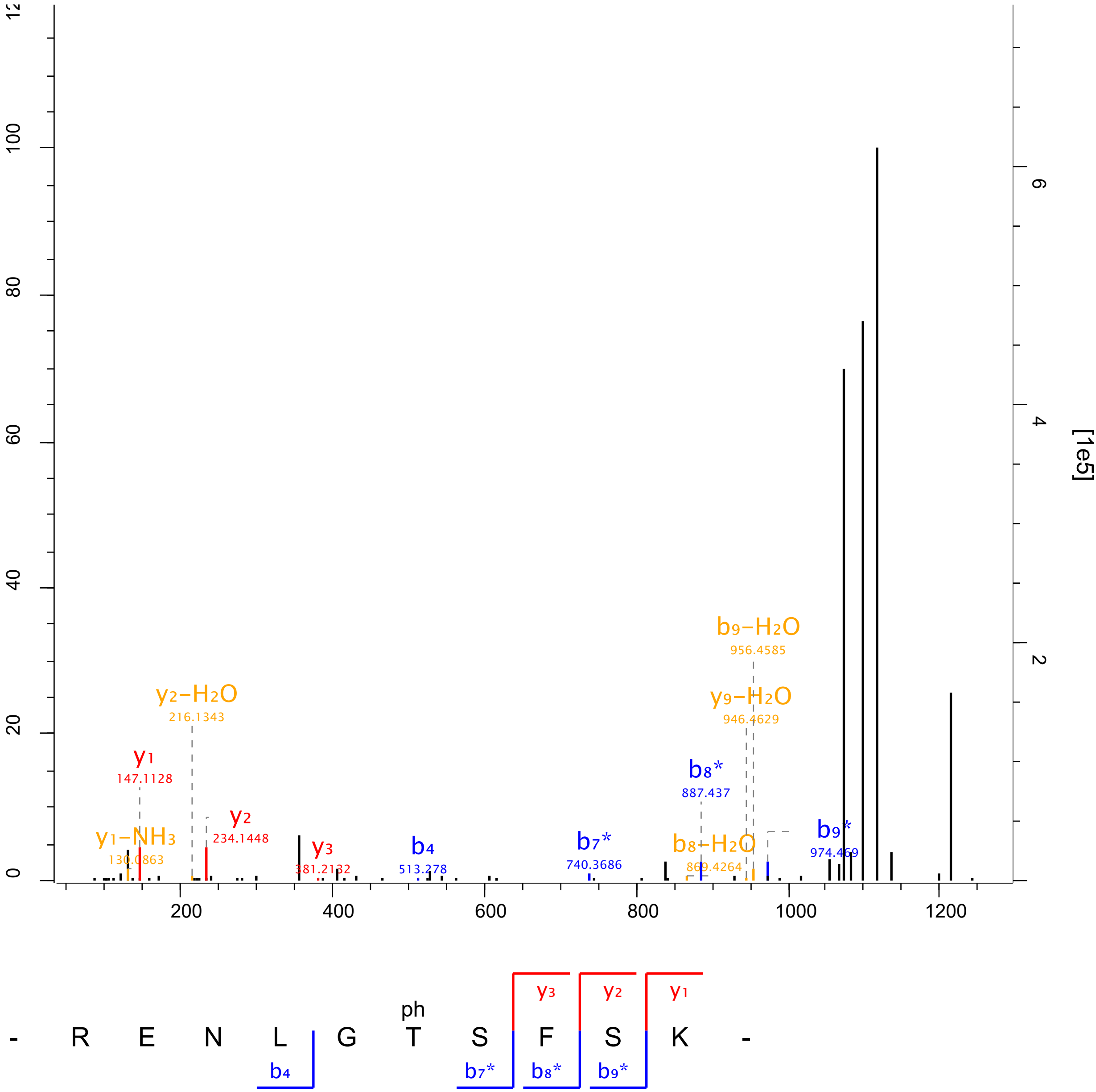

| Raw file | Scan | Method | Score | m/z |
|---|---|---|---|---|
| sys_02_2 | 6732 | FTMS; HCD | 130.01 | 480.71 |

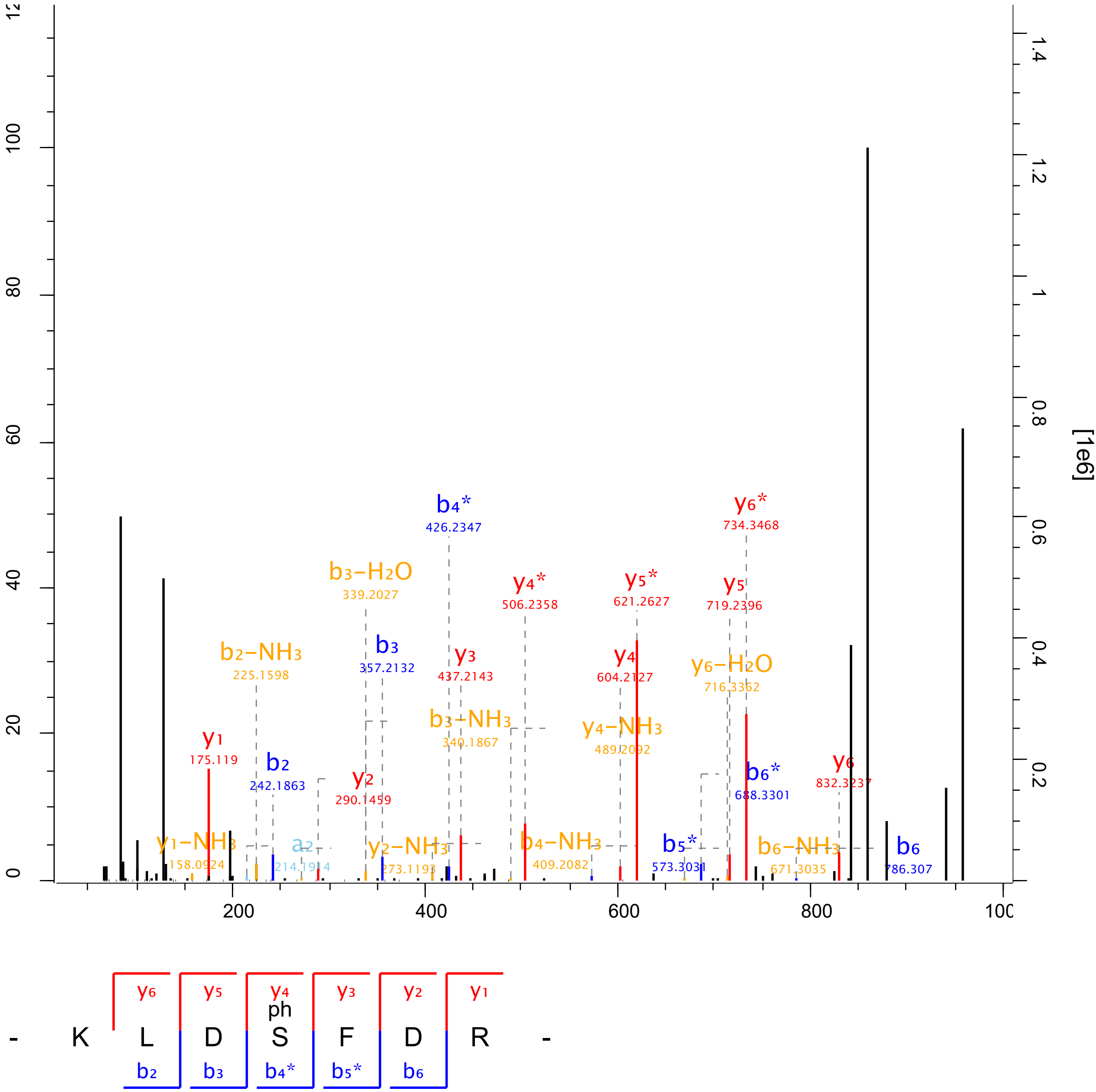

| Raw file | Scan | Method | Score | m/z |
|---|---|---|---|---|
| sys_02_2 | 6867 | FTMS; HCD | 107.06 | 559.74 |

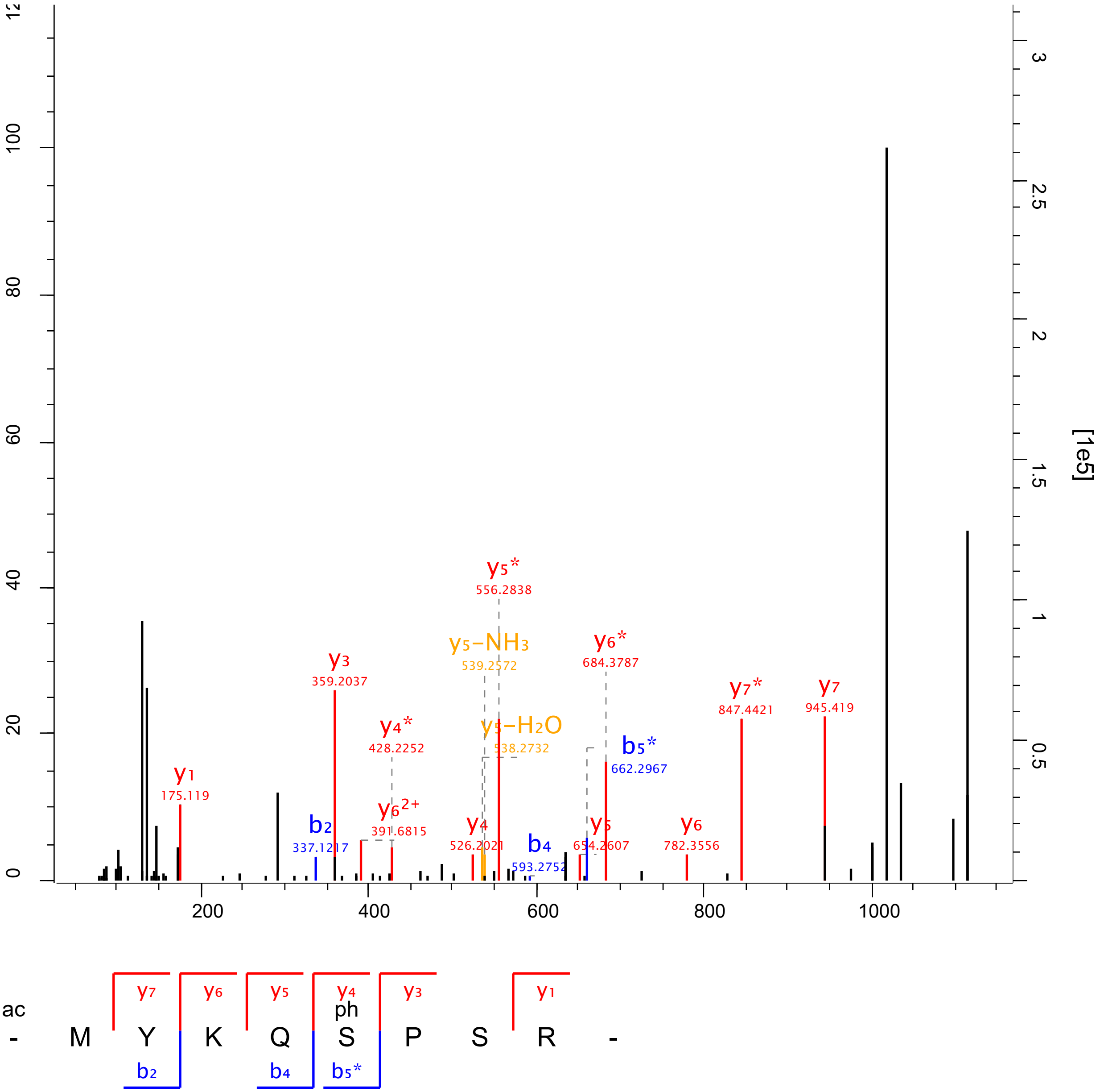

| Raw file | Scan | Method | Score | m/z |
|---|---|---|---|---|
| sys_02_2 | 7057 | FTMS; HCD | 45 | 529.22 |

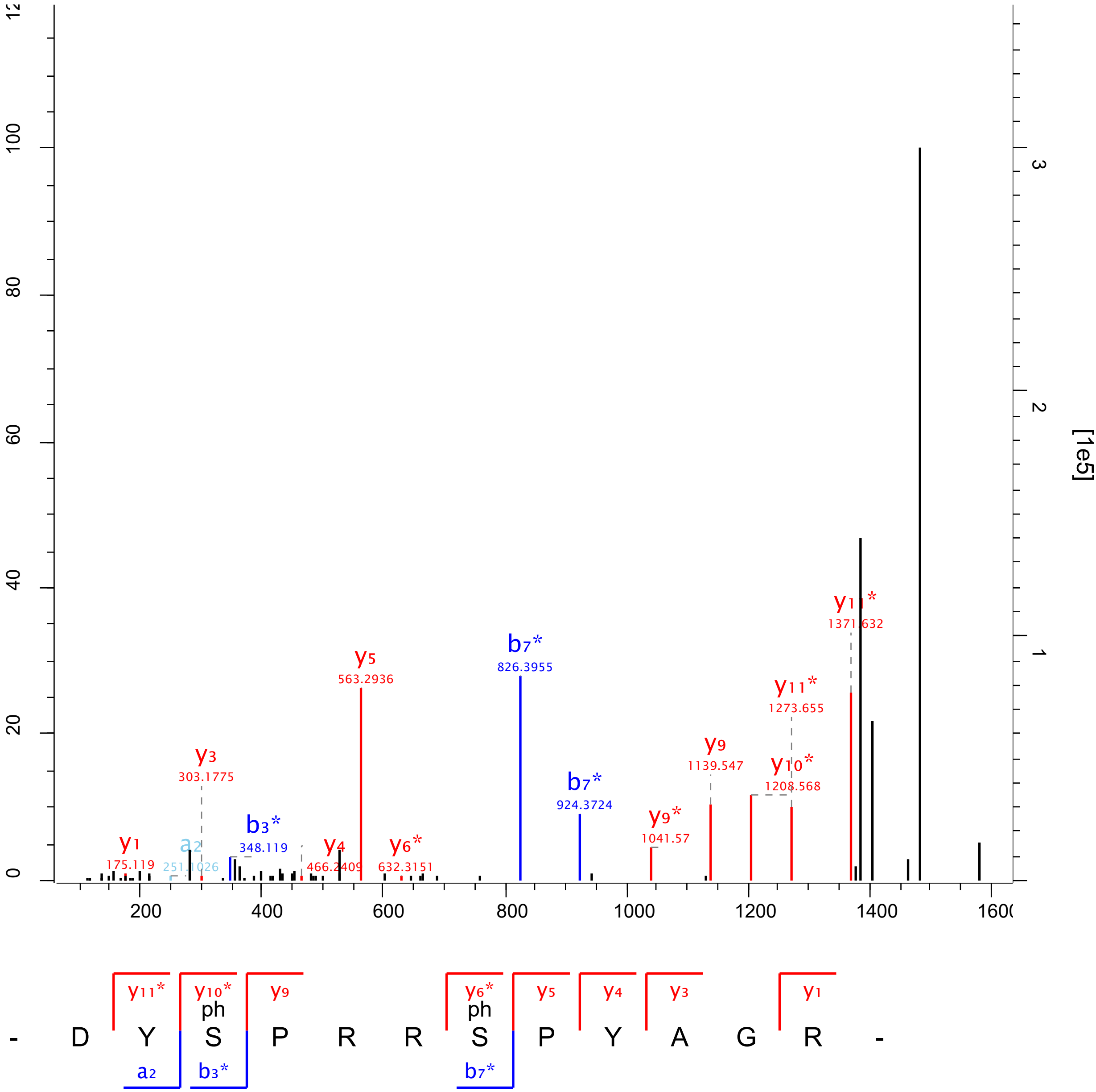

| Raw file | Scan | Method | Score | m/z |
|---|---|---|---|---|
| sys_02_2 | 7075 | FTMS; HCD | 89.47 | 724.29 |

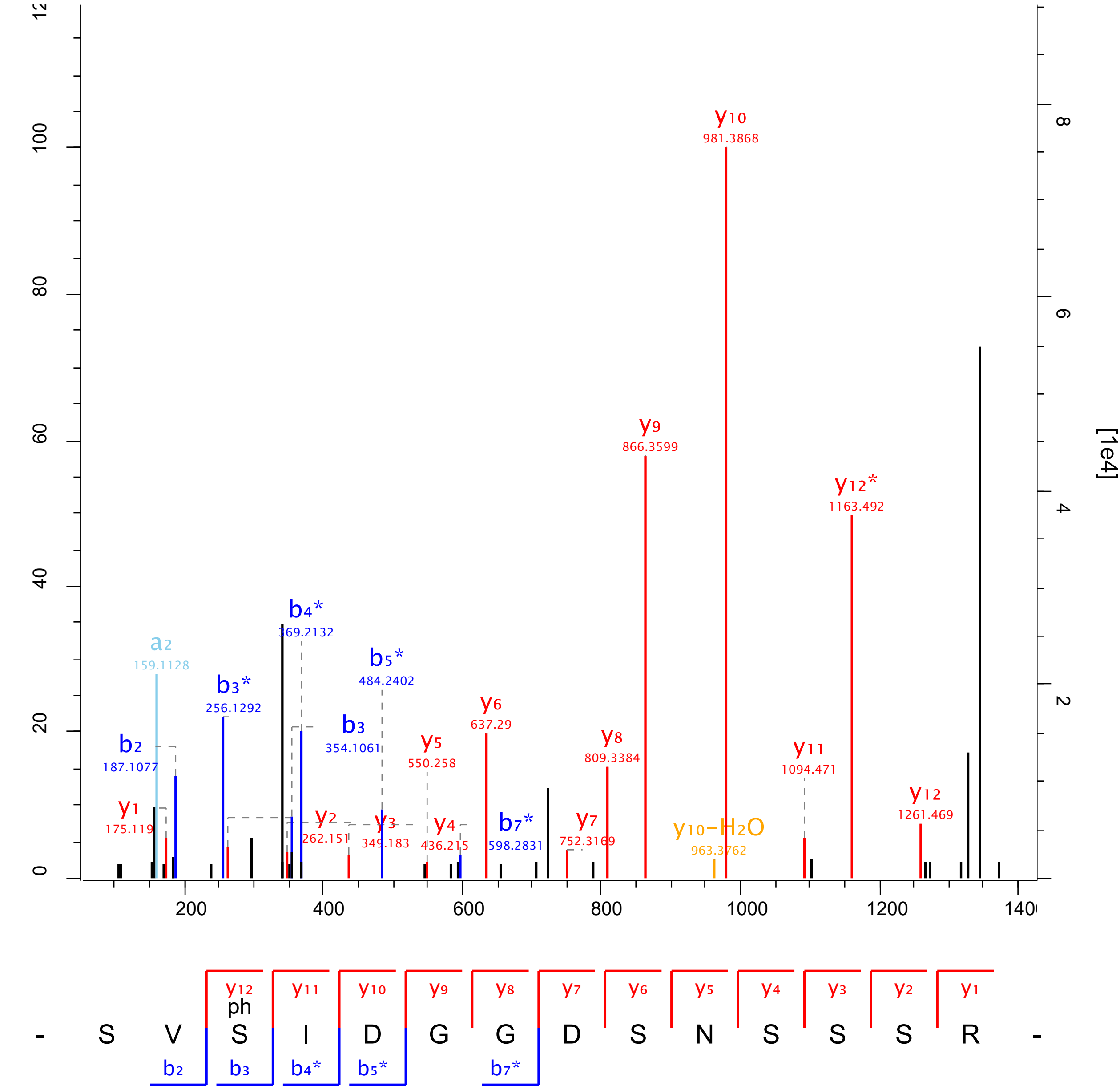

| Raw file | Scan | Method | Score | m/z |
|---|---|---|---|---|
| sys_02_2 | 7097 | FTMS; HCD | 67.65 | 639.93 |

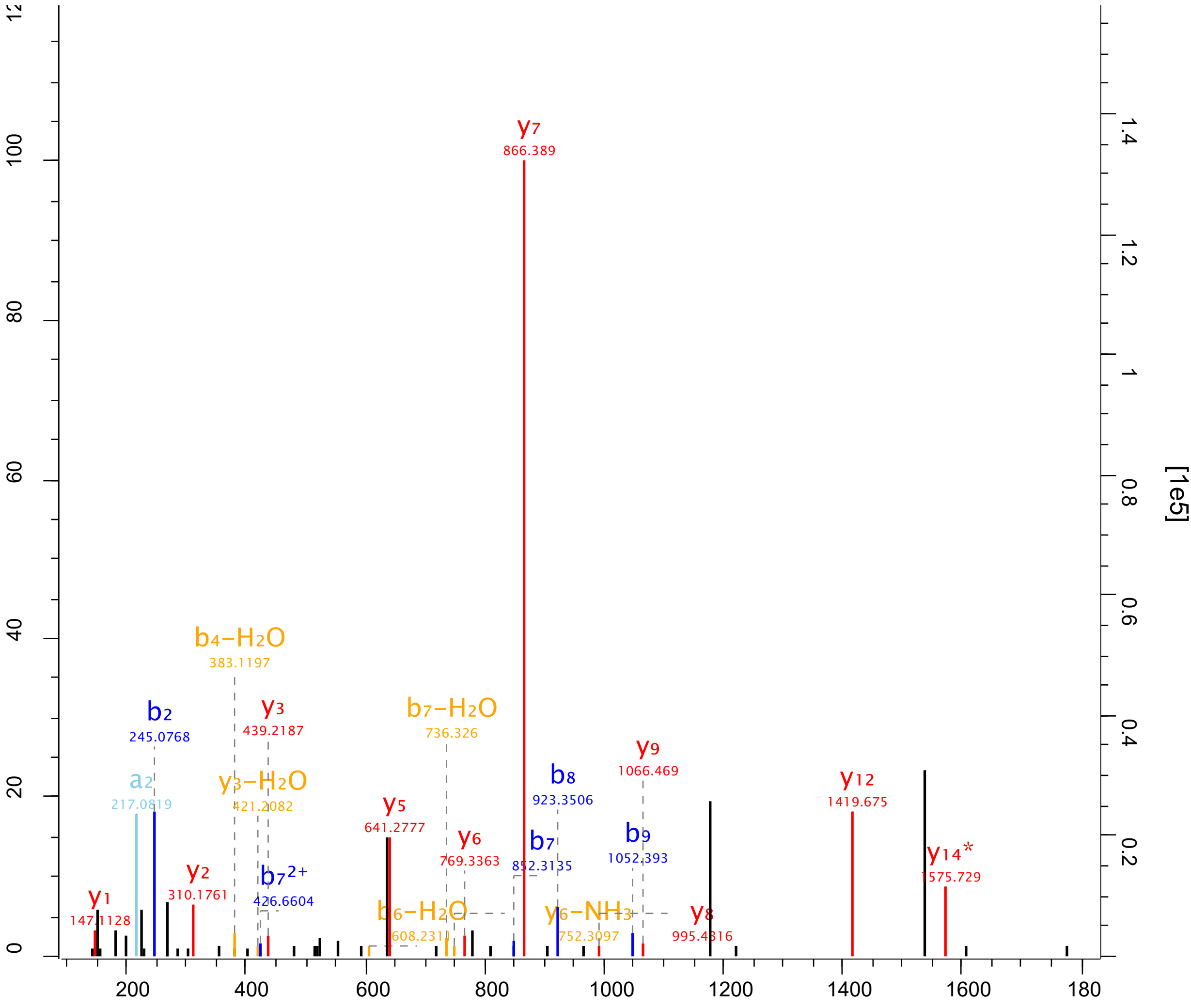


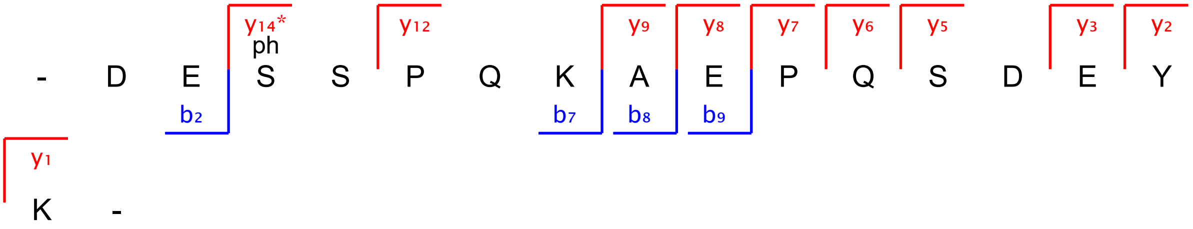

| Raw file | Scan | Method | Score | m/z |
|---|---|---|---|---|
| sys_02_2 | 7148 | FTMS; HCD | 106.23 | 580.9 |

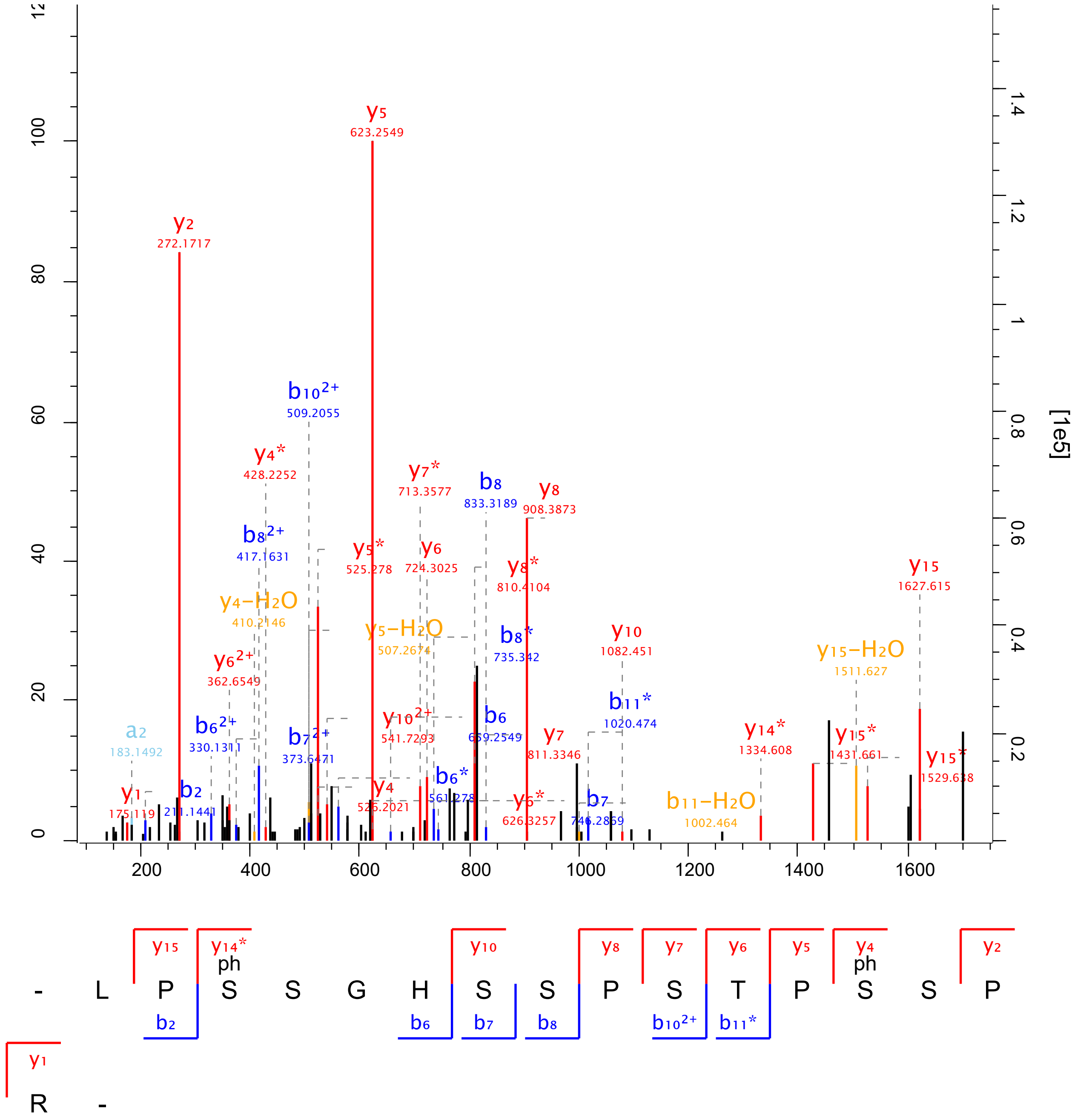

| Raw file | Scan | Method | Score | m/z |
|---|---|---|---|---|
| sys_02_2 | 7166 | FTMS; HCD | 255.93 | 827.86 |

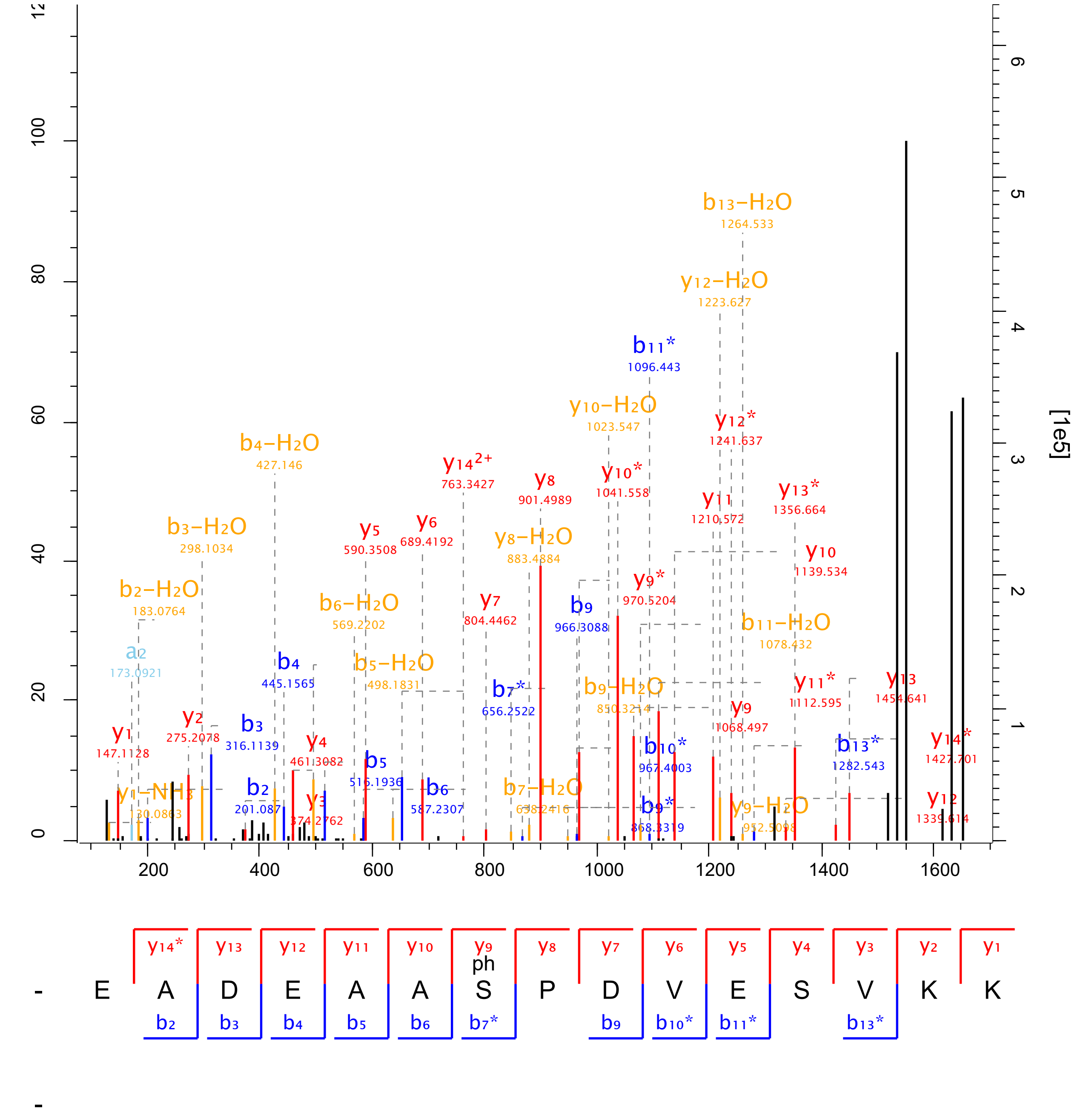

| Raw file | Scan | Method | Score | m/z |
|---|---|---|---|---|
| sys_02_2 | 7182 | FTMS; HCD | 82.63 | 616.62 |

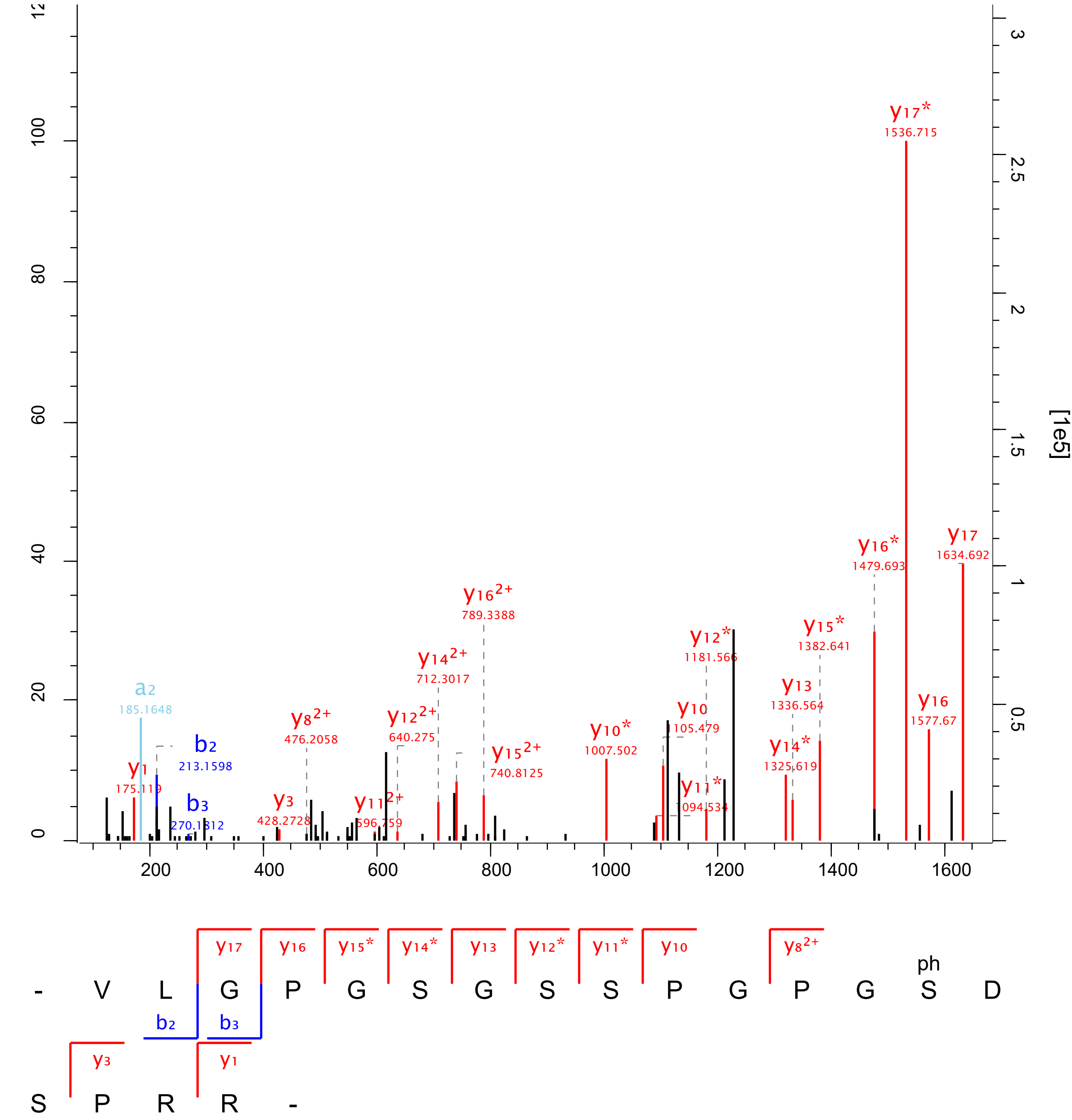

| Raw file | Scan | Method | Score | m/z |
|---|---|---|---|---|
| sys_02_2 | 7313 | FTMS; HCD | 50.46 | 429.85 |

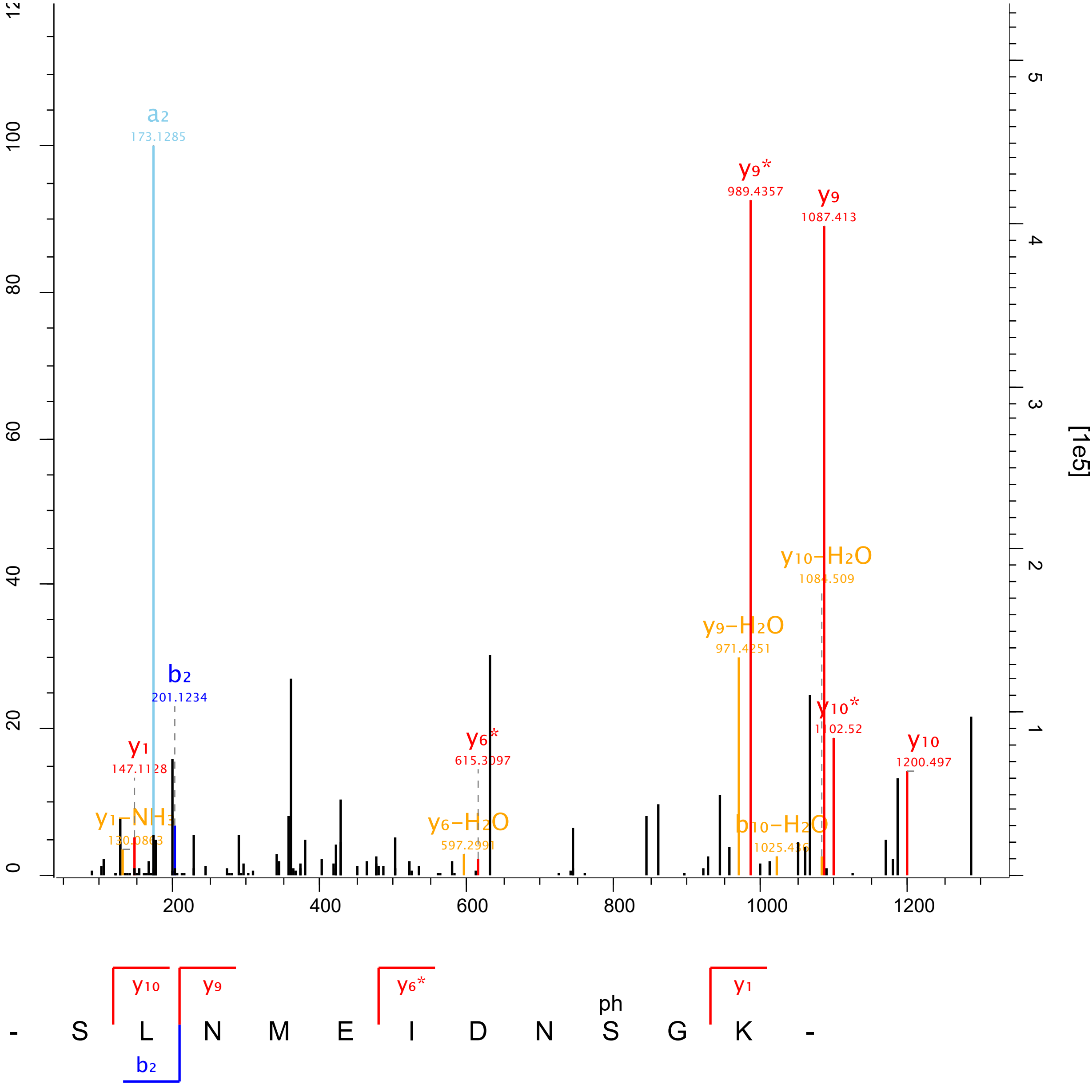

| Raw file | Scan | Method | Score | m/z |
|---|---|---|---|---|
| sys_02_2 | 7392 | FTMS; HCD | 182.86 | 565.27 |

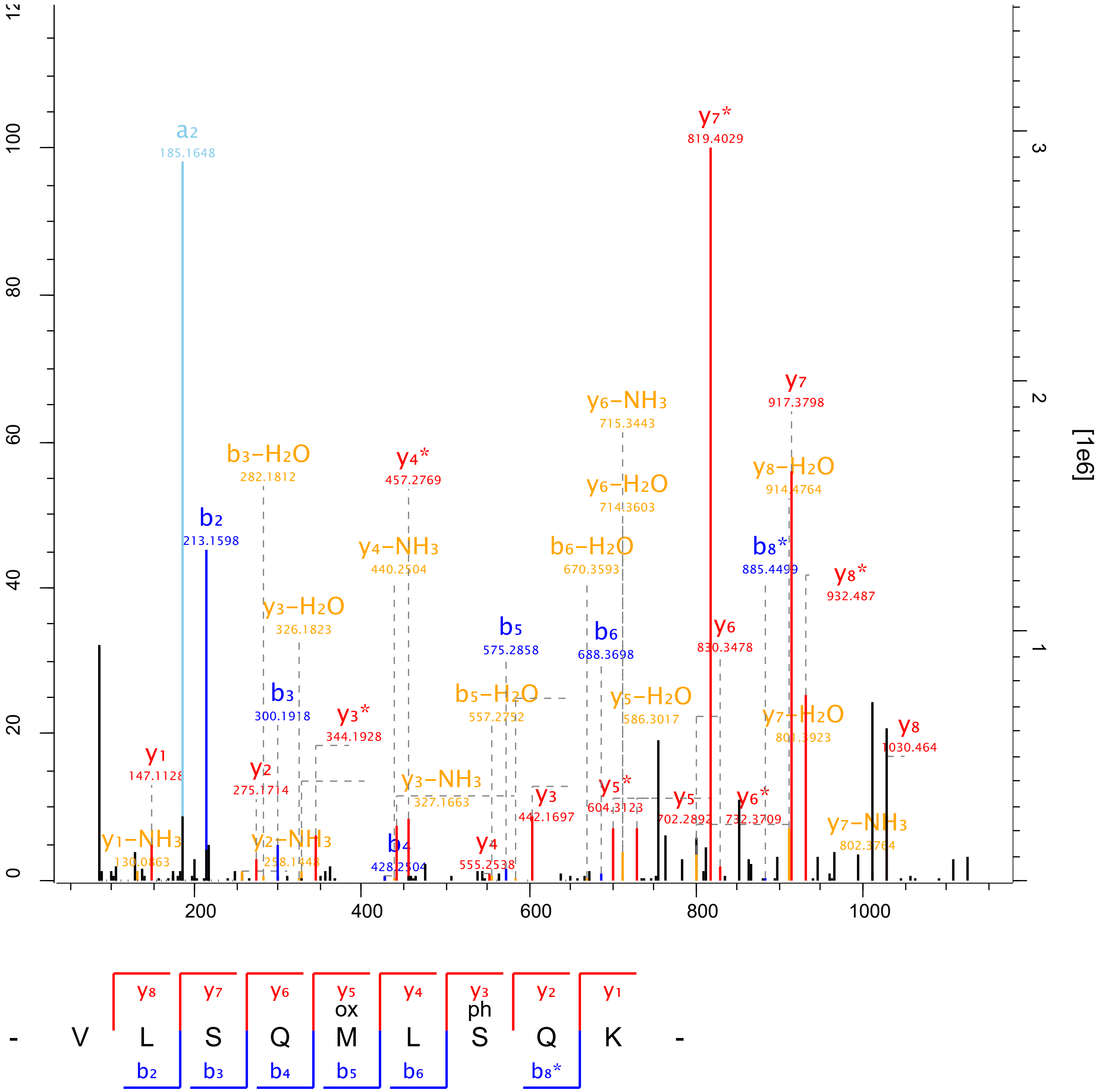

| Raw file | Scan | Method | Score | m/z |
|---|---|---|---|---|
| sys_02_2 | 7461 | FTMS; HCD | 189.51 | 572.76 |

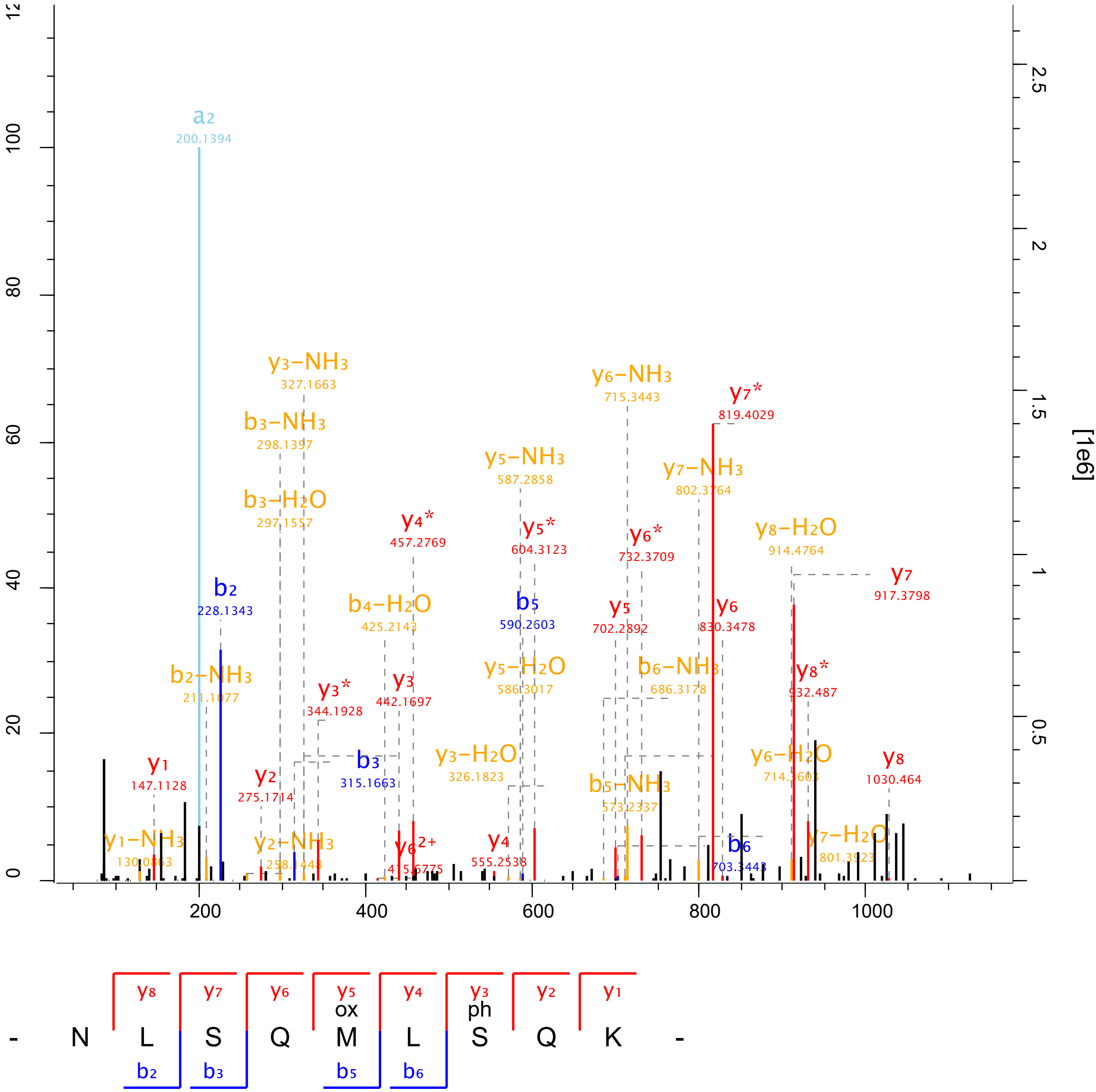

| Raw file | Scan | Method | Score | m/z |
|---|---|---|---|---|
| sys_02_2 | 7470 | FTMS; HCD | 83.54 | 768.79 |

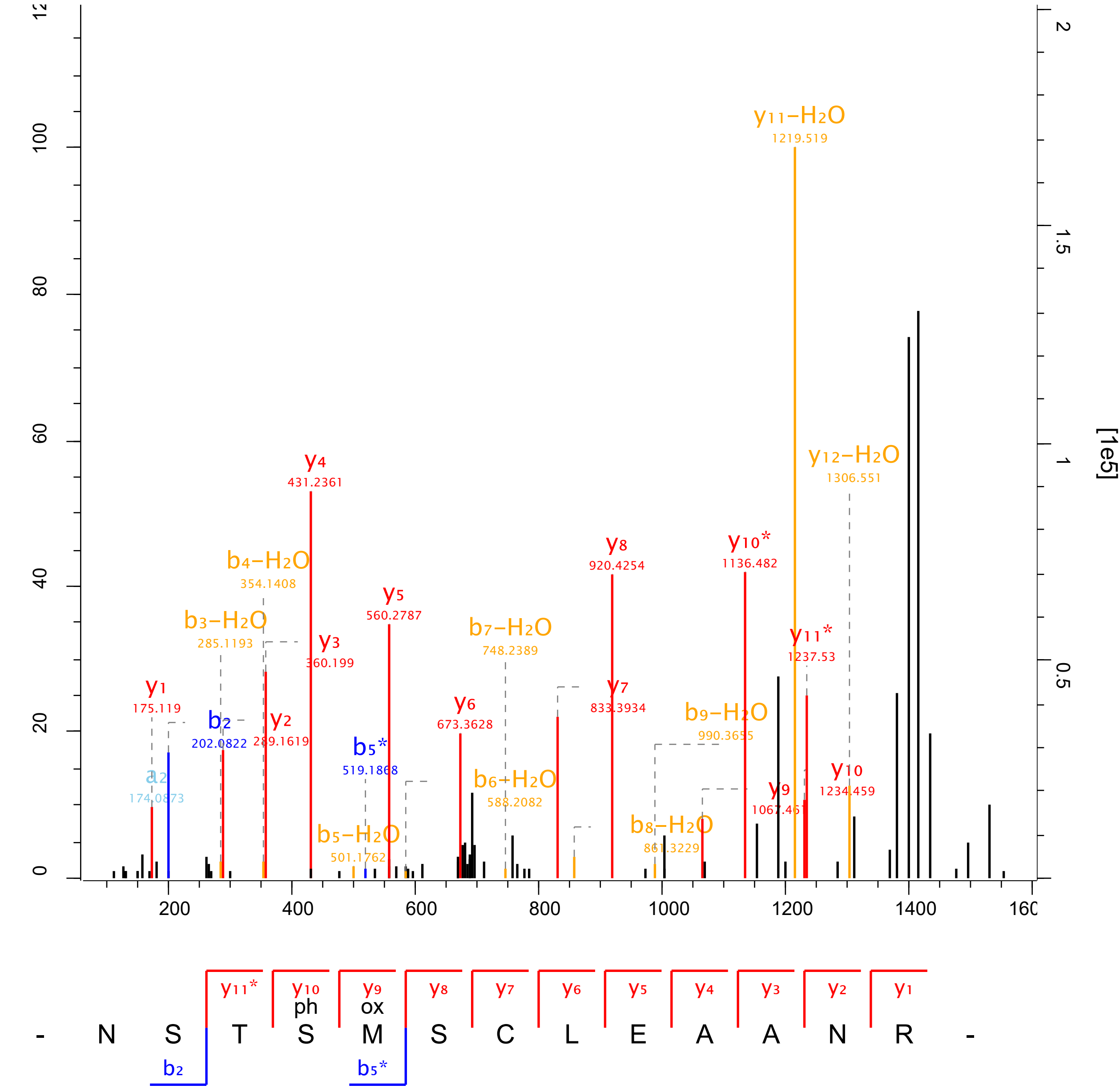

| Raw file | Scan | Method | Score | m/z |
| --- | --- | --- | --- | --- |
| sys_02_2 | 7514 | FTMS; HCD | 195.86 | 776.85 |

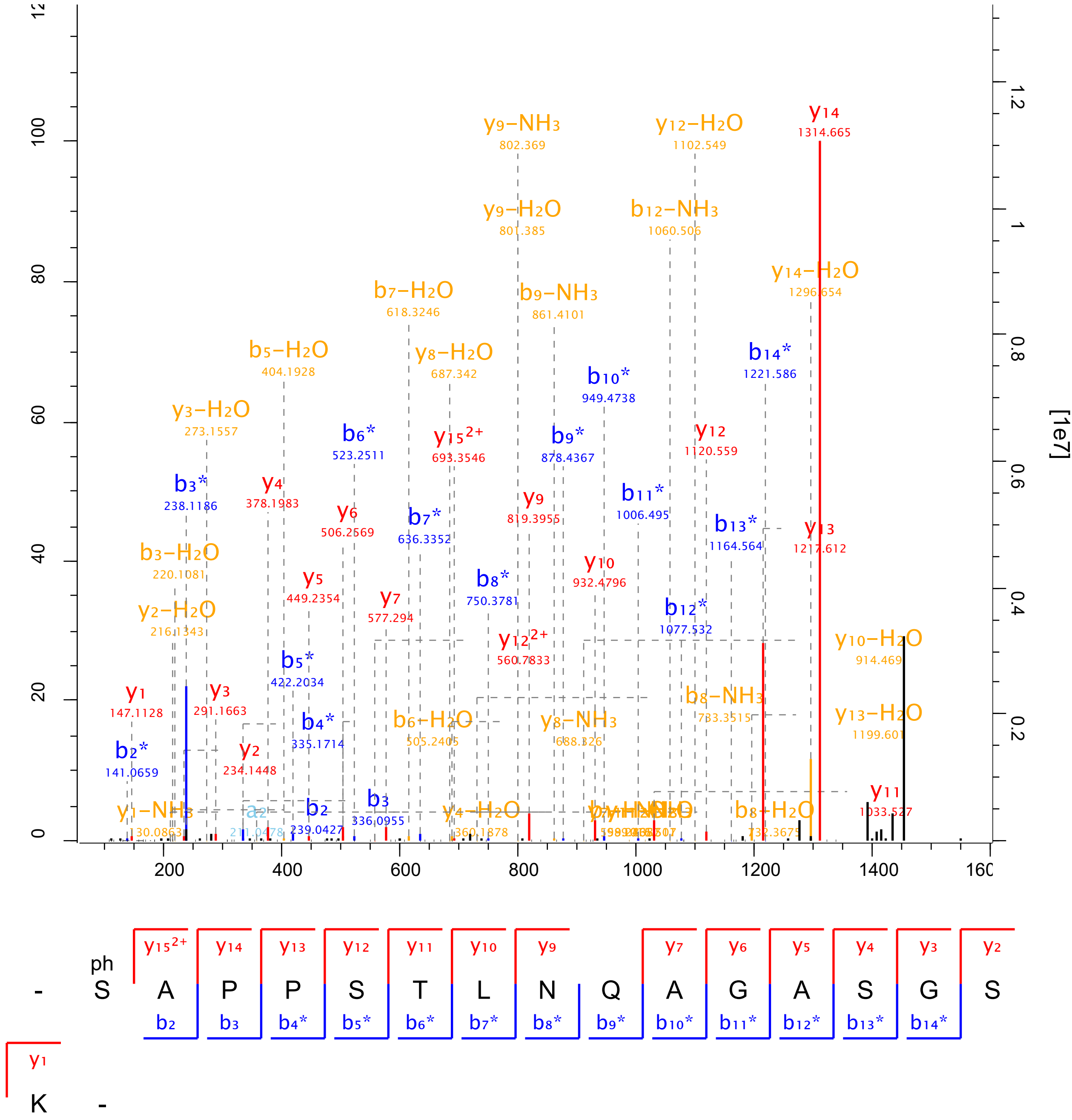

| Raw file | Scan | Method | Score | m/z |
|---|---|---|---|---|
| sys_02_2 | 7520 | FTMS; HCD | 86.29 | 535.72 |

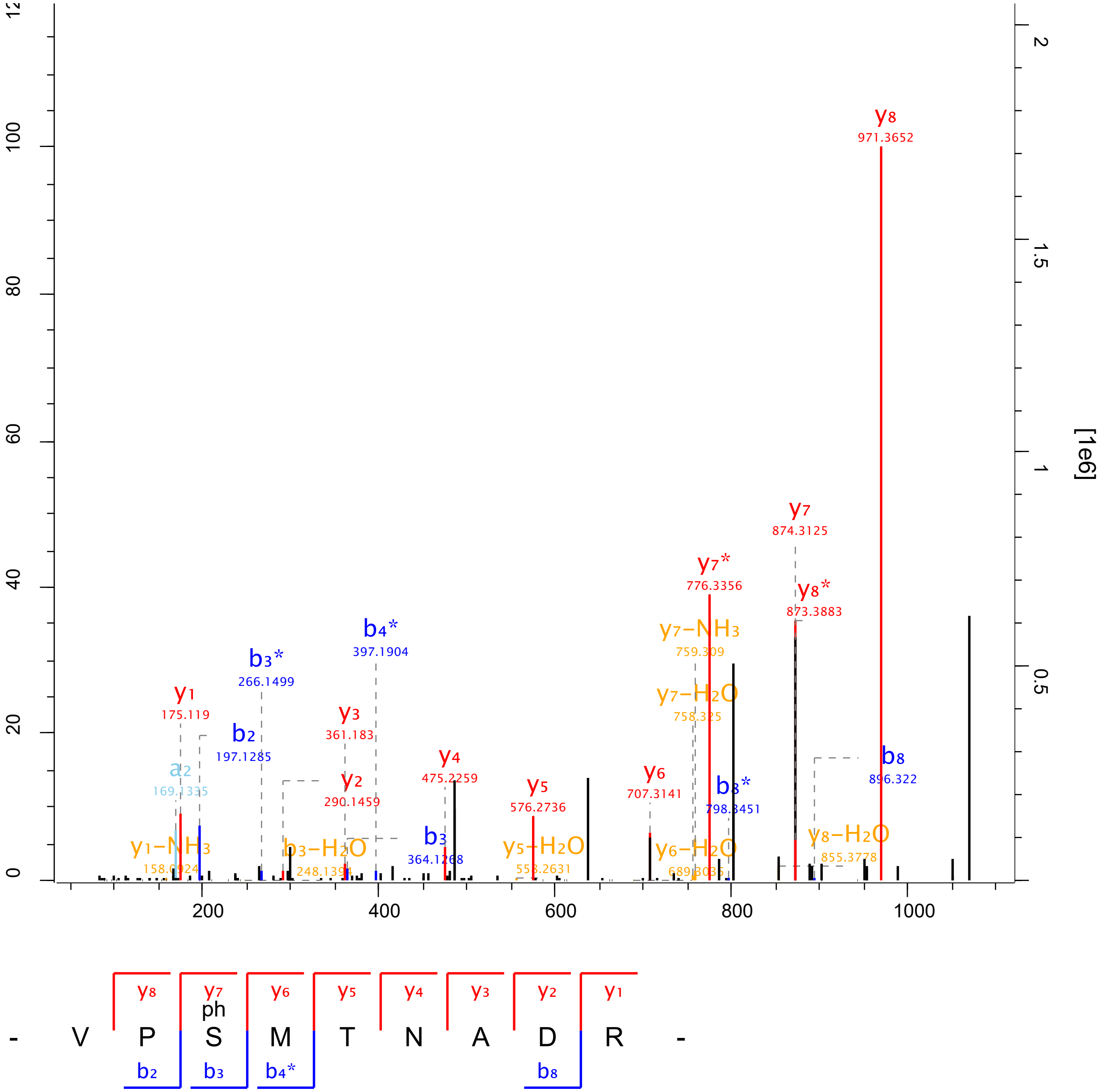

| Raw file | Scan | Method | Score | m/z |
|---|---|---|---|---|
| sys_02_2 | 7765 | FTMS; HCD | 102.21 | 648.94 |

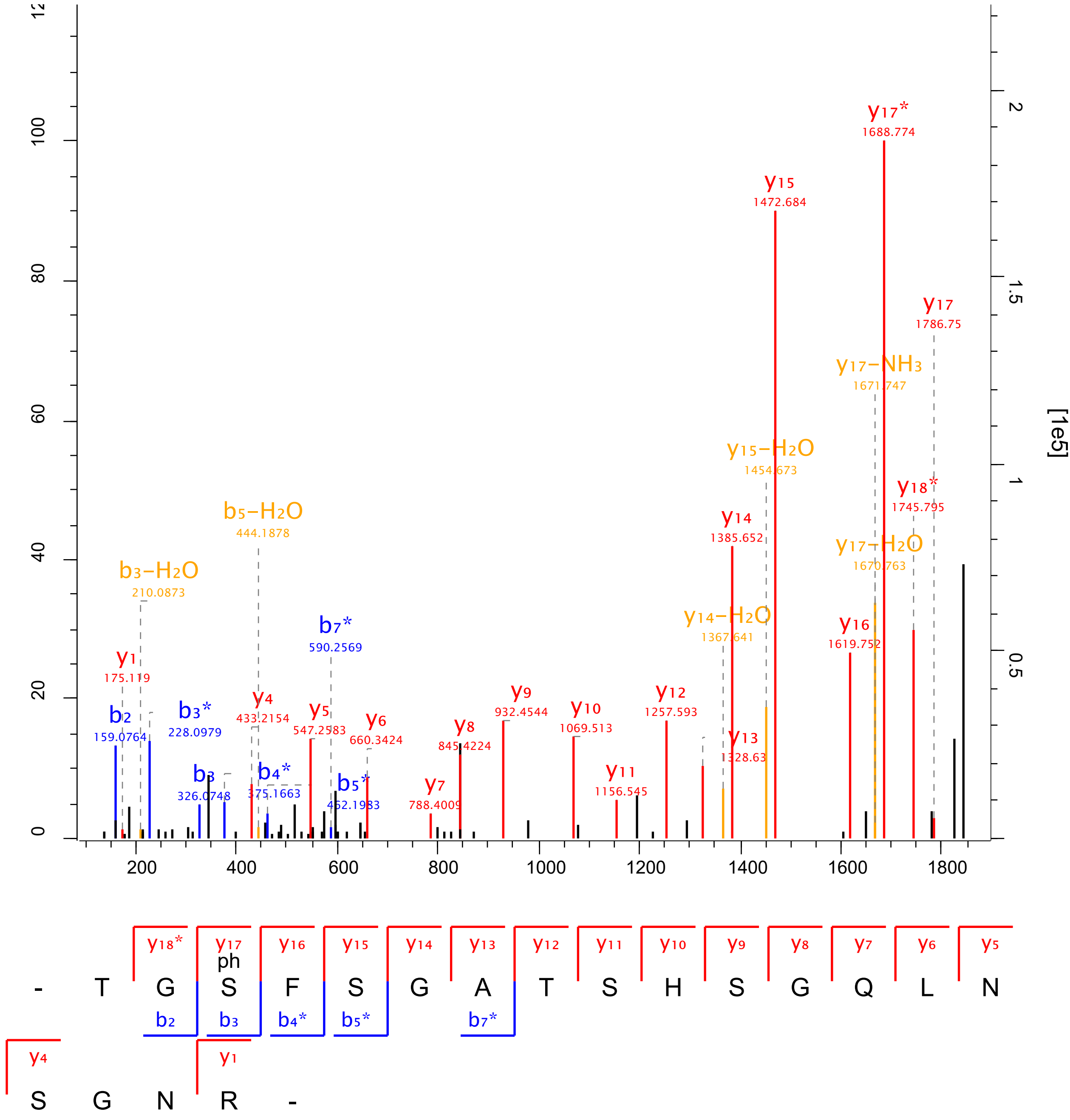

| Raw file | Scan | Method | Score | m/z |
|---|---|---|---|---|
| sys_02_2 | 7806 | FTMS; HCD | 83.4 | 656.26 |

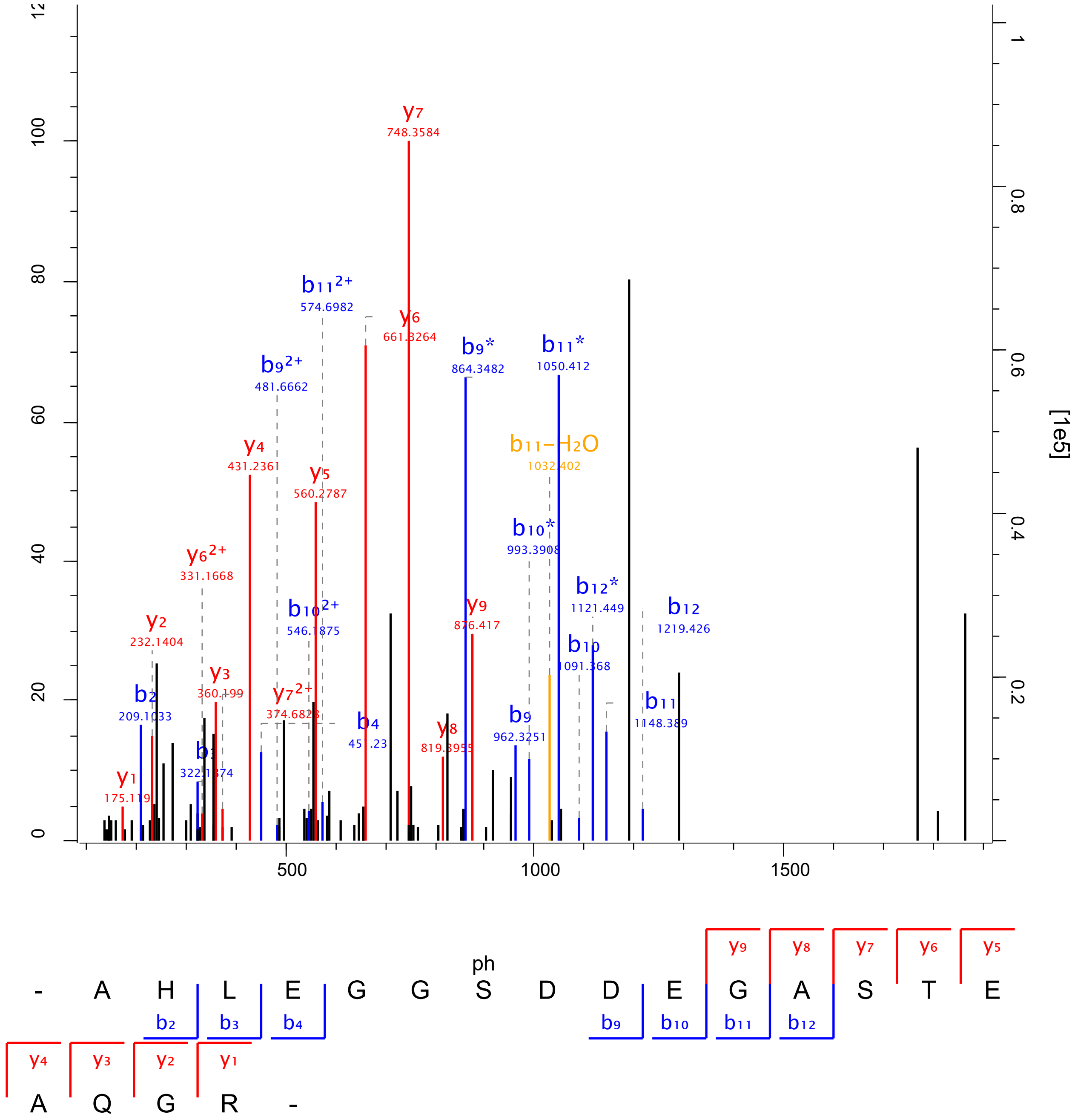

| Raw file | Scan | Method | Score | m/z |
|---|---|---|---|---|
| sys_02_2 | 7947 | FTMS; HCD | 41.54 | 478.56 |

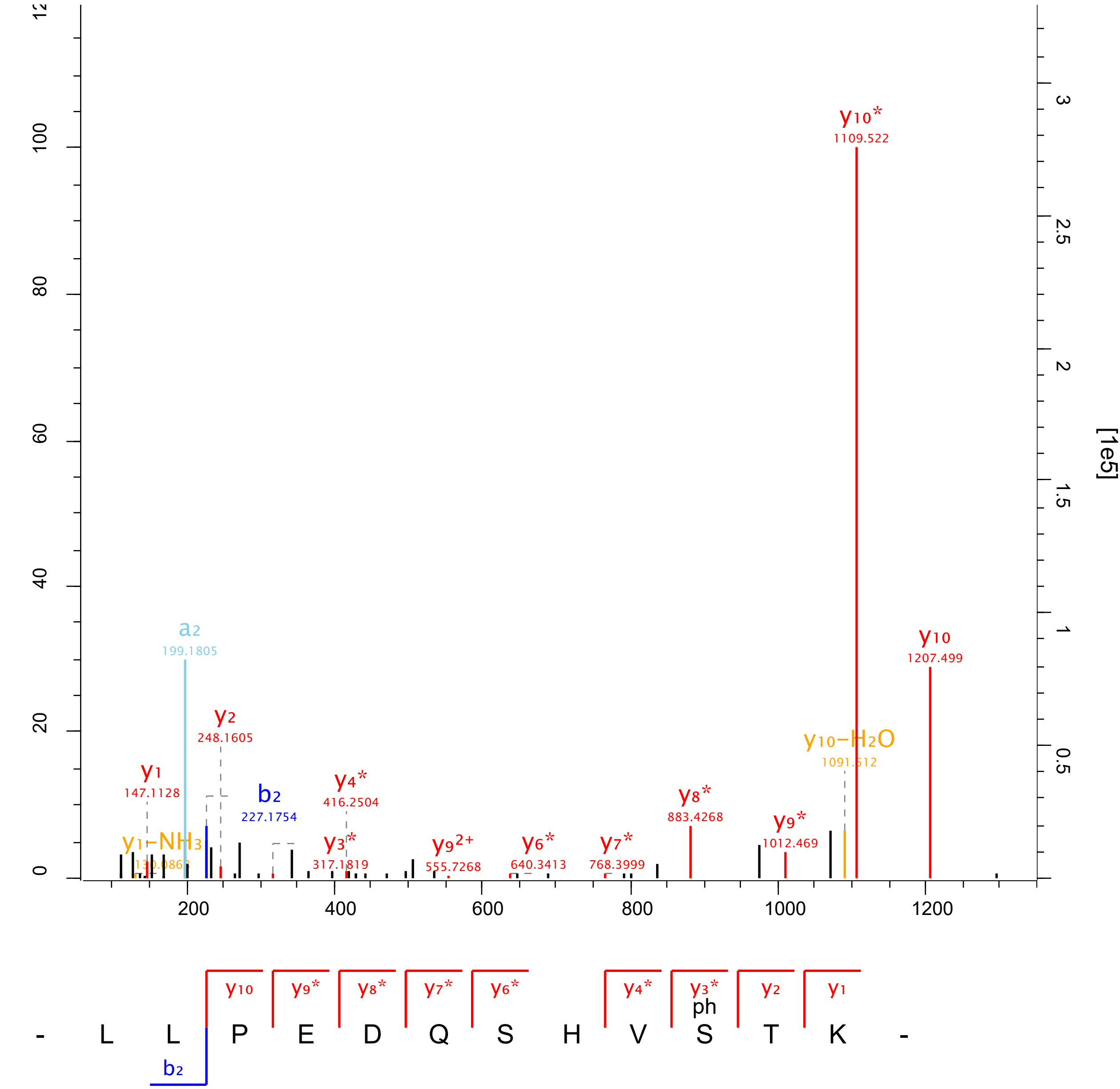

| Raw file | Scan | Method | Score | m/z |
|---|---|---|---|---|
| sys_02_2 | 8030 | FTMS; HCD | 46.32 | 450.87 |

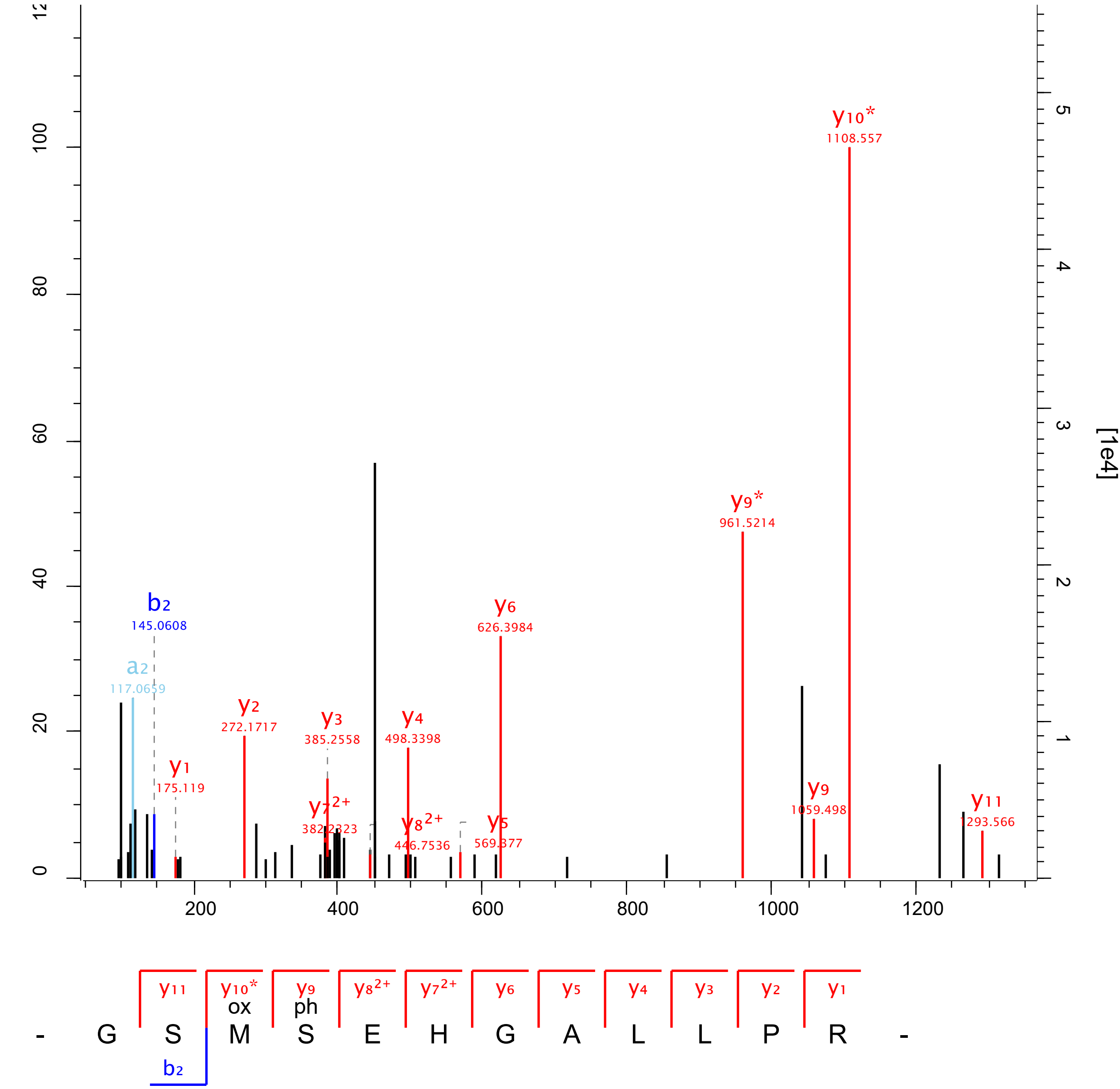

| Raw file | Scan | Method | Score | m/z |
|---|---|---|---|---|
| sys_02_2 | 8055 | FTMS; HCD | 66.27 | 552.75 |

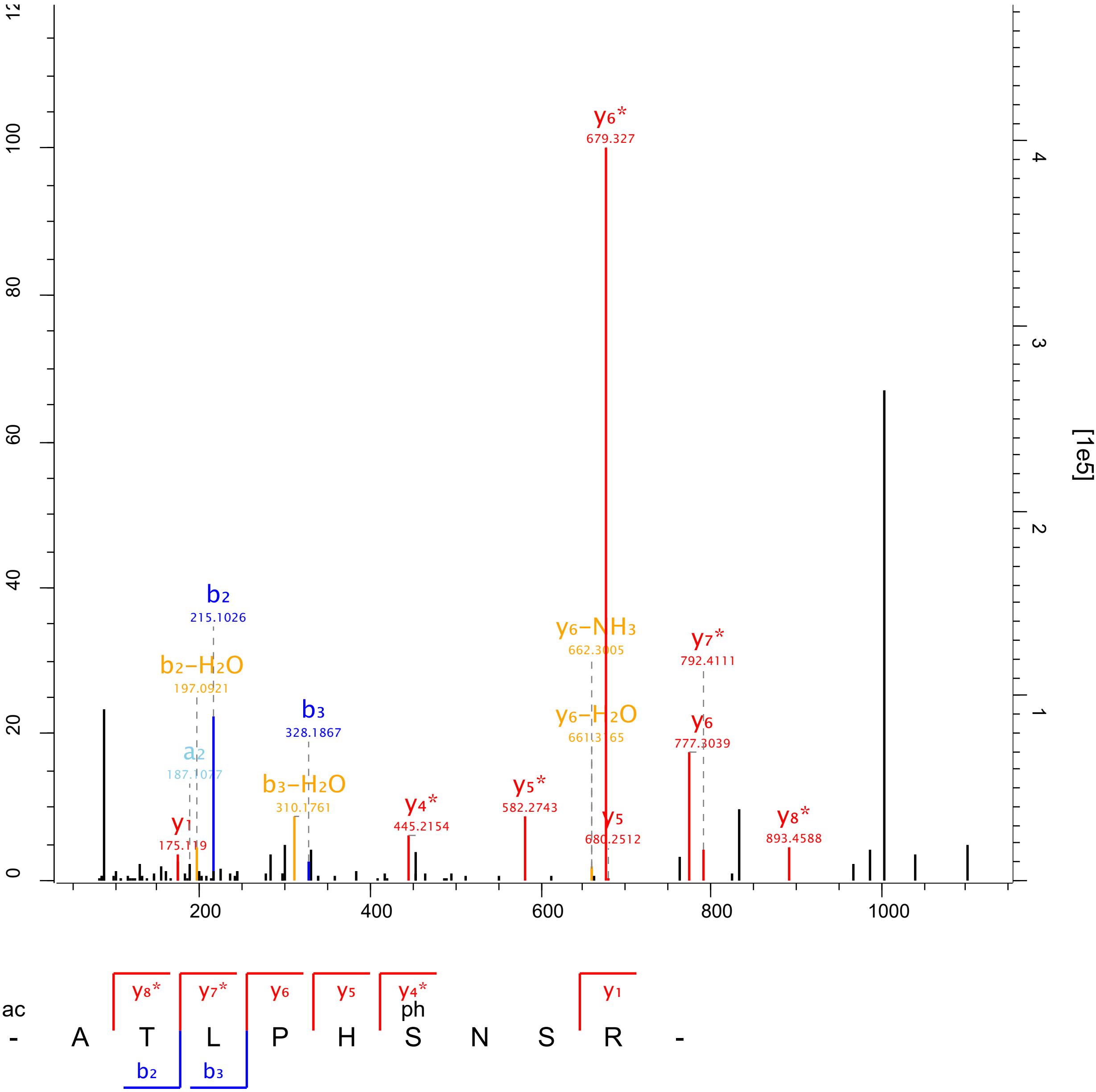

| Raw file | Scan | Method | Score | m/z |
|---|---|---|---|---|
| sys_02_2 | 8097 | FTMS; HCD | 104.26 | 481.23 |

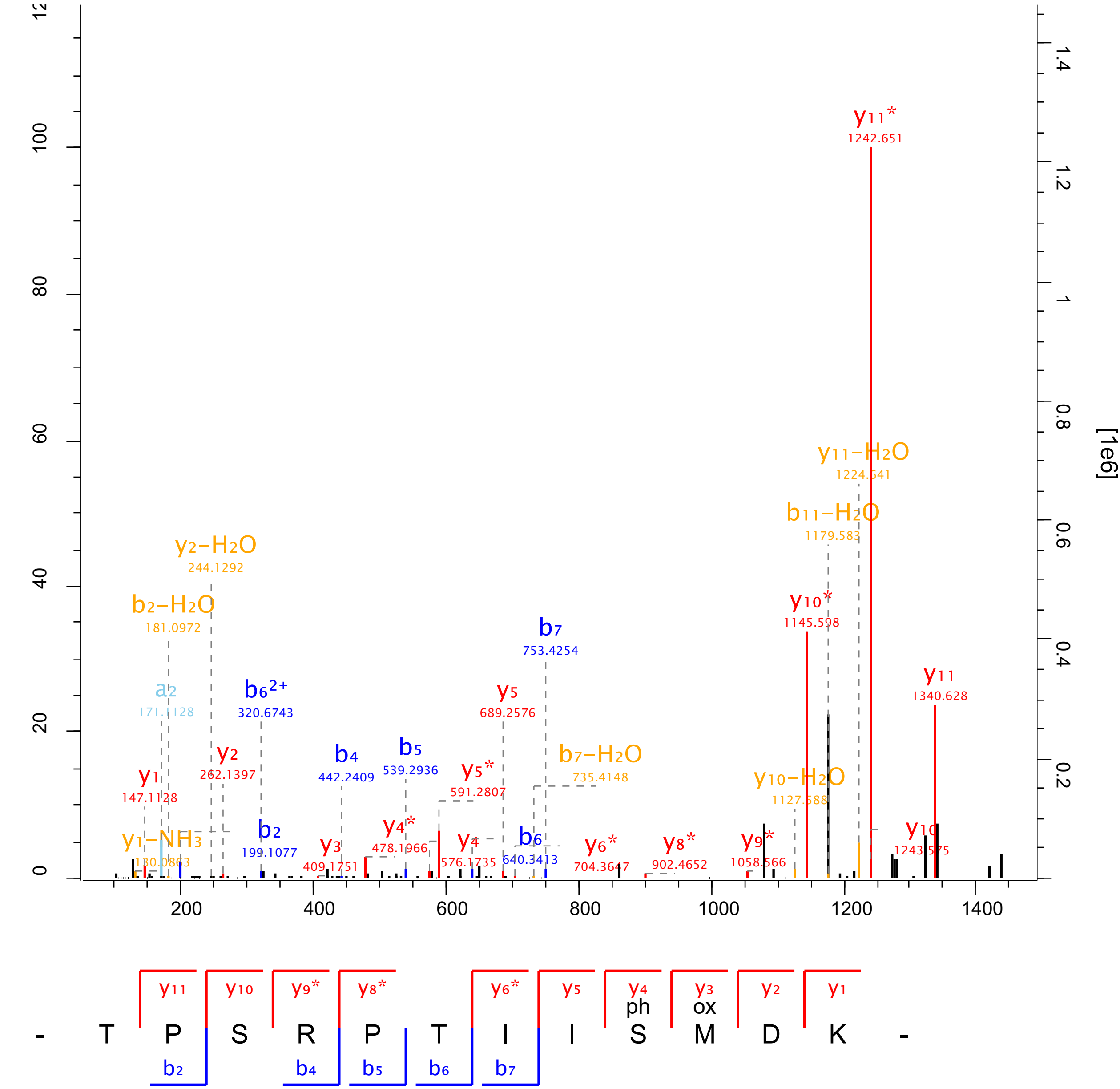

| Raw file | Scan | Method | Score | m/z |
| --- | --- | --- | --- | --- |
| sys_02_2 | 8101 | FTMS; HCD | 42.64 | 683.6 |

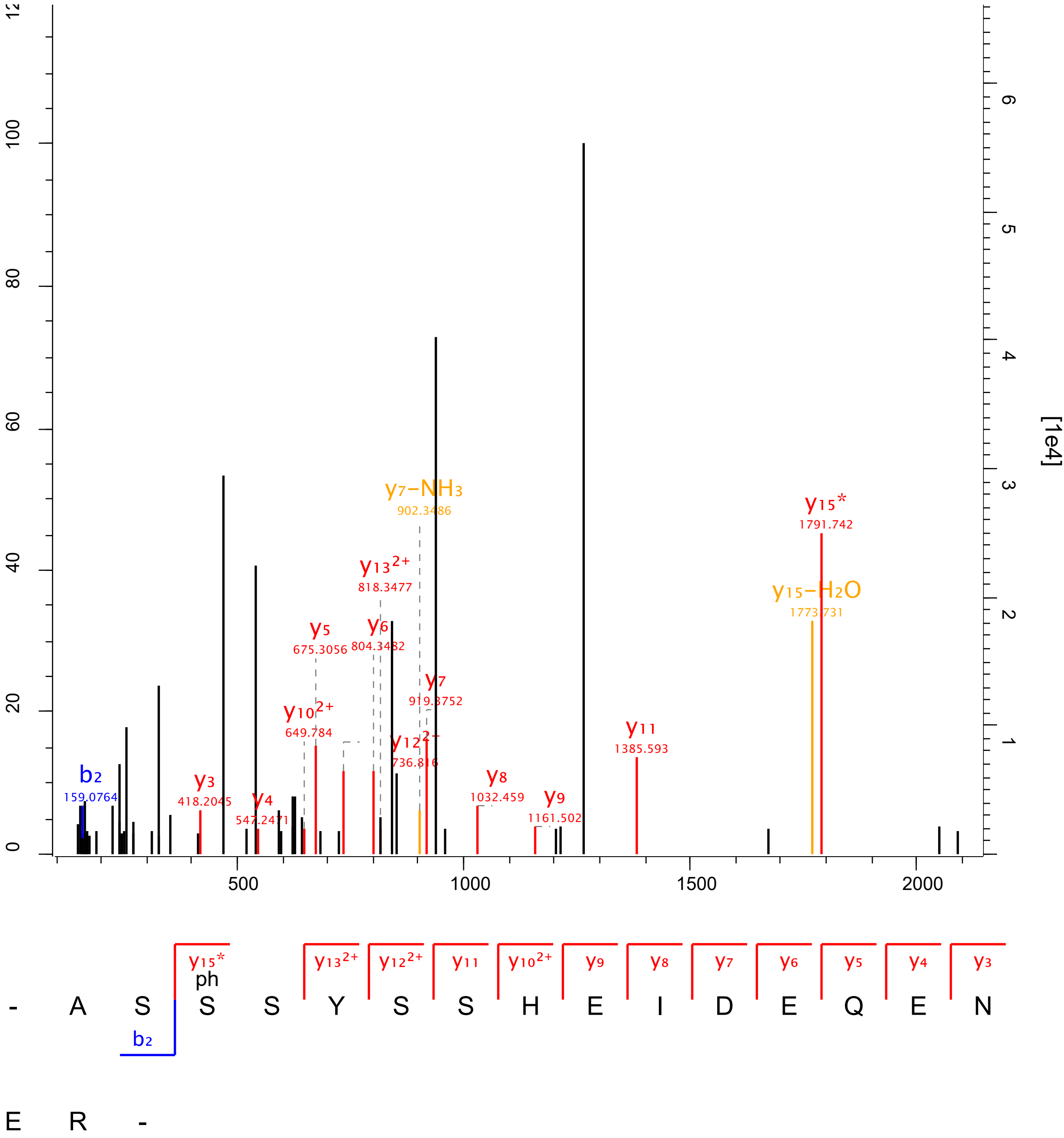

| Raw file | Scan | Method | Score | m/z |
| --- | --- | --- | --- | --- |
| sys_02_2 | 8104 | FTMS; HCD | 181.97 | 703.82 |

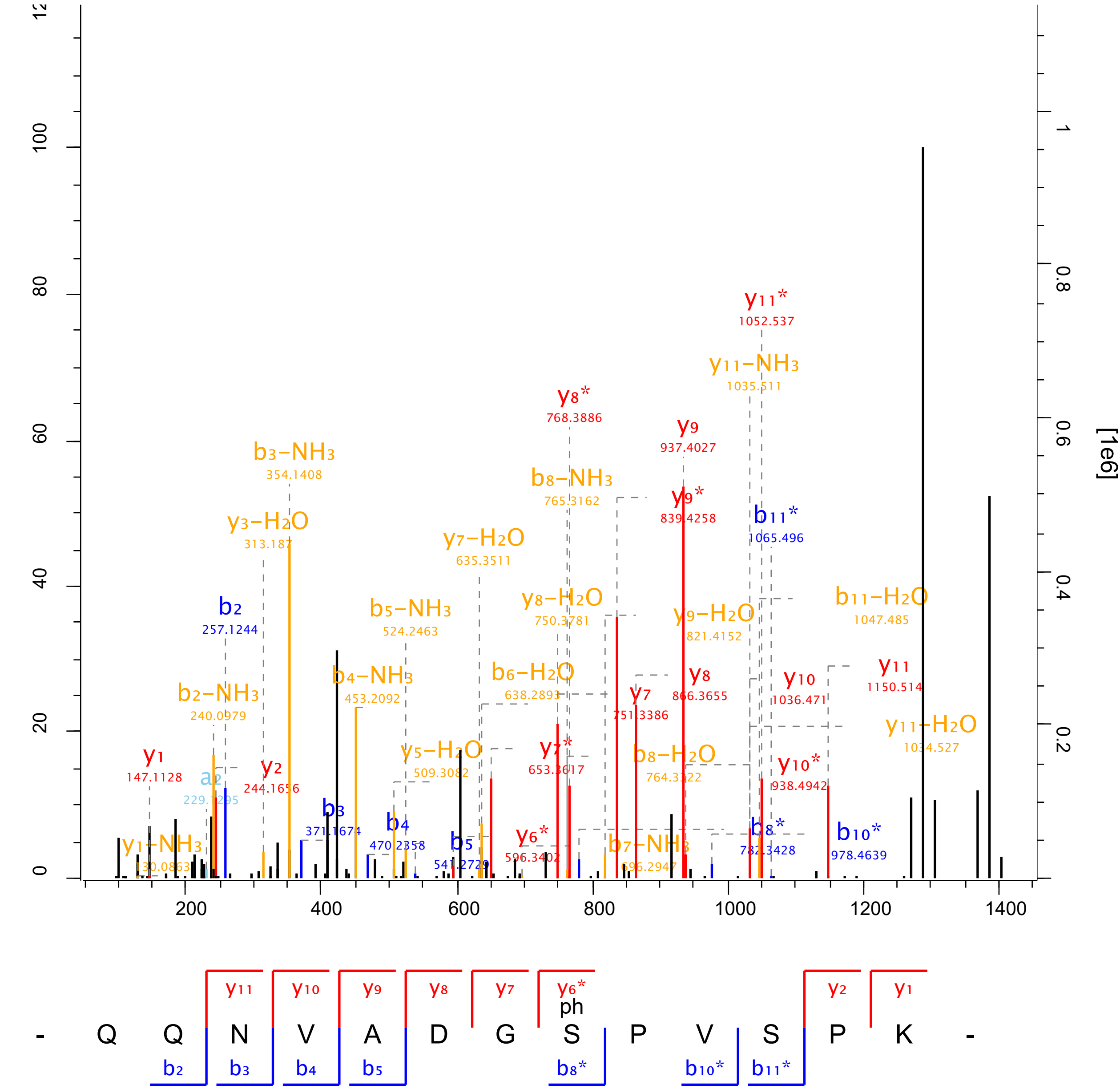

| Raw file | Scan | Method | Score | m/z |
|---|---|---|---|---|
| sys_02_2 | 8199 | FTMS; HCD | 115.56 | 796.86 |

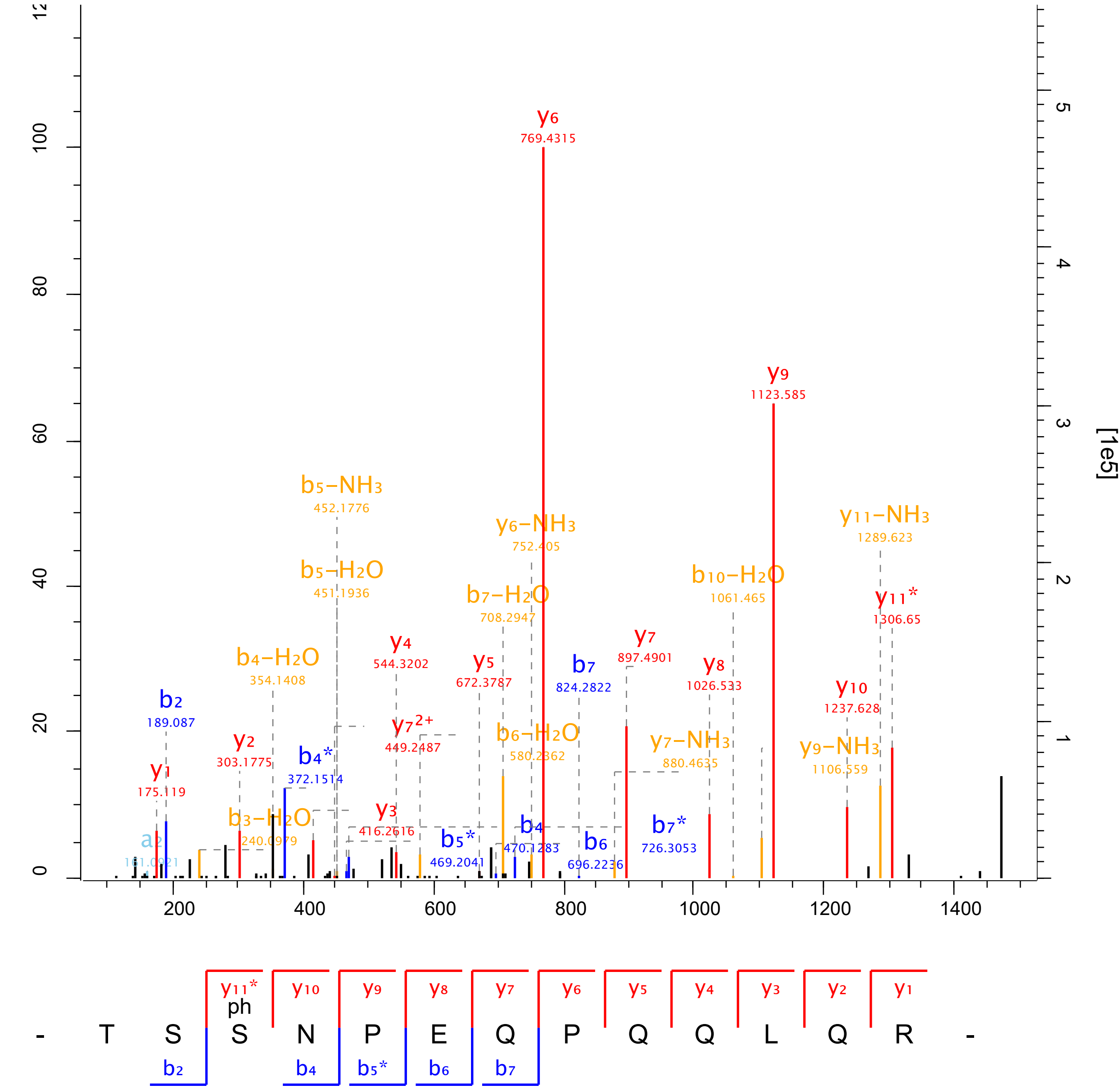

| Raw file | Scan | Method | Score | m/z |
|---|---|---|---|---|
| sys_02_2 | 8382 | FTMS; HCD | 46.16 | 505.57 |

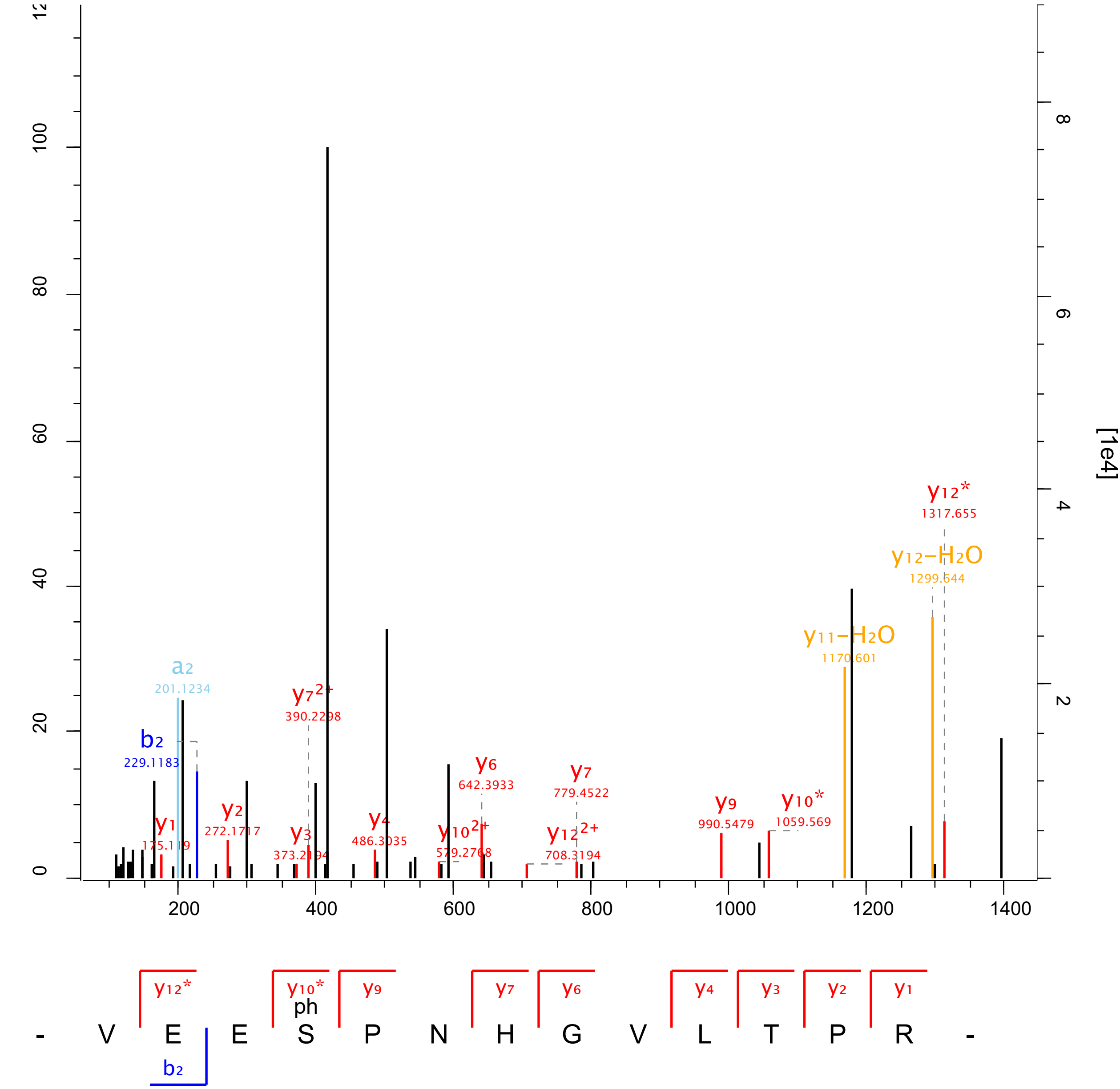

| Raw file | Scan | Method | Score | m/z |
|---|---|---|---|---|
| sys_02_2 | 8410 | FTMS; HCD | 233.49 | 743.82 |

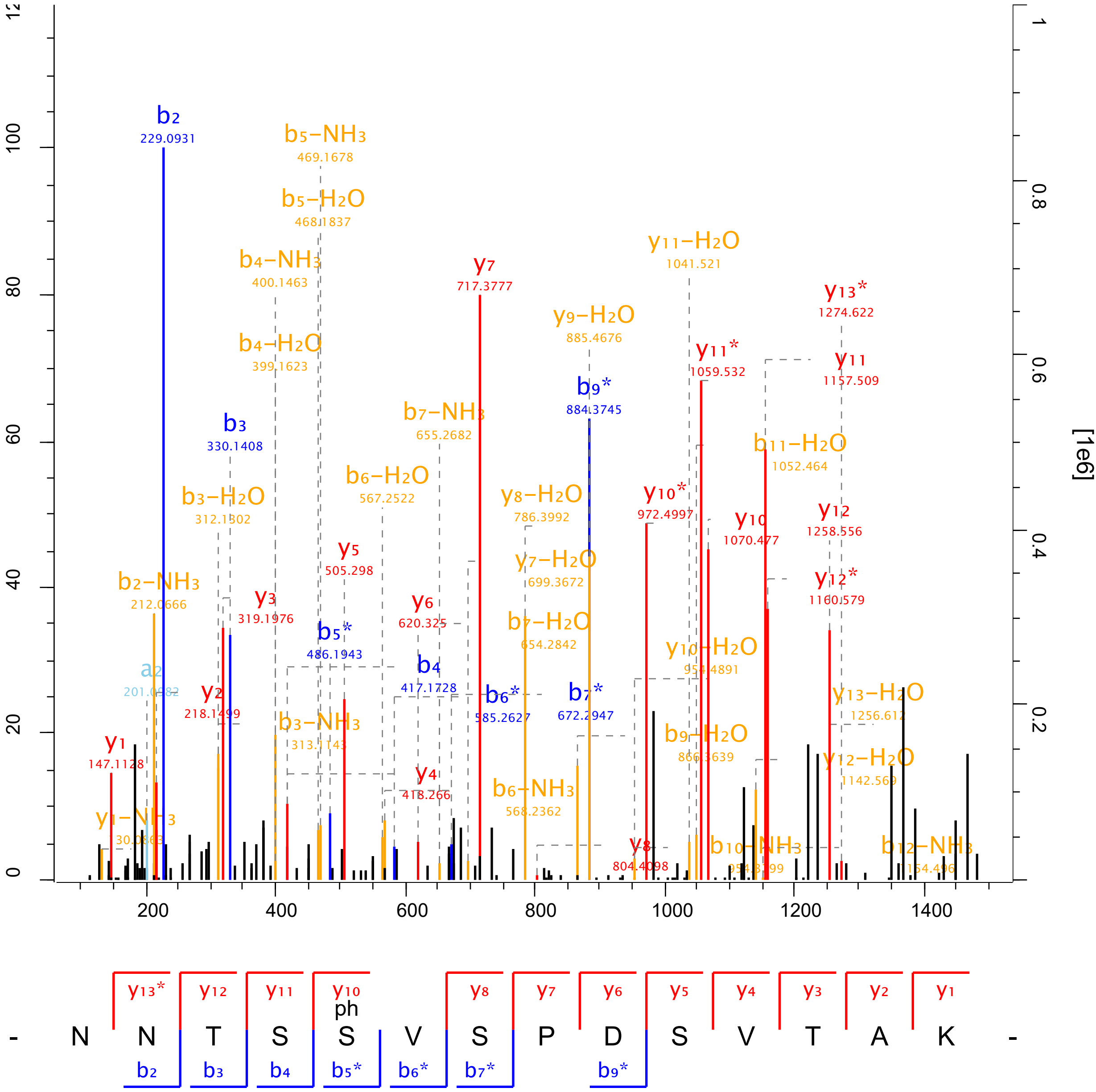

| Raw file | Scan | Method | Score | m/z |
|---|---|---|---|---|
| sys_02_2 | 8459 | FTMS; HCD | 132.65 | 681.64 |

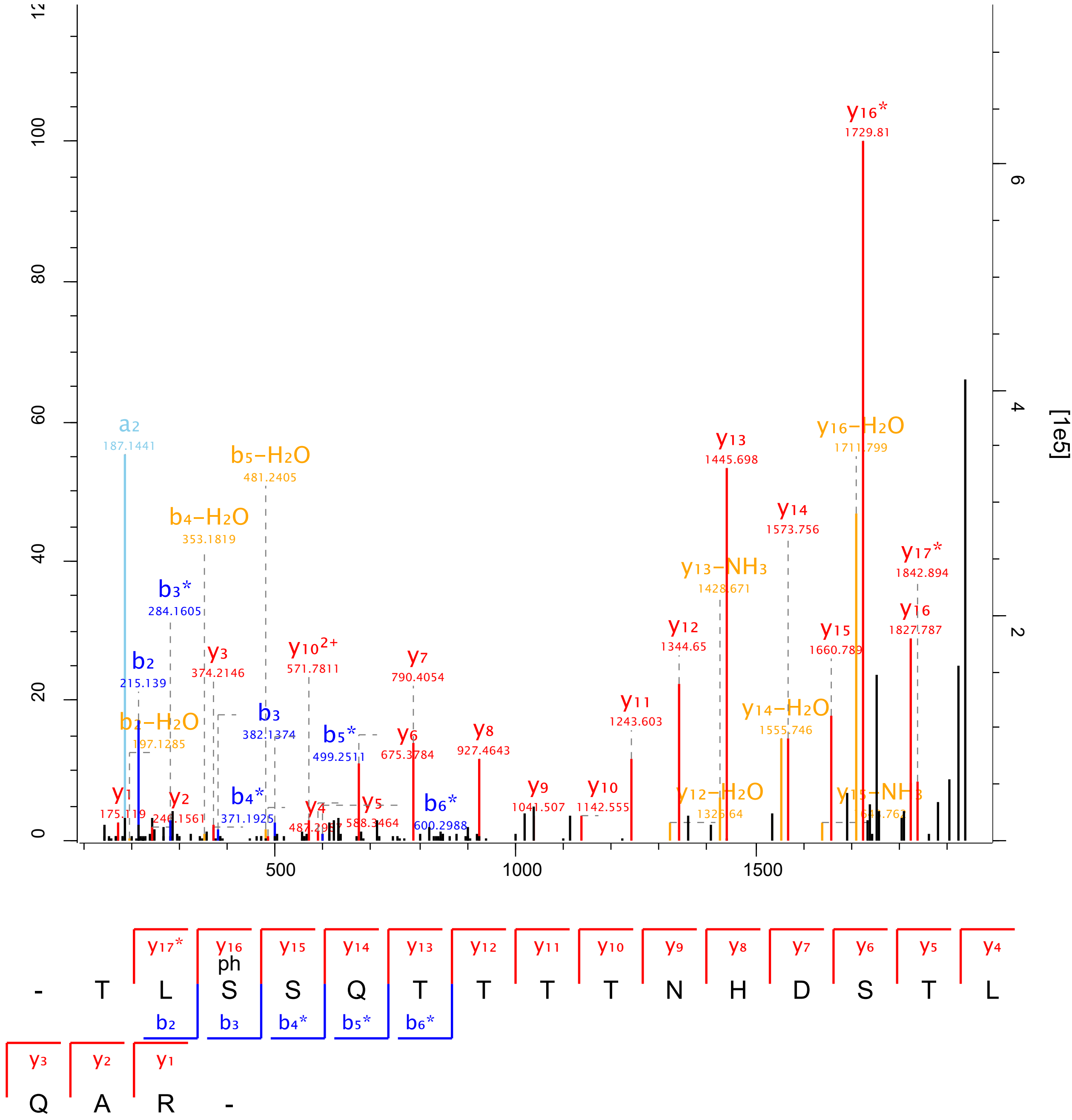

| Raw file | Scan | Method | Score | m/z |
|---|---|---|---|---|
| sys_02_2 | 8520 | FTMS; HCD | 206.62 | 670.34 |

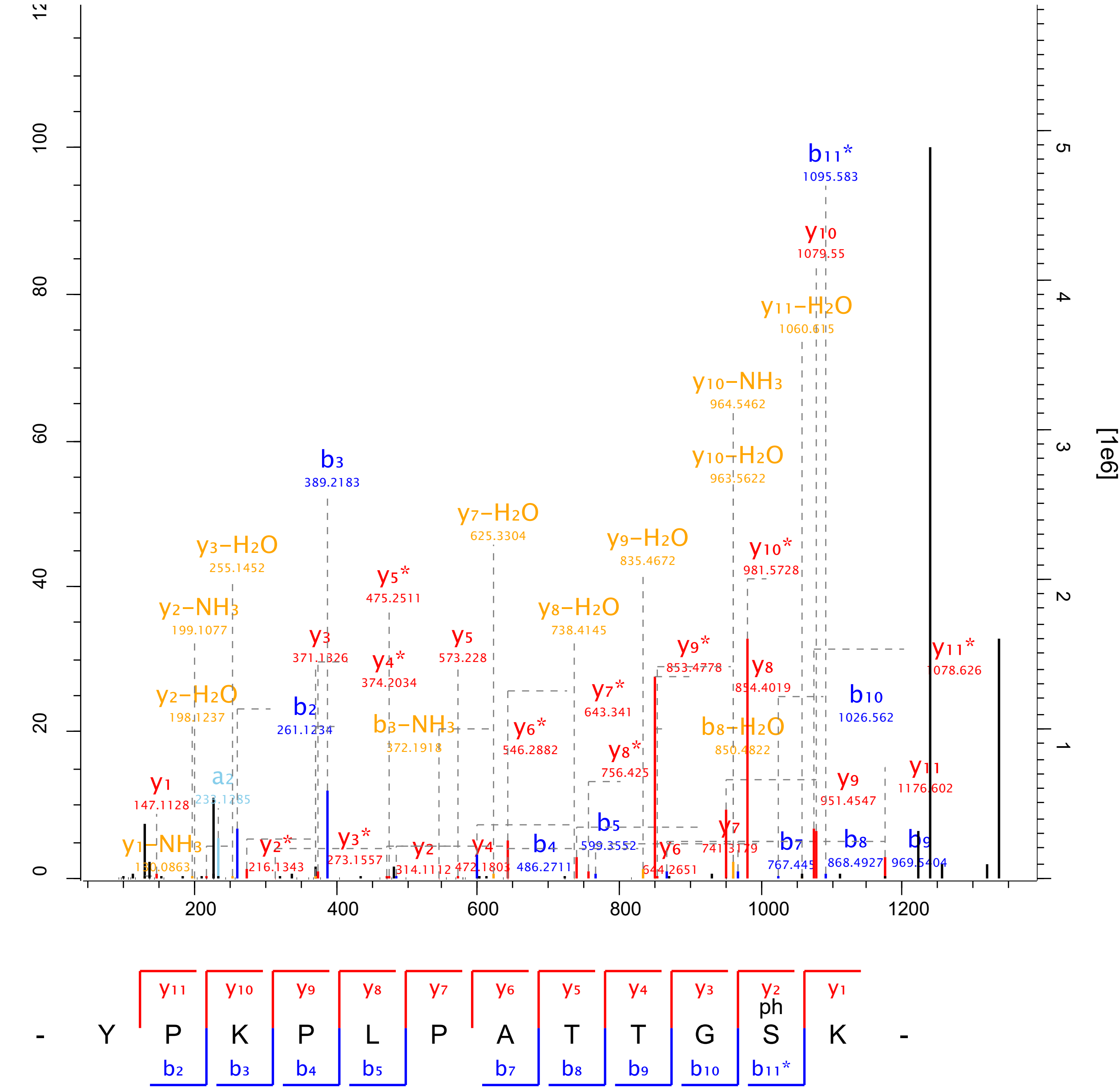

| Raw file | Scan | Method | Score | m/z |
|---|---|---|---|---|
| sys_02_2 | 8578 | FTMS; HCD | 41.7 | 480.24 |

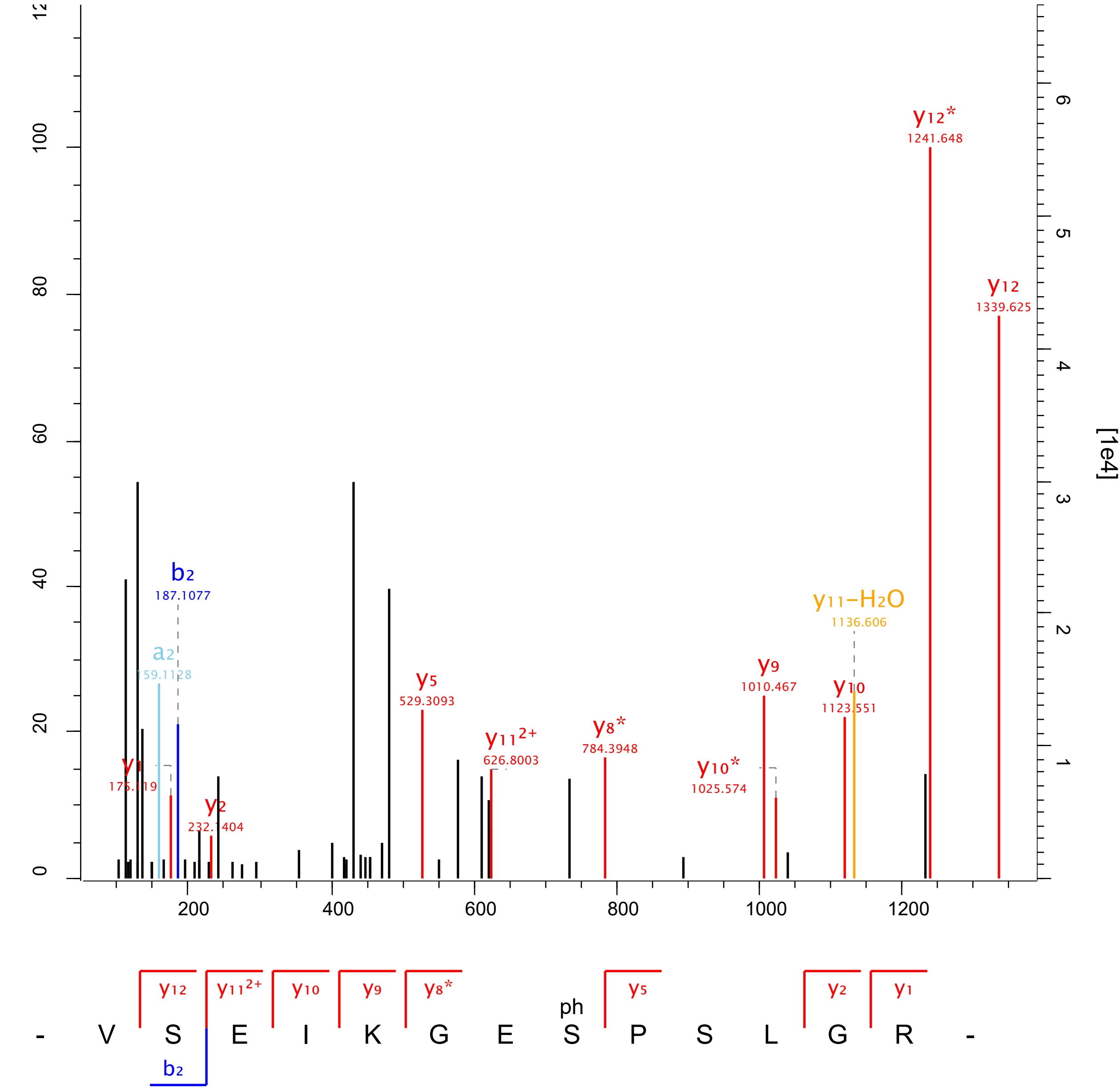

| Raw file | Scan | Method | Score | m/z |
| --- | --- | --- | --- | --- |
| sys_02_2 | 8660 | FTMS; HCD | 57.43 | 578.26 |

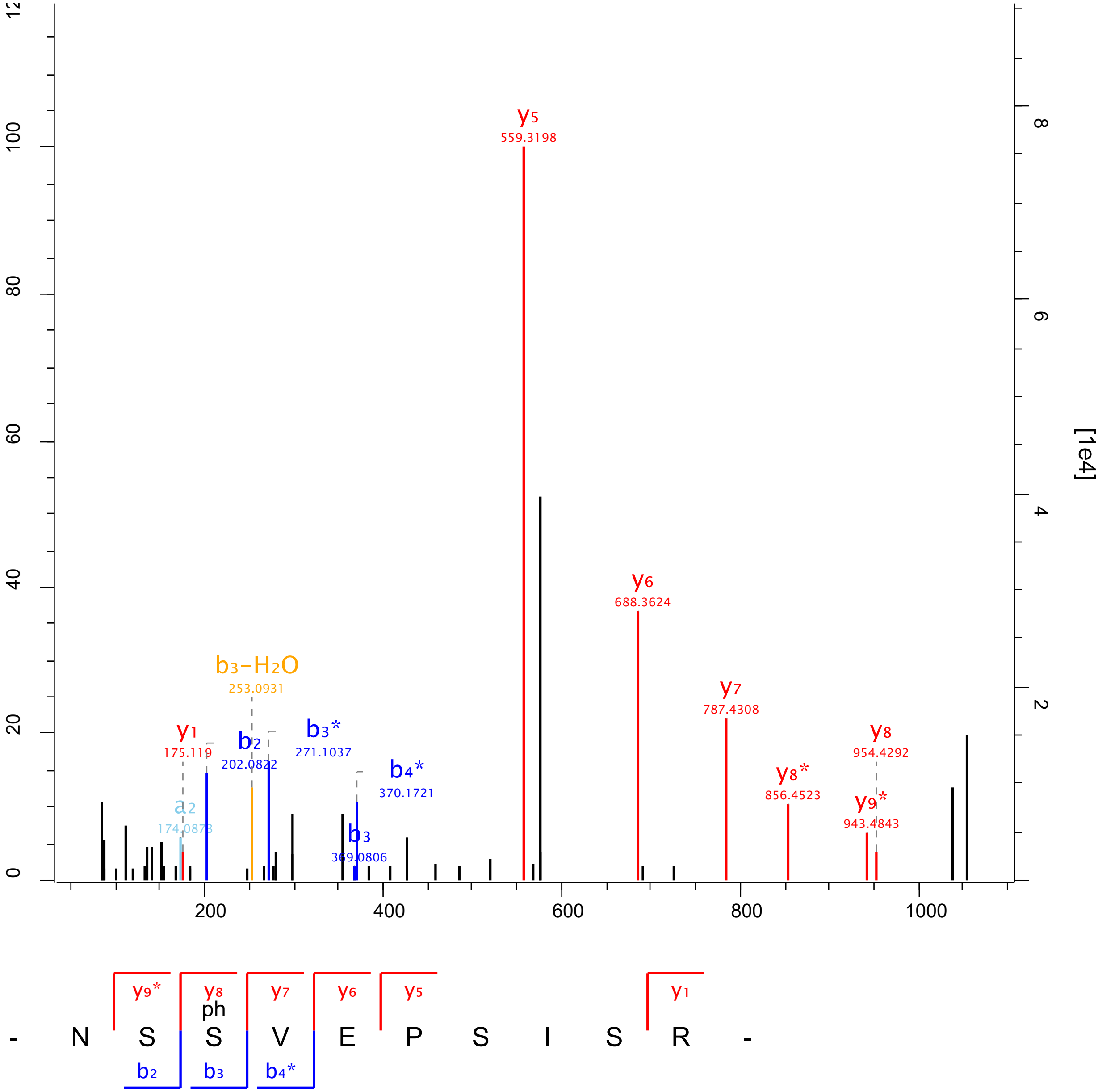

| Raw file | Scan | Method | Score | m/z |
|---|---|---|---|---|
| sys_02_2 | 8851 | FTMS; HCD | 104.84 | 780.35 |

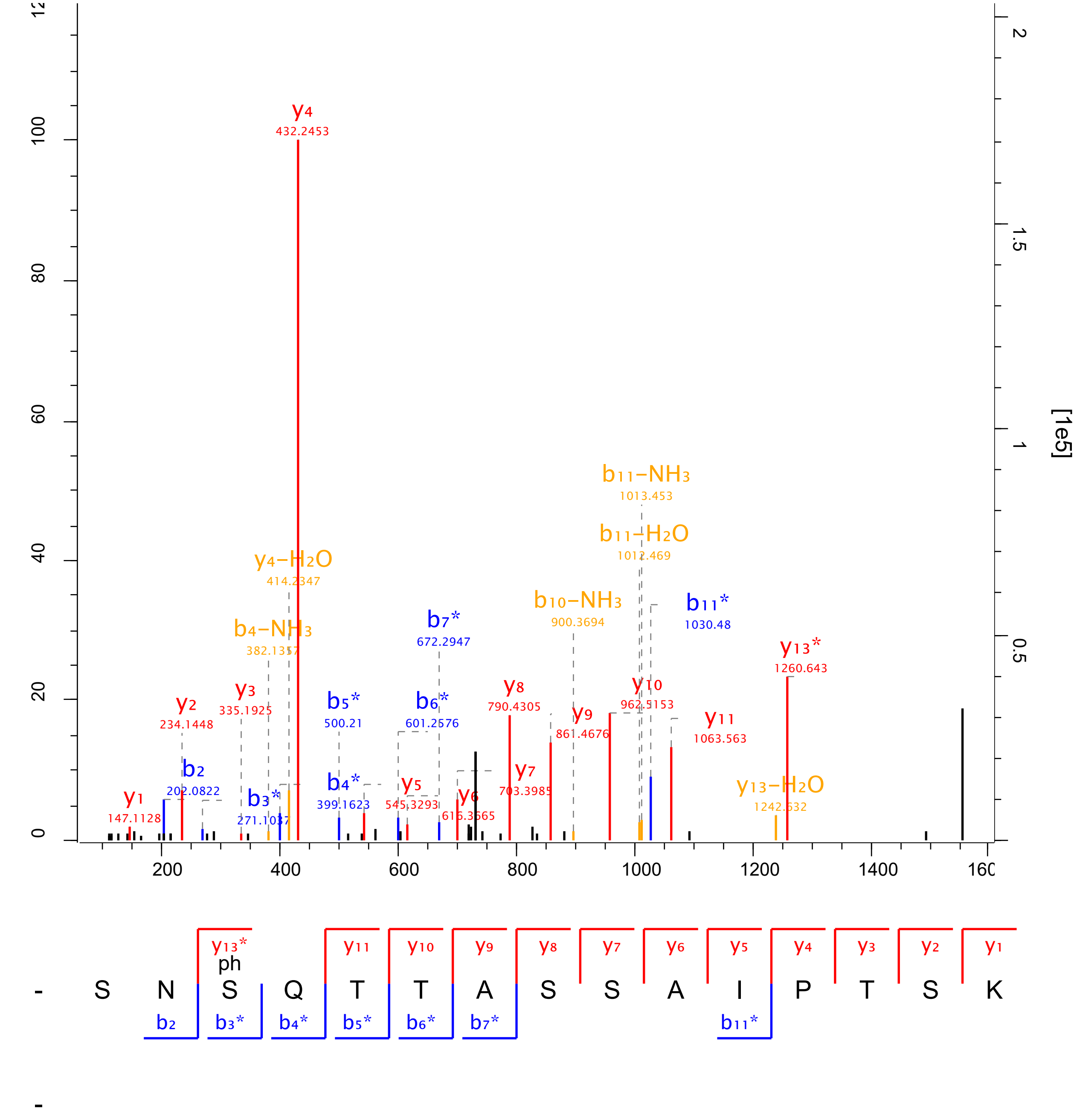

| Raw file | Scan | Method | Score | m/z |
|---|---|---|---|---|
| sys_02_2 | 8890 | FTMS; HCD | 66.27 | 502.22 |

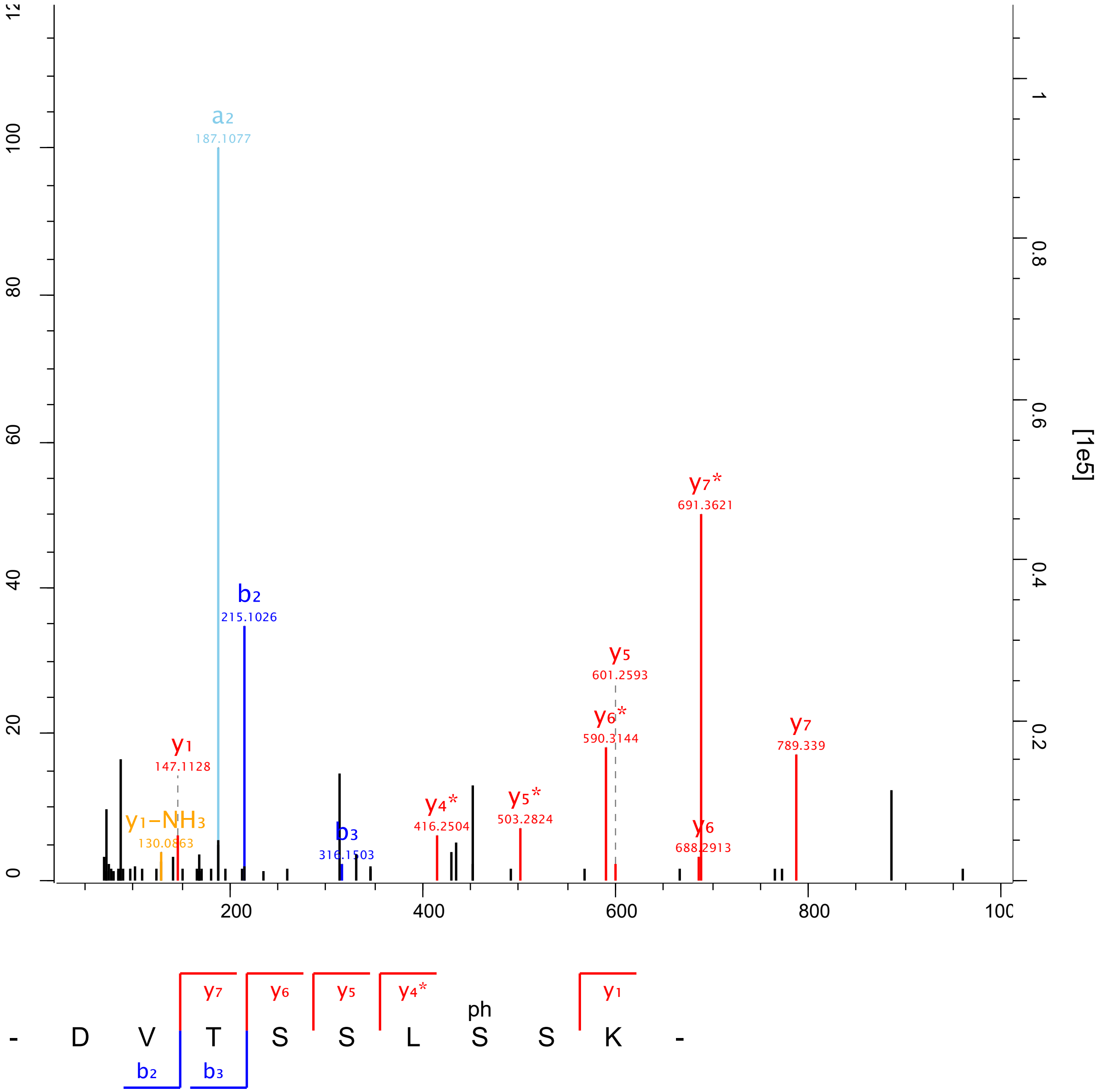

| Raw file | Scan | Method | Score | m/z |
|---|---|---|---|---|
| sys_02_2 | 8902 | FTMS; HCD | 76.23 | 456.73 |

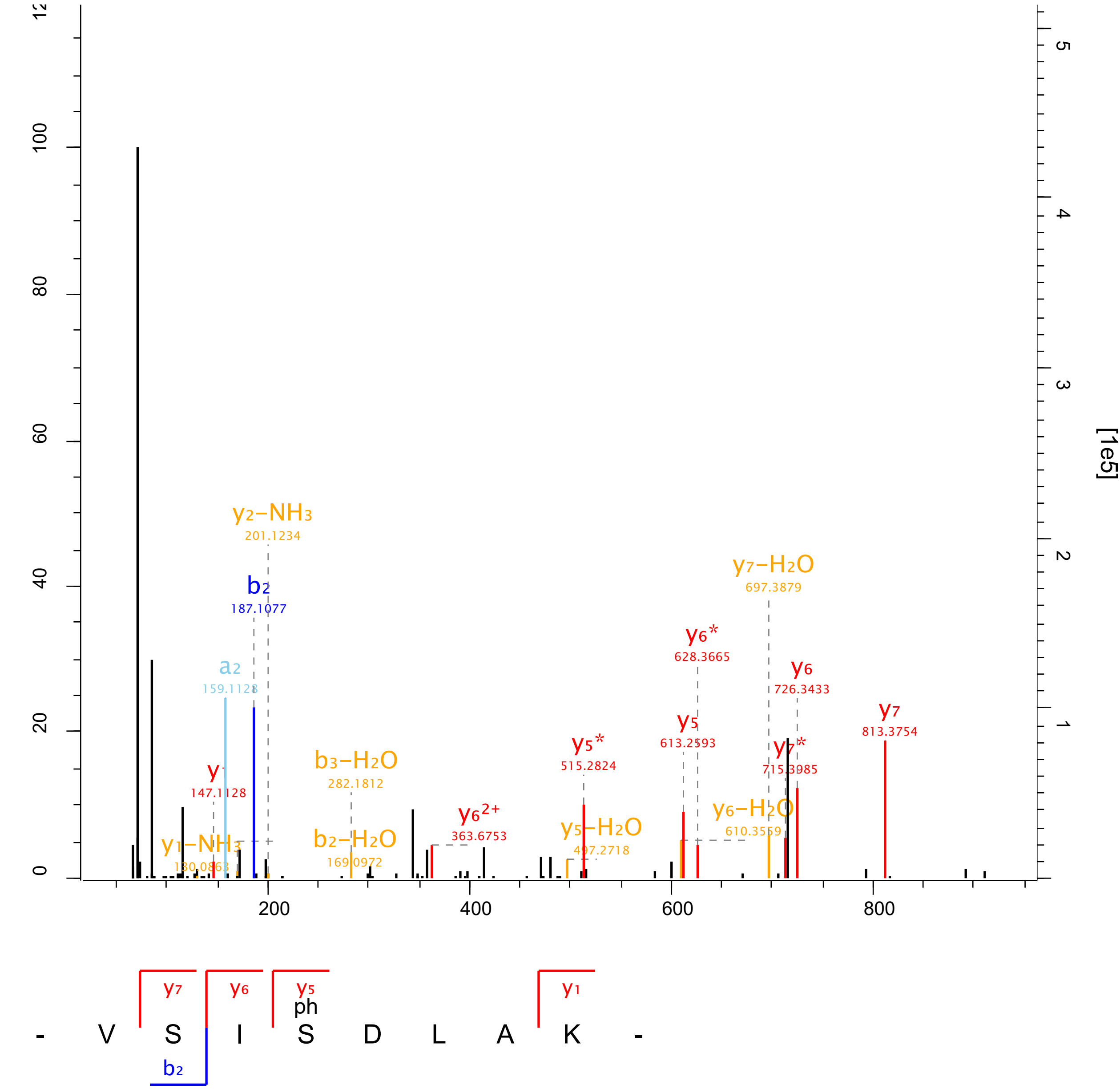

| Raw file | Scan | Method | Score | m/z |
| --- | --- | --- | --- | --- |
| sys_02_2 | 8969 | FTMS; HCD | 67.52 | 549.74 |

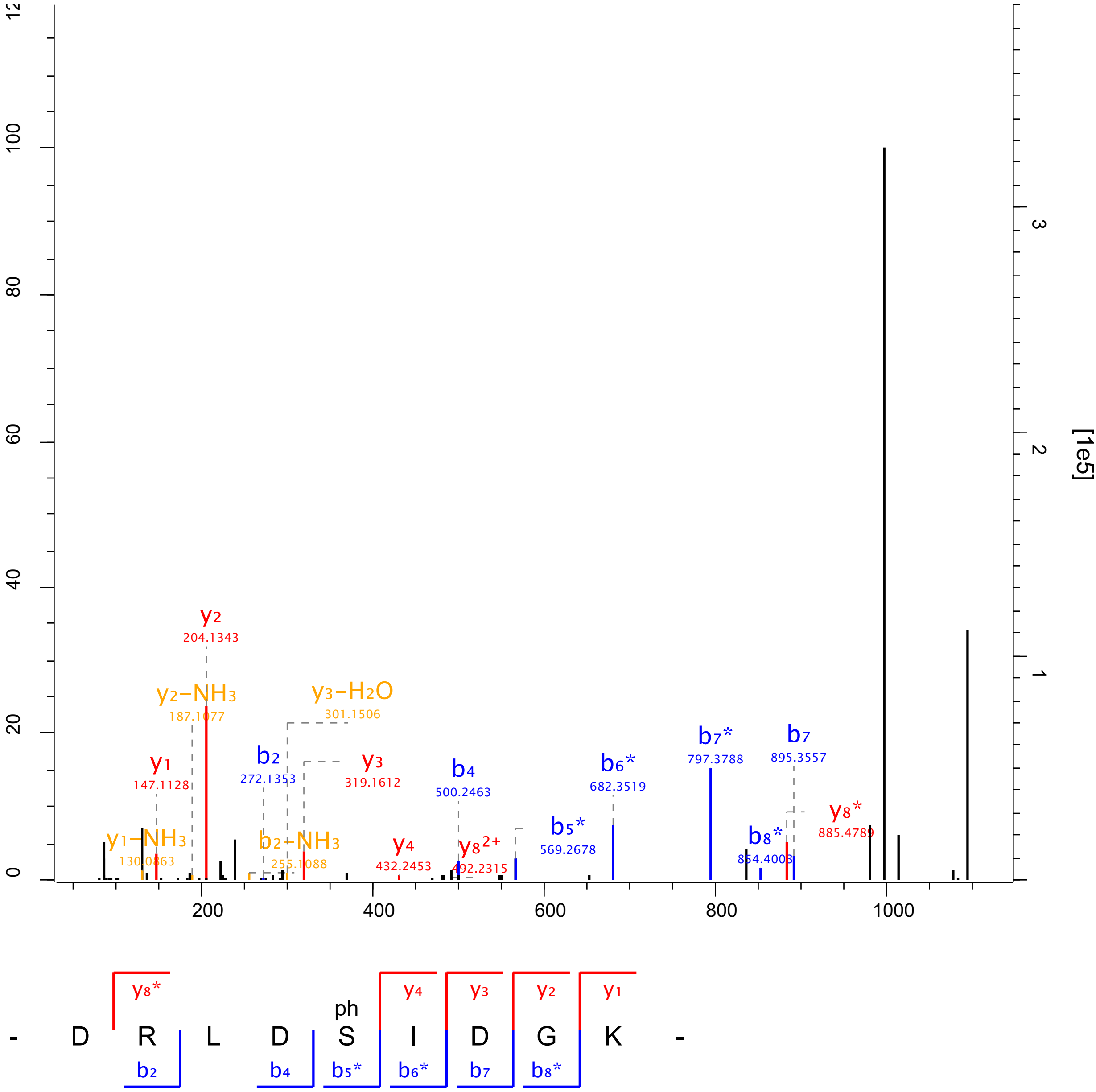

| Raw file | Scan | Method | Score | m/z |
|---|---|---|---|---|
| sys_02_2 | 8976 | FTMS; HCD | 233.79 | 750.81 |

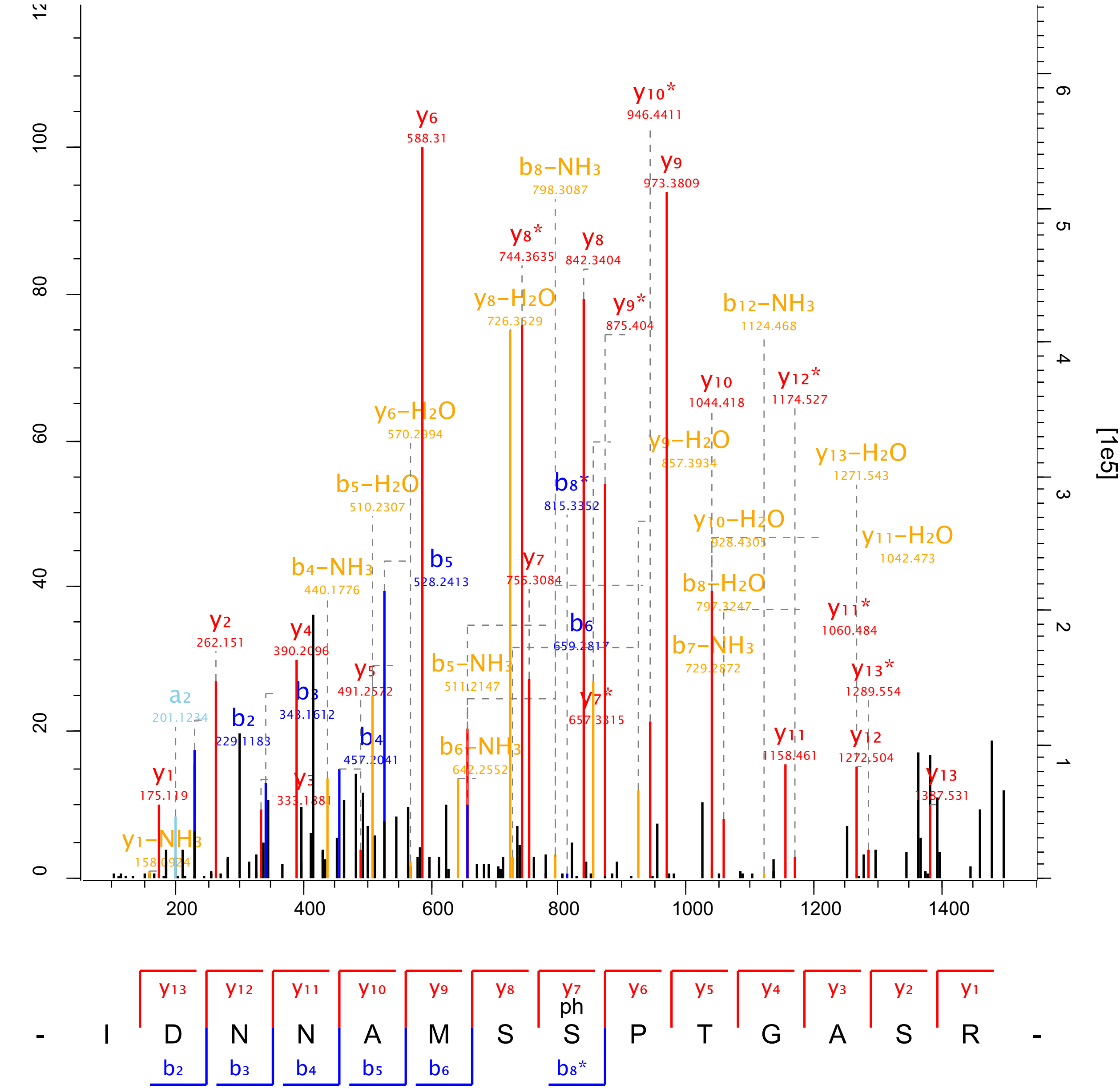

| Raw file | Scan | Method | Score | m/z |
|---|---|---|---|---|
| sys_02_2 | 8995 | FTMS; HCD | 123.72 | 688.28 |

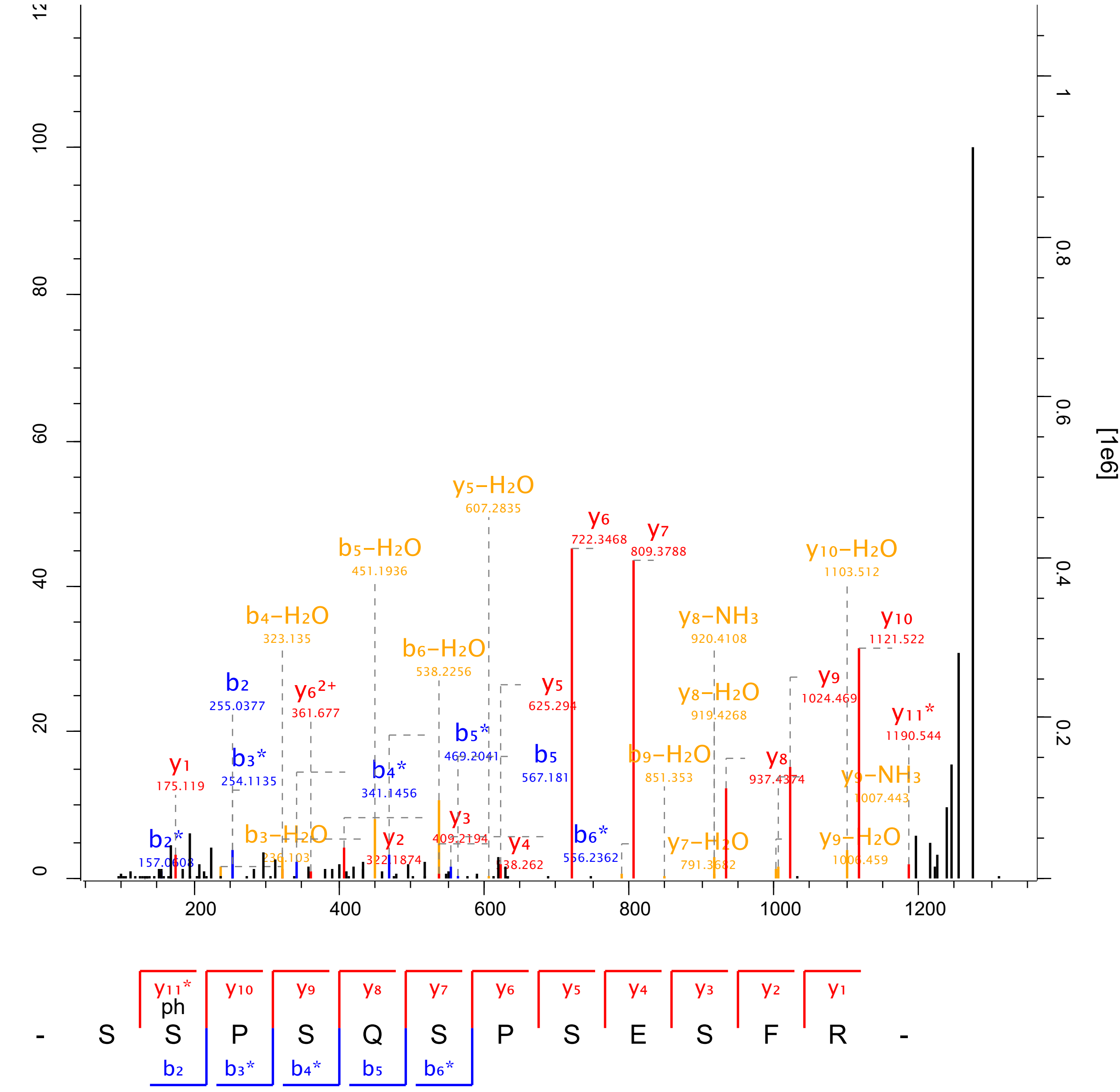

| Raw file | Scan | Method | Score | m/z |
|---|---|---|---|---|
| sys_02_2 | 9067 | FTMS; HCD | 47.19 | 493.72 |

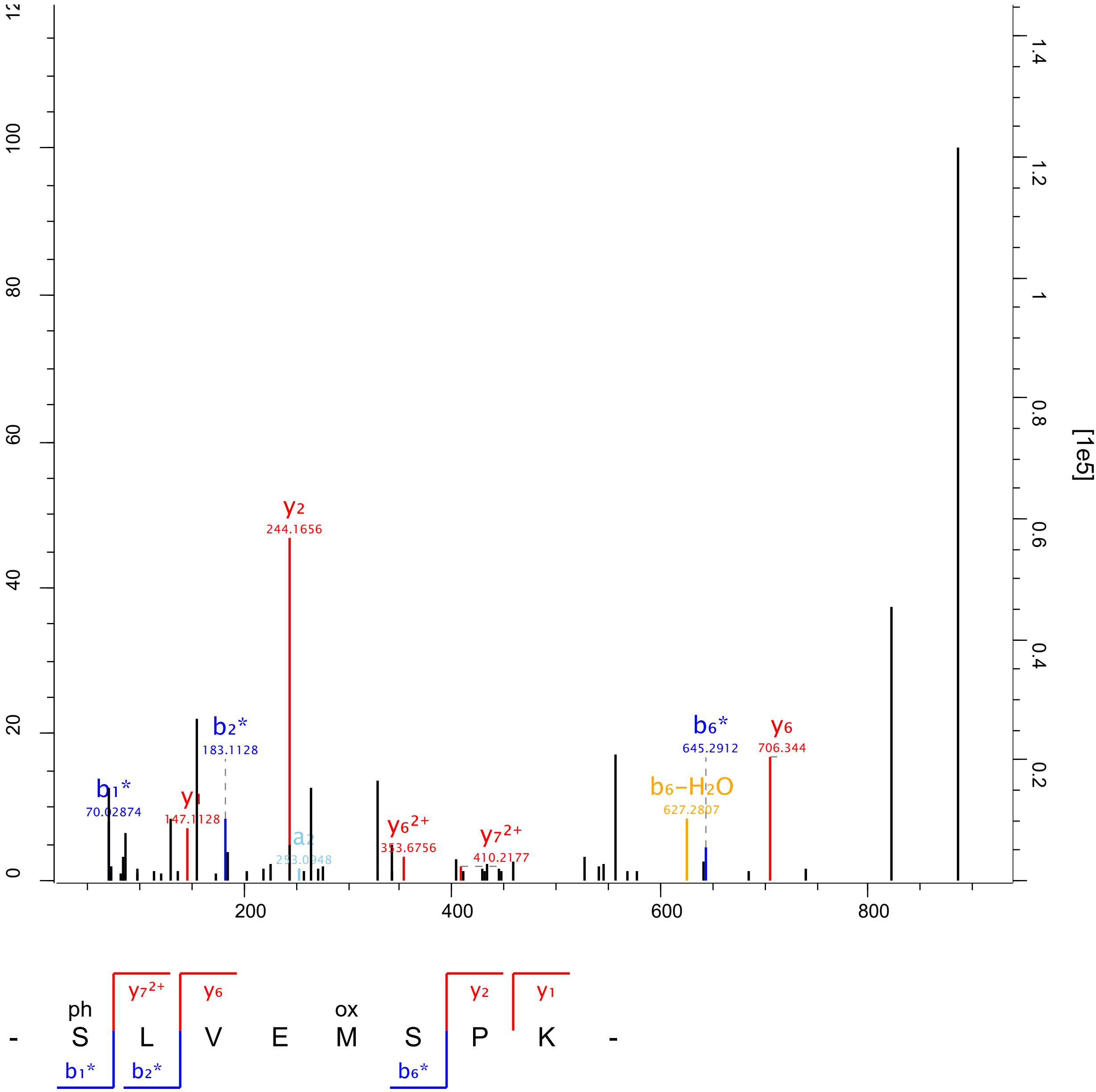

| Raw file | Scan | Method | Score | m/z |
|---|---|---|---|---|
| sys_02_2 | 9085 | FTMS; HCD | 49.59 | 731.77 |

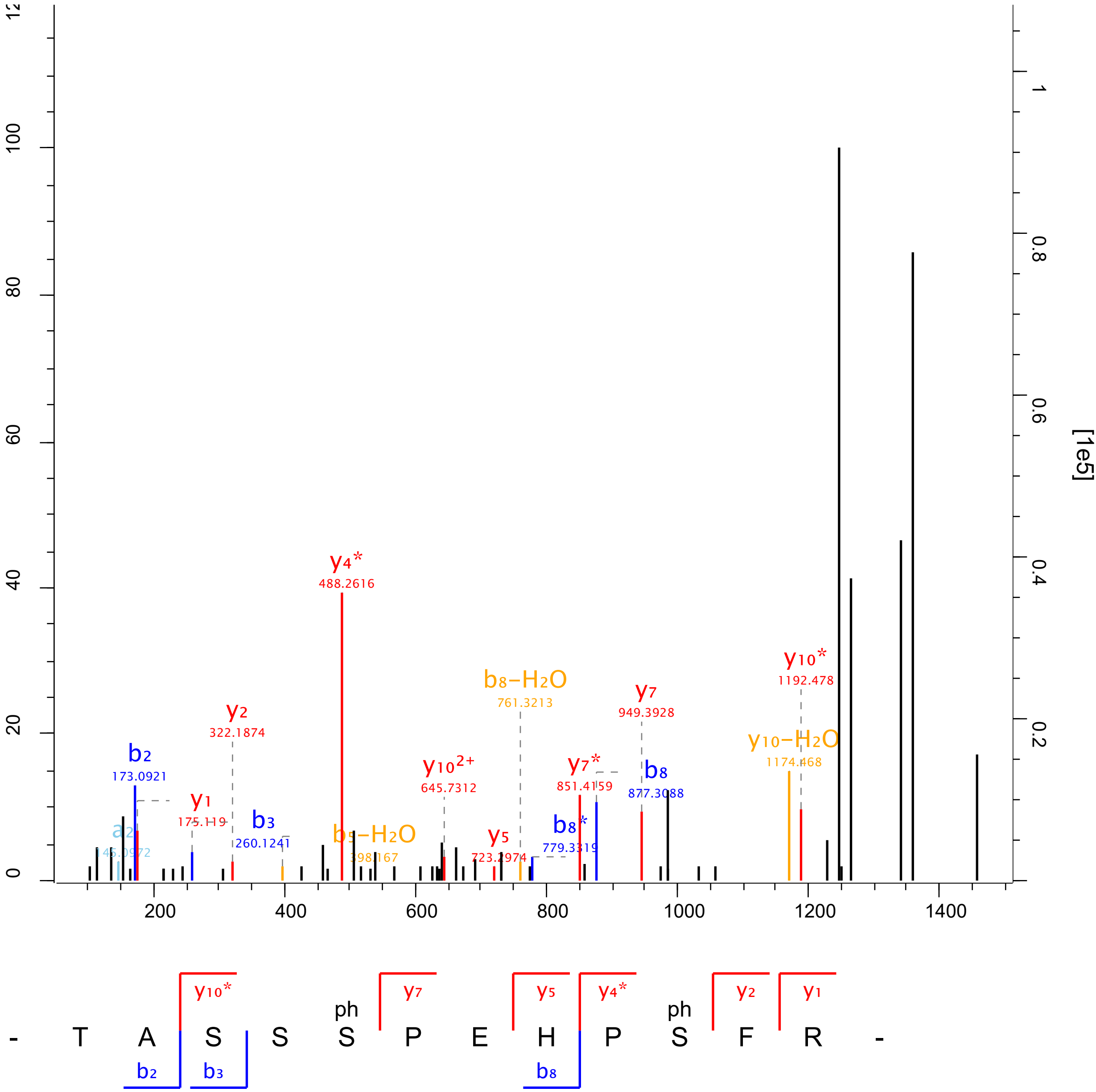

| Raw file | Scan | Method | Score | m/z |
|---|---|---|---|---|
| sys_02_2 | 9317 | FTMS; HCD | 51.73 | 415.54 |

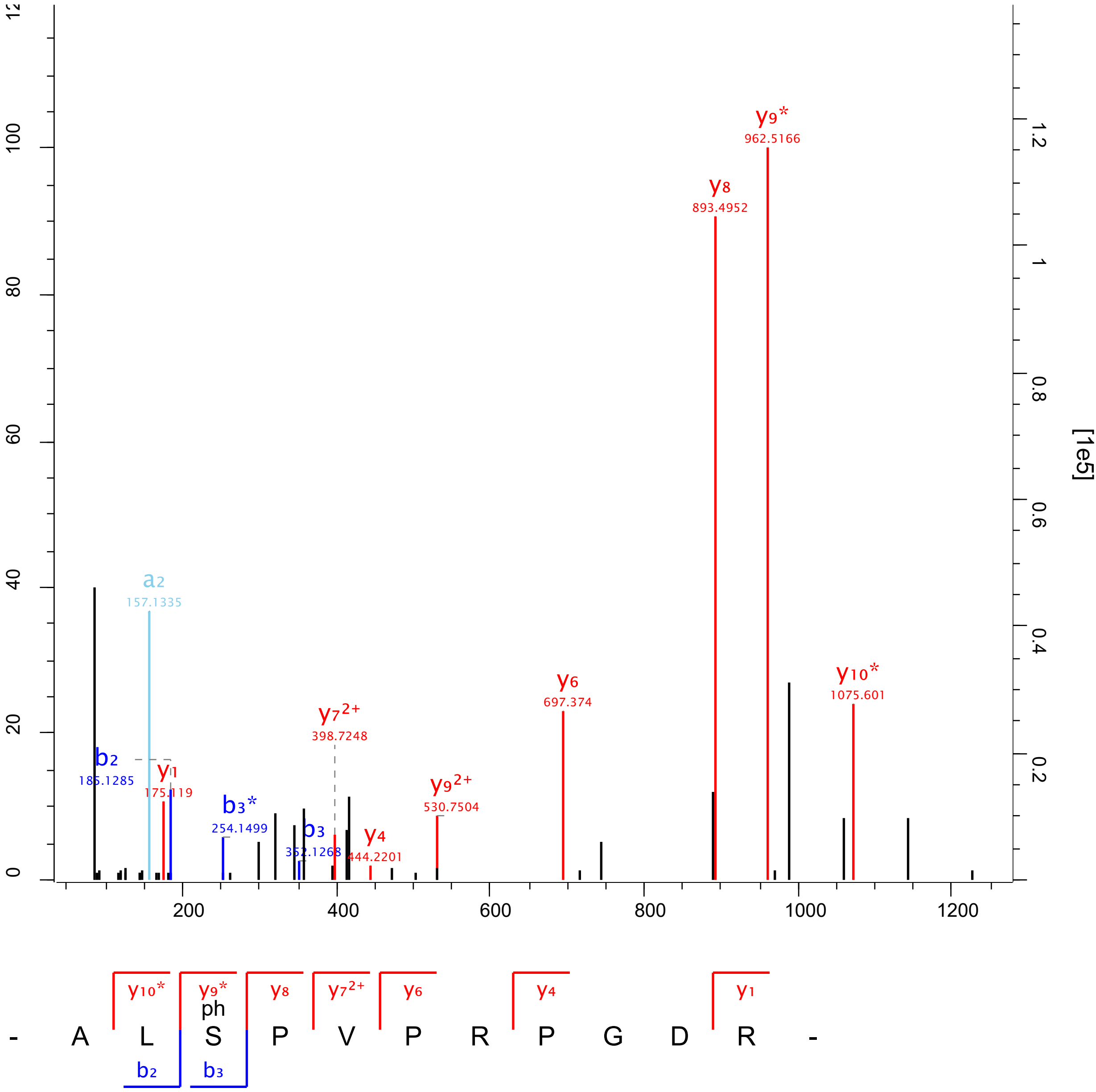

| Raw file | Scan | Method | Score | m/z |
|---|---|---|---|---|
| sys_02_2 | 9349 | FTMS; HCD | 62.09 | 655.29 |

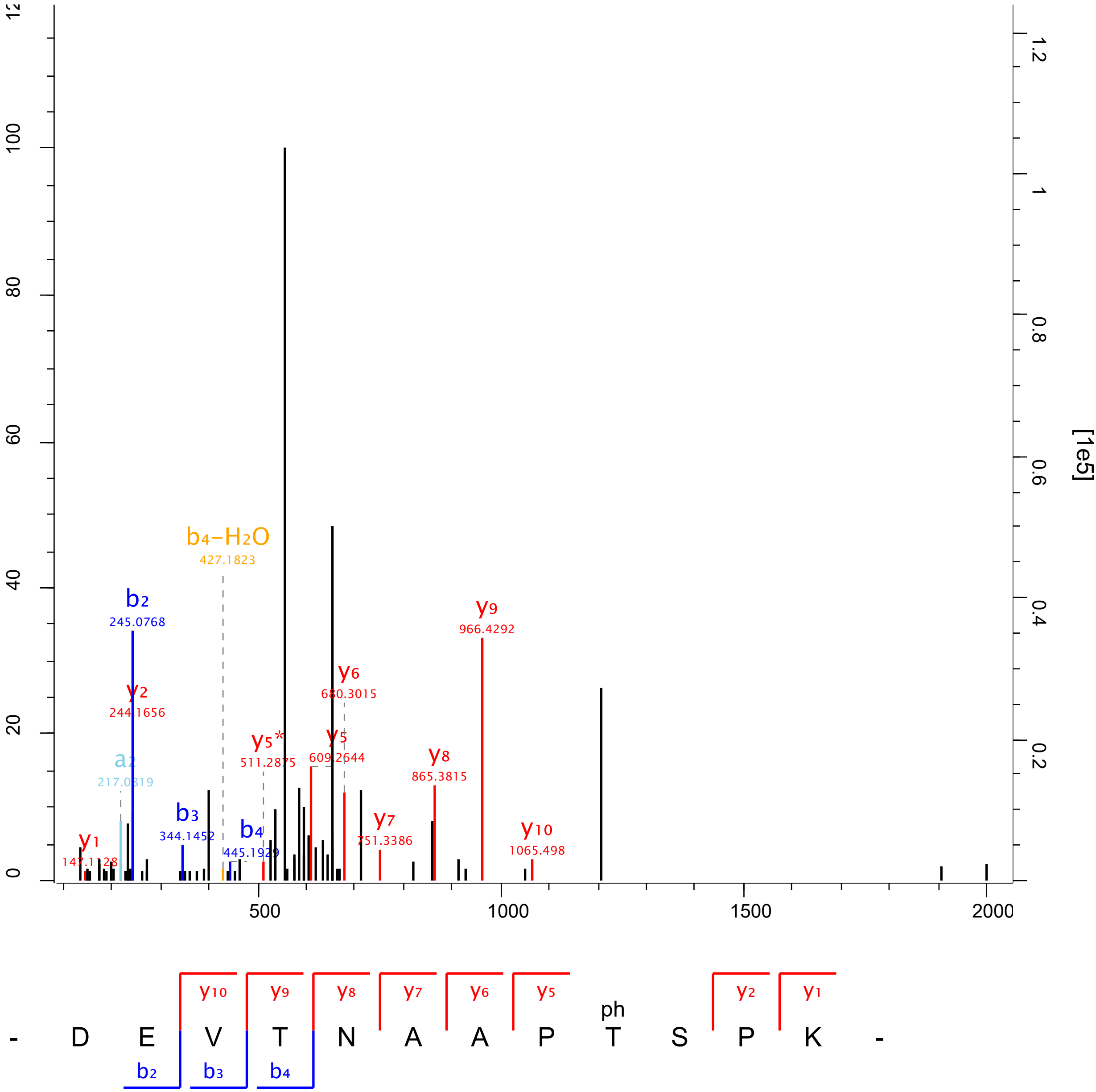

| Raw file | Scan | Method | Score | m/z |
|---|---|---|---|---|
| sys_02_2 | 9406 | FTMS; HCD | 197.36 | 570.75 |

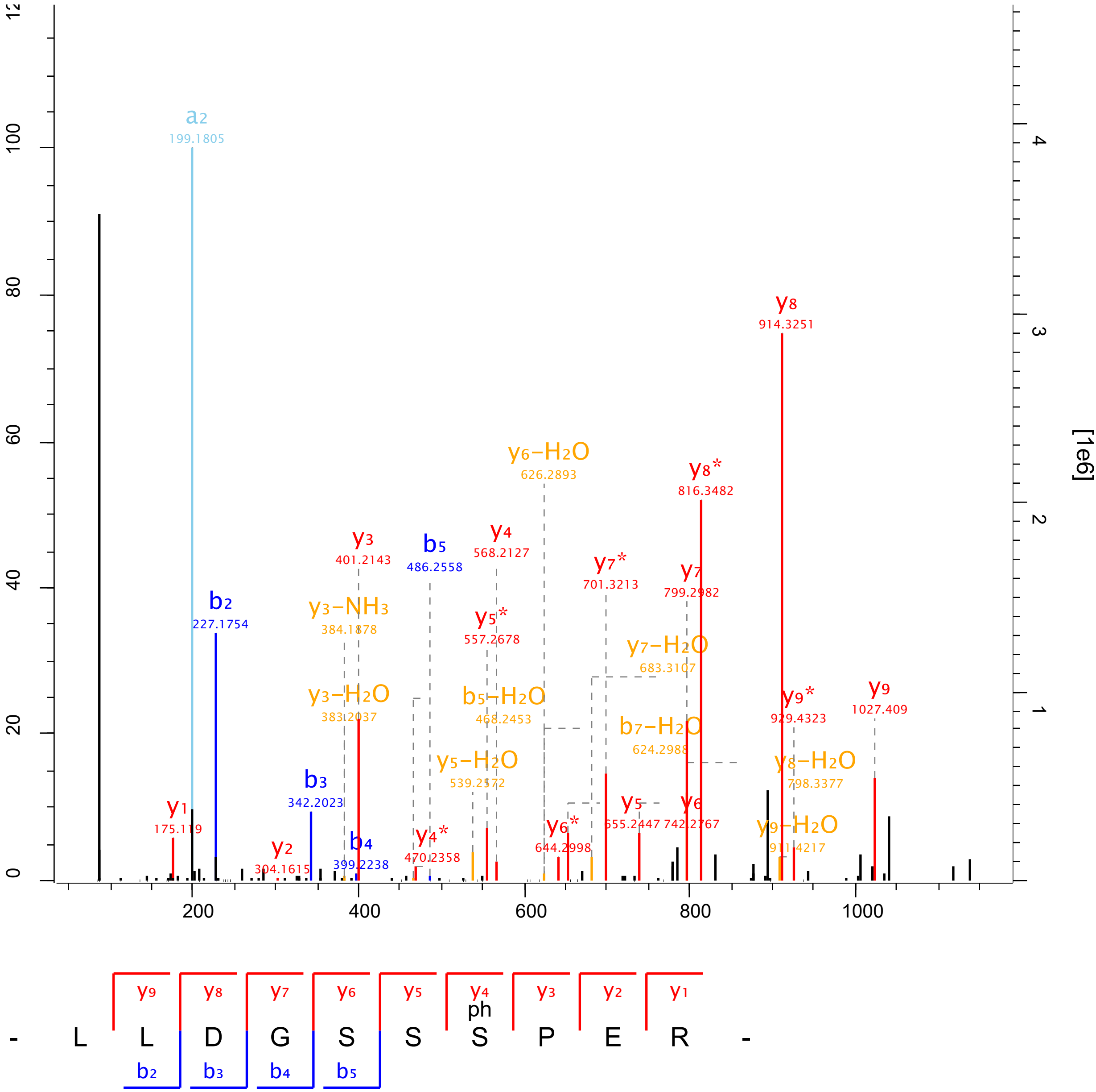

| Raw file | Scan | Method | Score | m/z |
|---|---|---|---|---|
| sys_02_2 | 9492 | FTMS; HCD | 41.54 | 615.26 |

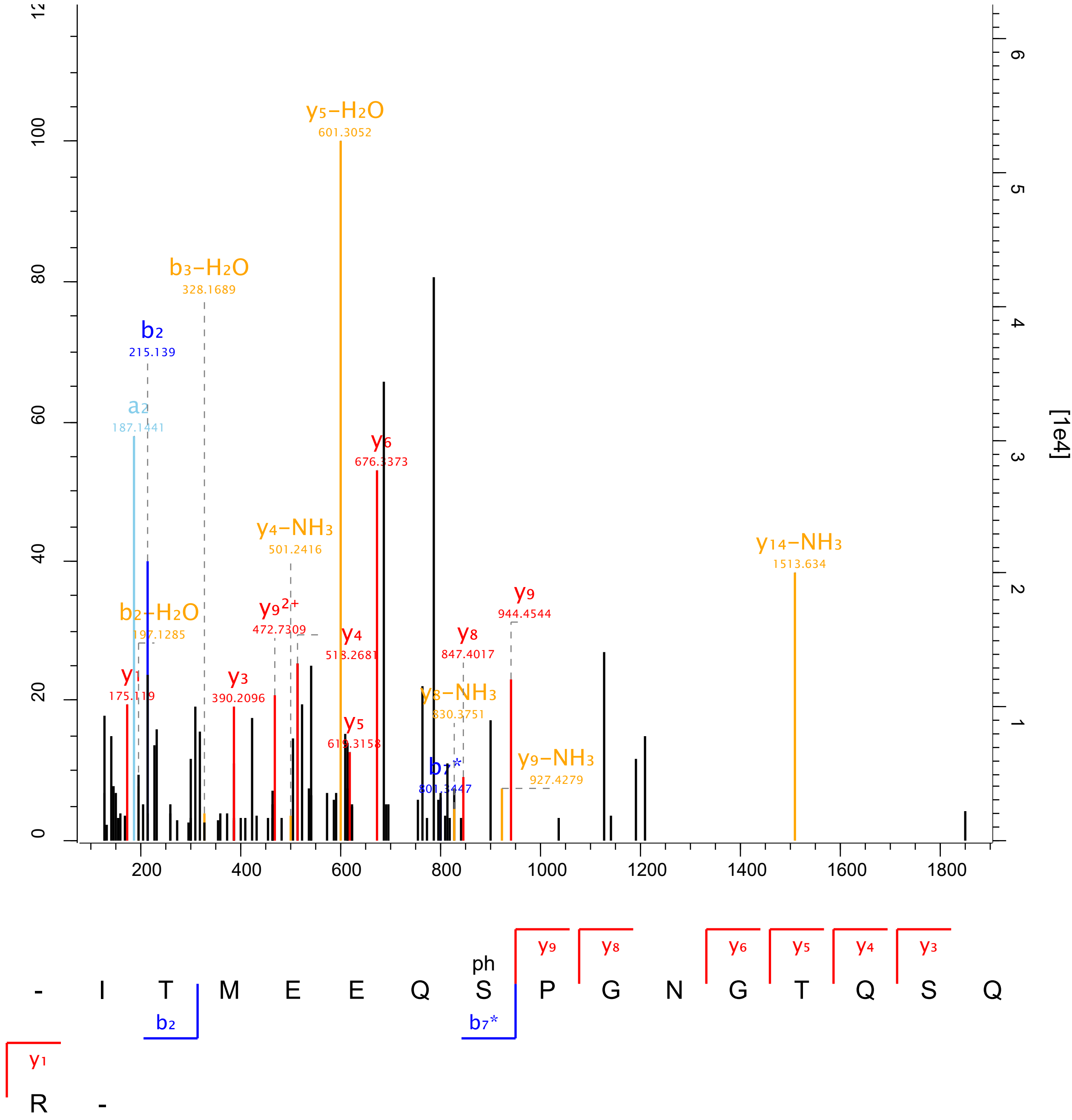

| Raw file | Scan | Method | Score | m/z |
| --- | --- | --- | --- | --- |
| sys_02_2 | 9554 | FTMS; HCD | 106.58 | 581.28 |

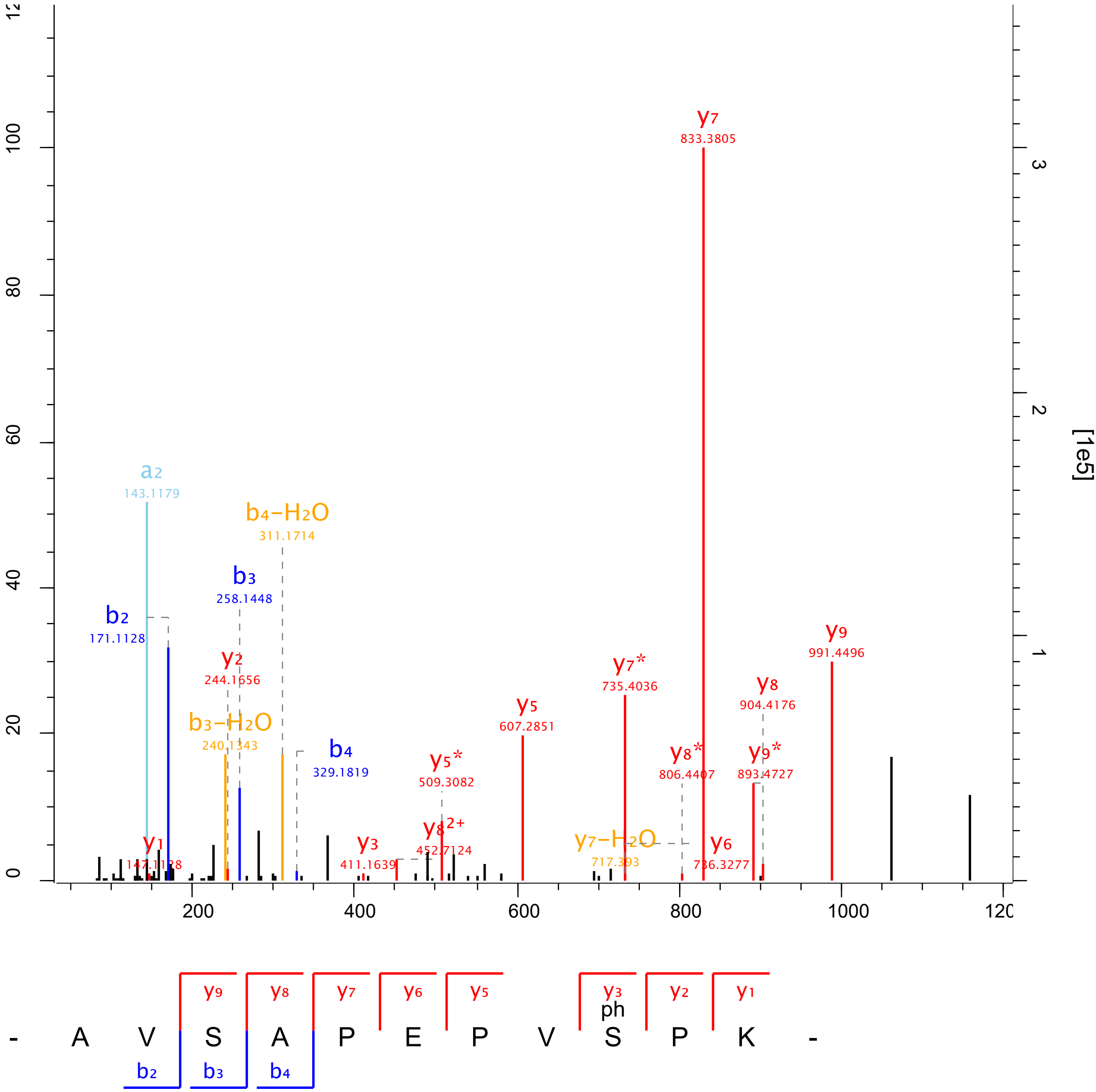

| Raw file | Scan | Method | Score | m/z |
|---|---|---|---|---|
| sys_02_2 | 9613 | FTMS; HCD | 170.38 | 786.34 |

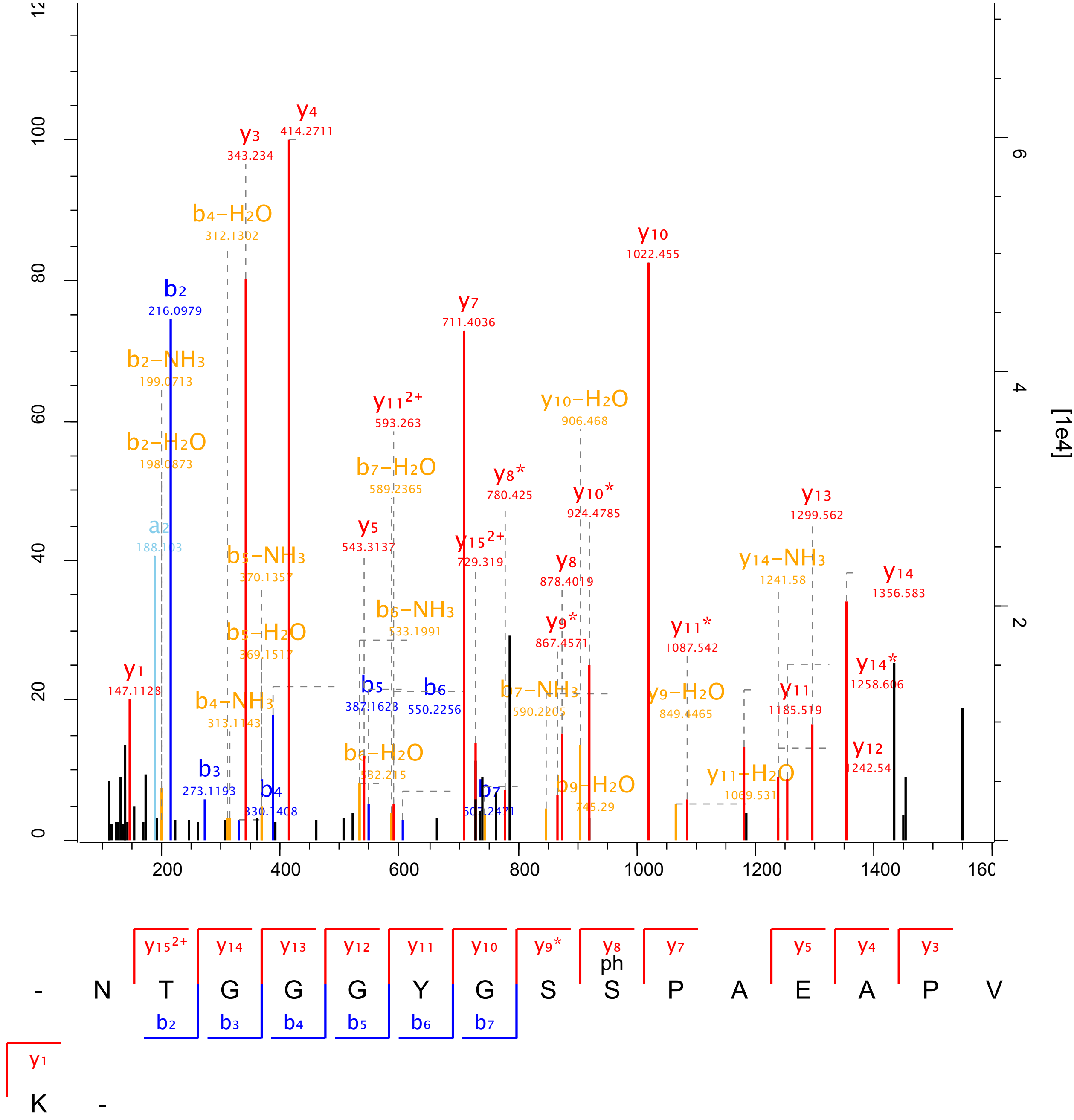

| Raw file | Scan | Method | Score | m/z |
| --- | --- | --- | --- | --- |
| sys_02_2 | 9679 | FTMS; HCD | 198.55 | 585.76 |

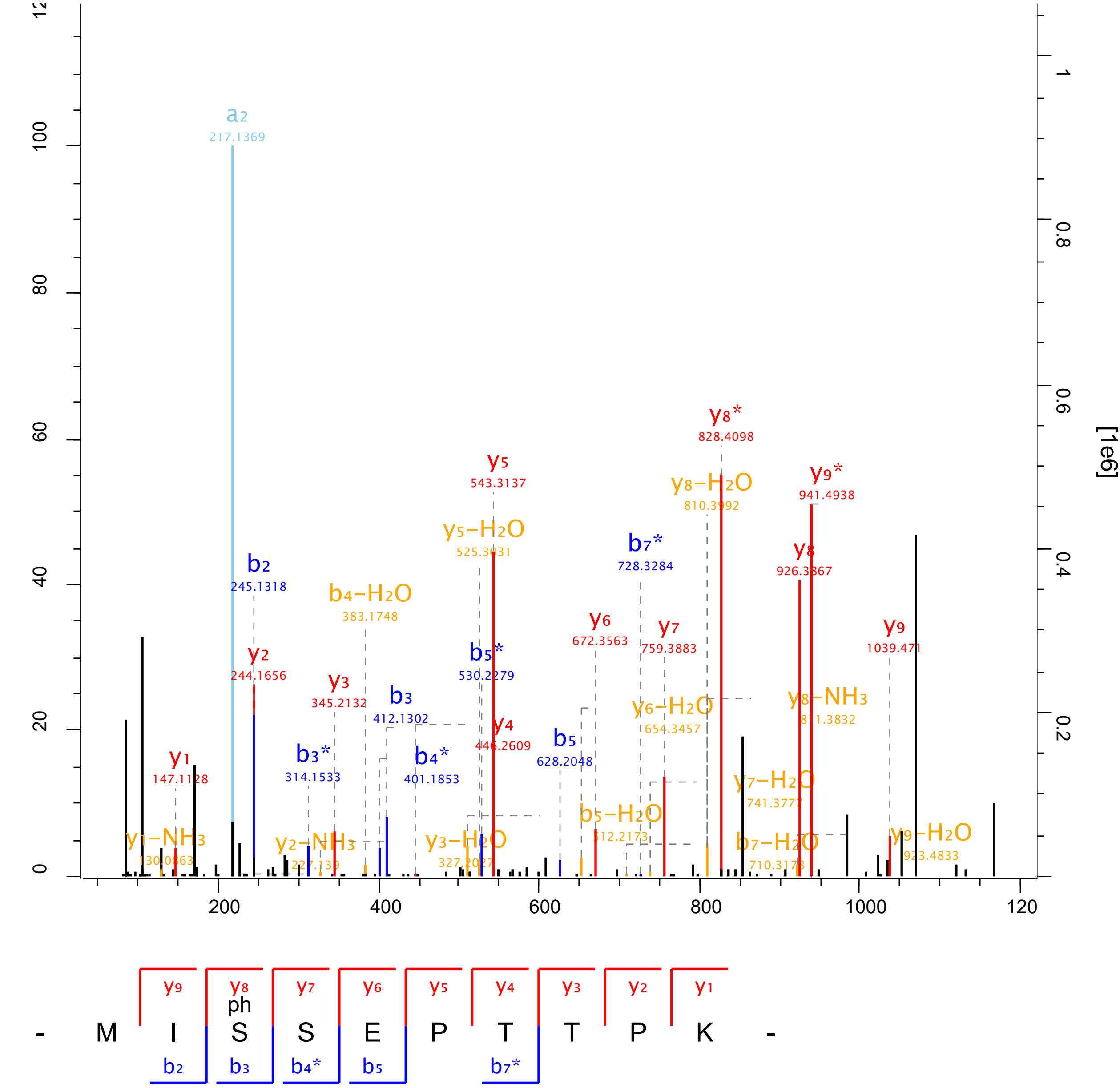

| Raw file | Scan | Method | Score | m/z |
|---|---|---|---|---|
| sys_02_2 | 9812 | FTMS; HCD | 94.49 | 449.57 |

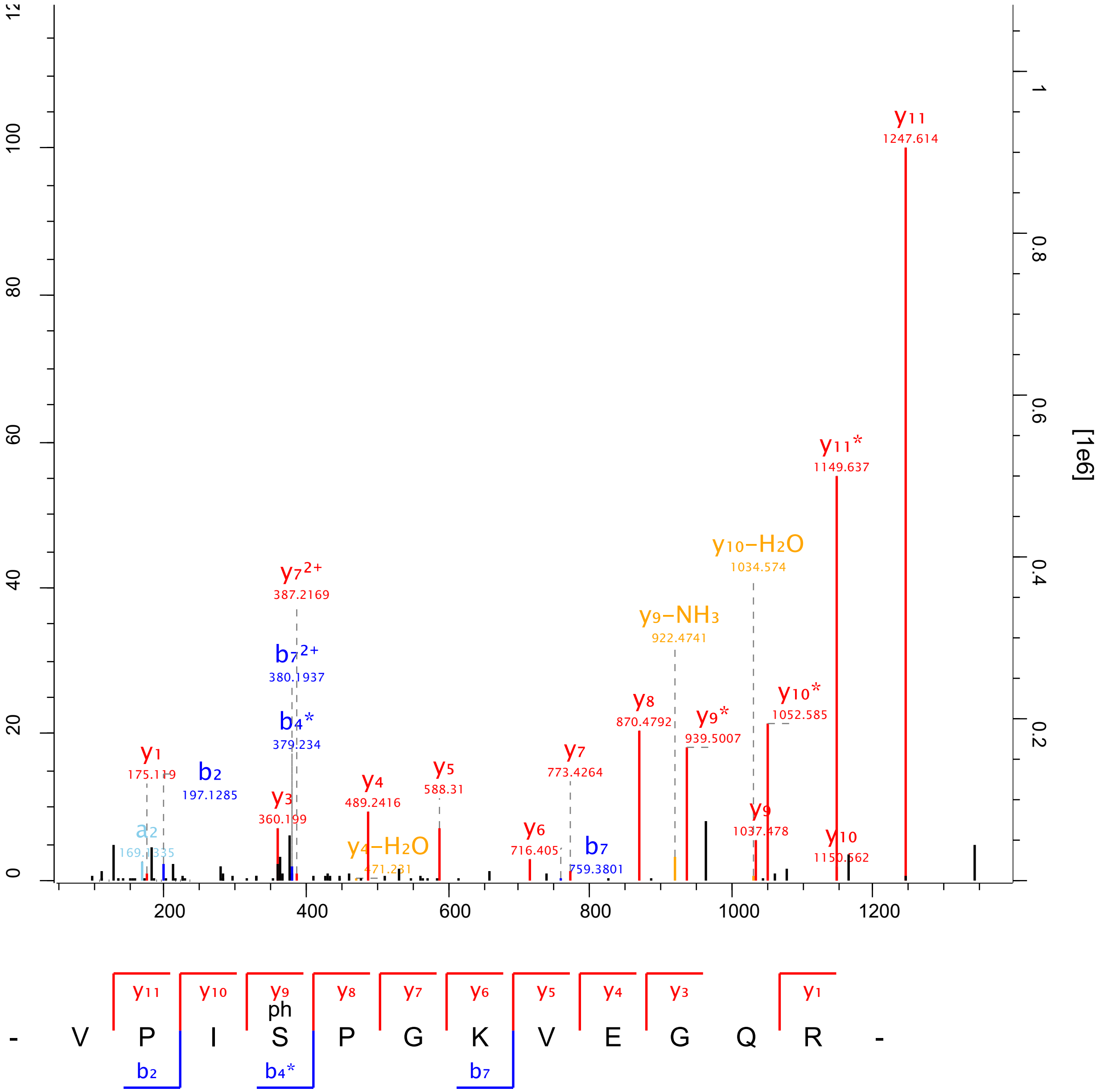

| Raw file | Scan | Method | Score | m/z |
|---|---|---|---|---|
| sys_02_2 | 9882 | FTMS; HCD | 85.54 | 692.3 |

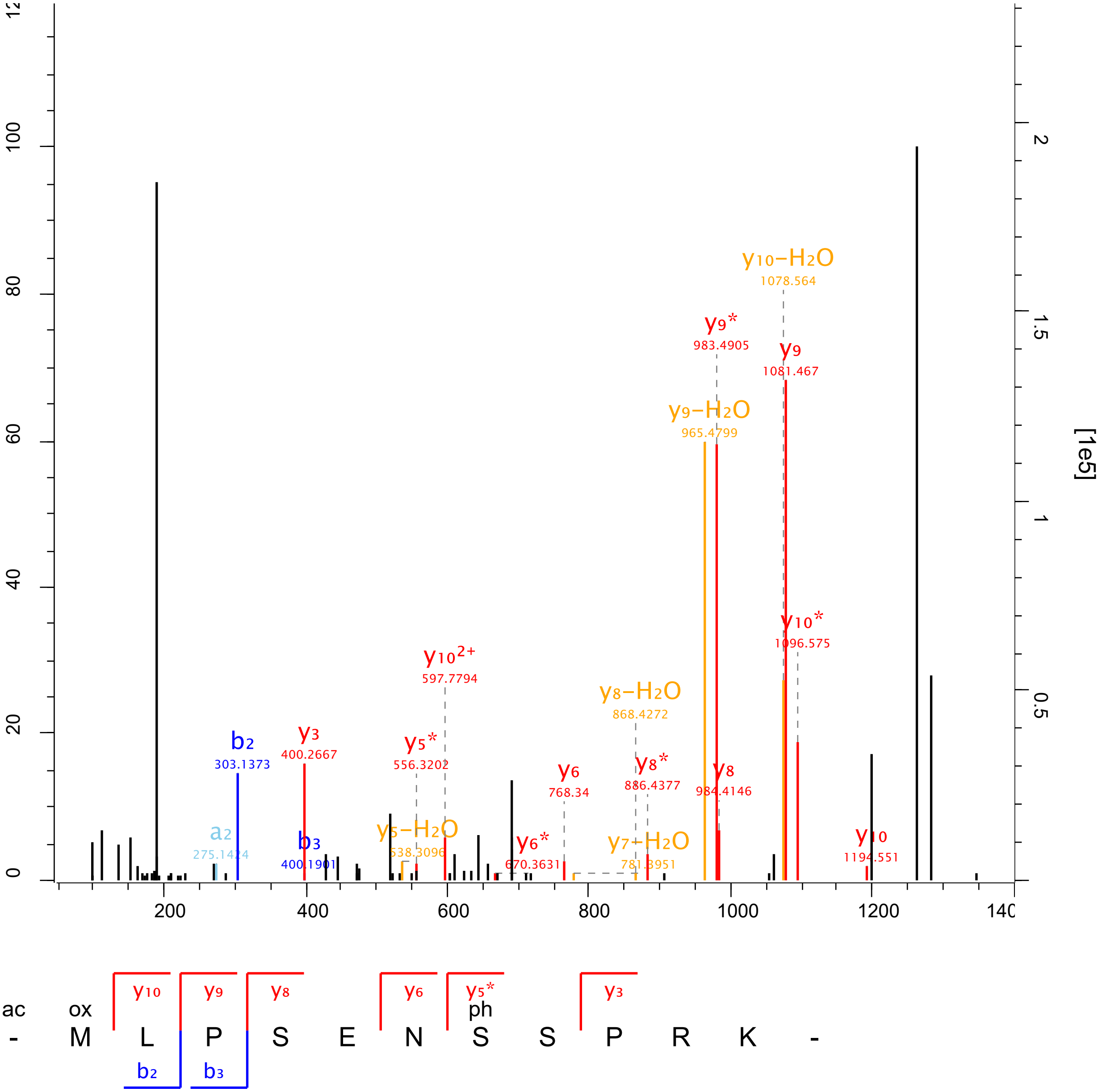

| Raw file | Scan | Method | Score | m/z |
|---|---|---|---|---|
| sys_02_2 | 9892 | FTMS; HCD | 100.88 | 465.72 |

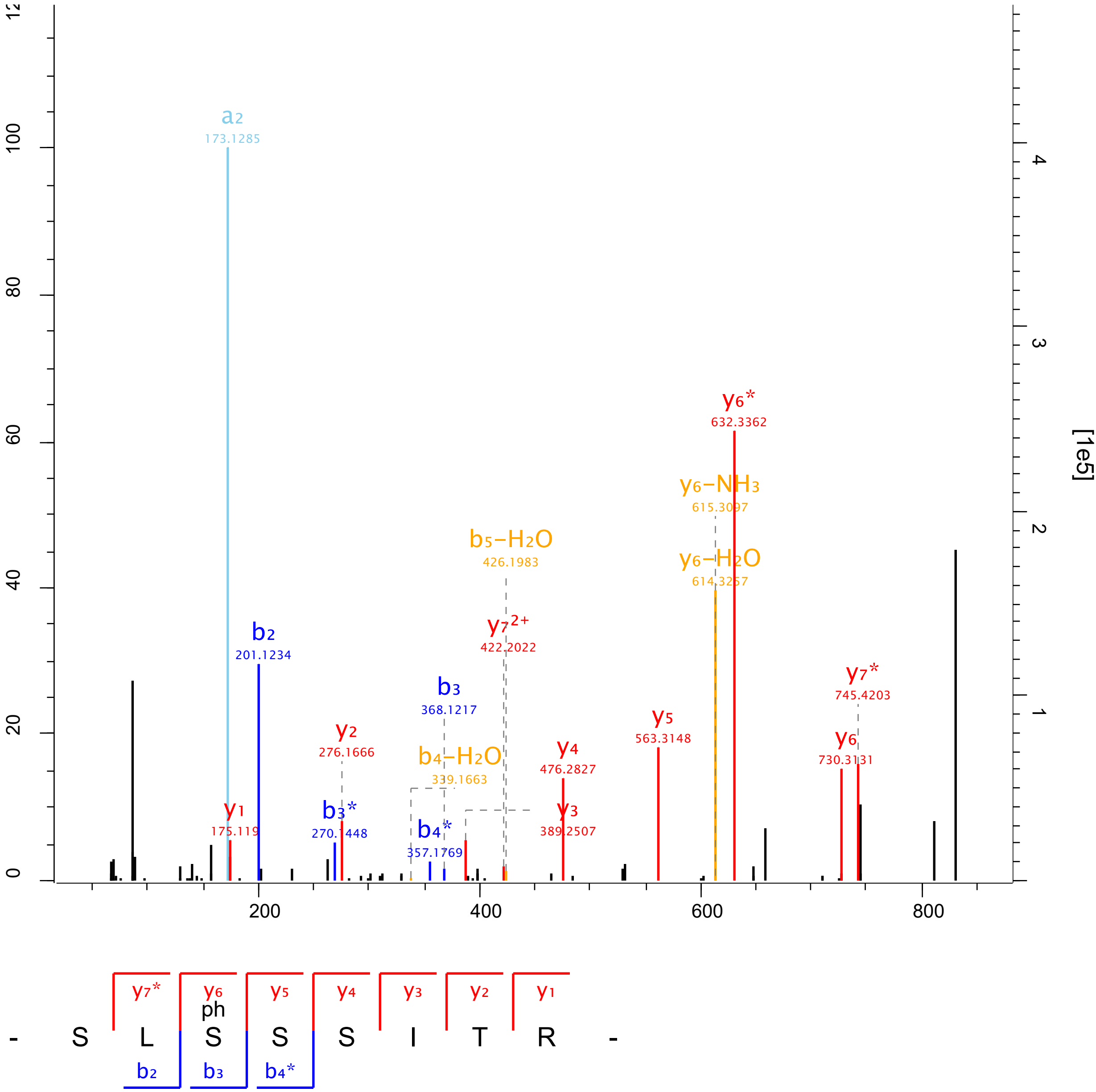

| Raw file | Scan | Method | Score | m/z |
|---|---|---|---|---|
| sys_02_3 | 10958 | FTMS; HCD | 113.47 | 677.31 |

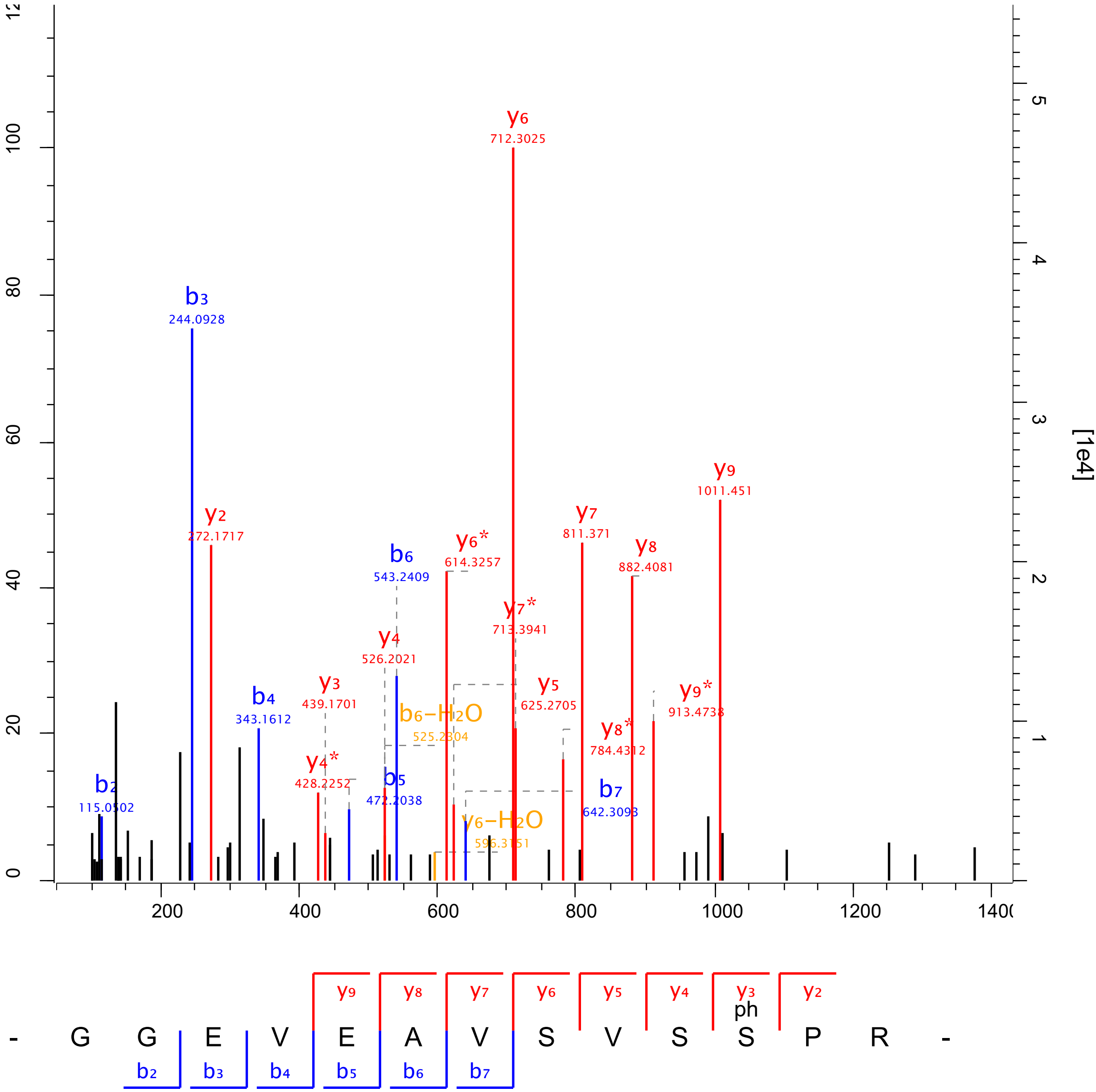

| Raw file | Scan | Method | Score | m/z |
|---|---|---|---|---|
| sys_02_3 | 14871 | FTMS; HCD | 47.77 | 707.33 |

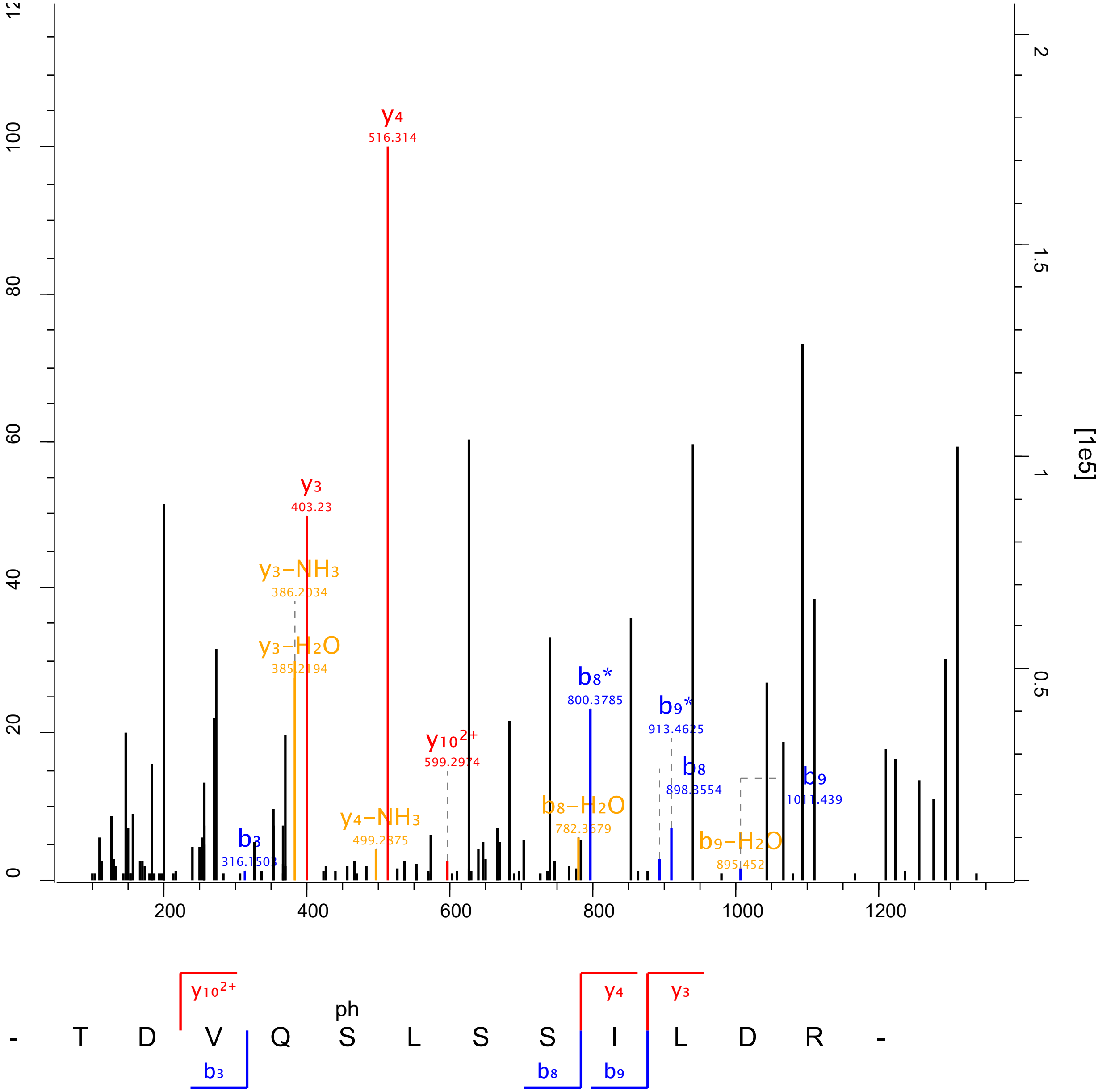

| Raw file | Scan | Method | Score | m/z |
|---|---|---|---|---|
| sys_02_3 | 18780 | FTMS; HCD | 43.3 | 567.25 |

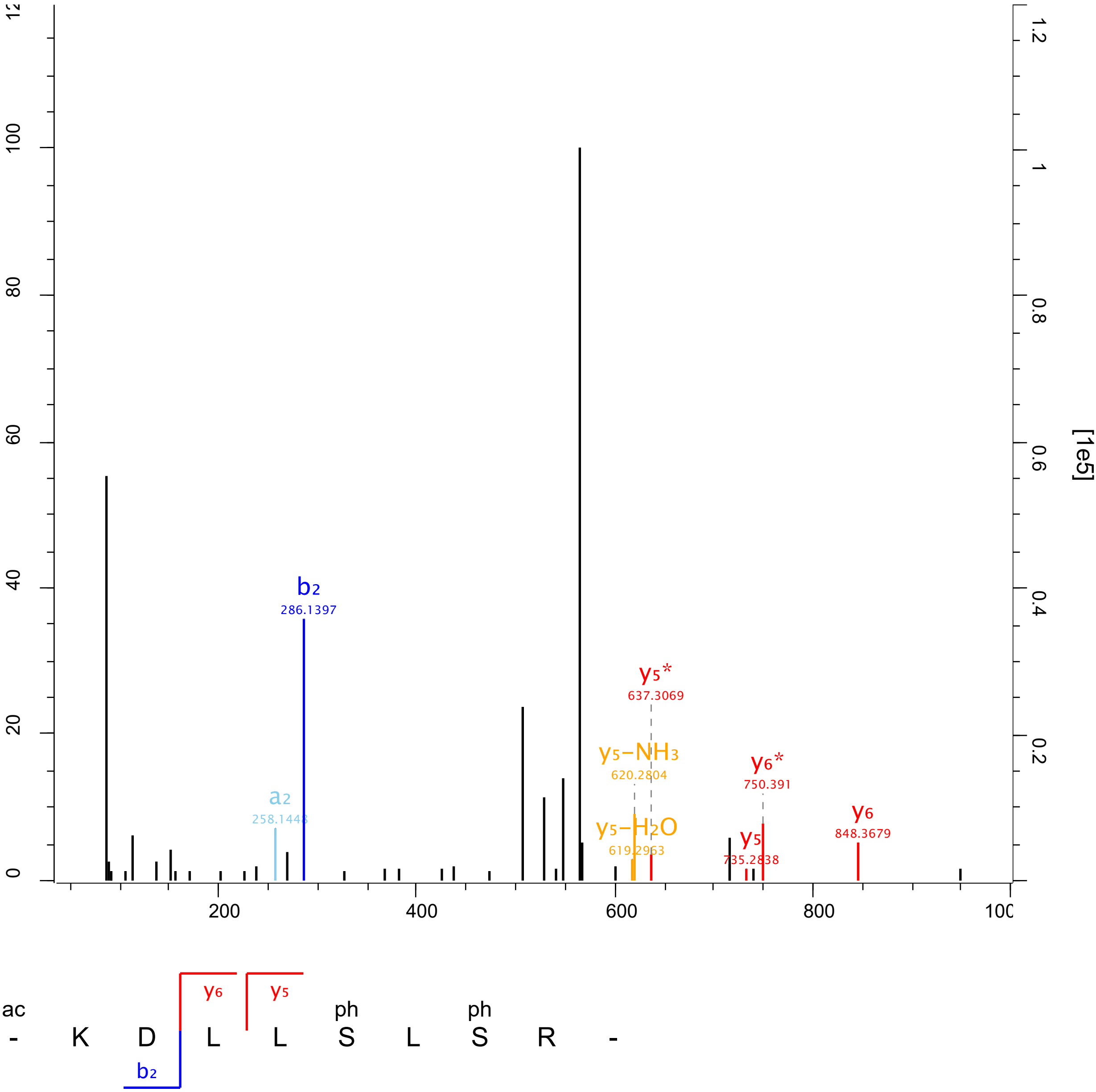

| Raw file | Scan | Method | Score | m/z |
|---|---|---|---|---|
| sys_02_3 | 19509 | FTMS; HCD | 45.43 | 548.24 |

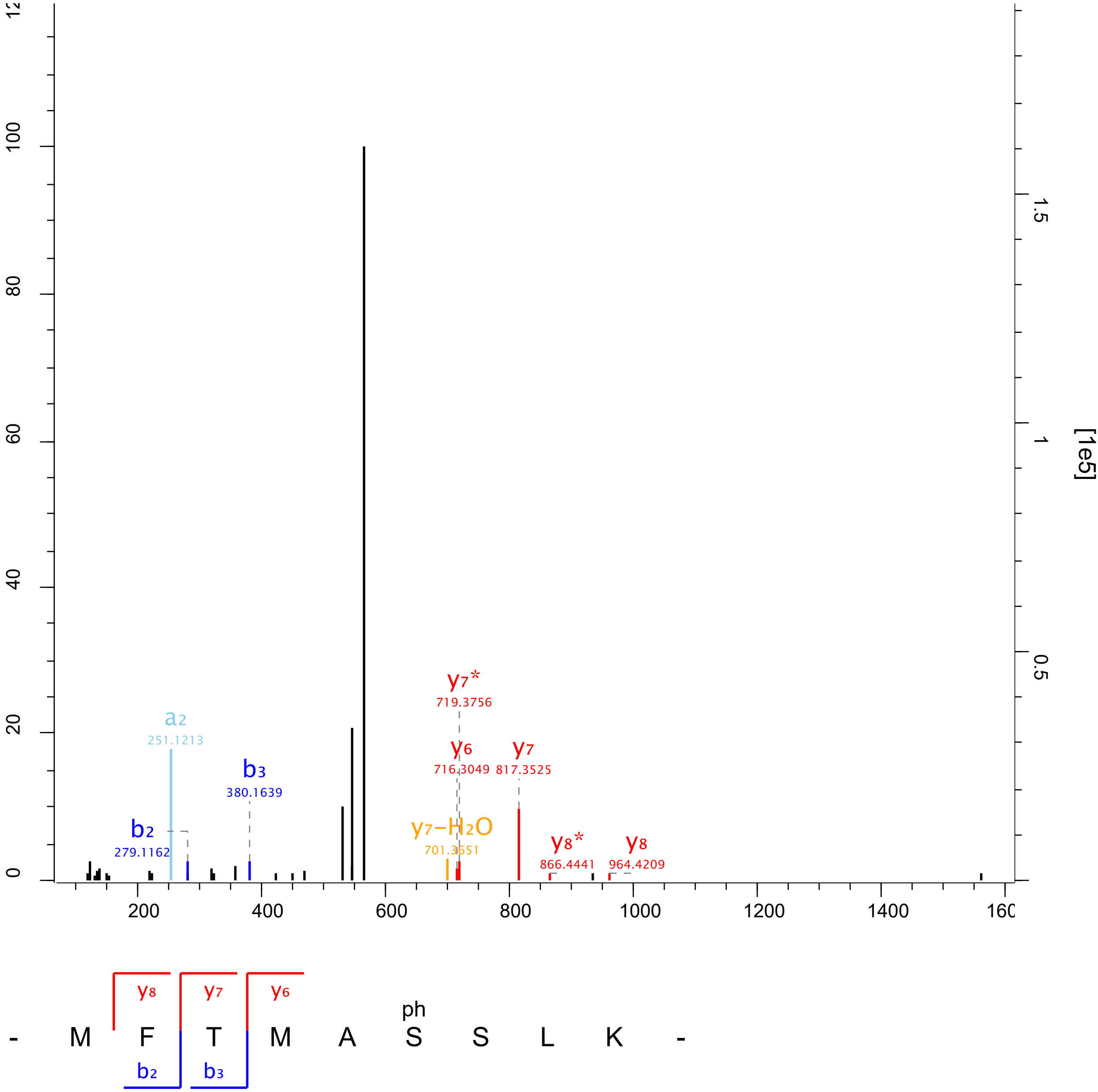

| Raw file | Scan | Method | Score | m/z |
| --- | --- | --- | --- | --- |
| sys_02_3 | 2240 | FTMS; HCD | 42.98 | 537.19 |

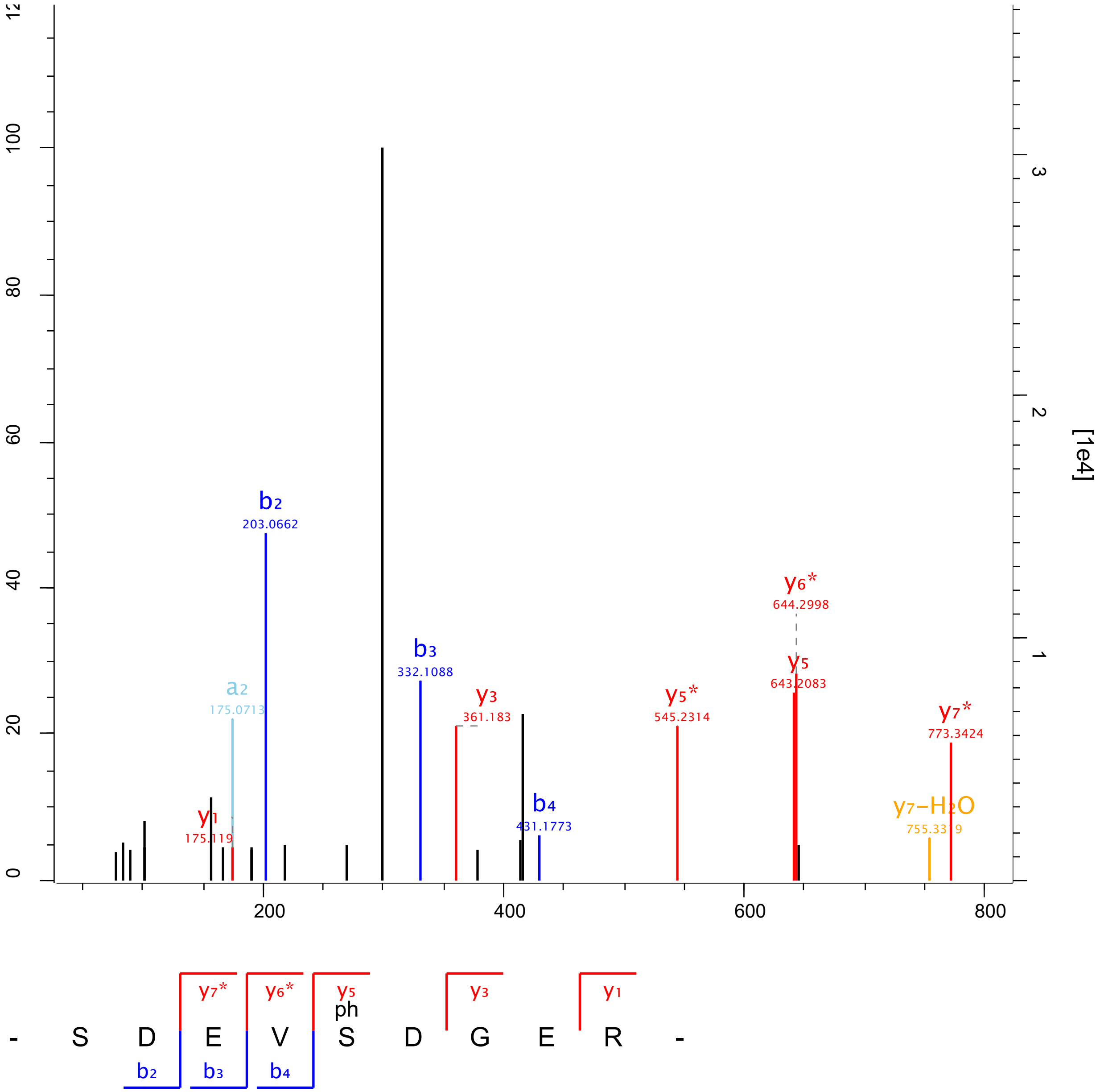

| Raw file | Scan | Method | Score | m/z |
|---|---|---|---|---|
| sys_02_3 | 23116 | FTMS; HCD | 154.1 | 570.29 |

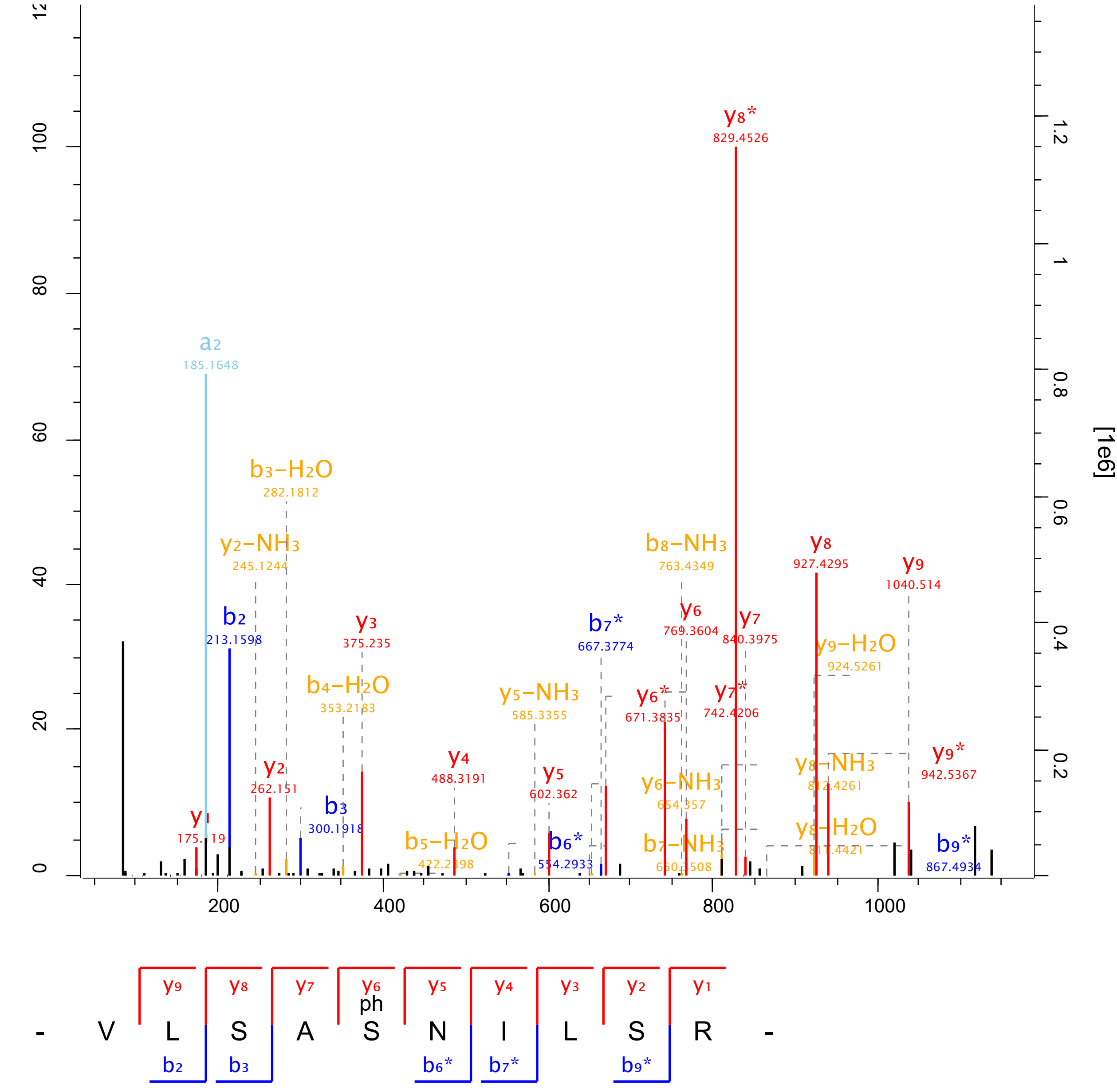

| Raw file | Scan | Method | Score | m/z |
|---|---|---|---|---|
| sys_02_3 | 23130 | FTMS; HCD | 41.26 | 918.42 |

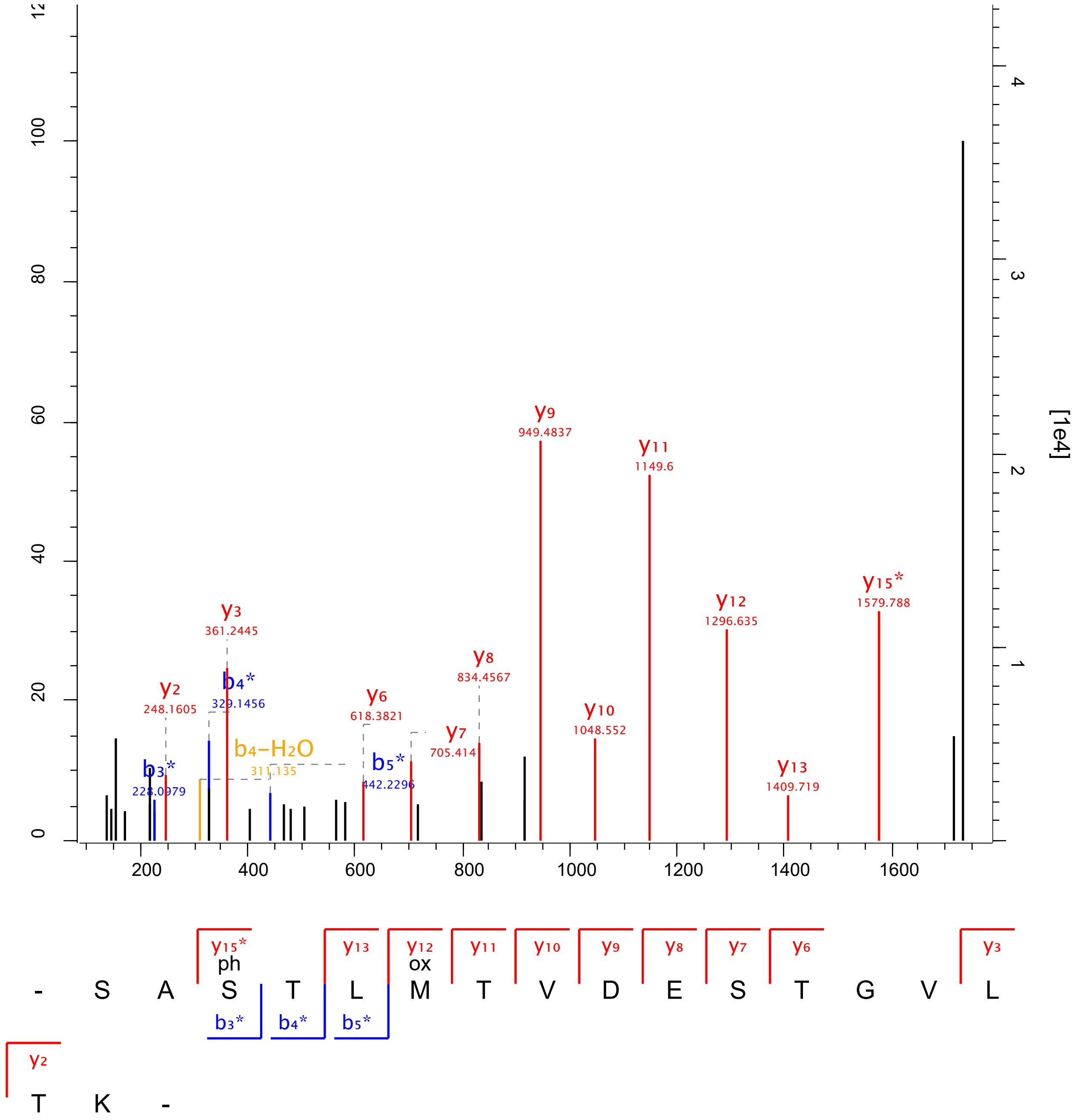

| Raw file | Scan | Method | Score | m/z |
|---|---|---|---|---|
| sys_02_3 | 2762 | FTMS; HCD | 56.26 | 441.74 |

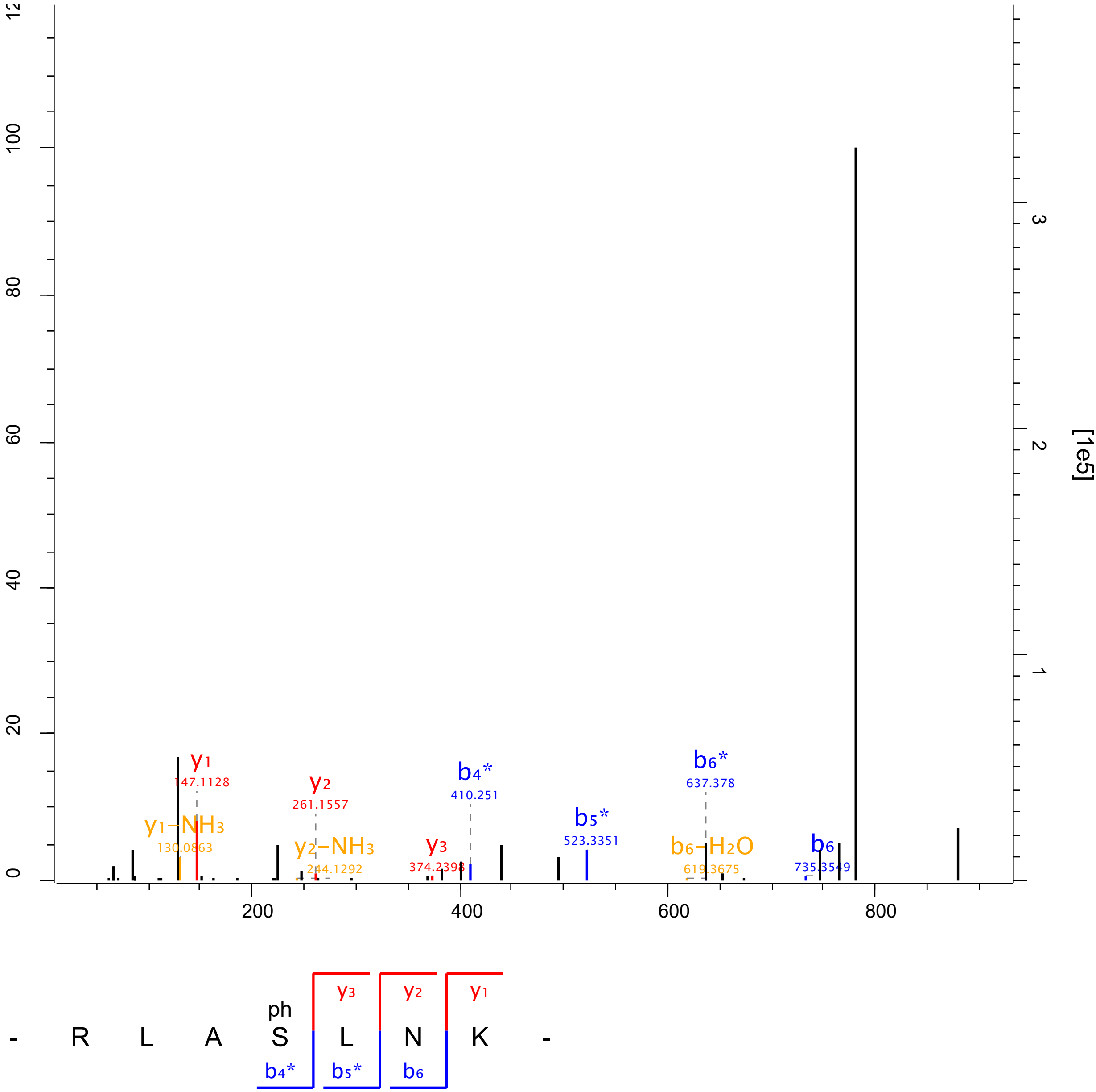

| Raw file | Scan | Method | Score | m/z |
|---|---|---|---|---|
| sys_02_3 | 3036 | FTMS; HCD | 48.57 | 556.89 |

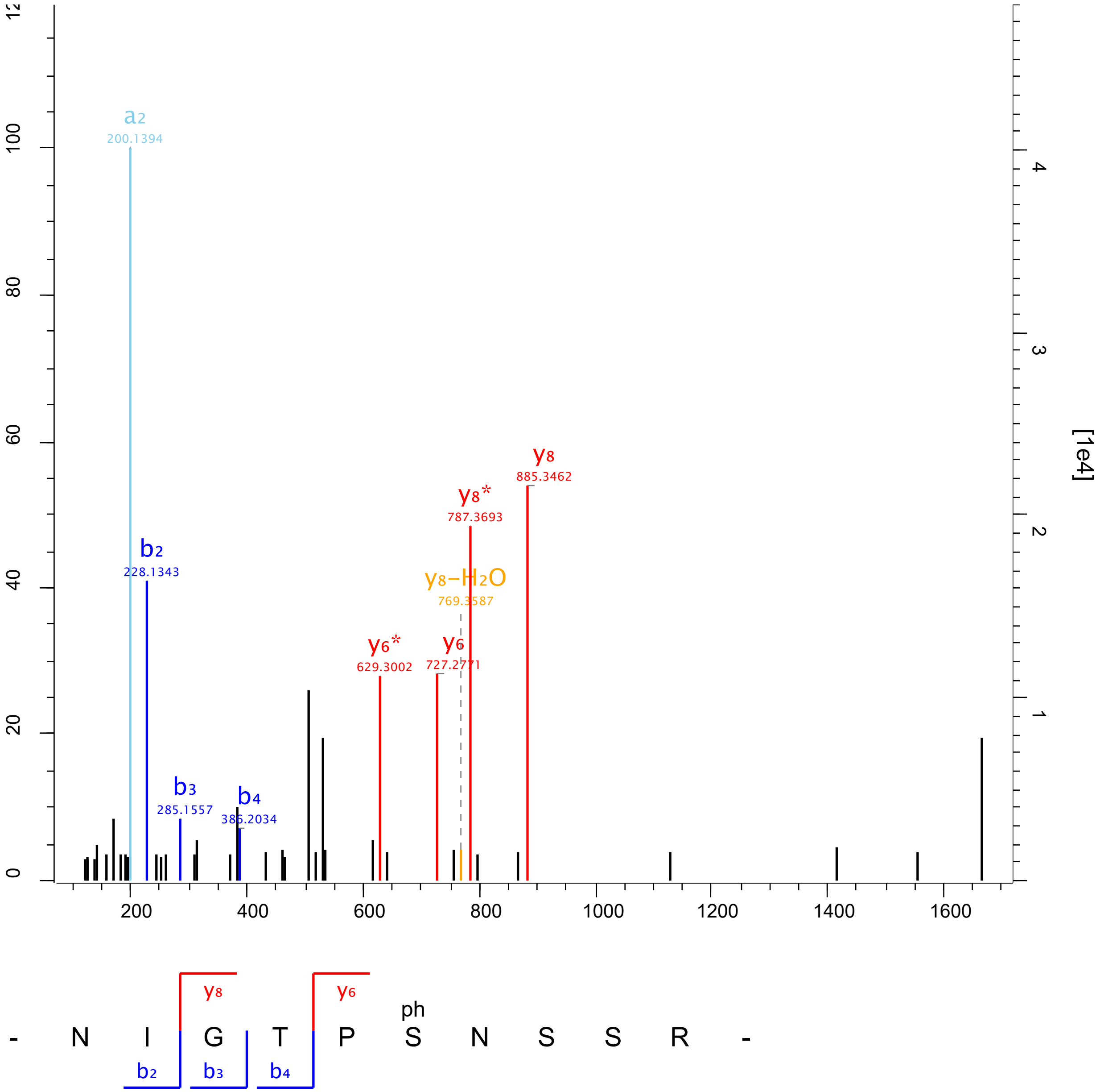

| Raw file | Scan | Method | Score | m/z |
| --- | --- | --- | --- | --- |
| sys_02_3 | 36058 | FTMS; HCD | 75.91 | 746.31 |

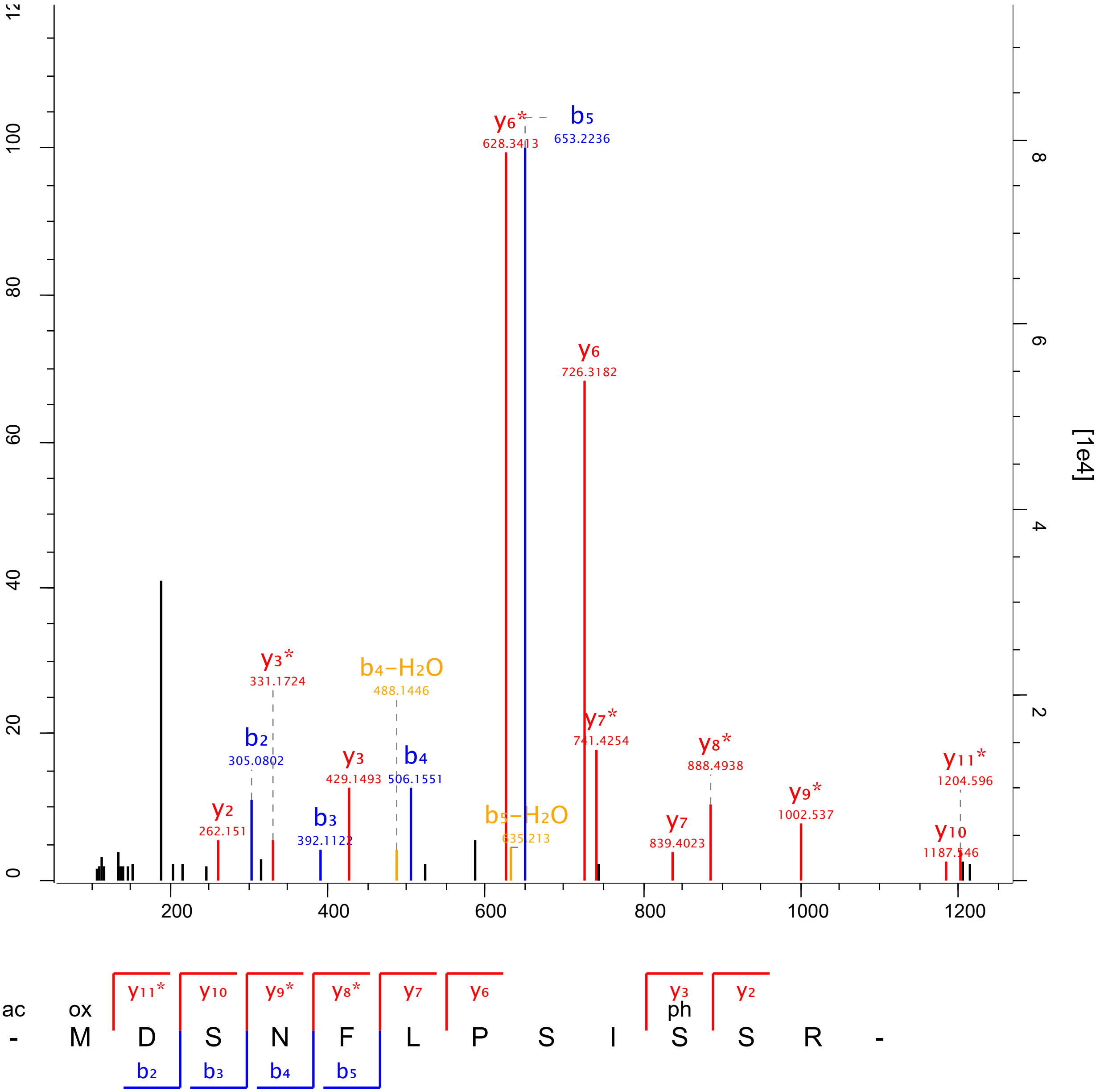

| Raw file | Scan | Method | Score | m/z |
|---|---|---|---|---|
| sys_02_3 | 3864 | FTMS; HCD | 144.73 | 638.73 |

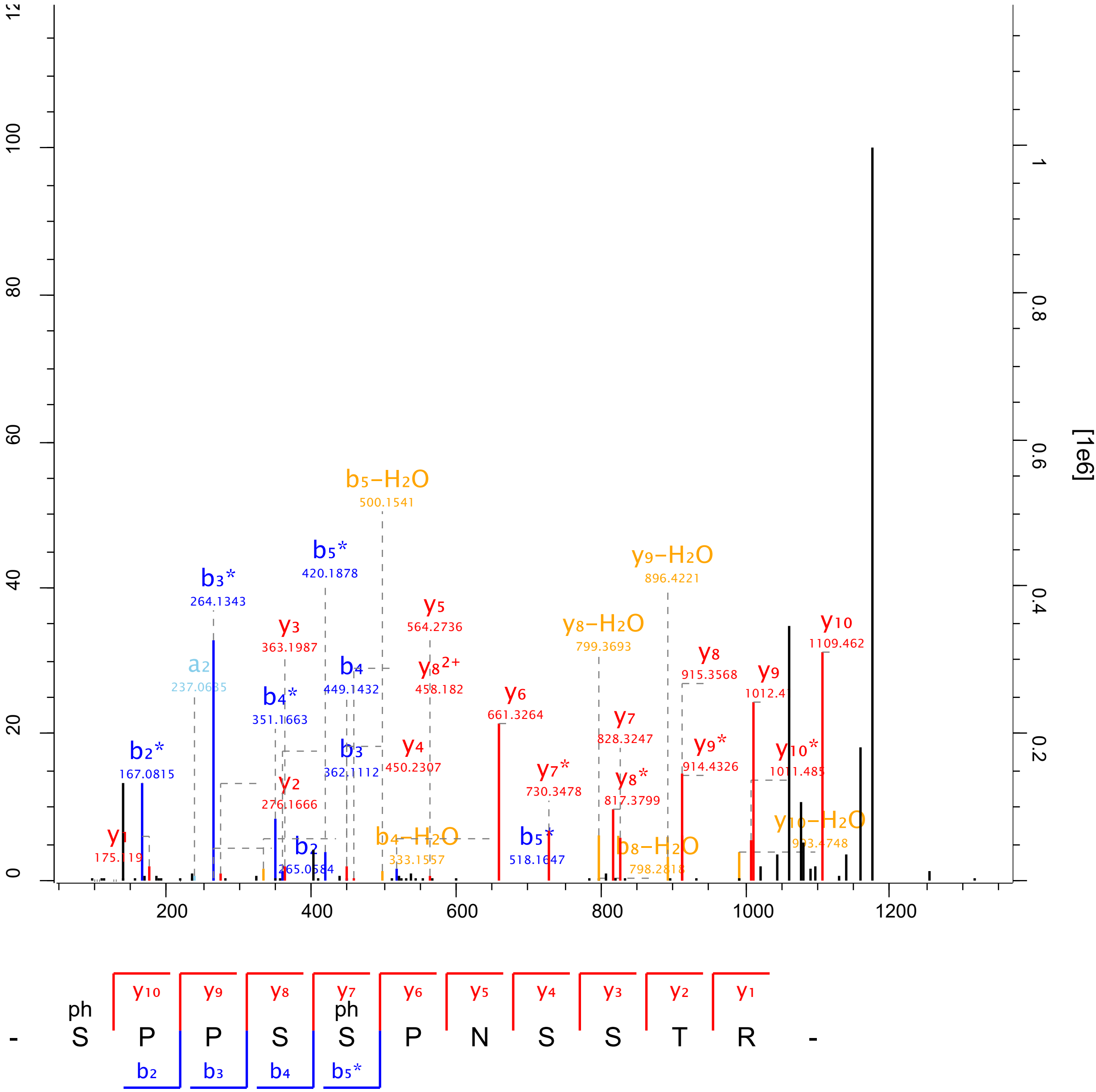

| Raw file | Scan | Method | Score | m/z |
|---|---|---|---|---|
| sys_02_3 | 4527 | FTMS; HCD | 138.22 | 457.22 |

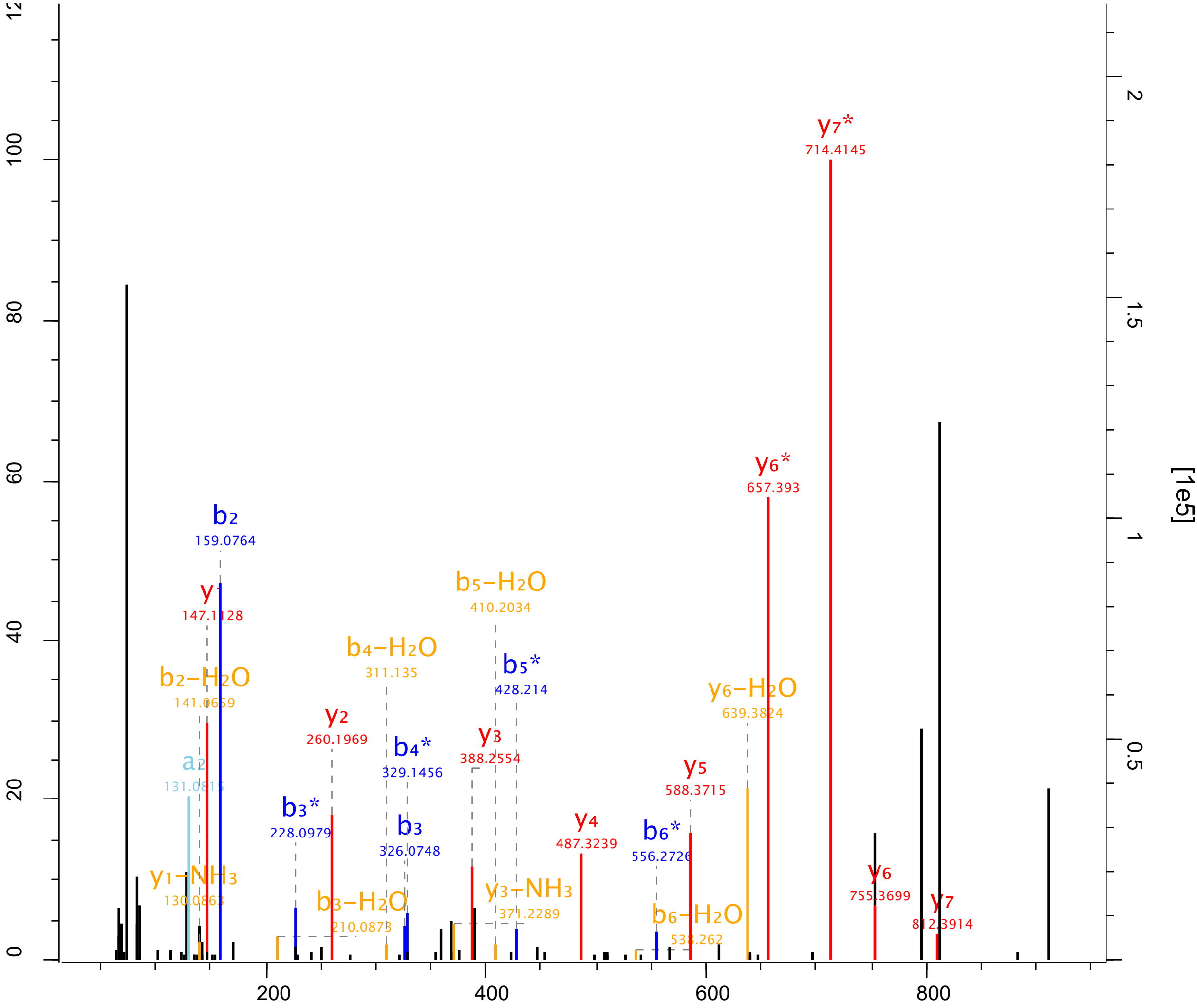


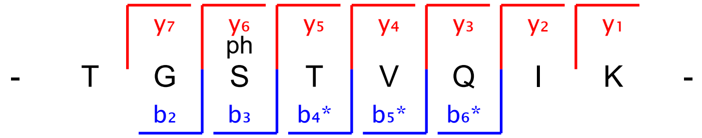

| Raw file | Scan | Method | Score | m/z |
|---|---|---|---|---|
| sys_02_3 | 5114 | FTMS; HCD | 92.25 | 492.74 |

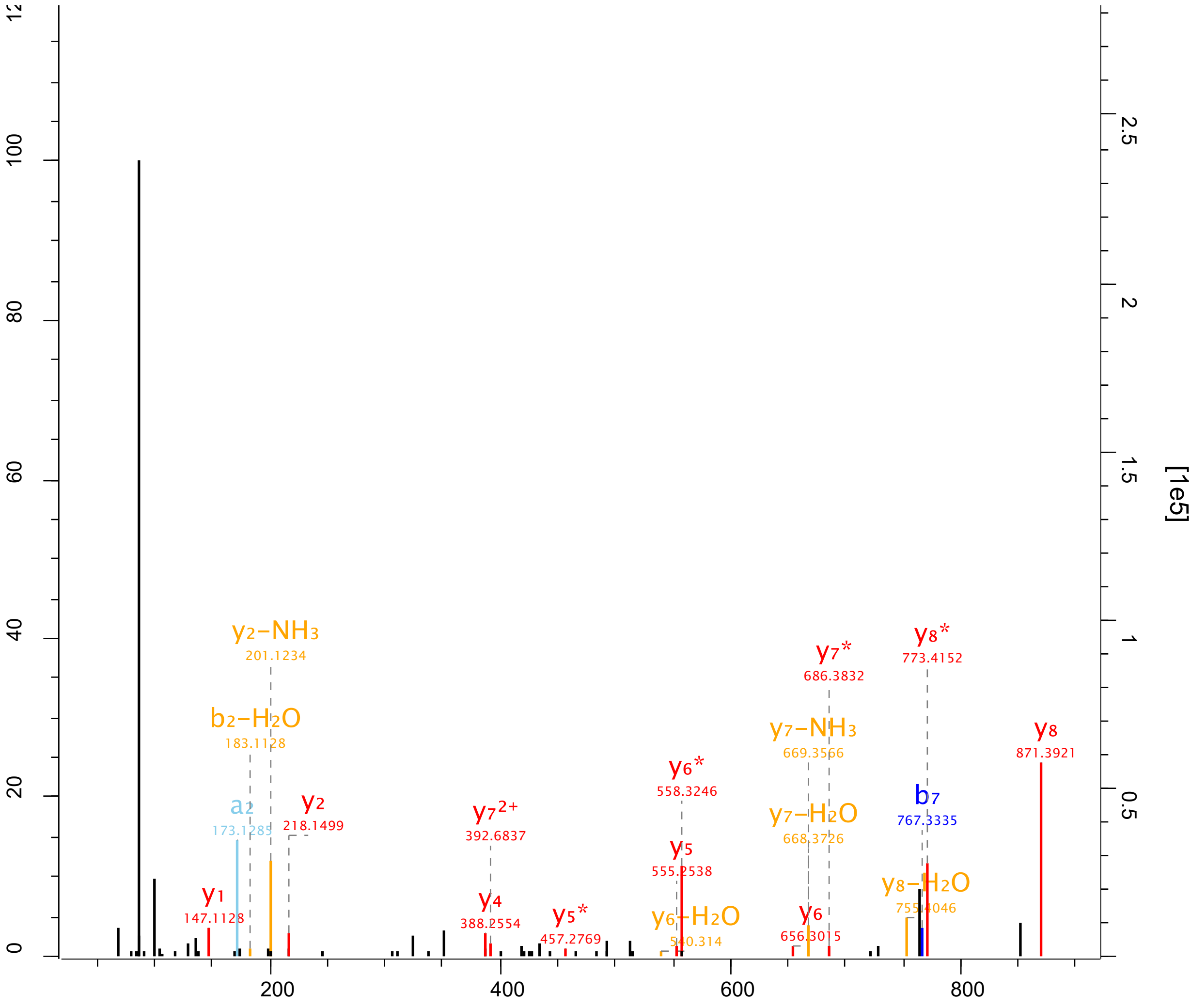


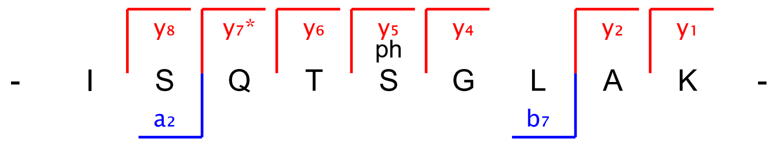

| Raw file | Scan | Method | Score | m/z |
|---|---|---|---|---|
| sys_02_3 | 7992 | FTMS; HCD | 78.32 | 630.78 |

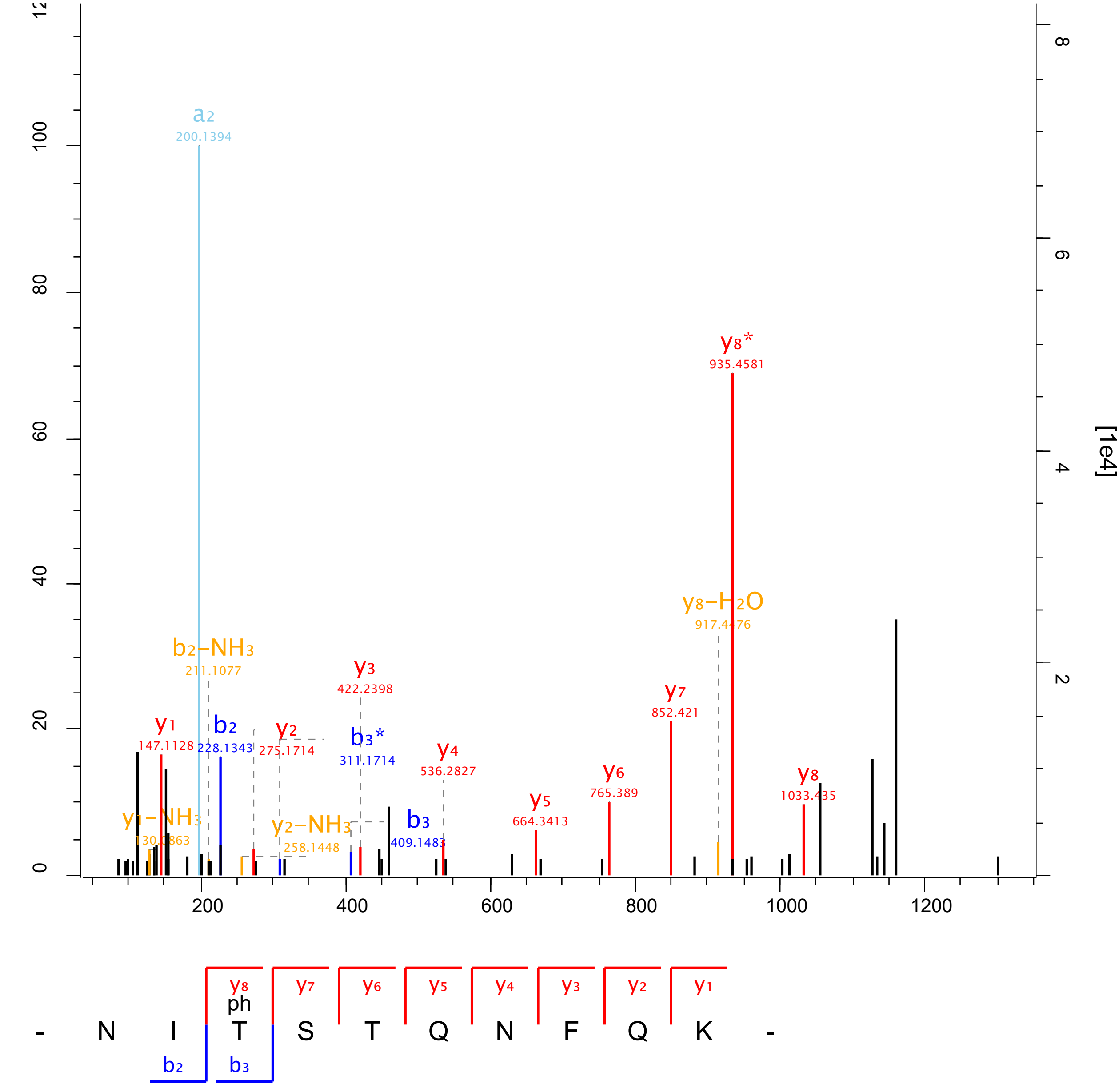

| Raw file | Scan | Method | Score | m/z |
|---|---|---|---|---|
| sys_02_3 | 8164 | FTMS; HCD | 60.26 | 564.23 |

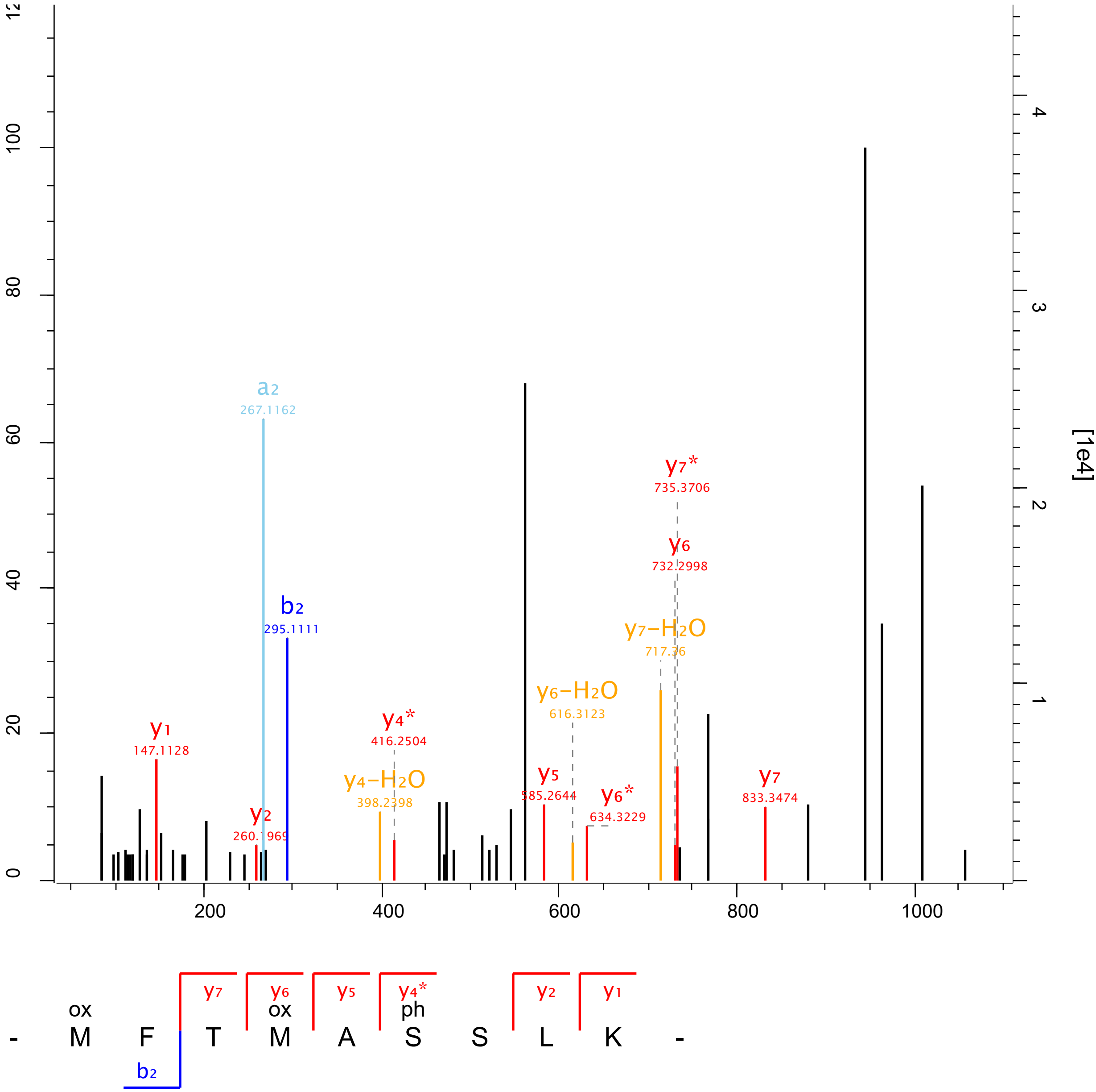

| Raw file | Scan | Method | Score | m/z |
|---|---|---|---|---|
| sys_05_1 | 13688 | FTMS; HCD | 136.24 | 584.22 |

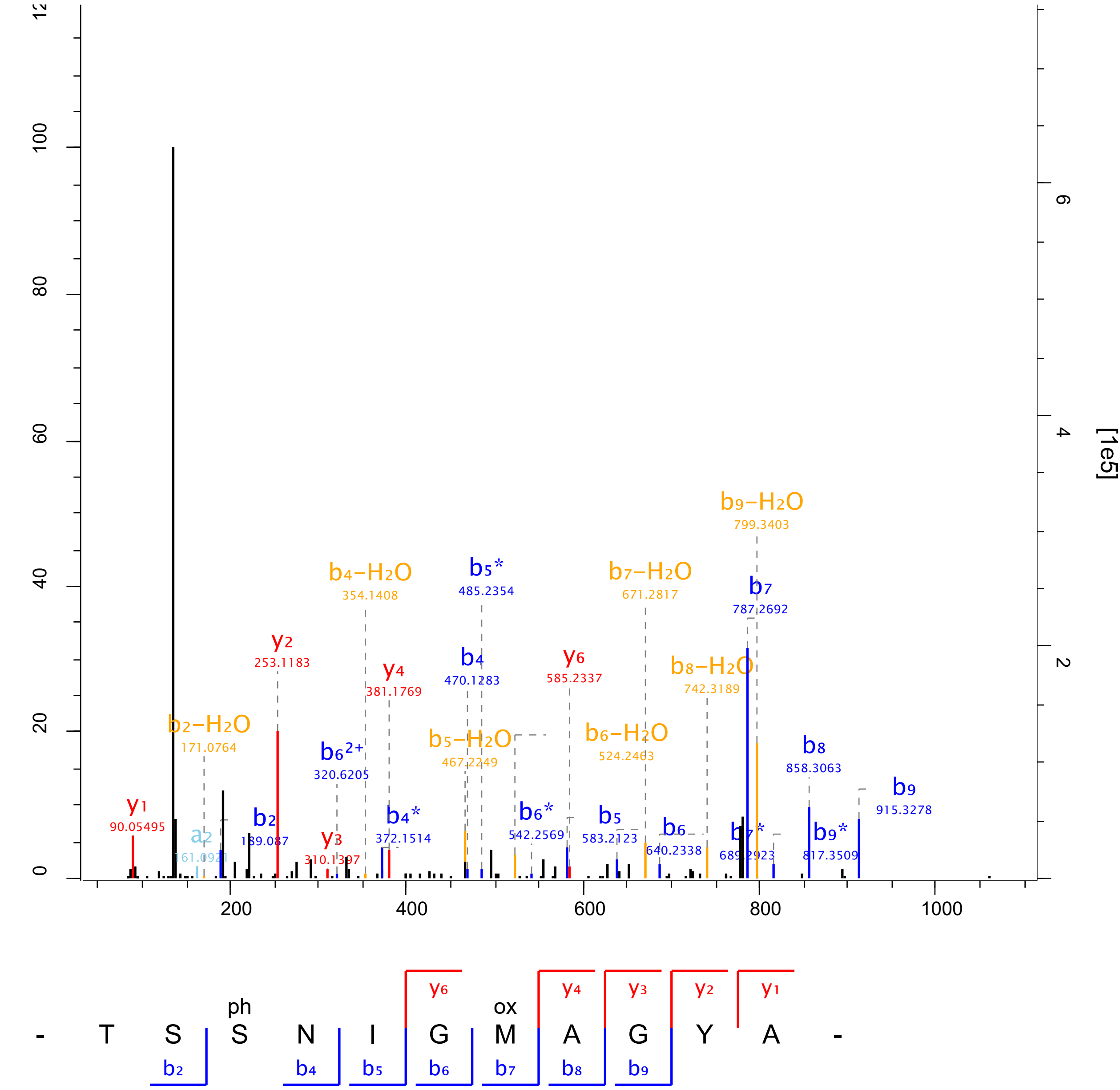

| Raw file | Scan | Method | Score | m/z |
| --- | --- | --- | --- | --- |
| sys_05_1 | 21449 | FTMS; HCD | 50.39 | 760.82 |

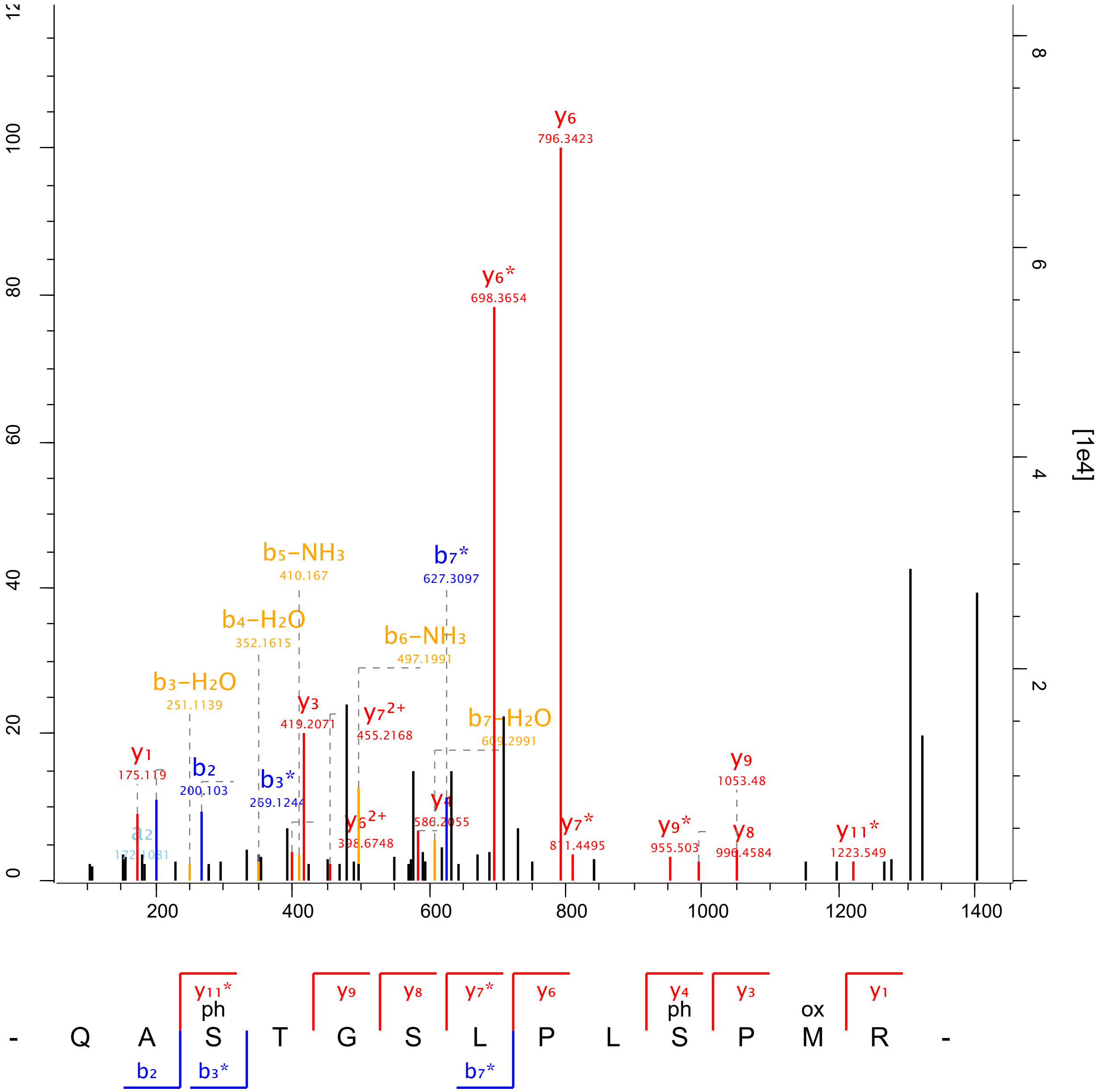

| Raw file | Scan | Method | Score | m/z |
|---|---|---|---|---|
| sys_05_1 | 23137 | FTMS; HCD | 89.01 | 623.95 |

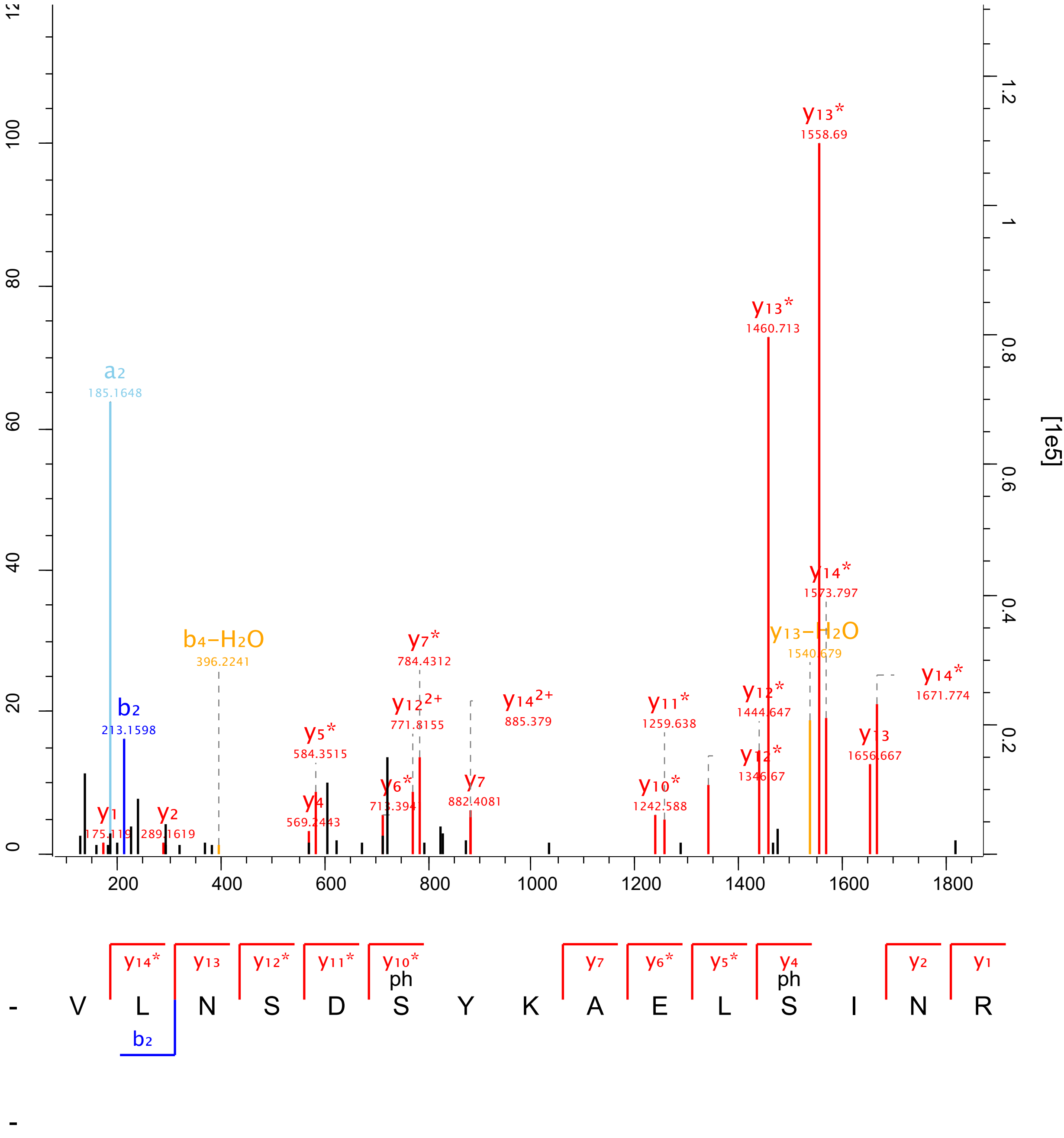

| Raw file | Scan | Method | Score | m/z |
| --- | --- | --- | --- | --- |
| sys_05_1 | 24556 | FTMS; HCD | 77.87 | 658.33 |

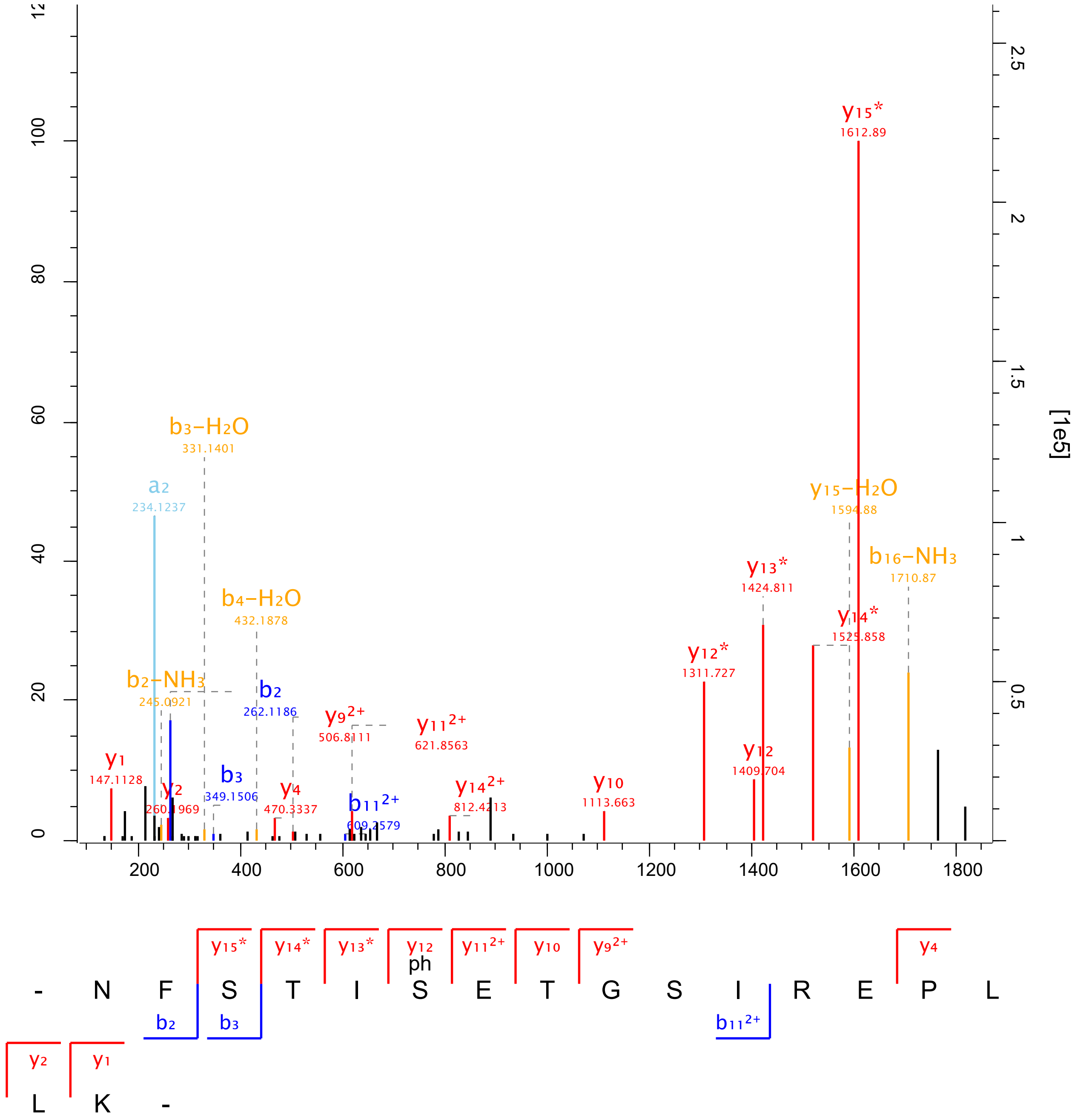

| Raw file | Scan | Method | Score | m/z |
|---|---|---|---|---|
| sys_05_1 | 2782 | FTMS; HCD | 75.82 | 448.21 |

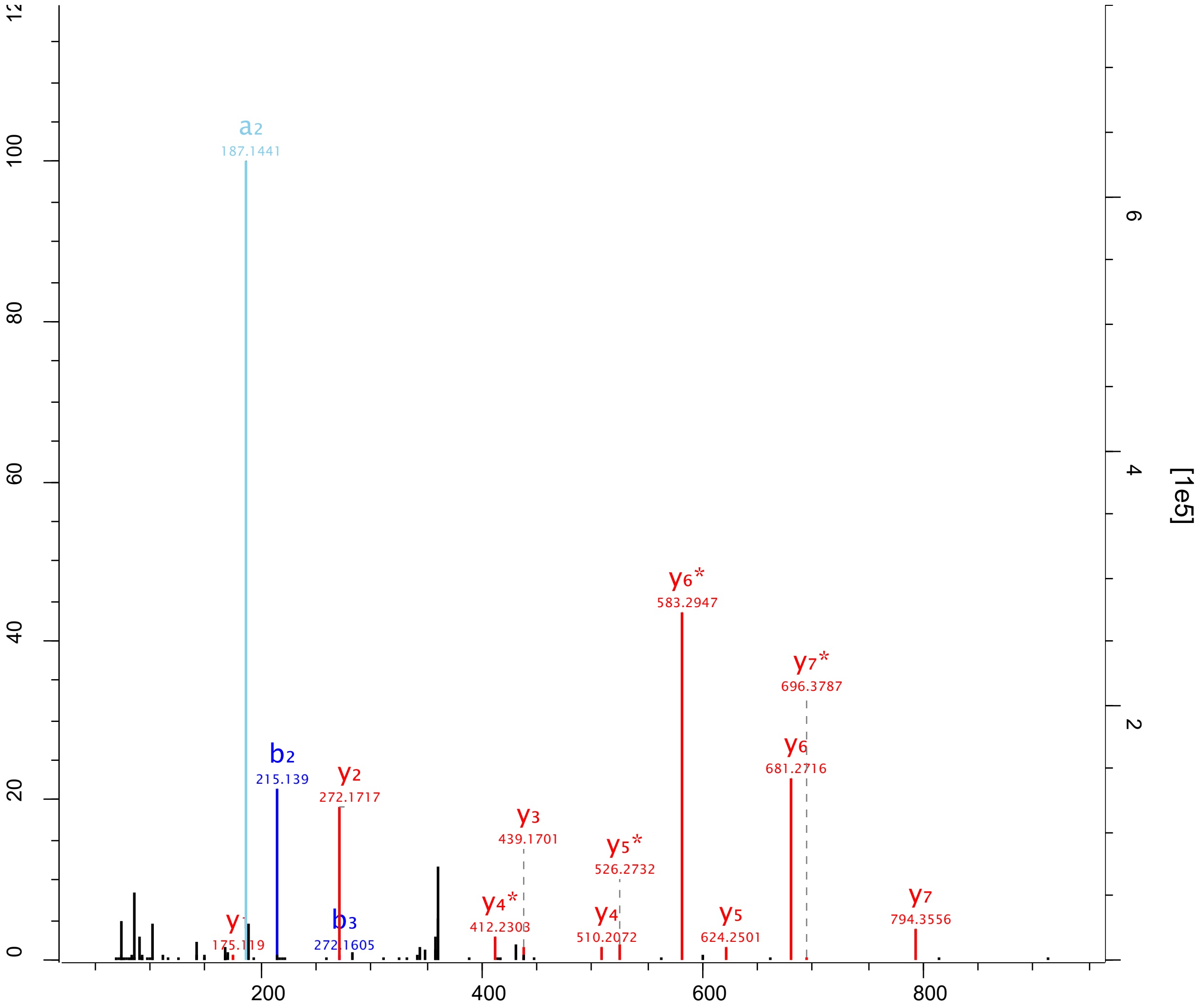


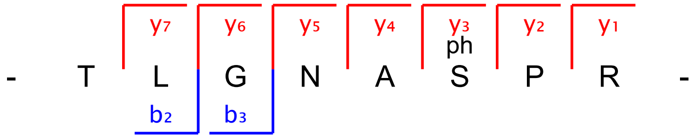

| Raw file | Scan | Method | Score | m/z |
| --- | --- | --- | --- | --- |
| sys_05_1 | 30797 | FTMS; HCD | 78.95 | 815.87 |

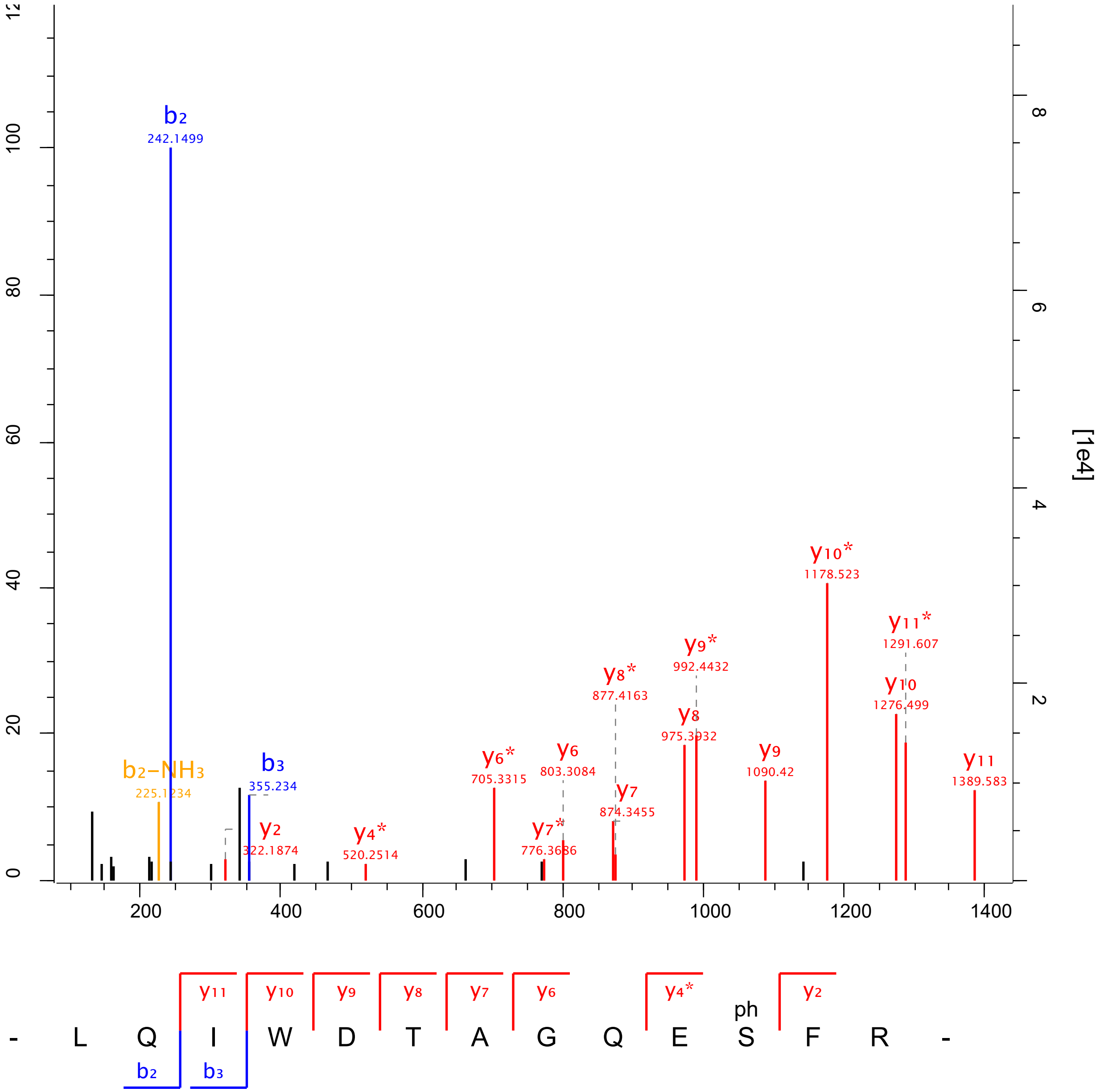

| Raw file | Scan | Method | Score | m/z |
| --- | --- | --- | --- | --- |
| sys_05_1 | 31096 | FTMS; HCD | 130.83 | 684.65 |

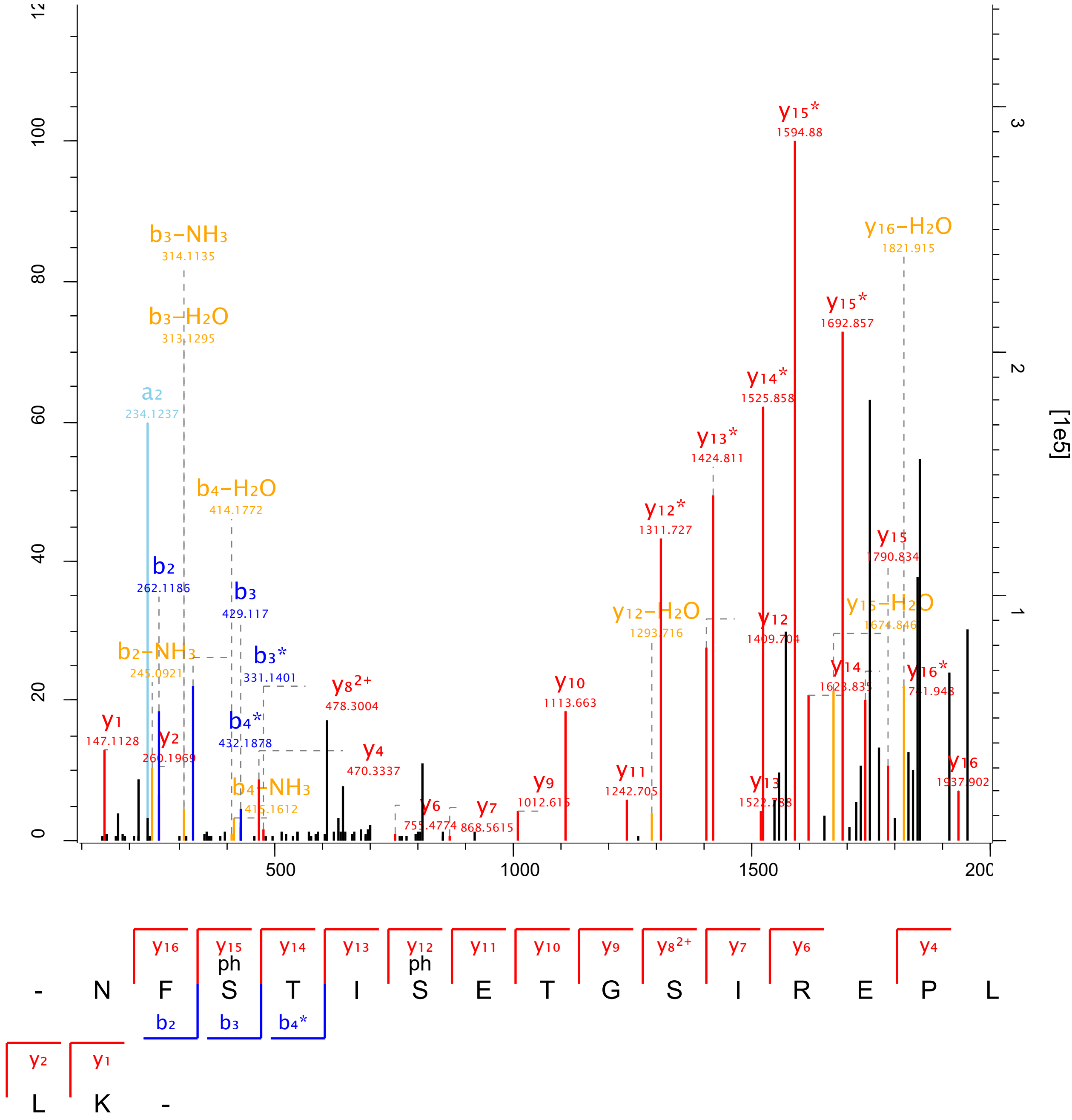

| Raw file | Scan | Method | Score | m/z |
|---|---|---|---|---|
| sys_05_1 | 32947 | FTMS; HCD | 48.29 | 814.36 |

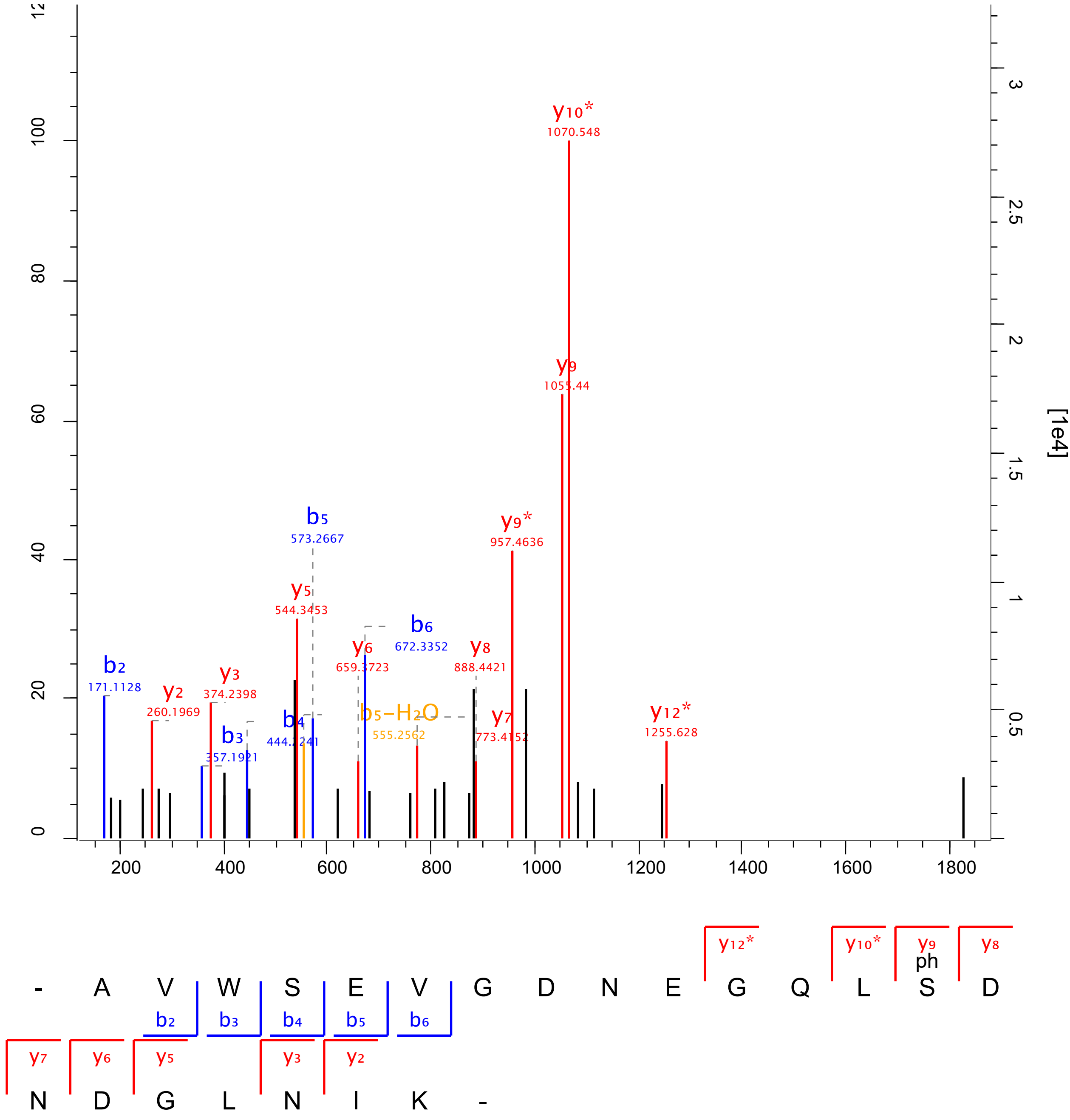

| Raw file | Scan | Method | Score | m/z |
|---|---|---|---|---|
| sys_05_1 | 3477 | FTMS; HCD | 113.38 | 485.7 |

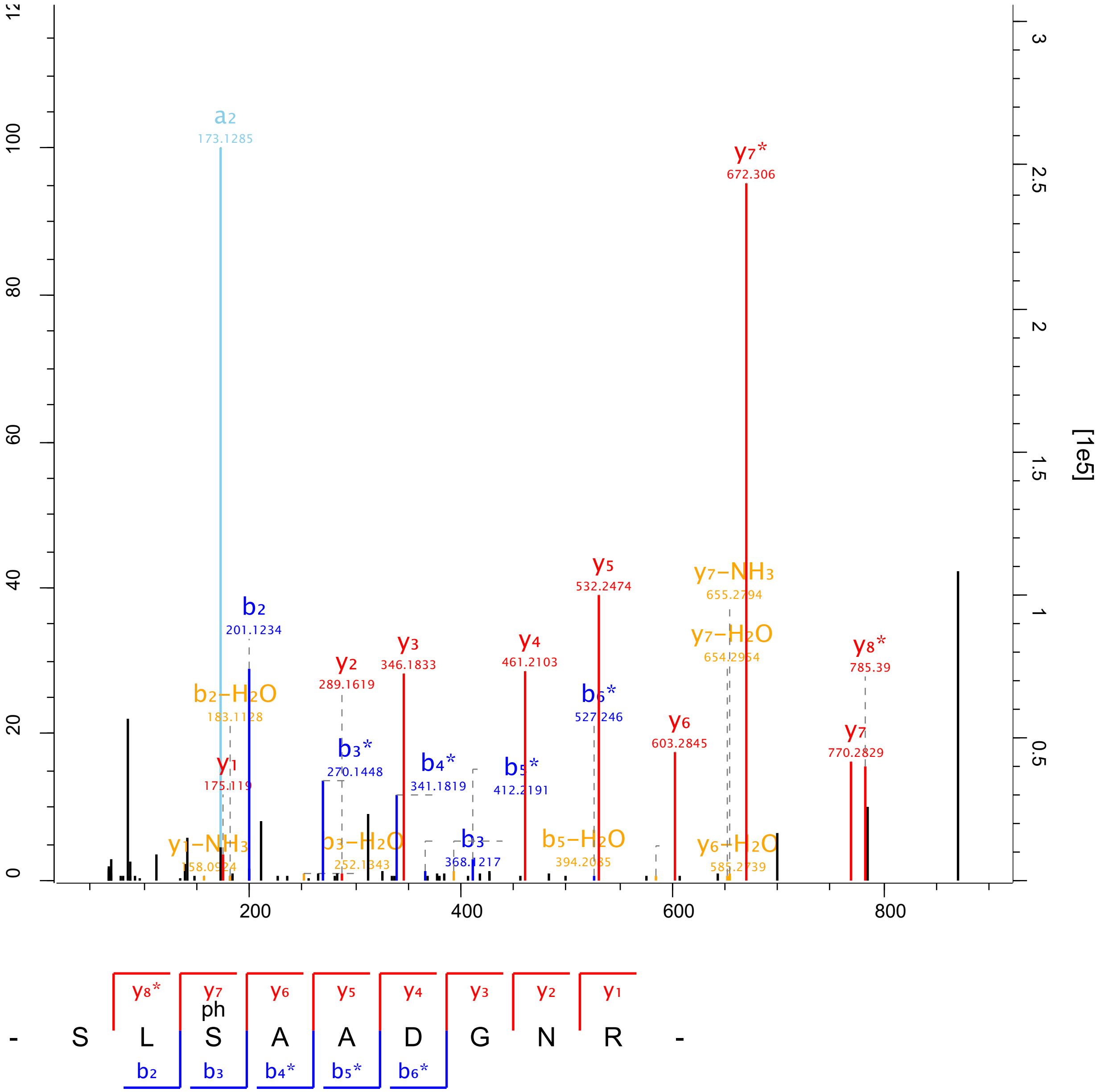

| Raw file | Scan | Method | Score | m/z |
|---|---|---|---|---|
| sys_05_1 | 39450 | FTMS; HCD | 146.78 | 736.8 |

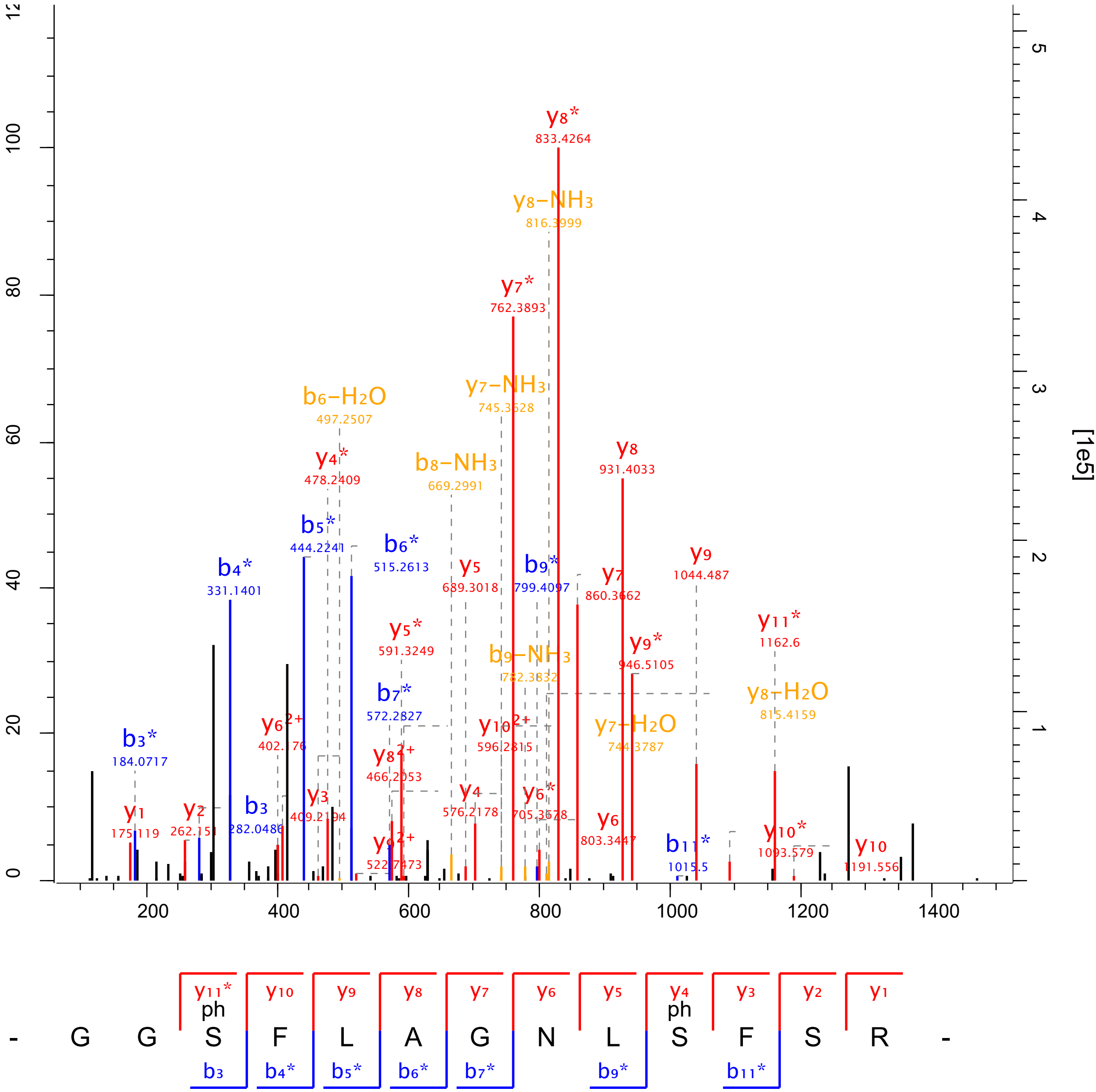

| Raw file | Scan | Method | Score | m/z |
|---|---|---|---|---|
| sys_05_1 | 3997 | FTMS; HCD | 107.11 | 461.7 |

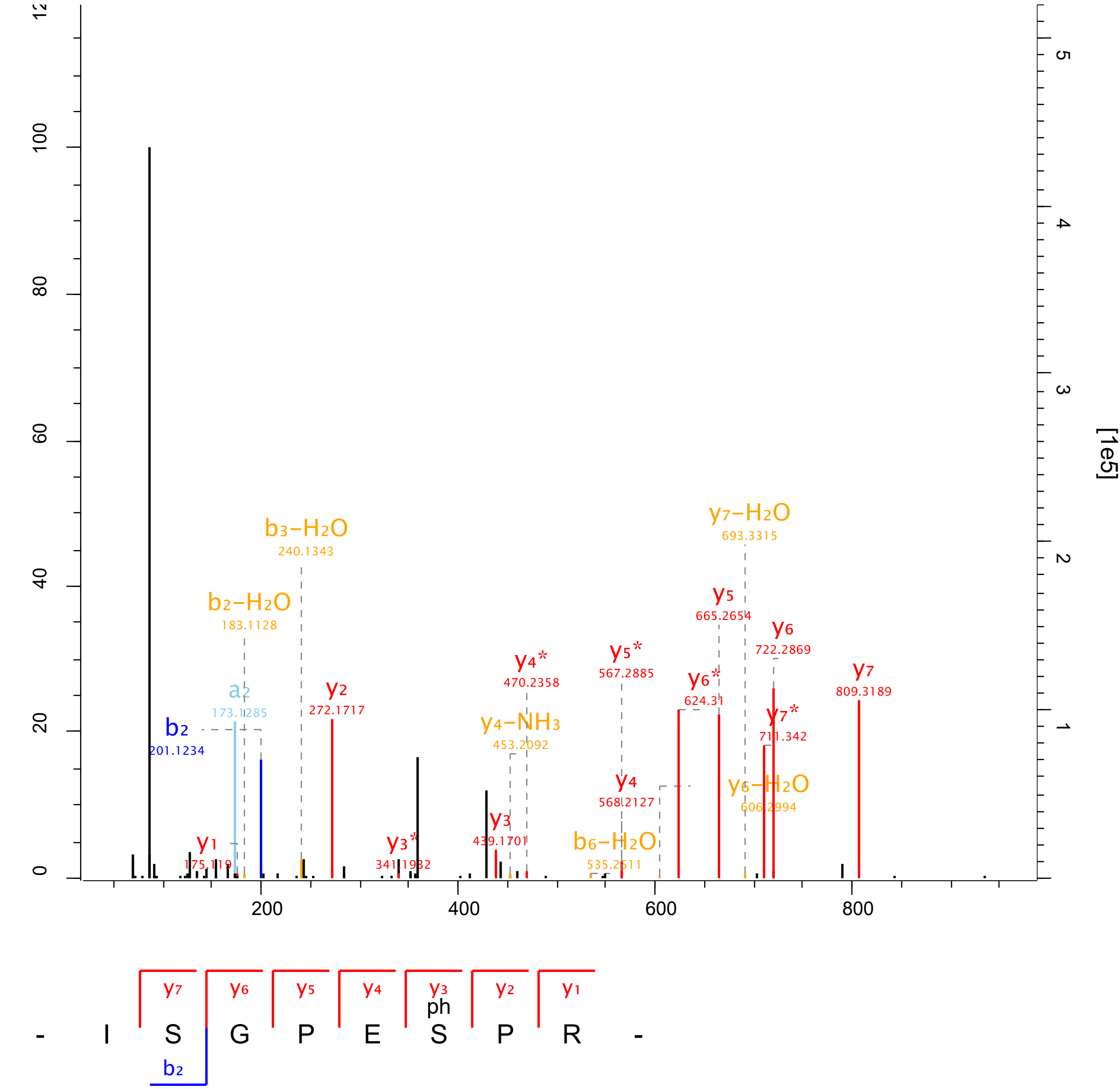

| Raw file | Scan | Method | Score | m/z |
|---|---|---|---|---|
| sys_05_1 | 40953 | FTMS; HCD | 77.75 | 725.34 |

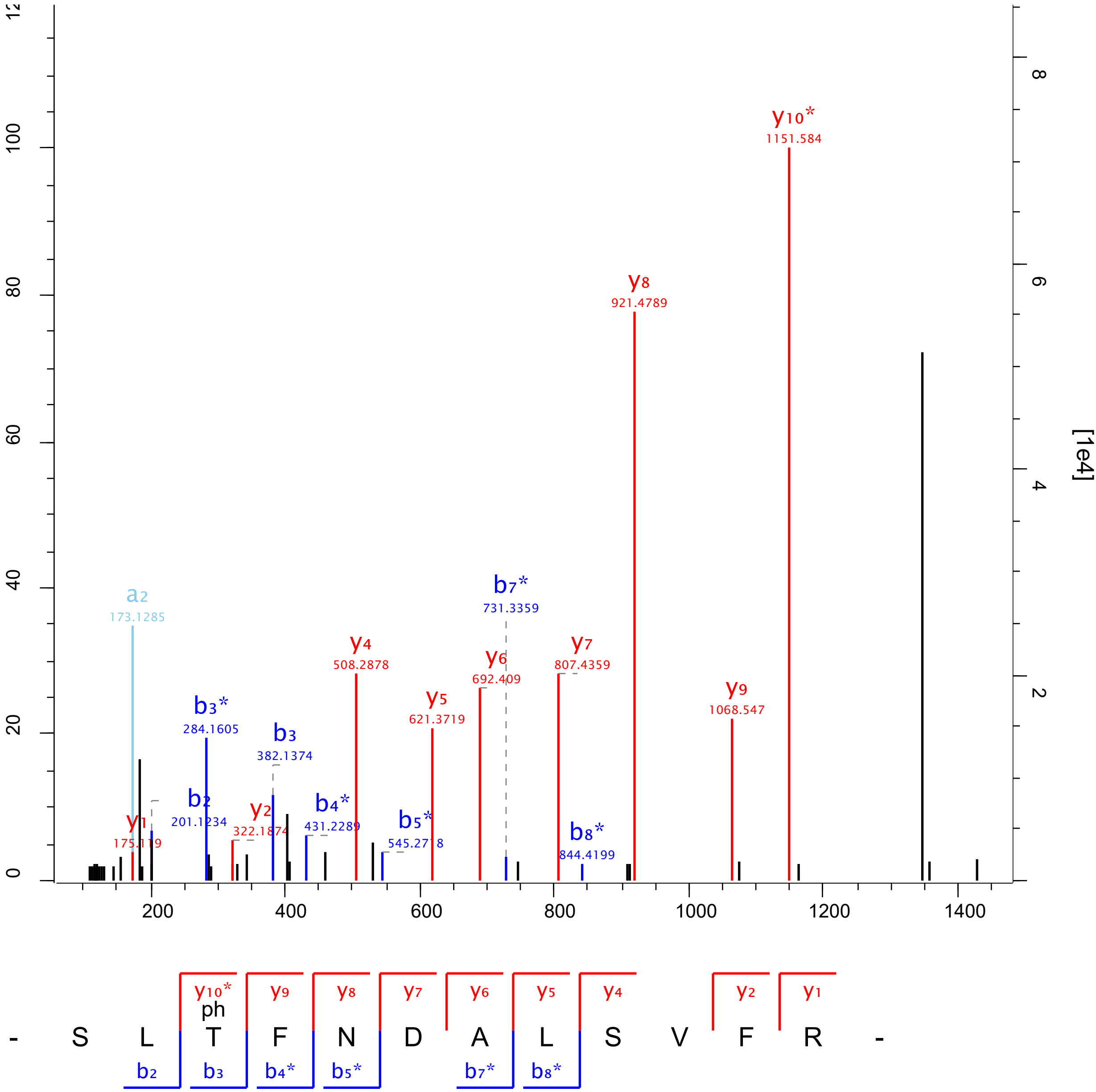

| Raw file | Scan | Method | Score | m/z |
| --- | --- | --- | --- | --- |
| sys_05_1 | 42484 | FTMS; HCD | 138.01 | 665.3 |

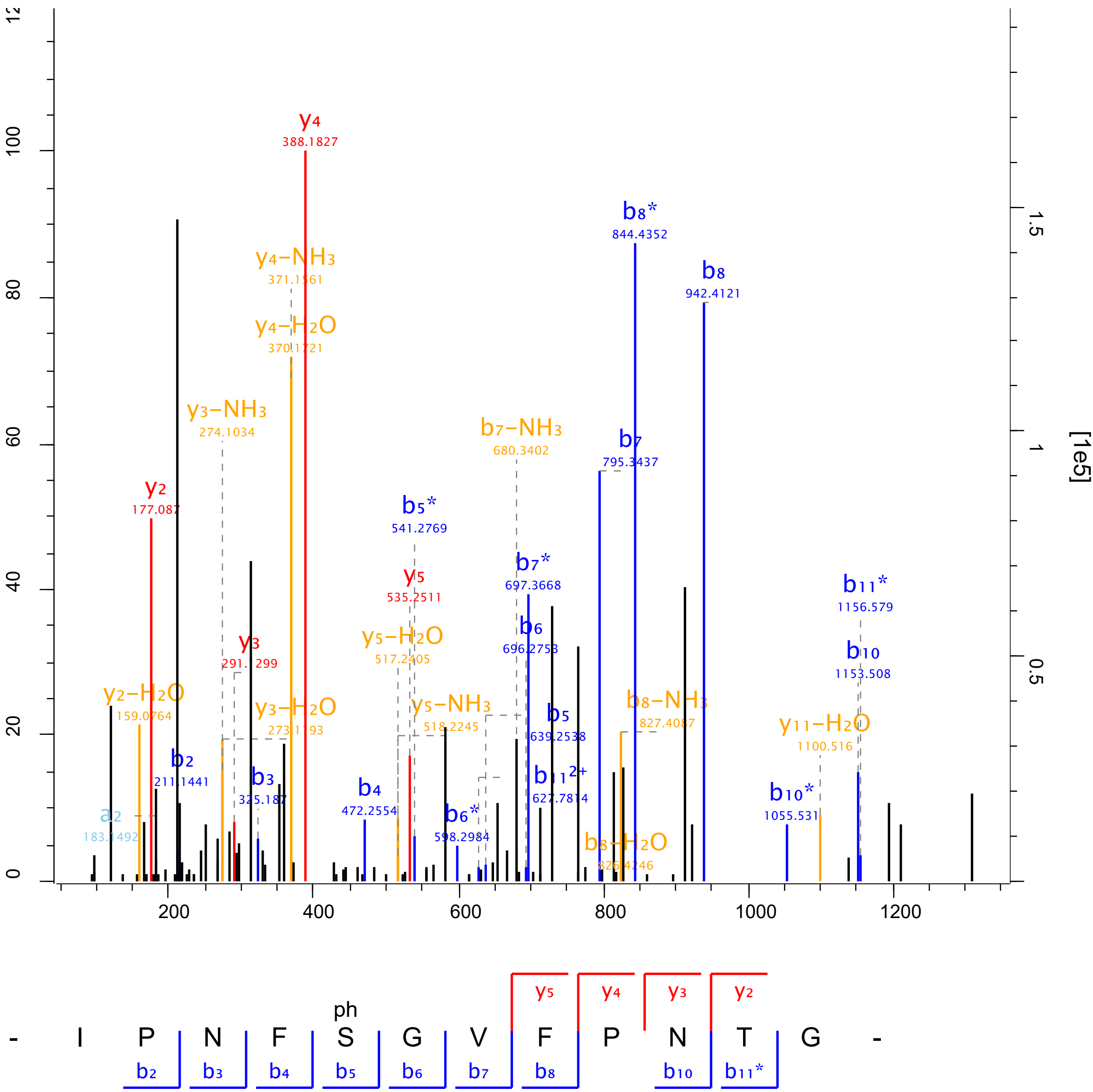

| Raw file | Scan | Method | Score | m/z |
|---|---|---|---|---|
| sys_05_1 | 43681 | FTMS; HCD | 75.91 | 699.81 |

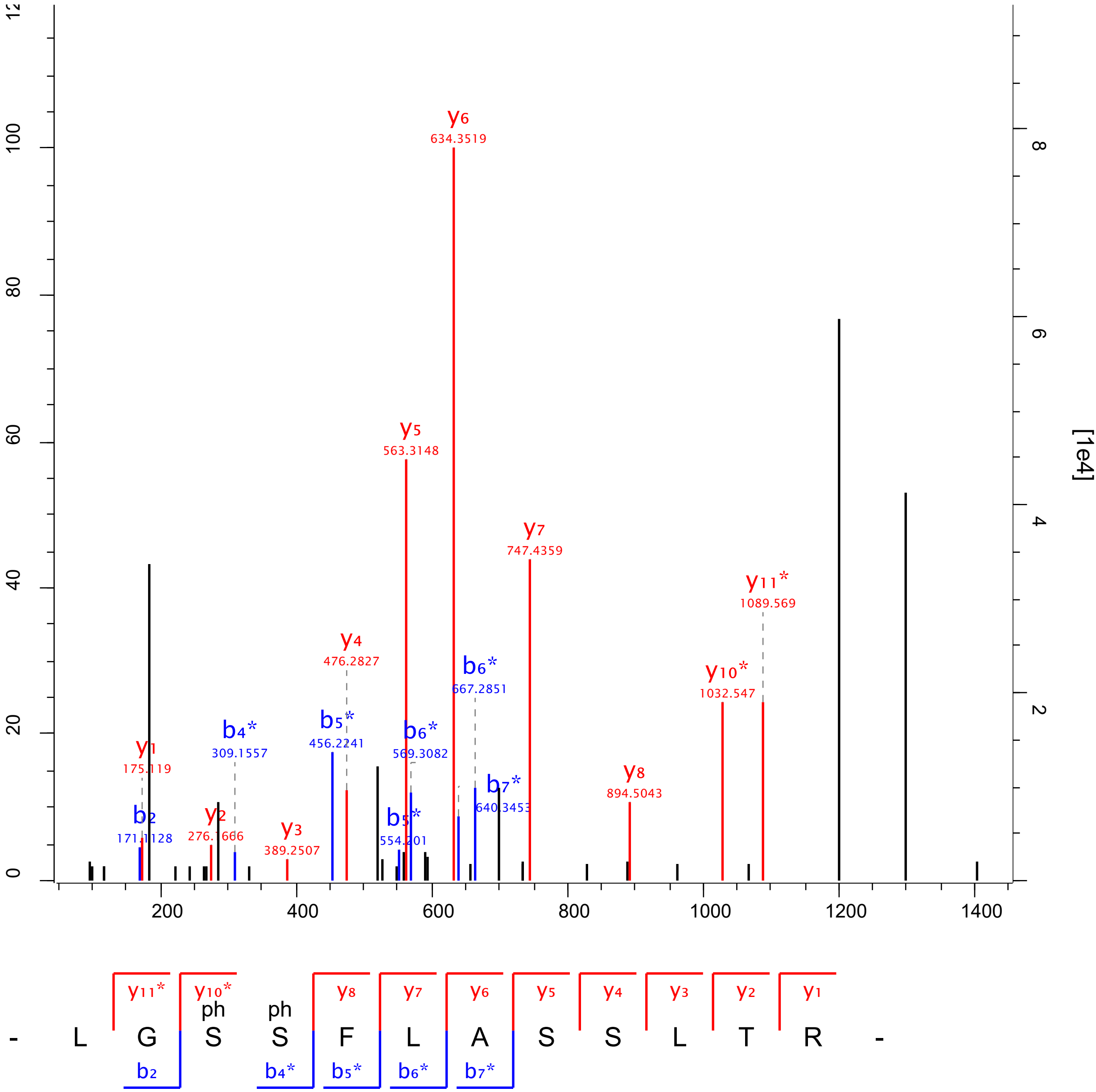

| Raw file | Scan | Method | Score | m/z |
| --- | --- | --- | --- | --- |
| sys_05_1 | 44181 | FTMS; HCD | 45.94 | 1008.8 |

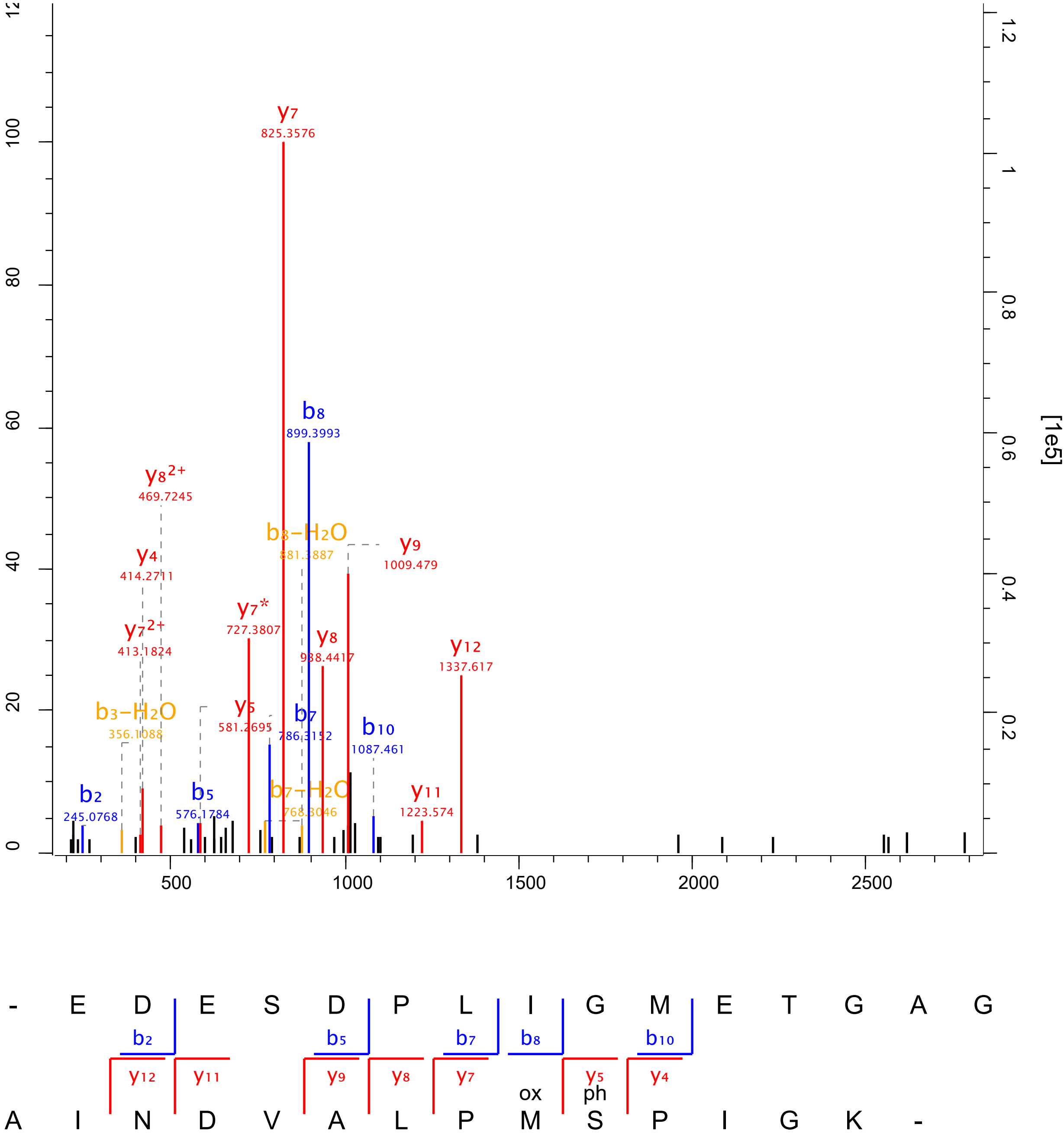

| Raw file | Scan | Method | Score | m/z |
|---|---|---|---|---|
| sys_05_1 | 45065 | FTMS; HCD | 58.43 | 827.38 |

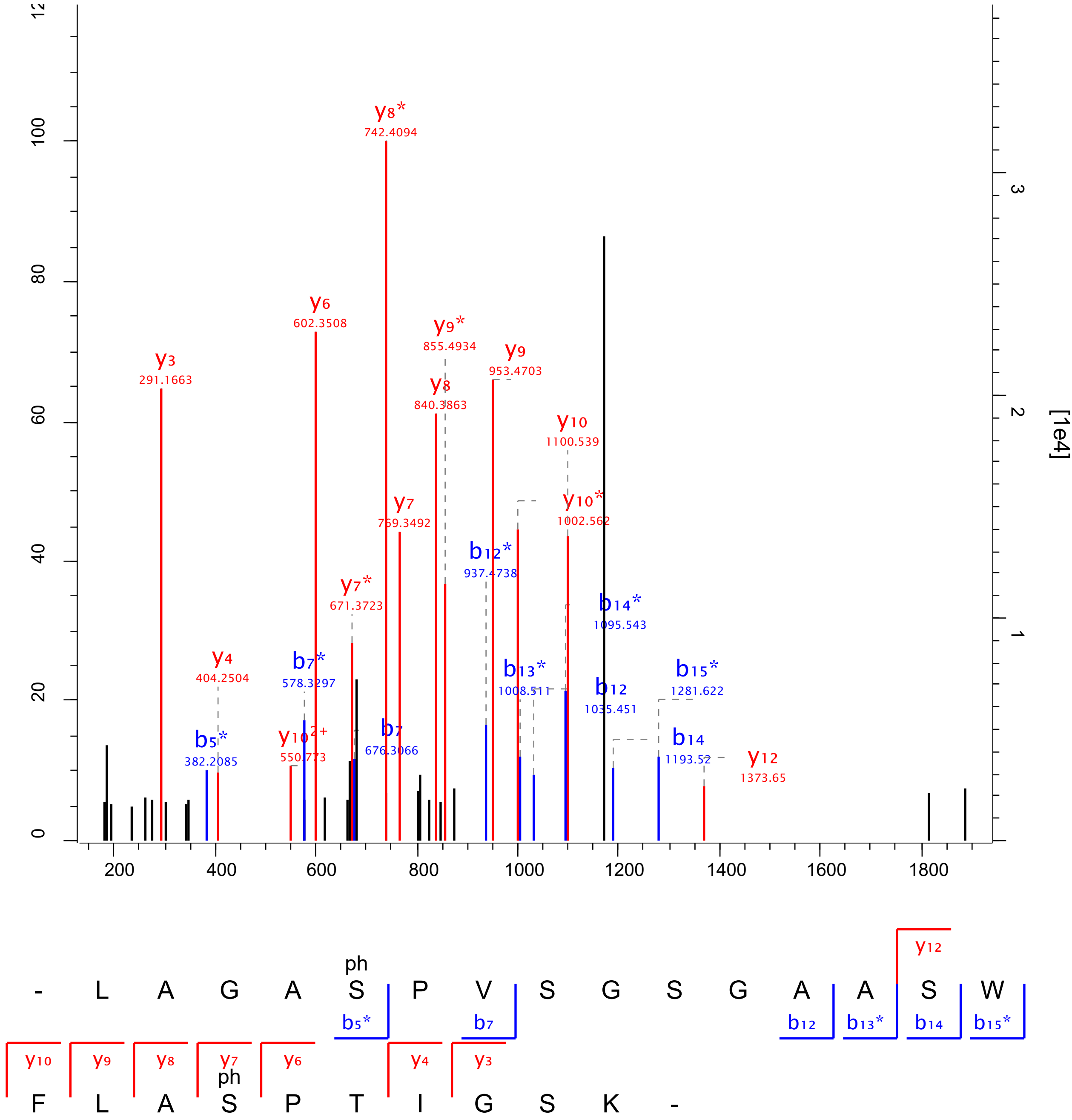

| Raw file | Scan | Method | Score | m/z |
| --- | --- | --- | --- | --- |
| sys_05_1 | 45327 | FTMS; HCD | 181.58 | 691.78 |

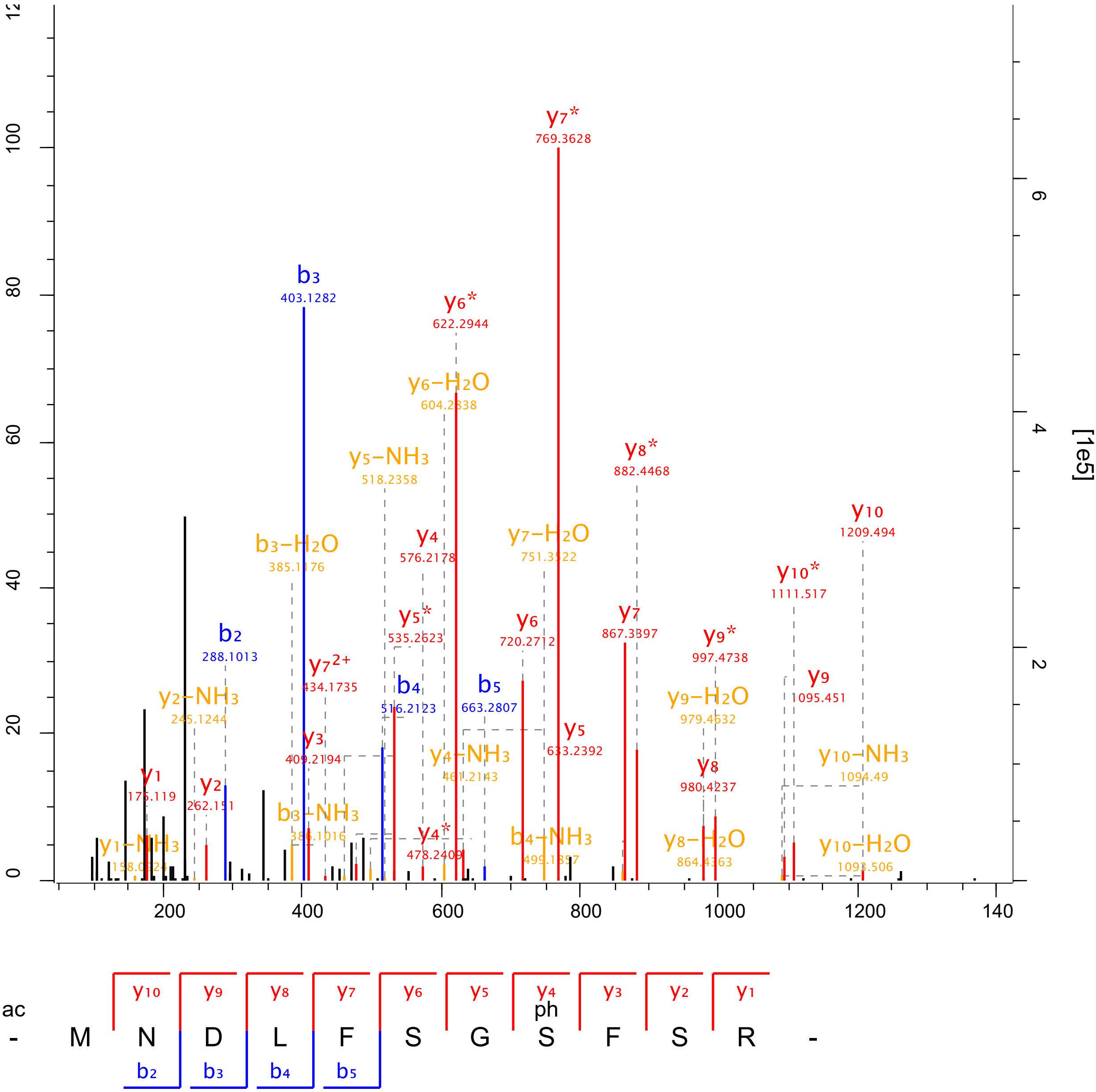

| Raw file | Scan | Method | Score | m/z |
| --- | --- | --- | --- | --- |
| sys_05_1 | 45334 | FTMS; HCD | 127.32 | 917.86 |

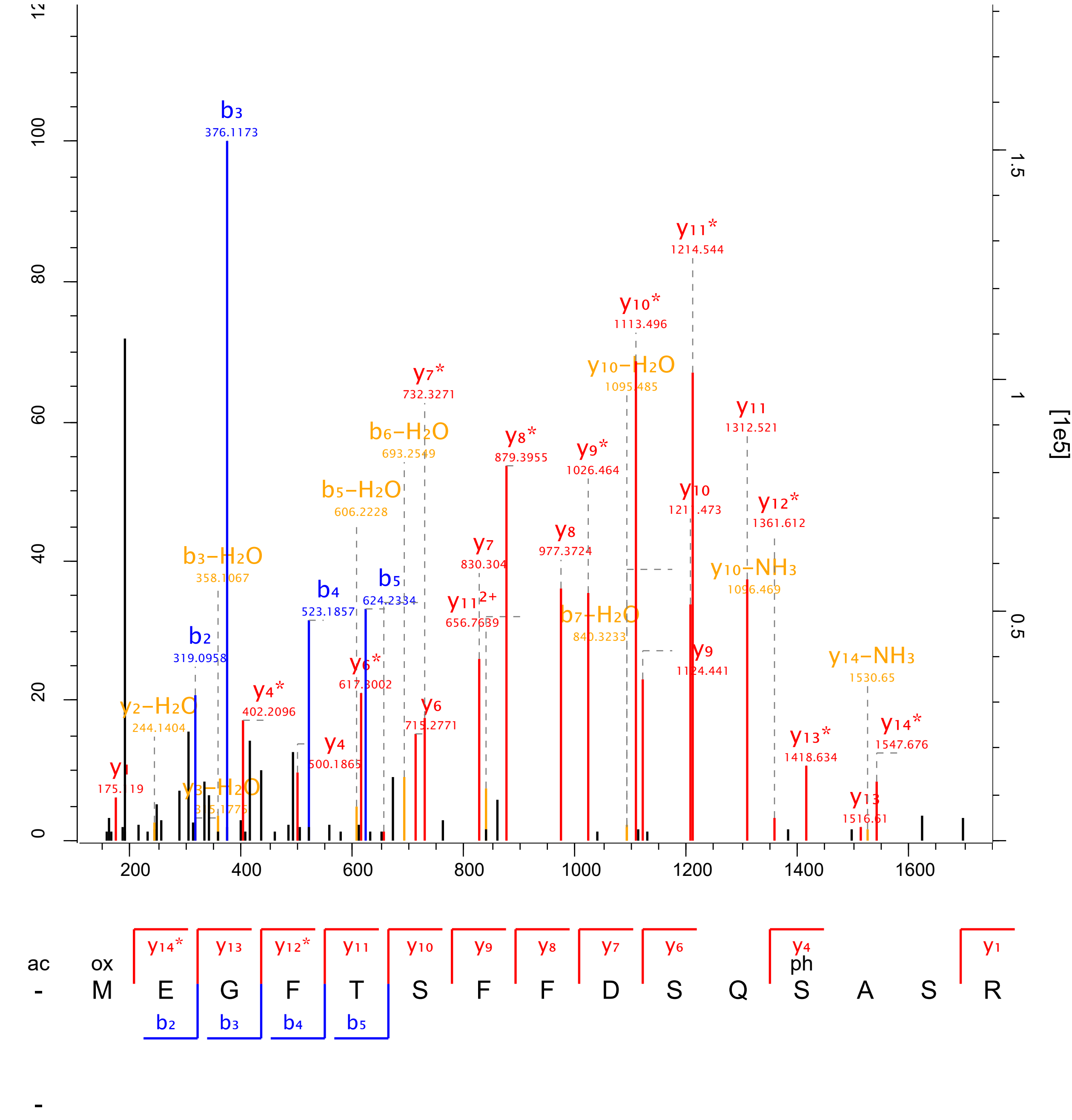

| Raw file | Scan | Method | Score | m/z |
|---|---|---|---|---|
| sys_05_1 | 7339 | FTMS; HCD | 92.54 | 427.71 |

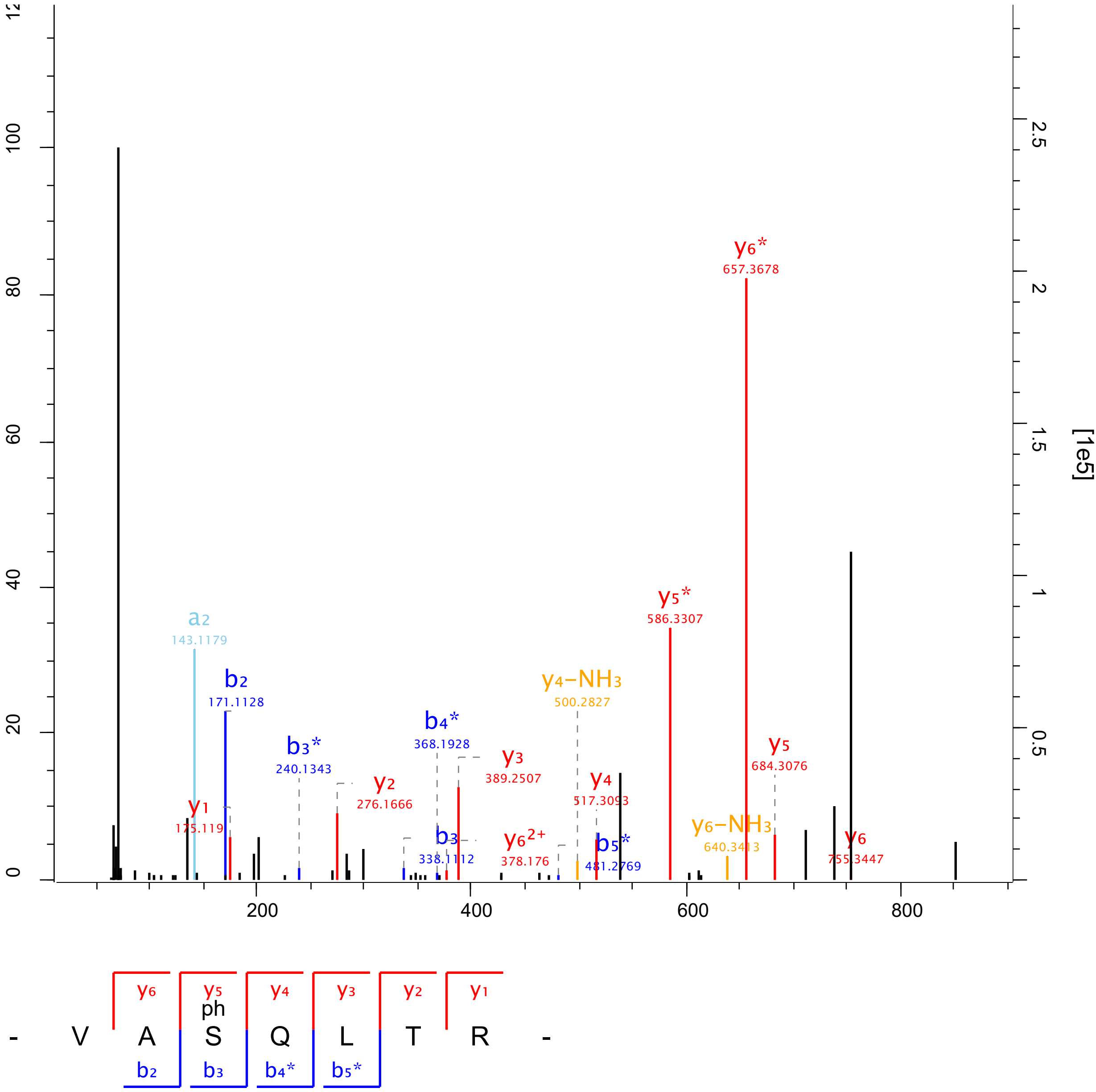

| Raw file | Scan | Method | Score | m/z |
| --- | --- | --- | --- | --- |
| sys_05_1 | 7348 | FTMS; HCD | 64.1 | 440.88 |

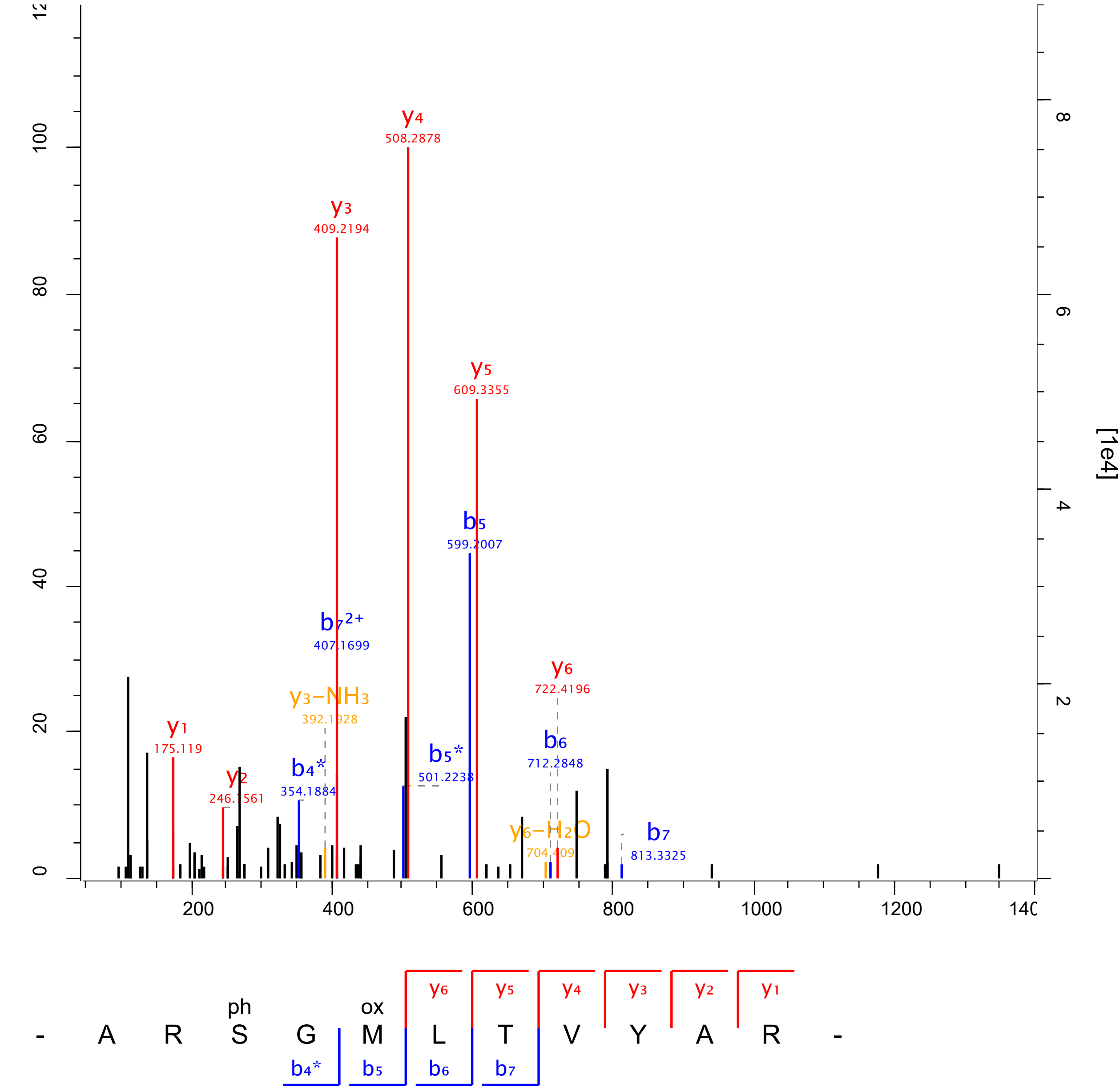

| Raw file | Scan | Method | Score | m/z |
|---|---|---|---|---|
| sys_05_1 | 8039 | FTMS; HCD | 174.44 | 633.74 |

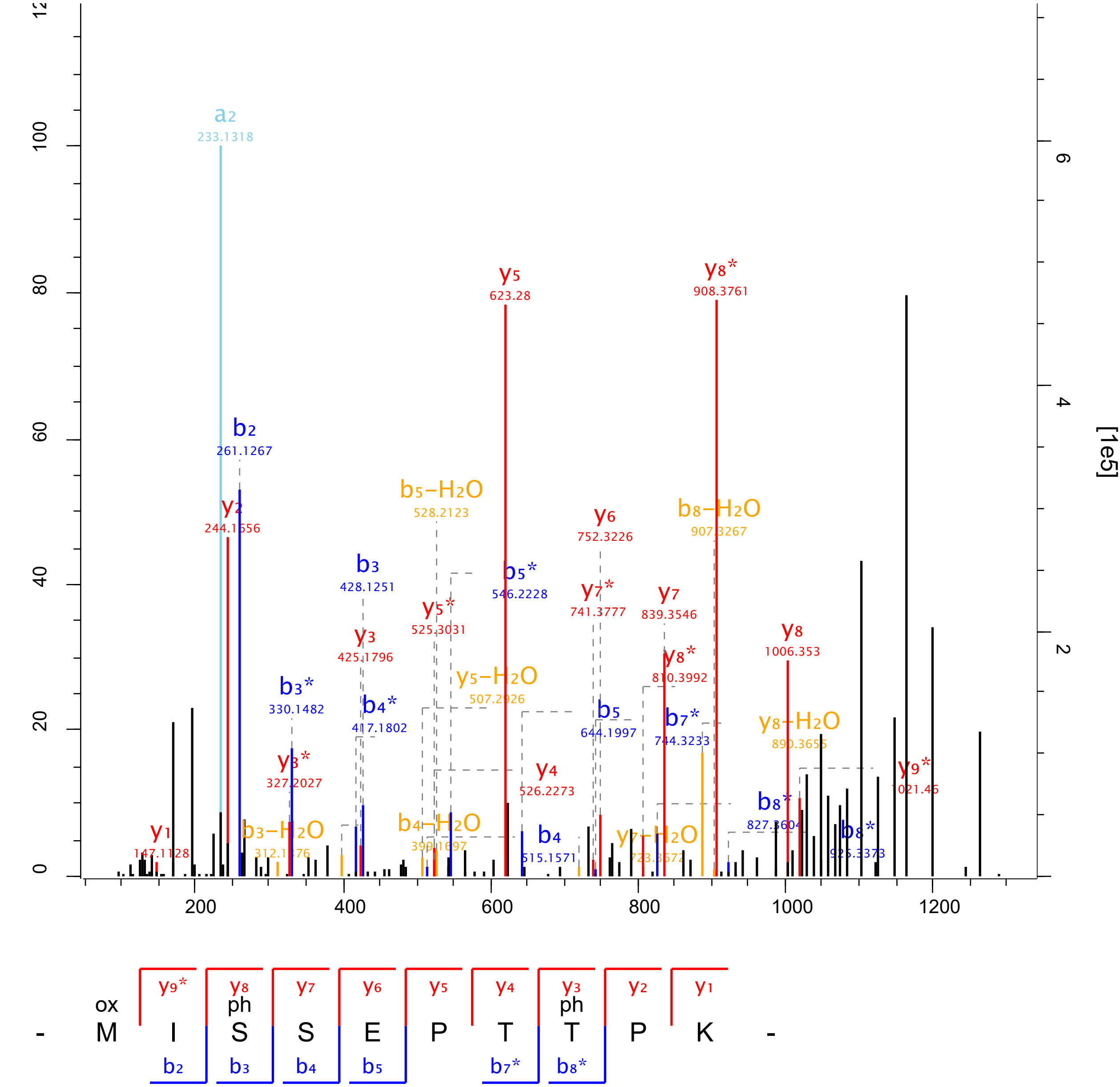

| Raw file | Scan | Method | Score | m/z |
| --- | --- | --- | --- | --- |
| sys_05_2 | 10087 | FTMS; HCD | 74.99 | 556.75 |

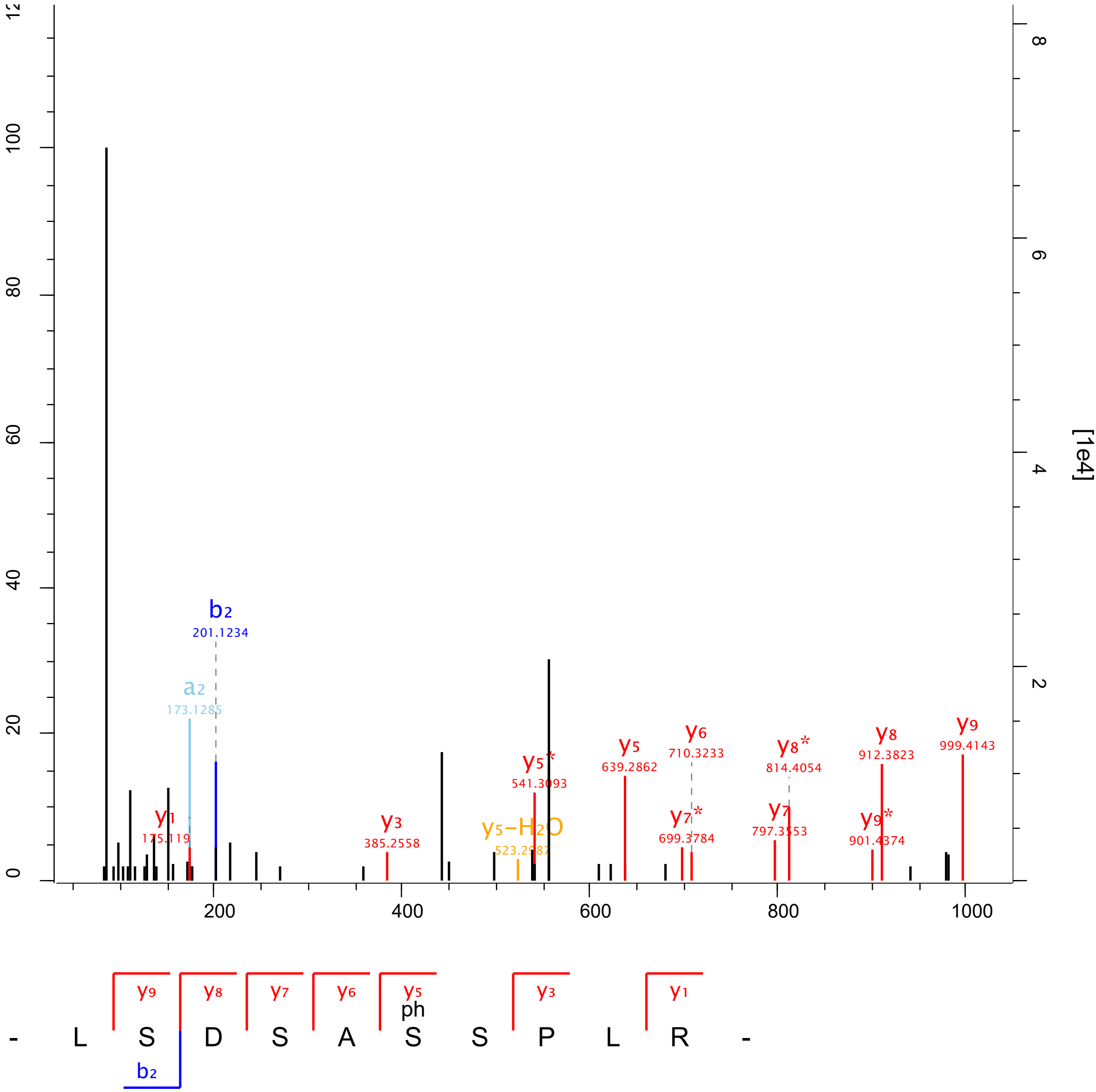

| Raw file | Scan | Method | Score | m/z |
|---|---|---|---|---|
| sys_05_2 | 10115 | FTMS; HCD | 114.97 | 428.7 |

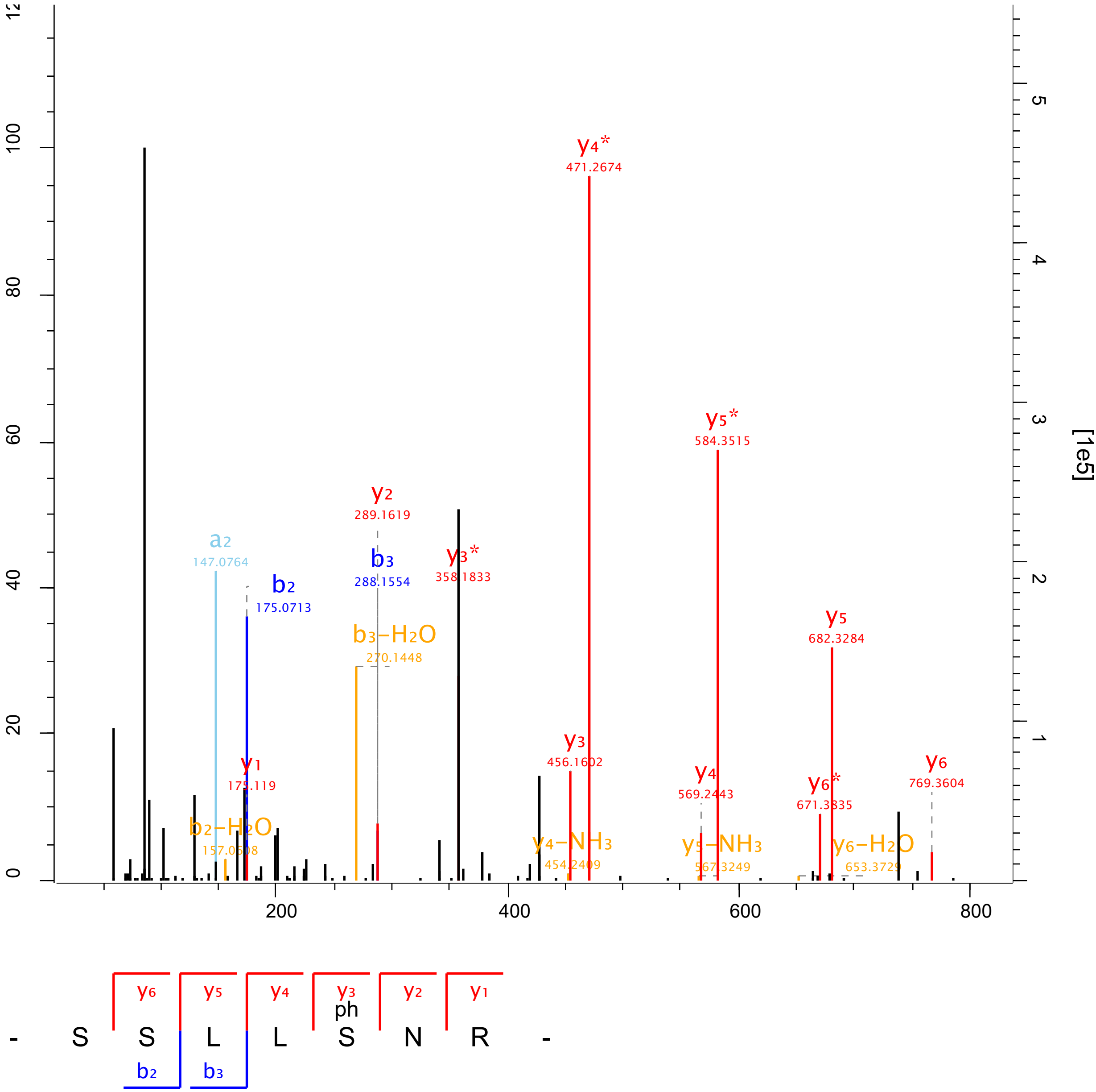

| Raw file | Scan | Method | Score | m/z |
| --- | --- | --- | --- | --- |
| sys_05_2 | 10137 | FTMS; HCD | 81.57 | 628.27 |

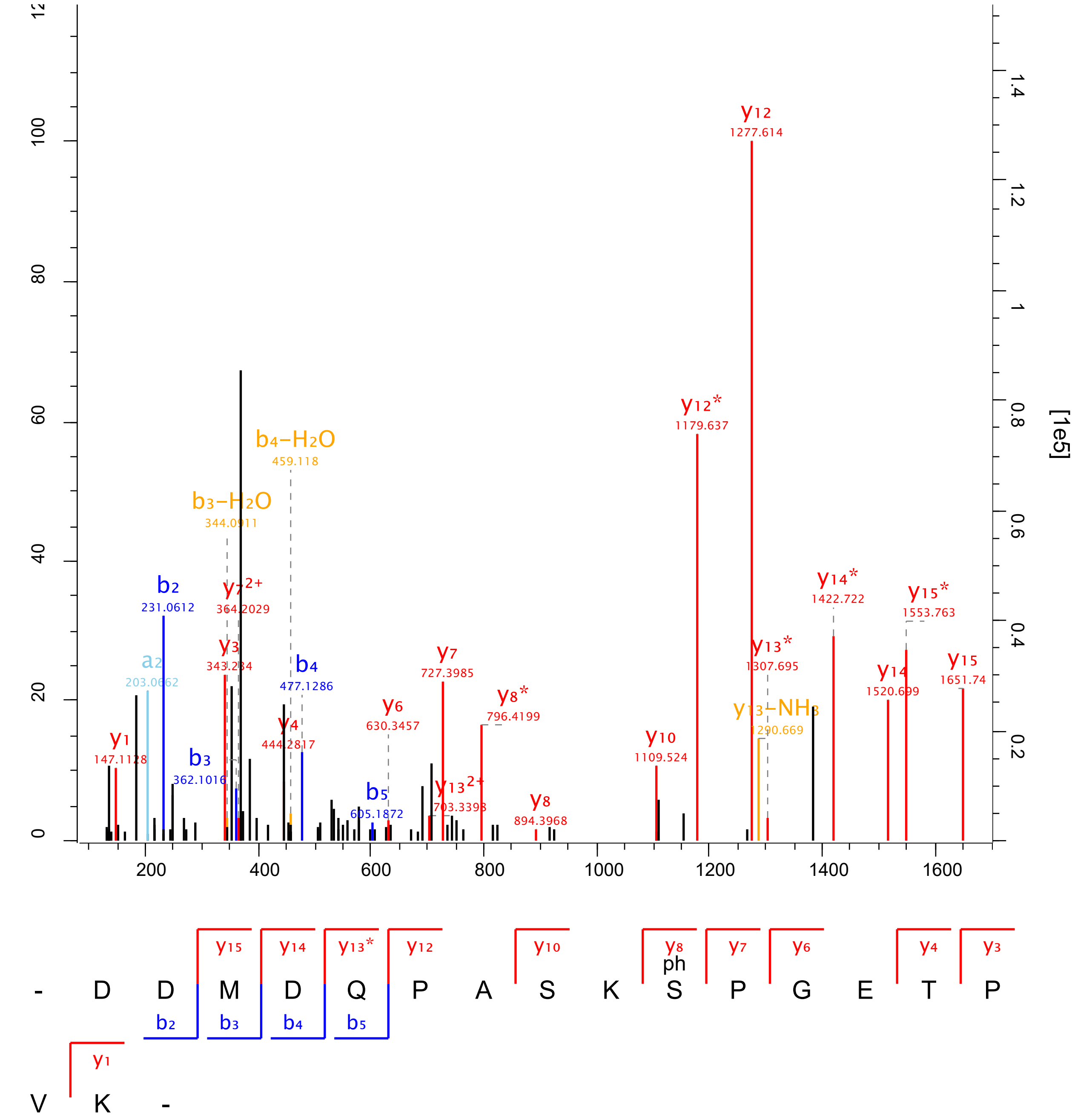

| Raw file | Scan | Method | Score | m/z |
|---|---|---|---|---|
| sys_05_2 | 10424 | FTMS; HCD | 96.11 | 427.72 |

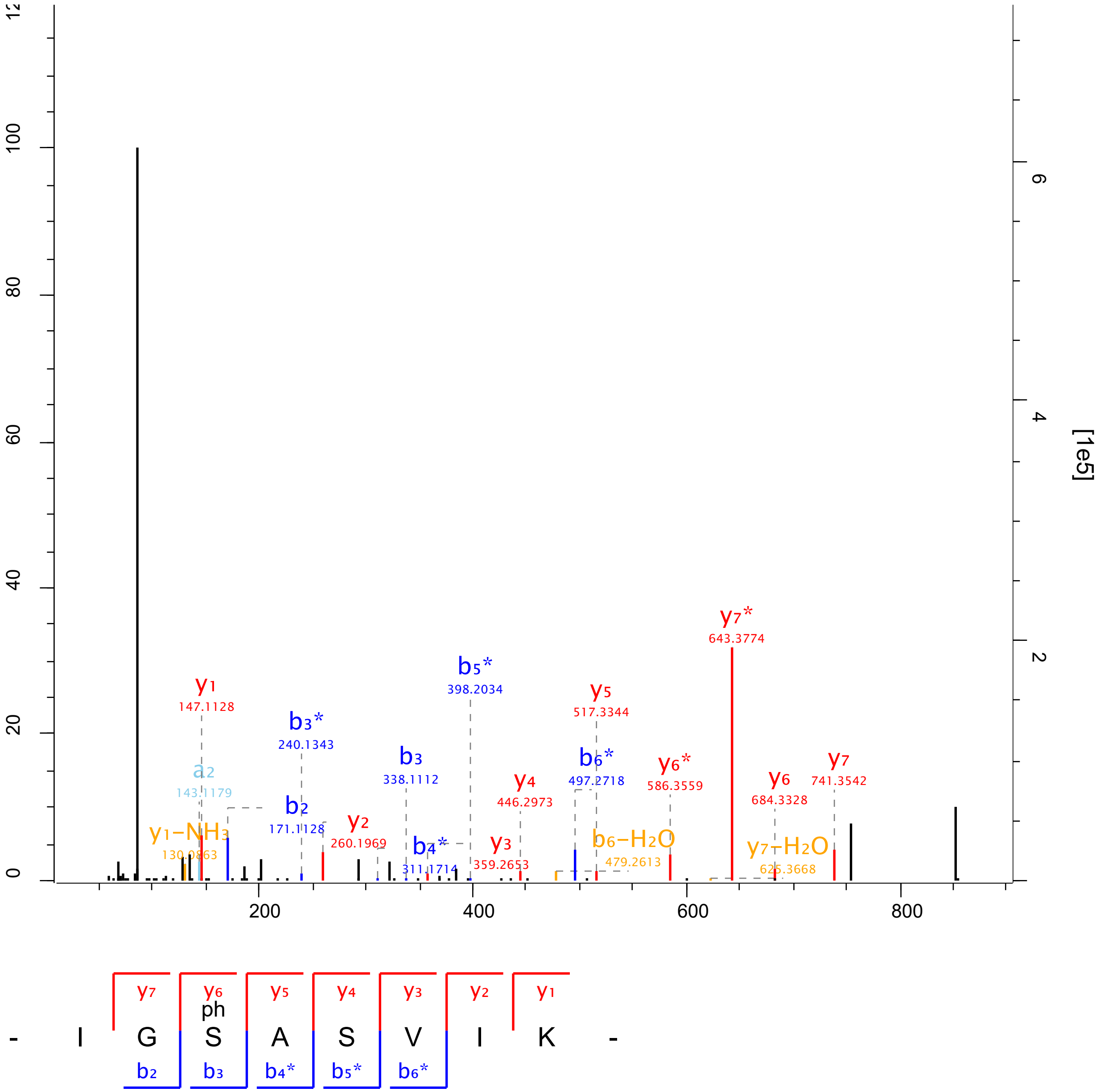

| Raw file | Scan | Method | Score | m/z |
| --- | --- | --- | --- | --- |
| sys_05_2 | 10455 | FTMS; HCD | 189.33 | 671.79 |

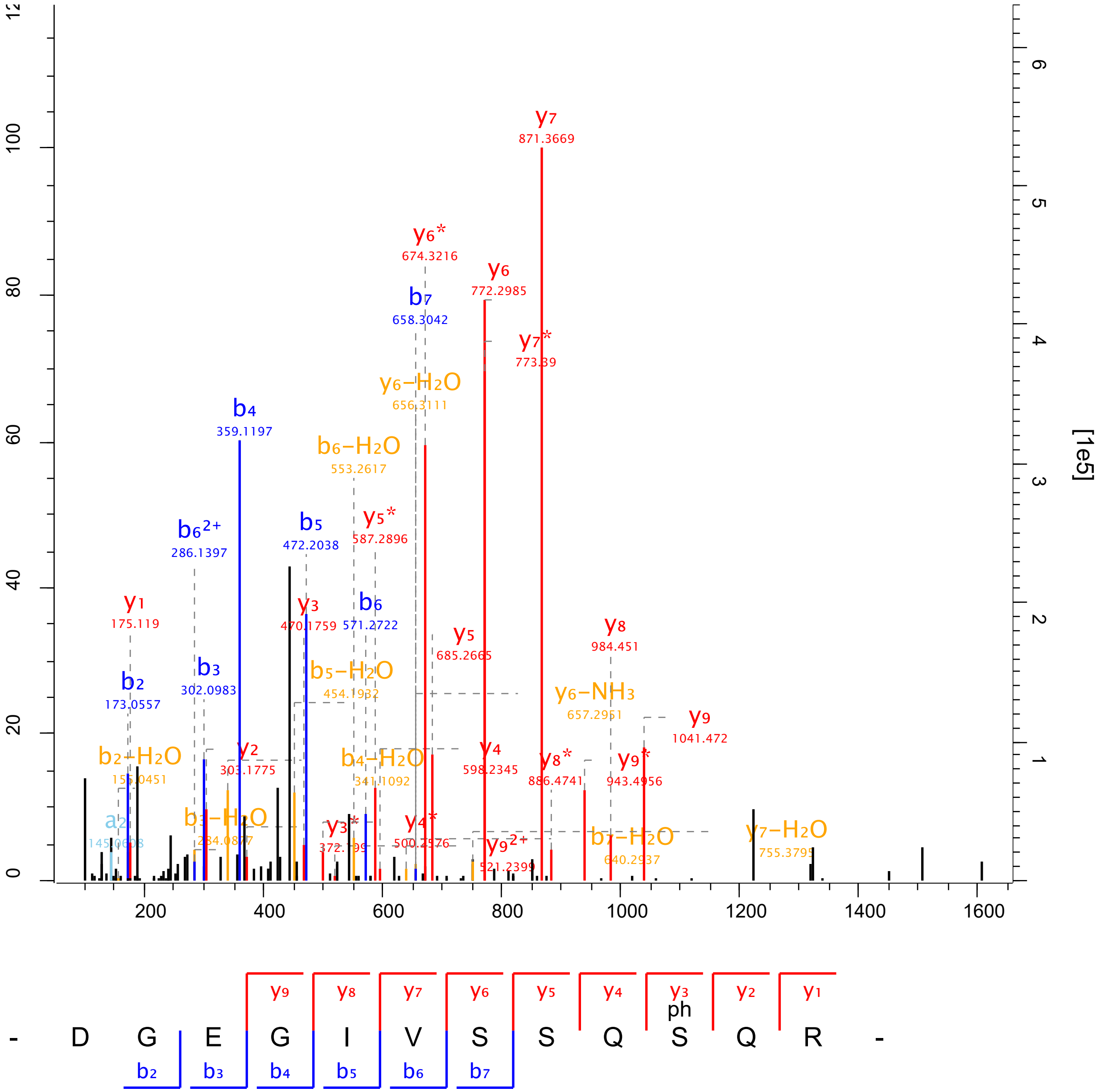

| Raw file | Scan | Method | Score | m/z |
|---|---|---|---|---|
| sys_05_2 | 10526 | FTMS; HCD | 275.14 | 935.84 |

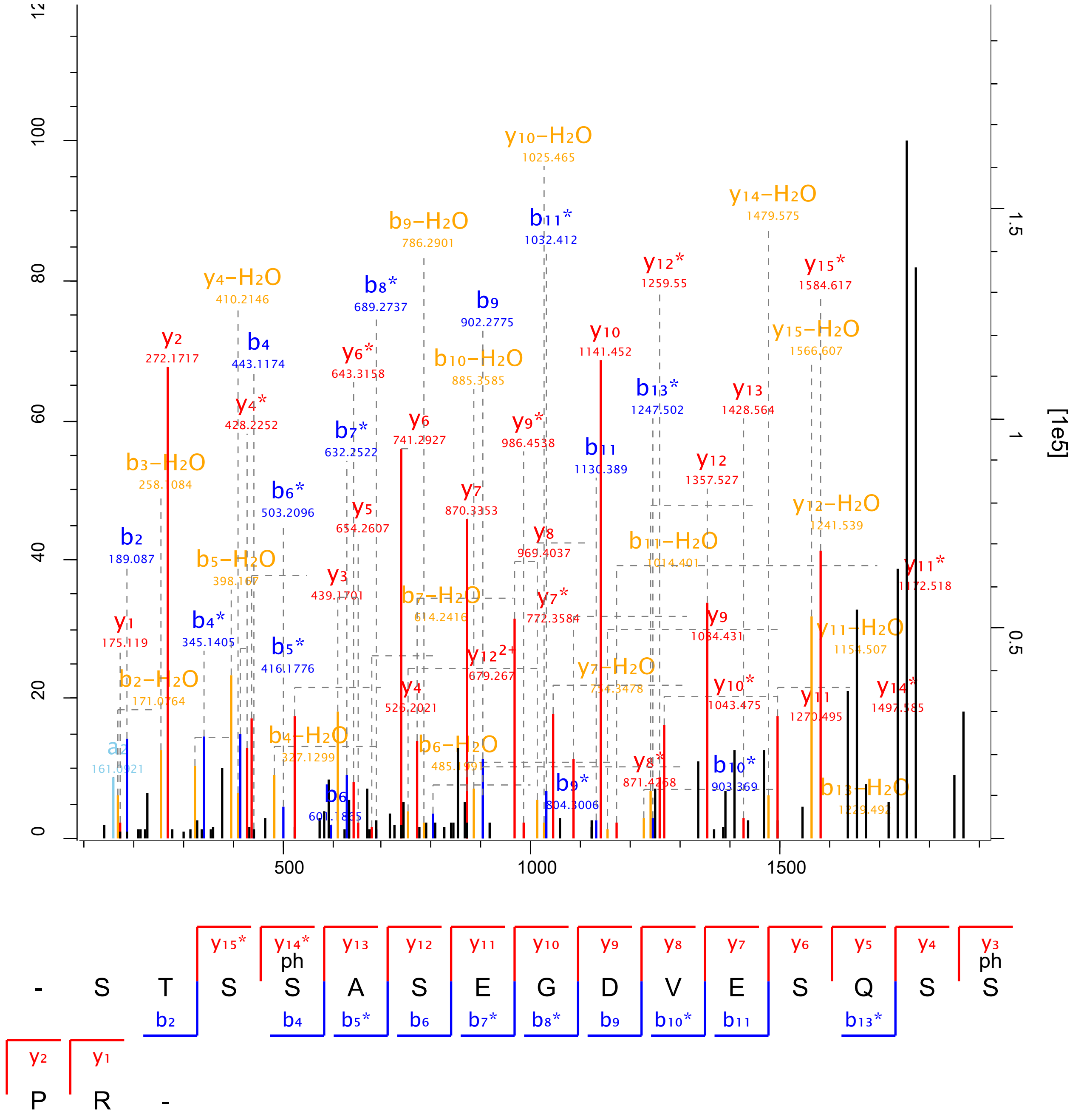

| Raw file | Scan | Method | Score | m/z |
|---|---|---|---|---|
| sys_05_2 | 10545 | FTMS; HCD | 184.58 | 495.72 |

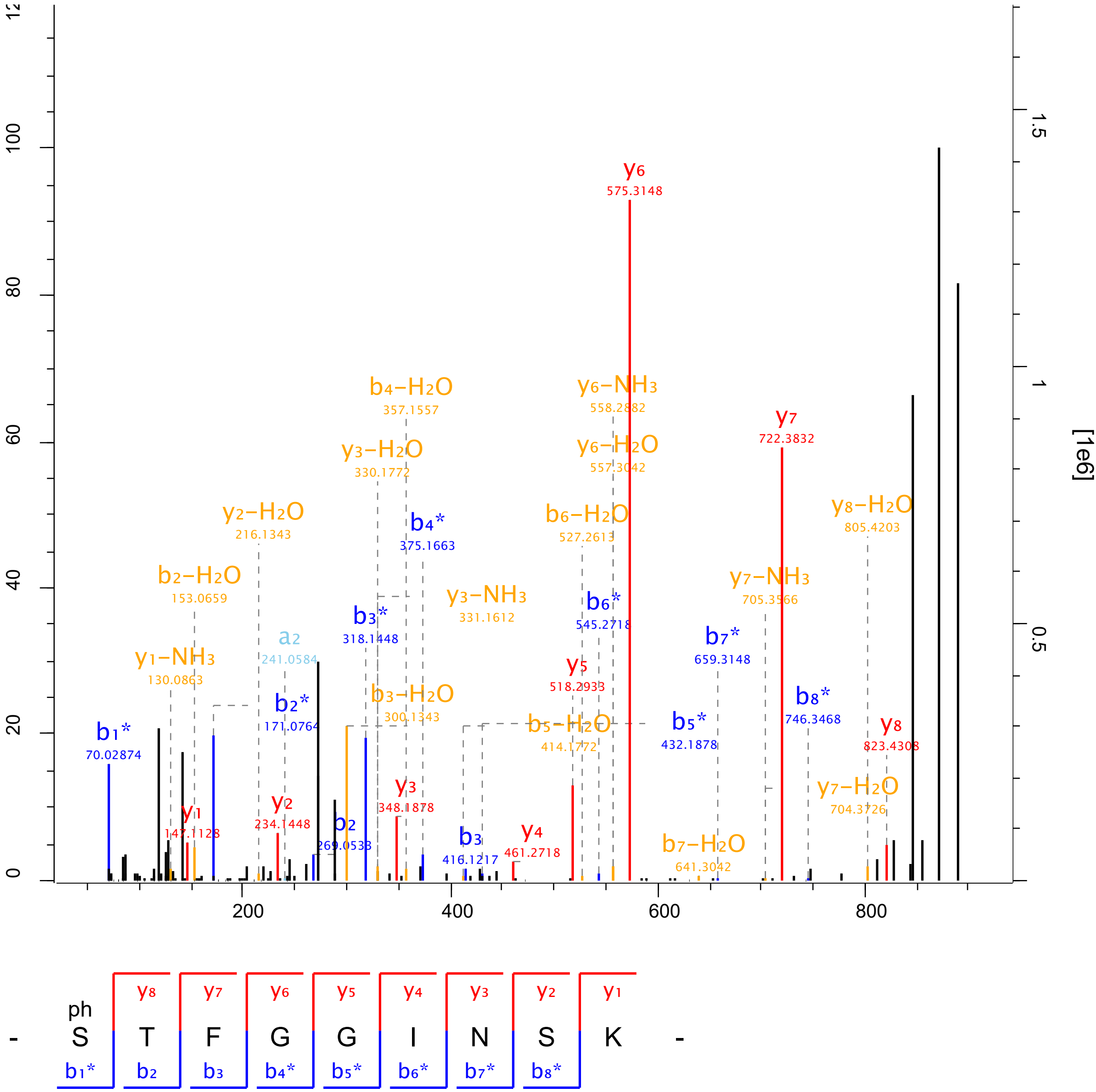

| Raw file | Scan | Method | Score | m/z |
| --- | --- | --- | --- | --- |
| sys_05_2 | 10652 | FTMS; HCD | 54.34 | 555.2 |

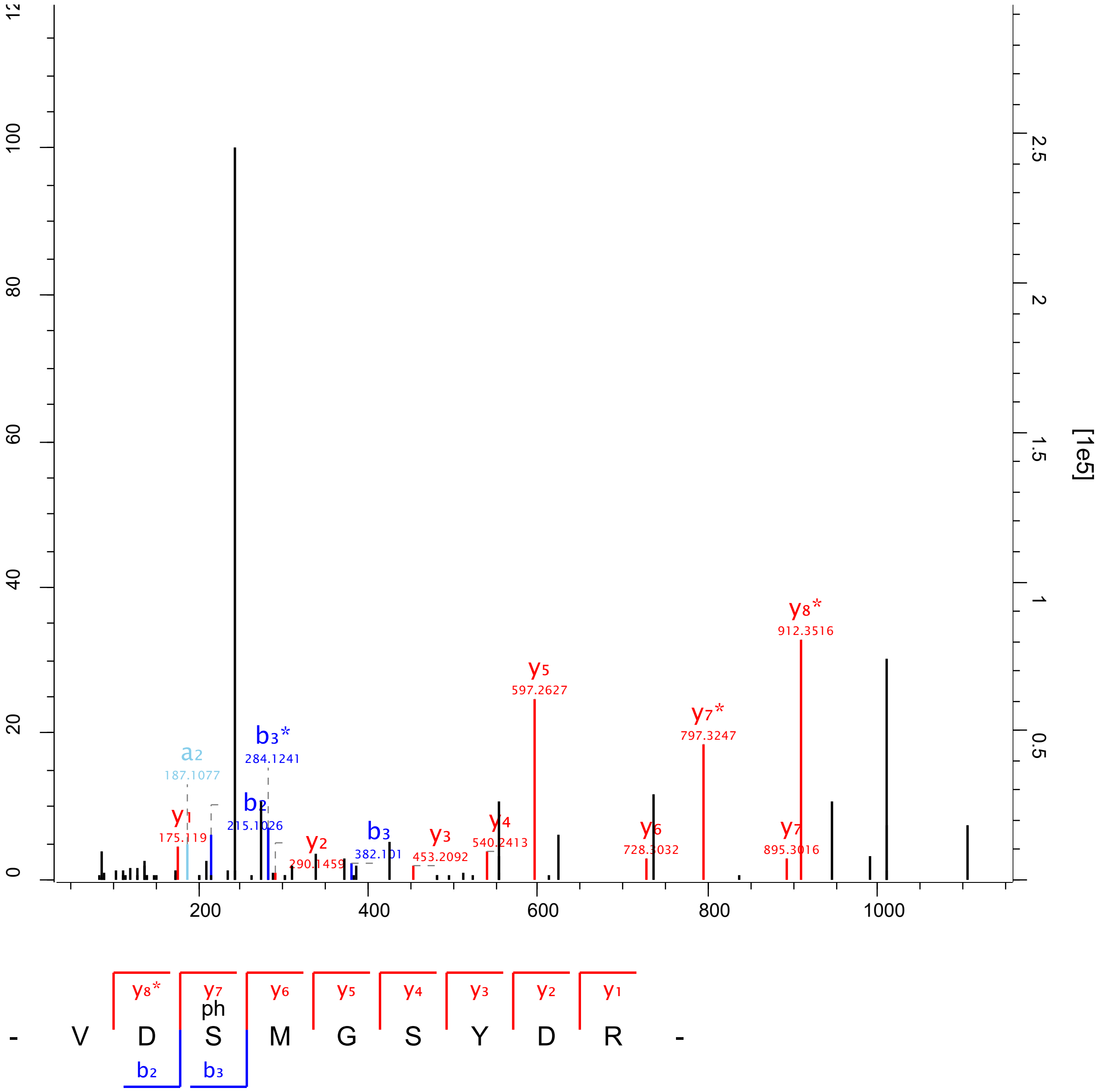

| Raw file | Scan | Method | Score | m/z |
| --- | --- | --- | --- | --- |
| sys_05_2 | 10685 | FTMS; HCD | 83.53 | 570.77 |

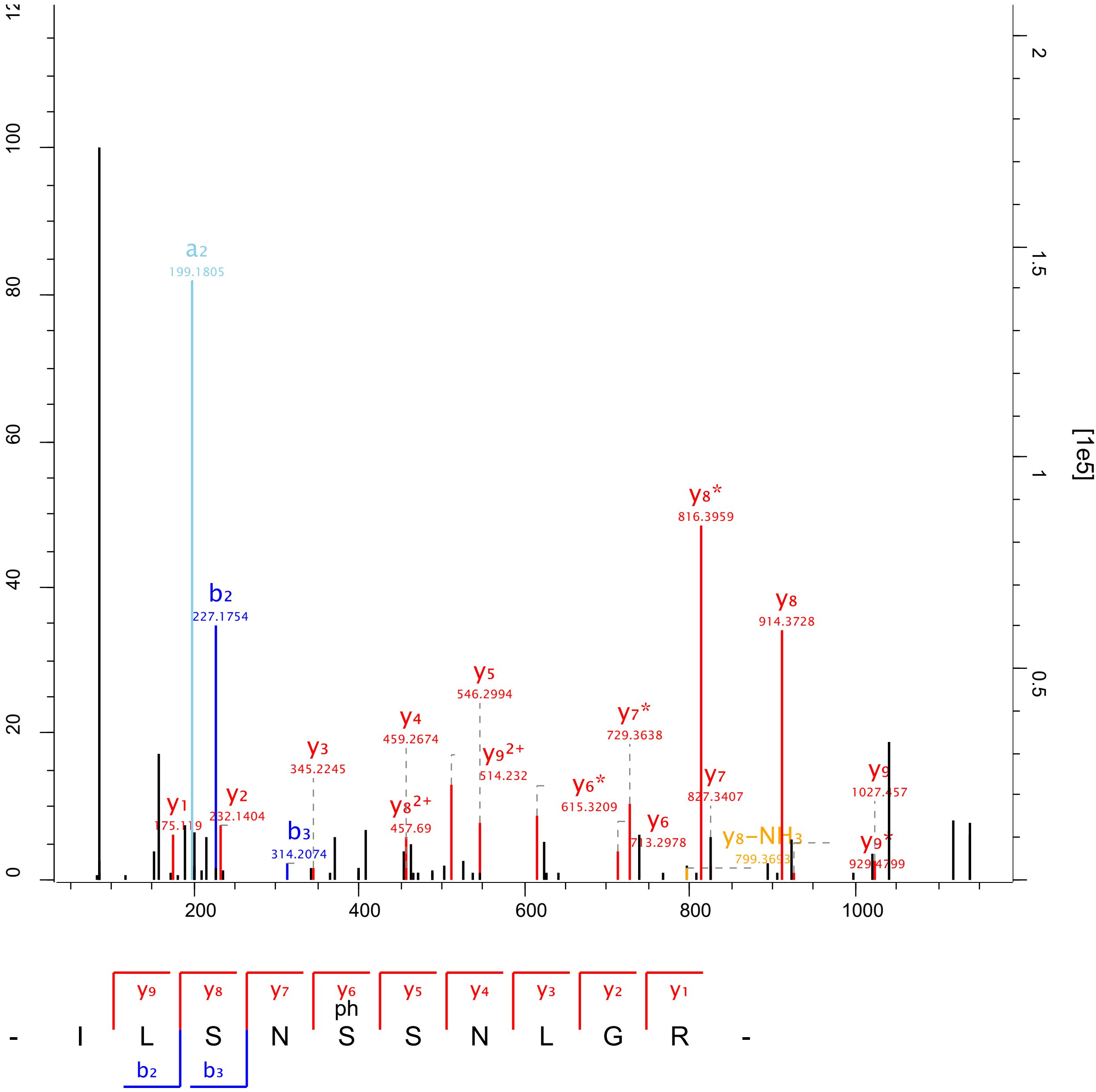

| Raw file | Scan | Method | Score | m/z |
| --- | --- | --- | --- | --- |
| sys_05_2 | 10875 | FTMS; HCD | 184.99 | 637.27 |

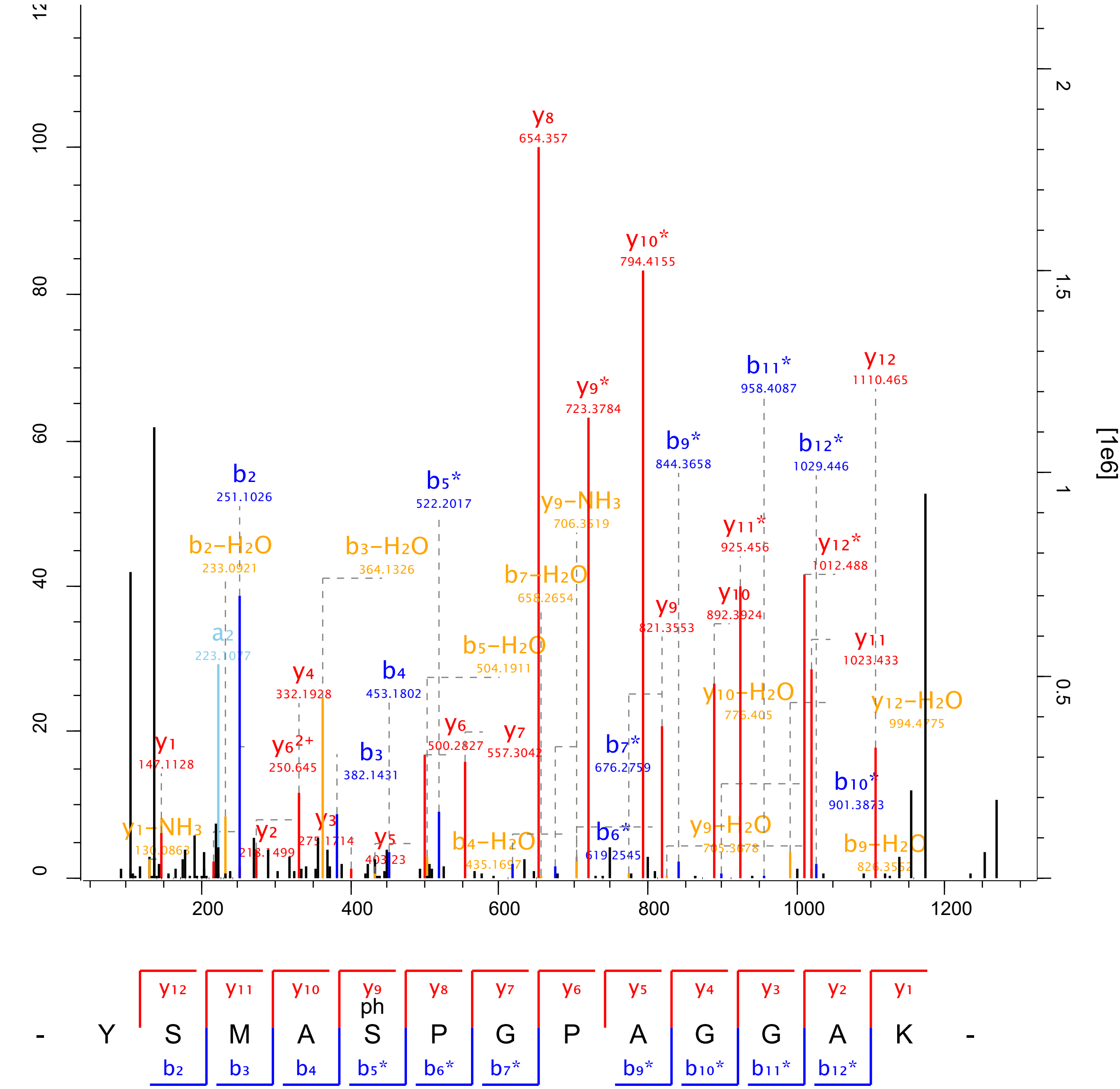

| Raw file | Scan | Method | Score | m/z |
|---|---|---|---|---|
| sys_05_2 | 11317 | FTMS; HCD | 62.16 | 598.73 |

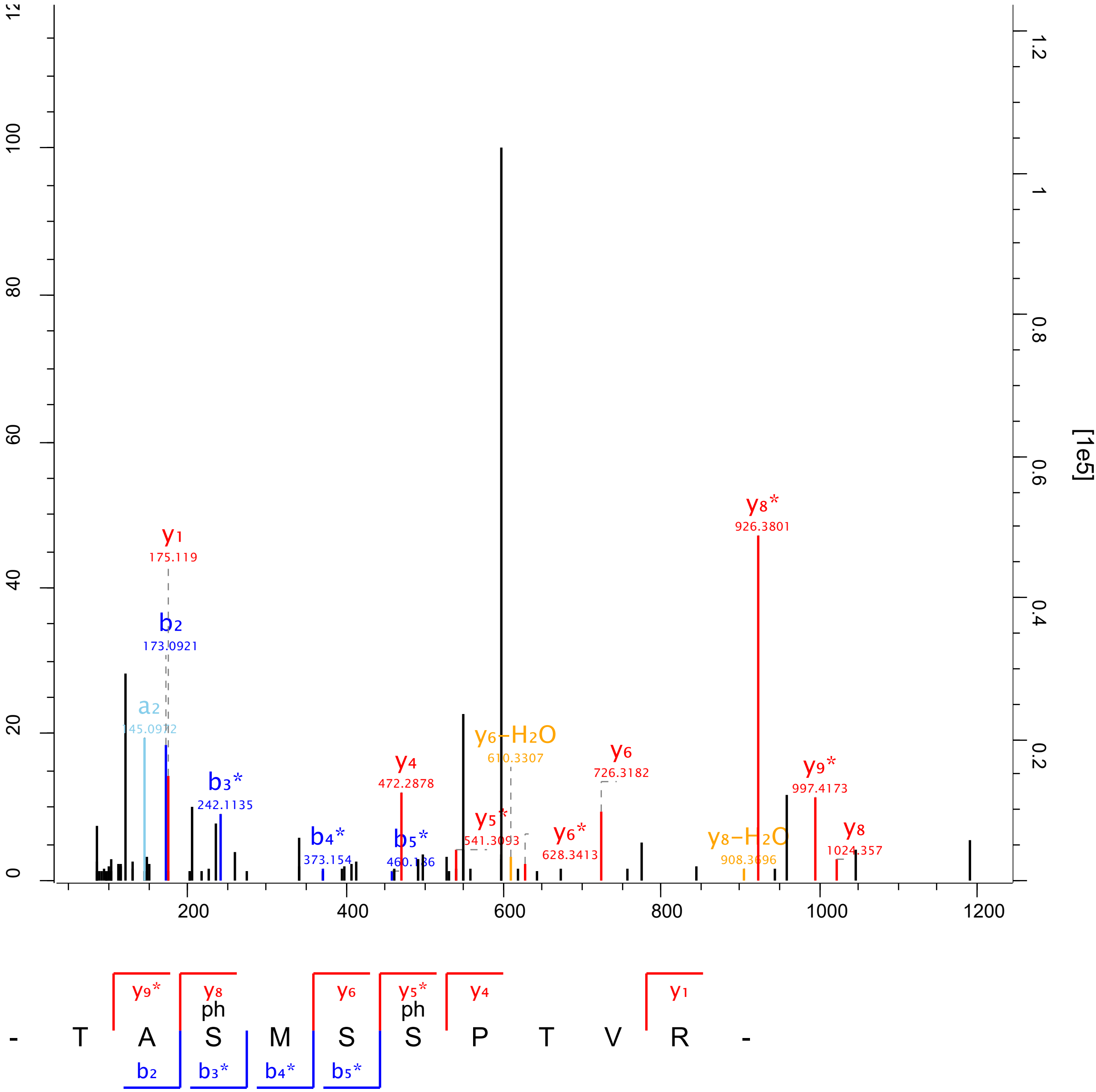

| Raw file | Scan | Method | Score | m/z |
|---|---|---|---|---|
| sys_05_2 | 11428 | FTMS; HCD | 229.59 | 584.22 |

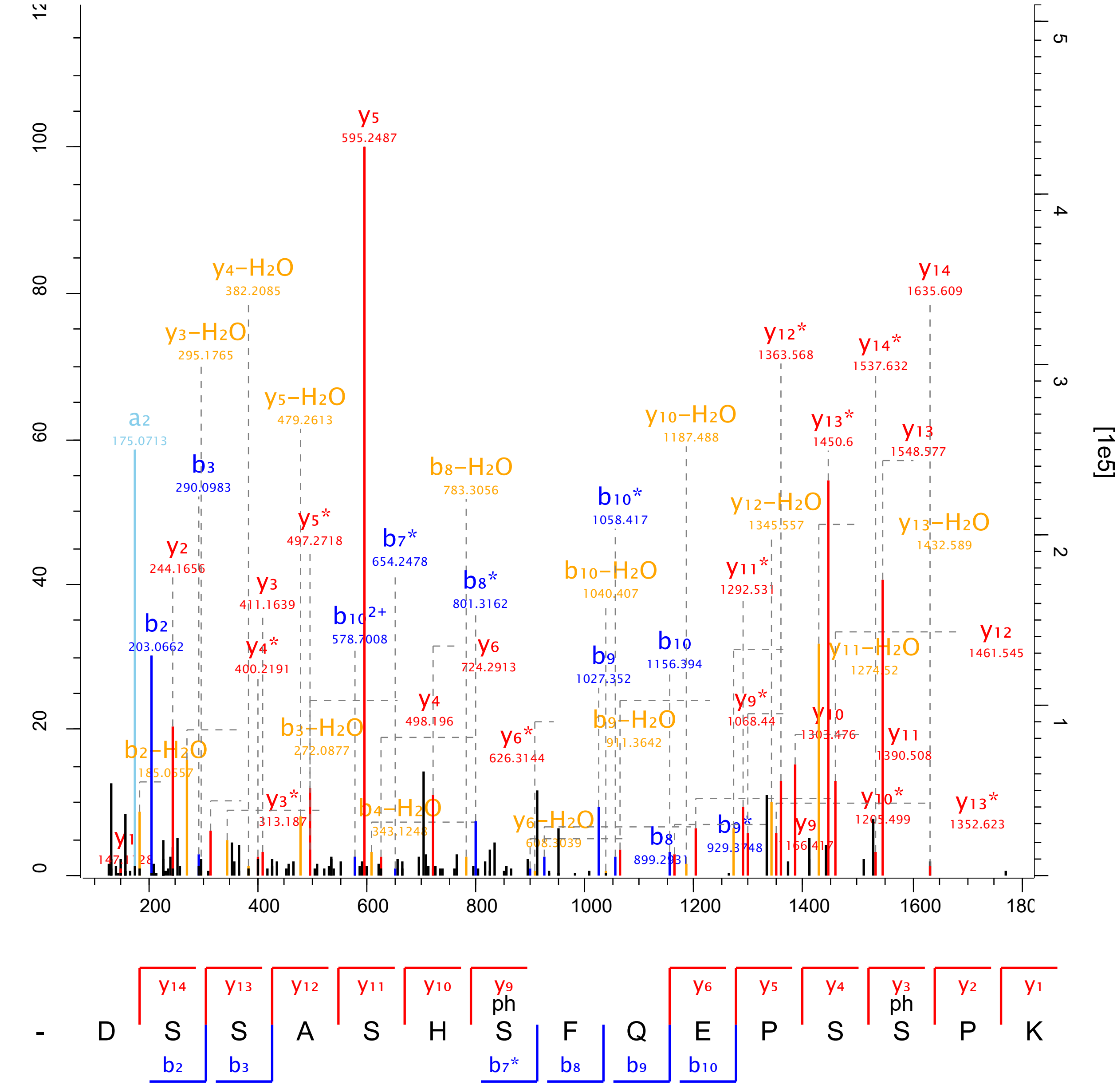

| Raw file | Scan | Method | Score | m/z |
|---|---|---|---|---|
| sys_05_2 | 11434 | FTMS; HCD | 196.16 | 530.25 |

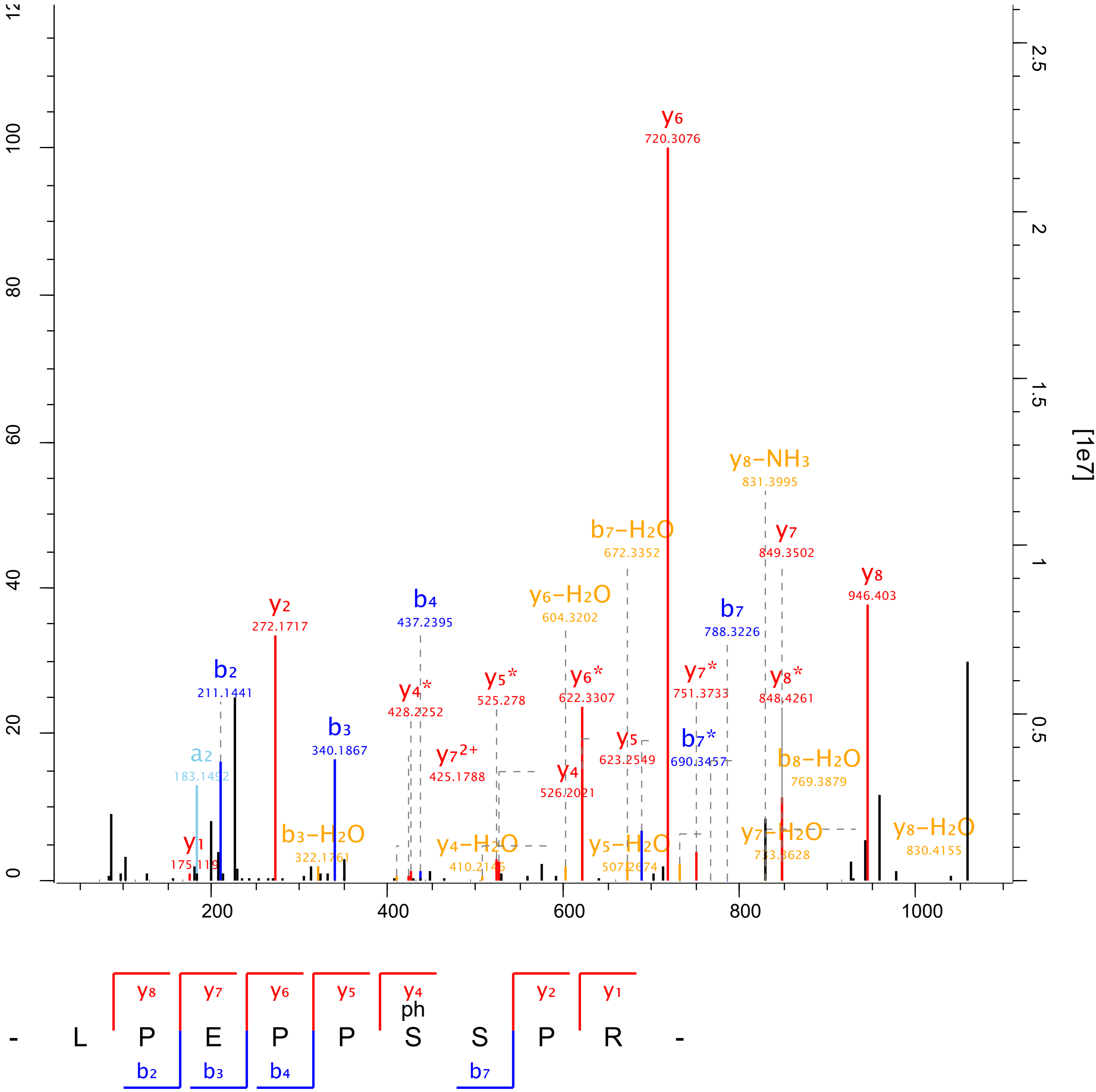

| Raw file | Scan | Method | Score | m/z |
|---|---|---|---|---|
| sys_05_2 | 11488 | FTMS; HCD | 220.25 | 560.24 |

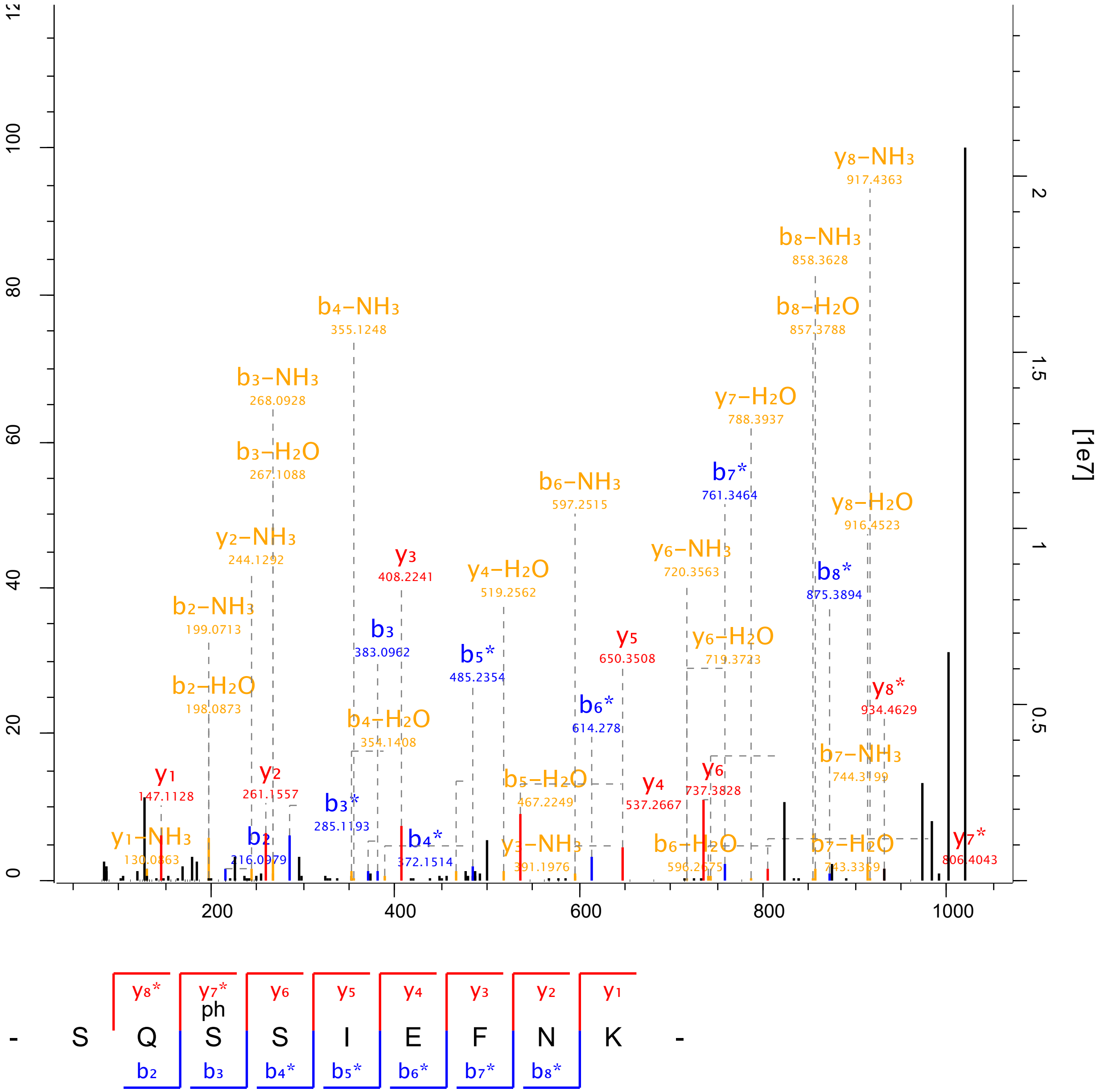

| Raw file | Scan | Method | Score | m/z |
|---|---|---|---|---|
| sys_05_2 | 11562 | FTMS; HCD | 155.96 | 701.79 |

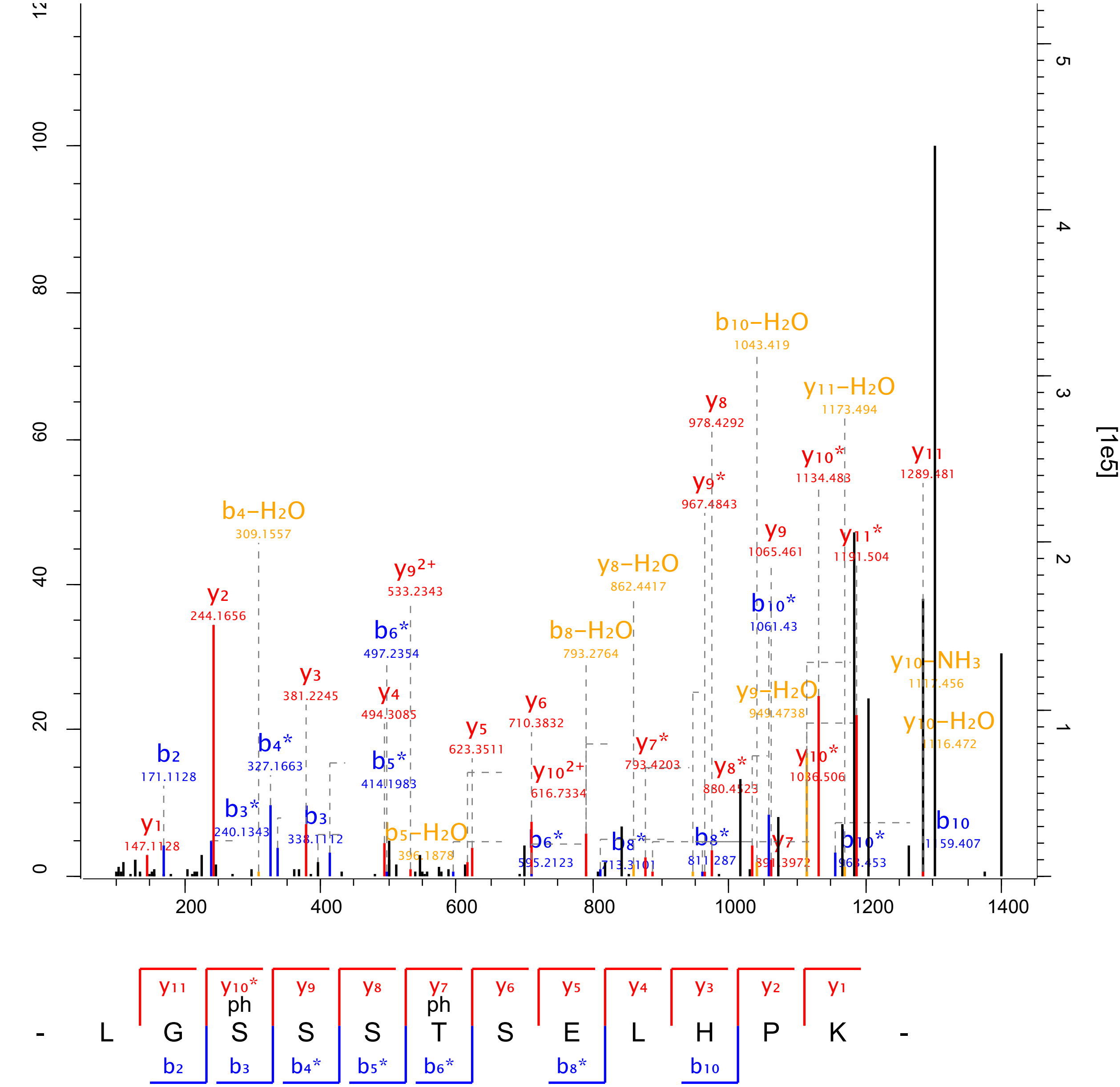

| Raw file | Scan | Method | Score | m/z |
|---|---|---|---|---|
| sys_05_2 | 11572 | FTMS; HCD | 59.91 | 535.24 |

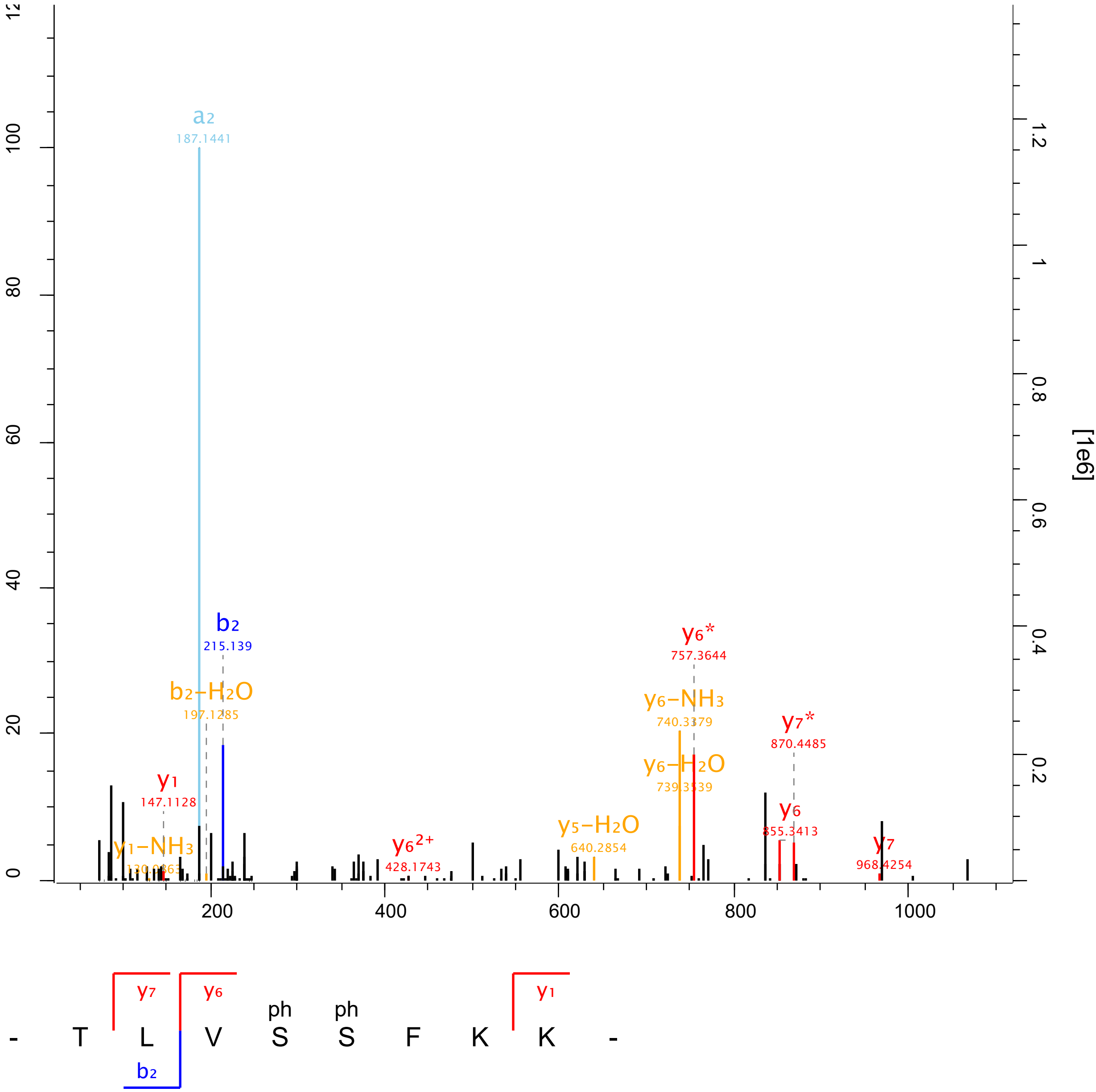

| Raw file | Scan | Method | Score | m/z |
| --- | --- | --- | --- | --- |
| sys_05_2 | 11623 | FTMS; HCD | 119.39 | 588.79 |

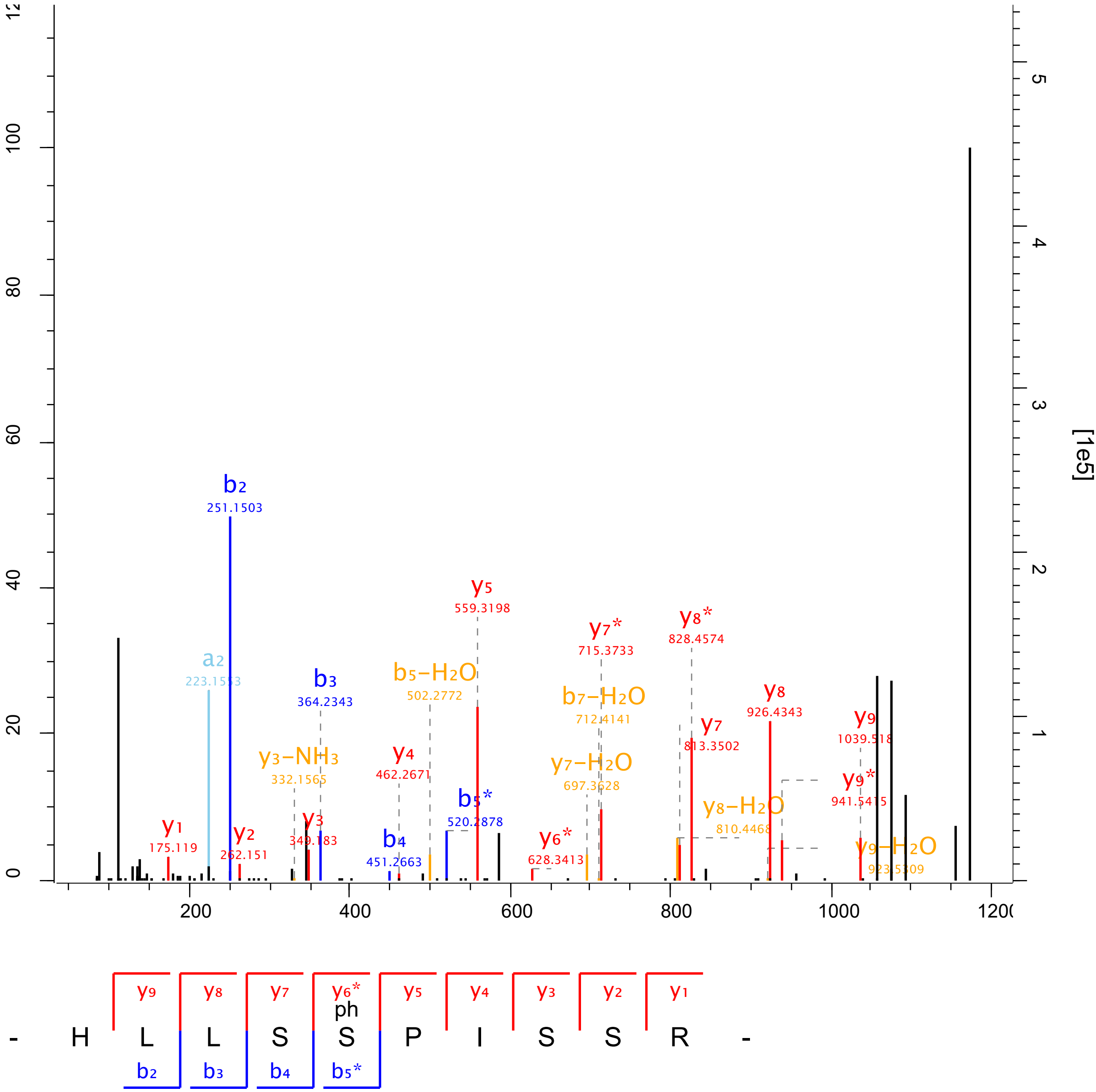

| Raw file | Scan | Method | Score | m/z |
|---|---|---|---|---|
| sys_05_2 | 11628 | FTMS; HCD | 81.16 | 685.82 |

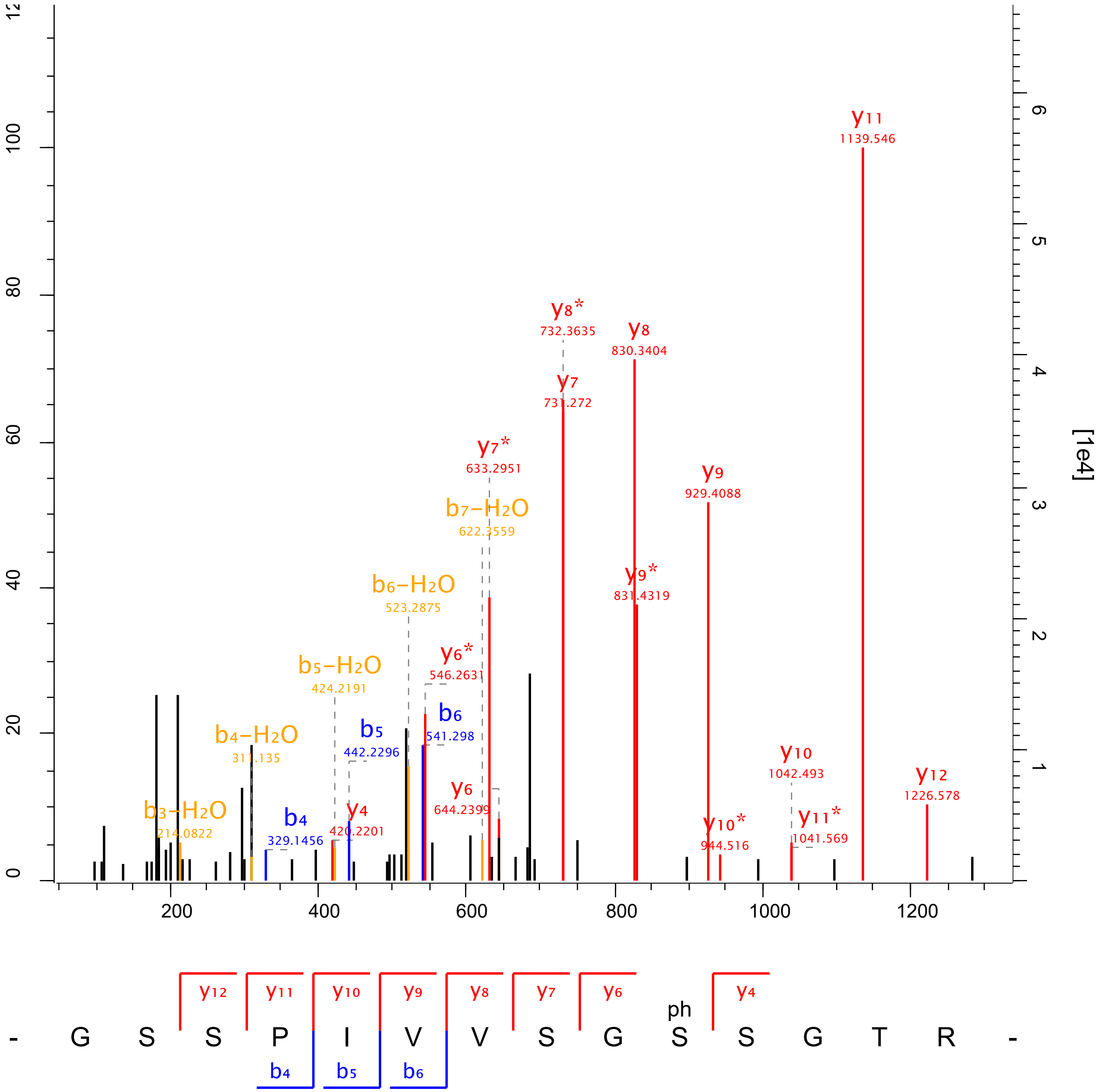

| Raw file | Scan | Method | Score | m/z |
| --- | --- | --- | --- | --- |
| sys_05_2 | 11721 | FTMS; HCD | 163.33 | 692.26 |

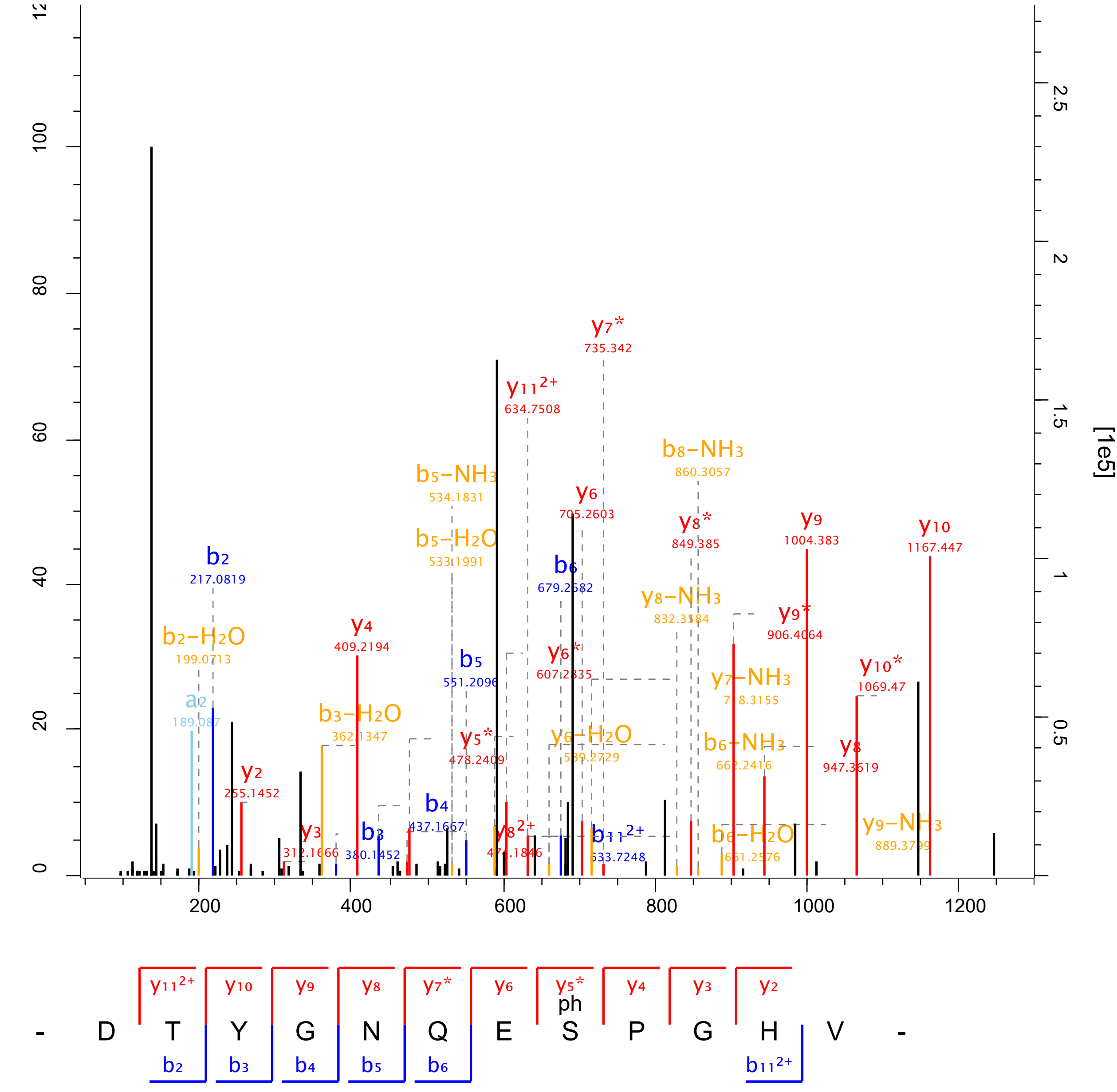

| Raw file | Scan | Method | Score | m/z |
|---|---|---|---|---|
| sys_05_2 | 11792 | FTMS; HCD | 132.84 | 810.78 |

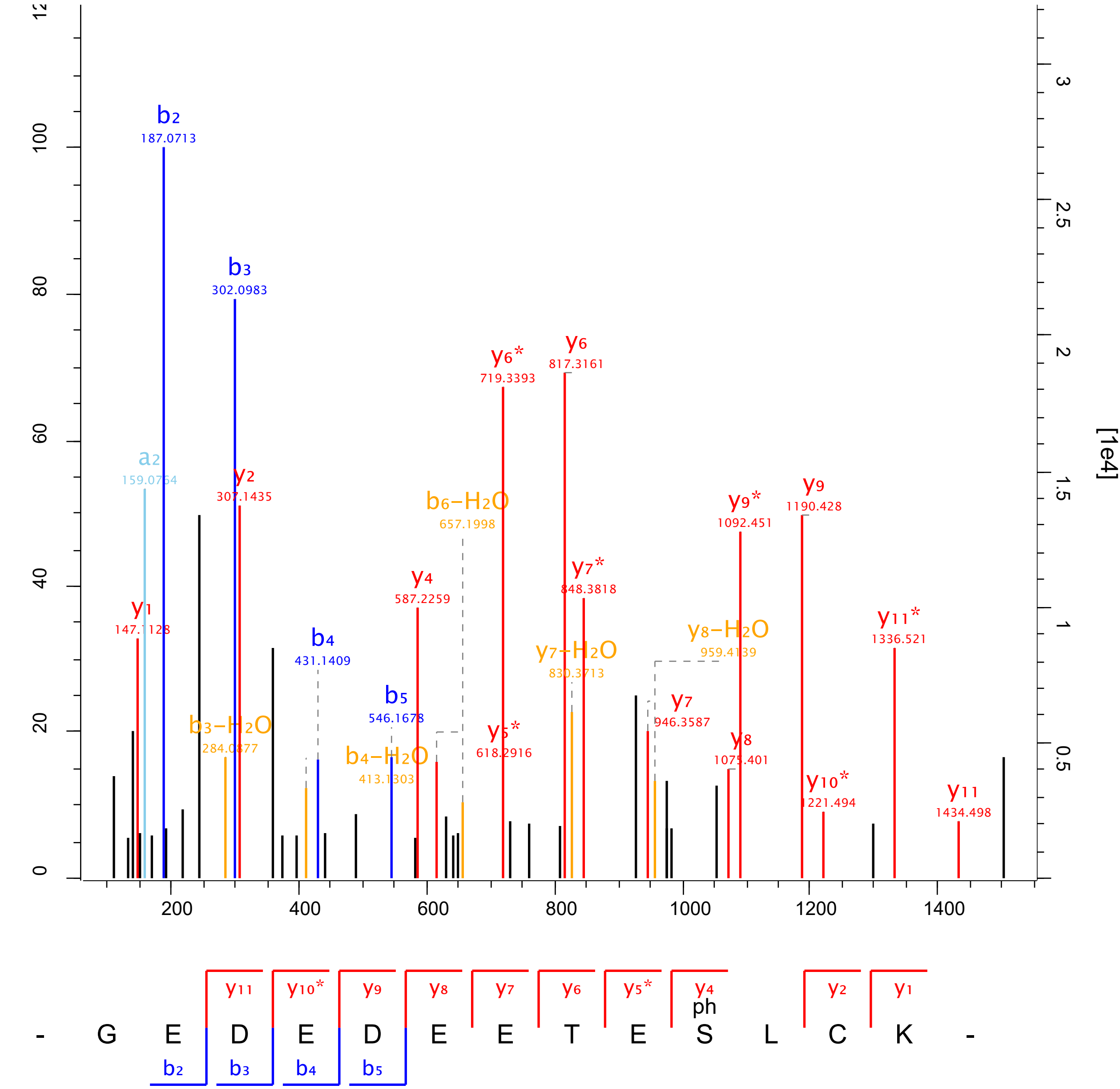

| Raw file | Scan | Method | Score | m/z |
|---|---|---|---|---|
| sys_05_2 | 11800 | FTMS; HCD | 74.24 | 544.24 |

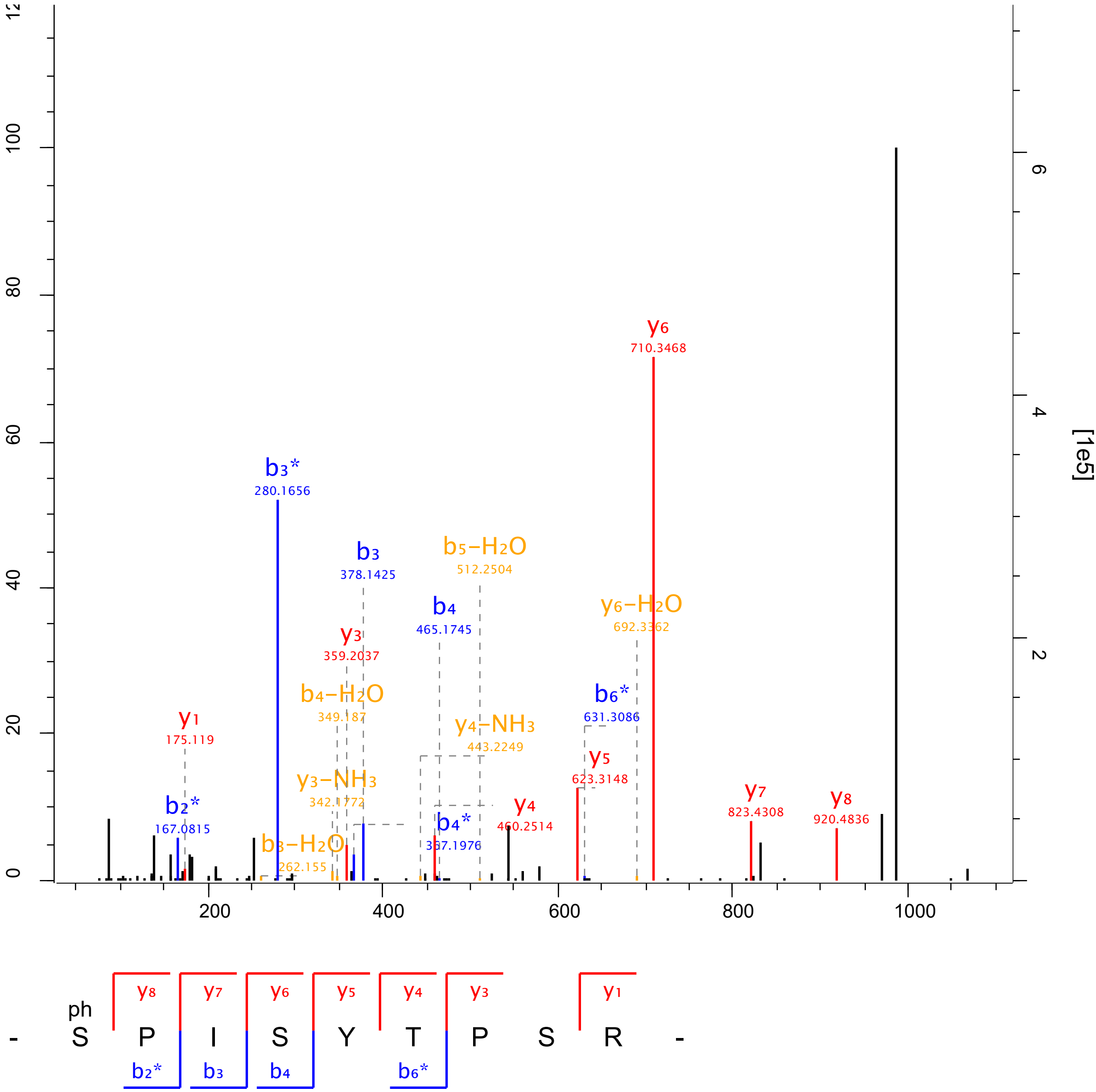

| Raw file | Scan | Method | Score | m/z |
|---|---|---|---|---|
| sys_05_2 | 12158 | FTMS; HCD | 76.94 | 505.72 |

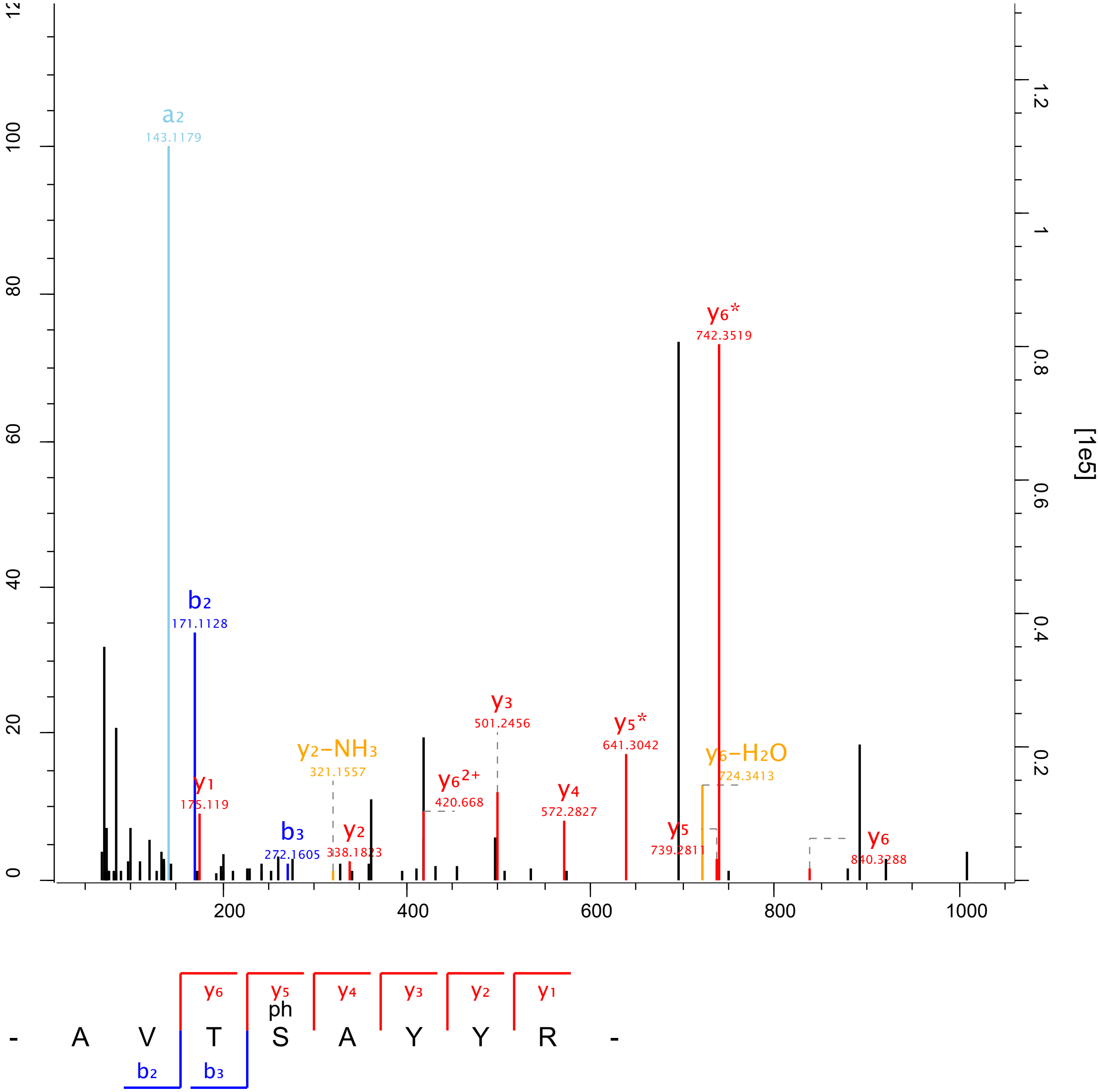

| Raw file | Scan | Method | Score | m/z |
| --- | --- | --- | --- | --- |
| sys_05_2 | 12330 | FTMS; HCD | 112.3 | 466.18 |

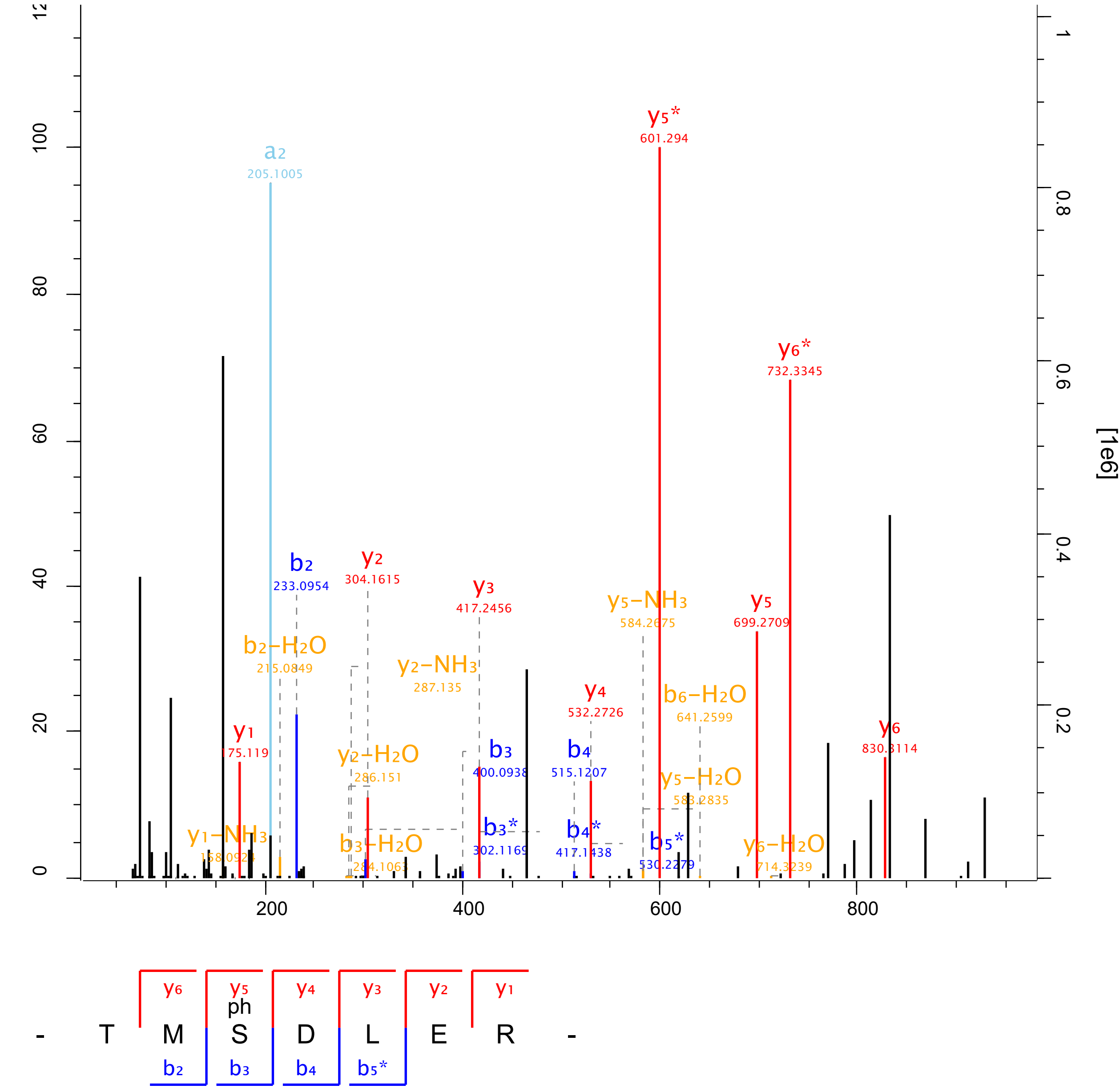

| Raw file | Scan | Method | Score | m/z |
|---|---|---|---|---|
| sys_05_2 | 12433 | FTMS; HCD | 73.44 | 507.23 |

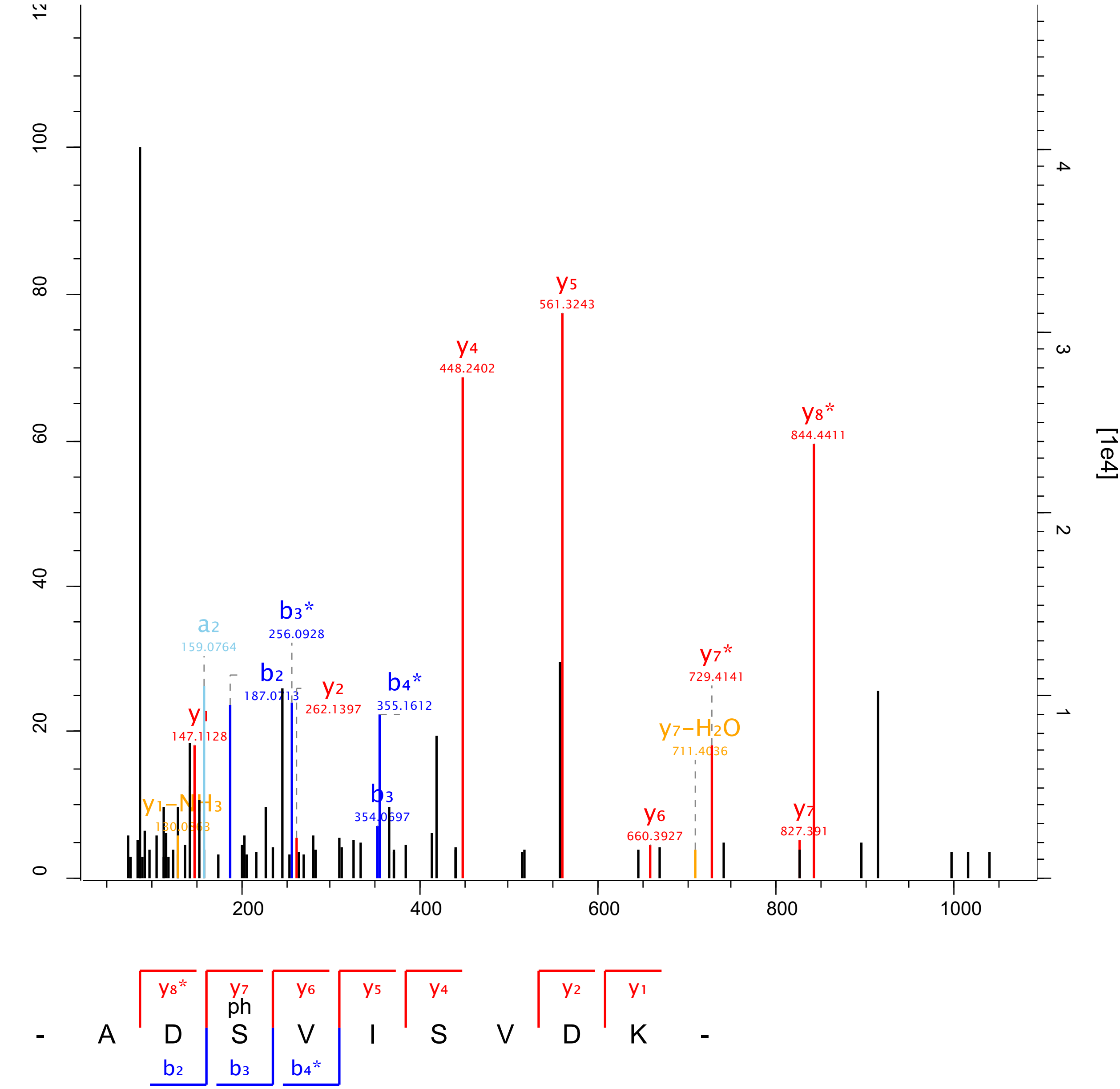

| Raw file | Scan | Method | Score | m/z |
|---|---|---|---|---|
| sys_05_2 | 12642 | FTMS; HCD | 145.9 | 436.7 |

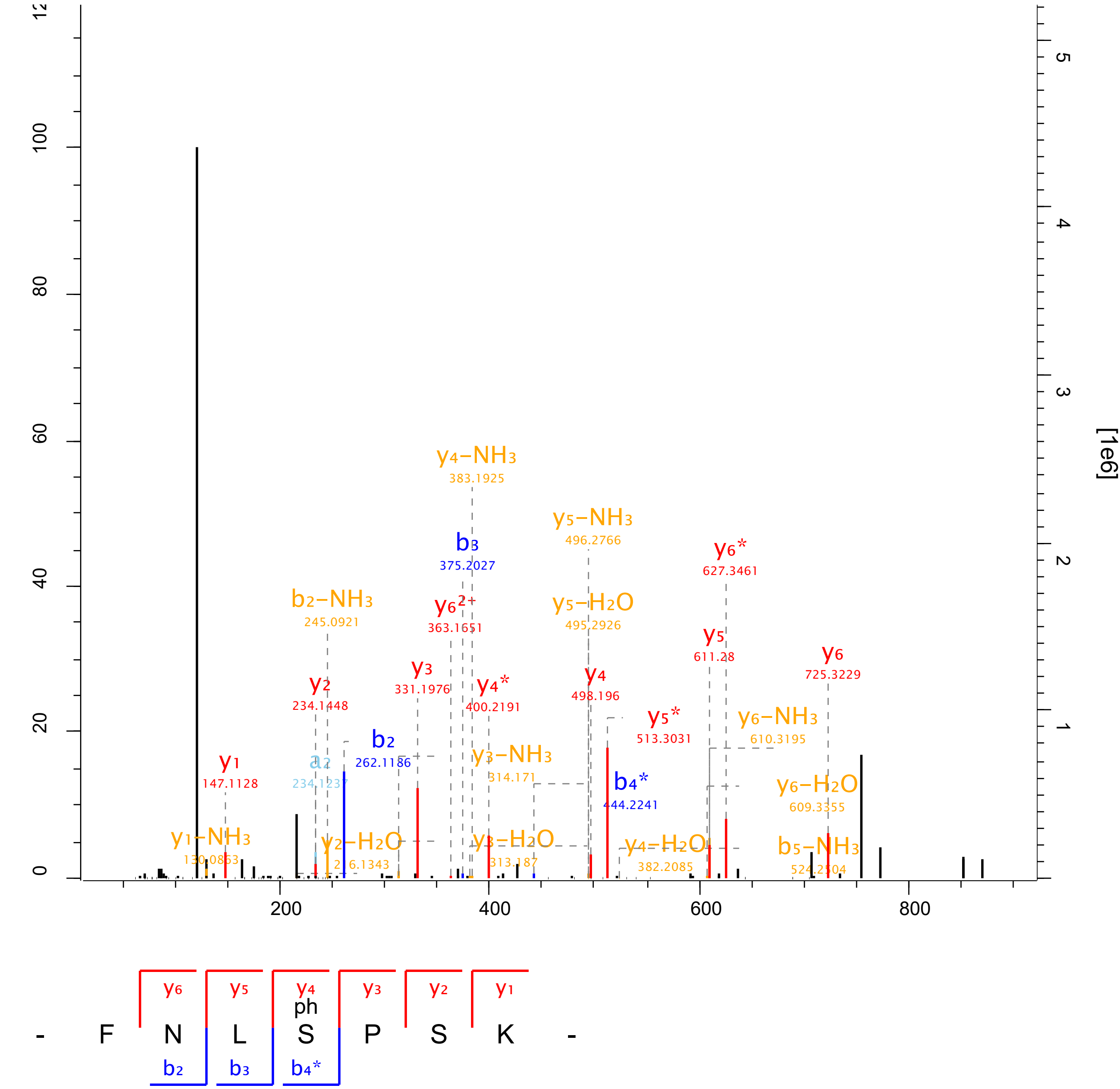

| Raw file | Scan | Method | Score | m/z |
| --- | --- | --- | --- | --- |
| sys_05_2 | 12954 | FTMS; HCD | 140.22 | 730.31 |

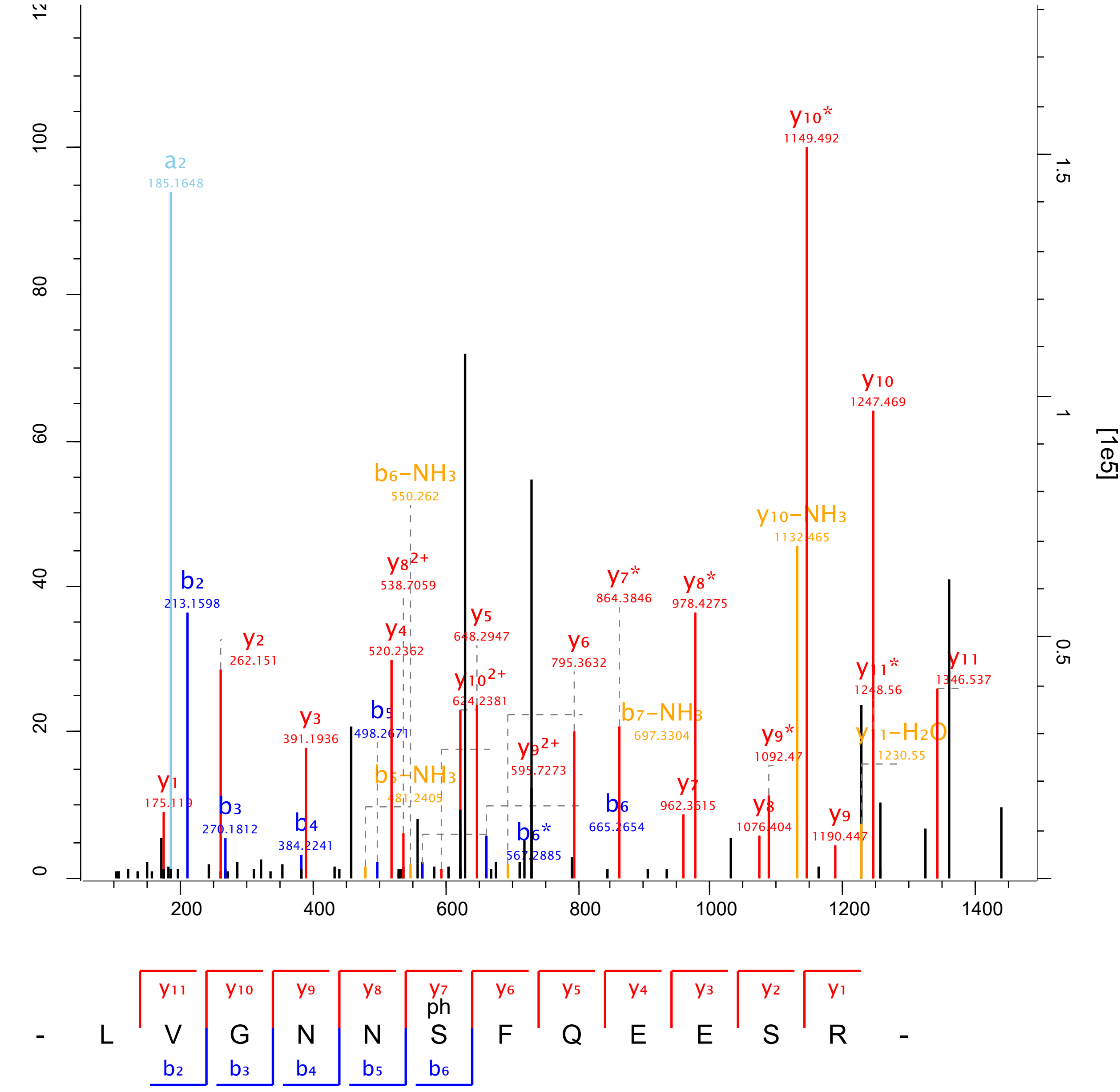

| Raw file | Scan | Method | Score | m/z |
|---|---|---|---|---|
| sys_05_2 | 12994 | FTMS; HCD | 109.07 | 632.81 |

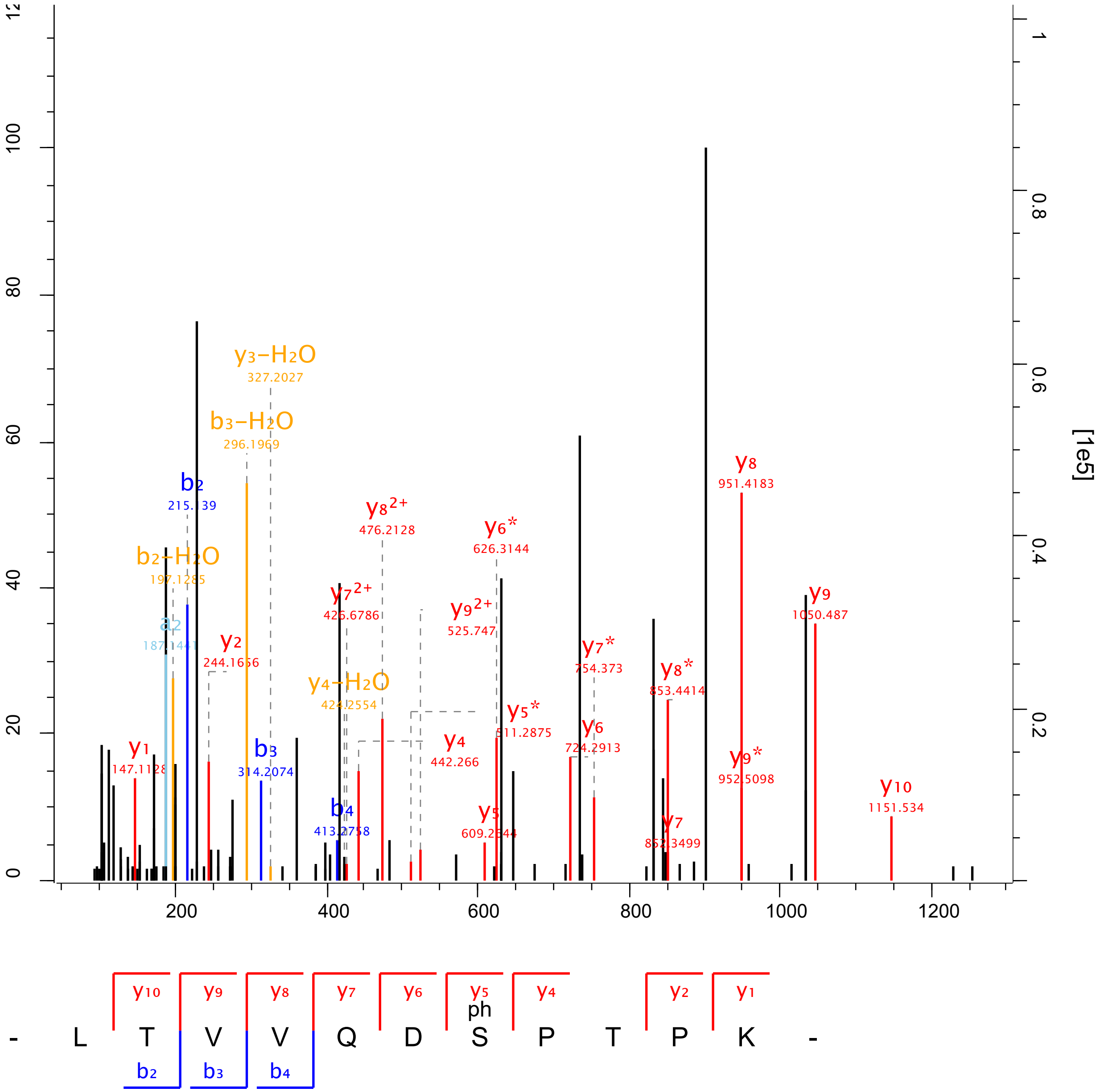

| Raw file | Scan | Method | Score | m/z |
|---|---|---|---|---|
| sys_05_2 | 13013 | FTMS; HCD | 118.37 | 809.35 |

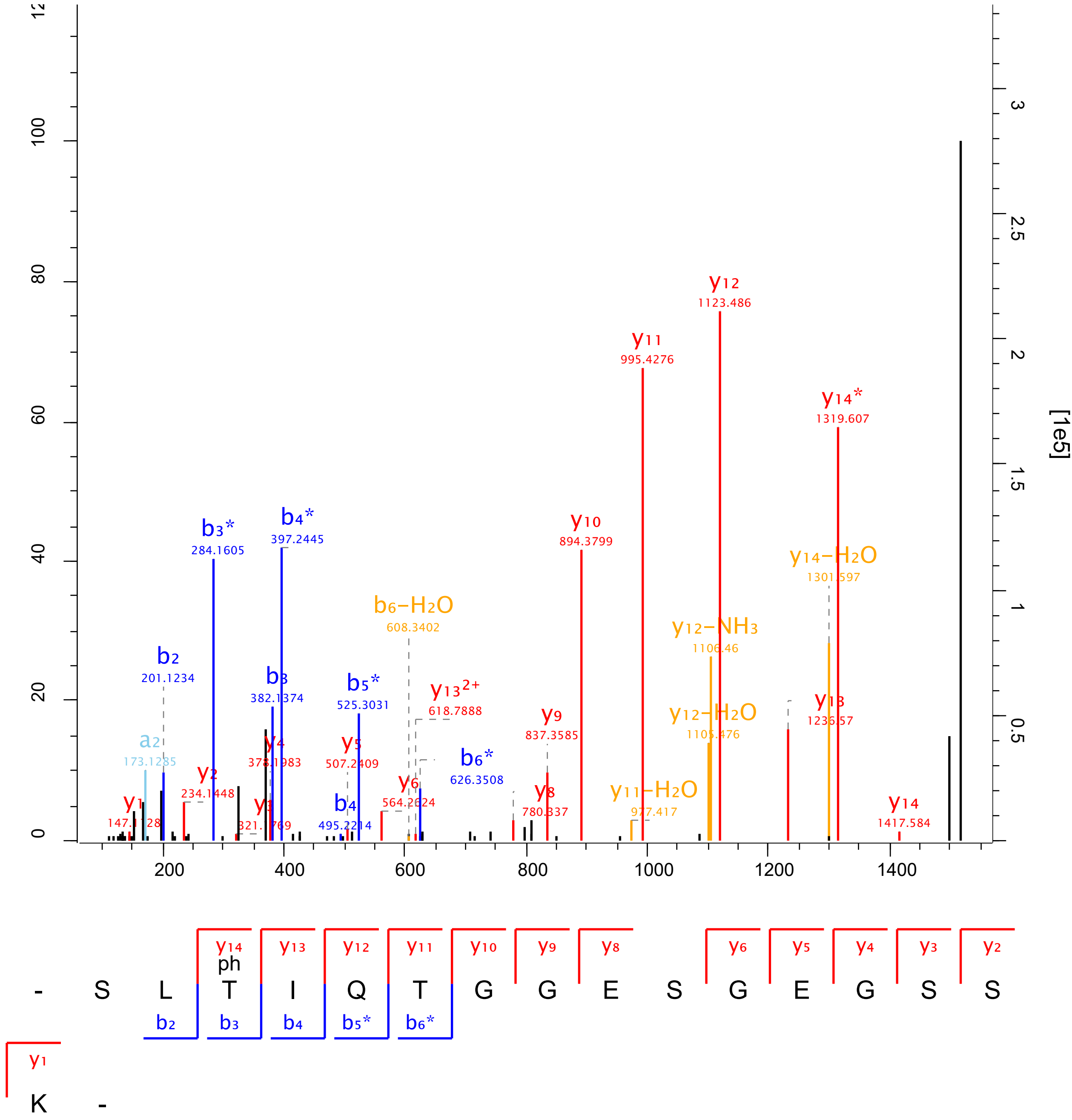

| Raw file | Scan | Method | Score | m/z |
|---|---|---|---|---|
| sys_05_2 | 13047 | FTMS; HCD | 117.2 | 531.73 |

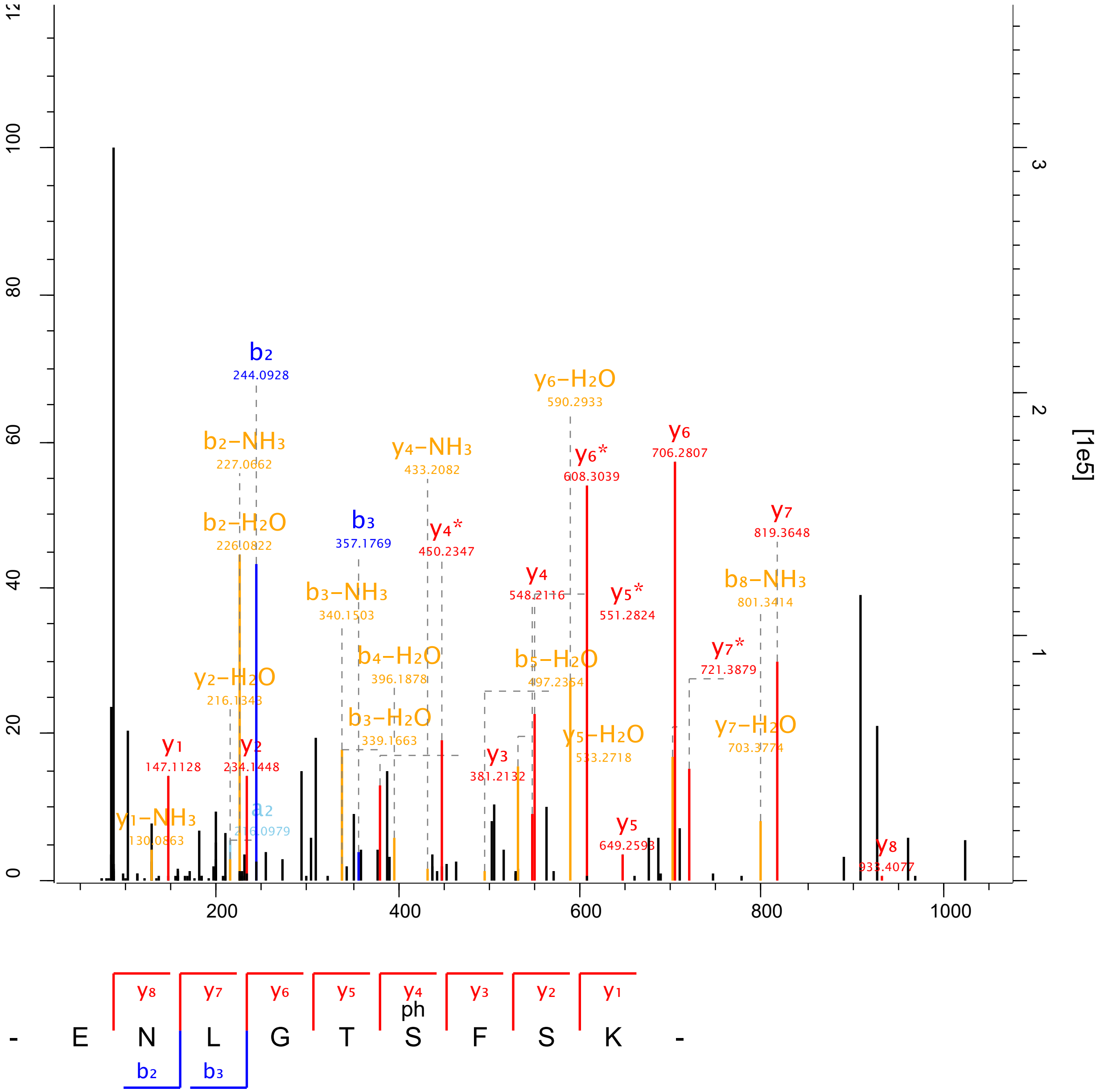

| Raw file | Scan | Method | Score | m/z |
|---|---|---|---|---|
| sys_05_2 | 13109 | FTMS; HCD | 188.81 | 563.24 |

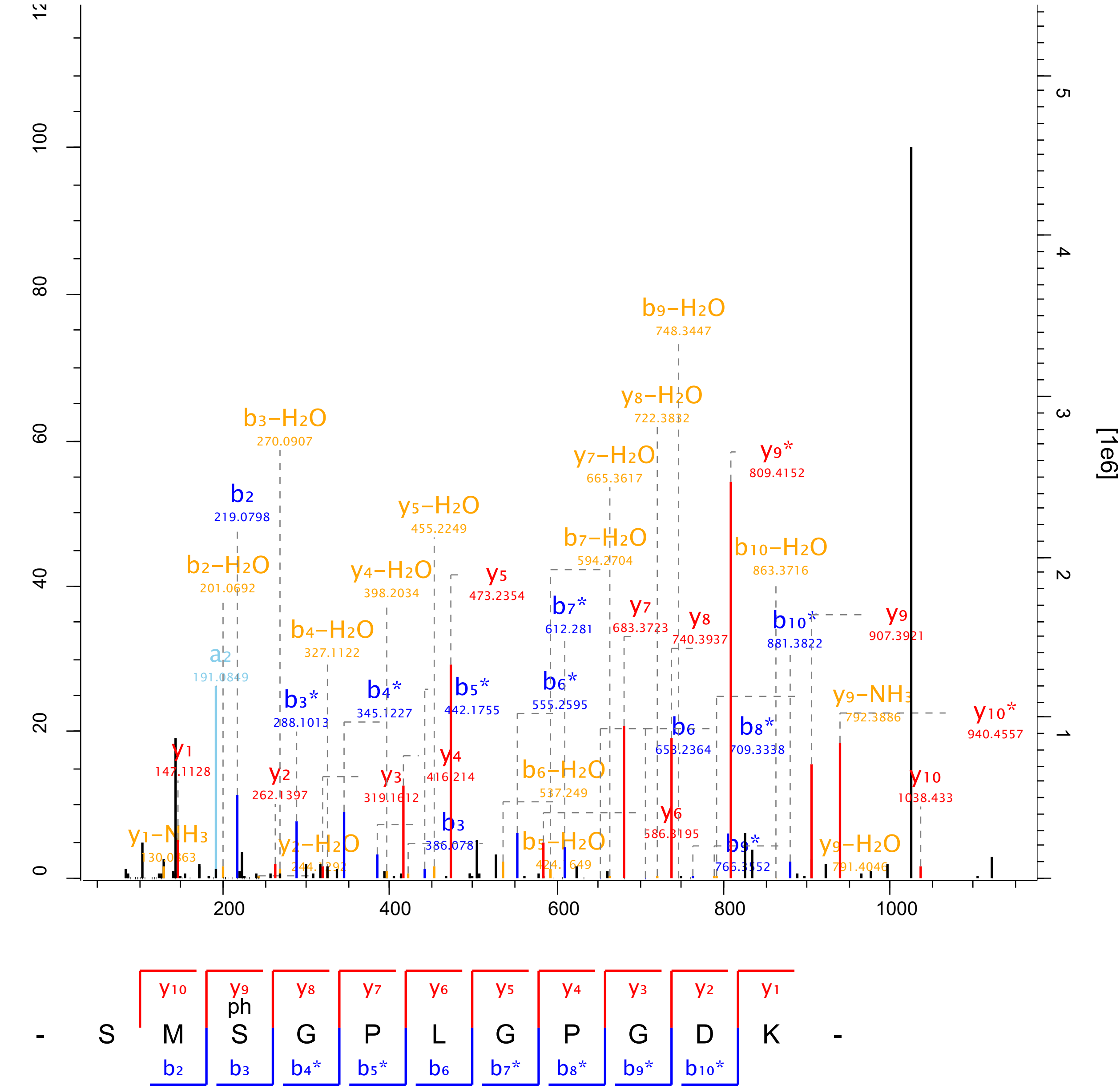

| Raw file | Scan | Method | Score | m/z |
|---|---|---|---|---|
| sys_05_2 | 13137 | FTMS; HCD | 206.64 | 849.39 |

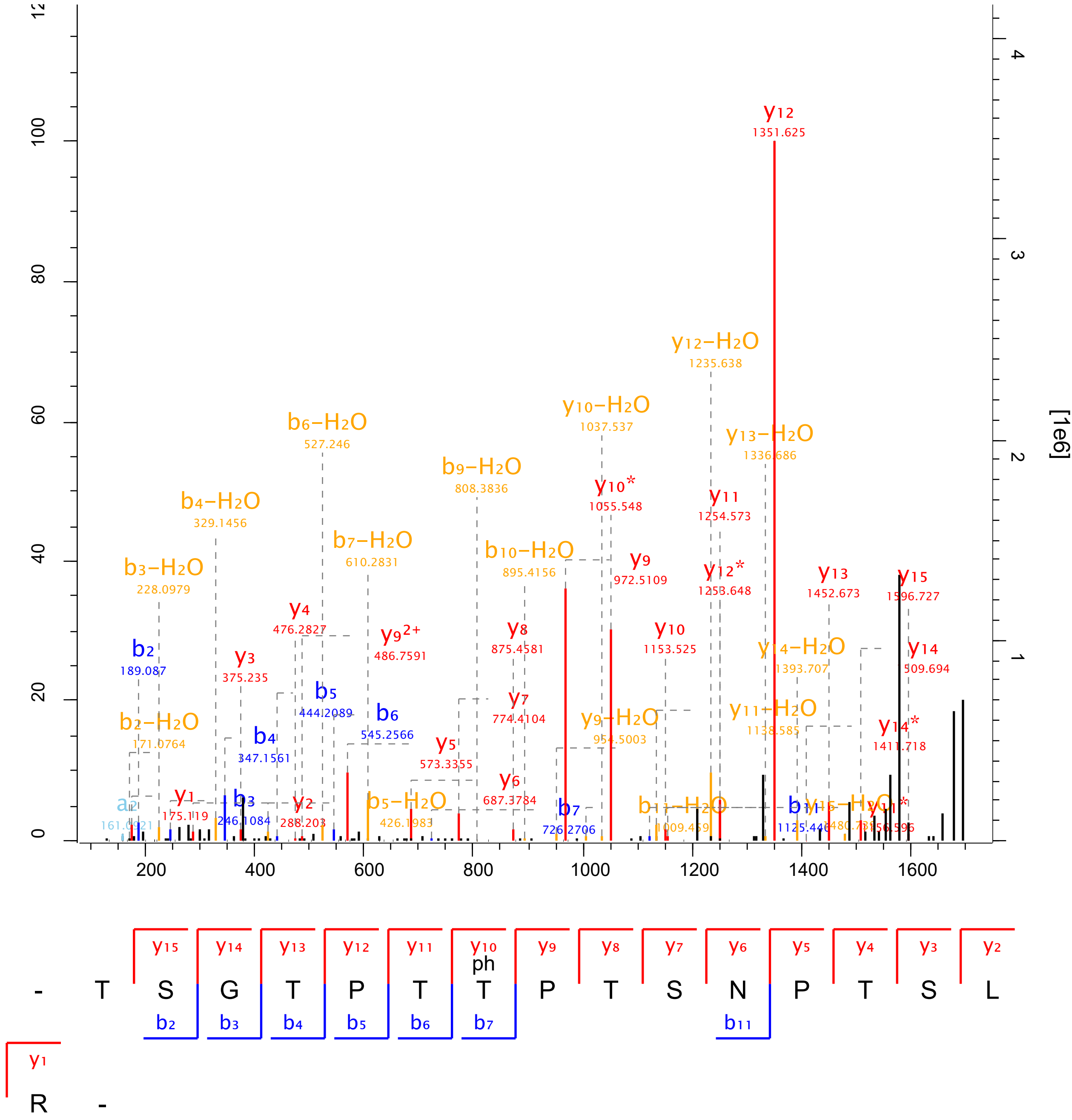

| Raw file | Scan | Method | Score | m/z |
|---|---|---|---|---|
| sys_05_2 | 13141 | FTMS; HCD | 55.75 | 760.77 |

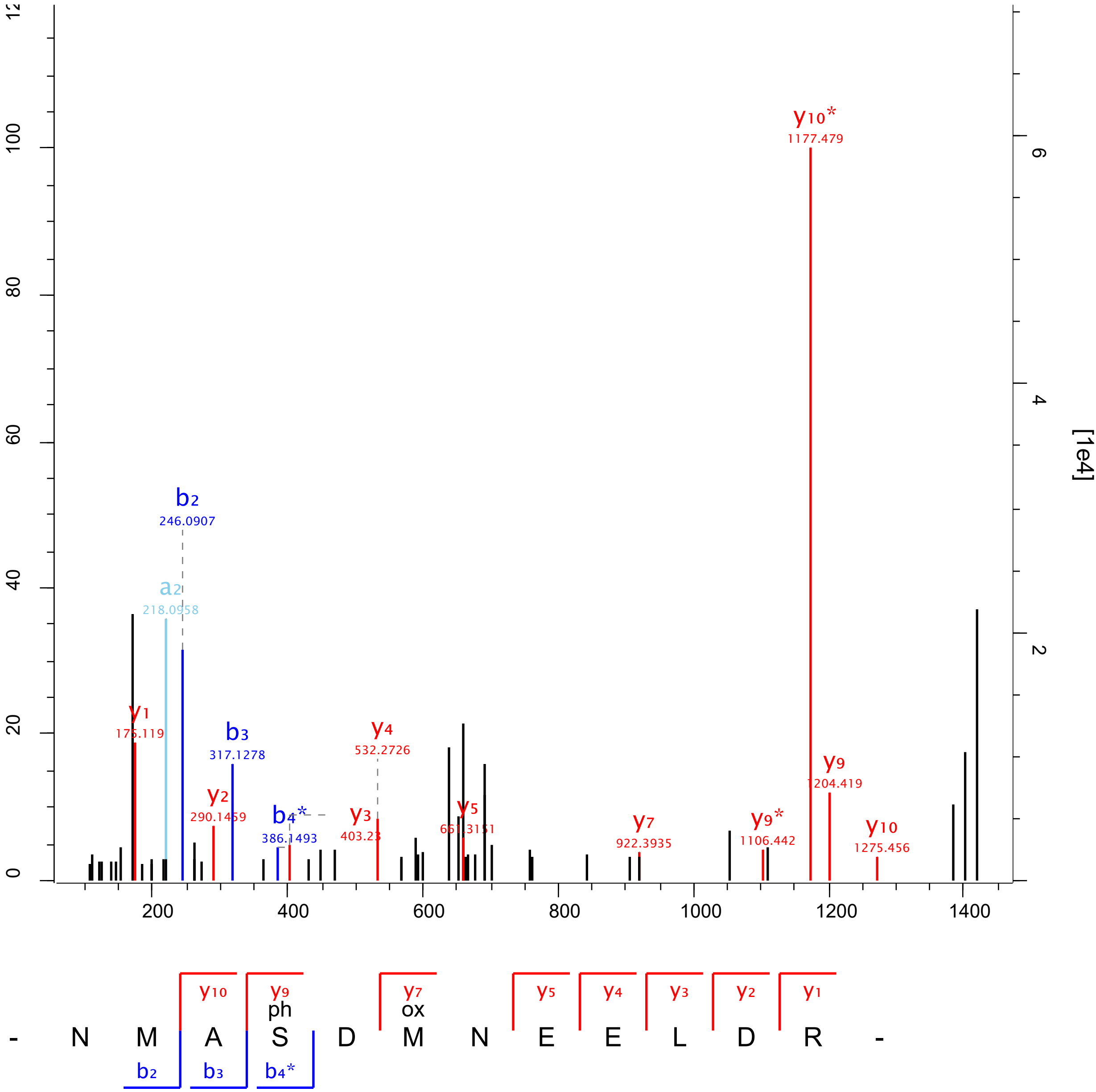

| Raw file | Scan | Method | Score | m/z |
|---|---|---|---|---|
| sys_05_2 | 13184 | FTMS; HCD | 142.1 | 572.24 |

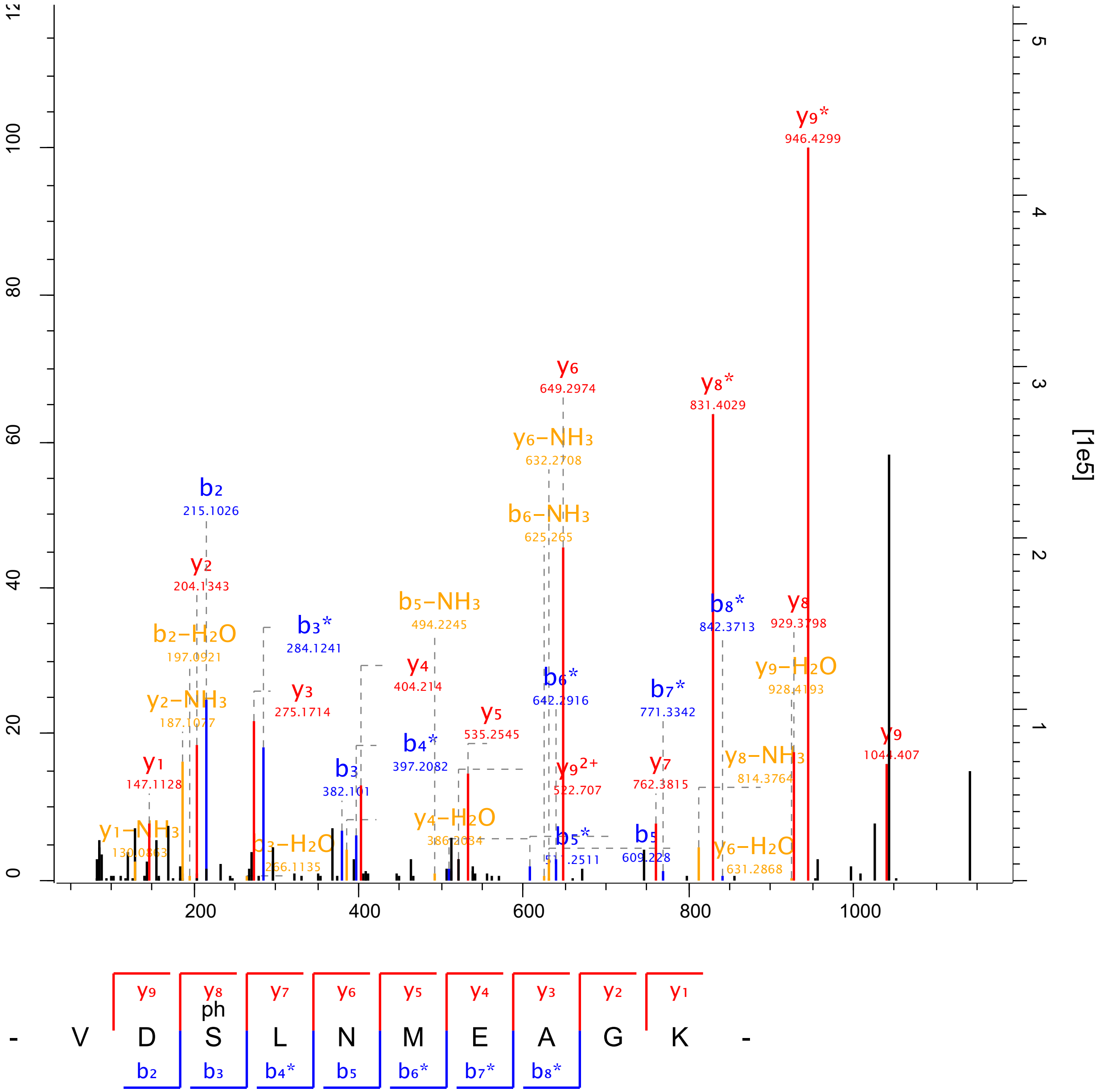

| Raw file | Scan | Method | Score | m/z |
|---|---|---|---|---|
| sys_05_2 | 13306 | FTMS; HCD | 113.65 | 645.78 |

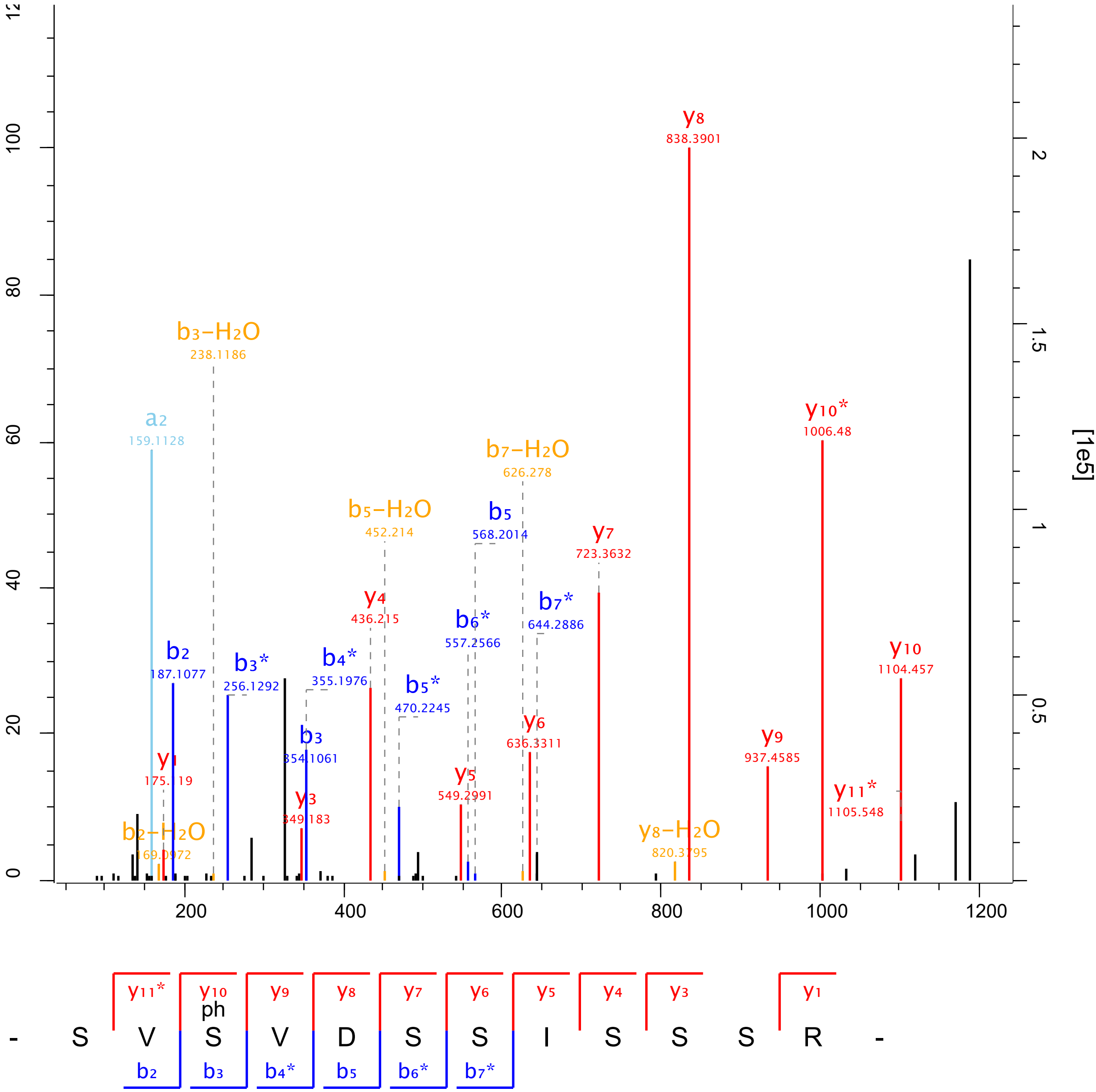

| Raw file | Scan | Method | Score | m/z |
|---|---|---|---|---|
| sys_05_2 | 13556 | FTMS; HCD | 81.52 | 500.22 |

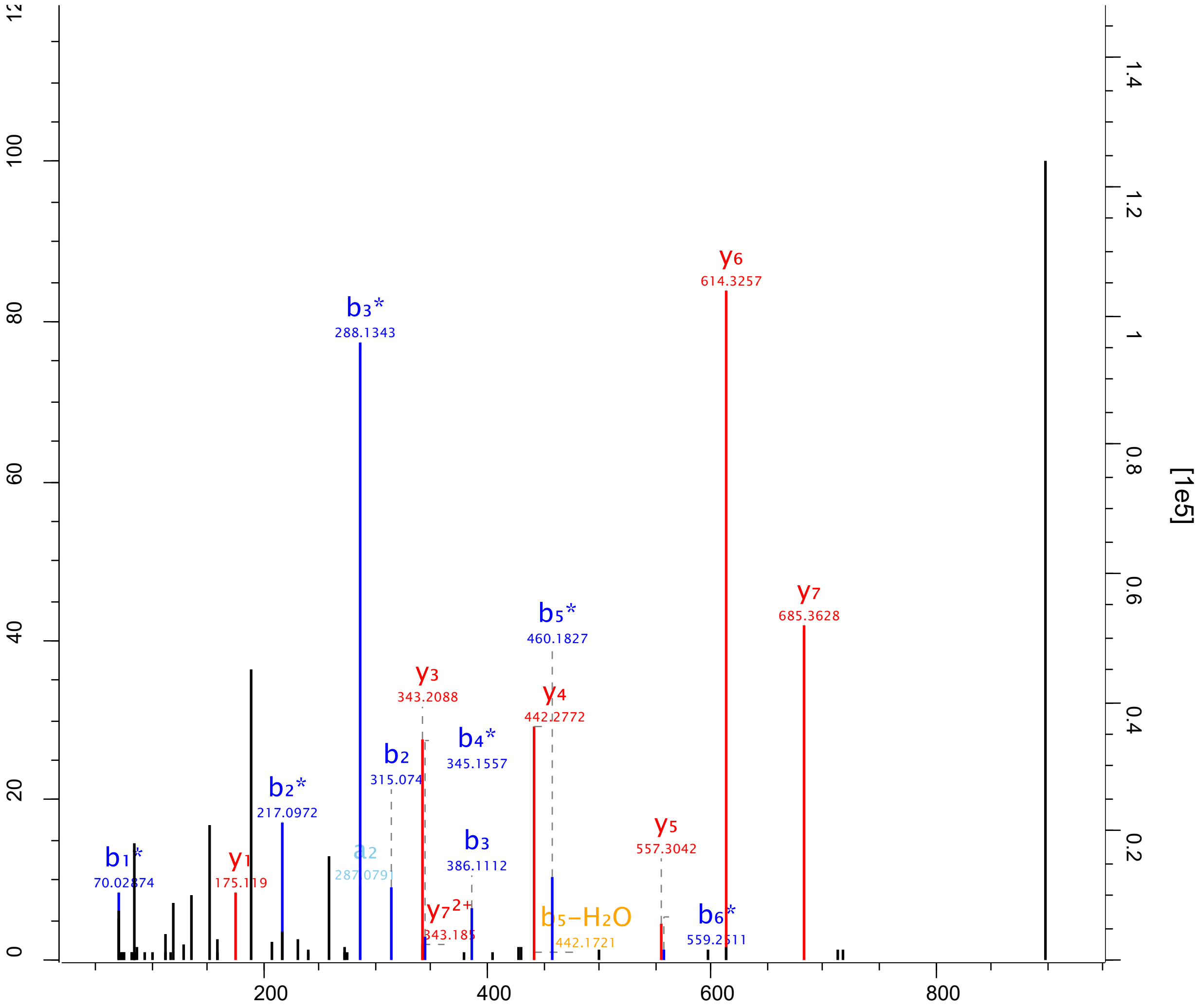


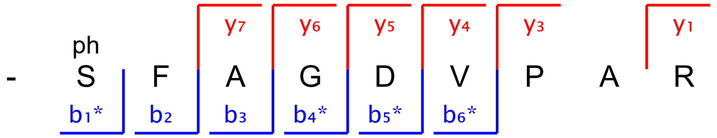

| Raw file | Scan | Method | Score | m/z |
| --- | --- | --- | --- | --- |
| sys_05_2 | 13614 | FTMS; HCD | 94.47 | 508.74 |

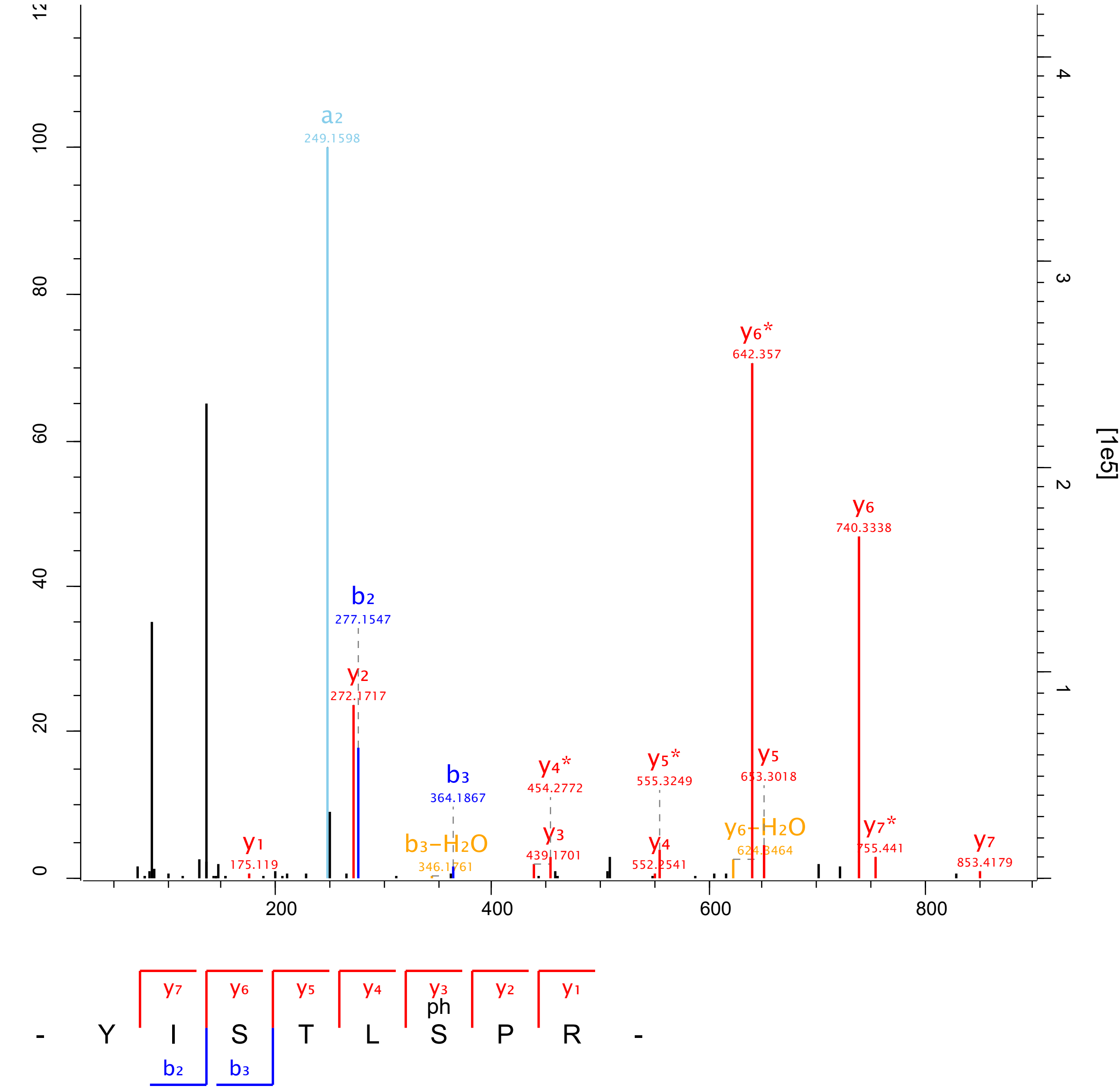

| Raw file | Scan | Method | Score | m/z |
|---|---|---|---|---|
| sys_05_2 | 13695 | FTMS; HCD | 116.73 | 892.87 |

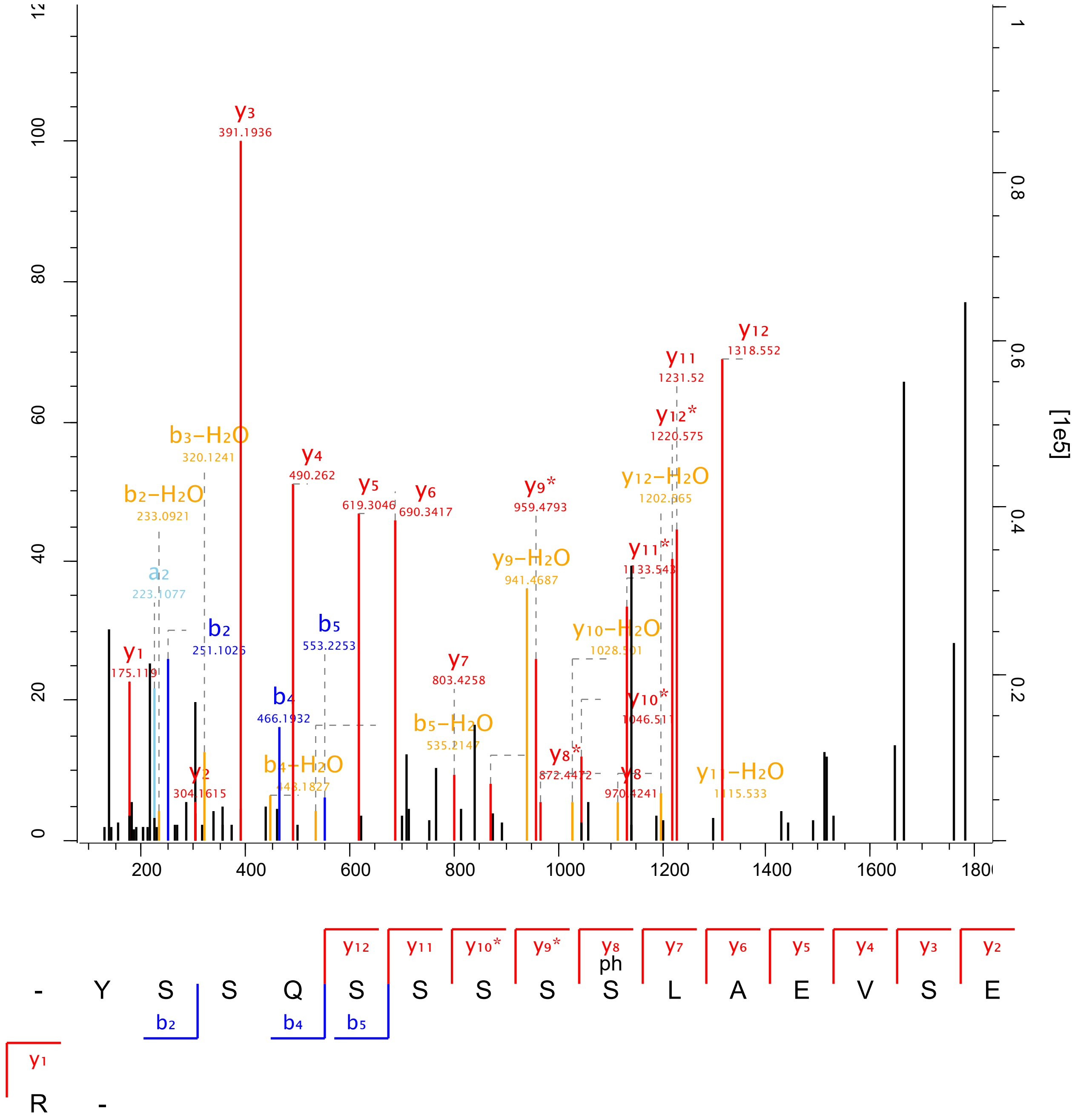

| Raw file | Scan | Method | Score | m/z |
|---|---|---|---|---|
| sys_05_2 | 13725 | FTMS; HCD | 108.77 | 786.33 |

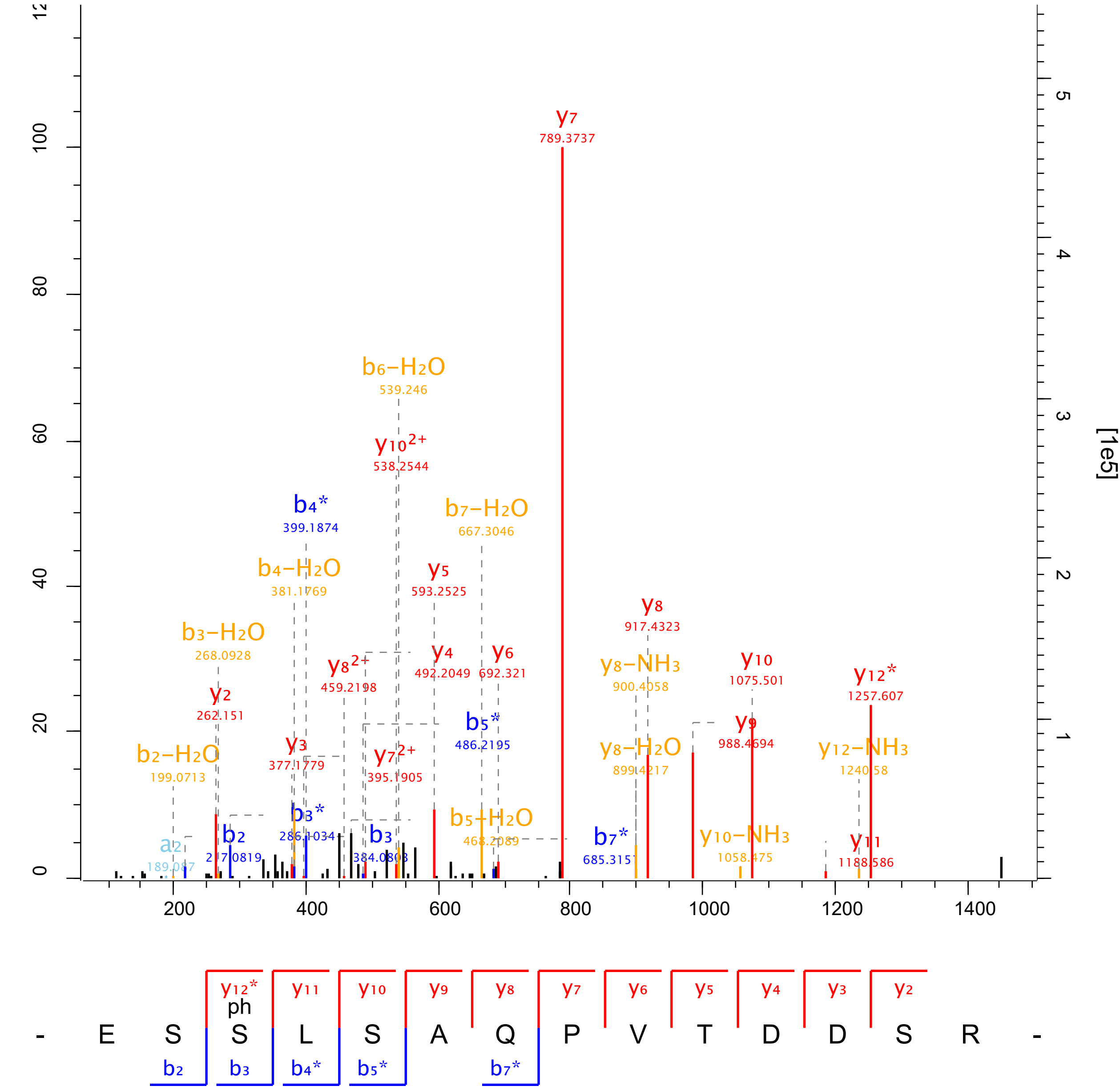

| Raw file | Scan | Method | Score | m/z |
|---|---|---|---|---|
| sys_05_2 | 13869 | FTMS; HCD | 107.09 | 528.24 |

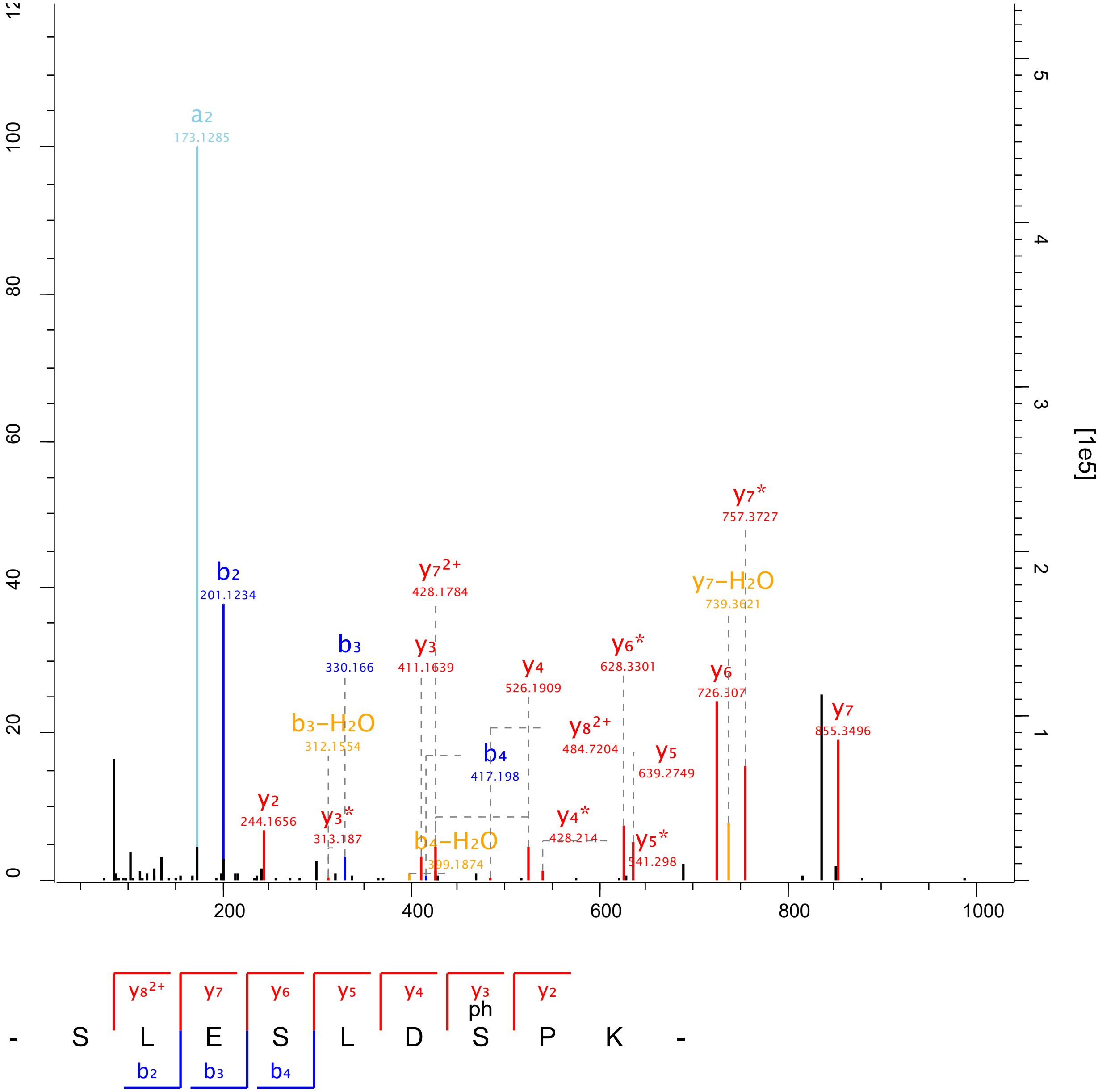

| Raw file | Scan | Method | Score | m/z |
|---|---|---|---|---|
| sys_05_2 | 13916 | FTMS; HCD | 47.71 | 586.24 |

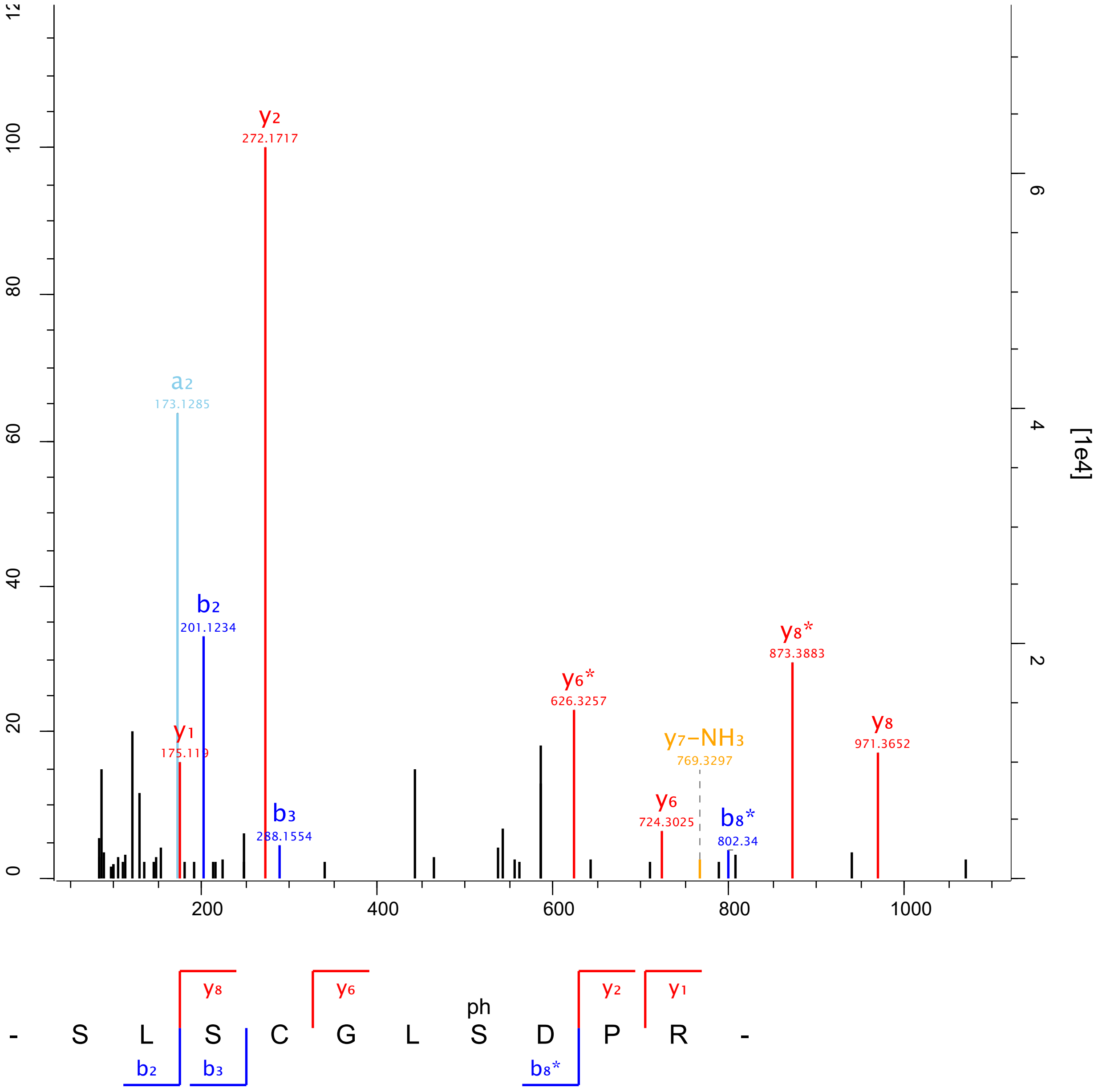

| Raw file | Scan | Method | Score | m/z |
| --- | --- | --- | --- | --- |
| sys_05_2 | 13946 | FTMS; HCD | 231.38 | 728.34 |

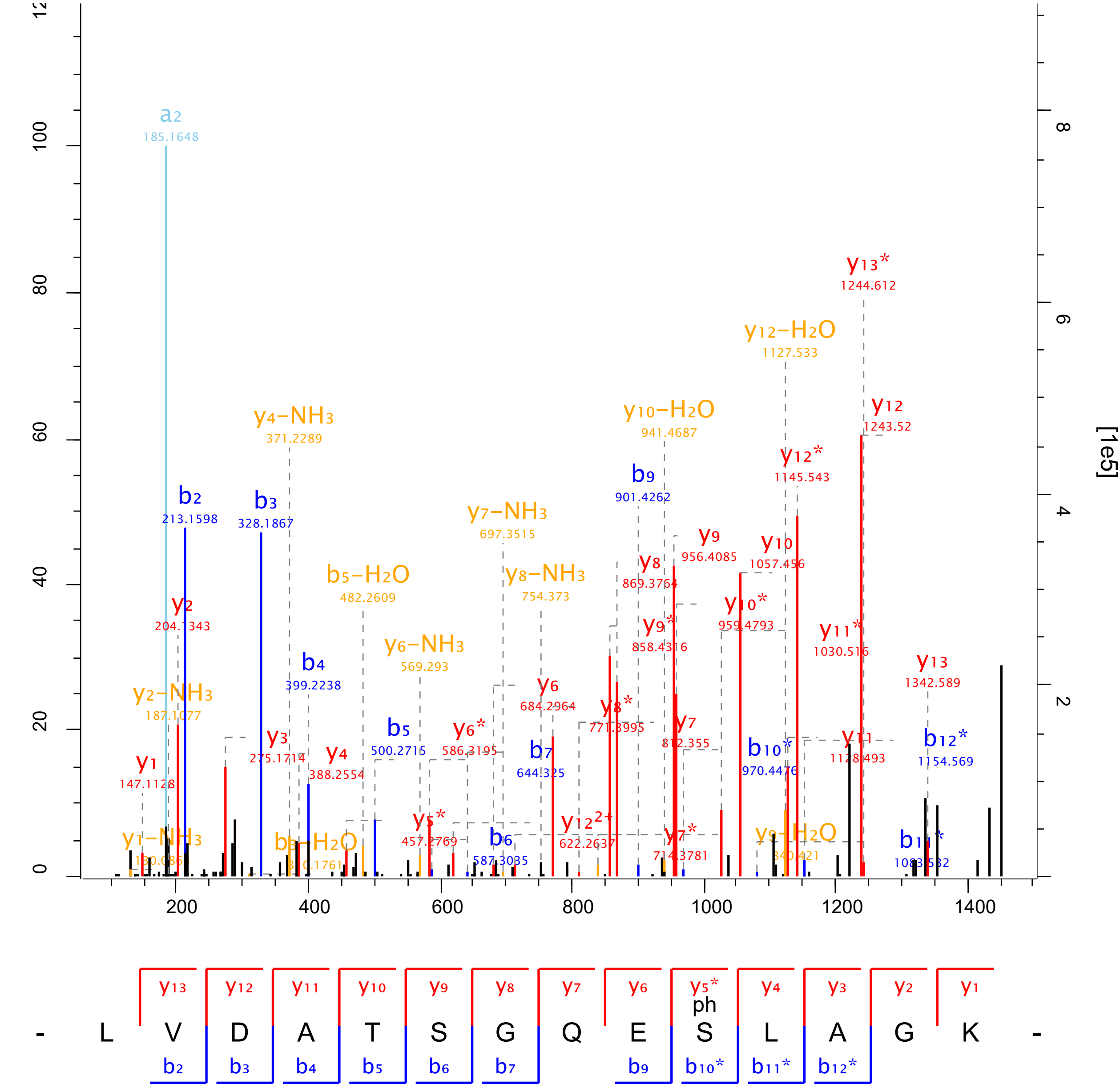

| Raw file | Scan | Method | Score | m/z |
|---|---|---|---|---|
| sys_05_2 | 13977 | FTMS; HCD | 225.3 | 861.35 |

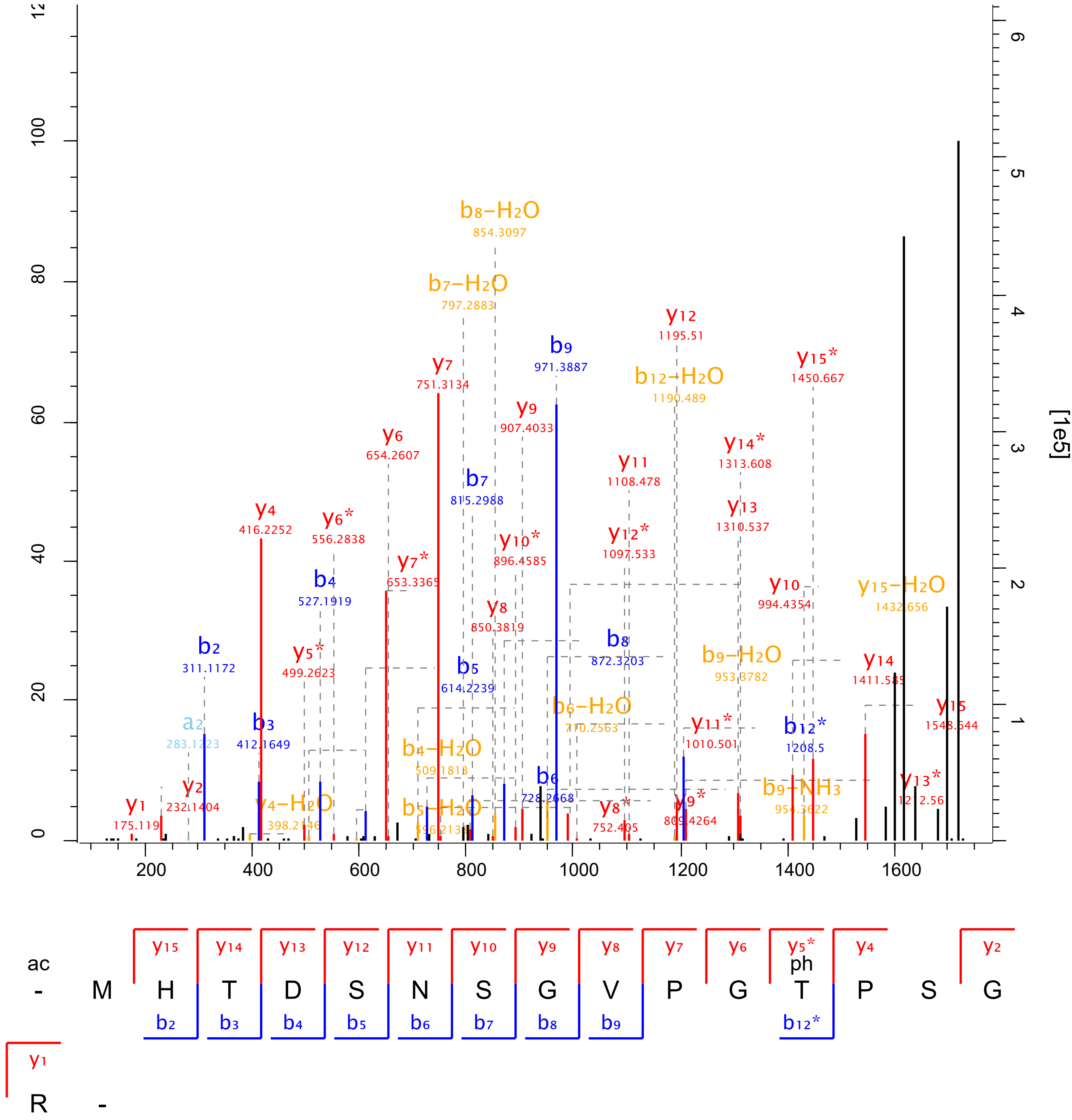

| Raw file | Scan | Method | Score | m/z |
|---|---|---|---|---|
| sys_05_2 | 13988 | FTMS; HCD | 186.08 | 662.27 |

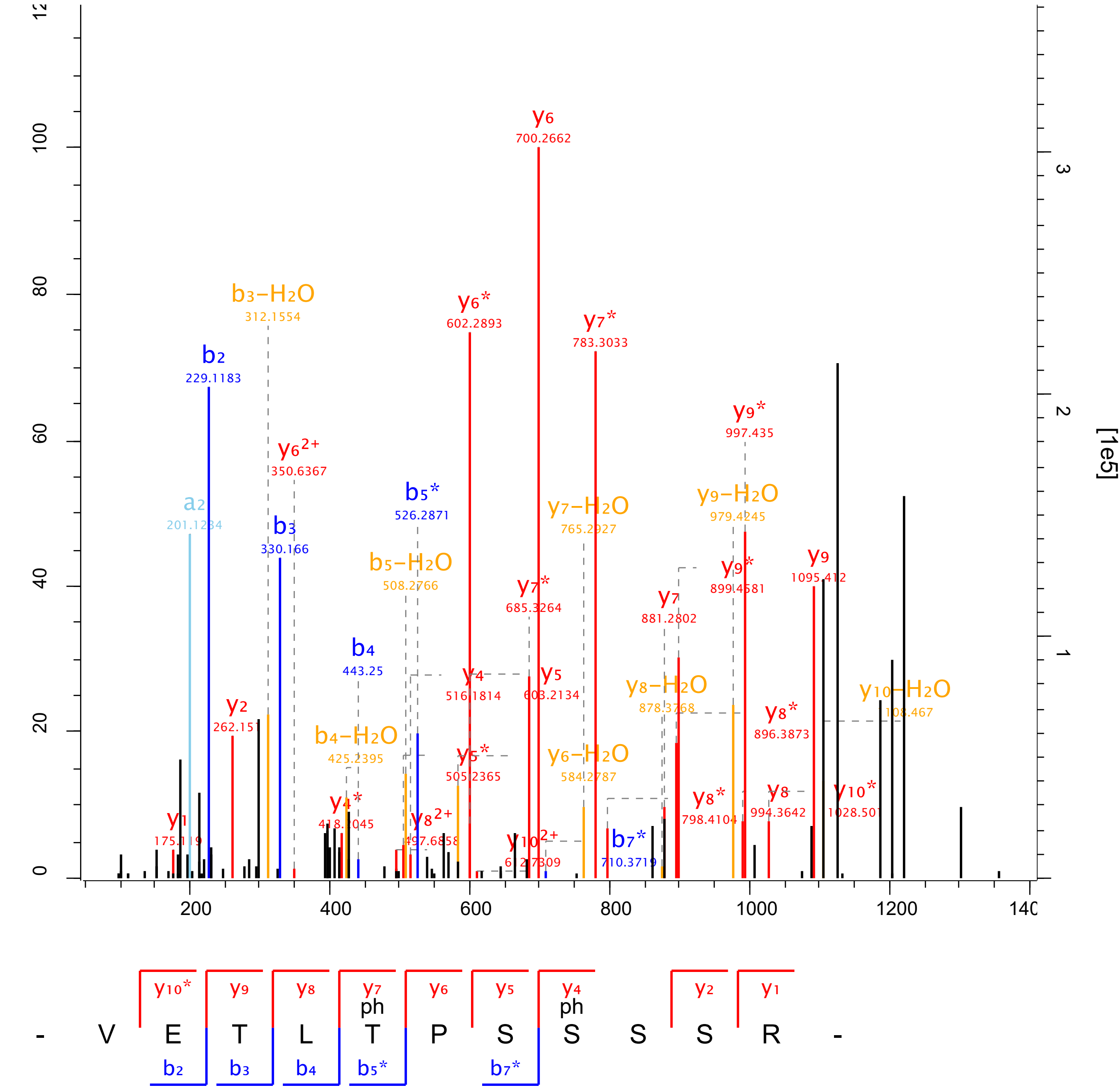

| Raw file | Scan | Method | Score | m/z |
| --- | --- | --- | --- | --- |
| sys_05_2 | 14017 | FTMS; HCD | 49.47 | 483.71 |

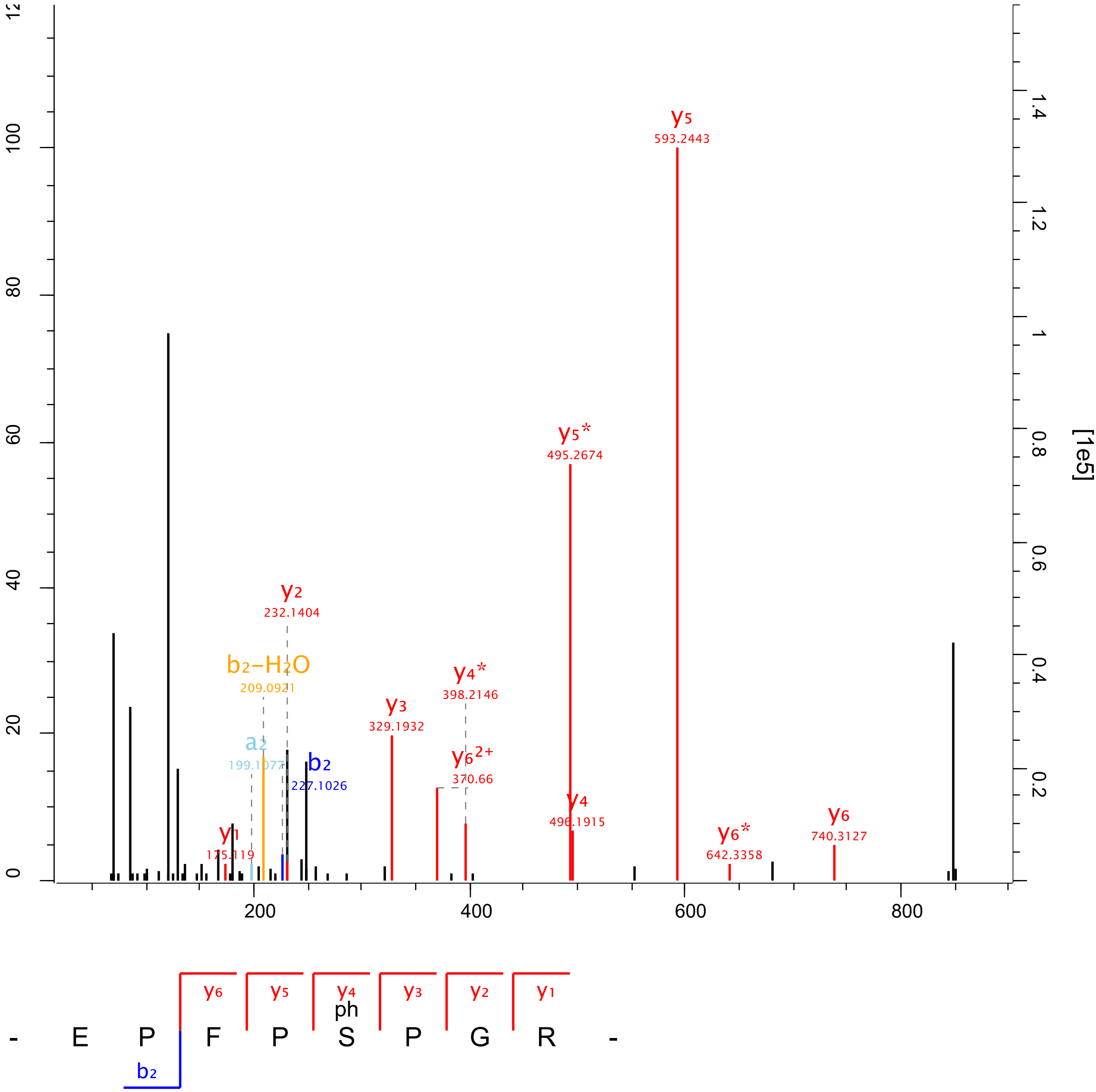

| Raw file | Scan | Method | Score | m/z |
|---|---|---|---|---|
| sys_05_2 | 14170 | FTMS; HCD | 192.46 | 693.31 |

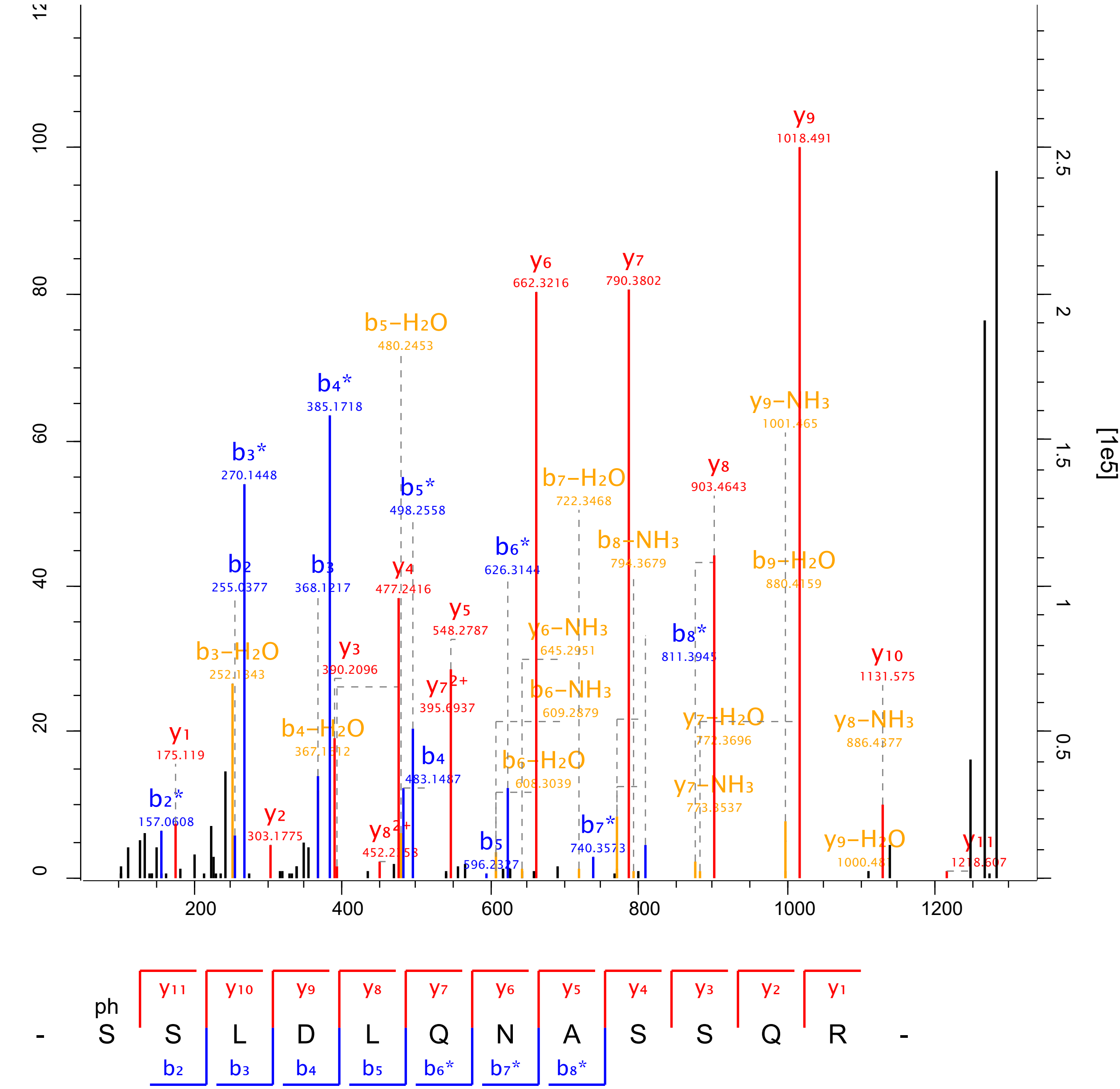

| Raw file | Scan | Method | Score | m/z |
|---|---|---|---|---|
| sys_05_2 | 14184 | FTMS; HCD | 71.18 | 510.69 |

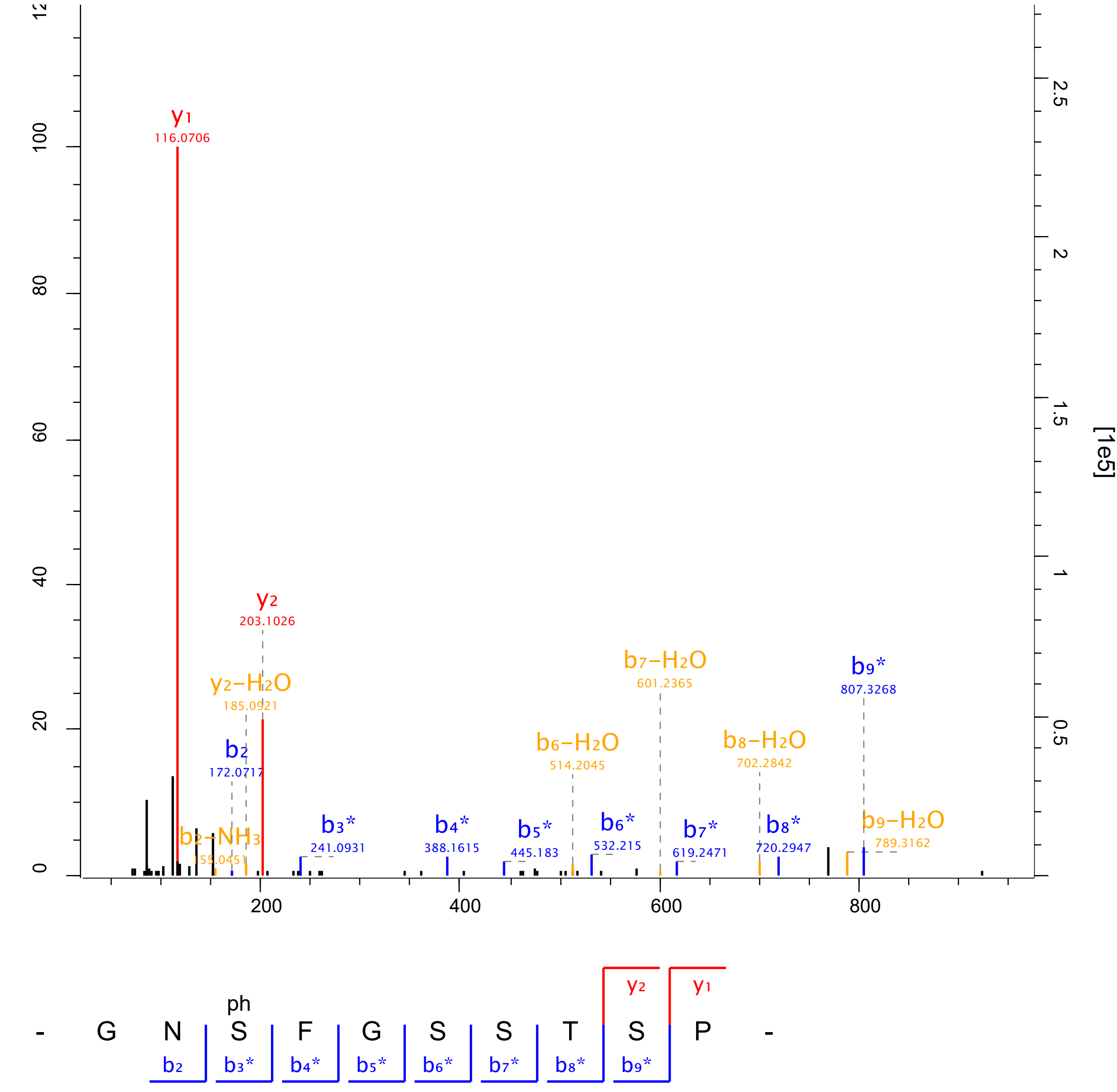

| Raw file | Scan | Method | Score | m/z |
|---|---|---|---|---|
| sys_05_2 | 14217 | FTMS; HCD | 103.88 | 758.83 |

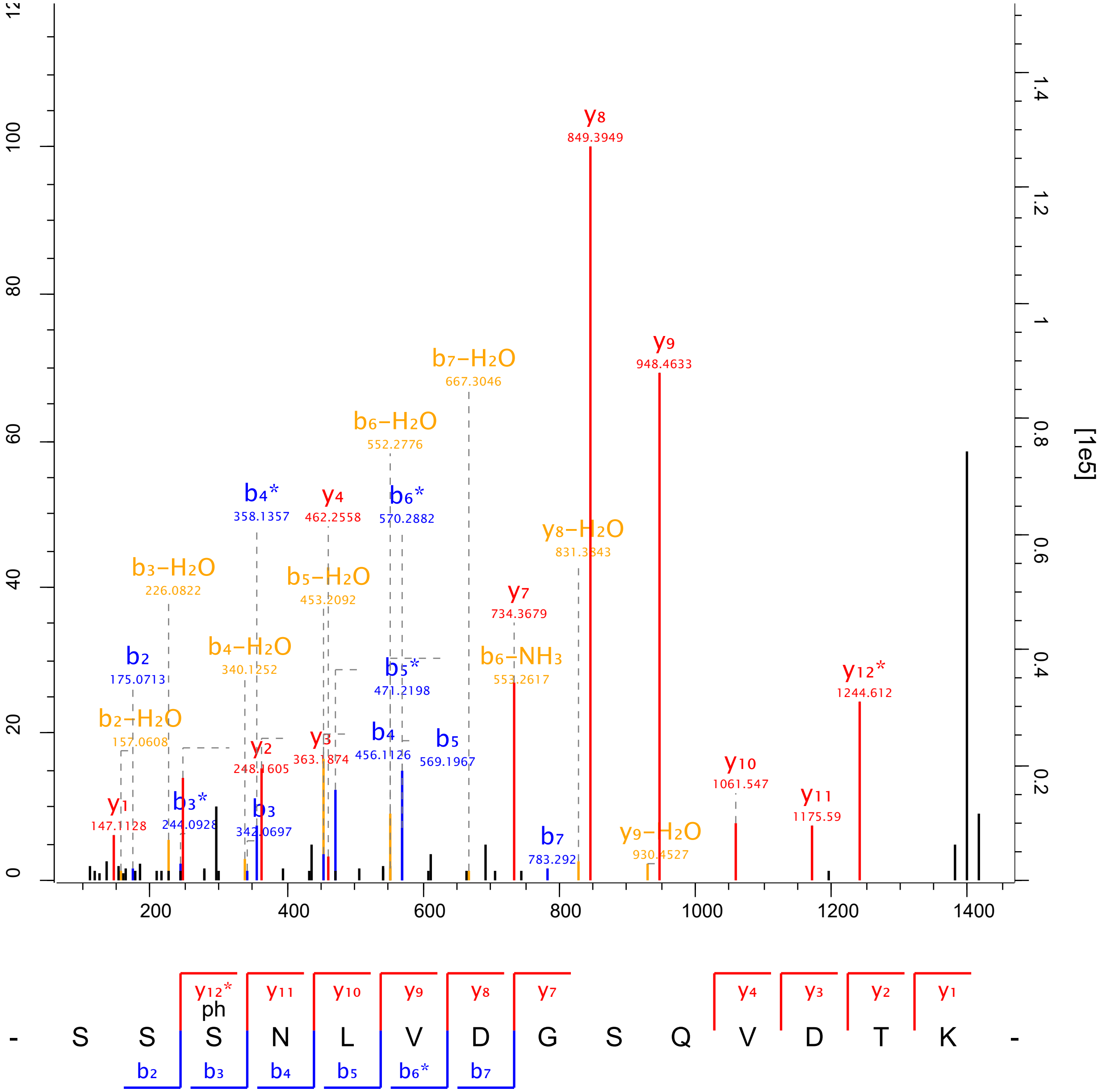

| Raw file | Scan | Method | Score | m/z |
| --- | --- | --- | --- | --- |
| sys_05_2 | 14279 | FTMS; HCD | 84.36 | 587.76 |

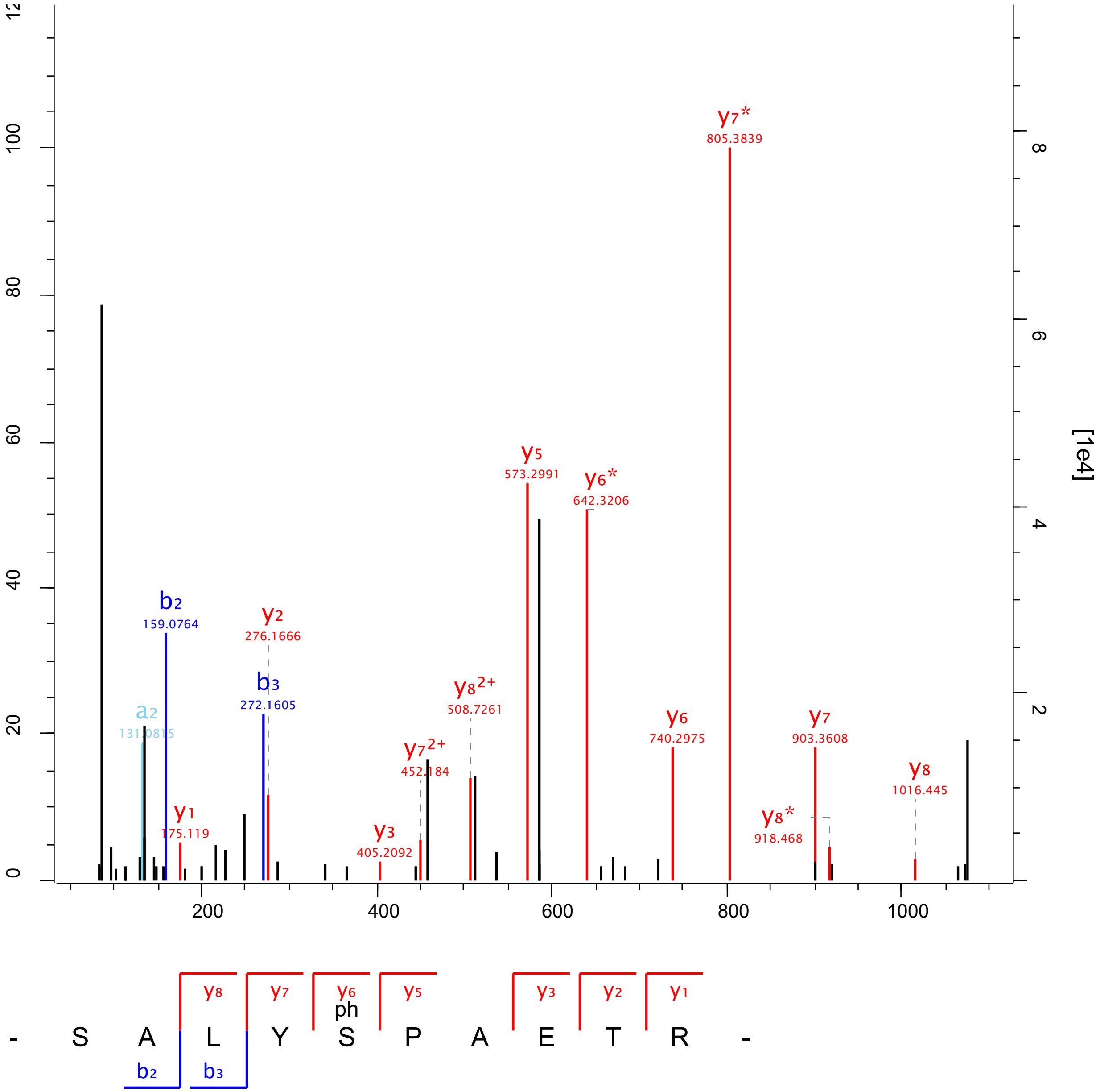

| Raw file | Scan | Method | Score | m/z |
|---|---|---|---|---|
| sys_05_2 | 14316 | FTMS; HCD | 108.47 | 521.73 |

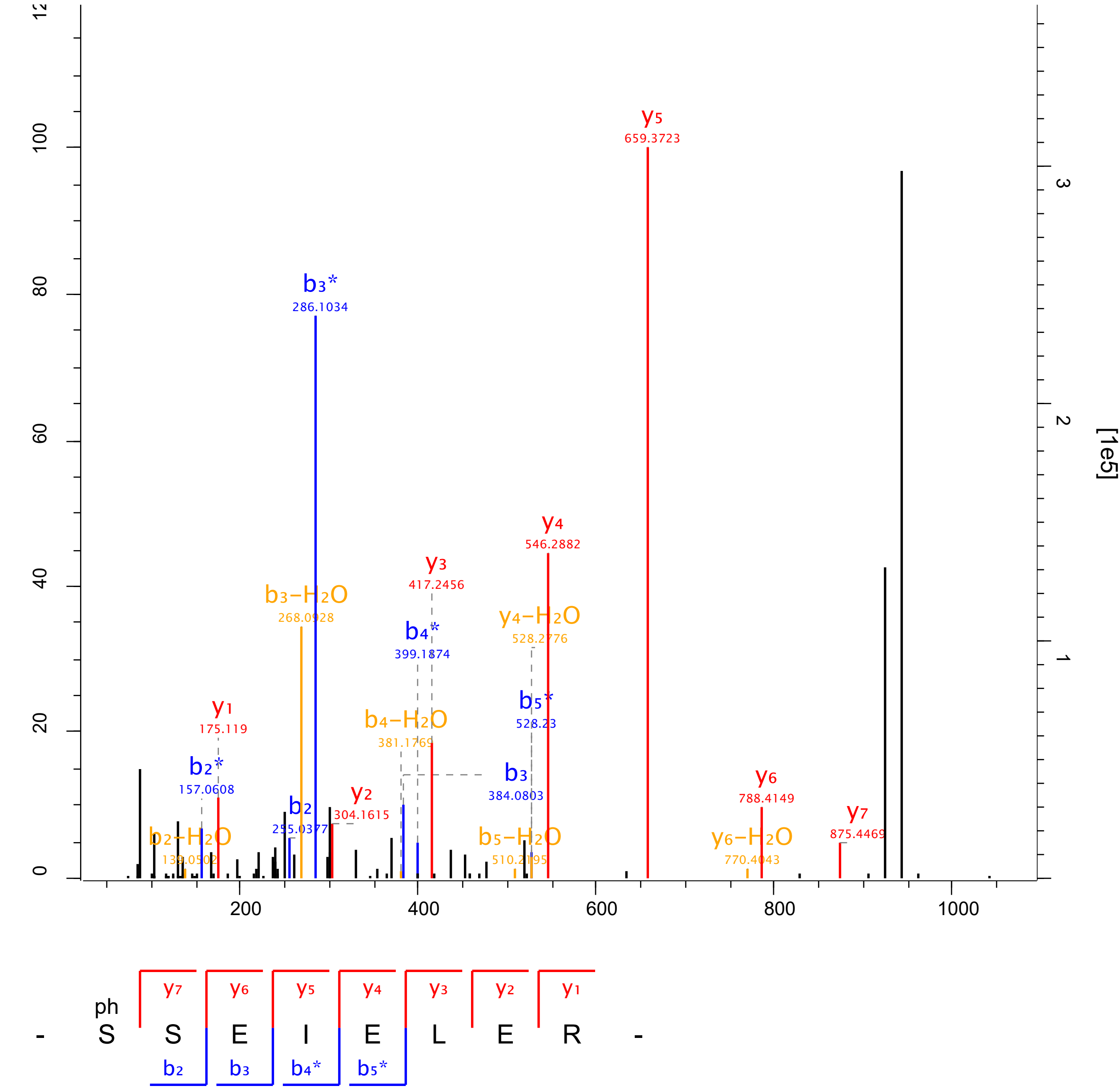

| Raw file | Scan | Method | Score | m/z |
|---|---|---|---|---|
| sys_05_2 | 14367 | FTMS; HCD | 152.94 | 723.82 |

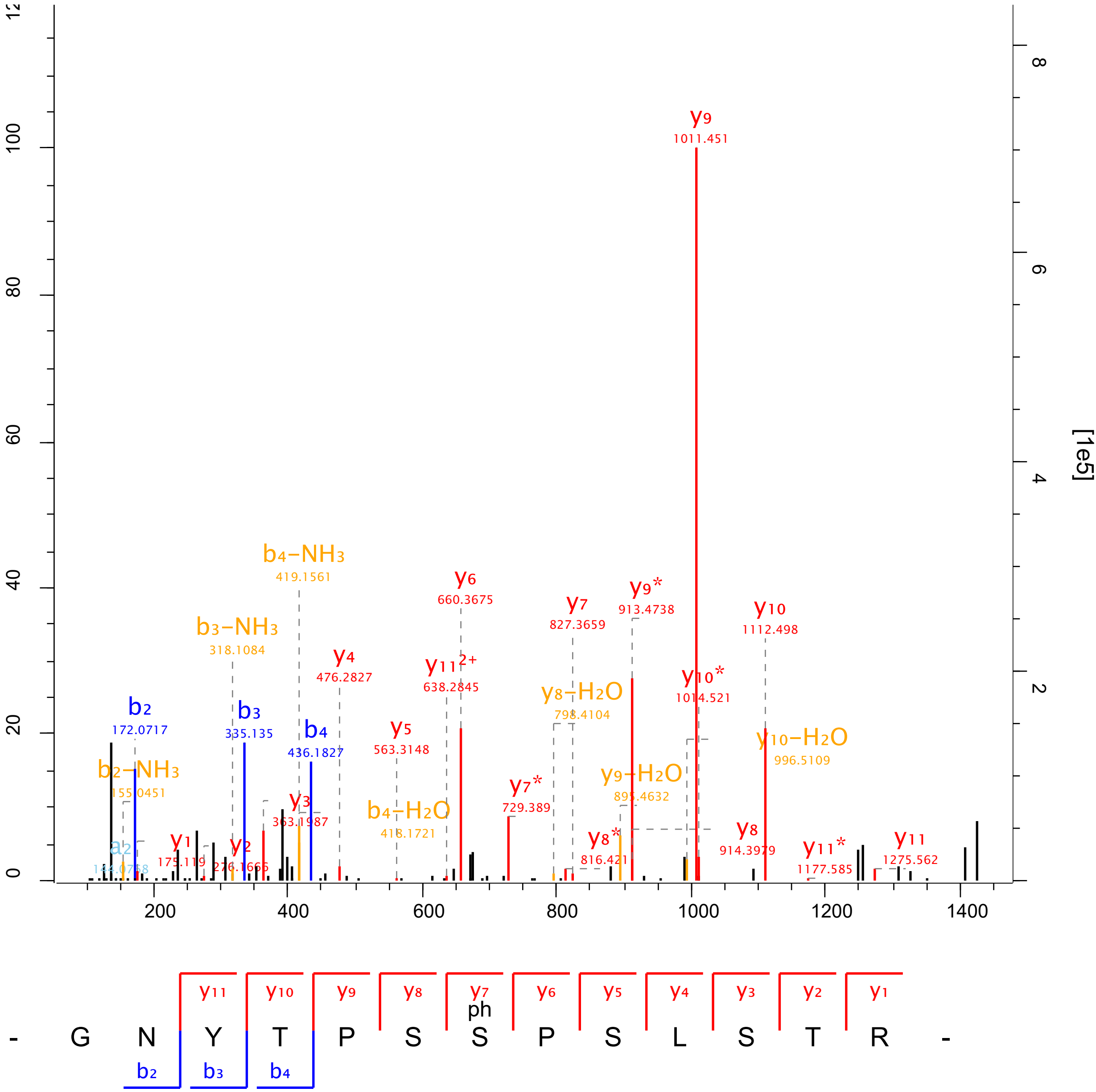

| Raw file | Scan | Method | Score | m/z |
| --- | --- | --- | --- | --- |
| sys_05_2 | 14385 | FTMS; HCD | 75.76 | 677.76 |

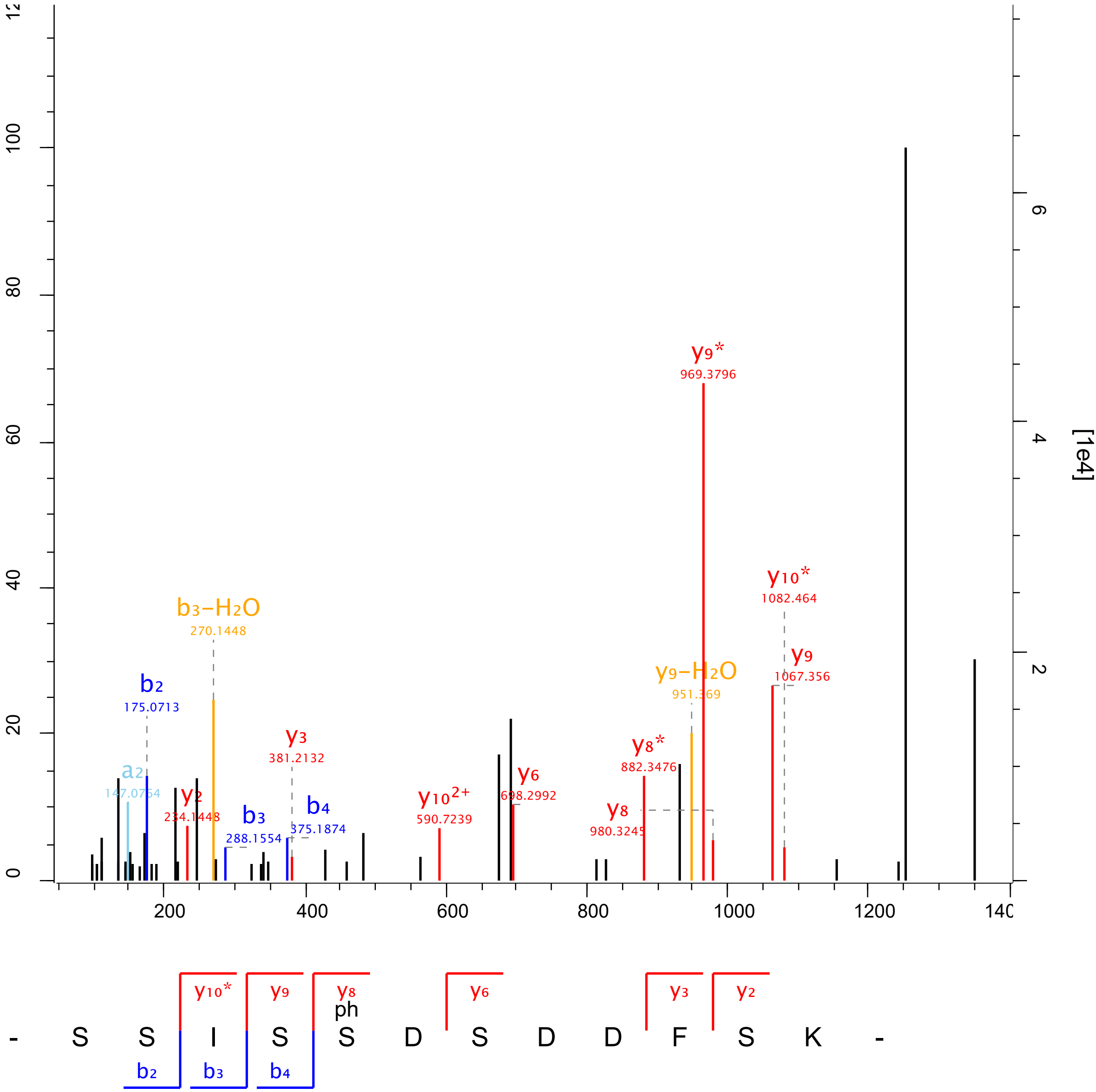

| Raw file | Scan | Method | Score | m/z |
|---|---|---|---|---|
| sys_05_2 | 14408 | FTMS; HCD | 51.77 | 631.76 |

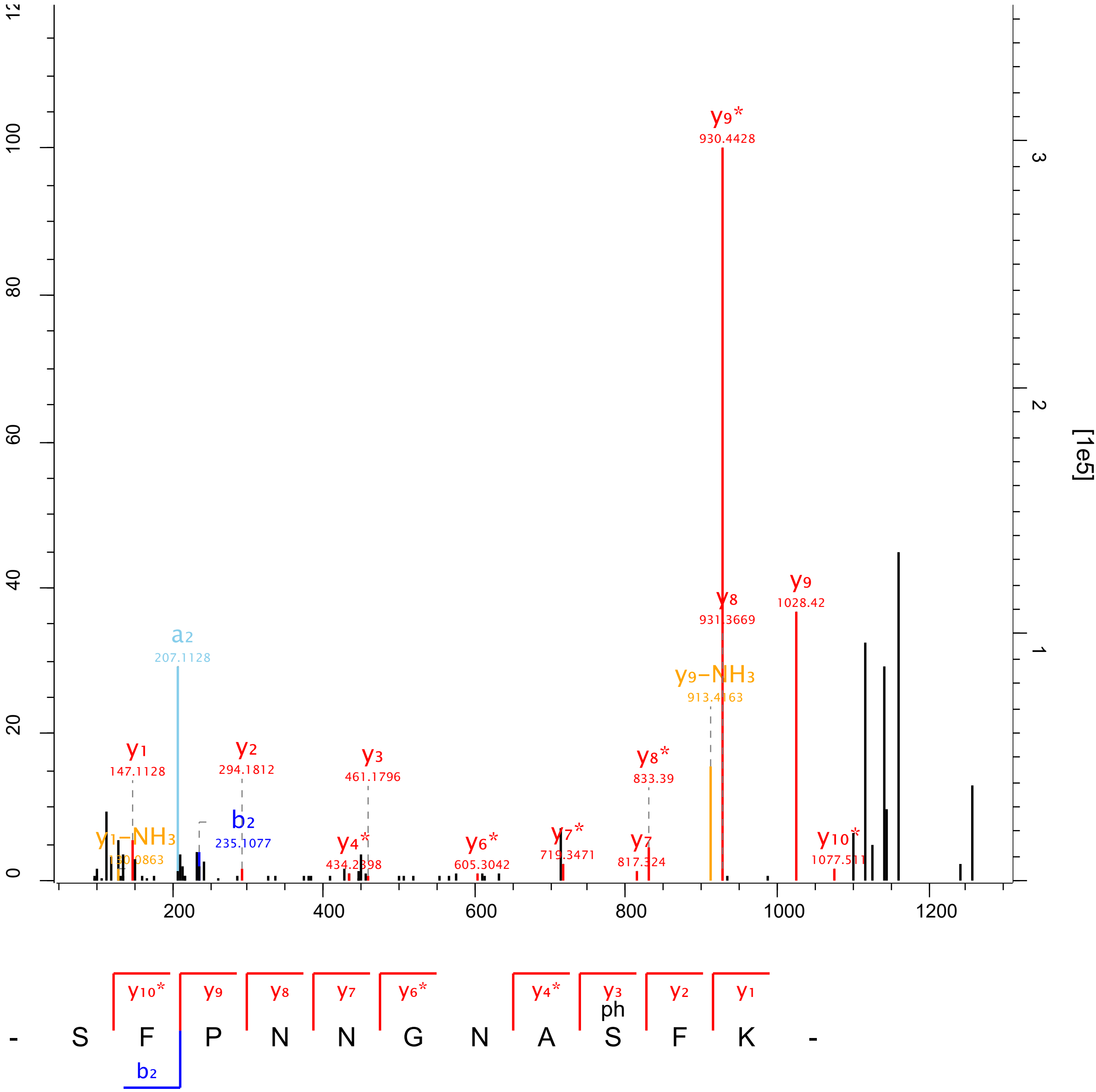

| Raw file | Scan | Method | Score | m/z |
|---|---|---|---|---|
| sys_05_2 | 14420 | FTMS; HCD | 197.41 | 711.81 |

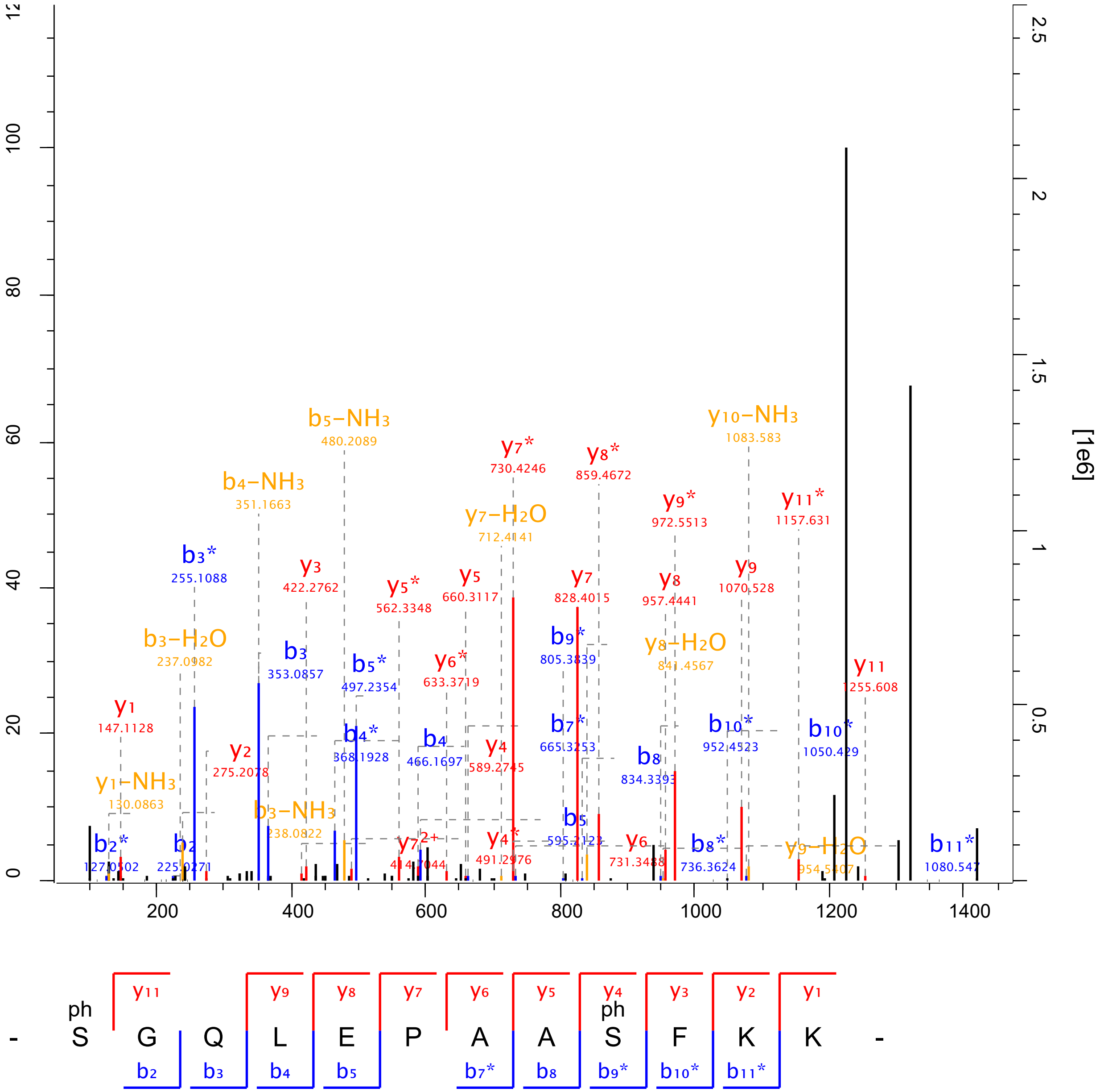

| Raw file | Scan | Method | Score | m/z |
| --- | --- | --- | --- | --- |
| sys_05_2 | 14445 | FTMS; HCD | 84.76 | 595.78 |

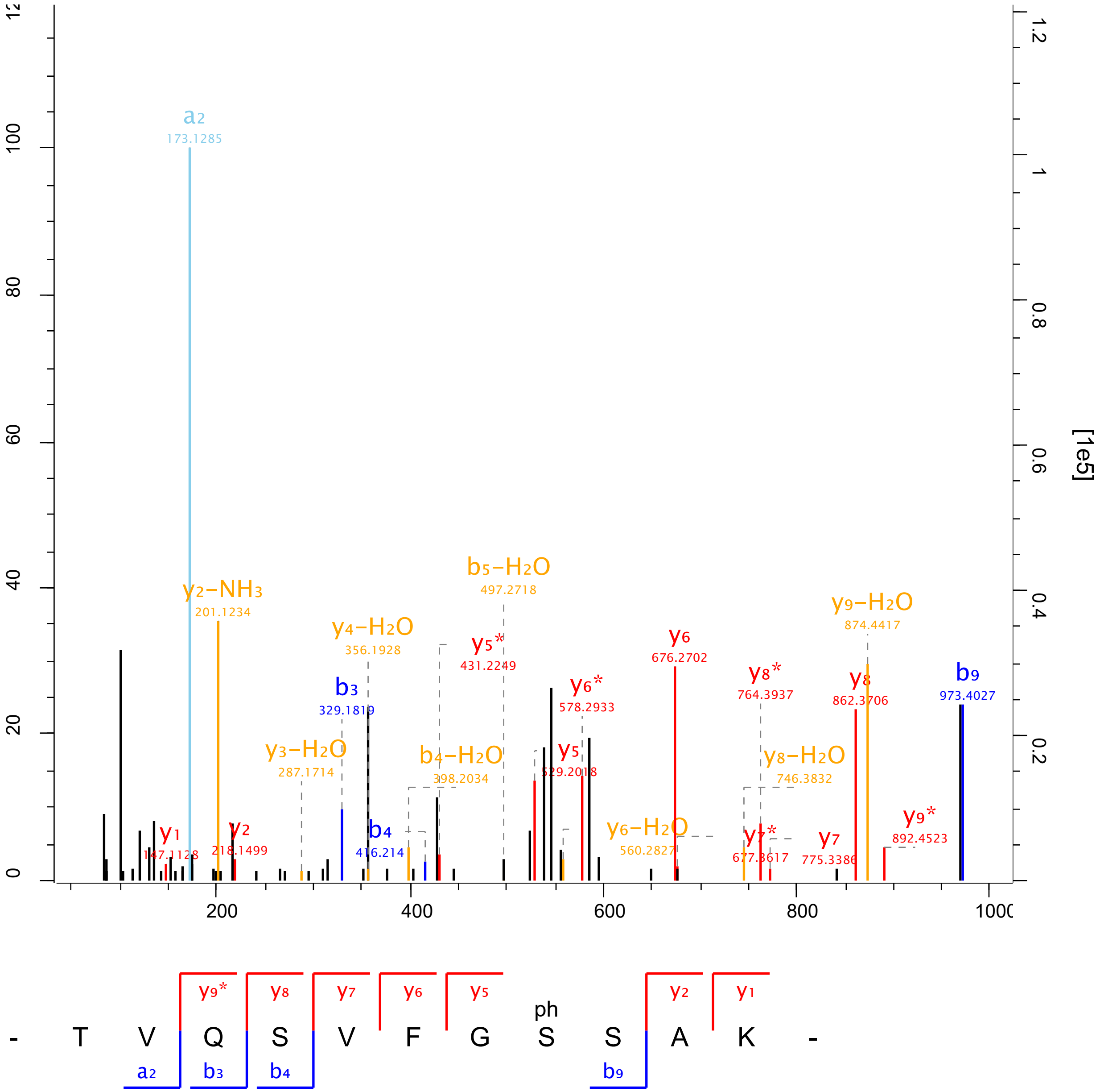

| Raw file | Scan | Method | Score | m/z |
|---|---|---|---|---|
| sys_05_2 | 14485 | FTMS; HCD | 97.69 | 601.26 |

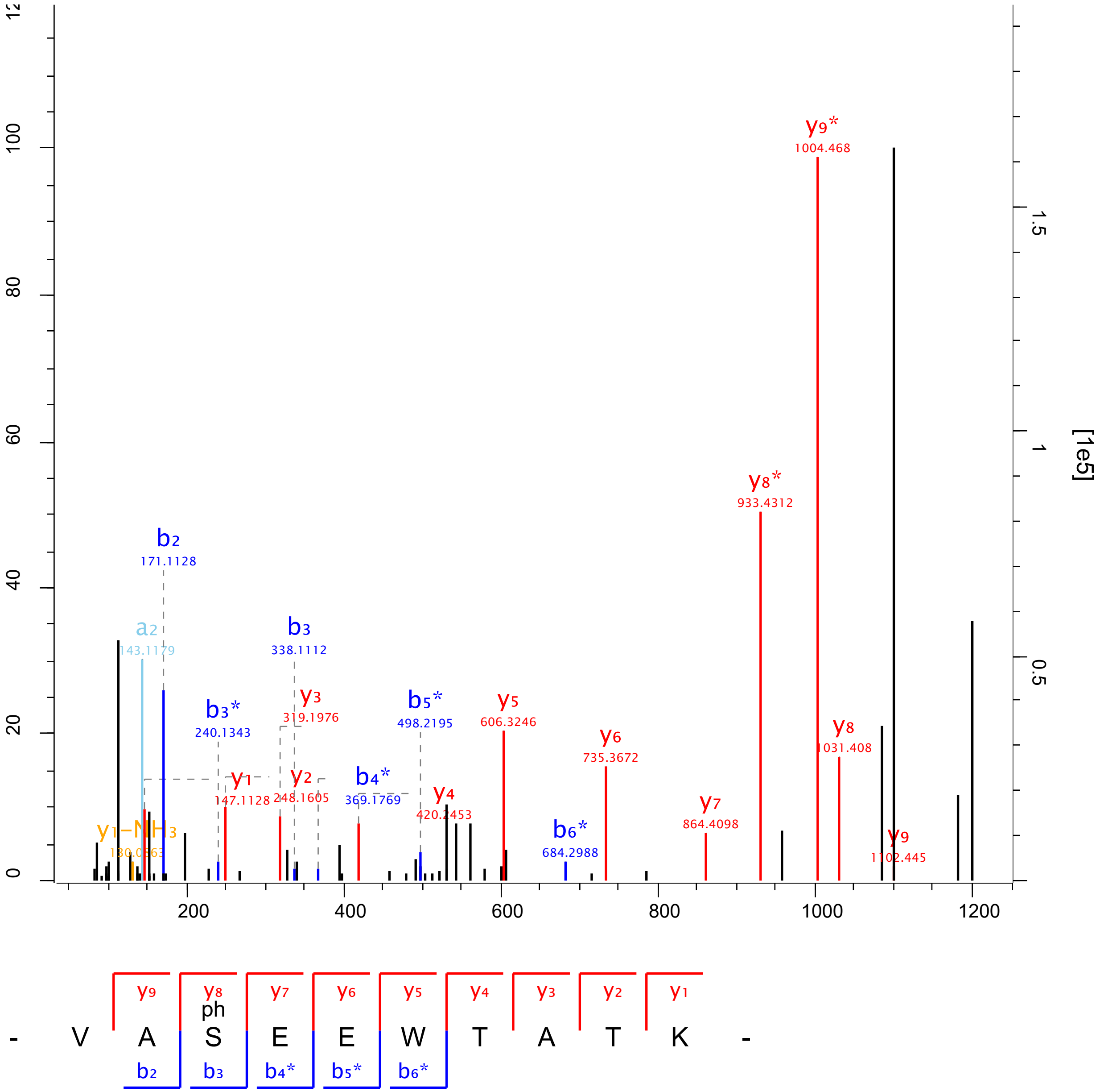

| Raw file | Scan | Method | Score | m/z |
| --- | --- | --- | --- | --- |
| sys_05_2 | 14530 | FTMS; HCD | 77.19 | 475.9 |

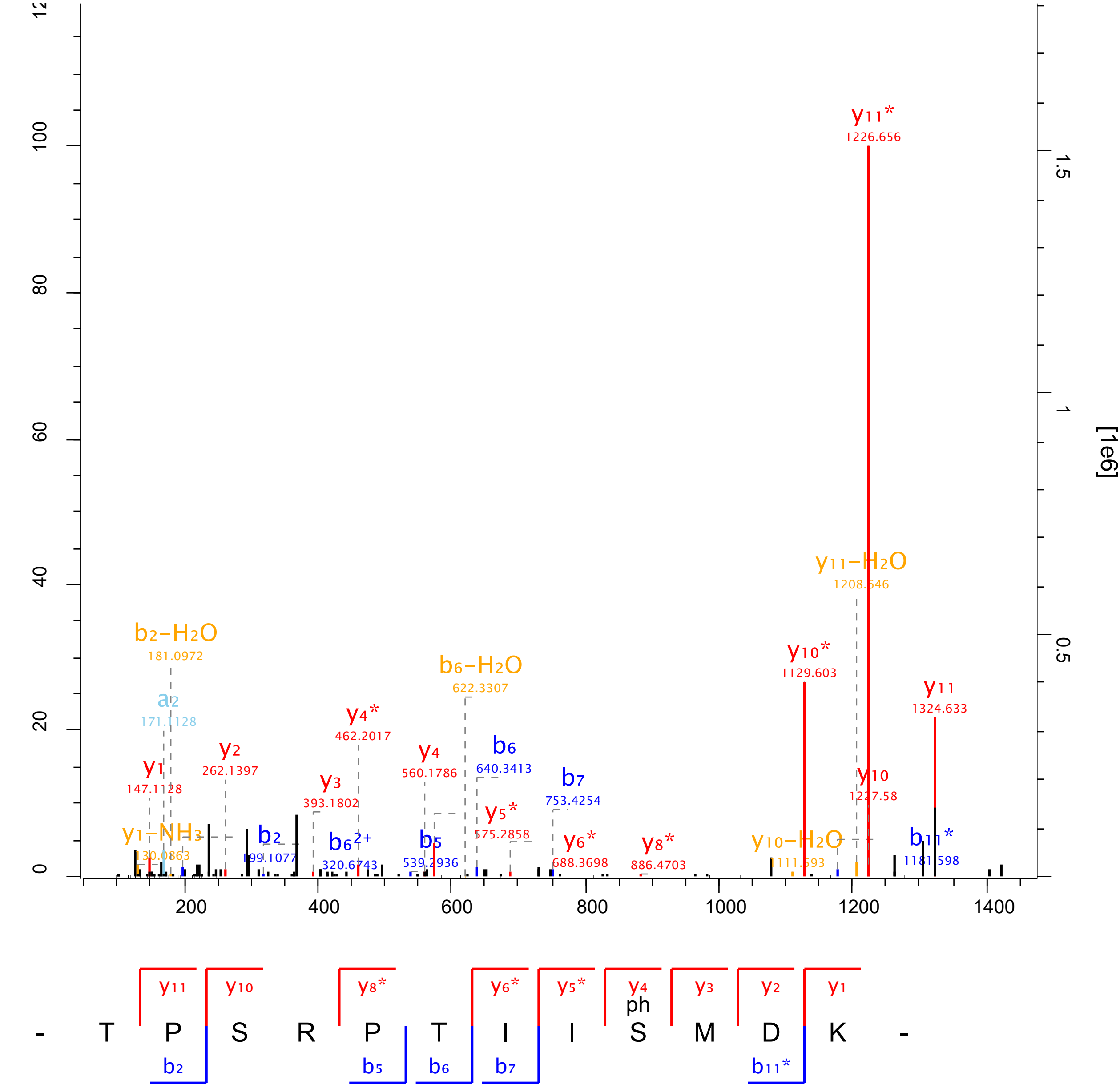

| Raw file | Scan | Method | Score | m/z |
|---|---|---|---|---|
| sys_05_2 | 14534 | FTMS; HCD | 192.46 | 625.74 |

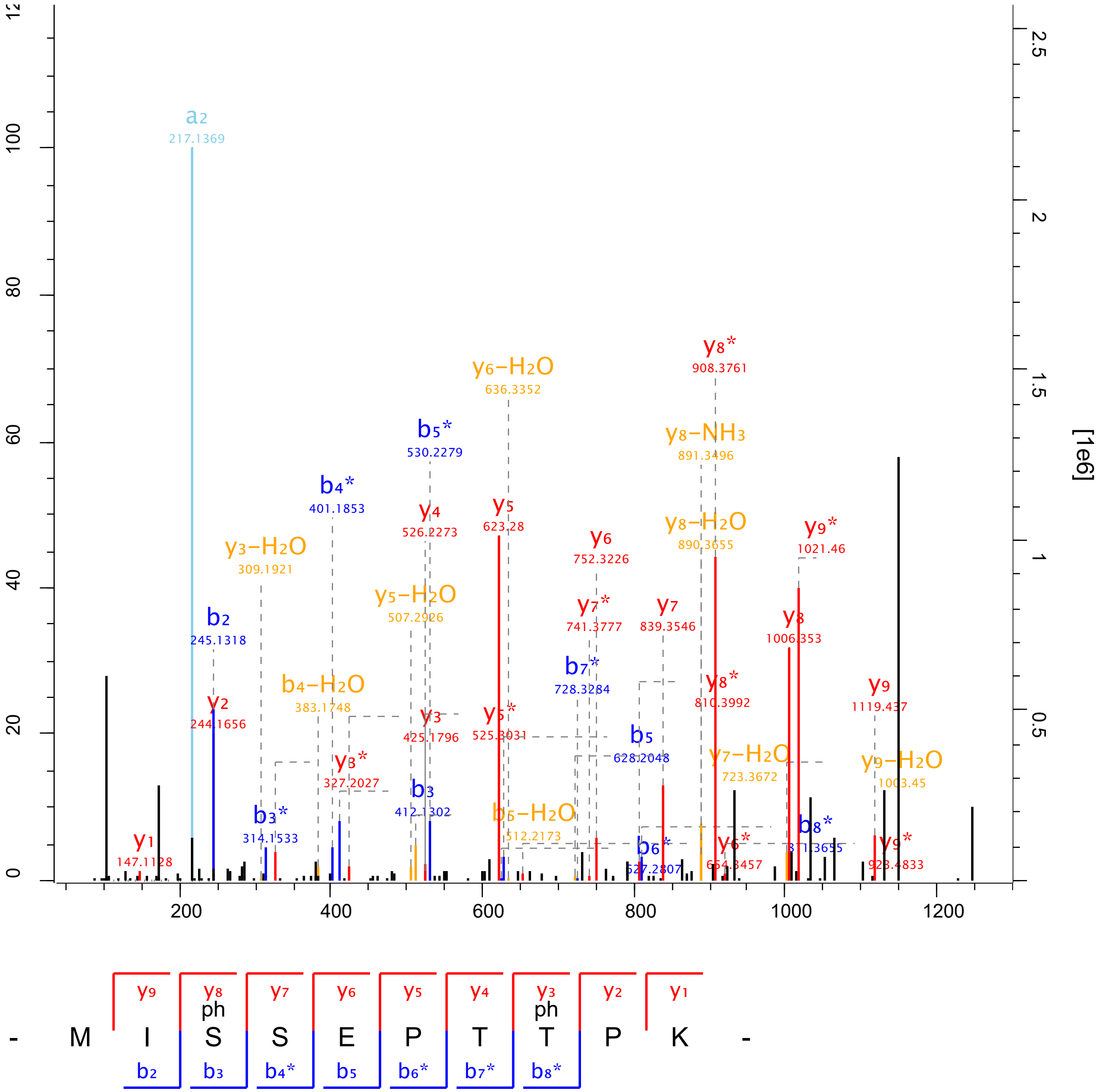

| Raw file | Scan | Method | Score | m/z |
|---|---|---|---|---|
| sys_05_2 | 14655 | FTMS; HCD | 49.99 | 517.9 |

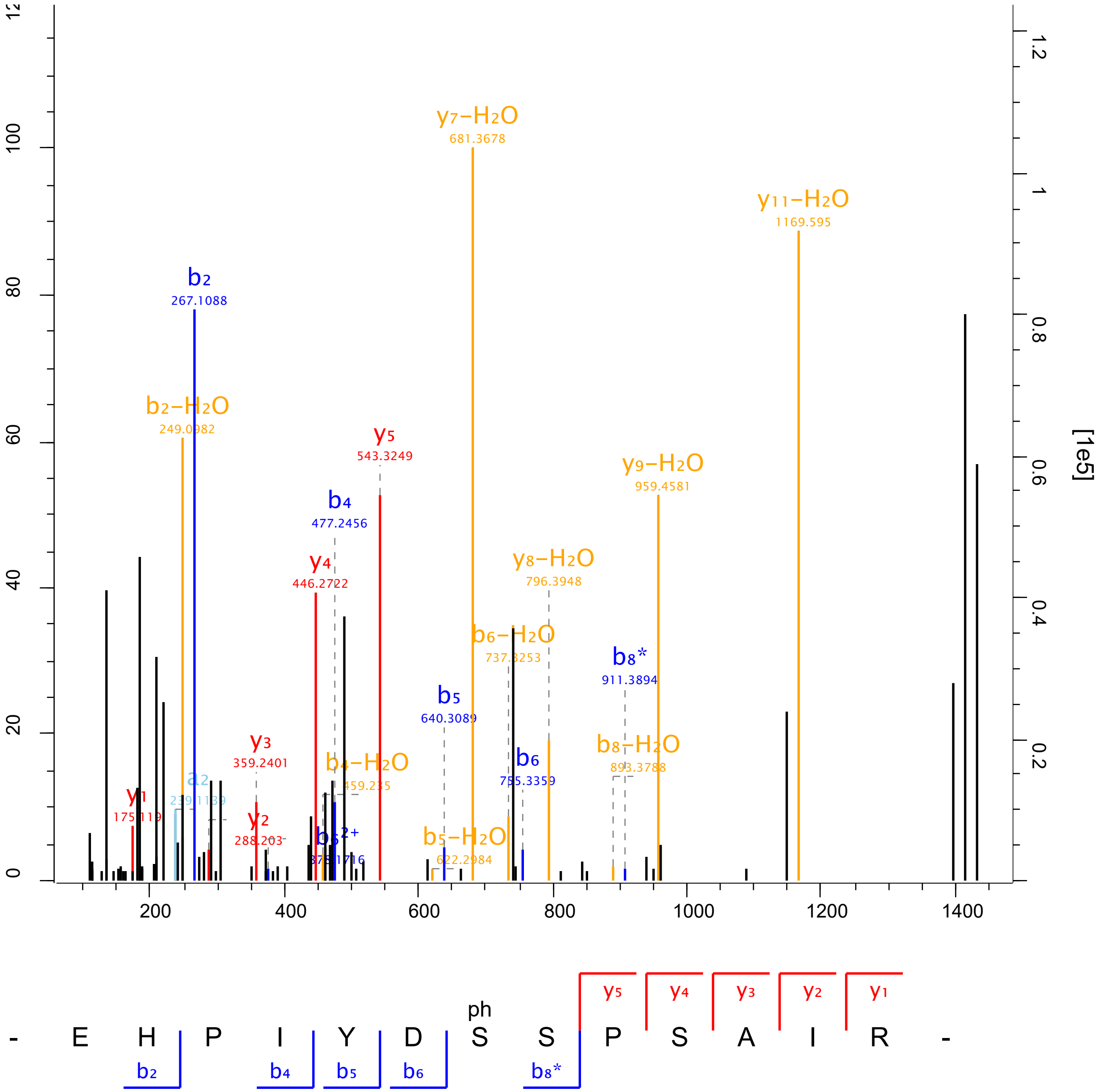

| Raw file | Scan | Method | Score | m/z |
| --- | --- | --- | --- | --- |
| sys_05_2 | 14733 | FTMS; HCD | 46.7 | 685.29 |

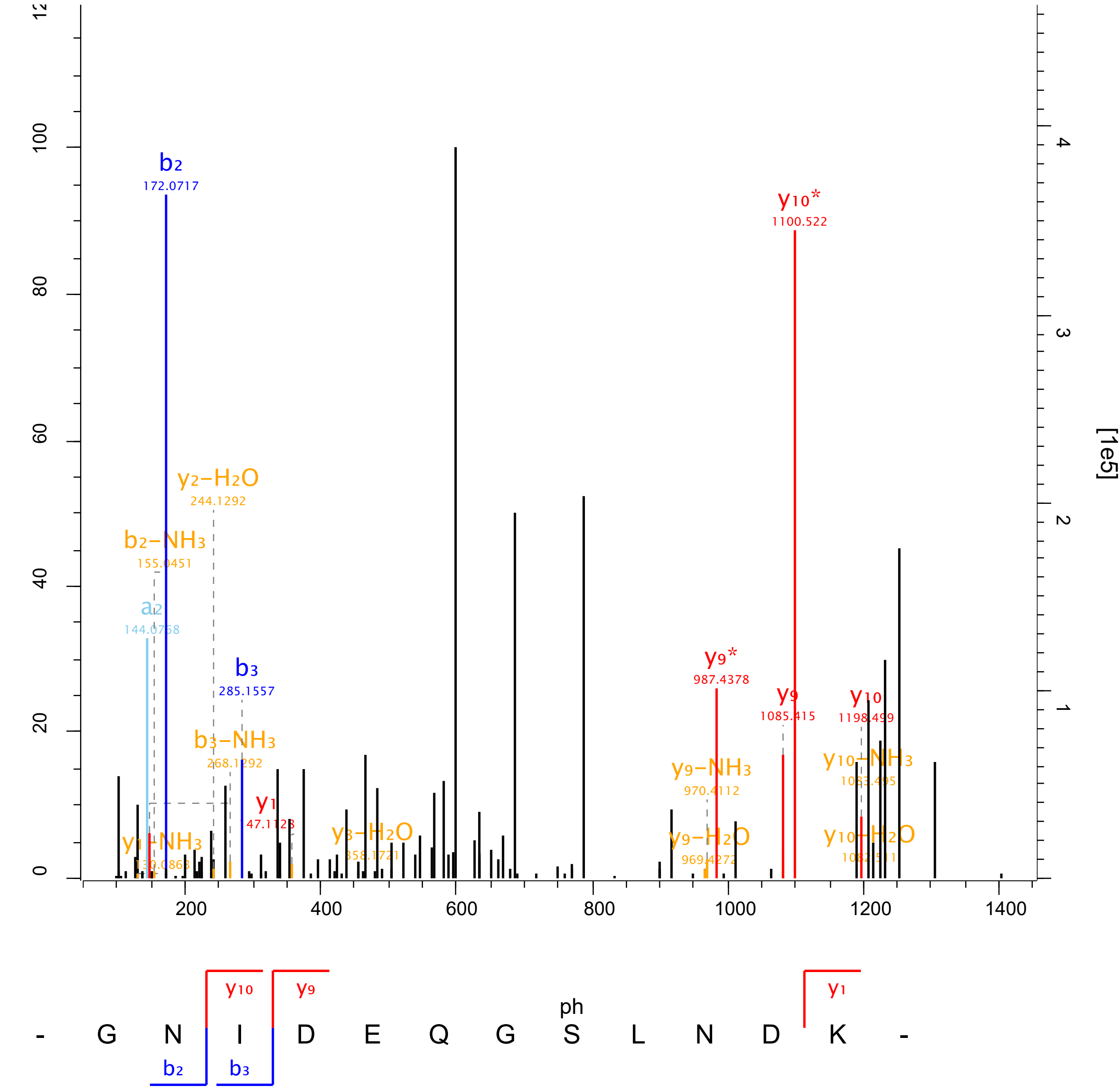

| Raw file | Scan | Method | Score | m/z |
|---|---|---|---|---|
| sys_05_2 | 14803 | FTMS; HCD | 119.17 | 682.29 |

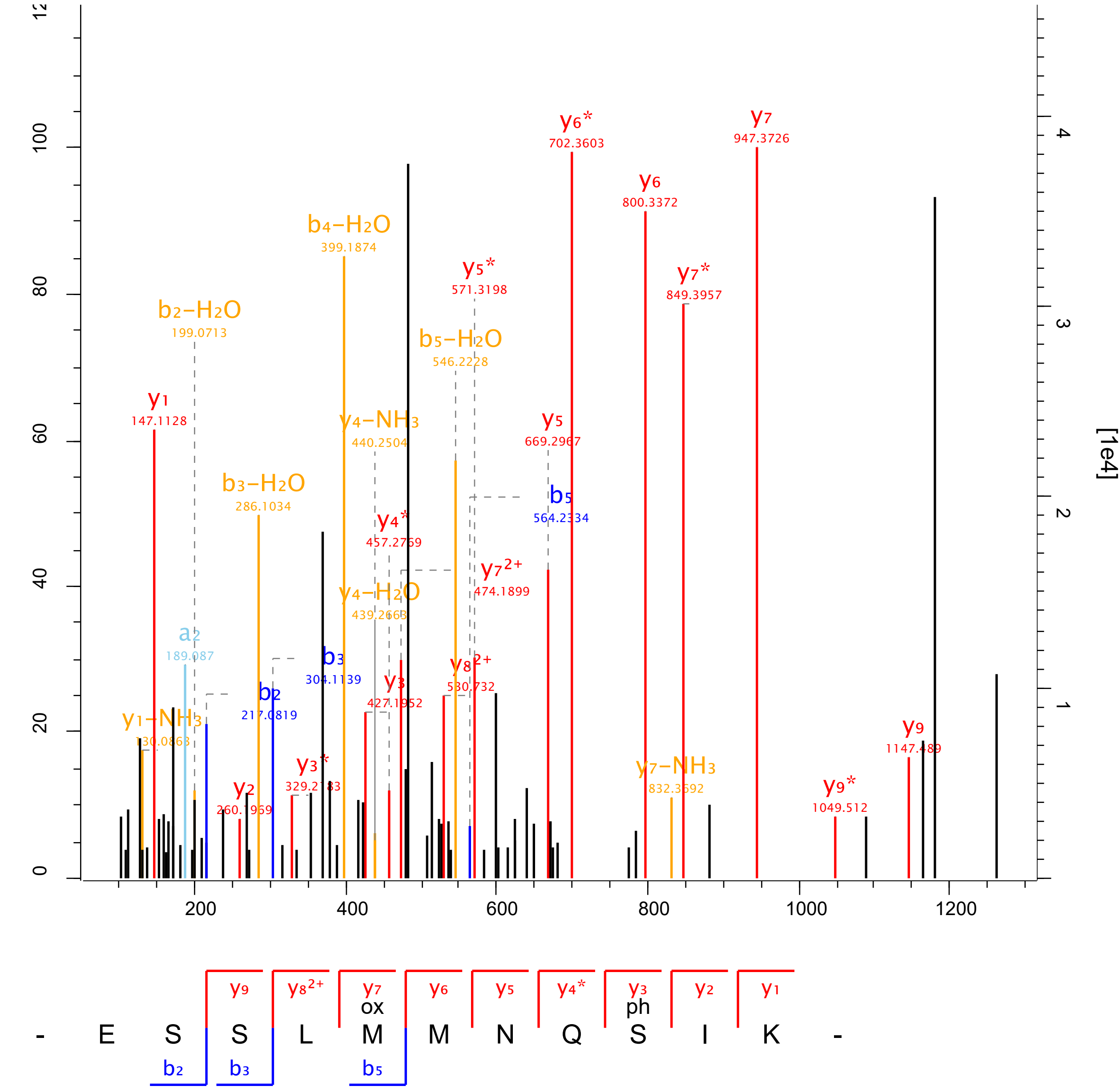

| Raw file | Scan | Method | Score | m/z |
|---|---|---|---|---|
| sys_05_2 | 14832 | FTMS; HCD | 61.96 | 731.83 |

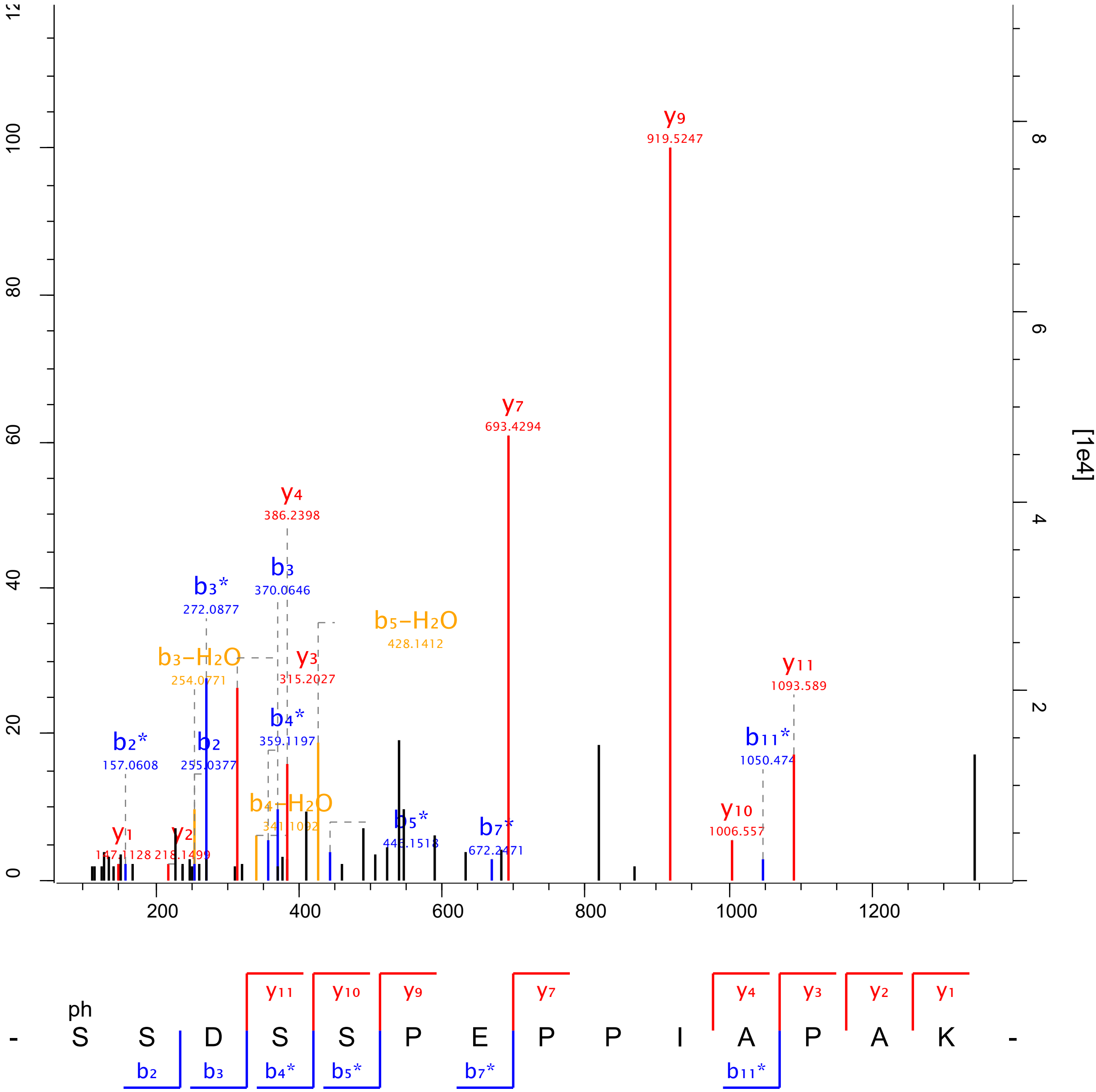

| Raw file | Scan | Method | Score | m/z |
|---|---|---|---|---|
| sys_05_2 | 14857 | FTMS; HCD | 57.2 | 555.78 |

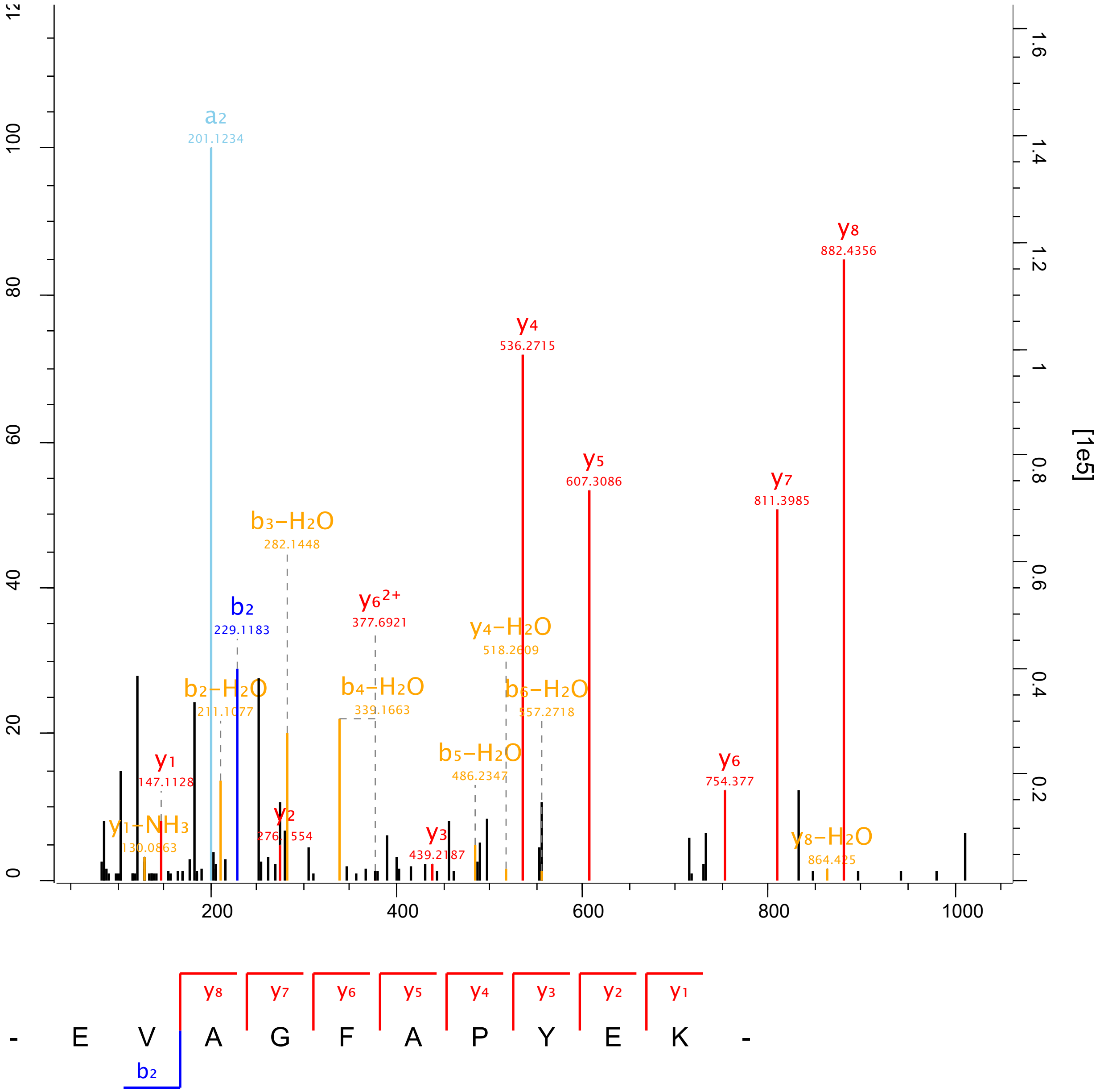

| Raw file | Scan | Method | Score | m/z |
|---|---|---|---|---|
| sys_05_2 | 14934 | FTMS; HCD | 108.71 | 606.93 |

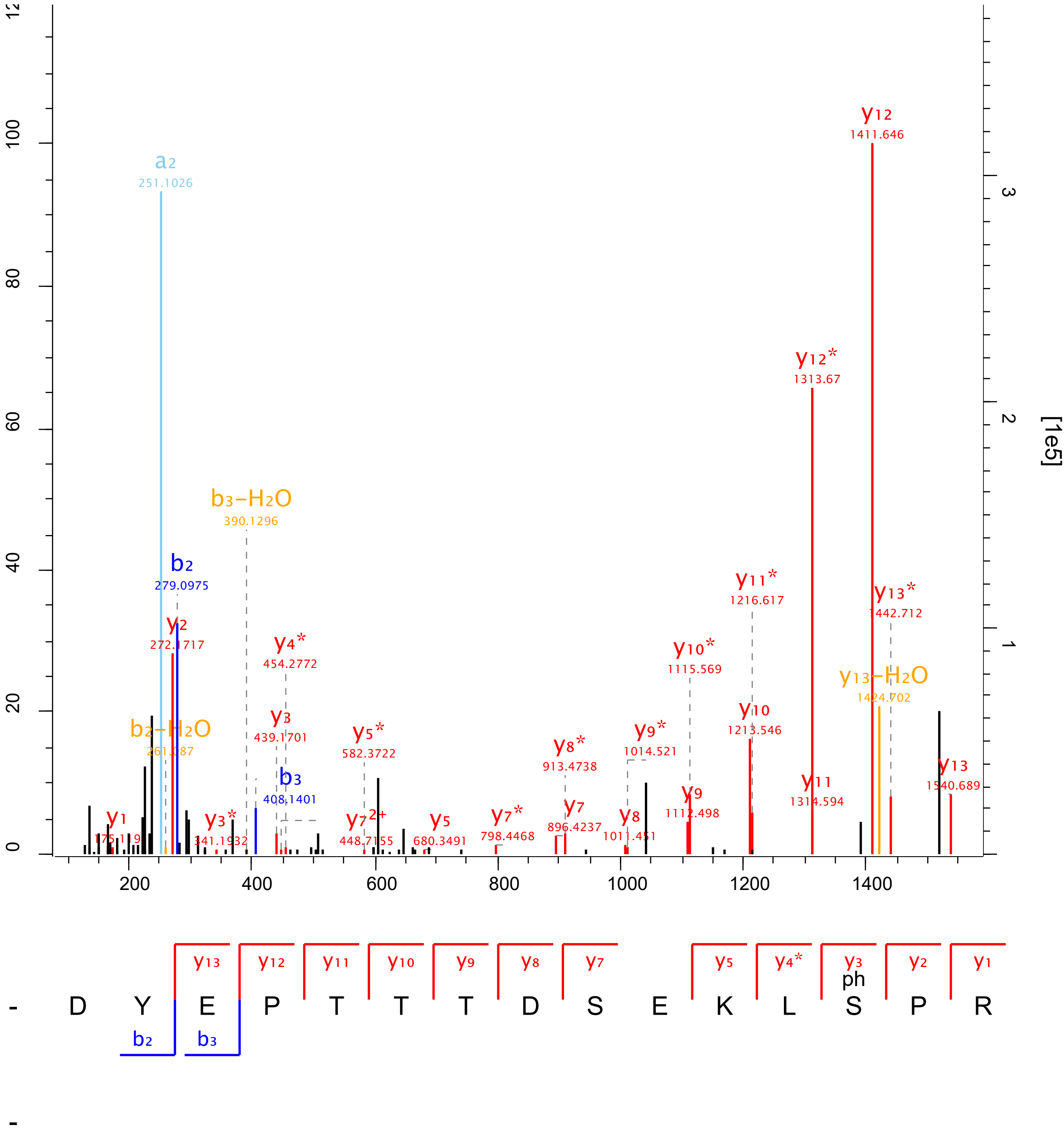

| Raw file | Scan | Method | Score | m/z |
|---|---|---|---|---|
| sys_05_2 | 1503 | FTMS; HCD | 66.19 | 566.74 |

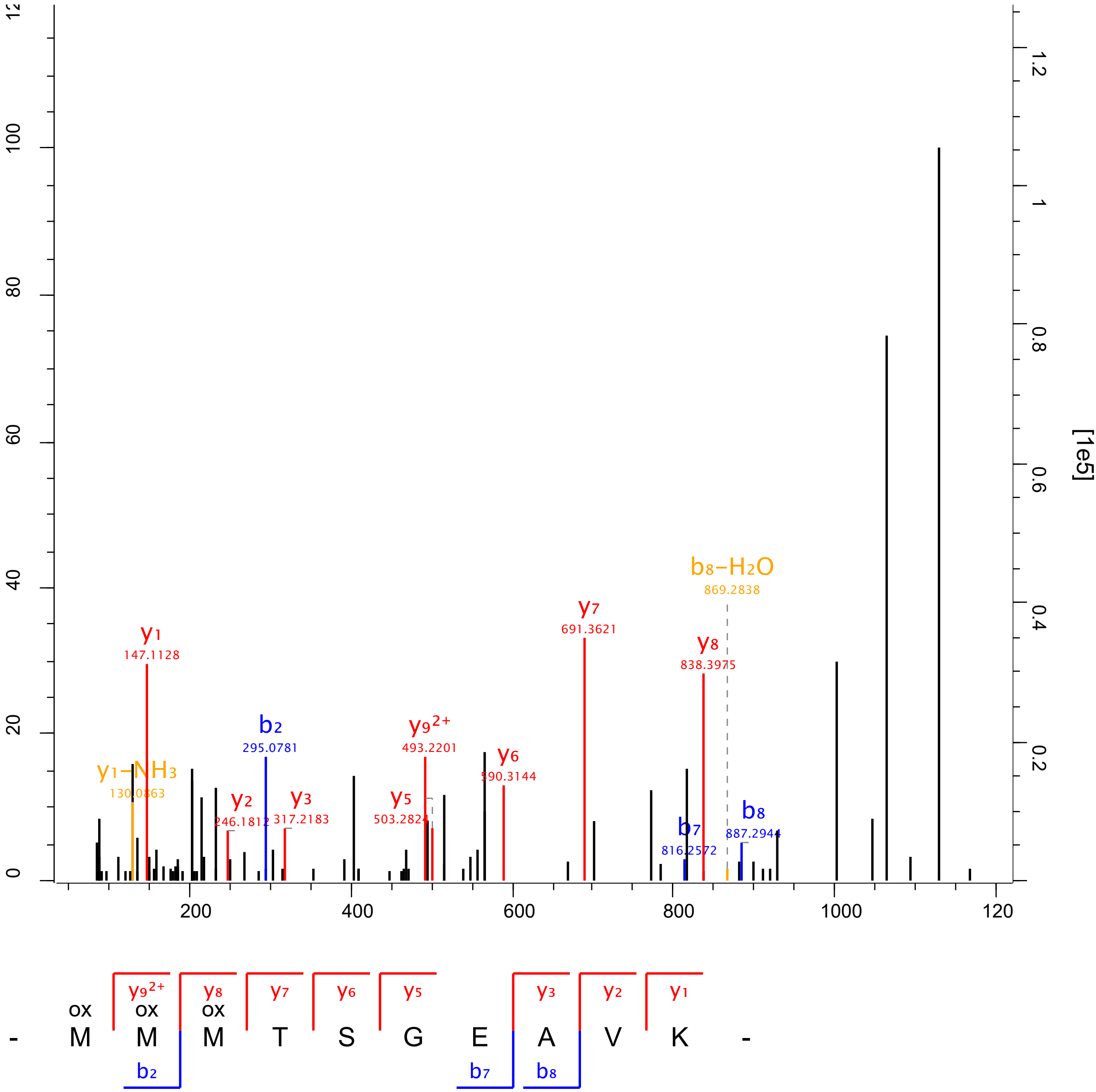

| Raw file | Scan | Method | Score | m/z |
|---|---|---|---|---|
| sys_05_2 | 15198 | FTMS; HCD | 120.32 | 840.88 |

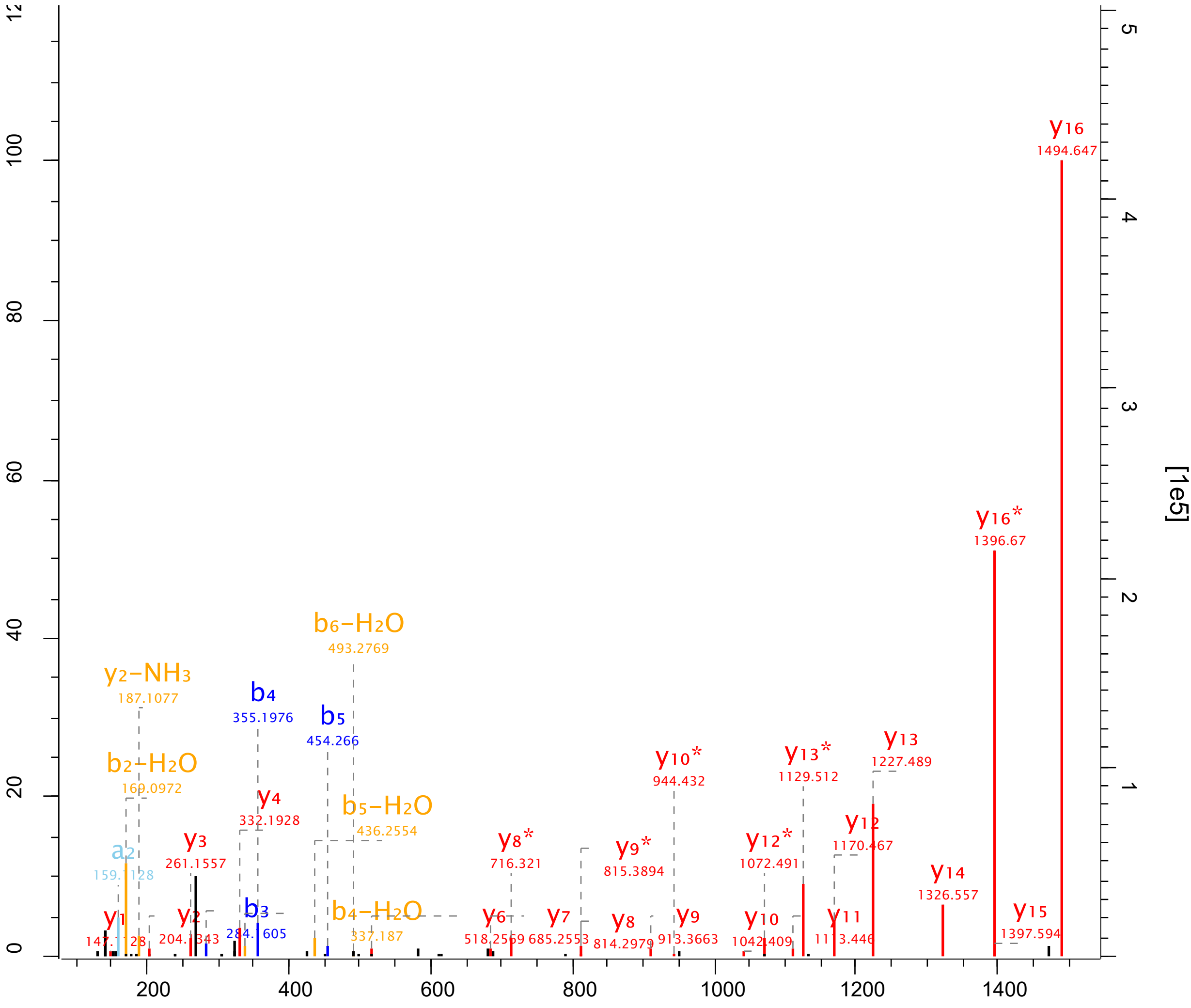


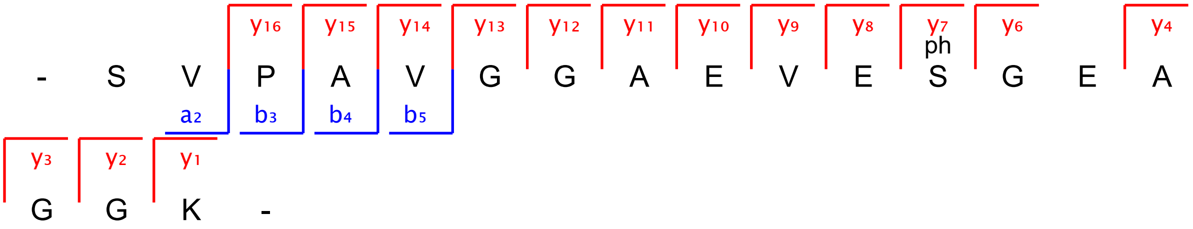

| Raw file | Scan | Method | Score | m/z |
|---|---|---|---|---|
| sys_05_2 | 15234 | FTMS; HCD | 54.26 | 540.73 |

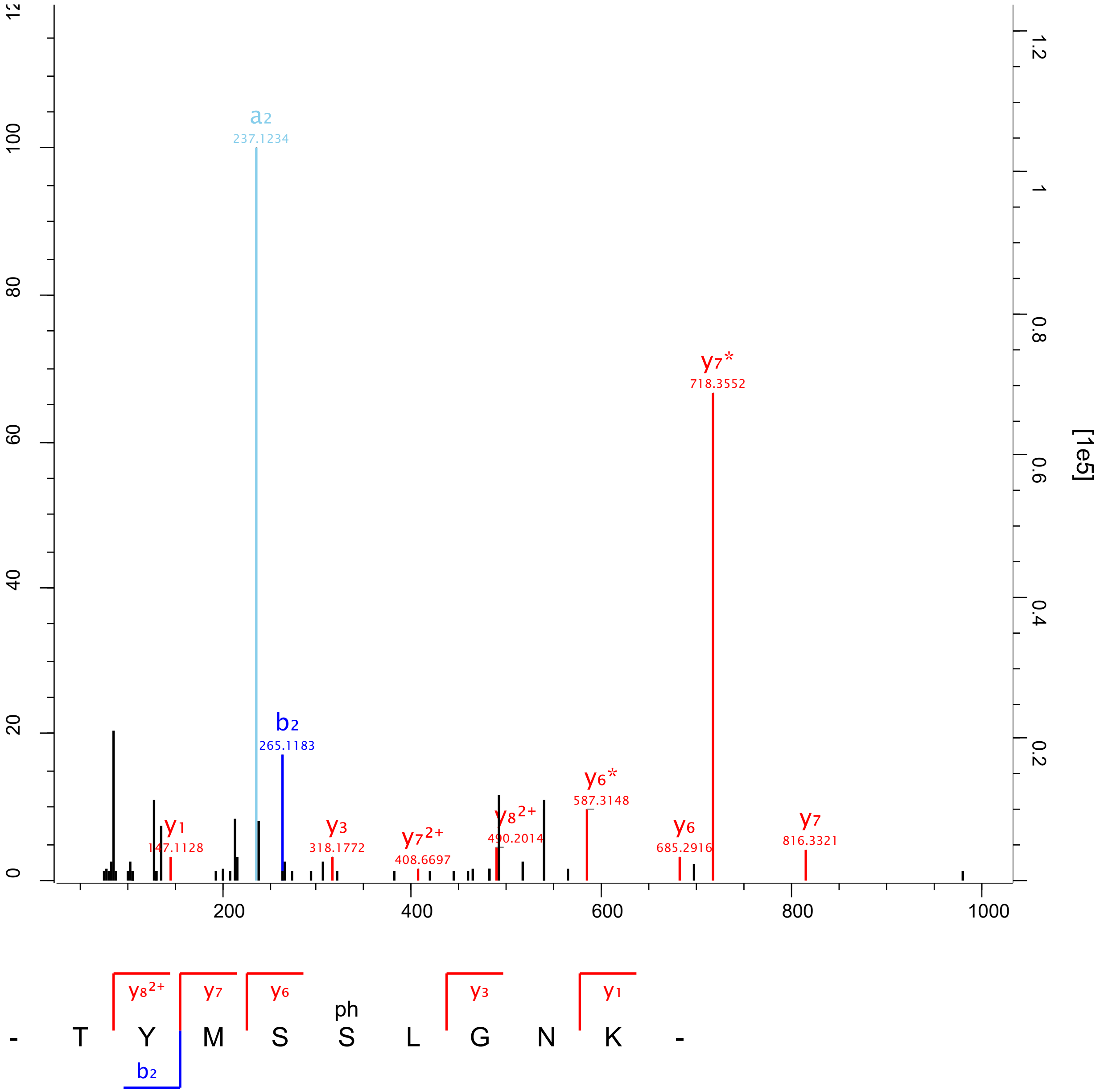

| Raw file | Scan | Method | Score | m/z |
|---|---|---|---|---|
| sys_05_2 | 15306 | FTMS; HCD | 139.11 | 880.84 |

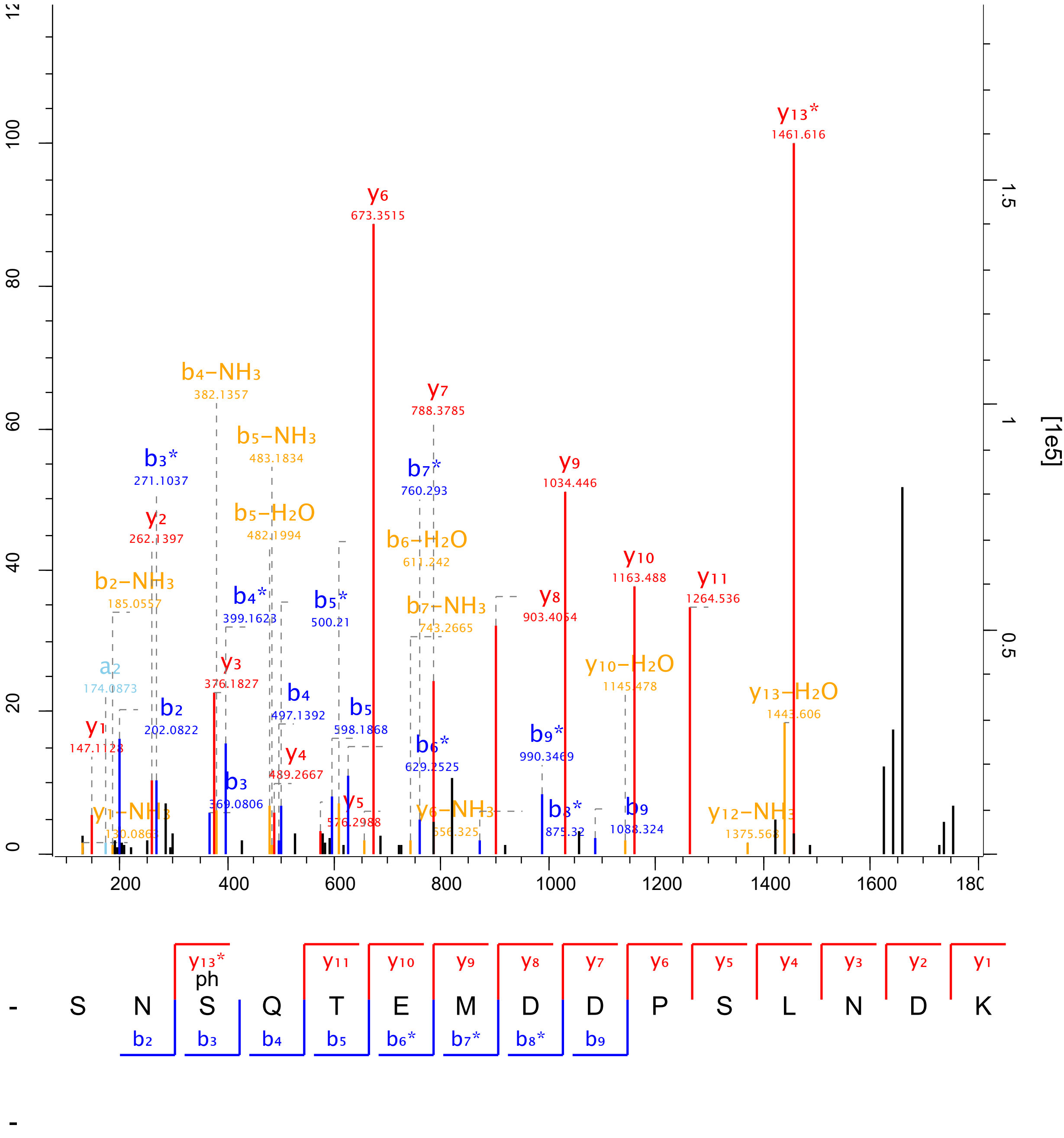

| Raw file | Scan | Method | Score | m/z |
| --- | --- | --- | --- | --- |
| sys_05_2 | 15355 | FTMS; HCD | 84.38 | 730.83 |

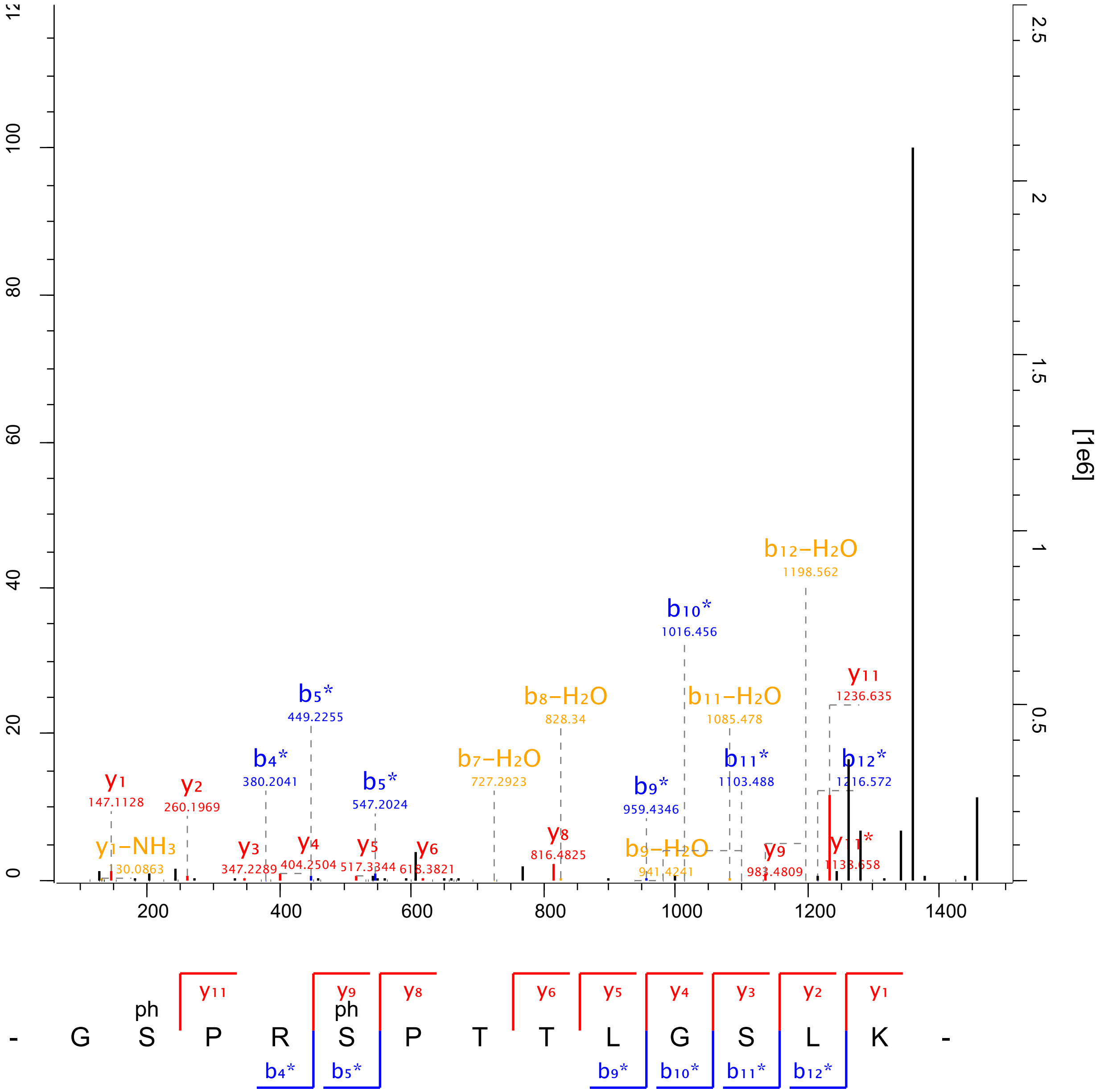

| Raw file | Scan | Method | Score | m/z |
|---|---|---|---|---|
| sys_05_2 | 15535 | FTMS; HCD | 50.87 | 619.29 |

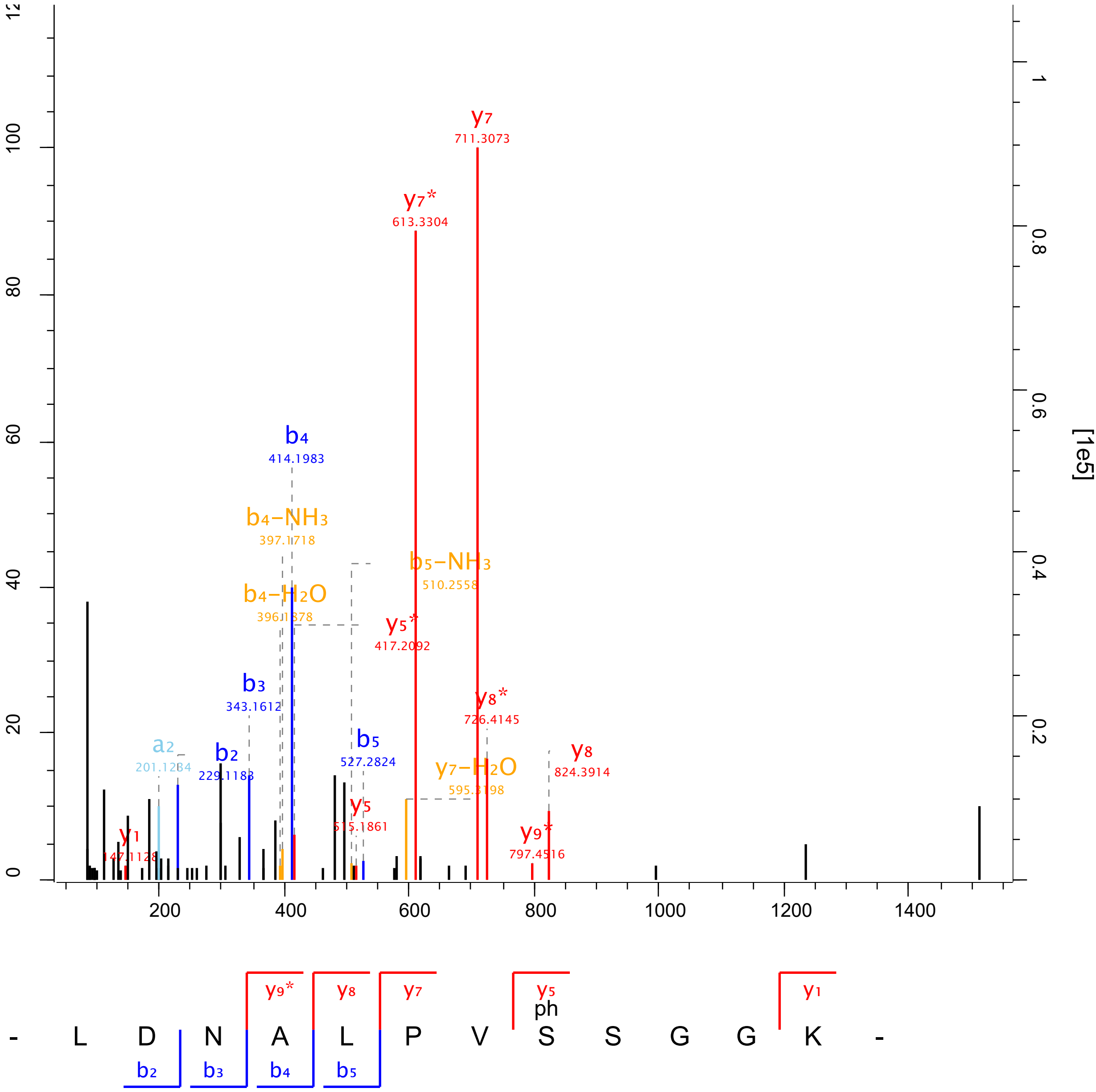

| Raw file | Scan | Method | Score | m/z |
|---|---|---|---|---|
| sys_05_2 | 15593 | FTMS; HCD | 40.49 | 700.31 |

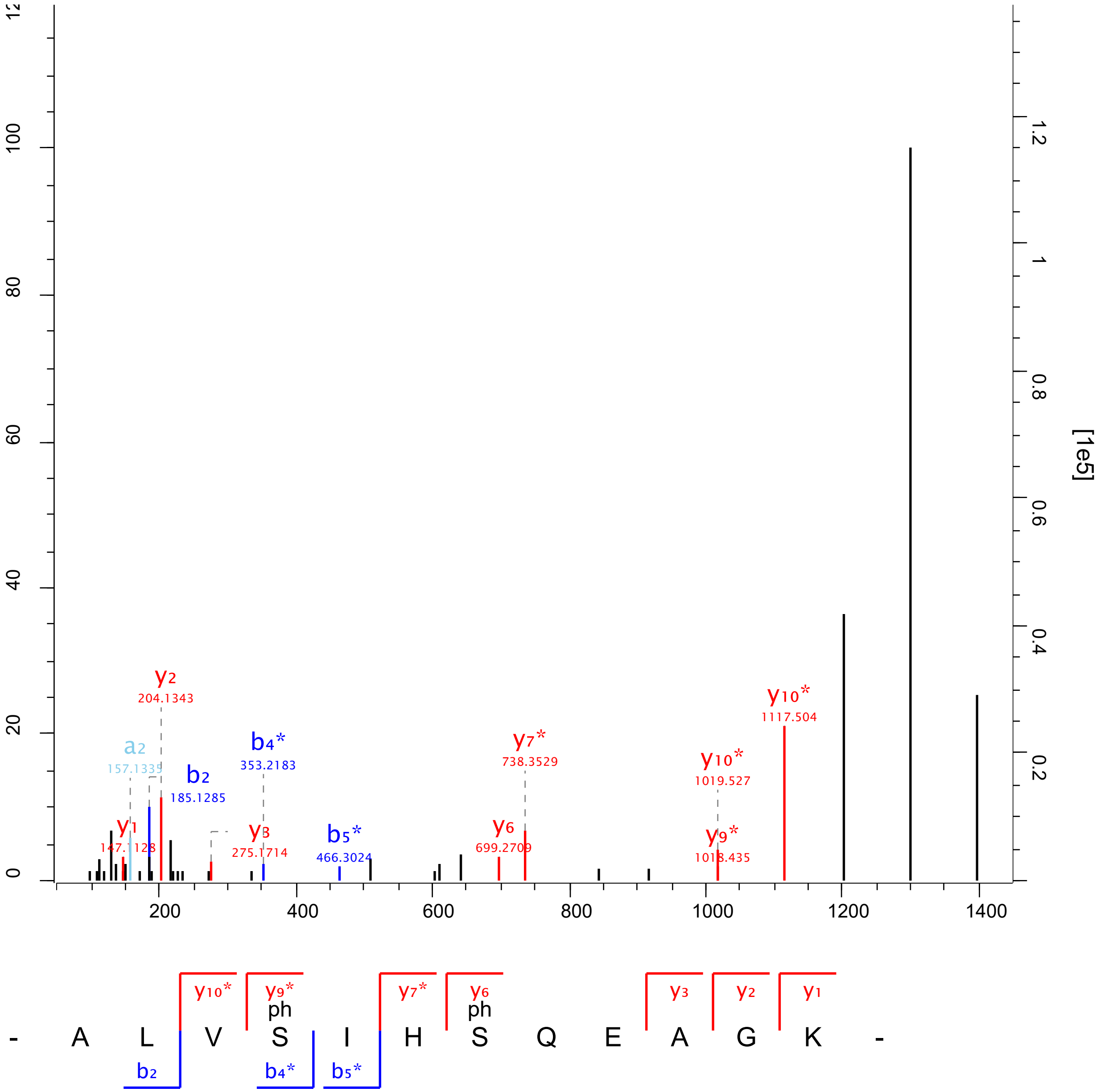

| Raw file | Scan | Method | Score | m/z |
|---|---|---|---|---|
| sys_05_2 | 15620 | FTMS; HCD | 74.14 | 638.34 |

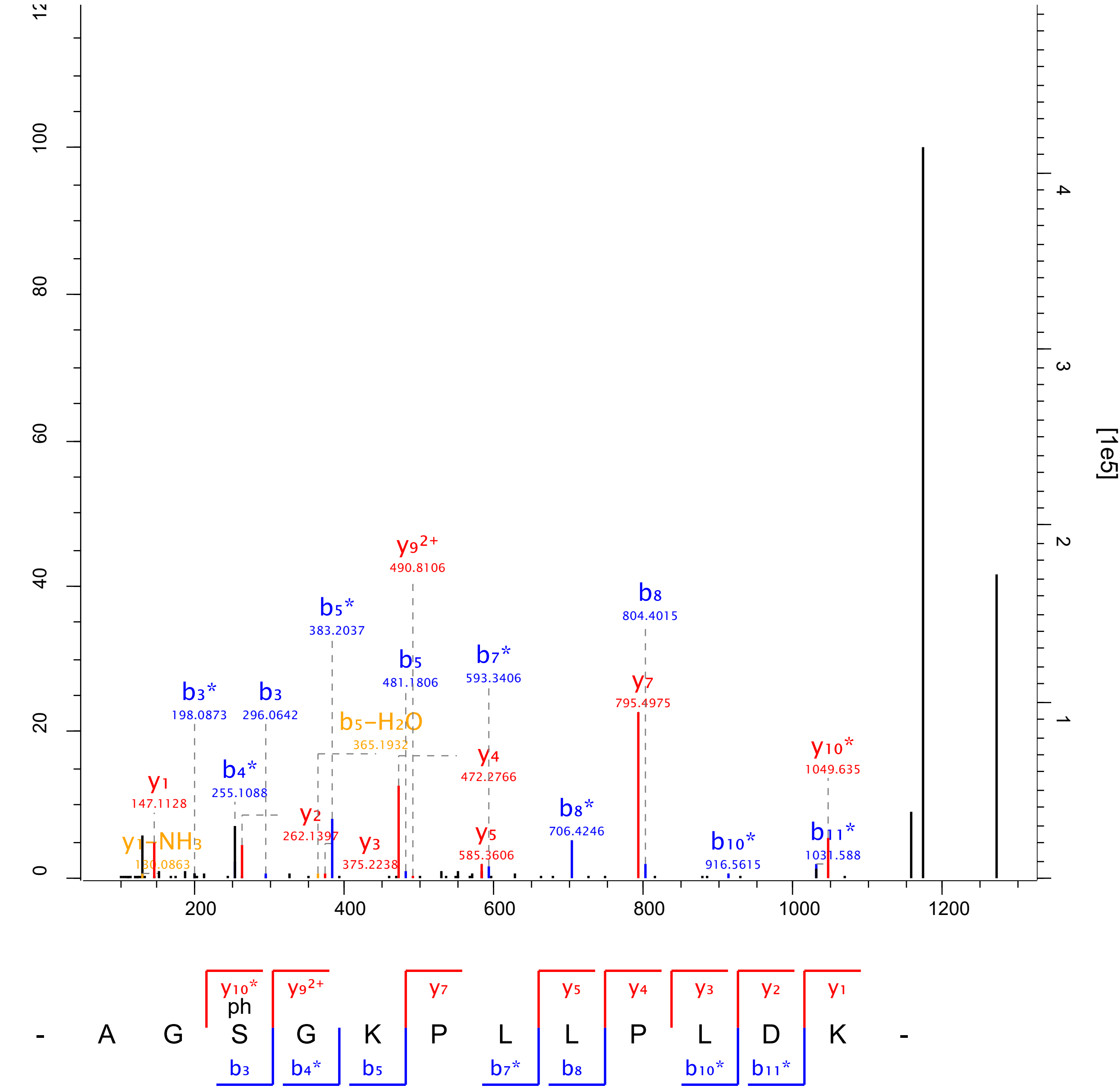

| Raw file | Scan | Method | Score | m/z |
|---|---|---|---|---|
| sys_05_2 | 15744 | FTMS; HCD | 59.2 | 482.24 |

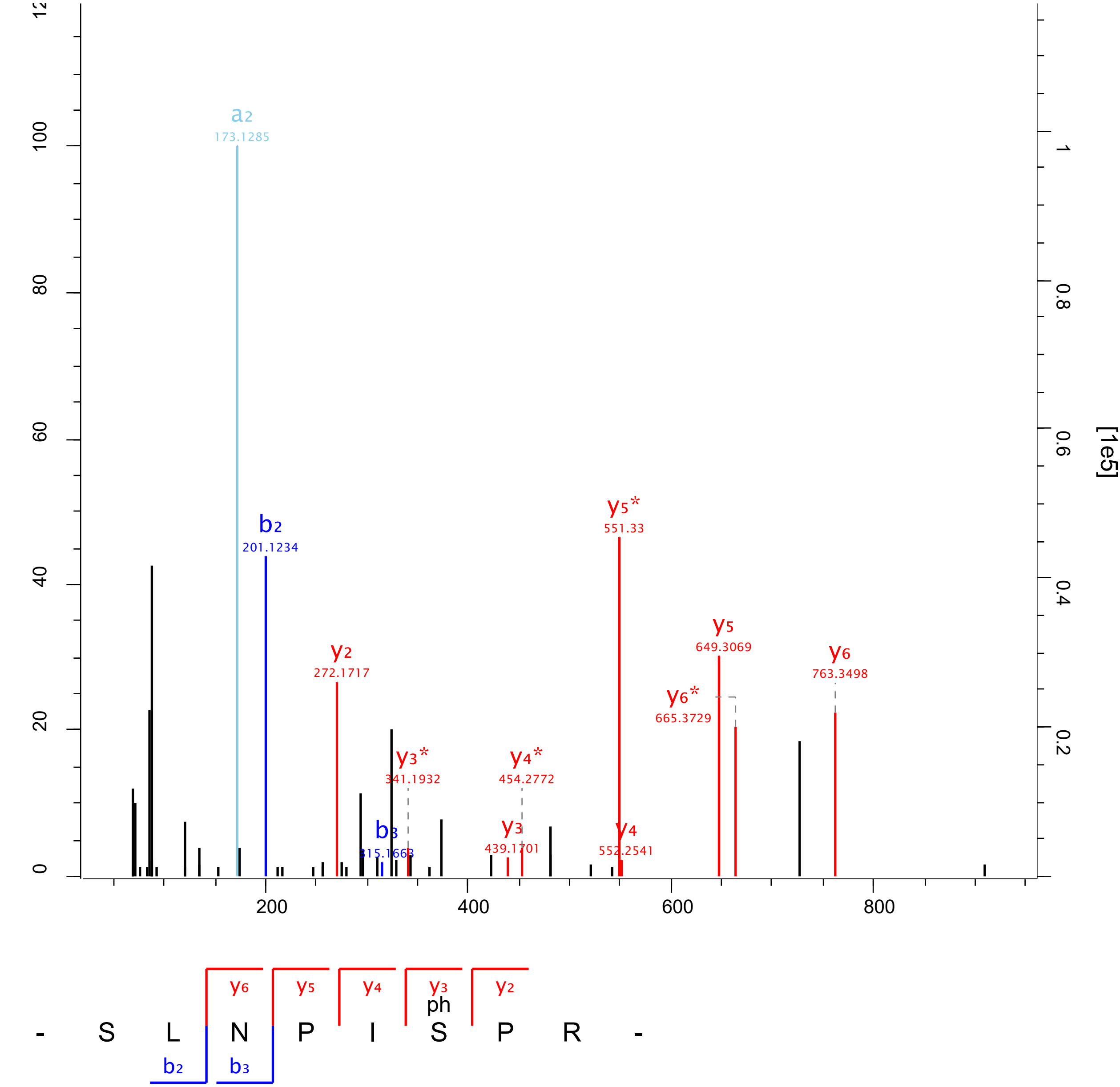

| Raw file | Scan | Method | Score | m/z |
|---|---|---|---|---|
| sys_05_2 | 15789 | FTMS; HCD | 64.64 | 512.26 |

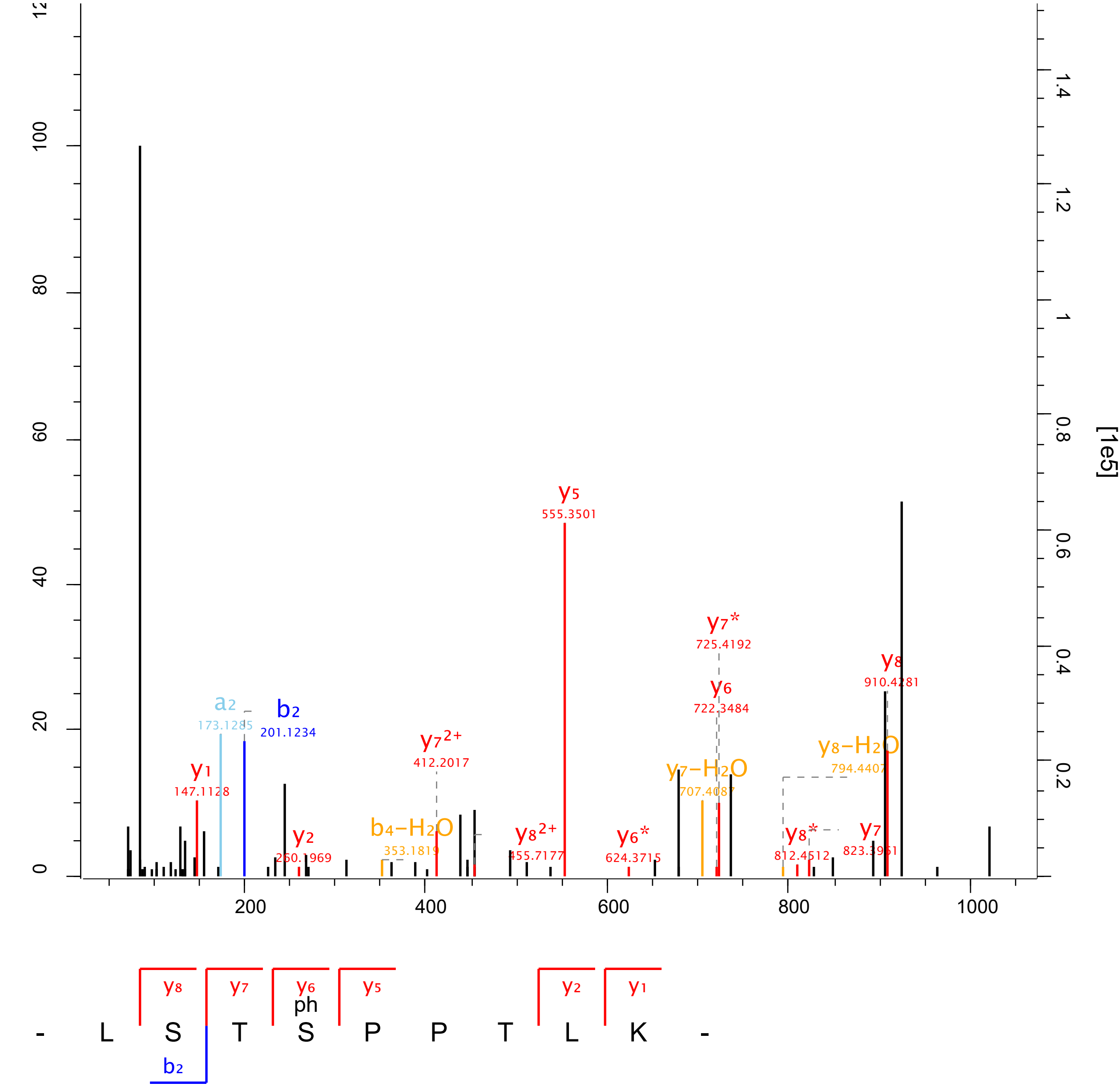

| Raw file | Scan | Method | Score | m/z |
| --- | --- | --- | --- | --- |
| sys_05_2 | 15849 | FTMS; HCD | 225.66 | 724.83 |

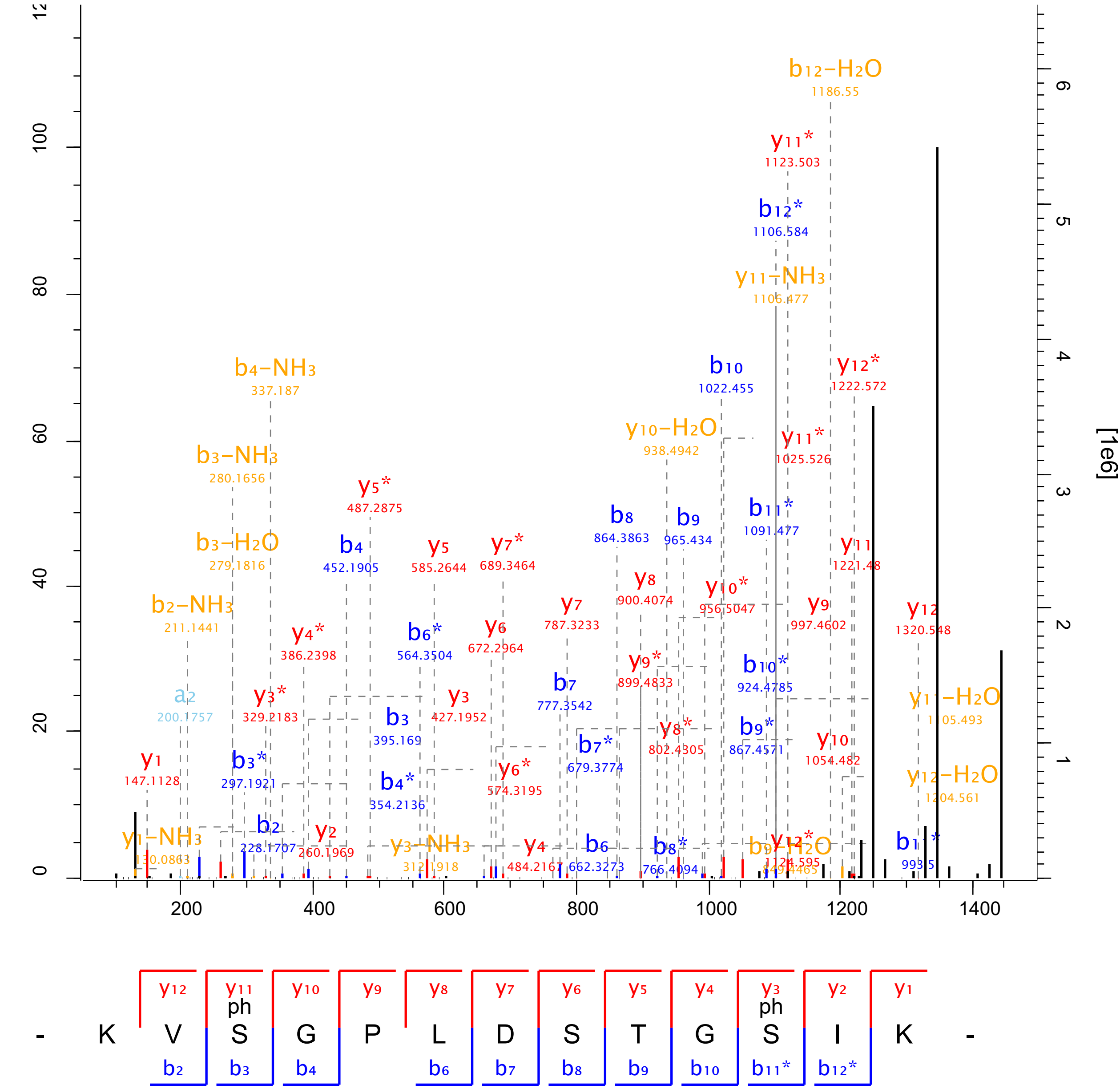

| Raw file | Scan | Method | Score | m/z |
|---|---|---|---|---|
| sys_05_2 | 15946 | FTMS; HCD | 67.65 | 480.23 |

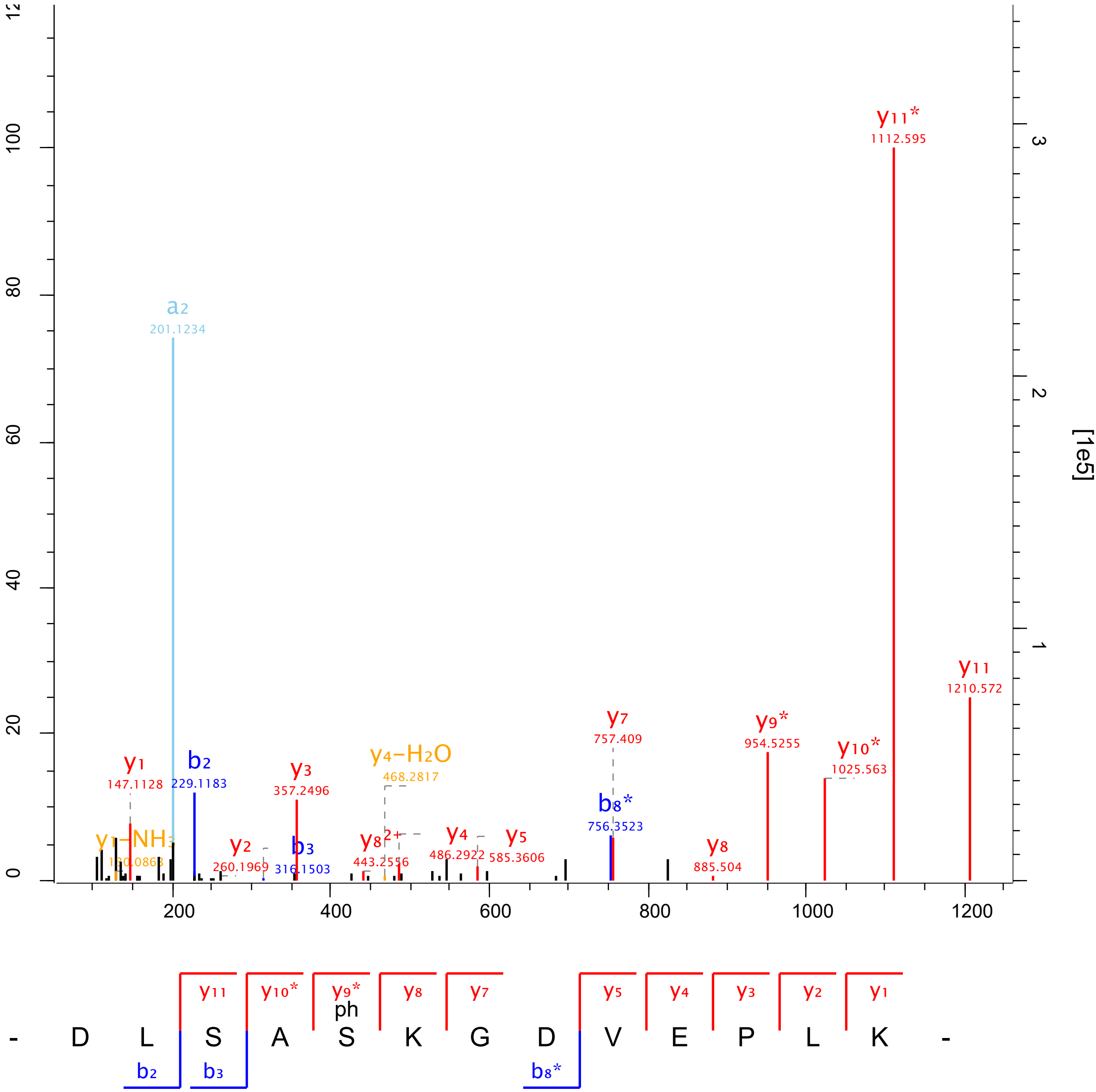

| Raw file | Scan | Method | Score | m/z |
|---|---|---|---|---|
| sys_05_2 | 16017 | FTMS; HCD | 111.52 | 542.26 |

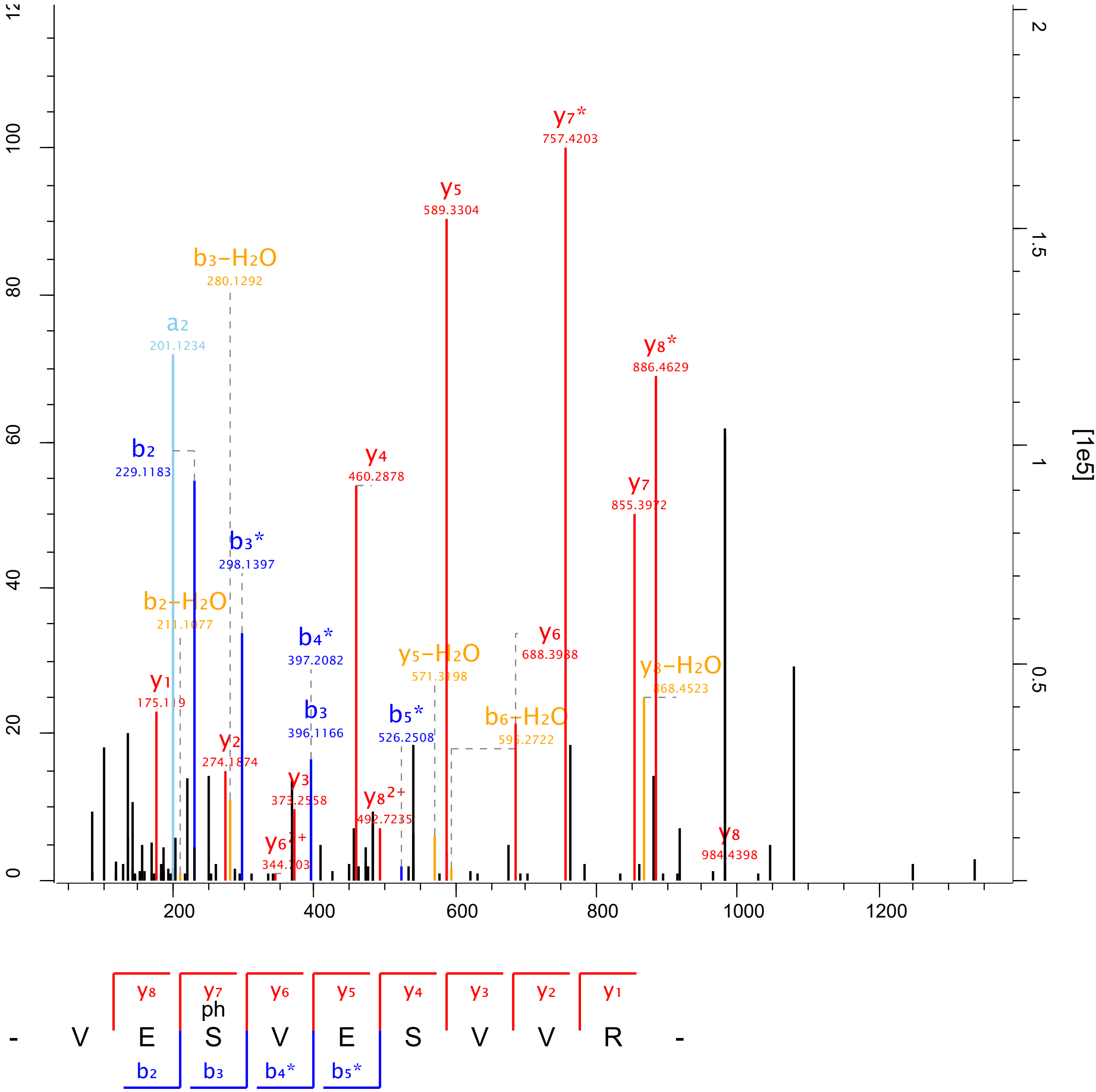

| Raw file | Scan | Method | Score | m/z |
| --- | --- | --- | --- | --- |
| sys_05_2 | 16035 | FTMS; HCD | 254.71 | 908.41 |

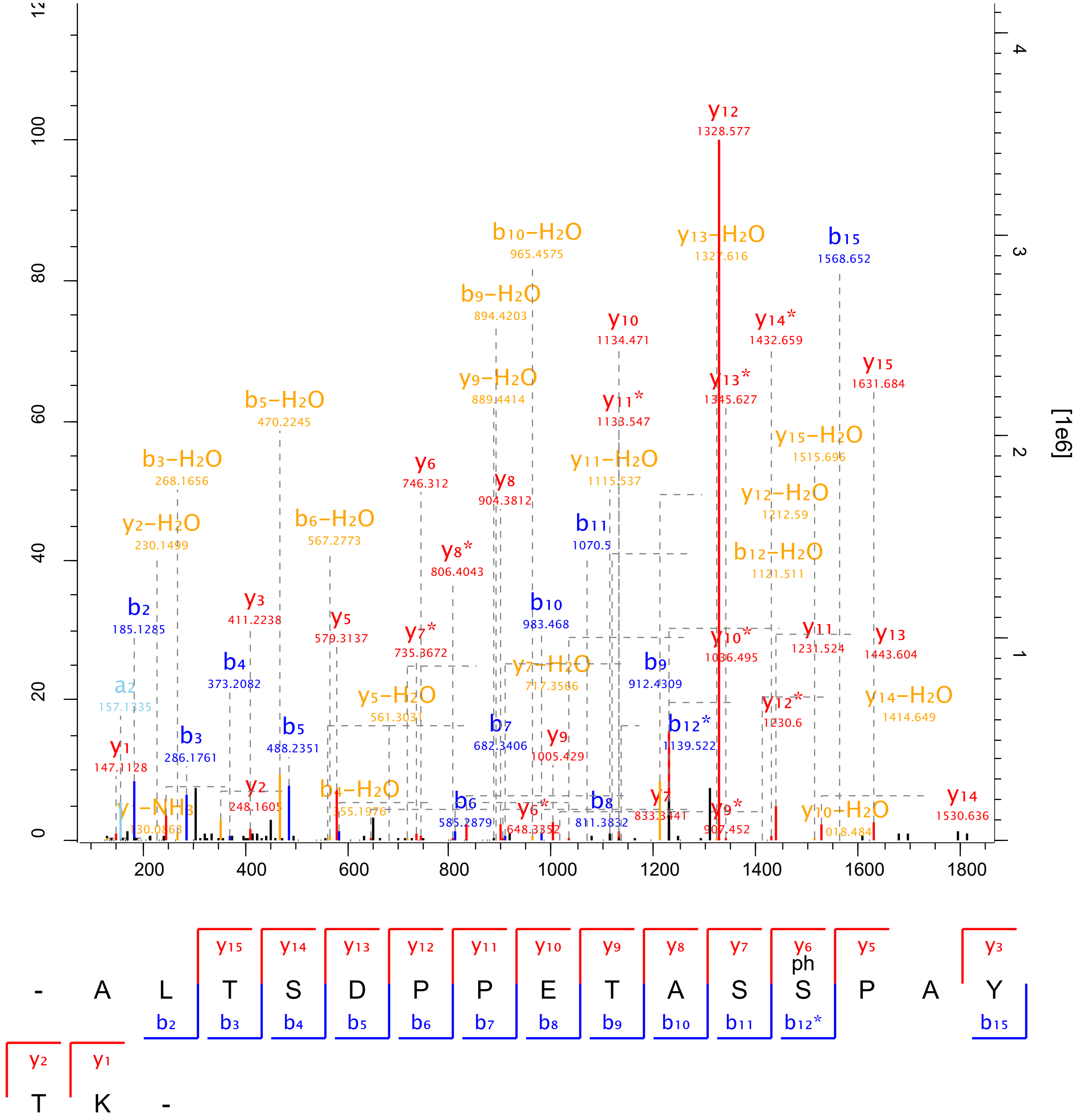

| Raw file | Scan | Method | Score | m/z |
|---|---|---|---|---|
| sys_05_2 | 16047 | FTMS; HCD | 103.9 | 630.25 |

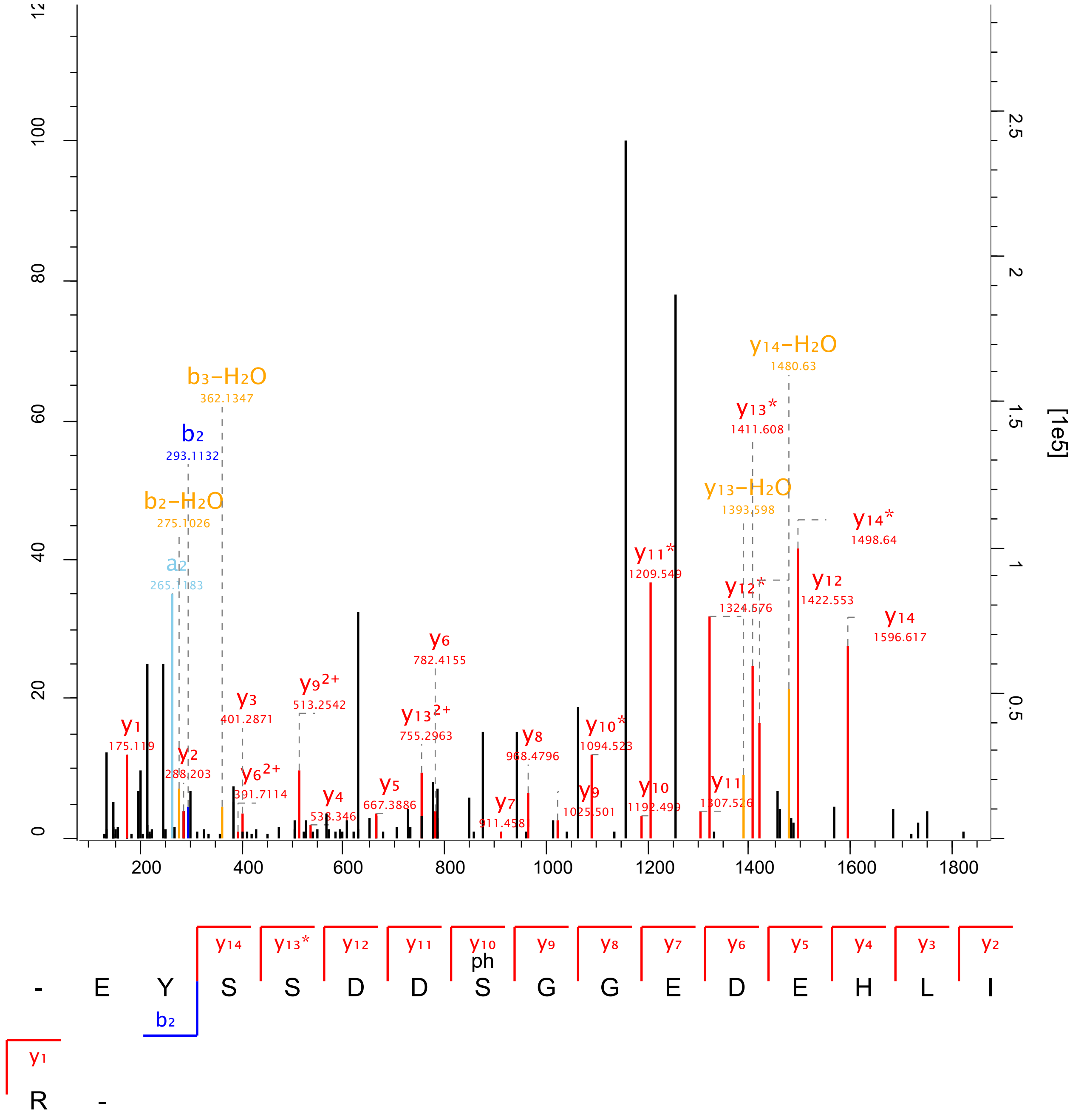

| Raw file | Scan | Method | Score | m/z |
|---|---|---|---|---|
| sys_05_2 | 16112 | FTMS; HCD | 42.32 | 653.62 |

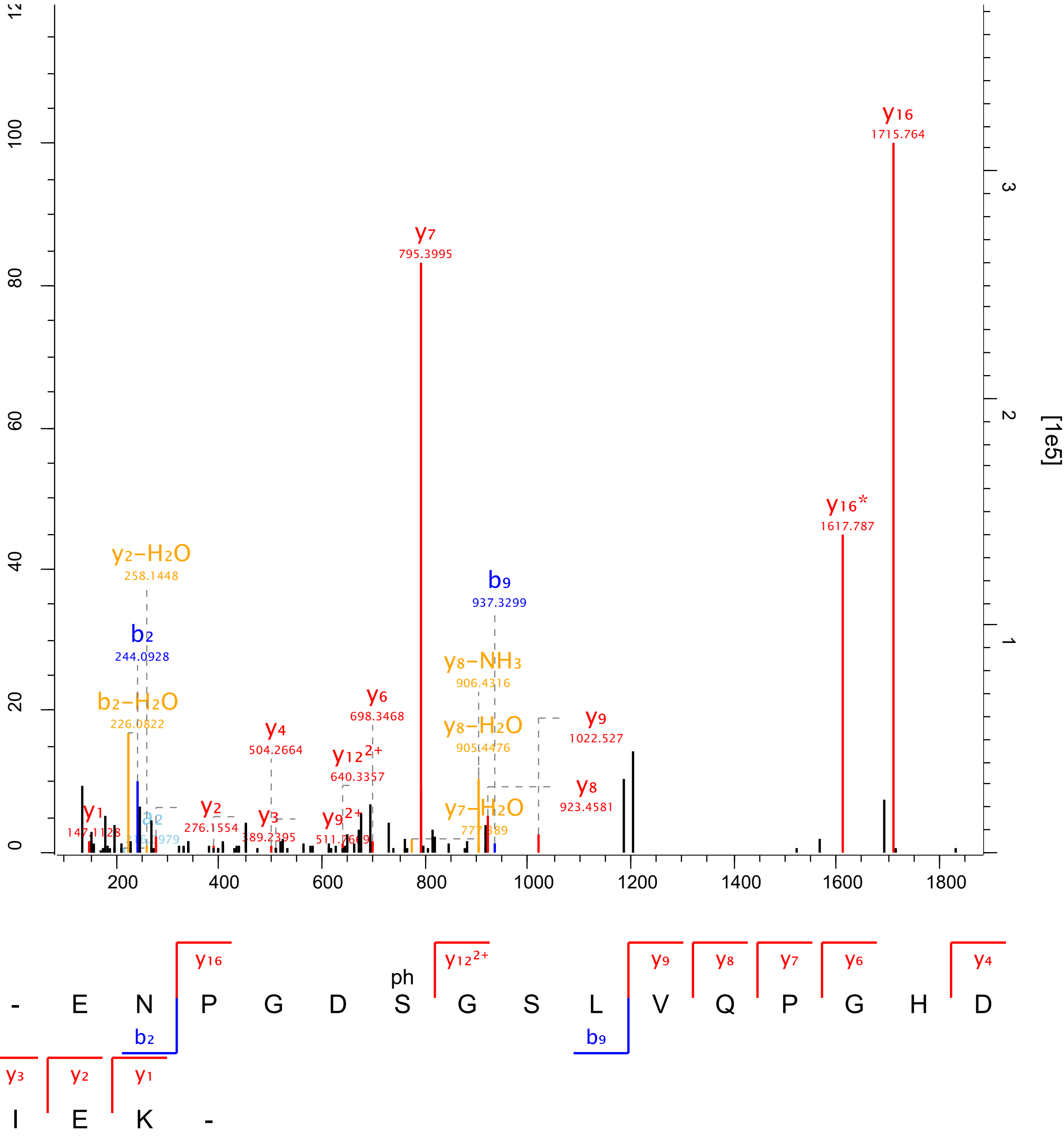

| Raw file | Scan | Method | Score | m/z |
| --- | --- | --- | --- | --- |
| sys_05_2 | 16131 | FTMS; HCD | 95.98 | 572.74 |

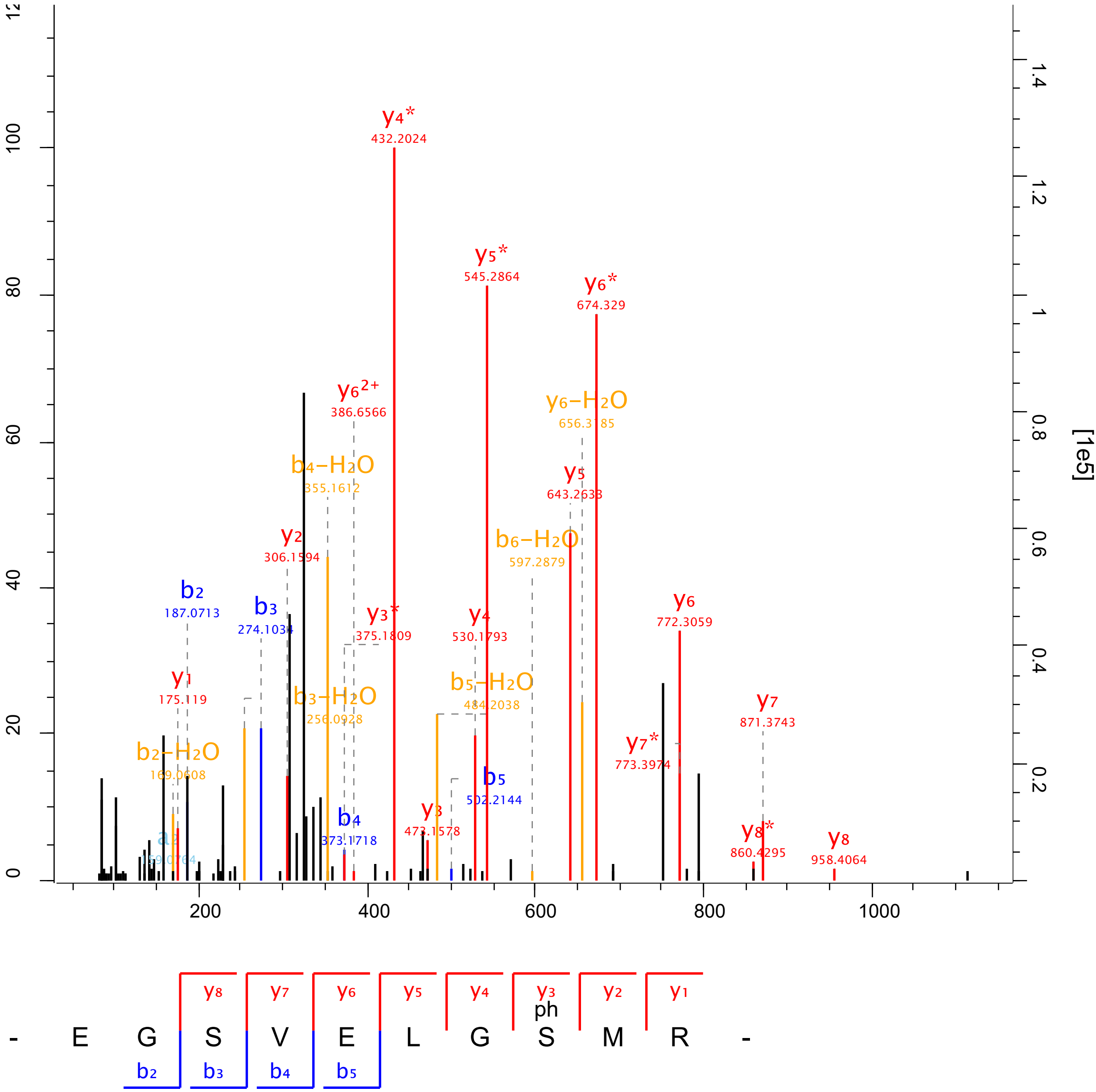

| Raw file | Scan | Method | Score | m/z |
| --- | --- | --- | --- | --- |
| sys_05_2 | 16235 | FTMS; HCD | 114.63 | 705.78 |

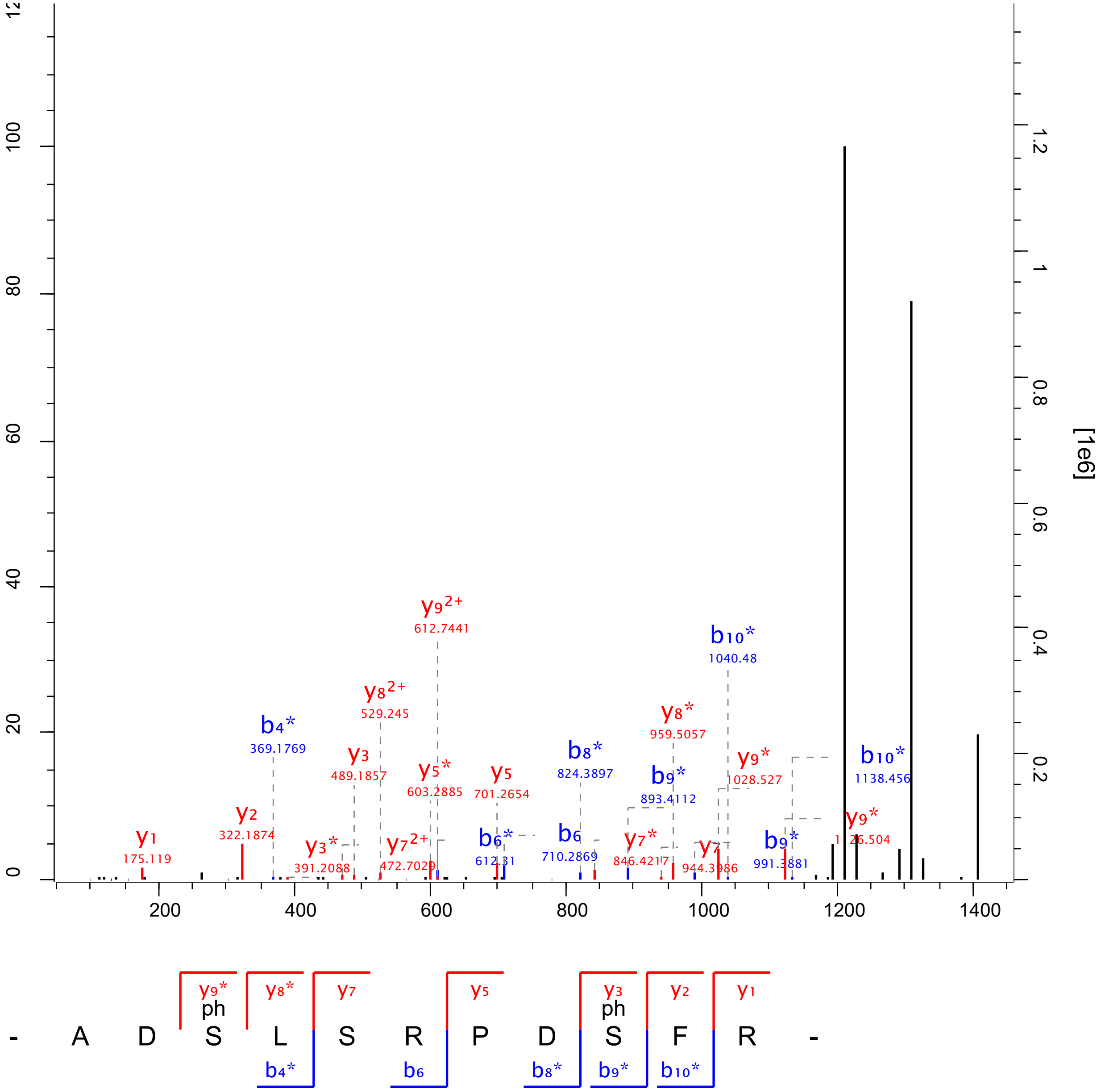

| Raw file | Scan | Method | Score | m/z |
|---|---|---|---|---|
| sys_05_2 | 16362 | FTMS; HCD | 100.04 | 741.31 |

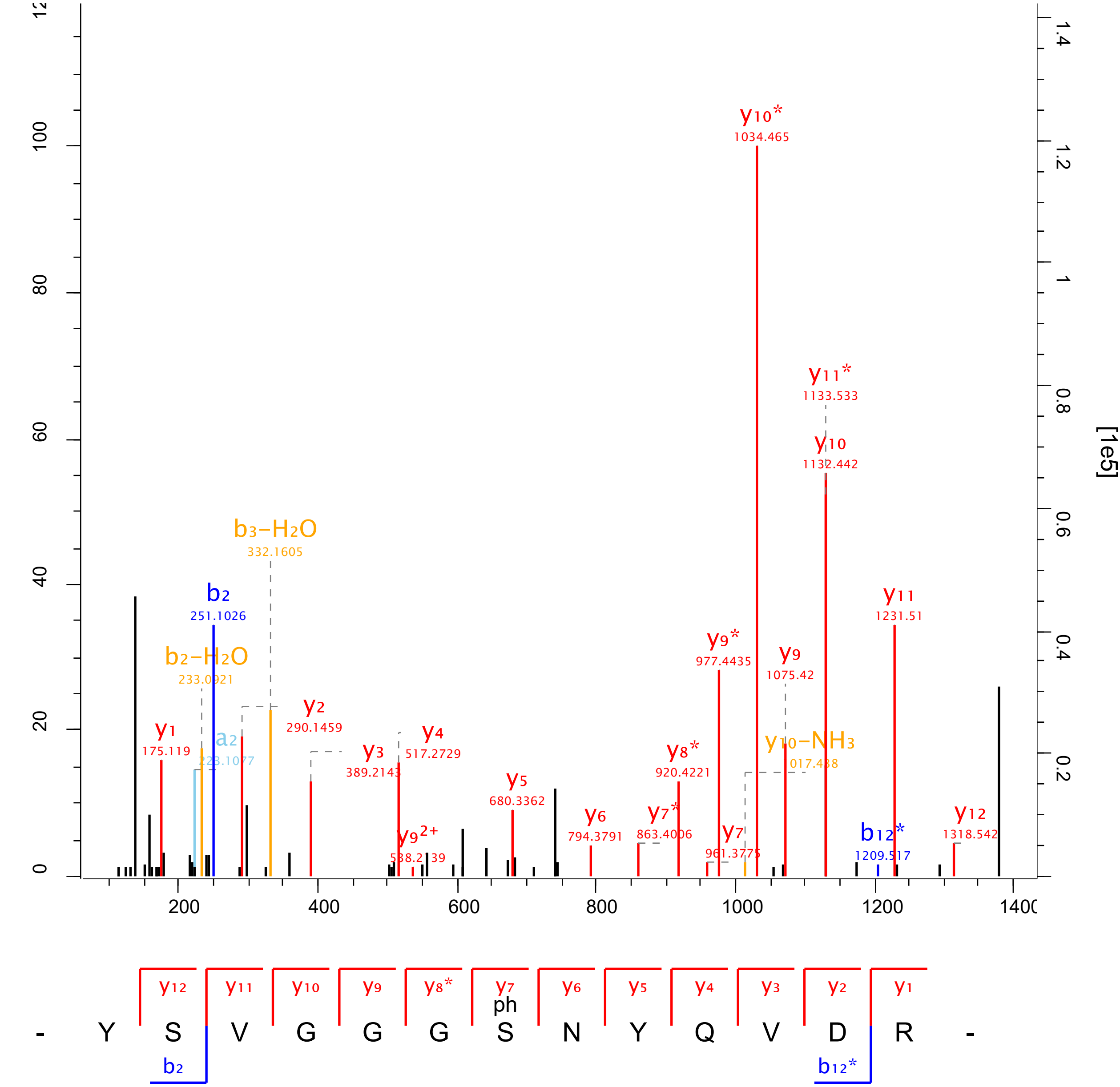

| Raw file | Scan | Method | Score | m/z |
|---|---|---|---|---|
| sys_05_2 | 16368 | FTMS; HCD | 259.77 | 782.85 |

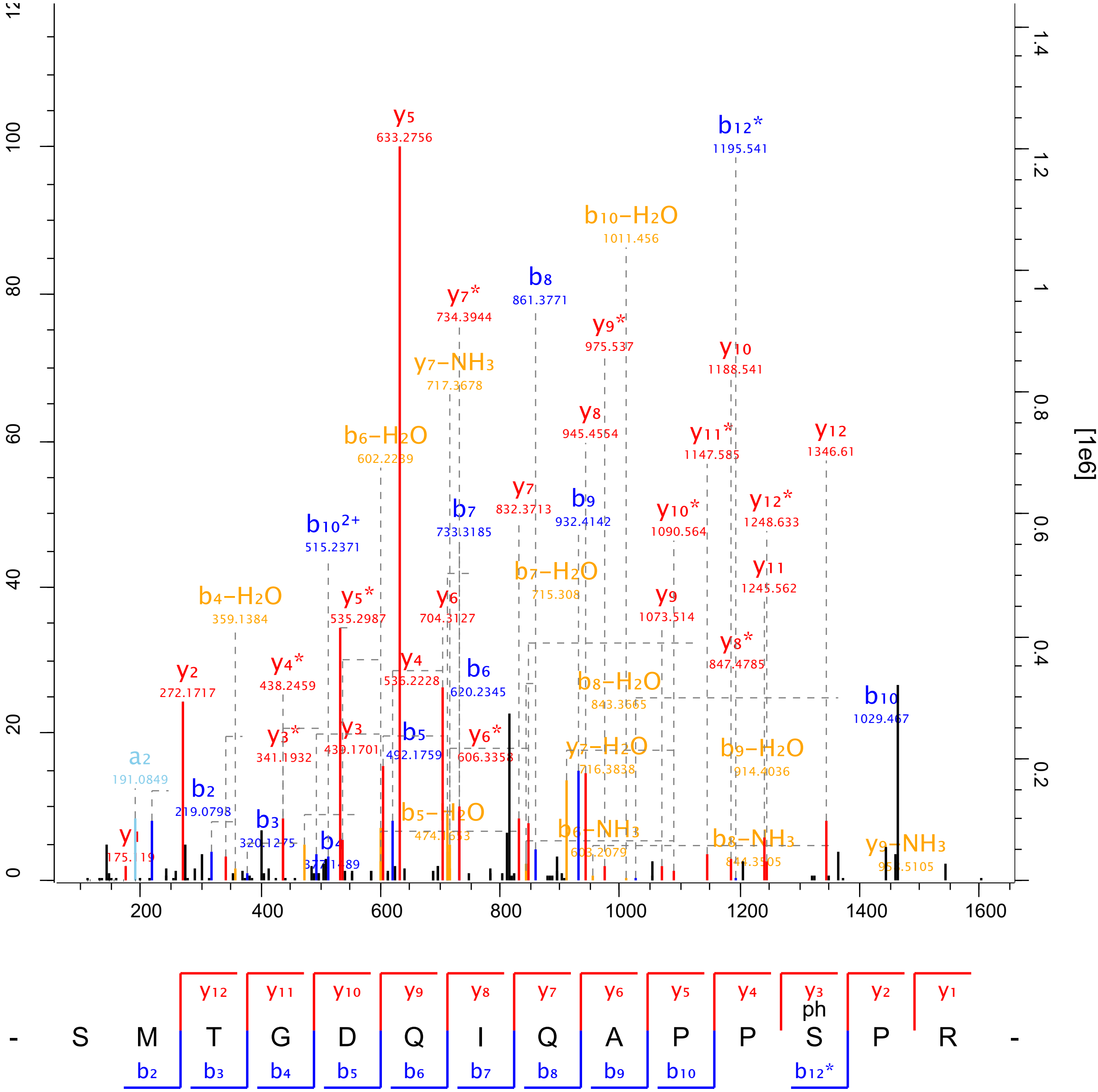

| Raw file | Scan | Method | Score | m/z |
|---|---|---|---|---|
| sys_05_2 | 16387 | FTMS; HCD | 77.18 | 607.73 |

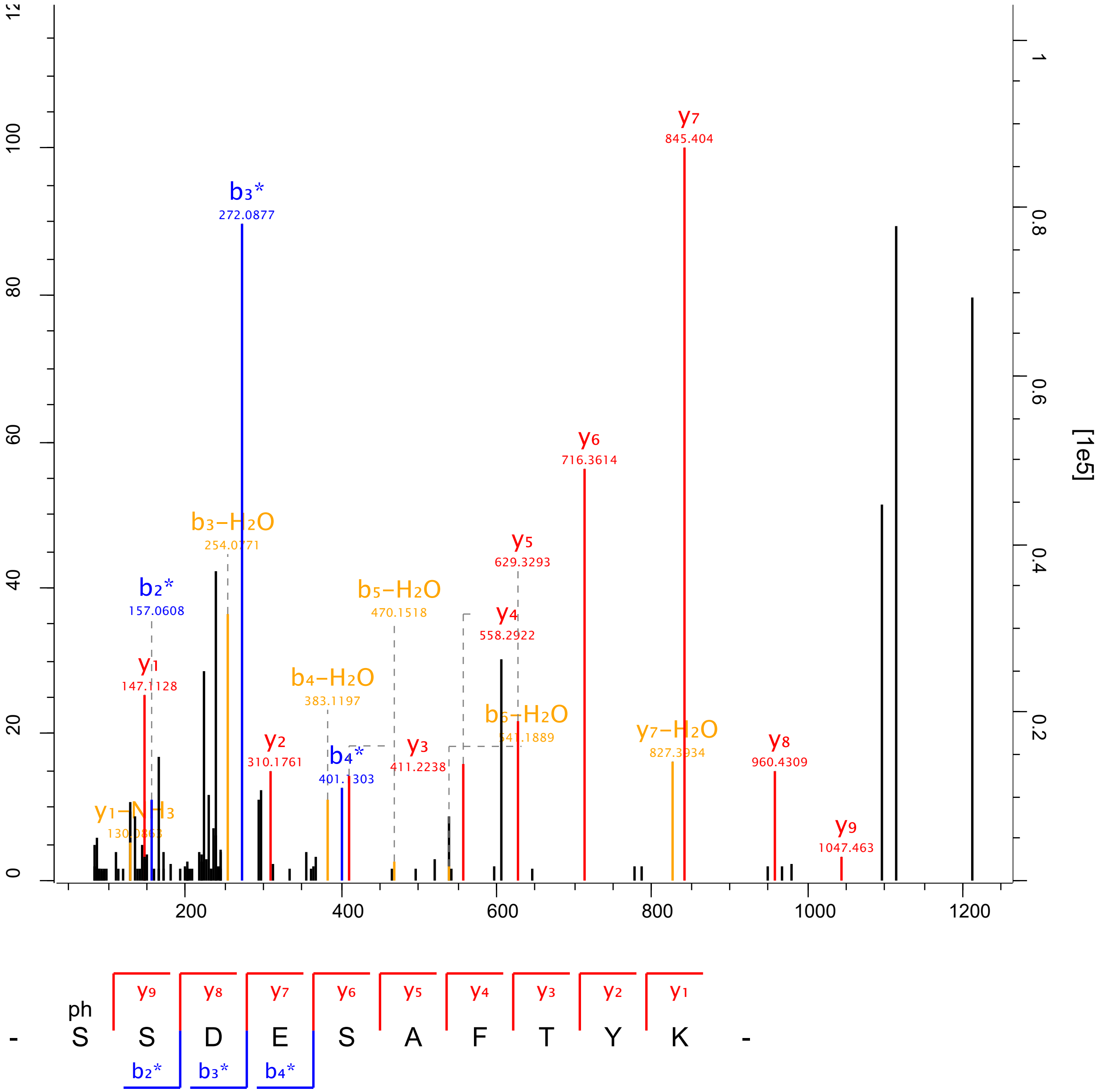

| Raw file | Scan | Method | Score | m/z |
|---|---|---|---|---|
| sys_05_2 | 16600 | FTMS; HCD | 124.42 | 633.81 |

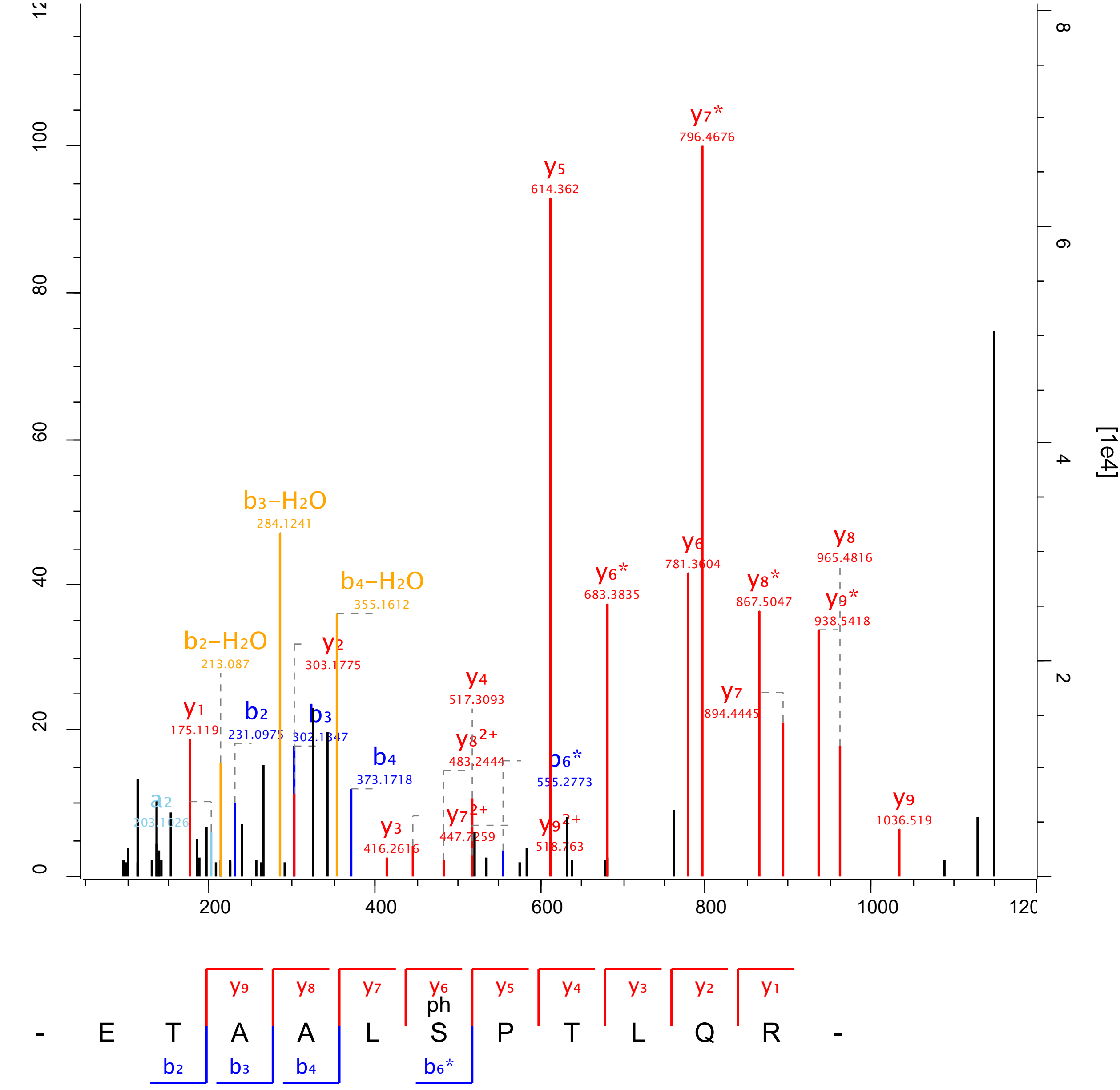

| Raw file | Scan | Method | Score | m/z |
|---|---|---|---|---|
| sys_05_2 | 16605 | FTMS; HCD | 214.58 | 643.79 |

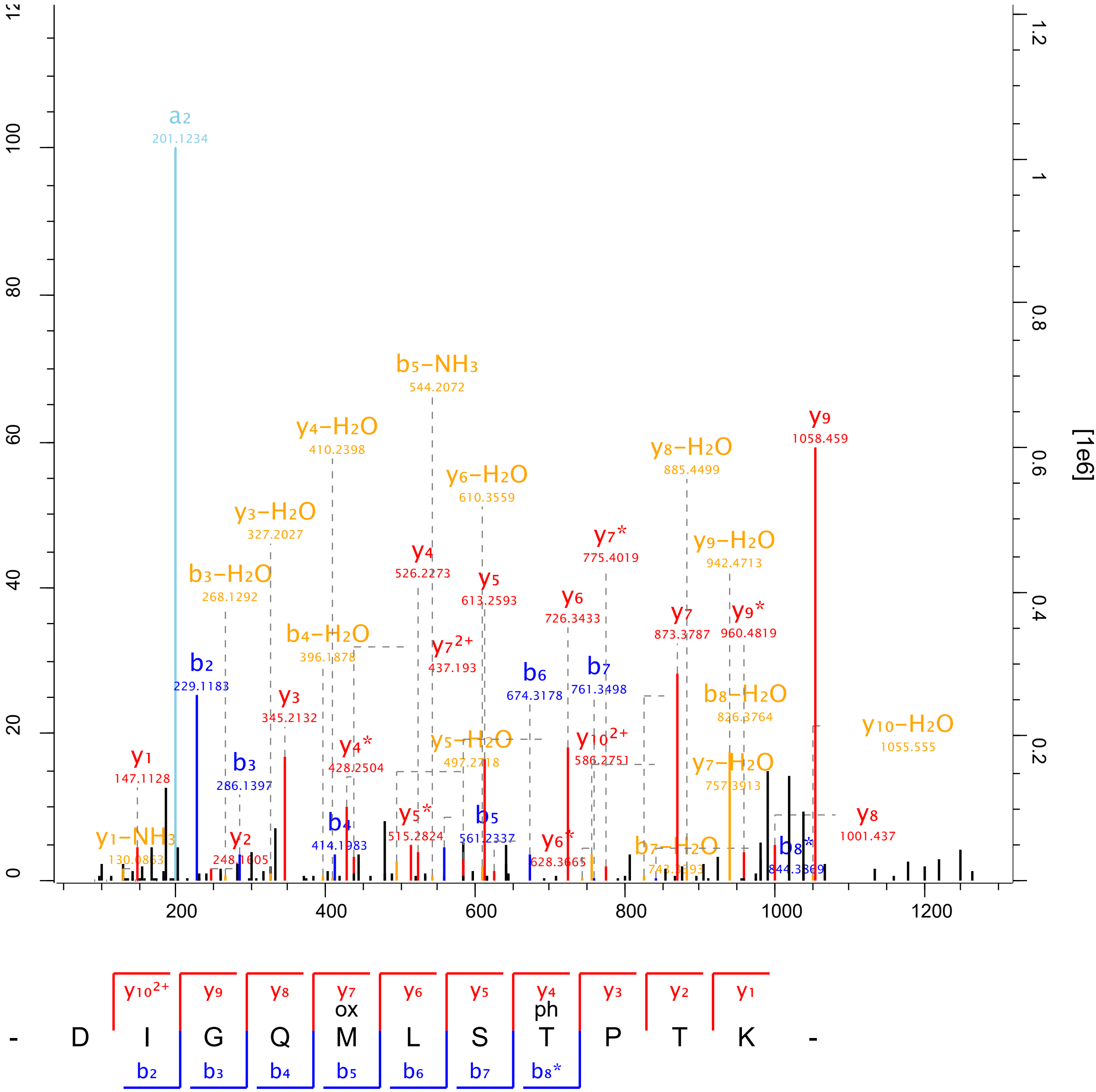

| Raw file | Scan | Method | Score | m/z |
|---|---|---|---|---|
| sys_05_2 | 16696 | FTMS; HCD | 162.29 | 692.81 |

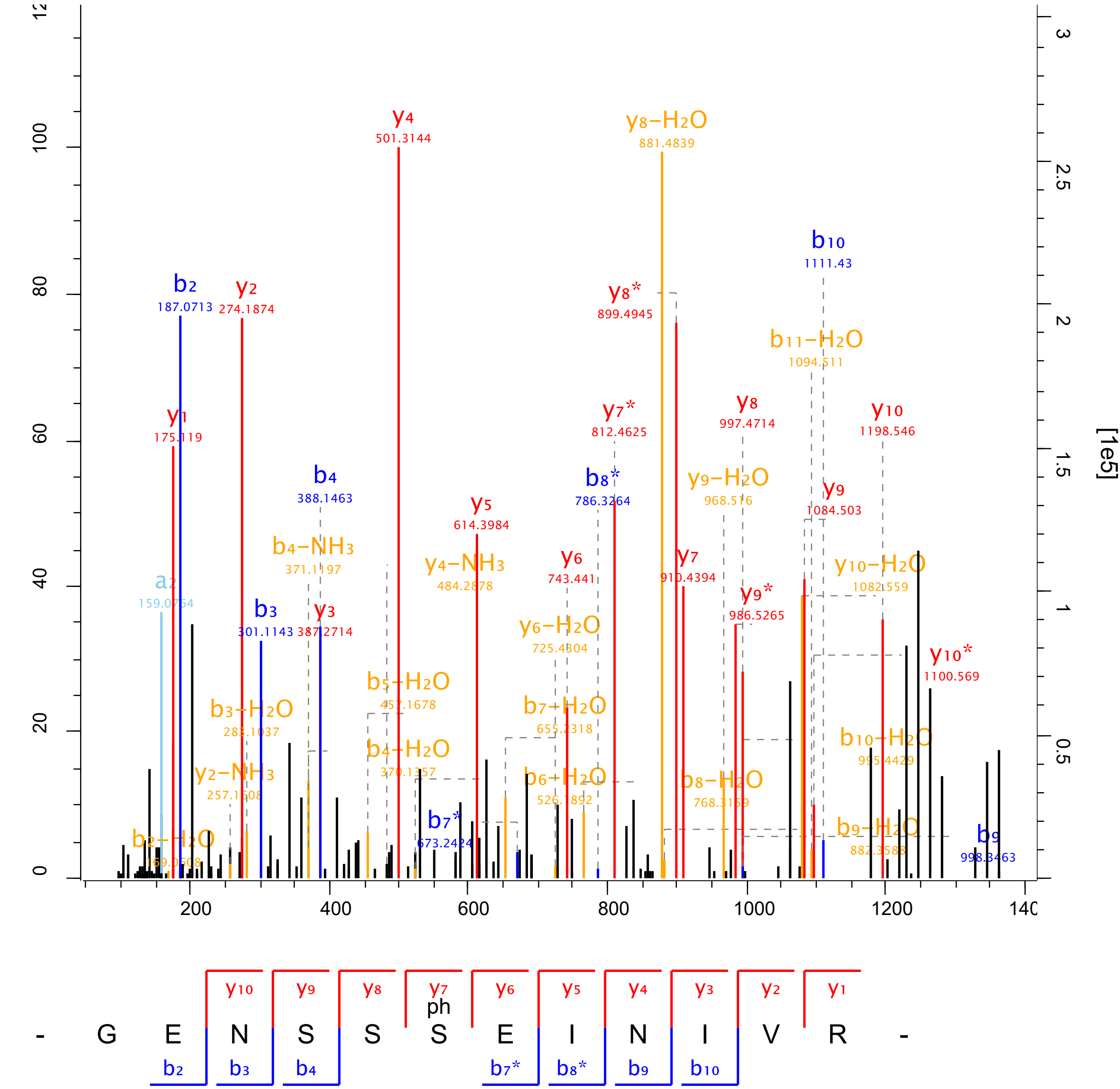

| Raw file | Scan | Method | Score | m/z |
| --- | --- | --- | --- | --- |
| sys_05_2 | 16794 | FTMS; HCD | 68.97 | 488.91 |

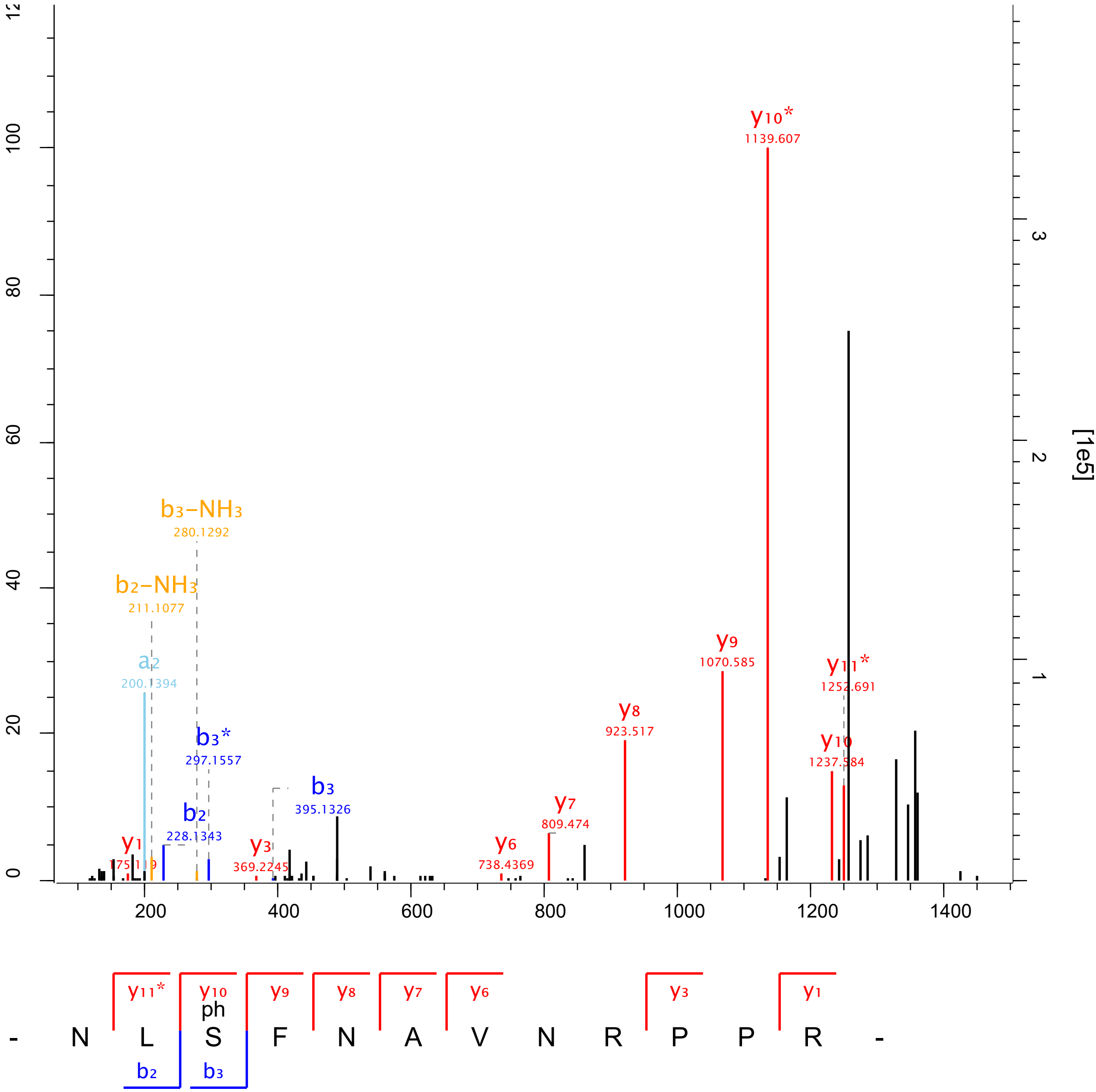

| Raw file | Scan | Method | Score | m/z |
|---|---|---|---|---|
| sys_05_2 | 16806 | FTMS; HCD | 82.65 | 578.22 |

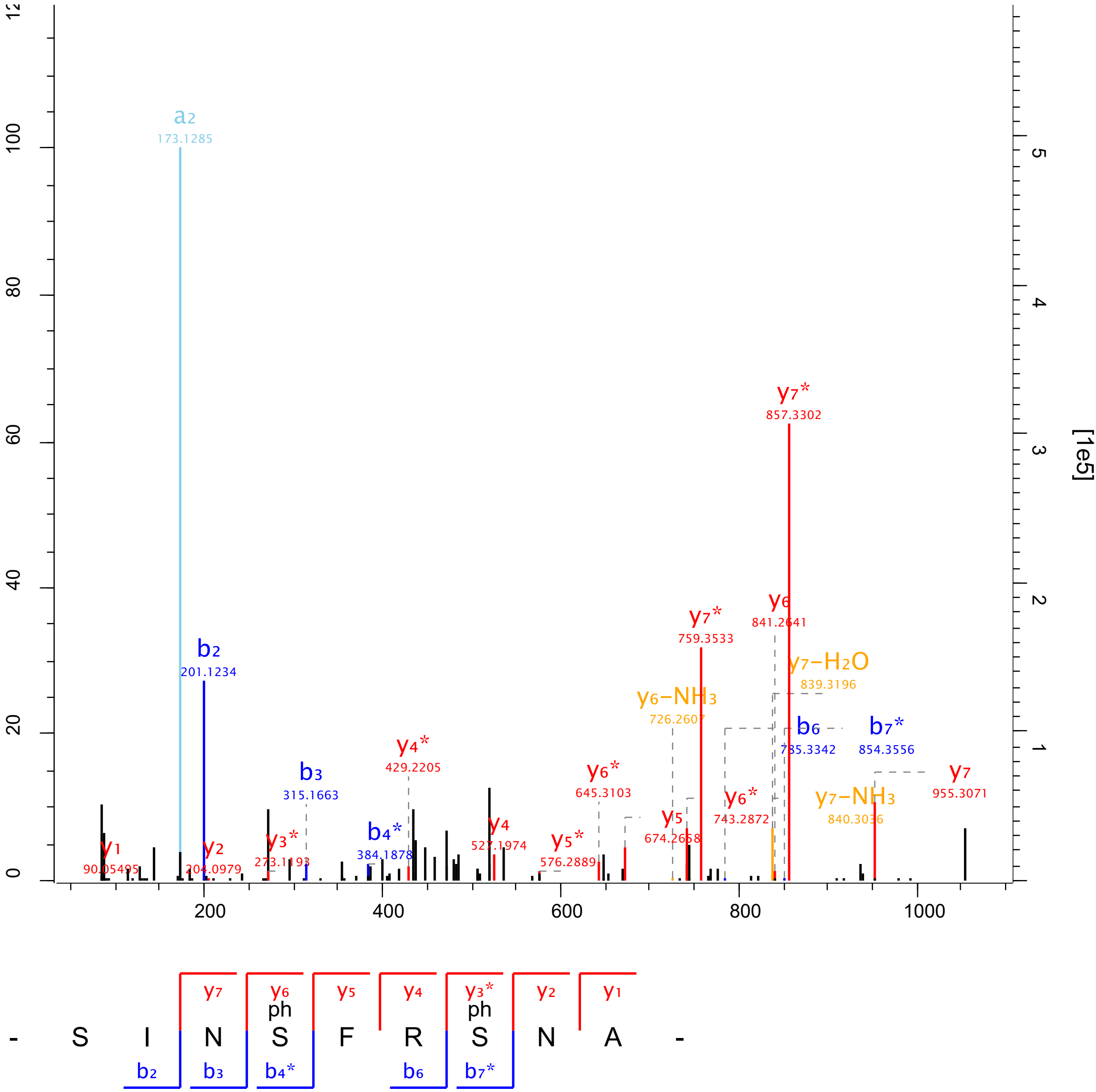

| Raw file | Scan | Method | Score | m/z |
| --- | --- | --- | --- | --- |
| sys_05_2 | 16886 | FTMS; HCD | 194.67 | 889.32 |

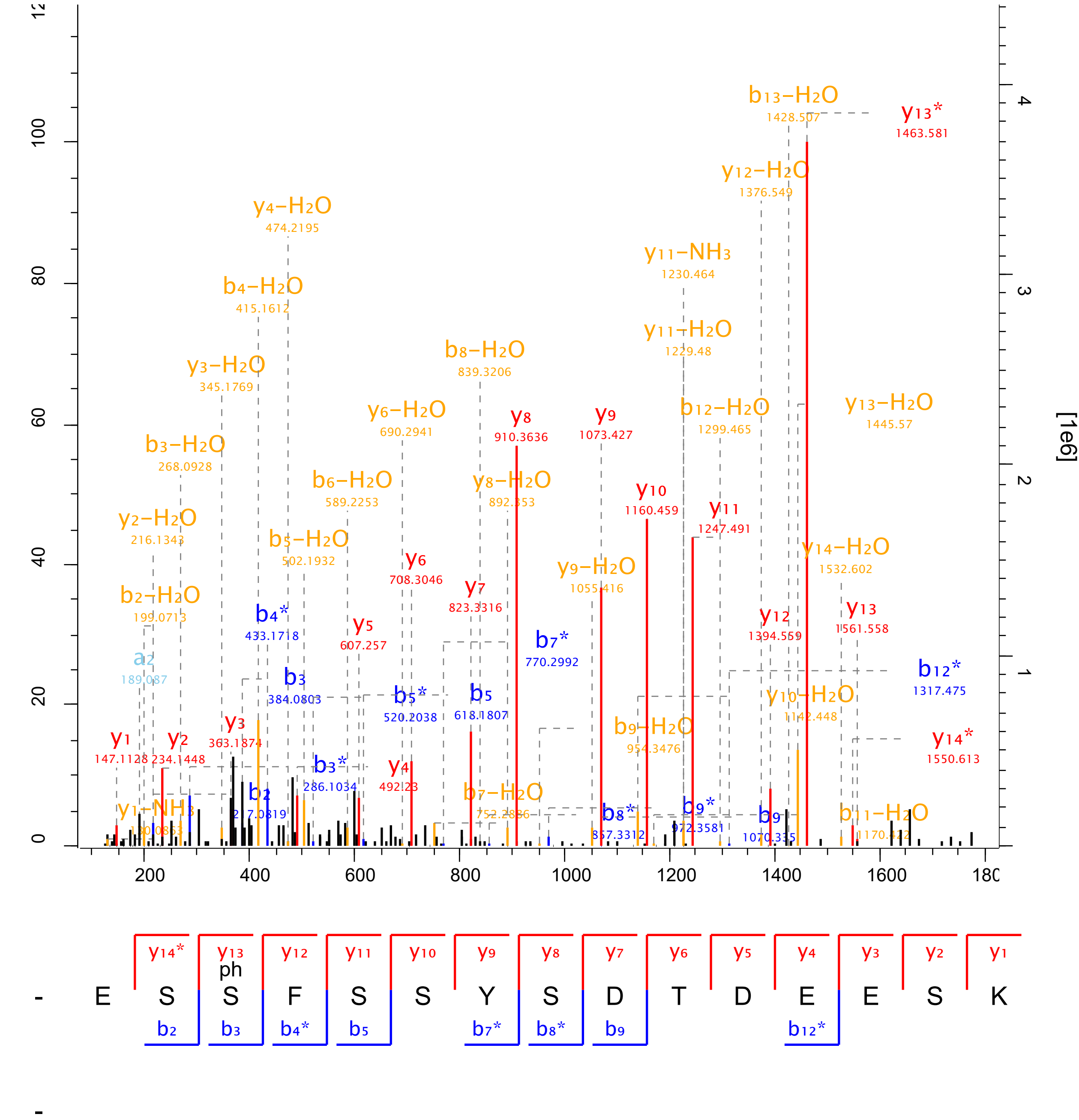

| Raw file | Scan | Method | Score | m/z |
|---|---|---|---|---|
| sys_05_2 | 16934 | FTMS; HCD | 50.9 | 530.58 |

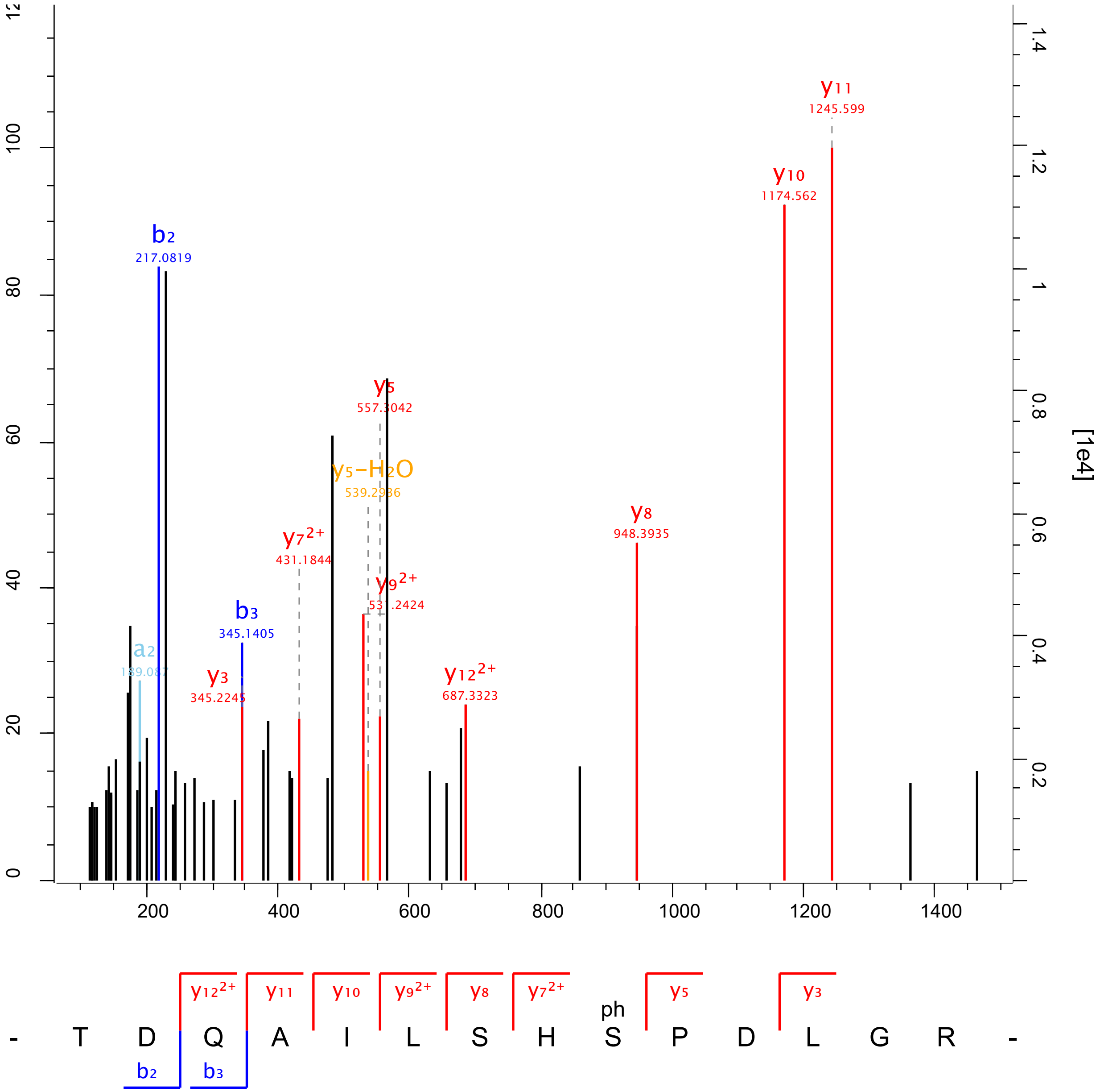

| Raw file | Scan | Method | Score | m/z |
|---|---|---|---|---|
| sys_05_2 | 16945 | FTMS; HCD | 171.26 | 729.82 |

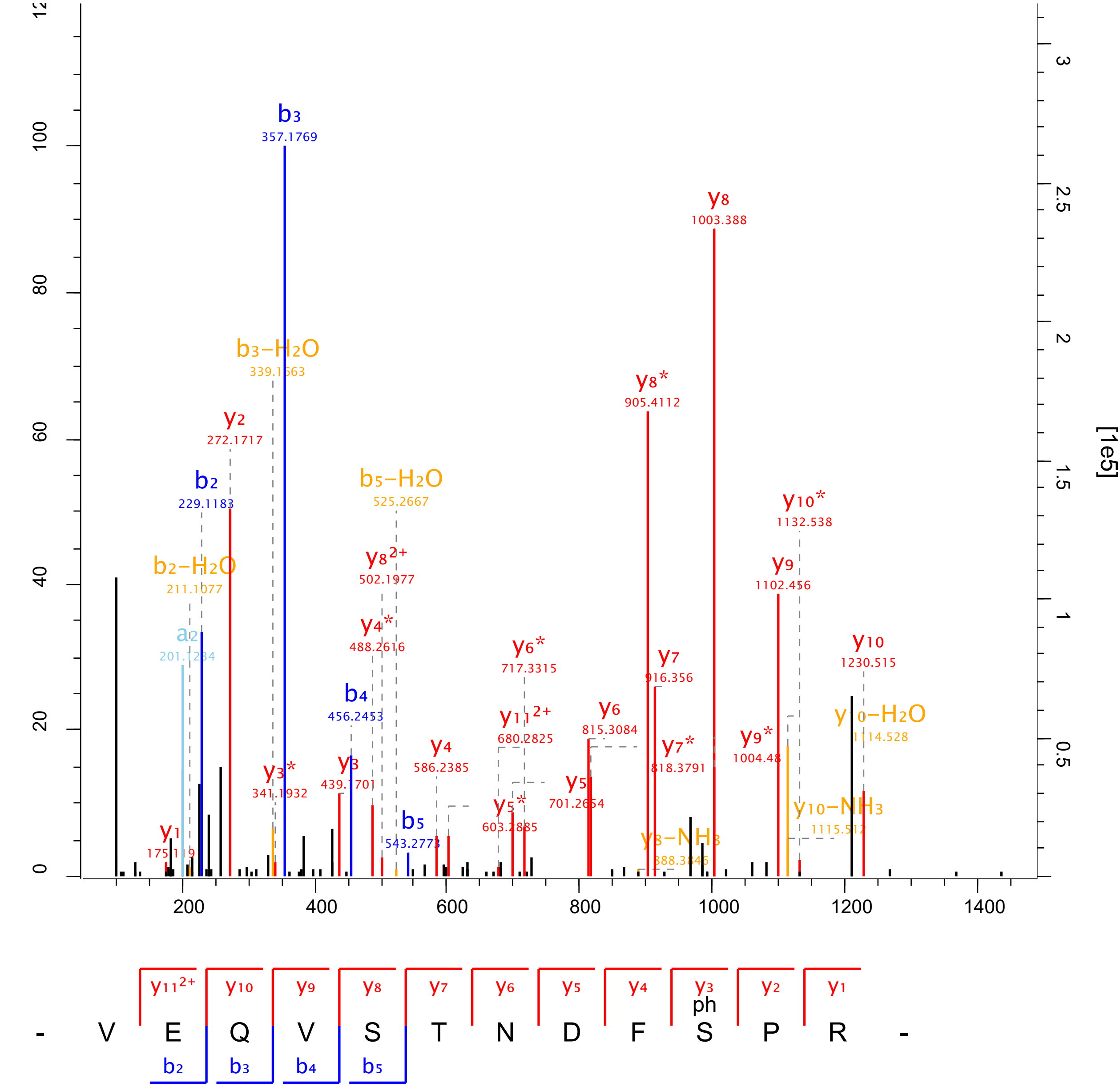

| Raw file | Scan | Method | Score | m/z |
|---|---|---|---|---|
| sys_05_2 | 16965 | FTMS; HCD | 49.45 | 571.27 |

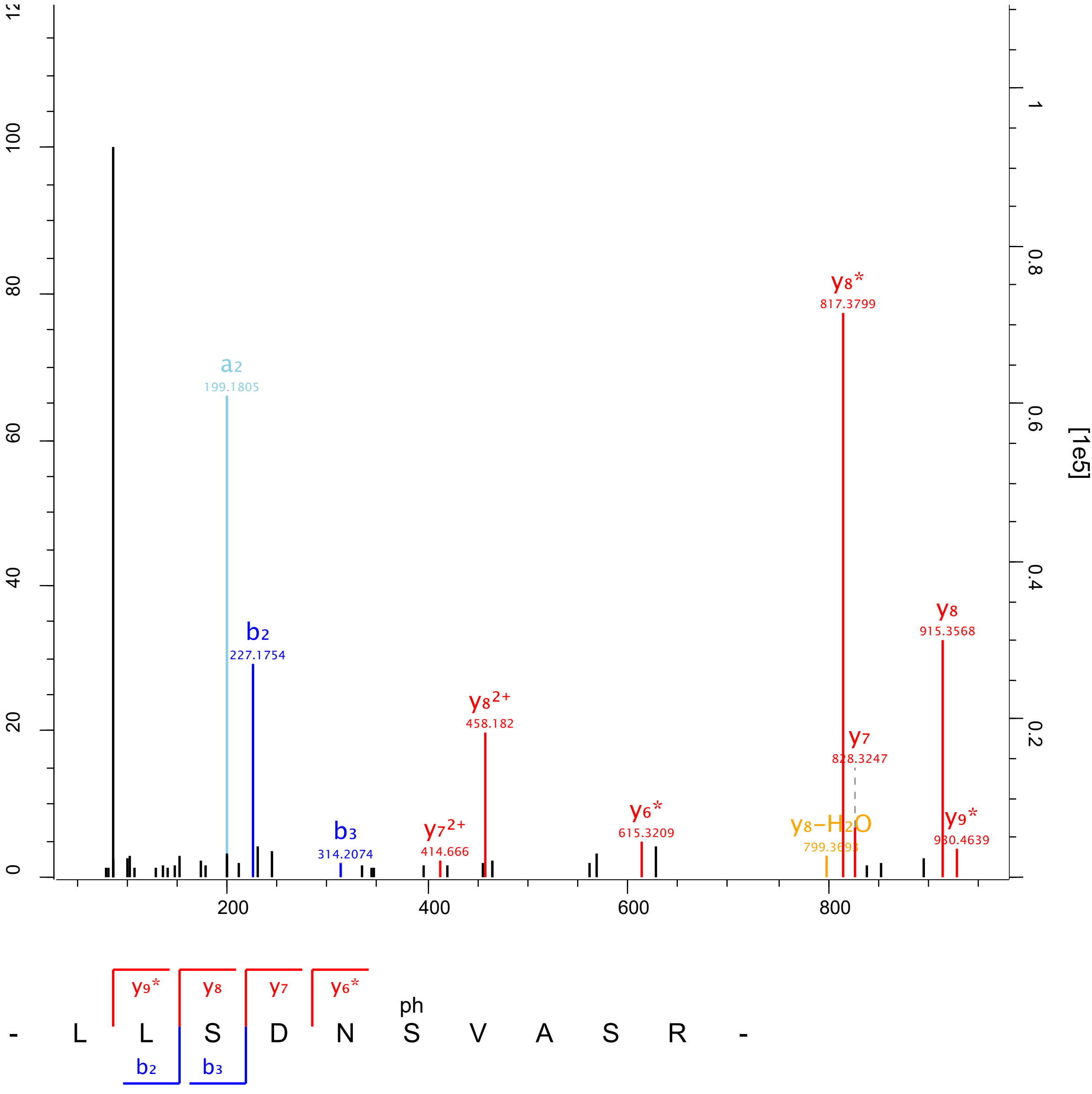

| Raw file | Scan | Method | Score | m/z |
|---|---|---|---|---|
| sys_05_2 | 16978 | FTMS; HCD | 197.38 | 762.33 |

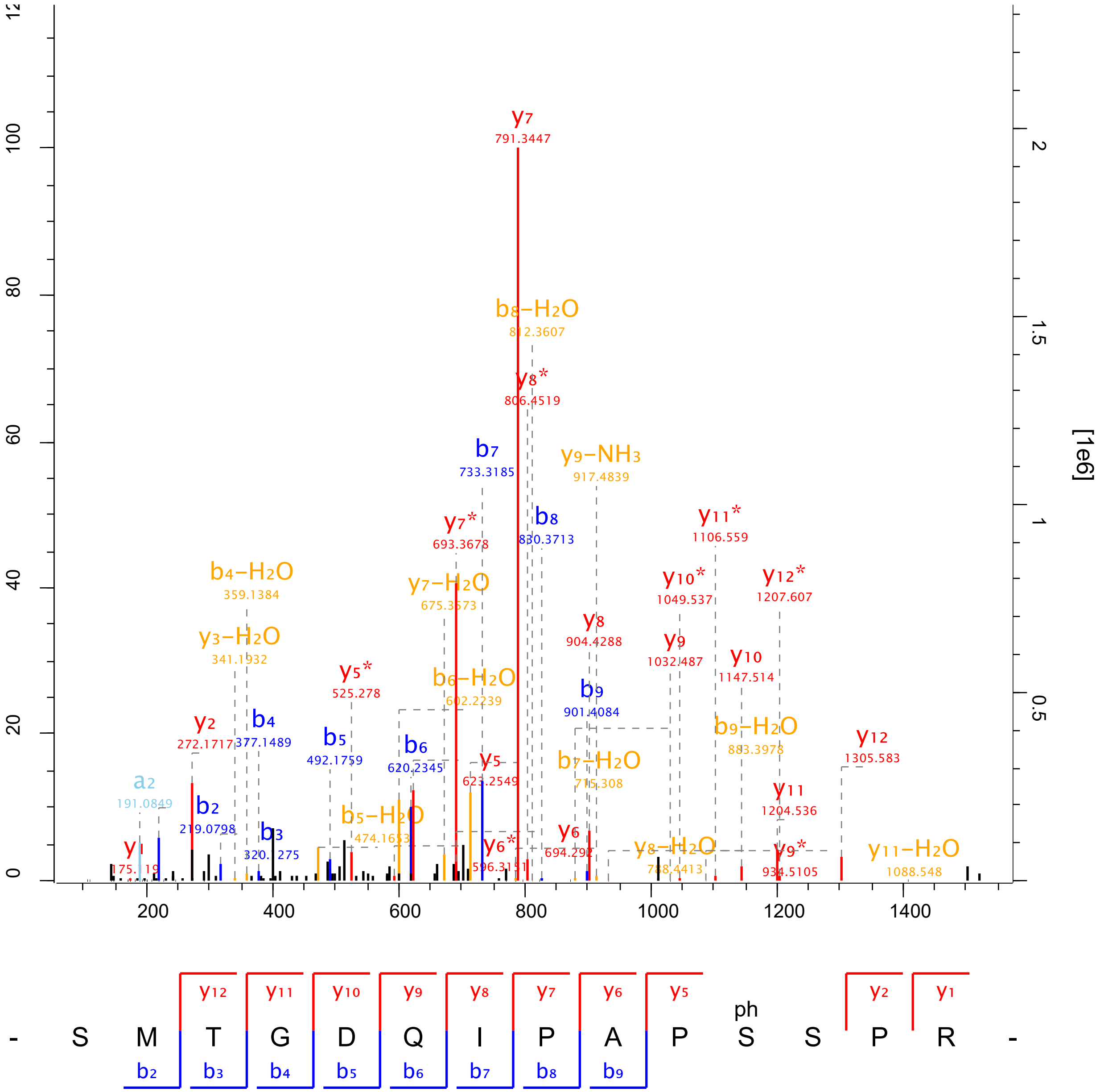

| Raw file | Scan | Method | Score | m/z |
|---|---|---|---|---|
| sys_05_2 | 17000 | FTMS; HCD | 144.73 | 644.77 |

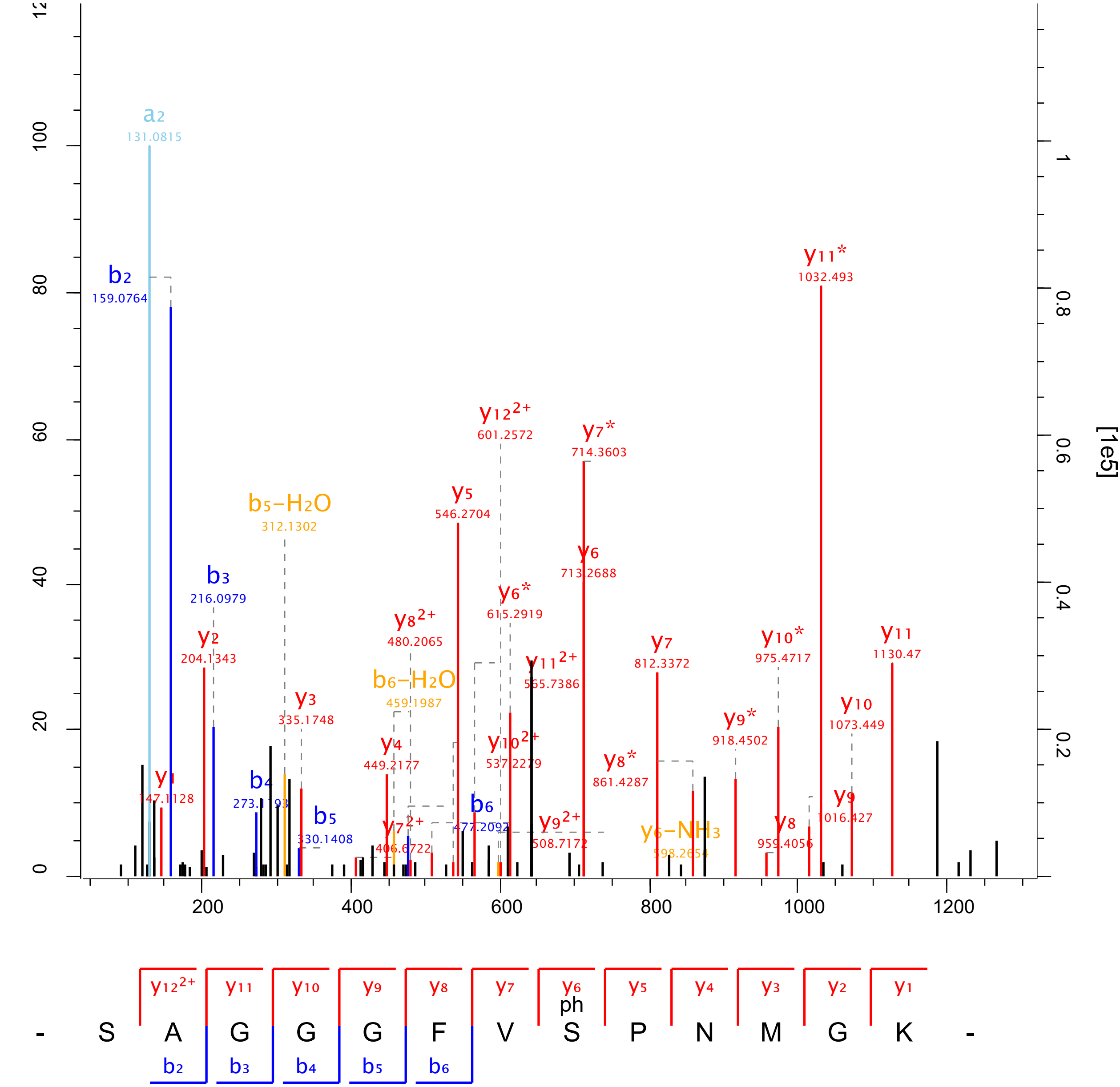

| Raw file | Scan | Method | Score | m/z |
|---|---|---|---|---|
| sys_05_2 | 17032 | FTMS; HCD | 186.74 | 851.35 |

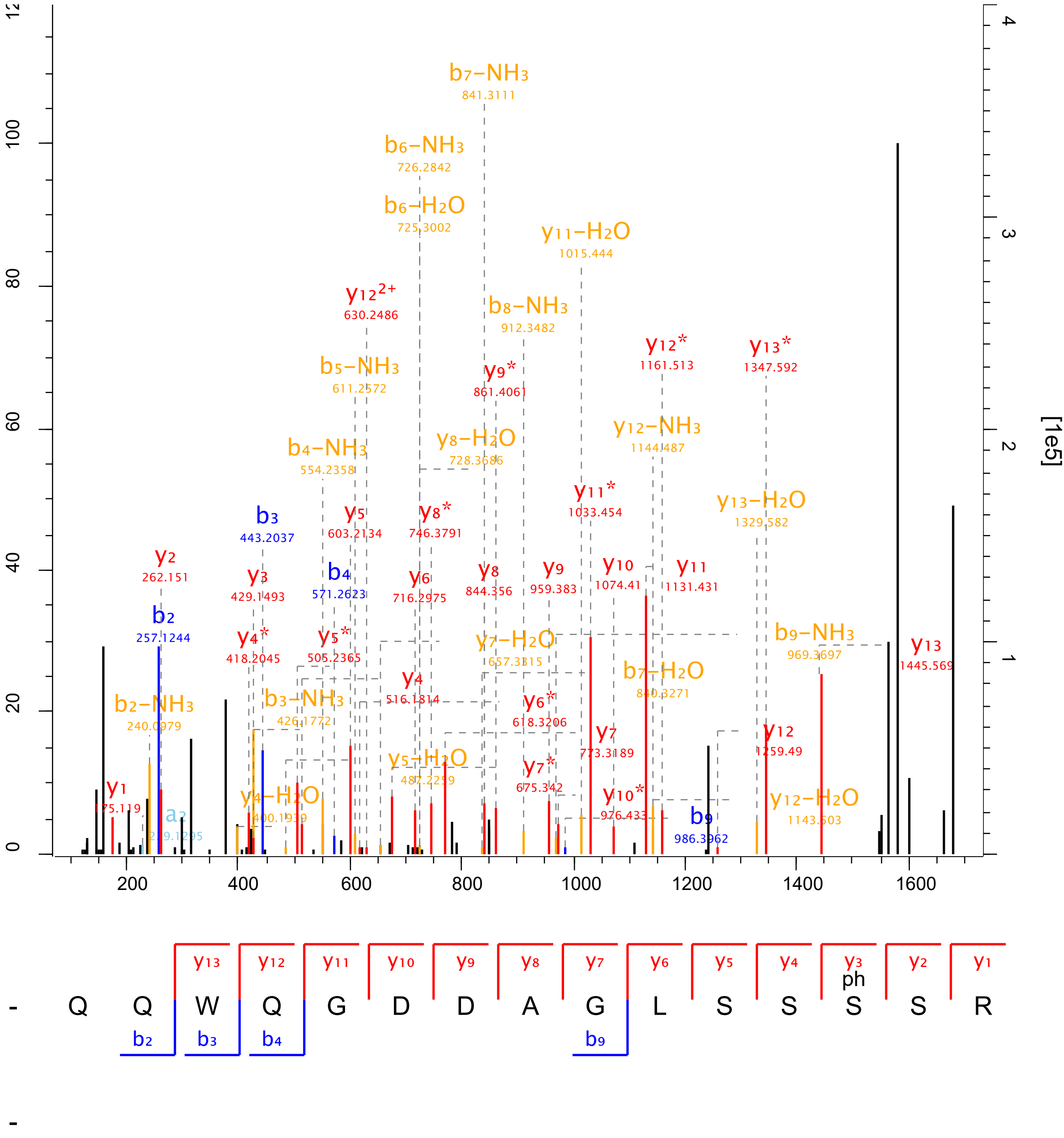

| Raw file | Scan | Method | Score | m/z |
|---|---|---|---|---|
| sys_05_2 | 17153 | FTMS; HCD | 64.82 | 495.71 |

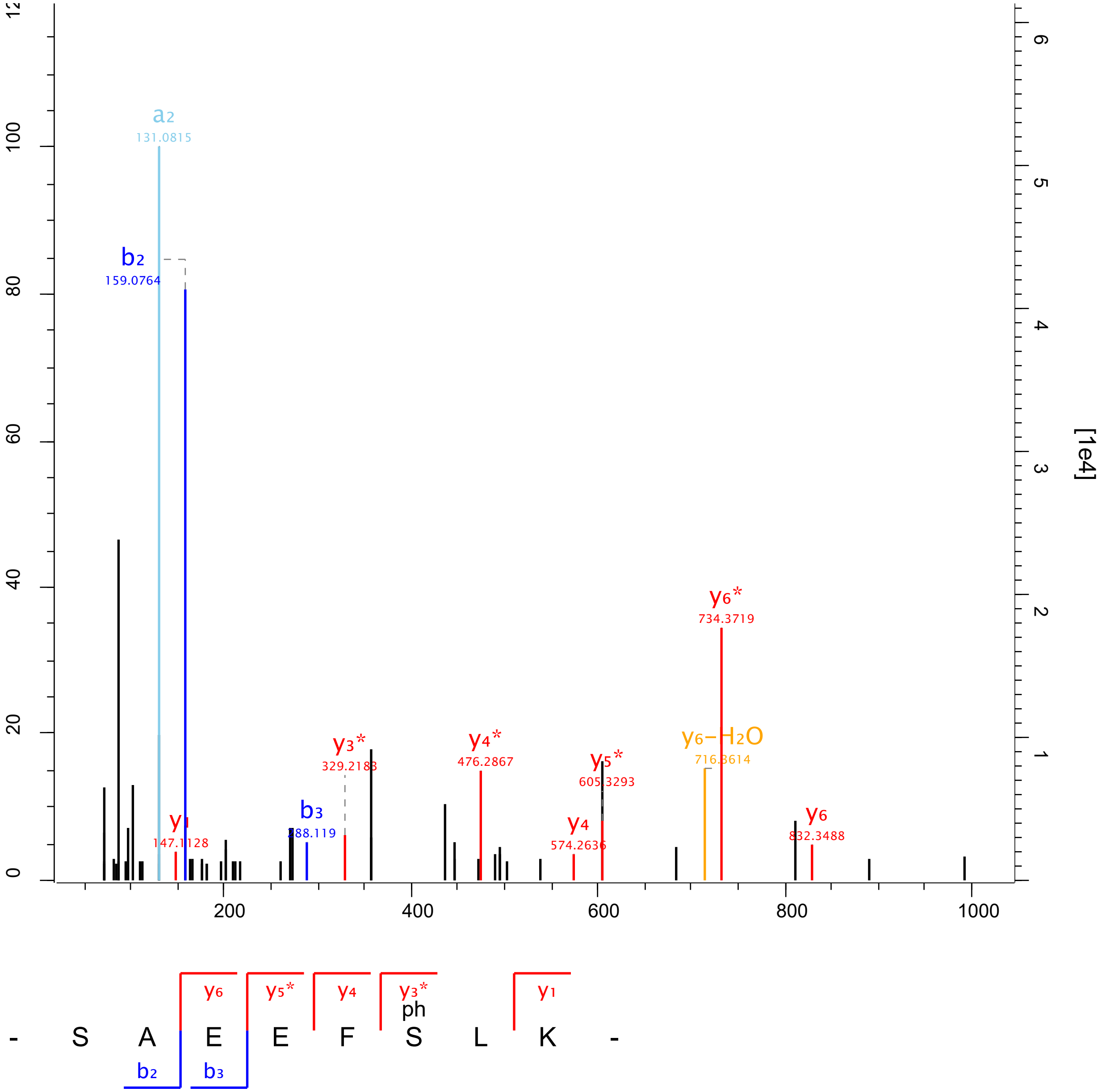

| Raw file | Scan | Method | Score | m/z |
|---|---|---|---|---|
| sys_05_2 | 17407 | FTMS; HCD | 115.88 | 856.4 |

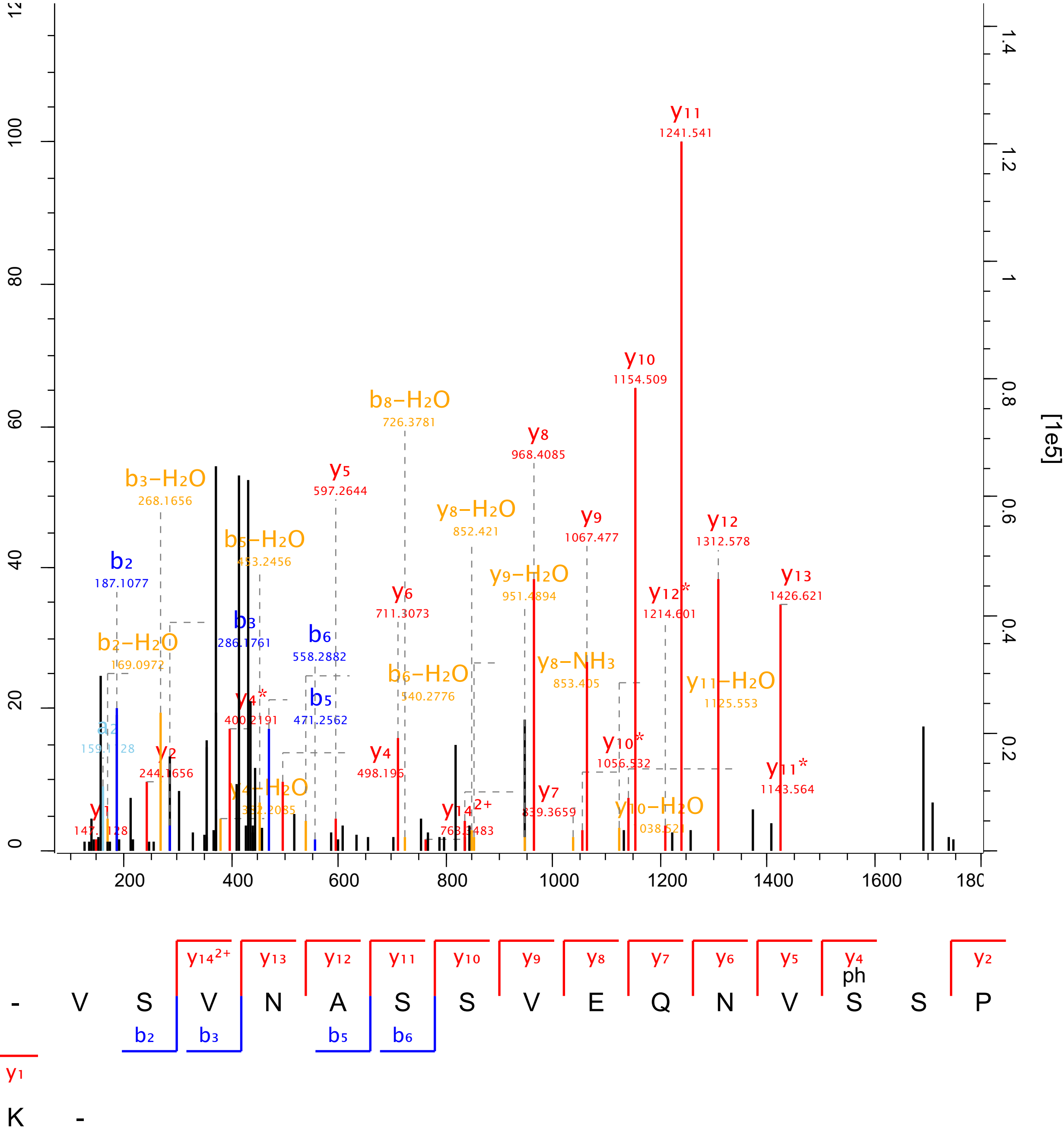

| Raw file | Scan | Method | Score | m/z |
|---|---|---|---|---|
| sys_05_2 | 17549 | FTMS; HCD | 163.79 | 782.84 |

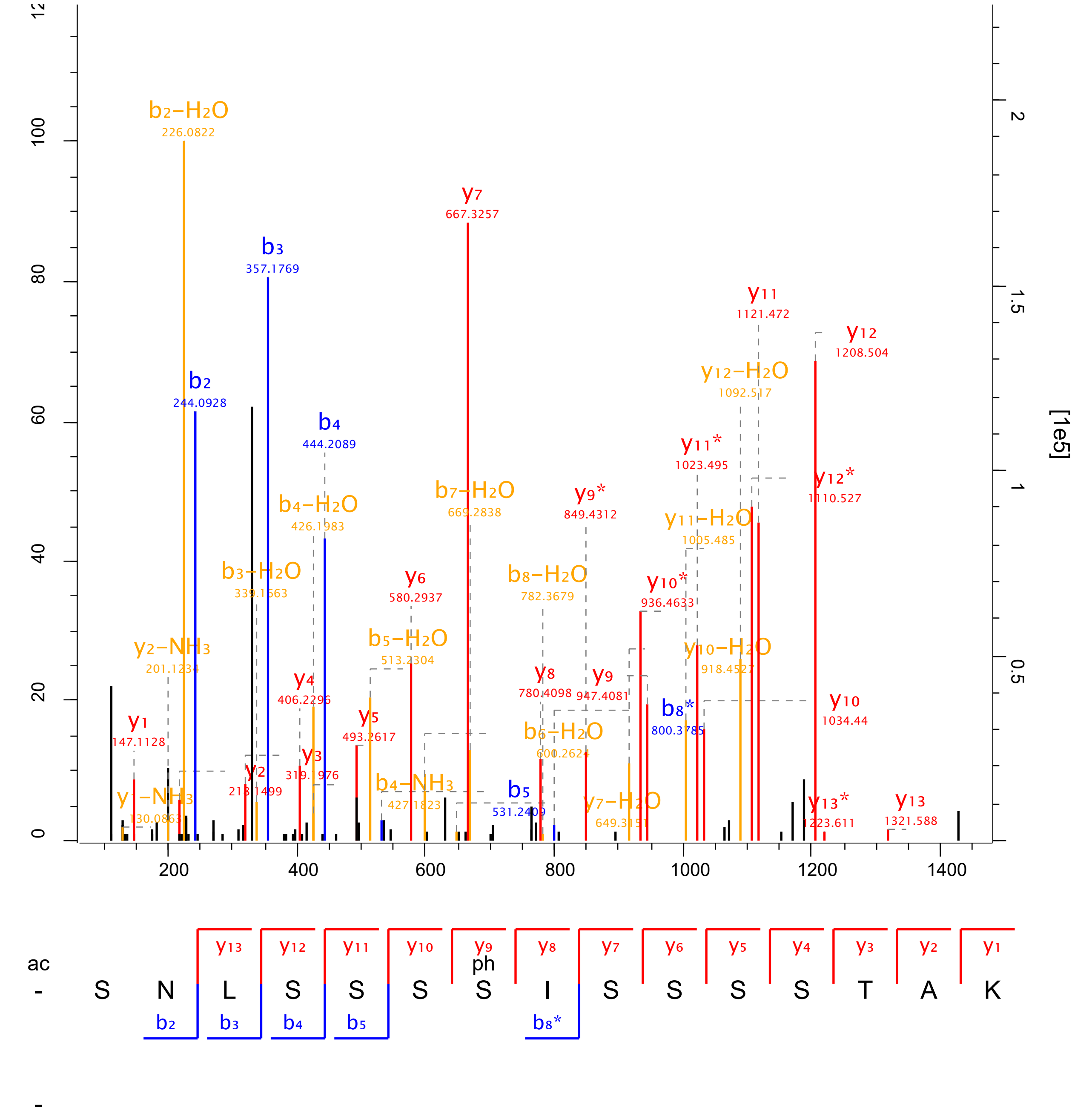

| Raw file | Scan | Method | Score | m/z |
| --- | --- | --- | --- | --- |
| sys_05_2 | 17614 | FTMS; HCD | 157.33 | 503.86 |

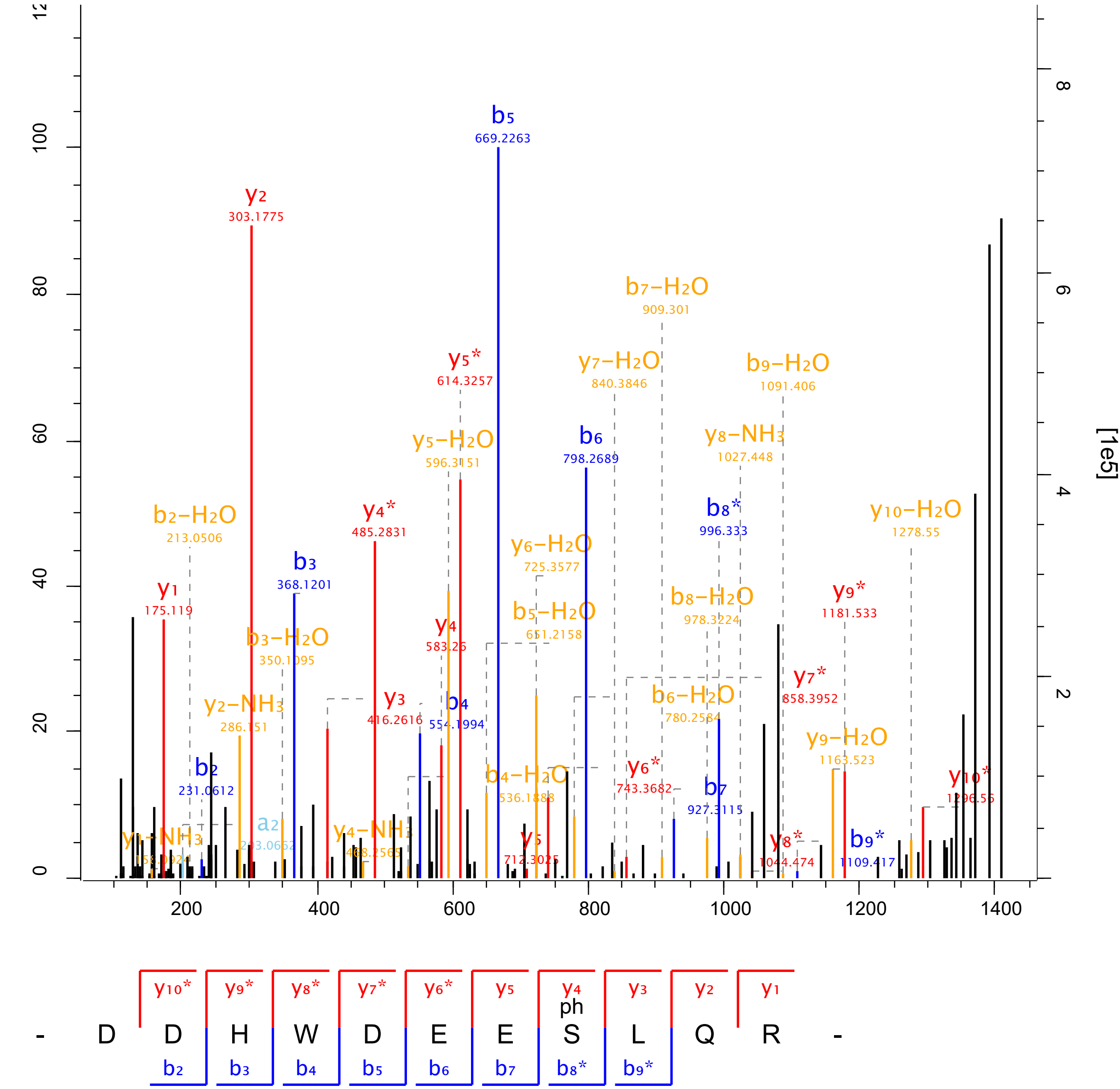

| Raw file | Scan | Method | Score | m/z |
|---|---|---|---|---|
| sys_05_2 | 17651 | FTMS; HCD | 171.79 | 794.35 |

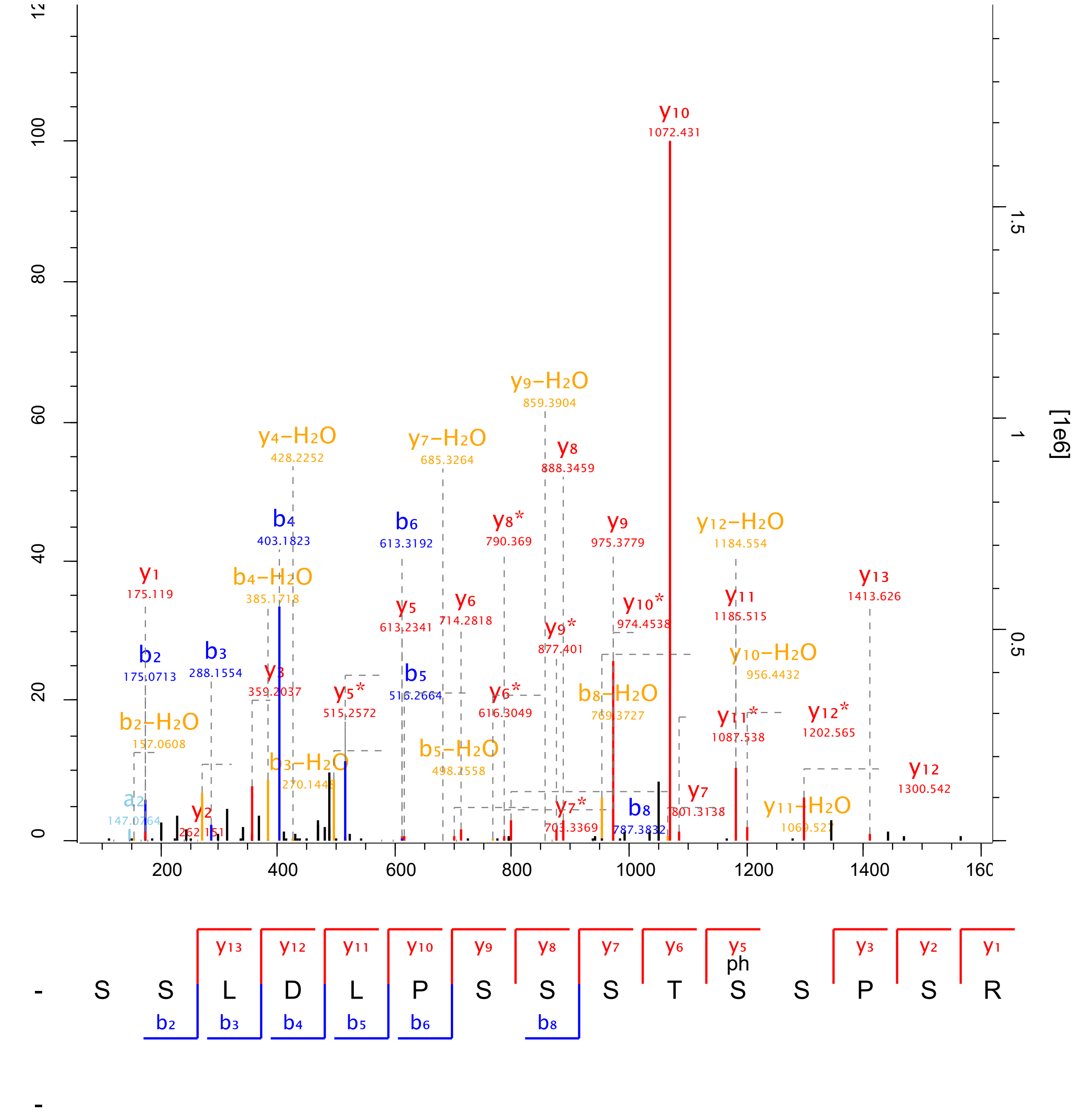

| Raw file | Scan | Method | Score | m/z |
|---|---|---|---|---|
| sys_05_2 | 17727 | FTMS; HCD | 58.08 | 744.8 |

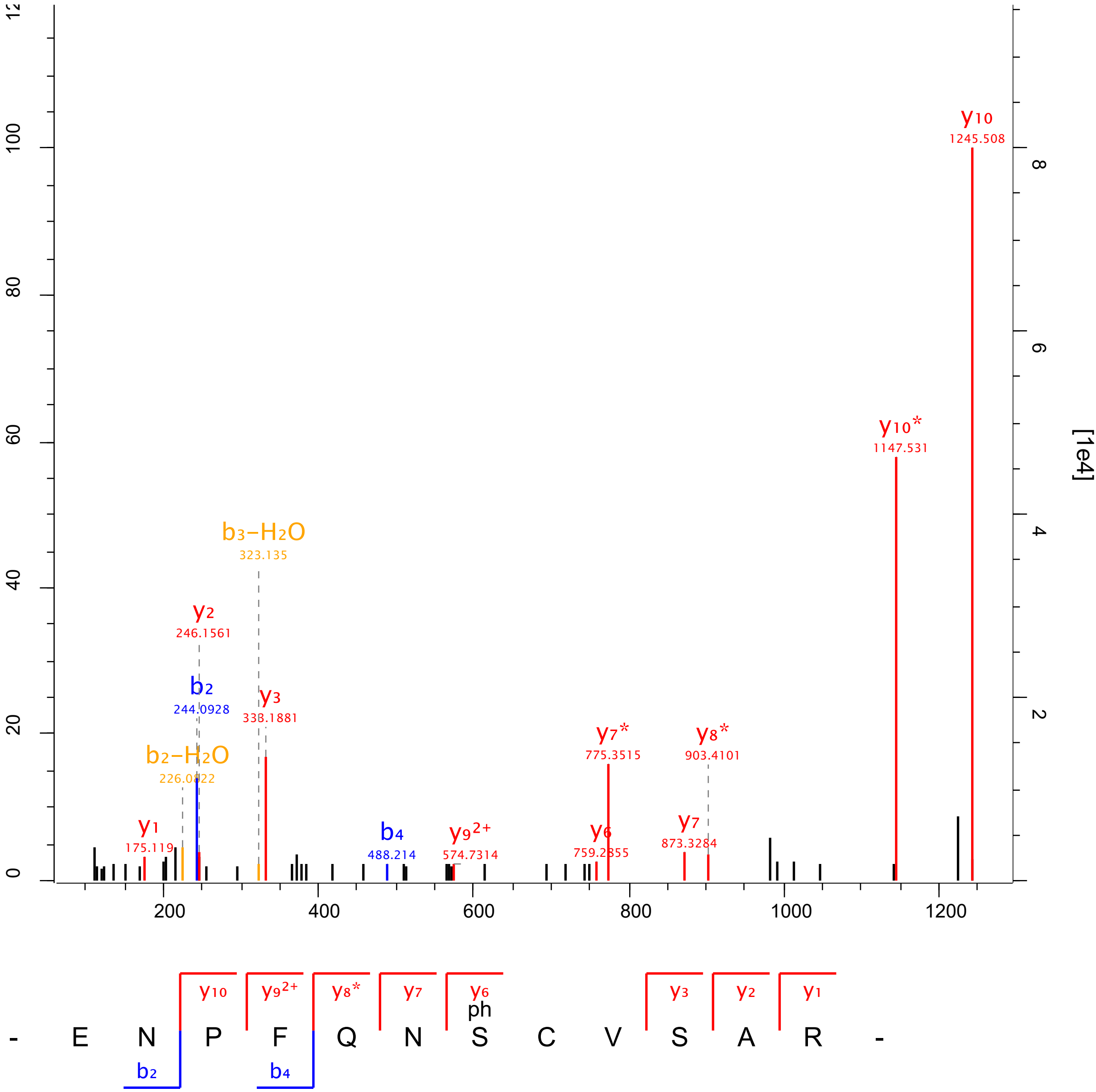

| Raw file | Scan | Method | Score | m/z |
| --- | --- | --- | --- | --- |
| sys_05_2 | 18324 | FTMS; HCD | 129.38 | 695.34 |

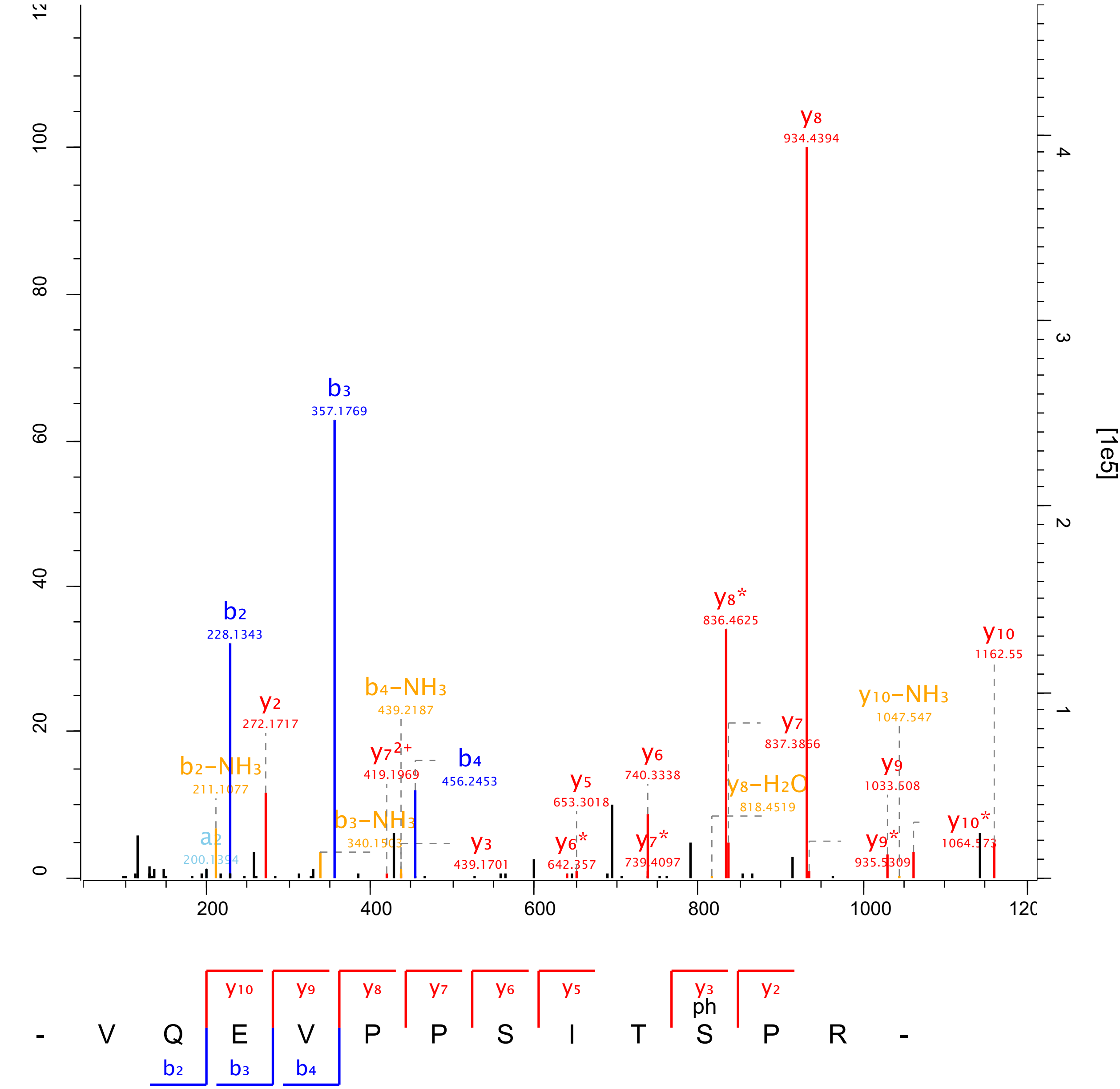

| Raw file | Scan | Method | Score | m/z |
|---|---|---|---|---|
| sys_05_2 | 18795 | FTMS; HCD | 213.72 | 564.76 |

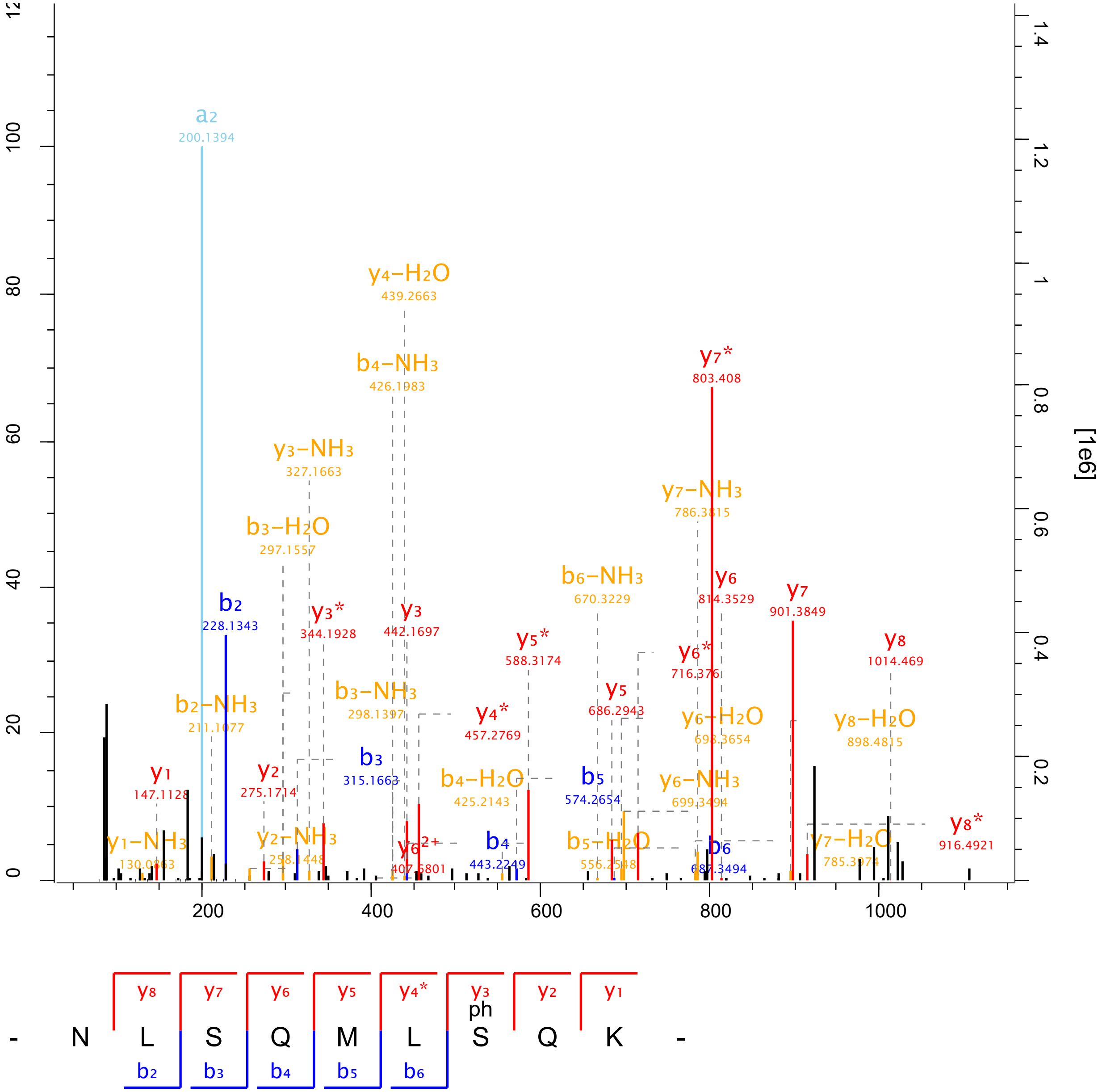

| Raw file | Scan | Method | Score | m/z |
|---|---|---|---|---|
| sys_05_2 | 18826 | FTMS; HCD | 128.76 | 454.52 |

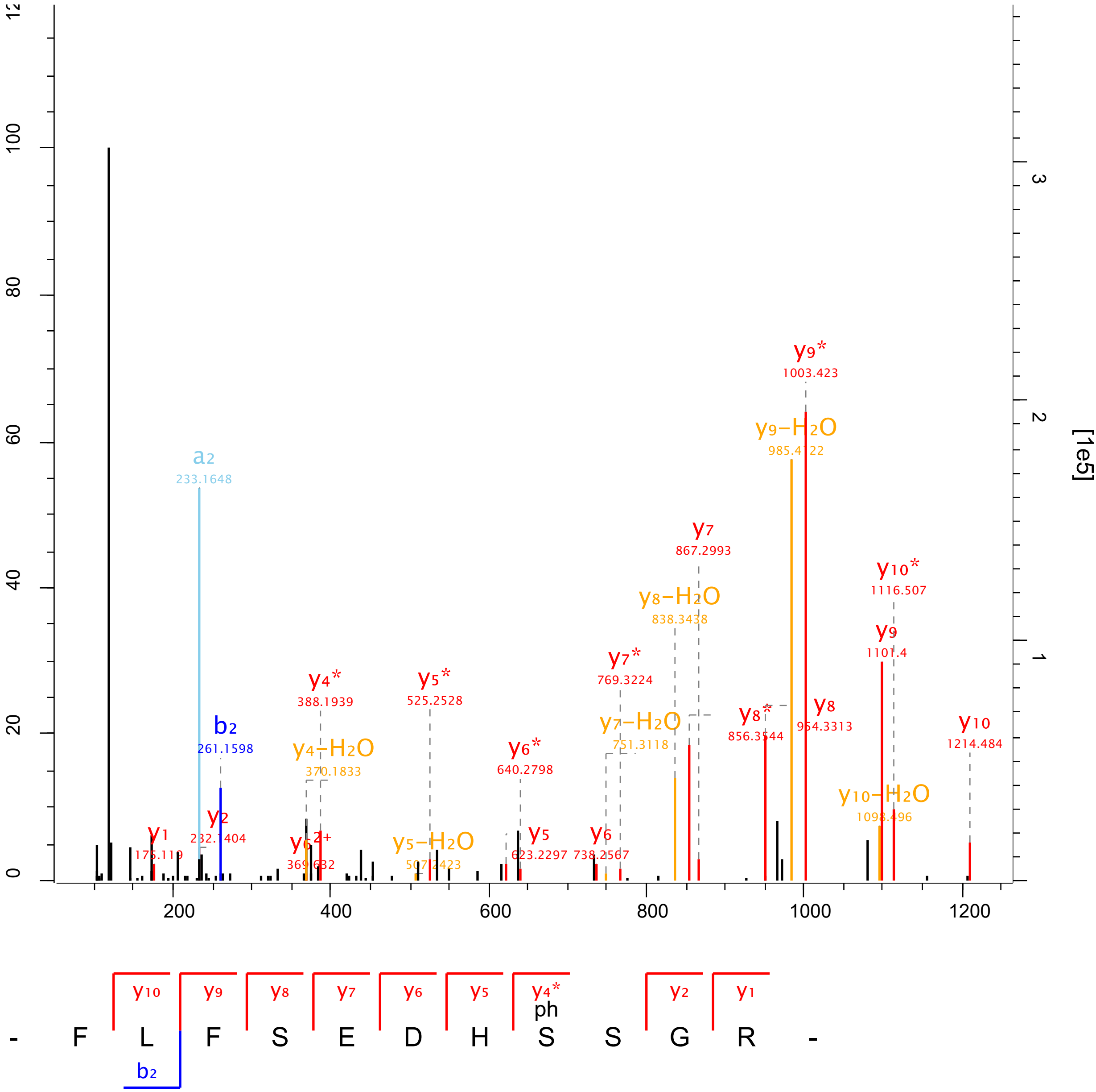

| Raw file | Scan | Method | Score | m/z |
|---|---|---|---|---|
| sys_05_2 | 19087 | FTMS; HCD | 55.45 | 840.87 |

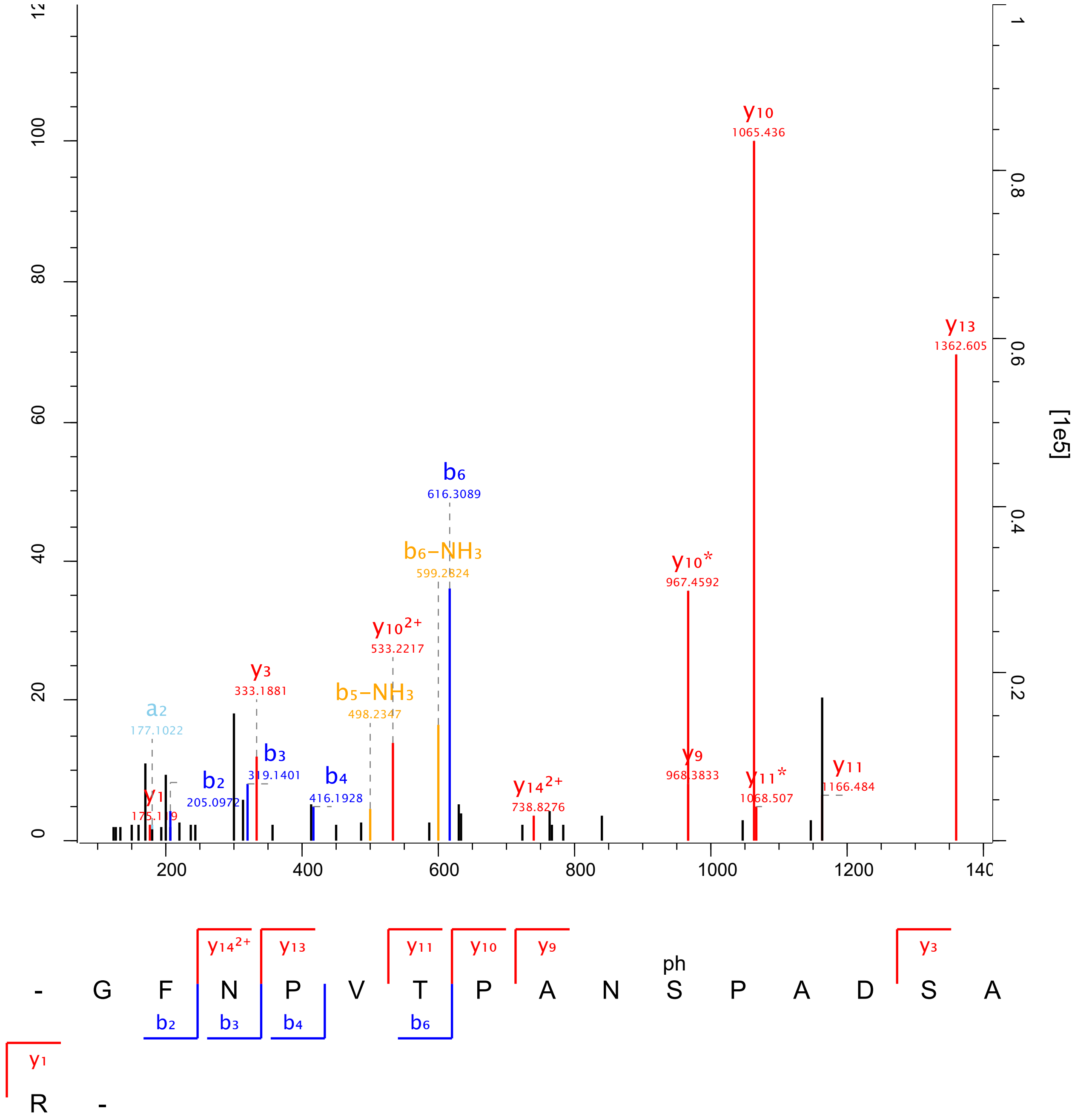

| Raw file | Scan | Method | Score | m/z |
|---|---|---|---|---|
| sys_05_2 | 19112 | FTMS; HCD | 97.27 | 643.3 |

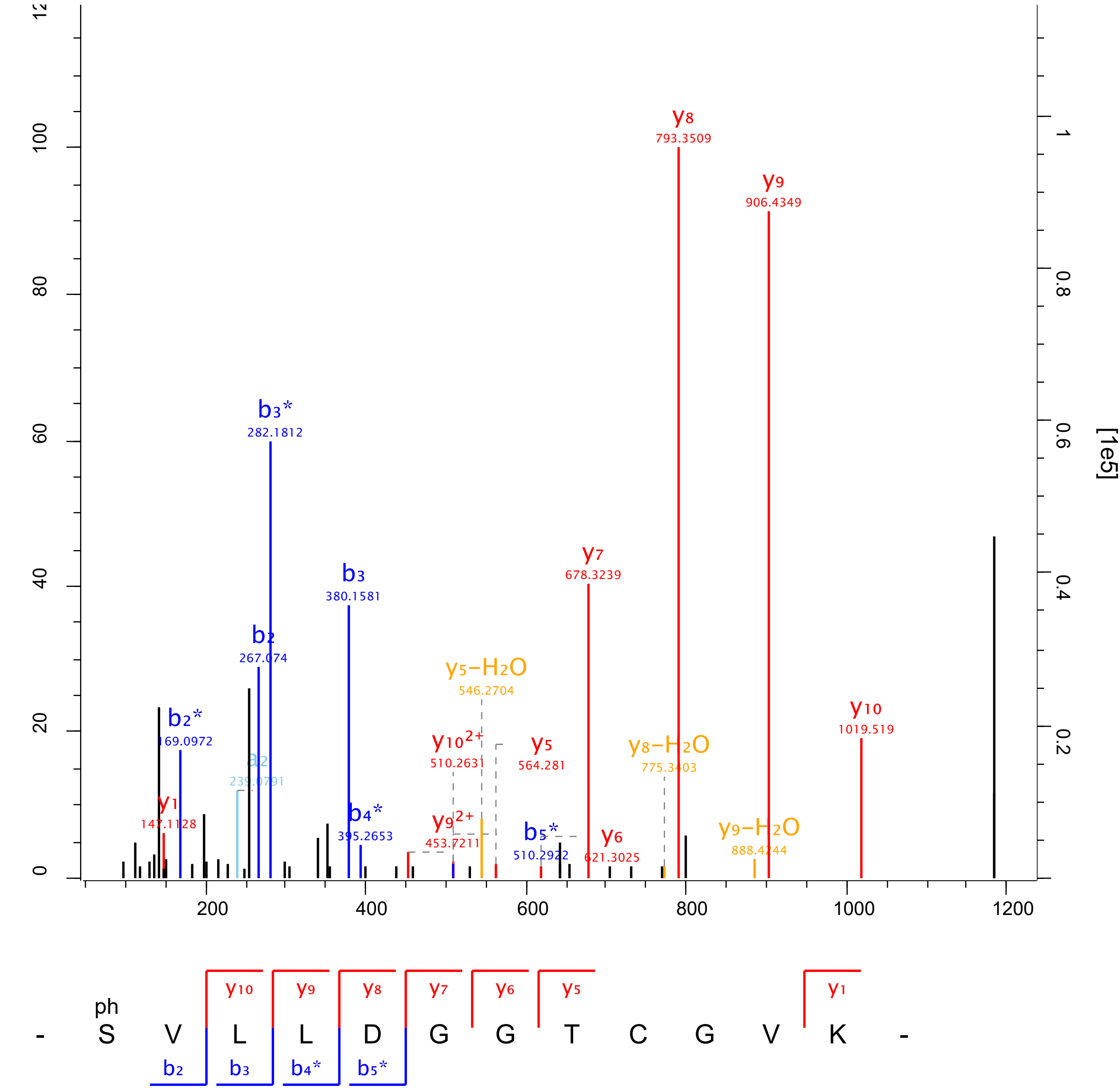

| Raw file | Scan | Method | Score | m/z |
|---|---|---|---|---|
| sys_05_2 | 19232 | FTMS; HCD | 135.73 | 519.2 |

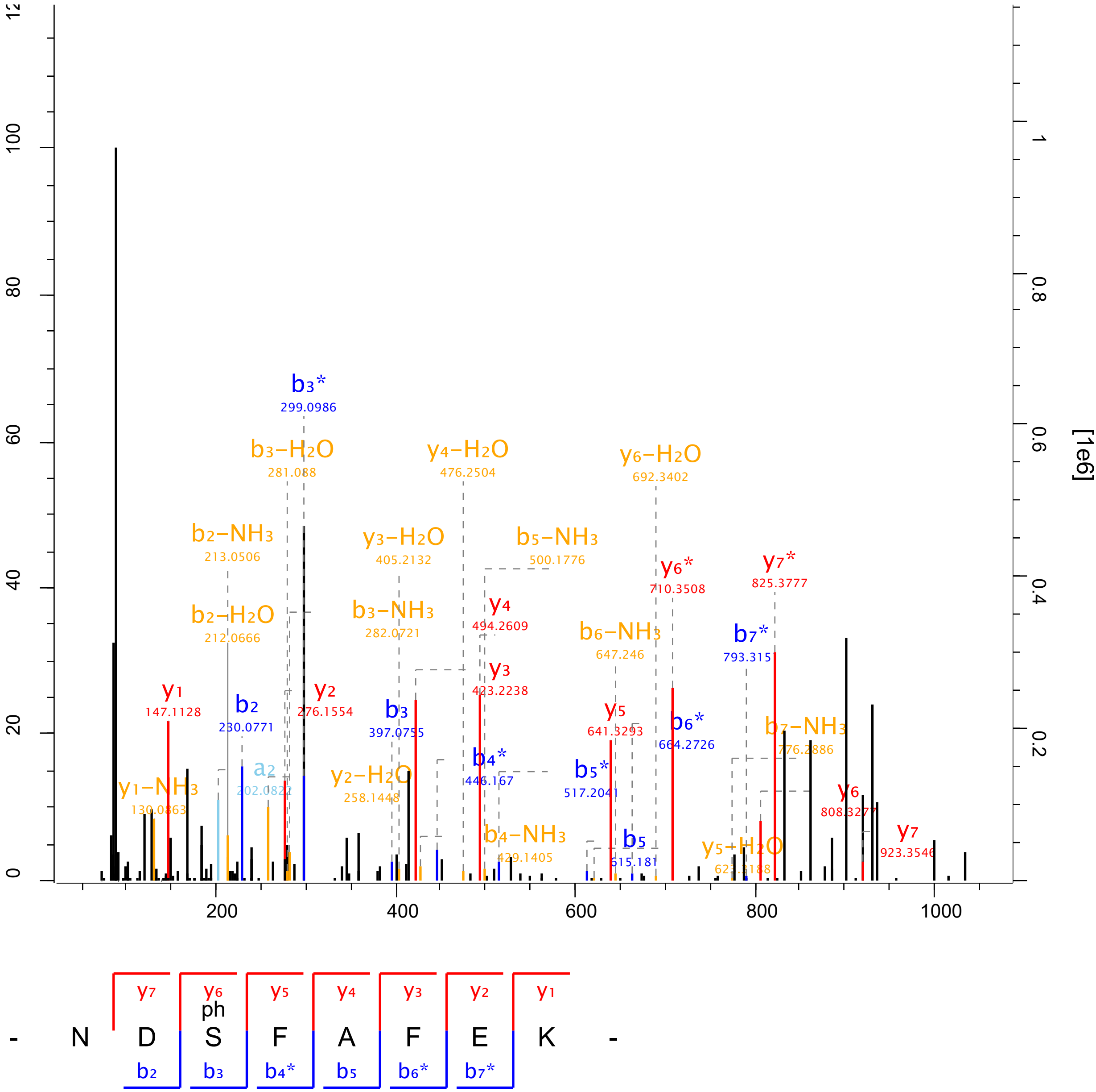

| Raw file | Scan | Method | Score | m/z |
|---|---|---|---|---|
| sys_05_2 | 19594 | FTMS; HCD | 177.32 | 914.88 |

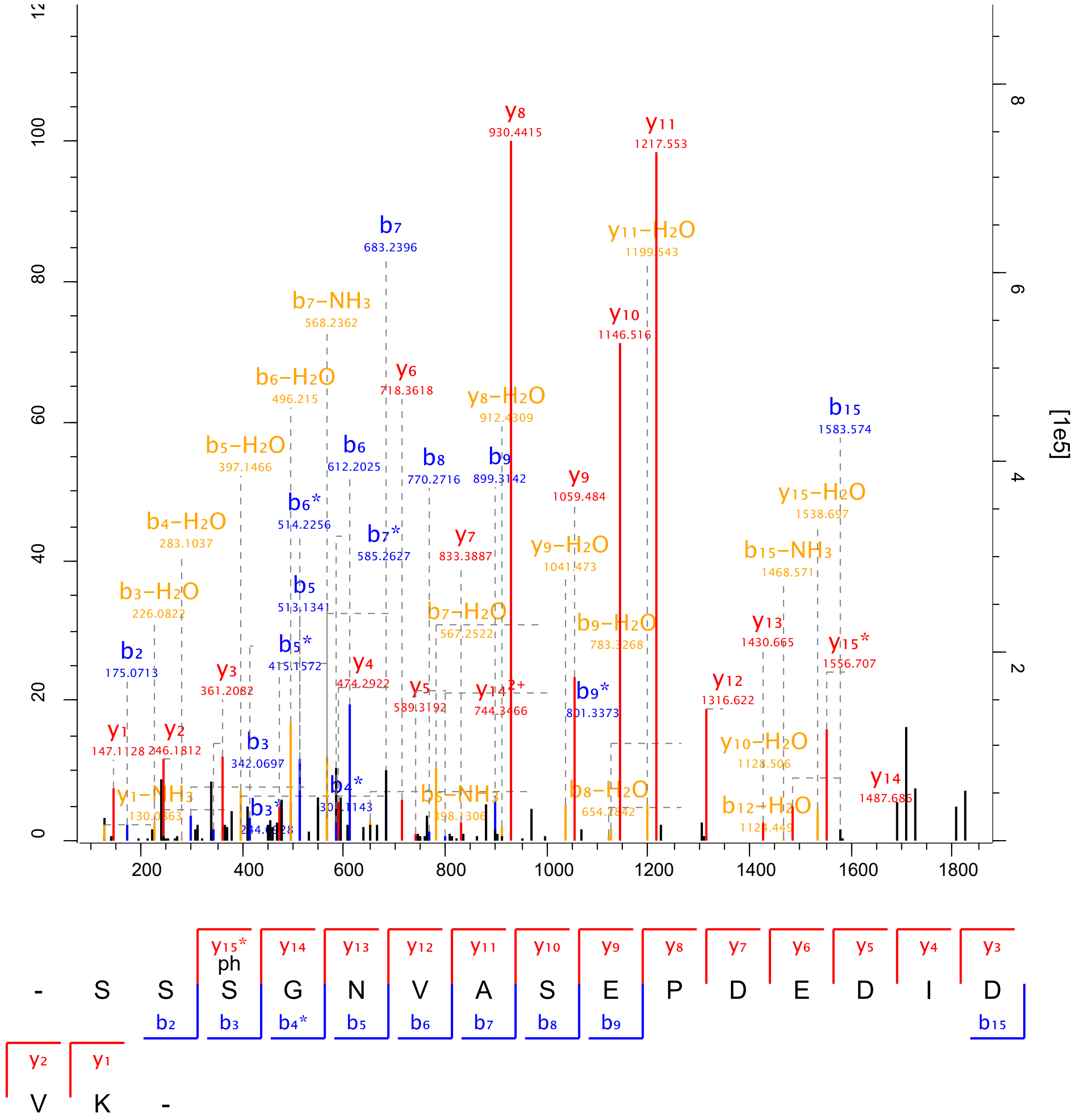

| Raw file | Scan | Method | Score | m/z |
|---|---|---|---|---|
| sys_05_2 | 19781 | FTMS; HCD | 110.44 | 795.85 |

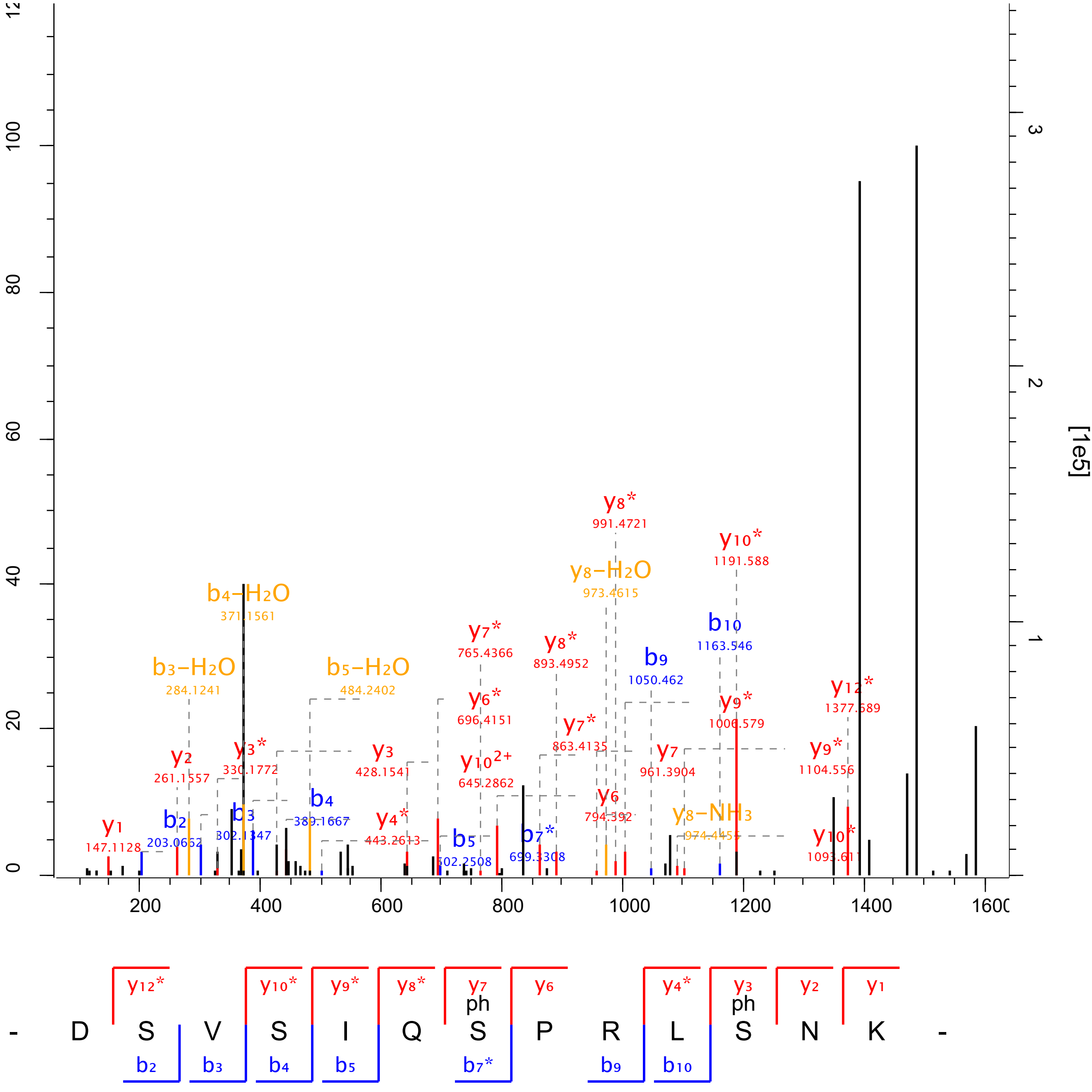

| Raw file | Scan | Method | Score | m/z |
| --- | --- | --- | --- | --- |
| sys_05_2 | 20126 | FTMS; HCD | 202.81 | 707.84 |

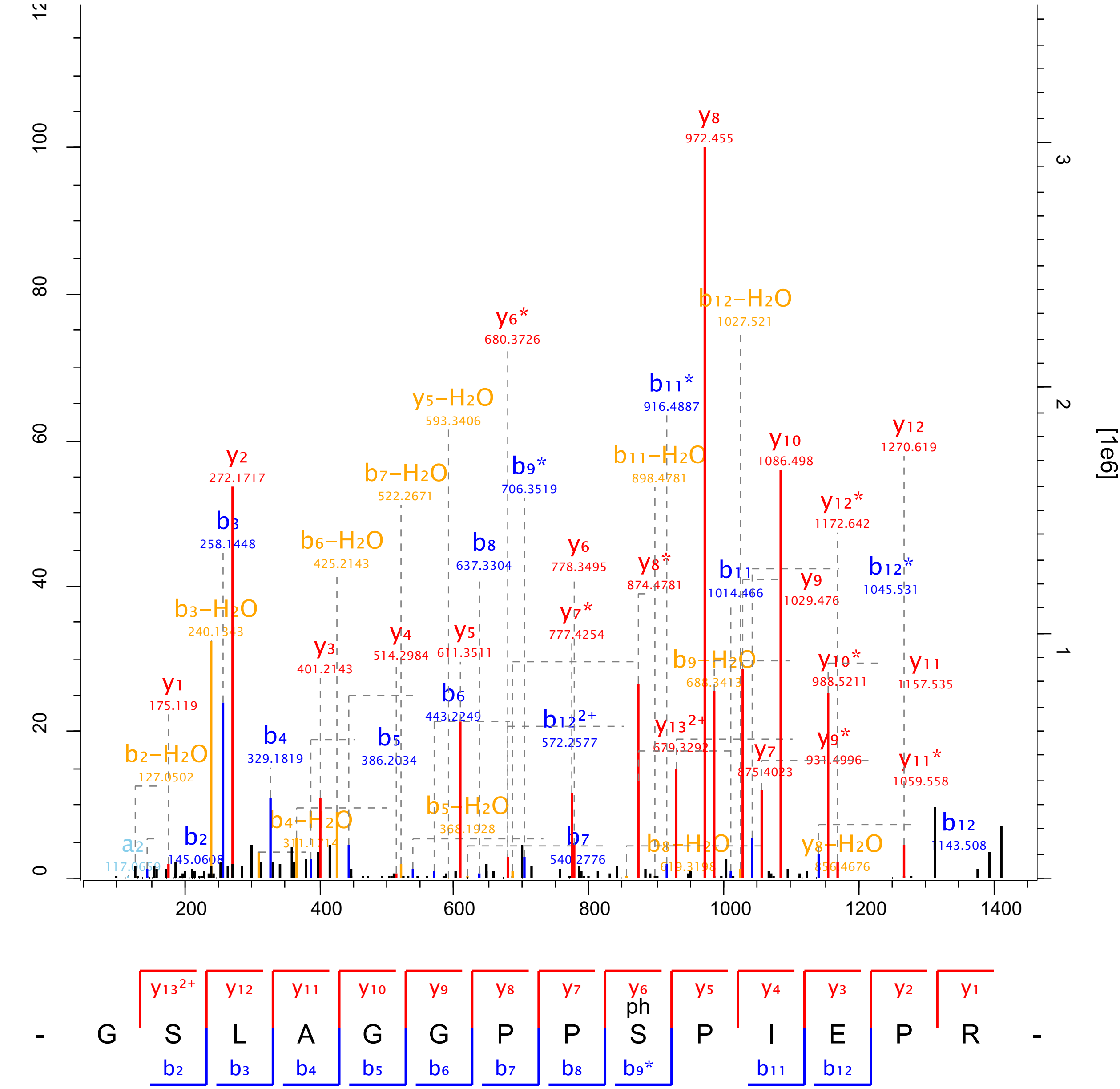

| Raw file | Scan | Method | Score | m/z |
| --- | --- | --- | --- | --- |
| sys_05_2 | 20341 | FTMS; HCD | 86.21 | 674.29 |

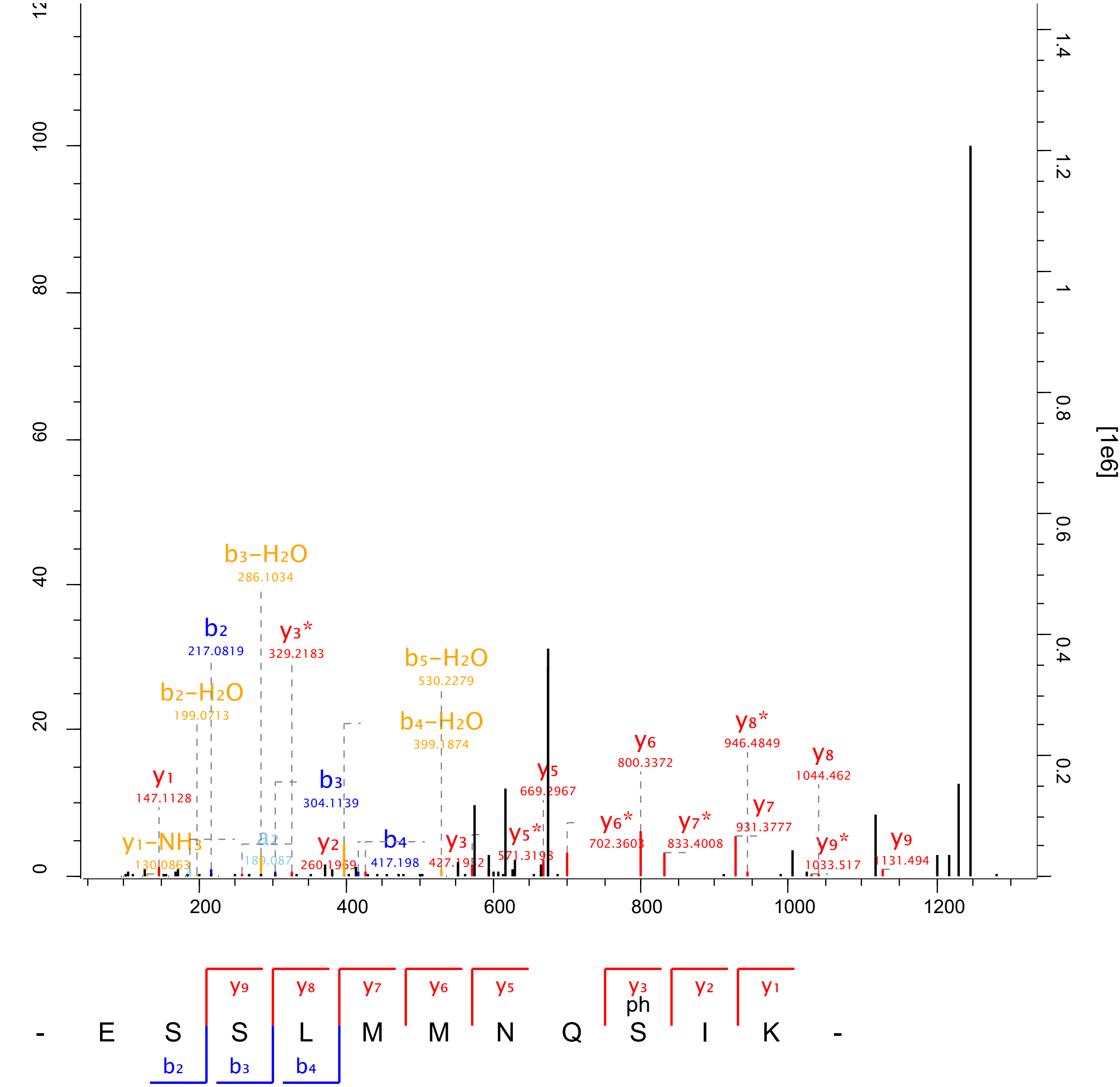

| Raw file | Scan | Method | Score | m/z |
|---|---|---|---|---|
| sys_05_2 | 20520 | FTMS; HCD | 93.65 | 433.23 |

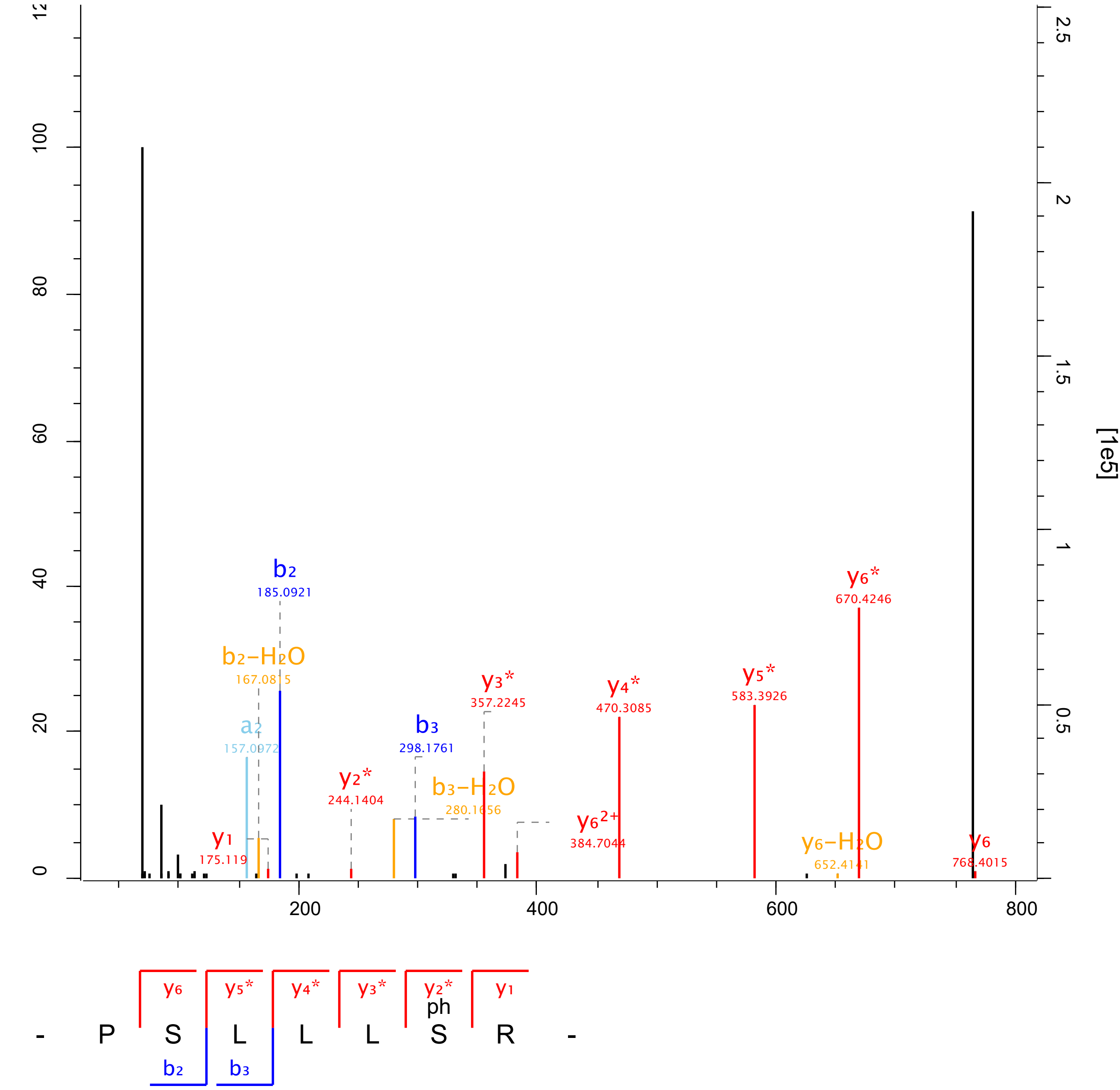

| Raw file | Scan | Method | Score | m/z |
| --- | --- | --- | --- | --- |
| sys_05_2 | 20773 | FTMS; HCD | 199.91 | 787.34 |

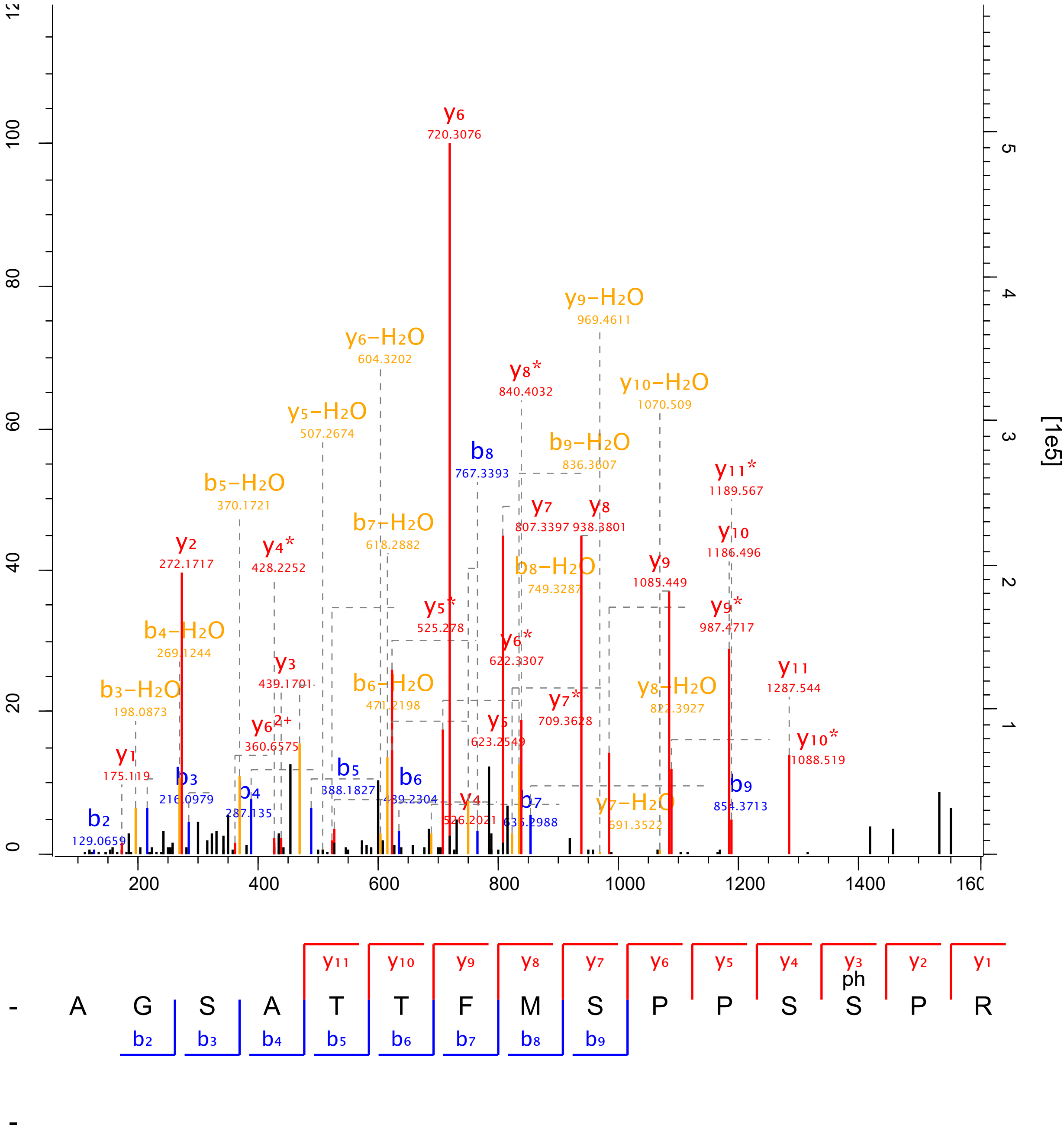

| Raw file | Scan | Method | Score | m/z |
| --- | --- | --- | --- | --- |
| sys_05_2 | 20939 | FTMS; HCD | 181.11 | 889.72 |

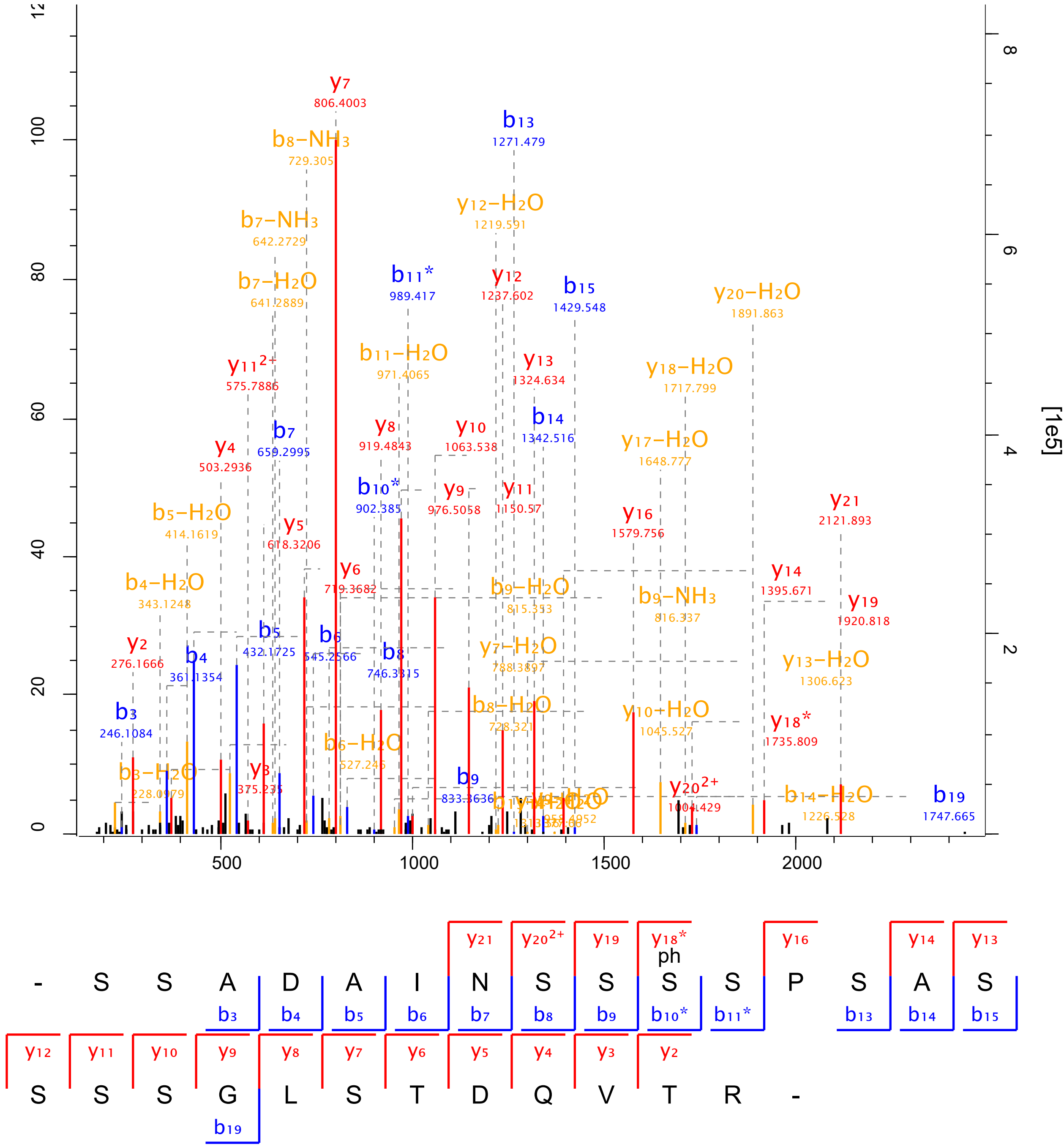

| Raw file | Scan | Method | Score | m/z |
| --- | --- | --- | --- | --- |
| sys_05_2 | 21084 | FTMS; HCD | 138.91 | 777.82 |

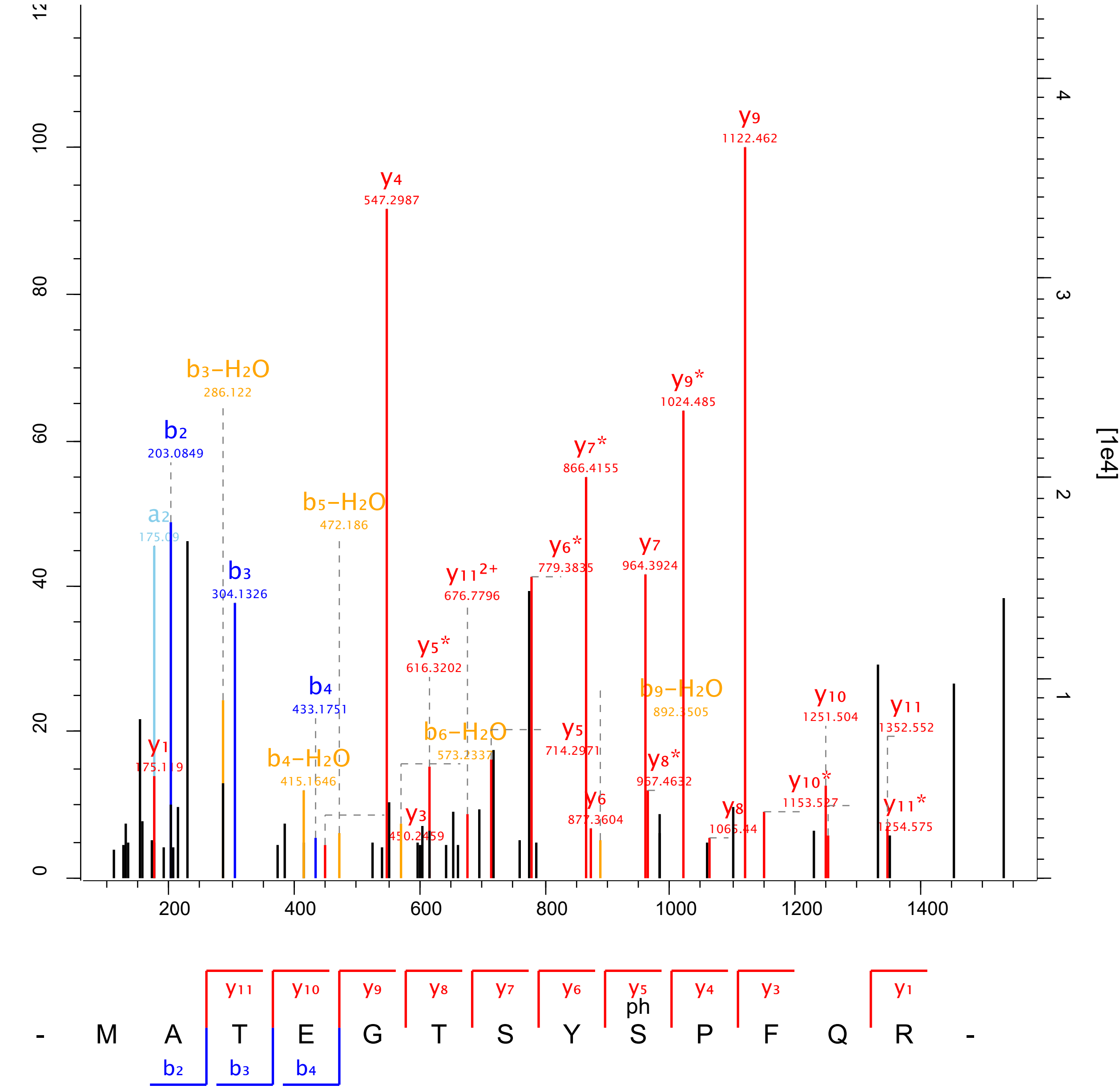

| Raw file | Scan | Method | Score | m/z |
| --- | --- | --- | --- | --- |
| sys_05_2 | 21250 | FTMS; HCD | 298 | 857.86 |

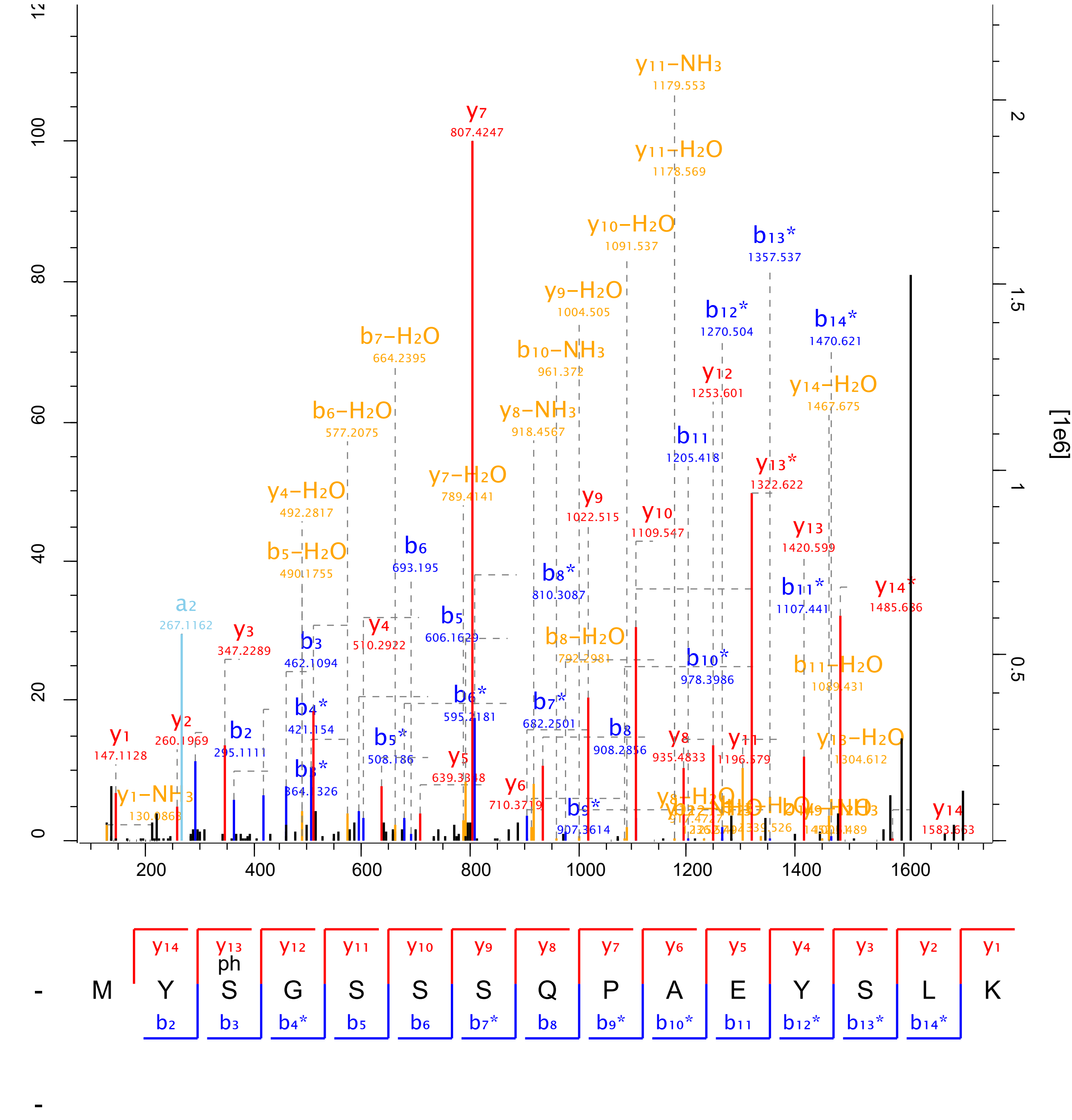

| Raw file | Scan | Method | Score | m/z |
|---|---|---|---|---|
| sys_05_2 | 21297 | FTMS; HCD | 76.82 | 932.35 |

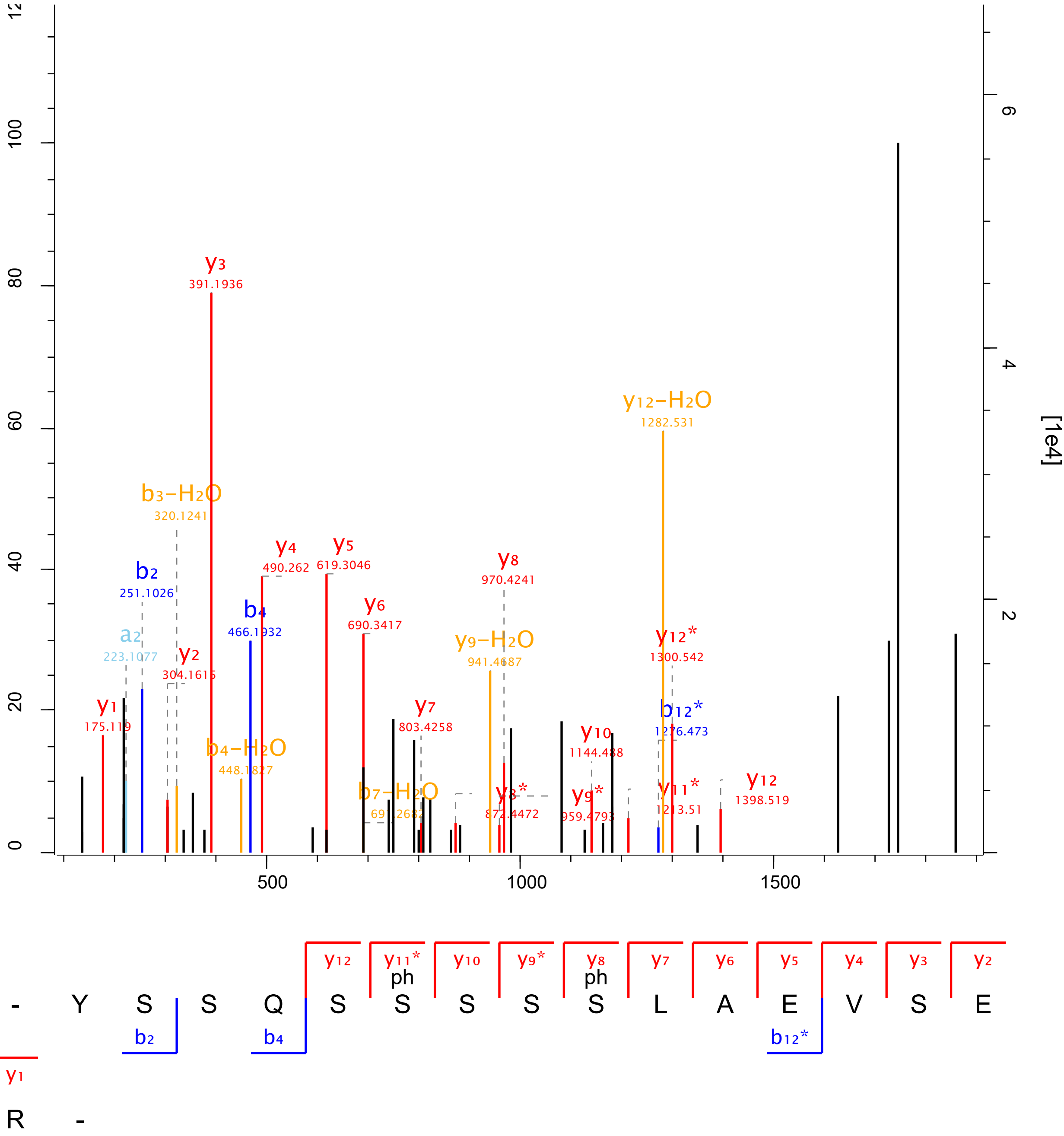

| Raw file | Scan | Method | Score | m/z |
| --- | --- | --- | --- | --- |
| sys_05_2 | 21658 | FTMS; HCD | 112.34 | 768.31 |

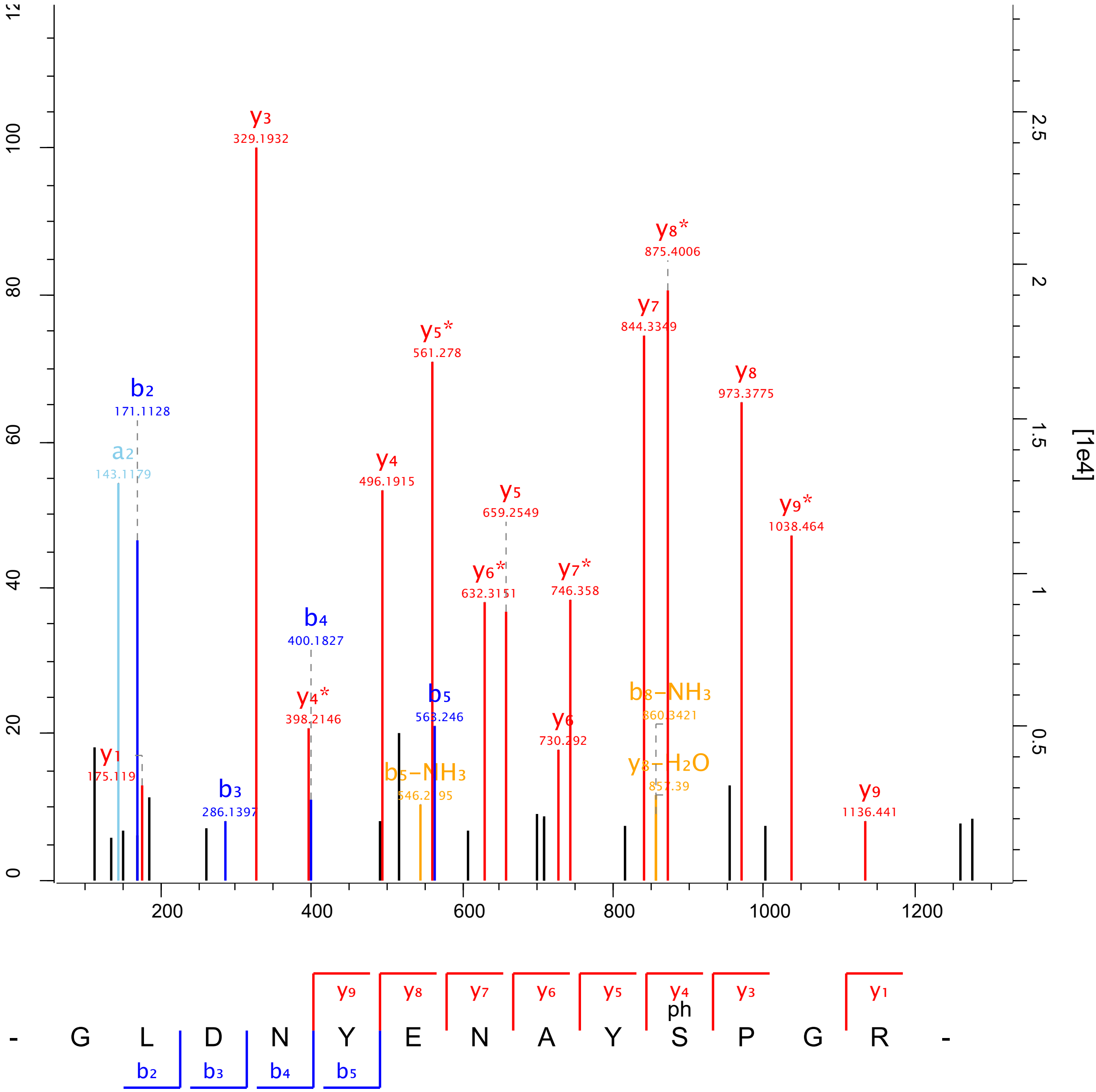

| Raw file | Scan | Method | Score | m/z |
|---|---|---|---|---|
| sys_05_2 | 21695 | FTMS; HCD | 49.19 | 617.27 |

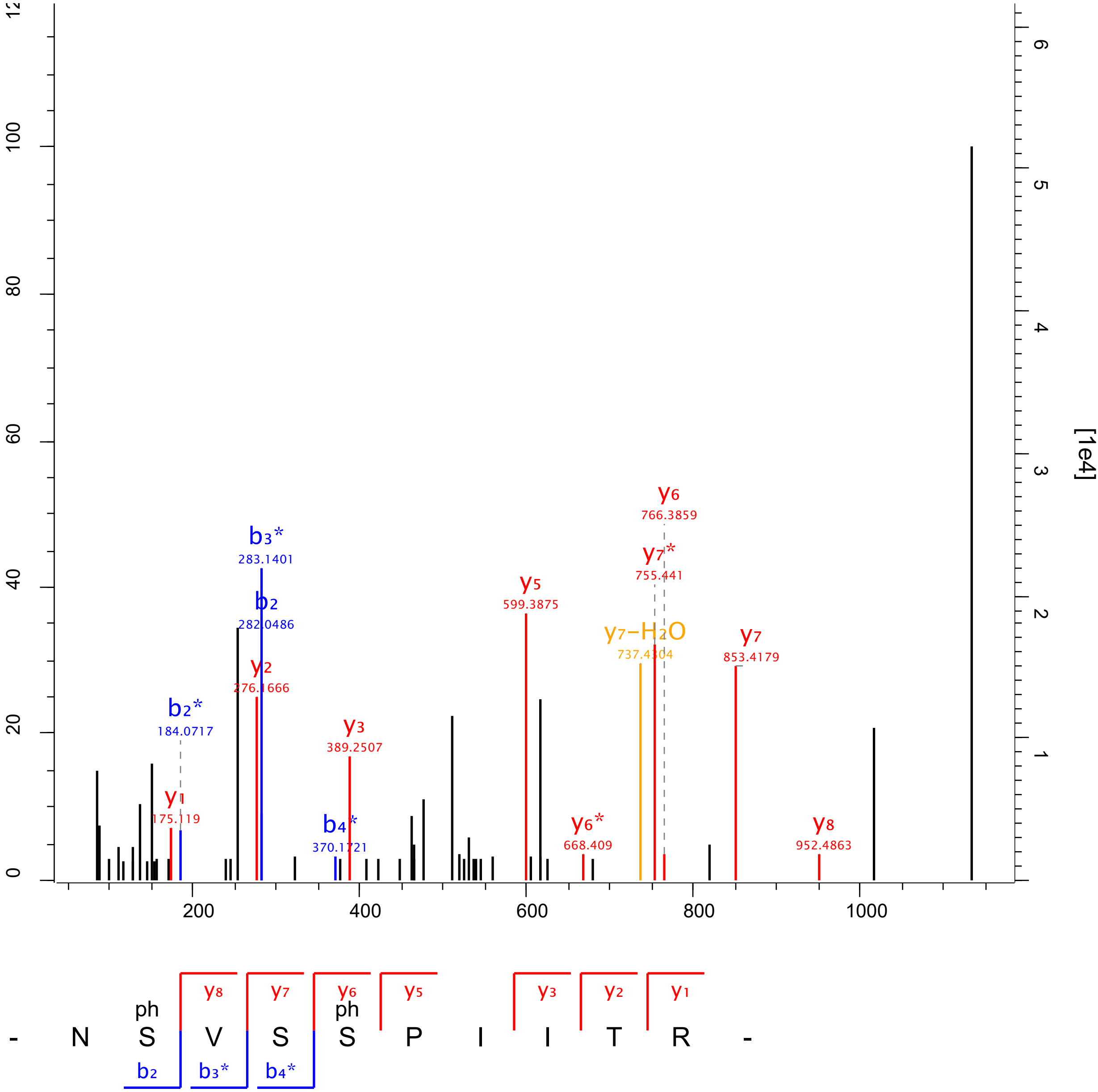

| Raw file | Scan | Method | Score | m/z |
|---|---|---|---|---|
| sys_05_2 | 21873 | FTMS; HCD | 136.66 | 804.86 |

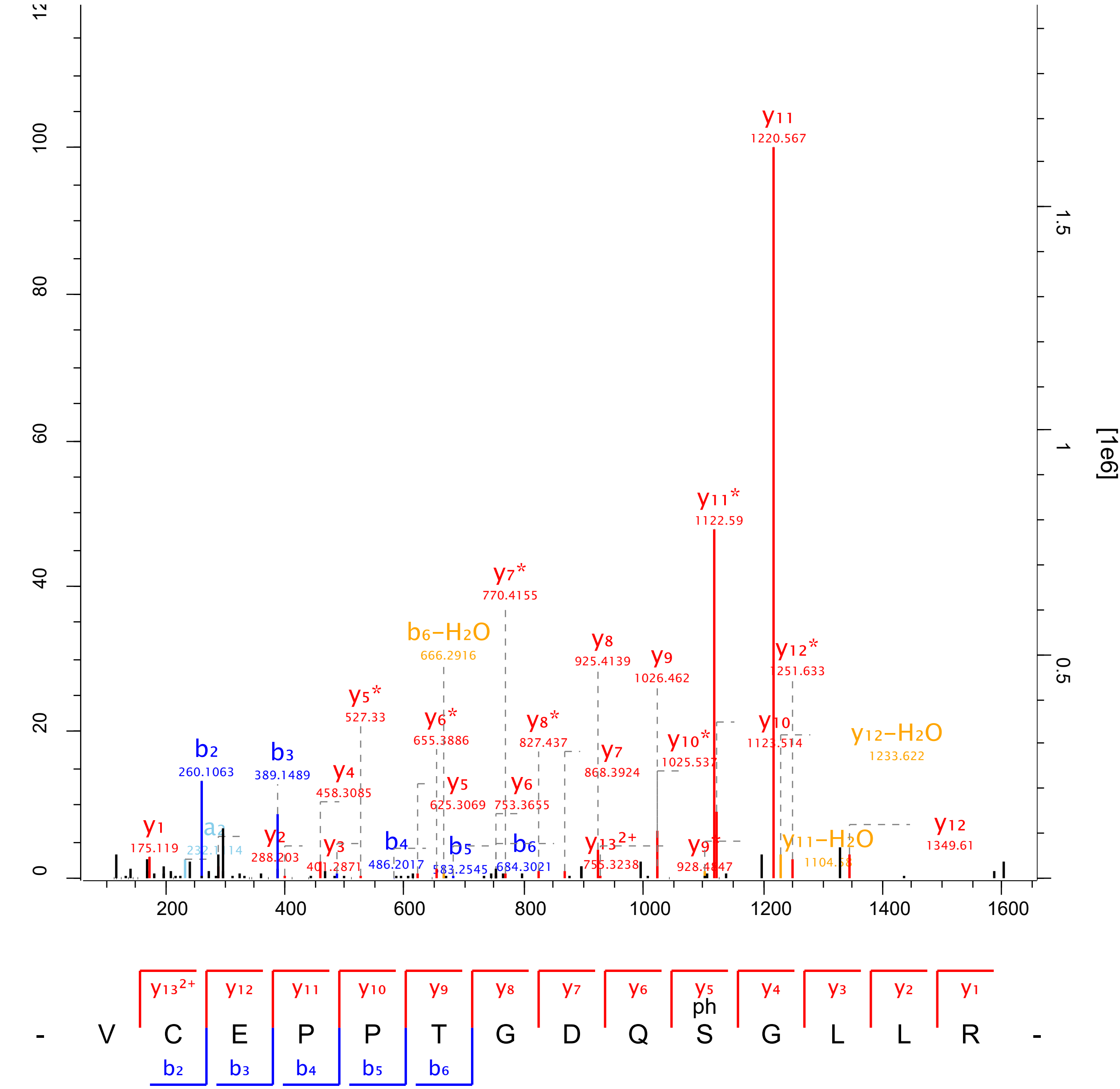

| Raw file | Scan | Method | Score | m/z |
| --- | --- | --- | --- | --- |
| sys_05_2 | 21880 | FTMS; HCD | 104.21 | 766.83 |

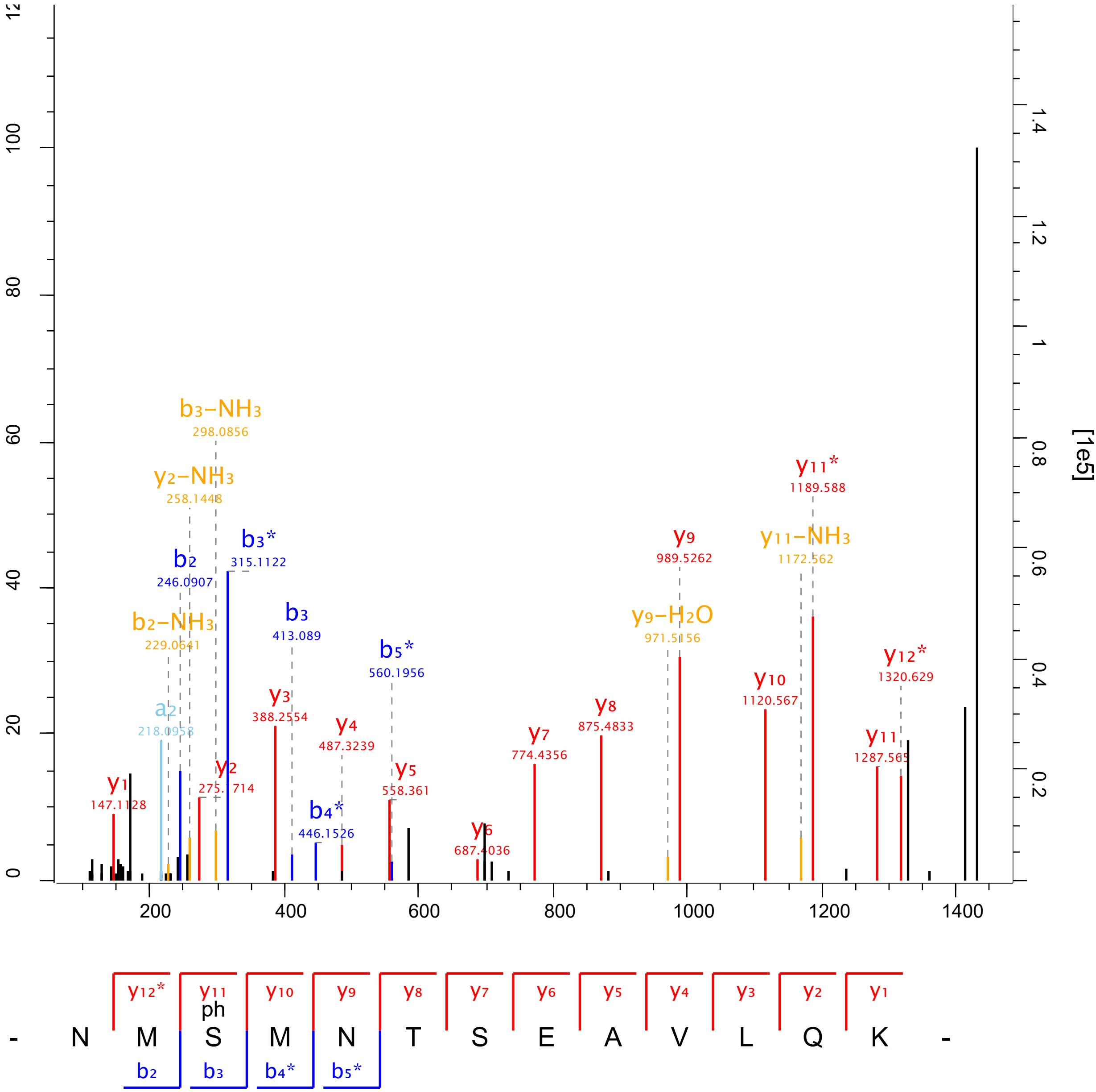

| Raw file | Scan | Method | Score | m/z |
| --- | --- | --- | --- | --- |
| sys_05_2 | 21887 | FTMS; HCD | 179.67 | 582.22 |

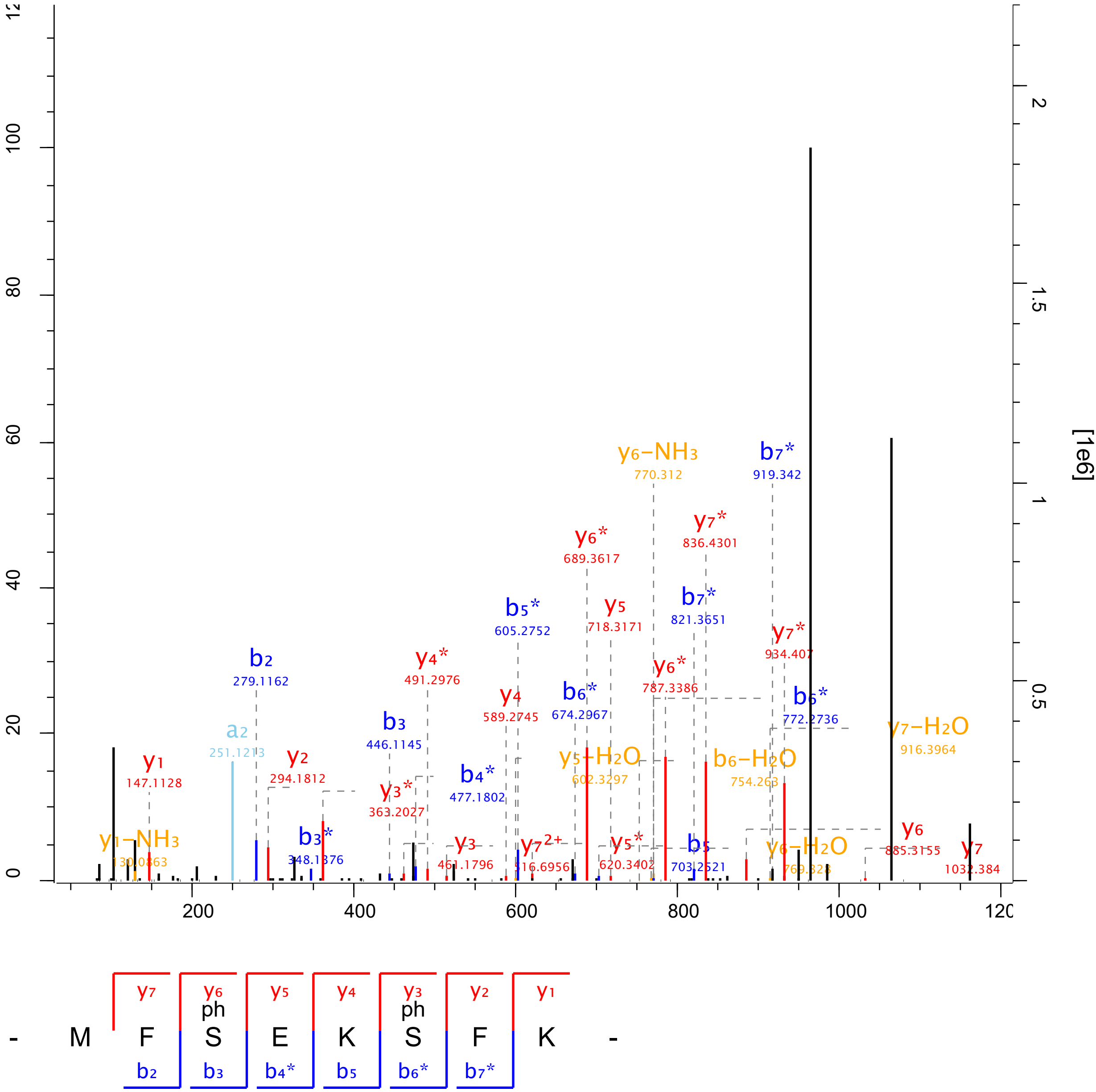

| Raw file | Scan | Method | Score | m/z |
| --- | --- | --- | --- | --- |
| sys_05_2 | 21922 | FTMS; HCD | 75.06 | 637.29 |

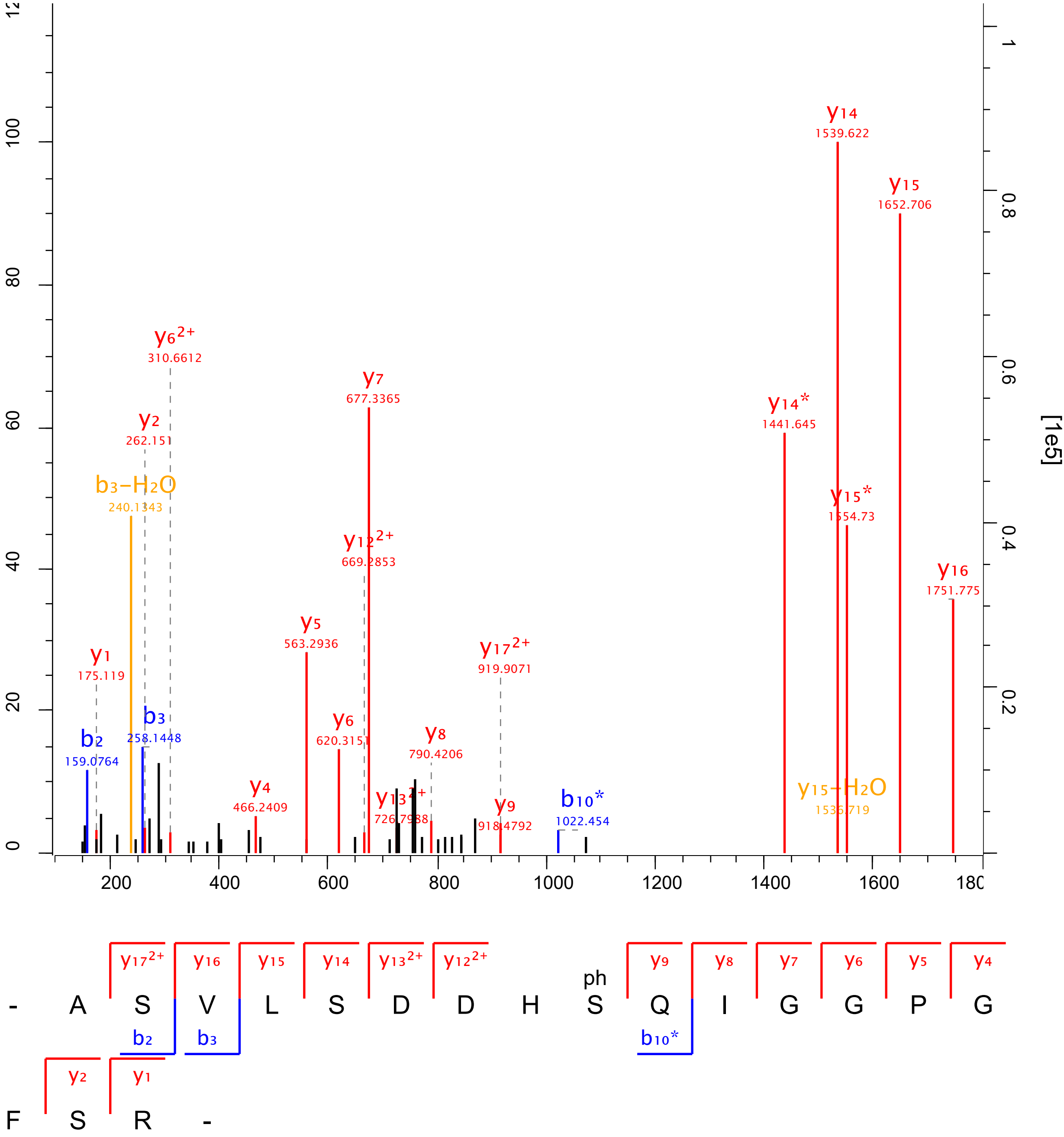

| Raw file | Scan | Method | Score | m/z |
|---|---|---|---|---|
| sys_05_2 | 22060 | FTMS; HCD | 255.83 | 669.62 |

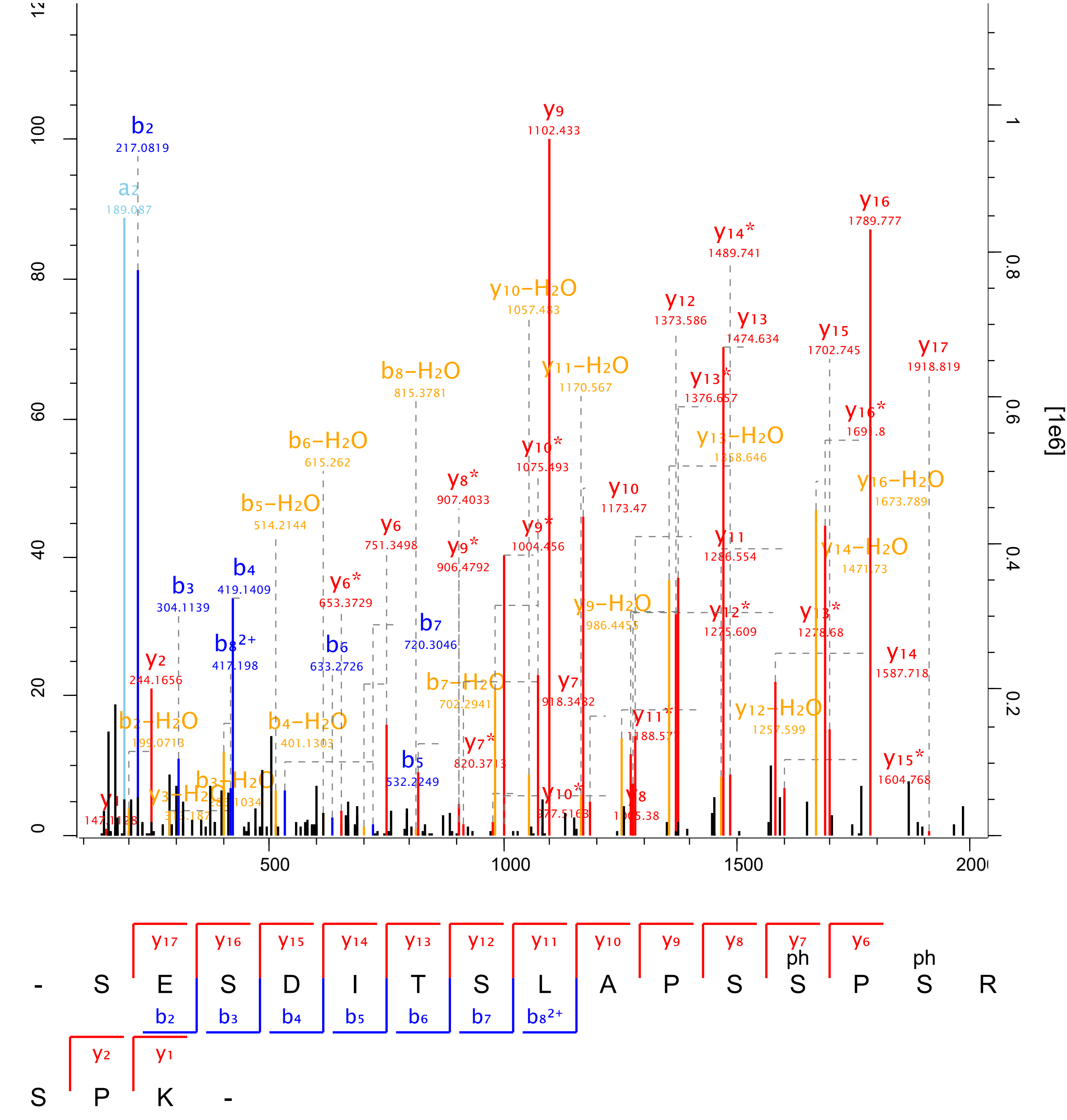

| Raw file | Scan | Method | Score | m/z |
|---|---|---|---|---|
| sys_05_2 | 22298 | FTMS; HCD | 67.08 | 524.72 |

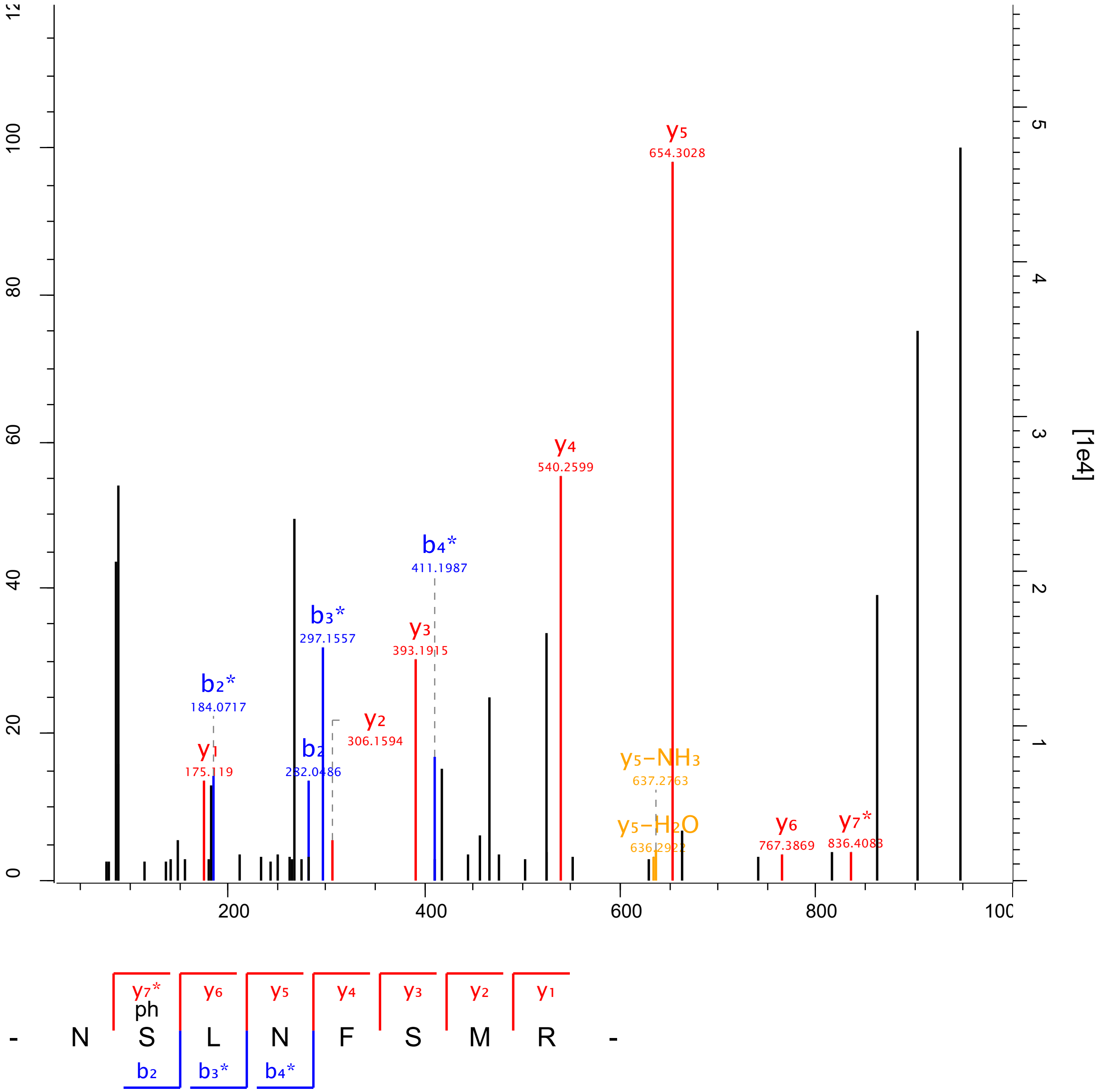

| Raw file | Scan | Method | Score | m/z |
|---|---|---|---|---|
| sys_05_2 | 22371 | FTMS; HCD | 147.64 | 782.34 |

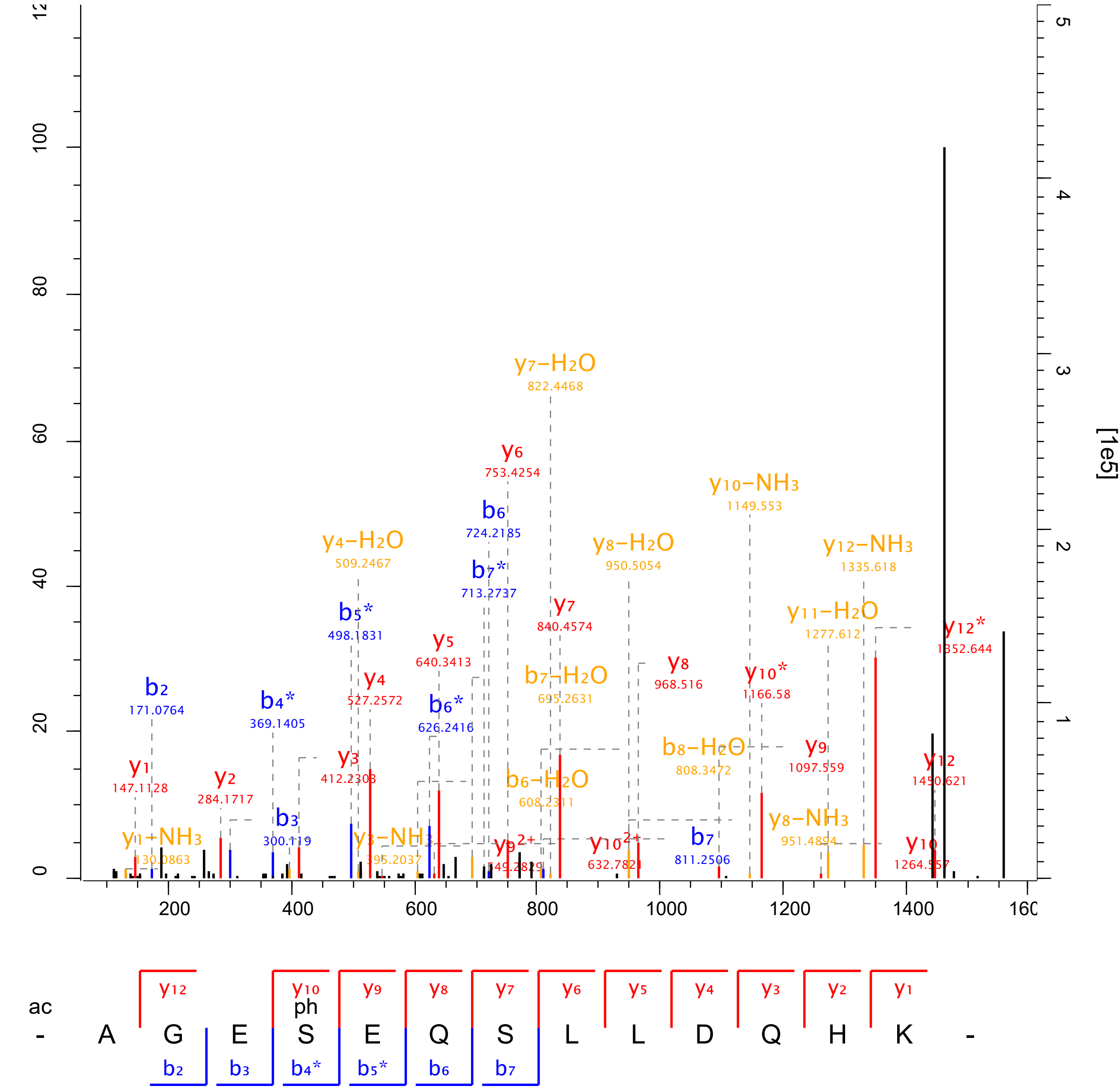

| Raw file | Scan | Method | Score | m/z |
|---|---|---|---|---|
| sys_05_2 | 22737 | FTMS; HCD | 113.65 | 685.82 |

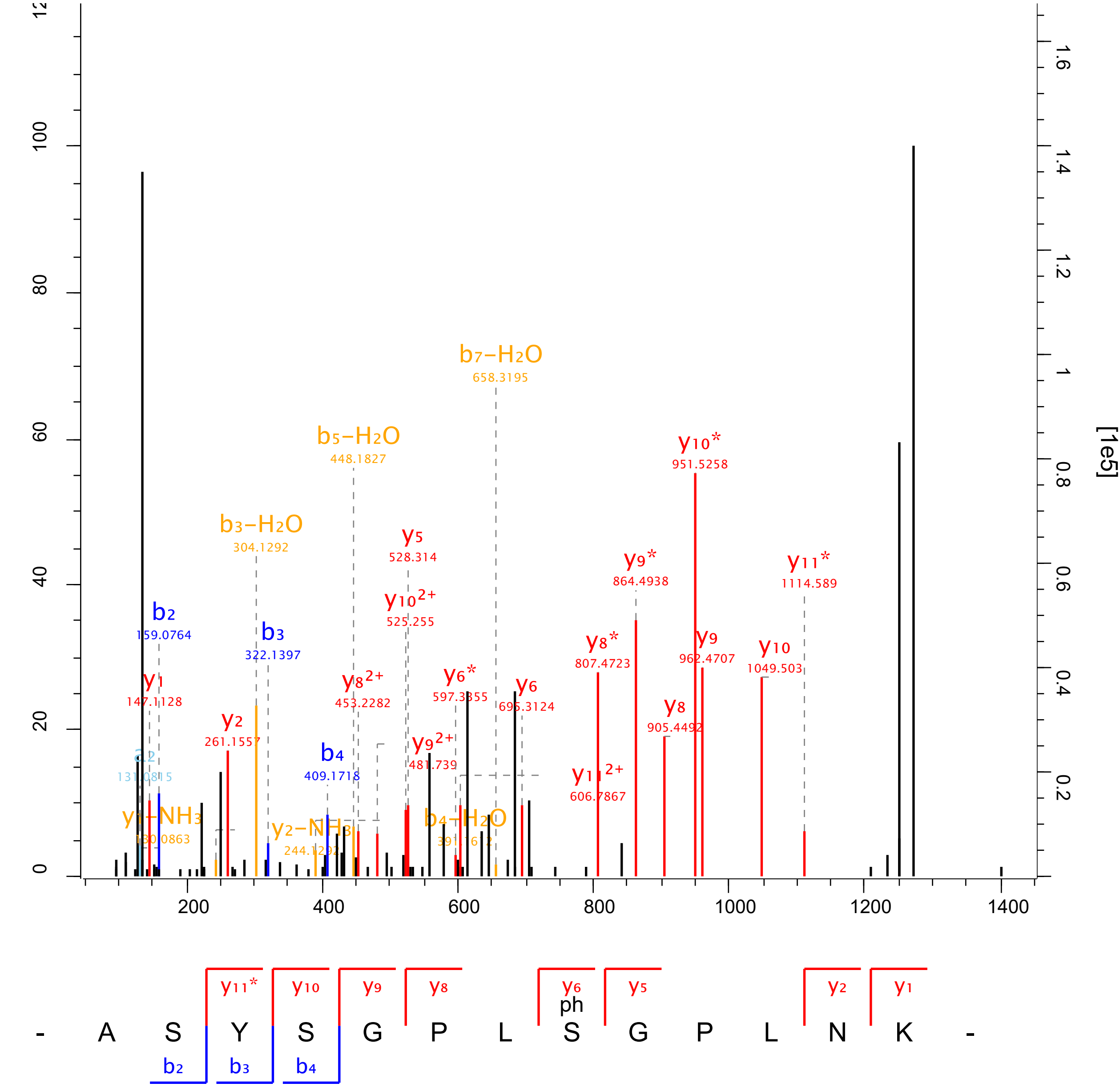

| Raw file | Scan | Method | Score | m/z |
|---|---|---|---|---|
| sys_05_2 | 23189 | FTMS; HCD | 104.84 | 697.29 |

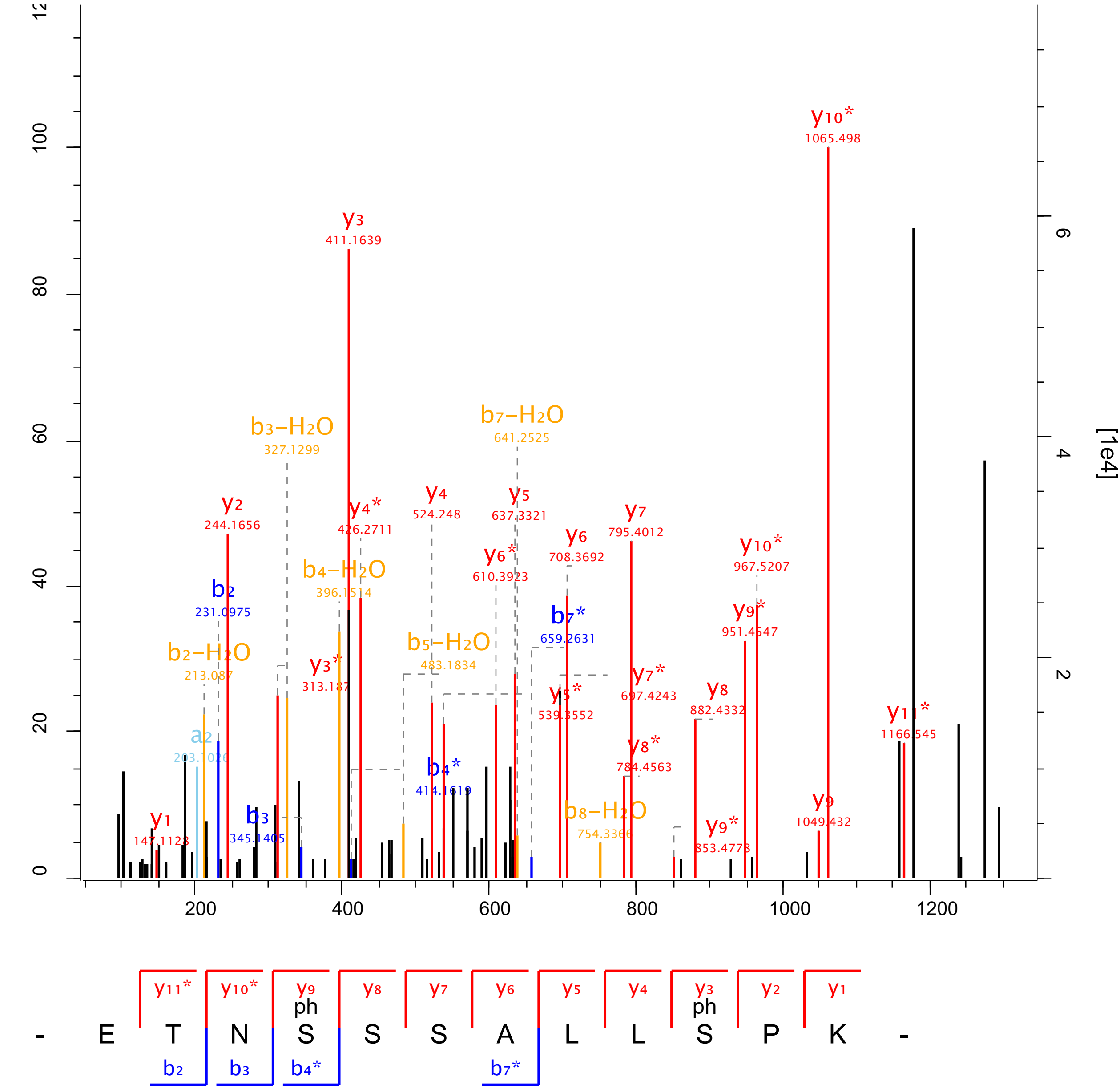

| Raw file | Scan | Method | Score | m/z |
| --- | --- | --- | --- | --- |
| sys_05_2 | 23202 | FTMS; HCD | 120.92 | 595.27 |

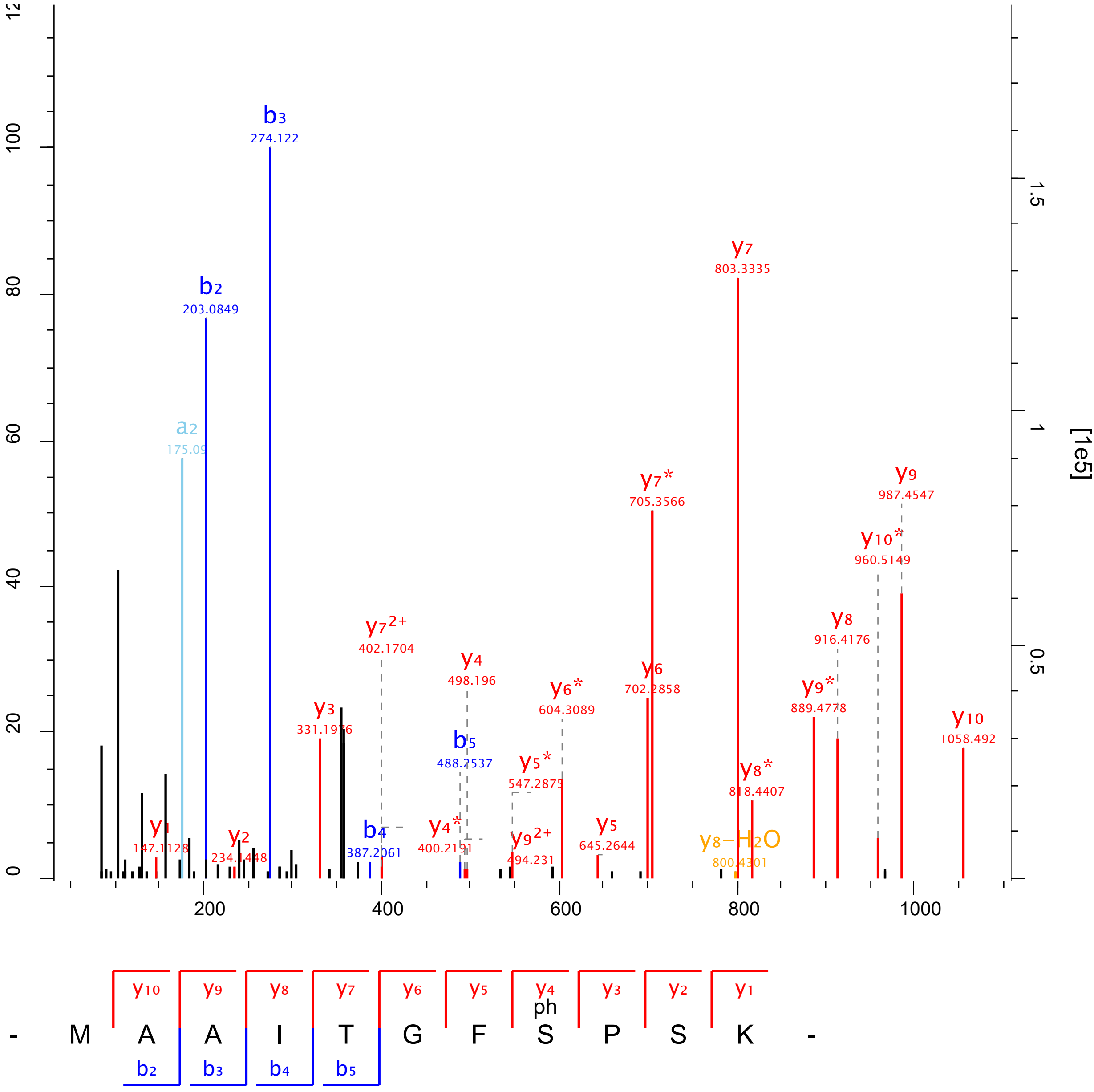

| Raw file | Scan | Method | Score | m/z |
|---|---|---|---|---|
| sys_05_2 | 23420 | FTMS; HCD | 141.82 | 639.27 |

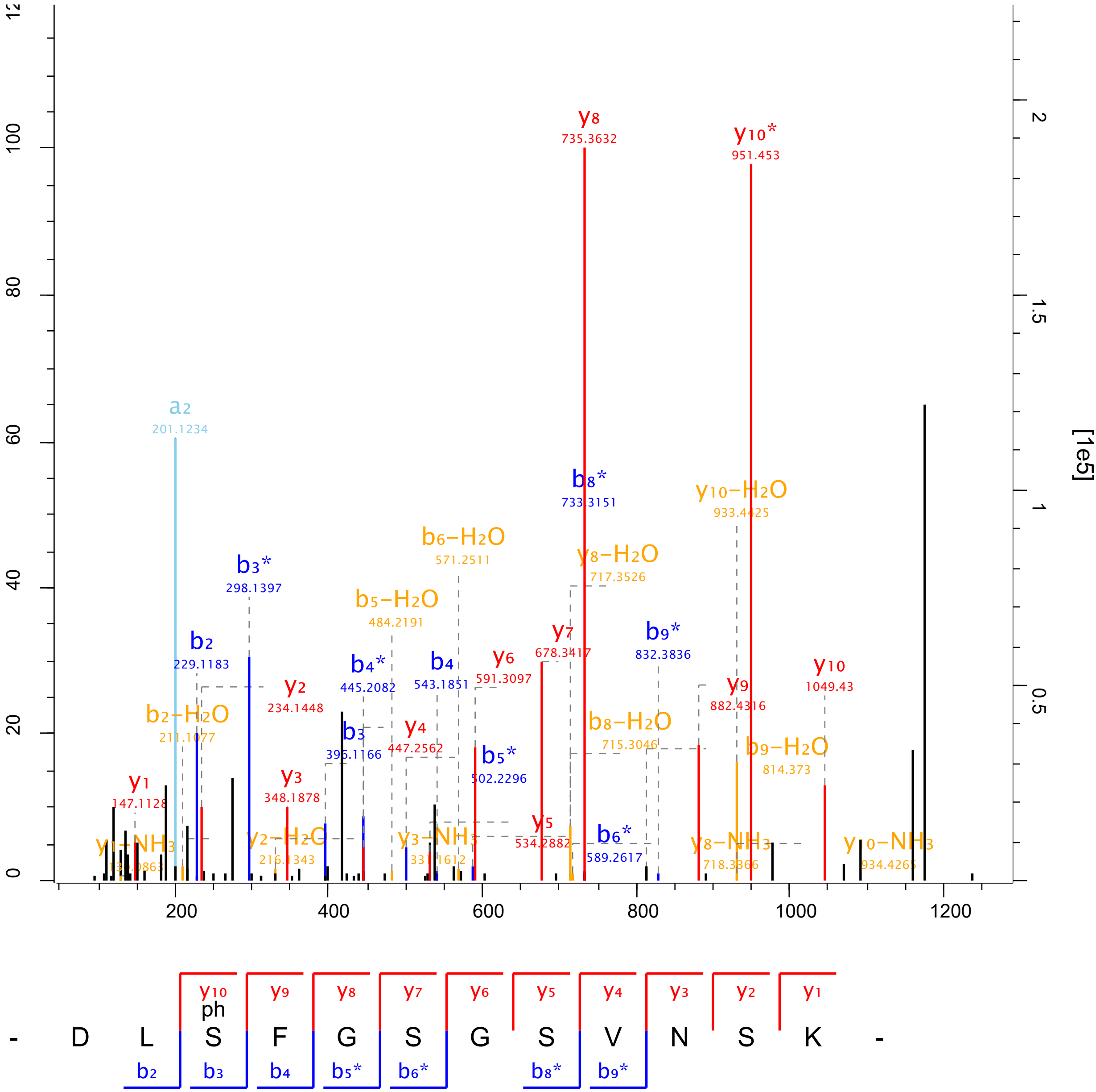

| Raw file | Scan | Method | Score | m/z |
|---|---|---|---|---|
| sys_05_2 | 23525 | FTMS; HCD | 176.46 | 738.32 |

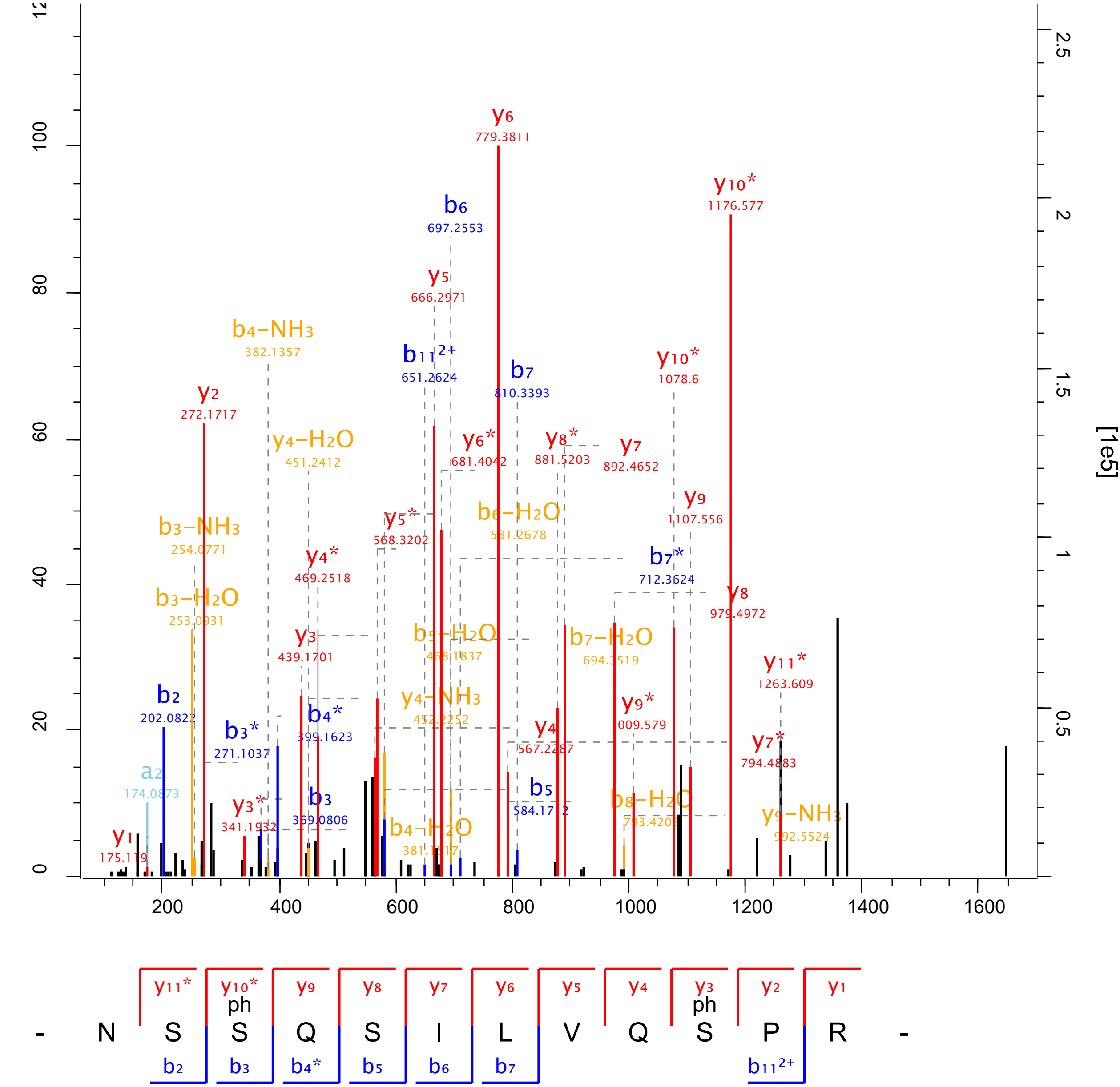

| Raw file | Scan | Method | Score | m/z |
|---|---|---|---|---|
| sys_05_2 | 23608 | FTMS; HCD | 359.08 | 913.35 |

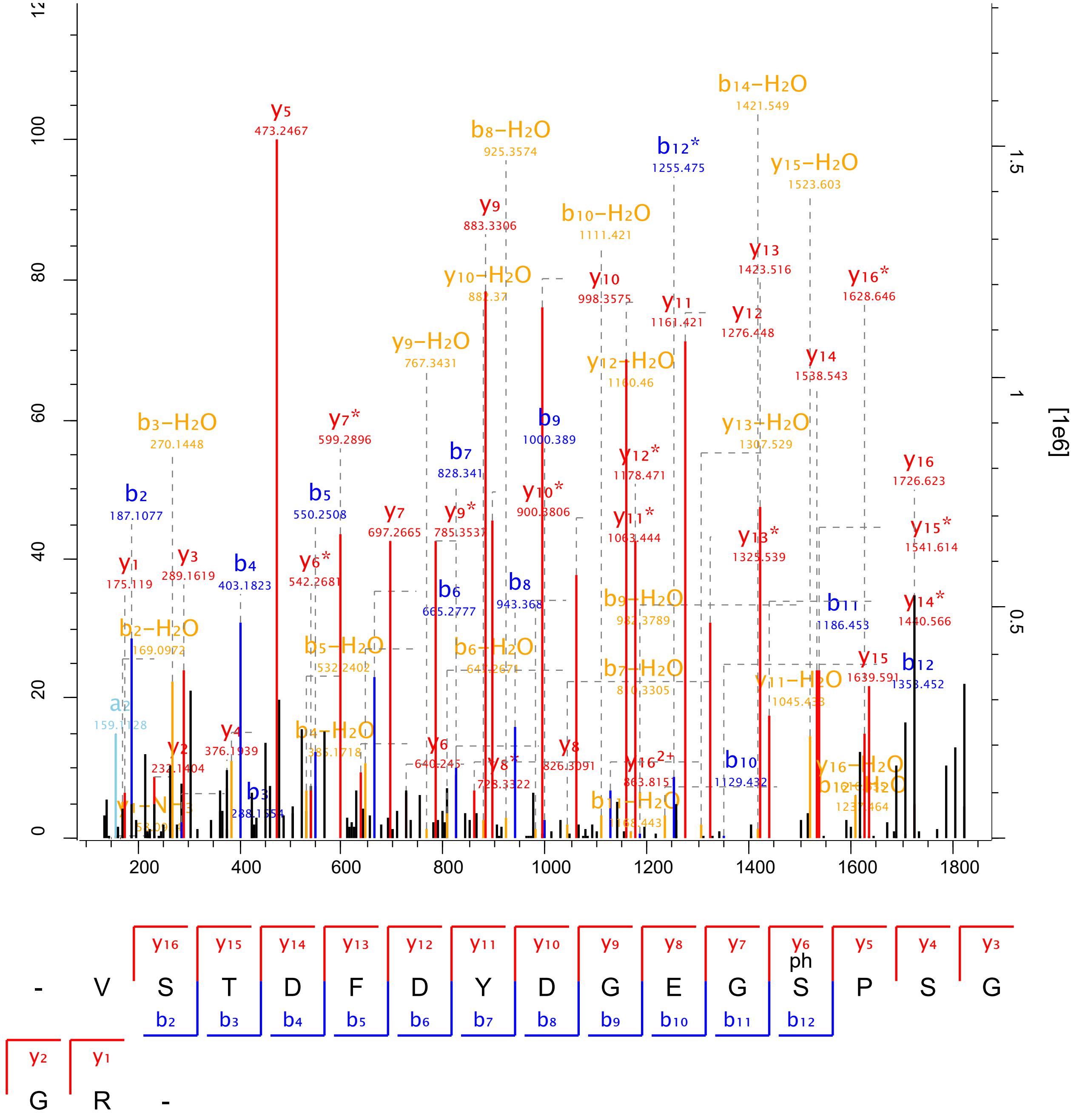

| Raw file | Scan | Method | Score | m/z |
|---|---|---|---|---|
| sys_05_2 | 23651 | FTMS; HCD | 71.88 | 681.81 |

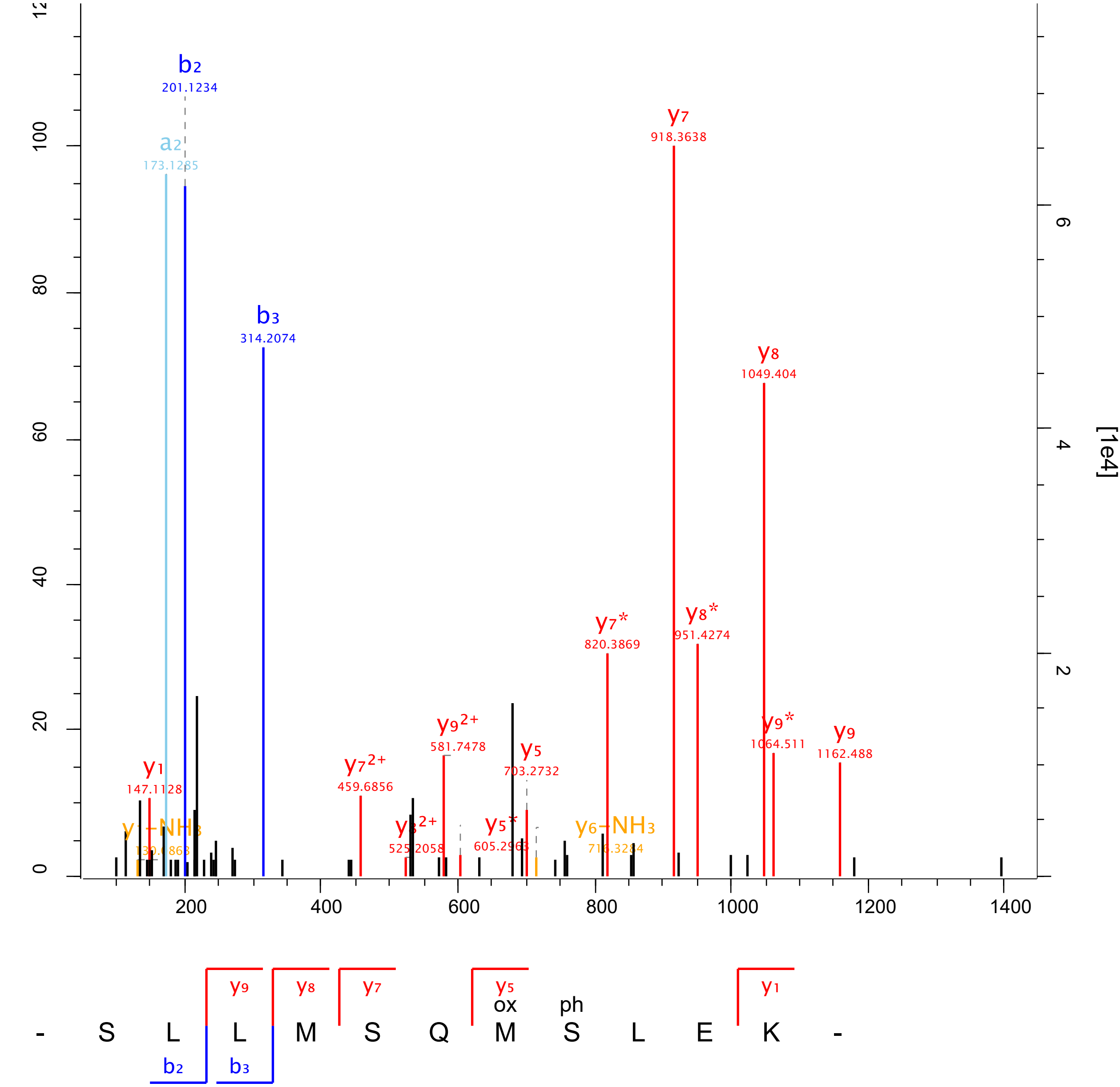

| Raw file | Scan | Method | Score | m/z |
|---|---|---|---|---|
| sys_05_2 | 23668 | FTMS; HCD | 177.57 | 539.24 |

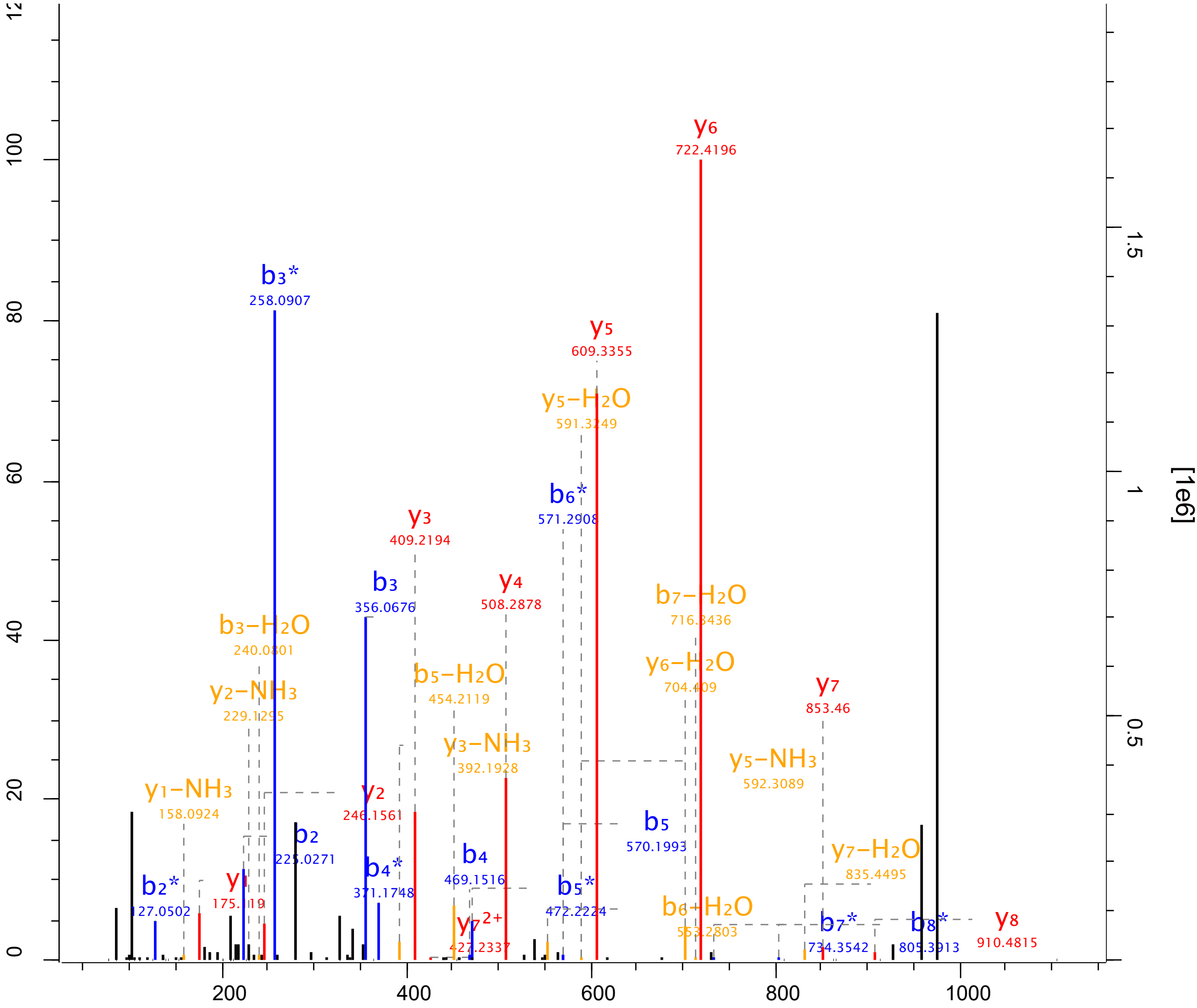


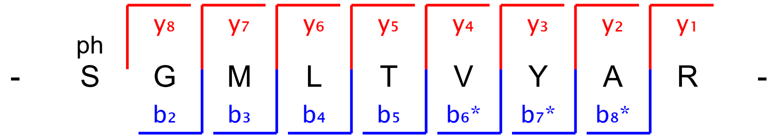

| Raw file | Scan | Method | Score | m/z |
| --- | --- | --- | --- | --- |
| sys_05_2 | 23760 | FTMS; HCD | 48.53 | 620.26 |

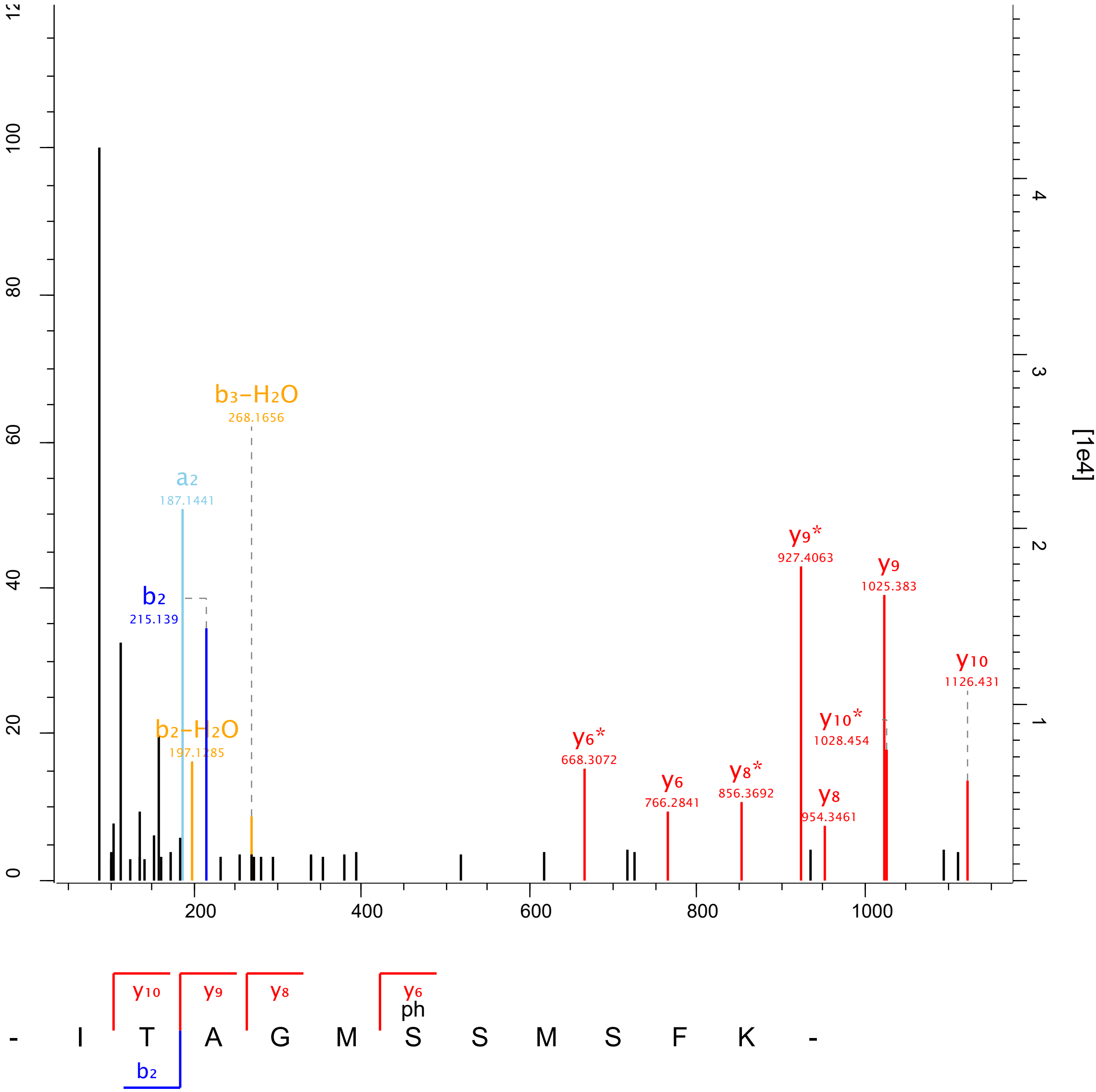

| Raw file | Scan | Method | Score | m/z |
|---|---|---|---|---|
| sys_05_2 | 23844 | FTMS; HCD | 159.26 | 775.38 |

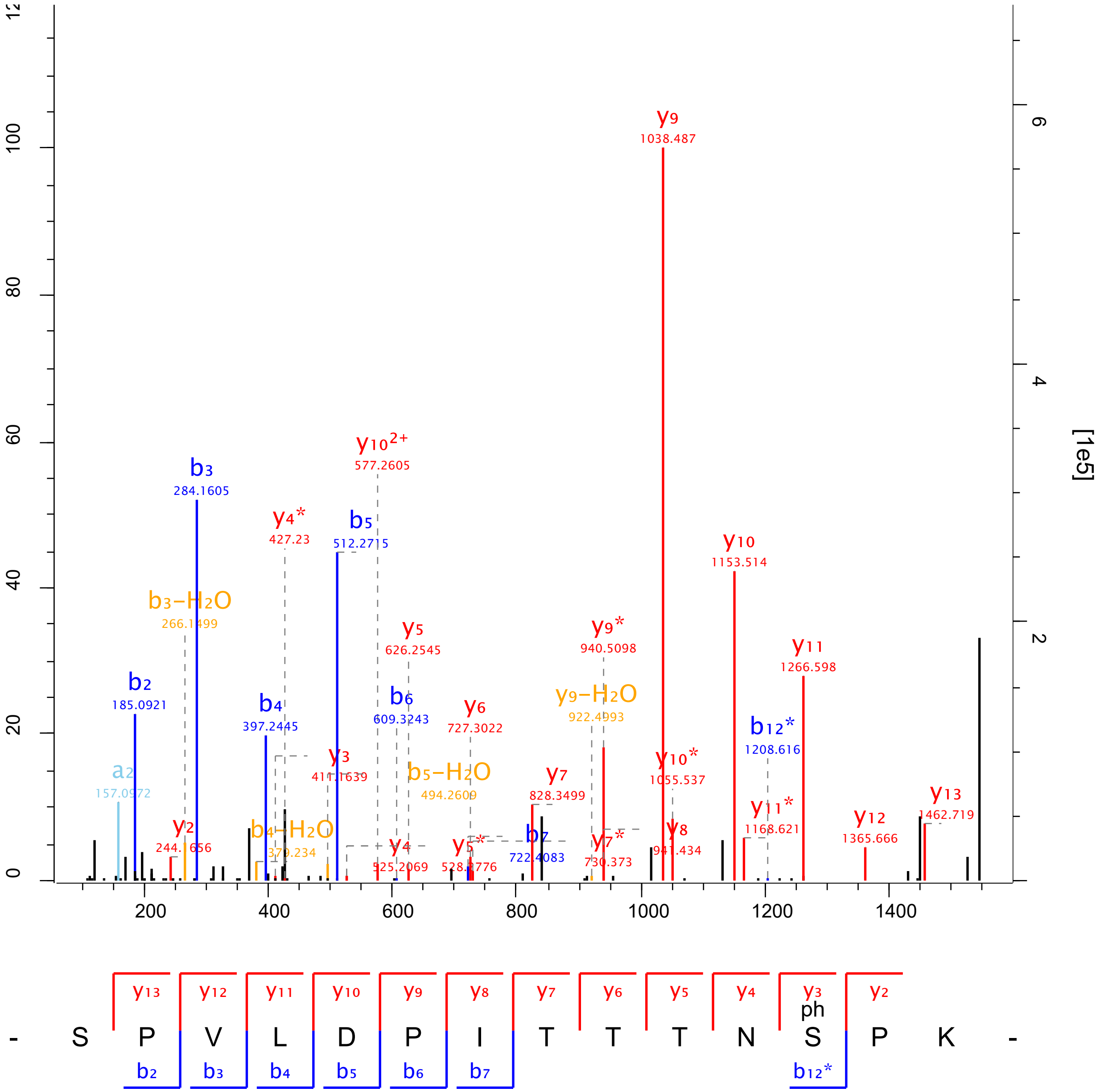

| Raw file | Scan | Method | Score | m/z |
|---|---|---|---|---|
| sys_05_2 | 23852 | FTMS; HCD | 138.05 | 843.89 |

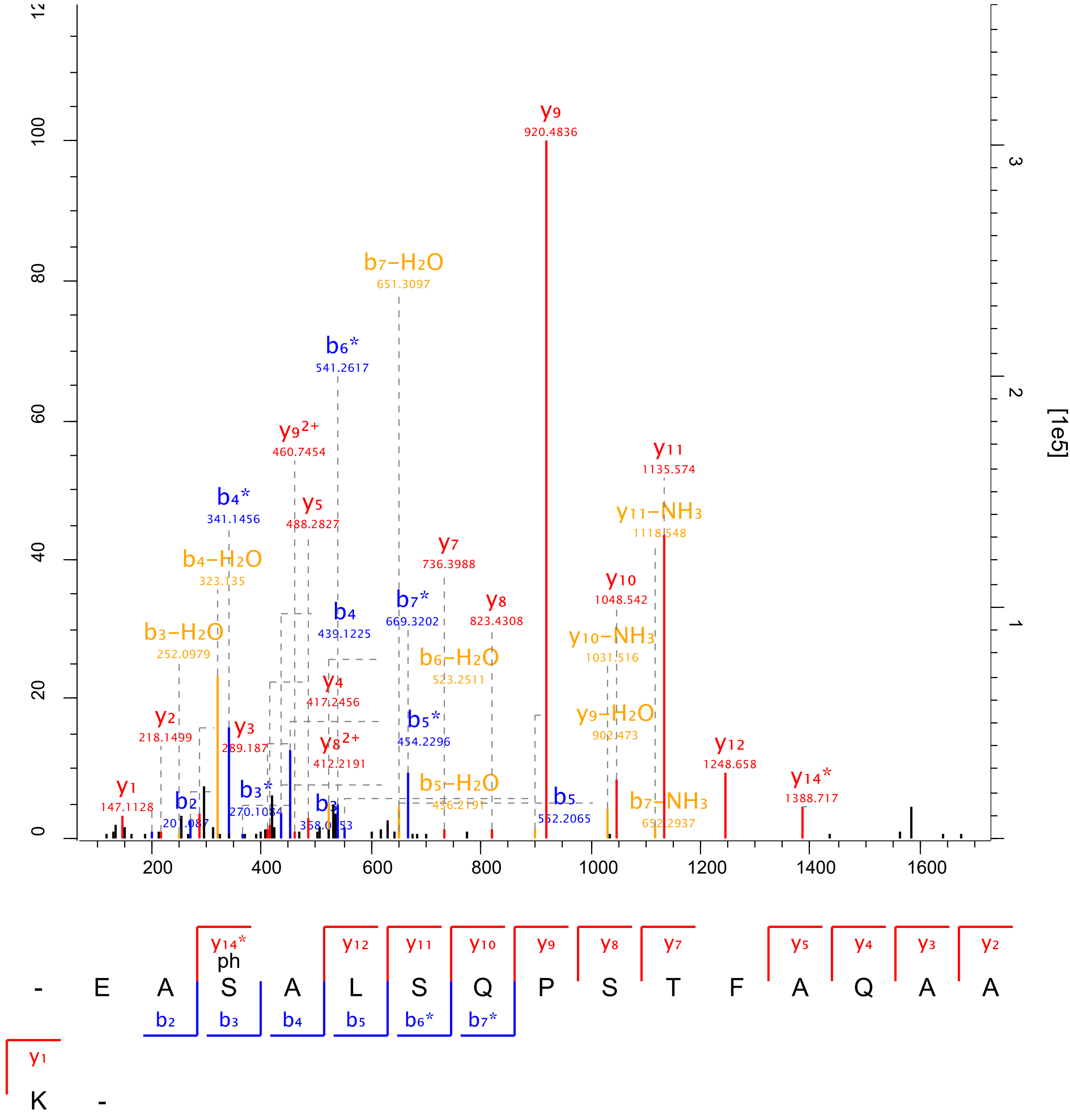

| Raw file | Scan | Method | Score | m/z |
| --- | --- | --- | --- | --- |
| sys_05_2 | 24230 | FTMS; HCD | 123.19 | 701.31 |

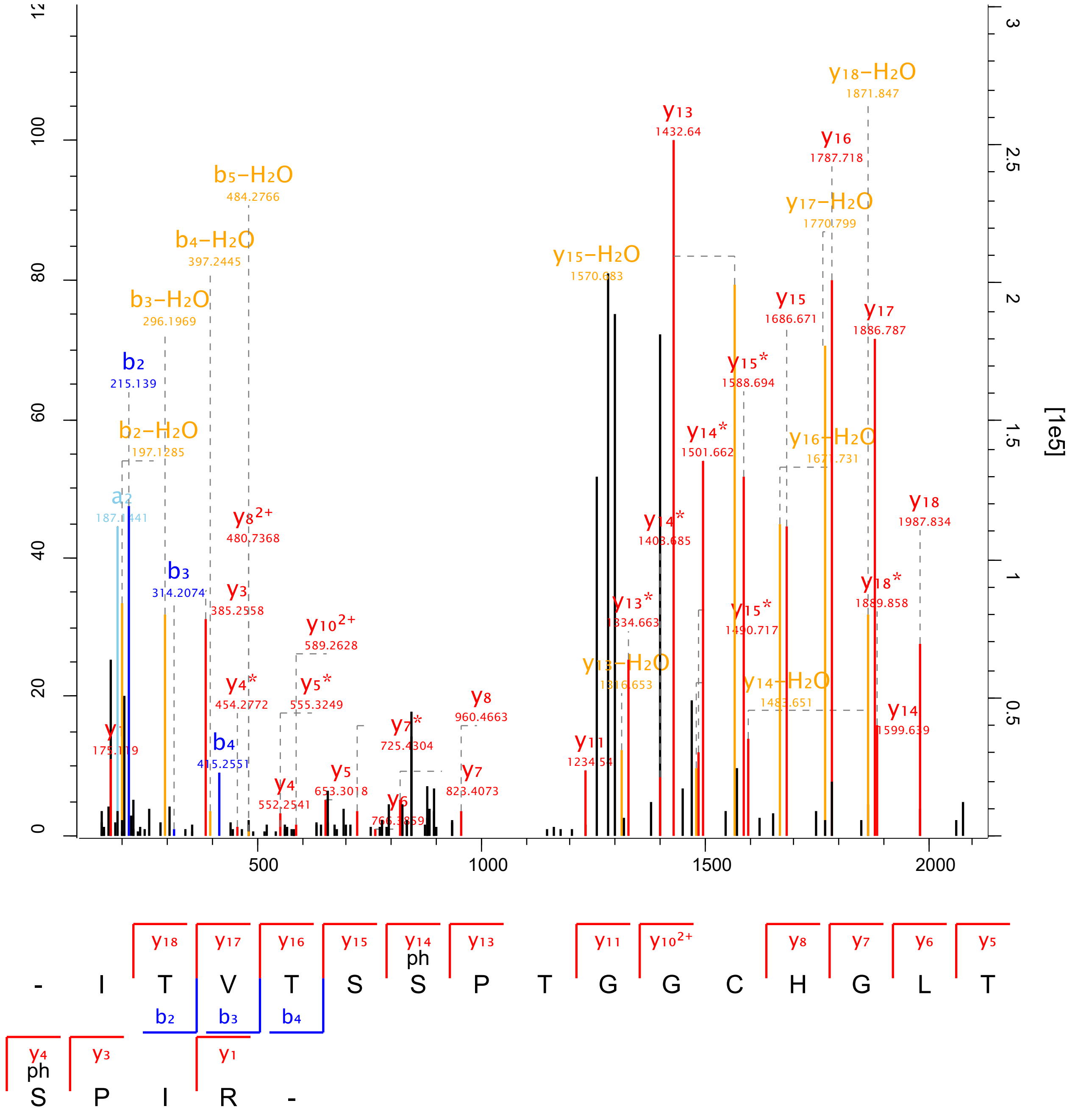

| Raw file | Scan | Method | Score | m/z |
| --- | --- | --- | --- | --- |
| sys_05_2 | 24402 | FTMS; HCD | 127.01 | 885.91 |

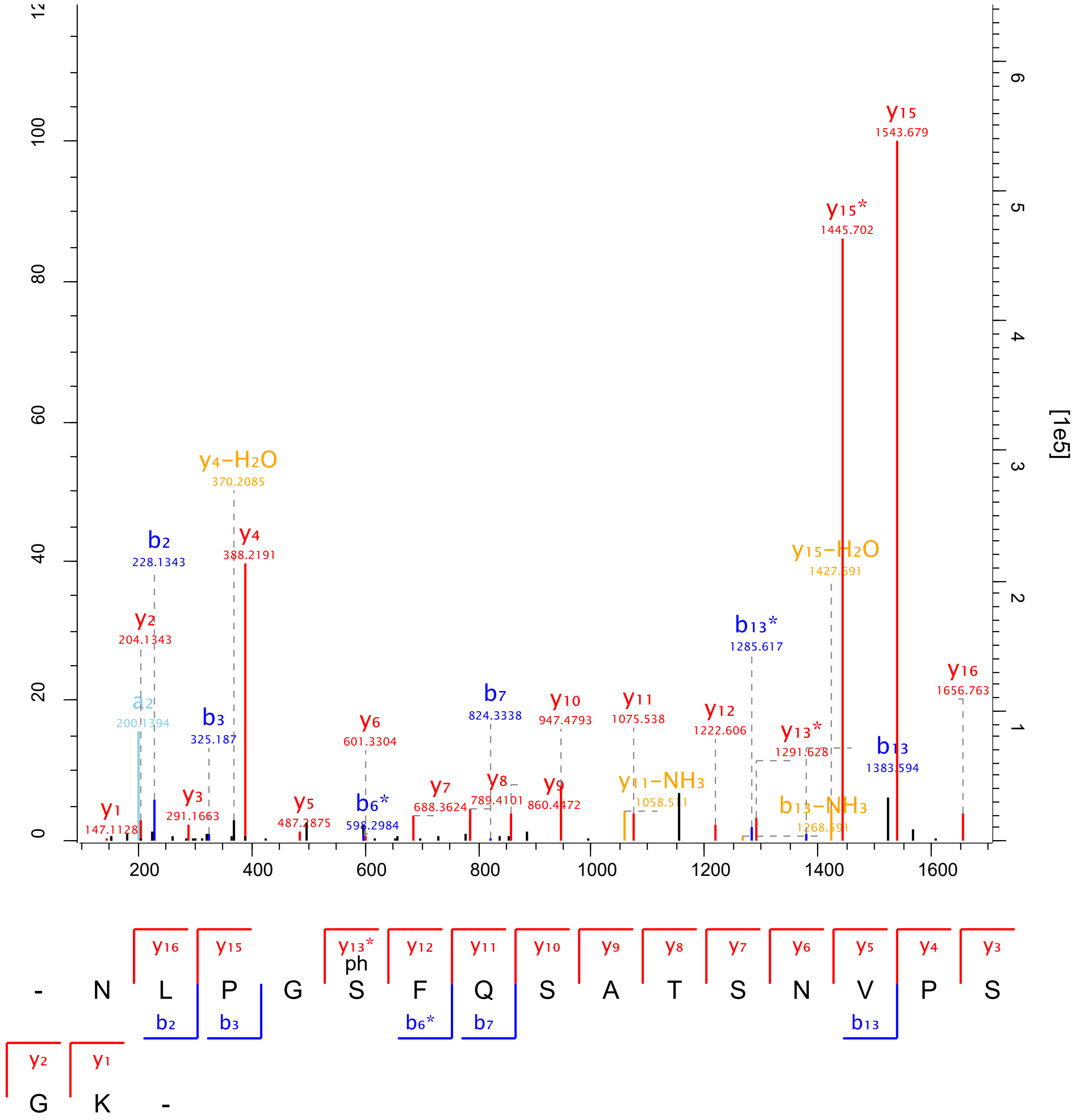

| Raw file | Scan | Method | Score | m/z |
| --- | --- | --- | --- | --- |
| sys_05_2 | 24423 | FTMS; HCD | 46.07 | 599.78 |

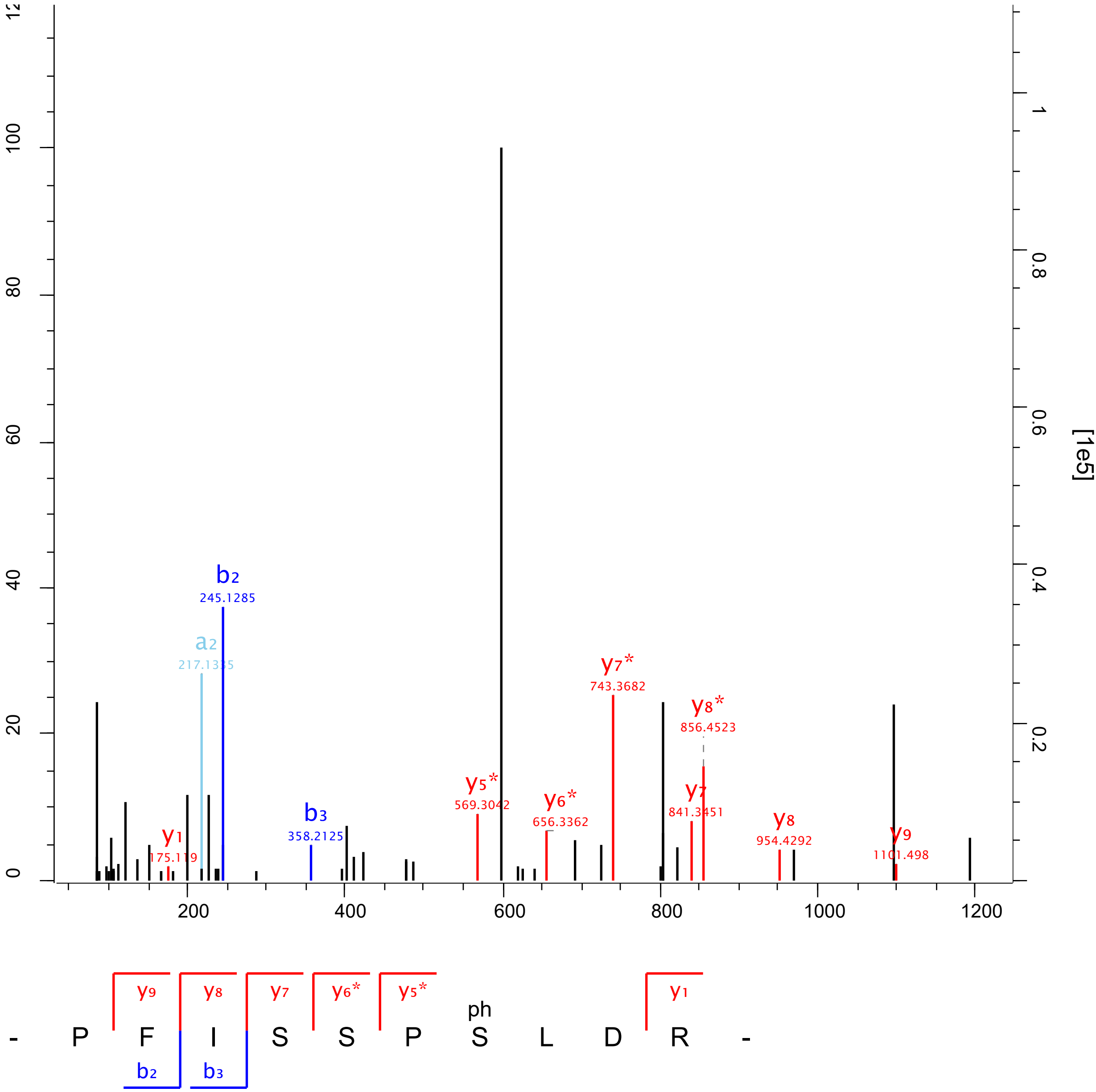

| Raw file | Scan | Method | Score | m/z |
|---|---|---|---|---|
| sys_05_2 | 24616 | FTMS; HCD | 224.42 | 627.3 |

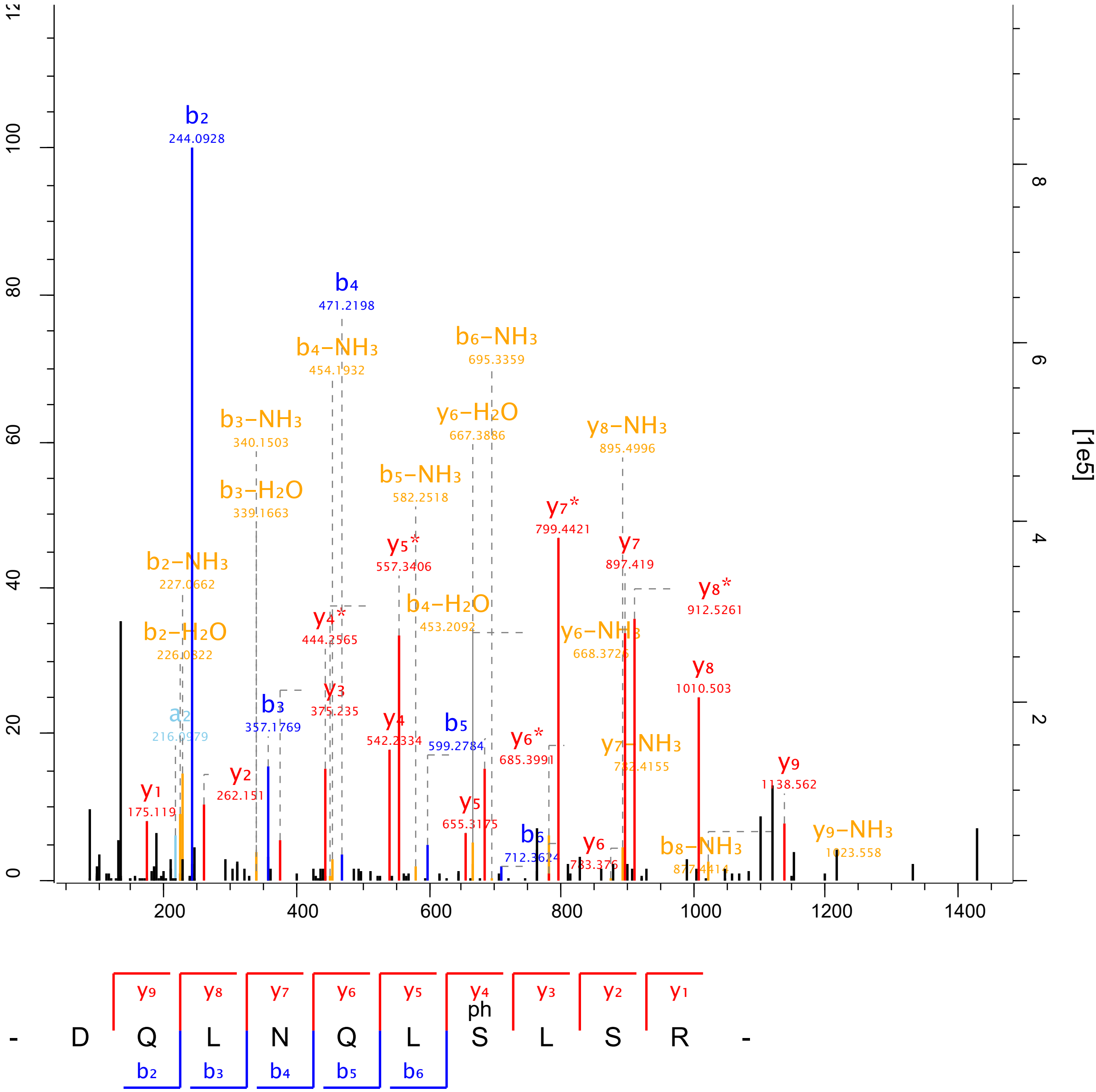

| Raw file | Scan | Method | Score | m/z |
| --- | --- | --- | --- | --- |
| sys_05_2 | 24877 | FTMS; HCD | 105.79 | 890.41 |

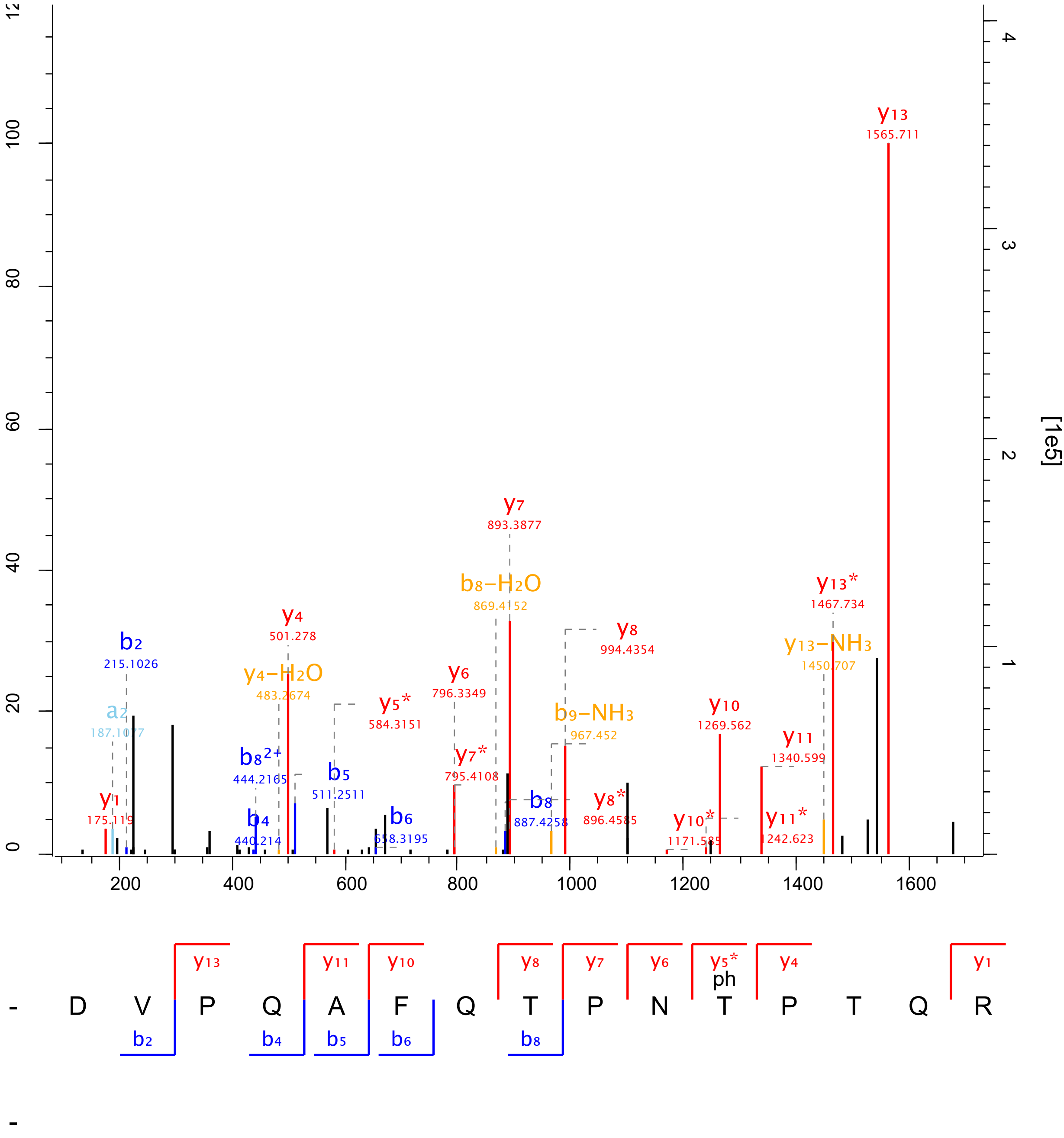

| Raw file | Scan | Method | Score | m/z |
|---|---|---|---|---|
| sys_05_2 | 24919 | FTMS; HCD | 121.82 | 766.32 |

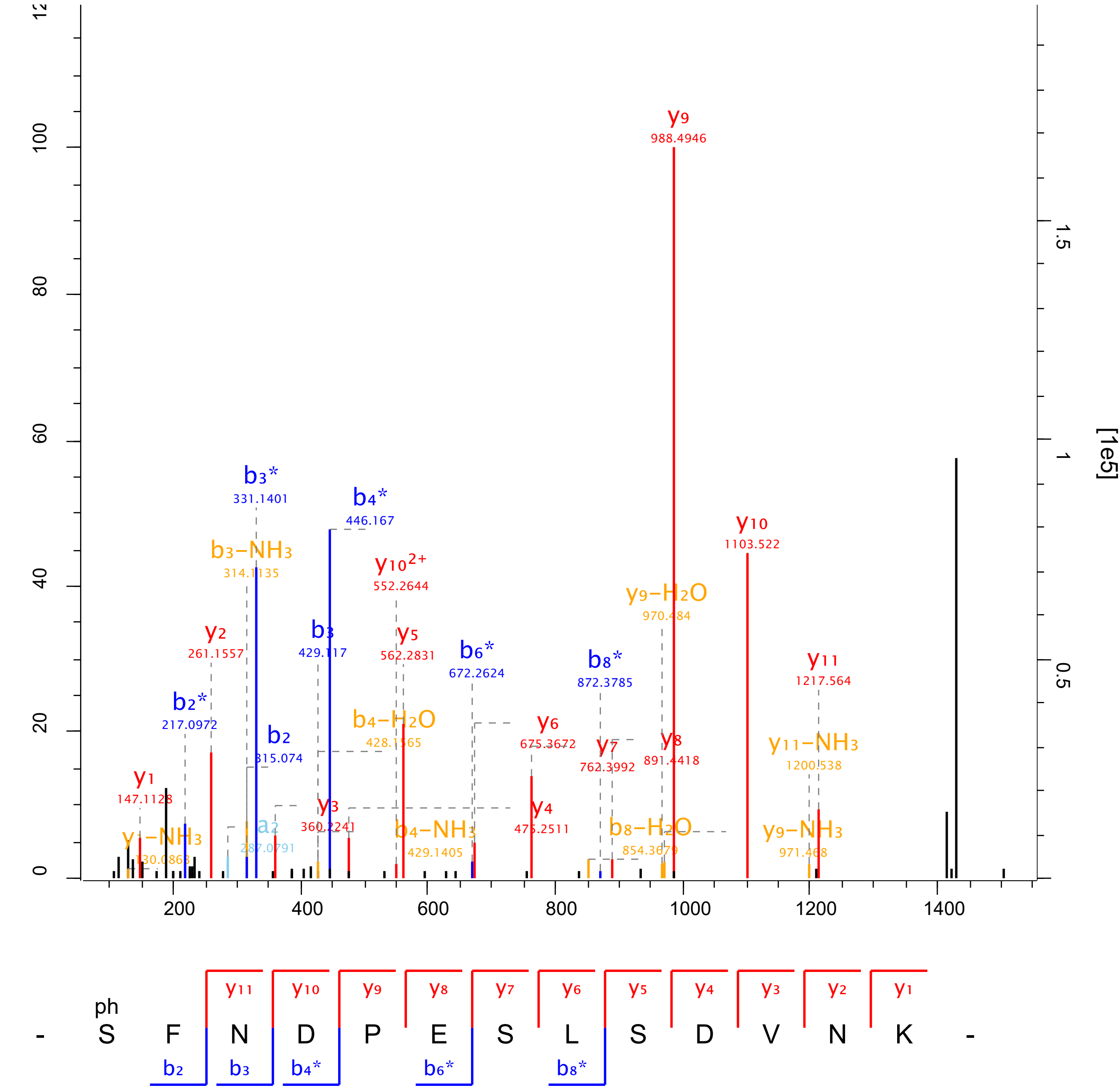

| Raw file | Scan | Method | Score | m/z |
|---|---|---|---|---|
| sys_05_2 | 25152 | FTMS; HCD | 66.81 | 697.86 |

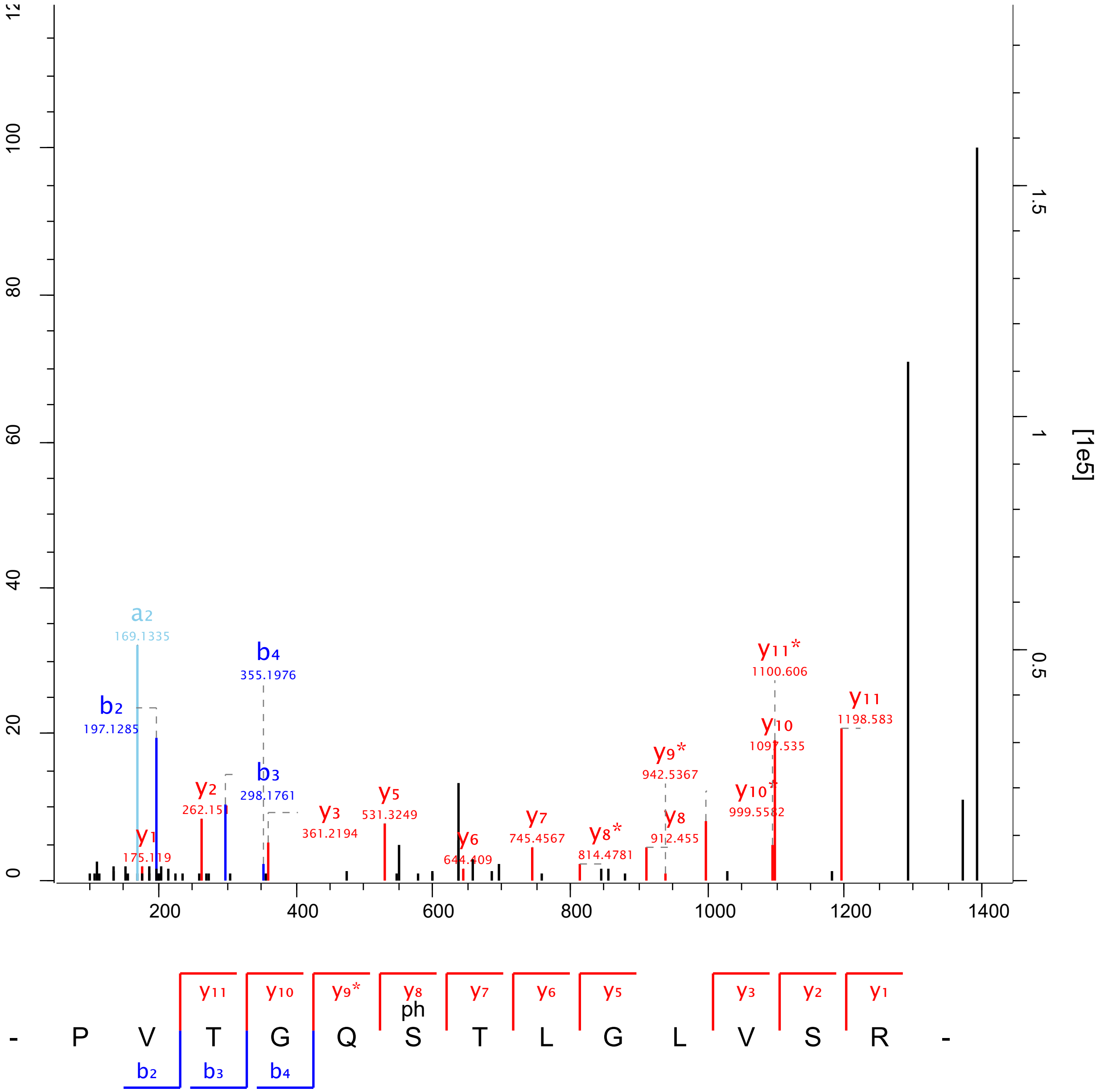

| Raw file | Scan | Method | Score | m/z |
| --- | --- | --- | --- | --- |
| sys_05_2 | 25263 | FTMS; HCD | 287.48 | 919.35 |

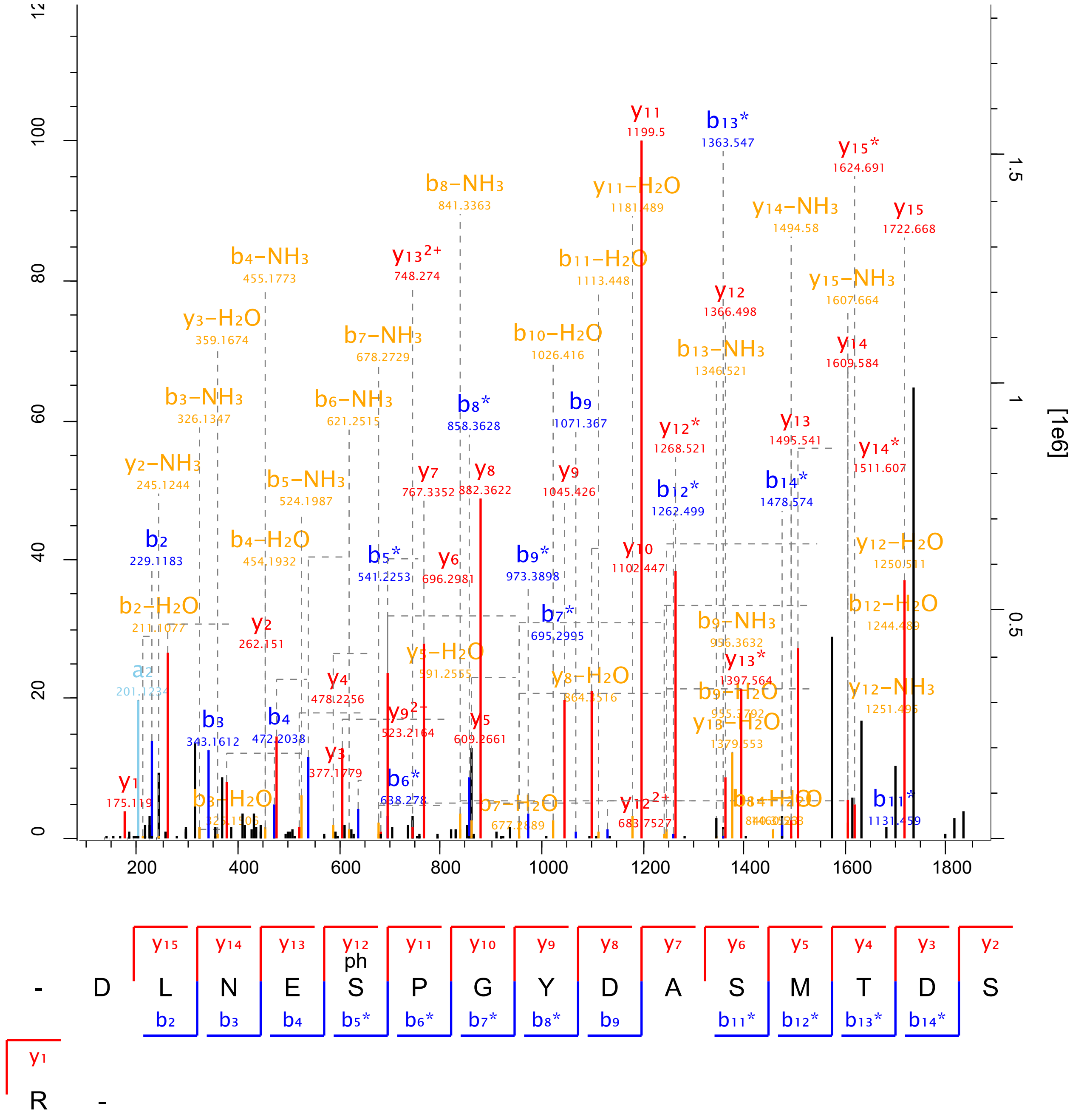

| Raw file | Scan | Method | Score | m/z |
|---|---|---|---|---|
| sys_05_2 | 25469 | FTMS; HCD | 41.62 | 708.66 |

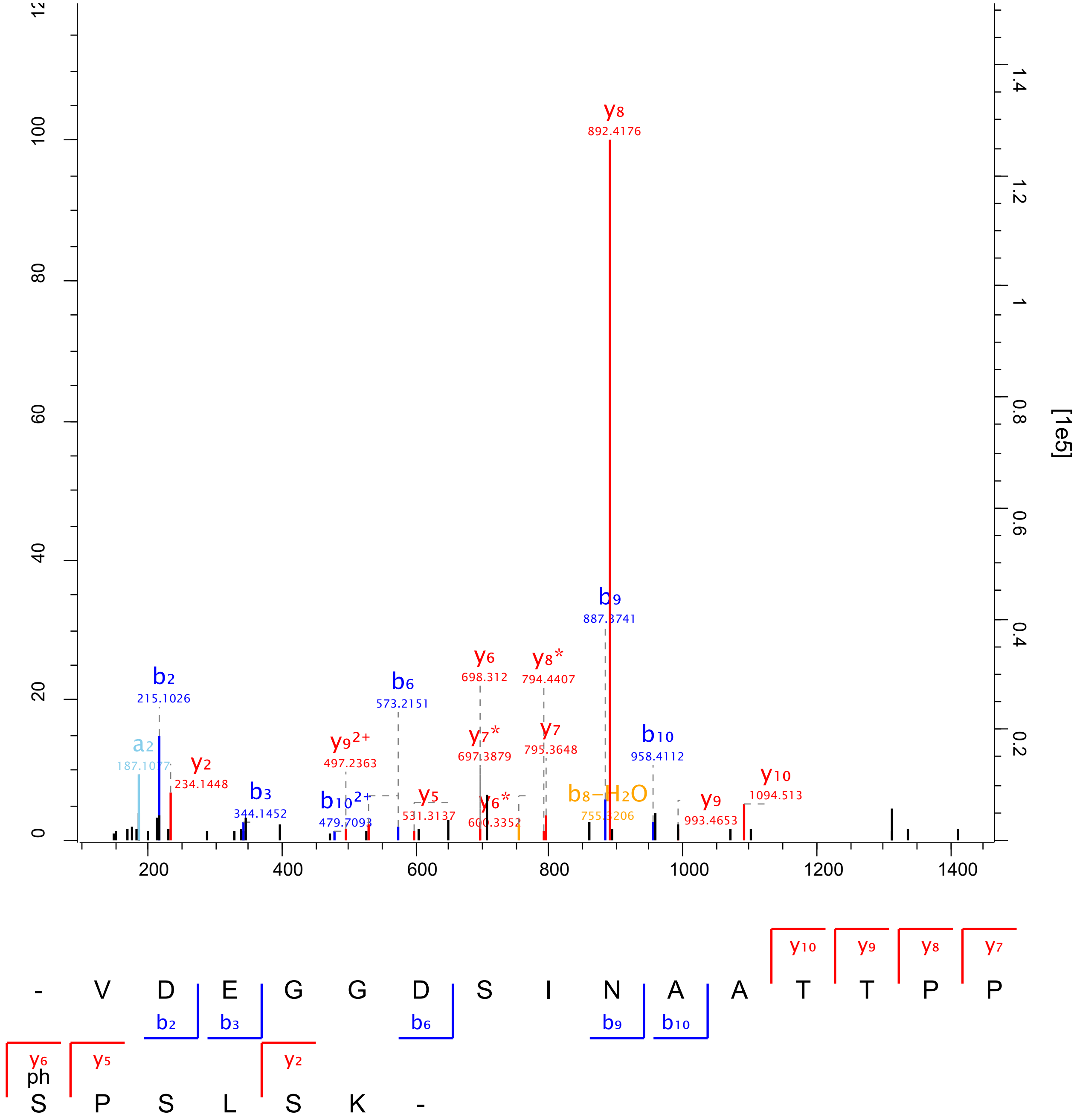

| Raw file | Scan | Method | Score | m/z |
|---|---|---|---|---|
| sys_05_2 | 25476 | FTMS; HCD | 198.67 | 749.36 |

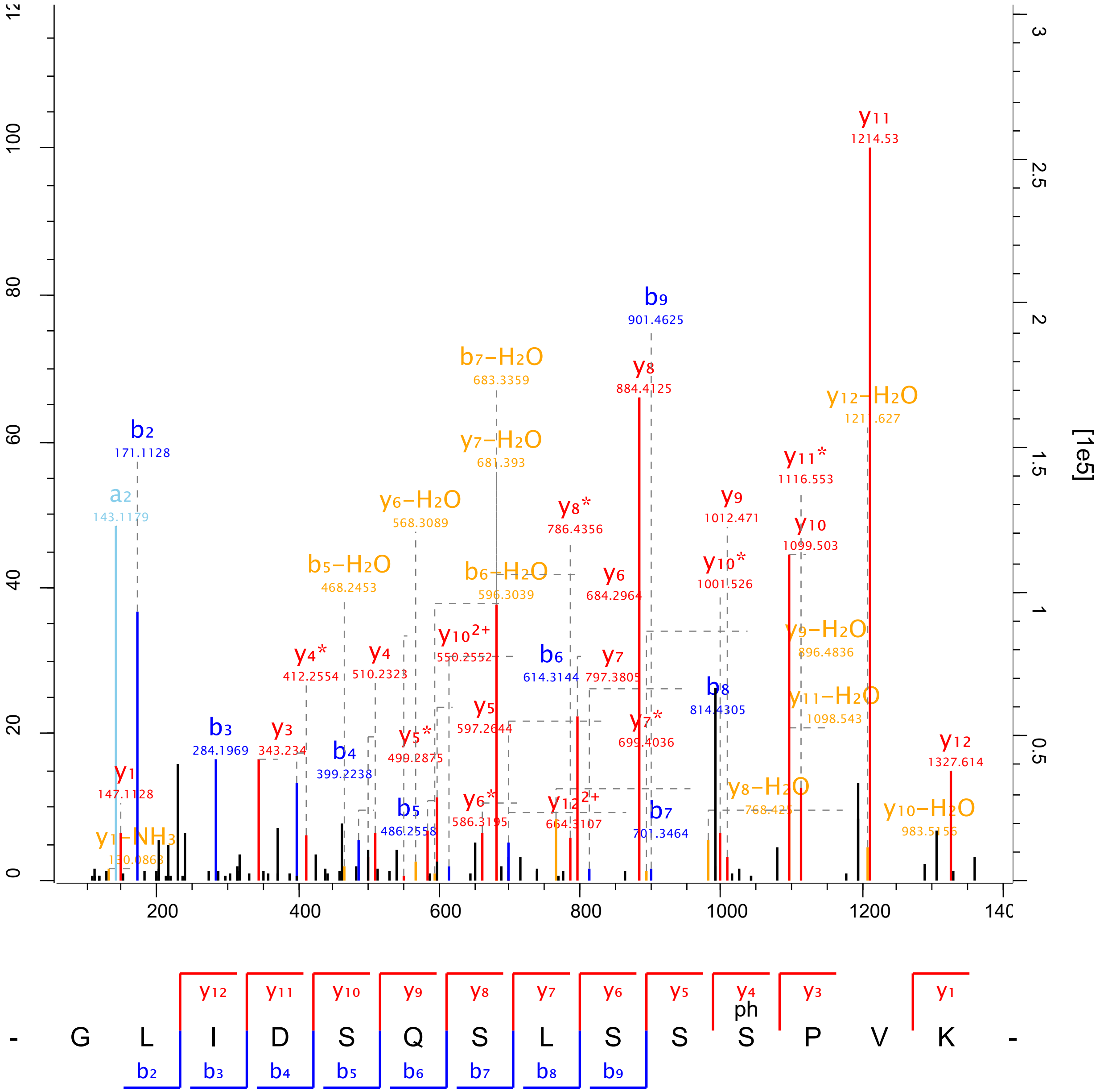

| Raw file | Scan | Method | Score | m/z |
| --- | --- | --- | --- | --- |
| sys_05_2 | 25726 | FTMS; HCD | 103.88 | 682.3 |

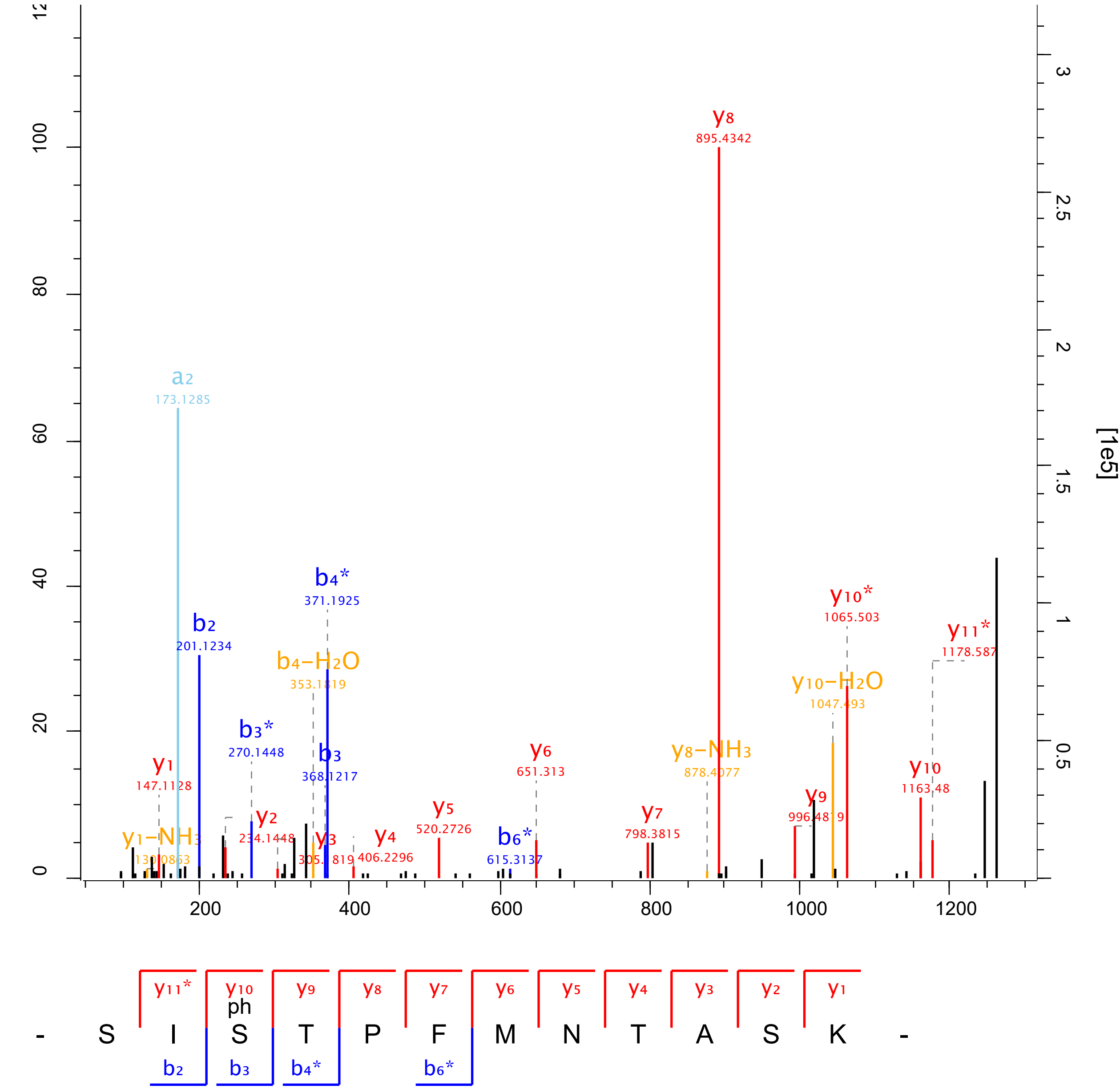

| Raw file | Scan | Method | Score | m/z |
|---|---|---|---|---|
| sys_05_2 | 25902 | FTMS; HCD | 154.72 | 564.76 |

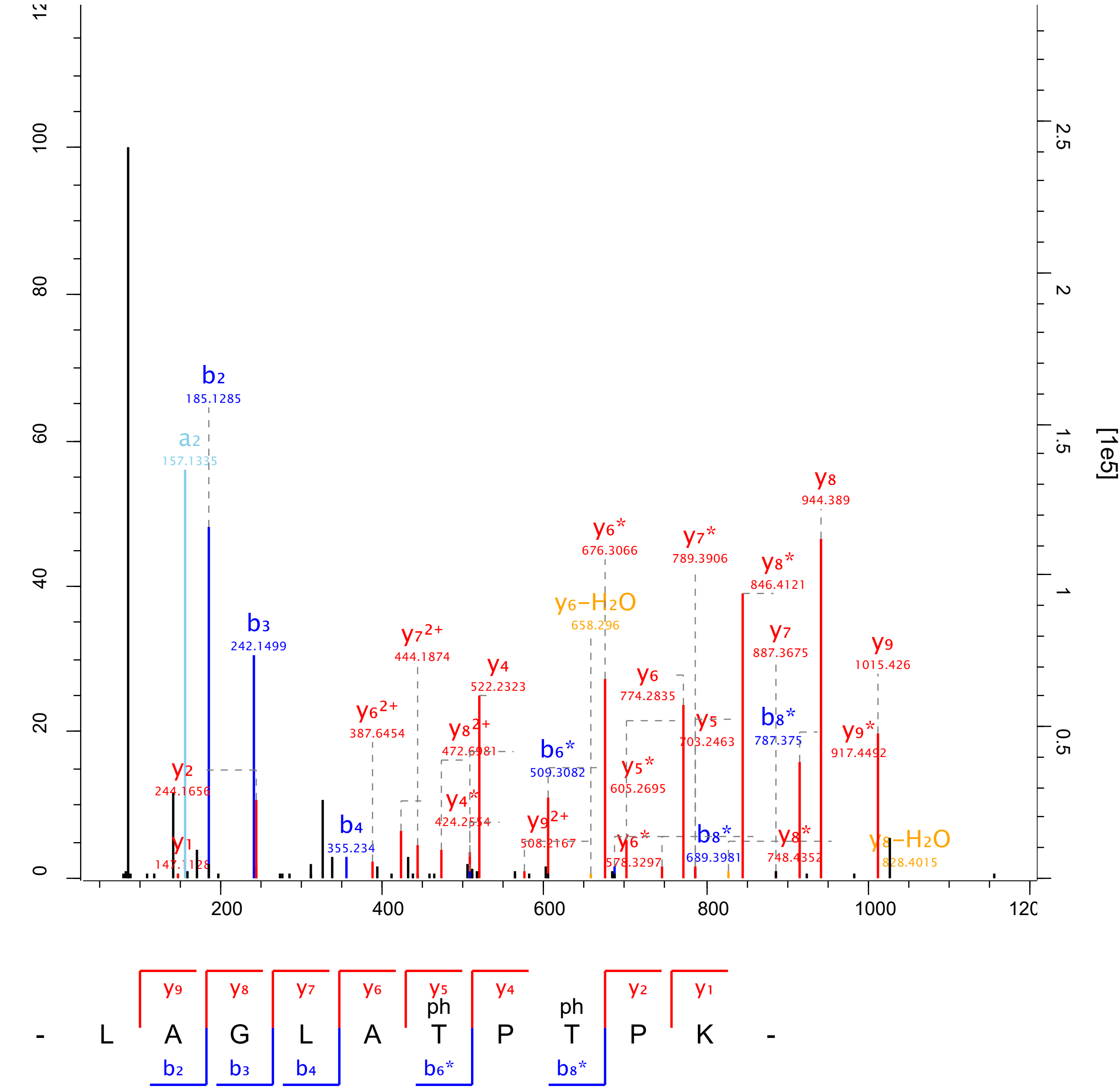

| Raw file | Scan | Method | Score | m/z |
|---|---|---|---|---|
| sys_05_2 | 25944 | FTMS; HCD | 96.77 | 647.76 |

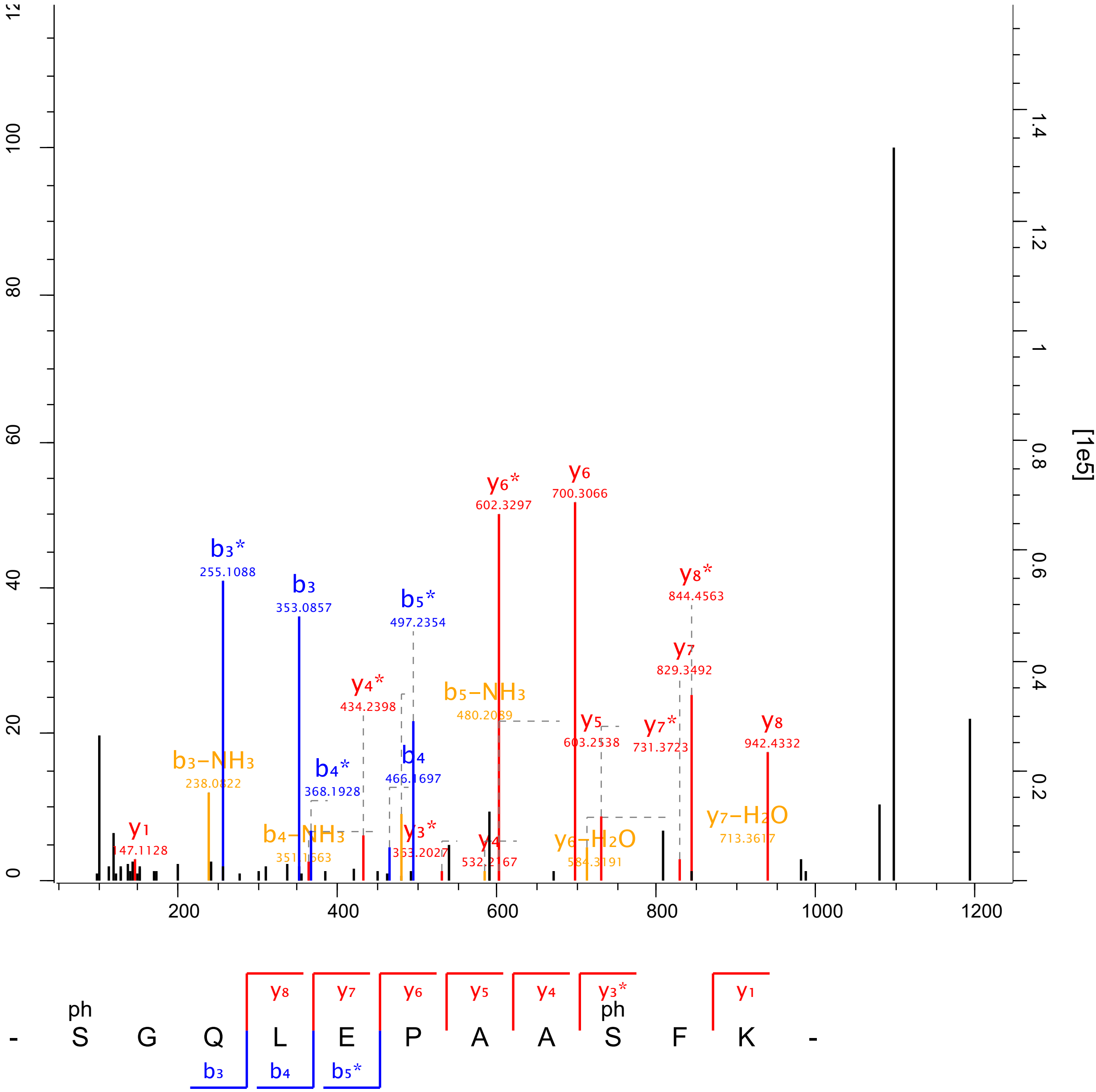

| Raw file | Scan | Method | Score | m/z |
|---|---|---|---|---|
| sys_05_2 | 26024 | FTMS; HCD | 67.58 | 967.93 |

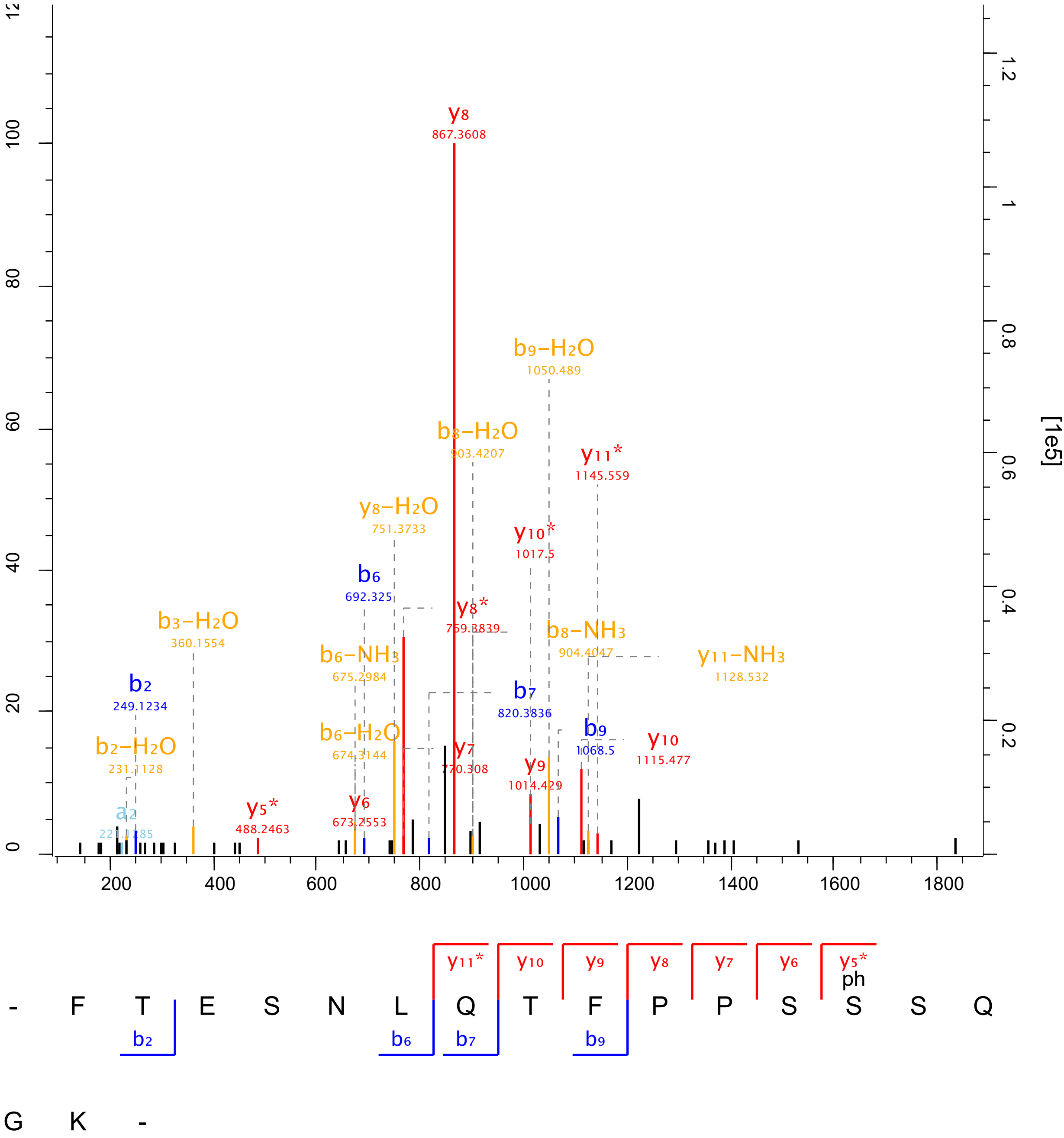

| Raw file | Scan | Method | Score | m/z |
|---|---|---|---|---|
| sys_05_2 | 26246 | FTMS; HCD | 251.02 | 555.78 |

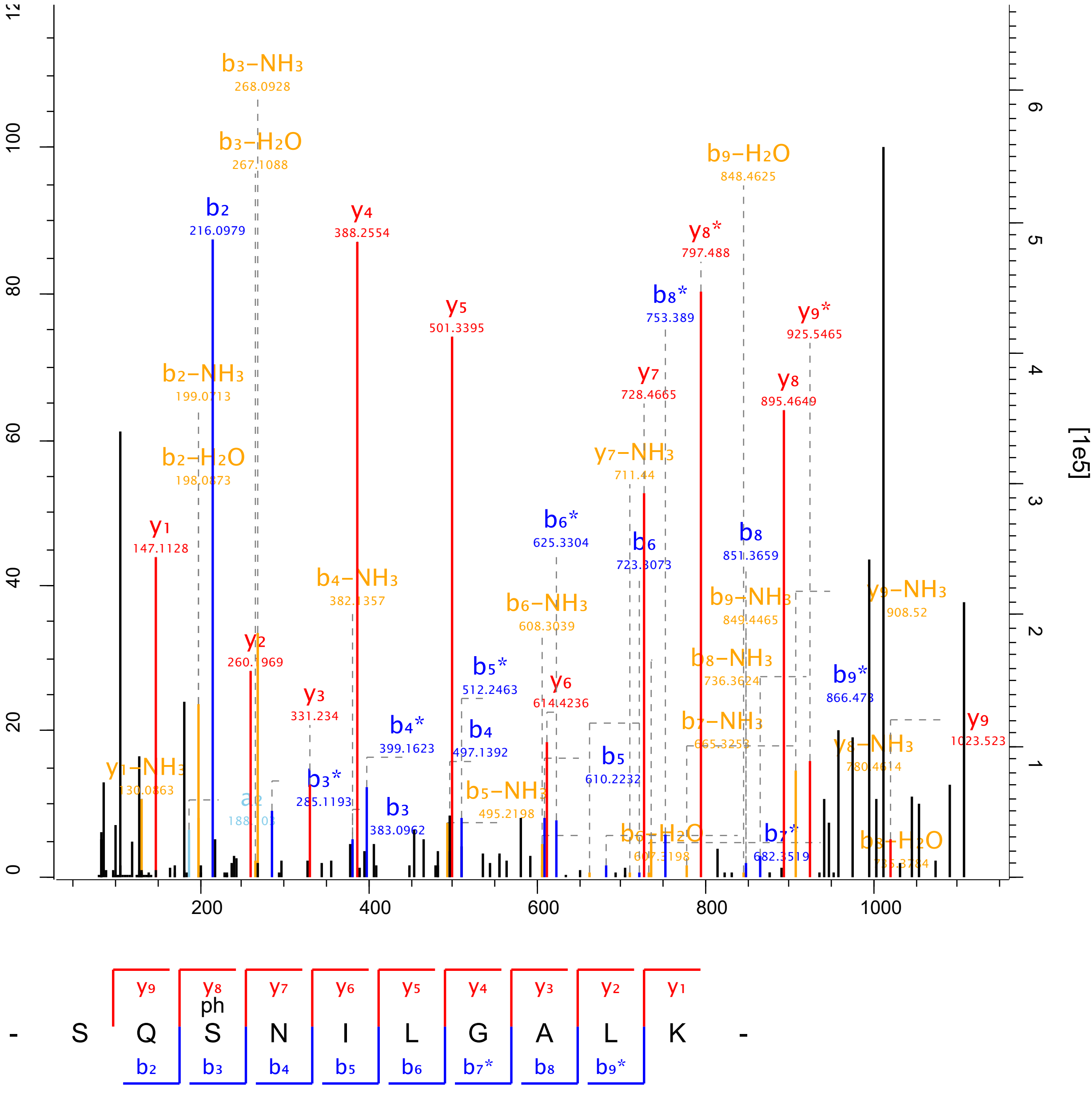

| Raw file | Scan | Method | Score | m/z |
|---|---|---|---|---|
| sys_05_2 | 26258 | FTMS; HCD | 247.5 | 832.36 |

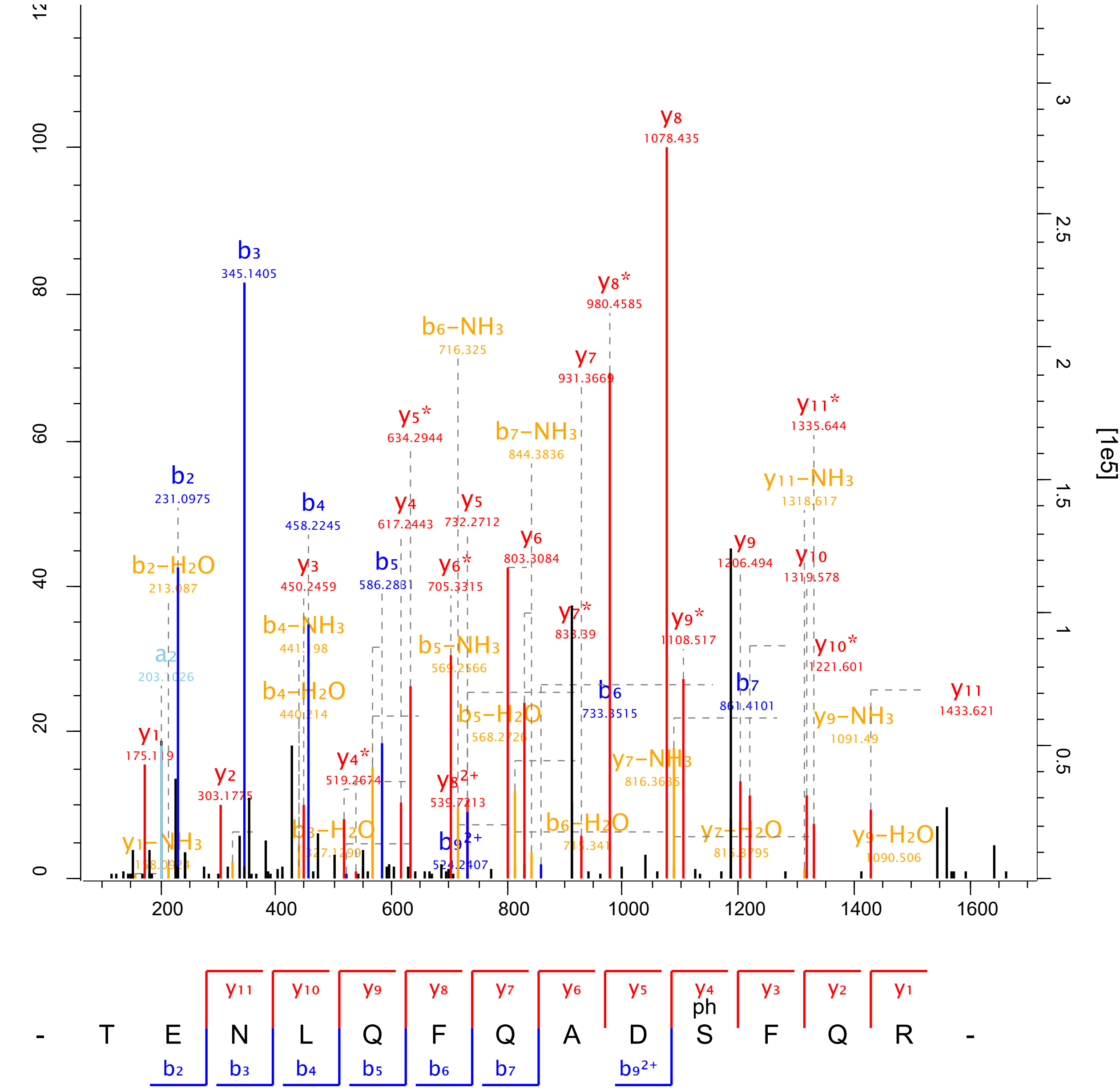

| Raw file | Scan | Method | Score | m/z |
|---|---|---|---|---|
| sys_05_2 | 26297 | FTMS; HCD | 127.37 | 526.2 |

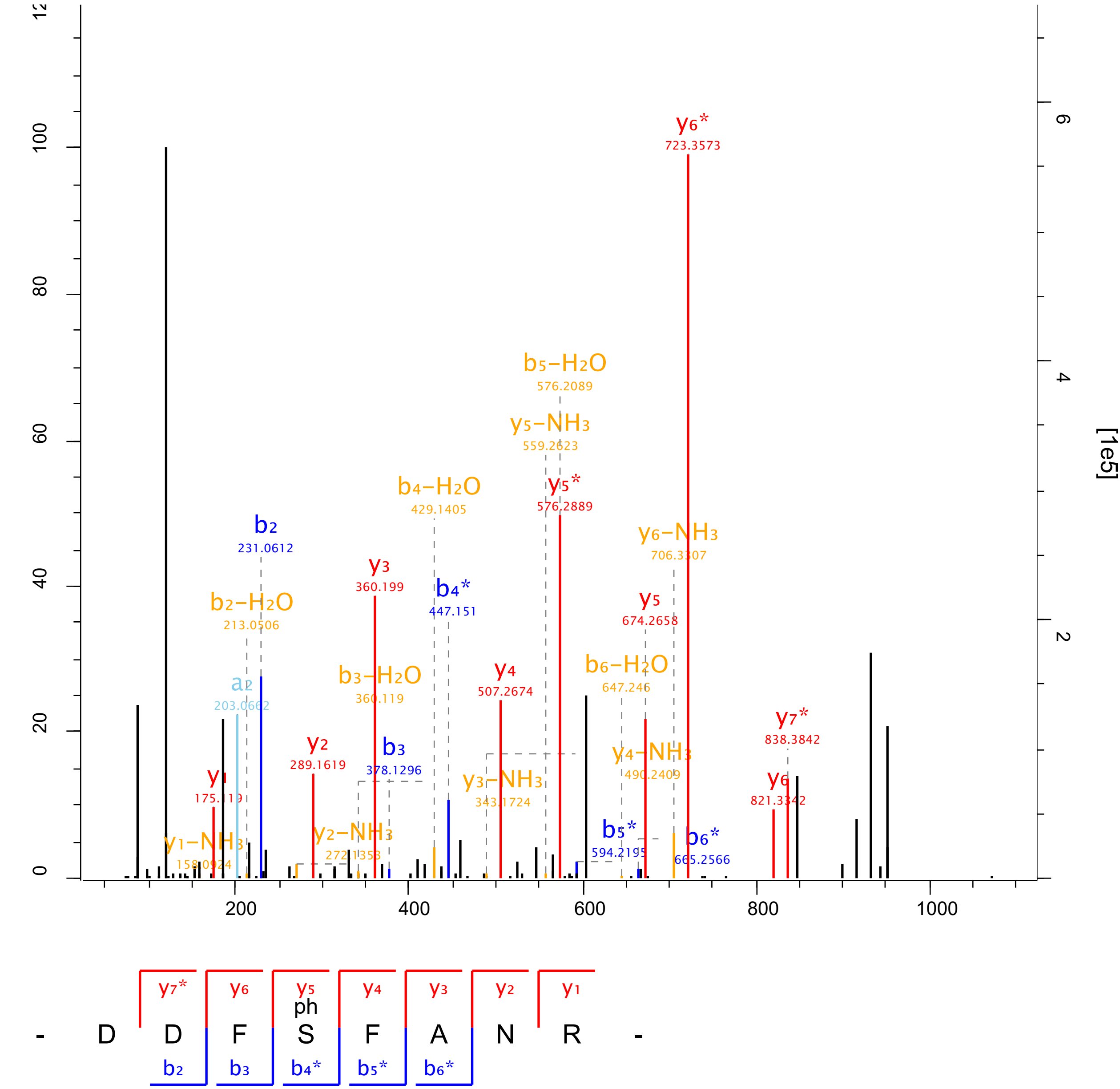

| Raw file | Scan | Method | Score | m/z |
|---|---|---|---|---|
| sys_05_2 | 26304 | FTMS; HCD | 180.86 | 889.84 |

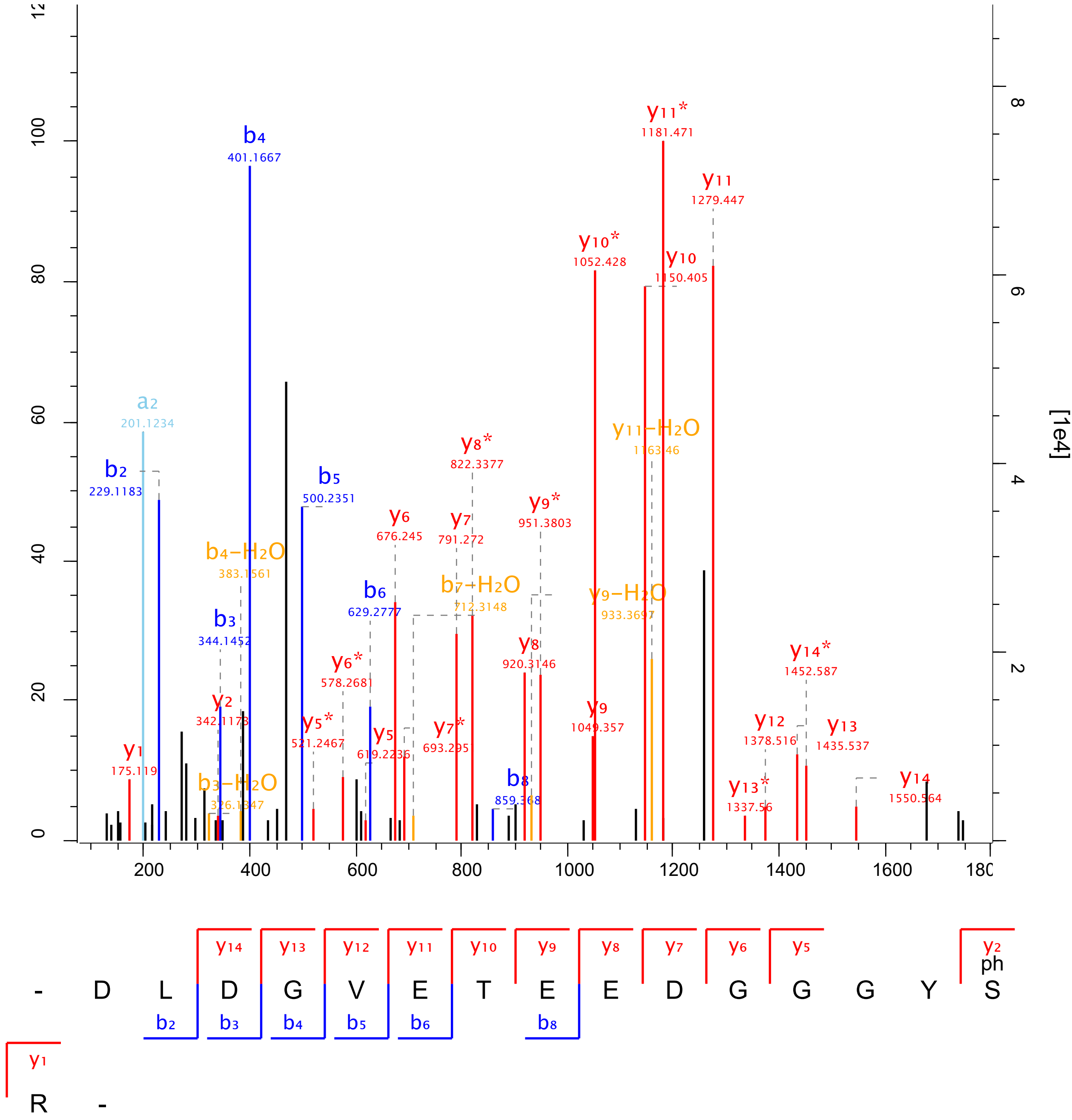

| Raw file | Scan | Method | Score | m/z |
|---|---|---|---|---|
| sys_05_2 | 26667 | FTMS; HCD | 42 | 583.75 |

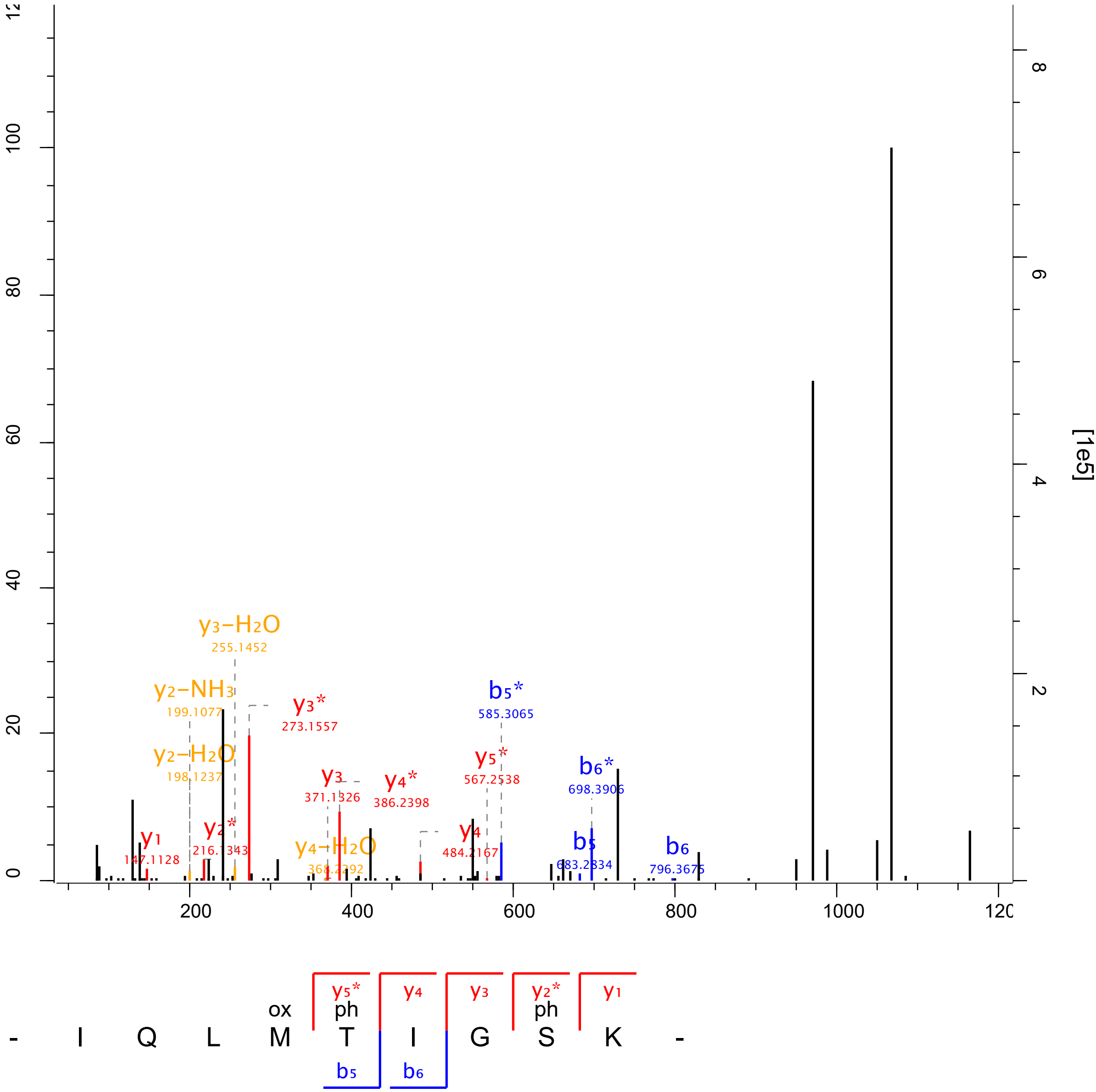

| Raw file | Scan | Method | Score | m/z |
| --- | --- | --- | --- | --- |
| sys_05_2 | 26767 | FTMS; HCD | 97.64 | 478.71 |

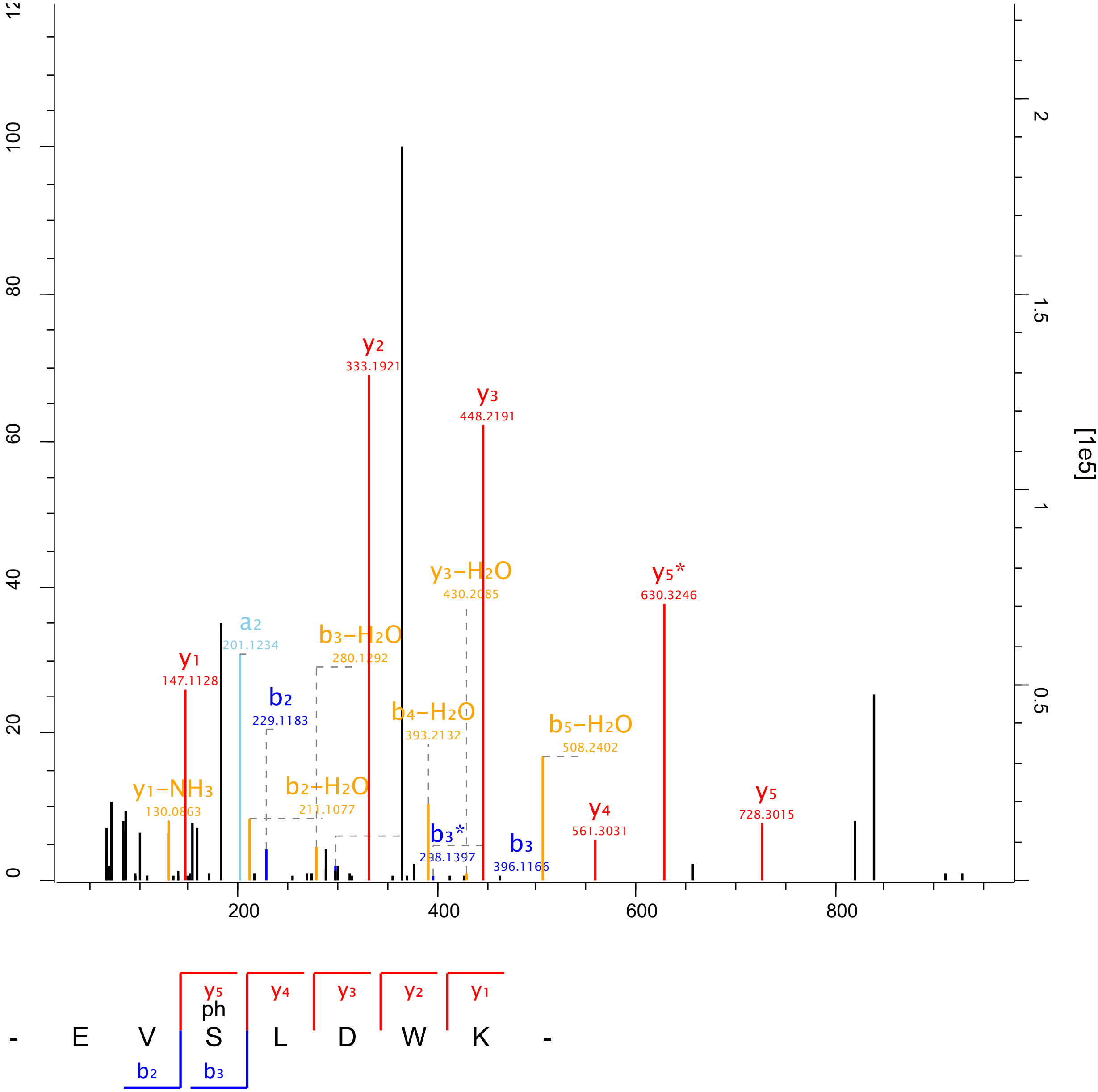

| Raw file | Scan | Method | Score | m/z |
|---|---|---|---|---|
| sys_05_2 | 26806 | FTMS; HCD | 40.28 | 671.83 |

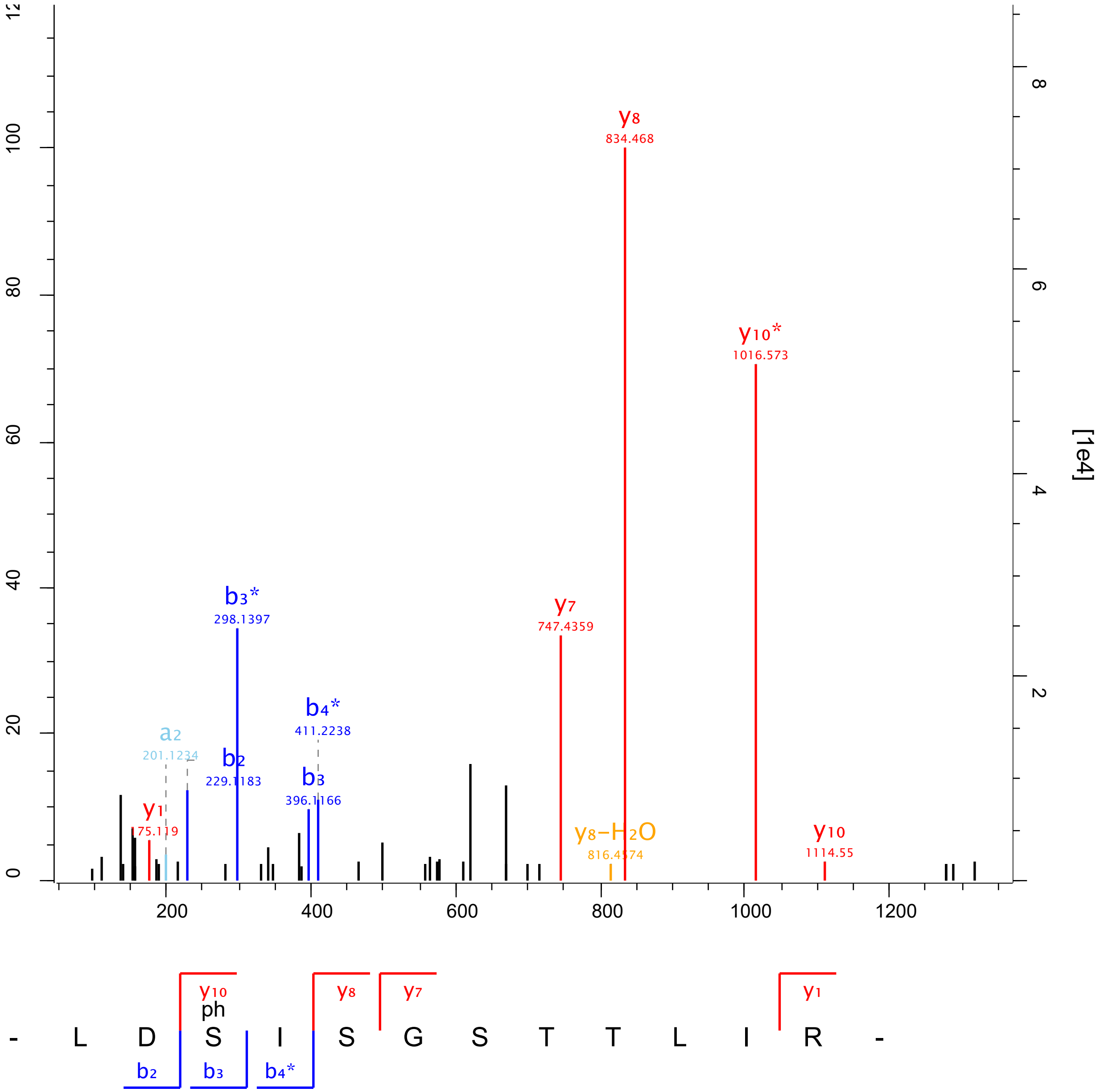

| Raw file | Scan | Method | Score | m/z |
|---|---|---|---|---|
| sys_05_2 | 26856 | FTMS; HCD | 106.38 | 752.78 |

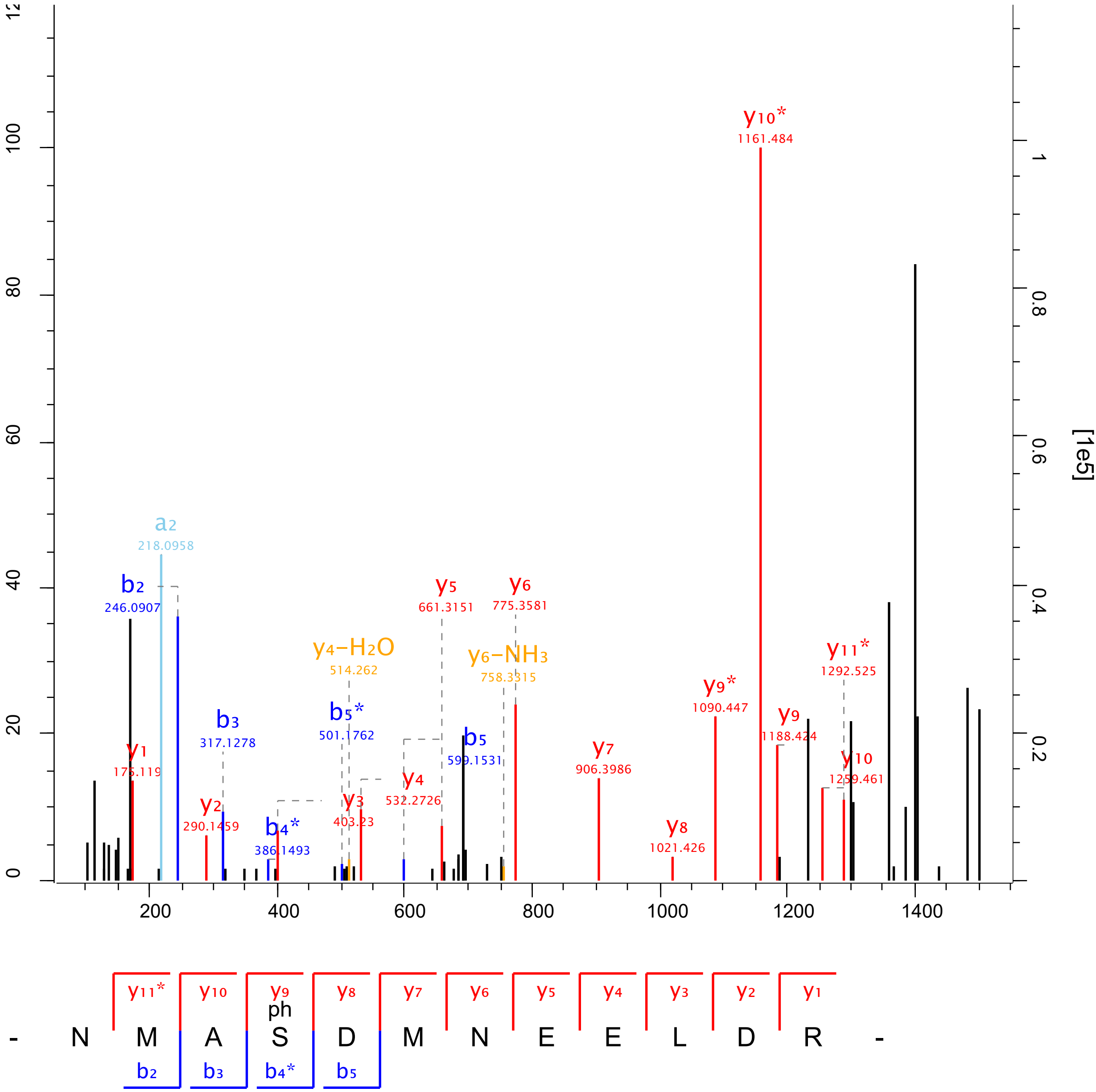

| Raw file | Scan | Method | Score | m/z |
|---|---|---|---|---|
| sys_05_2 | 26898 | FTMS; HCD | 118.54 | 871.4 |

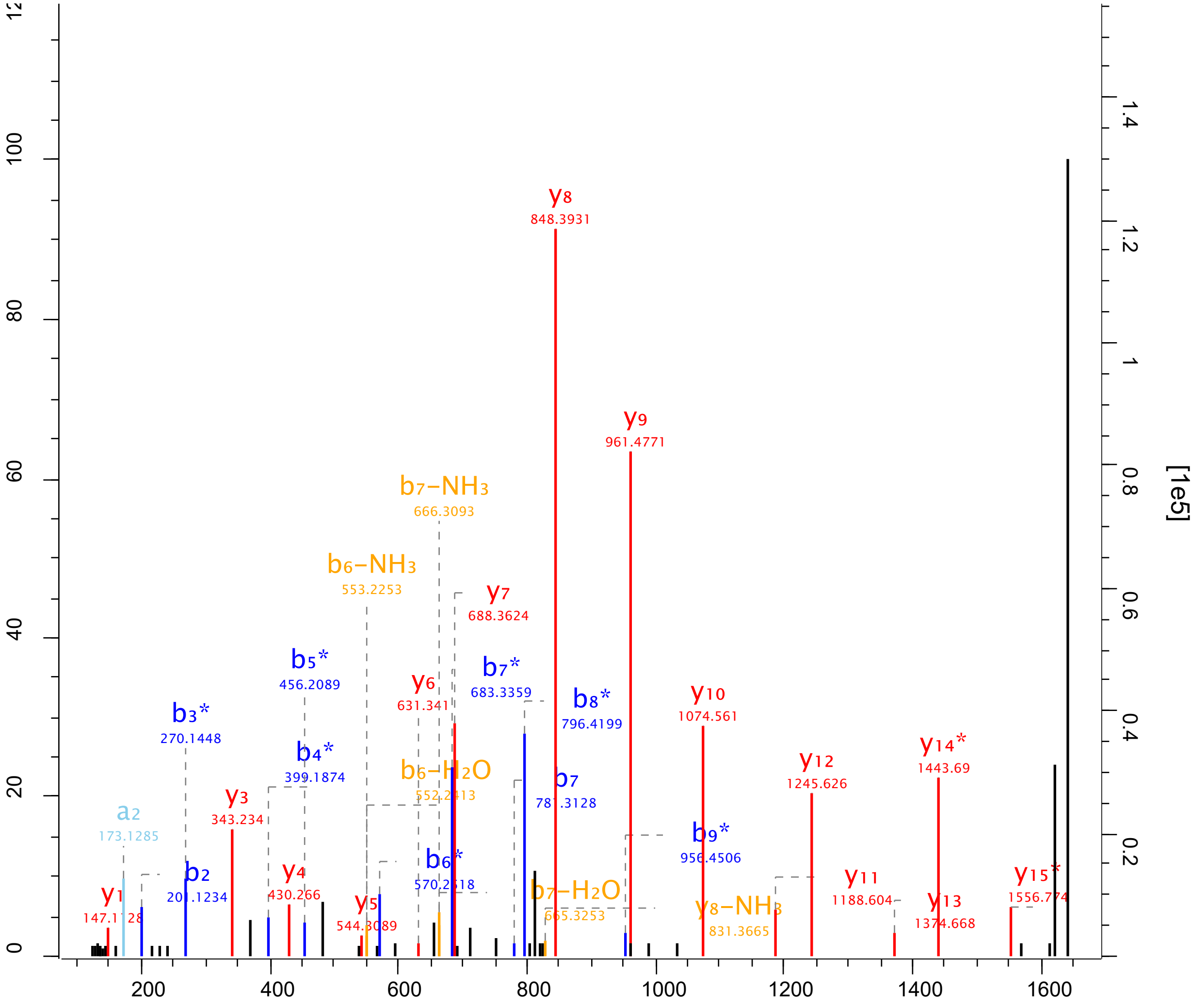


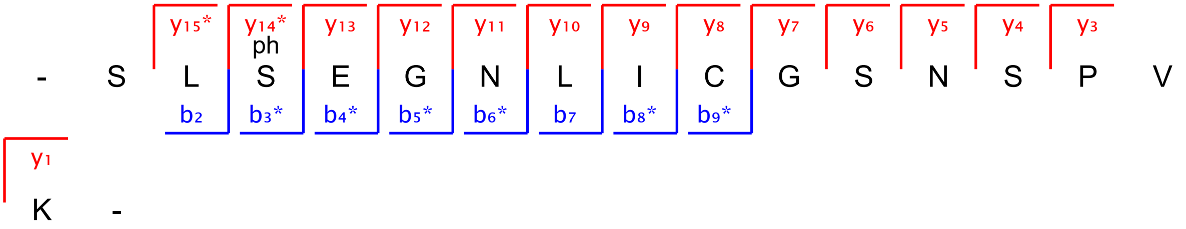

| Raw file | Scan | Method | Score | m/z |
| --- | --- | --- | --- | --- |
| sys_05_2 | 27043 | FTMS; HCD | 204.09 | 1151.45 |

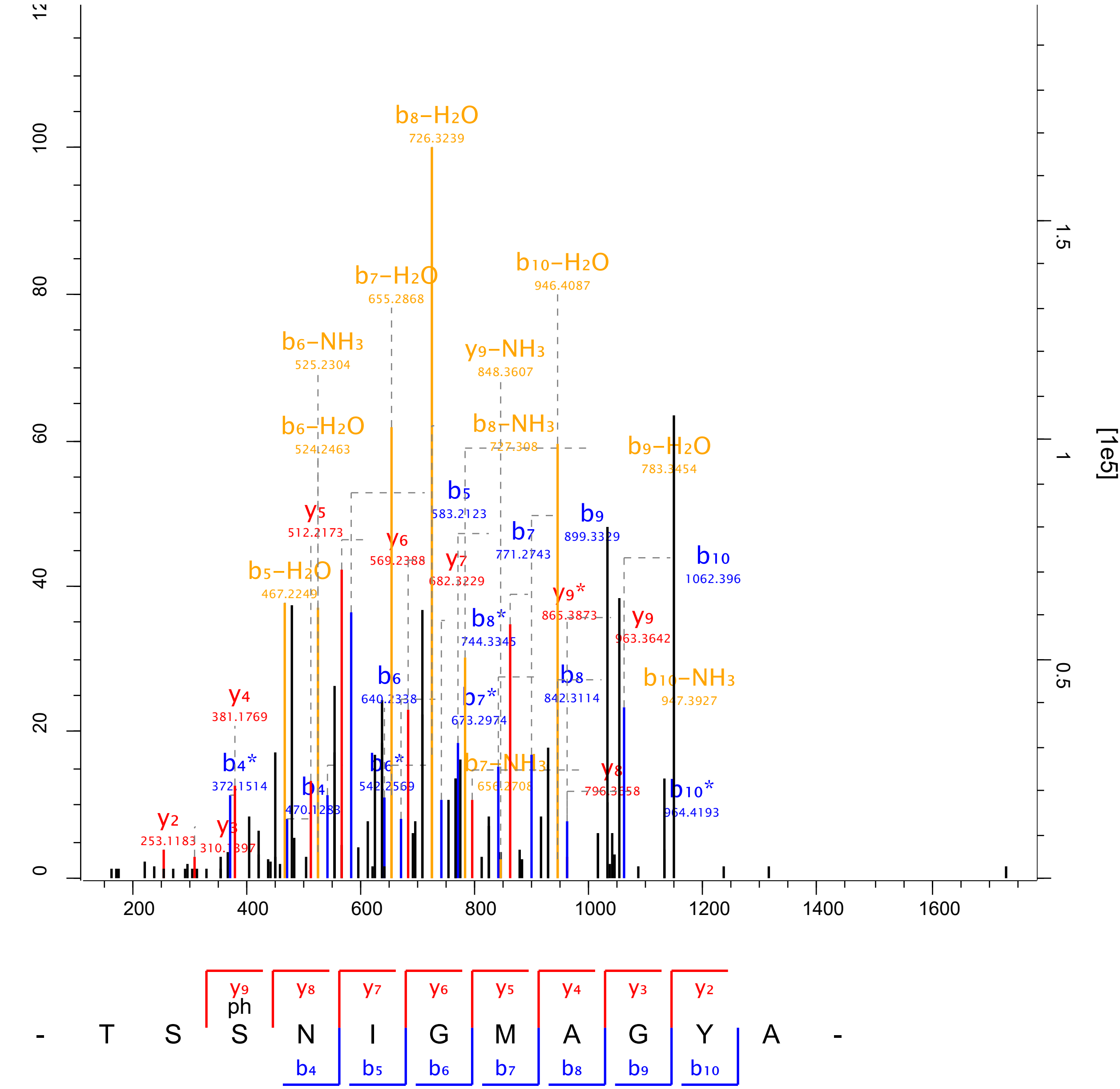

| Raw file | Scan | Method | Score | m/z |
| --- | --- | --- | --- | --- |
| sys_05_2 | 27224 | FTMS; HCD | 199.41 | 934.92 |

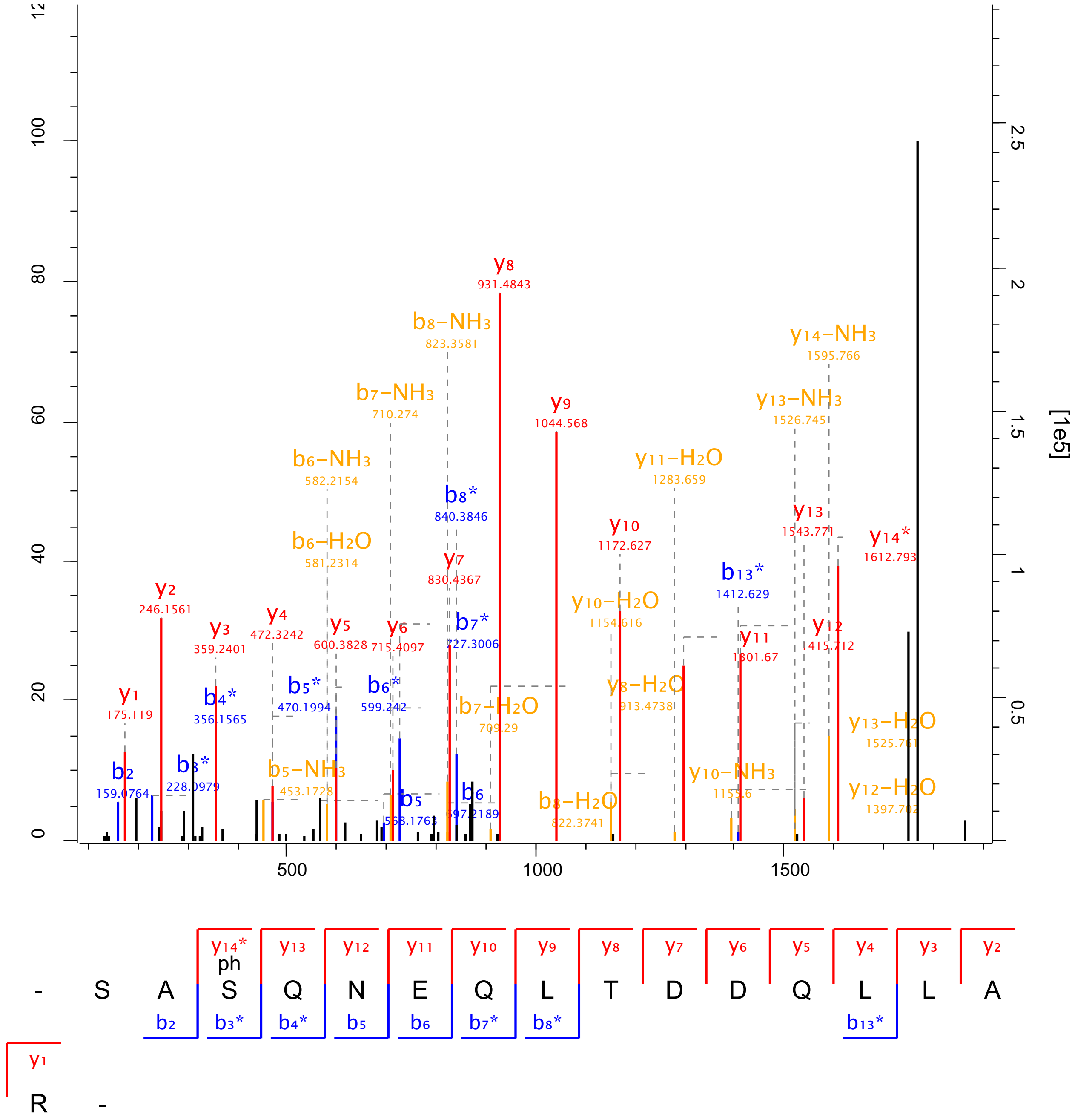

| Raw file | Scan | Method | Score | m/z |
|---|---|---|---|---|
| sys_05_2 | 27318 | FTMS; HCD | 220.06 | 635.79 |

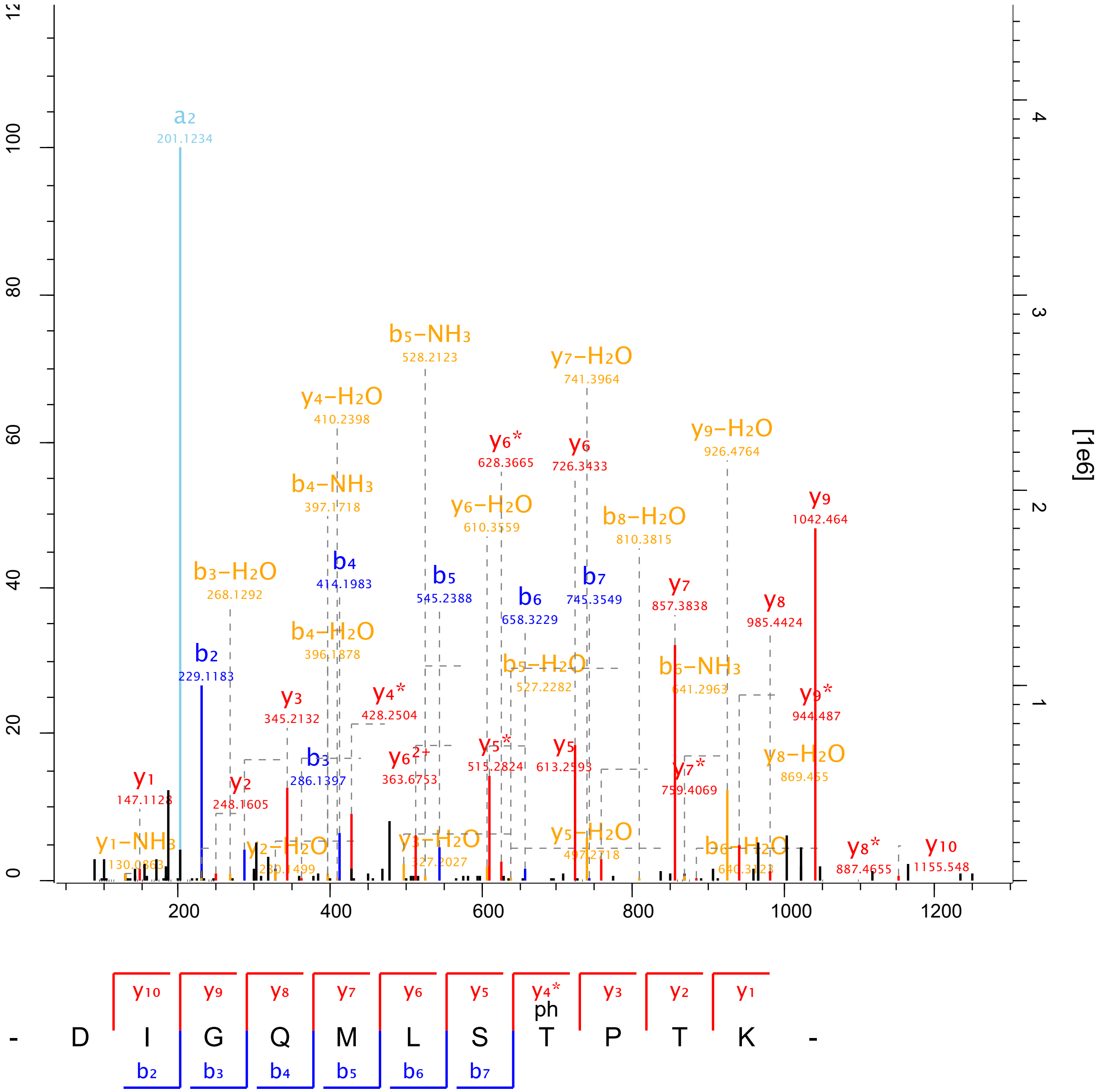

| Raw file | Scan | Method | Score | m/z |
|---|---|---|---|---|
| sys_05_2 | 27838 | FTMS; HCD | 99.27 | 912.43 |

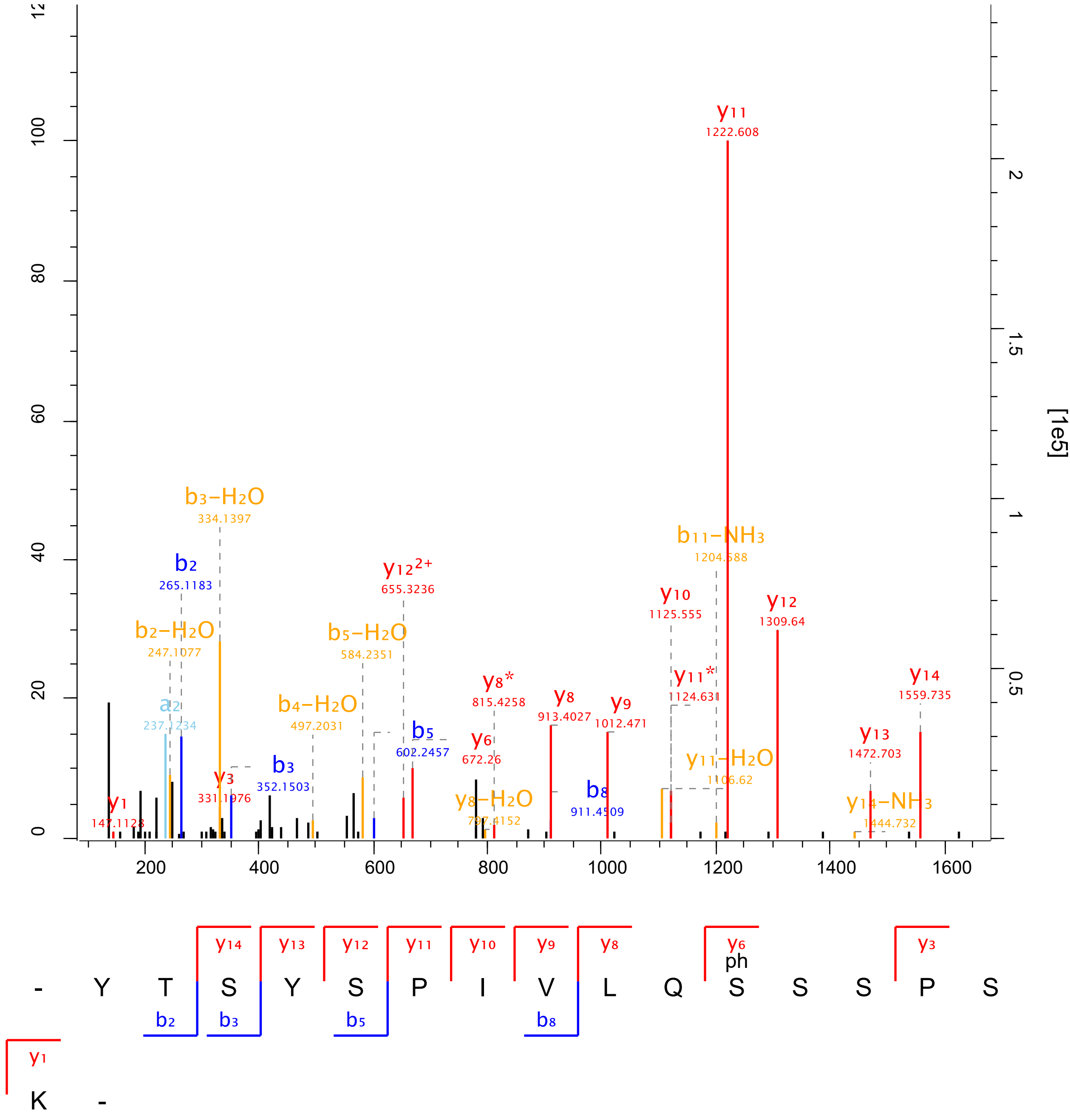

| Raw file | Scan | Method | Score | m/z |
|---|---|---|---|---|
| sys_05_2 | 27987 | FTMS; HCD | 122.33 | 606.77 |

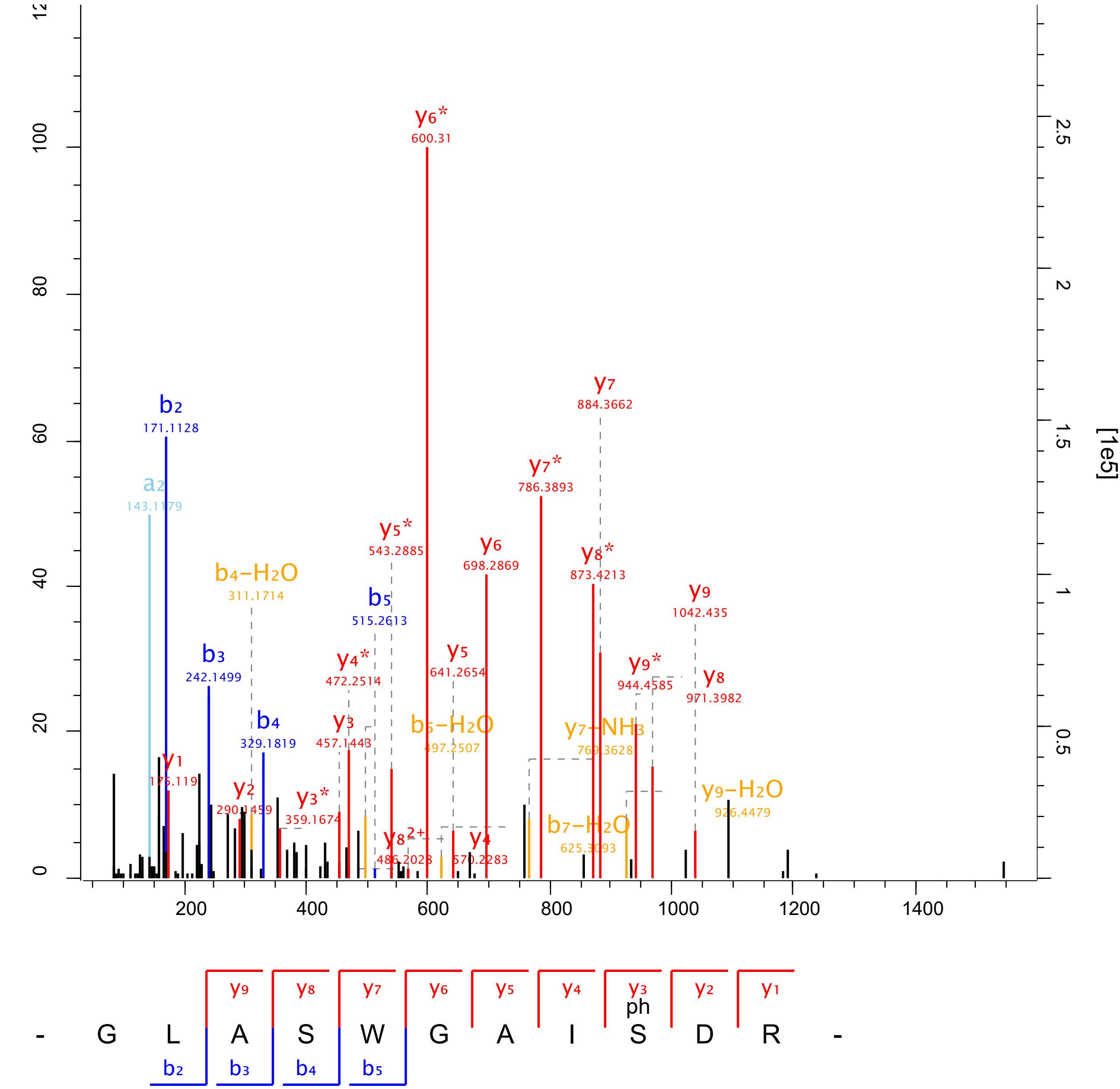

| Raw file | Scan | Method | Score | m/z |
|---|---|---|---|---|
| sys_05_2 | 28580 | FTMS; HCD | 207.66 | 686.79 |

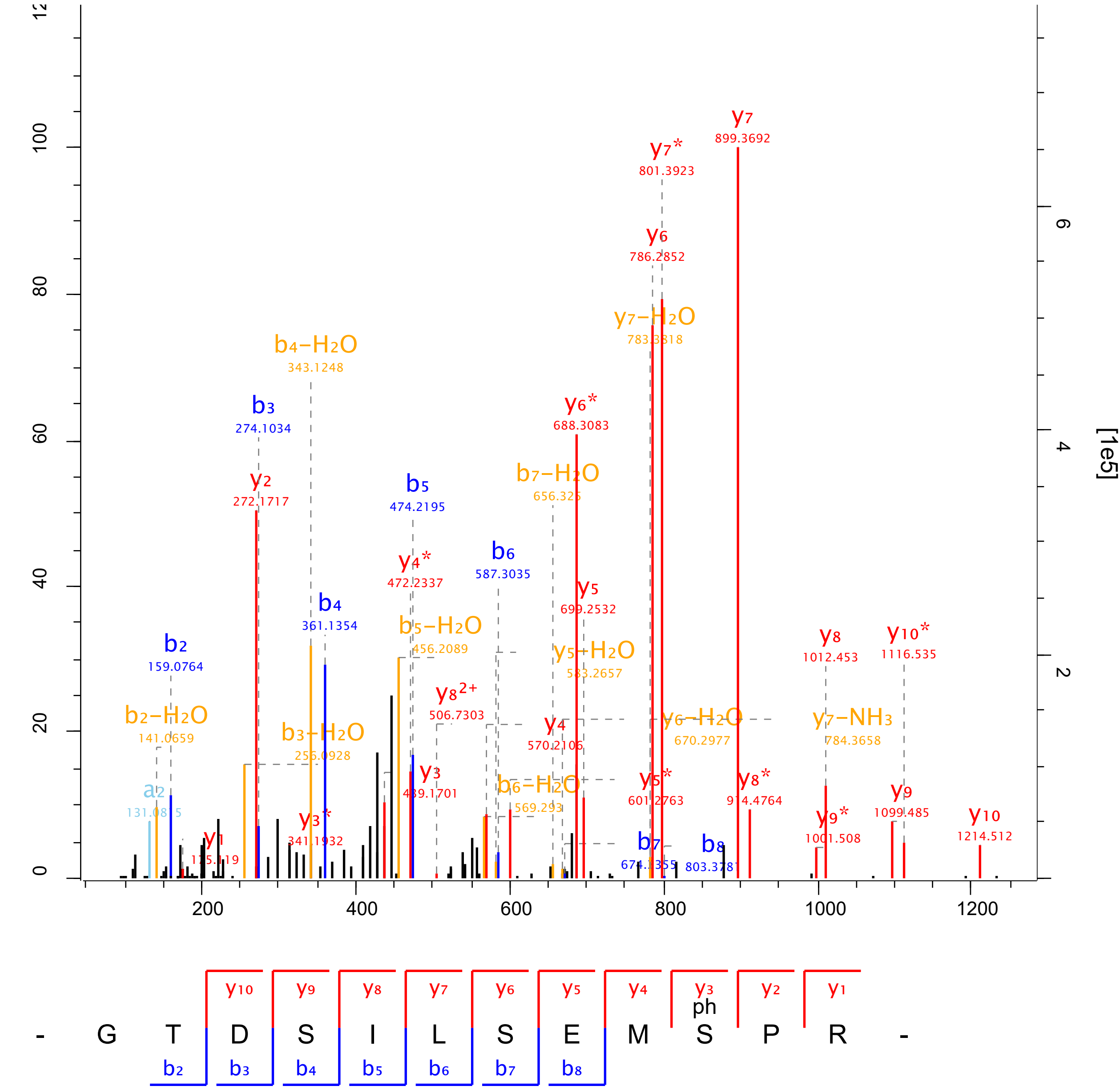

| Raw file | Scan | Method | Score | m/z |
|---|---|---|---|---|
| sys_05_2 | 28619 | FTMS; HCD | 52.5 | 607.75 |

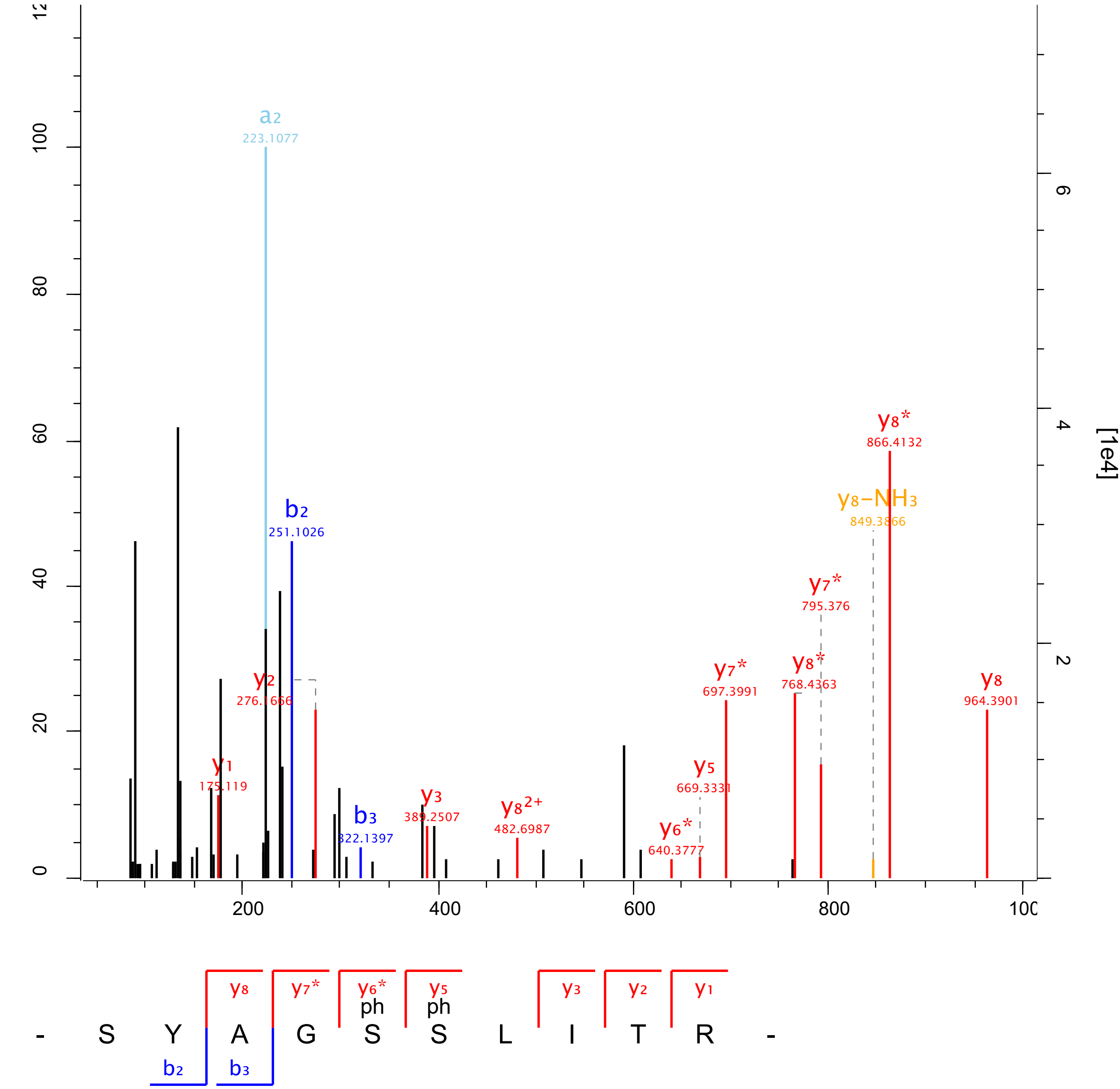

| Raw file | Scan | Method | Score | m/z |
|---|---|---|---|---|
| sys_05_2 | 28630 | FTMS; HCD | 99.32 | 894.38 |

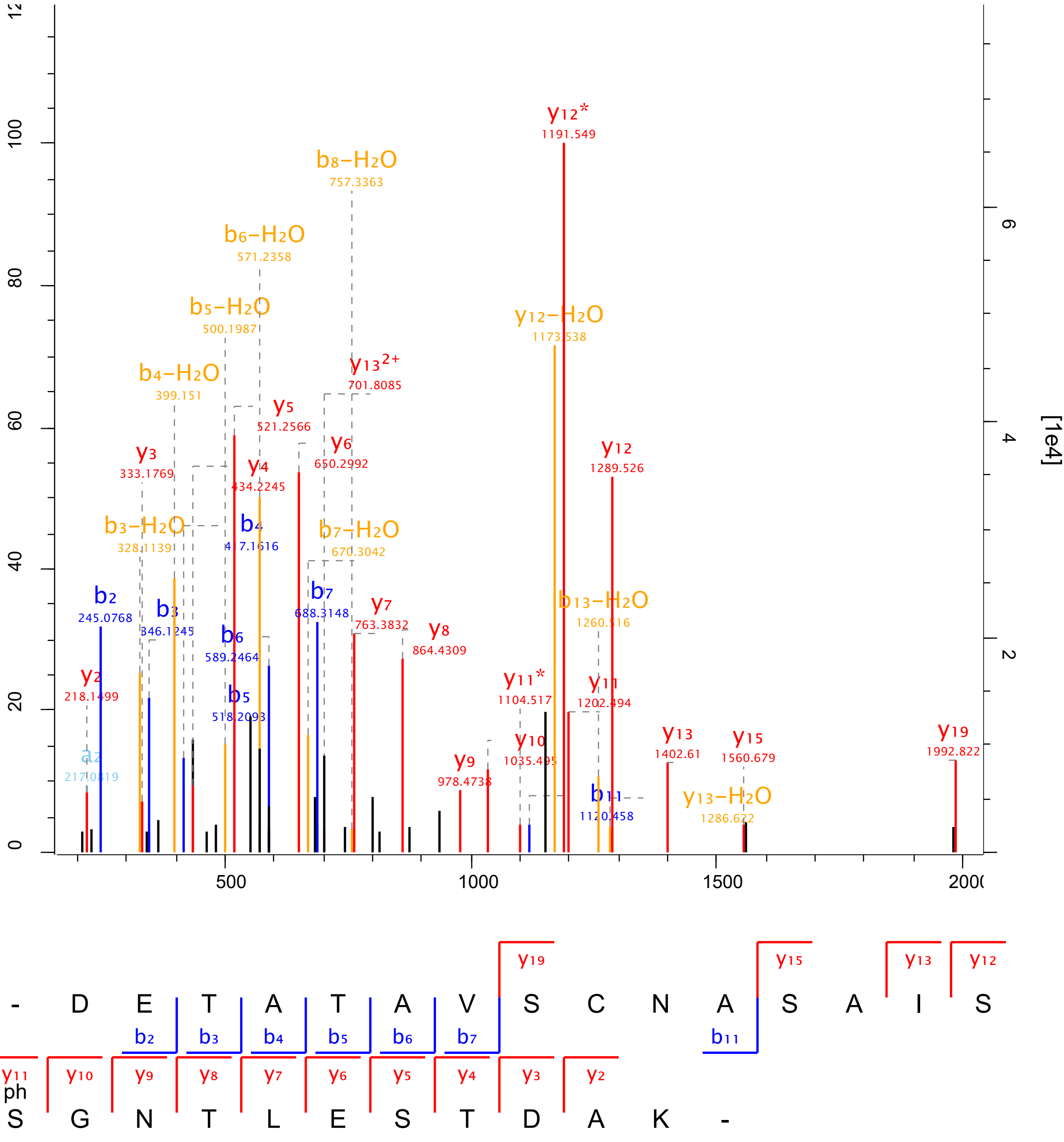

| Raw file | Scan | Method | Score | m/z |
|---|---|---|---|---|
| sys_05_2 | 28908 | FTMS; HCD | 274.91 | 827.32 |

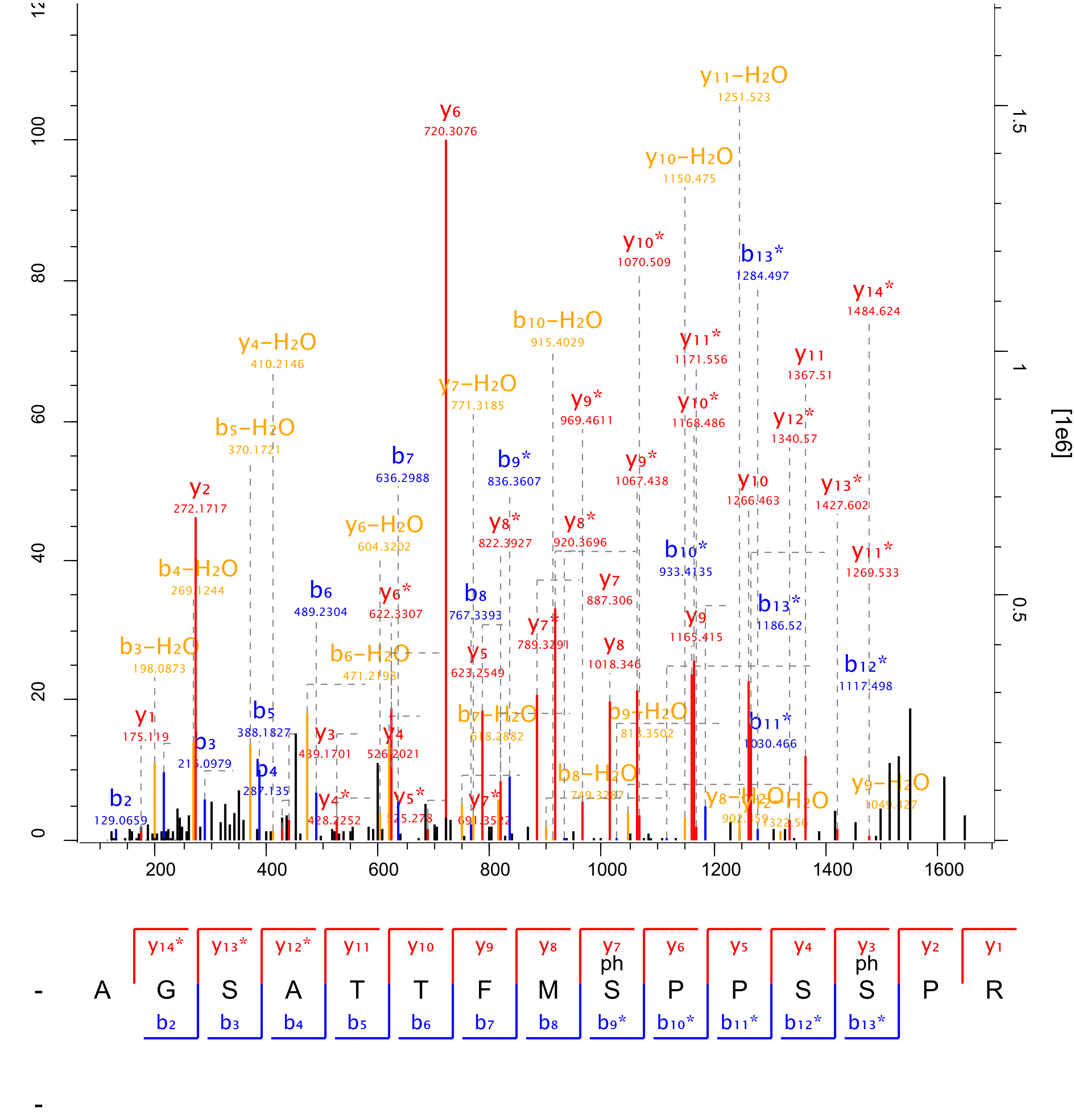

| Raw file | Scan | Method | Score | m/z |
|---|---|---|---|---|
| sys_05_2 | 28958 | FTMS; HCD | 50.9 | 643.79 |

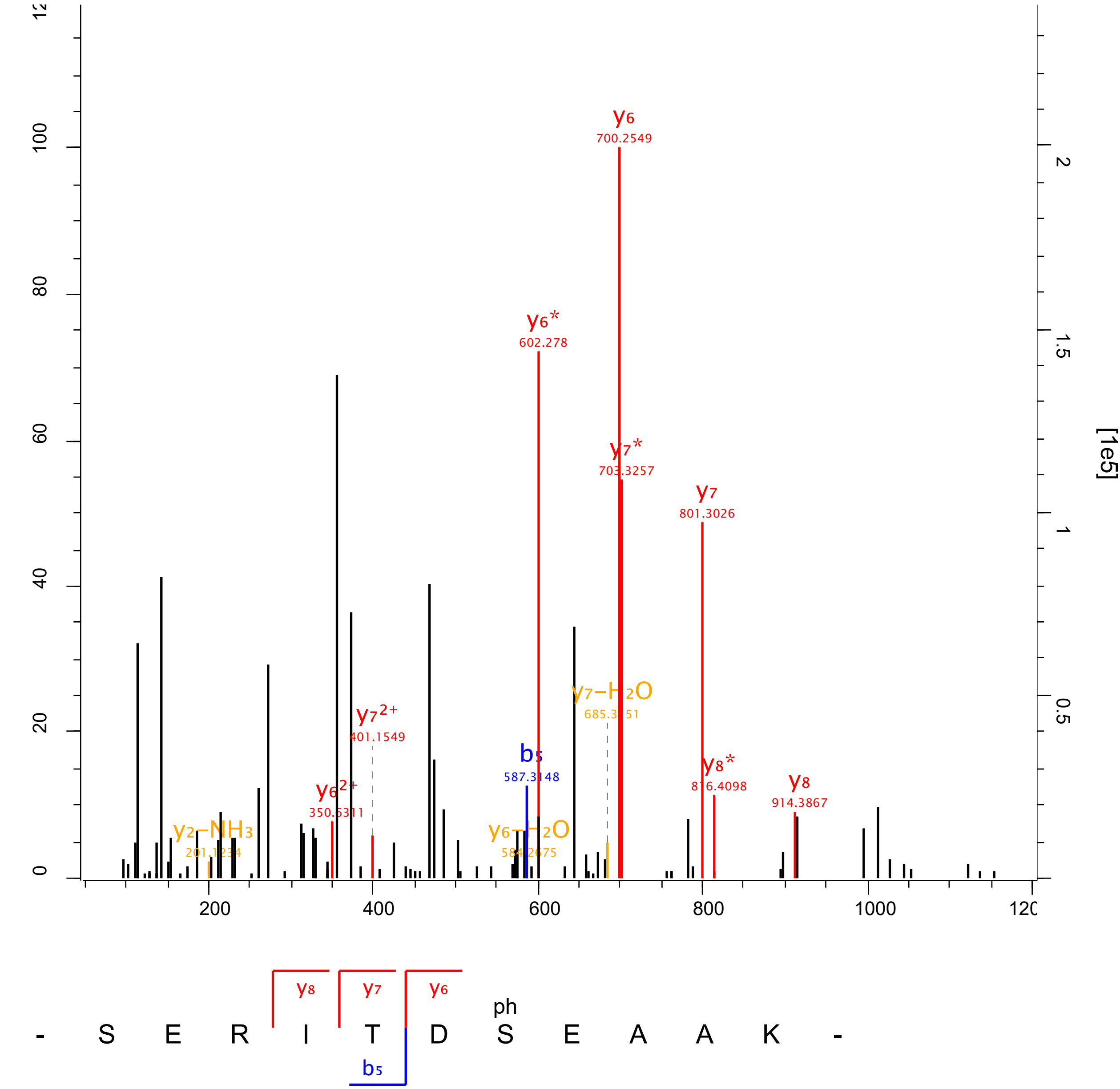

| Raw file | Scan | Method | Score | m/z |
| --- | --- | --- | --- | --- |
| sys_05_2 | 29072 | FTMS; HCD | 92.44 | 804.35 |

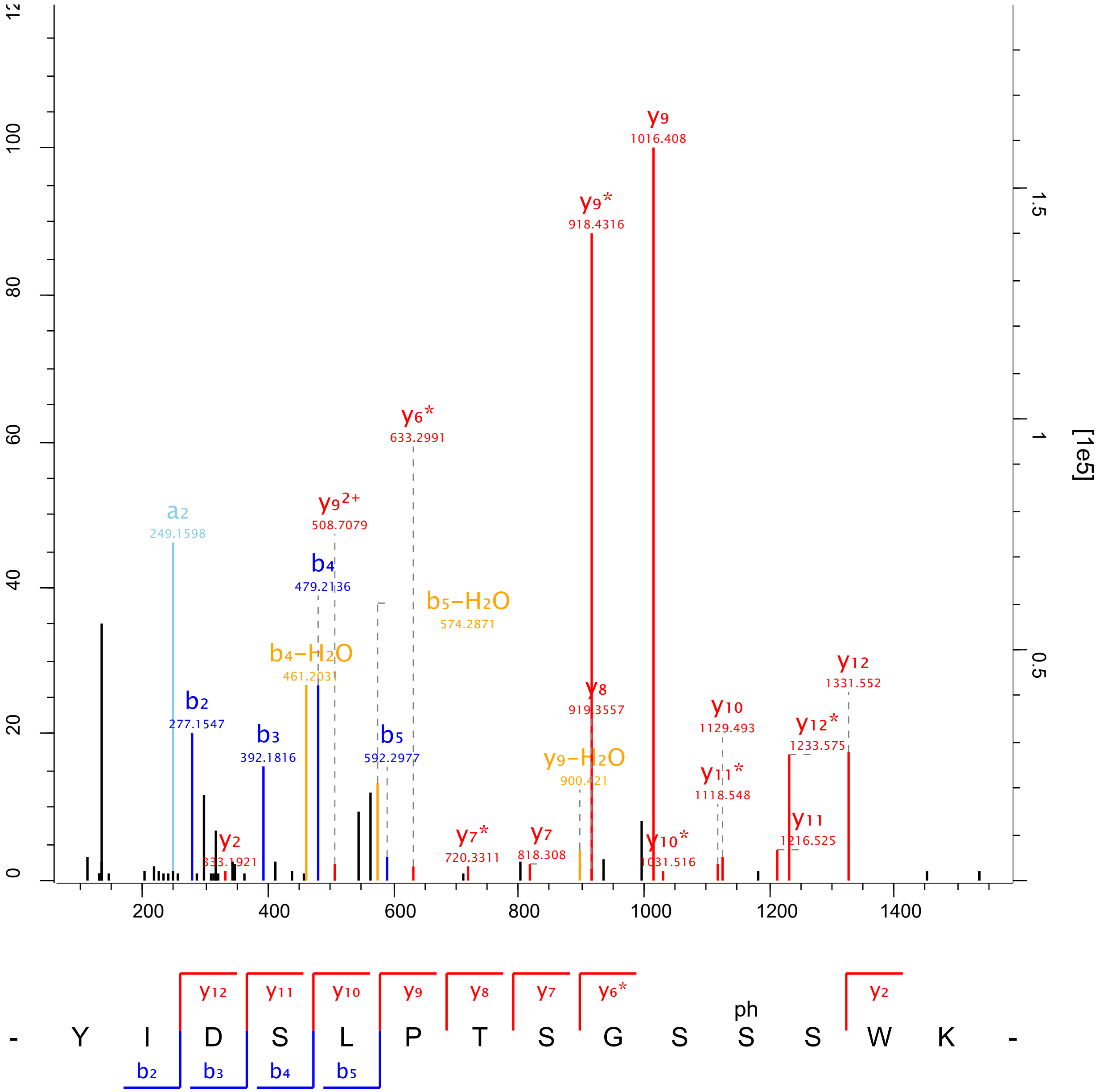

| Raw file | Scan | Method | Score | m/z |
|---|---|---|---|---|
| sys_05_2 | 29132 | FTMS; HCD | 104.88 | 867.39 |

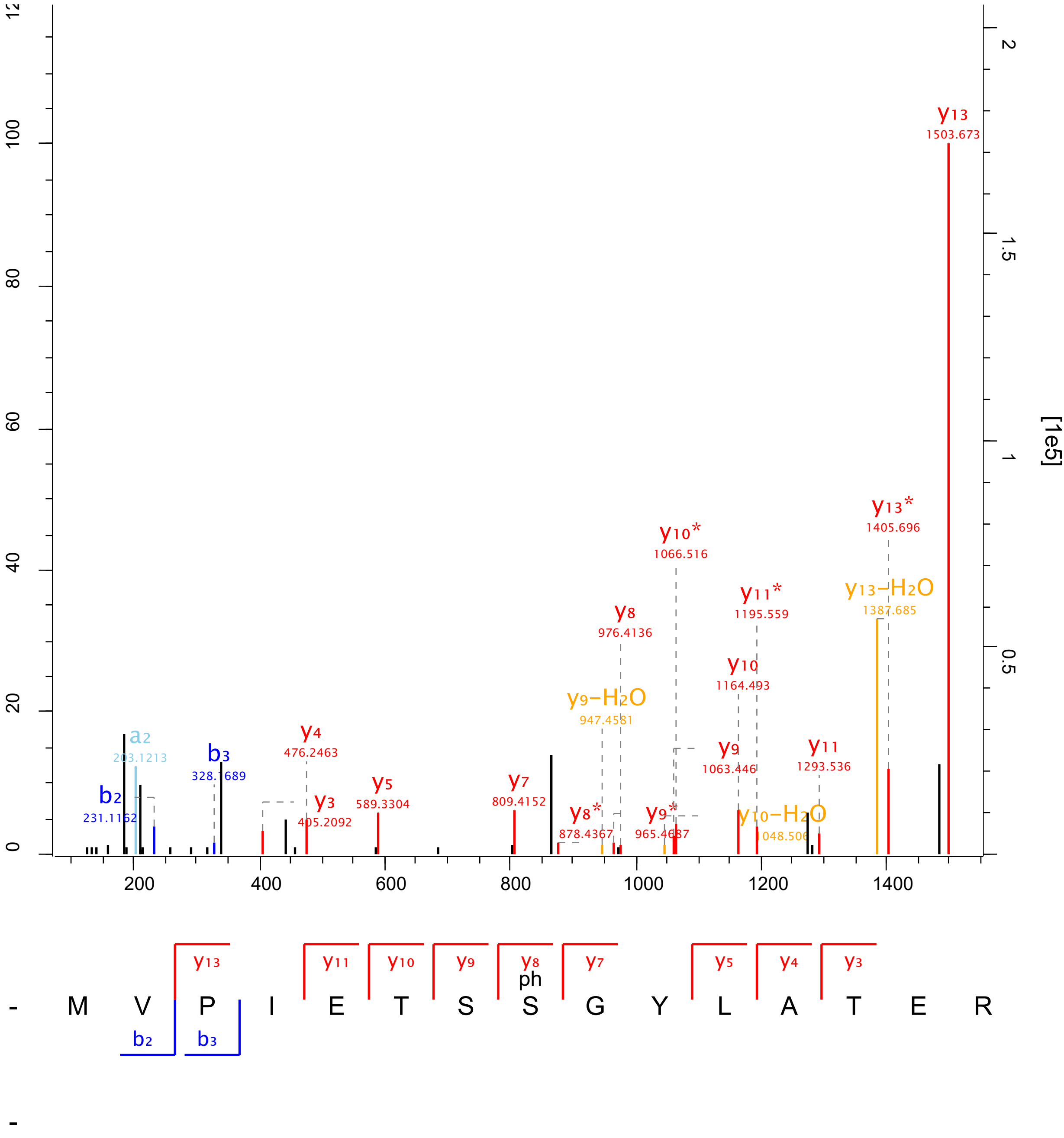

| Raw file | Scan | Method | Score | m/z |
|---|---|---|---|---|
| sys_05_2 | 29141 | FTMS; HCD | 131.18 | 854.9 |

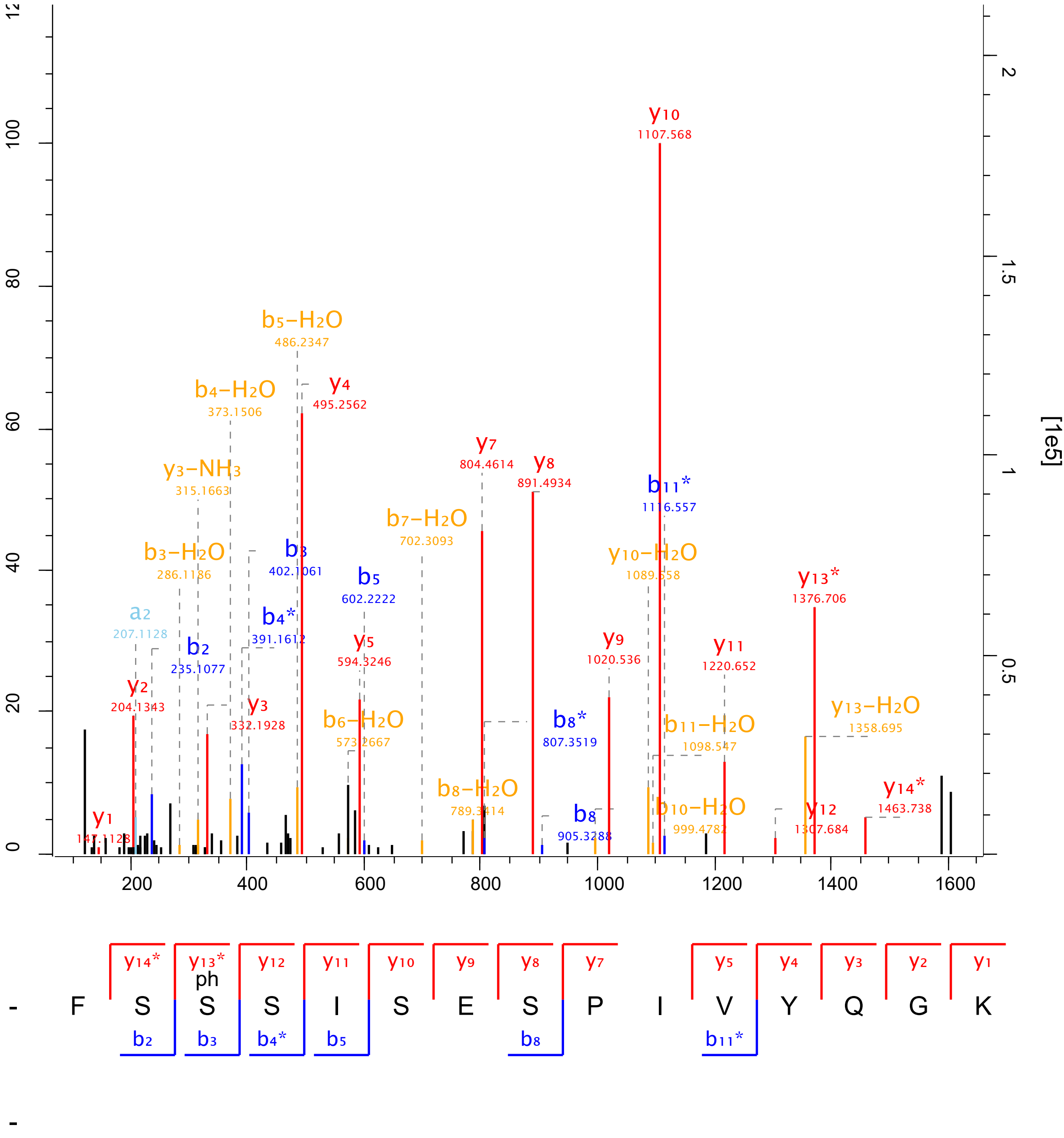

| Raw file | Scan | Method | Score | m/z |
|---|---|---|---|---|
| sys_05_2 | 29684 | FTMS; HCD | 79.36 | 1089.5 |

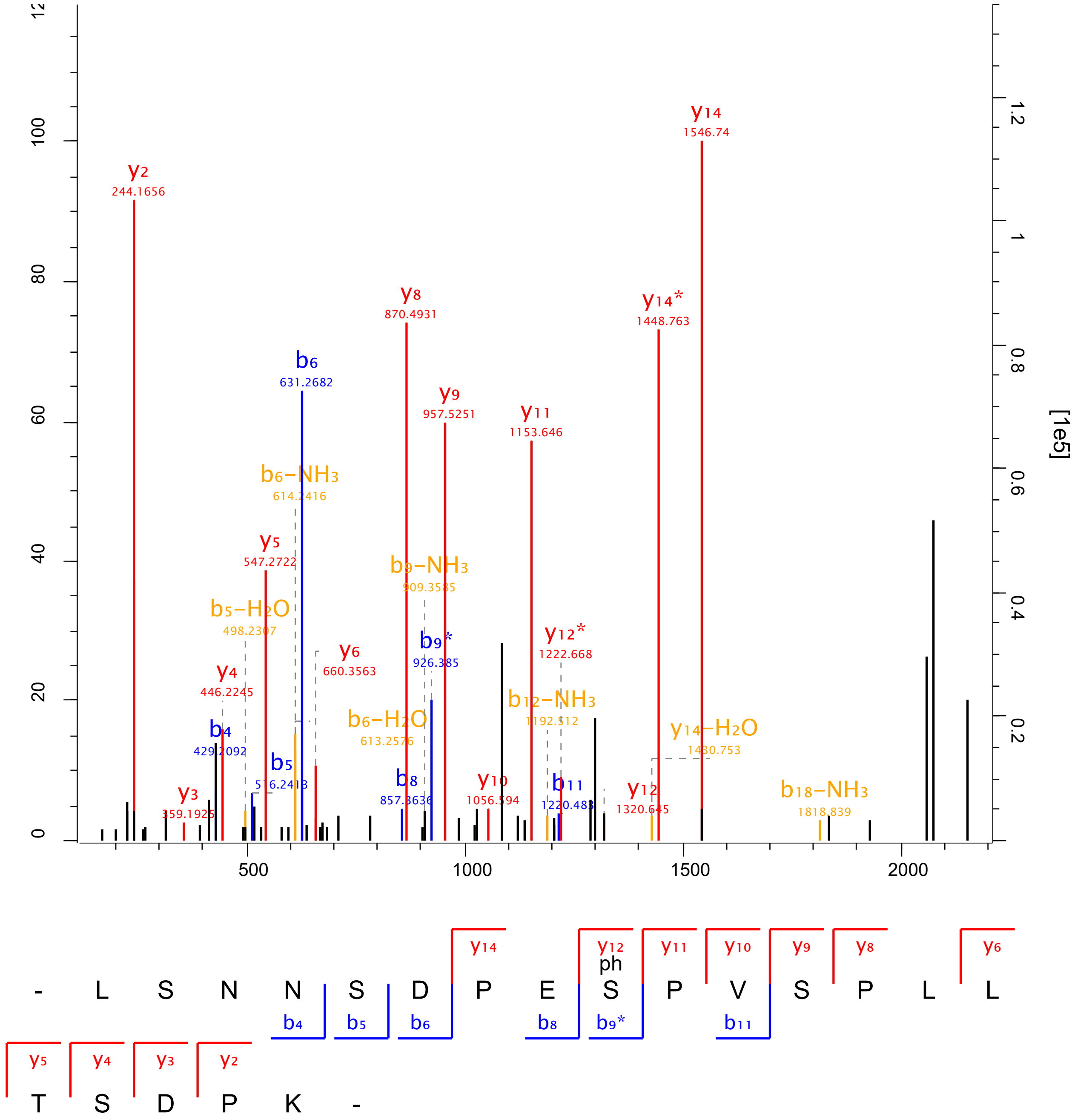

| Raw file | Scan | Method | Score | m/z |
|---|---|---|---|---|
| sys_05_2 | 30131 | FTMS; HCD | 217.89 | 871.36 |

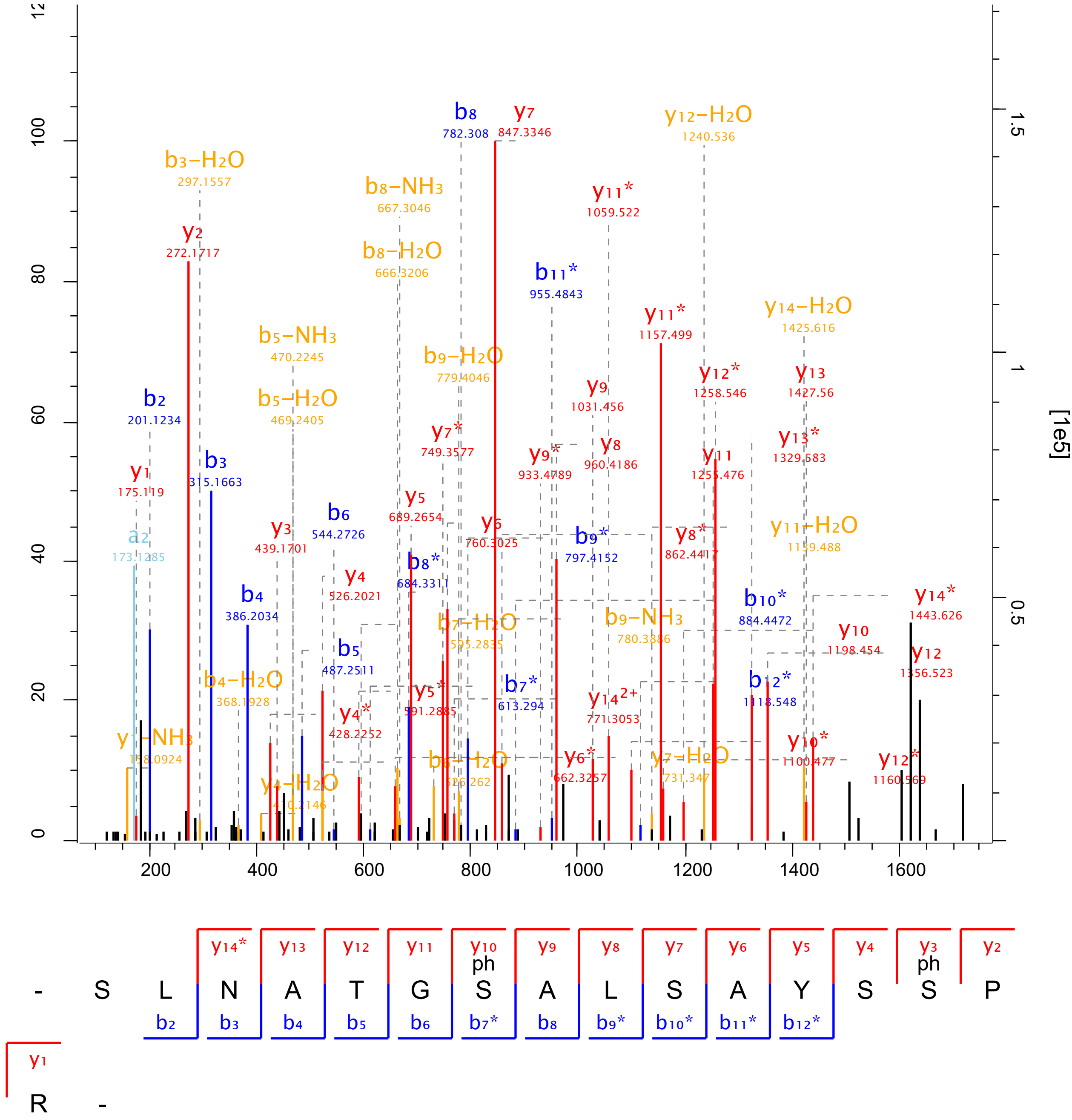

| Raw file | Scan | Method | Score | m/z |
|---|---|---|---|---|
| sys_05_2 | 30162 | FTMS; HCD | 181.87 | 912.39 |

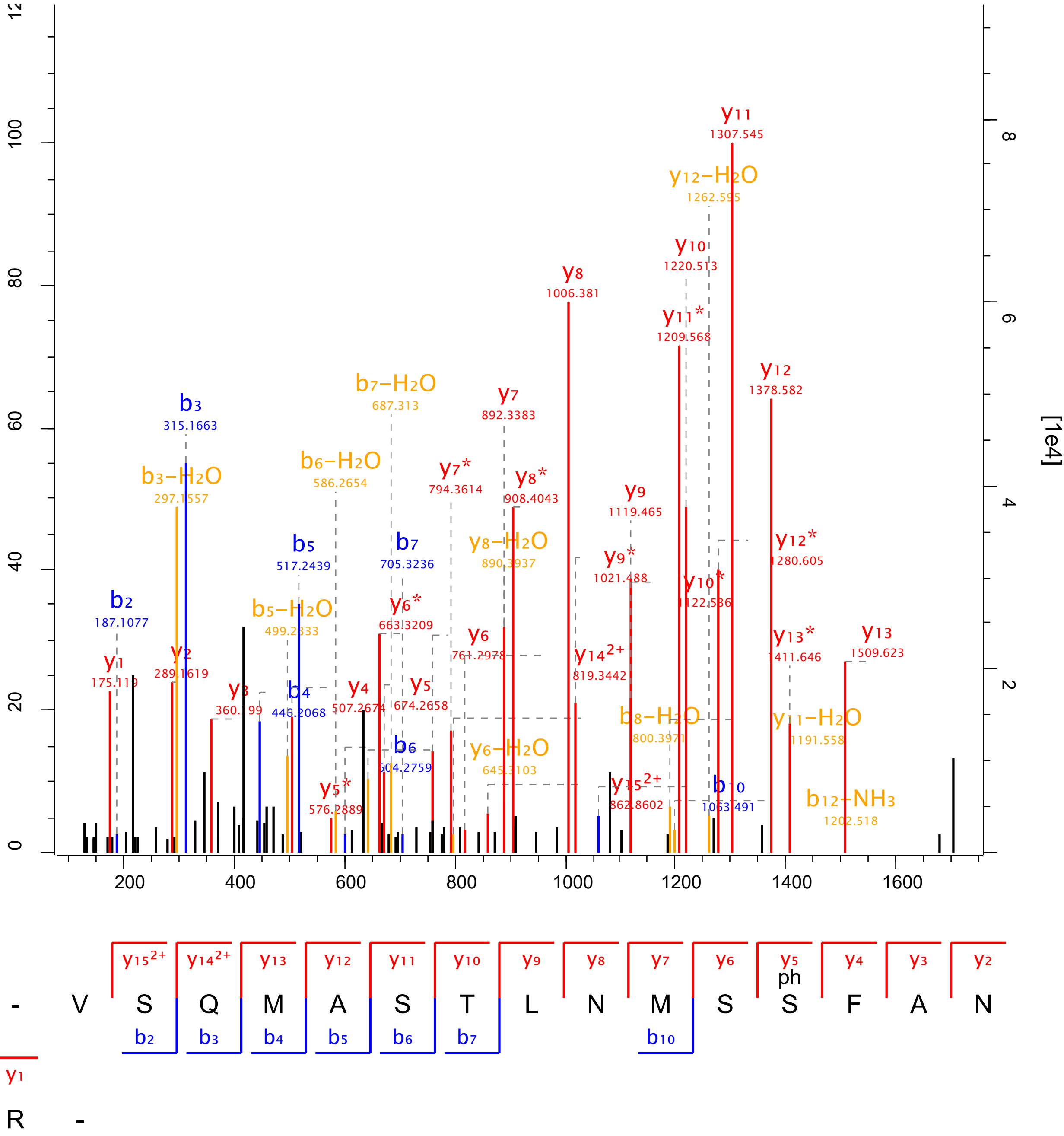

| Raw file | Scan | Method | Score | m/z |
| --- | --- | --- | --- | --- |
| sys_05_2 | 30283 | FTMS; HCD | 42.31 | 745.87 |

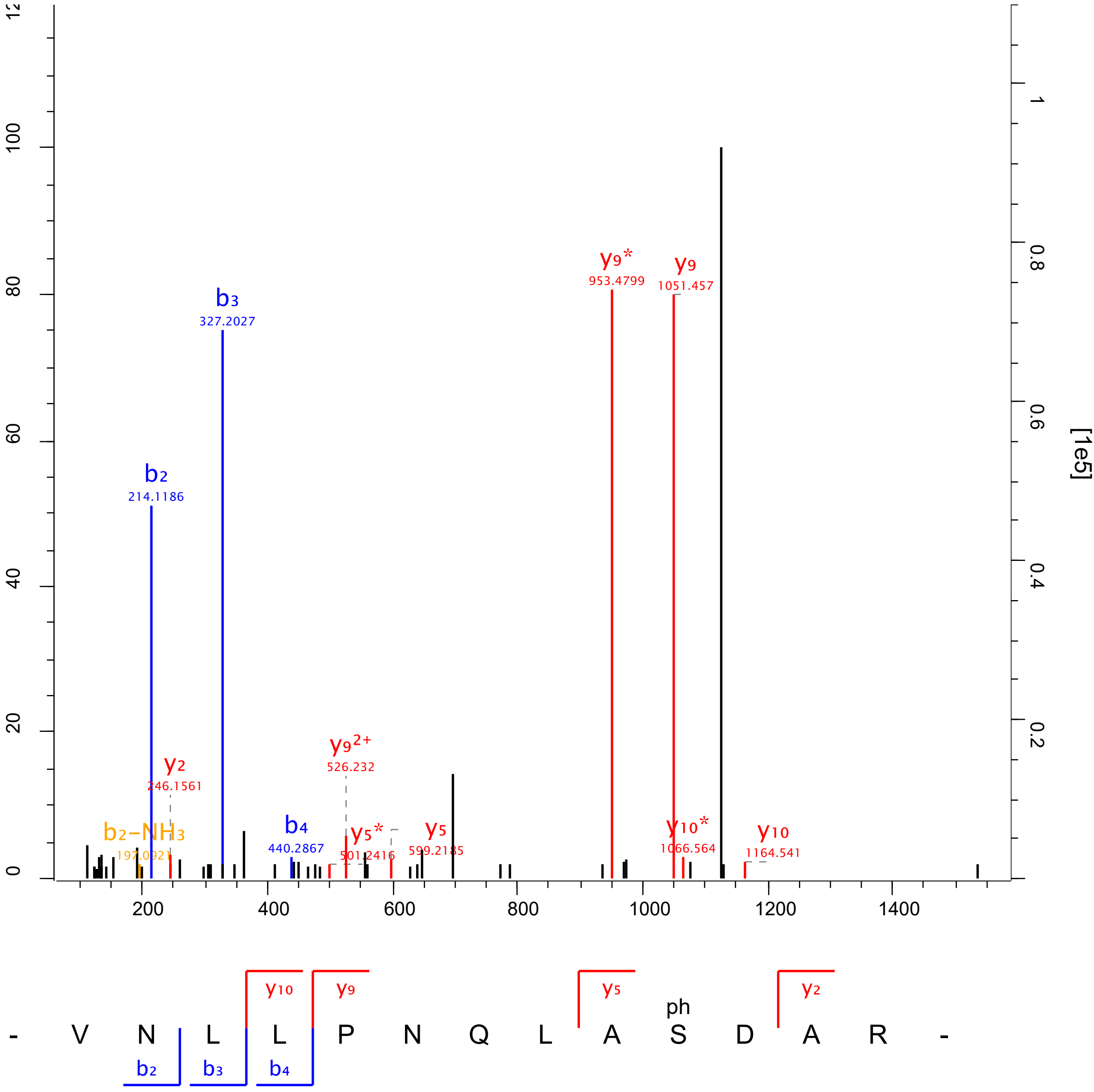

| Raw file | Scan | Method | Score | m/z |
|---|---|---|---|---|
| sys_05_2 | 30398 | FTMS; HCD | 72.66 | 663.27 |

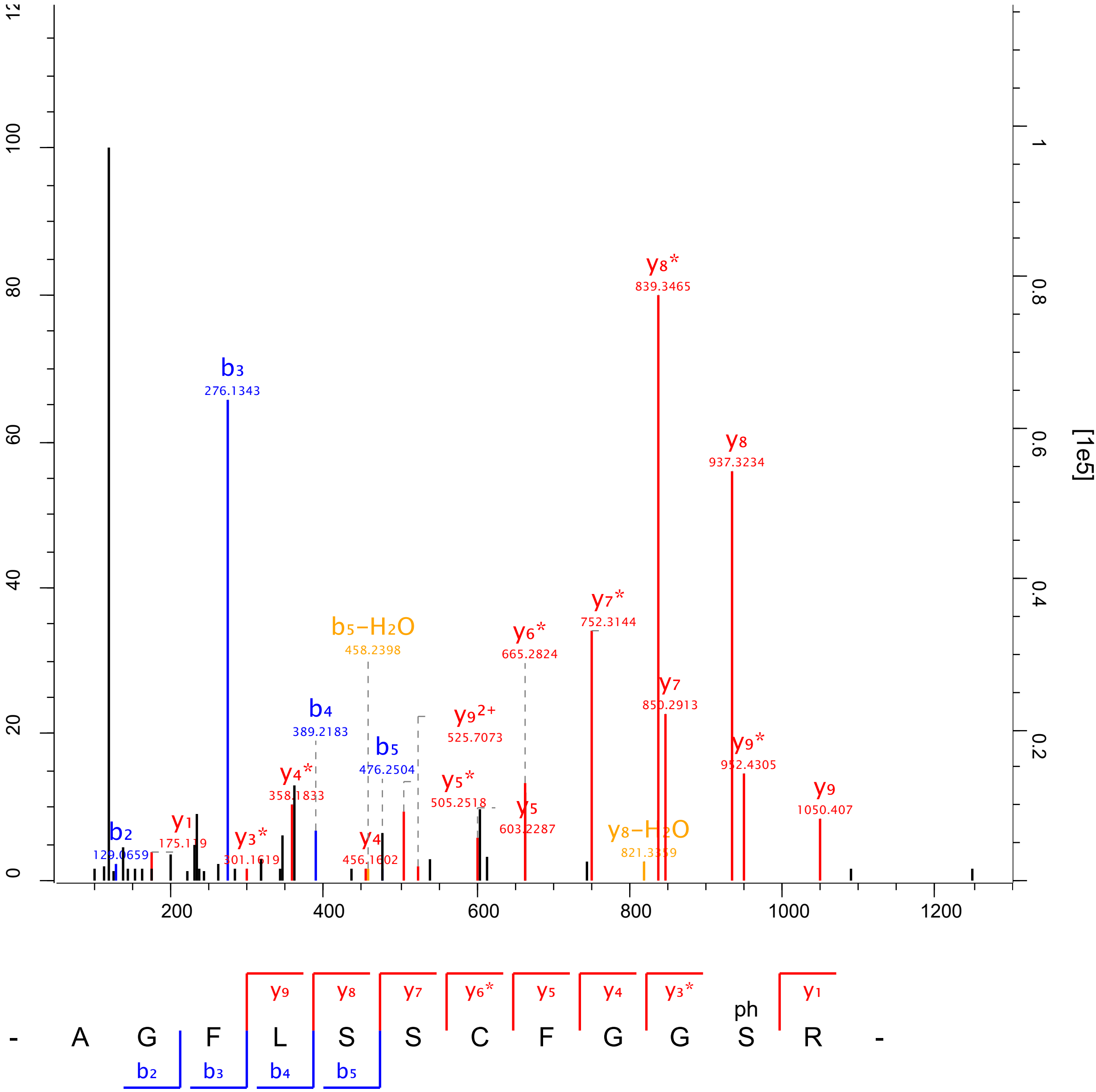

| Raw file | Scan | Method | Score | m/z |
| --- | --- | --- | --- | --- |
| sys_05_2 | 30514 | FTMS; HCD | 162.59 | 730.33 |

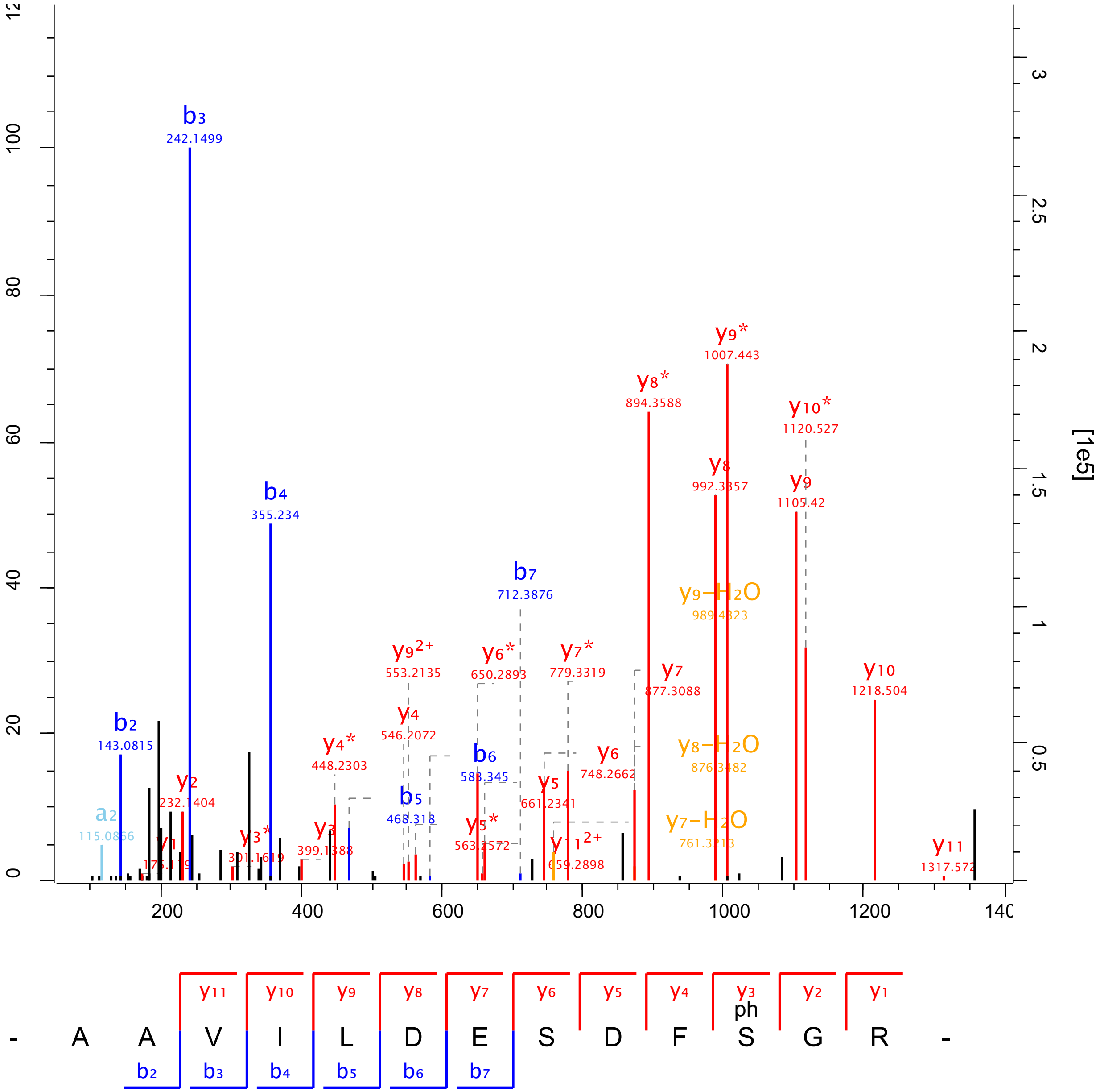

| Raw file | Scan | Method | Score | m/z |
| --- | --- | --- | --- | --- |
| sys_05_2 | 30700 | FTMS; HCD | 52.58 | 635.79 |

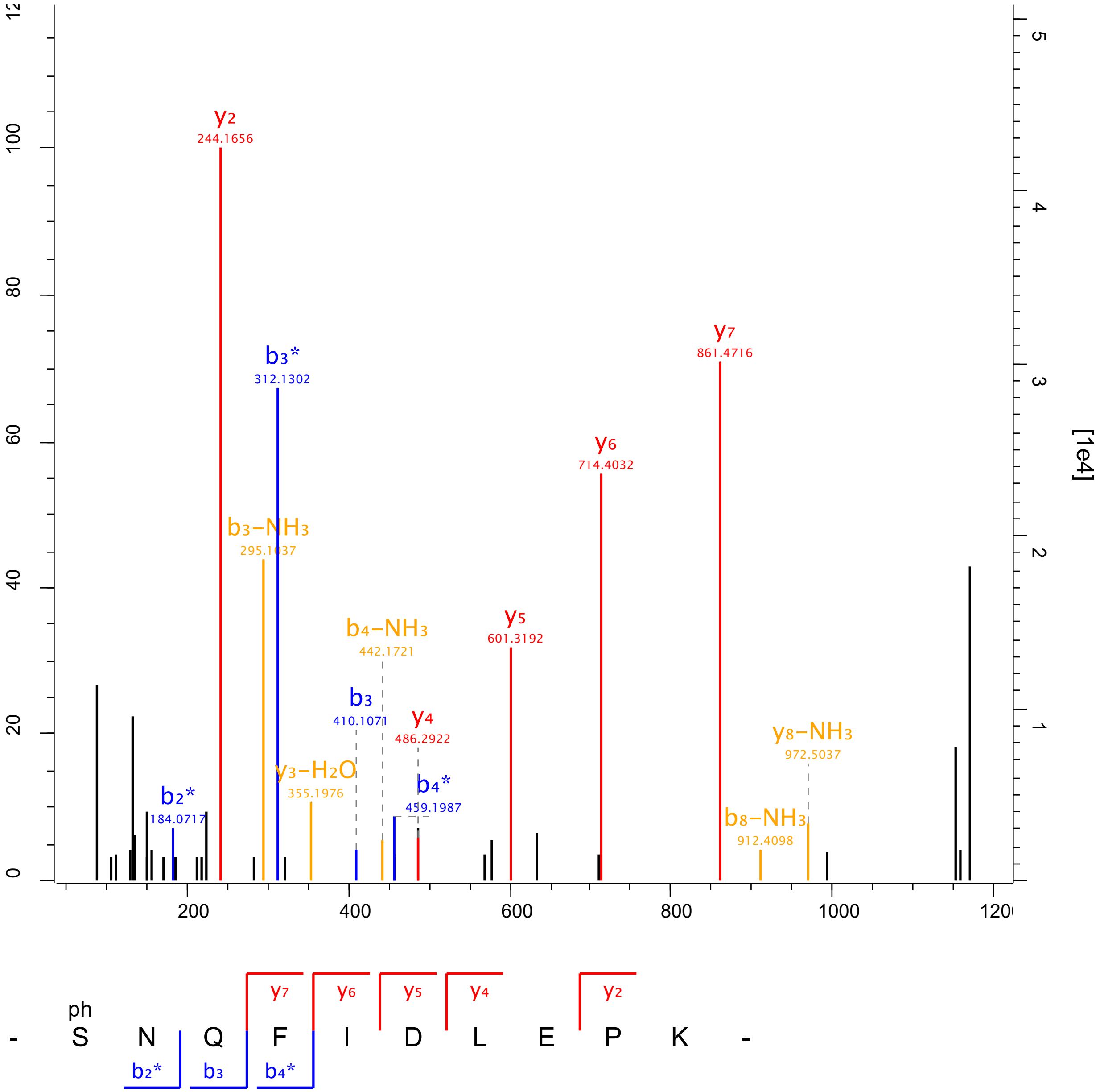

| Raw file | Scan | Method | Score | m/z |
|---|---|---|---|---|
| sys_05_2 | 30743 | FTMS; HCD | 89.3 | 574.78 |

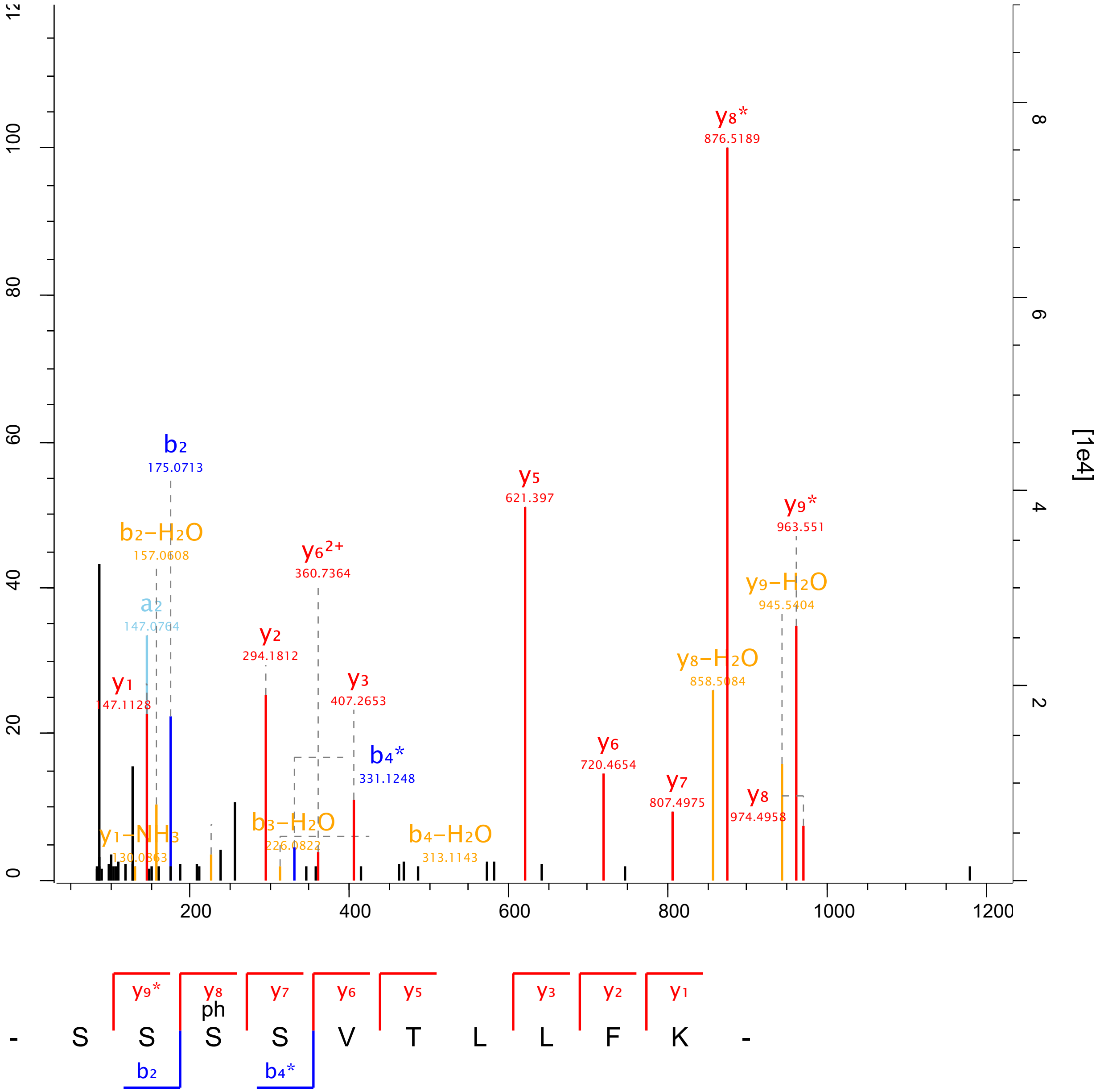

| Raw file | Scan | Method | Score | m/z |
|---|---|---|---|---|
| sys_05_2 | 30774 | FTMS; HCD | 84.95 | 931.41 |

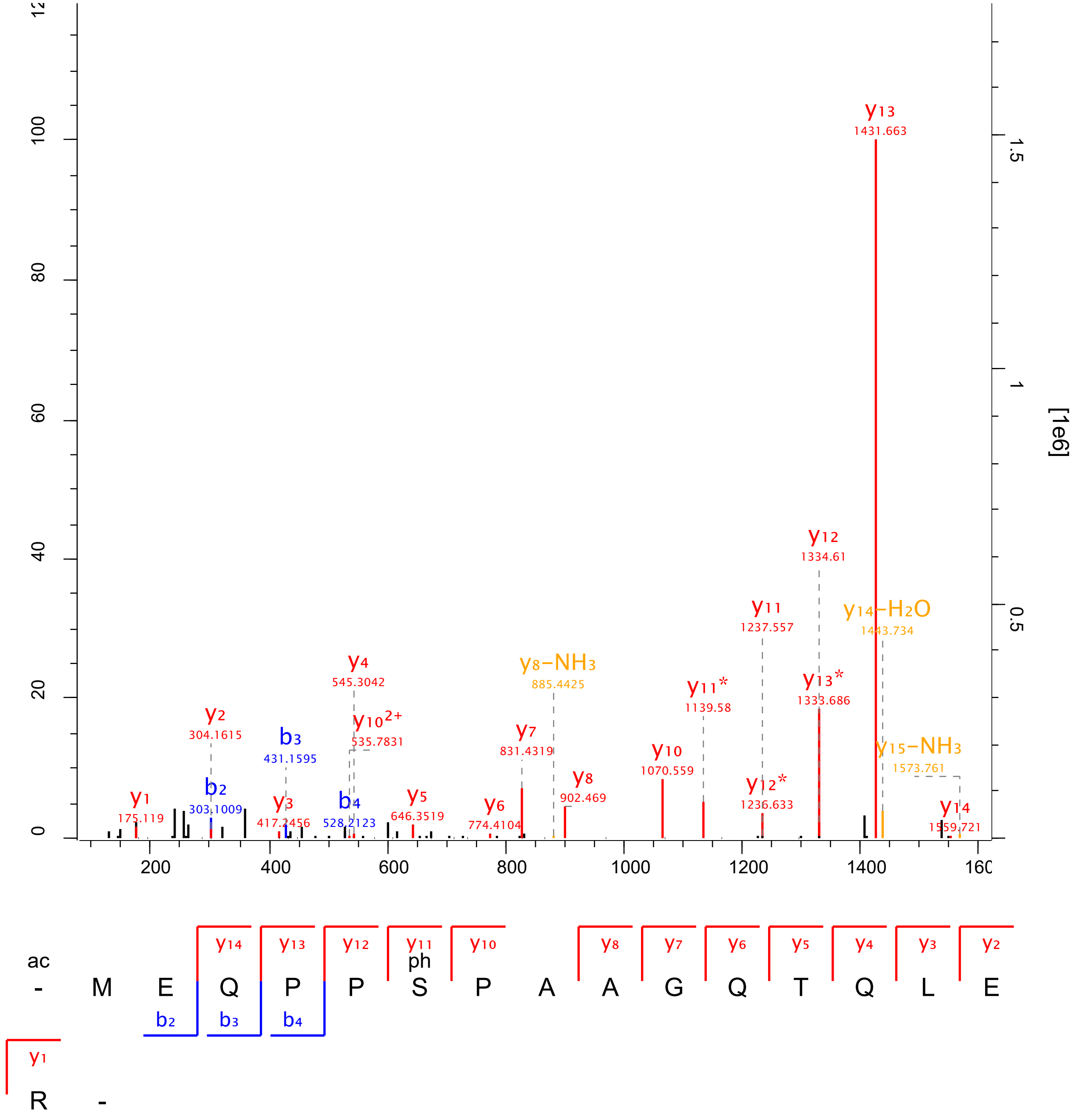

| Raw file | Scan | Method | Score | m/z |
|---|---|---|---|---|
| sys_05_2 | 3081 | FTMS; HCD | 115.91 | 628.27 |

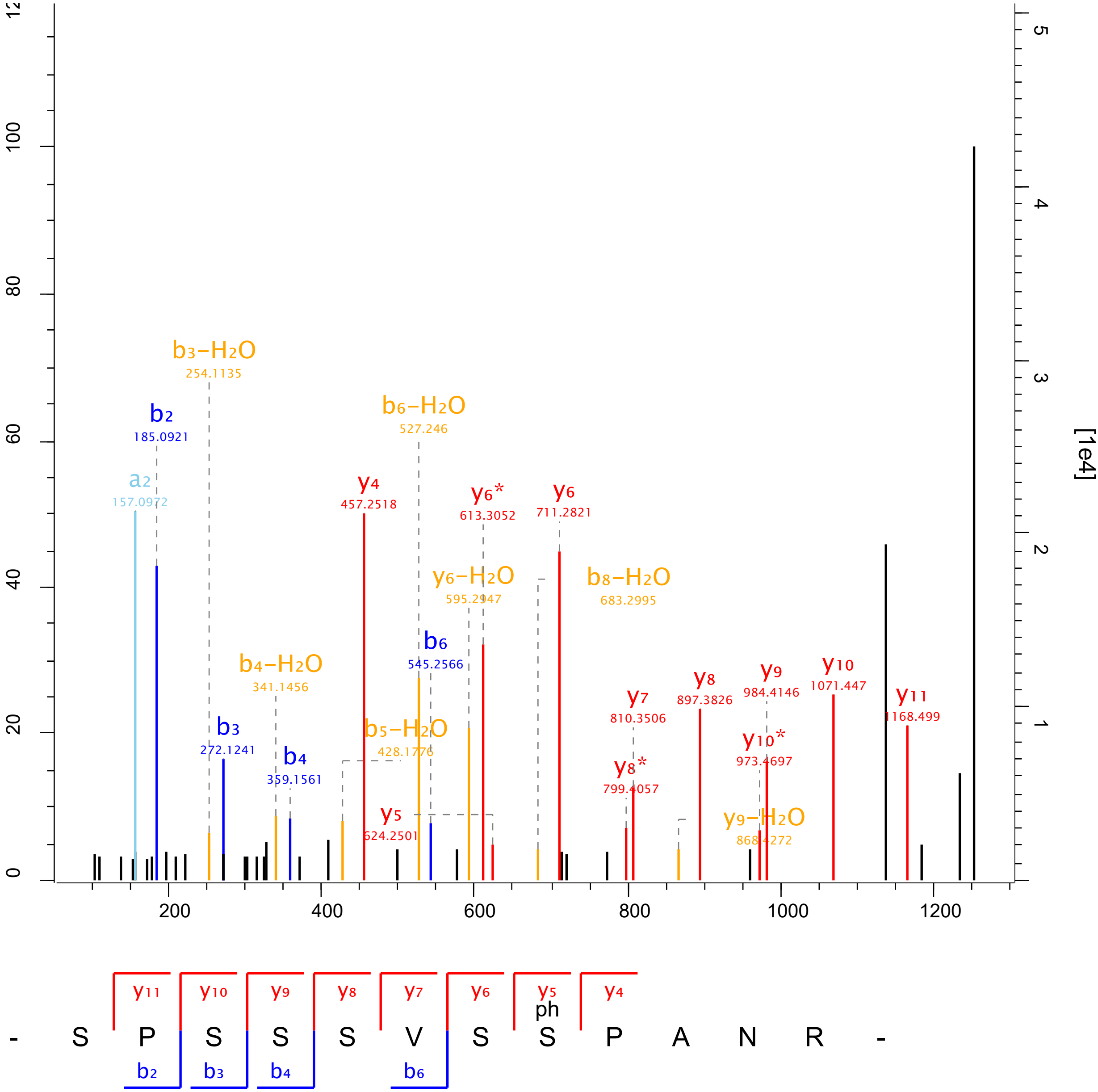

| Raw file | Scan | Method | Score | m/z |
|---|---|---|---|---|
| sys_05_2 | 30905 | FTMS; HCD | 54.09 | 779.89 |

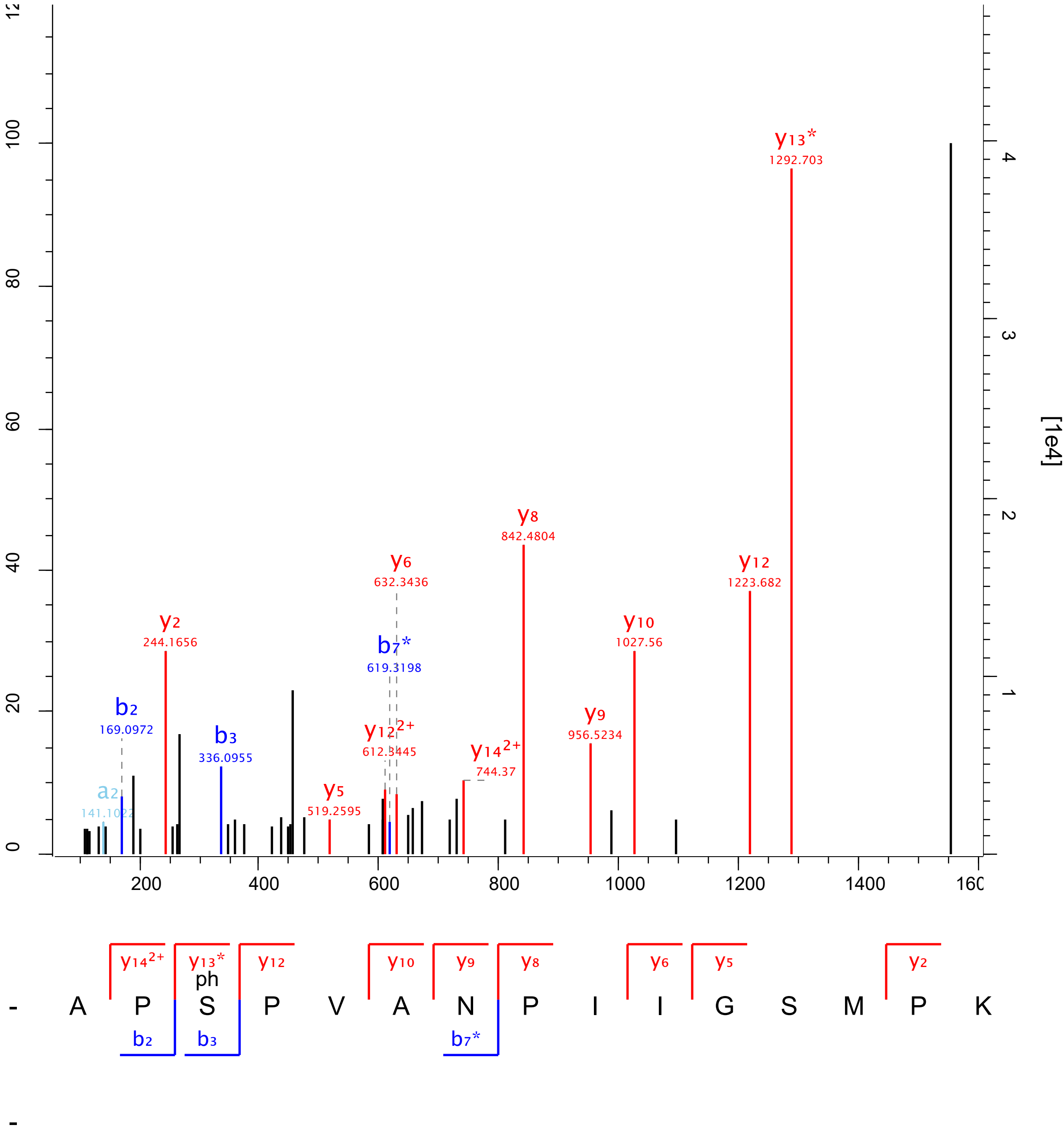

| Raw file | Scan | Method | Score | m/z |
|---|---|---|---|---|
| sys_05_2 | 31025 | FTMS; HCD | 248.67 | 698.82 |

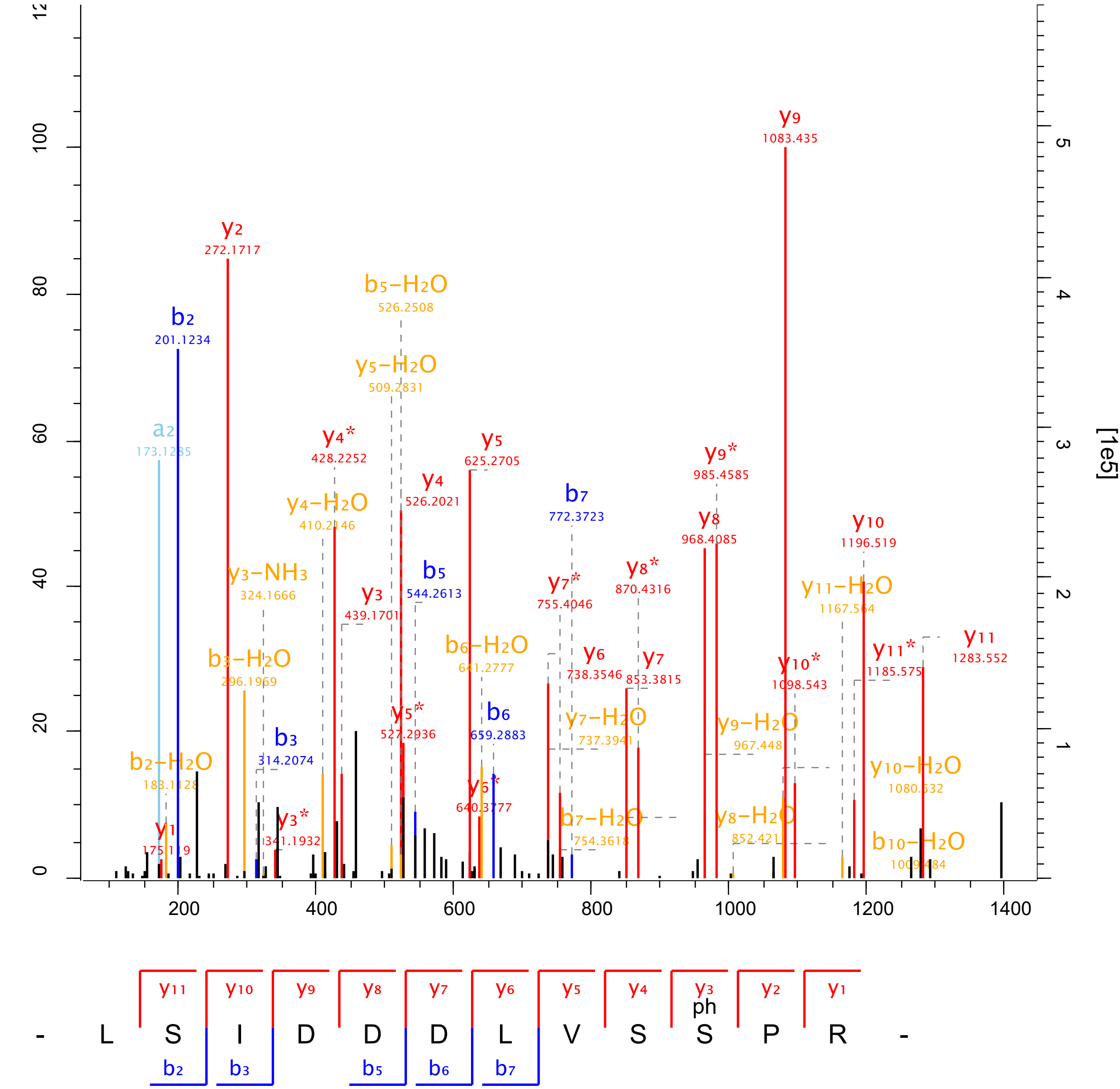

| Raw file | Scan | Method | Score | m/z |
| --- | --- | --- | --- | --- |
| sys_05_2 | 31109 | FTMS; HCD | 84.75 | 673.81 |

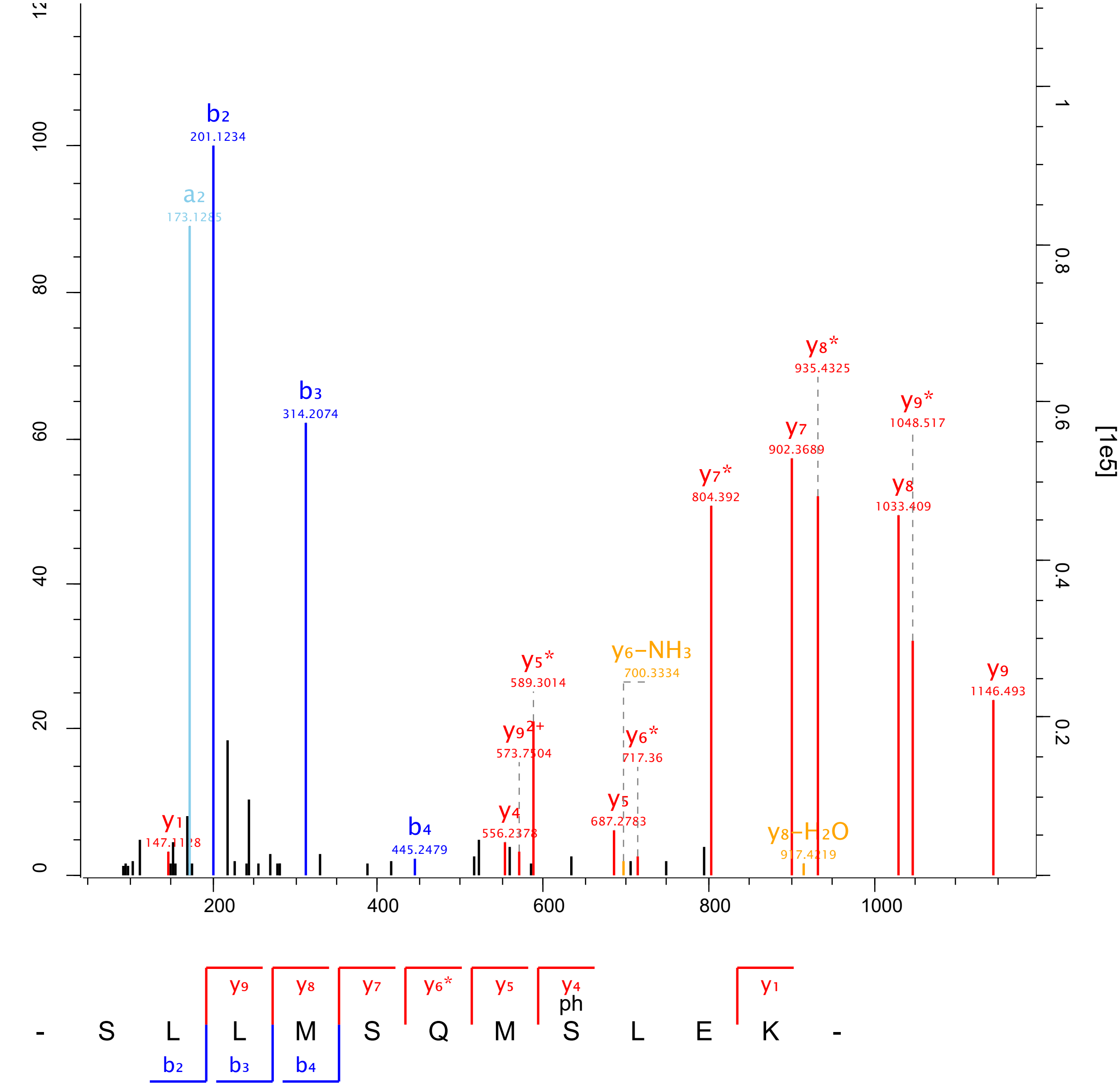

| Raw file | Scan | Method | Score | m/z |
|---|---|---|---|---|
| sys_05_2 | 31487 | FTMS; HCD | 109.79 | 667.26 |

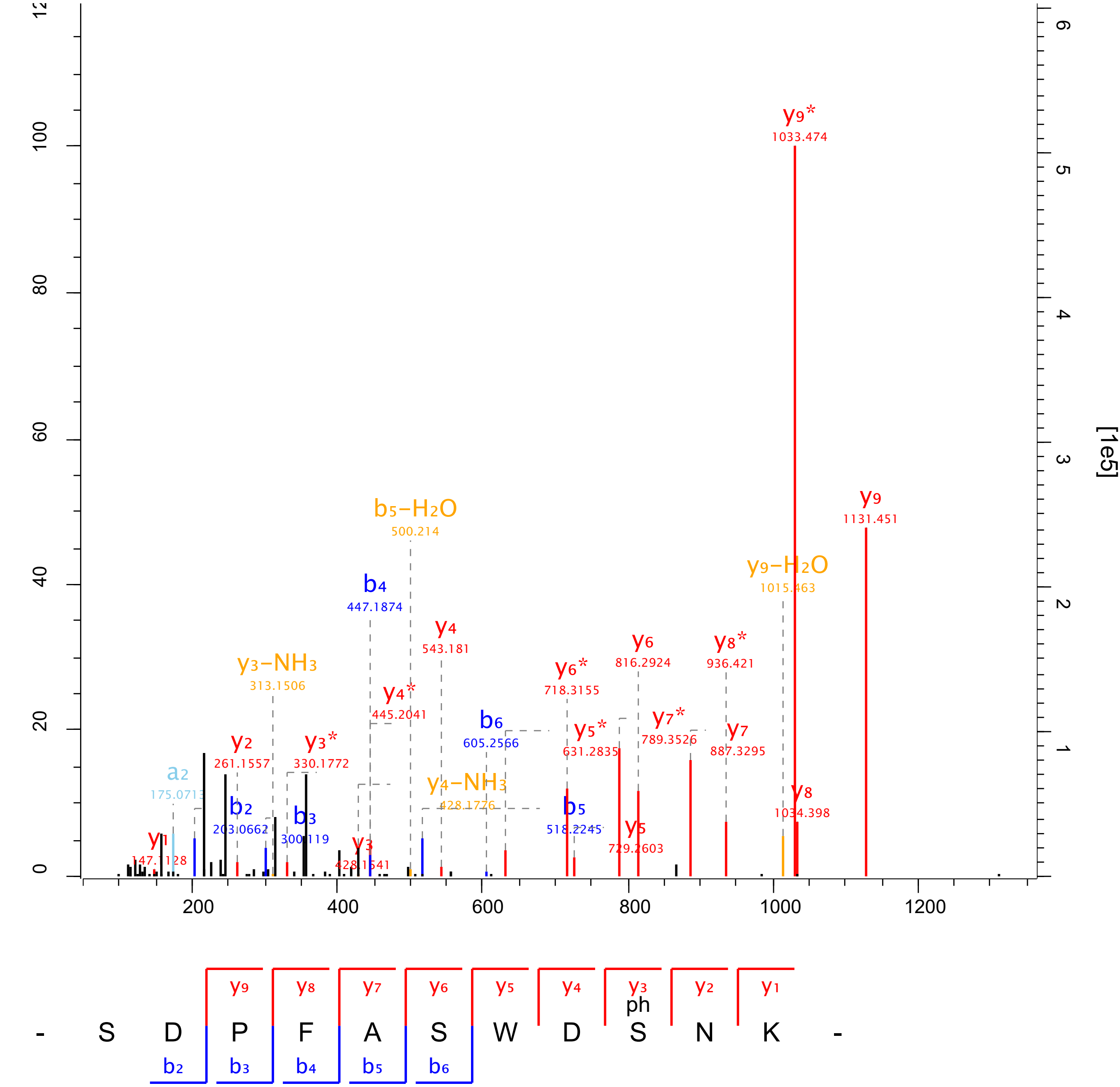

| Raw file | Scan | Method | Score | m/z |
|---|---|---|---|---|
| sys_05_2 | 31591 | FTMS; HCD | 47.55 | 732.83 |

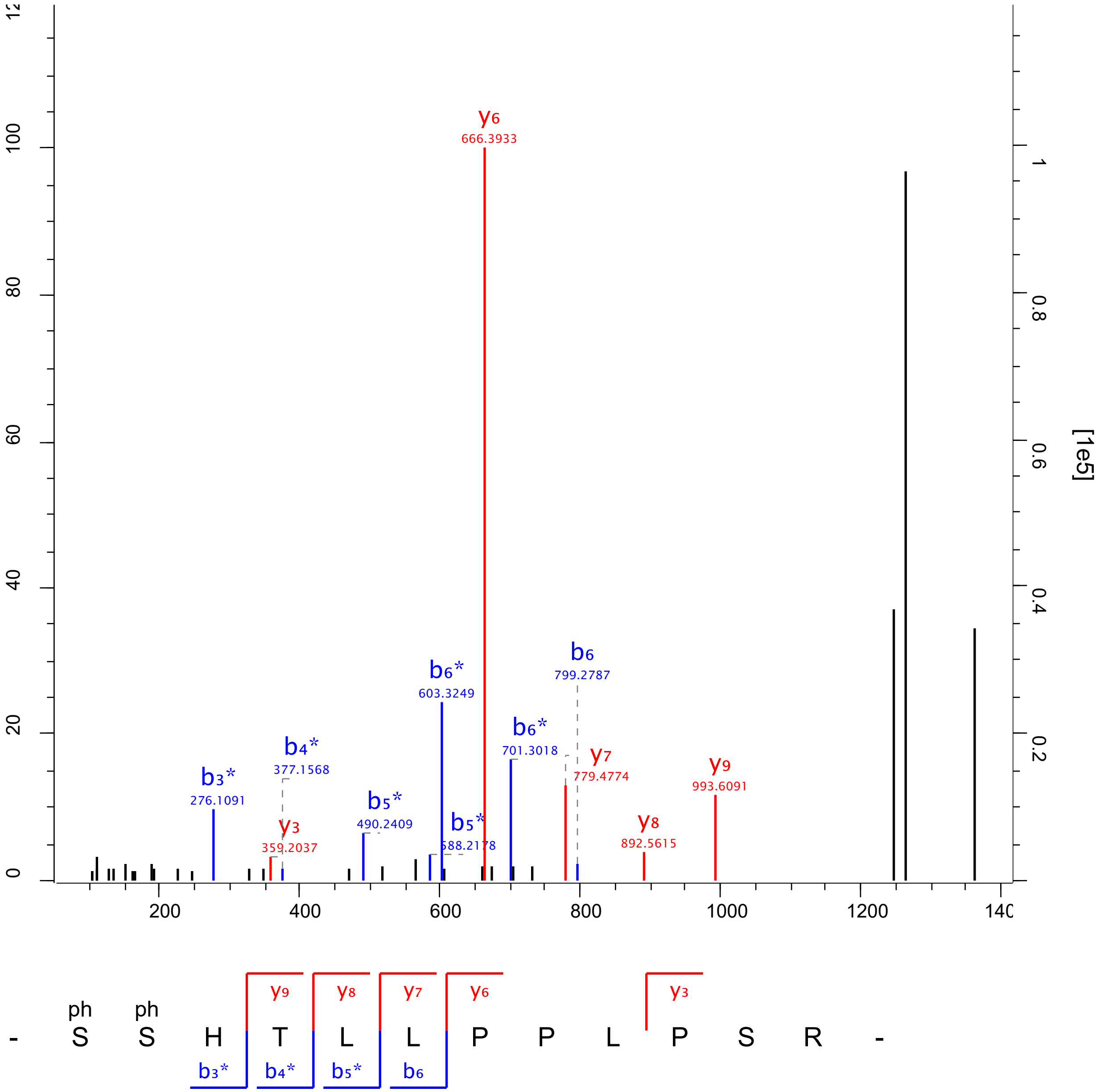

| Raw file | Scan | Method | Score | m/z |
| --- | --- | --- | --- | --- |
| sys_05_2 | 31802 | FTMS; HCD | 66.02 | 664.3 |

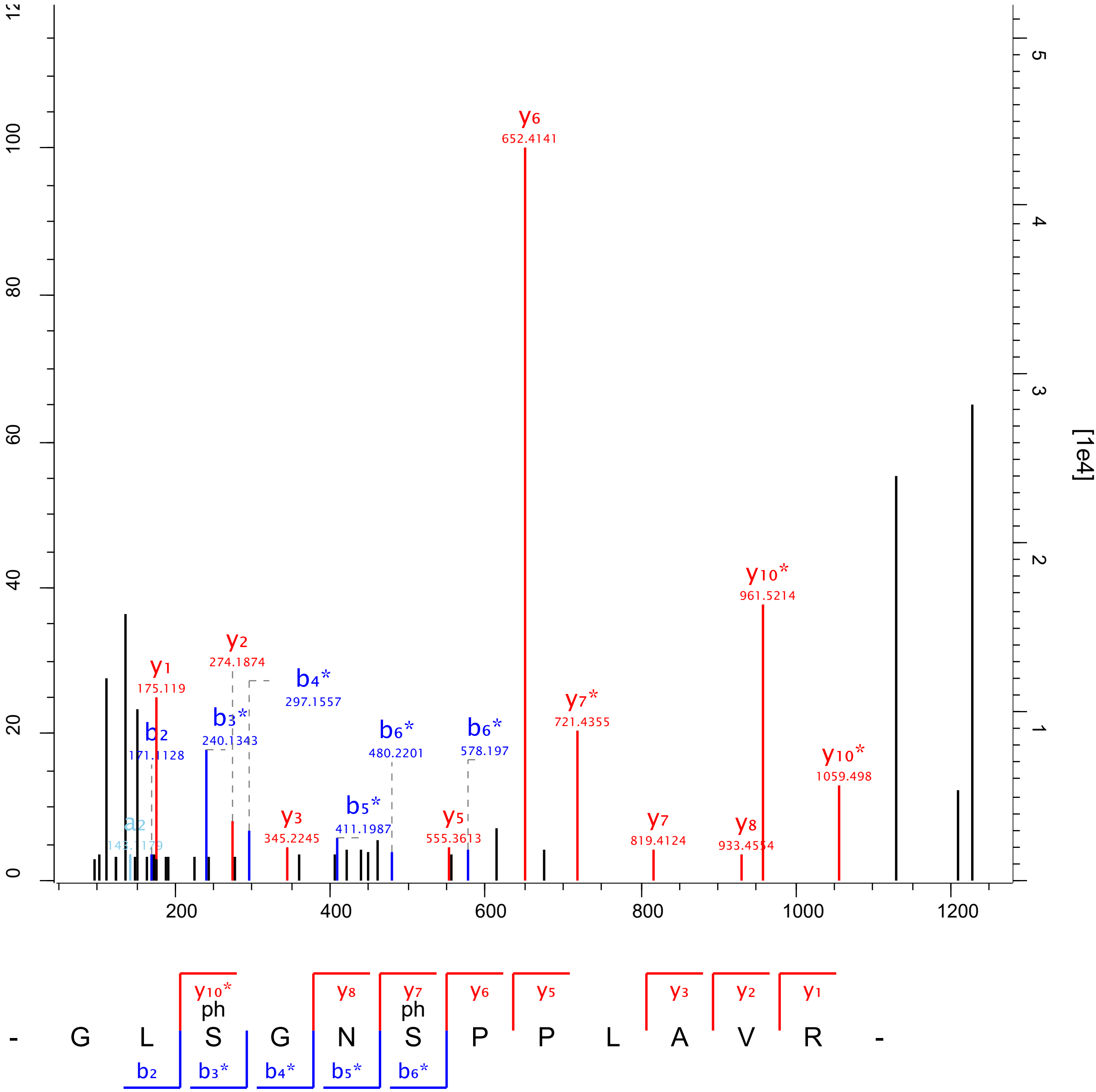

| Raw file | Scan | Method | Score | m/z |
|---|---|---|---|---|
| sys_05_2 | 31849 | FTMS; HCD | 160.81 | 696.34 |

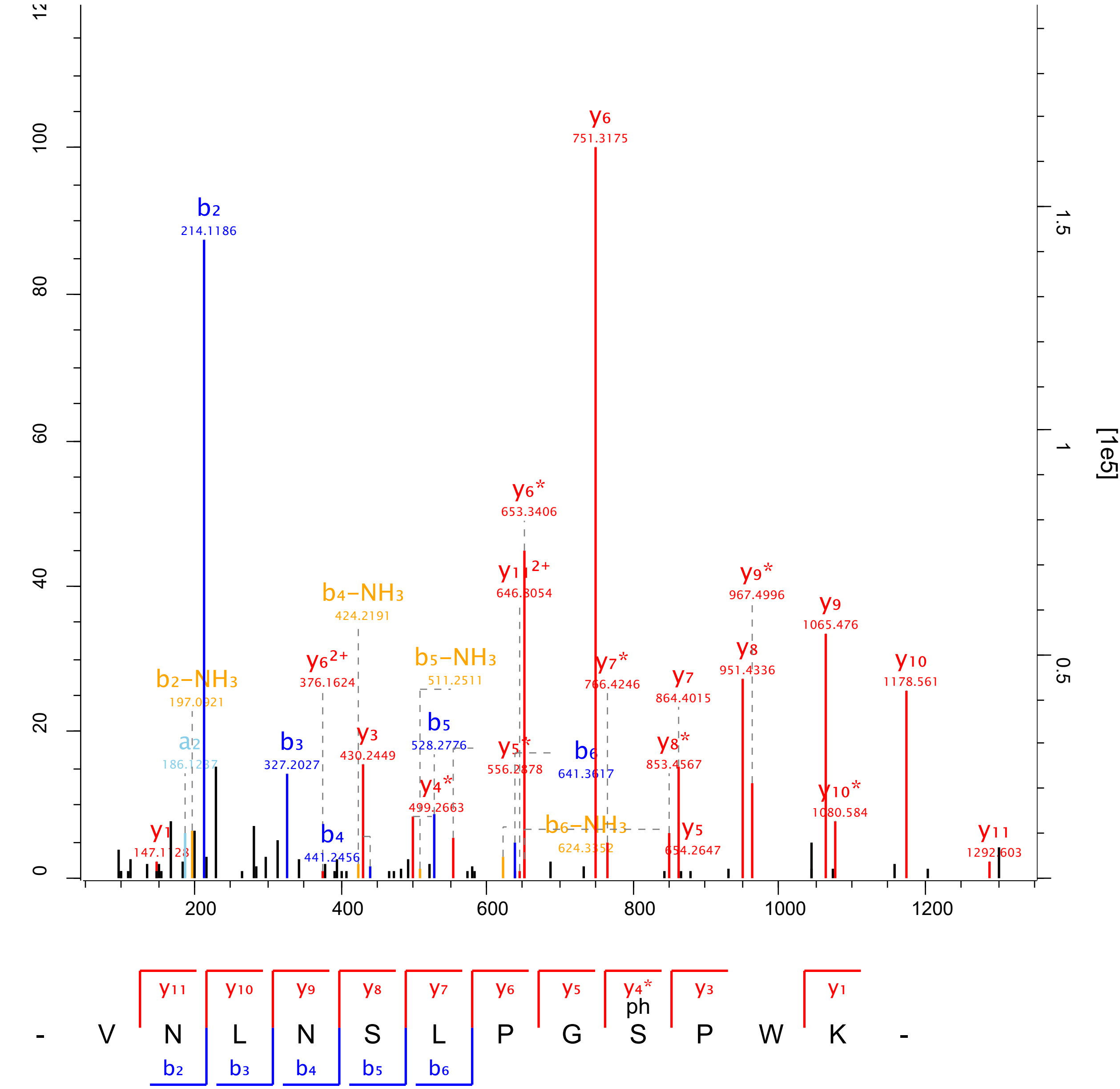

| Raw file | Scan | Method | Score | m/z |
| --- | --- | --- | --- | --- |
| sys_05_2 | 31871 | FTMS; HCD | 62.2 | 813.86 |

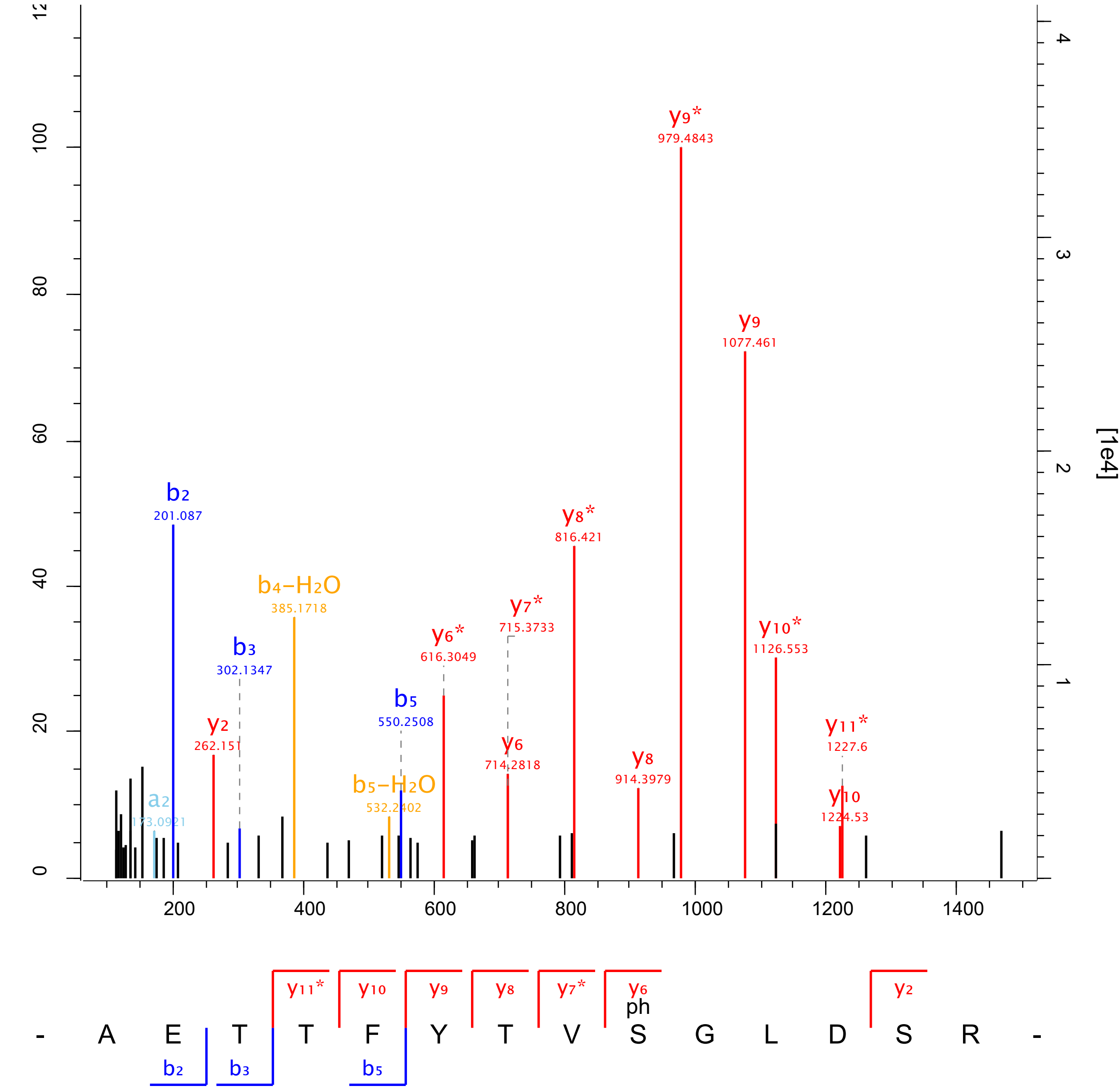

| Raw file | Scan | Method | Score | m/z |
|---|---|---|---|---|
| sys_05_2 | 32079 | FTMS; HCD | 72.68 | 677.98 |

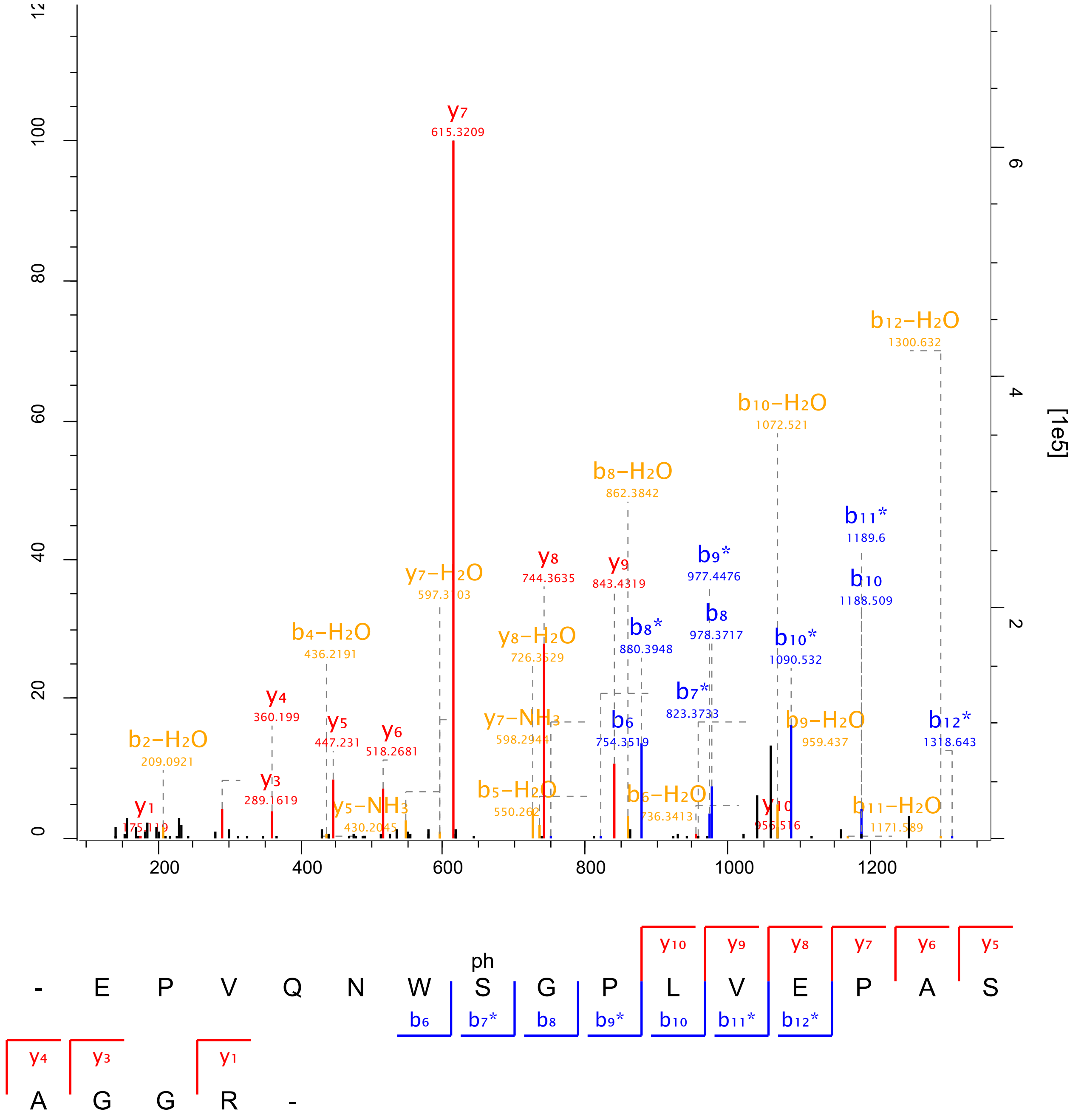

| Raw file | Scan | Method | Score | m/z |
| --- | --- | --- | --- | --- |
| sys_05_2 | 32203 | FTMS; HCD | 223.93 | 802.34 |

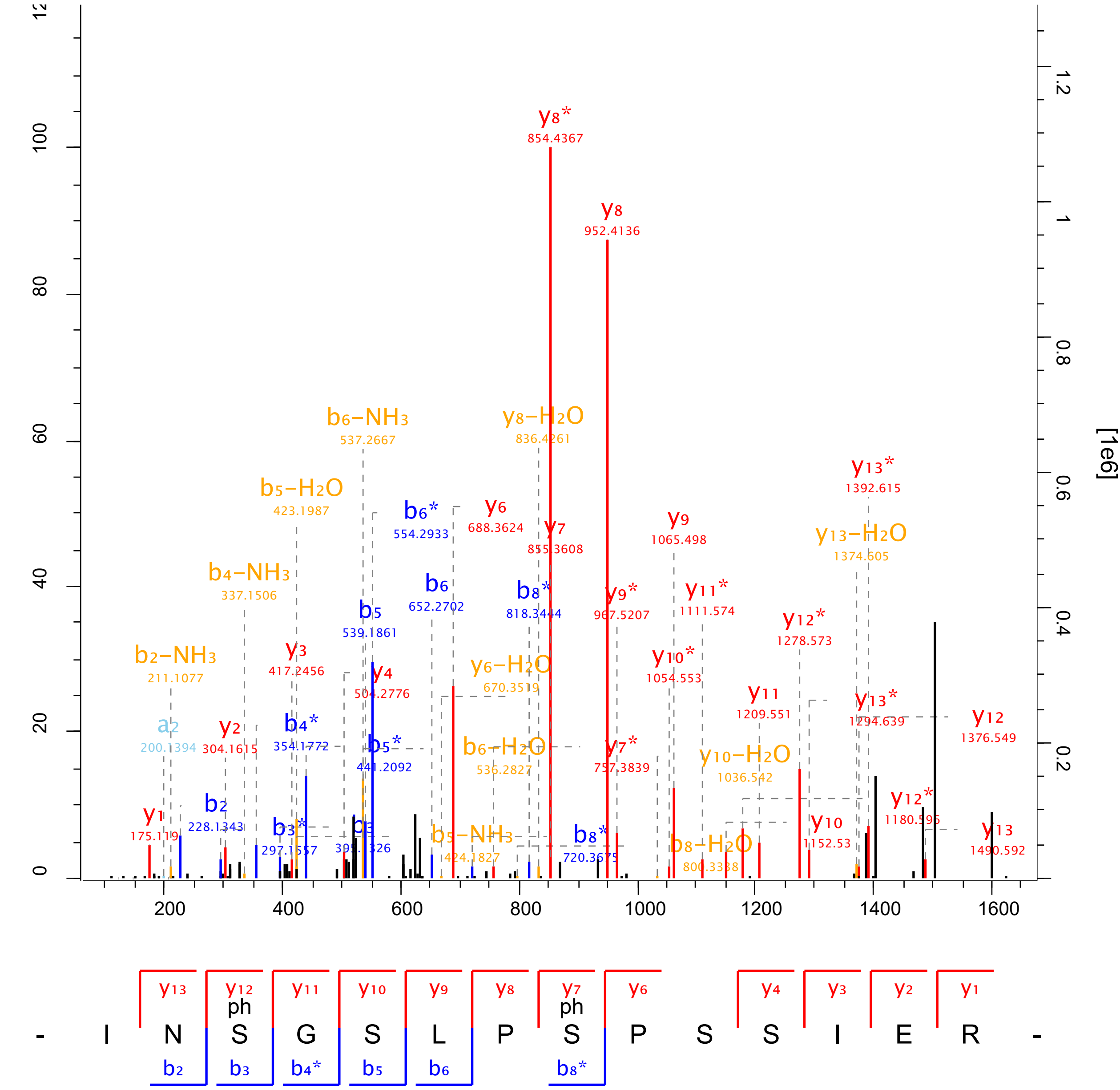

| Raw file | Scan | Method | Score | m/z |
|---|---|---|---|---|
| sys_05_2 | 32263 | FTMS; HCD | 215.51 | 722.33 |

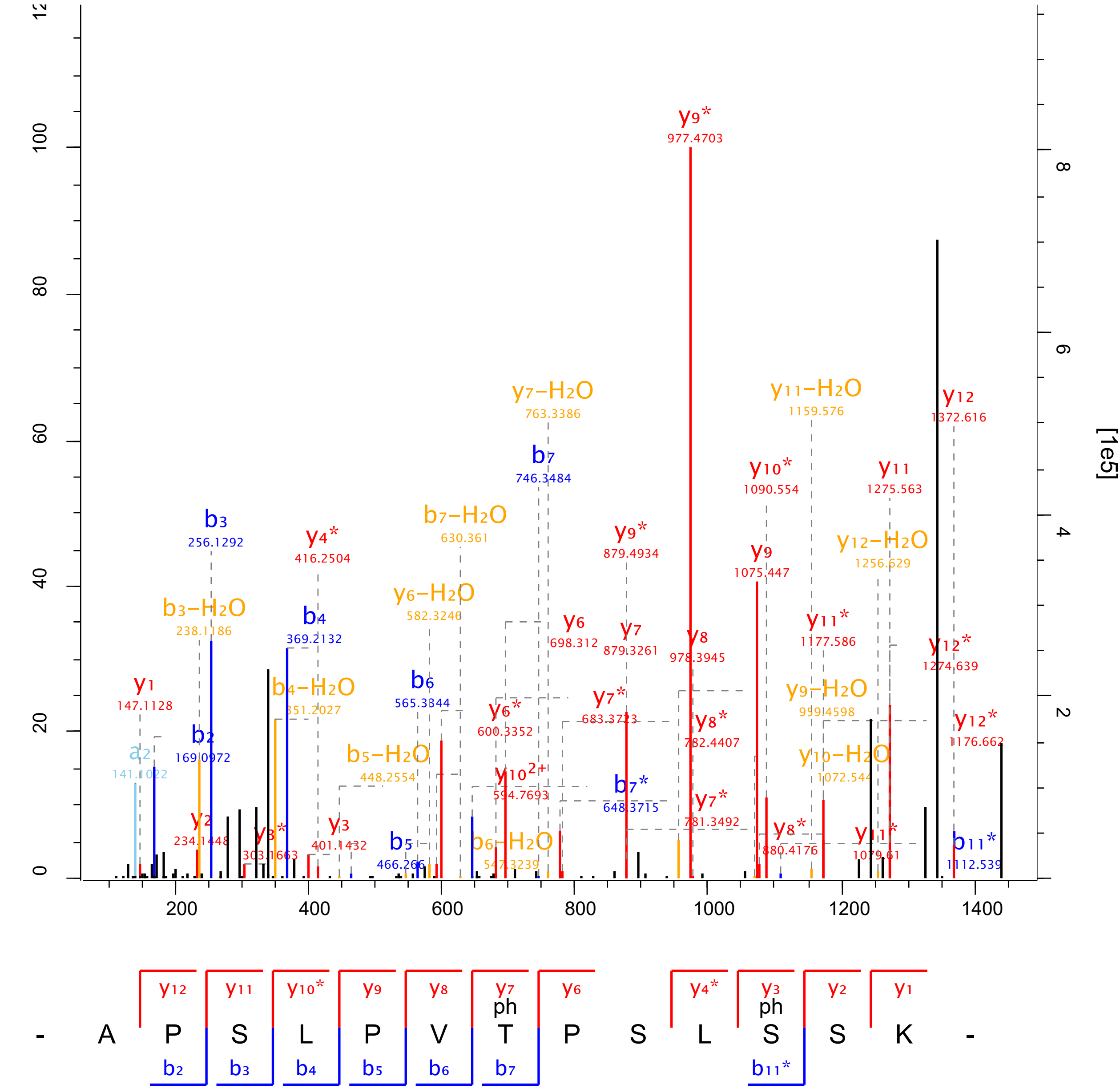

| Raw file | Scan | Method | Score | m/z |
|---|---|---|---|---|
| sys_05_2 | 32348 | FTMS; HCD | 125.41 | 719.82 |

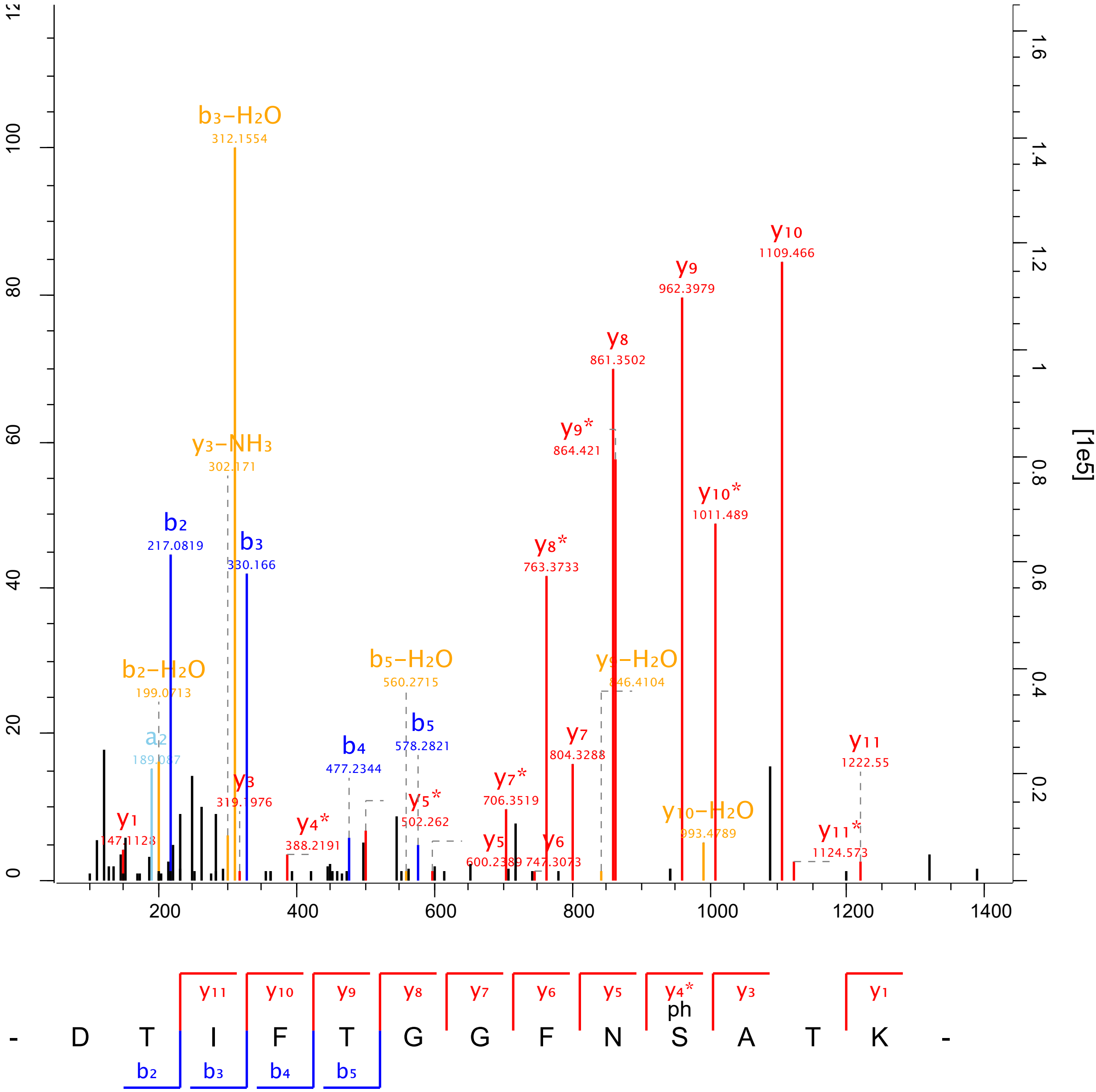

| Raw file | Scan | Method | Score | m/z |
|---|---|---|---|---|
| sys_05_2 | 32375 | FTMS; HCD | 44.44 | 696.81 |

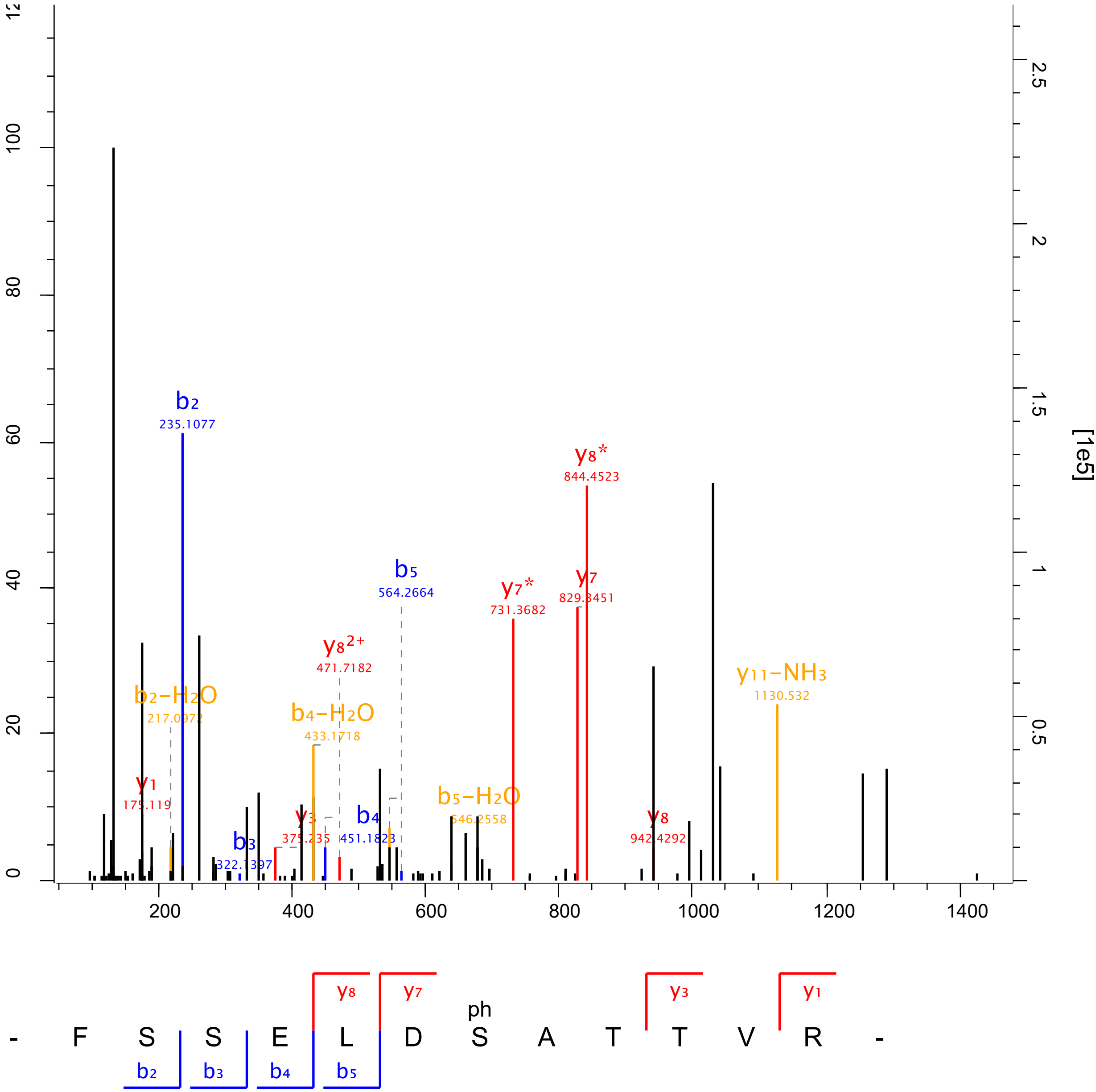

| Raw file | Scan | Method | Score | m/z |
|---|---|---|---|---|
| sys_05_2 | 32652 | FTMS; HCD | 65.02 | 958.92 |

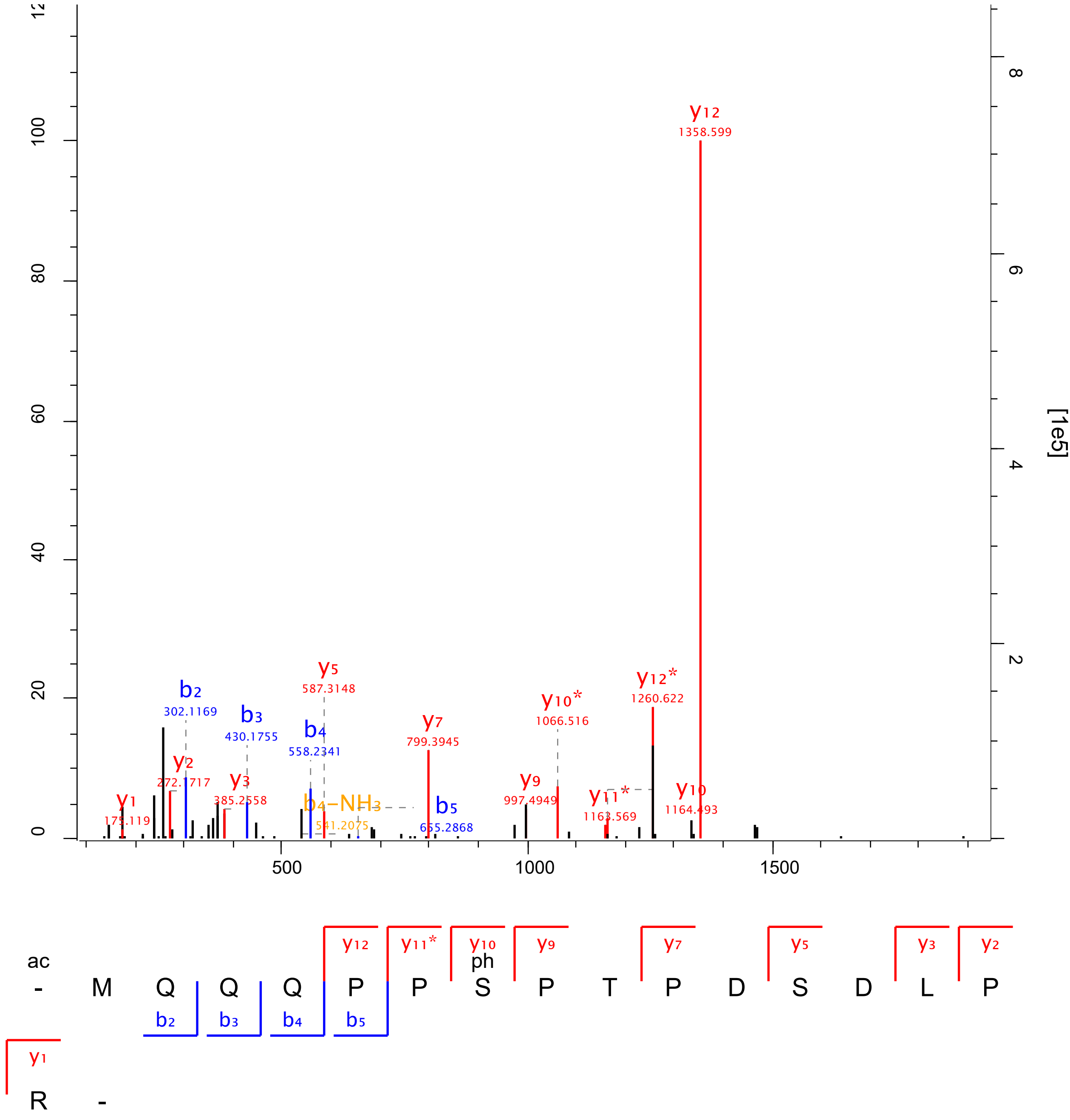

| Raw file | Scan | Method | Score | m/z |
|---|---|---|---|---|
| sys_05_2 | 32749 | FTMS; HCD | 54.76 | 789.35 |

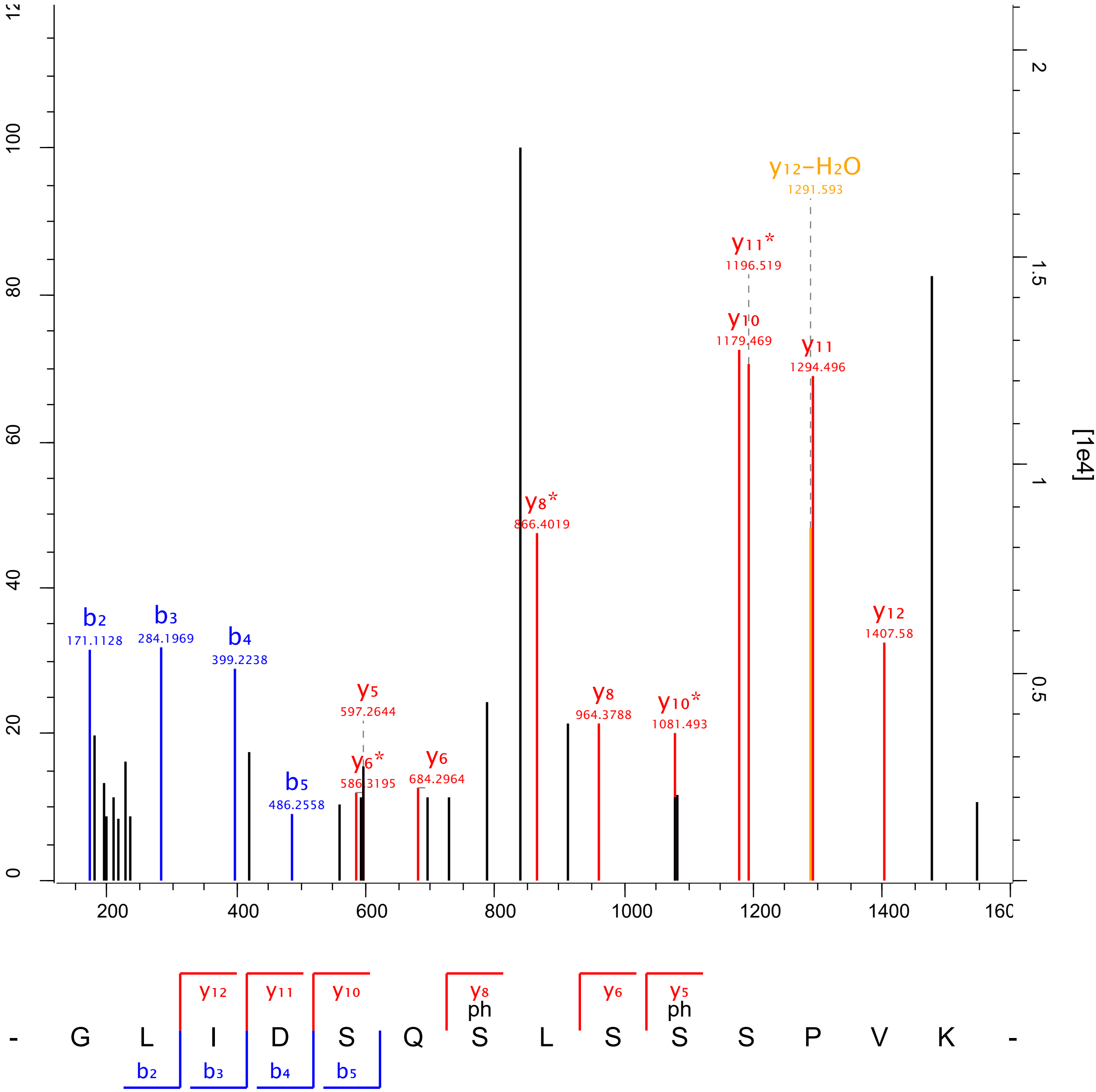

| Raw file | Scan | Method | Score | m/z |
|---|---|---|---|---|
| sys_05_2 | 32799 | FTMS; HCD | 110.44 | 729.84 |

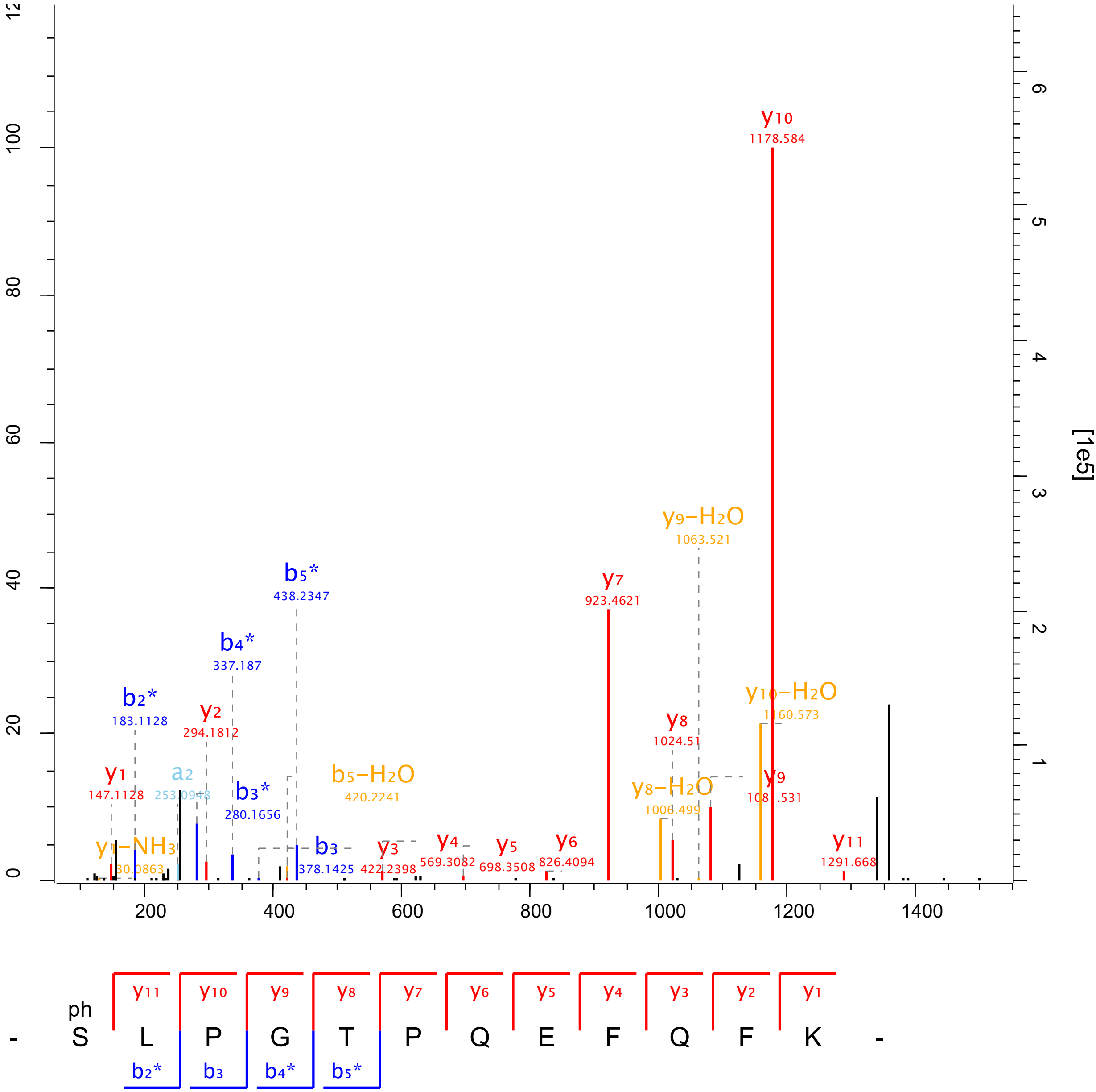

| Raw file | Scan | Method | Score | m/z |
|---|---|---|---|---|
| sys_05_2 | 3295 | FTMS; HCD | 52.58 | 633.79 |

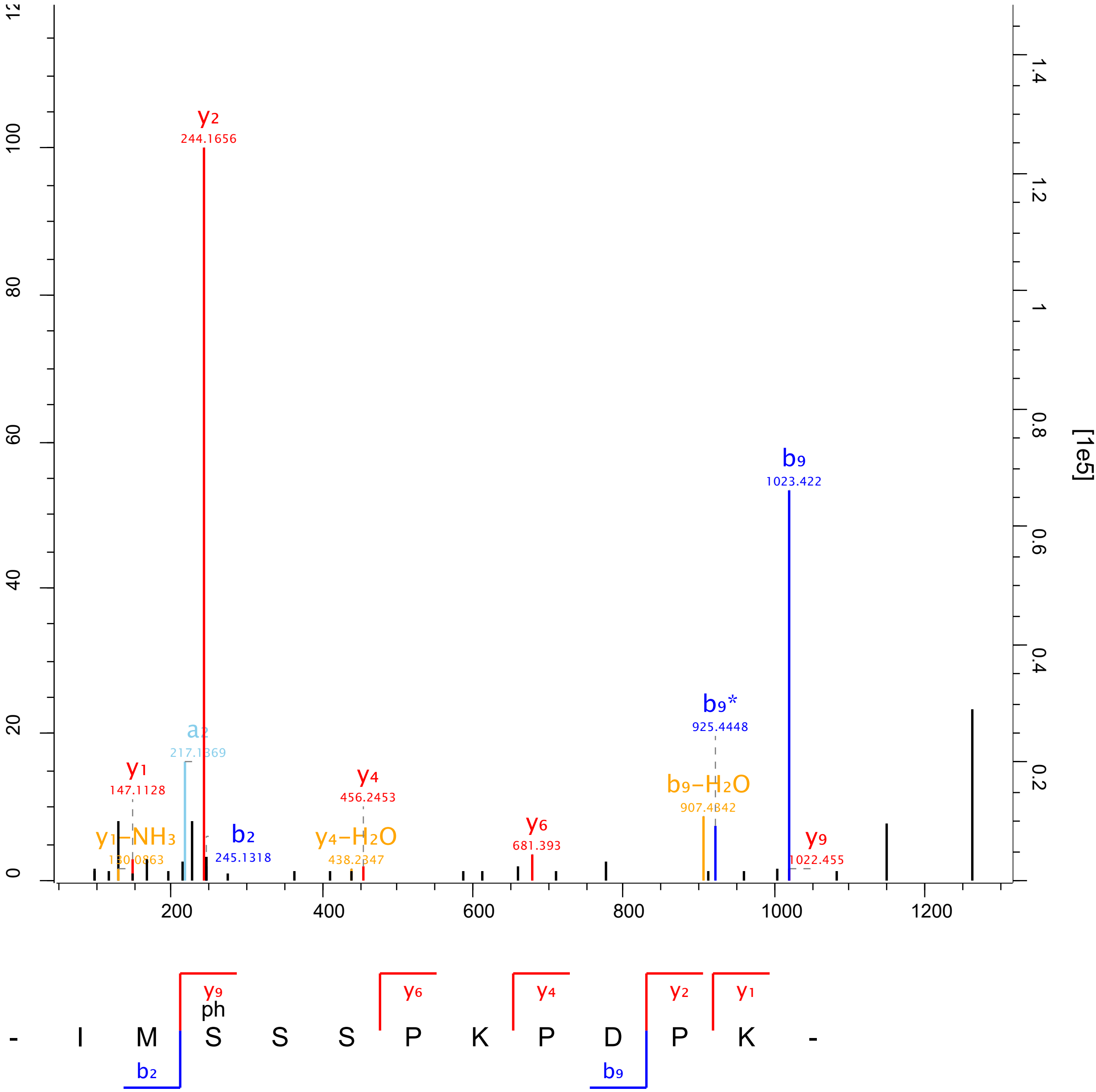

| Raw file | Scan | Method | Score | m/z |
|---|---|---|---|---|
| sys_05_2 | 33063 | FTMS; HCD | 105.79 | 752.82 |

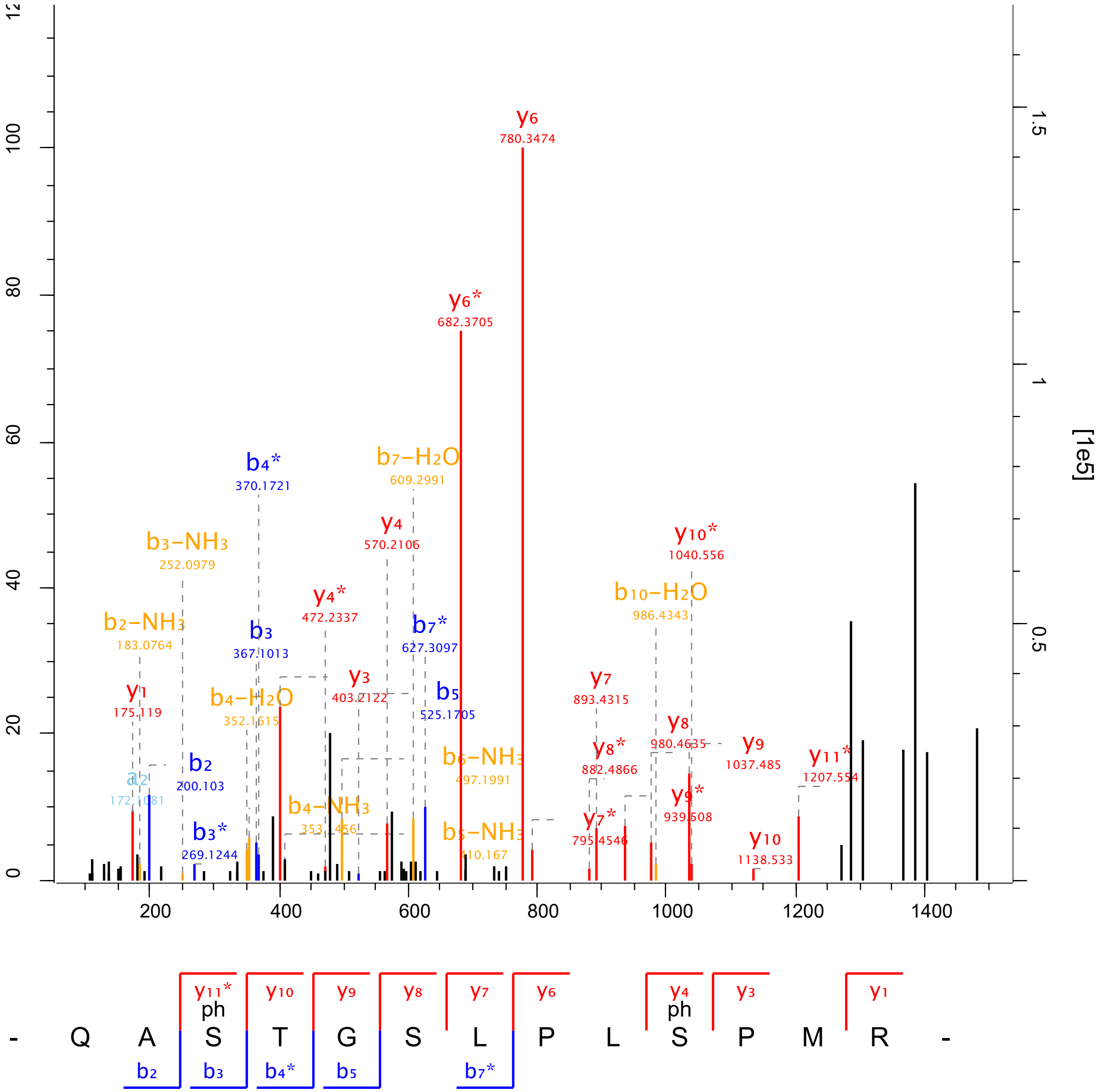

| Raw file | Scan | Method | Score | m/z |
|---|---|---|---|---|
| sys_05_2 | 33199 | FTMS; HCD | 163.18 | 1023.42 |

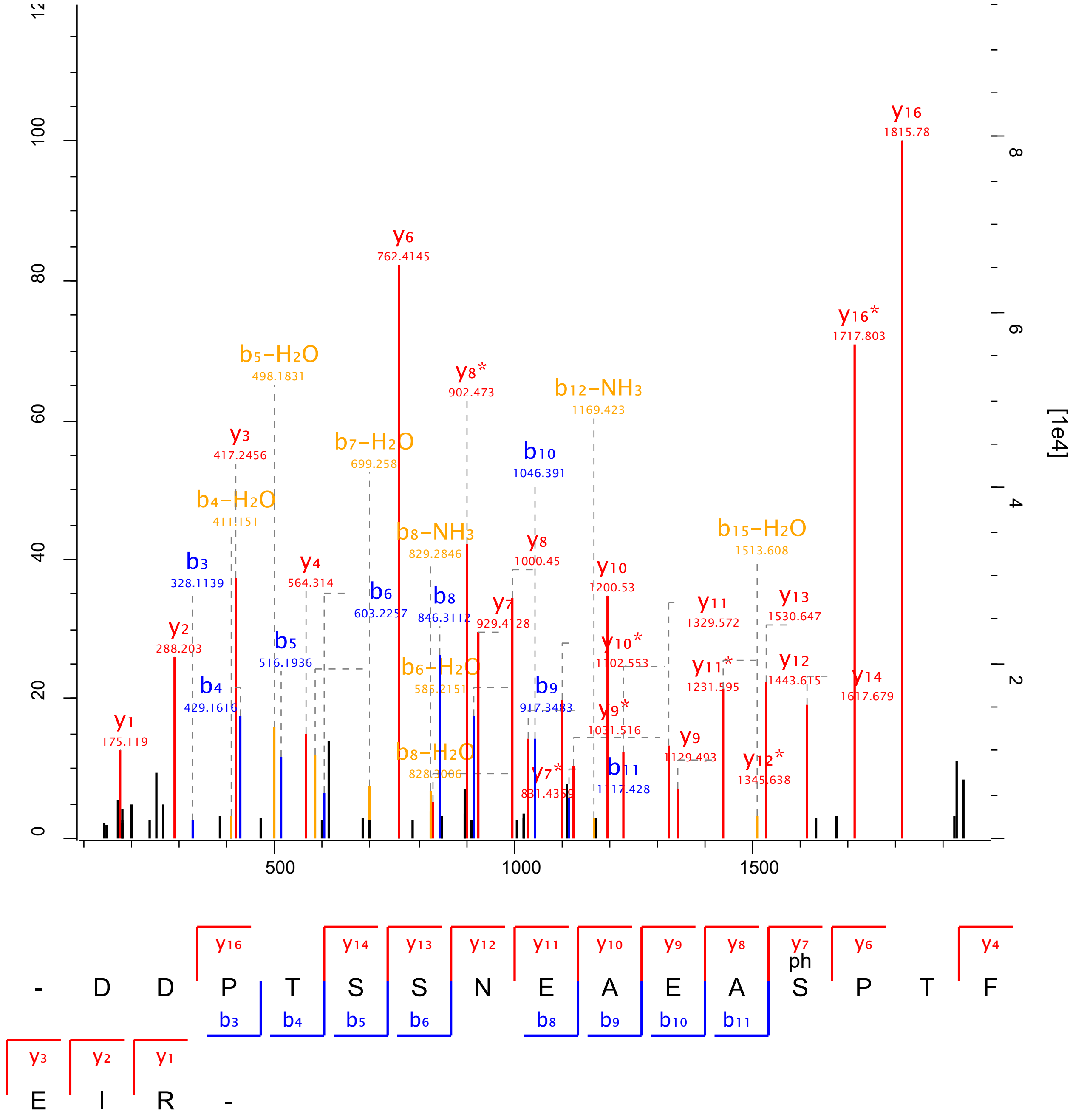

| Raw file | Scan | Method | Score | m/z |
|---|---|---|---|---|
| sys_05_2 | 33301 | FTMS; HCD | 218.2 | 742.34 |

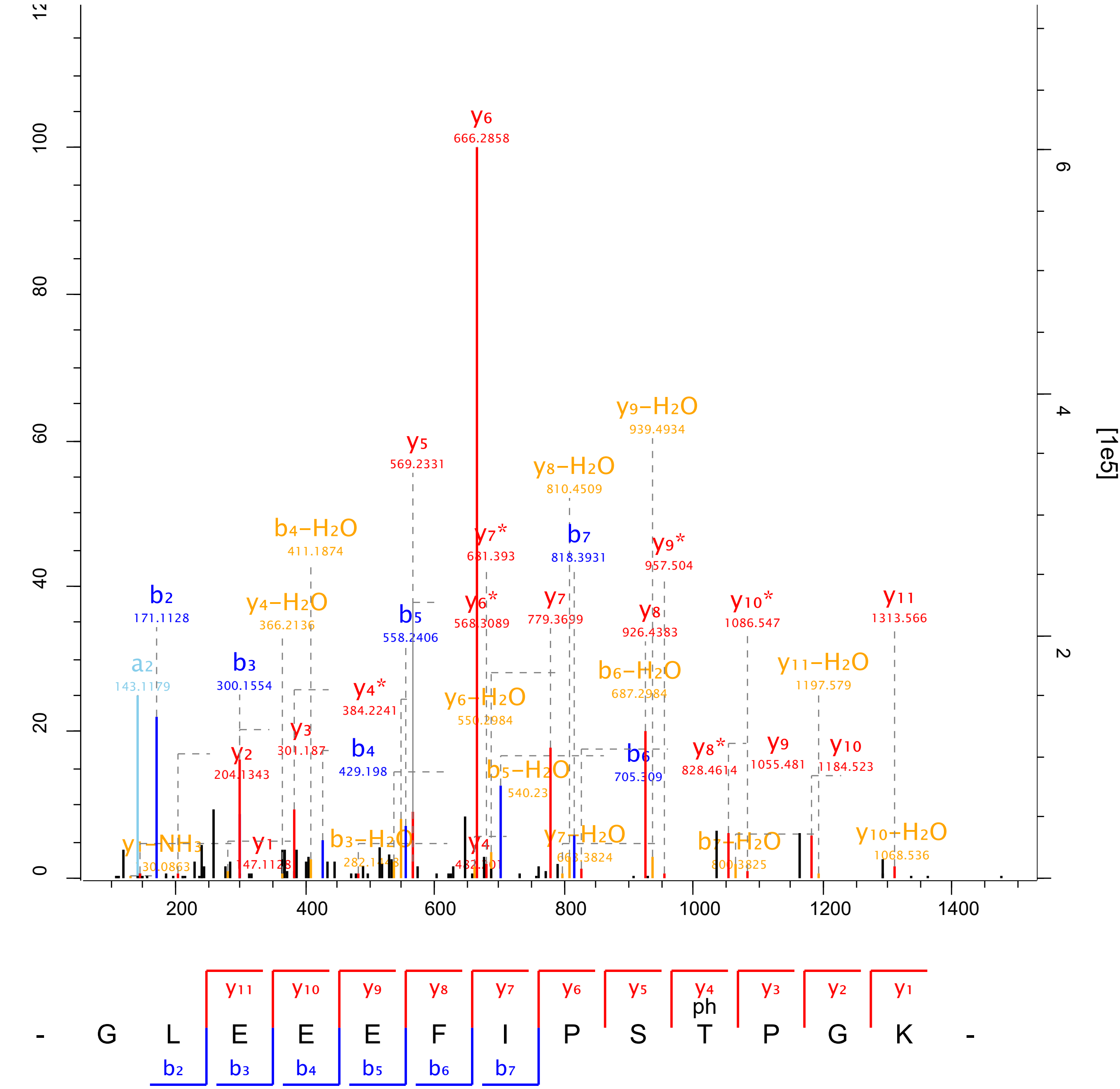

| Raw file | Scan | Method | Score | m/z |
|---|---|---|---|---|
| sys_05_2 | 33521 | FTMS; HCD | 47.04 | 727.29 |

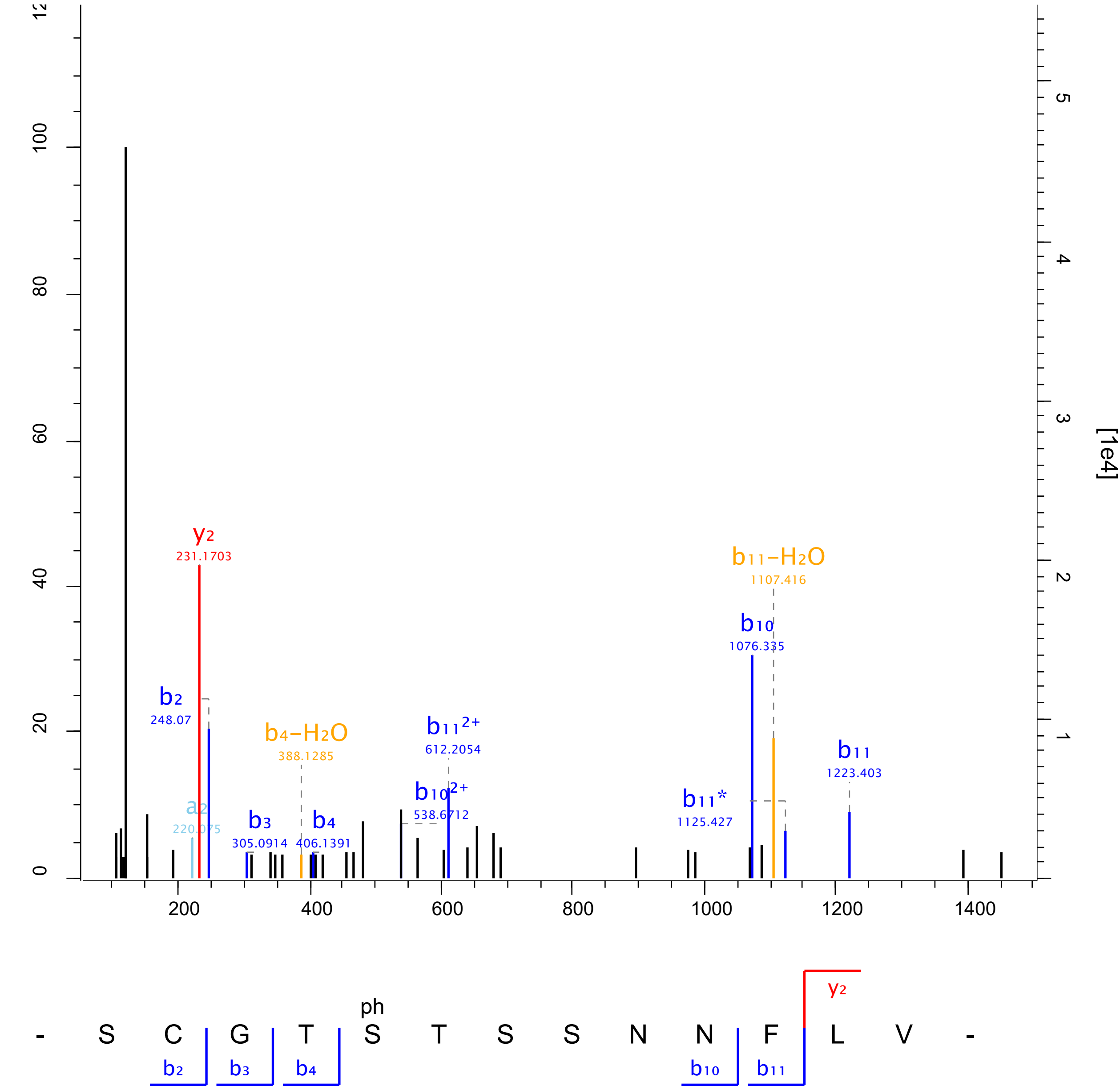

| Raw file | Scan | Method | Score | m/z |
| --- | --- | --- | --- | --- |
| sys_05_2 | 33540 | FTMS; HCD | 104.26 | 741.35 |

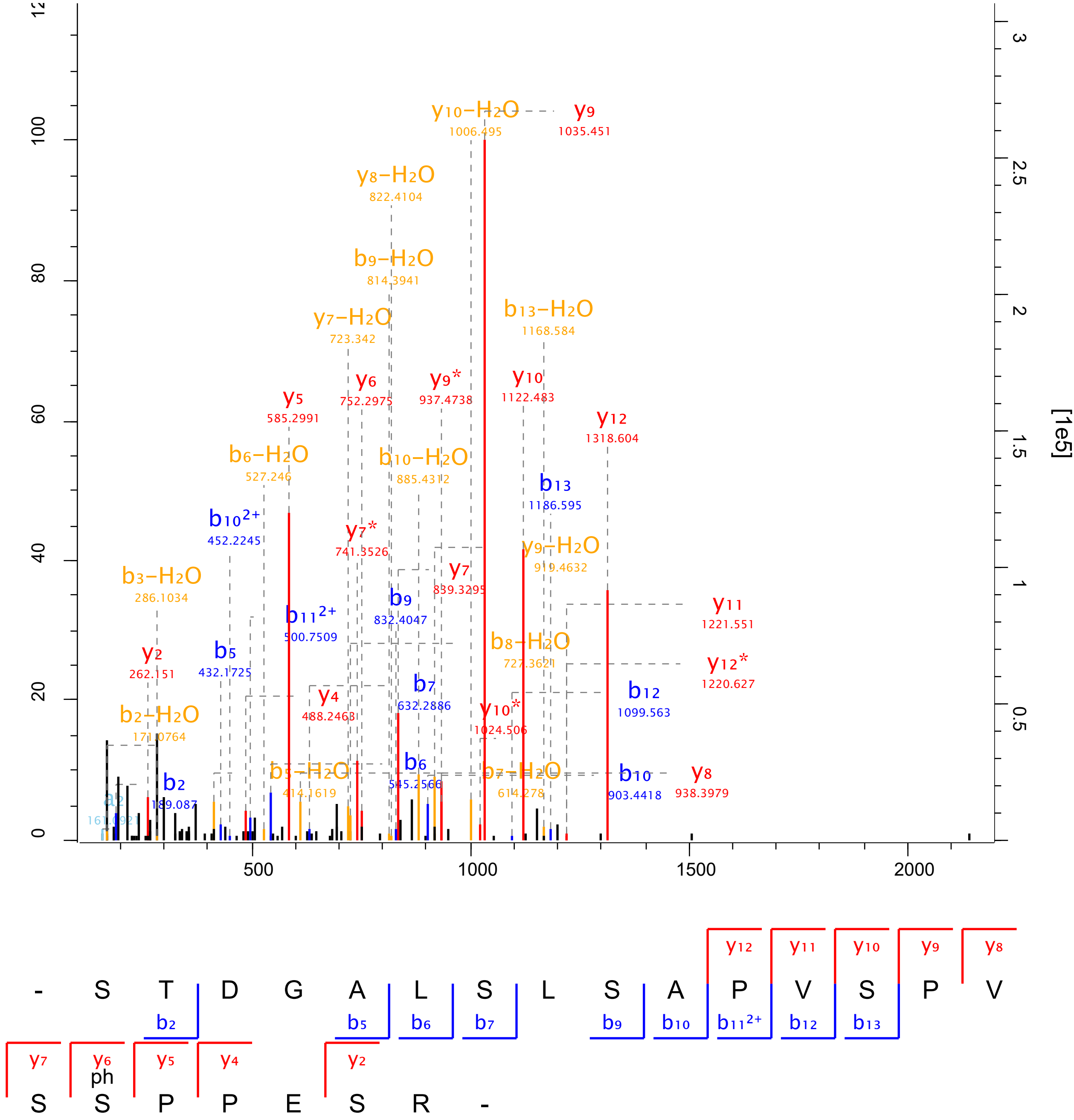

| Raw file | Scan | Method | Score | m/z |
| --- | --- | --- | --- | --- |
| sys_05_2 | 33583 | FTMS; HCD | 41.7 | 757.82 |

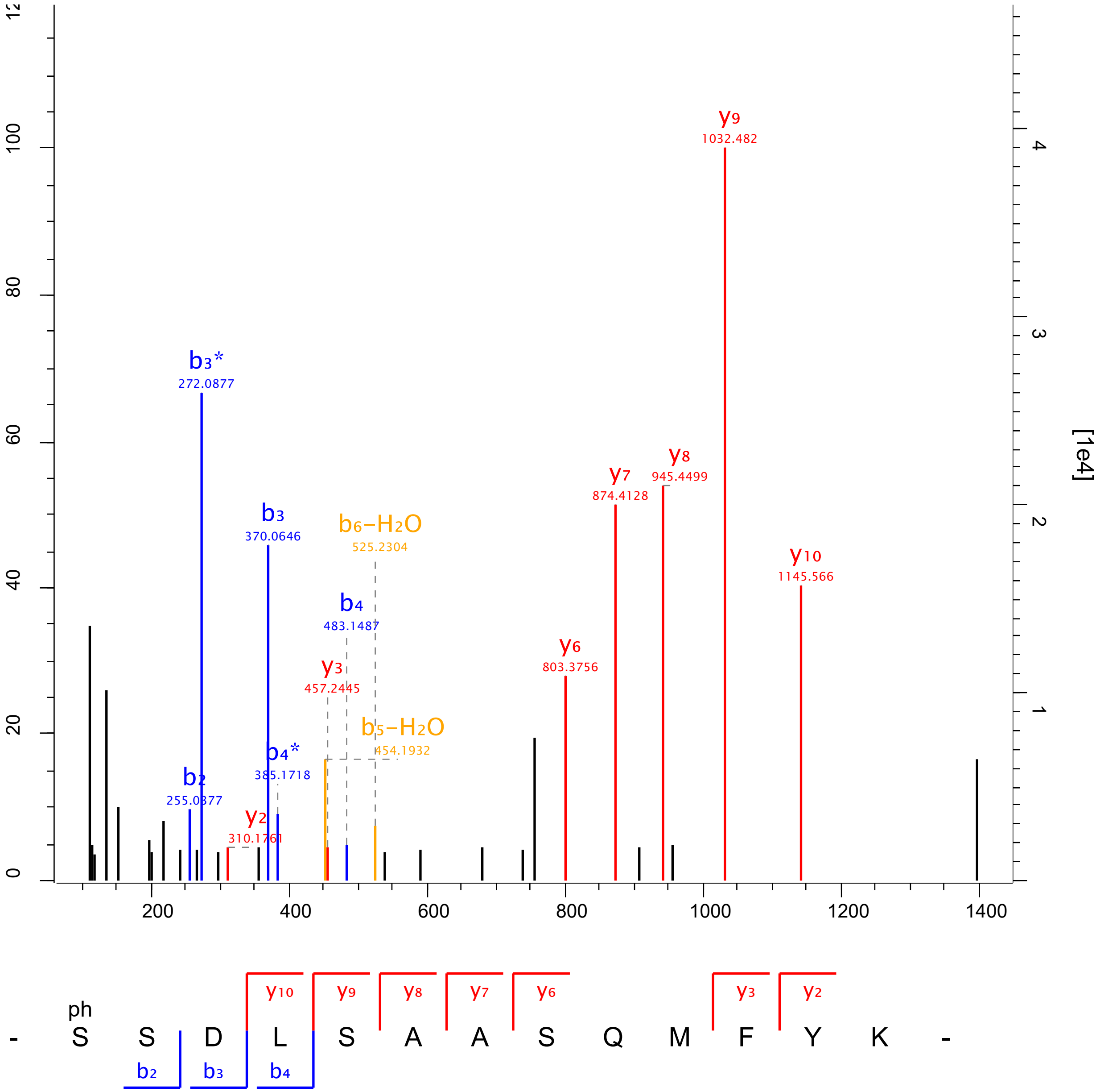

| Raw file | Scan | Method | Score | m/z |
|---|---|---|---|---|
| sys_05_2 | 34587 | FTMS; HCD | 106.67 | 580.73 |

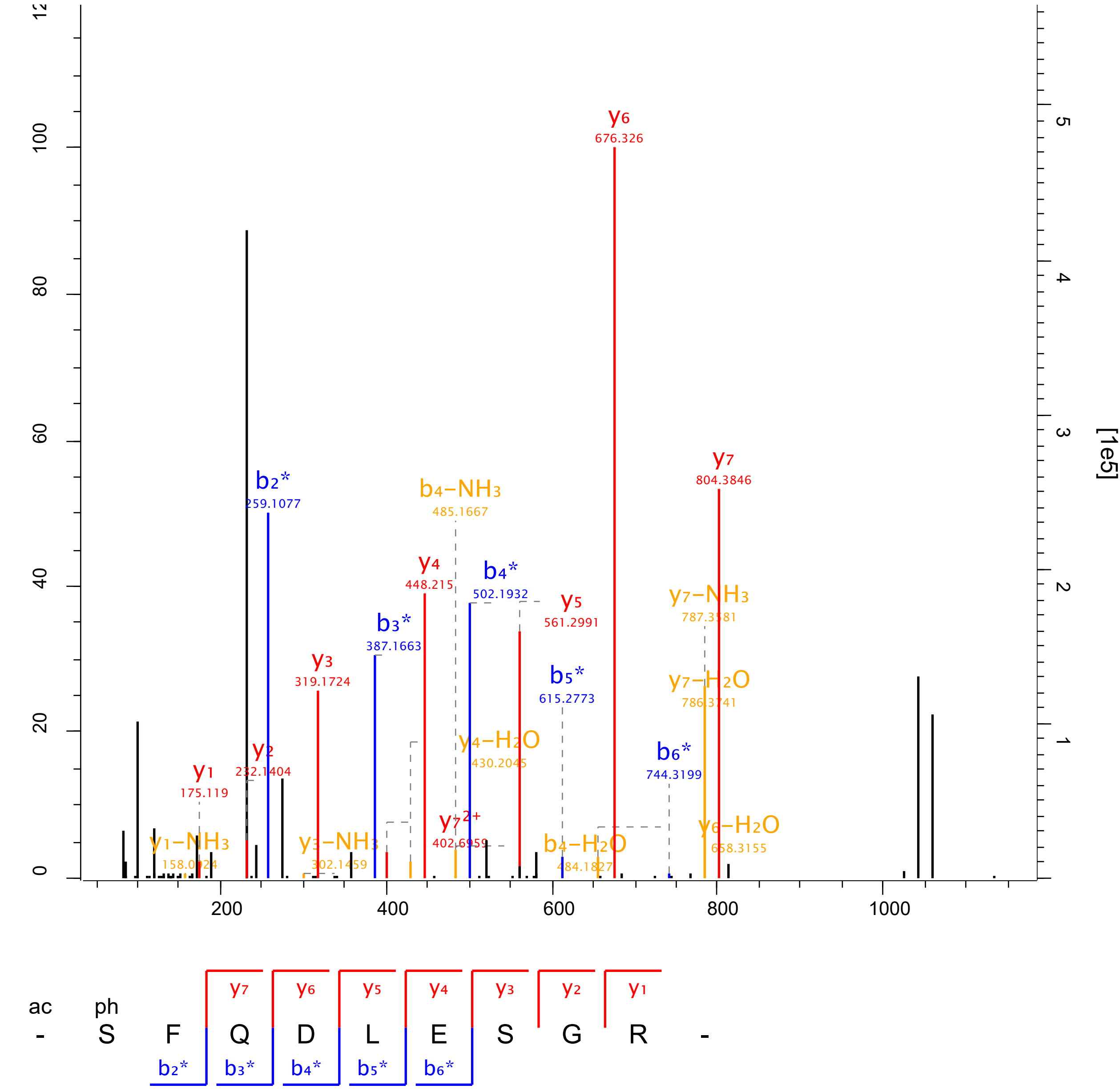

| Raw file | Scan | Method | Score | m/z |
|---|---|---|---|---|
| sys_05_2 | 34708 | FTMS; HCD | 247.91 | 731.78 |

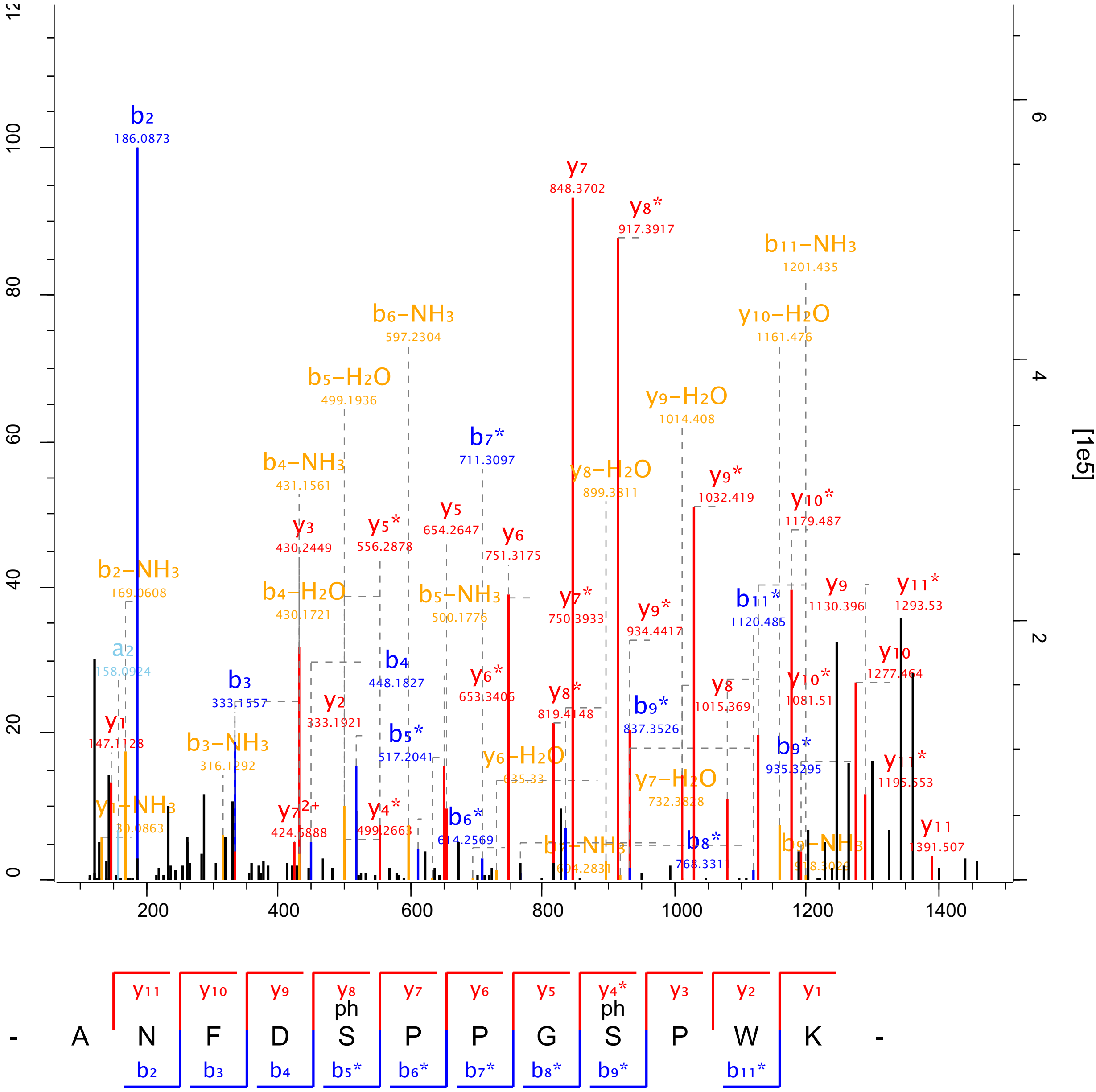

| Raw file | Scan | Method | Score | m/z |
| --- | --- | --- | --- | --- |
| sys_05_2 | 35017 | FTMS; HCD | 68.68 | 662.33 |

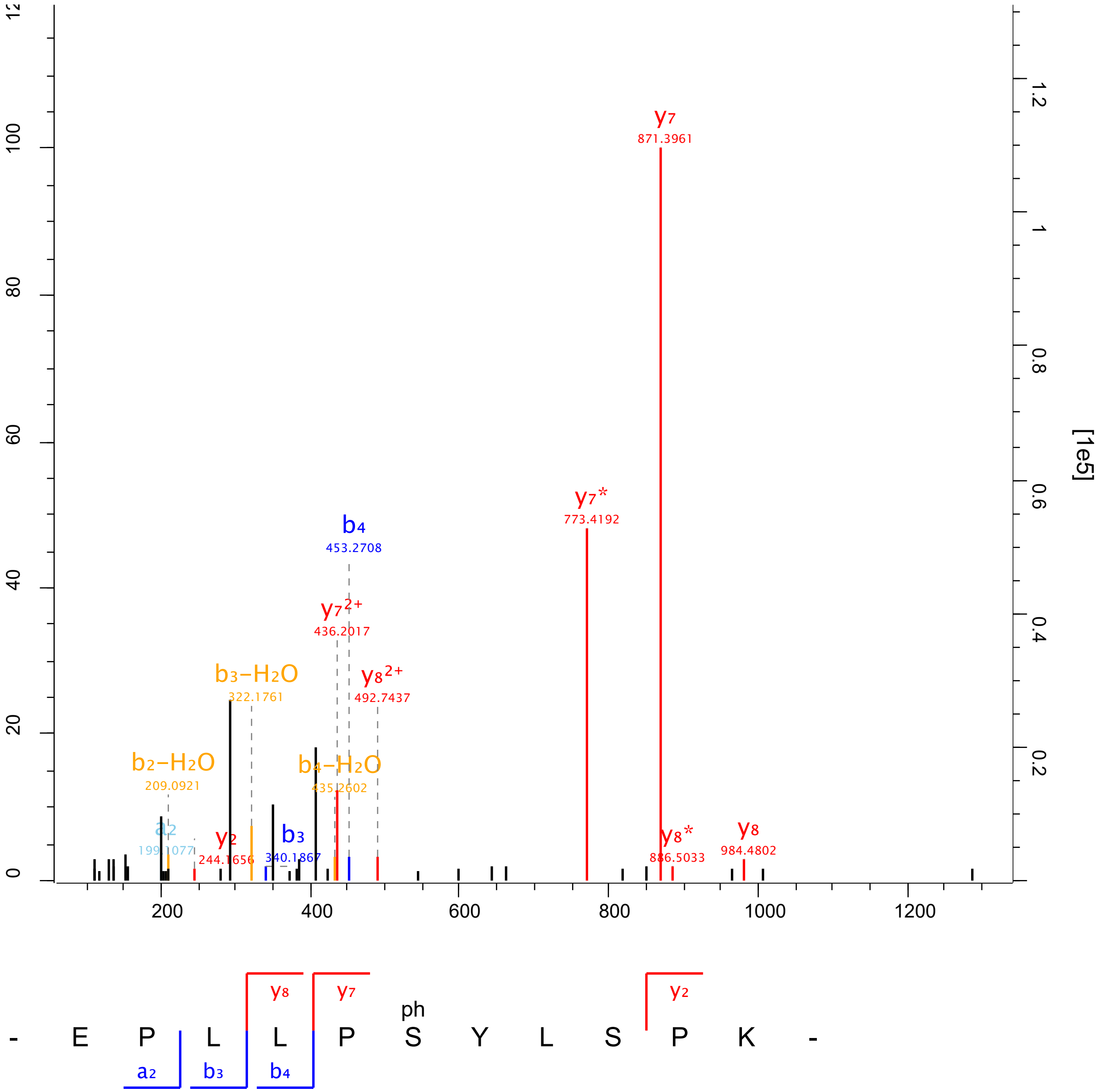

| Raw file | Scan | Method | Score | m/z |
|---|---|---|---|---|
| sys_05_2 | 35388 | FTMS; HCD | 185.66 | 597.26 |

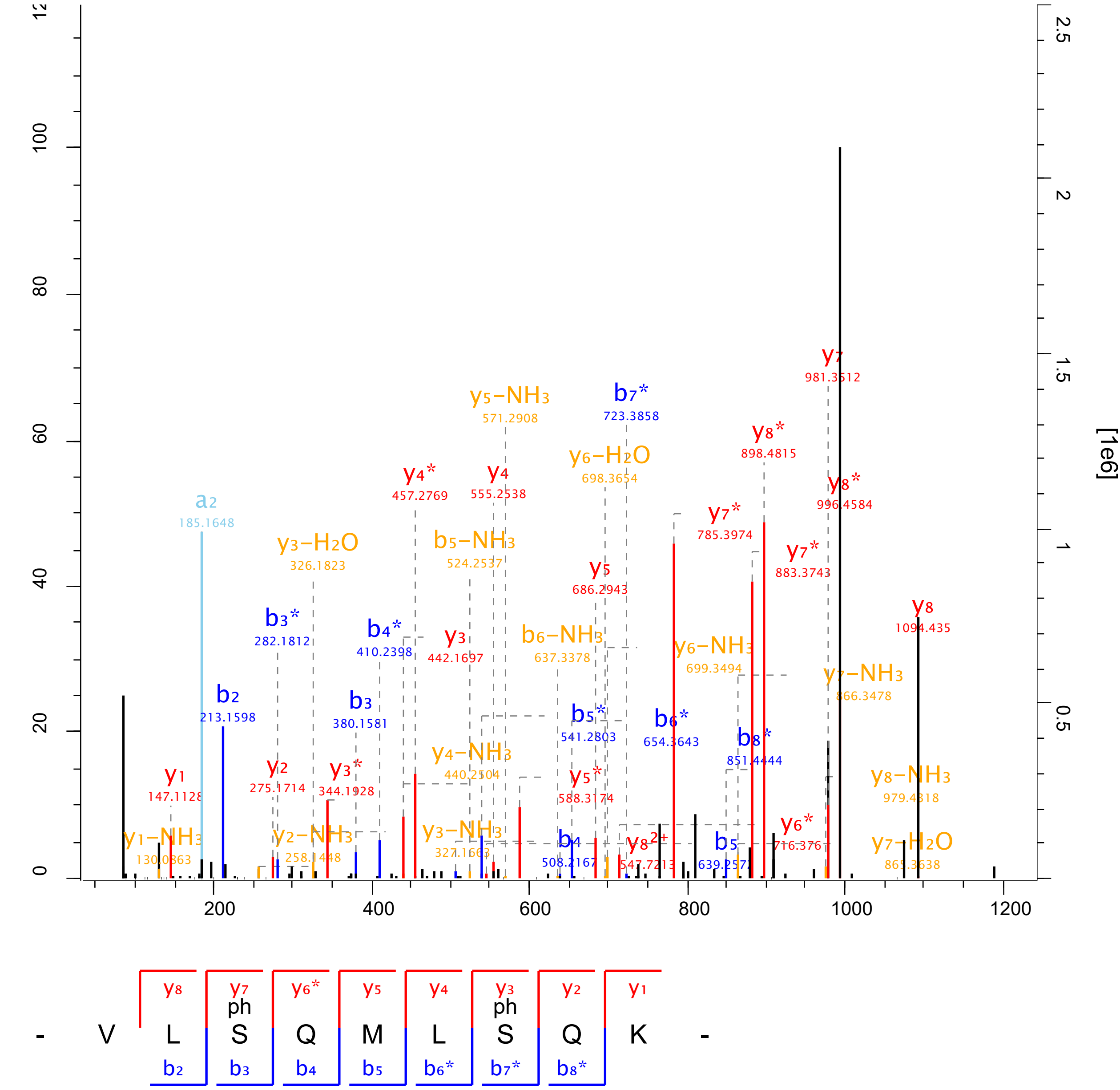

| Raw file | Scan | Method | Score | m/z |
| --- | --- | --- | --- | --- |
| sys_05_2 | 3551 | FTMS; HCD | 80.69 | 606.25 |

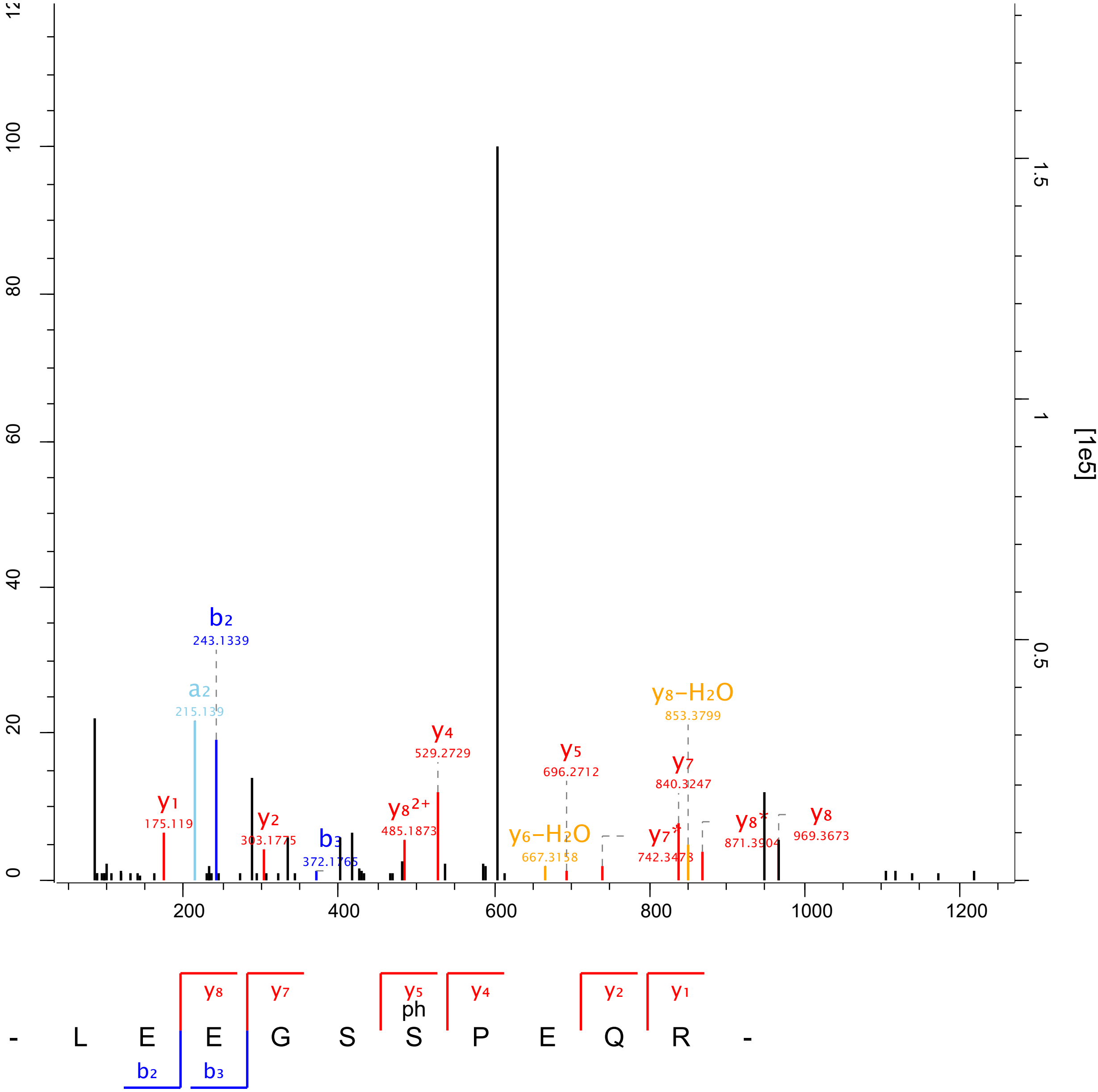

| Raw file | Scan | Method | Score | m/z |
|---|---|---|---|---|
| sys_05_2 | 3575 | FTMS; HCD | 61.24 | 545.73 |

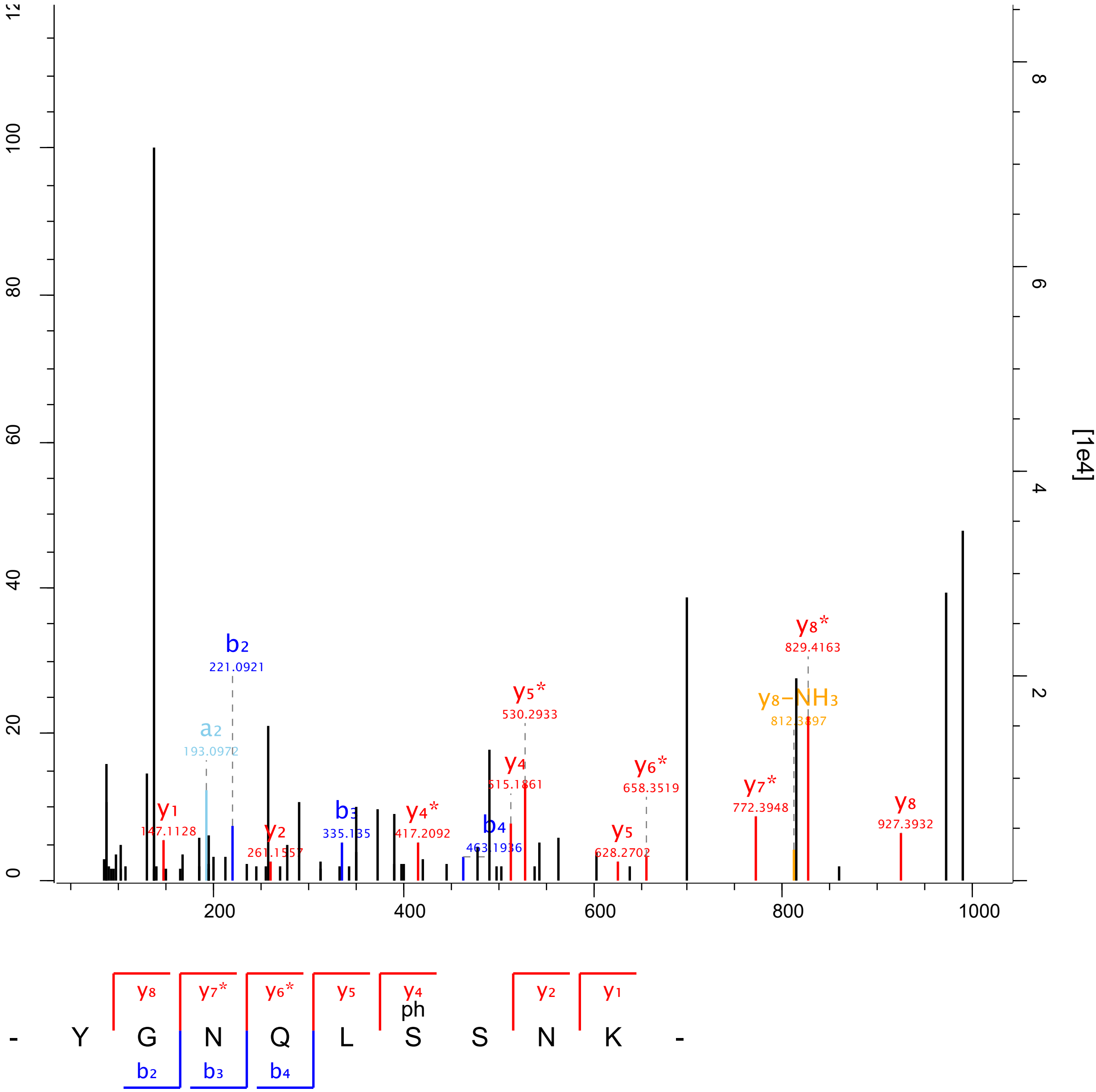

| Raw file | Scan | Method | Score | m/z |
| --- | --- | --- | --- | --- |
| sys_05_2 | 35983 | FTMS; HCD | 68.58 | 946.92 |

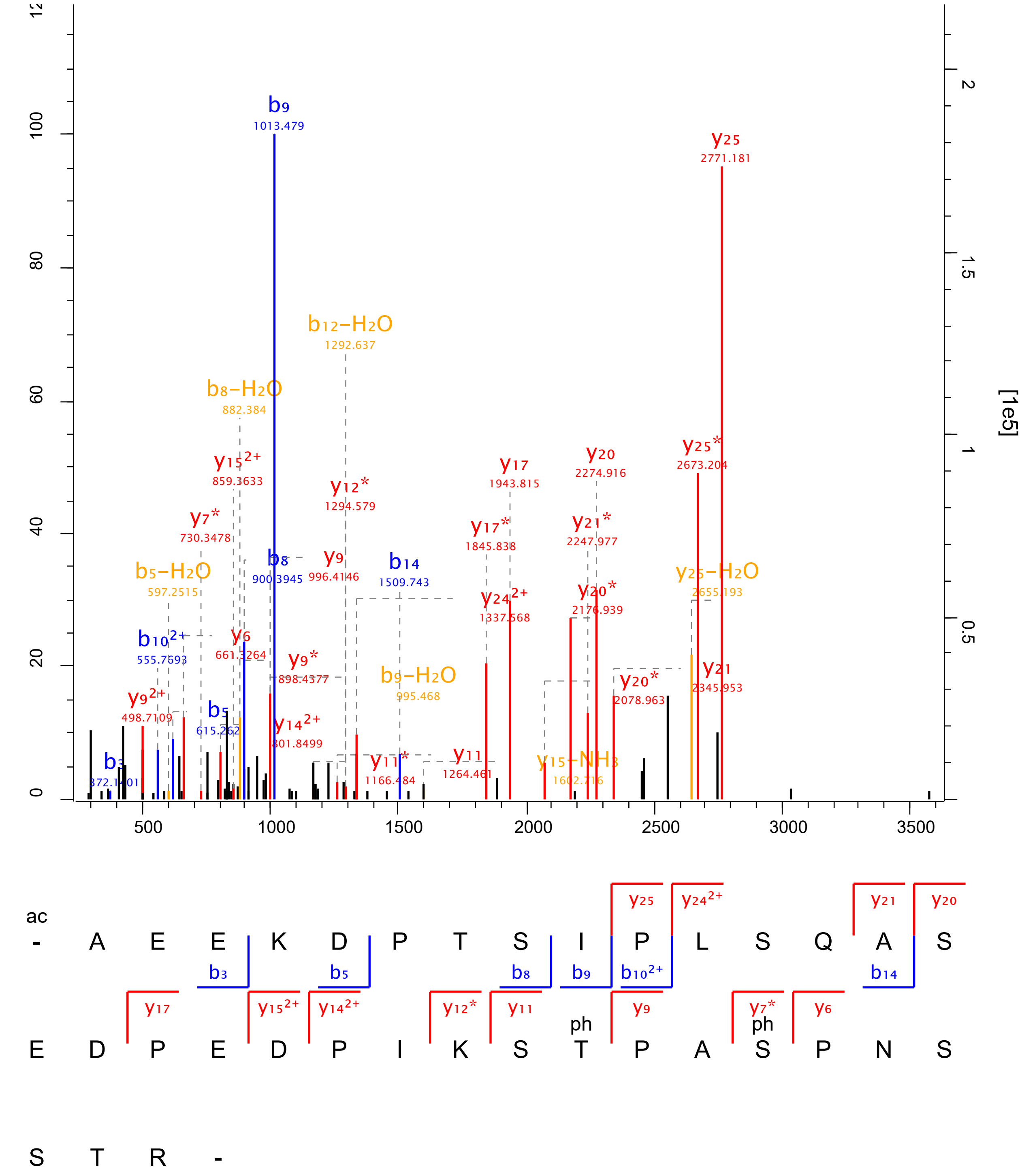

| Raw file | Scan | Method | Score | m/z |
|---|---|---|---|---|
| sys_05_2 | 36133 | FTMS; HCD | 170.09 | 799.37 |

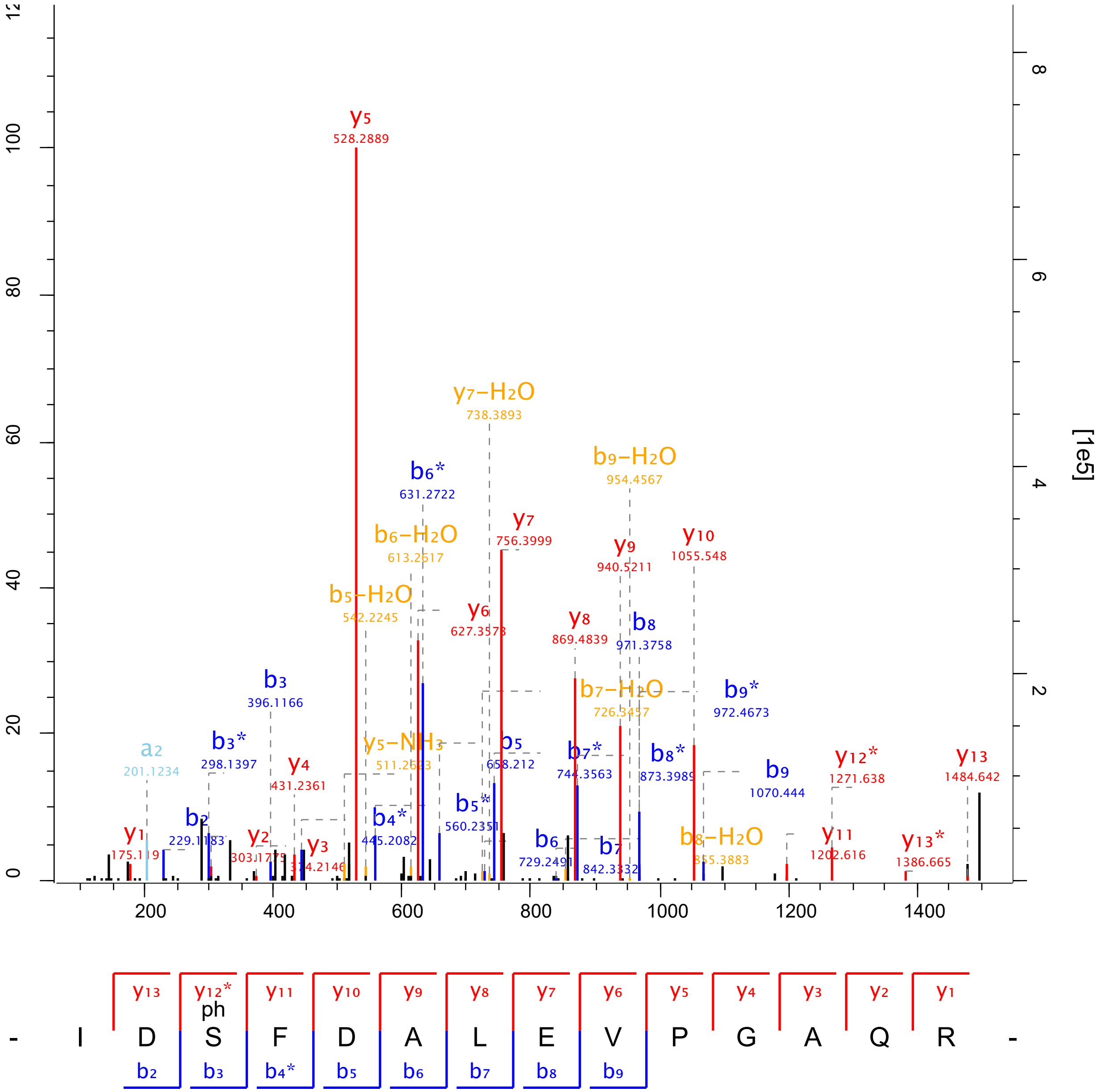

| Raw file | Scan | Method | Score | m/z |
|---|---|---|---|---|
| sys_05_2 | 3714 | FTMS; HCD | 83.88 | 507.22 |

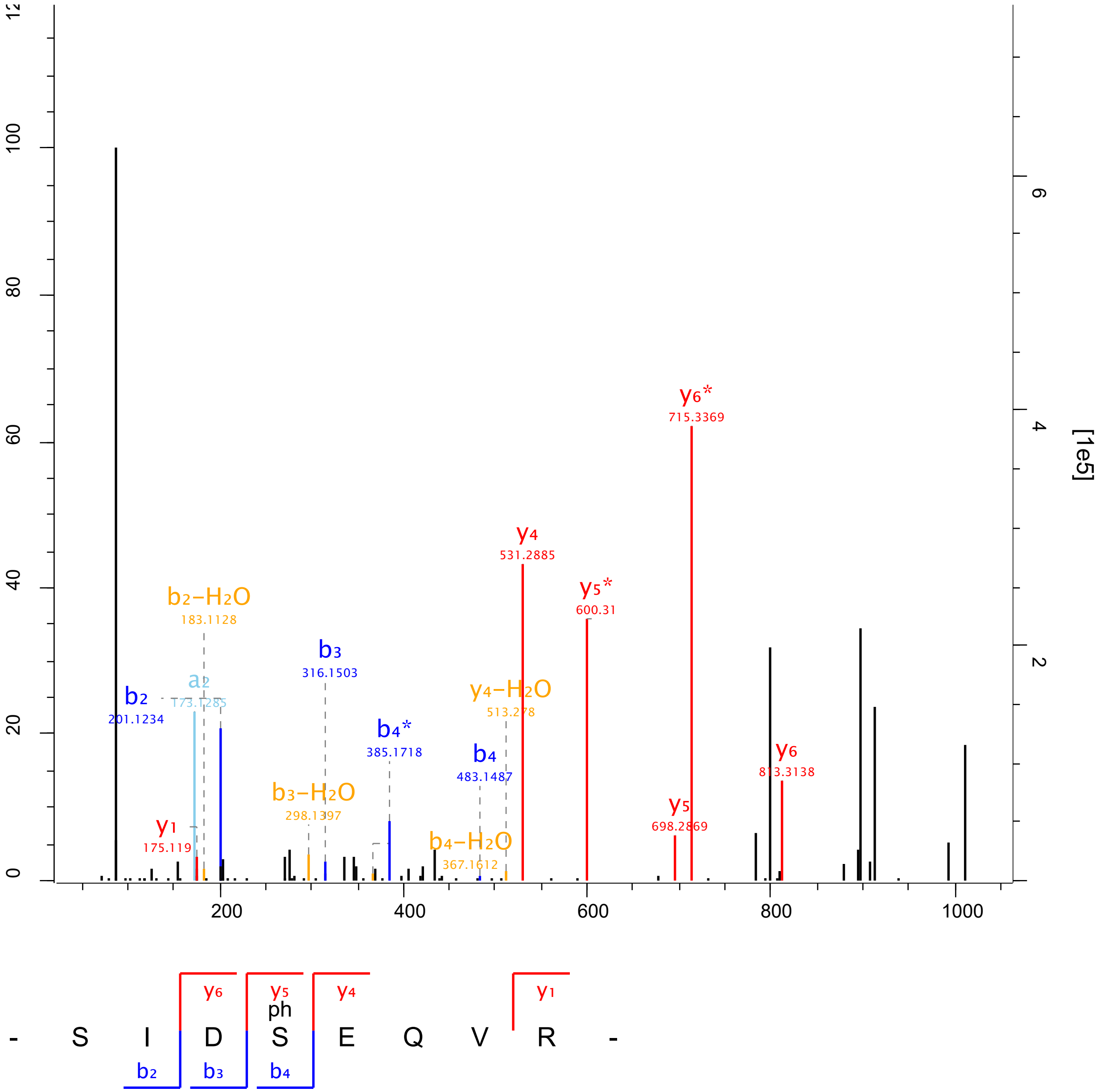

| Raw file | Scan | Method | Score | m/z |
|---|---|---|---|---|
| sys_05_2 | 38471 | FTMS; HCD | 47.97 | 550.7 |

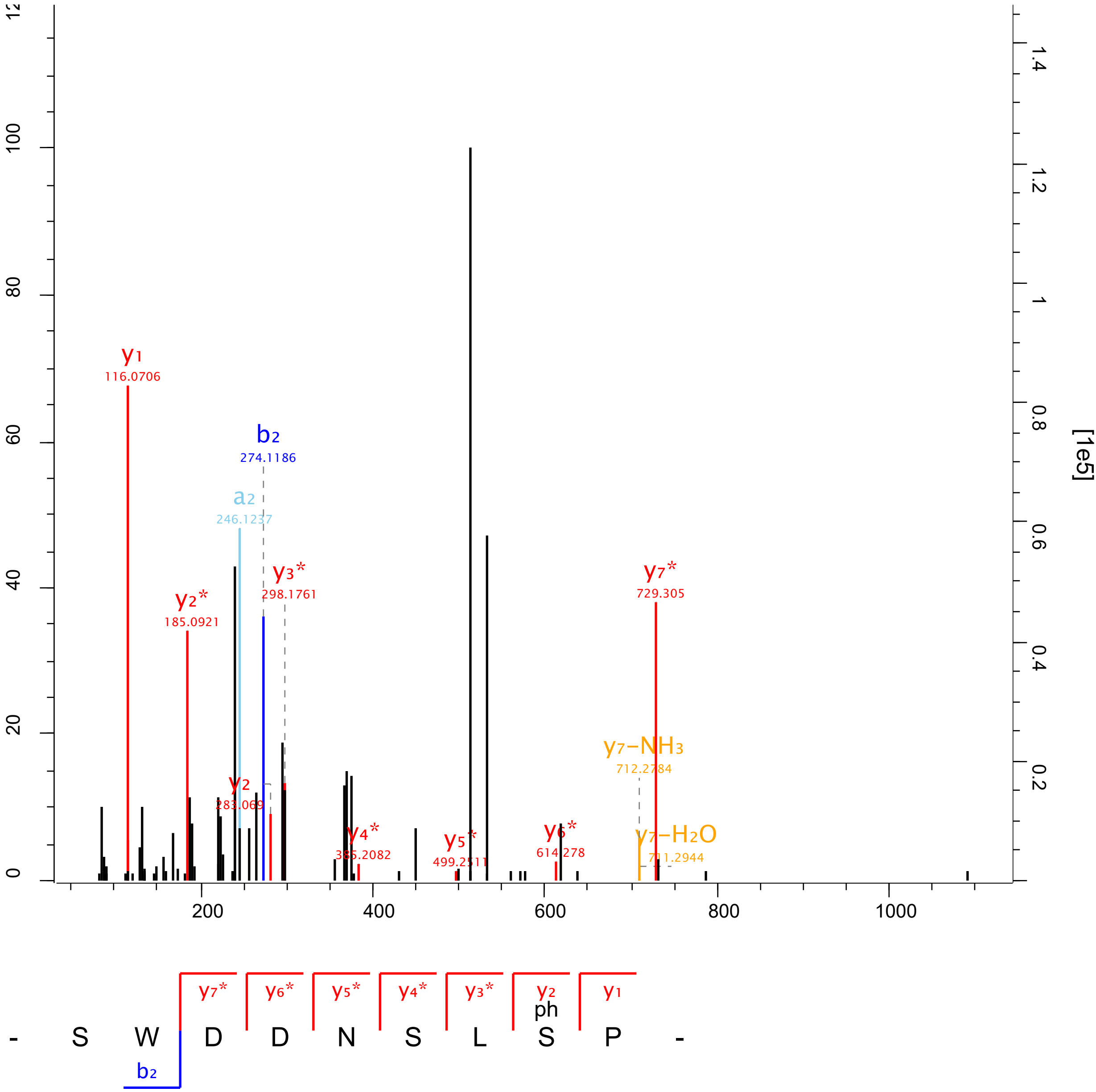

| Raw file | Scan | Method | Score | m/z |
|---|---|---|---|---|
| sys_05_2 | 38501 | FTMS; HCD | 313.45 | 819.86 |

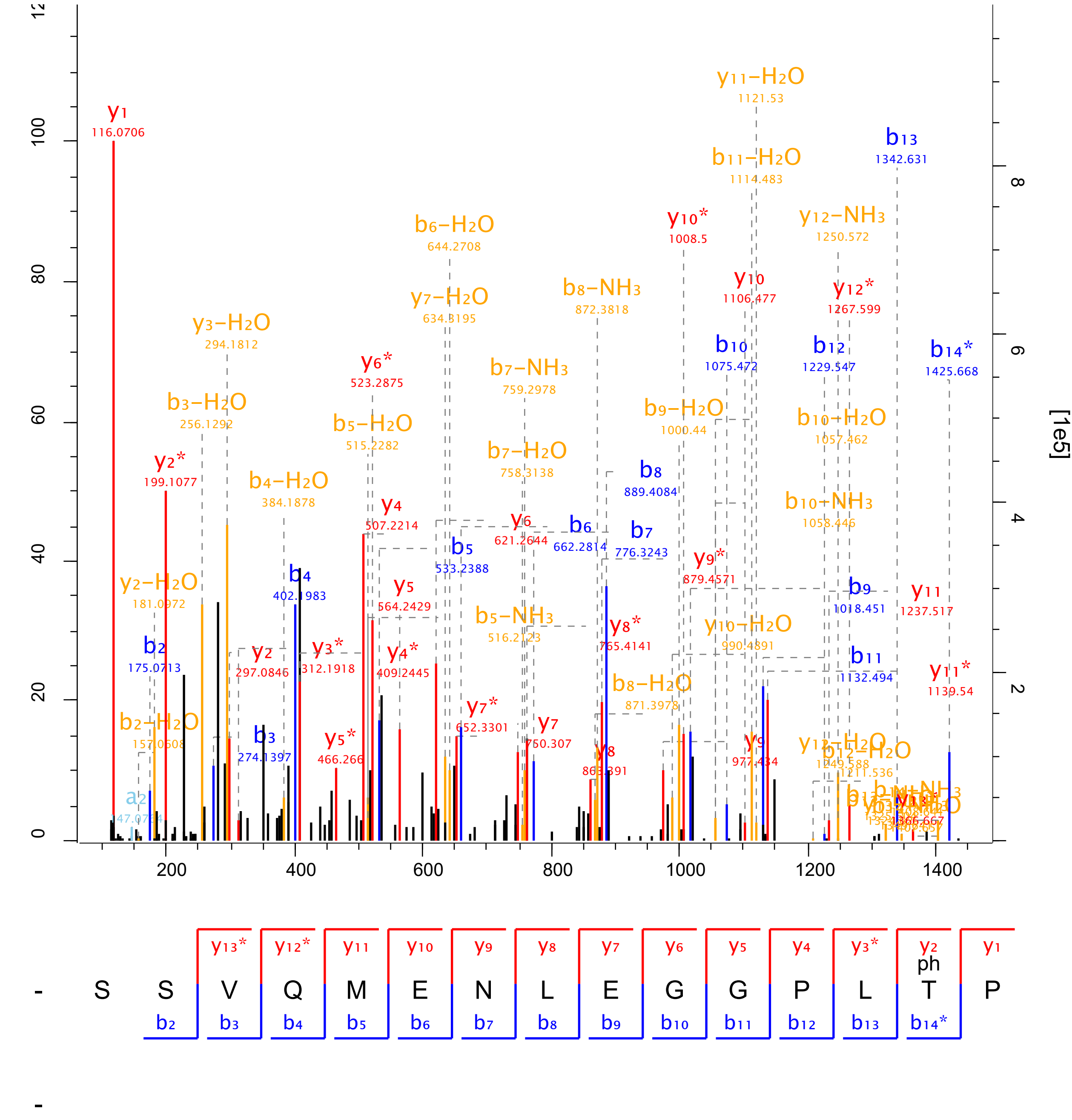

| Raw file | Scan | Method | Score | m/z |
|---|---|---|---|---|
| sys_05_2 | 3904 | FTMS; HCD | 86.77 | 548.74 |

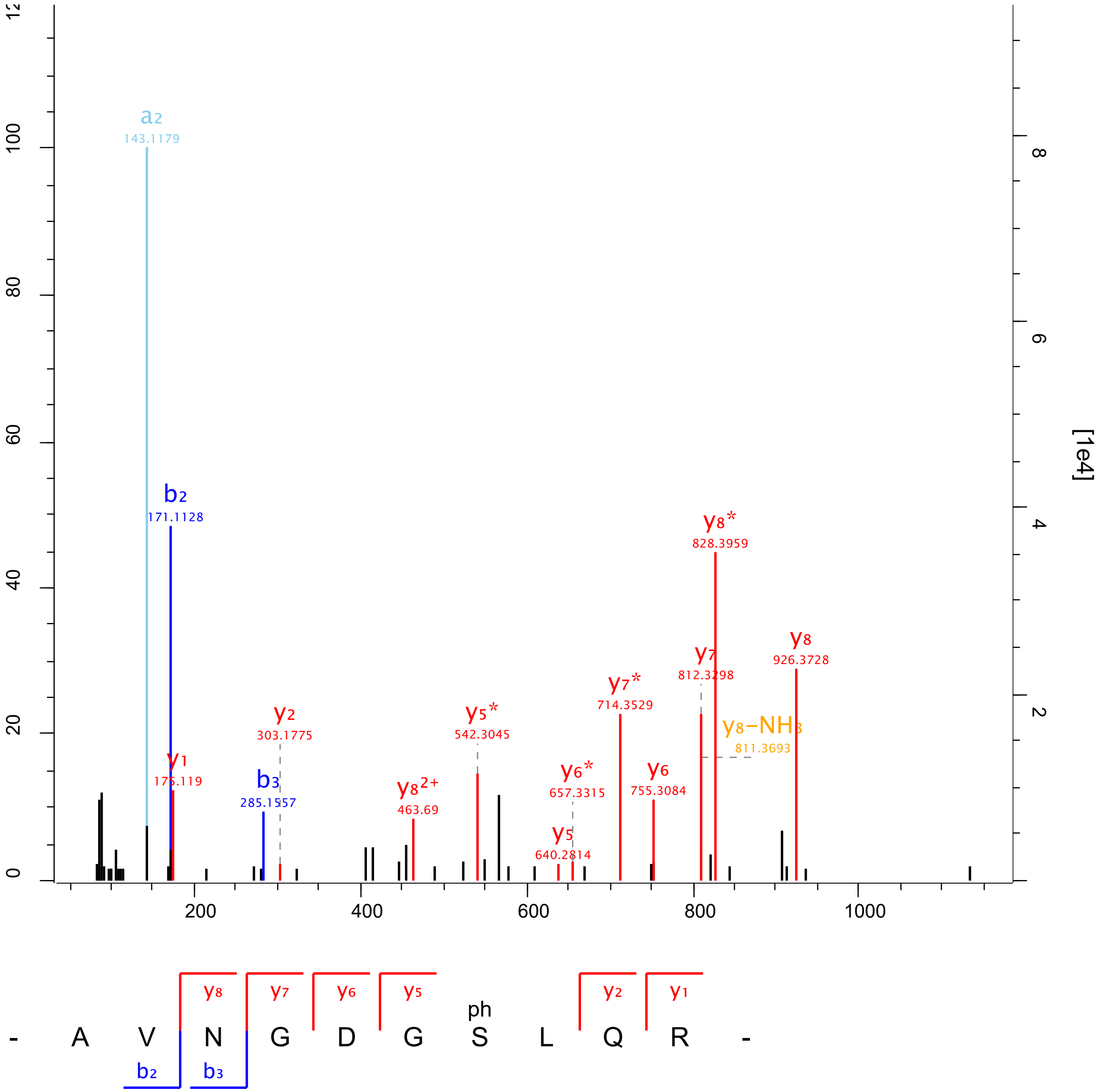

| Raw file | Scan | Method | Score | m/z |
| --- | --- | --- | --- | --- |
| sys_05_2 | 39396 | FTMS; HCD | 60.47 | 704.64 |

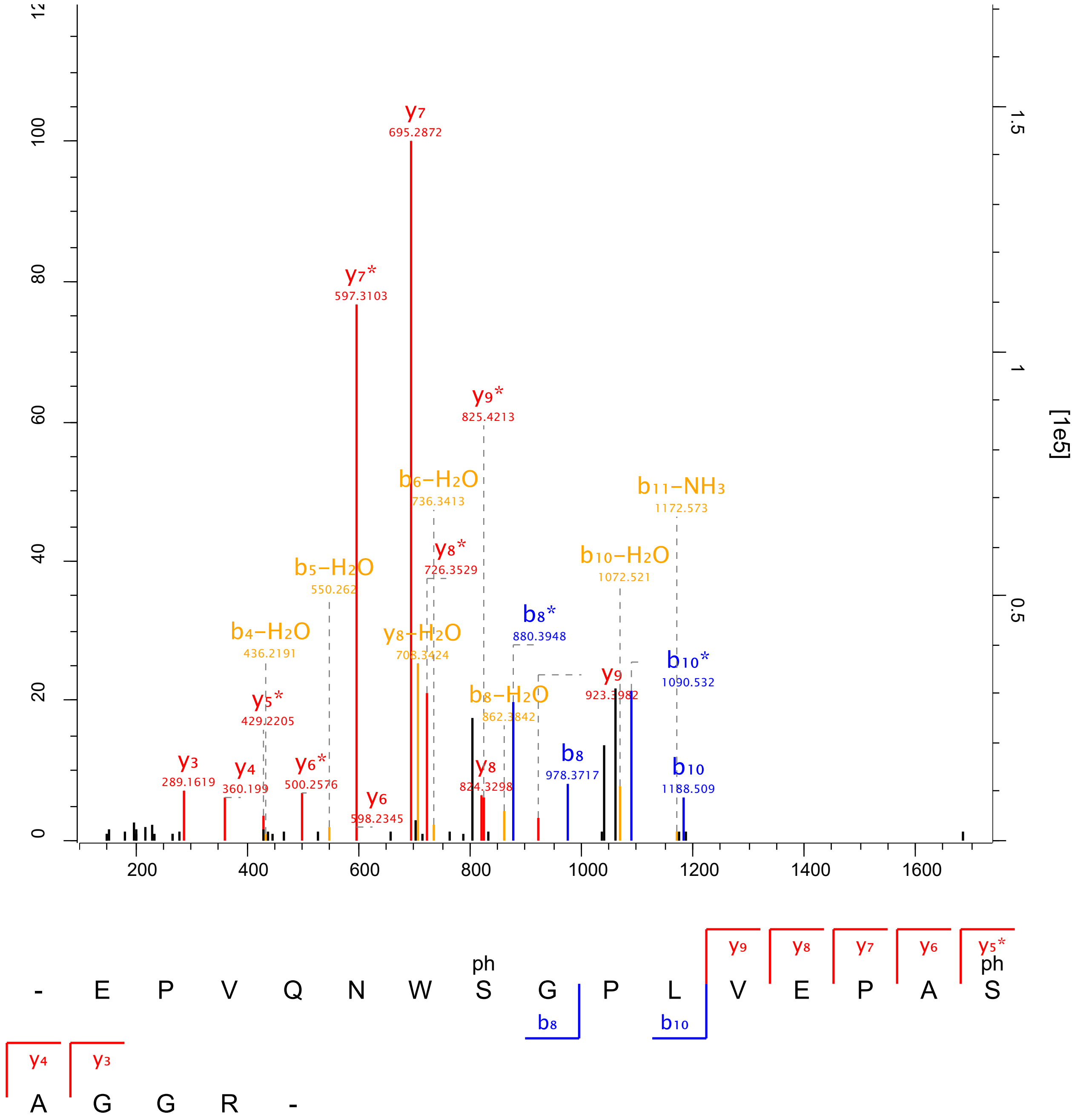

| Raw file | Scan | Method | Score | m/z |
| --- | --- | --- | --- | --- |
| sys_05_2 | 39835 | FTMS; HCD | 104.67 | 856.35 |

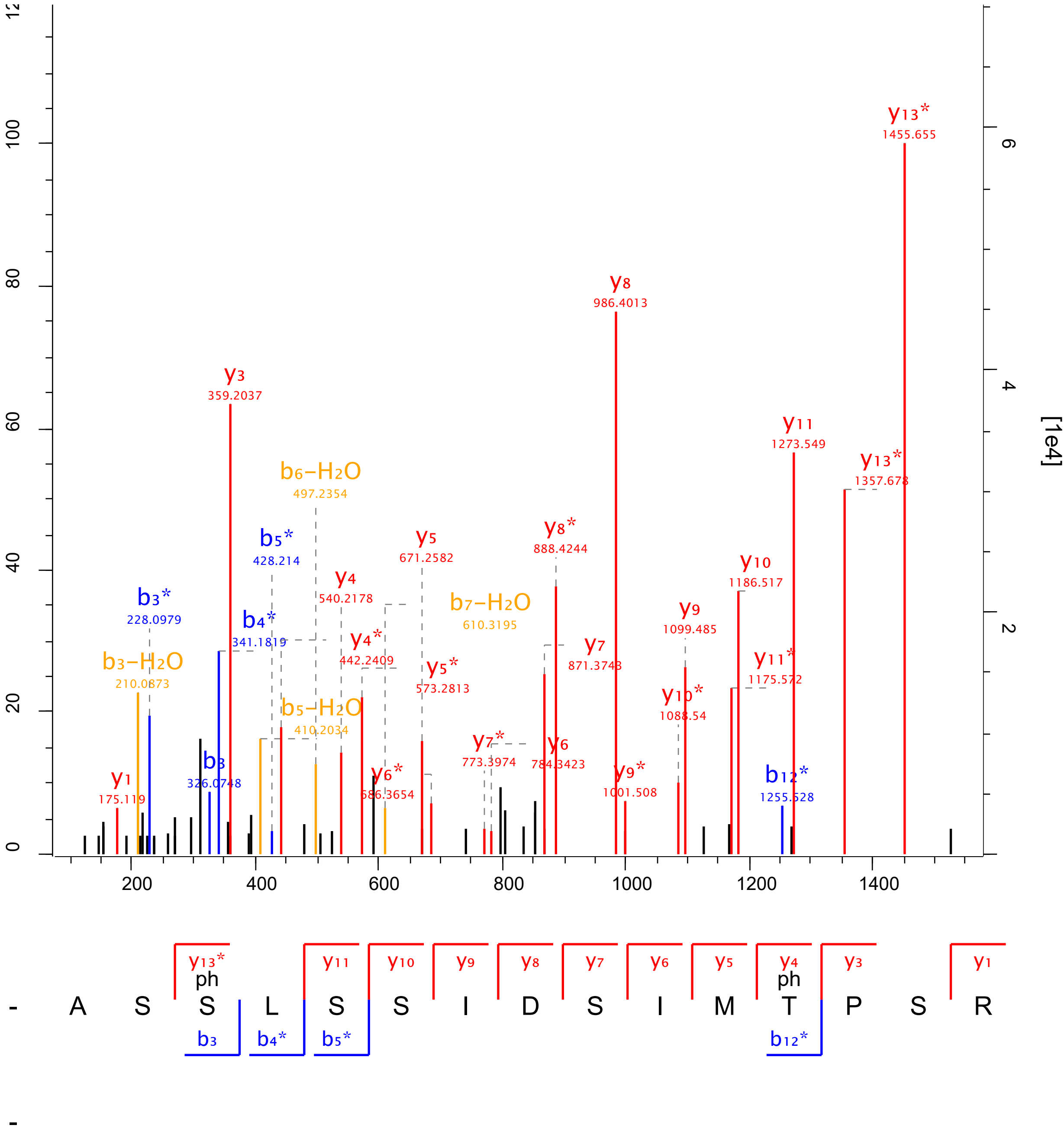

| Raw file | Scan | Method | Score | m/z |
| --- | --- | --- | --- | --- |
| sys_05_2 | 40238 | FTMS; HCD | 110.97 | 732.32 |

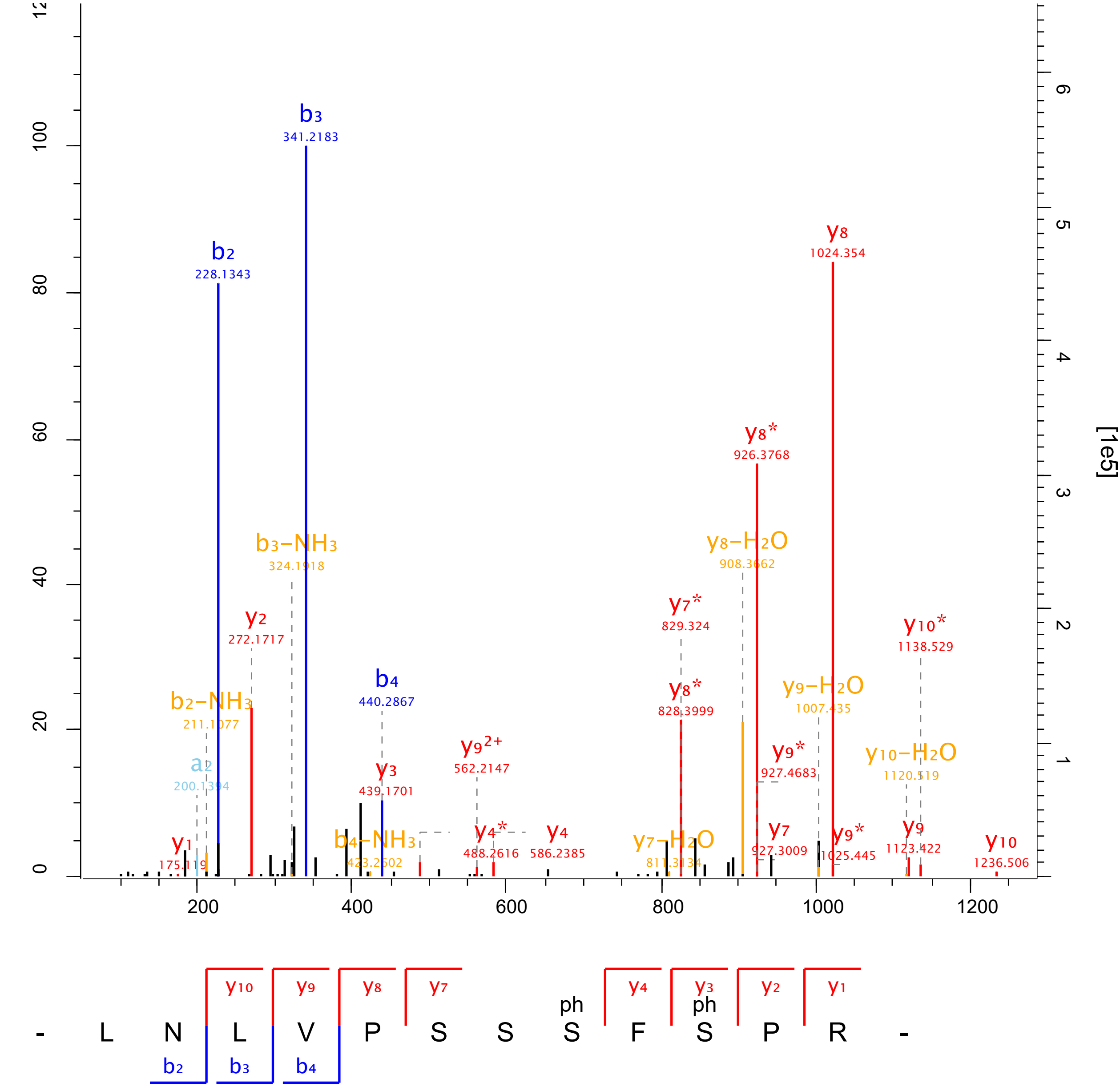

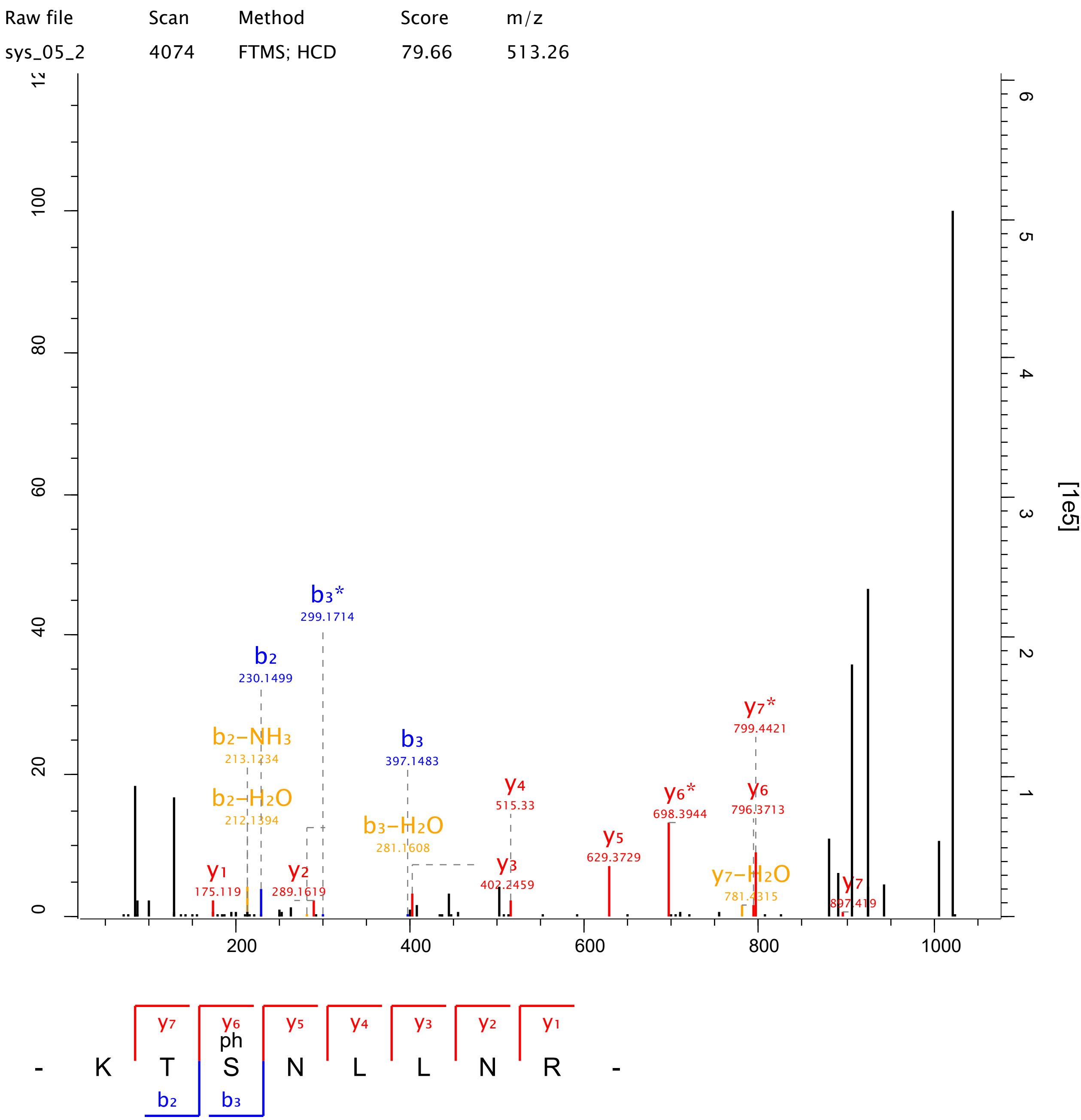

Raw file
Scan
Method
Score
m/z
sys_05_2
4074
FTMS; HCD
79.66
513.26
b3*
299.1714
b2
230.1499
b2−NH3
213.1234
b2−H2O
212.1394
b3
397.1483
b3−H2O
281.1608
y1
175.119
y2
289.1619
y3
402.2459
y4
515.33
y5
629.3729
y6*
698.3944
y6
796.3713
y7*
799.4421
y7−H2O
781.4315
y7
897.419
[1e5]
- K T S N L L N R -
ph
y7 y6 y5 y4 y3 y2 y1
b2 b3

| Raw file | Scan | Method | Score | m/z |
|---|---|---|---|---|
| sys_05_2 | 41352 | FTMS; HCD | 117.18 | 691.31 |

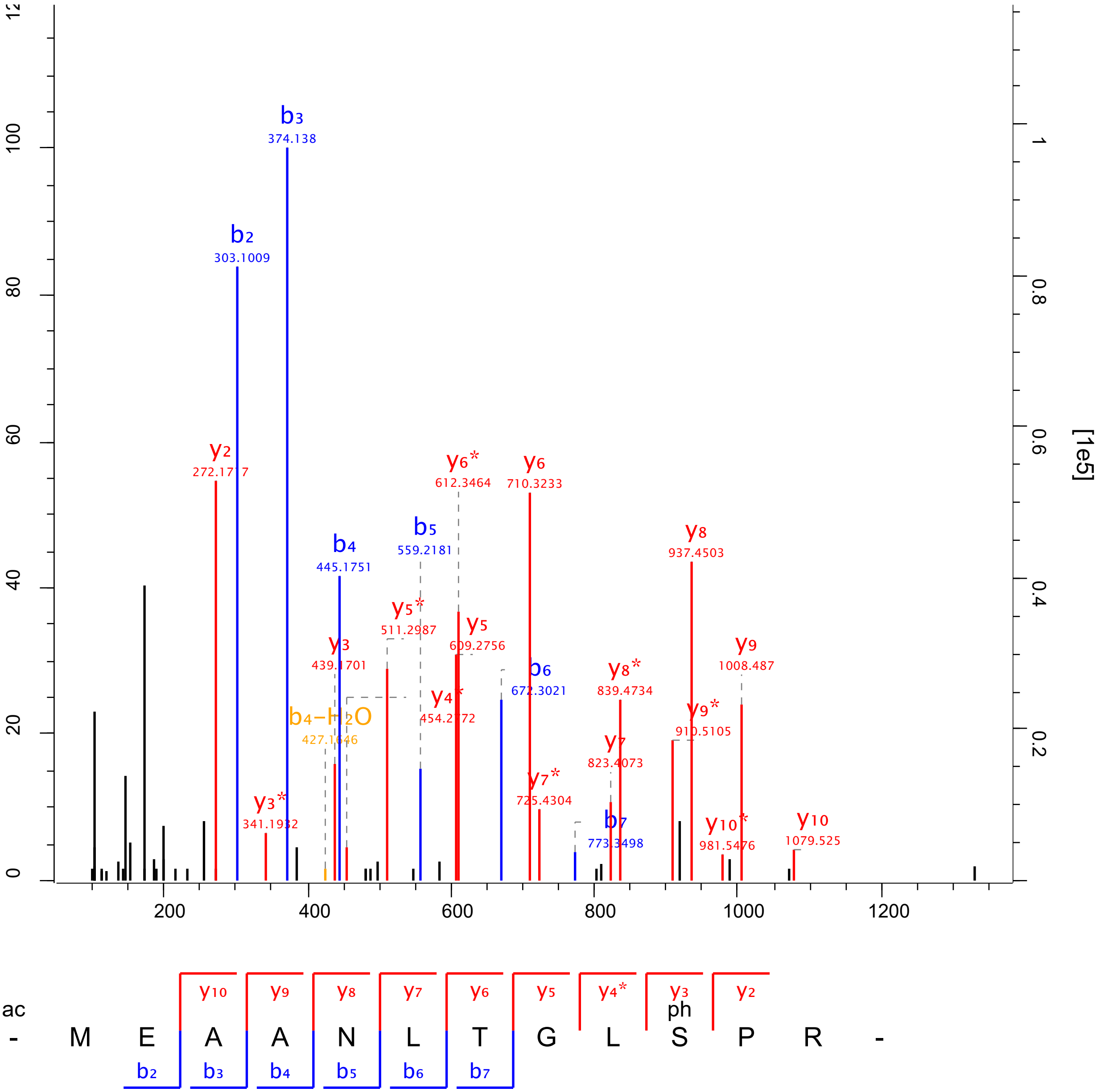

| Raw file | Scan | Method | Score | m/z |
|---|---|---|---|---|
| sys_05_2 | 41402 | FTMS; HCD | 155.1 | 809.88 |

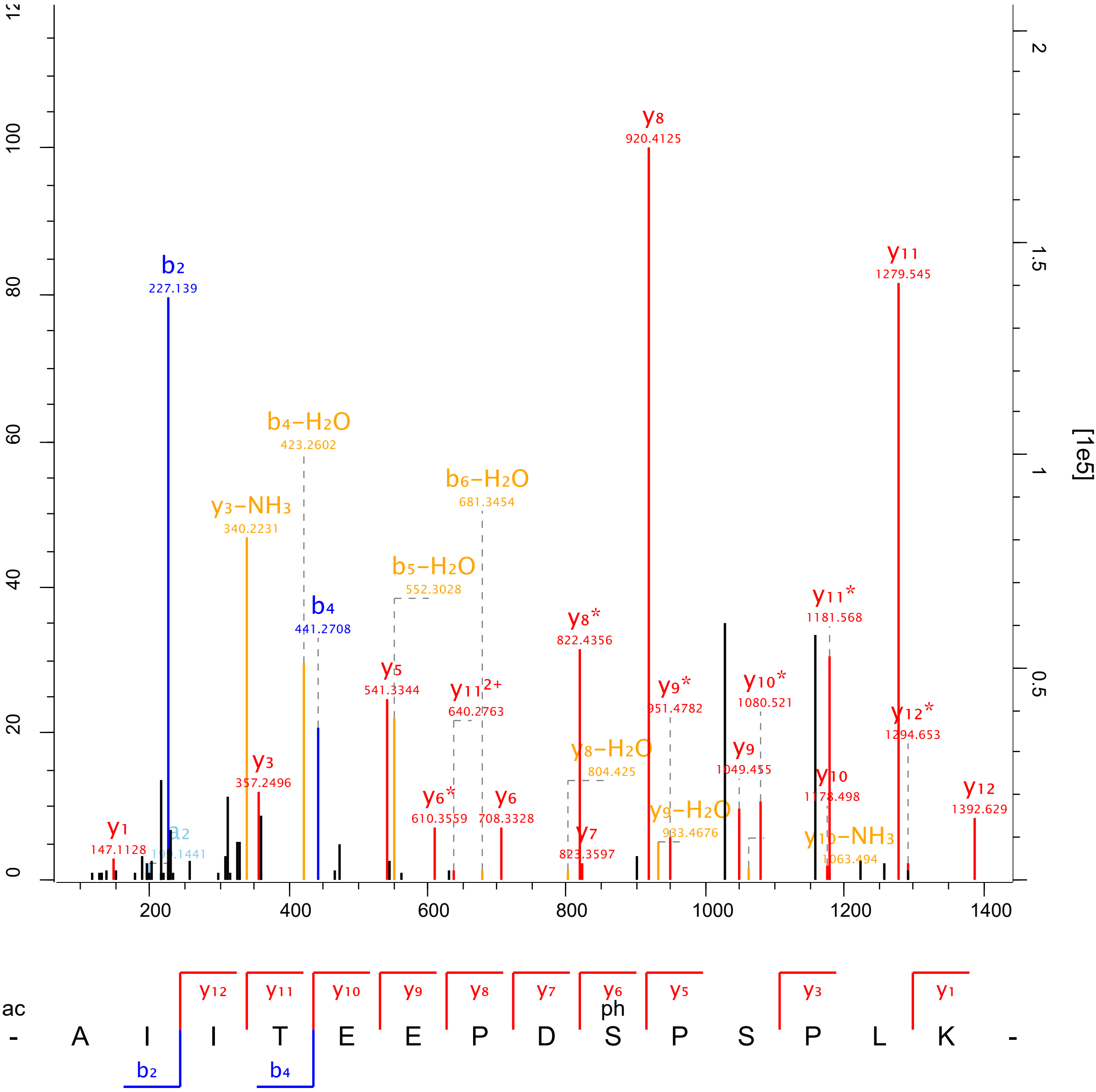

| Raw file | Scan | Method | Score | m/z |
|---|---|---|---|---|
| sys_05_2 | 42234 | FTMS; HCD | 108.75 | 790.86 |

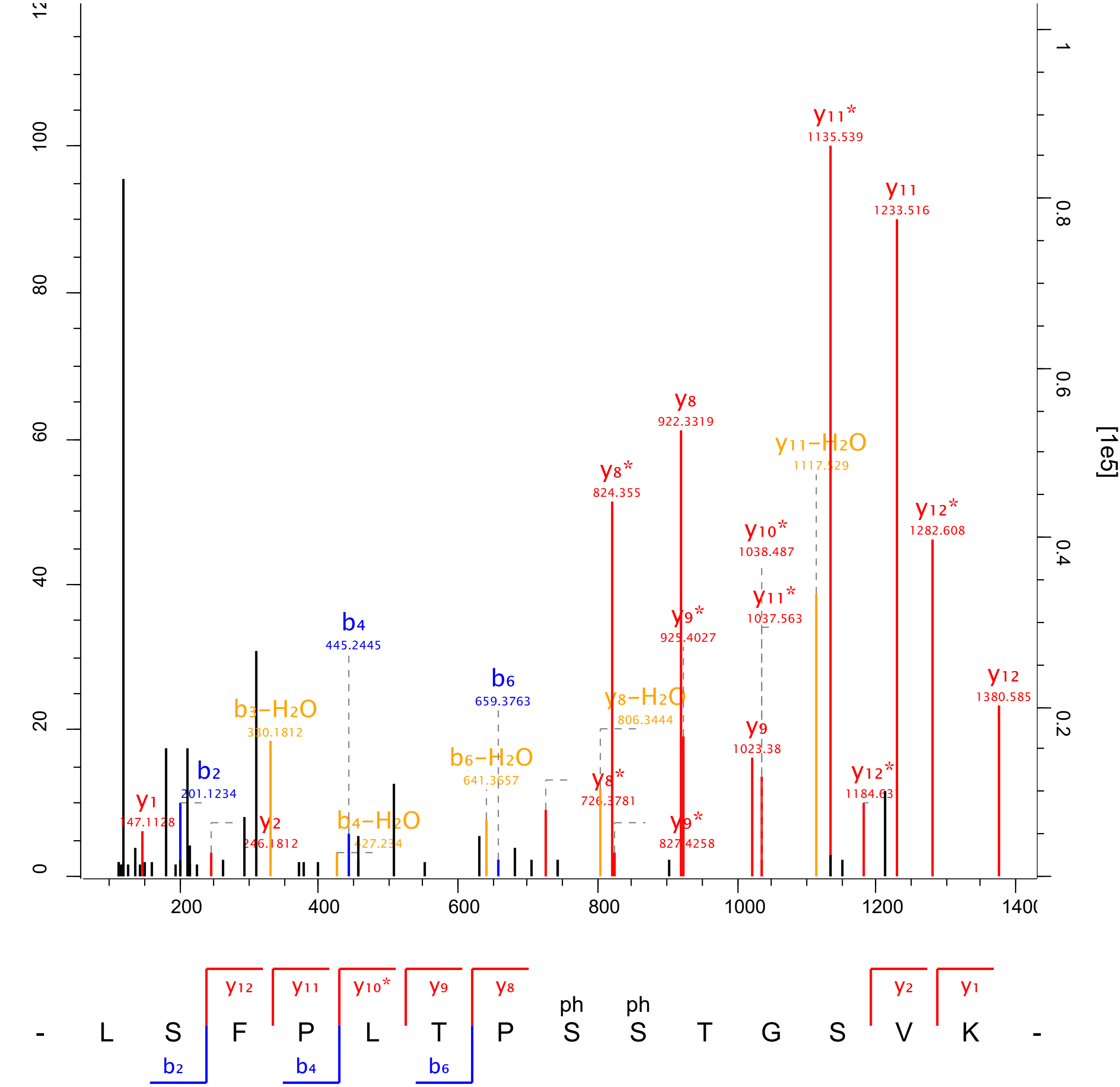

| Raw file | Scan | Method | Score | m/z |
| --- | --- | --- | --- | --- |
| sys_05_2 | 42536 | FTMS; HCD | 273.25 | 993.42 |

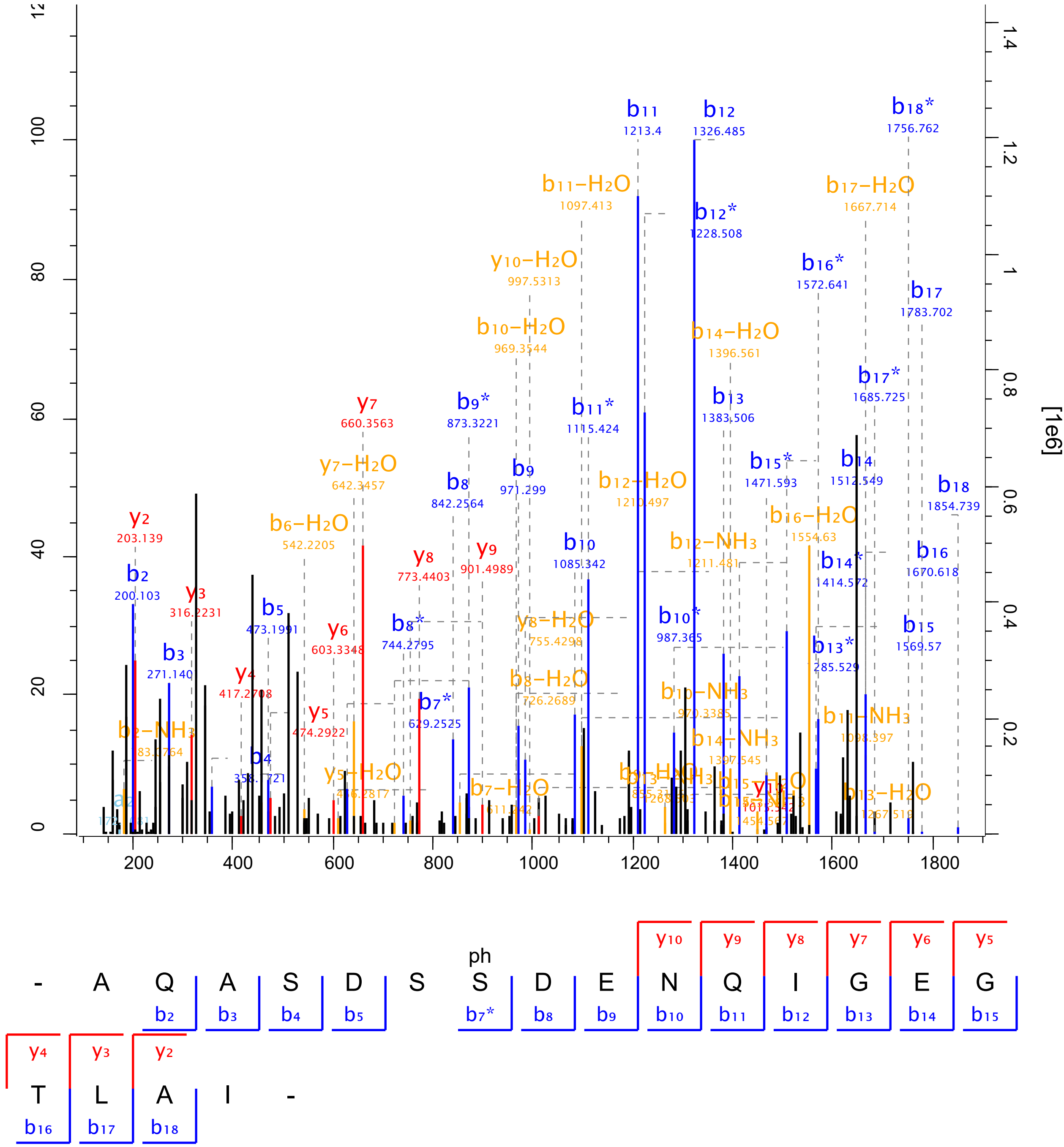

| Raw file | Scan | Method | Score | m/z |
|---|---|---|---|---|
| sys_05_2 | 42764 | FTMS; HCD | 264.01 | 835.87 |

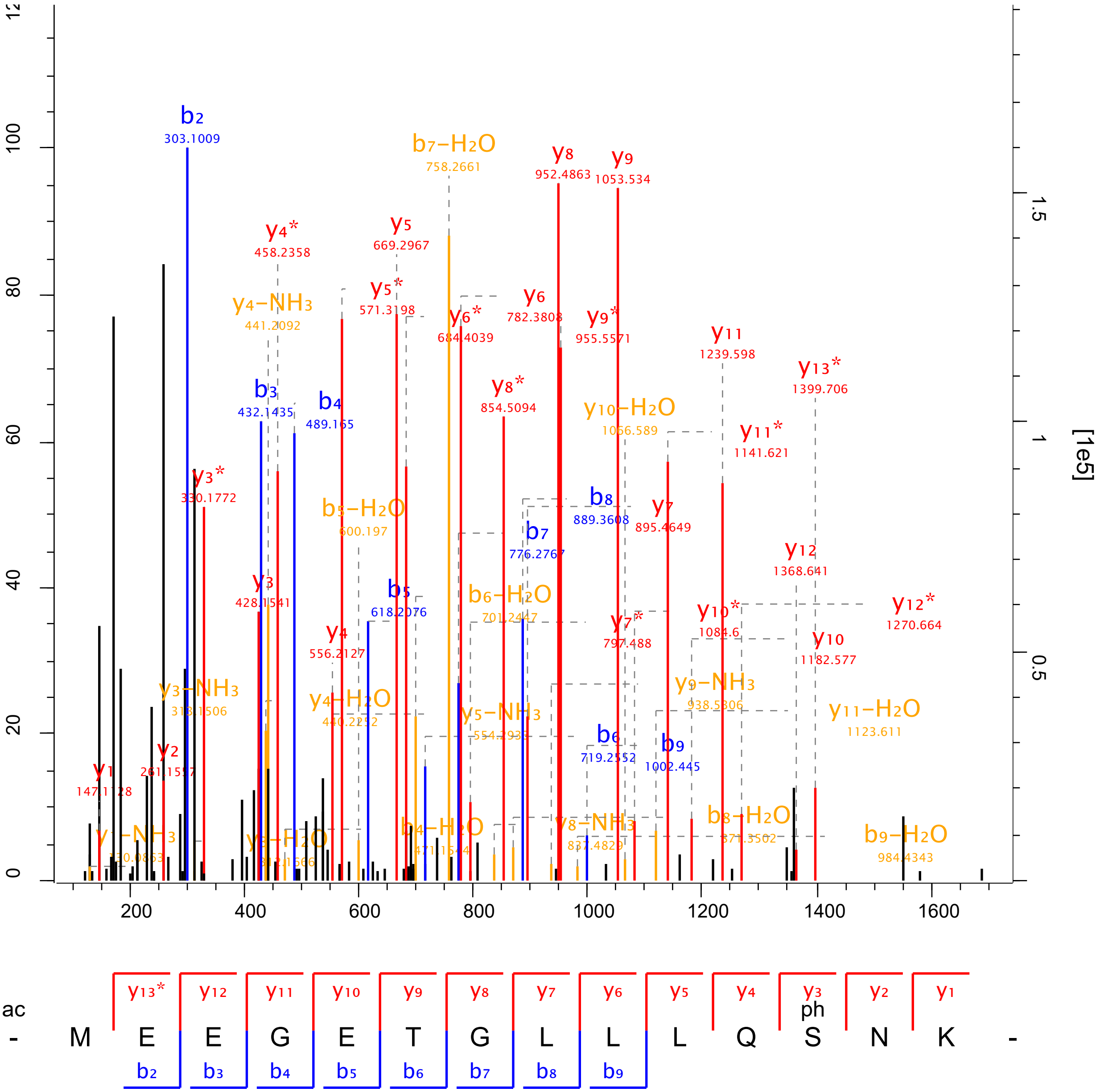

| Raw file | Scan | Method | Score | m/z |
| --- | --- | --- | --- | --- |
| sys_05_2 | 4327 | FTMS; HCD | 167.89 | 585.26 |

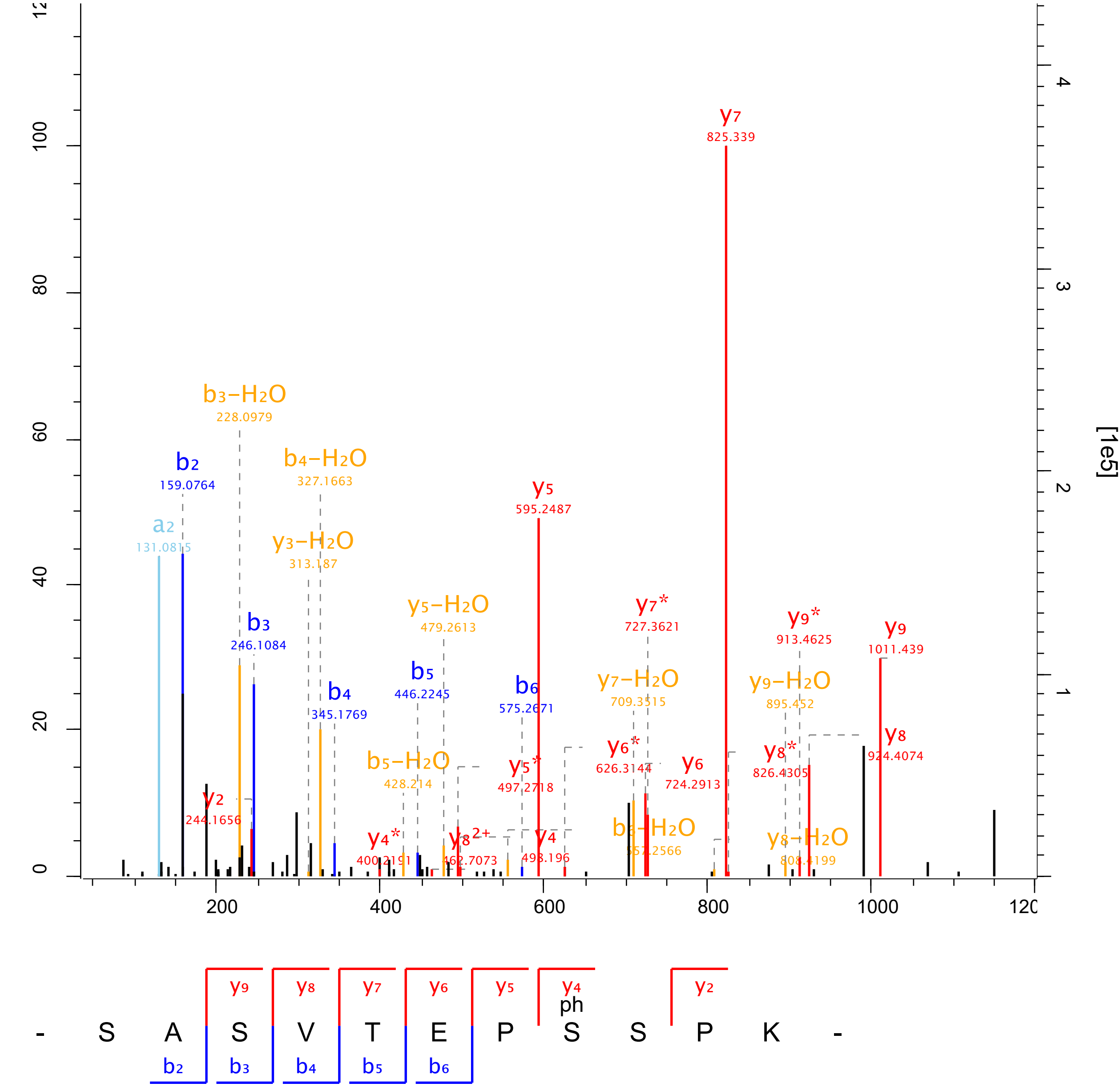

| Raw file | Scan | Method | Score | m/z |
|---|---|---|---|---|
| sys_05_2 | 43787 | FTMS; HCD | 48.9 | 722.31 |

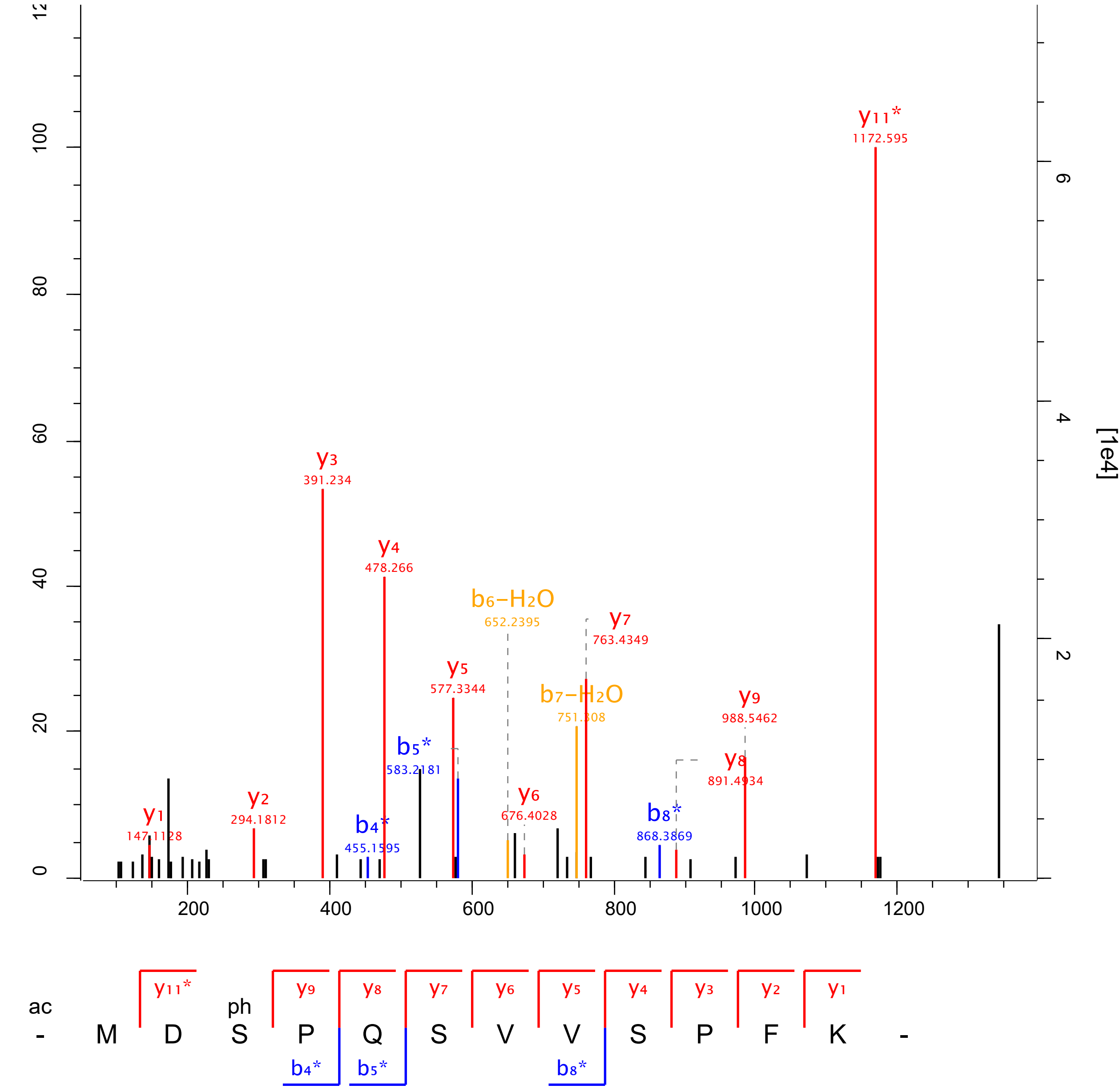

| Raw file | Scan | Method | Score | m/z |
|---|---|---|---|---|
| sys_05_2 | 45695 | FTMS; HCD | 130.01 | 534.76 |

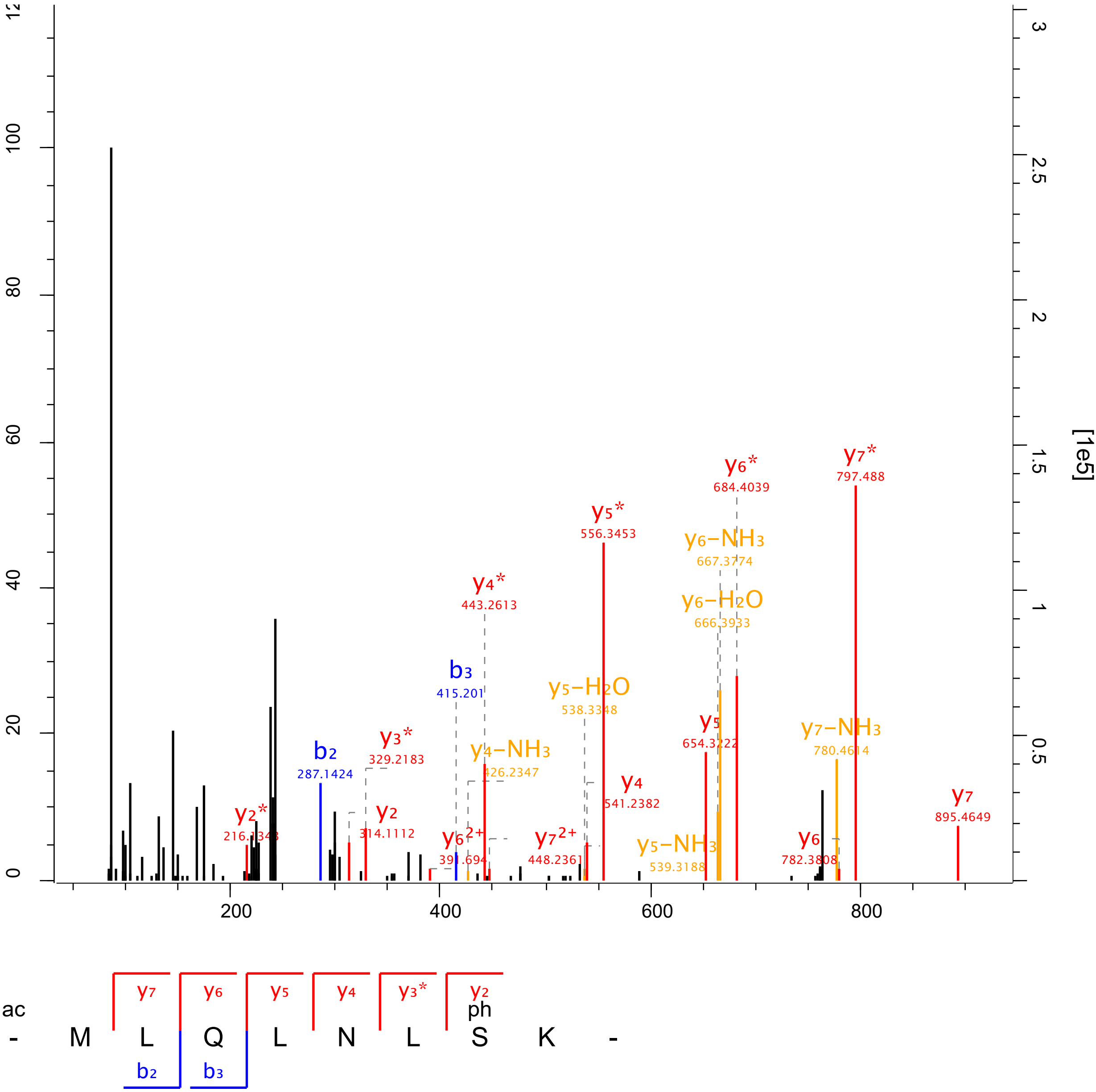

| Raw file | Scan | Method | Score | m/z |
|---|---|---|---|---|
| sys_05_2 | 4607 | FTMS; HCD | 91.62 | 557.76 |

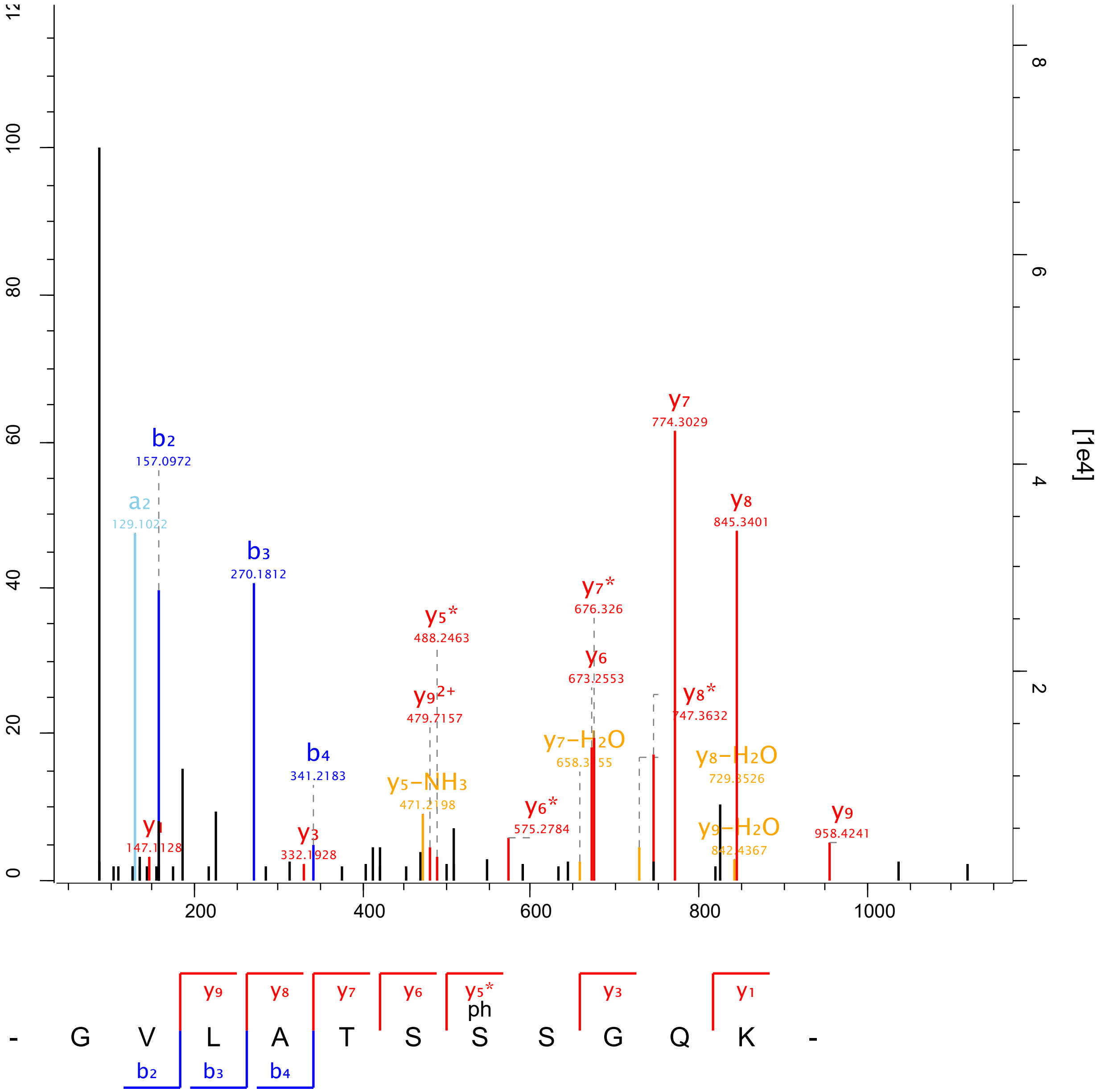

| Raw file | Scan | Method | Score | m/z |
|---|---|---|---|---|
| sys_05_2 | 4614 | FTMS; HCD | 44.93 | 632.76 |

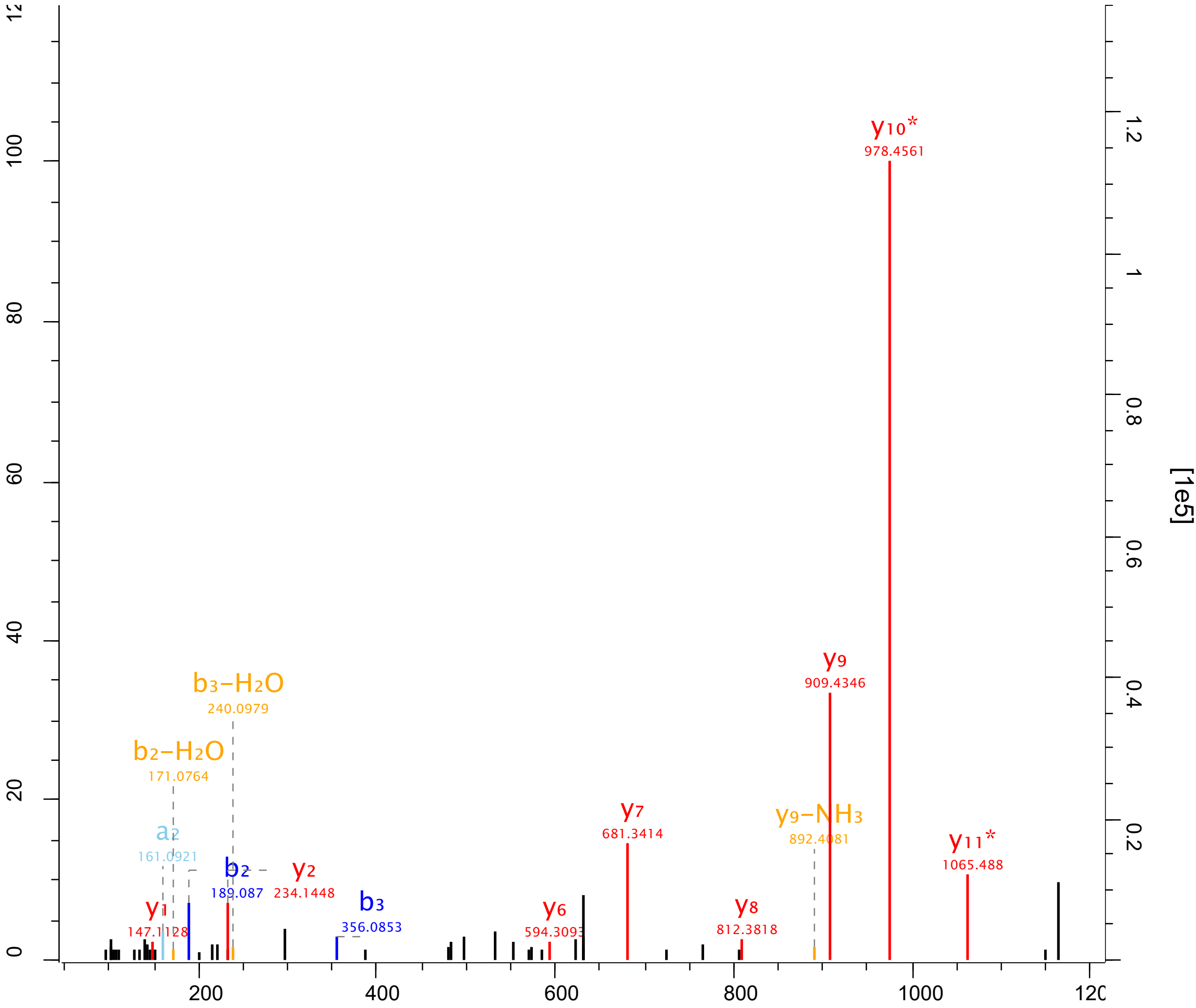


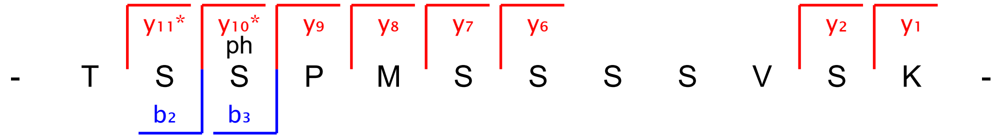

| Raw file | Scan | Method | Score | m/z |
|---|---|---|---|---|
| sys_05_2 | 4635 | FTMS; HCD | 140.78 | 506.73 |

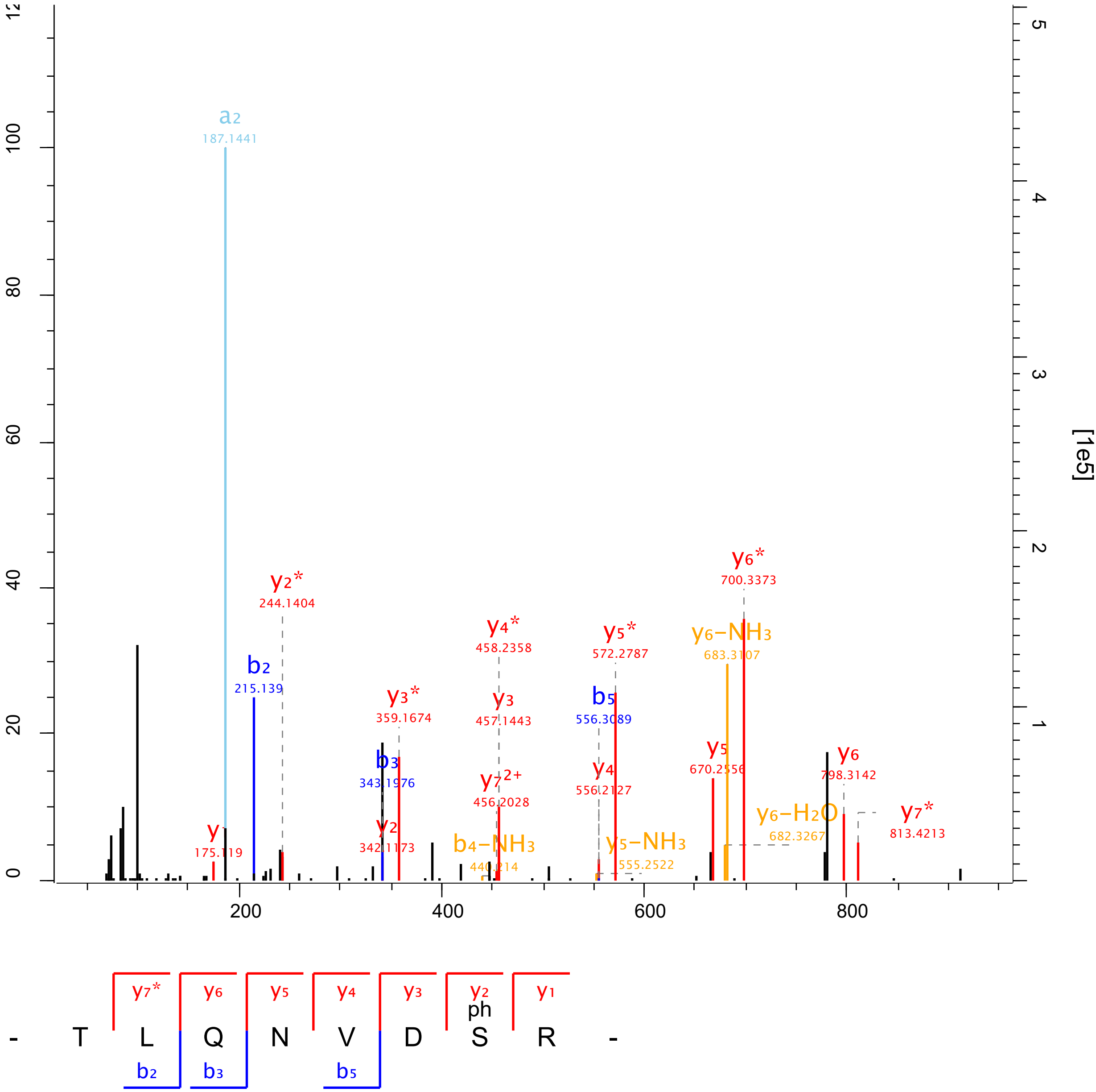

| Raw file | Scan | Method | Score | m/z |
|---|---|---|---|---|
| sys_05_2 | 4649 | FTMS; HCD | 108.47 | 485.72 |

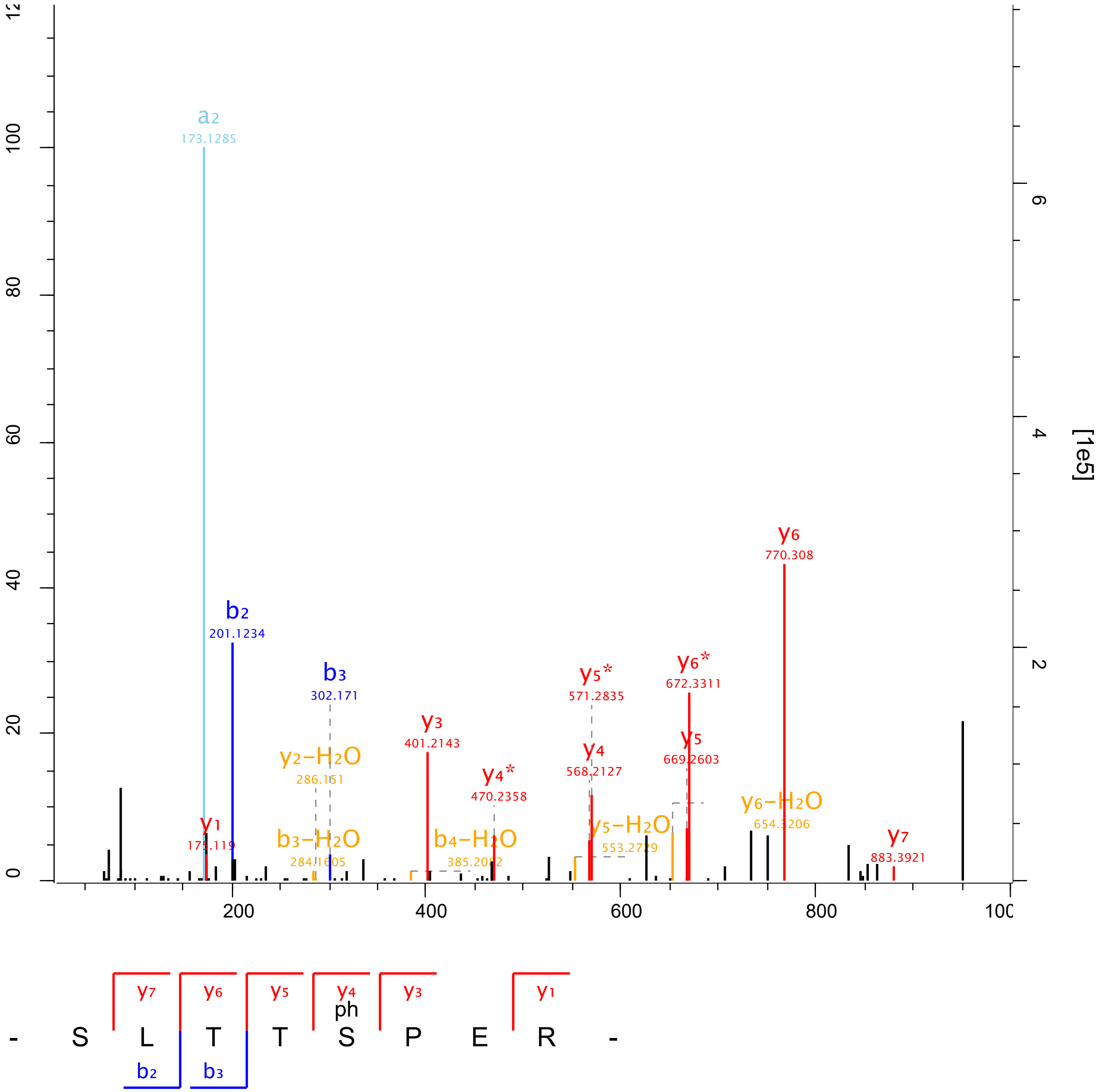

| Raw file | Scan | Method | Score | m/z |
|---|---|---|---|---|
| sys_05_2 | 4729 | FTMS; HCD | 203.76 | 658.28 |

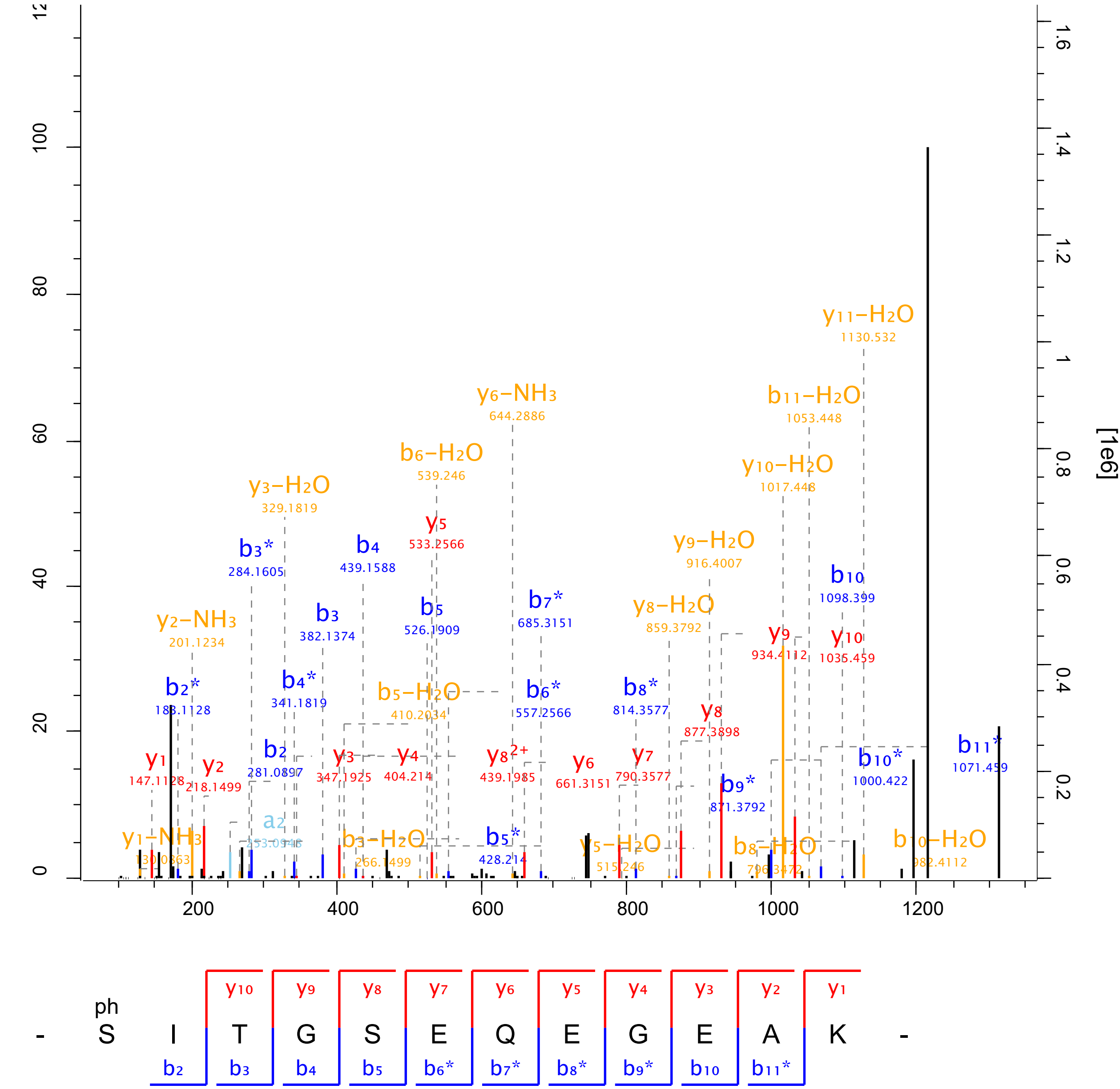

| Raw file | Scan | Method | Score | m/z |
|---|---|---|---|---|
| sys_05_2 | 4834 | FTMS; HCD | 68.68 | 599.74 |

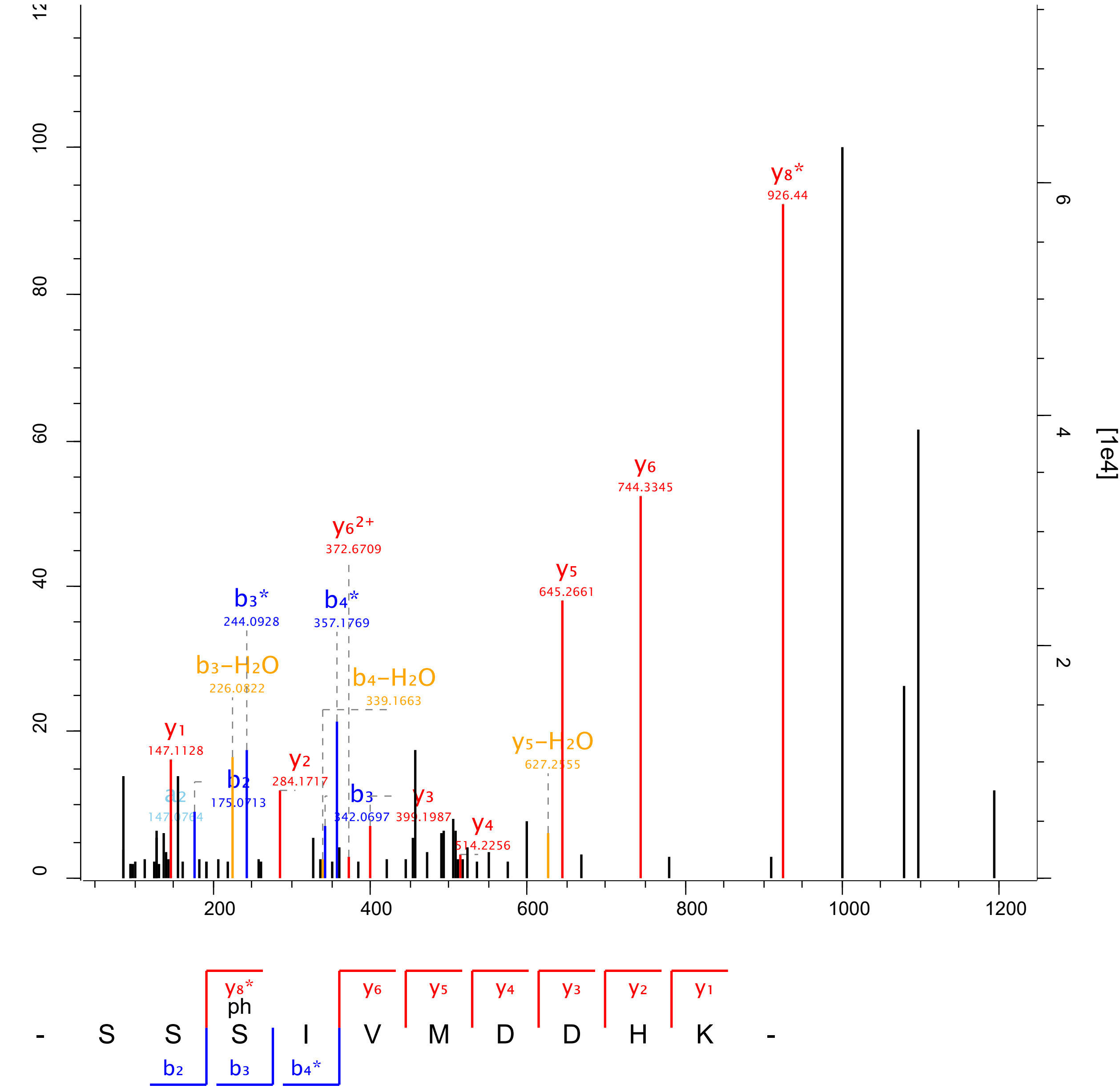

| Raw file | Scan | Method | Score | m/z |
|---|---|---|---|---|
| sys_05_2 | 4870 | FTMS; HCD | 179.6 | 507.72 |

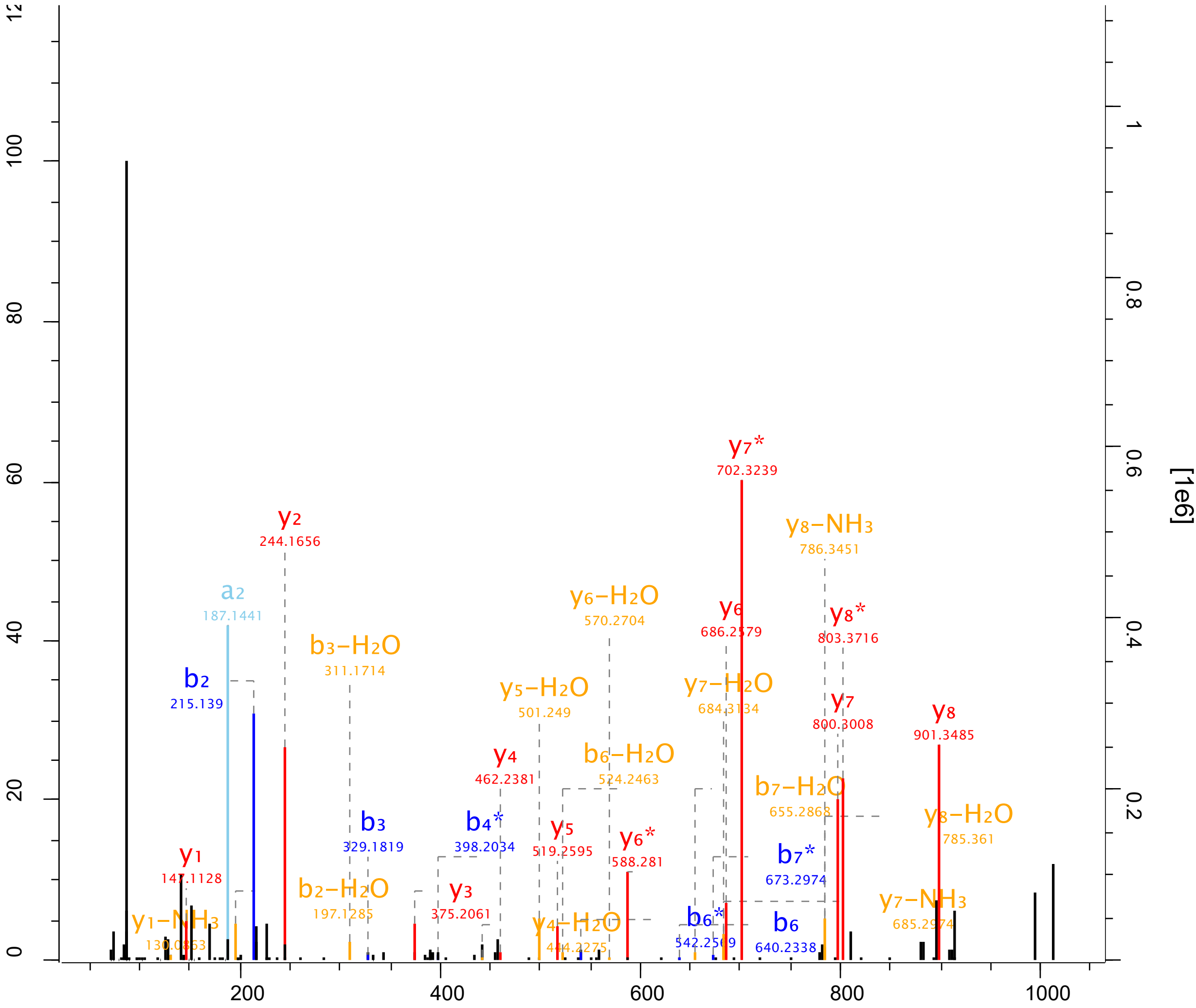


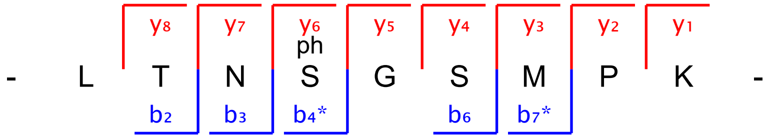

| Raw file | Scan | Method | Score | m/z |
|---|---|---|---|---|
| sys_05_2 | 49176 | FTMS; HCD | 111.57 | 994.41 |

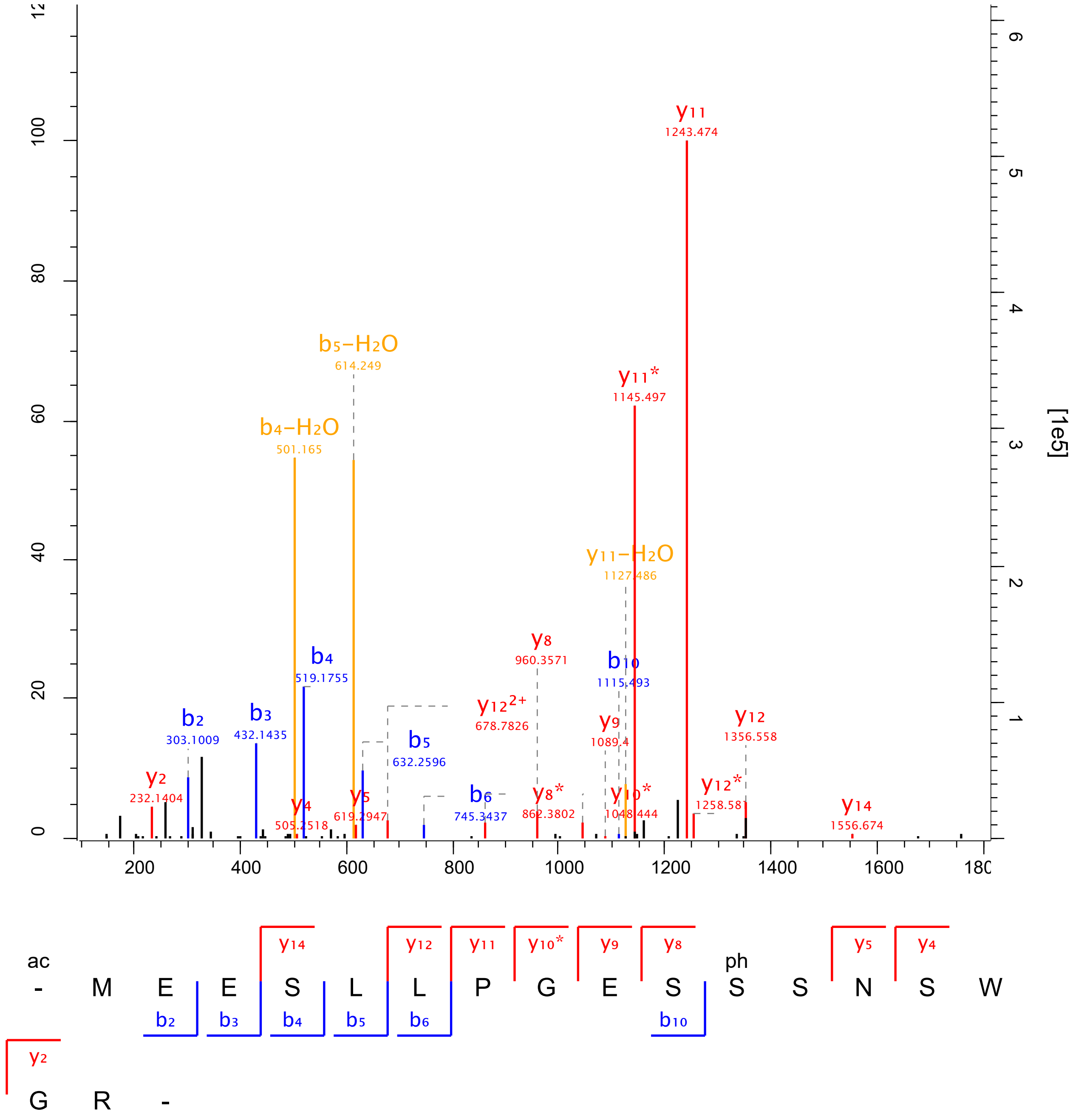

| Raw file | Scan | Method | Score | m/z |
|---|---|---|---|---|
| sys_05_2 | 49177 | FTMS; HCD | 50.3 | 738.31 |

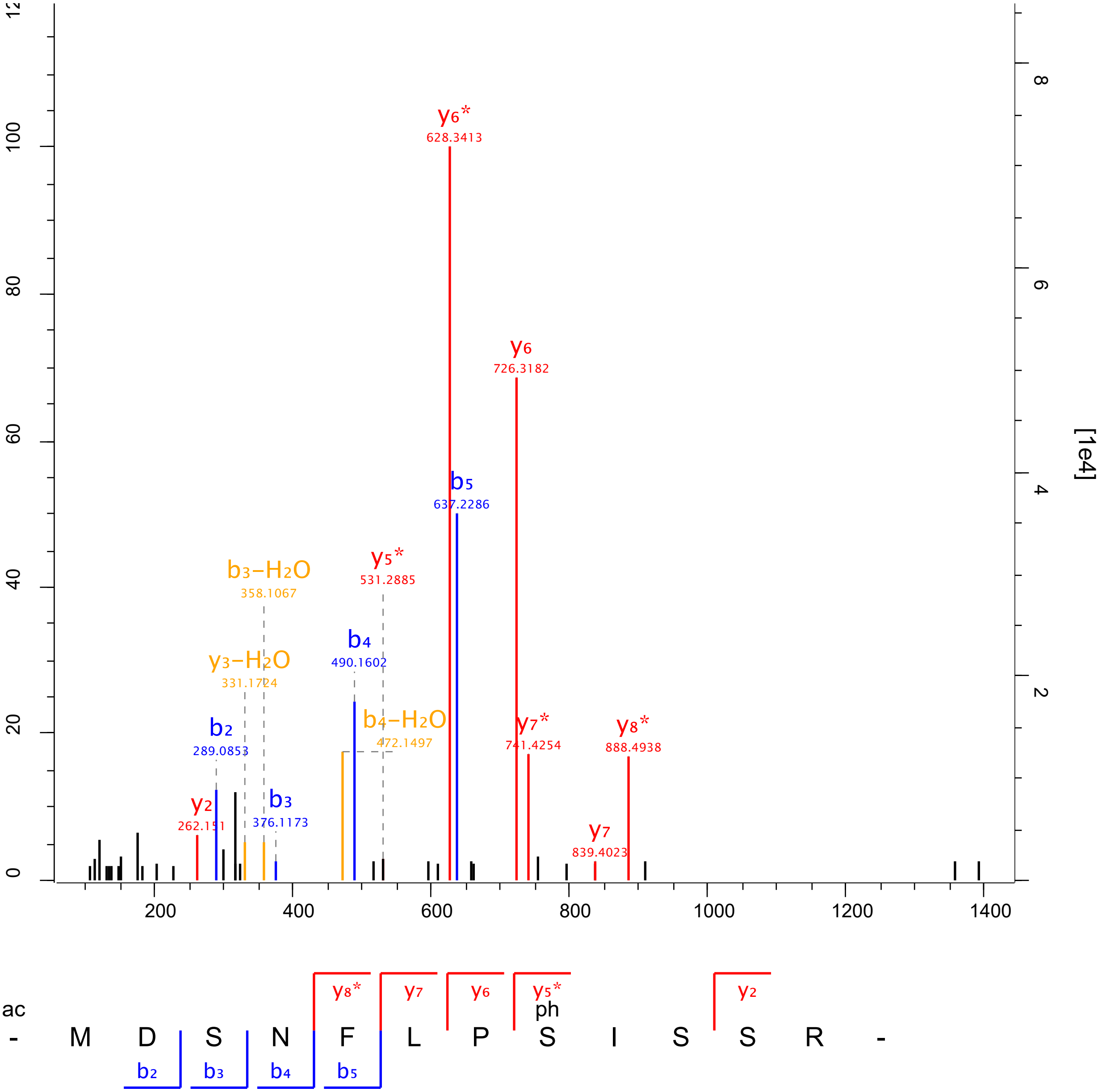

| Raw file | Scan | Method | Score | m/z |
|---|---|---|---|---|
| sys_05_2 | 5204 | FTMS; HCD | 67.33 | 554.73 |

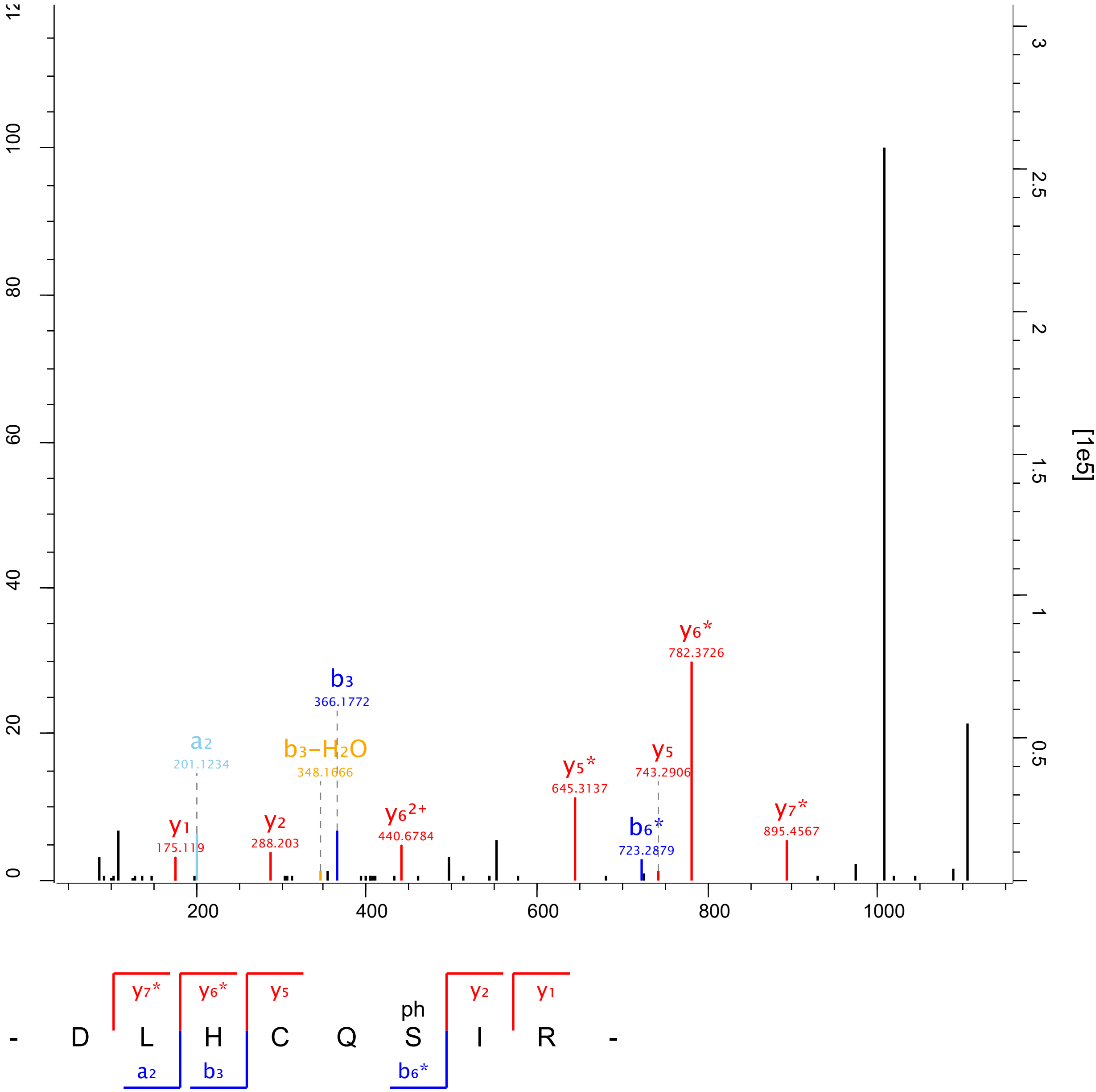

| Raw file | Scan | Method | Score | m/z |
|---|---|---|---|---|
| sys_05_2 | 5273 | FTMS; HCD | 68.66 | 508.75 |

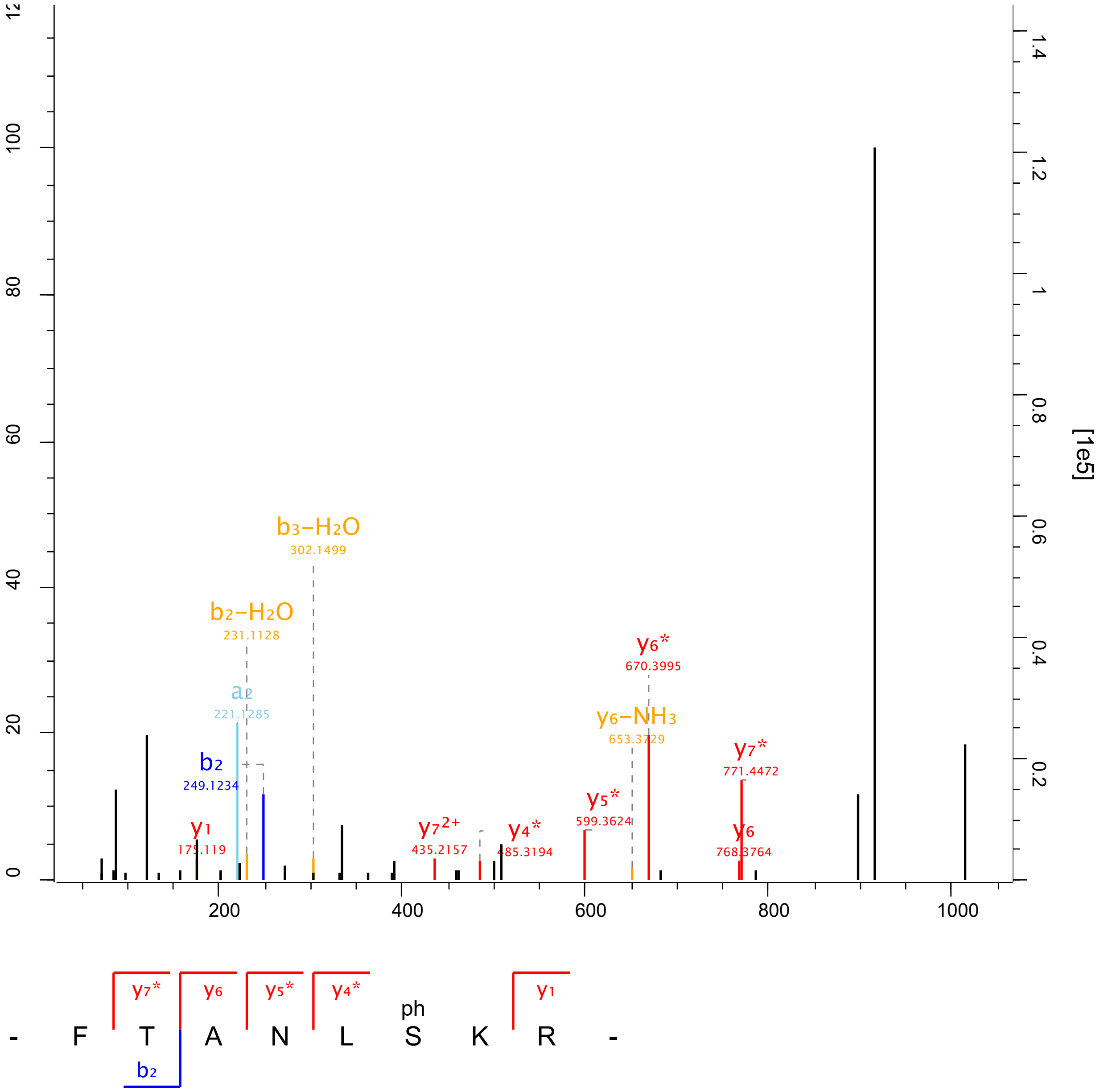

| Raw file | Scan | Method | Score | m/z |
|---|---|---|---|---|
| sys_05_2 | 5406 | FTMS; HCD | 107.35 | 684.28 |

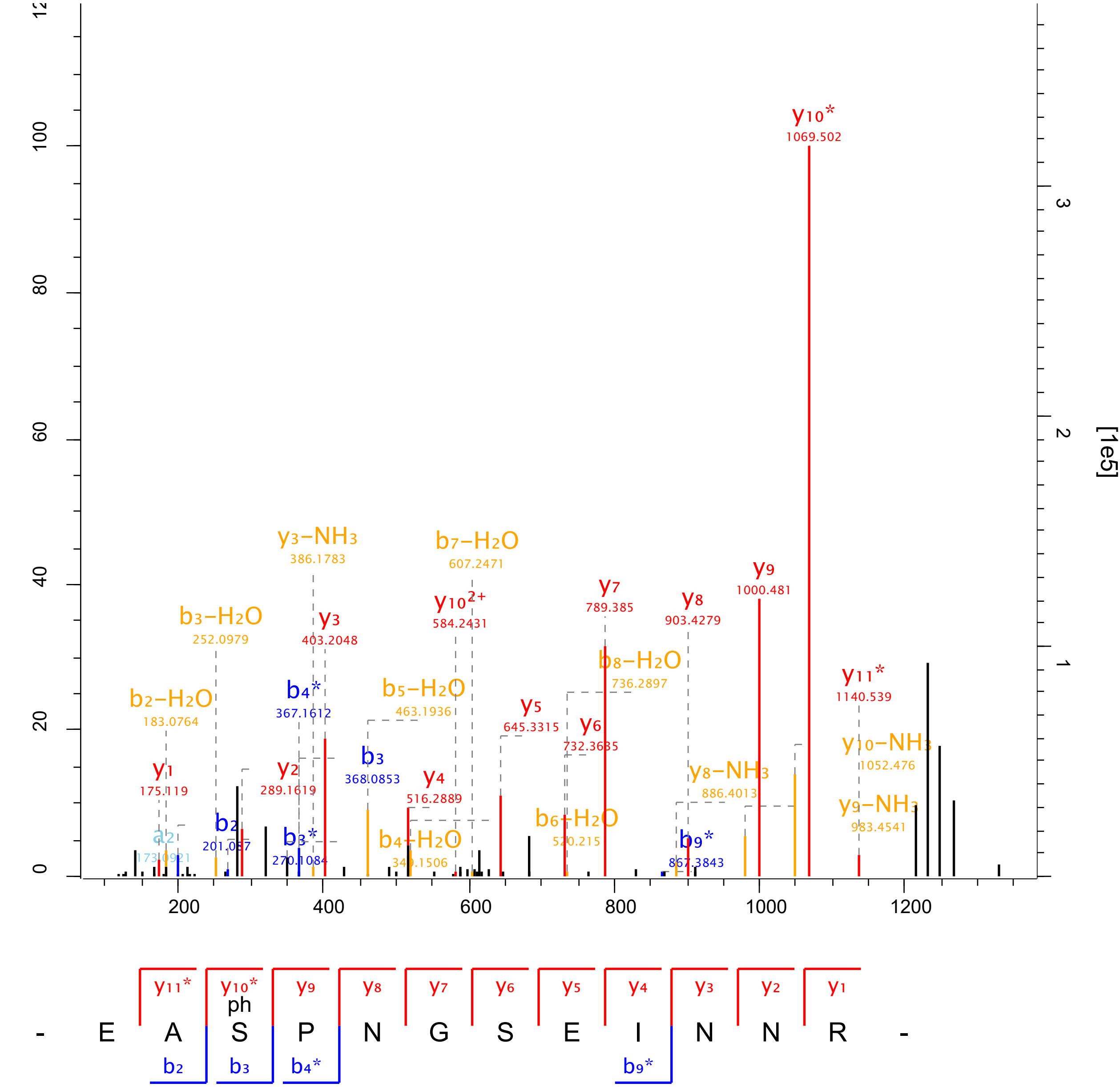

| Raw file | Scan | Method | Score | m/z |
|---|---|---|---|---|
| sys_05_2 | 5611 | FTMS; HCD | 189.09 | 829.32 |

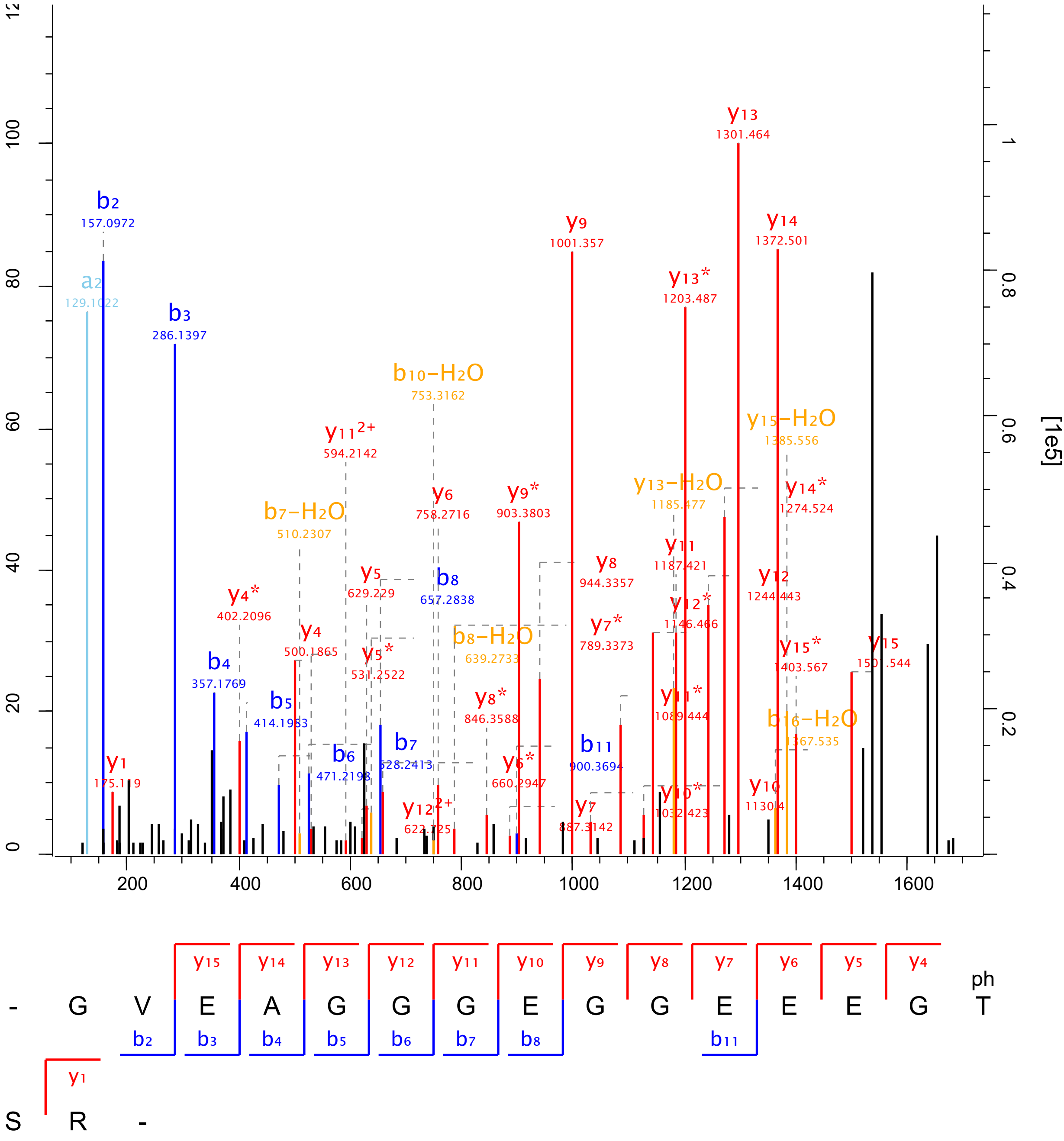

| Raw file | Scan | Method | Score | m/z |
|---|---|---|---|---|
| sys_05_2 | 5630 | FTMS; HCD | 68.54 | 452.69 |

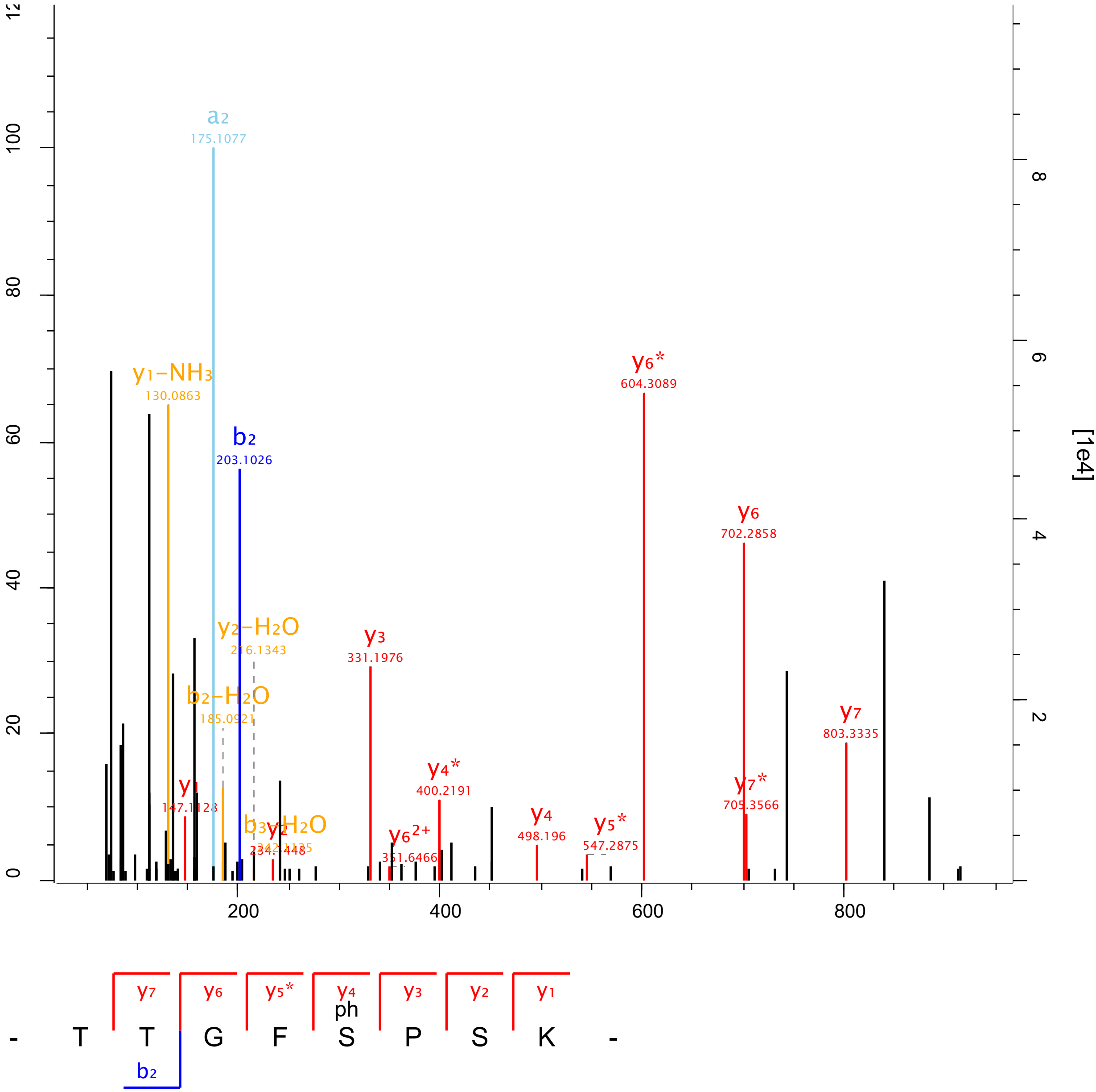

| Raw file | Scan | Method | Score | m/z |
| --- | --- | --- | --- | --- |
| sys_05_2 | 5681 | FTMS; HCD | 311.16 | 895.86 |

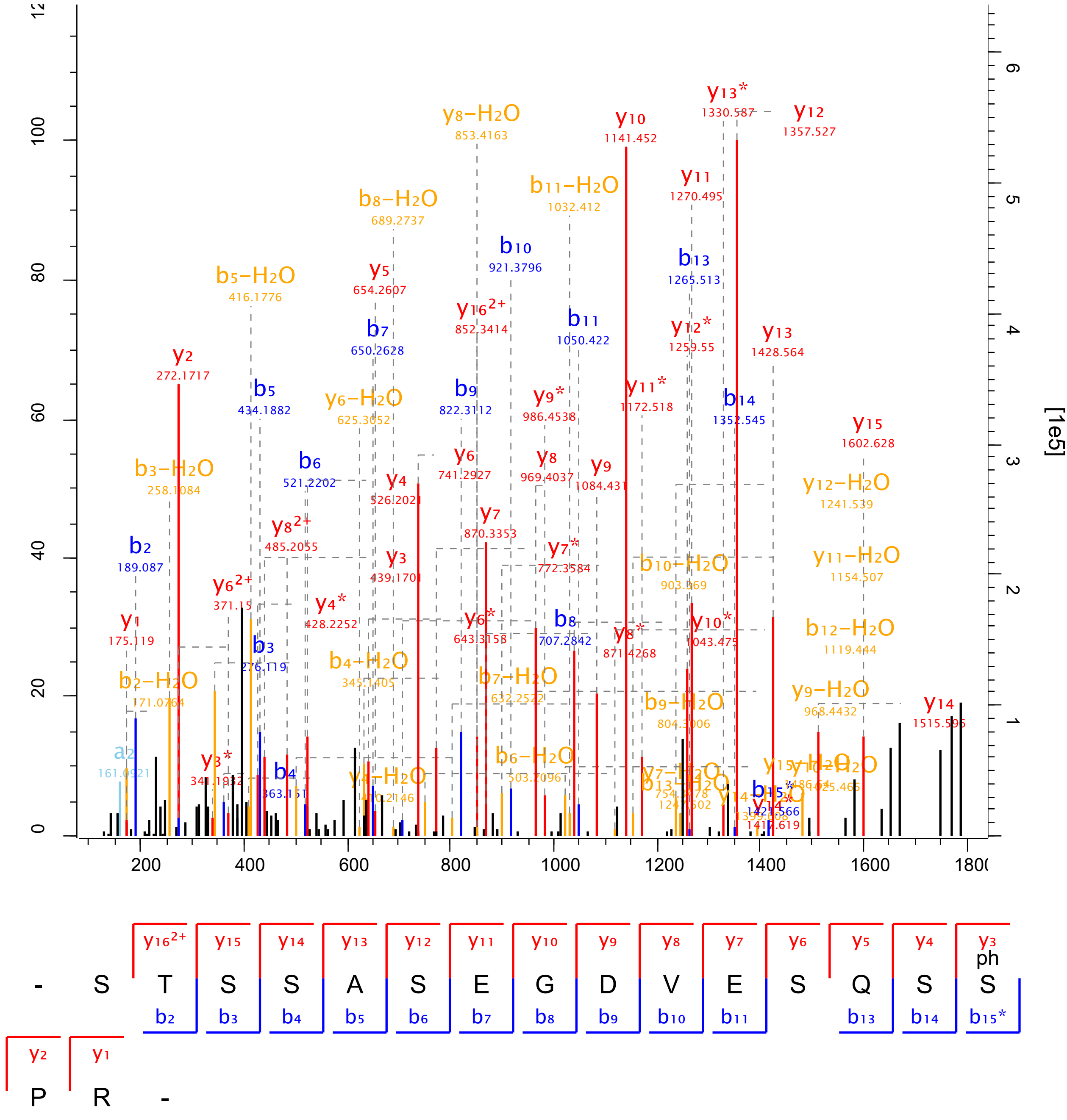

| Raw file | Scan | Method | Score | m/z |
|---|---|---|---|---|
| sys_05_2 | 5694 | FTMS; HCD | 194.58 | 769.81 |

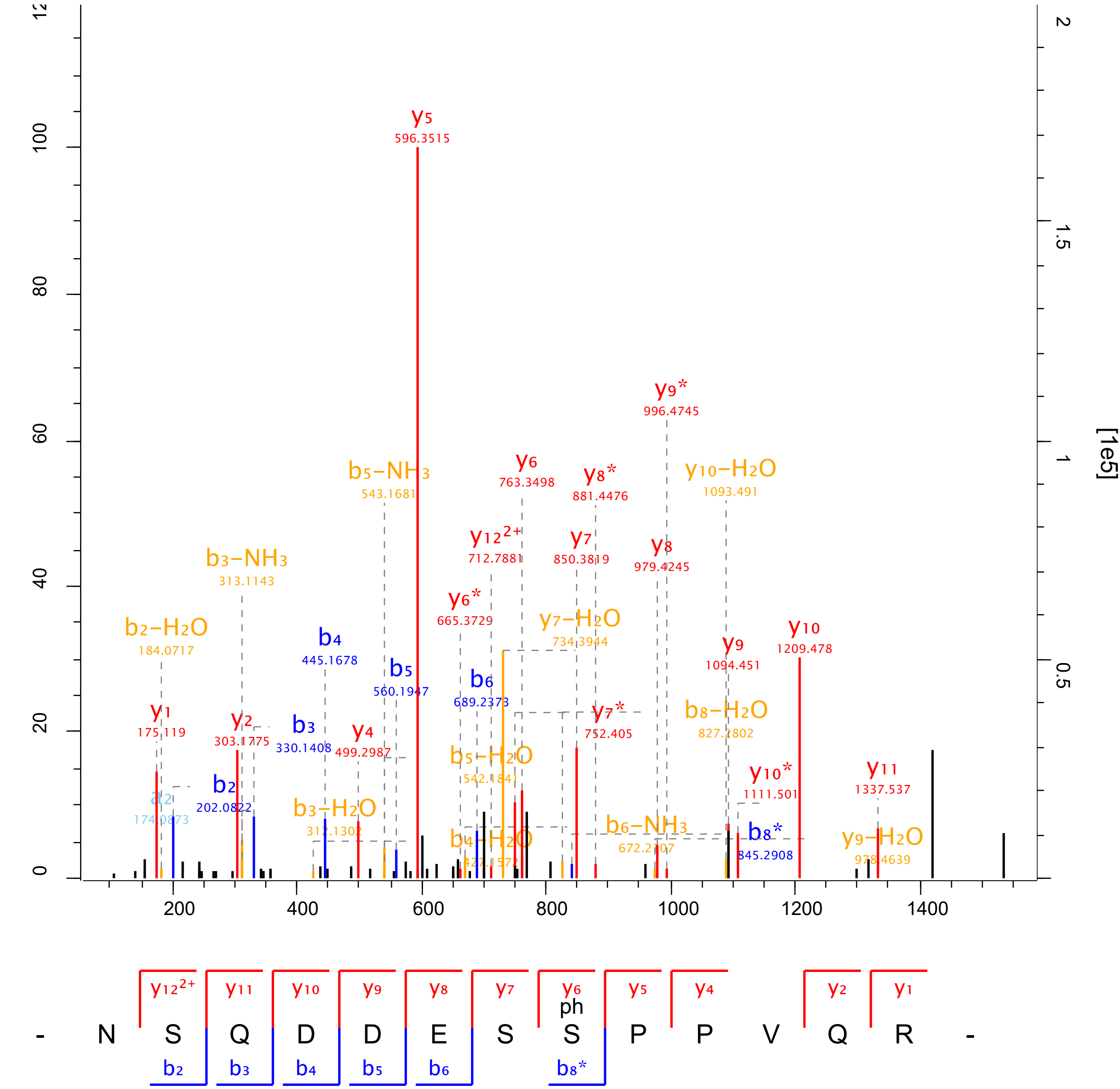

| Raw file | Scan | Method | Score | m/z |
| --- | --- | --- | --- | --- |
| sys_05_2 | 5859 | FTMS; HCD | 206.63 | 684.3 |

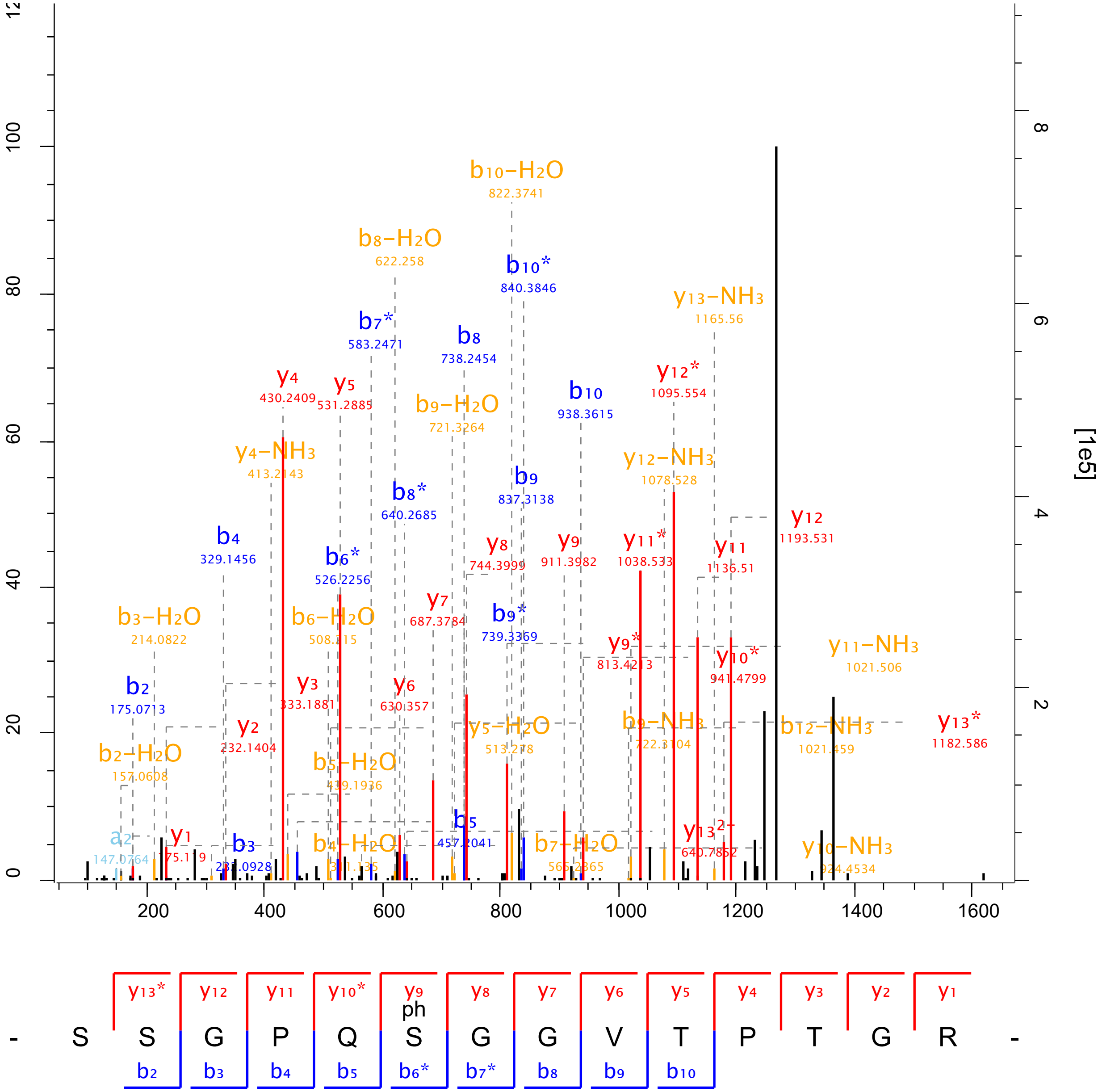

| Raw file | Scan | Method | Score | m/z |
|---|---|---|---|---|
| sys_05_2 | 6060 | FTMS; HCD | 123.76 | 618.77 |

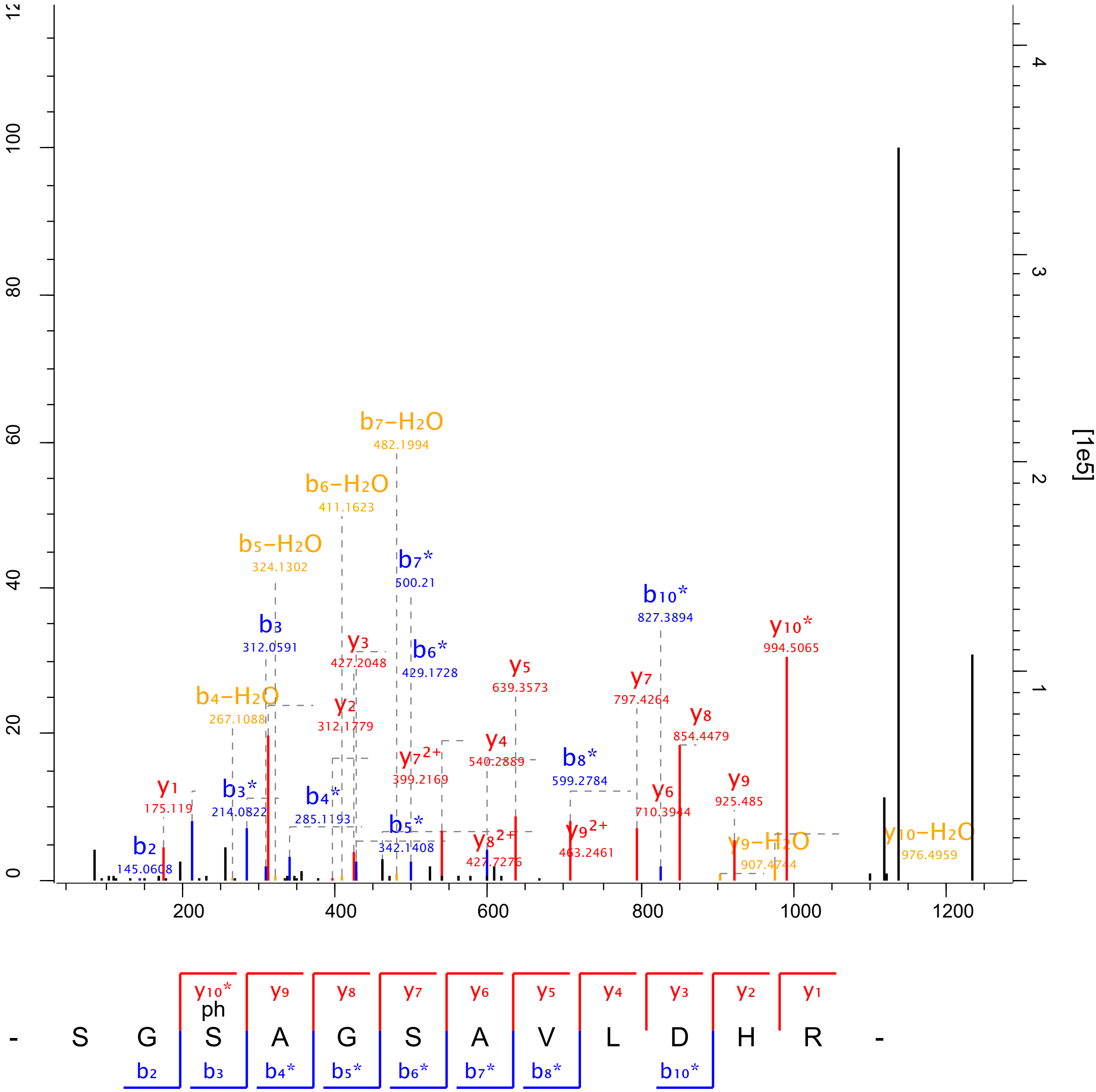

| Raw file | Scan | Method | Score | m/z |
|---|---|---|---|---|
| sys_05_2 | 6073 | FTMS; HCD | 151.18 | 676.6 |

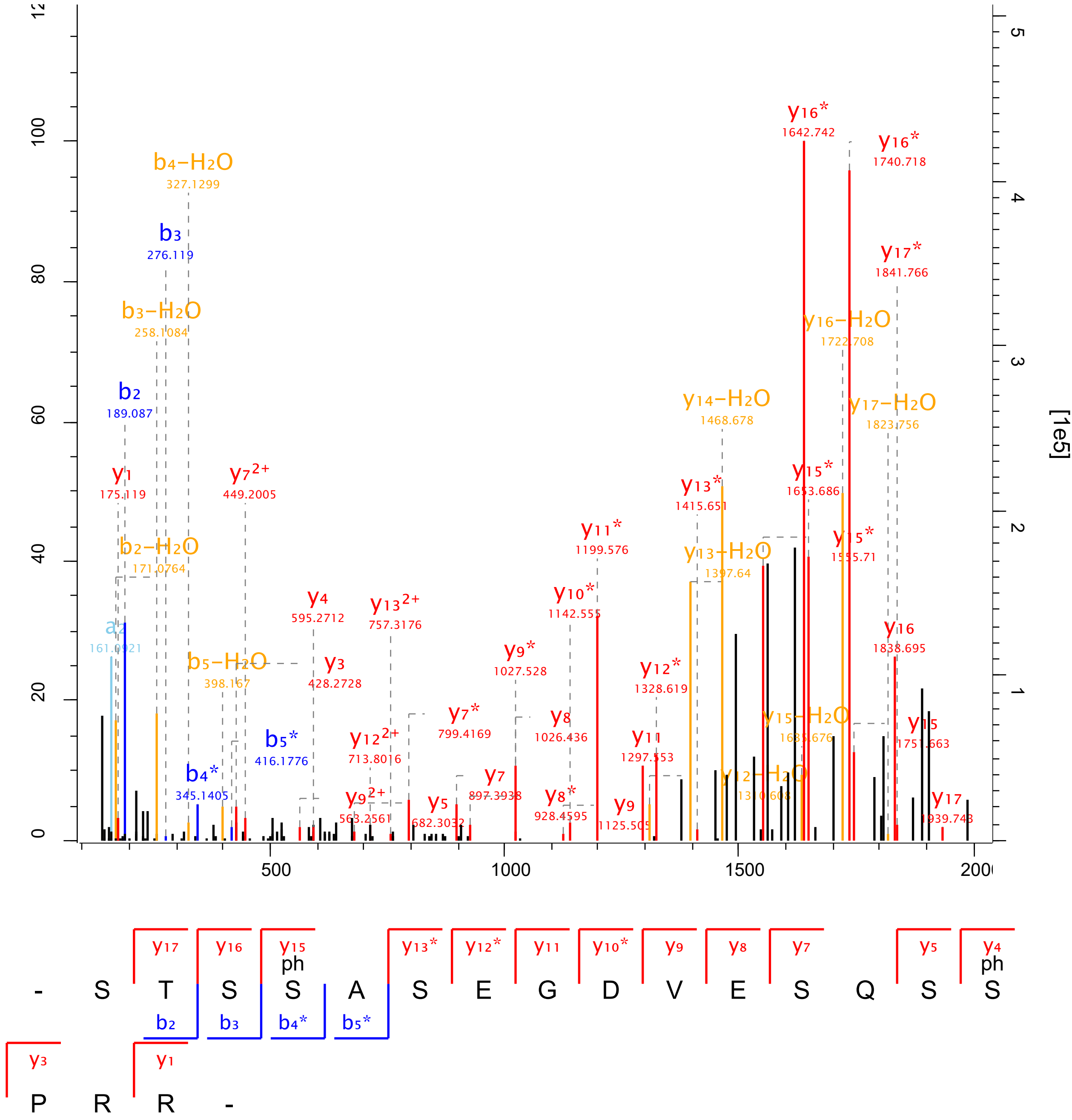

| Raw file | Scan | Method | Score | m/z |
|---|---|---|---|---|
| sys_05_2 | 6121 | FTMS; HCD | 85.68 | 431.71 |

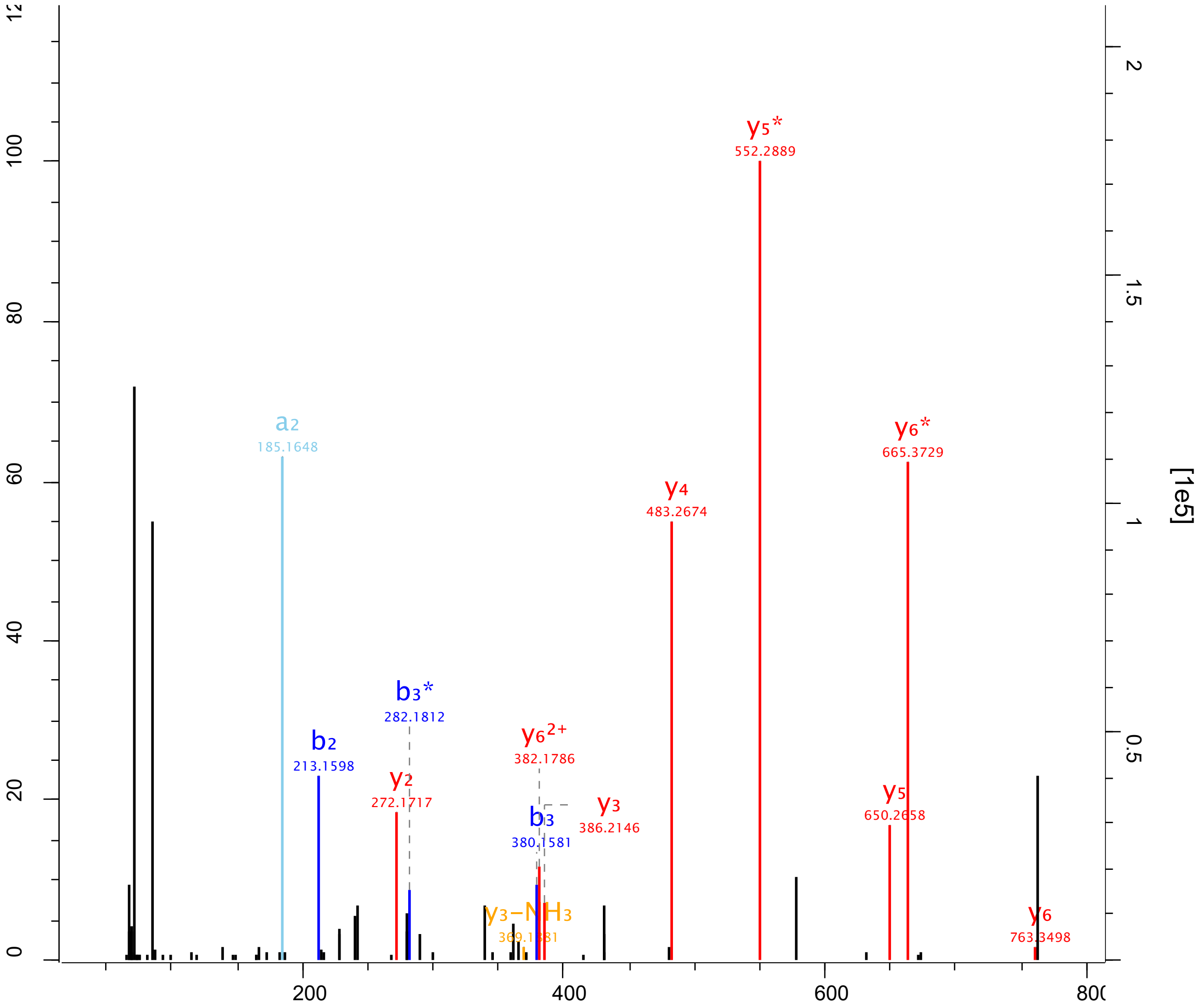


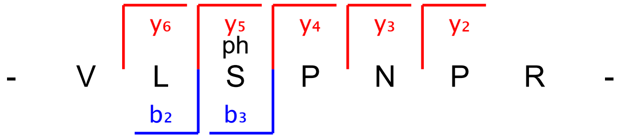

| Raw file | Scan | Method | Score | m/z |
|---|---|---|---|---|
| sys_05_2 | 6151 | FTMS; HCD | 144.97 | 739.83 |

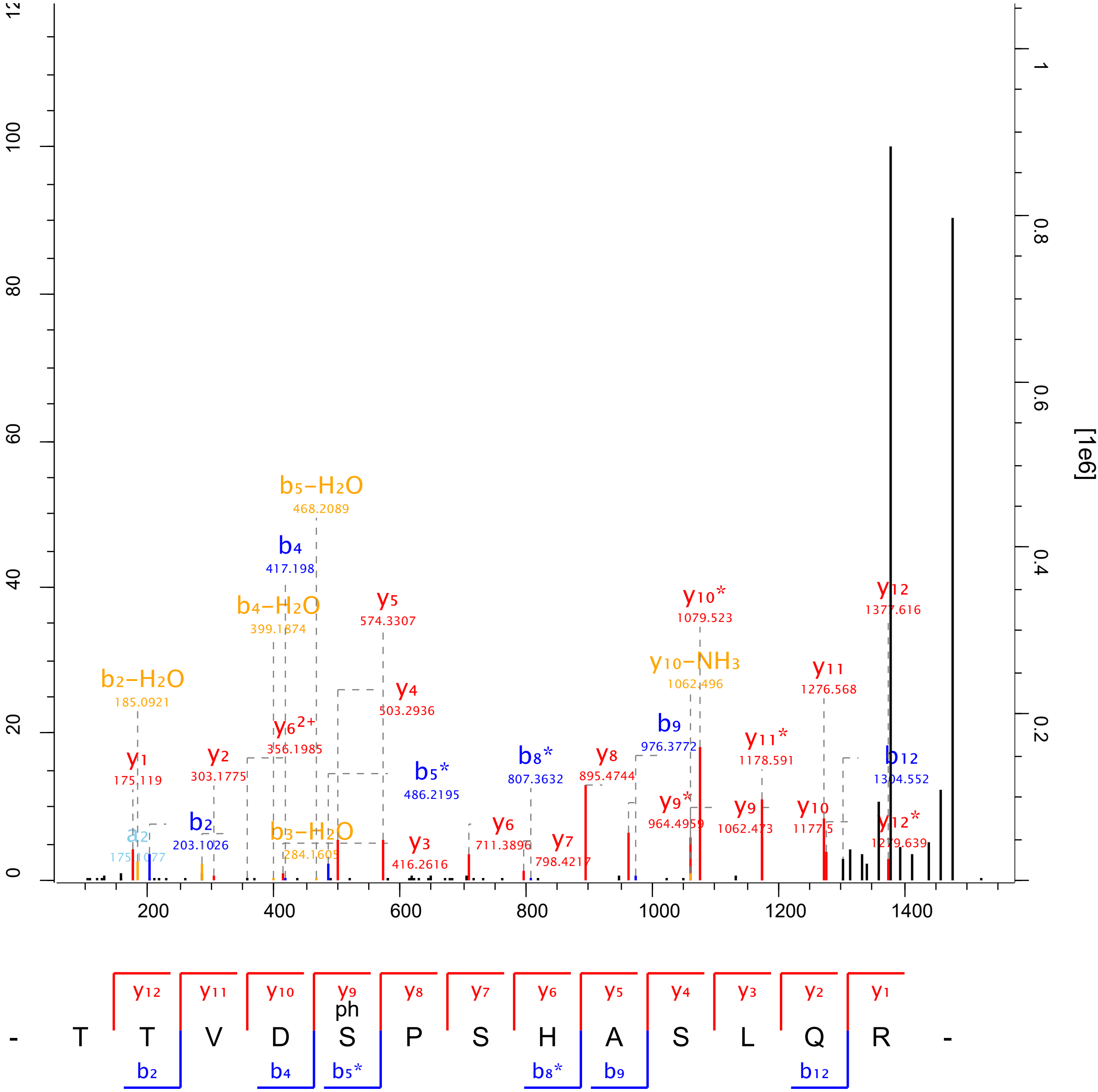

| Raw file | Scan | Method | Score | m/z |
|---|---|---|---|---|
| sys_05_2 | 6228 | FTMS; HCD | 95.48 | 585.77 |

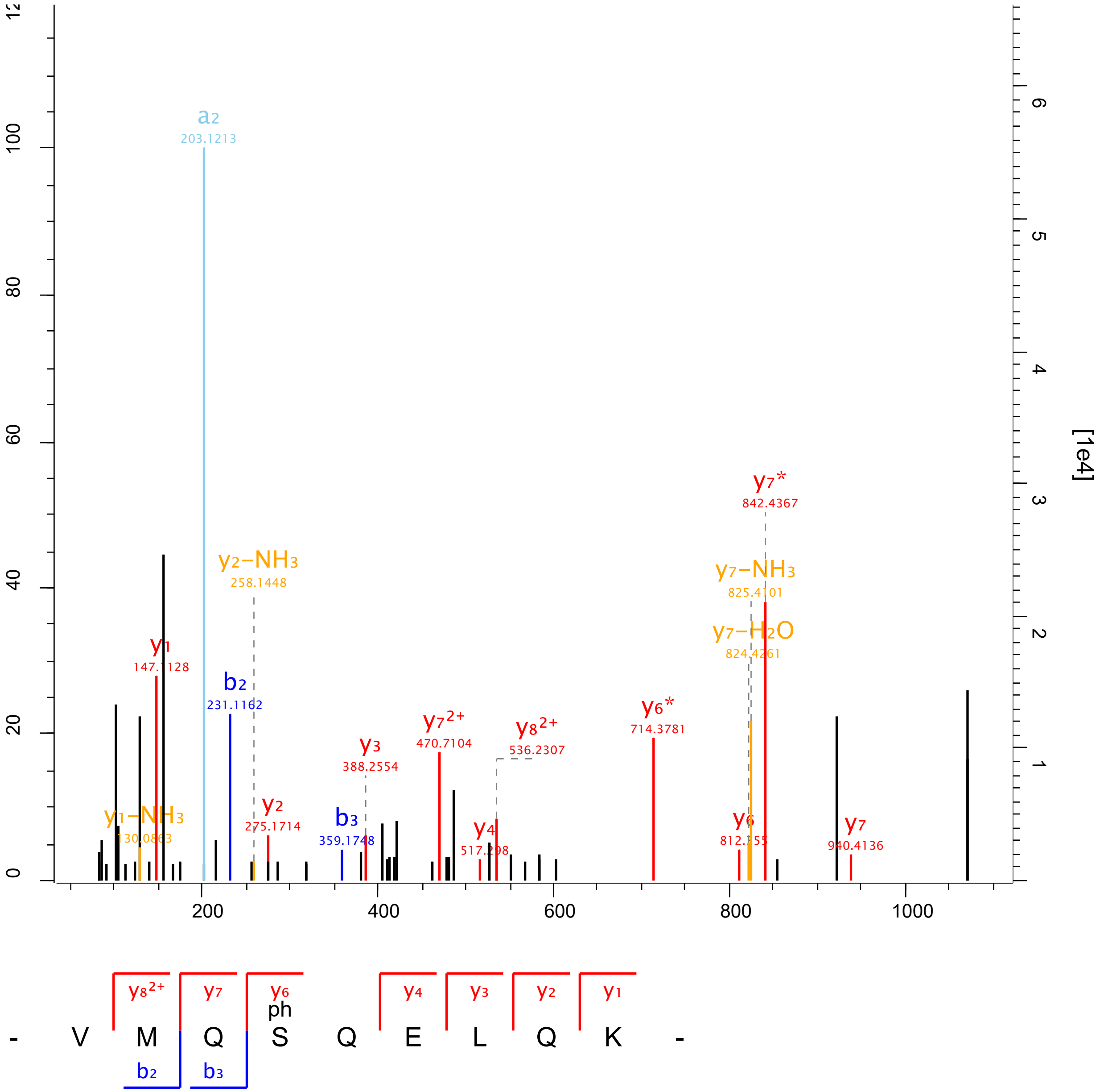

| Raw file | Scan | Method | Score | m/z |
|---|---|---|---|---|
| sys_05_2 | 6346 | FTMS; HCD | 204.63 | 736.8 |

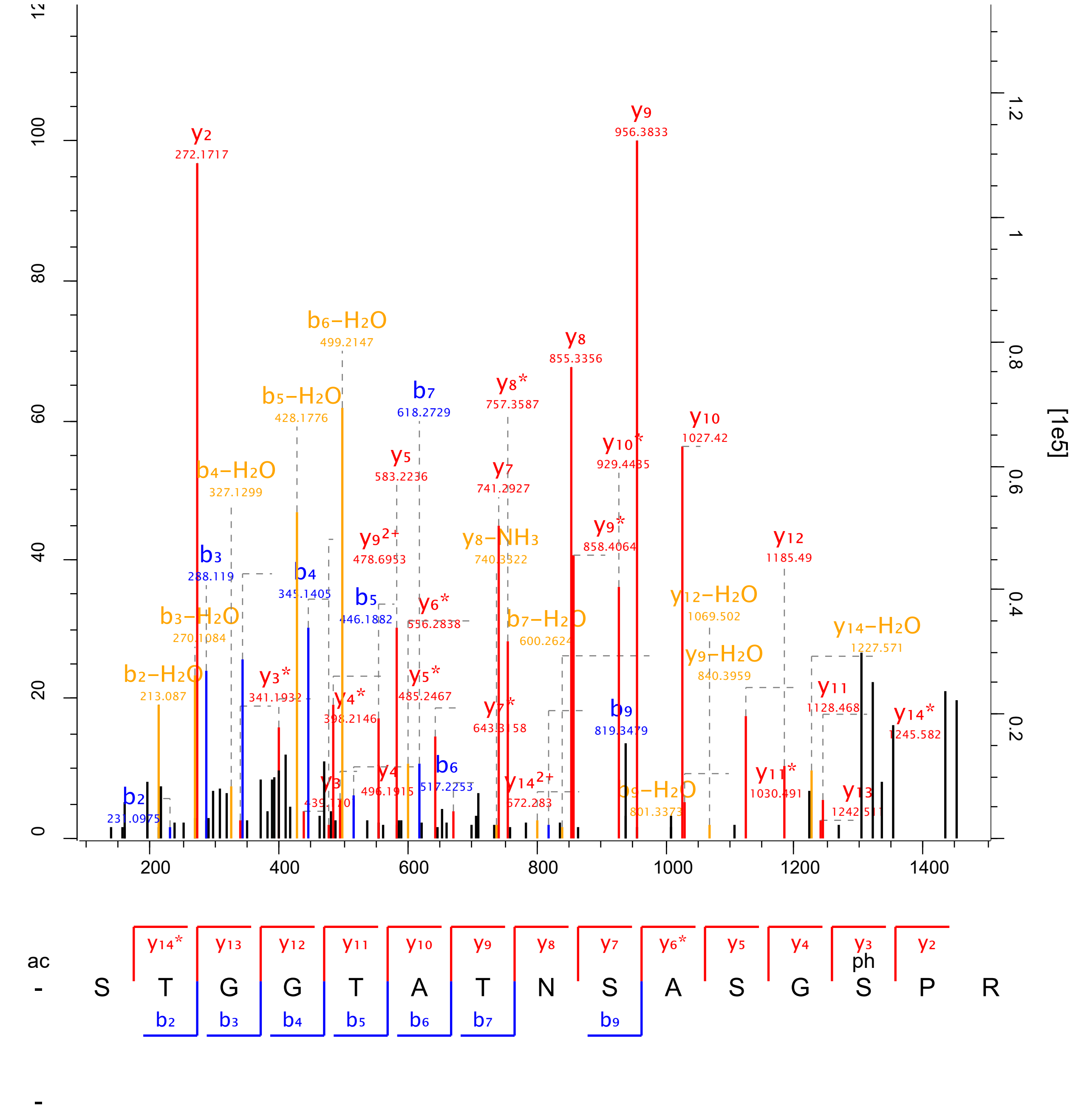

| Raw file | Scan | Method | Score | m/z |
| --- | --- | --- | --- | --- |
| sys_05_2 | 6385 | FTMS; HCD | 127.32 | 622.29 |

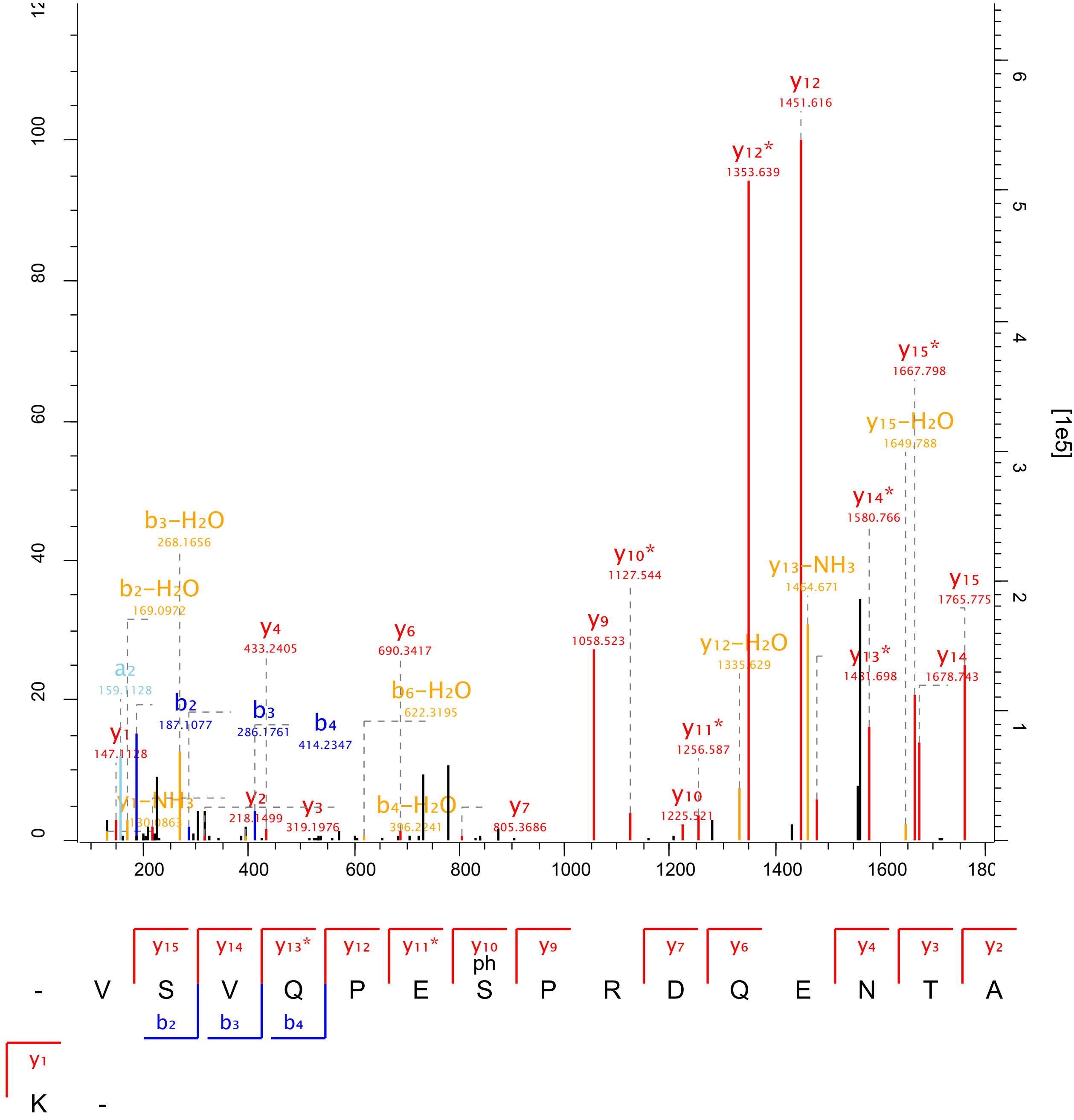

| Raw file | Scan | Method | Score | m/z |
|---|---|---|---|---|
| sys_05_2 | 6423 | FTMS; HCD | 140.93 | 515.88 |

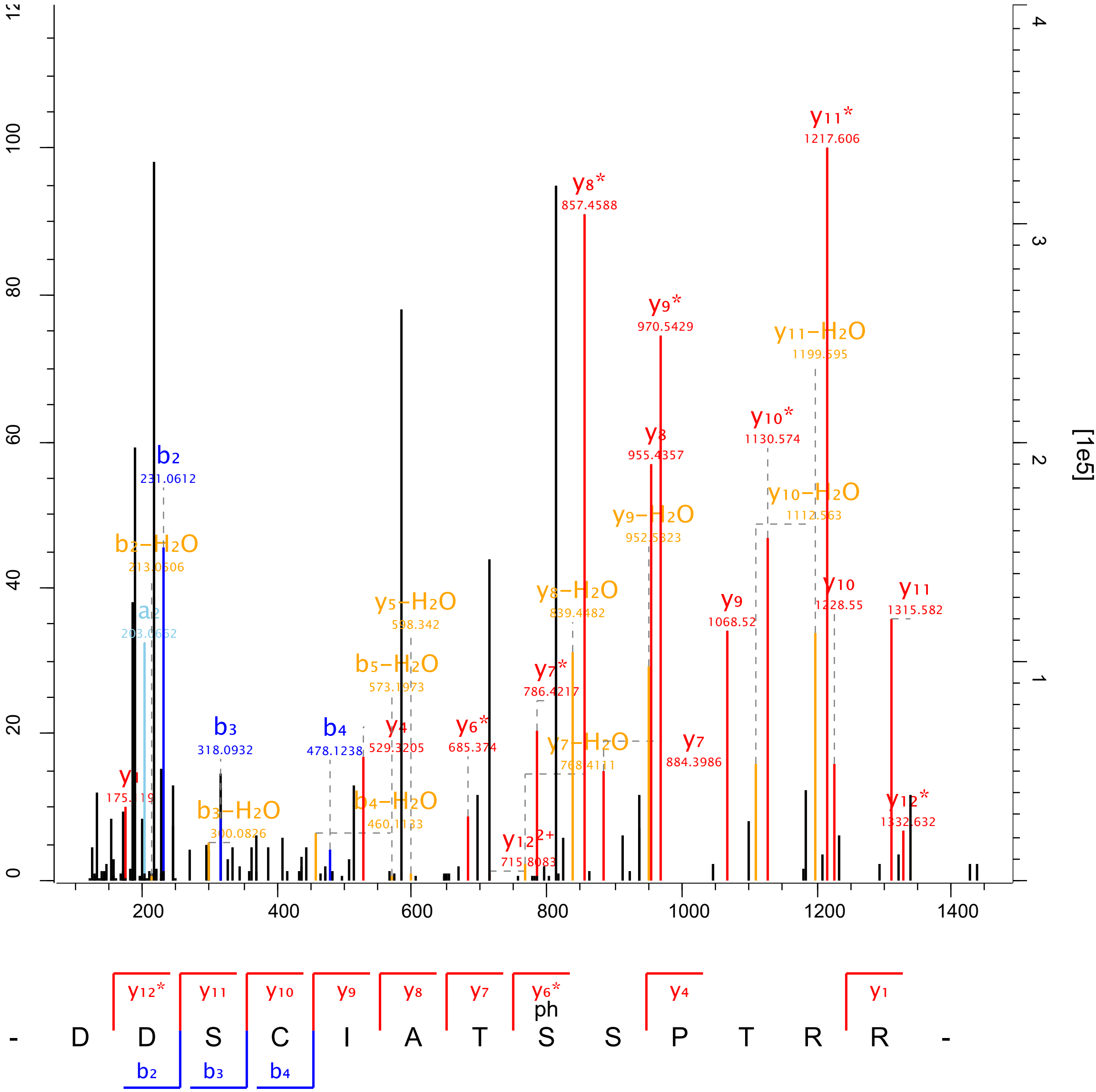

| Raw file | Scan | Method | Score | m/z |
|---|---|---|---|---|
| sys_05_2 | 6453 | FTMS; HCD | 137.45 | 789.3 |

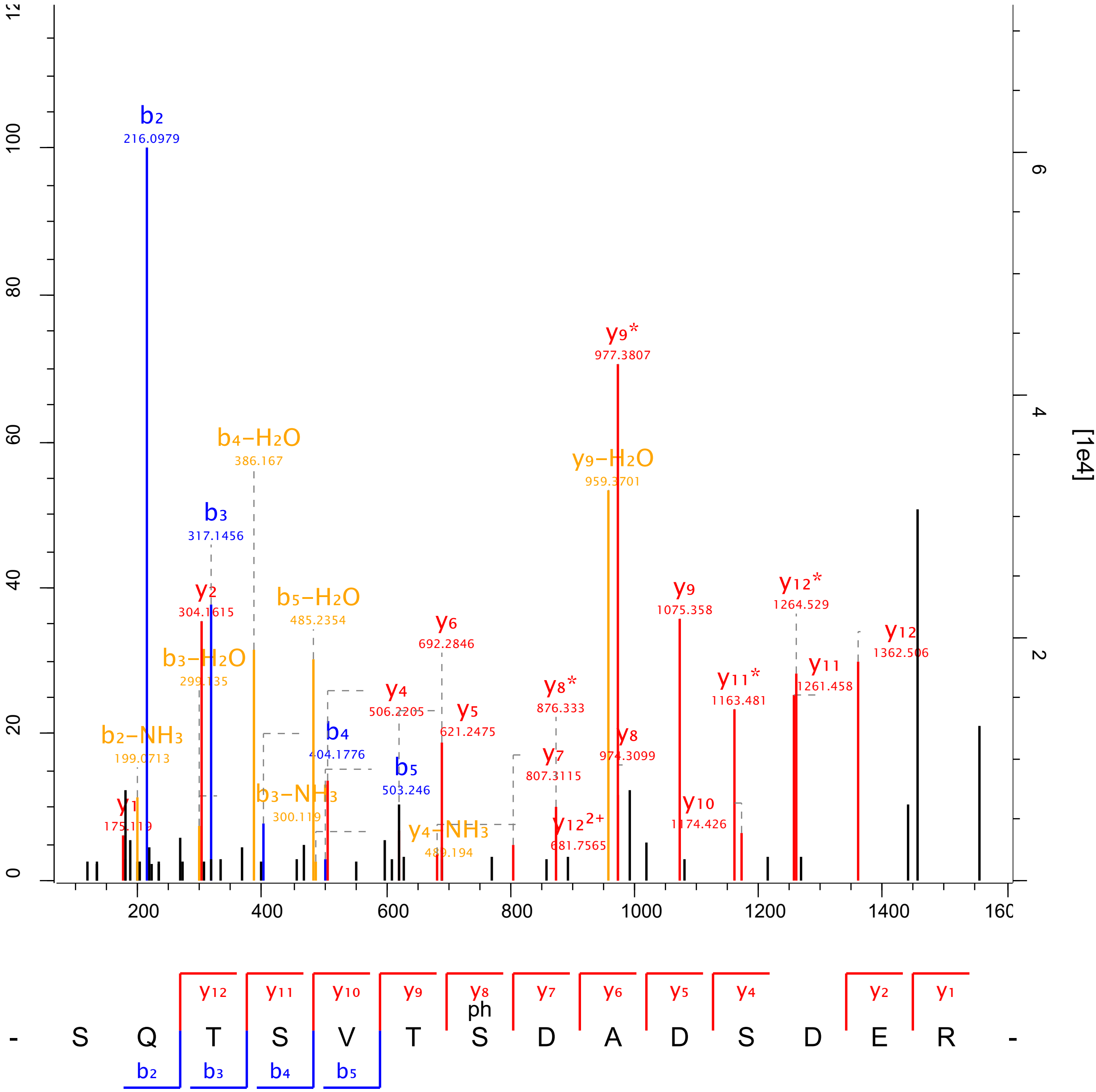

| Raw file | Scan | Method | Score | m/z |
| --- | --- | --- | --- | --- |
| sys_05_2 | 6524 | FTMS; HCD | 95.85 | 585.76 |

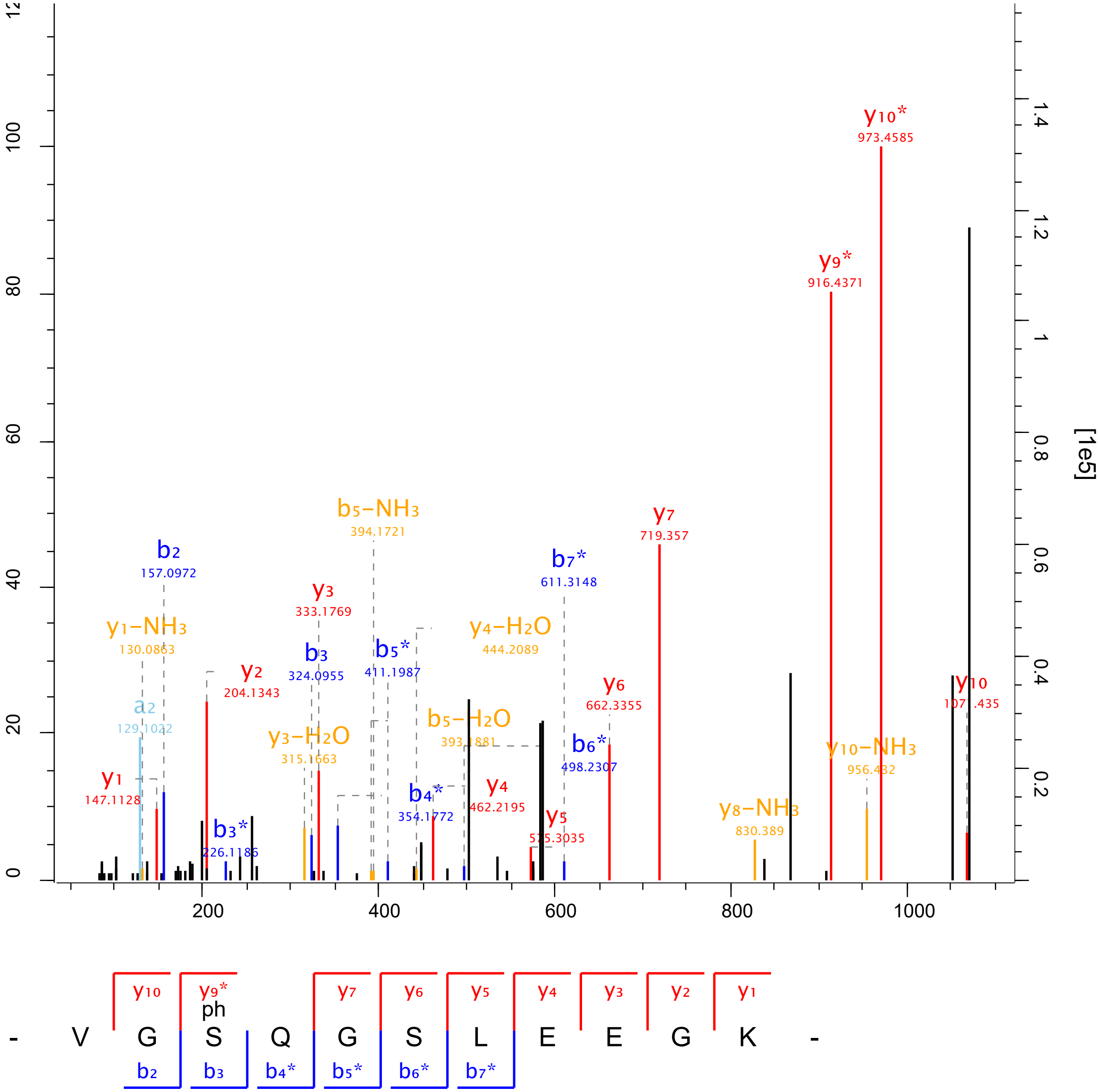

| Raw file | Scan | Method | Score | m/z |
|---|---|---|---|---|
| sys_05_2 | 6539 | FTMS; HCD | 200.93 | 456.17 |

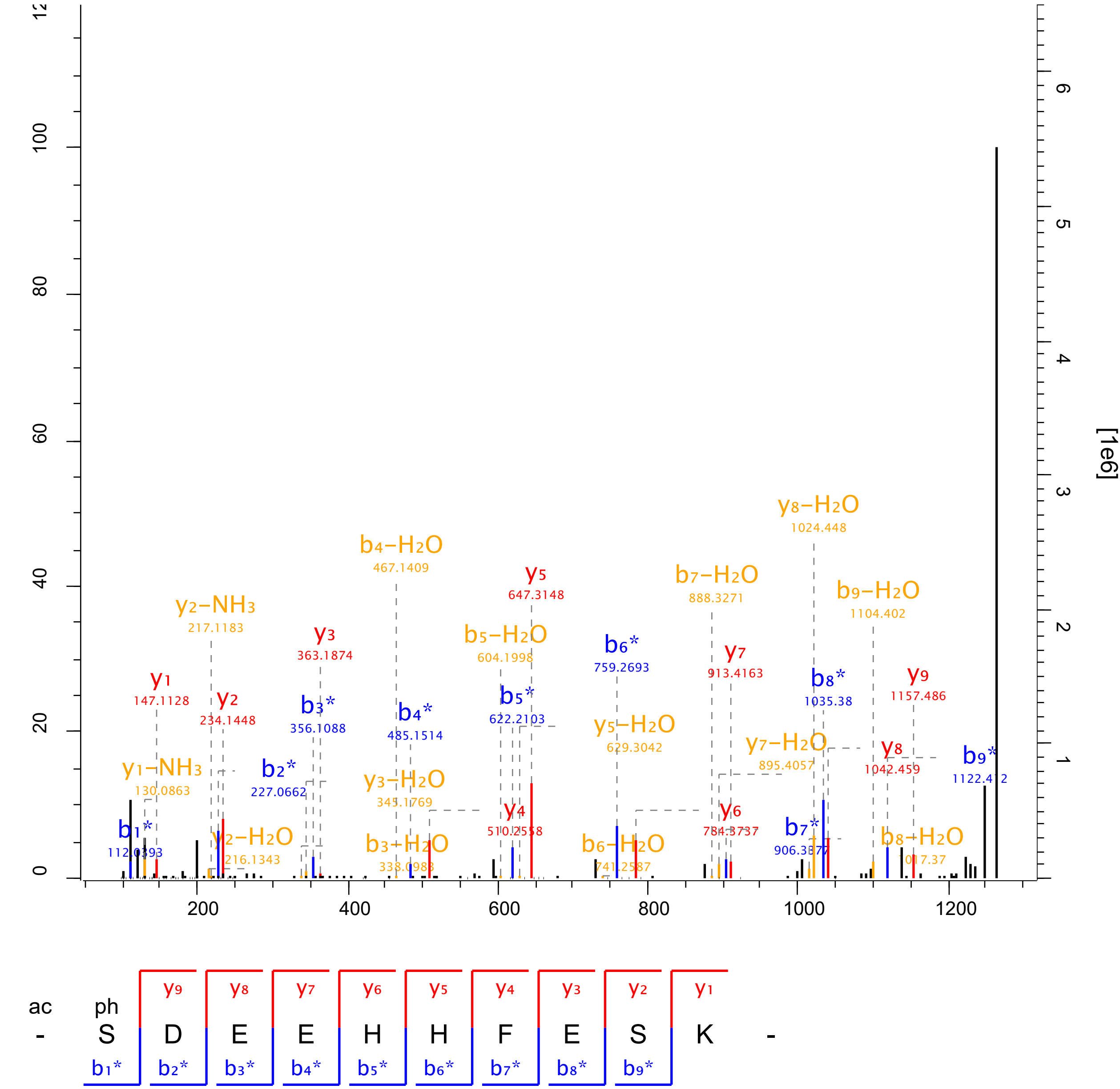

| Raw file | Scan | Method | Score | m/z |
| --- | --- | --- | --- | --- |
| sys_05_2 | 6588 | FTMS; HCD | 133.89 | 503.22 |

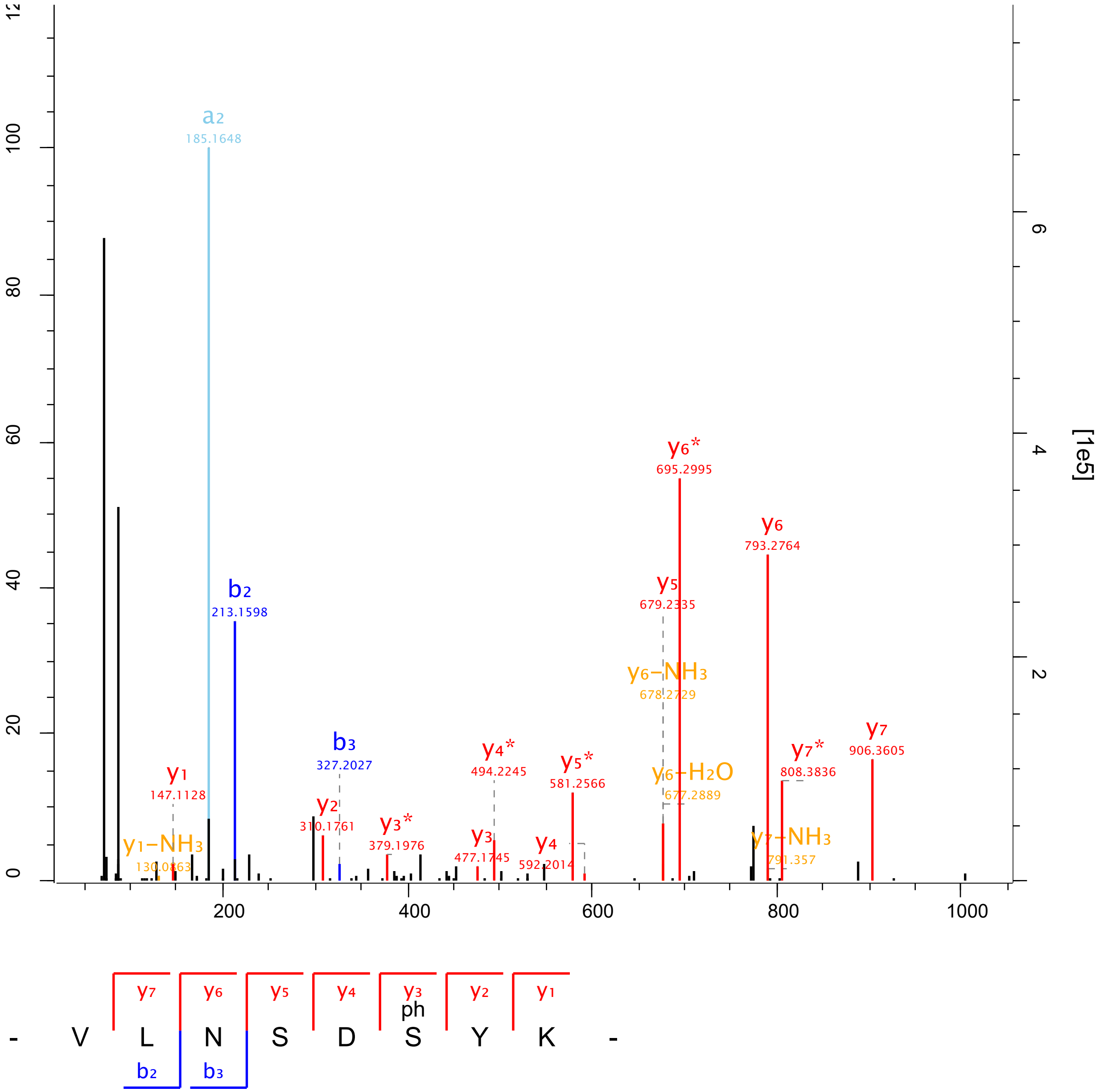

| Raw file | Scan | Method | Score | m/z |
|---|---|---|---|---|
| sys_05_2 | 6614 | FTMS; HCD | 140.11 | 434.7 |

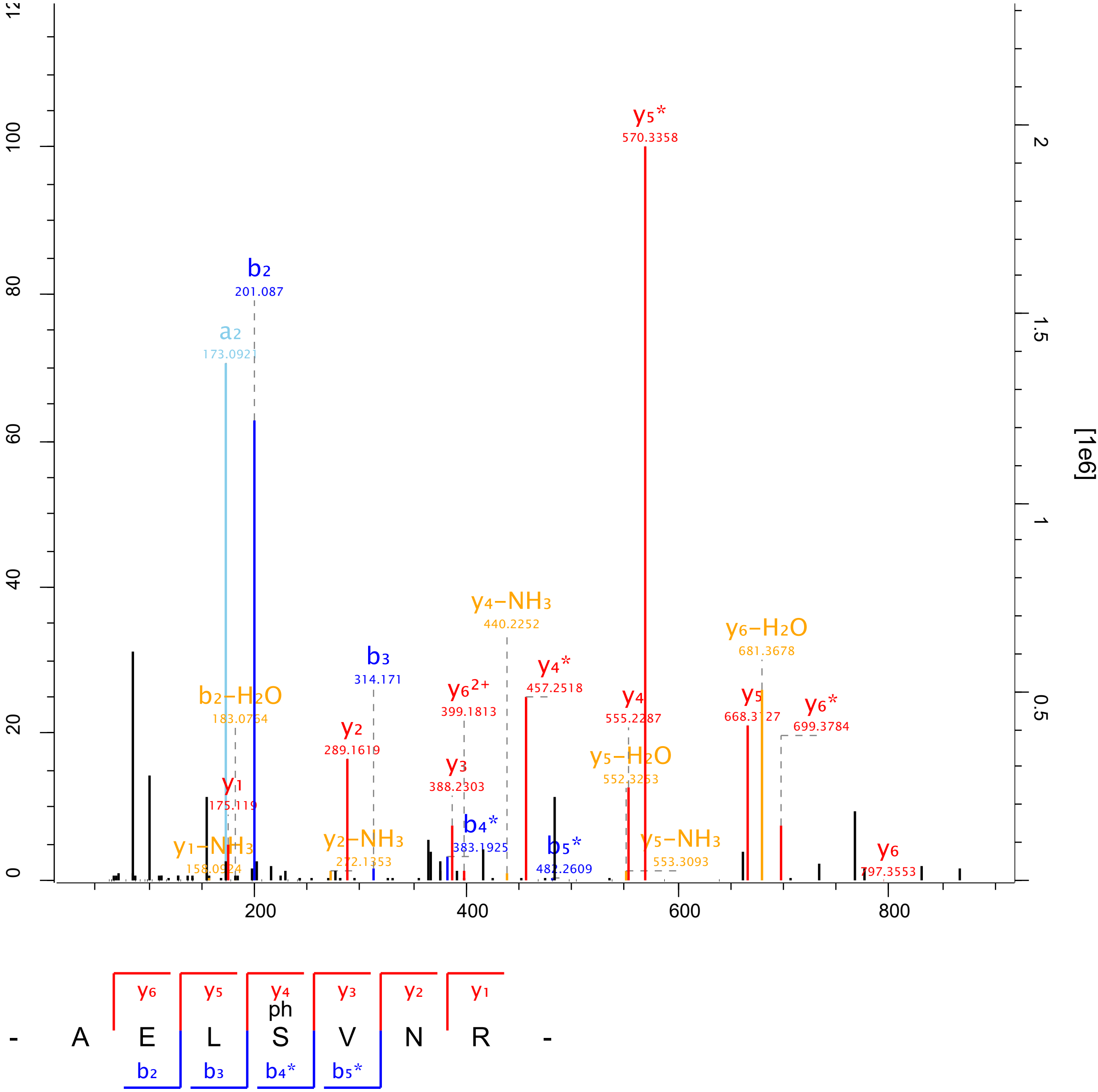

| Raw file | Scan | Method | Score | m/z |
|---|---|---|---|---|
| sys_05_2 | 6624 | FTMS; HCD | 177.63 | 548.28 |

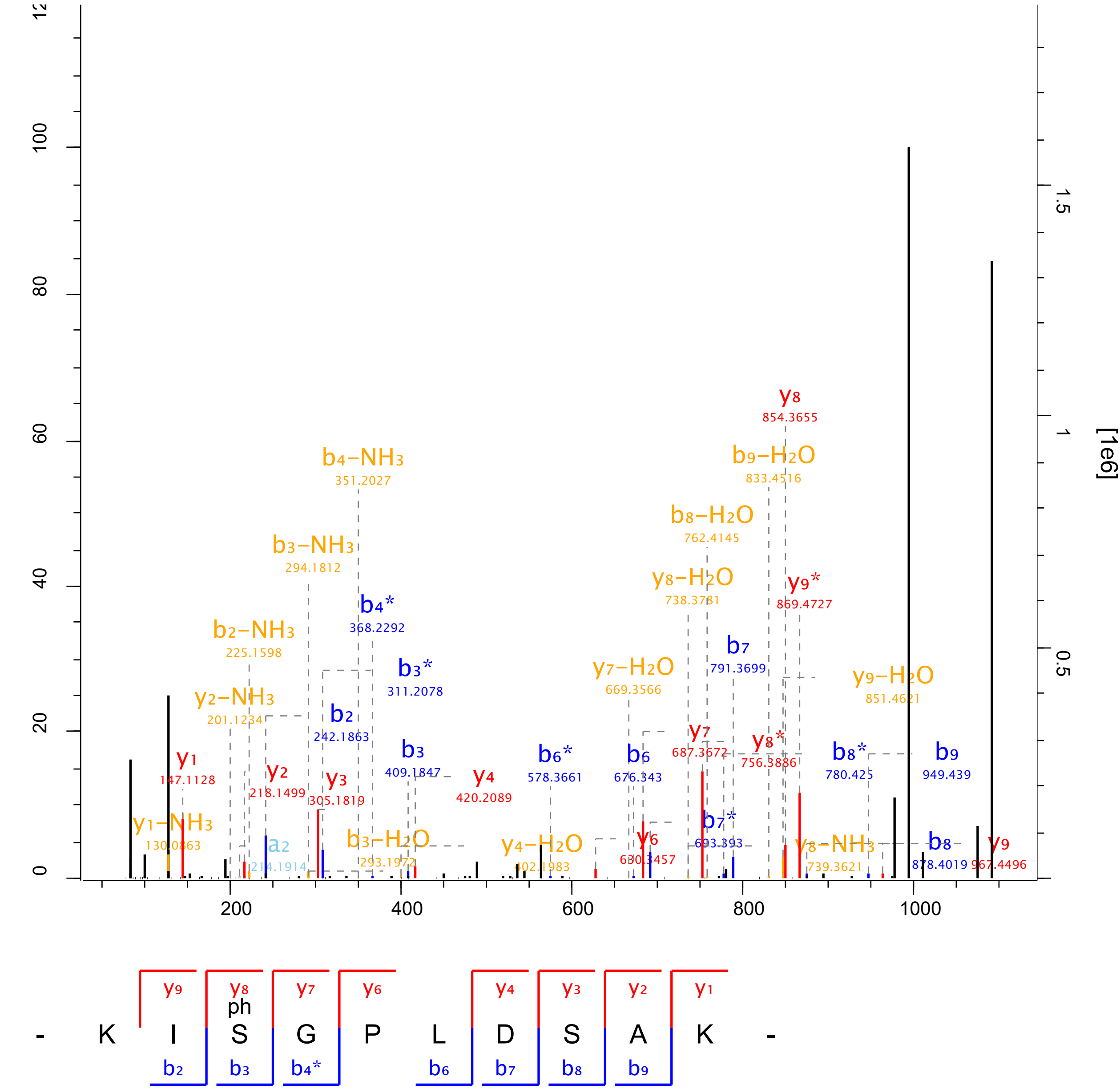

| Raw file | Scan | Method | Score | m/z |
|---|---|---|---|---|
| sys_05_2 | 6638 | FTMS; HCD | 85.54 | 579.26 |

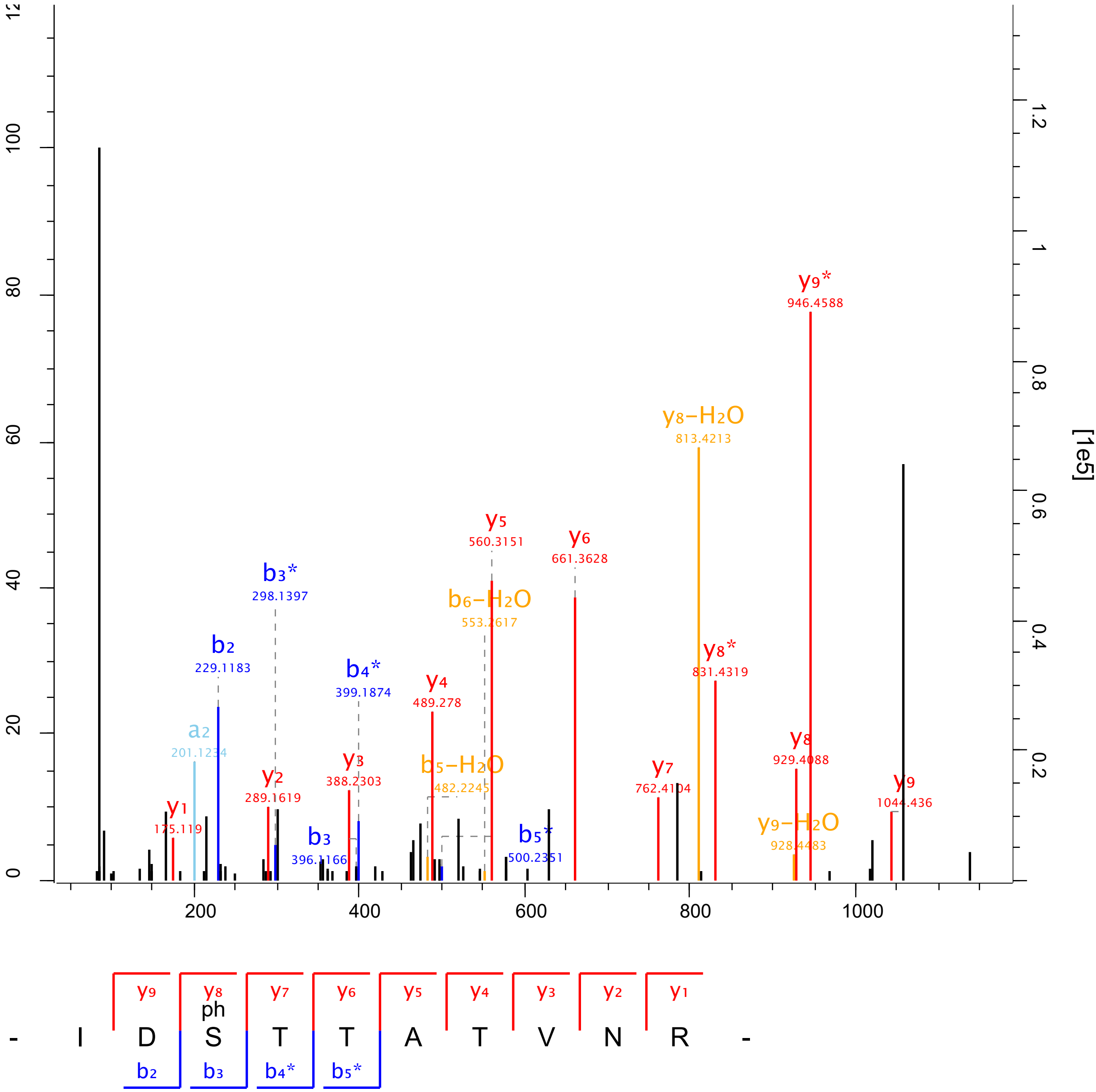

| Raw file | Scan | Method | Score | m/z |
|---|---|---|---|---|
| sys_05_2 | 6780 | FTMS; HCD | 43.37 | 706.3 |

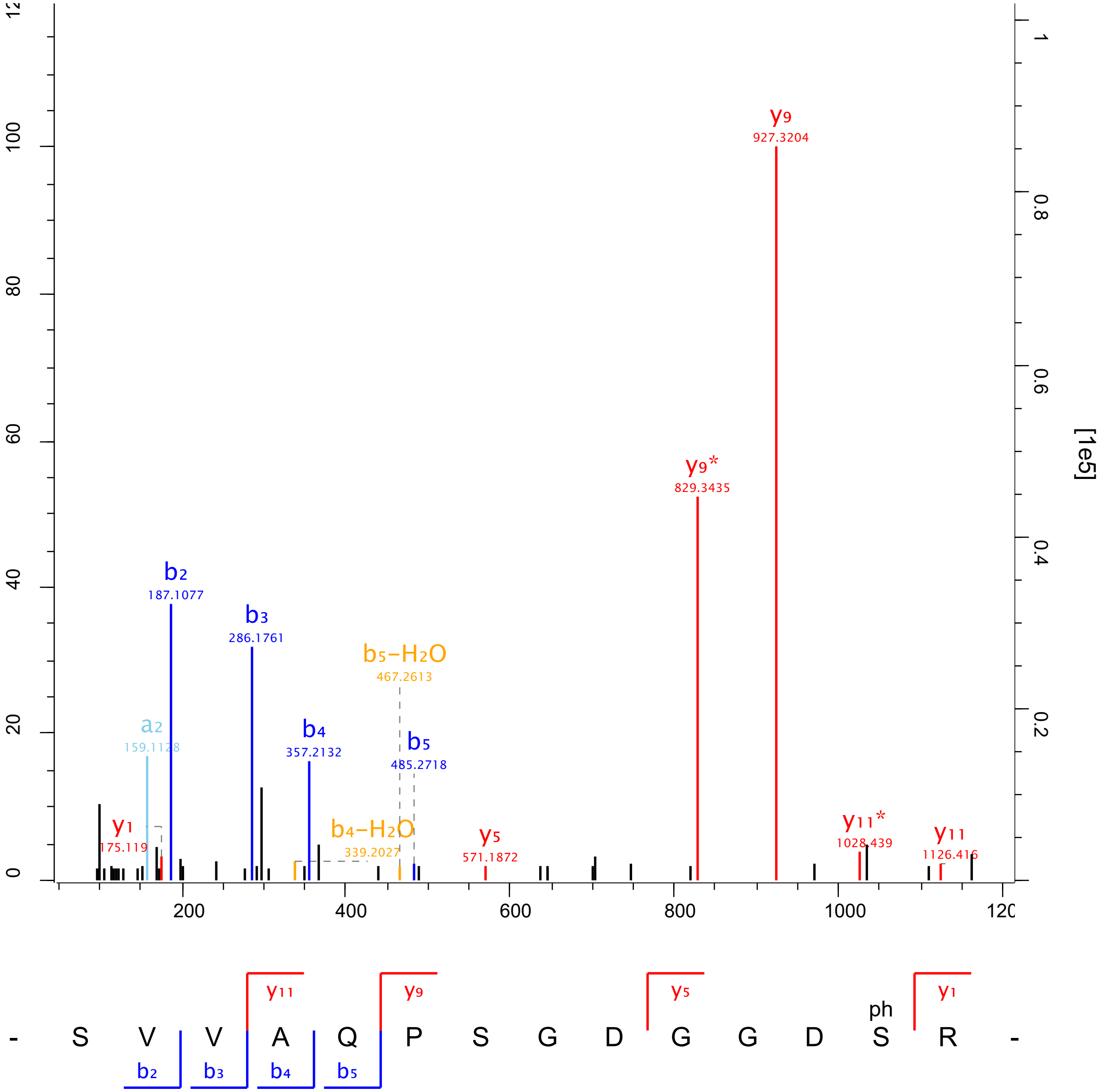

| Raw file | Scan | Method | Score | m/z |
|---|---|---|---|---|
| sys_05_2 | 6827 | FTMS; HCD | 119.77 | 595.24 |

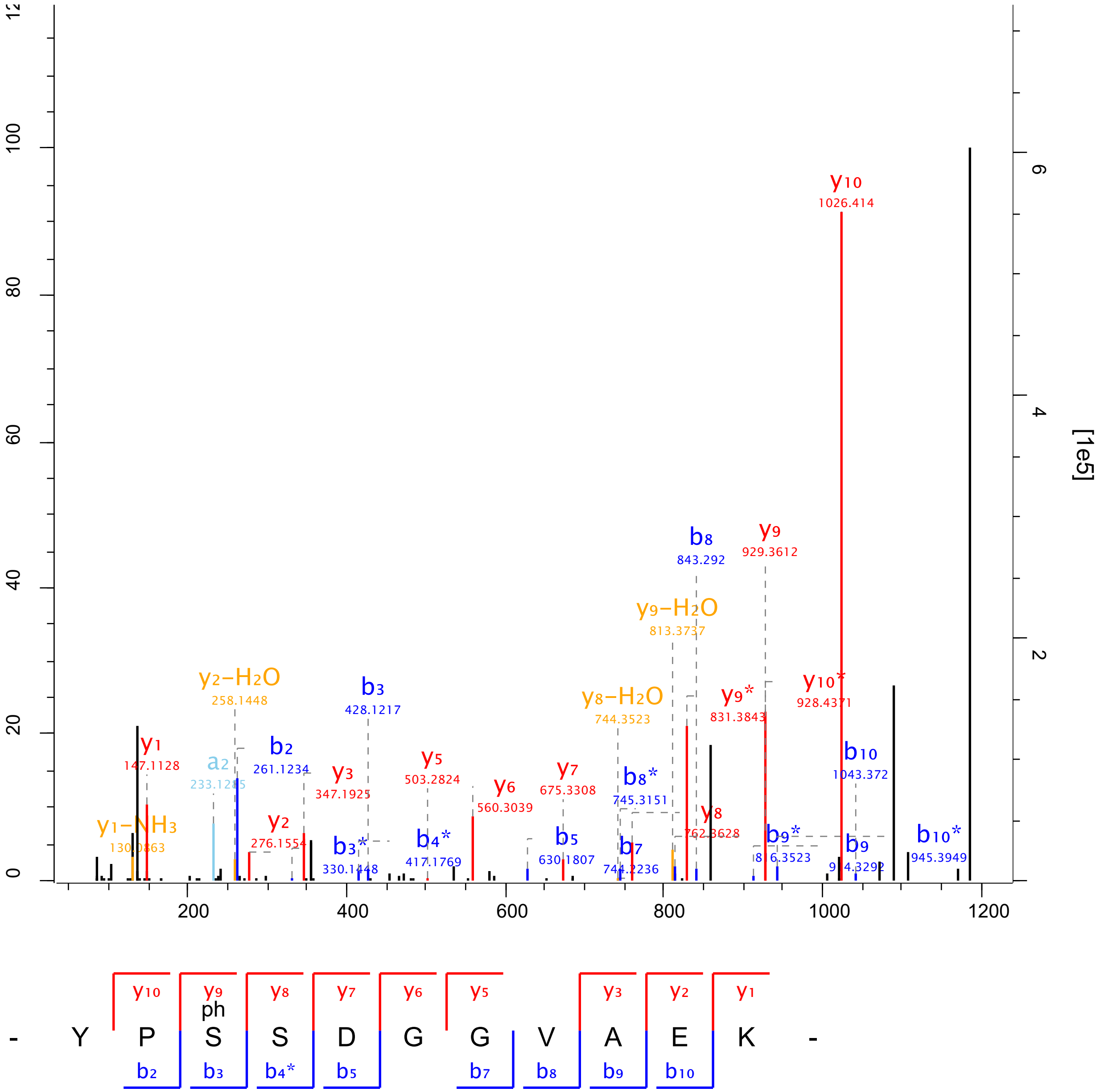

| Raw file | Scan | Method | Score | m/z |
|---|---|---|---|---|
| sys_05_2 | 6918 | FTMS; HCD | 120.53 | 627.25 |

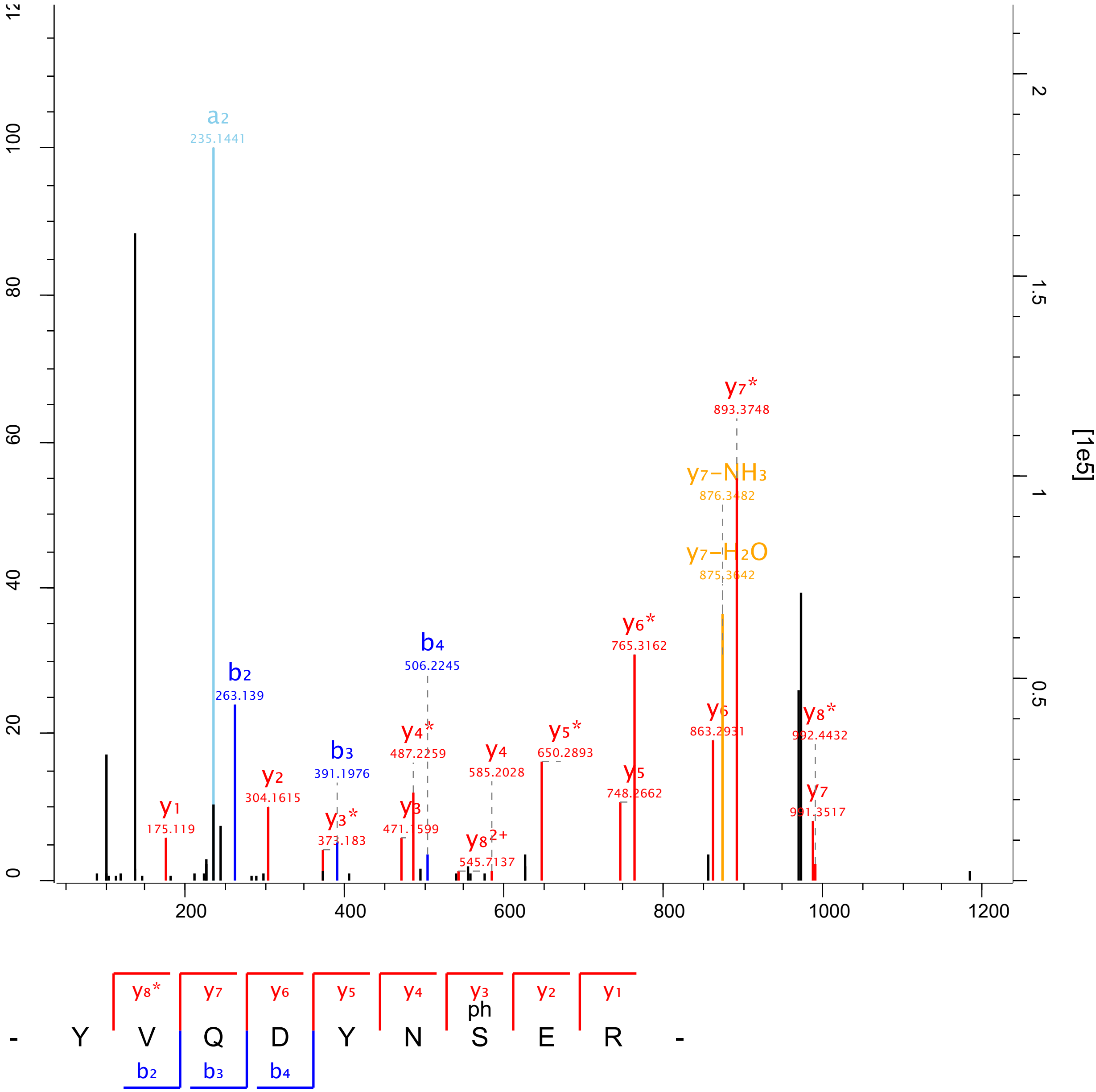

| Raw file | Scan | Method | Score | m/z |
| --- | --- | --- | --- | --- |
| sys_05_2 | 6972 | FTMS; HCD | 192.39 | 661.8 |

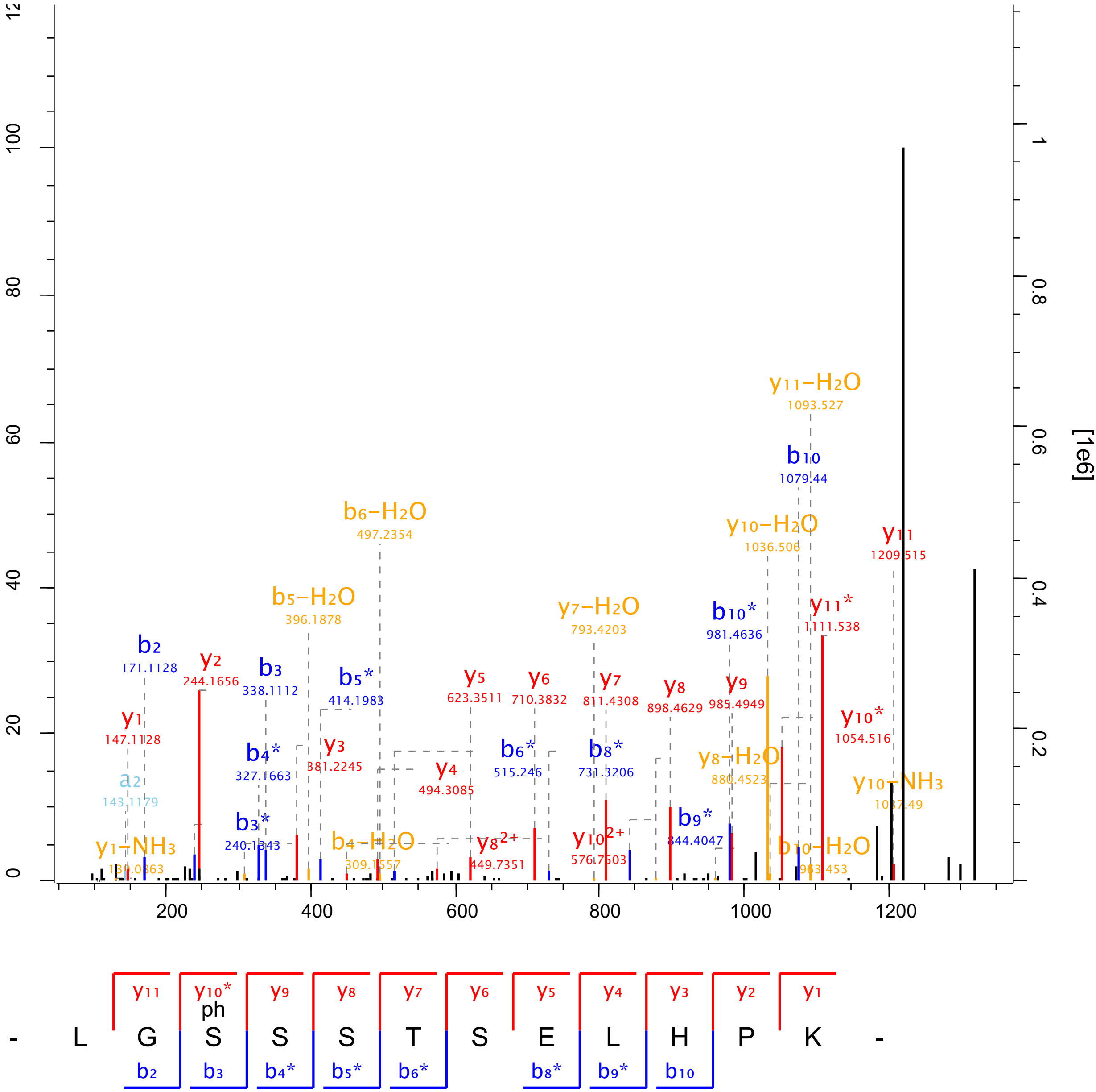

| Raw file | Scan | Method | Score | m/z |
| --- | --- | --- | --- | --- |
| sys_05_2 | 7017 | FTMS; HCD | 84.51 | 476.71 |

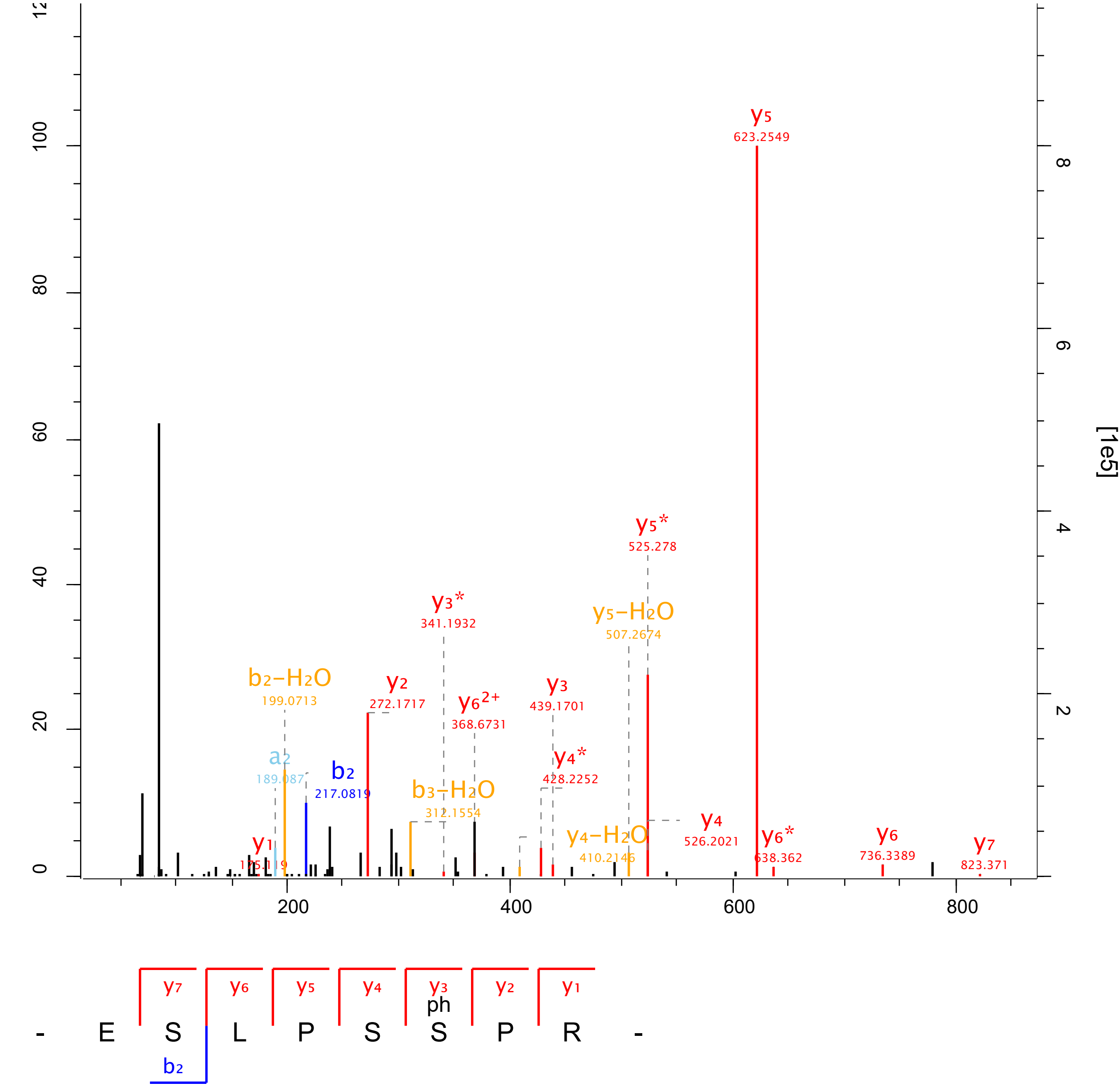

| Raw file | Scan | Method | Score | m/z |
|---|---|---|---|---|
| sys_05_2 | 7133 | FTMS; HCD | 129.53 | 566.73 |

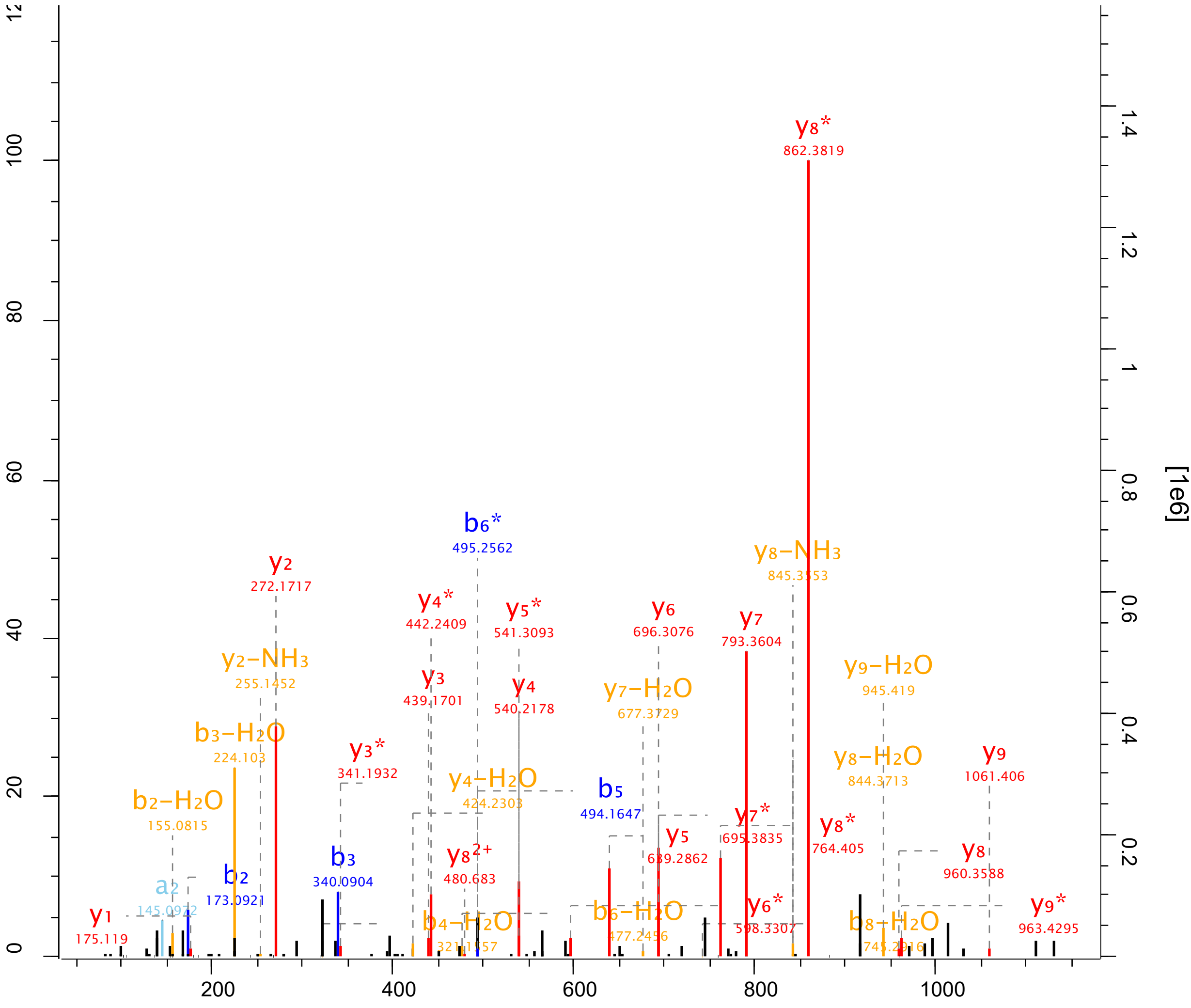


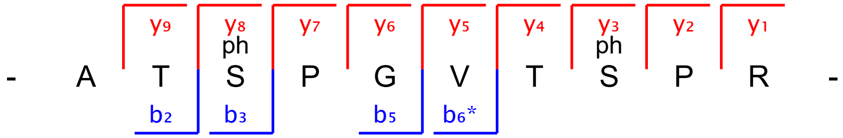

| Raw file | Scan | Method | Score | m/z |
|---|---|---|---|---|
| sys_05_2 | 7161 | FTMS; HCD | 92.01 | 762.65 |

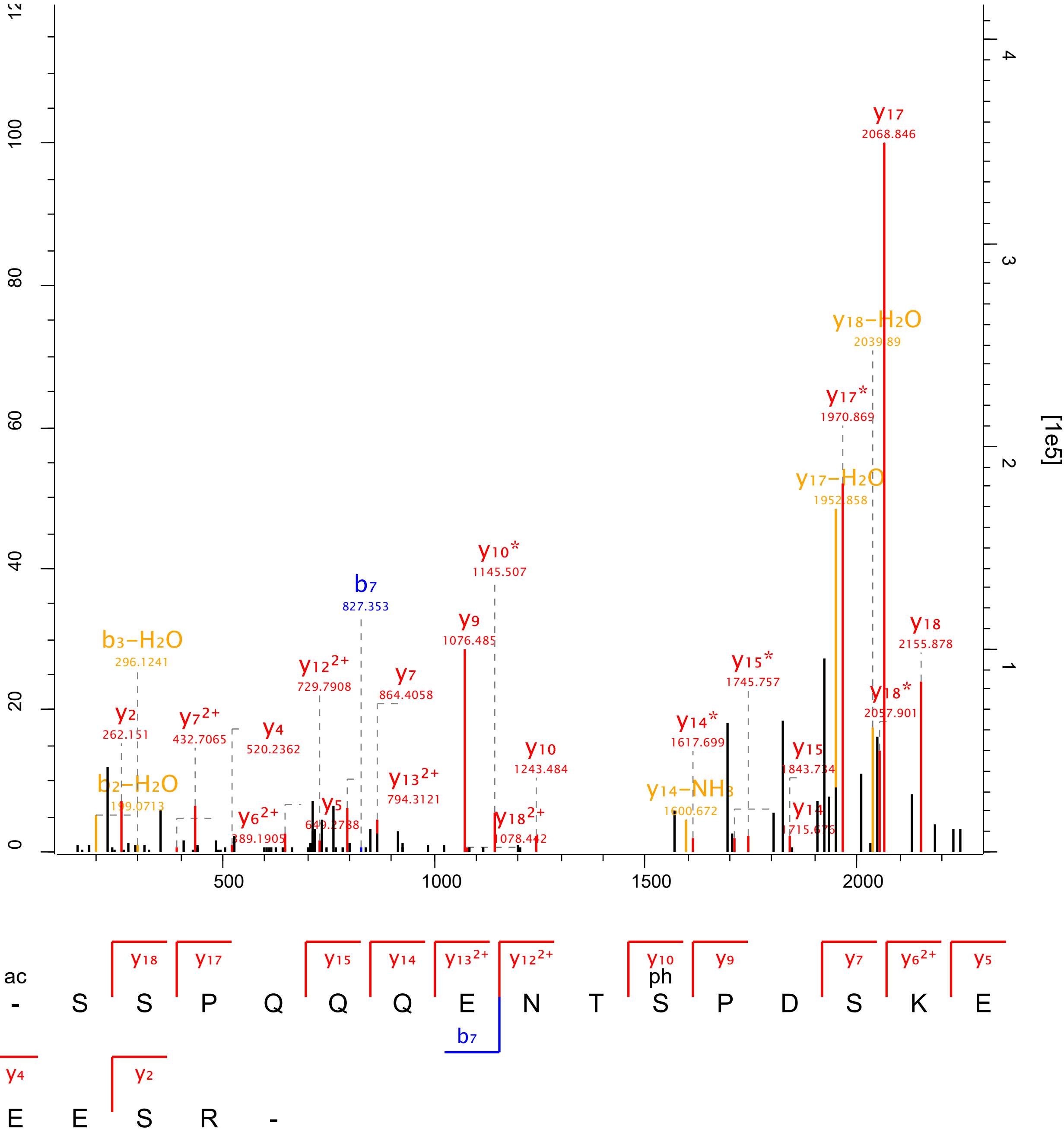

| Raw file | Scan | Method | Score | m/z |
| --- | --- | --- | --- | --- |
| sys_05_2 | 7209 | FTMS; HCD | 56.57 | 646.8 |

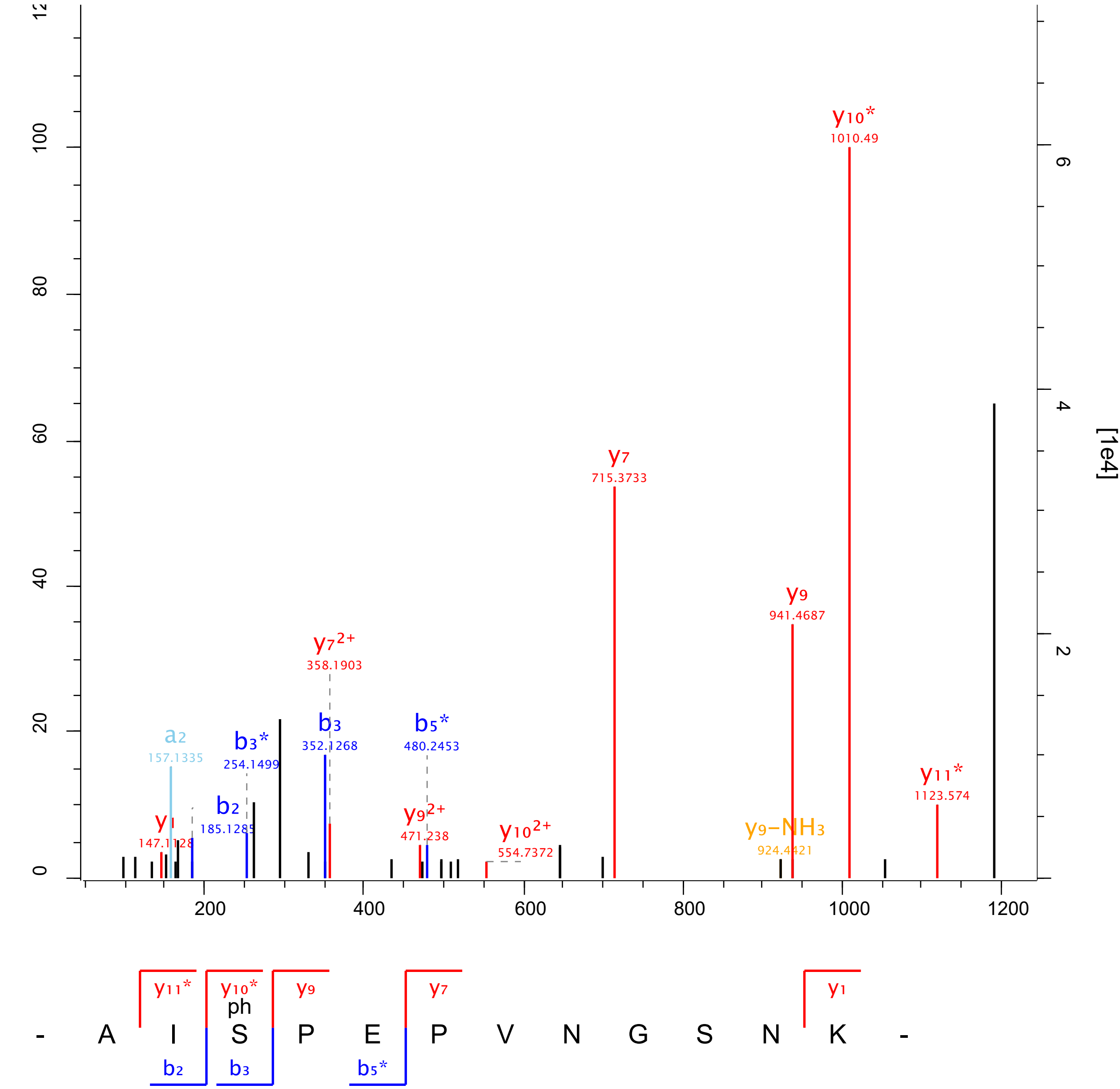

| Raw file | Scan | Method | Score | m/z |
|---|---|---|---|---|
| sys_05_2 | 7267 | FTMS; HCD | 90.83 | 576.79 |

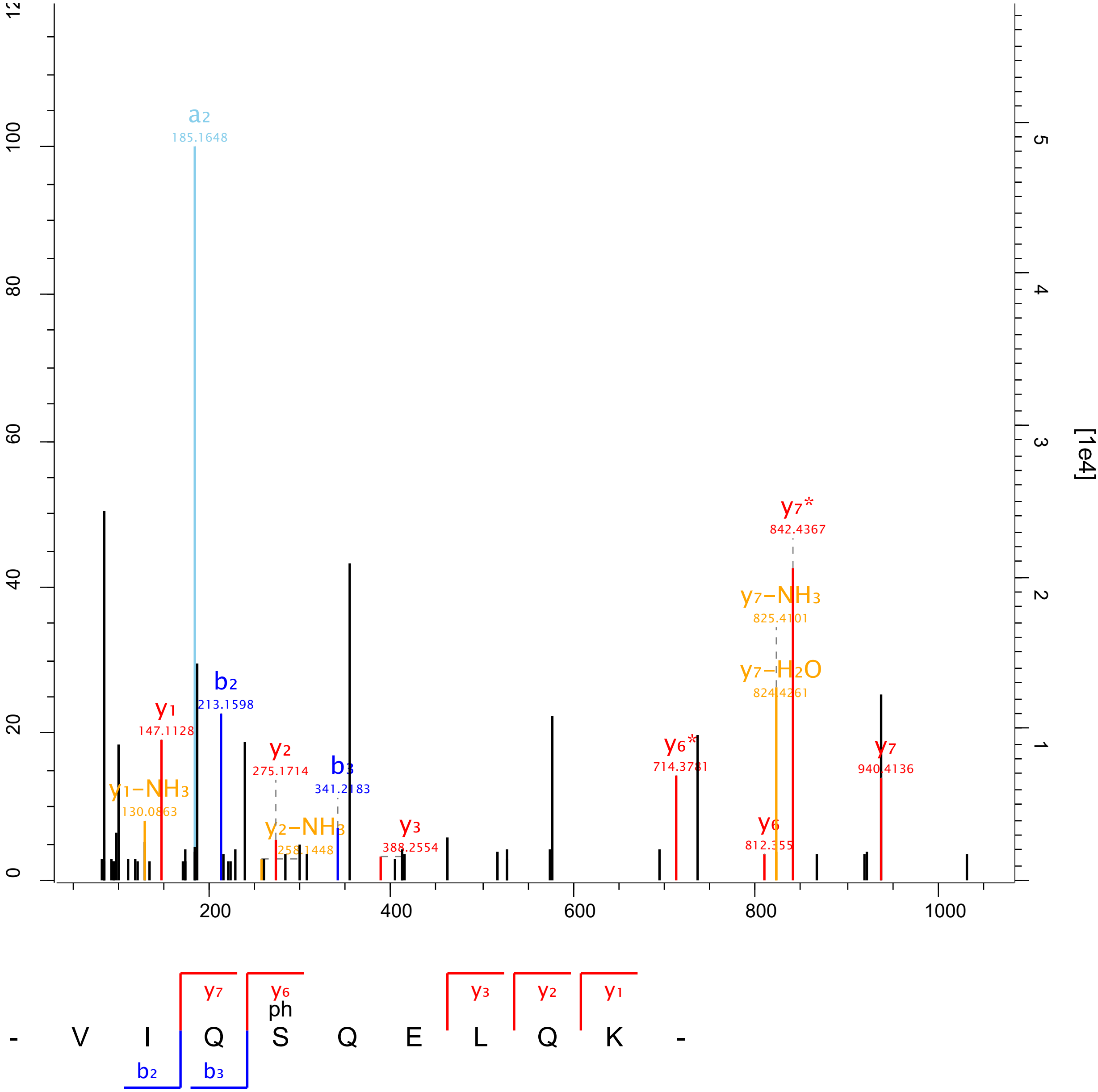

| Raw file | Scan | Method | Score | m/z |
|---|---|---|---|---|
| sys_05_2 | 7308 | FTMS; HCD | 93.37 | 839.83 |

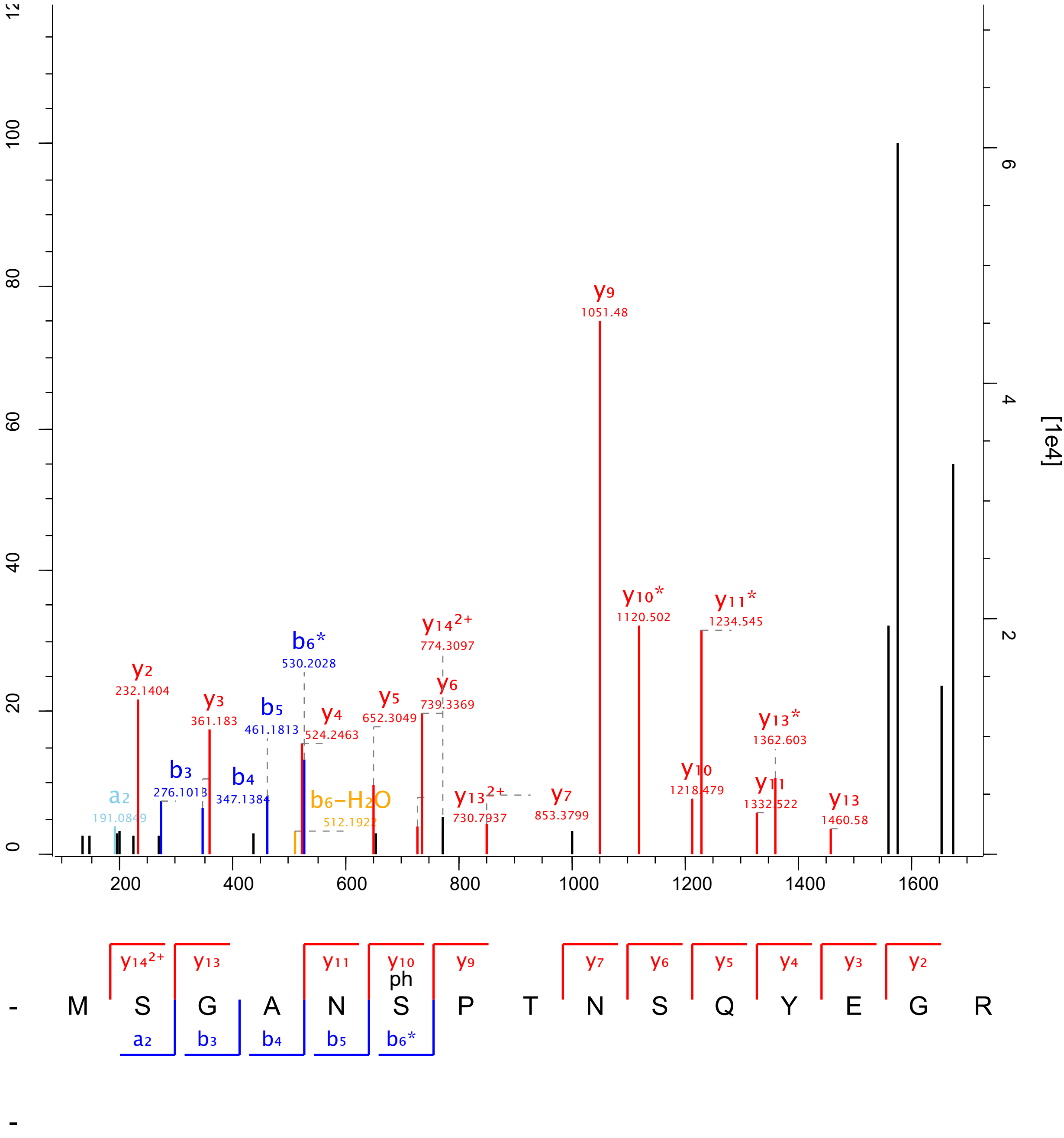

| Raw file | Scan | Method | Score | m/z |
| --- | --- | --- | --- | --- |
| sys_05_2 | 7373 | FTMS; HCD | 116.37 | 412.22 |

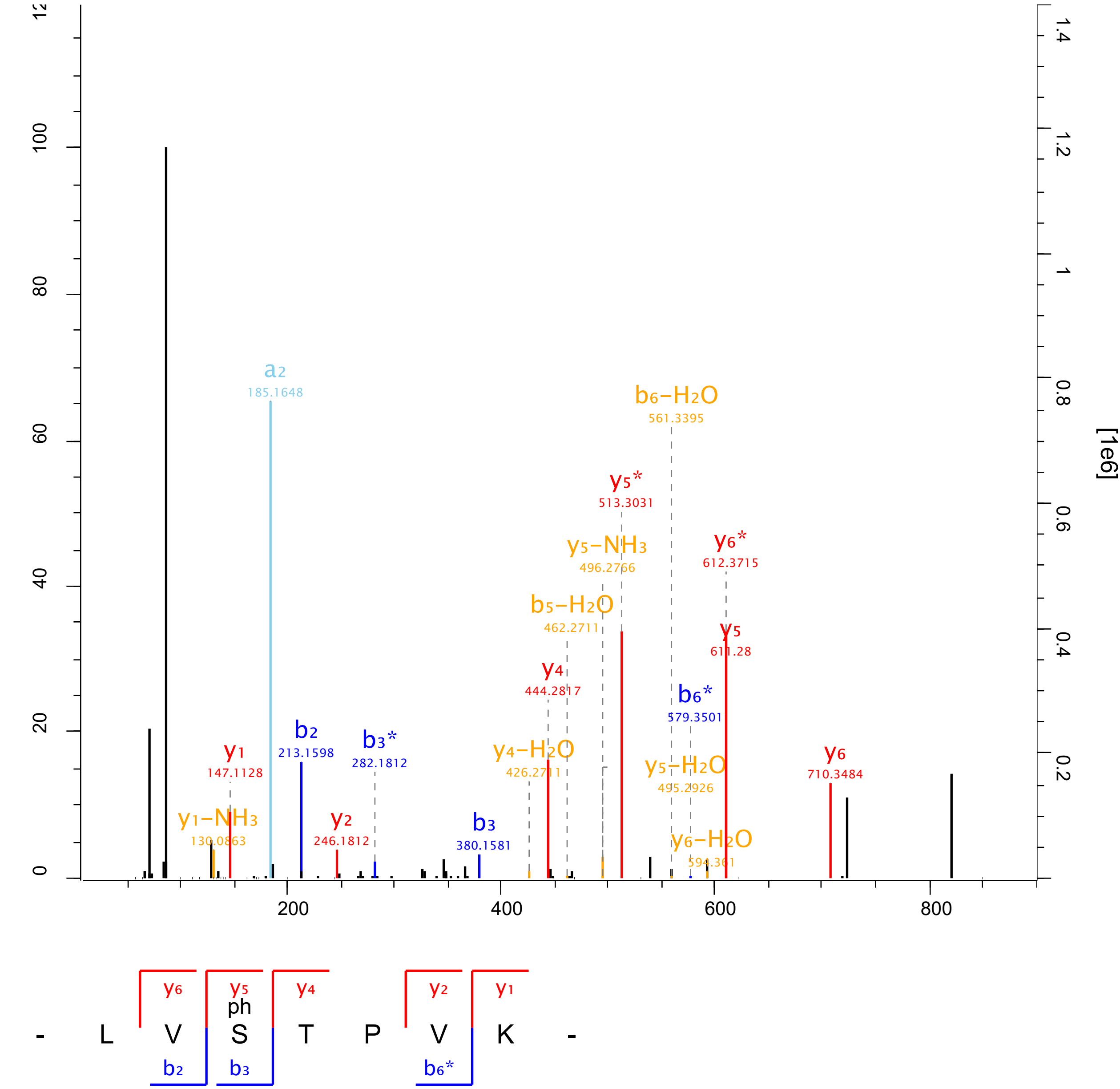

| Raw file | Scan | Method | Score | m/z |
| --- | --- | --- | --- | --- |
| sys_05_2 | 7474 | FTMS; HCD | 280.5 | 663.27 |

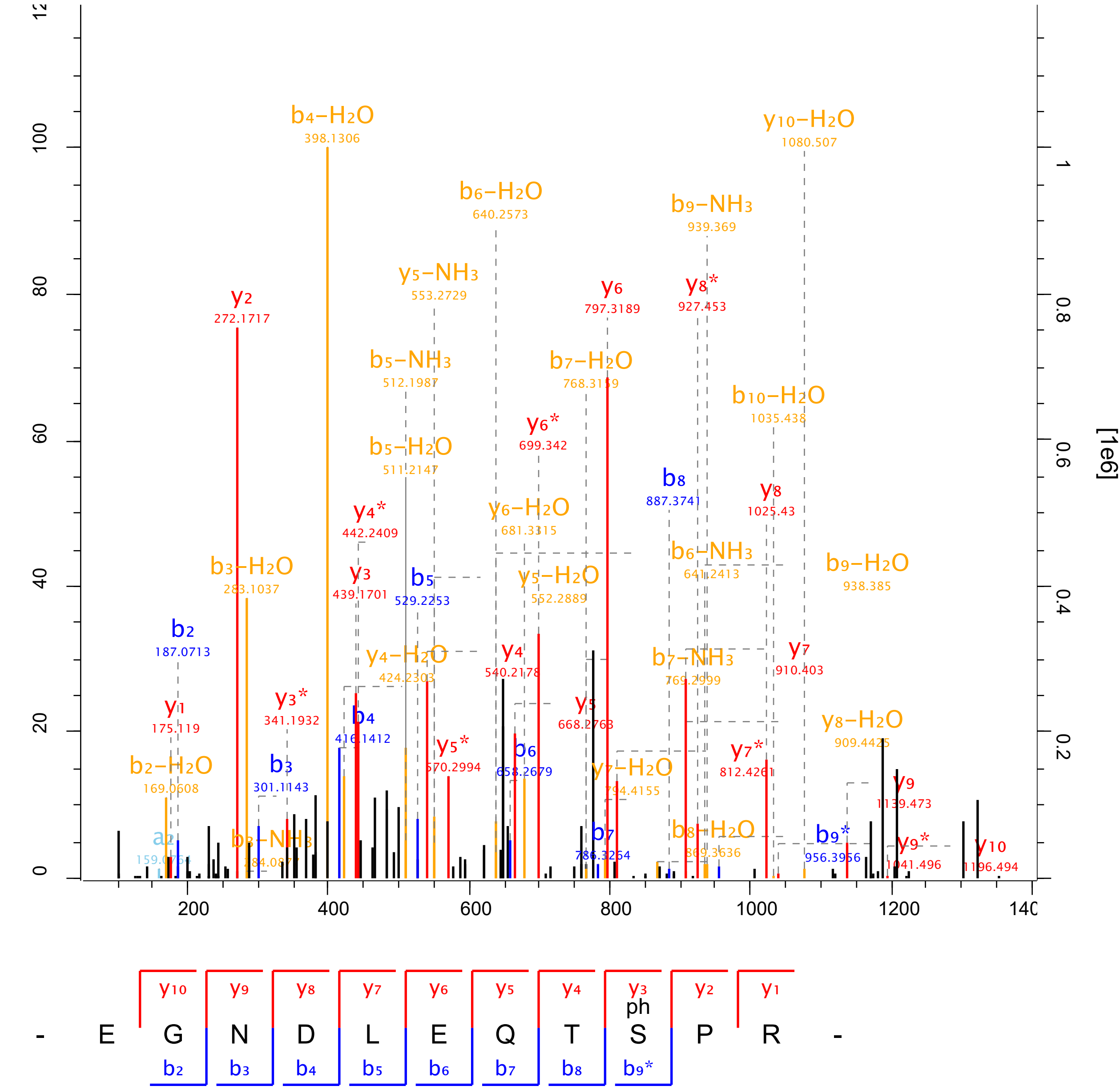

| Raw file | Scan | Method | Score | m/z |
|---|---|---|---|---|
| sys_05_2 | 7489 | FTMS; HCD | 69.23 | 591.26 |

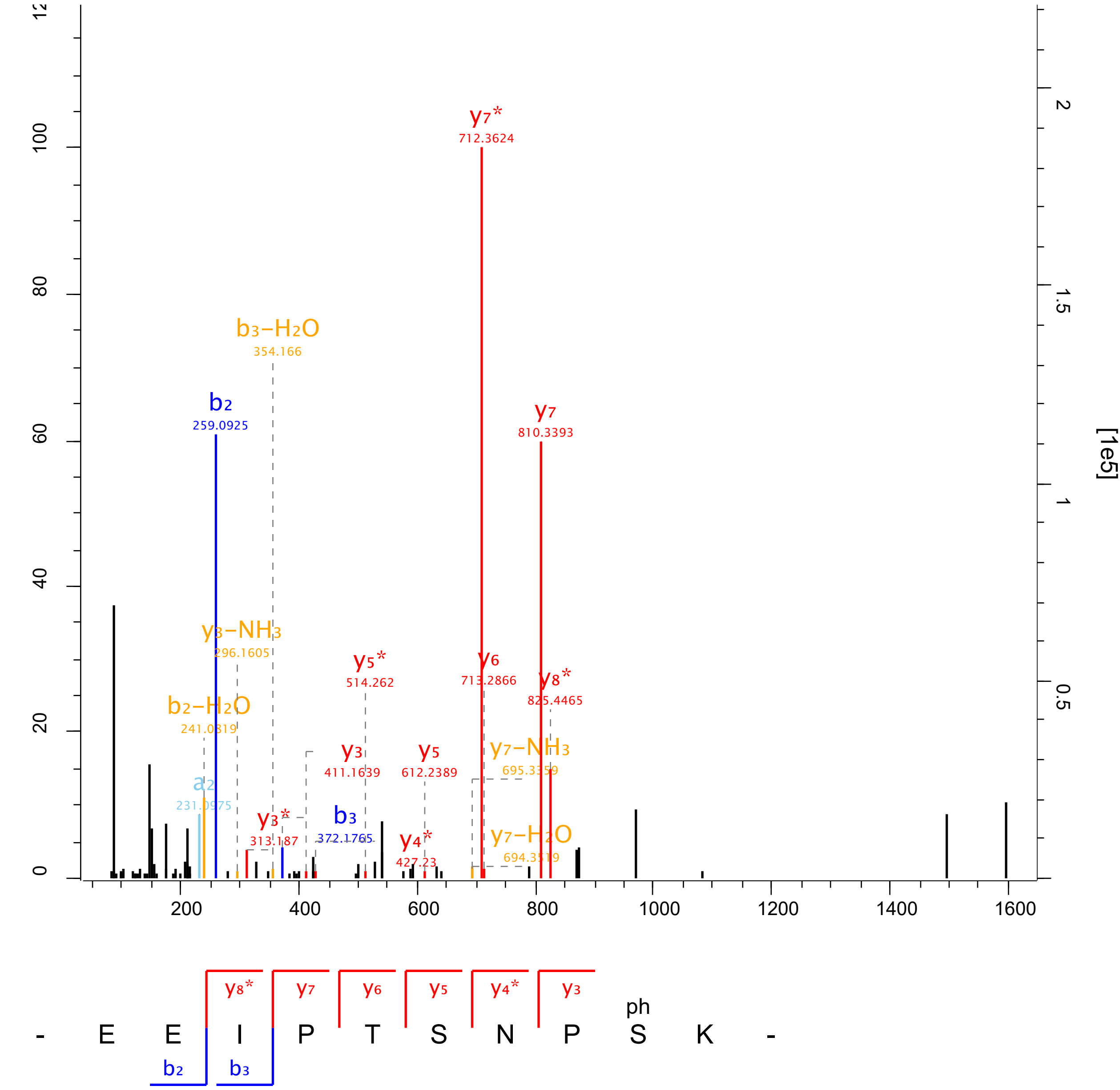

| Raw file | Scan | Method | Score | m/z |
|---|---|---|---|---|
| sys_05_2 | 7502 | FTMS; HCD | 82.07 | 558.74 |

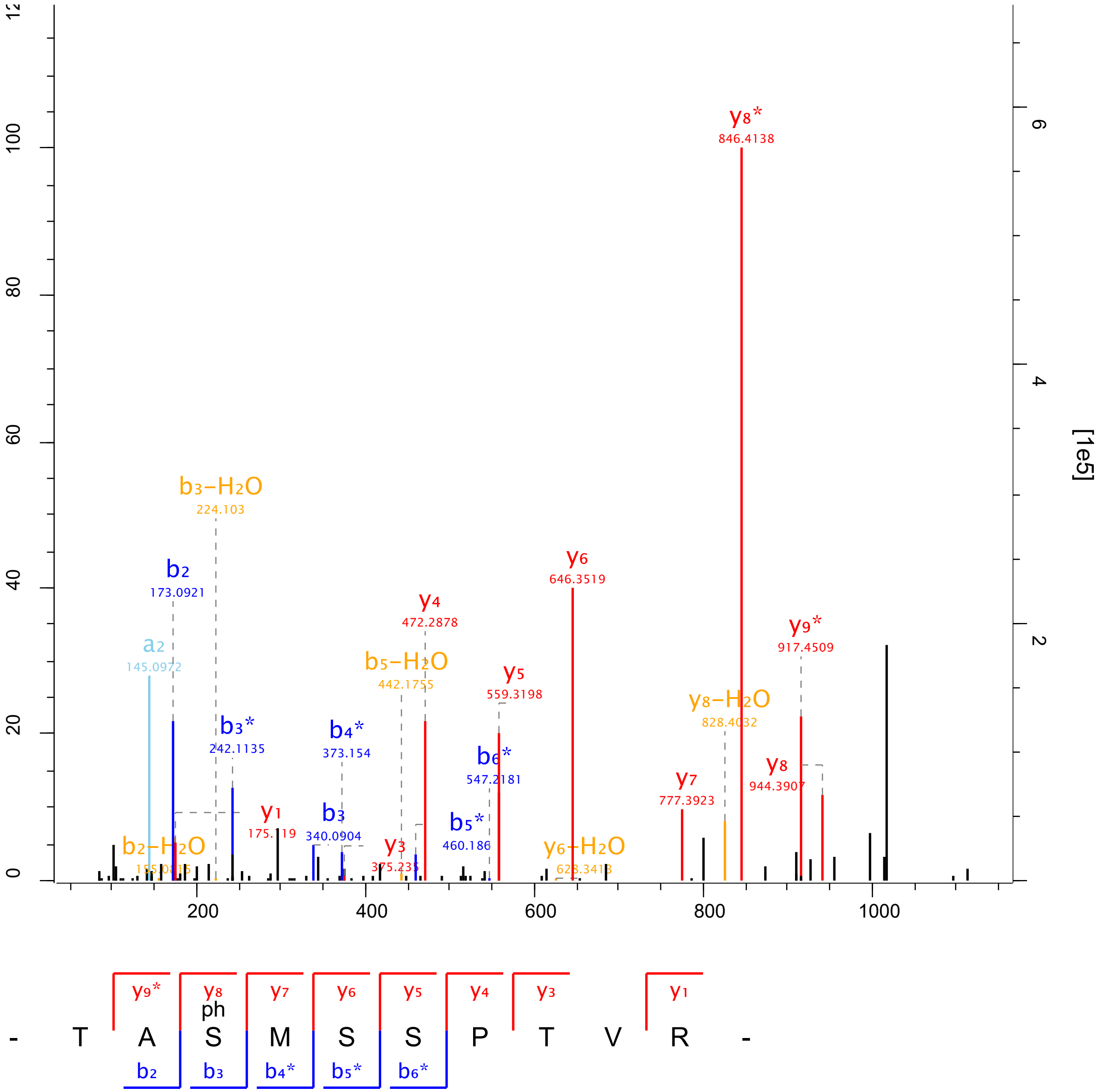

| Raw file | Scan | Method | Score | m/z |
| --- | --- | --- | --- | --- |
| sys_05_2 | 7587 | FTMS; HCD | 109.1 | 573.75 |

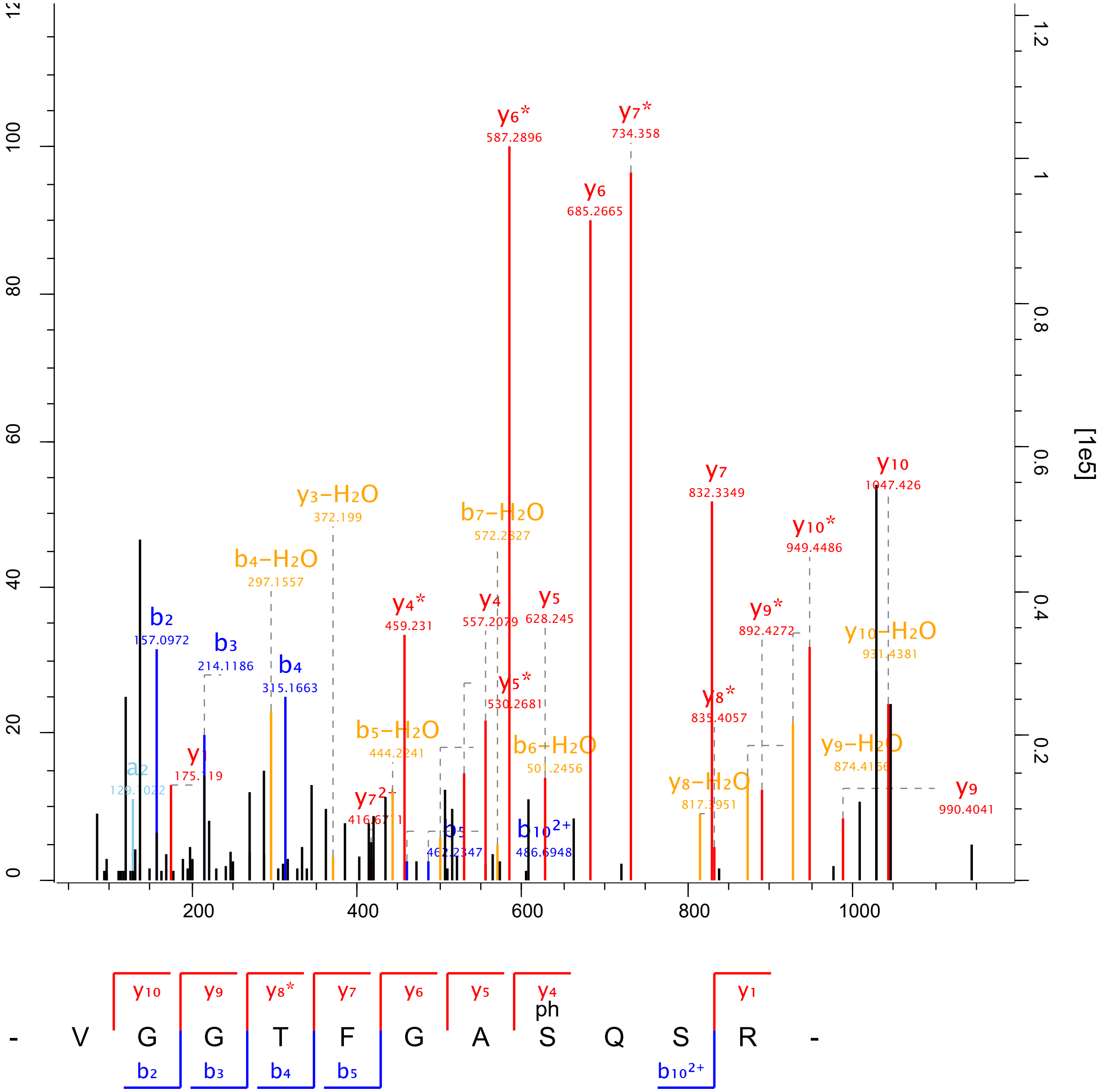

| Raw file | Scan | Method | Score | m/z |
|---|---|---|---|---|
| sys_05_2 | 7605 | FTMS; HCD | 123.19 | 627.24 |

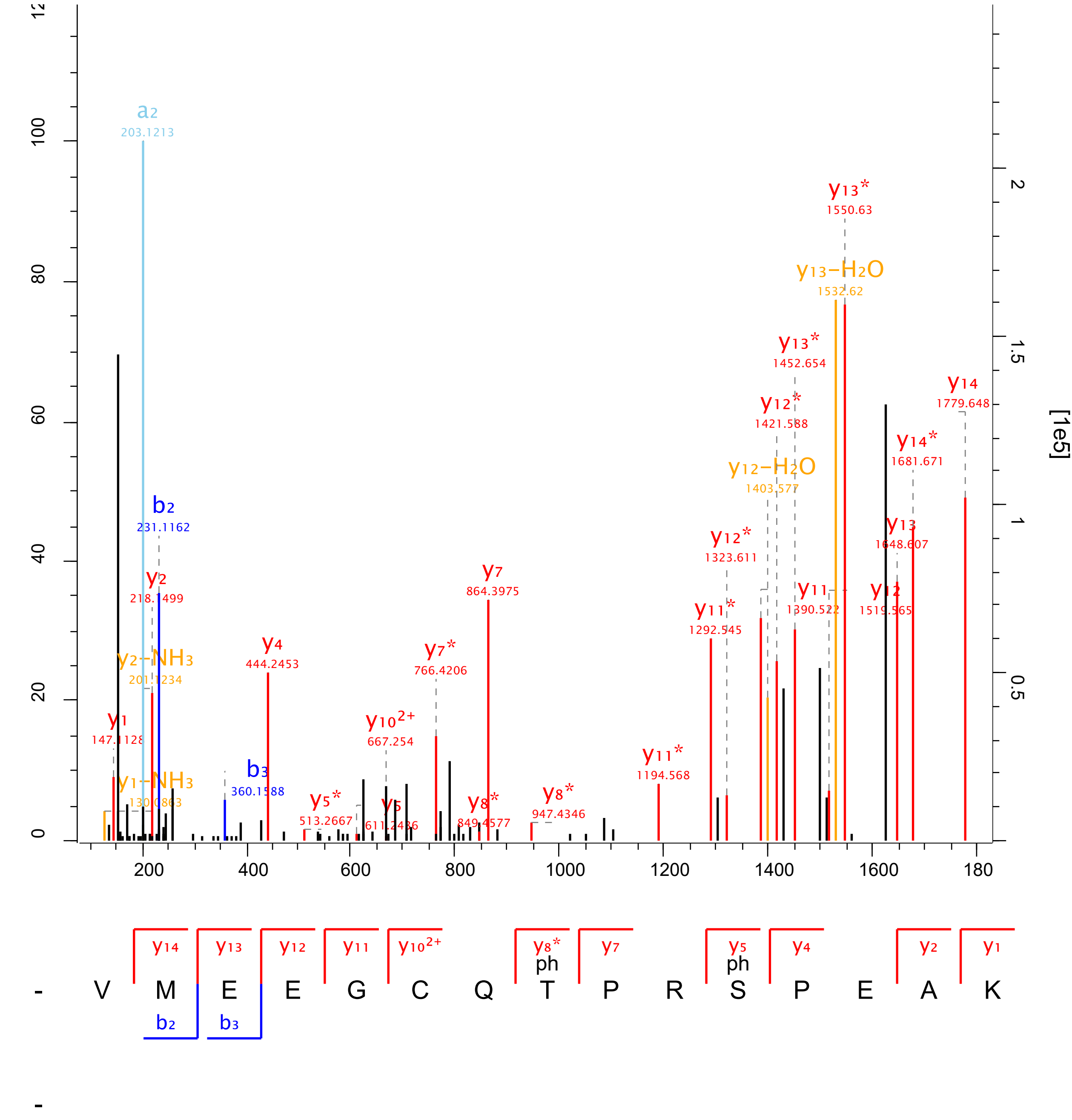

| Raw file | Scan | Method | Score | m/z |
| --- | --- | --- | --- | --- |
| sys_05_2 | 7753 | FTMS; HCD | 228.38 | 557.56 |

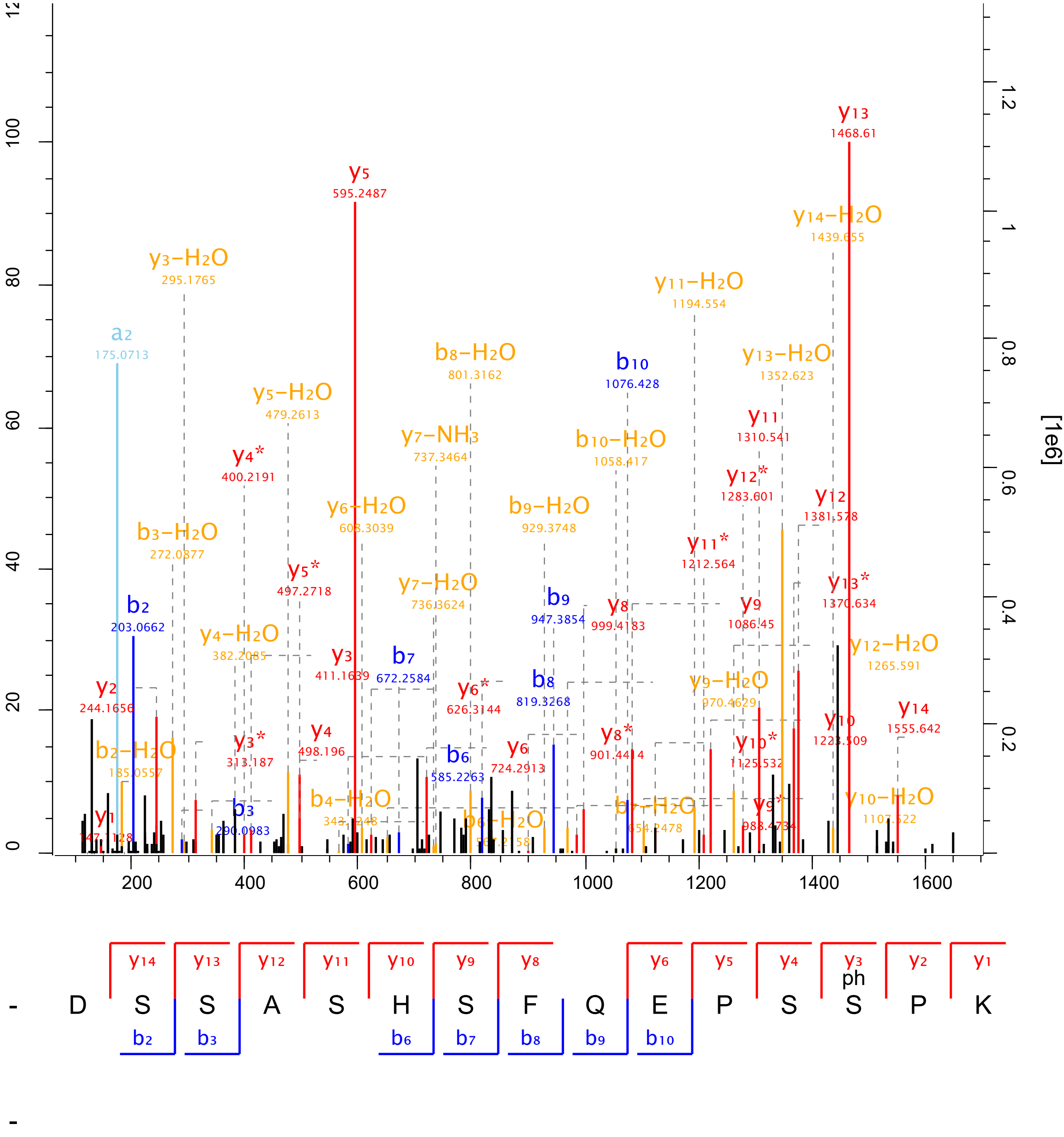

| Raw file | Scan | Method | Score | m/z |
| --- | --- | --- | --- | --- |
| sys_05_2 | 7769 | FTMS; HCD | 66.02 | 526.74 |

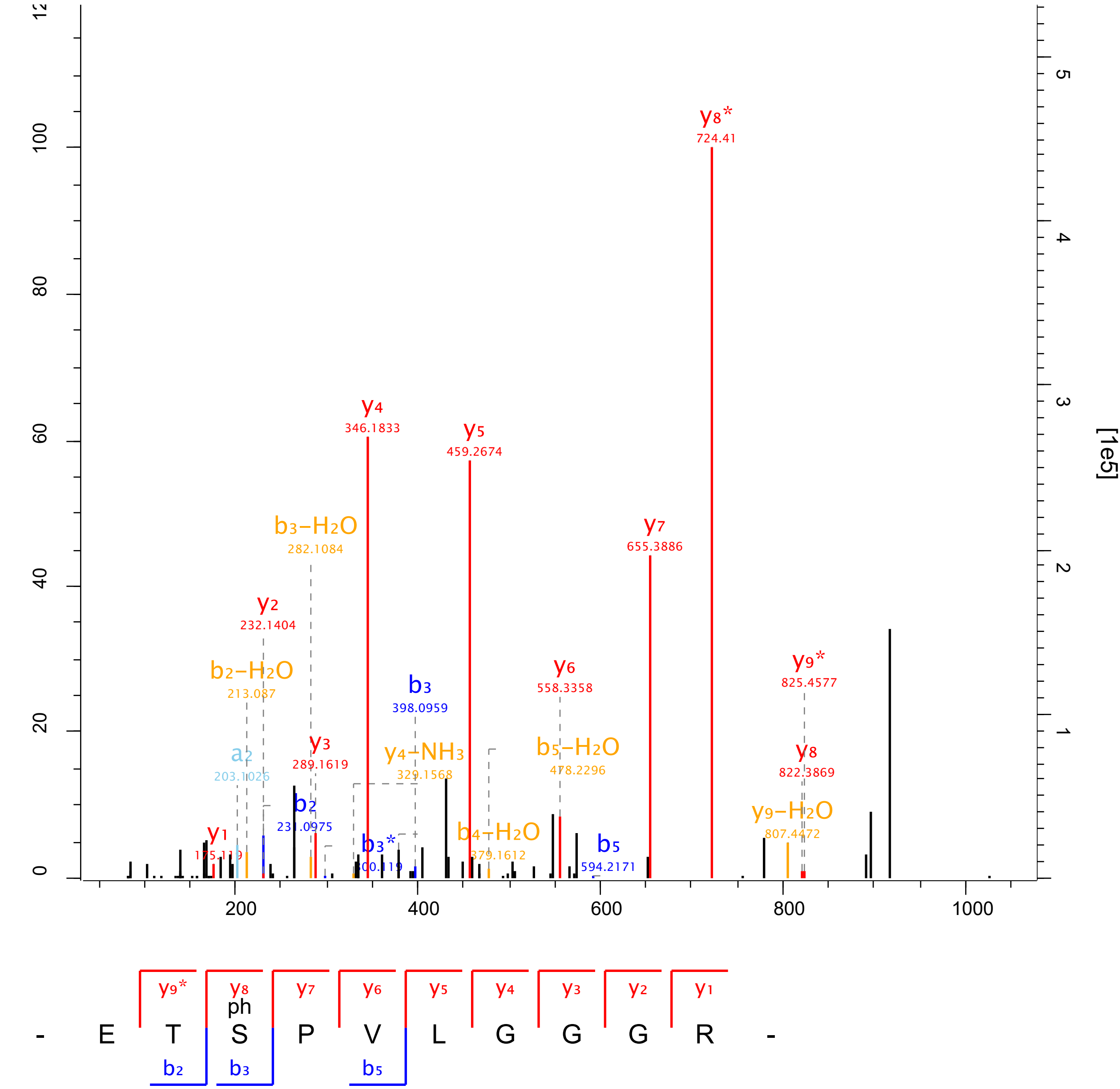

| Raw file | Scan | Method | Score | m/z |
|---|---|---|---|---|
| sys_05_2 | 7897 | FTMS; HCD | 97.16 | 390.52 |

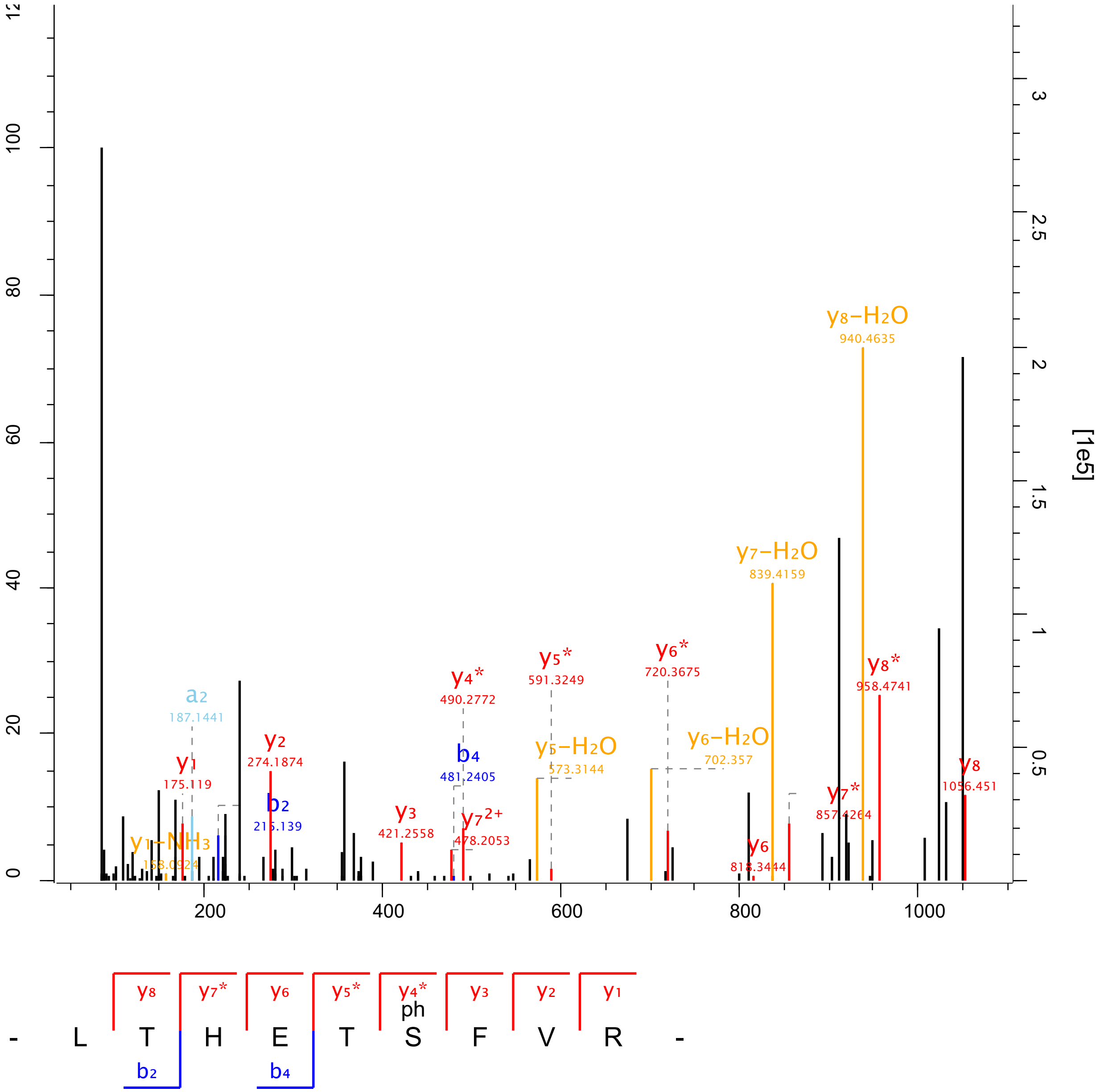

| Raw file | Scan | Method | Score | m/z |
|---|---|---|---|---|
| sys_05_2 | 8021 | FTMS; HCD | 46.32 | 440.55 |

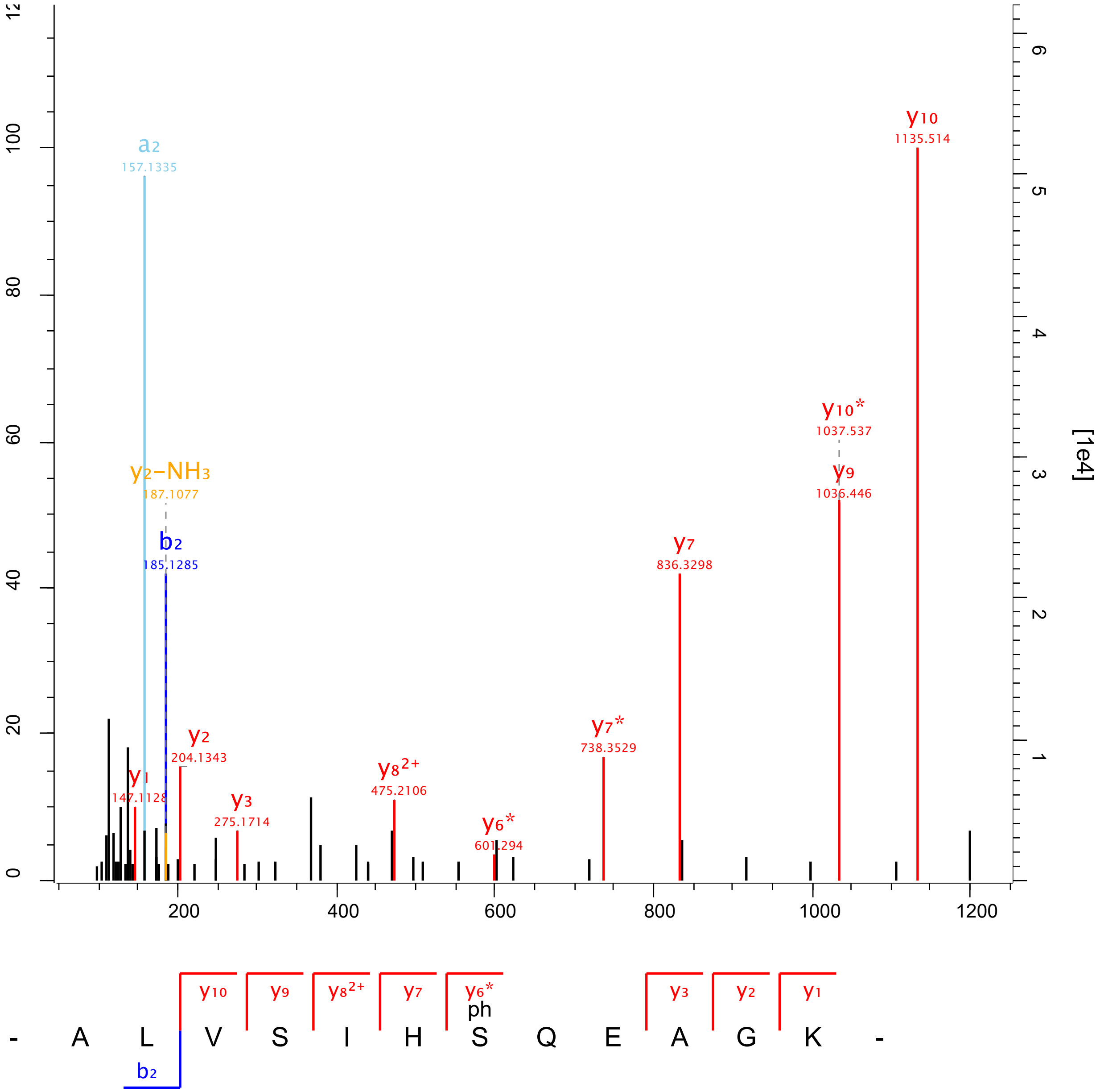

| Raw file | Scan | Method | Score | m/z |
|---|---|---|---|---|
| sys_05_2 | 8192 | FTMS; HCD | 194.19 | 726.79 |

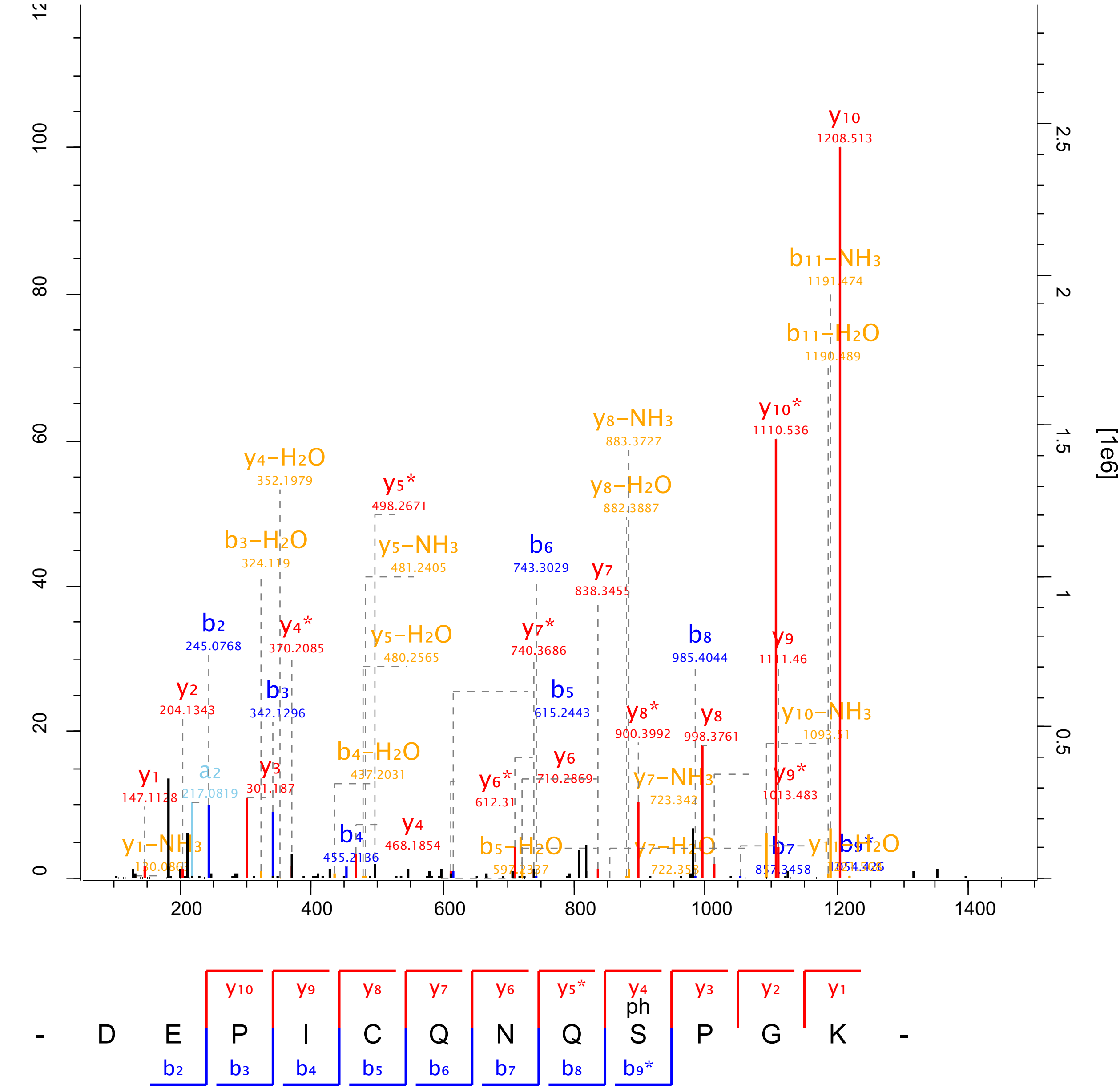

| Raw file | Scan | Method | Score | m/z |
| --- | --- | --- | --- | --- |
| sys_05_2 | 8483 | FTMS; HCD | 90.56 | 678.77 |

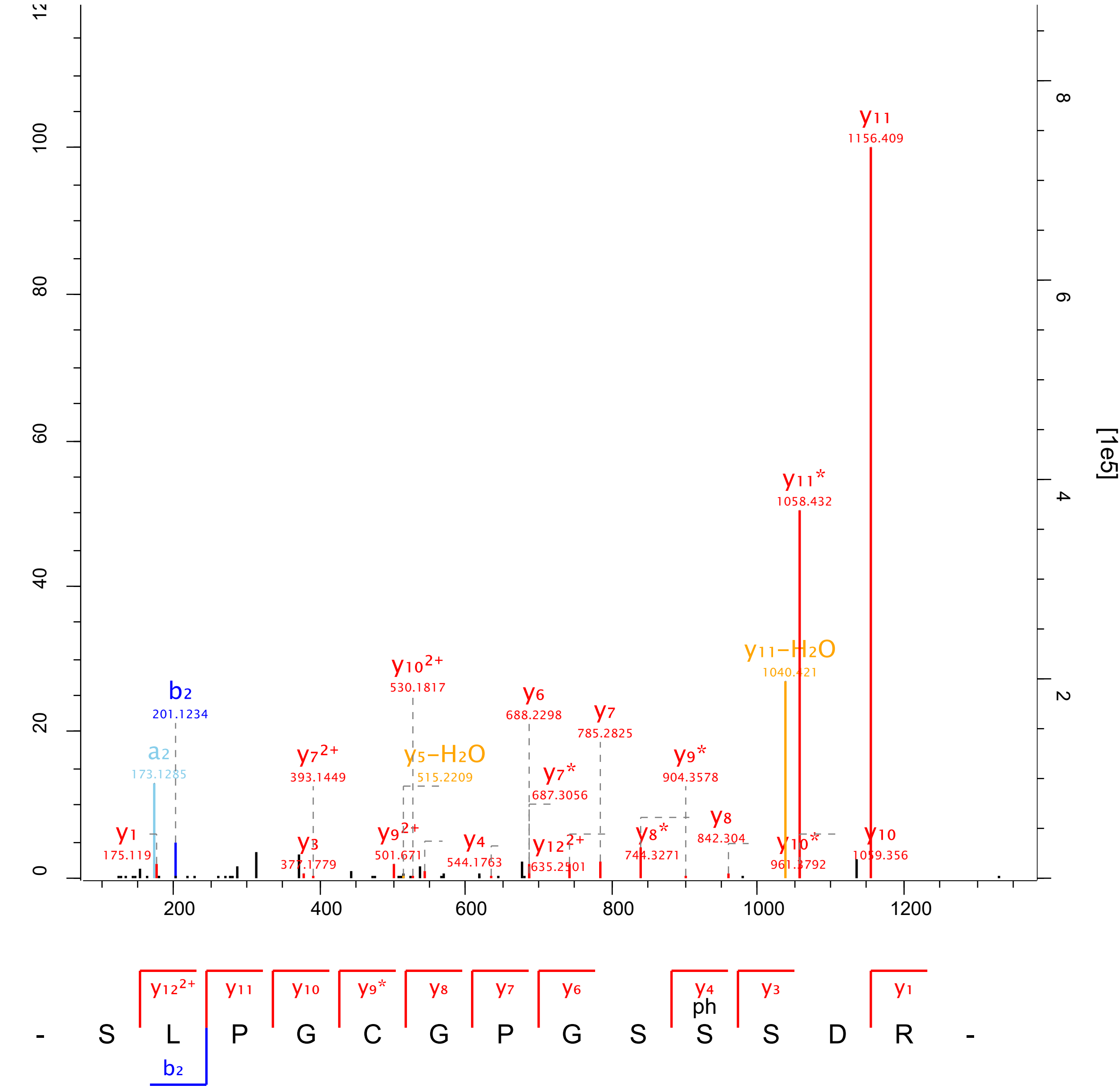

| Raw file | Scan | Method | Score | m/z |
|---|---|---|---|---|
| sys_05_2 | 8666 | FTMS; HCD | 103.88 | 640.29 |

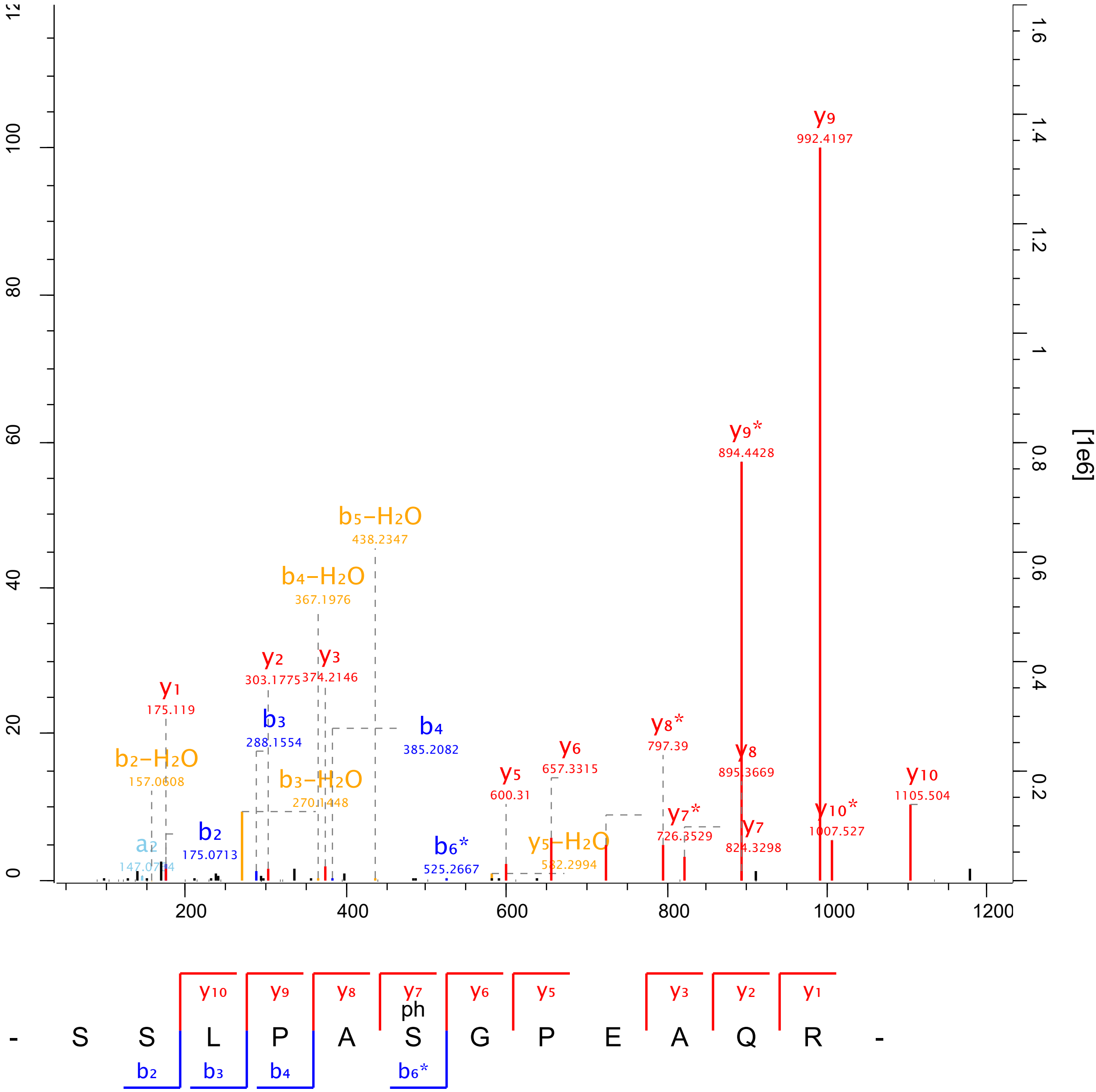

| Raw file | Scan | Method | Score | m/z |
|---|---|---|---|---|
| sys_05_2 | 8807 | FTMS; HCD | 68.16 | 541.57 |

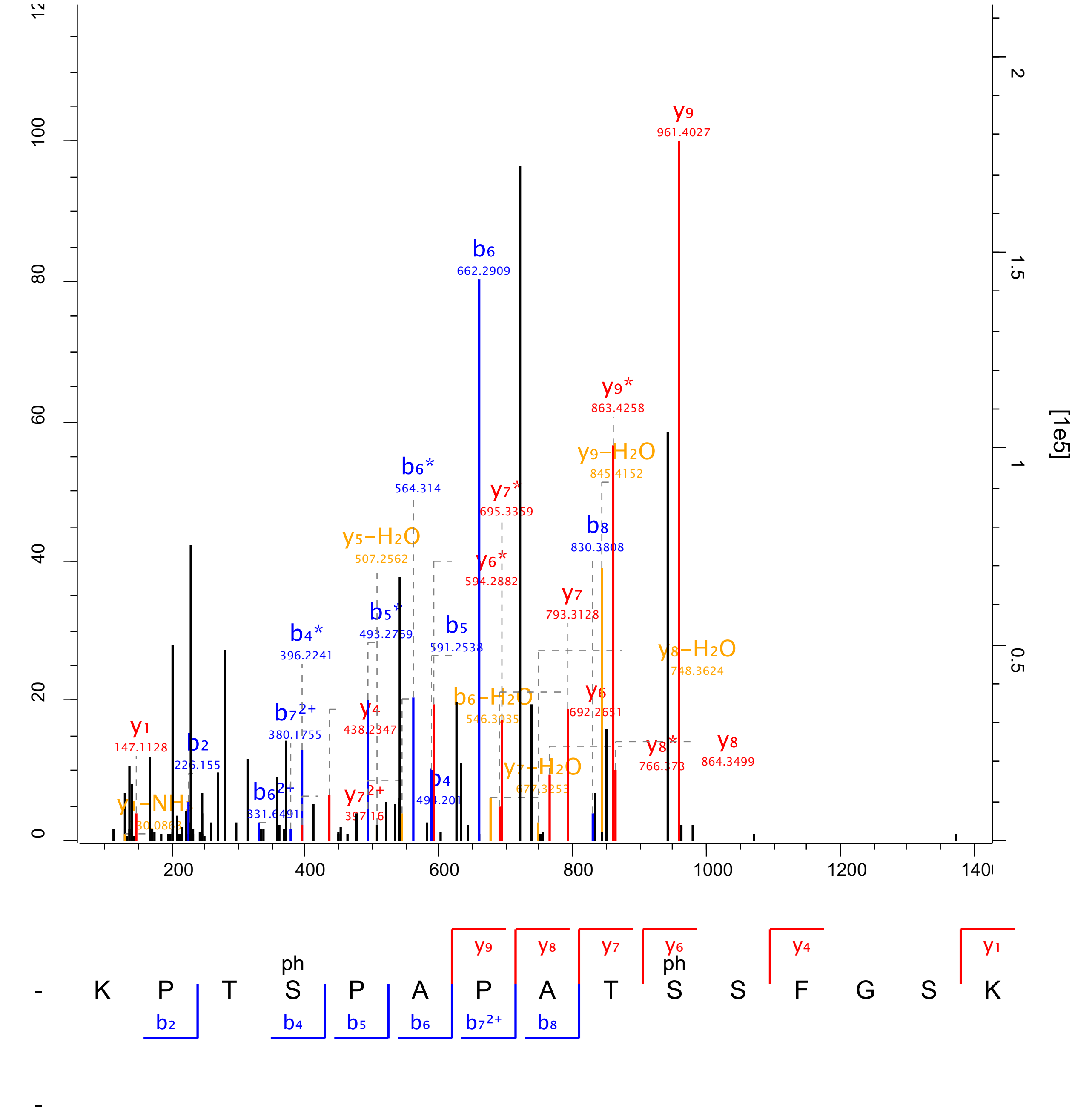

| Raw file | Scan | Method | Score | m/z |
|---|---|---|---|---|
| sys_05_2 | 8887 | FTMS; HCD | 91.62 | 580.74 |

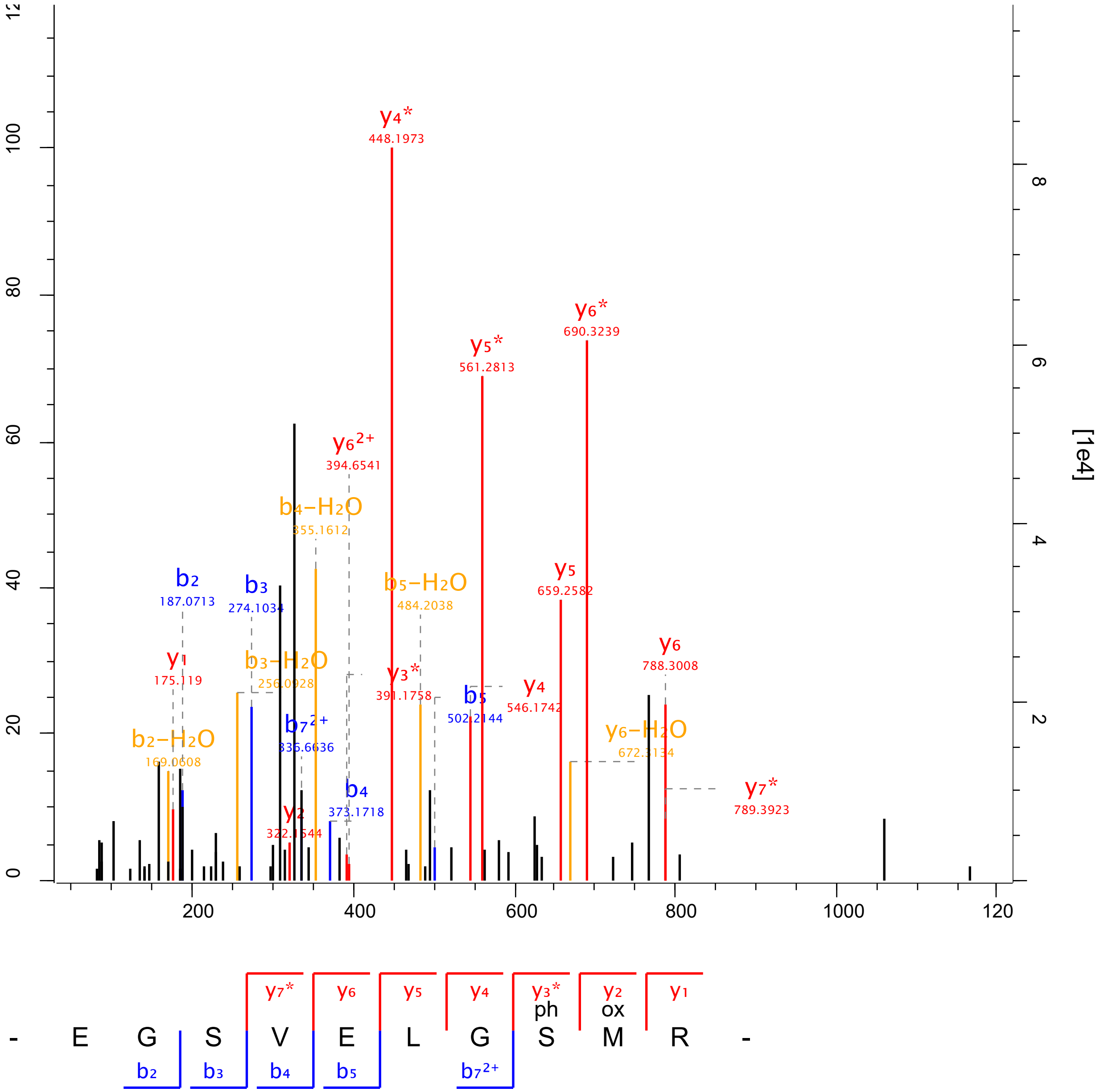

| Raw file | Scan | Method | Score | m/z |
| --- | --- | --- | --- | --- |
| sys_05_2 | 8974 | FTMS; HCD | 287.16 | 826.89 |

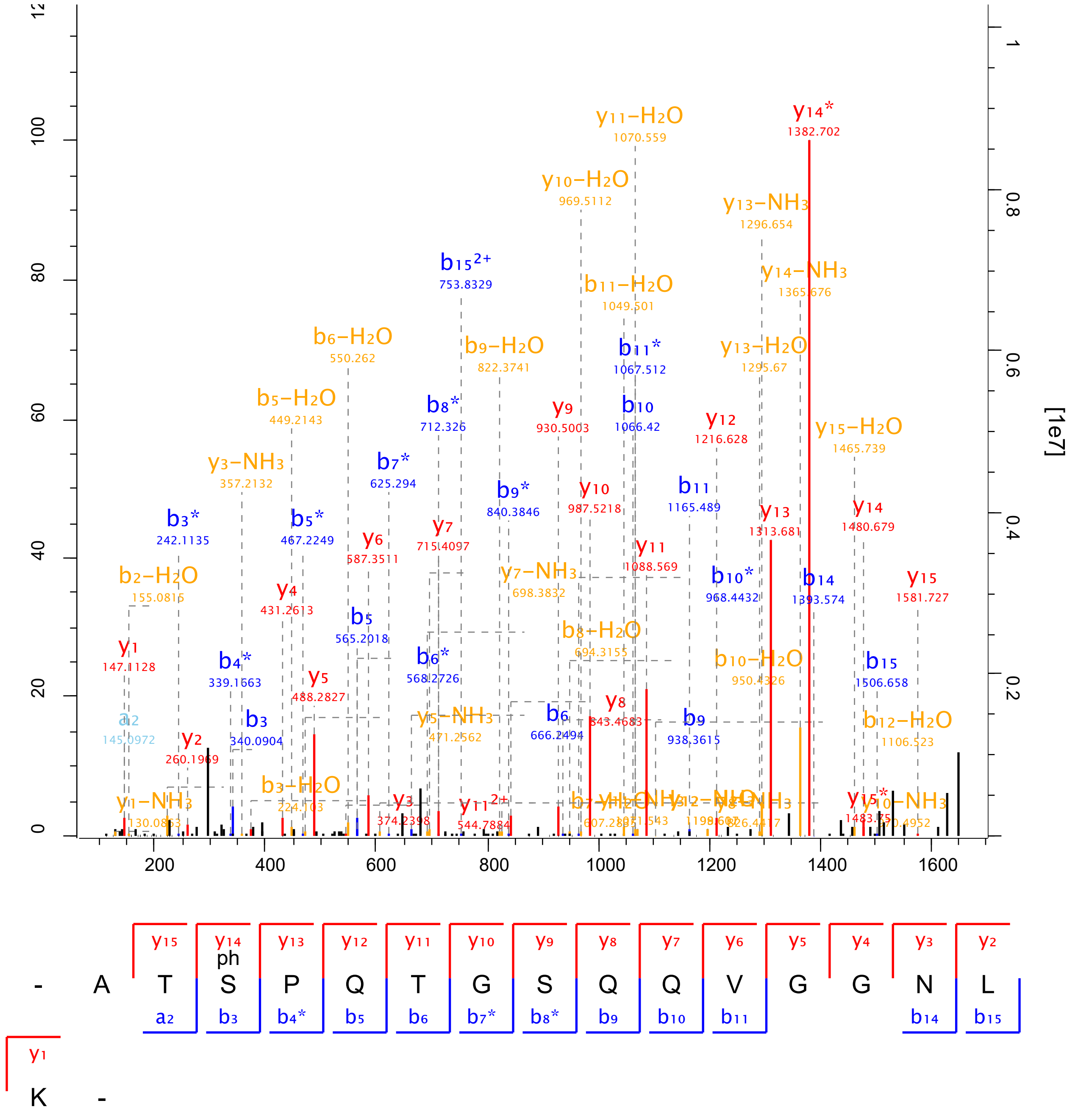

| Raw file | Scan | Method | Score | m/z |
|---|---|---|---|---|
| sys_05_2 | 9095 | FTMS; HCD | 74.78 | 572.6 |

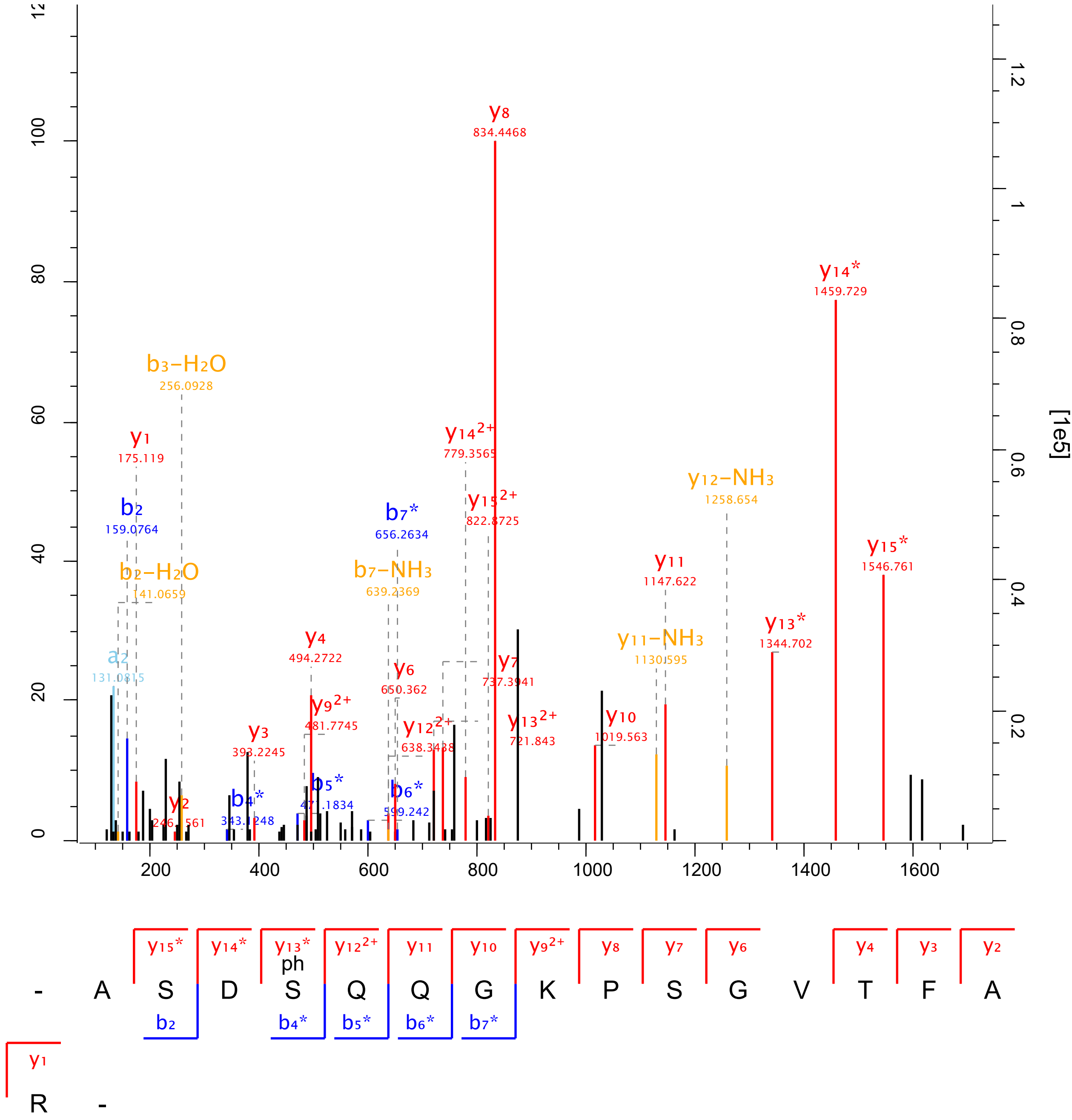

| Raw file | Scan | Method | Score | m/z |
|---|---|---|---|---|
| sys_05_2 | 9241 | FTMS; HCD | 181.35 | 701.79 |

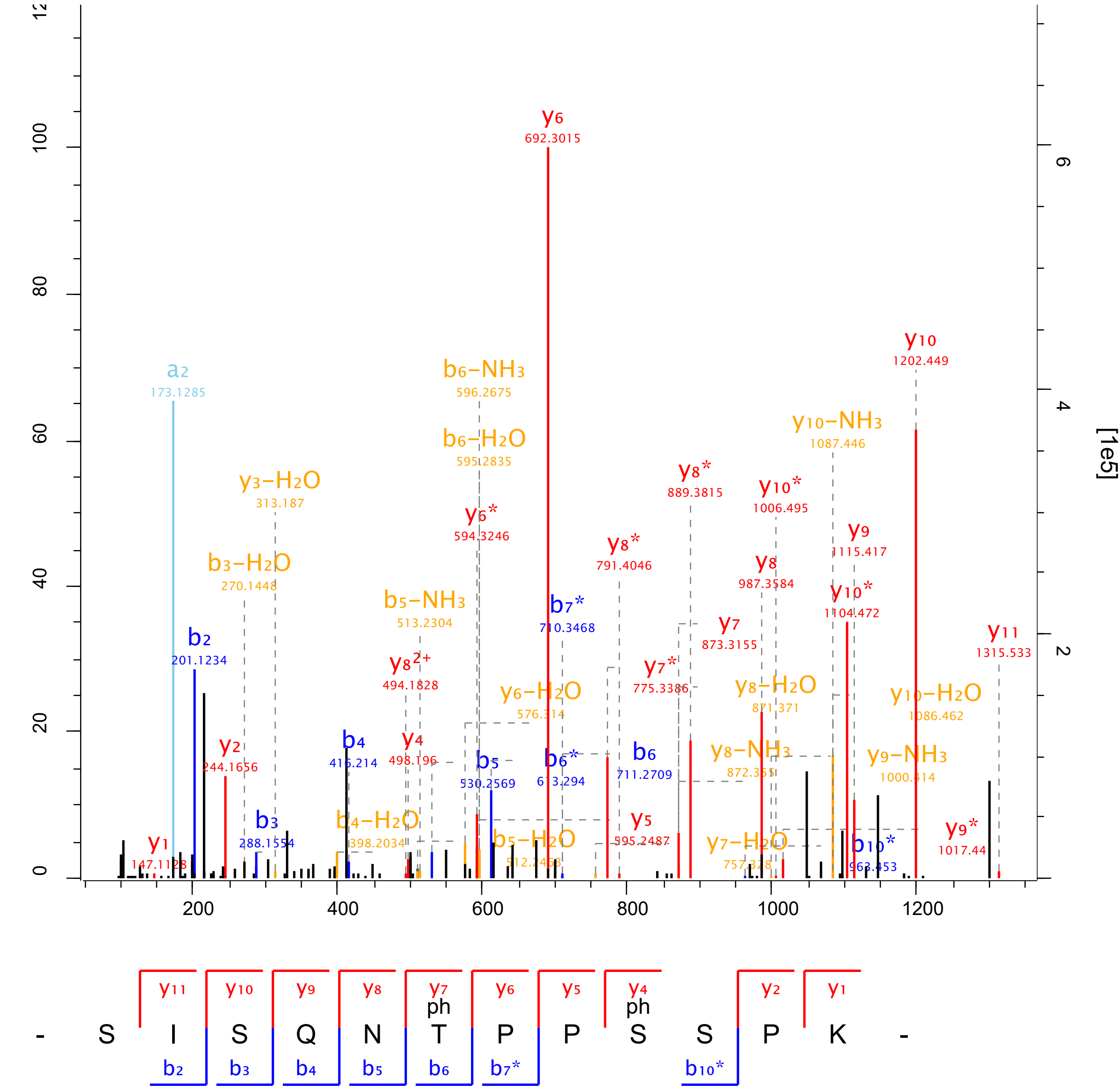

| Raw file | Scan | Method | Score | m/z |
| --- | --- | --- | --- | --- |
| sys_05_2 | 9347 | FTMS; HCD | 79.49 | 676.74 |

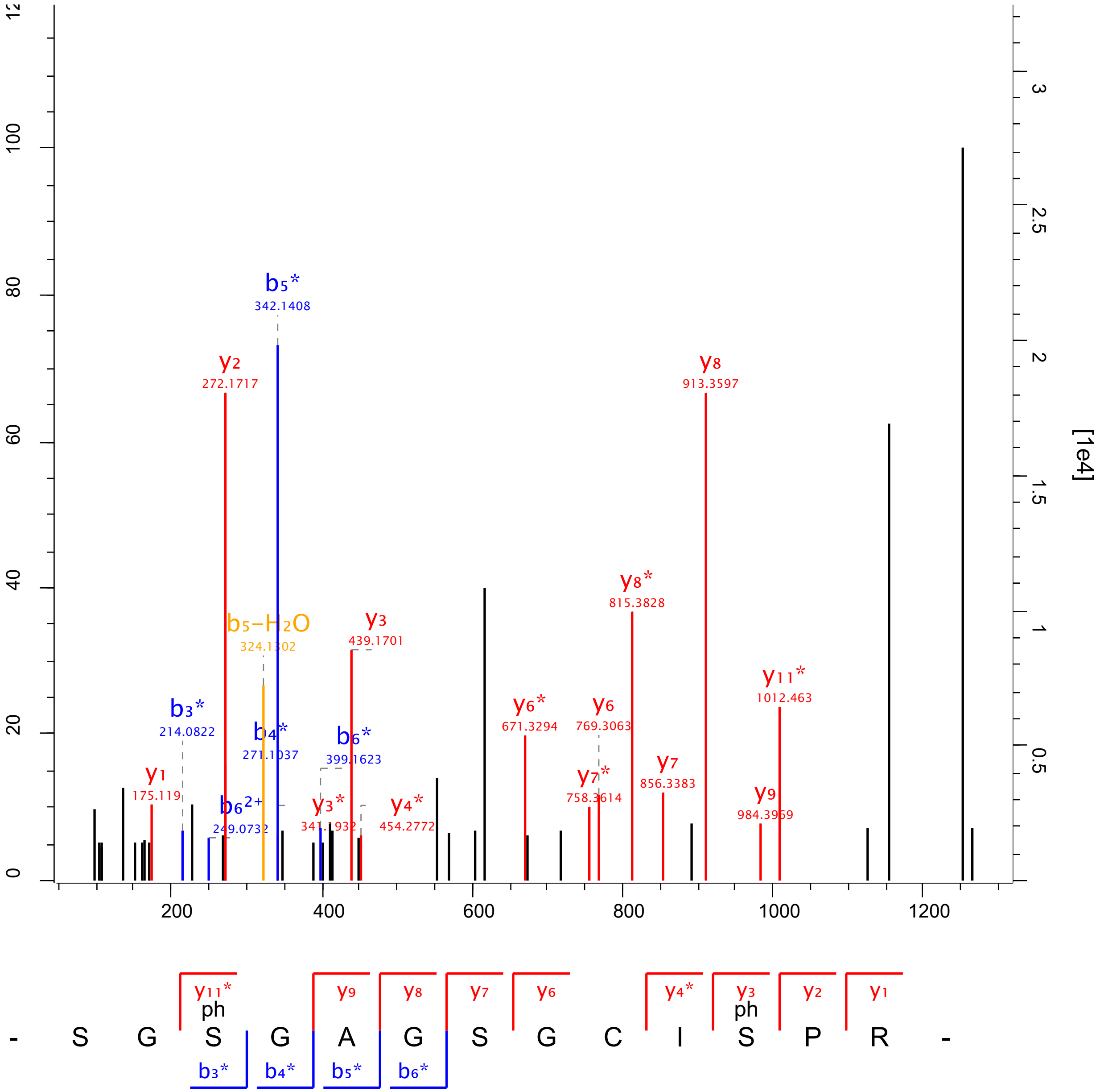

| Raw file | Scan | Method | Score | m/z |
| --- | --- | --- | --- | --- |
| sys_05_2 | 9355 | FTMS; HCD | 63.09 | 511.69 |

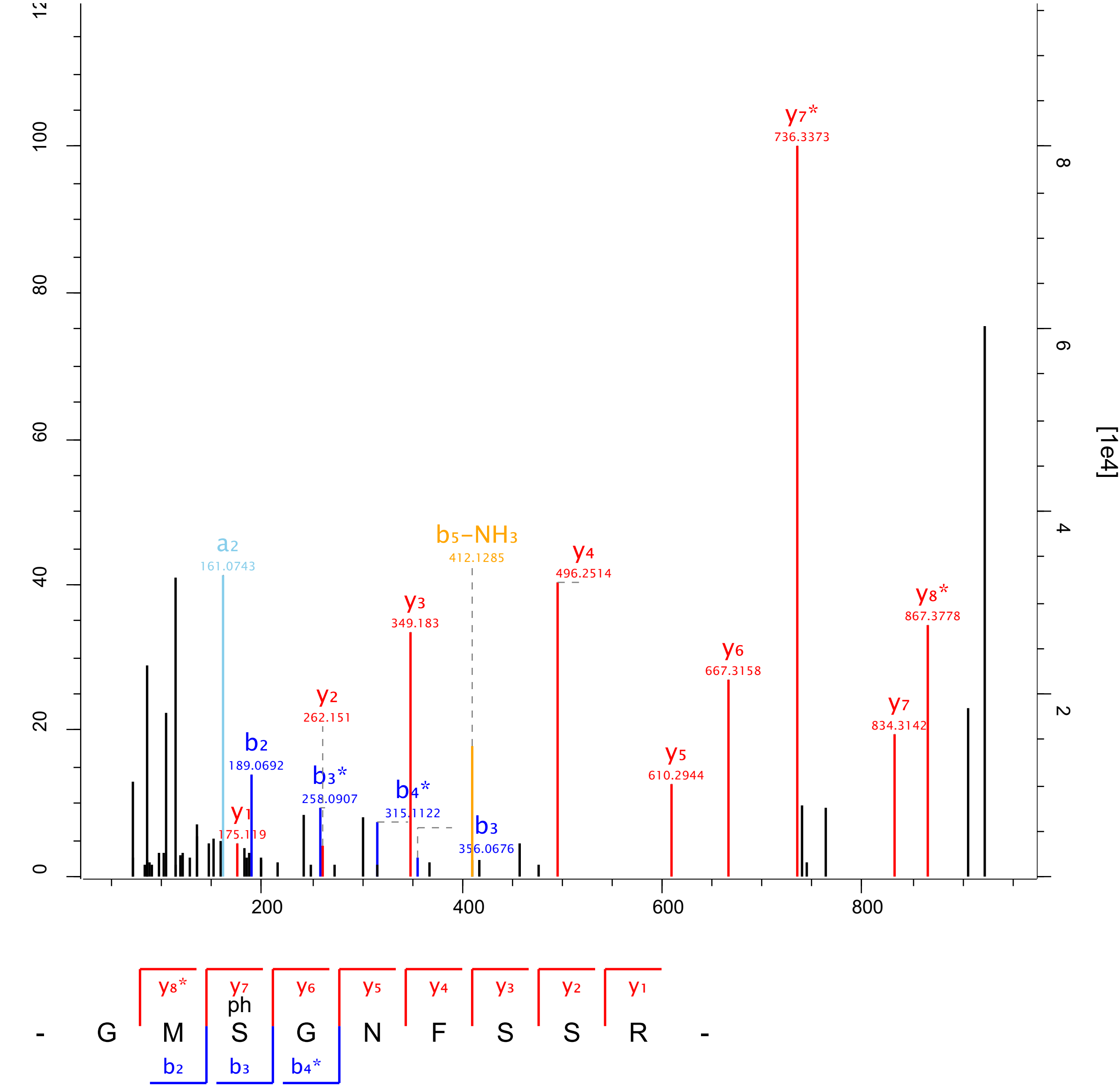

| Raw file | Scan | Method | Score | m/z |
| --- | --- | --- | --- | --- |
| sys_05_2 | 9458 | FTMS; HCD | 40.97 | 561.22 |

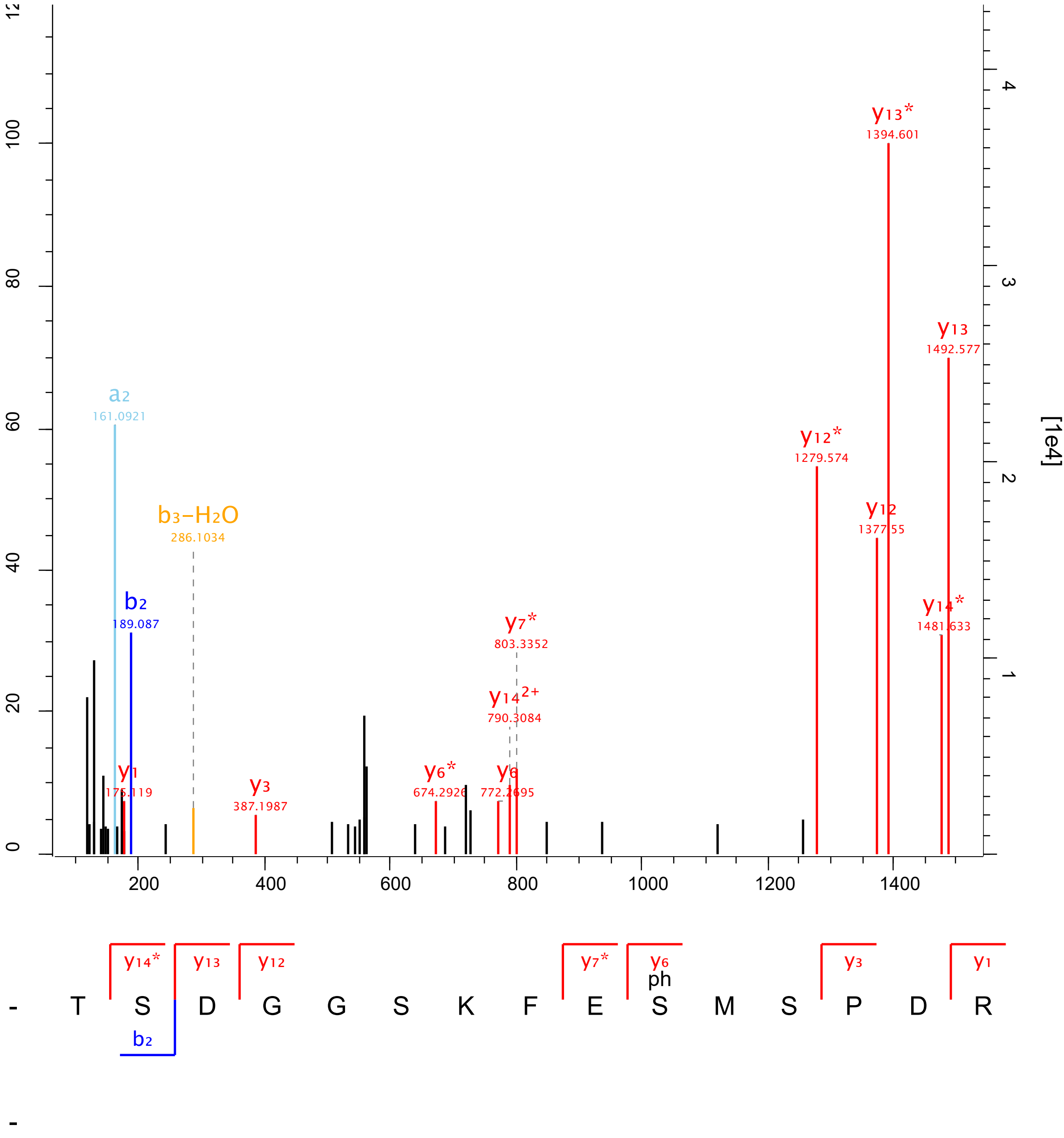

| Raw file | Scan | Method | Score | m/z |
|---|---|---|---|---|
| sys_05_2 | 9492 | FTMS; HCD | 102.73 | 547.7 |

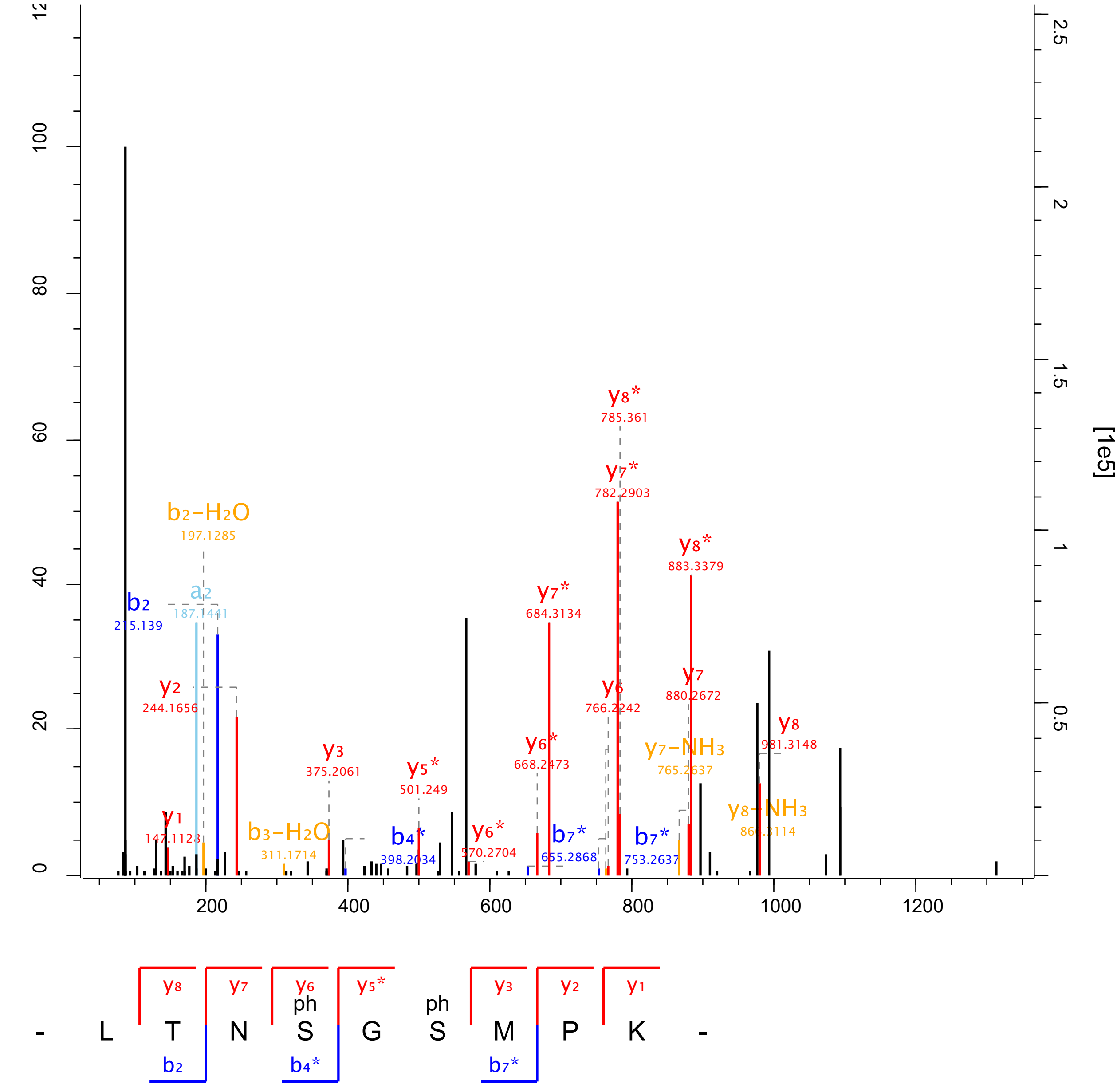

| Raw file | Scan | Method | Score | m/z |
|---|---|---|---|---|
| sys_05_2 | 9617 | FTMS; HCD | 127.83 | 528.73 |

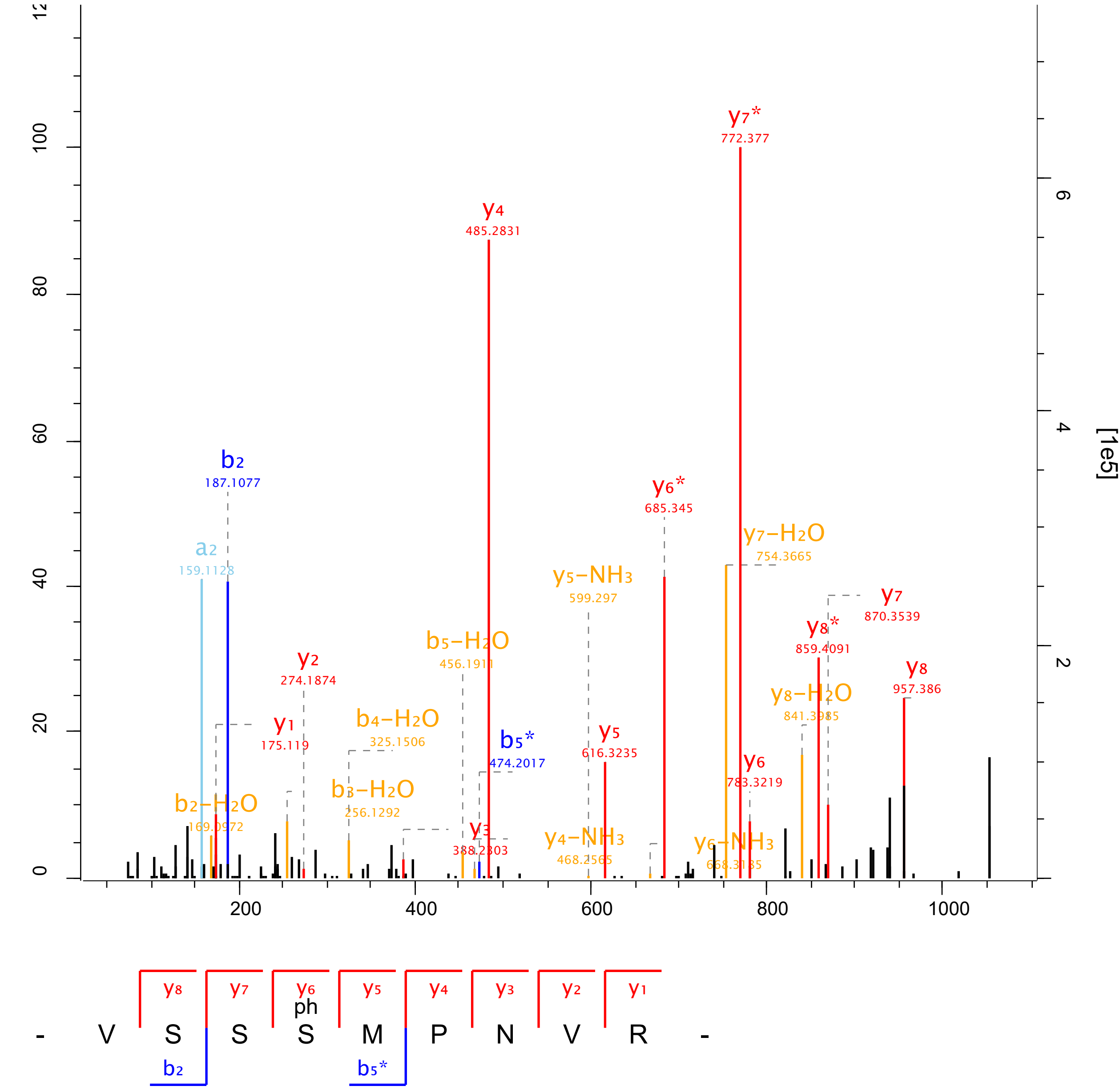

| Raw file | Scan | Method | Score | m/z |
| --- | --- | --- | --- | --- |
| sys_05_2 | 9646 | FTMS; HCD | 43.5 | 492.21 |

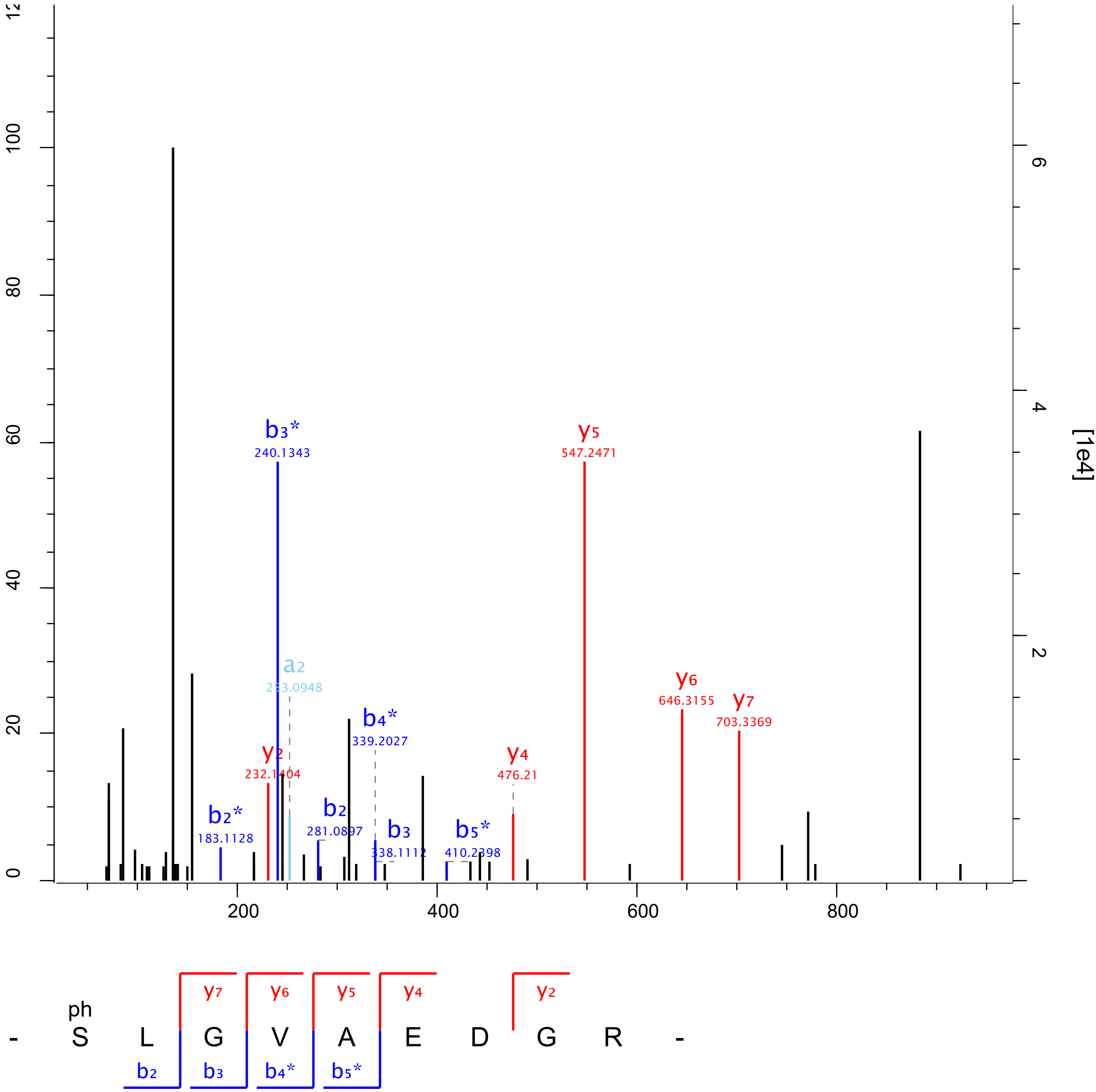

| Raw file | Scan | Method | Score | m/z |
|---|---|---|---|---|
| sys_05_2 | 9769 | FTMS; HCD | 59.42 | 506.73 |

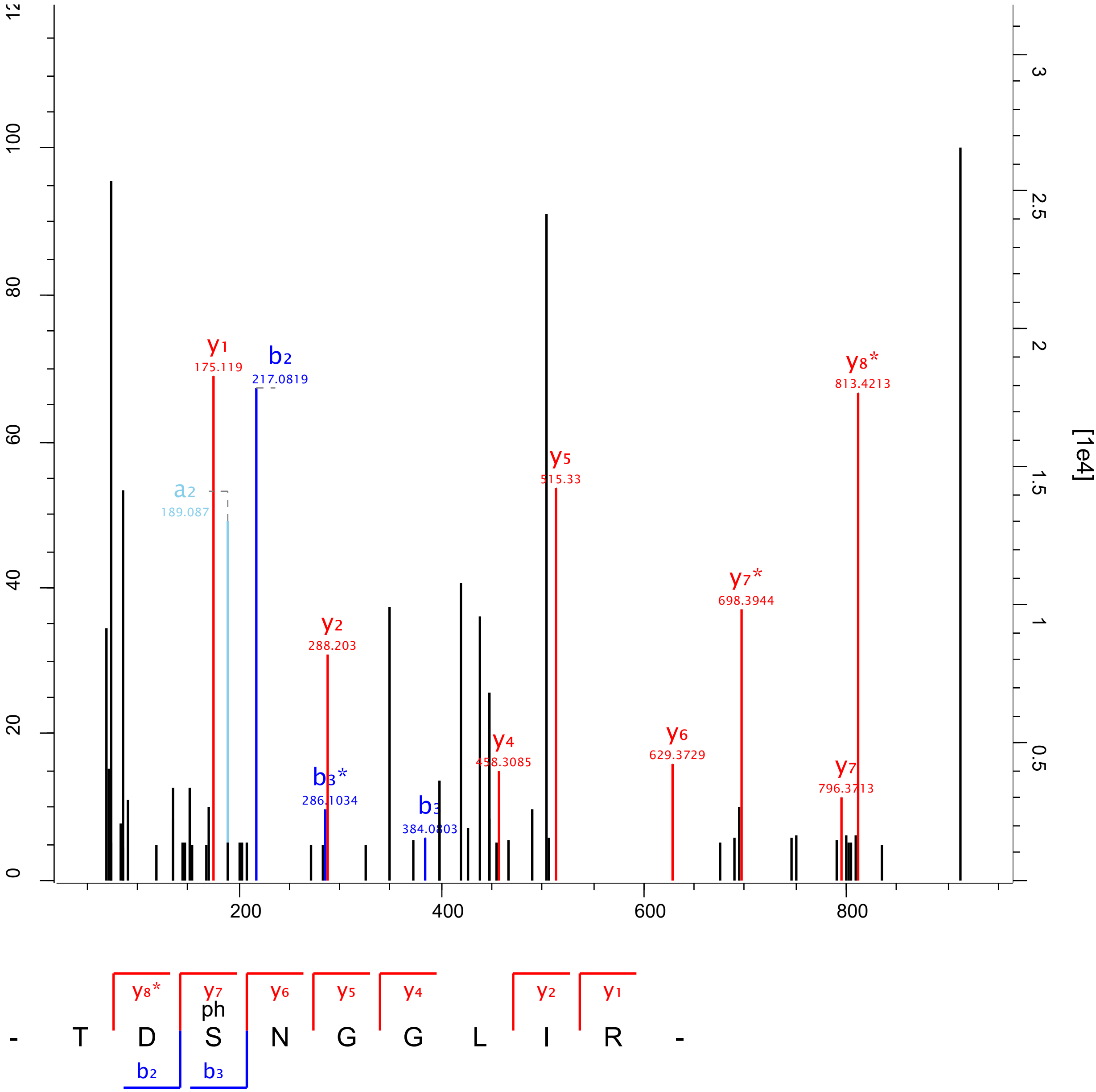

| Raw file | Scan | Method | Score | m/z |
|---|---|---|---|---|
| sys_05_2 | 9902 | FTMS; HCD | 246.8 | 676.3 |

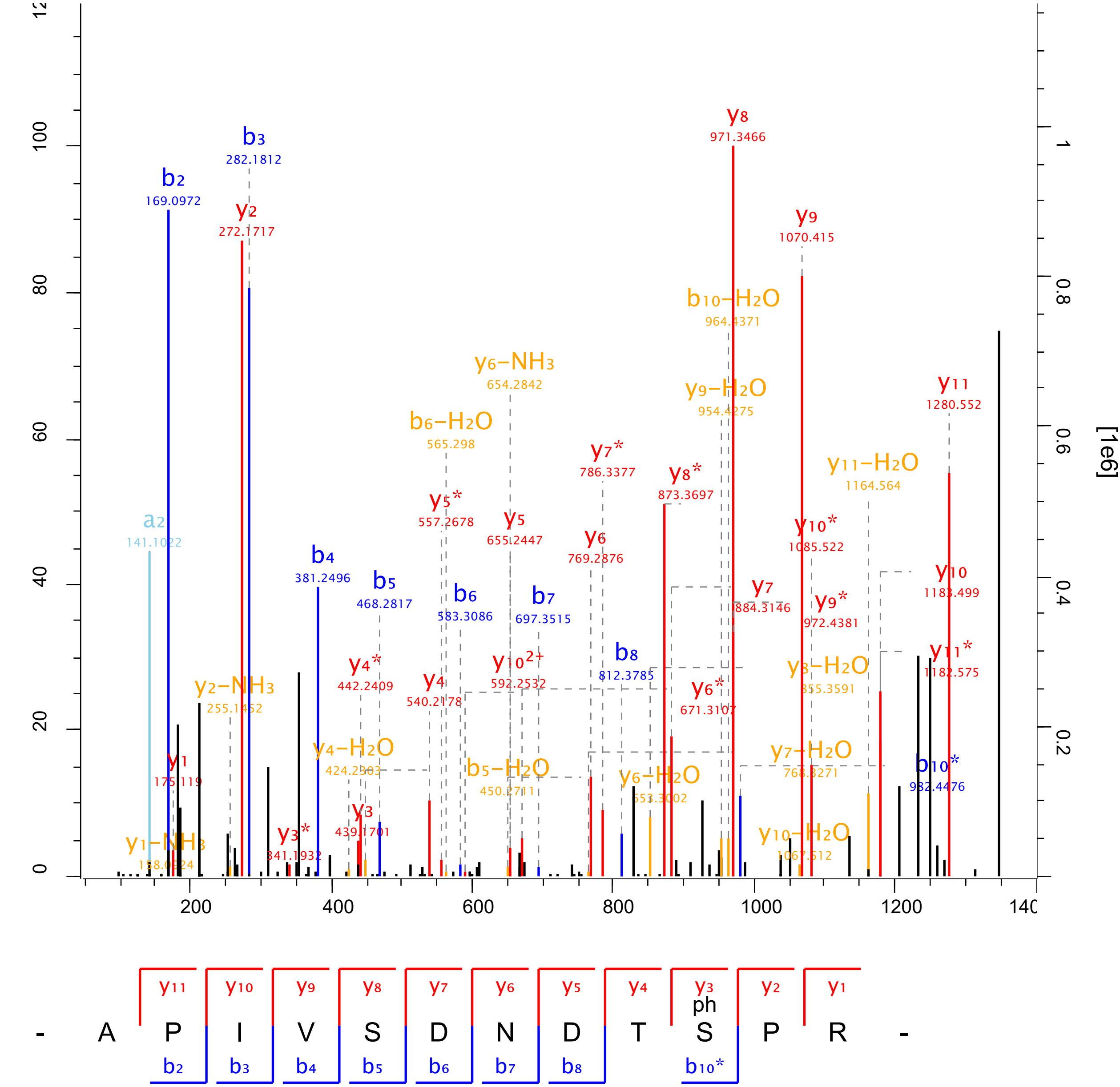

| Raw file | Scan | Method | Score | m/z |
|---|---|---|---|---|
| sys_05_2 | 9997 | FTMS; HCD | 41.7 | 785.32 |

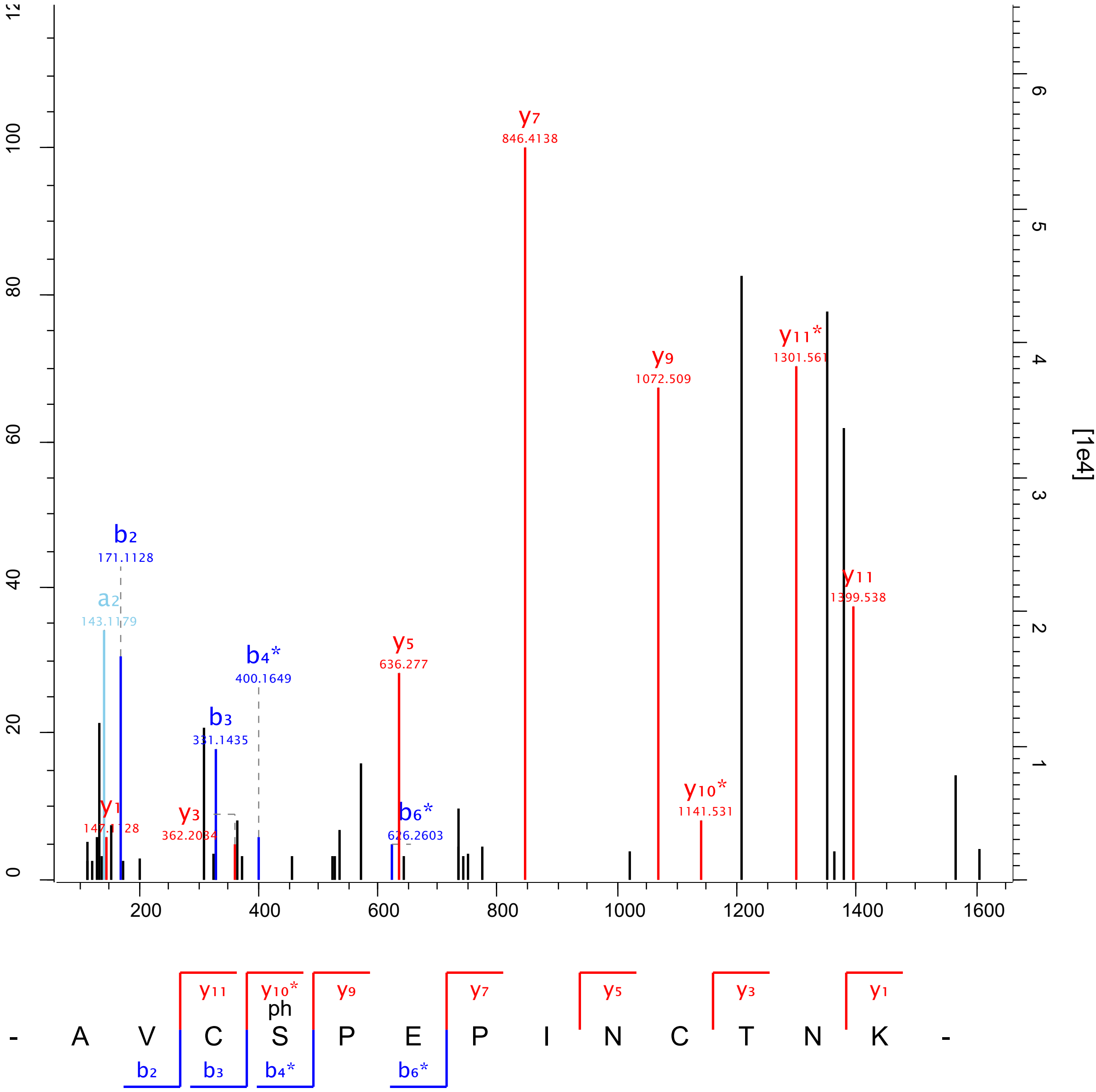

| Raw file | Scan | Method | Score | m/z |
| --- | --- | --- | --- | --- |
| sys_05_3 | 13608 | FTMS; HCD | 93.35 | 503.77 |

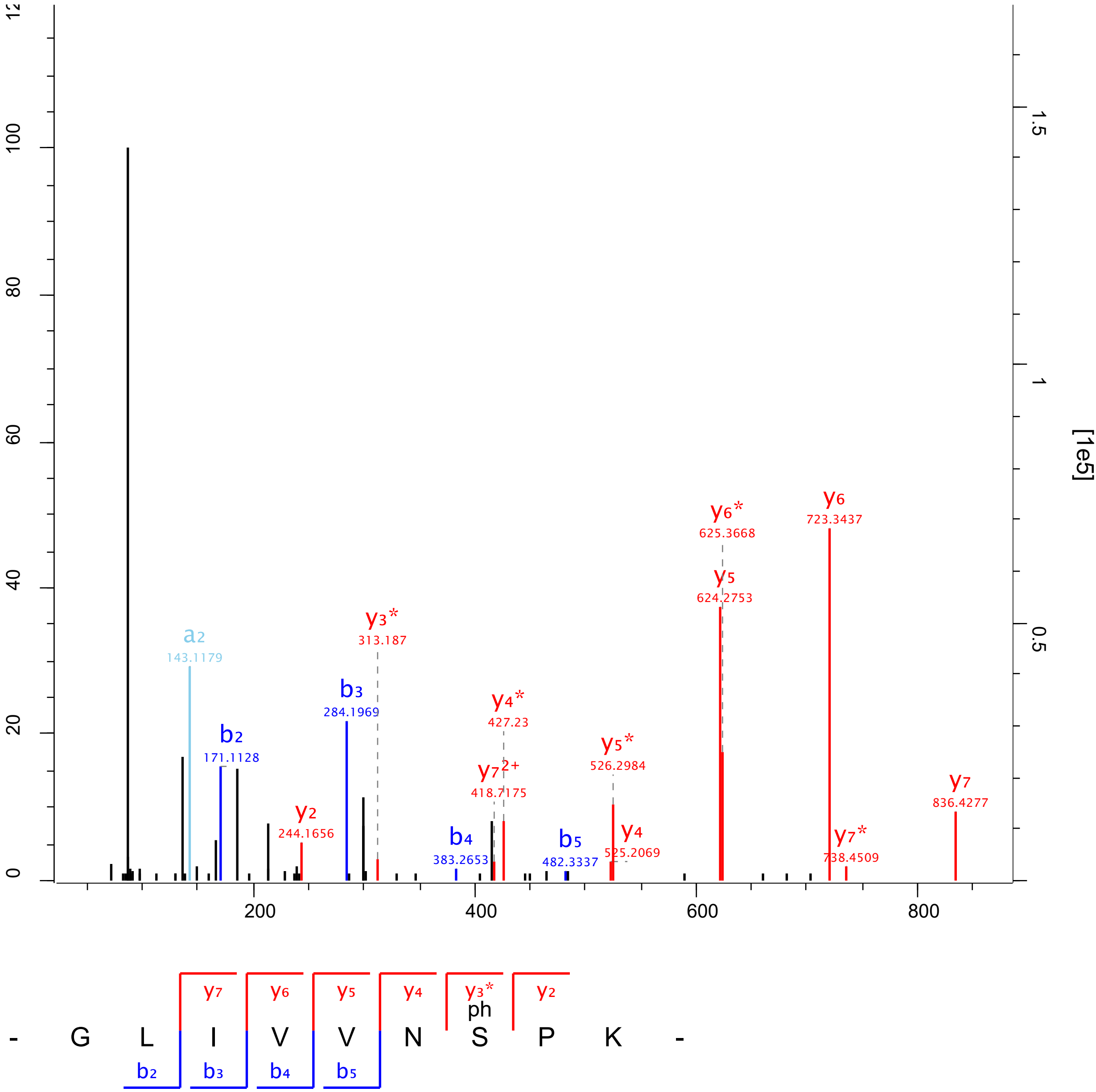

| Raw file | Scan | Method | Score | m/z |
| --- | --- | --- | --- | --- |
| sys_05_3 | 4381 | FTMS; HCD | 172.06 | 634.26 |

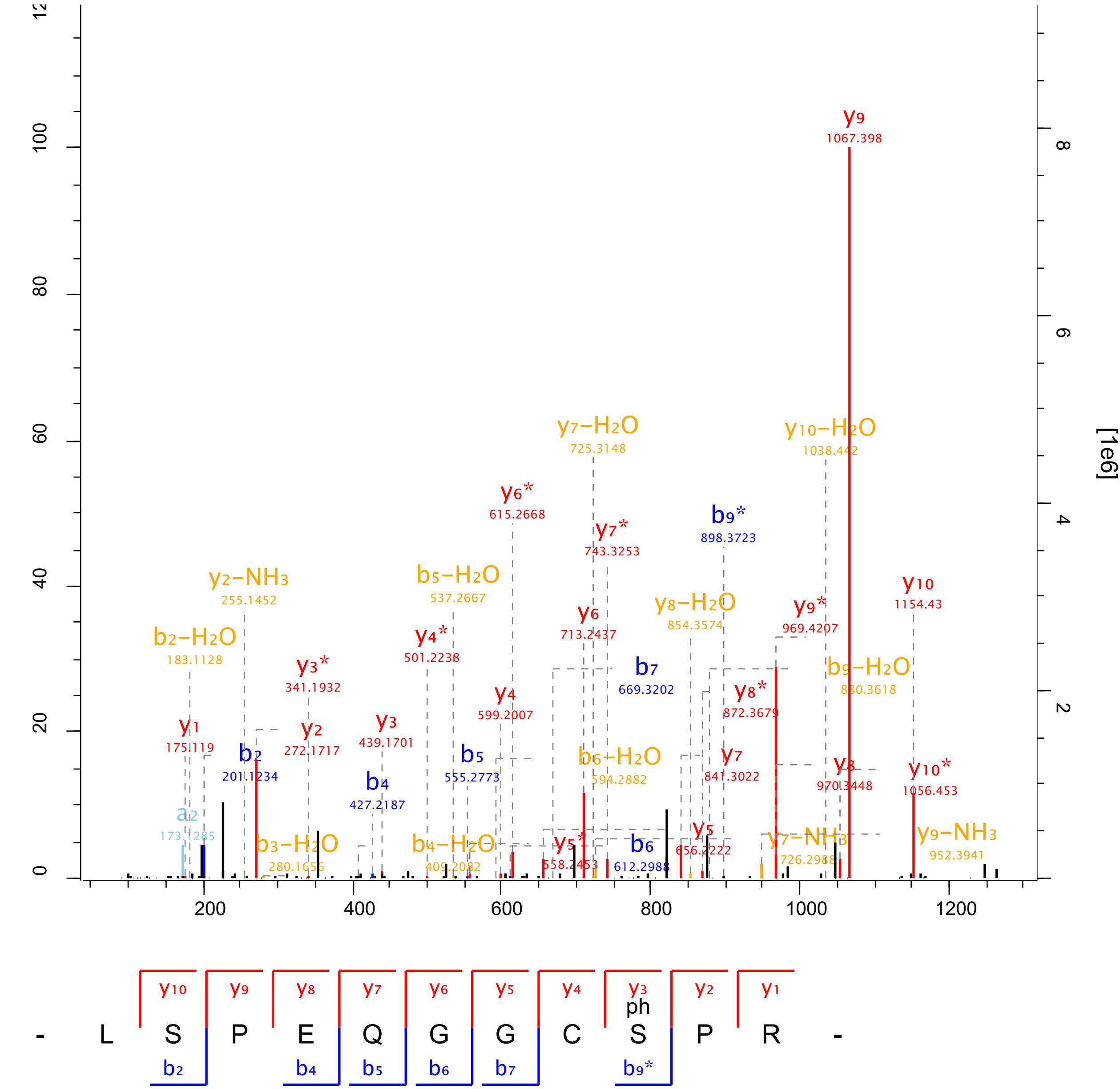

| Raw file | Scan | Method | Score | m/z |
|---|---|---|---|---|
| sys_05_3 | 8633 | FTMS; HCD | 57.79 | 443.69 |

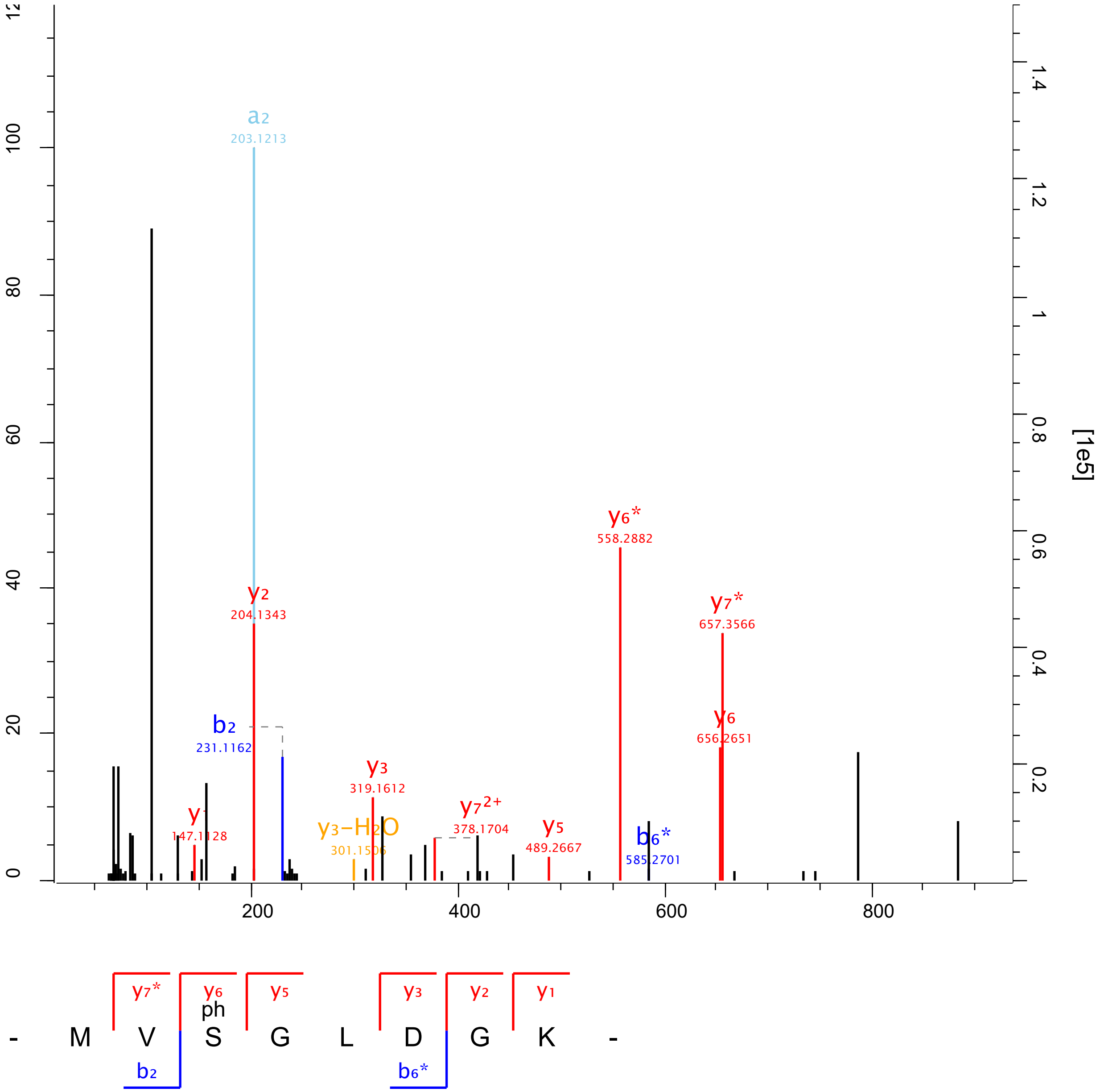

| Raw file | Scan | Method | Score | m/z |
|---|---|---|---|---|
| sys_15_1 | 10064 | FTMS; HCD | 109.44 | 684.28 |

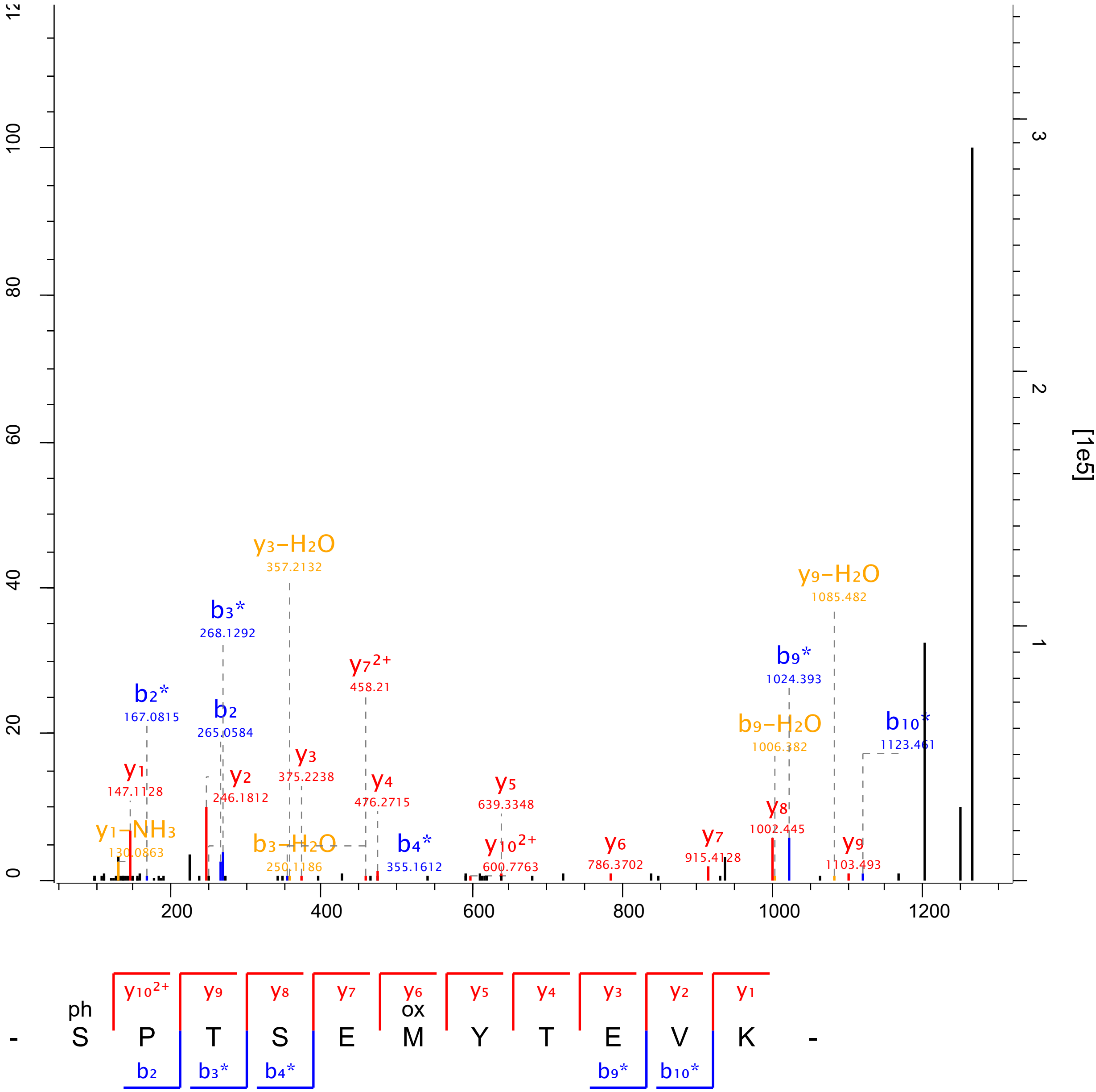

| Raw file | Scan | Method | Score | m/z |
| --- | --- | --- | --- | --- |
| sys_15_1 | 10104 | FTMS; HCD | 111.95 | 421.7 |

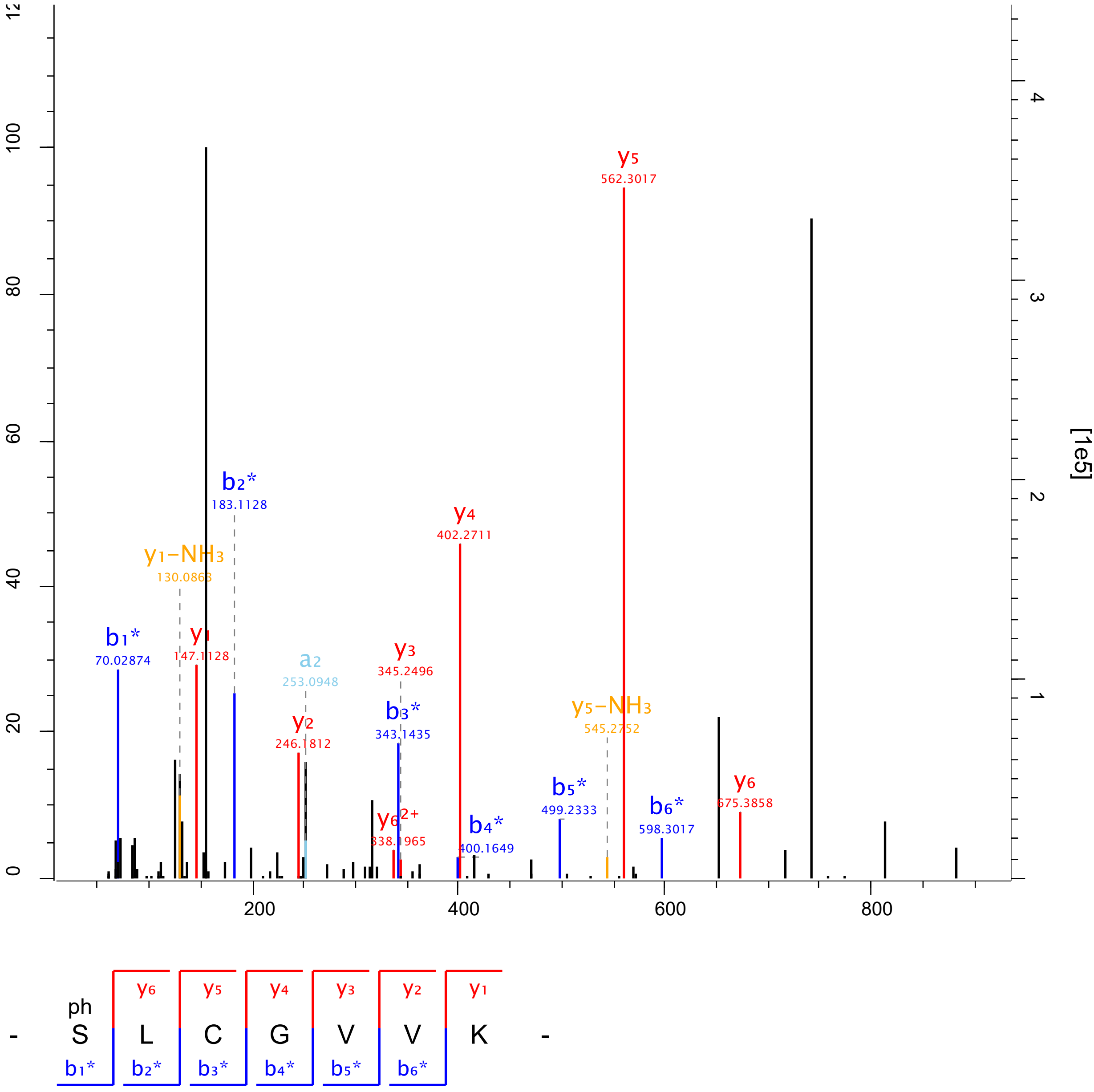

| Raw file | Scan | Method | Score | m/z |
| --- | --- | --- | --- | --- |
| sys_15_1 | 10119 | FTMS; HCD | 117.02 | 517.71 |

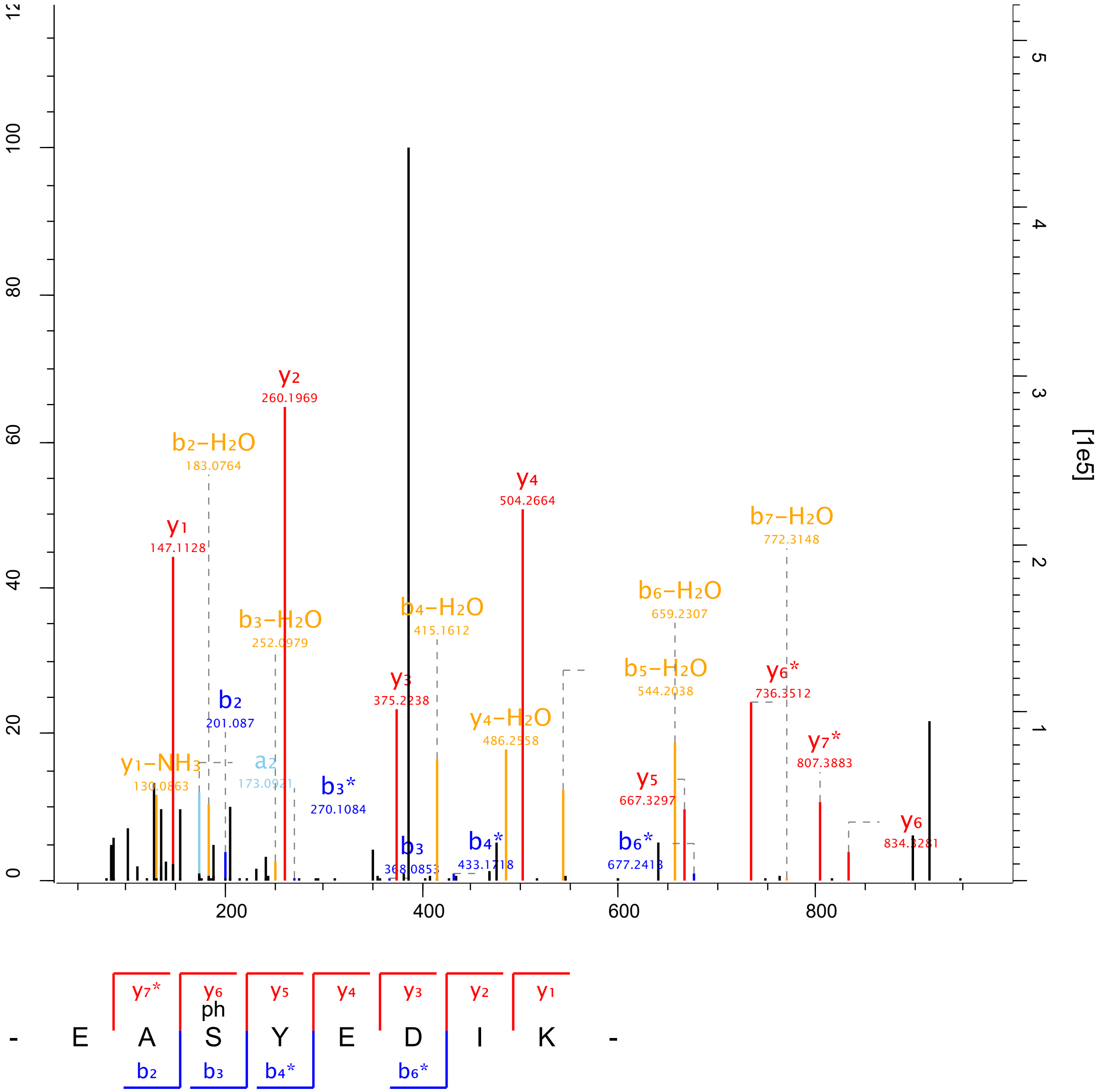

| Raw file | Scan | Method | Score | m/z |
|---|---|---|---|---|
| sys_15_1 | 10136 | FTMS; HCD | 139.81 | 853.3 |

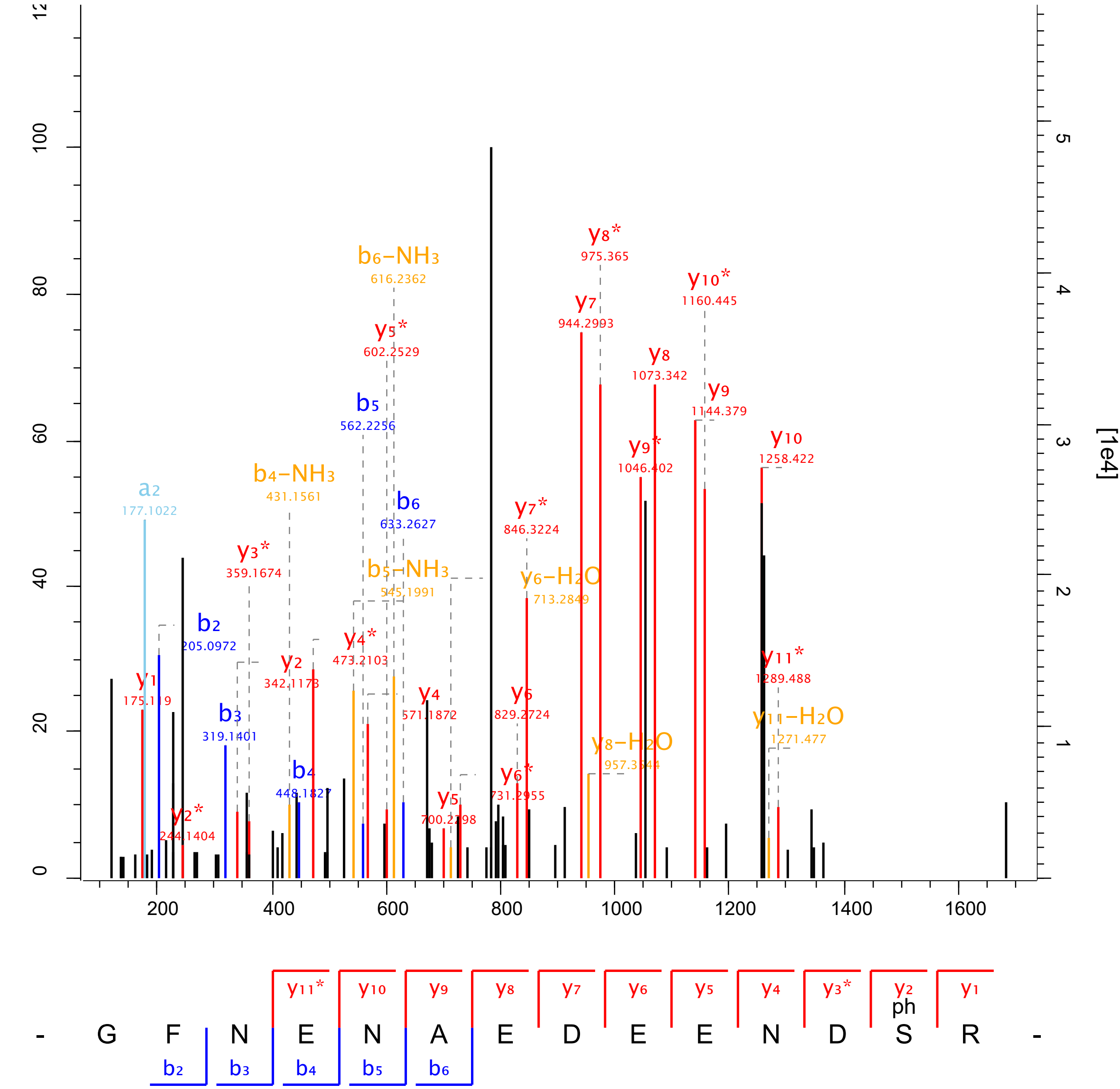

| Raw file | Scan | Method | Score | m/z |
|---|---|---|---|---|
| sys_15_1 | 10175 | FTMS; HCD | 281.96 | 834.34 |

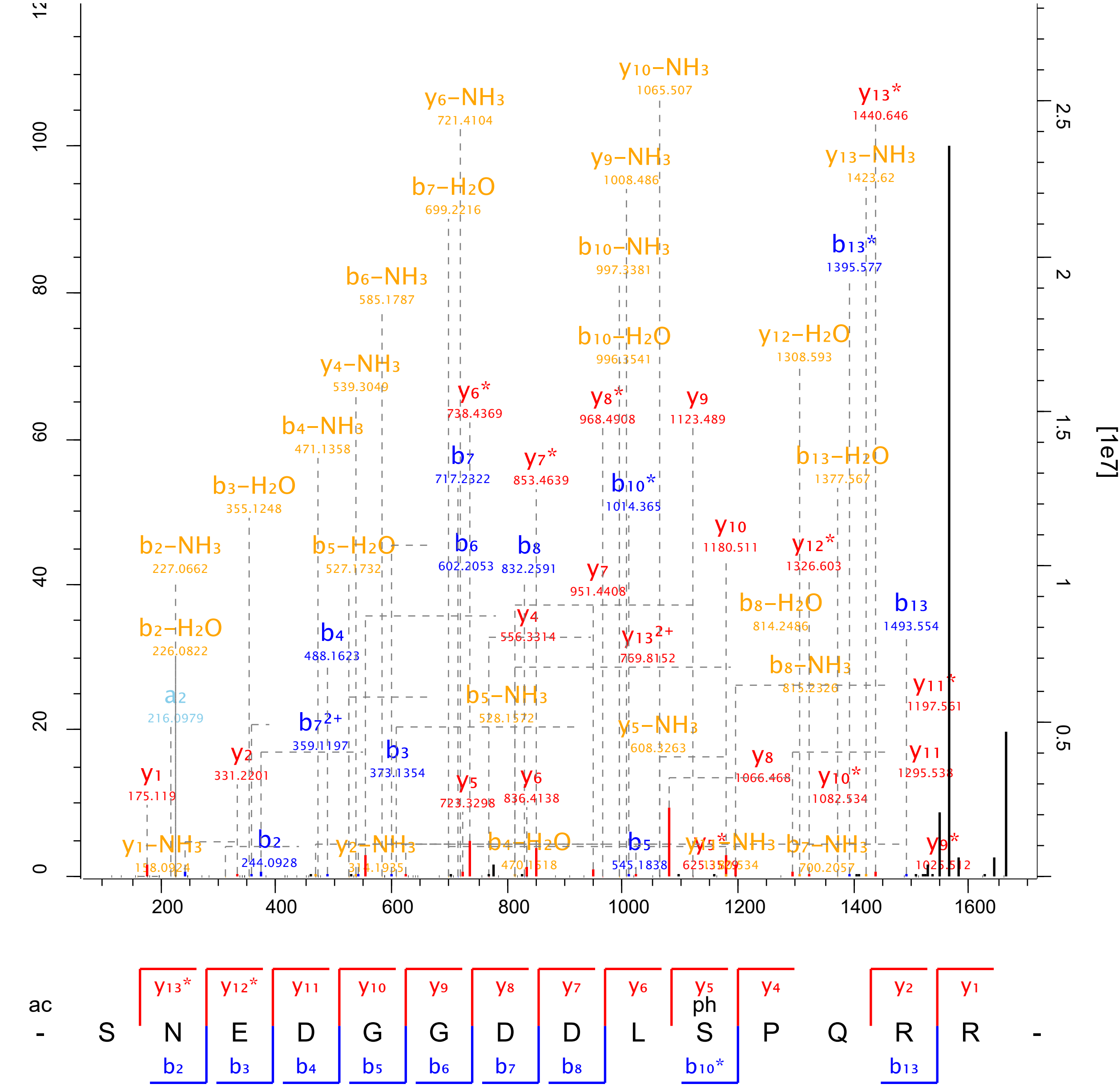

| Raw file | Scan | Method | Score | m/z |
|---|---|---|---|---|
| sys_15_1 | 10178 | FTMS; HCD | 70.44 | 540.75 |

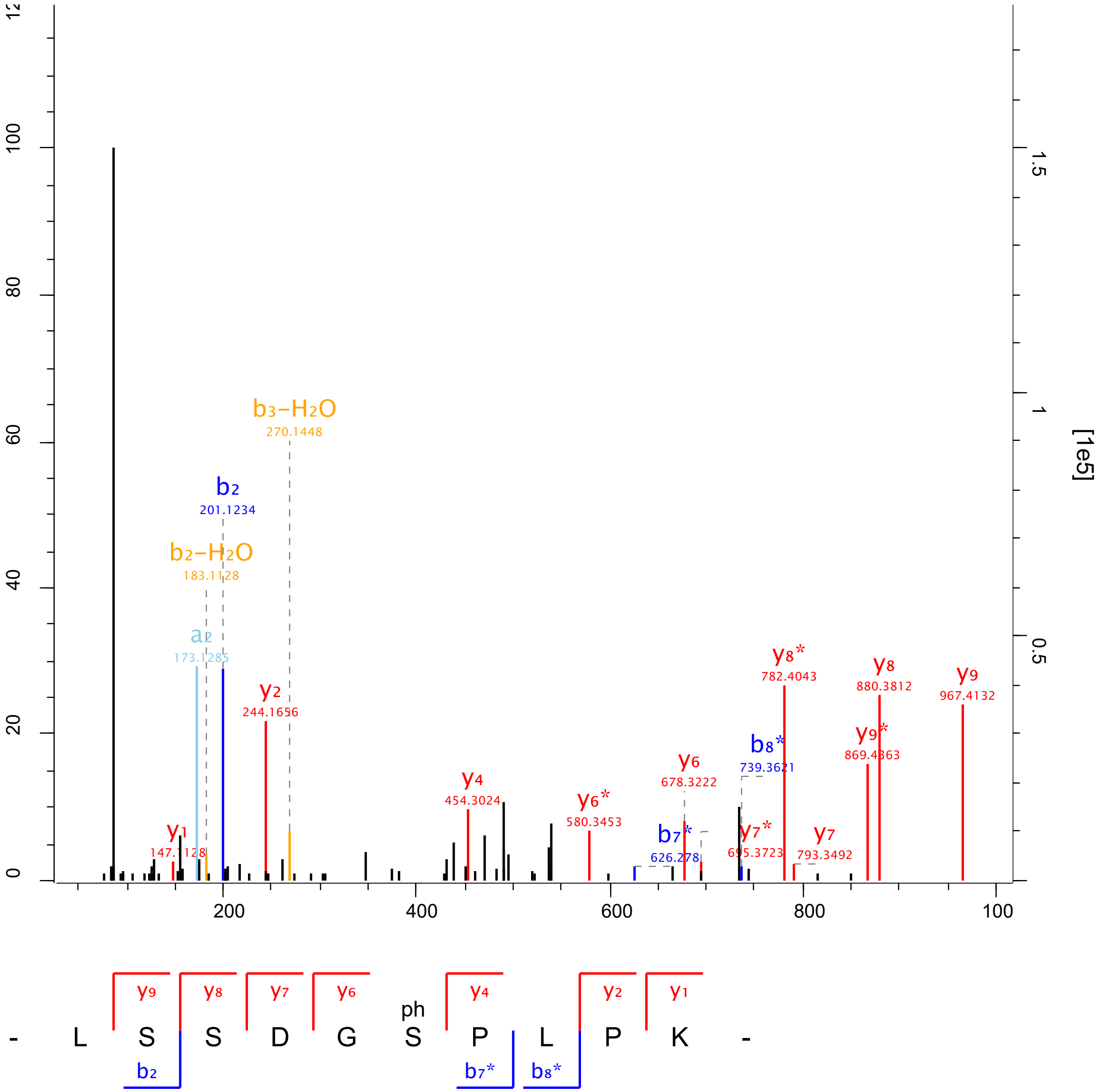

| Raw file | Scan | Method | Score | m/z |
|---|---|---|---|---|
| sys_15_1 | 10190 | FTMS; HCD | 81.85 | 768.26 |

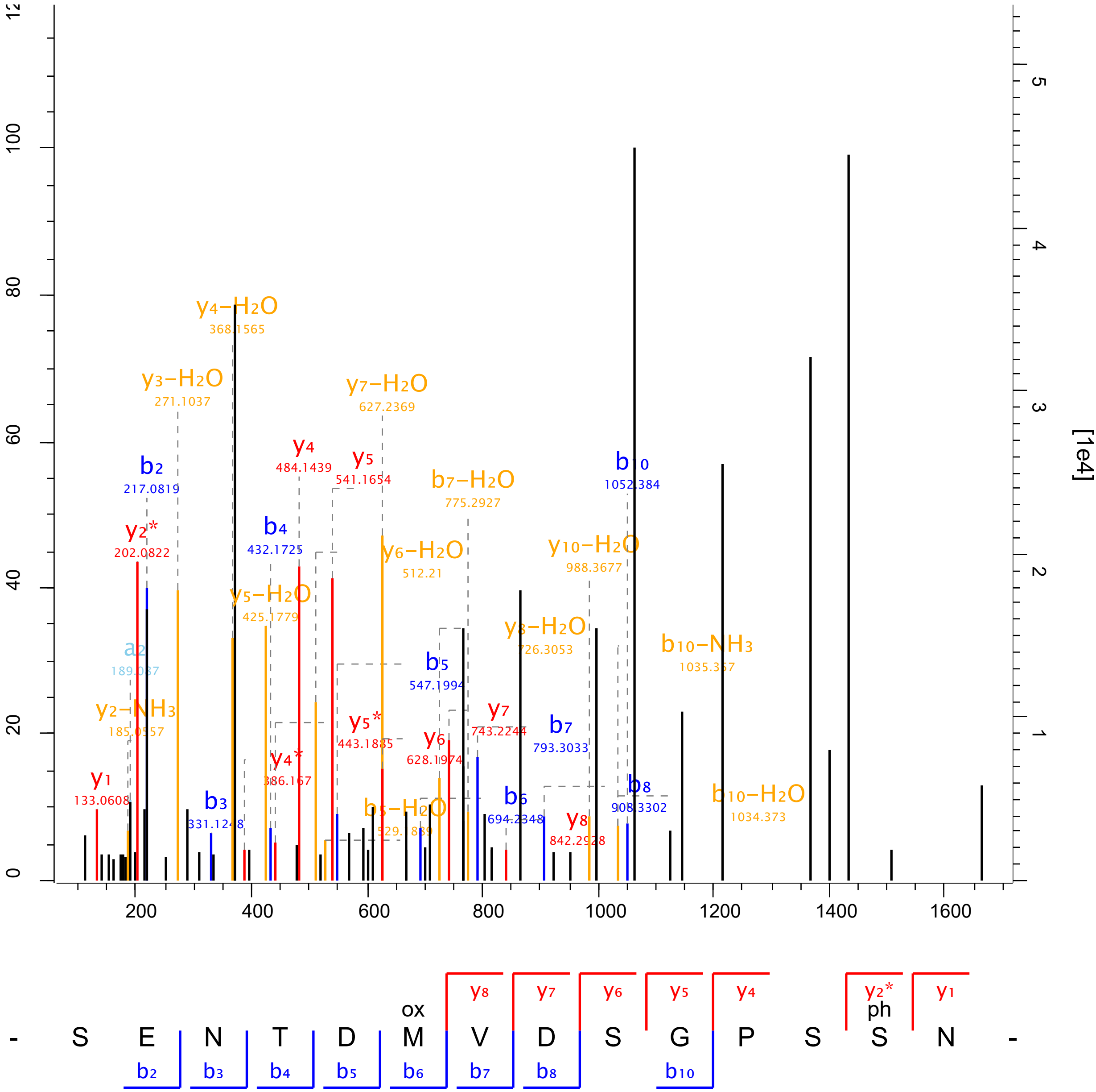

| Raw file | Scan | Method | Score | m/z |
| --- | --- | --- | --- | --- |
| sys_15_1 | 10226 | FTMS; HCD | 214.89 | 713.83 |

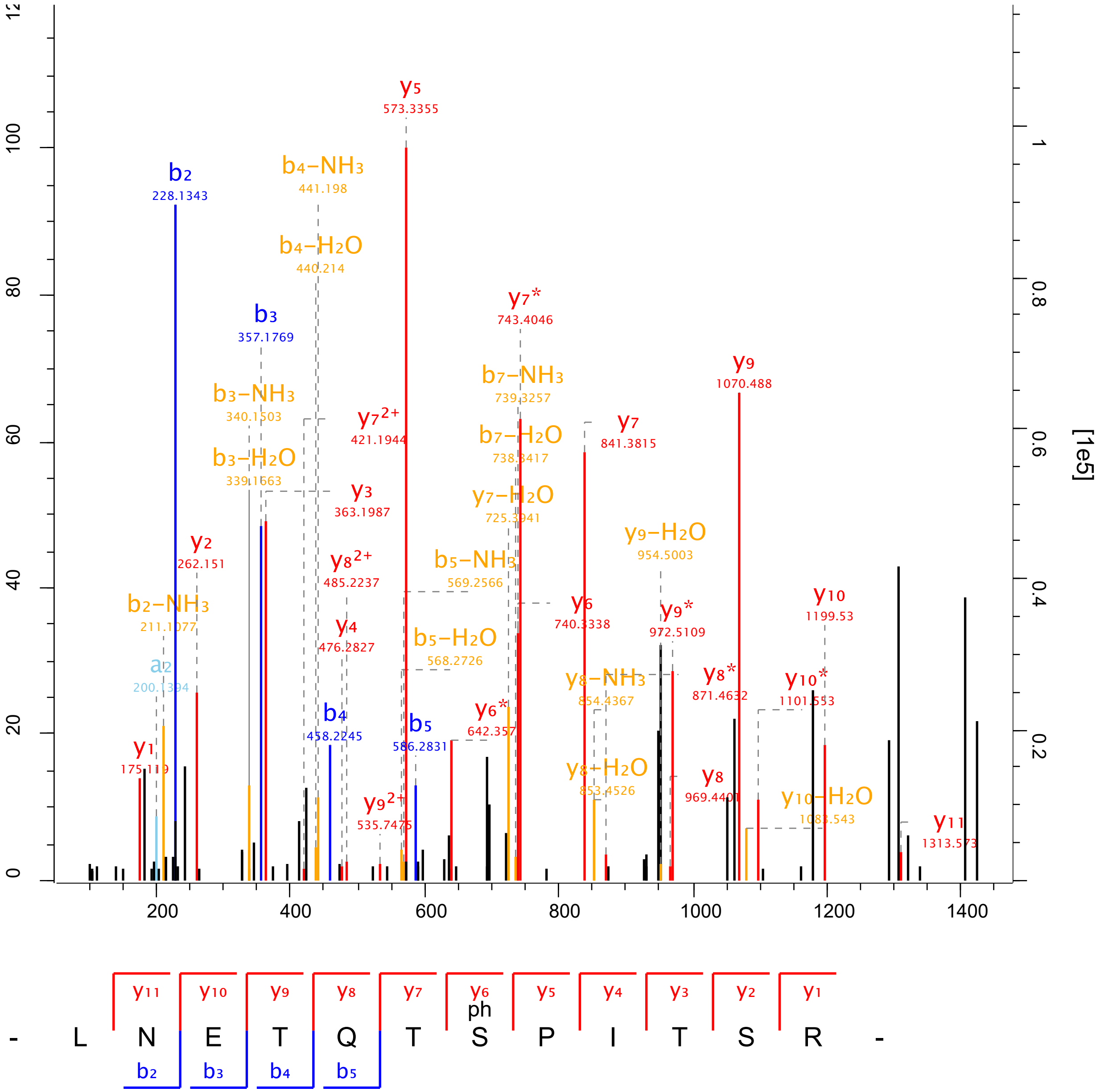

| Raw file | Scan | Method | Score | m/z |
|---|---|---|---|---|
| sys_15_1 | 10242 | FTMS; HCD | 101.2 | 645.8 |

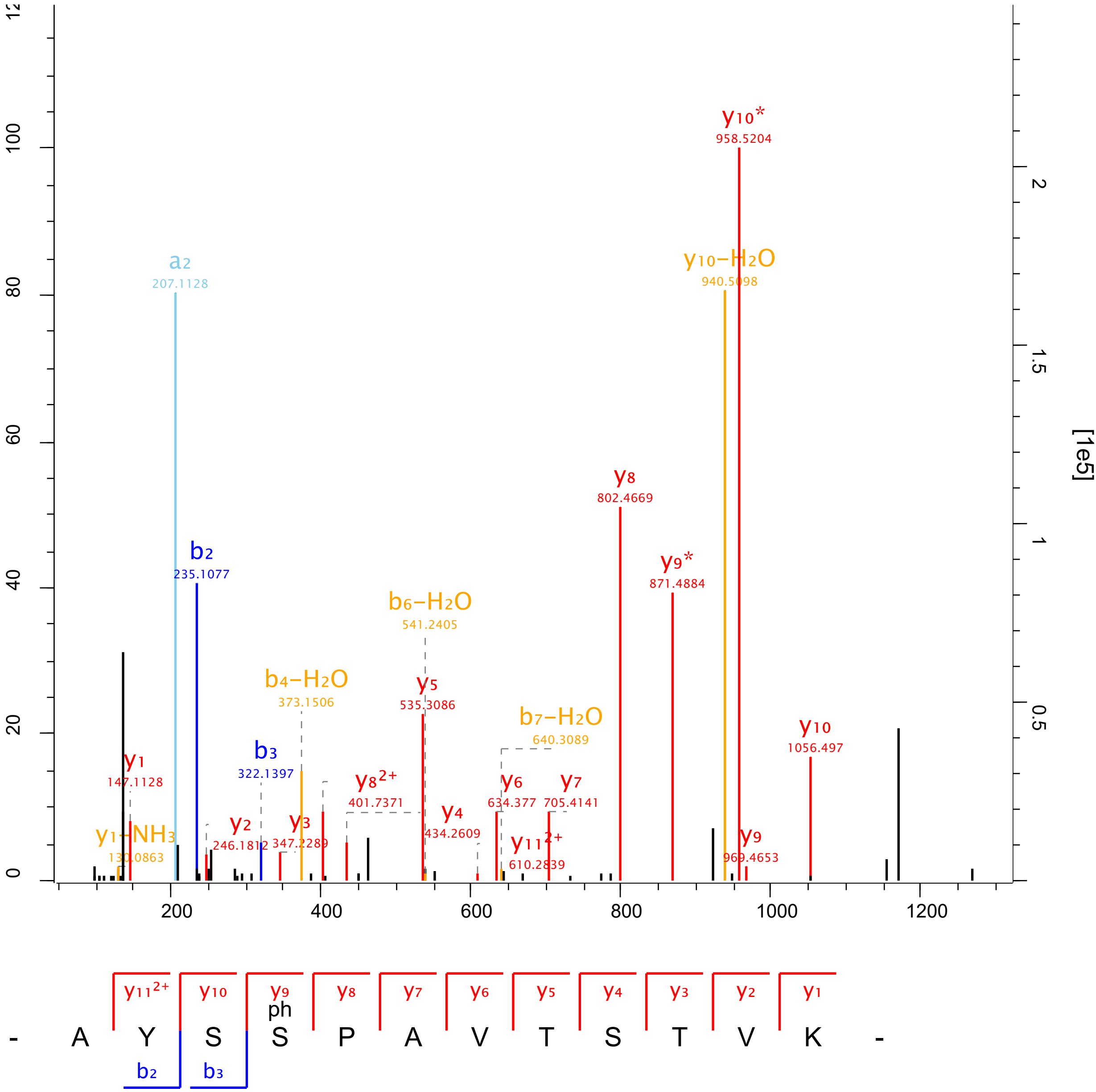

| Raw file | Scan | Method | Score | m/z |
| --- | --- | --- | --- | --- |
| sys_15_1 | 10313 | FTMS; HCD | 110.12 | 672.3 |

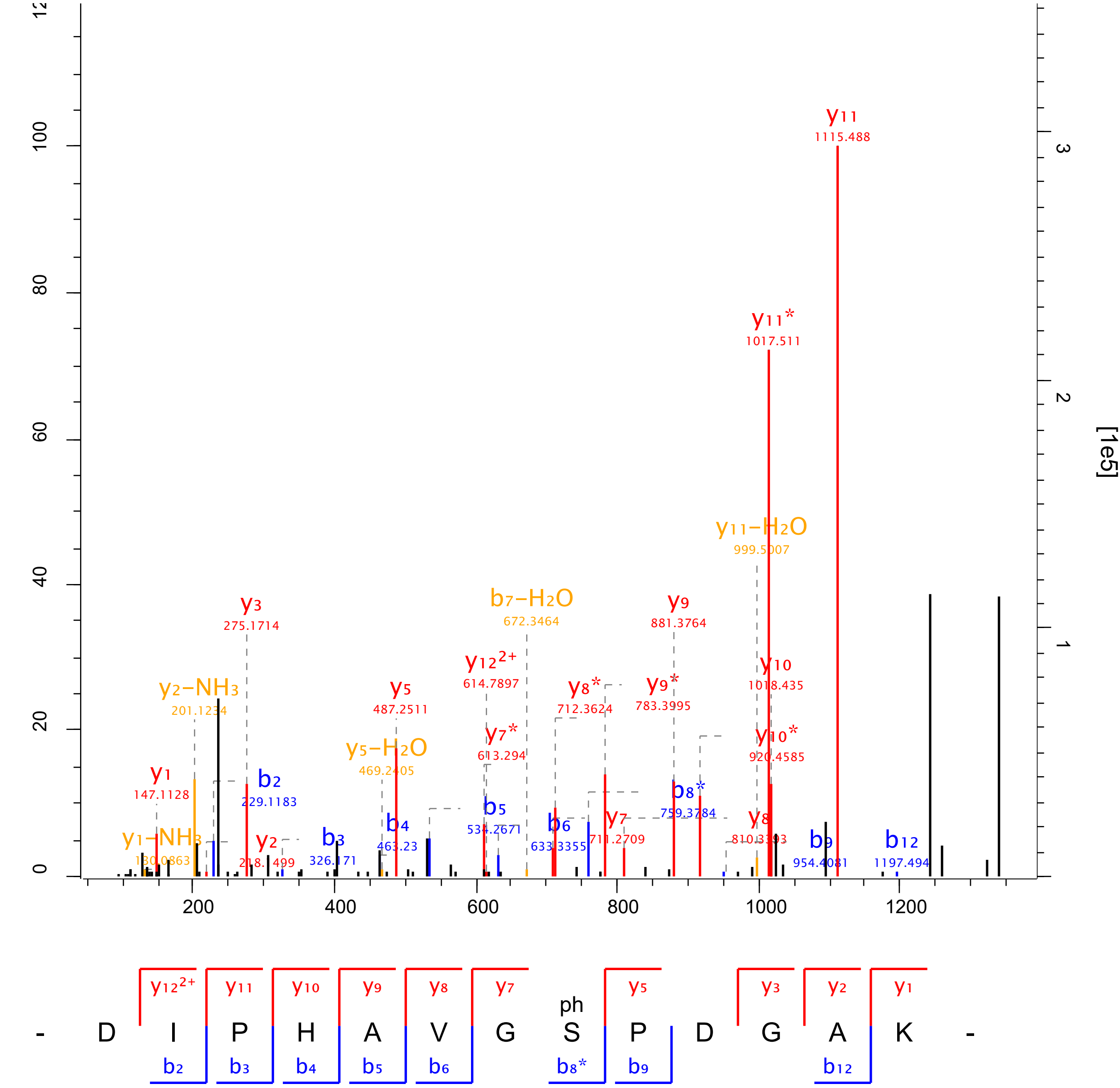

| Raw file | Scan | Method | Score | m/z |
| --- | --- | --- | --- | --- |
| sys_15_1 | 10314 | FTMS; HCD | 73.5 | 516.71 |

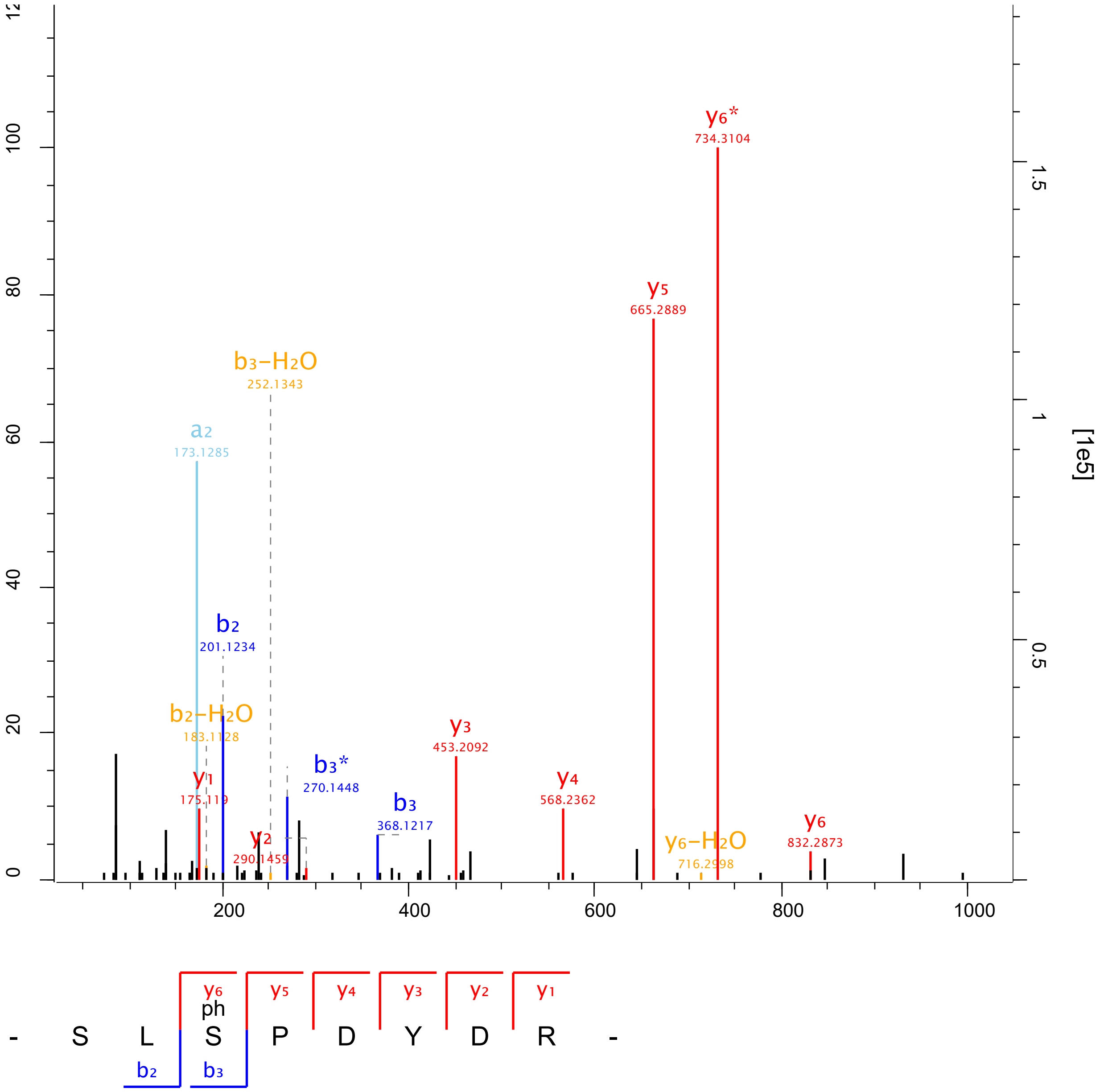

| Raw file | Scan | Method | Score | m/z |
| --- | --- | --- | --- | --- |
| sys_15_1 | 10355 | FTMS; HCD | 46.07 | 586.72 |

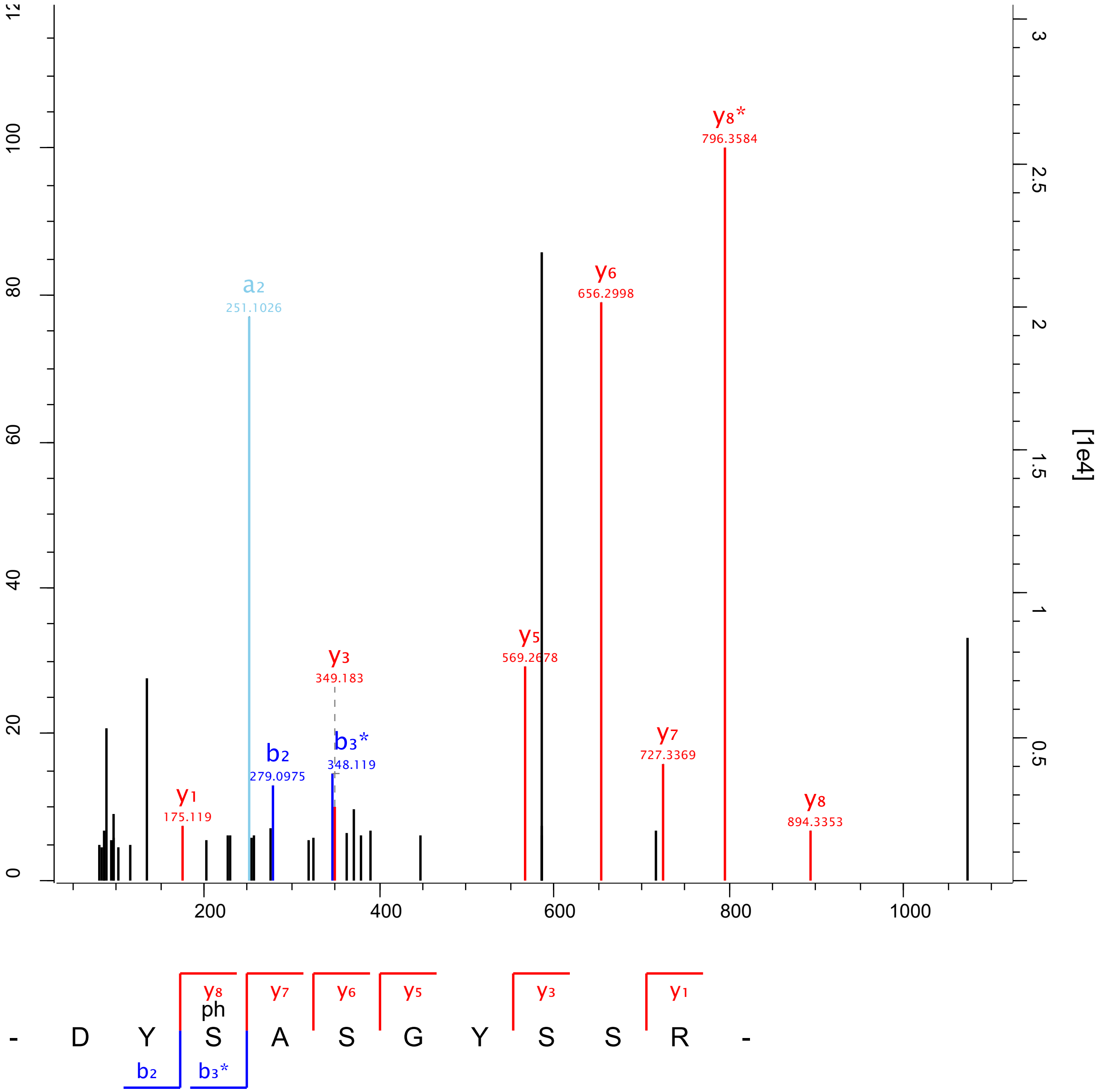

| Raw file | Scan | Method | Score | m/z |
| --- | --- | --- | --- | --- |
| sys_15_1 | 10454 | FTMS; HCD | 41.45 | 600.24 |

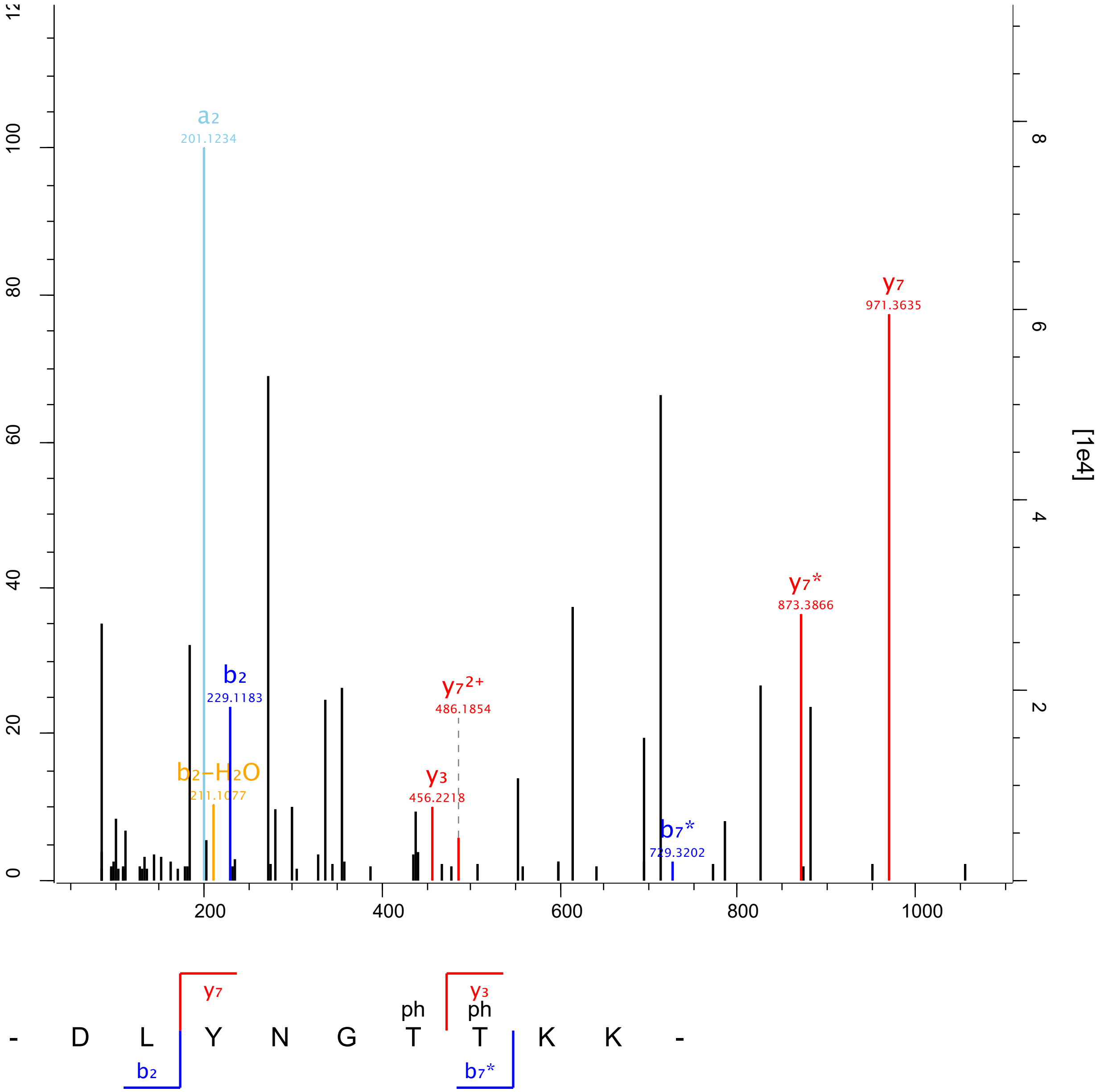

| Raw file | Scan | Method | Score | m/z |
|---|---|---|---|---|
| sys_15_1 | 10480 | FTMS; HCD | 205.31 | 557.76 |

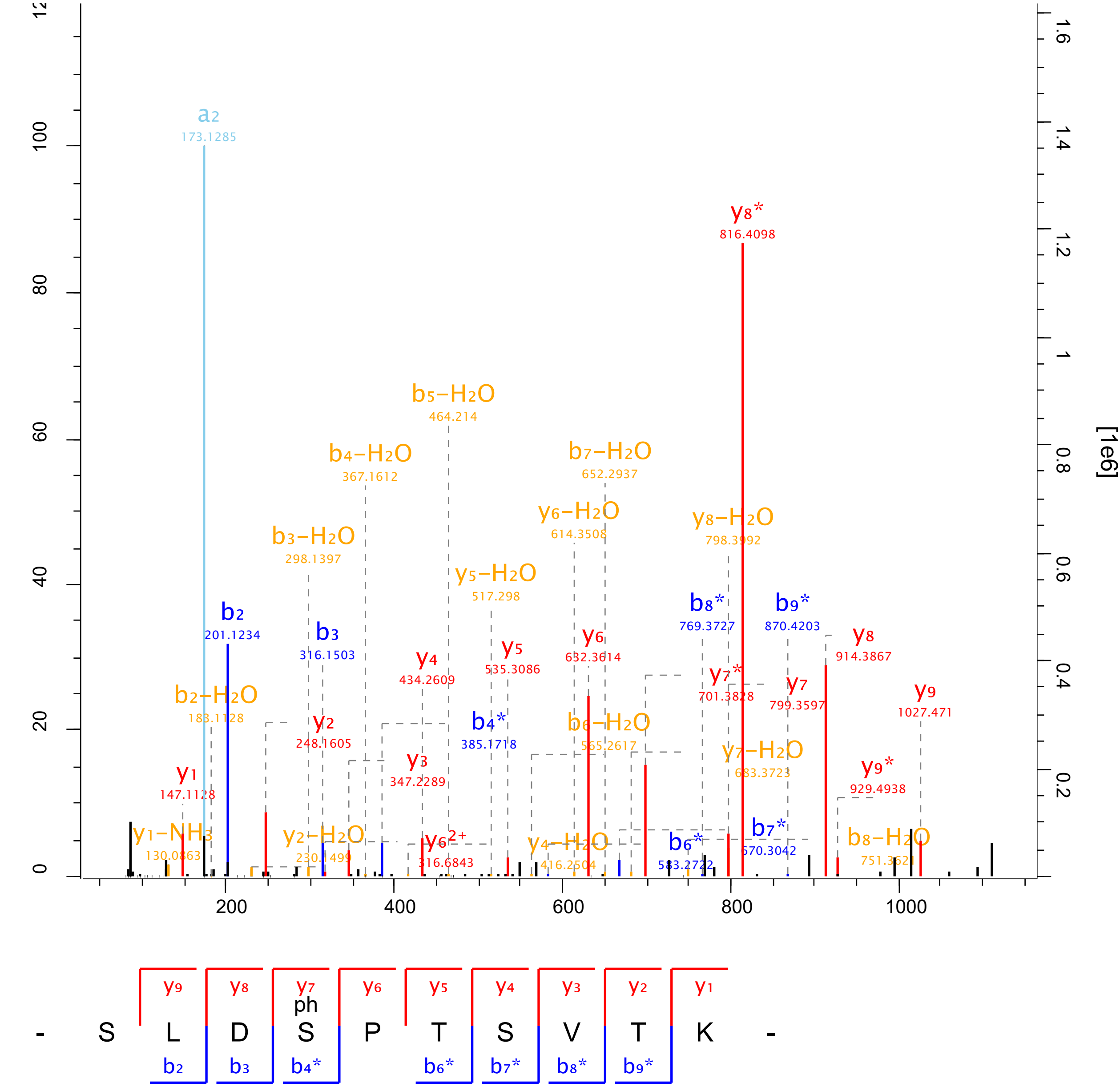

| Raw file | Scan | Method | Score | m/z |
|---|---|---|---|---|
| sys_15_1 | 10614 | FTMS; HCD | 40.14 | 887.38 |

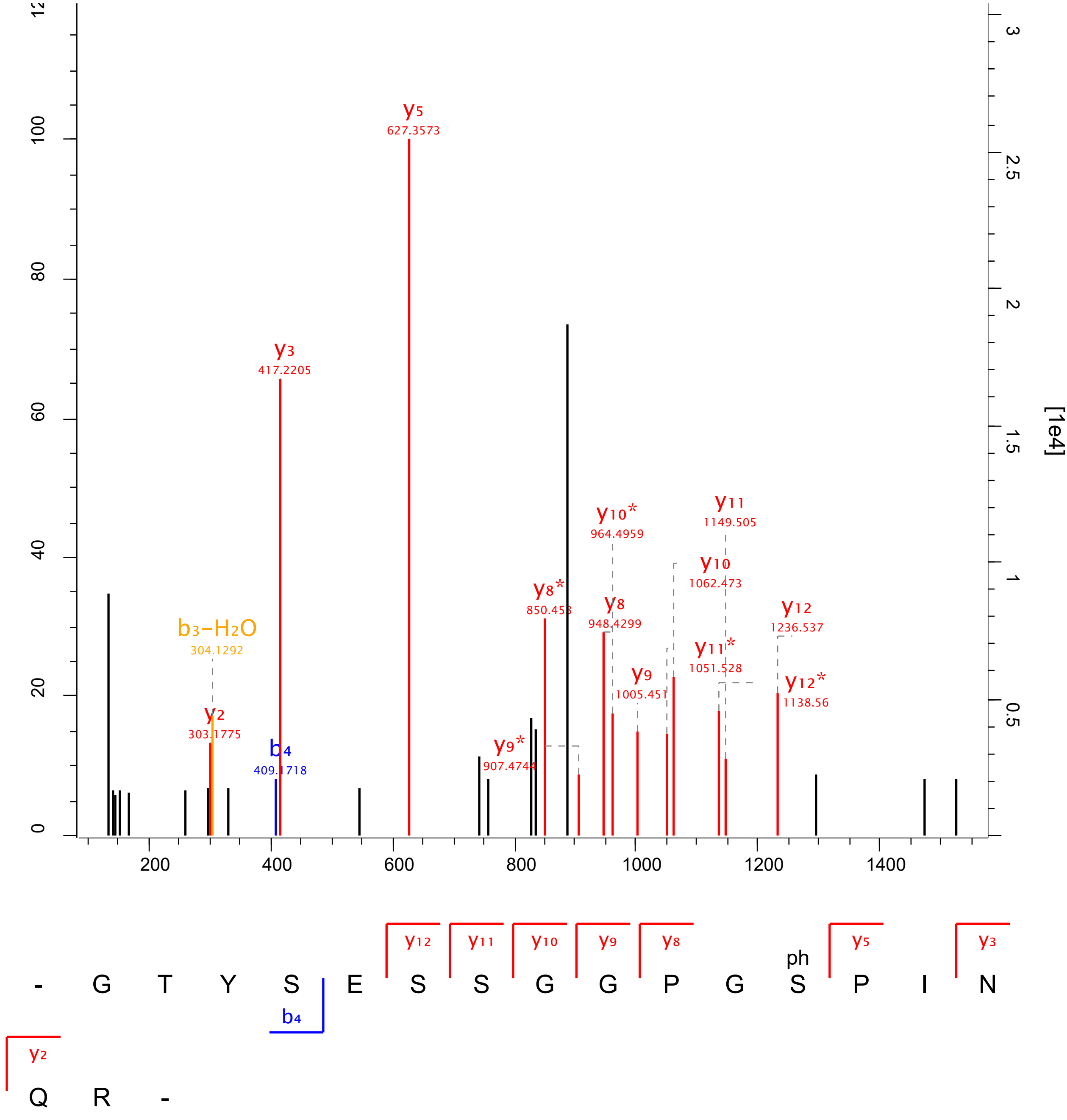

| Raw file | Scan | Method | Score | m/z |
|---|---|---|---|---|
| sys_15_1 | 10712 | FTMS; HCD | 93.24 | 620.26 |

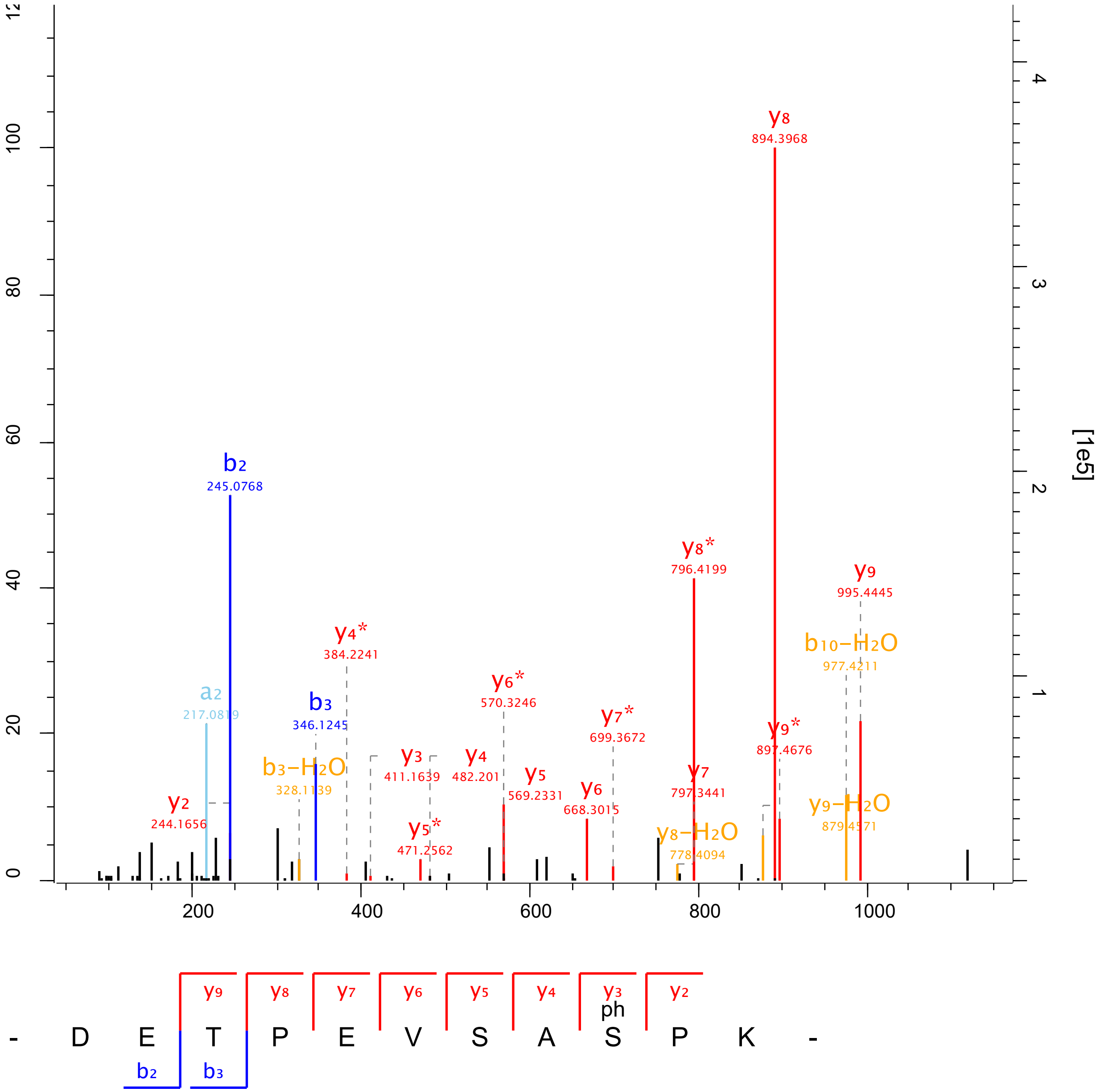

| Raw file | Scan | Method | Score | m/z |
|---|---|---|---|---|
| sys_15_1 | 10777 | FTMS; HCD | 59.12 | 616.78 |

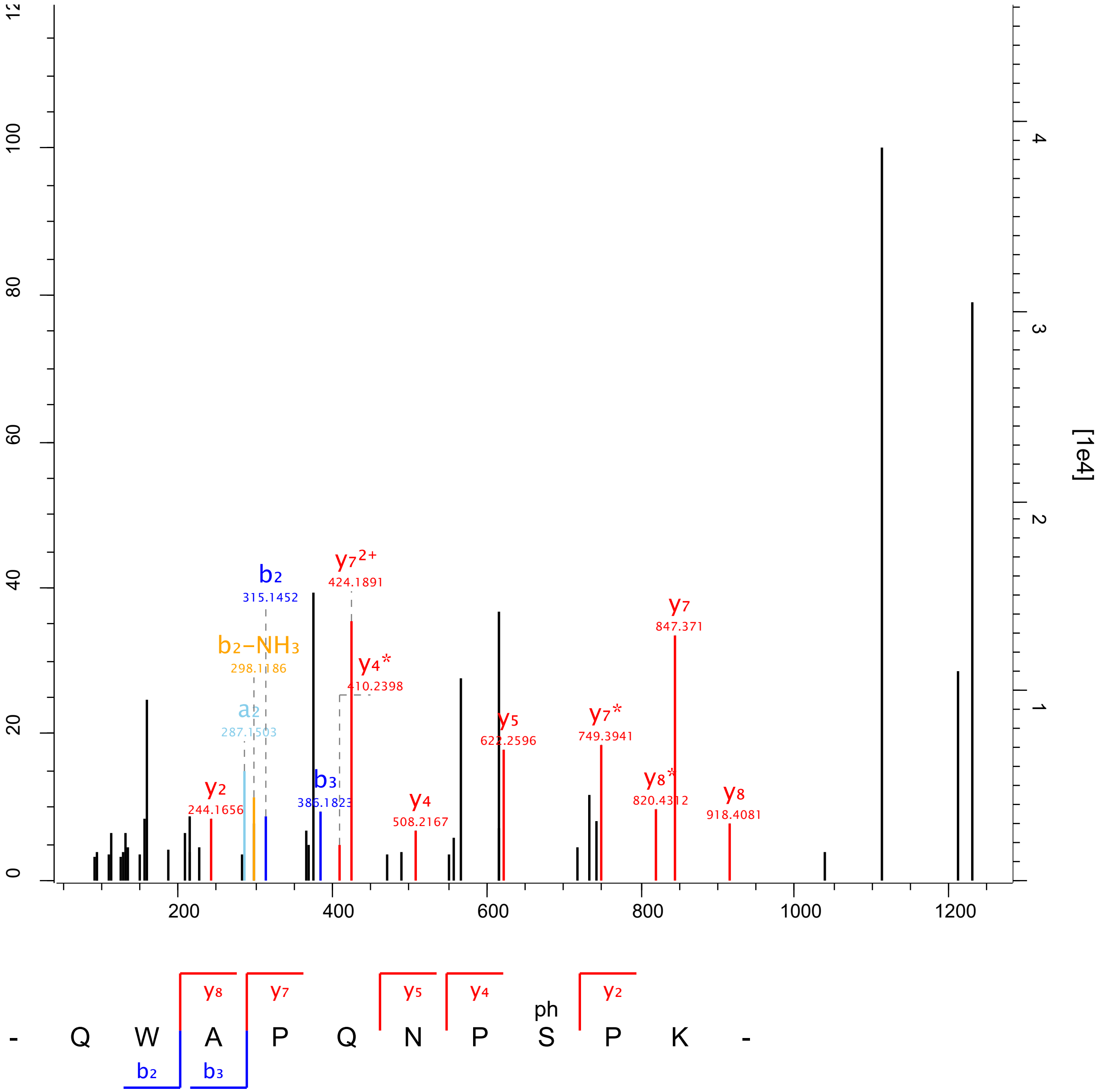

| Raw file | Scan | Method | Score | m/z |
| --- | --- | --- | --- | --- |
| sys_15_1 | 10780 | FTMS; HCD | 106.12 | 750.64 |

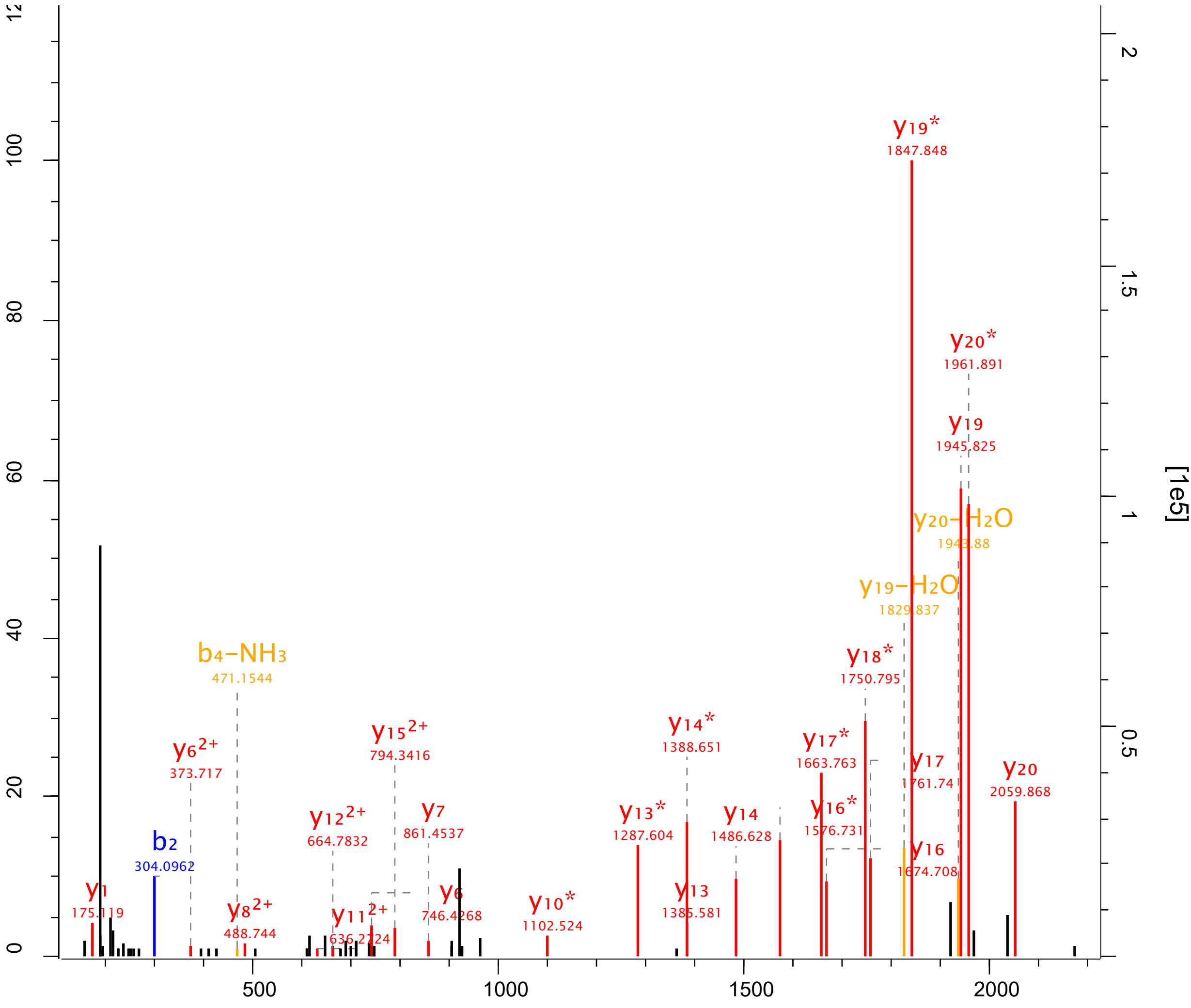


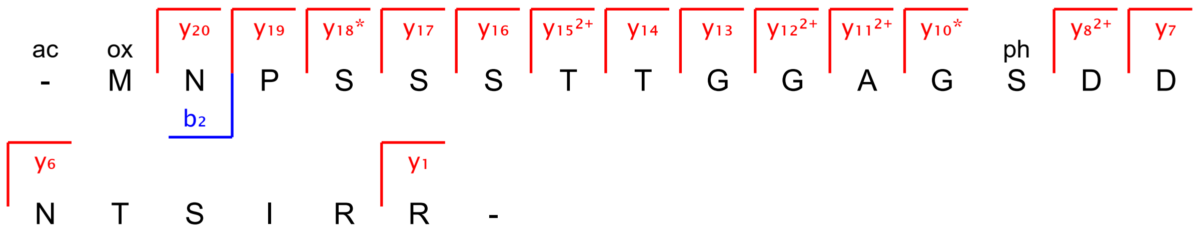

| Raw file | Scan | Method | Score | m/z |
| --- | --- | --- | --- | --- |
| sys_15_1 | 10830 | FTMS; HCD | 48.4 | 525.21 |

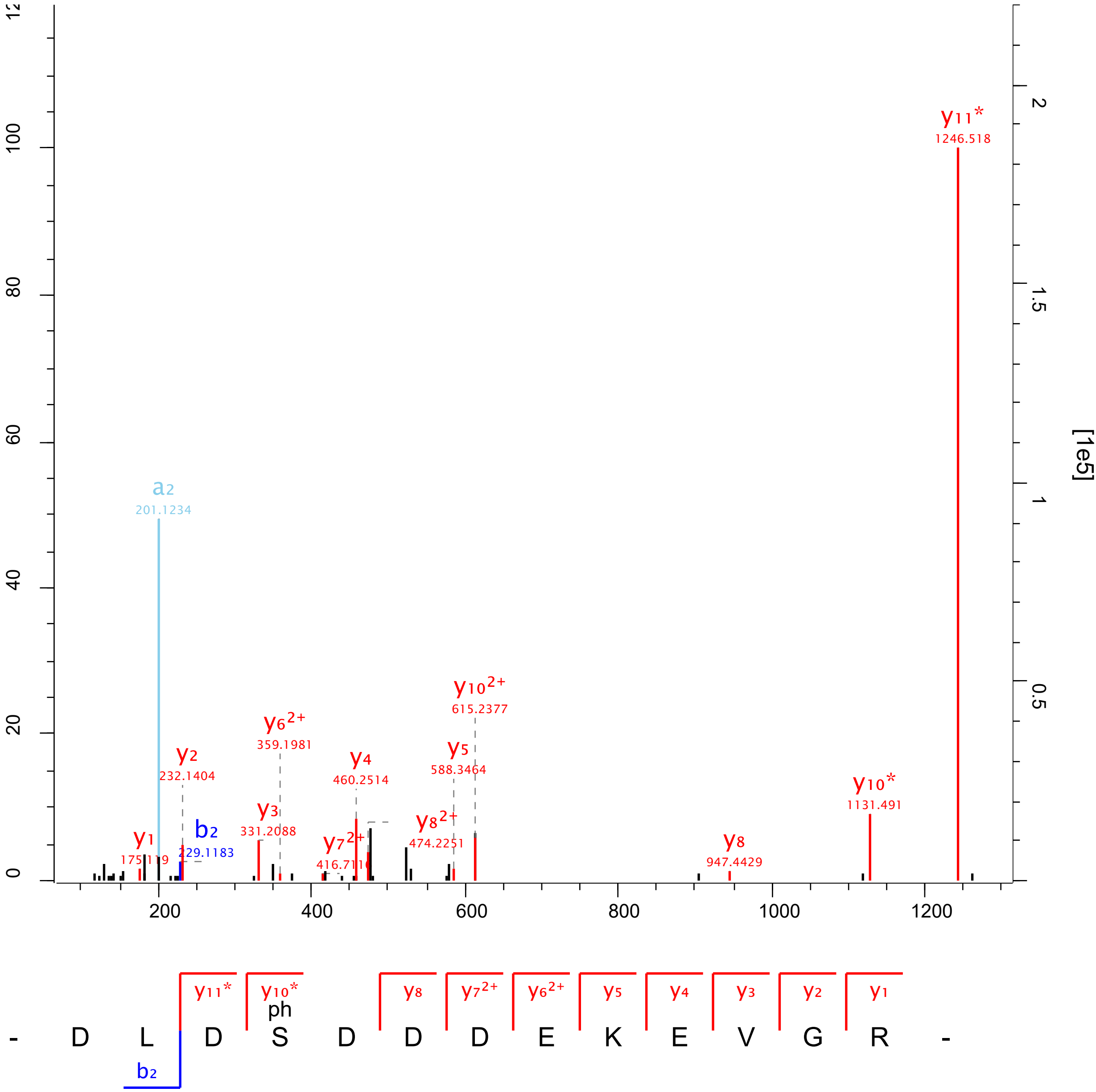

| Raw file | Scan | Method | Score | m/z |
| --- | --- | --- | --- | --- |
| sys_15_1 | 10865 | FTMS; HCD | 89.48 | 753.81 |

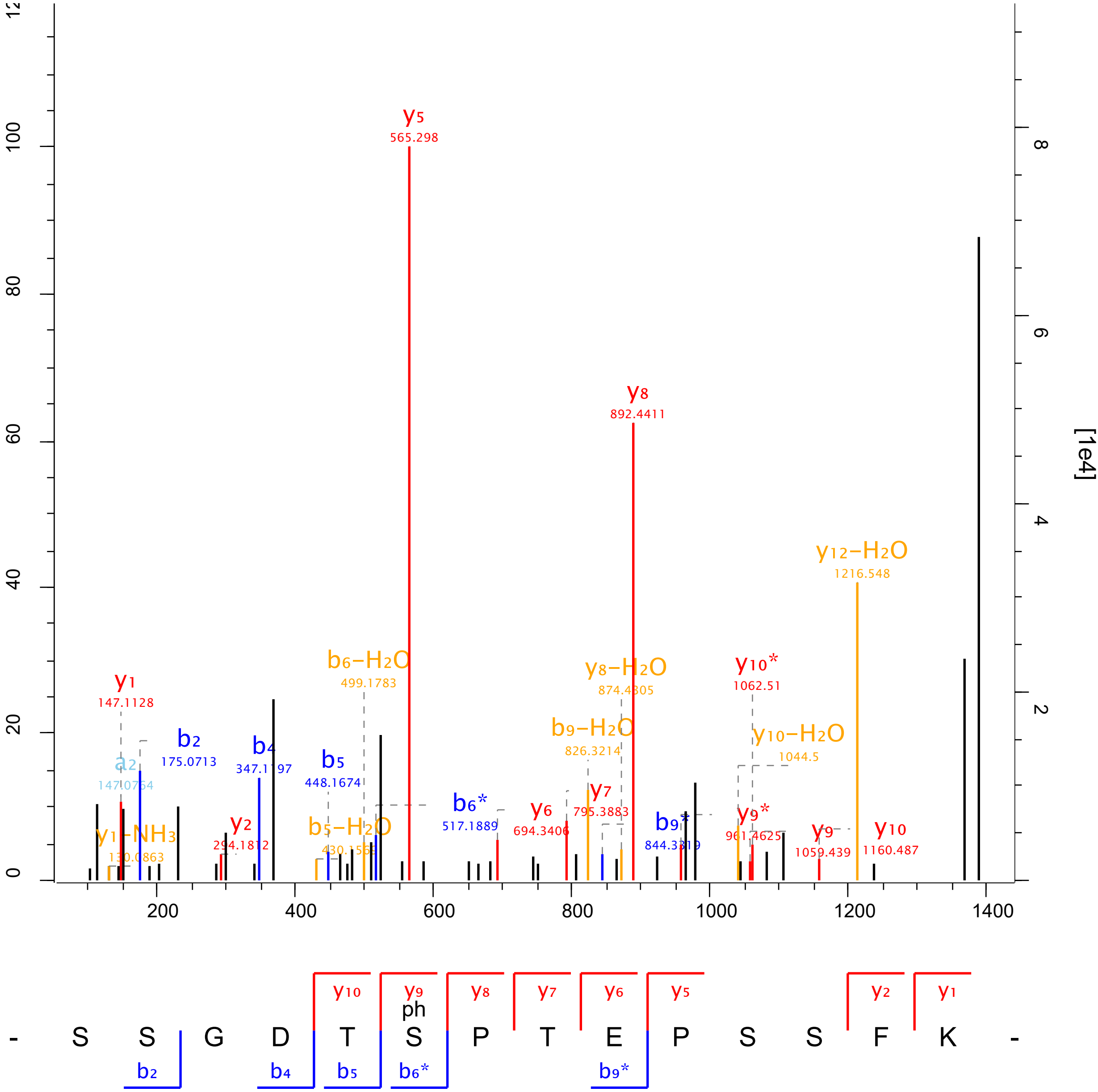

| Raw file | Scan | Method | Score | m/z |
|---|---|---|---|---|
| sys_15_1 | 10870 | FTMS; HCD | 133.91 | 519.22 |

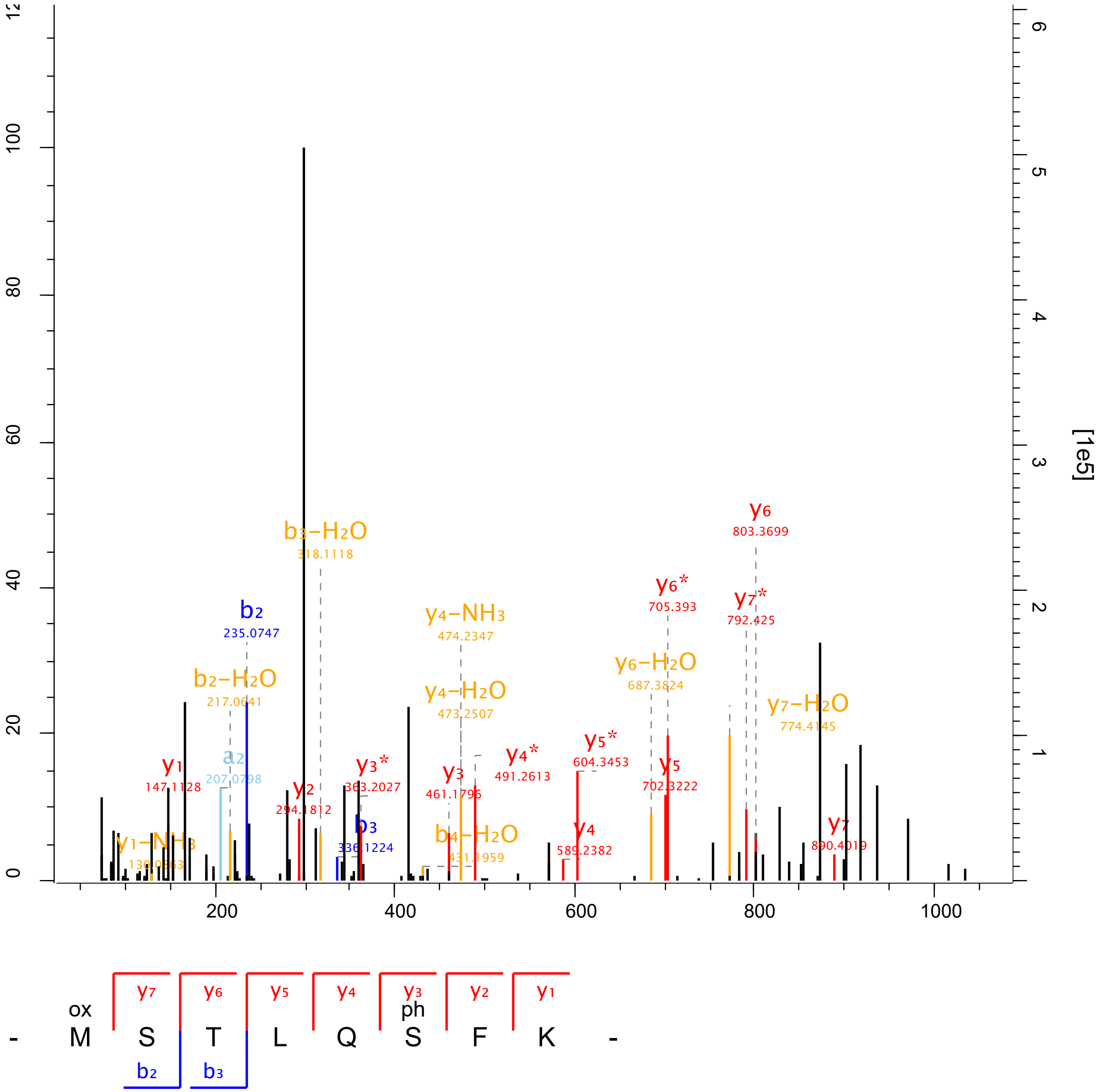

| Raw file | Scan | Method | Score | m/z |
|---|---|---|---|---|
| sys_15_1 | 10878 | FTMS; HCD | 139.74 | 573.76 |

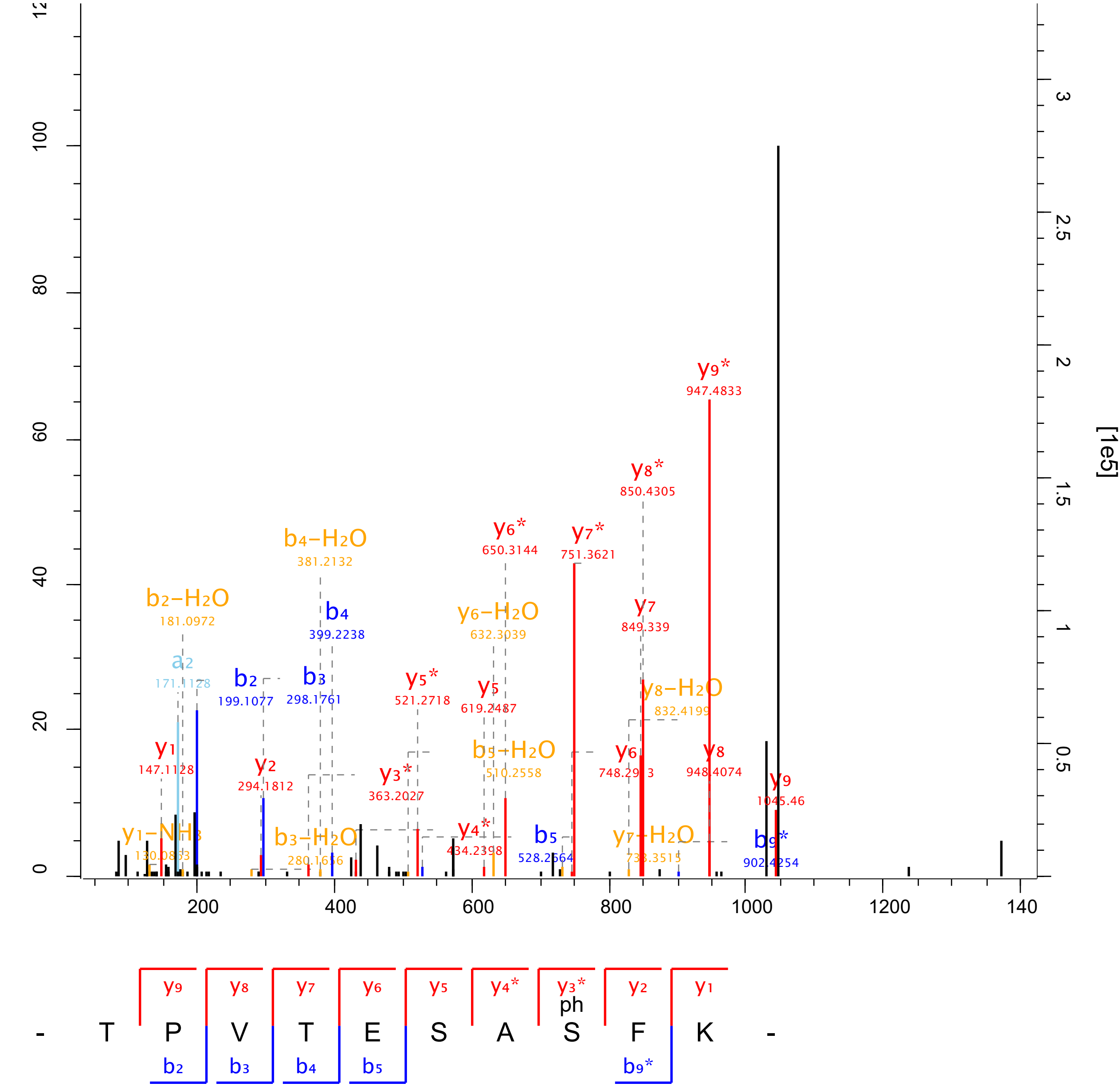

| Raw file | Scan | Method | Score | m/z |
|---|---|---|---|---|
| sys_15_1 | 10949 | FTMS; HCD | 86.49 | 678.81 |

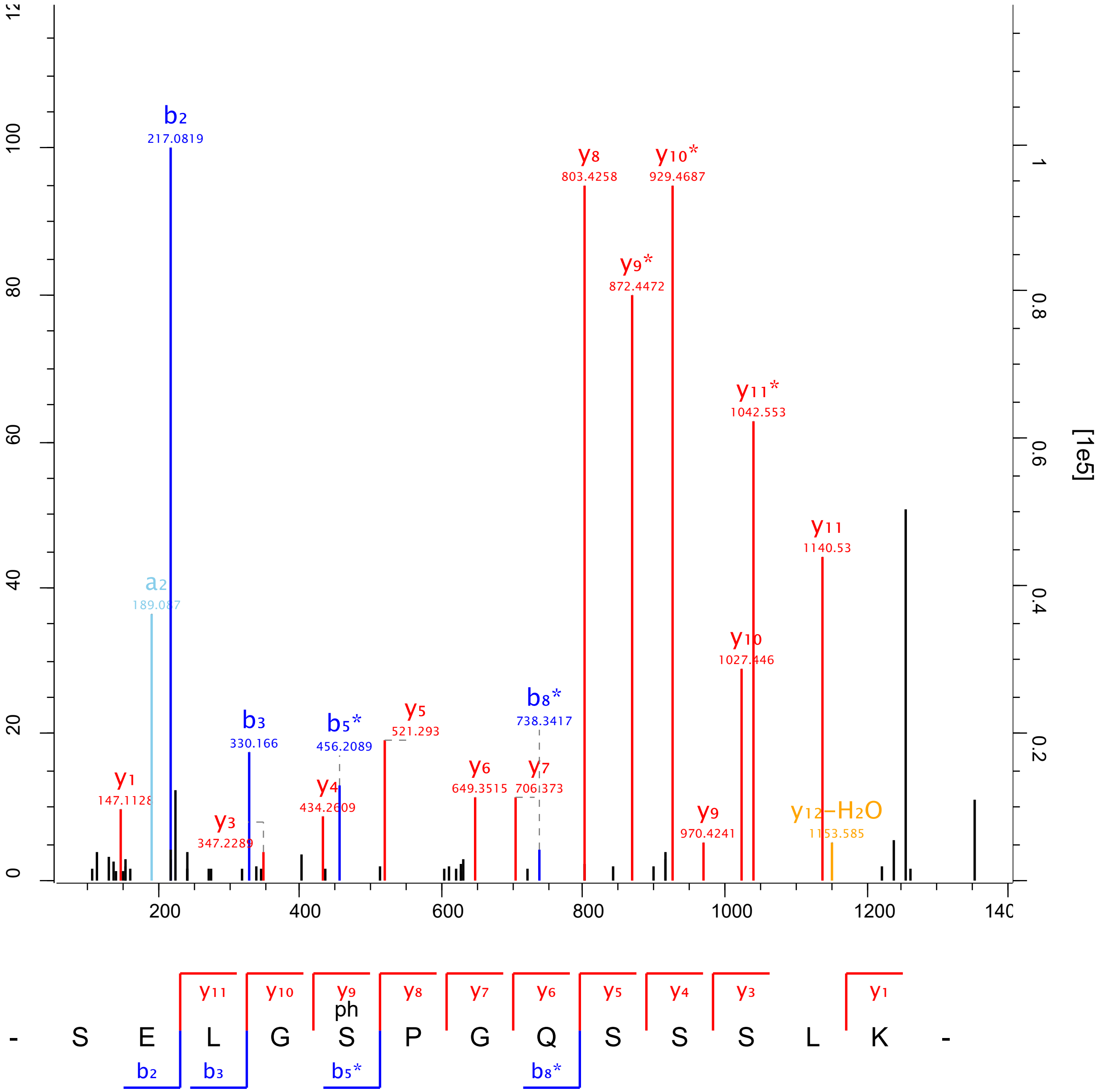

| Raw file | Scan | Method | Score | m/z |
|---|---|---|---|---|
| sys_15_1 | 10962 | FTMS; HCD | 97.21 | 658.77 |

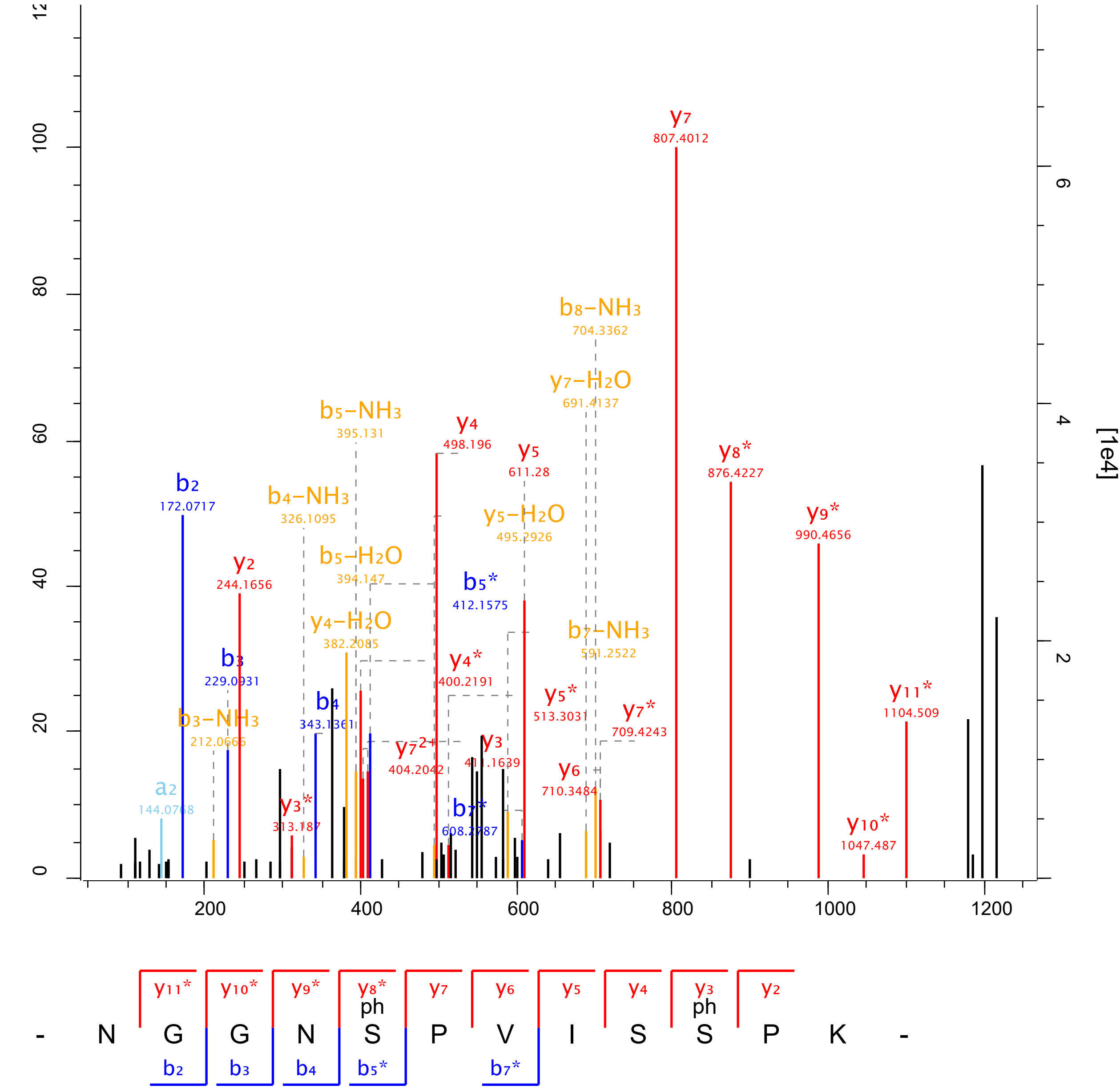

| Raw file | Scan | Method | Score | m/z |
|---|---|---|---|---|
| sys_15_1 | 11046 | FTMS; HCD | 91.59 | 753.82 |

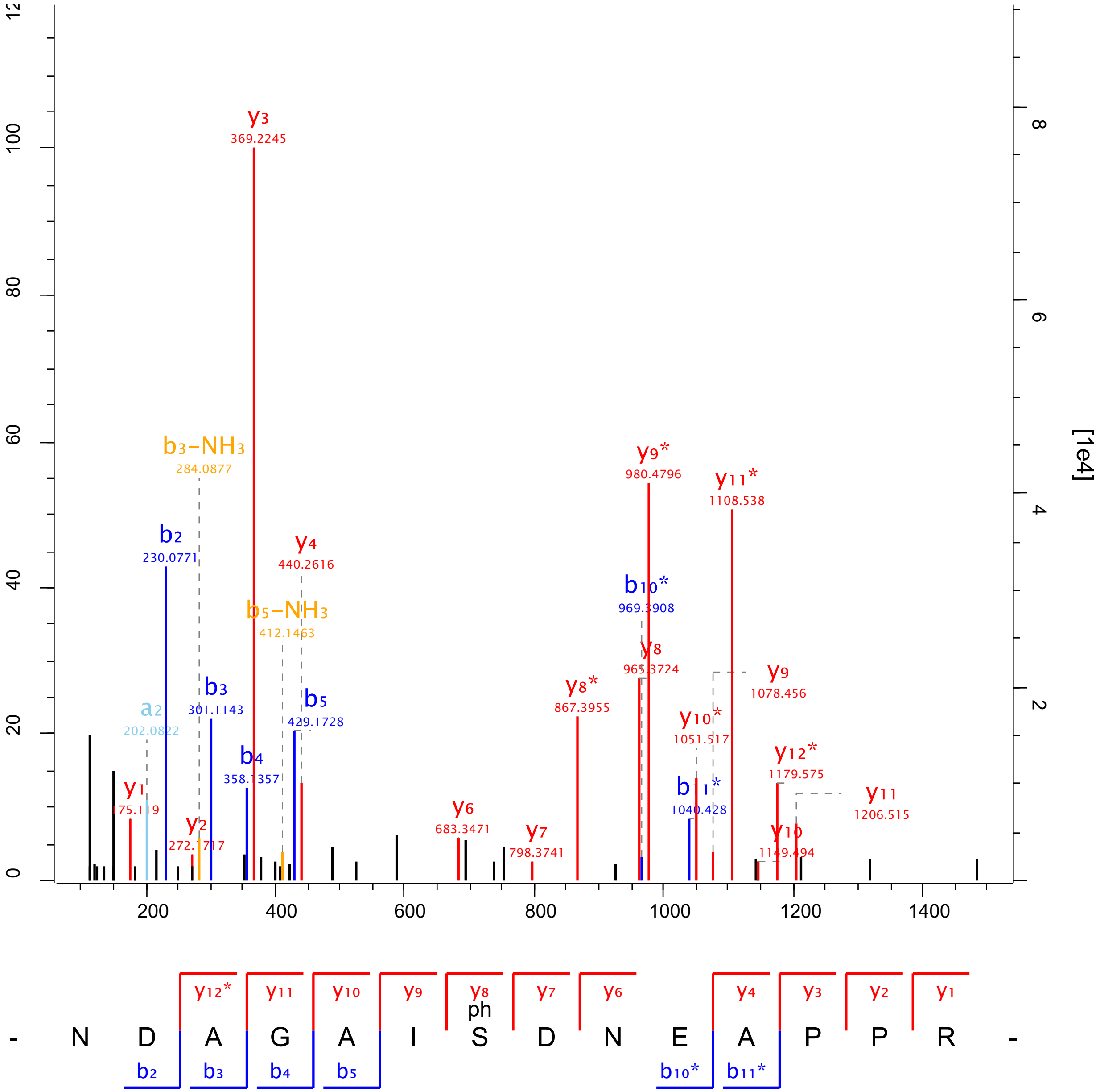

| Raw file | Scan | Method | Score | m/z |
| --- | --- | --- | --- | --- |
| sys_15_1 | 11068 | FTMS; HCD | 136.33 | 608.24 |

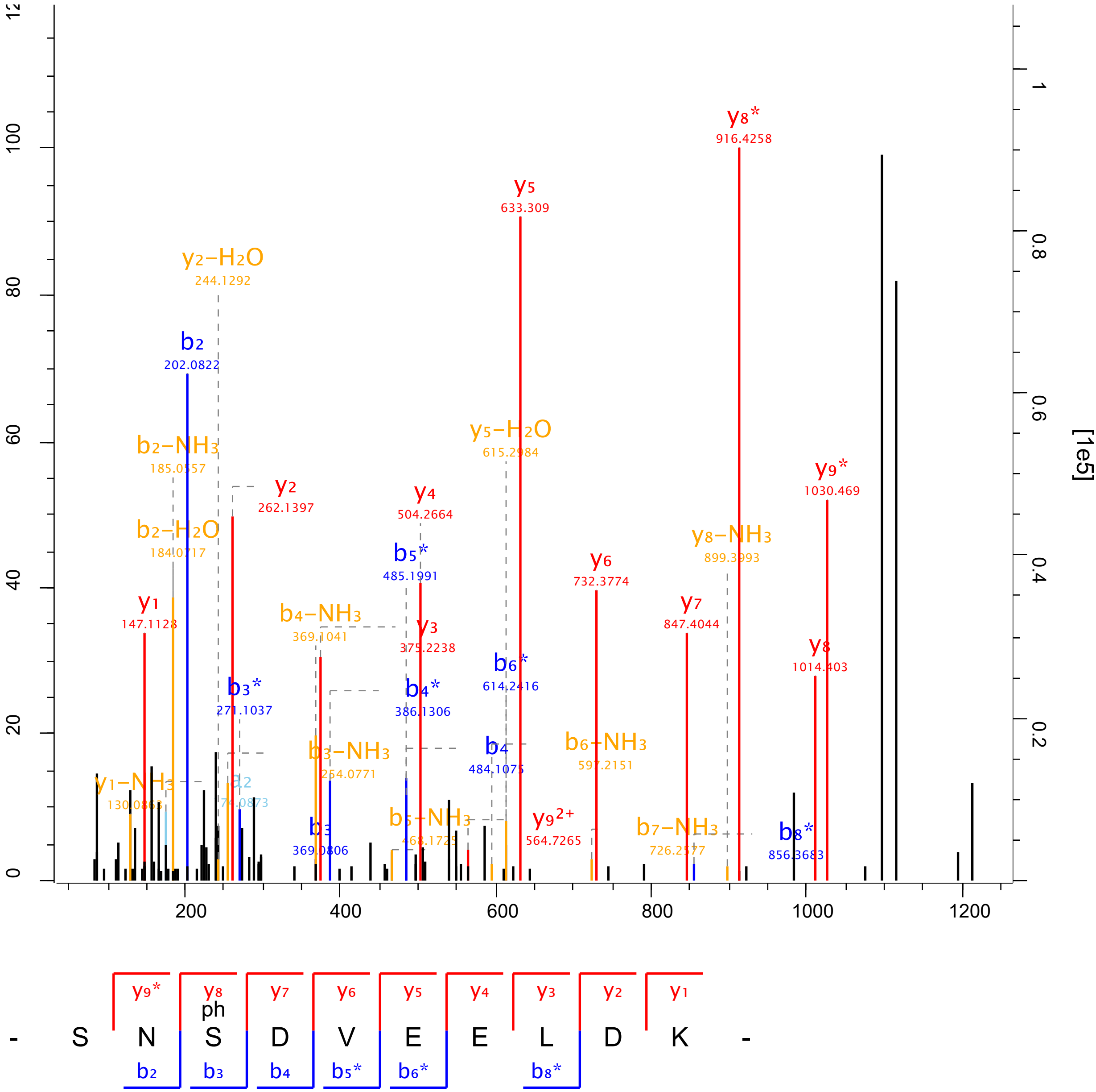

| Raw file | Scan | Method | Score | m/z |
| --- | --- | --- | --- | --- |
| sys_15_1 | 11078 | FTMS; HCD | 98.06 | 556.24 |

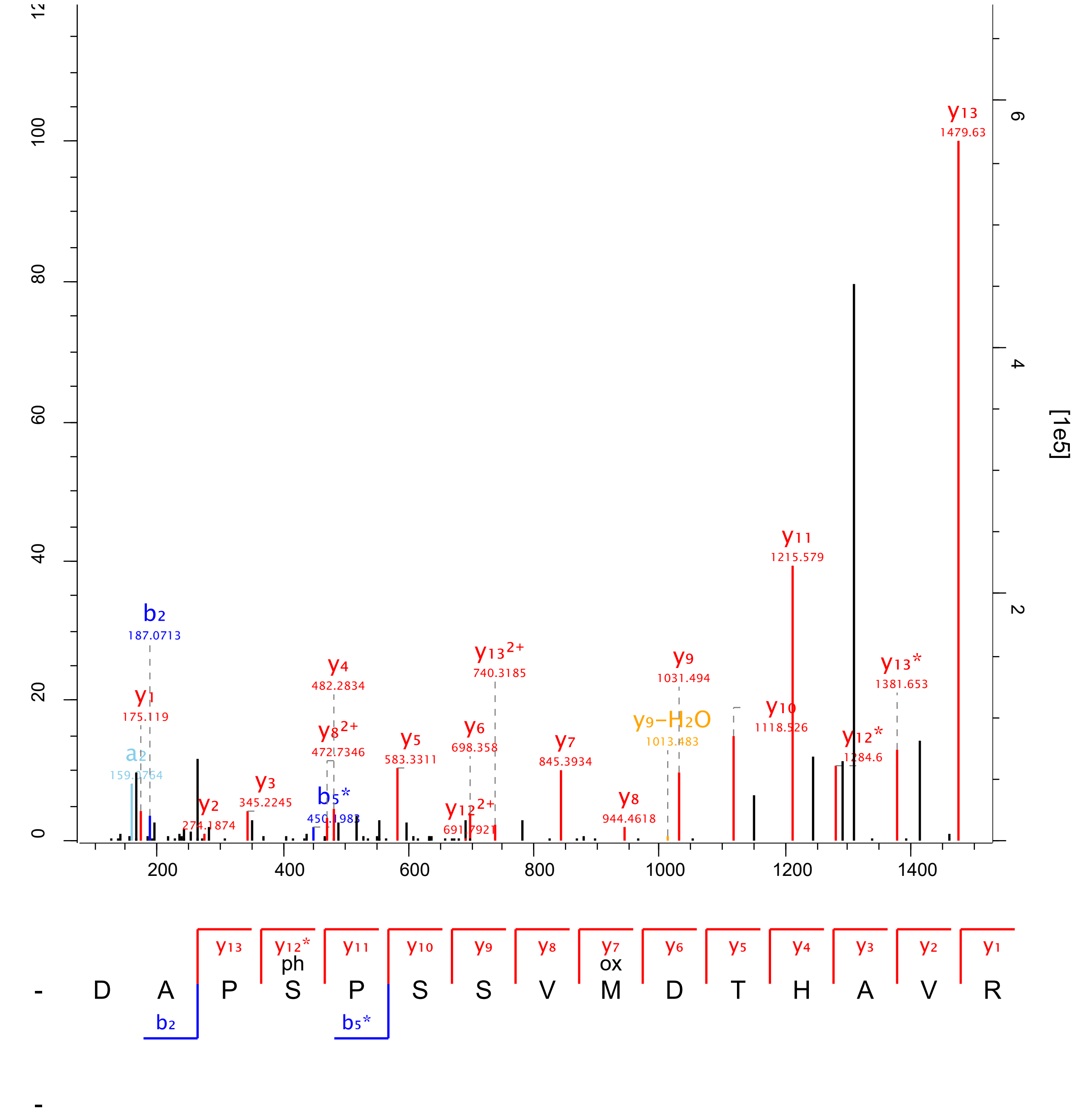

| Raw file | Scan | Method | Score | m/z |
|---|---|---|---|---|
| sys_15_1 | 11081 | FTMS; HCD | 186.56 | 681.81 |

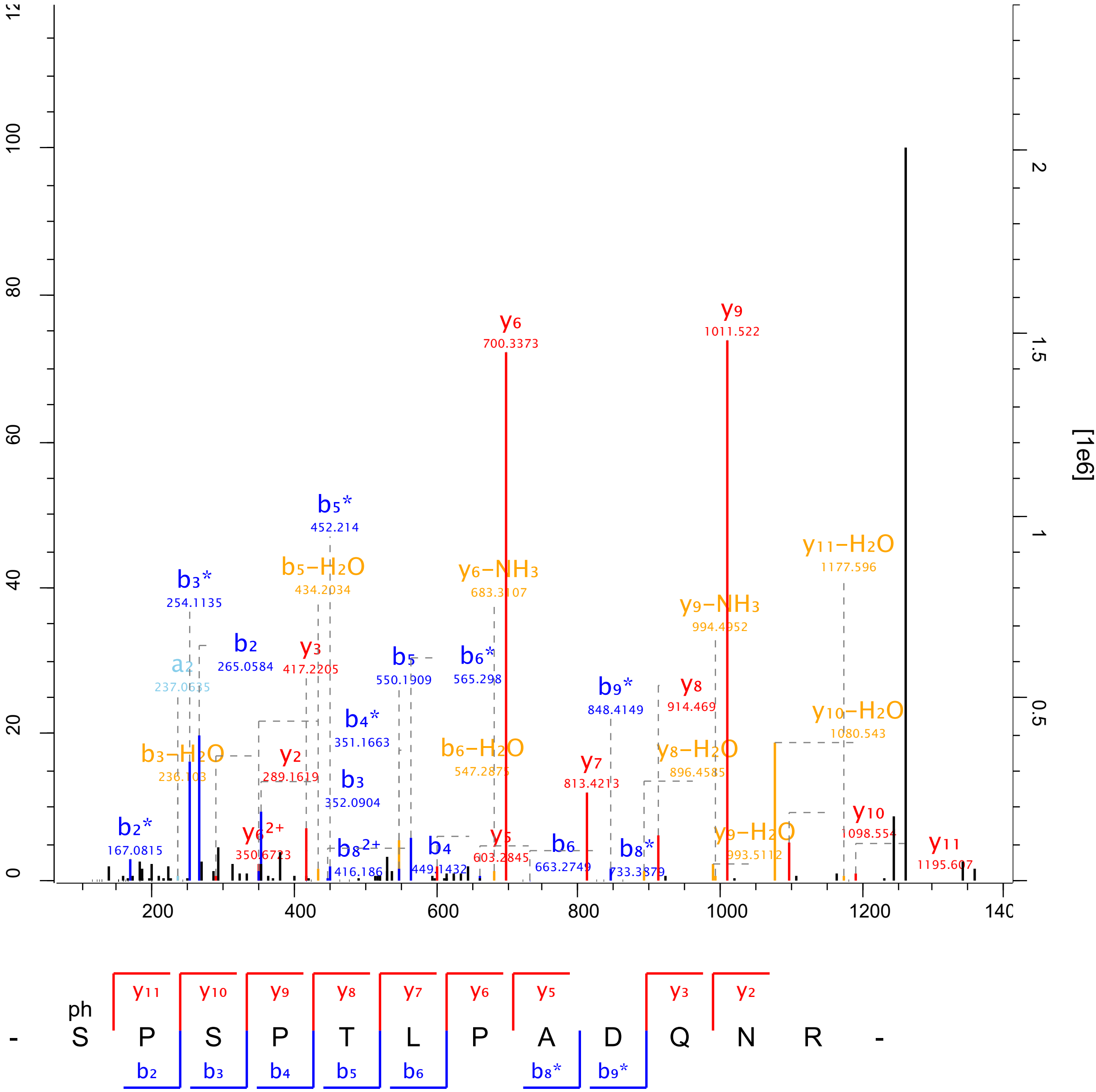

| Raw file | Scan | Method | Score | m/z |
|---|---|---|---|---|
| sys_15_1 | 11082 | FTMS; HCD | 78.9 | 720.33 |

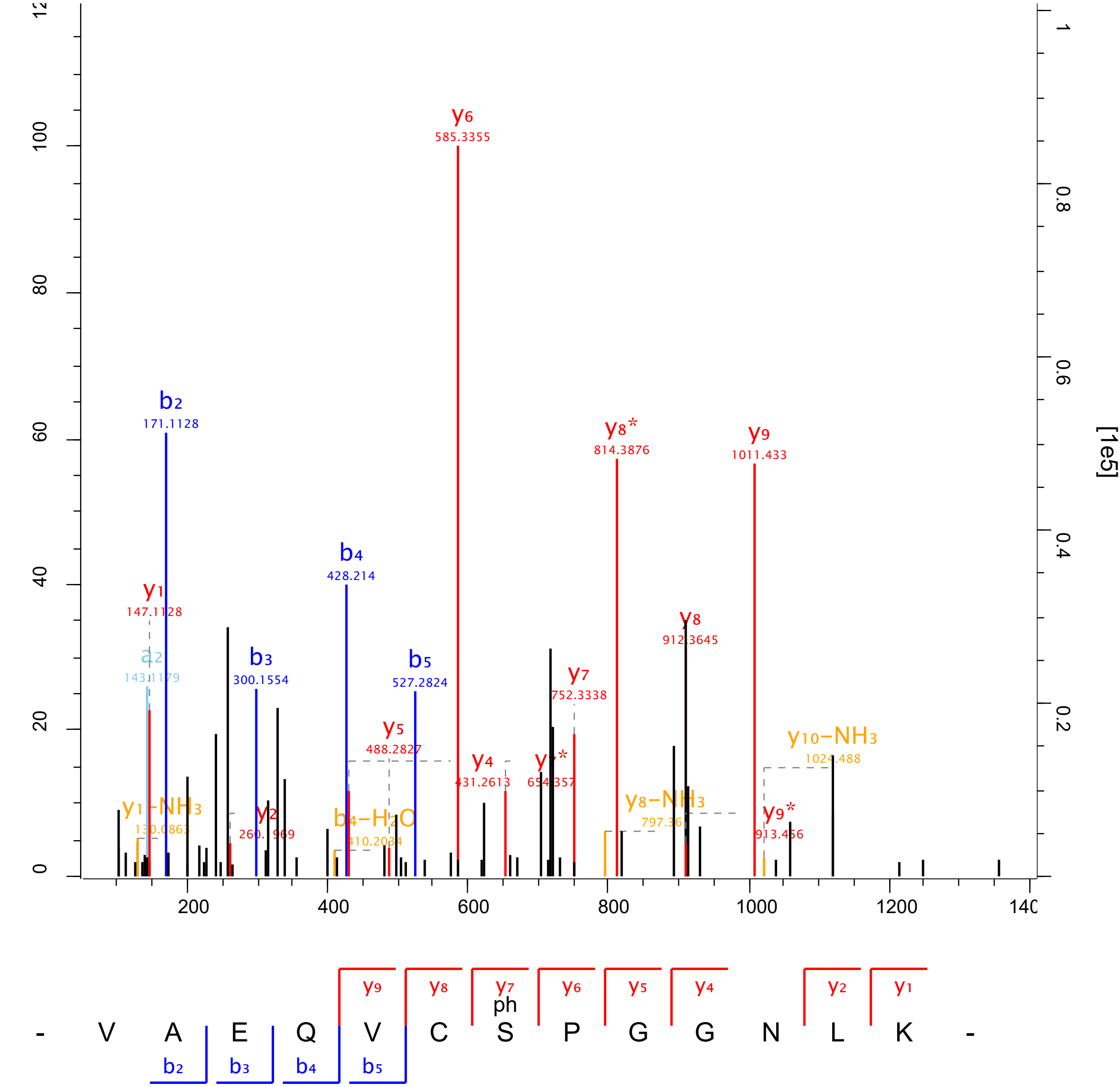

| Raw file | Scan | Method | Score | m/z |
|---|---|---|---|---|
| sys_15_1 | 11115 | FTMS; HCD | 58.78 | 495.22 |

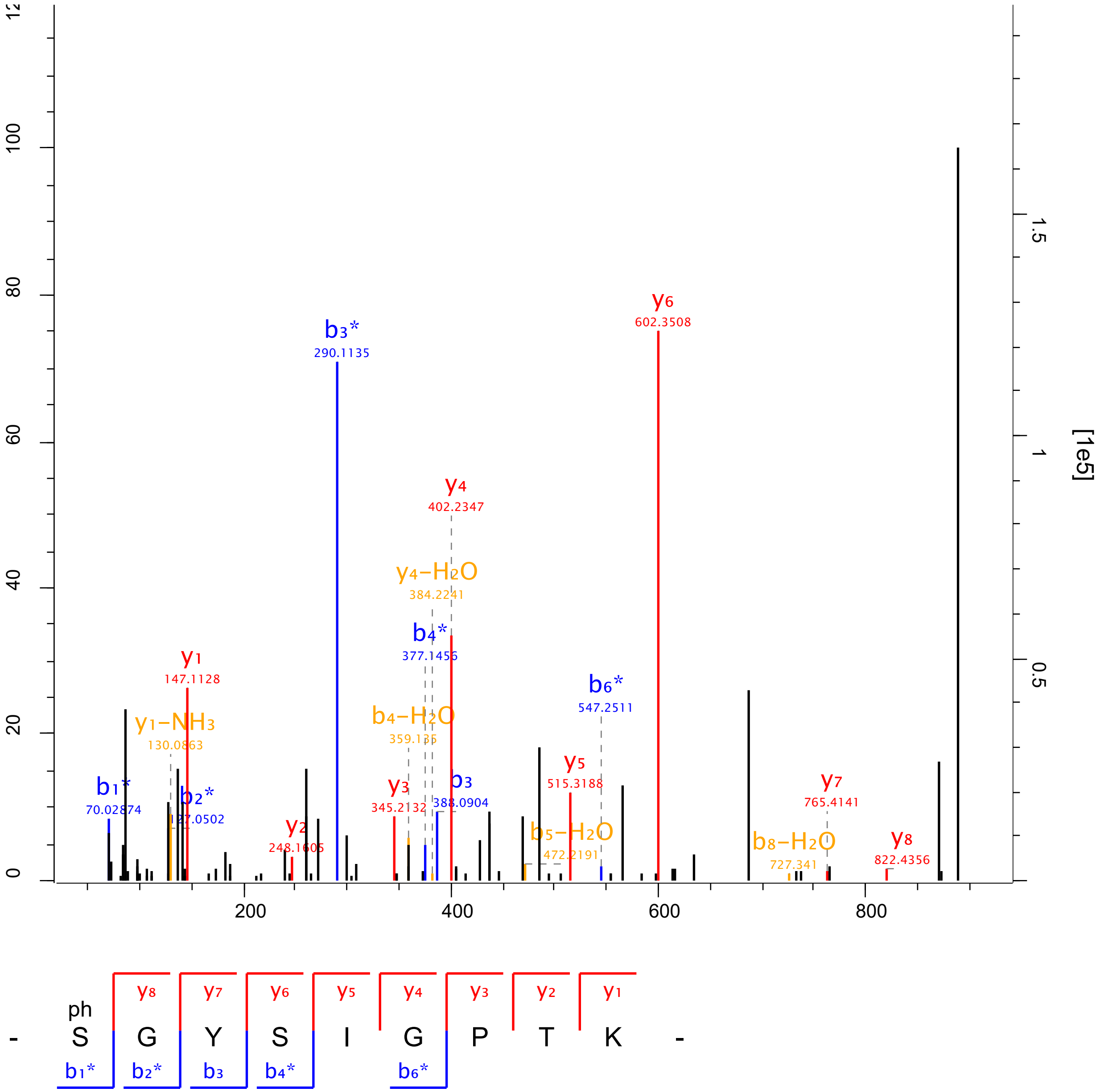

| Raw file | Scan | Method | Score | m/z |
|---|---|---|---|---|
| sys_15_1 | 11218 | FTMS; HCD | 89.23 | 650.27 |

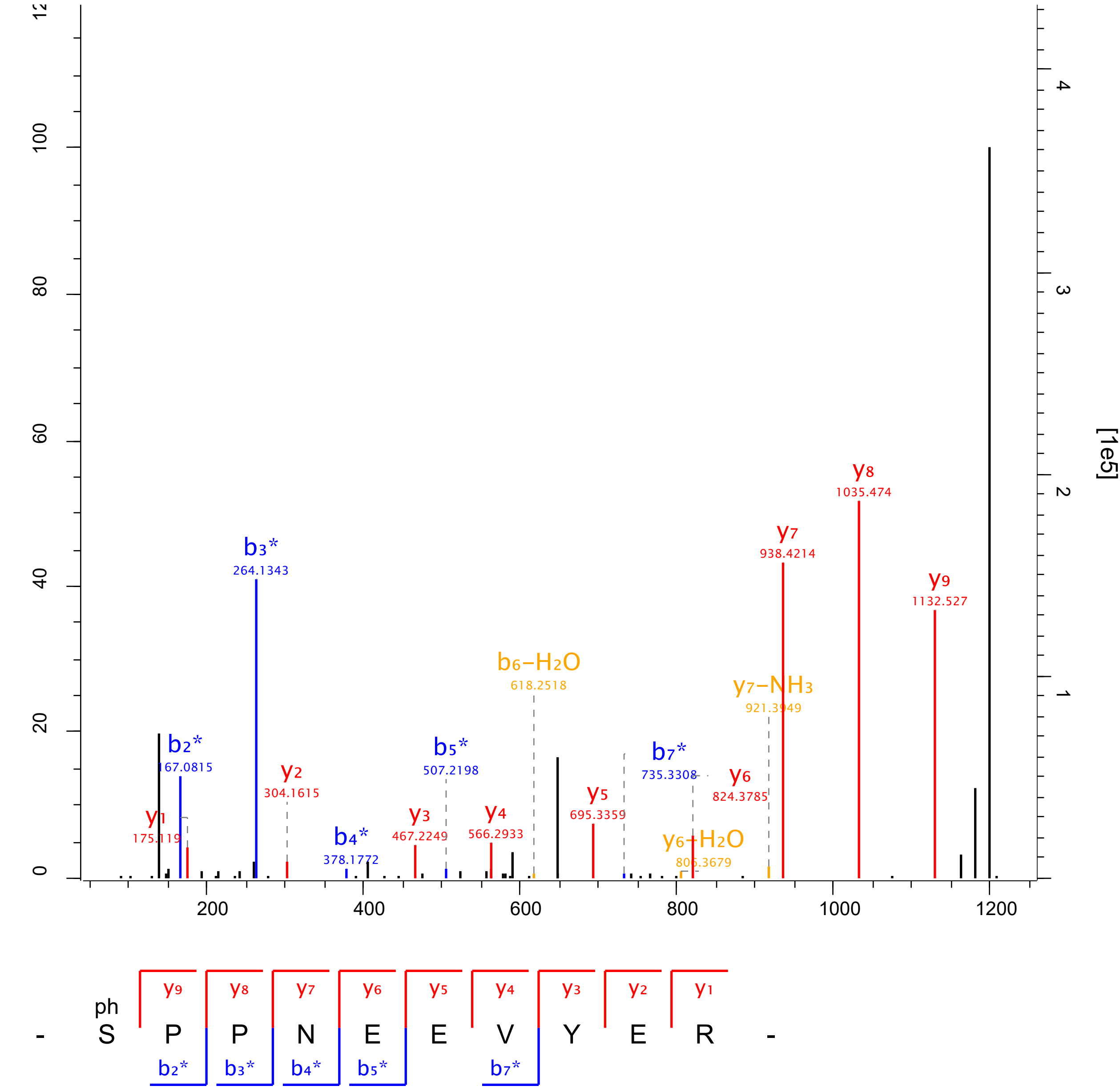

| Raw file | Scan | Method | Score | m/z |
|---|---|---|---|---|
| sys_15_1 | 11314 | FTMS; HCD | 93.35 | 575.74 |

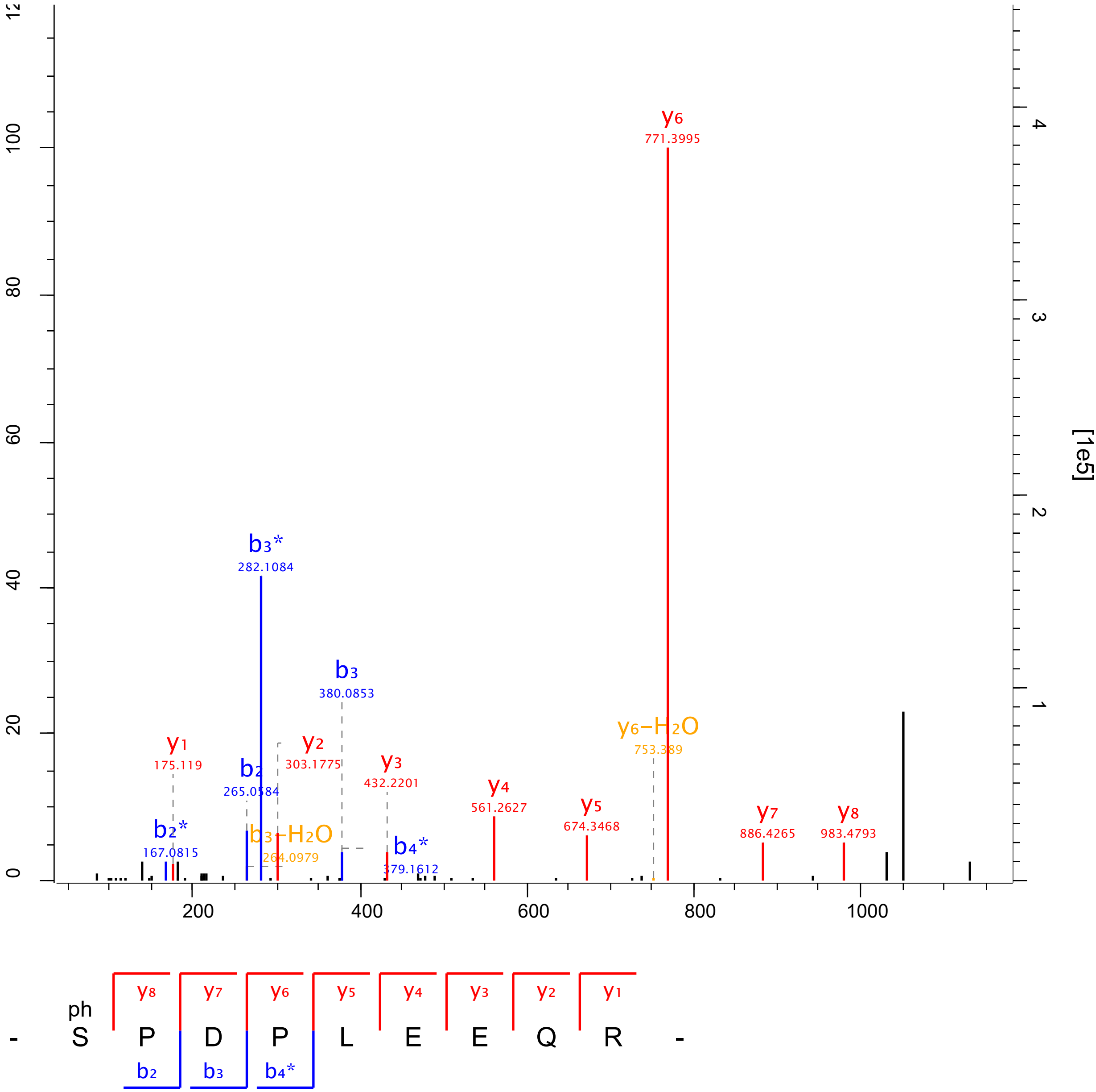

| Raw file | Scan | Method | Score | m/z |
|---|---|---|---|---|
| sys_15_1 | 11322 | FTMS; HCD | 76.36 | 627.78 |

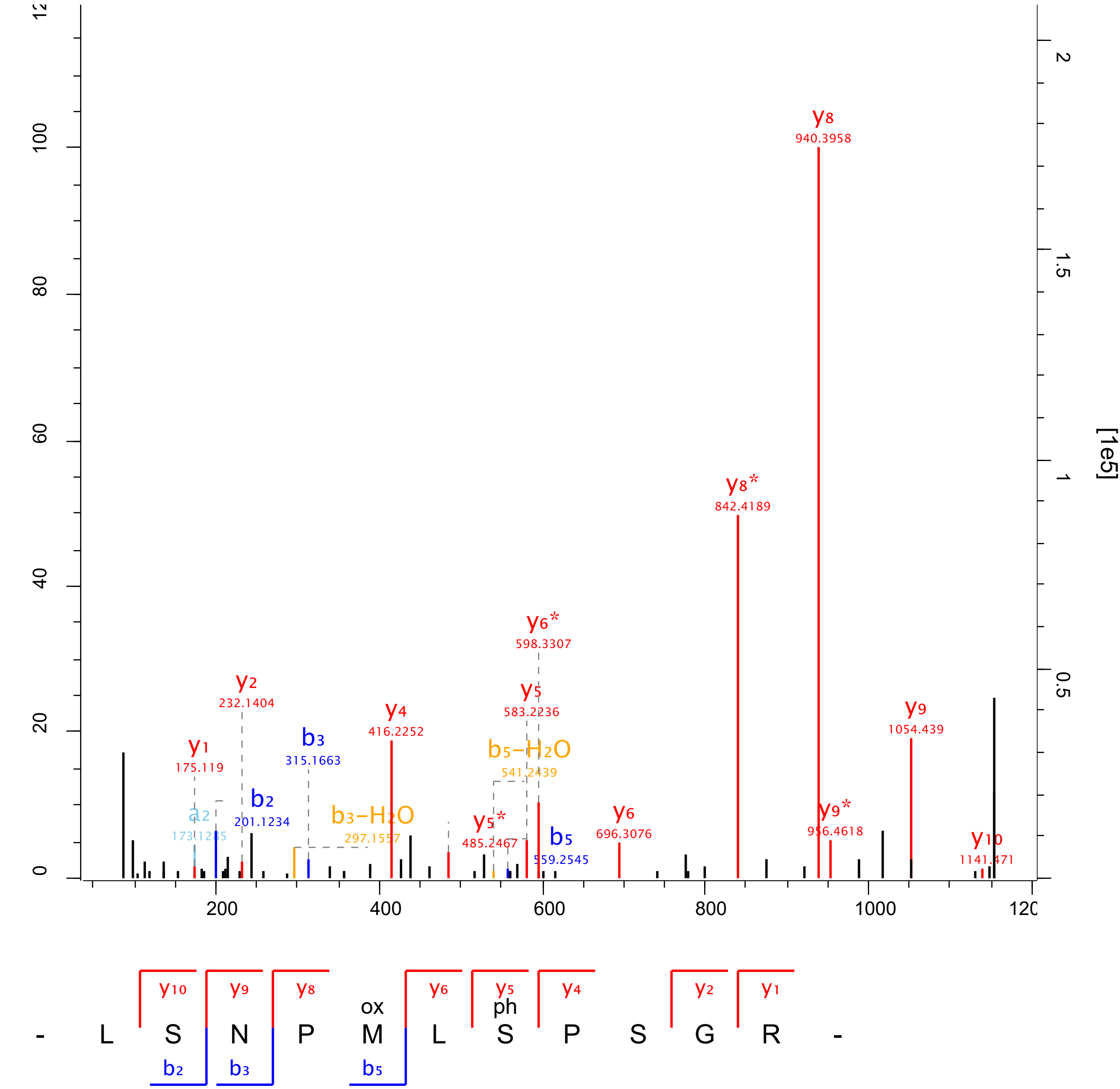

| Raw file | Scan | Method | Score | m/z |
|---|---|---|---|---|
| sys_15_1 | 11357 | FTMS; HCD | 53.45 | 748.33 |

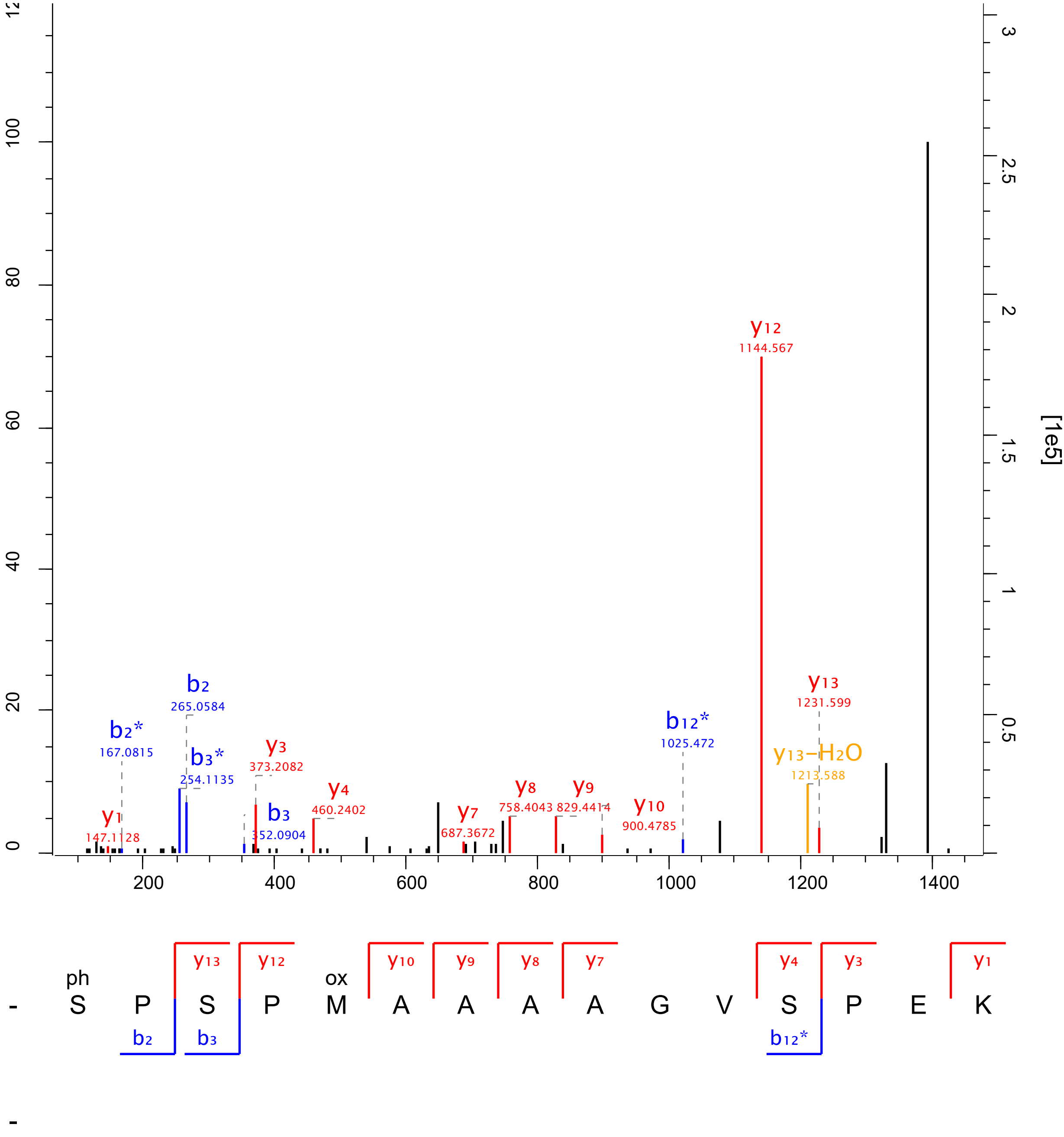

| Raw file | Scan | Method | Score | m/z |
|---|---|---|---|---|
| sys_15_1 | 11360 | FTMS; HCD | 51.77 | 727.3 |

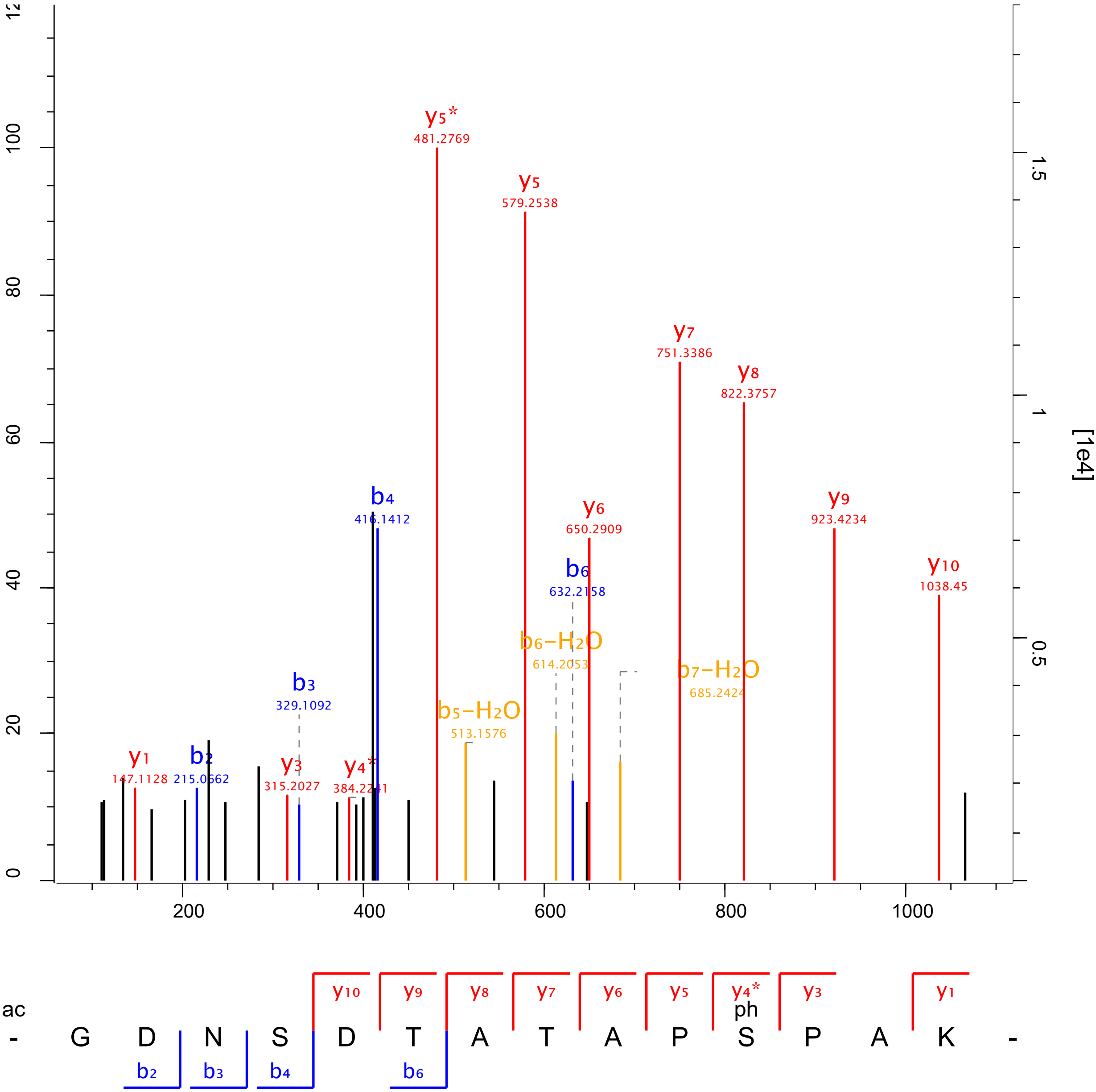

| Raw file | Scan | Method | Score | m/z |
| --- | --- | --- | --- | --- |
| sys_15_1 | 11429 | FTMS; HCD | 198.77 | 788.85 |

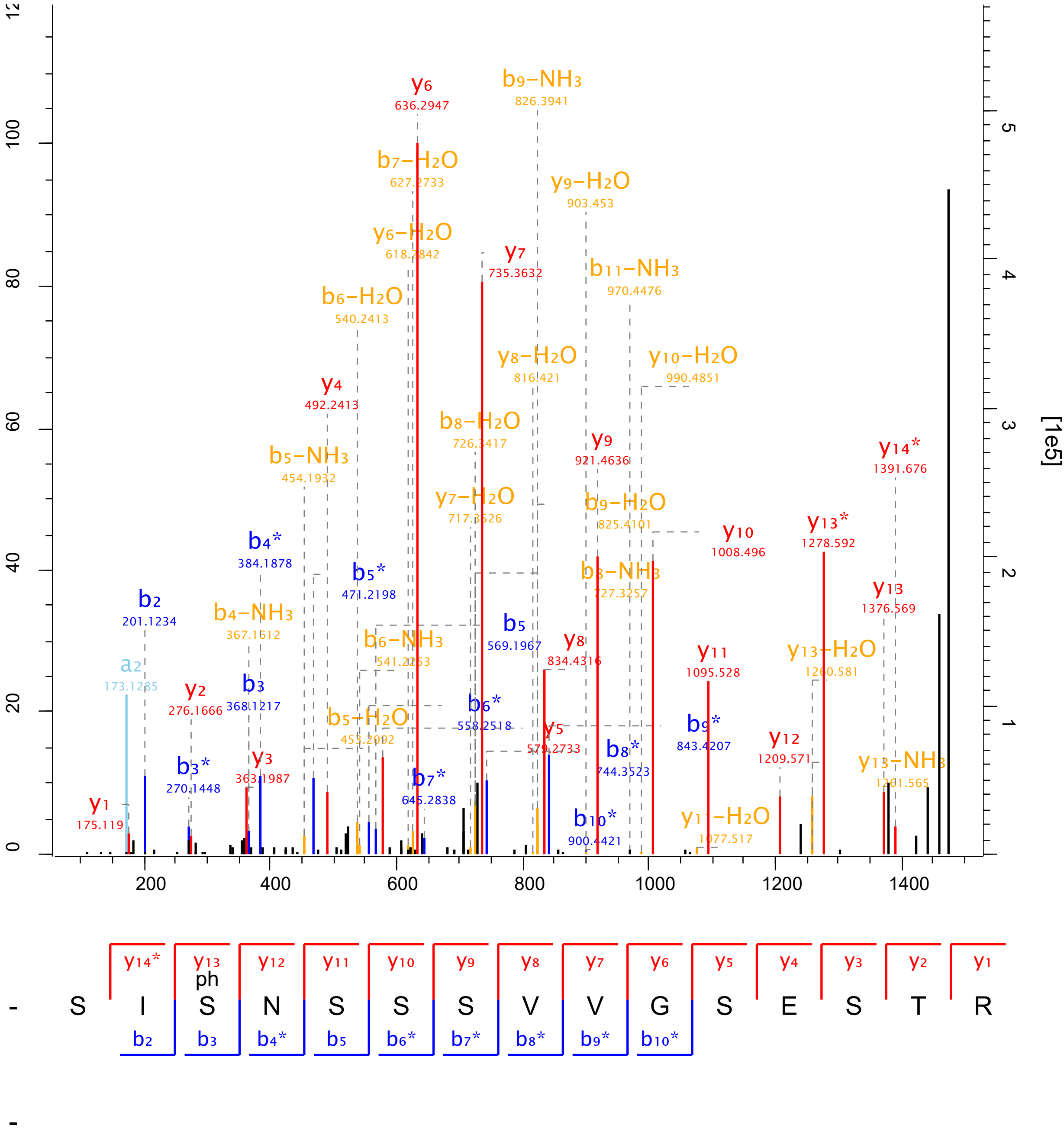

| Raw file | Scan | Method | Score | m/z |
| --- | --- | --- | --- | --- |
| sys_15_1 | 11688 | FTMS; HCD | 126.26 | 898.33 |

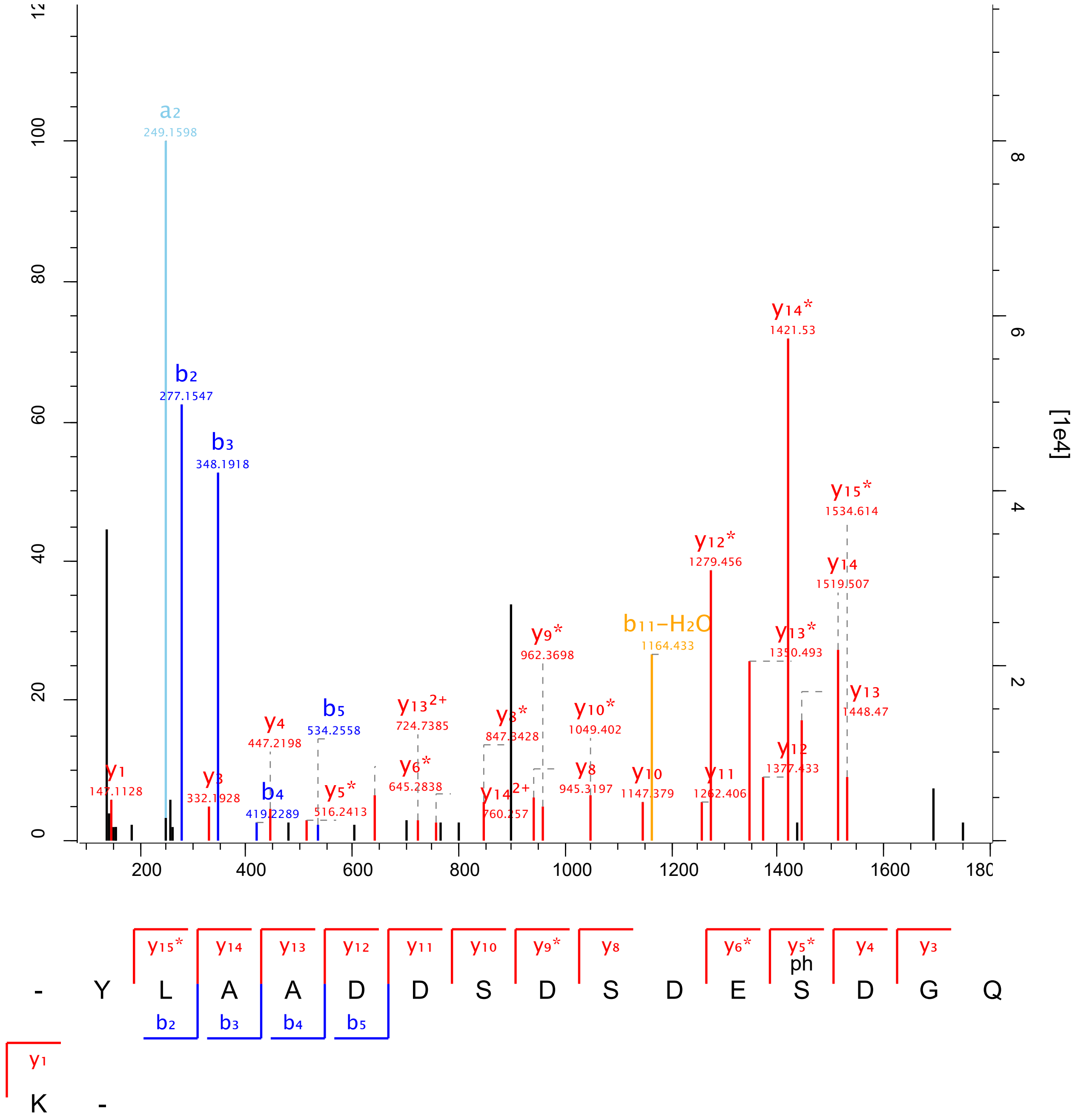

| Raw file | Scan | Method | Score | m/z |
|---|---|---|---|---|
| sys_15_1 | 11860 | FTMS; HCD | 107.09 | 617.77 |

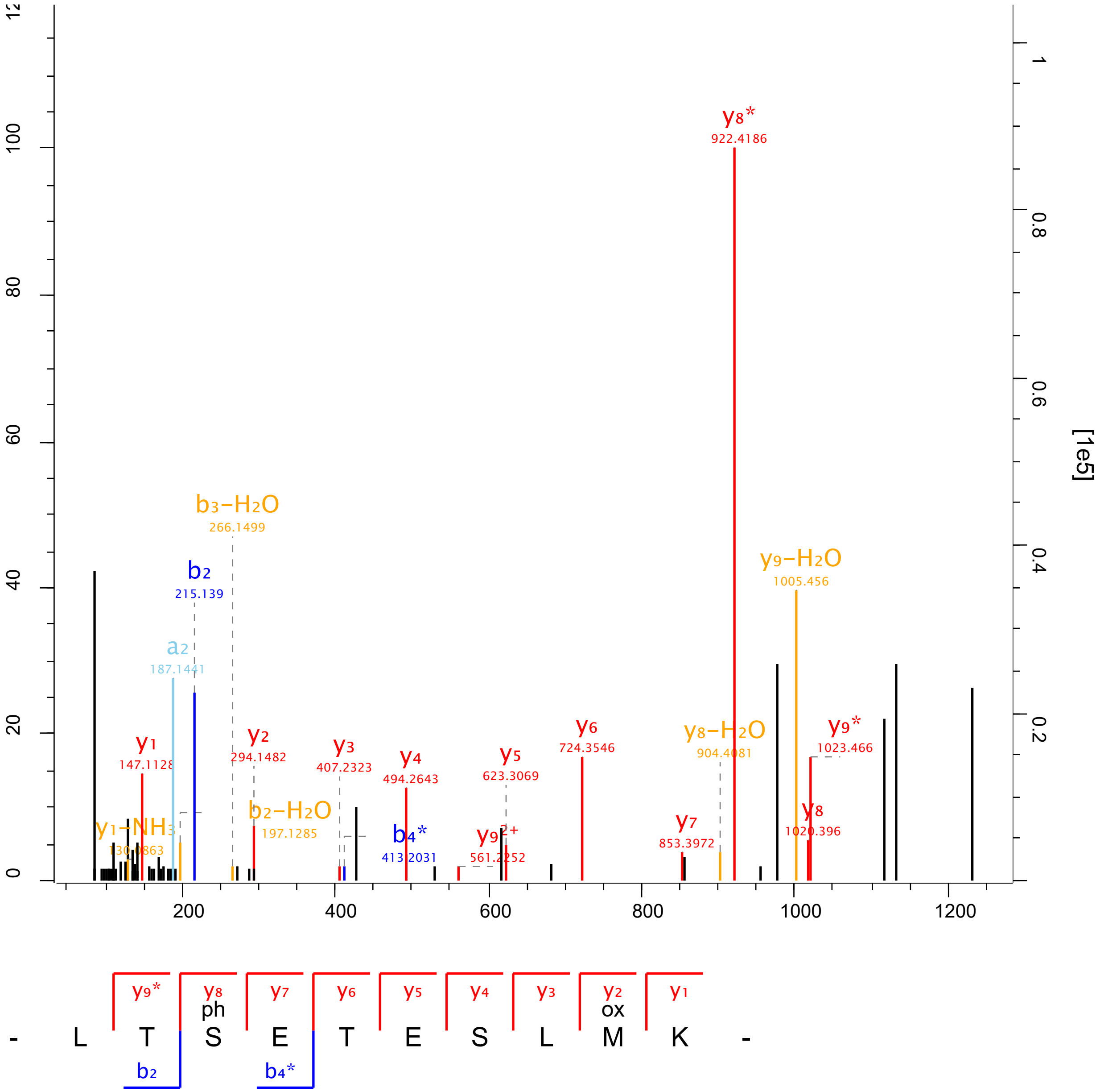

| Raw file | Scan | Method | Score | m/z |
|---|---|---|---|---|
| sys_15_1 | 11872 | FTMS; HCD | 269.61 | 790.84 |

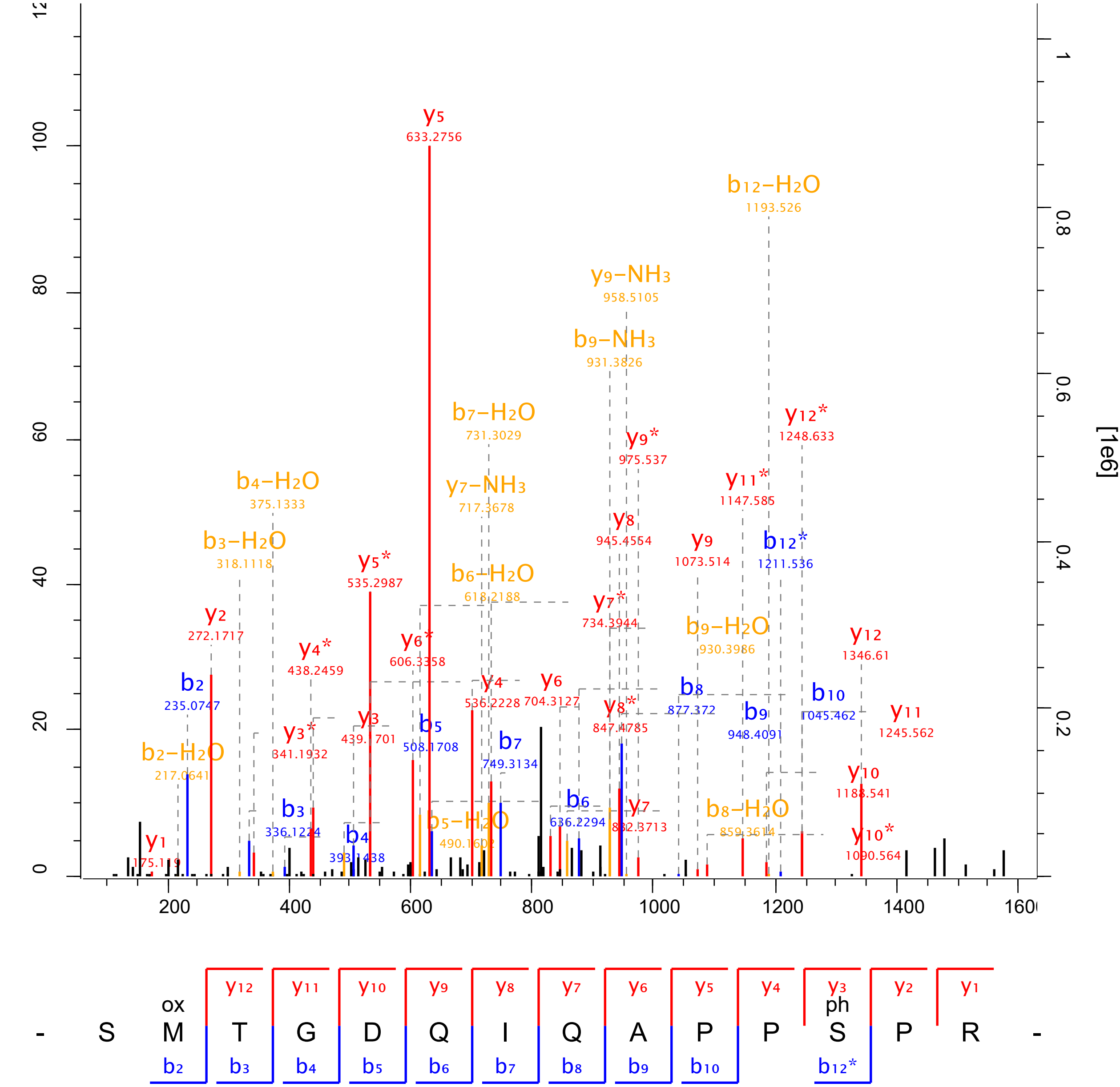

| Raw file | Scan | Method | Score | m/z |
| --- | --- | --- | --- | --- |
| sys_15_1 | 11873 | FTMS; HCD | 112.44 | 595.75 |

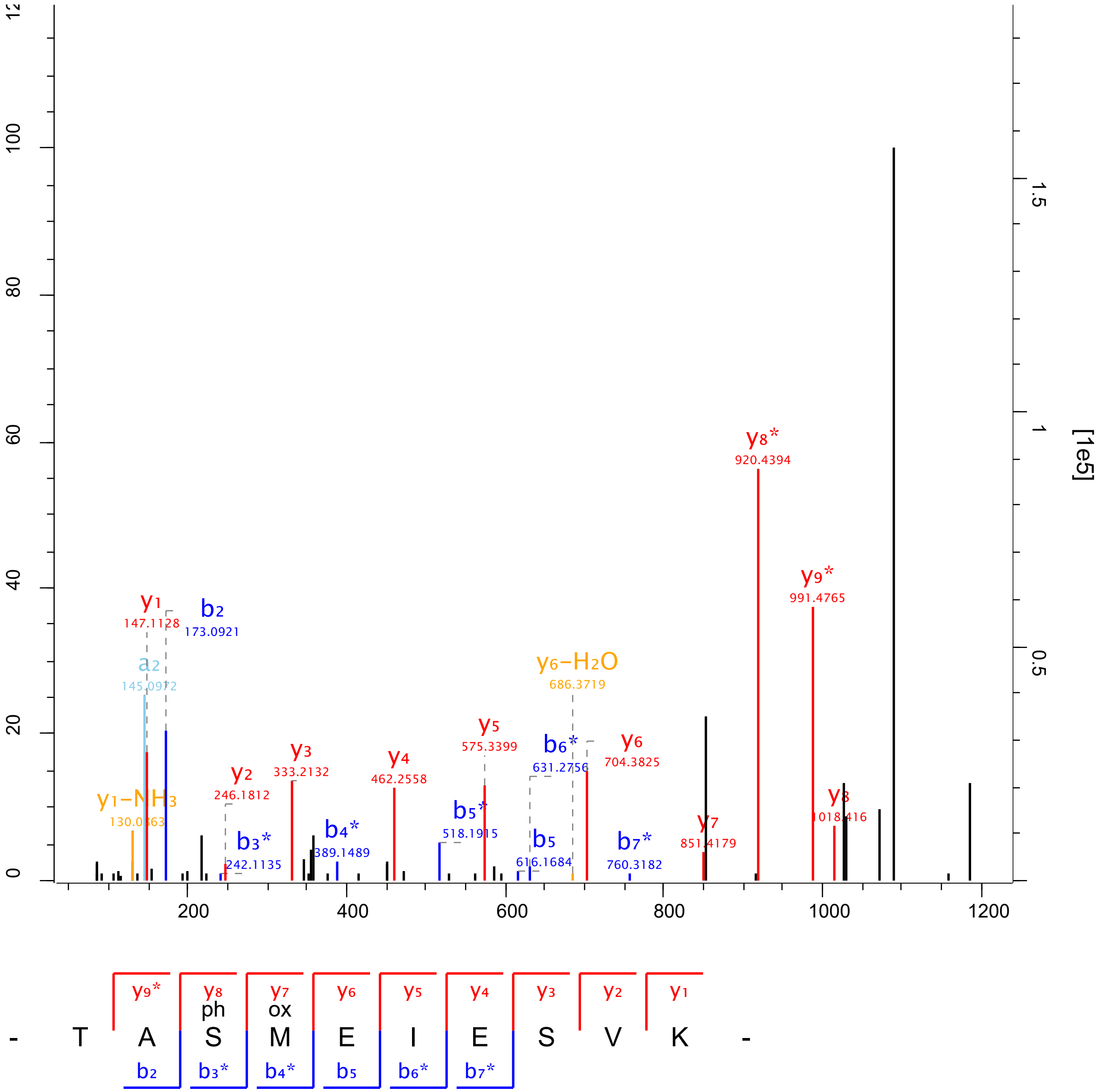

| Raw file | Scan | Method | Score | m/z |
|---|---|---|---|---|
| sys_15_1 | 11896 | FTMS; HCD | 251 | 829.35 |

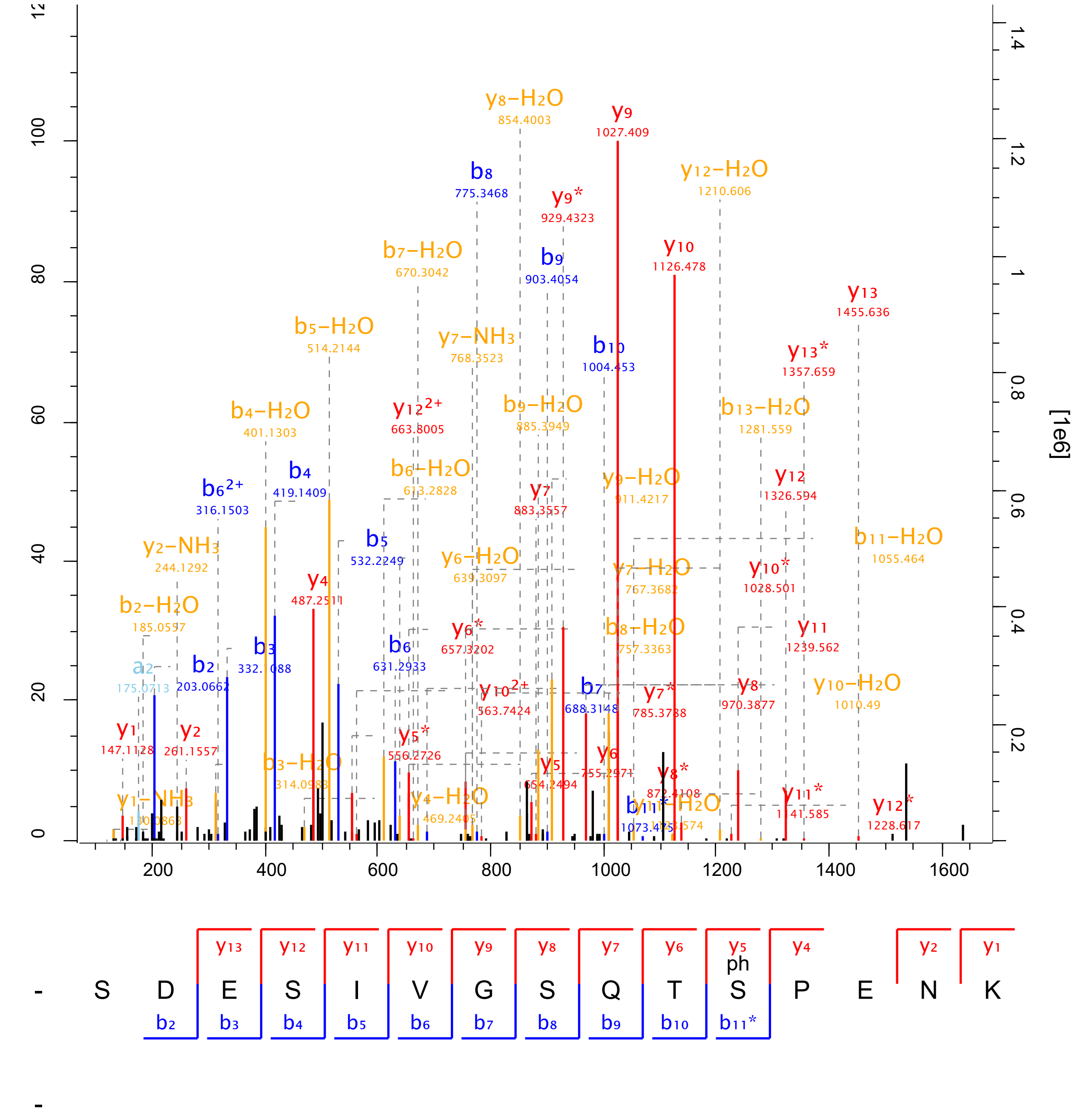

| Raw file | Scan | Method | Score | m/z |
| --- | --- | --- | --- | --- |
| sys_15_1 | 12078 | FTMS; HCD | 107.65 | 675.81 |

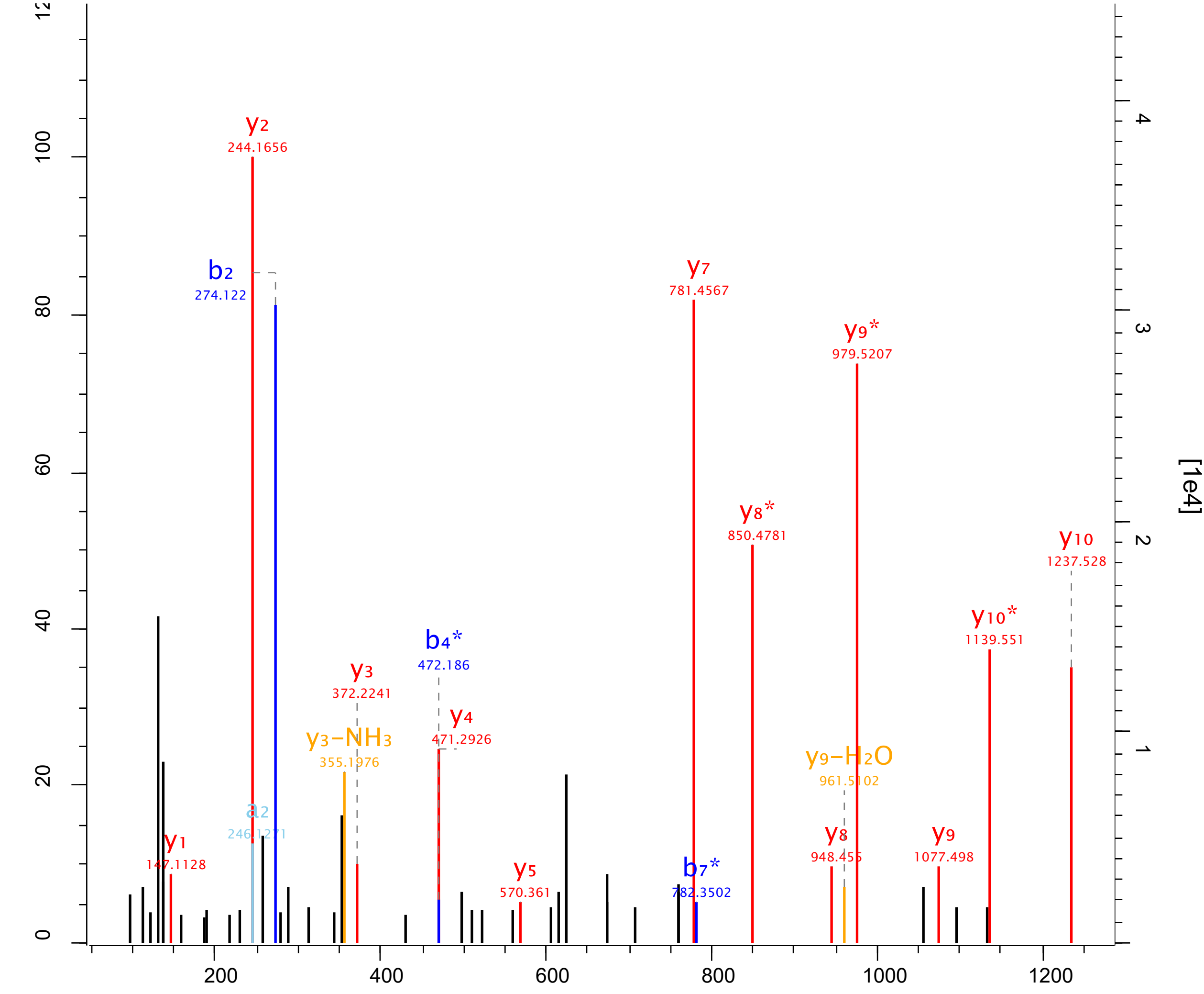


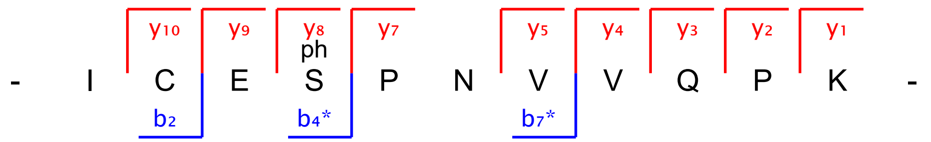

| Raw file | Scan | Method | Score | m/z |
|---|---|---|---|---|
| sys_15_1 | 12117 | FTMS; HCD | 190.1 | 693.81 |

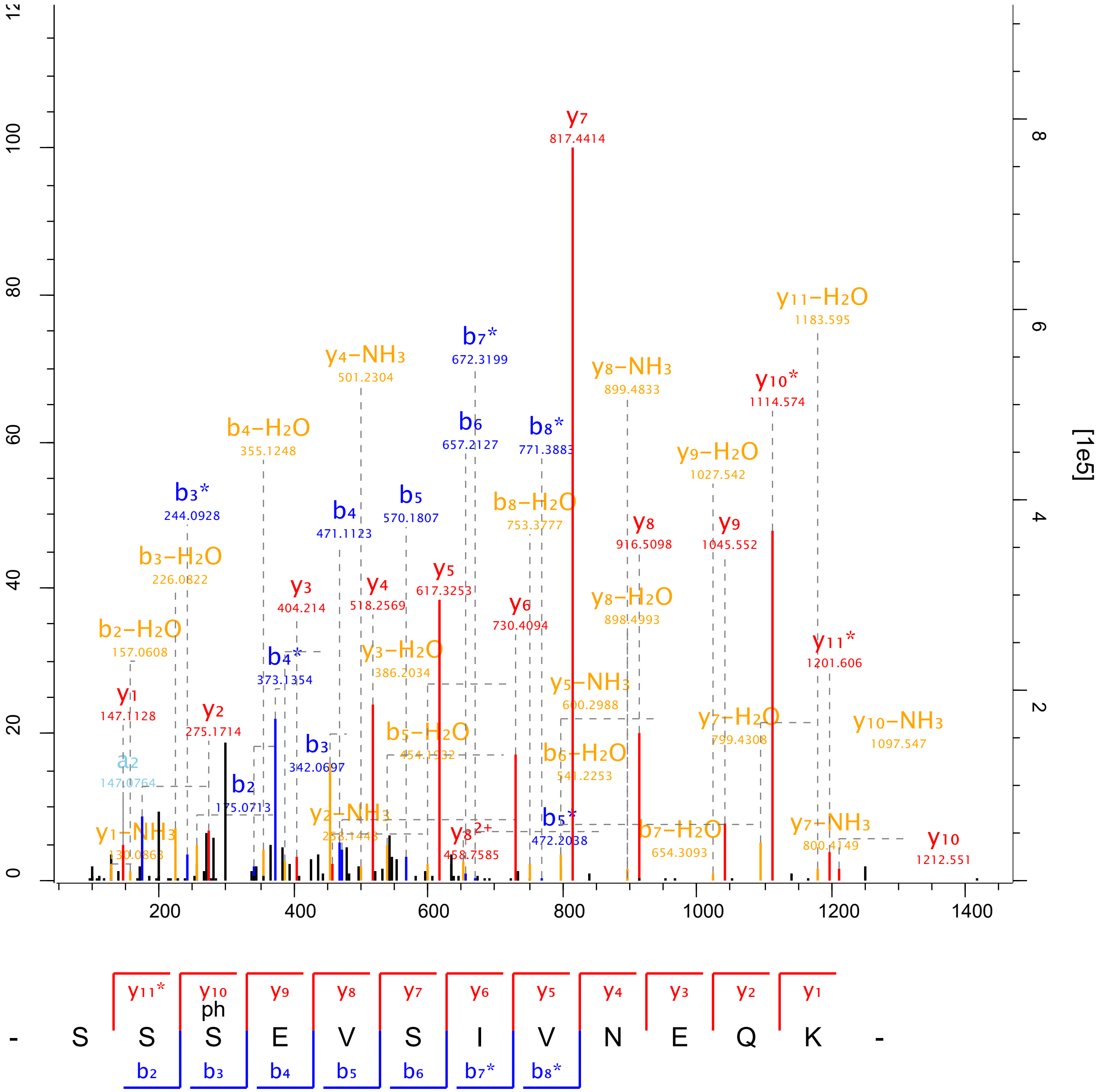

| Raw file | Scan | Method | Score | m/z |
|---|---|---|---|---|
| sys_15_1 | 12148 | FTMS; HCD | 116.19 | 556.26 |

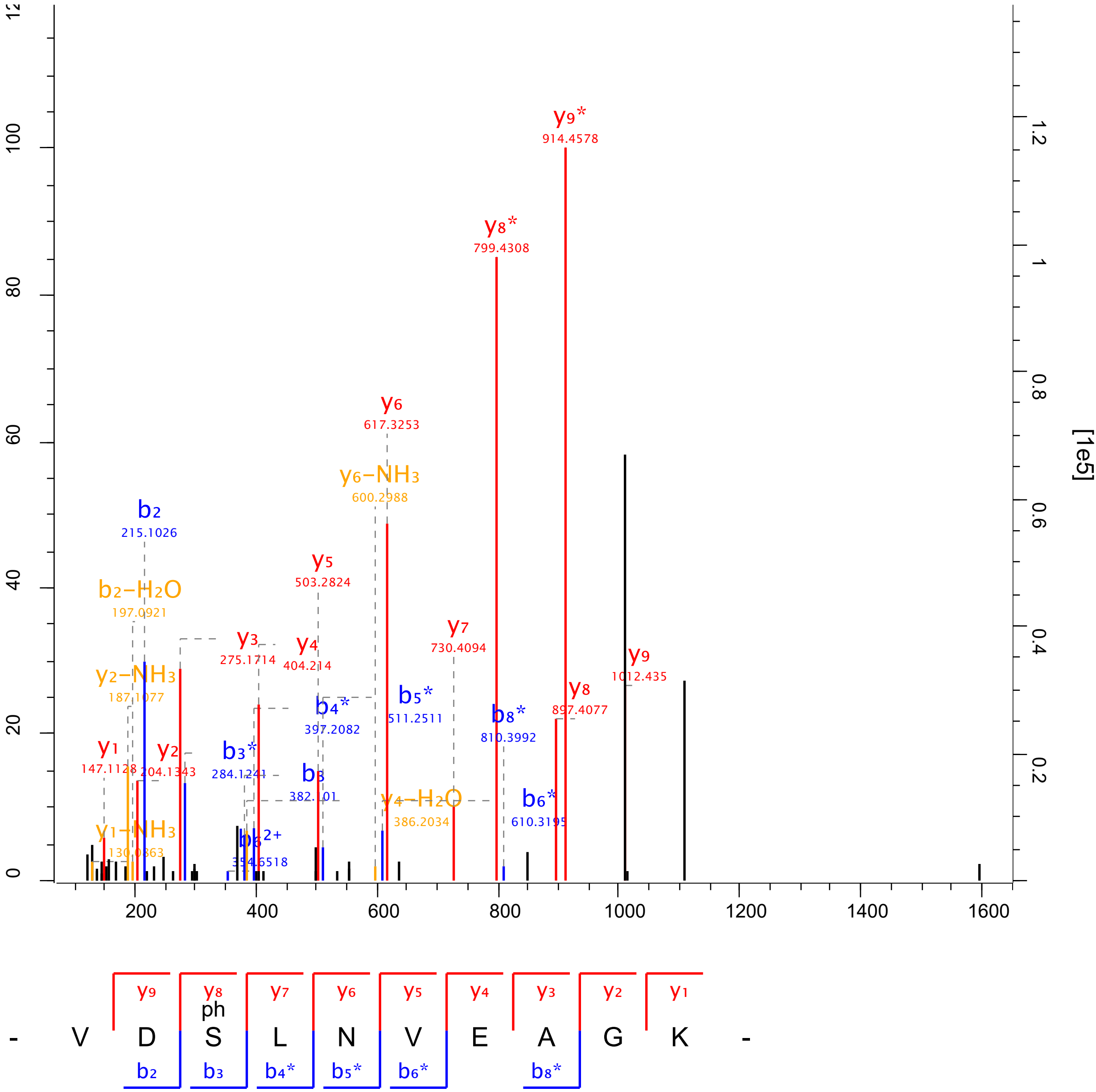

| Raw file | Scan | Method | Score | m/z |
|---|---|---|---|---|
| sys_15_1 | 12158 | FTMS; HCD | 99.28 | 455.72 |

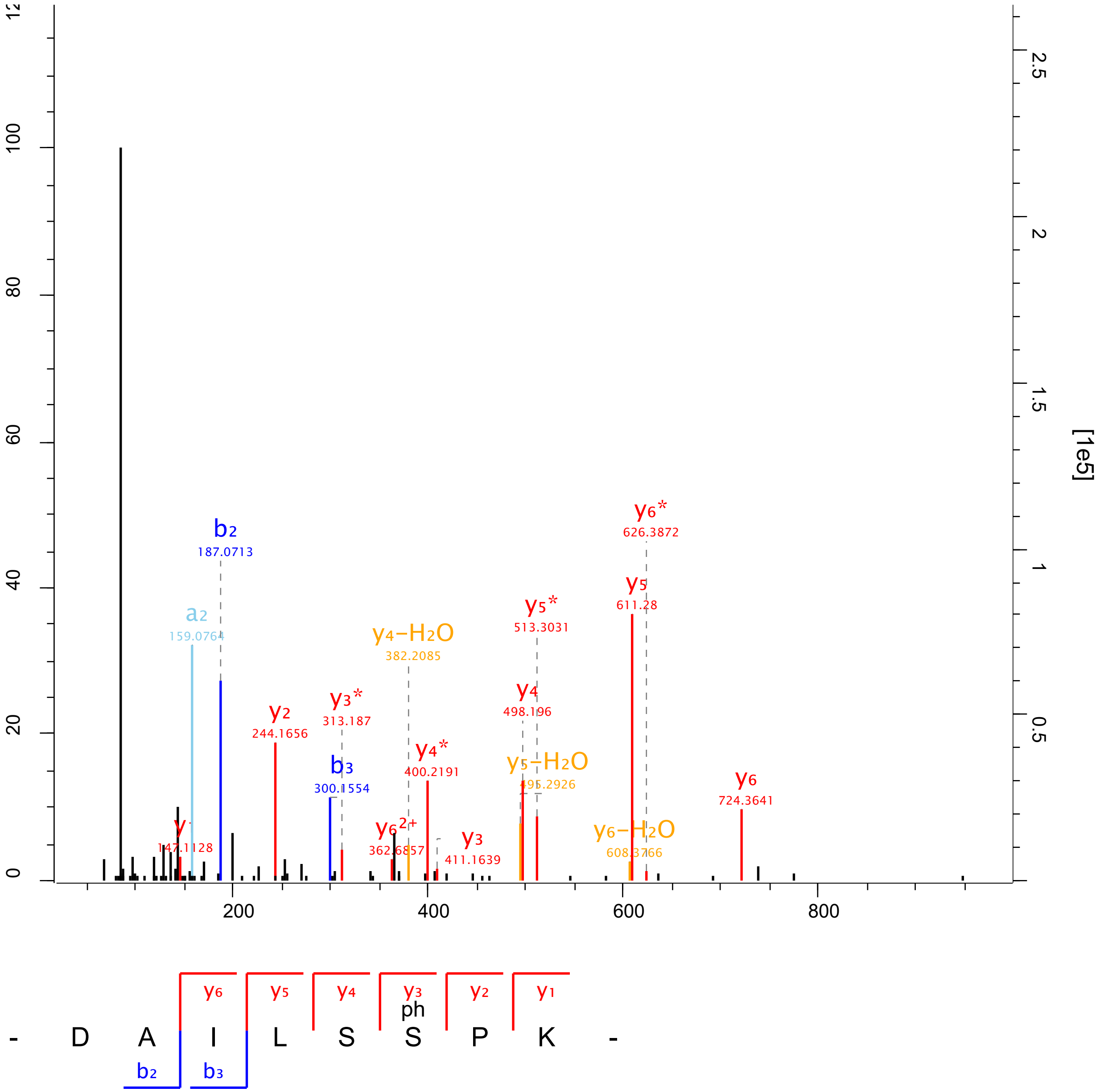

| Raw file | Scan | Method | Score | m/z |
| --- | --- | --- | --- | --- |
| sys_15_1 | 12159 | FTMS; HCD | 109.04 | 849.3 |

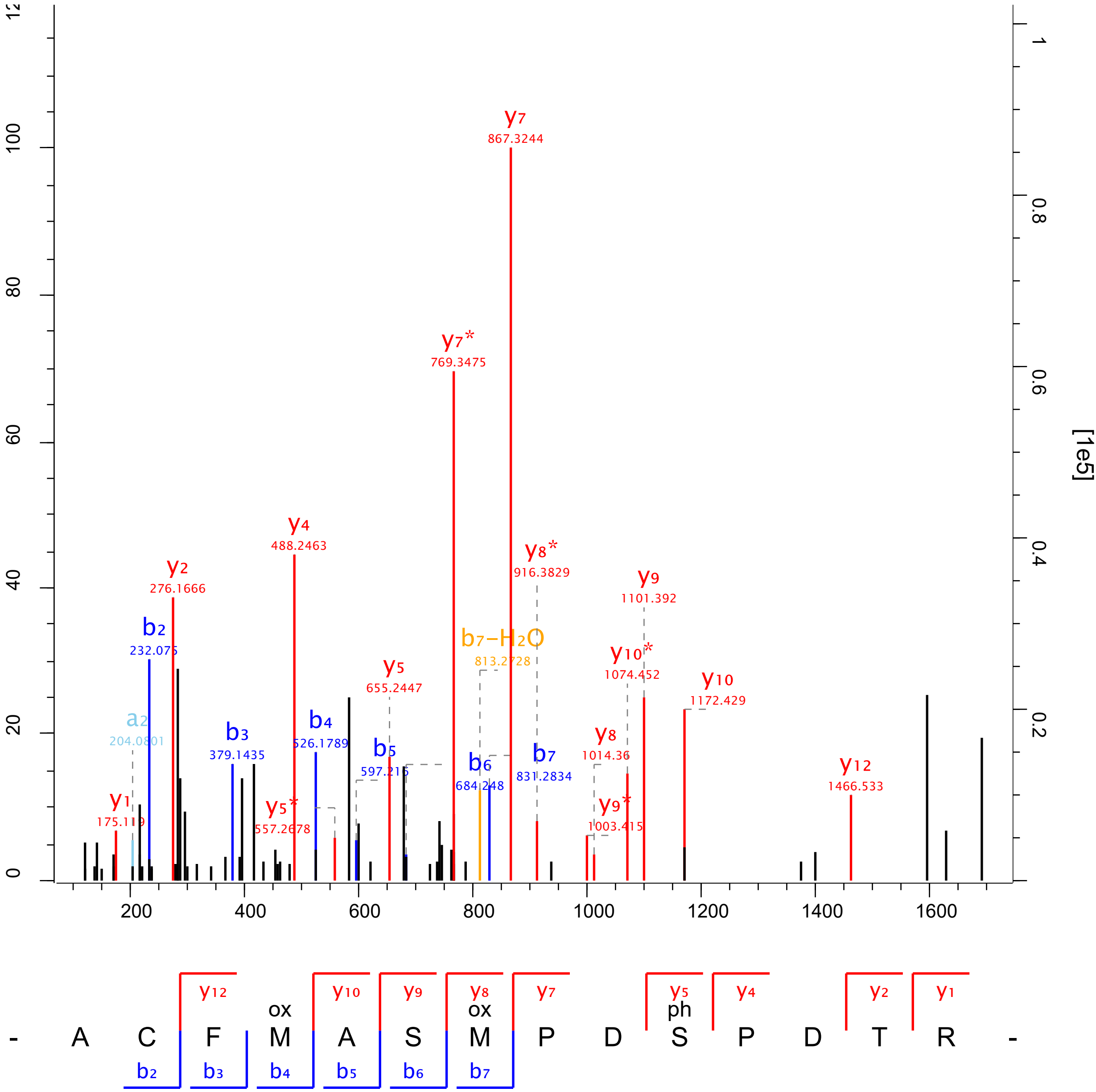

| Raw file | Scan | Method | Score | m/z |
| --- | --- | --- | --- | --- |
| sys_15_1 | 12182 | FTMS; HCD | 50.04 | 432.7 |

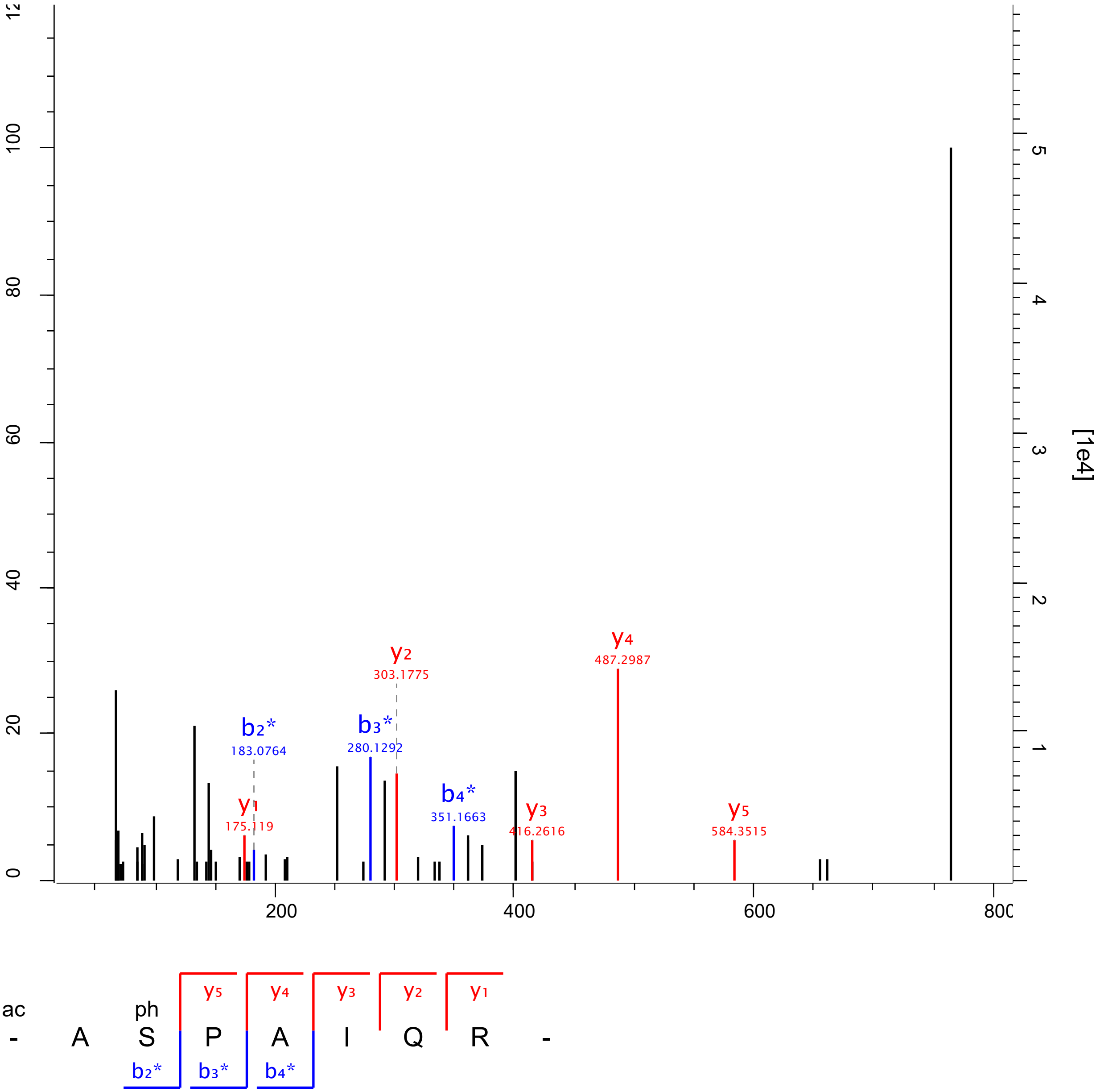

| Raw file | Scan | Method | Score | m/z |
|---|---|---|---|---|
| sys_15_1 | 12236 | FTMS; HCD | 104.43 | 477.23 |

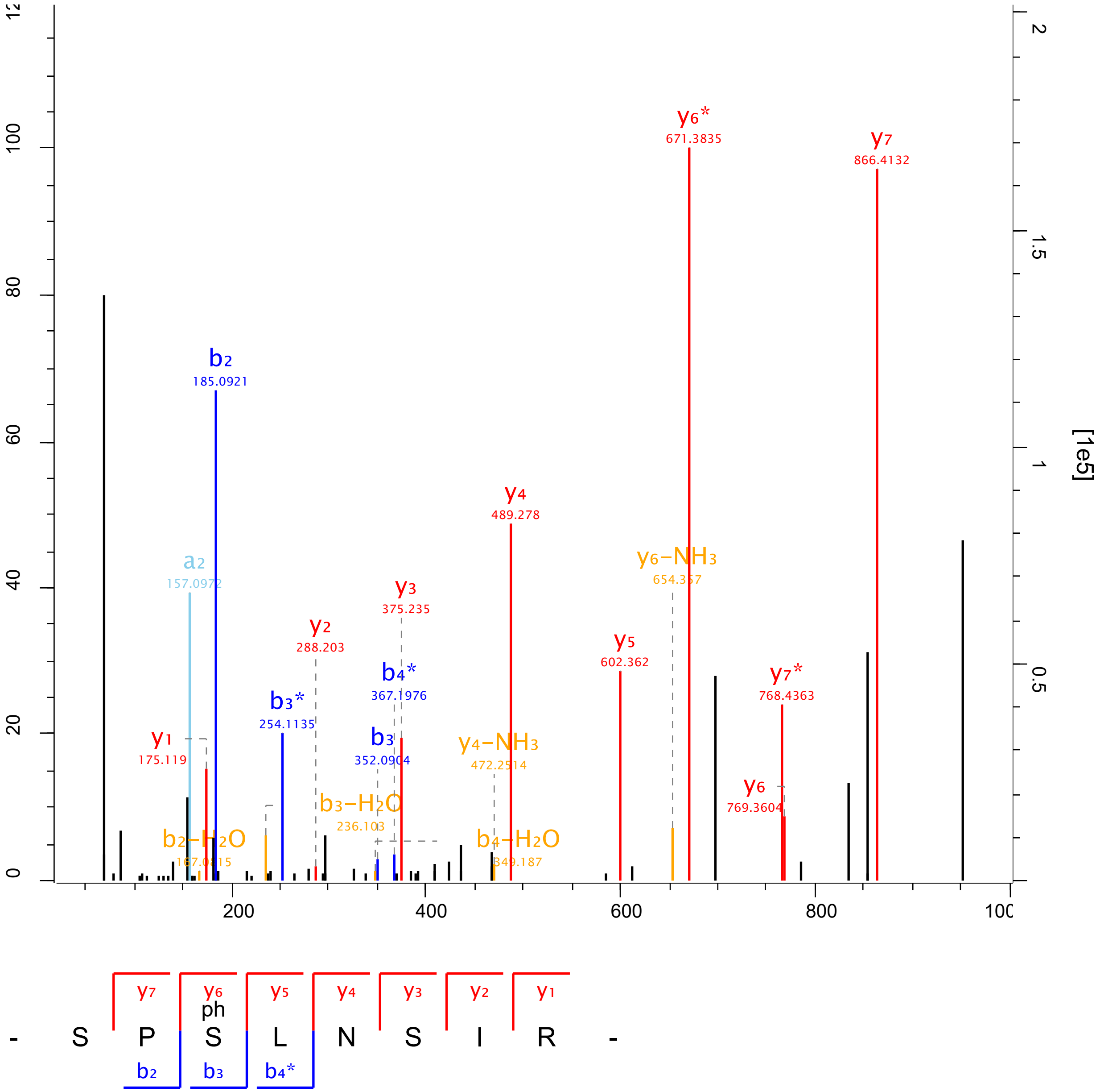

| Raw file | Scan | Method | Score | m/z |
|---|---|---|---|---|
| sys_15_1 | 12327 | FTMS; HCD | 137.64 | 841.87 |

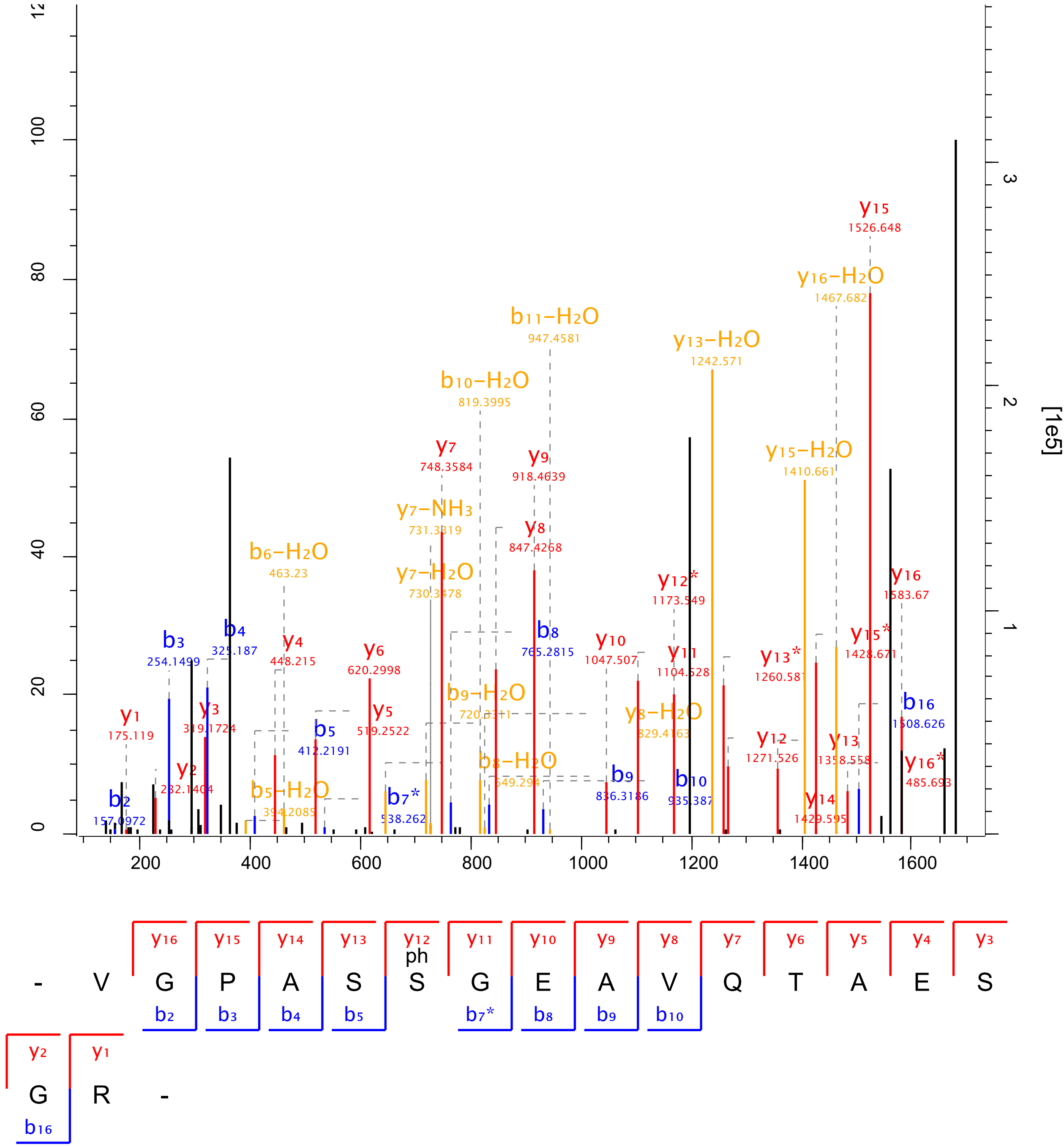

| Raw file | Scan | Method | Score | m/z |
|---|---|---|---|---|
| sys_15_1 | 12357 | FTMS; HCD | 69.7 | 949.06 |

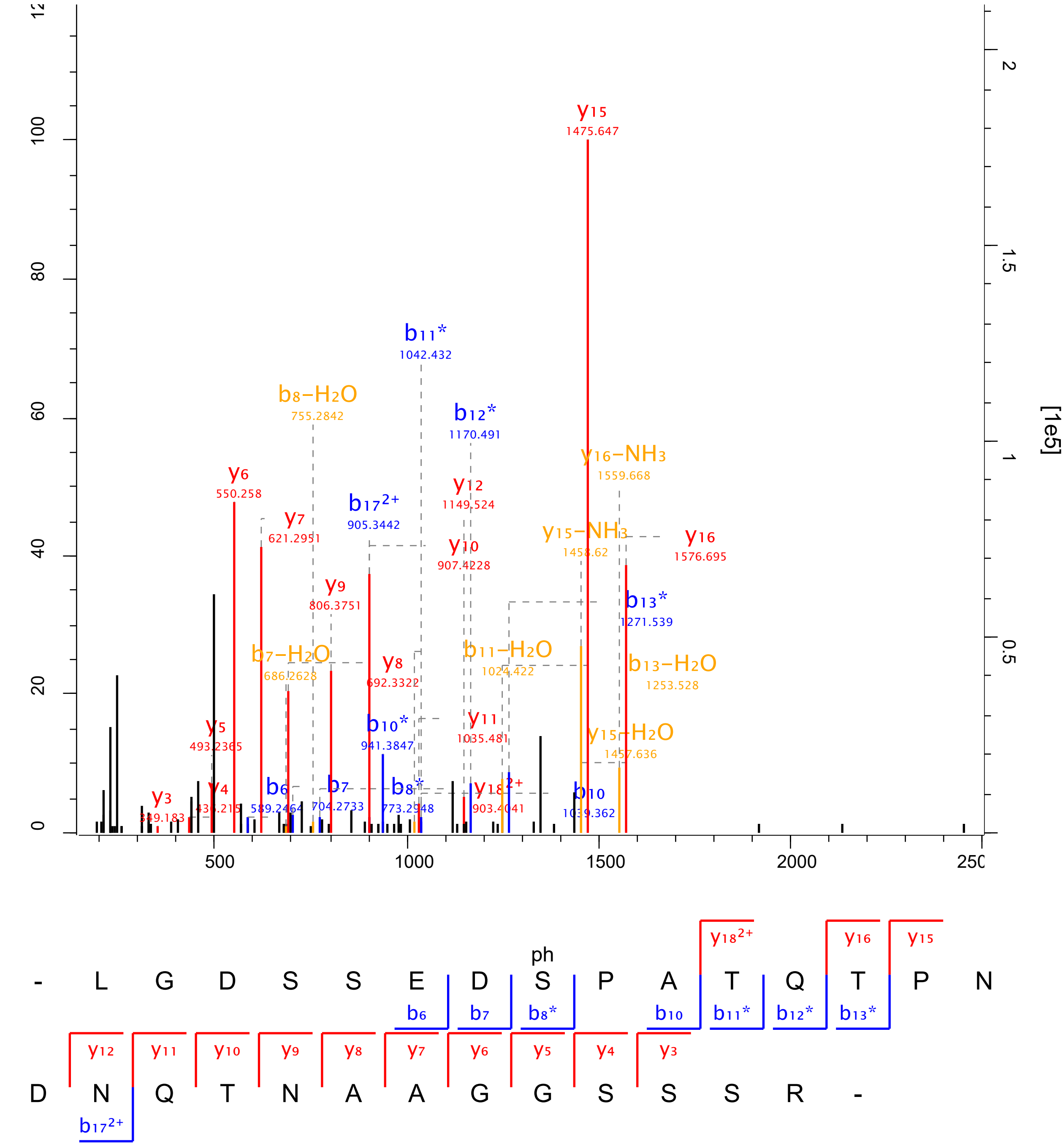

| Raw file | Scan | Method | Score | m/z |
|---|---|---|---|---|
| sys_15_1 | 12376 | FTMS; HCD | 186.45 | 833.82 |

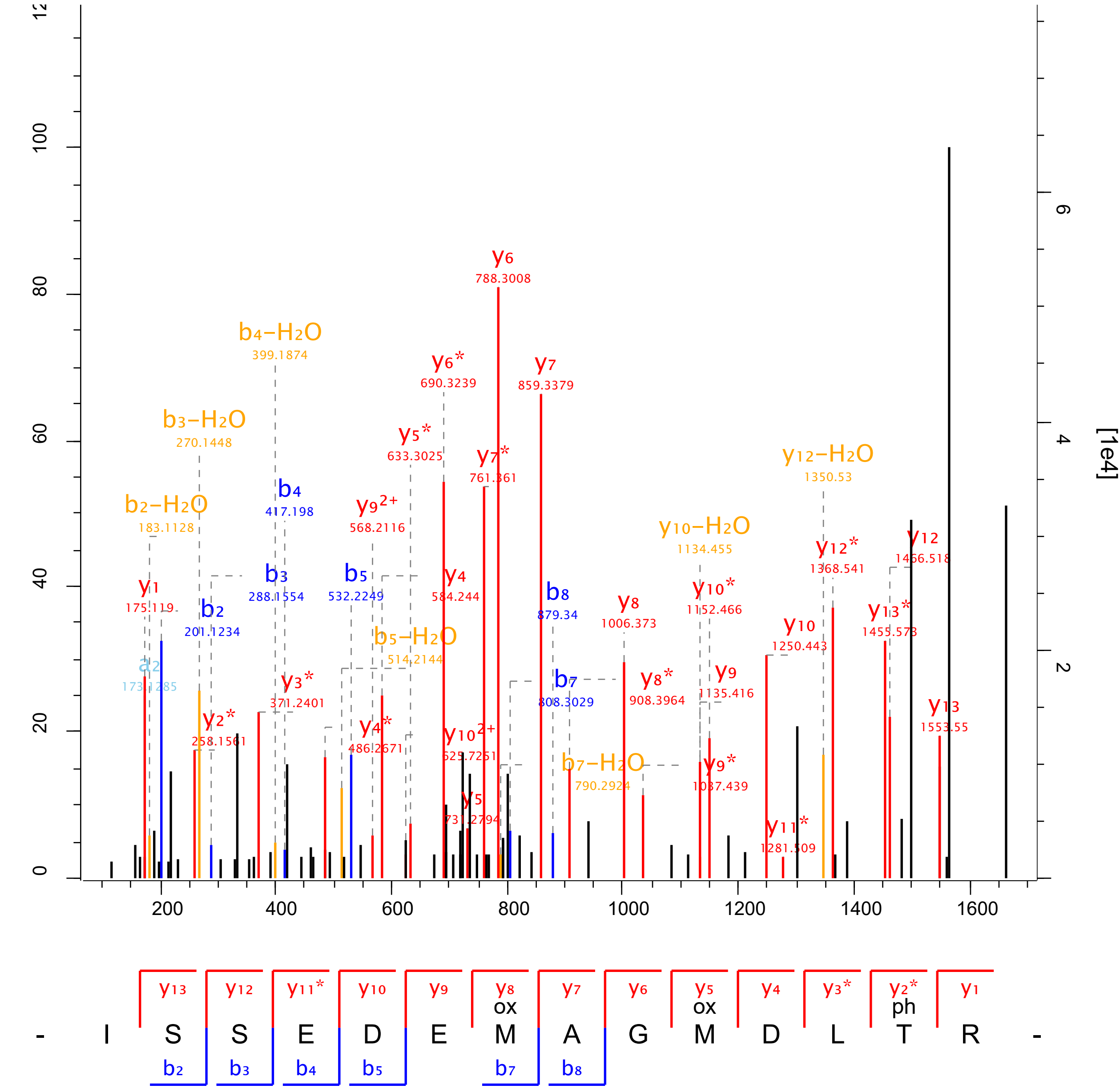

| Raw file | Scan | Method | Score | m/z |
|---|---|---|---|---|
| sys_15_1 | 12564 | FTMS; HCD | 108.63 | 702.28 |

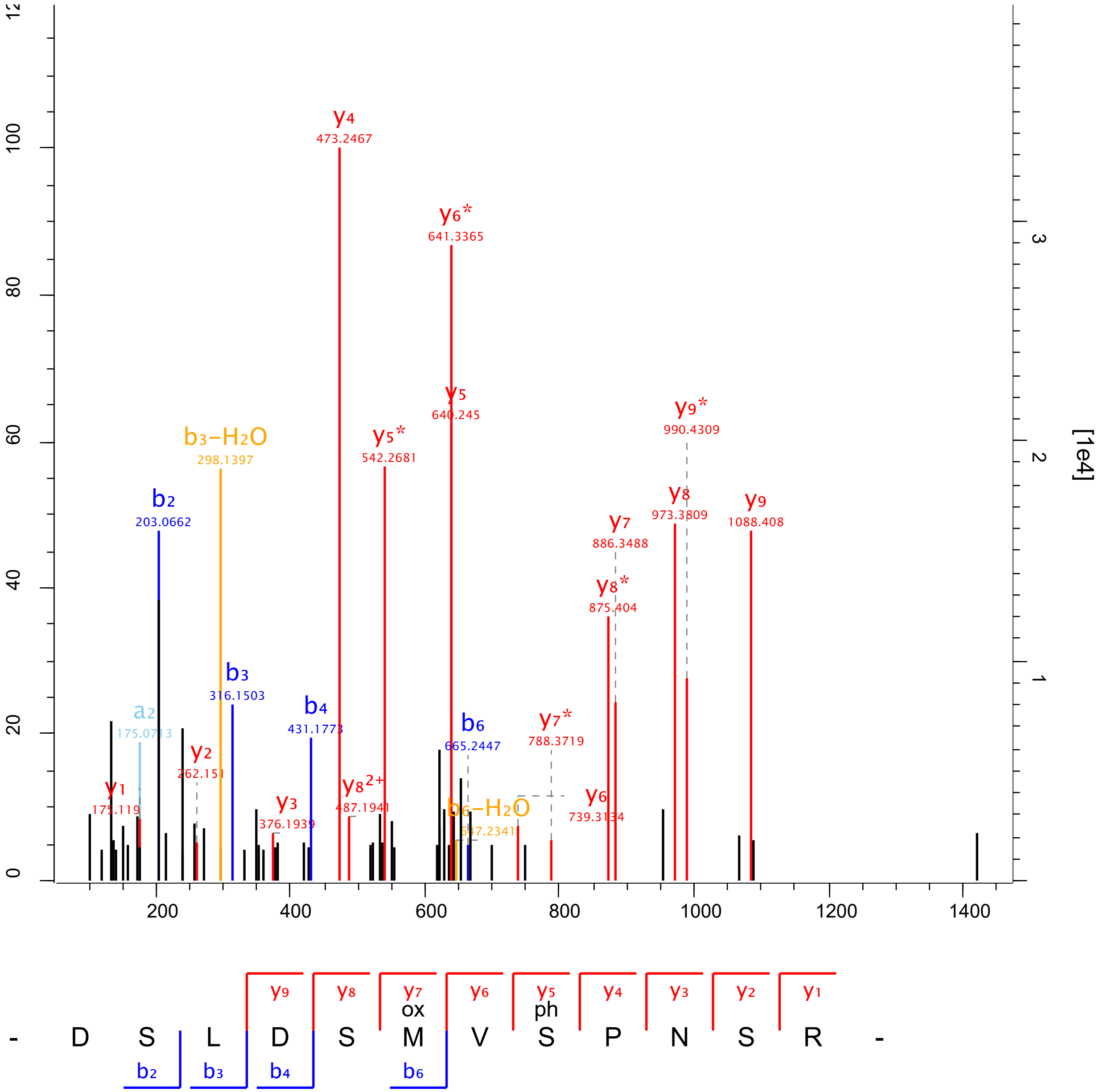

| Raw file | Scan | Method | Score | m/z |
|---|---|---|---|---|
| sys_15_1 | 12592 | FTMS; HCD | 203.93 | 776.32 |

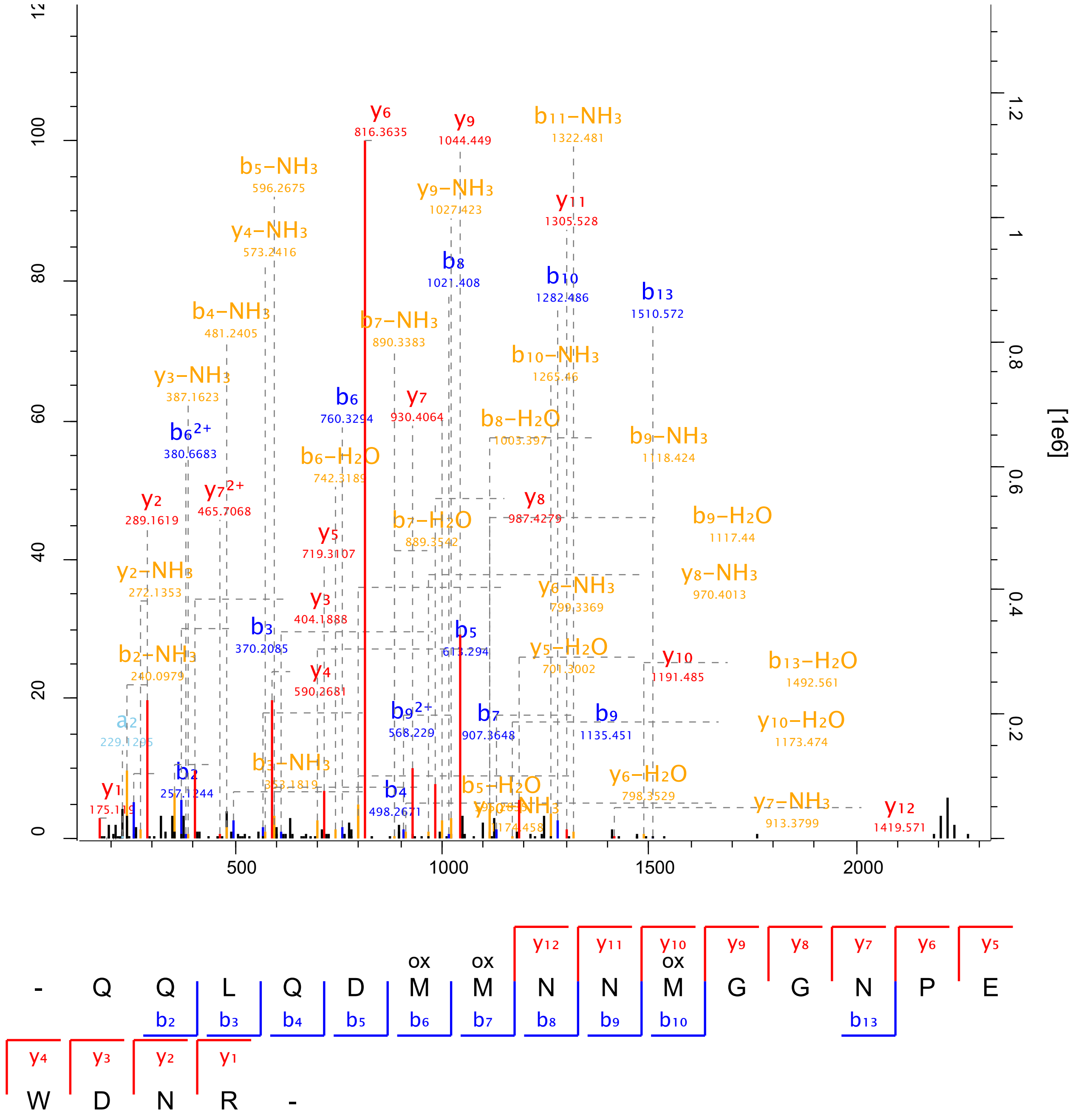

| Raw file | Scan | Method | Score | m/z |
| --- | --- | --- | --- | --- |
| sys_15_1 | 12713 | FTMS; HCD | 54.27 | 532.72 |

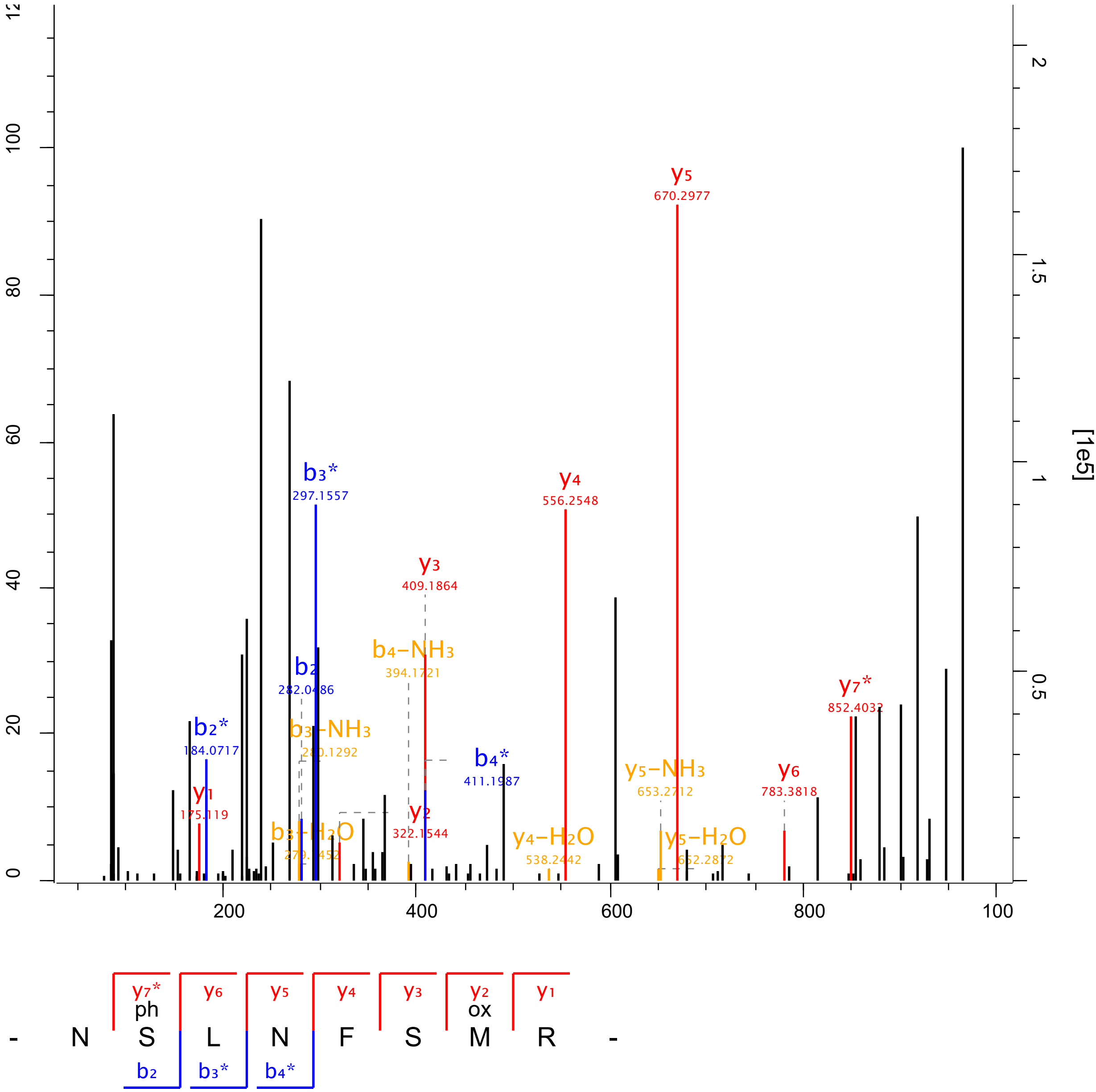

| Raw file | Scan | Method | Score | m/z |
|---|---|---|---|---|
| sys_15_1 | 12735 | FTMS; HCD | 92.65 | 635.27 |

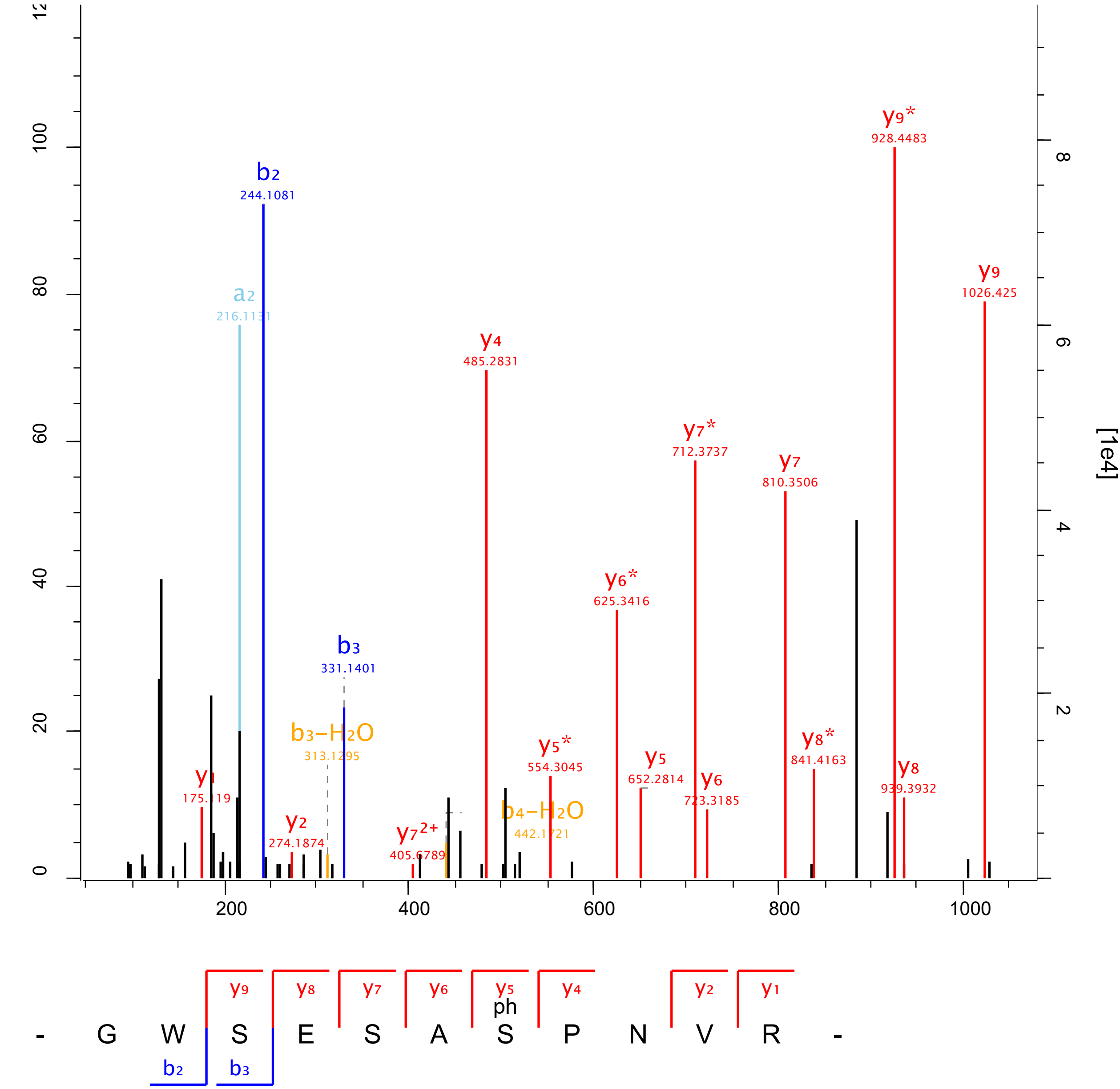

| Raw file | Scan | Method | Score | m/z |
| --- | --- | --- | --- | --- |
| sys_15_1 | 12905 | FTMS; HCD | 157.3 | 671.63 |

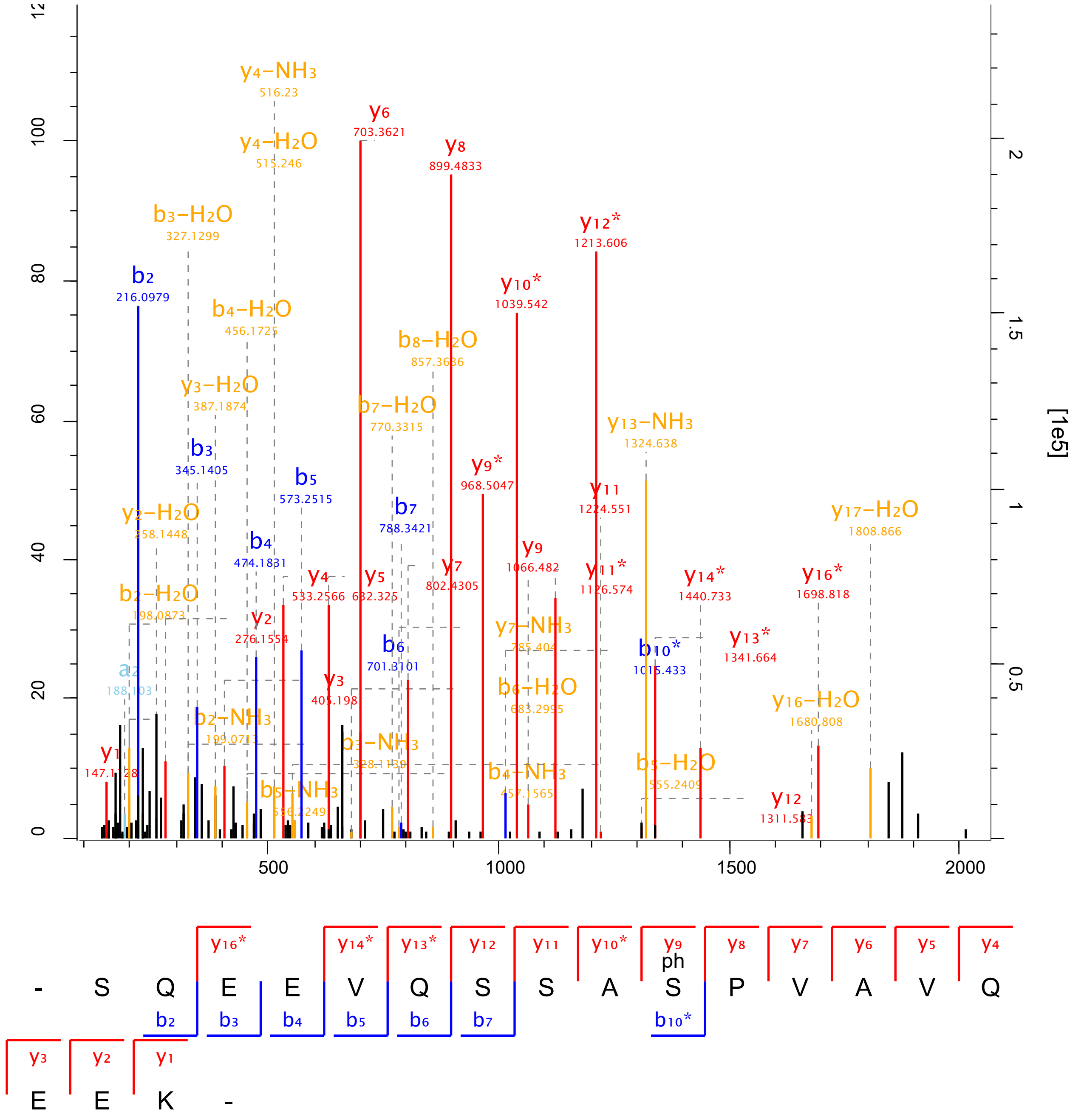

| Raw file | Scan | Method | Score | m/z |
|---|---|---|---|---|
| sys_15_1 | 12913 | FTMS; HCD | 48.22 | 737.29 |

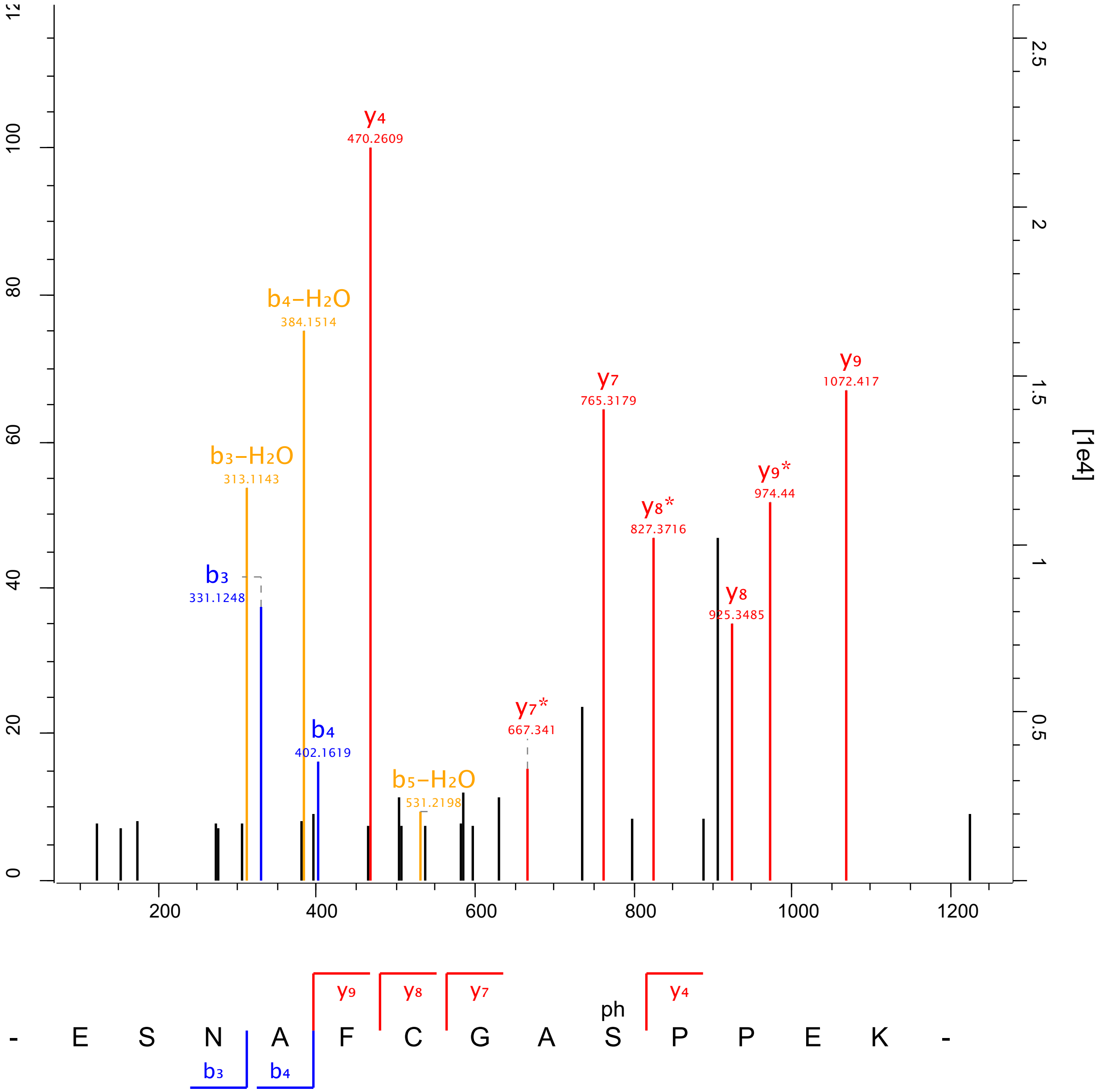

| Raw file | Scan | Method | Score | m/z |
|---|---|---|---|---|
| sys_15_1 | 12918 | FTMS; HCD | 40.94 | 608.8 |

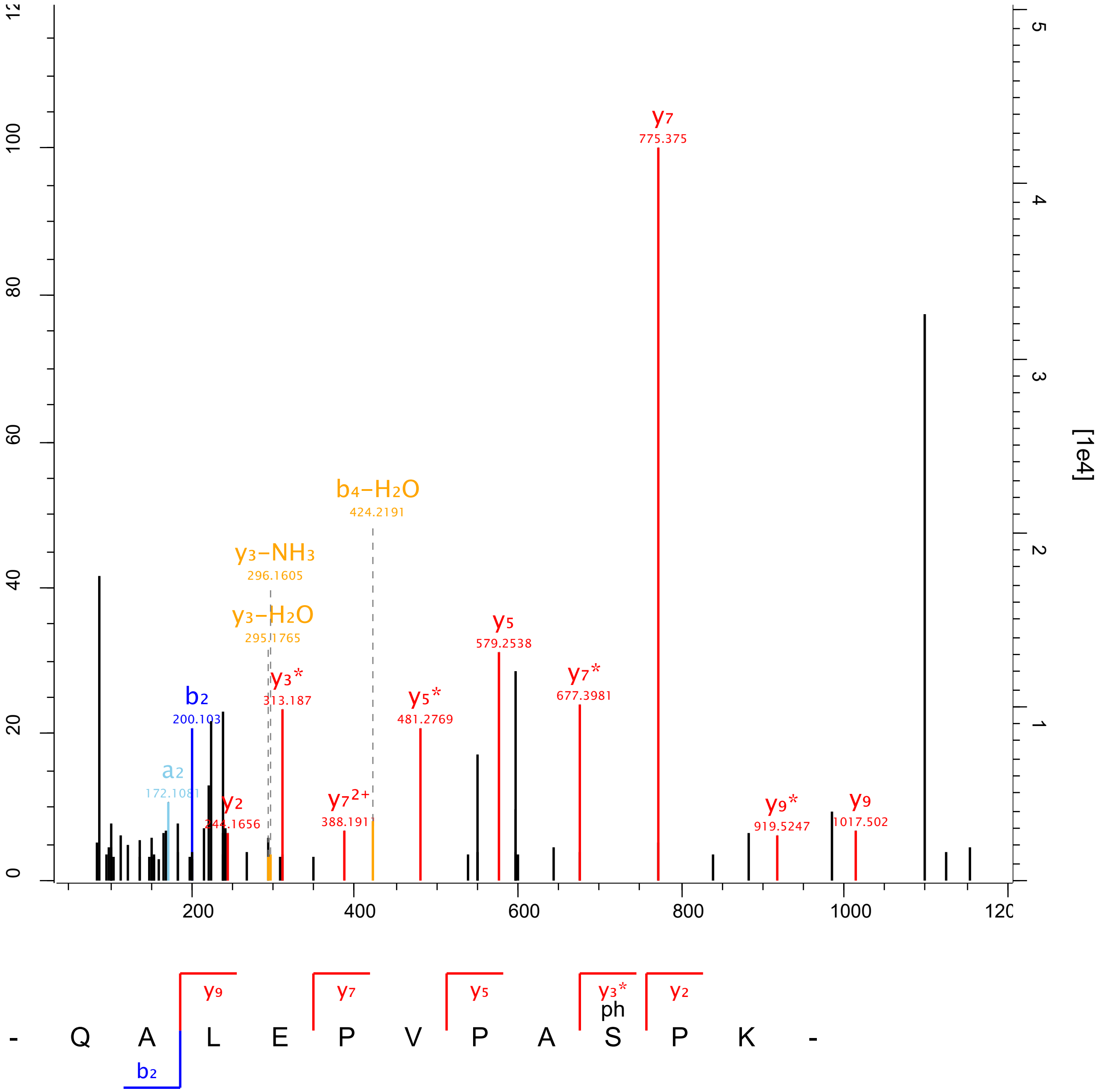

| Raw file | Scan | Method | Score | m/z |
|---|---|---|---|---|
| sys_15_1 | 12998 | FTMS; HCD | 146.86 | 728.78 |

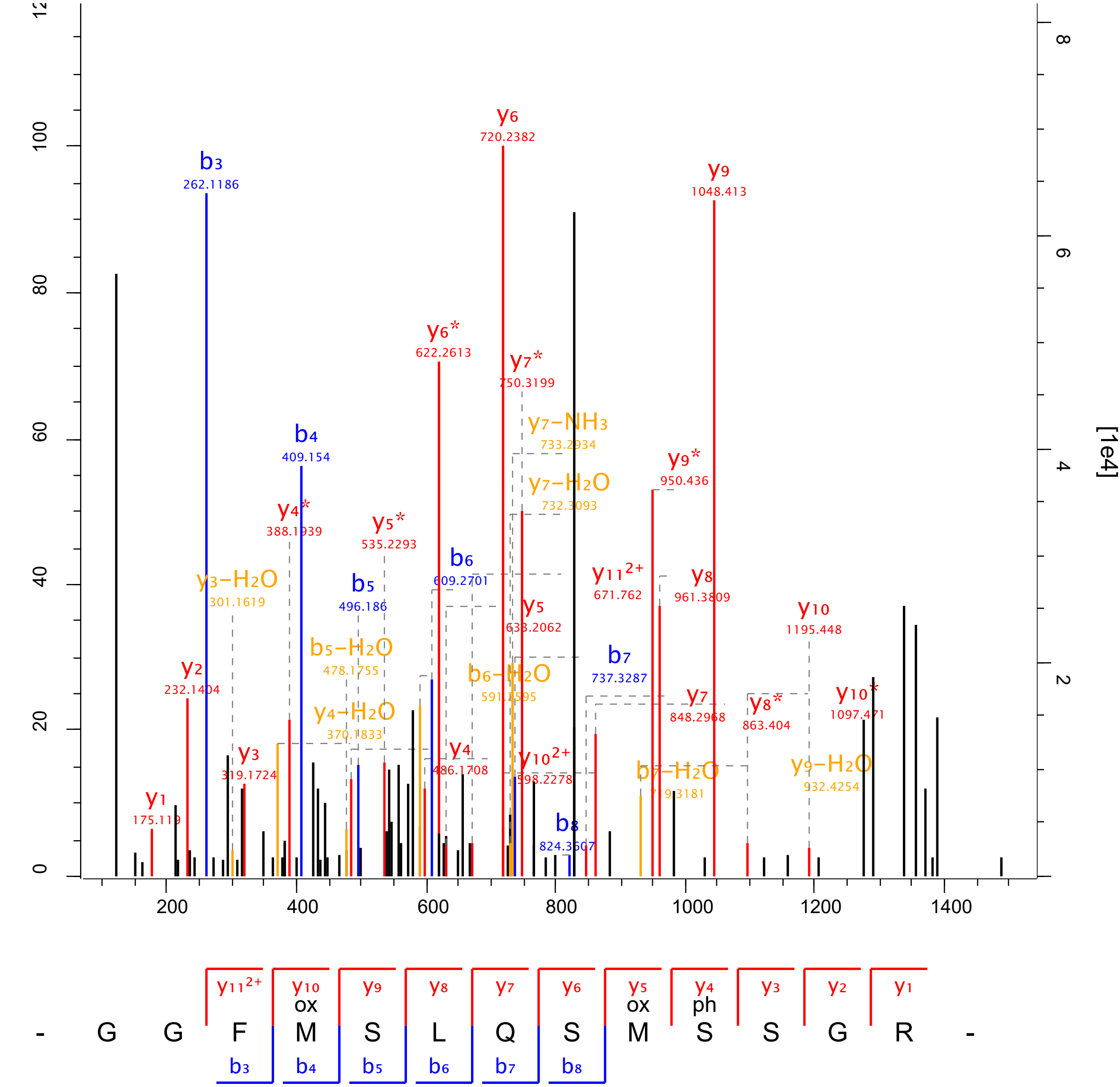

| Raw file | Scan | Method | Score | m/z |
| --- | --- | --- | --- | --- |
| sys_15_1 | 13068 | FTMS; HCD | 79.66 | 418.7 |

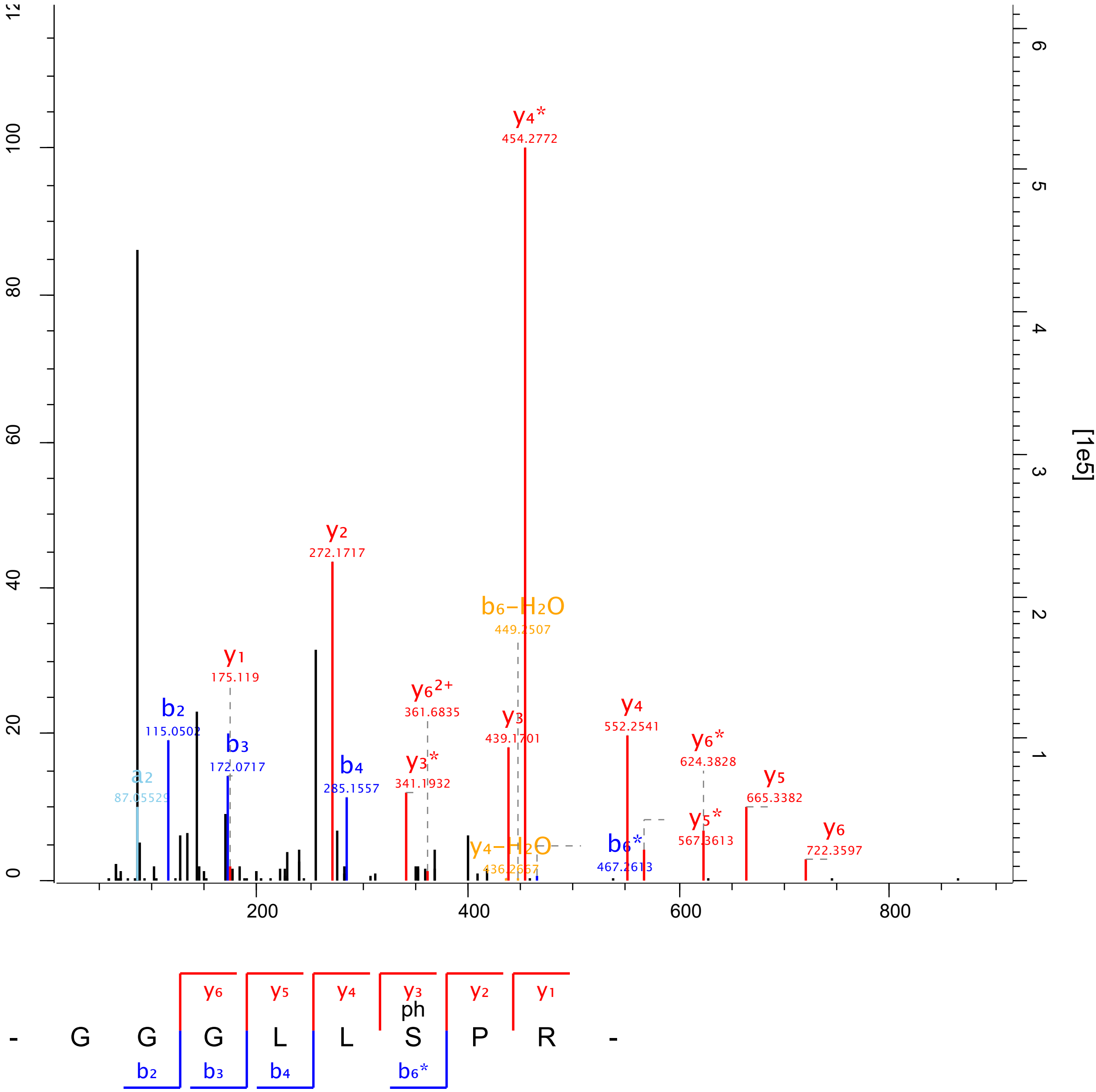

| Raw file | Scan | Method | Score | m/z |
|---|---|---|---|---|
| sys_15_1 | 13098 | FTMS; HCD | 45.92 | 830.36 |

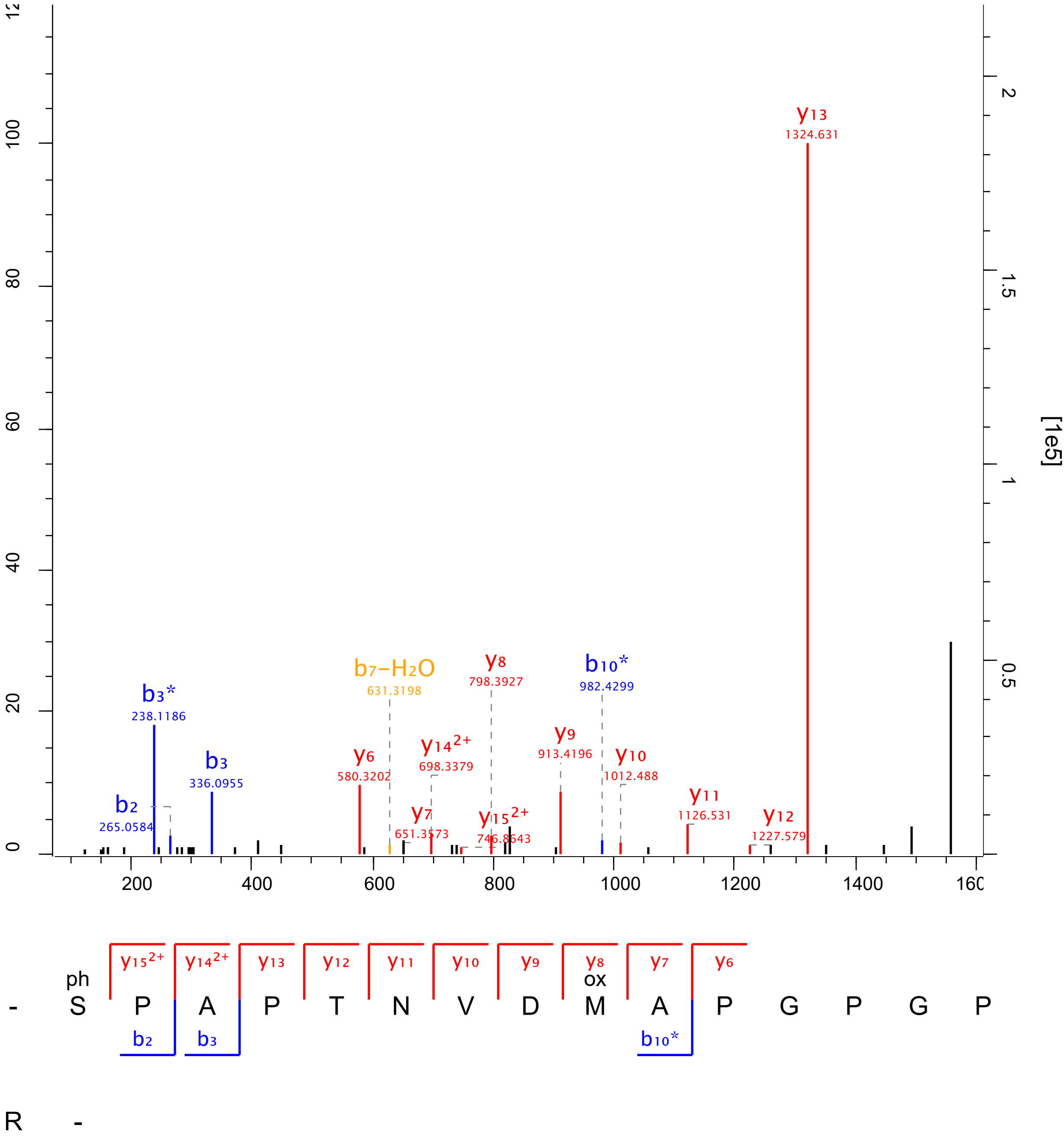

| Raw file | Scan | Method | Score | m/z |
|---|---|---|---|---|
| sys_15_1 | 13100 | FTMS; HCD | 149.3 | 724.8 |

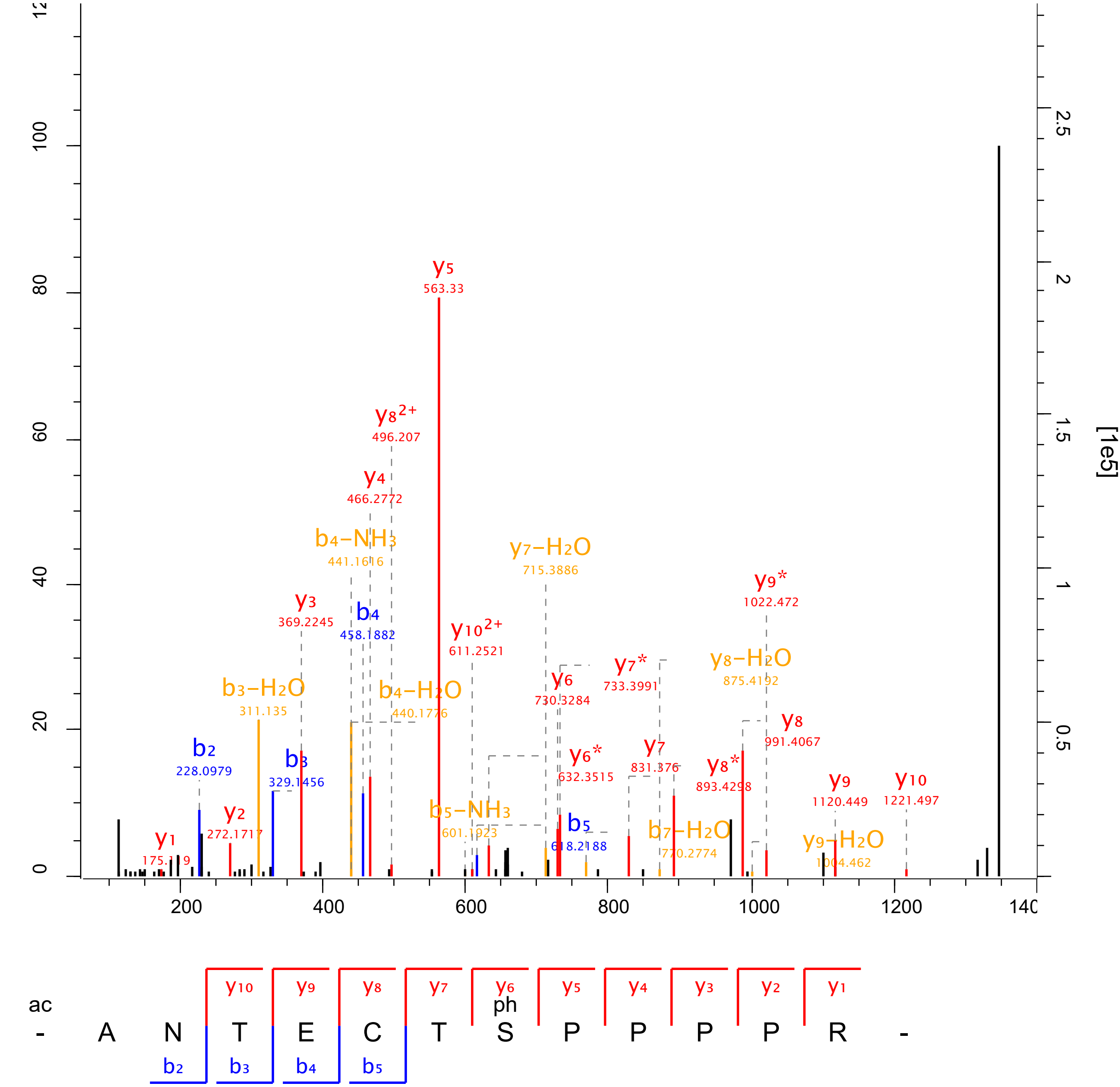

| Raw file | Scan | Method | Score | m/z |
|---|---|---|---|---|
| sys_15_1 | 13106 | FTMS; HCD | 104.2 | 730.82 |

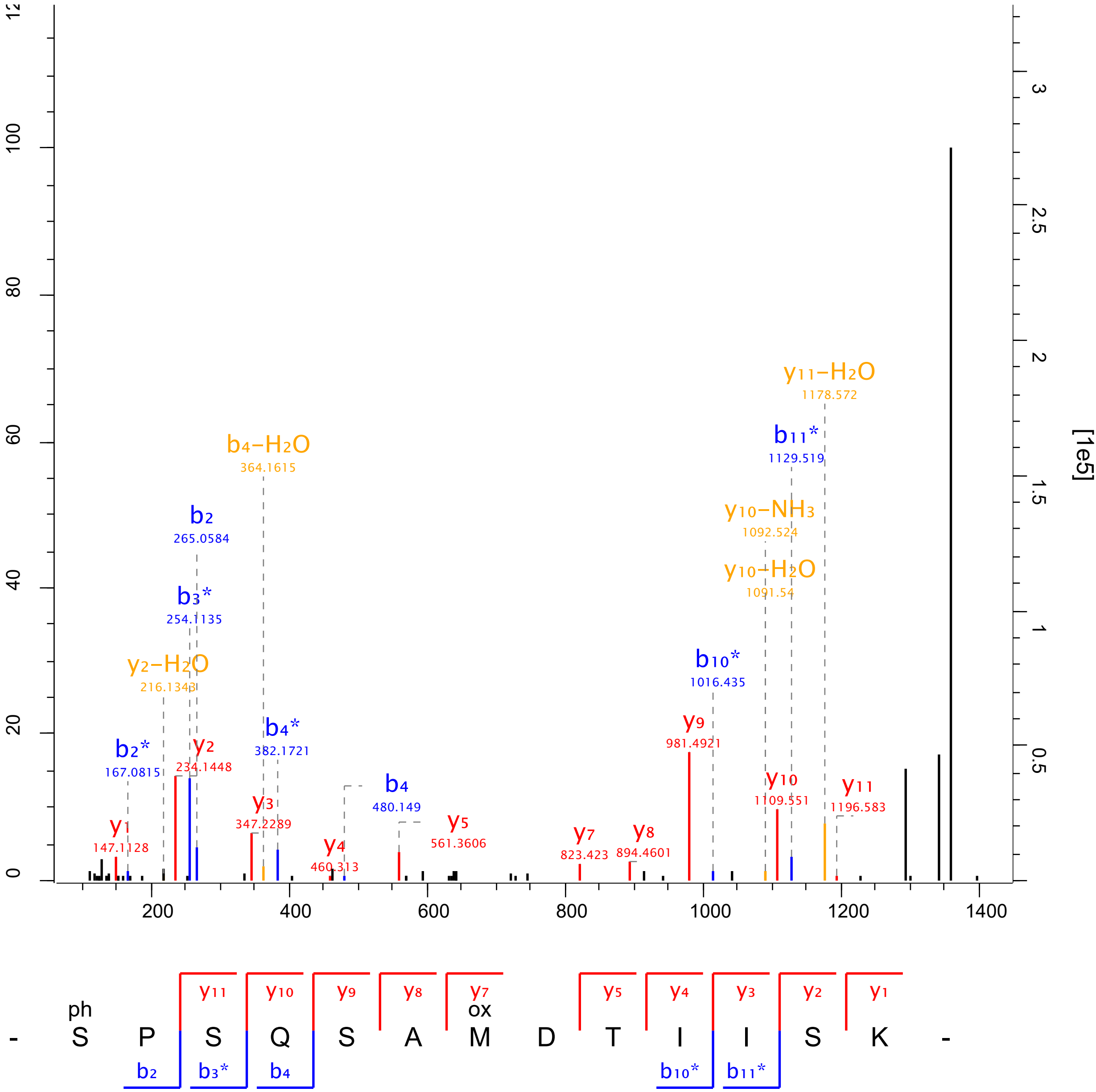

| Raw file | Scan | Method | Score | m/z |
|---|---|---|---|---|
| sys_15_1 | 13210 | FTMS; HCD | 55.35 | 526.21 |

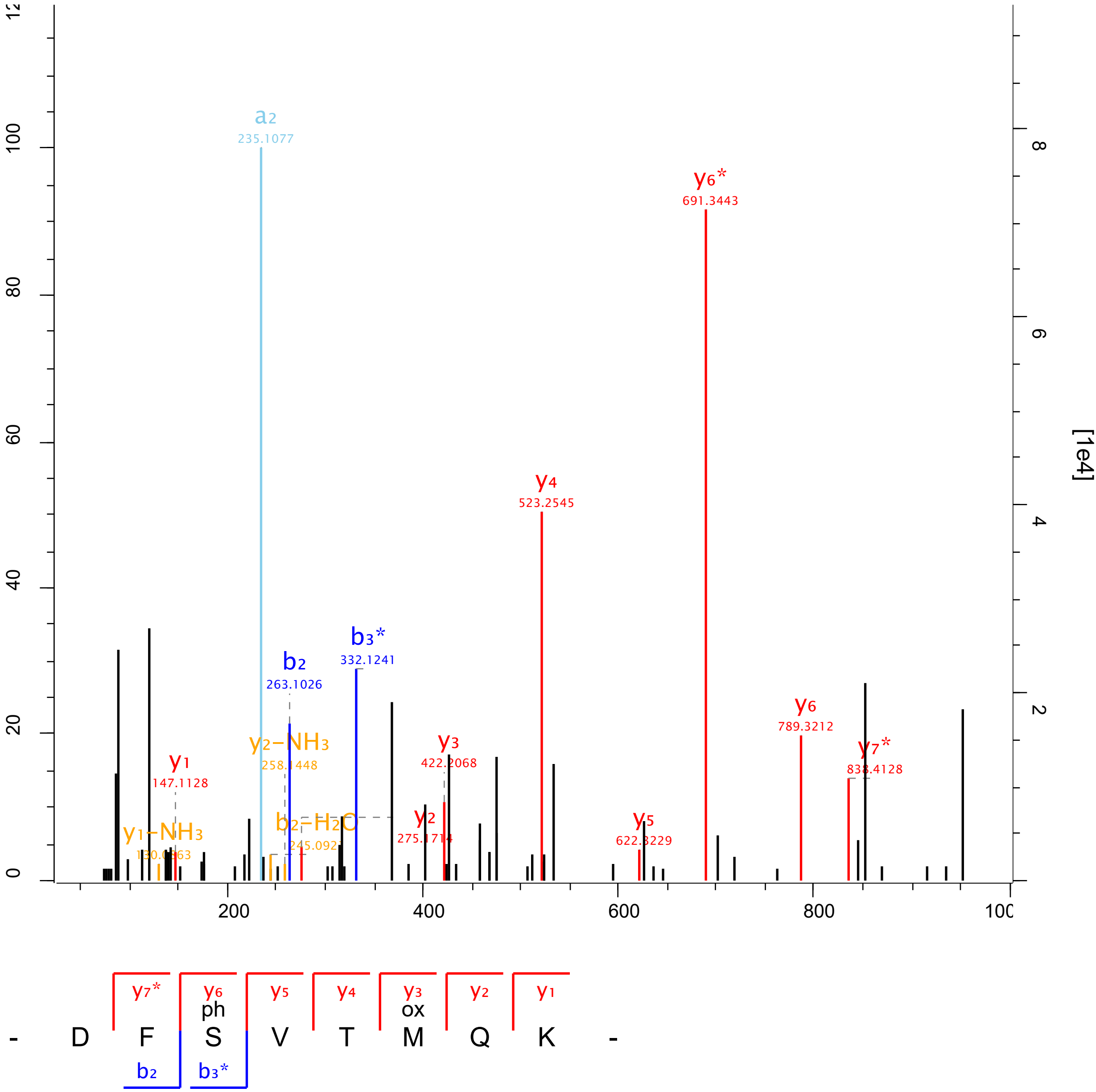

| Raw file | Scan | Method | Score | m/z |
|---|---|---|---|---|
| sys_15_1 | 13283 | FTMS; HCD | 179.5 | 577.28 |

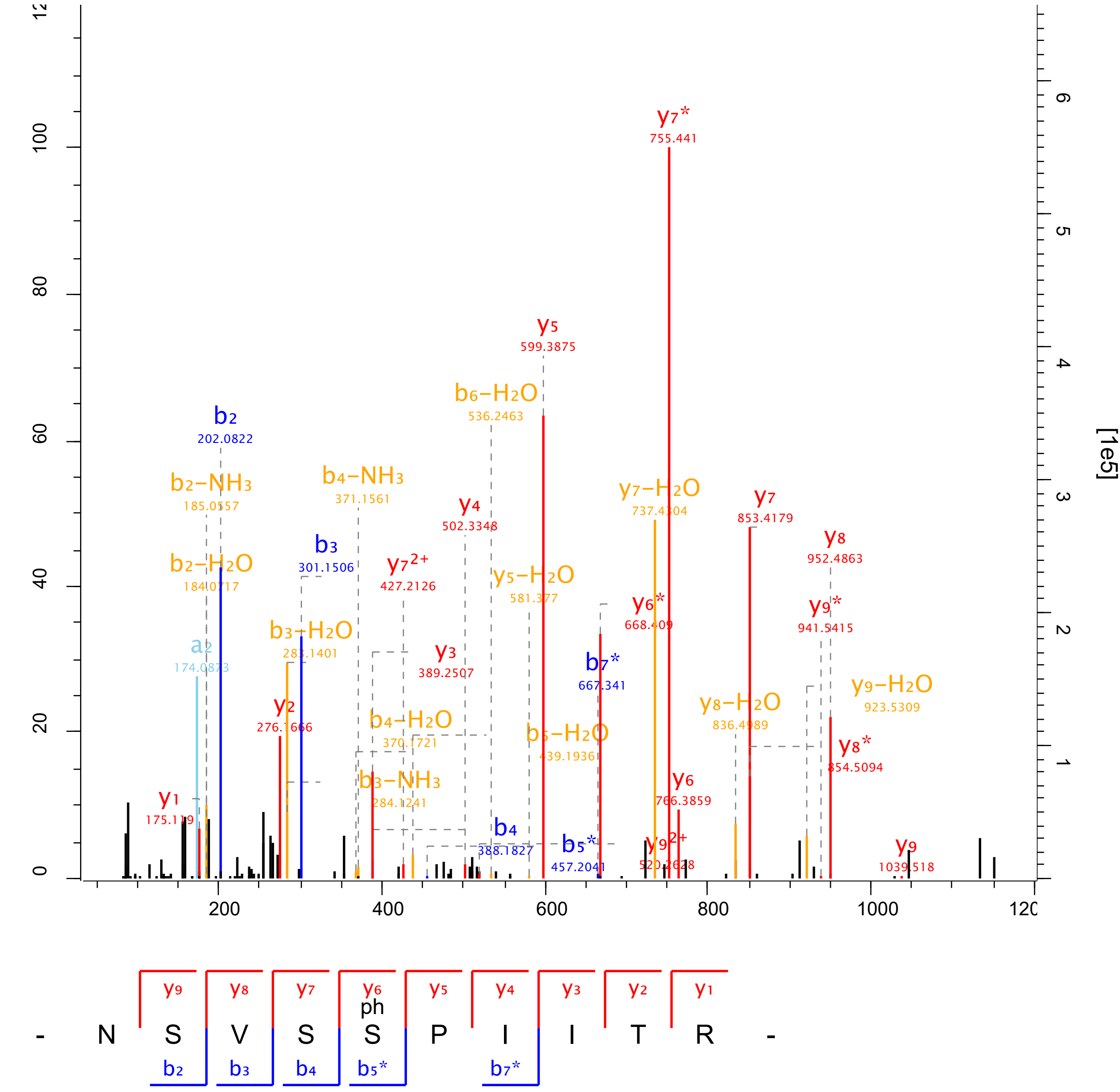

| Raw file | Scan | Method | Score | m/z |
| --- | --- | --- | --- | --- |
| sys_15_1 | 13304 | FTMS; HCD | 100.93 | 535.25 |

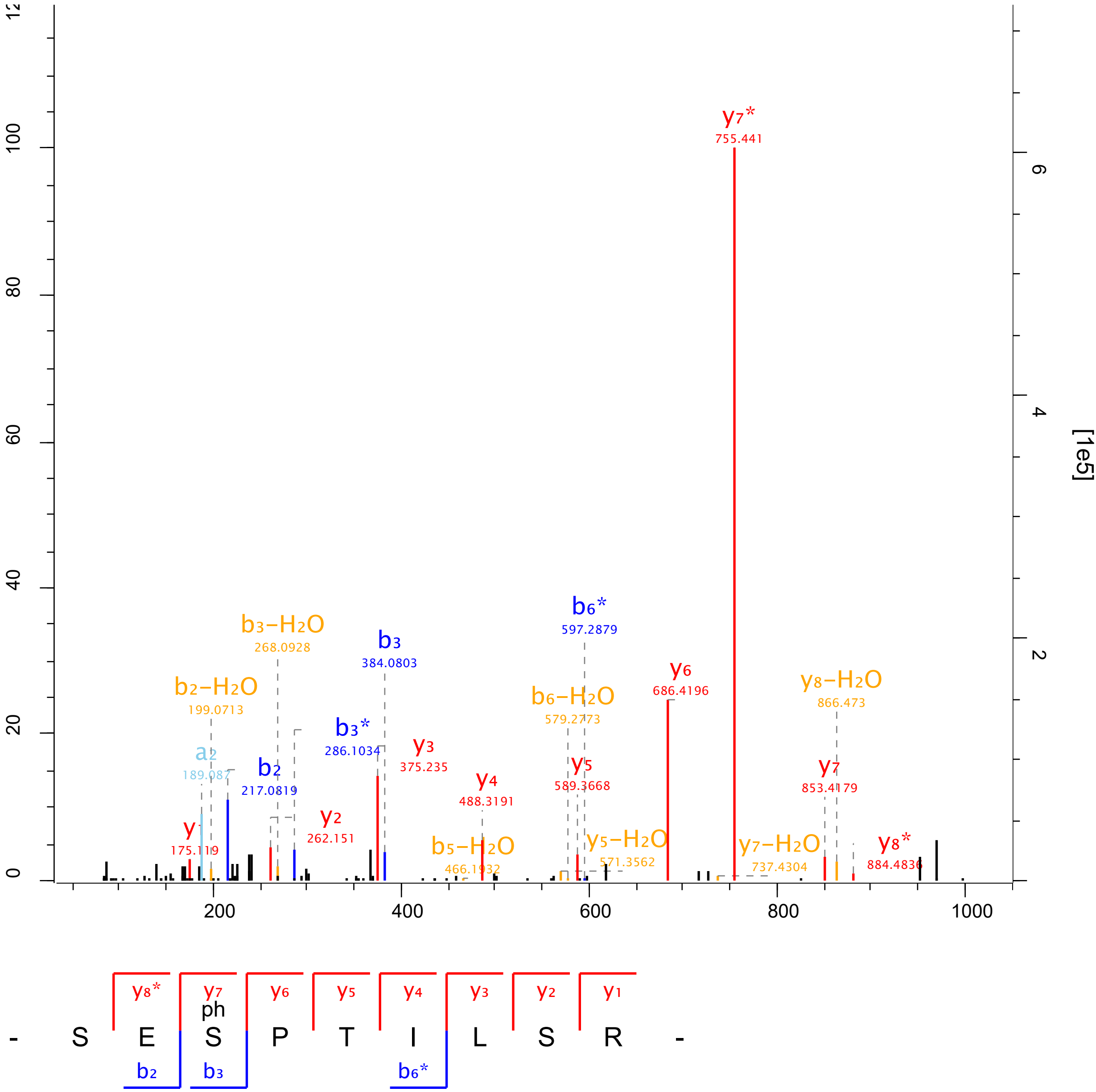

| Raw file | Scan | Method | Score | m/z |
|---|---|---|---|---|
| sys_15_1 | 13322 | FTMS; HCD | 146.15 | 721.27 |

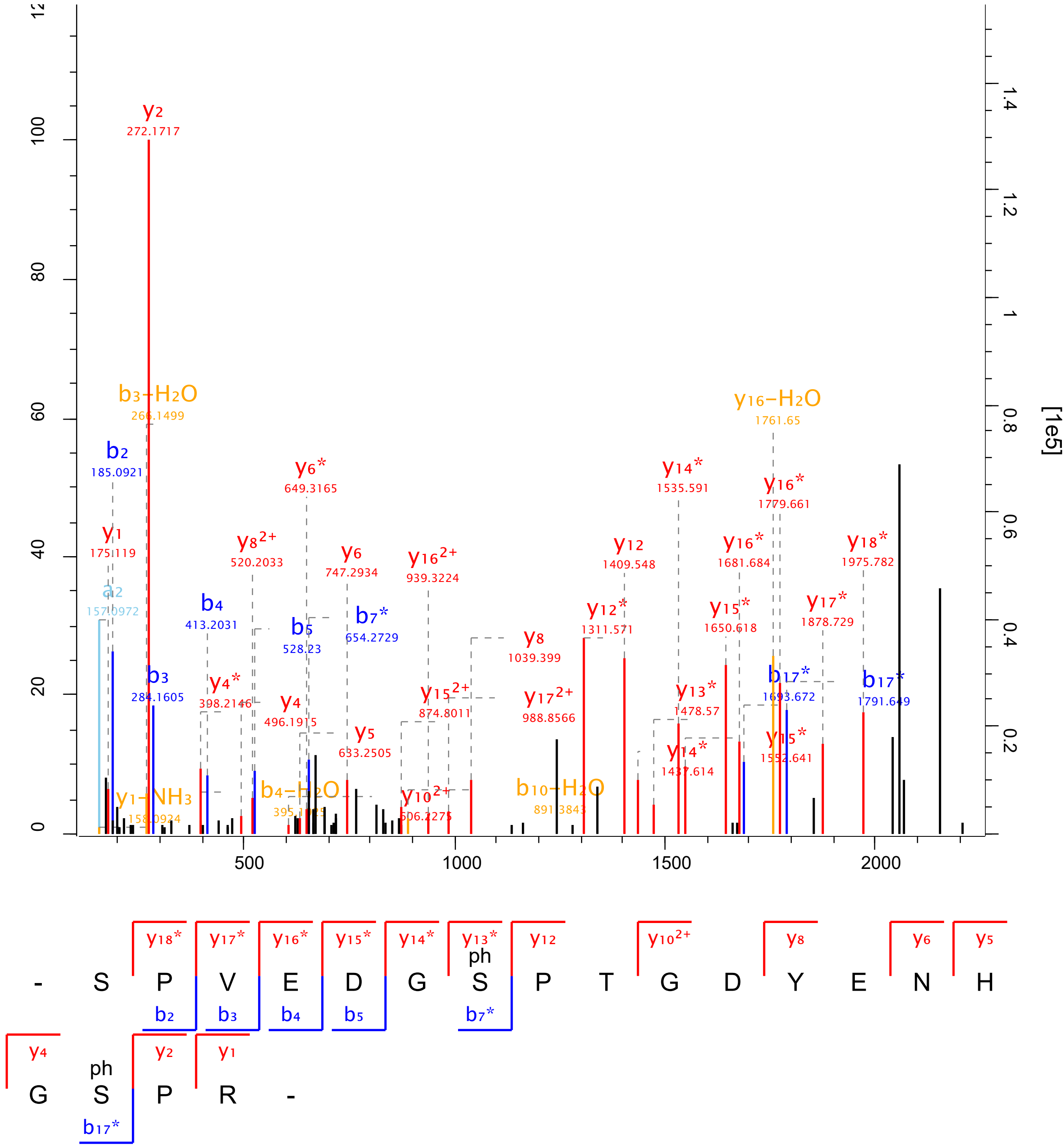

| Raw file | Scan | Method | Score | m/z |
|---|---|---|---|---|
| sys_15_1 | 13388 | FTMS; HCD | 92.38 | 683.29 |

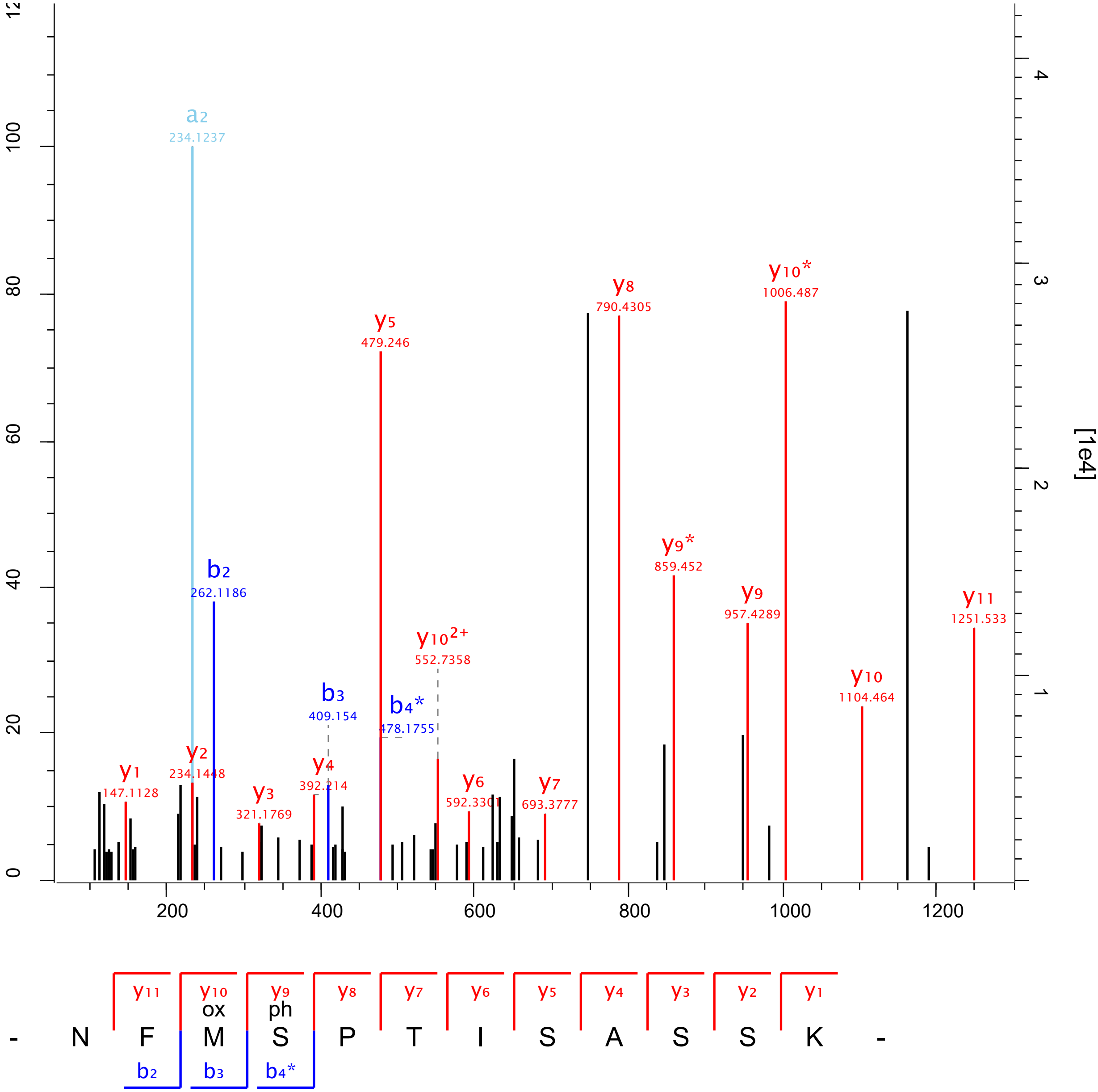

| Raw file | Scan | Method | Score | m/z |
|---|---|---|---|---|
| sys_15_1 | 13479 | FTMS; HCD | 126.8 | 898.4 |

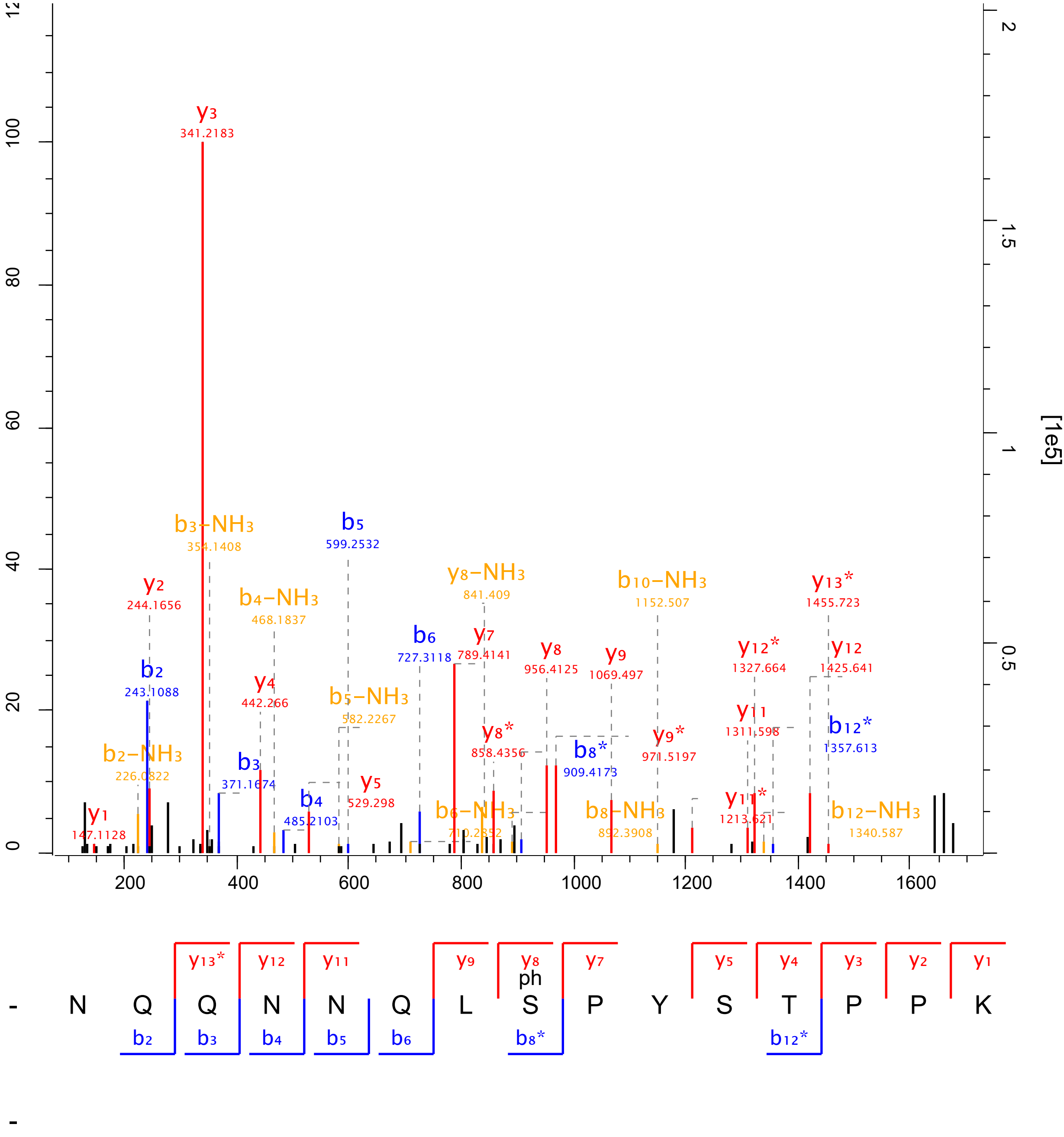

| Raw file | Scan | Method | Score | m/z |
|---|---|---|---|---|
| sys_15_1 | 13716 | FTMS; HCD | 50.15 | 488.23 |

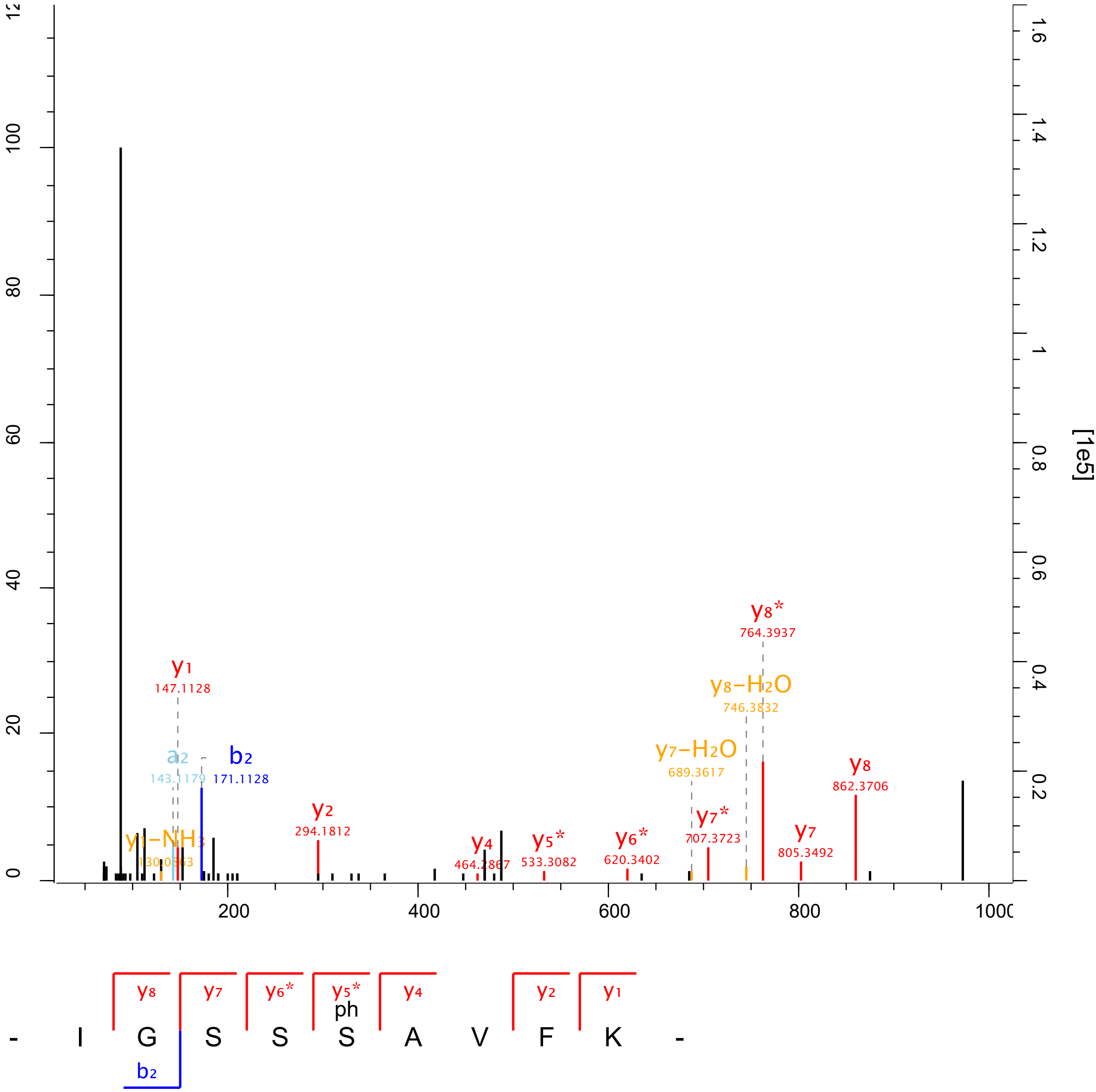

| Raw file | Scan | Method | Score | m/z |
| --- | --- | --- | --- | --- |
| sys_15_1 | 13820 | FTMS; HCD | 144.8 | 662.24 |

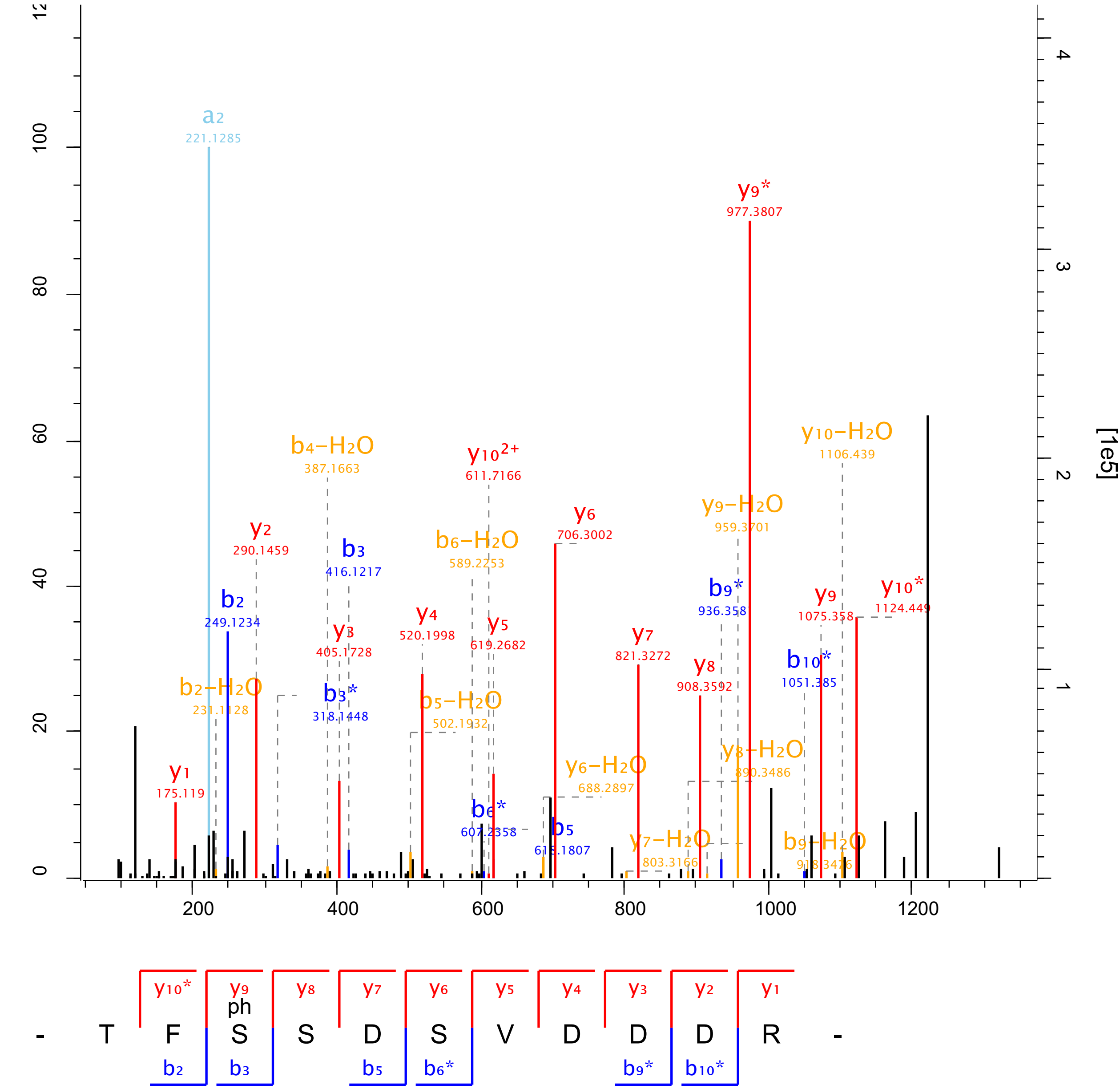

| Raw file | Scan | Method | Score | m/z |
|---|---|---|---|---|
| sys_15_1 | 13846 | FTMS; HCD | 97.77 | 577.27 |

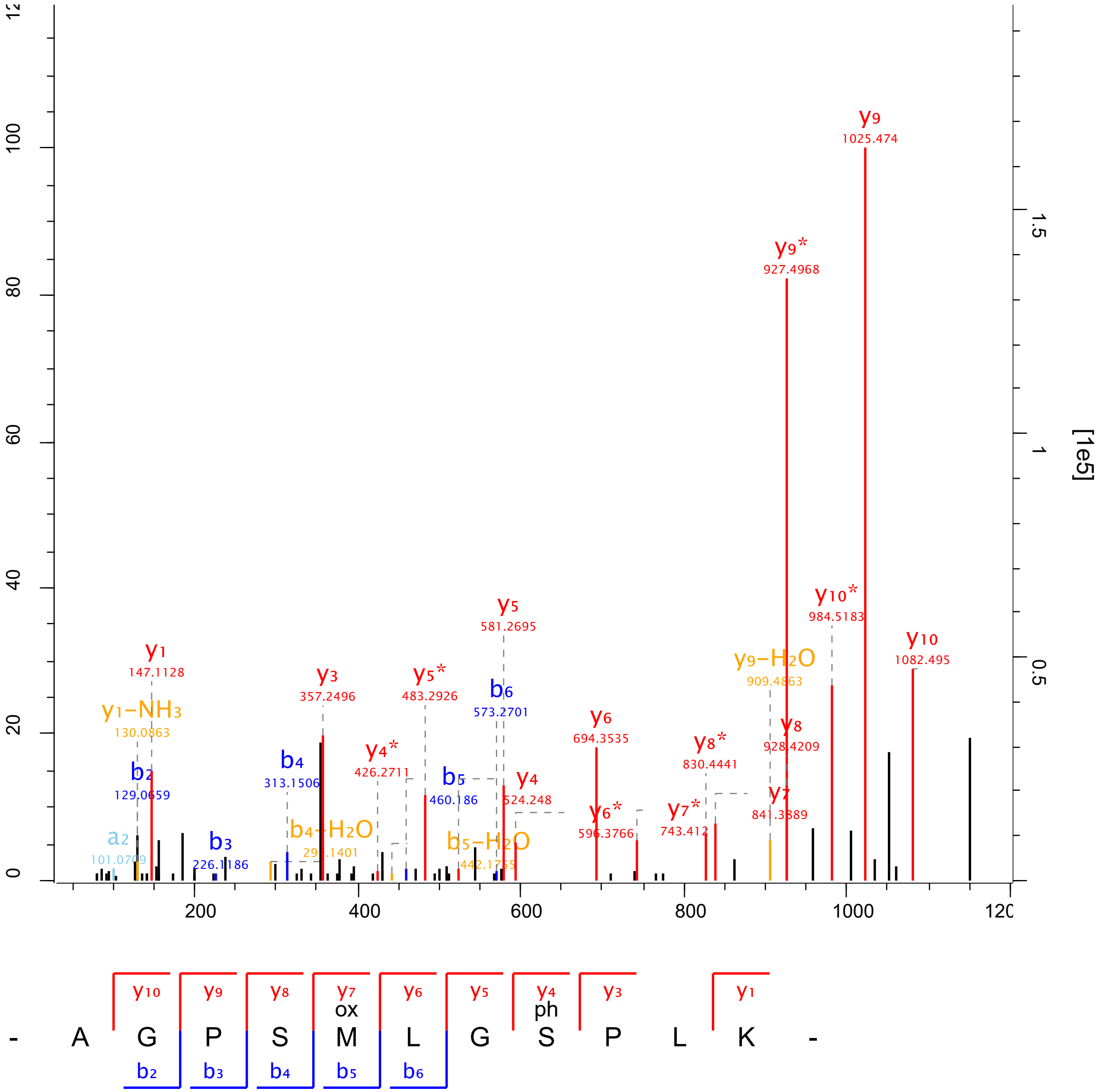

| Raw file | Scan | Method | Score | m/z |
| --- | --- | --- | --- | --- |
| sys_15_1 | 13874 | FTMS; HCD | 139.7 | 579.76 |

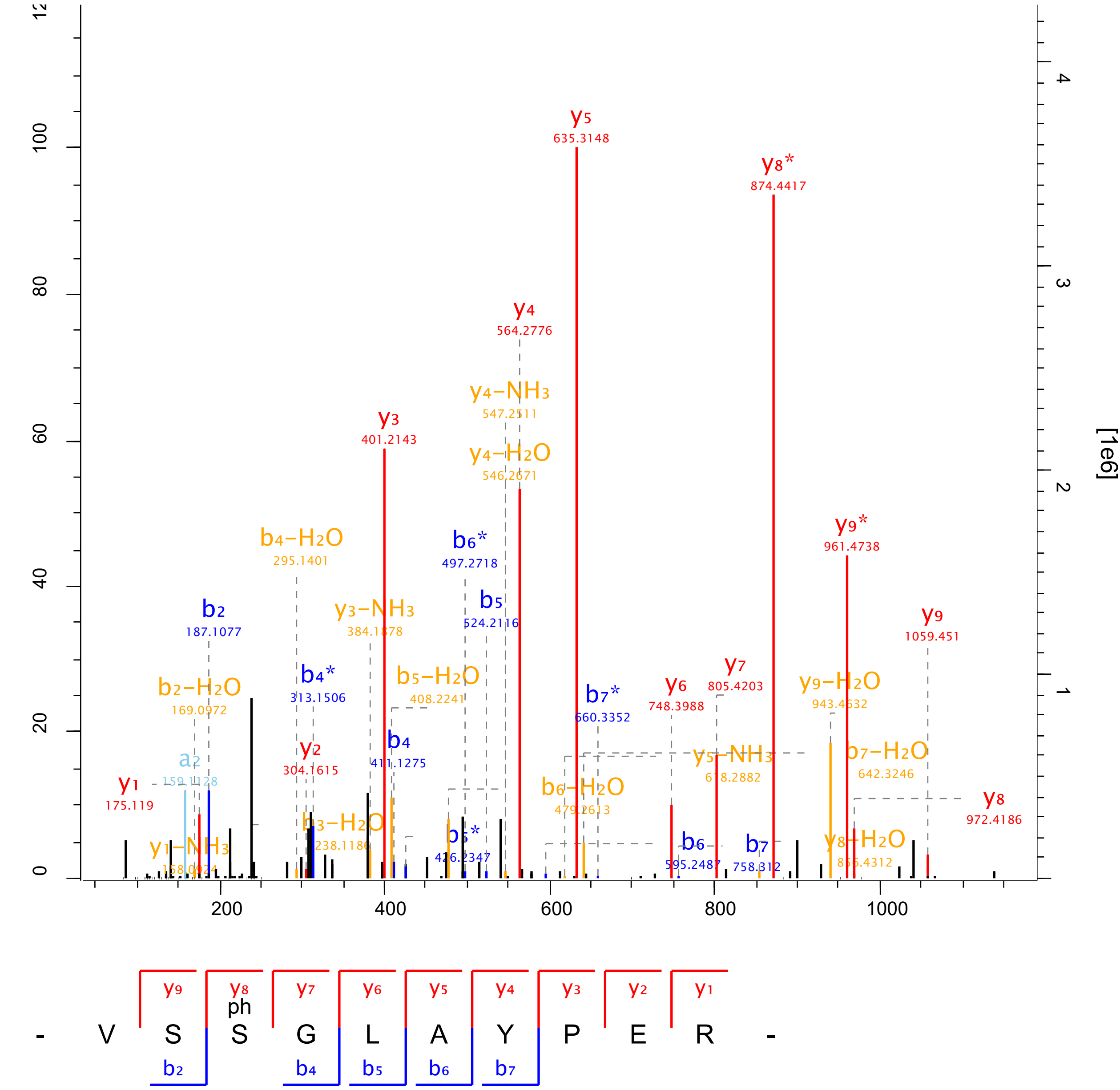

| Raw file | Scan | Method | Score | m/z |
| --- | --- | --- | --- | --- |
| sys_15_1 | 13875 | FTMS; HCD | 325.18 | 946.36 |

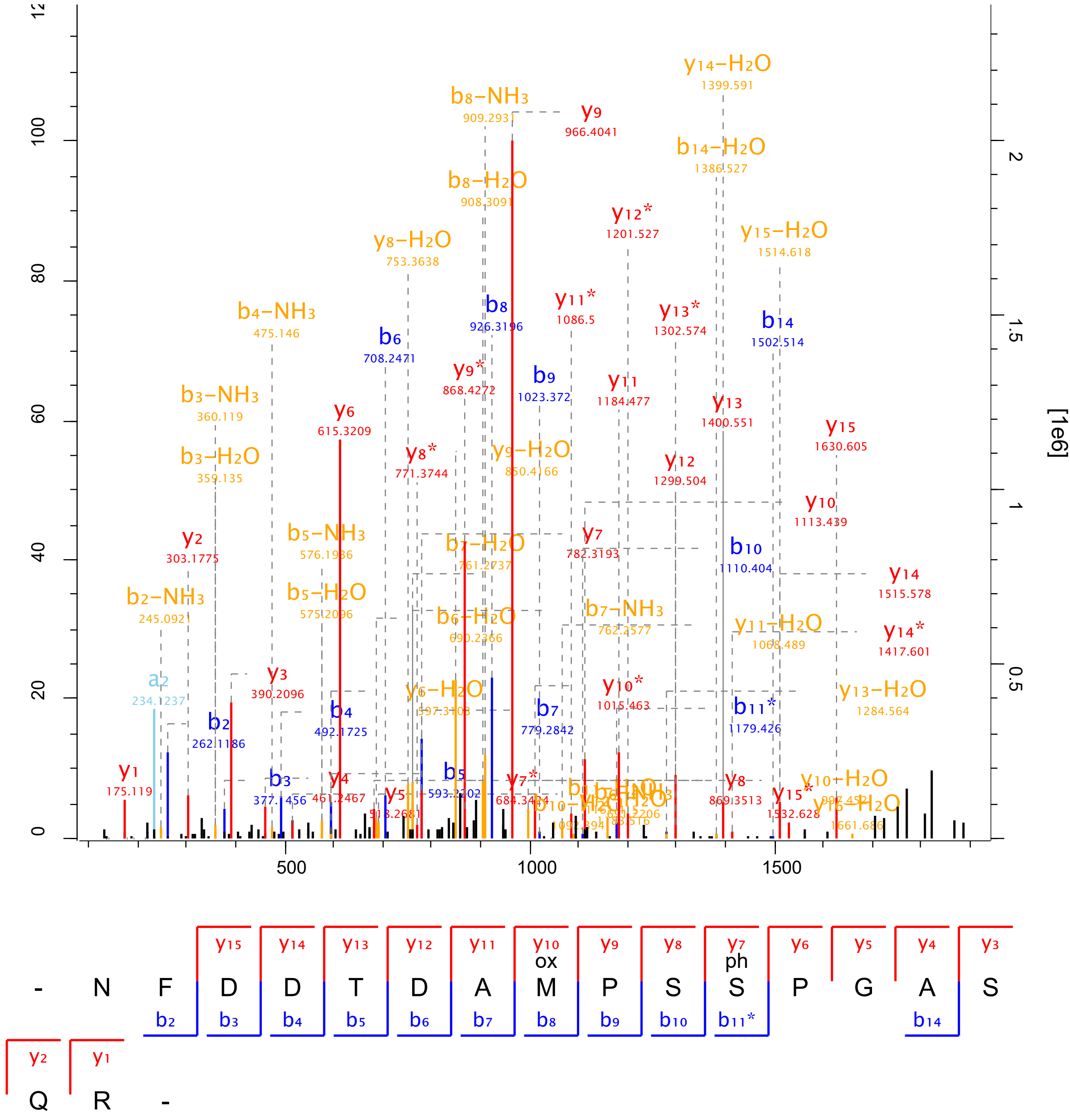

| Raw file | Scan | Method | Score | m/z |
| --- | --- | --- | --- | --- |
| sys_15_1 | 13948 | FTMS; HCD | 50.18 | 803.33 |

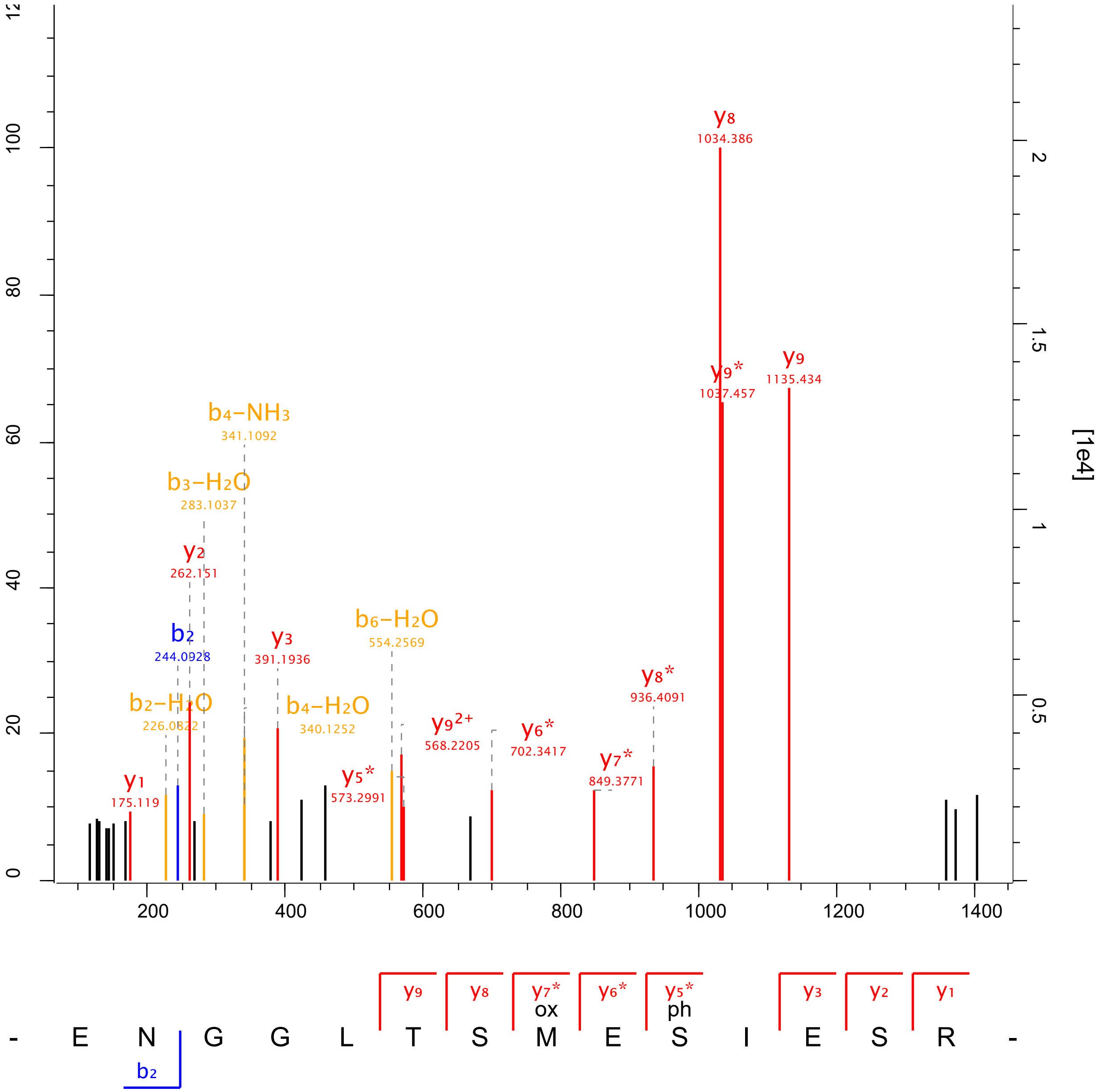

| Raw file | Scan | Method | Score | m/z |
| --- | --- | --- | --- | --- |
| sys_15_1 | 14016 | FTMS; HCD | 45.37 | 559.69 |

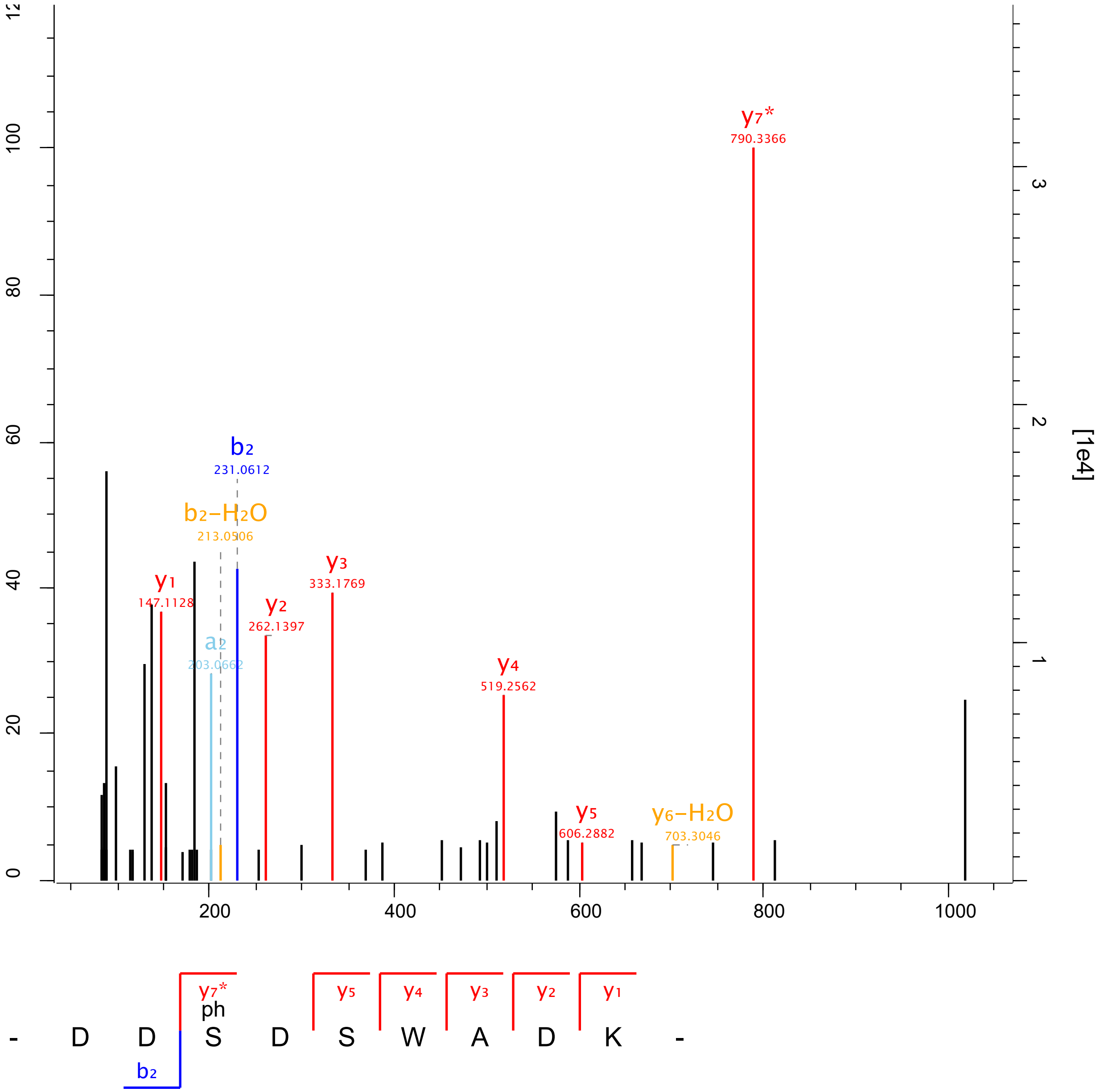

| Raw file | Scan | Method | Score | m/z |
|---|---|---|---|---|
| sys_15_1 | 14039 | FTMS; HCD | 95.57 | 610.79 |

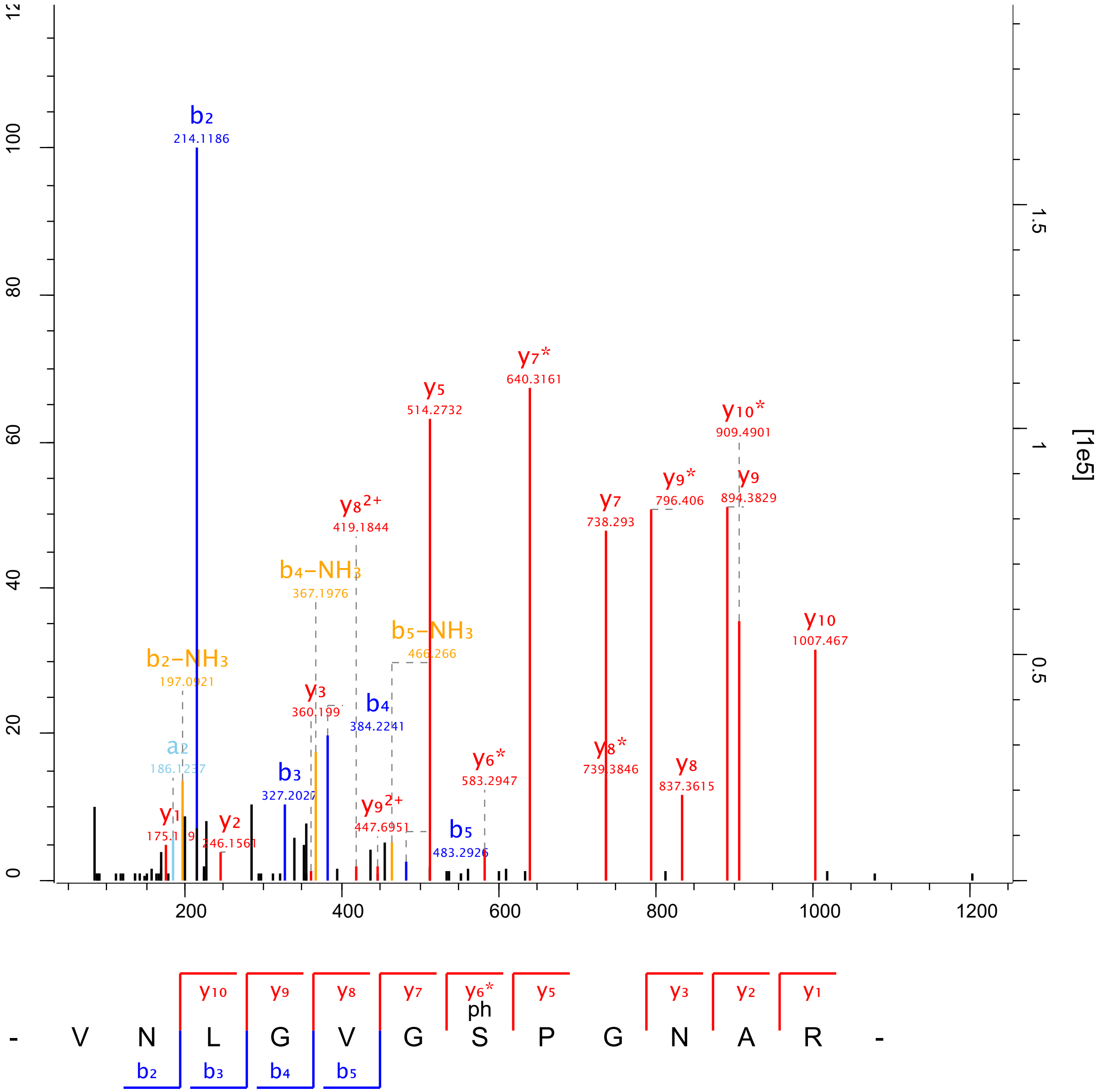

| Raw file | Scan | Method | Score | m/z |
|---|---|---|---|---|
| sys_15_1 | 14217 | FTMS; HCD | 136.14 | 547.27 |

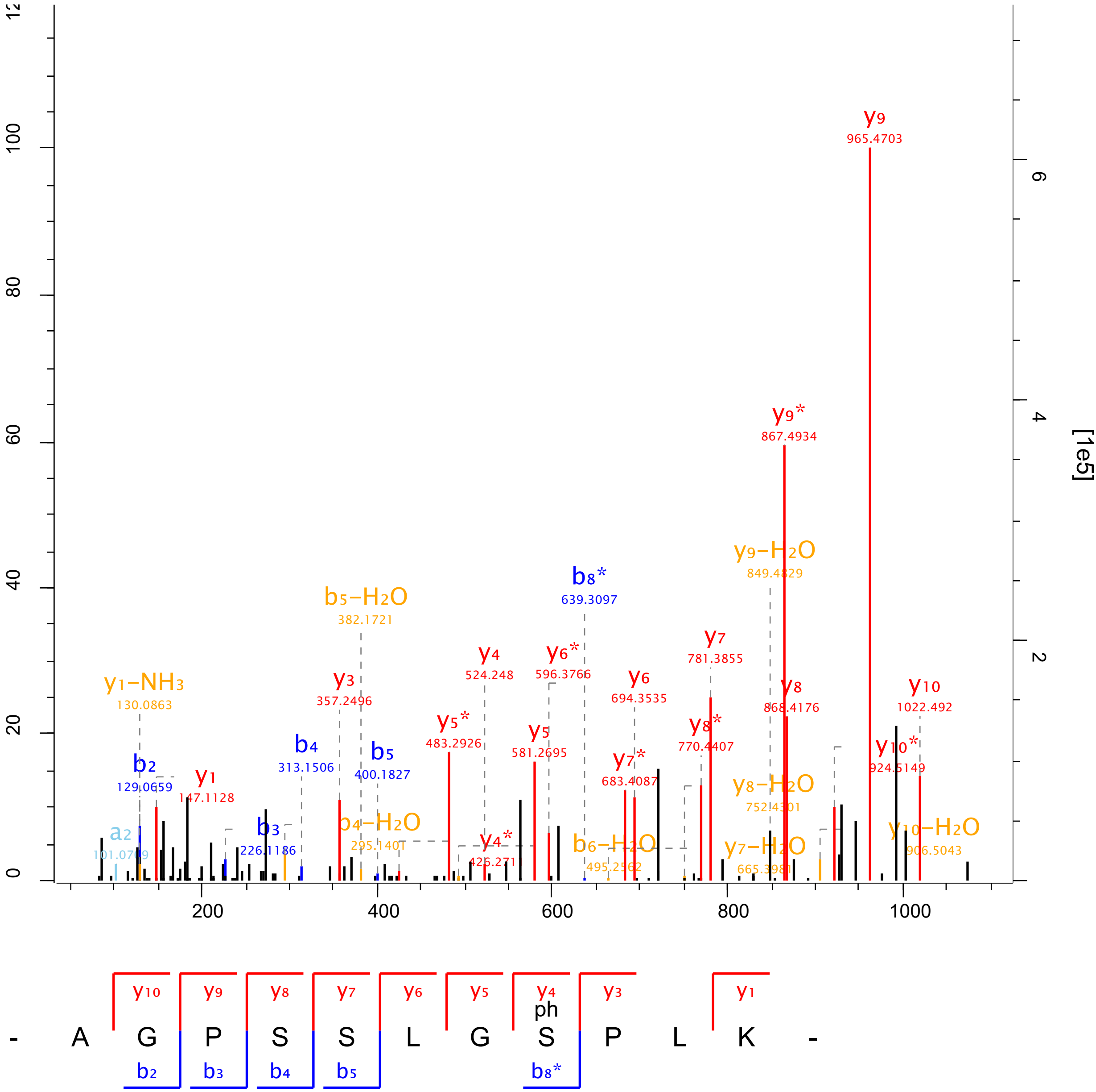

| Raw file | Scan | Method | Score | m/z |
|---|---|---|---|---|
| sys_15_1 | 14263 | FTMS; HCD | 248.23 | 891.87 |

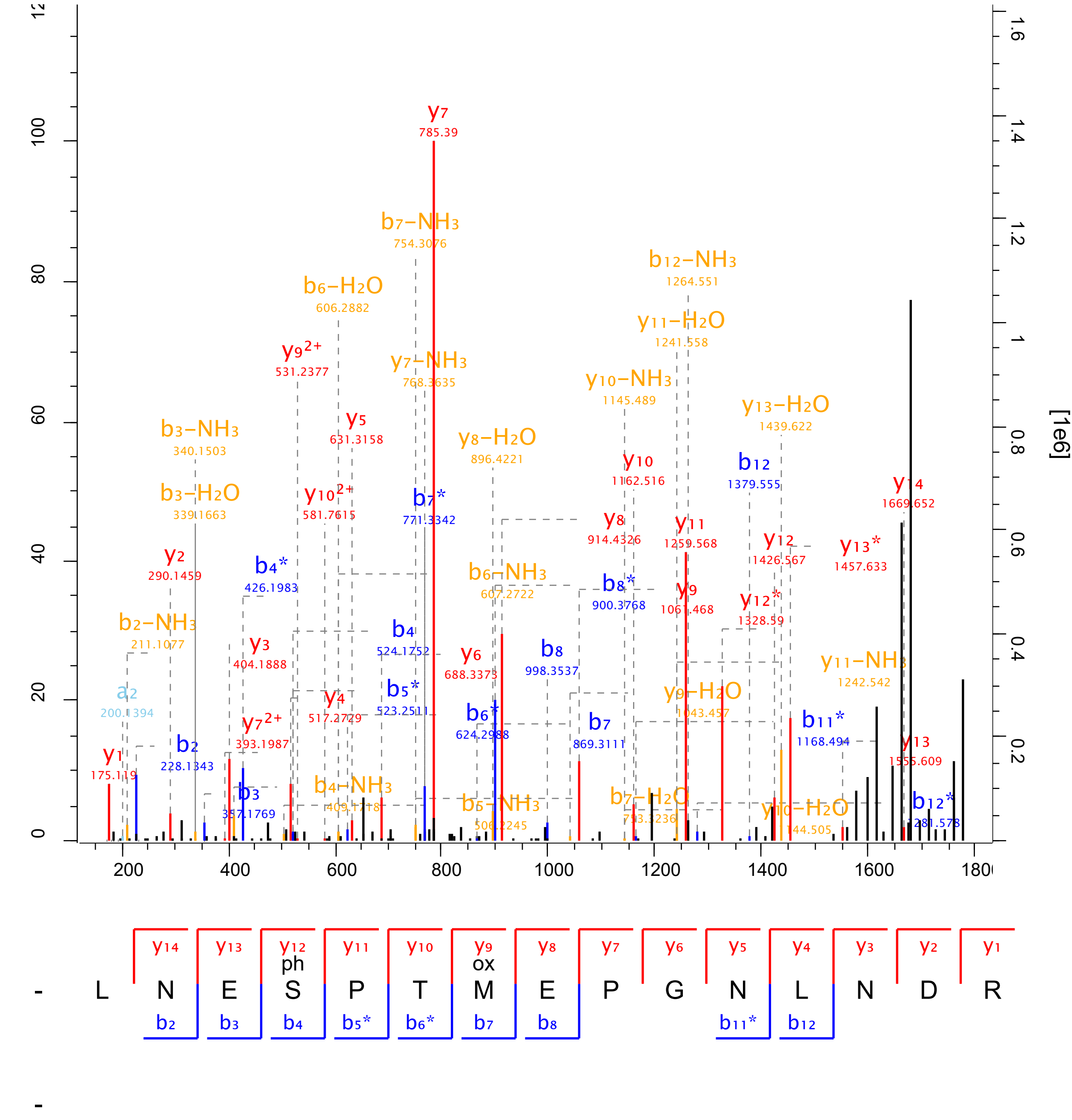

| Raw file | Scan | Method | Score | m/z |
|---|---|---|---|---|
| sys_15_1 | 14334 | FTMS; HCD | 136.43 | 665.28 |

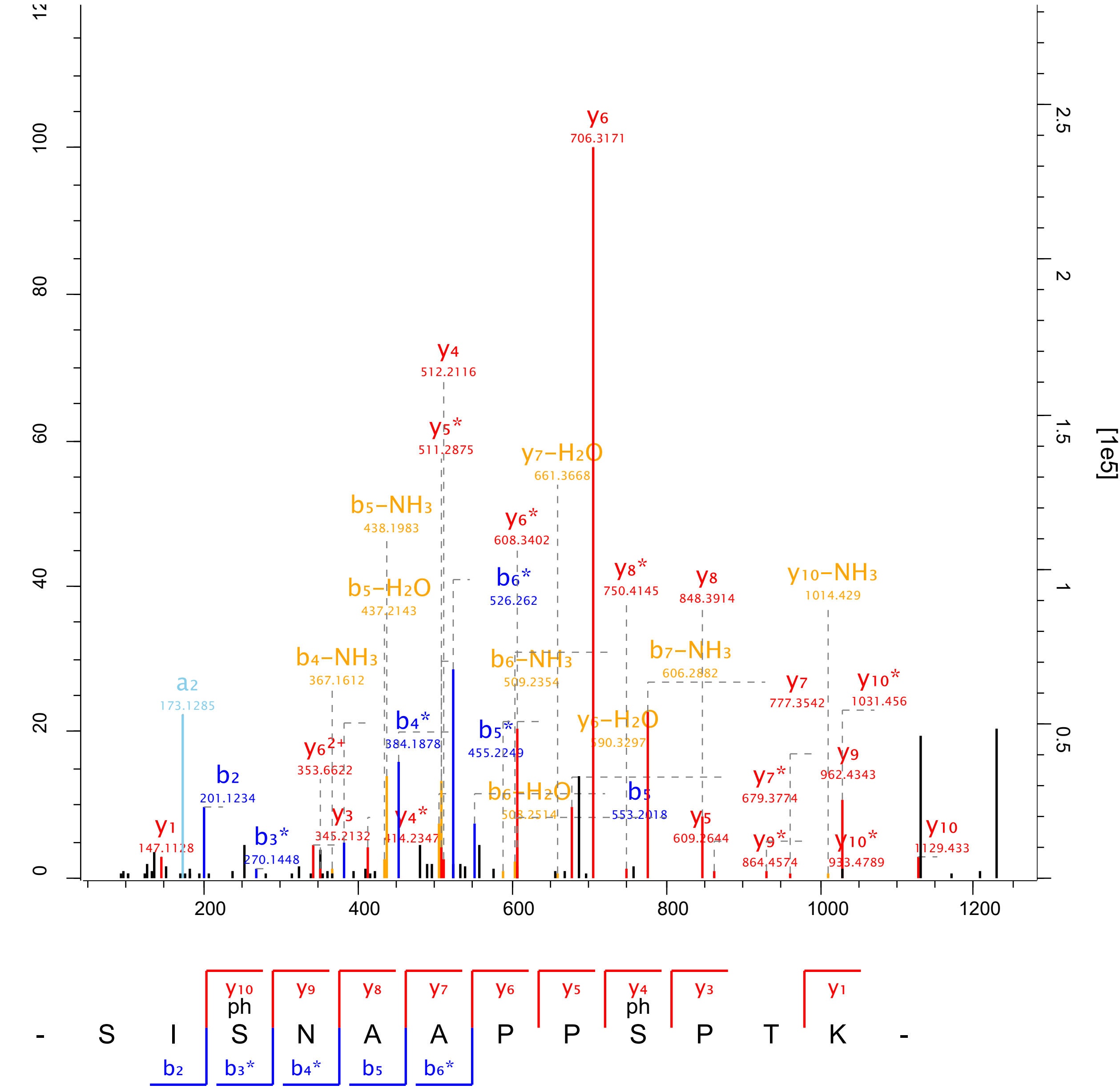

| Raw file | Scan | Method | Score | m/z |
|---|---|---|---|---|
| sys_15_1 | 14339 | FTMS; HCD | 167.18 | 742.28 |

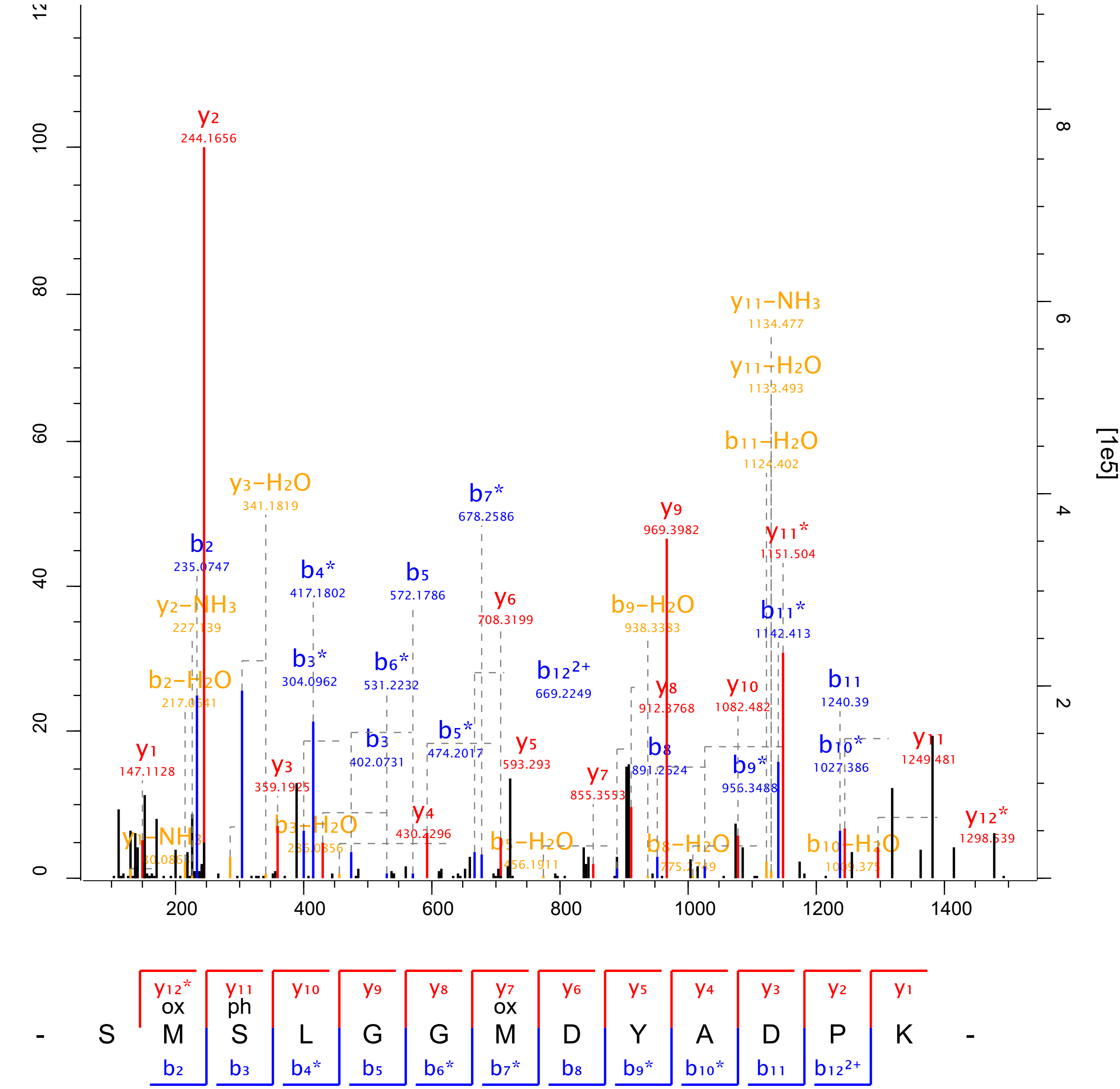

| Raw file | Scan | Method | Score | m/z |
|---|---|---|---|---|
| sys_15_1 | 14353 | FTMS; HCD | 267.38 | 728.82 |

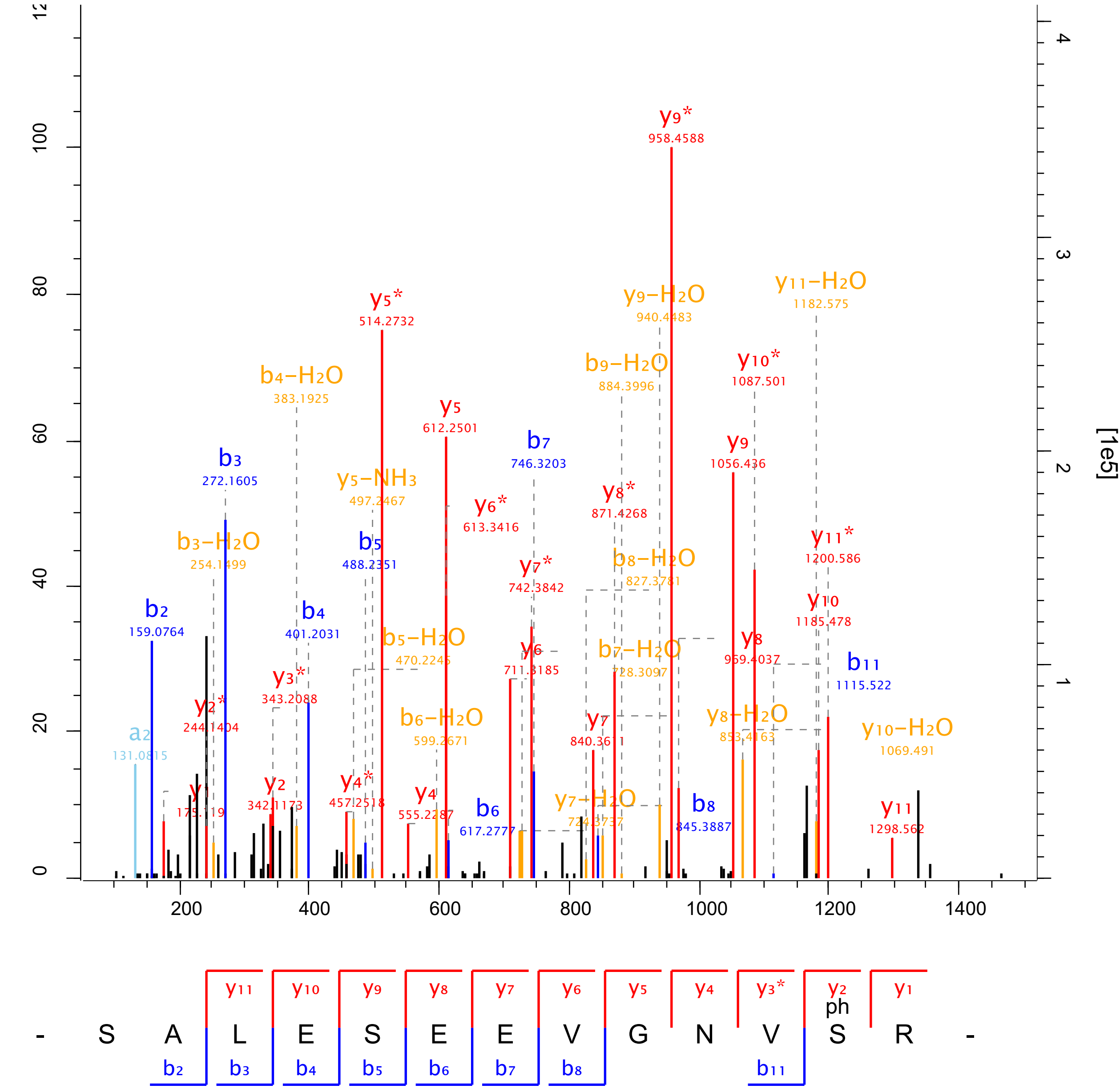

| Raw file | Scan | Method | Score | m/z |
|---|---|---|---|---|
| sys_15_1 | 14357 | FTMS; HCD | 77.22 | 816.8 |

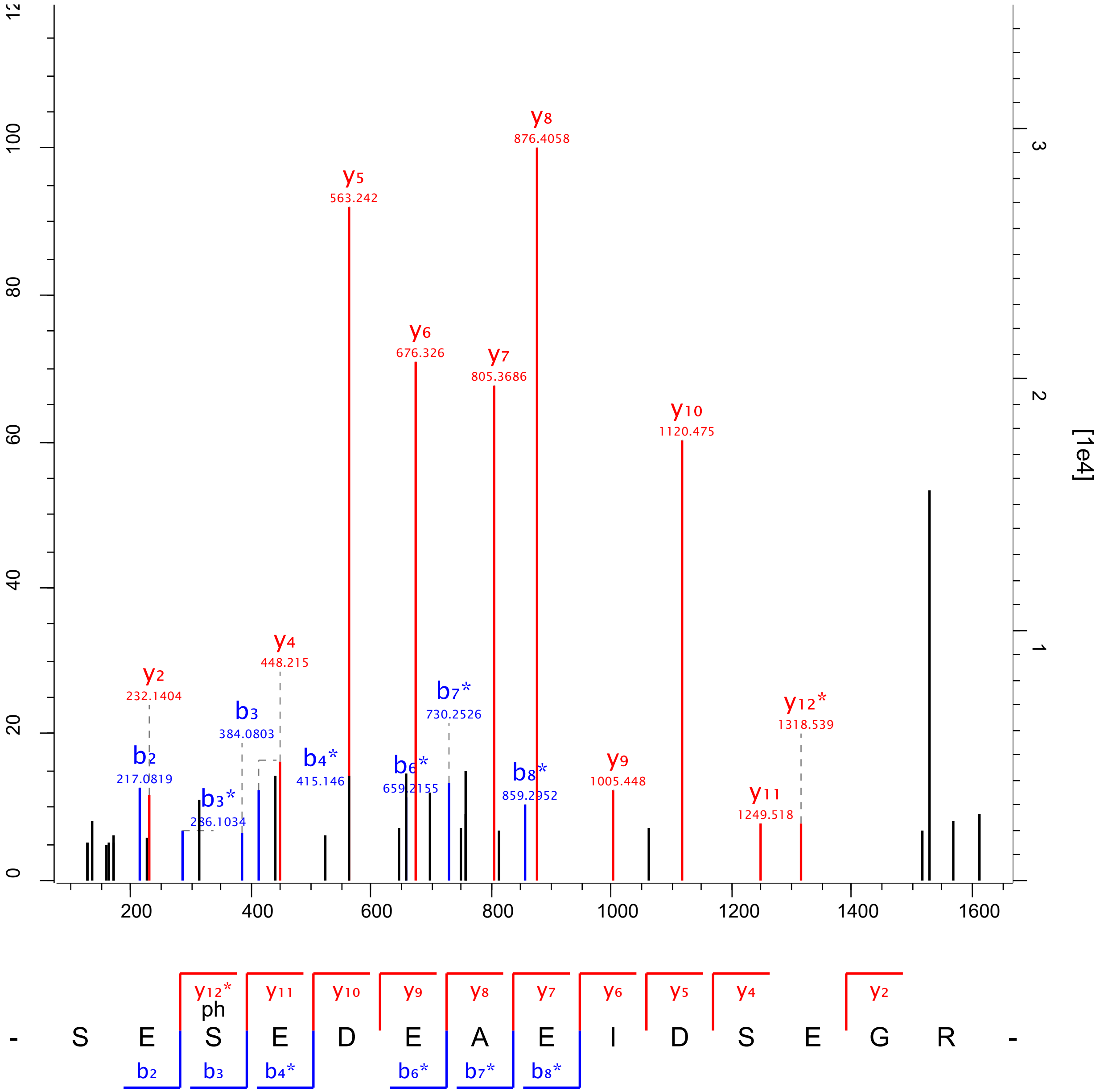

| Raw file | Scan | Method | Score | m/z |
|---|---|---|---|---|
| sys_15_1 | 14392 | FTMS; HCD | 72.9 | 755.29 |

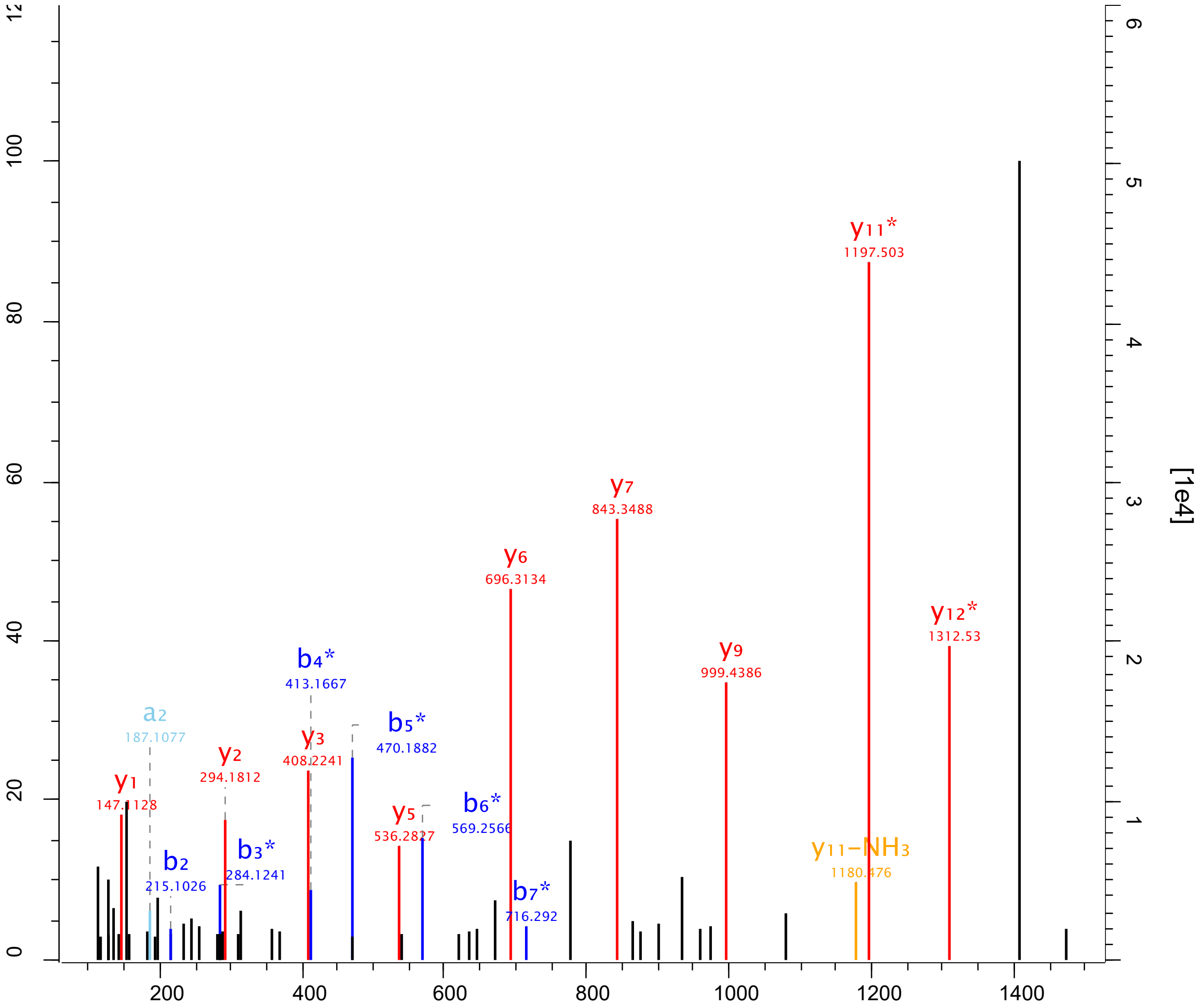


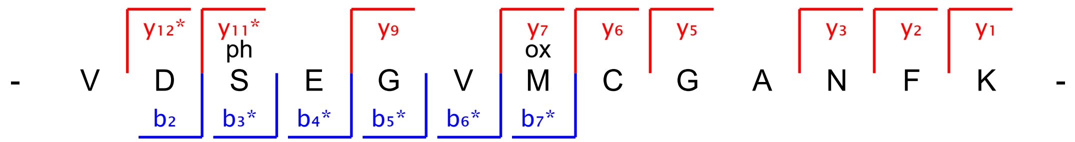

| Raw file | Scan | Method | Score | m/z |
|---|---|---|---|---|
| sys_15_1 | 14453 | FTMS; HCD | 237.5 | 928.39 |

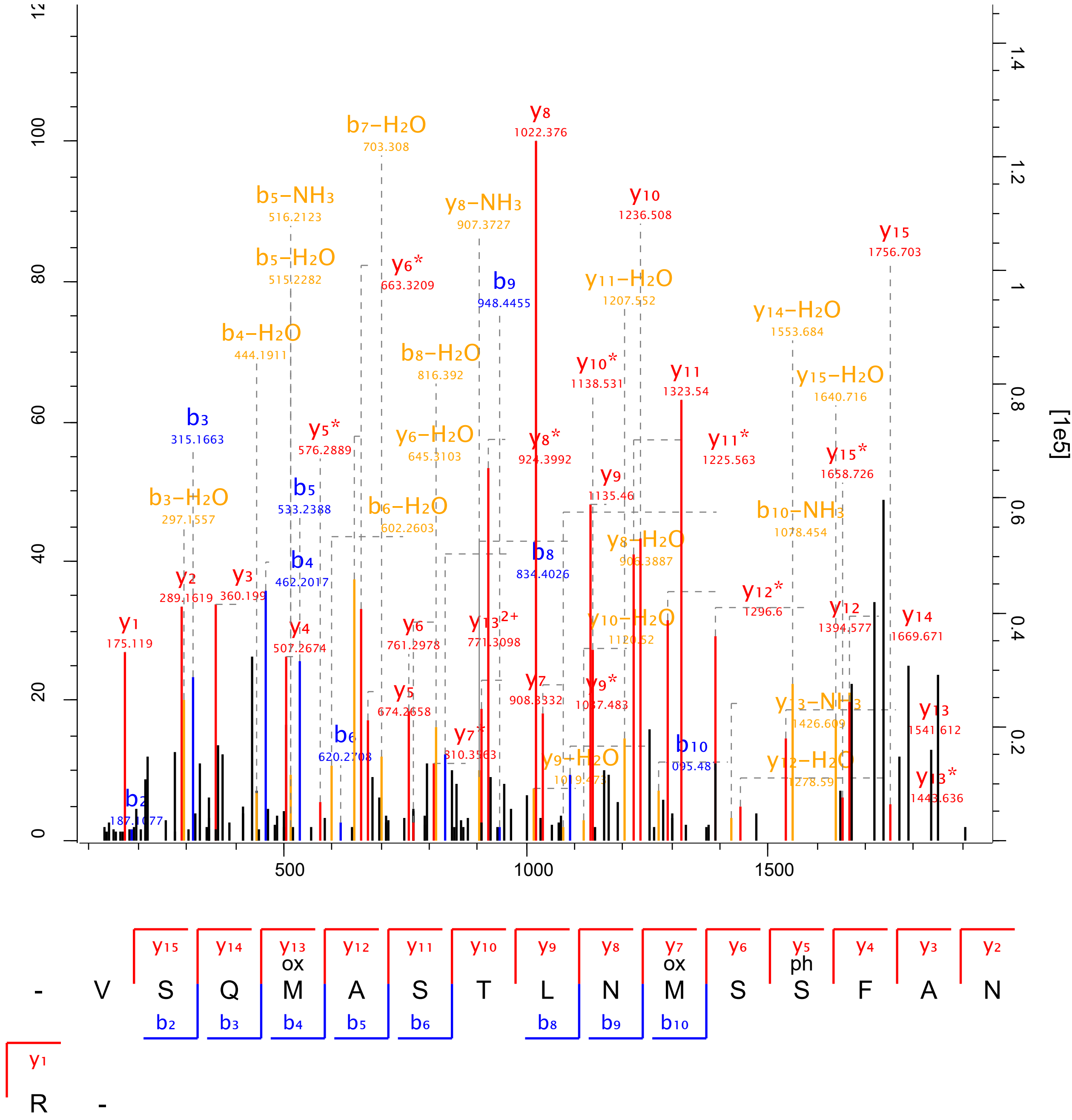

| Raw file | Scan | Method | Score | m/z |
|---|---|---|---|---|
| sys_15_1 | 14462 | FTMS; HCD | 119.39 | 556.75 |

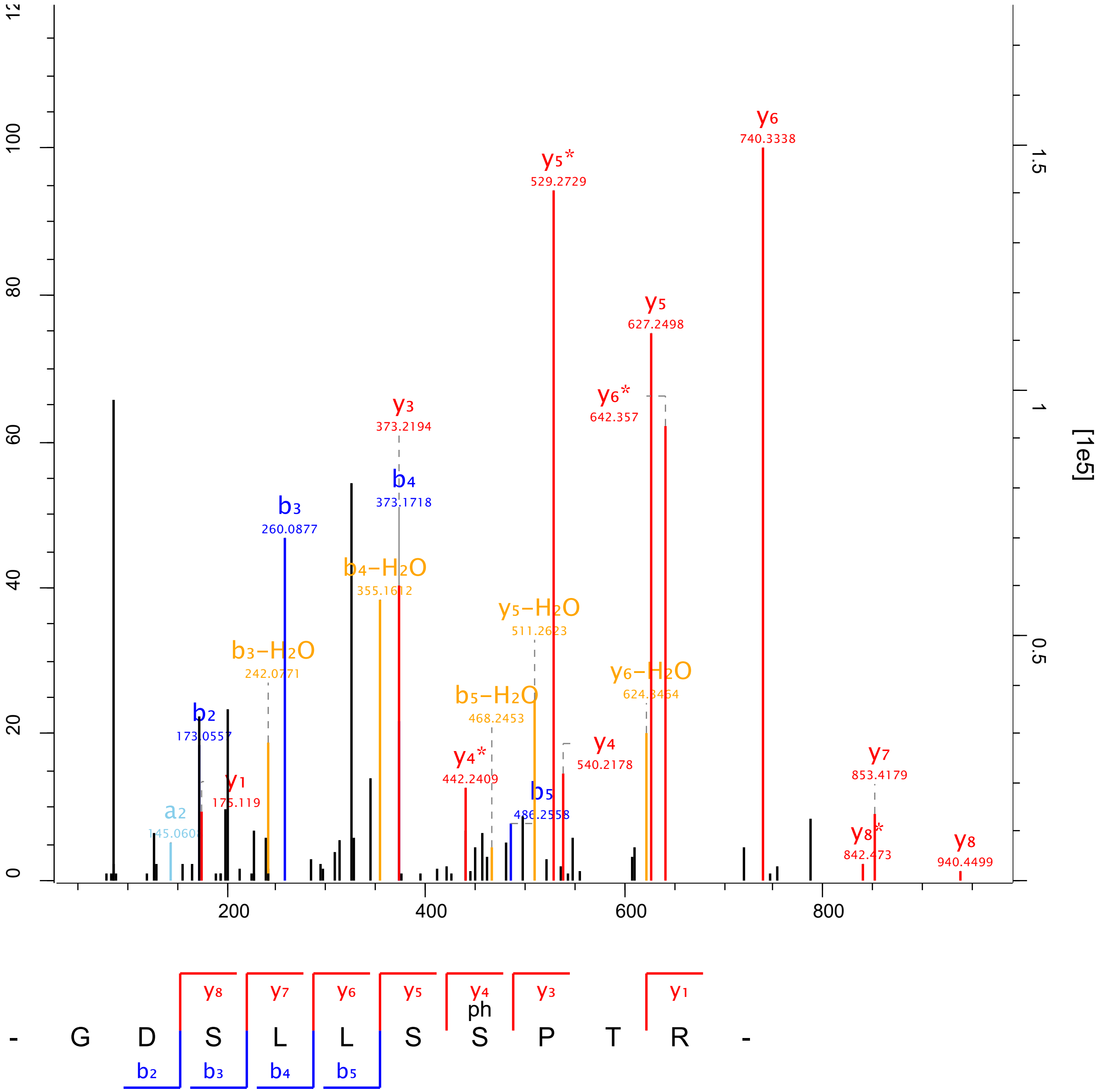

| Raw file | Scan | Method | Score | m/z |
|---|---|---|---|---|
| sys_15_1 | 14488 | FTMS; HCD | 242.71 | 1012.43 |

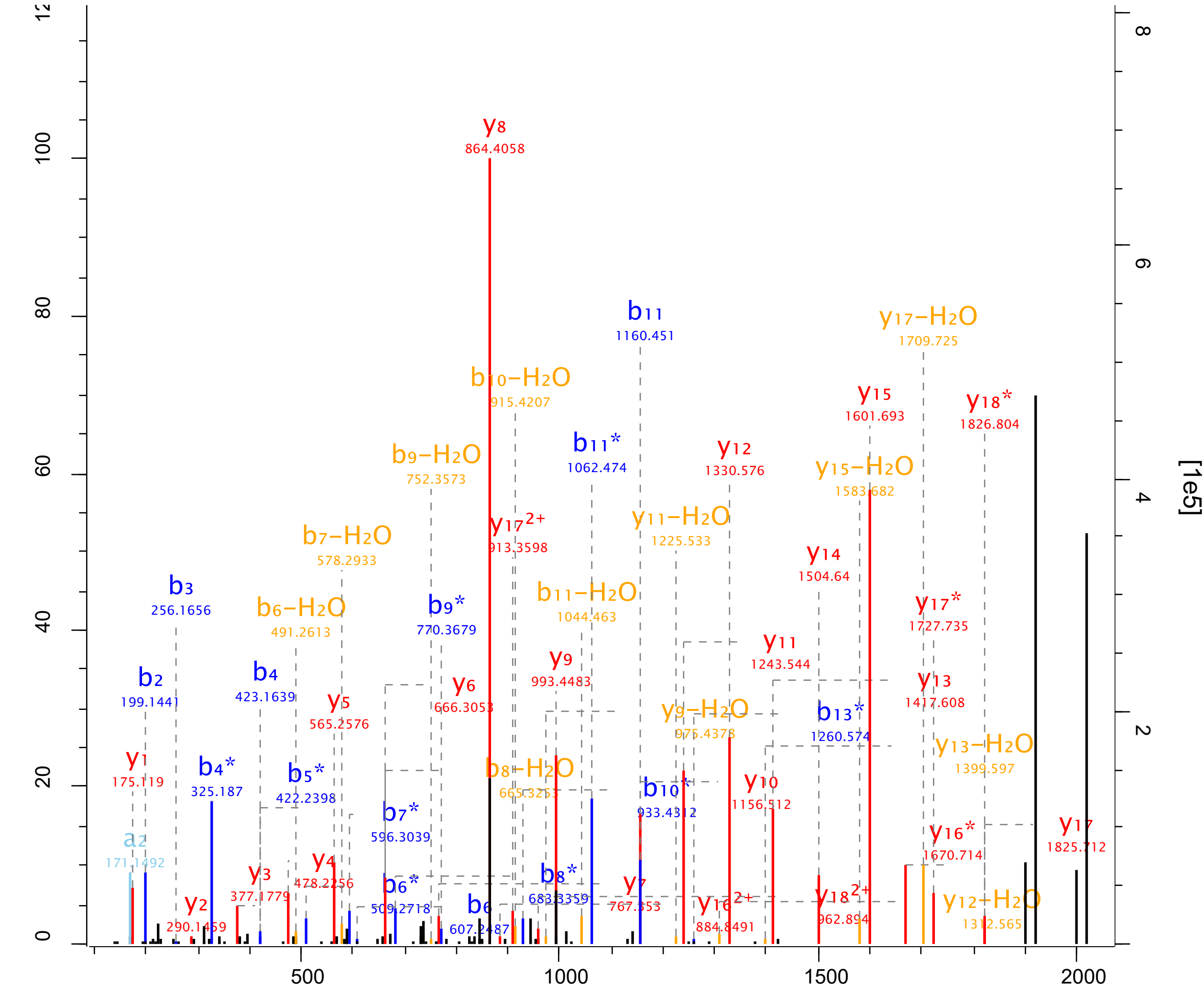


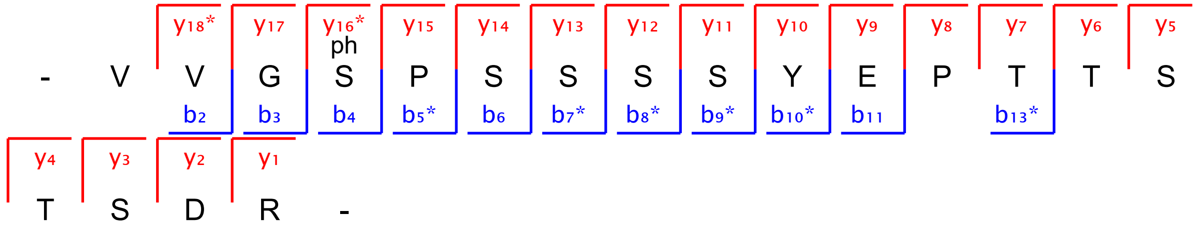

| Raw file | Scan | Method | Score | m/z |
| --- | --- | --- | --- | --- |
| sys_15_1 | 14562 | FTMS; HCD | 67.65 | 806.36 |

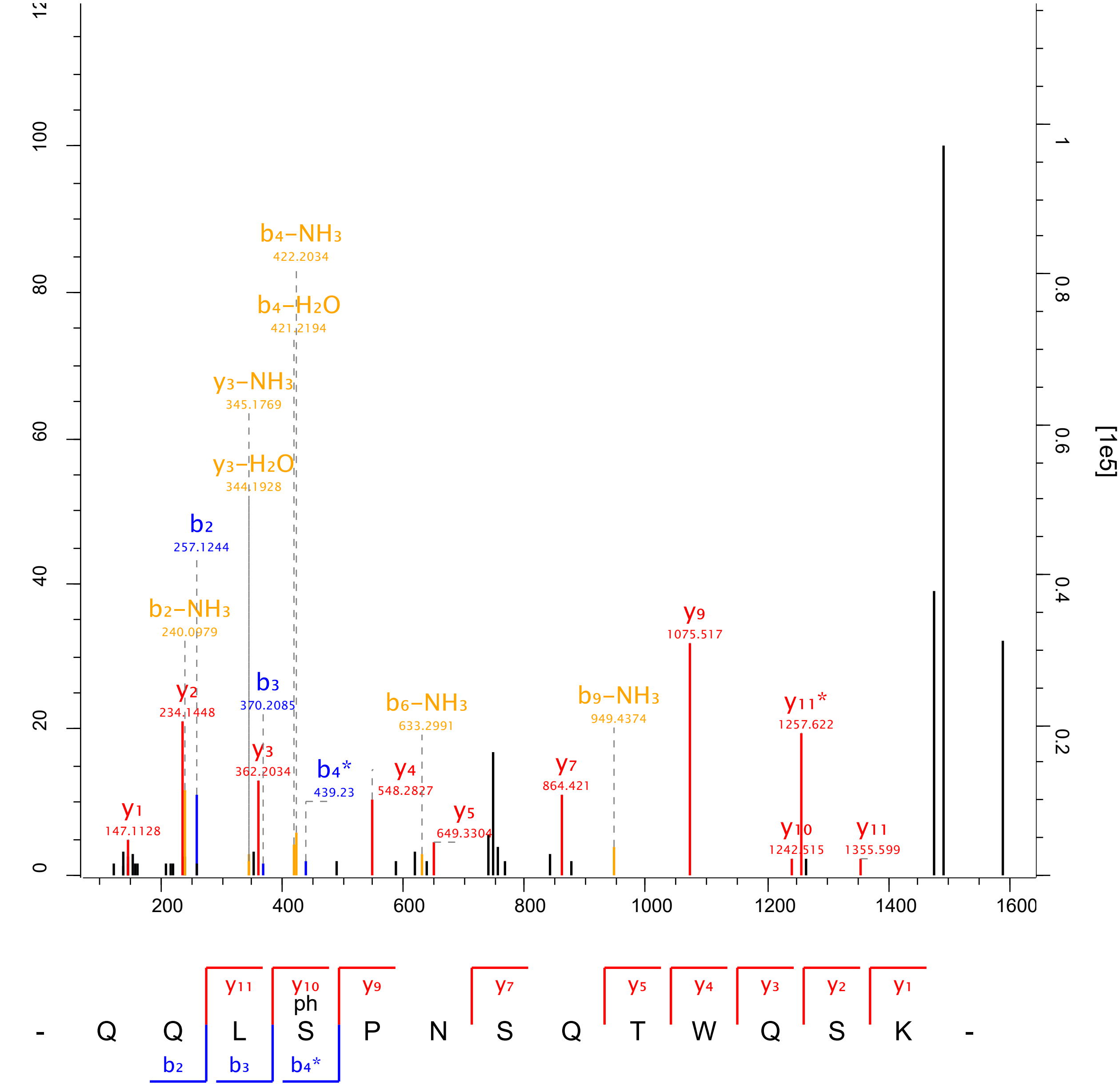

| Raw file | Scan | Method | Score | m/z |
|---|---|---|---|---|
| sys_15_1 | 14567 | FTMS; HCD | 80.01 | 565.73 |

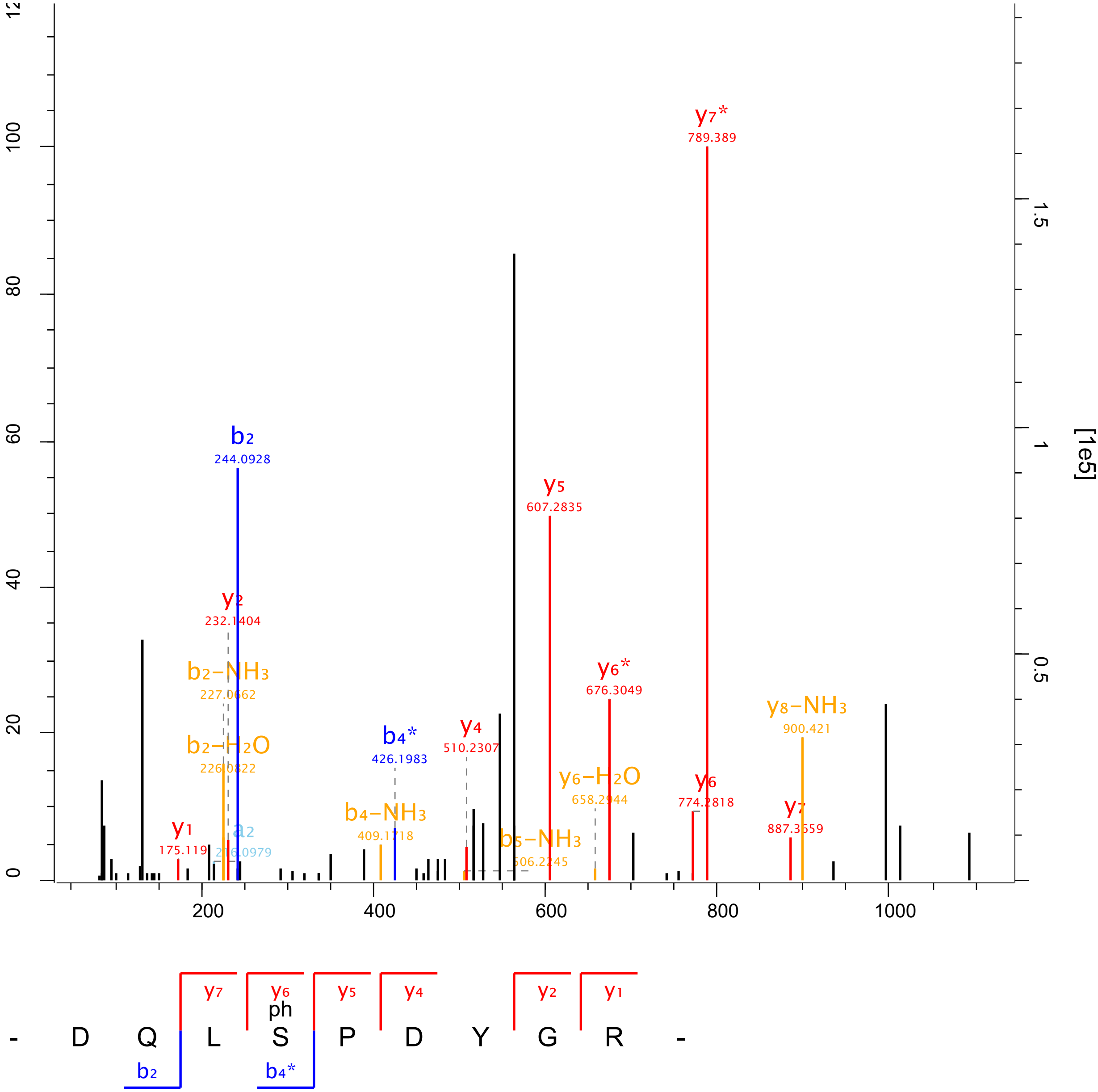

| Raw file | Scan | Method | Score | m/z |
|---|---|---|---|---|
| sys_15_1 | 14675 | FTMS; HCD | 254.72 | 725.29 |

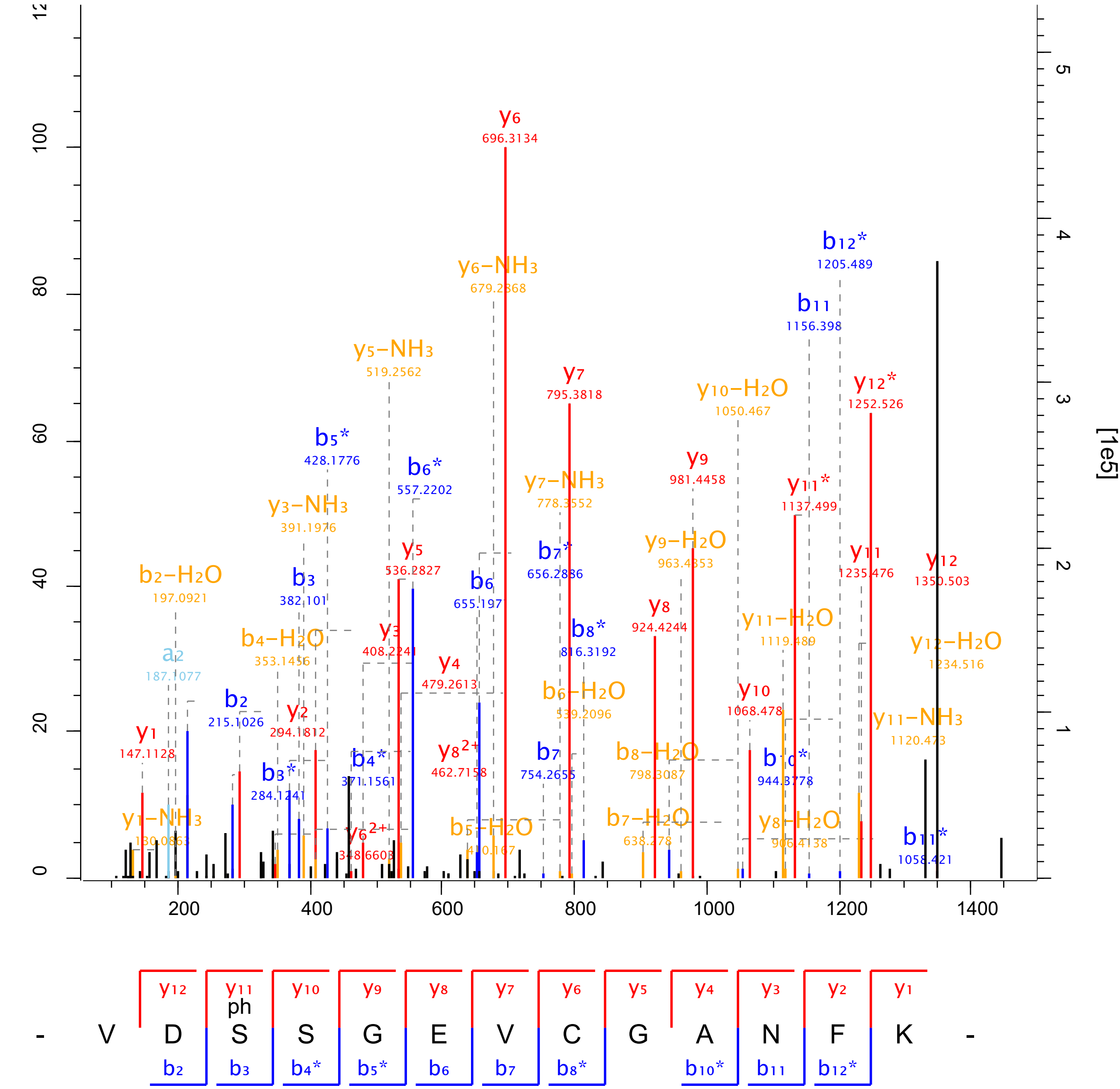

| Raw file | Scan | Method | Score | m/z |
| --- | --- | --- | --- | --- |
| sys_15_1 | 14695 | FTMS; HCD | 41.51 | 960.94 |

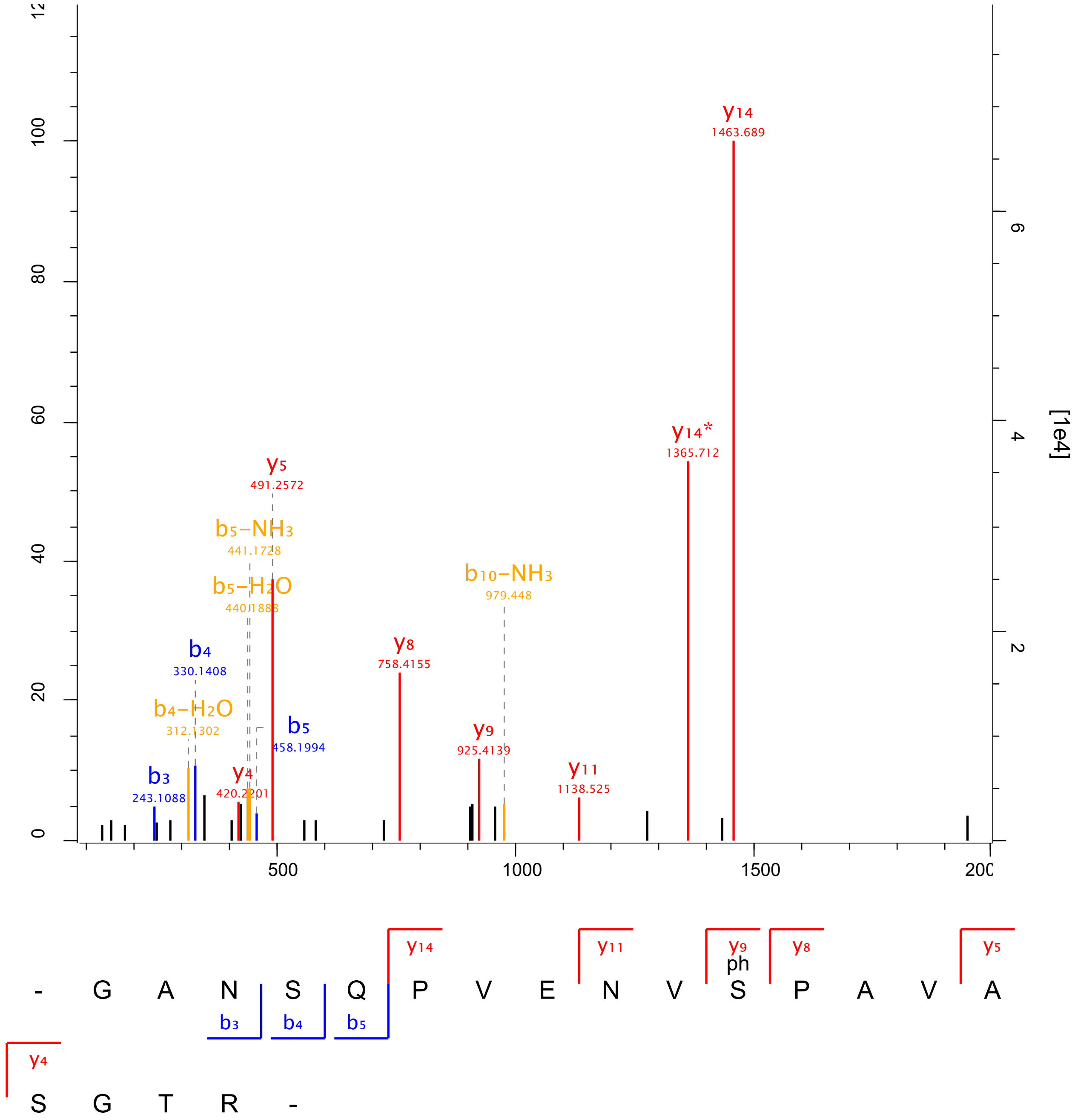

| Raw file | Scan | Method | Score | m/z |
|---|---|---|---|---|
| sys_15_1 | 14850 | FTMS; HCD | 94.09 | 679.27 |

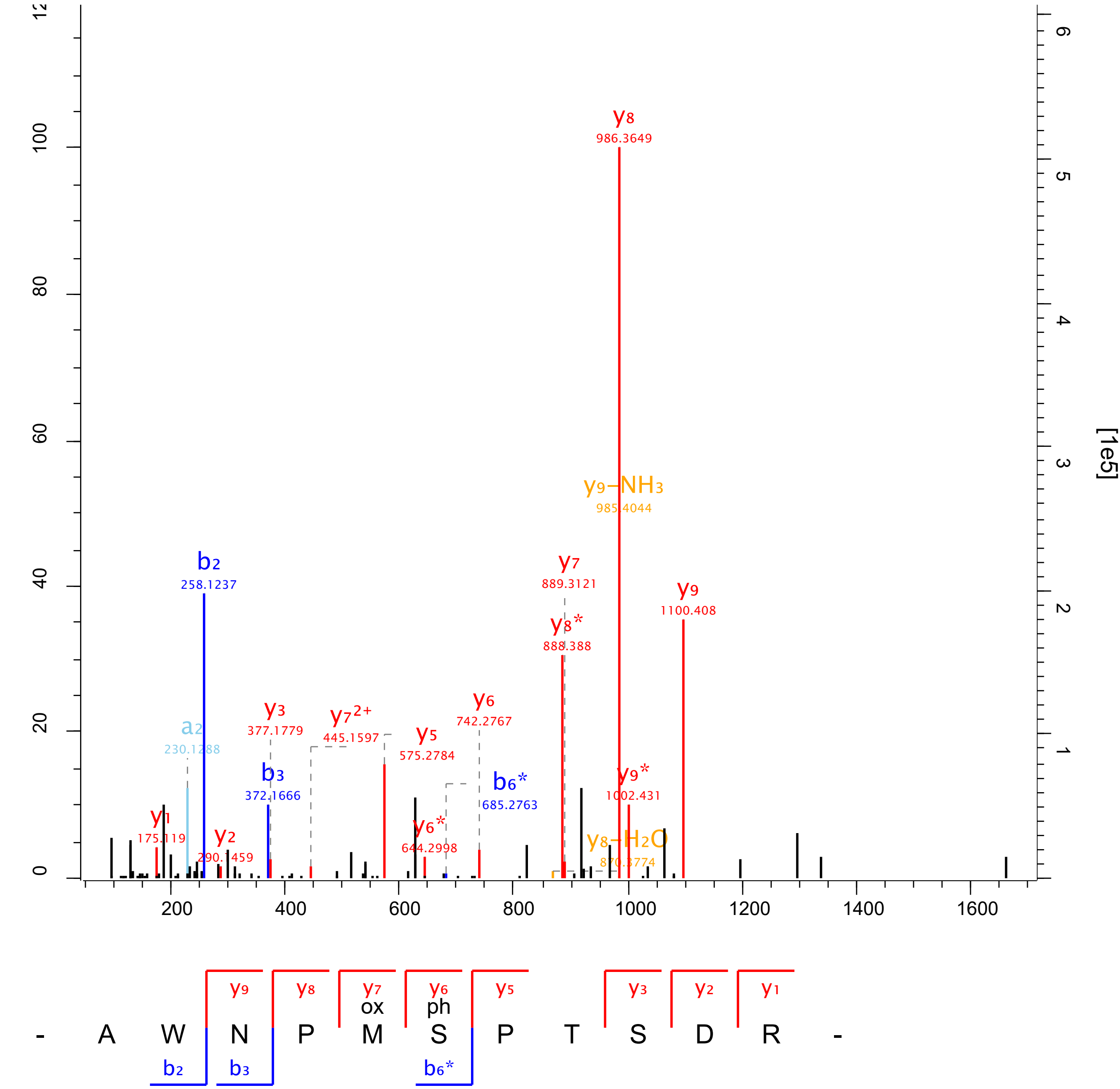

| Raw file | Scan | Method | Score | m/z |
|---|---|---|---|---|
| sys_15_1 | 14894 | FTMS; HCD | 118.2 | 670.3 |

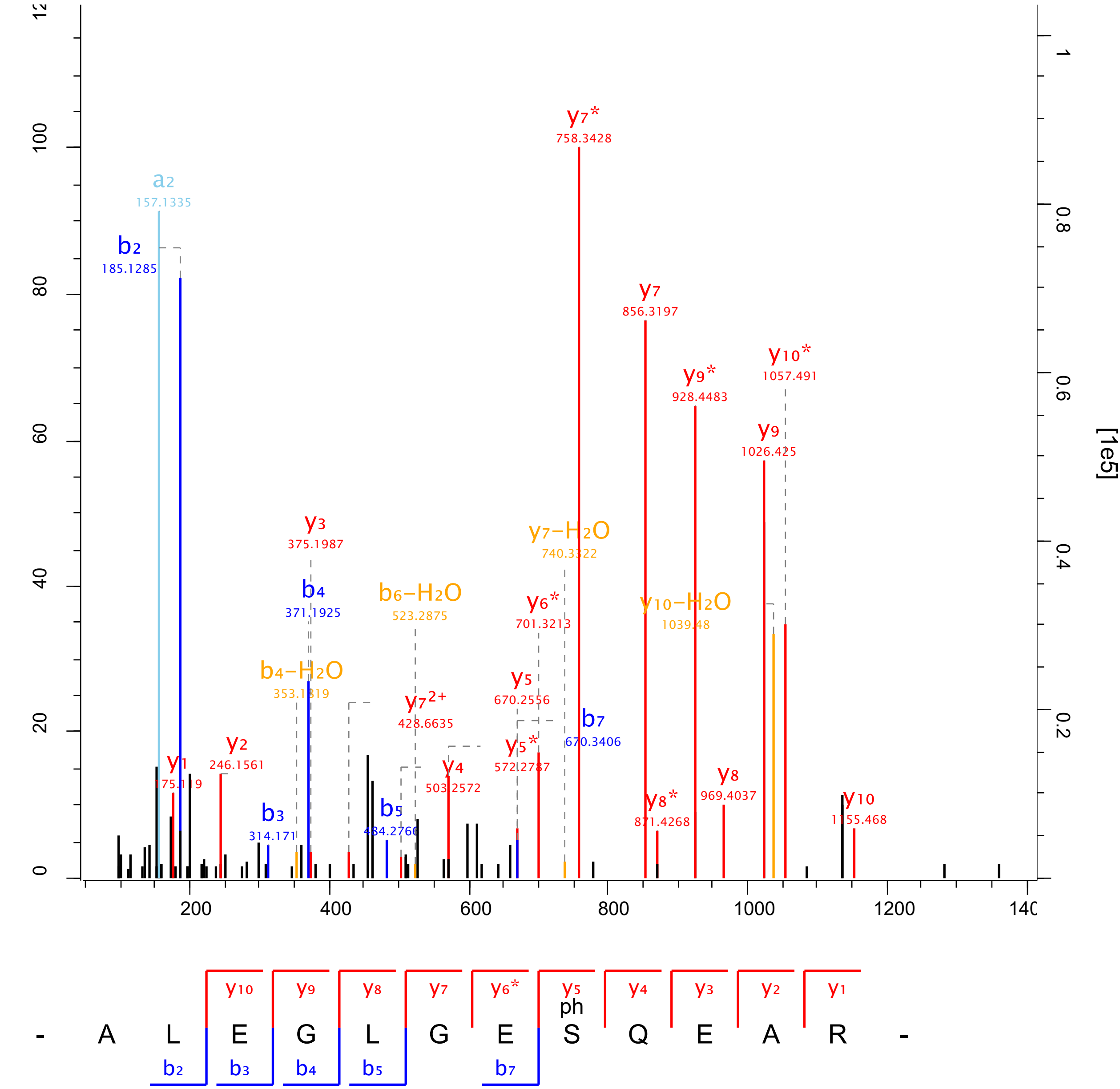

| Raw file | Scan | Method | Score | m/z |
|---|---|---|---|---|
| sys_15_1 | 14948 | FTMS; HCD | 111.4 | 926.4 |

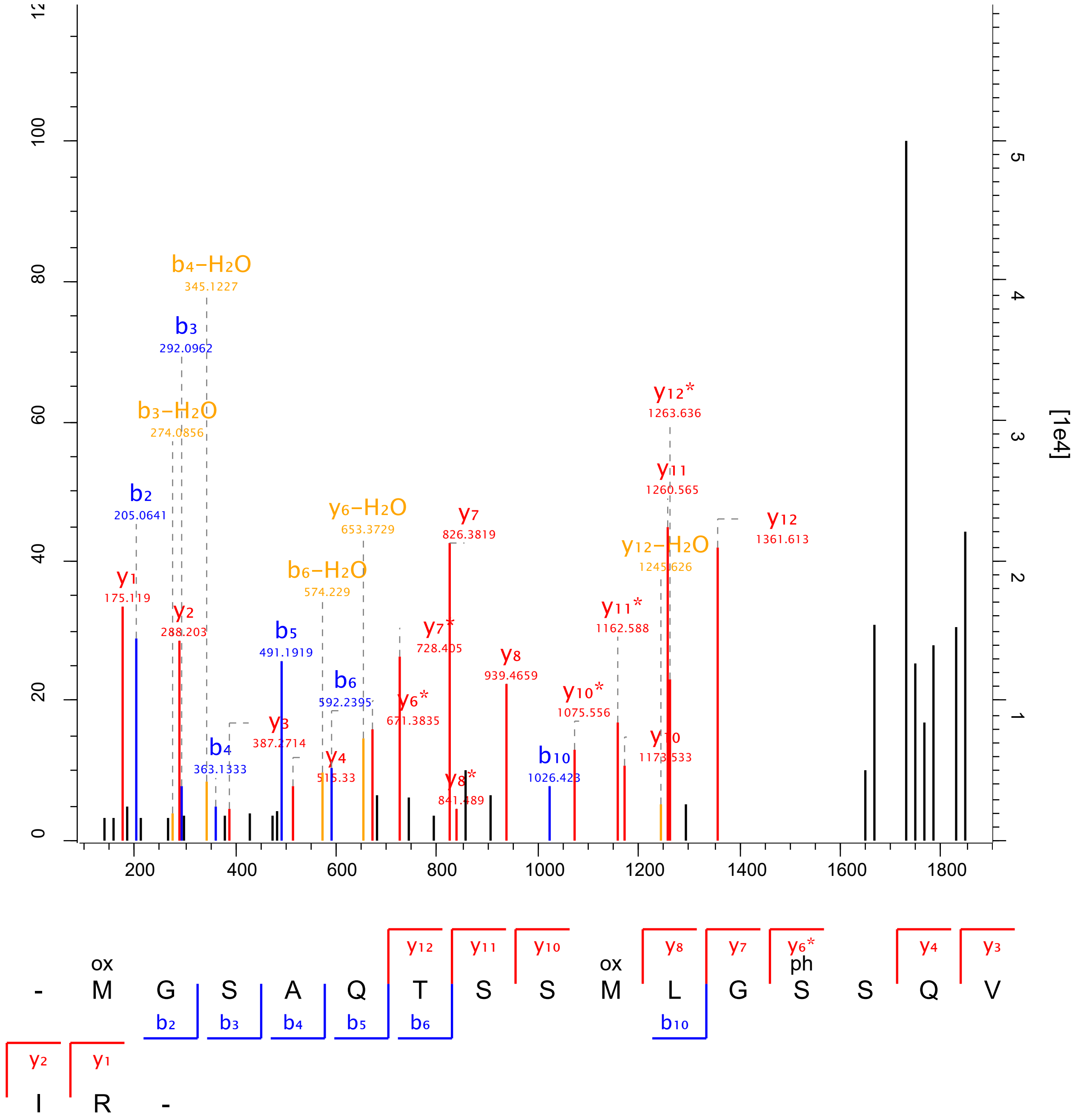

| Raw file | Scan | Method | Score | m/z |
|---|---|---|---|---|
| sys_15_1 | 14991 | FTMS; HCD | 162.59 | 751.33 |

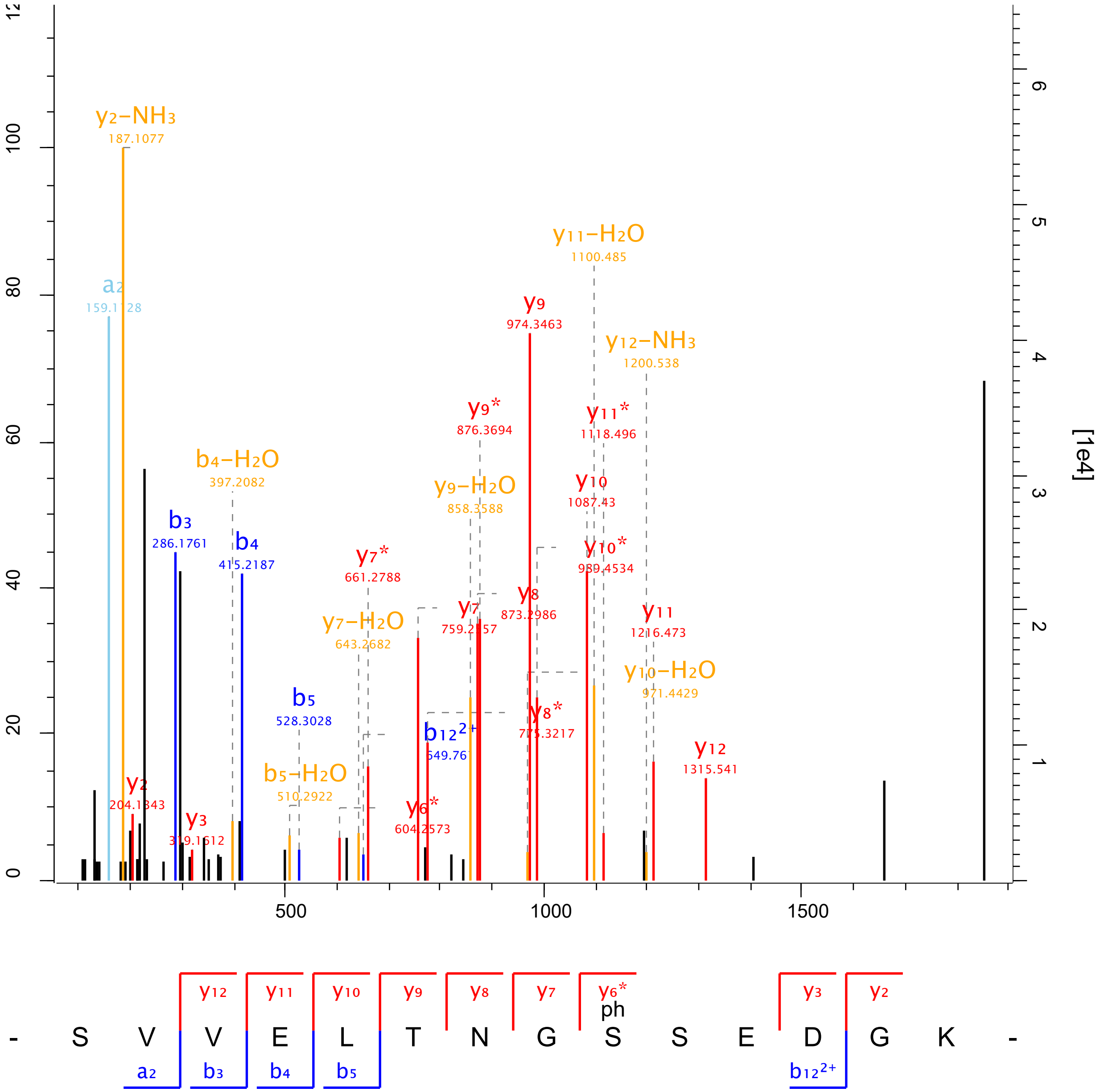

| Raw file | Scan | Method | Score | m/z |
|---|---|---|---|---|
| sys_15_1 | 15005 | FTMS; HCD | 49.3 | 634.79 |

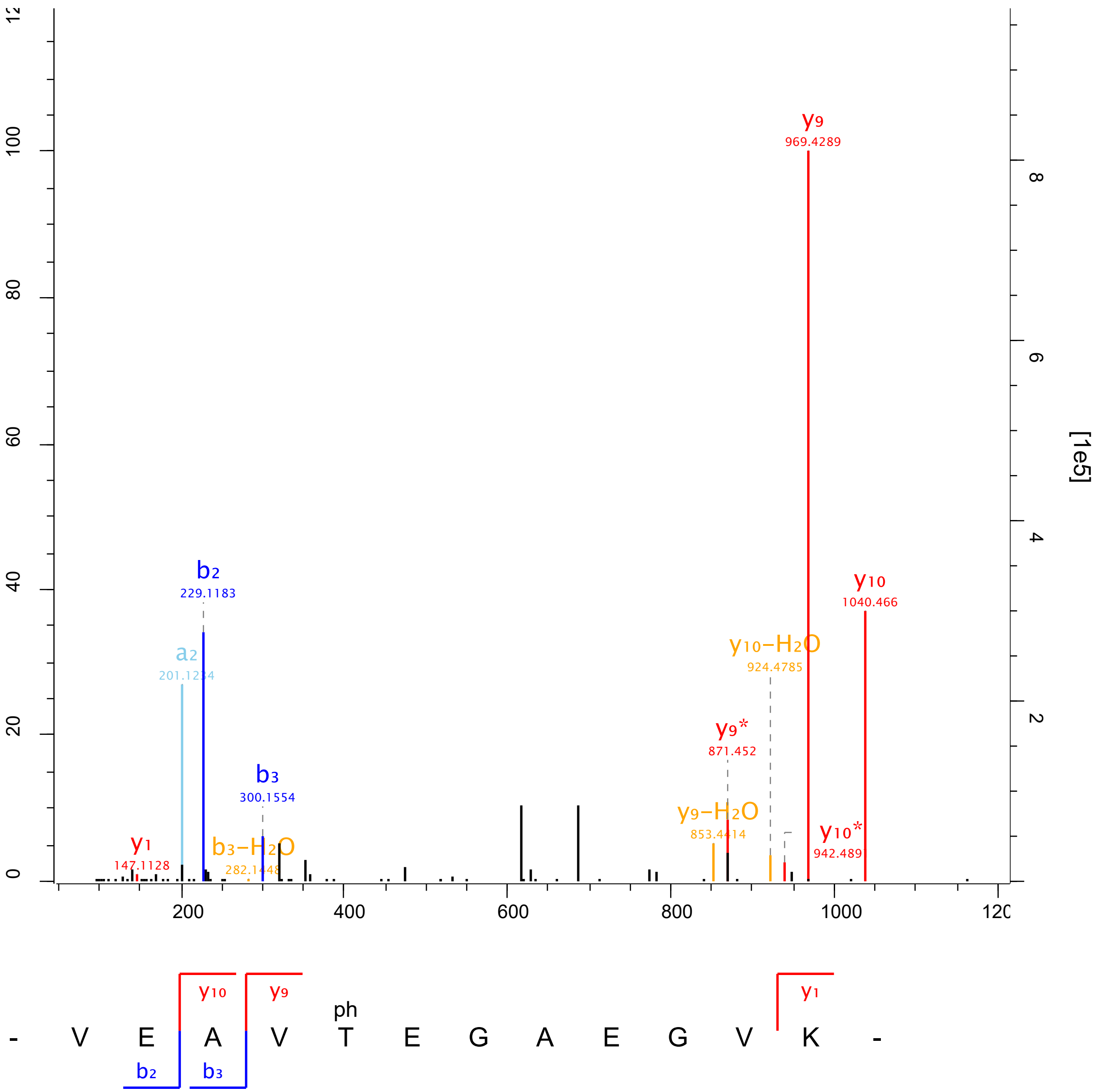

| Raw file | Scan | Method | Score | m/z |
|---|---|---|---|---|
| sys_15_1 | 15127 | FTMS; HCD | 63.69 | 877.35 |

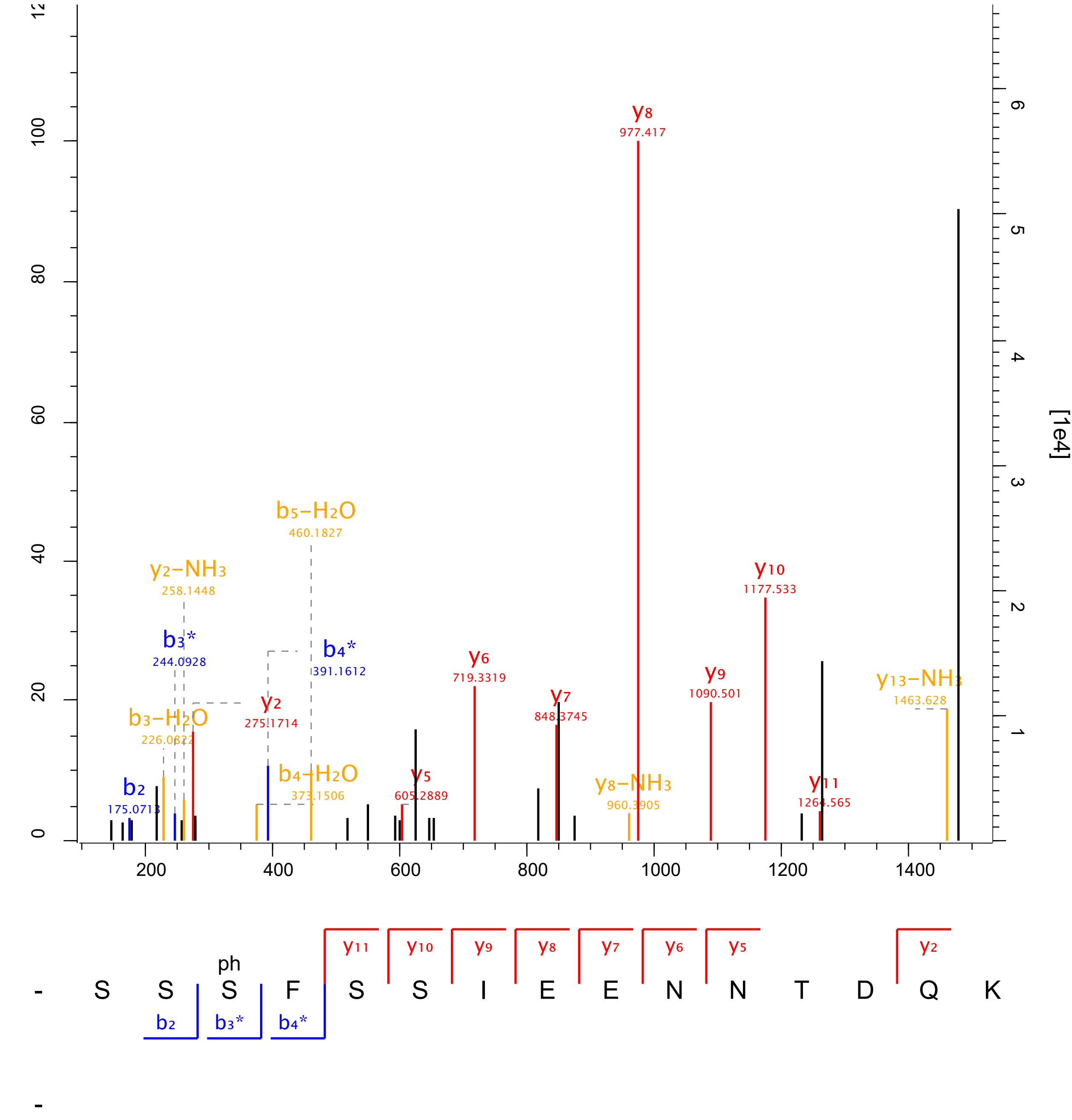

| Raw file | Scan | Method | Score | m/z |
|---|---|---|---|---|
| sys_15_1 | 15152 | FTMS; HCD | 96.25 | 701.28 |

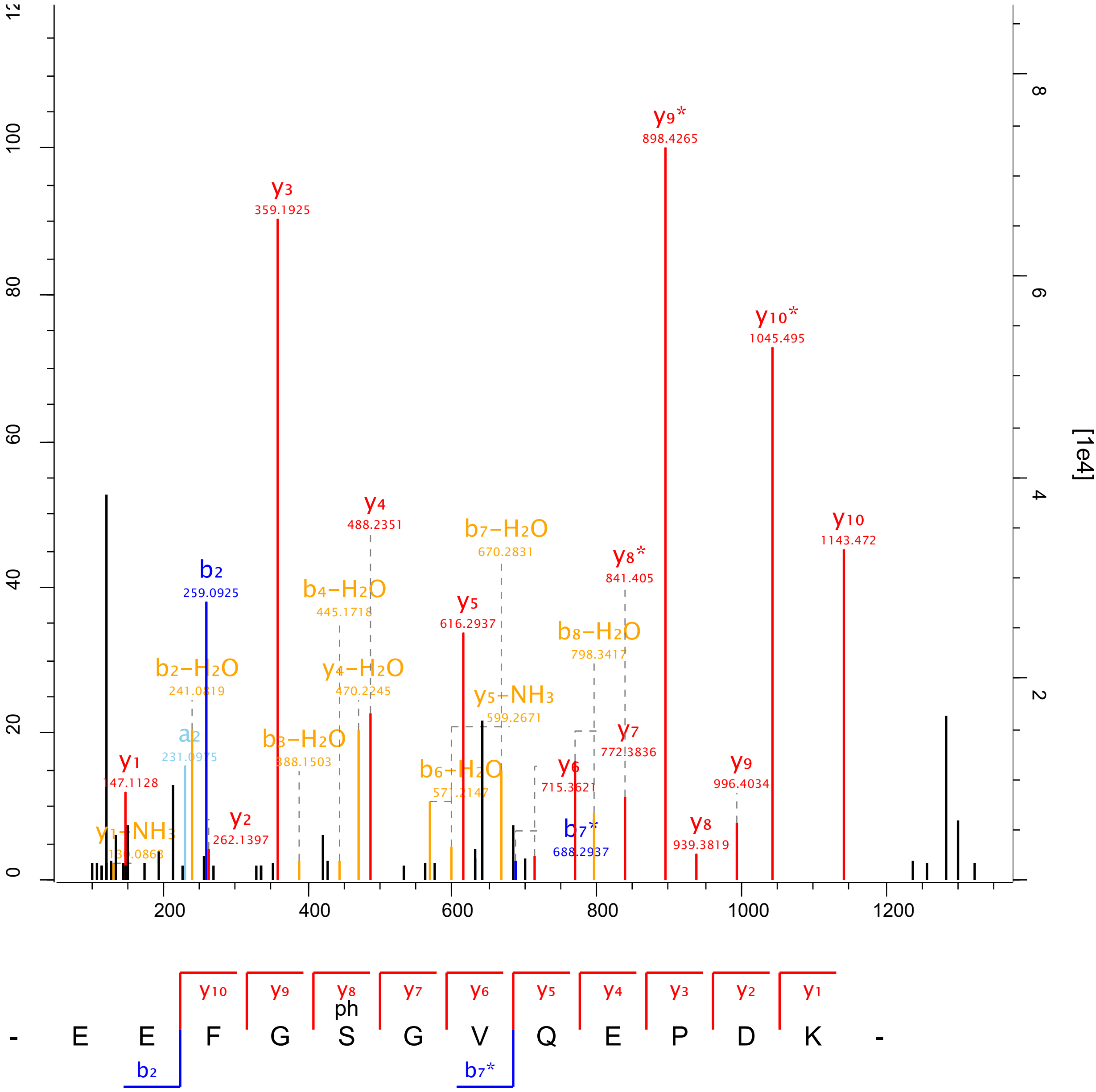

| Raw file | Scan | Method | Score | m/z |
|---|---|---|---|---|
| sys_15_1 | 15390 | FTMS; HCD | 288.25 | 840.79 |

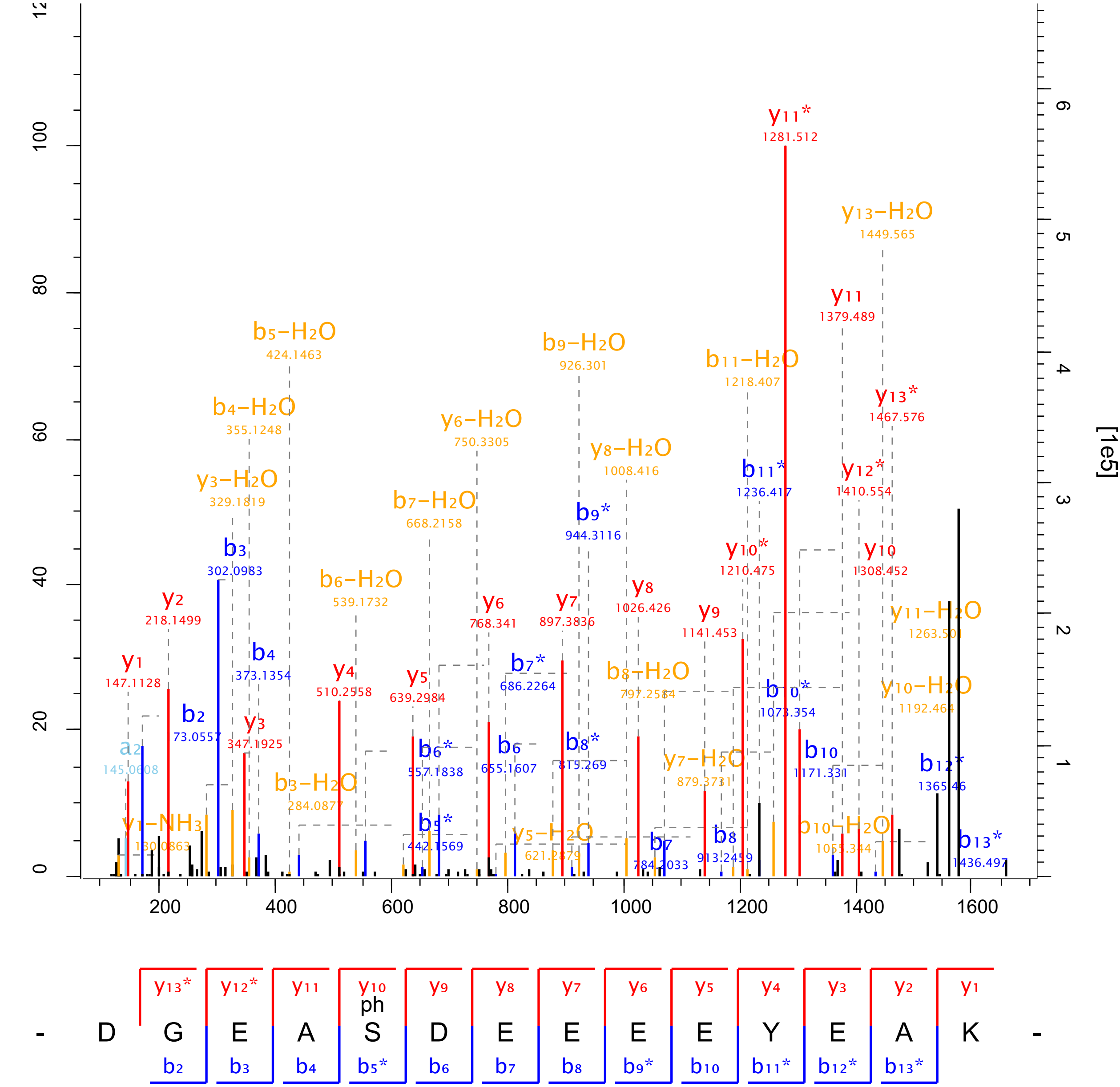

| Raw file | Scan | Method | Score | m/z |
|---|---|---|---|---|
| sys_15_1 | 15393 | FTMS; HCD | 134.14 | 669.73 |

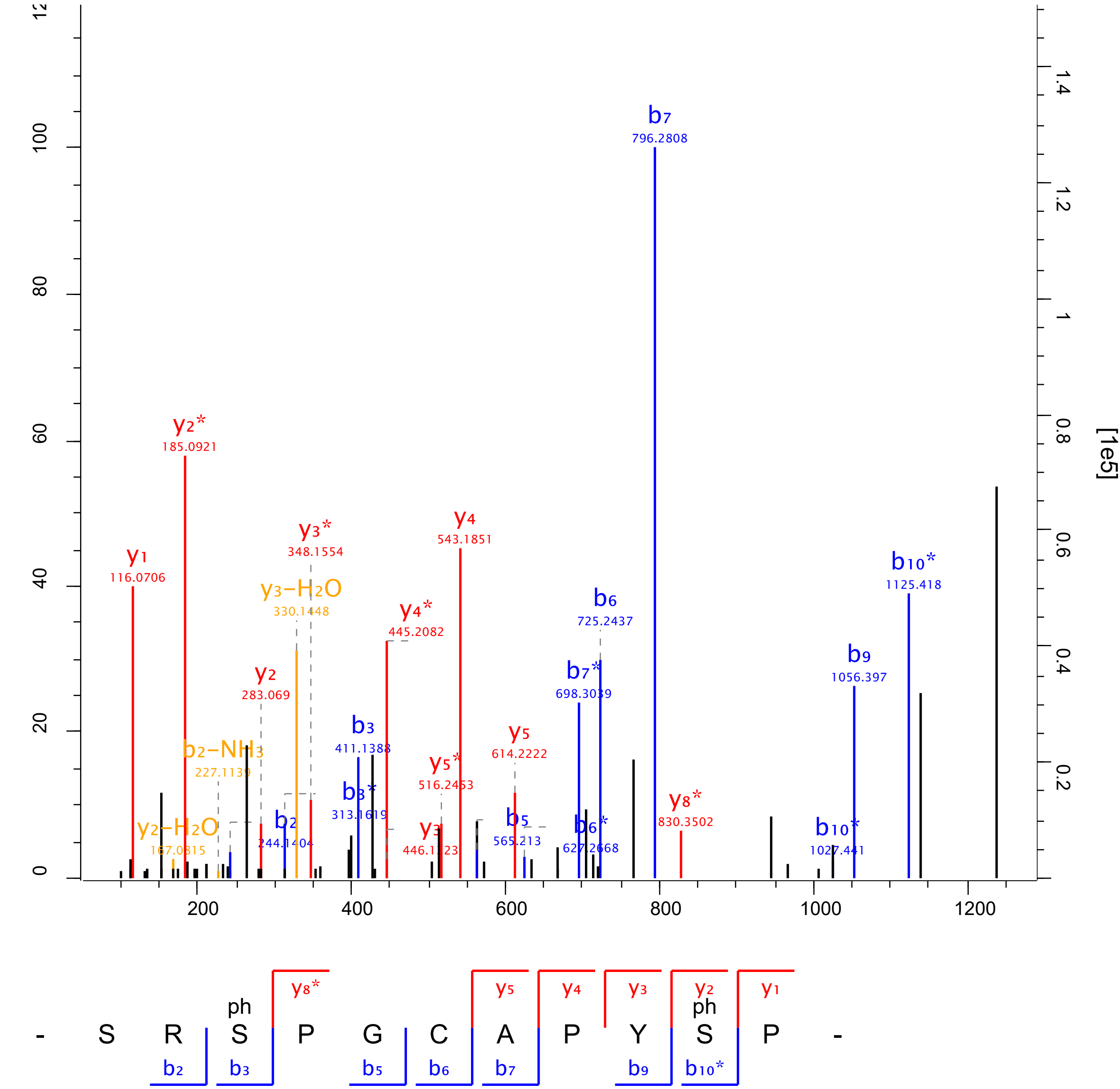

| Raw file | Scan | Method | Score | m/z |
| --- | --- | --- | --- | --- |
| sys_15_1 | 15435 | FTMS; HCD | 102.52 | 676.3 |

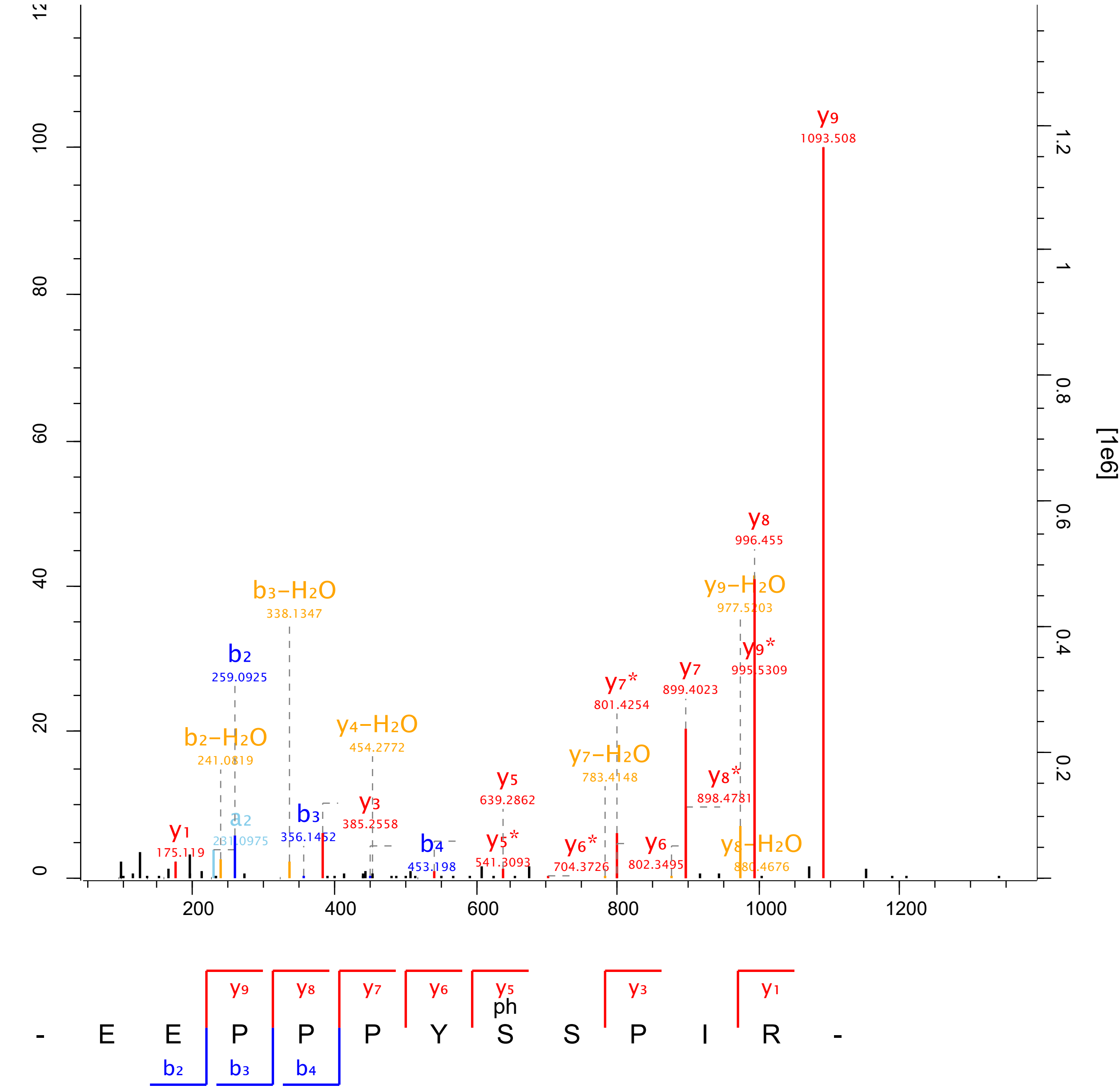

| Raw file | Scan | Method | Score | m/z |
|---|---|---|---|---|
| sys_15_1 | 15475 | FTMS; HCD | 68.48 | 664.27 |

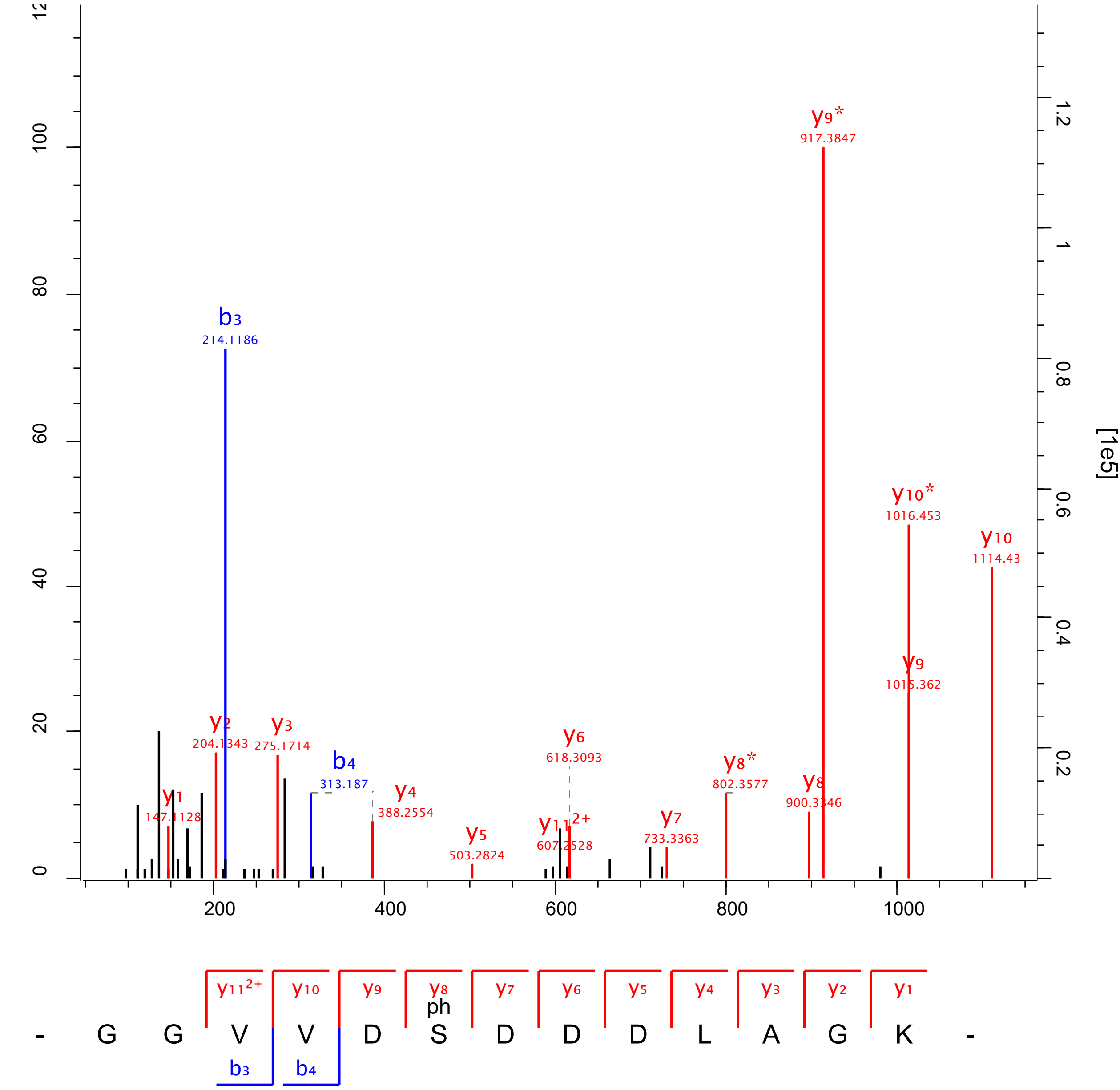

| Raw file | Scan | Method | Score | m/z |
|---|---|---|---|---|
| sys_15_1 | 15551 | FTMS; HCD | 66.15 | 646.25 |

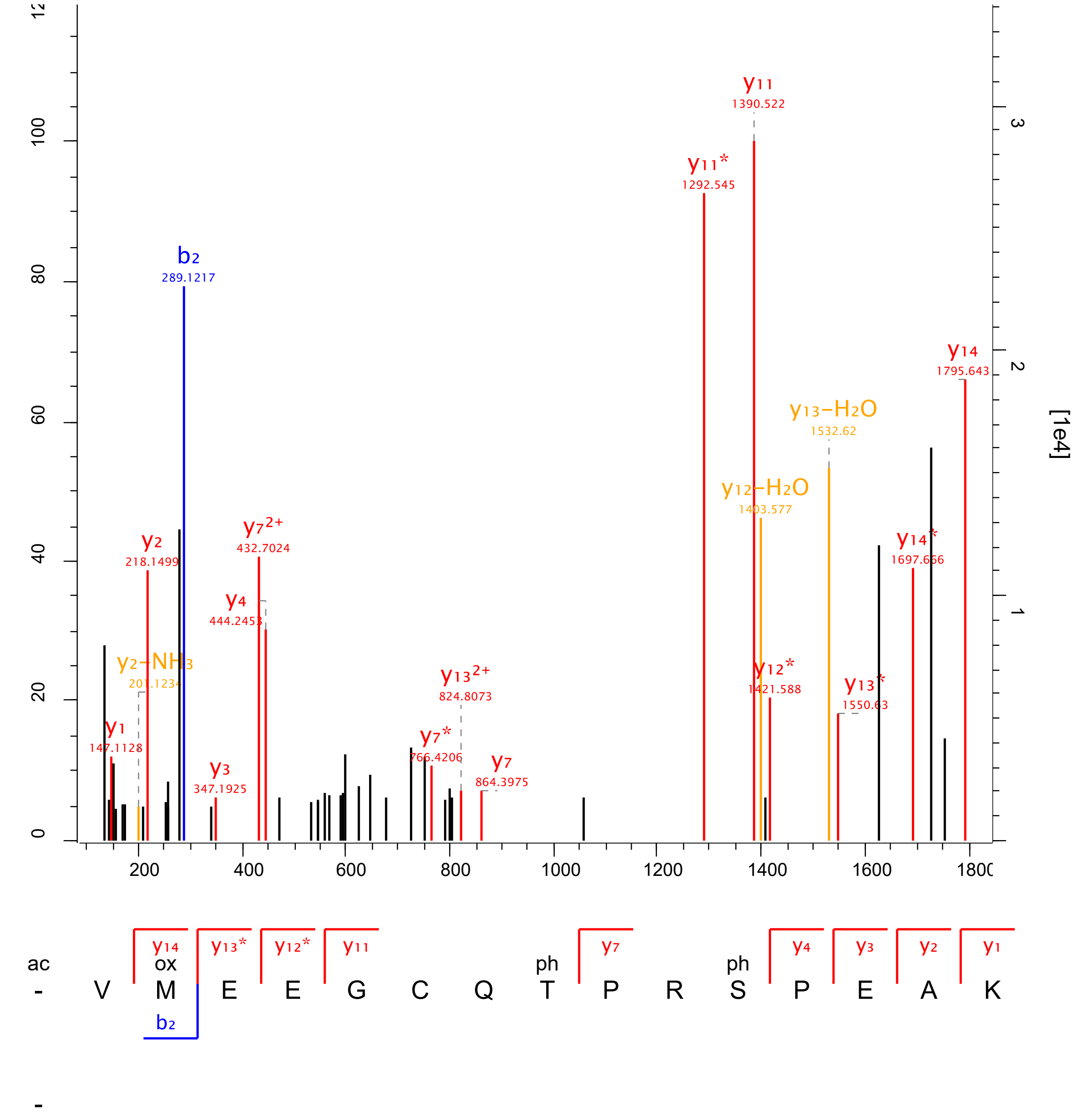

| Raw file | Scan | Method | Score | m/z |
| --- | --- | --- | --- | --- |
| sys_15_1 | 1558 | FTMS; HCD | 81.95 | 421.17 |

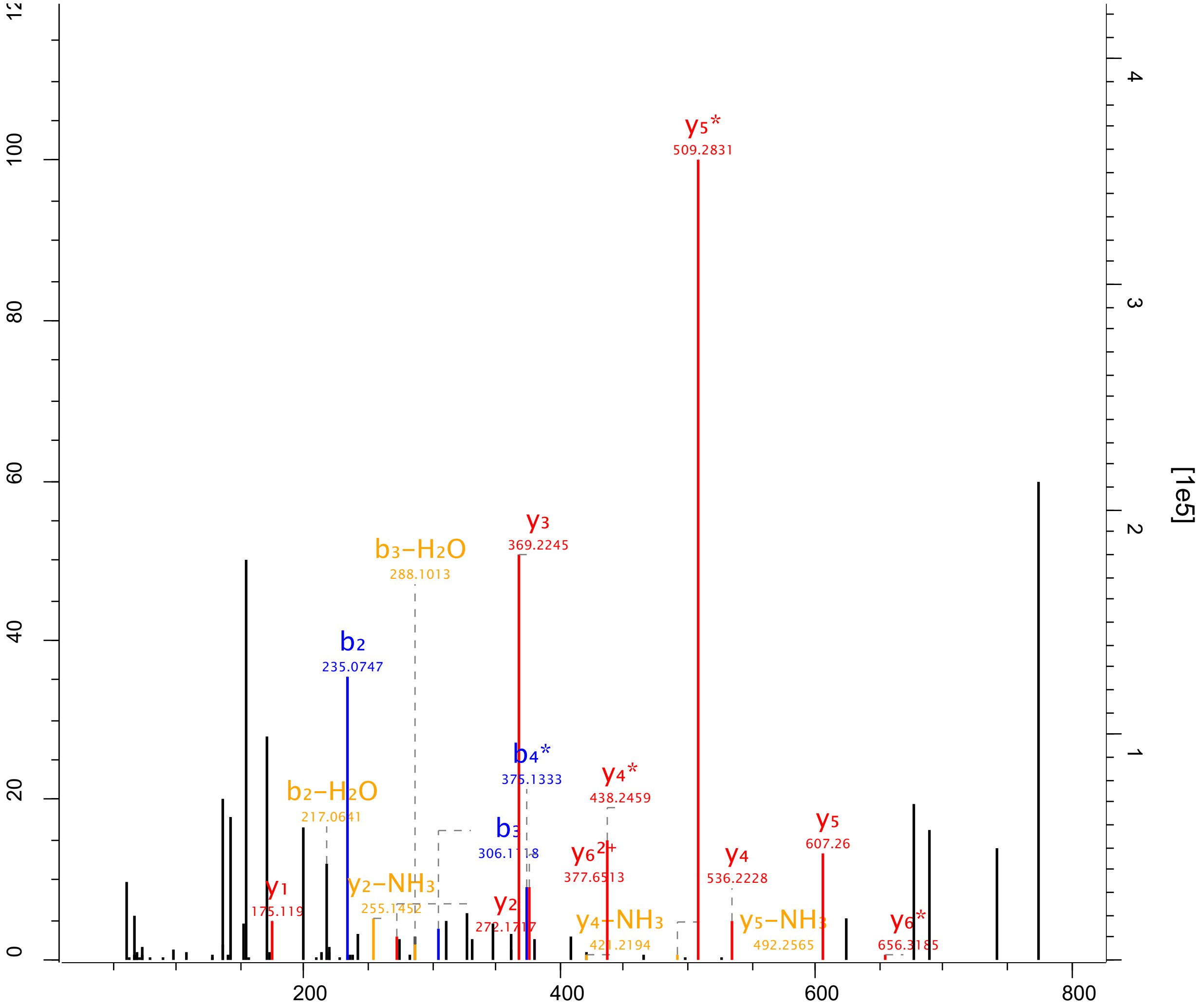


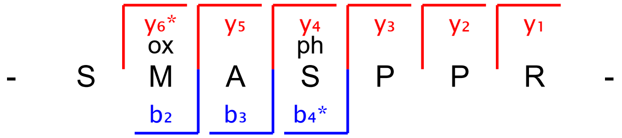

| Raw file | Scan | Method | Score | m/z |
|---|---|---|---|---|
| sys_15_1 | 15622 | FTMS; HCD | 54.34 | 593.75 |

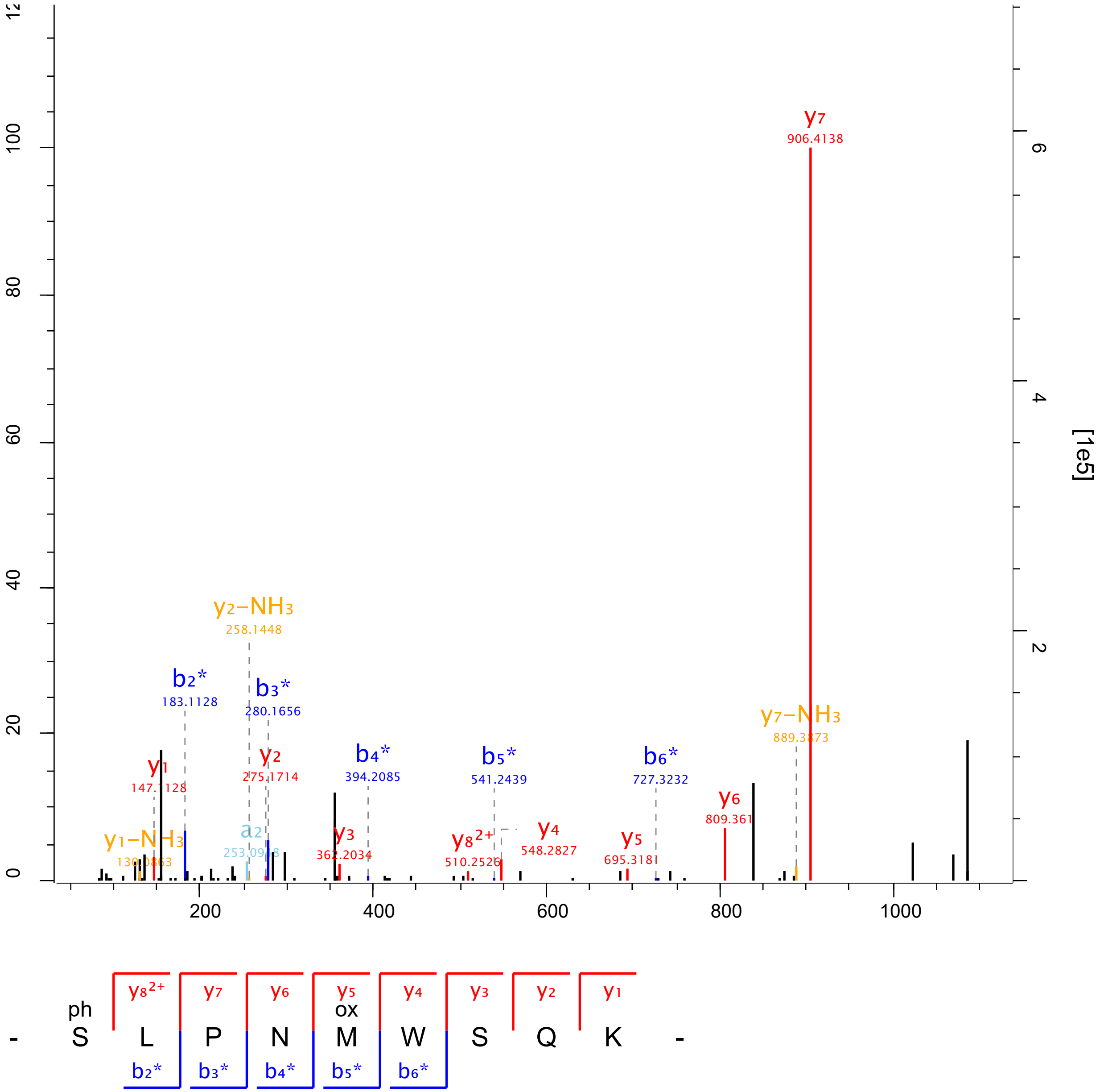

| Raw file | Scan | Method | Score | m/z |
| --- | --- | --- | --- | --- |
| sys_15_1 | 15645 | FTMS; HCD | 48.22 | 728.81 |

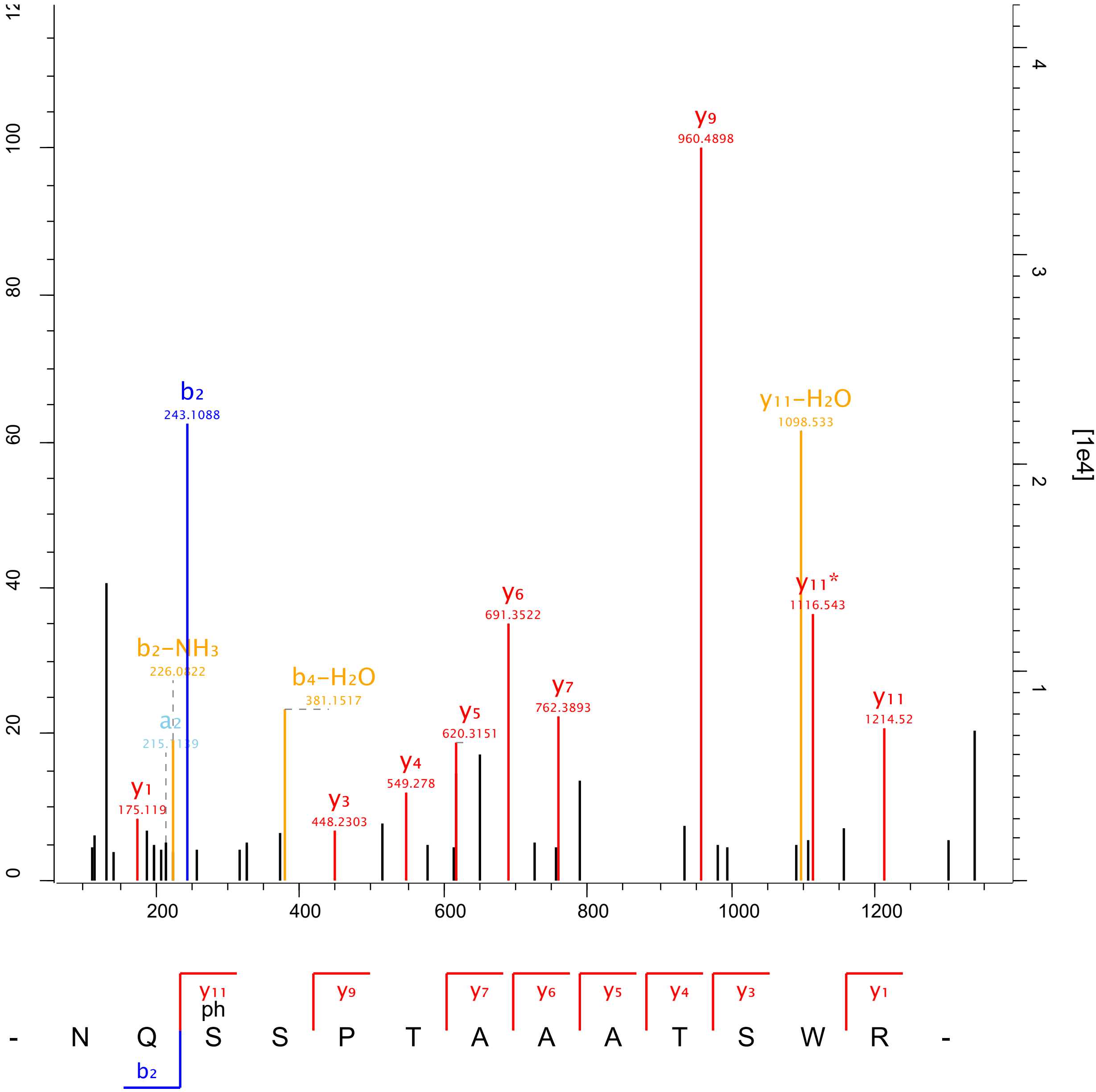

| Raw file | Scan | Method | Score | m/z |
|---|---|---|---|---|
| sys_15_1 | 15684 | FTMS; HCD | 118.05 | 1013.46 |

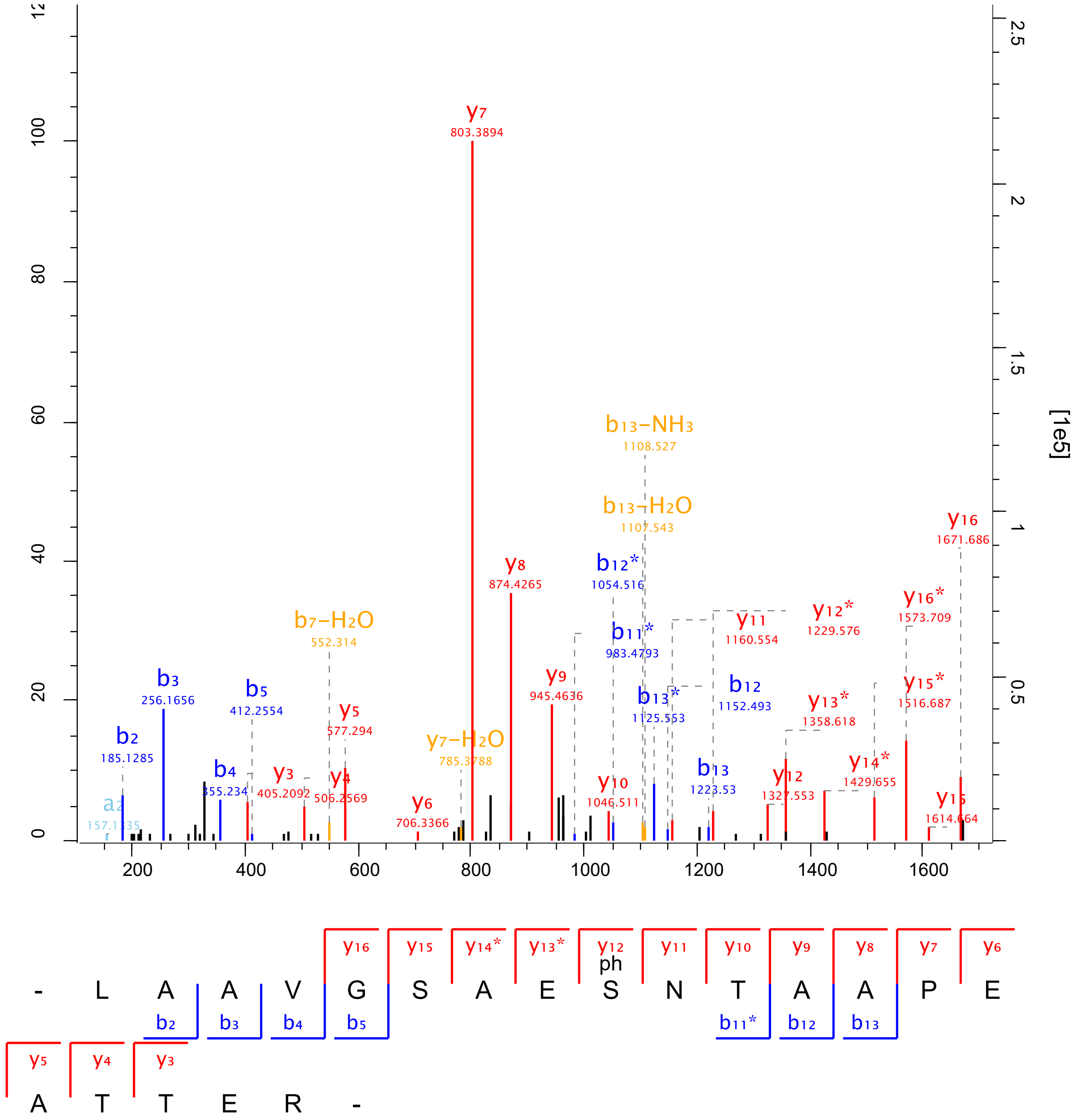

| Raw file | Scan | Method | Score | m/z |
|---|---|---|---|---|
| sys_15_1 | 15880 | FTMS; HCD | 133.47 | 620.24 |

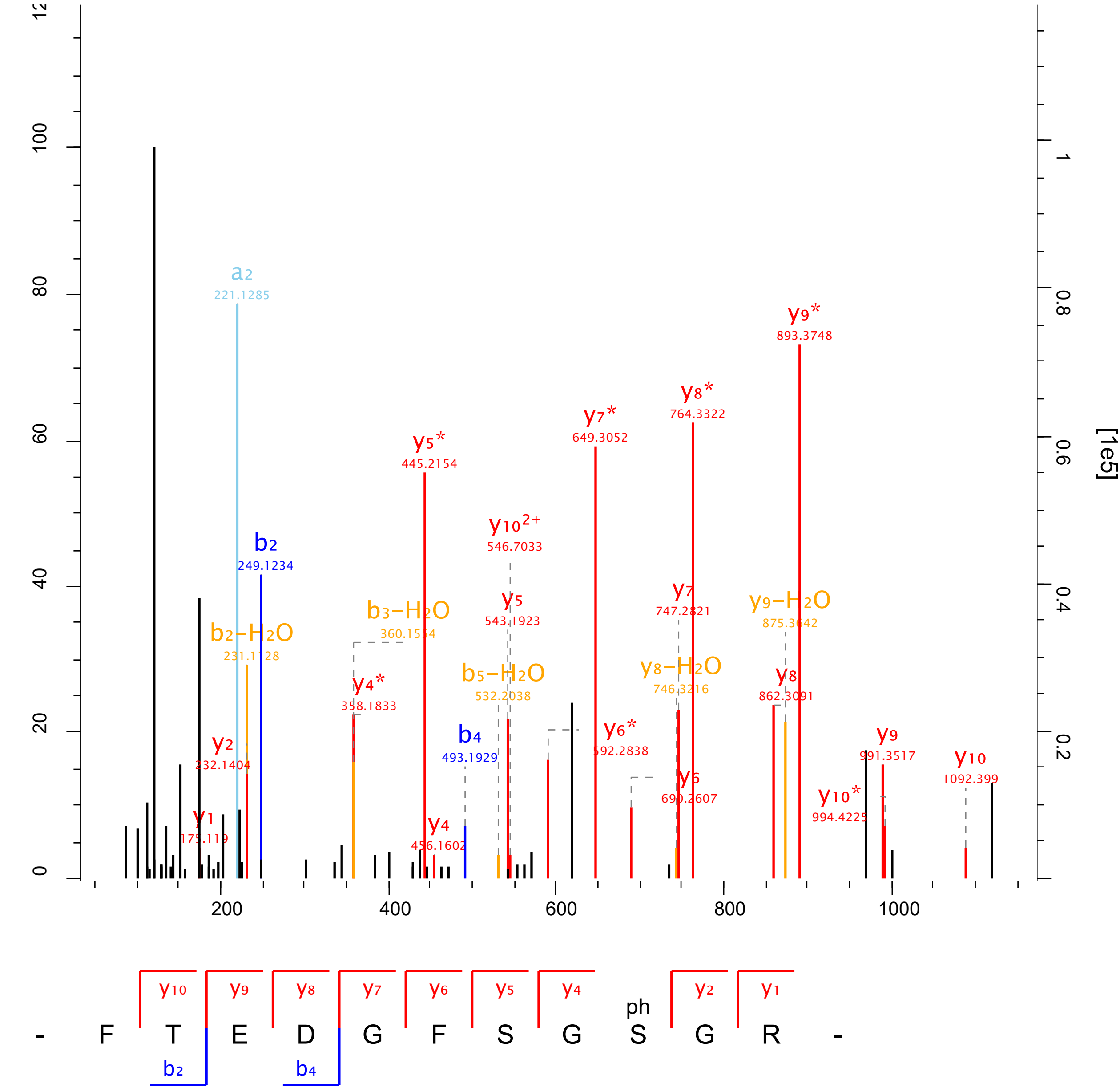

| Raw file | Scan | Method | Score | m/z |
|---|---|---|---|---|
| sys_15_1 | 15968 | FTMS; HCD | 61.28 | 624.93 |

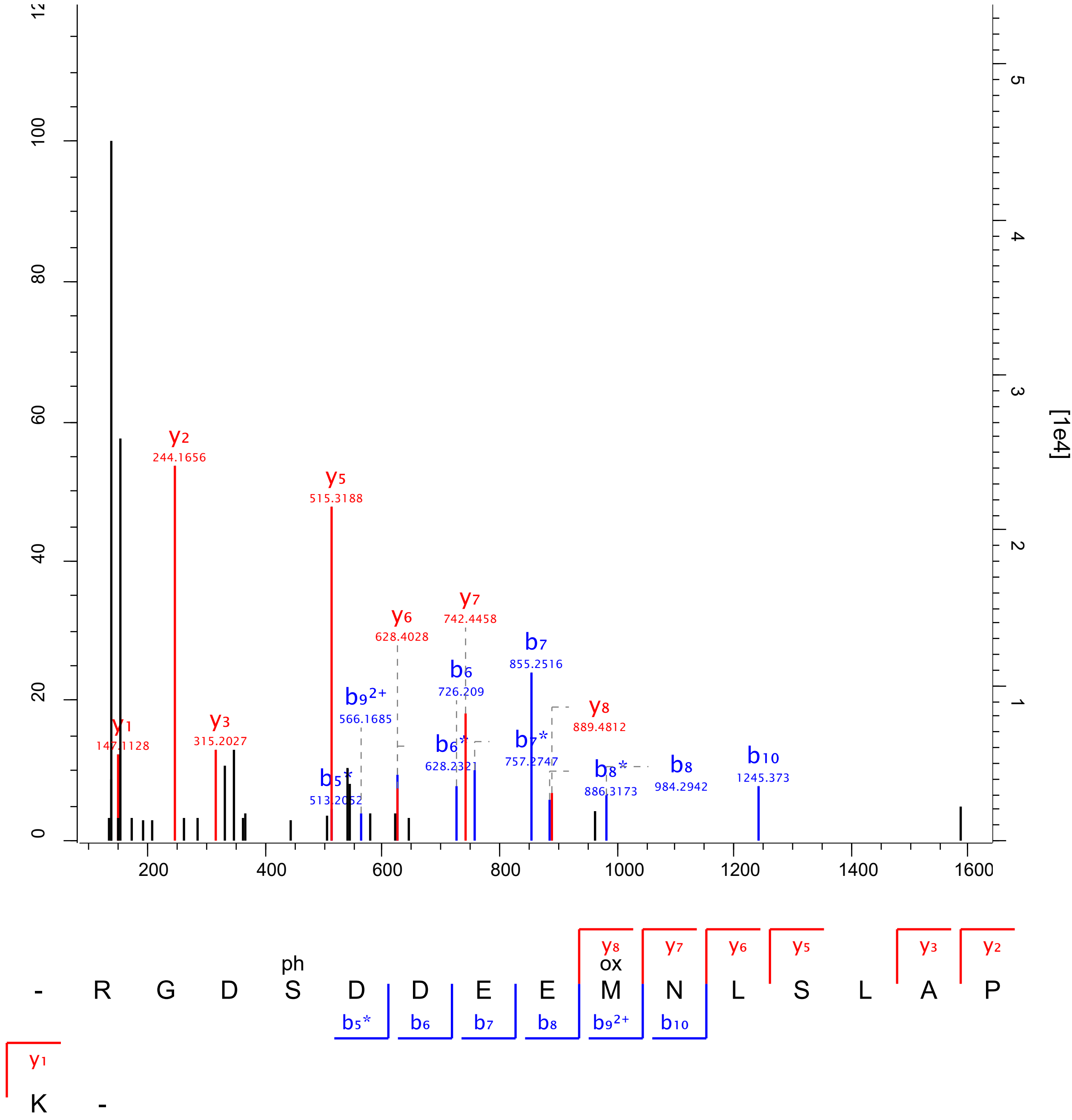

| Raw file | Scan | Method | Score | m/z |
| --- | --- | --- | --- | --- |
| sys_15_1 | 16033 | FTMS; HCD | 75.09 | 681.28 |

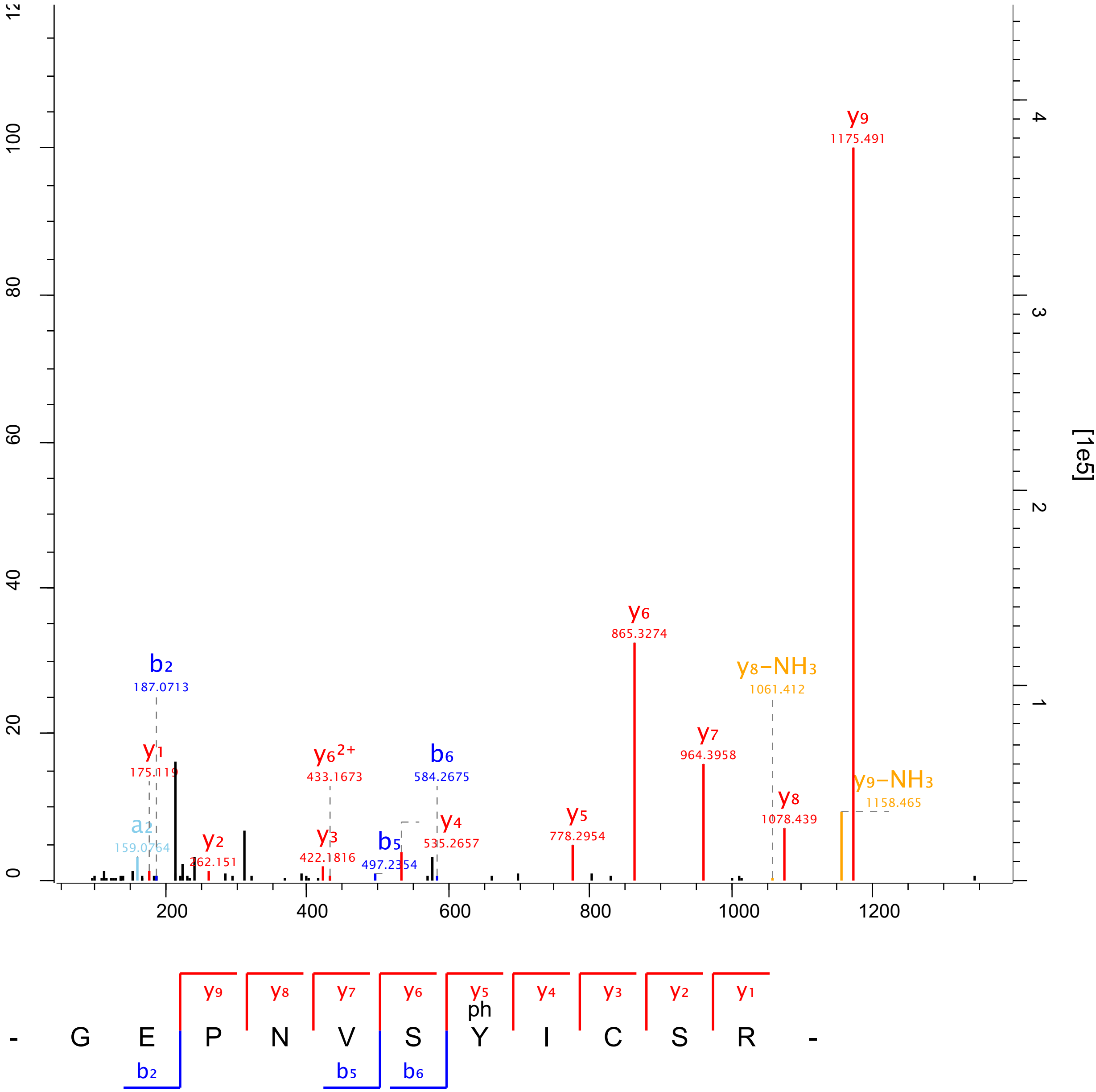

| Raw file | Scan | Method | Score | m/z |
|---|---|---|---|---|
| sys_15_1 | 16217 | FTMS; HCD | 185.86 | 723.29 |

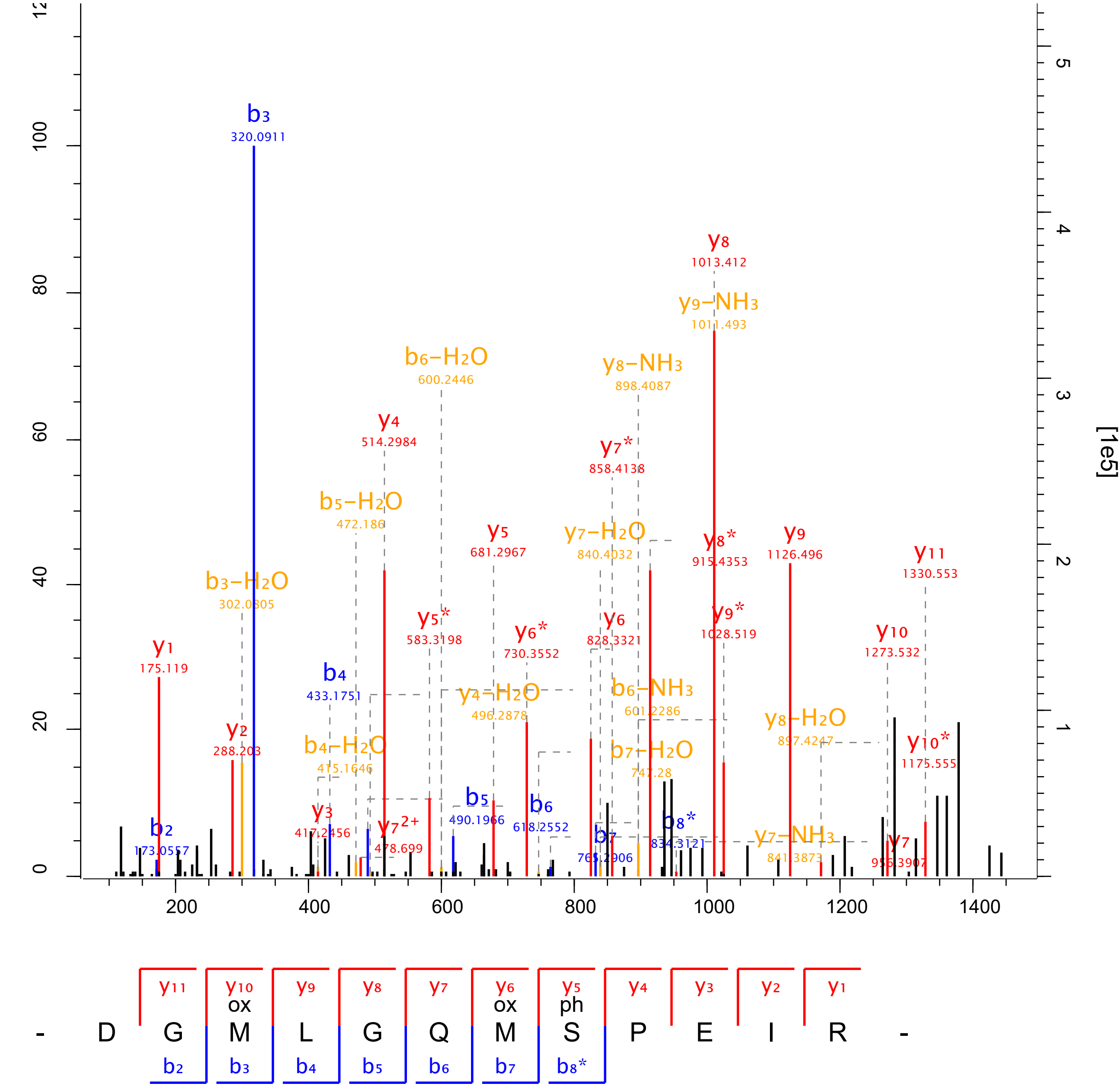

| Raw file | Scan | Method | Score | m/z |
| --- | --- | --- | --- | --- |
| sys_15_1 | 16236 | FTMS; HCD | 92.46 | 780.67 |

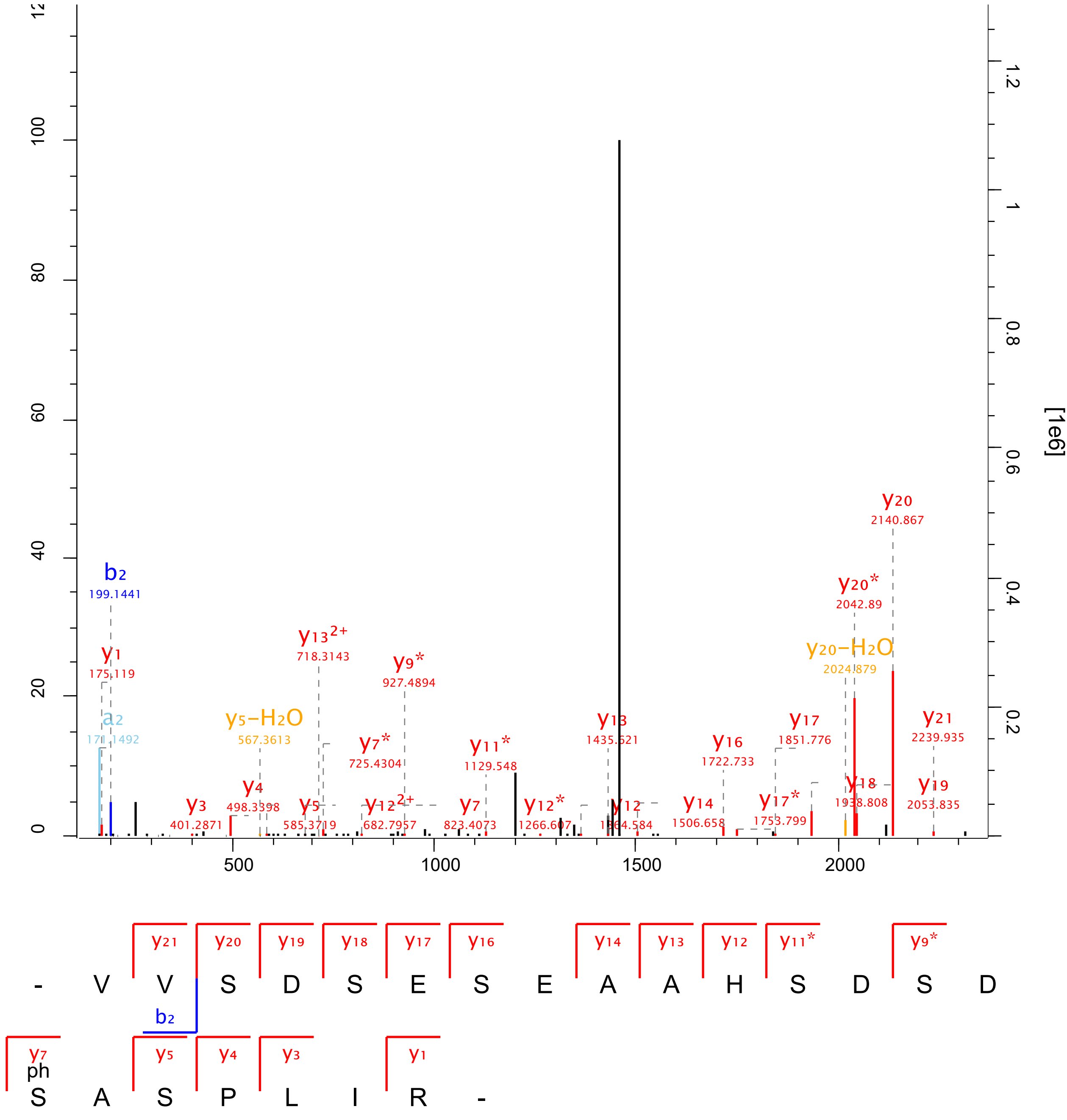

| Raw file | Scan | Method | Score | m/z |
|---|---|---|---|---|
| sys_15_1 | 16367 | FTMS; HCD | 152.34 | 686.29 |

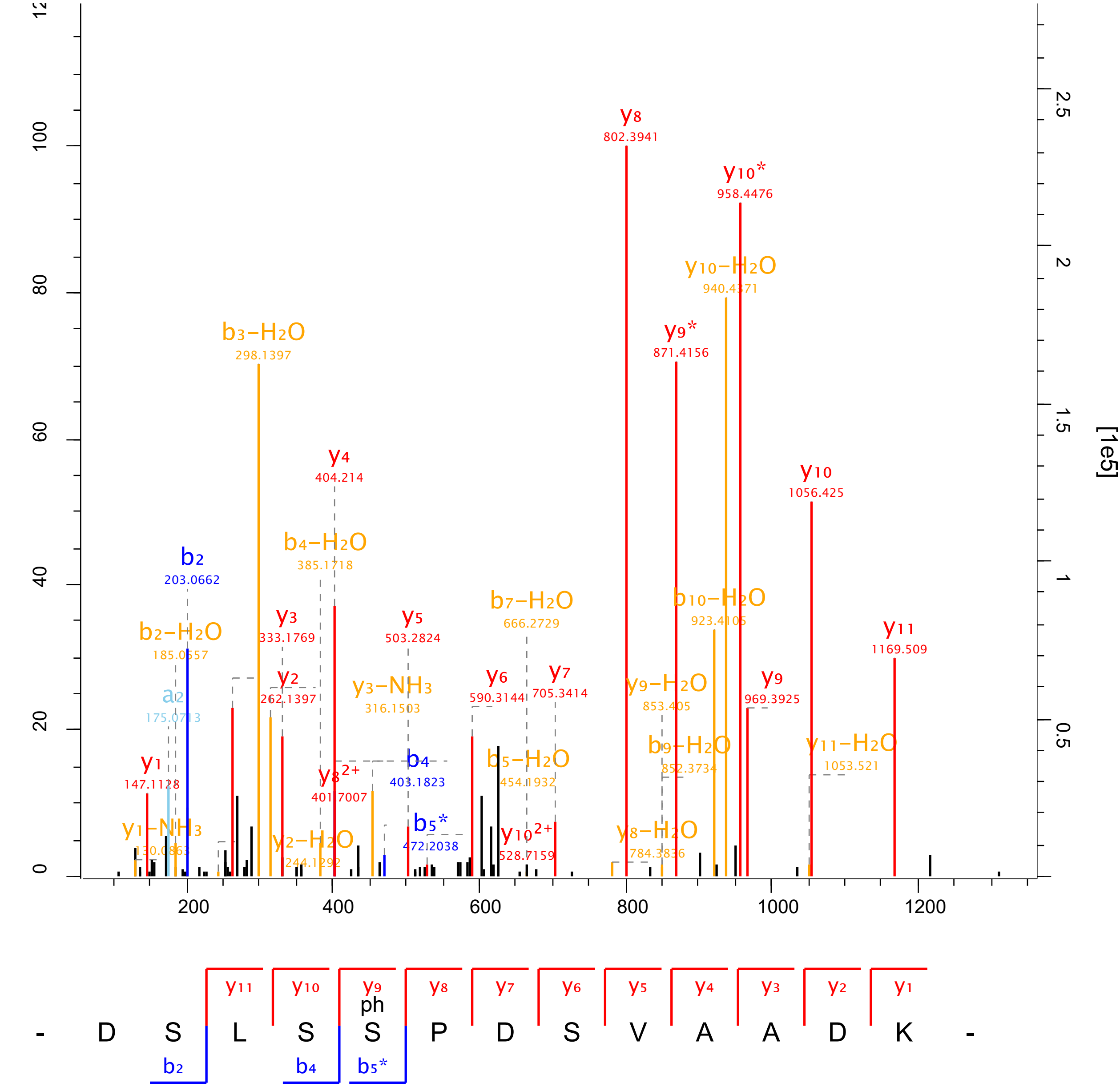

| Raw file | Scan | Method | Score | m/z |
| --- | --- | --- | --- | --- |
| sys_15_1 | 16386 | FTMS; HCD | 60.16 | 615.27 |

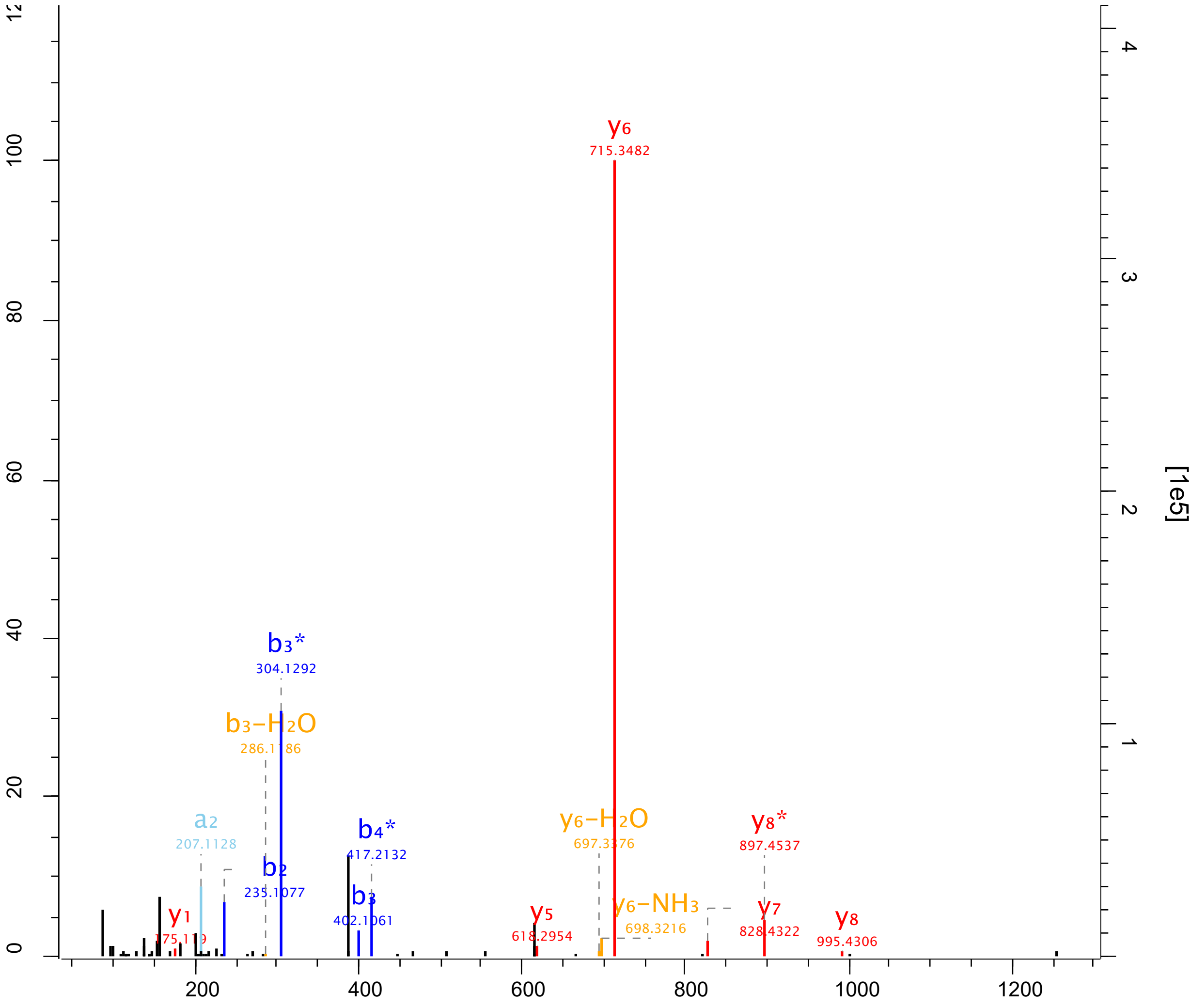


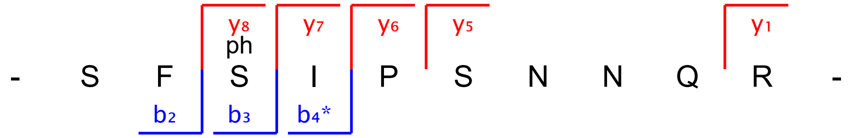

| Raw file | Scan | Method | Score | m/z |
|---|---|---|---|---|
| sys_15_1 | 16413 | FTMS; HCD | 47.55 | 698.24 |

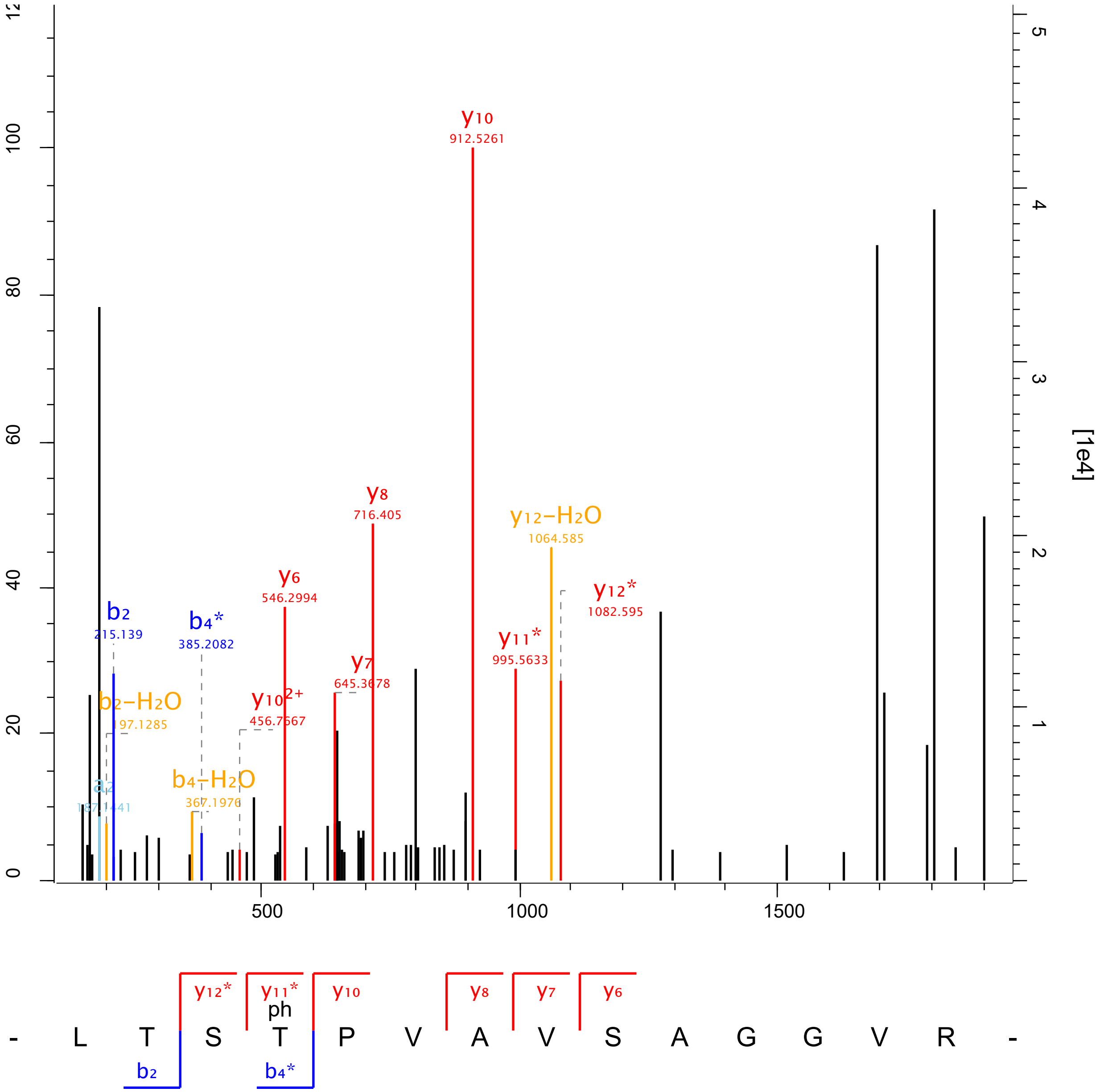

| Raw file | Scan | Method | Score | m/z |
| --- | --- | --- | --- | --- |
| sys_15_1 | 16442 | FTMS; HCD | 182.22 | 873.36 |

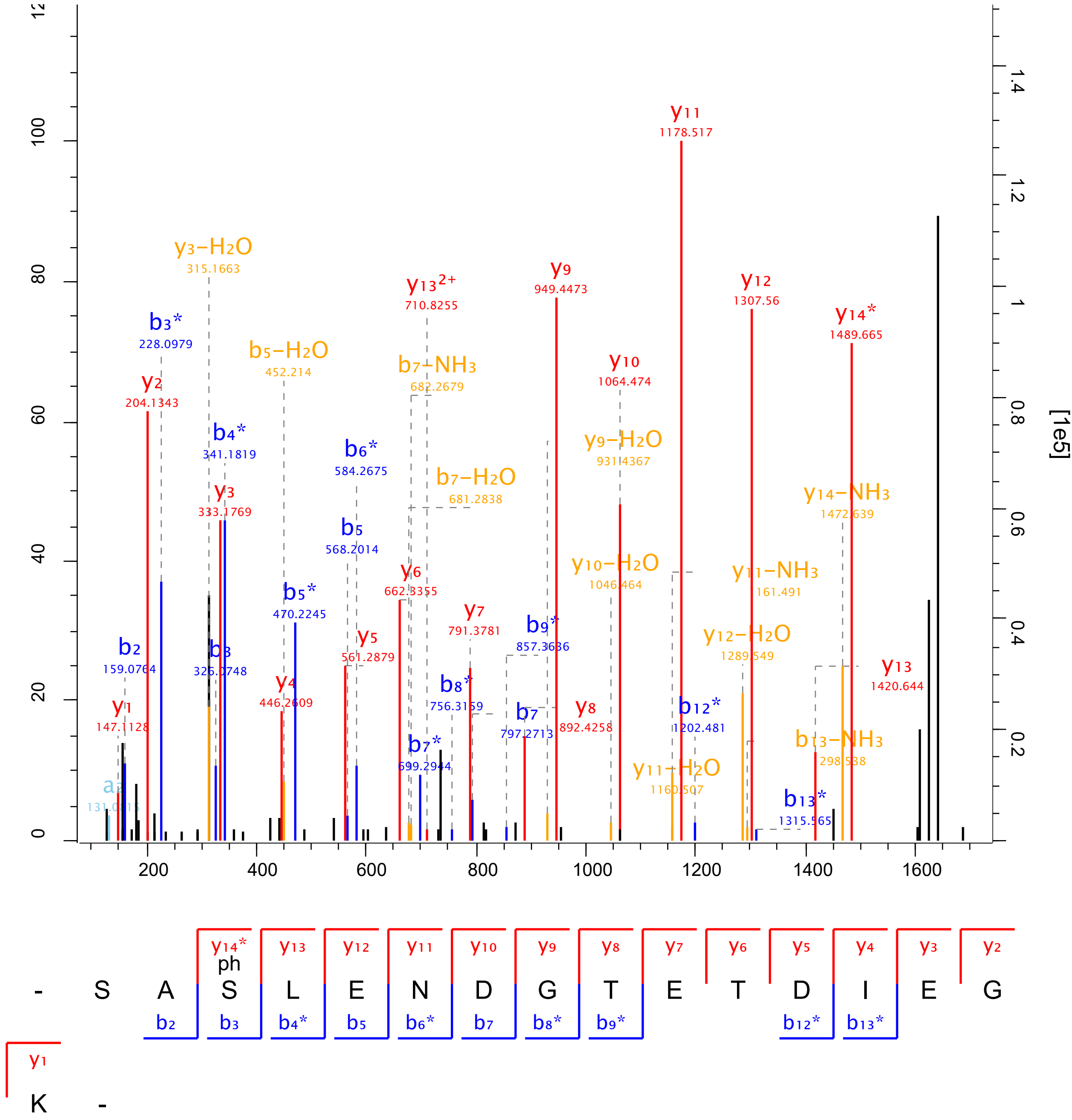

| Raw file | Scan | Method | Score | m/z |
|---|---|---|---|---|
| sys_15_1 | 16502 | FTMS; HCD | 116.87 | 720.83 |

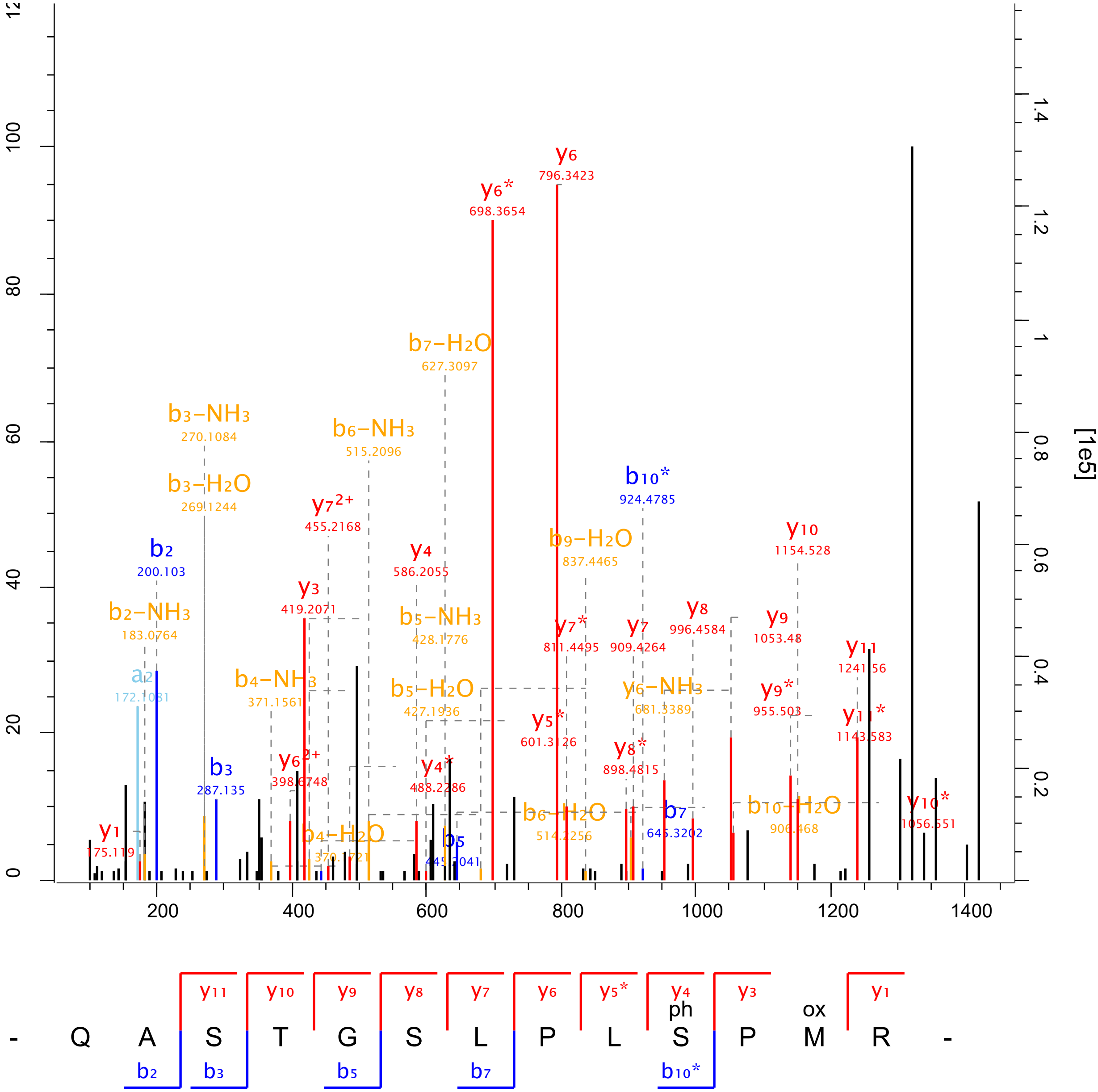

| Raw file | Scan | Method | Score | m/z |
|---|---|---|---|---|
| sys_15_1 | 16533 | FTMS; HCD | 80.31 | 540.76 |

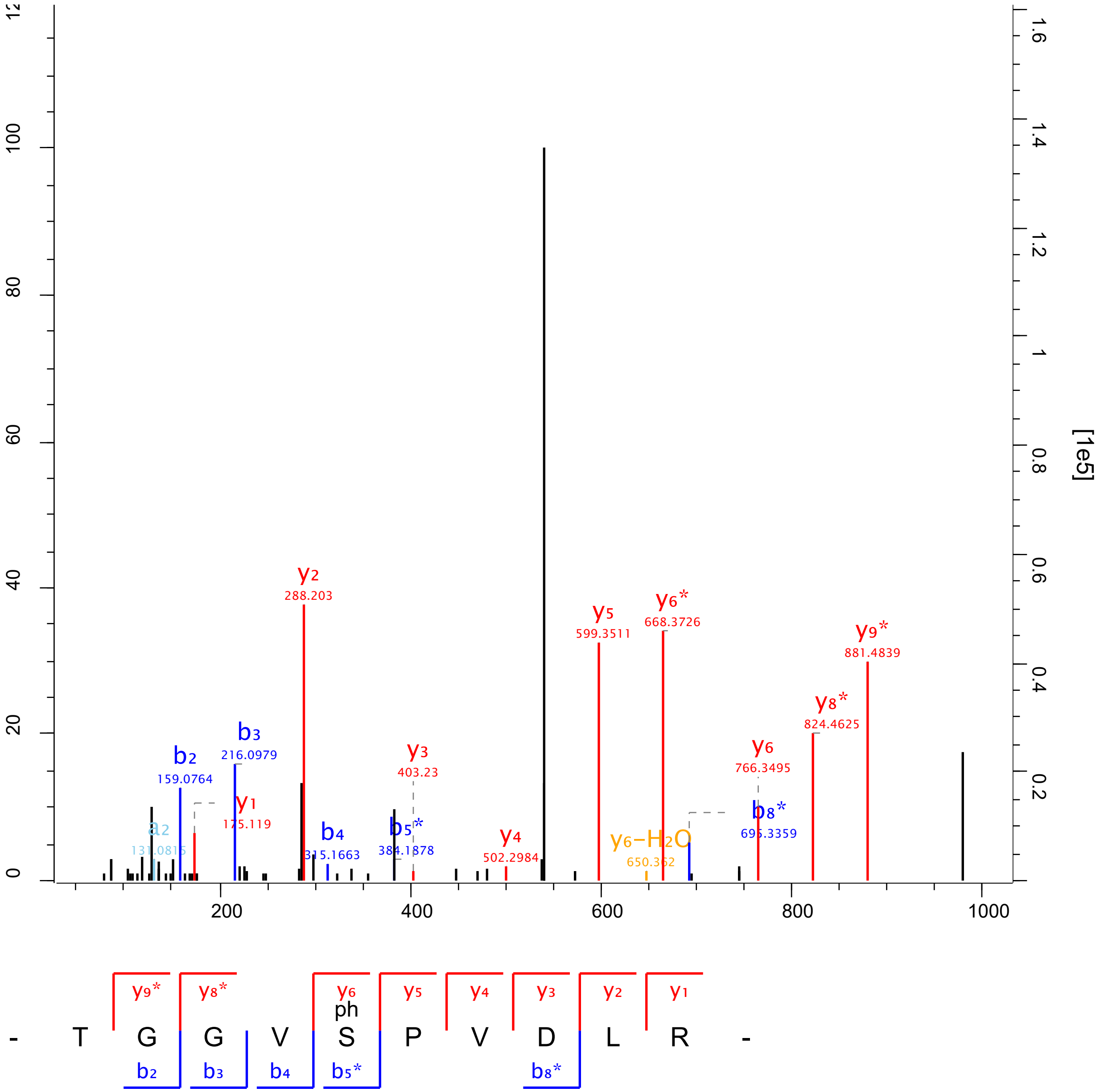

| Raw file | Scan | Method | Score | m/z |
| --- | --- | --- | --- | --- |
| sys_15_1 | 16536 | FTMS; HCD | 142.85 | 656.64 |

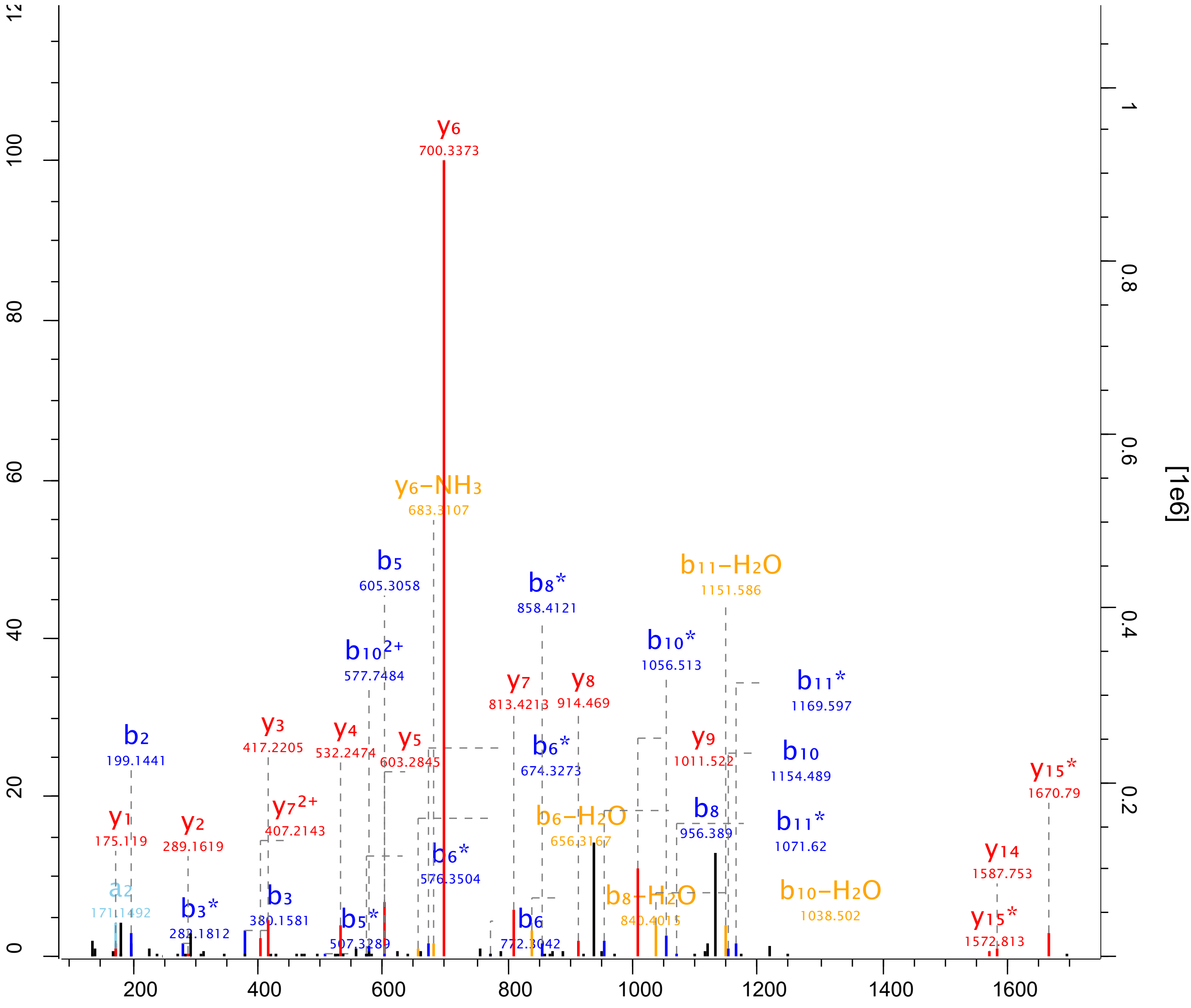


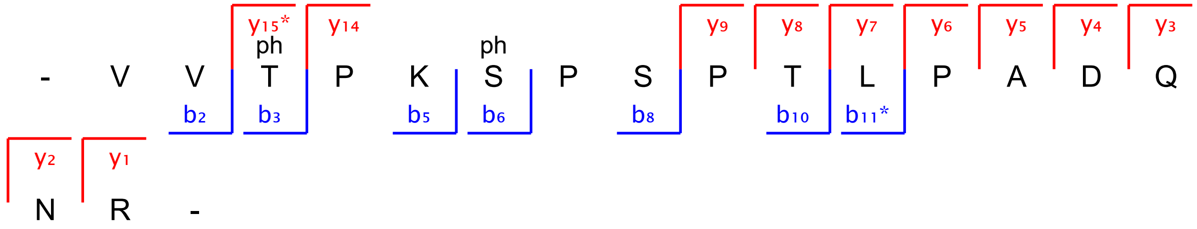

| Raw file | Scan | Method | Score | m/z |
|---|---|---|---|---|
| sys_15_1 | 16584 | FTMS; HCD | 92.94 | 635.28 |

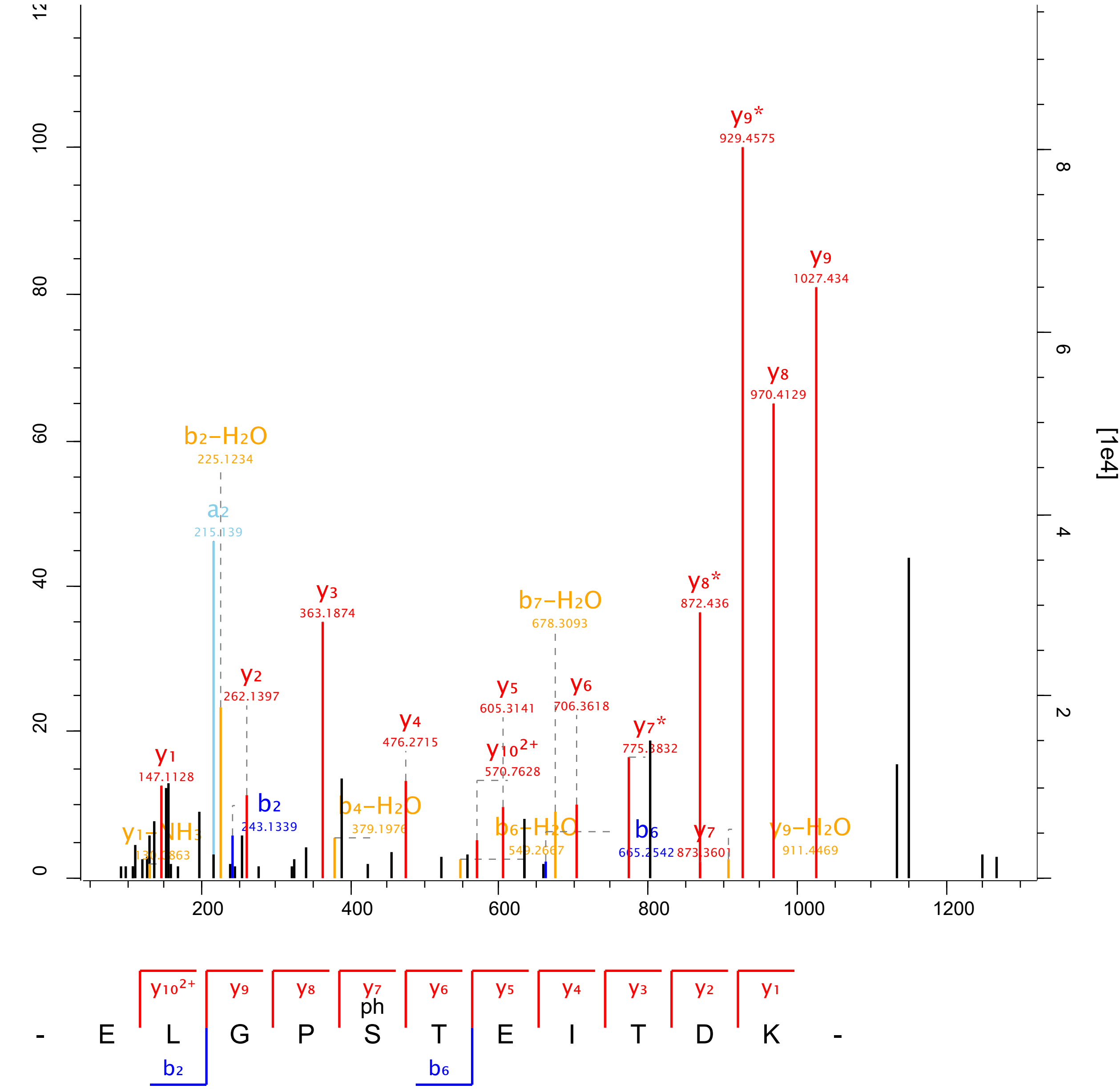

| Raw file | Scan | Method | Score | m/z |
|---|---|---|---|---|
| sys_15_1 | 16585 | FTMS; HCD | 42.86 | 732.67 |

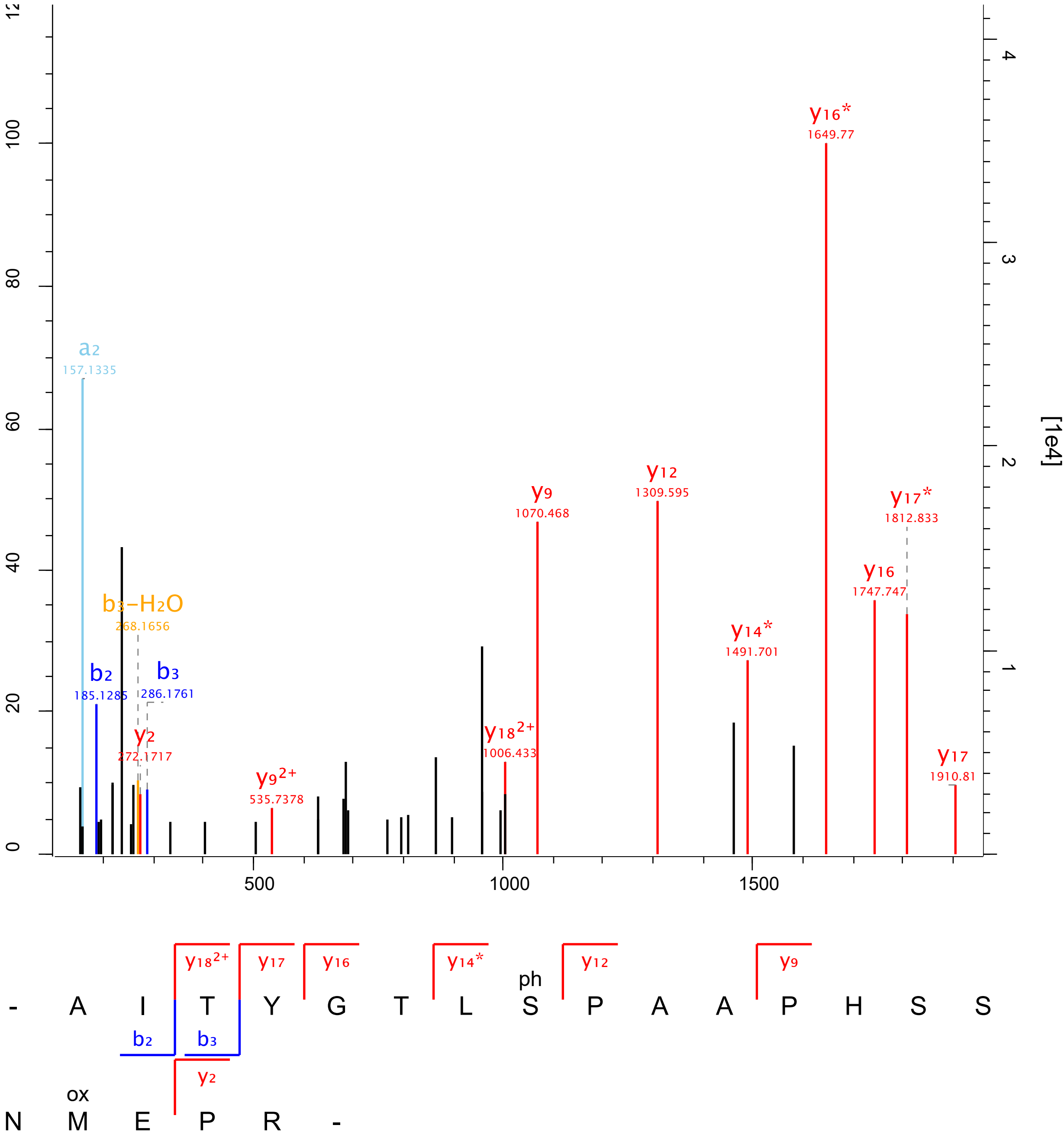

| Raw file | Scan | Method | Score | m/z |
| --- | --- | --- | --- | --- |
| sys_15_1 | 16667 | FTMS; HCD | 107.78 | 883.87 |

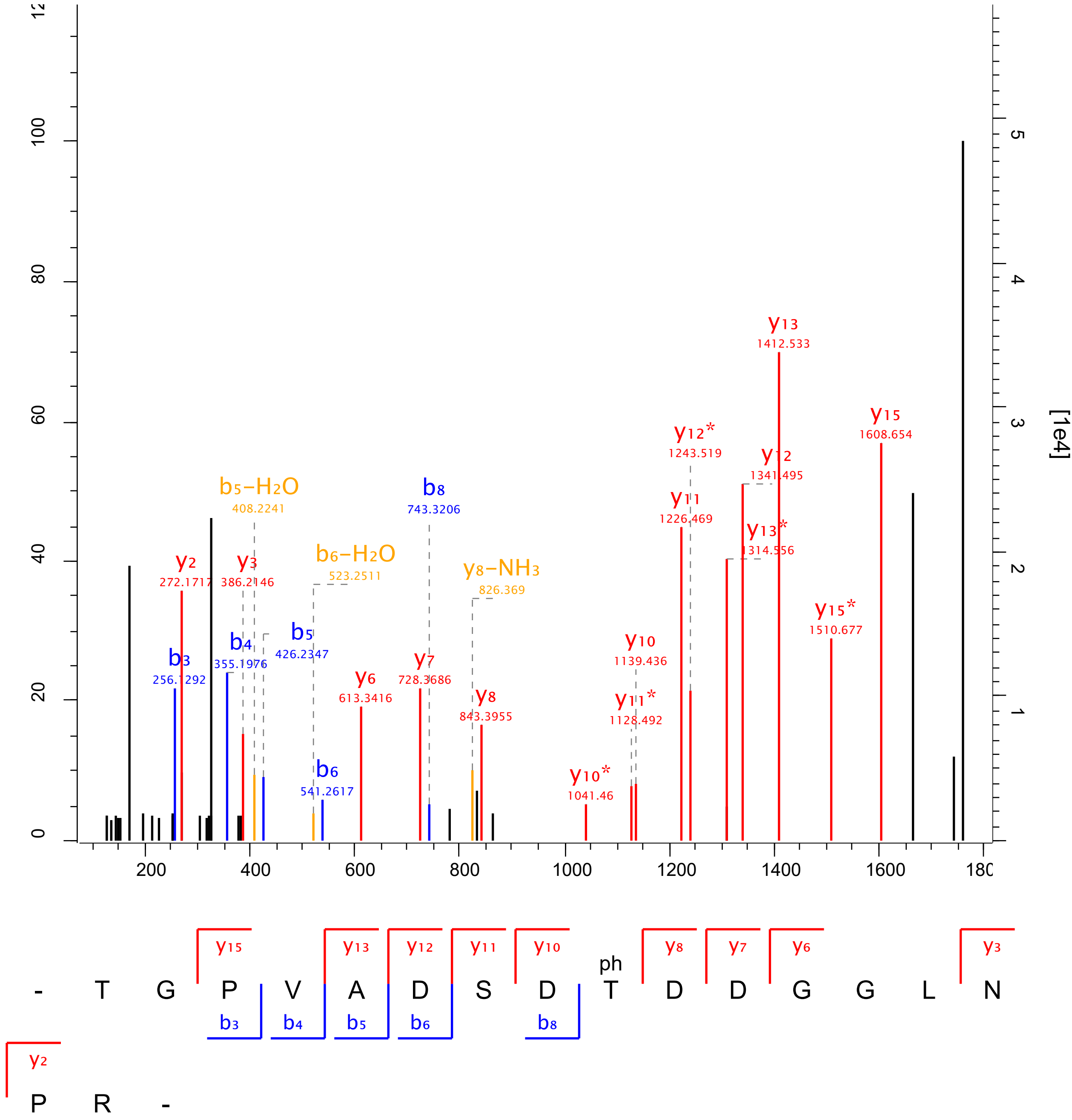

| Raw file | Scan | Method | Score | m/z |
| --- | --- | --- | --- | --- |
| sys_15_1 | 16679 | FTMS; HCD | 151.96 | 841.34 |

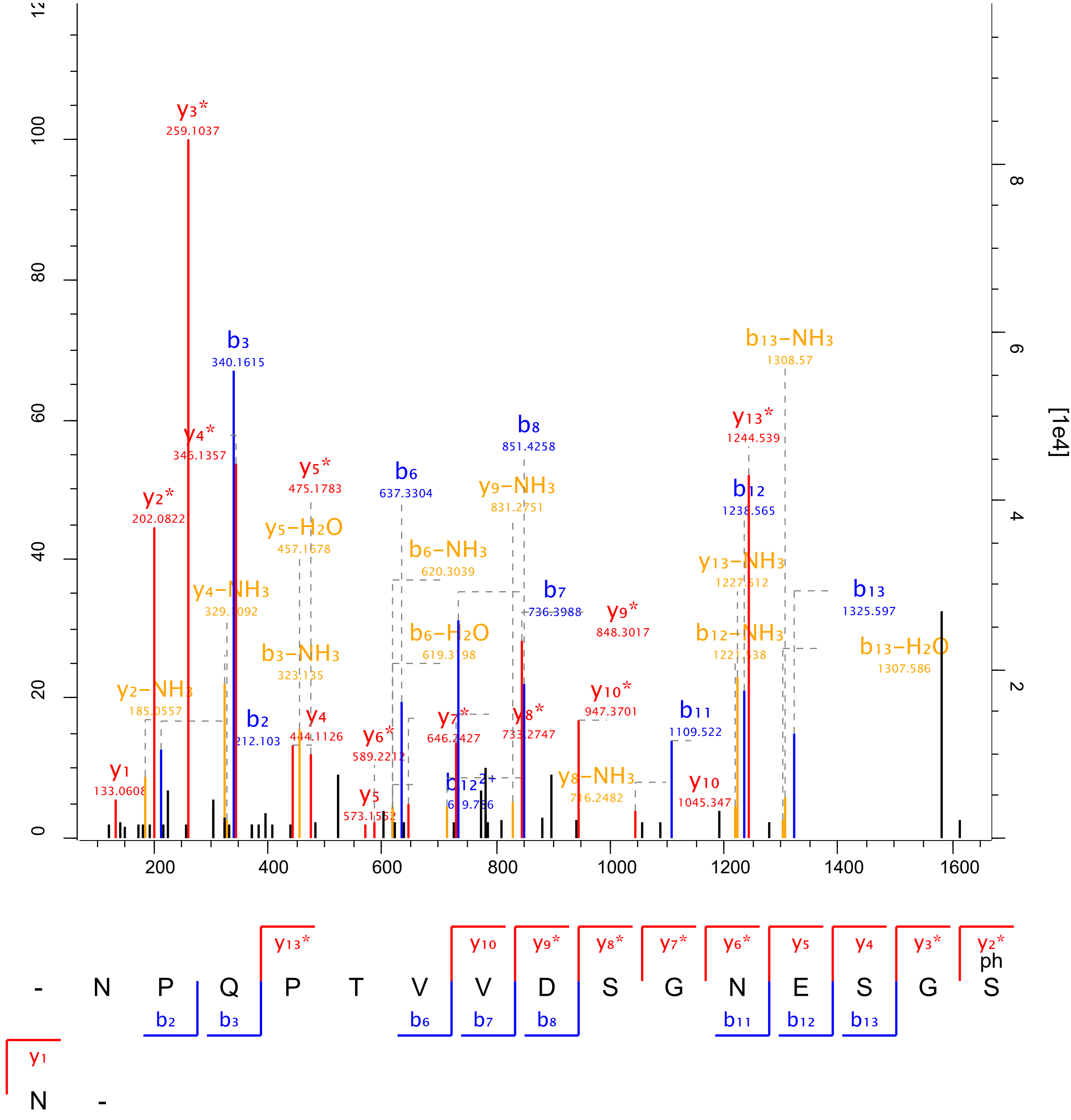

| Raw file | Scan | Method | Score | m/z |
|---|---|---|---|---|
| sys_15_1 | 16695 | FTMS; HCD | 45 | 746.86 |

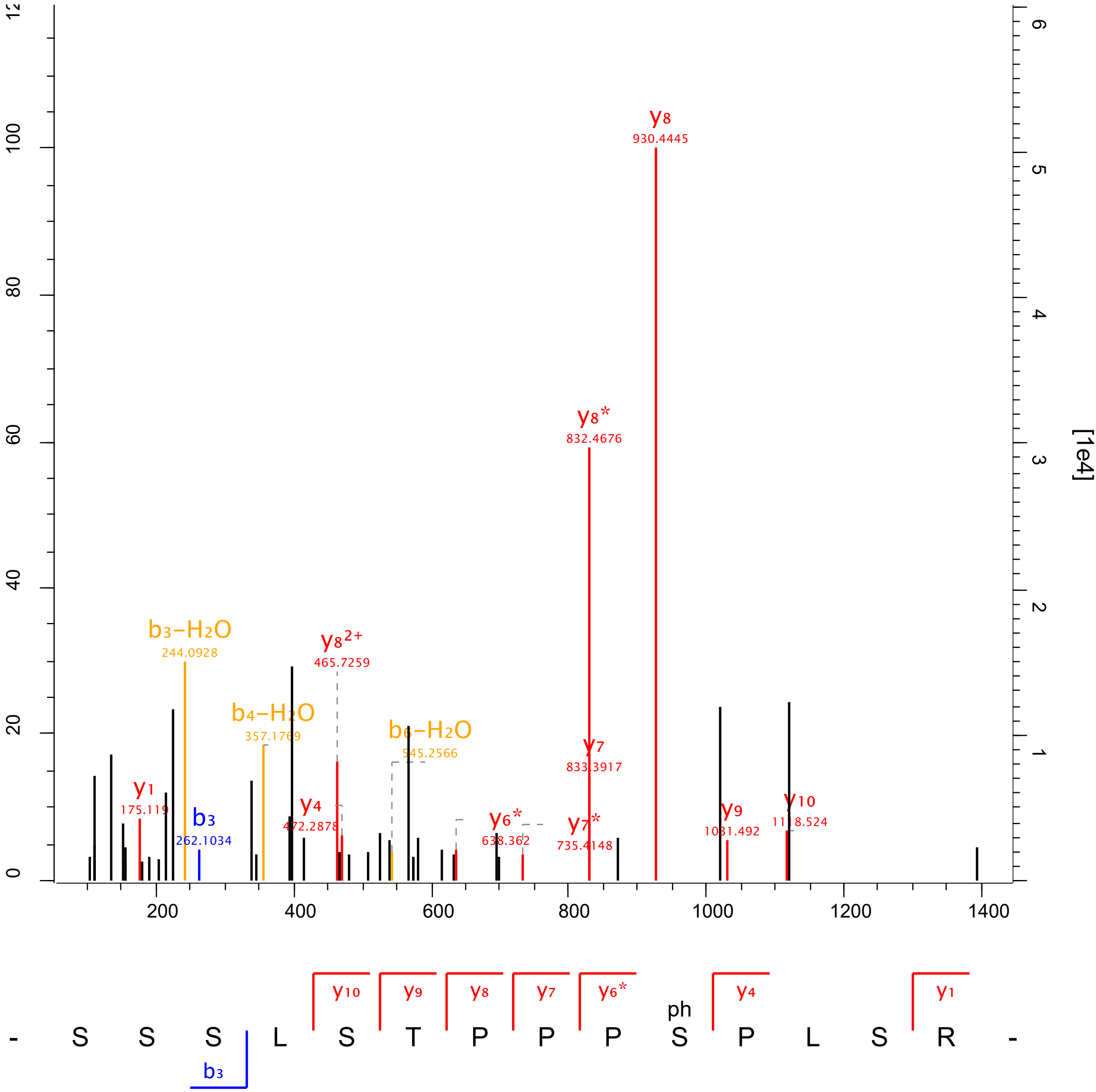

| Raw file | Scan | Method | Score | m/z |
| --- | --- | --- | --- | --- |
| sys_15_1 | 16709 | FTMS; HCD | 117.08 | 603.27 |

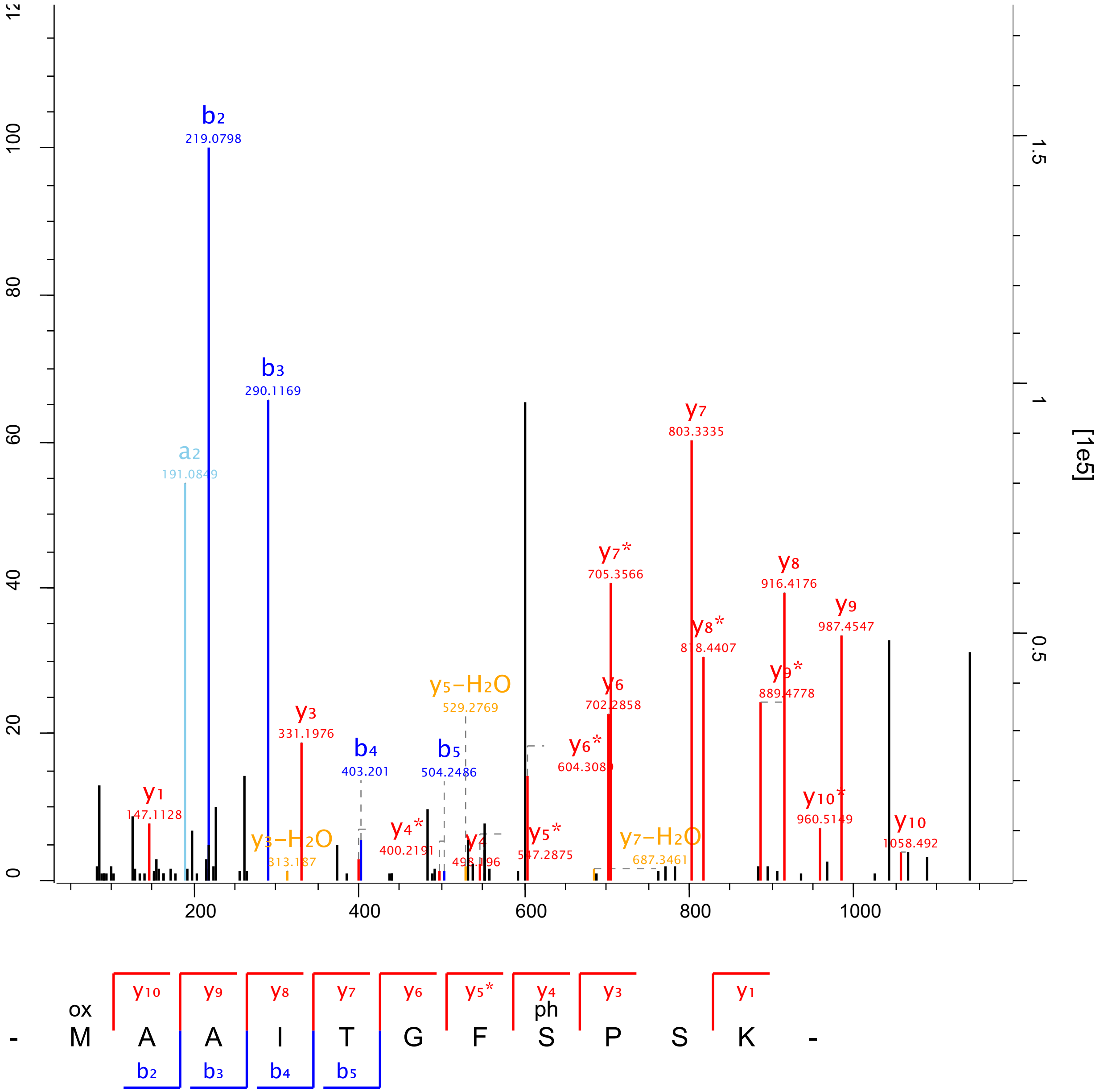

| Raw file | Scan | Method | Score | m/z |
| --- | --- | --- | --- | --- |
| sys_15_1 | 16718 | FTMS; HCD | 46.07 | 785.31 |

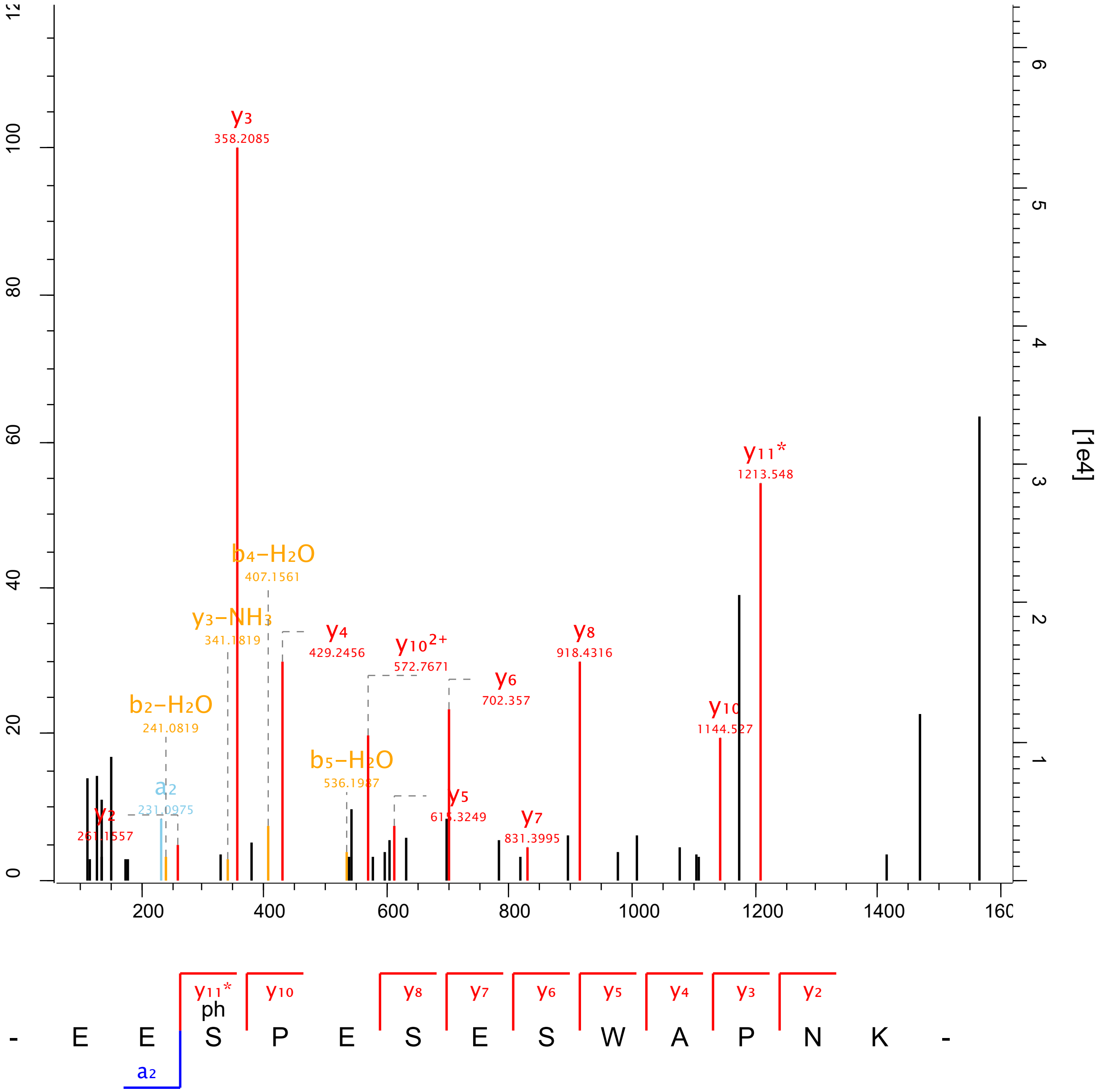

| Raw file | Scan | Method | Score | m/z |
| --- | --- | --- | --- | --- |
| sys_15_1 | 16829 | FTMS; HCD | 107.09 | 542.23 |

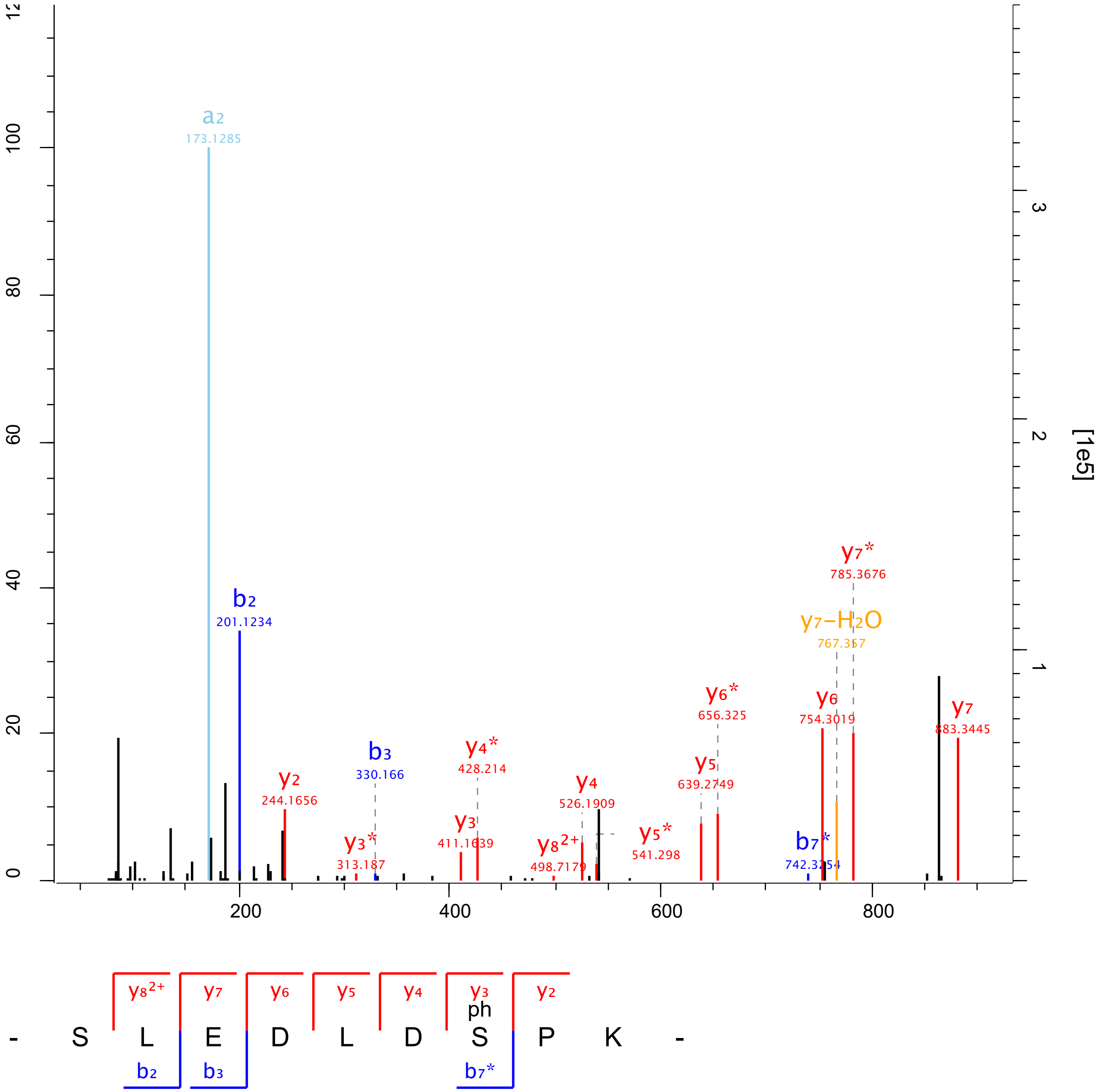

| Raw file | Scan | Method | Score | m/z |
|---|---|---|---|---|
| sys_15_1 | 16844 | FTMS; HCD | 44.93 | 512.25 |

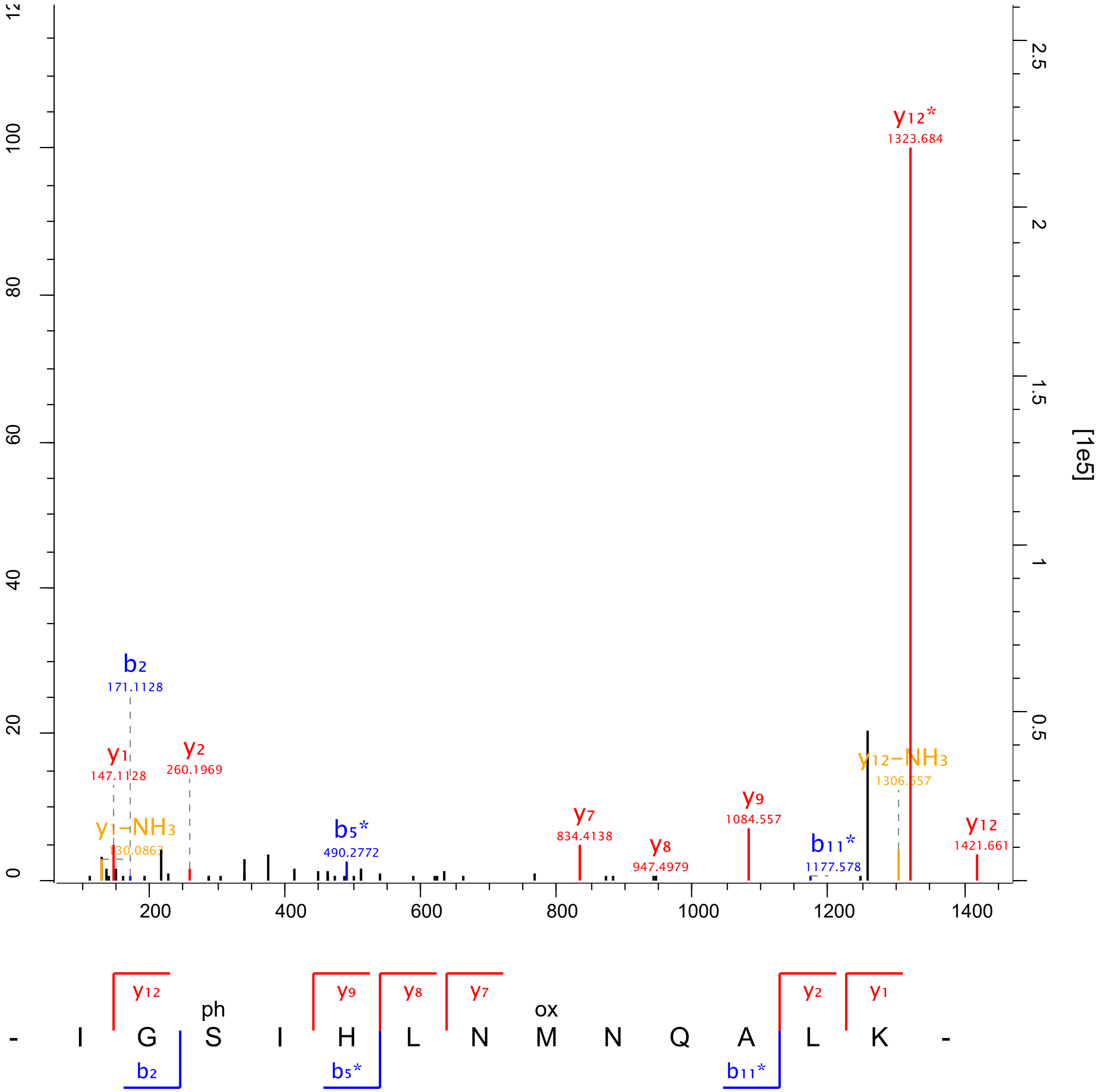

| Raw file | Scan | Method | Score | m/z |
|---|---|---|---|---|
| sys_15_1 | 1685 | FTMS; HCD | 91.55 | 365.18 |

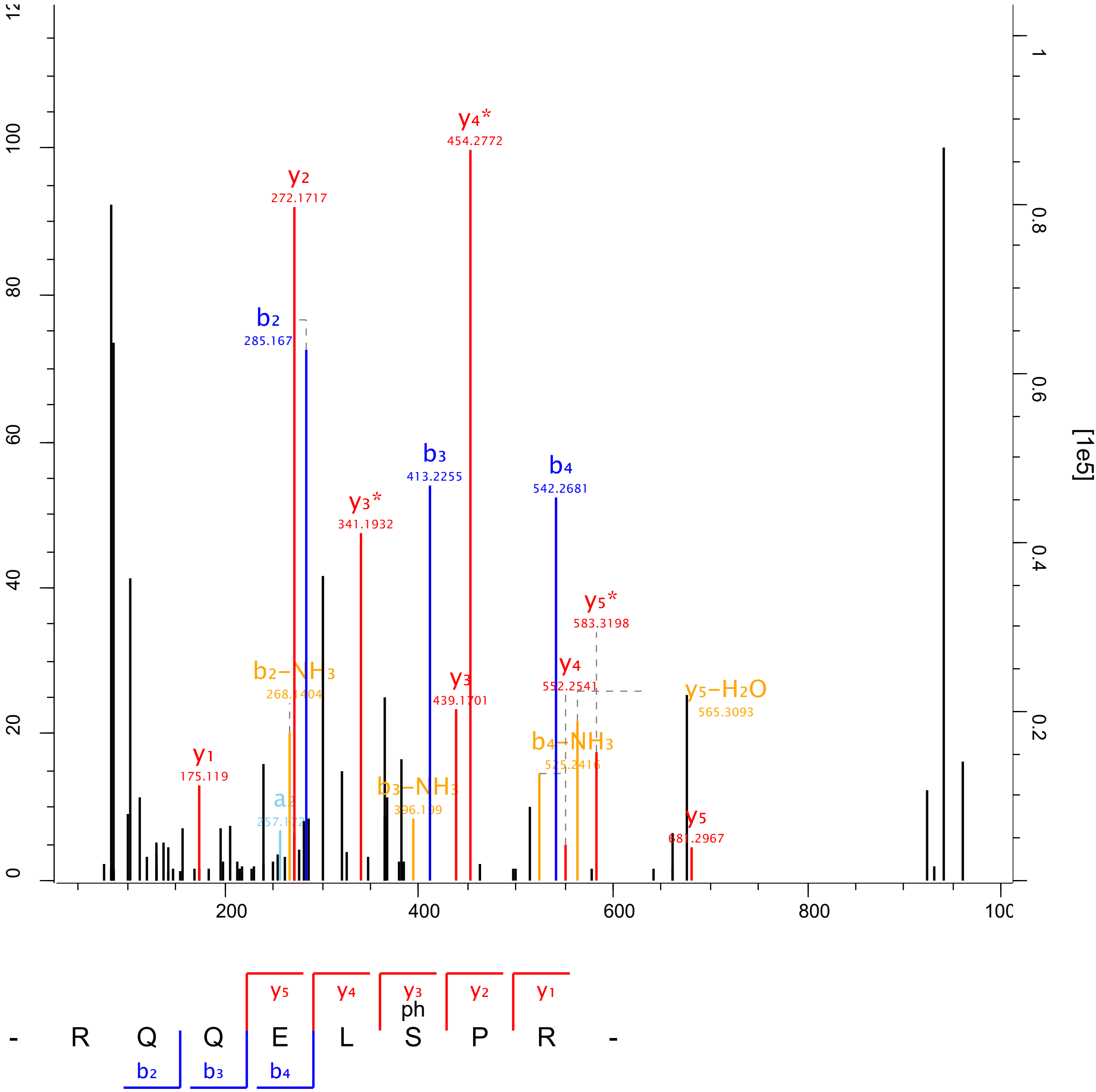

| Raw file | Scan | Method | Score | m/z |
| --- | --- | --- | --- | --- |
| sys_15_1 | 16850 | FTMS; HCD | 44.56 | 651.81 |

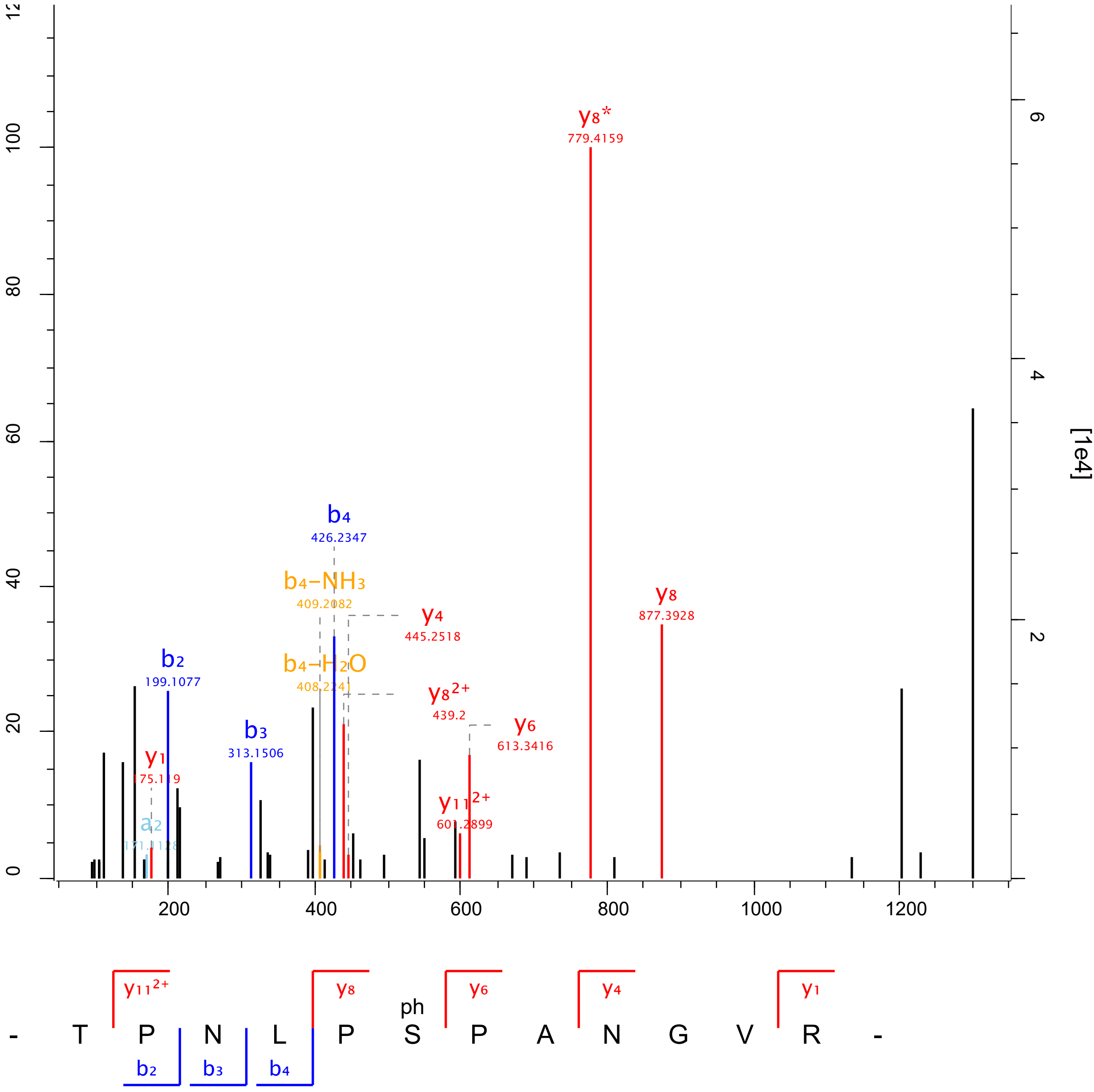

| Raw file | Scan | Method | Score | m/z |
| --- | --- | --- | --- | --- |
| sys_15_1 | 16904 | FTMS; HCD | 162.81 | 693.32 |

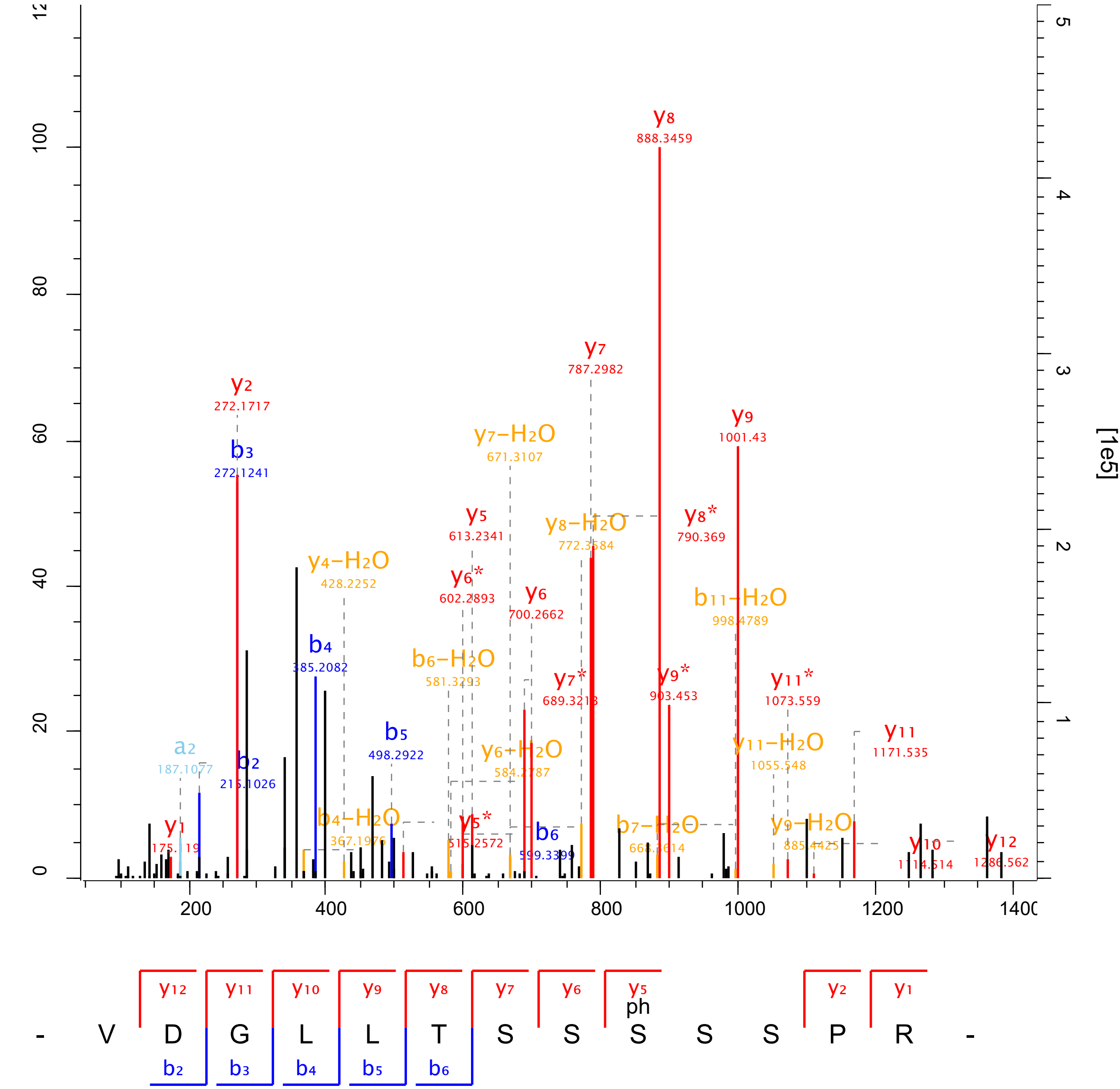

| Raw file | Scan | Method | Score | m/z |
| --- | --- | --- | --- | --- |
| sys_15_1 | 17036 | FTMS; HCD | 234.74 | 926.71 |

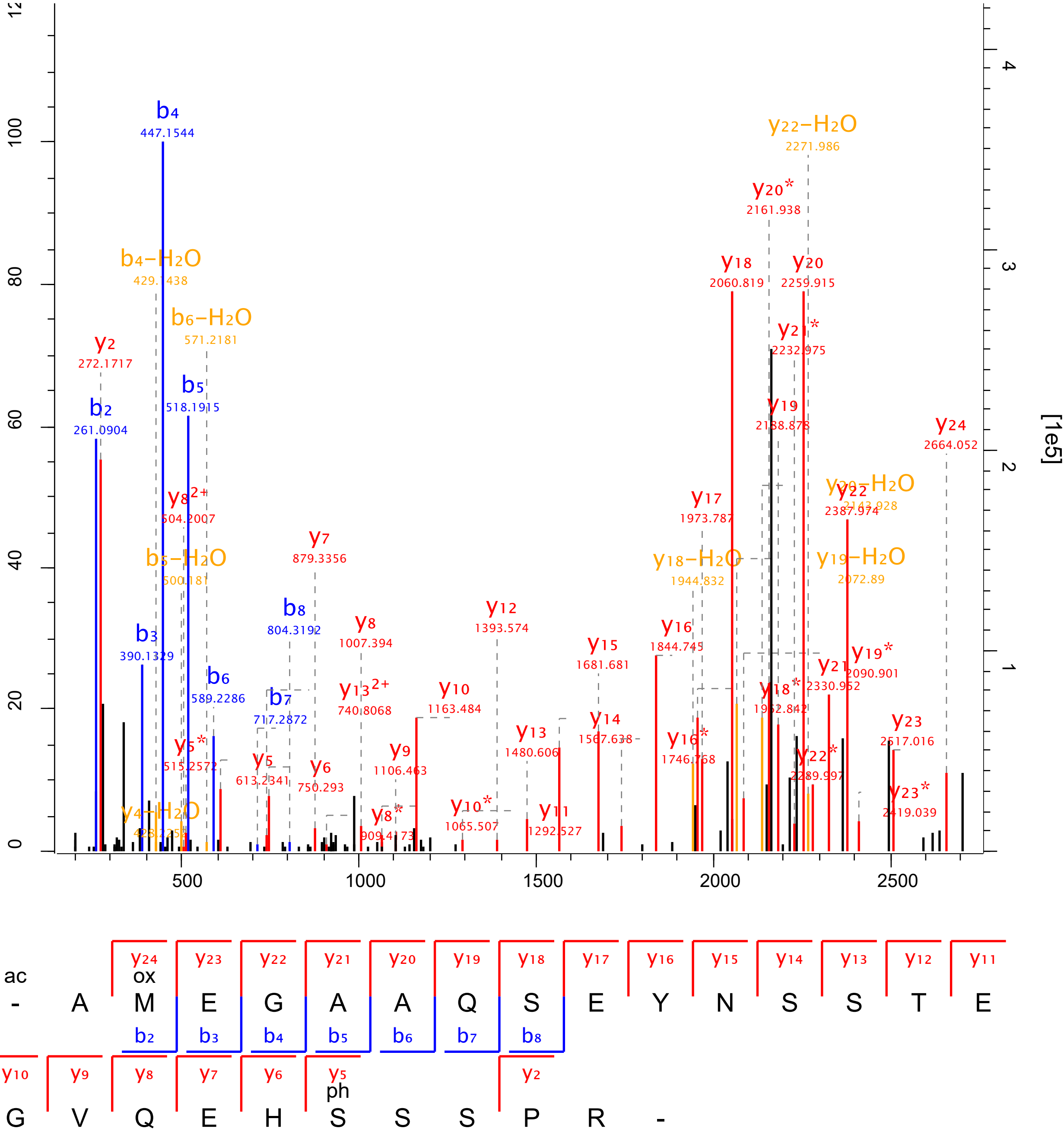

| Raw file | Scan | Method | Score | m/z |
| --- | --- | --- | --- | --- |
| sys_15_1 | 17048 | FTMS; HCD | 135.14 | 689.8 |

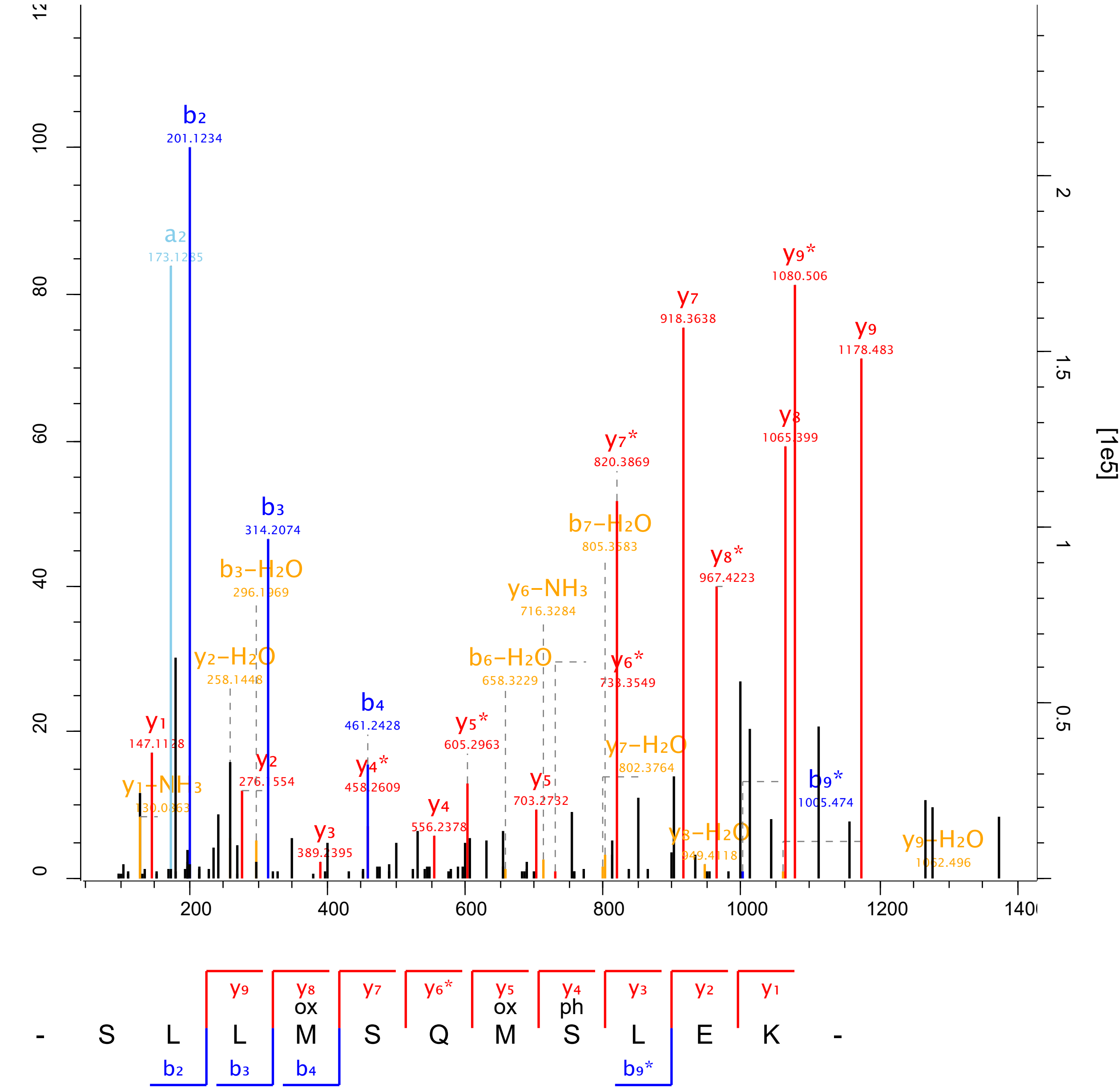

| Raw file | Scan | Method | Score | m/z |
|---|---|---|---|---|
| sys_15_1 | 17081 | FTMS; HCD | 89.23 | 642.8 |

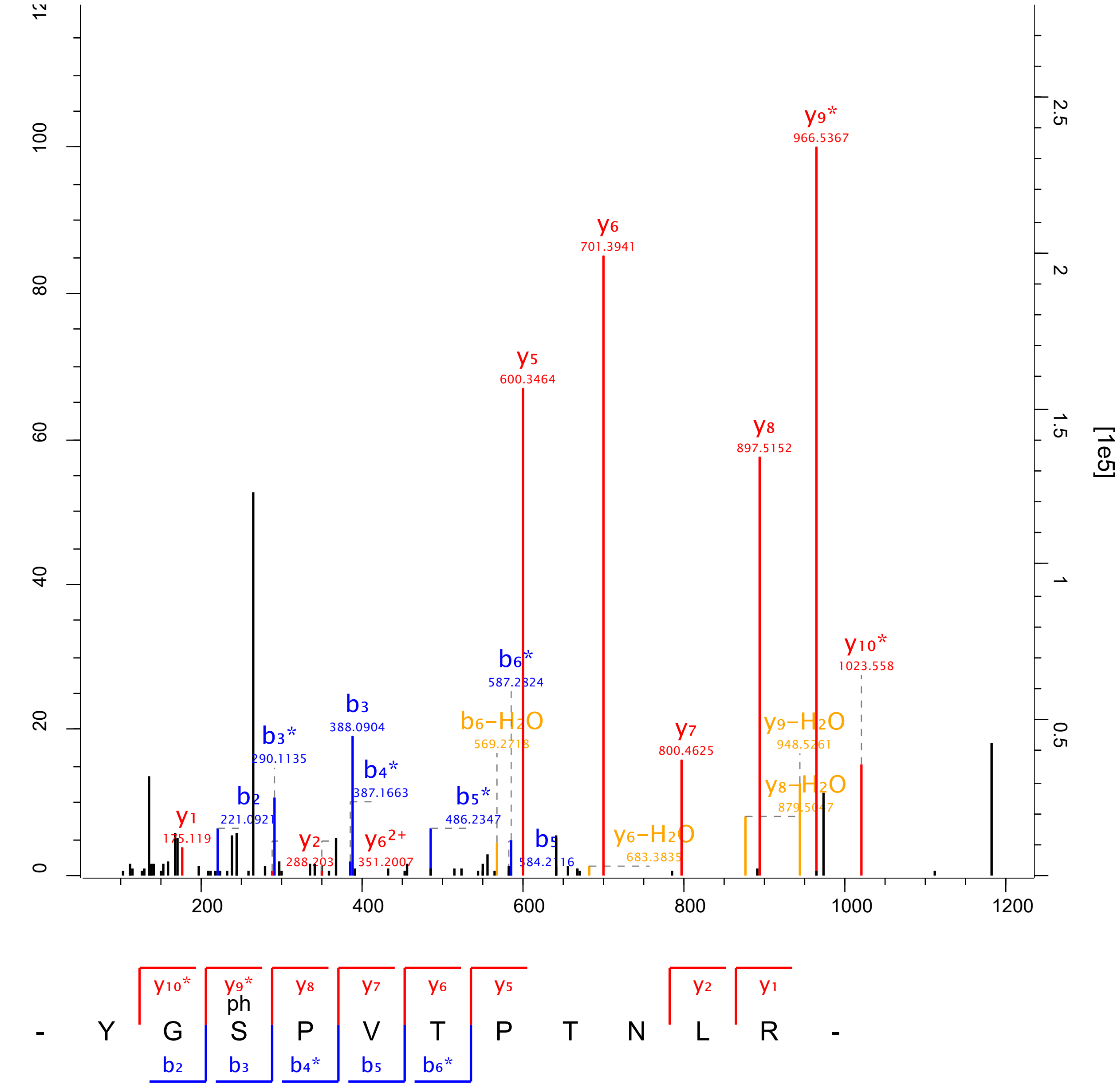

| Raw file | Scan | Method | Score | m/z |
|---|---|---|---|---|
| sys_15_1 | 17228 | FTMS; HCD | 43.84 | 979.92 |

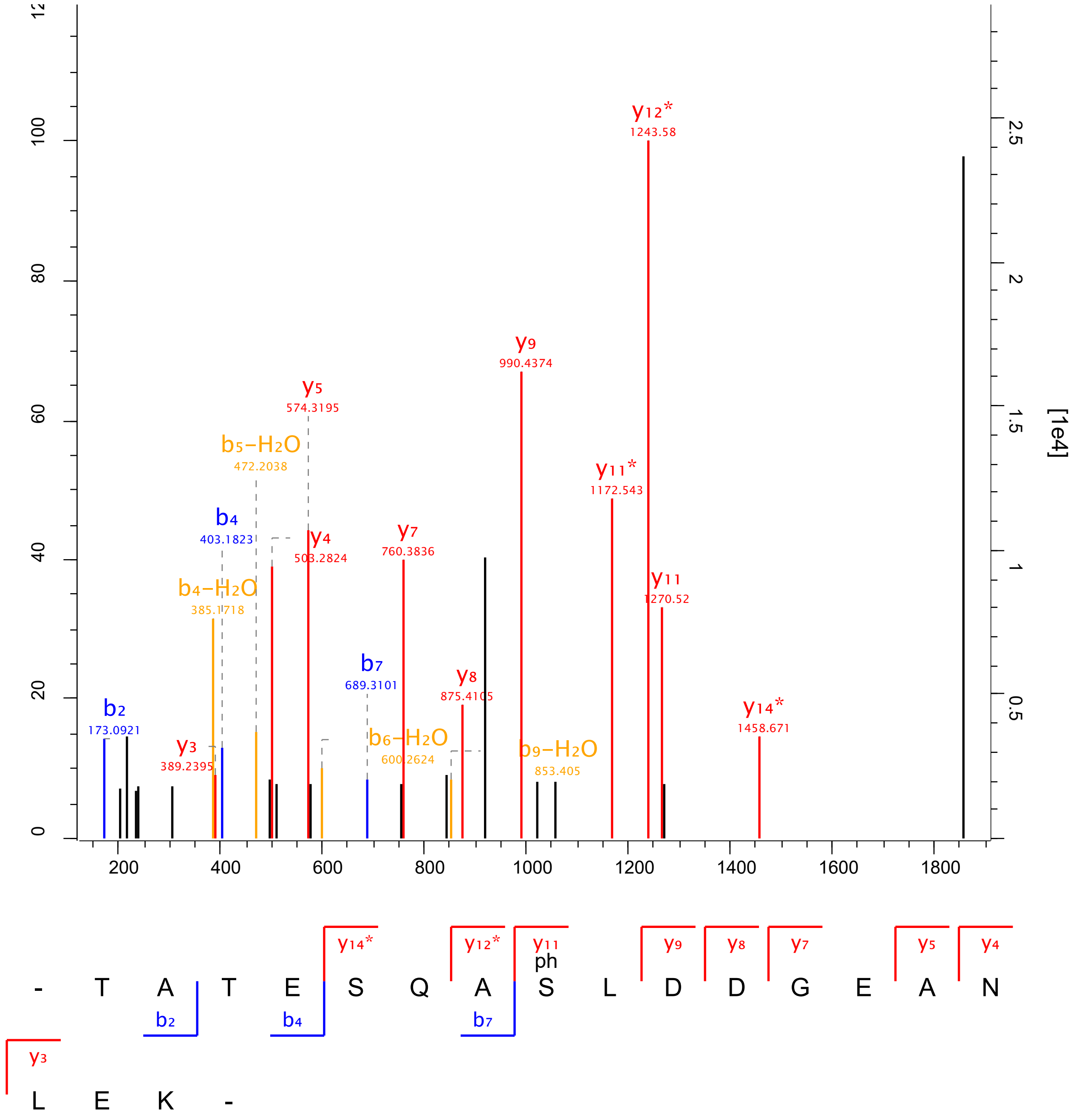

| Raw file | Scan | Method | Score | m/z |
| --- | --- | --- | --- | --- |
| sys_15_1 | 17289 | FTMS; HCD | 100.97 | 839.84 |

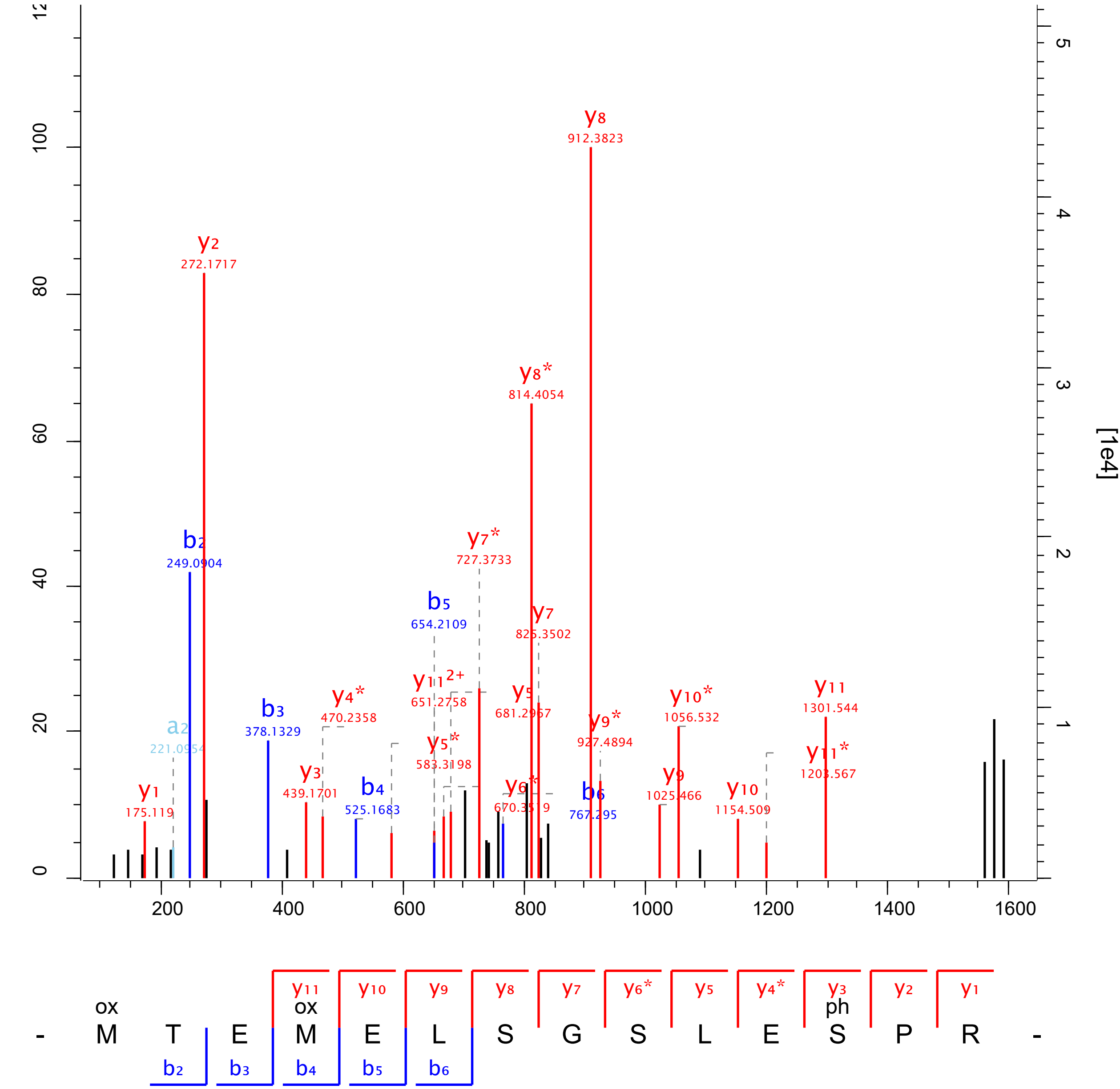

| Raw file | Scan | Method | Score | m/z |
|---|---|---|---|---|
| sys_15_1 | 17319 | FTMS; HCD | 178.15 | 1046.91 |

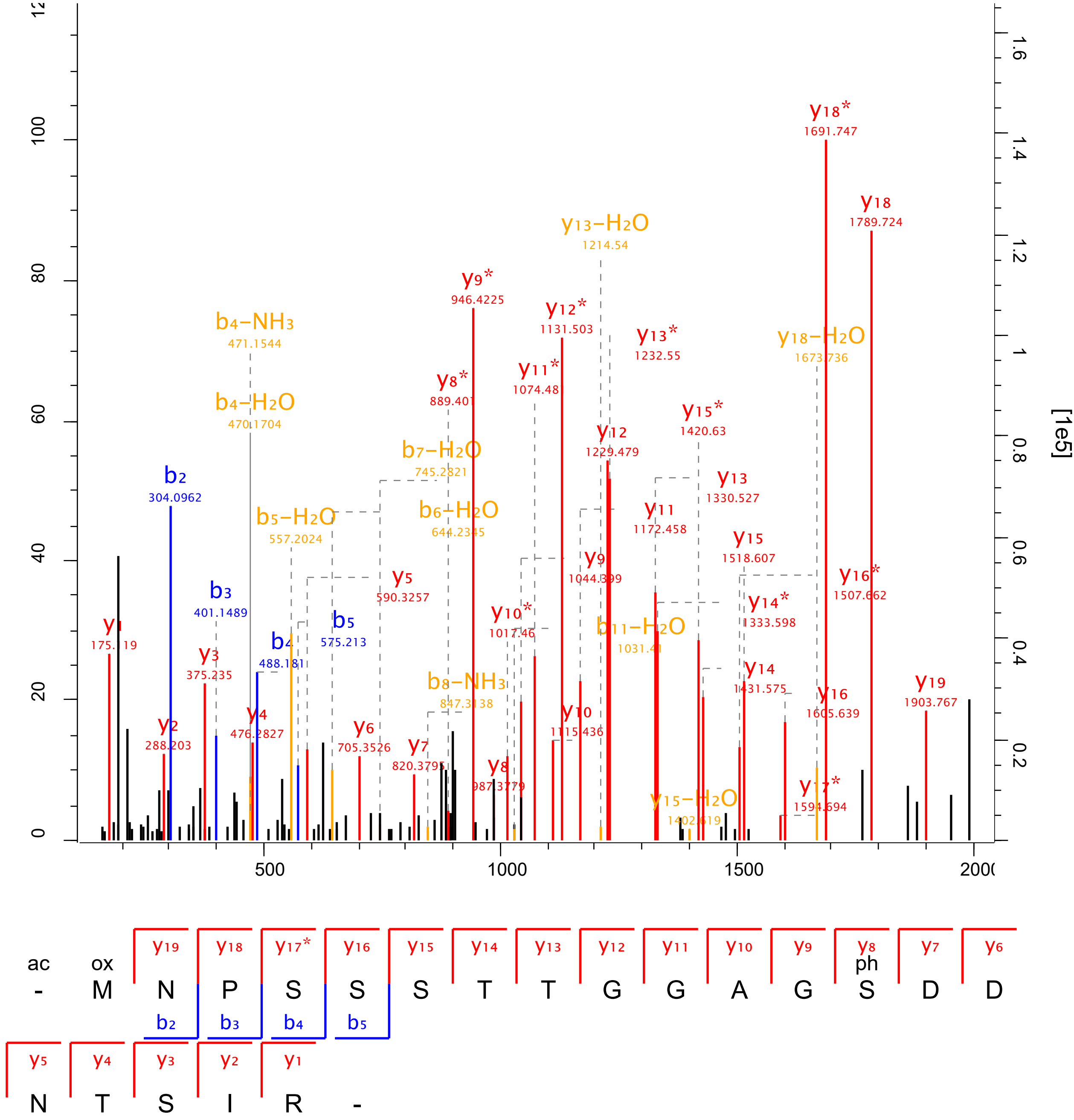

| Raw file | Scan | Method | Score | m/z |
|---|---|---|---|---|
| sys_15_1 | 17338 | FTMS; HCD | 195.92 | 872.35 |

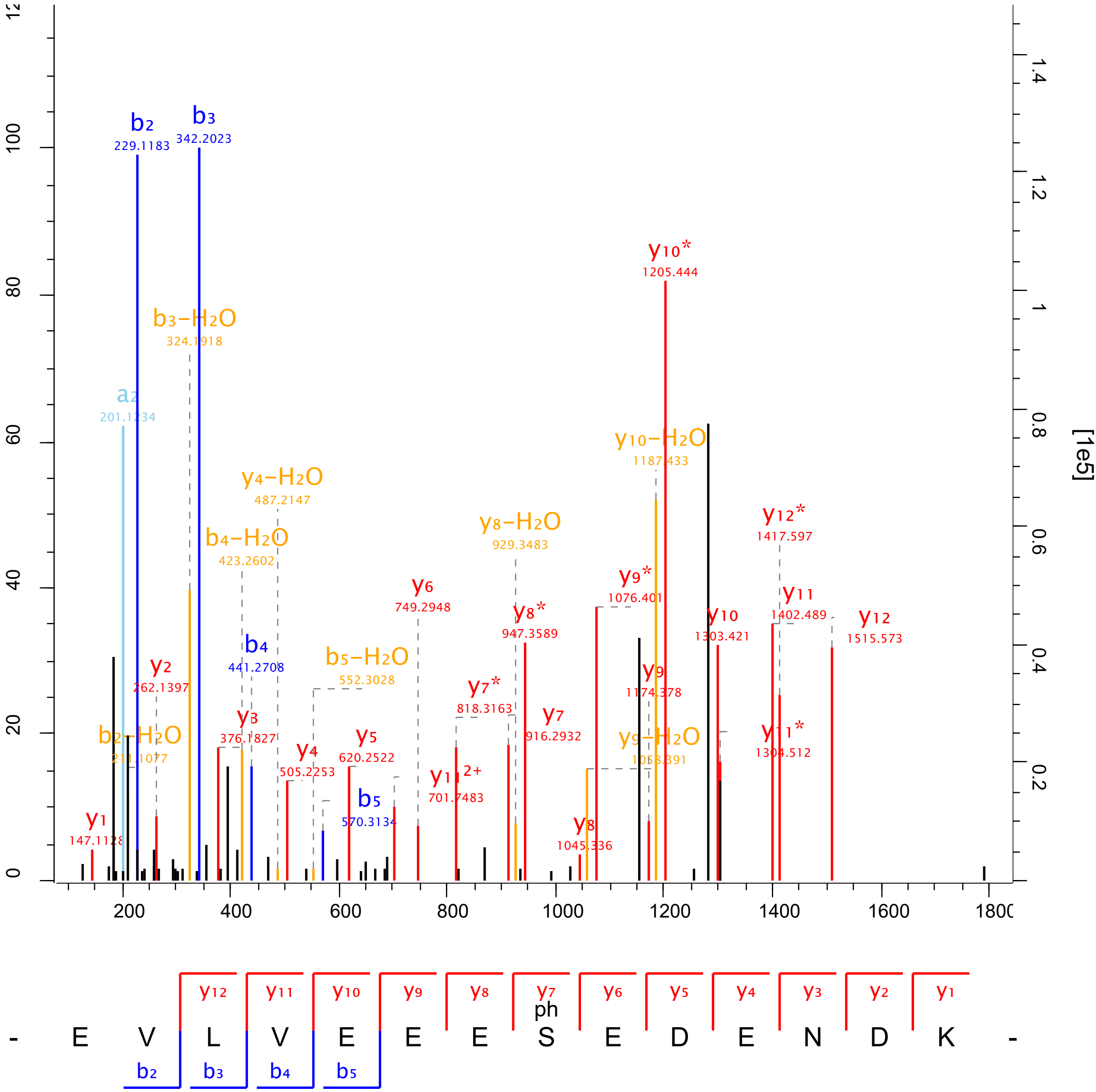

| Raw file | Scan | Method | Score | m/z |
| --- | --- | --- | --- | --- |
| sys_15_1 | 17496 | FTMS; HCD | 172.13 | 576.26 |

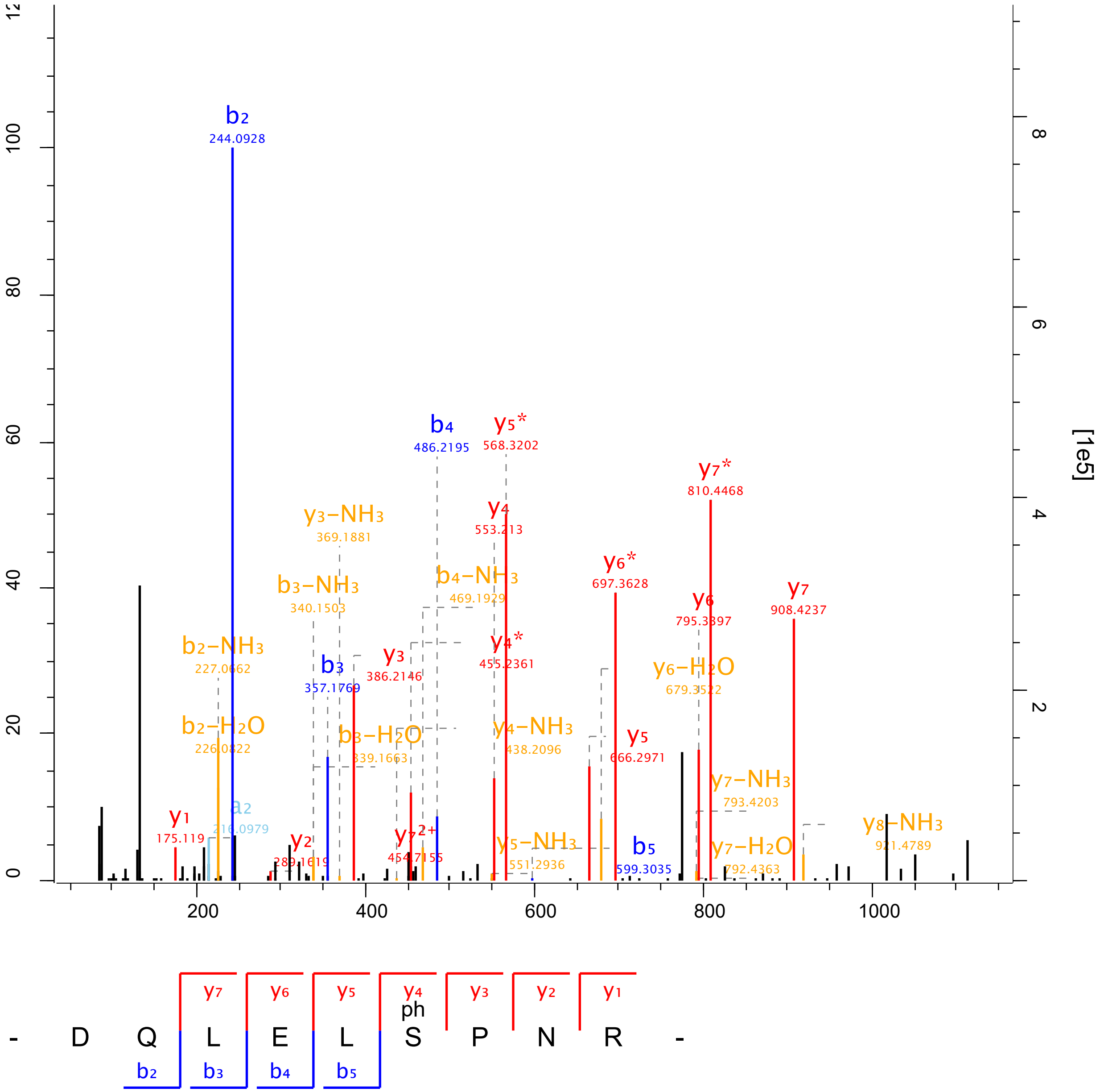

| Raw file | Scan | Method | Score | m/z |
|---|---|---|---|---|
| sys_15_1 | 17509 | FTMS; HCD | 69.98 | 848.86 |

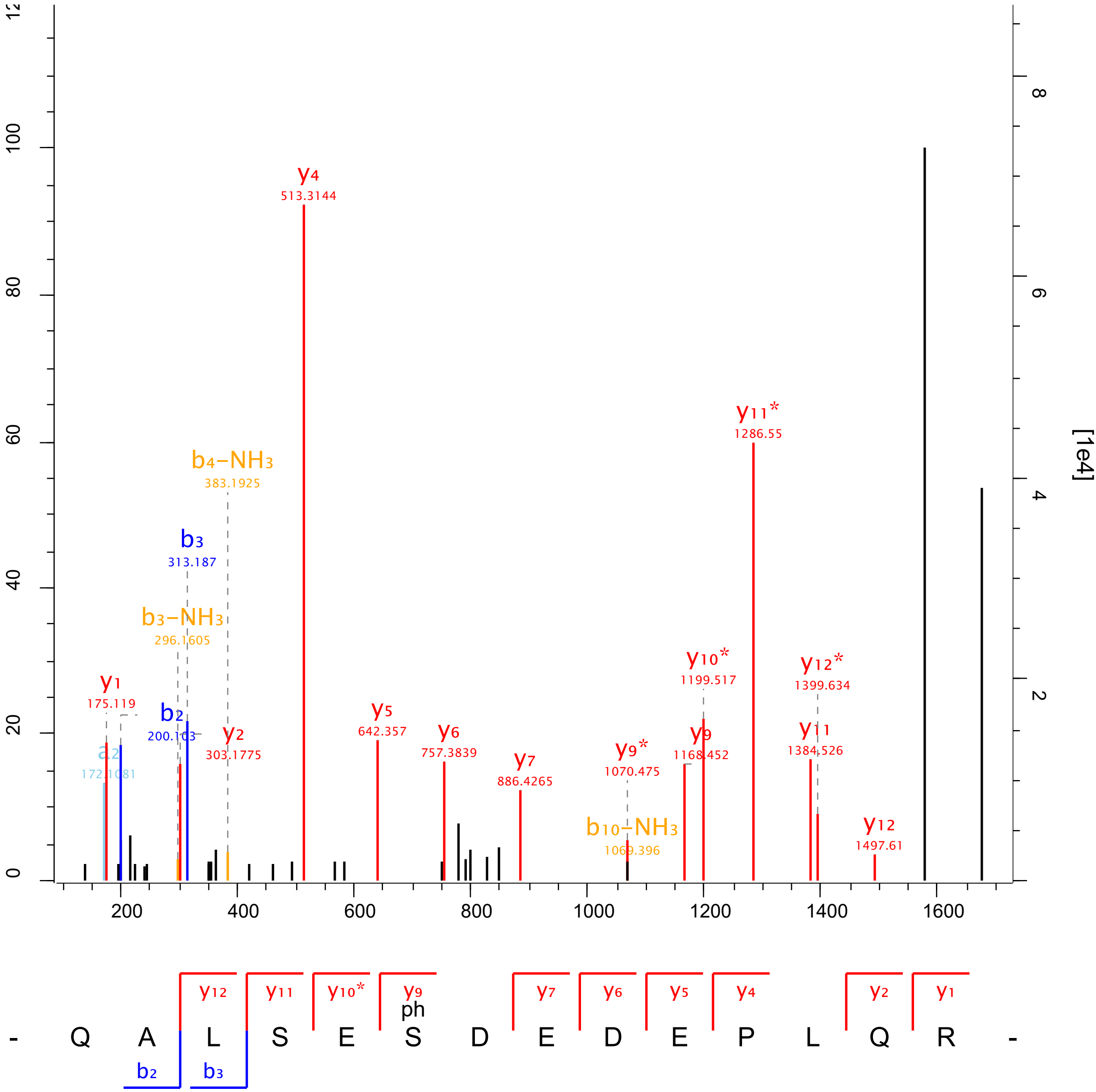

| Raw file | Scan | Method | Score | m/z |
|---|---|---|---|---|
| sys_15_1 | 17529 | FTMS; HCD | 148.96 | 580.75 |

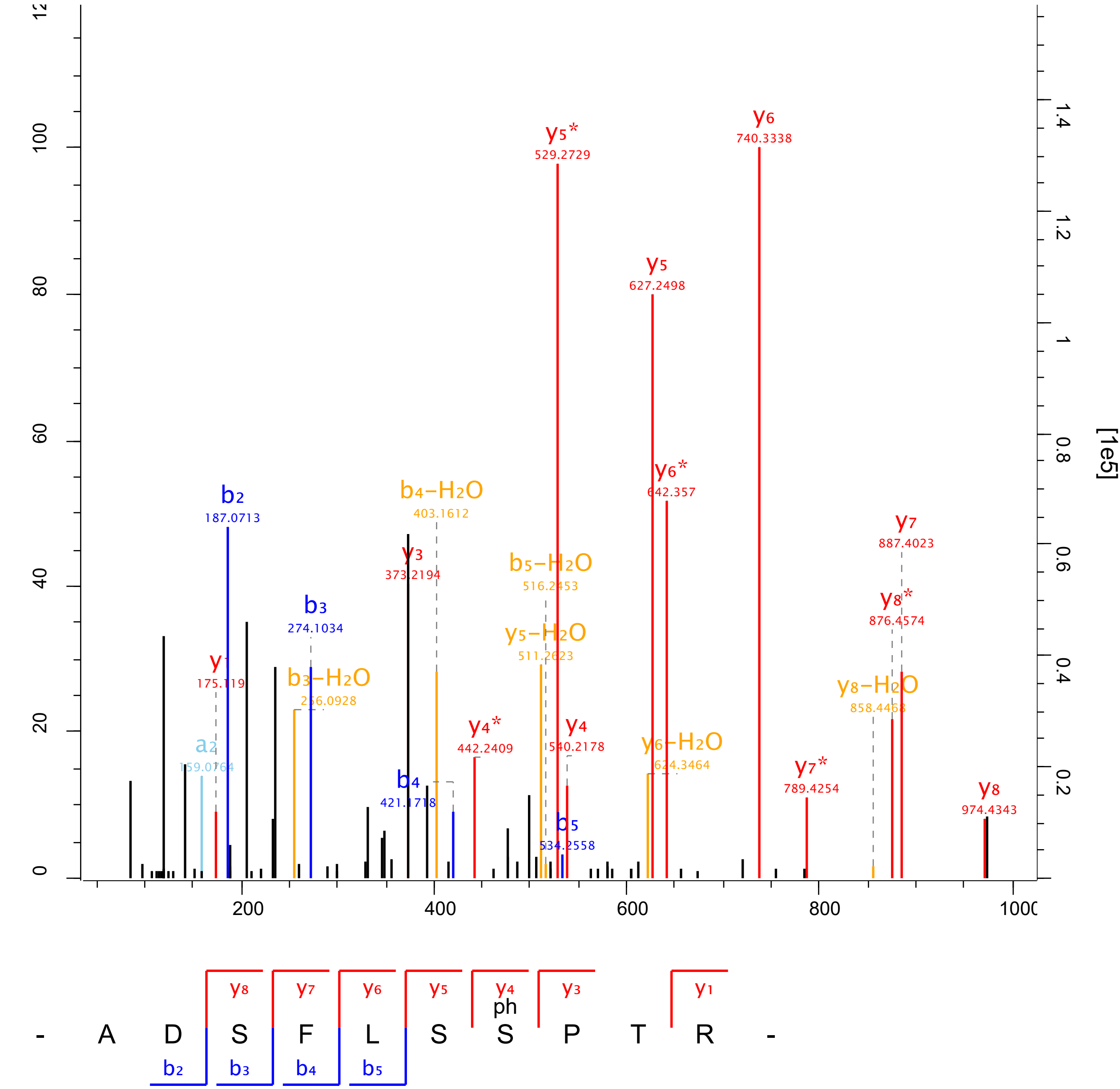

| Raw file | Scan | Method | Score | m/z |
| --- | --- | --- | --- | --- |
| sys_15_1 | 17623 | FTMS; HCD | 111.34 | 697.83 |

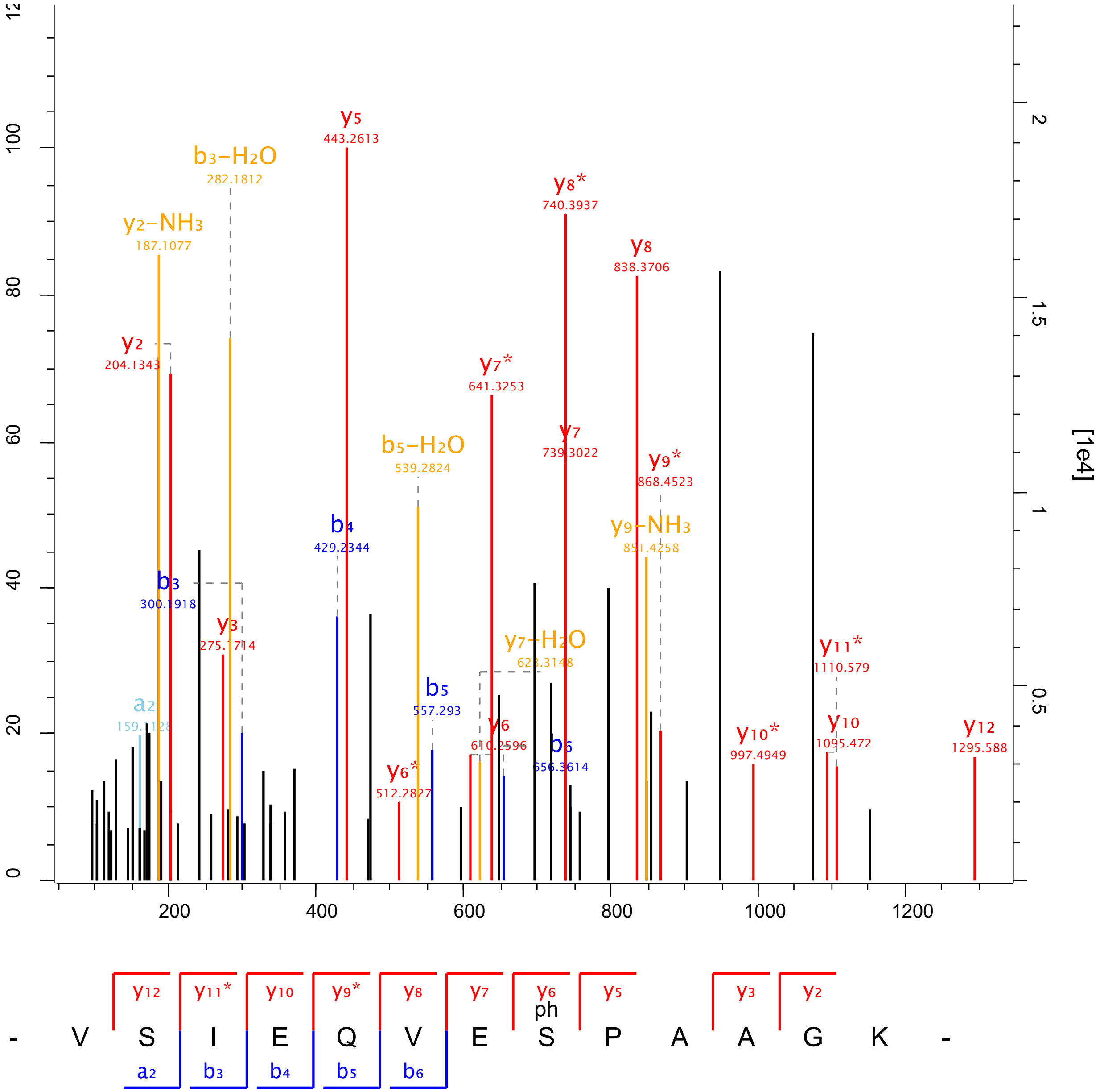

| Raw file | Scan | Method | Score | m/z |
|---|---|---|---|---|
| sys_15_1 | 17648 | FTMS; HCD | 79.68 | 591.93 |

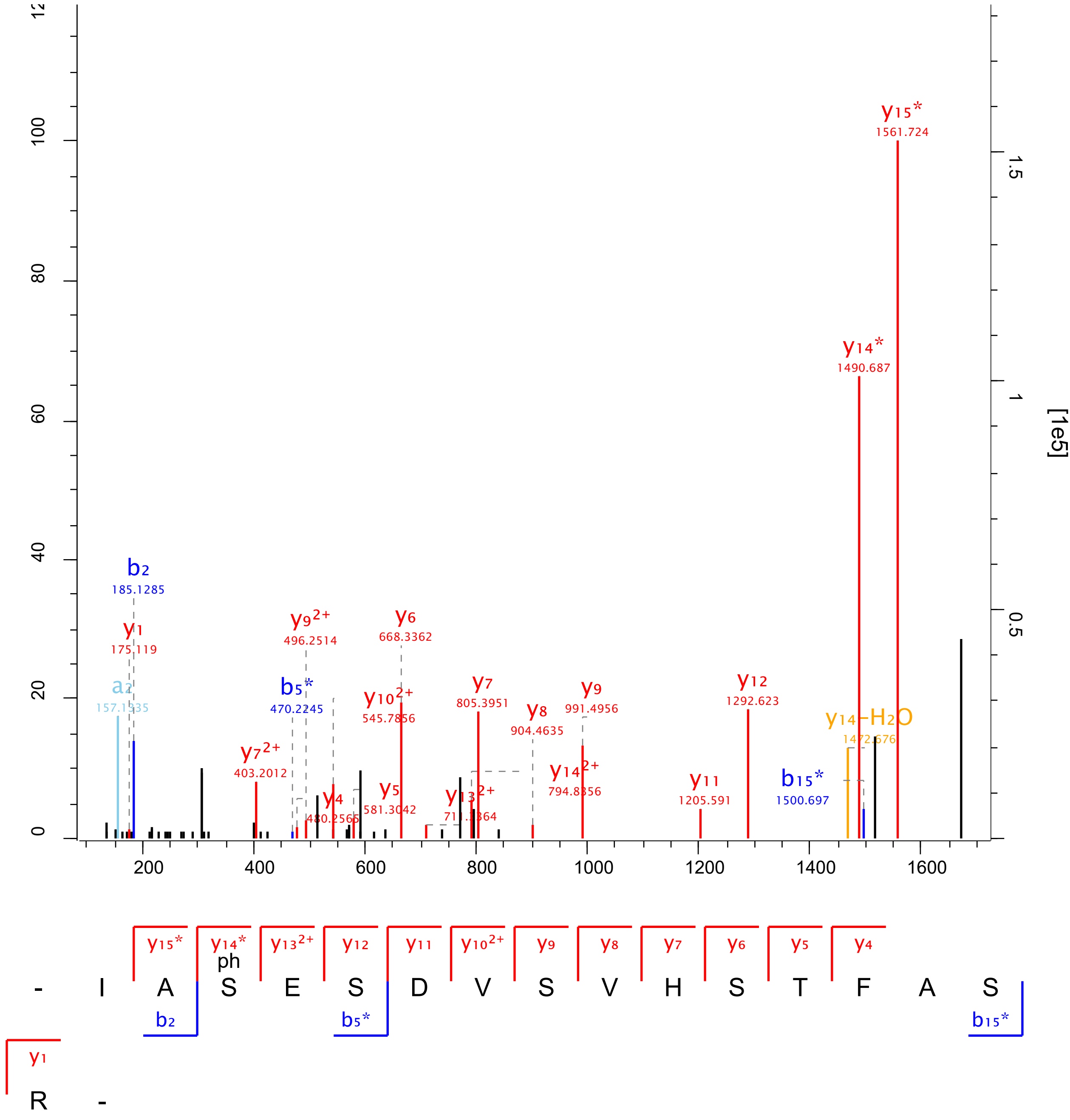

| Raw file | Scan | Method | Score | m/z |
|---|---|---|---|---|
| sys_15_1 | 17651 | FTMS; HCD | 97.69 | 598.27 |

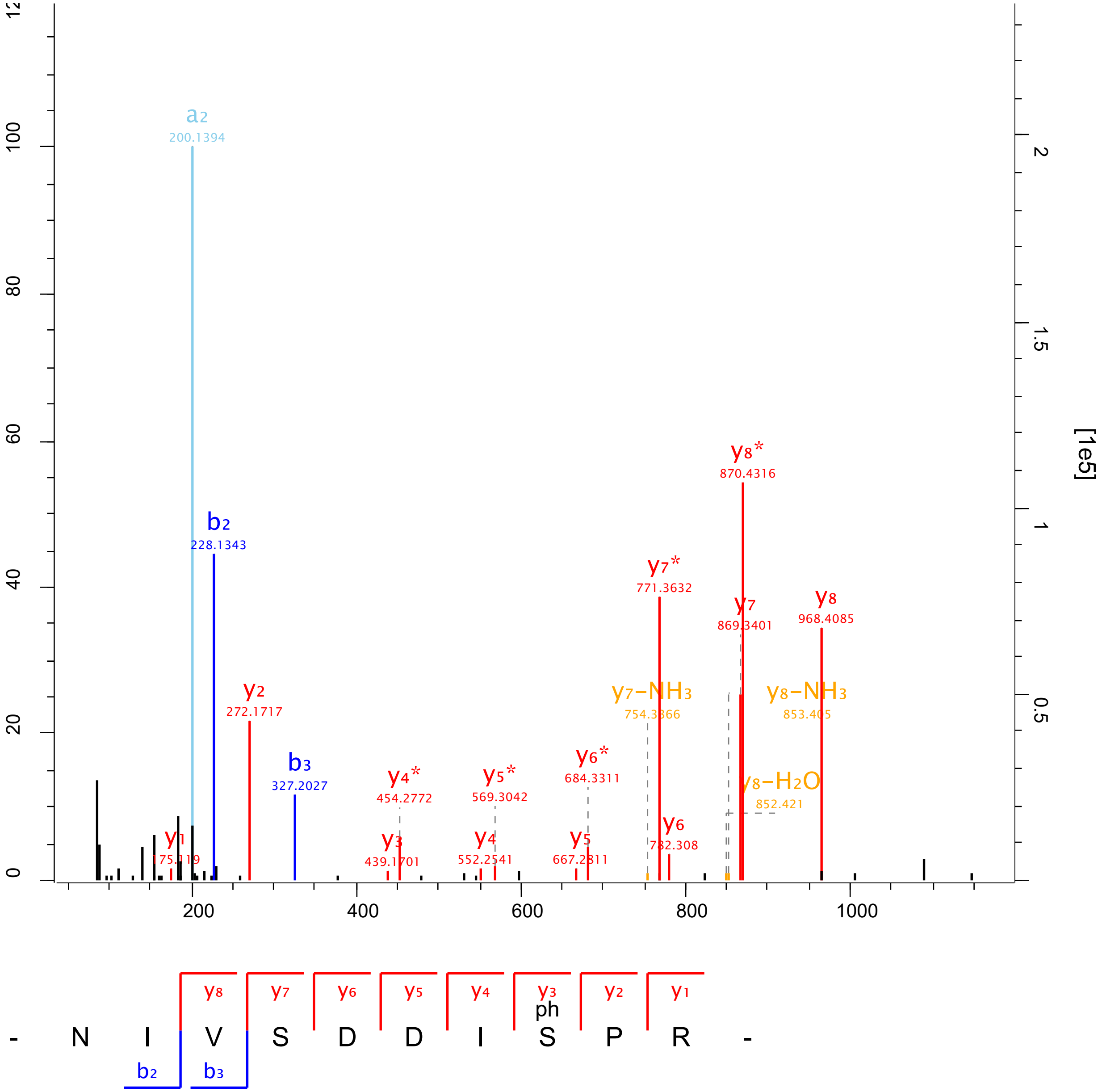

| Raw file | Scan | Method | Score | m/z |
| --- | --- | --- | --- | --- |
| sys_15_1 | 17772 | FTMS; HCD | 91.81 | 657.74 |

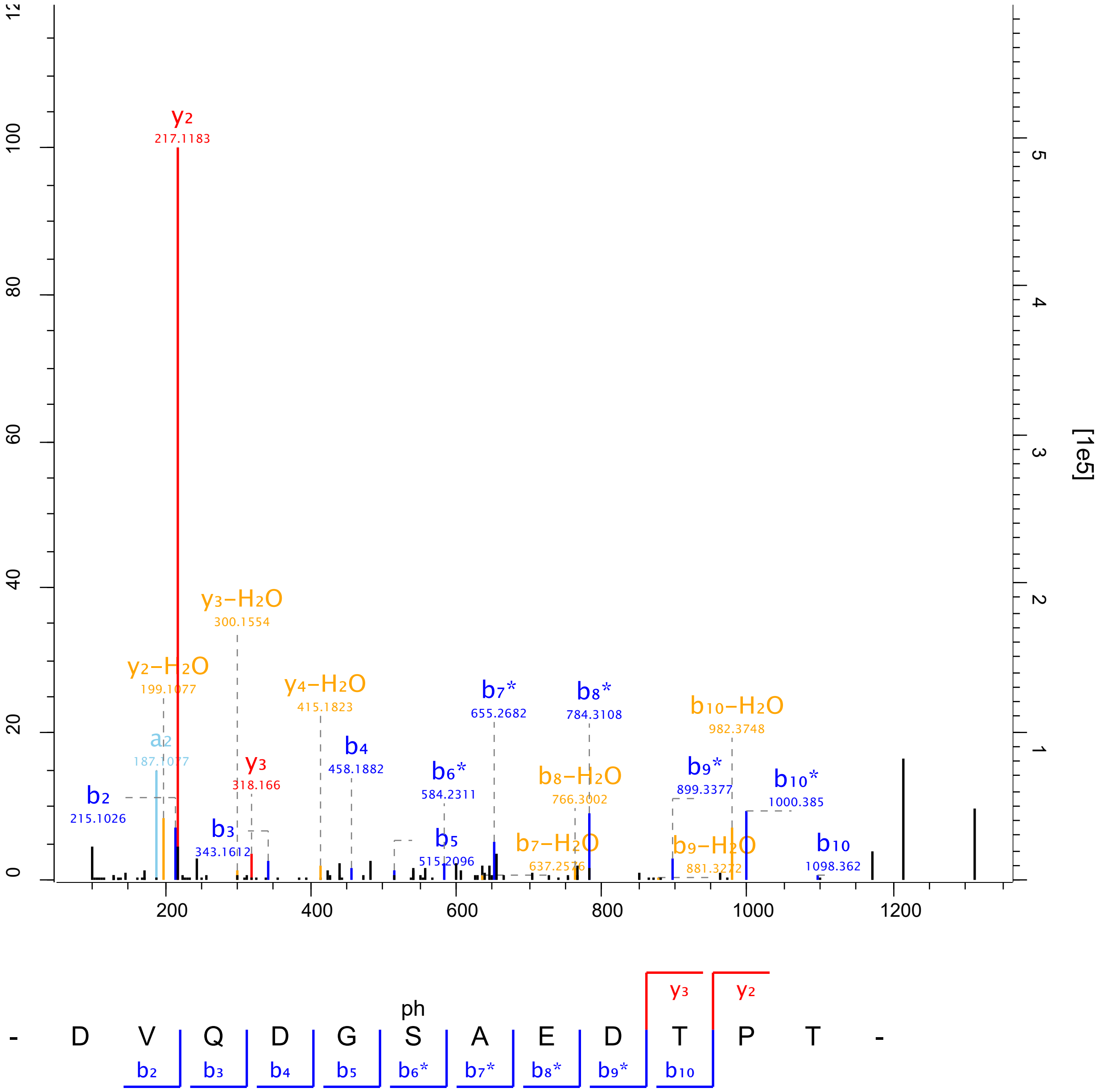

| Raw file | Scan | Method | Score | m/z |
|---|---|---|---|---|
| sys_15_1 | 17862 | FTMS; HCD | 69.7 | 885.86 |

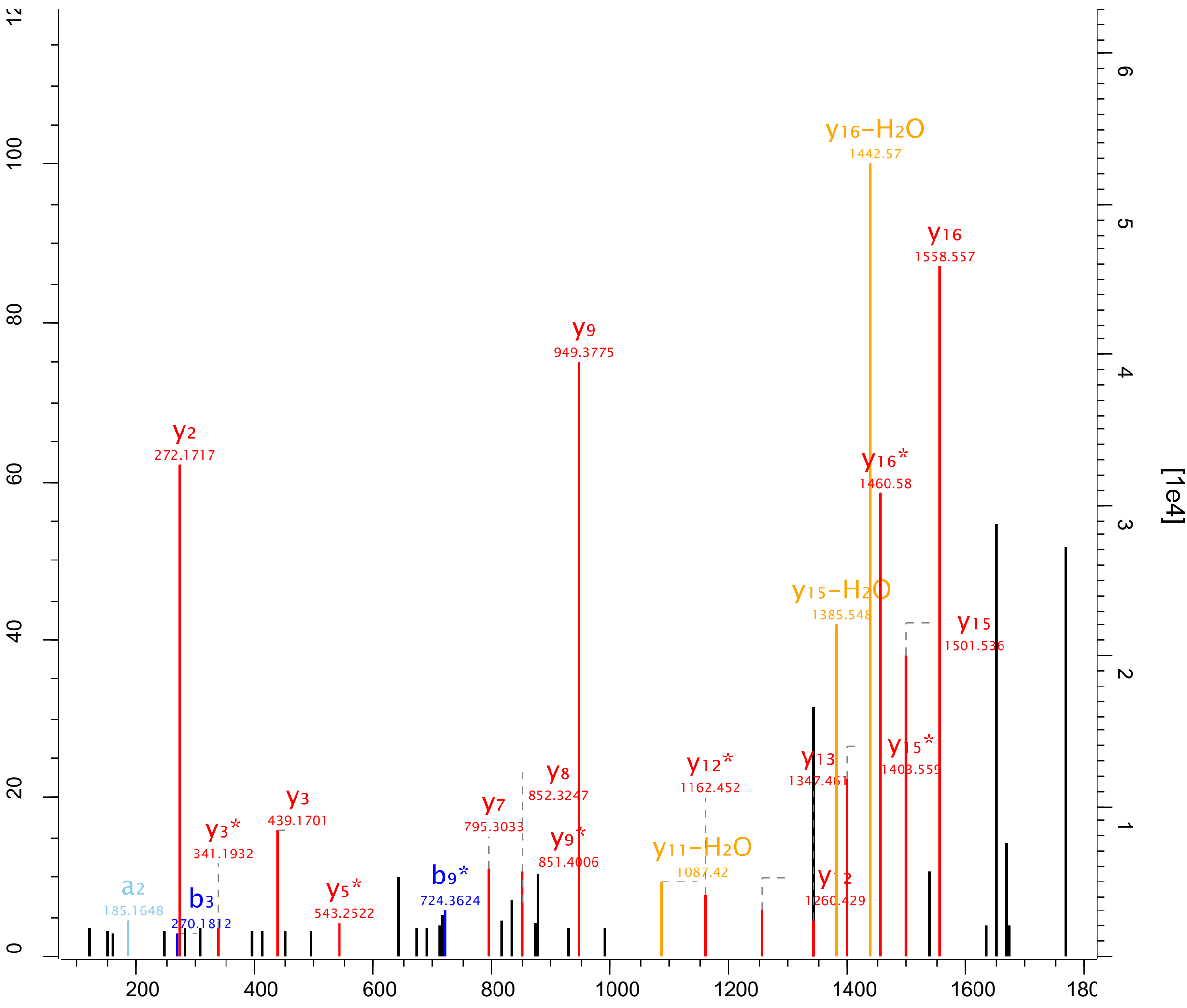


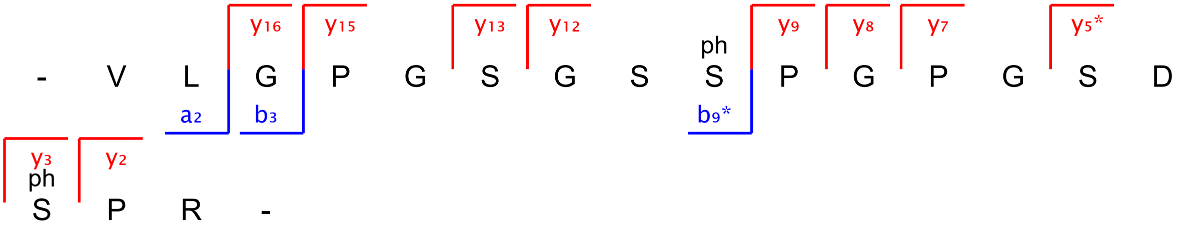

| Raw file | Scan | Method | Score | m/z |
| --- | --- | --- | --- | --- |
| sys_15_1 | 17892 | FTMS; HCD | 62.62 | 795.01 |

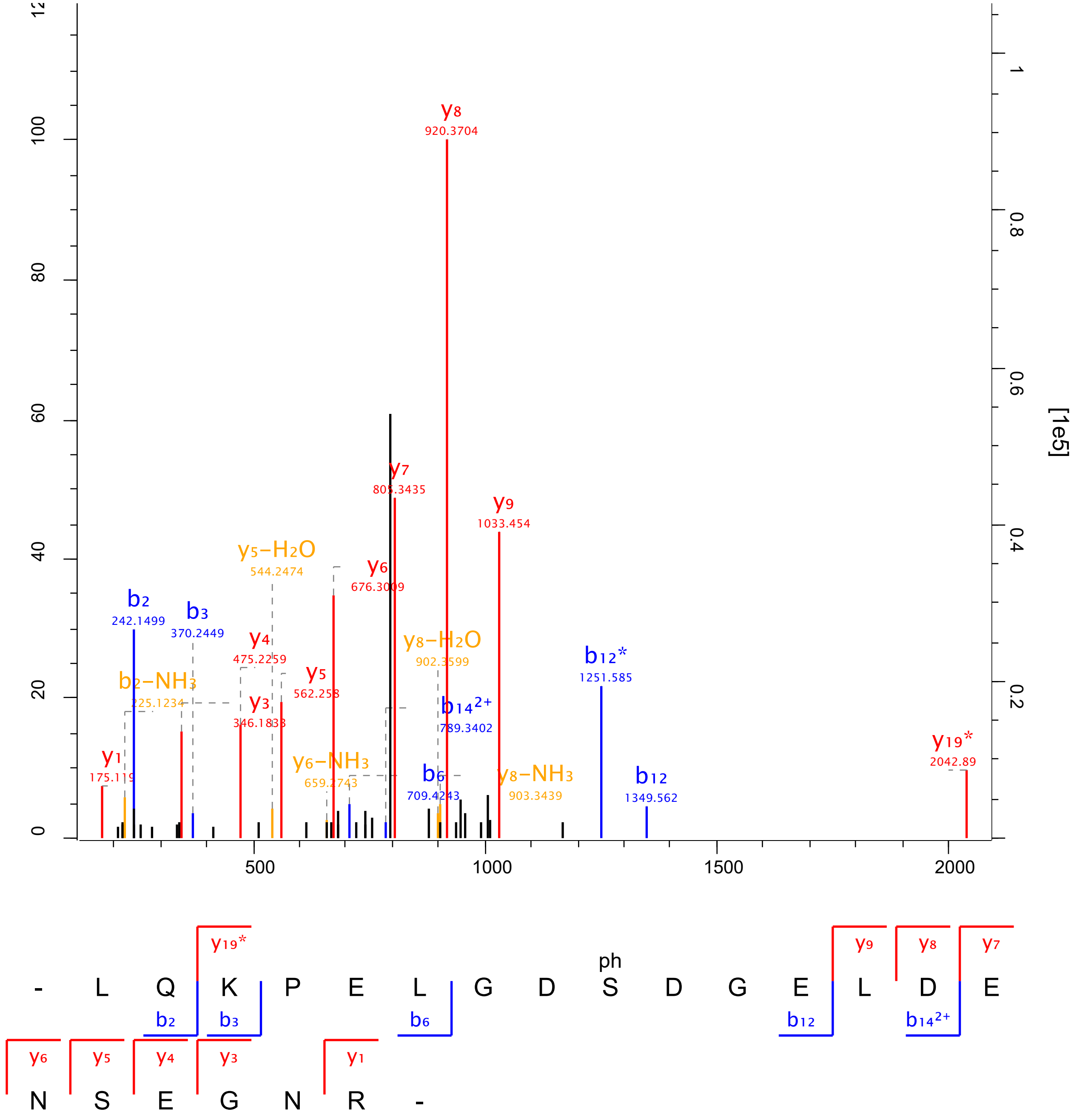

| Raw file | Scan | Method | Score | m/z |
|---|---|---|---|---|
| sys_15_1 | 17953 | FTMS; HCD | 125.79 | 888.9 |

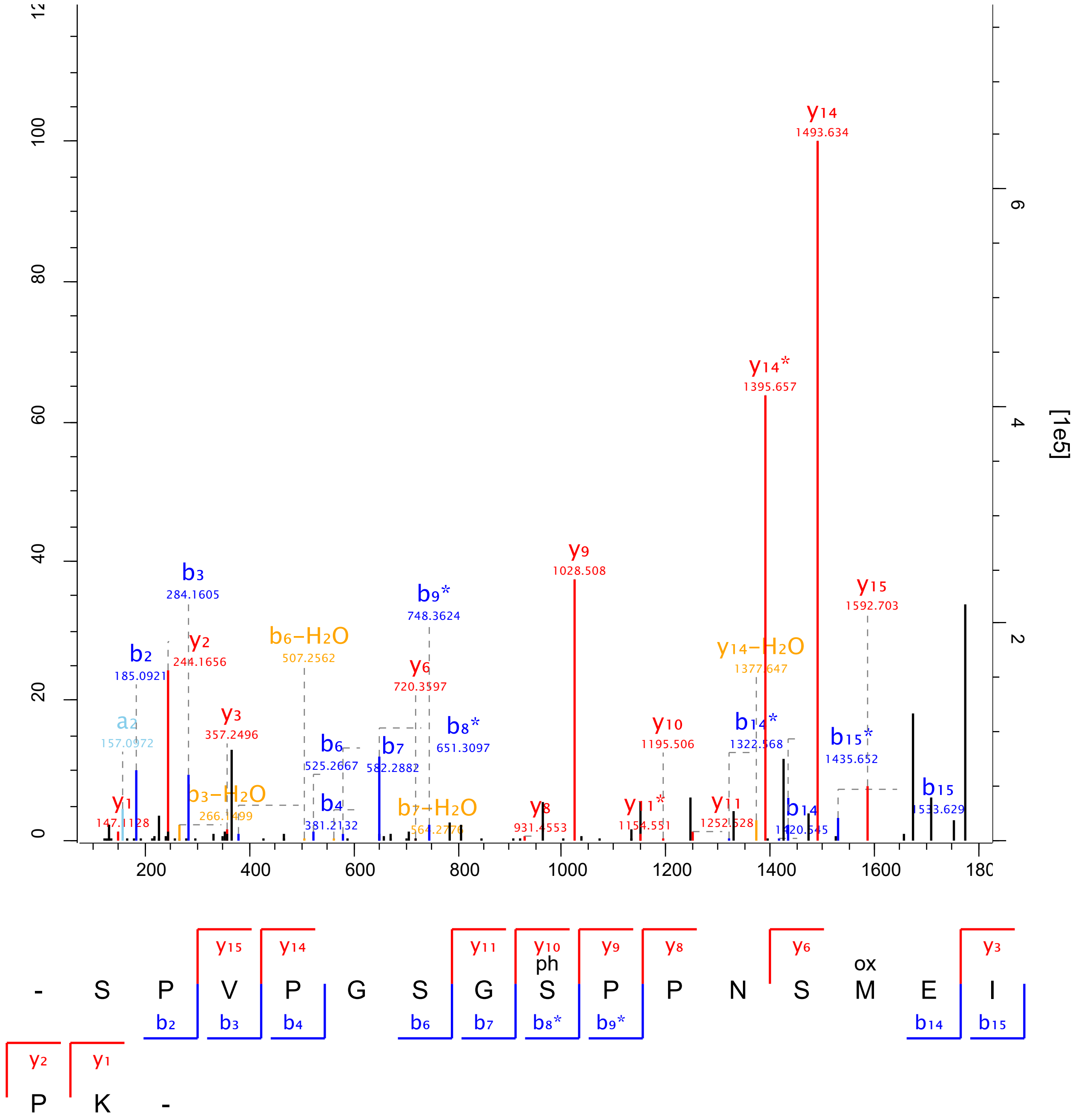

| Raw file | Scan | Method | Score | m/z |
| --- | --- | --- | --- | --- |
| sys_15_1 | 18008 | FTMS; HCD | 51.94 | 563.78 |

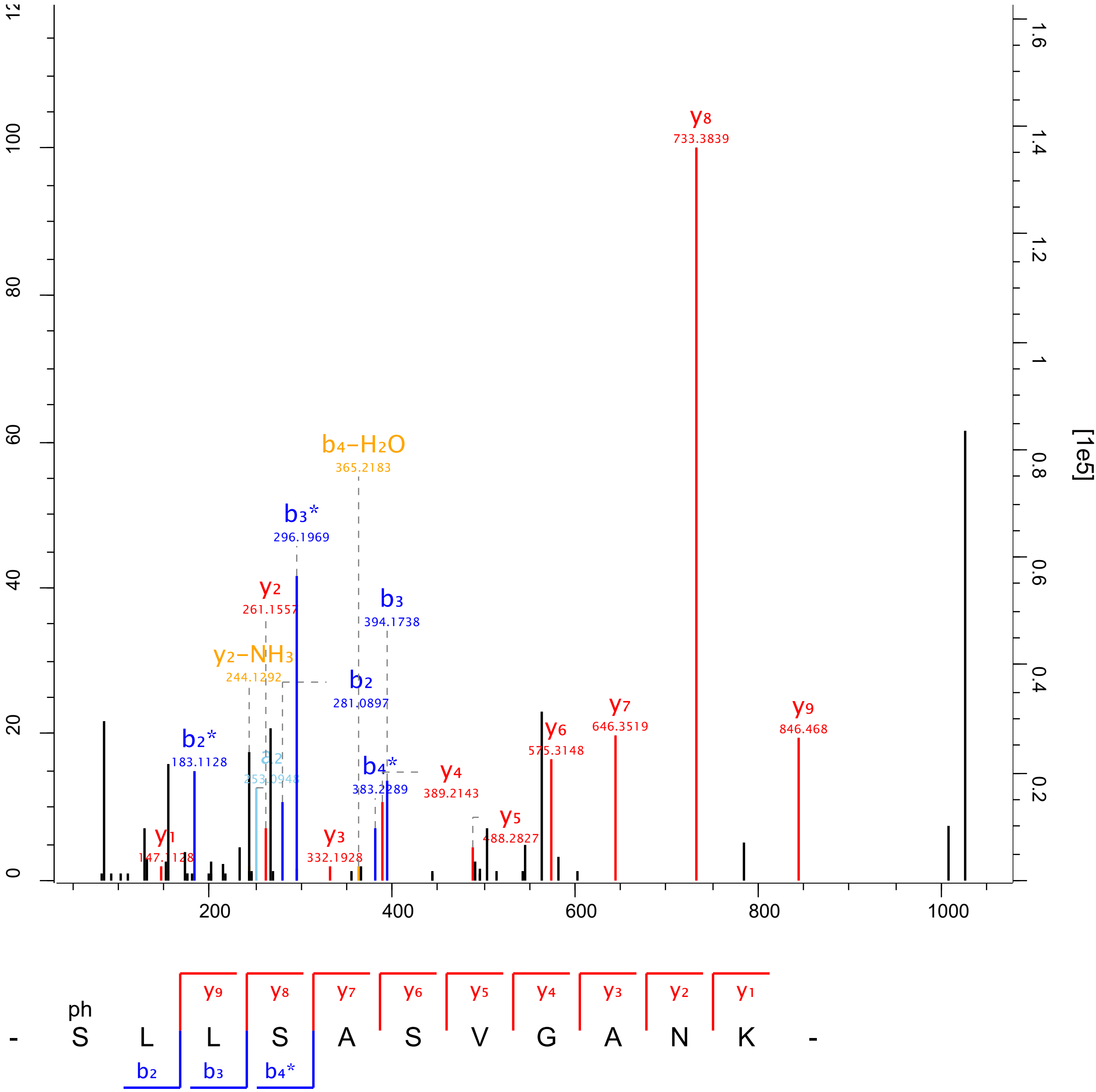

| Raw file | Scan | Method | Score | m/z |
|---|---|---|---|---|
| sys_15_1 | 18070 | FTMS; HCD | 211.45 | 705.82 |

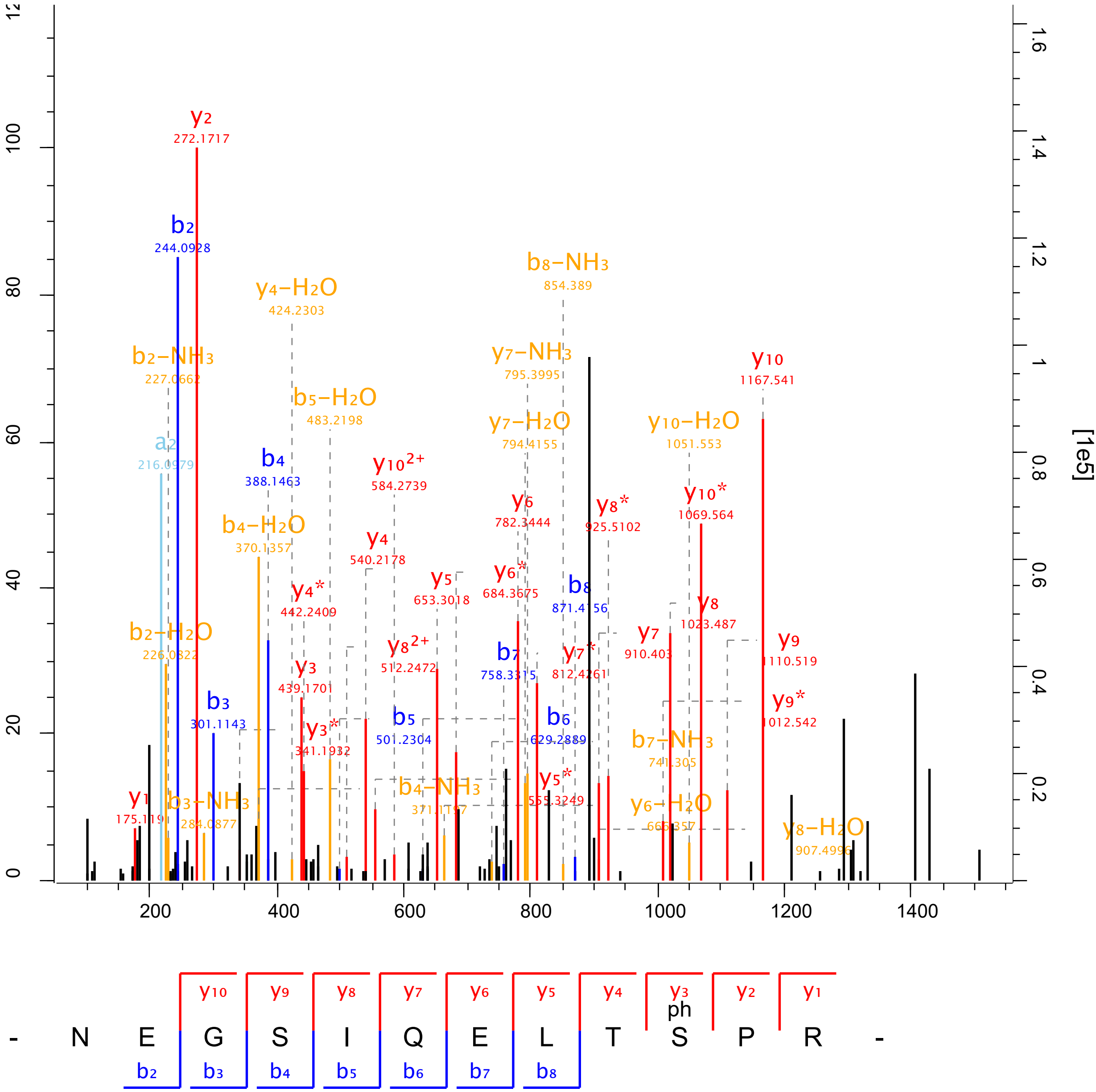

| Raw file | Scan | Method | Score | m/z |
|---|---|---|---|---|
| sys_15_1 | 1810 | FTMS; HCD | 56.57 | 641.79 |

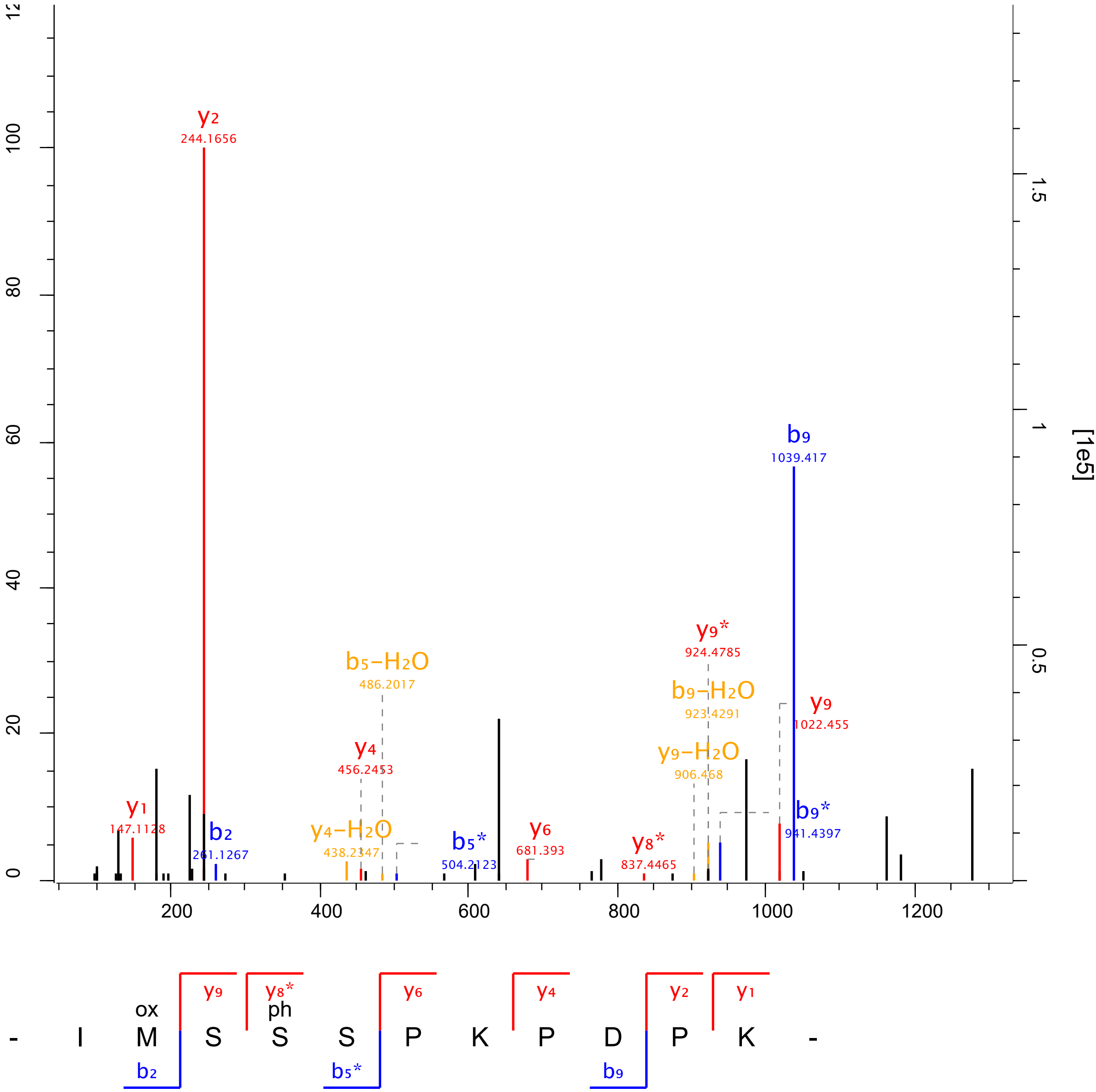

| Raw file | Scan | Method | Score | m/z |
|---|---|---|---|---|
| sys_15_1 | 18134 | FTMS; HCD | 135.8 | 527.73 |

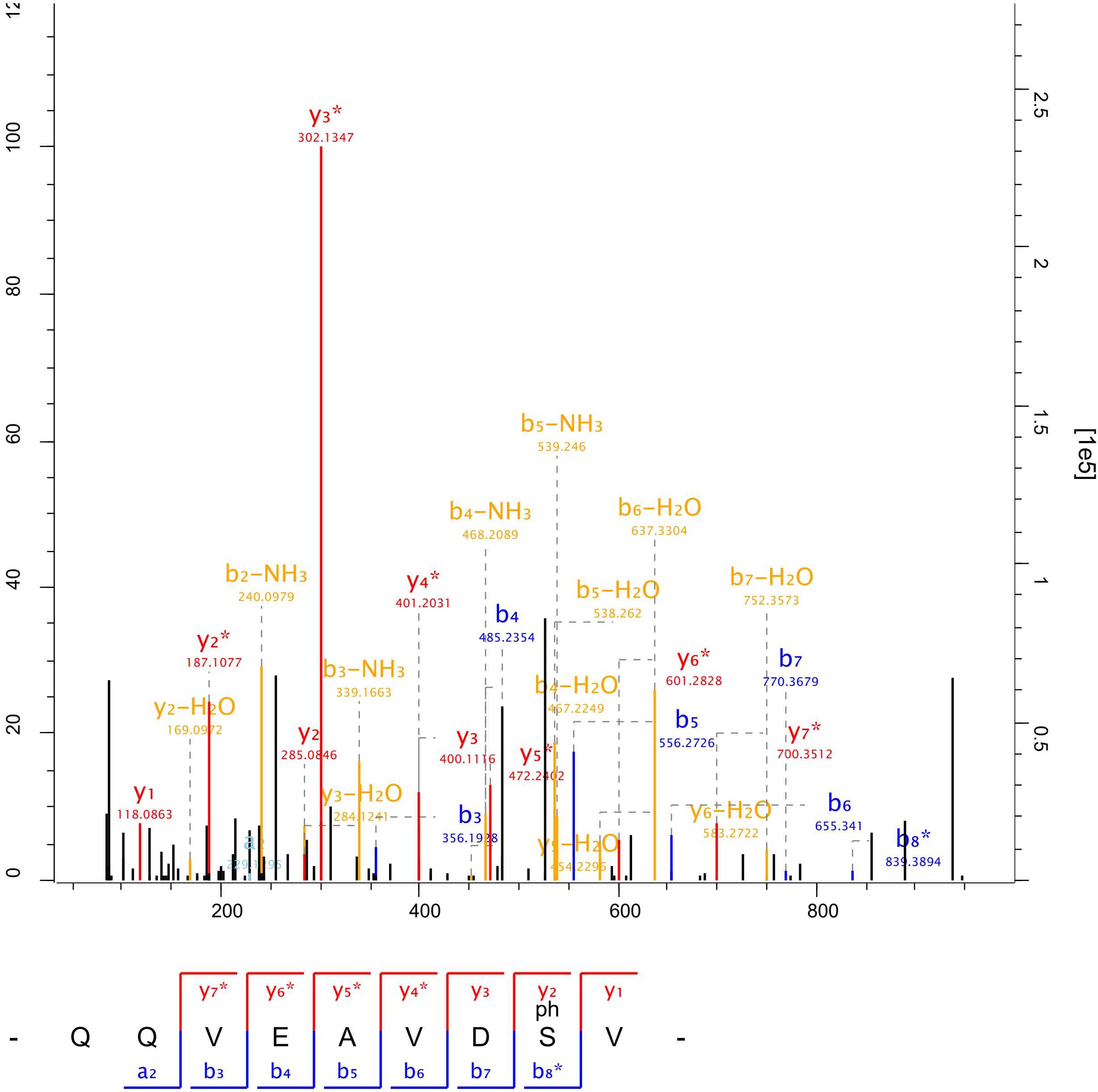

| Raw file | Scan | Method | Score | m/z |
|---|---|---|---|---|
| sys_15_1 | 18194 | FTMS; HCD | 50.86 | 904.38 |

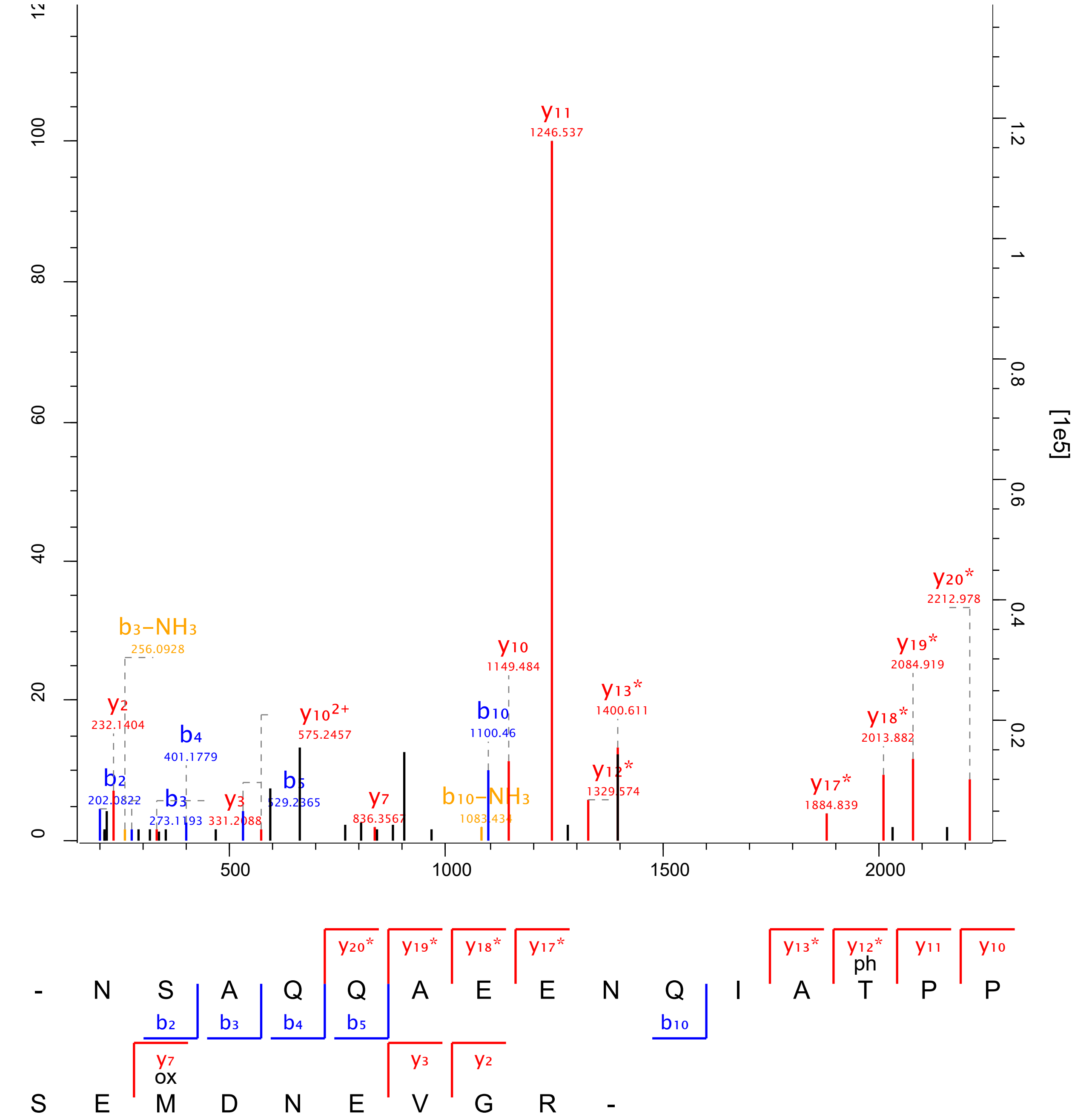

| Raw file | Scan | Method | Score | m/z |
|---|---|---|---|---|
| sys_15_1 | 18273 | FTMS; HCD | 159.66 | 557.27 |

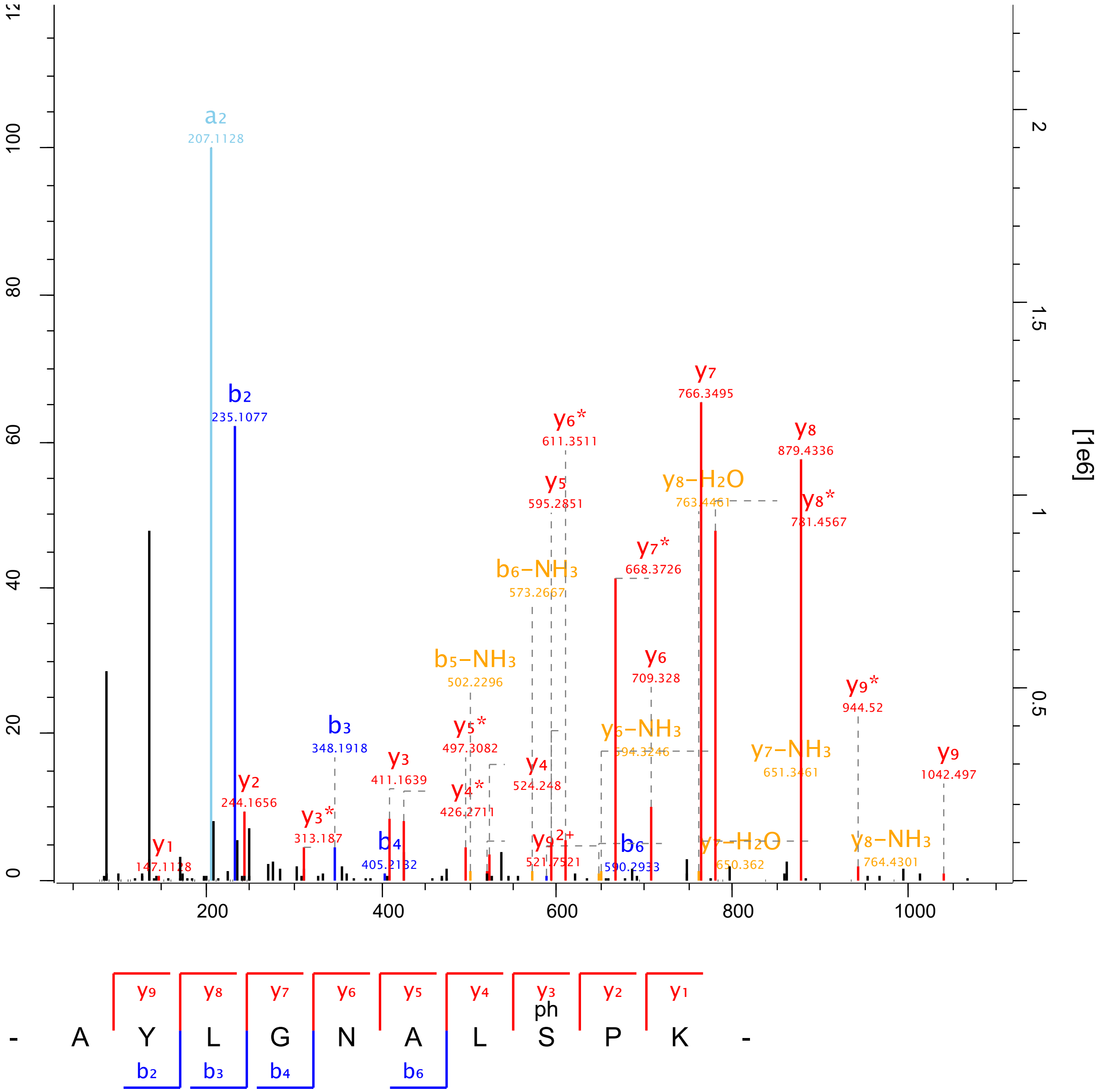

| Raw file | Scan | Method | Score | m/z |
|---|---|---|---|---|
| sys_15_1 | 18285 | FTMS; HCD | 64.12 | 553.73 |

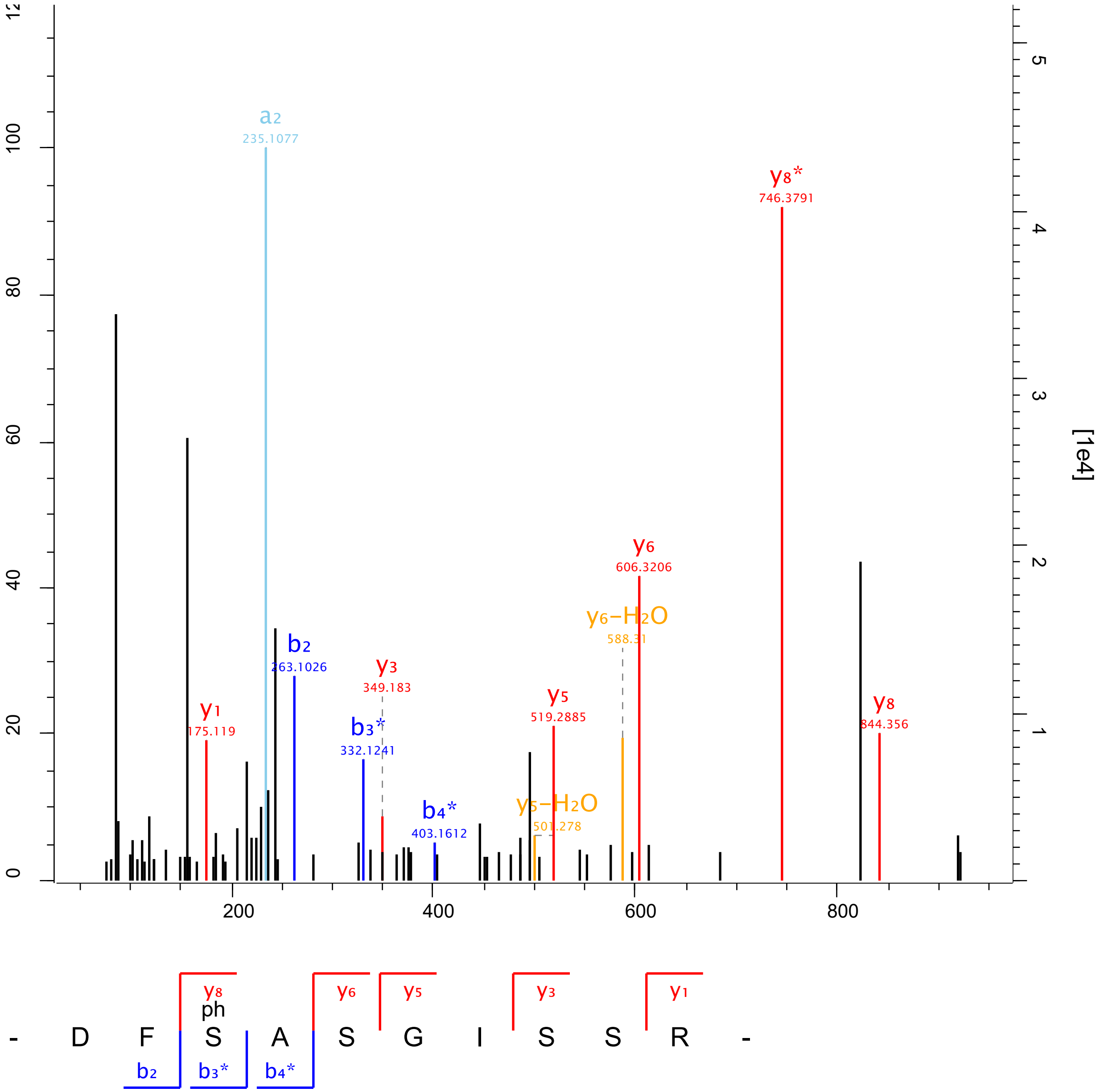

| Raw file | Scan | Method | Score | m/z |
| --- | --- | --- | --- | --- |
| sys_15_1 | 18288 | FTMS; HCD | 261.76 | 761.88 |

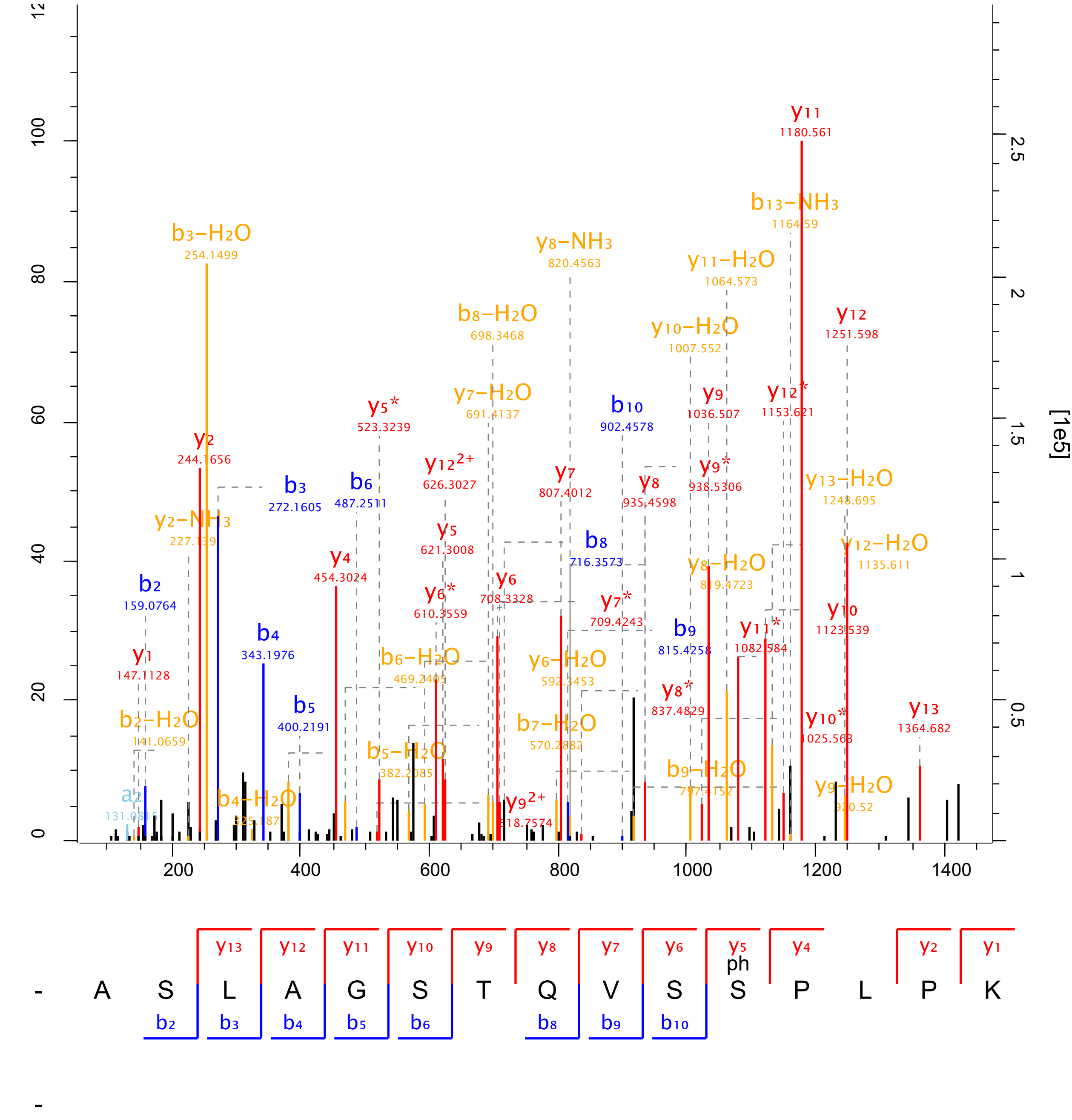

| Raw file | Scan | Method | Score | m/z |
|---|---|---|---|---|
| sys_15_1 | 18394 | FTMS; HCD | 41.8 | 950.41 |

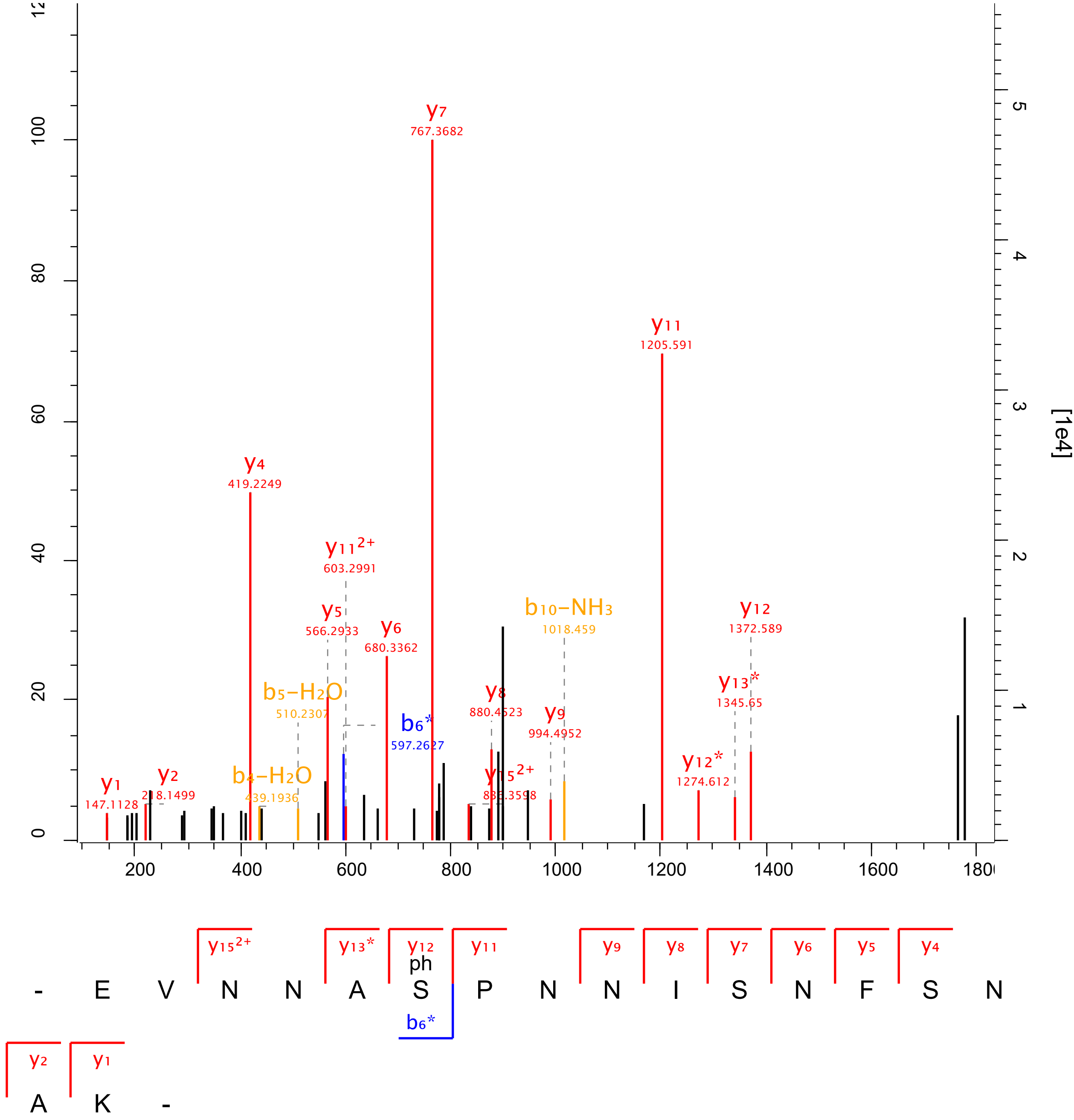

| Raw file | Scan | Method | Score | m/z |
|---|---|---|---|---|
| sys_15_1 | 18410 | FTMS; HCD | 42.95 | 511.57 |

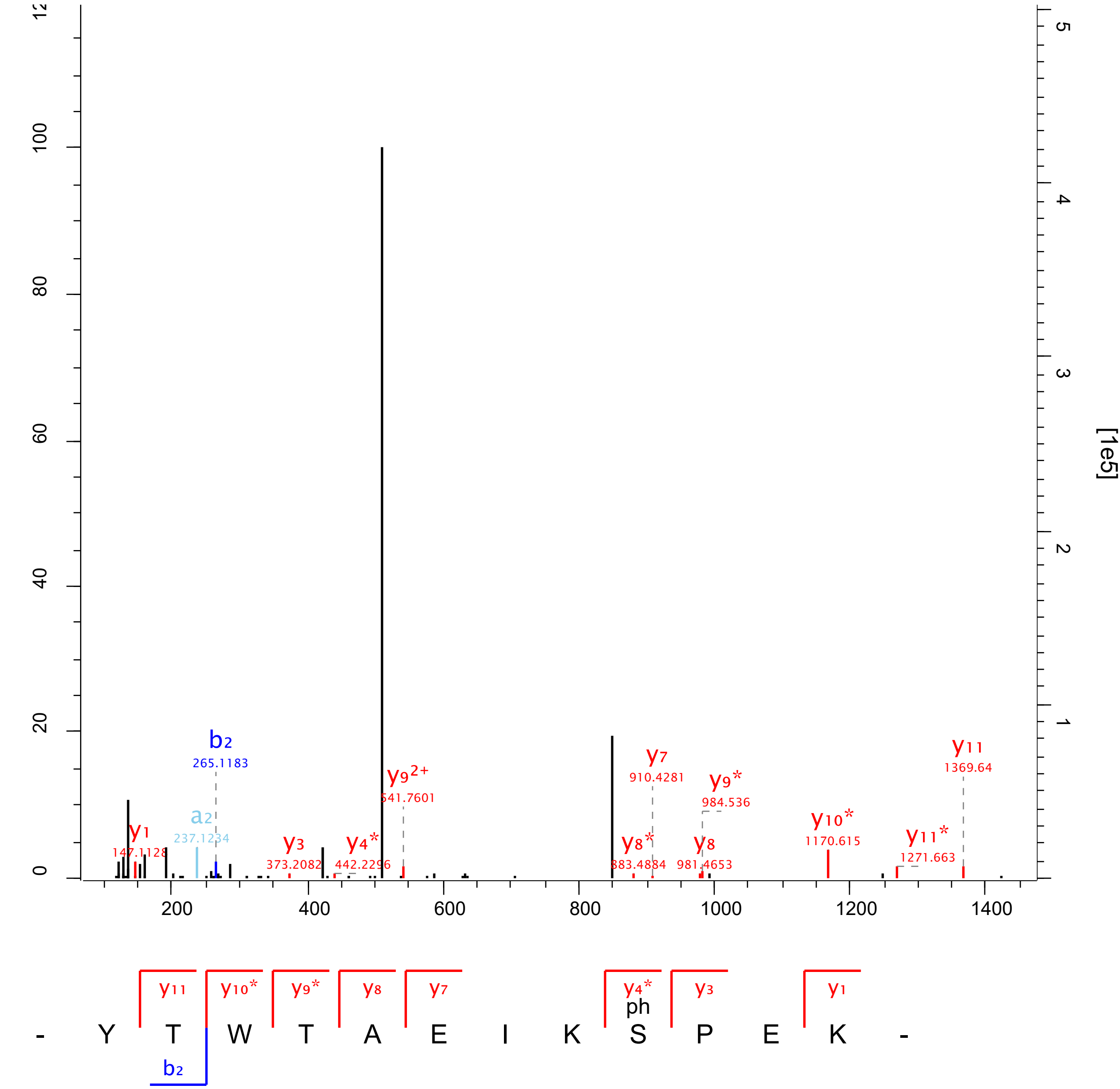

| Raw file | Scan | Method | Score | m/z |
| --- | --- | --- | --- | --- |
| sys_15_1 | 18432 | FTMS; HCD | 105.65 | 552.78 |

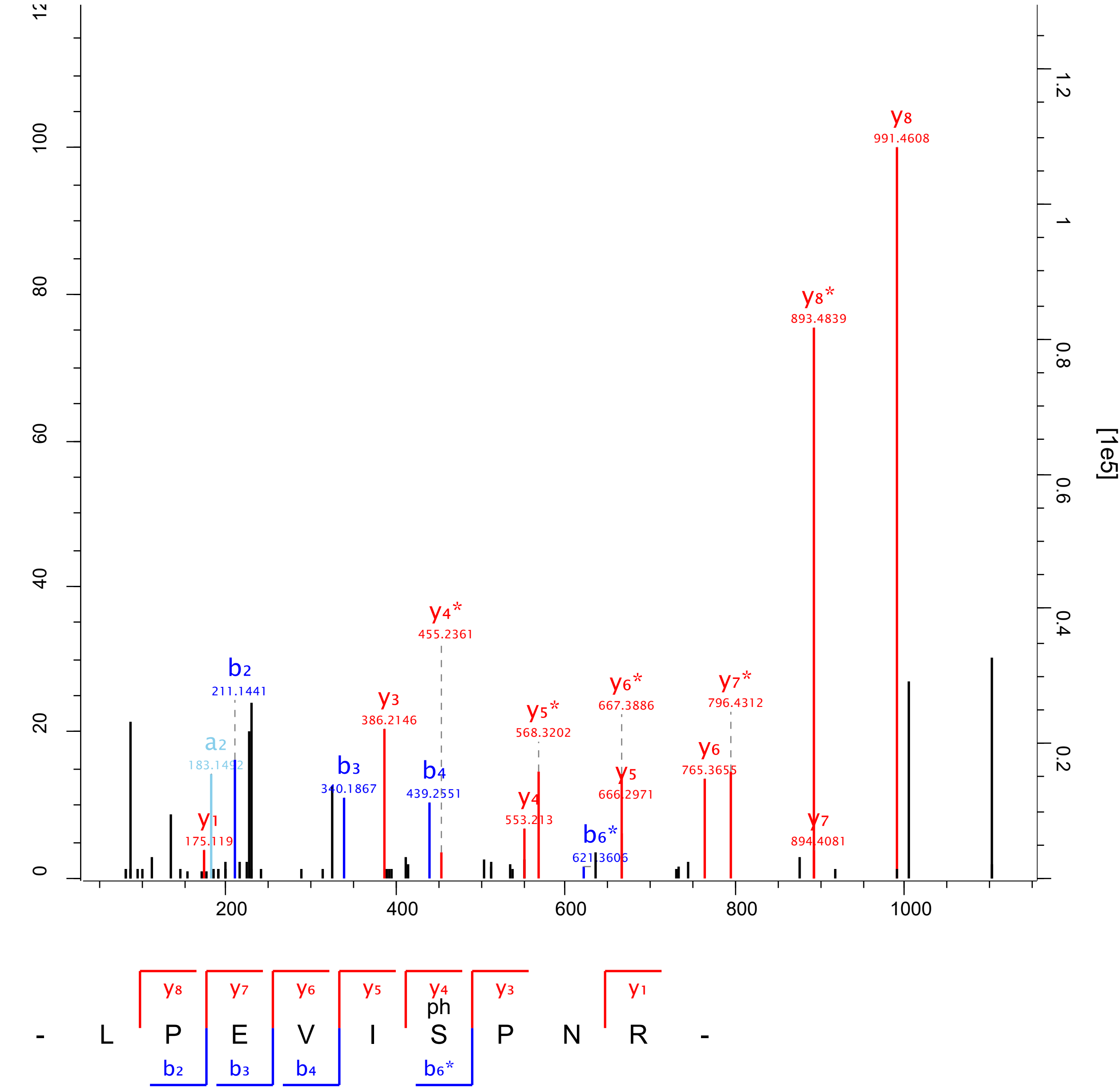

| Raw file | Scan | Method | Score | m/z |
| --- | --- | --- | --- | --- |
| sys_15_1 | 18537 | FTMS; HCD | 120.4 | 889.37 |

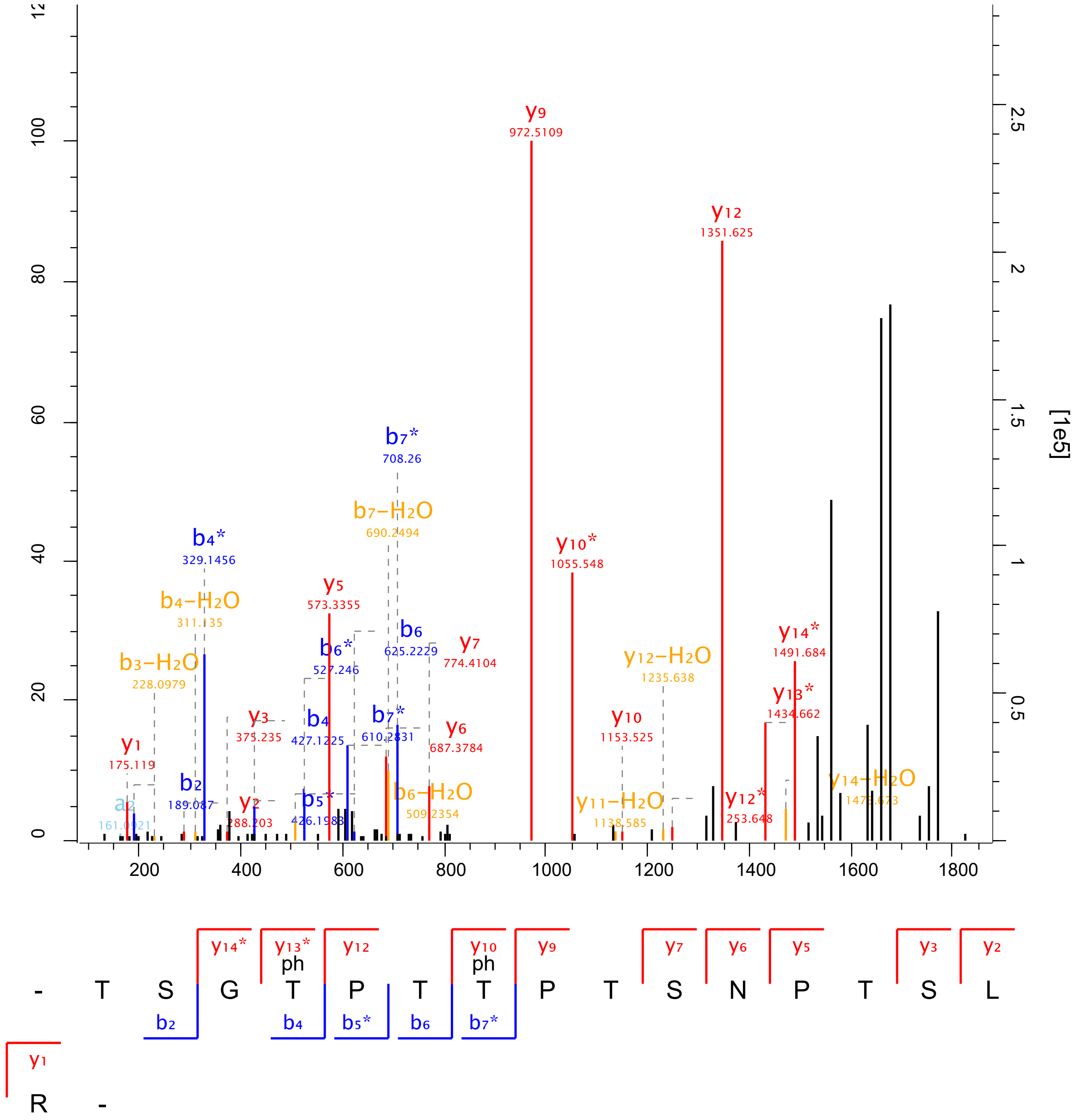

| Raw file | Scan | Method | Score | m/z |
|---|---|---|---|---|
| sys_15_1 | 18559 | FTMS; HCD | 86.54 | 828.83 |

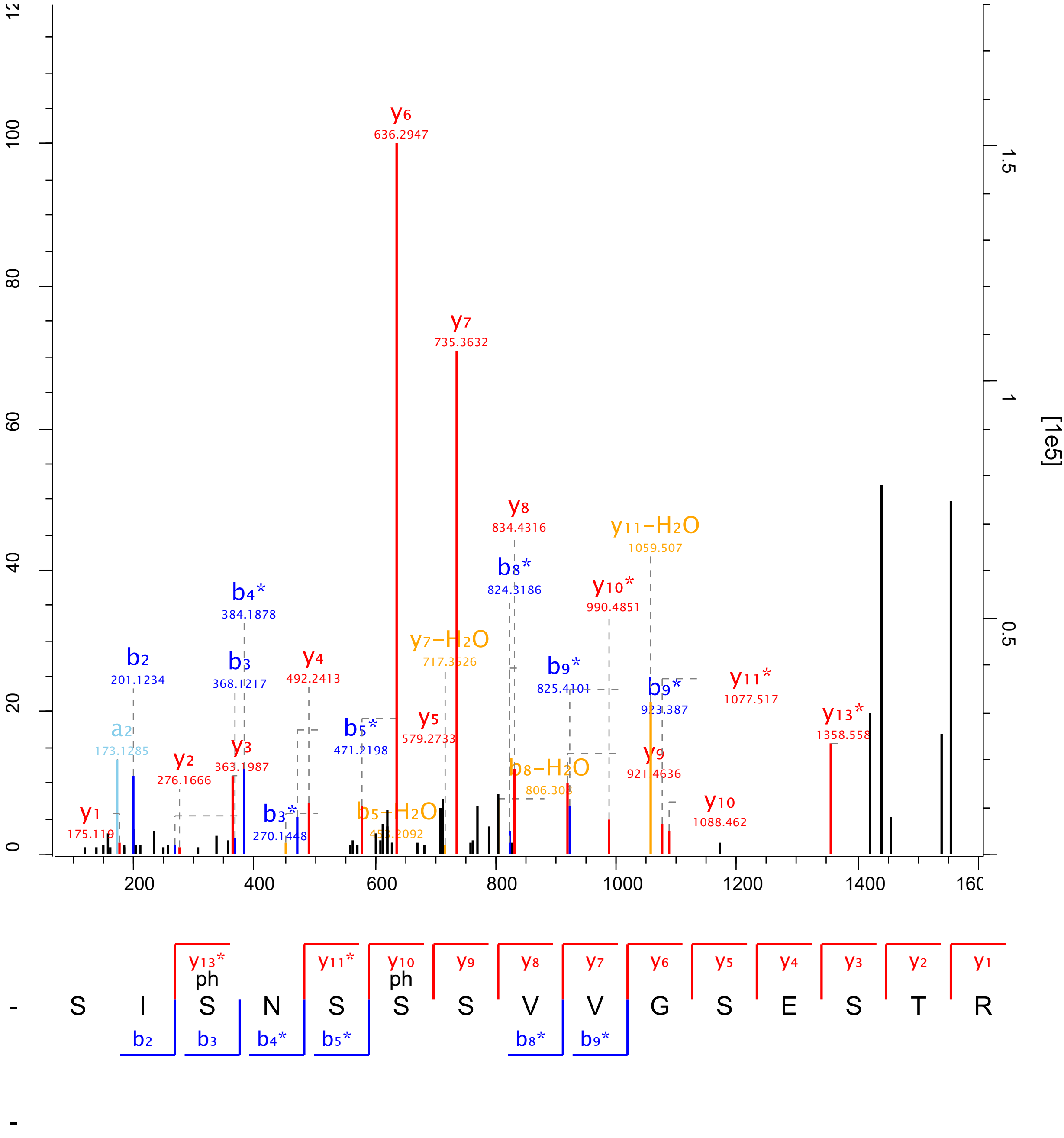

| Raw file | Scan | Method | Score | m/z |
|---|---|---|---|---|
| sys_15_1 | 18582 | FTMS; HCD | 53.24 | 481.75 |

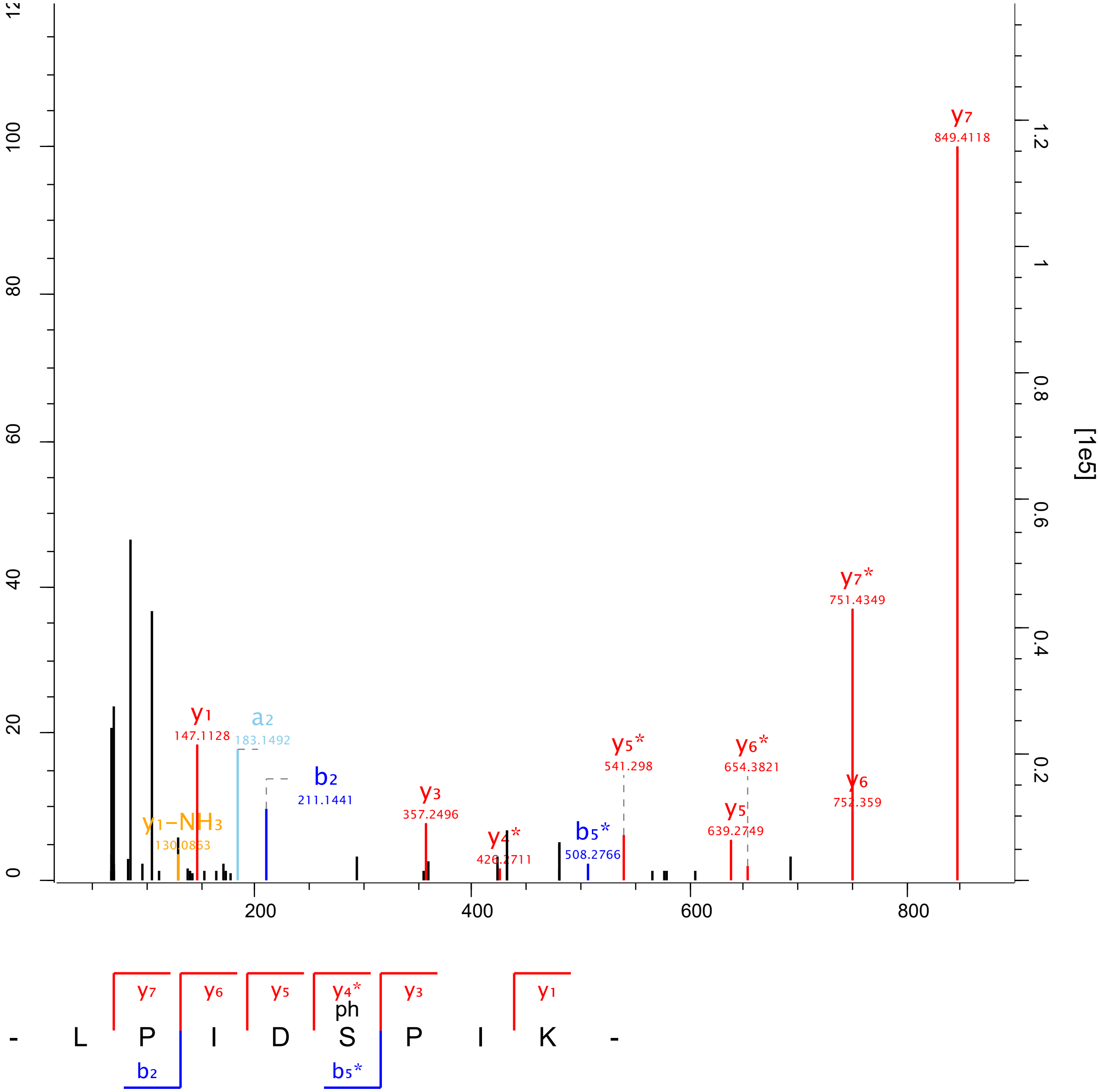

| Raw file | Scan | Method | Score | m/z |
| --- | --- | --- | --- | --- |
| sys_15_1 | 18646 | FTMS; HCD | 286.81 | 835.32 |

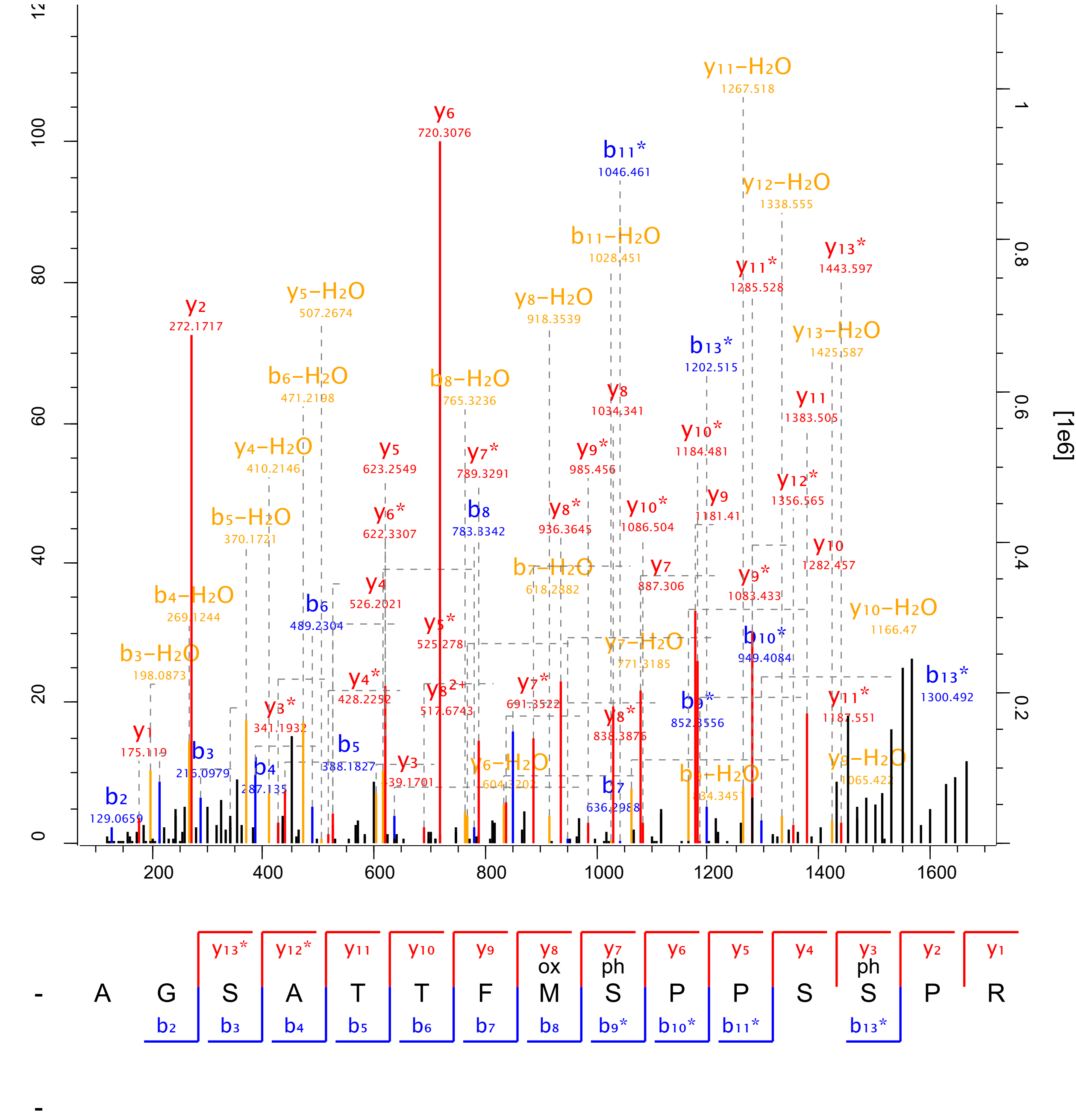

| Raw file | Scan | Method | Score | m/z |
|---|---|---|---|---|
| sys_15_1 | 18746 | FTMS; HCD | 46.42 | 1003.38 |

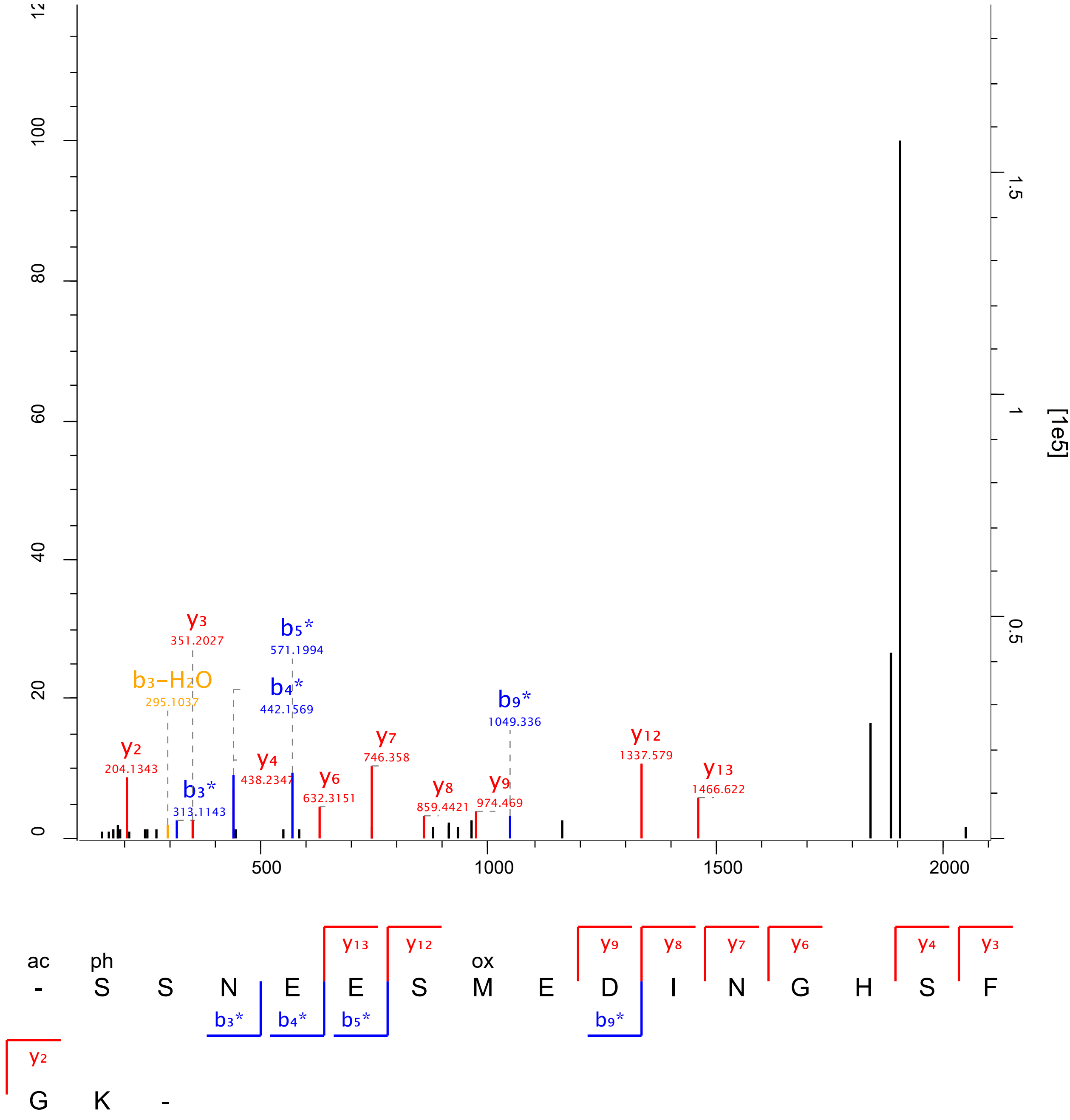

| Raw file | Scan | Method | Score | m/z |
|---|---|---|---|---|
| sys_15_1 | 18776 | FTMS; HCD | 107.35 | 708.28 |

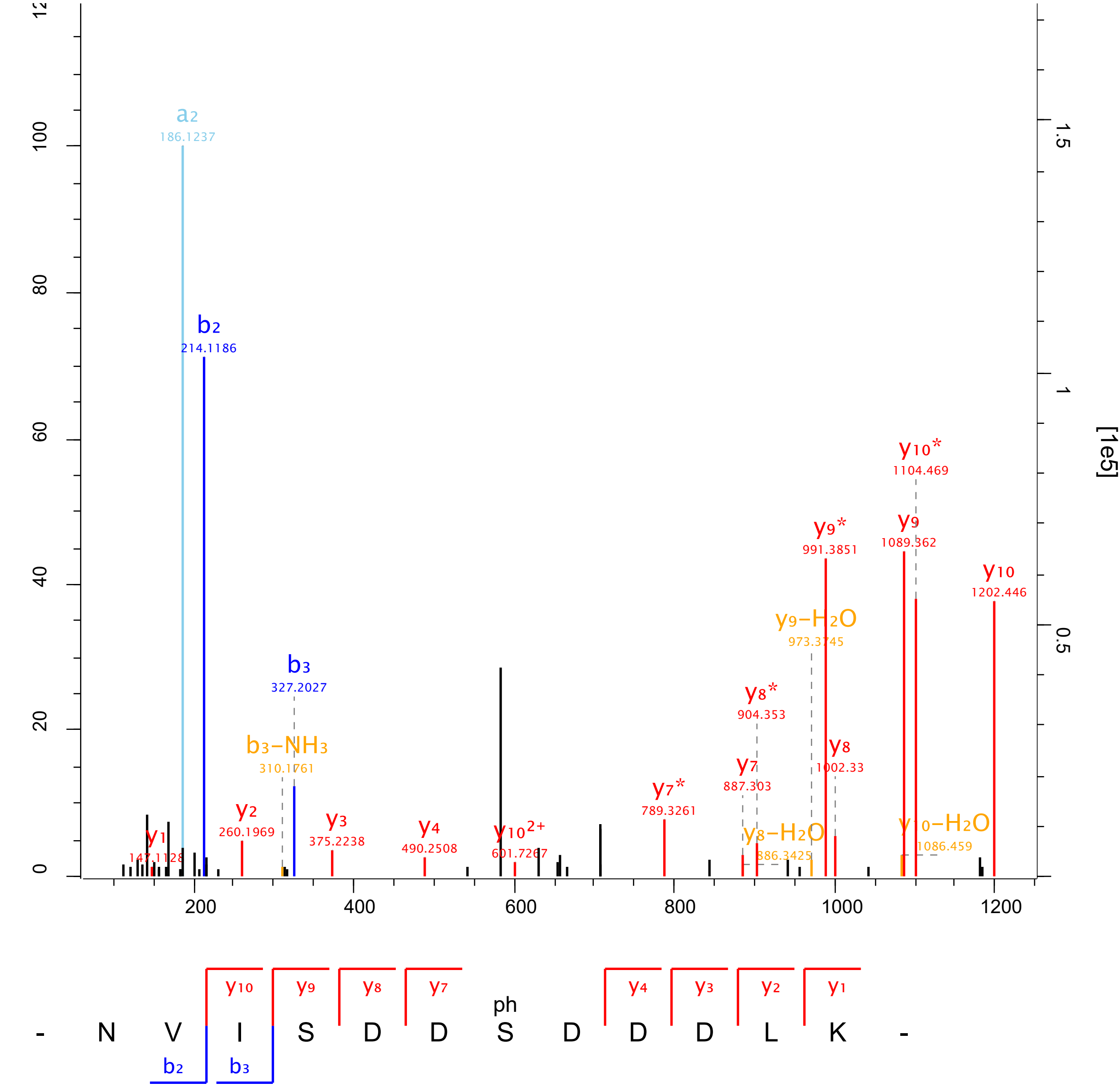

| Raw file | Scan | Method | Score | m/z |
|---|---|---|---|---|
| sys_15_1 | 18830 | FTMS; HCD | 193.45 | 652.27 |

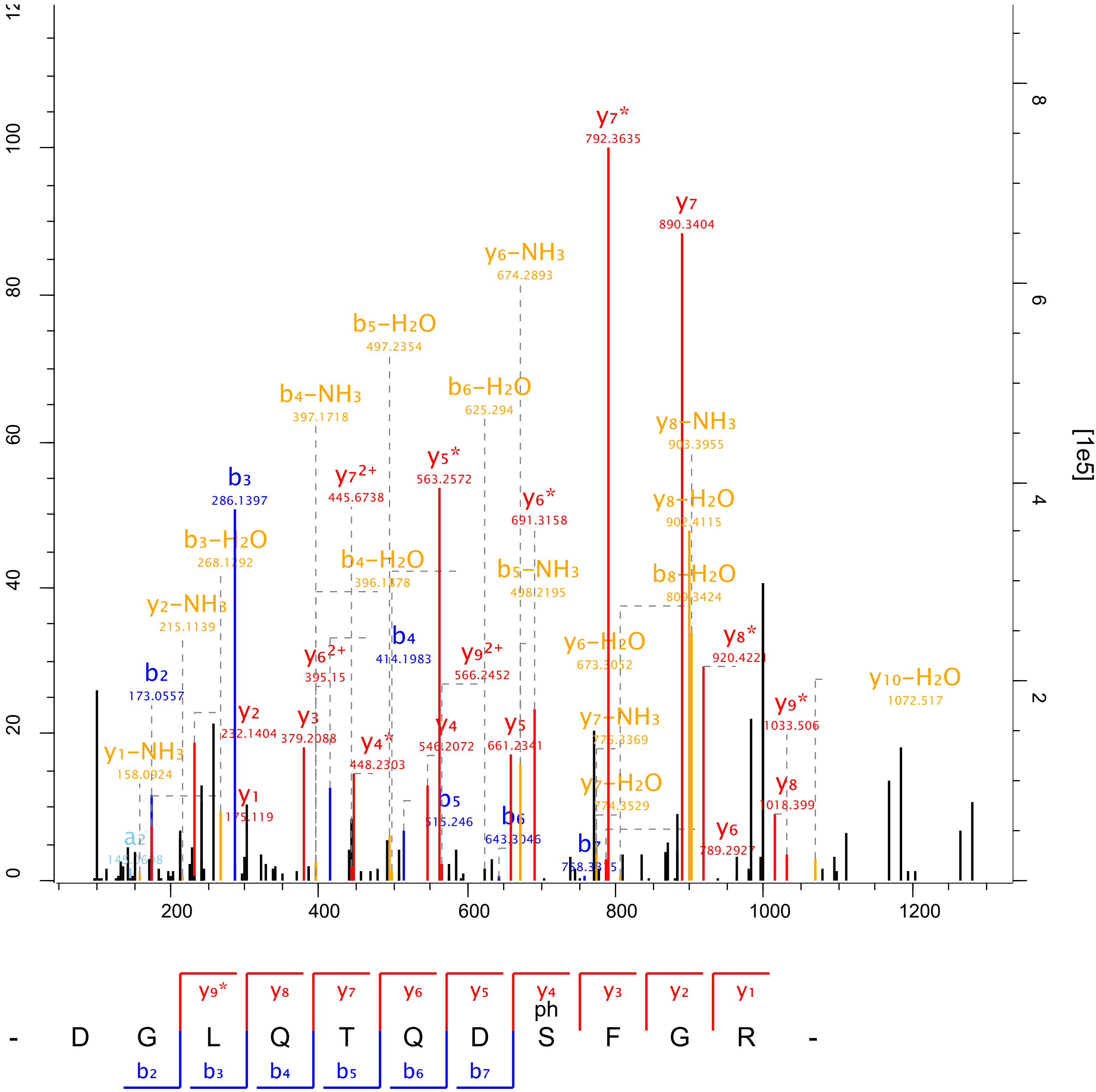

| Raw file | Scan | Method | Score | m/z |
|---|---|---|---|---|
| sys_15_1 | 18840 | FTMS; HCD | 103.83 | 667.29 |

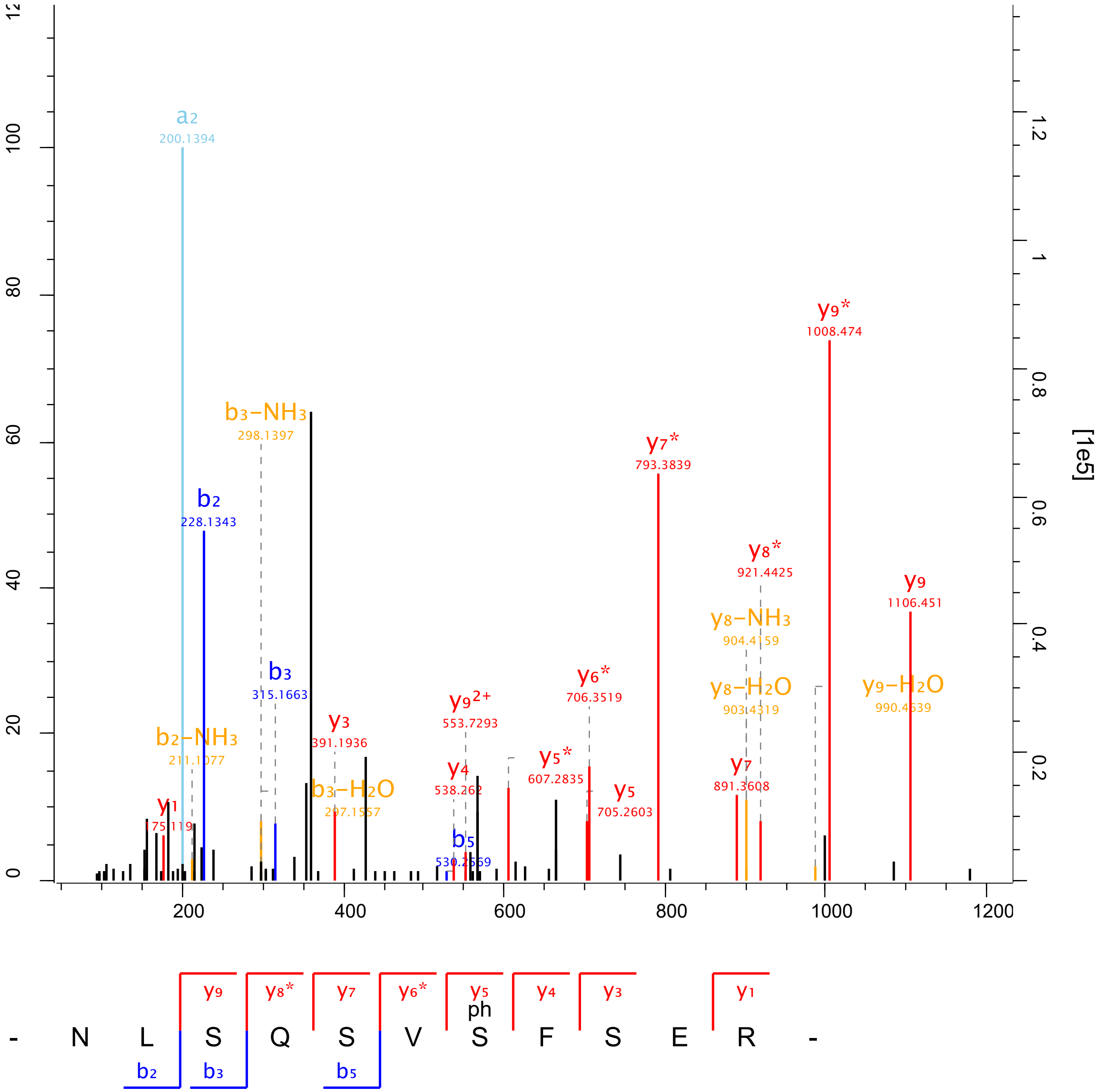

| Raw file | Scan | Method | Score | m/z |
|---|---|---|---|---|
| sys_15_1 | 18844 | FTMS; HCD | 140.77 | 624.31 |

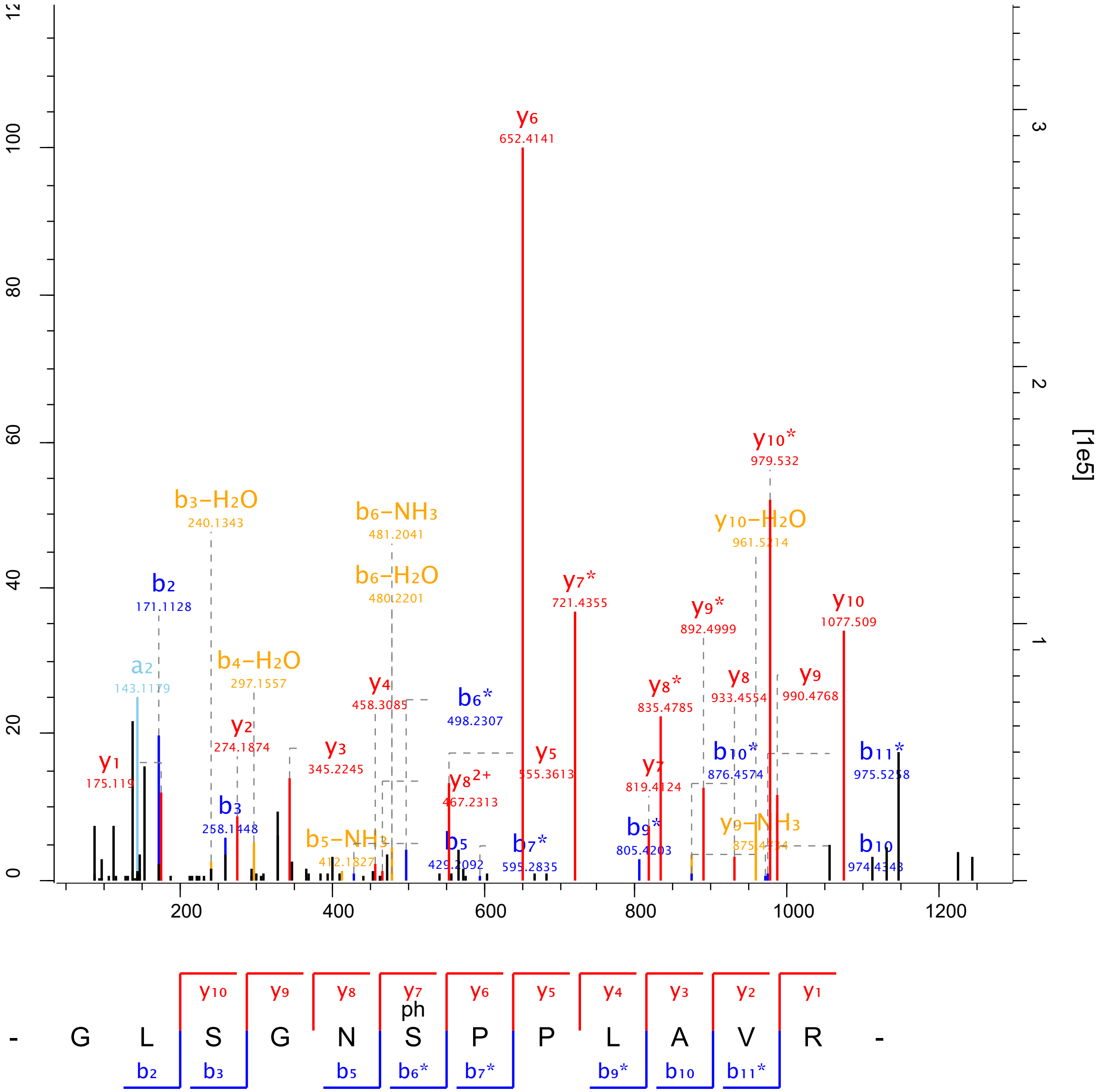

| Raw file | Scan | Method | Score | m/z |
|---|---|---|---|---|
| sys_15_1 | 18956 | FTMS; HCD | 136.92 | 739.65 |

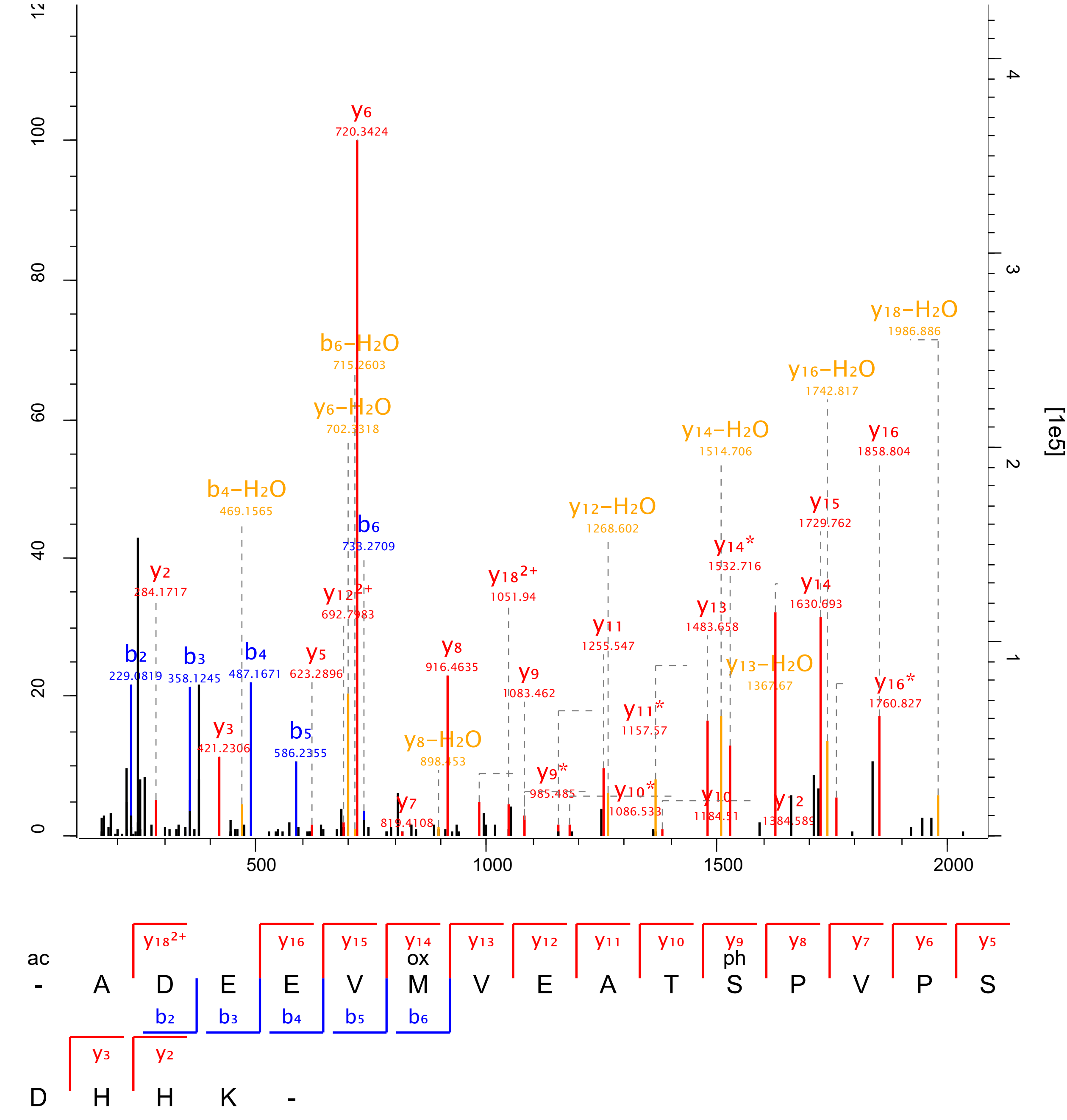

| Raw file | Scan | Method | Score | m/z |
|---|---|---|---|---|
| sys_15_1 | 19027 | FTMS; HCD | 51.29 | 542.76 |

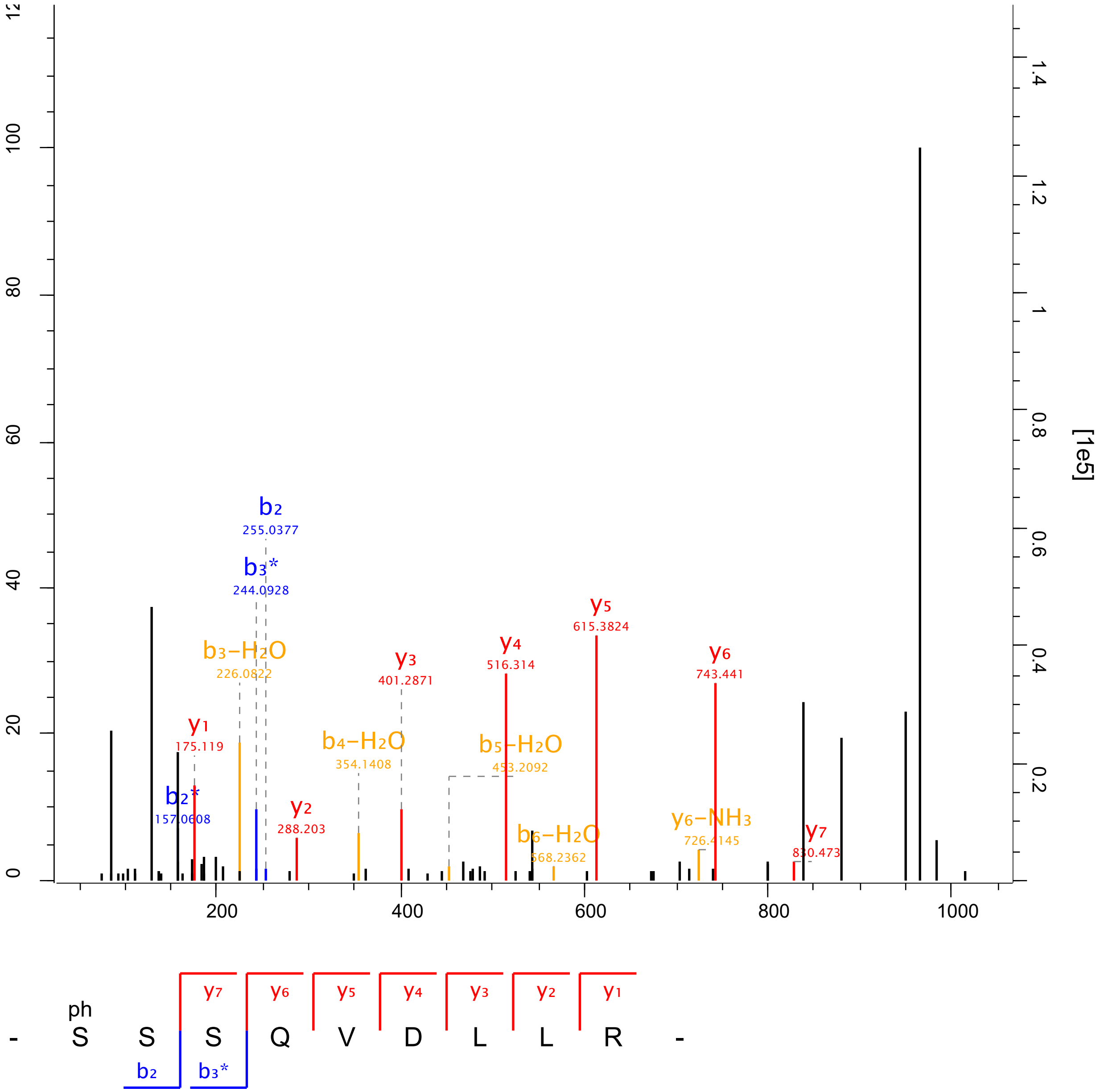

| Raw file | Scan | Method | Score | m/z |
| --- | --- | --- | --- | --- |
| sys_15_1 | 19043 | FTMS; HCD | 219.41 | 645.32 |

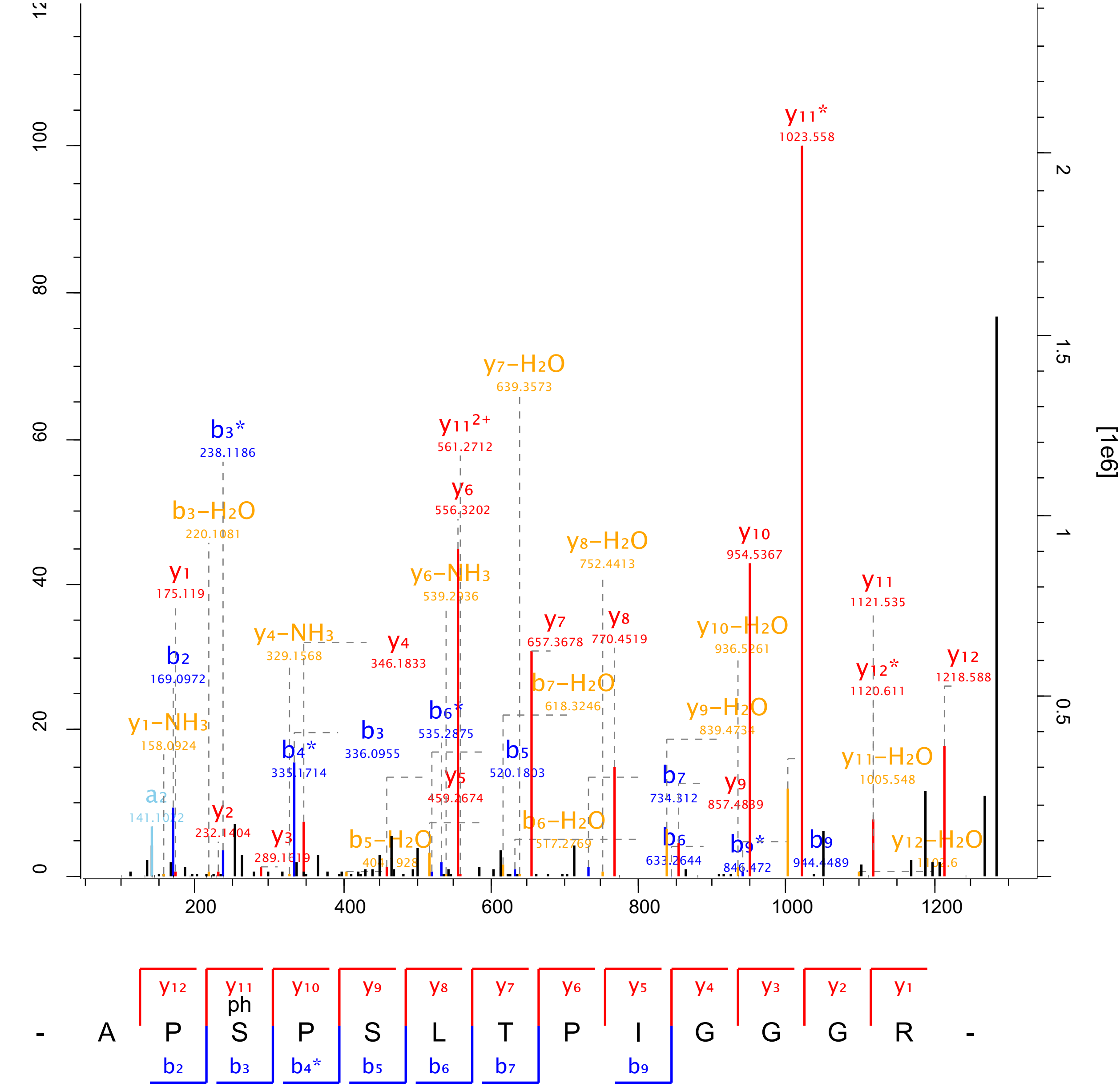

| Raw file | Scan | Method | Score | m/z |
|---|---|---|---|---|
| sys_15_1 | 19057 | FTMS; HCD | 70.2 | 503.91 |

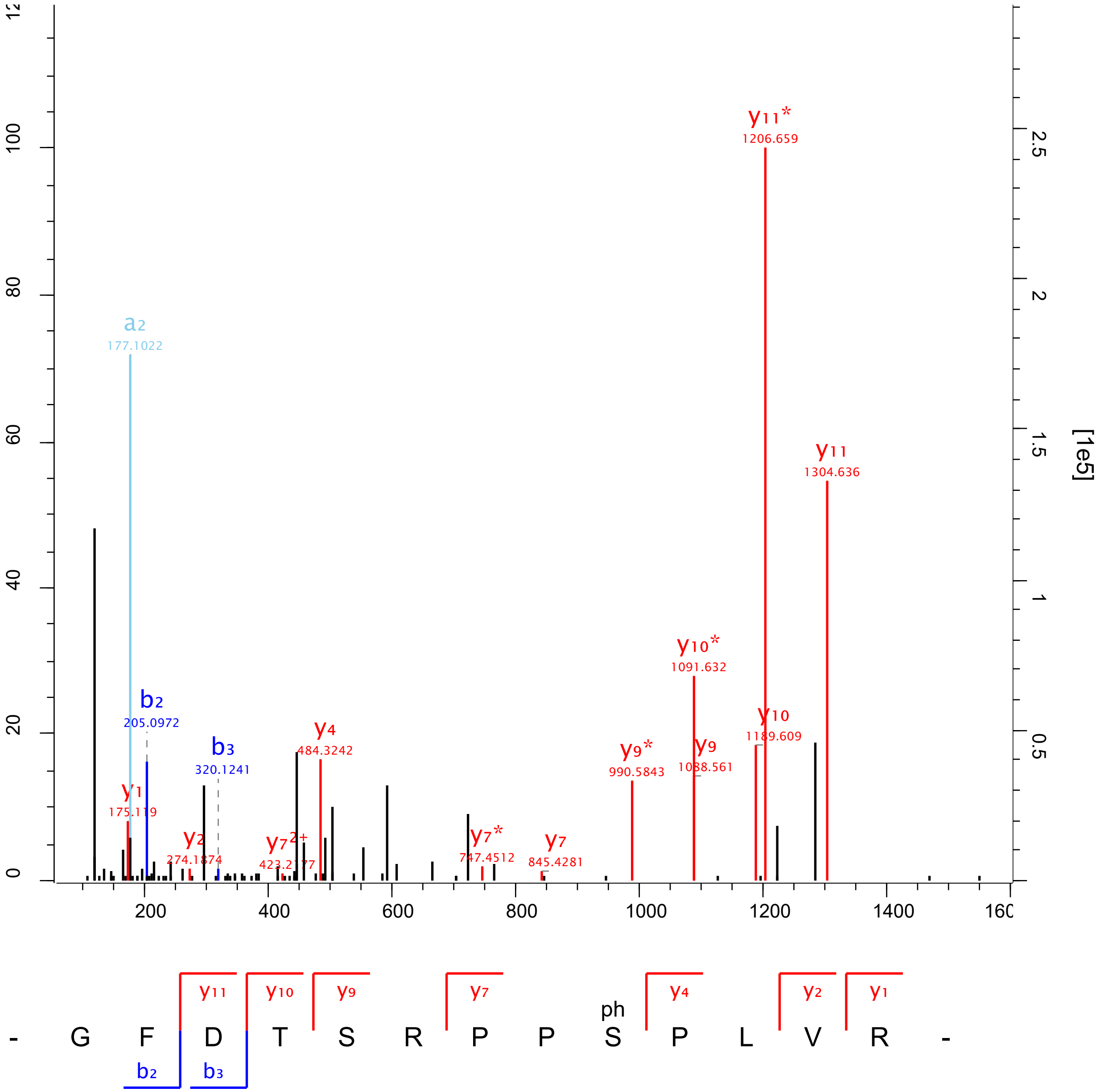

| Raw file | Scan | Method | Score | m/z |
| --- | --- | --- | --- | --- |
| sys_15_1 | 19091 | FTMS; HCD | 109.07 | 626.81 |

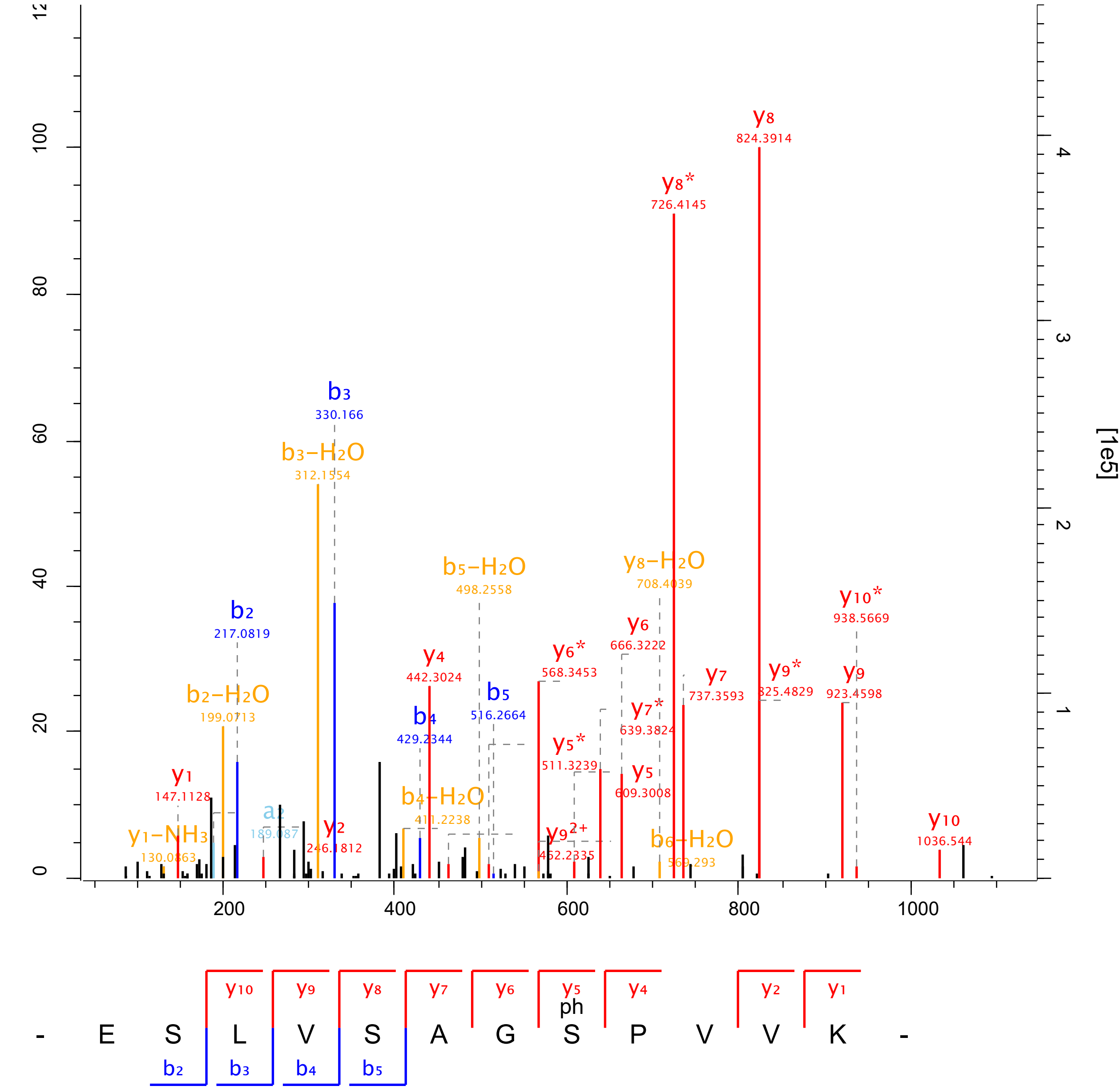

| Raw file | Scan | Method | Score | m/z |
| --- | --- | --- | --- | --- |
| sys_15_1 | 19198 | FTMS; HCD | 63.22 | 670.01 |

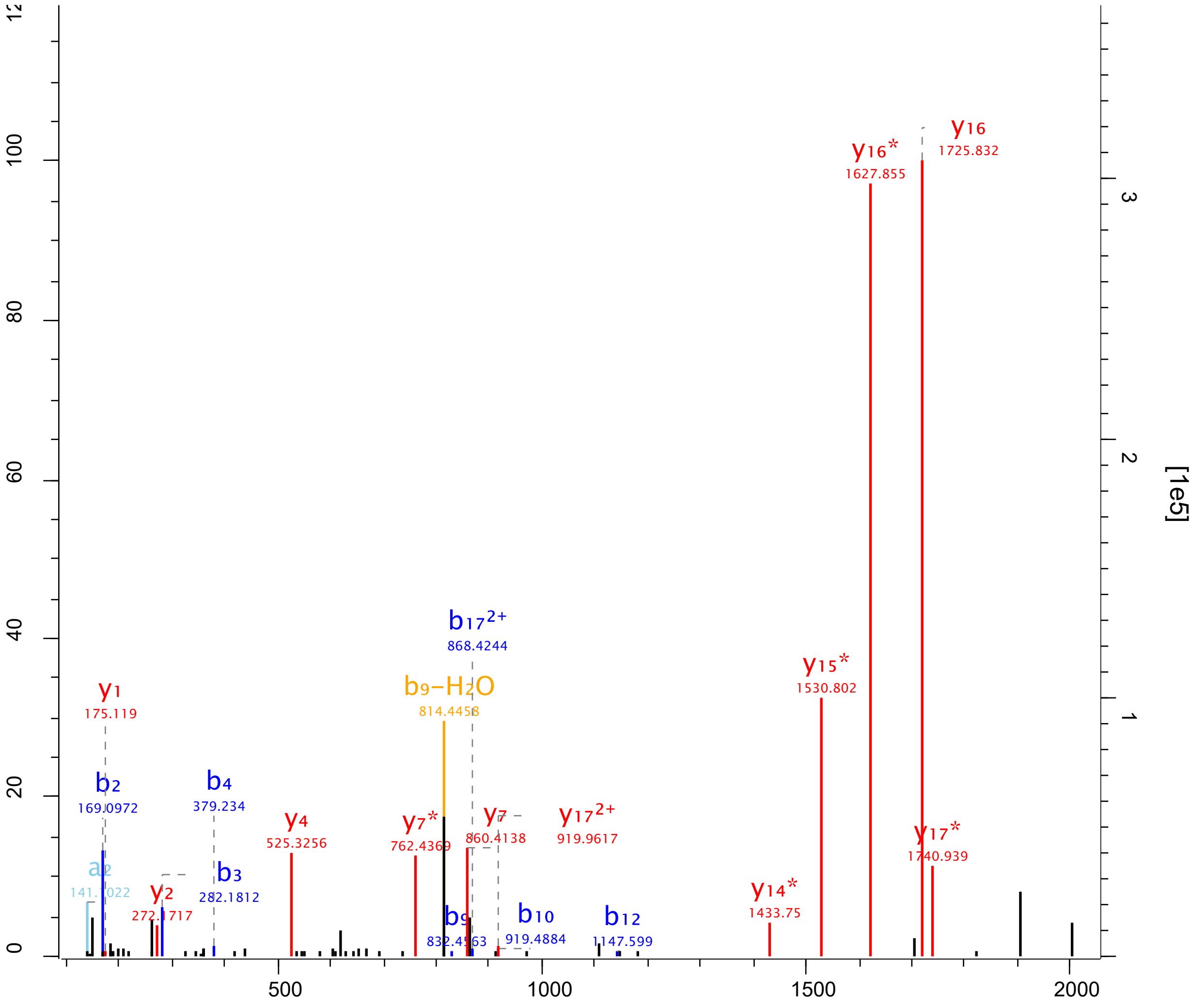


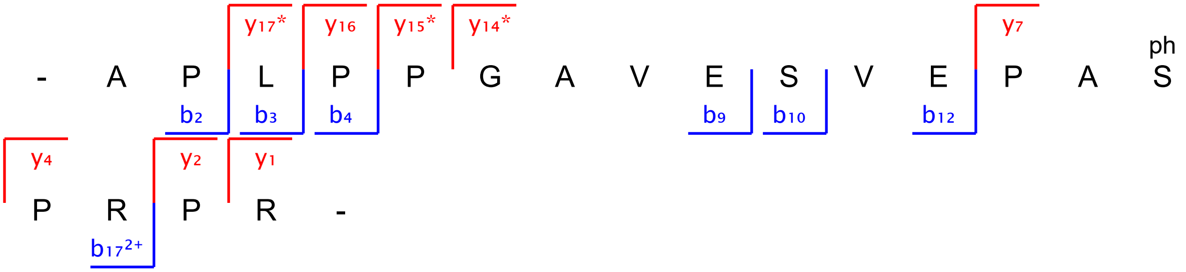

| Raw file | Scan | Method | Score | m/z |
|---|---|---|---|---|
| sys_15_1 | 19204 | FTMS; HCD | 110.26 | 544.57 |

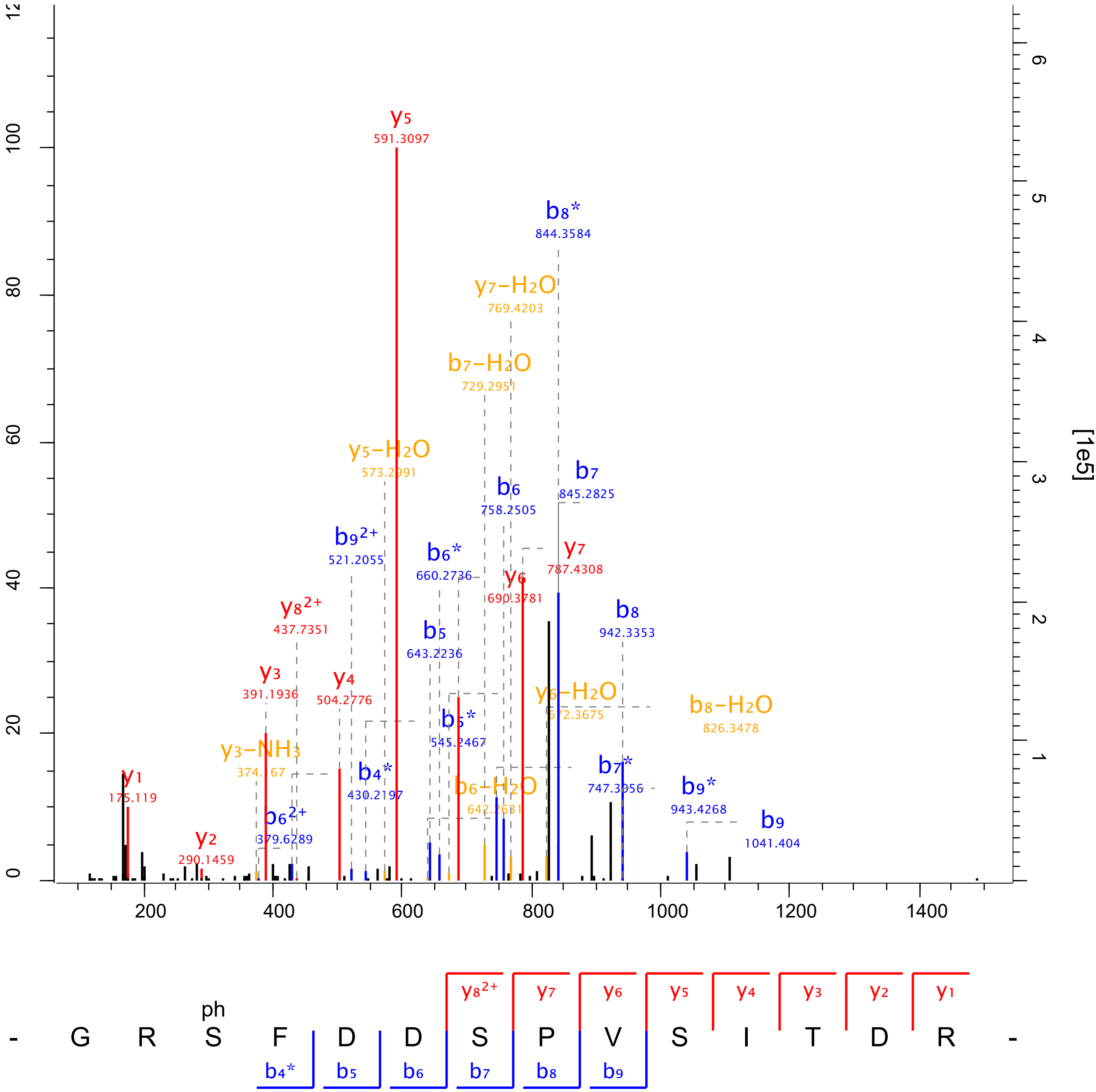

| Raw file | Scan | Method | Score | m/z |
|---|---|---|---|---|
| sys_15_1 | 19254 | FTMS; HCD | 63.98 | 822.28 |

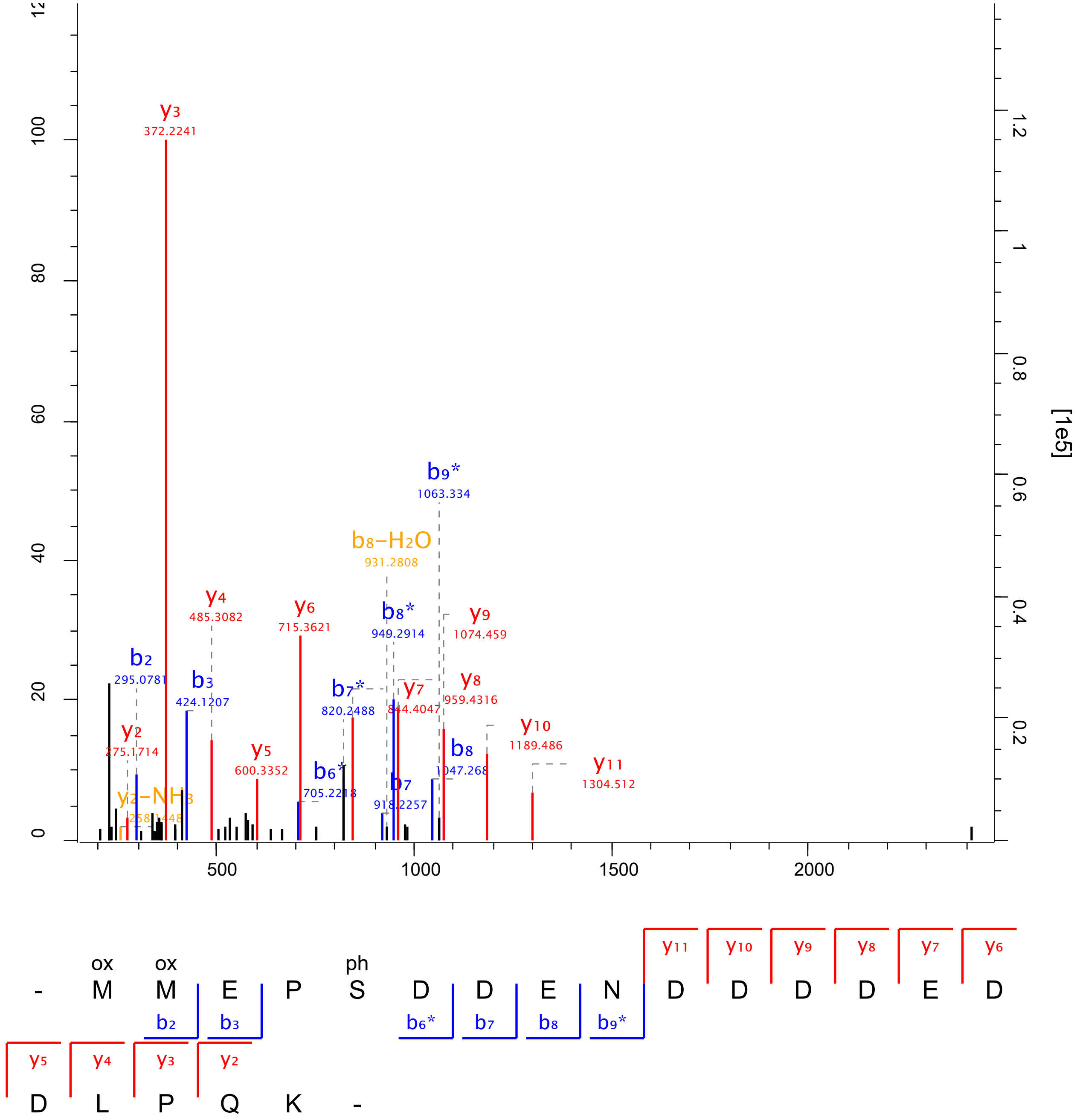

| Raw file | Scan | Method | Score | m/z |
| --- | --- | --- | --- | --- |
| sys_15_1 | 19271 | FTMS; HCD | 194.24 | 1031.42 |

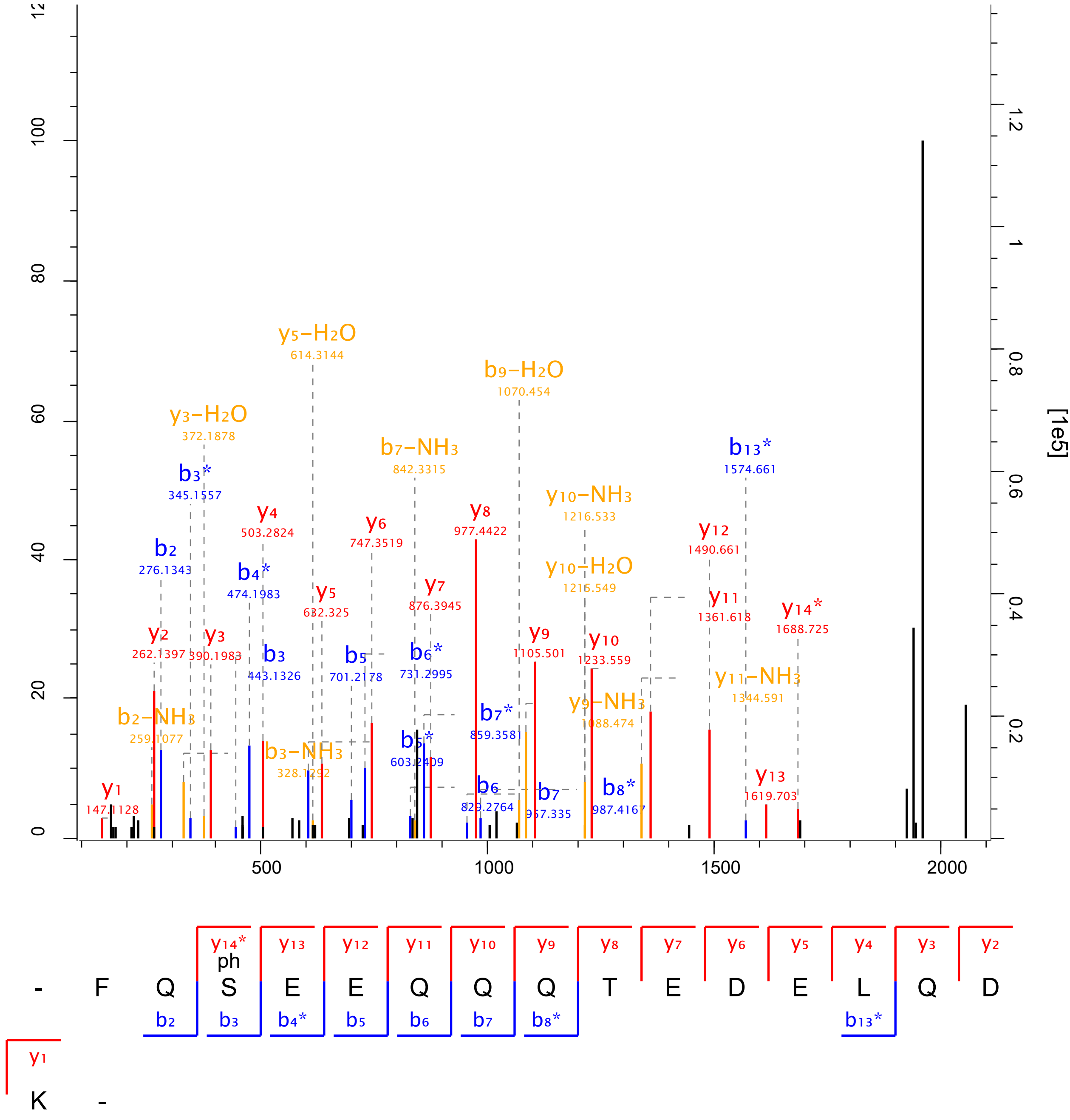

| Raw file | Scan | Method | Score | m/z |
|---|---|---|---|---|
| sys_15_1 | 19278 | FTMS; HCD | 166.3 | 769.37 |

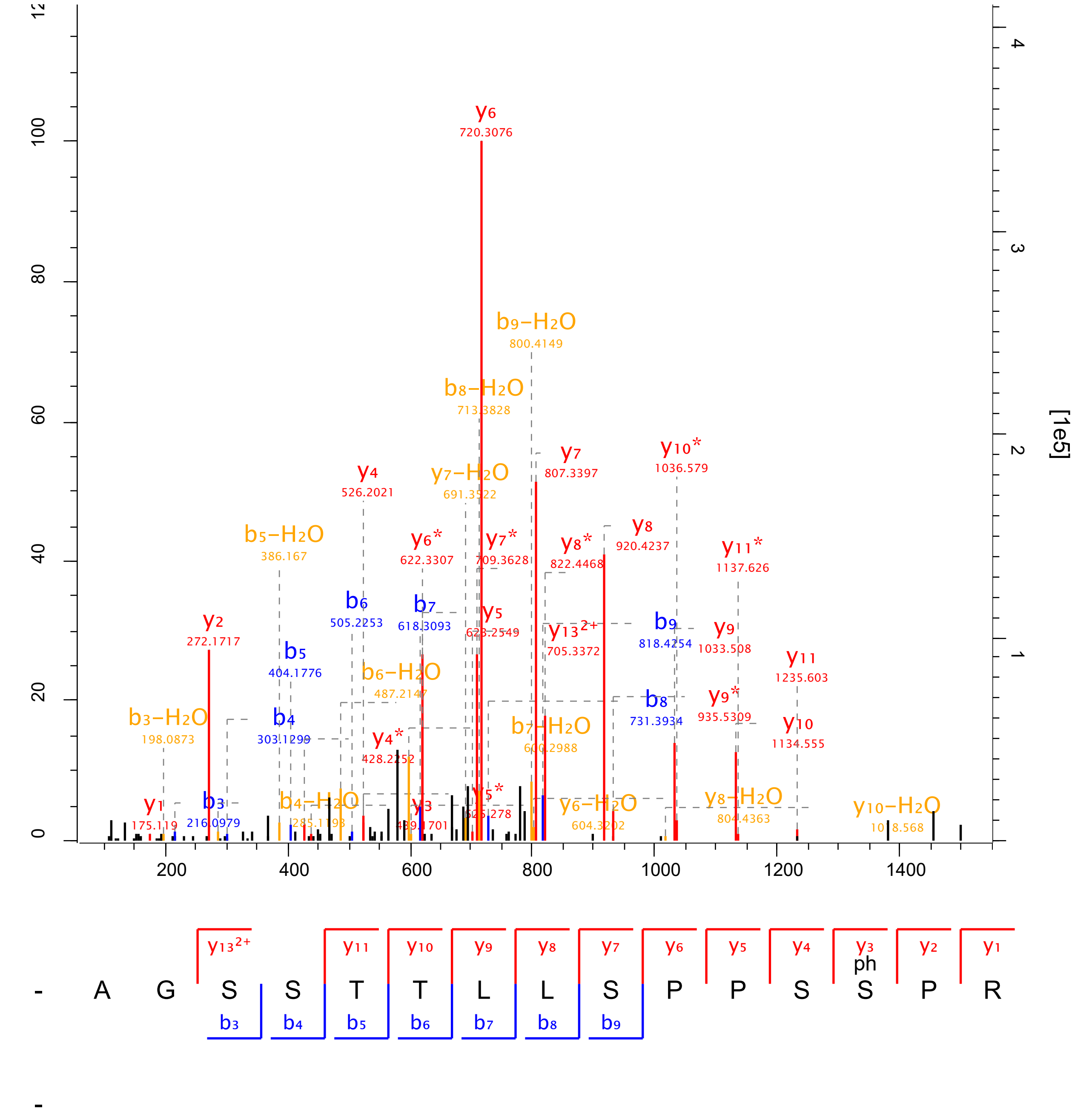

| Raw file | Scan | Method | Score | m/z |
|---|---|---|---|---|
| sys_15_1 | 19283 | FTMS; HCD | 47.55 | 703.28 |

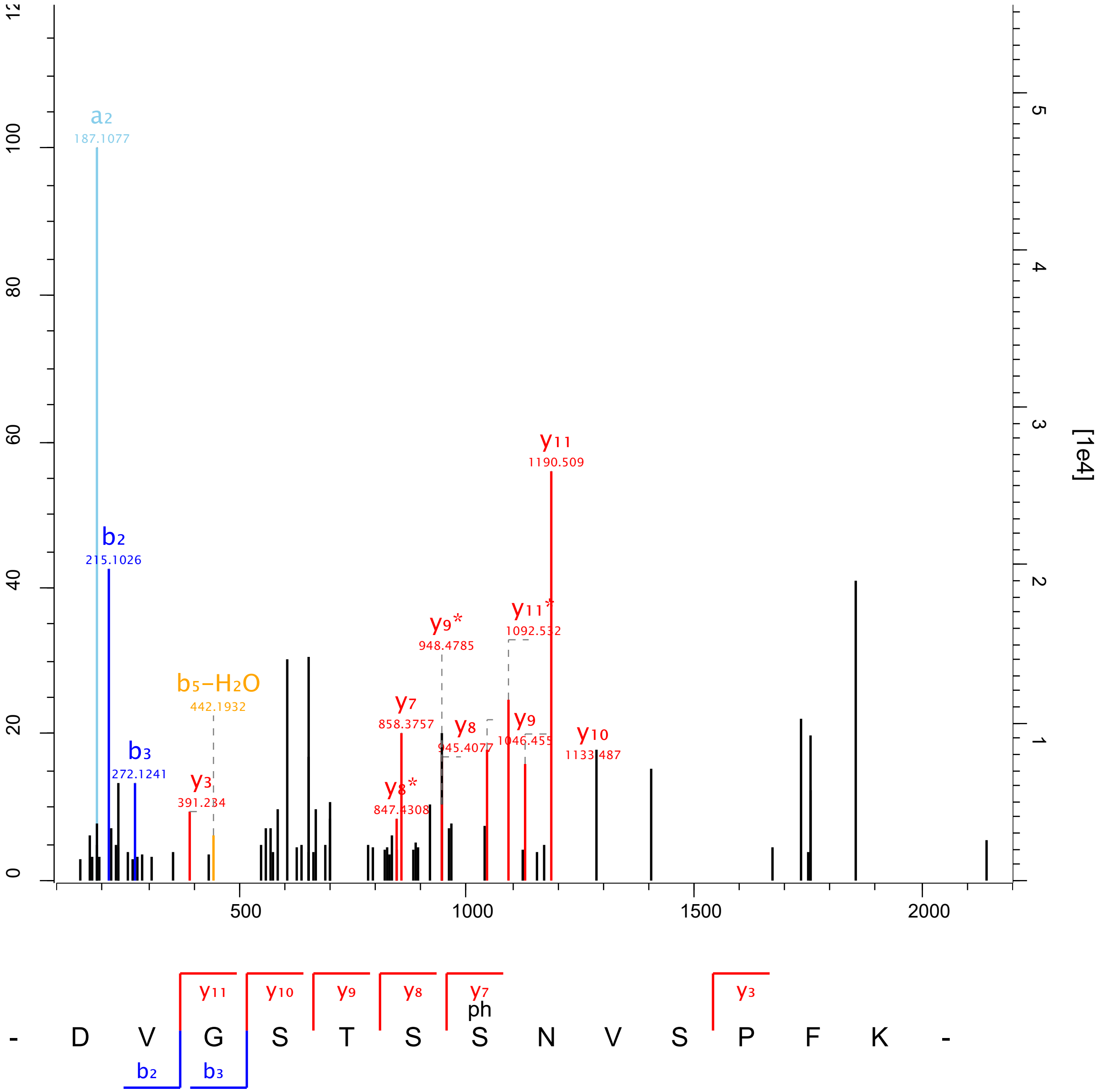

| Raw file | Scan | Method | Score | m/z |
| --- | --- | --- | --- | --- |
| sys_15_1 | 19329 | FTMS; HCD | 133.16 | 650.26 |

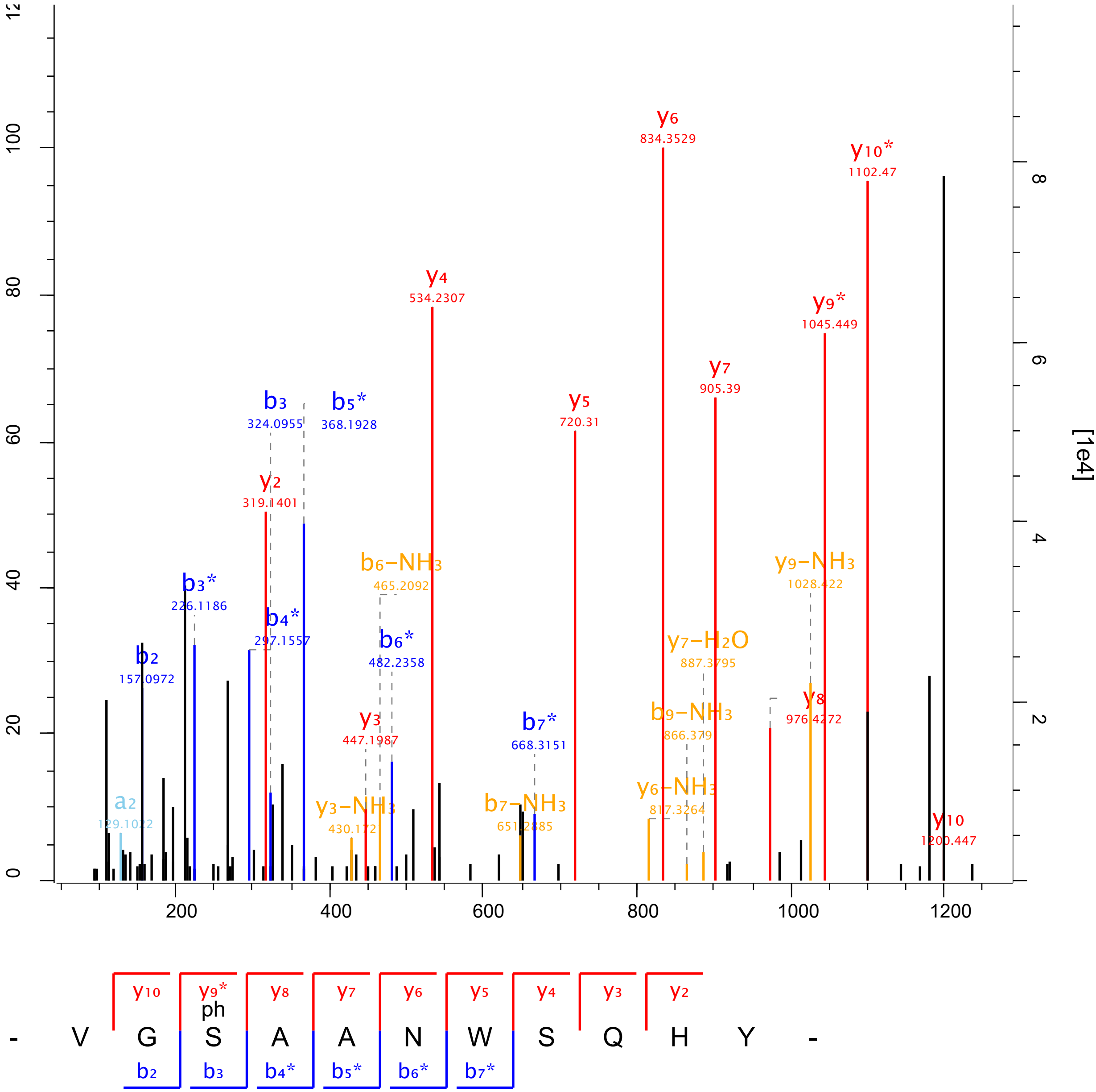

| Raw file | Scan | Method | Score | m/z |
| --- | --- | --- | --- | --- |
| sys_15_1 | 1935 | FTMS; HCD | 57.59 | 437.18 |

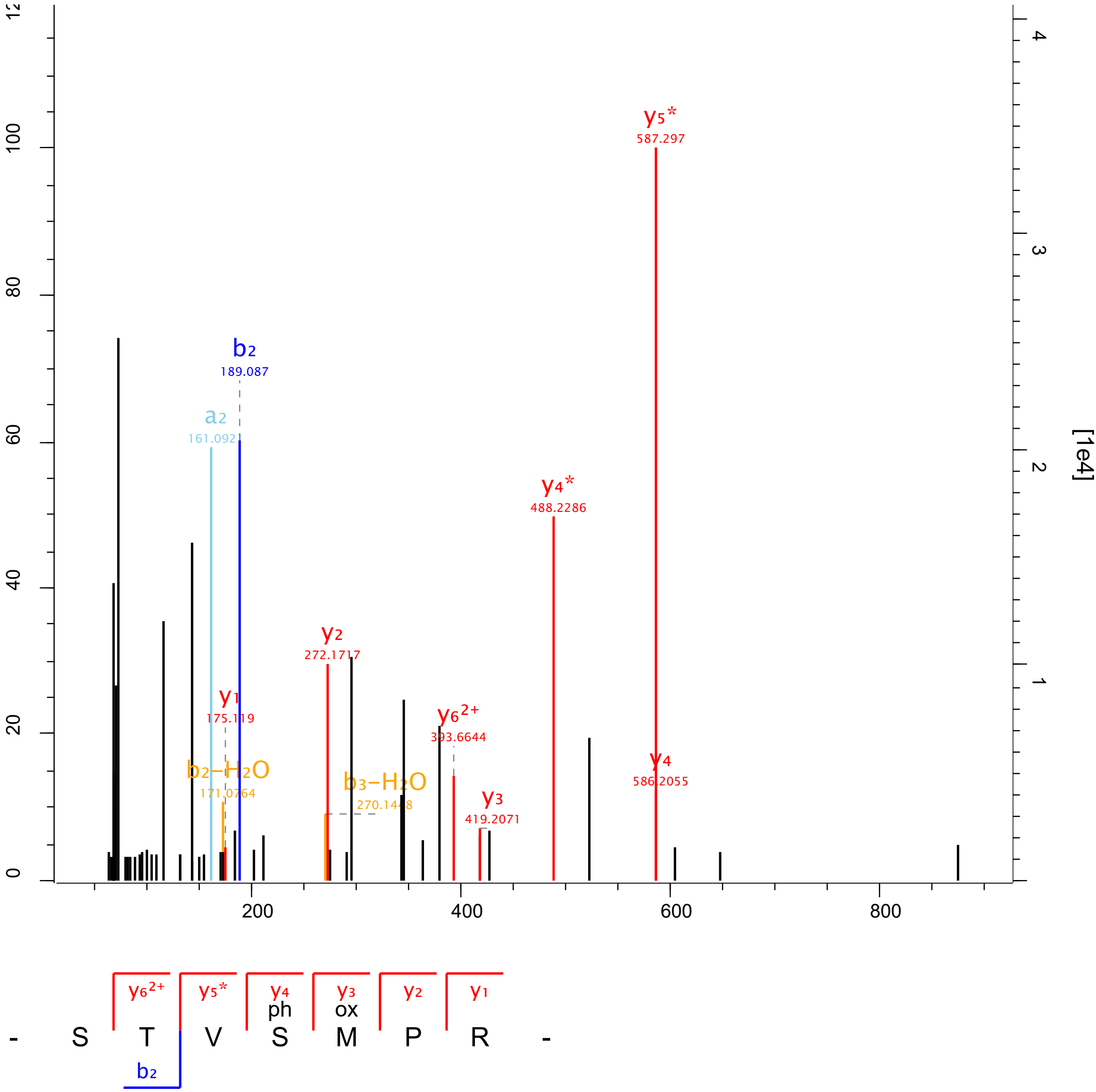

| Raw file | Scan | Method | Score | m/z |
|---|---|---|---|---|
| sys_15_1 | 19441 | FTMS; HCD | 41.24 | 628.25 |

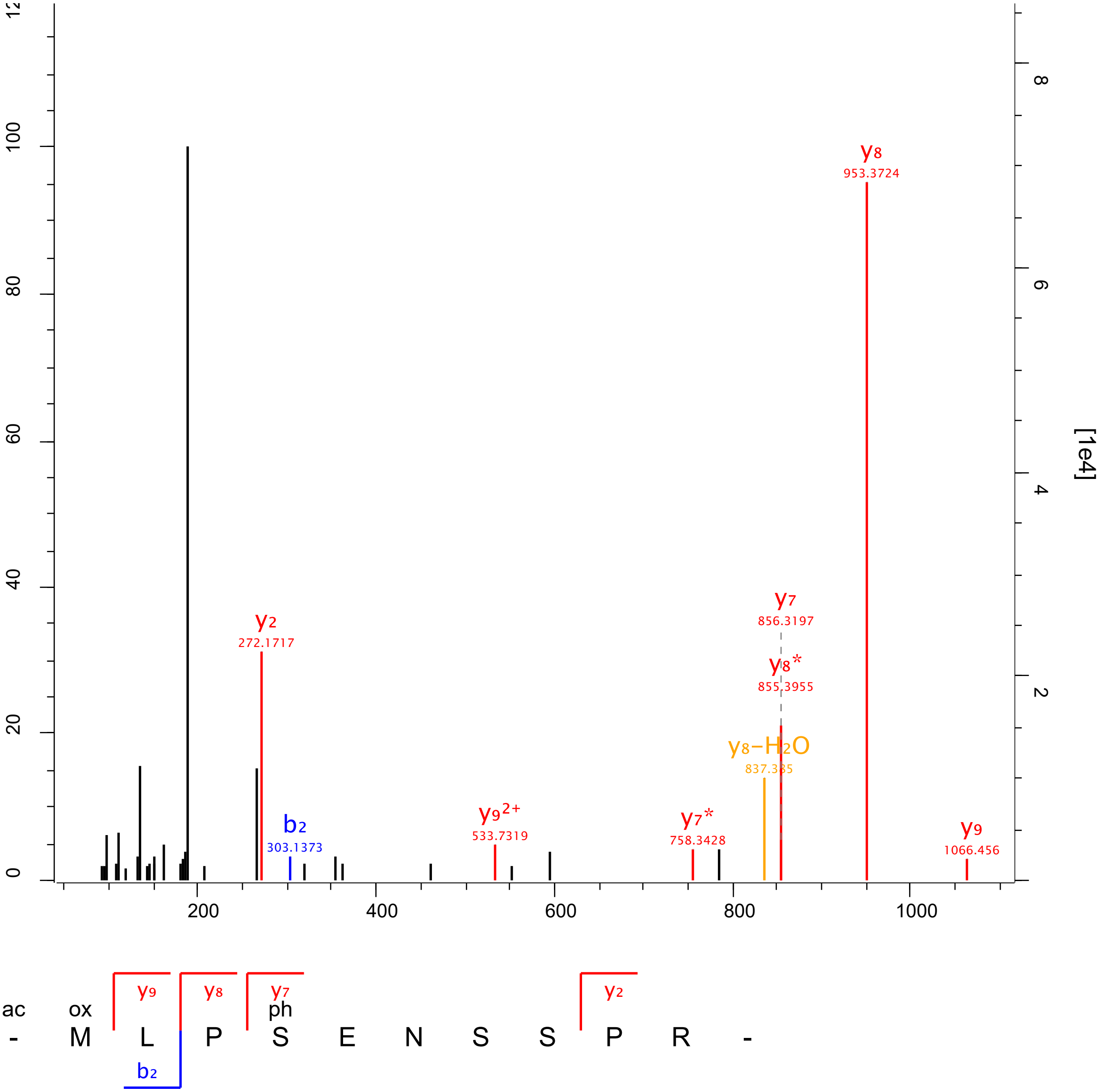

| Raw file | Scan | Method | Score | m/z |
|---|---|---|---|---|
| sys_15_1 | 19541 | FTMS; HCD | 63.91 | 740.29 |

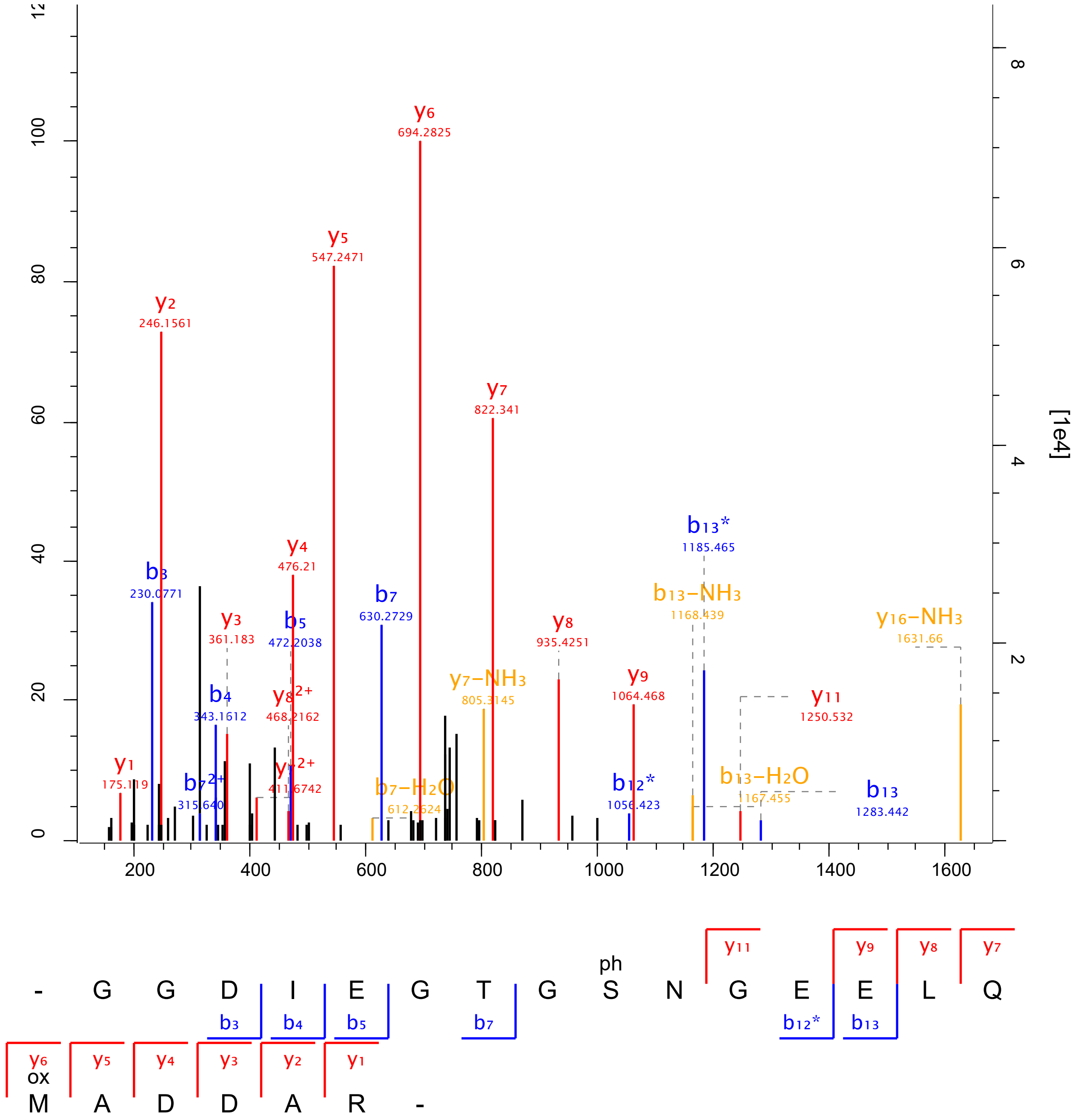

| Raw file | Scan | Method | Score | m/z |
|---|---|---|---|---|
| sys_15_1 | 19597 | FTMS; HCD | 166.05 | 780.97 |

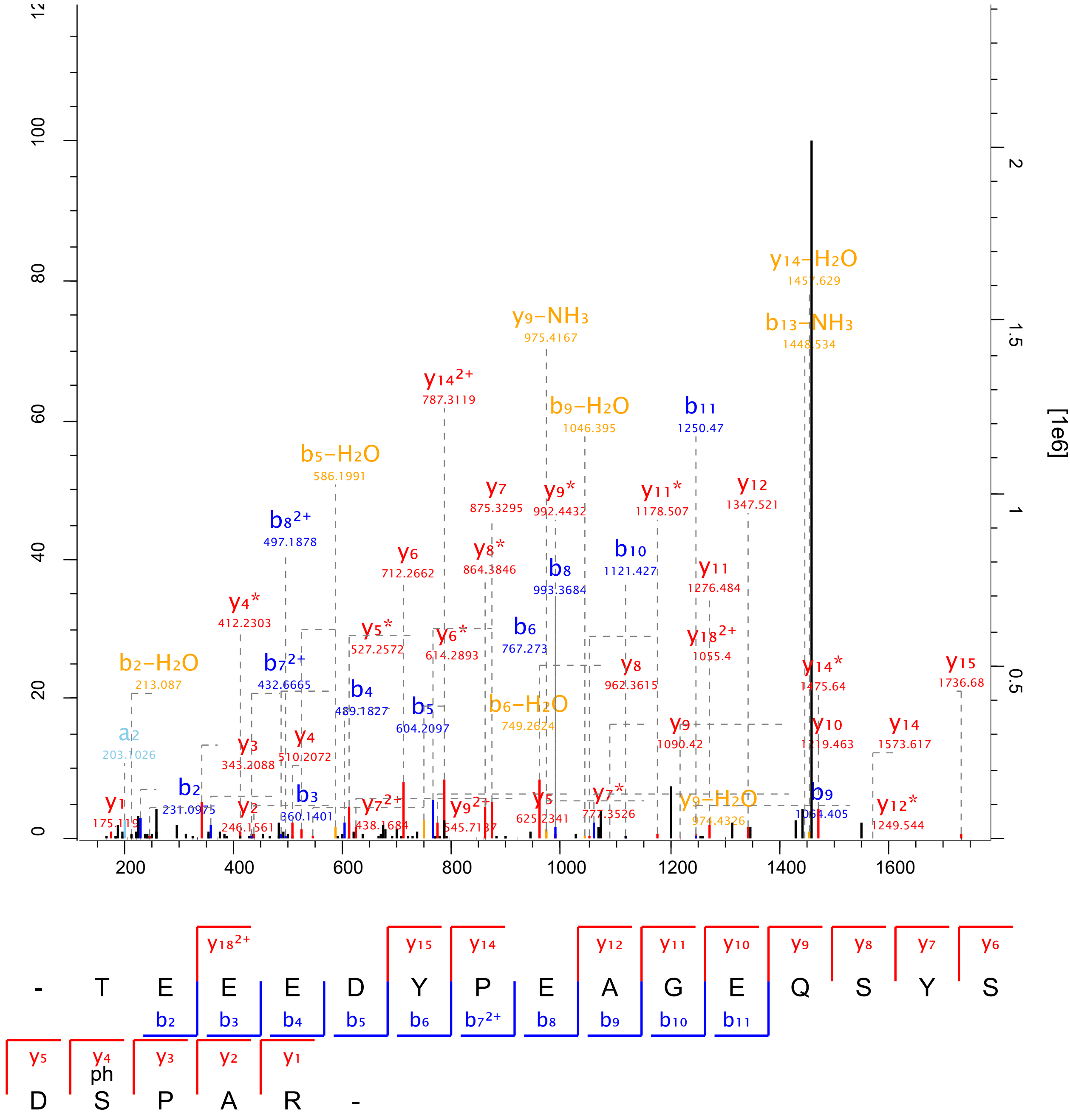

| Raw file | Scan | Method | Score | m/z |
|---|---|---|---|---|
| sys_15_1 | 19668 | FTMS; HCD | 93.63 | 959.38 |

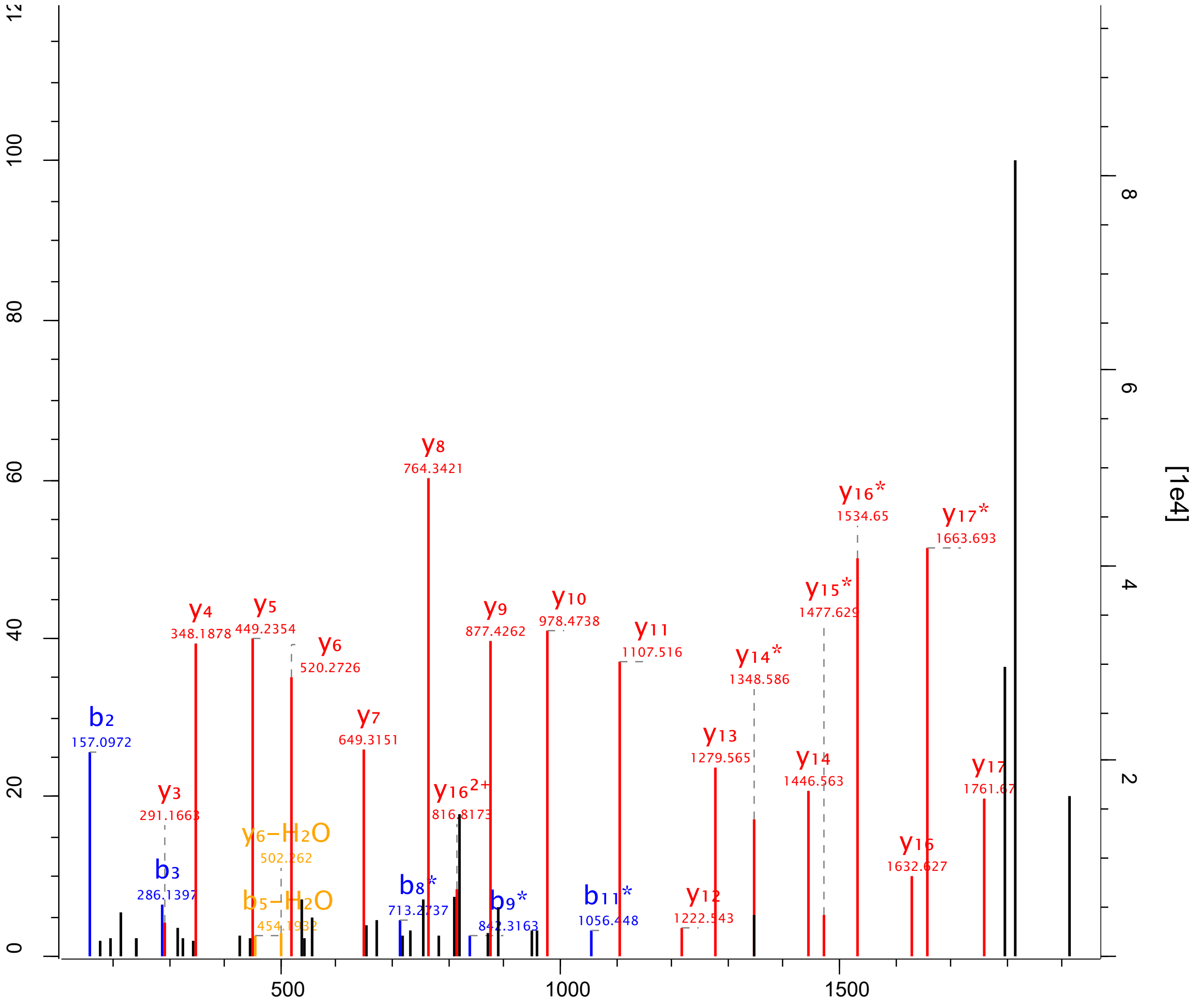


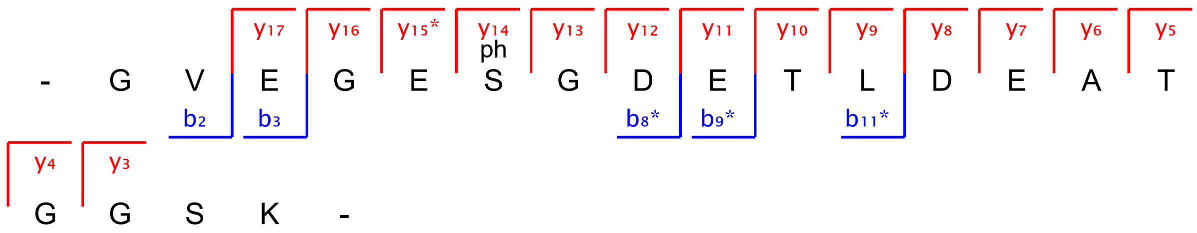

| Raw file | Scan | Method | Score | m/z |
| --- | --- | --- | --- | --- |
| sys_15_1 | 1967 | FTMS; HCD | 51.28 | 487.2 |

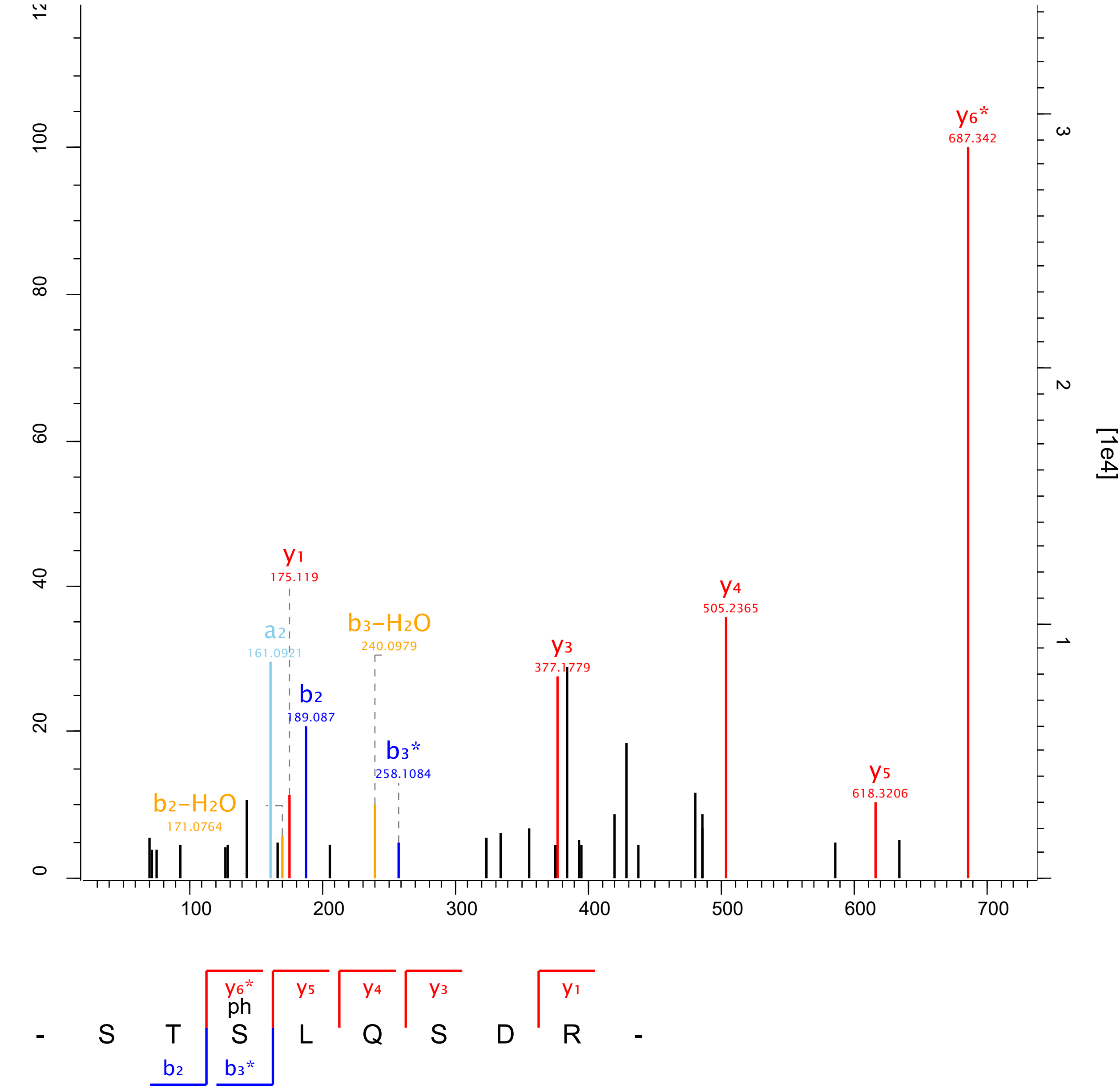

| Raw file | Scan | Method | Score | m/z |
|---|---|---|---|---|
| sys_15_1 | 19786 | FTMS; HCD | 79.84 | 549.91 |

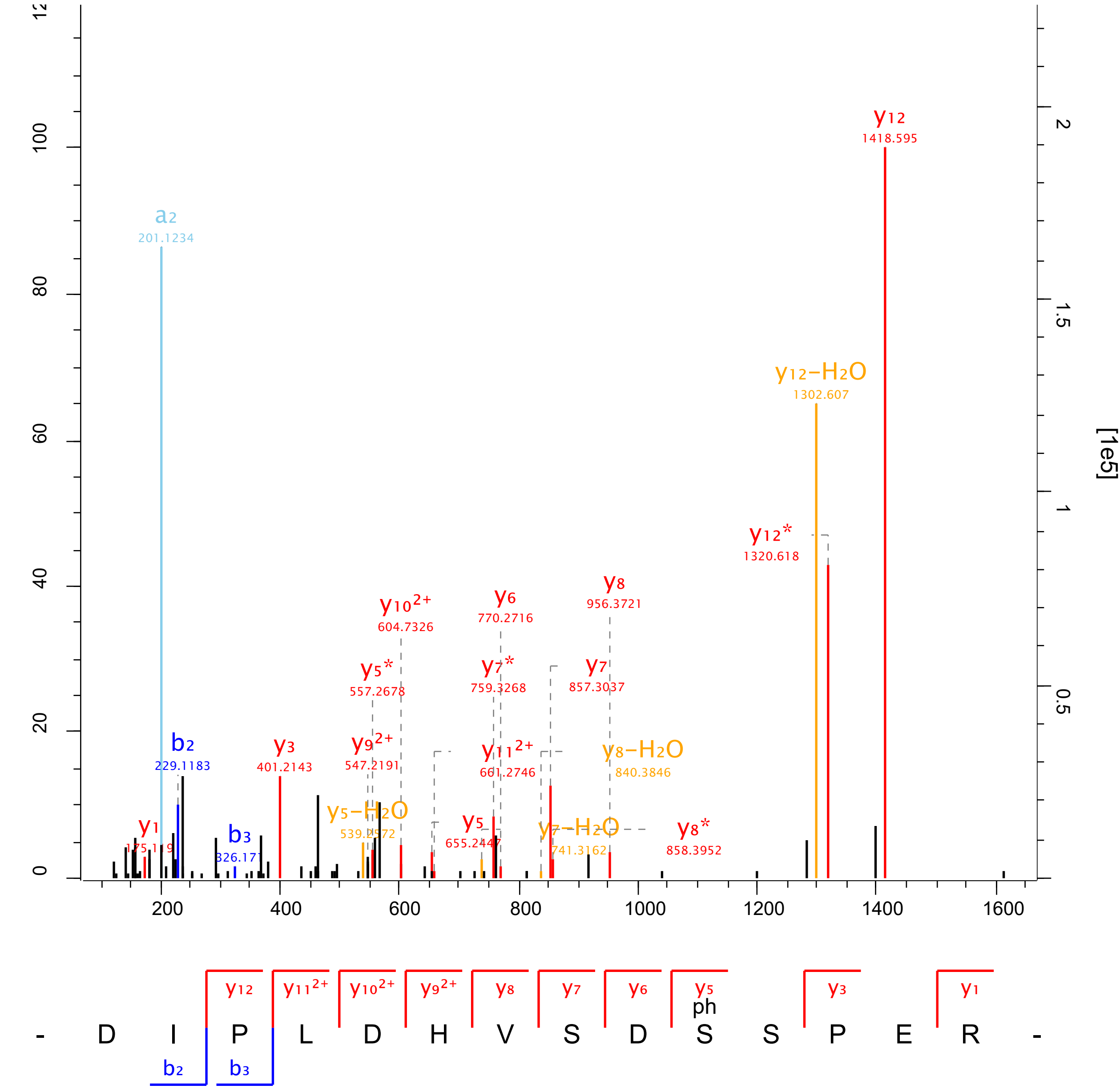

| Raw file | Scan | Method | Score | m/z |
|---|---|---|---|---|
| sys_15_1 | 1984 | FTMS; HCD | 69.72 | 546.23 |

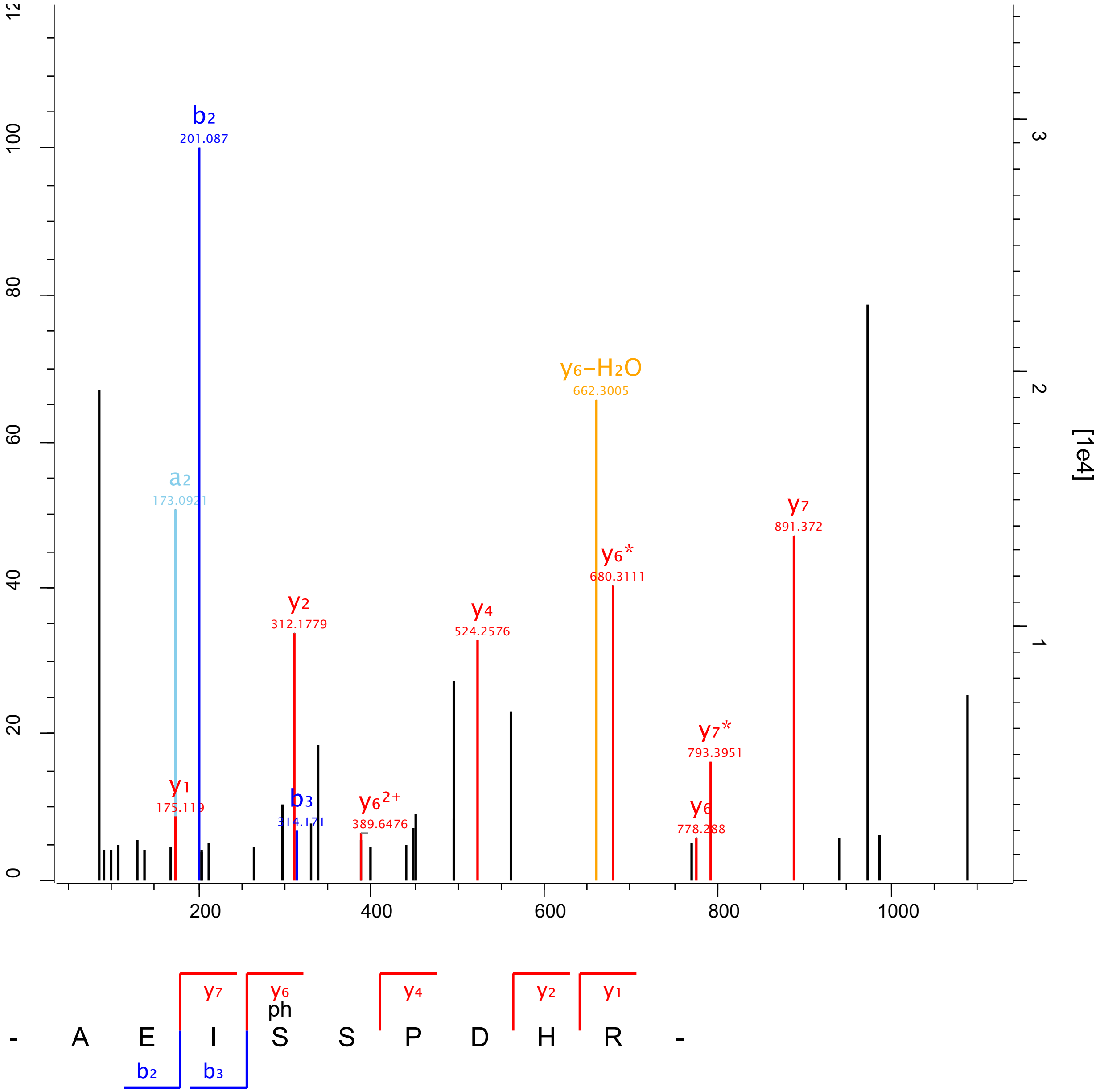

| Raw file | Scan | Method | Score | m/z |
|---|---|---|---|---|
| sys_15_1 | 19935 | FTMS; HCD | 52.49 | 515.74 |

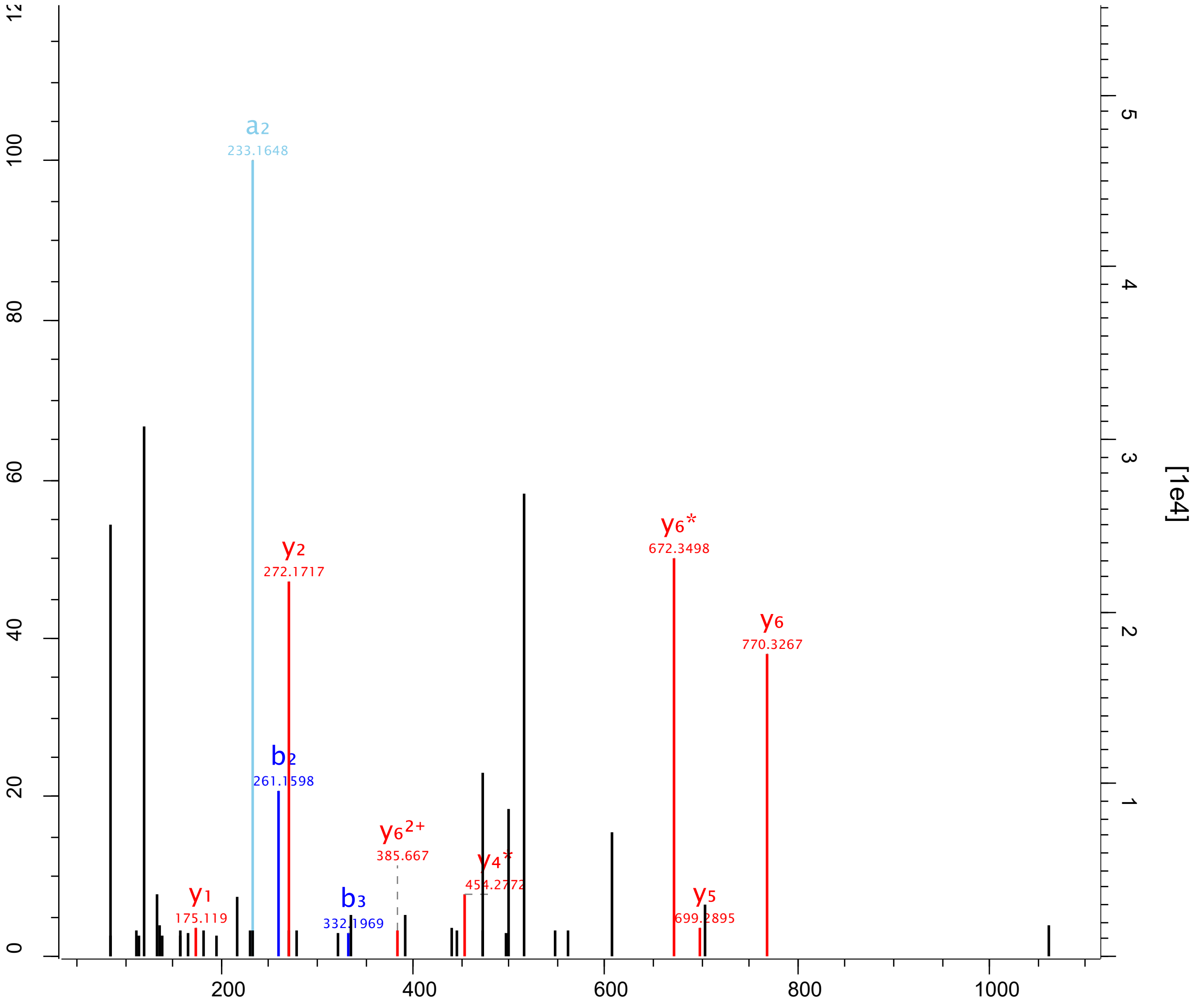


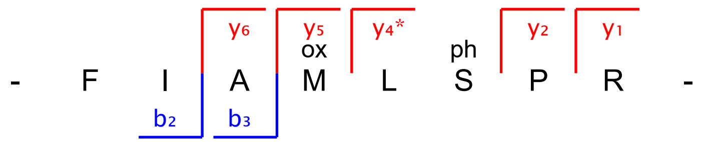

| Raw file | Scan | Method | Score | m/z |
|---|---|---|---|---|
| sys_15_1 | 19985 | FTMS; HCD | 80.07 | 645.28 |

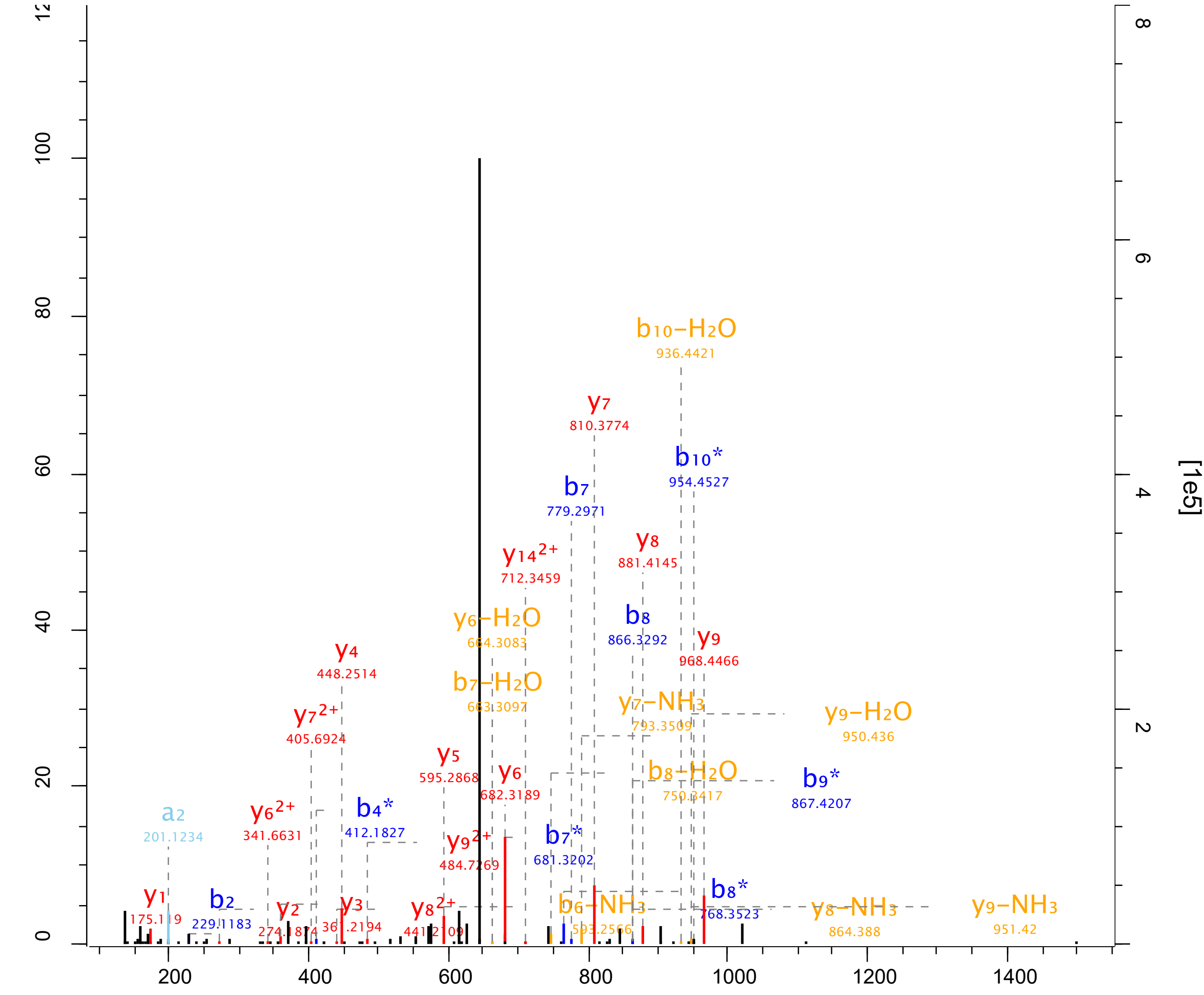


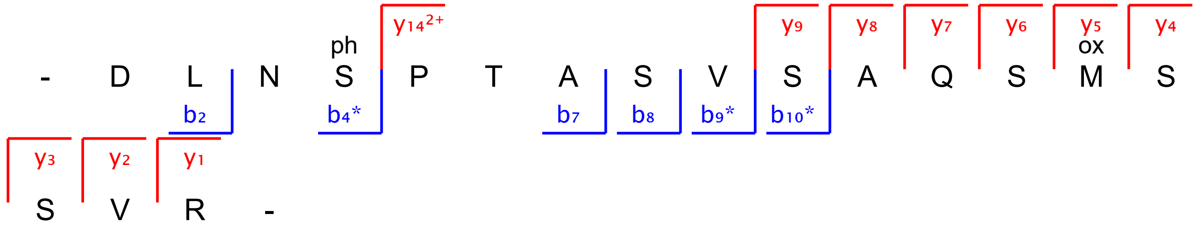

| Raw file | Scan | Method | Score | m/z |
| --- | --- | --- | --- | --- |
| sys_15_1 | 20120 | FTMS; HCD | 69.33 | 727.33 |

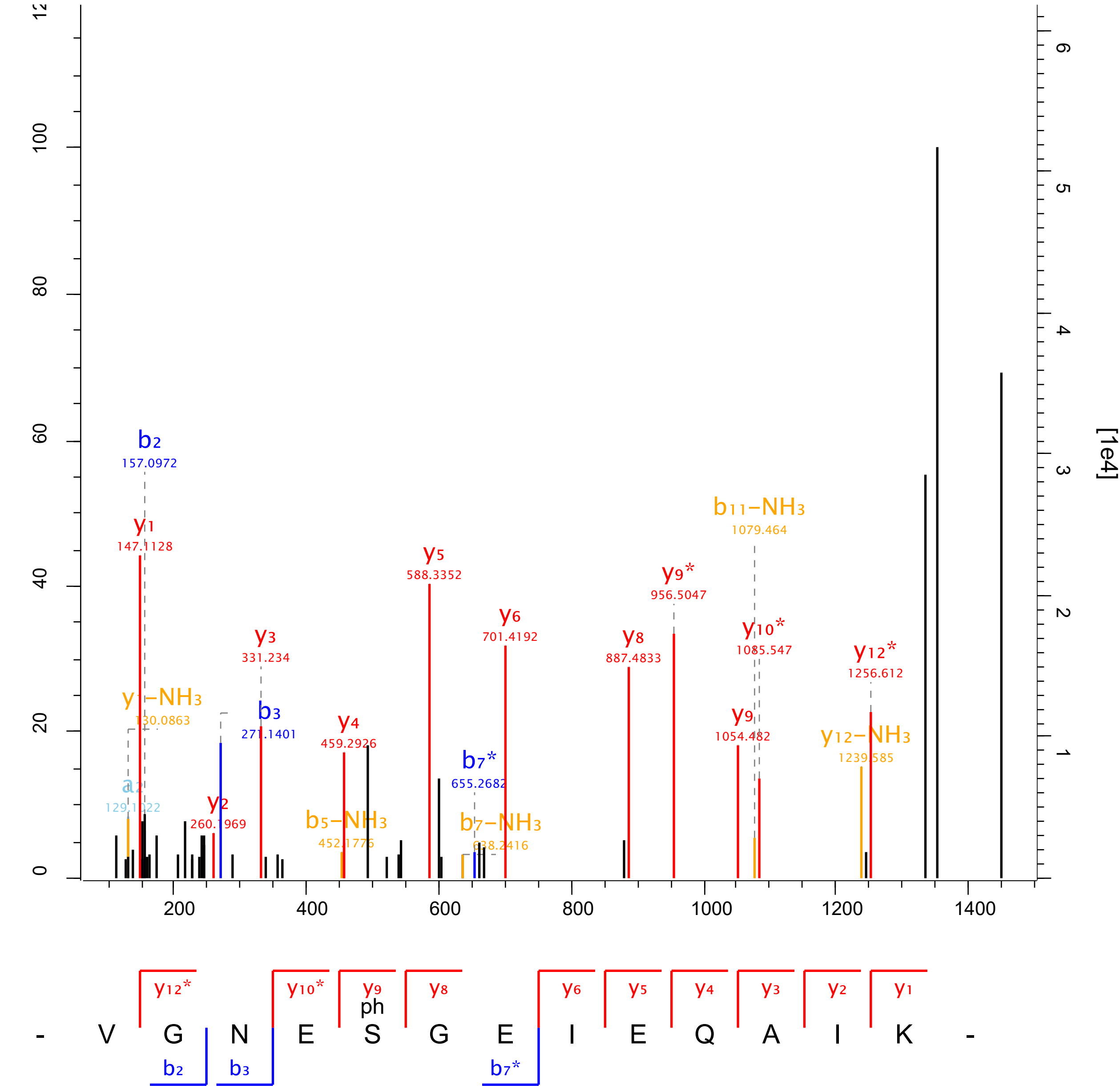

| Raw file | Scan | Method | Score | m/z |
|---|---|---|---|---|
| sys_15_1 | 20130 | FTMS; HCD | 52.17 | 768.31 |

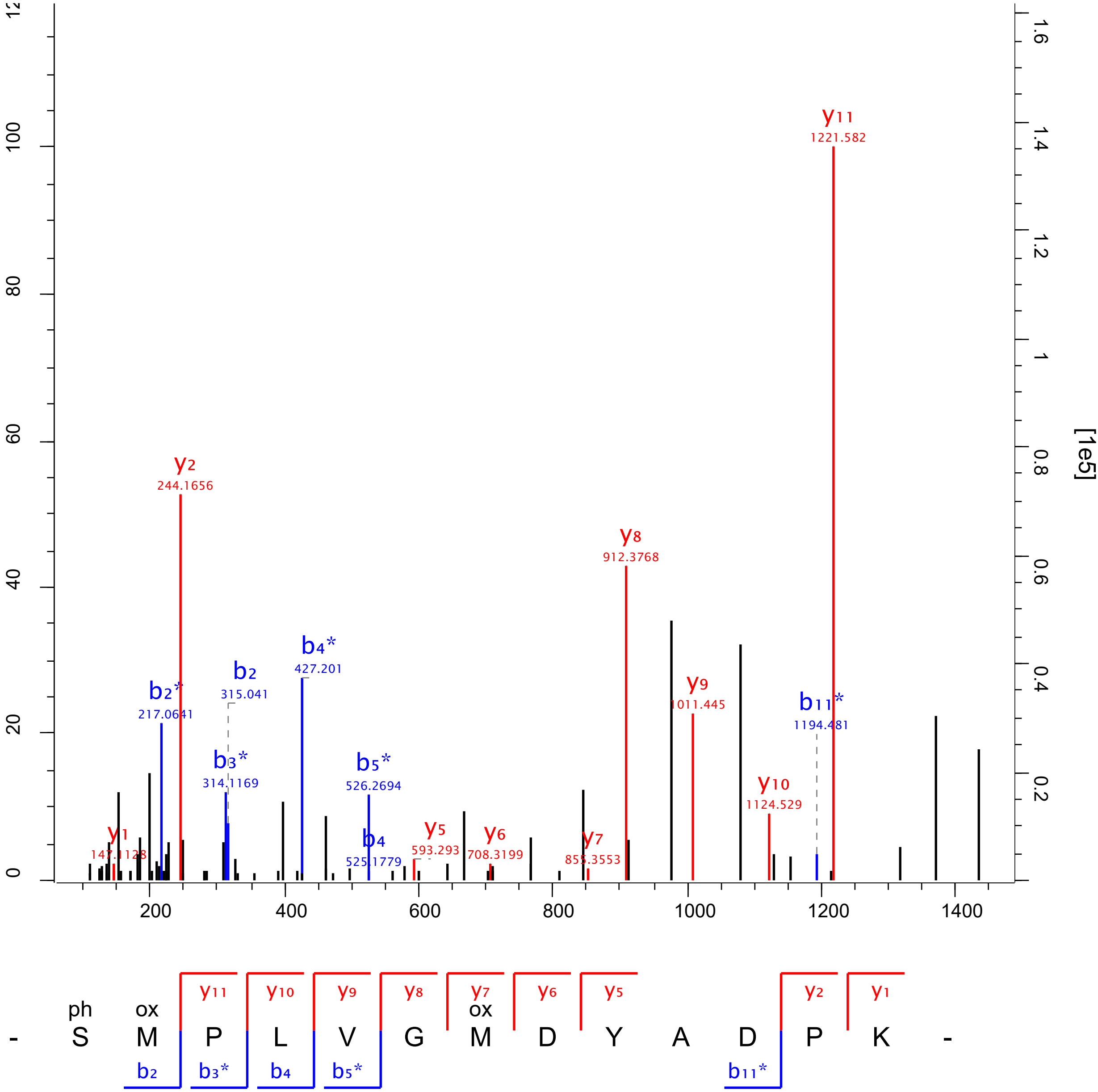

| Raw file | Scan | Method | Score | m/z |
| --- | --- | --- | --- | --- |
| sys_15_1 | 20134 | FTMS; HCD | 58.32 | 439.86 |

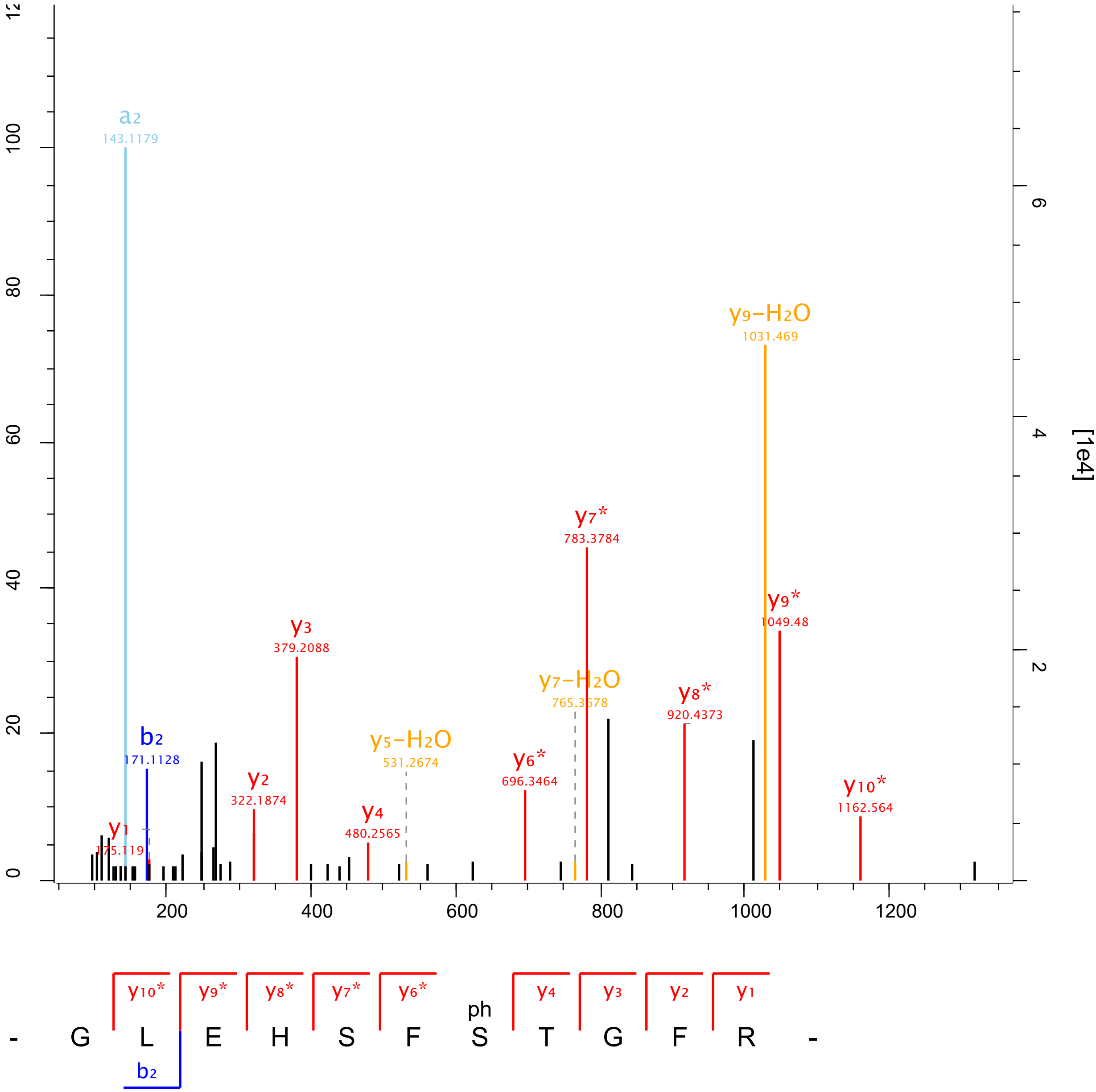

| Raw file | Scan | Method | Score | m/z |
| --- | --- | --- | --- | --- |
| sys_15_1 | 20154 | FTMS; HCD | 105.79 | 822.87 |

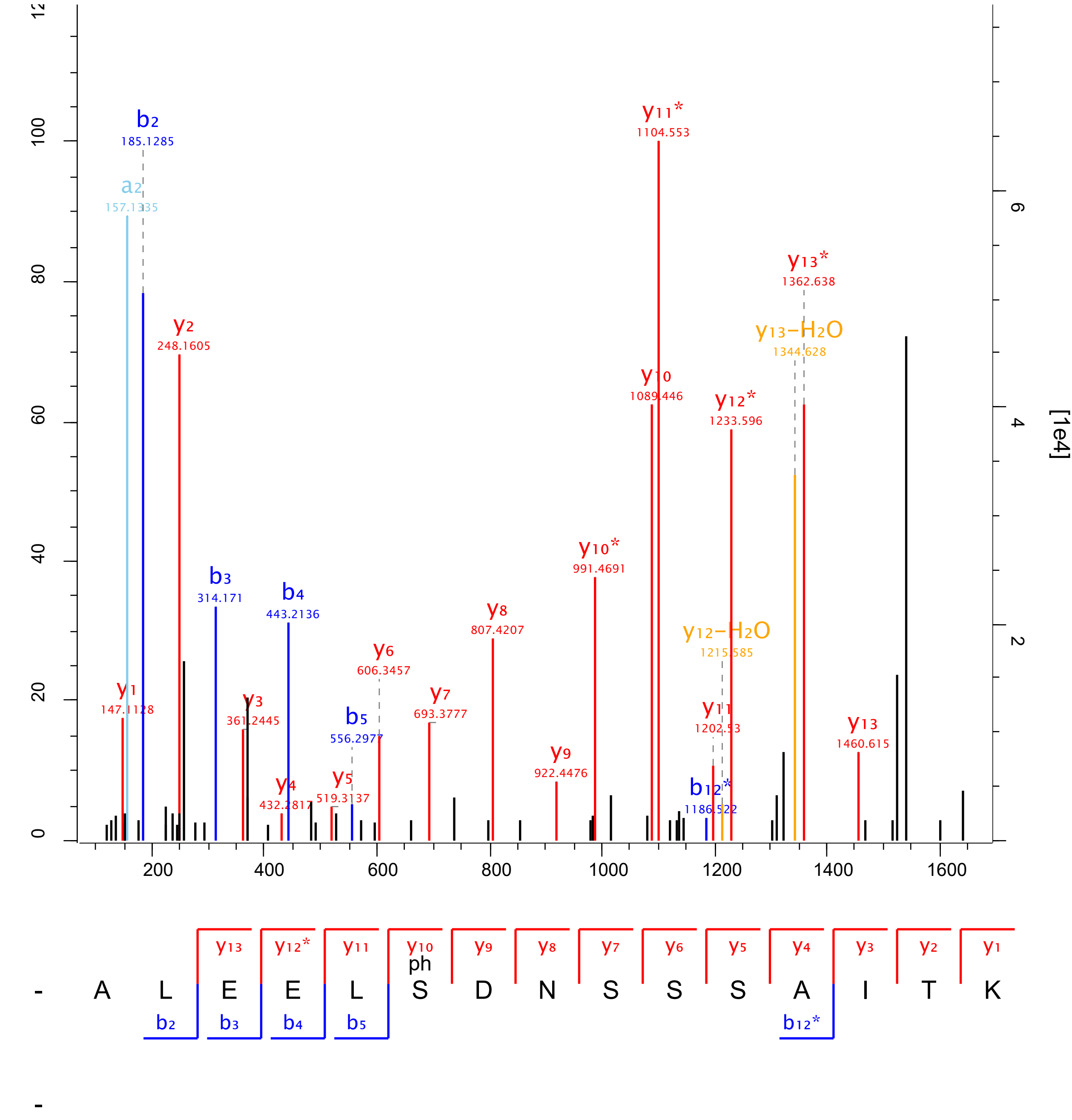

| Raw file | Scan | Method | Score | m/z |
| --- | --- | --- | --- | --- |
| sys_15_1 | 20564 | FTMS; HCD | 162.81 | 632.25 |

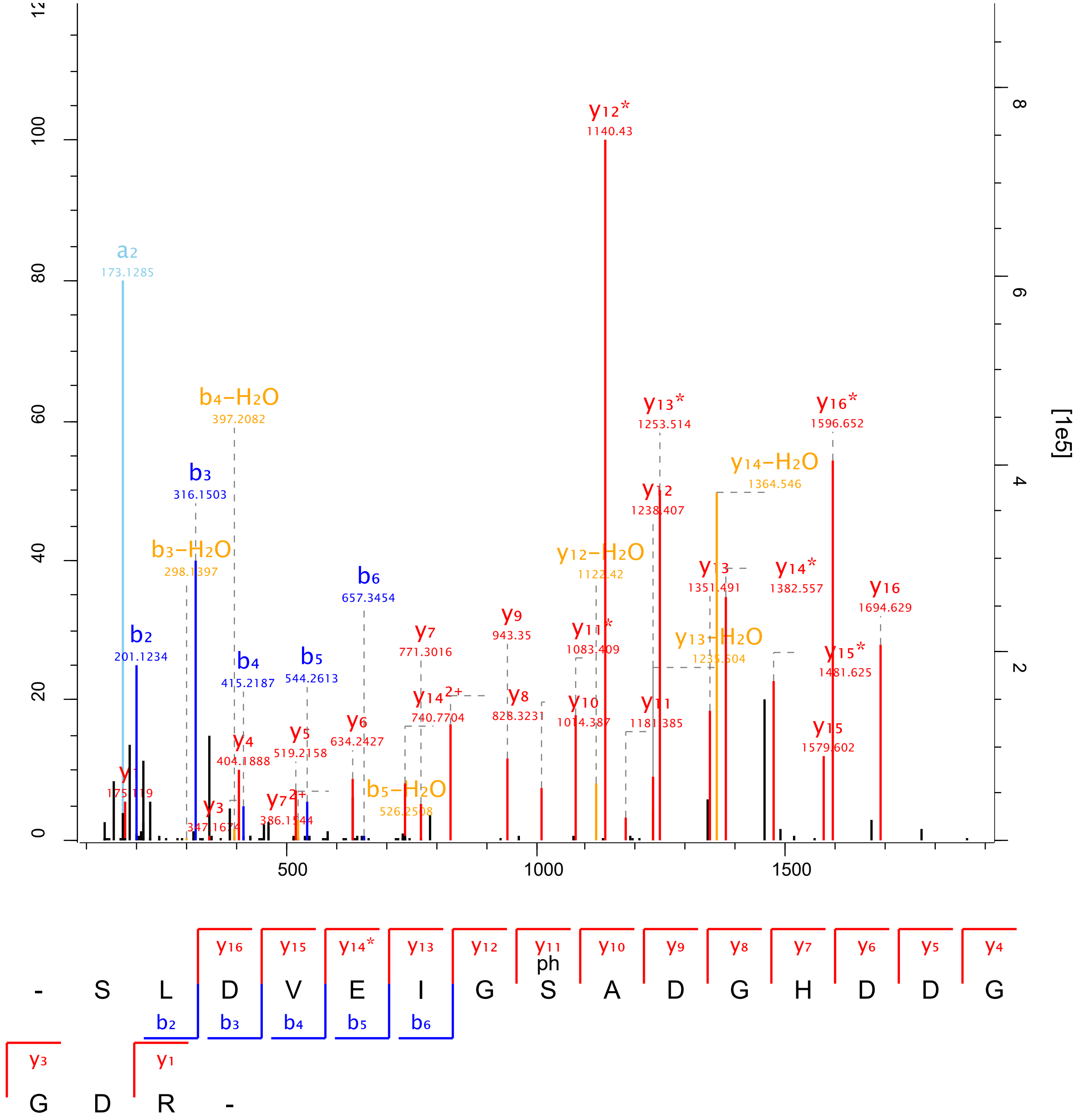

| Raw file | Scan | Method | Score | m/z |
|---|---|---|---|---|
| sys_15_1 | 20596 | FTMS; HCD | 119.74 | 688.29 |

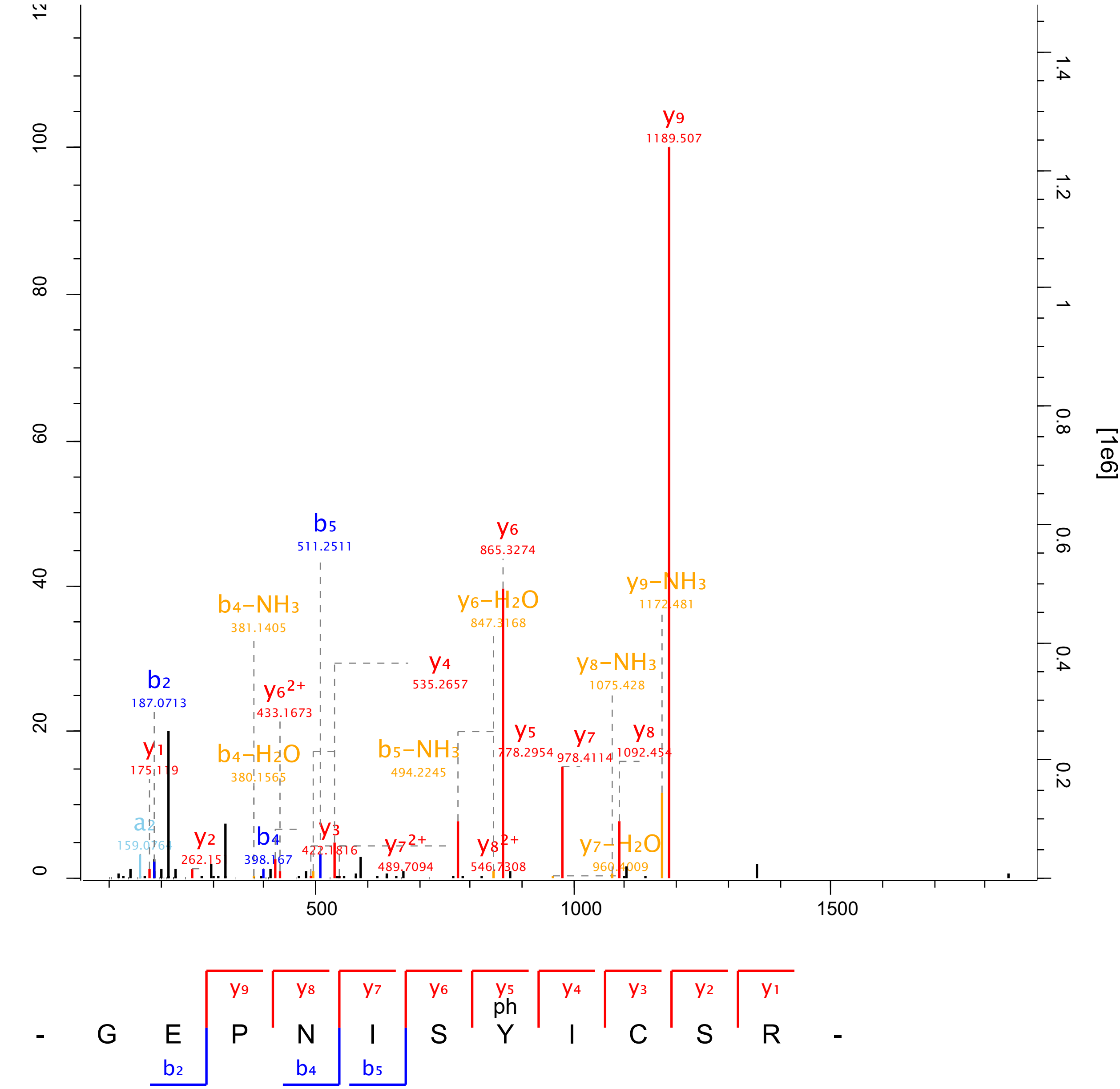

| Raw file | Scan | Method | Score | m/z |
| --- | --- | --- | --- | --- |
| sys_15_1 | 20659 | FTMS; HCD | 88.34 | 517.27 |

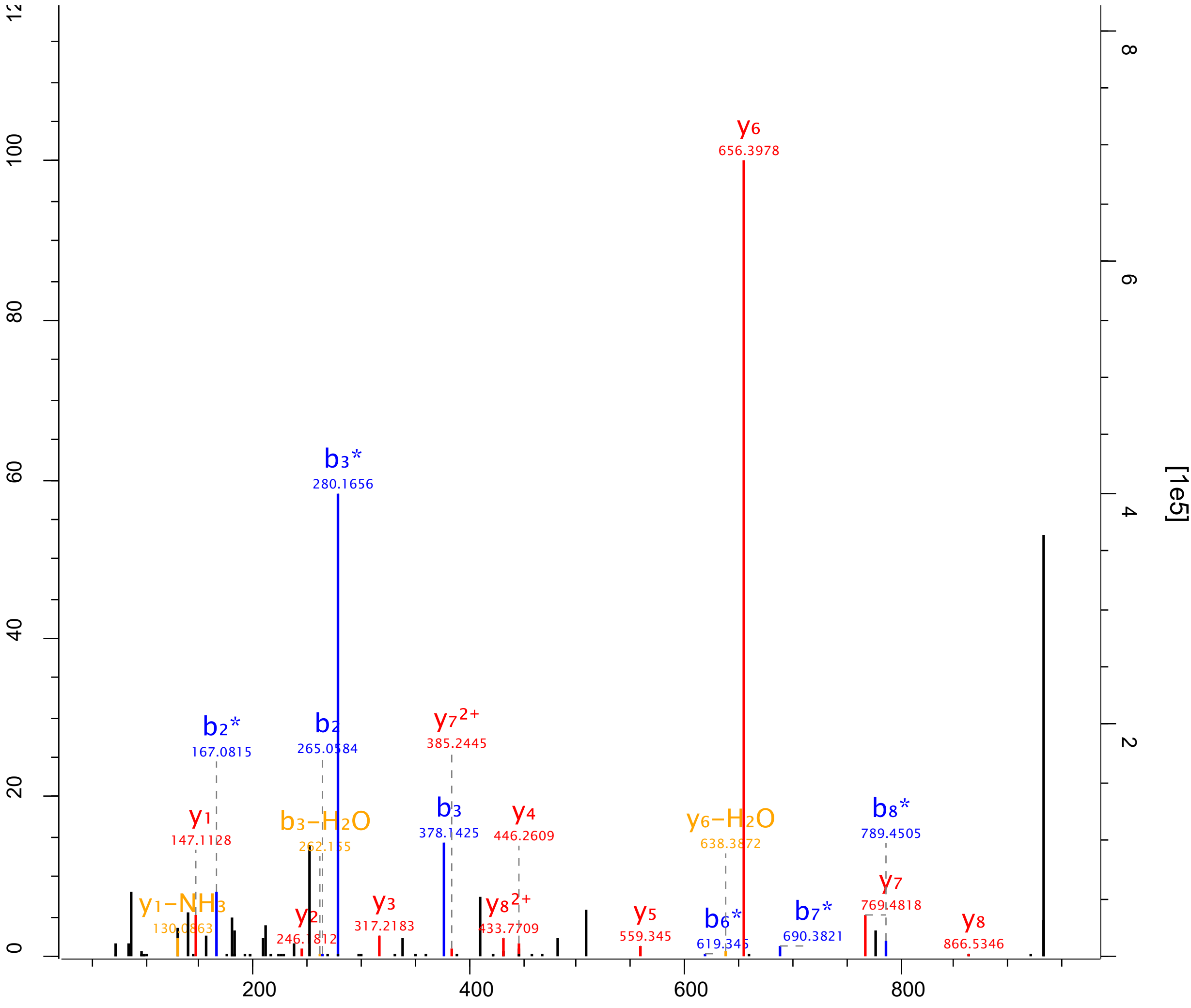


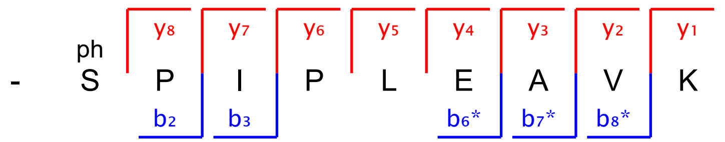

| Raw file | Scan | Method | Score | m/z |
|---|---|---|---|---|
| sys_15_1 | 20683 | FTMS; HCD | 189.77 | 806.88 |

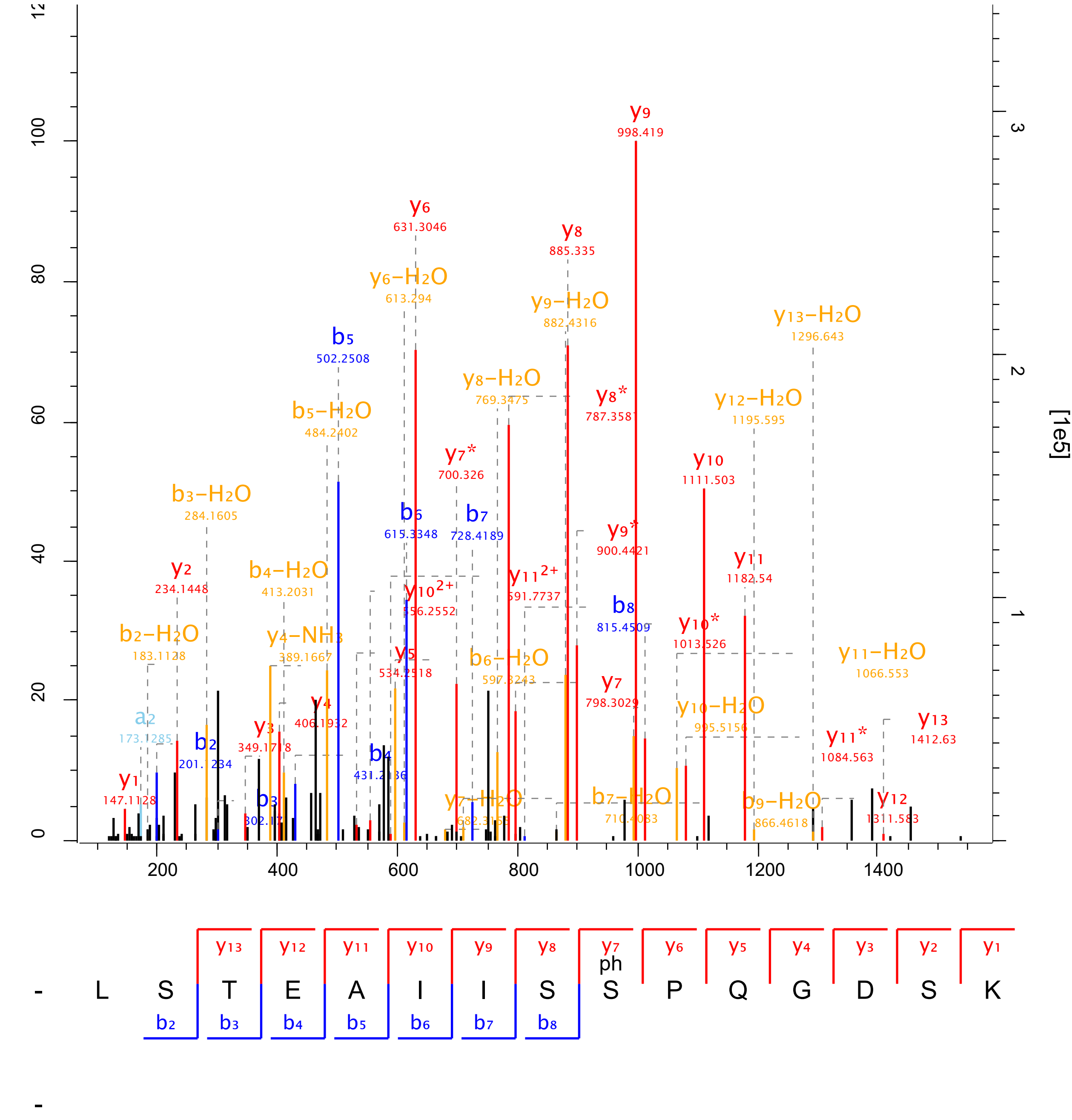

| Raw file | Scan | Method | Score | m/z |
|---|---|---|---|---|
| sys_15_1 | 20702 | FTMS; HCD | 263.55 | 894.36 |

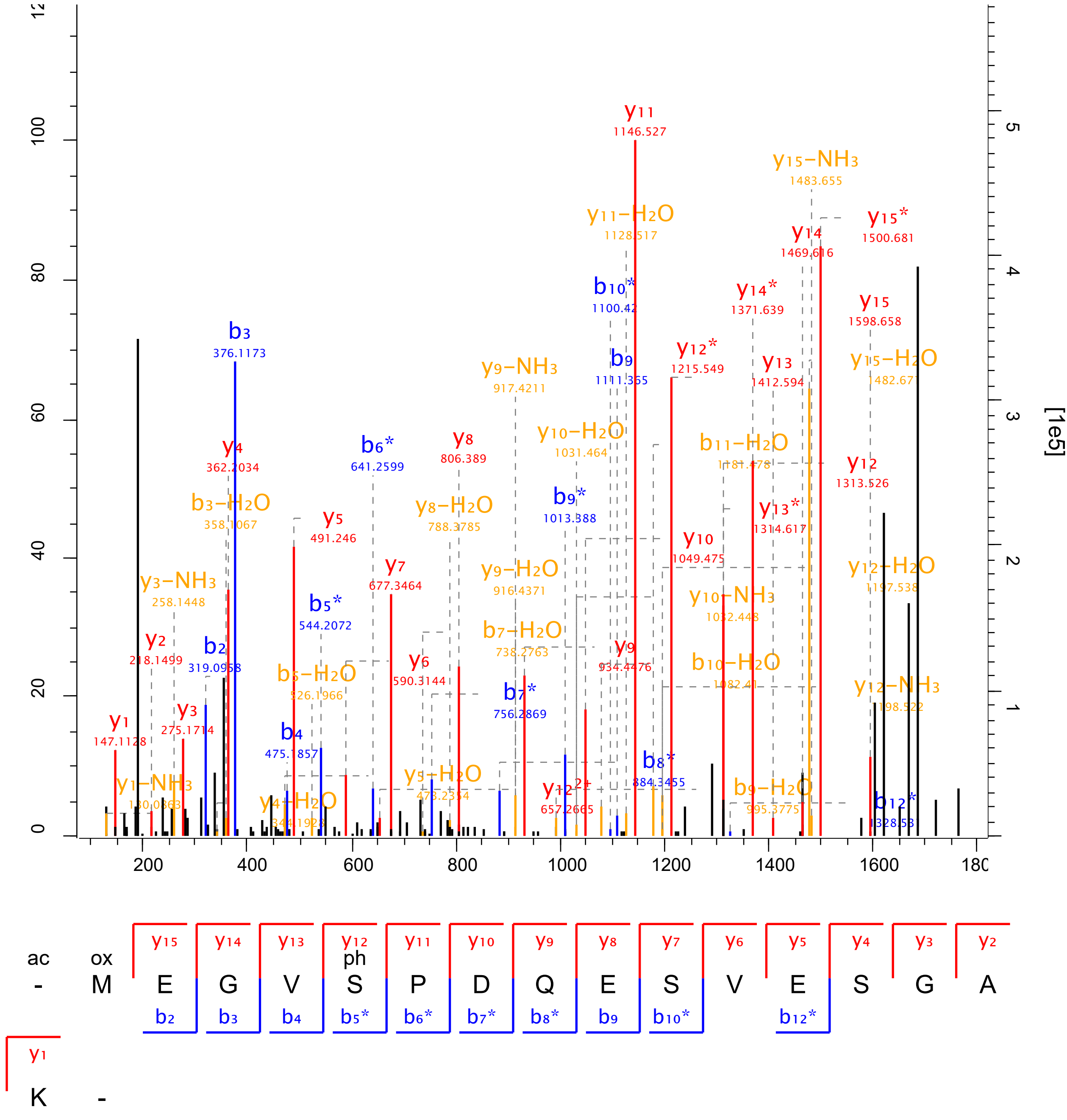

| Raw file | Scan | Method | Score | m/z |
| --- | --- | --- | --- | --- |
| sys_15_1 | 20782 | FTMS; HCD | 101.6 | 694.63 |

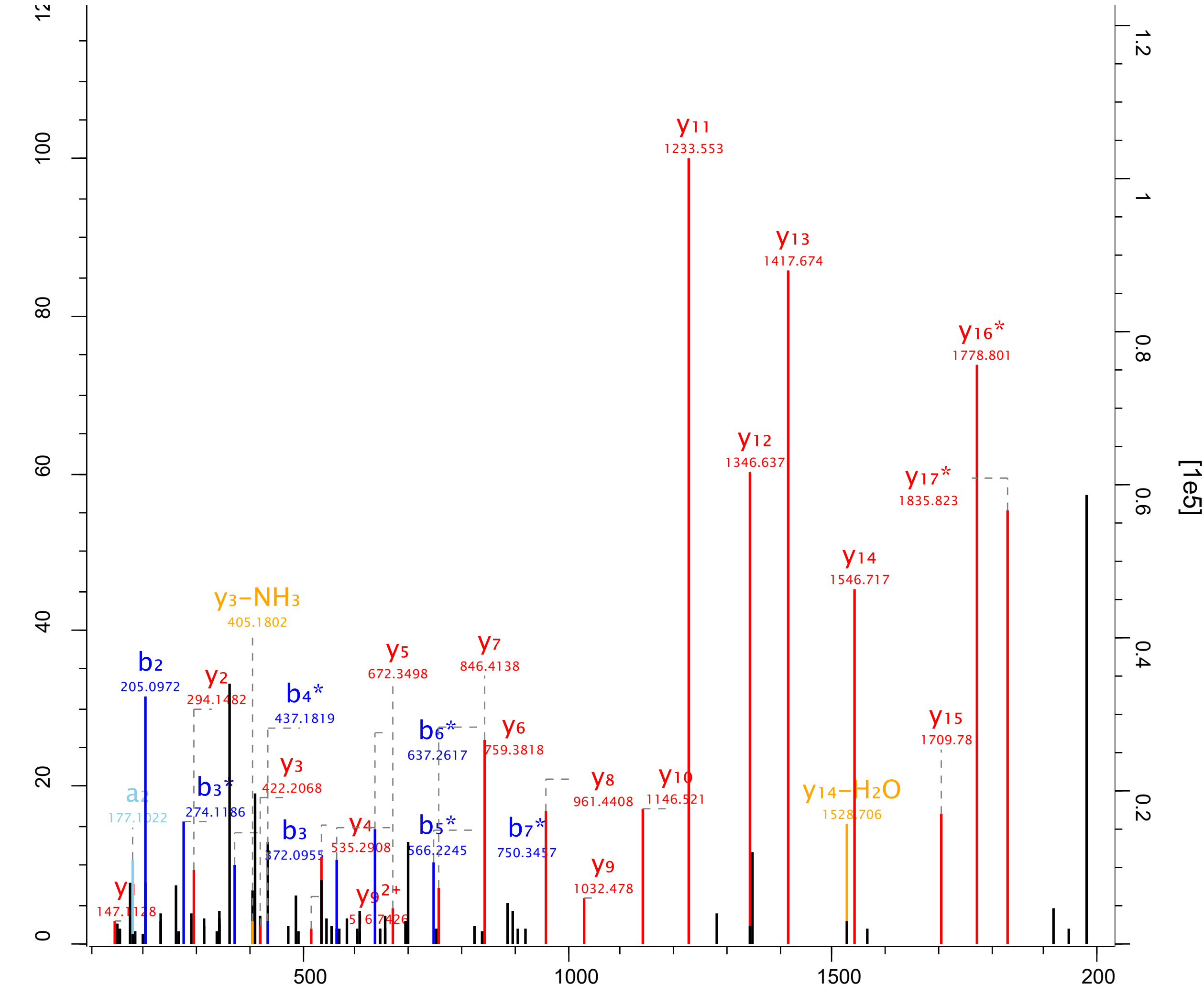


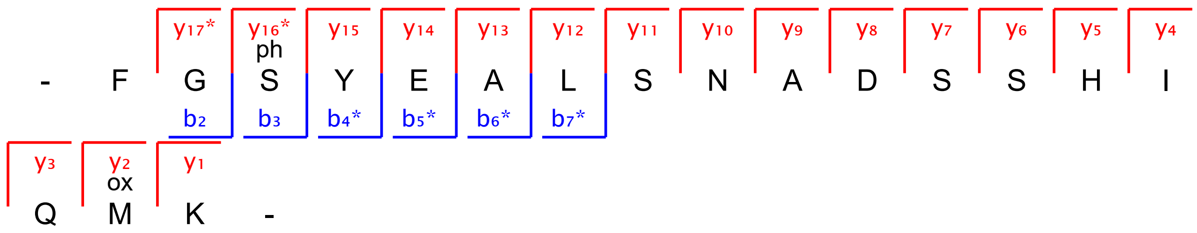

| Raw file | Scan | Method | Score | m/z |
| --- | --- | --- | --- | --- |
| sys_15_1 | 20799 | FTMS; HCD | 75.54 | 883.73 |

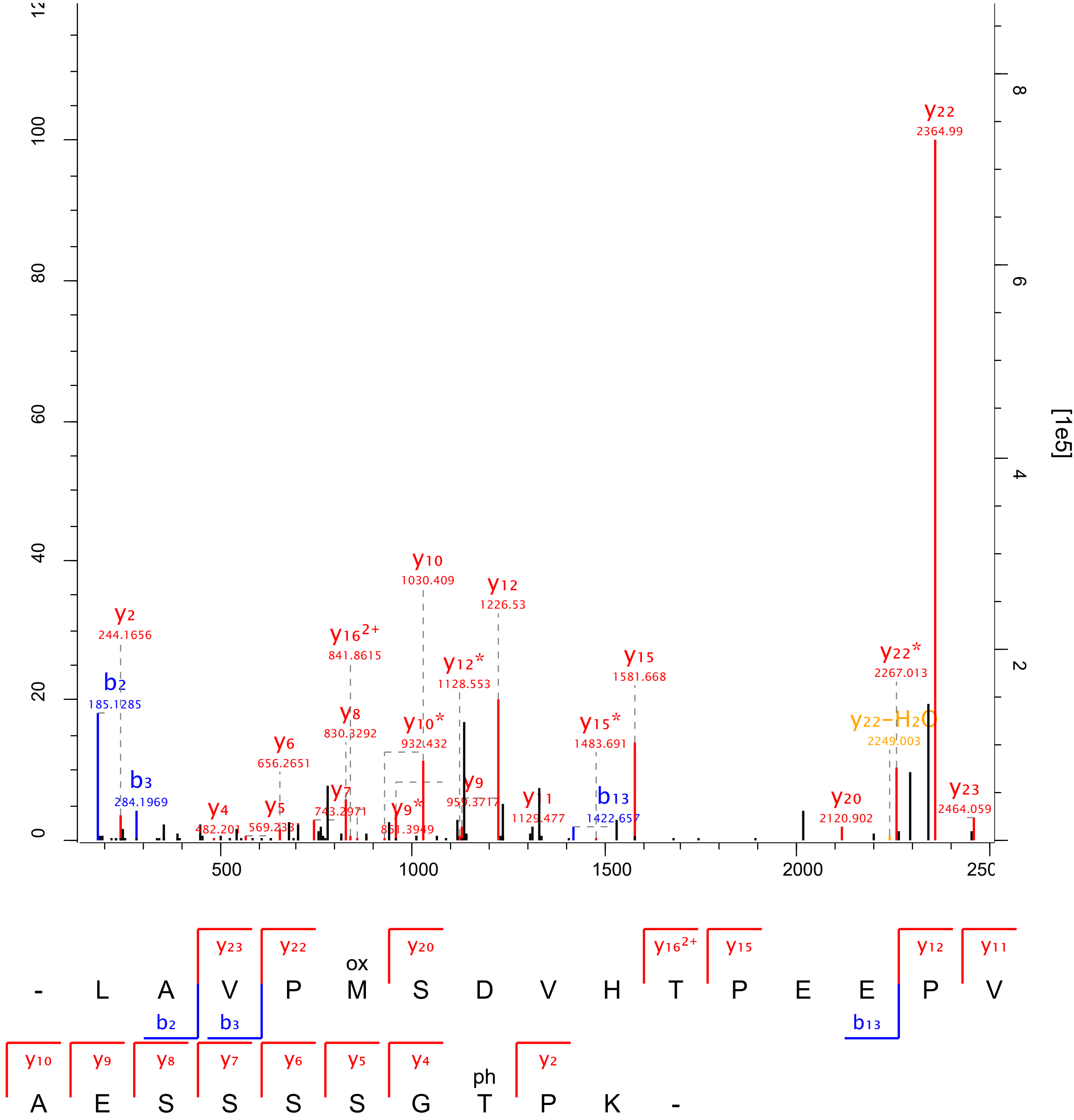

| Raw file | Scan | Method | Score | m/z |
|---|---|---|---|---|
| sys_15_1 | 20955 | FTMS; HCD | 62.09 | 615.76 |

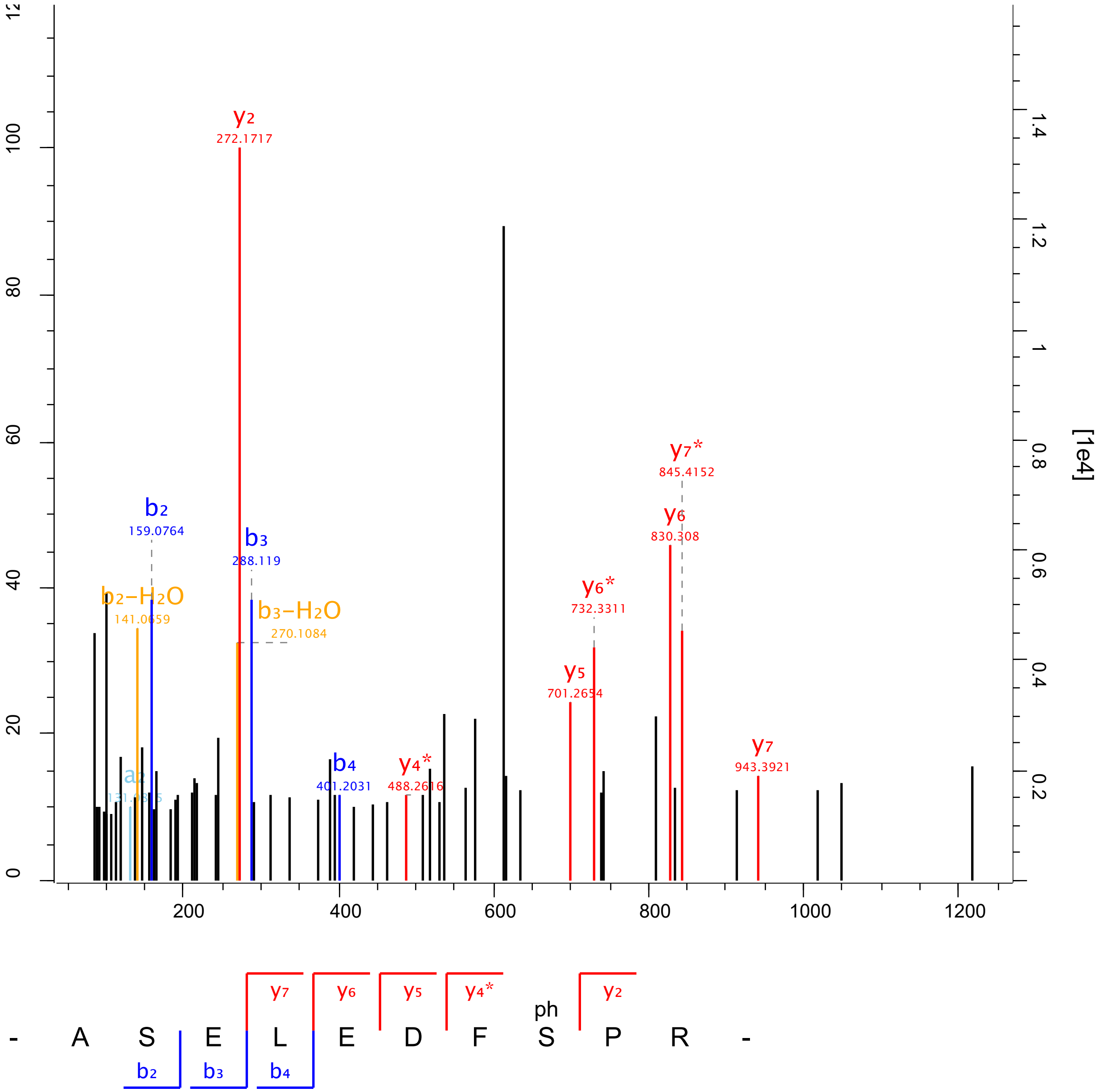

| Raw file | Scan | Method | Score | m/z |
|---|---|---|---|---|
| sys_15_1 | 21058 | FTMS; HCD | 71.12 | 847.37 |

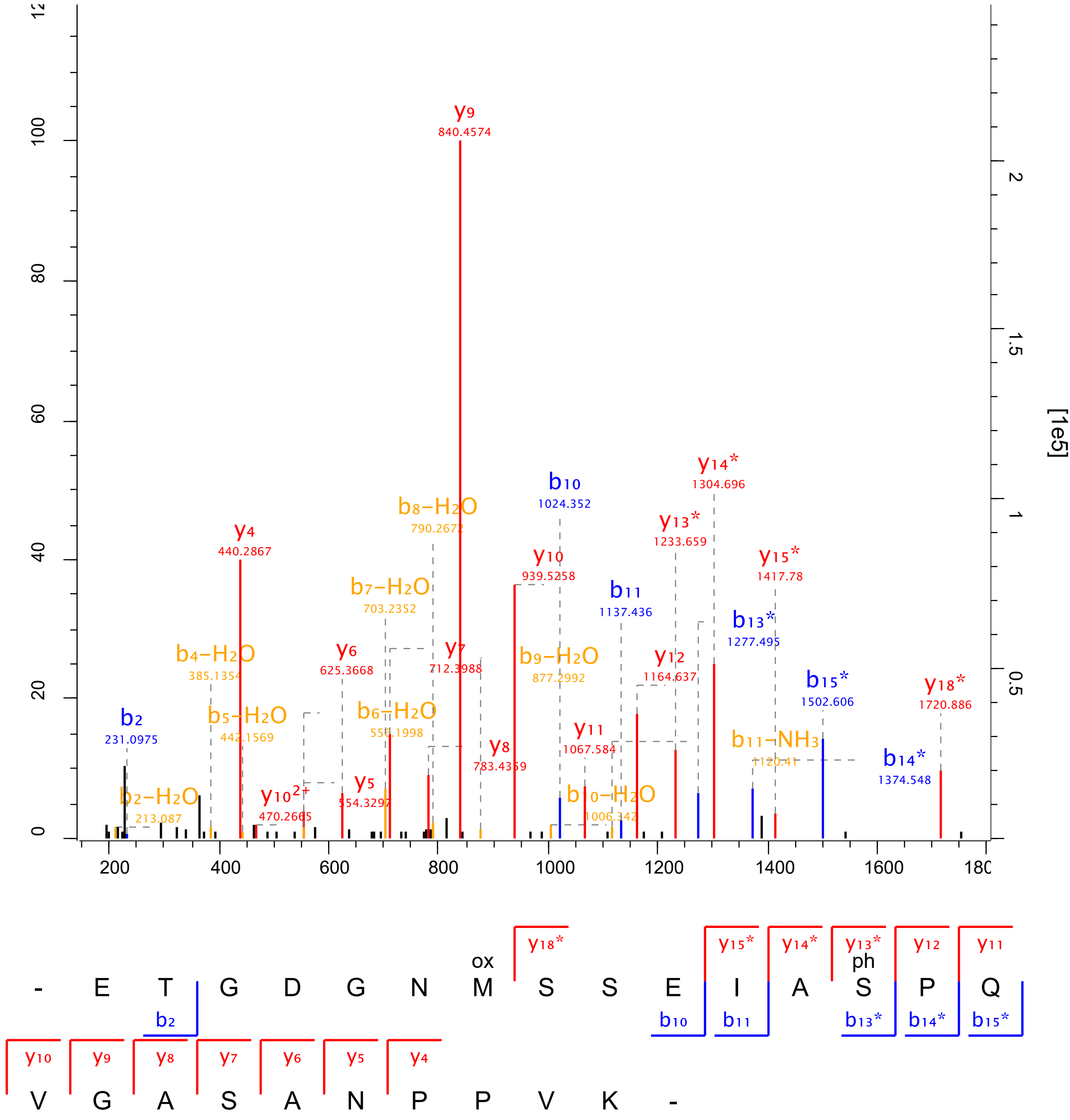

| Raw file | Scan | Method | Score | m/z |
|---|---|---|---|---|
| sys_15_1 | 21078 | FTMS; HCD | 148.15 | 871.91 |

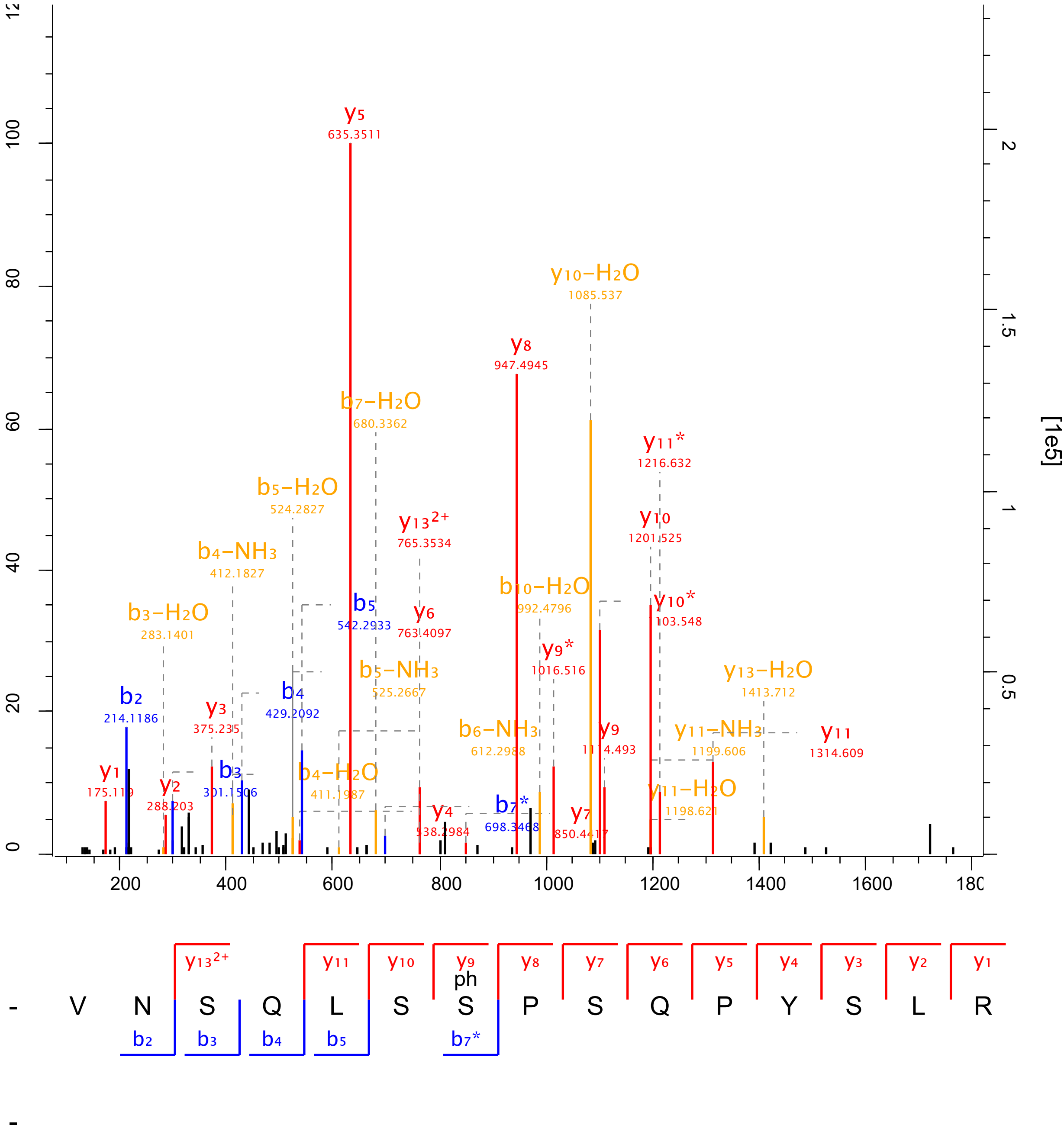

| Raw file | Scan | Method | Score | m/z |
|---|---|---|---|---|
| sys_15_1 | 21530 | FTMS; HCD | 279.97 | 831.38 |

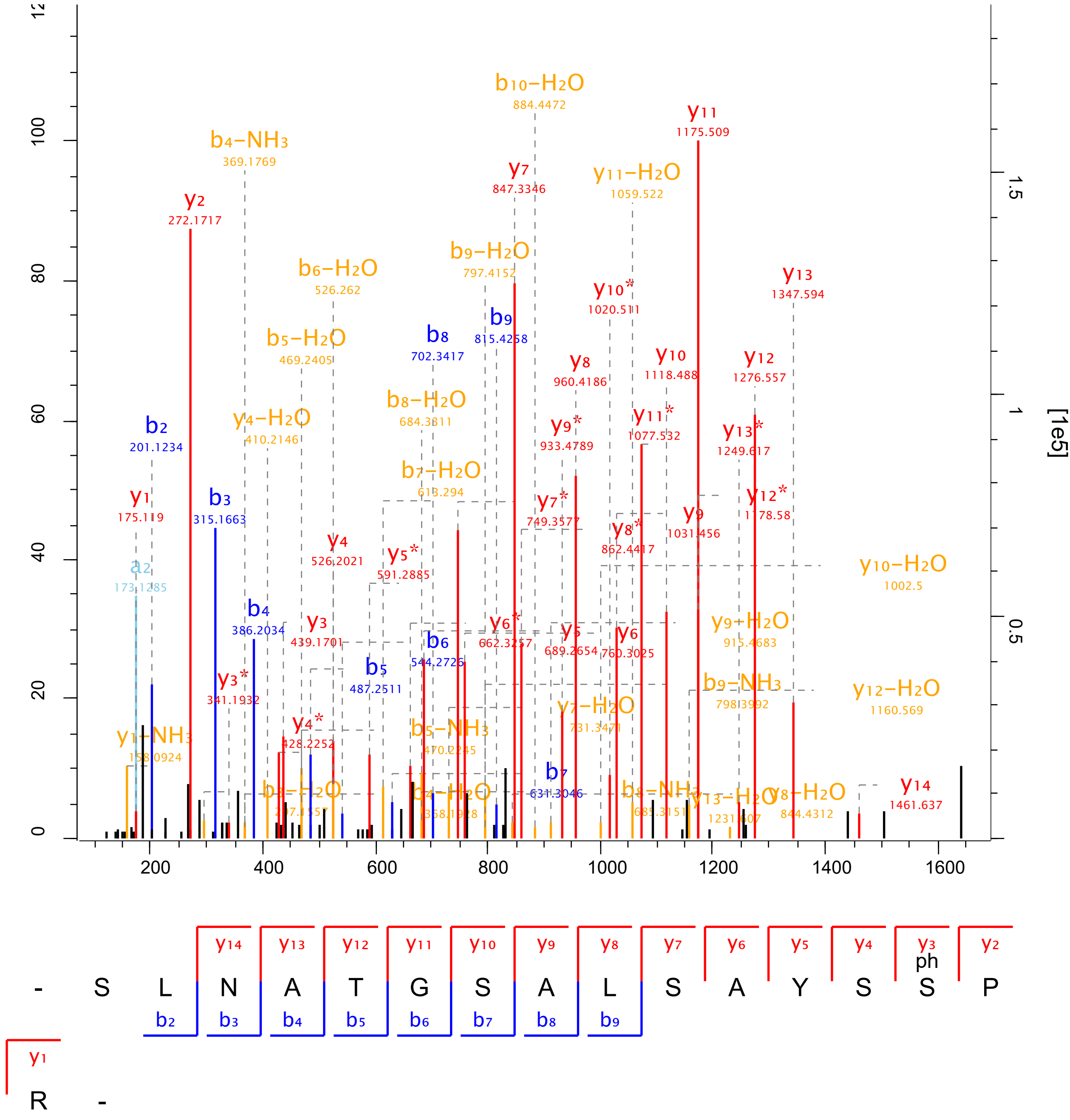

| Raw file | Scan | Method | Score | m/z |
|---|---|---|---|---|
| sys_15_1 | 21543 | FTMS; HCD | 164.2 | 851.38 |

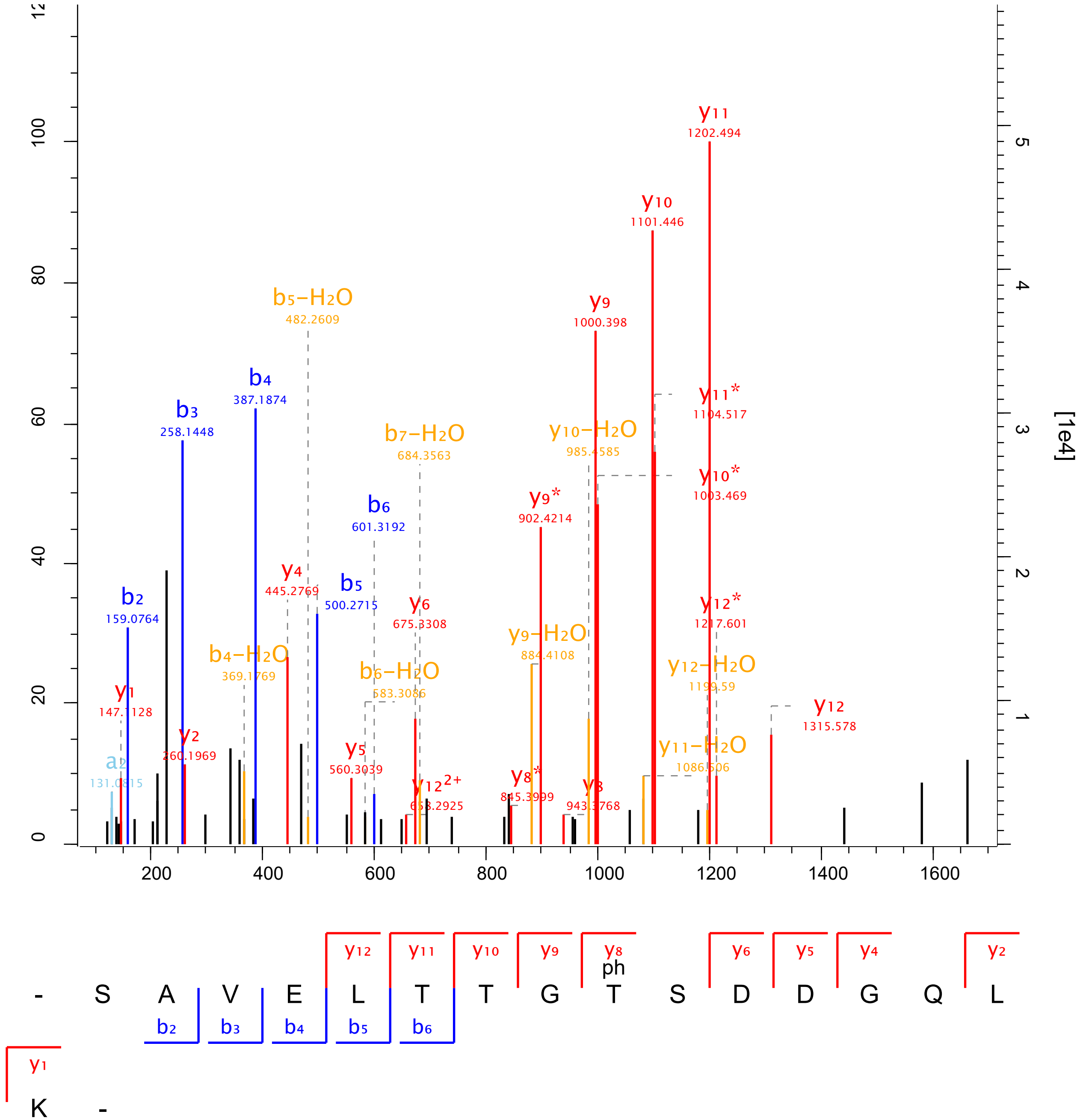


- S A V E L T T G T S D D G Q L

K -

(y12, y11, y10, y9, y8, y6, y5, y4, y2, y1; b2, b3, b4, b5, b6; ph on T)

| Raw file | Scan | Method | Score | m/z |
|---|---|---|---|---|
| sys_15_1 | 21660 | FTMS; HCD | 104.84 | 865.87 |

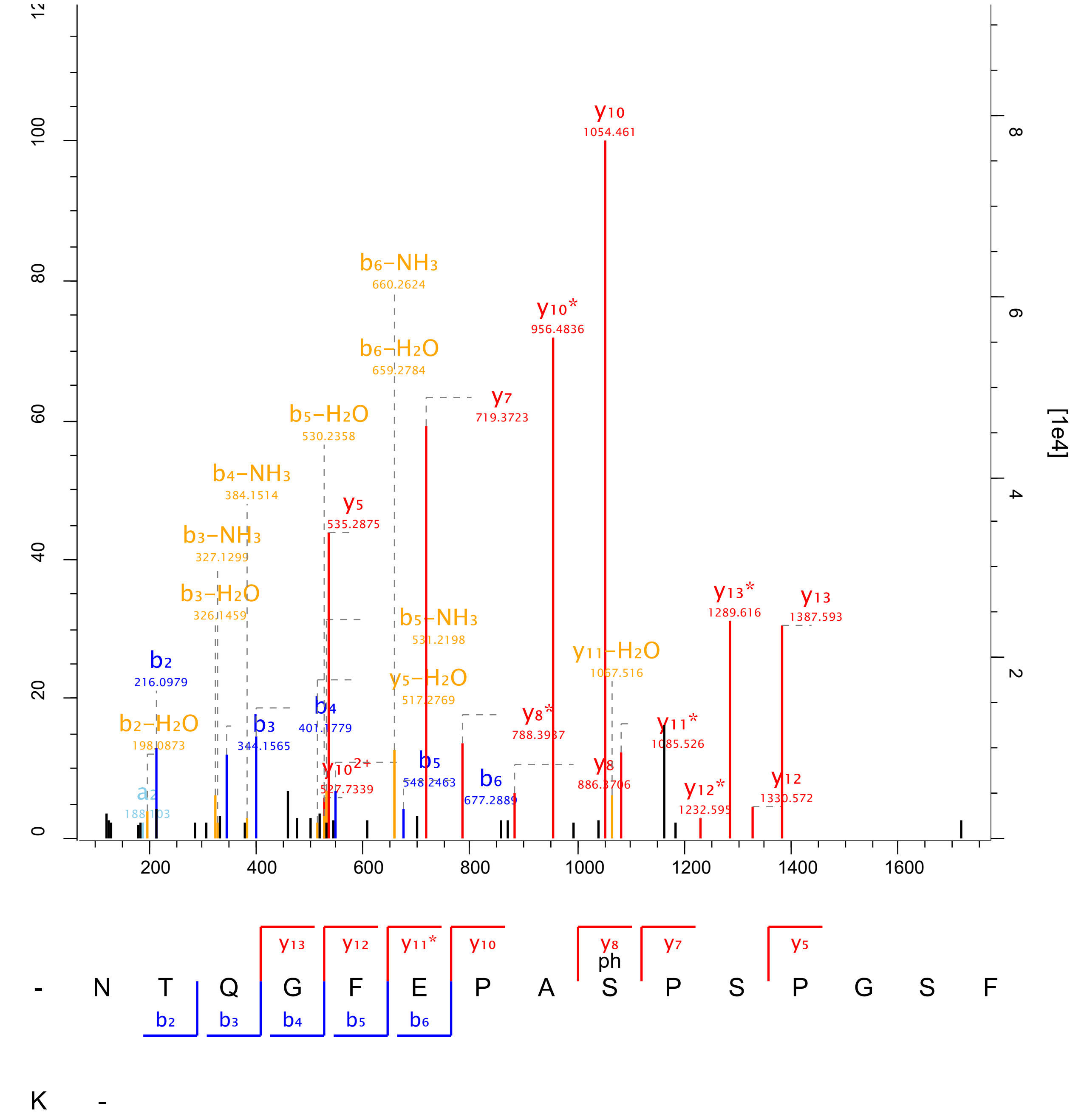

| Raw file | Scan | Method | Score | m/z |
| --- | --- | --- | --- | --- |
| sys_15_1 | 21760 | FTMS; HCD | 94.36 | 501.23 |

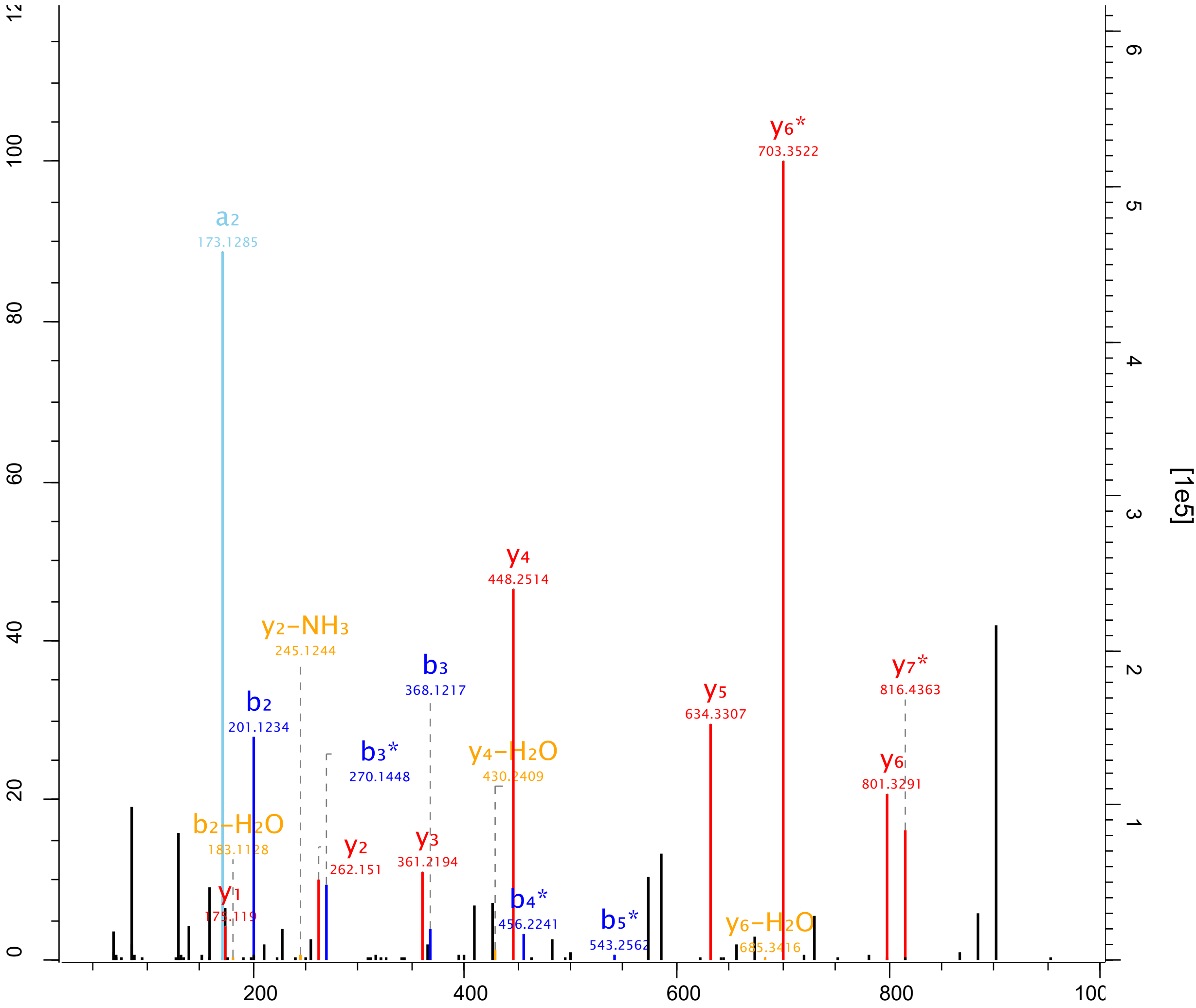


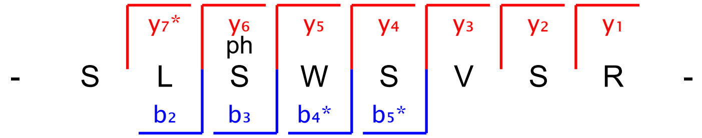

| Raw file | Scan | Method | Score | m/z |
| --- | --- | --- | --- | --- |
| sys_15_1 | 21772 | FTMS; HCD | 64.71 | 707.3 |

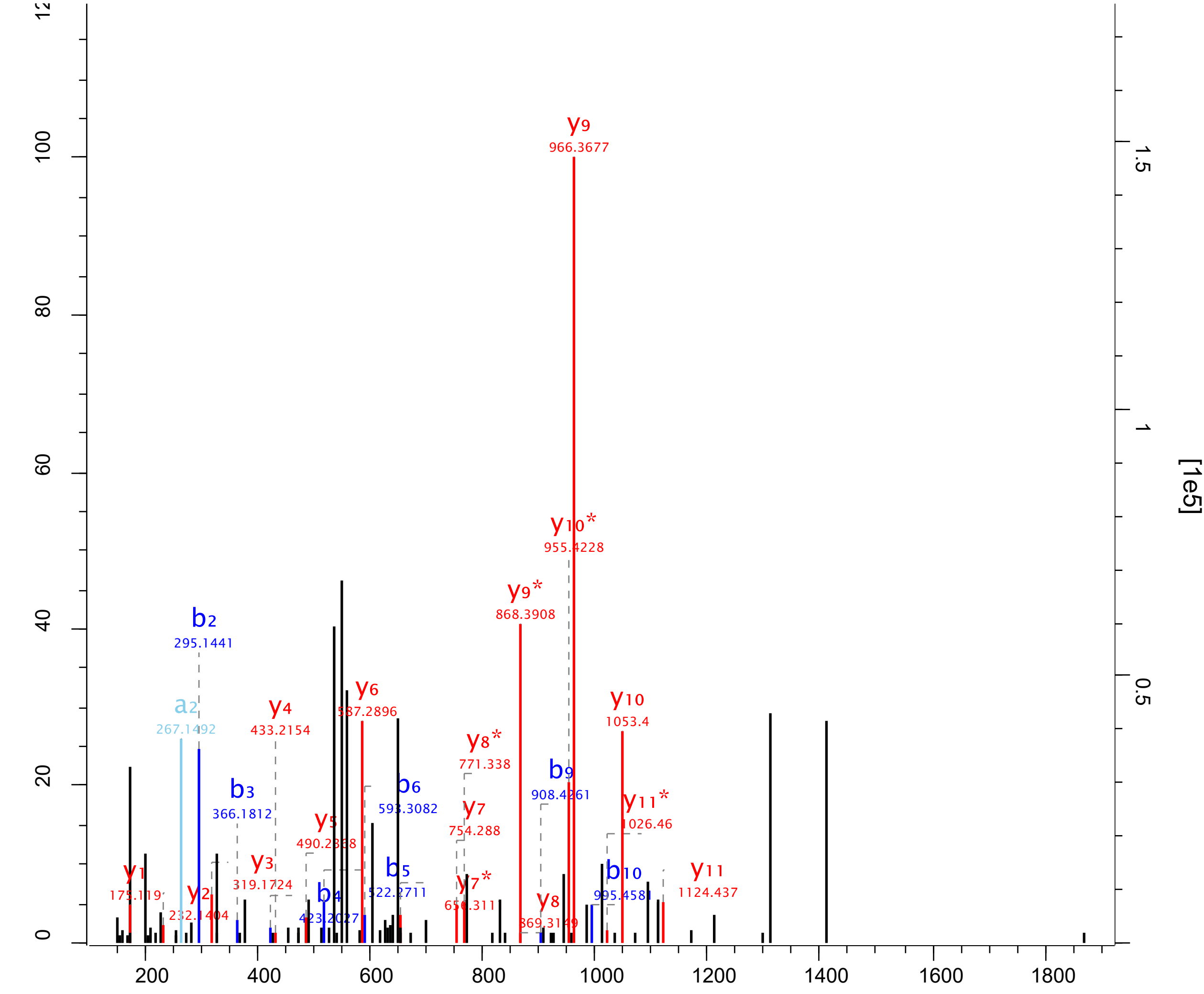


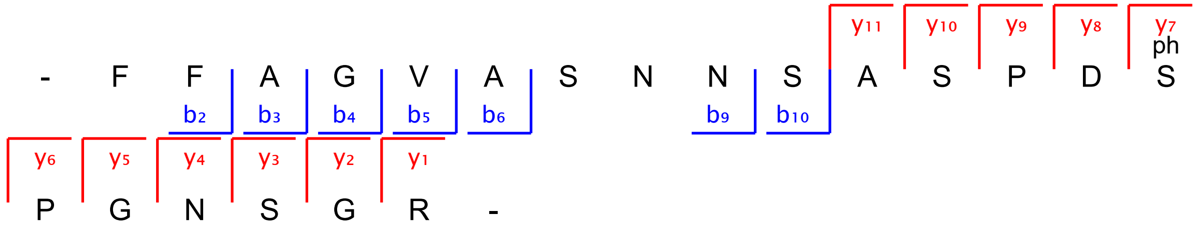

| Raw file | Scan | Method | Score | m/z |
|---|---|---|---|---|
| sys_15_1 | 21780 | FTMS; HCD | 165.4 | 679.98 |

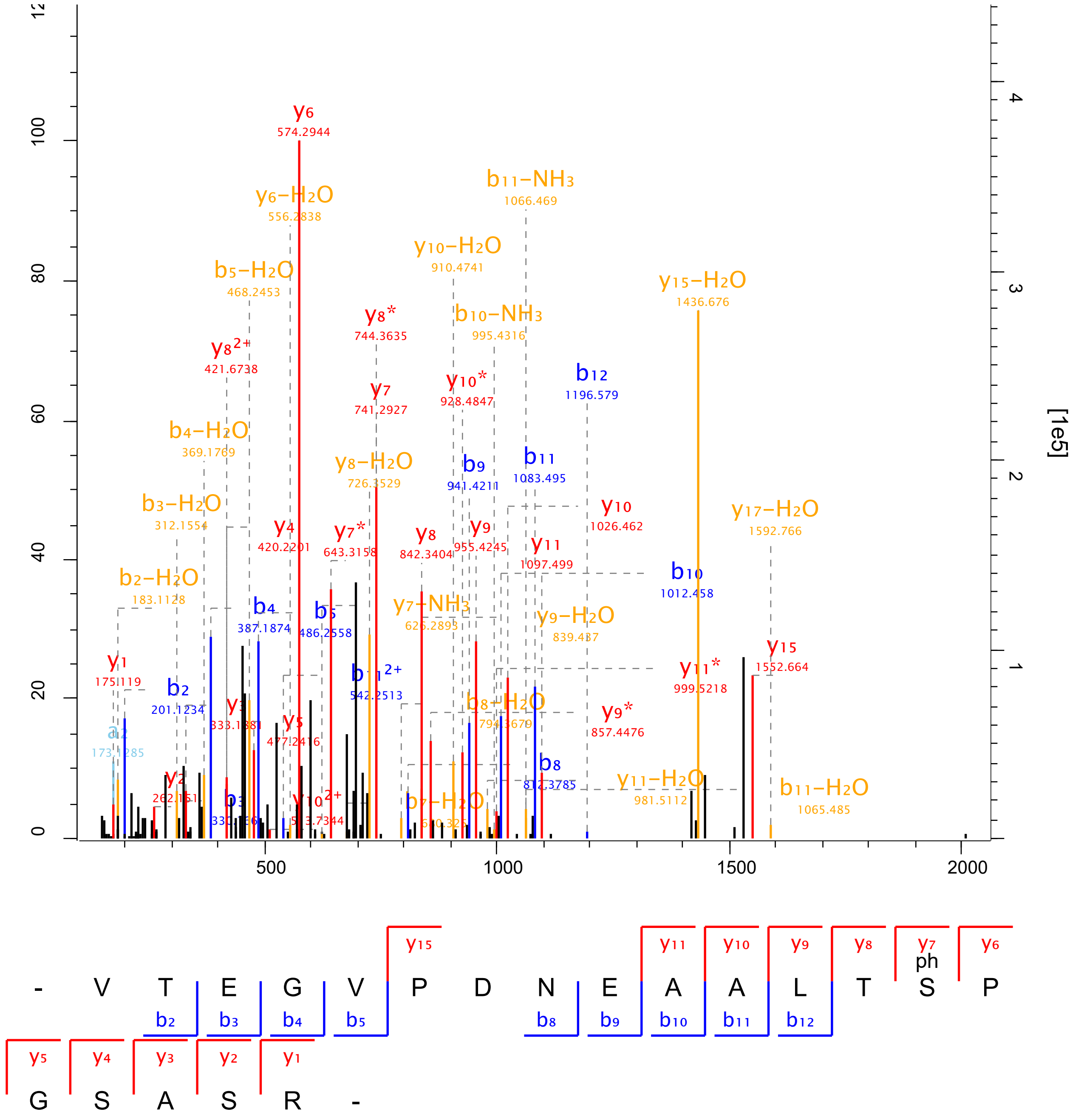

| Raw file | Scan | Method | Score | m/z |
| --- | --- | --- | --- | --- |
| sys_15_1 | 21863 | FTMS; HCD | 108.63 | 750.82 |

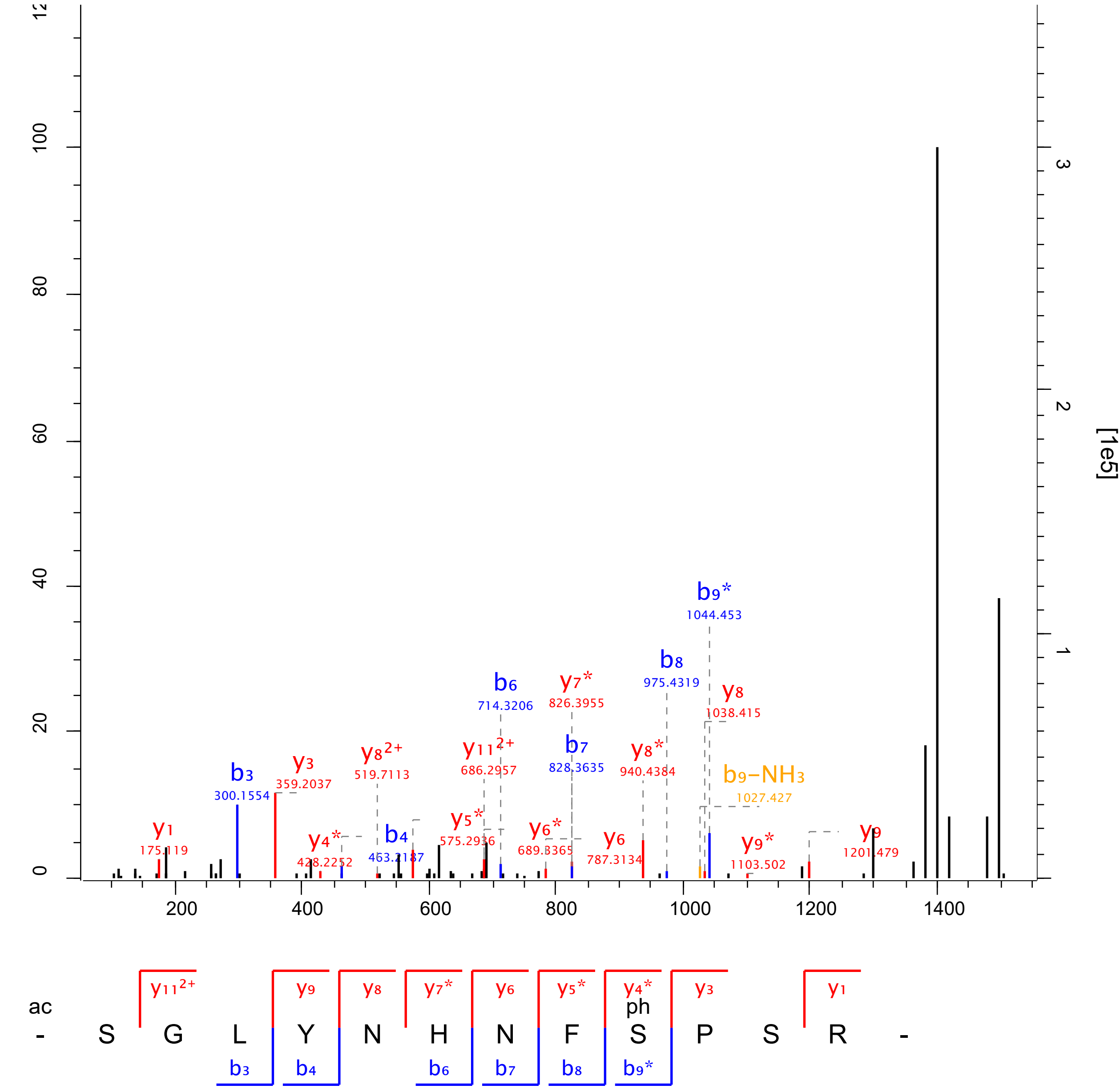

| Raw file | Scan | Method | Score | m/z |
| --- | --- | --- | --- | --- |
| sys_15_1 | 21888 | FTMS; HCD | 70.09 | 587.29 |

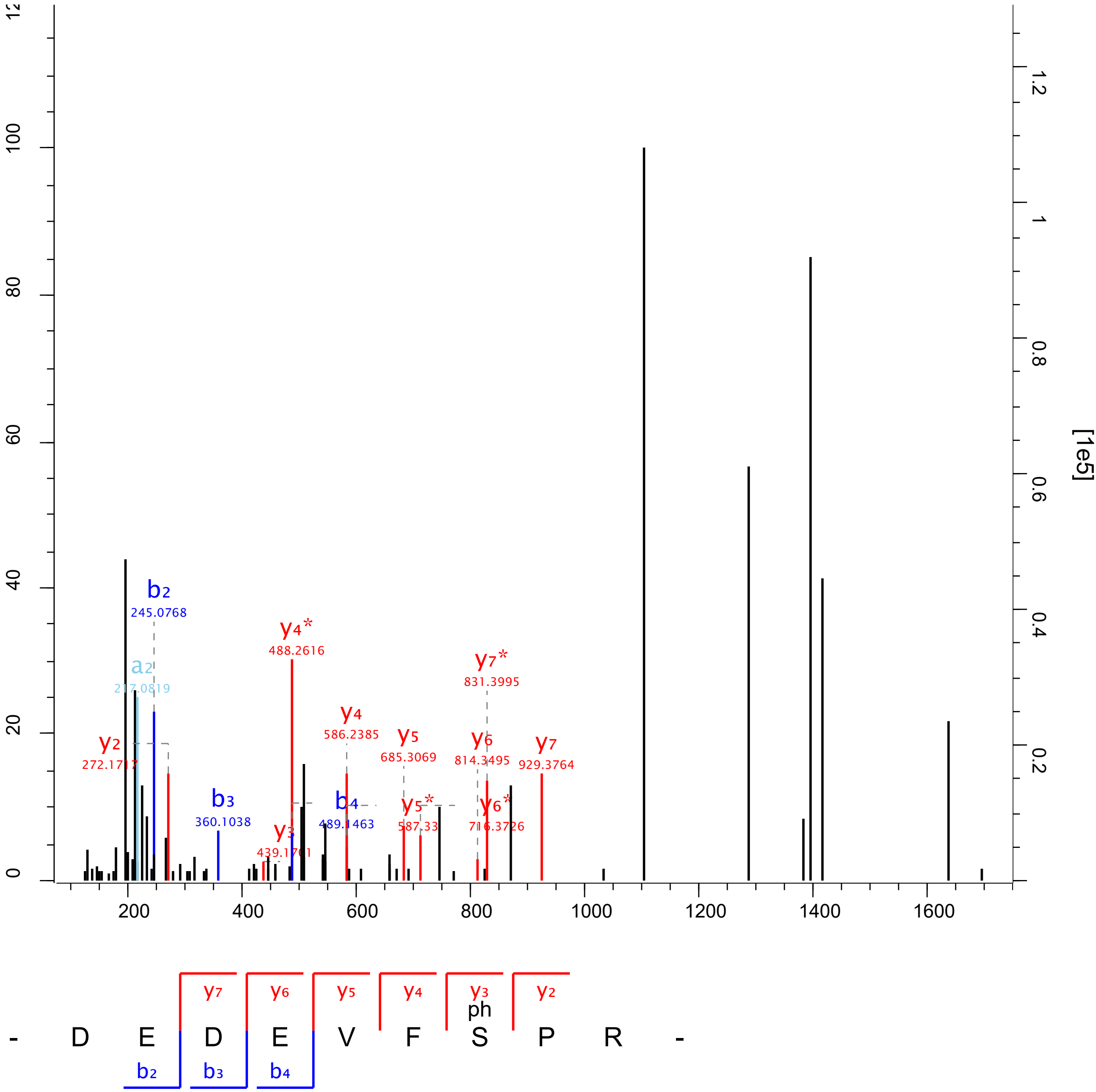

| Raw file | Scan | Method | Score | m/z |
| --- | --- | --- | --- | --- |
| sys_15_1 | 21920 | FTMS; HCD | 58.32 | 761.29 |

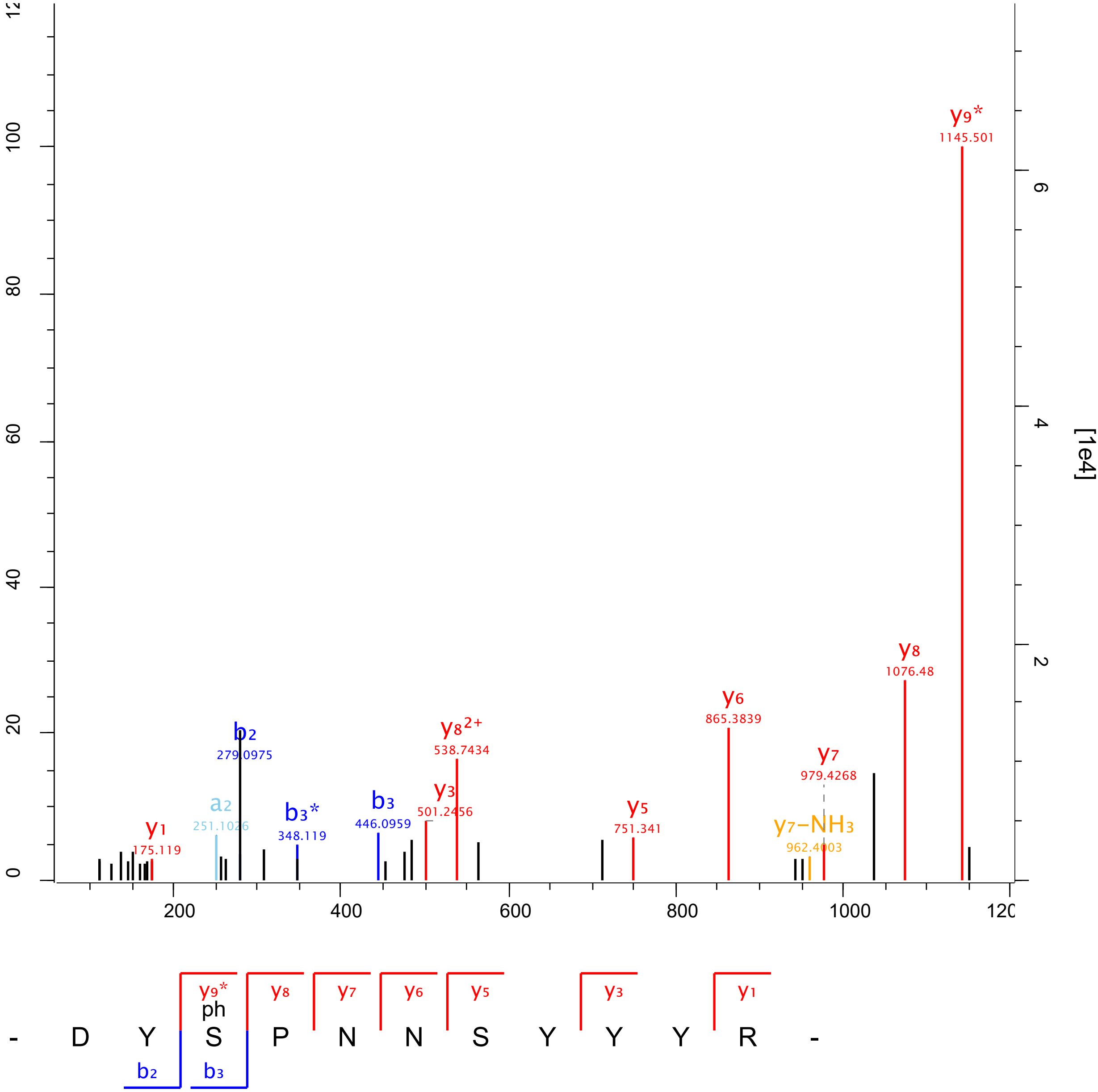

| Raw file | Scan | Method | Score | m/z |
|---|---|---|---|---|
| sys_15_1 | 21946 | FTMS; HCD | 96.21 | 796.35 |

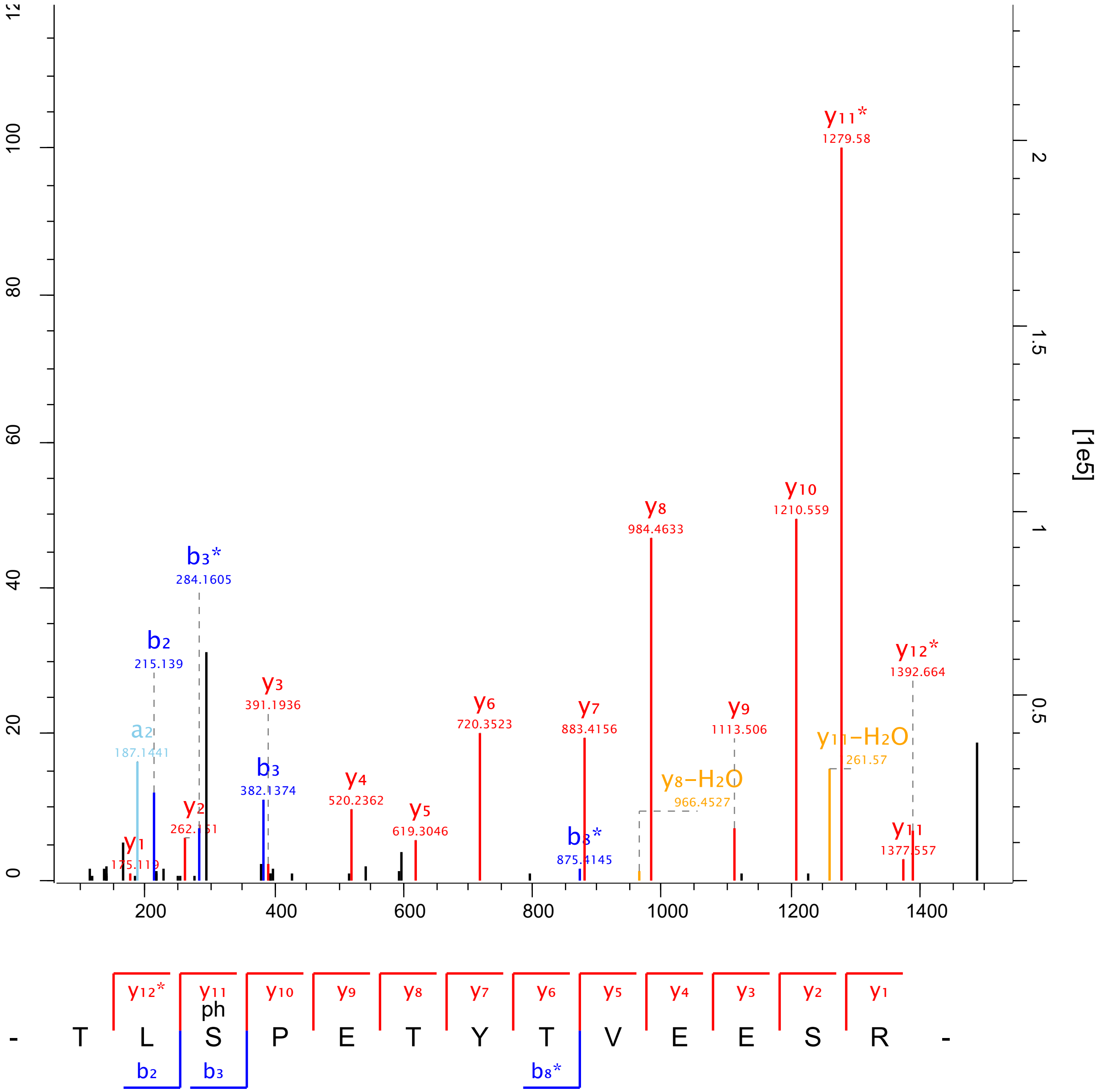

| Raw file | Scan | Method | Score | m/z |
| --- | --- | --- | --- | --- |
| sys_15_1 | 22049 | FTMS; HCD | 69.72 | 480.73 |

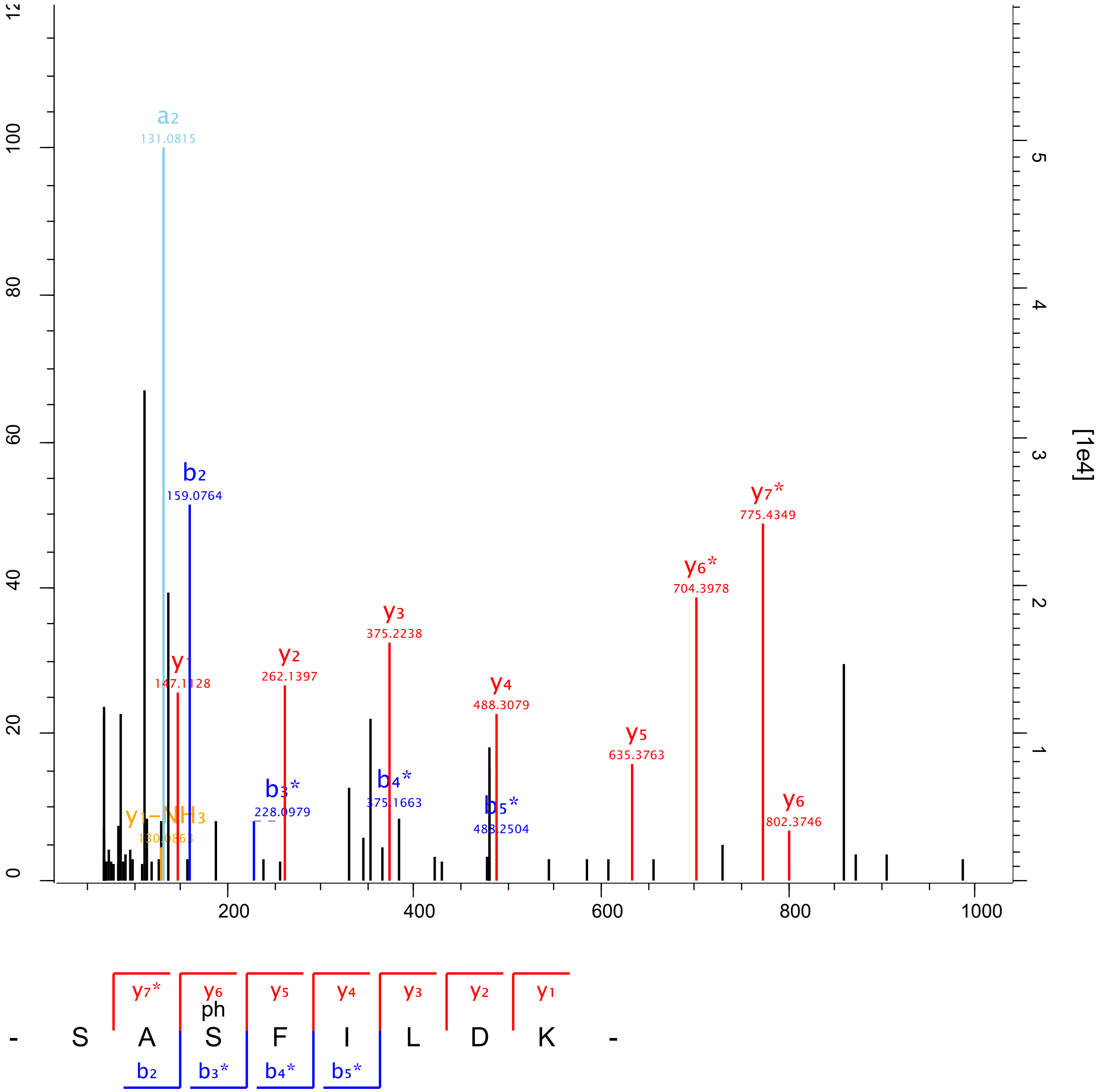

| Raw file | Scan | Method | Score | m/z |
| --- | --- | --- | --- | --- |
| sys_15_1 | 22102 | FTMS; HCD | 107.99 | 649.82 |

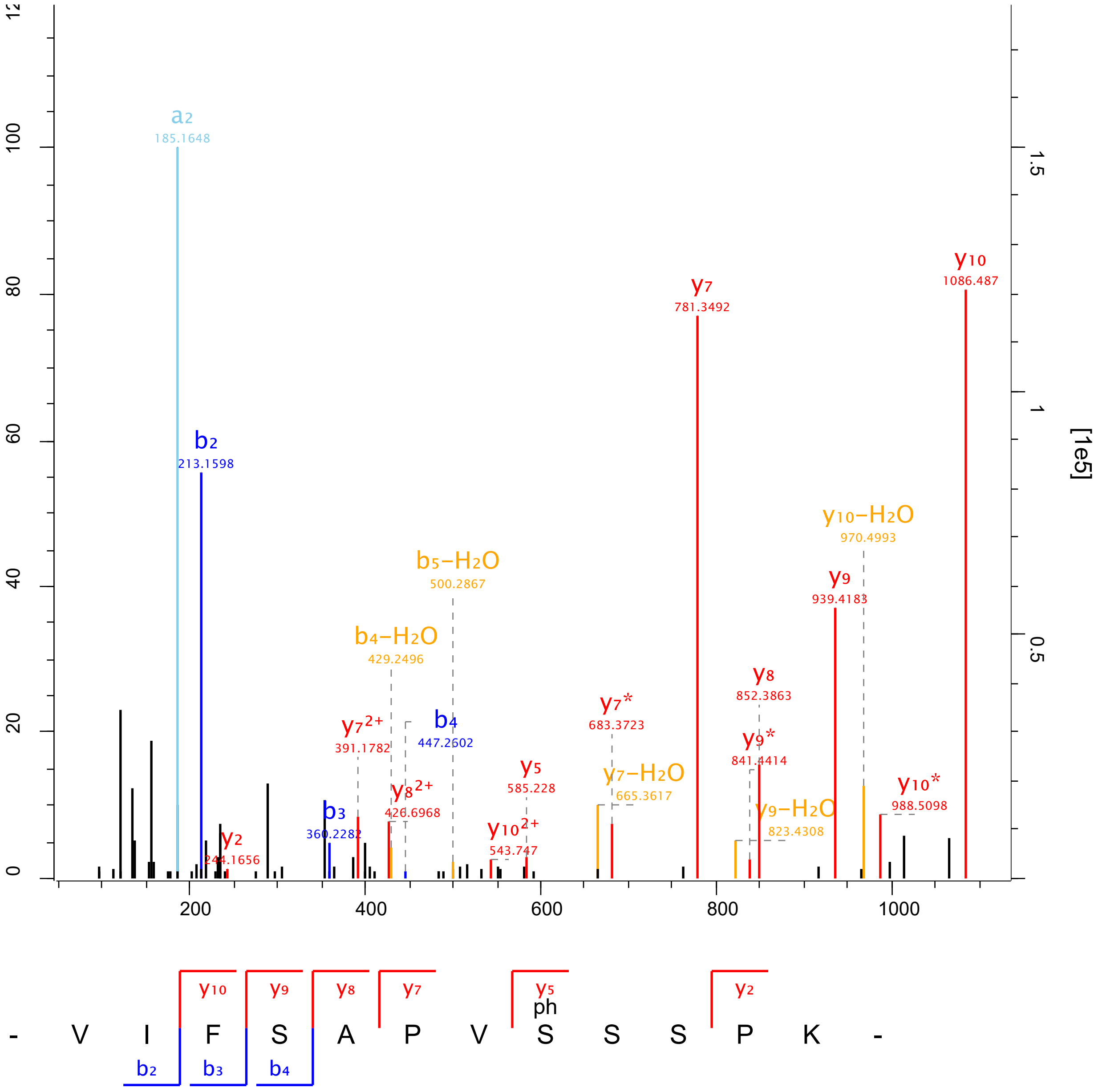

| Raw file | Scan | Method | Score | m/z |
| --- | --- | --- | --- | --- |
| sys_15_1 | 22126 | FTMS; HCD | 86.8 | 1005.38 |

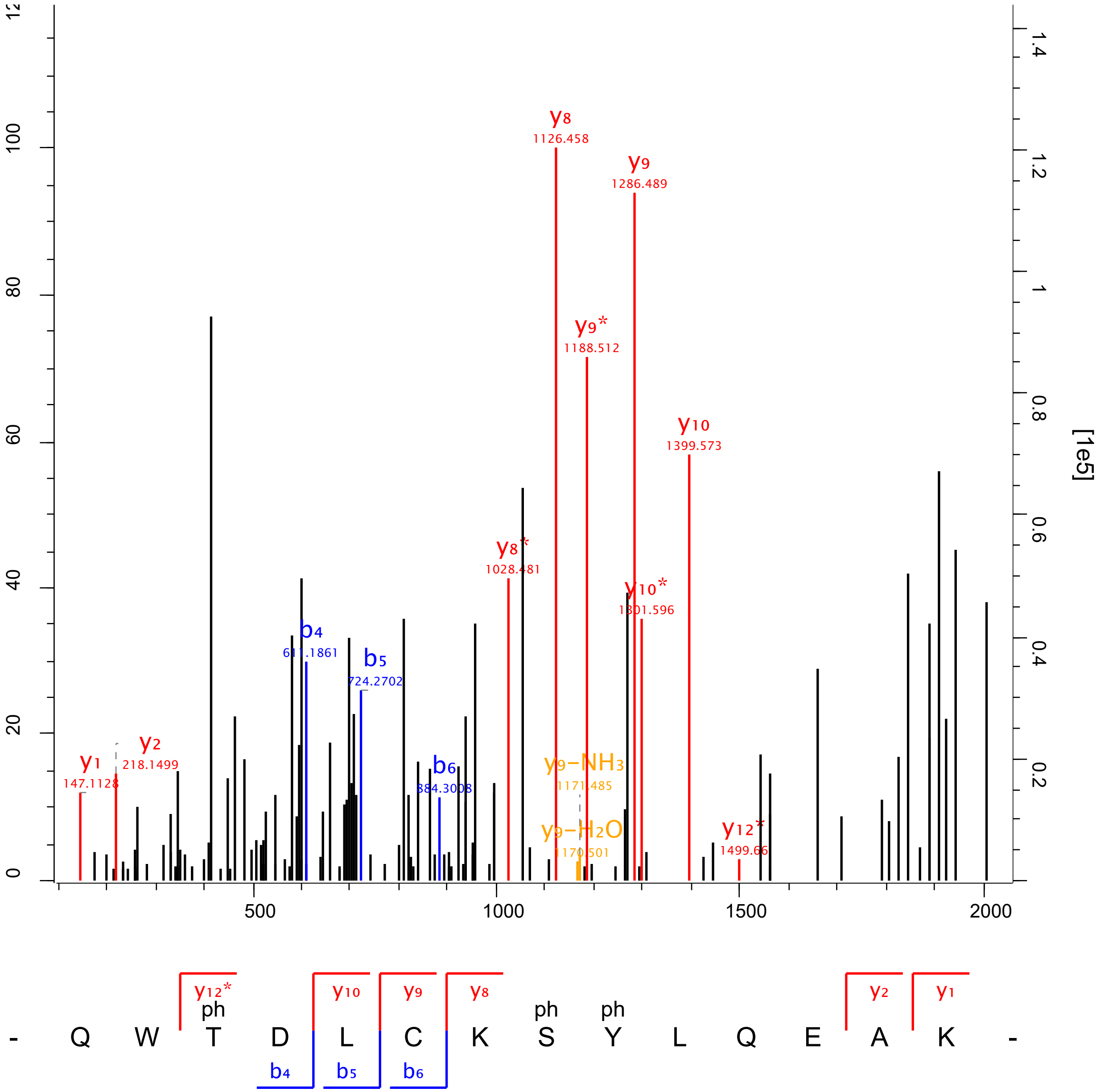

| Raw file | Scan | Method | Score | m/z |
|---|---|---|---|---|
| sys_15_1 | 22130 | FTMS; HCD | 174.78 | 810.89 |

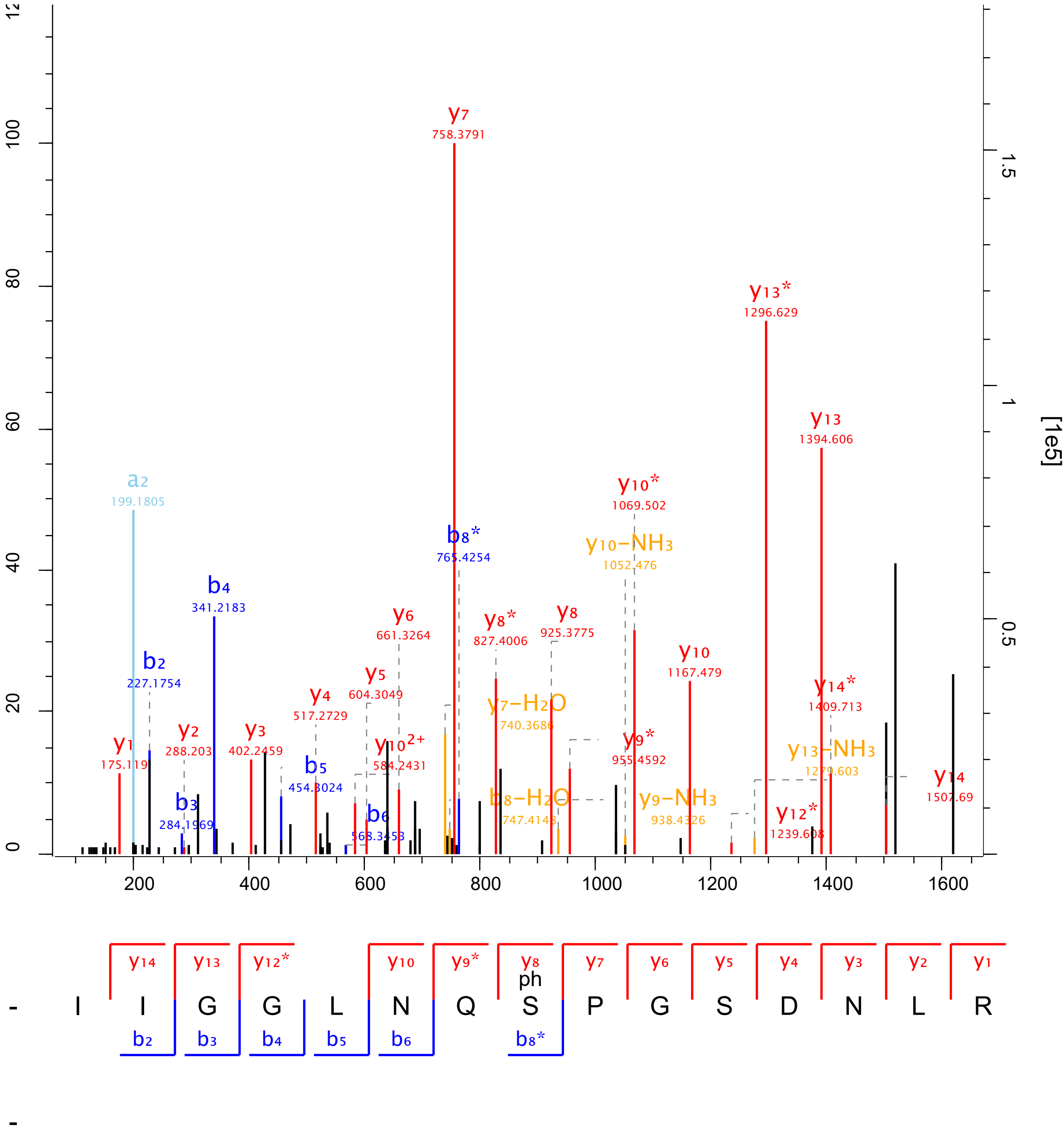

| Raw file | Scan | Method | Score | m/z |
| --- | --- | --- | --- | --- |
| sys_15_1 | 22140 | FTMS; HCD | 55.01 | 599.6 |

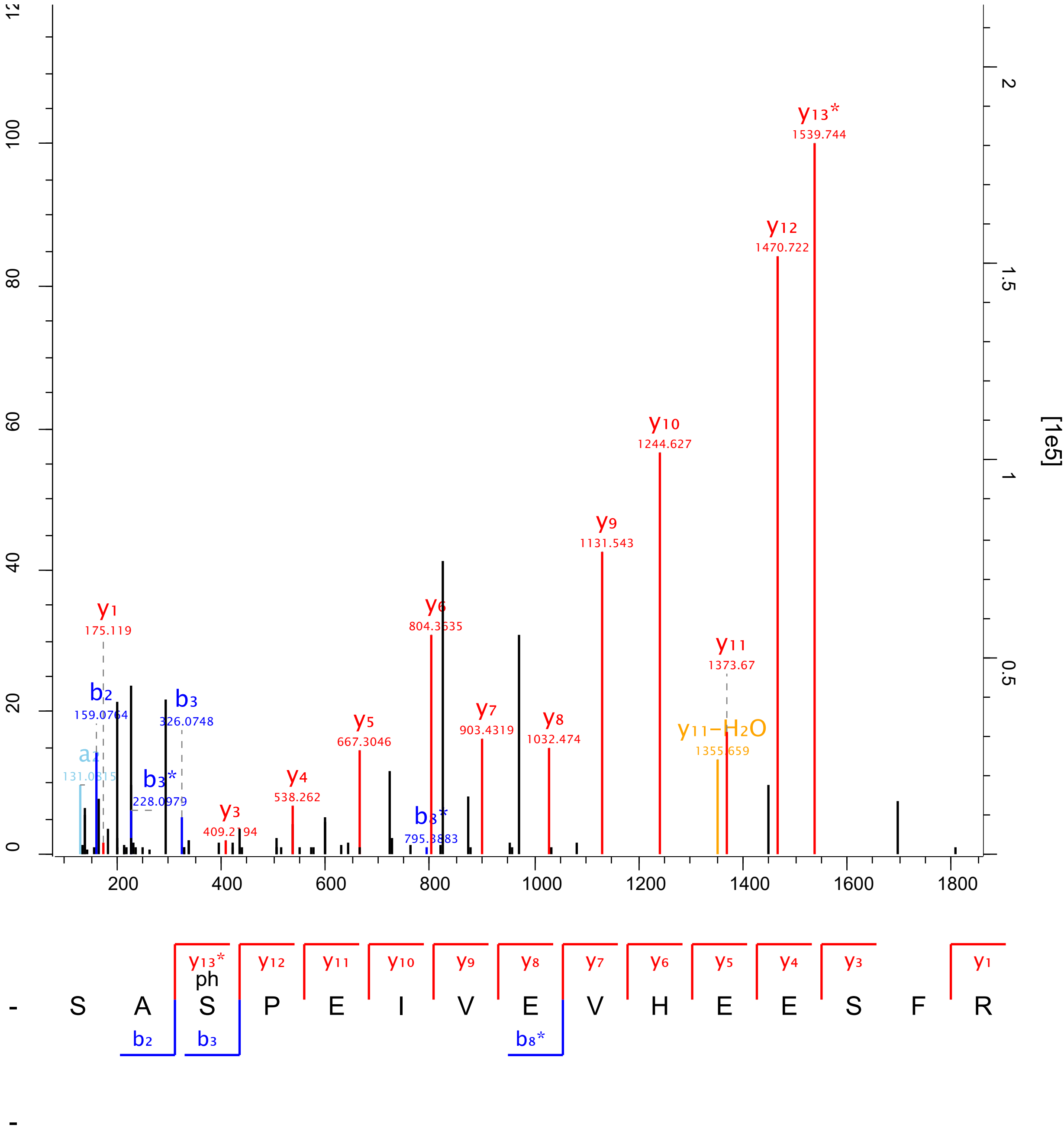

| Raw file | Scan | Method | Score | m/z |
| --- | --- | --- | --- | --- |
| sys_15_1 | 22159 | FTMS; HCD | 70.45 | 909.39 |

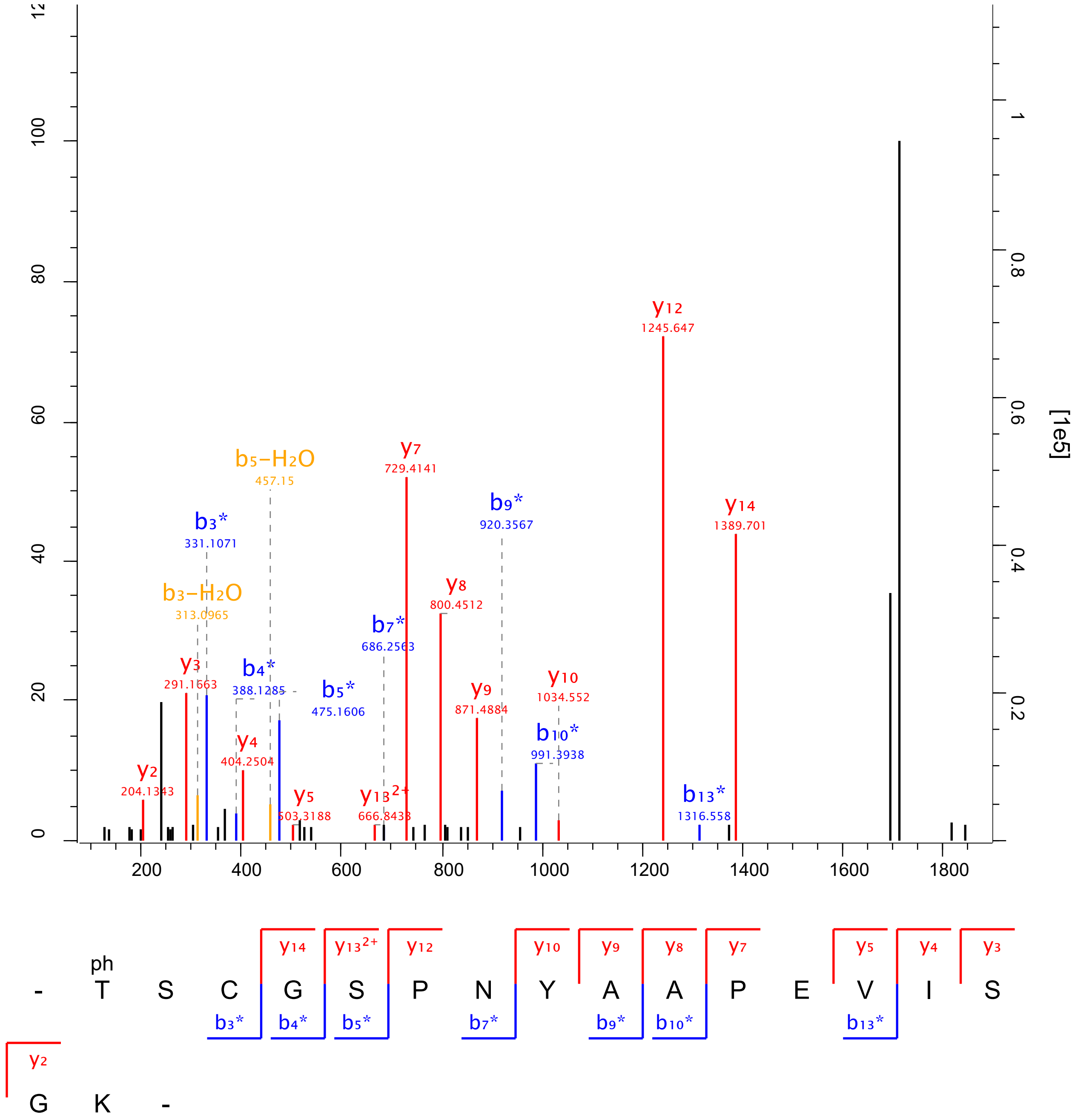

| Raw file | Scan | Method | Score | m/z |
|---|---|---|---|---|
| sys_15_1 | 22245 | FTMS; HCD | 150.34 | 723.33 |

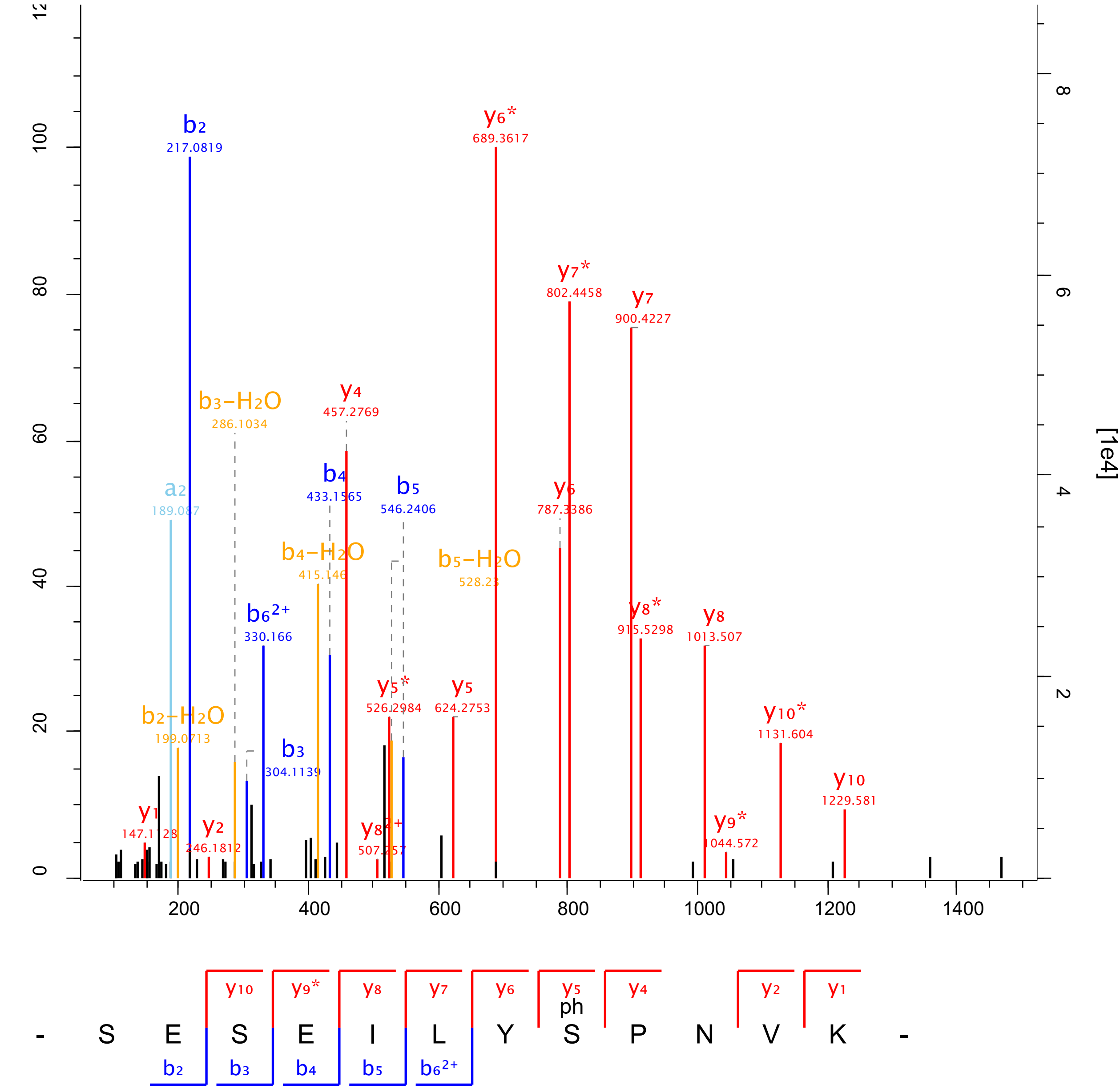

| Raw file | Scan | Method | Score | m/z |
| --- | --- | --- | --- | --- |
| sys_15_1 | 22365 | FTMS; HCD | 134.52 | 827.82 |

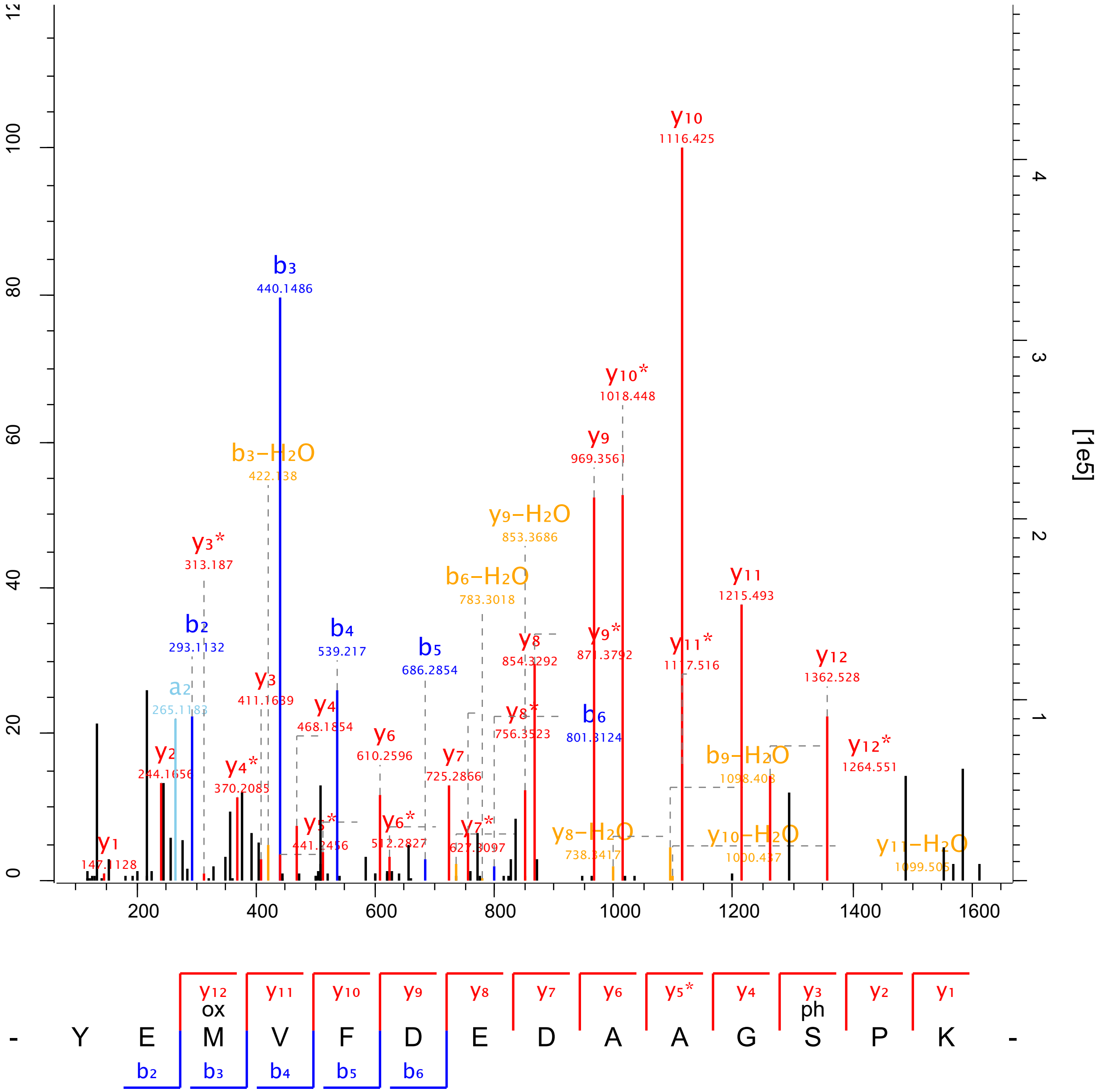

| Raw file | Scan | Method | Score | m/z |
| --- | --- | --- | --- | --- |
| sys_15_1 | 2250 | FTMS; HCD | 48.57 | 405.5 |

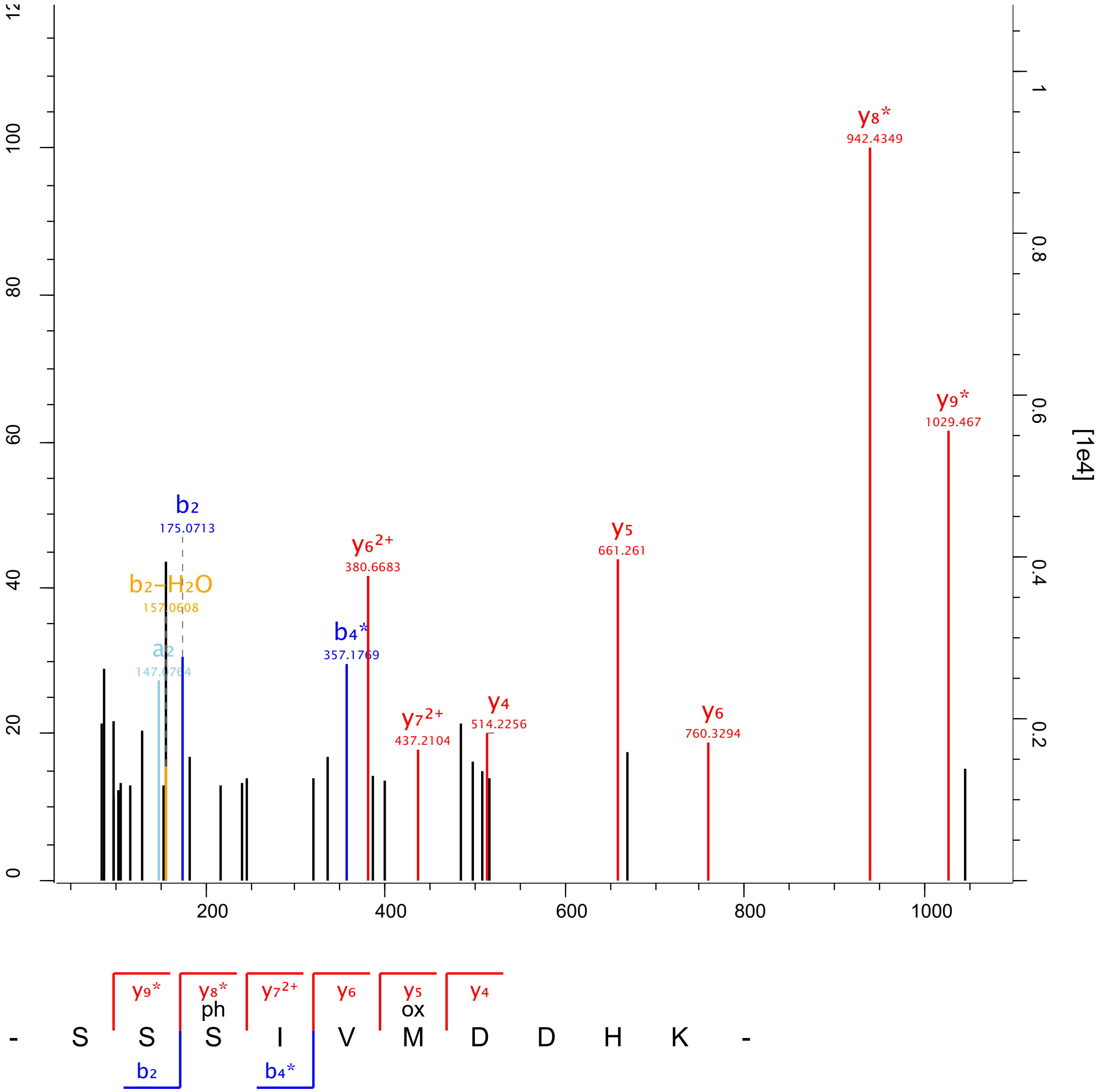

| Raw file | Scan | Method | Score | m/z |
|---|---|---|---|---|
| sys_15_1 | 22527 | FTMS; HCD | 59.71 | 636.8 |

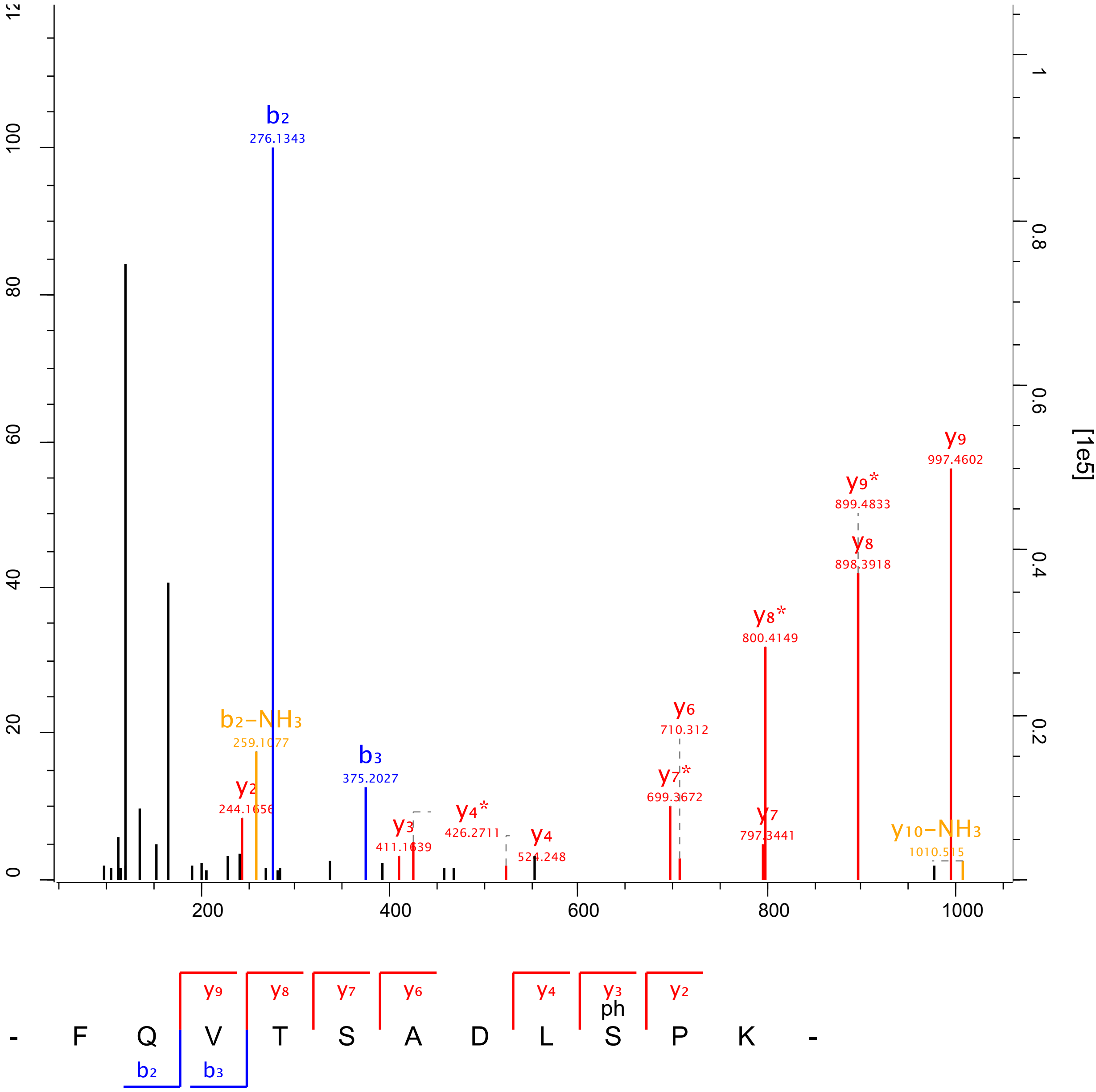

| Raw file | Scan | Method | Score | m/z |
| --- | --- | --- | --- | --- |
| sys_15_1 | 22543 | FTMS; HCD | 63.19 | 436.91 |

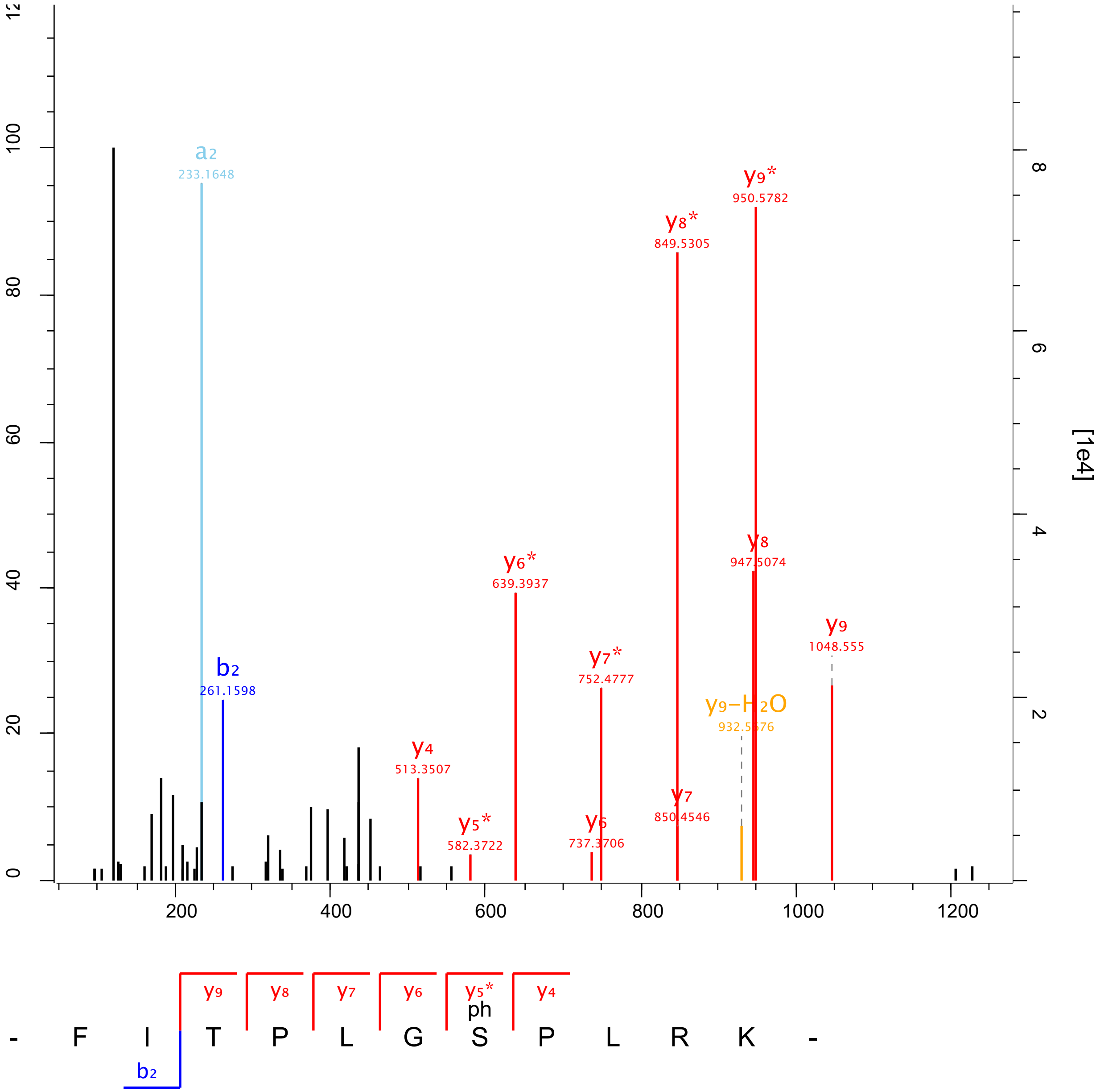

| Raw file | Scan | Method | Score | m/z |
|---|---|---|---|---|
| sys_15_1 | 22596 | FTMS; HCD | 42.47 | 581.74 |

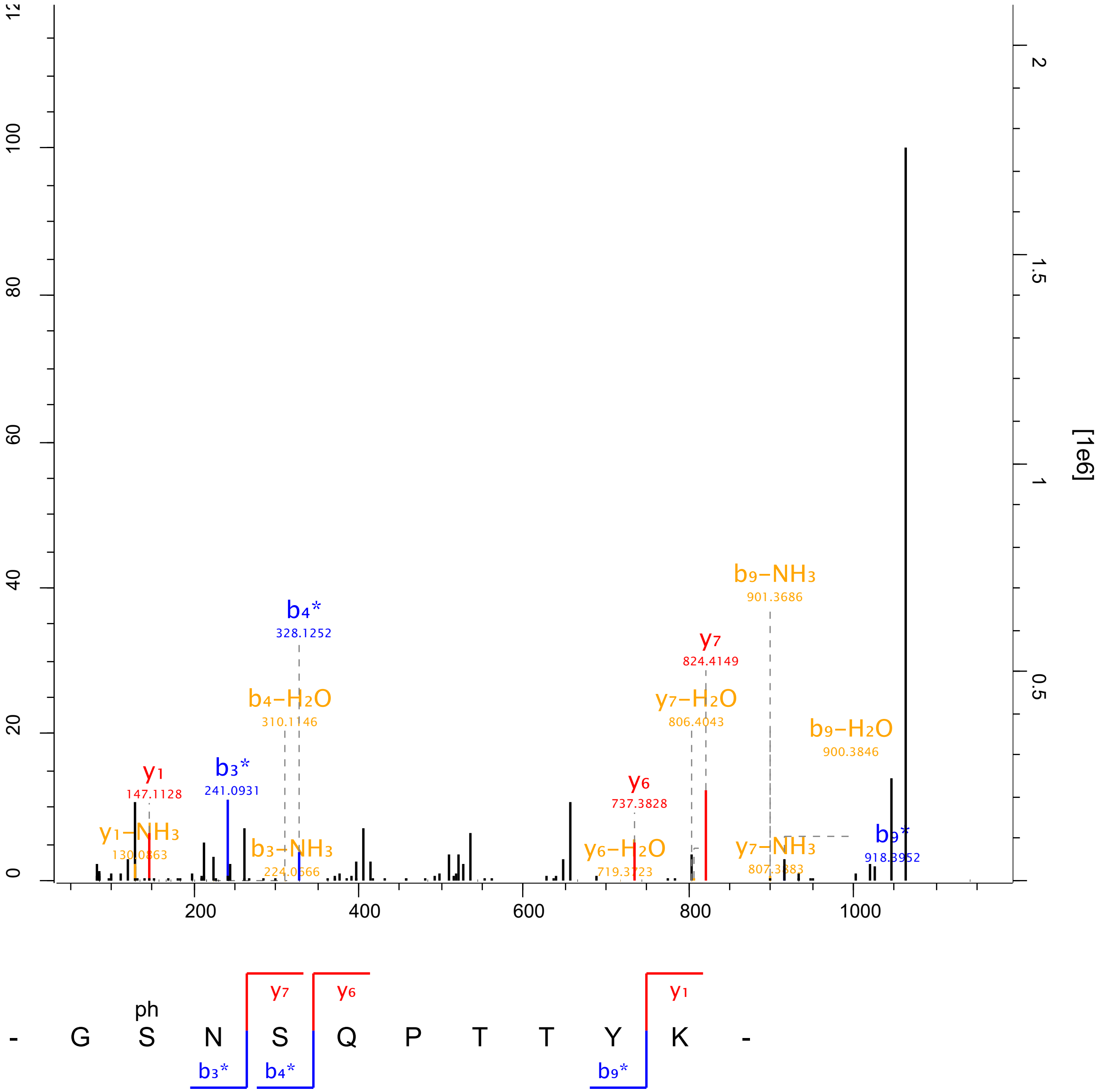

| Raw file | Scan | Method | Score | m/z |
| --- | --- | --- | --- | --- |
| sys_15_1 | 22703 | FTMS; HCD | 151.42 | 626.96 |

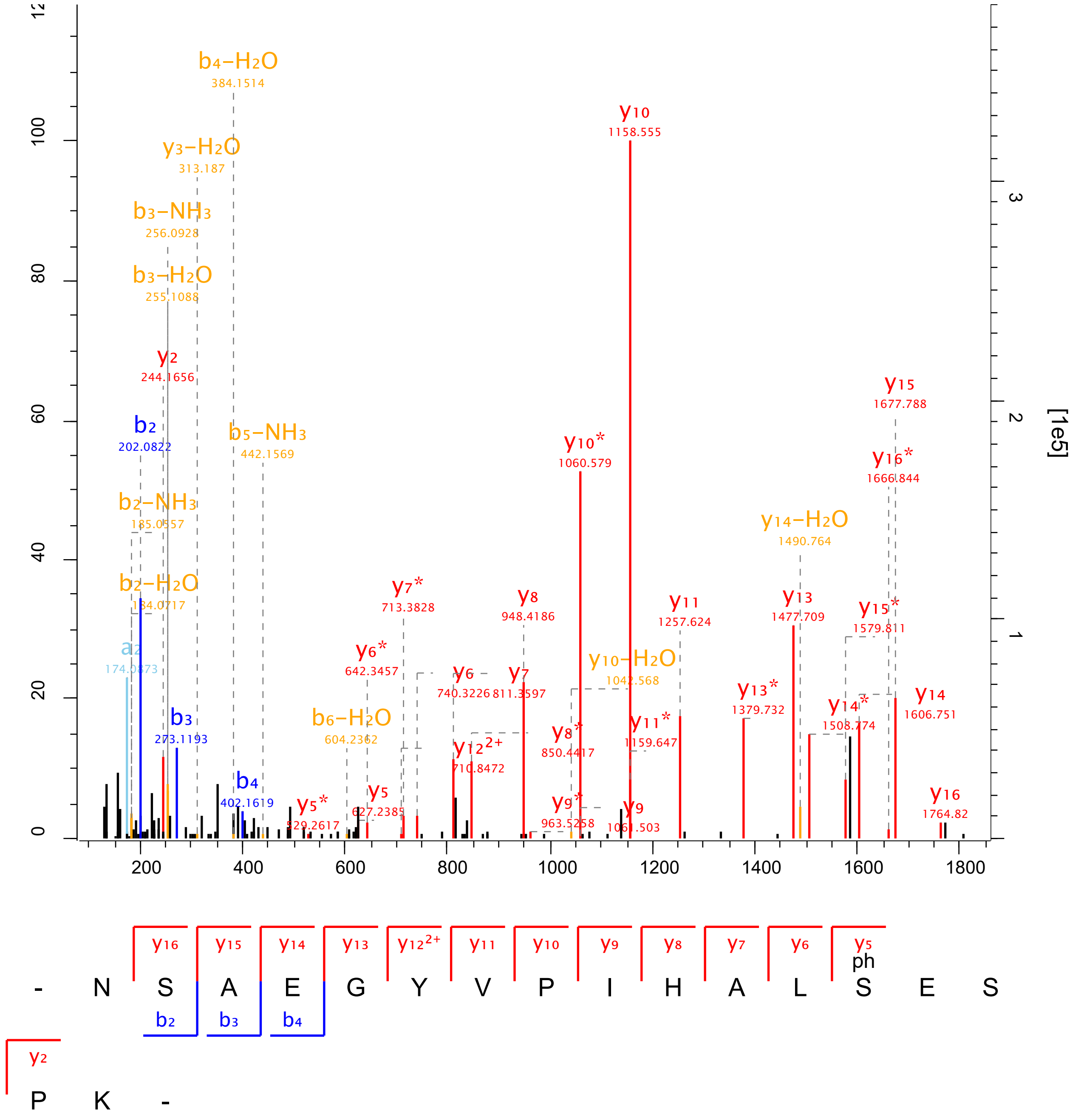

| Raw file | Scan | Method | Score | m/z |
|---|---|---|---|---|
| sys_15_1 | 22916 | FTMS; HCD | 71.65 | 735.98 |

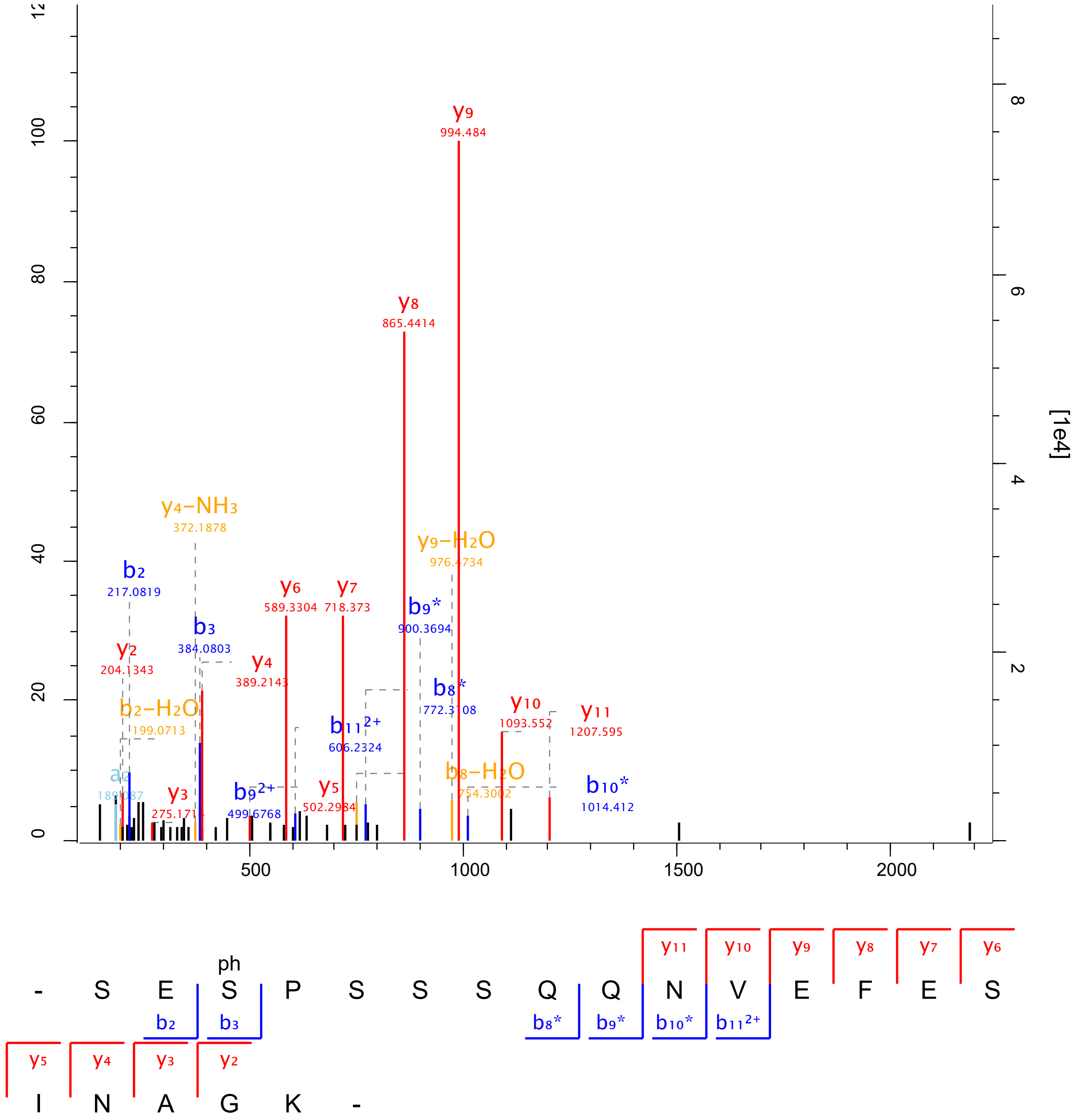

| Raw file | Scan | Method | Score | m/z |
|---|---|---|---|---|
| sys_15_1 | 22976 | FTMS; HCD | 68.41 | 618.63 |

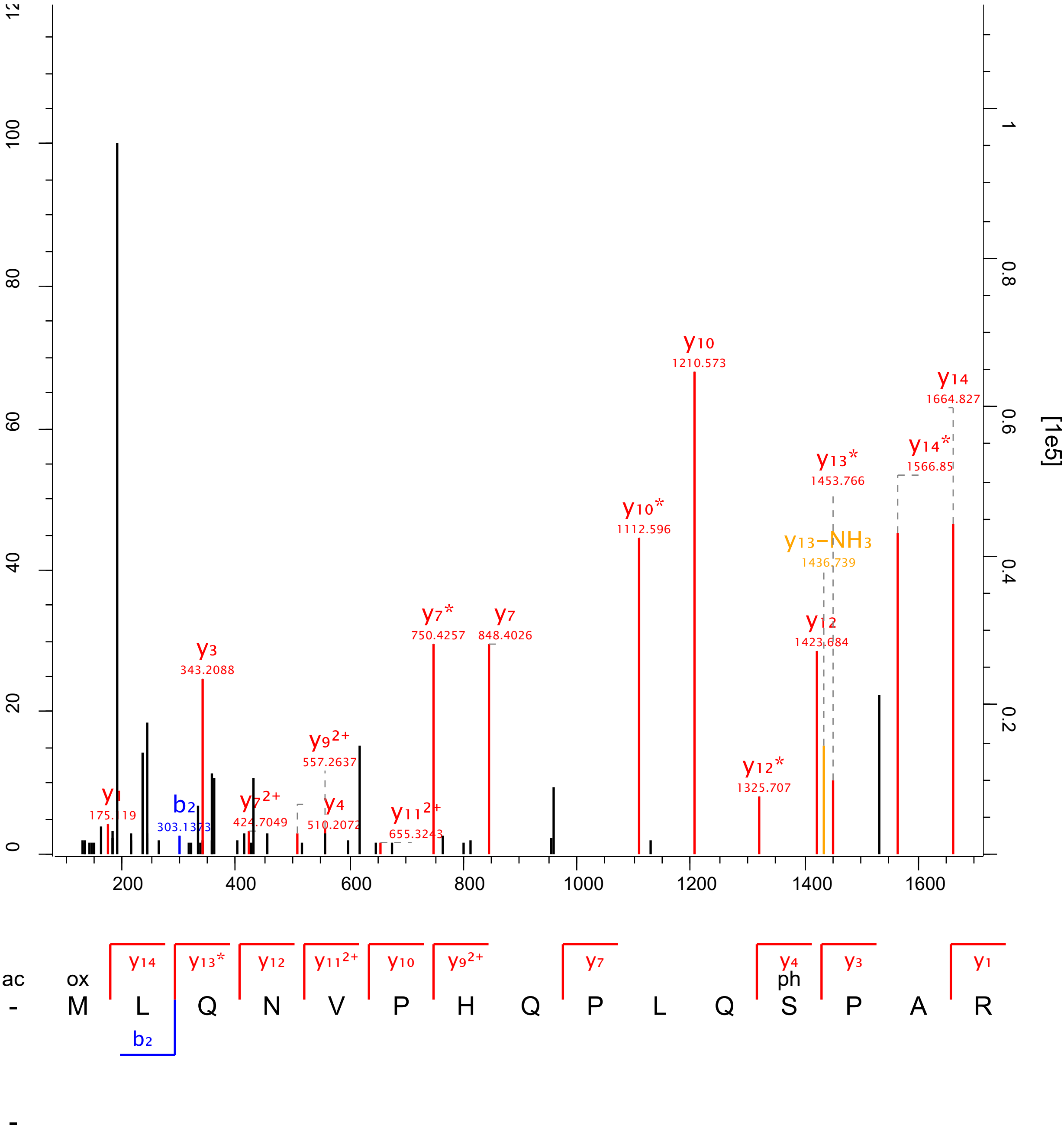

| Raw file | Scan | Method | Score | m/z |
|---|---|---|---|---|
| sys_15_1 | 23027 | FTMS; HCD | 145.46 | 509.73 |

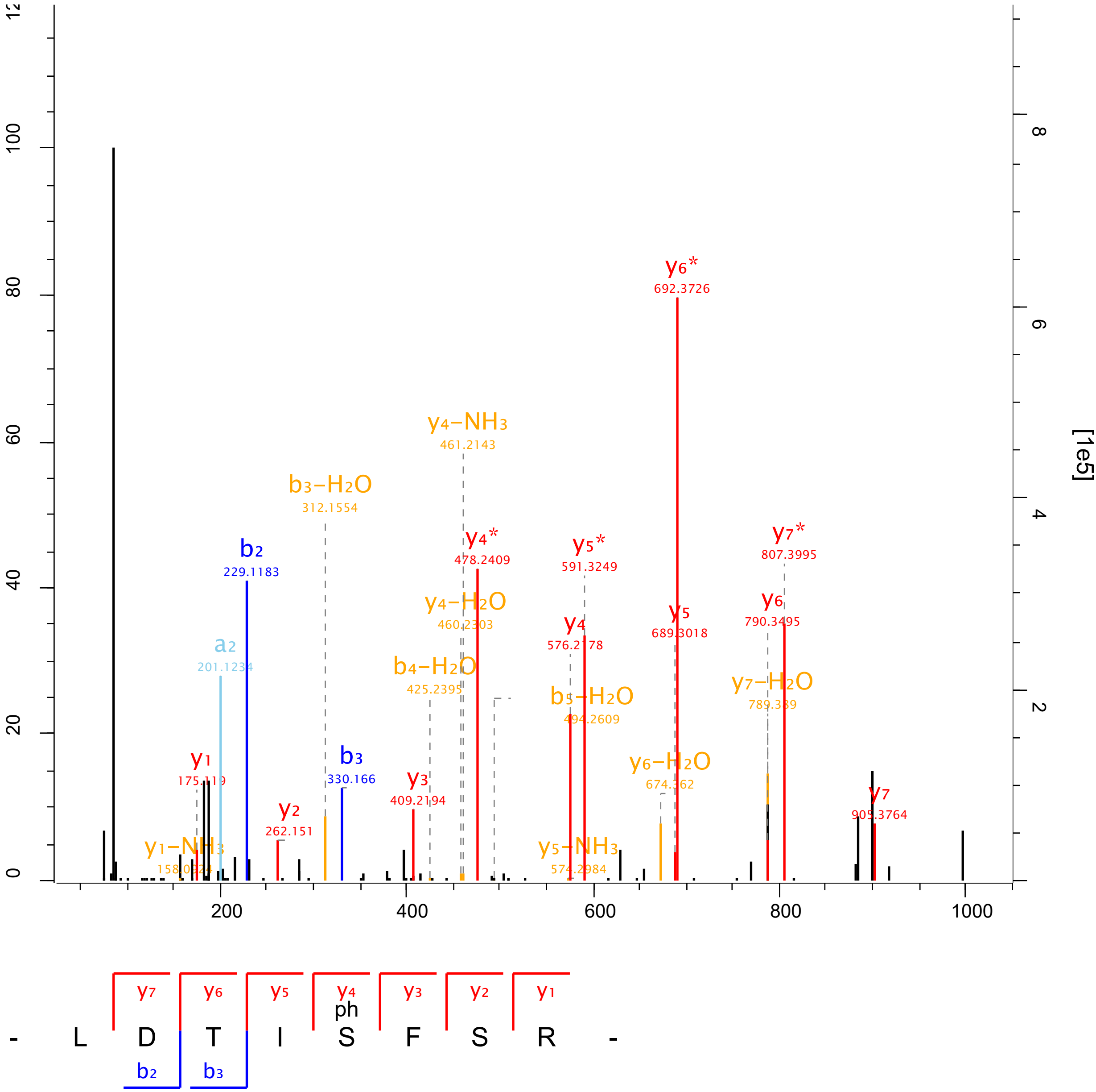

| Raw file | Scan | Method | Score | m/z |
|---|---|---|---|---|
| sys_15_1 | 23150 | FTMS; HCD | 224.86 | 827.89 |

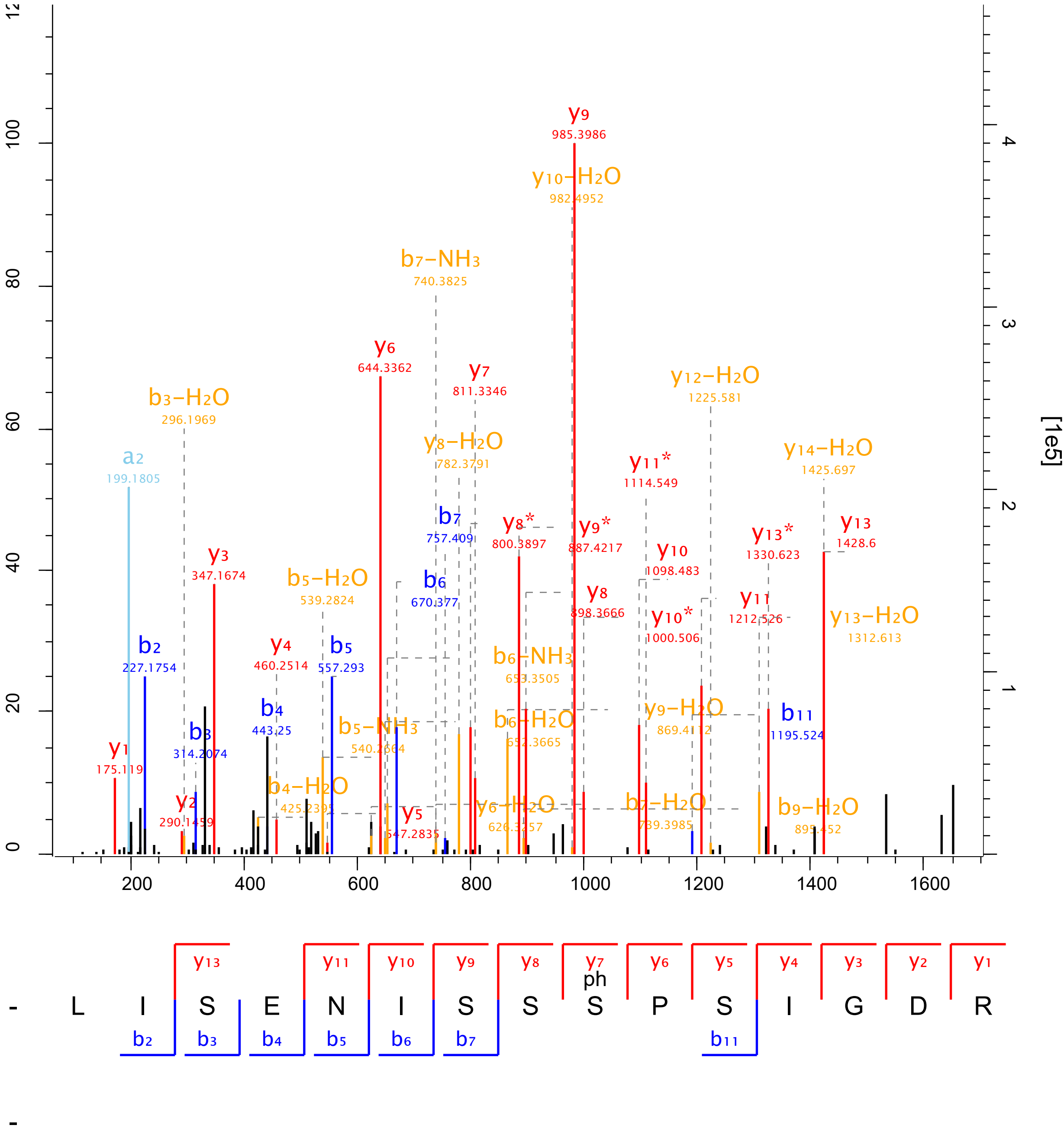

| Raw file | Scan | Method | Score | m/z |
|---|---|---|---|---|
| sys_15_1 | 2320 | FTMS; HCD | 85.55 | 555.7 |

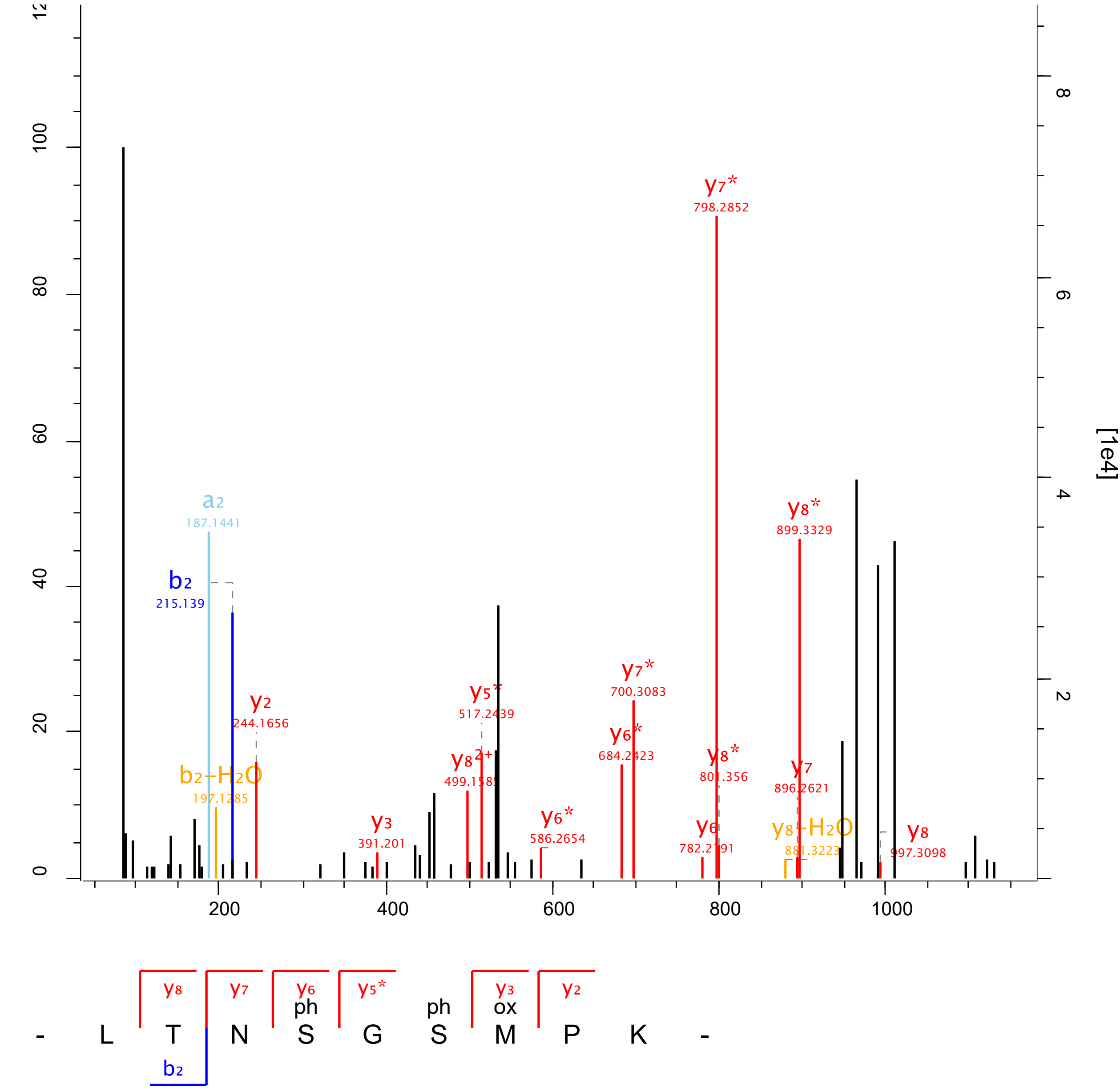

| Raw file | Scan | Method | Score | m/z |
| --- | --- | --- | --- | --- |
| sys_15_1 | 23221 | FTMS; HCD | 127.3 | 768.32 |

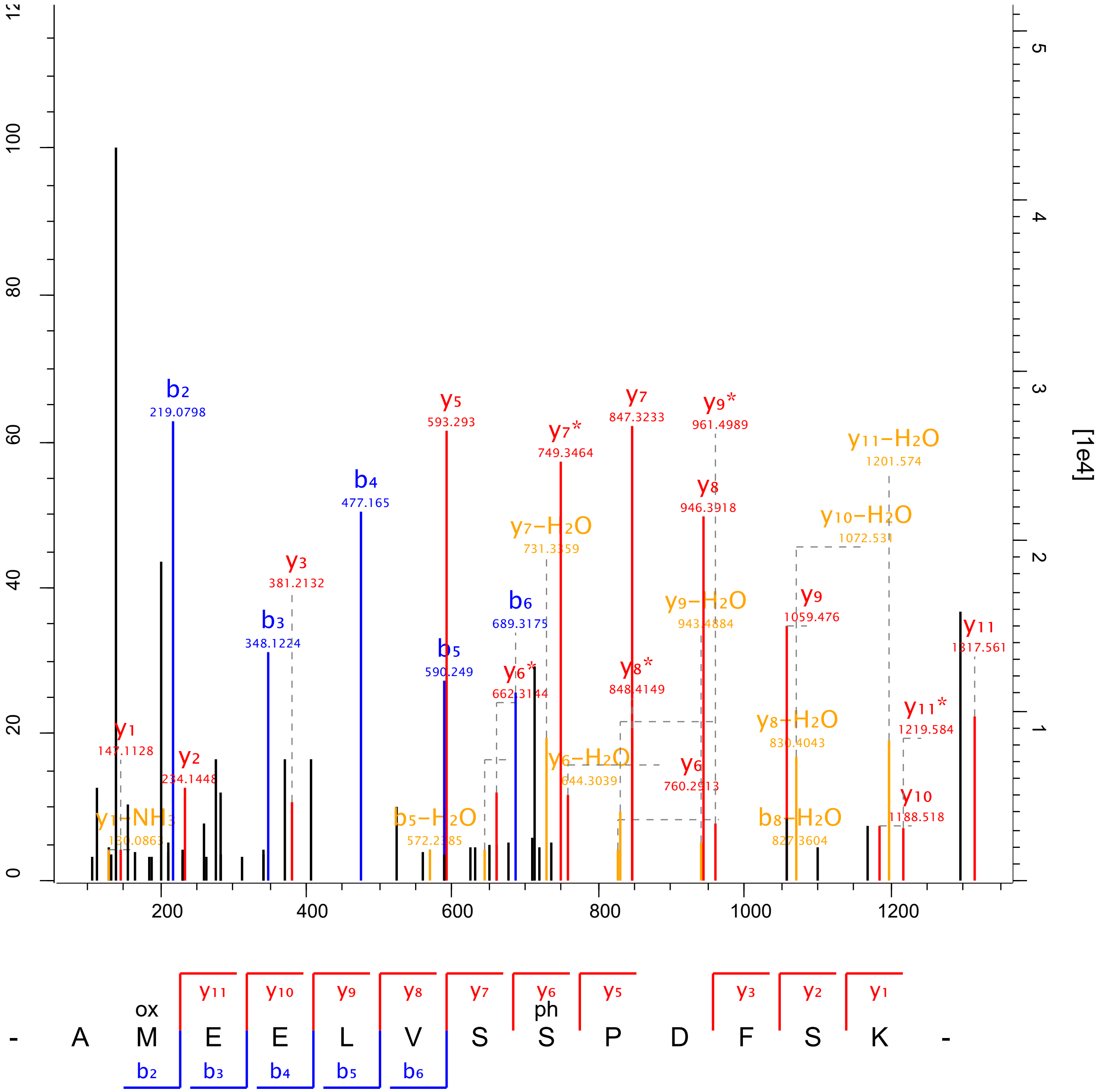

| Raw file | Scan | Method | Score | m/z |
|---|---|---|---|---|
| sys_15_1 | 23270 | FTMS; HCD | 52.86 | 999.93 |

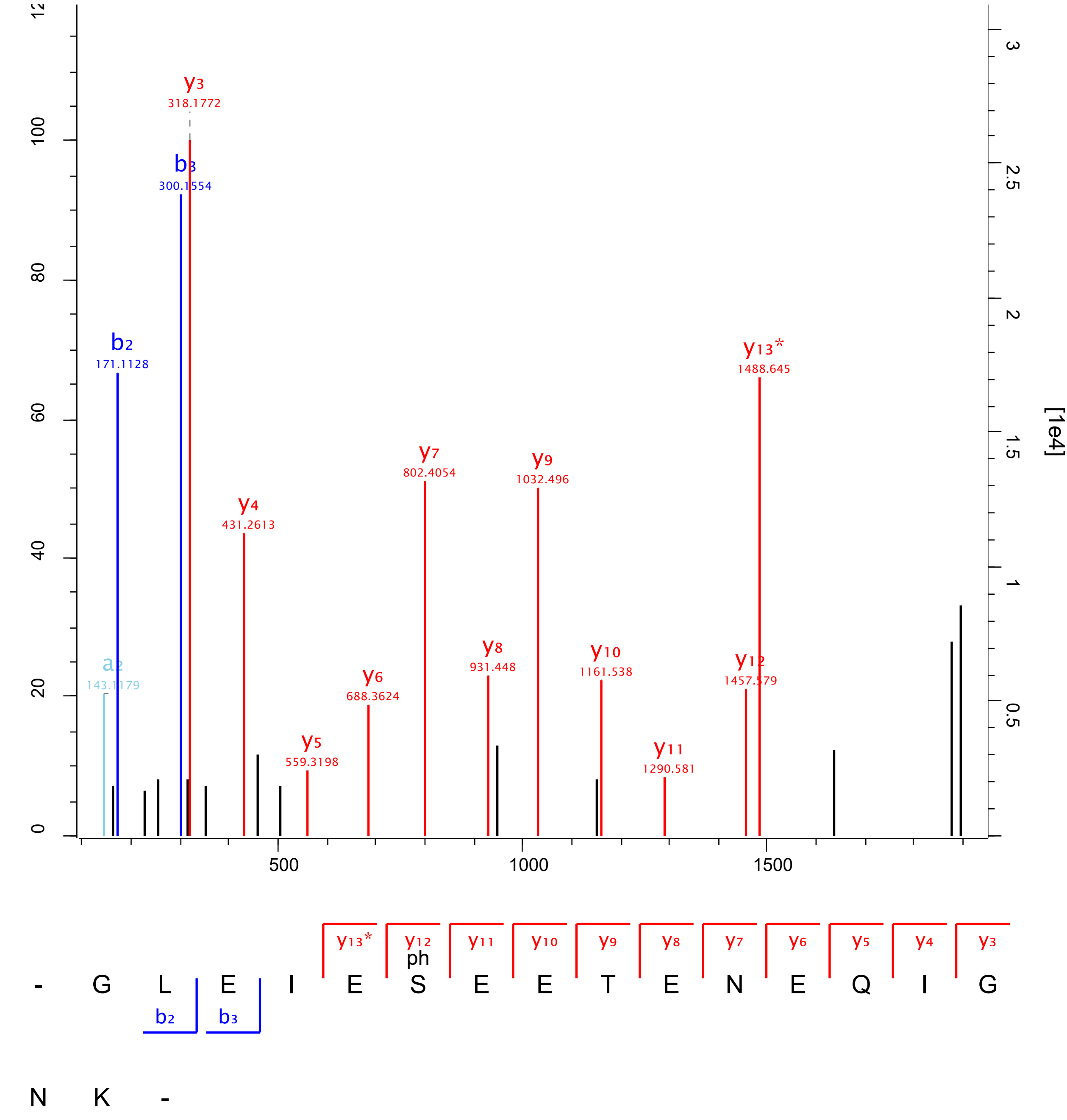

| Raw file | Scan | Method | Score | m/z |
|---|---|---|---|---|
| sys_15_1 | 23342 | FTMS; HCD | 263.1 | 955.41 |

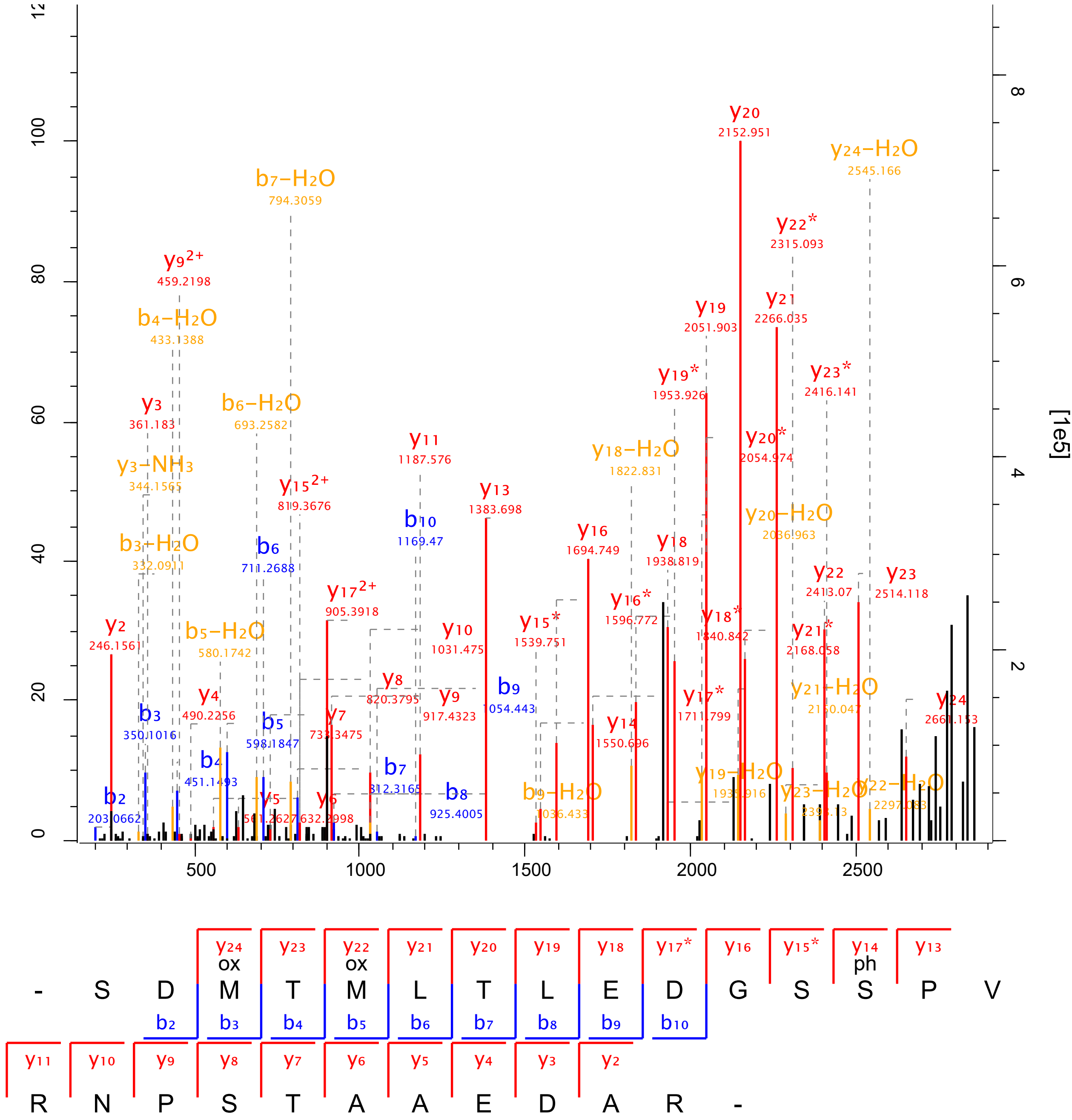

| Raw file | Scan | Method | Score | m/z |
| --- | --- | --- | --- | --- |
| sys_15_1 | 23369 | FTMS; HCD | 100.22 | 576.31 |

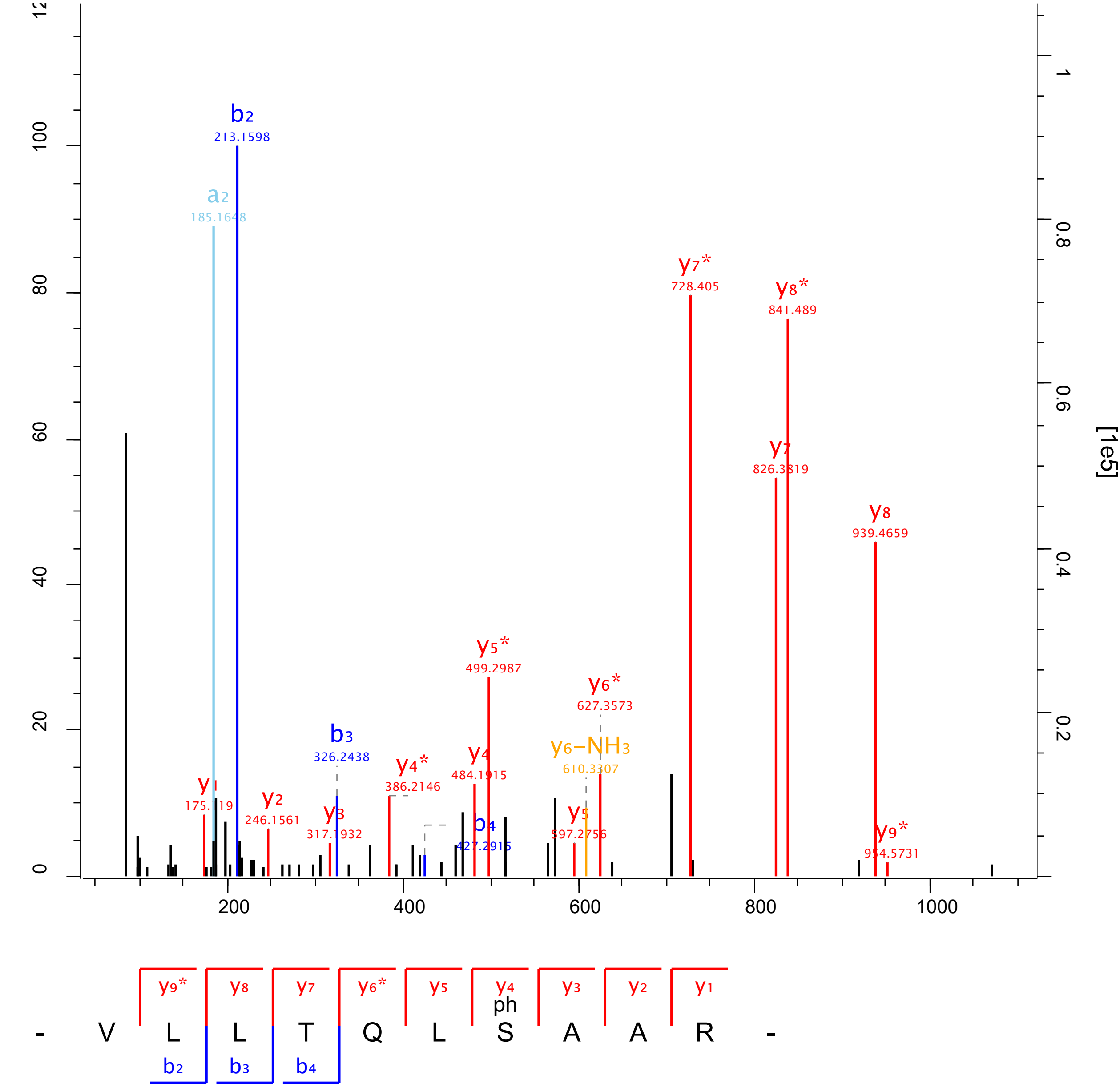

| Raw file | Scan | Method | Score | m/z |
|---|---|---|---|---|
| sys_15_1 | 23406 | FTMS; HCD | 51.57 | 632.63 |

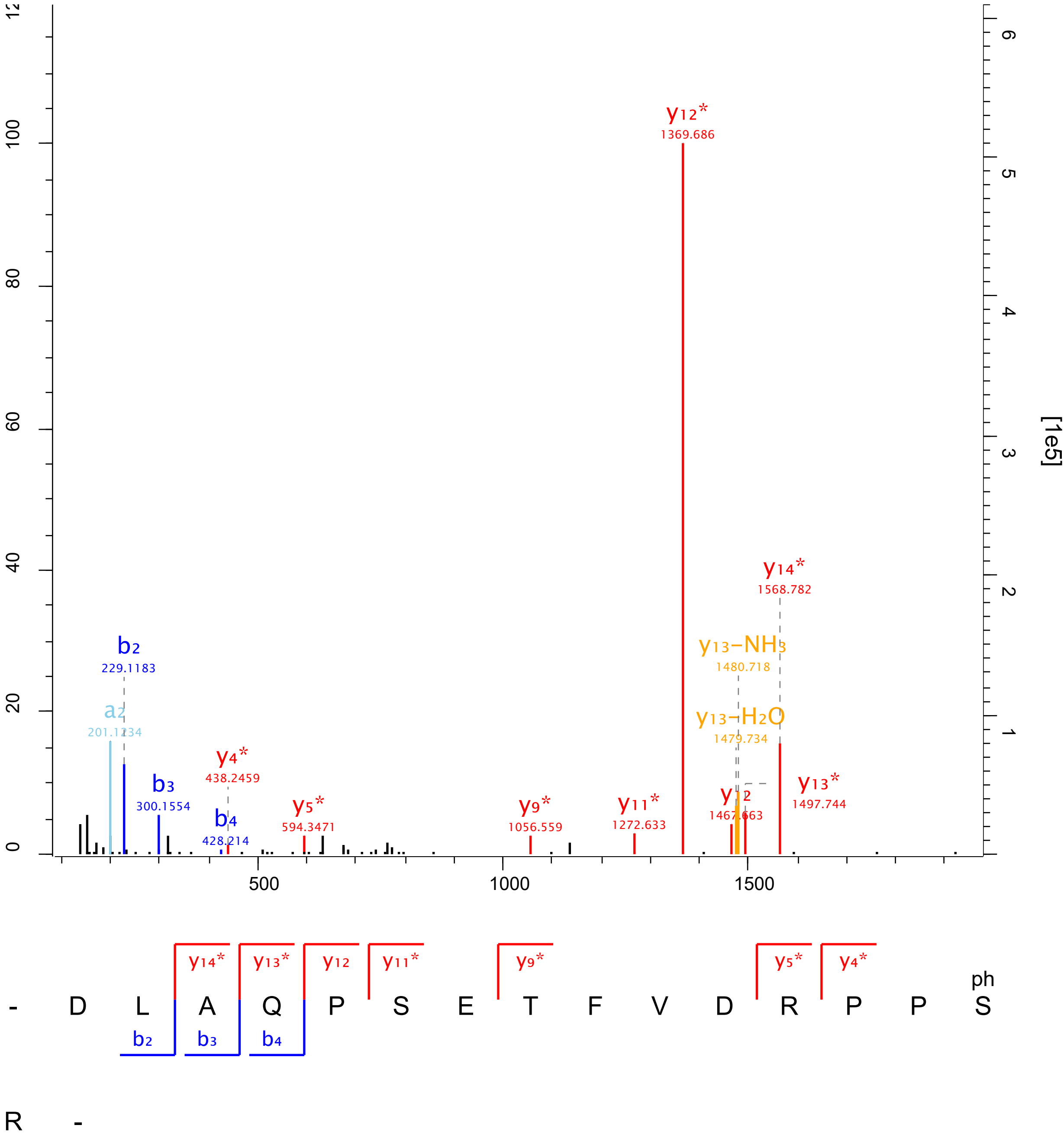

| Raw file | Scan | Method | Score | m/z |
| --- | --- | --- | --- | --- |
| sys_15_1 | 23436 | FTMS; HCD | 43.9 | 524.25 |

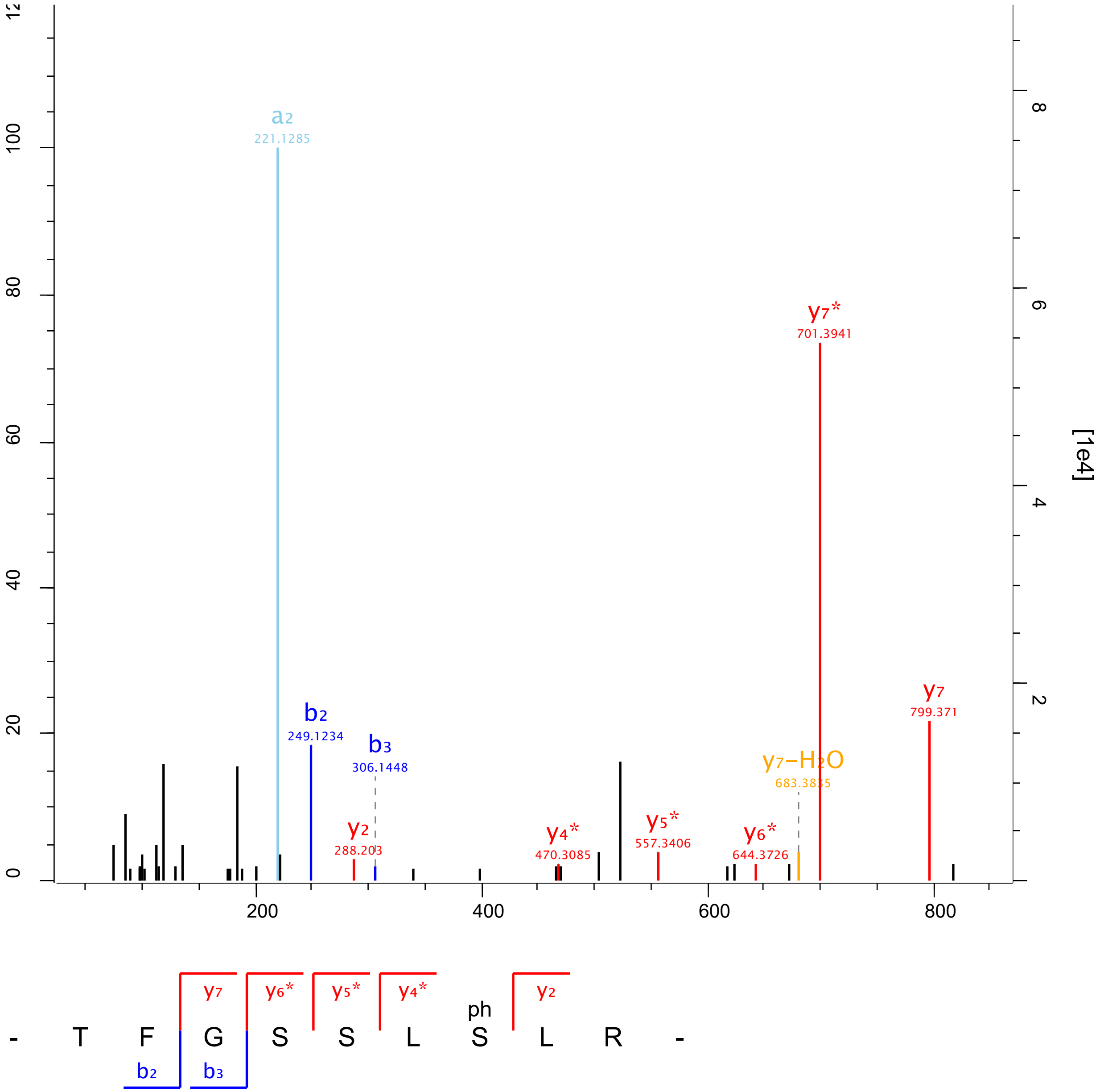

| Raw file | Scan | Method | Score | m/z |
|---|---|---|---|---|
| sys_15_1 | 23453 | FTMS; HCD | 139.3 | 559.27 |

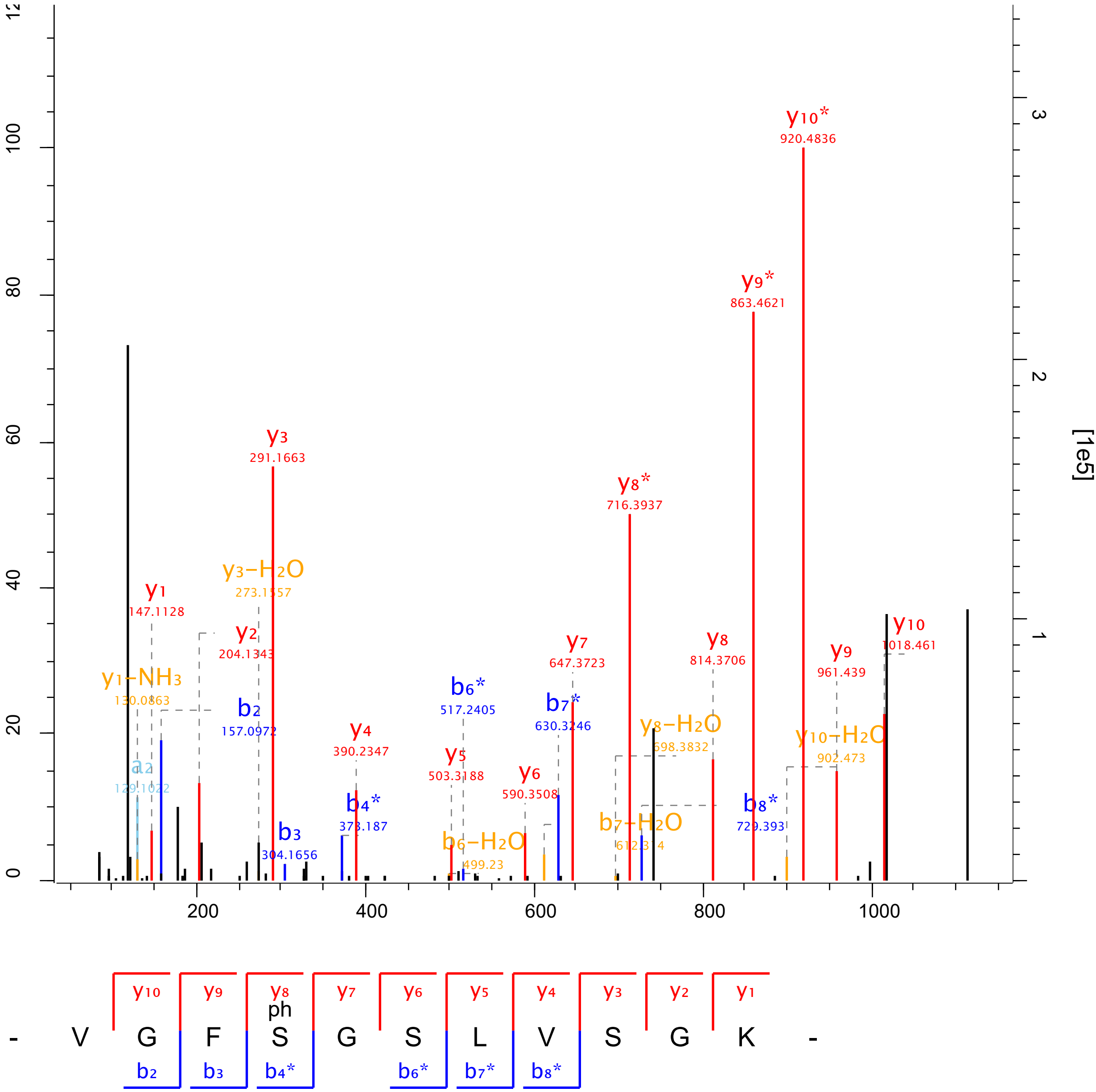

| Raw file | Scan | Method | Score | m/z |
| --- | --- | --- | --- | --- |
| sys_15_1 | 23465 | FTMS; HCD | 77.44 | 748.04 |

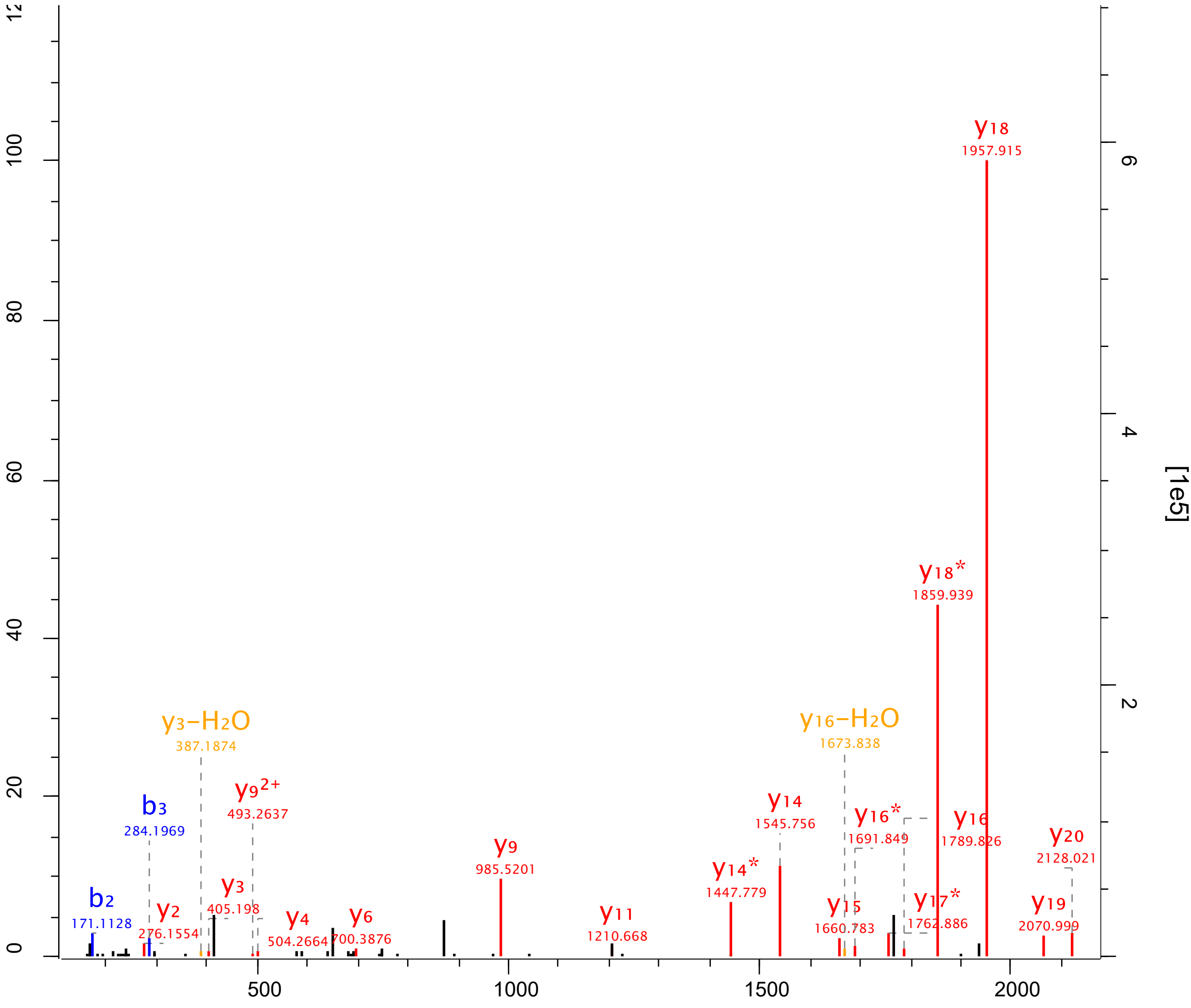


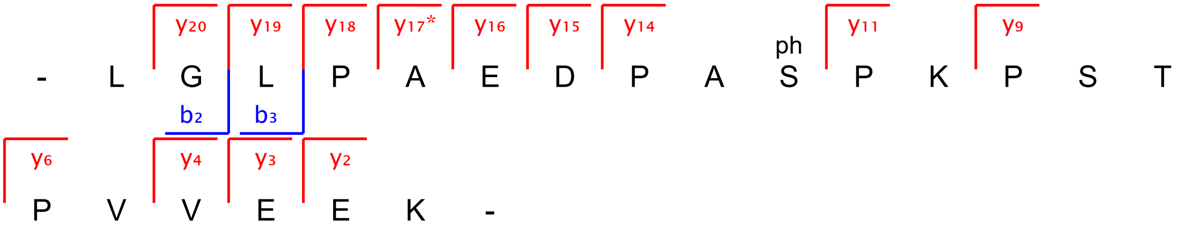

| Raw file | Scan | Method | Score | m/z |
|---|---|---|---|---|
| sys_15_1 | 23501 | FTMS; HCD | 100.22 | 581.76 |

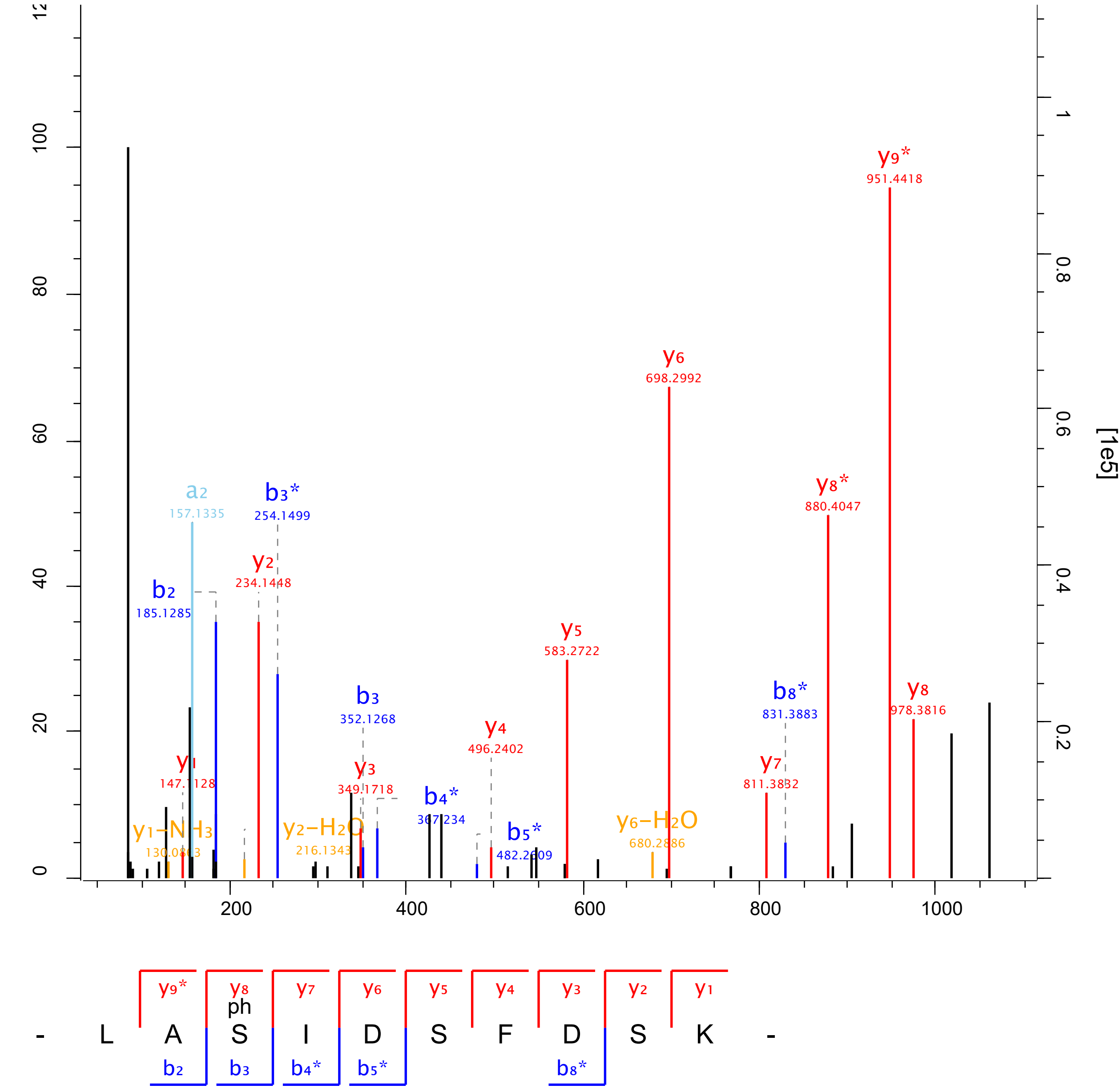

| Raw file | Scan | Method | Score | m/z |
|---|---|---|---|---|
| sys_15_1 | 23512 | FTMS; HCD | 134.57 | 532.73 |

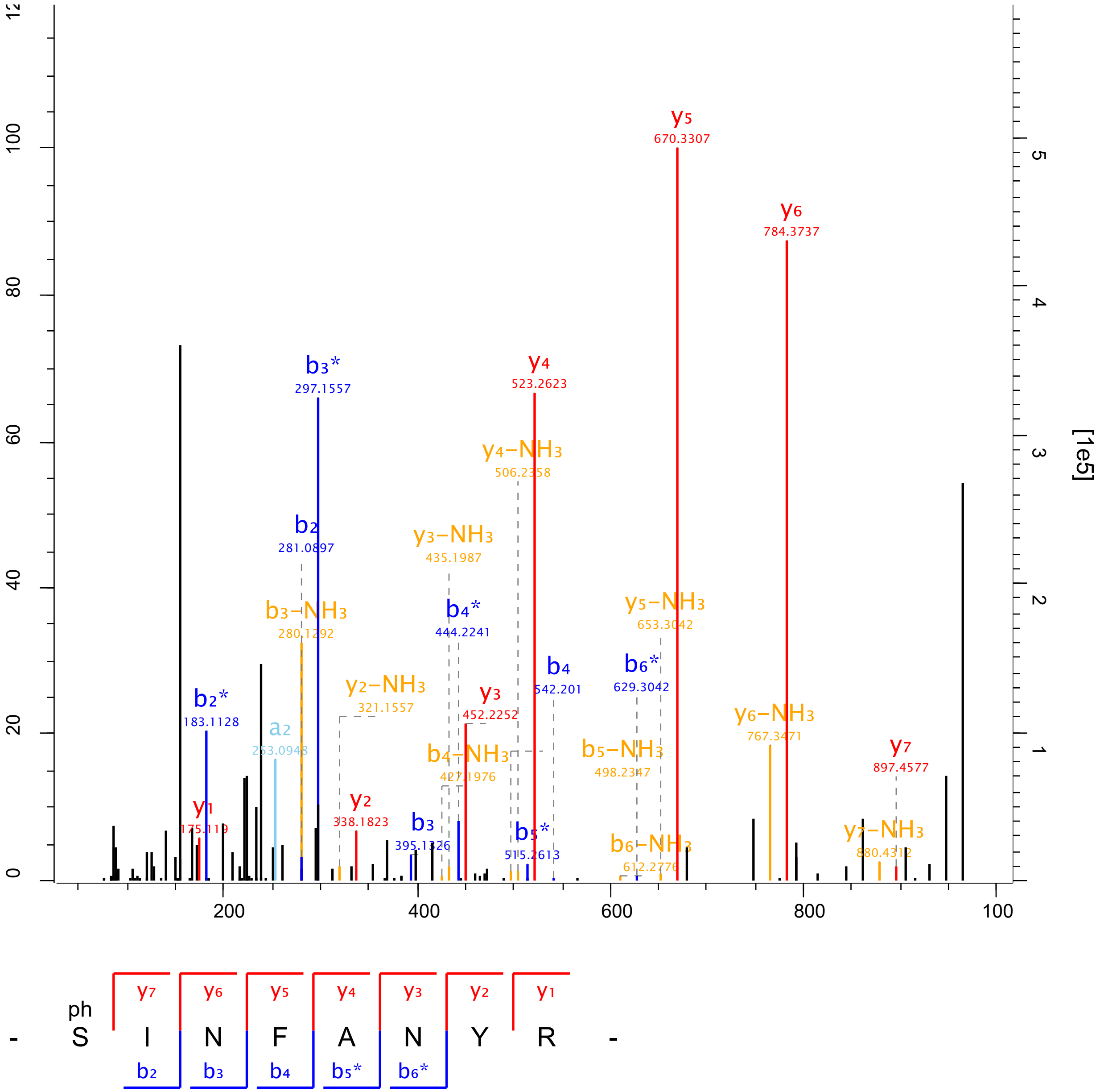

| Raw file | Scan | Method | Score | m/z |
|---|---|---|---|---|
| sys_15_1 | 23579 | FTMS; HCD | 100.23 | 707.79 |

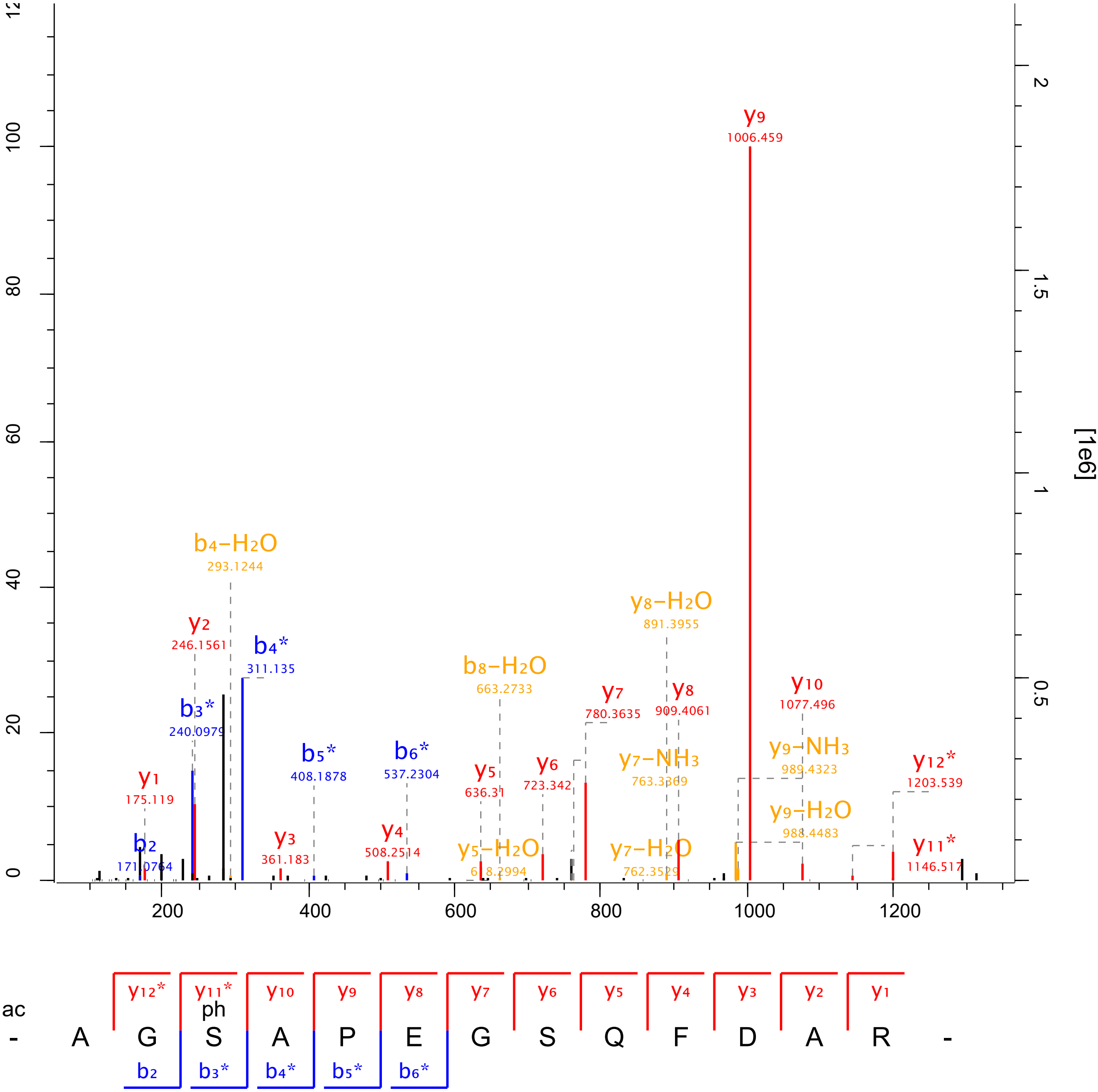

| Raw file | Scan | Method | Score | m/z |
|---|---|---|---|---|
| sys_15_1 | 23587 | FTMS; HCD | 106.93 | 496.21 |

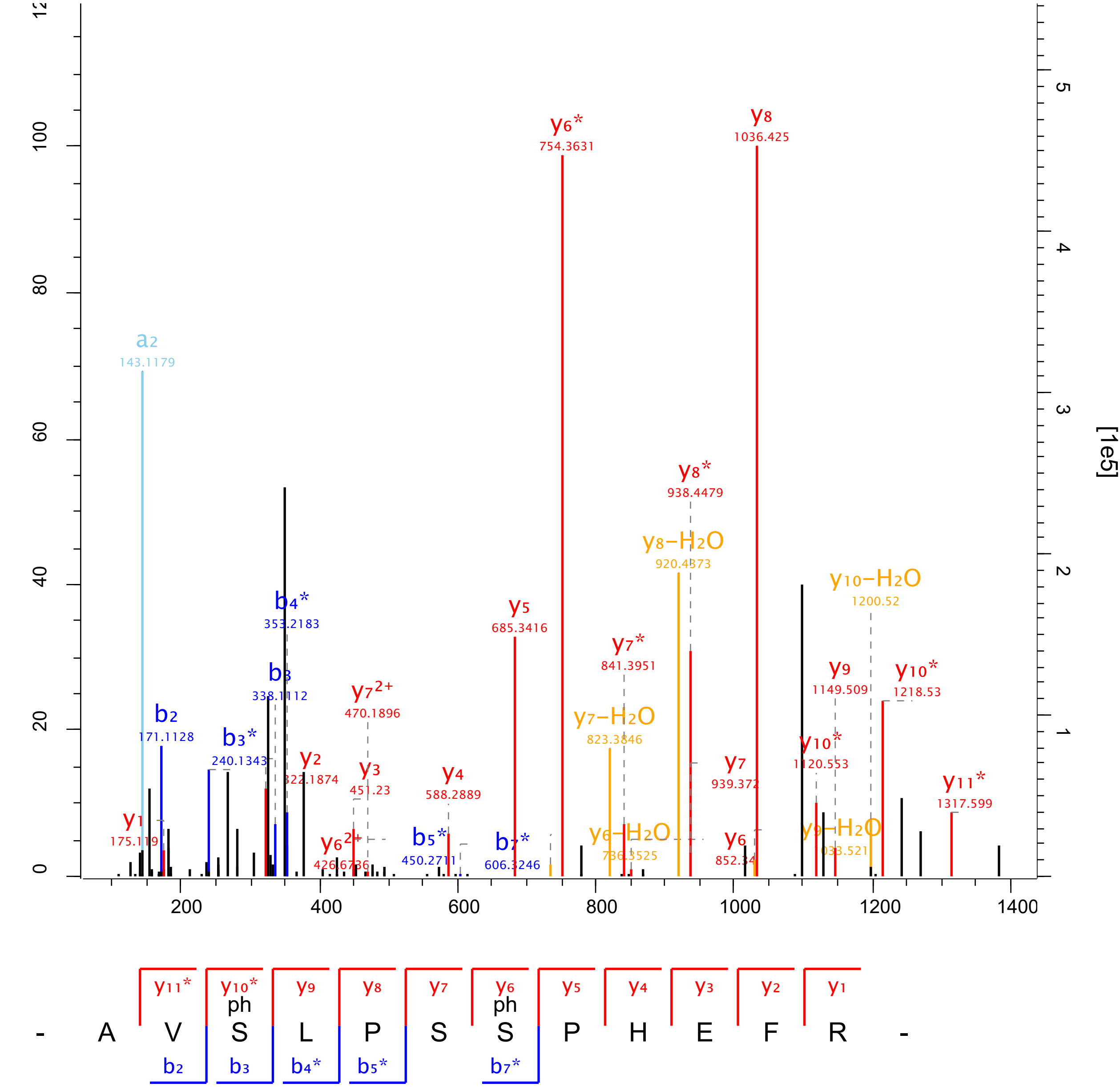

| Raw file | Scan | Method | Score | m/z |
| --- | --- | --- | --- | --- |
| sys_15_1 | 23727 | FTMS; HCD | 209.43 | 784.84 |

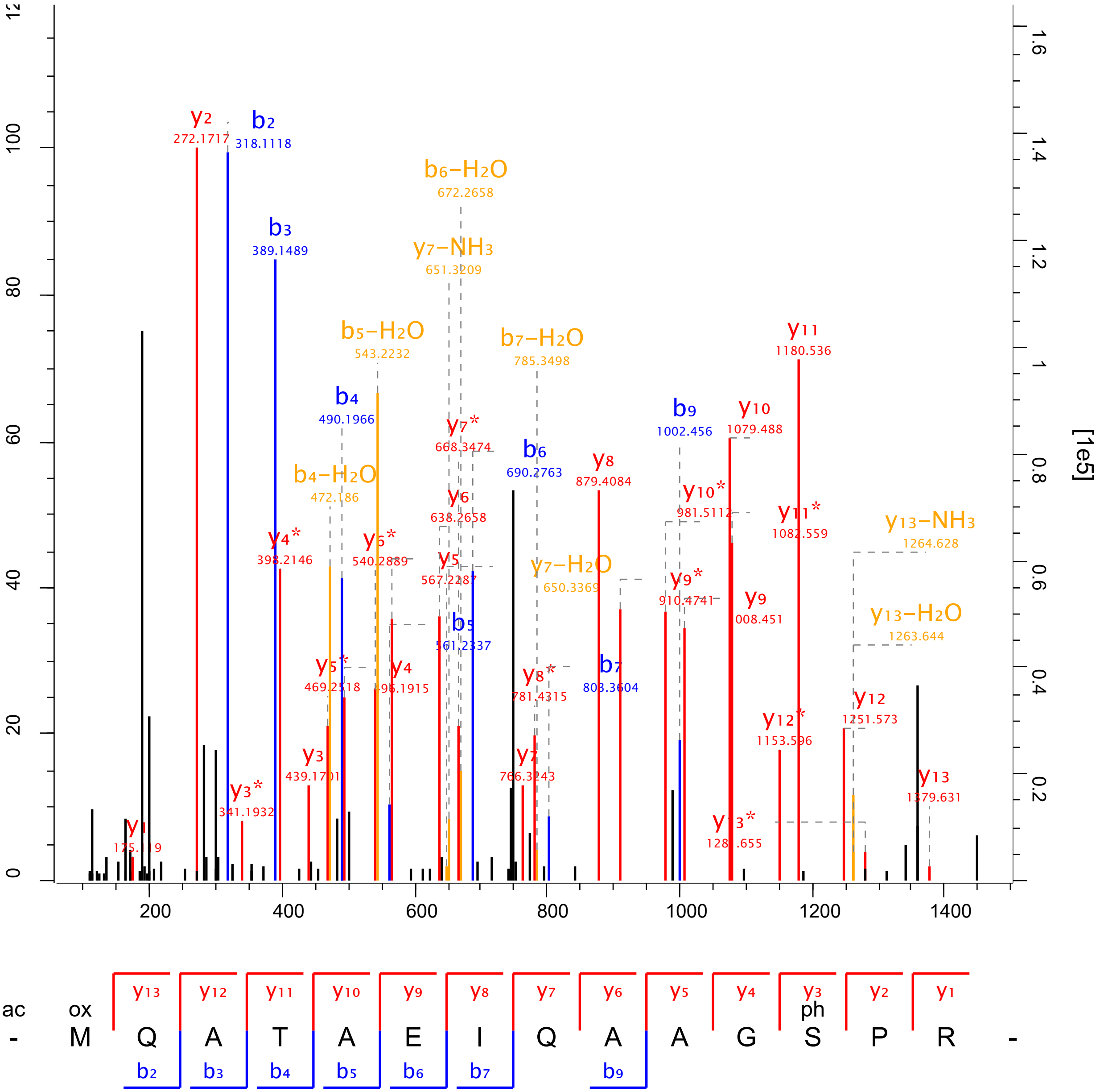

| Raw file | Scan | Method | Score | m/z |
|---|---|---|---|---|
| sys_15_1 | 23736 | FTMS; HCD | 166.29 | 925.88 |

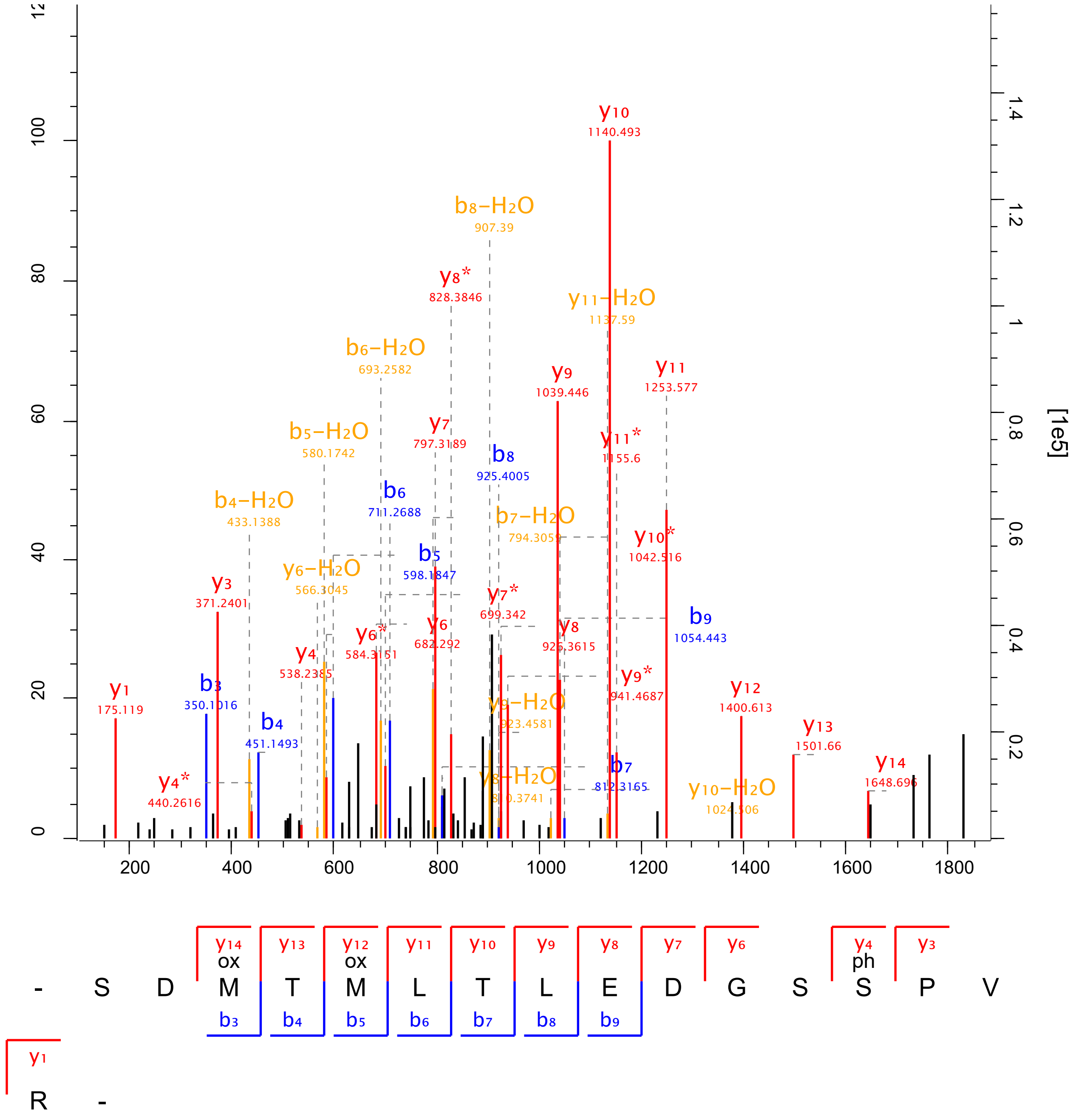

| Raw file | Scan | Method | Score | m/z |
|---|---|---|---|---|
| sys_15_1 | 23777 | FTMS; HCD | 41.09 | 756.33 |

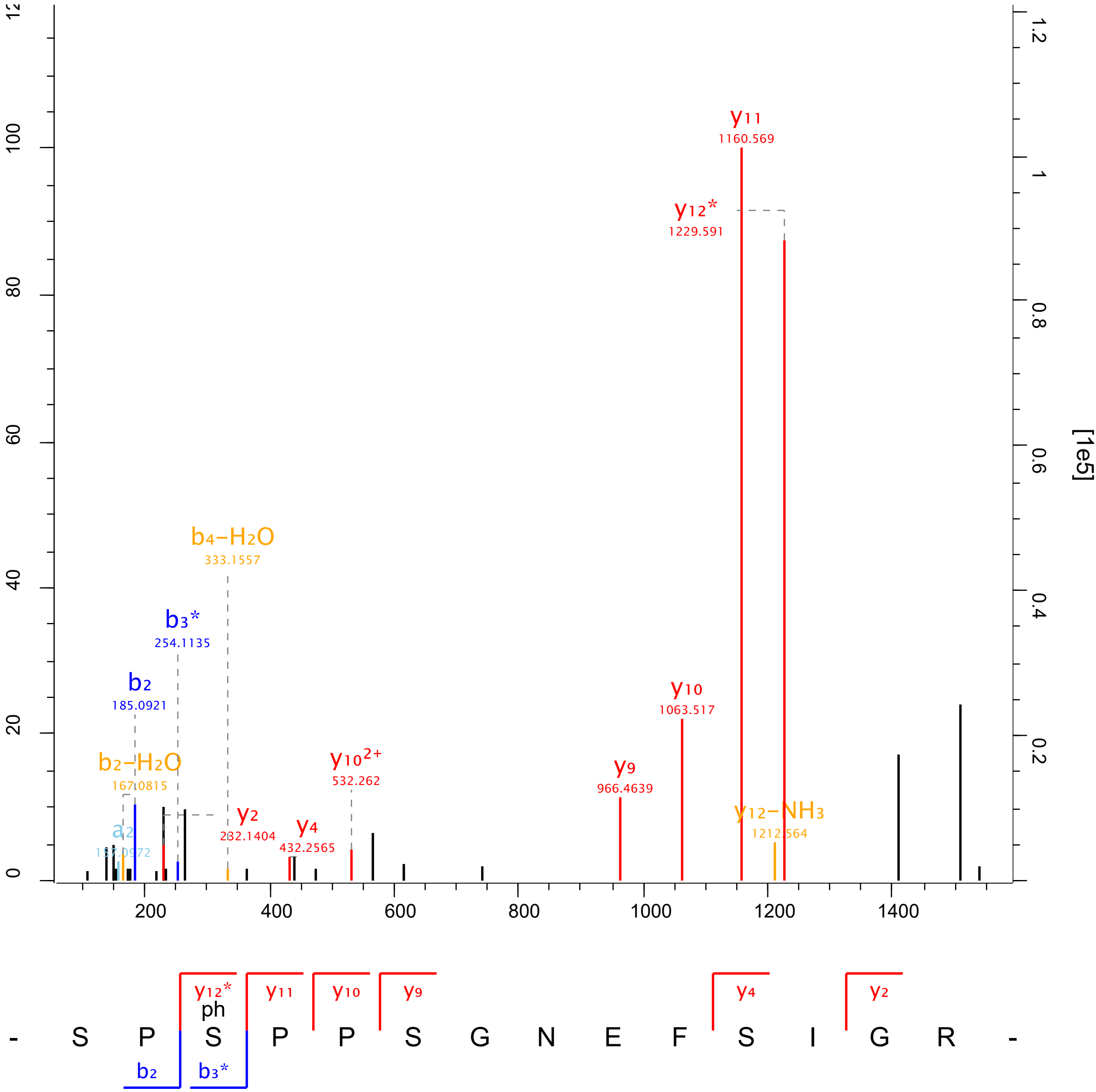

| Raw file | Scan | Method | Score | m/z |
| --- | --- | --- | --- | --- |
| sys_15_1 | 23780 | FTMS; HCD | 58.04 | 585.79 |

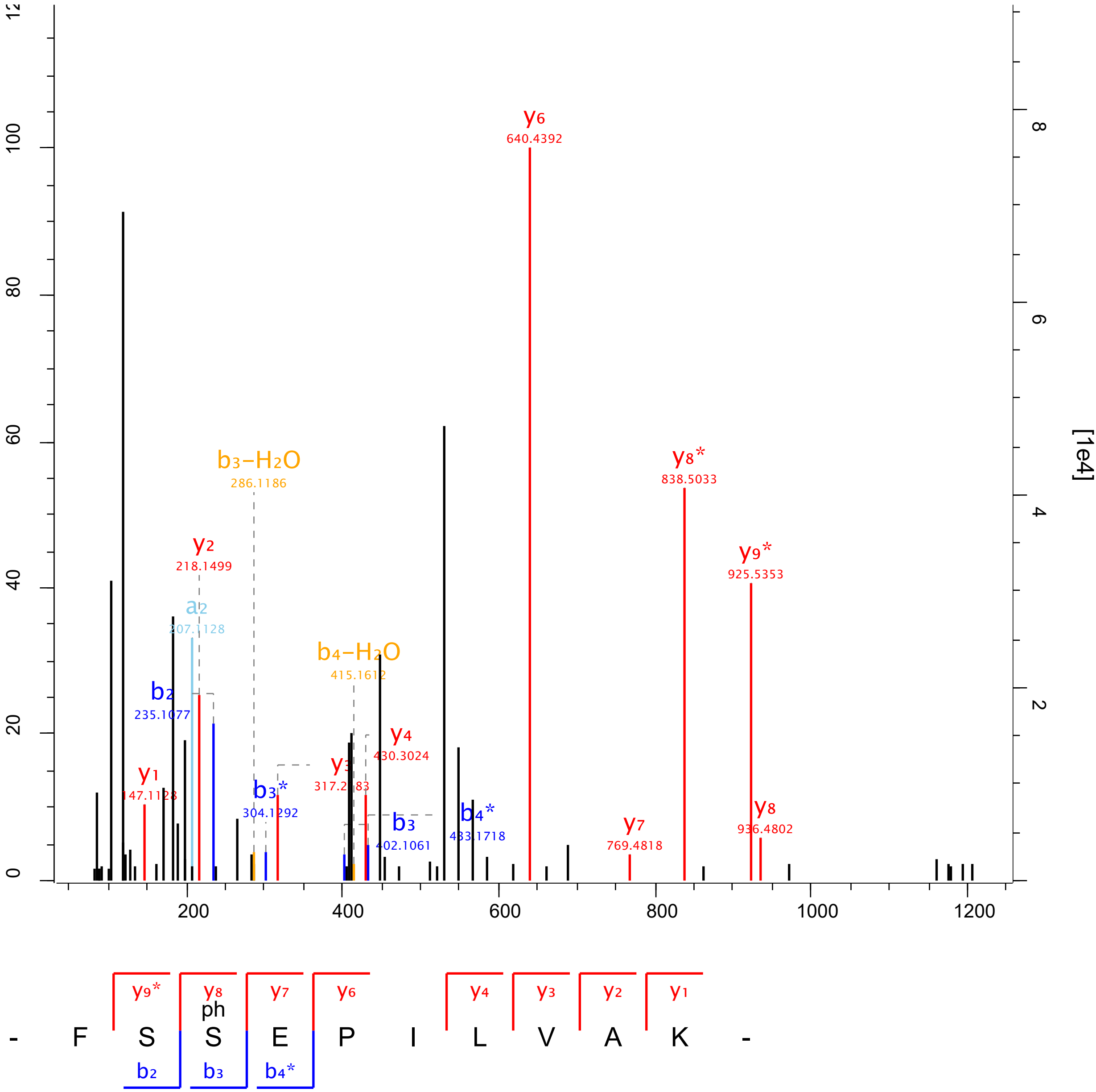

| Raw file | Scan | Method | Score | m/z |
| --- | --- | --- | --- | --- |
| sys_15_1 | 23894 | FTMS; HCD | 53.45 | 607.79 |

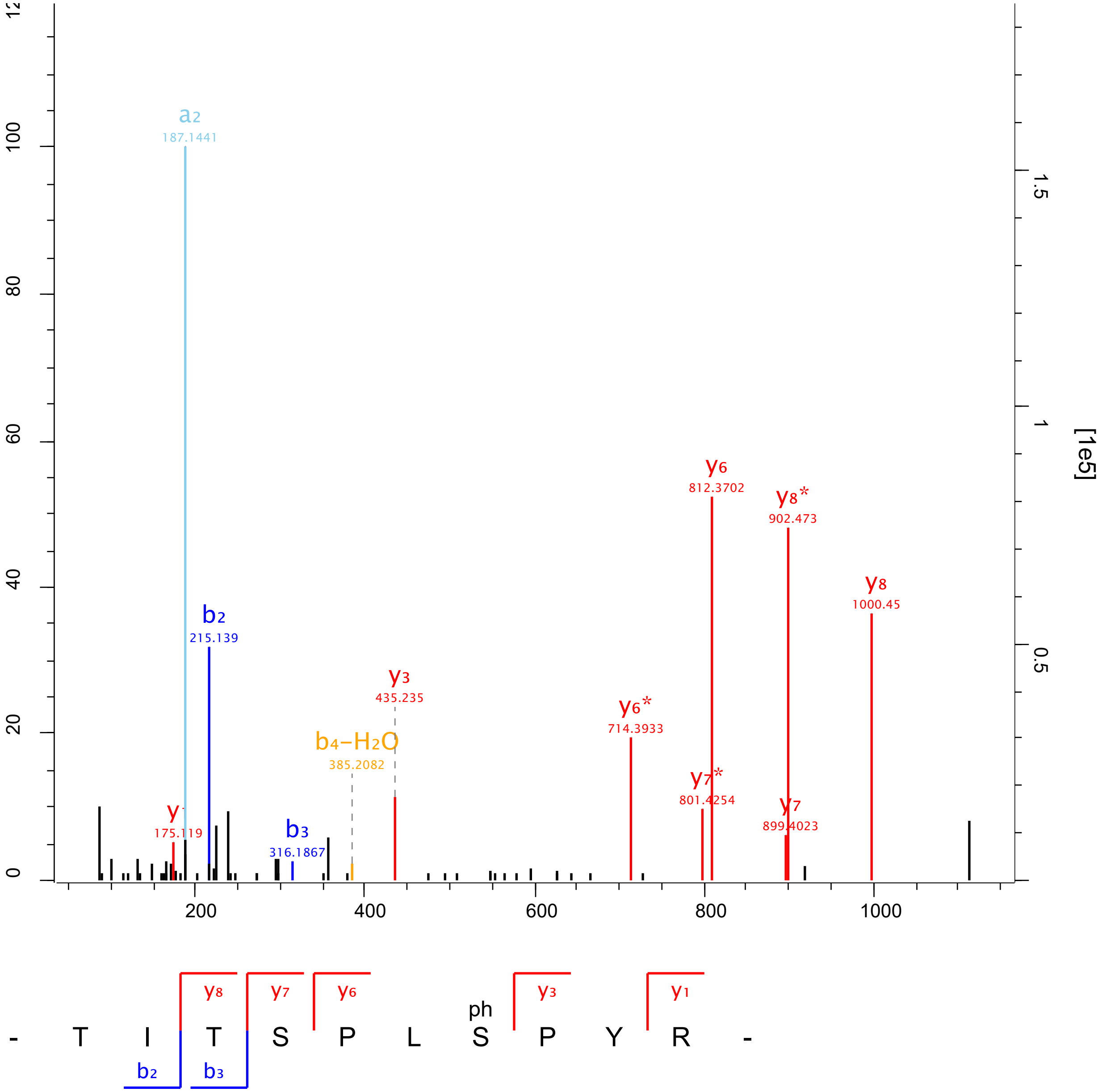

| Raw file | Scan | Method | Score | m/z |
|---|---|---|---|---|
| sys_15_1 | 24073 | FTMS; HCD | 57.53 | 558.62 |

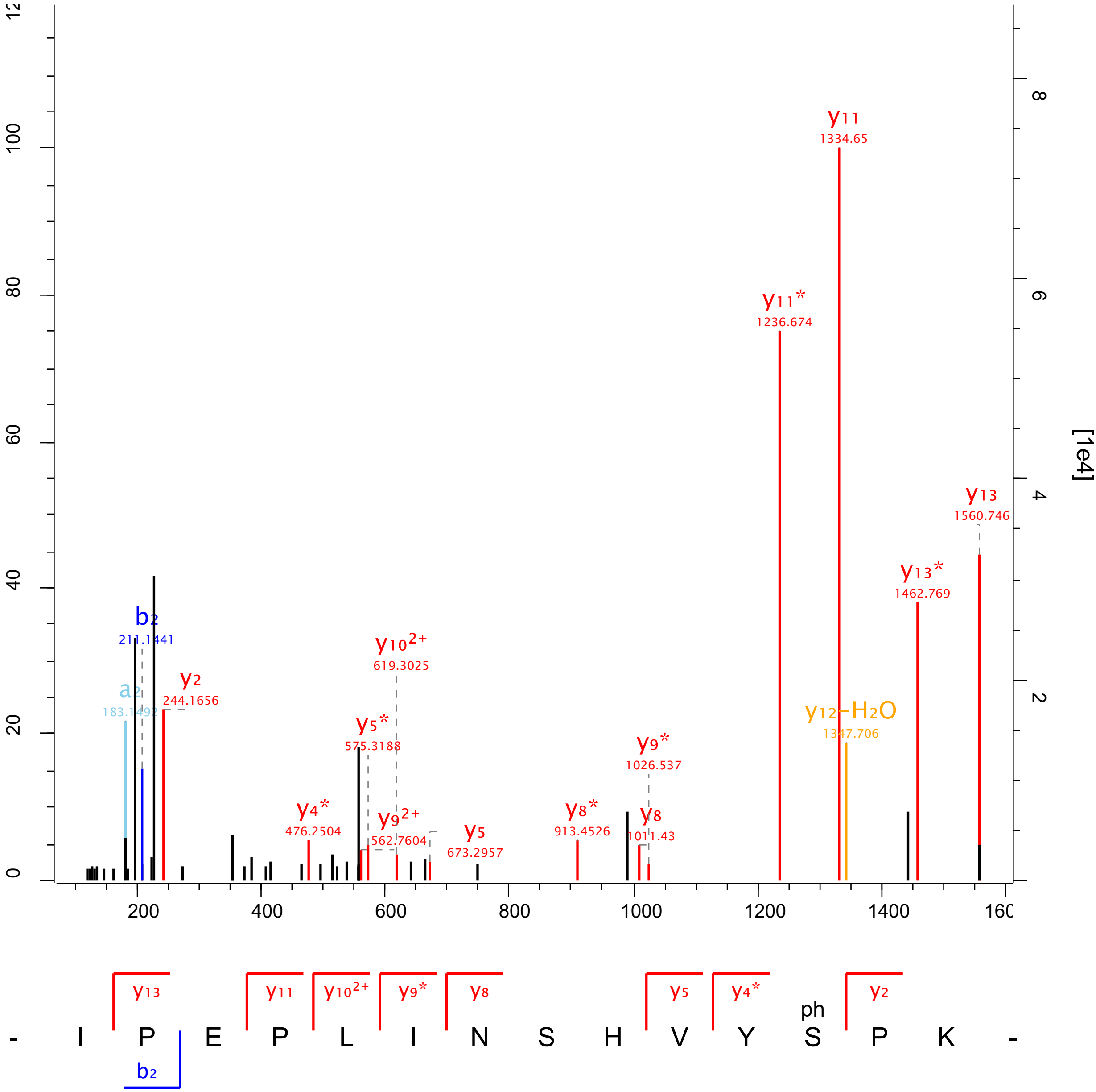

| Raw file | Scan | Method | Score | m/z |
| --- | --- | --- | --- | --- |
| sys_15_1 | 24104 | FTMS; HCD | 95.67 | 775.03 |

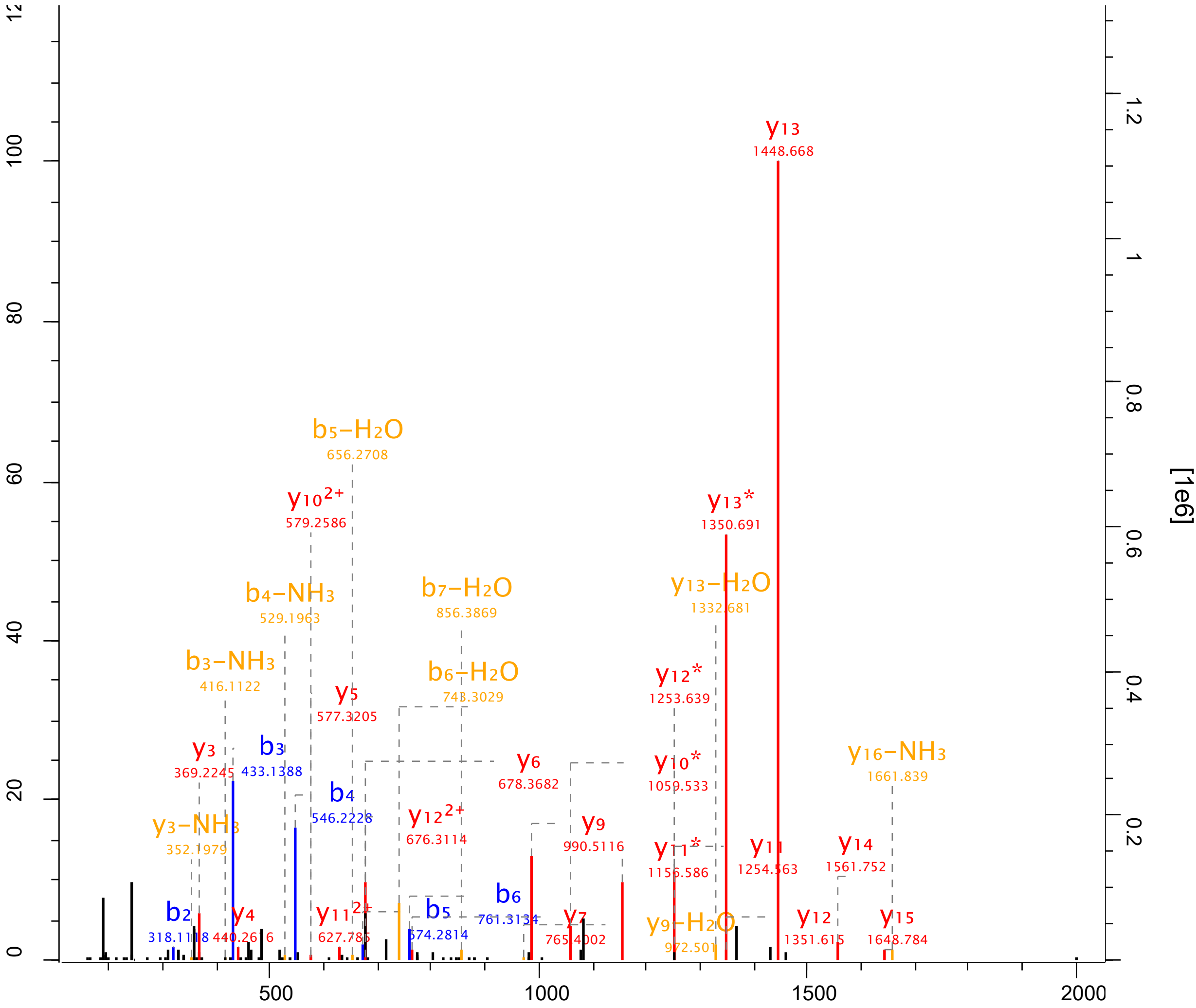


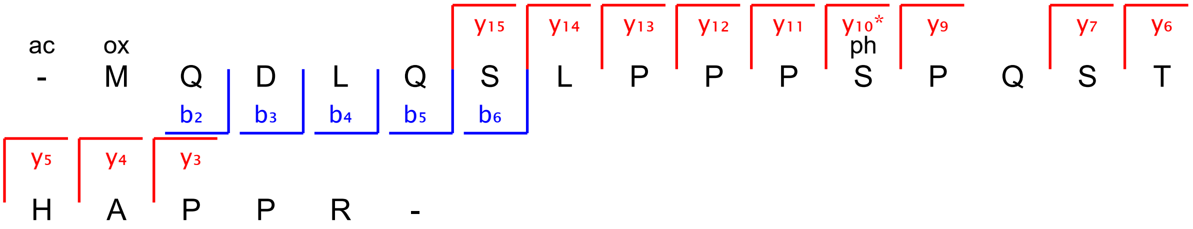

| Raw file | Scan | Method | Score | m/z |
|---|---|---|---|---|
| sys_15_1 | 24237 | FTMS; HCD | 57.29 | 749.33 |

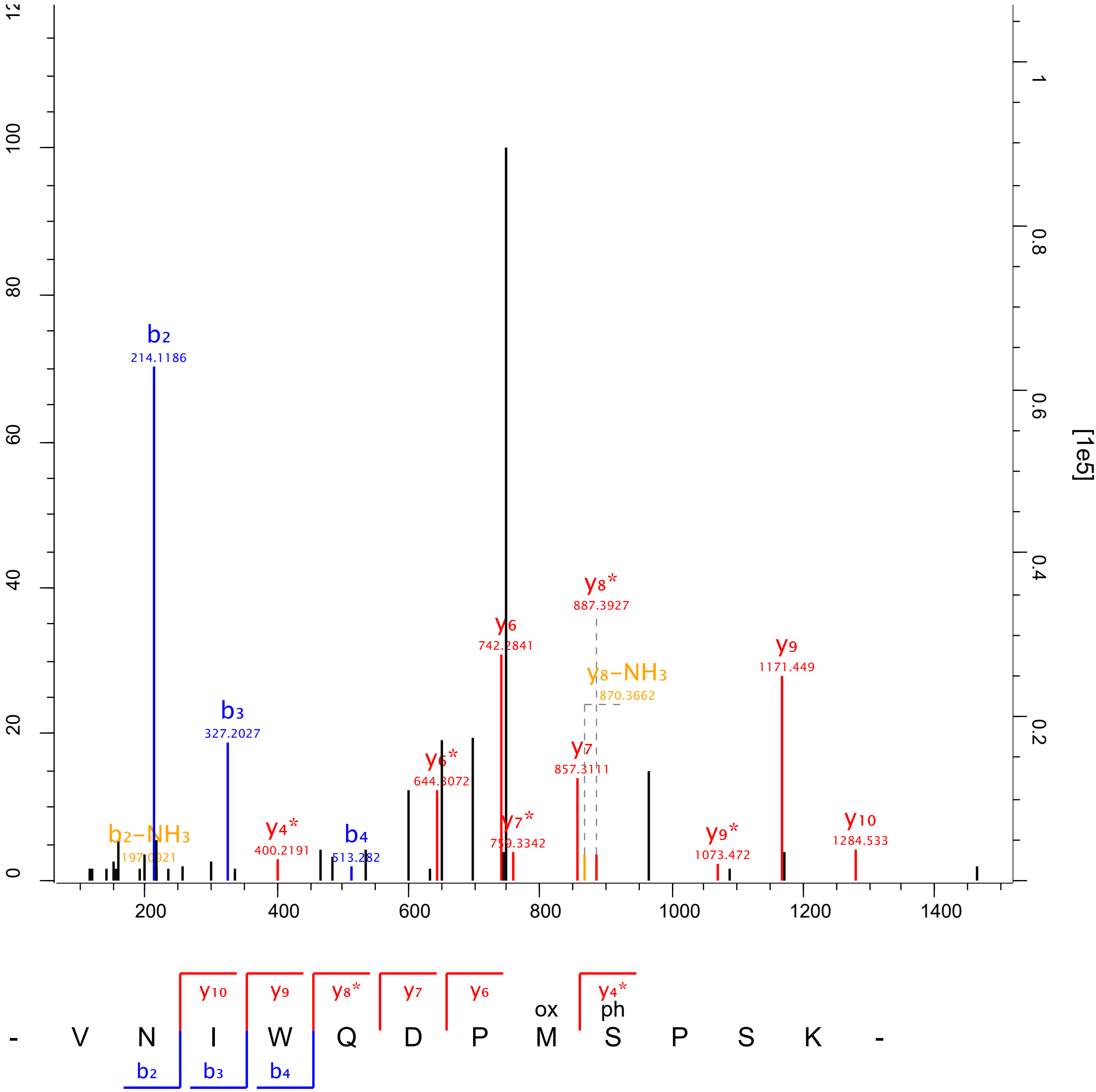

| Raw file | Scan | Method | Score | m/z |
|---|---|---|---|---|
| sys_15_1 | 24322 | FTMS; HCD | 78.81 | 674.8 |

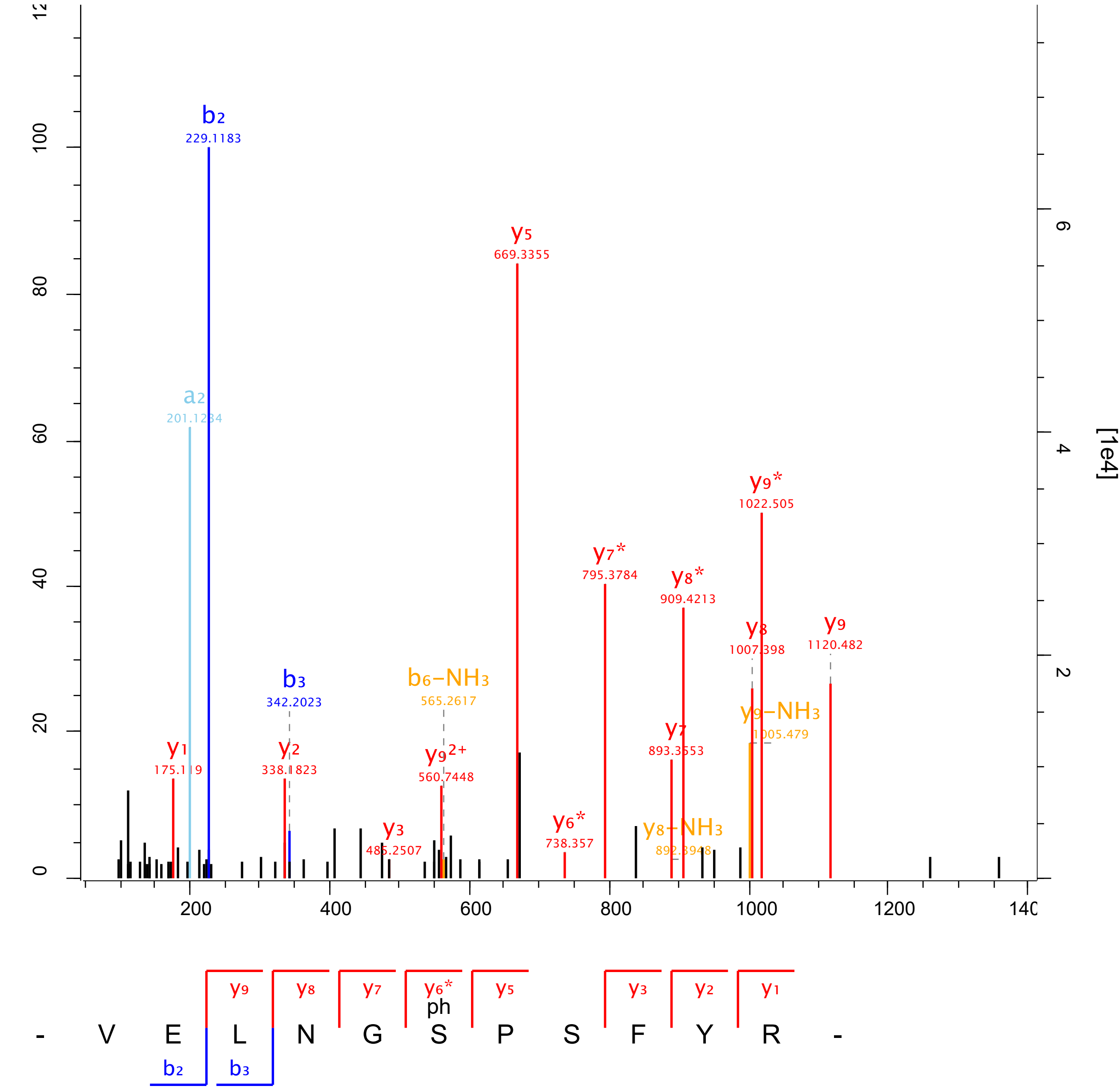

| Raw file | Scan | Method | Score | m/z |
|---|---|---|---|---|
| sys_15_1 | 24333 | FTMS; HCD | 65.23 | 794.84 |

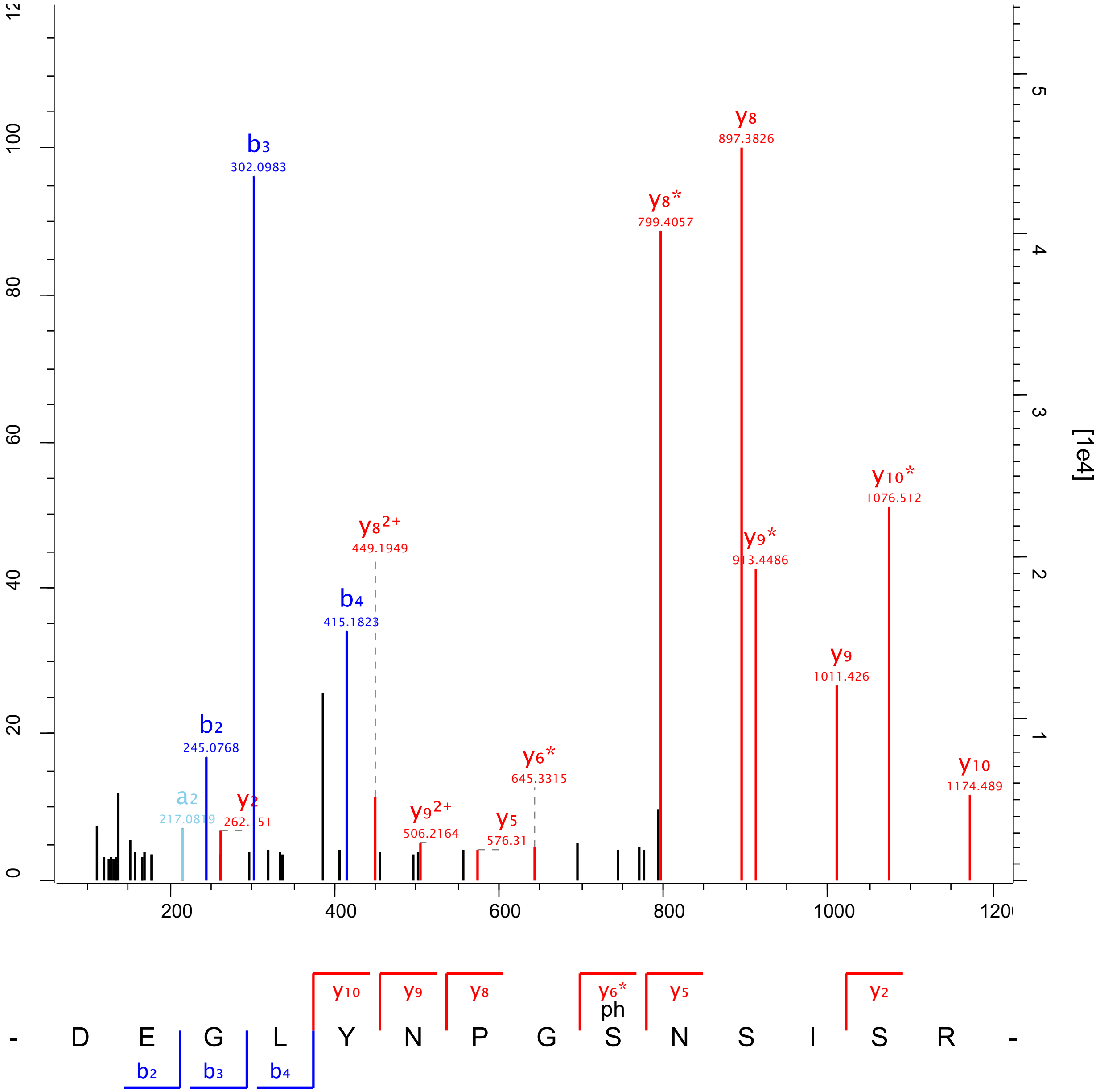

| Raw file | Scan | Method | Score | m/z |
| --- | --- | --- | --- | --- |
| sys_15_1 | 24374 | FTMS; HCD | 71.88 | 803.69 |

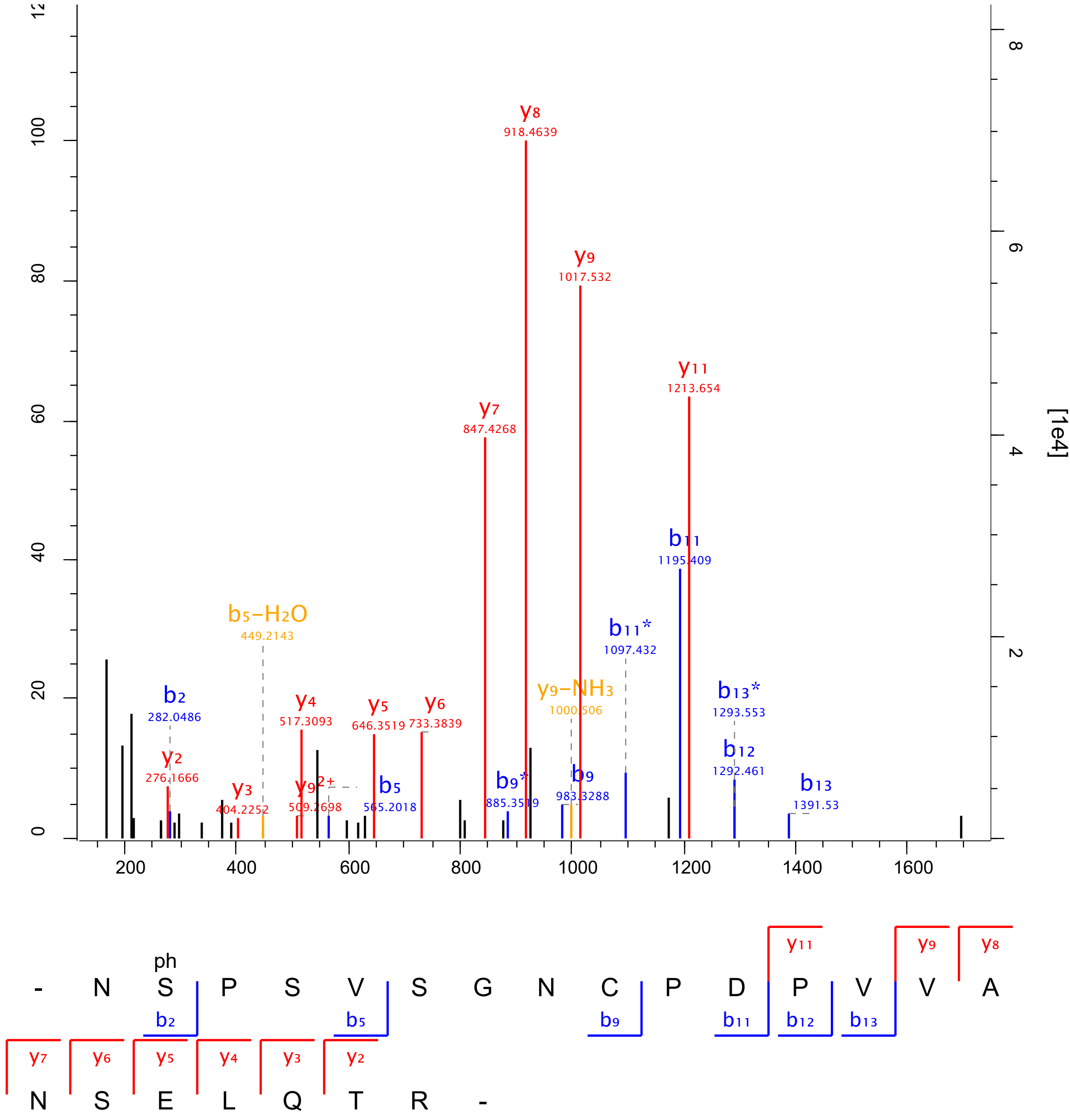

| Raw file | Scan | Method | Score | m/z |
|---|---|---|---|---|
| sys_15_1 | 24464 | FTMS; HCD | 238.88 | 713.84 |

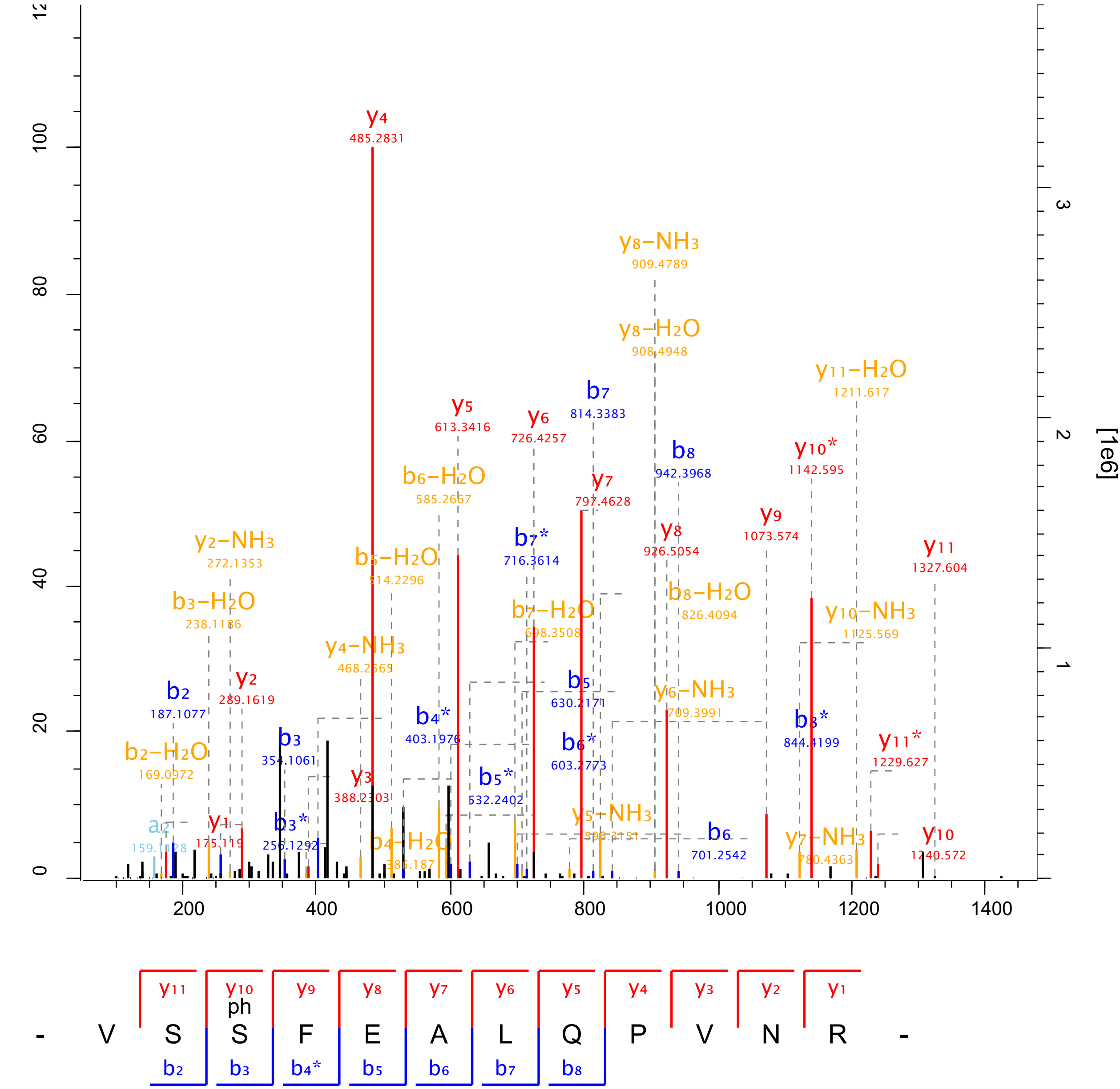

| Raw file | Scan | Method | Score | m/z |
|---|---|---|---|---|
| sys_15_1 | 24523 | FTMS; HCD | 127.83 | 740.27 |

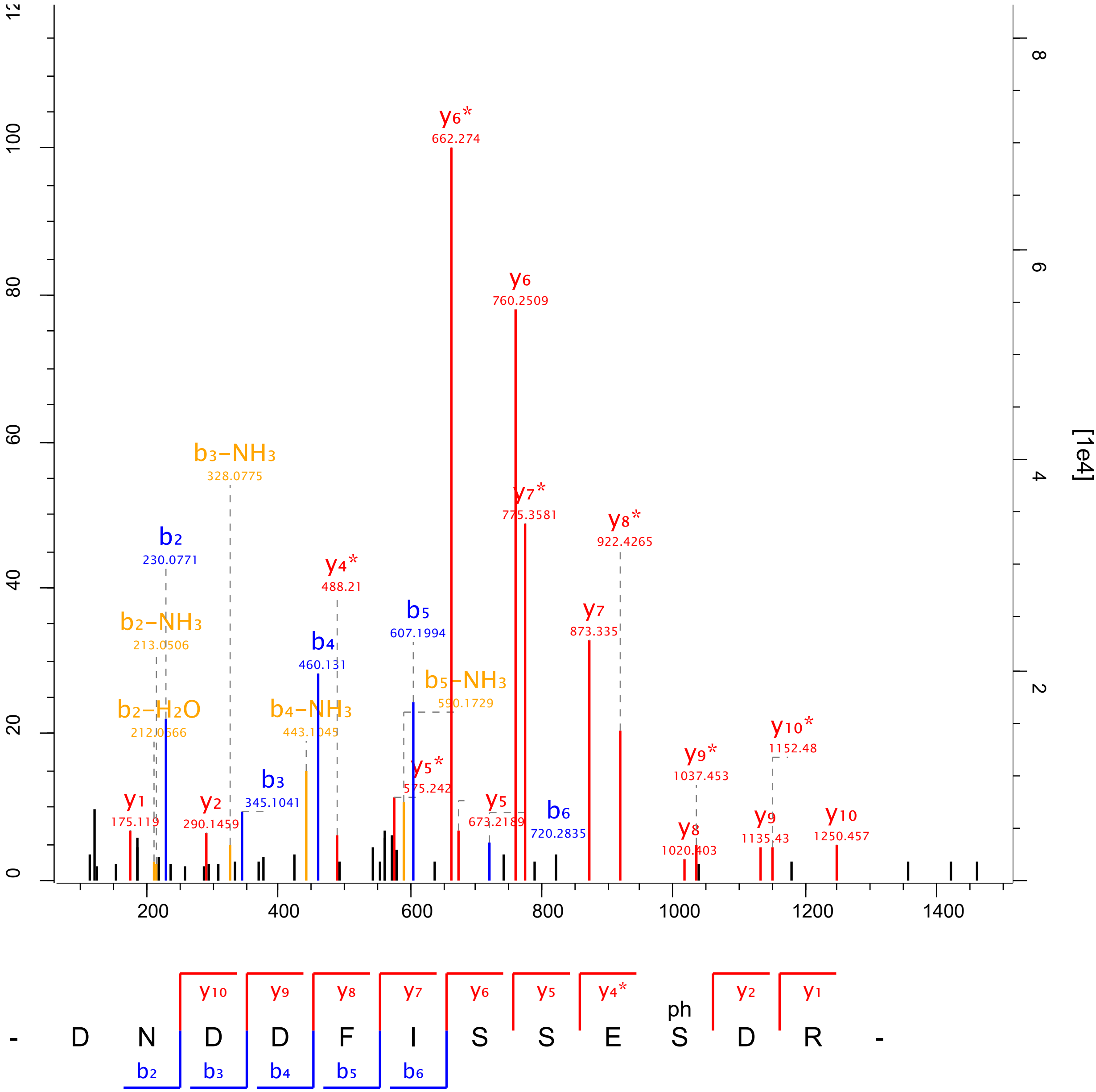

| Raw file | Scan | Method | Score | m/z |
|---|---|---|---|---|
| sys_15_1 | 24677 | FTMS; HCD | 46.42 | 743.68 |

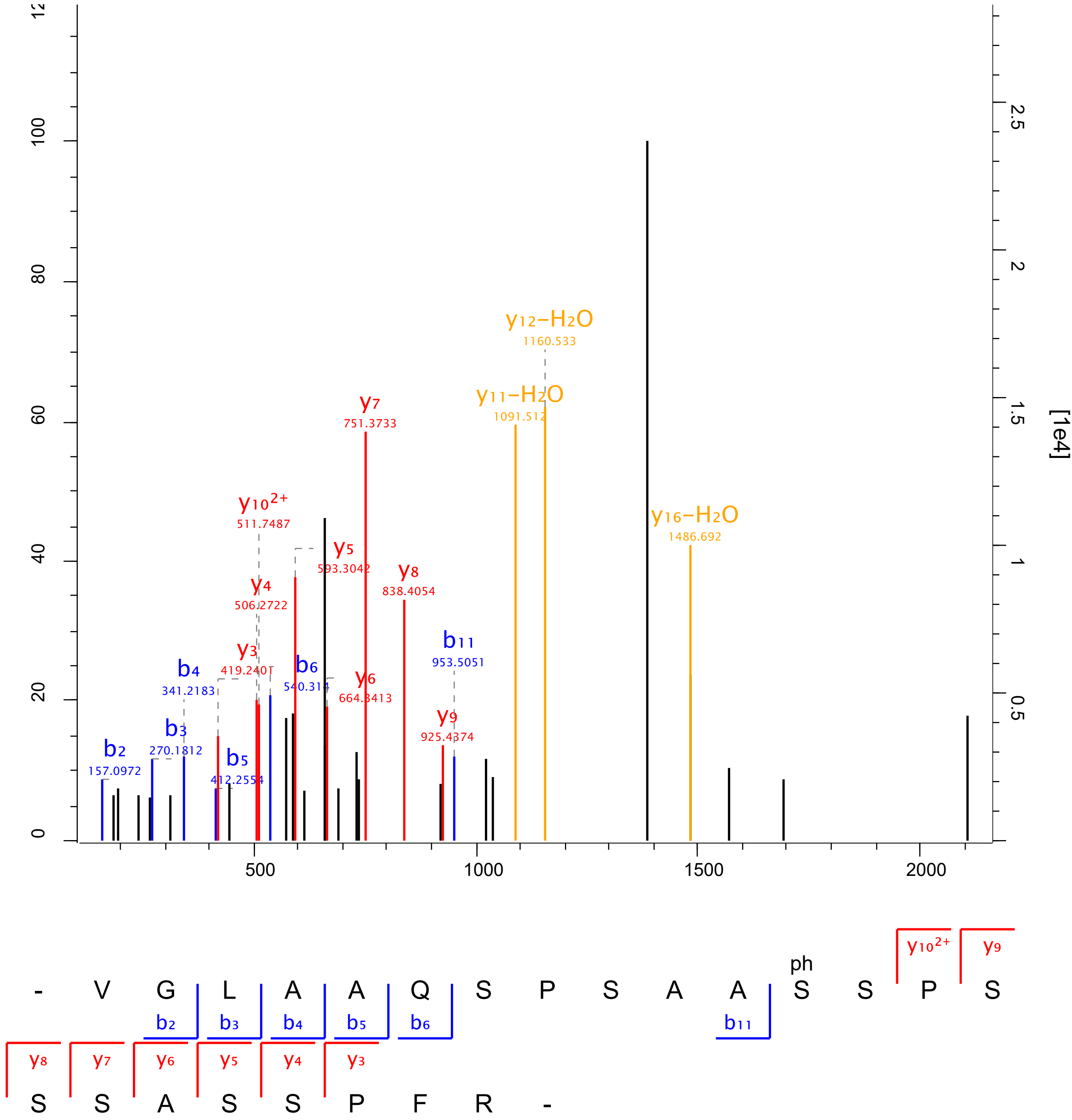

| Raw file | Scan | Method | Score | m/z |
|---|---|---|---|---|
| sys_15_1 | 24740 | FTMS; HCD | 116.08 | 832.02 |

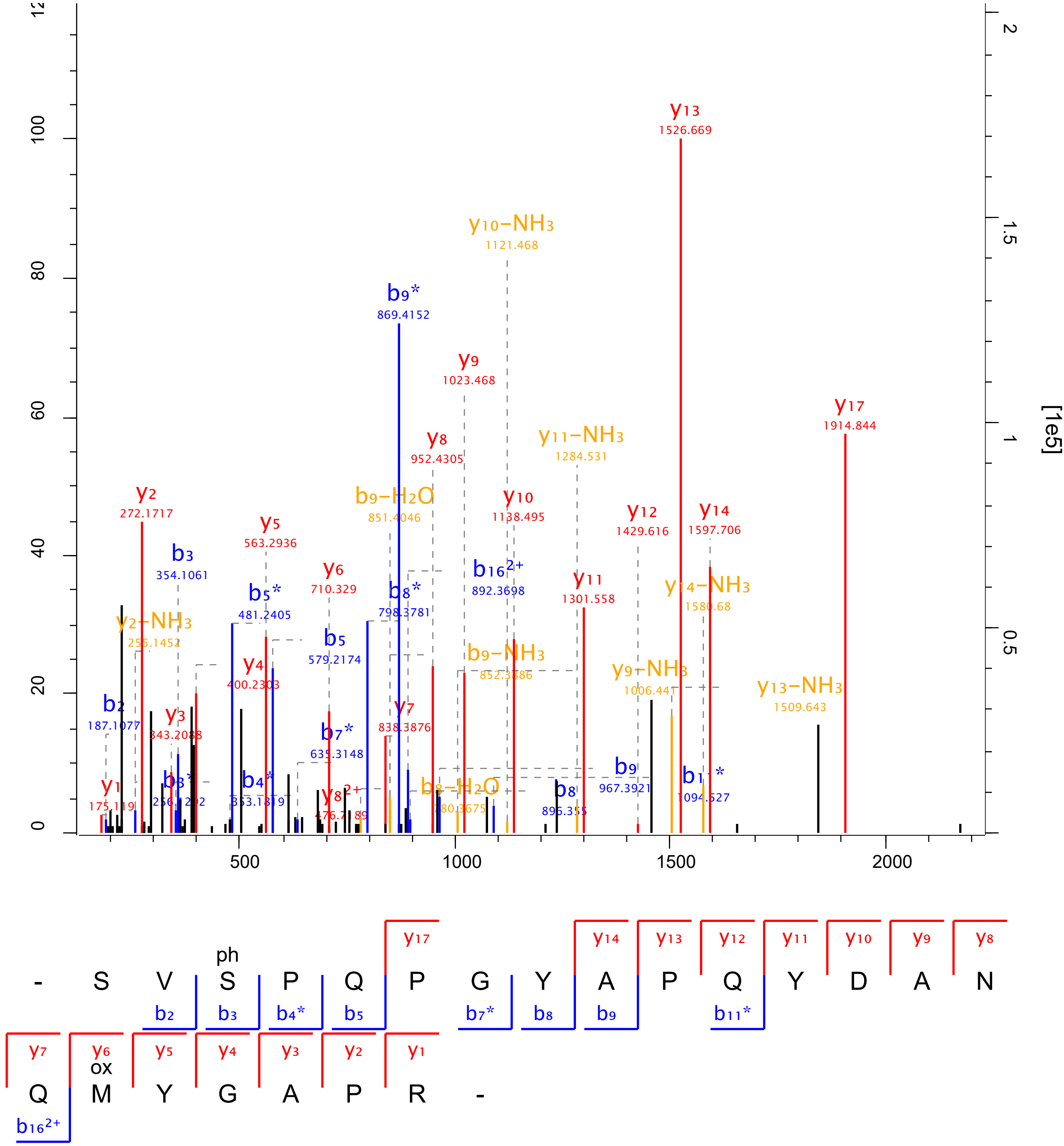

| Raw file | Scan | Method | Score | m/z |
|---|---|---|---|---|
| sys_15_1 | 24741 | FTMS; HCD | 99.09 | 895.88 |

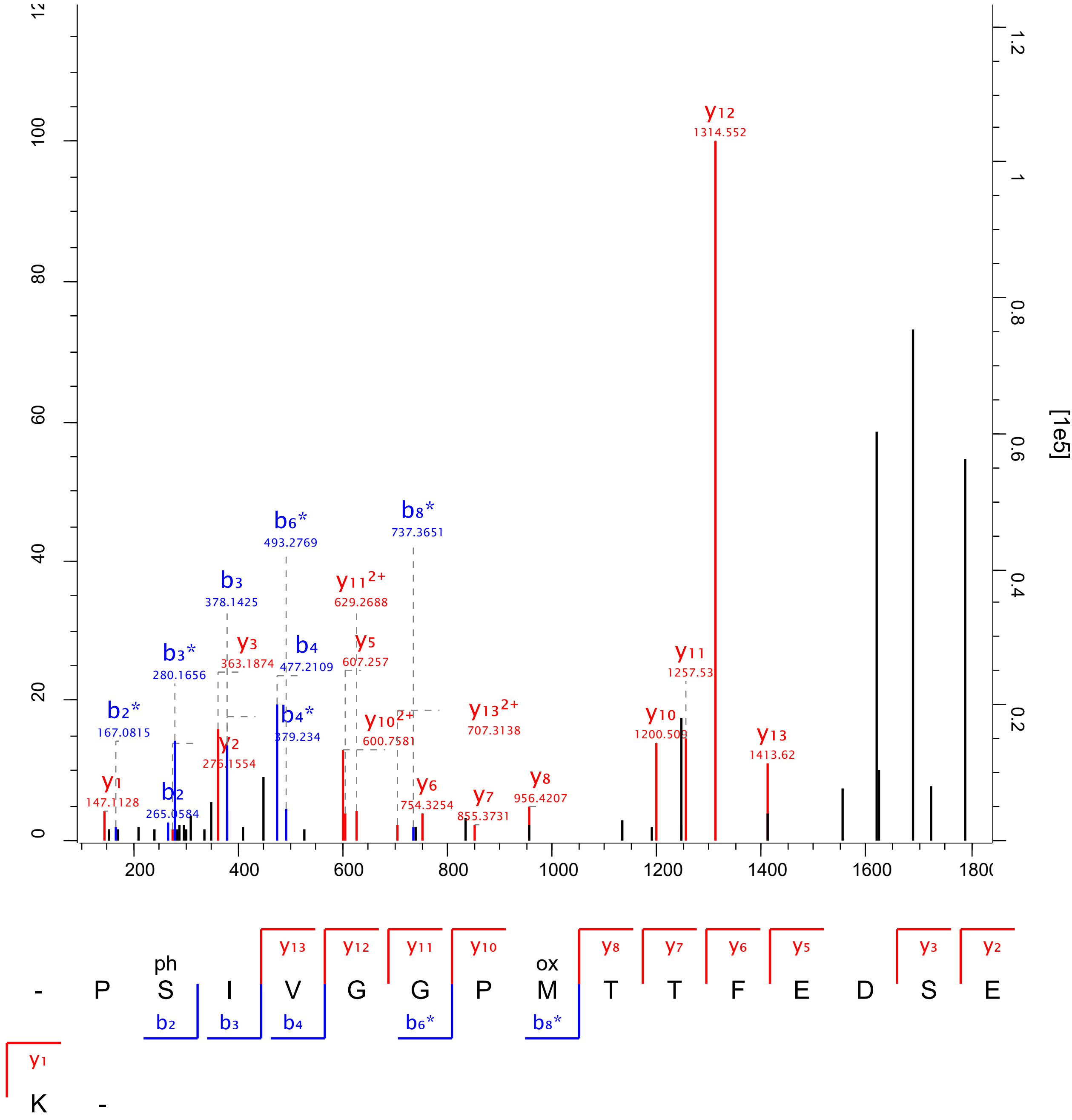

| Raw file | Scan | Method | Score | m/z |
|---|---|---|---|---|
| sys_15_1 | 24756 | FTMS; HCD | 73.29 | 550.61 |

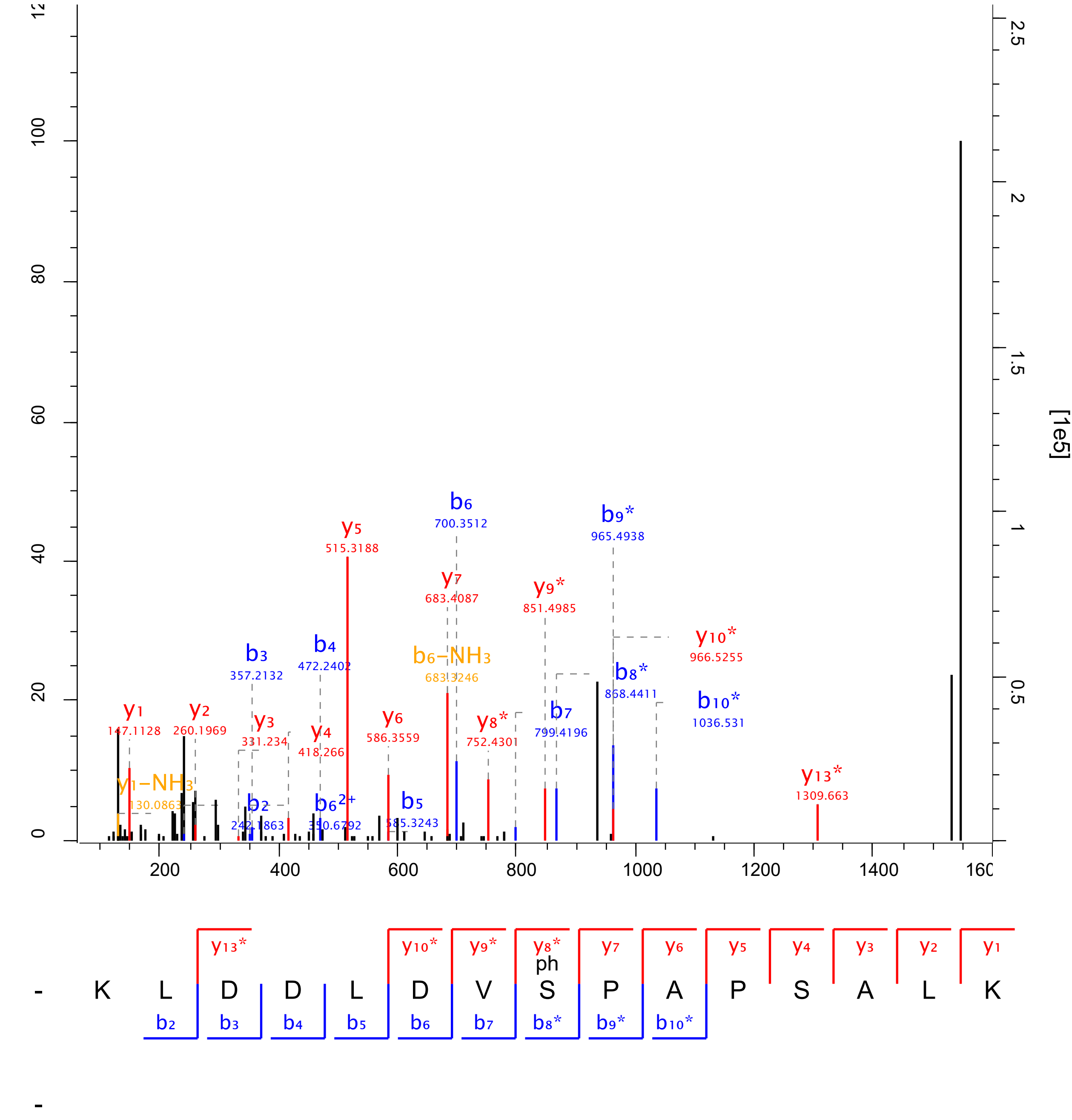

| Raw file | Scan | Method | Score | m/z |
| --- | --- | --- | --- | --- |
| sys_15_1 | 24885 | FTMS; HCD | 66.89 | 875.39 |

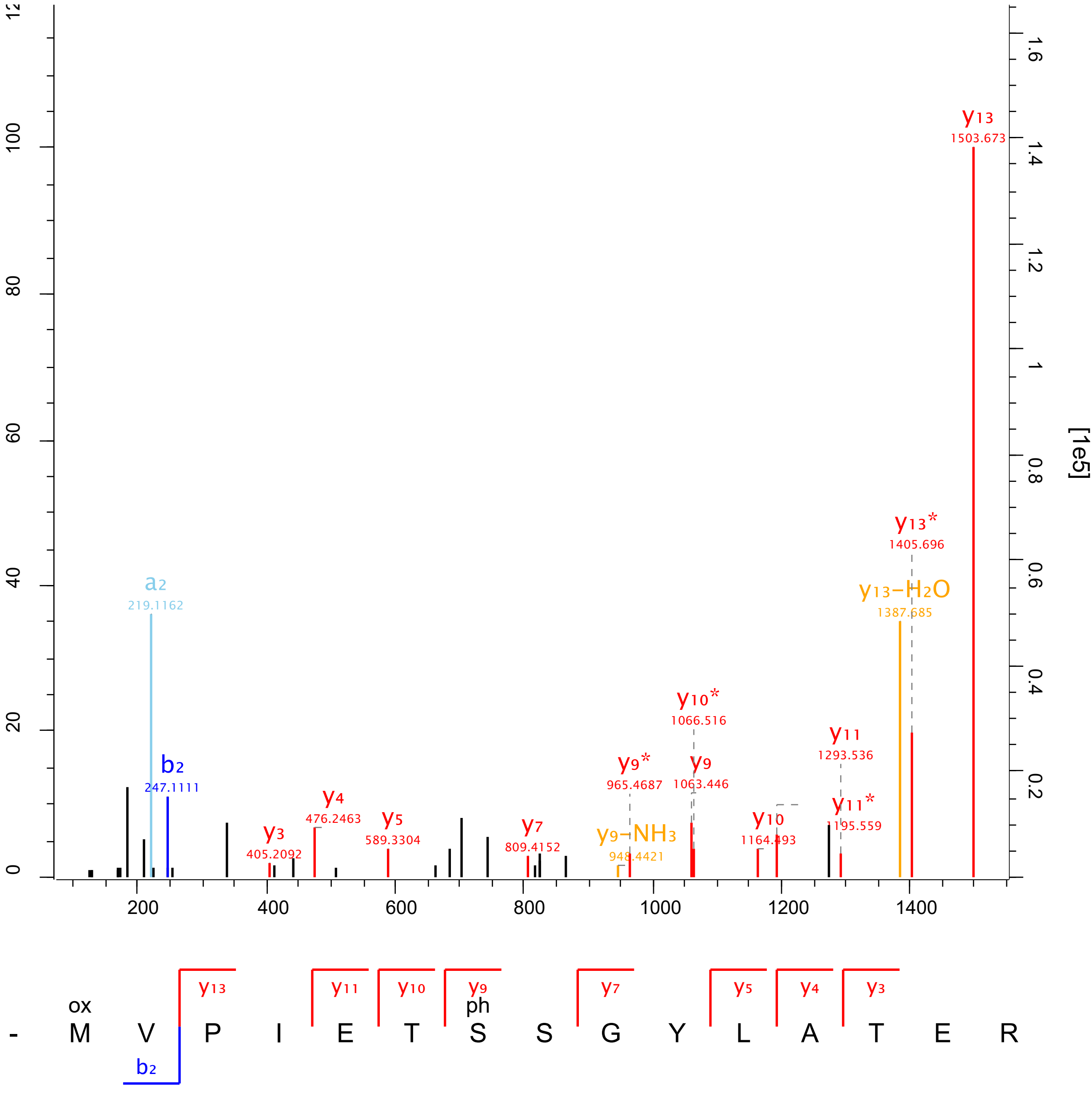

| Raw file | Scan | Method | Score | m/z |
| --- | --- | --- | --- | --- |
| sys_15_1 | 24947 | FTMS; HCD | 100.11 | 696.28 |

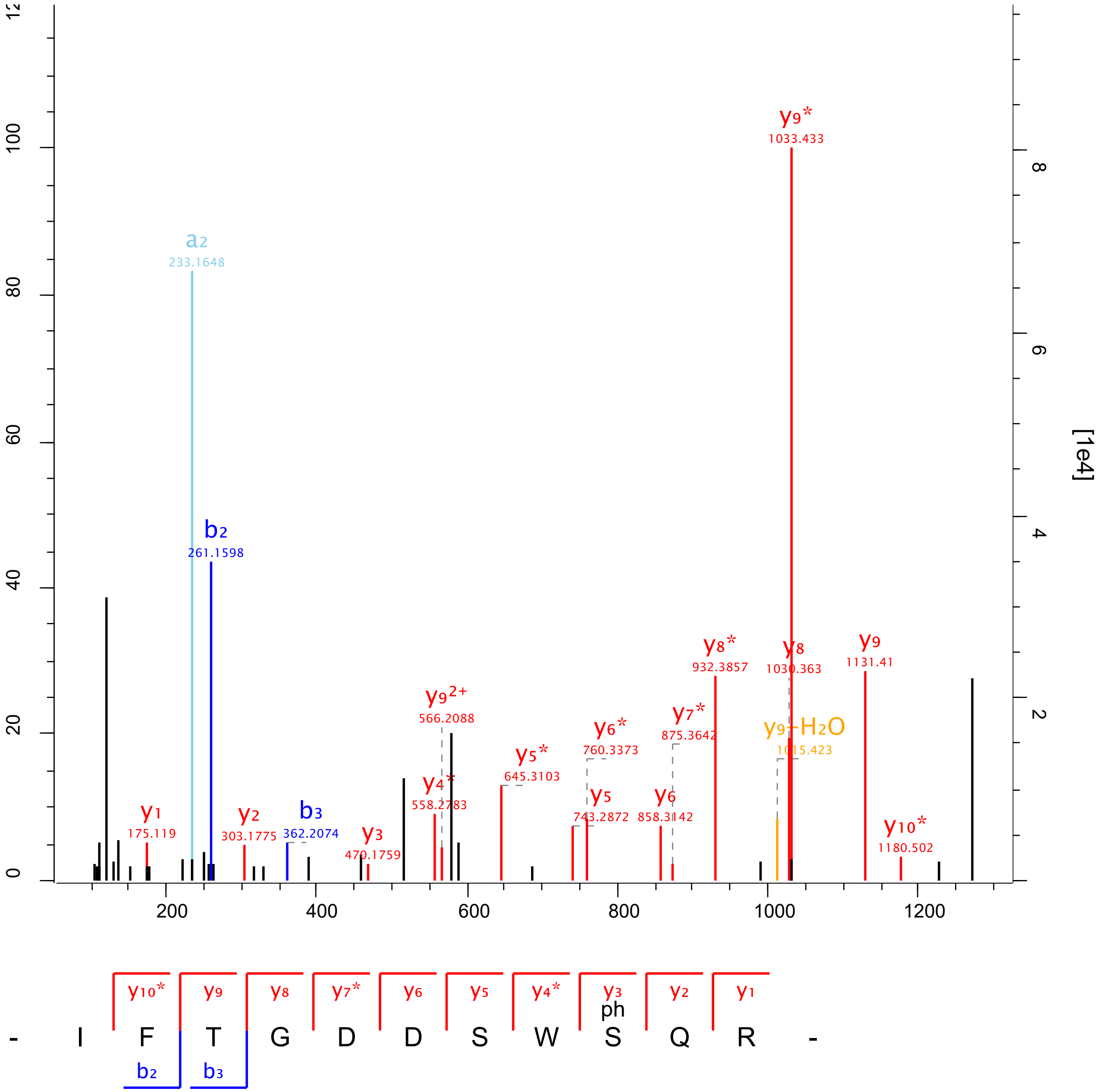

| Raw file | Scan | Method | Score | m/z |
| --- | --- | --- | --- | --- |
| sys_15_1 | 25000 | FTMS; HCD | 42.03 | 642.31 |

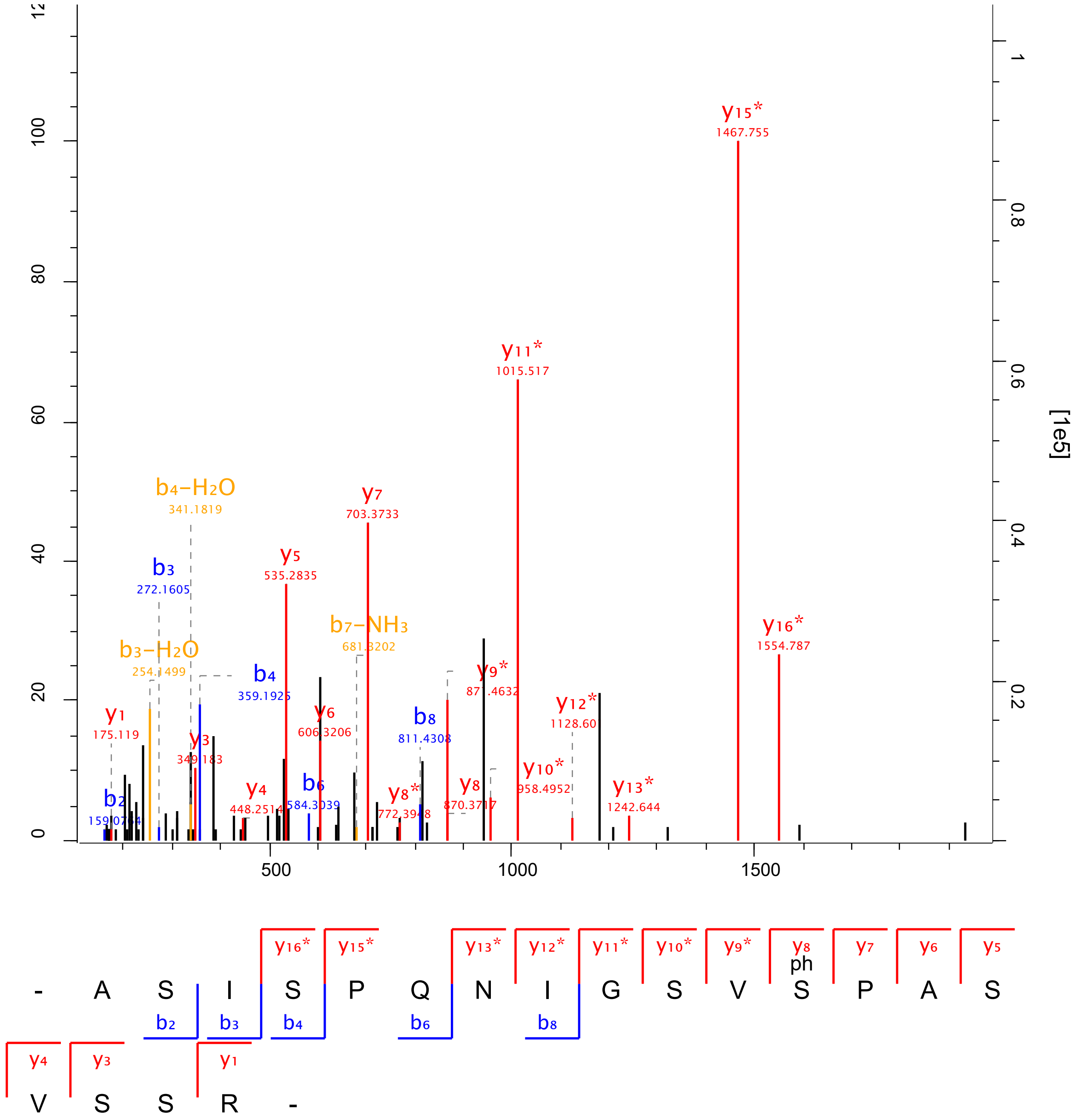

| Raw file | Scan | Method | Score | m/z |
|---|---|---|---|---|
| sys_15_1 | 25112 | FTMS; HCD | 58.95 | 691.63 |

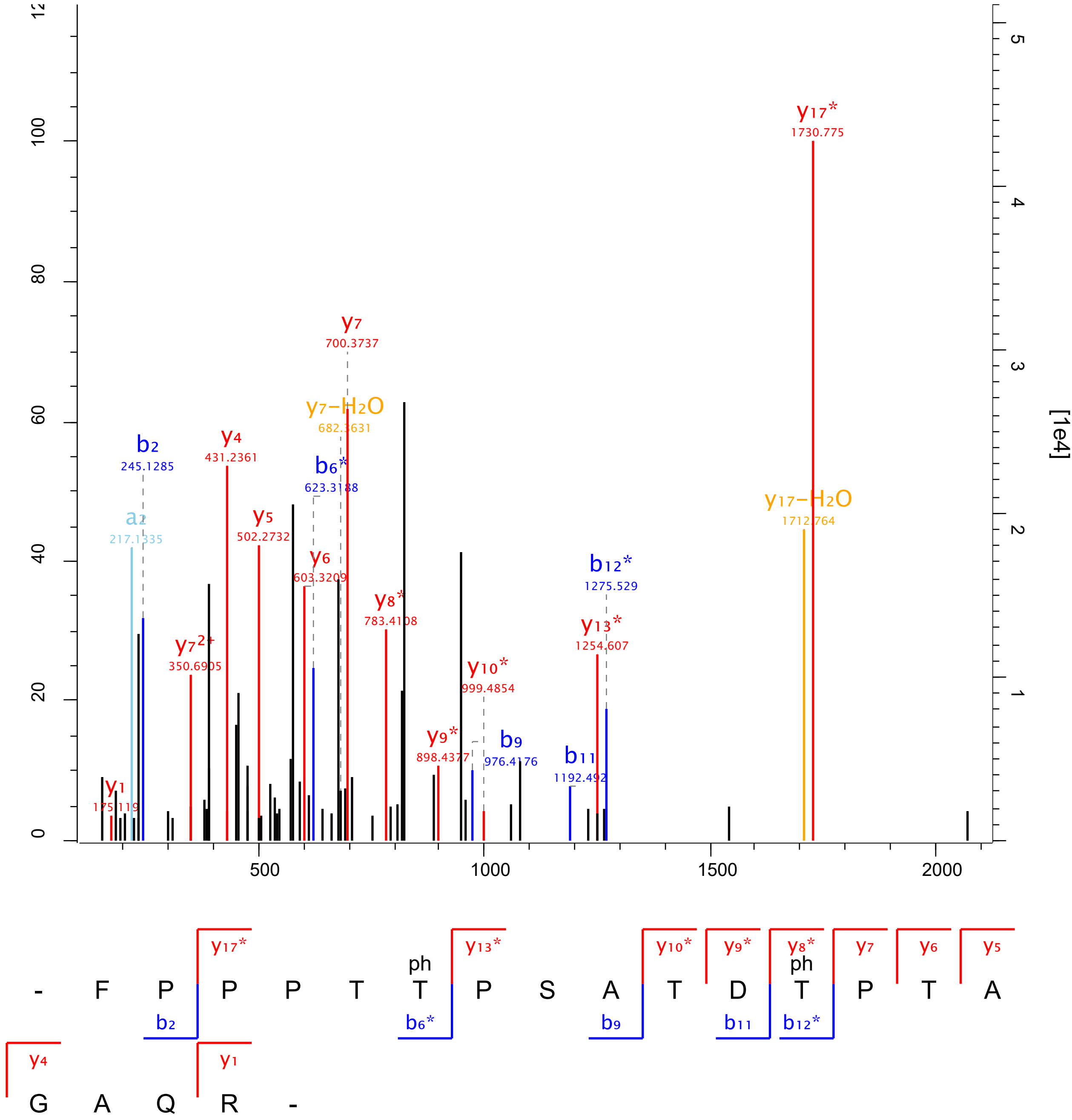

| Raw file | Scan | Method | Score | m/z |
|---|---|---|---|---|
| sys_15_1 | 25171 | FTMS; HCD | 288.96 | 949.39 |

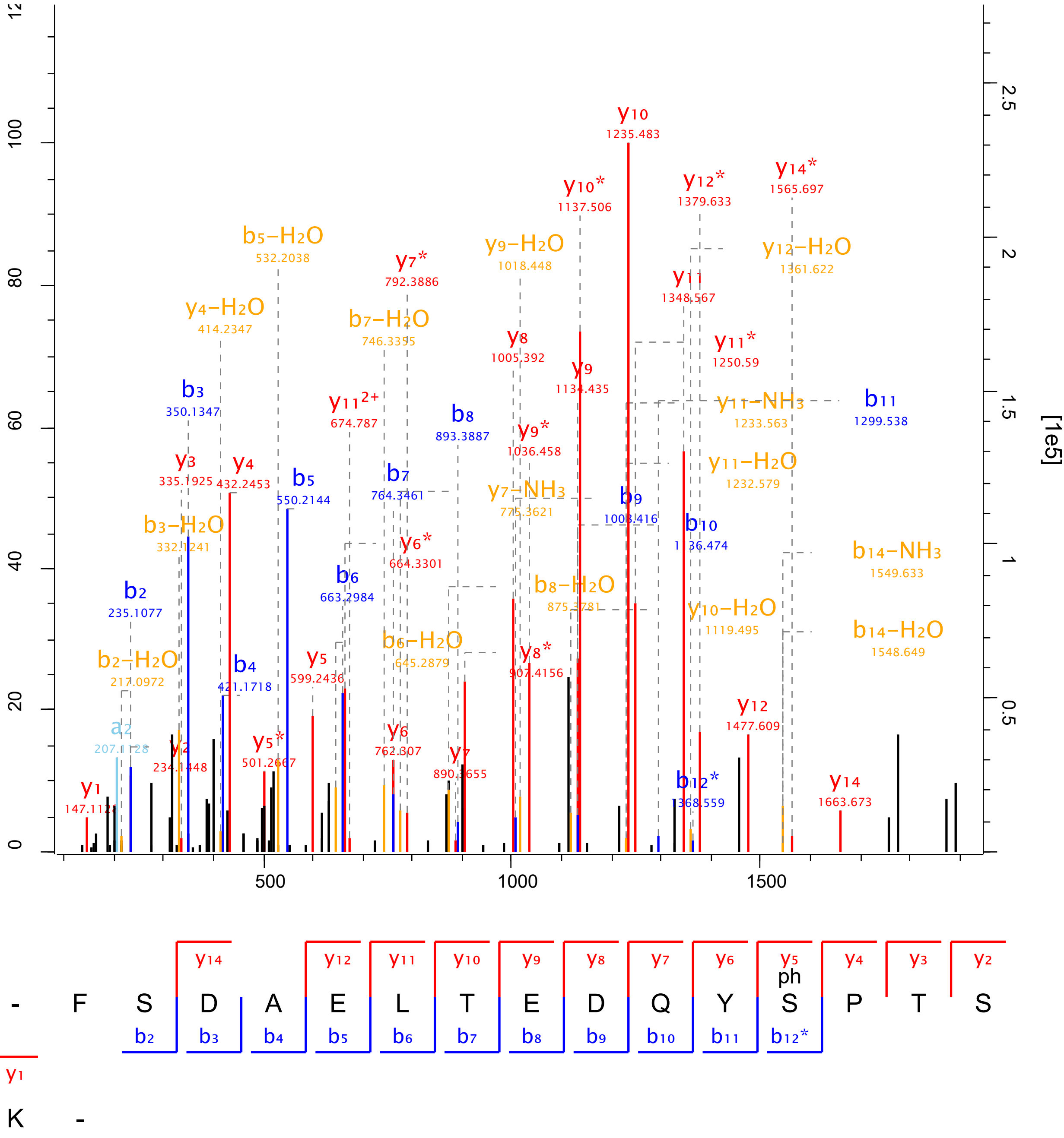

| Raw file | Scan | Method | Score | m/z |
|---|---|---|---|---|
| sys_15_1 | 25239 | FTMS; HCD | 159.18 | 643.81 |

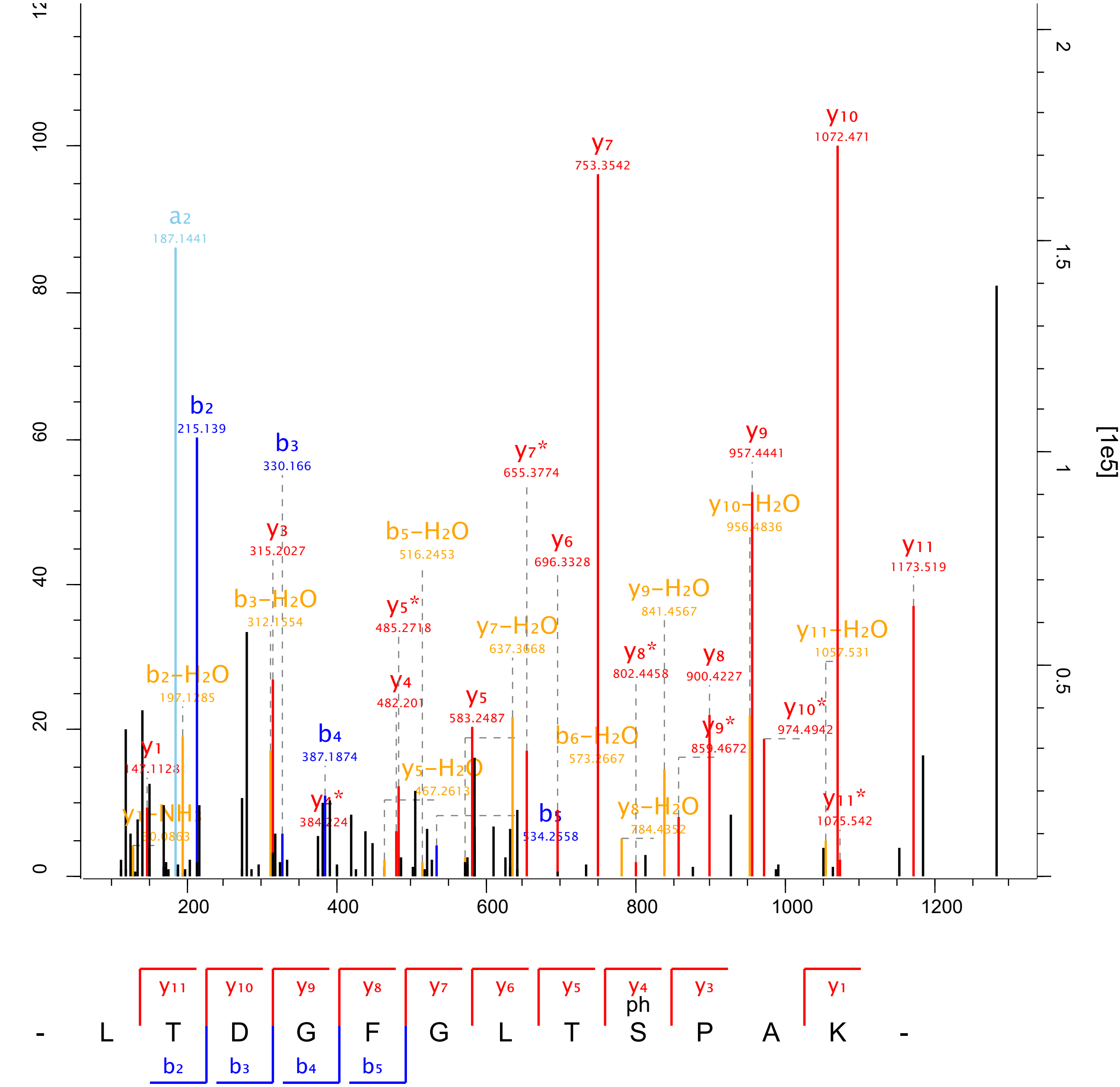

| Raw file | Scan | Method | Score | m/z |
| --- | --- | --- | --- | --- |
| sys_15_1 | 25246 | FTMS; HCD | 106.93 | 748.84 |

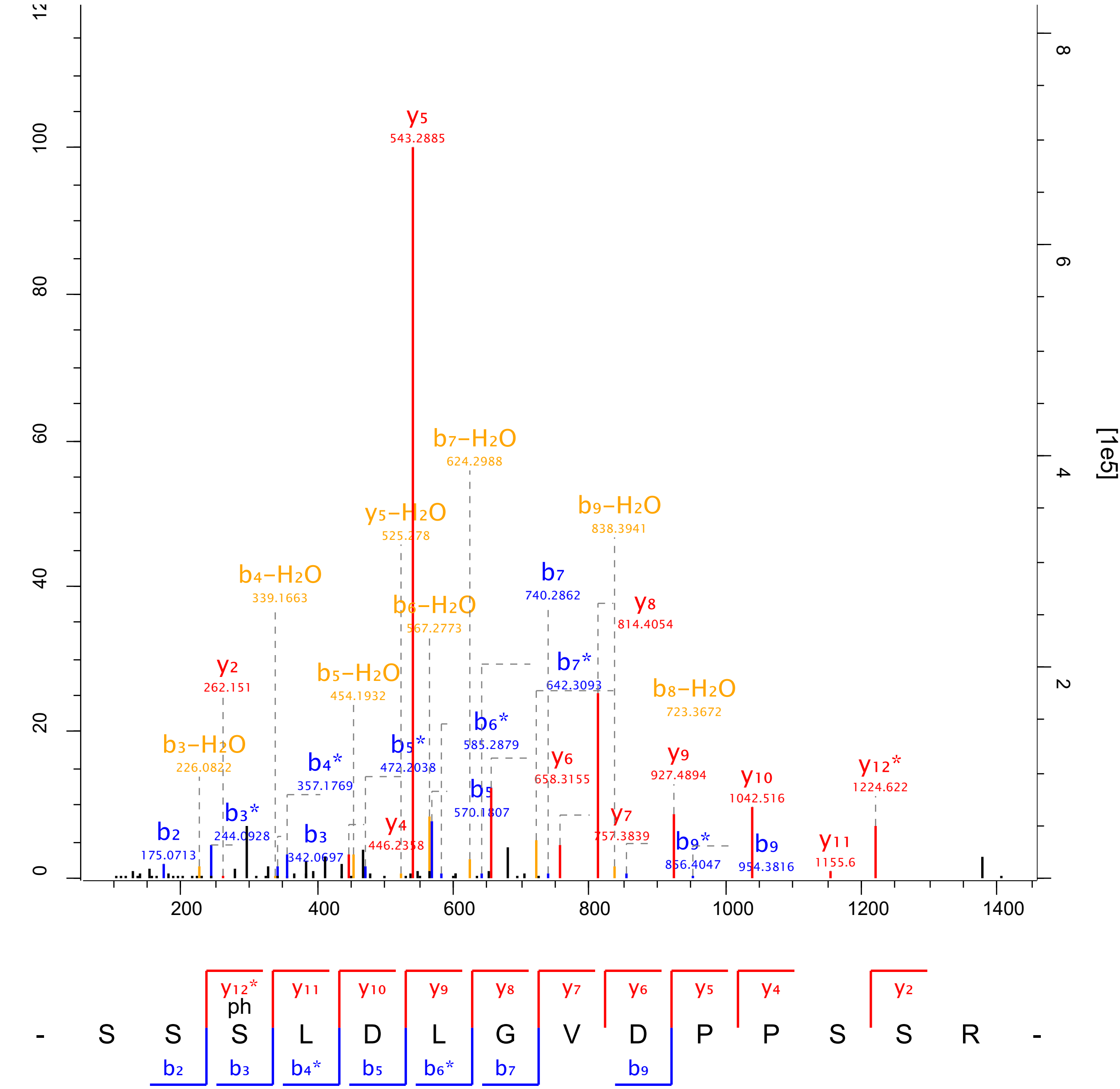

| Raw file | Scan | Method | Score | m/z |
|---|---|---|---|---|
| sys_15_1 | 25285 | FTMS; HCD | 101.53 | 814.85 |

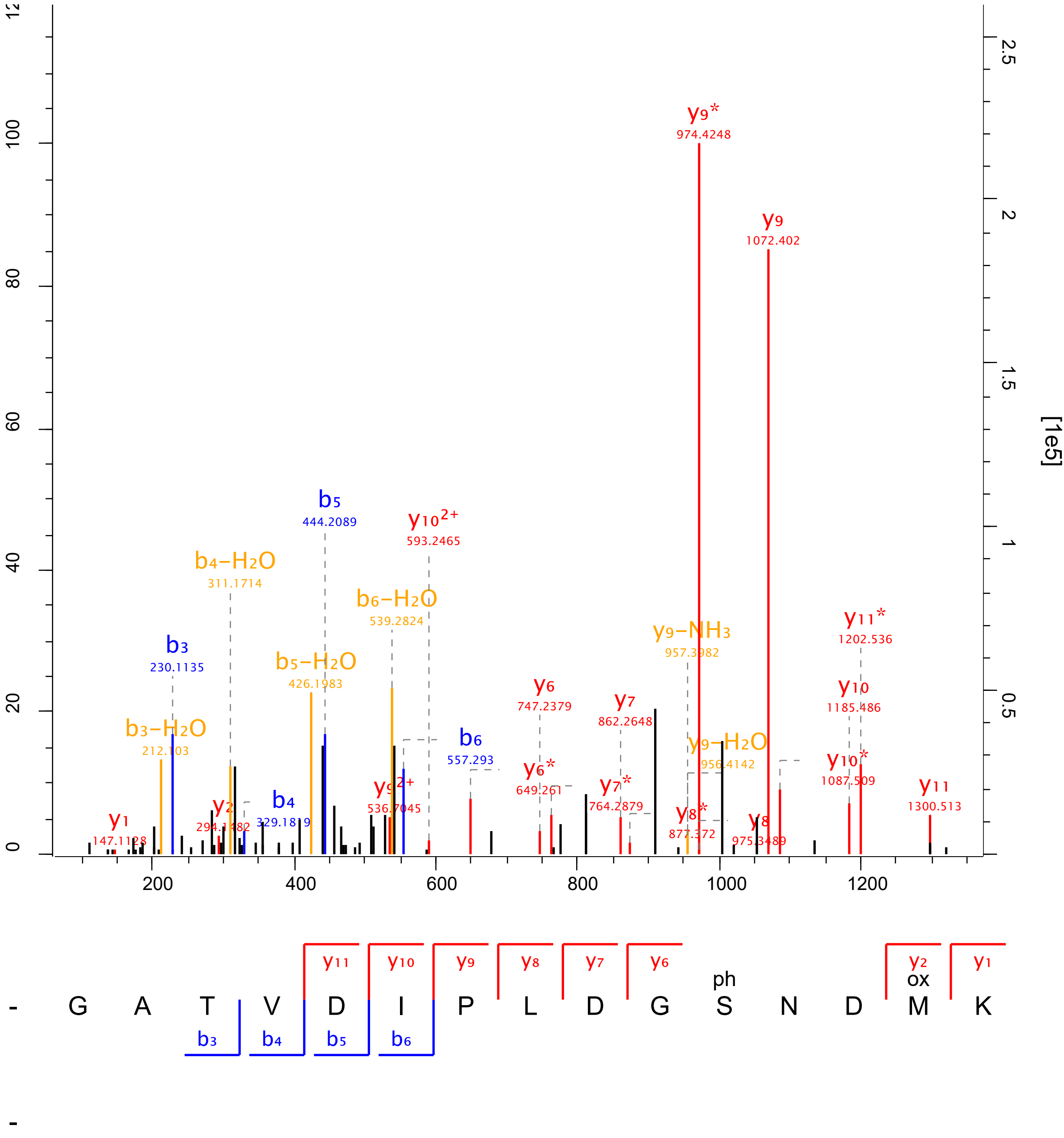

| Raw file | Scan | Method | Score | m/z |
| --- | --- | --- | --- | --- |
| sys_15_1 | 25499 | FTMS; HCD | 82.89 | 1159.97 |

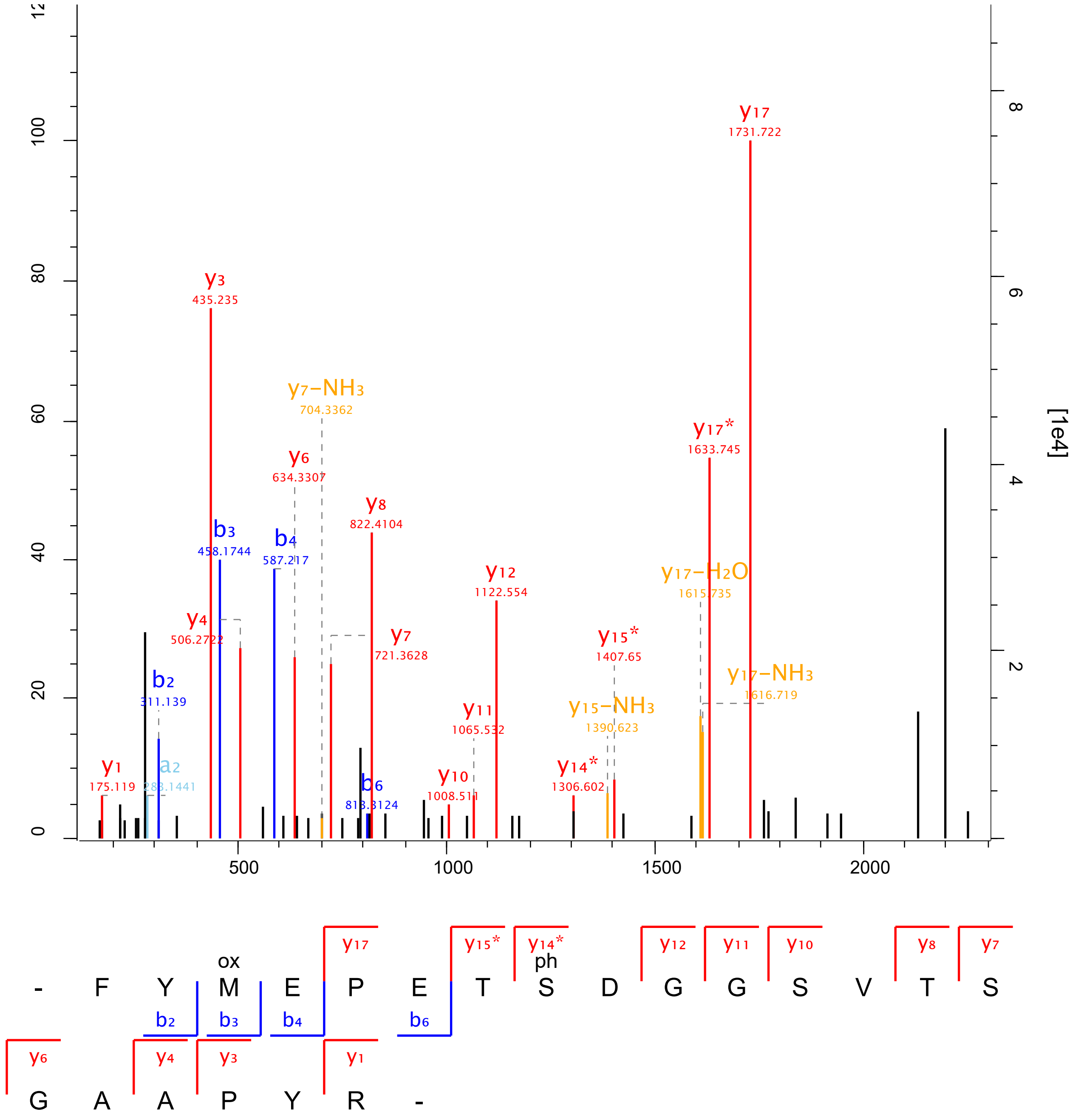

| Raw file | Scan | Method | Score | m/z |
|---|---|---|---|---|
| sys_15_1 | 25504 | FTMS; HCD | 110.97 | 765.81 |

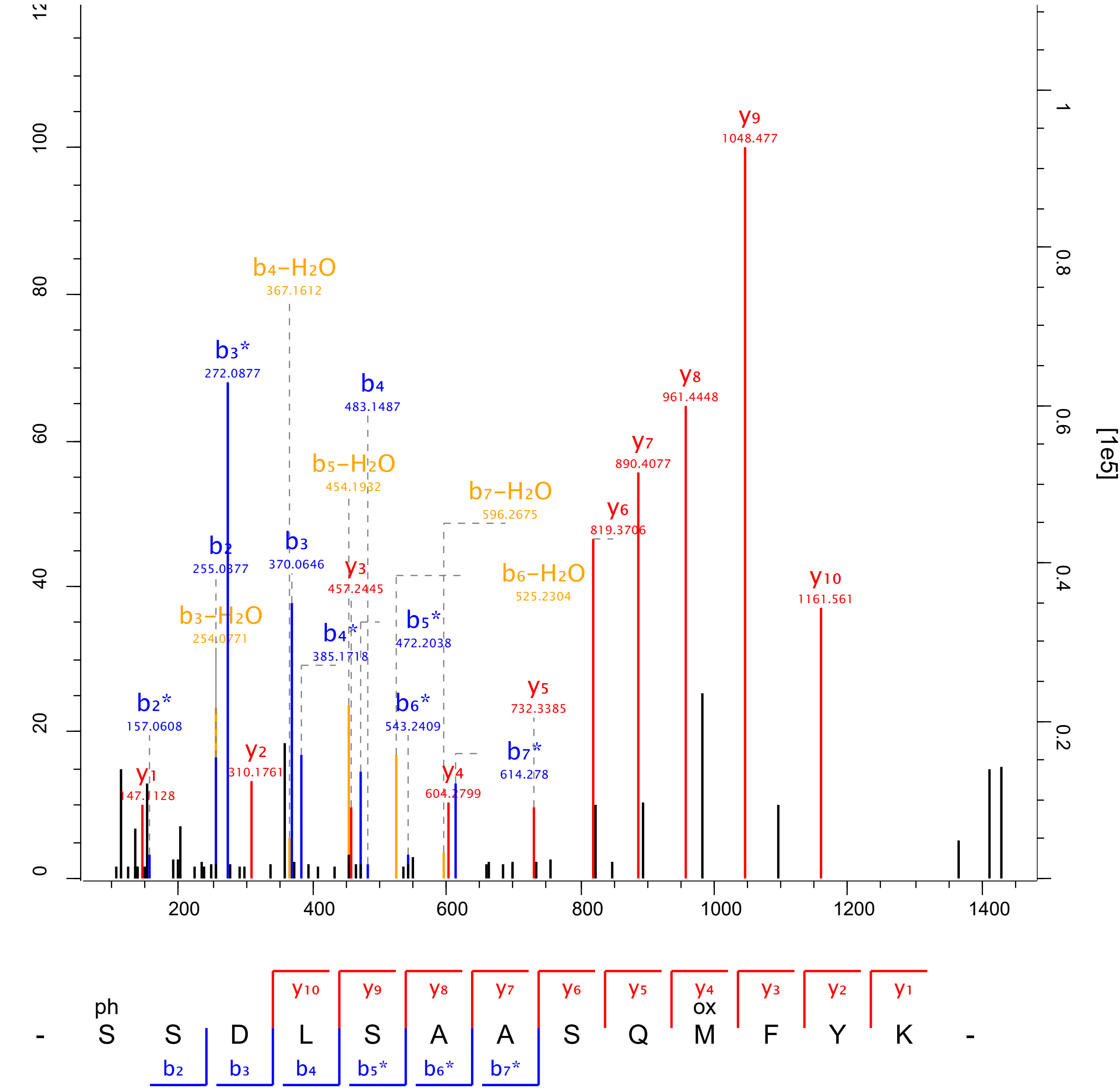

| Raw file | Scan | Method | Score | m/z |
| --- | --- | --- | --- | --- |
| sys_15_1 | 25545 | FTMS; HCD | 190.86 | 937.4 |

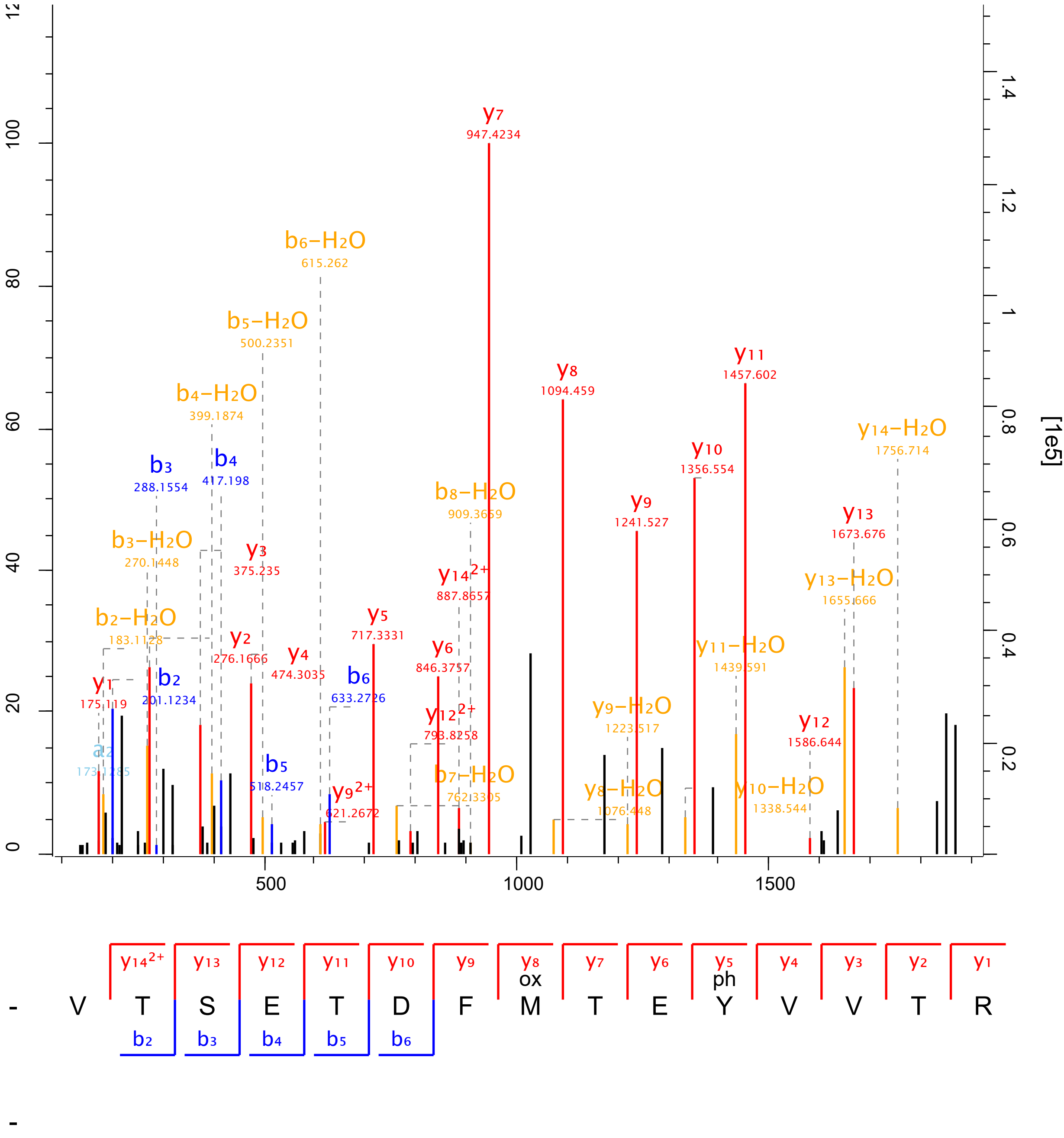

| Raw file | Scan | Method | Score | m/z |
| --- | --- | --- | --- | --- |
| sys_15_1 | 25788 | FTMS; HCD | 43.68 | 599.77 |

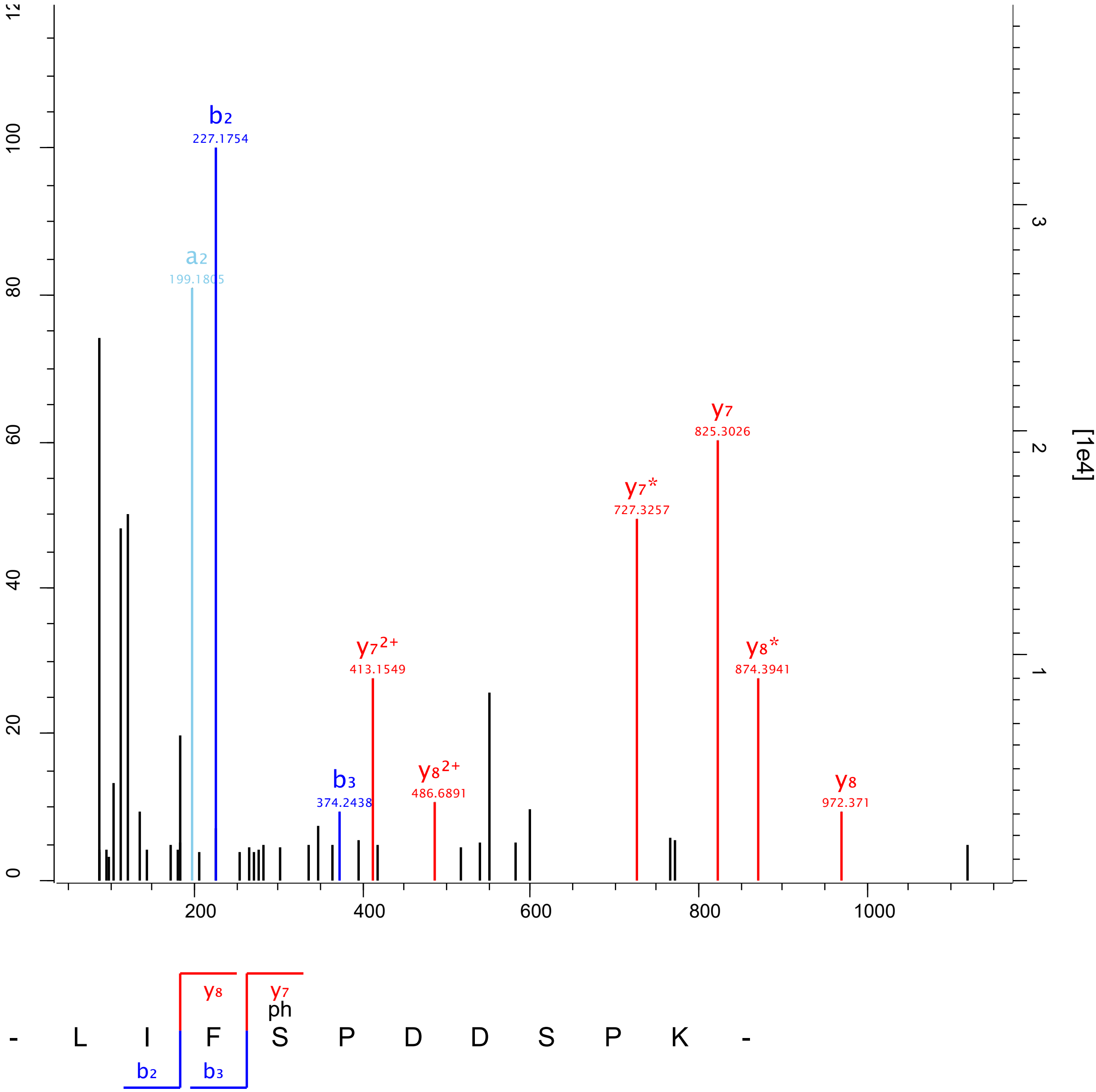

| Raw file | Scan | Method | Score | m/z |
| --- | --- | --- | --- | --- |
| sys_15_1 | 25982 | FTMS; HCD | 118.48 | 729.85 |

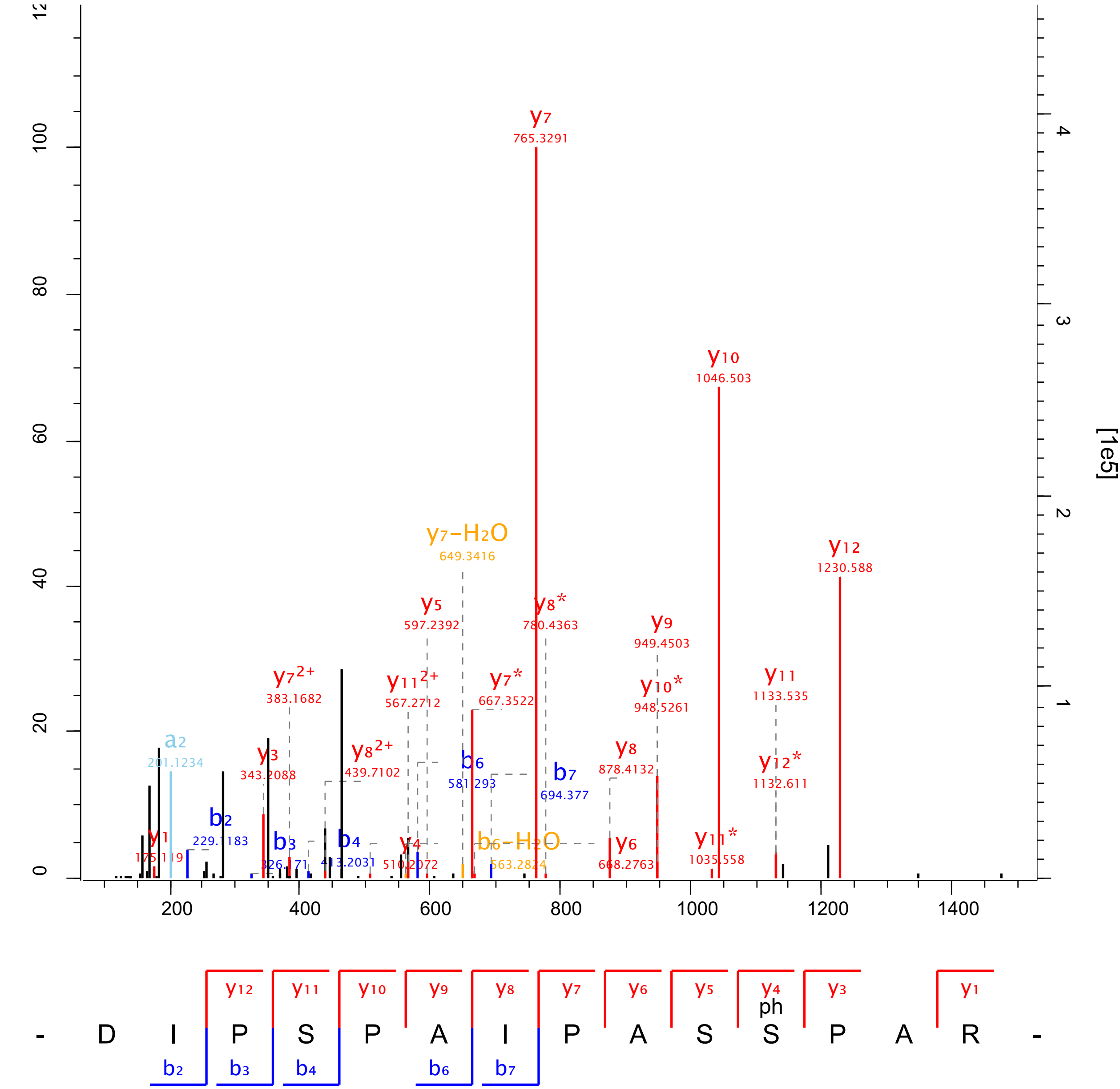

| Raw file | Scan | Method | Score | m/z |
| --- | --- | --- | --- | --- |
| sys_15_1 | 26076 | FTMS; HCD | 80.96 | 692.81 |

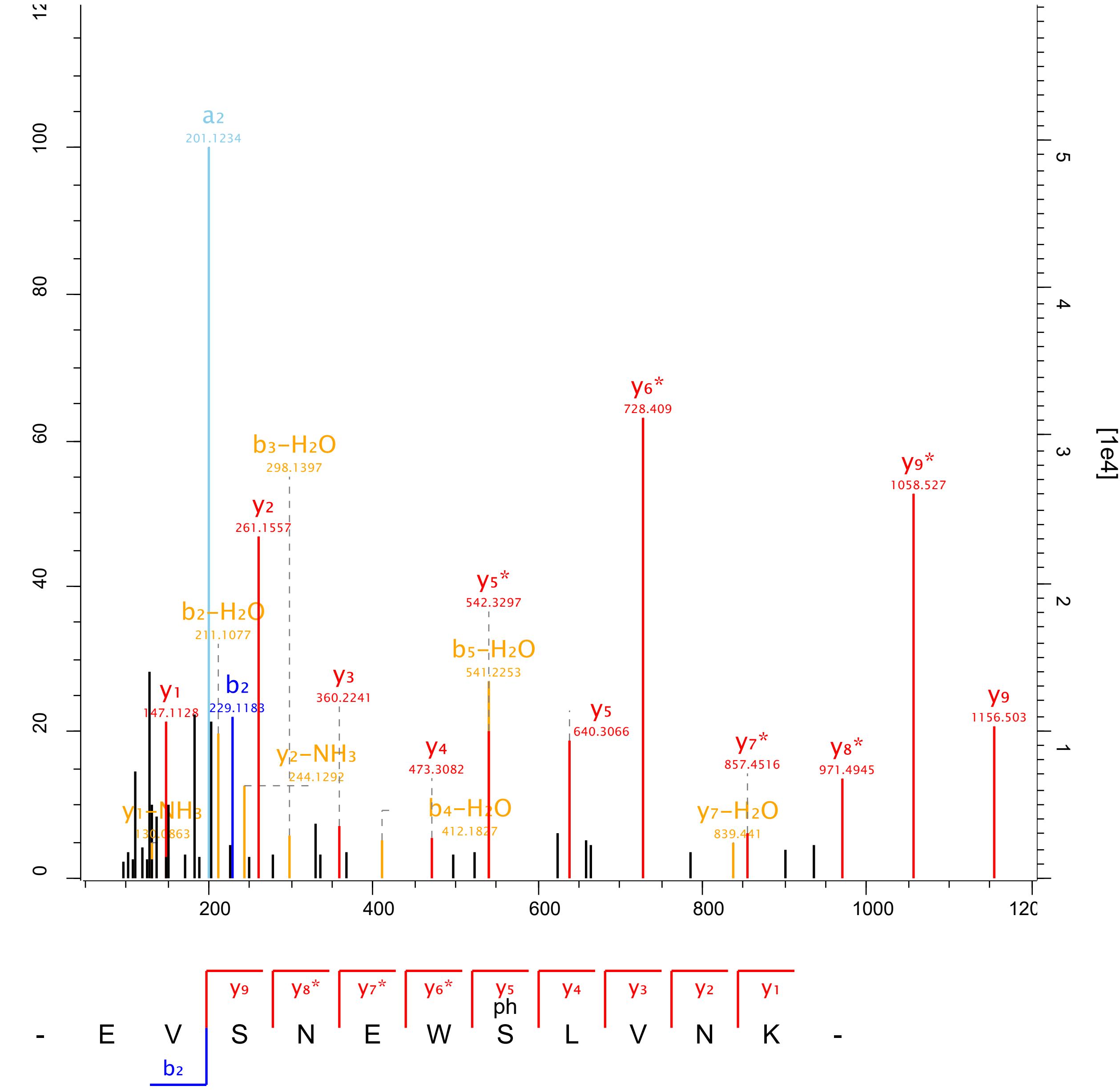

| Raw file | Scan | Method | Score | m/z |
|---|---|---|---|---|
| sys_15_1 | 26147 | FTMS; HCD | 61.11 | 544.23 |

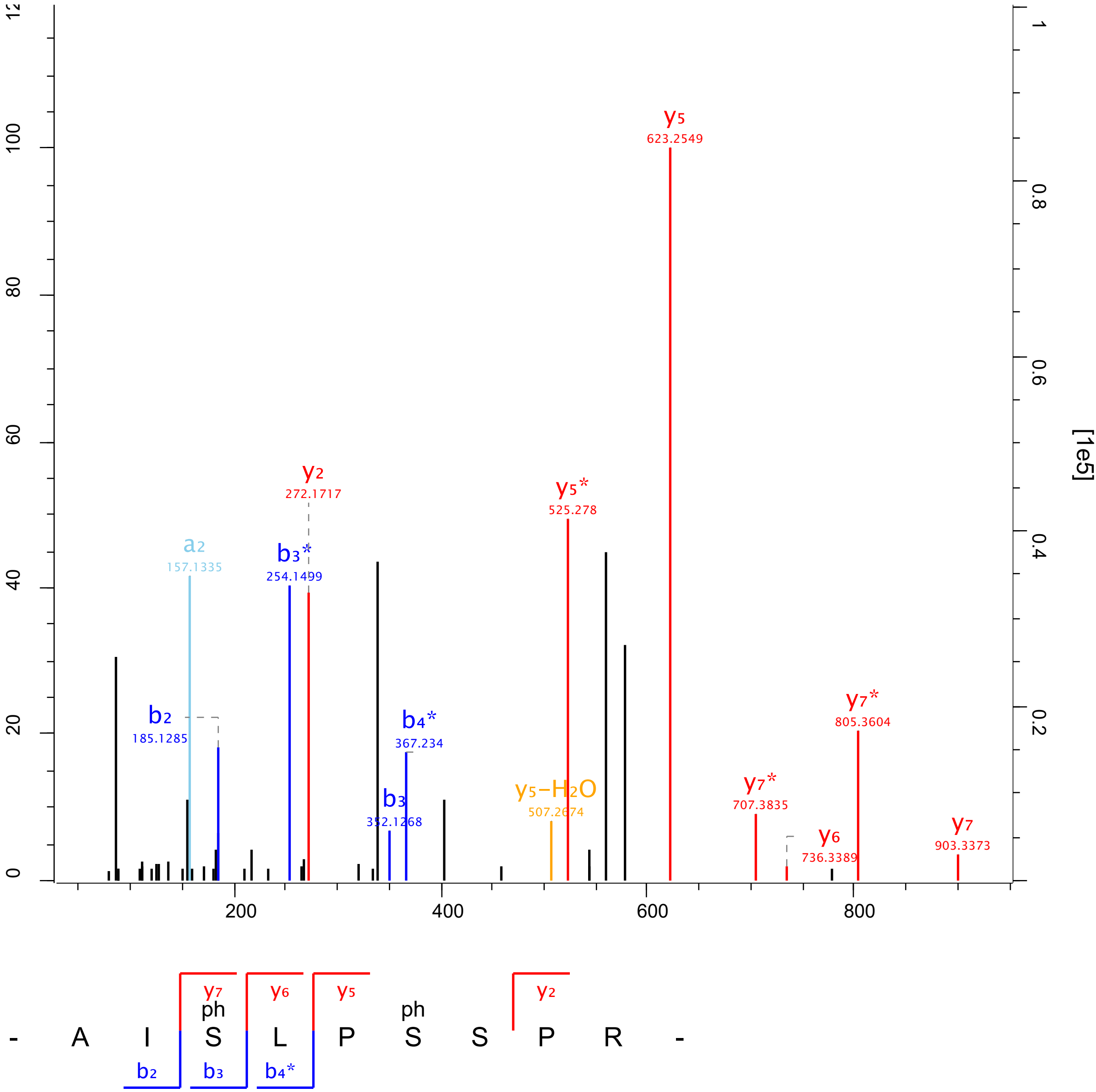

| Raw file | Scan | Method | Score | m/z |
|---|---|---|---|---|
| sys_15_1 | 26149 | FTMS; HCD | 144.72 | 951.87 |

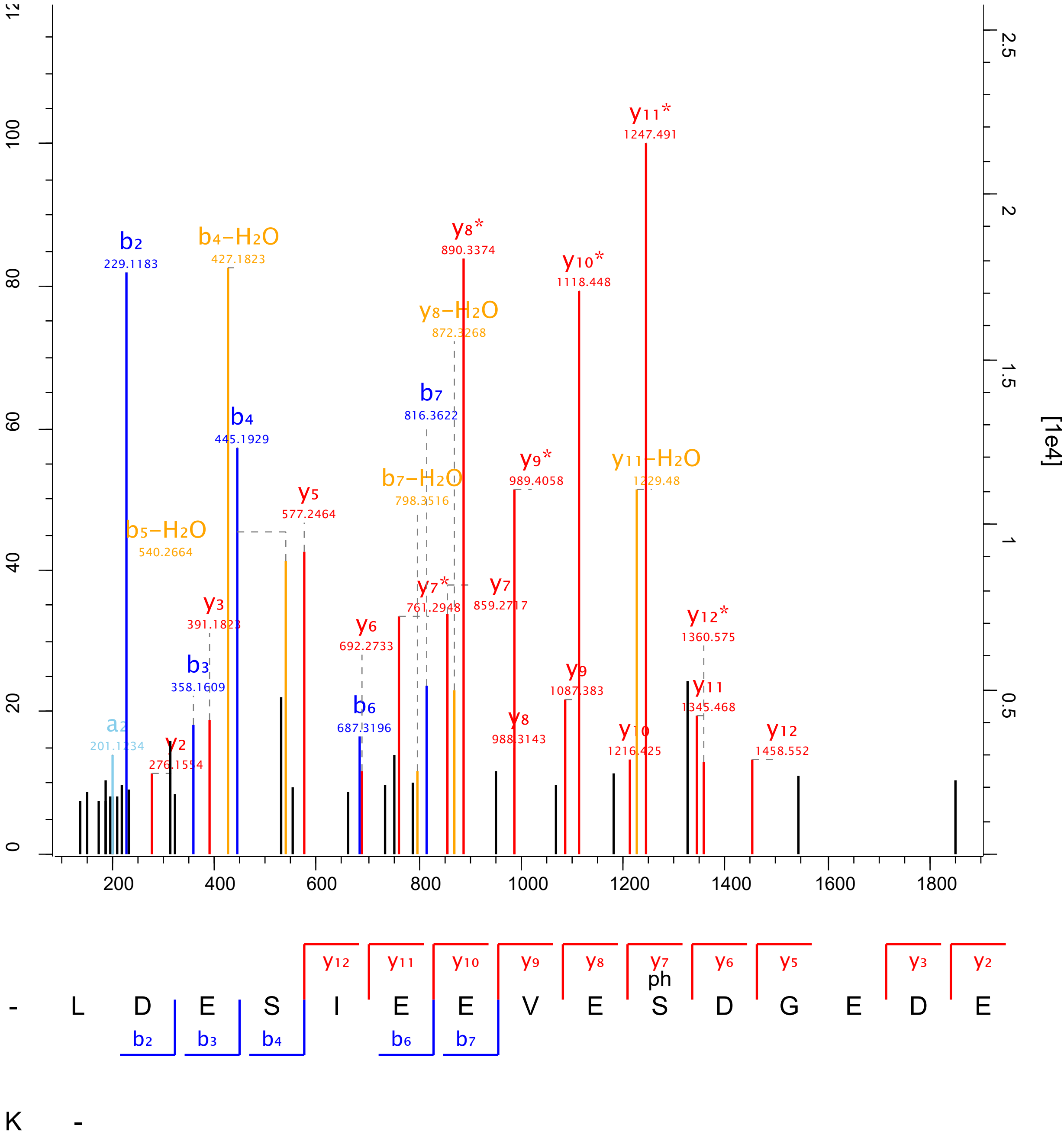

| Raw file | Scan | Method | Score | m/z |
|---|---|---|---|---|
| sys_15_1 | 26250 | FTMS; HCD | 150.36 | 623.97 |

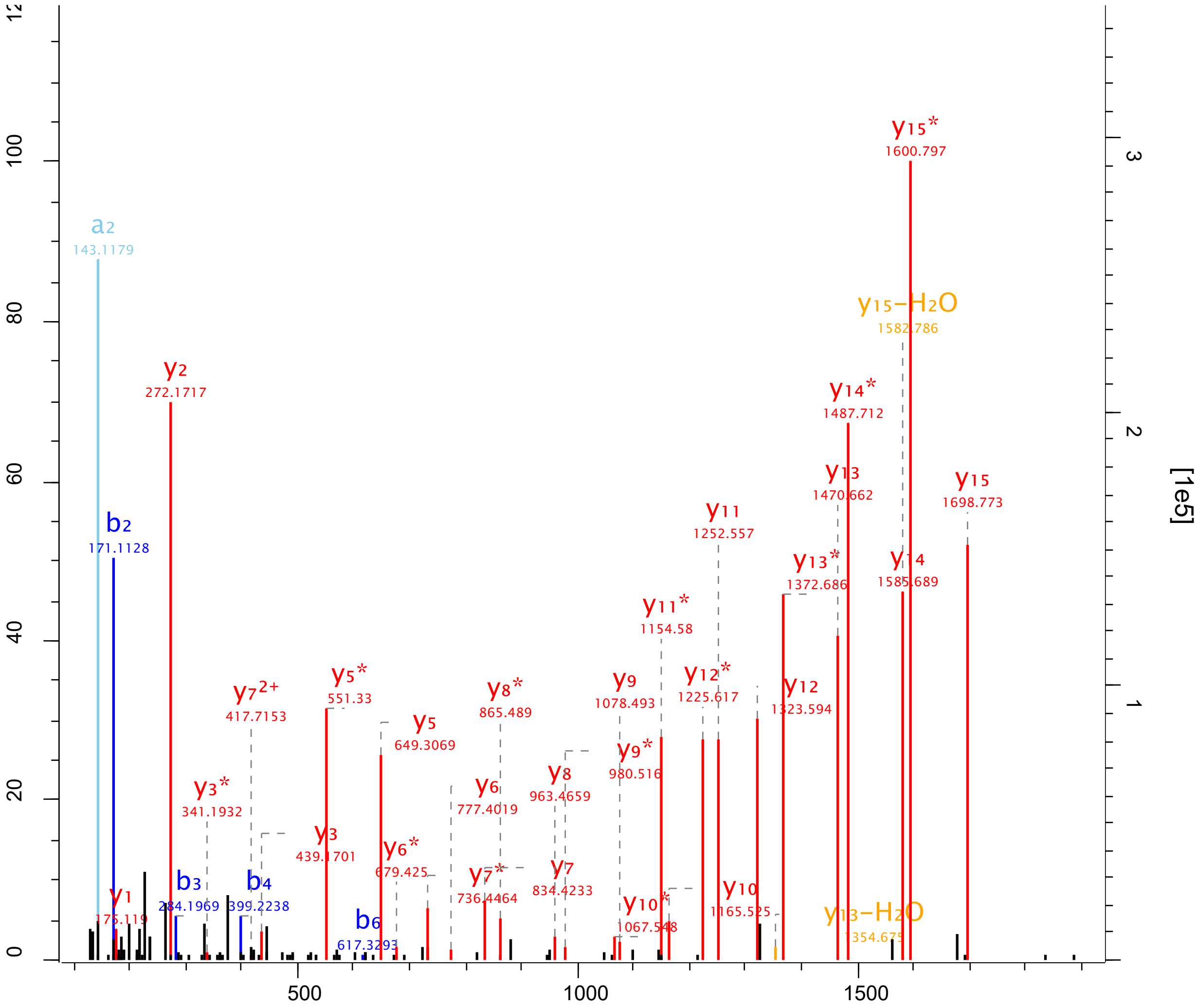


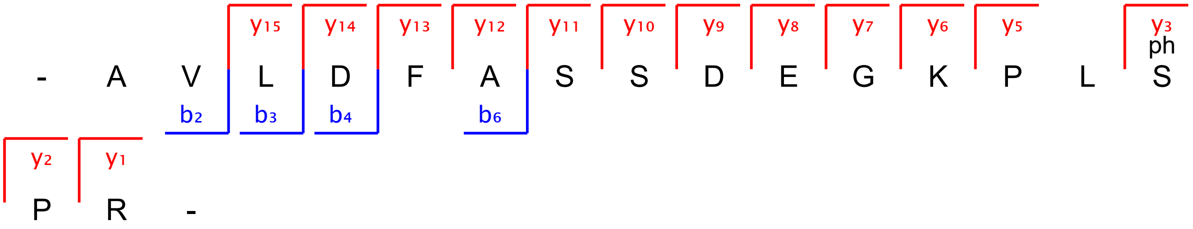

| Raw file | Scan | Method | Score | m/z |
|---|---|---|---|---|
| sys_15_1 | 26314 | FTMS; HCD | 161.68 | 1039.82 |

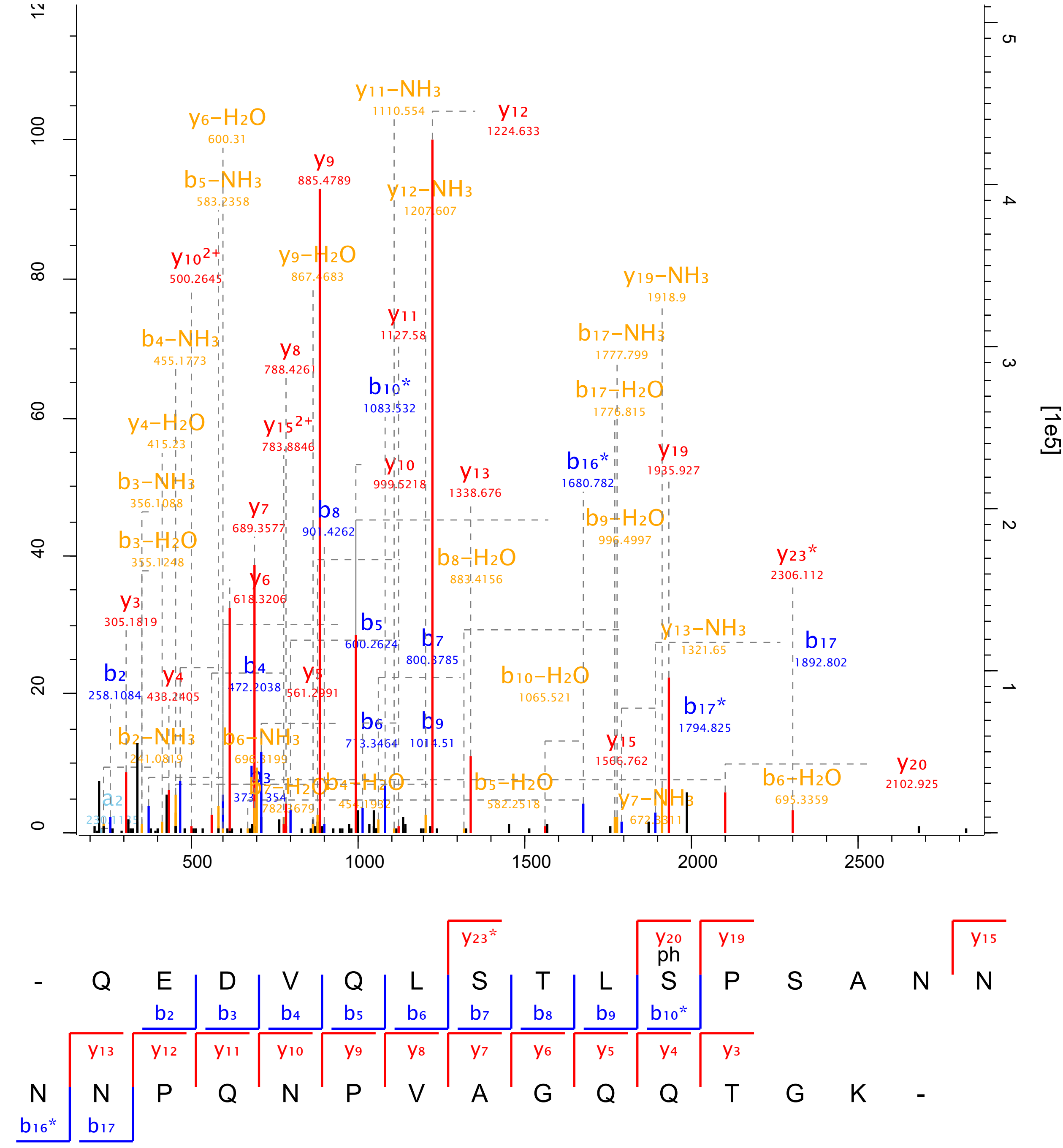

| Raw file | Scan | Method | Score | m/z |
| --- | --- | --- | --- | --- |
| sys_15_1 | 26419 | FTMS; HCD | 165.3 | 801.04 |

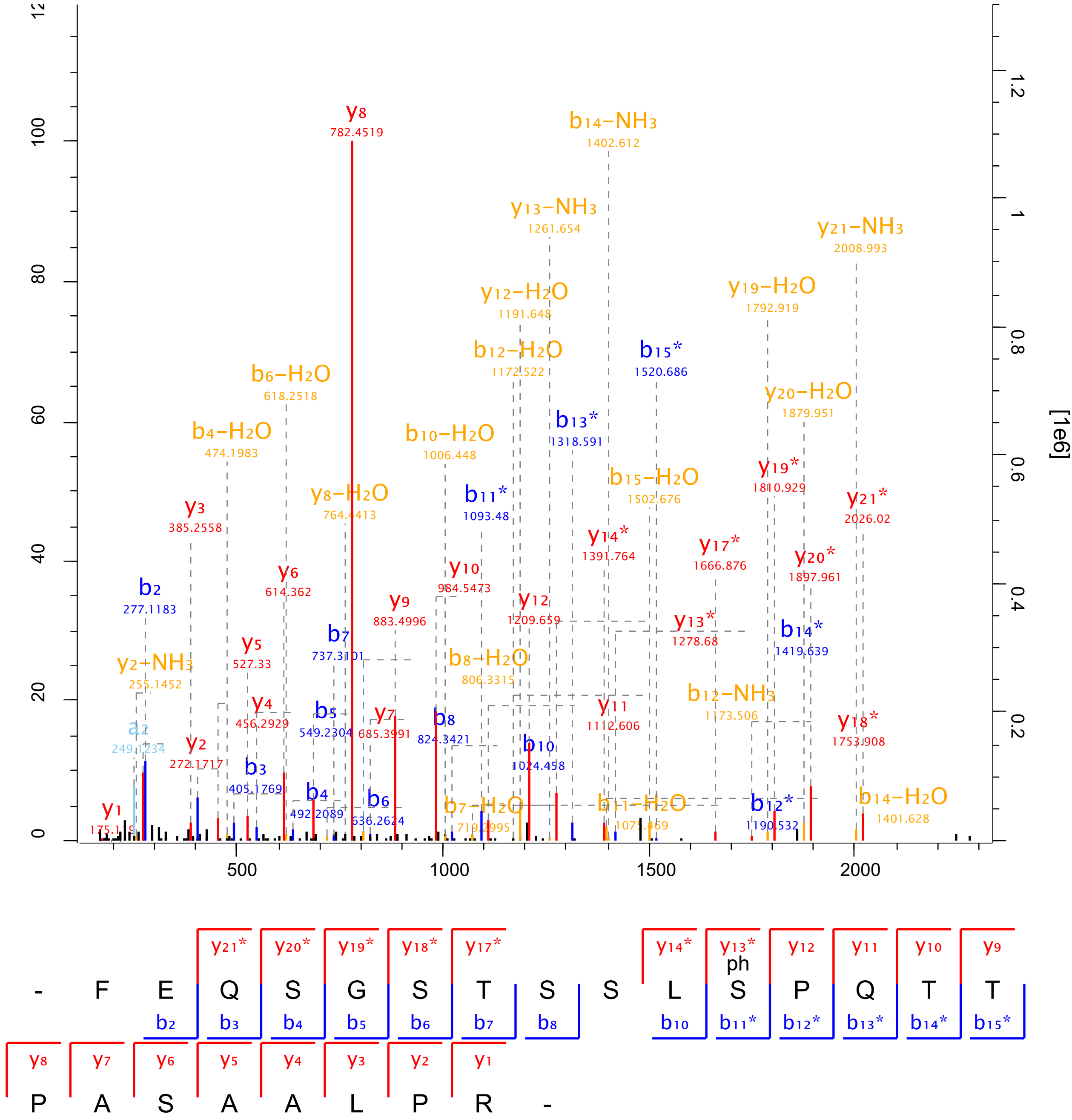

| Raw file | Scan | Method | Score | m/z |
| --- | --- | --- | --- | --- |
| sys_15_1 | 26467 | FTMS; HCD | 63.68 | 714.81 |

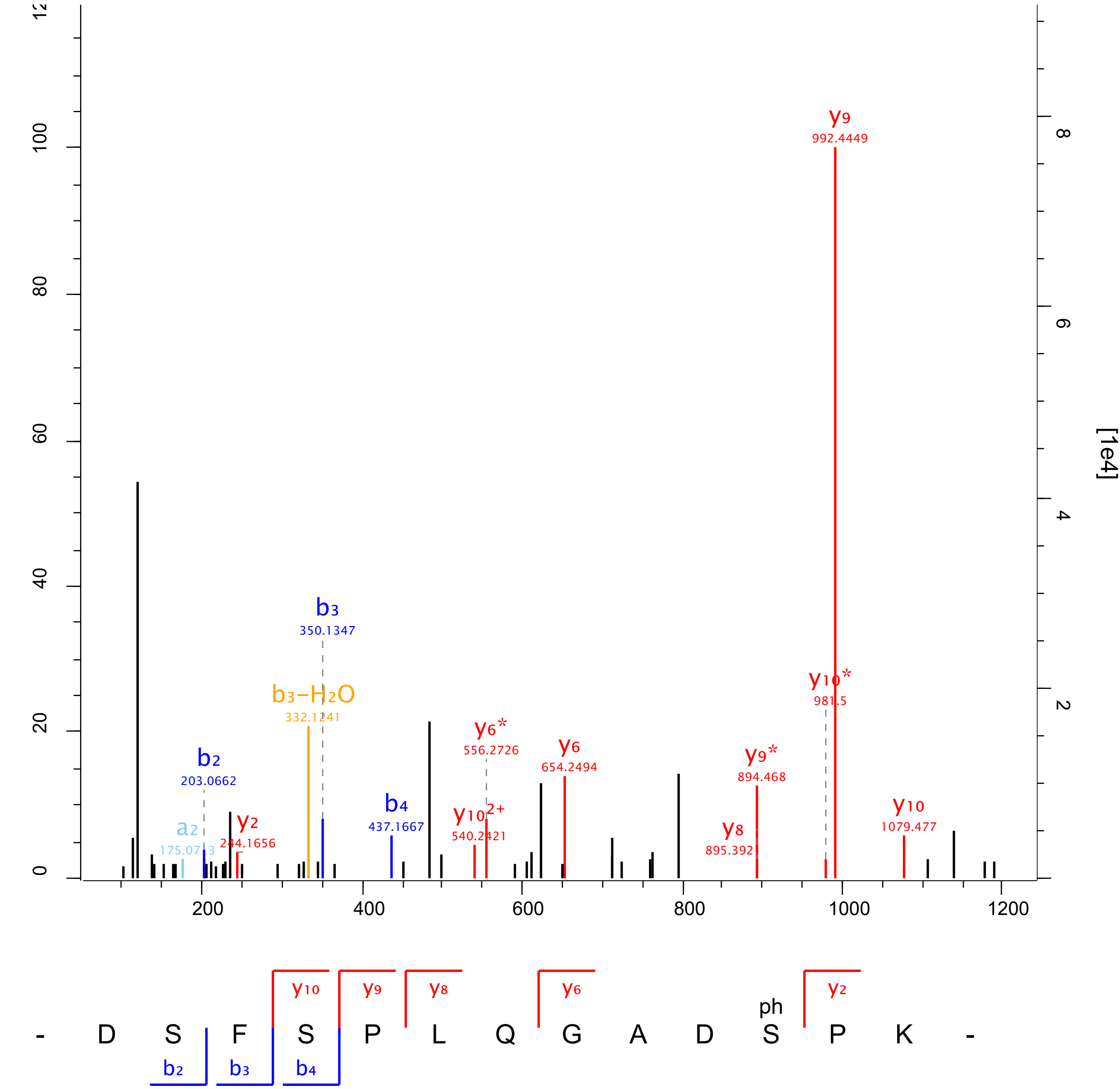

| Raw file | Scan | Method | Score | m/z |
| --- | --- | --- | --- | --- |
| sys_15_1 | 2650 | FTMS; HCD | 198.98 | 697.78 |

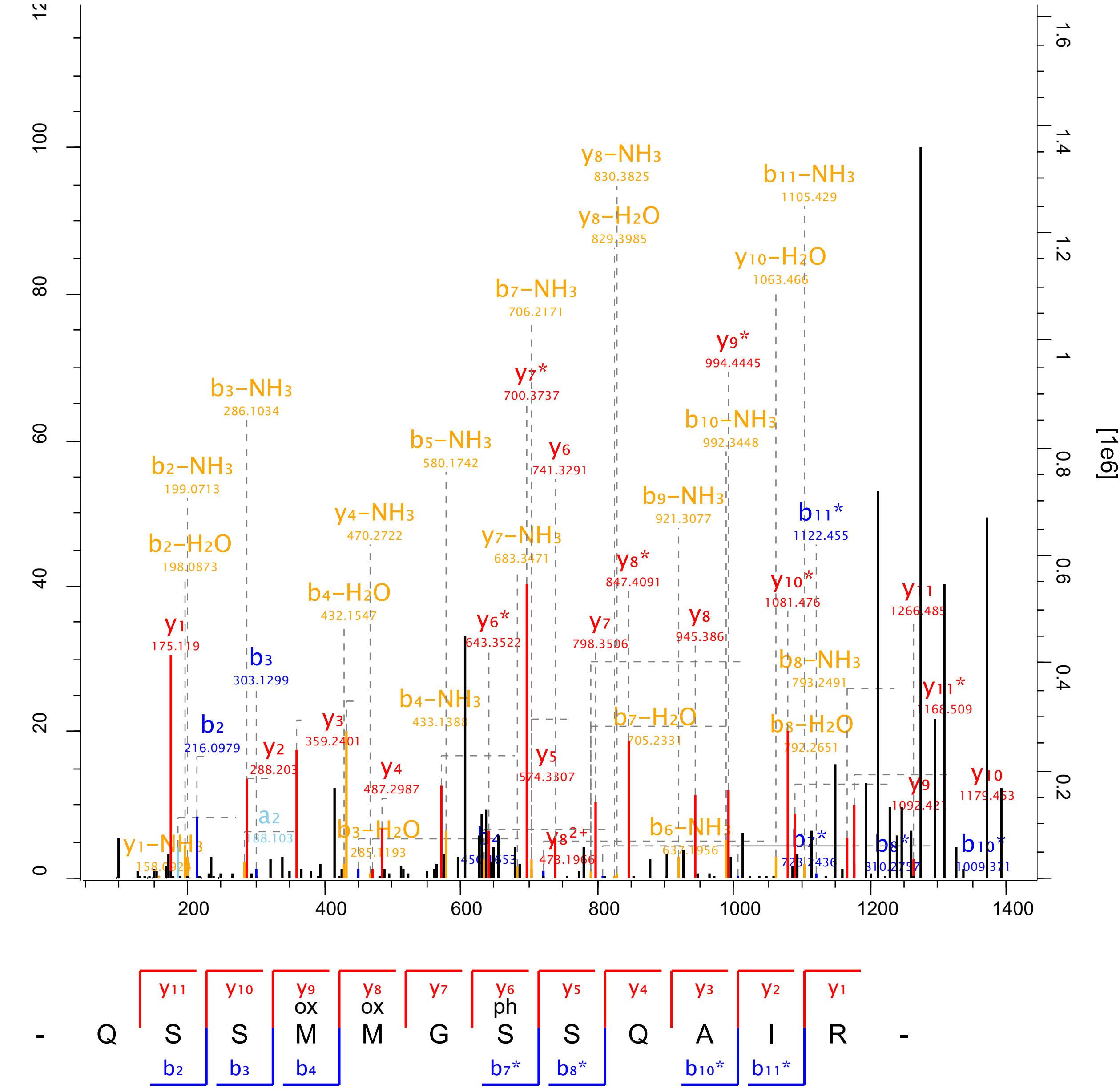

| Raw file | Scan | Method | Score | m/z |
| --- | --- | --- | --- | --- |
| sys_15_1 | 26518 | FTMS; HCD | 88.54 | 749 |

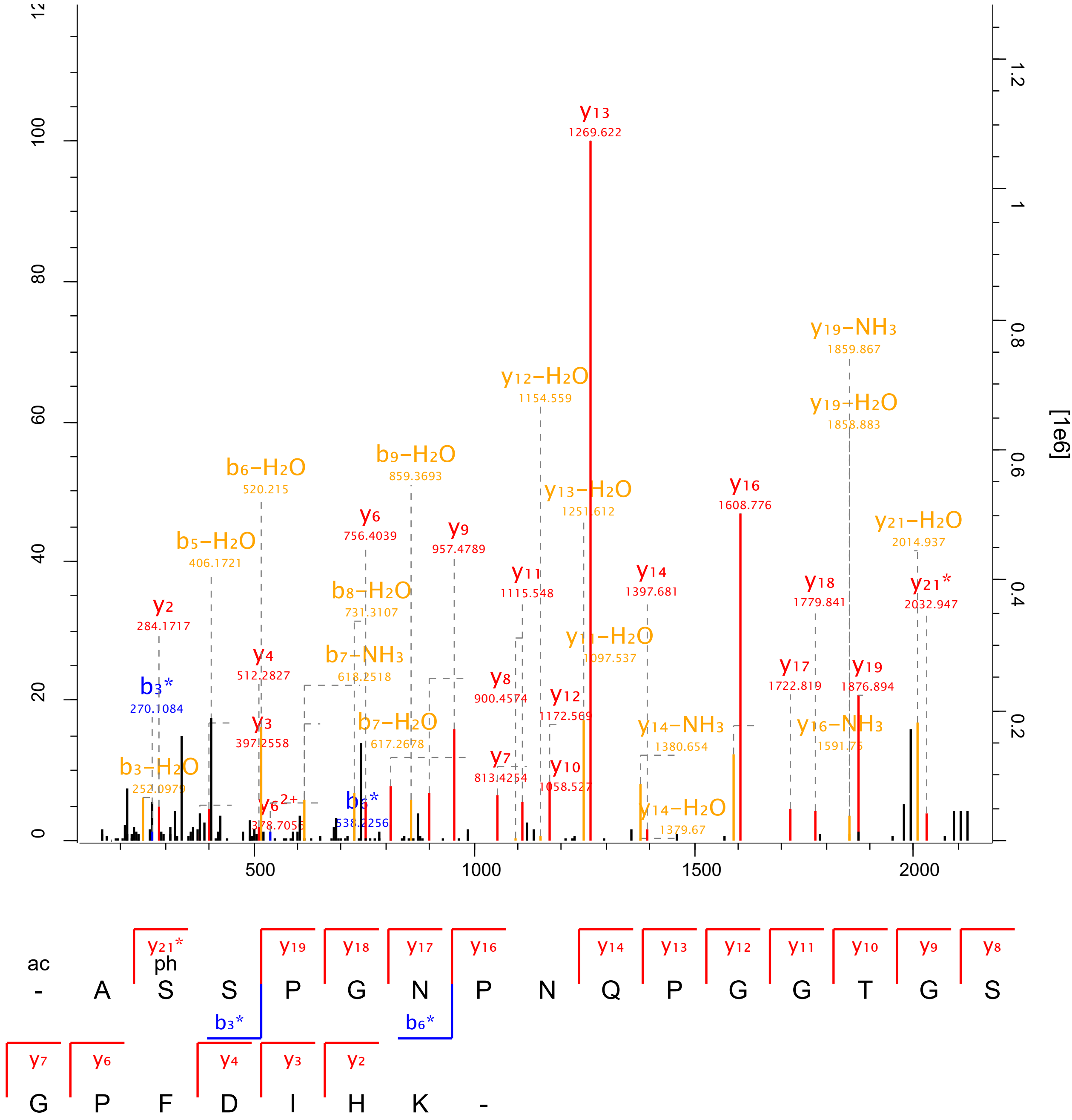

| Raw file | Scan | Method | Score | m/z |
|---|---|---|---|---|
| sys_15_1 | 26601 | FTMS; HCD | 139.86 | 607.76 |

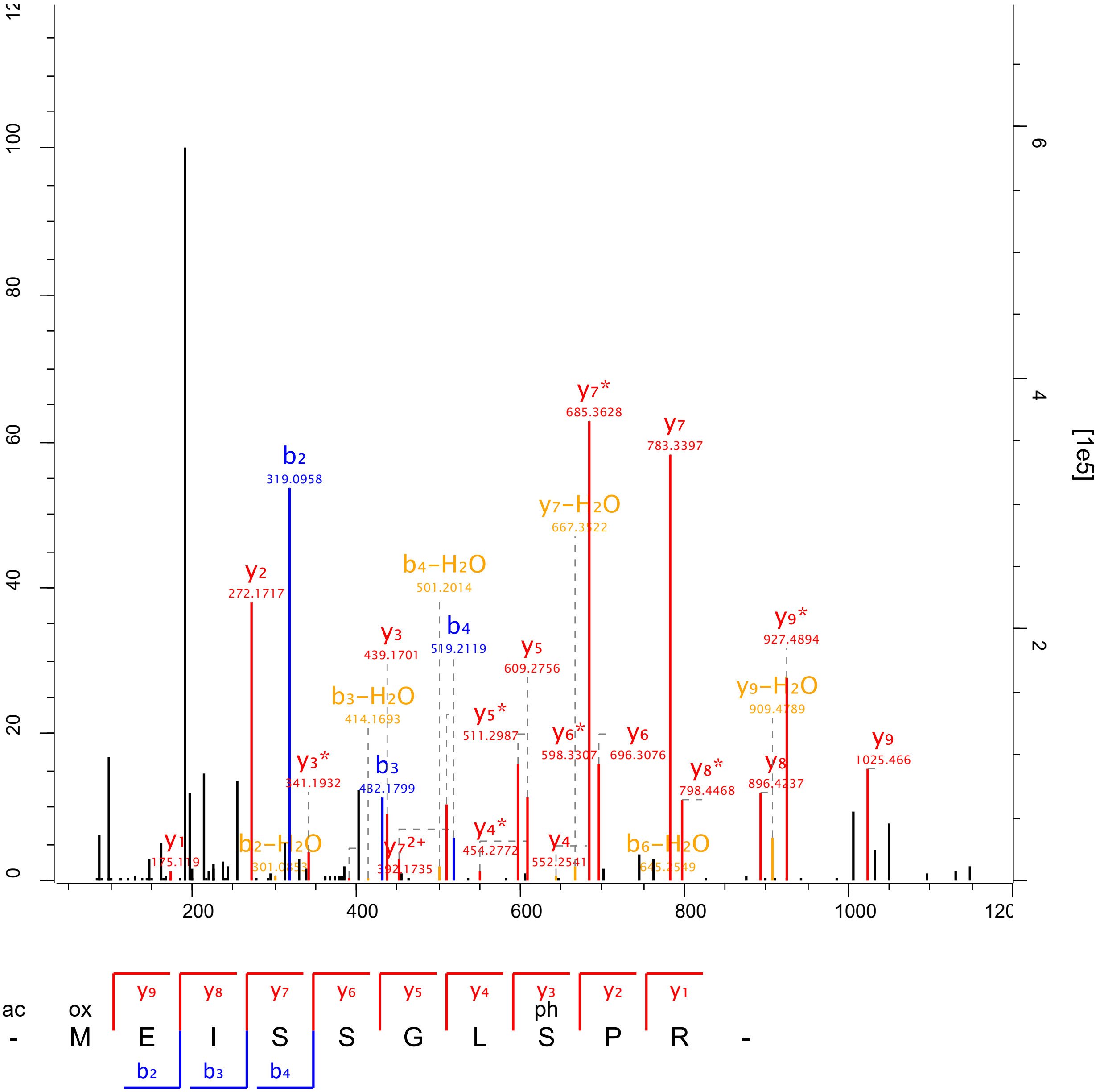

| Raw file | Scan | Method | Score | m/z |
|---|---|---|---|---|
| sys_15_1 | 26741 | FTMS; HCD | 100.97 | 871.87 |

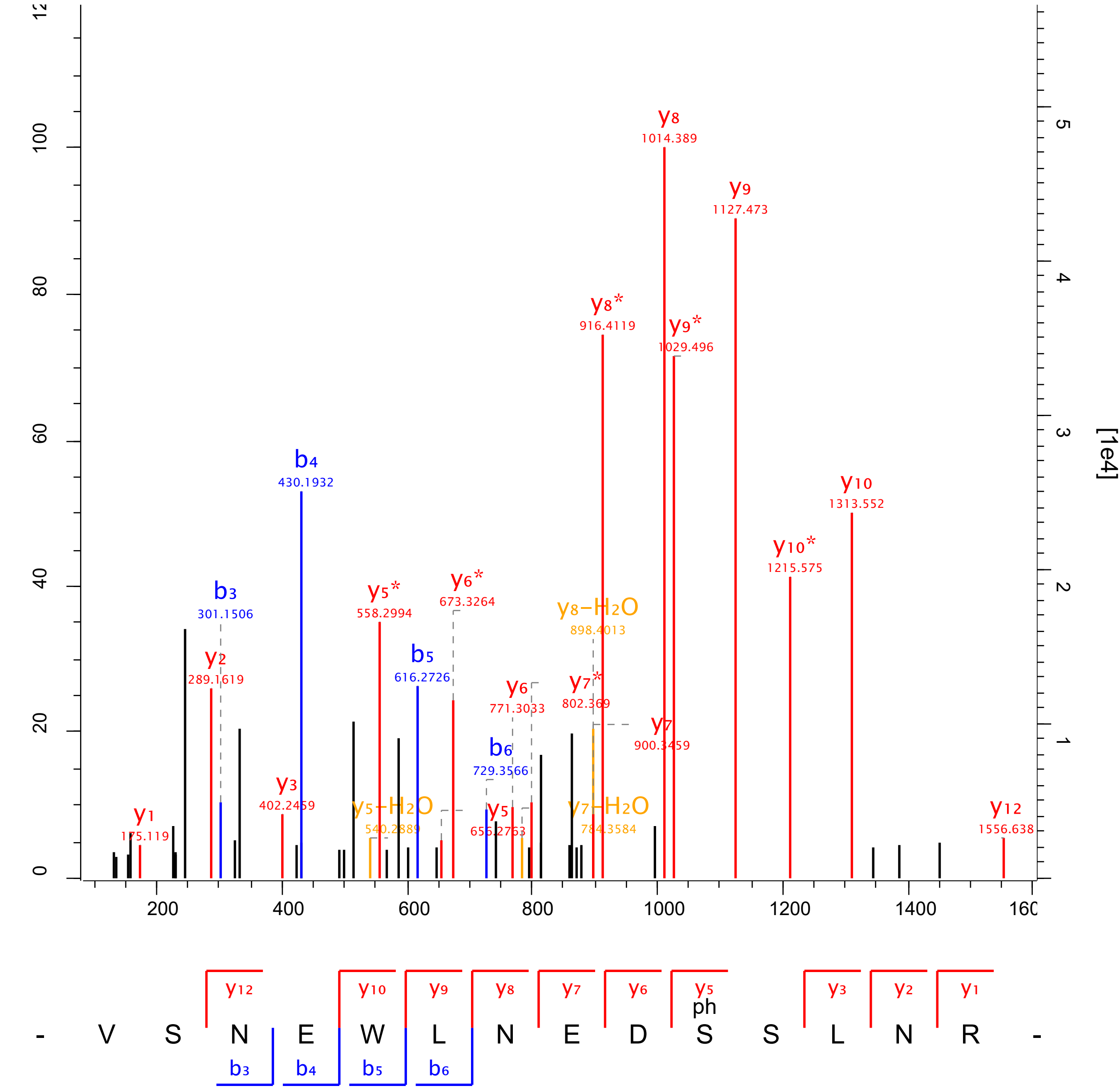

| Raw file | Scan | Method | Score | m/z |
| --- | --- | --- | --- | --- |
| sys_15_1 | 26783 | FTMS; HCD | 112.96 | 755.79 |

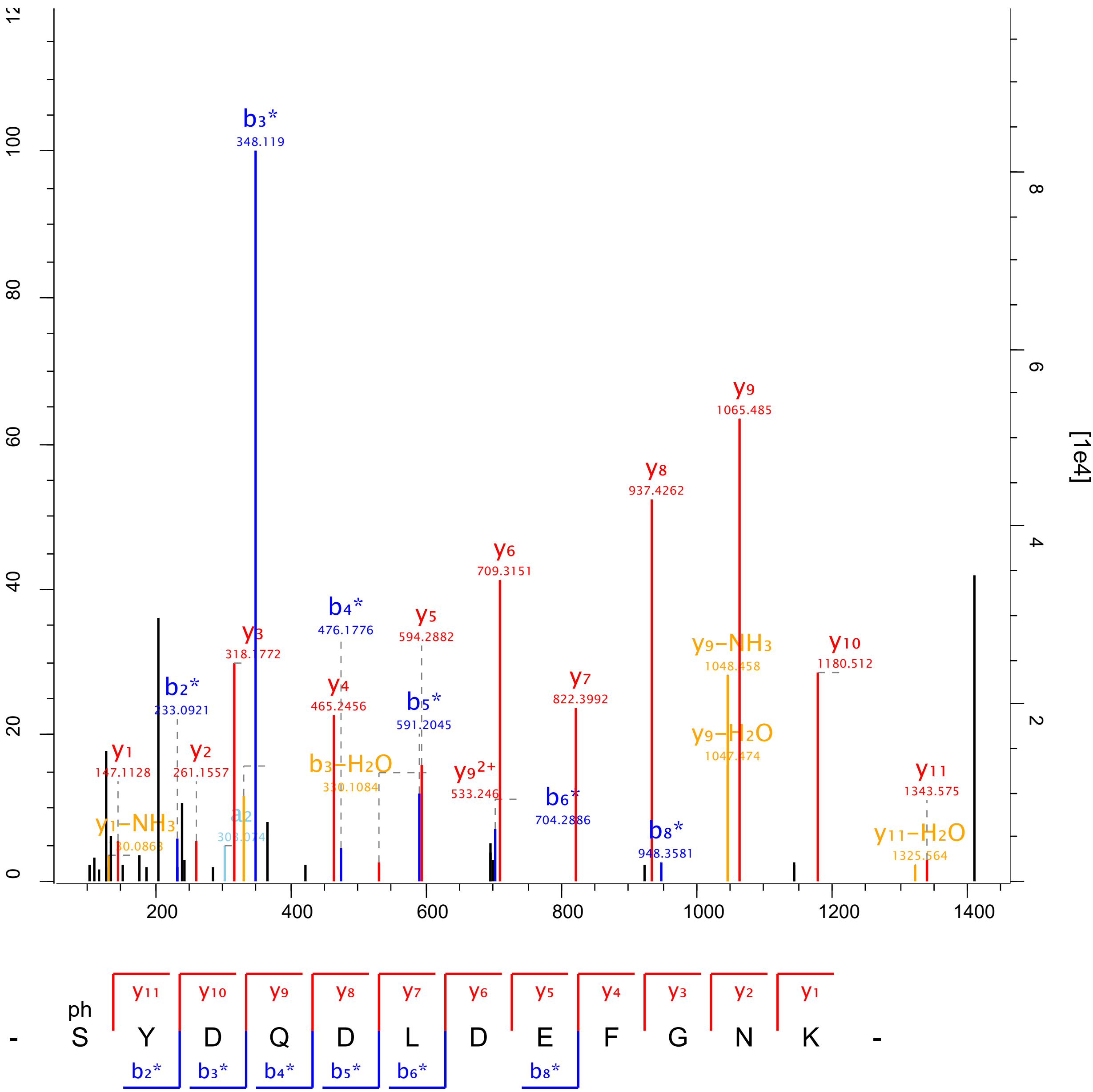

| Raw file | Scan | Method | Score | m/z |
|---|---|---|---|---|
| sys_15_1 | 26808 | FTMS; HCD | 118.15 | 966.41 |

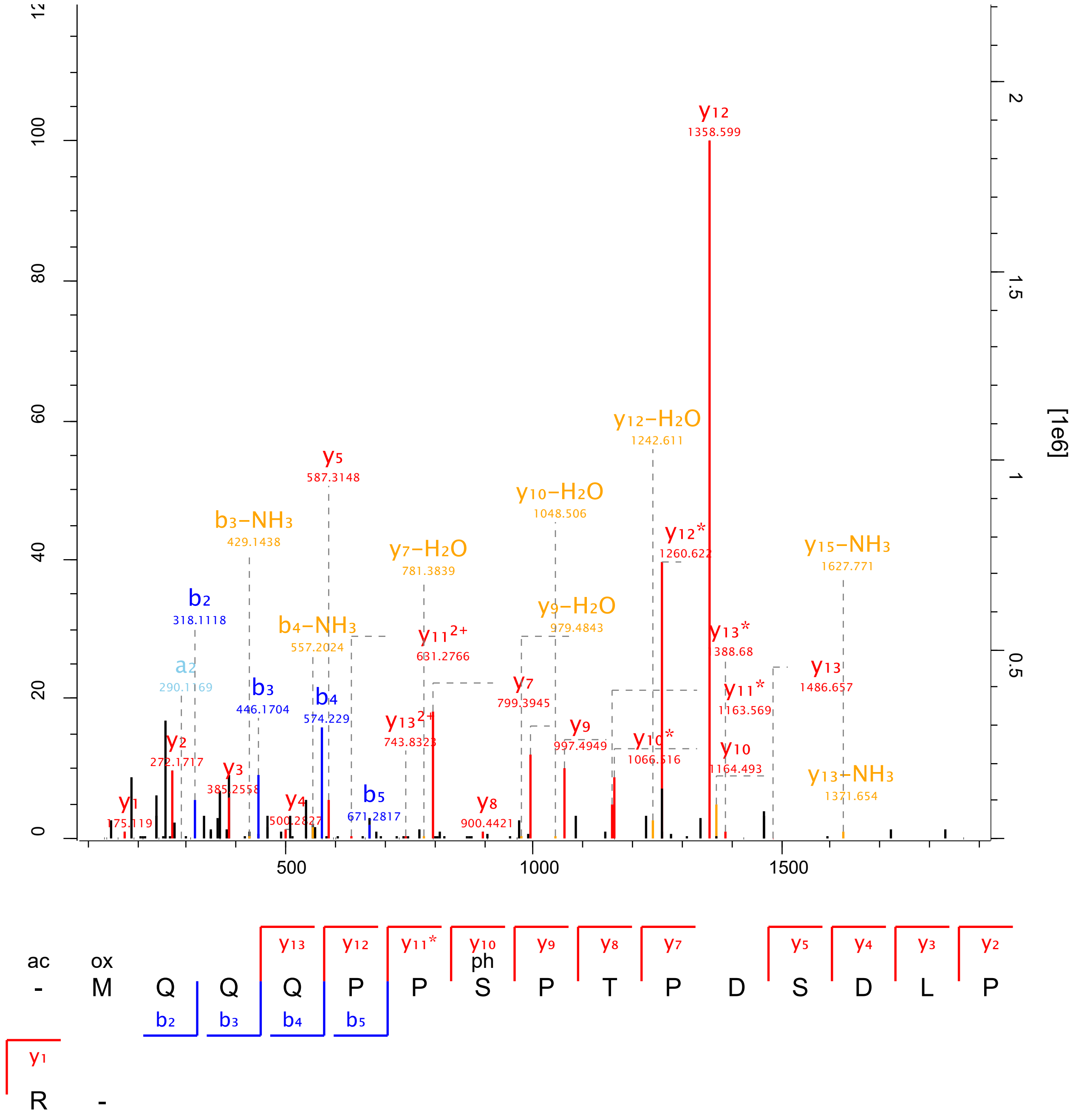

| Raw file | Scan | Method | Score | m/z |
| --- | --- | --- | --- | --- |
| sys_15_1 | 26823 | FTMS; HCD | 52.36 | 761.03 |

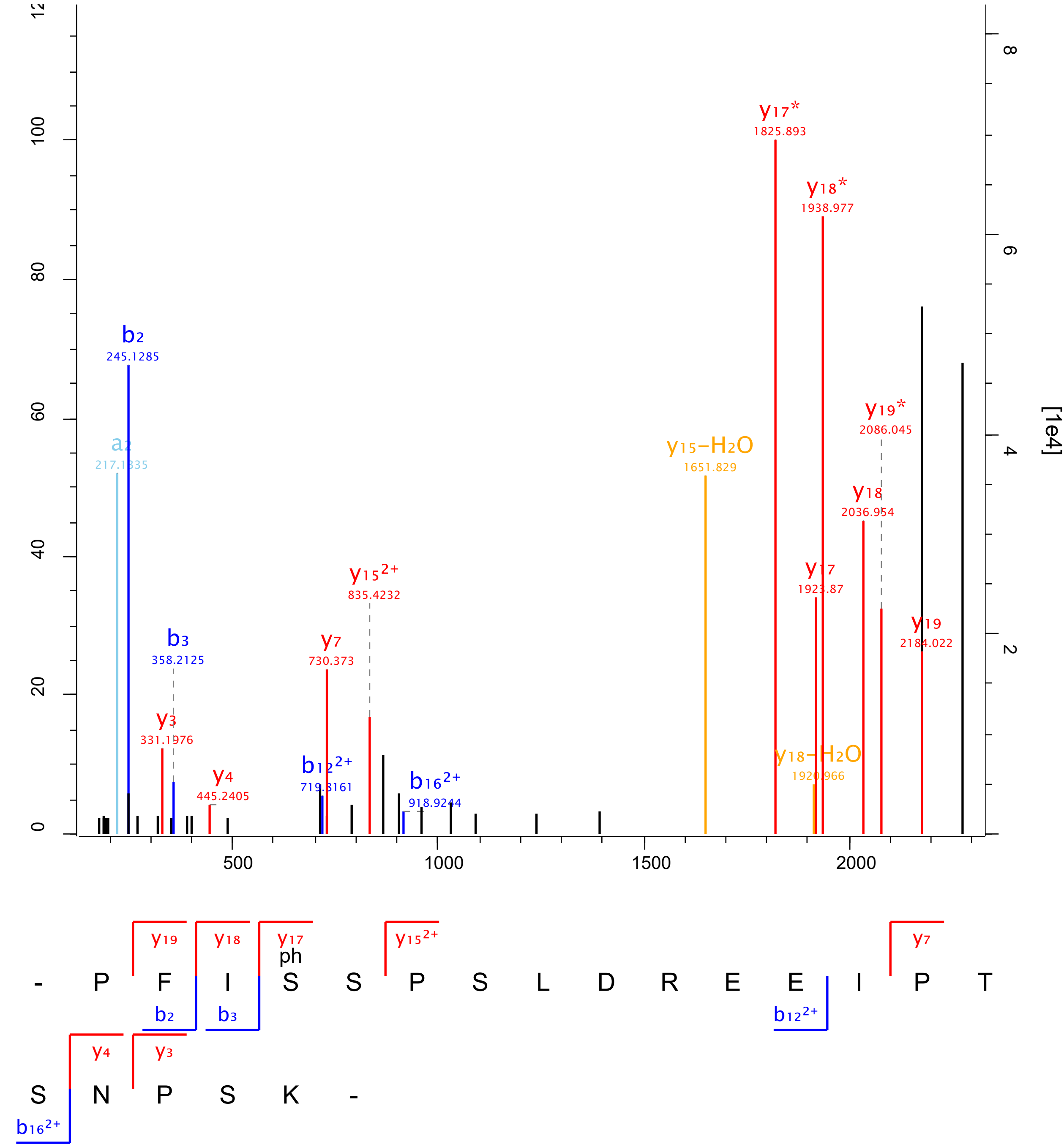

| Raw file | Scan | Method | Score | m/z |
|---|---|---|---|---|
| sys_15_1 | 26825 | FTMS; HCD | 86.8 | 847.39 |

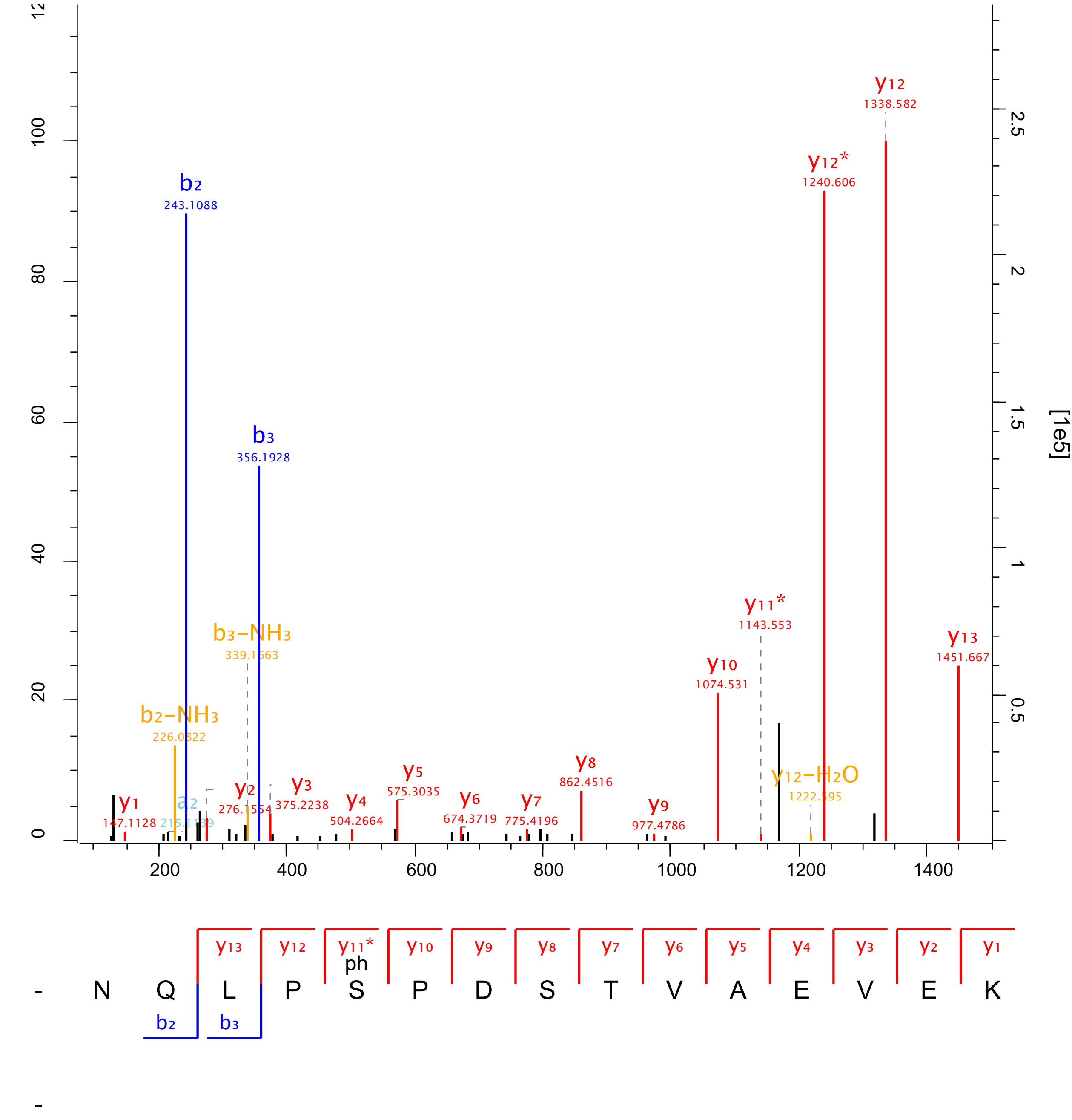

| Raw file | Scan | Method | Score | m/z |
|---|---|---|---|---|
| sys_15_1 | 26888 | FTMS; HCD | 42.35 | 913.74 |

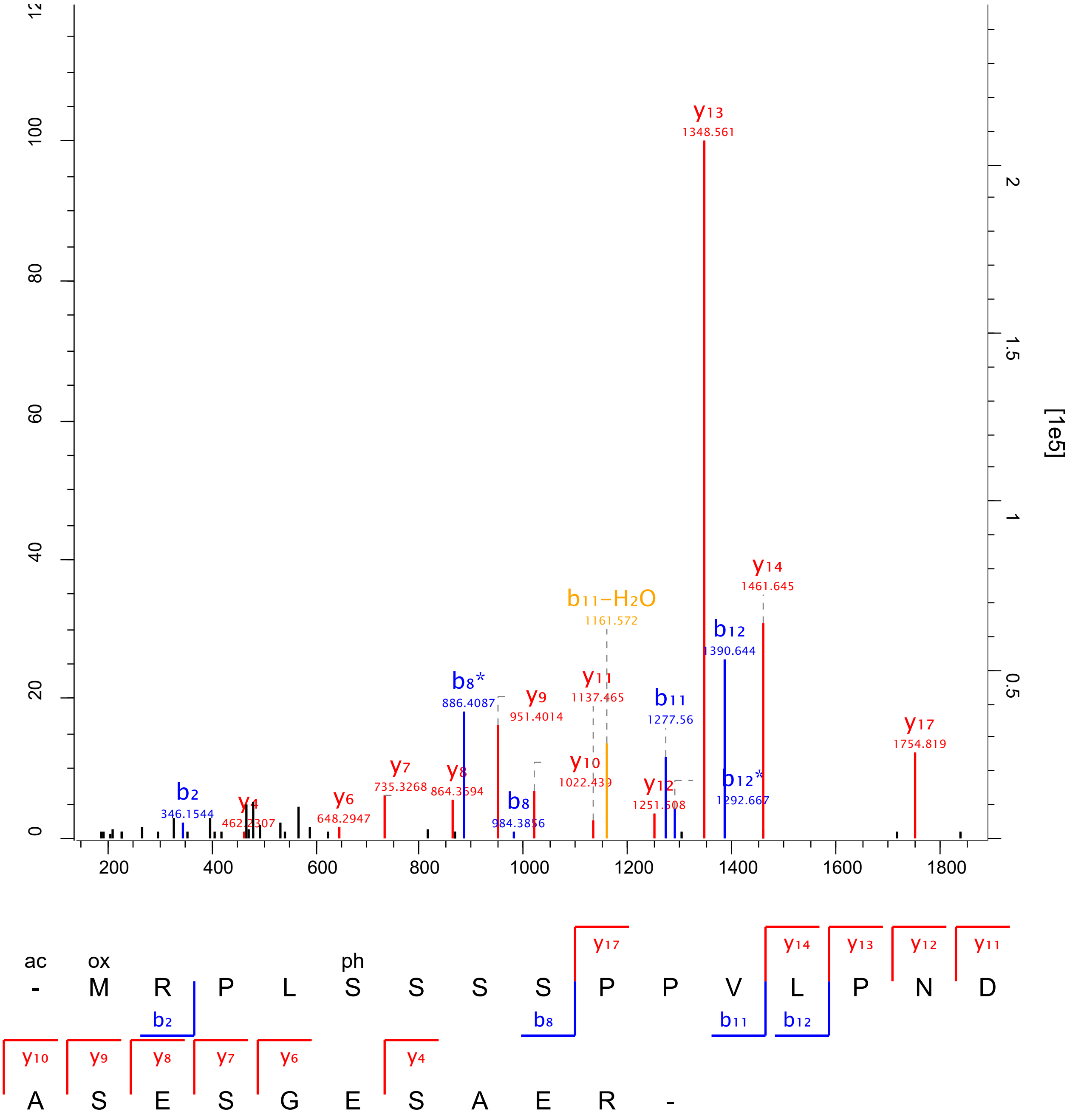

| Raw file | Scan | Method | Score | m/z |
| --- | --- | --- | --- | --- |
| sys_15_1 | 26921 | FTMS; HCD | 56.51 | 728.36 |

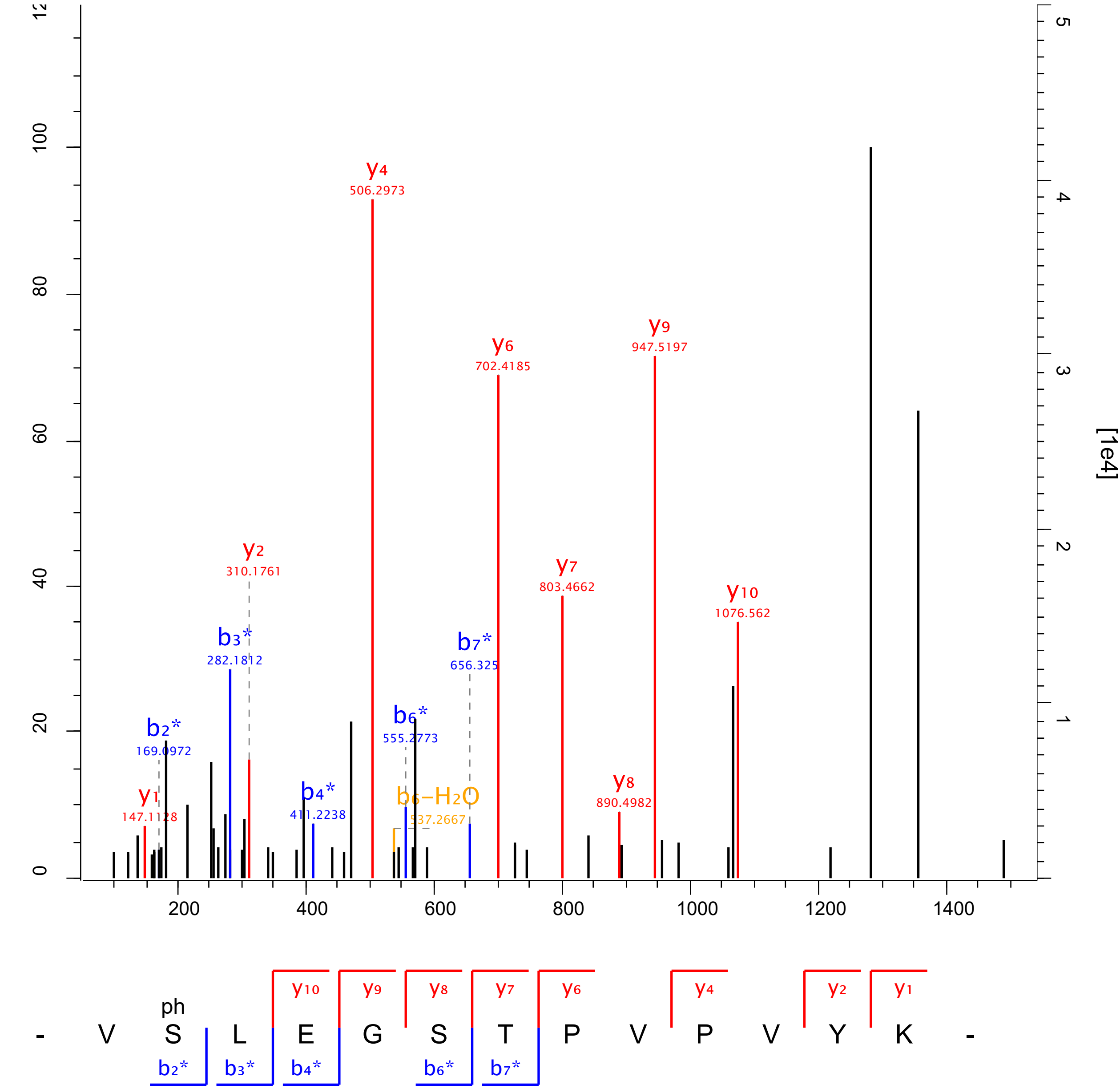

| Raw file | Scan | Method | Score | m/z |
|---|---|---|---|---|
| sys_15_1 | 2696 | FTMS; HCD | 170.74 | 599.76 |

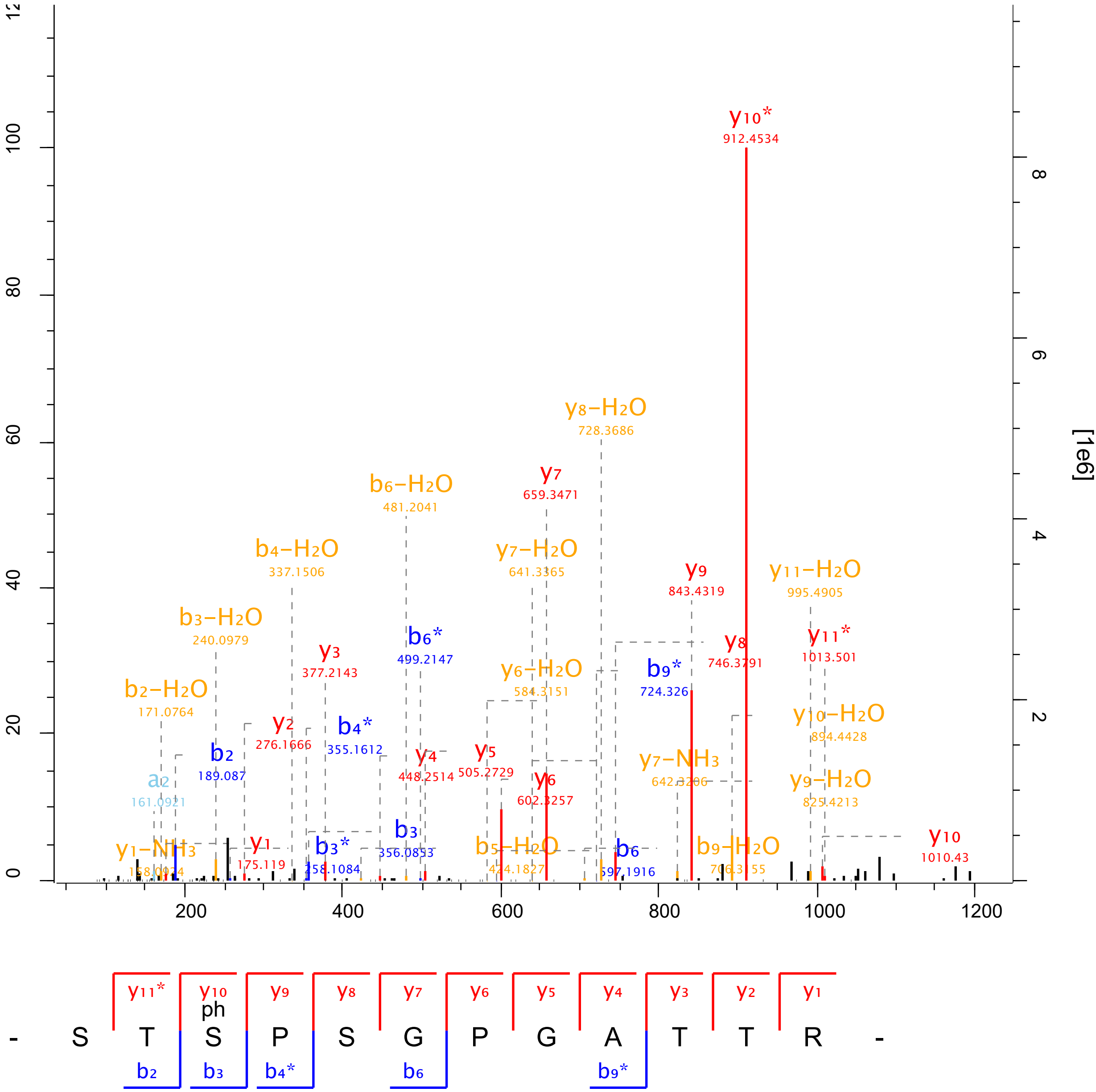

| Raw file | Scan | Method | Score | m/z |
|---|---|---|---|---|
| sys_15_1 | 26962 | FTMS; HCD | 242.64 | 774.65 |

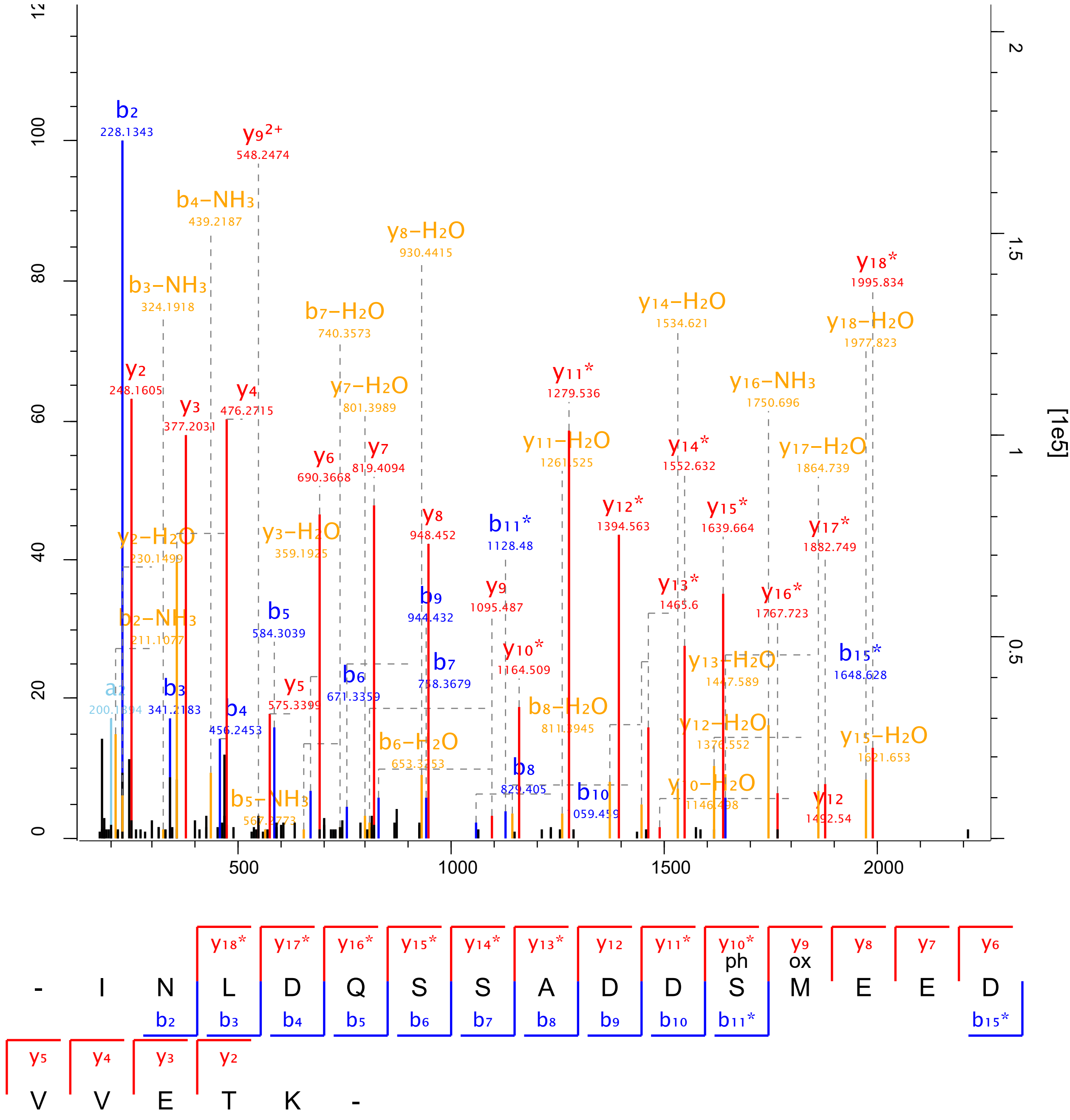

| Raw file | Scan | Method | Score | m/z |
| --- | --- | --- | --- | --- |
| sys_15_1 | 26965 | FTMS; HCD | 84.36 | 625.97 |

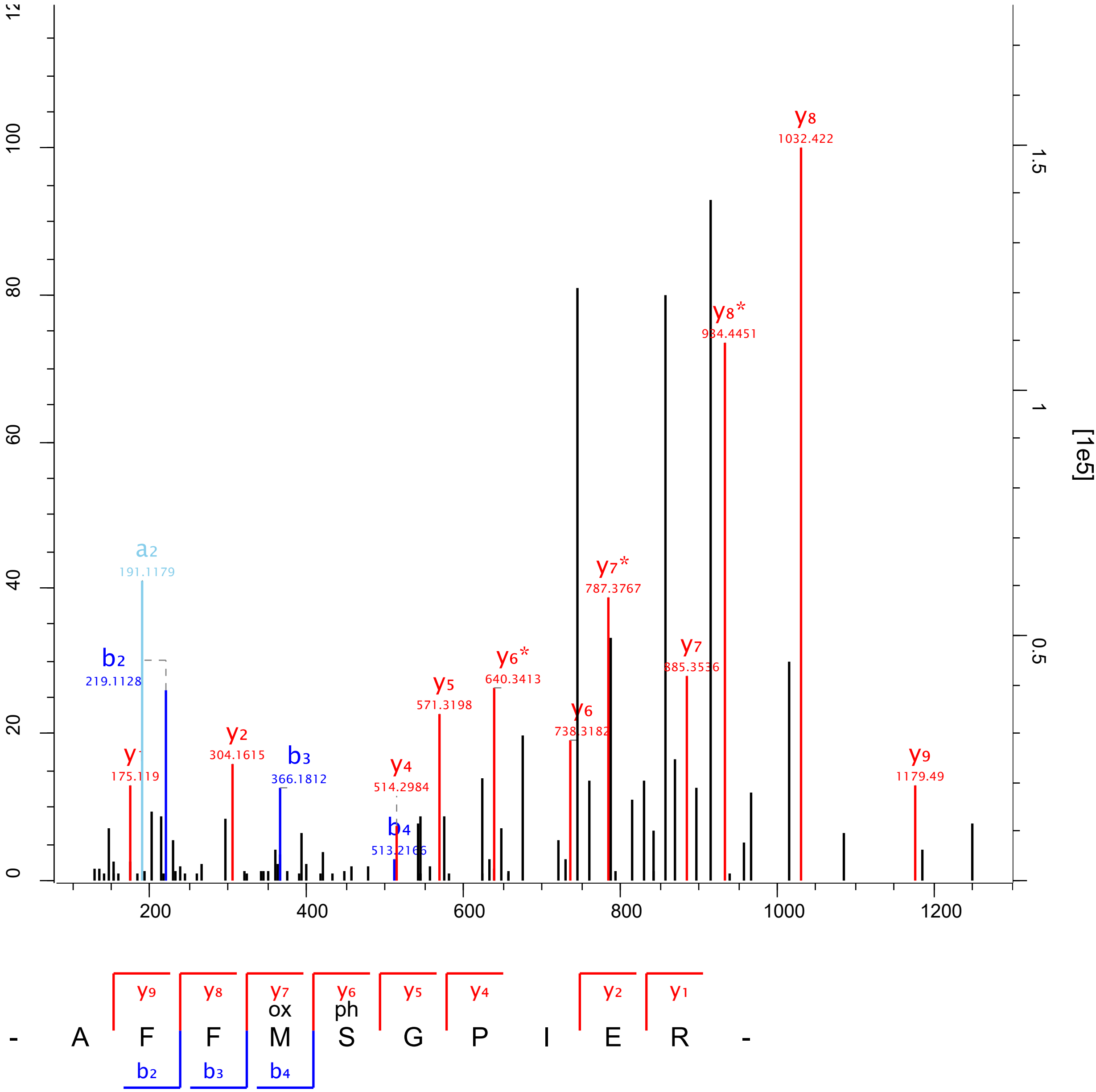

| Raw file | Scan | Method | Score | m/z |
|---|---|---|---|---|
| sys_15_1 | 2704 | FTMS; HCD | 70.41 | 447.7 |

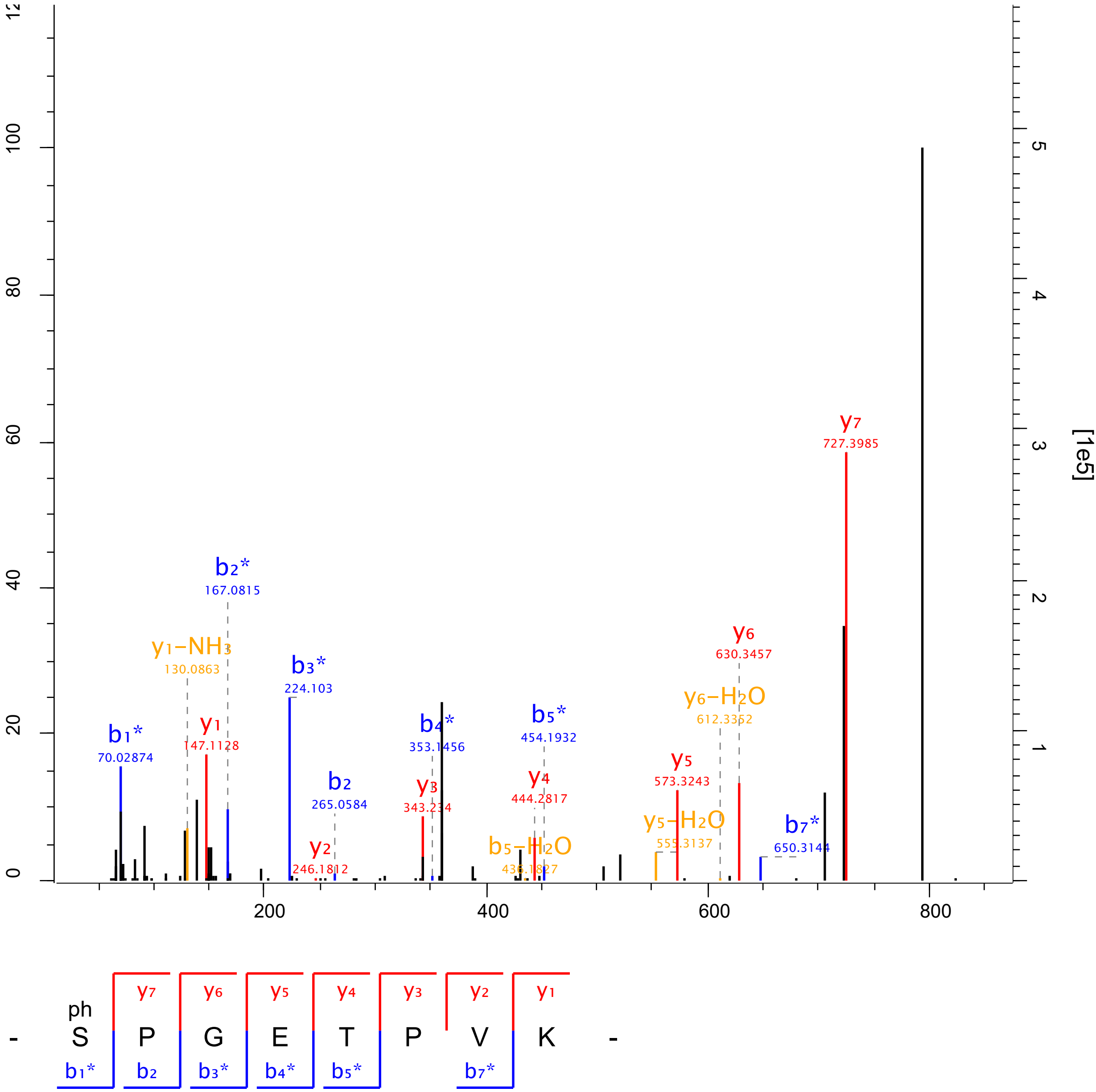

| Raw file | Scan | Method | Score | m/z |
|---|---|---|---|---|
| sys_15_1 | 27054 | FTMS; HCD | 86.54 | 941.93 |

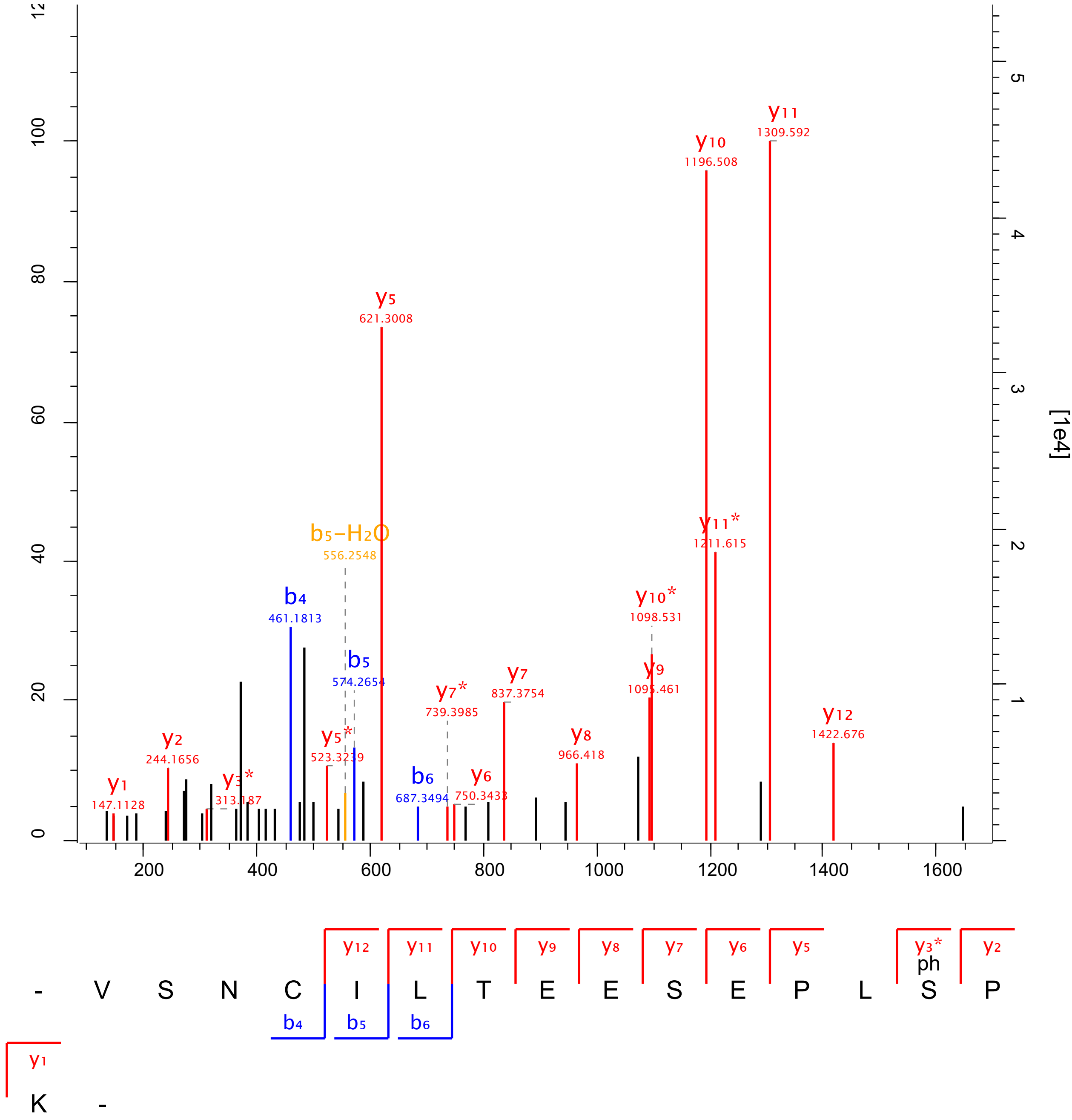

| Raw file | Scan | Method | Score | m/z |
|---|---|---|---|---|
| sys_15_1 | 27078 | FTMS; HCD | 51.03 | 724.88 |

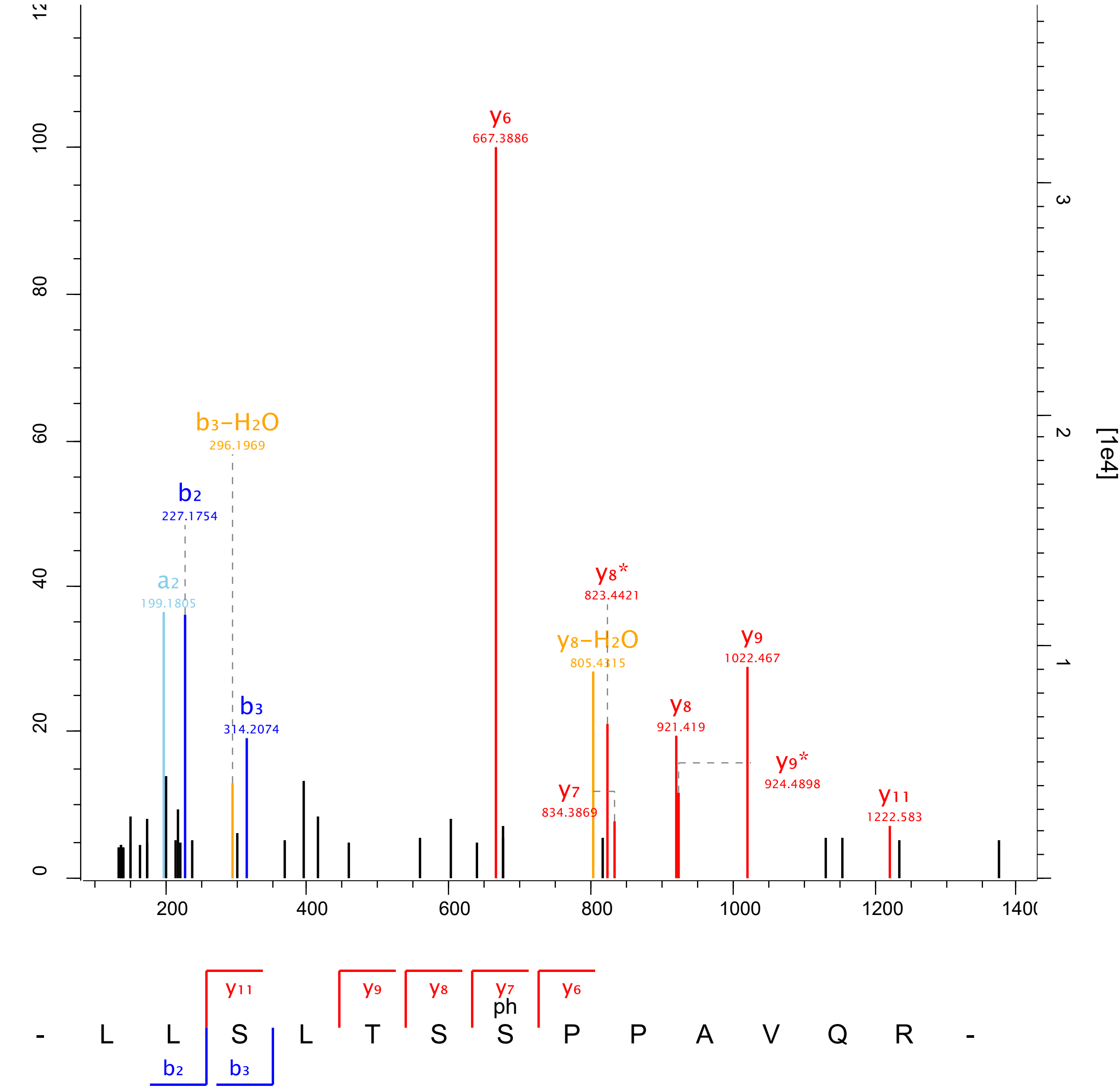

| Raw file | Scan | Method | Score | m/z |
| --- | --- | --- | --- | --- |
| sys_15_1 | 27132 | FTMS; HCD | 178.35 | 784.68 |

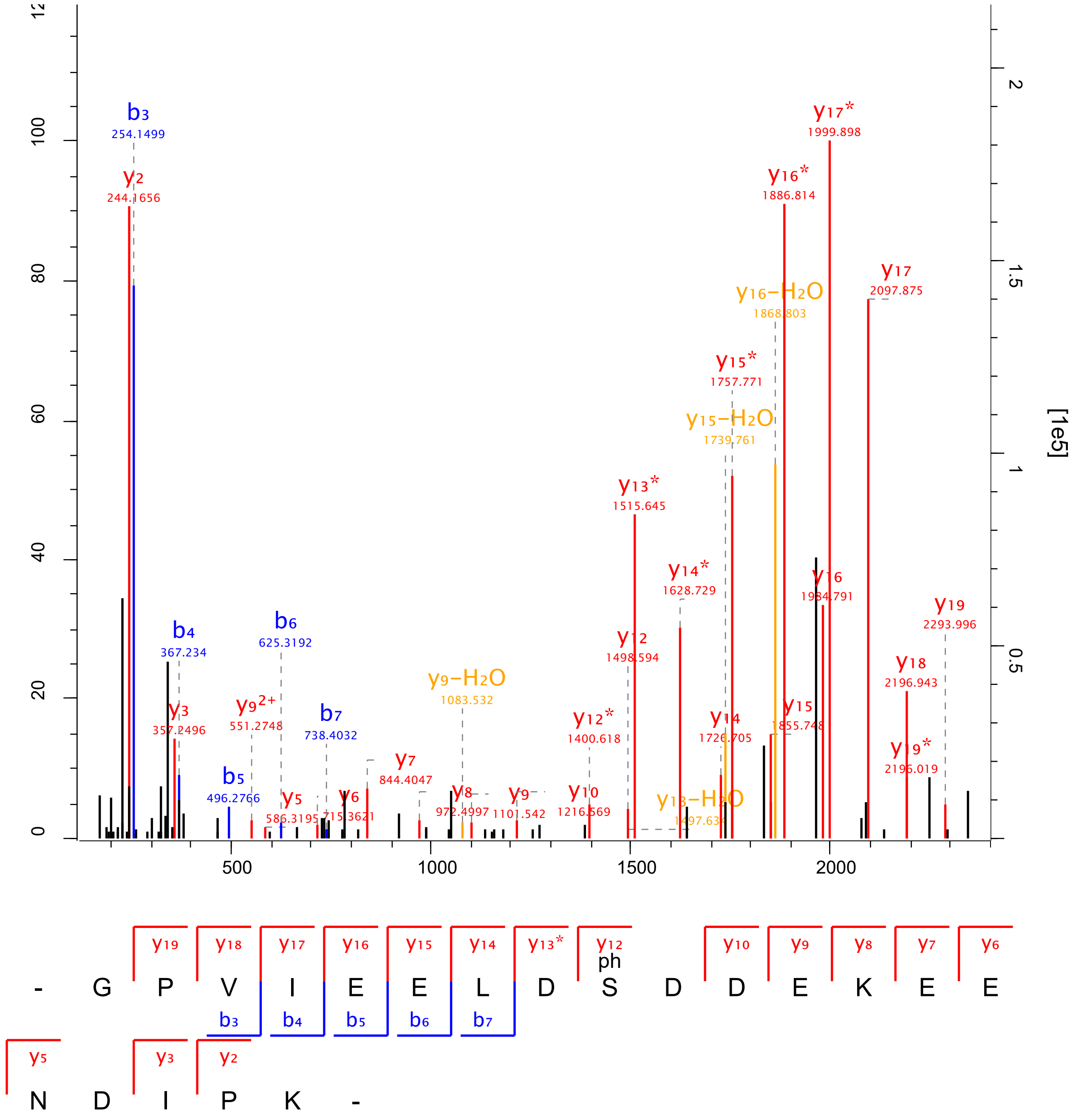

| Raw file | Scan | Method | Score | m/z |
| --- | --- | --- | --- | --- |
| sys_15_1 | 27308 | FTMS; HCD | 86.56 | 626.62 |

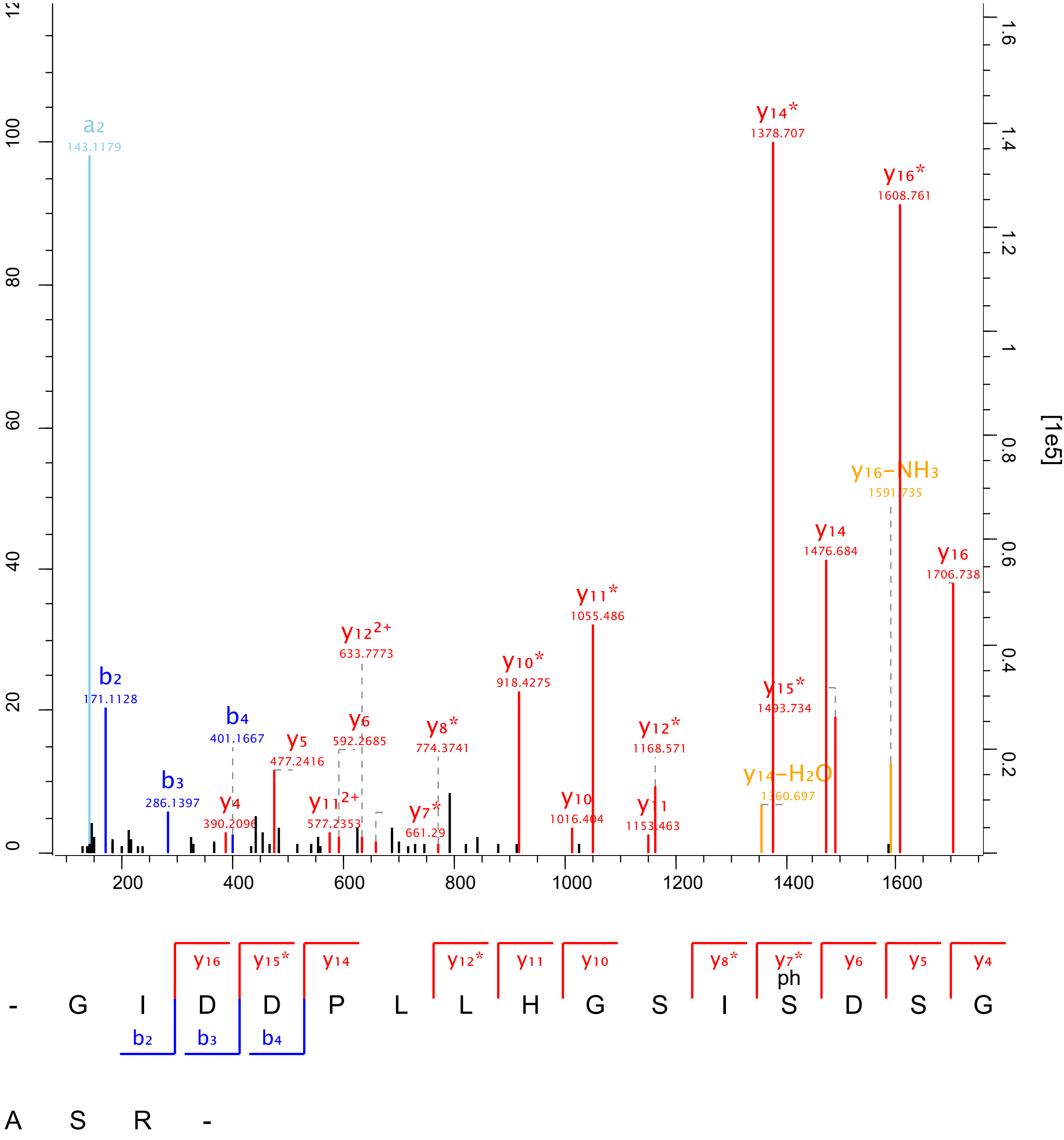

| Raw file | Scan | Method | Score | m/z |
| --- | --- | --- | --- | --- |
| sys_15_1 | 27335 | FTMS; HCD | 40.86 | 1031.08 |

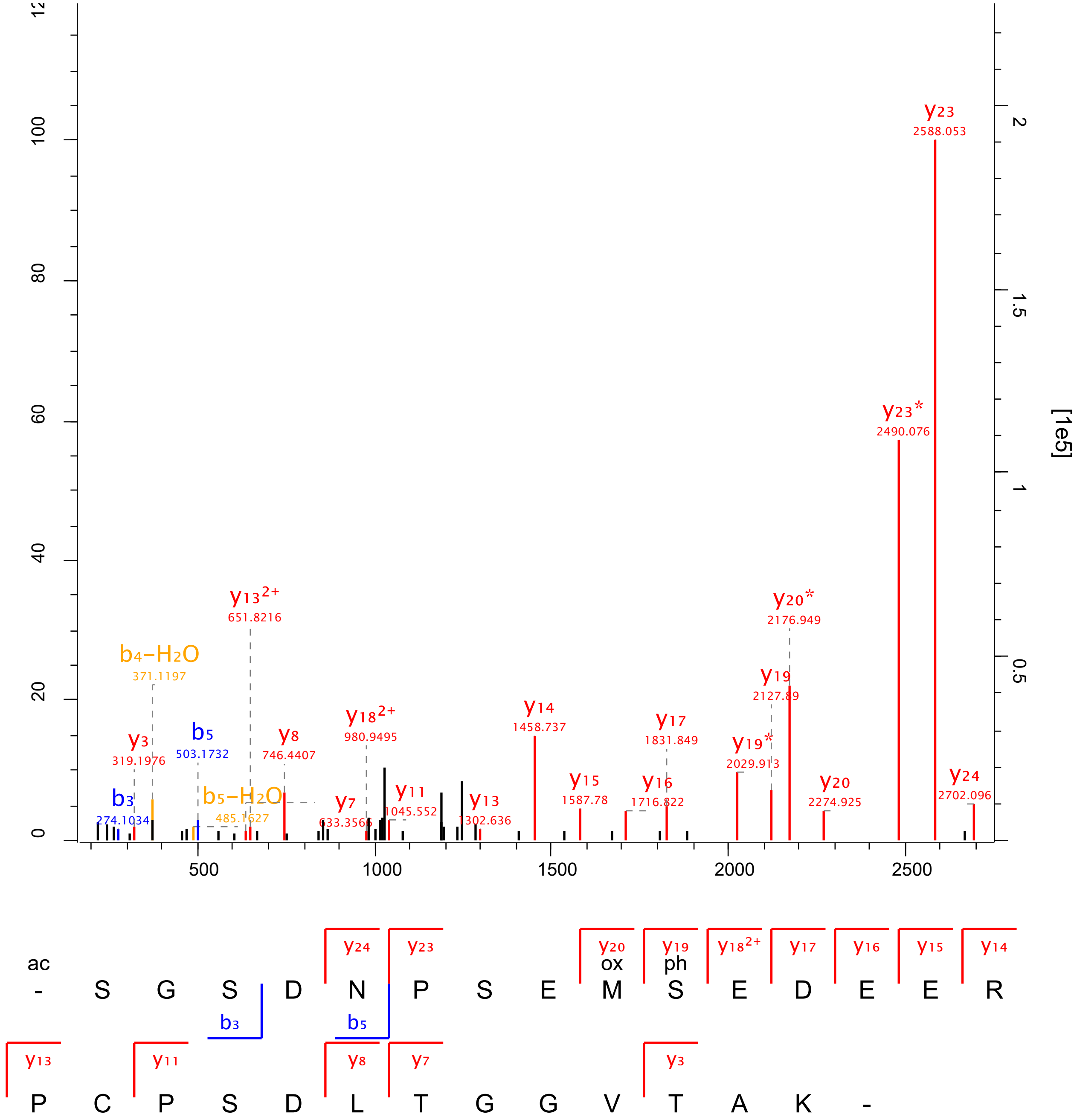

| Raw file | Scan | Method | Score | m/z |
|---|---|---|---|---|
| sys_15_1 | 2737 | FTMS; HCD | 59.07 | 497.69 |

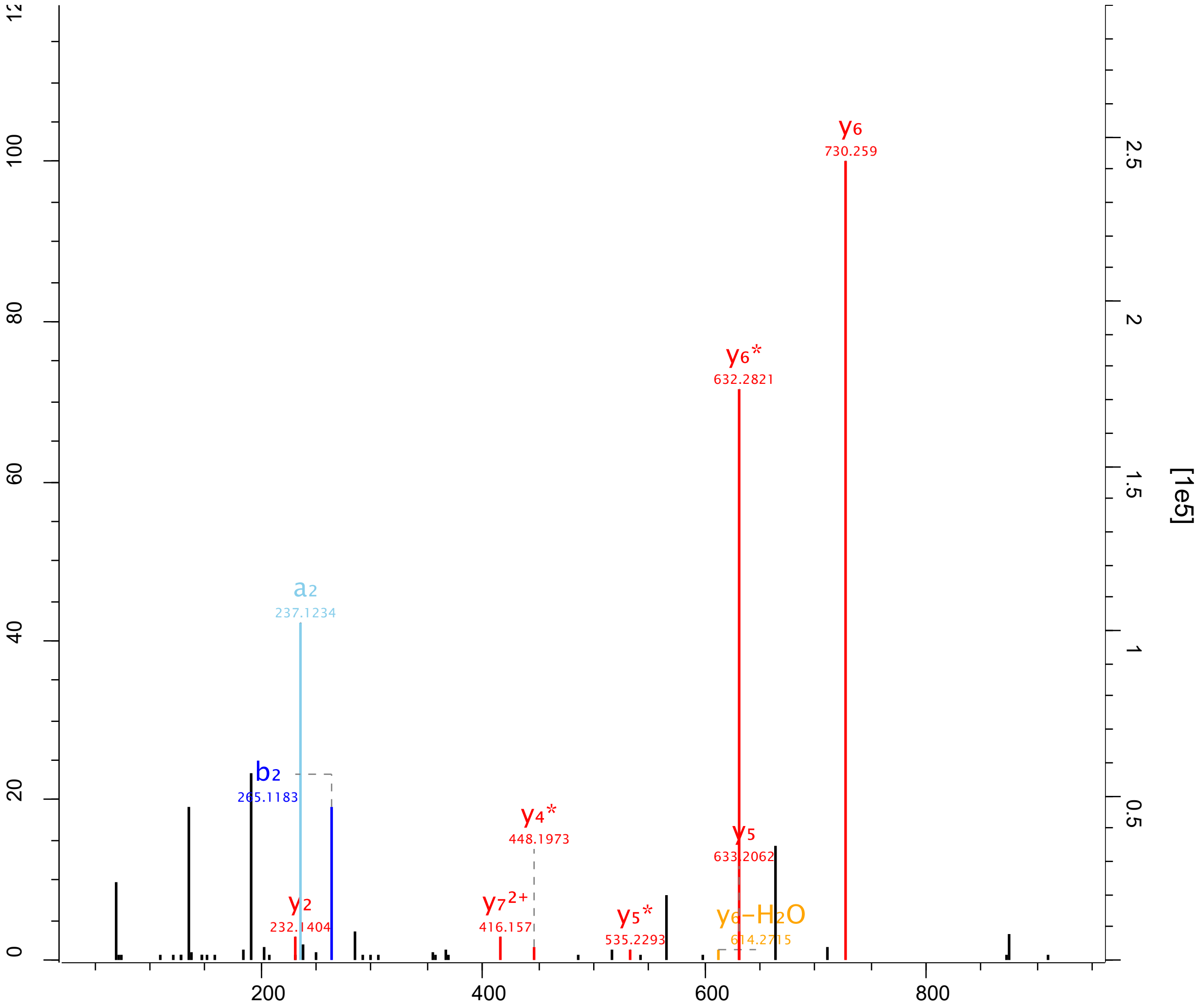


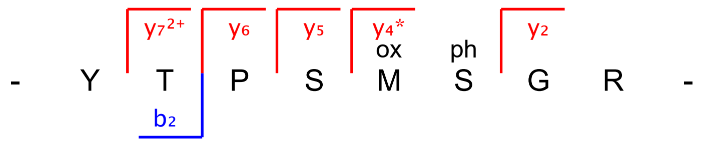

| Raw file | Scan | Method | Score | m/z |
|---|---|---|---|---|
| sys_15_1 | 27442 | FTMS; HCD | 94.38 | 709.68 |

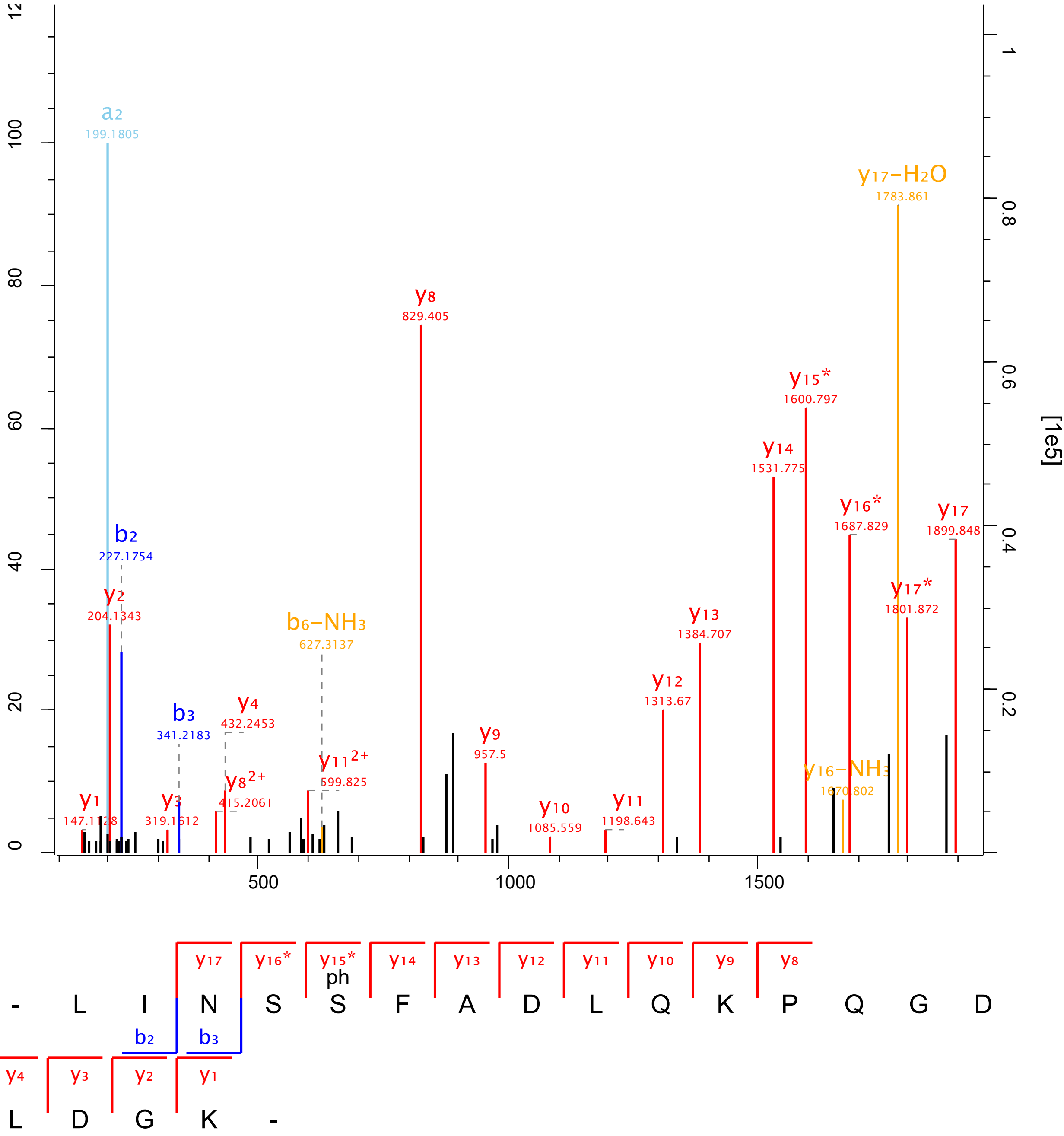

| Raw file | Scan | Method | Score | m/z |
| --- | --- | --- | --- | --- |
| sys_15_1 | 27502 | FTMS; HCD | 149.71 | 896.89 |

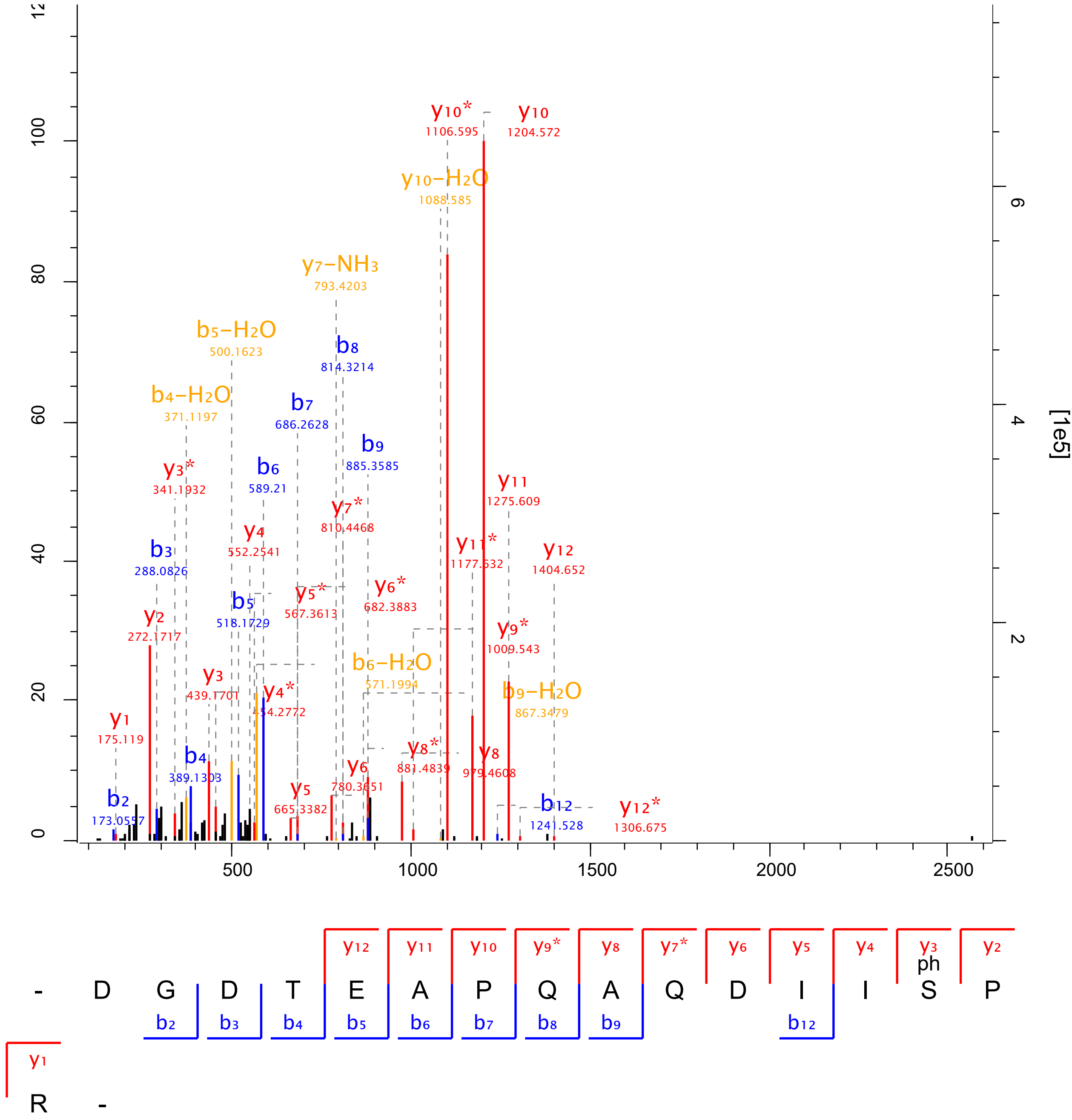

| Raw file | Scan | Method | Score | m/z |
|---|---|---|---|---|
| sys_15_1 | 27569 | FTMS; HCD | 172.98 | 954.93 |

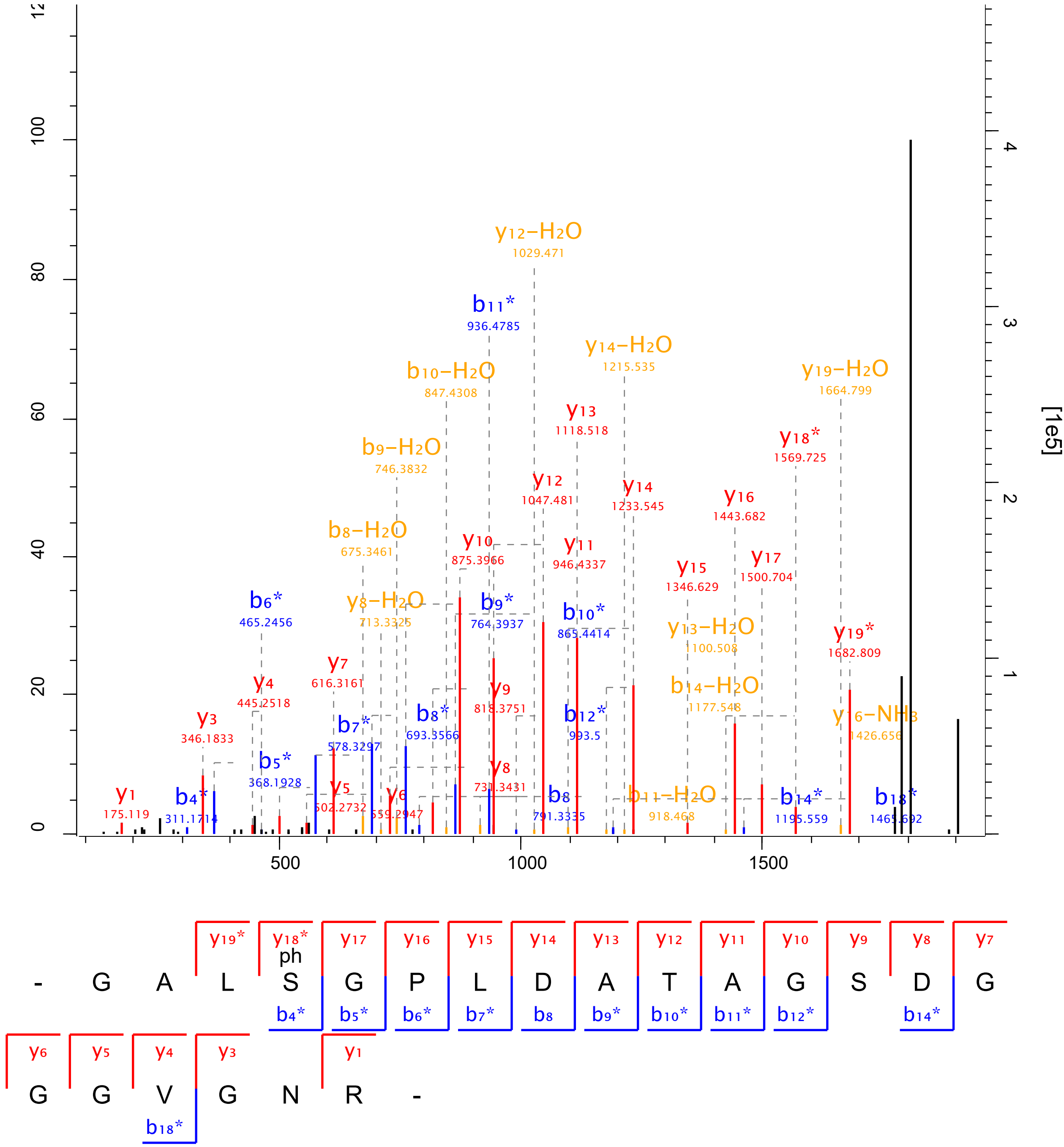

| Raw file | Scan | Method | Score | m/z |
|---|---|---|---|---|
| sys_15_1 | 27604 | FTMS; HCD | 82.07 | 596.77 |

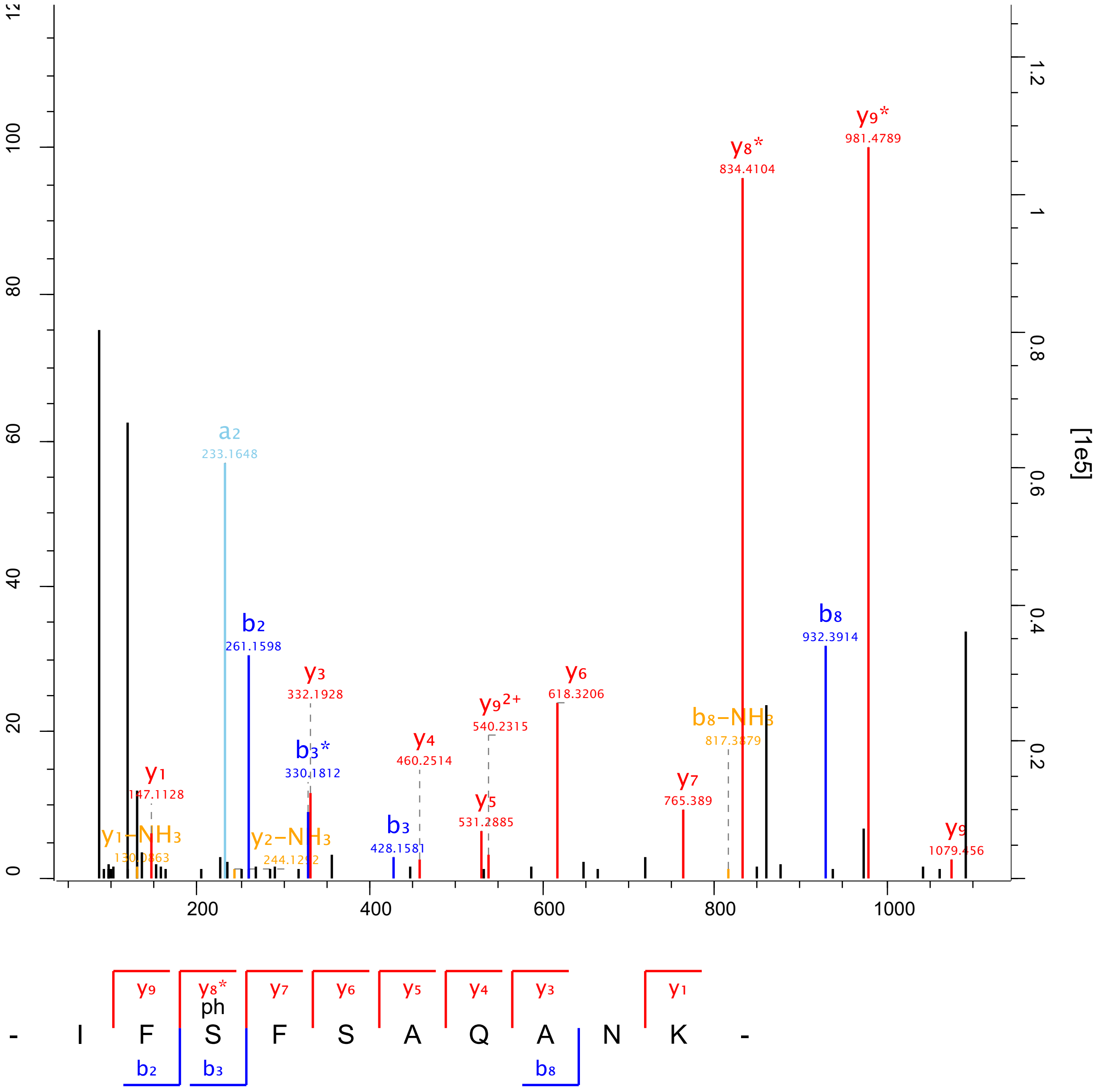

| Raw file | Scan | Method | Score | m/z |
| --- | --- | --- | --- | --- |
| sys_15_1 | 27623 | FTMS; HCD | 139.78 | 444.71 |

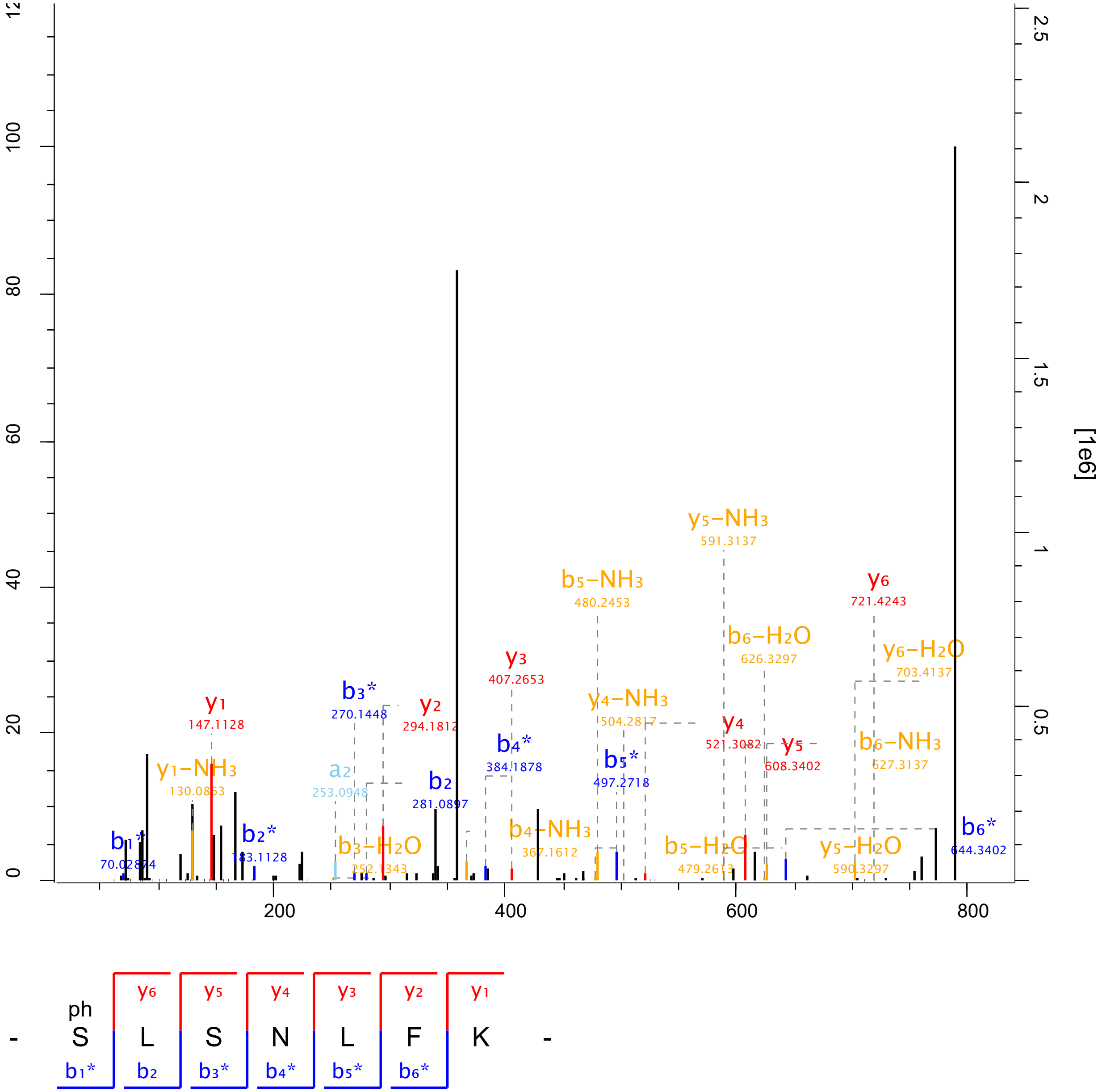

| Raw file | Scan | Method | Score | m/z |
|---|---|---|---|---|
| sys_15_1 | 27796 | FTMS; HCD | 60.49 | 693.82 |

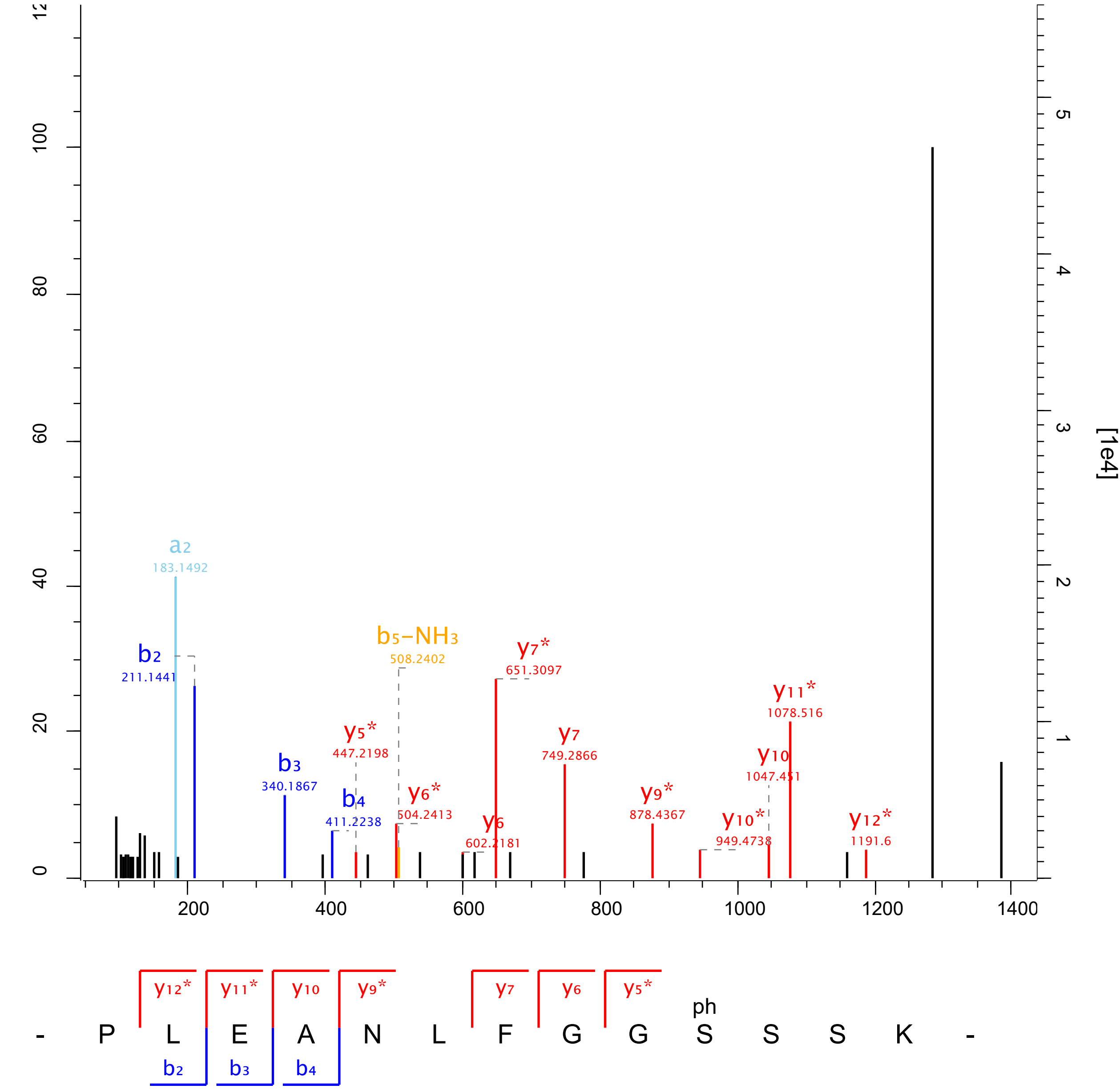

| Raw file | Scan | Method | Score | m/z |
|---|---|---|---|---|
| sys_15_1 | 27899 | FTMS; HCD | 110.26 | 961.91 |

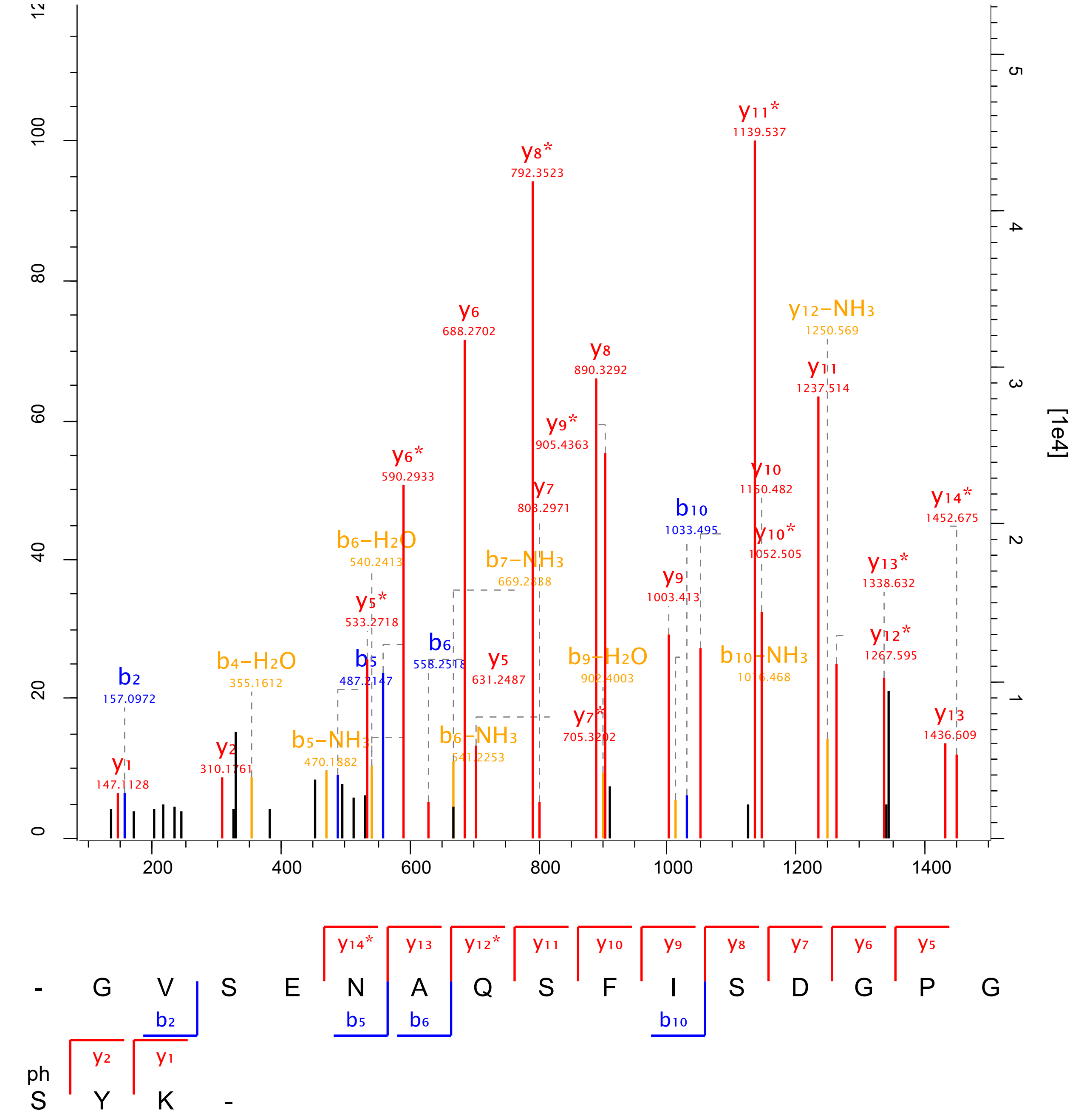

| Raw file | Scan | Method | Score | m/z |
|---|---|---|---|---|
| sys_15_1 | 27936 | FTMS; HCD | 77.22 | 740.02 |

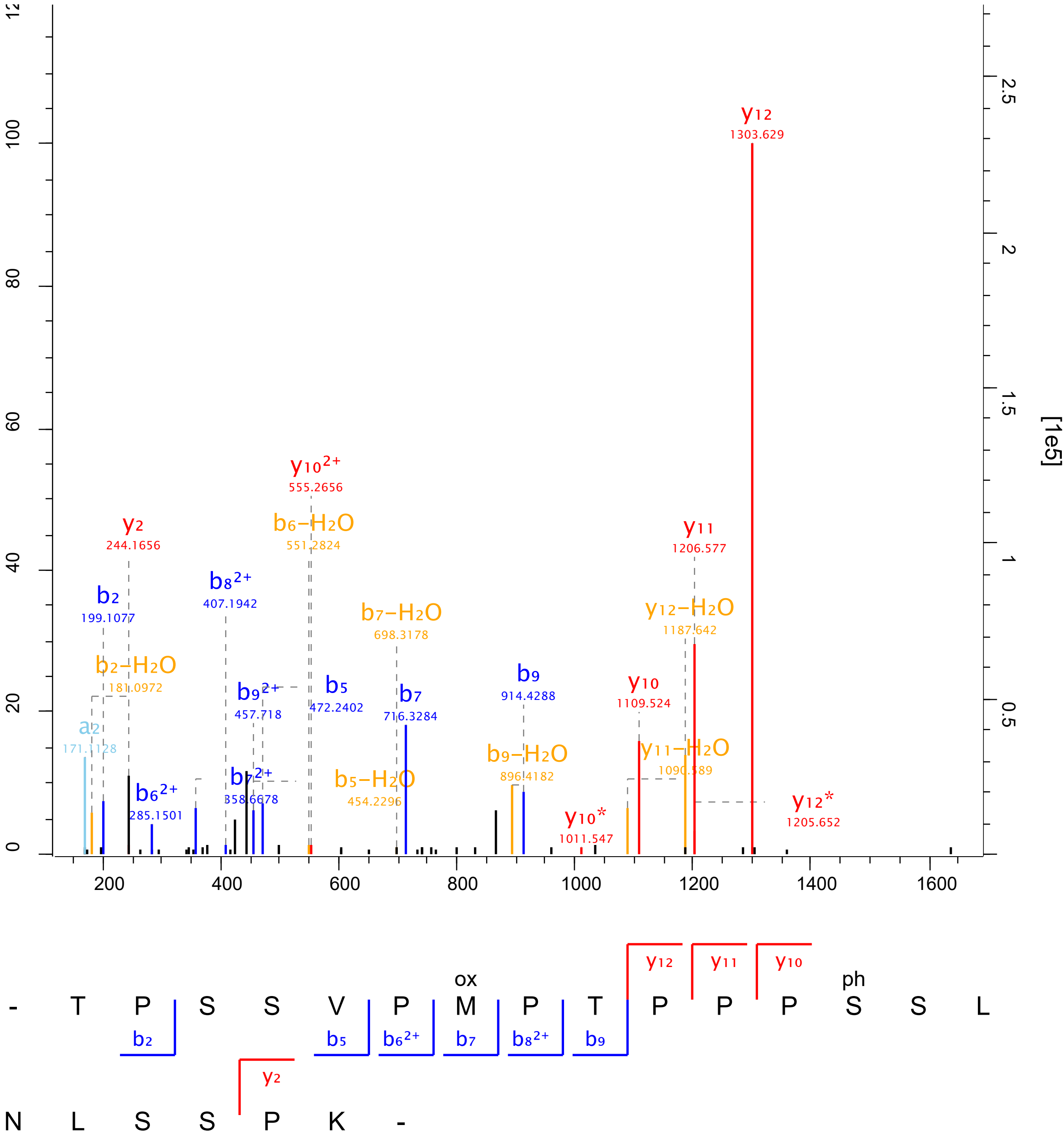

| Raw file | Scan | Method | Score | m/z |
|---|---|---|---|---|
| sys_15_1 | 27985 | FTMS; HCD | 49.66 | 667.34 |

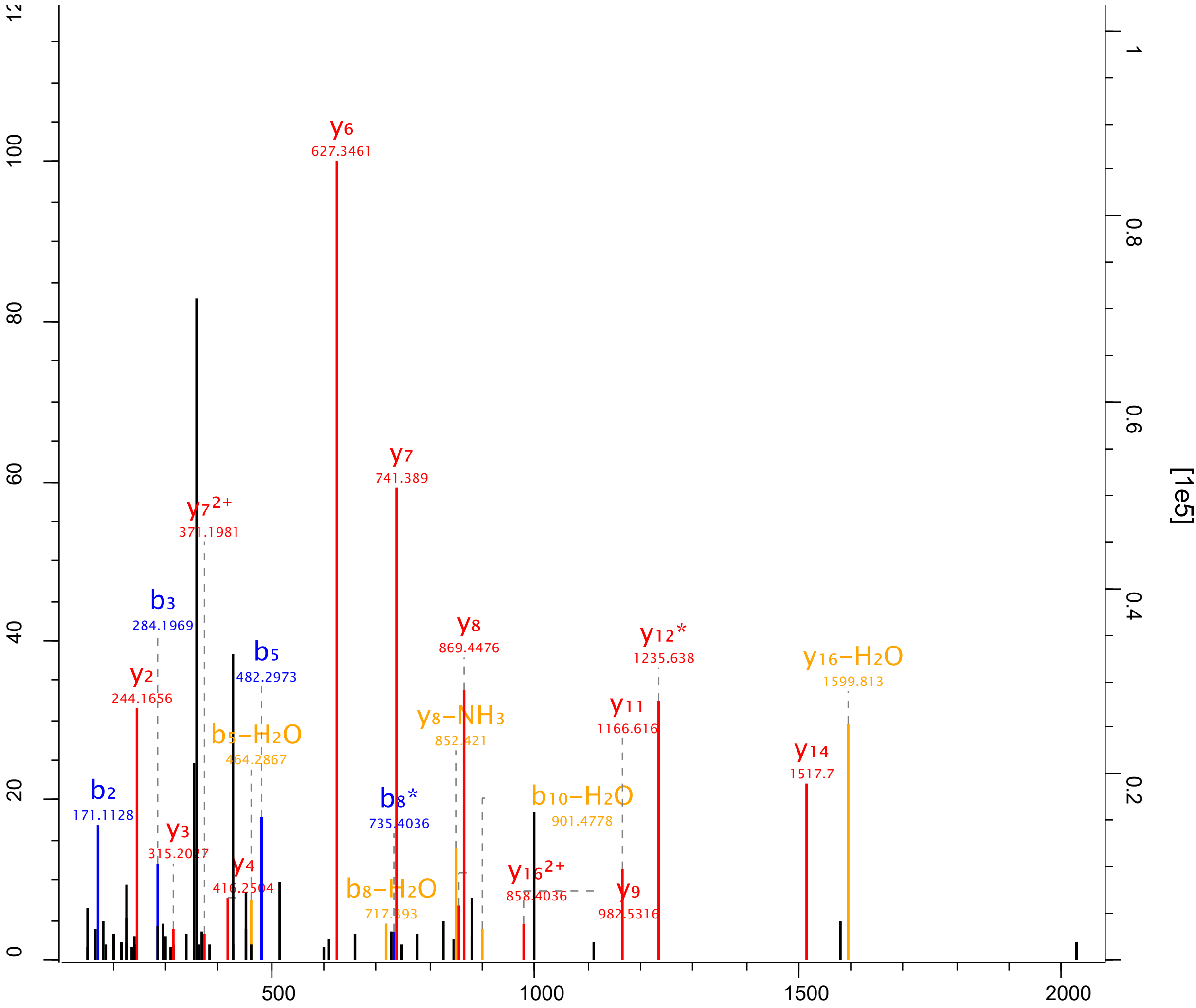


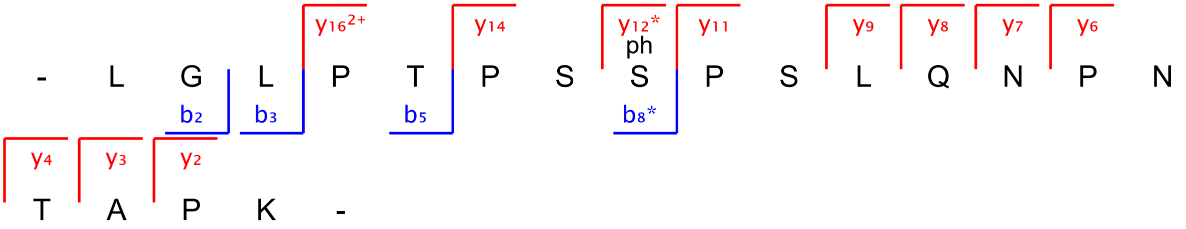

| Raw file | Scan | Method | Score | m/z |
|---|---|---|---|---|
| sys_15_1 | 28118 | FTMS; HCD | 105.46 | 587.26 |

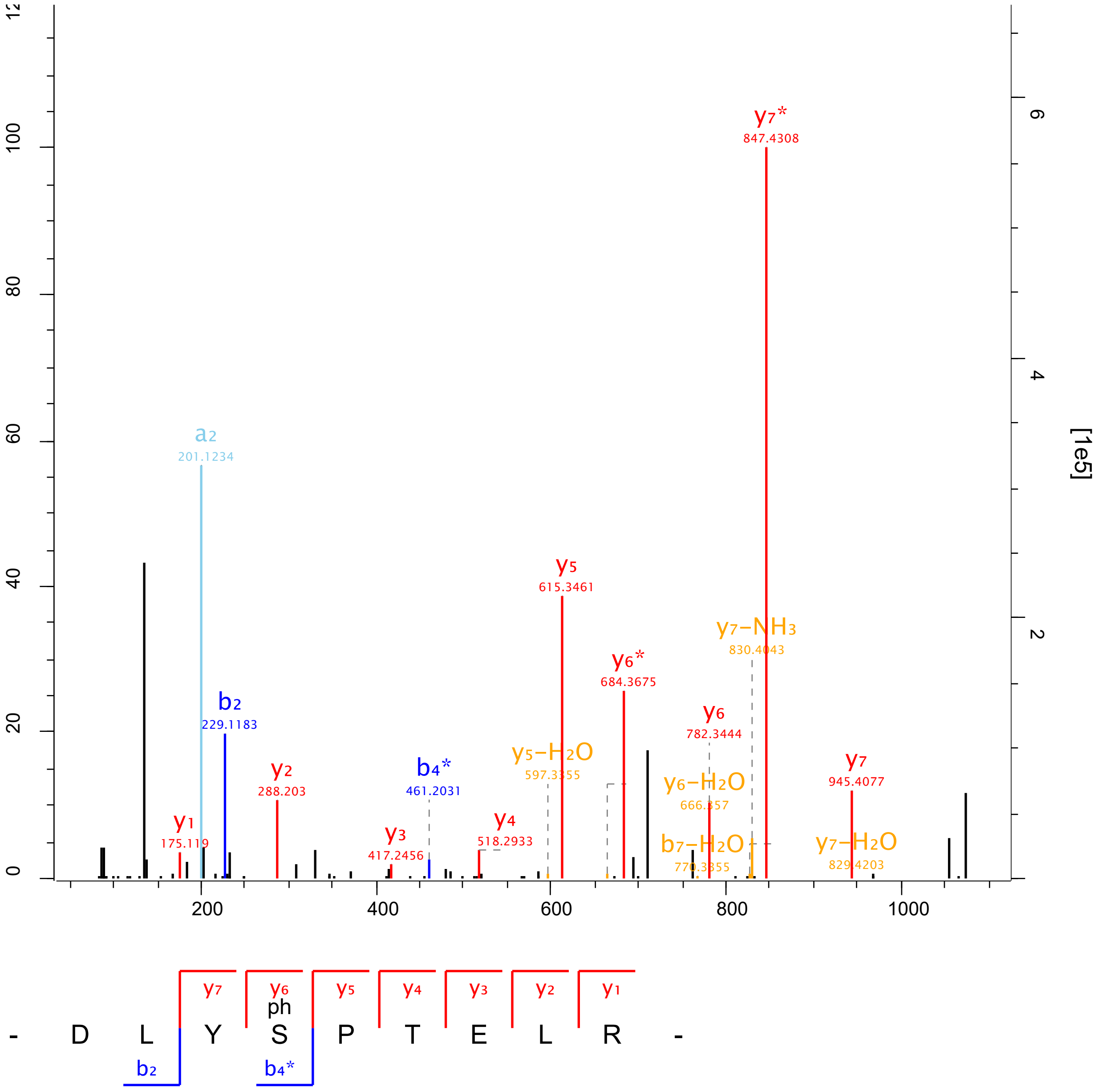

| Raw file | Scan | Method | Score | m/z |
|---|---|---|---|---|
| sys_15_1 | 28139 | FTMS; HCD | 50.5 | 792.35 |

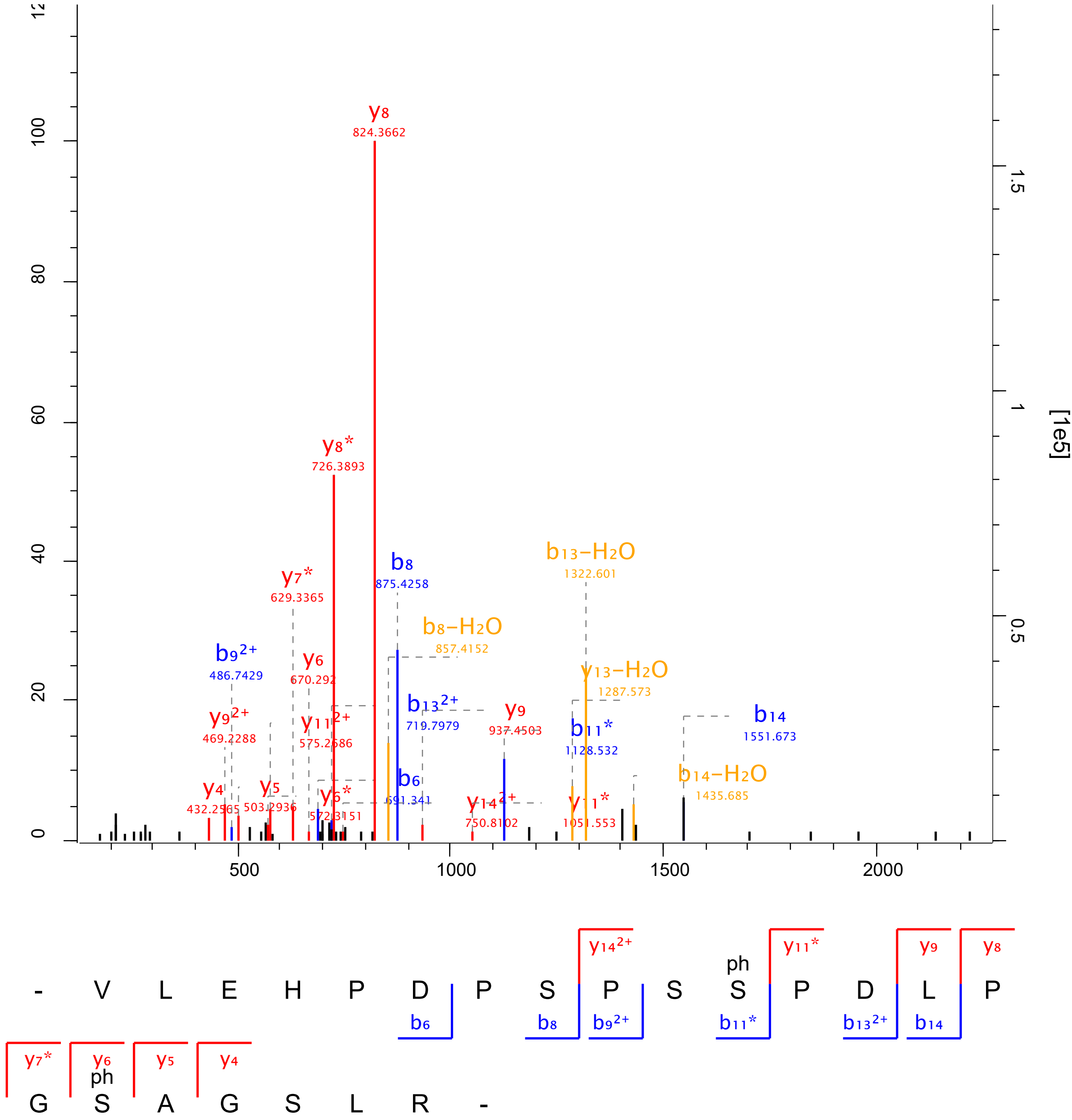

| Raw file | Scan | Method | Score | m/z |
|---|---|---|---|---|
| sys_15_1 | 28290 | FTMS; HCD | 204.17 | 709.8 |

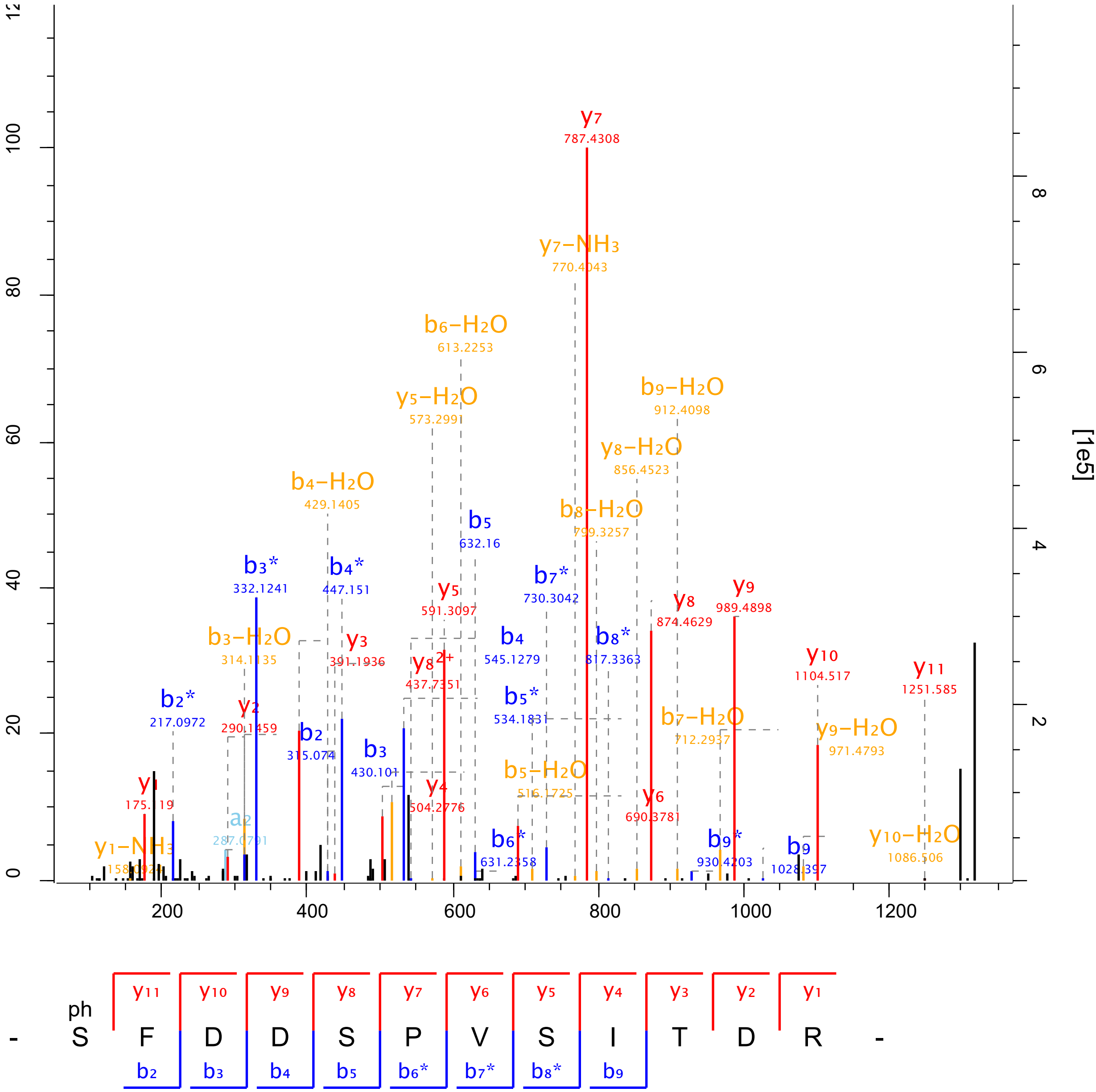

| Raw file | Scan | Method | Score | m/z |
| --- | --- | --- | --- | --- |
| sys_15_1 | 28328 | FTMS; HCD | 131.35 | 694.81 |

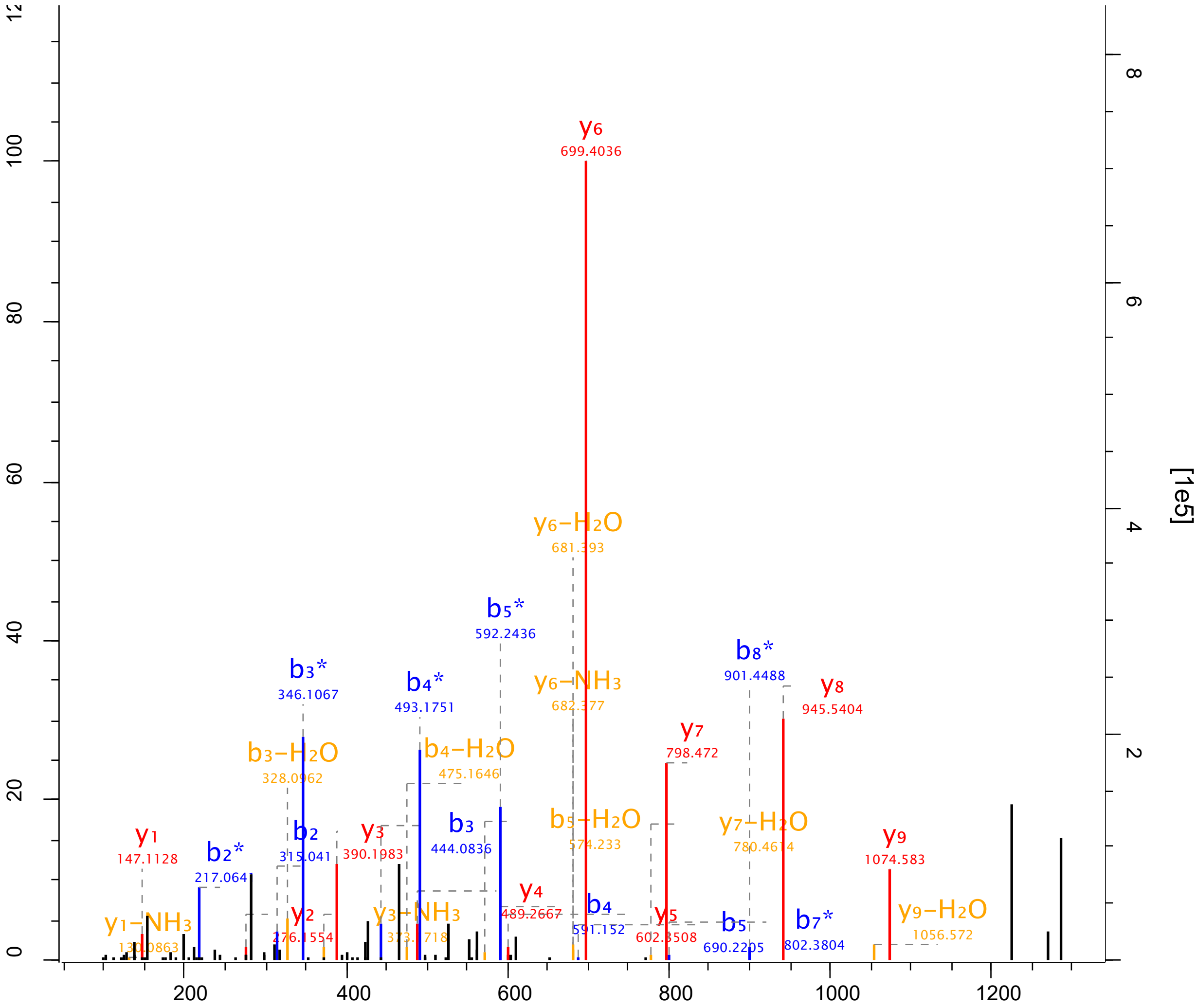


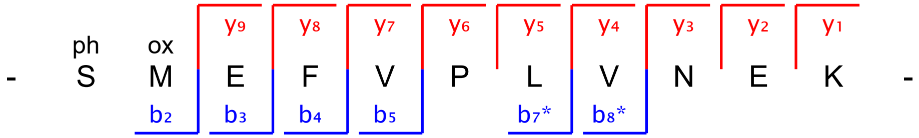

| Raw file | Scan | Method | Score | m/z |
|---|---|---|---|---|
| sys_15_1 | 28403 | FTMS; HCD | 173.09 | 801.38 |

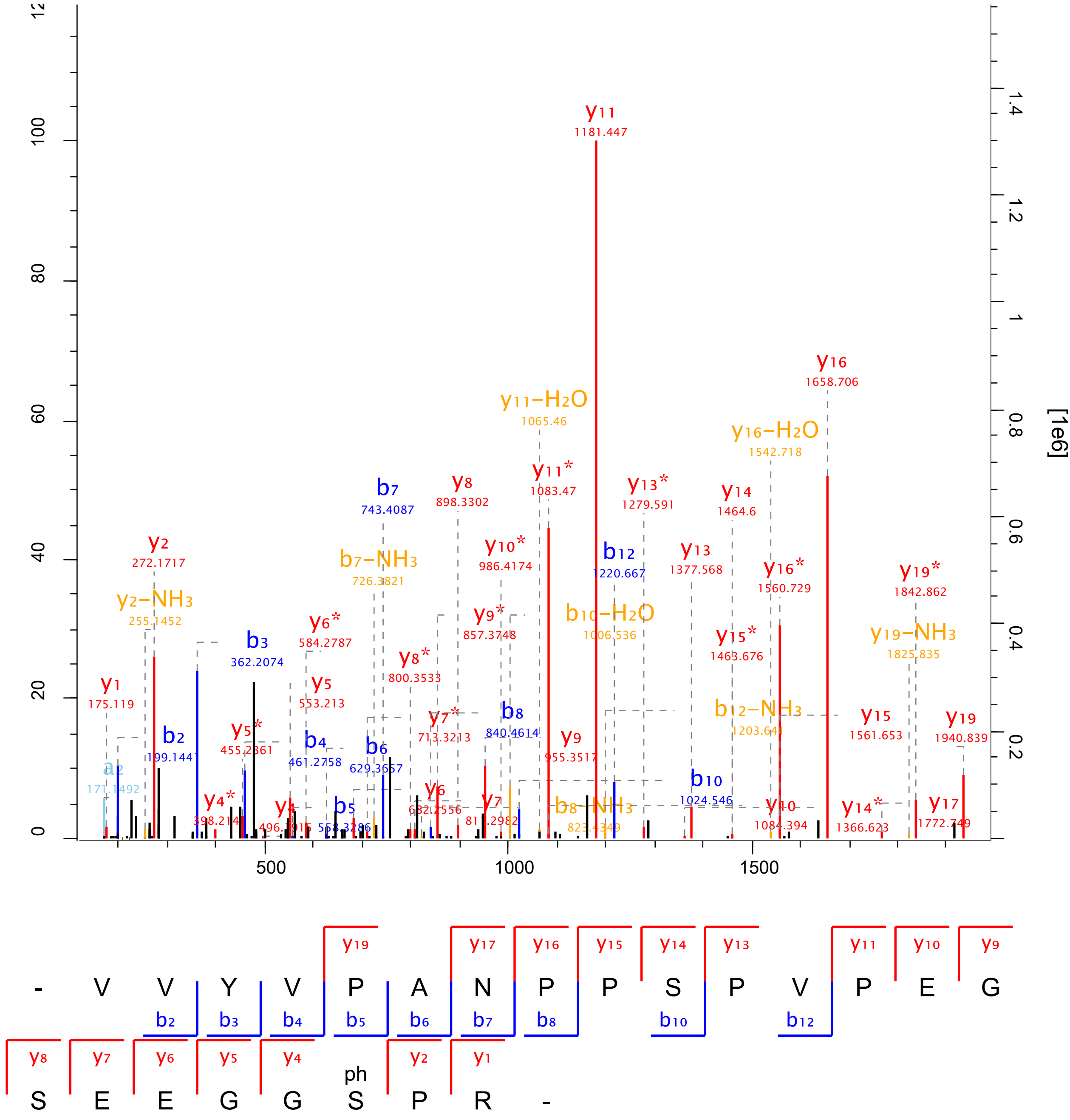

| Raw file | Scan | Method | Score | m/z |
|---|---|---|---|---|
| sys_15_1 | 28460 | FTMS; HCD | 109.79 | 735.33 |

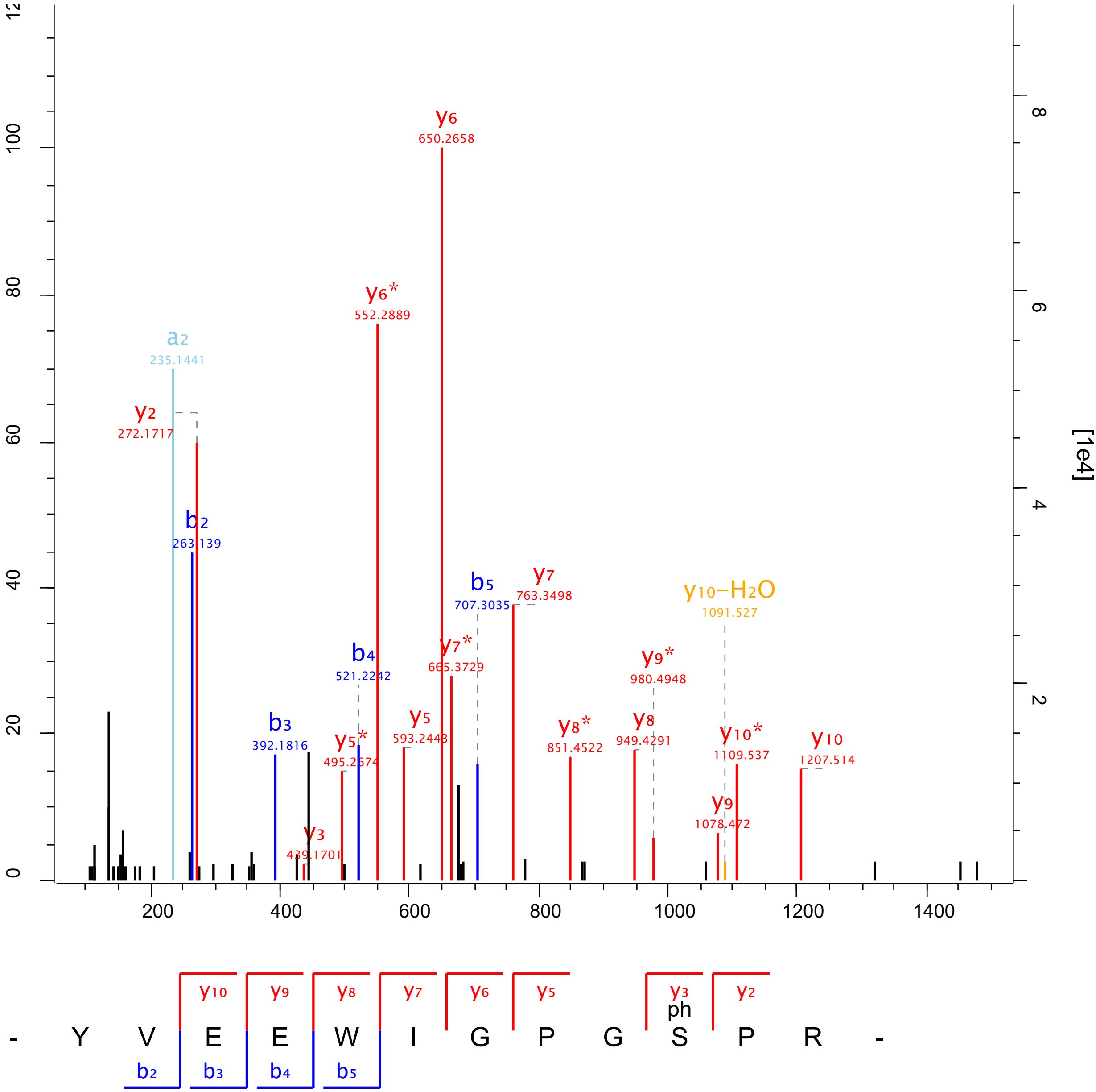

| Raw file | Scan | Method | Score | m/z |
|---|---|---|---|---|
| sys_15_1 | 28491 | FTMS; HCD | 123.29 | 781.31 |

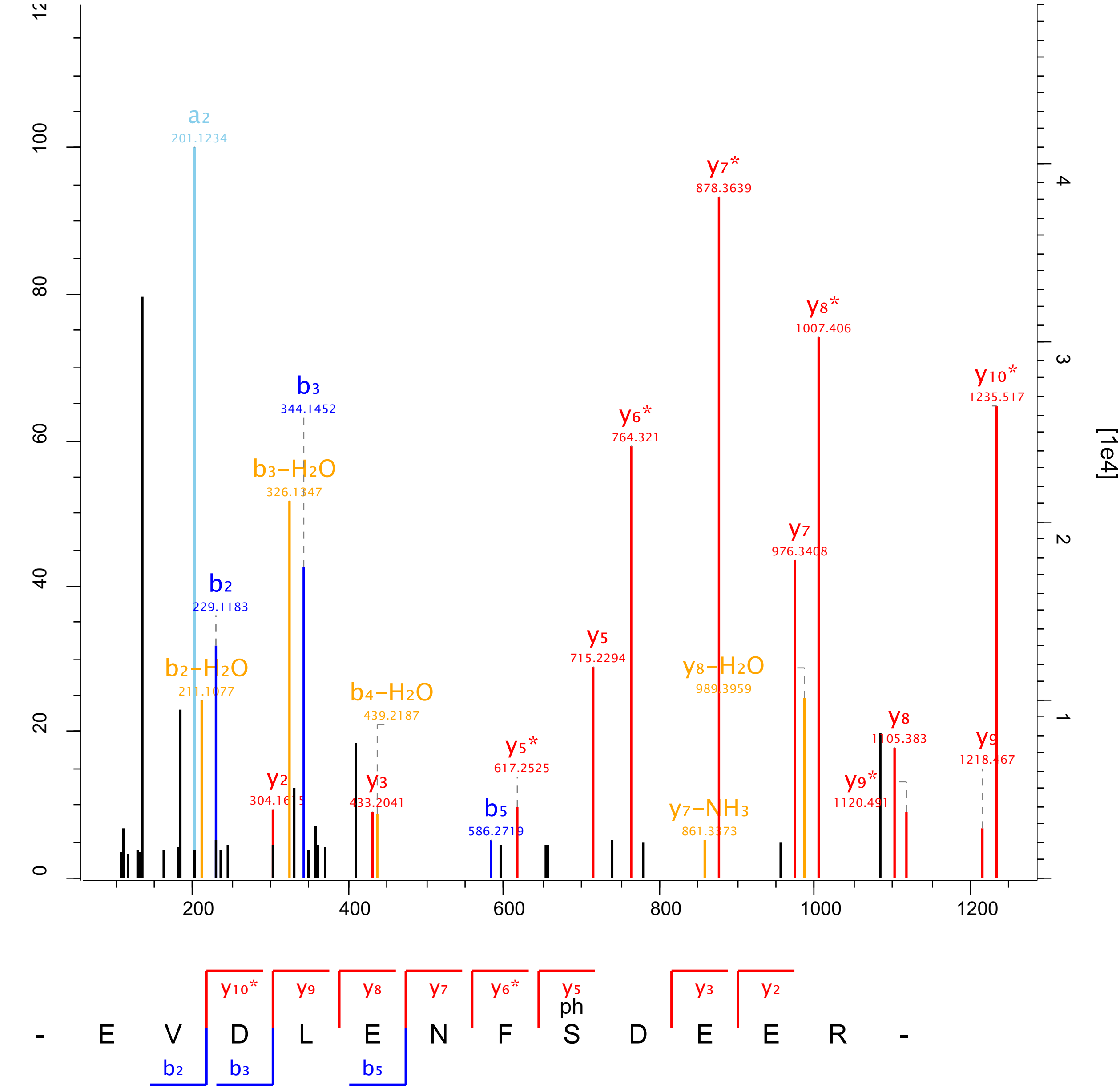

| Raw file | Scan | Method | Score | m/z |
| --- | --- | --- | --- | --- |
| sys_15_1 | 28549 | FTMS; HCD | 97.71 | 757.87 |

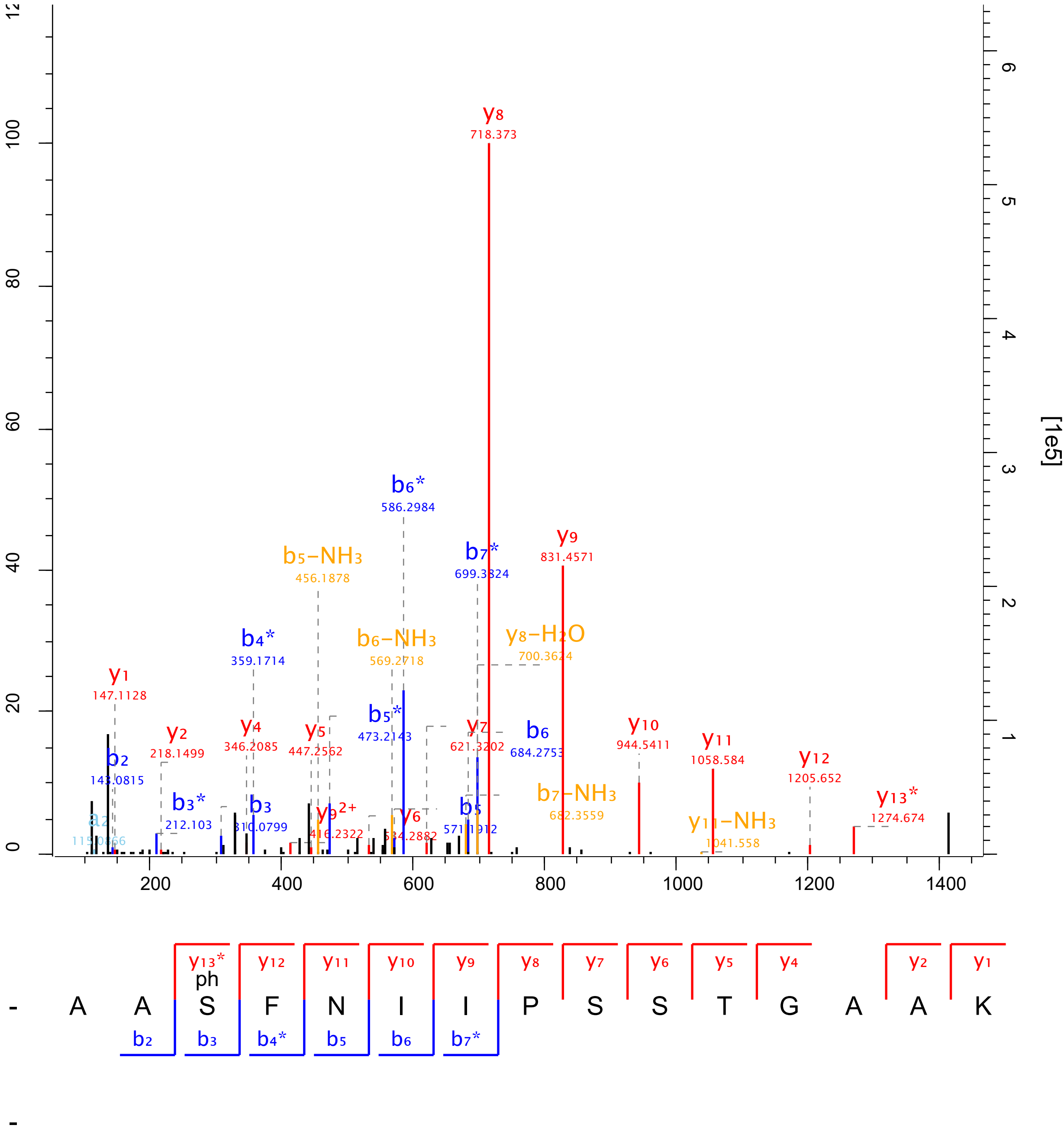

| Raw file | Scan | Method | Score | m/z |
|---|---|---|---|---|
| sys_15_1 | 28886 | FTMS; HCD | 77.07 | 797.83 |

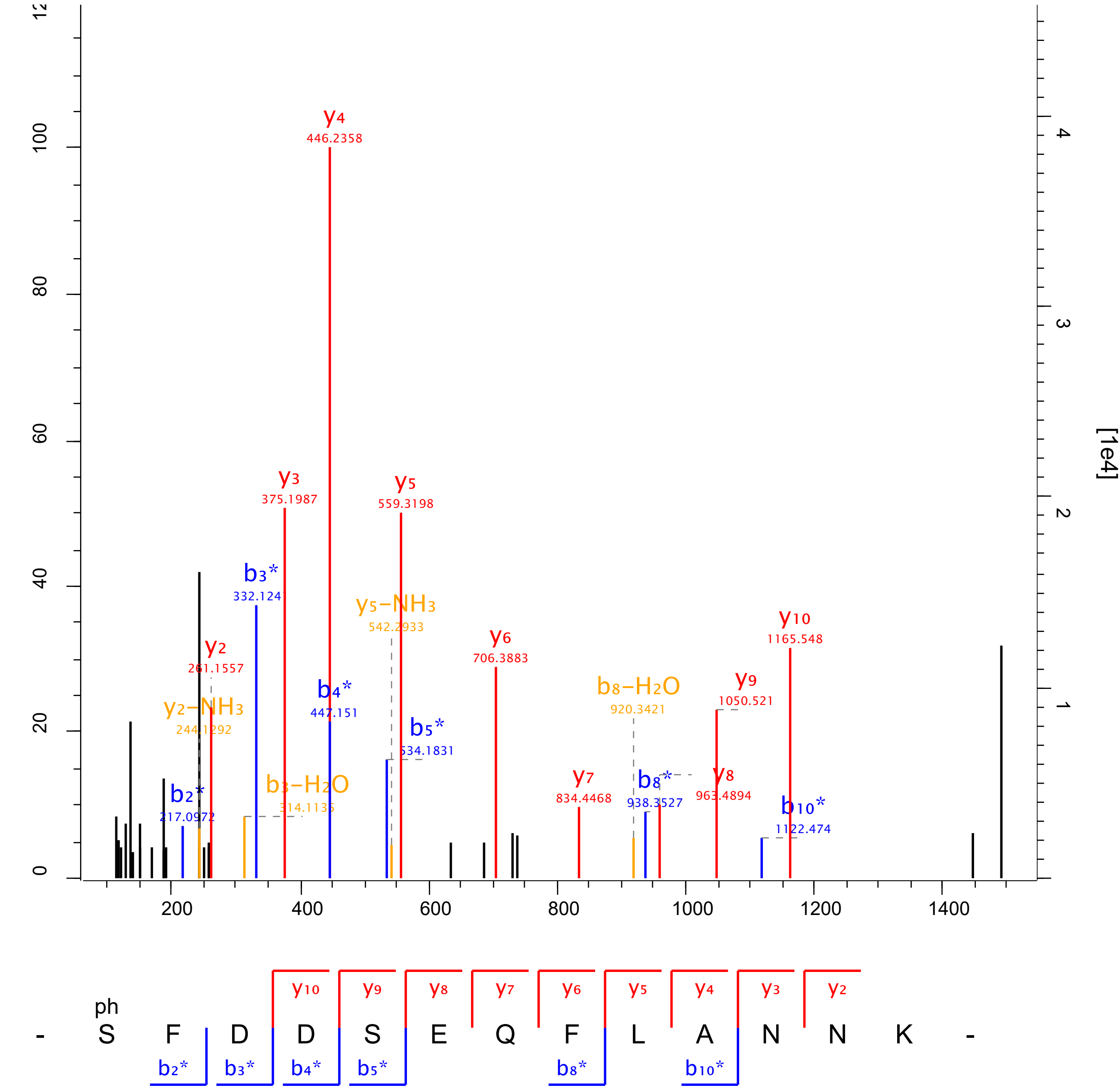

| Raw file | Scan | Method | Score | m/z |
| --- | --- | --- | --- | --- |
| sys_15_1 | 28929 | FTMS; HCD | 108.57 | 764.05 |

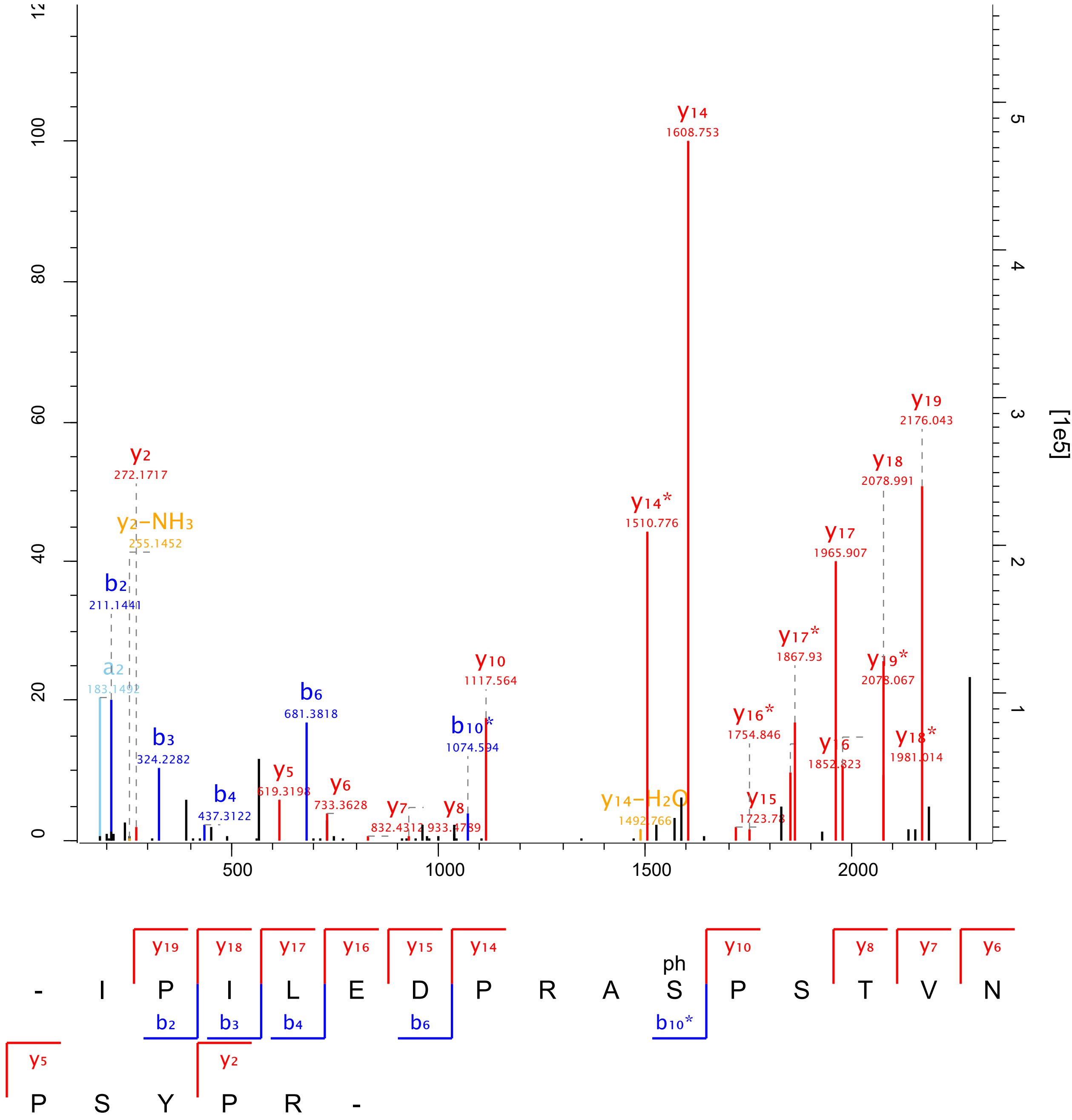

| Raw file | Scan | Method | Score | m/z |
|---|---|---|---|---|
| sys_15_1 | 28942 | FTMS; HCD | 76.76 | 678.77 |

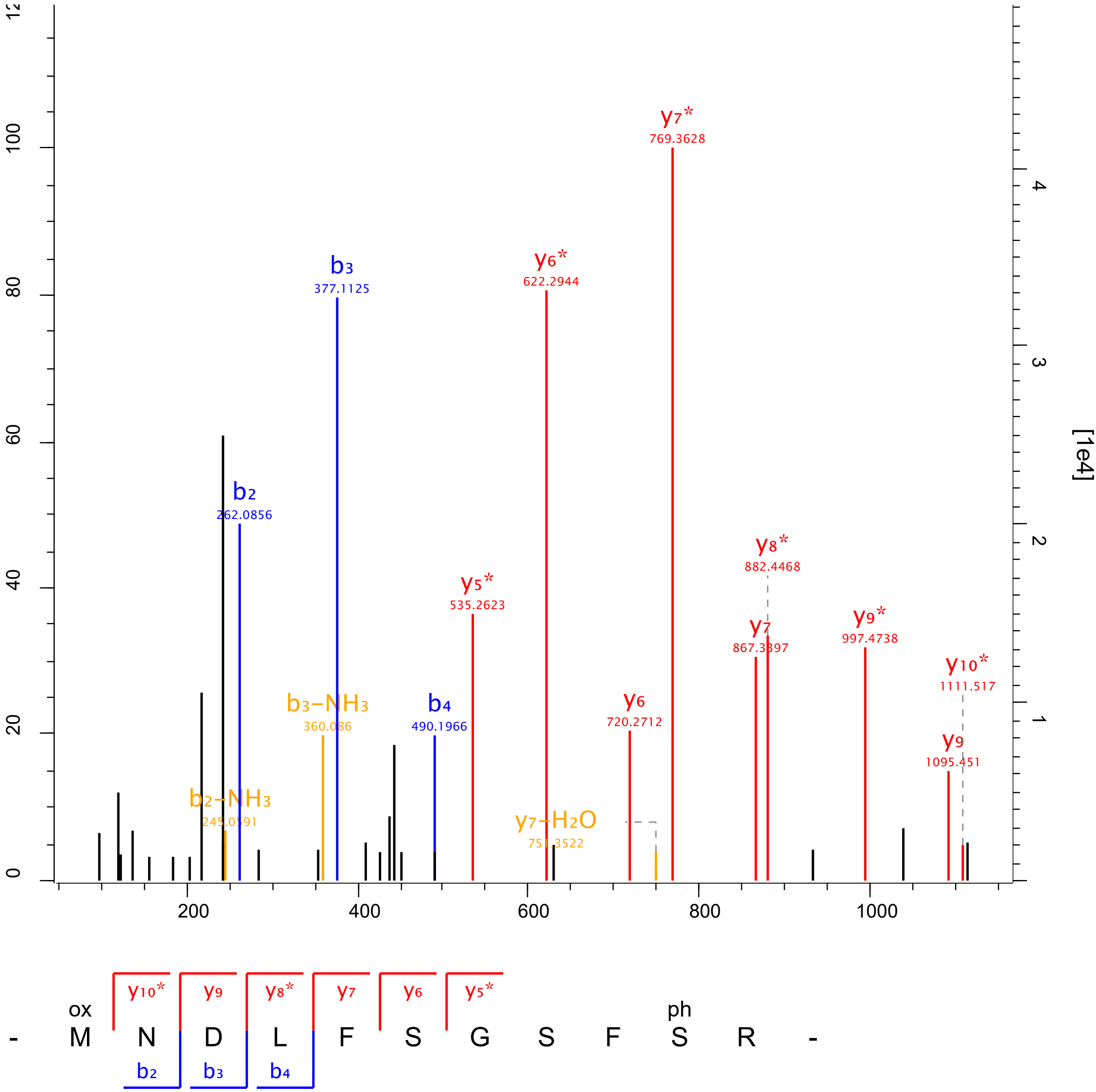

| Raw file | Scan | Method | Score | m/z |
|---|---|---|---|---|
| sys_15_1 | 29114 | FTMS; HCD | 48.57 | 548.28 |

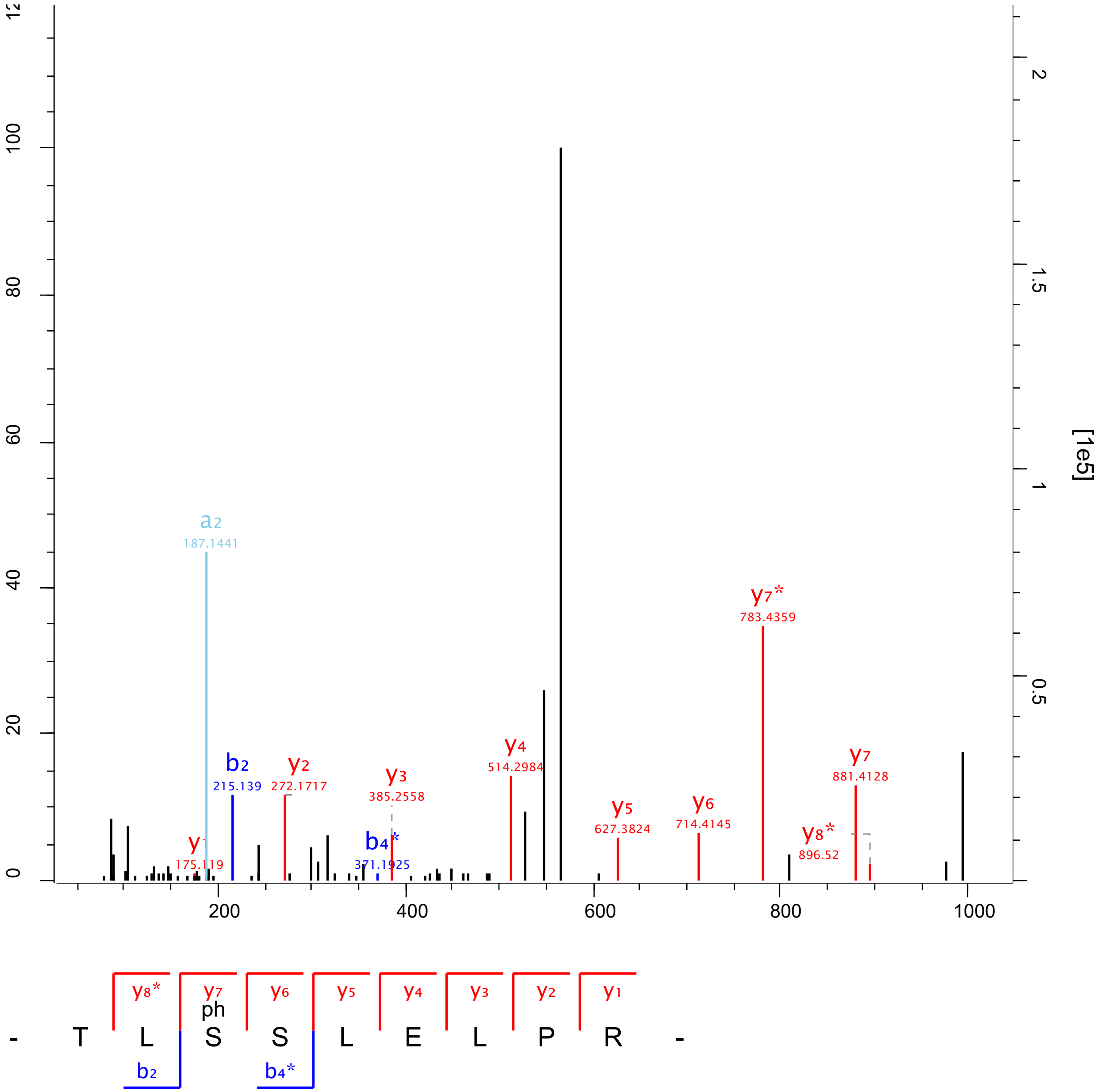

| Raw file | Scan | Method | Score | m/z |
|---|---|---|---|---|
| sys_15_1 | 29185 | FTMS; HCD | 50.3 | 710.3 |

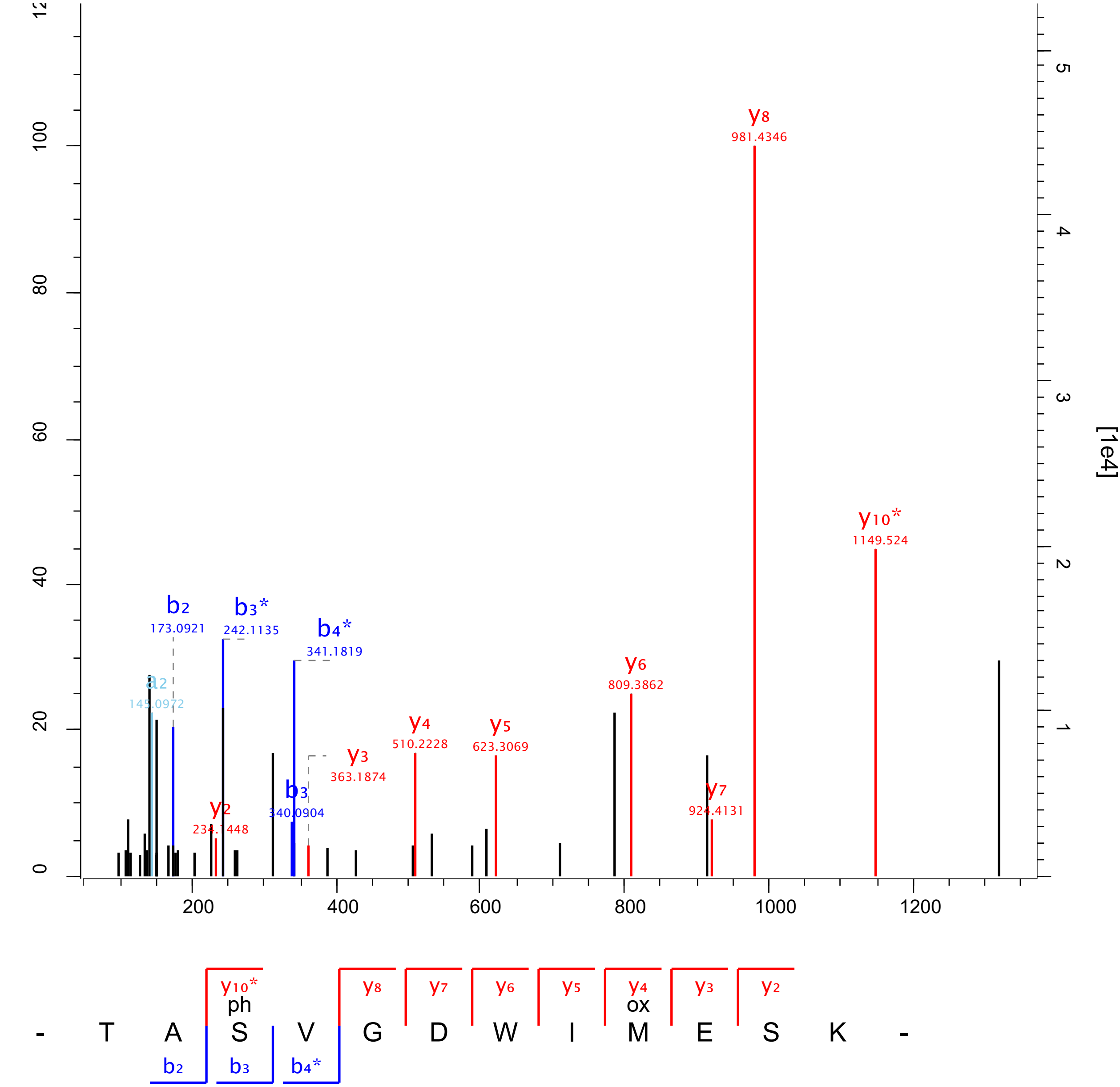

| Raw file | Scan | Method | Score | m/z |
| --- | --- | --- | --- | --- |
| sys_15_1 | 29224 | FTMS; HCD | 191.93 | 887.88 |

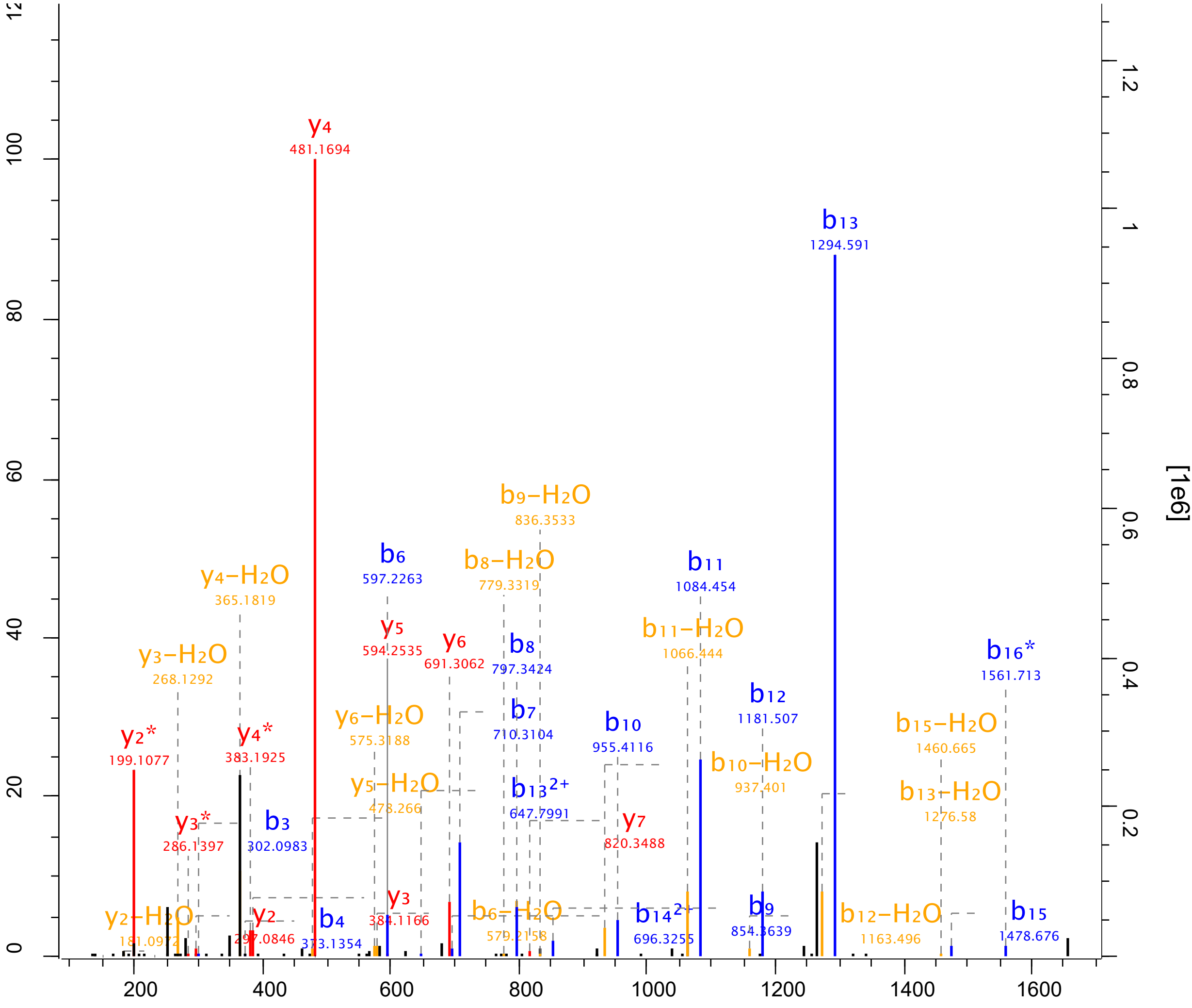

| Raw file | Scan | Method | Score | m/z |
| --- | --- | --- | --- | --- |
| sys_15_1 | 2931 | FTMS; HCD | 42.03 | 711.32 |

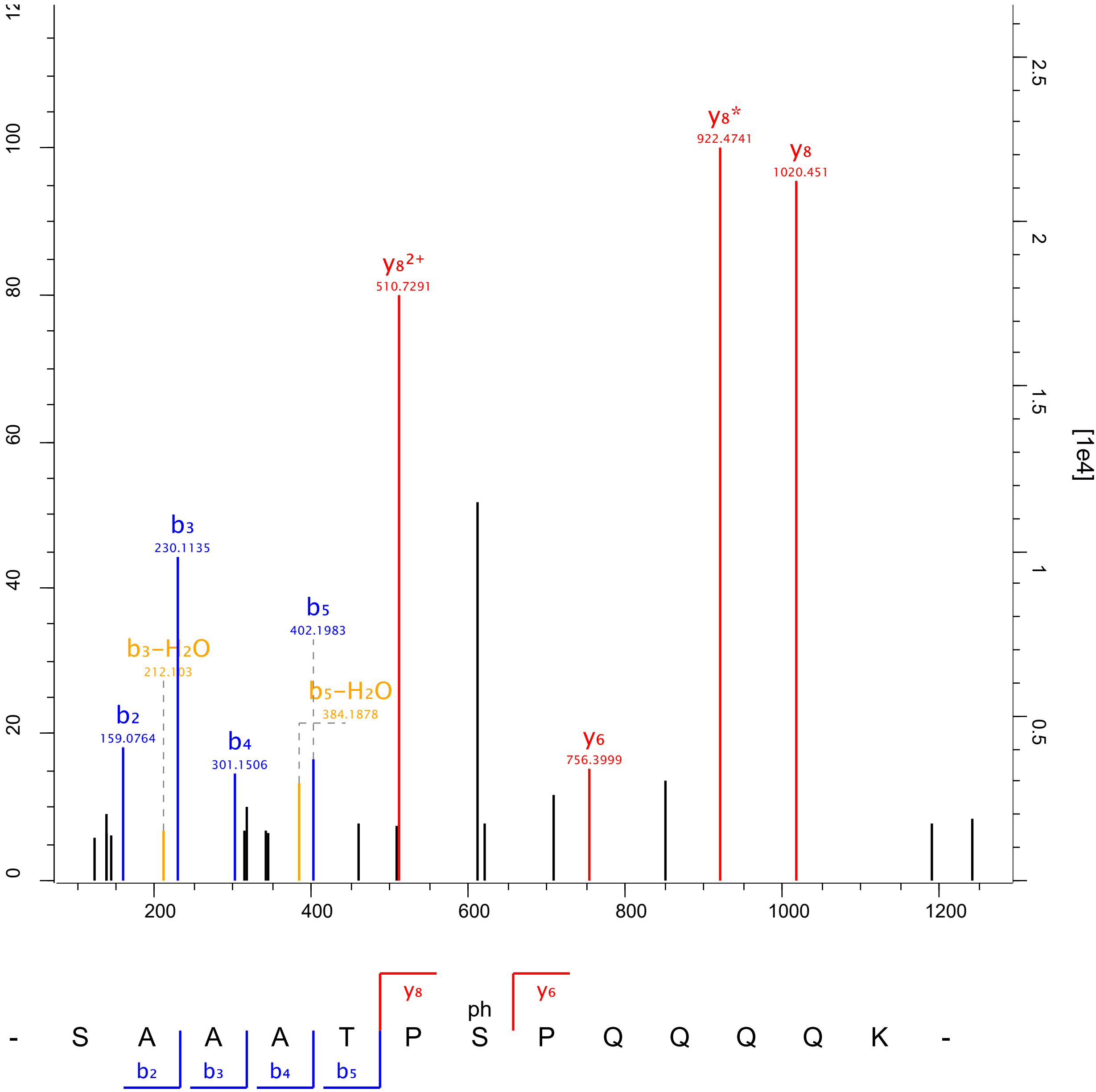

| Raw file | Scan | Method | Score | m/z |
| --- | --- | --- | --- | --- |
| sys_15_1 | 29353 | FTMS; HCD | 44.34 | 723.96 |

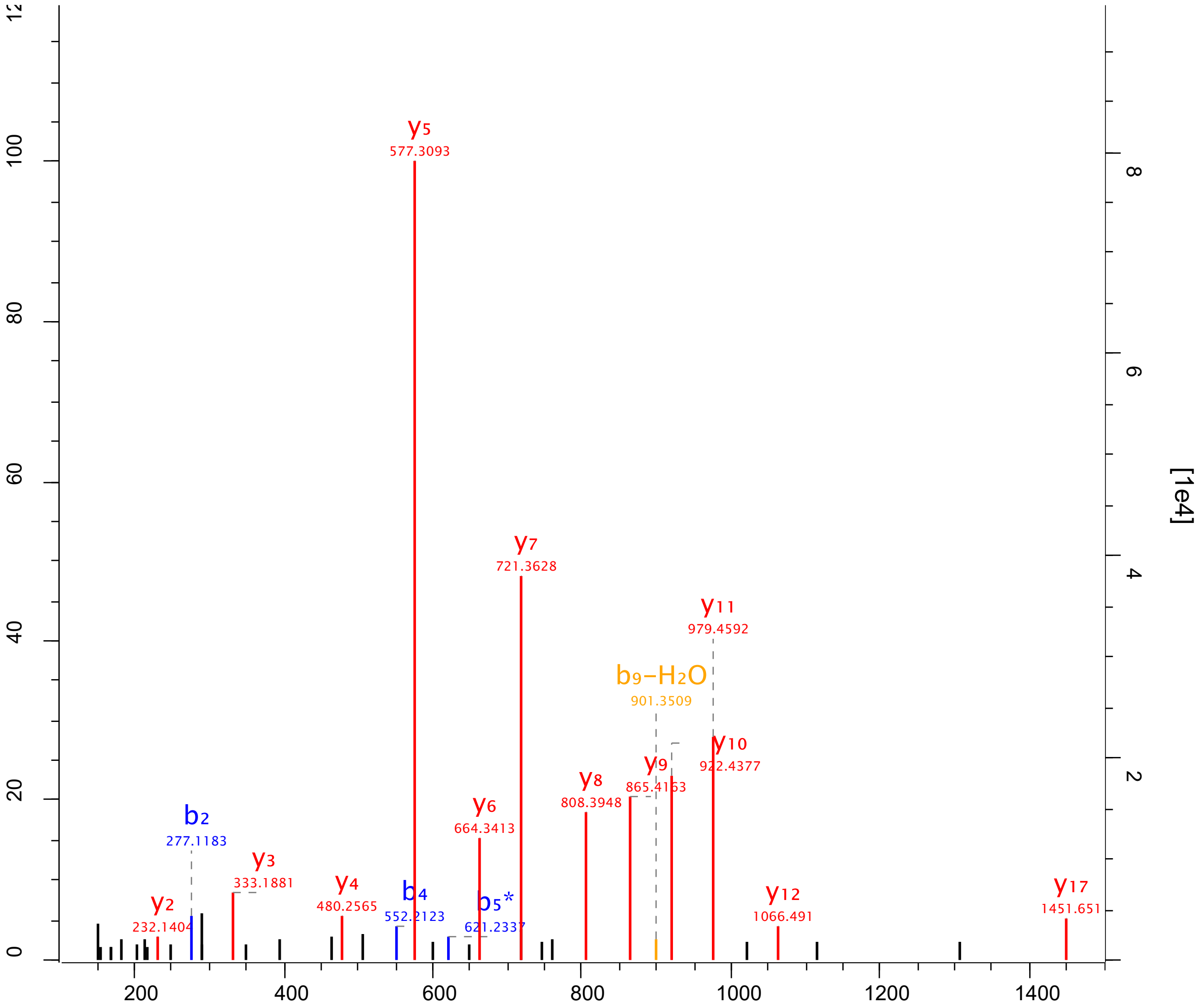


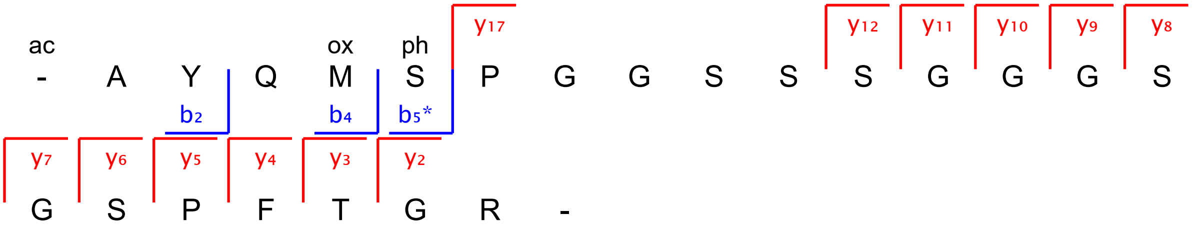

| Raw file | Scan | Method | Score | m/z |
|---|---|---|---|---|
| sys_15_1 | 2937 | FTMS; HCD | 43.76 | 659.27 |

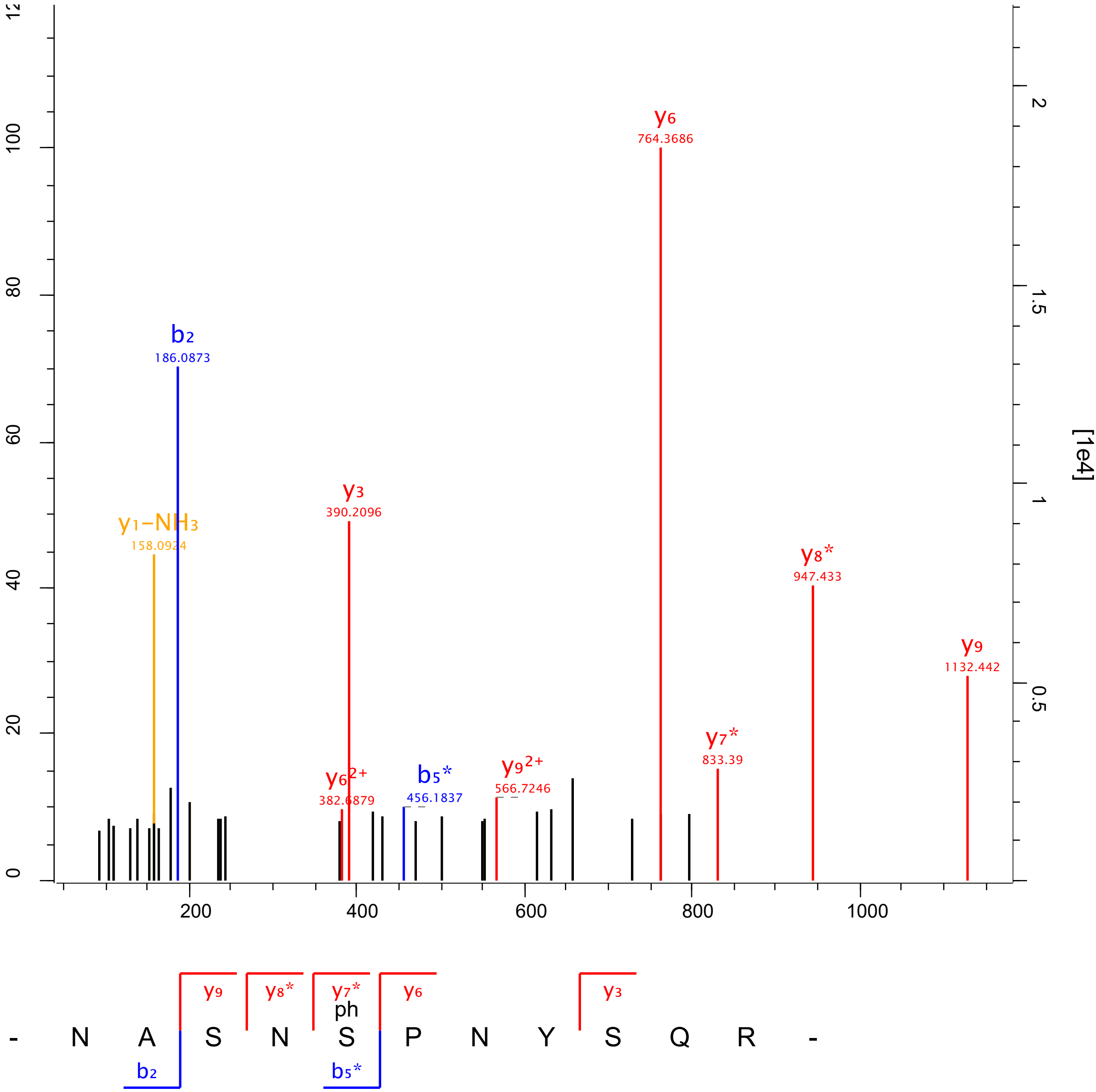

| Raw file | Scan | Method | Score | m/z |
|---|---|---|---|---|
| sys_15_1 | 2946 | FTMS; HCD | 86.34 | 498.73 |

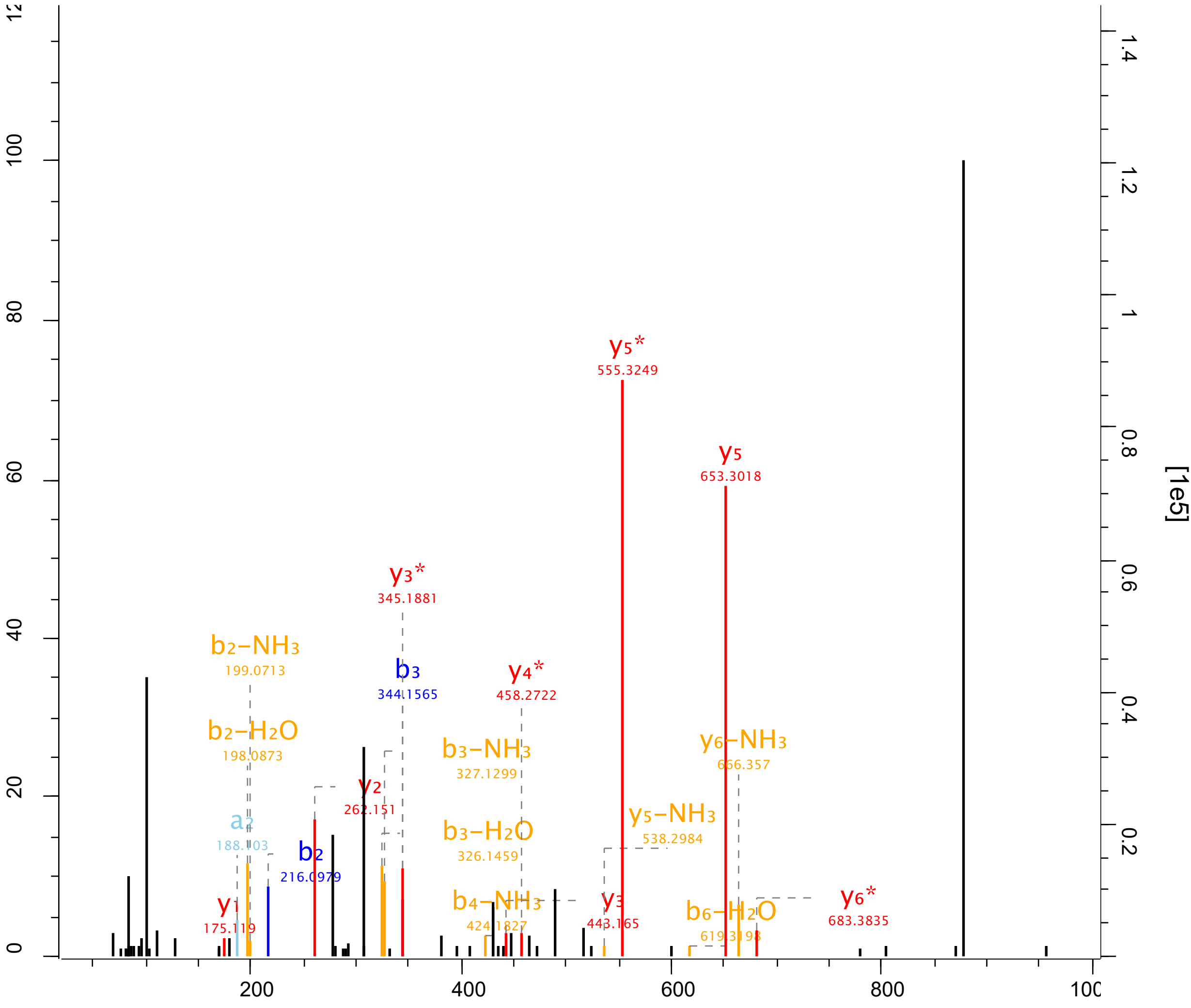


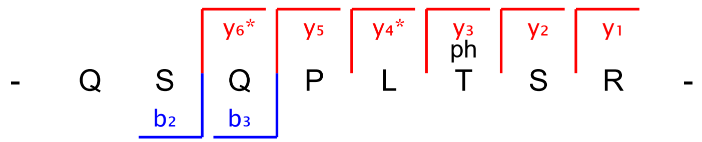

| Raw file | Scan | Method | Score | m/z |
|---|---|---|---|---|
| sys_15_1 | 29469 | FTMS; HCD | 159.76 | 1079.98 |

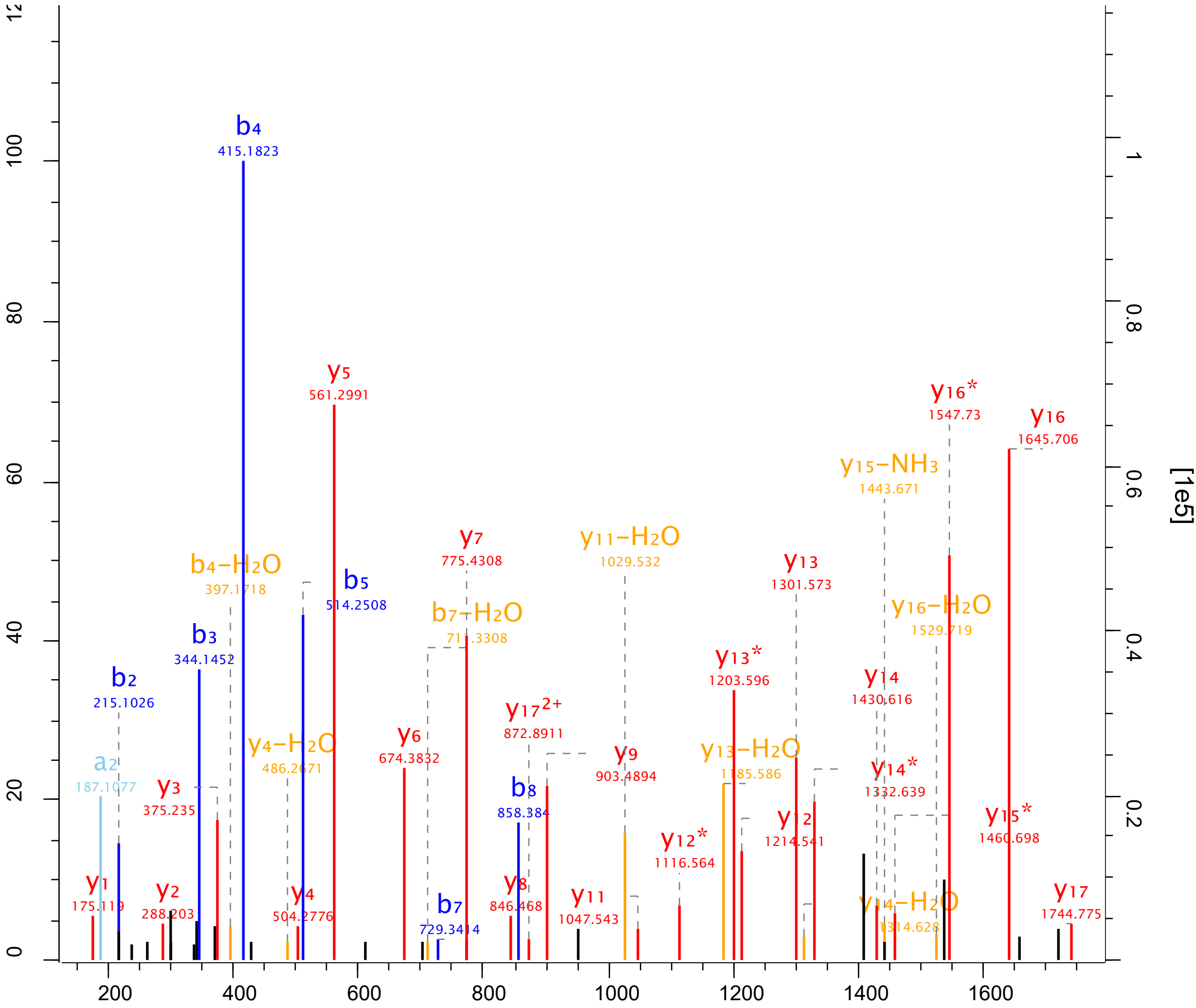


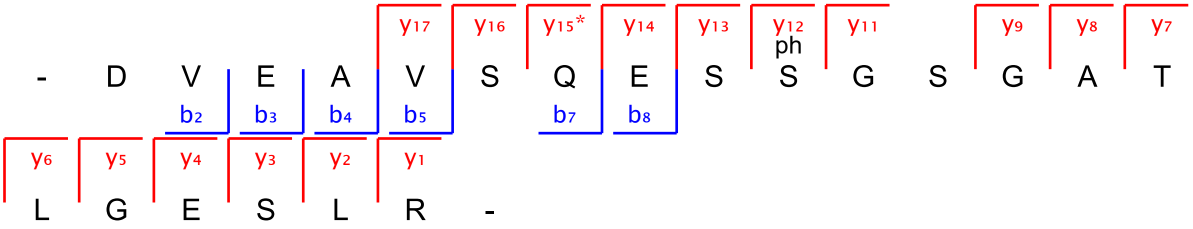

| Raw file | Scan | Method | Score | m/z |
|---|---|---|---|---|
| sys_15_1 | 29561 | FTMS; HCD | 72.99 | 728.69 |

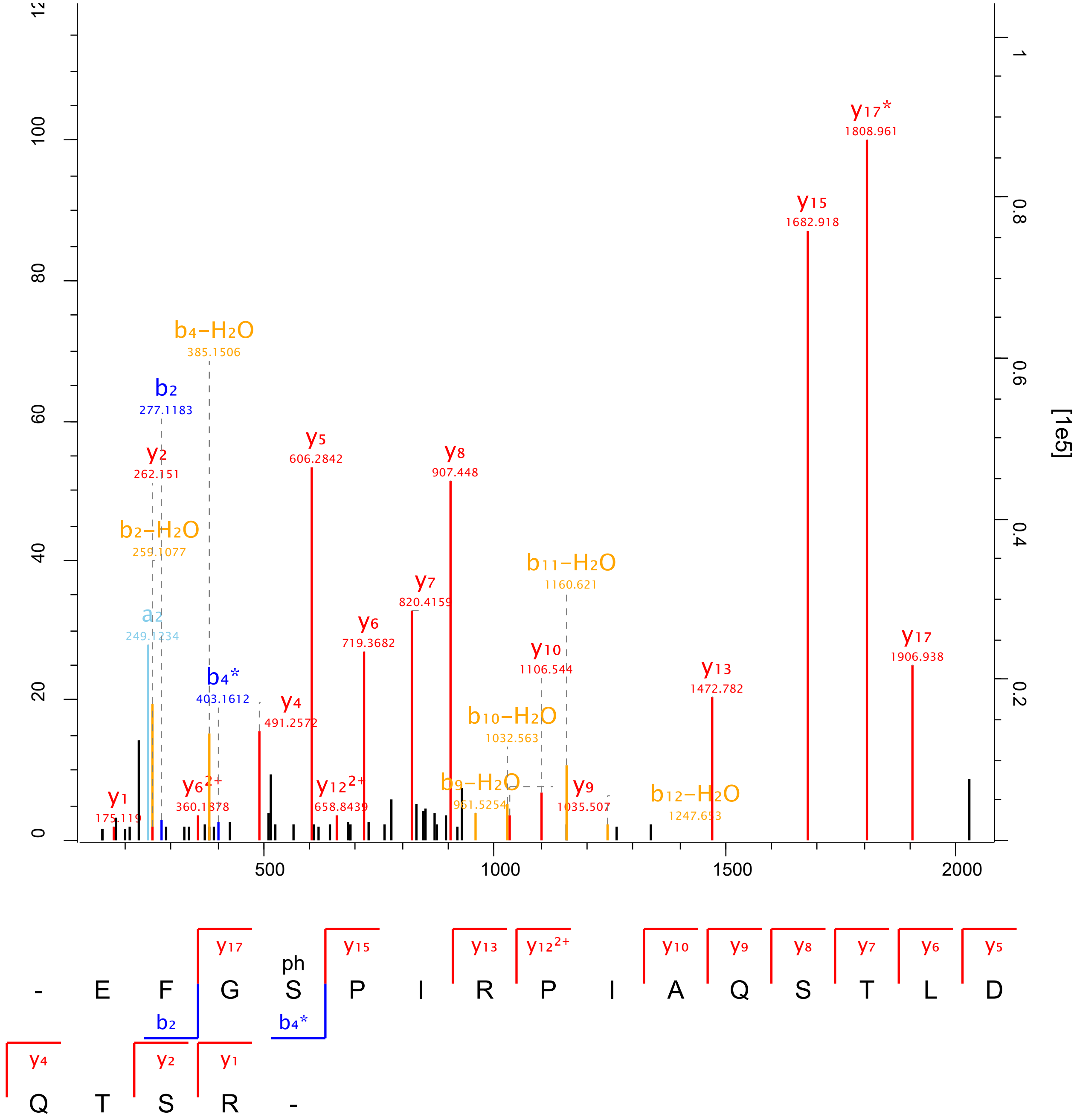

| Raw file | Scan | Method | Score | m/z |
|---|---|---|---|---|
| sys_15_1 | 2965 | FTMS; HCD | 50.35 | 506.71 |

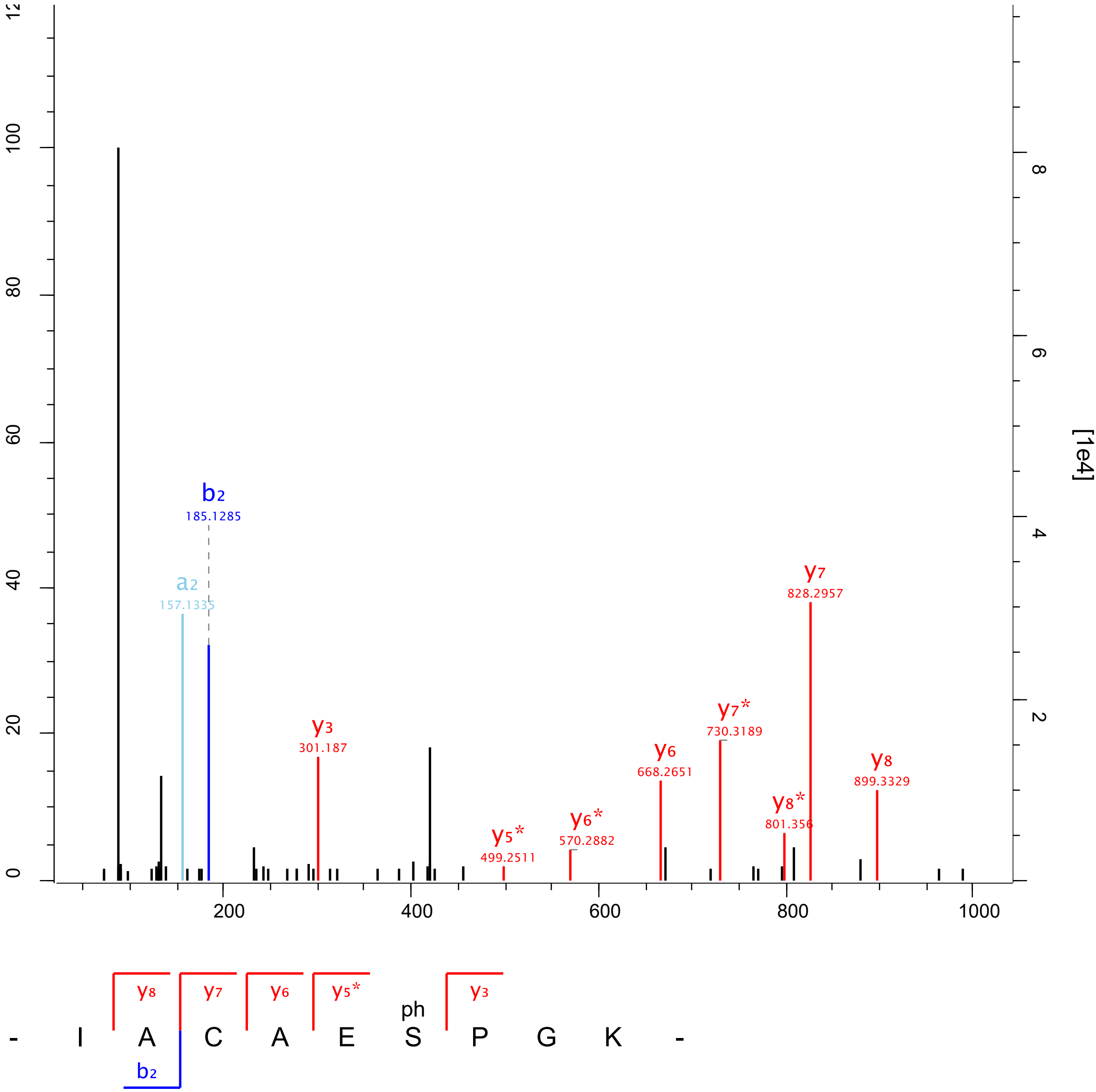

| Raw file | Scan | Method | Score | m/z |
|---|---|---|---|---|
| sys_15_1 | 29675 | FTMS; HCD | 51.73 | 583.25 |

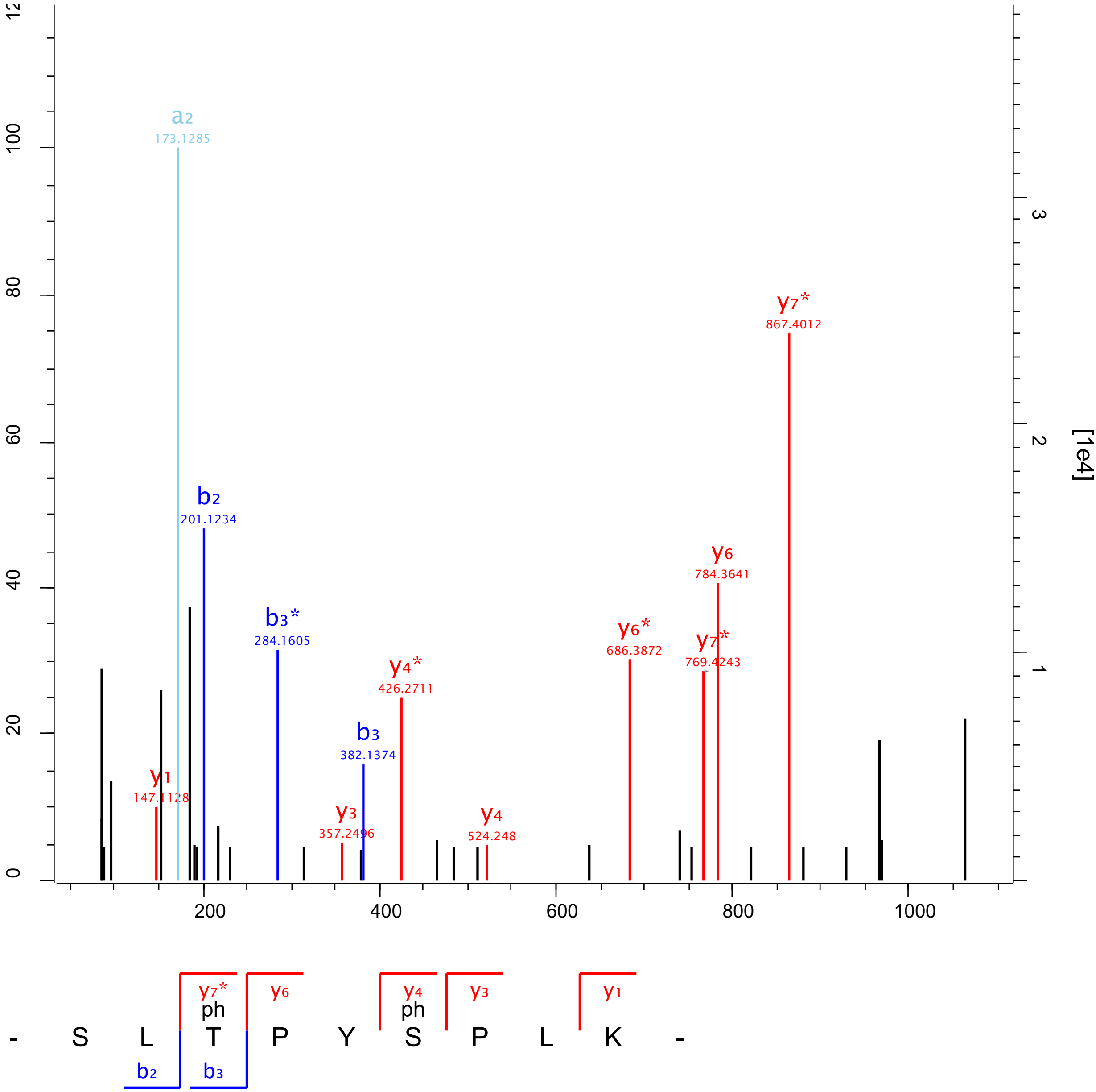

| Raw file | Scan | Method | Score | m/z |
|---|---|---|---|---|
| sys_15_1 | 29855 | FTMS; HCD | 42.06 | 657.64 |

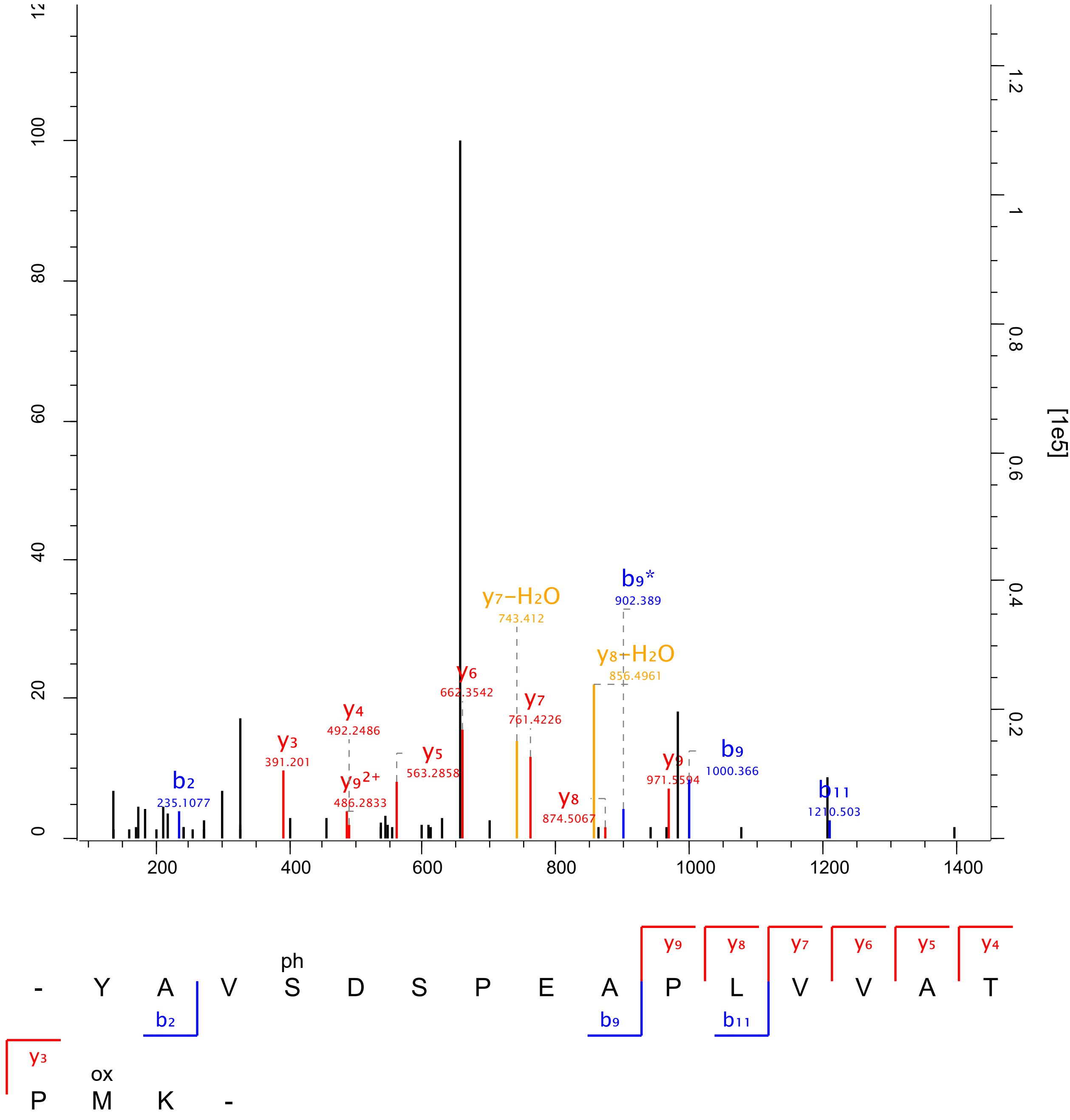

| Raw file | Scan | Method | Score | m/z |
|---|---|---|---|---|
| sys_15_1 | 29863 | FTMS; HCD | 113.4 | 758.33 |

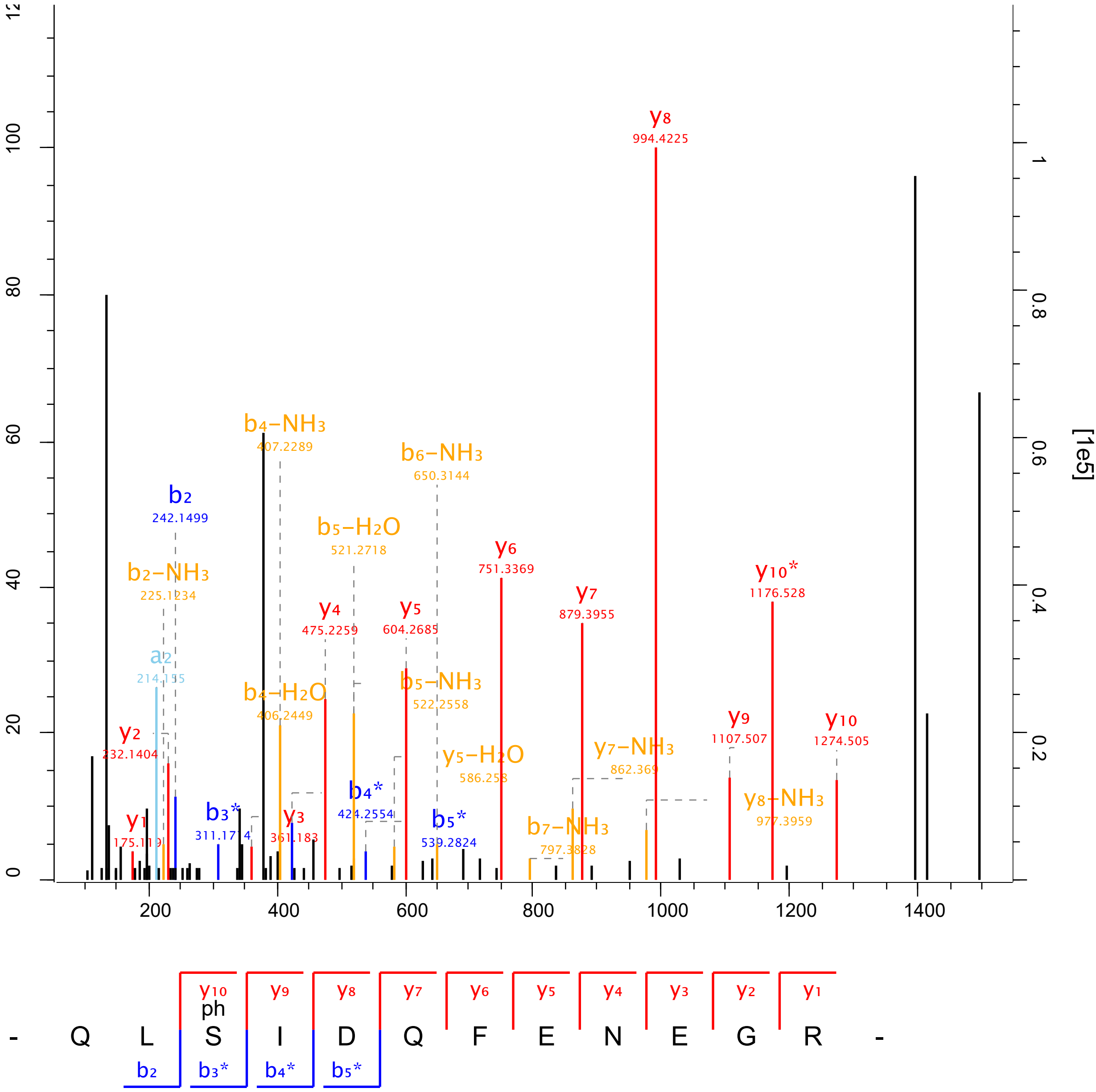

| Raw file | Scan | Method | Score | m/z |
|---|---|---|---|---|
| sys_15_1 | 29954 | FTMS; HCD | 166.28 | 854.89 |

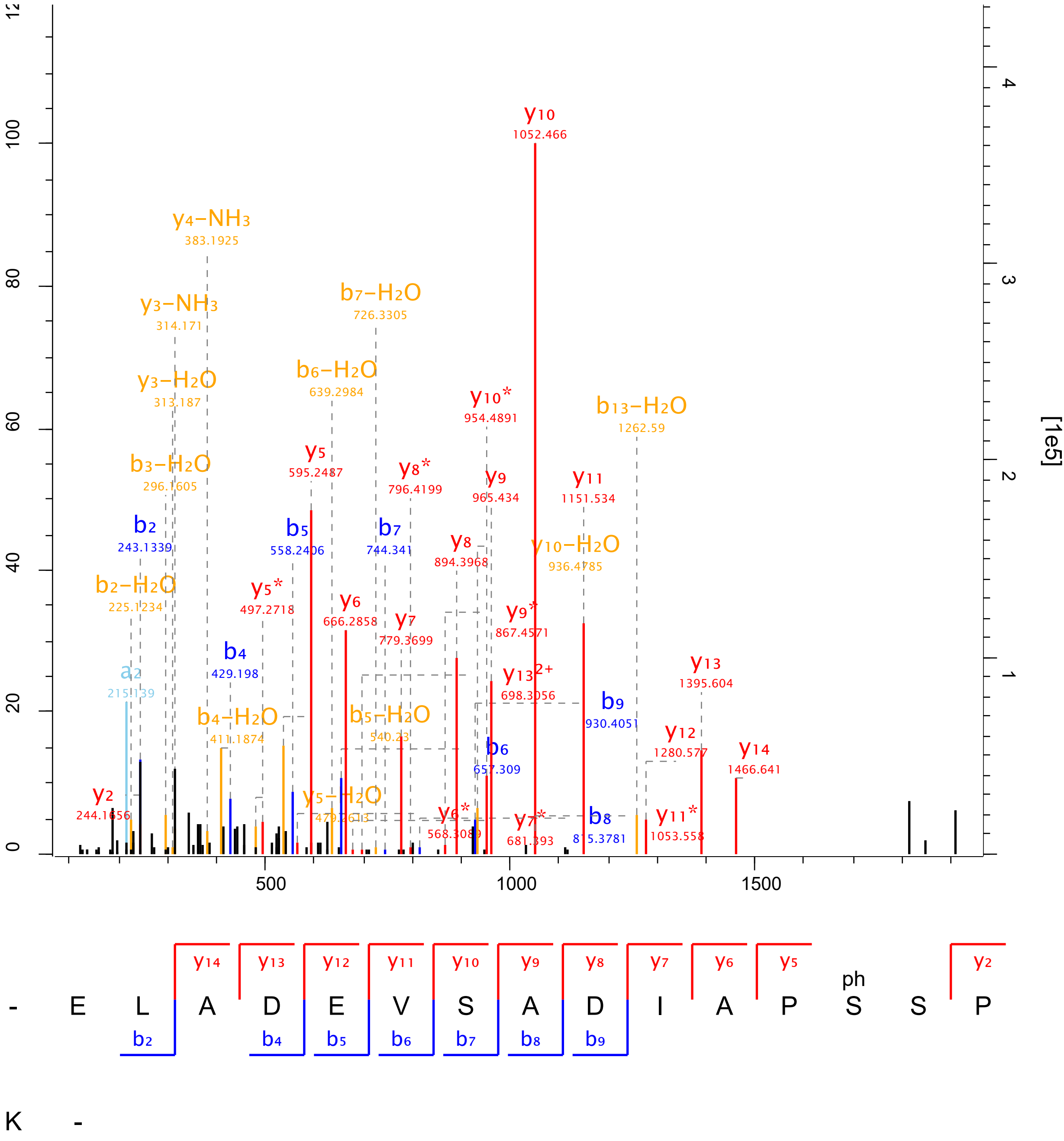

| Raw file | Scan | Method | Score | m/z |
|---|---|---|---|---|
| sys_15_1 | 29977 | FTMS; HCD | 98.16 | 549.23 |

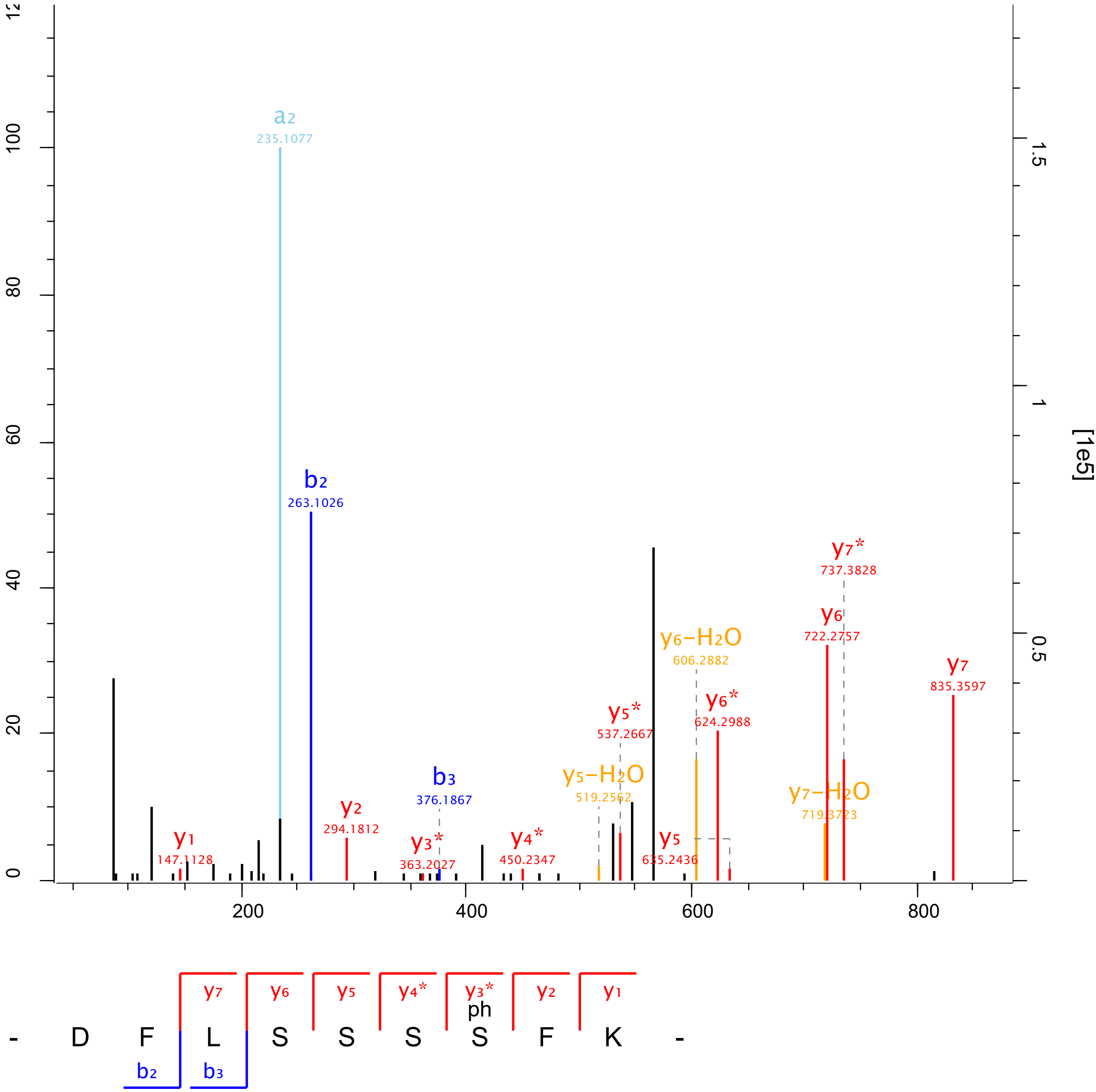

| Raw file | Scan | Method | Score | m/z |
|---|---|---|---|---|
| sys_15_1 | 3002 | FTMS; HCD | 93.78 | 871.78 |

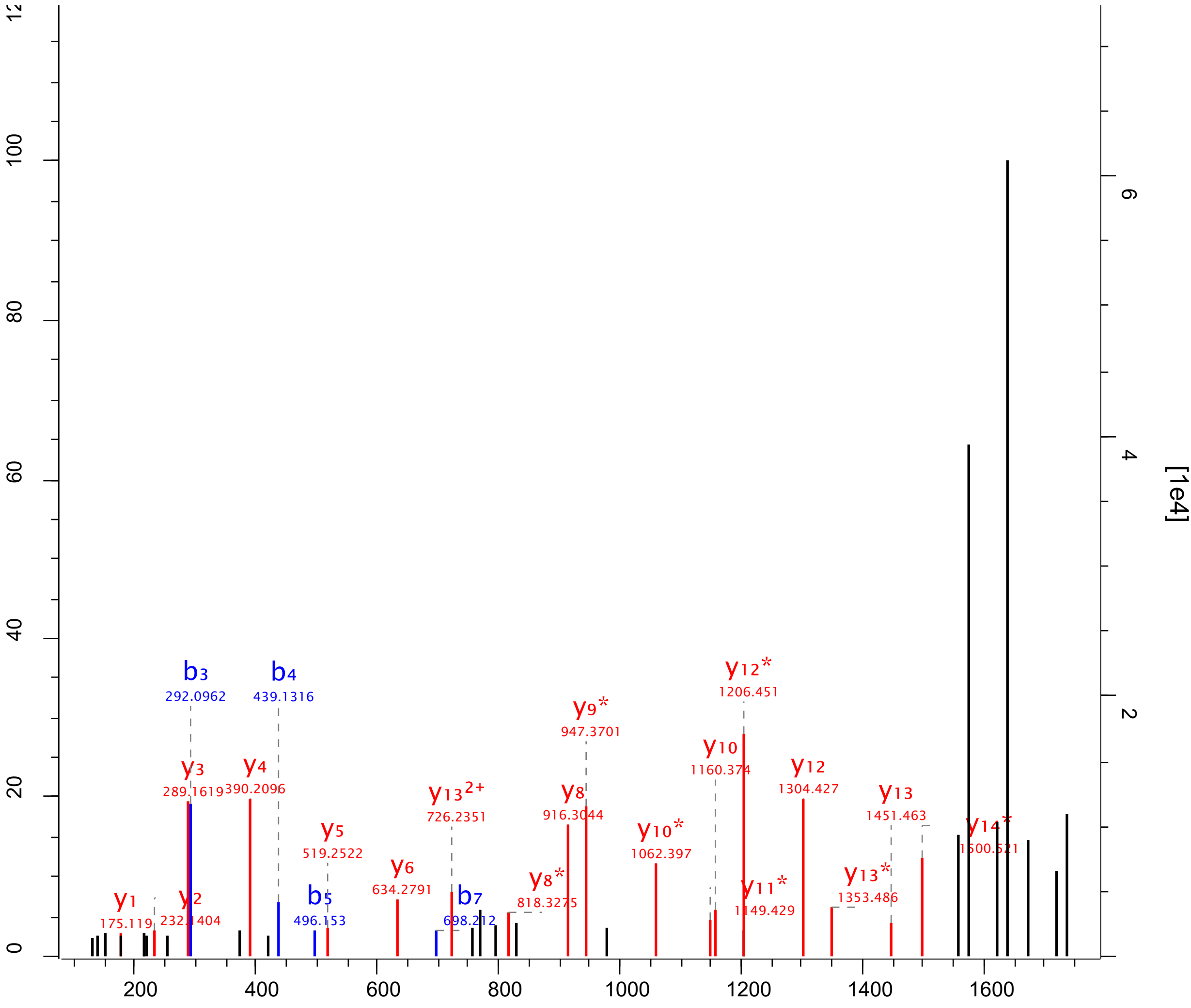


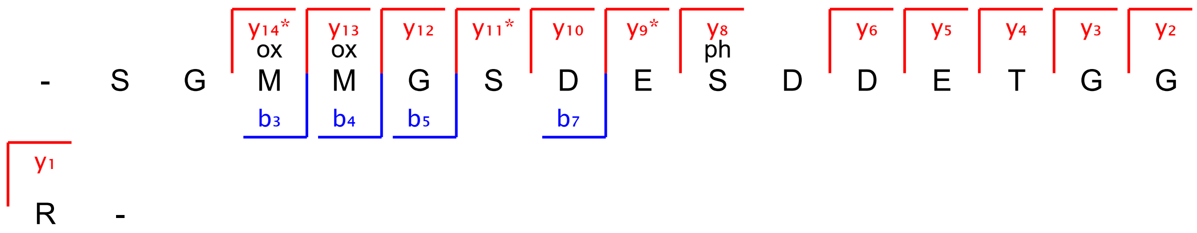

| Raw file | Scan | Method | Score | m/z |
| --- | --- | --- | --- | --- |
| sys_15_1 | 30040 | FTMS; HCD | 50.75 | 724.65 |

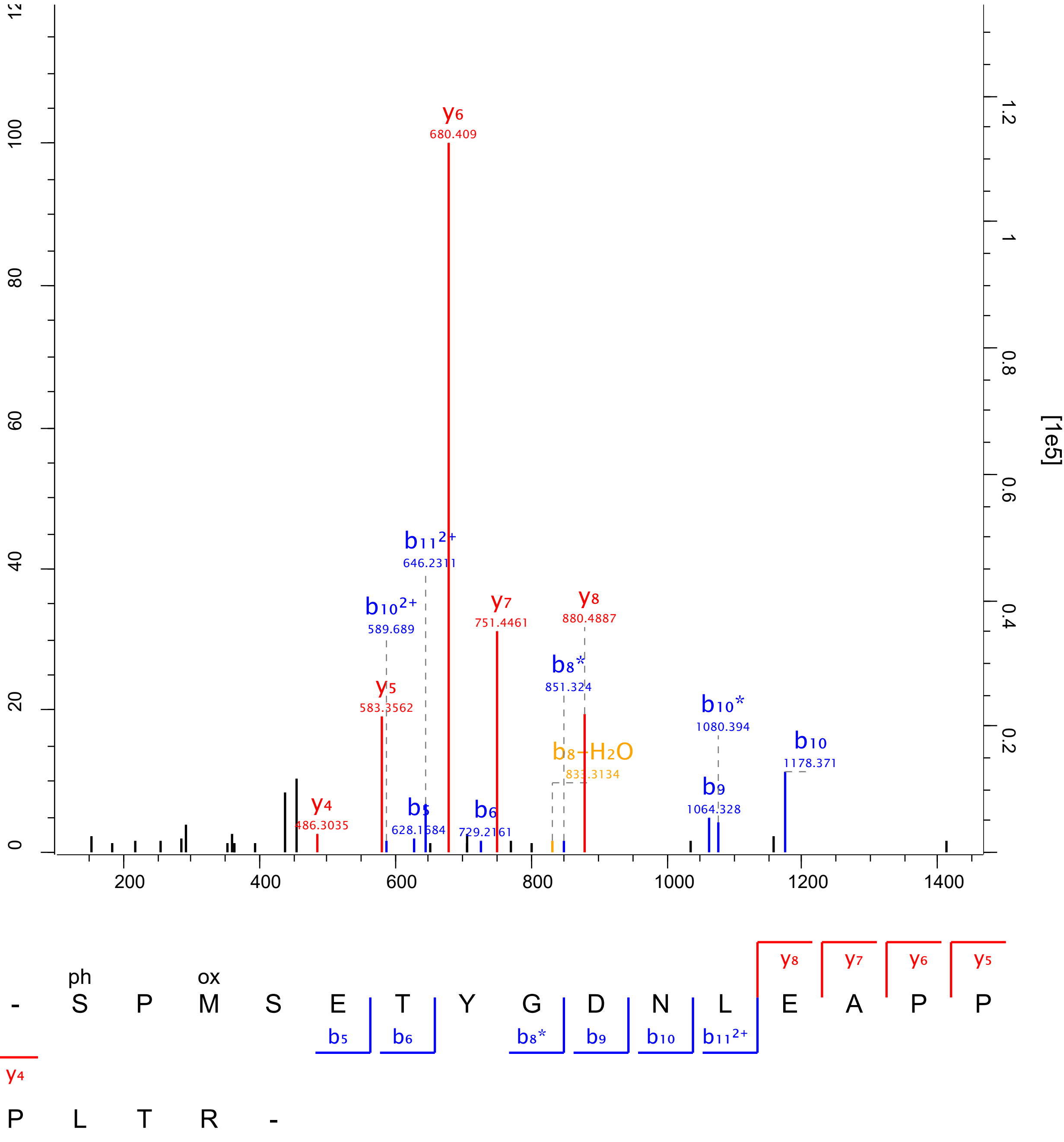

| Raw file | Scan | Method | Score | m/z |
|---|---|---|---|---|
| sys_15_1 | 3006 | FTMS; HCD | 121.82 | 769.76 |

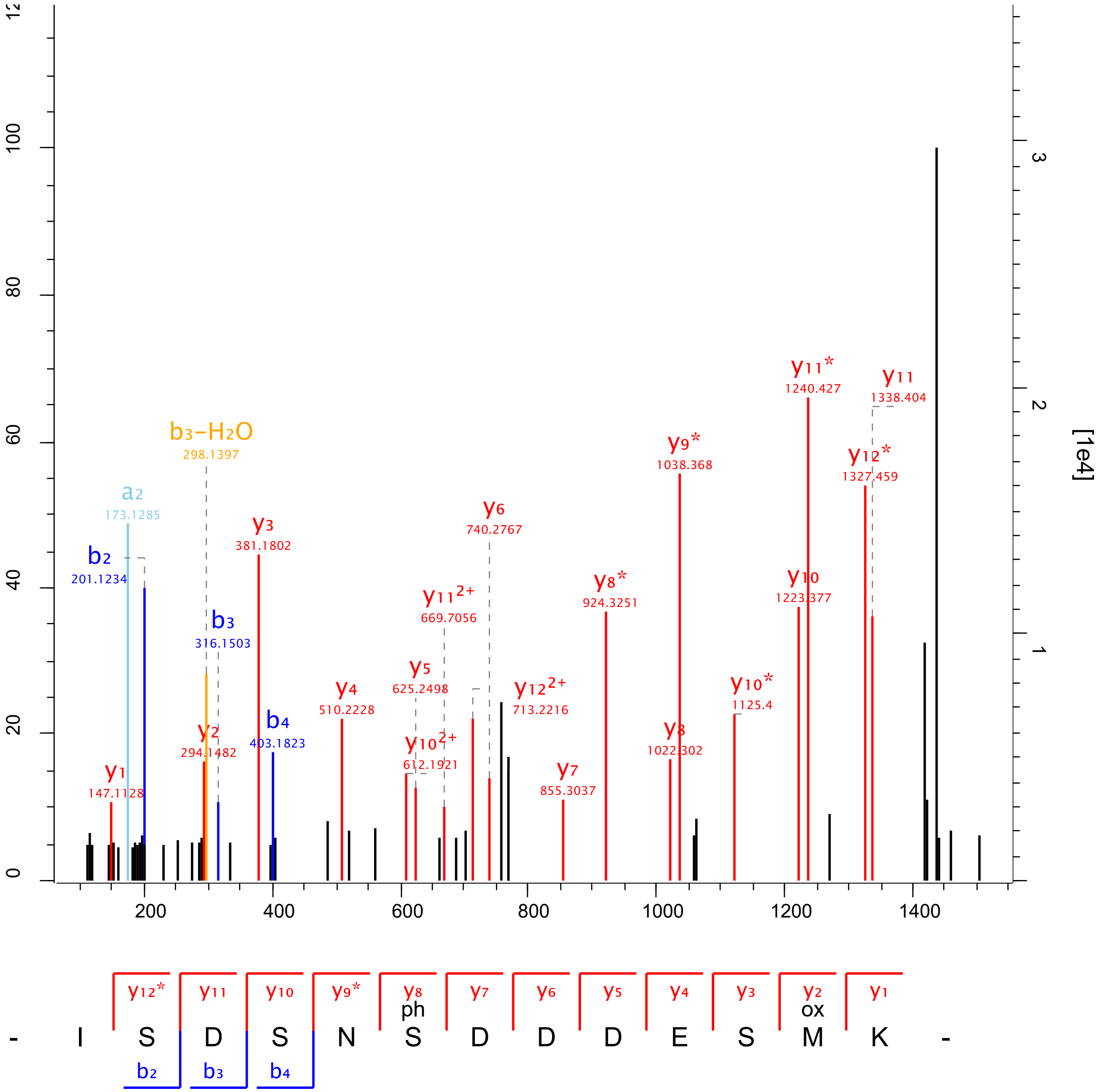

| Raw file | Scan | Method | Score | m/z |
|---|---|---|---|---|
| sys_15_1 | 30146 | FTMS; HCD | 285.5 | 827.85 |

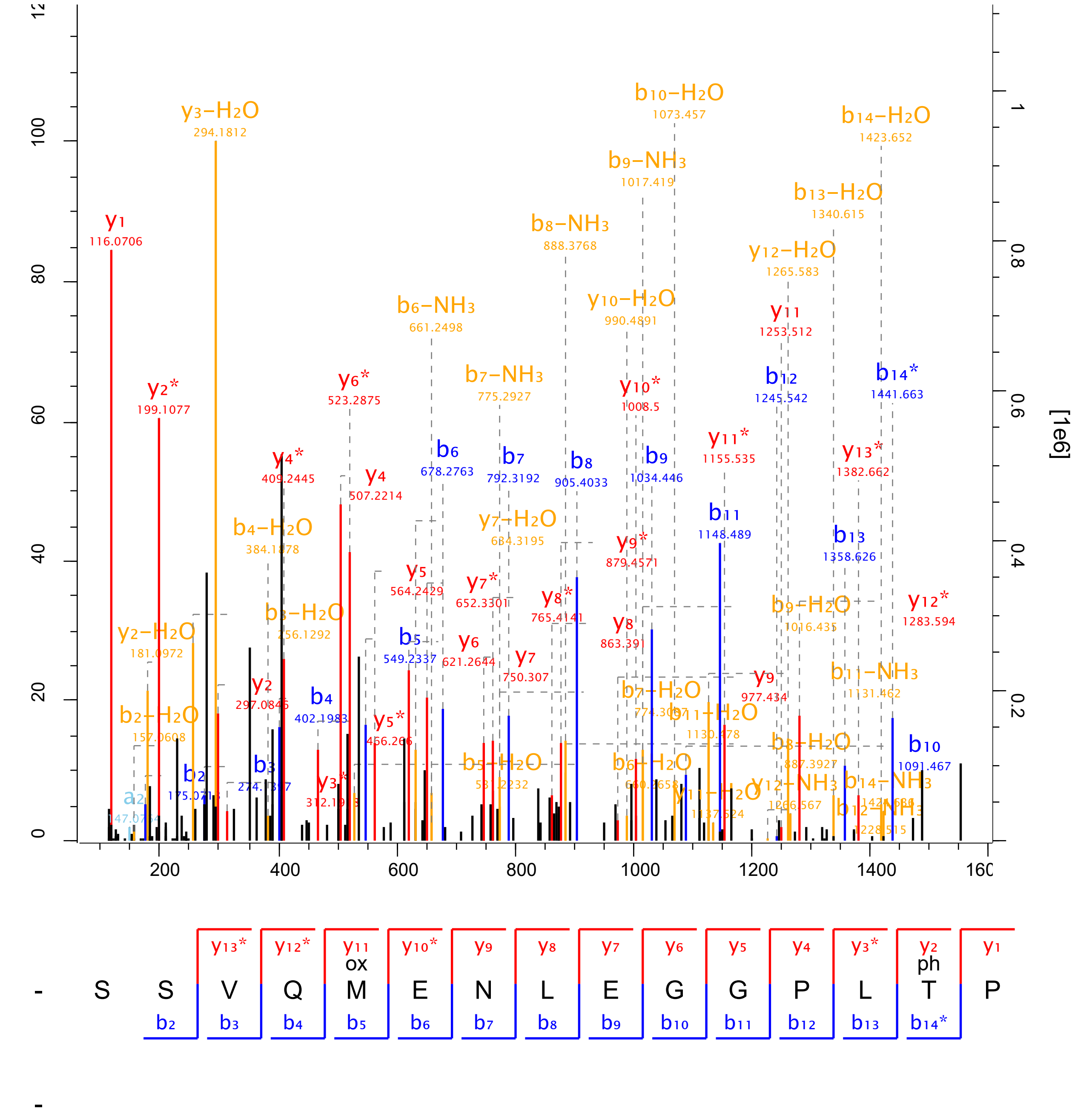

| Raw file | Scan | Method | Score | m/z |
|---|---|---|---|---|
| sys_15_1 | 3021 | FTMS; HCD | 46.84 | 593.75 |

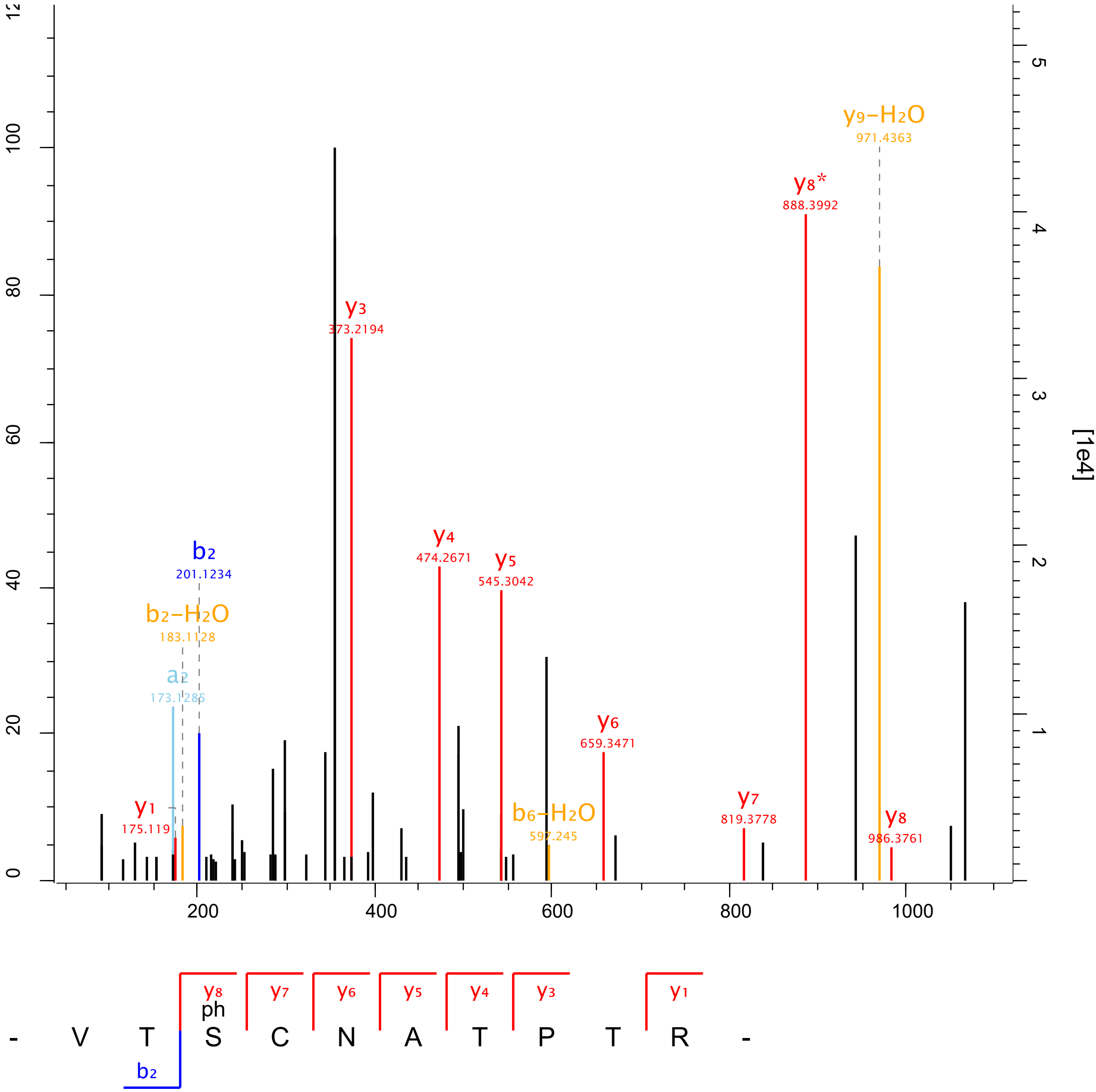

| Raw file | Scan | Method | Score | m/z |
|---|---|---|---|---|
| sys_15_1 | 30232 | FTMS; HCD | 141.43 | 805.67 |

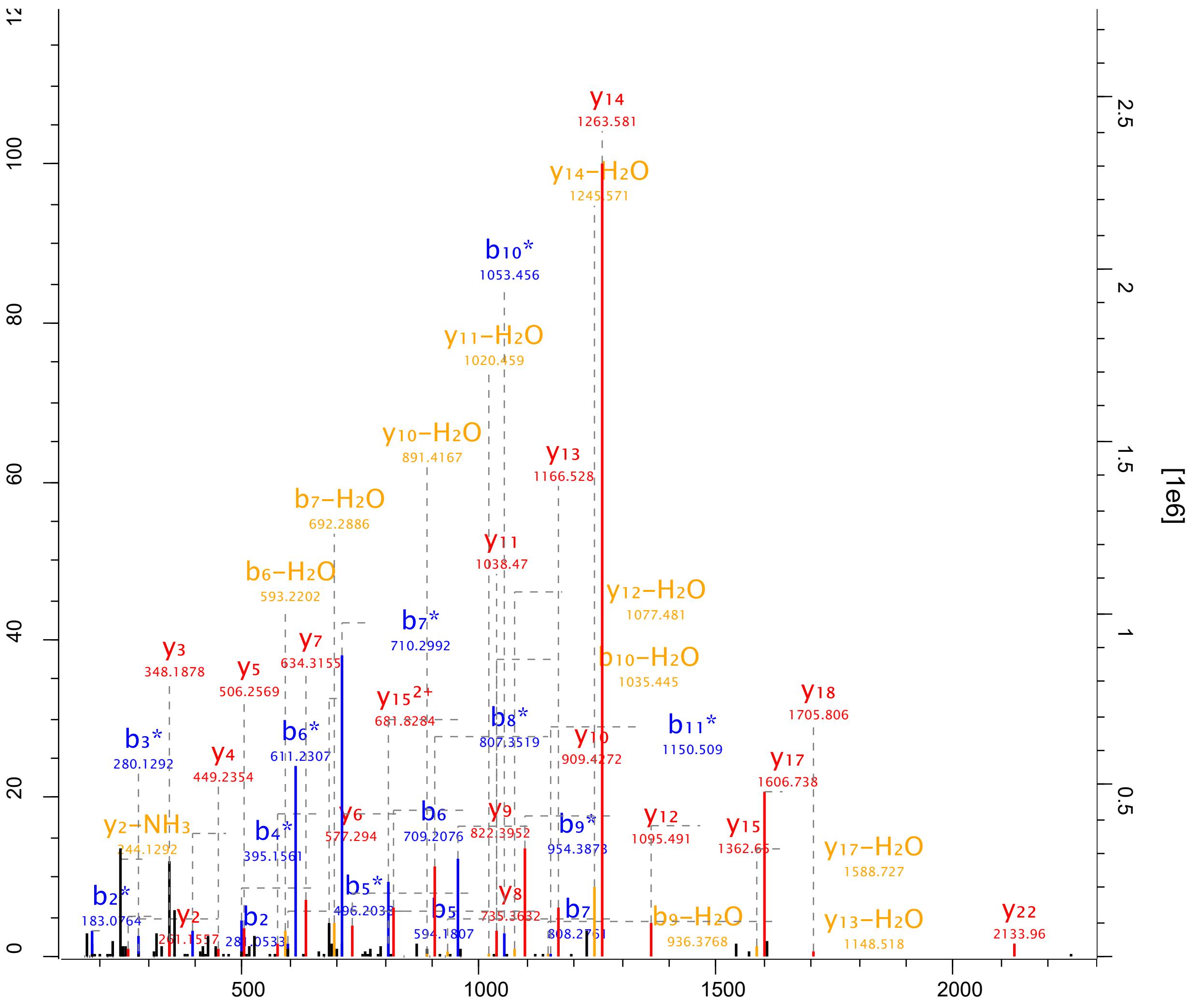


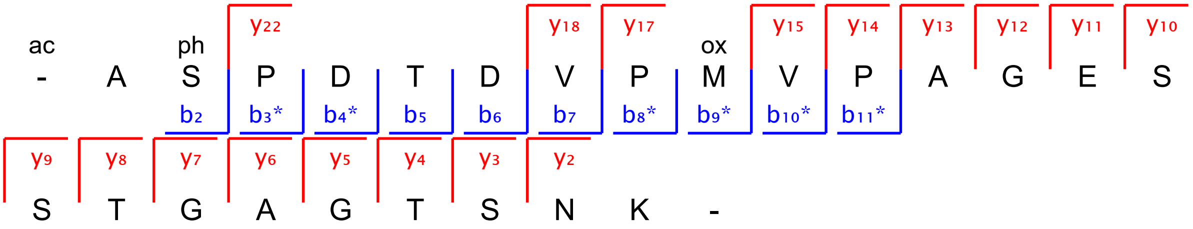

| Raw file | Scan | Method | Score | m/z |
|---|---|---|---|---|
| sys_15_1 | 30420 | FTMS; HCD | 84.16 | 653.31 |

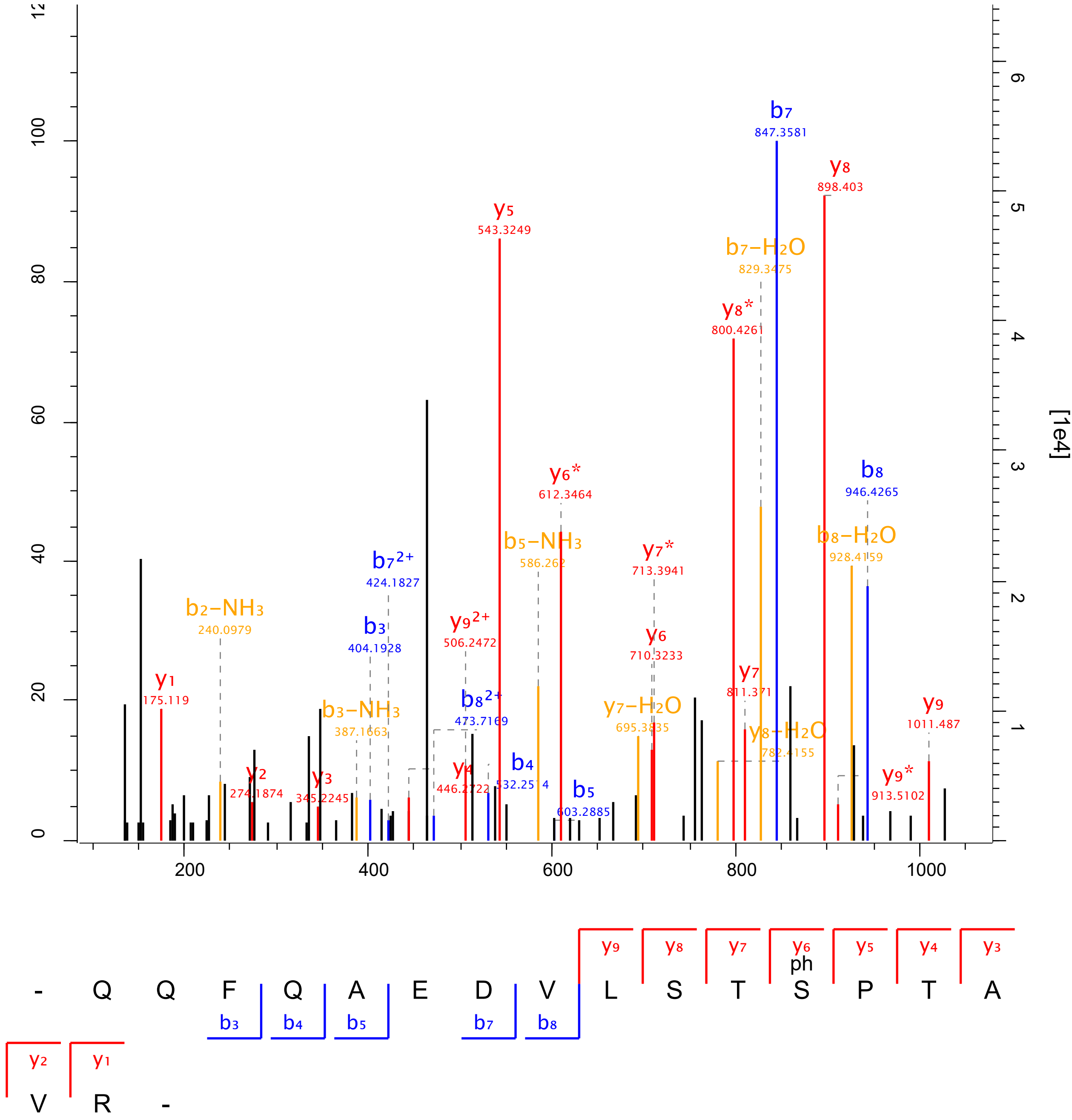

| Raw file | Scan | Method | Score | m/z |
|---|---|---|---|---|
| sys_15_1 | 30426 | FTMS; HCD | 87.35 | 948.9 |

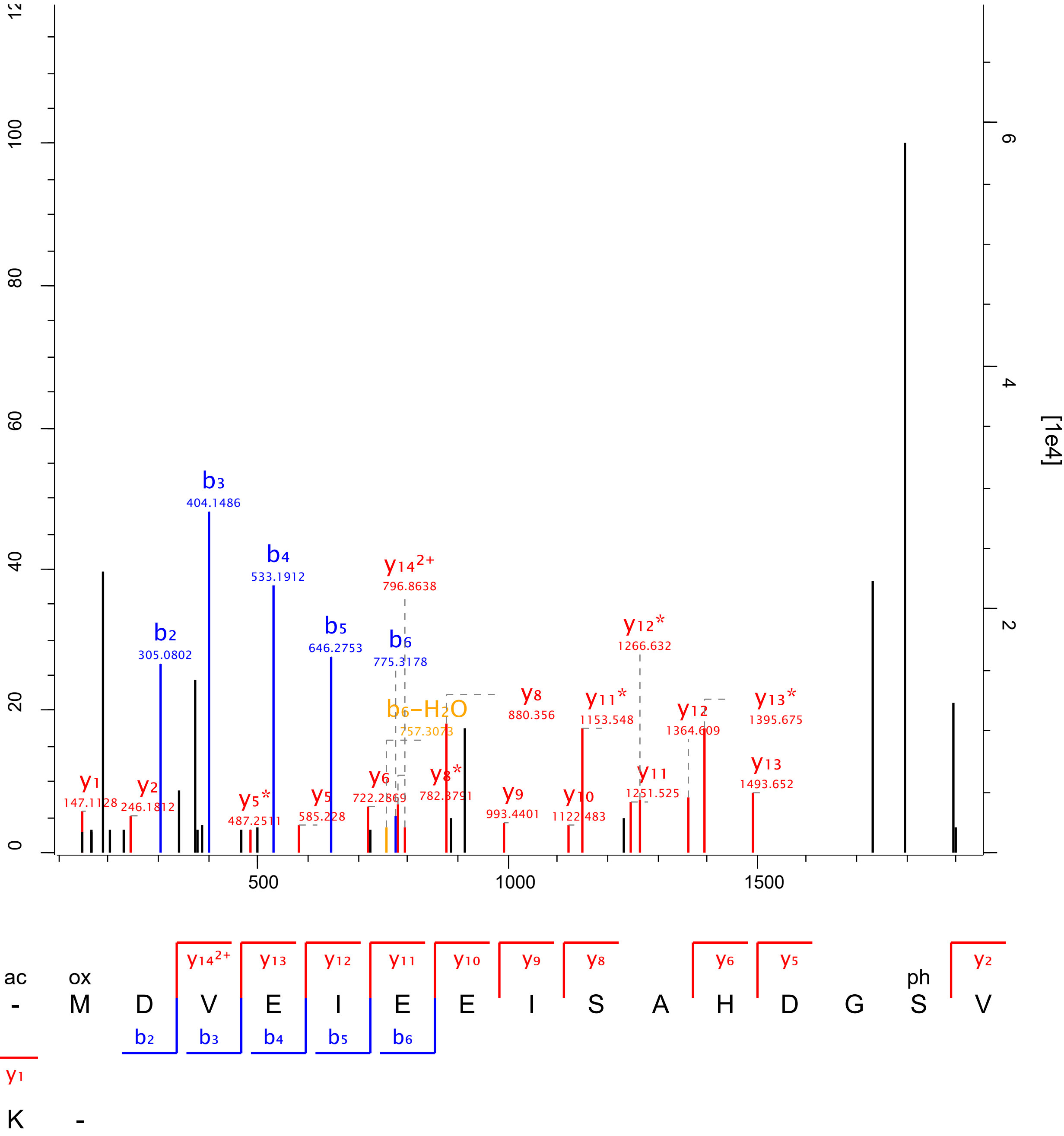

| Raw file | Scan | Method | Score | m/z |
|---|---|---|---|---|
| sys_15_1 | 30464 | FTMS; HCD | 55.71 | 926.41 |

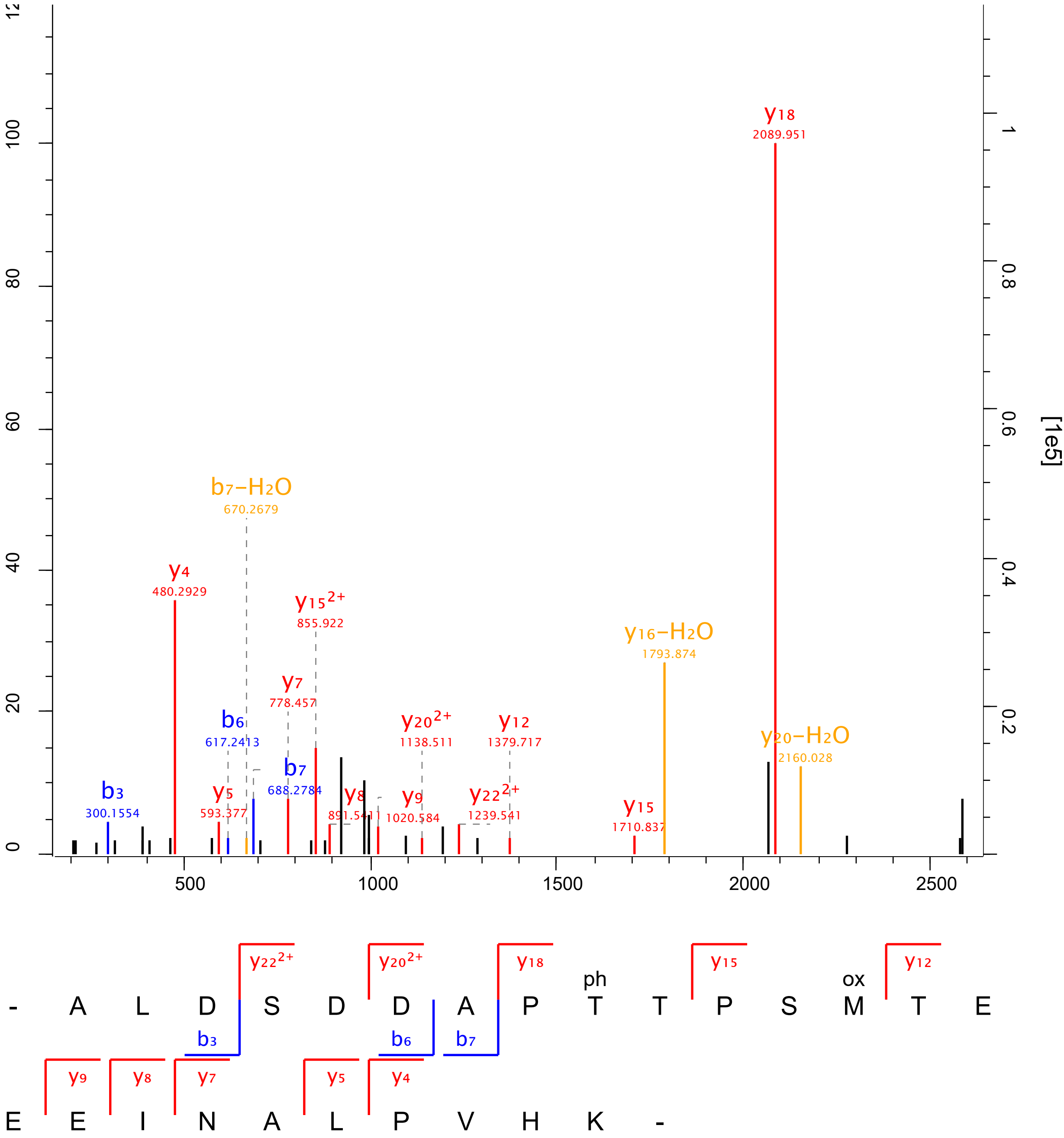

| Raw file | Scan | Method | Score | m/z |
|---|---|---|---|---|
| sys_15_1 | 30524 | FTMS; HCD | 91.62 | 755.87 |

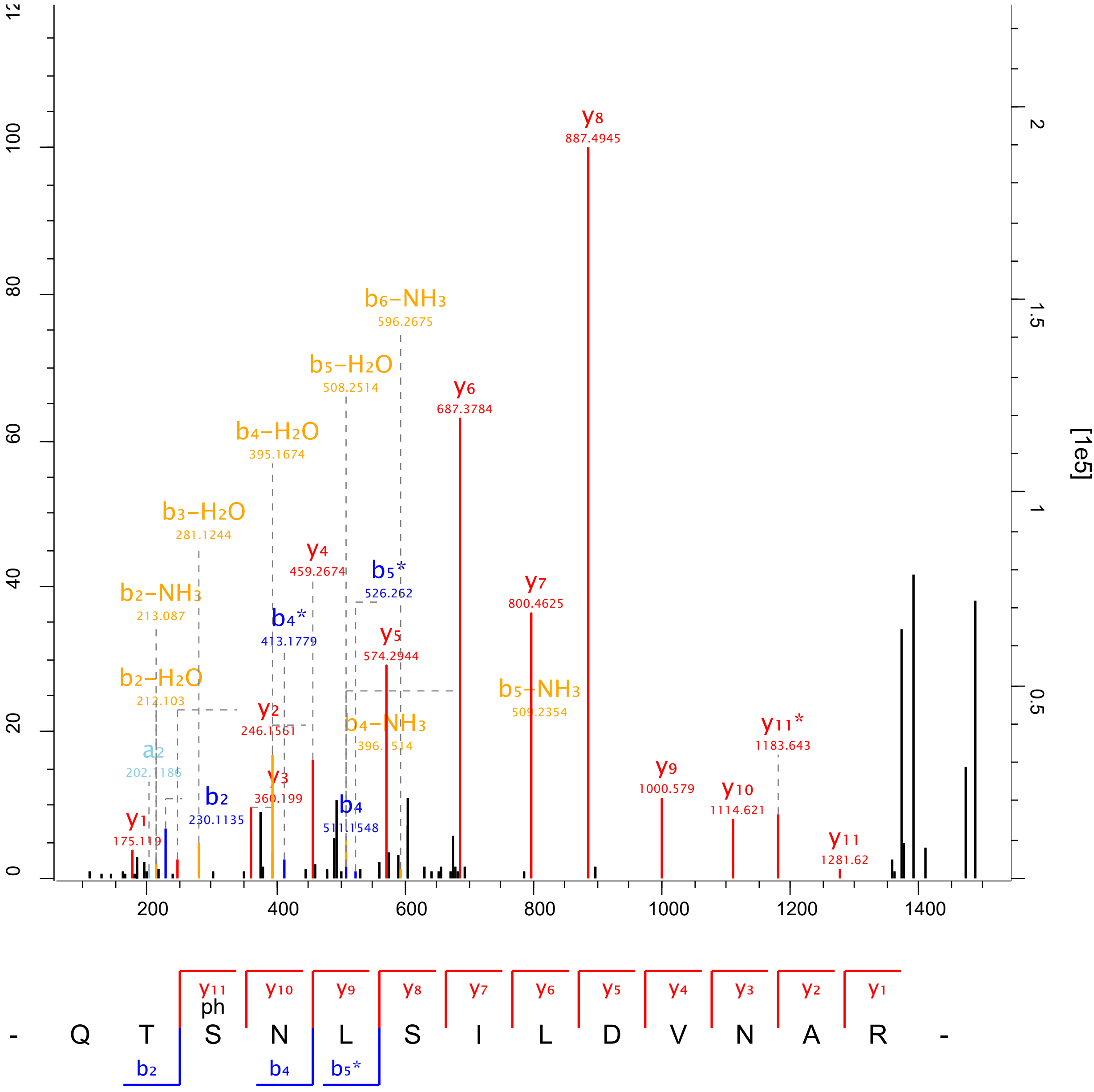

| Raw file | Scan | Method | Score | m/z |
|---|---|---|---|---|
| sys_15_1 | 30549 | FTMS; HCD | 114.46 | 901.91 |

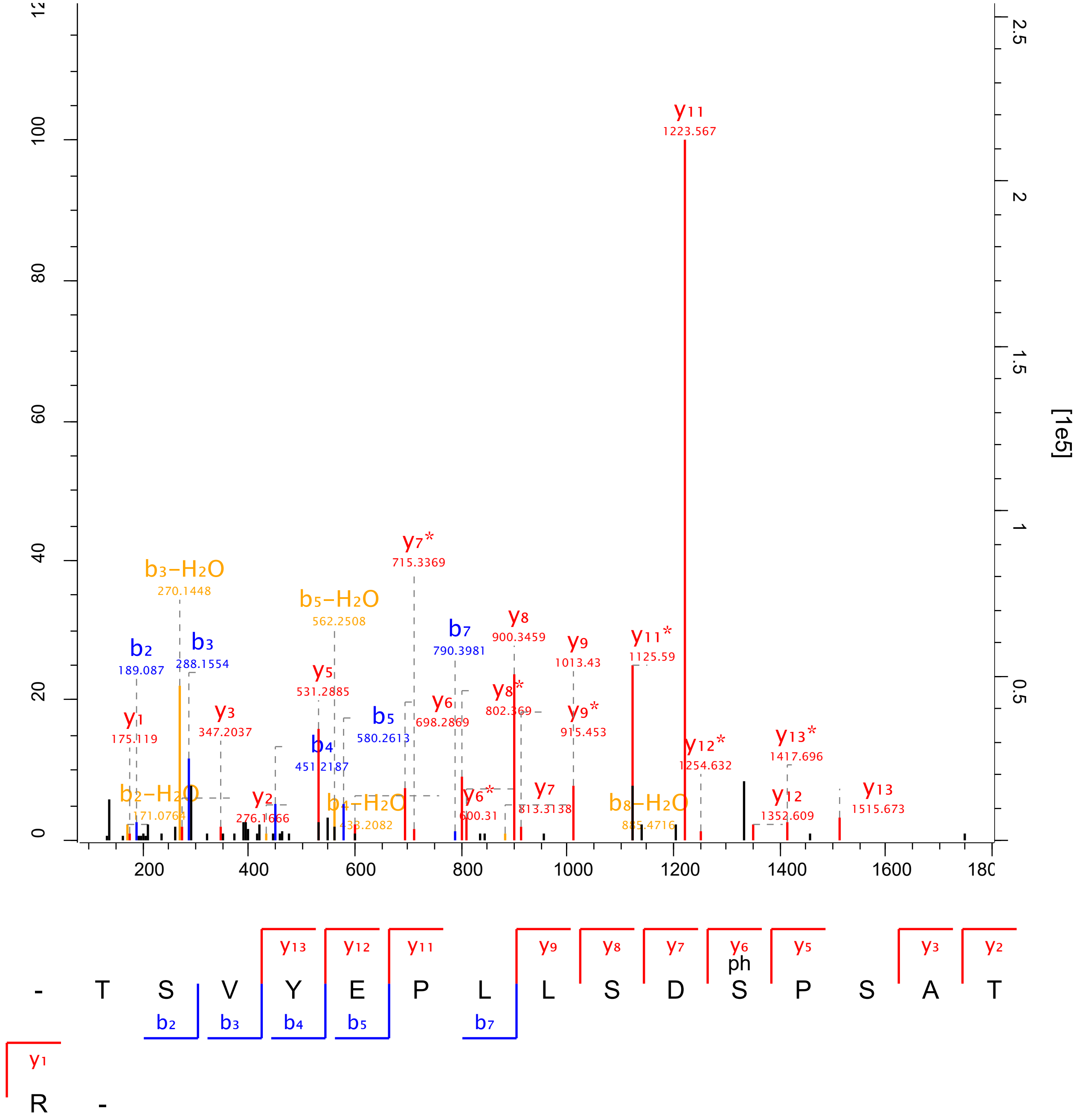

| Raw file | Scan | Method | Score | m/z |
| --- | --- | --- | --- | --- |
| sys_15_1 | 30603 | FTMS; HCD | 46.88 | 607.78 |

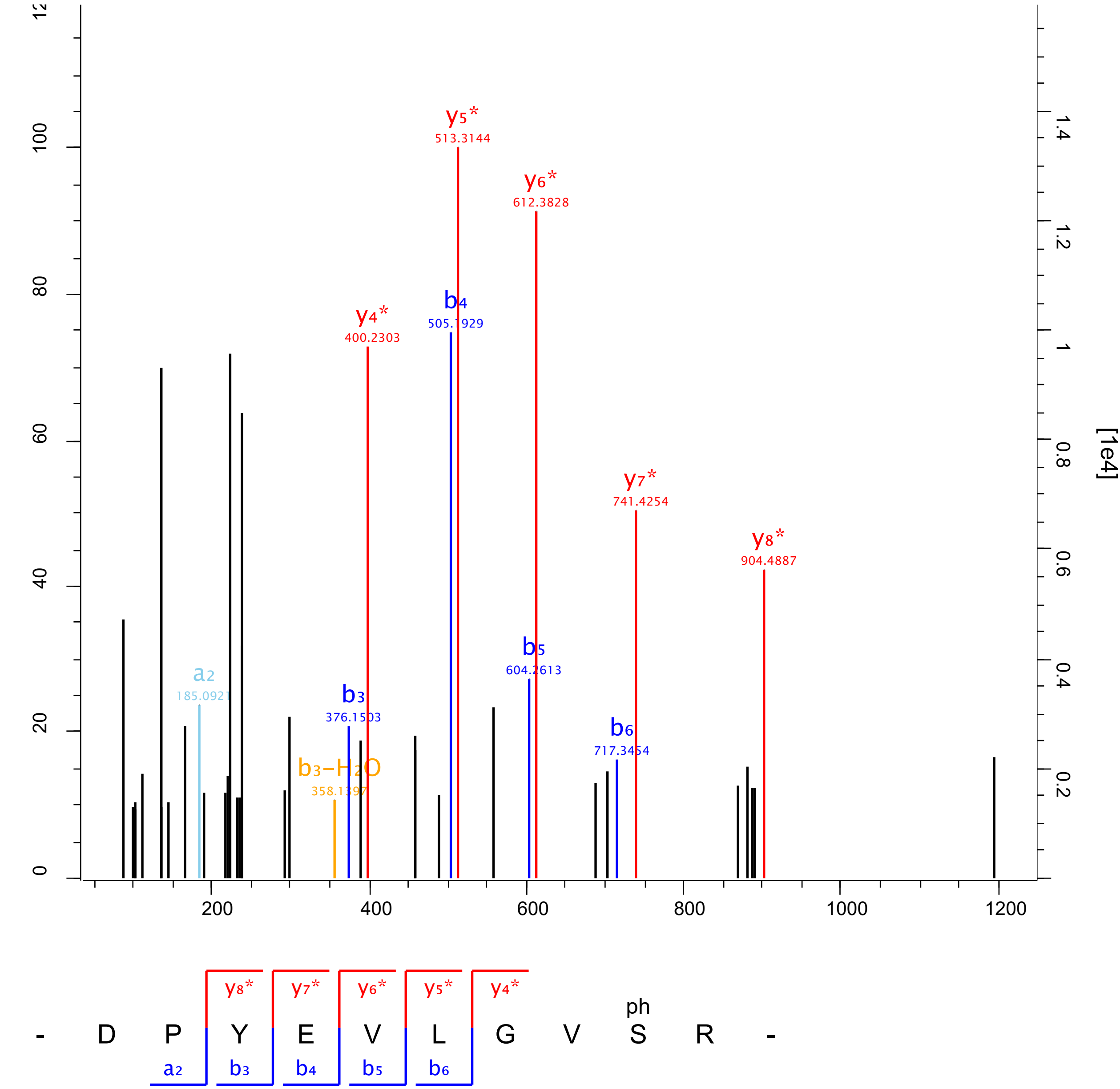

| Raw file | Scan | Method | Score | m/z |
|---|---|---|---|---|
| sys_15_1 | 30651 | FTMS; HCD | 208.28 | 642.29 |

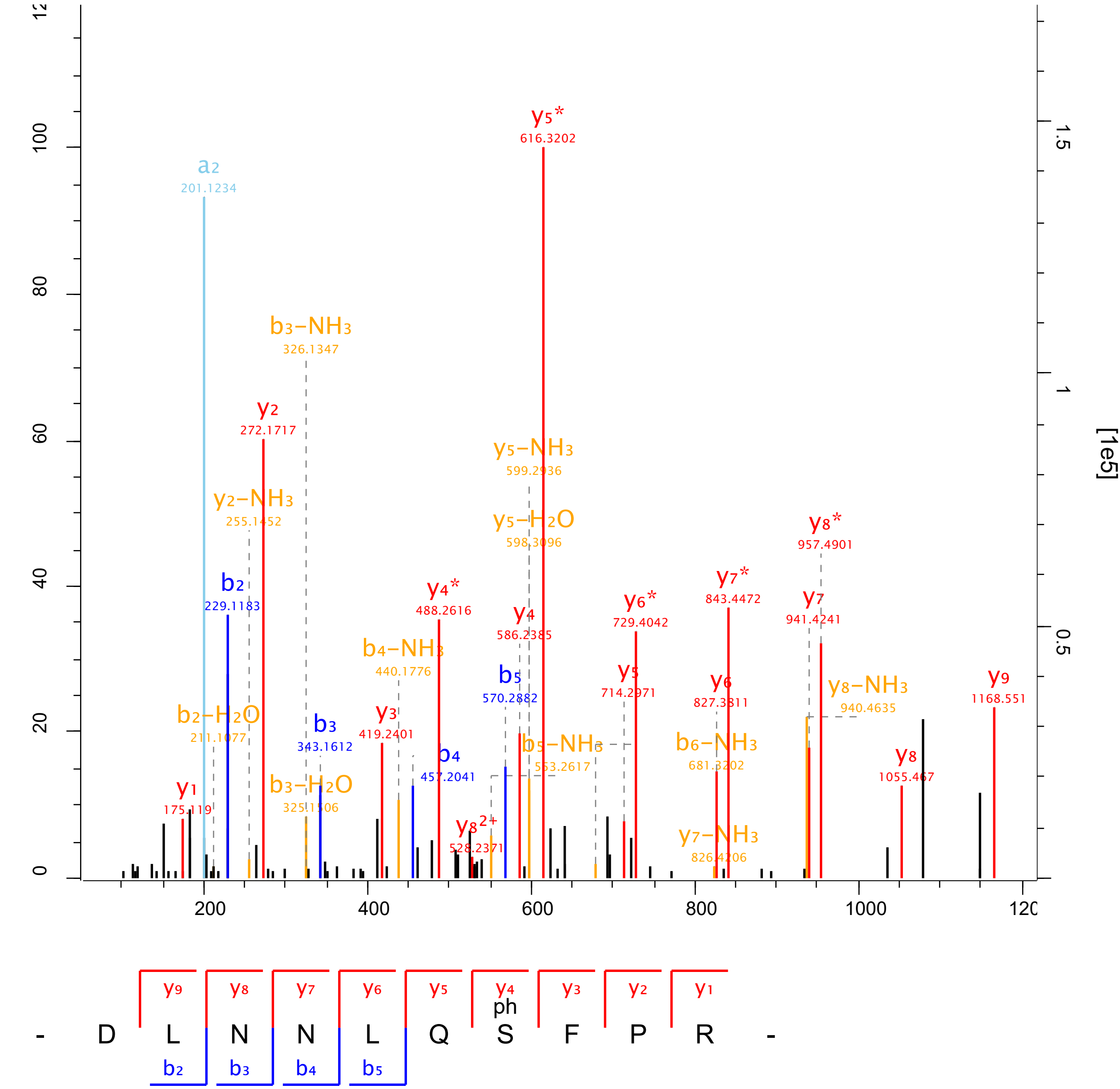

| Raw file | Scan | Method | Score | m/z |
|---|---|---|---|---|
| sys_15_1 | 30662 | FTMS; HCD | 181.4 | 924.35 |

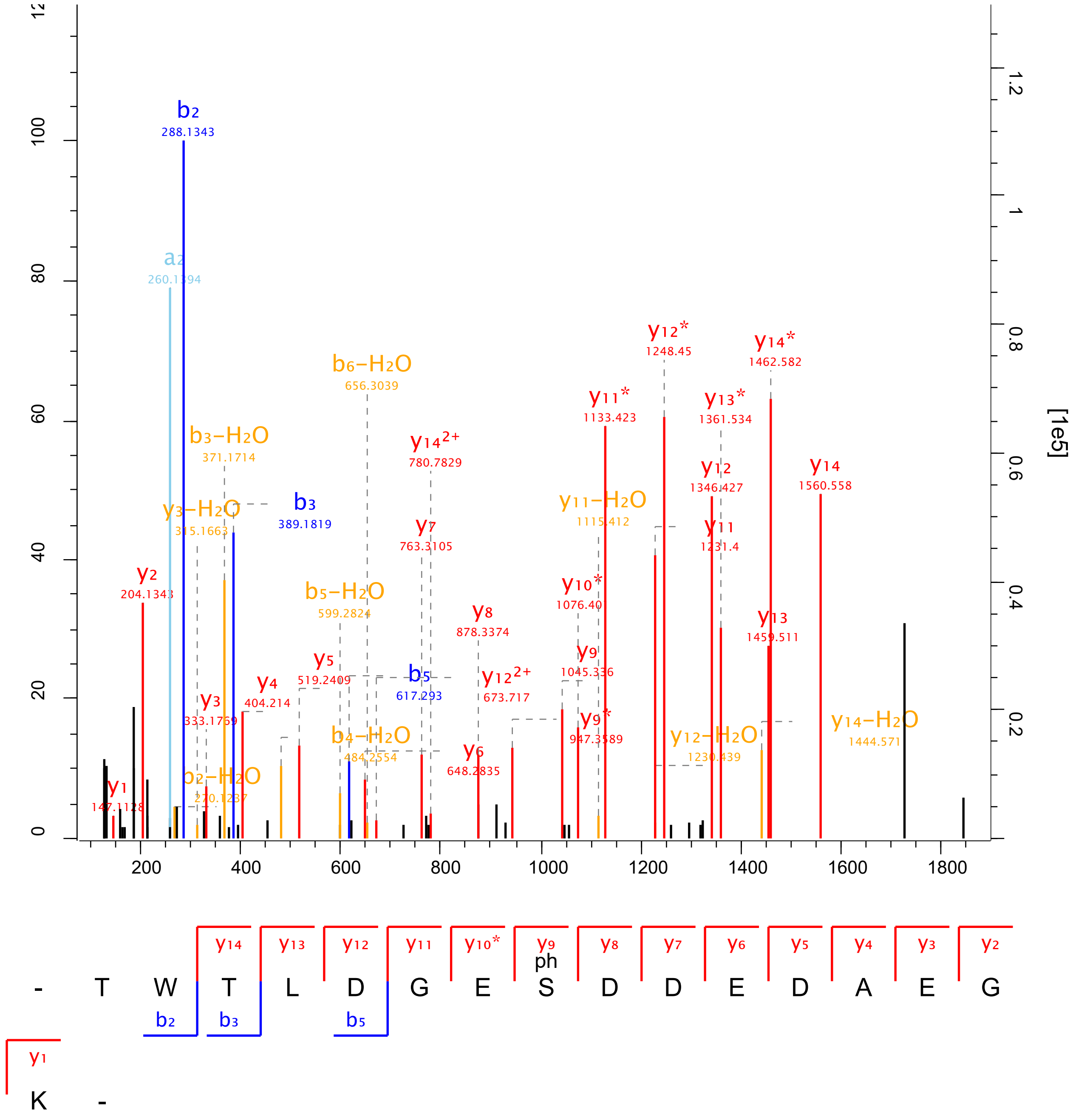

| Raw file | Scan | Method | Score | m/z |
|---|---|---|---|---|
| sys_15_1 | 30668 | FTMS; HCD | 91.96 | 545.76 |

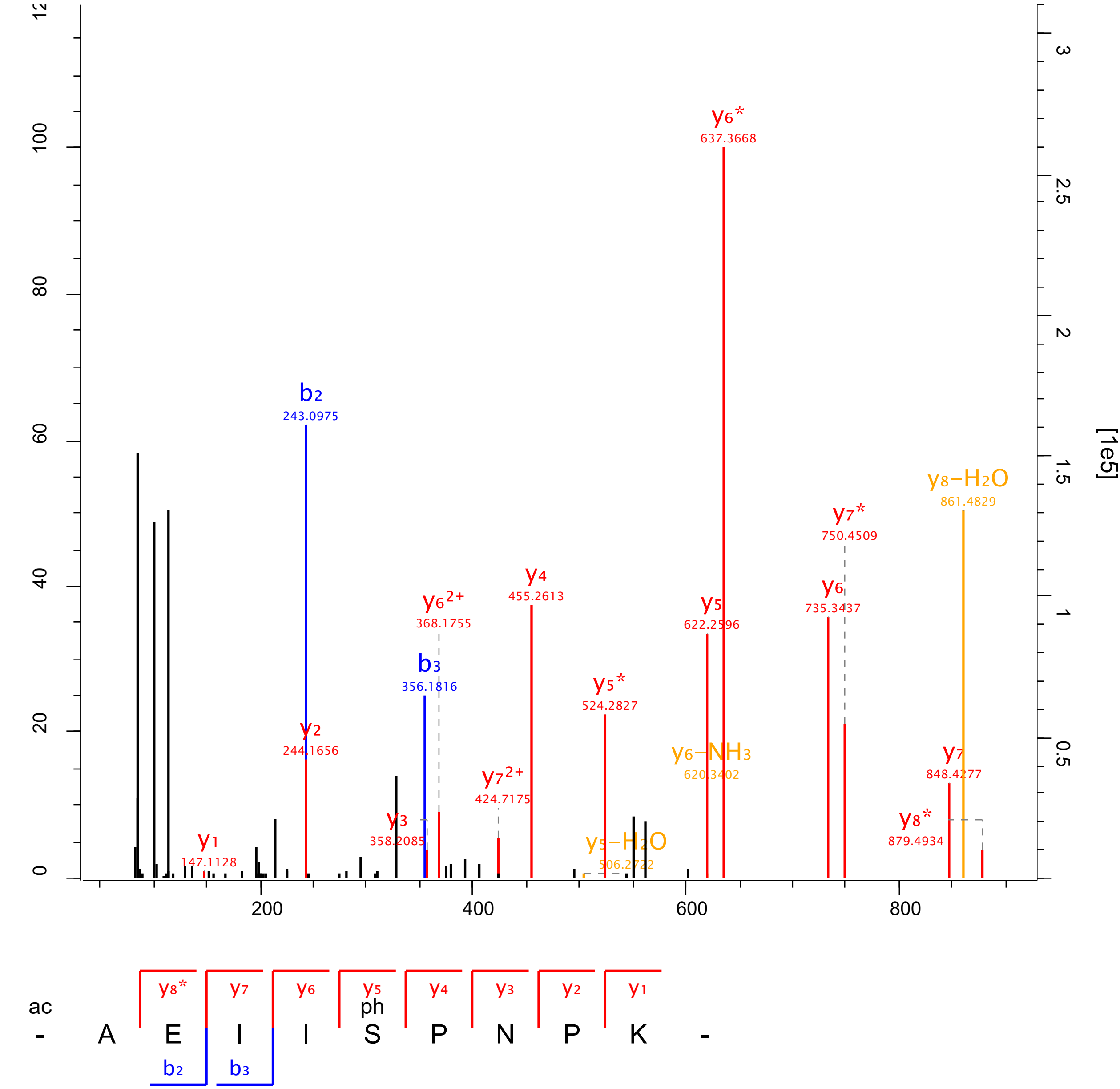

| Raw file | Scan | Method | Score | m/z |
| --- | --- | --- | --- | --- |
| sys_15_1 | 30685 | FTMS; HCD | 64.22 | 583.81 |

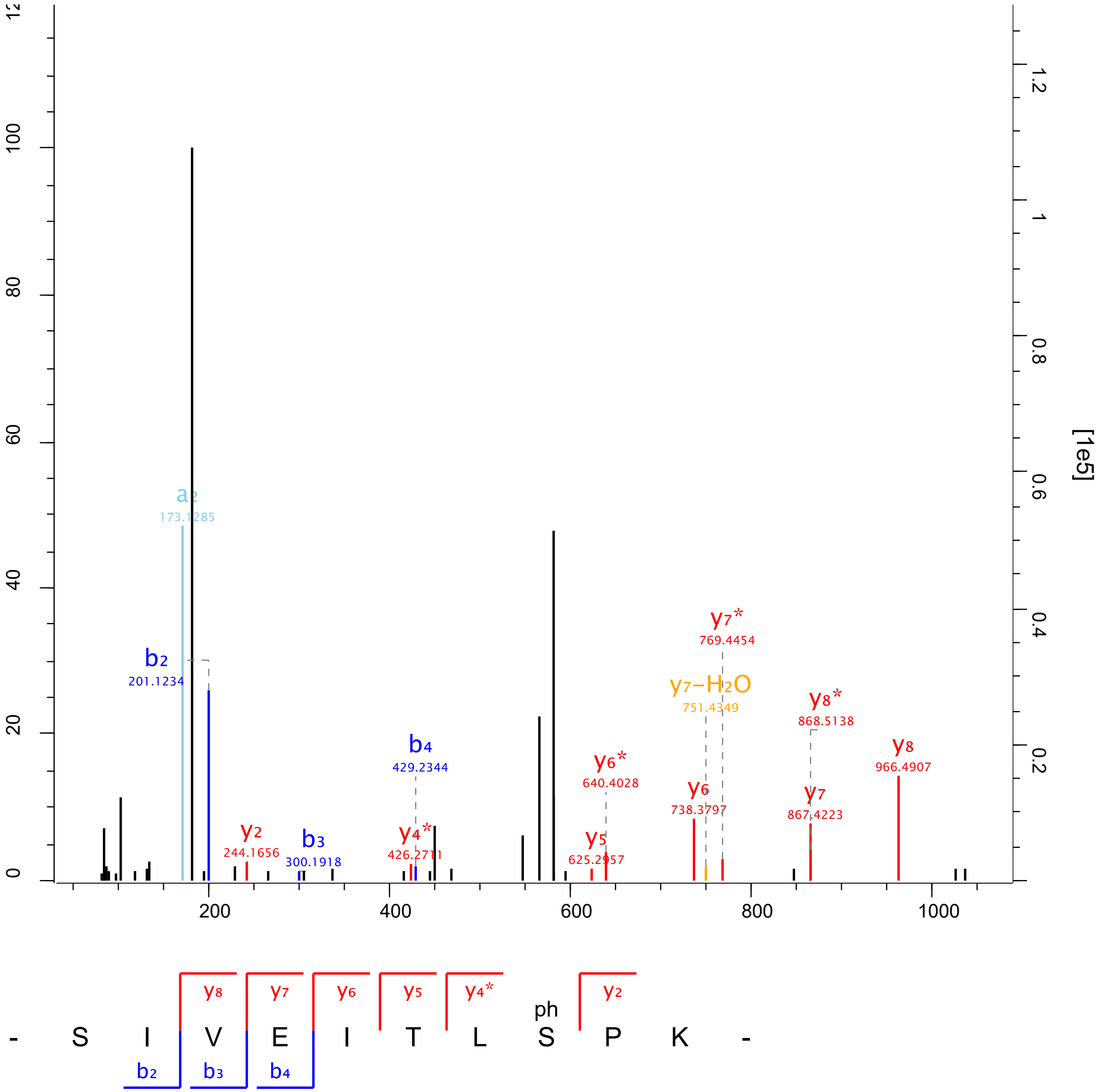

| Raw file | Scan | Method | Score | m/z |
| --- | --- | --- | --- | --- |
| sys_15_1 | 30778 | FTMS; HCD | 136.64 | 851.37 |

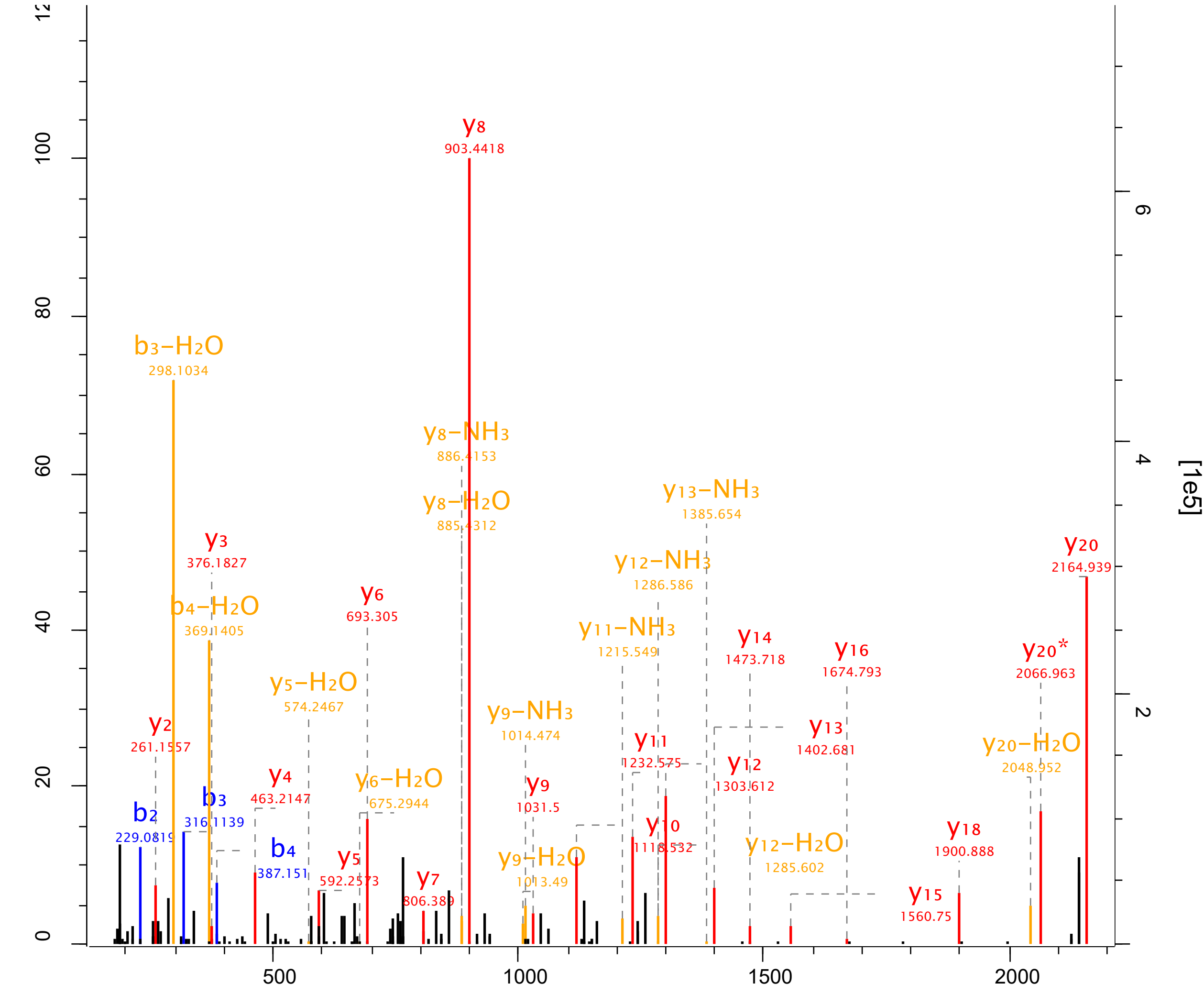


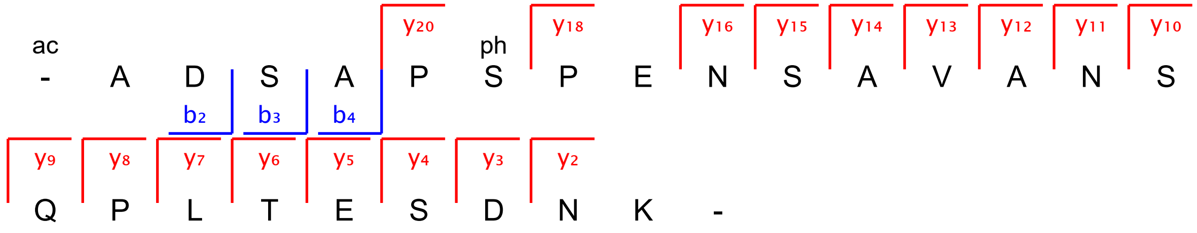

| Raw file | Scan | Method | Score | m/z |
|---|---|---|---|---|
| sys_15_1 | 30825 | FTMS; HCD | 154.69 | 773.33 |

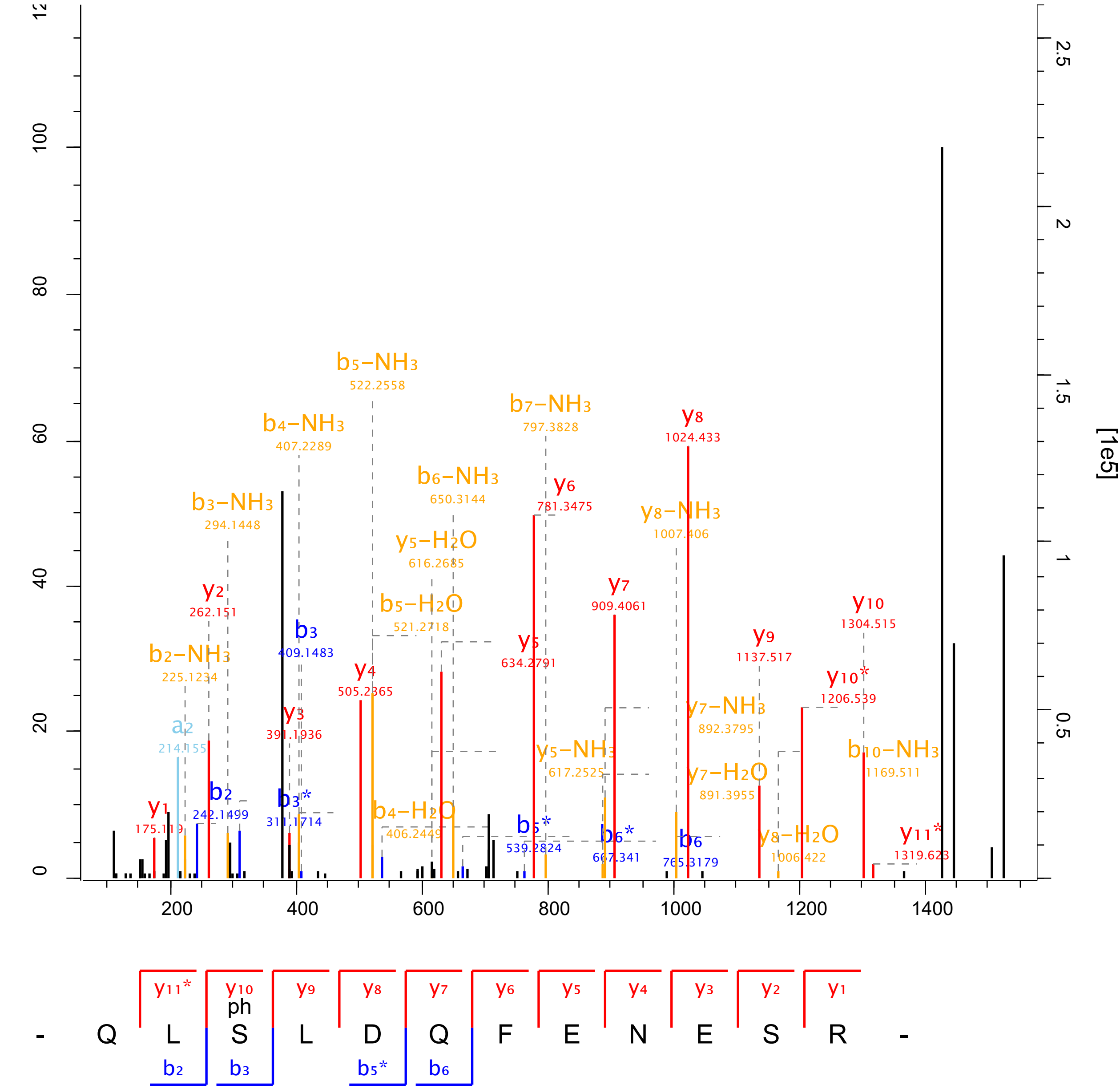

| Raw file | Scan | Method | Score | m/z |
|---|---|---|---|---|
| sys_15_1 | 30853 | FTMS; HCD | 140.73 | 864.35 |

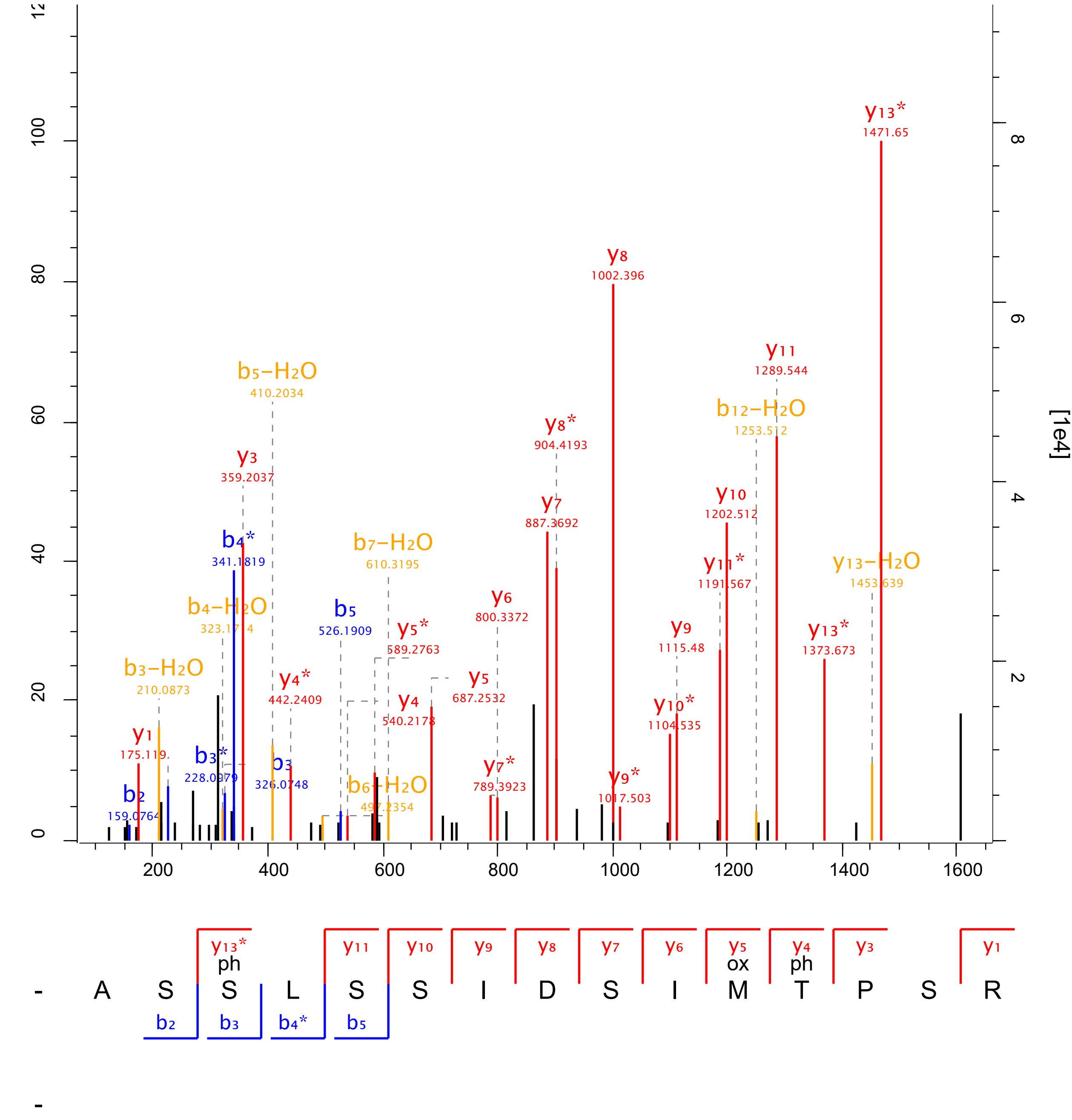

| Raw file | Scan | Method | Score | m/z |
| --- | --- | --- | --- | --- |
| sys_15_1 | 30880 | FTMS; HCD | 275.69 | 977.38 |

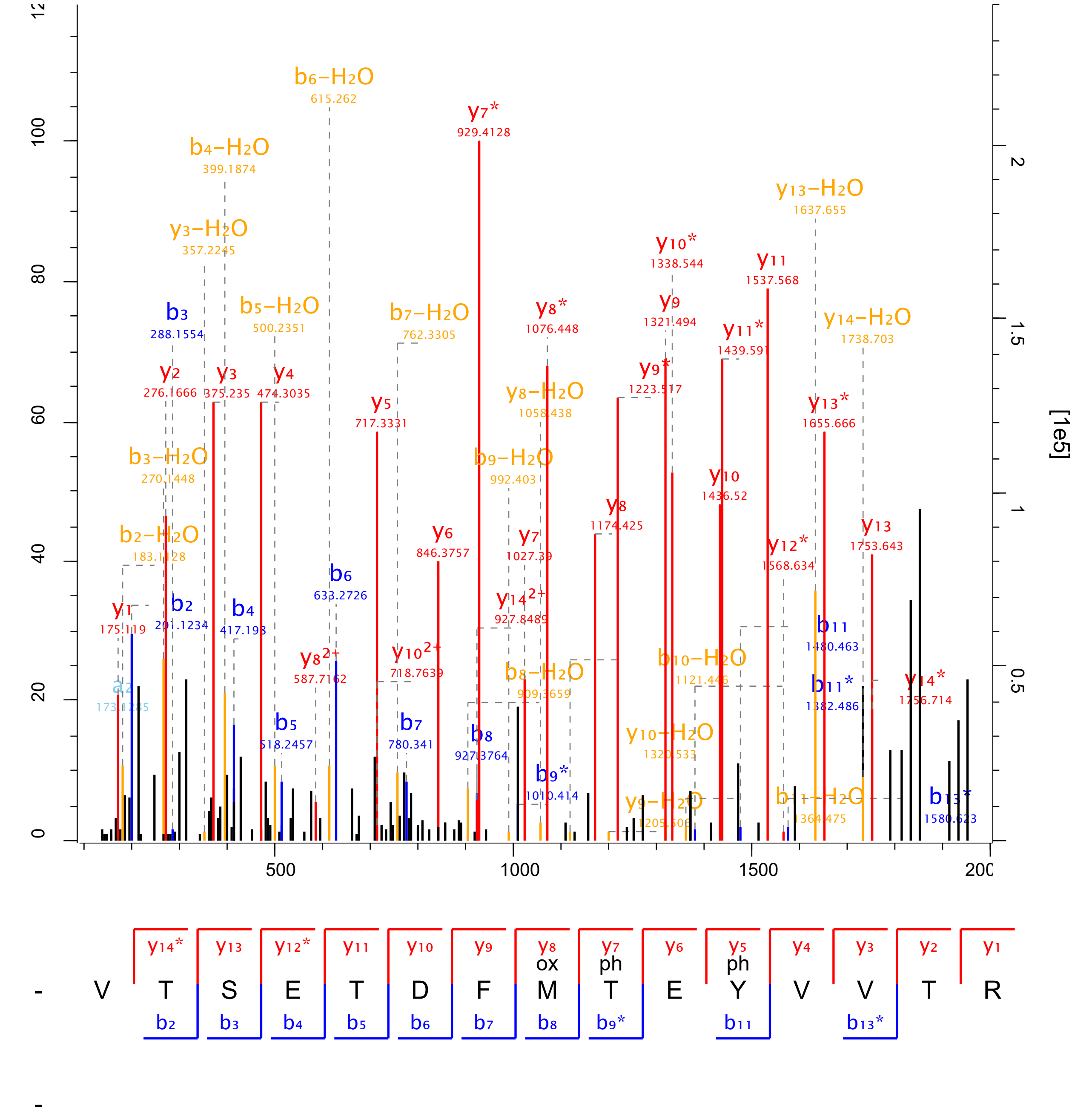

| Raw file | Scan | Method | Score | m/z |
|---|---|---|---|---|
| sys_15_1 | 3111 | FTMS; HCD | 66.06 | 451.69 |

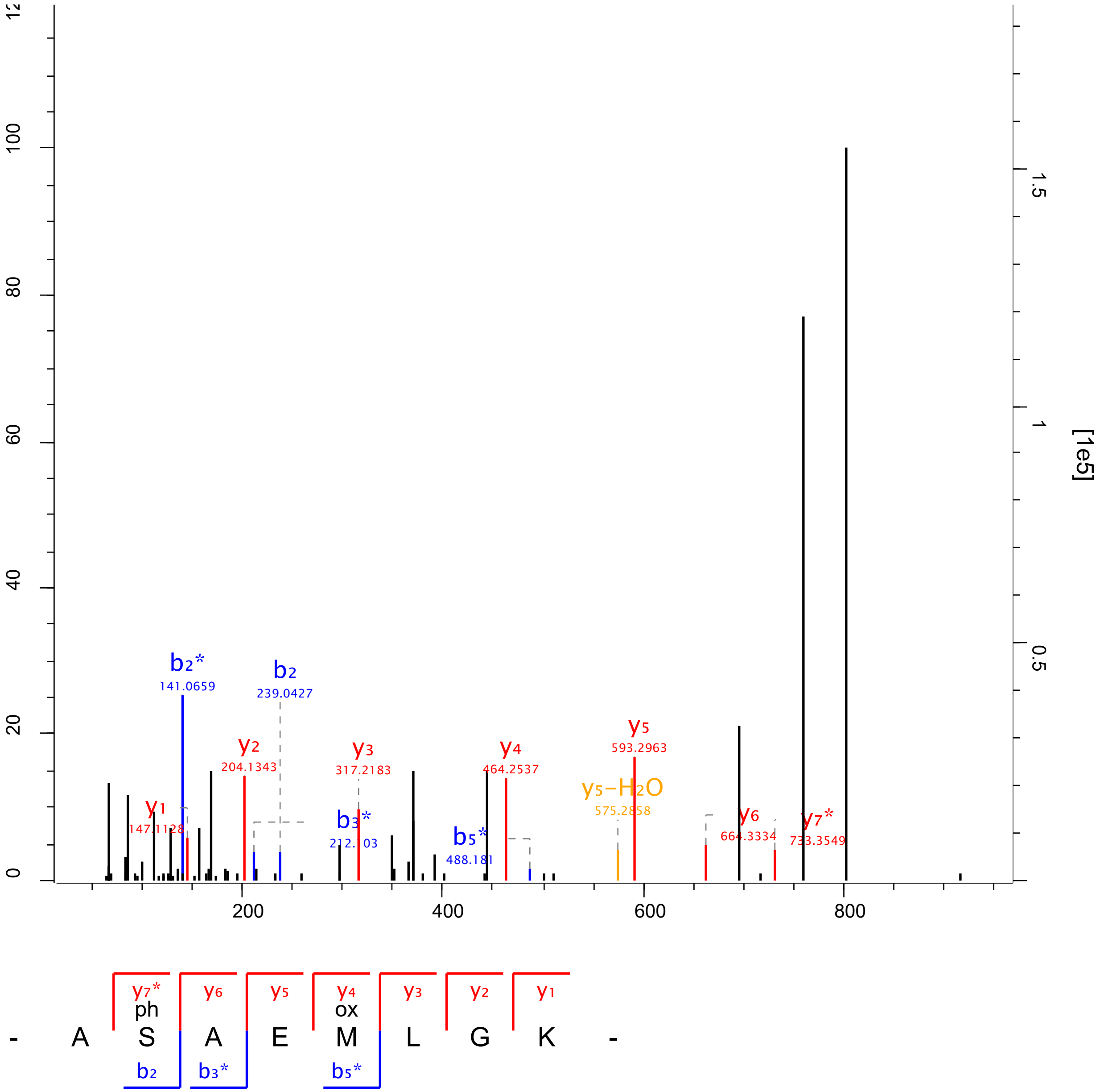

| Raw file | Scan | Method | Score | m/z |
| --- | --- | --- | --- | --- |
| sys_15_1 | 31129 | FTMS; HCD | 152.4 | 674.27 |

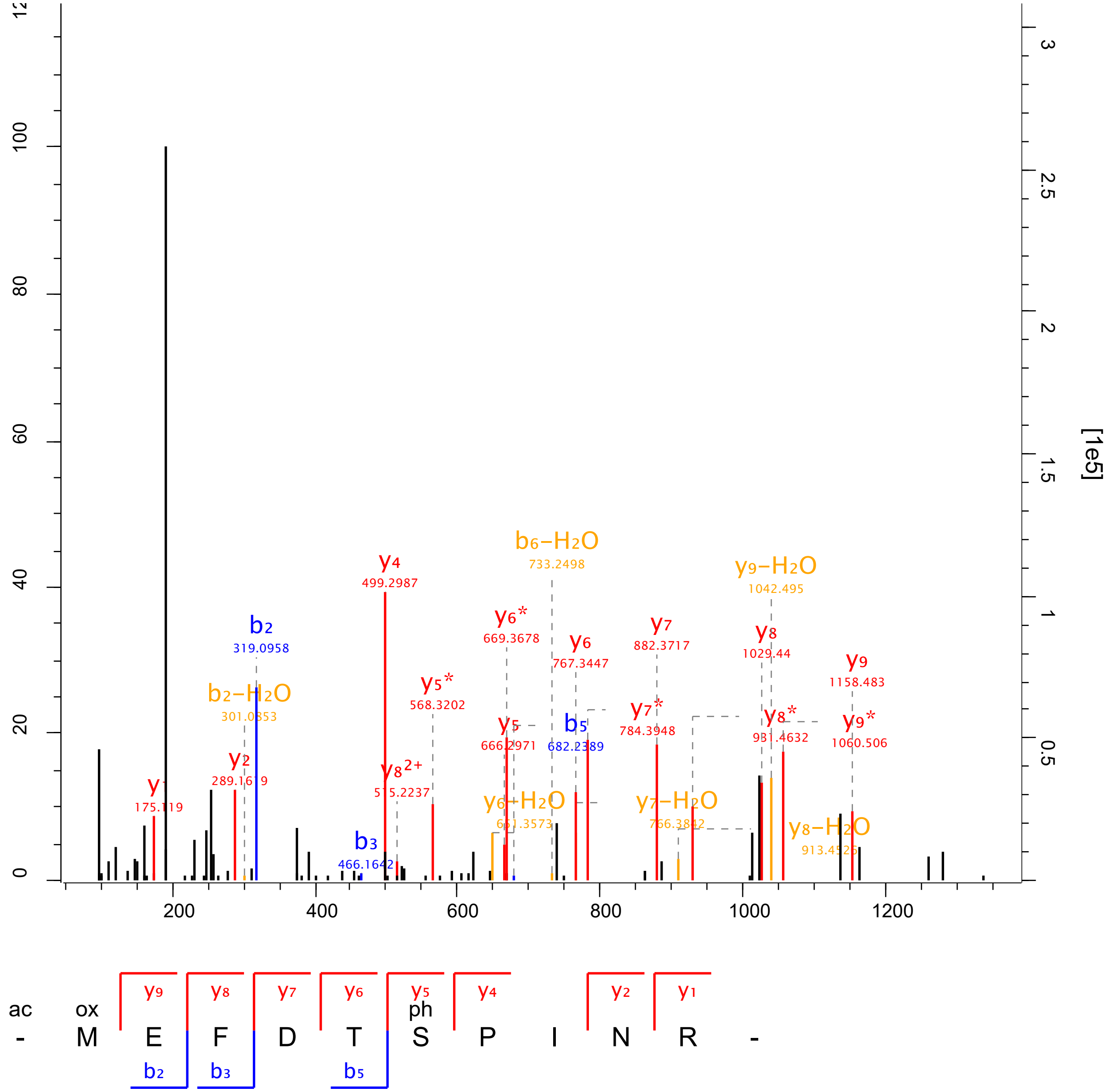

| Raw file | Scan | Method | Score | m/z |
| --- | --- | --- | --- | --- |
| sys_15_1 | 31200 | FTMS; HCD | 139.9 | 820.9 |

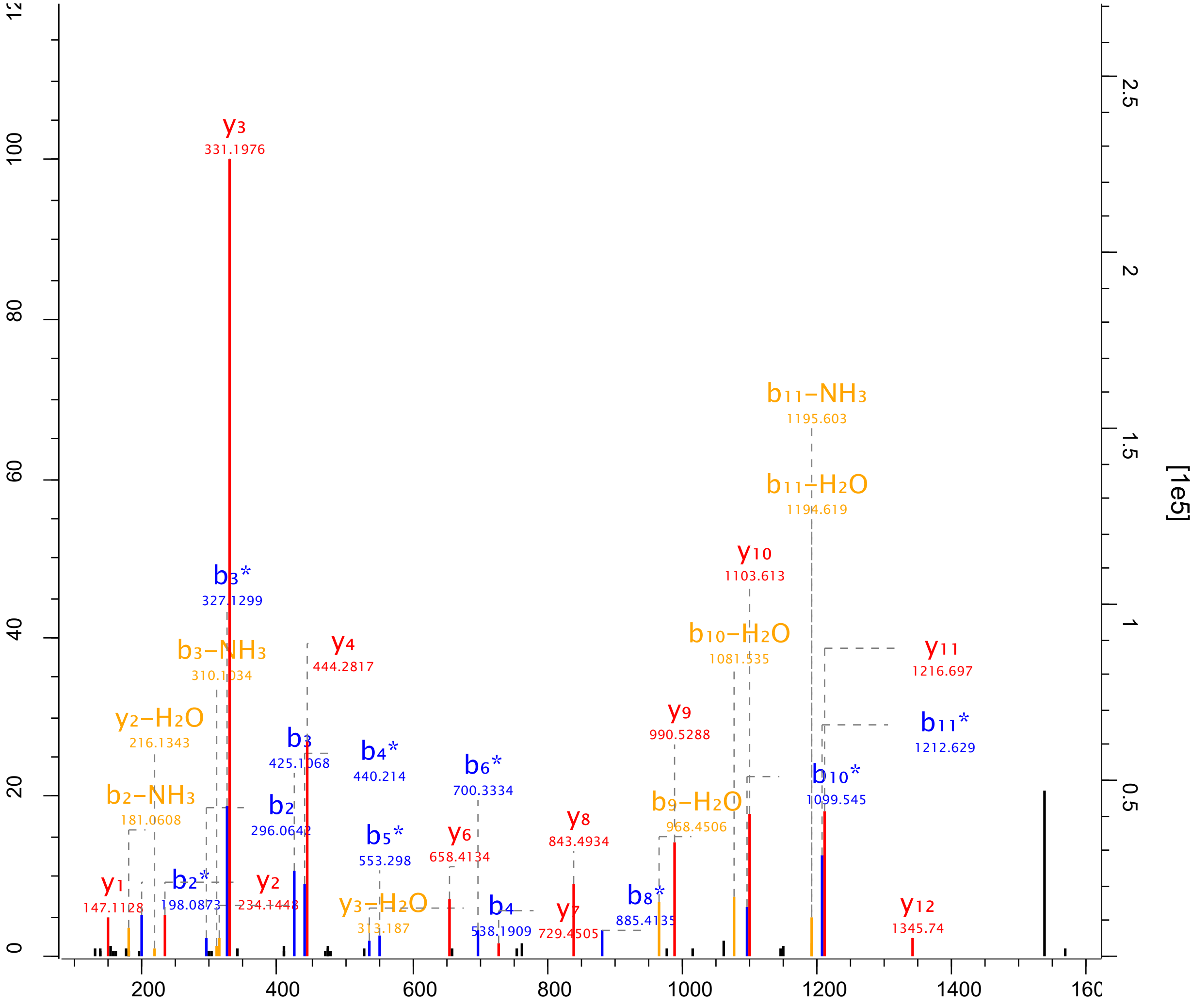


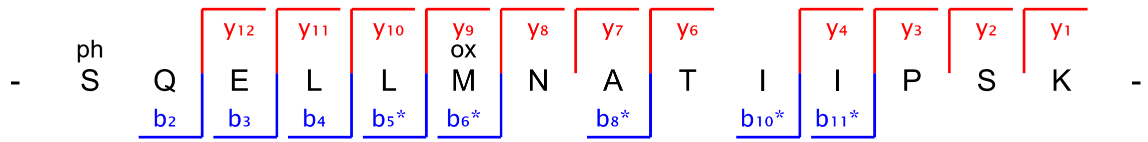

| Raw file | Scan | Method | Score | m/z |
|---|---|---|---|---|
| sys_15_1 | 31232 | FTMS; HCD | 91.32 | 803.42 |

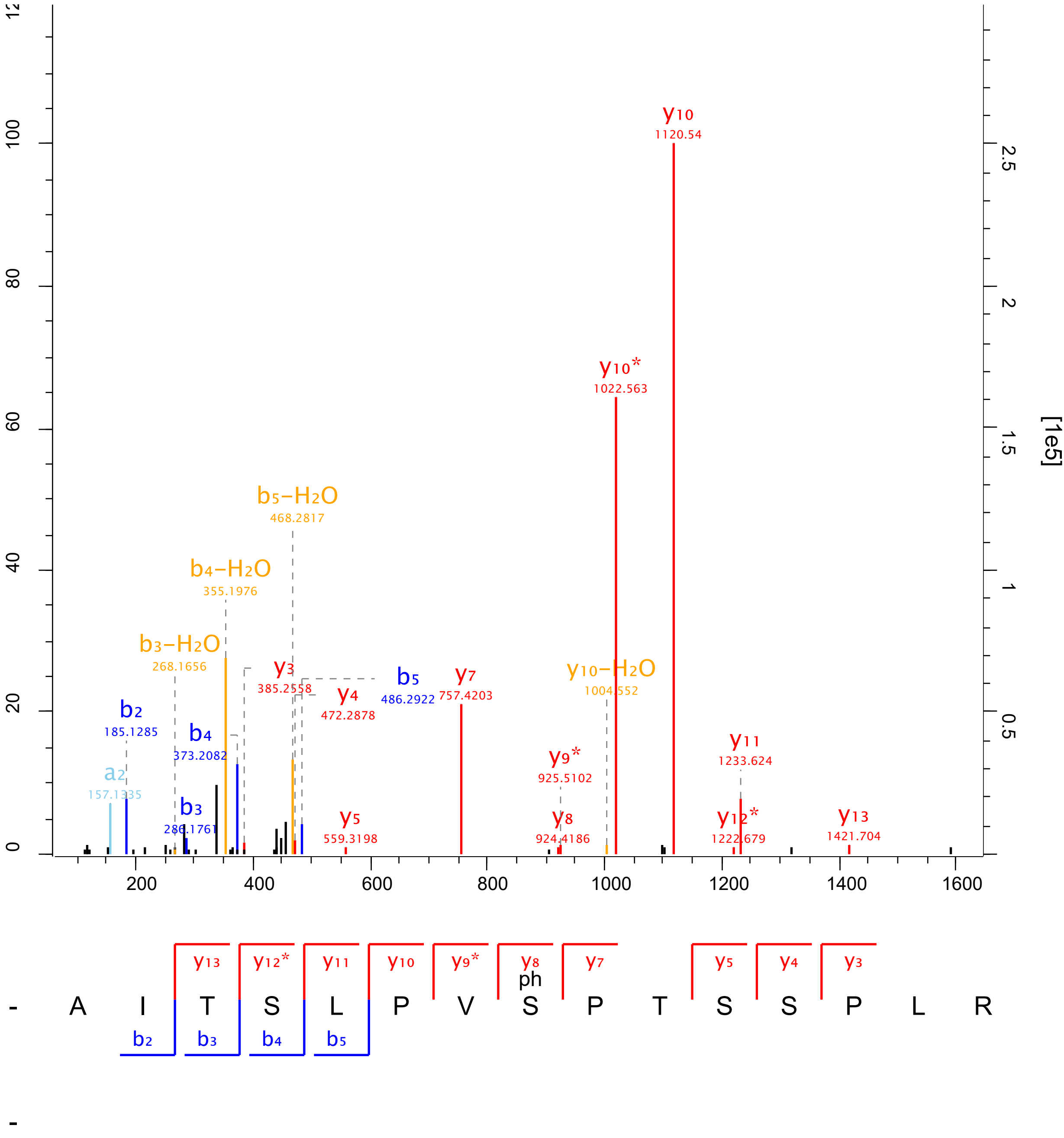

| Raw file | Scan | Method | Score | m/z |
| --- | --- | --- | --- | --- |
| sys_15_1 | 31373 | FTMS; HCD | 41.45 | 639.31 |

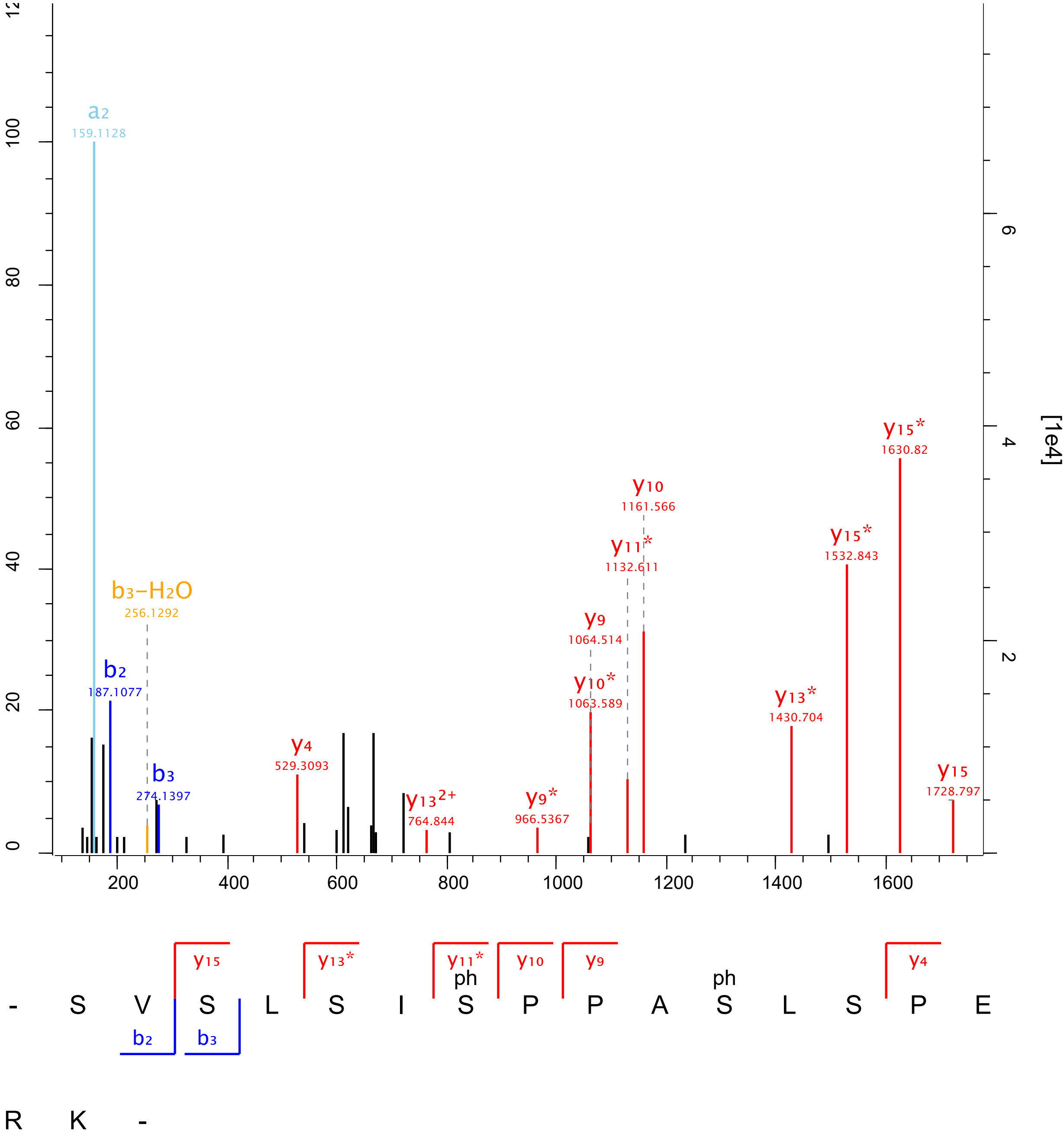

| Raw file | Scan | Method | Score | m/z |
|---|---|---|---|---|
| sys_15_1 | 31482 | FTMS; HCD | 227.95 | 920.88 |

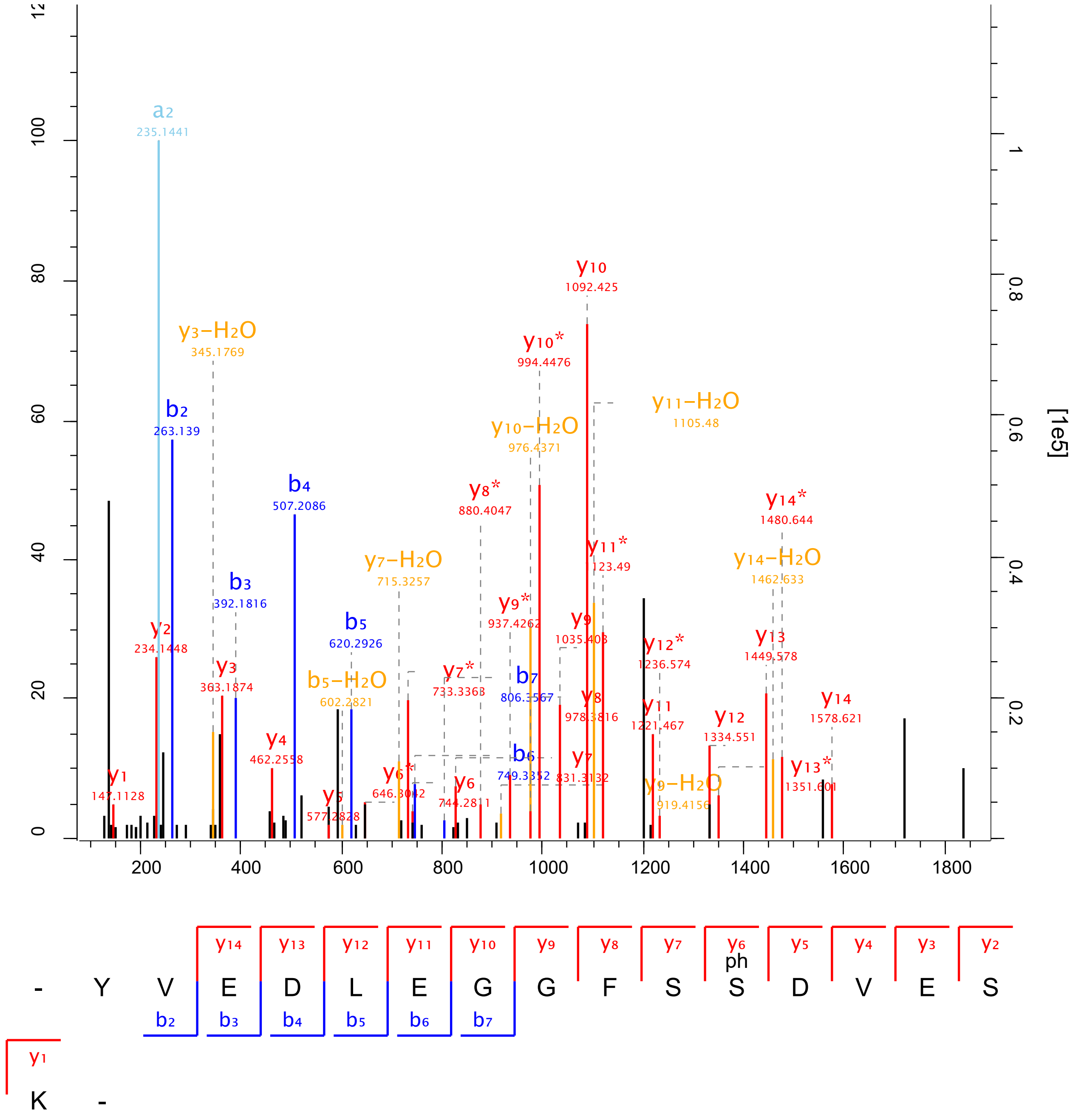

| Raw file | Scan | Method | Score | m/z |
|---|---|---|---|---|
| sys_15_1 | 31524 | FTMS; HCD | 58.95 | 751.01 |

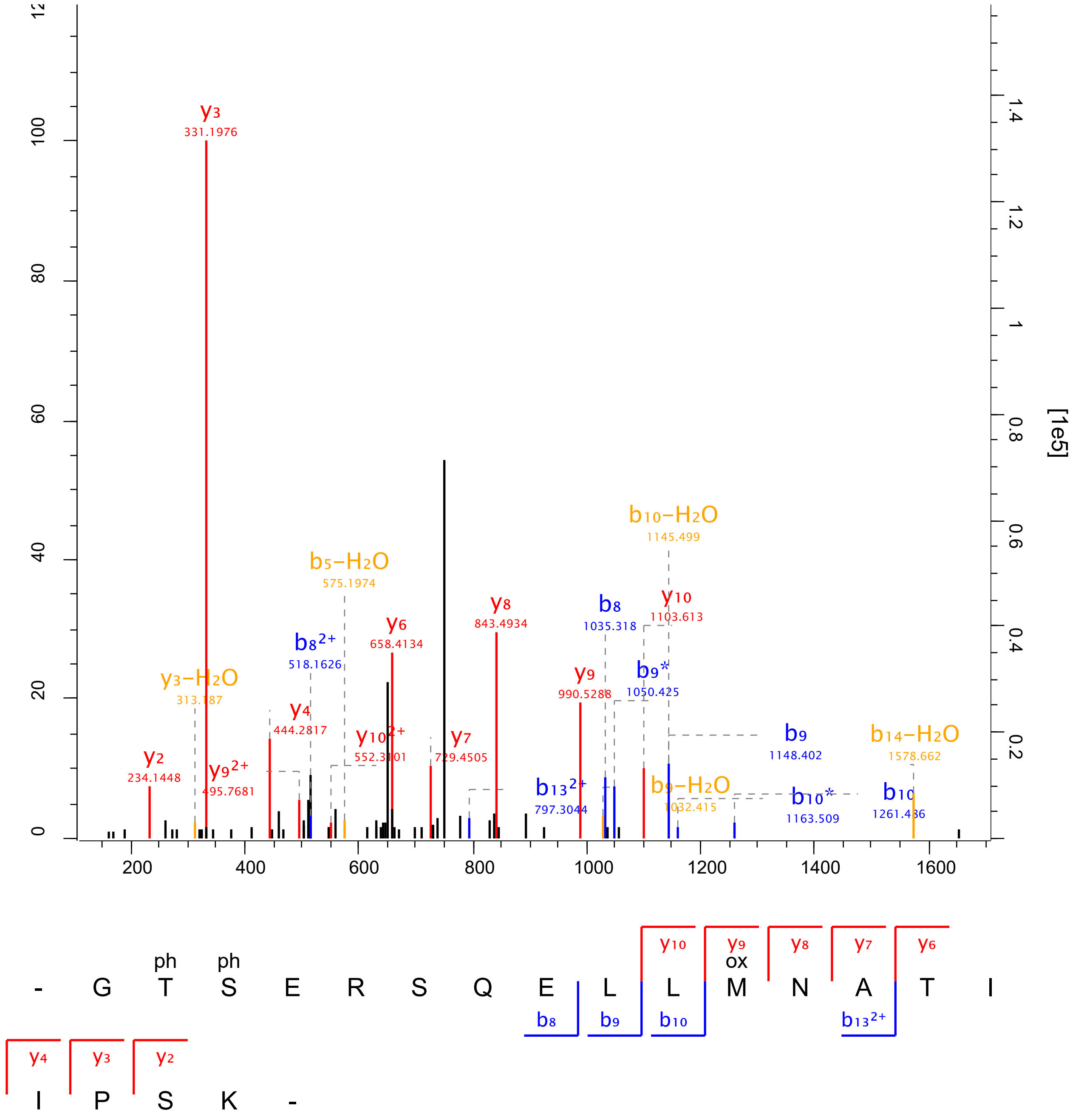

| Raw file | Scan | Method | Score | m/z |
|---|---|---|---|---|
| sys_15_1 | 31552 | FTMS; HCD | 41.65 | 668.97 |

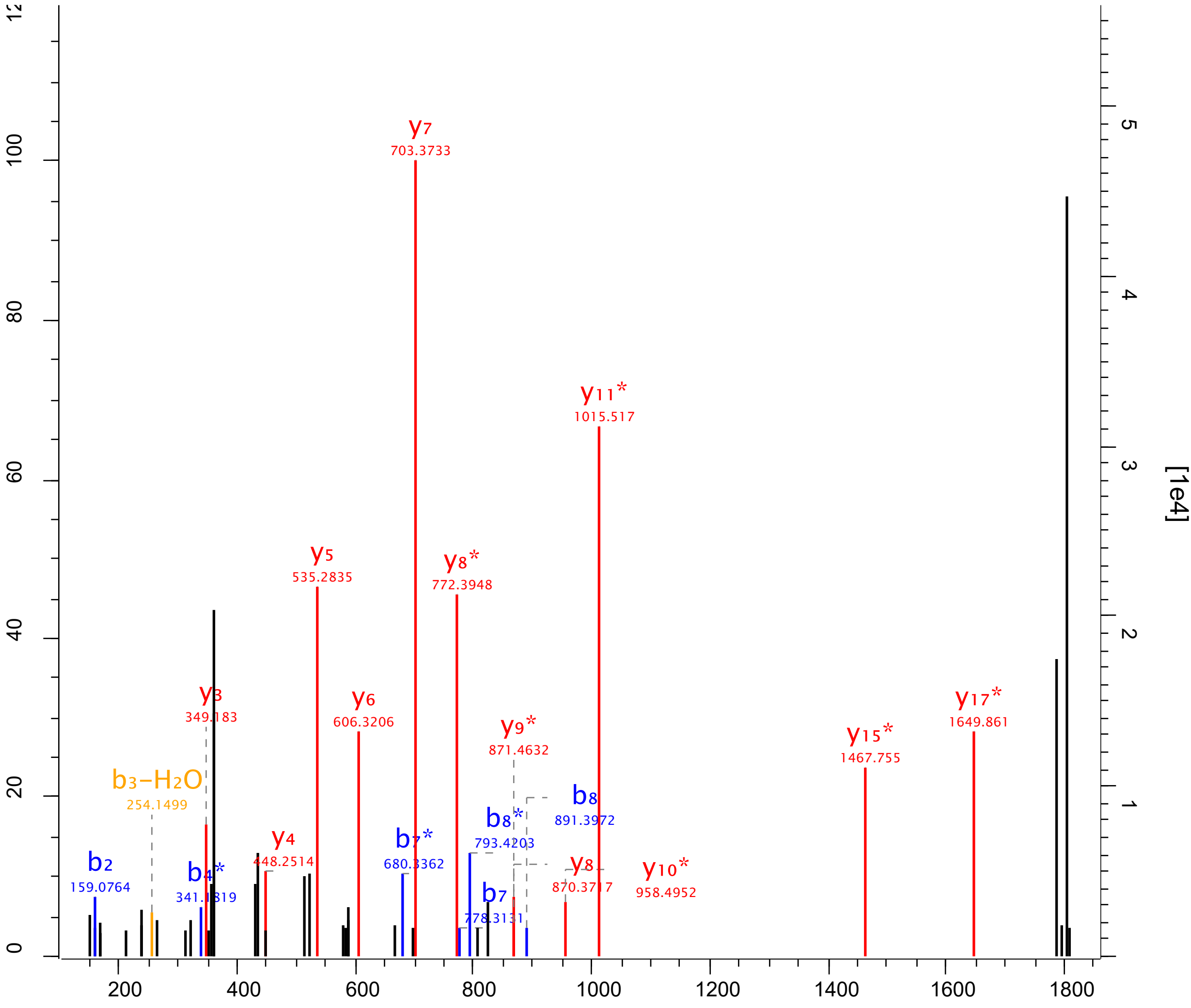


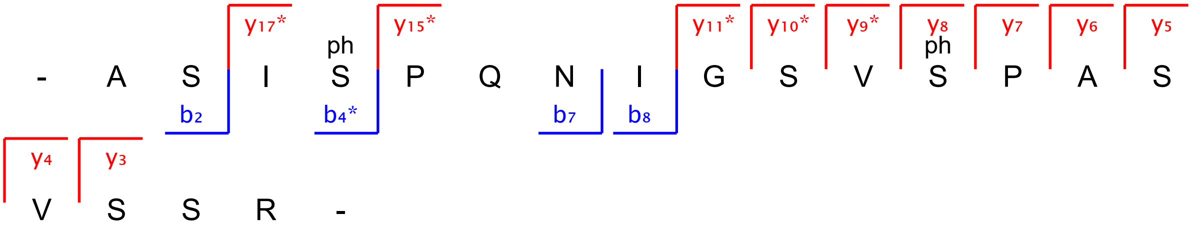

| Raw file | Scan | Method | Score | m/z |
|---|---|---|---|---|
| sys_15_1 | 31651 | FTMS; HCD | 100.97 | 904.39 |

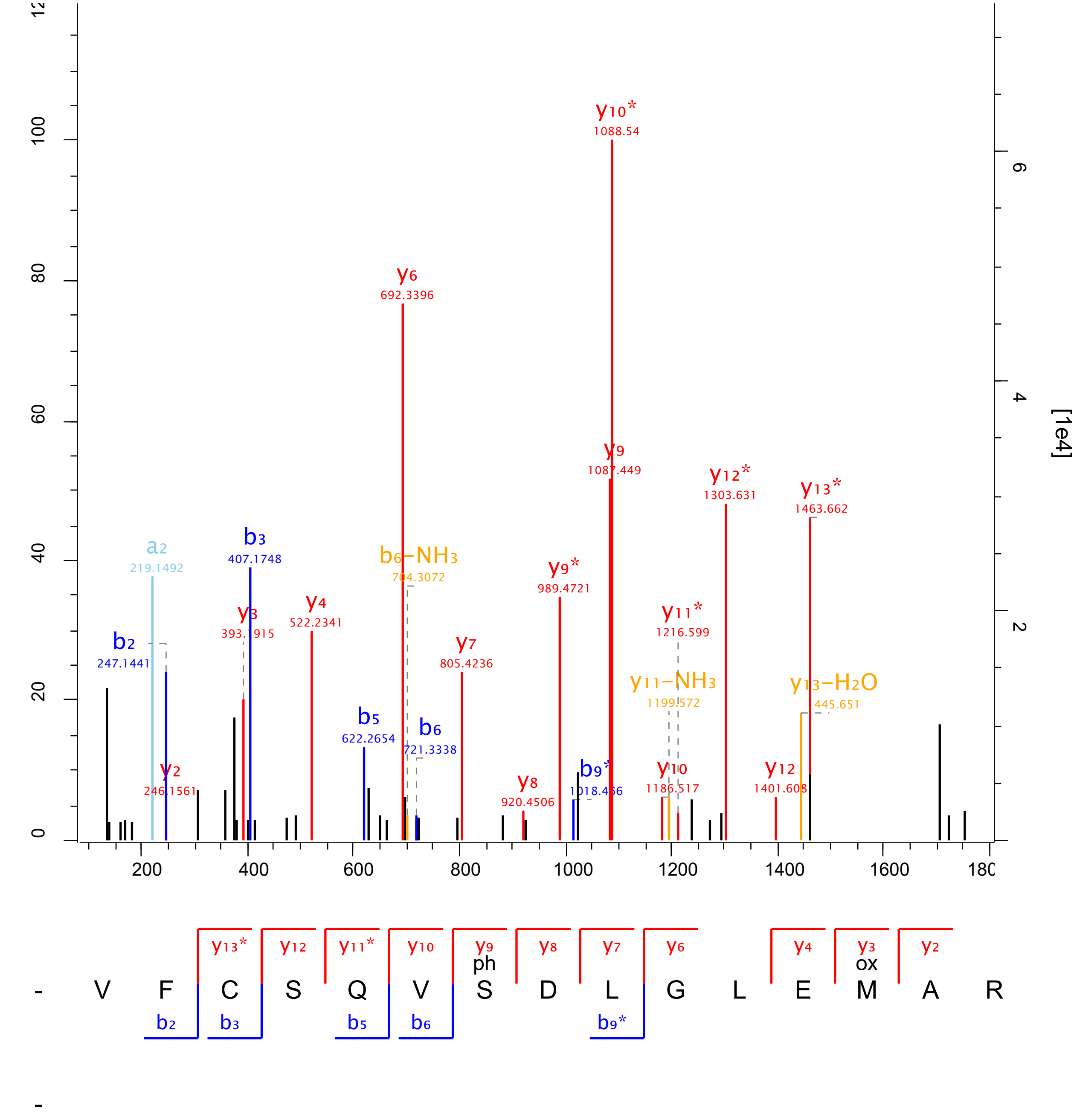

| Raw file | Scan | Method | Score | m/z |
|---|---|---|---|---|
| sys_15_1 | 3181 | FTMS; HCD | 68.97 | 453.54 |

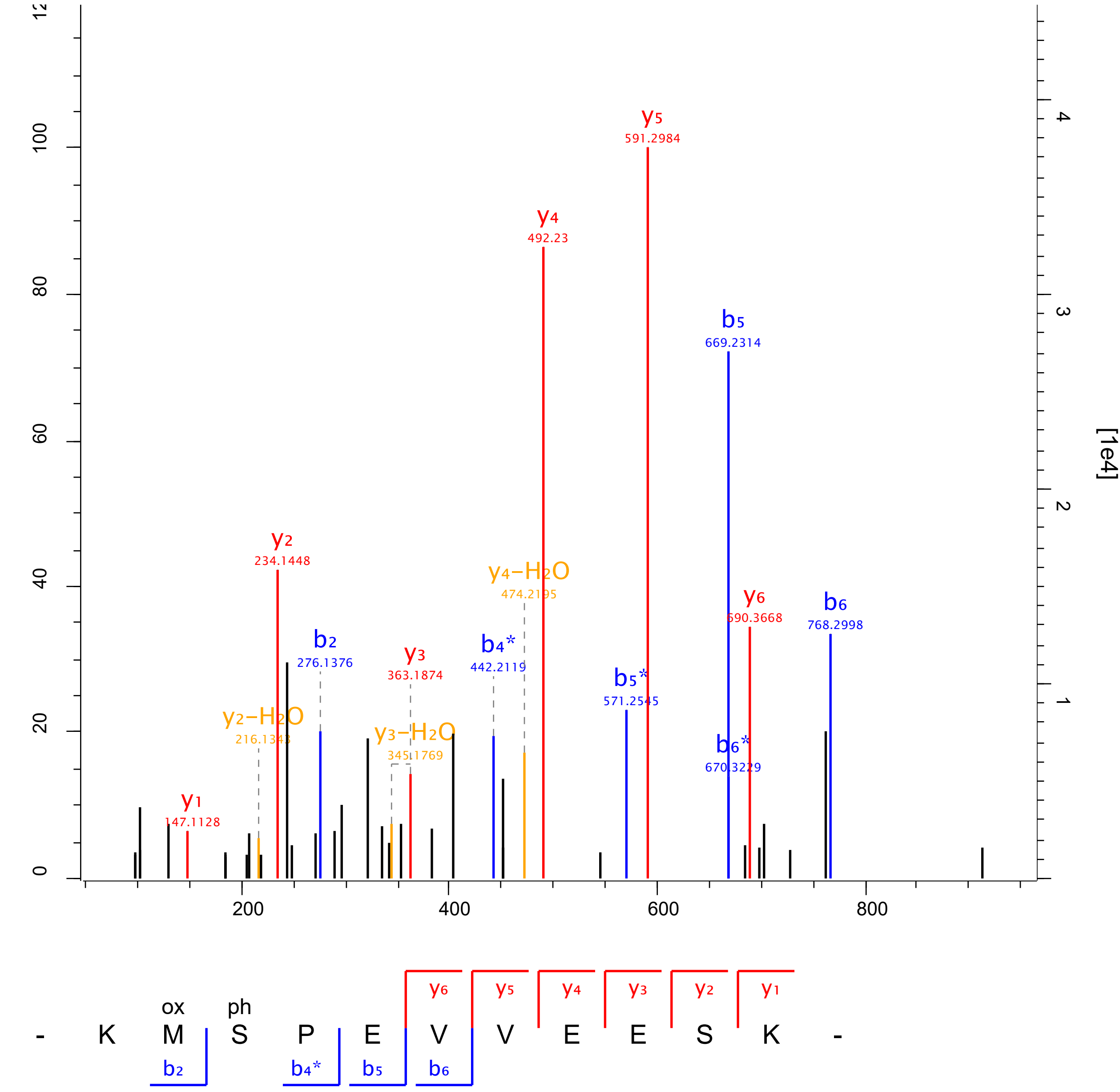

| Raw file | Scan | Method | Score | m/z |
| --- | --- | --- | --- | --- |
| sys_15_1 | 31811 | FTMS; HCD | 171.47 | 1005.38 |

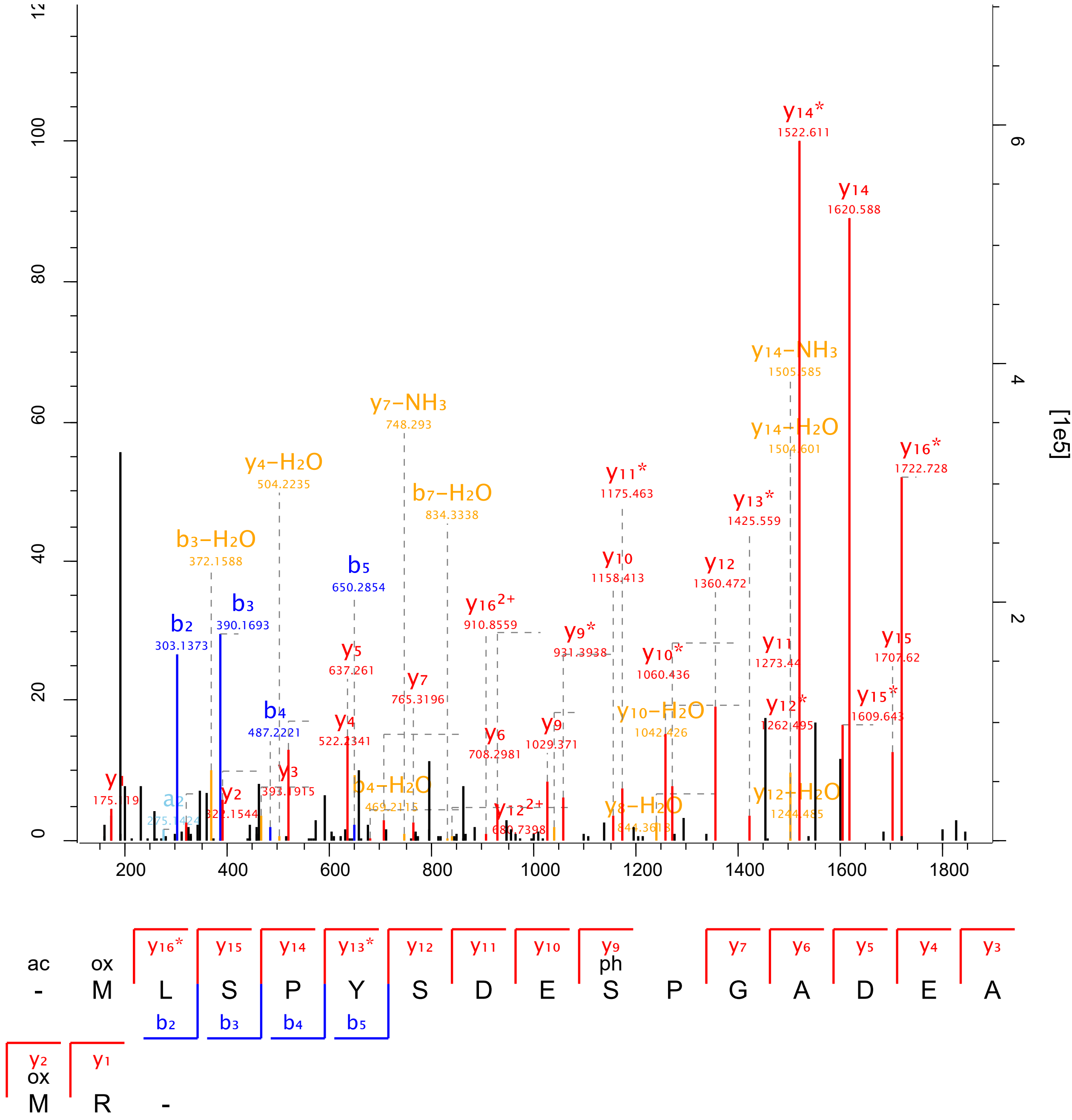

| Raw file | Scan | Method | Score | m/z |
|---|---|---|---|---|
| sys_15_1 | 32081 | FTMS; HCD | 112.26 | 941.77 |

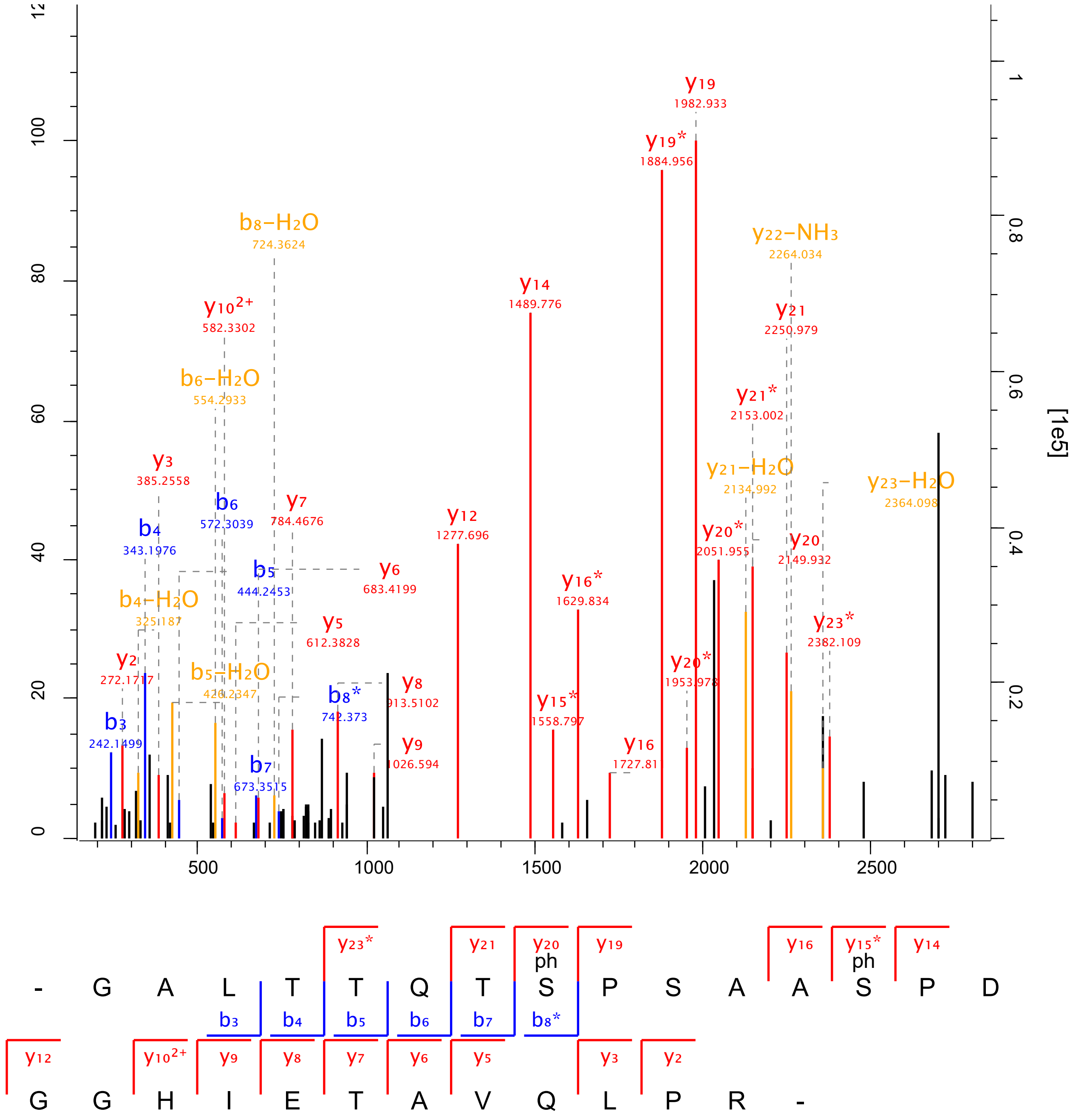

| Raw file | Scan | Method | Score | m/z |
|---|---|---|---|---|
| sys_15_1 | 32091 | FTMS; HCD | 49.99 | 916.41 |

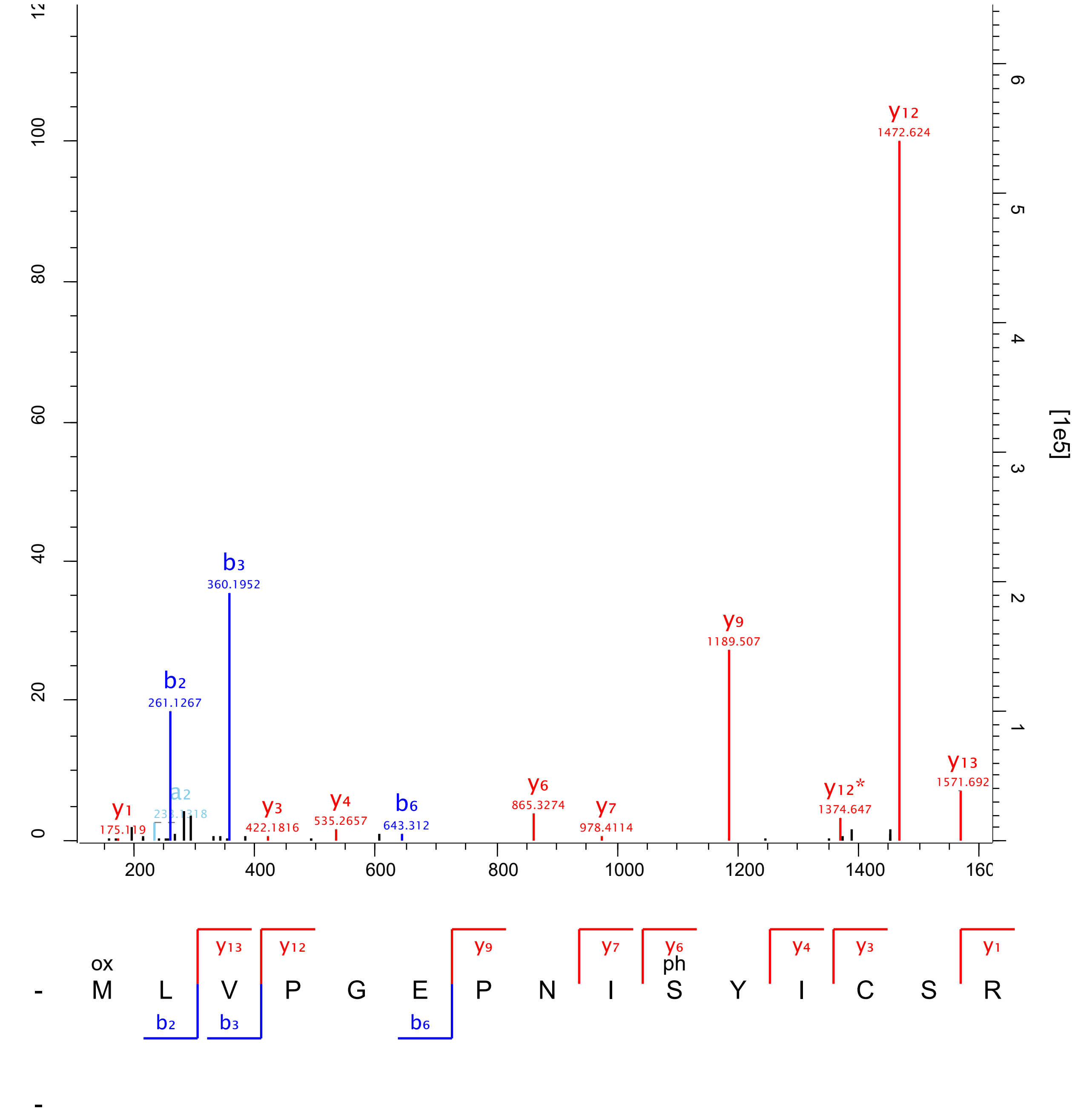

| Raw file | Scan | Method | Score | m/z |
|---|---|---|---|---|
| sys_15_1 | 32151 | FTMS; HCD | 109 | 632.78 |

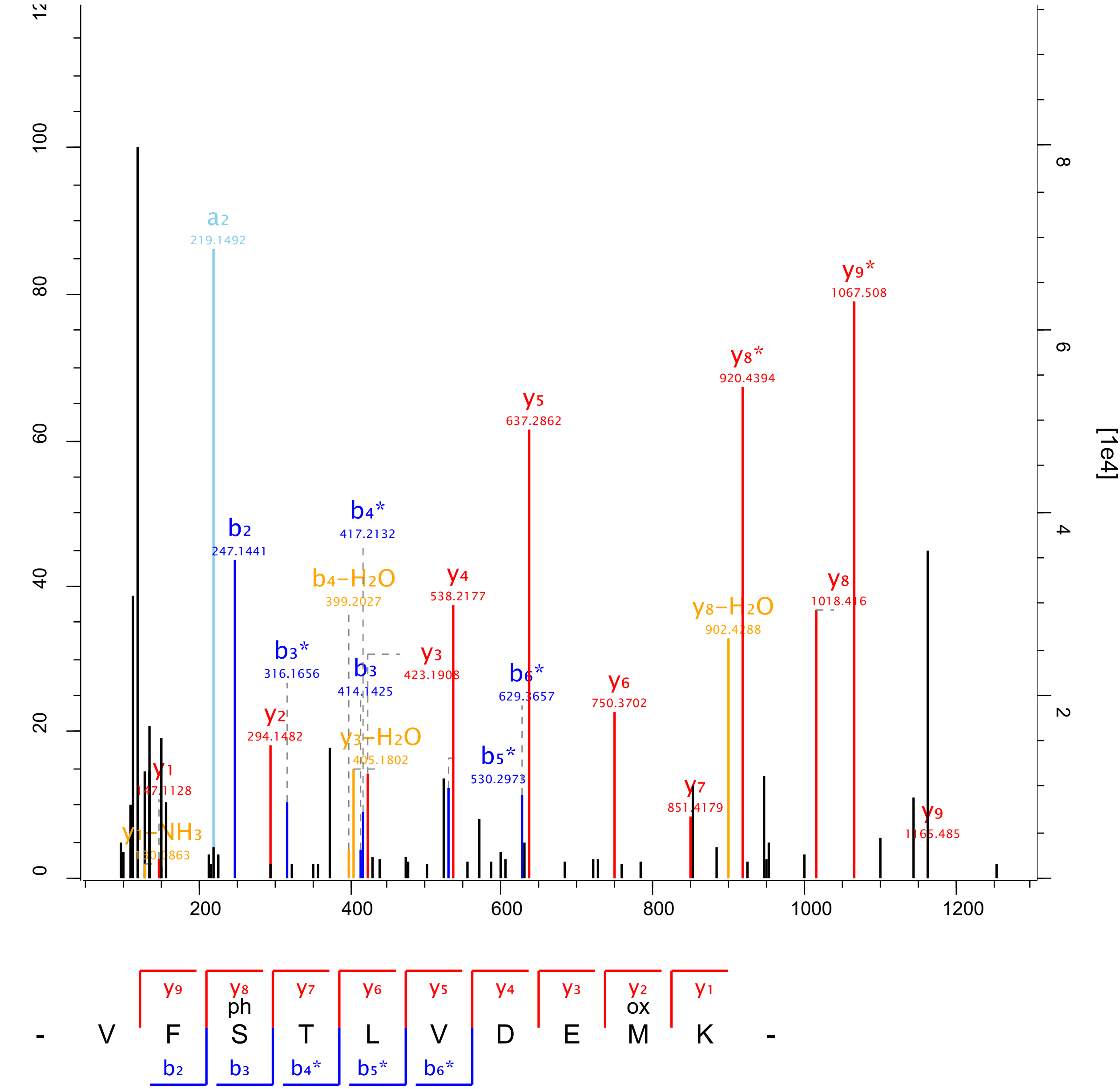

| Raw file | Scan | Method | Score | m/z |
|---|---|---|---|---|
| sys_15_1 | 32255 | FTMS; HCD | 41.91 | 688.65 |

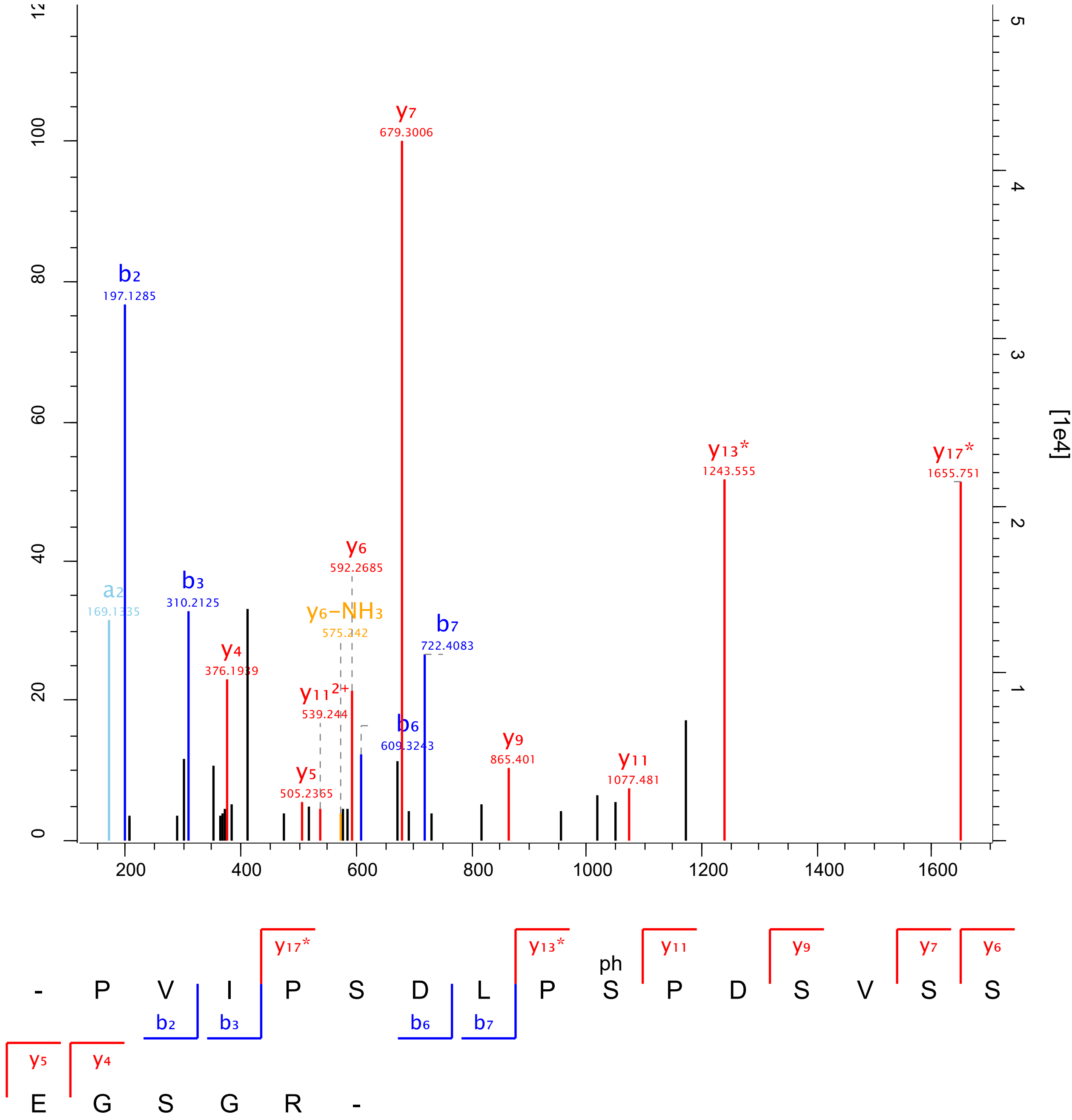

| Raw file | Scan | Method | Score | m/z |
| --- | --- | --- | --- | --- |
| sys_15_1 | 32257 | FTMS; HCD | 166.6 | 853.43 |

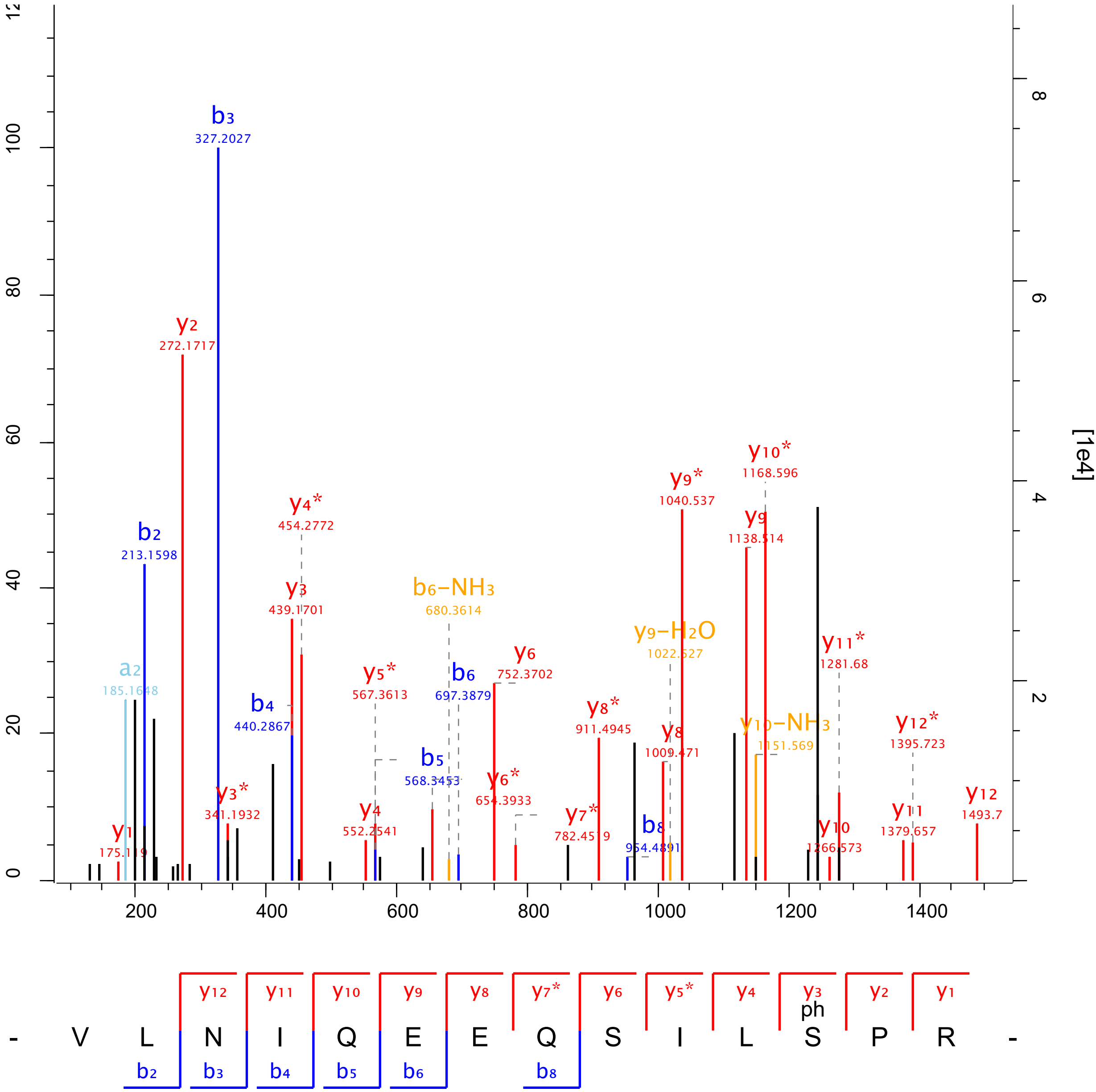

| Raw file | Scan | Method | Score | m/z |
|---|---|---|---|---|
| sys_15_1 | 32262 | FTMS; HCD | 45.58 | 741.65 |

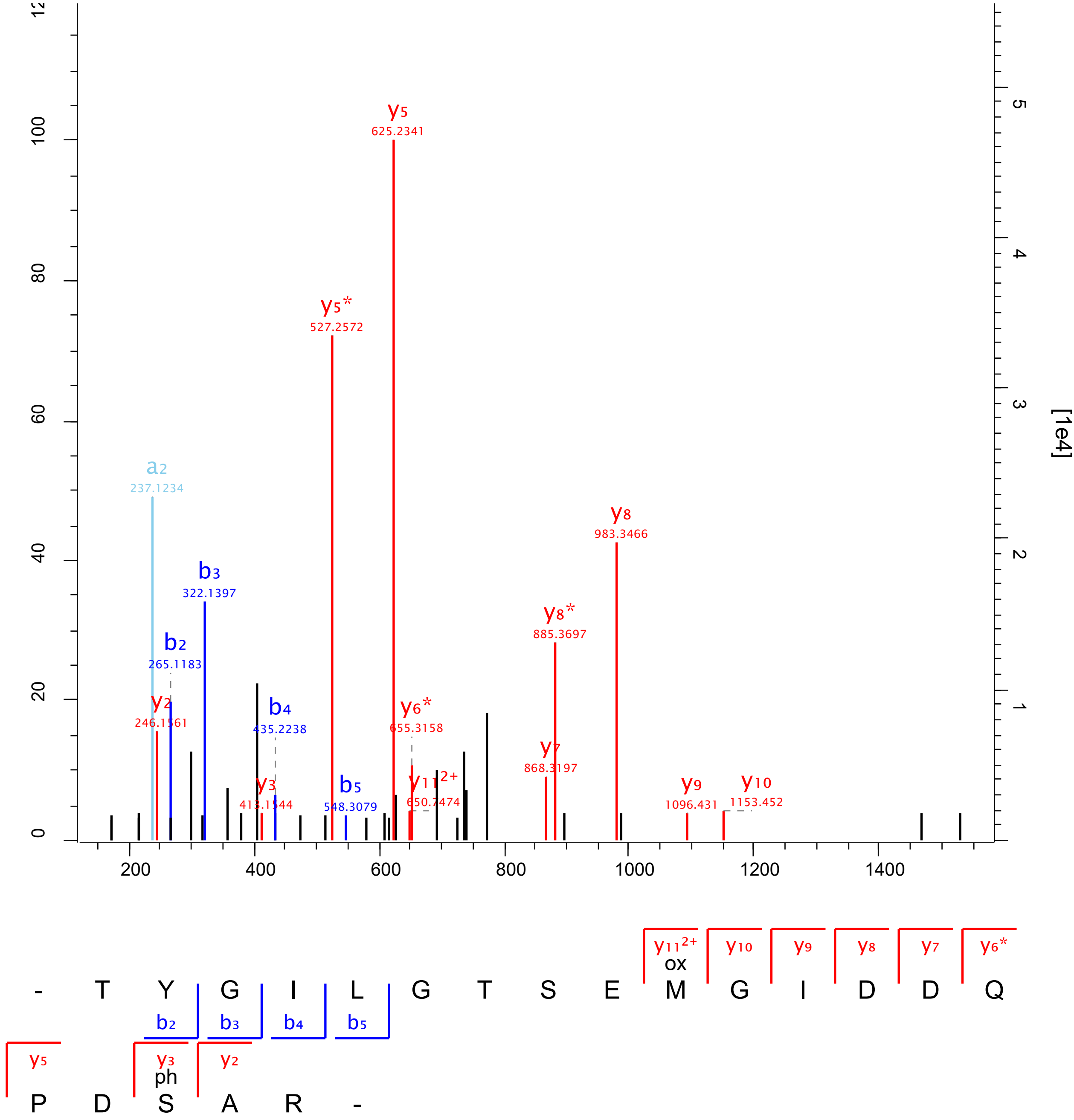

| Raw file | Scan | Method | Score | m/z |
| --- | --- | --- | --- | --- |
| sys_15_1 | 32282 | FTMS; HCD | 110.33 | 1108.45 |

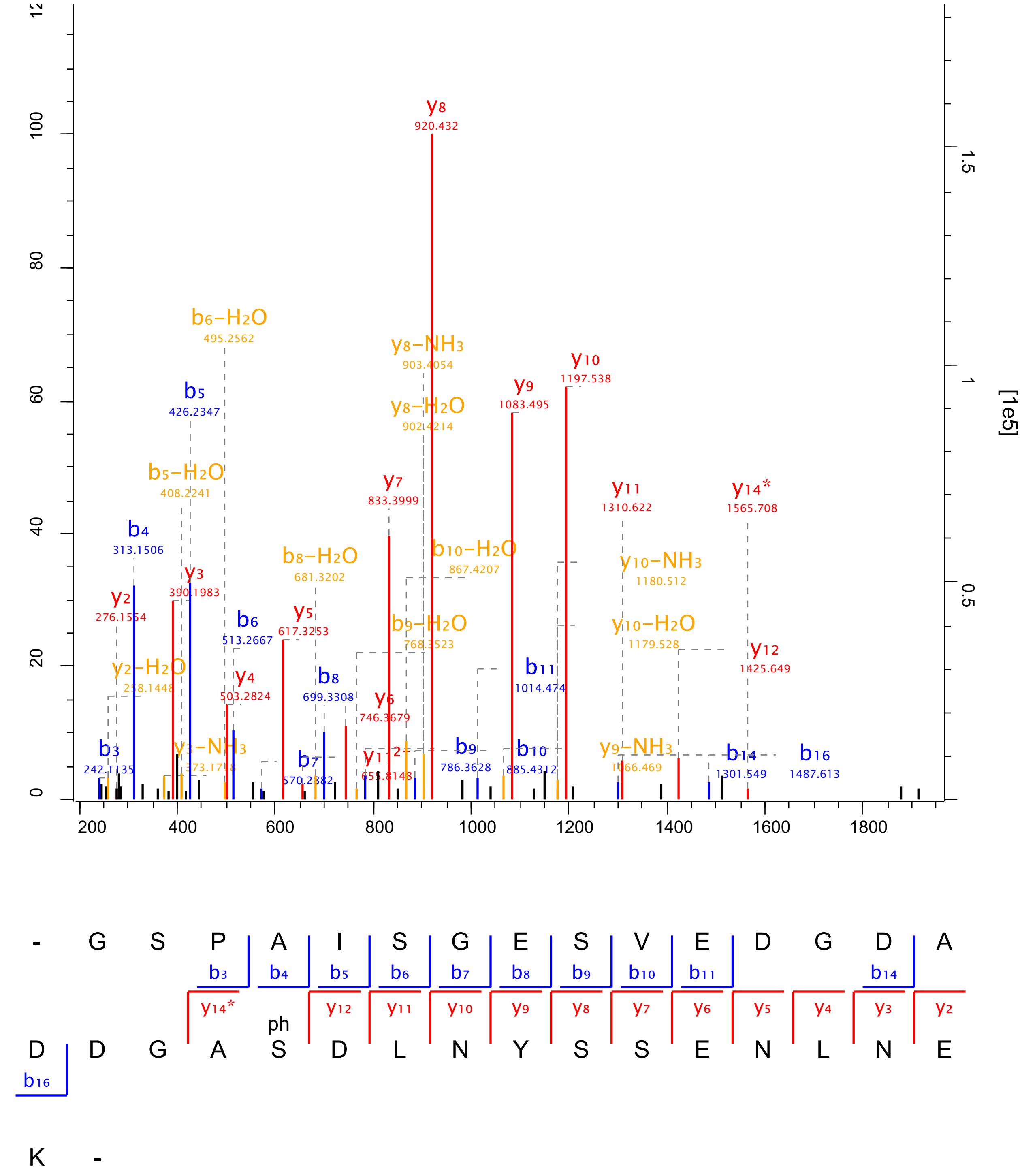

| Raw file | Scan | Method | Score | m/z |
|---|---|---|---|---|
| sys_15_1 | 32313 | FTMS; HCD | 56.69 | 957.76 |

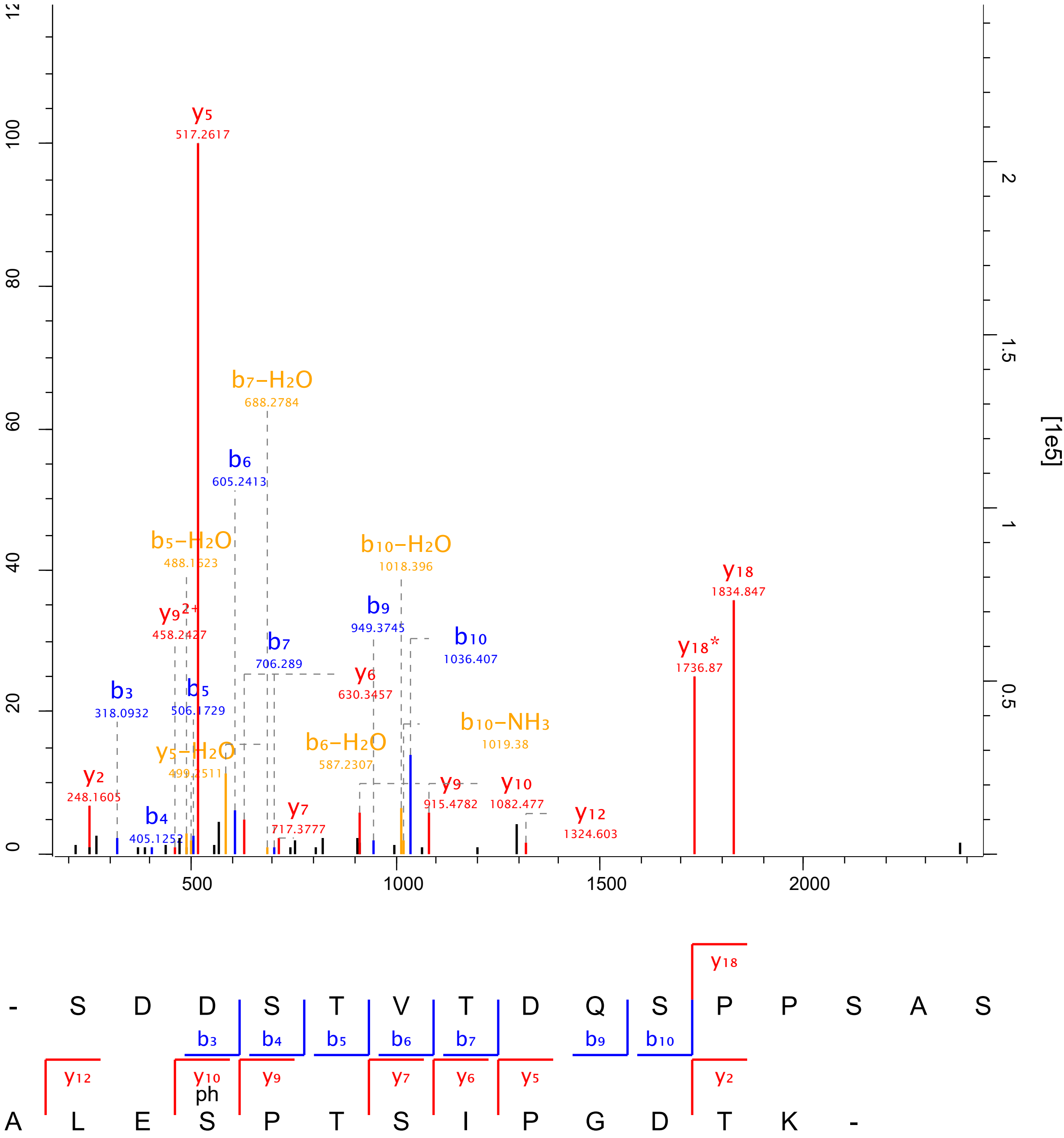

| Raw file | Scan | Method | Score | m/z |
| --- | --- | --- | --- | --- |
| sys_15_1 | 32356 | FTMS; HCD | 101.41 | 755.33 |

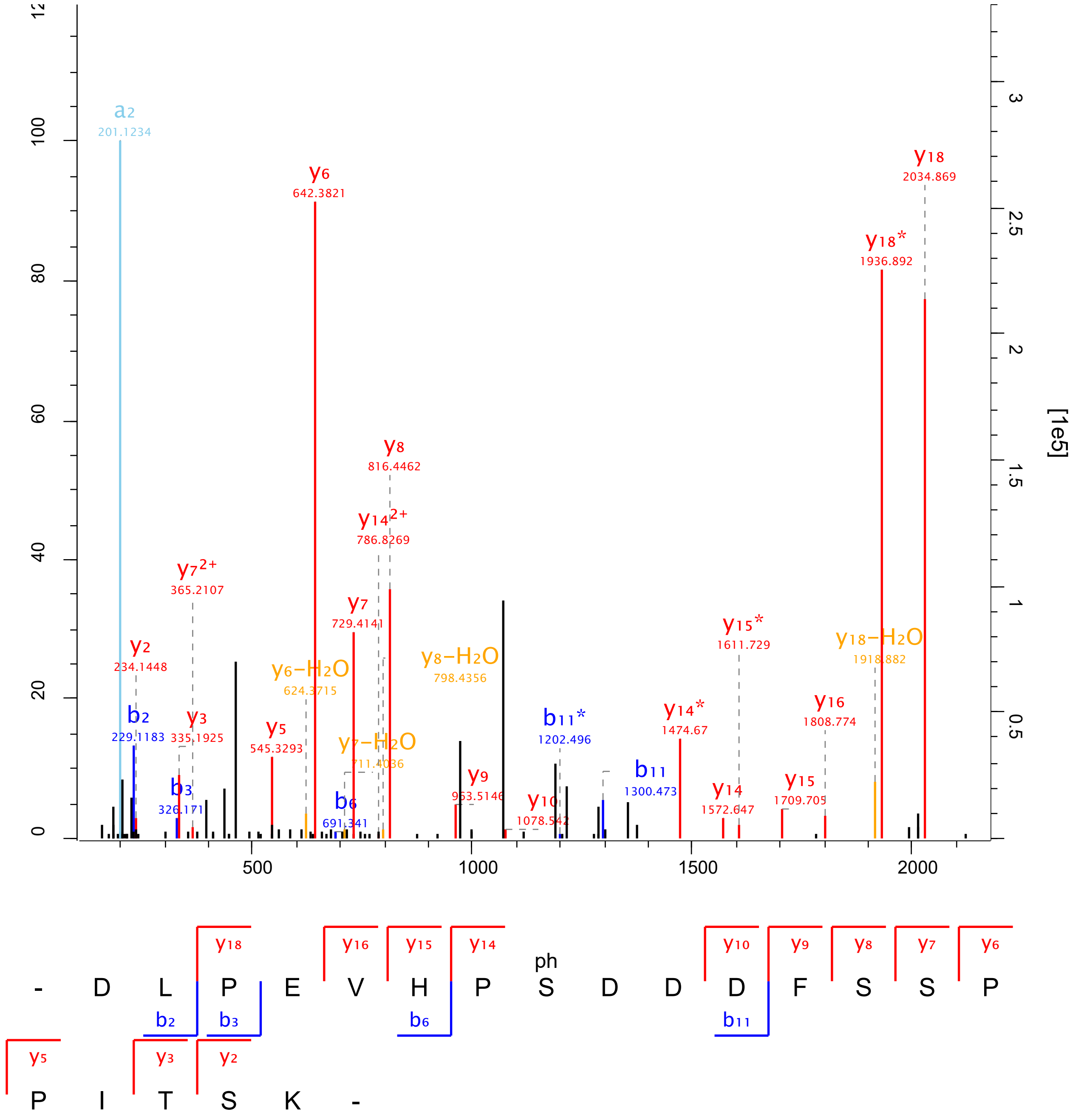

| Raw file | Scan | Method | Score | m/z |
|---|---|---|---|---|
| sys_15_1 | 32391 | FTMS; HCD | 71.38 | 769.84 |

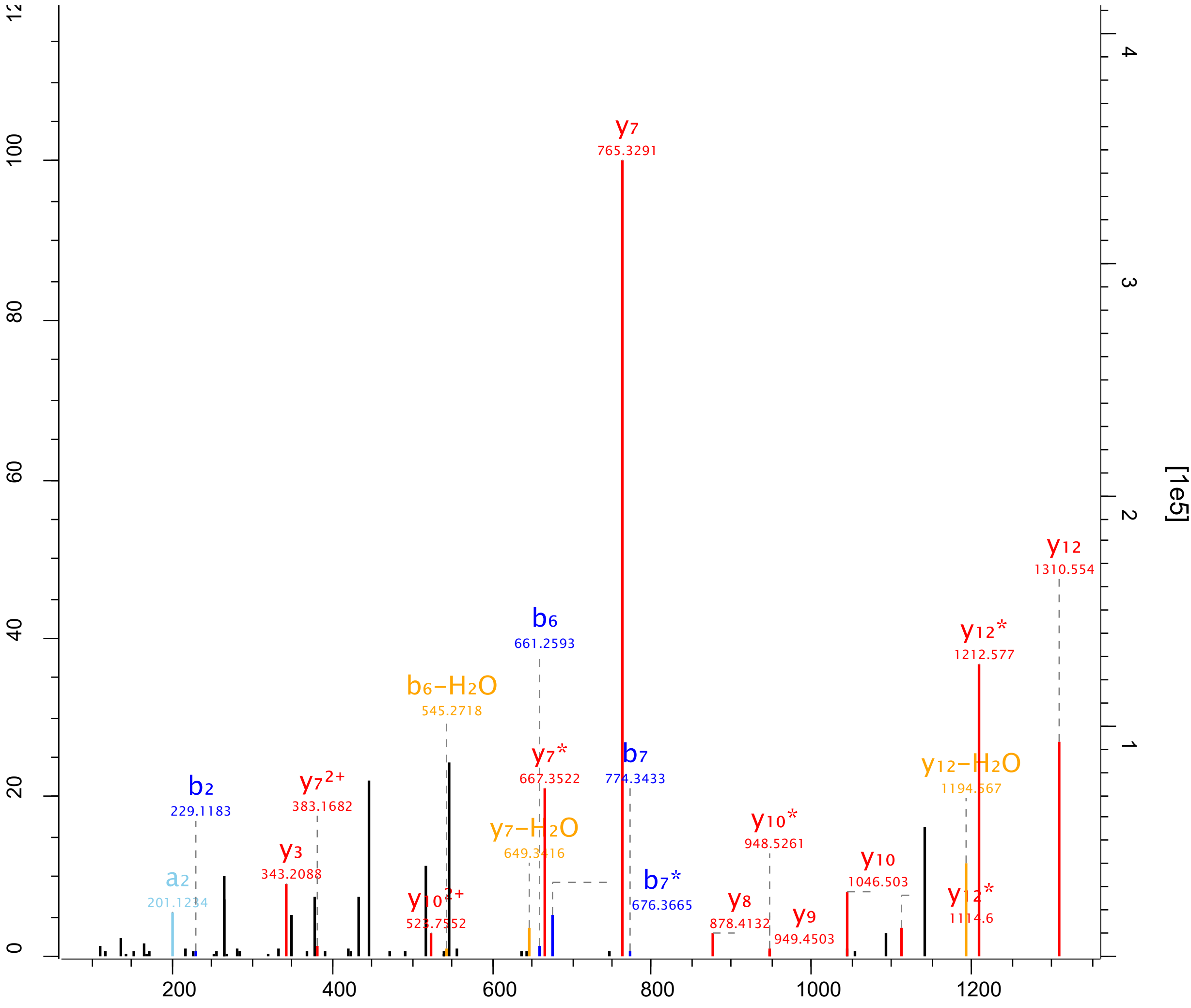


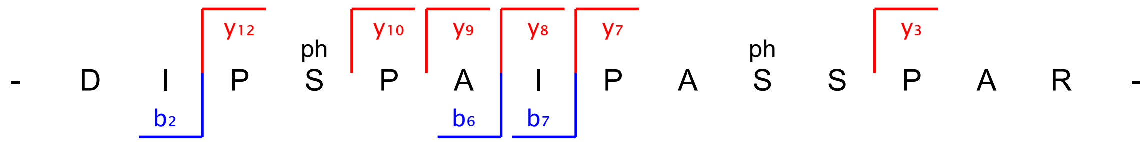

| Raw file | Scan | Method | Score | m/z |
|---|---|---|---|---|
| sys_15_1 | 32427 | FTMS; HCD | 159.81 | 699.31 |

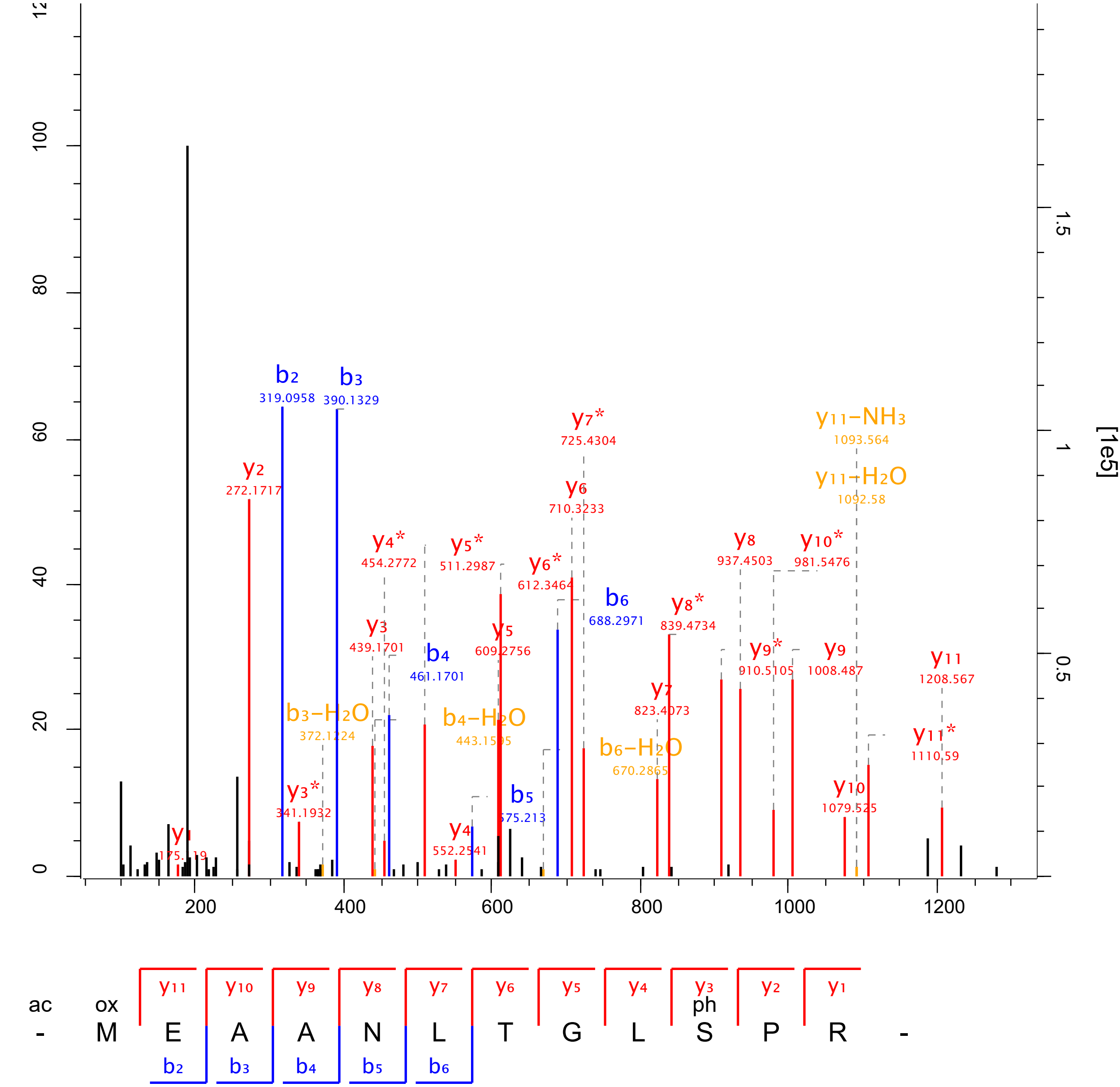

| Raw file | Scan | Method | Score | m/z |
|---|---|---|---|---|
| sys_15_1 | 32593 | FTMS; HCD | 142.94 | 869.4 |

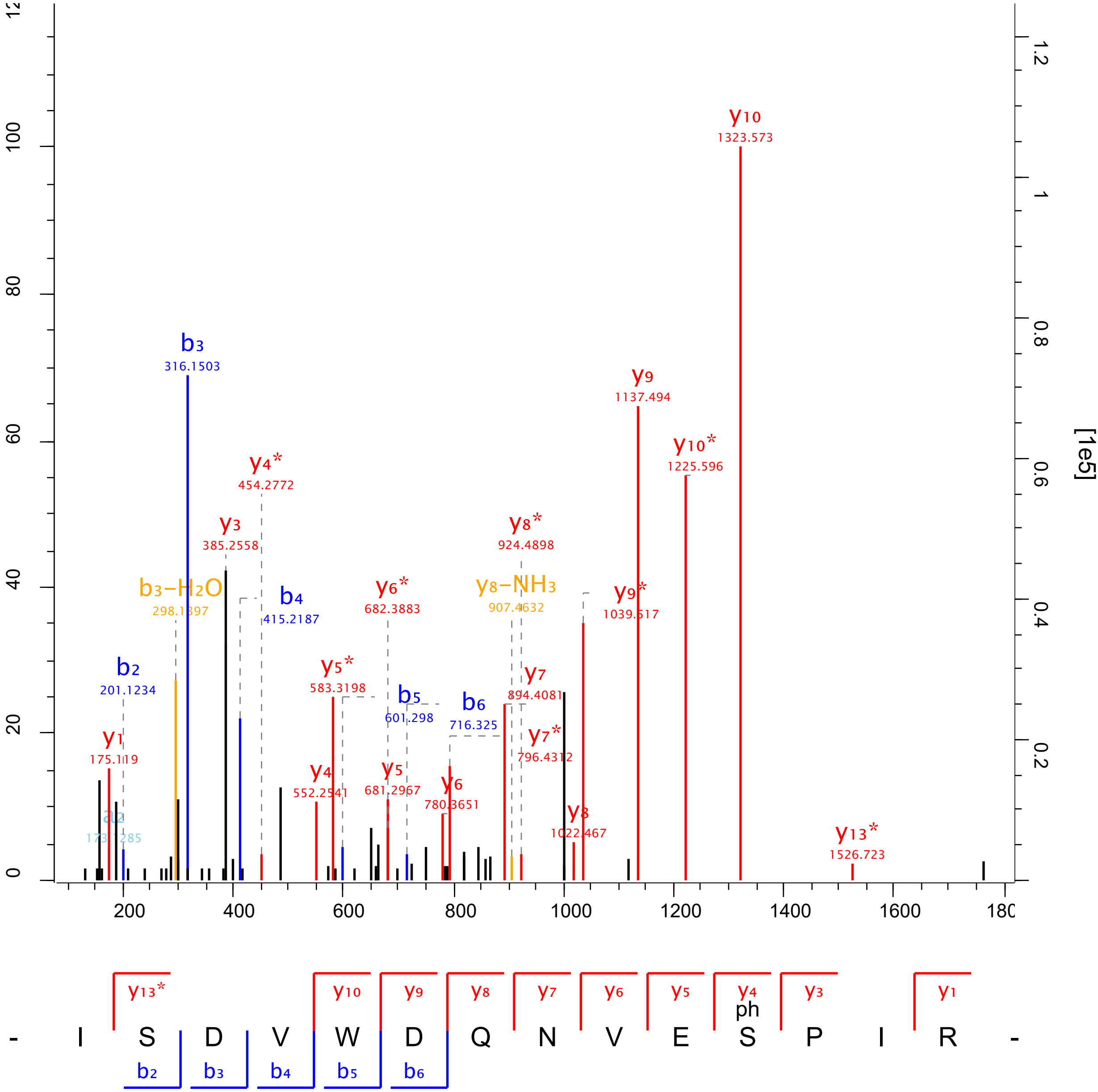

| Raw file | Scan | Method | Score | m/z |
| --- | --- | --- | --- | --- |
| sys_15_1 | 32823 | FTMS; HCD | 108.72 | 876.41 |

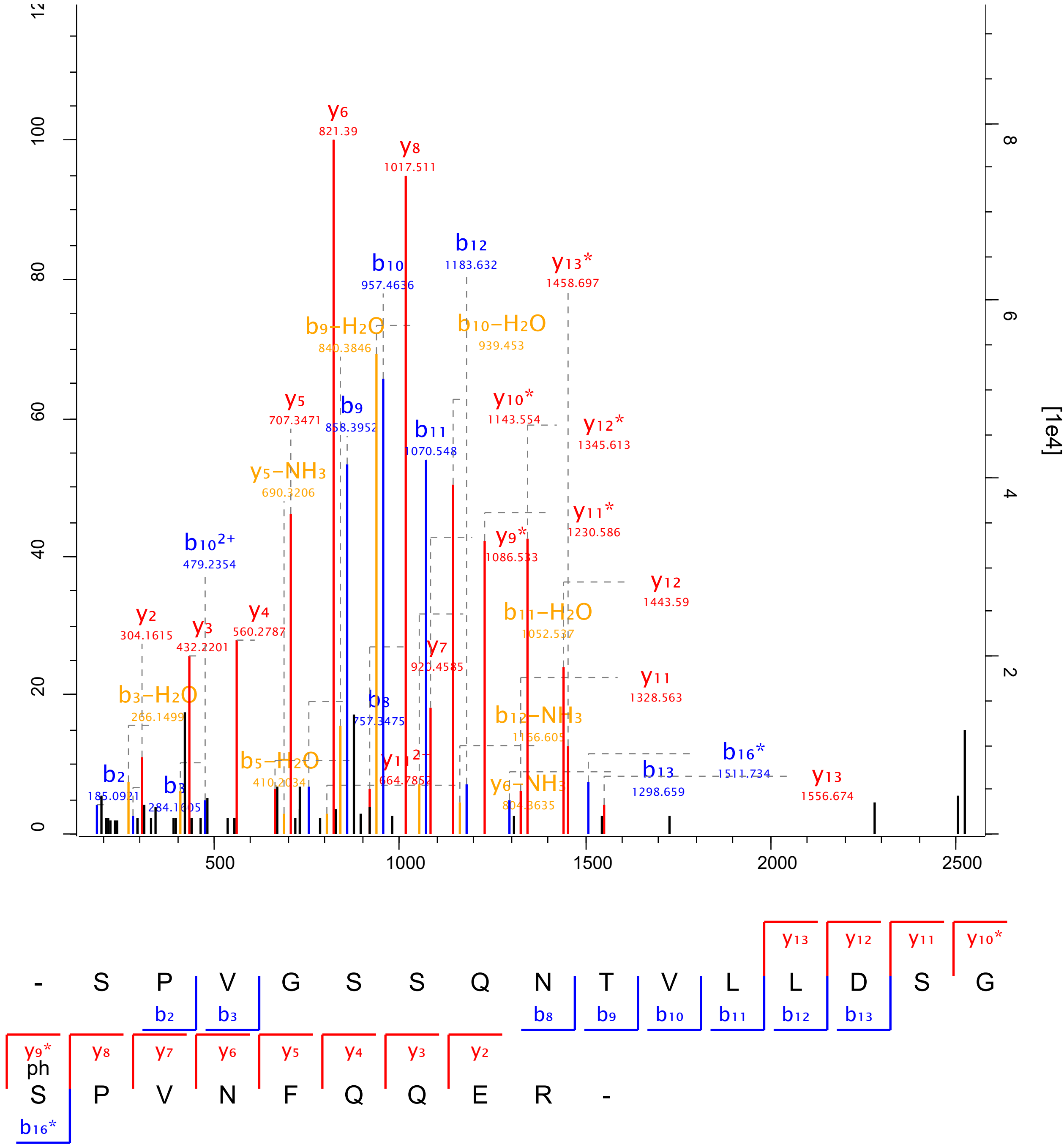

| Raw file | Scan | Method | Score | m/z |
| --- | --- | --- | --- | --- |
| sys_15_1 | 32848 | FTMS; HCD | 67.14 | 881.37 |

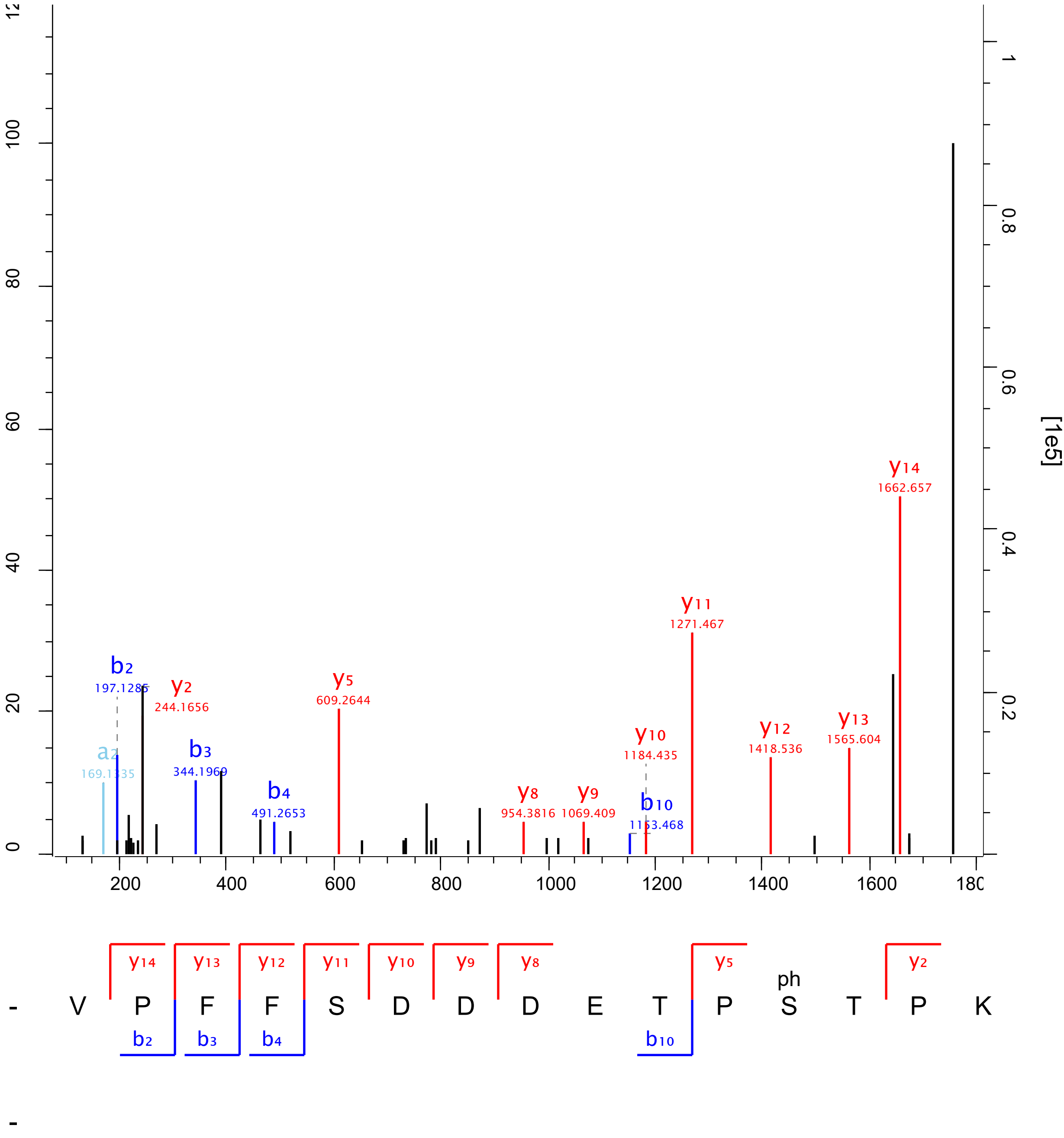

| Raw file | Scan | Method | Score | m/z |
|---|---|---|---|---|
| sys_15_1 | 32999 | FTMS; HCD | 88.28 | 815.83 |

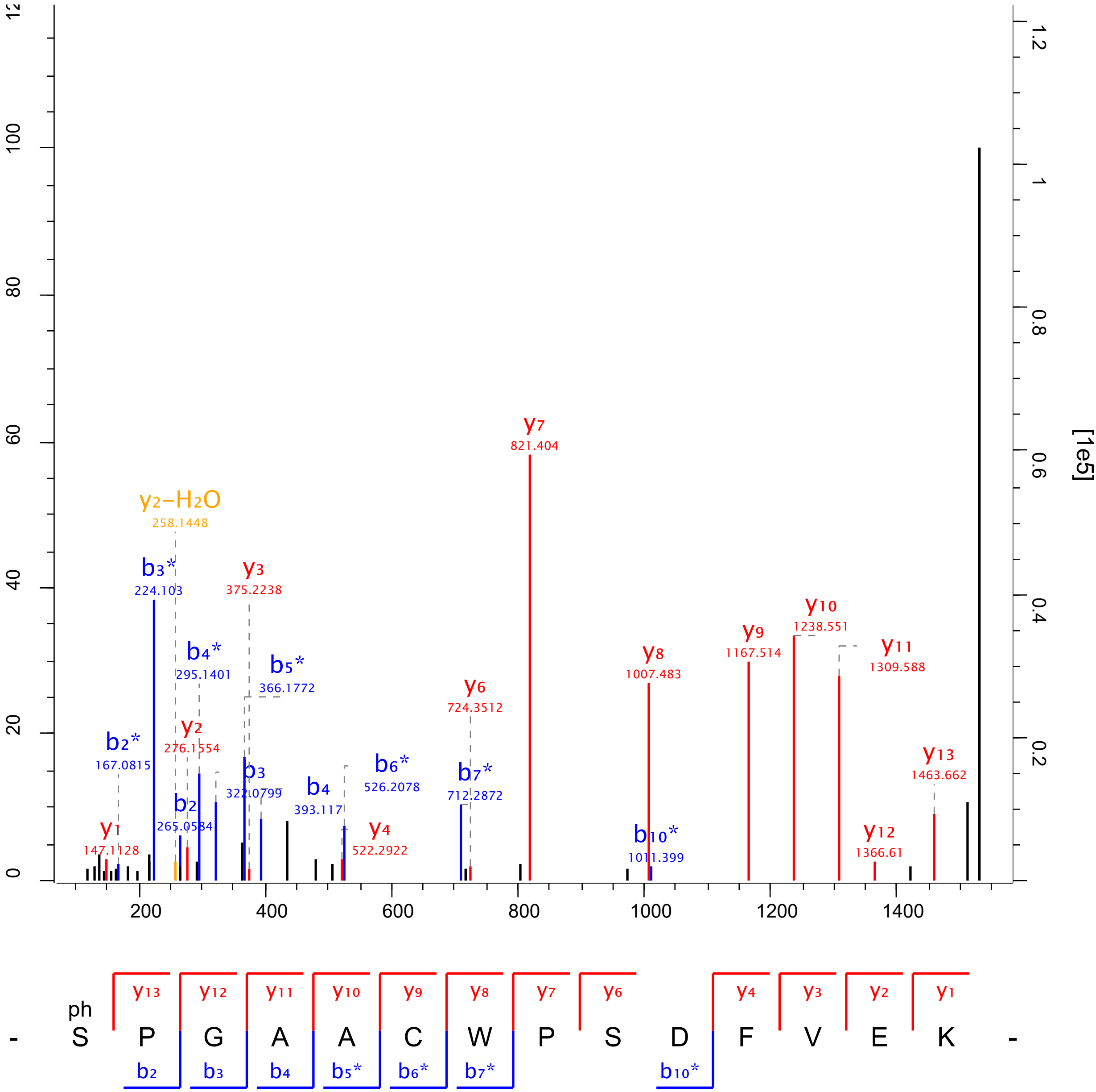

| Raw file | Scan | Method | Score | m/z |
|---|---|---|---|---|
| sys_15_1 | 3321 | FTMS; HCD | 103.01 | 670.3 |

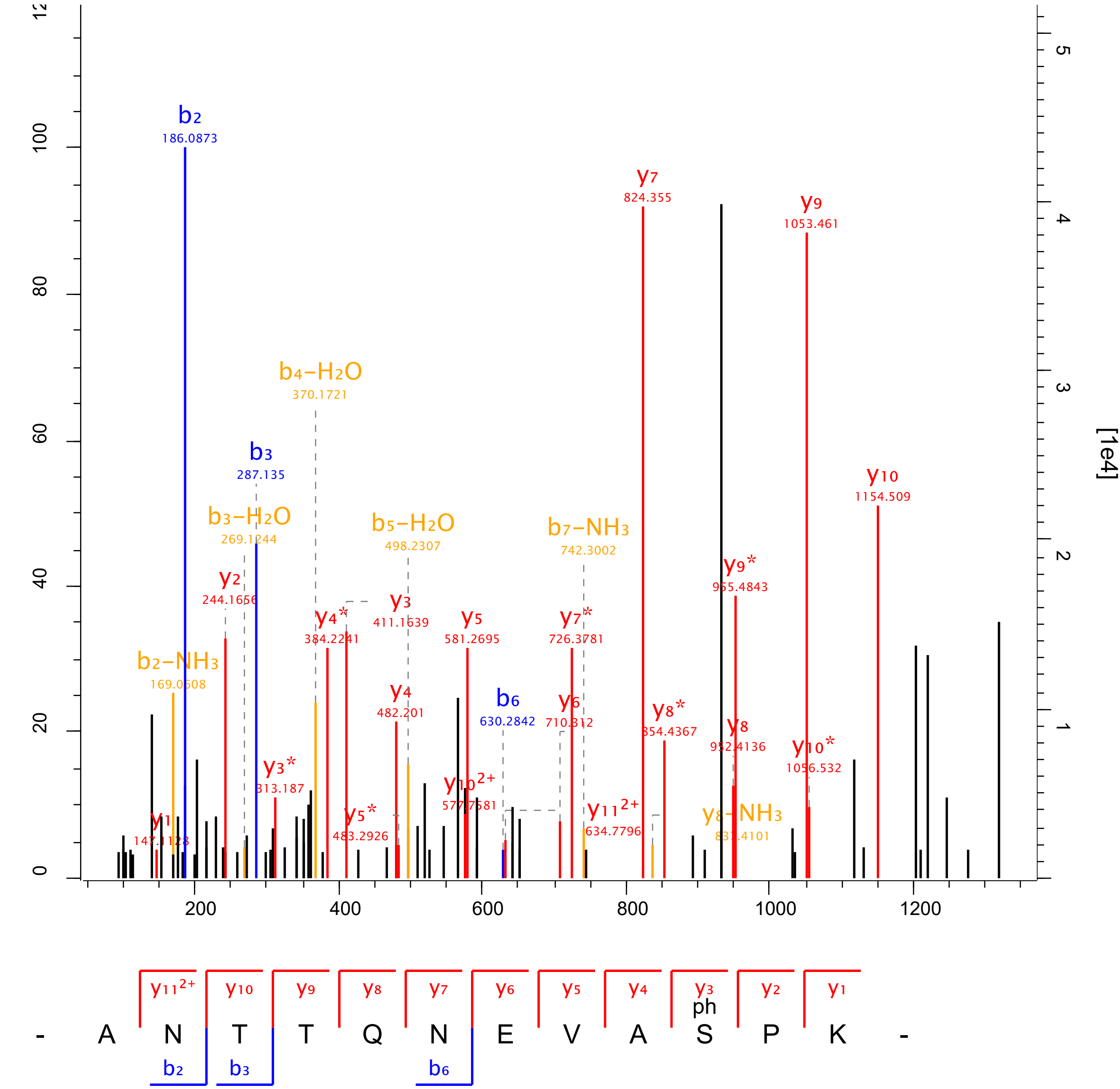

| Raw file | Scan | Method | Score | m/z |
|---|---|---|---|---|
| sys_15_1 | 33364 | FTMS; HCD | 250.94 | 818.37 |

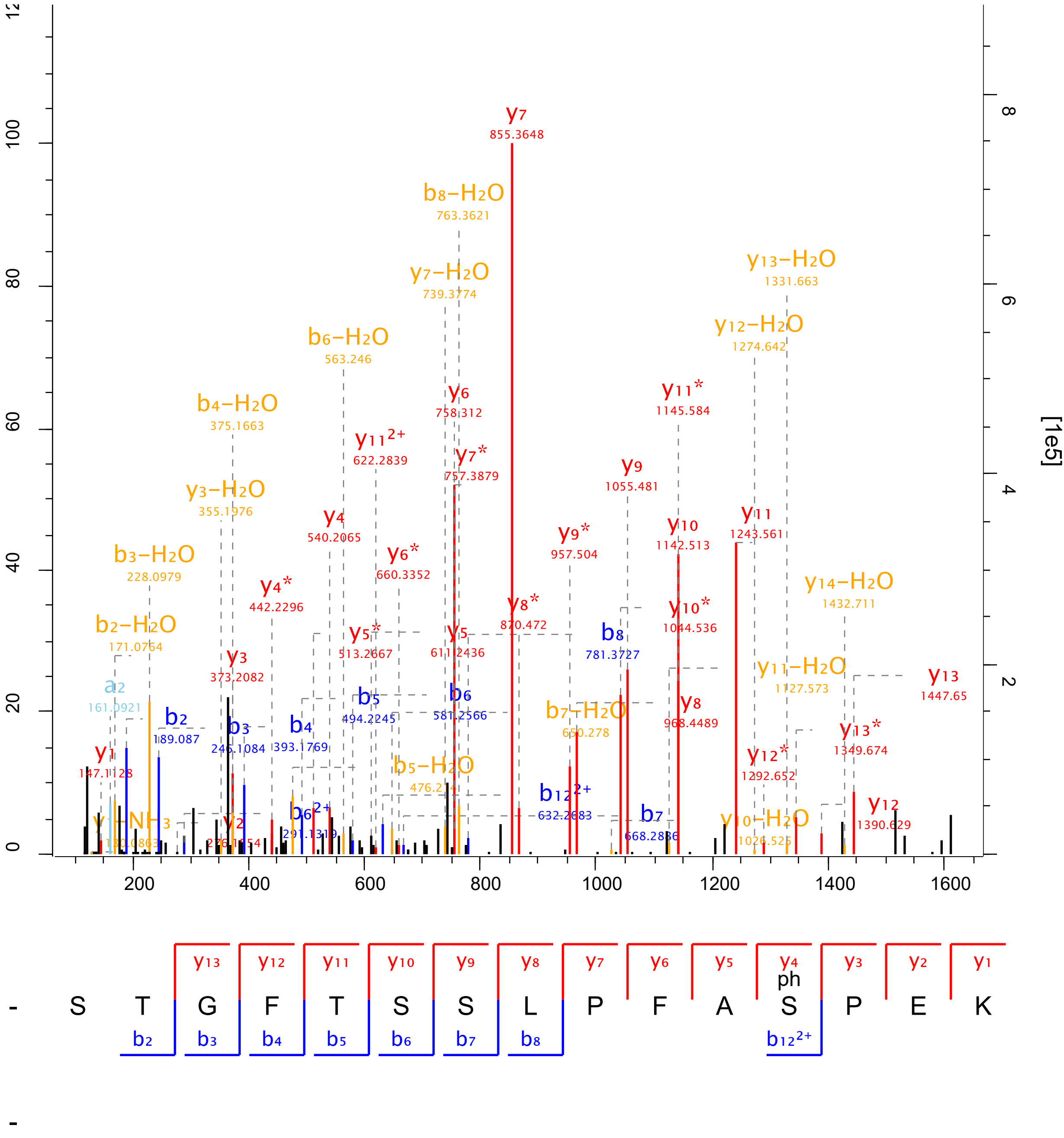

| Raw file | Scan | Method | Score | m/z |
| --- | --- | --- | --- | --- |
| sys_15_1 | 3348 | FTMS; HCD | 87.12 | 474.18 |

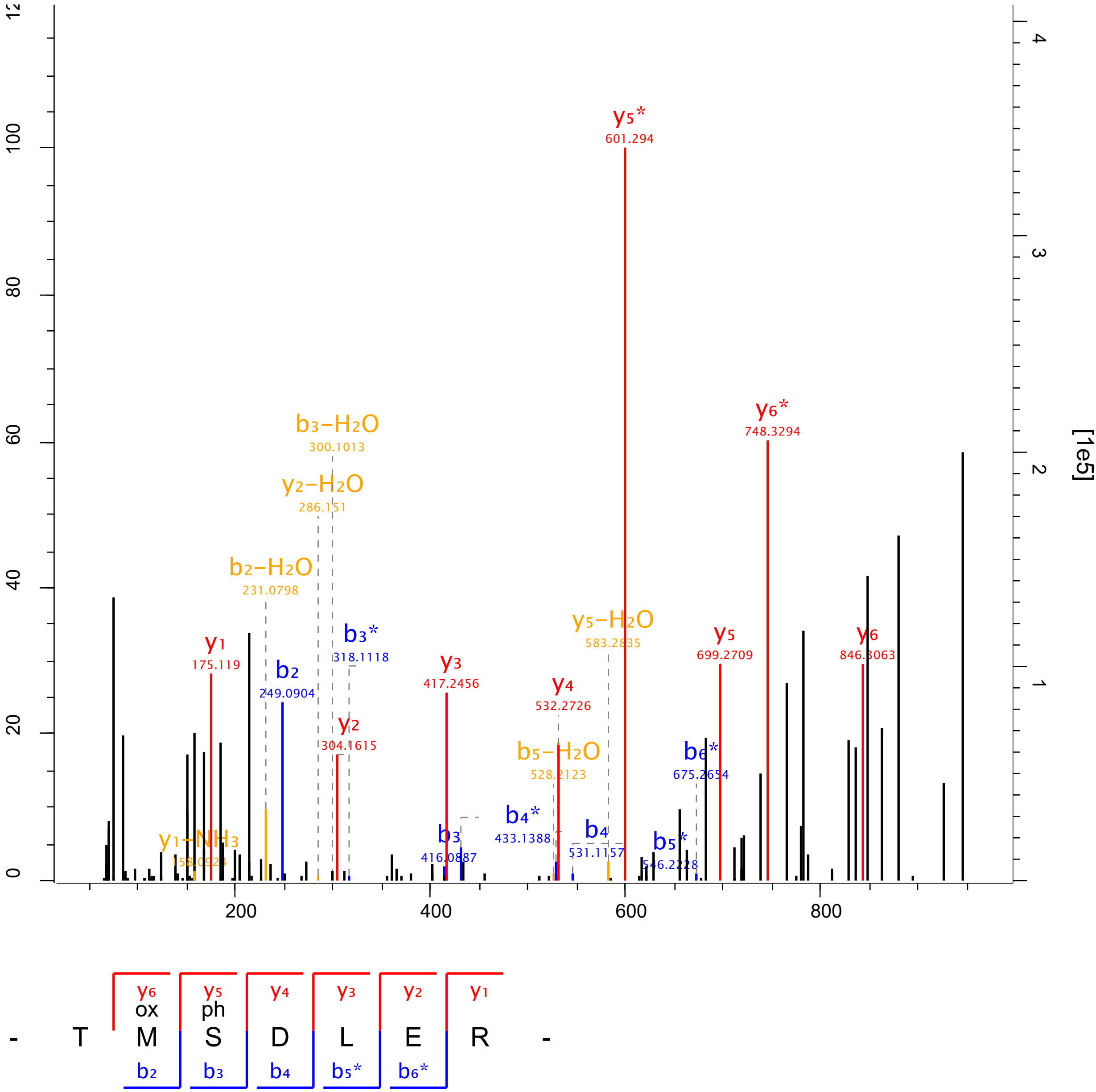

| Raw file | Scan | Method | Score | m/z |
|---|---|---|---|---|
| sys_15_1 | 3351 | FTMS; HCD | 122.33 | 593.76 |

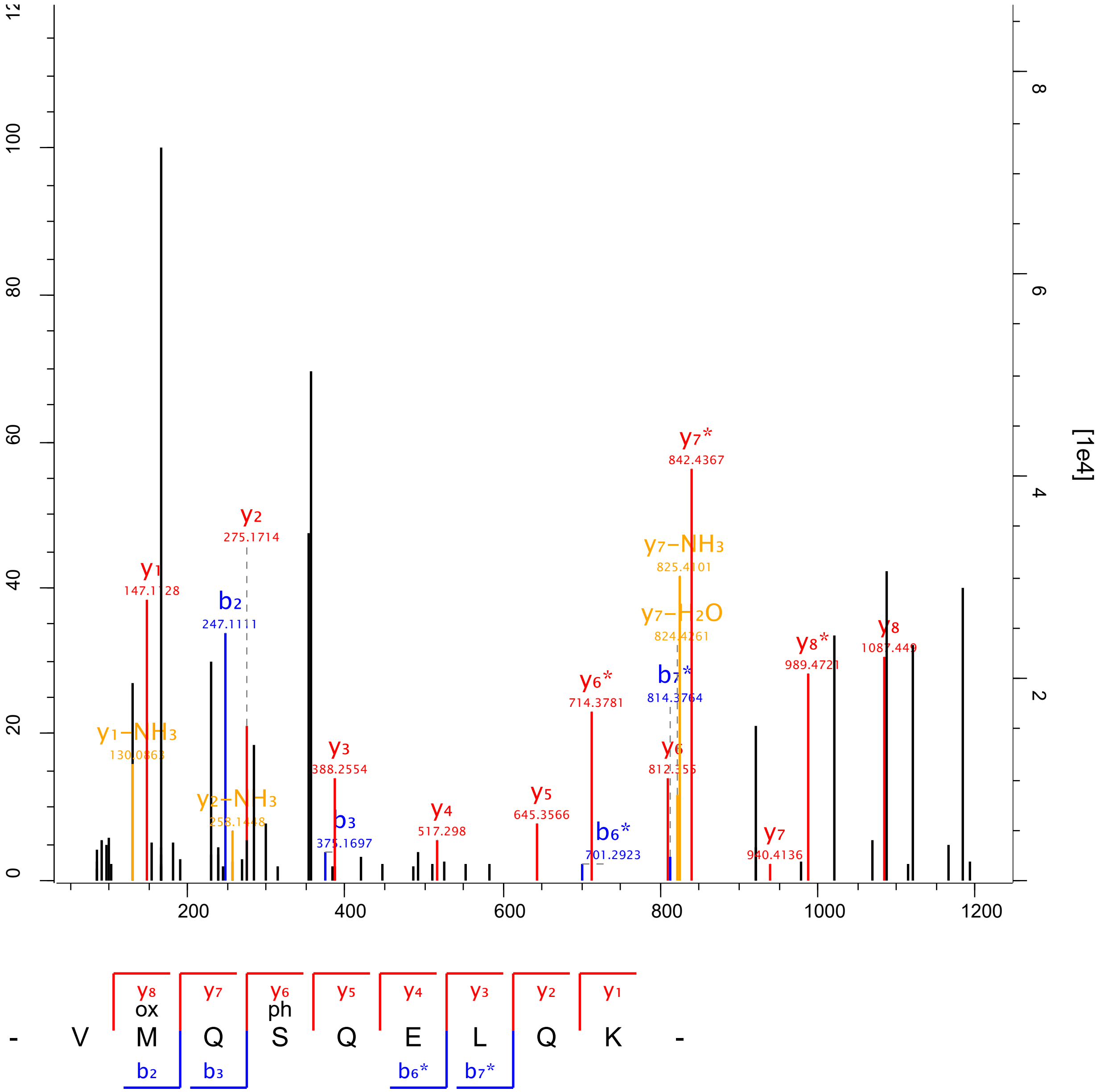

| Raw file | Scan | Method | Score | m/z |
|---|---|---|---|---|
| sys_15_1 | 33585 | FTMS; HCD | 46.07 | 642.78 |

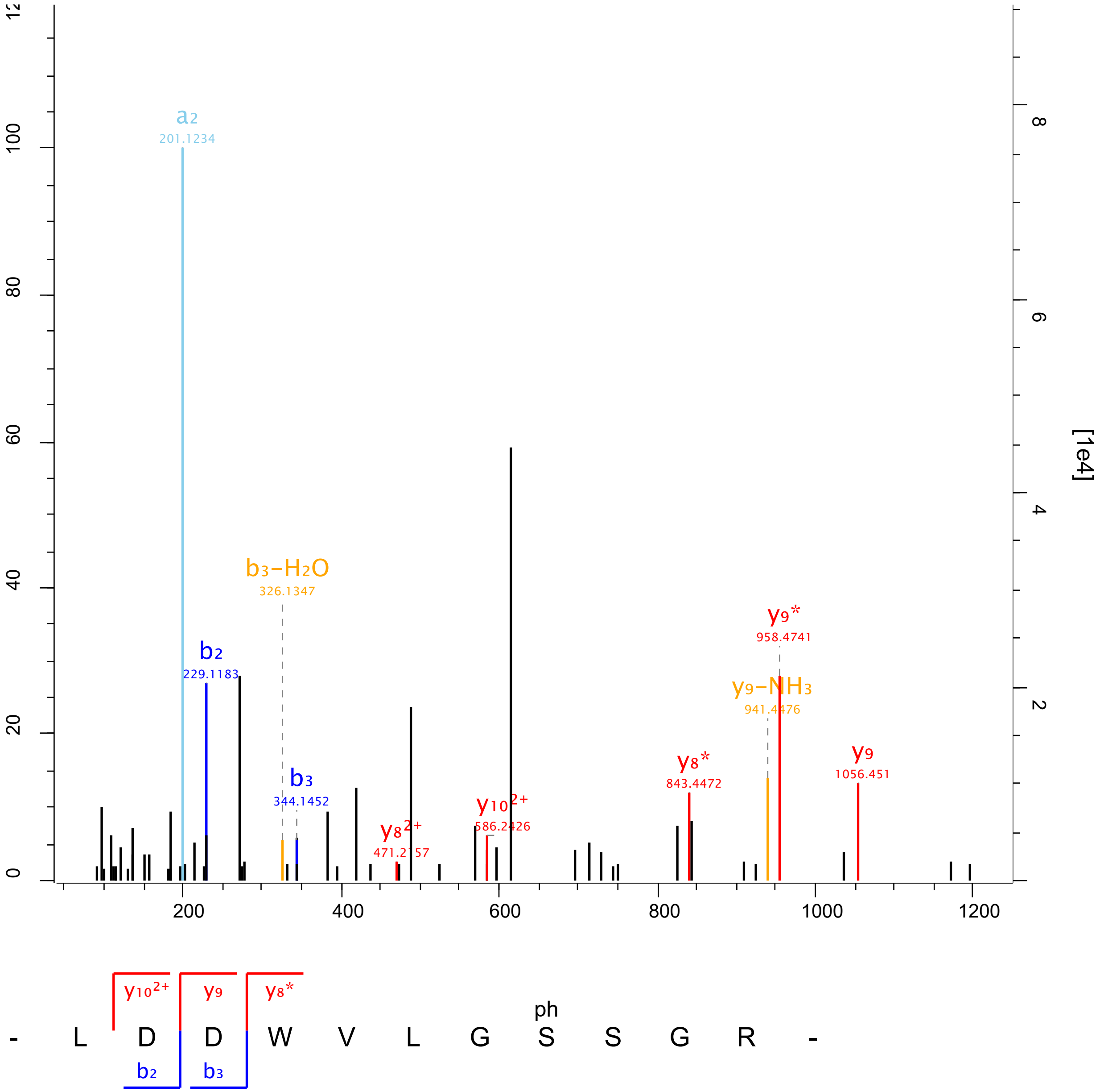

| Raw file | Scan | Method | Score | m/z |
| --- | --- | --- | --- | --- |
| sys_15_1 | 3379 | FTMS; HCD | 62.17 | 533.73 |

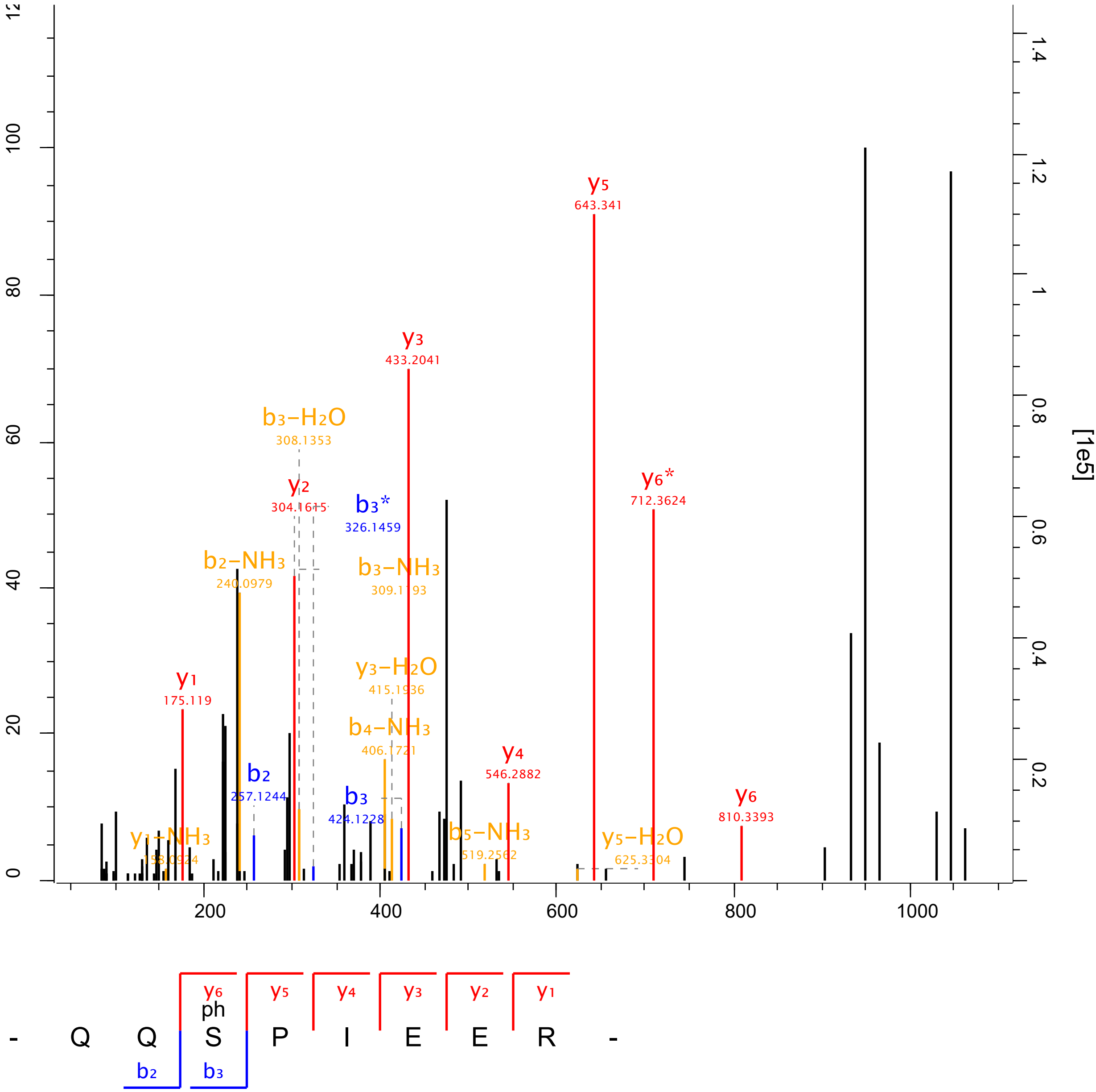

| Raw file | Scan | Method | Score | m/z |
|---|---|---|---|---|
| sys_15_1 | 34031 | FTMS; HCD | 117 | 804.36 |

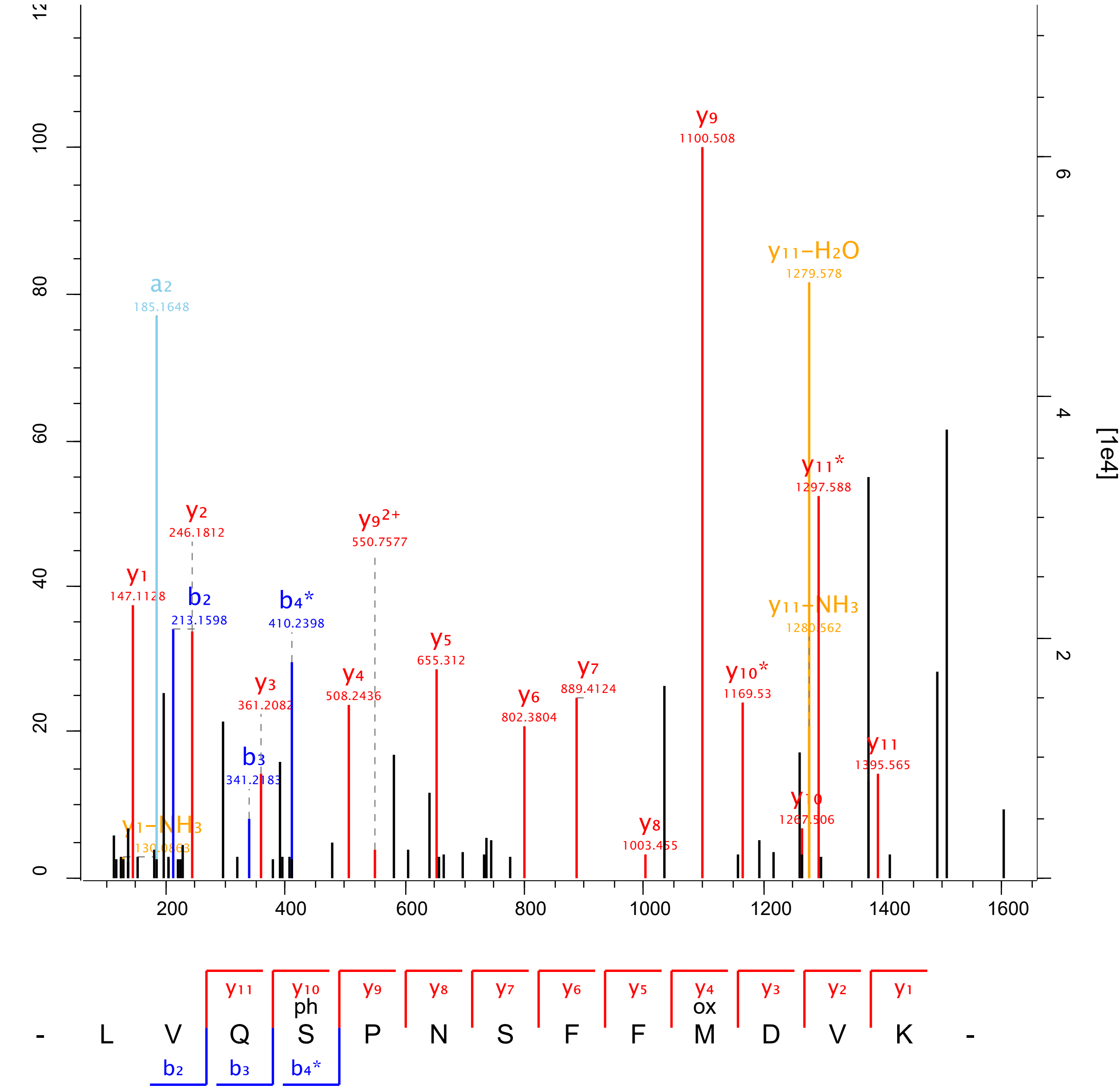

| Raw file | Scan | Method | Score | m/z |
|---|---|---|---|---|
| sys_15_1 | 3430 | FTMS; HCD | 64.71 | 435.19 |

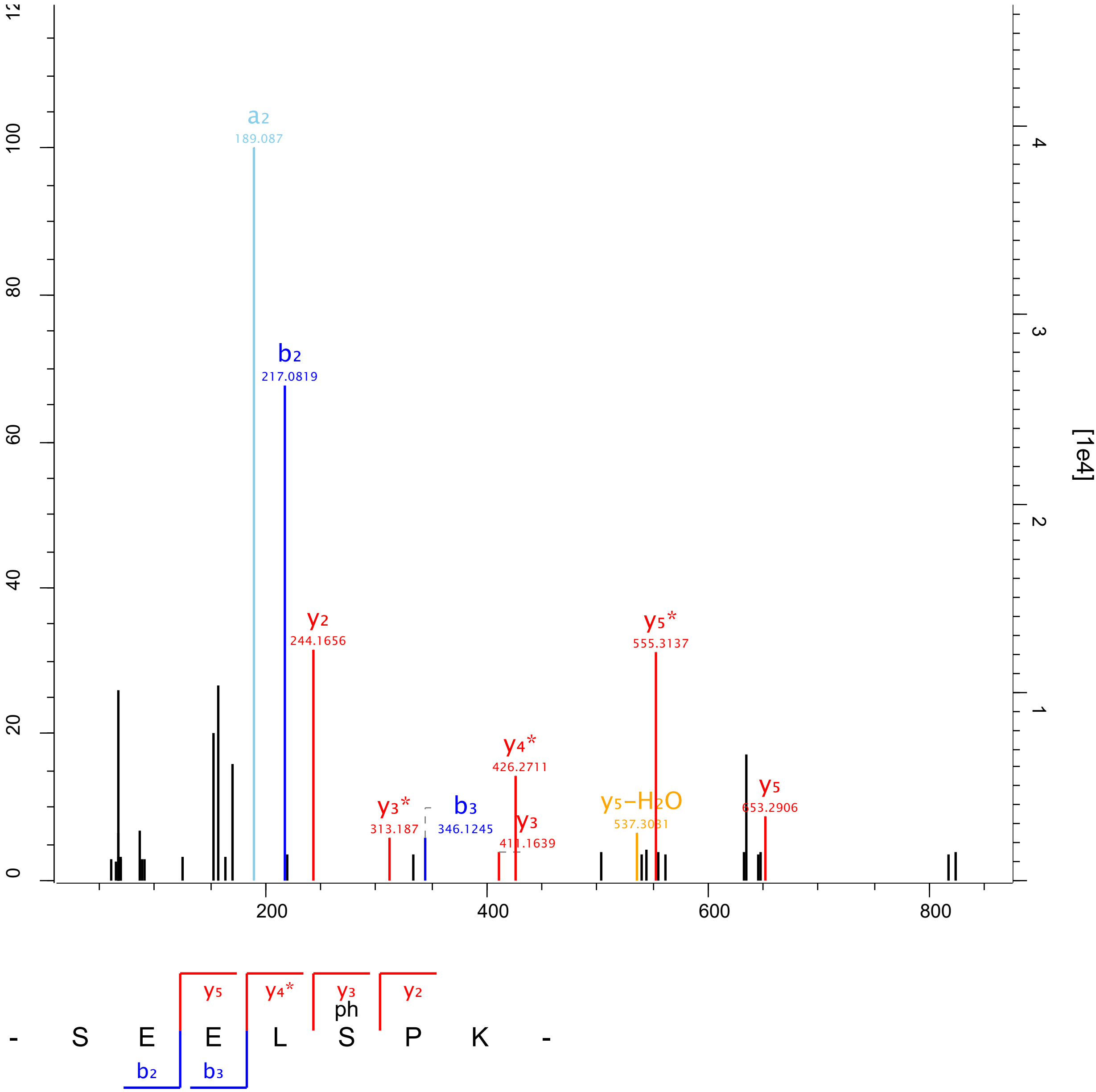

| Raw file | Scan | Method | Score | m/z |
| --- | --- | --- | --- | --- |
| sys_15_1 | 34334 | FTMS; HCD | 62.05 | 881.69 |

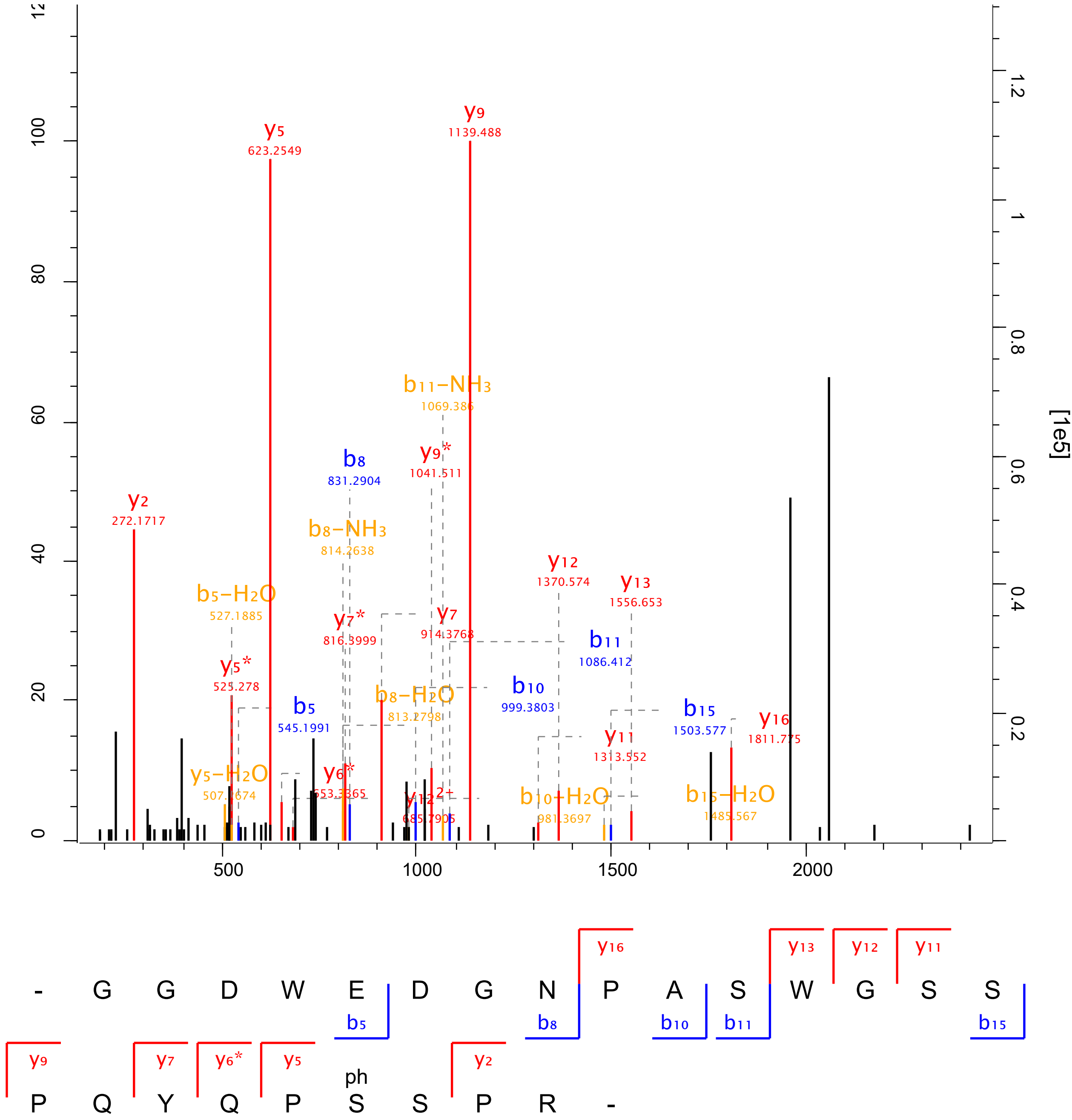

| Raw file | Scan | Method | Score | m/z |
|---|---|---|---|---|
| sys_15_1 | 34340 | FTMS; HCD | 79.69 | 1104.43 |

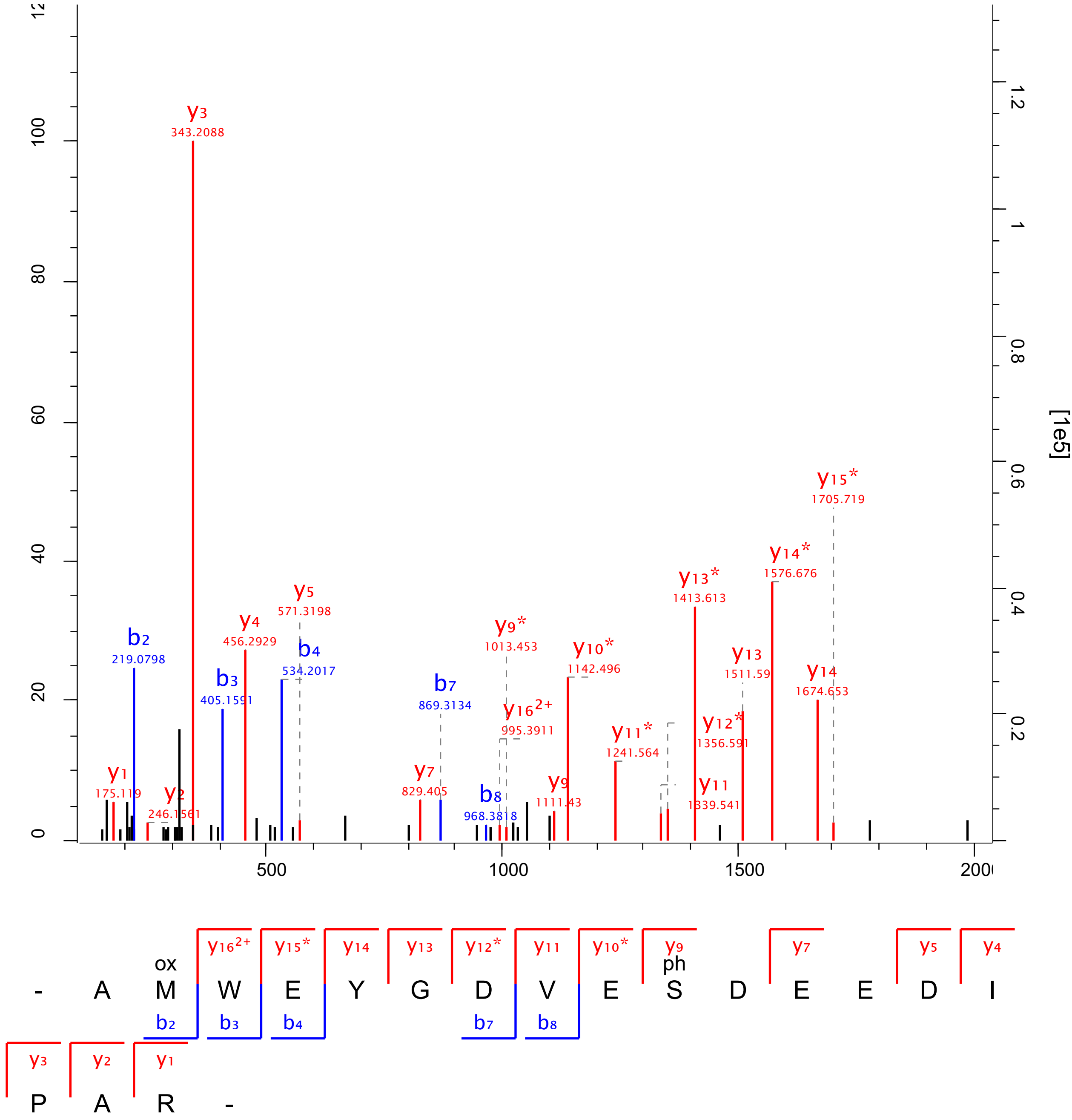

| Raw file | Scan | Method | Score | m/z |
|---|---|---|---|---|
| sys_15_1 | 34631 | FTMS; HCD | 42.71 | 884.07 |

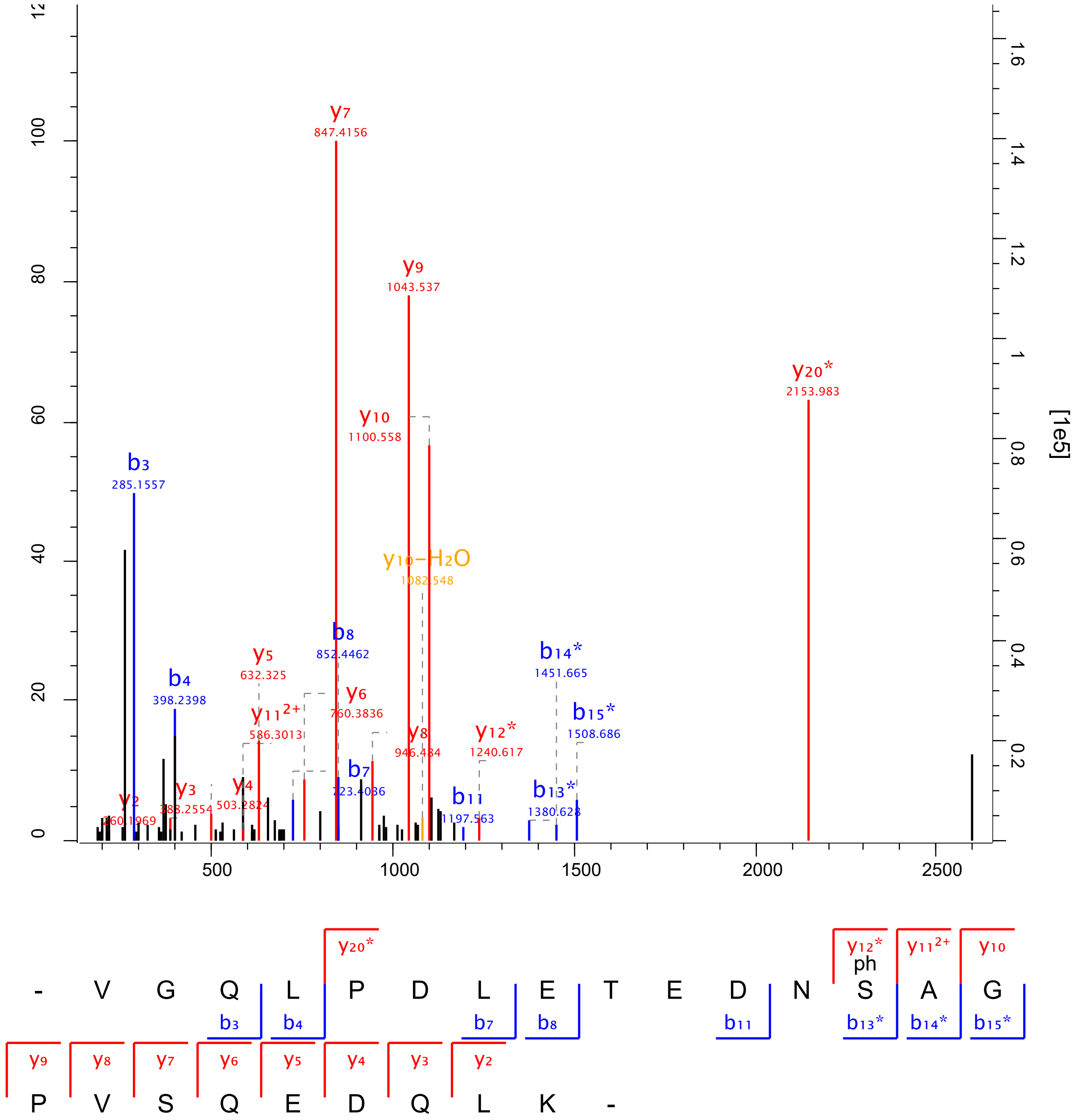

| Raw file | Scan | Method | Score | m/z |
| --- | --- | --- | --- | --- |
| sys_15_1 | 34723 | FTMS; HCD | 40.68 | 1030.13 |

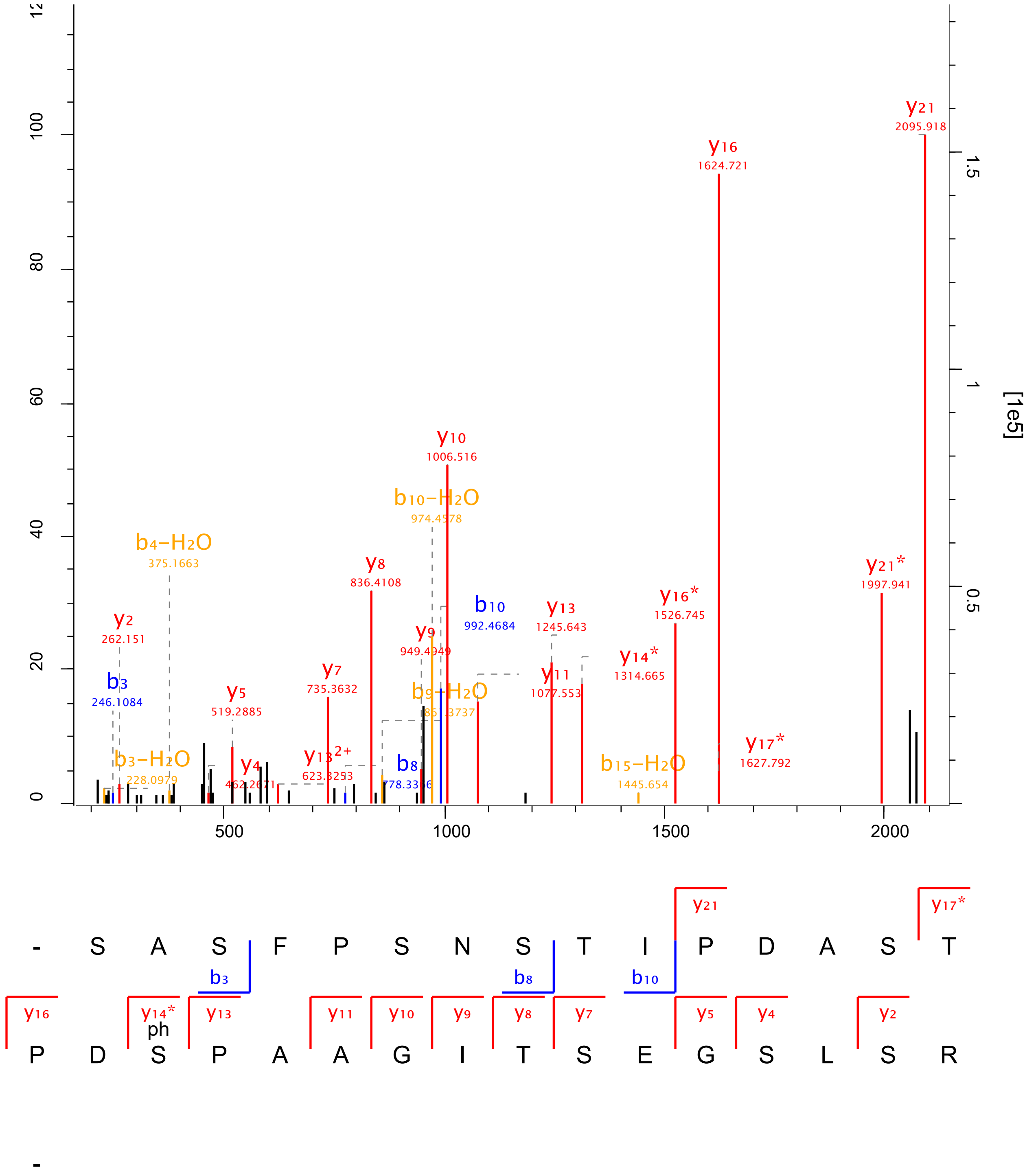

| Raw file | Scan | Method | Score | m/z |
|---|---|---|---|---|
| sys_15_1 | 34929 | FTMS; HCD | 182.76 | 819.88 |

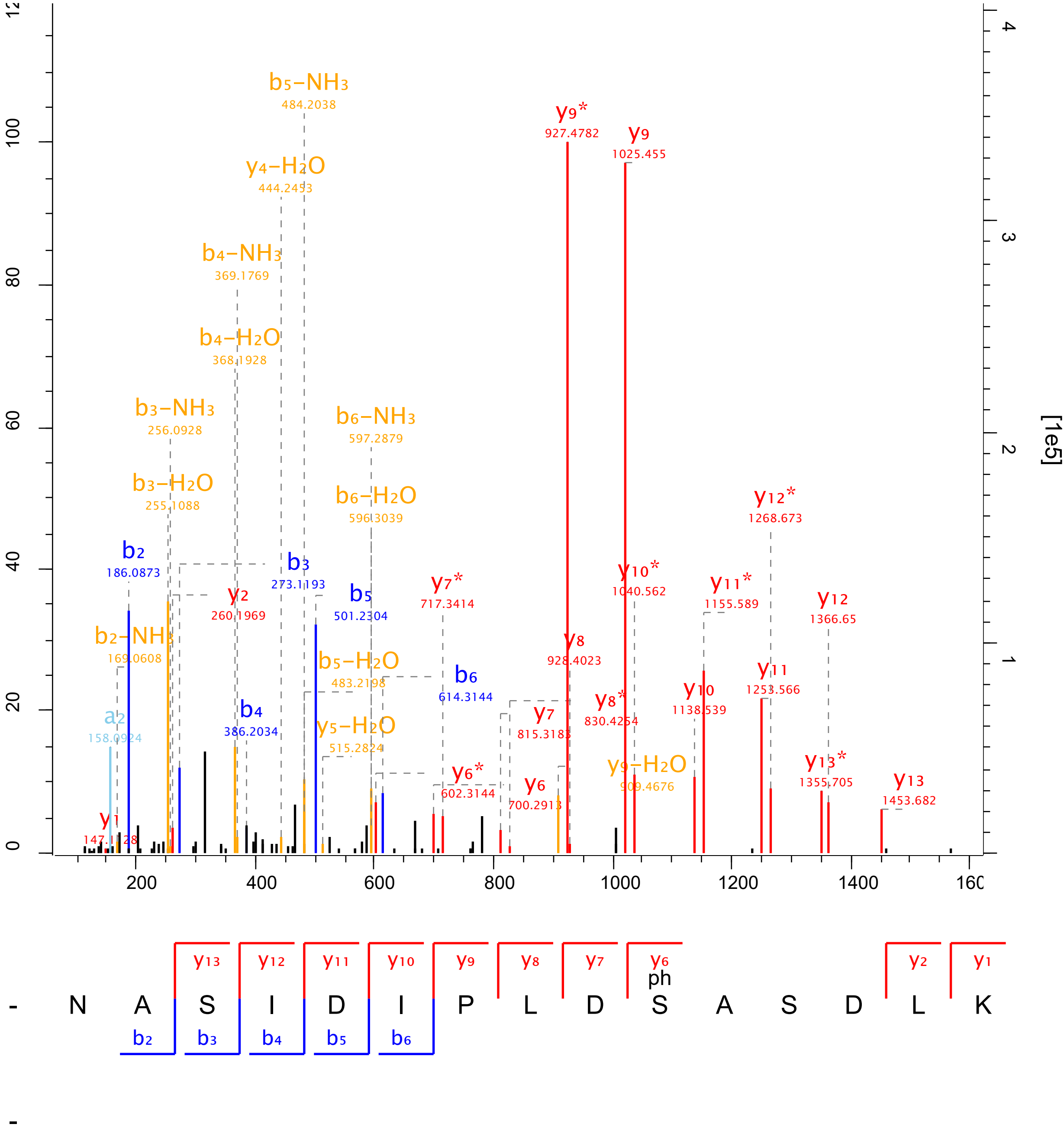

| Raw file | Scan | Method | Score | m/z |
|---|---|---|---|---|
| sys_15_1 | 34951 | FTMS; HCD | 134.64 | 906.4 |

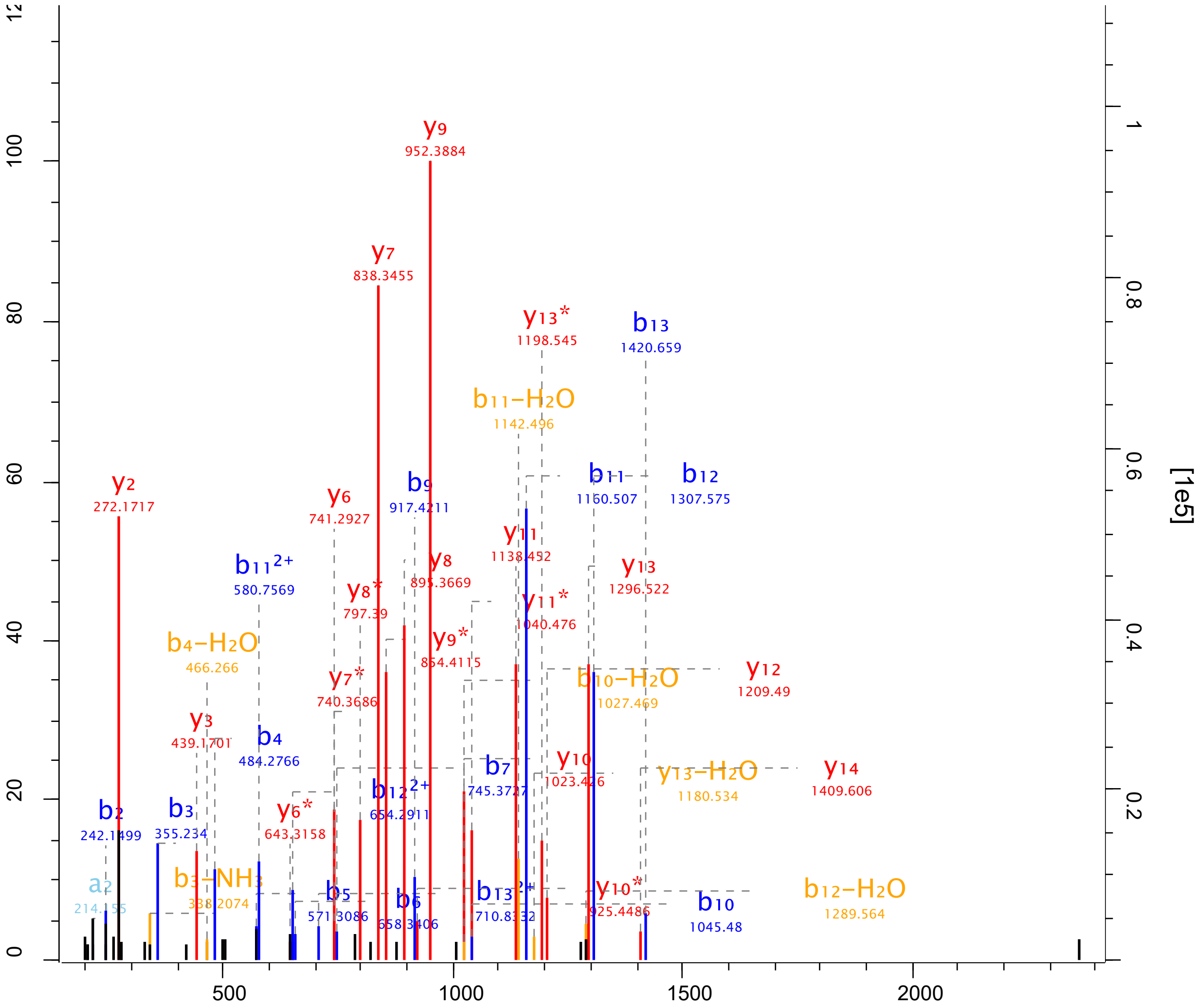


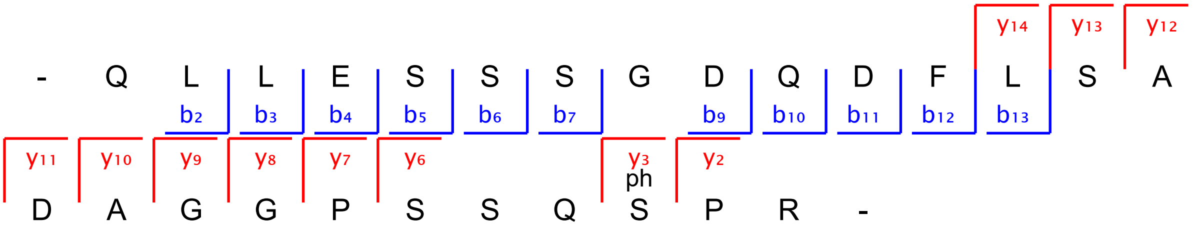

| Raw file | Scan | Method | Score | m/z |
| --- | --- | --- | --- | --- |
| sys_15_1 | 35311 | FTMS; HCD | 50.83 | 842.91 |

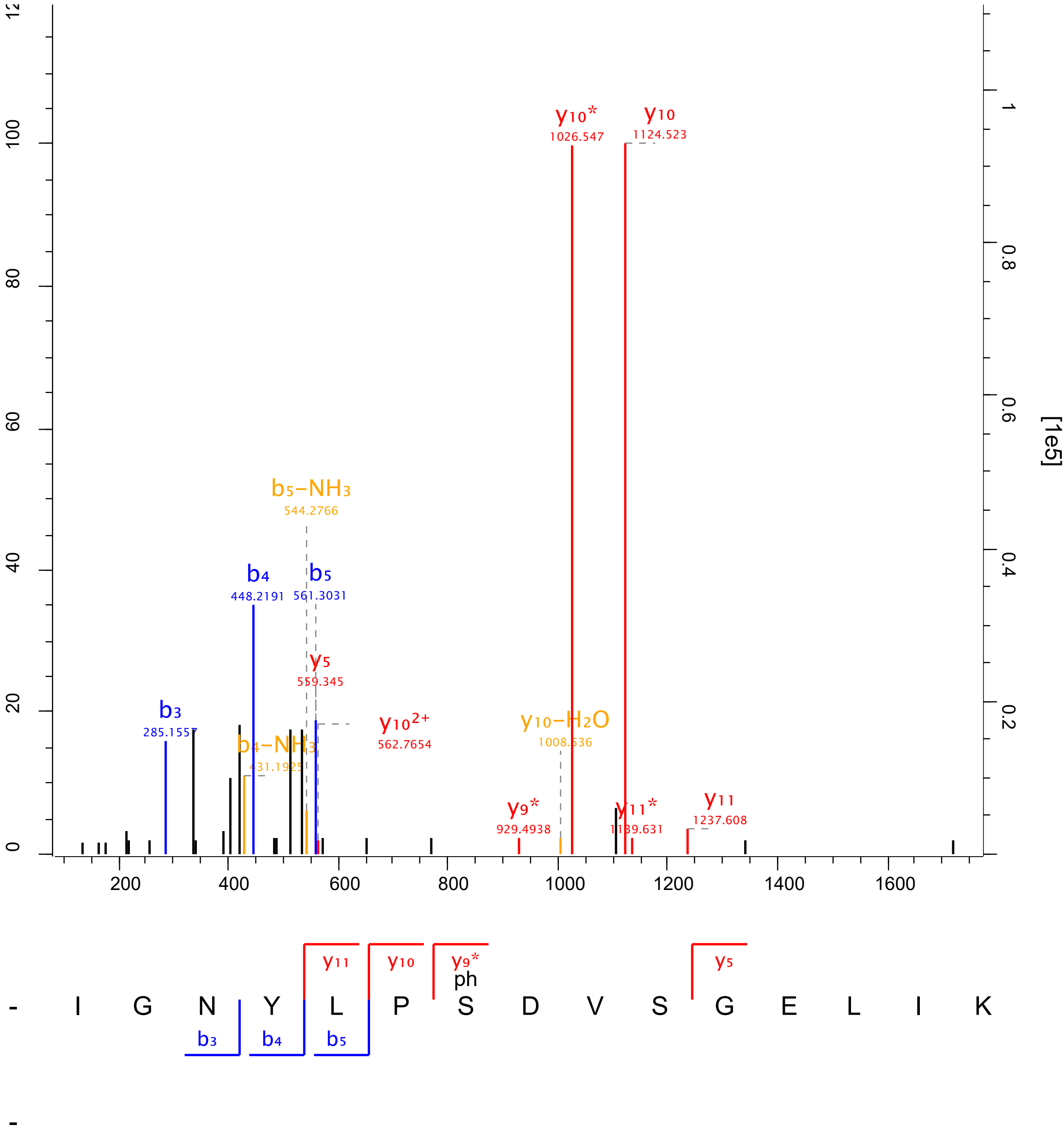

| Raw file | Scan | Method | Score | m/z |
|---|---|---|---|---|
| sys_15_1 | 35426 | FTMS; HCD | 109.65 | 926.92 |

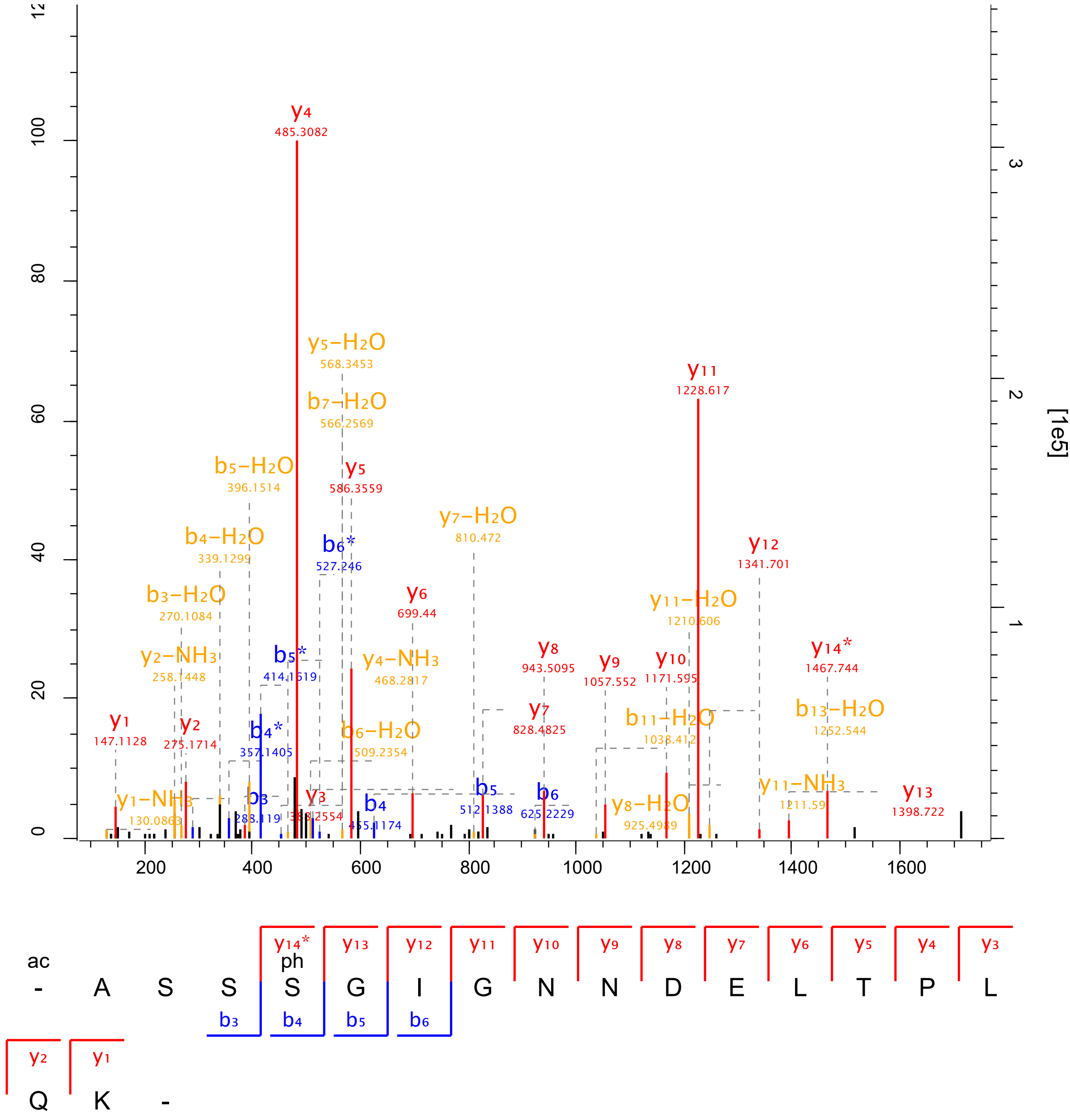

| Raw file | Scan | Method | Score | m/z |
|---|---|---|---|---|
| sys_15_1 | 35427 | FTMS; HCD | 57.43 | 647.78 |

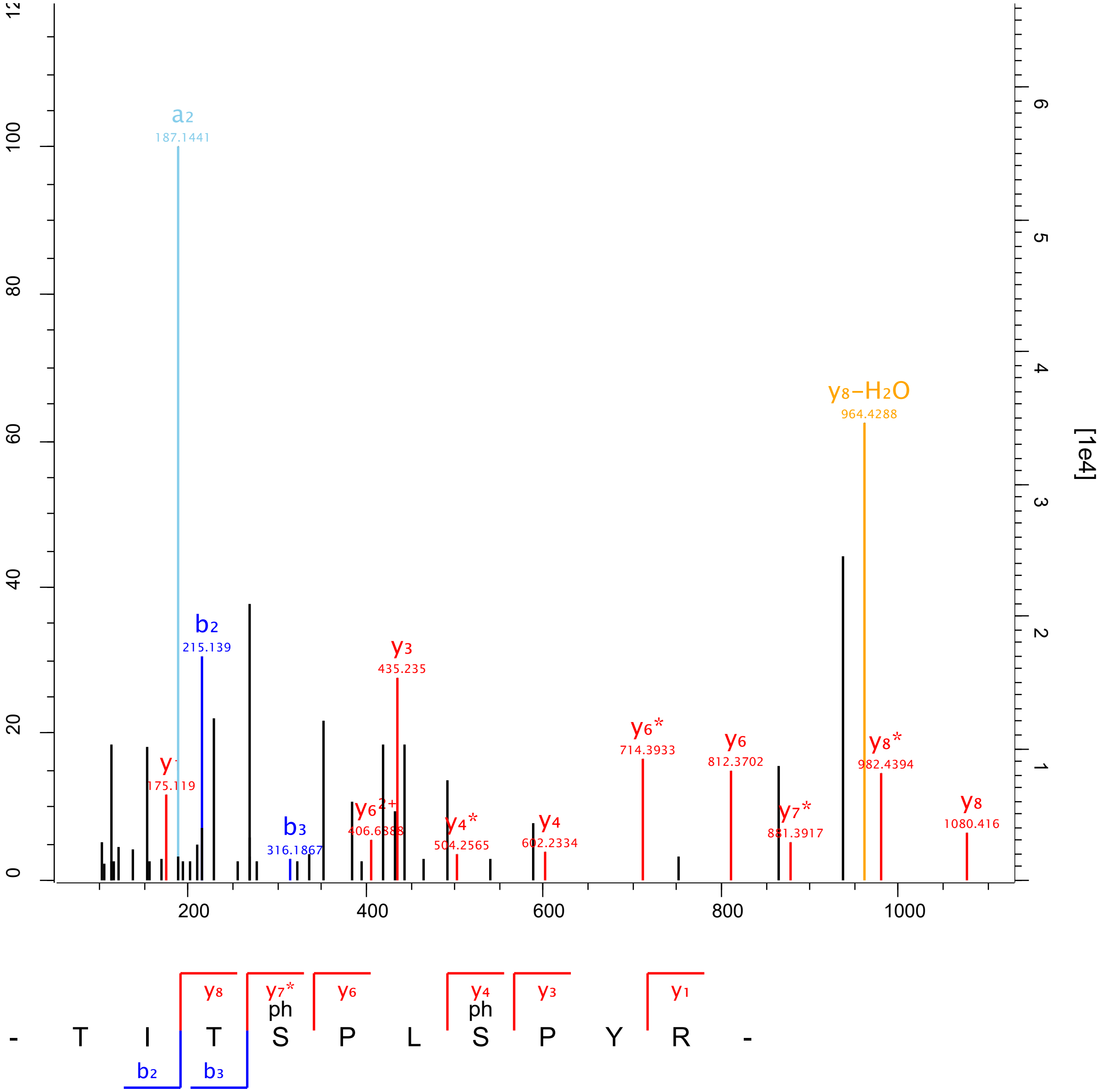

| Raw file | Scan | Method | Score | m/z |
|---|---|---|---|---|
| sys_15_1 | 35597 | FTMS; HCD | 159.46 | 1010.92 |

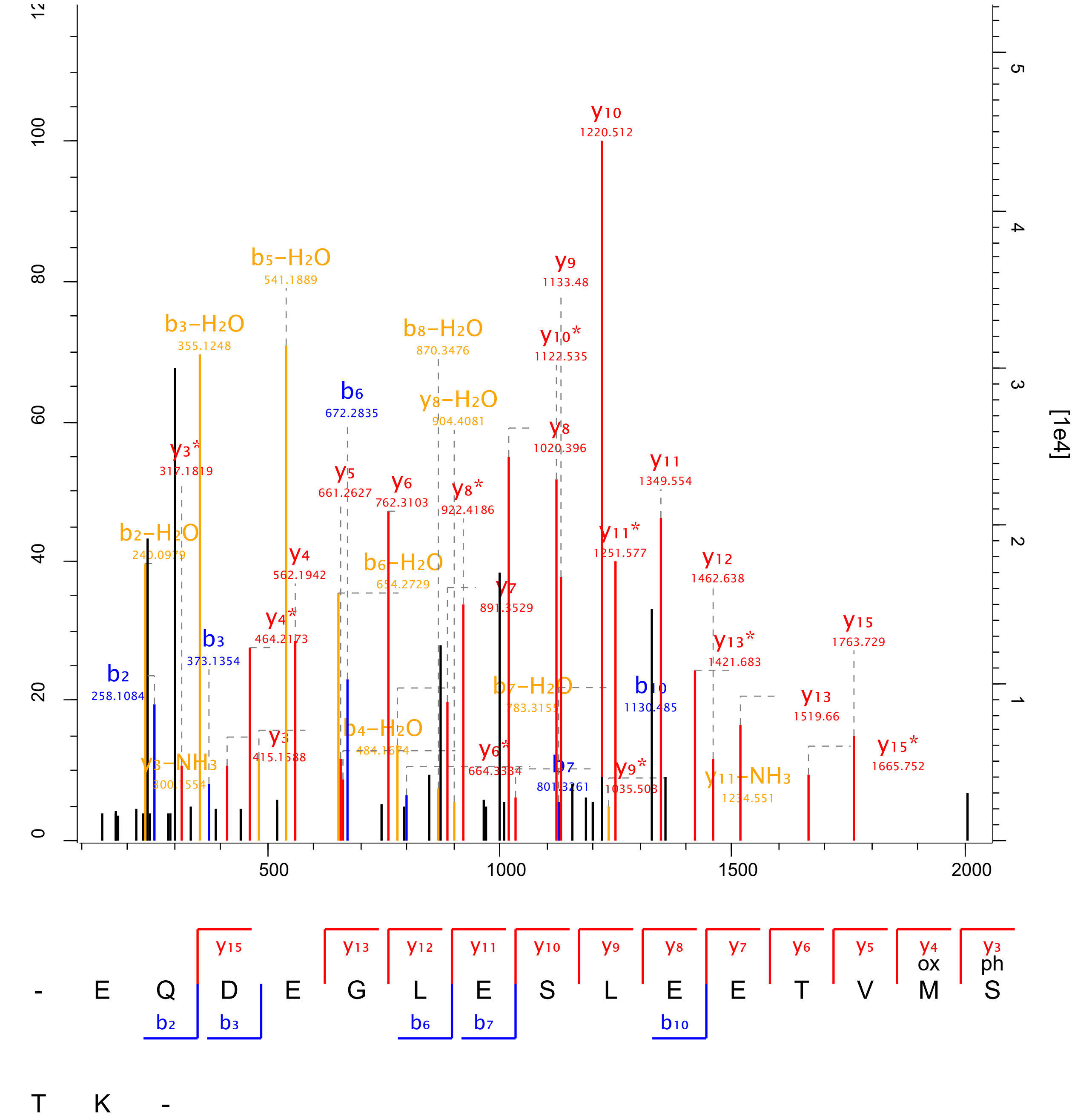

| Raw file | Scan | Method | Score | m/z |
| --- | --- | --- | --- | --- |
| sys_15_1 | 3560 | FTMS; HCD | 271.97 | 556.89 |

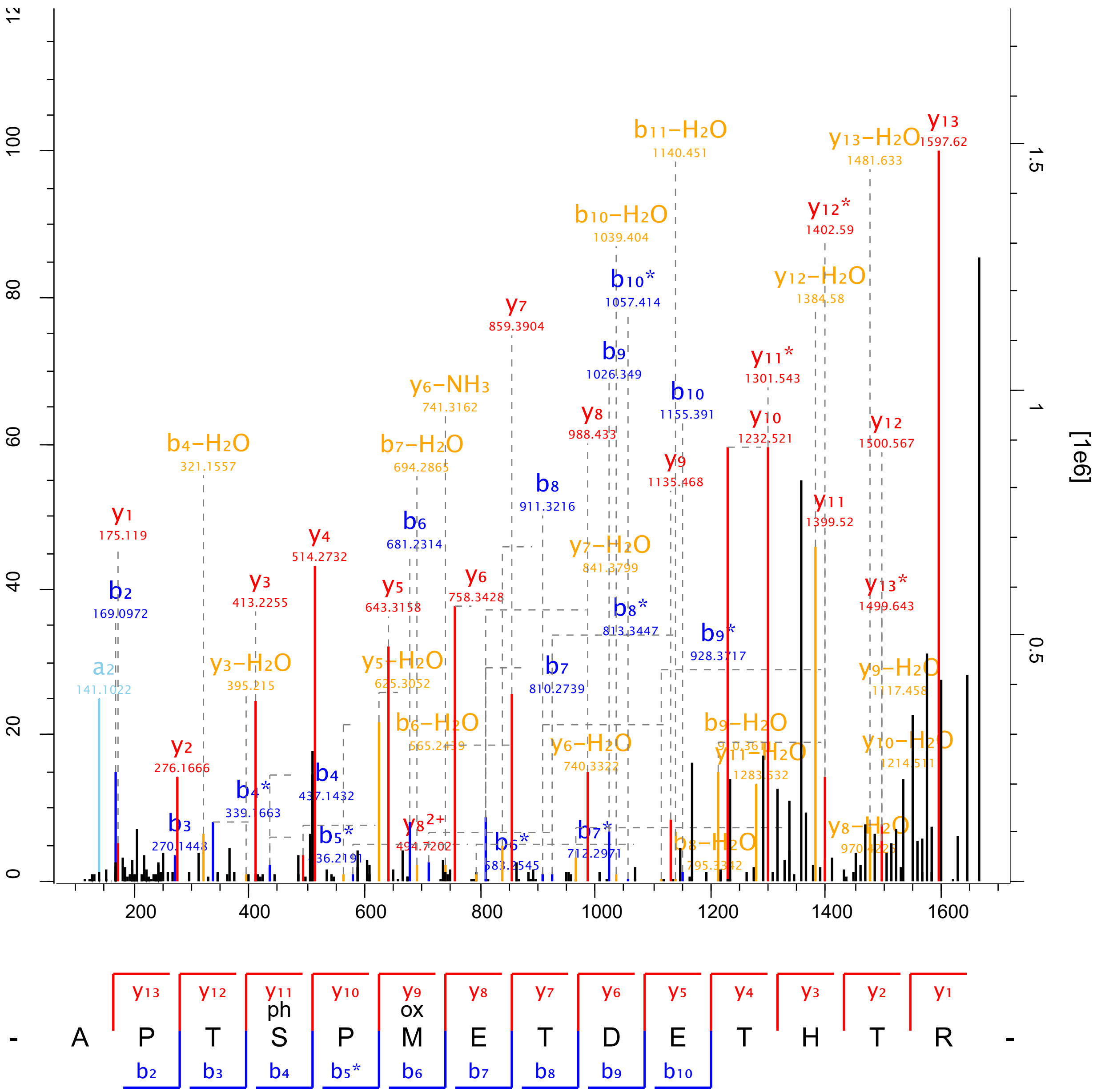

| Raw file | Scan | Method | Score | m/z |
| --- | --- | --- | --- | --- |
| sys_15_1 | 35600 | FTMS; HCD | 210.98 | 882.42 |

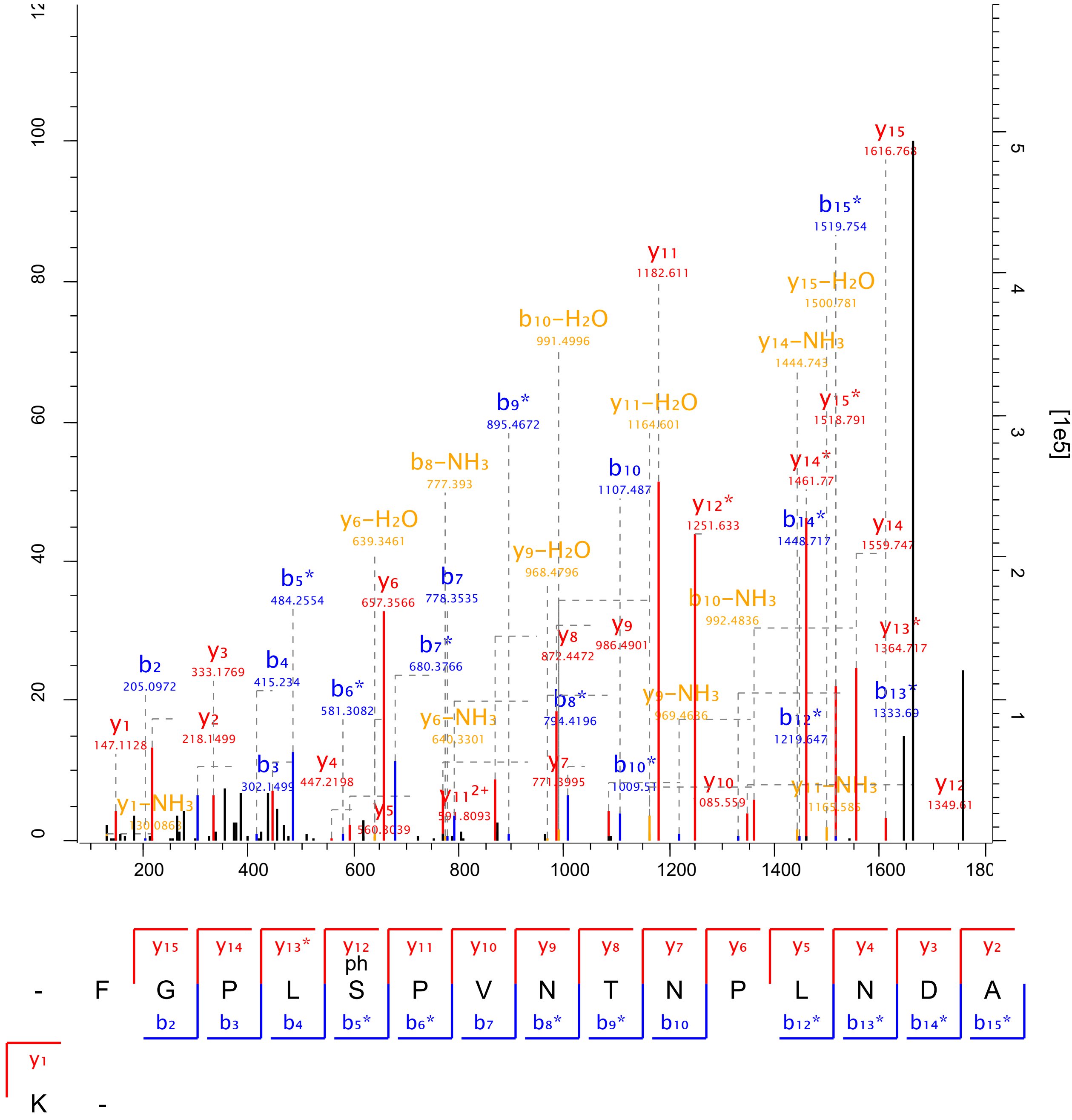

| Raw file | Scan | Method | Score | m/z |
| --- | --- | --- | --- | --- |
| sys_15_1 | 35649 | FTMS; HCD | 43.79 | 975.44 |

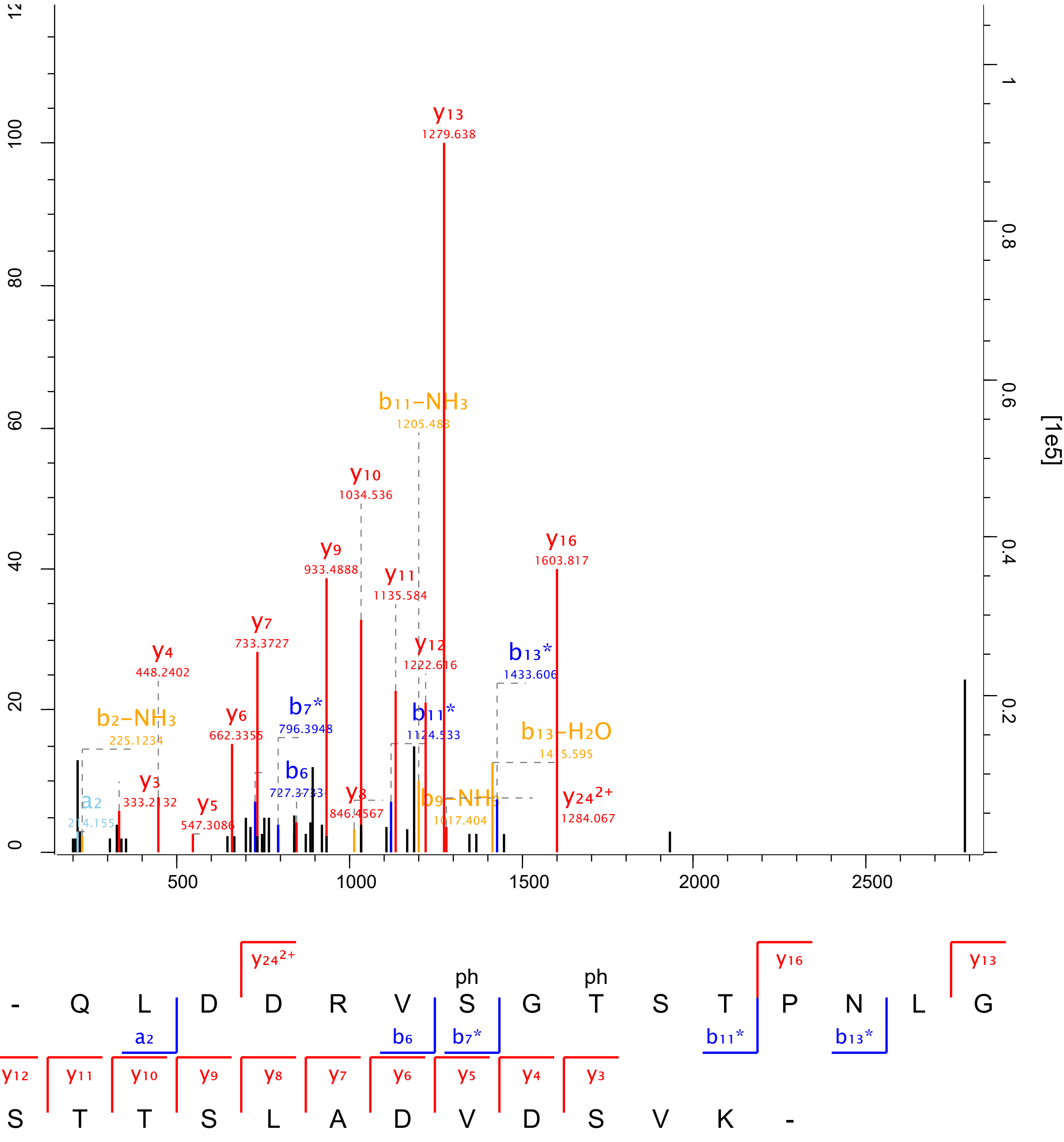

| Raw file | Scan | Method | Score | m/z |
| --- | --- | --- | --- | --- |
| sys_15_1 | 35985 | FTMS; HCD | 56.57 | 656.87 |

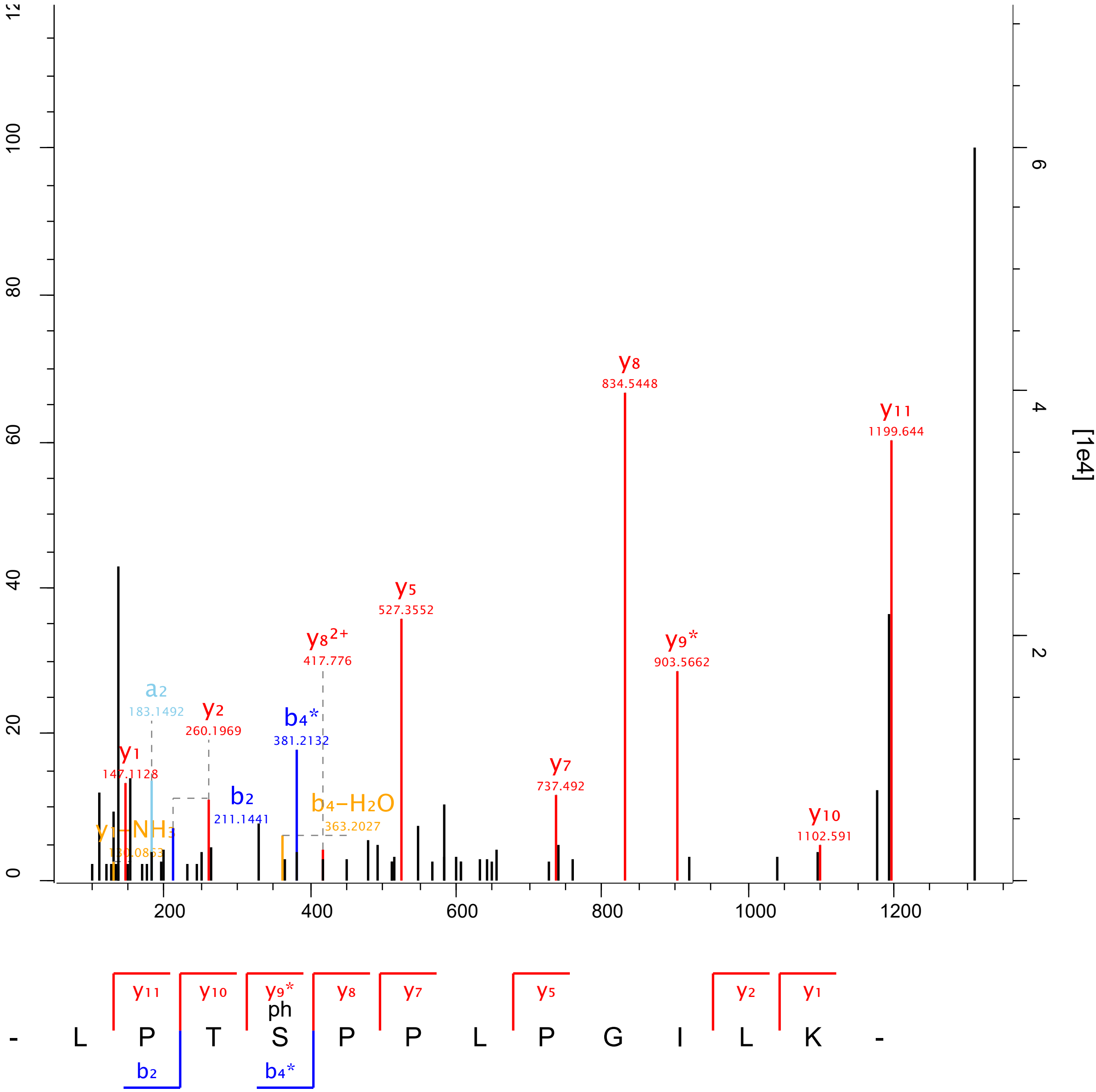

| Raw file | Scan | Method | Score | m/z |
| --- | --- | --- | --- | --- |
| sys_15_1 | 36049 | FTMS; HCD | 74.78 | 923.86 |

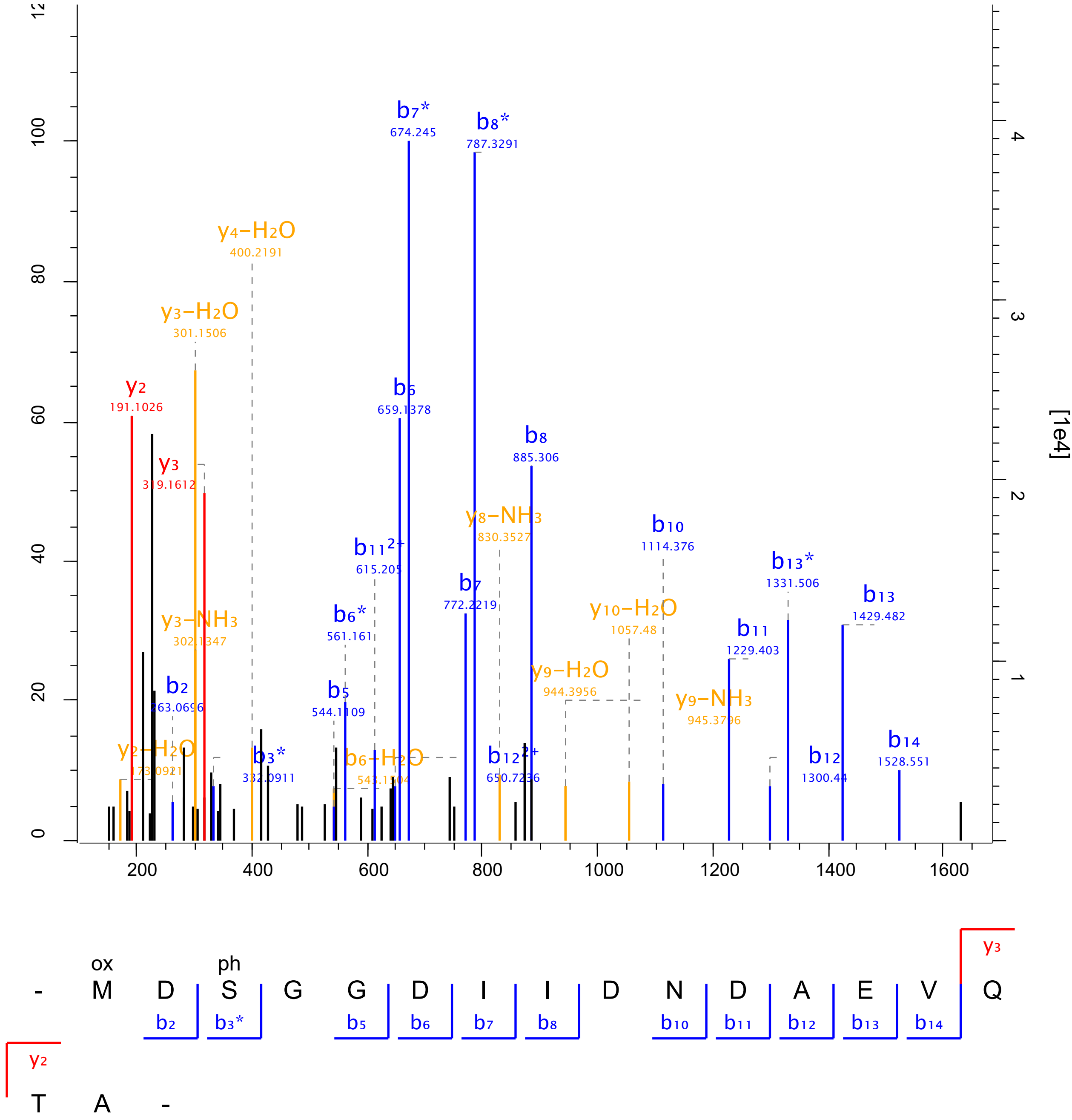

| Raw file | Scan | Method | Score | m/z |
|---|---|---|---|---|
| sys_15_1 | 36225 | FTMS; HCD | 50.36 | 750.65 |

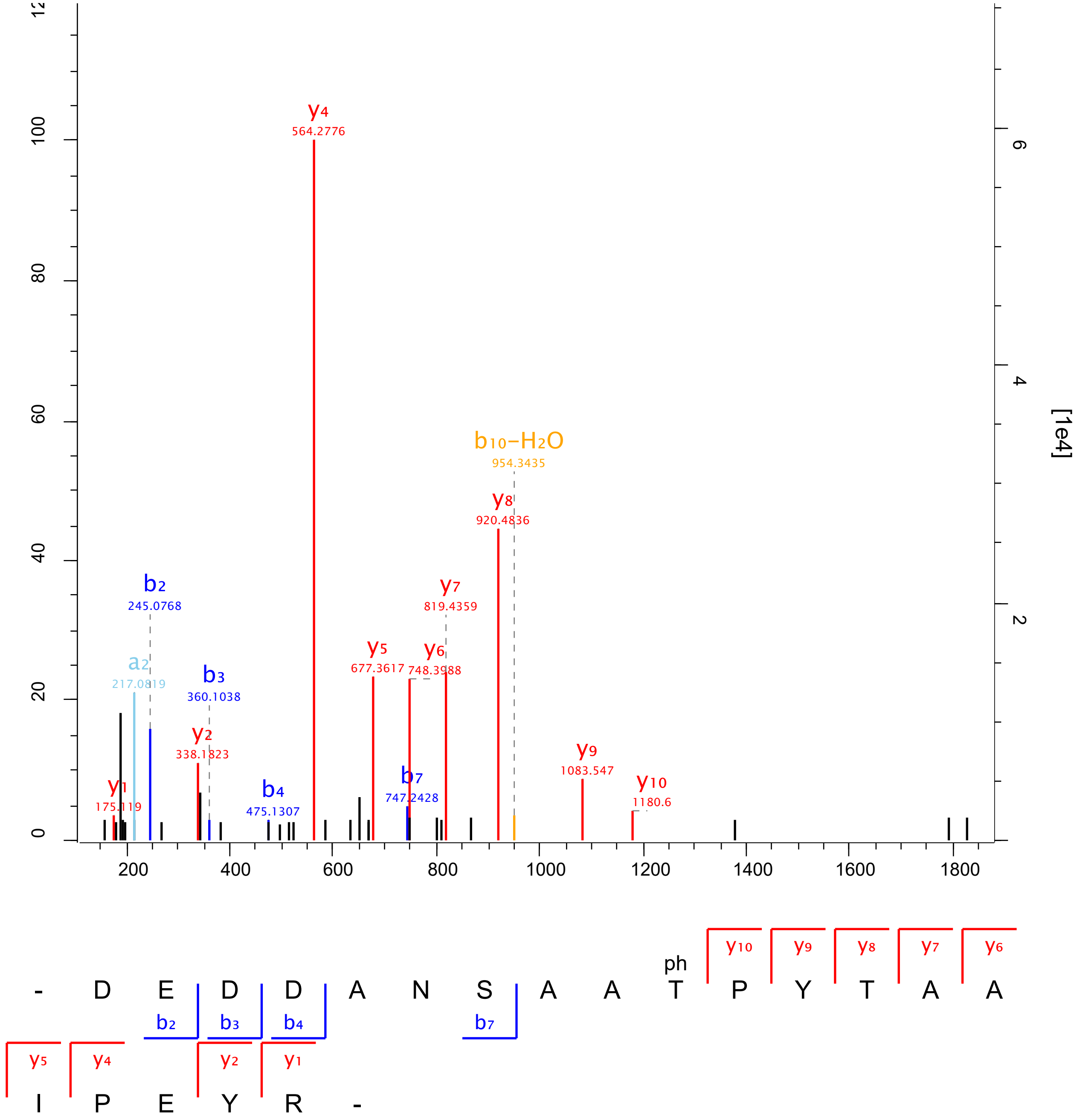

| Raw file | Scan | Method | Score | m/z |
|---|---|---|---|---|
| sys_15_1 | 36288 | FTMS; HCD | 79.51 | 823.06 |

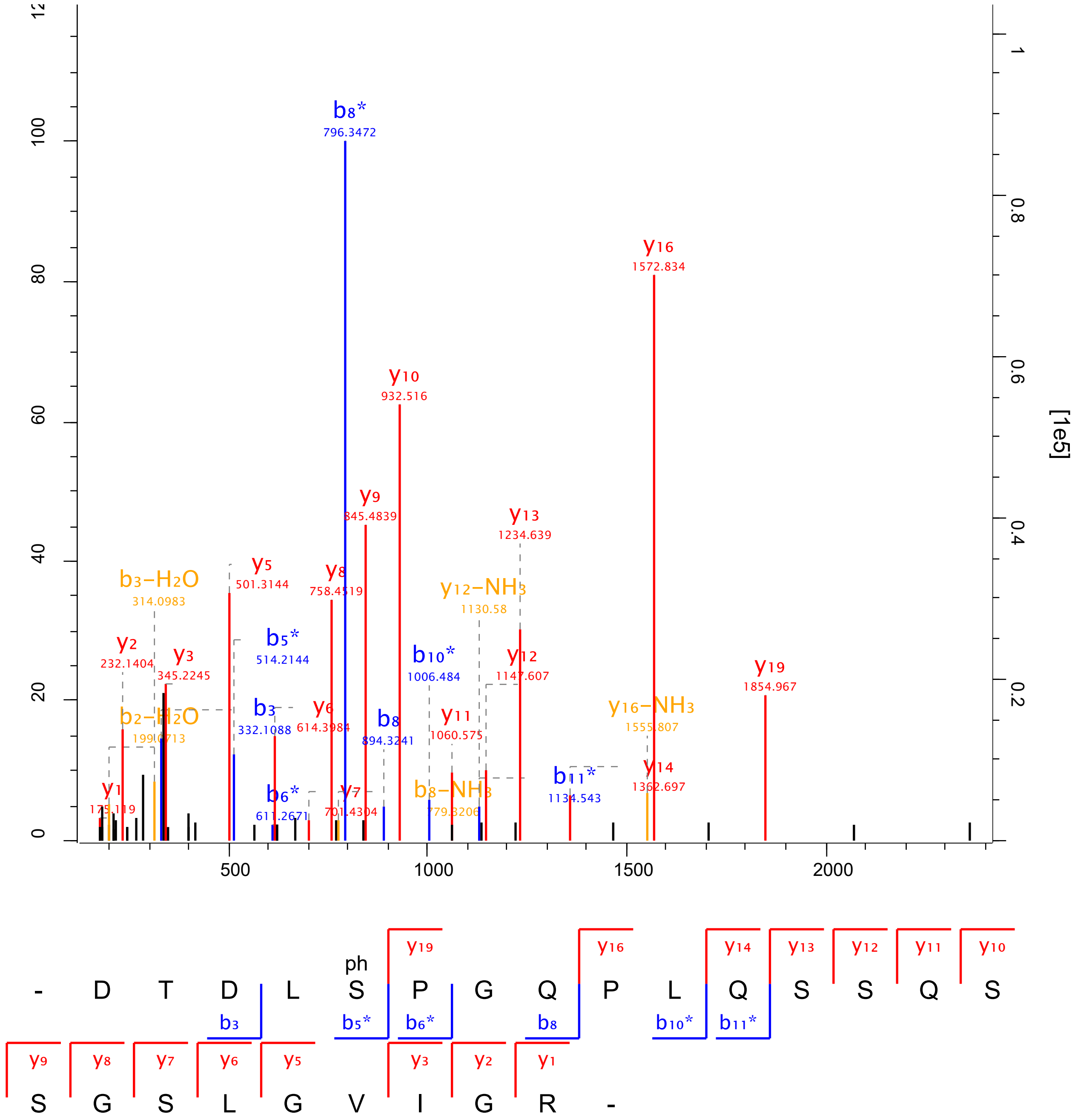

| Raw file | Scan | Method | Score | m/z |
| --- | --- | --- | --- | --- |
| sys_15_1 | 36340 | FTMS; HCD | 144.7 | 542.76 |

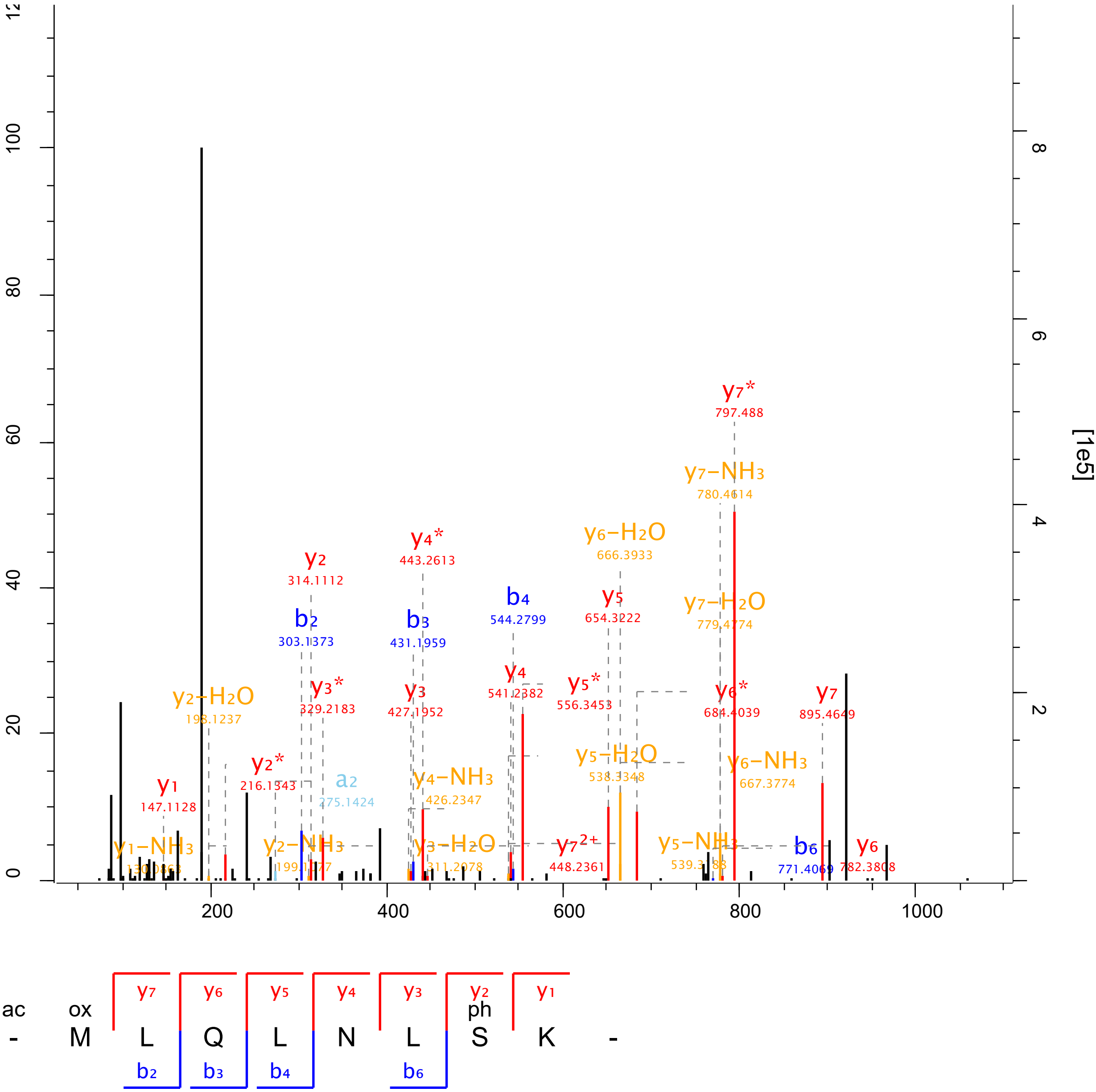

| Raw file | Scan | Method | Score | m/z |
|---|---|---|---|---|
| sys_15_1 | 36373 | FTMS; HCD | 55.13 | 848.4 |

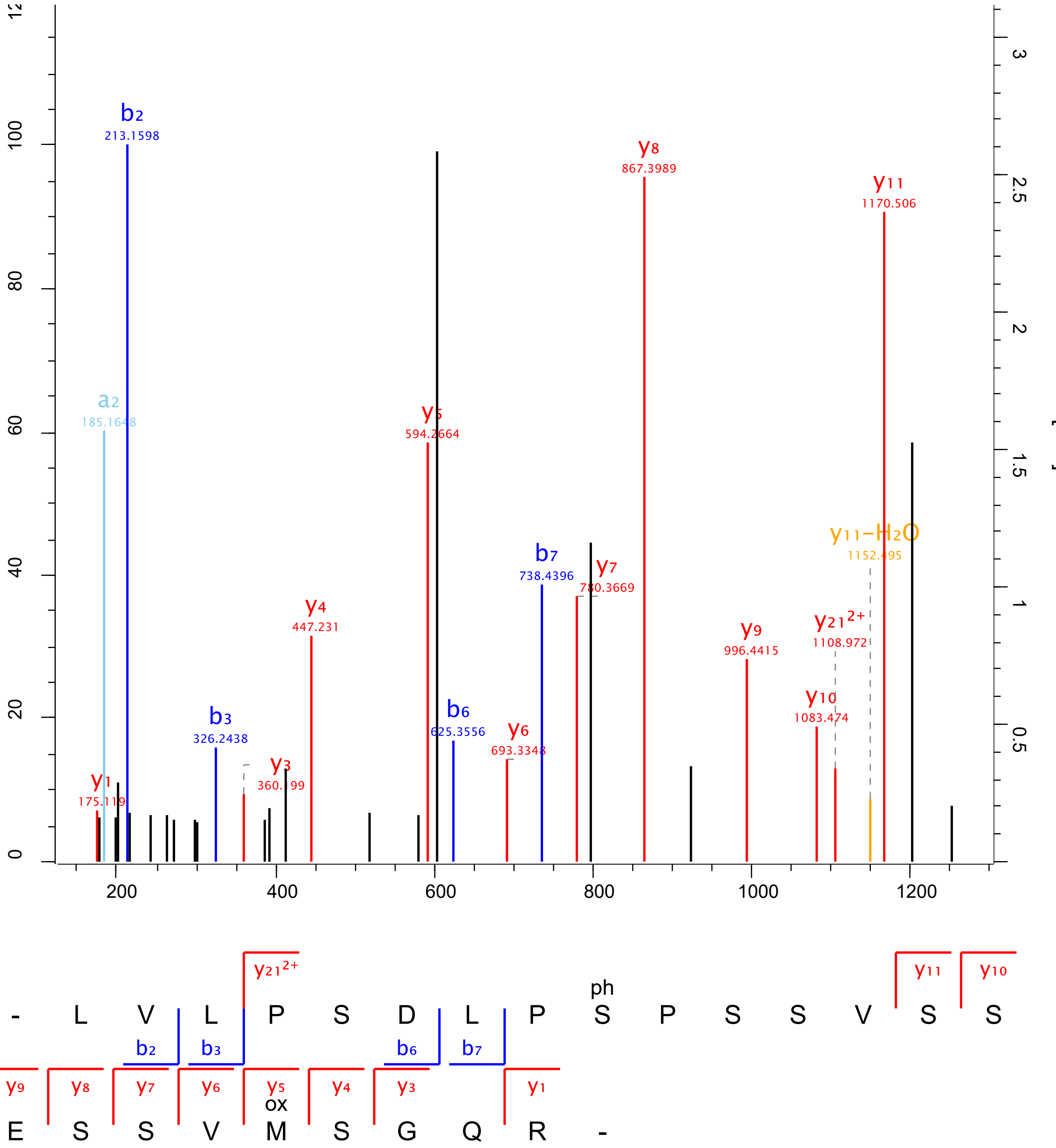

| Raw file | Scan | Method | Score | m/z |
|---|---|---|---|---|
| sys_15_1 | 3644 | FTMS; HCD | 115.29 | 536.73 |

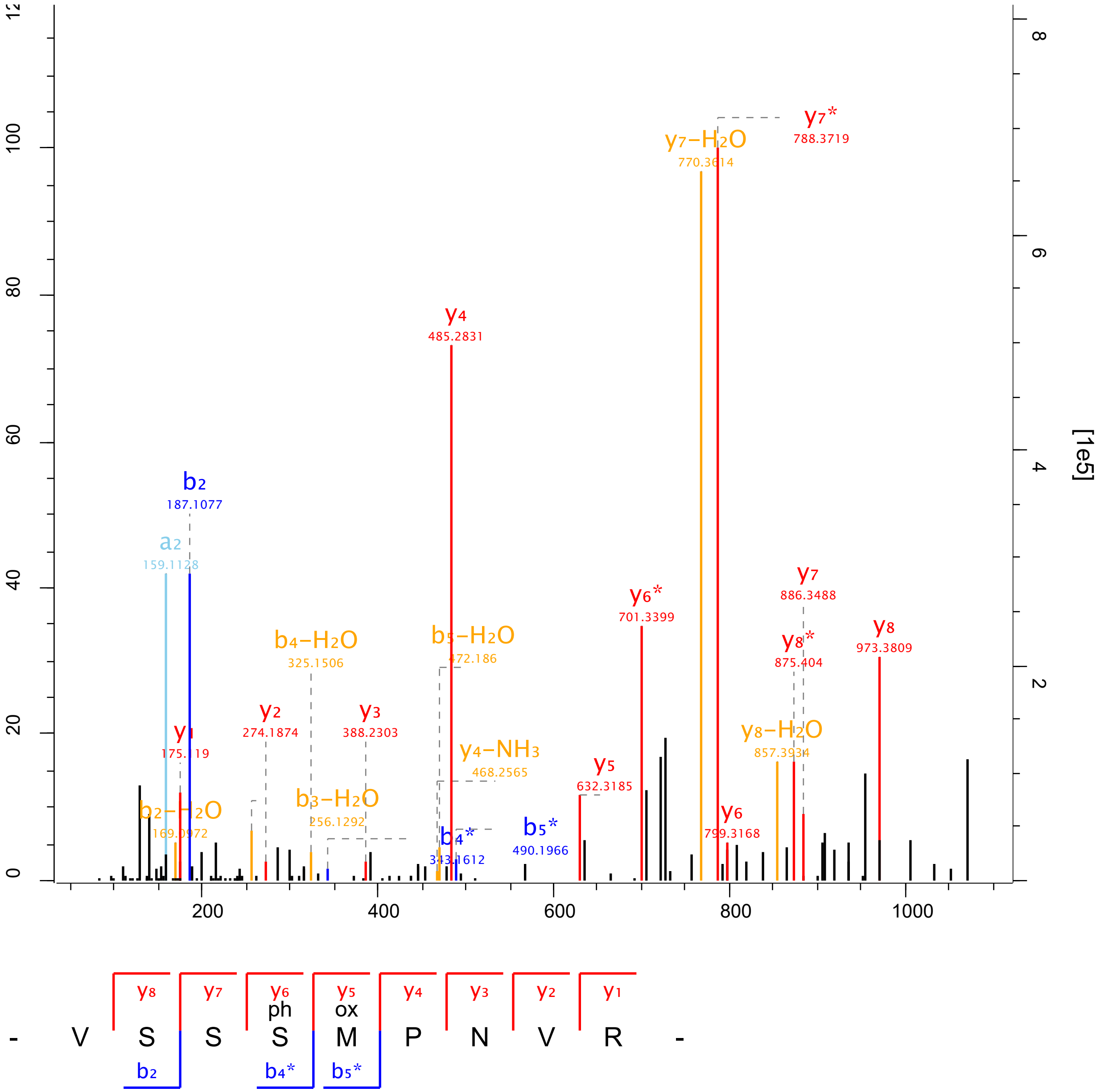

| Raw file | Scan | Method | Score | m/z |
|---|---|---|---|---|
| sys_15_1 | 36442 | FTMS; HCD | 55.59 | 956.45 |

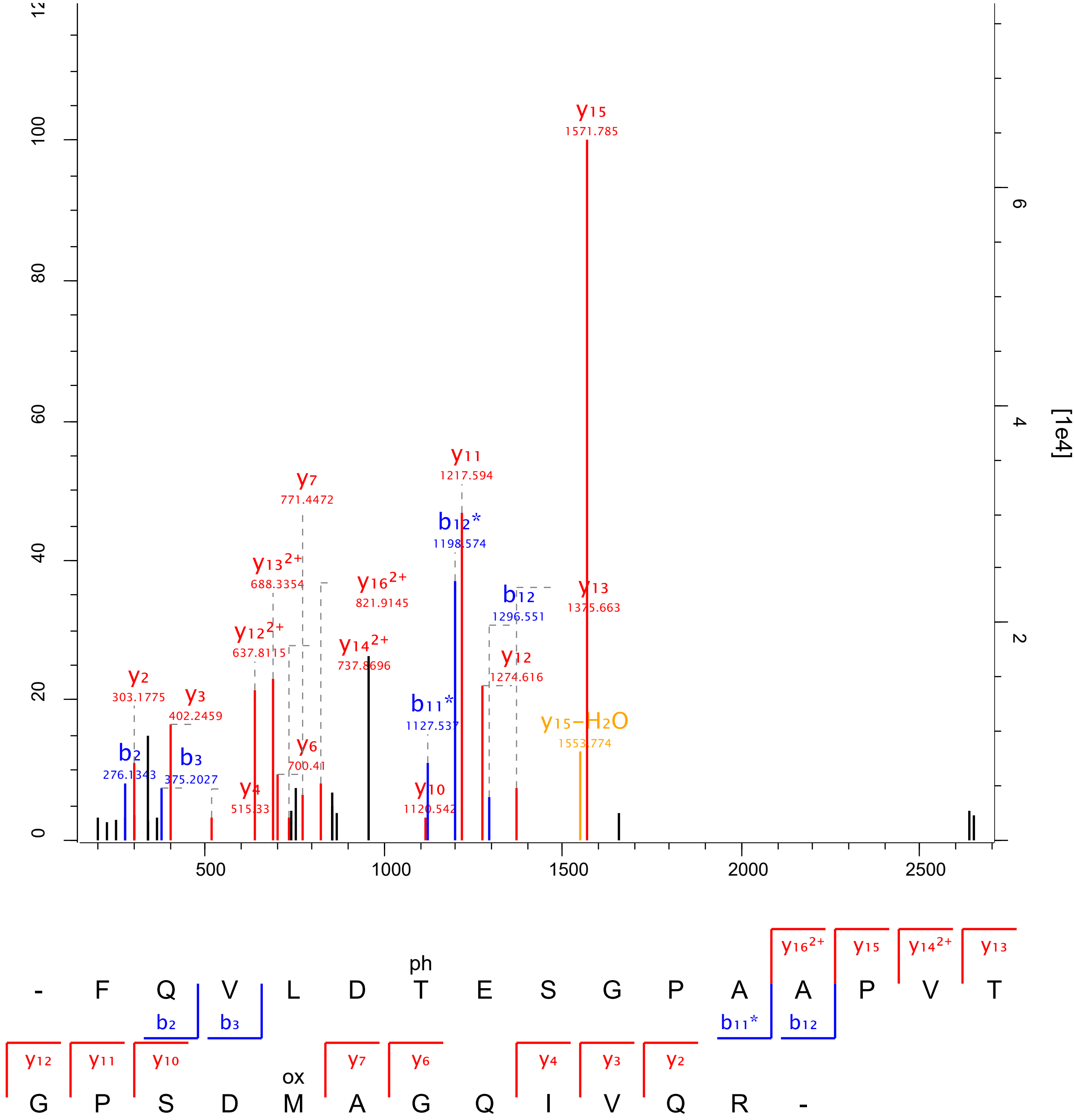

| Raw file | Scan | Method | Score | m/z |
|---|---|---|---|---|
| sys_15_1 | 36610 | FTMS; HCD | 102.65 | 722.82 |

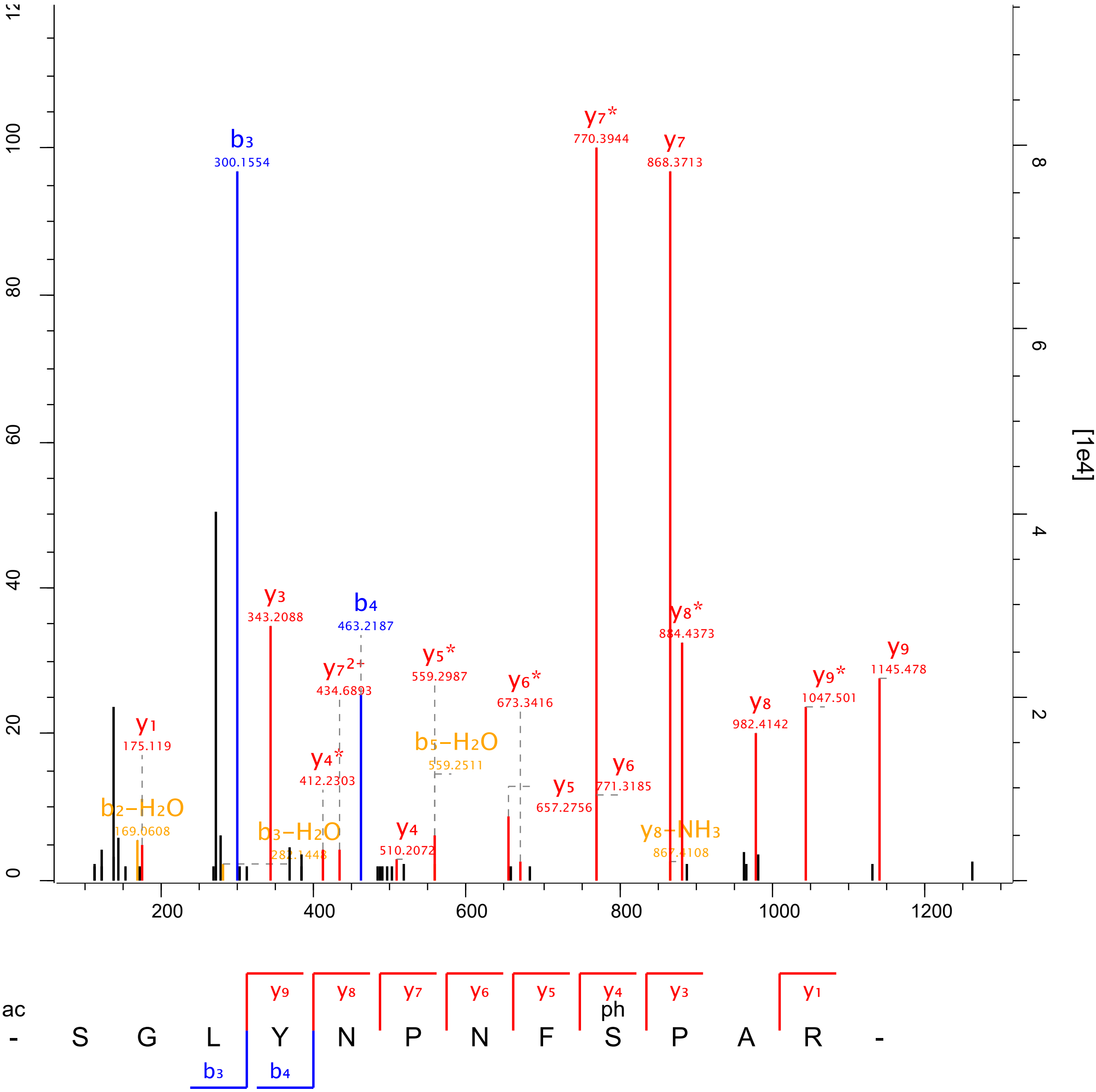

| Raw file | Scan | Method | Score | m/z |
|---|---|---|---|---|
| sys_15_1 | 3663 | FTMS; HCD | 76.36 | 583.75 |

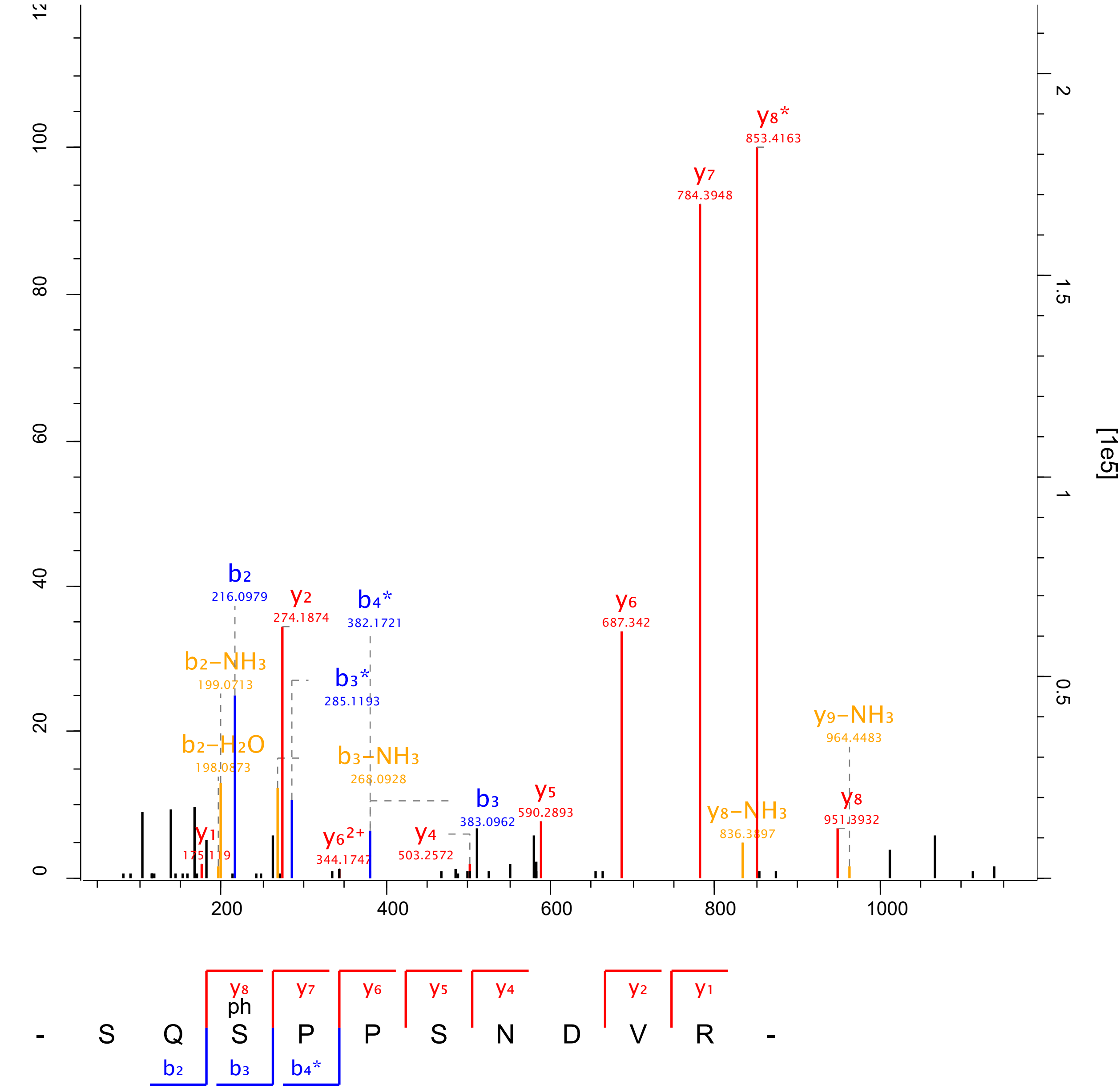

| Raw file | Scan | Method | Score | m/z |
| --- | --- | --- | --- | --- |
| sys_15_1 | 36782 | FTMS; HCD | 85.36 | 778.89 |

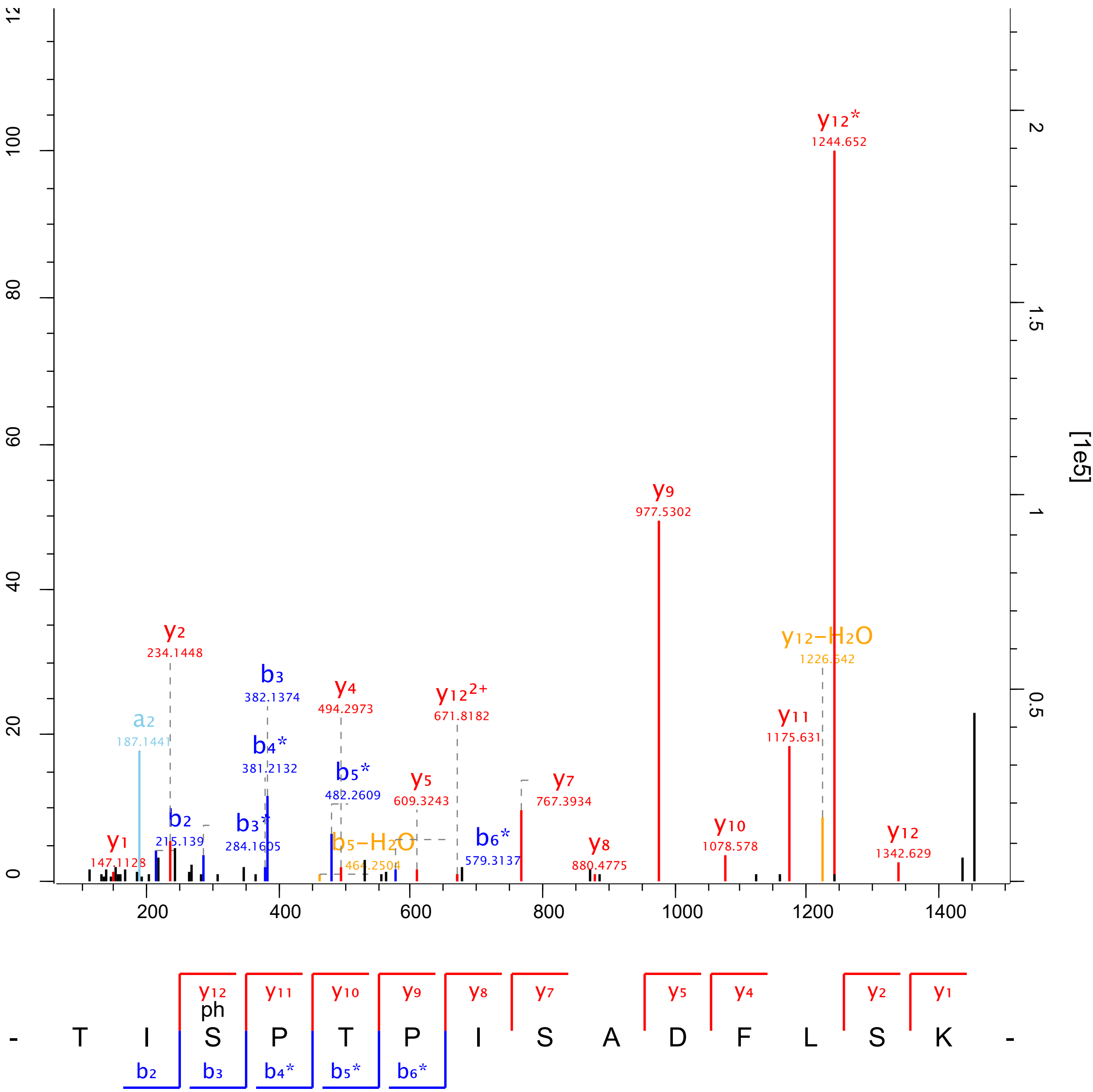

| Raw file | Scan | Method | Score | m/z |
| --- | --- | --- | --- | --- |
| sys_15_1 | 36785 | FTMS; HCD | 227.49 | 889.69 |

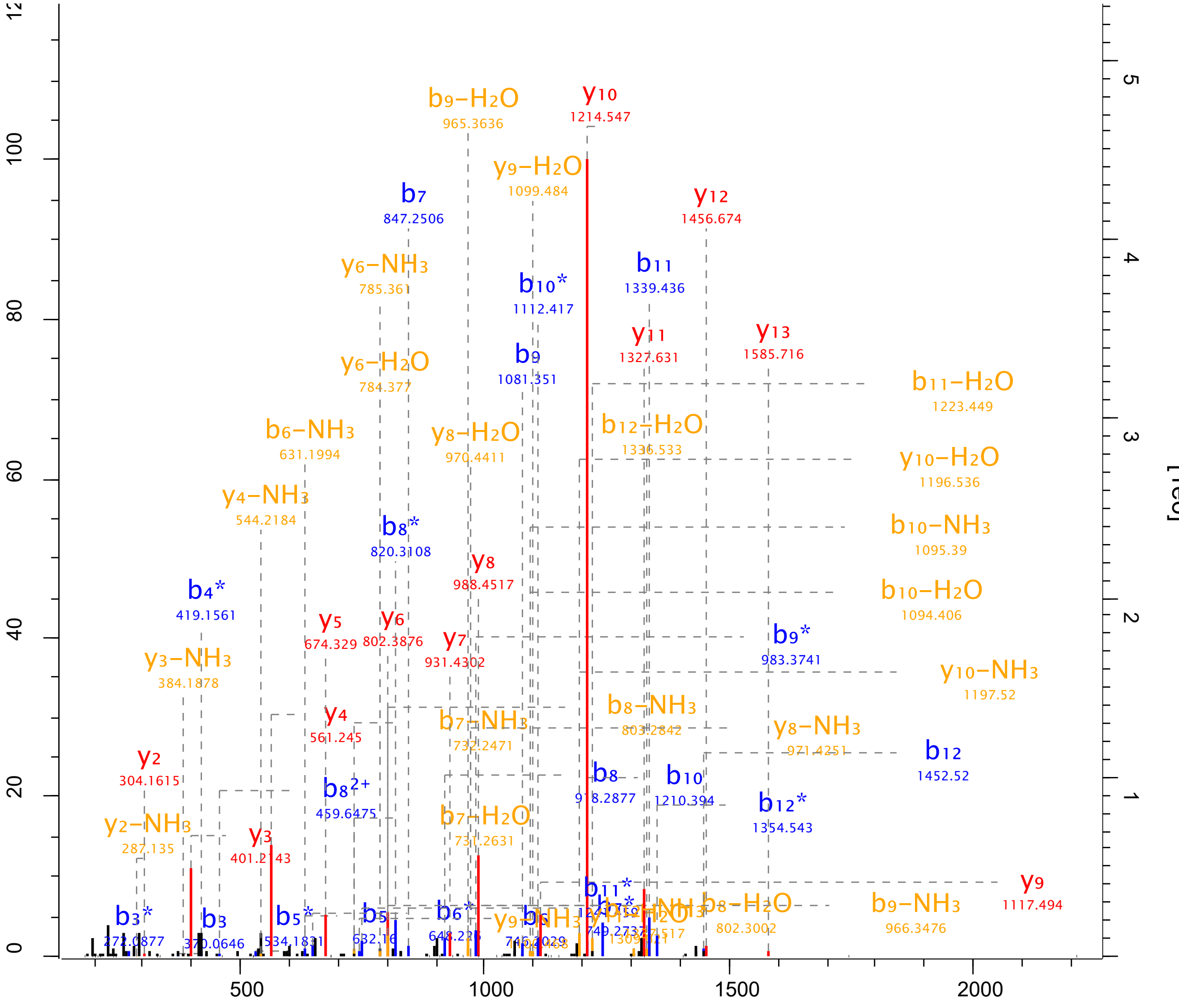


| | | | | | | | | | | | | | | |
| --- | --- | --- | --- | --- | --- | --- | --- | --- | --- | --- | --- | --- | --- | --- |
| | | | | | | | | | | y13 | y12 | y11 | y10 | y9 | y8 |
| | ph | | | | | | | | | | | | | | |
| - | S | D | S | F | D | N | T | A | Y | E | E | L | P | E | G |
| | | | b3 | b4* | b5 | b6 | b7 | b8 | b9 | b10 | b11 | b12 | | | |
| y7 | y6 | y5 | y4 | y3 | y2 | | | | | | | | | | |
| E | Q | I | C | P | E | R | - | | | | | | | | |

| Raw file | Scan | Method | Score | m/z |
| --- | --- | --- | --- | --- |
| sys_15_1 | 36852 | FTMS; HCD | 104.18 | 858.39 |

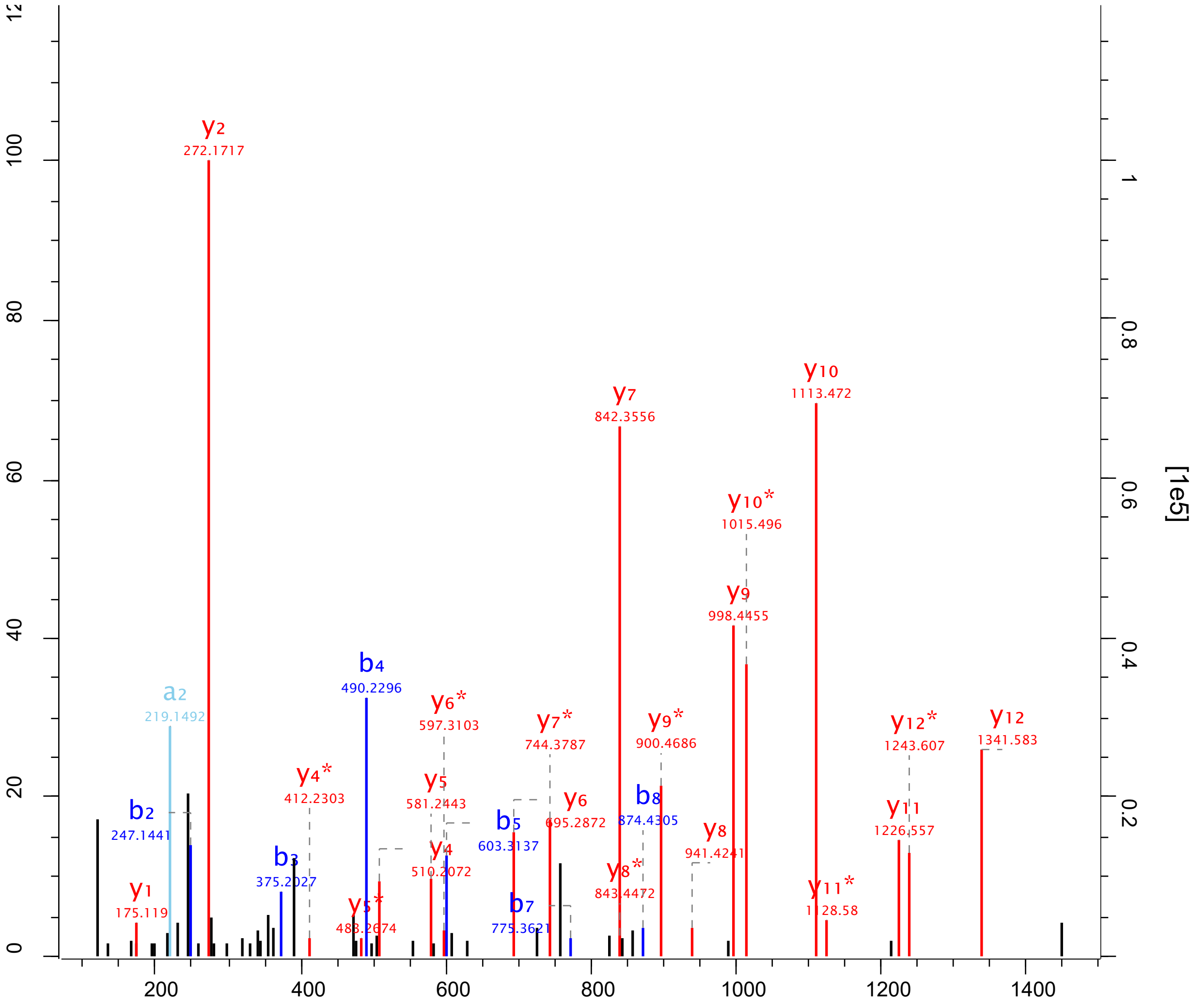


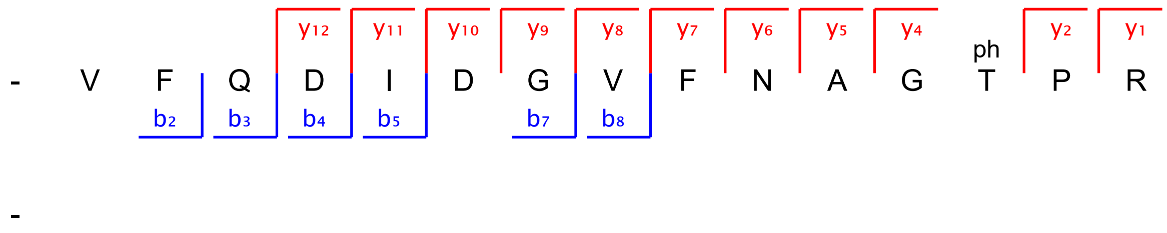

| Raw file | Scan | Method | Score | m/z |
|---|---|---|---|---|
| sys_15_1 | 36975 | FTMS; HCD | 137.73 | 843.4 |

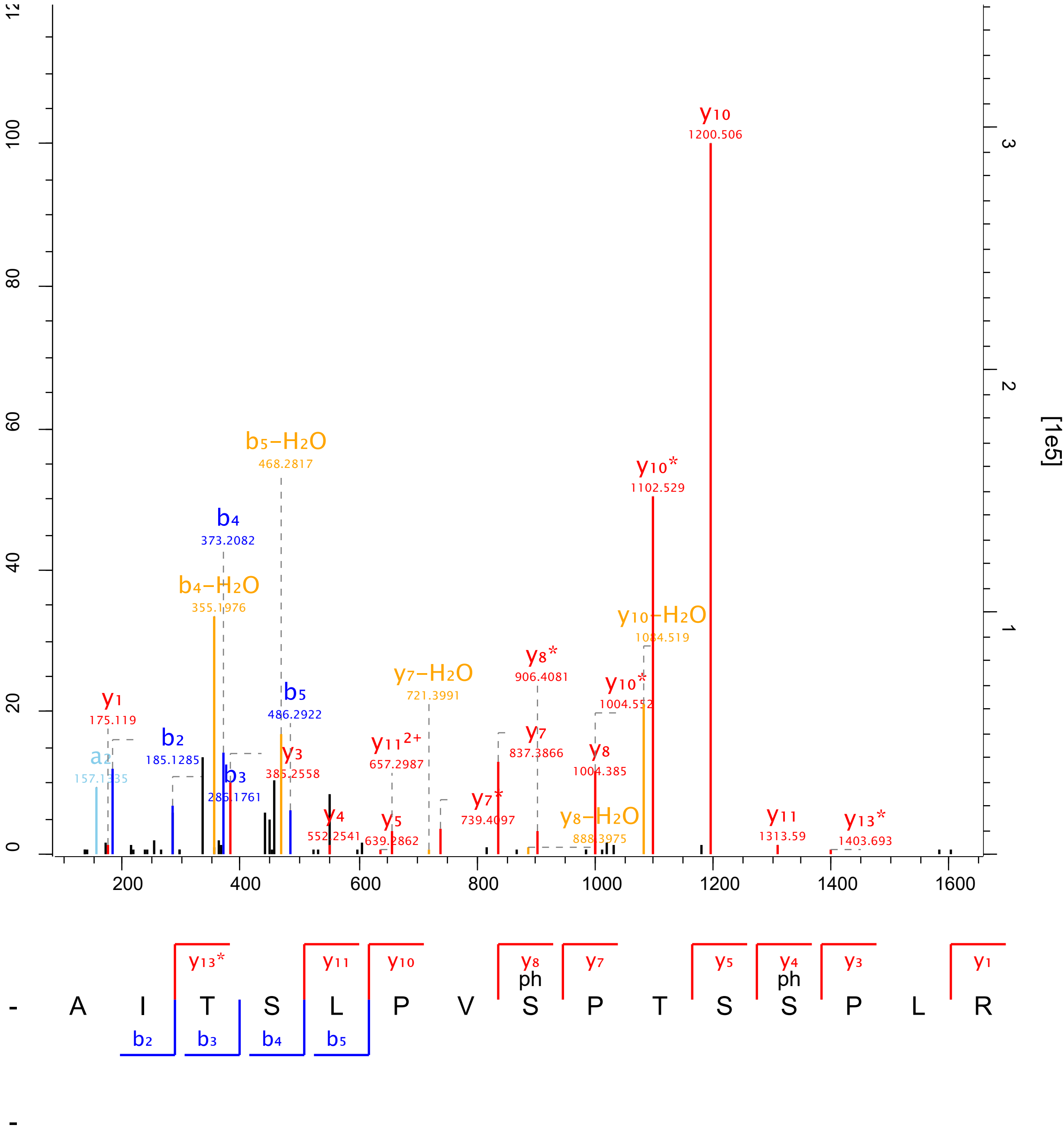

| Raw file | Scan | Method | Score | m/z |
|---|---|---|---|---|
| sys_15_1 | 3705 | FTMS; HCD | 50.35 | 476.72 |

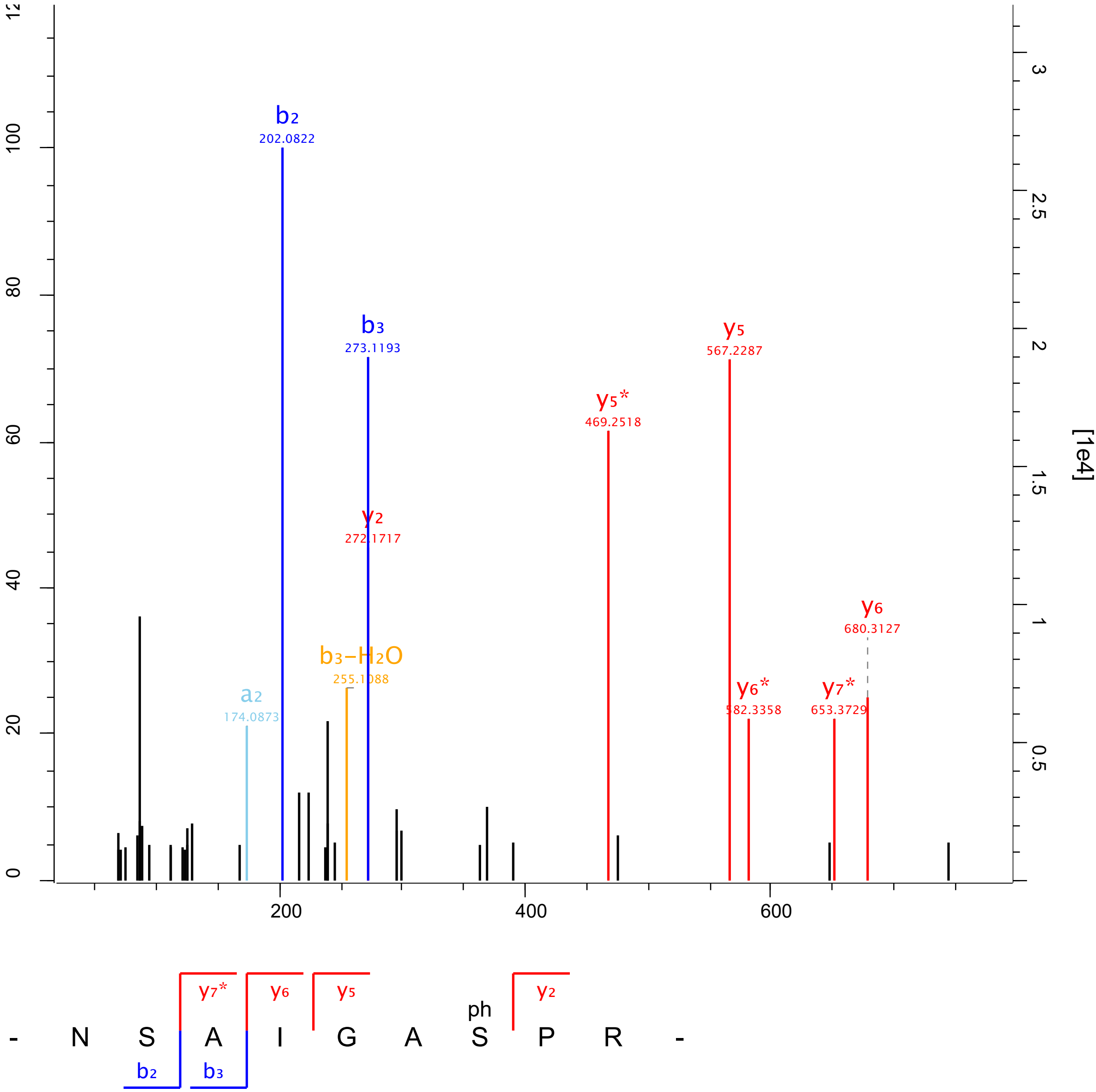

| Raw file | Scan | Method | Score | m/z |
|---|---|---|---|---|
| sys_15_1 | 37280 | FTMS; HCD | 141.45 | 986 |

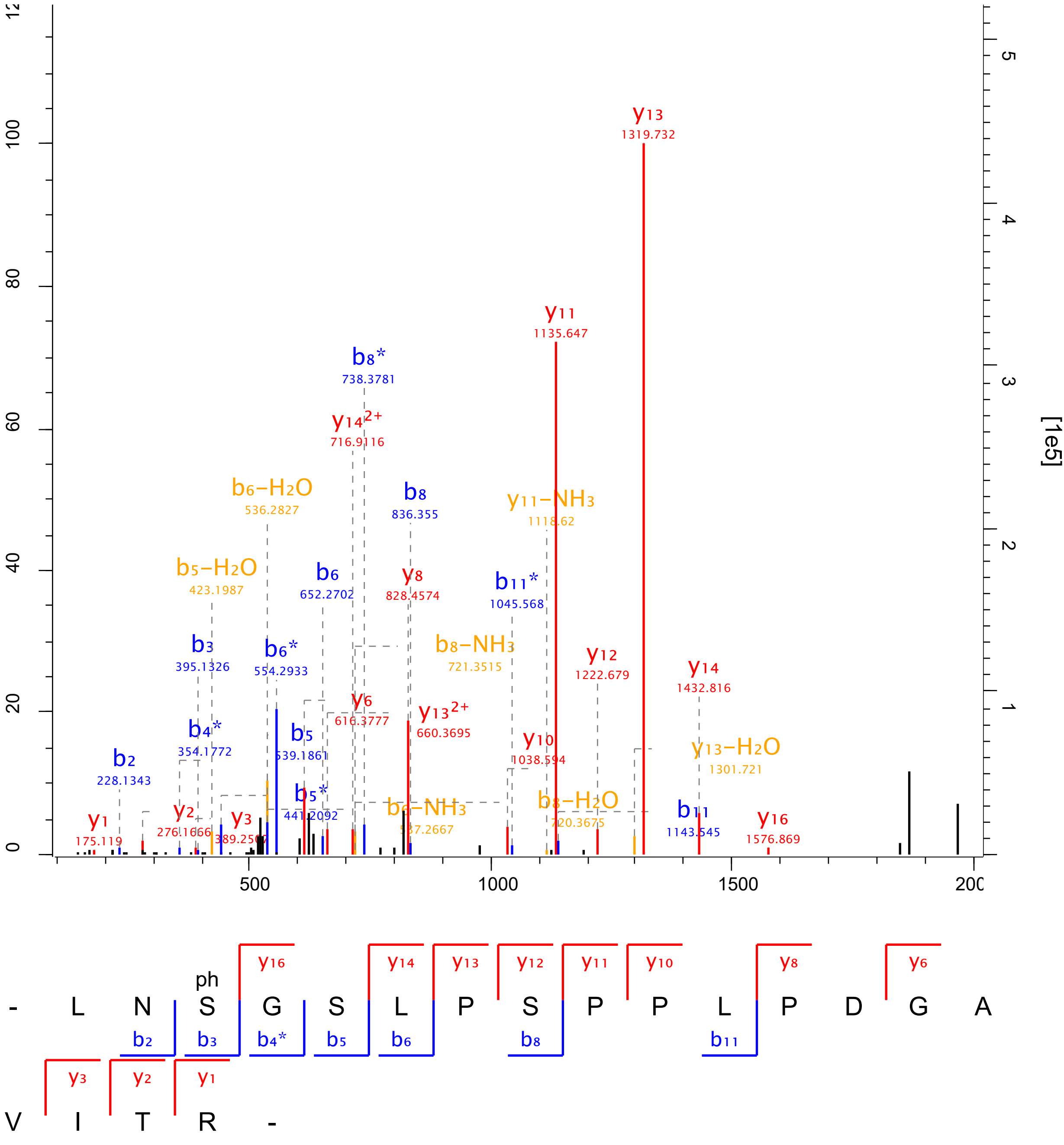

| Raw file | Scan | Method | Score | m/z |
|---|---|---|---|---|
| sys_15_1 | 37388 | FTMS; HCD | 171.46 | 809.36 |

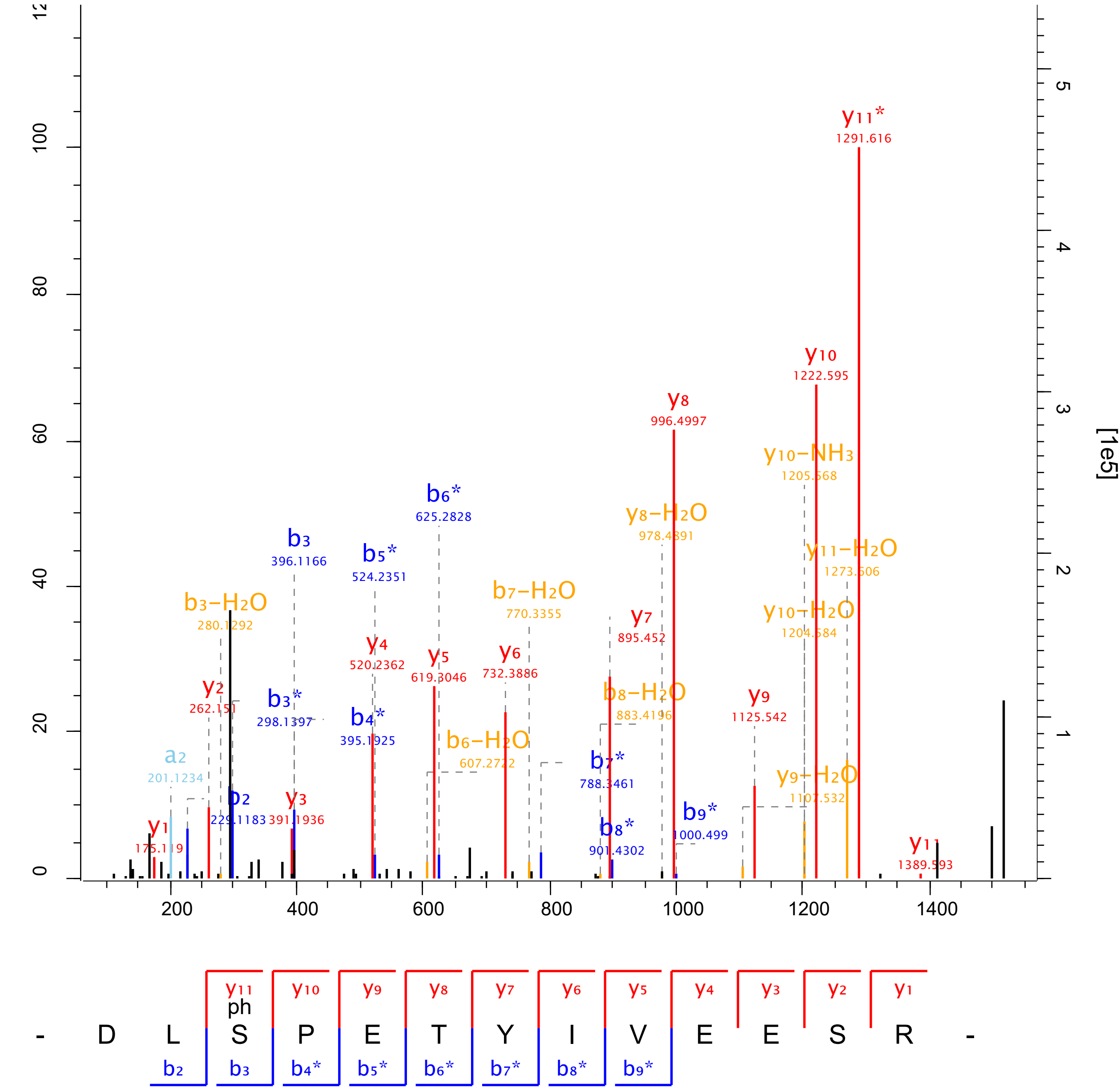

| Raw file | Scan | Method | Score | m/z |
| --- | --- | --- | --- | --- |
| sys_15_1 | 37469 | FTMS; HCD | 129.05 | 797.89 |

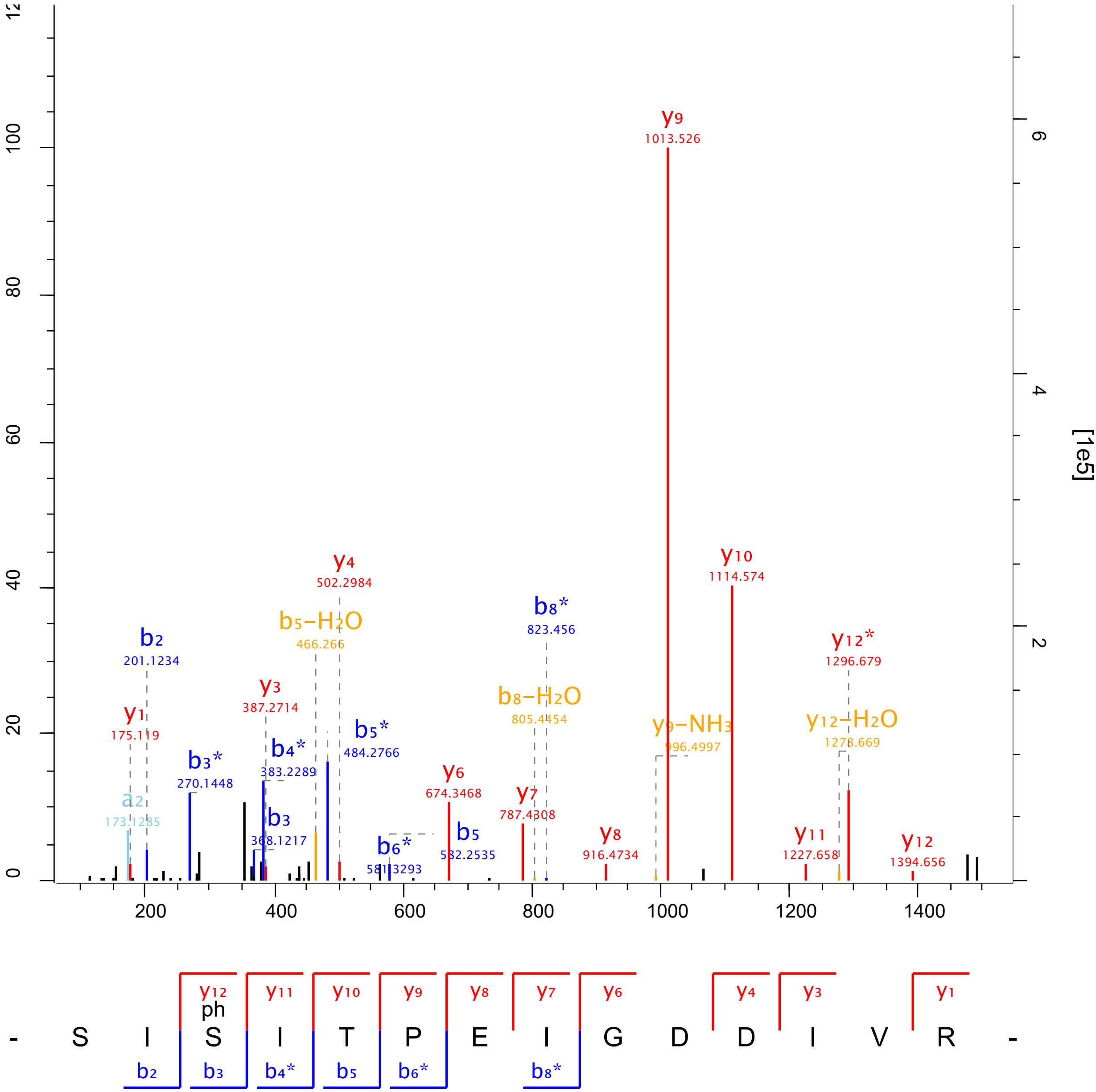

| Raw file | Scan | Method | Score | m/z |
|---|---|---|---|---|
| sys_15_1 | 37610 | FTMS; HCD | 106.35 | 730.31 |

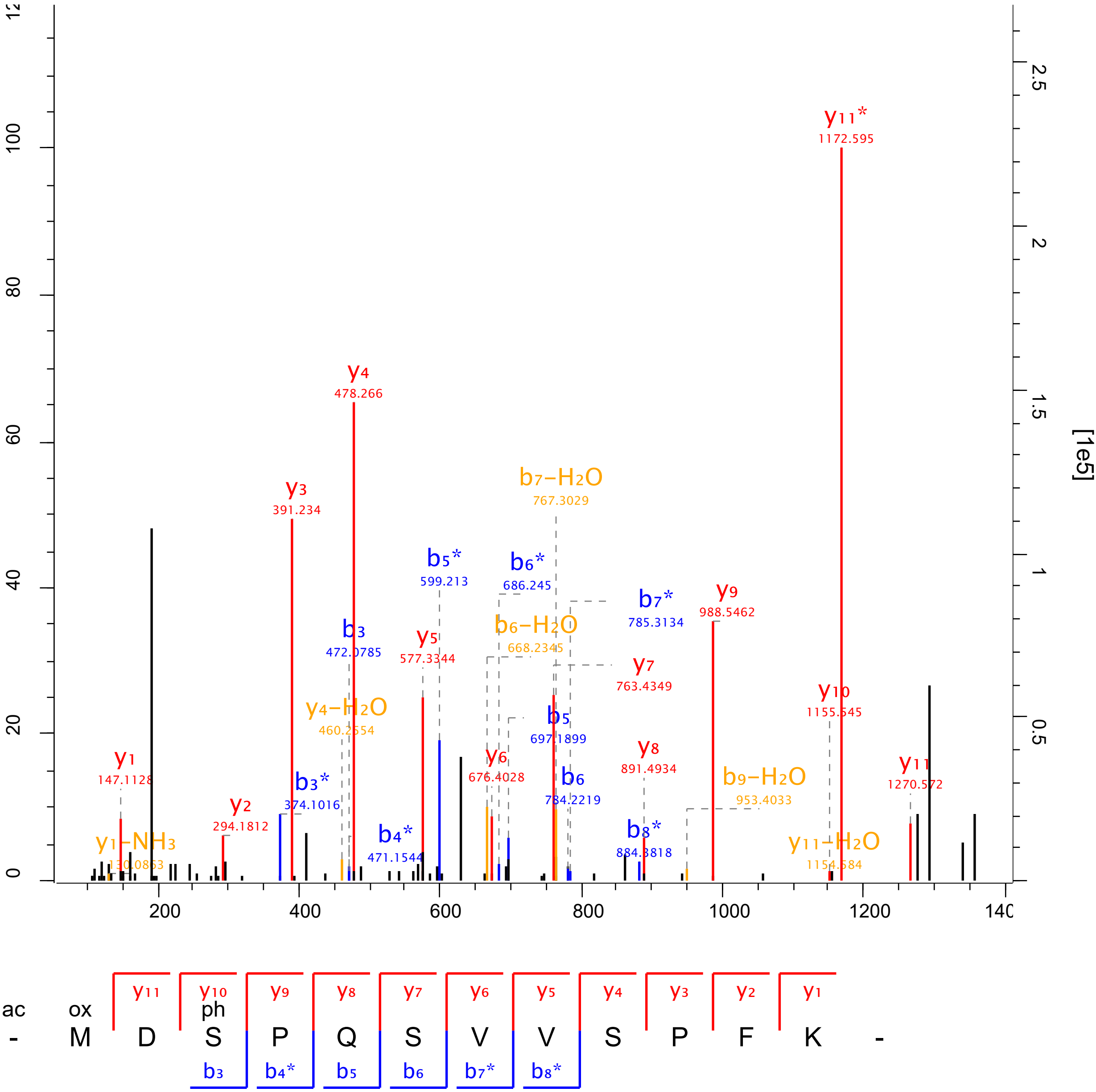

| Raw file | Scan | Method | Score | m/z |
|---|---|---|---|---|
| sys_15_1 | 3780 | FTMS; HCD | 107.5 | 641.79 |

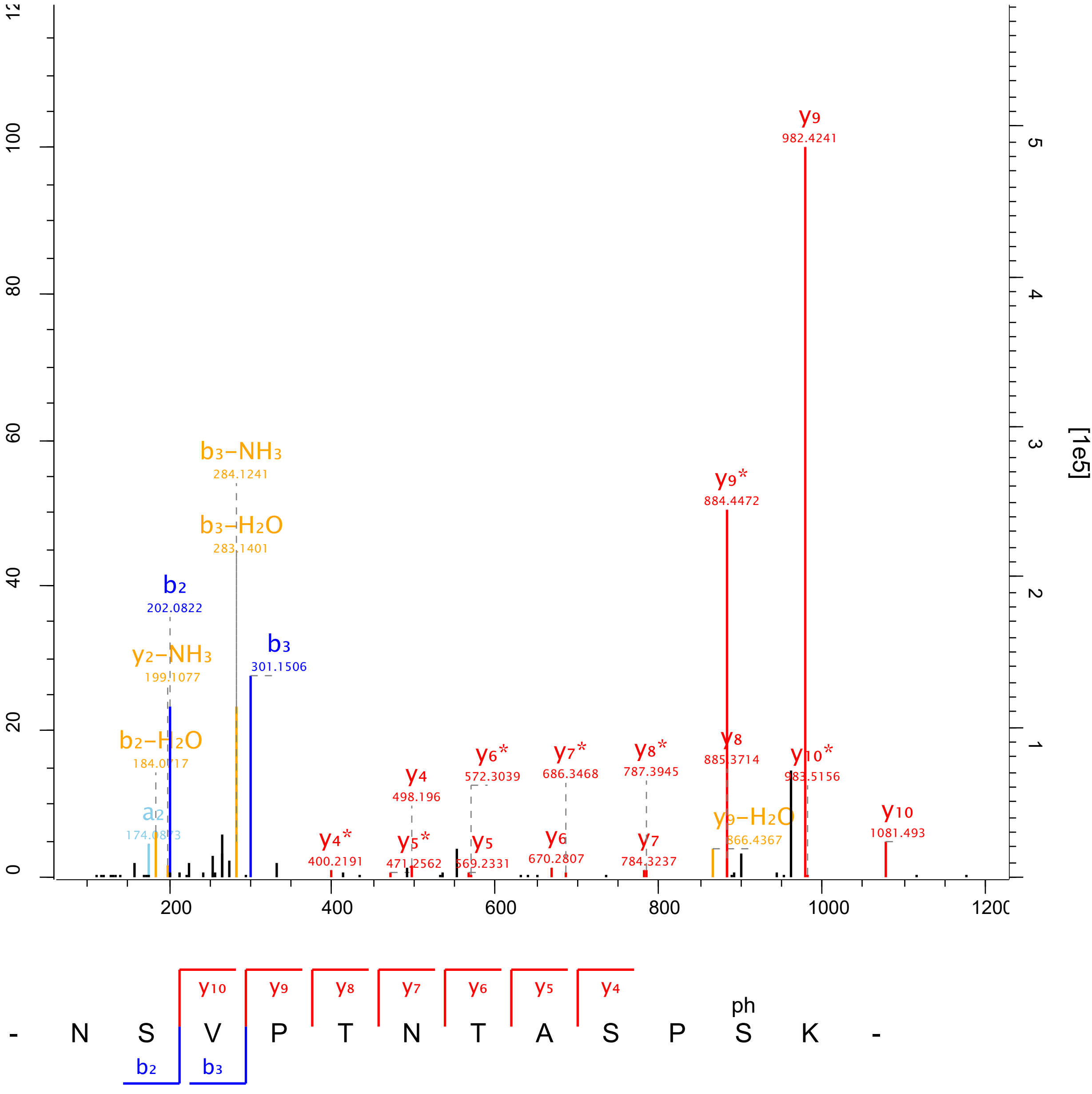

| Raw file | Scan | Method | Score | m/z |
|---|---|---|---|---|
| sys_15_1 | 37883 | FTMS; HCD | 95.2 | 958.93 |

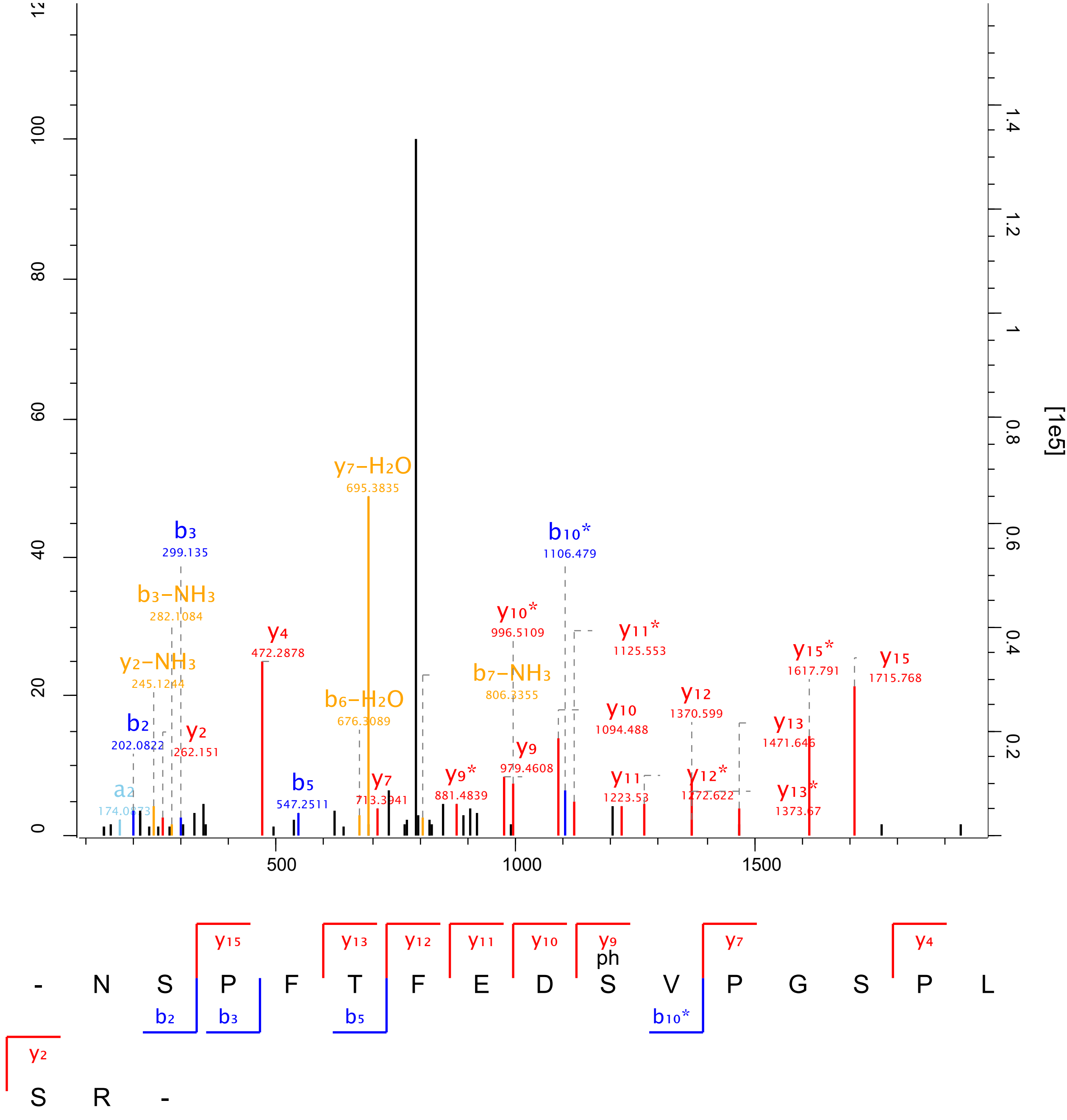

| Raw file | Scan | Method | Score | m/z |
| --- | --- | --- | --- | --- |
| sys_15_1 | 38118 | FTMS; HCD | 142.26 | 876.84 |

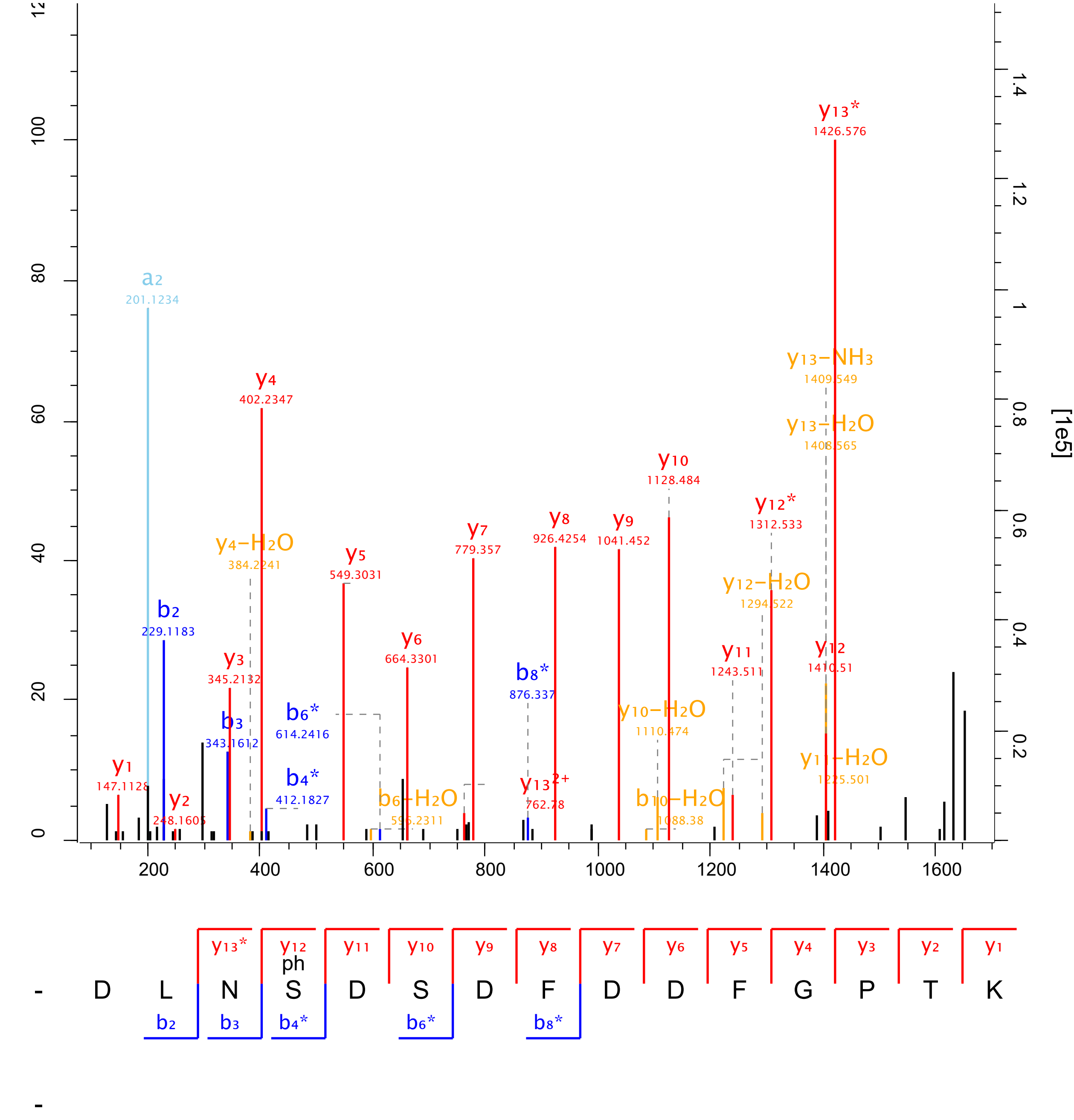

| Raw file | Scan | Method | Score | m/z |
|---|---|---|---|---|
| sys_15_1 | 3815 | FTMS; HCD | 226.99 | 870.37 |

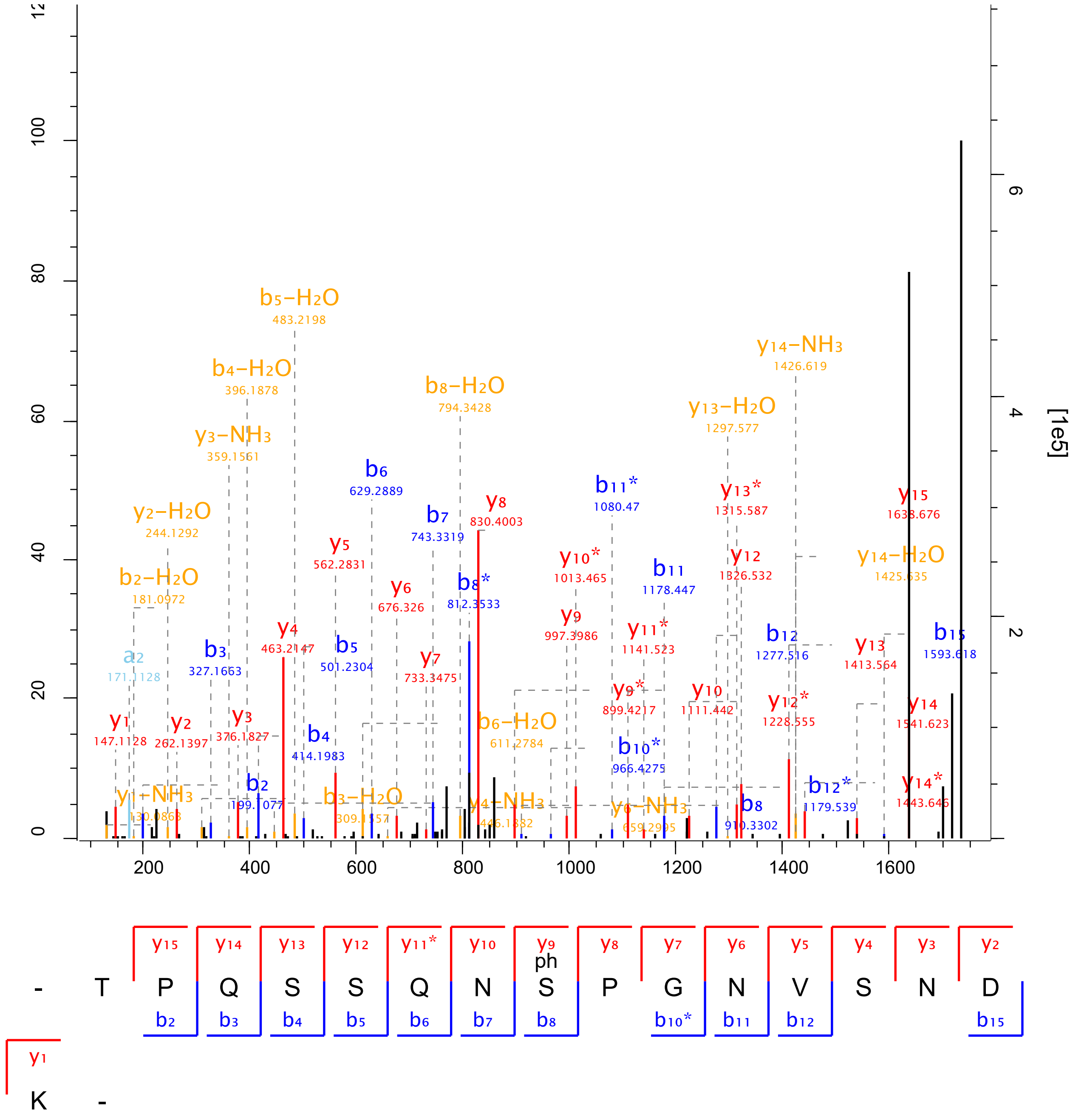

| Raw file | Scan | Method | Score | m/z |
| --- | --- | --- | --- | --- |
| sys_15_1 | 38231 | FTMS; HCD | 131.33 | 718.35 |

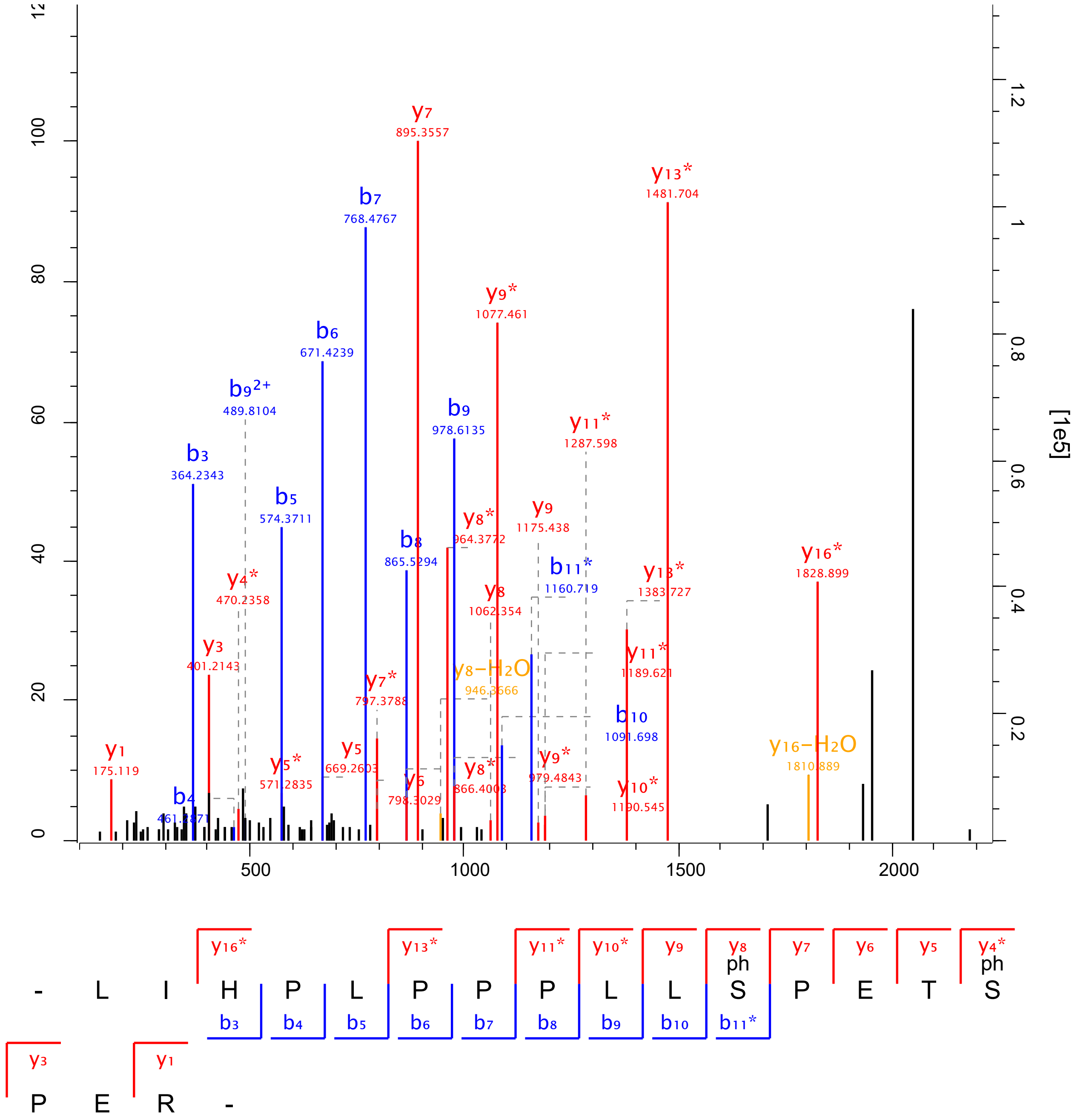

| Raw file | Scan | Method | Score | m/z |
|---|---|---|---|---|
| sys_15_1 | 3835 | FTMS; HCD | 58.69 | 802.78 |

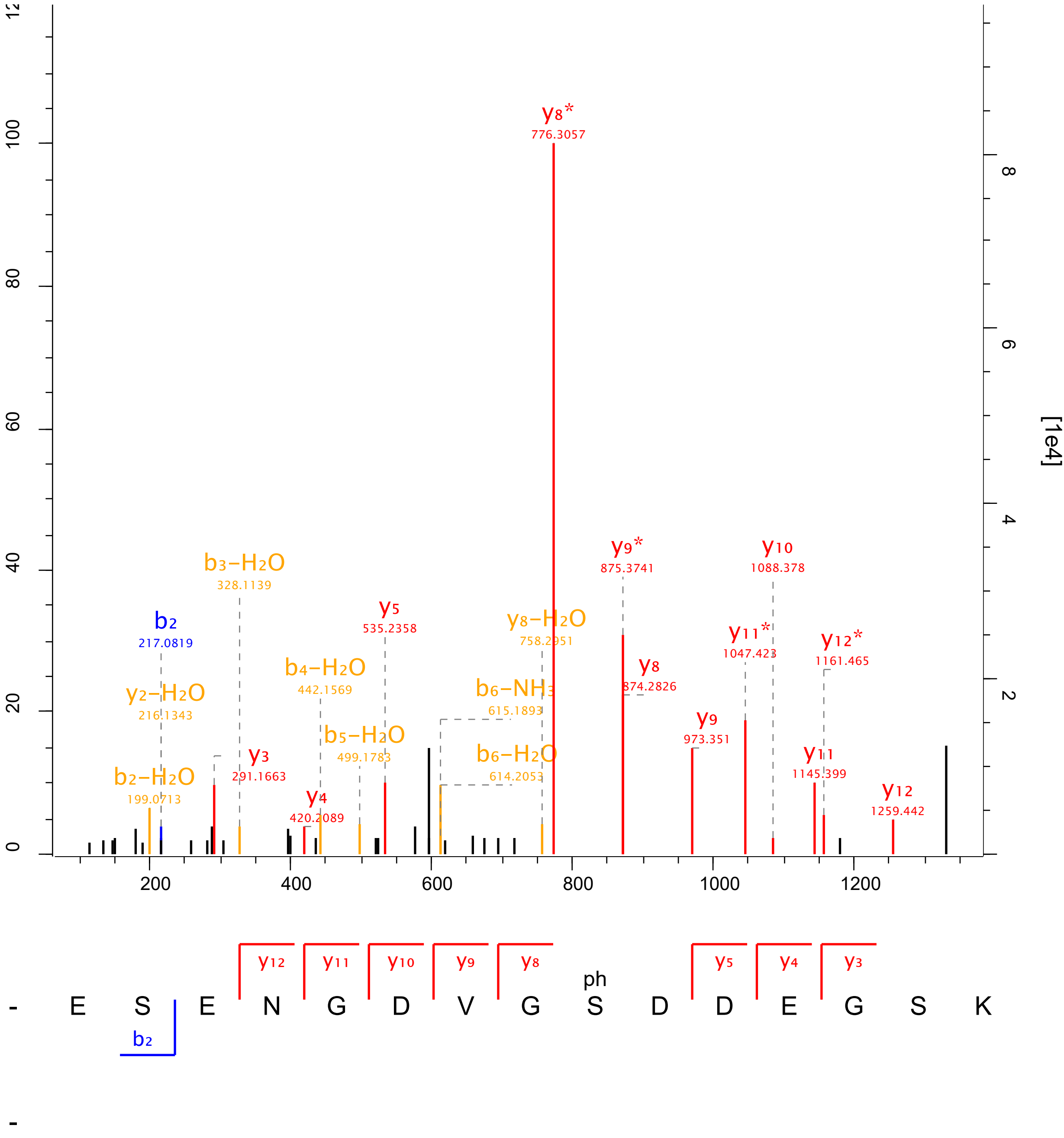

| Raw file | Scan | Method | Score | m/z |
|---|---|---|---|---|
| sys_15_1 | 38425 | FTMS; HCD | 159.14 | 1085.99 |

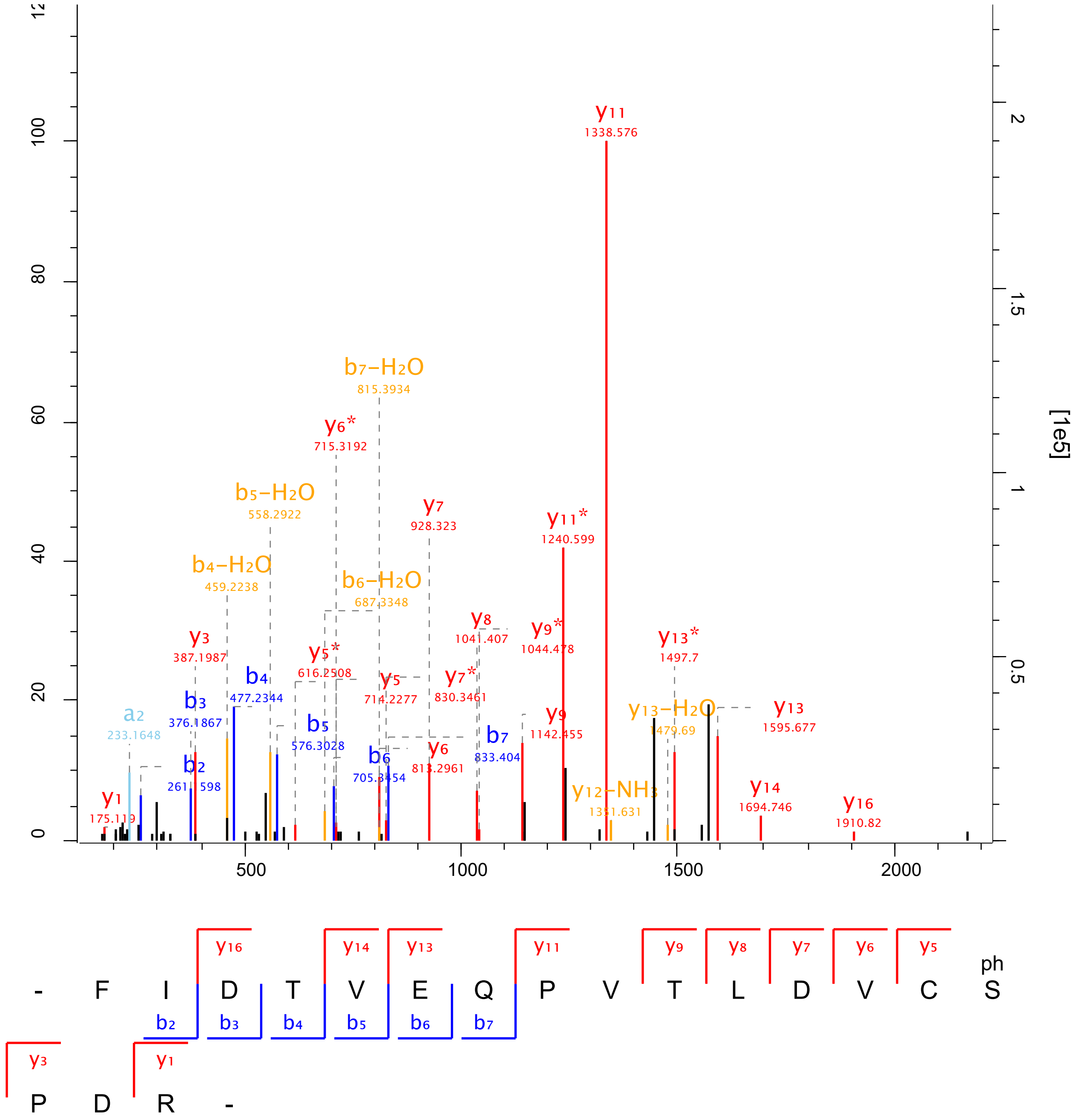

| Raw file | Scan | Method | Score | m/z |
|---|---|---|---|---|
| sys_15_1 | 38484 | FTMS; HCD | 124.51 | 1138.46 |

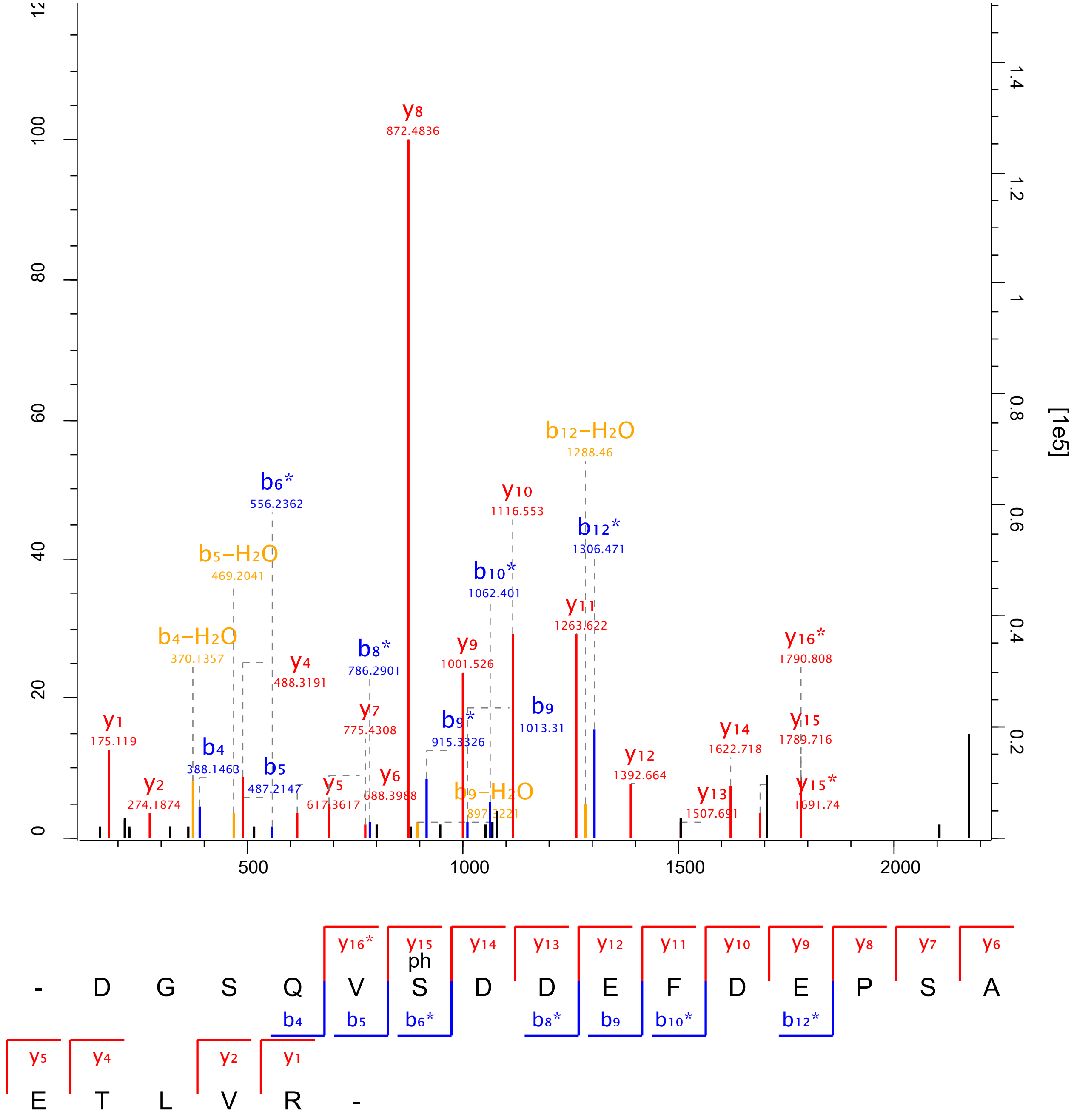

| Raw file | Scan | Method | Score | m/z |
| --- | --- | --- | --- | --- |
| sys_15_1 | 3851 | FTMS; HCD | 88.71 | 594.76 |

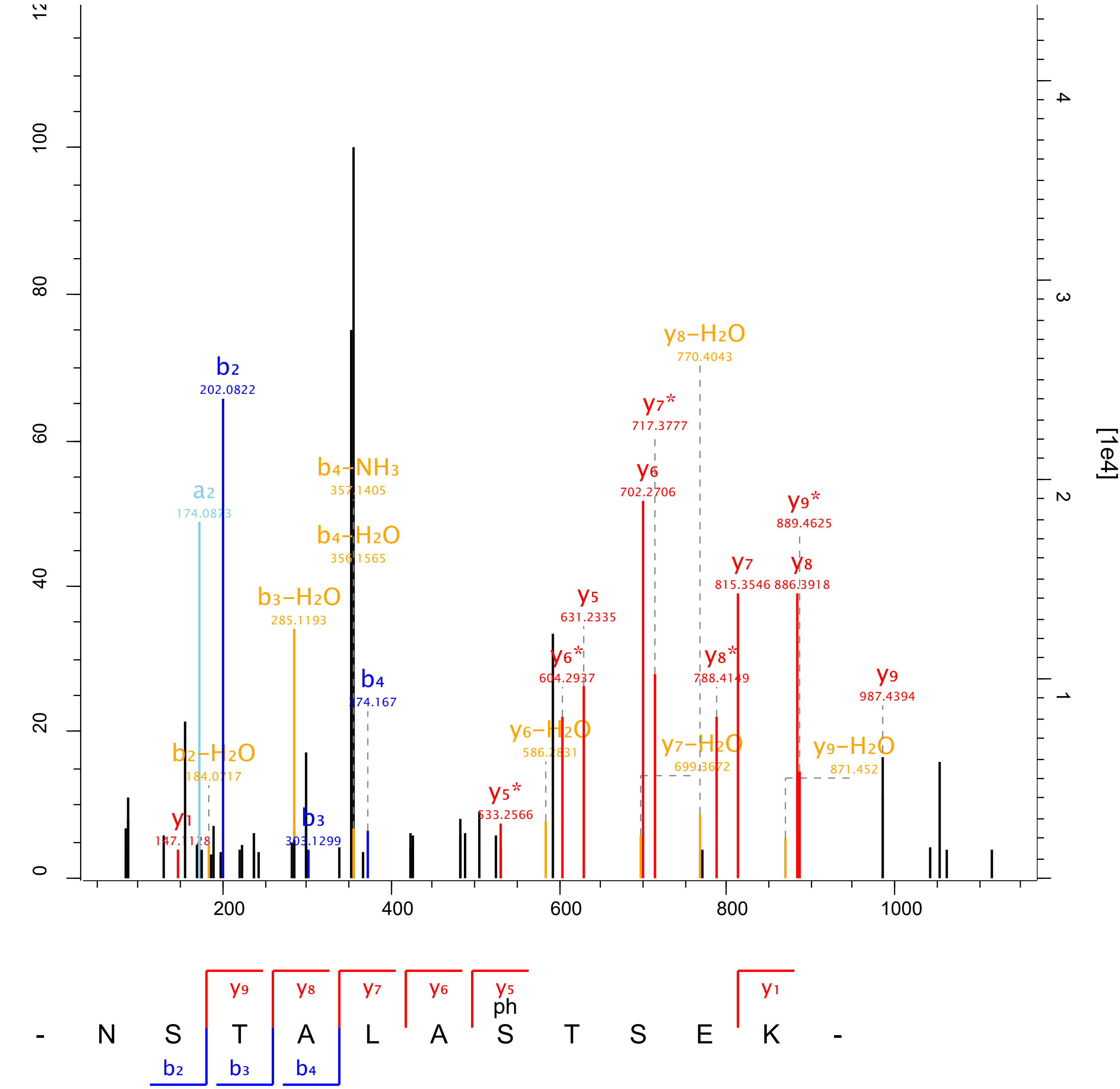

| Raw file | Scan | Method | Score | m/z |
| --- | --- | --- | --- | --- |
| sys_15_1 | 38596 | FTMS; HCD | 111.01 | 887.03 |

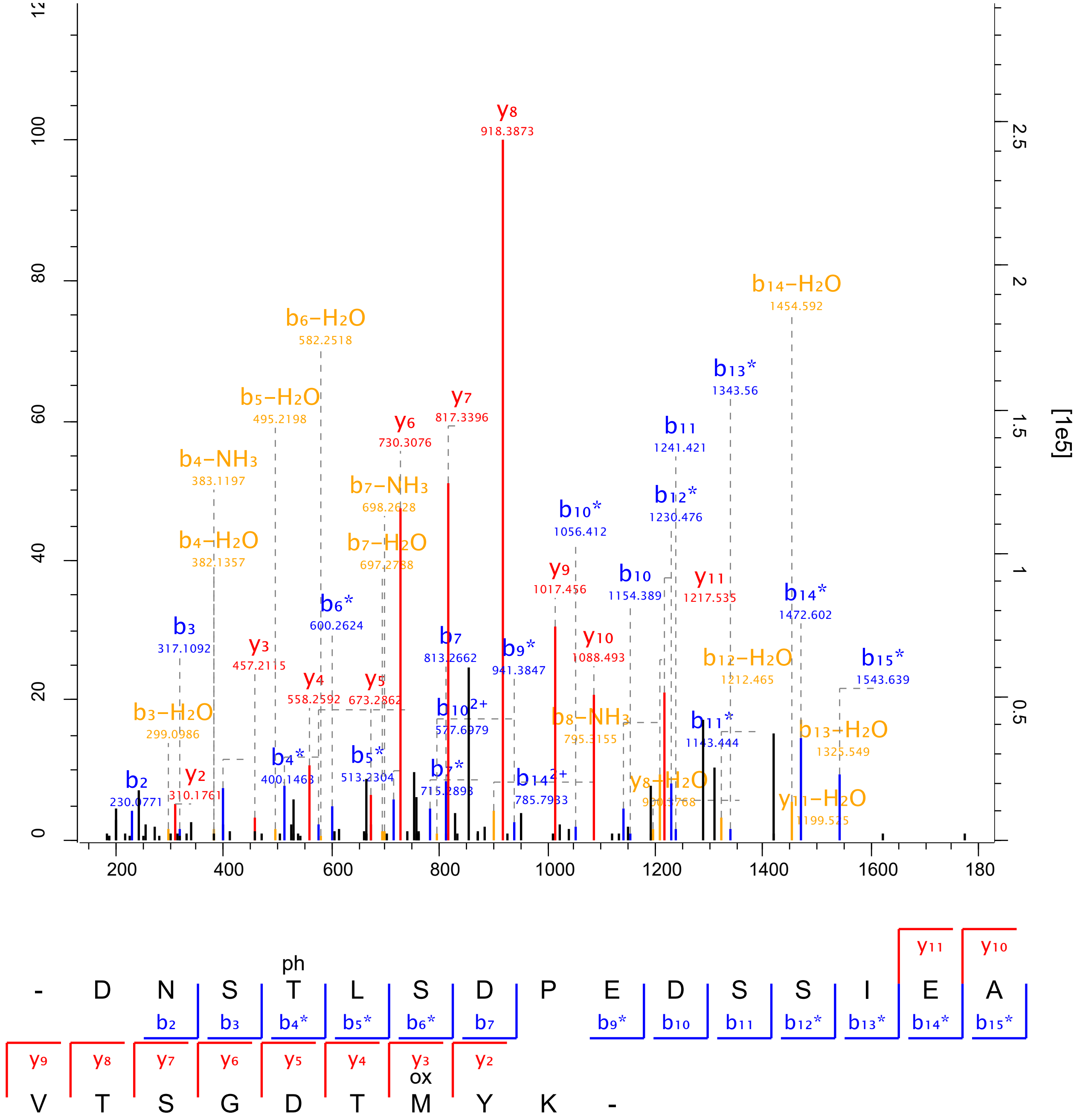

| Raw file | Scan | Method | Score | m/z |
|---|---|---|---|---|
| sys_15_1 | 38621 | FTMS; HCD | 203.19 | 902.03 |

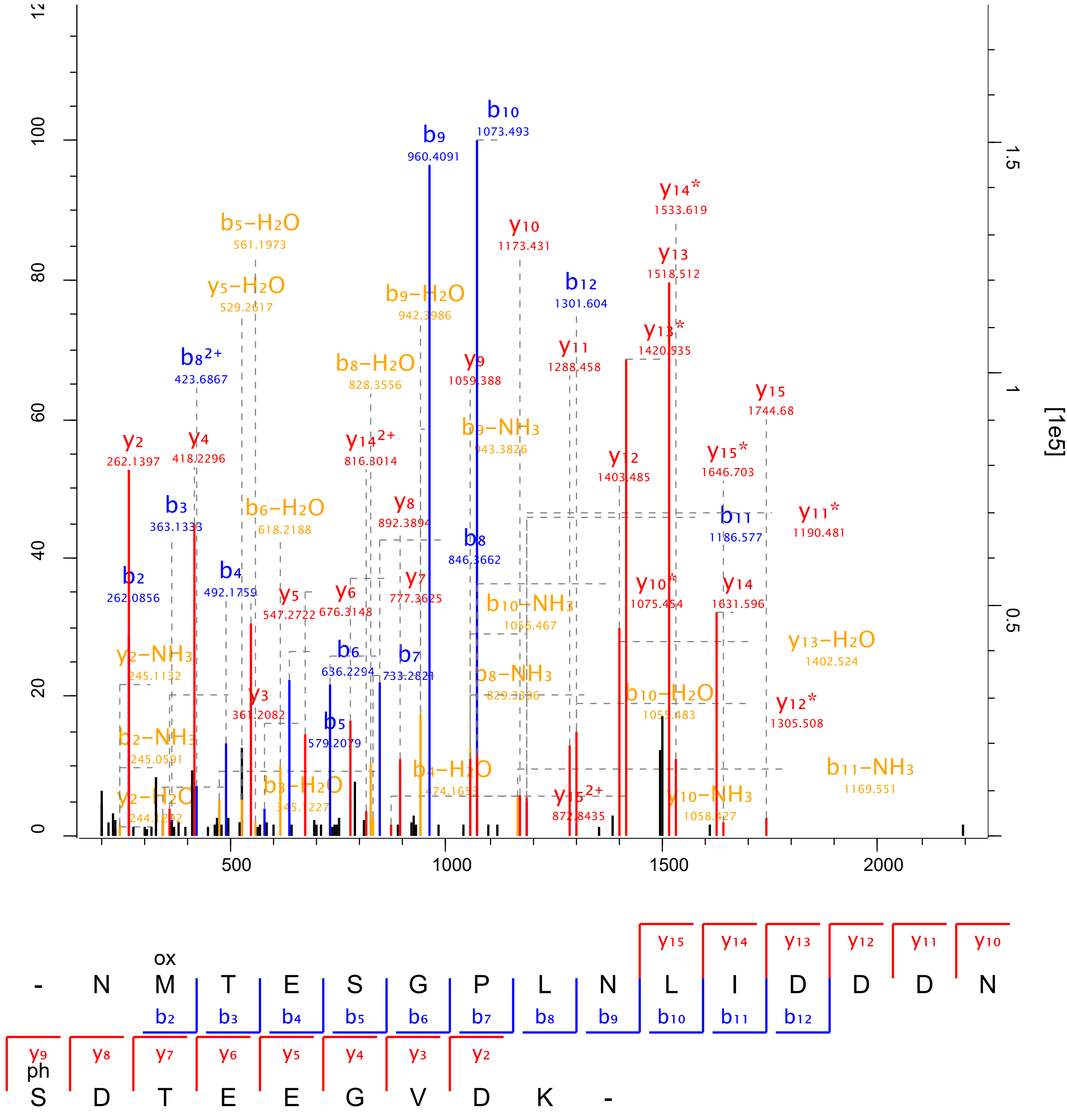

| Raw file | Scan | Method | Score | m/z |
|---|---|---|---|---|
| sys_15_1 | 38623 | FTMS; HCD | 205.95 | 805.87 |

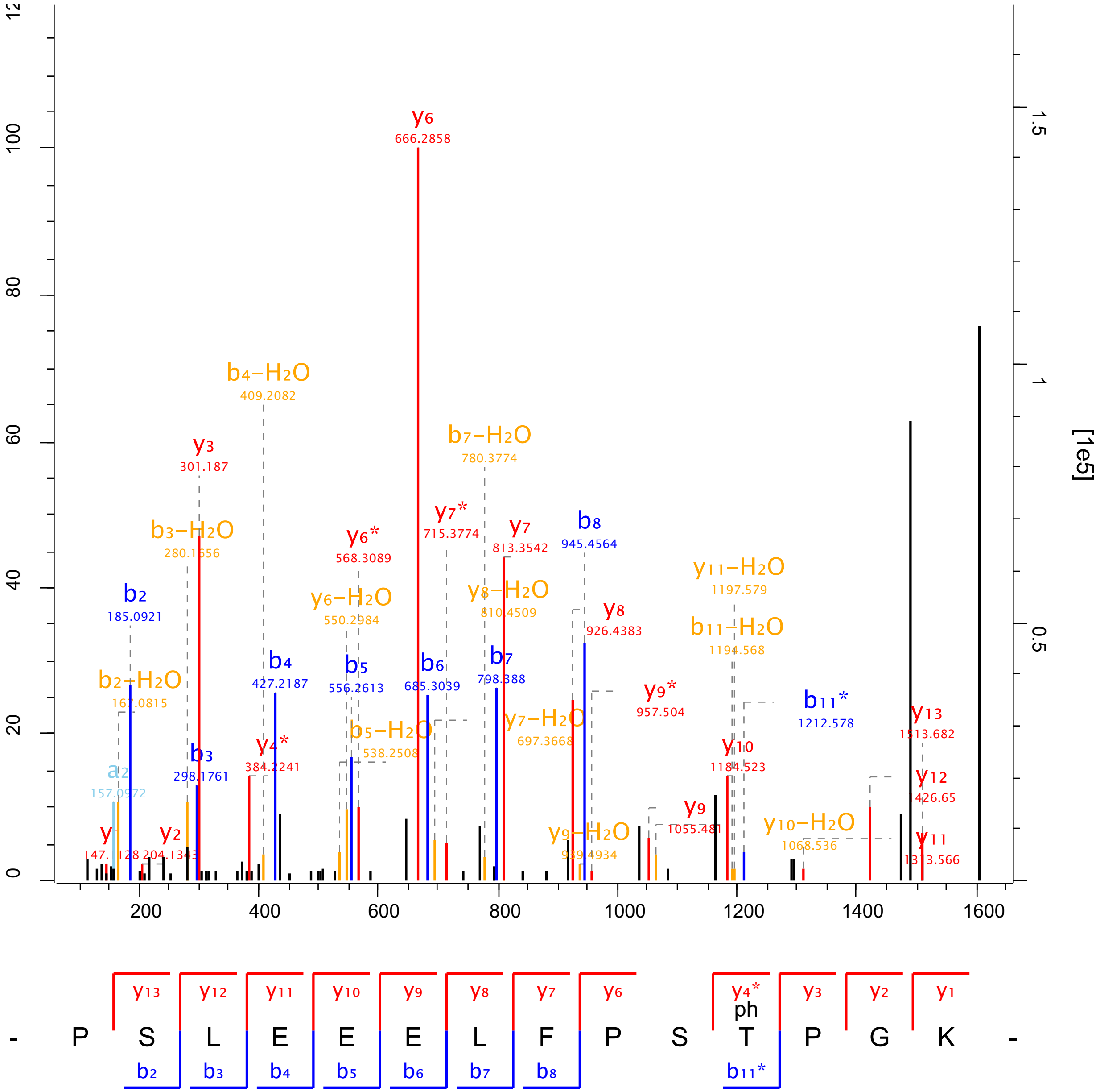

| Raw file | Scan | Method | Score | m/z |
| --- | --- | --- | --- | --- |
| sys_15_1 | 38872 | FTMS; HCD | 98.3 | 761.03 |

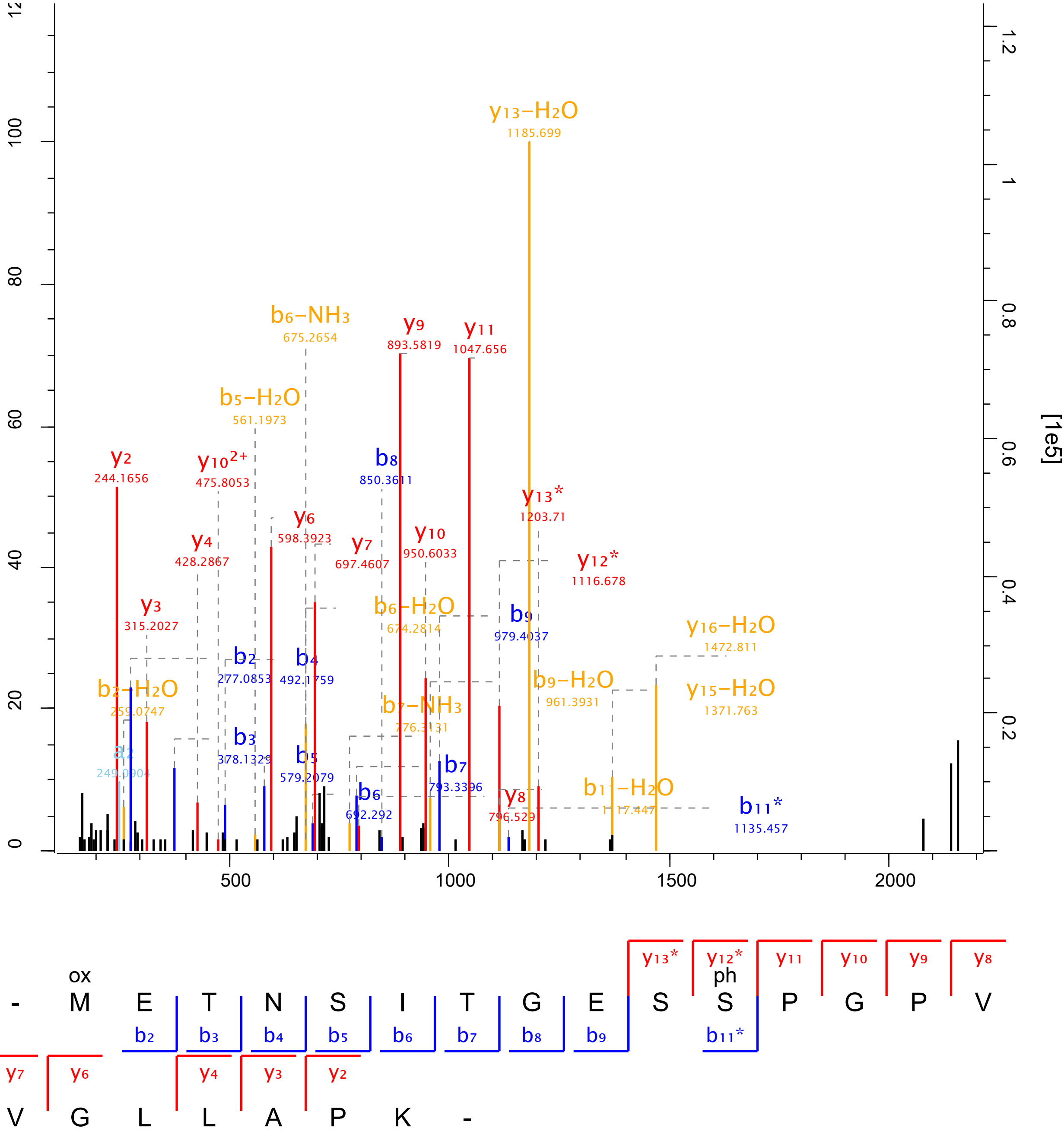

| Raw file | Scan | Method | Score | m/z |
|---|---|---|---|---|
| sys_15_1 | 3888 | FTMS; HCD | 61.78 | 462.69 |

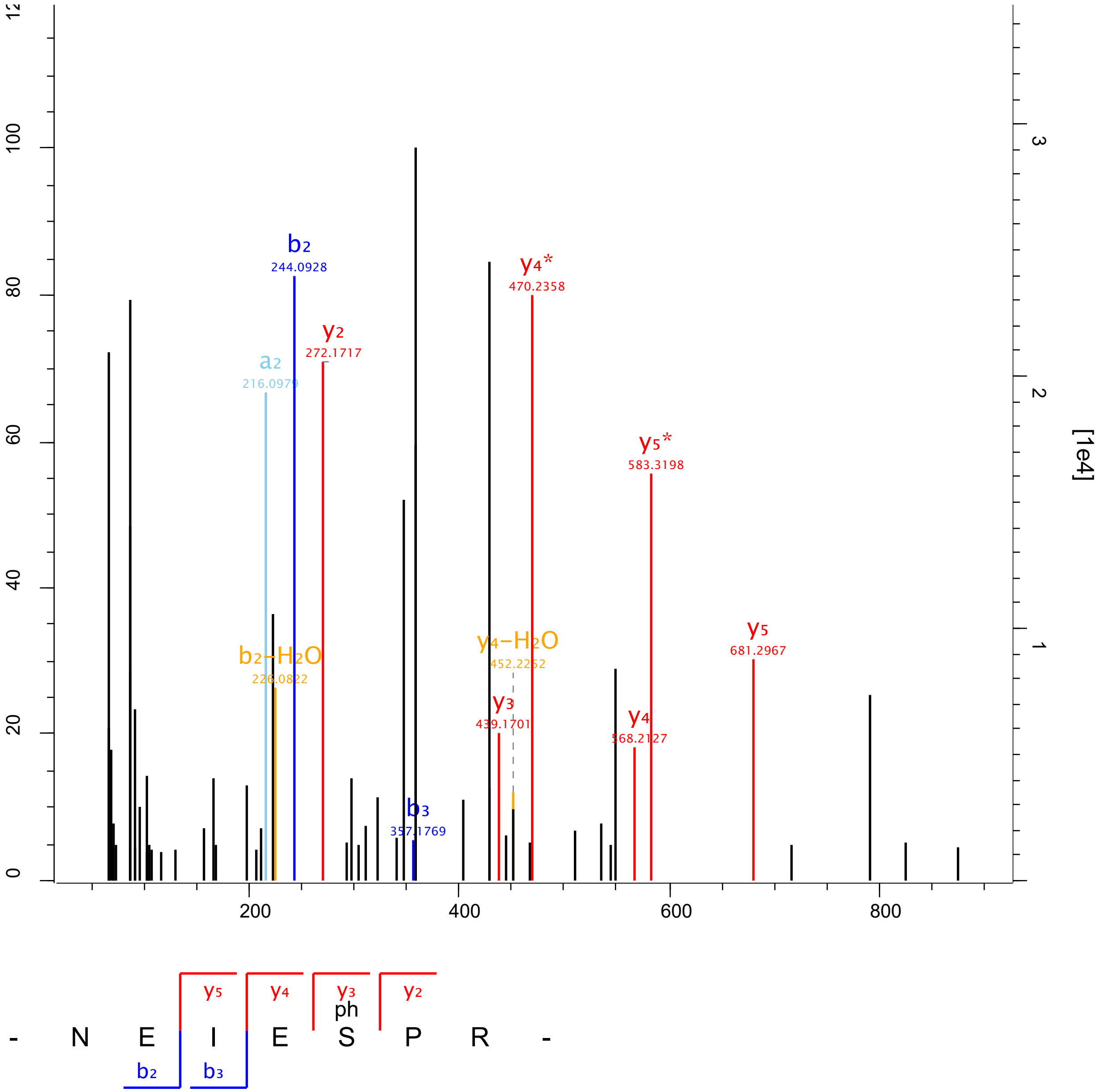

| Raw file | Scan | Method | Score | m/z |
| --- | --- | --- | --- | --- |
| sys_15_1 | 39109 | FTMS; HCD | 45.92 | 837.88 |

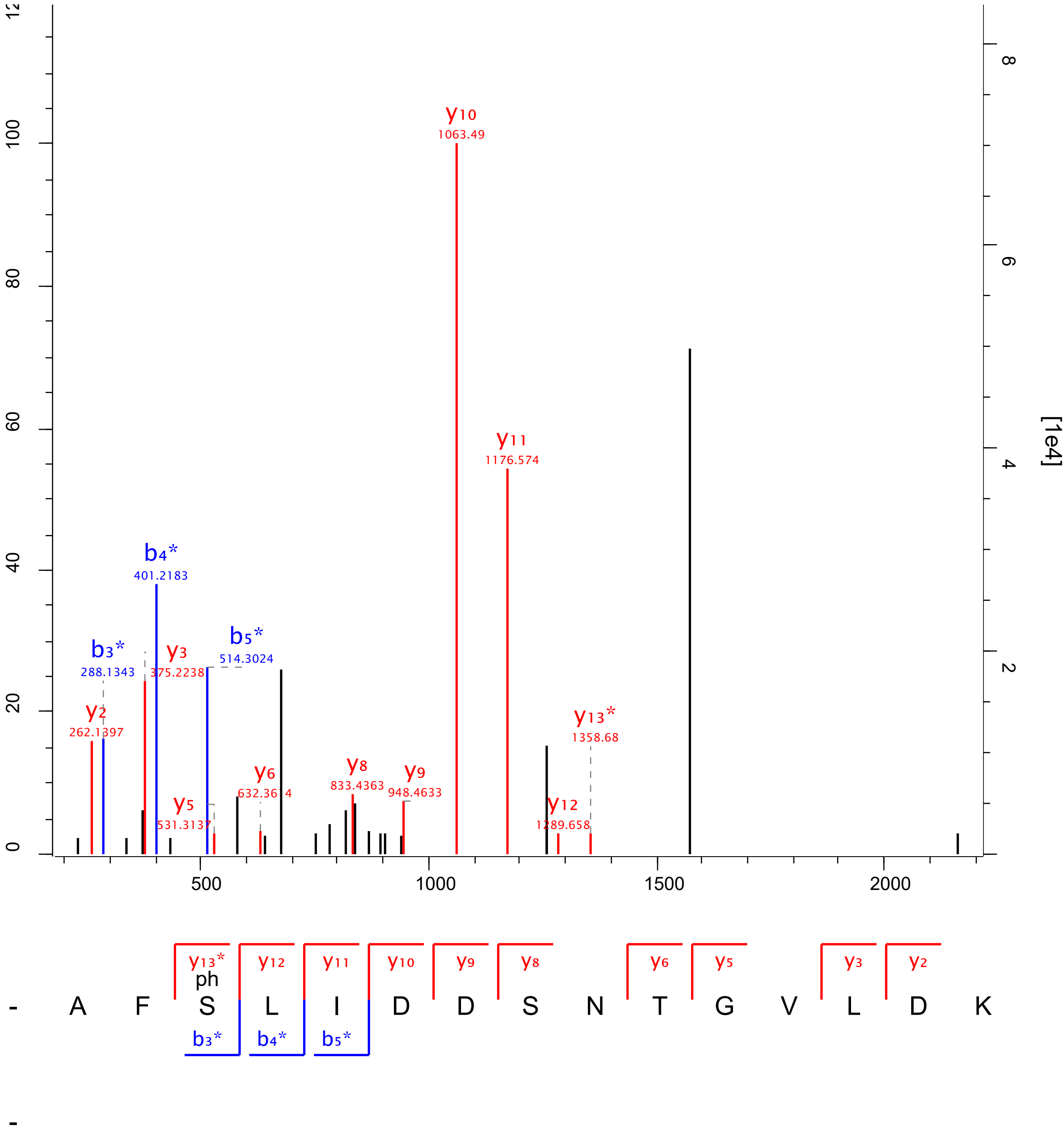

| Raw file | Scan | Method | Score | m/z |
|---|---|---|---|---|
| sys_15_1 | 39289 | FTMS; HCD | 64.7 | 838.42 |

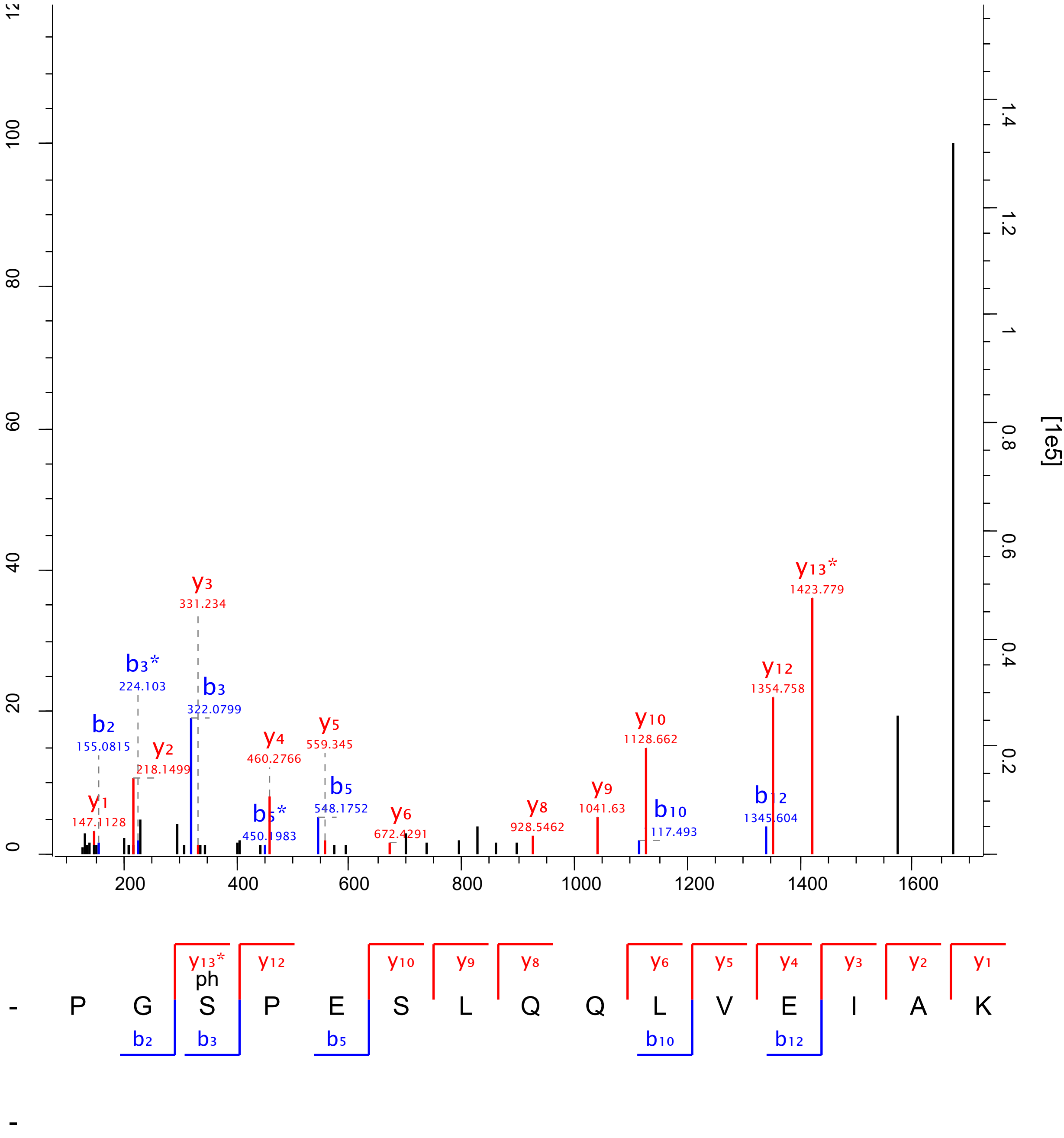

| Raw file | Scan | Method | Score | m/z |
|---|---|---|---|---|
| sys_15_1 | 39425 | FTMS; HCD | 218.12 | 747.88 |

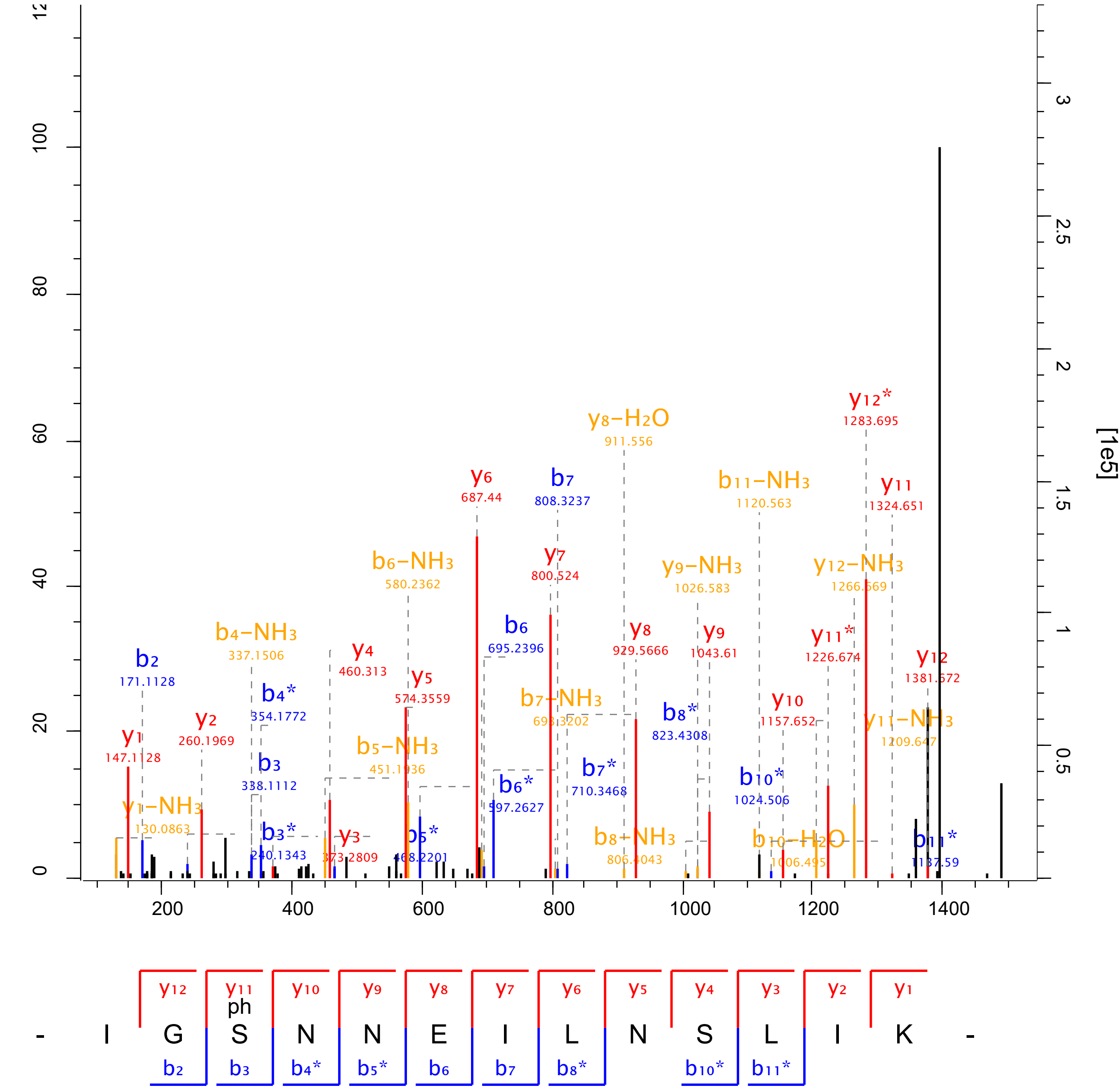

| Raw file | Scan | Method | Score | m/z |
|---|---|---|---|---|
| sys_15_1 | 3966 | FTMS; HCD | 120.77 | 532.69 |

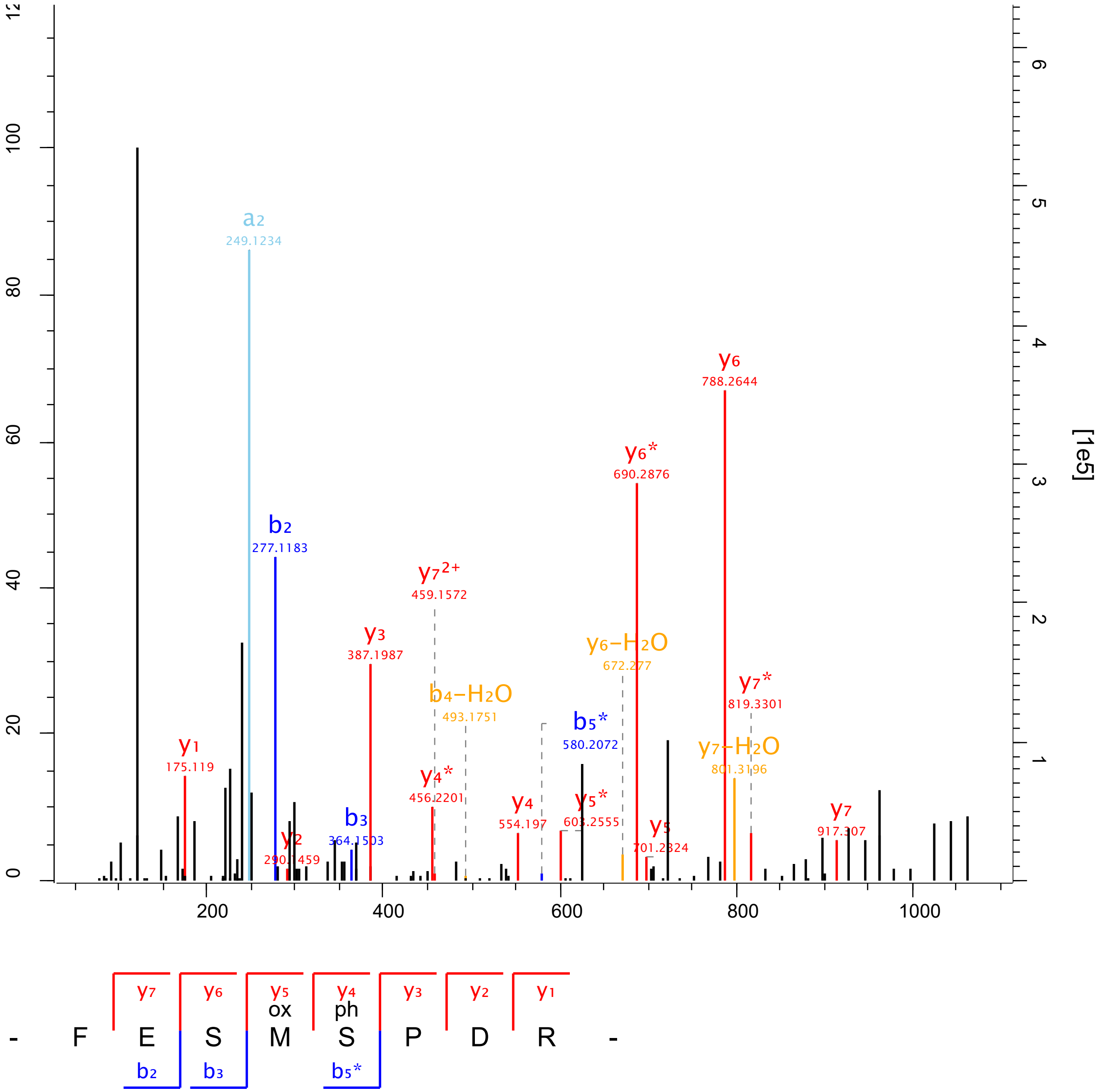

| Raw file | Scan | Method | Score | m/z |
|---|---|---|---|---|
| sys_15_1 | 39821 | FTMS; HCD | 43.7 | 747.31 |

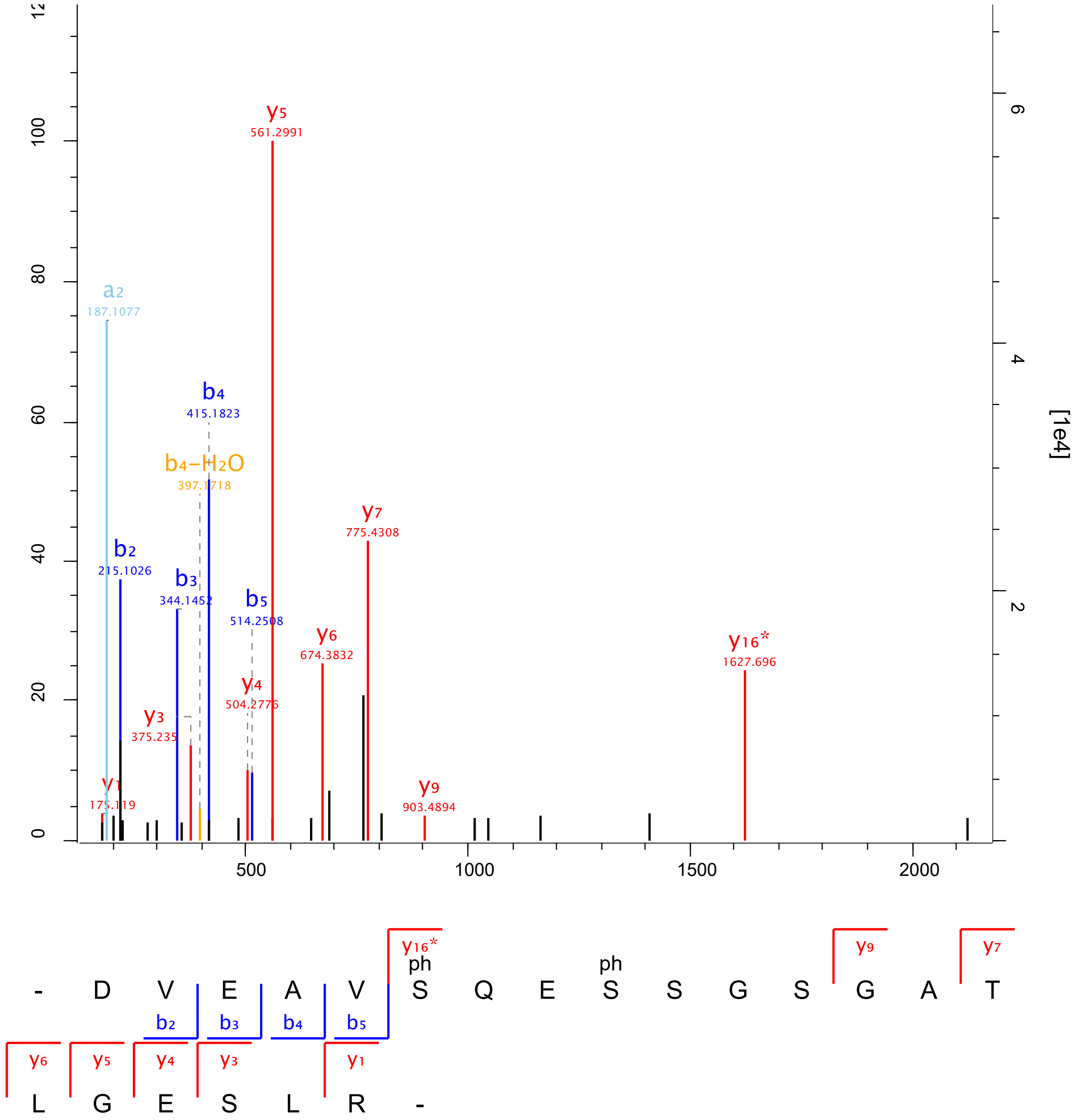

| Raw file | Scan | Method | Score | m/z |
|---|---|---|---|---|
| sys_15_1 | 39897 | FTMS; HCD | 166.47 | 869.87 |

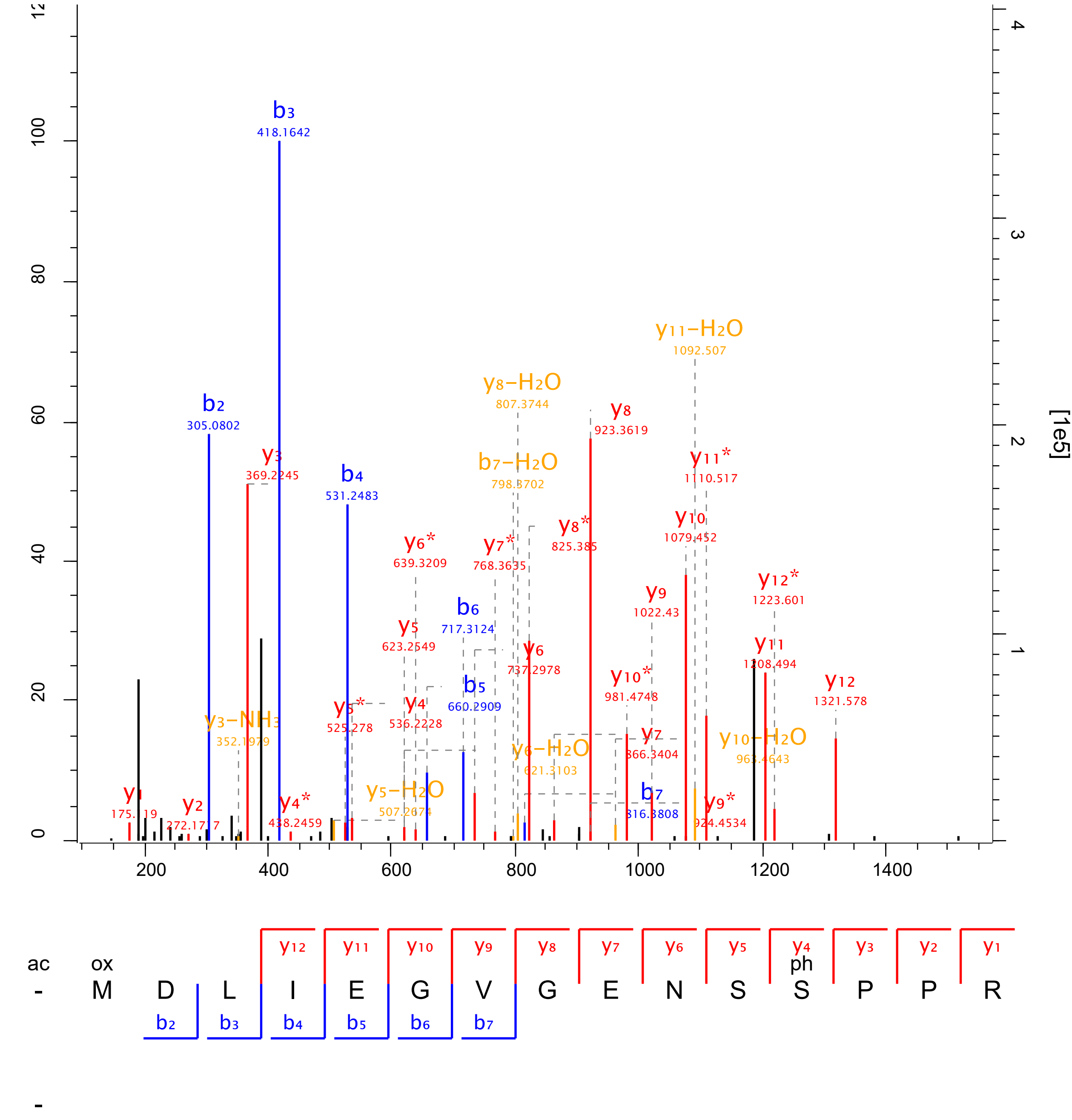

| Raw file | Scan | Method | Score | m/z |
|---|---|---|---|---|
| sys_15_1 | 4012 | FTMS; HCD | 51.73 | 556.18 |

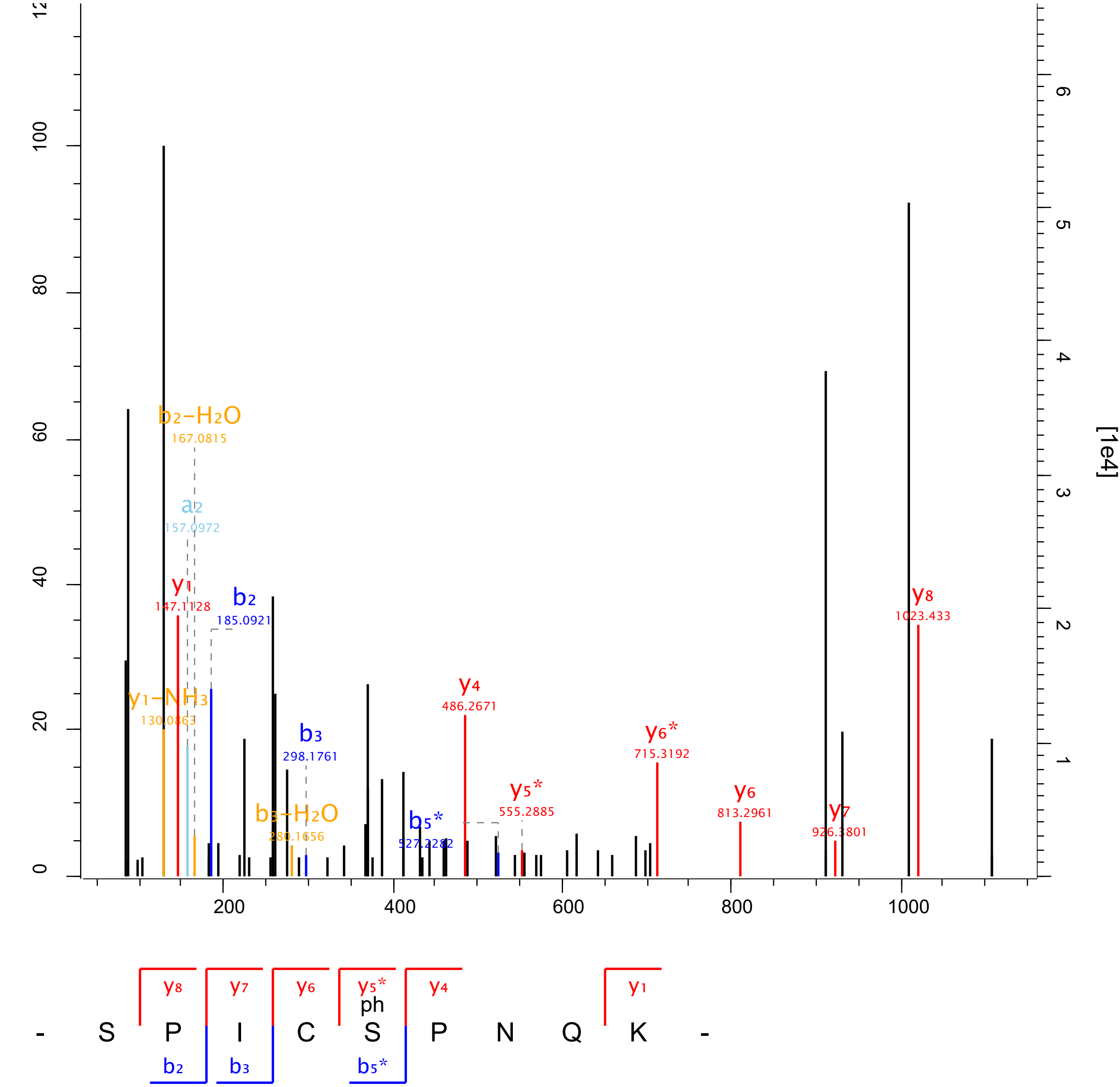

| Raw file | Scan | Method | Score | m/z |
|---|---|---|---|---|
| sys_15_1 | 40251 | FTMS; HCD | 162.87 | 930.74 |

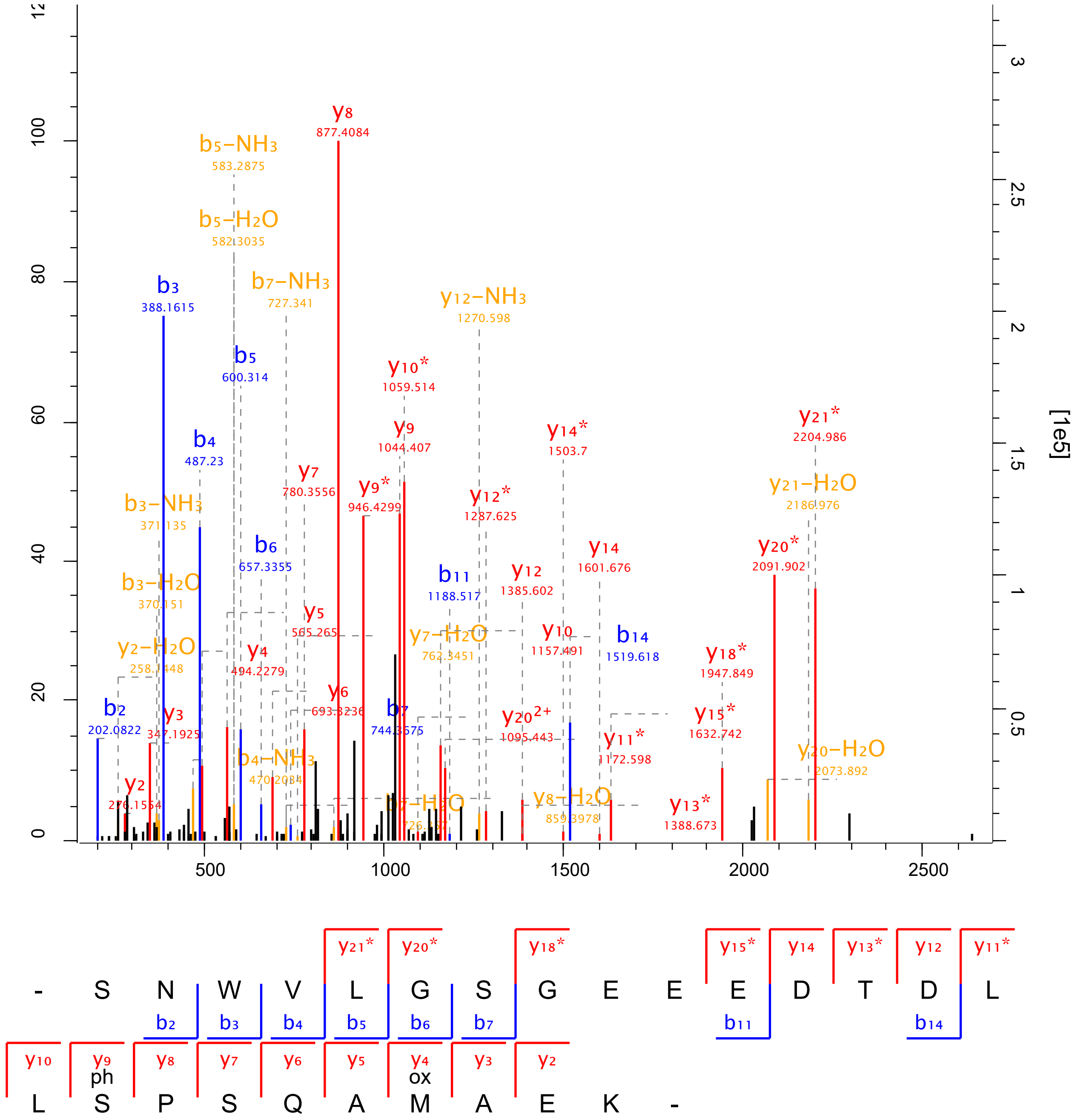

| Raw file | Scan | Method | Score | m/z |
| --- | --- | --- | --- | --- |
| sys_15_1 | 4031 | FTMS; HCD | 160.44 | 578.74 |

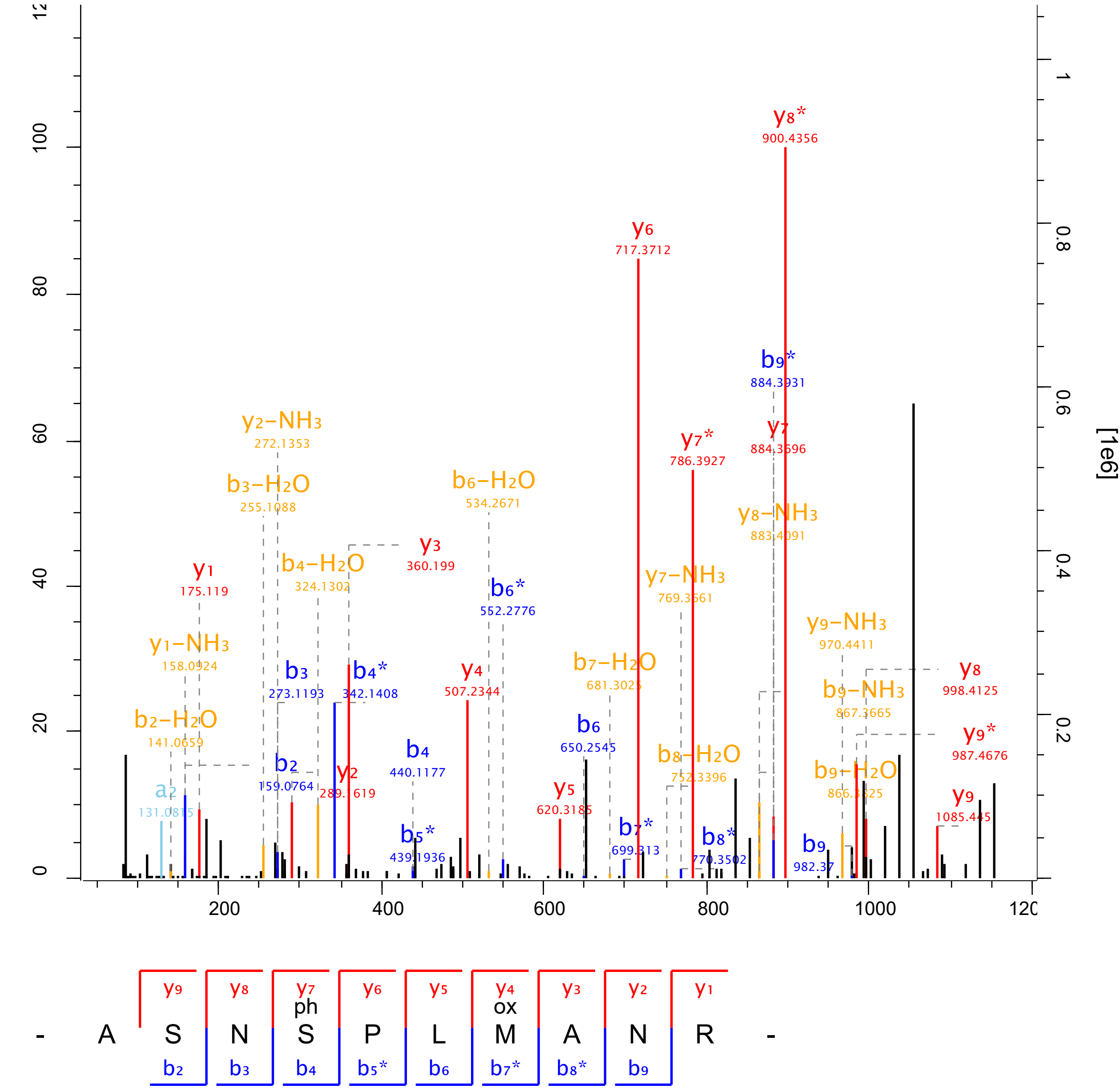

| Raw file | Scan | Method | Score | m/z |
|---|---|---|---|---|
| sys_15_1 | 40941 | FTMS; HCD | 42 | 740.36 |

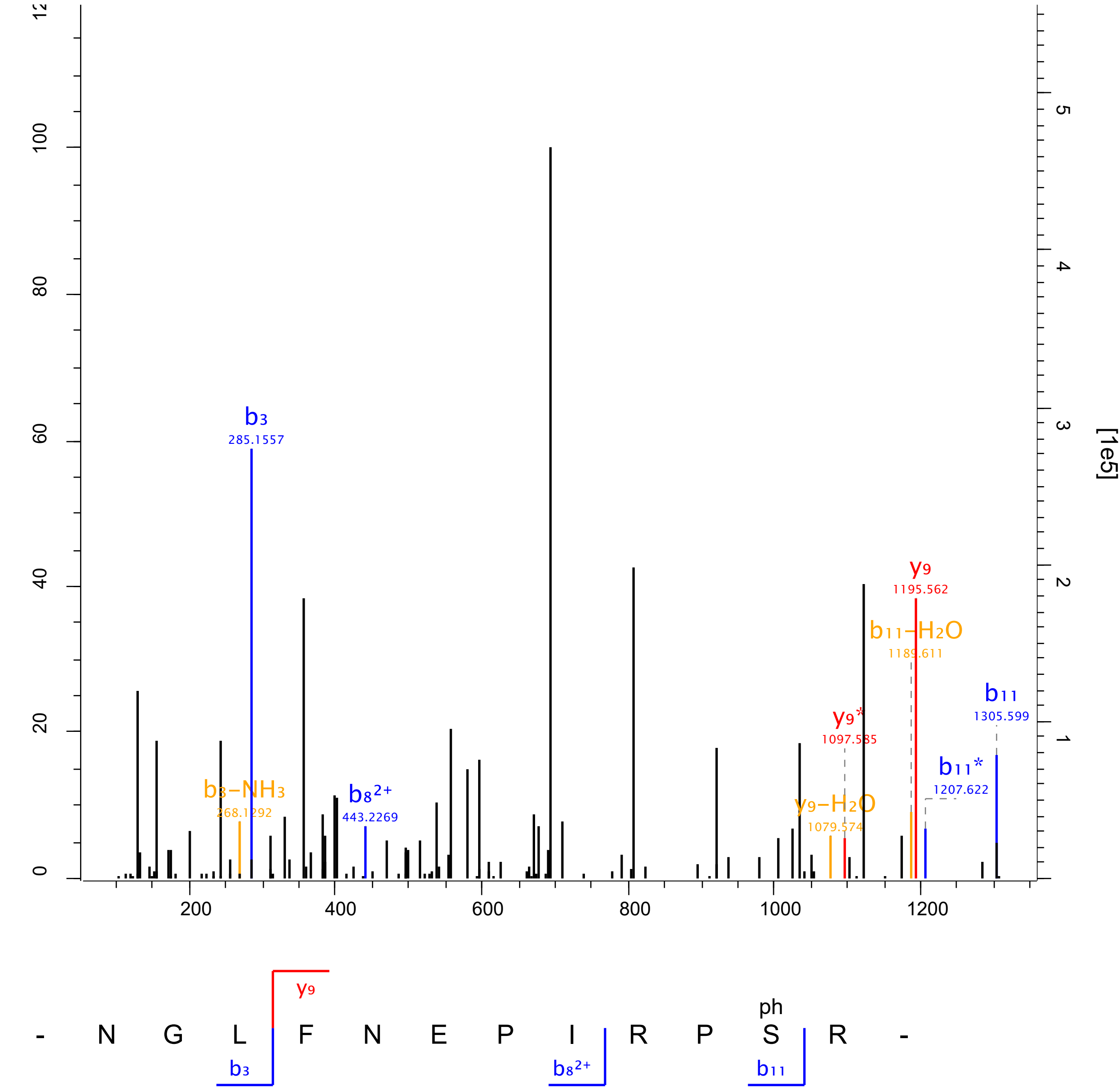

| Raw file | Scan | Method | Score | m/z |
|---|---|---|---|---|
| sys_15_1 | 41018 | FTMS; HCD | 154.48 | 847.4 |

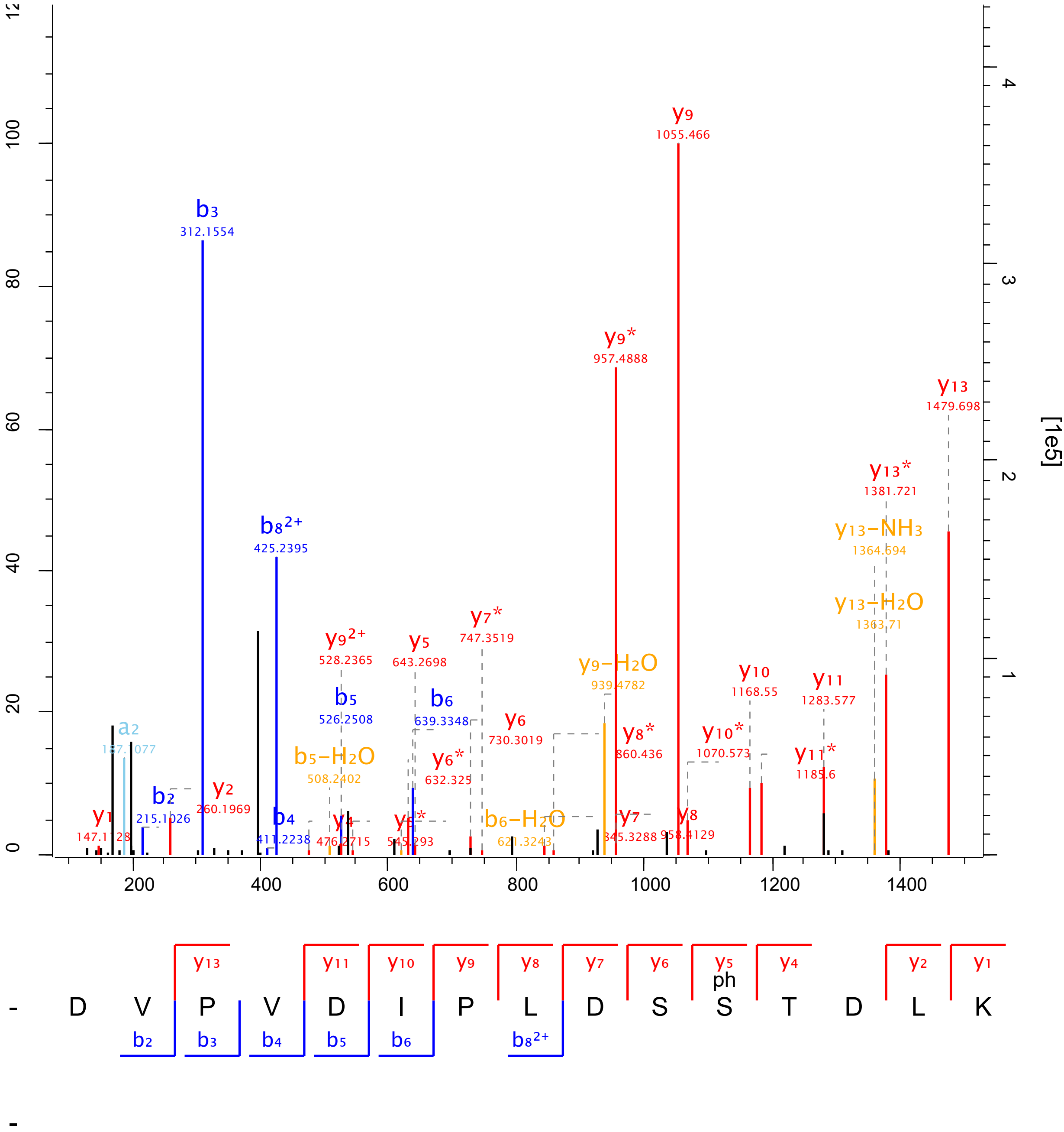

| Raw file | Scan | Method | Score | m/z |
|---|---|---|---|---|
| sys_15_1 | 41058 | FTMS; HCD | 72.43 | 816.35 |

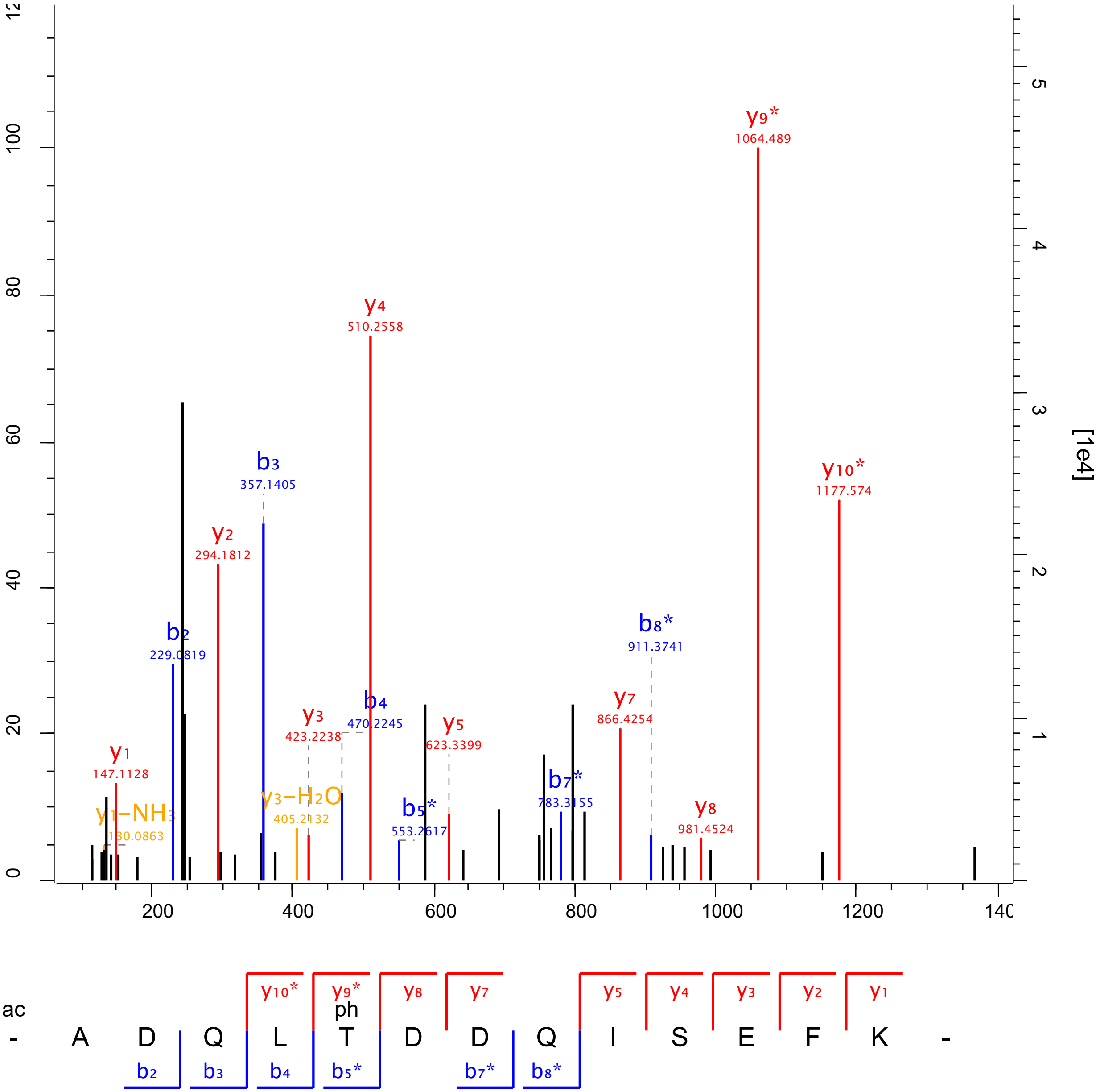

| Raw file | Scan | Method | Score | m/z |
| --- | --- | --- | --- | --- |
| sys_15_1 | 41078 | FTMS; HCD | 96.79 | 800.41 |

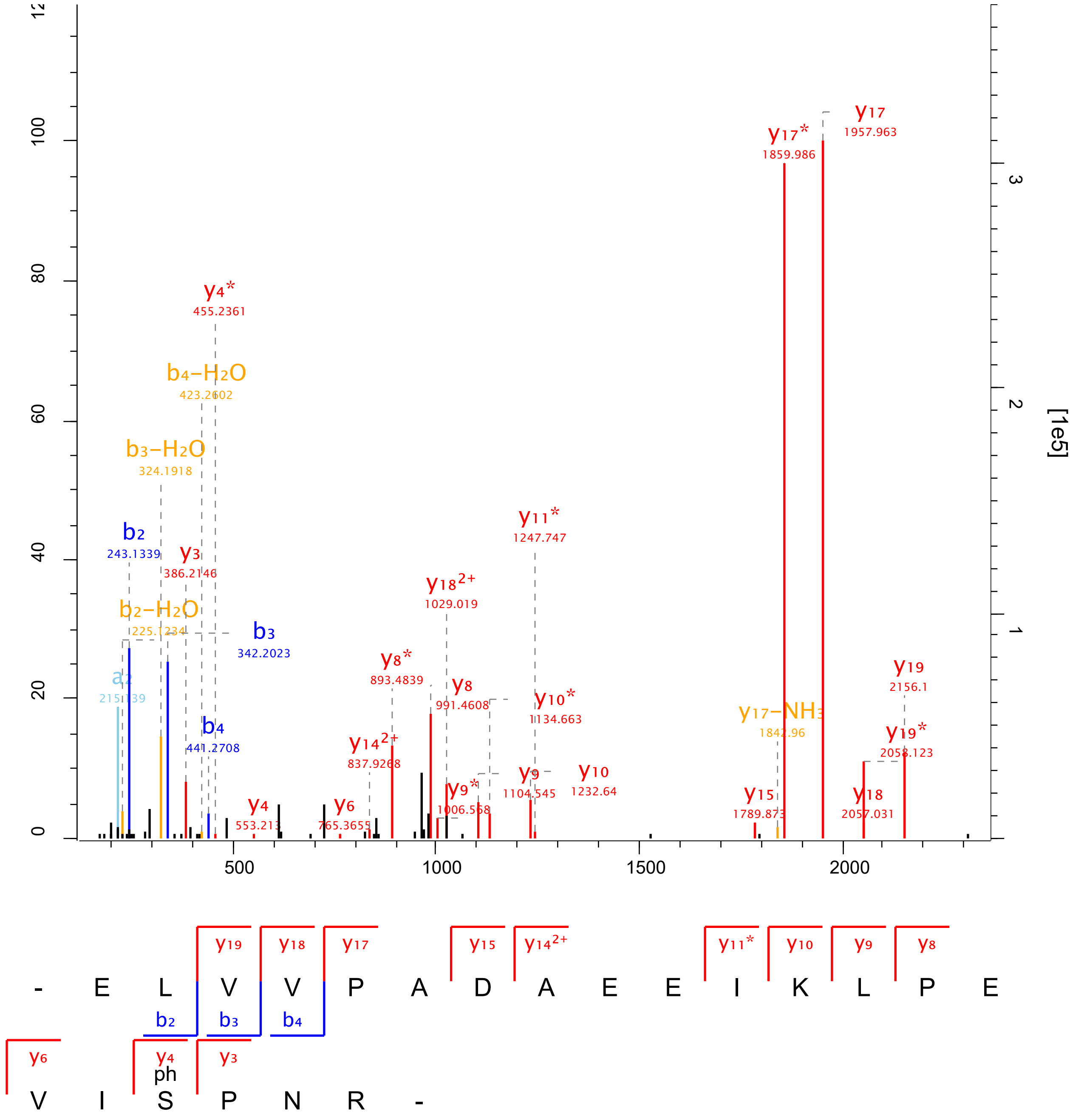

| Raw file | Scan | Method | Score | m/z |
|---|---|---|---|---|
| sys_15_1 | 41706 | FTMS; HCD | 68.58 | 1056.79 |

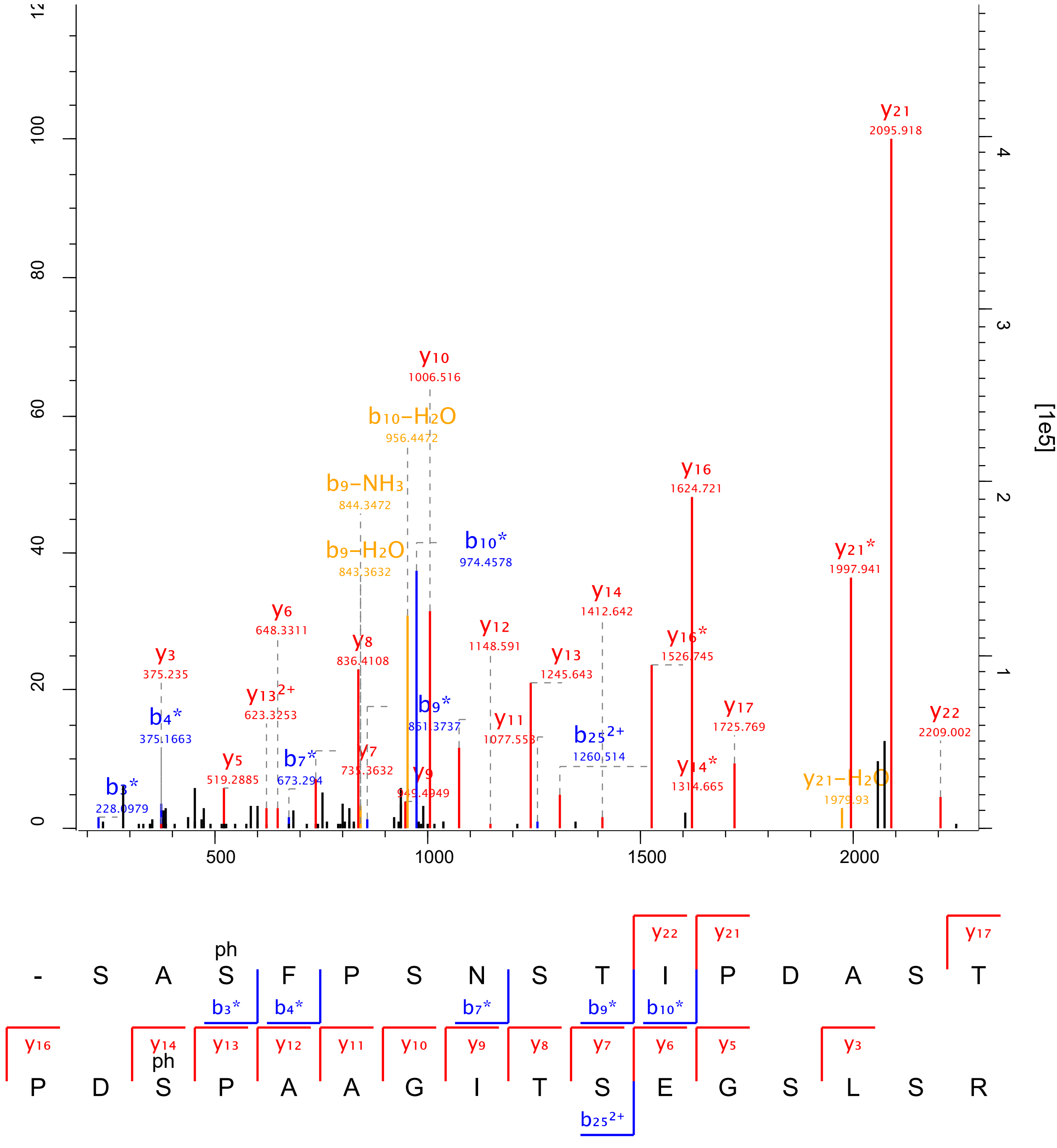

| Raw file | Scan | Method | Score | m/z |
| --- | --- | --- | --- | --- |
| sys_15_1 | 41892 | FTMS; HCD | 107.21 | 978 |

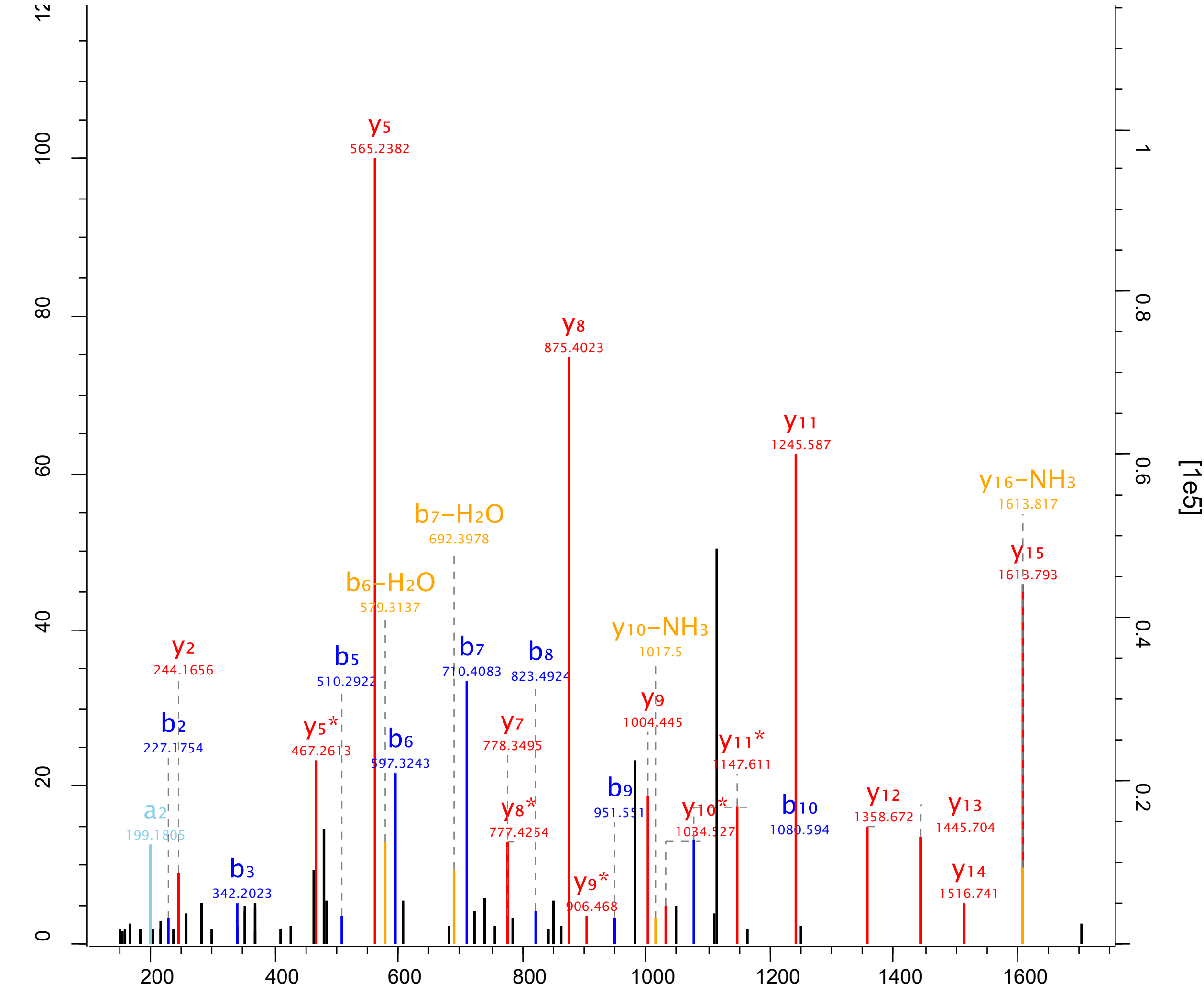


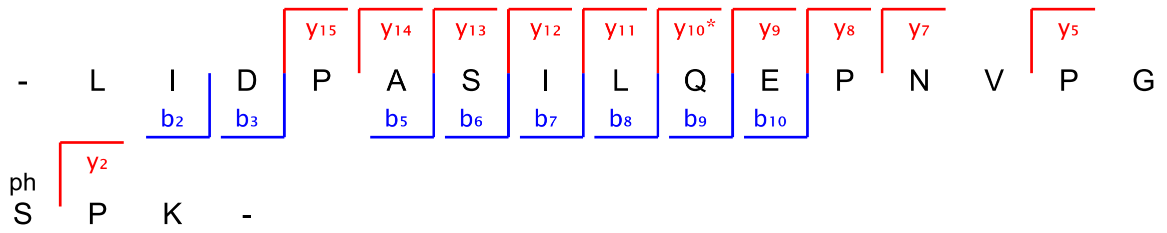

| Raw file | Scan | Method | Score | m/z |
|---|---|---|---|---|
| sys_15_1 | 4205 | FTMS; HCD | 102.73 | 570.76 |

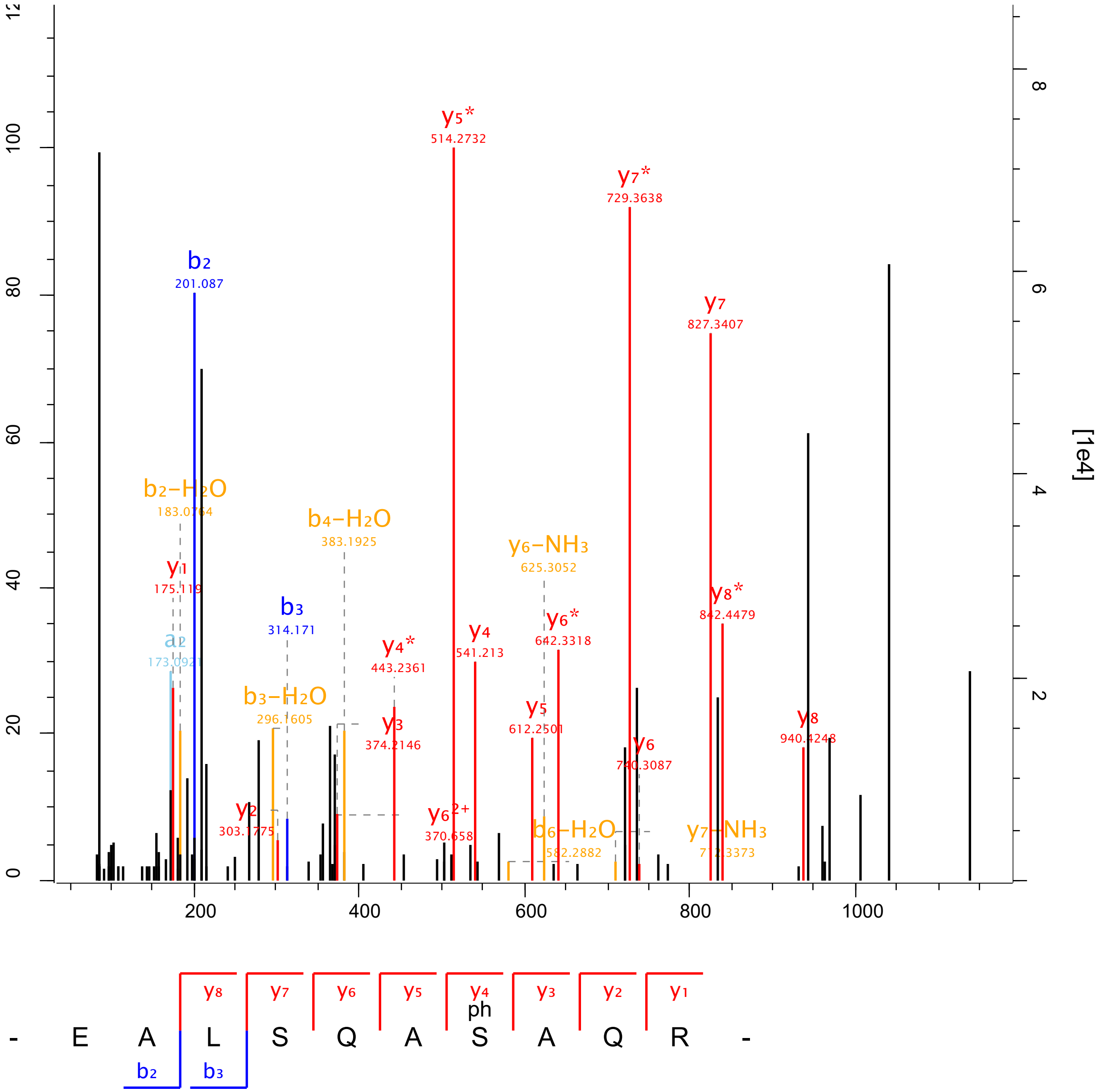

| Raw file | Scan | Method | Score | m/z |
|---|---|---|---|---|
| sys_15_1 | 4227 | FTMS; HCD | 68.04 | 580.72 |

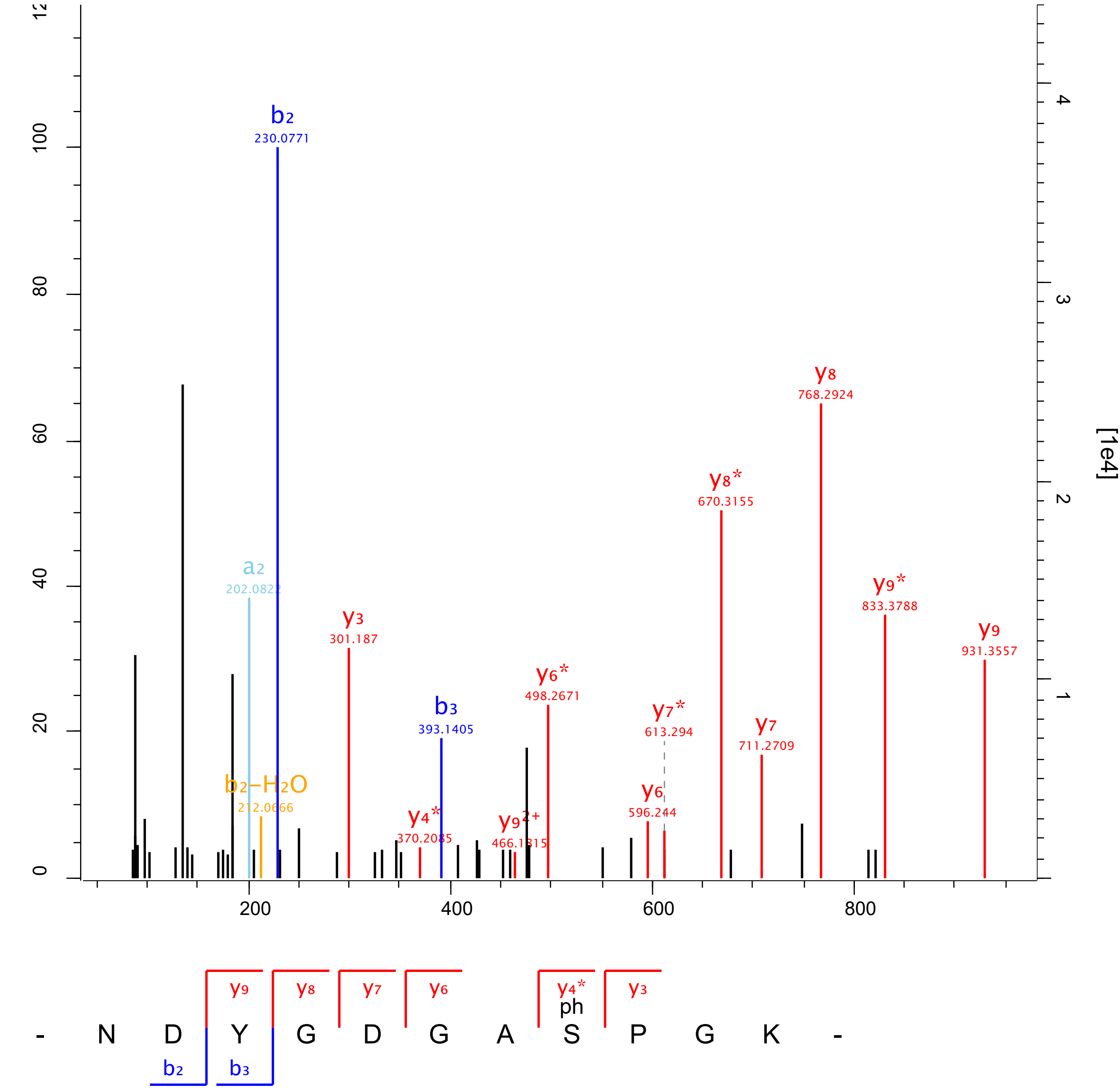

| Raw file | Scan | Method | Score | m/z |
|---|---|---|---|---|
| sys_15_1 | 42328 | FTMS; HCD | 134.26 | 771.75 |

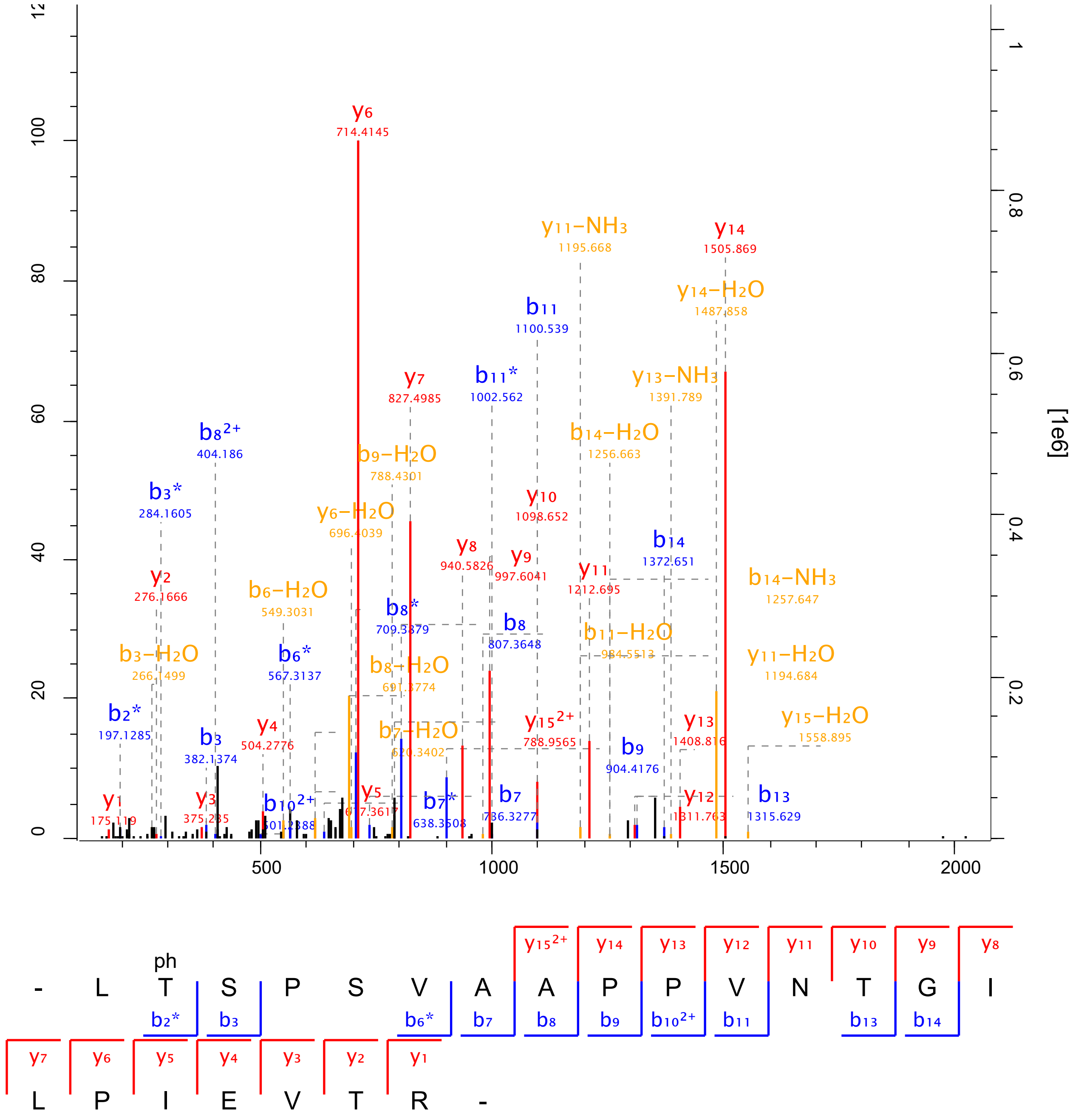

| Raw file | Scan | Method | Score | m/z |
|---|---|---|---|---|
| sys_15_1 | 4244 | FTMS; HCD | 49.3 | 549.23 |

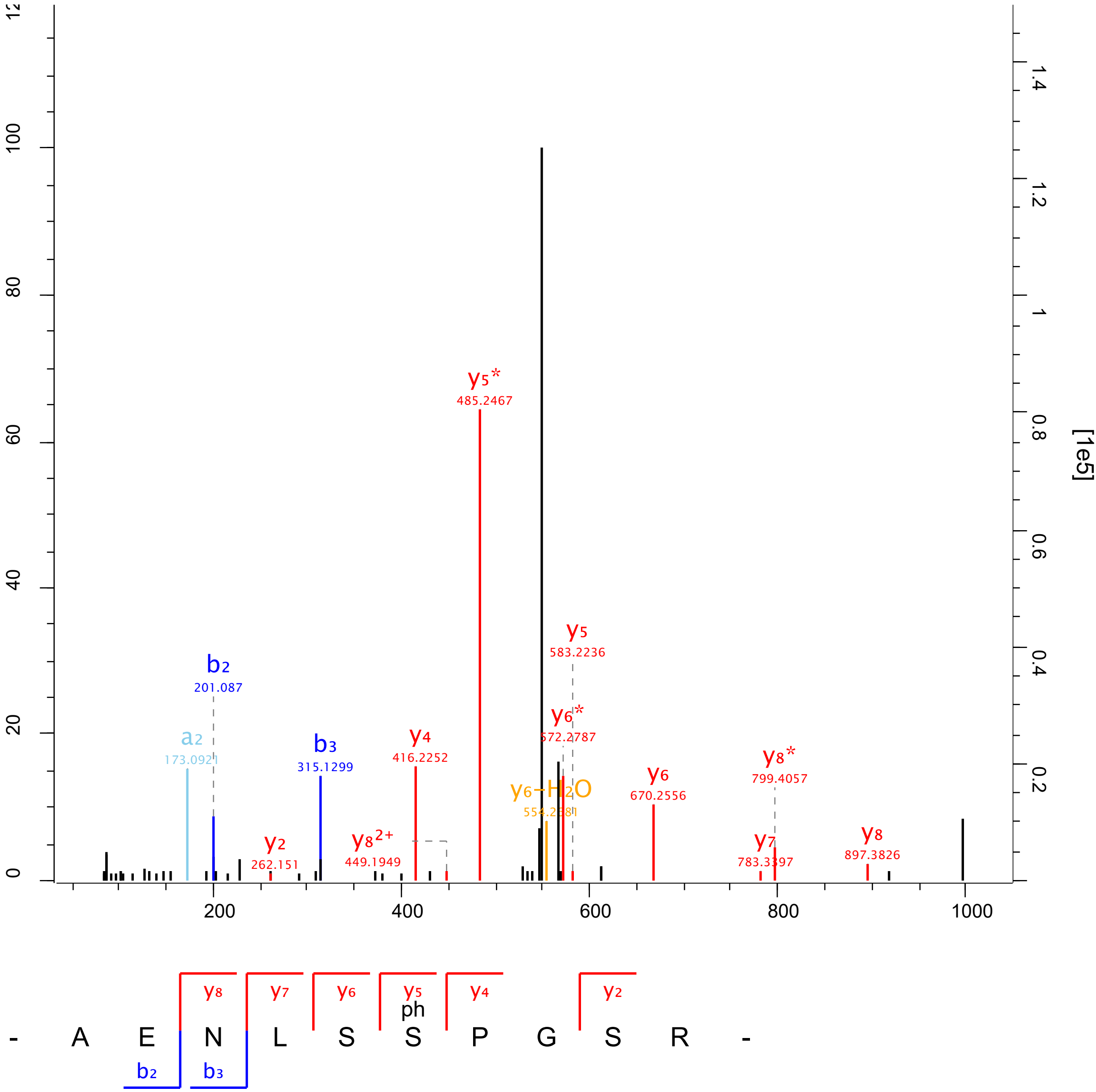

| Raw file | Scan | Method | Score | m/z |
|---|---|---|---|---|
| sys_15_1 | 4256 | FTMS; HCD | 43.59 | 688.26 |

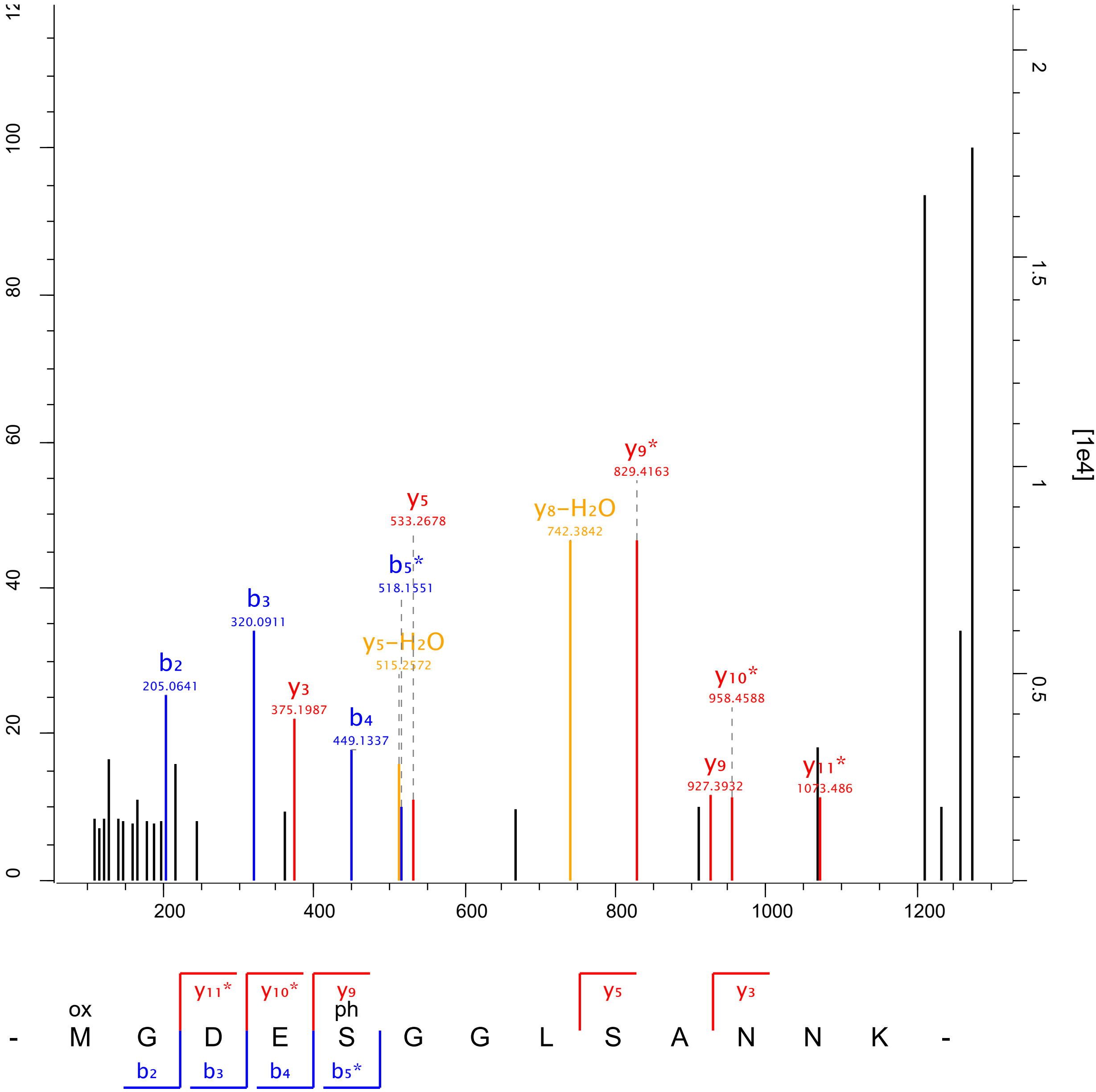

| Raw file | Scan | Method | Score | m/z |
|---|---|---|---|---|
| sys_15_1 | 42613 | FTMS; HCD | 63.14 | 710.01 |

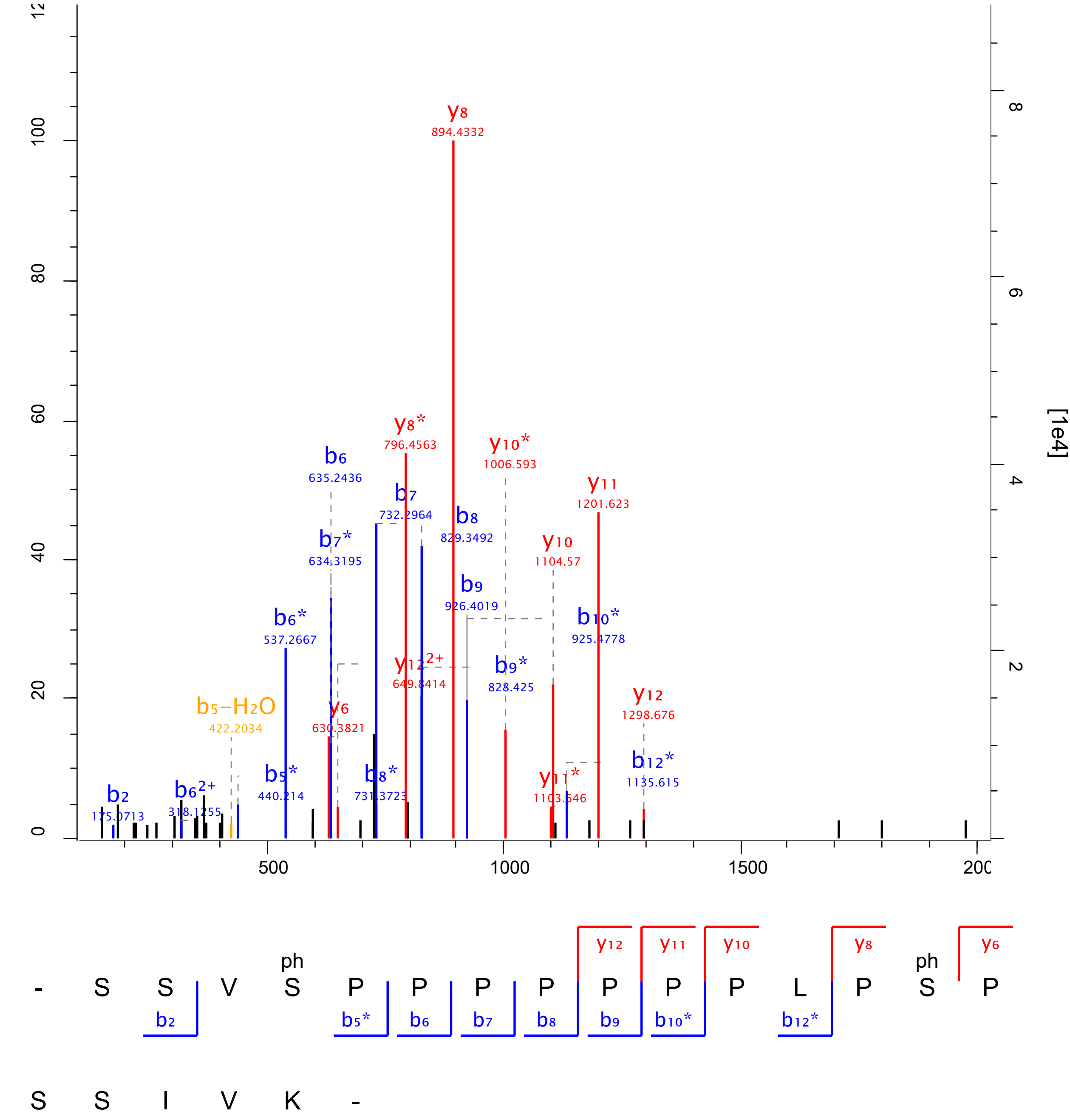

| Raw file | Scan | Method | Score | m/z |
|---|---|---|---|---|
| sys_15_1 | 4294 | FTMS; HCD | 81.79 | 645.76 |

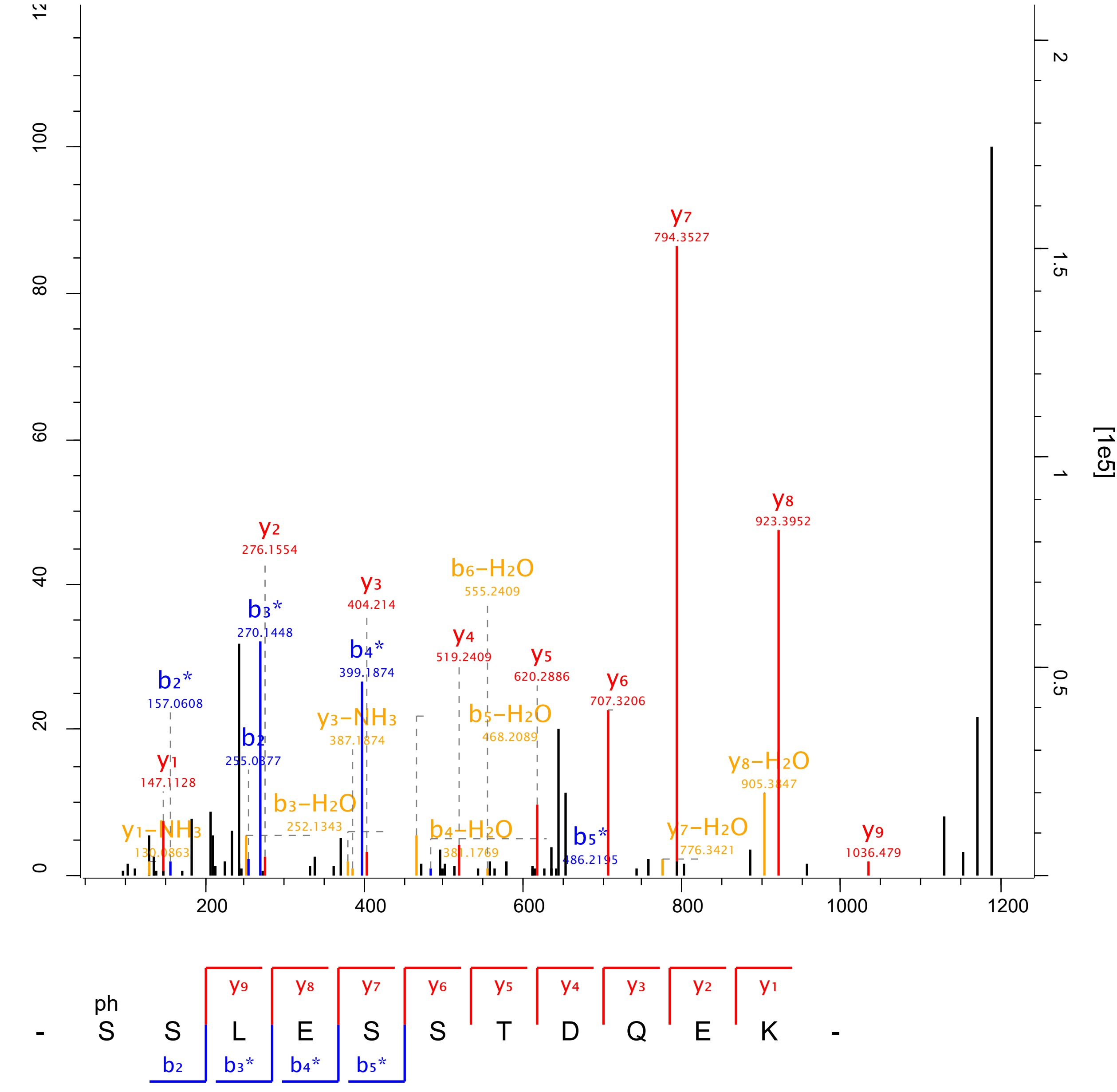

| Raw file | Scan | Method | Score | m/z |
|---|---|---|---|---|
| sys_15_1 | 42954 | FTMS; HCD | 74.3 | 770.83 |

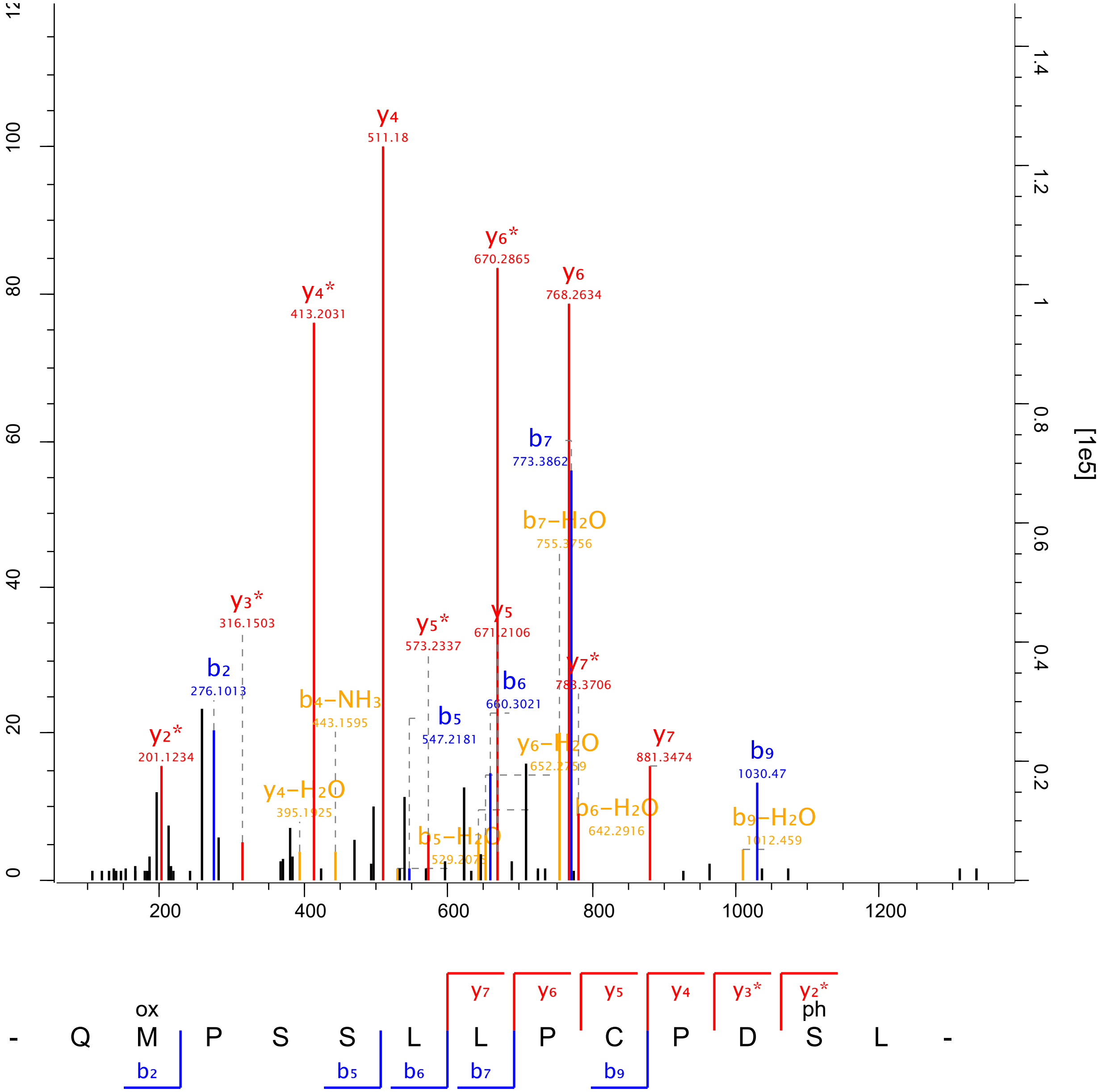

| Raw file | Scan | Method | Score | m/z |
|---|---|---|---|---|
| sys_15_1 | 43087 | FTMS; HCD | 178.32 | 1002.41 |

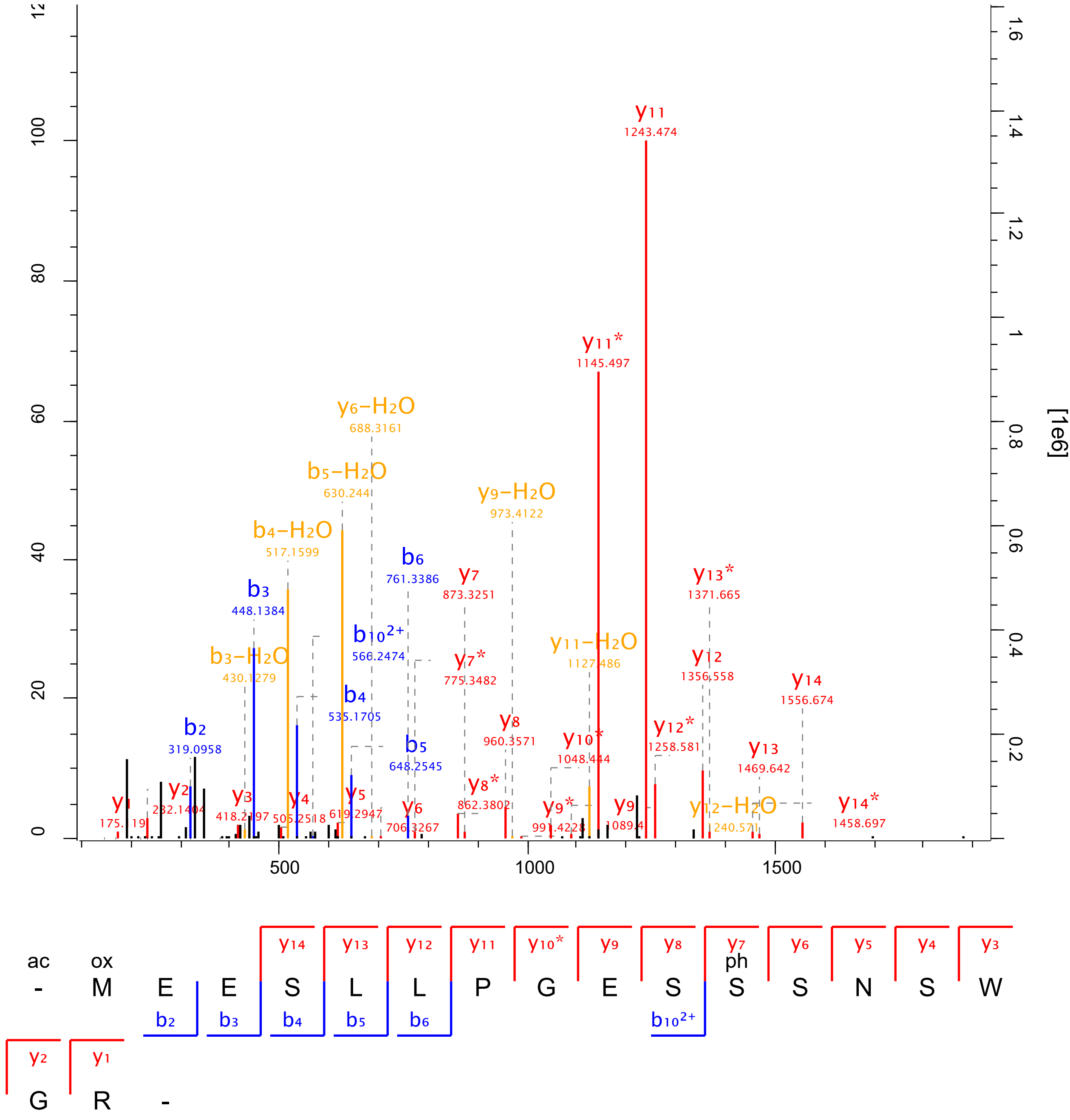

| Raw file | Scan | Method | Score | m/z |
|---|---|---|---|---|
| sys_15_1 | 4332 | FTMS; HCD | 80.69 | 563.2 |

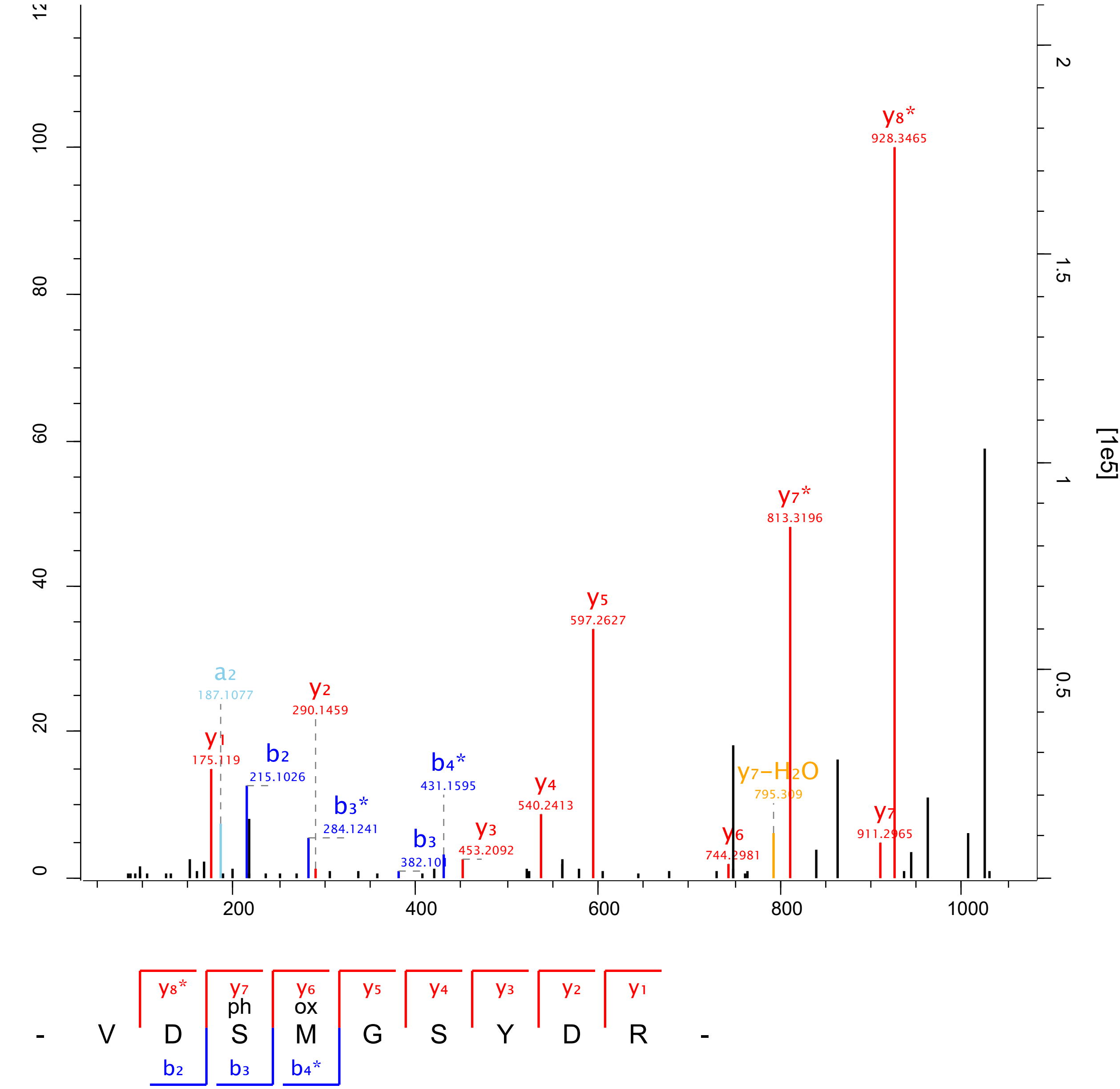

| Raw file | Scan | Method | Score | m/z |
|---|---|---|---|---|
| sys_15_1 | 4338 | FTMS; HCD | 90.61 | 548.74 |

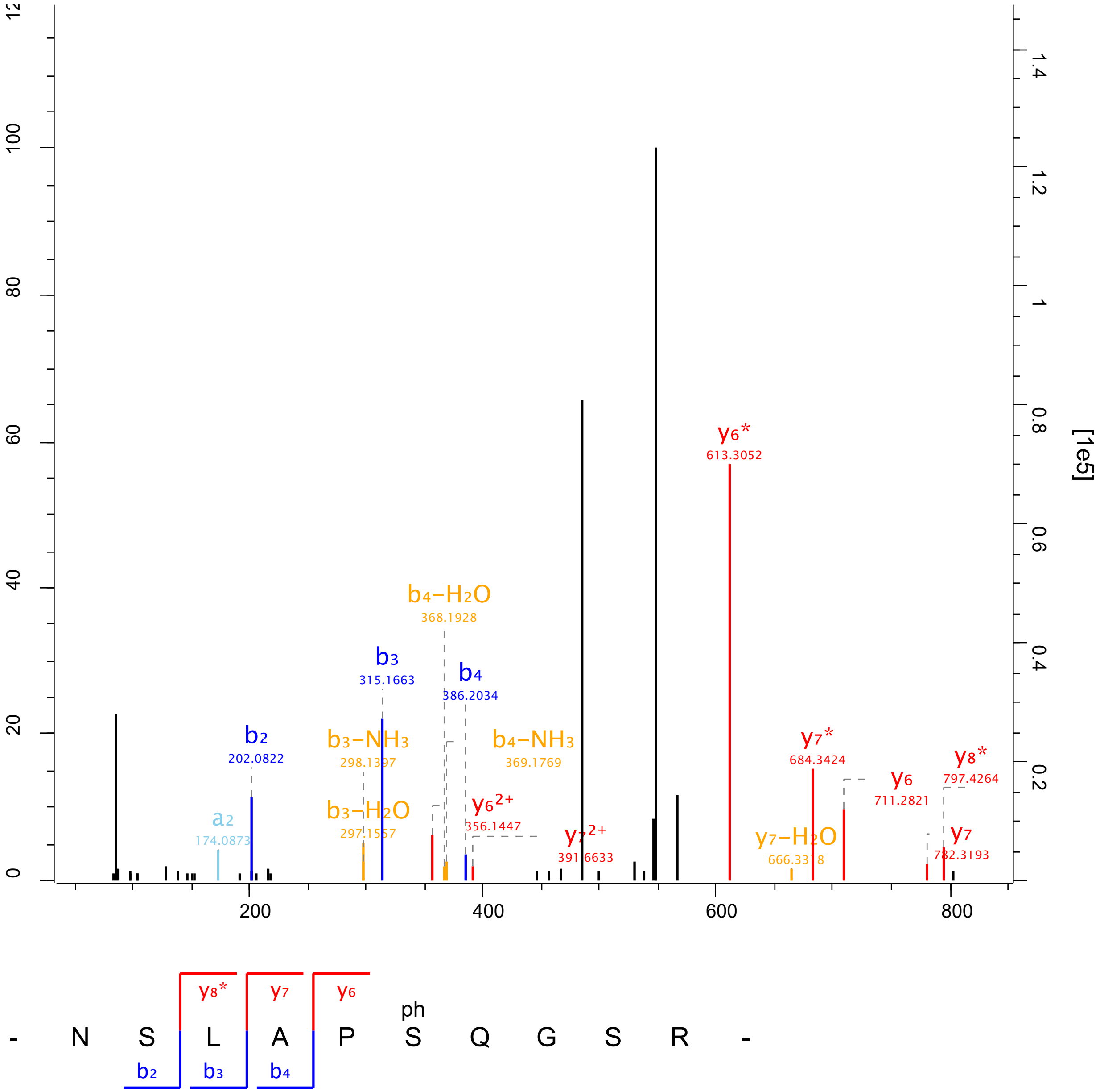

| Raw file | Scan | Method | Score | m/z |
| --- | --- | --- | --- | --- |
| sys_15_1 | 43423 | FTMS; HCD | 142.45 | 847.06 |

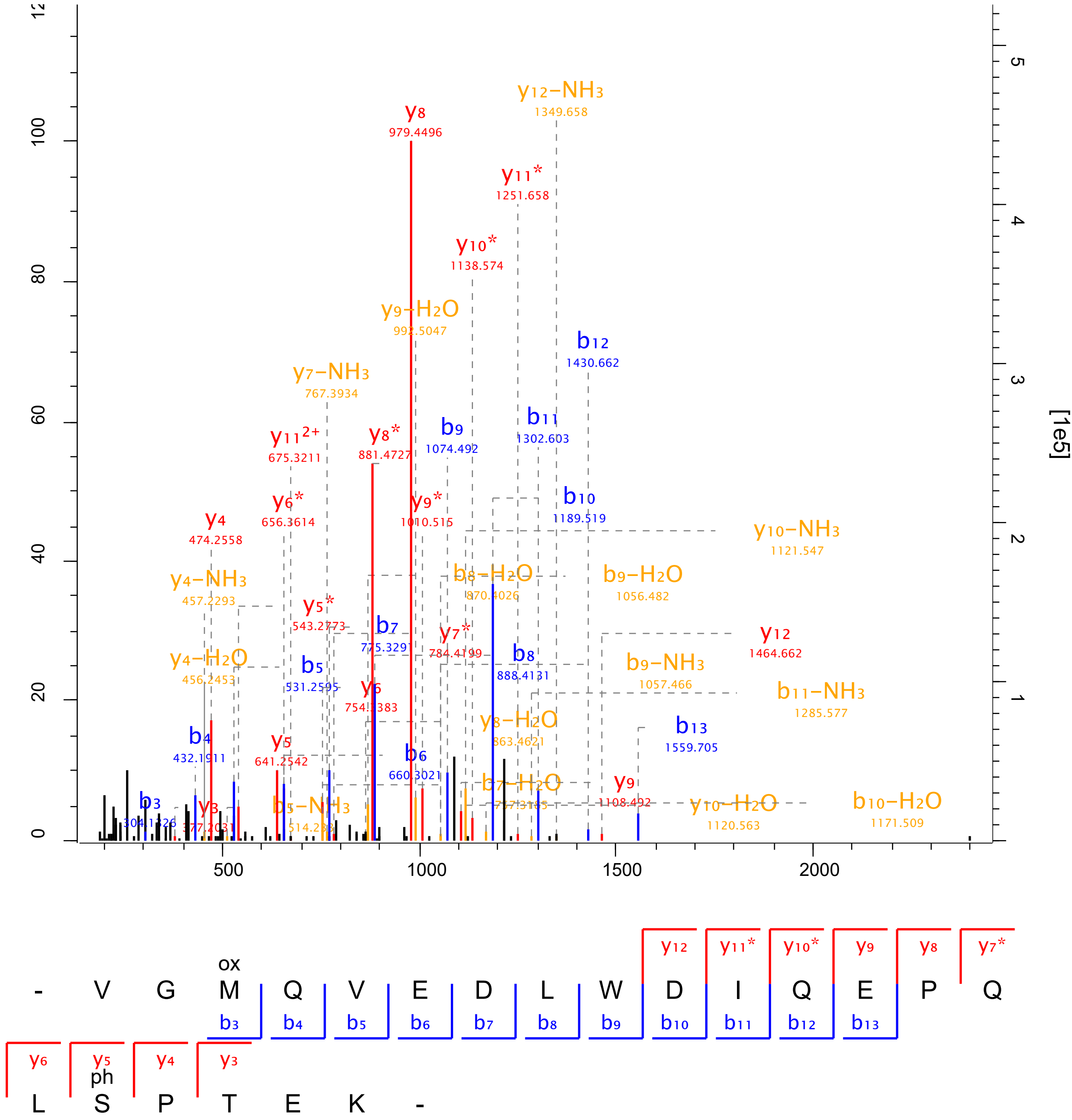

| Raw file | Scan | Method | Score | m/z |
|---|---|---|---|---|
| sys_15_1 | 44236 | FTMS; HCD | 120.84 | 801.76 |

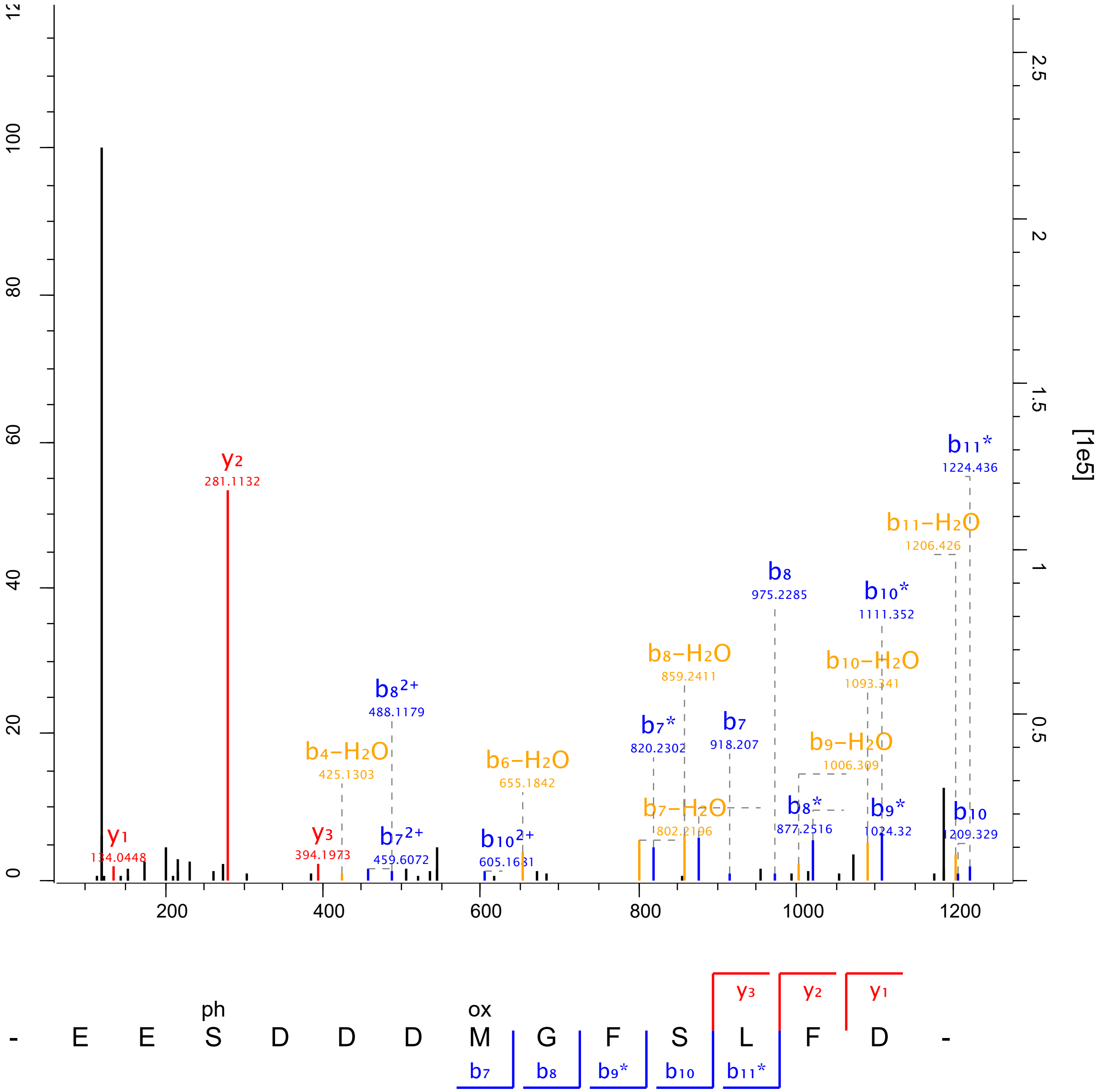

| Raw file | Scan | Method | Score | m/z |
|---|---|---|---|---|
| sys_15_1 | 44495 | FTMS; HCD | 157.5 | 1014.13 |

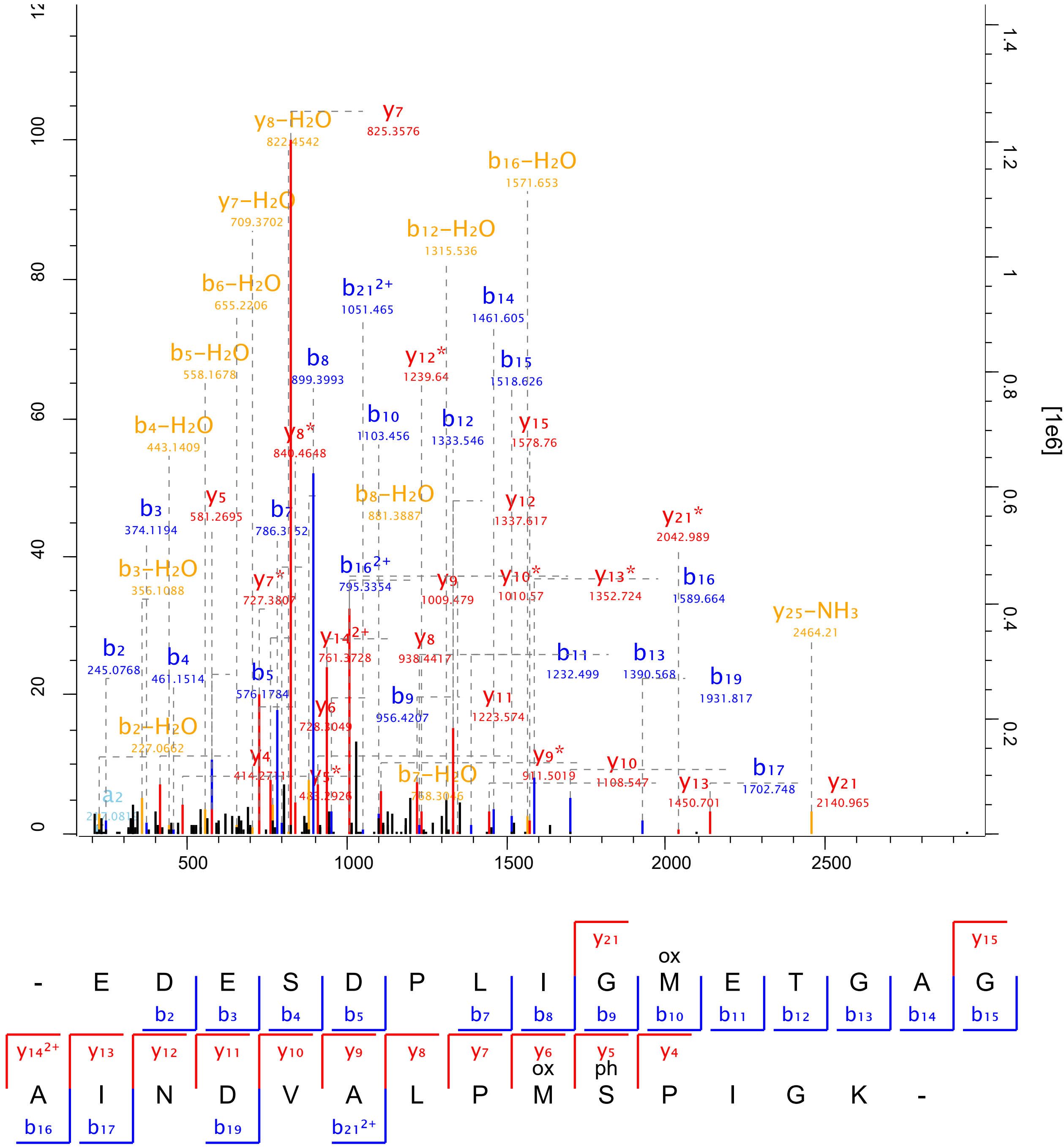

| Raw file | Scan | Method | Score | m/z |
|---|---|---|---|---|
| sys_15_1 | 4501 | FTMS; HCD | 101.71 | 543.21 |

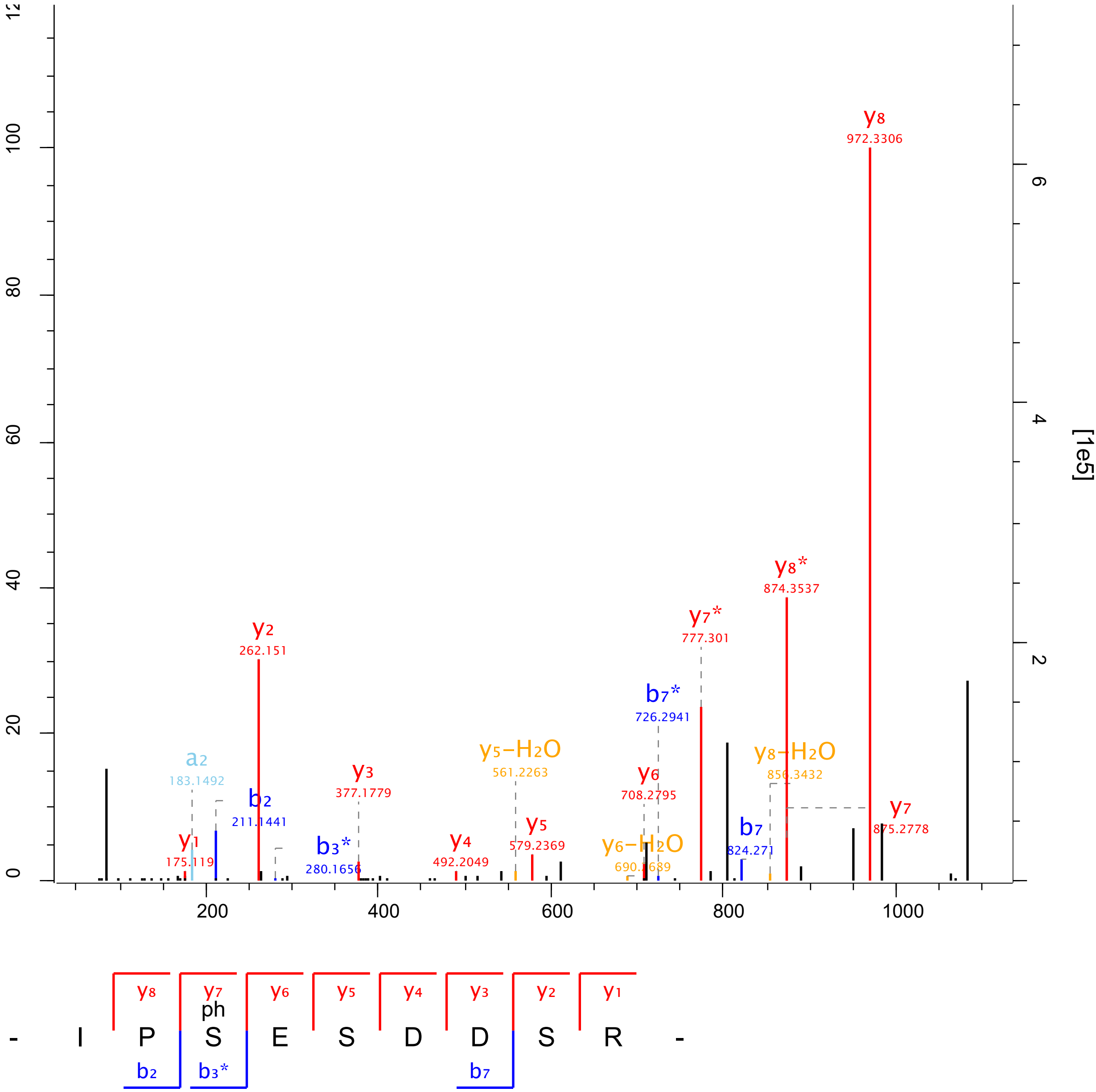

| Raw file | Scan | Method | Score | m/z |
|---|---|---|---|---|
| sys_15_1 | 4553 | FTMS; HCD | 77.18 | 414.84 |

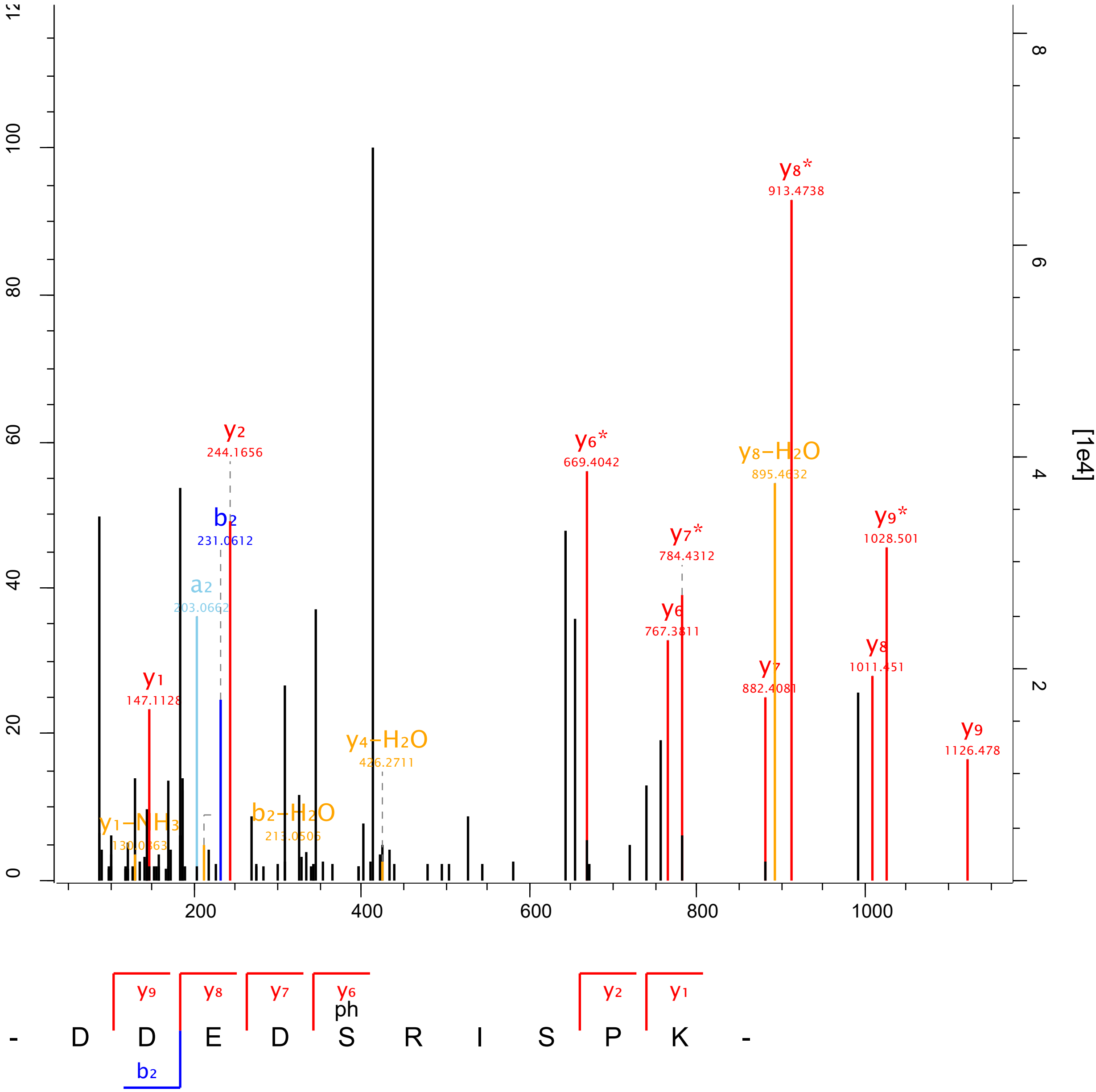

| Raw file | Scan | Method | Score | m/z |
|---|---|---|---|---|
| sys_15_1 | 45702 | FTMS; HCD | 106.86 | 1044.39 |

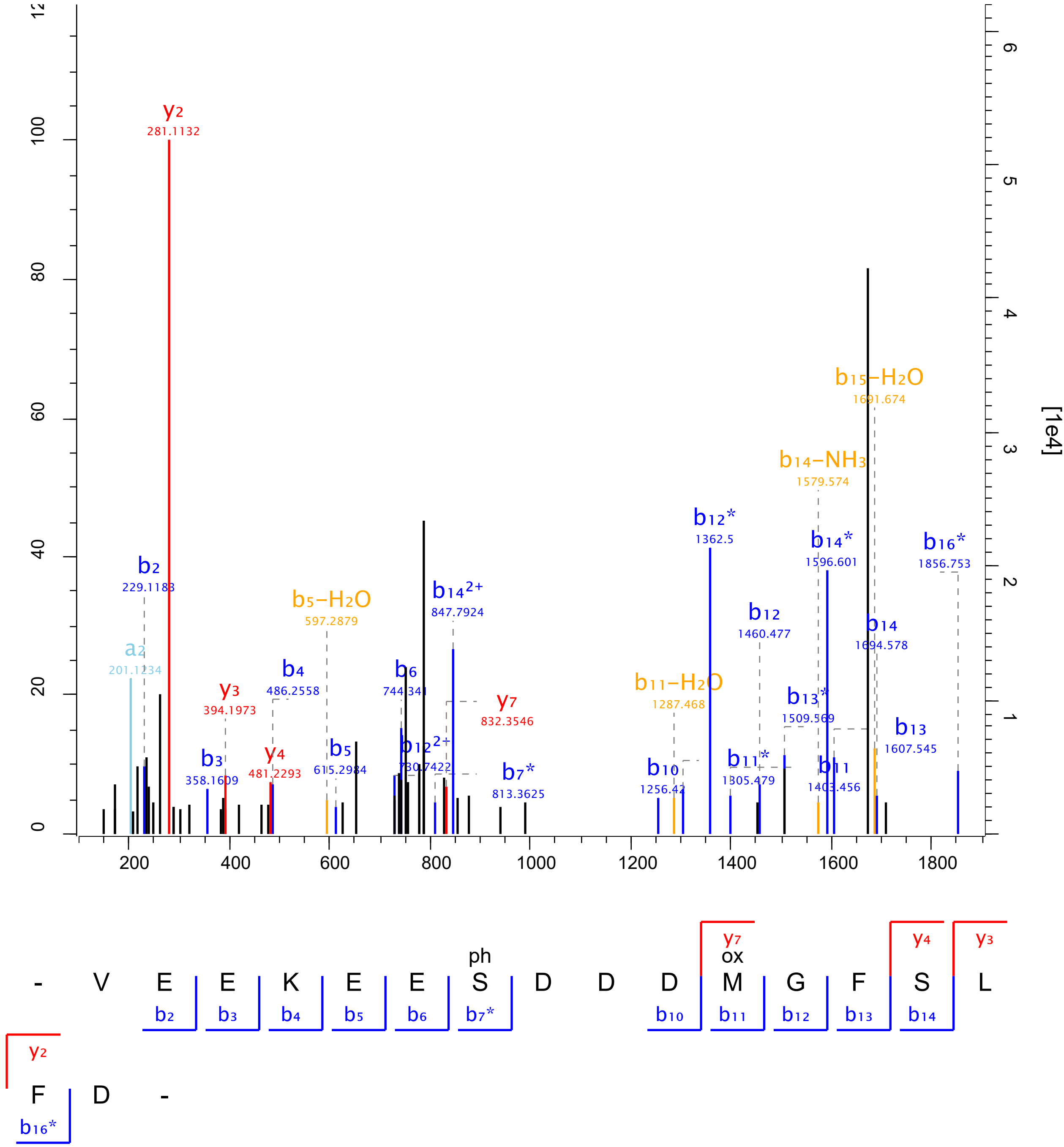

| Raw file | Scan | Method | Score | m/z |
|---|---|---|---|---|
| sys_15_1 | 46699 | FTMS; HCD | 56.72 | 903.89 |

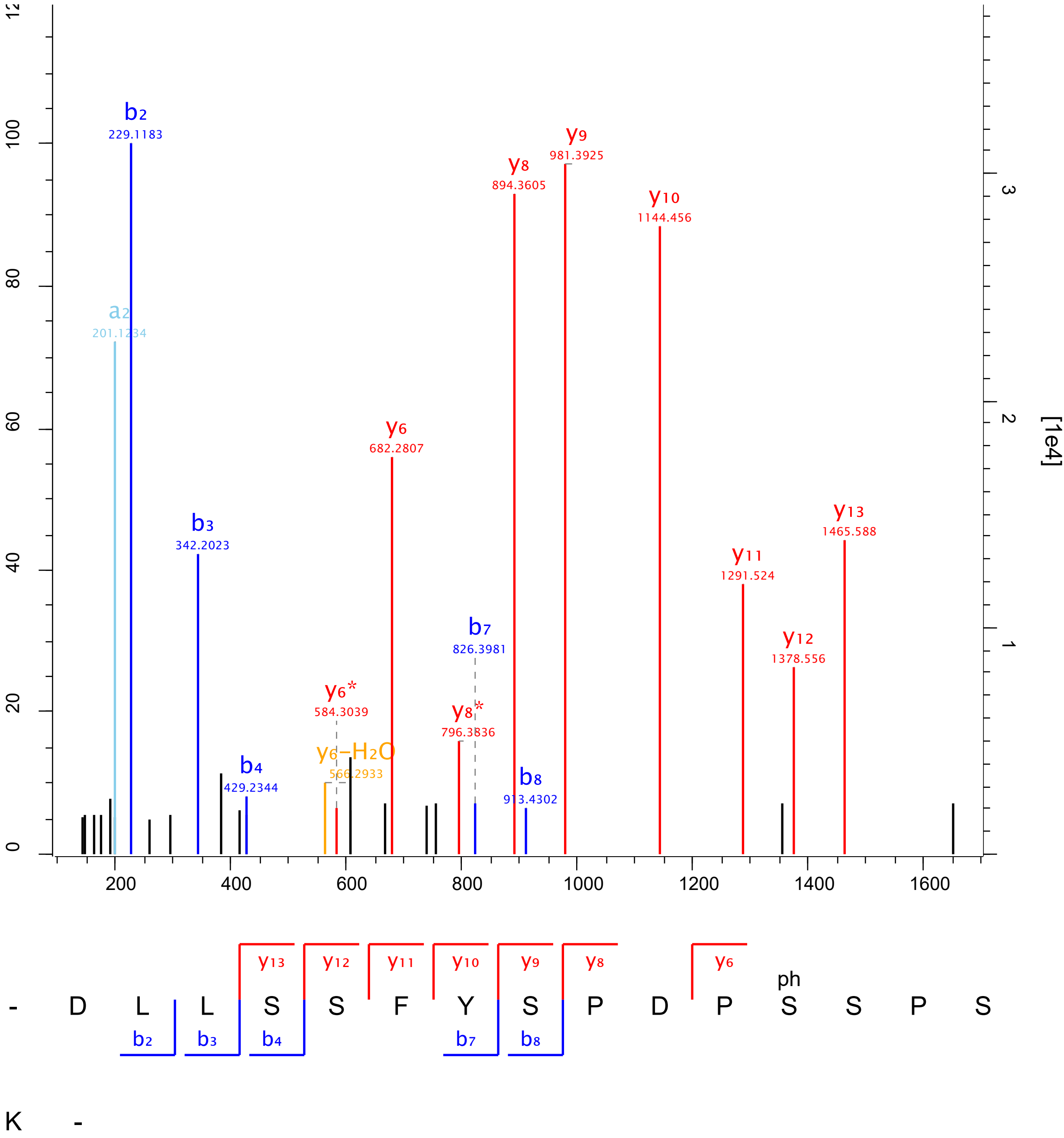

| Raw file | Scan | Method | Score | m/z |
|---|---|---|---|---|
| sys_15_1 | 46904 | FTMS; HCD | 52.17 | 824.9 |

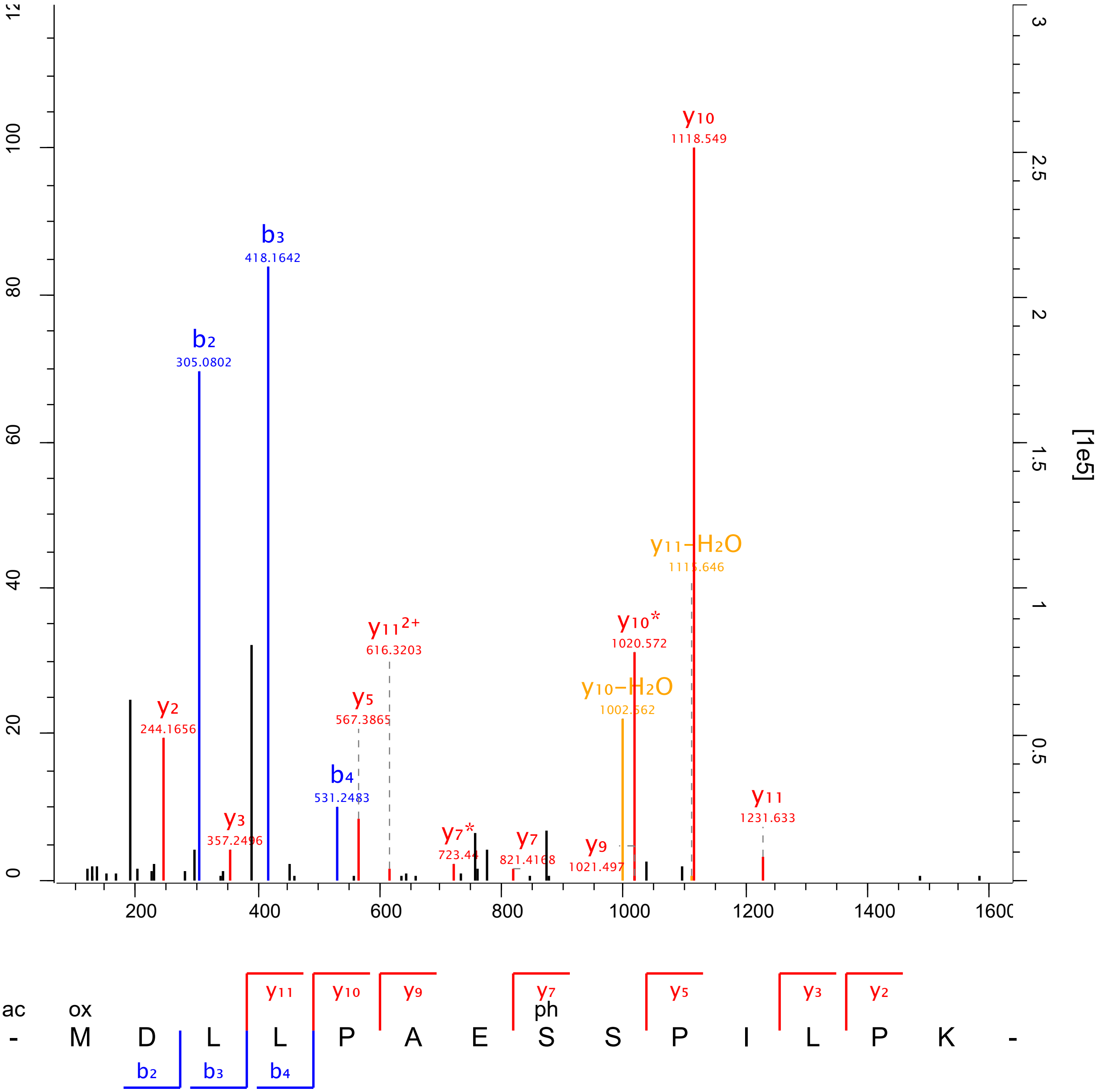

| Raw file | Scan | Method | Score | m/z |
|---|---|---|---|---|
| sys_15_1 | 46923 | FTMS; HCD | 190.55 | 885.89 |

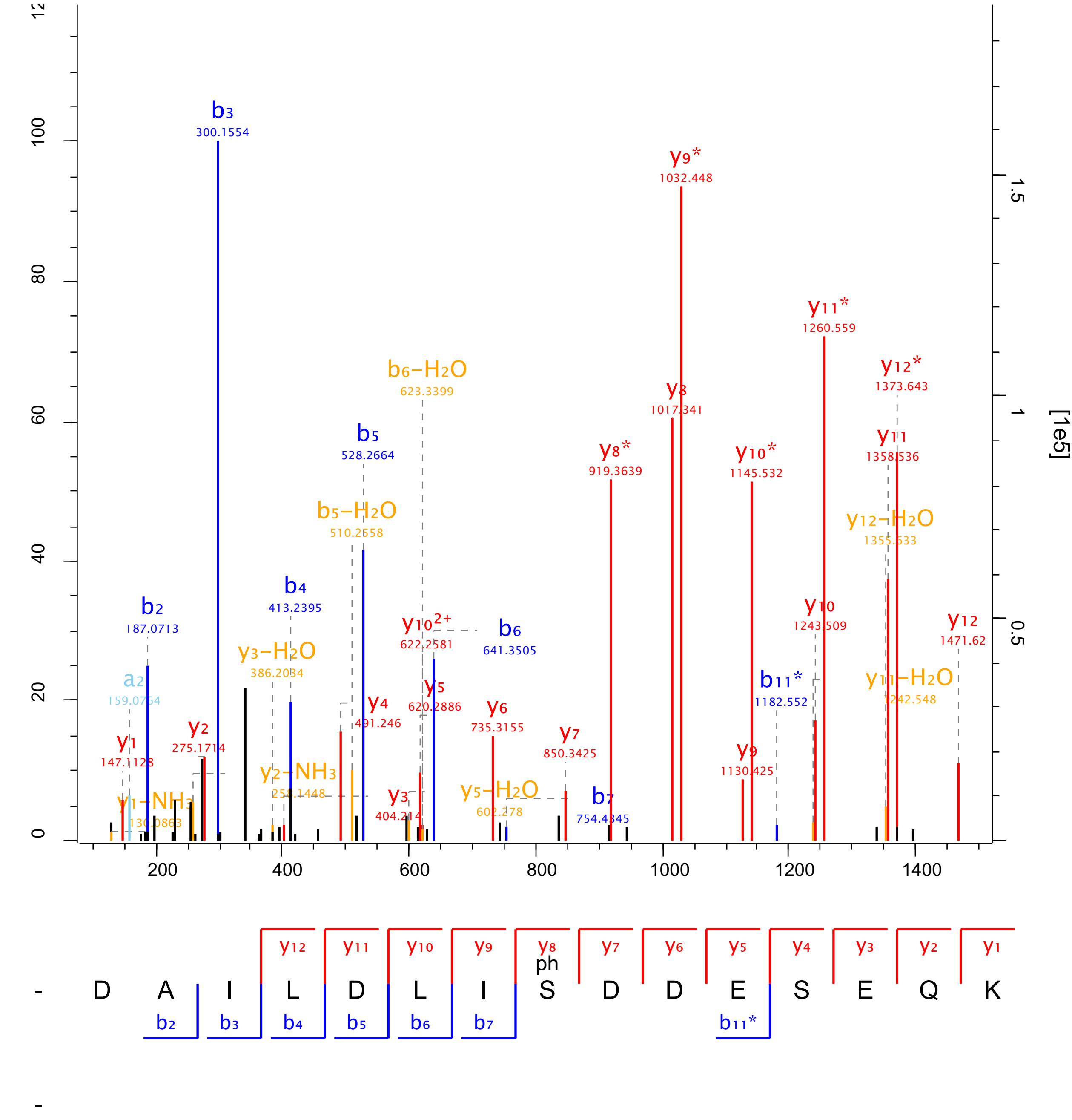

| Raw file | Scan | Method | Score | m/z |
|---|---|---|---|---|
| sys_15_1 | 4711 | FTMS; HCD | 188.48 | 886.85 |

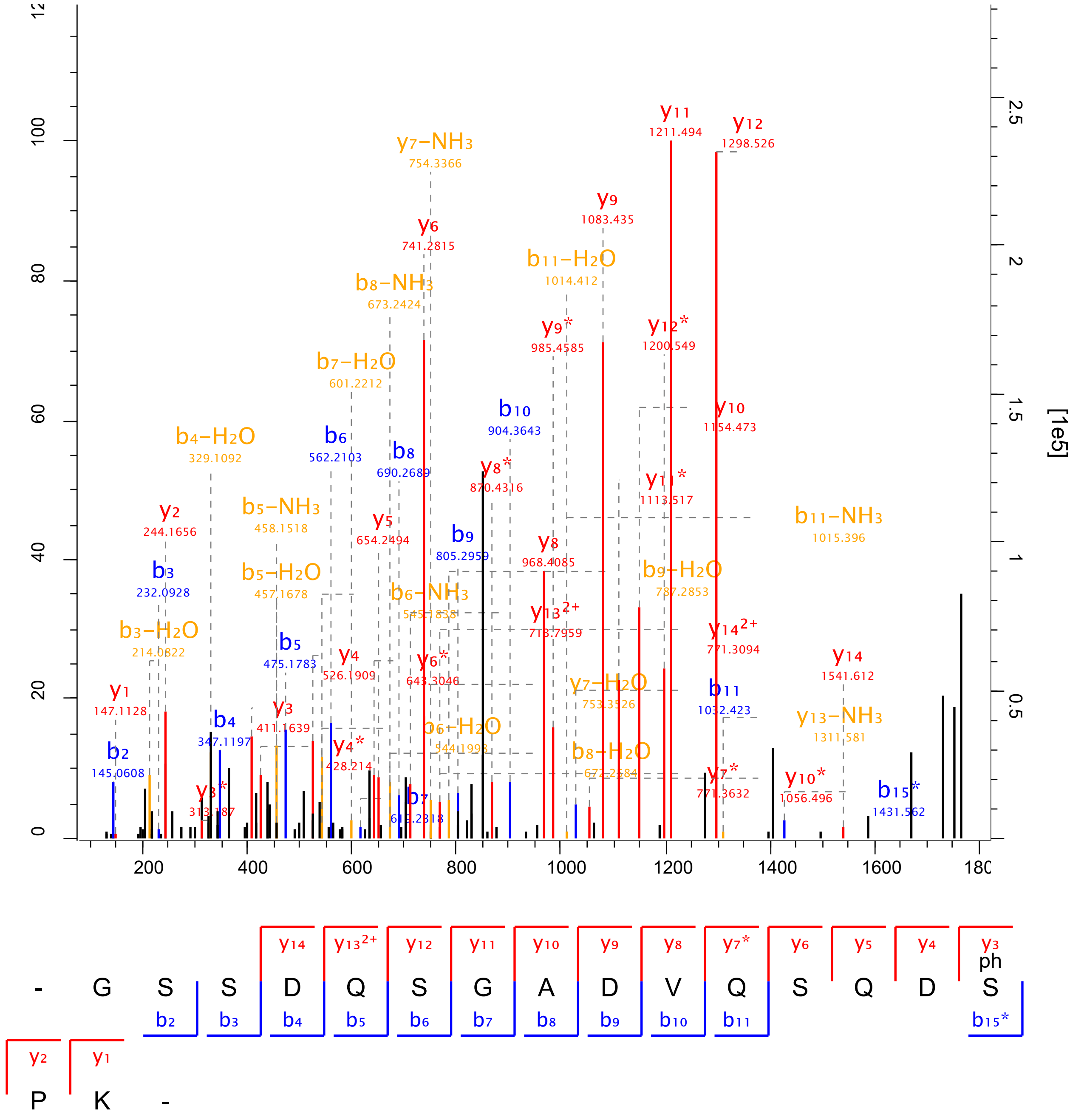

| Raw file | Scan | Method | Score | m/z |
|---|---|---|---|---|
| sys_15_1 | 4723 | FTMS; HCD | 63.3 | 620.26 |

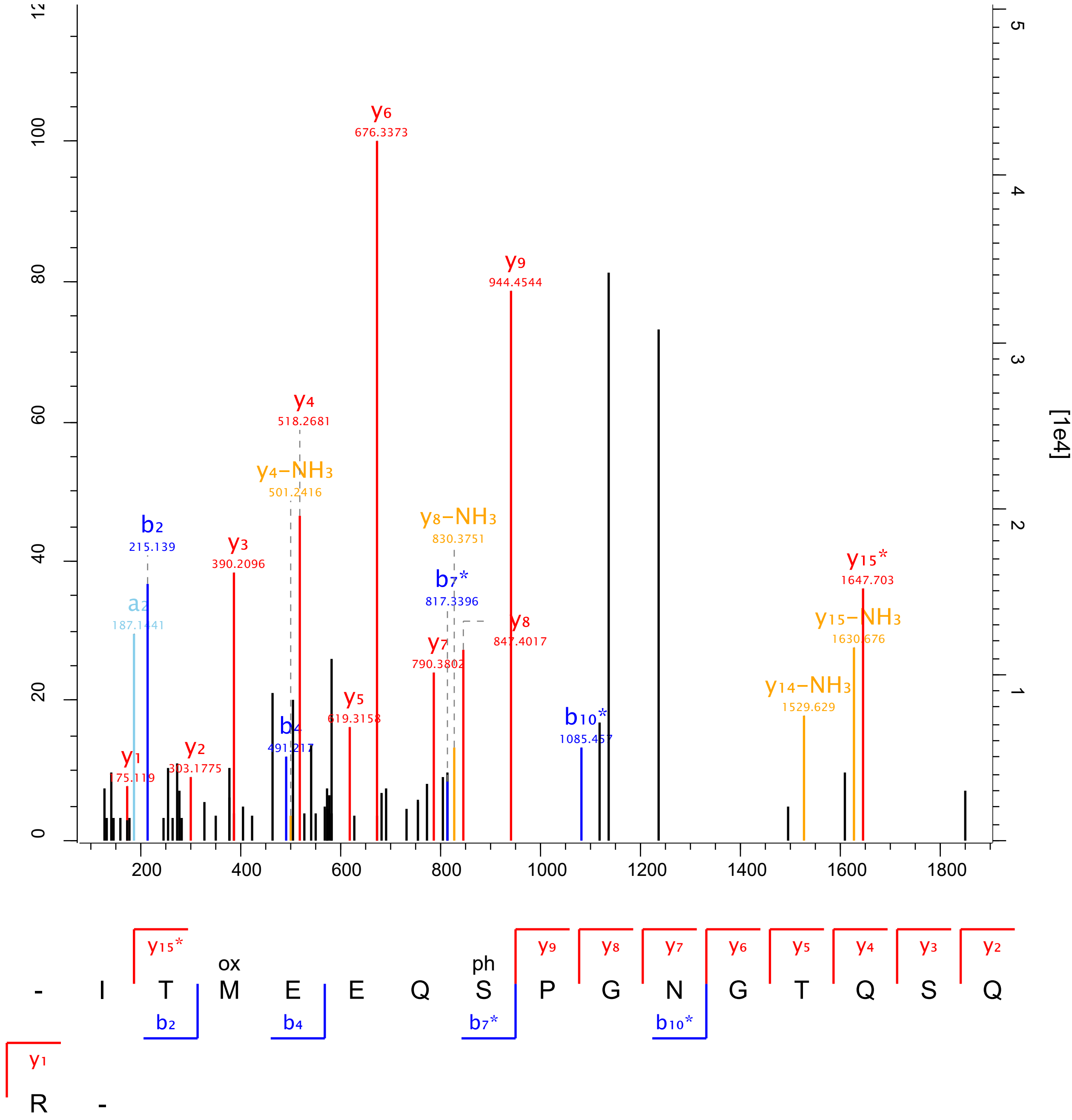

| Raw file | Scan | Method | Score | m/z |
| --- | --- | --- | --- | --- |
| sys_15_1 | 47430 | FTMS; HCD | 67.11 | 701.68 |

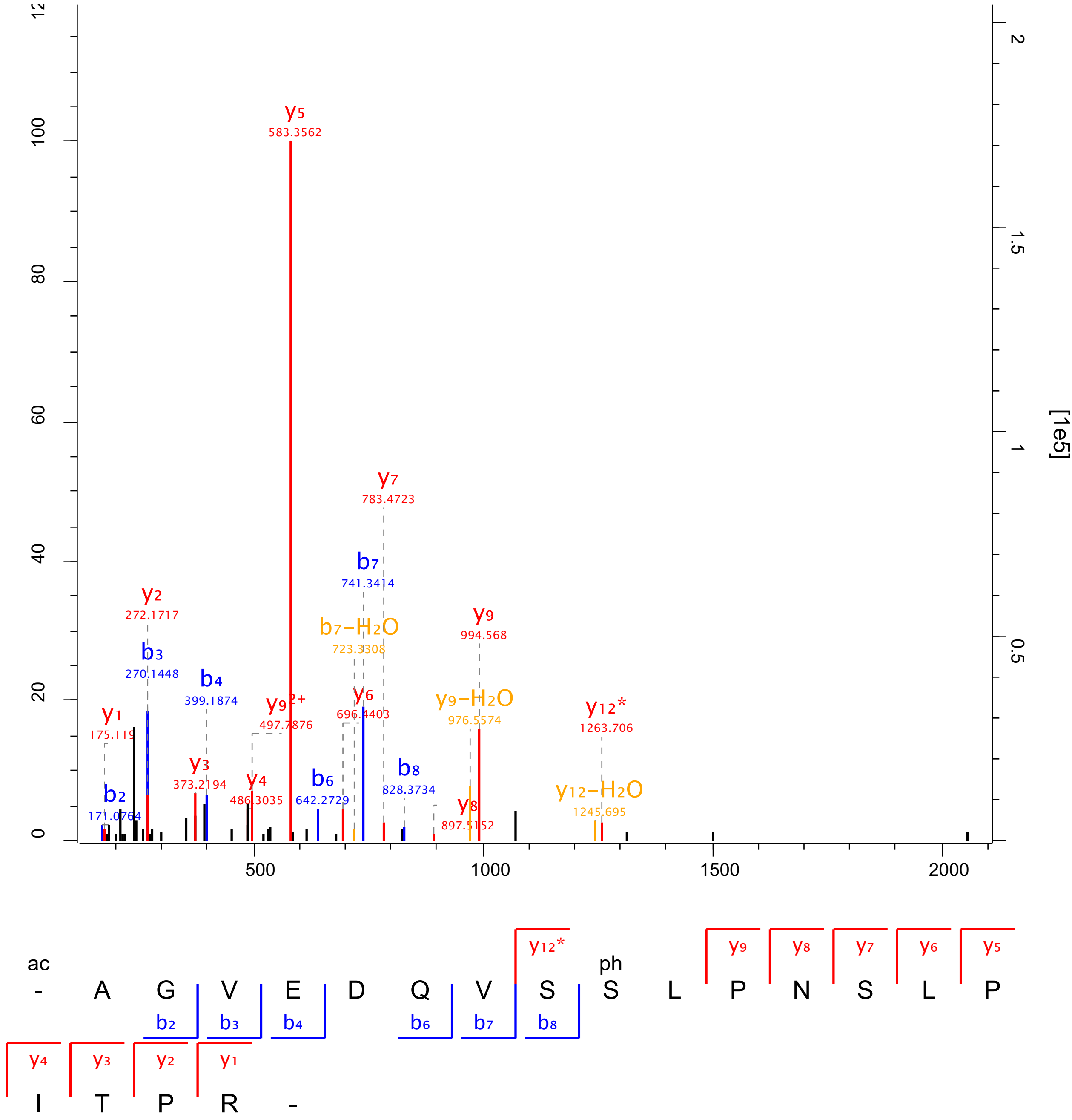

| Raw file | Scan | Method | Score | m/z |
|---|---|---|---|---|
| sys_15_1 | 47474 | FTMS; HCD | 134.04 | 1043.38 |

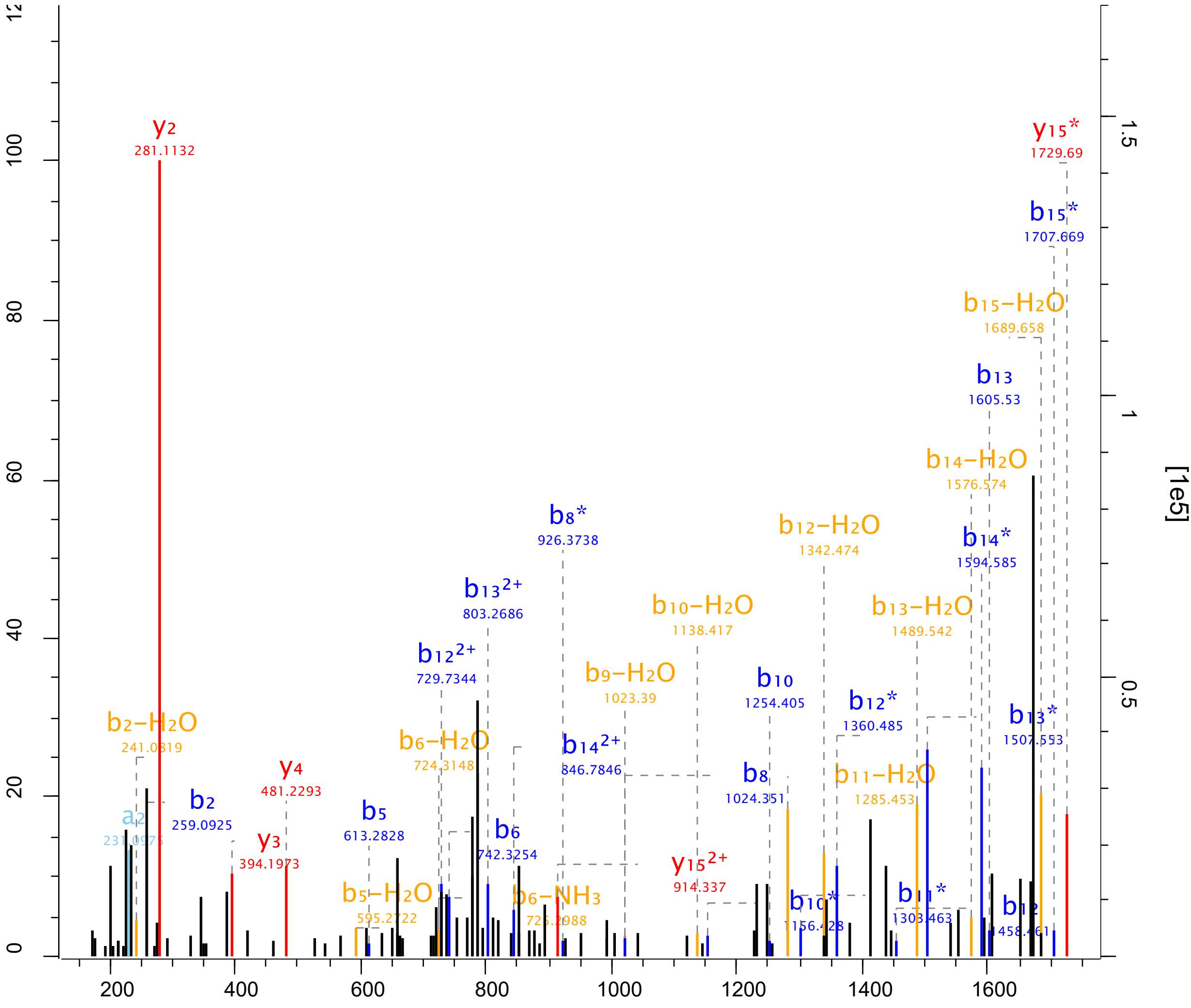


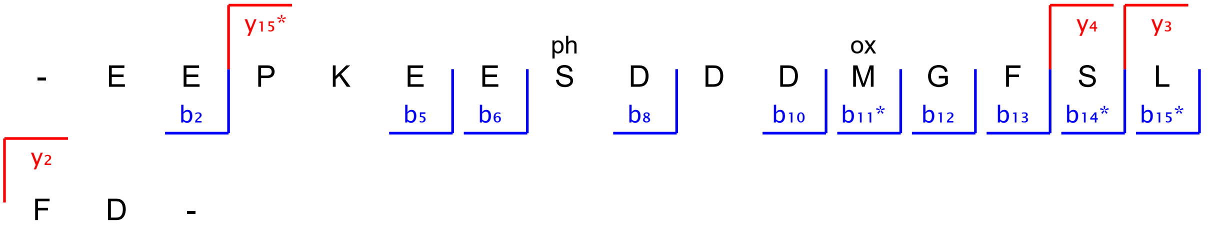

| Raw file | Scan | Method | Score | m/z |
|---|---|---|---|---|
| sys_15_1 | 4748 | FTMS; HCD | 78.33 | 668.78 |

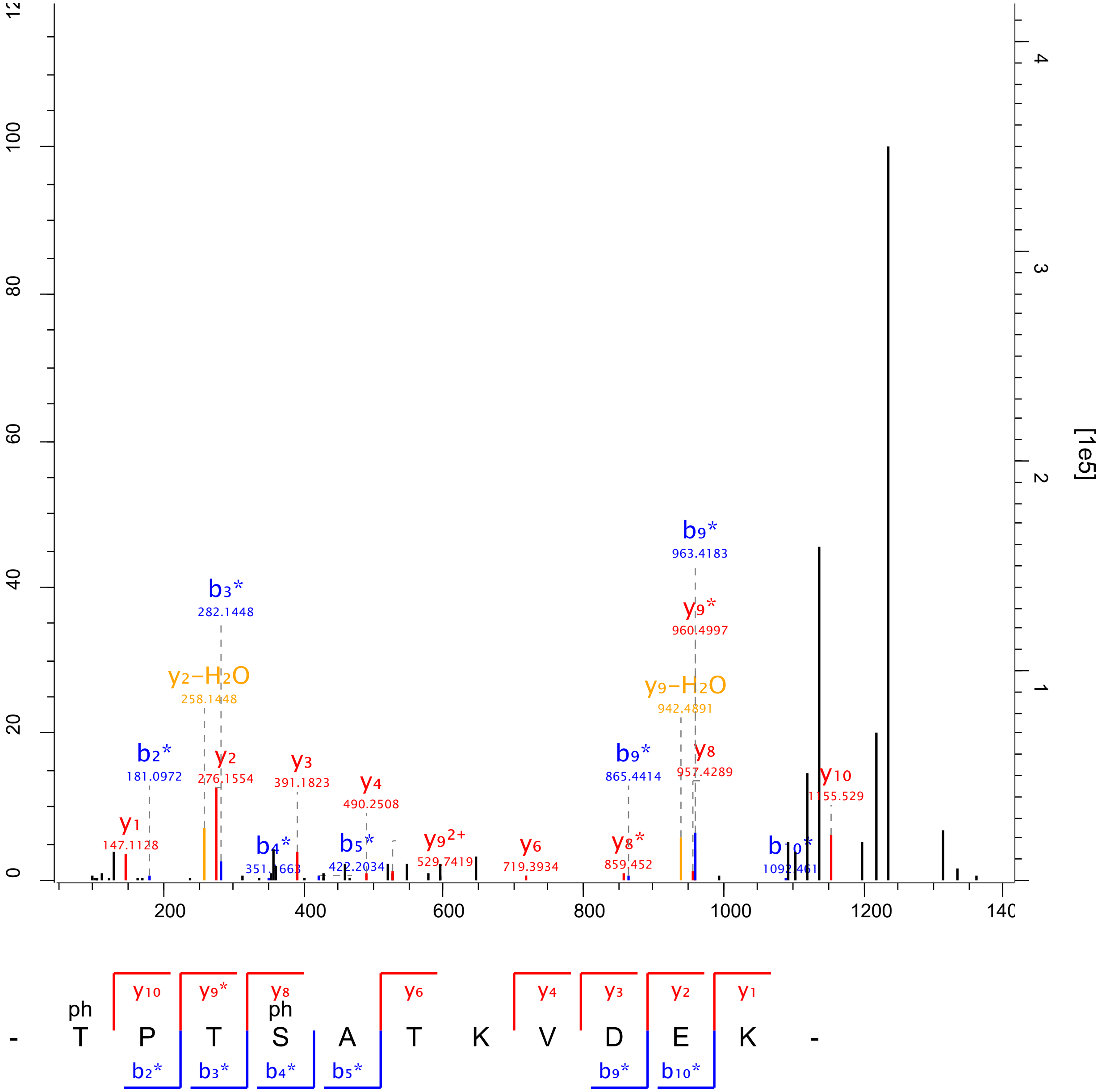

| Raw file | Scan | Method | Score | m/z |
|---|---|---|---|---|
| sys_15_1 | 48276 | FTMS; HCD | 149.44 | 944.42 |

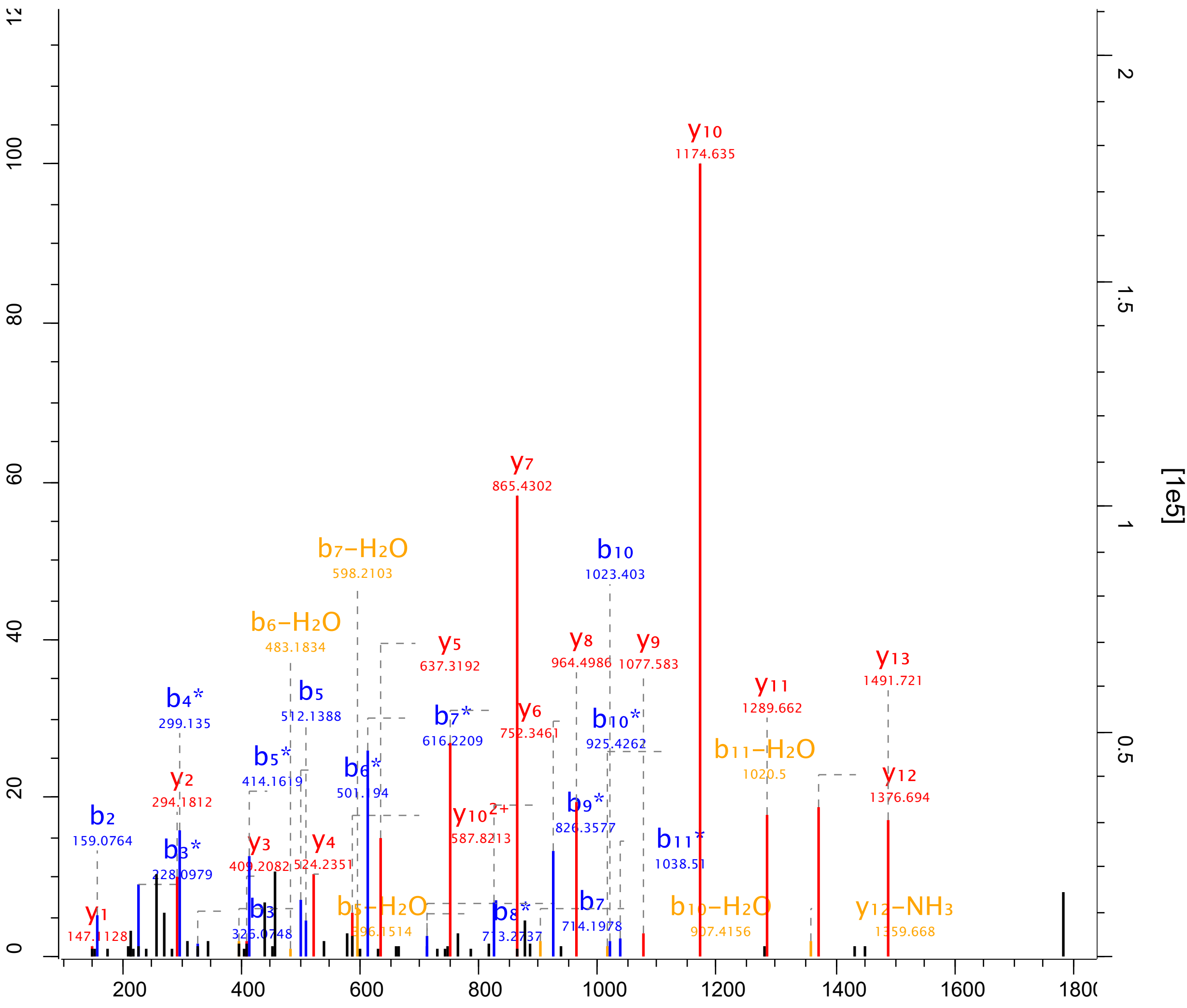


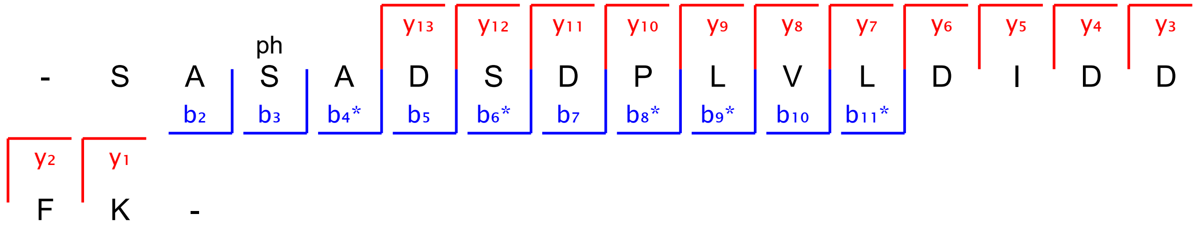

| Raw file | Scan | Method | Score | m/z |
|---|---|---|---|---|
| sys_15_1 | 4858 | FTMS; HCD | 66.27 | 604.77 |

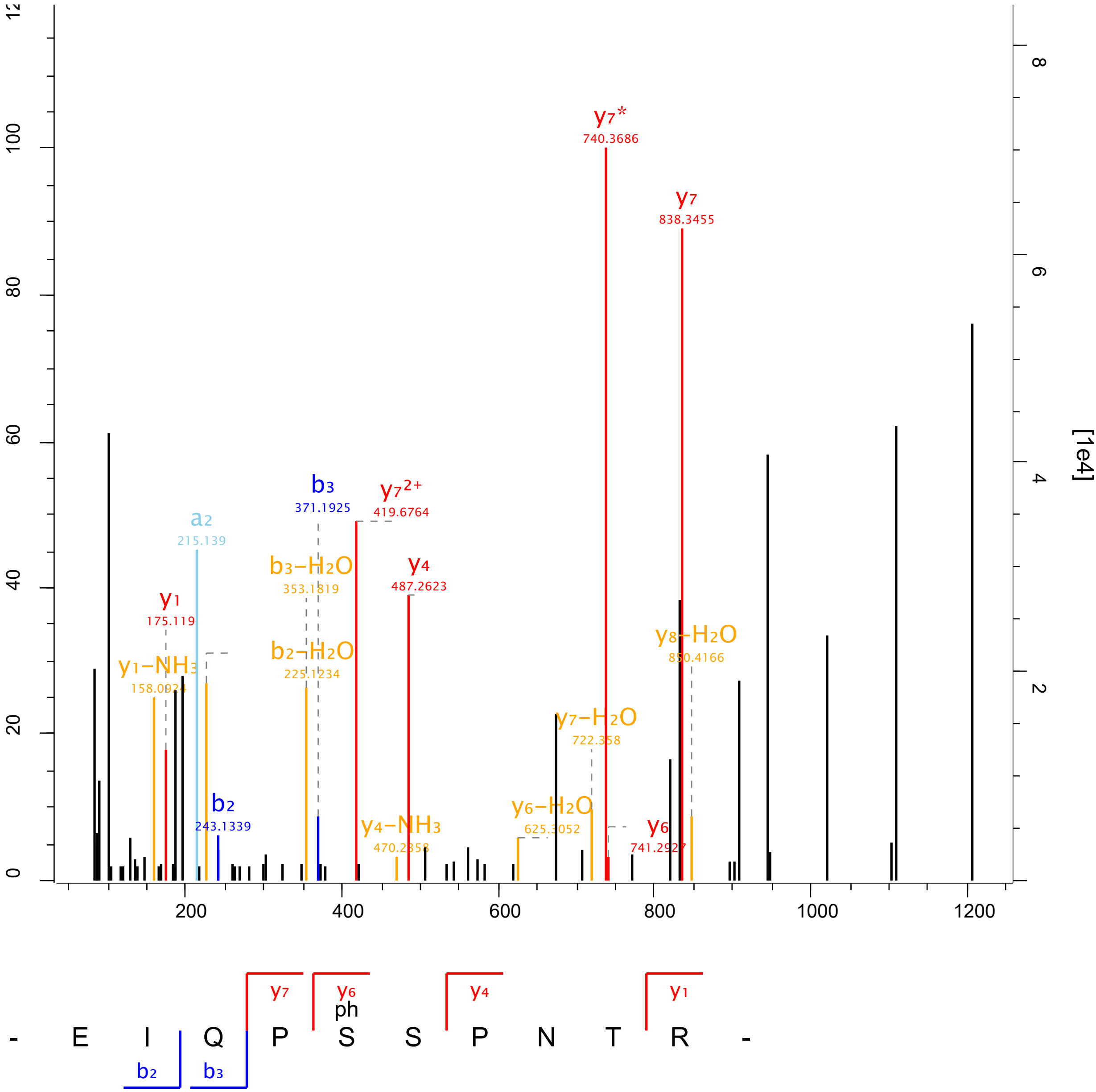

| Raw file | Scan | Method | Score | m/z |
| --- | --- | --- | --- | --- |
| sys_15_1 | 48906 | FTMS; HCD | 112.26 | 776.38 |

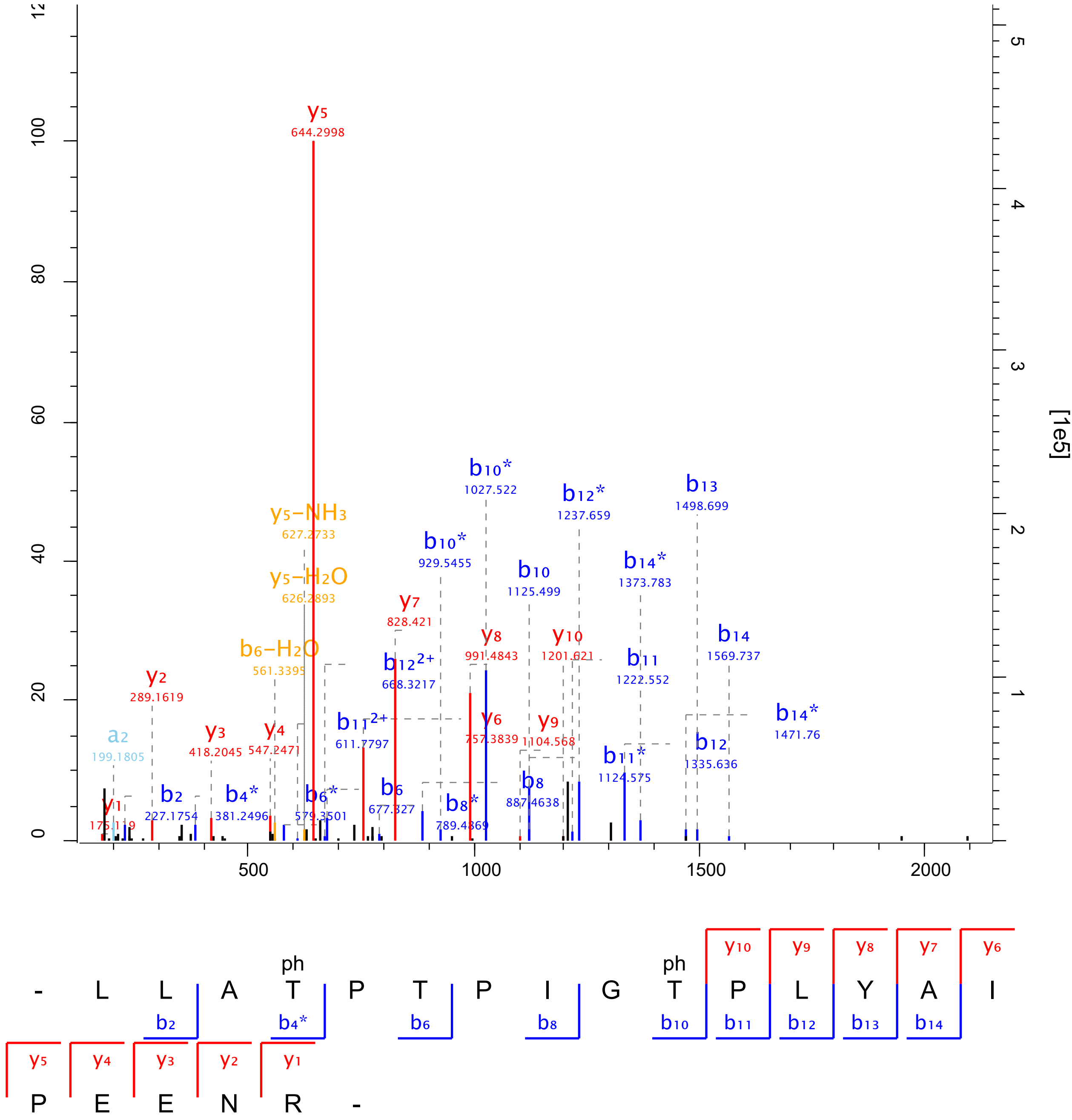

| Raw file | Scan | Method | Score | m/z |
|---|---|---|---|---|
| sys_15_1 | 48919 | FTMS; HCD | 77.08 | 889.44 |

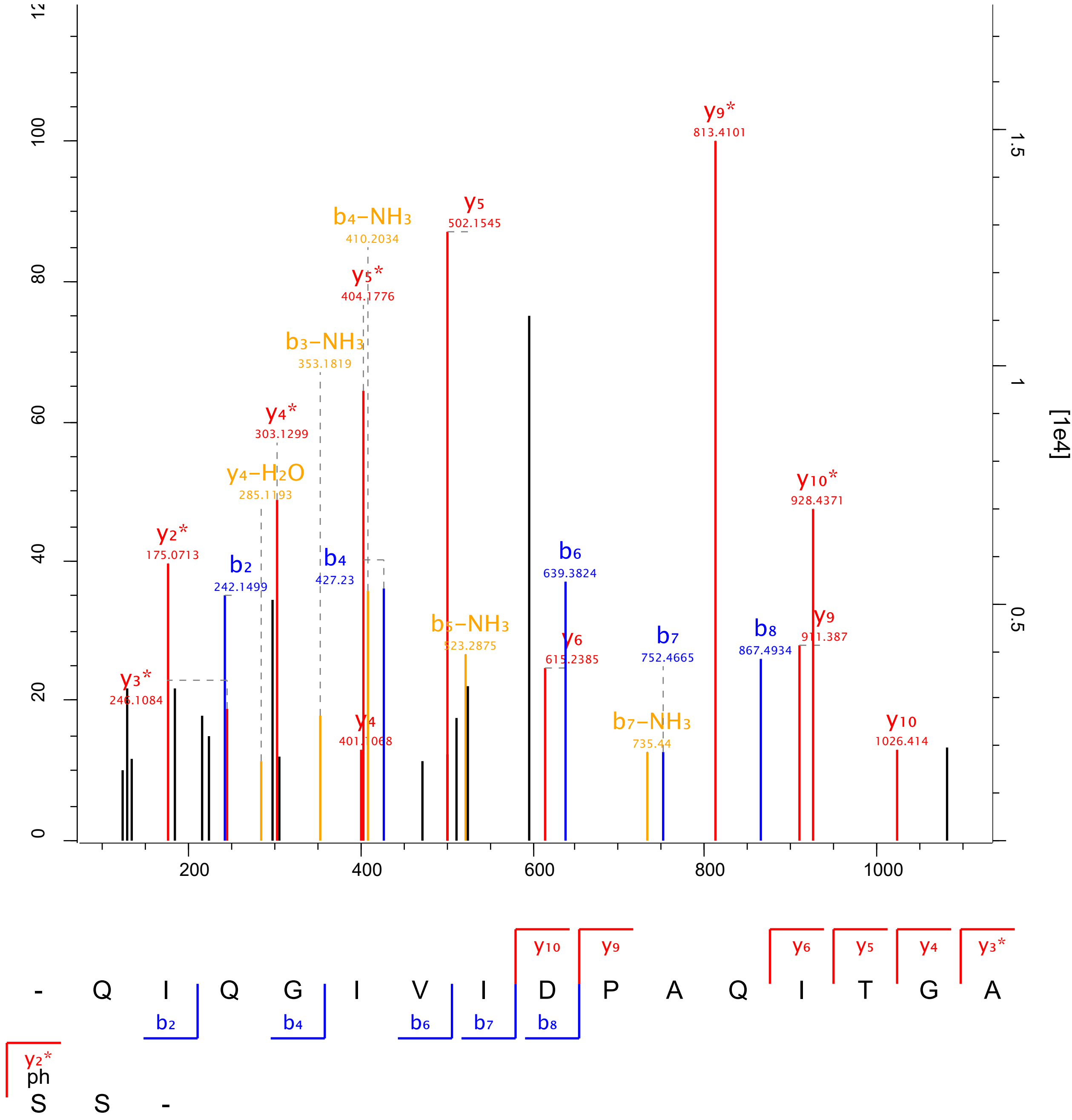

| Raw file | Scan | Method | Score | m/z |
|---|---|---|---|---|
| sys_15_1 | 49508 | FTMS; HCD | 46.89 | 853.39 |

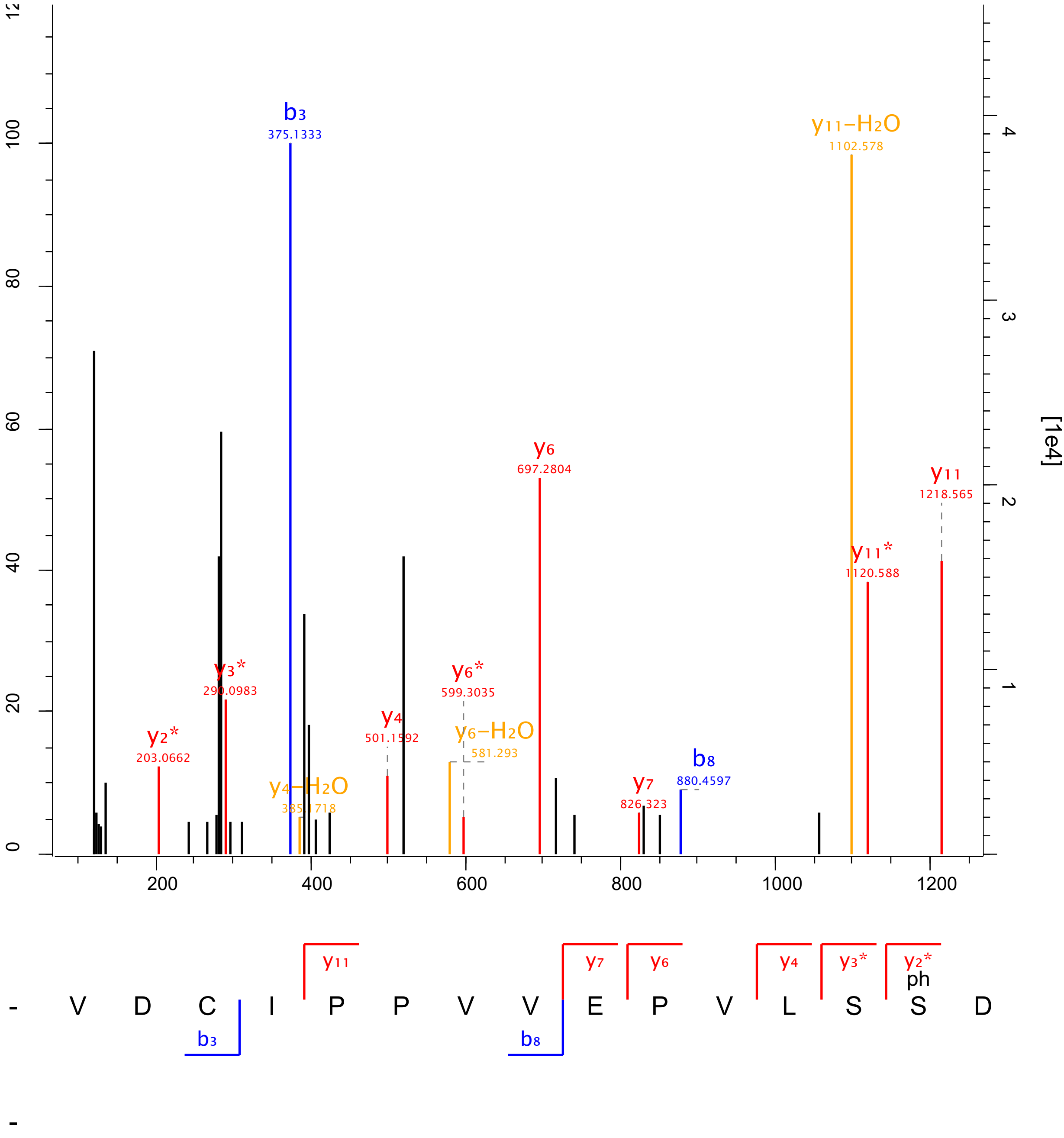

| Raw file | Scan | Method | Score | m/z |
|---|---|---|---|---|
| sys_15_1 | 4959 | FTMS; HCD | 166.65 | 629.28 |

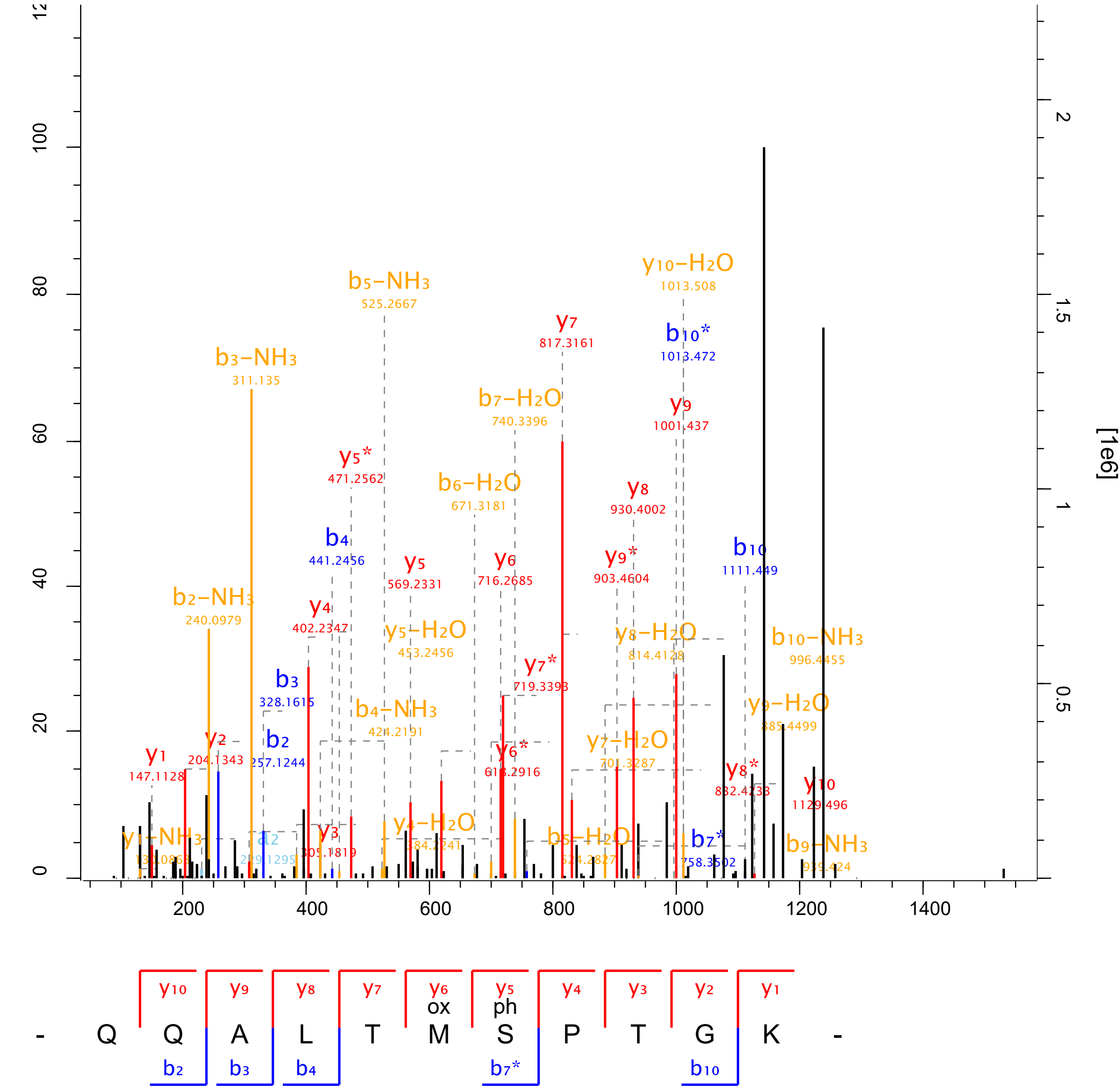

| Raw file | Scan | Method | Score | m/z |
|---|---|---|---|---|
| sys_15_1 | 50021 | FTMS; HCD | 44.79 | 969.41 |

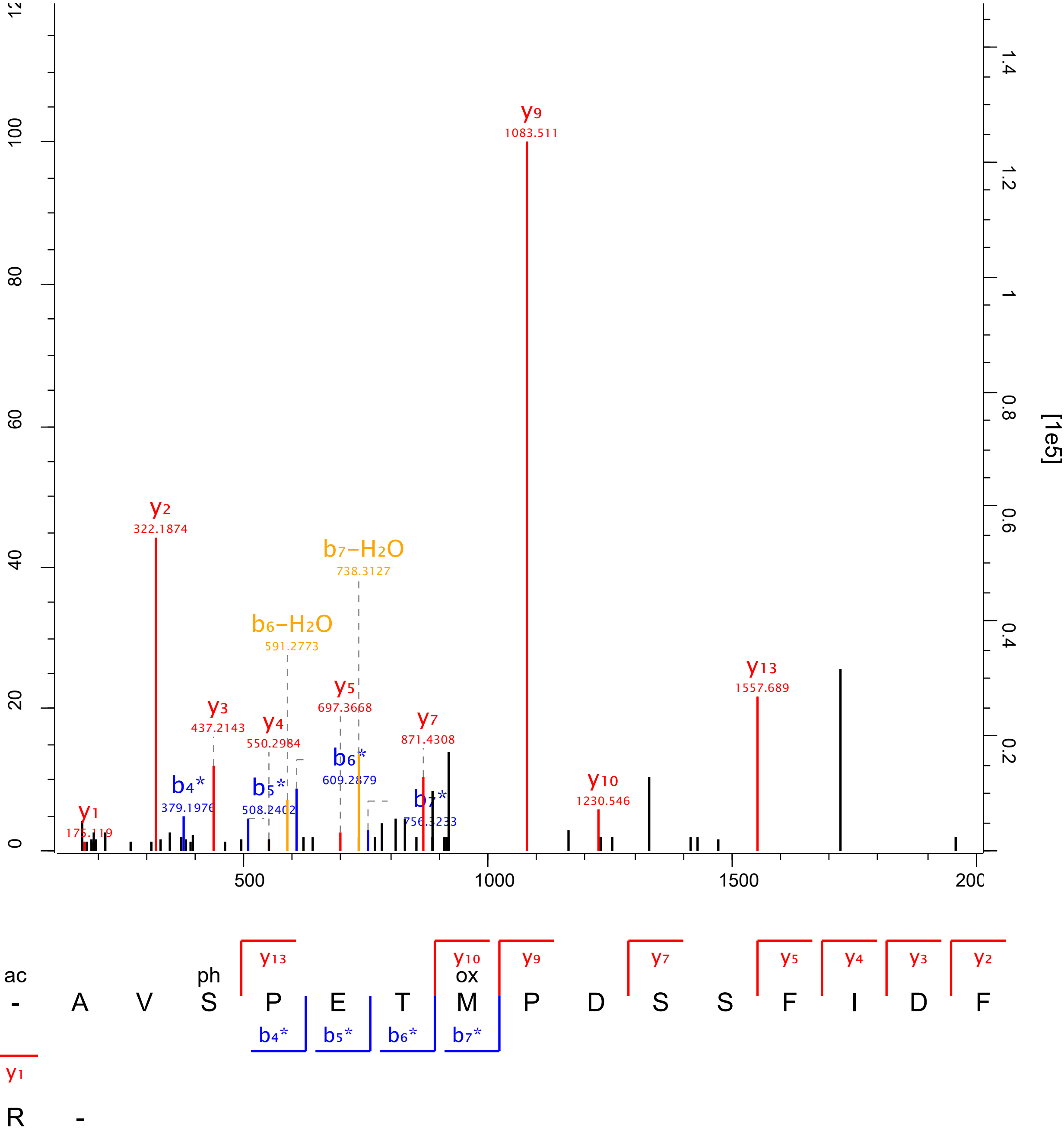

| Raw file | Scan | Method | Score | m/z |
| --- | --- | --- | --- | --- |
| sys_15_1 | 5021 | FTMS; HCD | 131.41 | 632.24 |

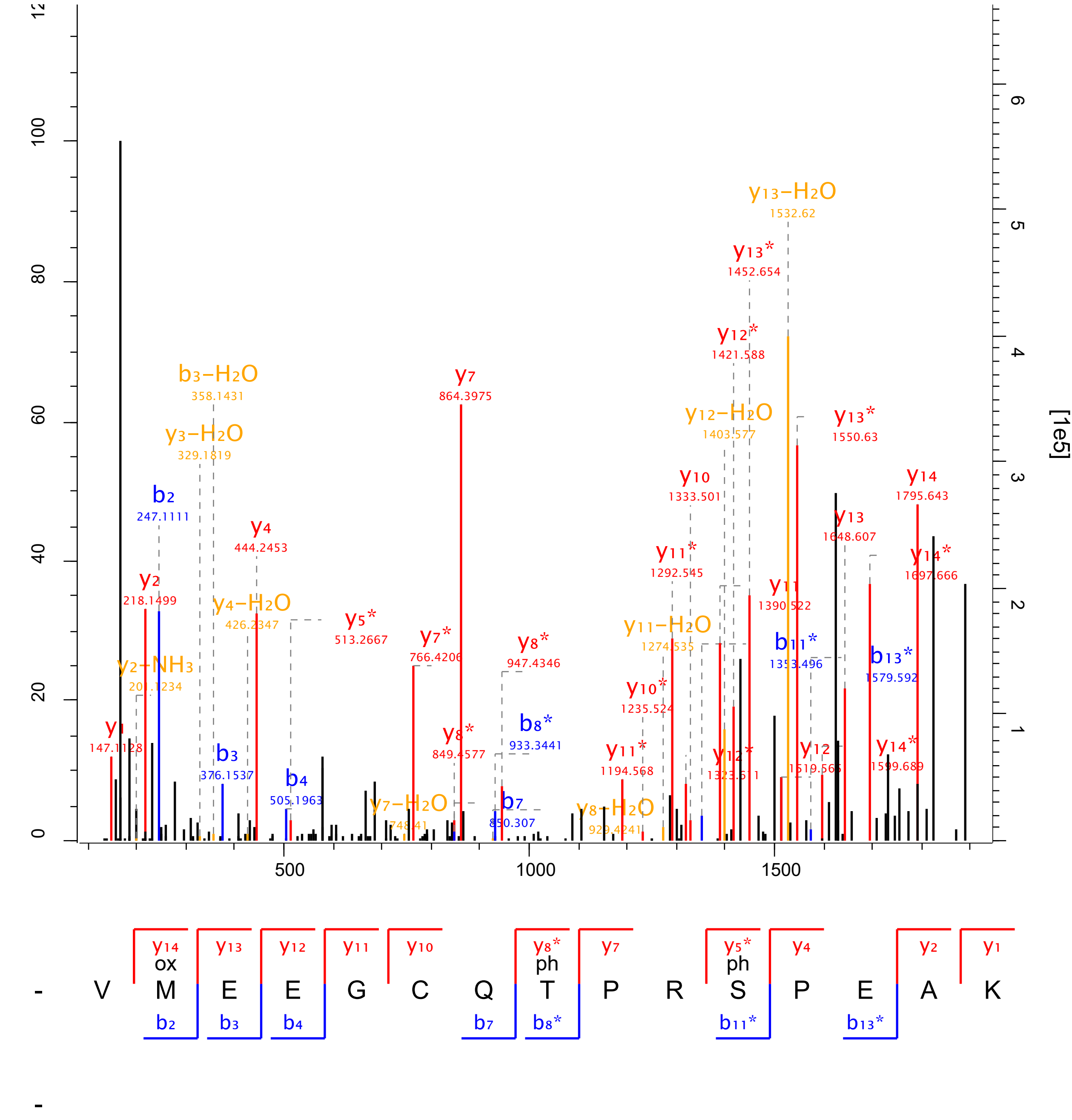

| Raw file | Scan | Method | Score | m/z |
| --- | --- | --- | --- | --- |
| sys_15_1 | 5055 | FTMS; HCD | 66.27 | 582.72 |

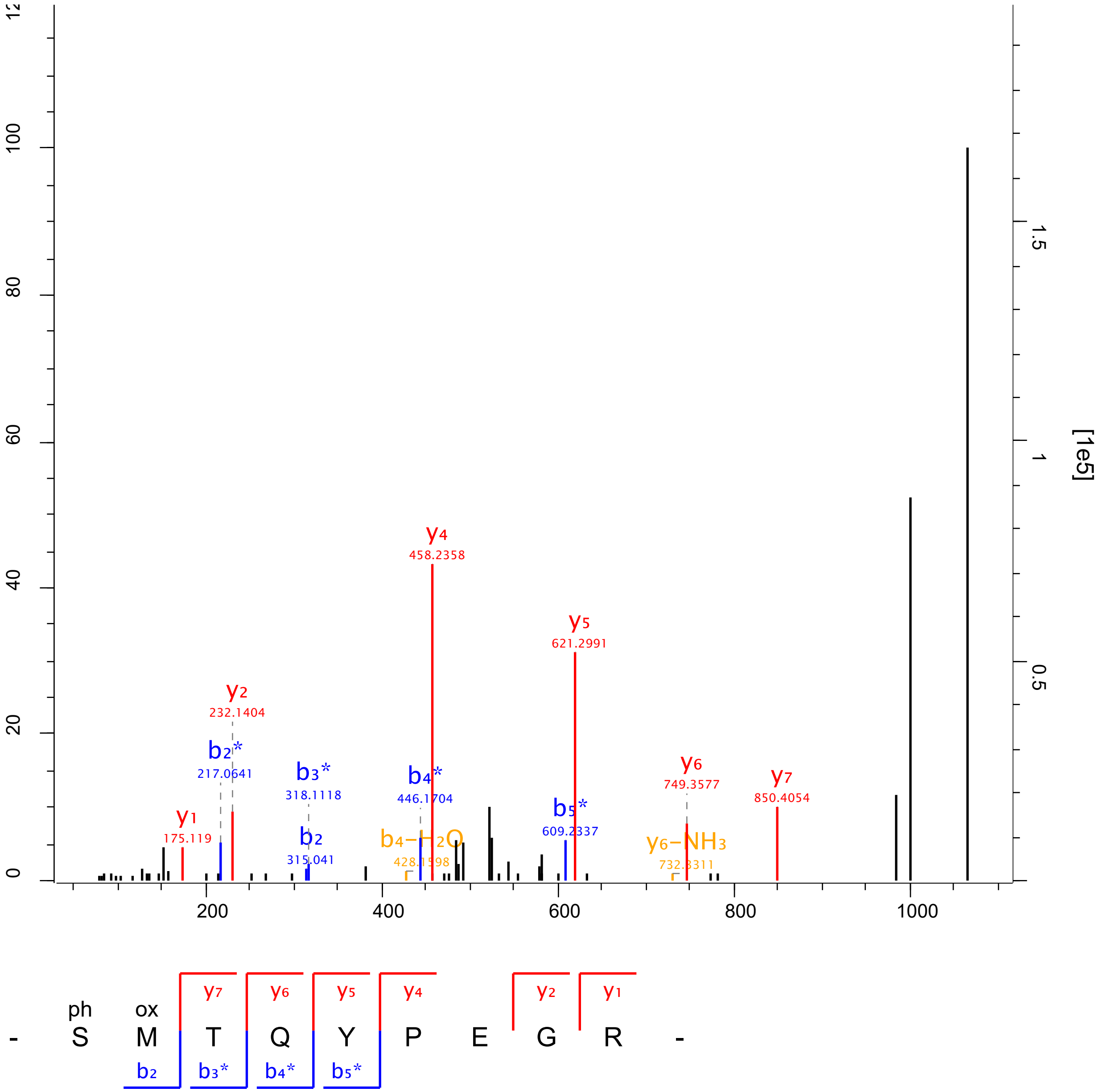

| Raw file | Scan | Method | Score | m/z |
|---|---|---|---|---|
| sys_15_1 | 50600 | FTMS; HCD | 108.08 | 1018.01 |

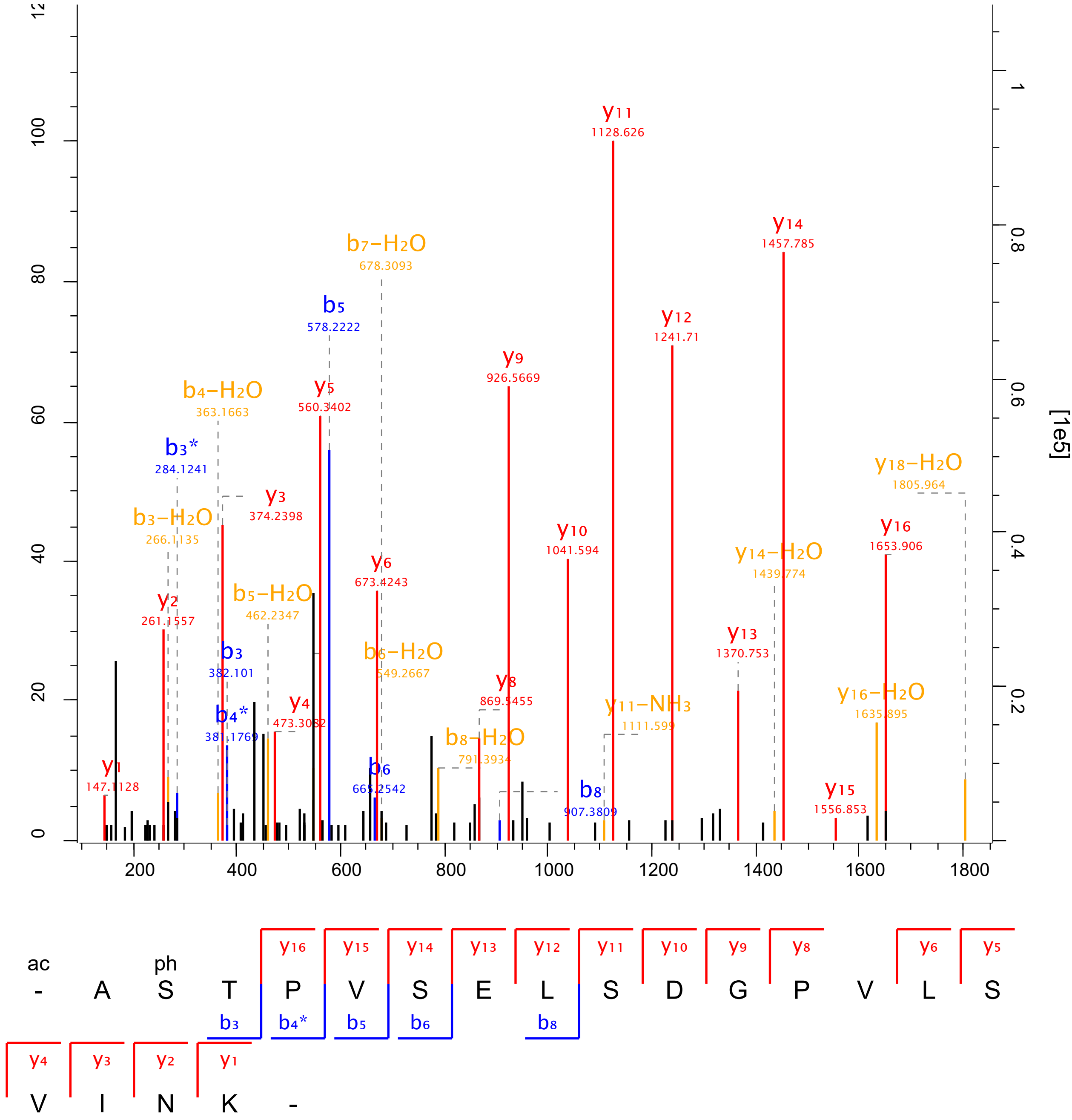

| Raw file | Scan | Method | Score | m/z |
|---|---|---|---|---|
| sys_15_1 | 5077 | FTMS; HCD | 80.71 | 461.72 |

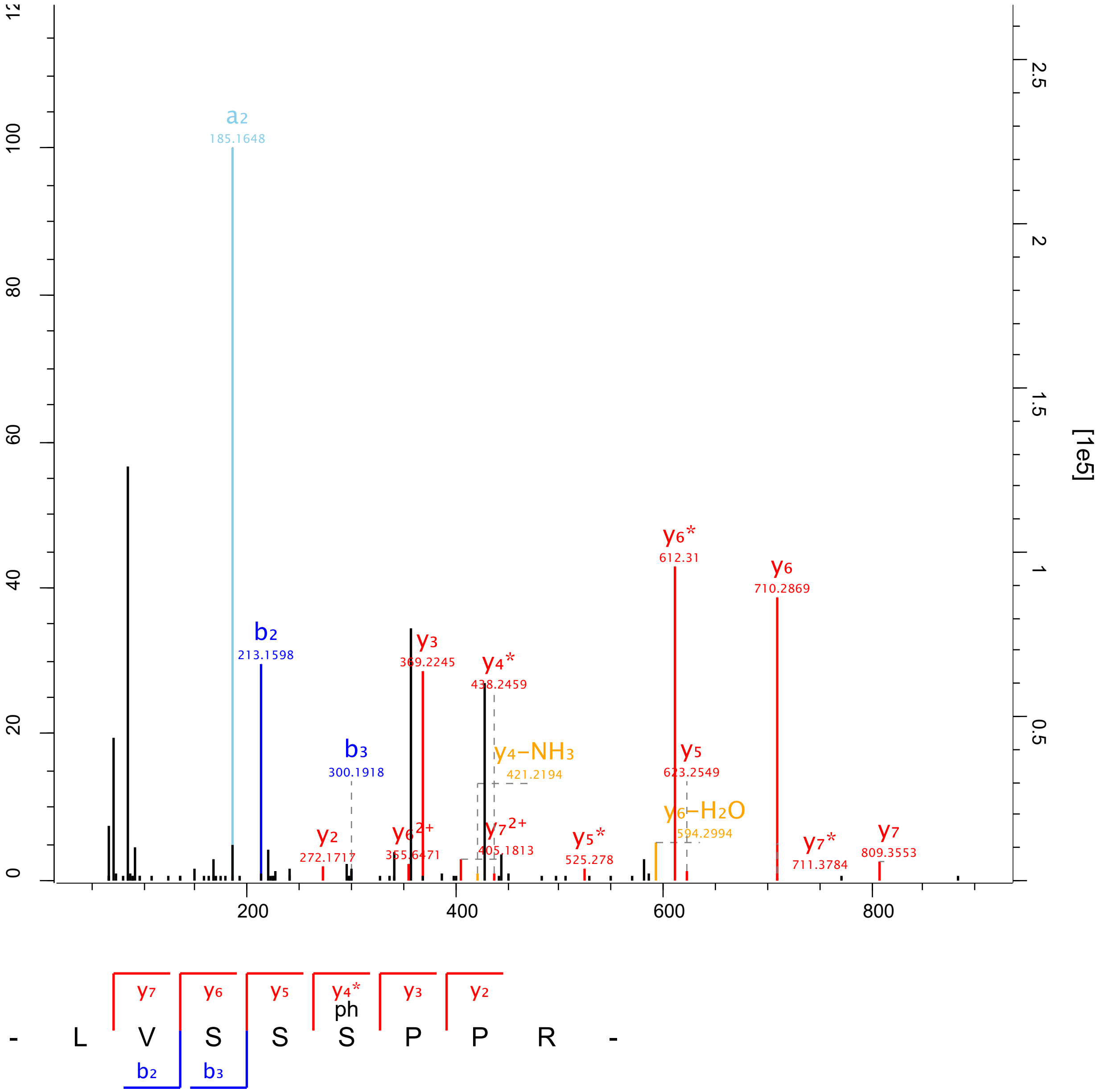

| Raw file | Scan | Method | Score | m/z |
| --- | --- | --- | --- | --- |
| sys_15_1 | 5107 | FTMS; HCD | 108.74 | 472.7 |

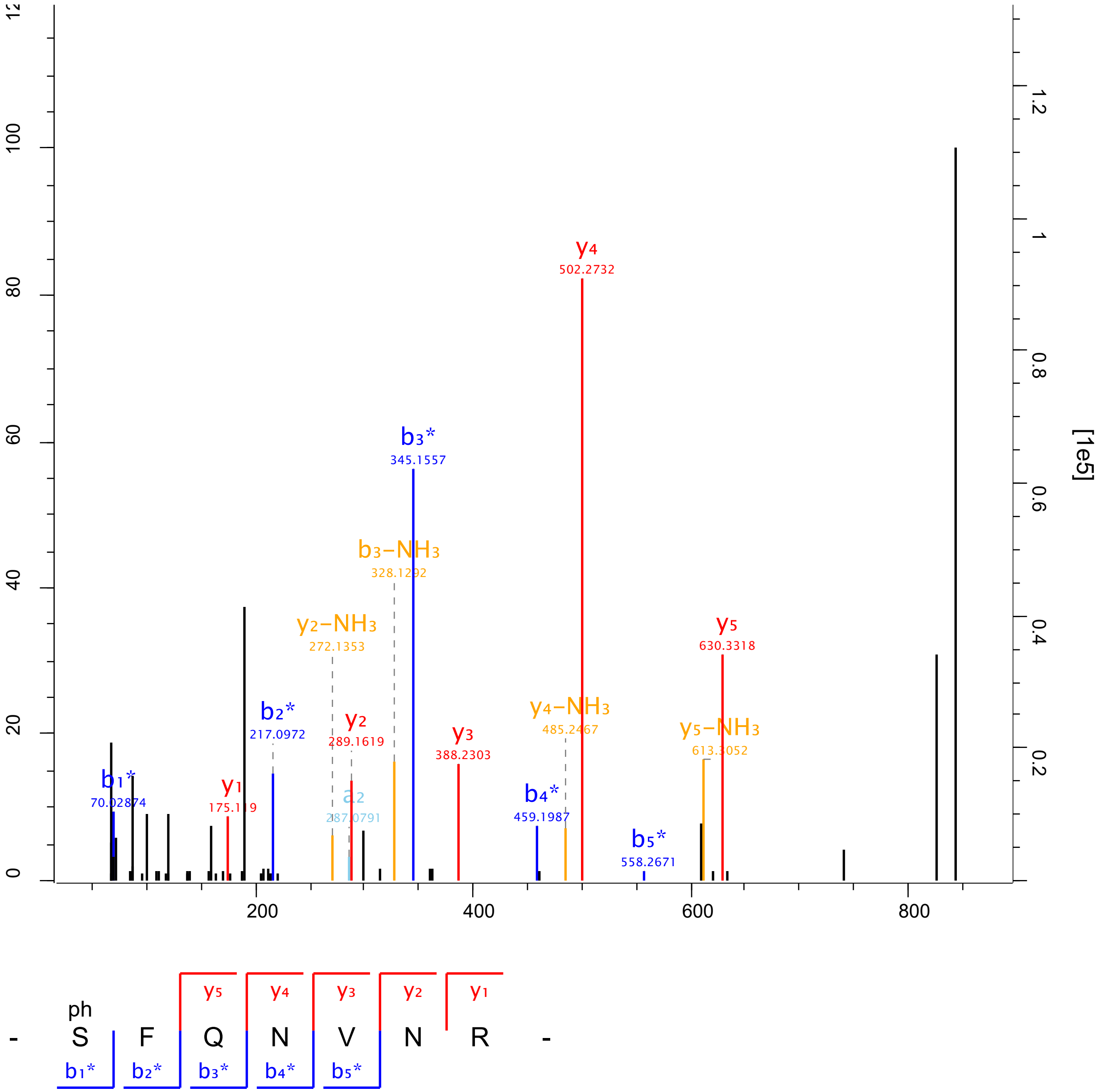

| Raw file | Scan | Method | Score | m/z |
| --- | --- | --- | --- | --- |
| sys_15_1 | 5137 | FTMS; HCD | 64.1 | 451.69 |

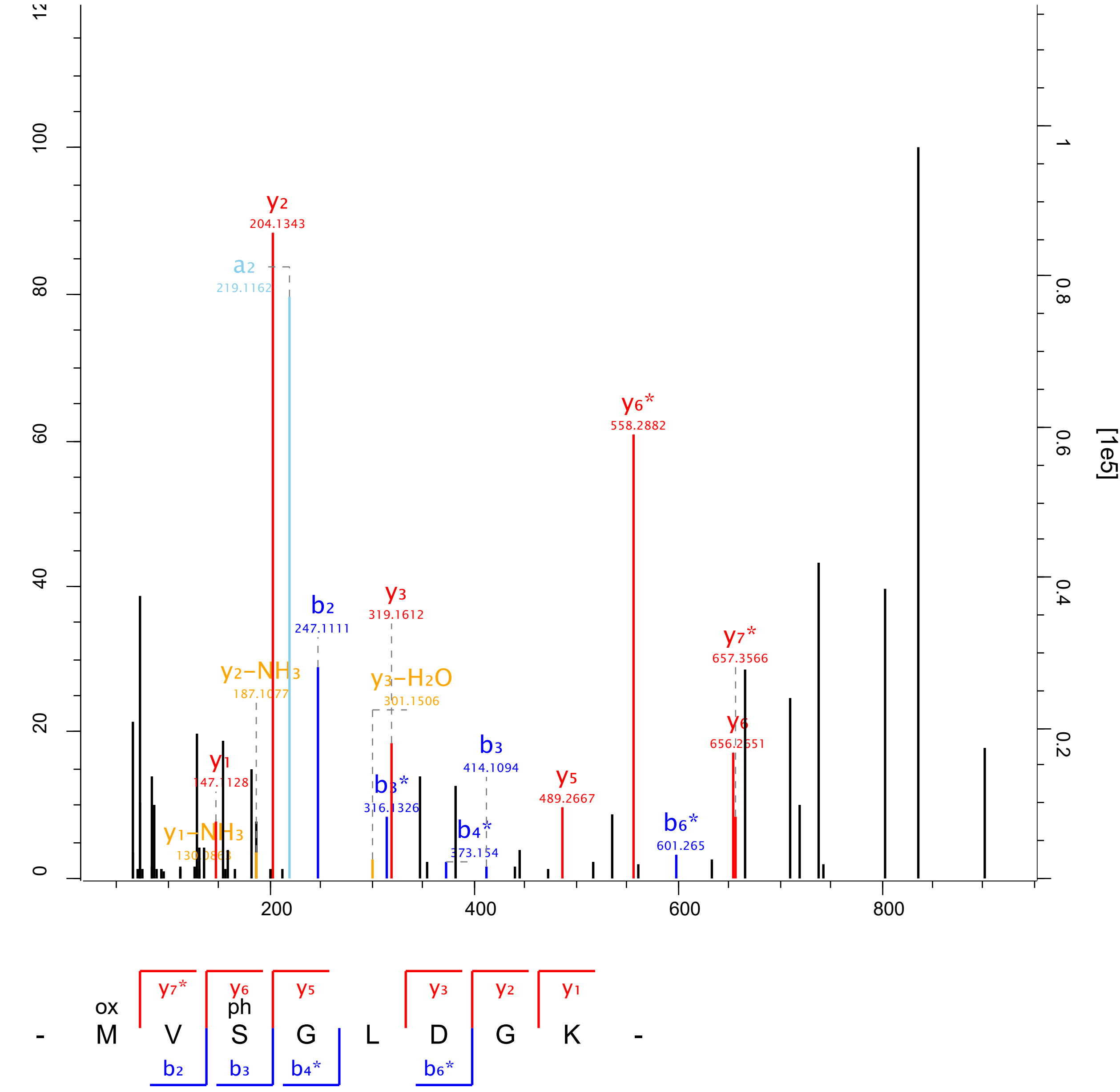

| Raw file | Scan | Method | Score | m/z |
|---|---|---|---|---|
| sys_15_1 | 5155 | FTMS; HCD | 83.05 | 532.26 |

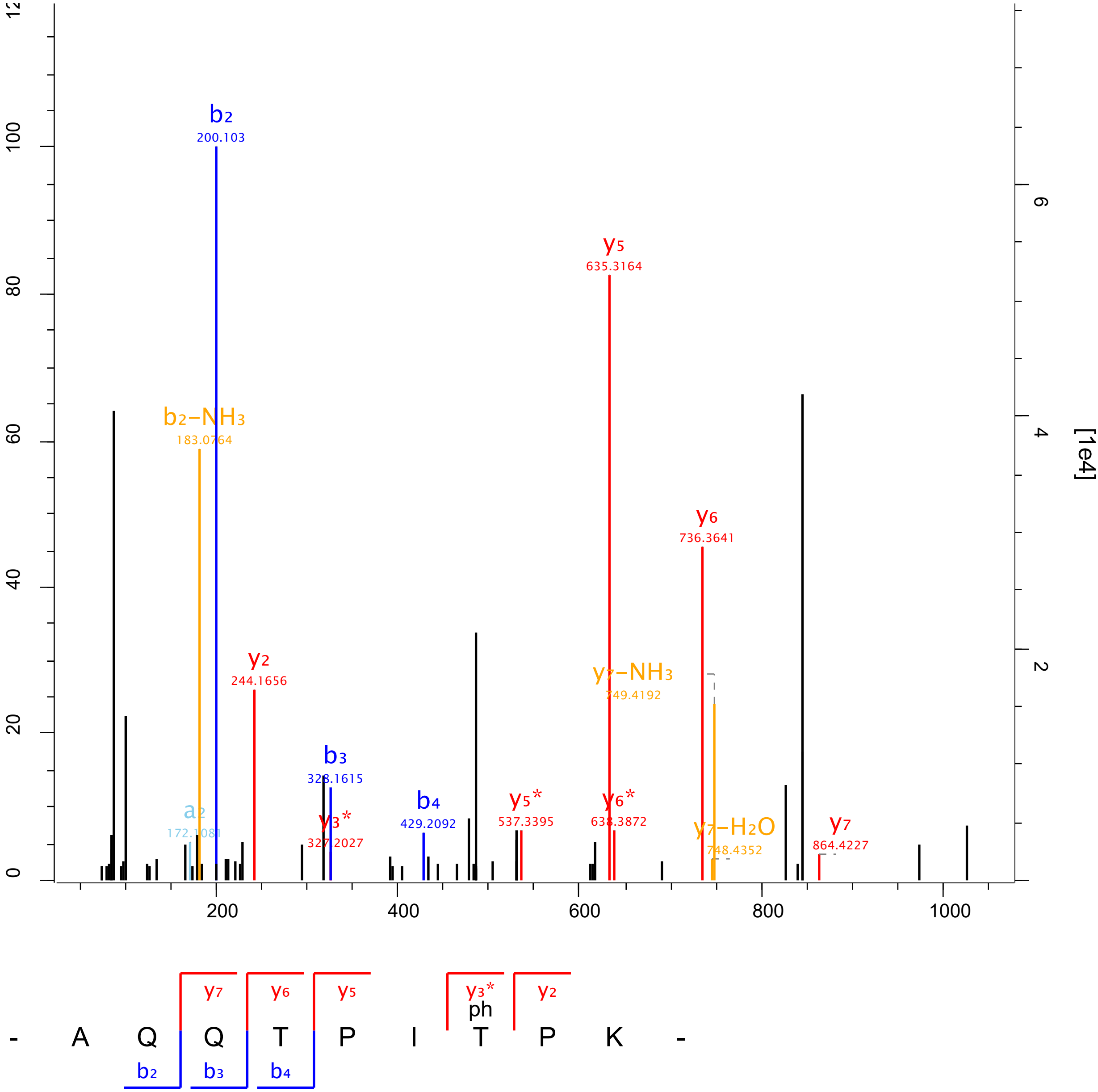

| Raw file | Scan | Method | Score | m/z |
|---|---|---|---|---|
| sys_15_1 | 5170 | FTMS; HCD | 93.48 | 656.25 |

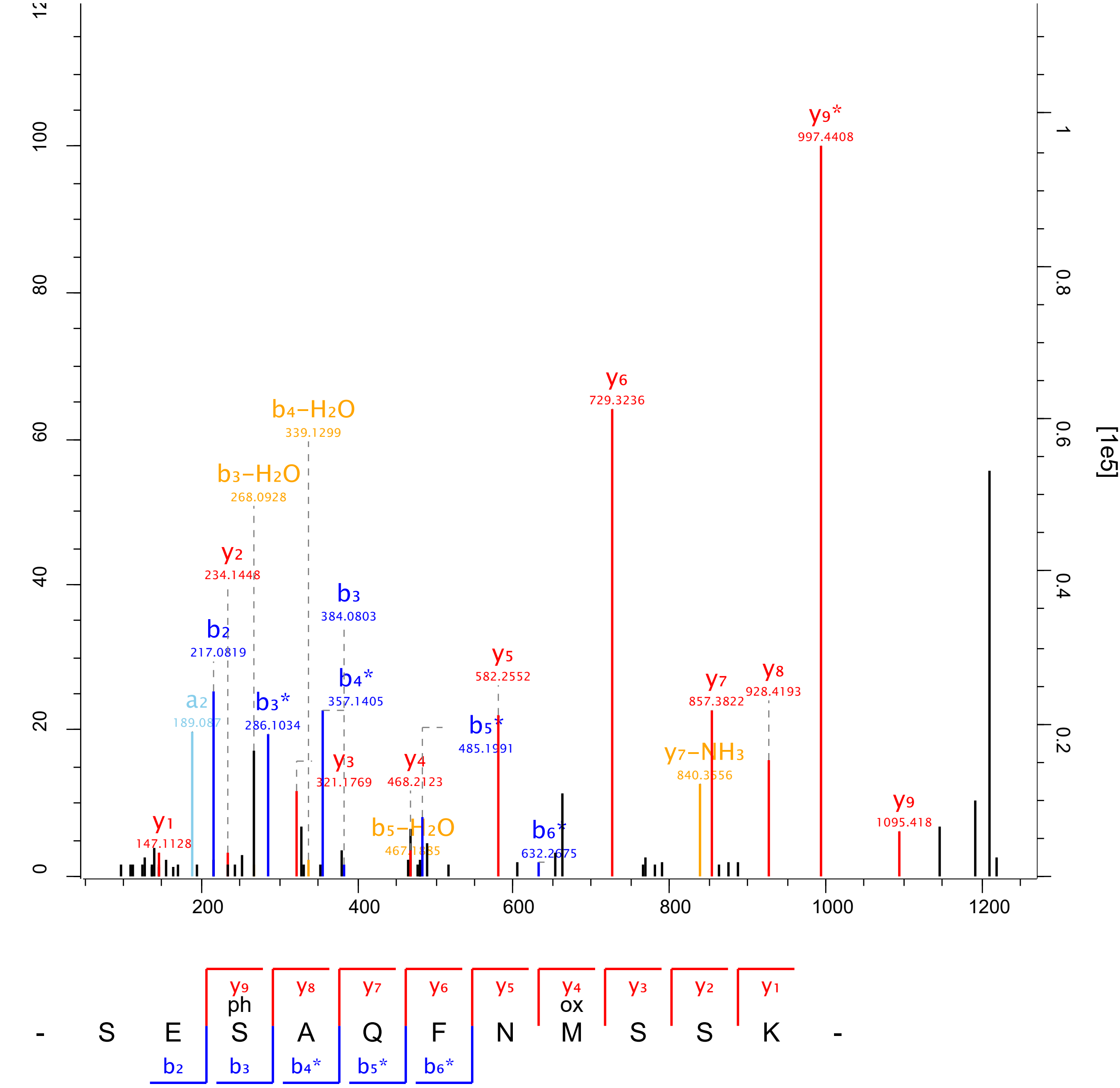

| Raw file | Scan | Method | Score | m/z |
|---|---|---|---|---|
| sys_15_1 | 5203 | FTMS; HCD | 86.77 | 545.23 |

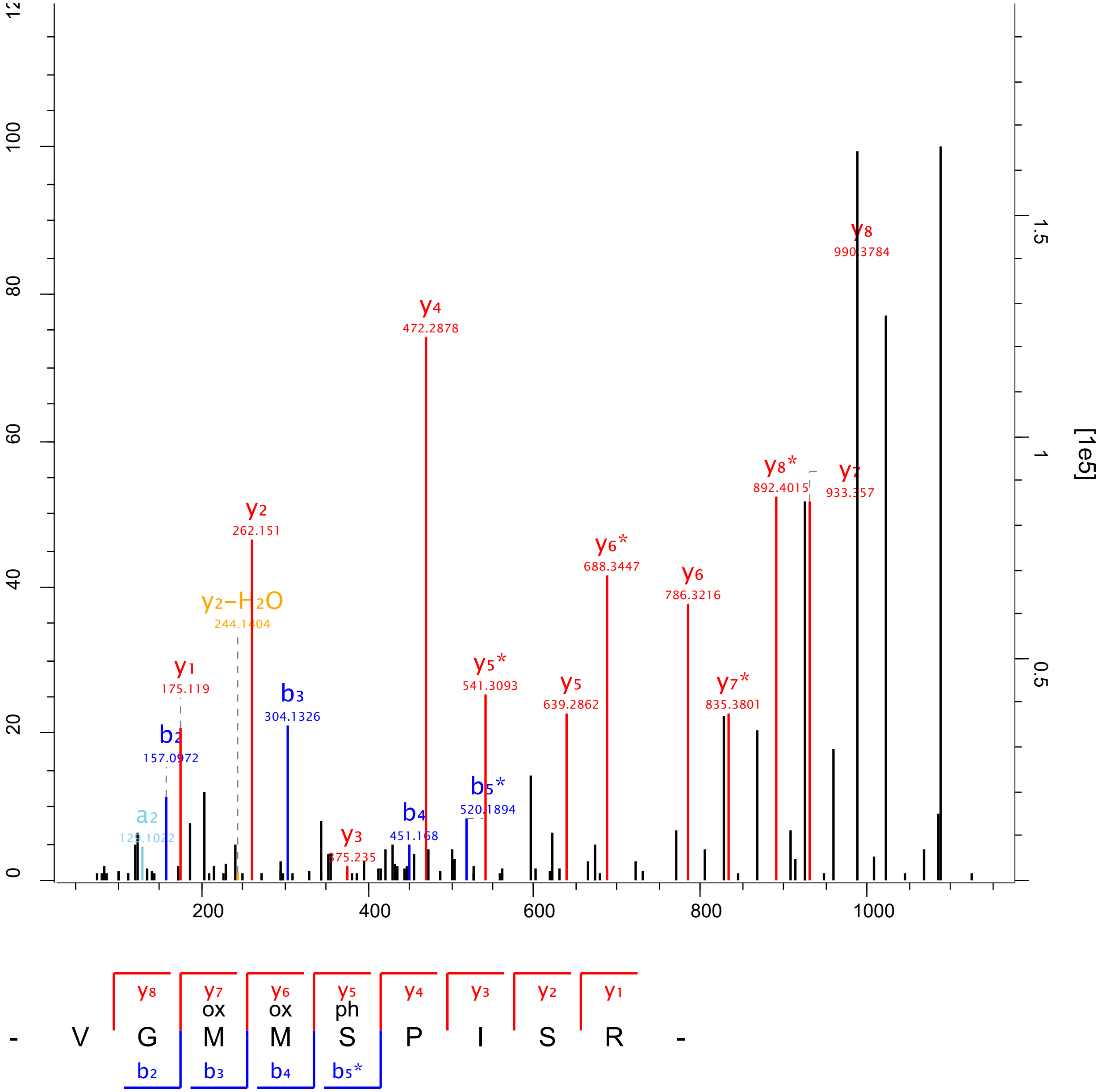

| Raw file | Scan | Method | Score | m/z |
| --- | --- | --- | --- | --- |
| sys_15_1 | 5233 | FTMS; HCD | 102.52 | 659.25 |

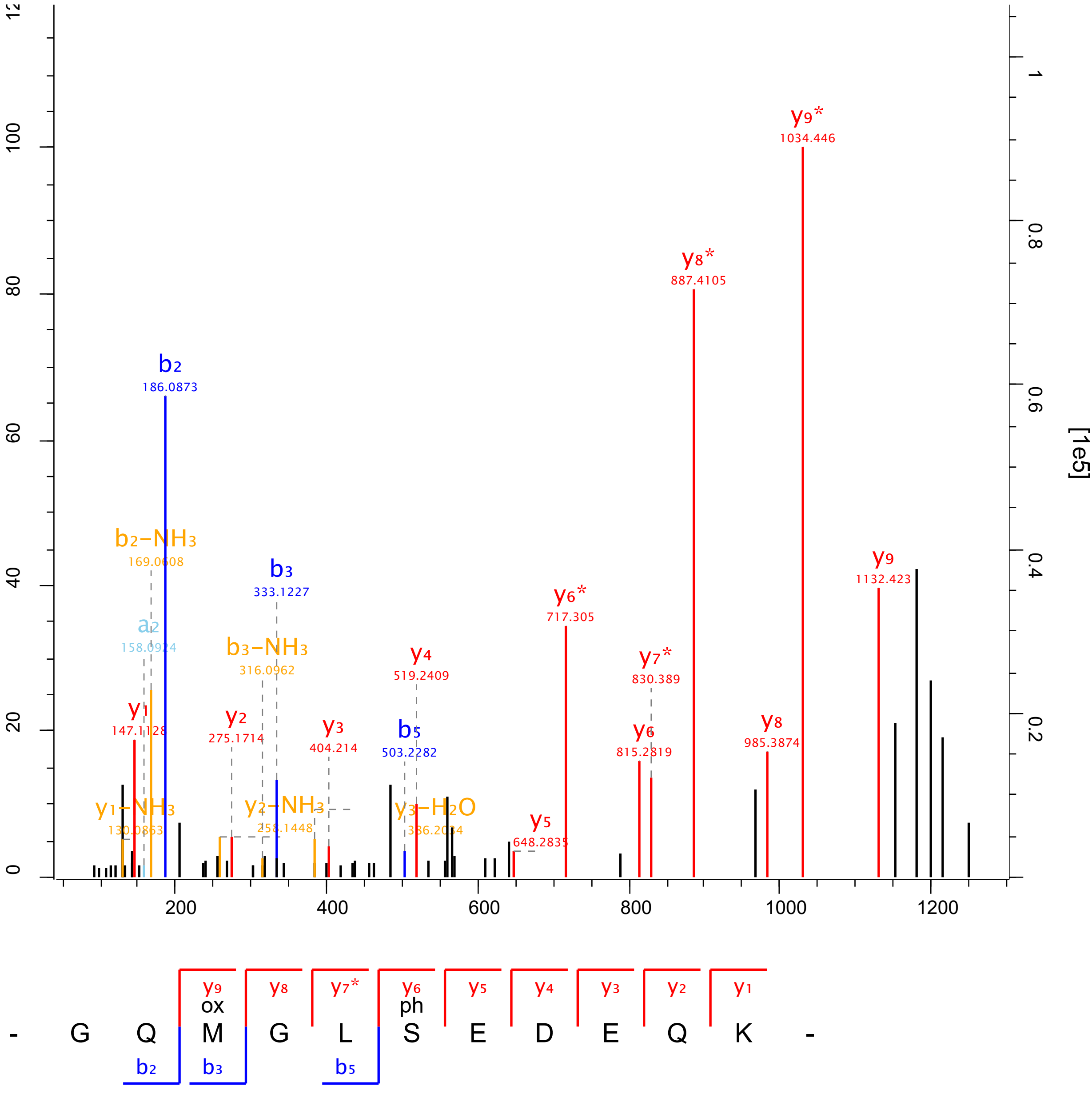

| Raw file | Scan | Method | Score | m/z |
|---|---|---|---|---|
| sys_15_1 | 5280 | FTMS; HCD | 63.3 | 847.83 |

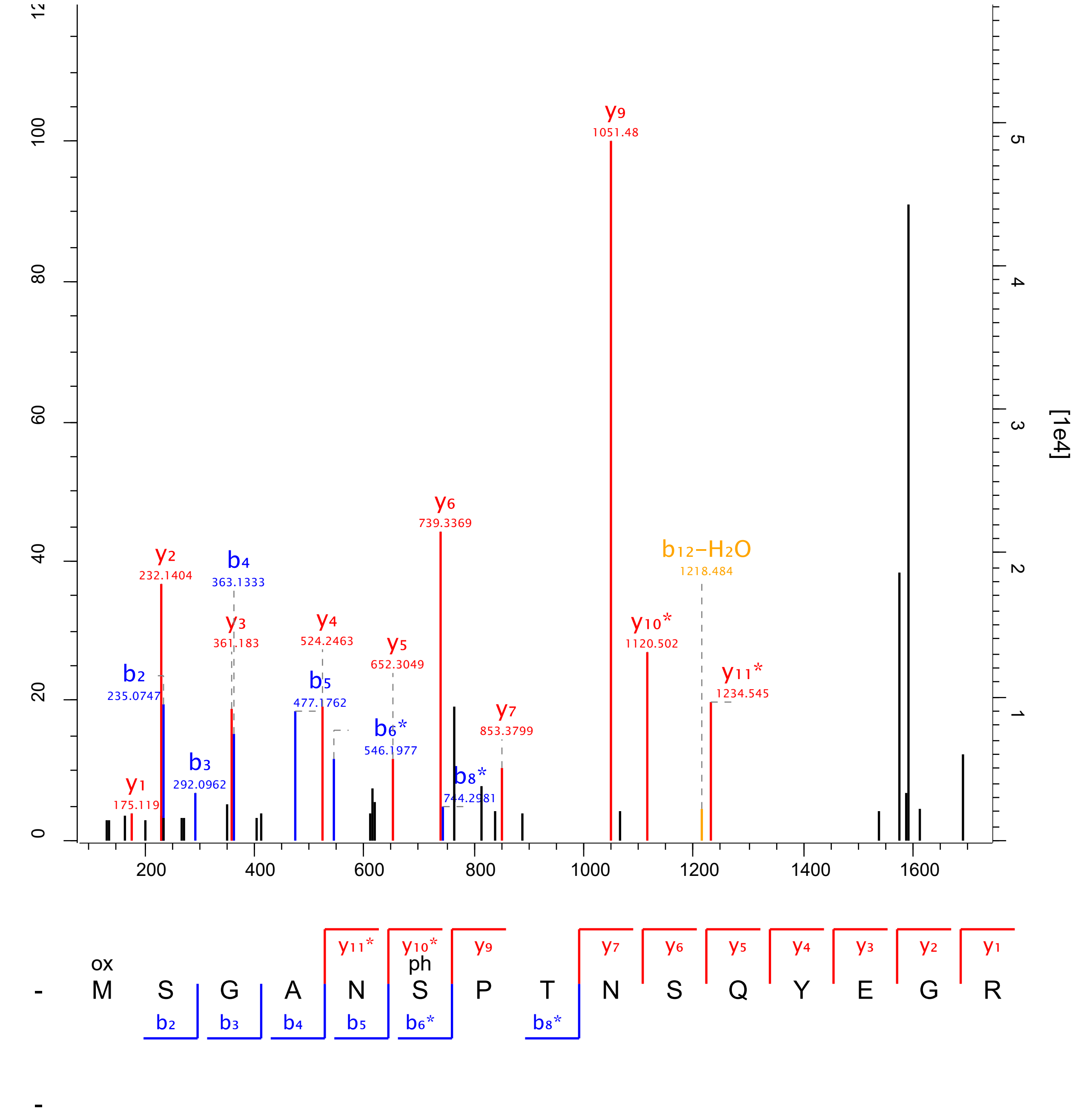

| Raw file | Scan | Method | Score | m/z |
|---|---|---|---|---|
| sys_15_1 | 5382 | FTMS; HCD | 65.72 | 489.7 |

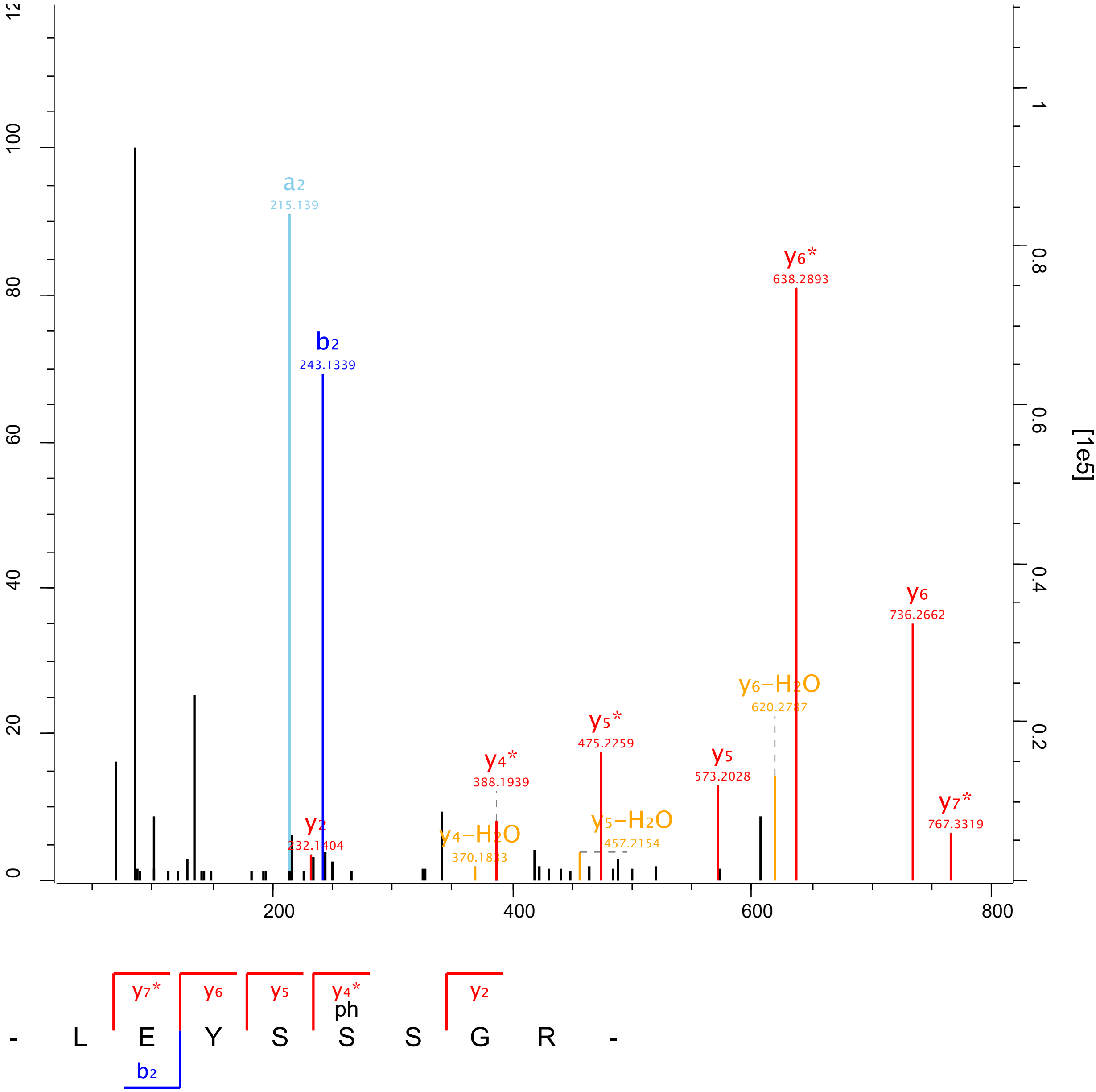

| Raw file | Scan | Method | Score | m/z |
|---|---|---|---|---|
| sys_15_1 | 5390 | FTMS; HCD | 61.44 | 606.72 |

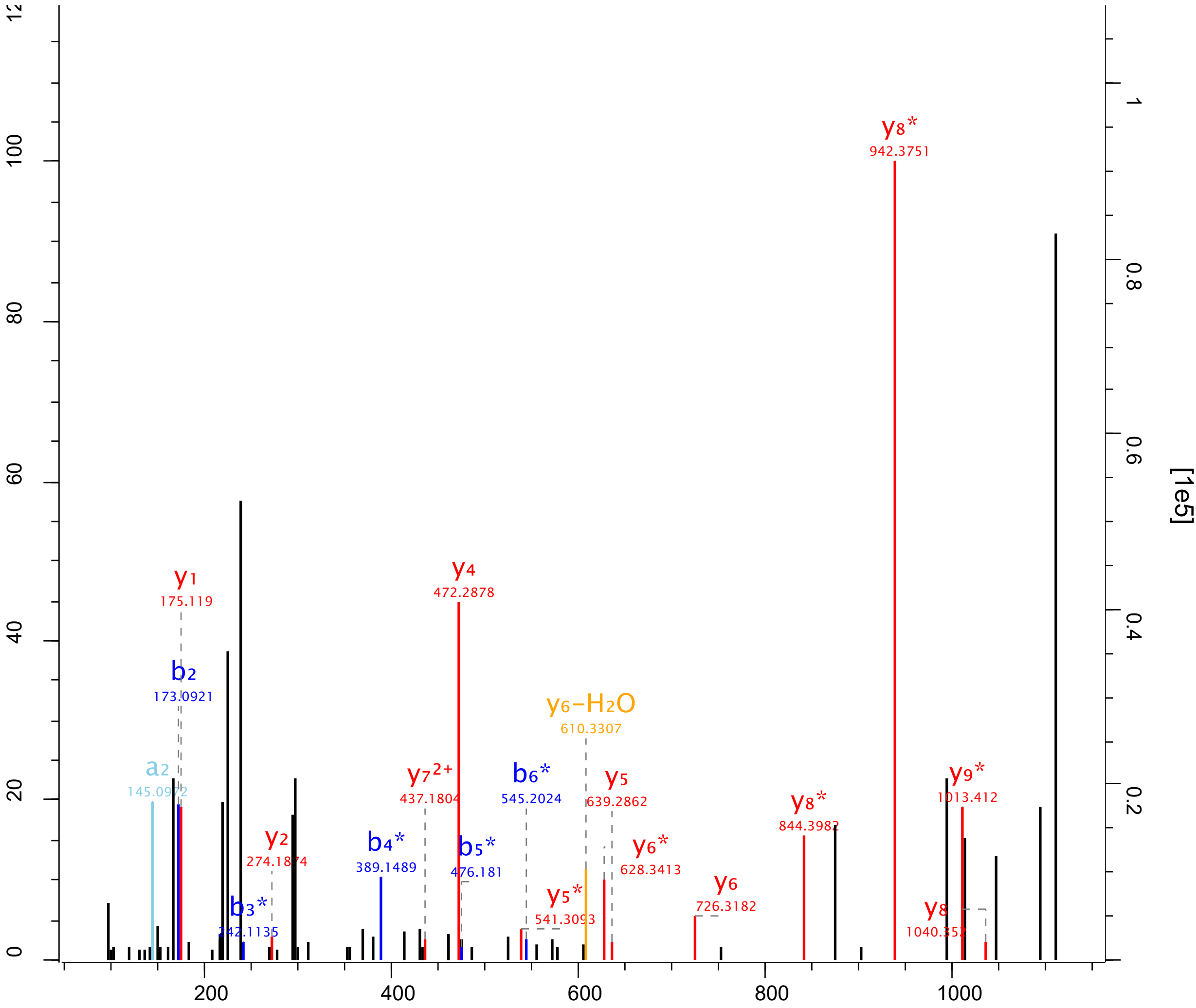


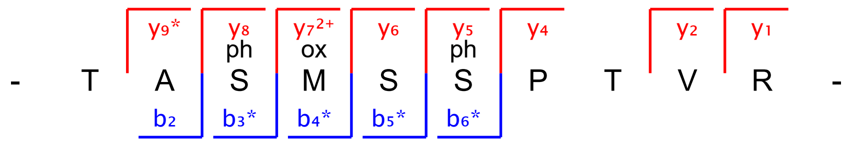

| Raw file | Scan | Method | Score | m/z |
|---|---|---|---|---|
| sys_15_1 | 5415 | FTMS; HCD | 71.56 | 529.23 |

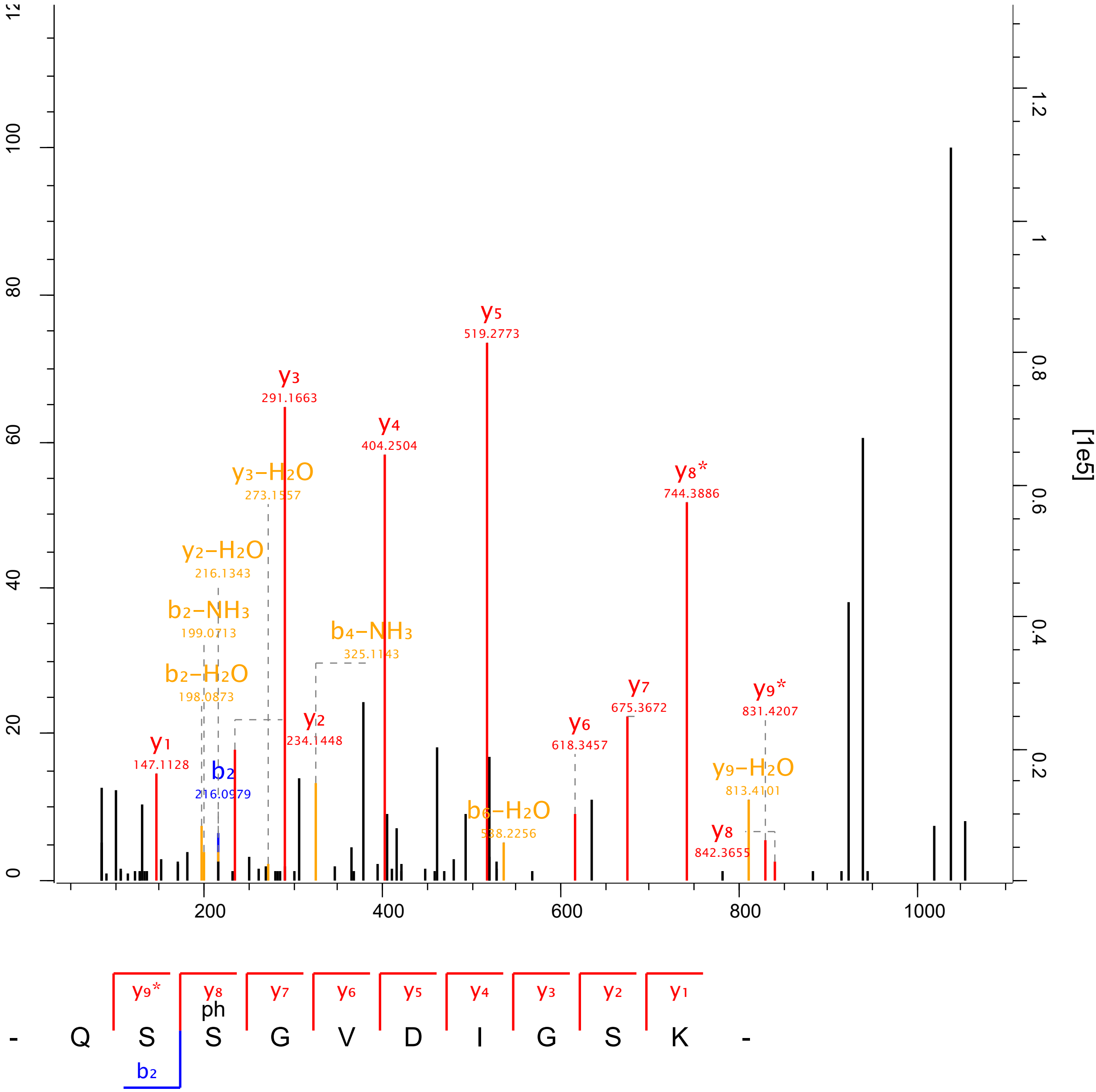

| Raw file | Scan | Method | Score | m/z |
| --- | --- | --- | --- | --- |
| sys_15_1 | 5445 | FTMS; HCD | 107.06 | 457.71 |

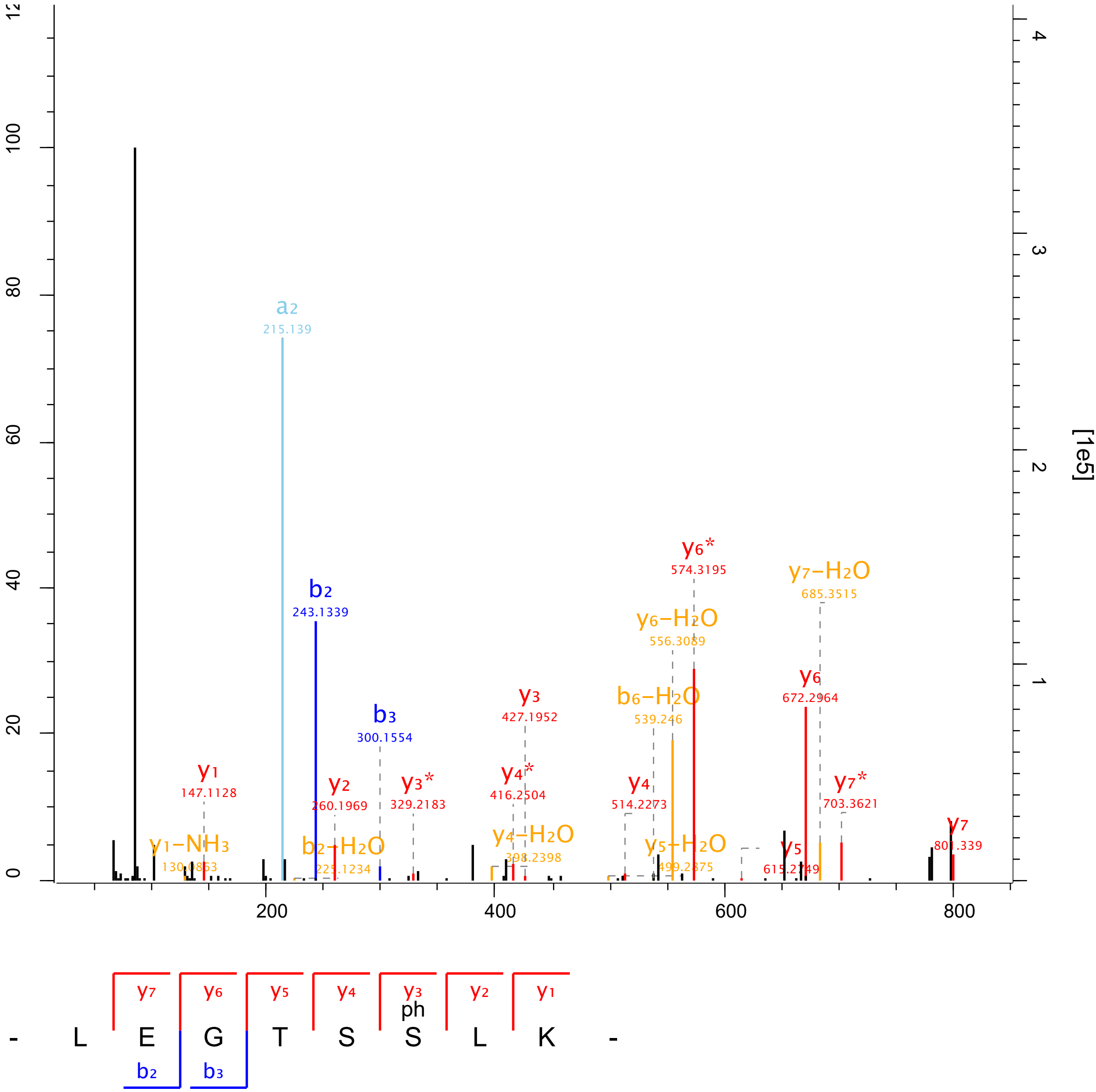

| Raw file | Scan | Method | Score | m/z |
|---|---|---|---|---|
| sys_15_1 | 5463 | FTMS; HCD | 103.67 | 541.24 |

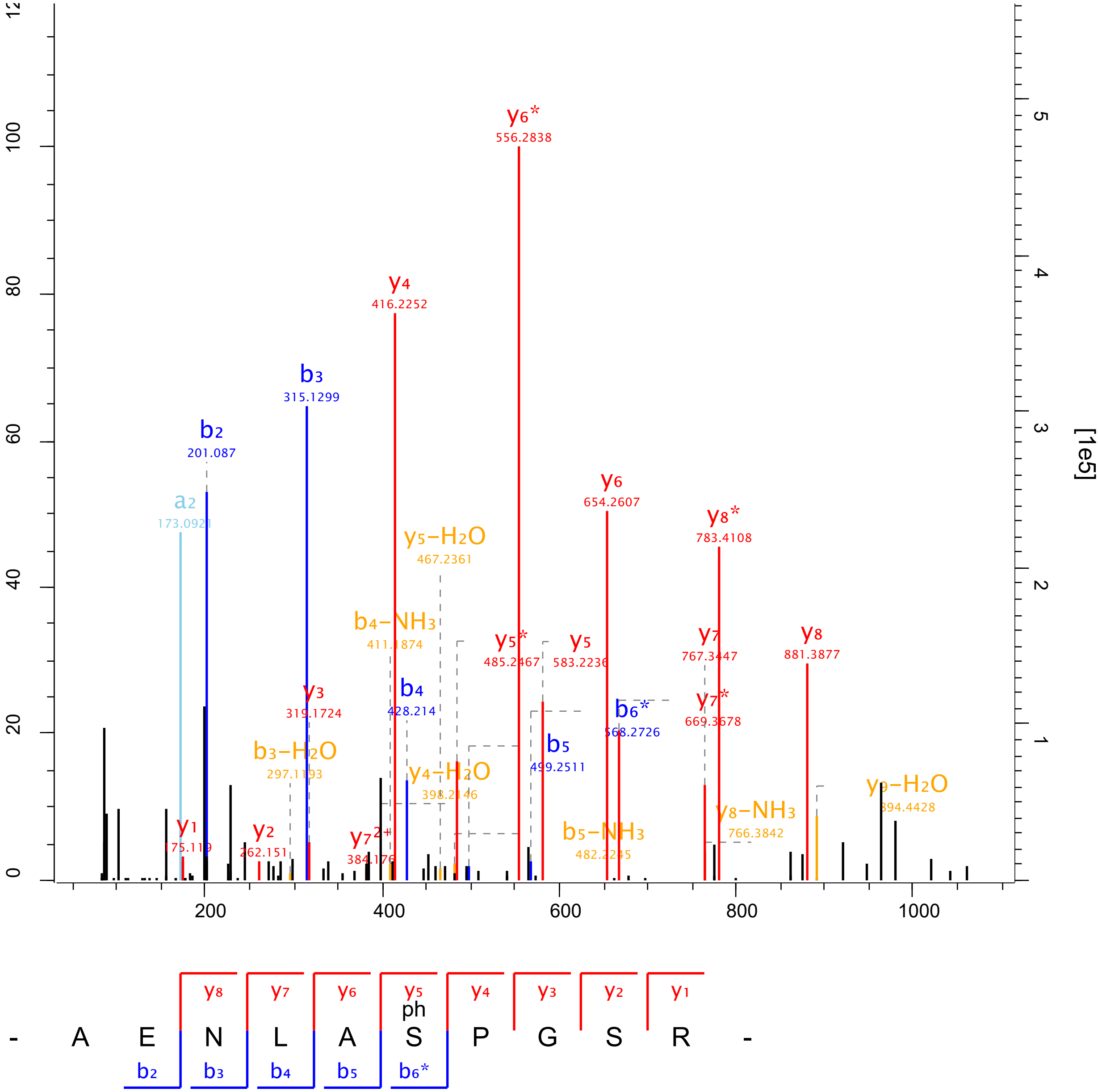

| Raw file | Scan | Method | Score | m/z |
|---|---|---|---|---|
| sys_15_1 | 5653 | FTMS; HCD | 150.22 | 578.24 |

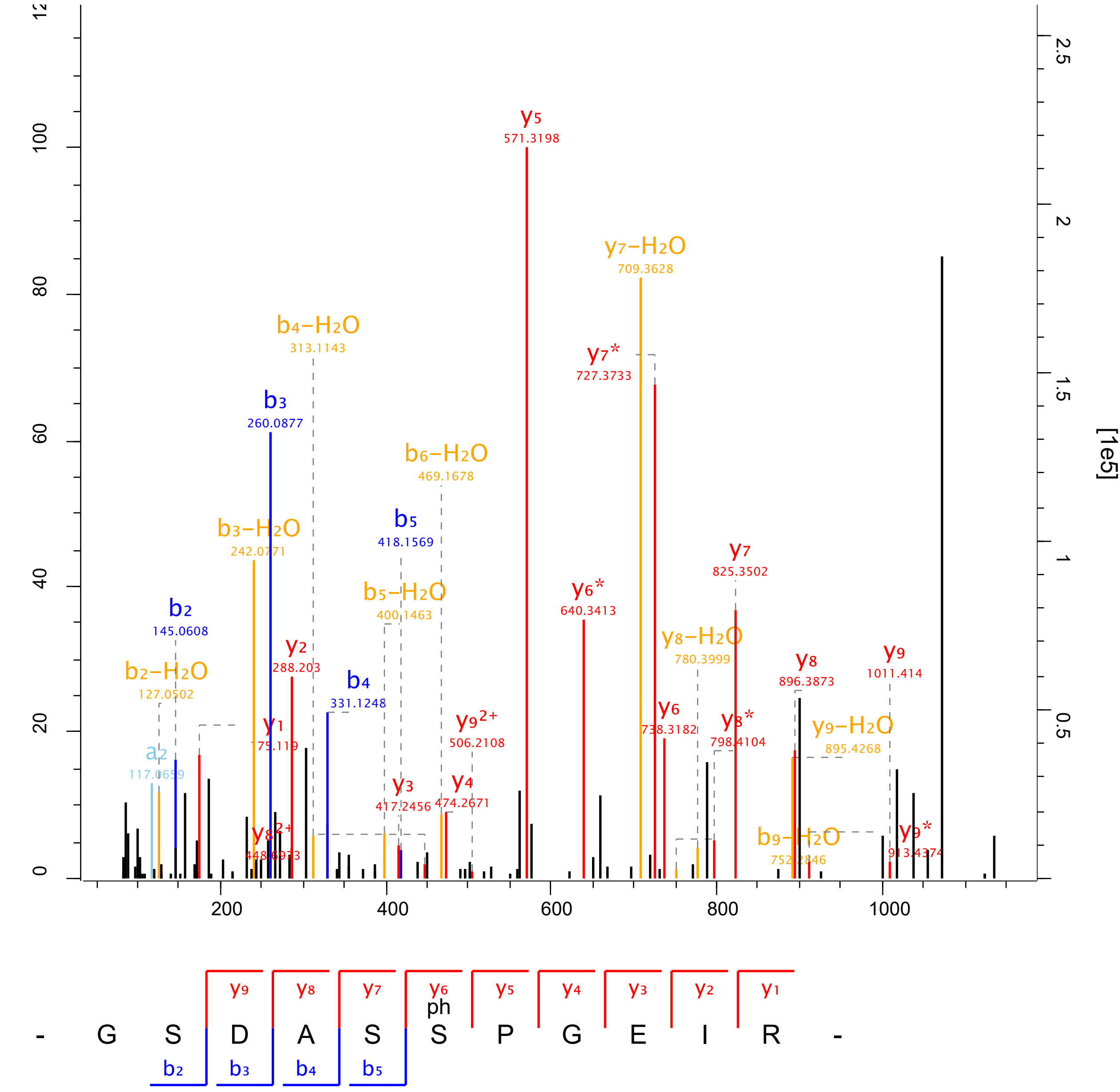

| Raw file | Scan | Method | Score | m/z |
| --- | --- | --- | --- | --- |
| sys_15_1 | 5754 | FTMS; HCD | 92.87 | 462.7 |

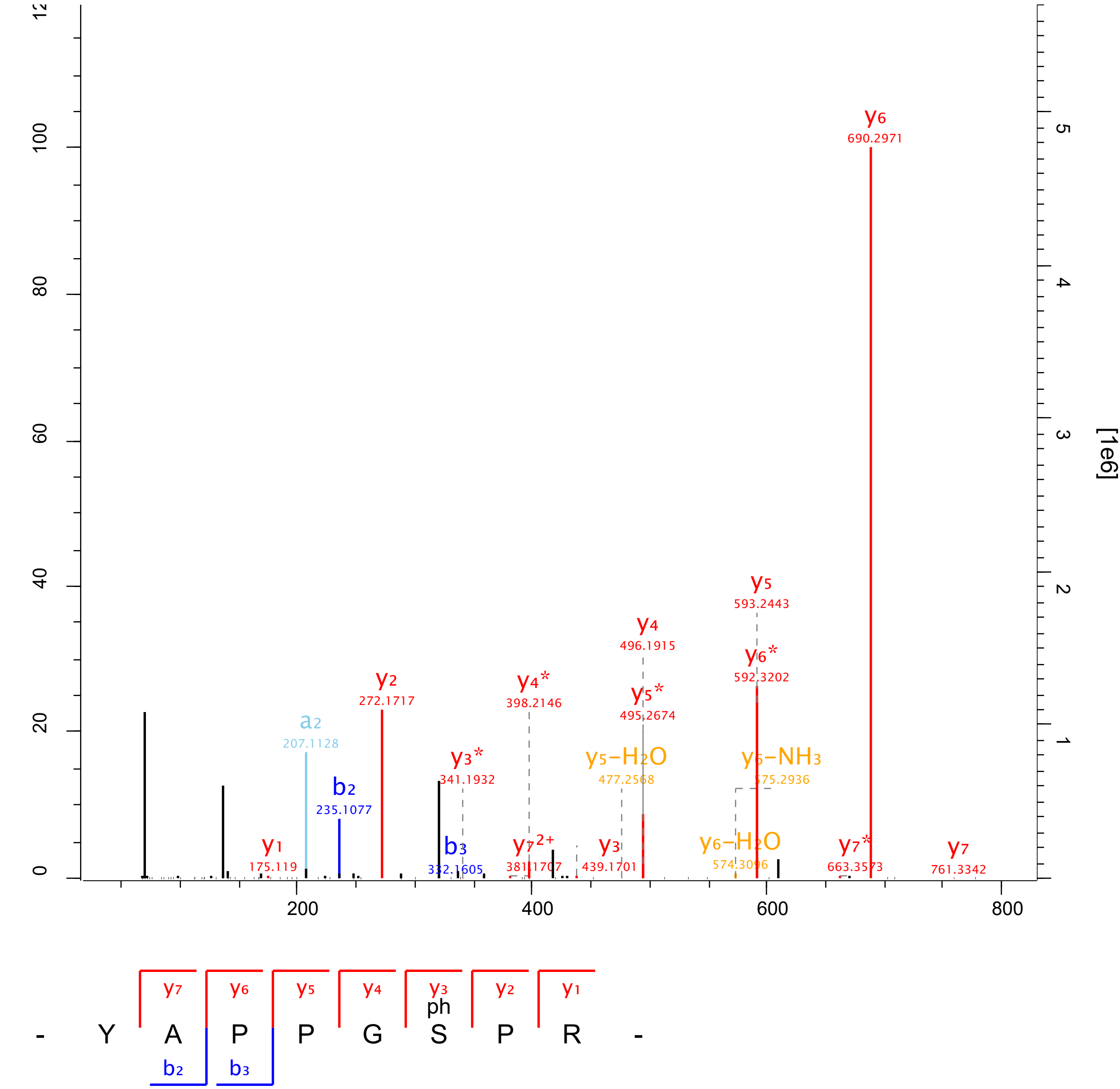

| Raw file | Scan | Method | Score | m/z |
| --- | --- | --- | --- | --- |
| sys_15_1 | 5758 | FTMS; HCD | 112.36 | 496.69 |

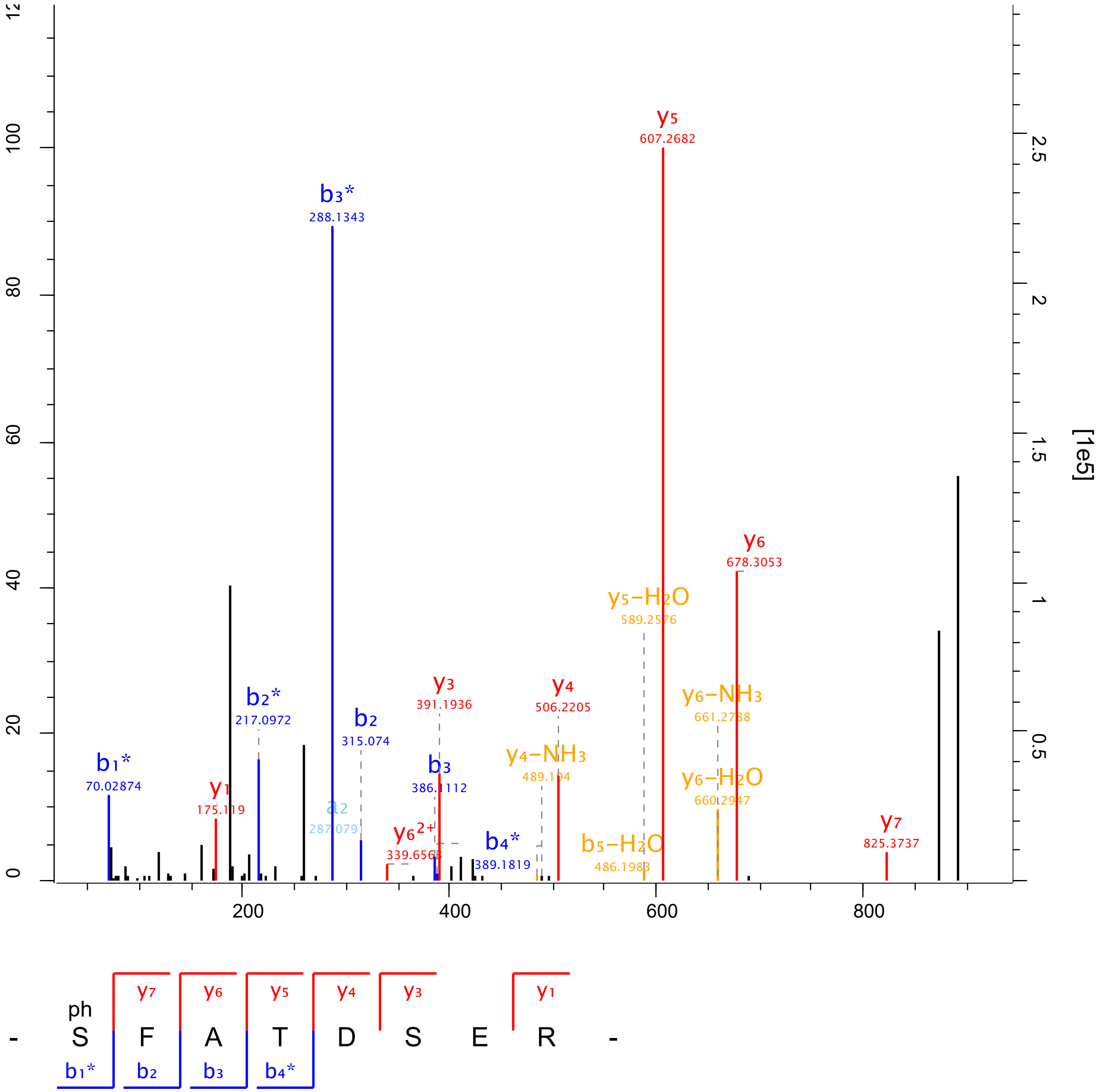

| Raw file | Scan | Method | Score | m/z |
|---|---|---|---|---|
| sys_15_1 | 5771 | FTMS; HCD | 143.25 | 525.26 |

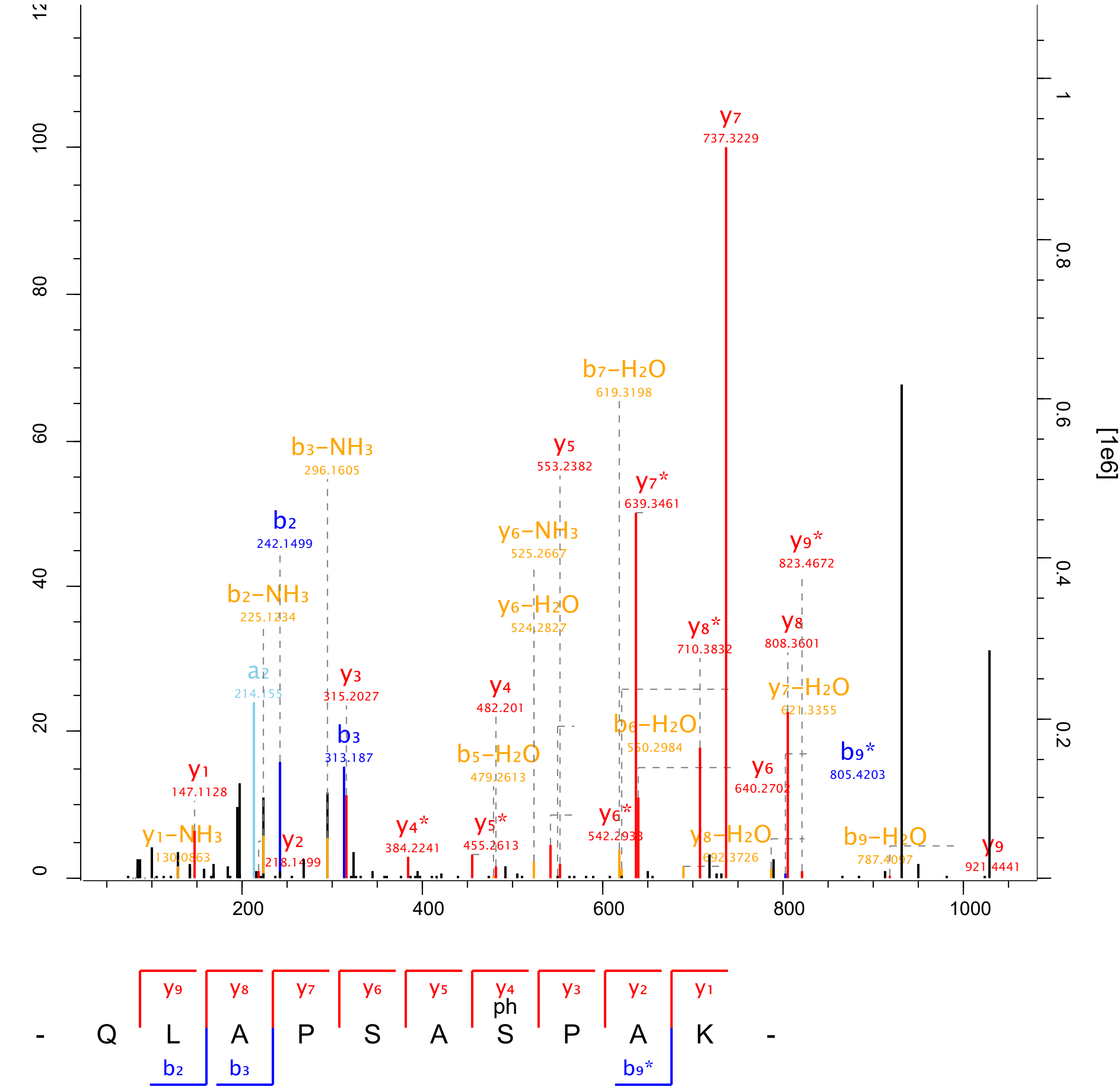

| Raw file | Scan | Method | Score | m/z |
|---|---|---|---|---|
| sys_15_1 | 5791 | FTMS; HCD | 216.01 | 635.76 |

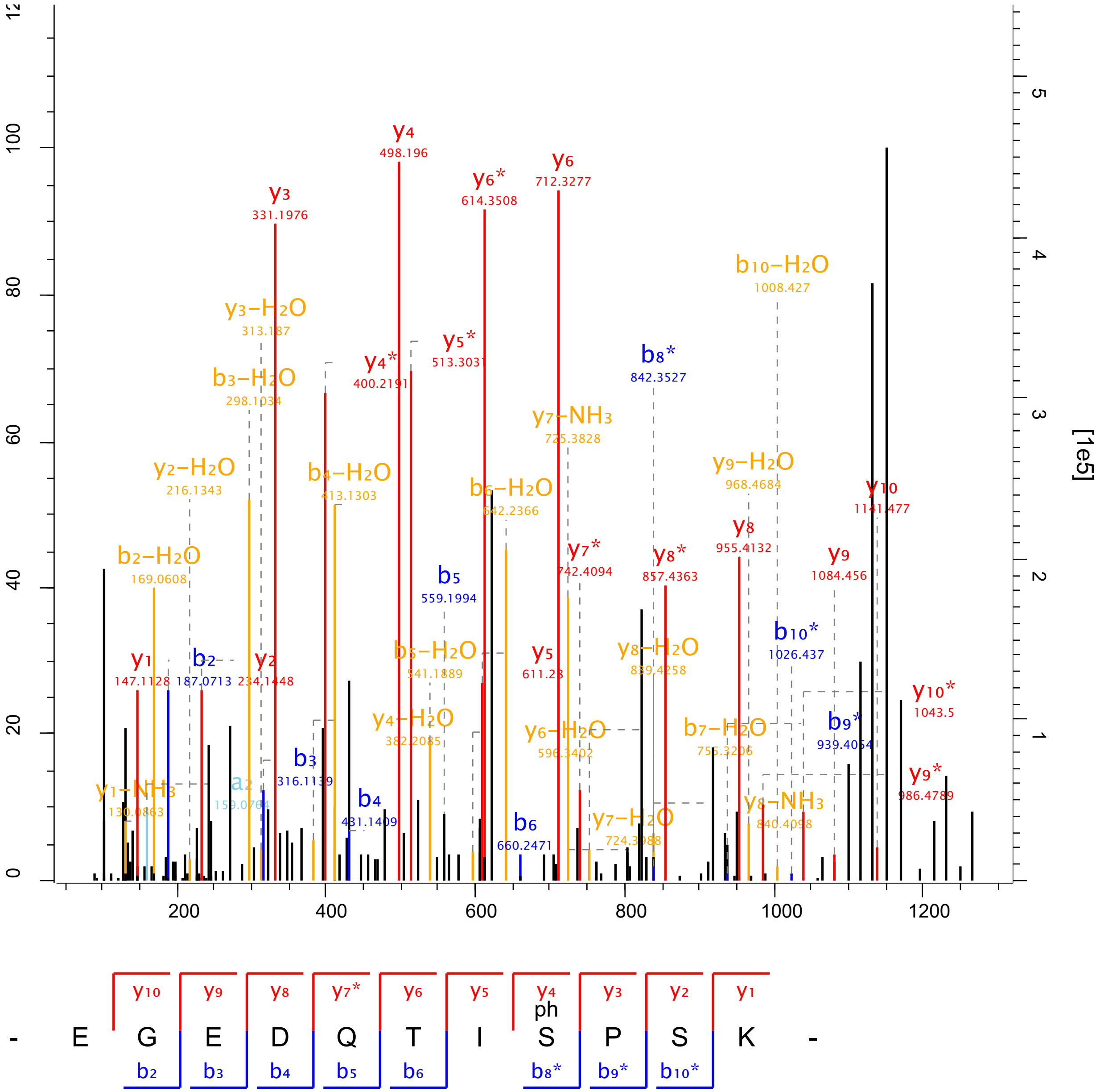

| Raw file | Scan | Method | Score | m/z |
|---|---|---|---|---|
| sys_15_1 | 5796 | FTMS; HCD | 158.47 | 580.24 |

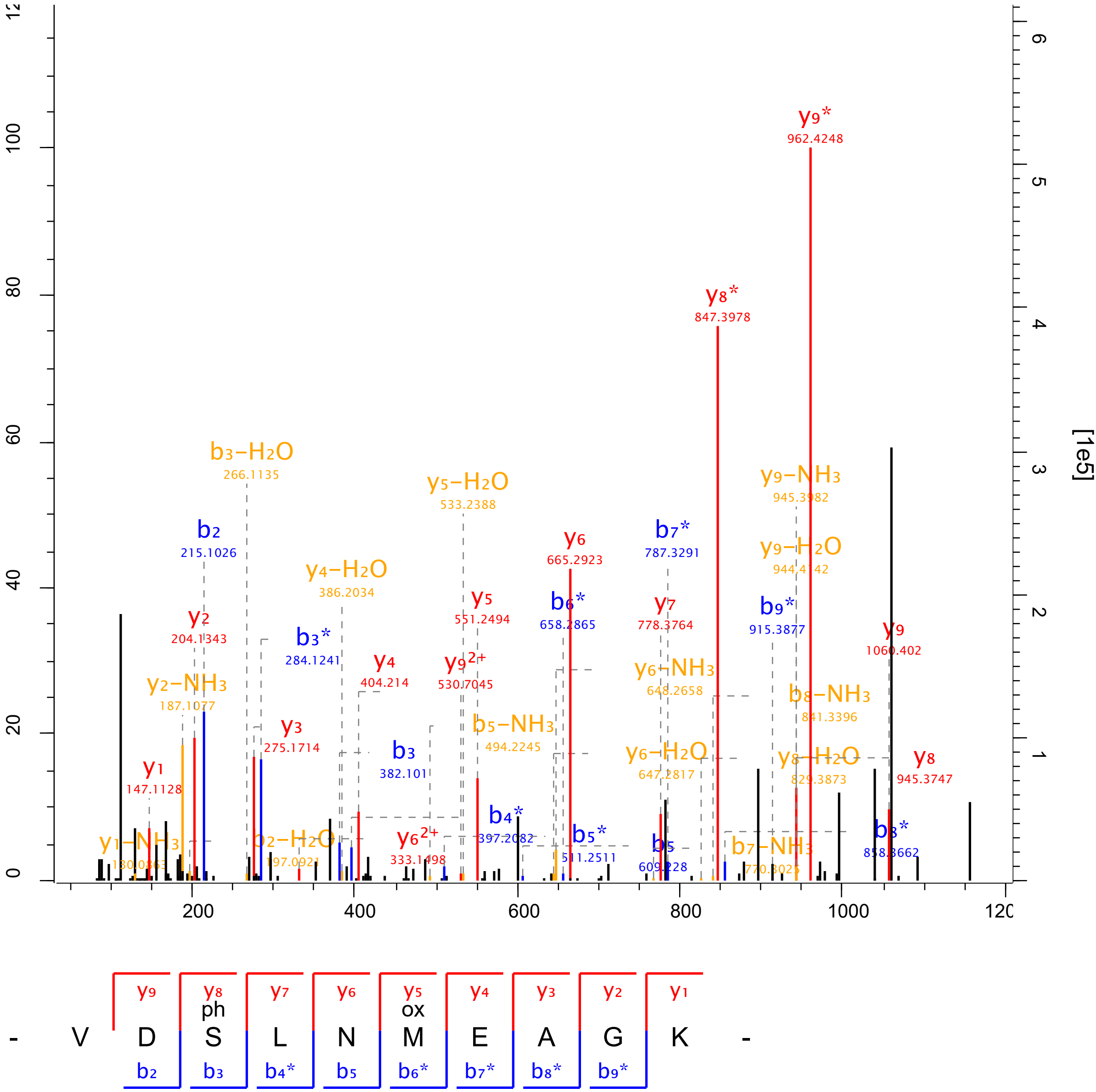

| Raw file | Scan | Method | Score | m/z |
|---|---|---|---|---|
| sys_15_1 | 5819 | FTMS; HCD | 45.83 | 473.19 |

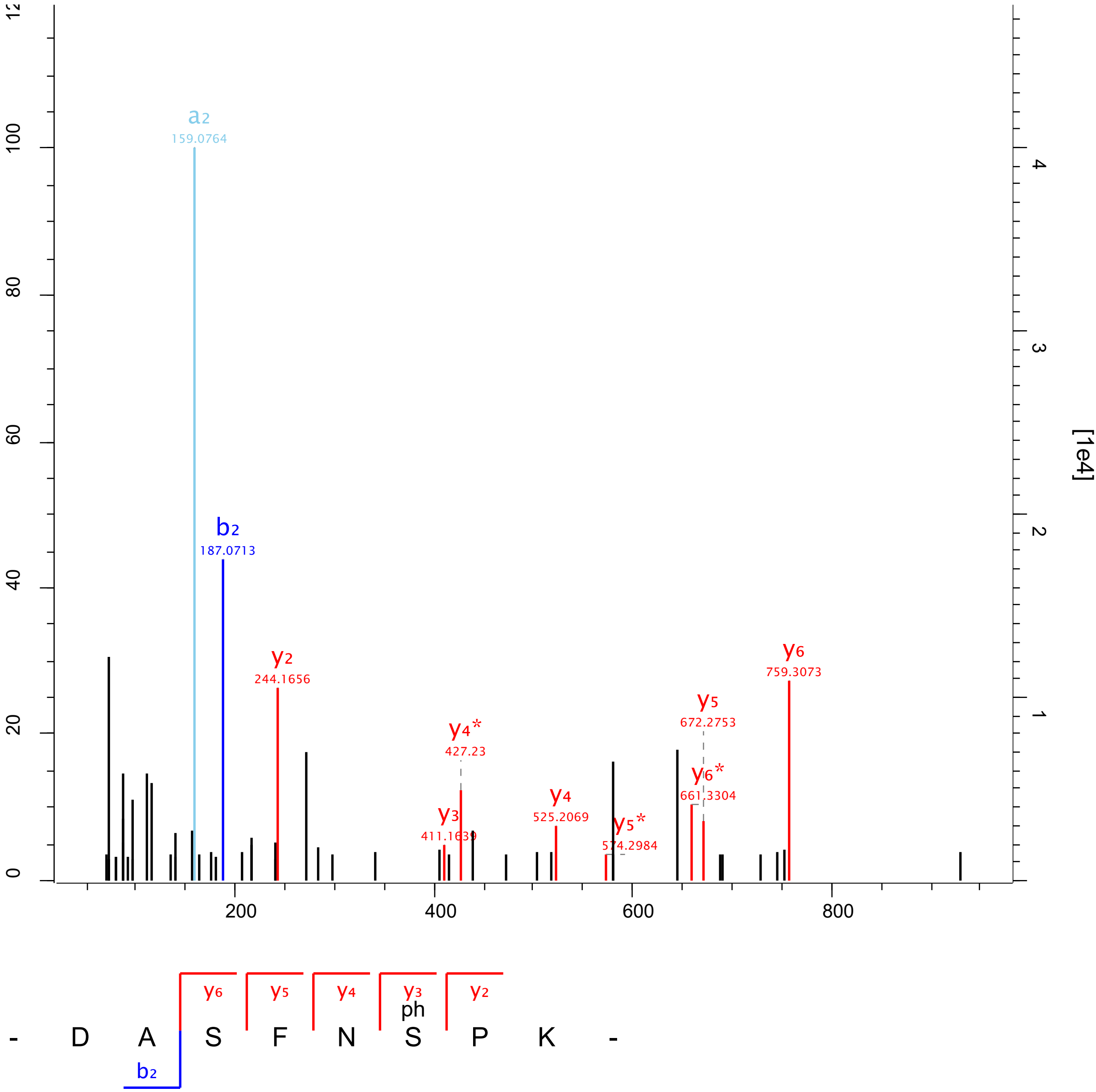

| Raw file | Scan | Method | Score | m/z |
|---|---|---|---|---|
| sys_15_1 | 5841 | FTMS; HCD | 96.17 | 459.7 |

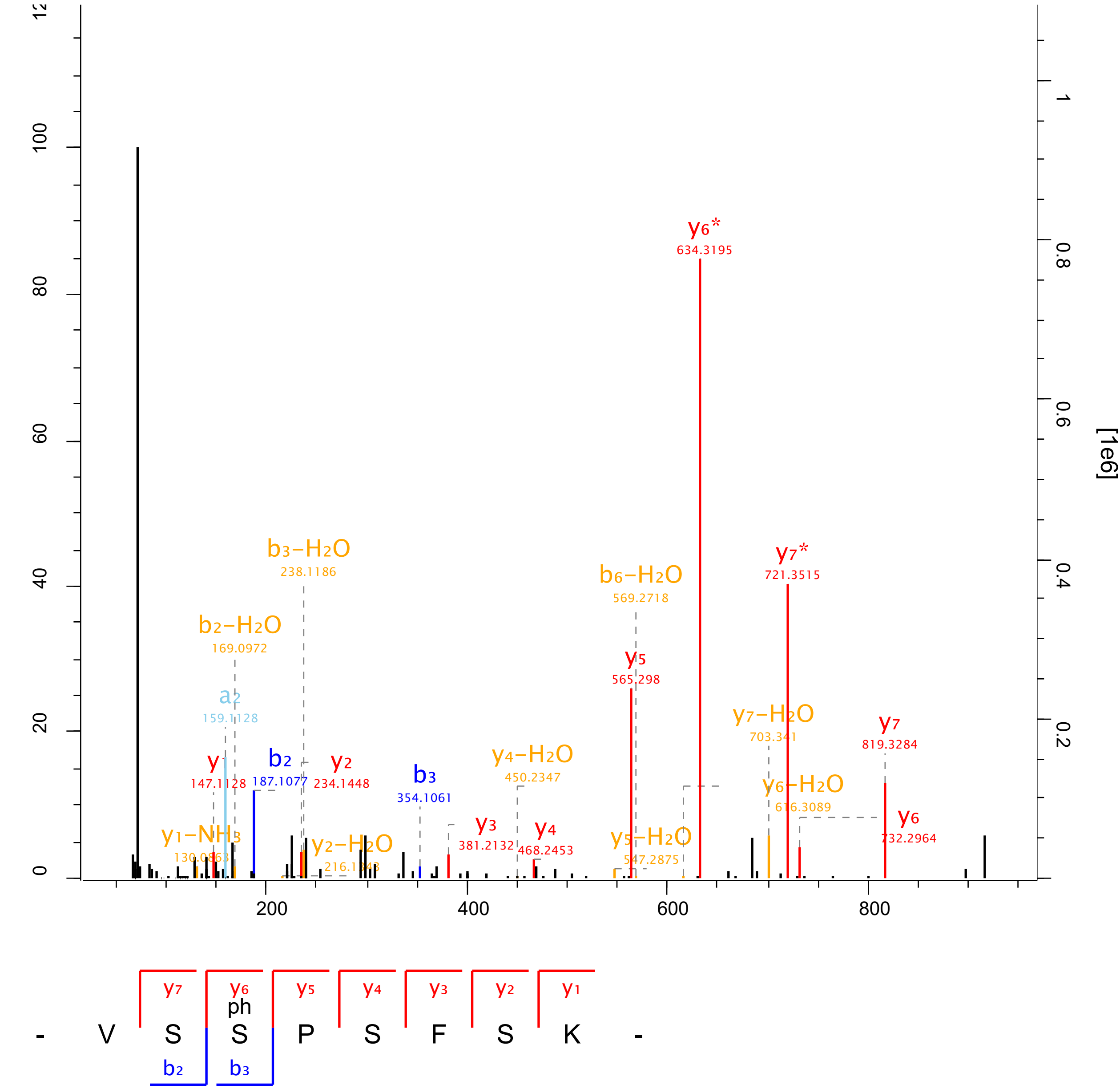

| Raw file | Scan | Method | Score | m/z |
|---|---|---|---|---|
| sys_15_1 | 5895 | FTMS; HCD | 50.9 | 657.76 |

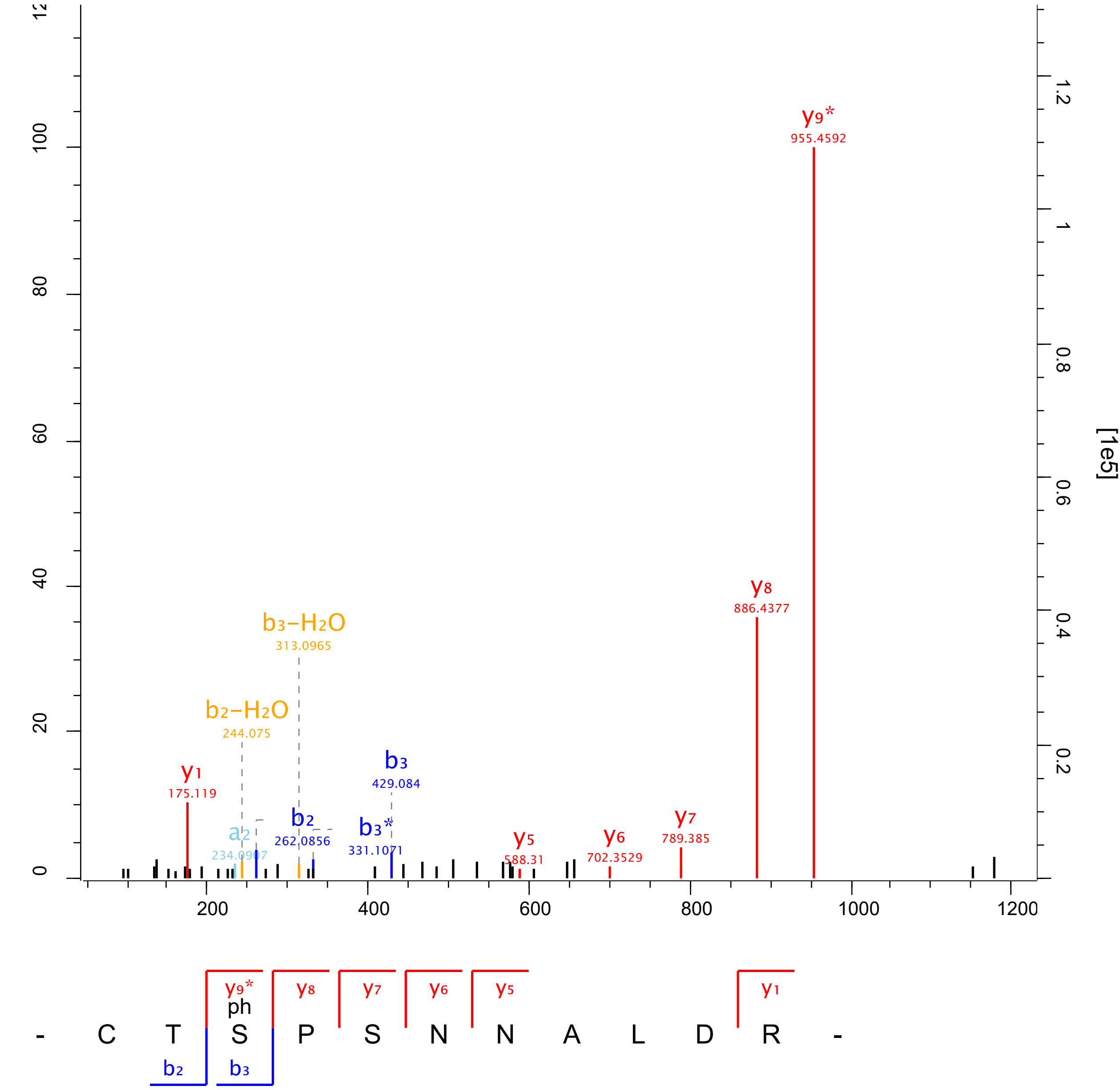

| Raw file | Scan | Method | Score | m/z |
|---|---|---|---|---|
| sys_15_1 | 5980 | FTMS; HCD | 51.45 | 508.73 |

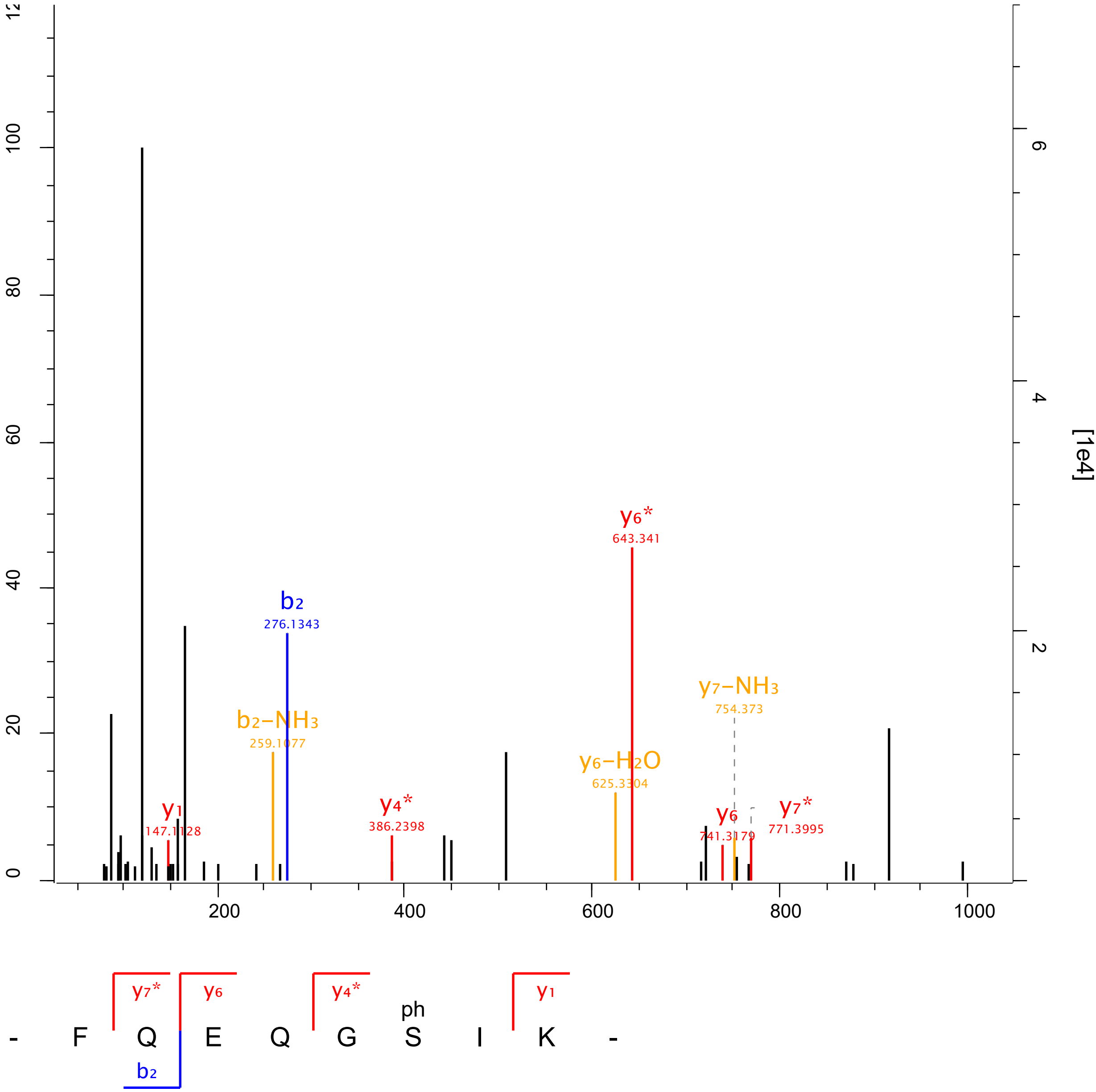

| Raw file | Scan | Method | Score | m/z |
| --- | --- | --- | --- | --- |
| sys_15_1 | 6010 | FTMS; HCD | 123.26 | 532.21 |

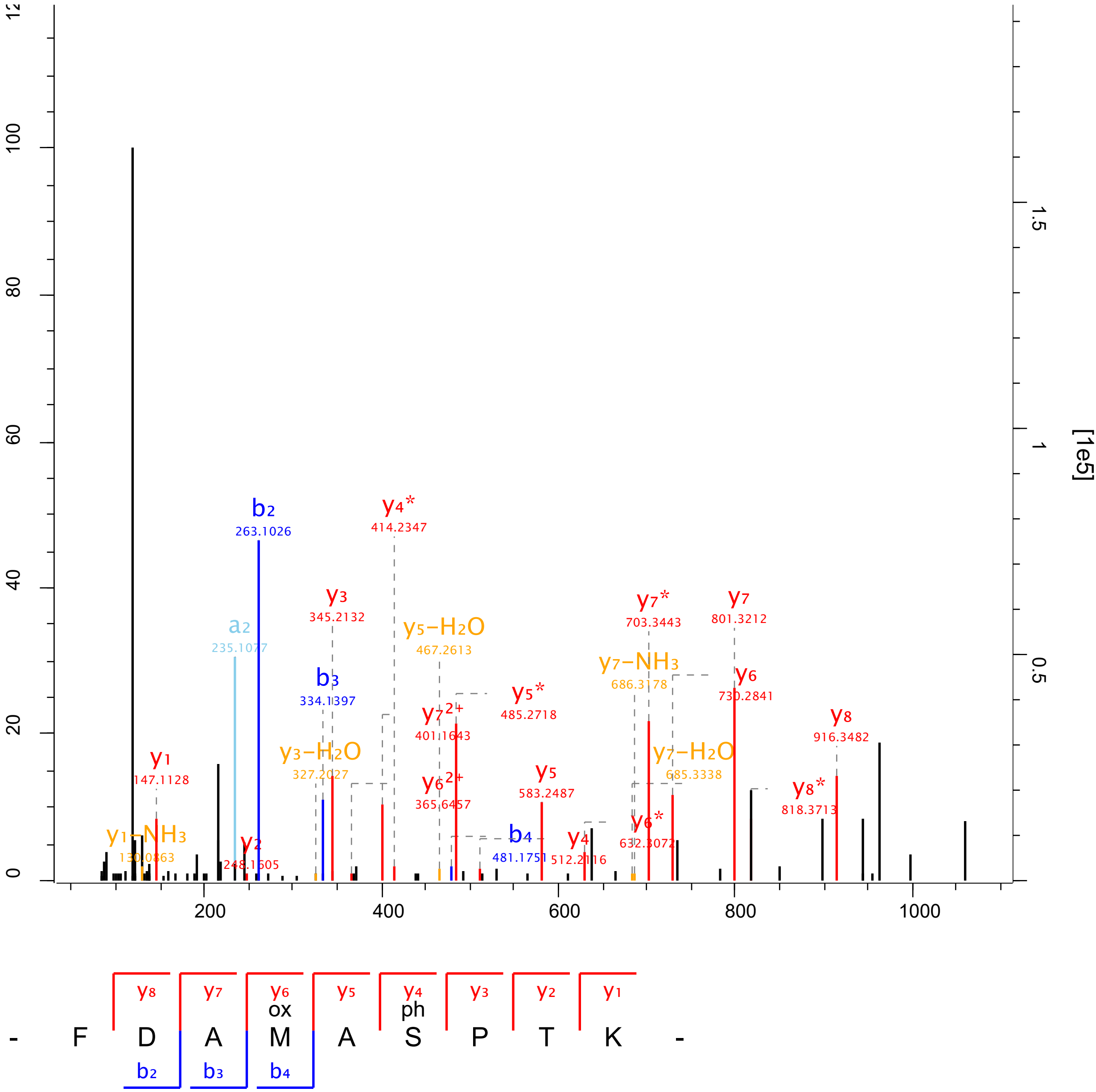

| Raw file | Scan | Method | Score | m/z |
| --- | --- | --- | --- | --- |
| sys_15_1 | 6053 | FTMS; HCD | 49.45 | 545.71 |

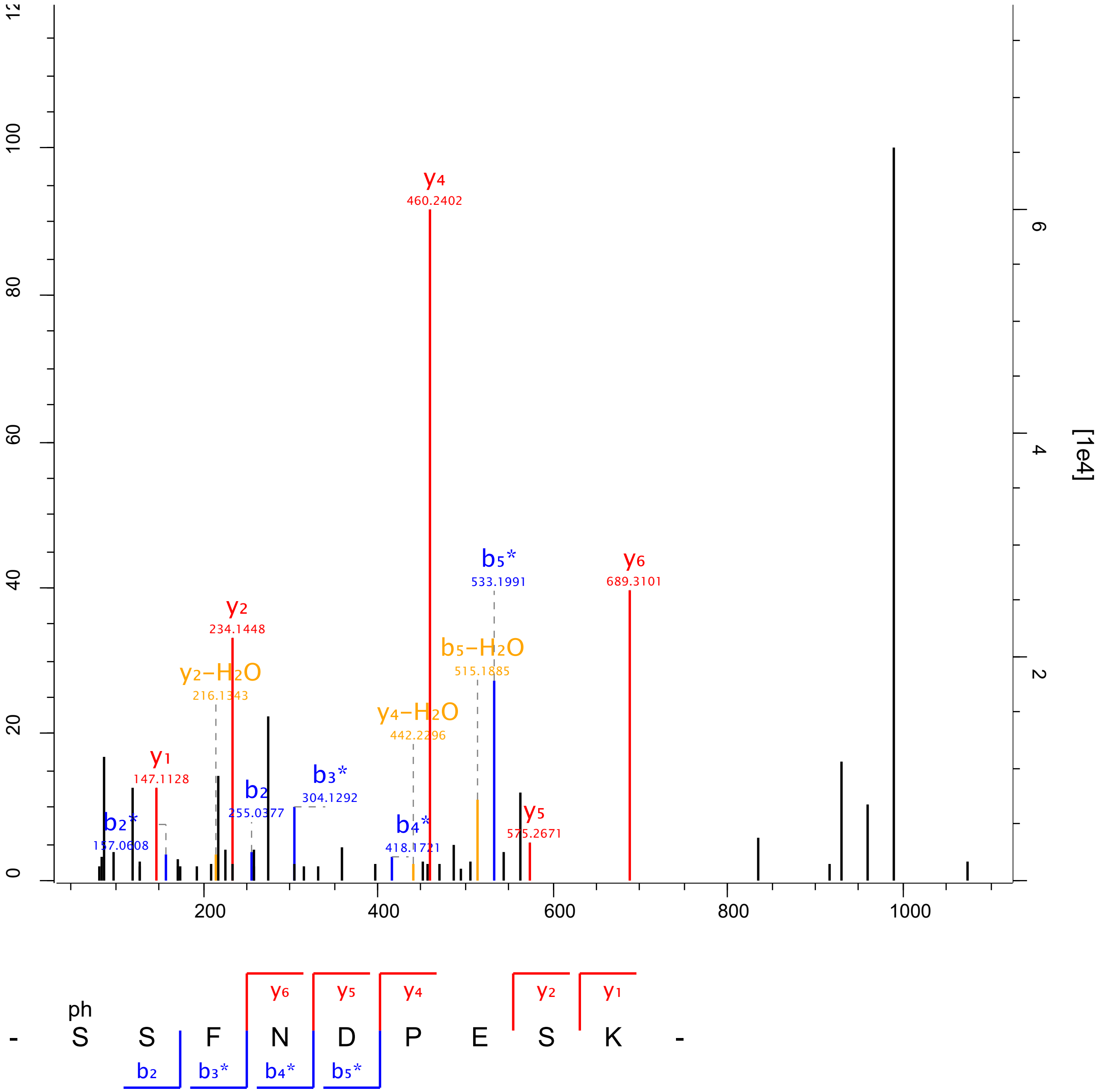

| Raw file | Scan | Method | Score | m/z |
|---|---|---|---|---|
| sys_15_1 | 6138 | FTMS; HCD | 92.25 | 576.21 |

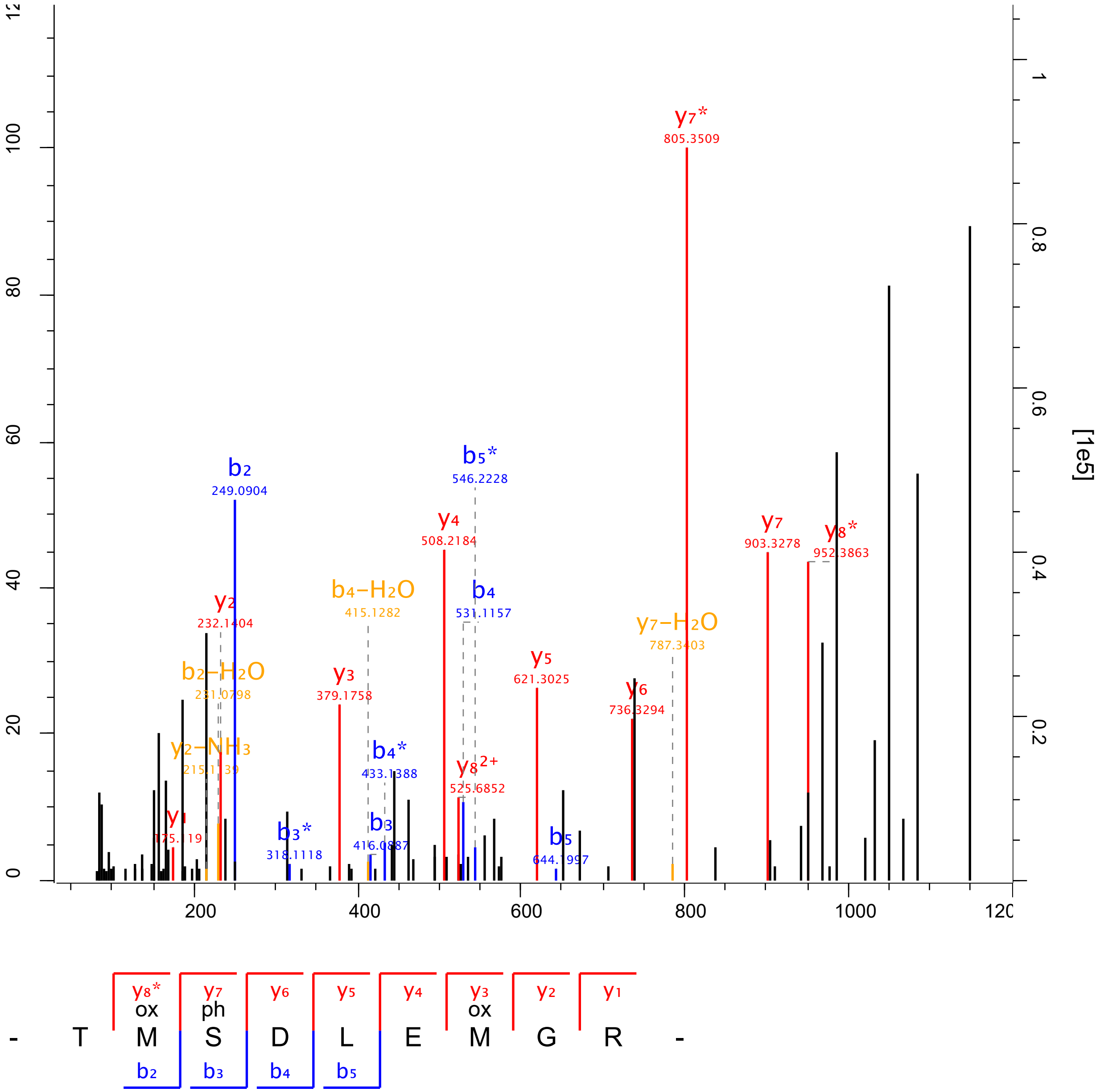

| Raw file | Scan | Method | Score | m/z |
|---|---|---|---|---|
| sys_15_1 | 6191 | FTMS; HCD | 71.35 | 584.21 |

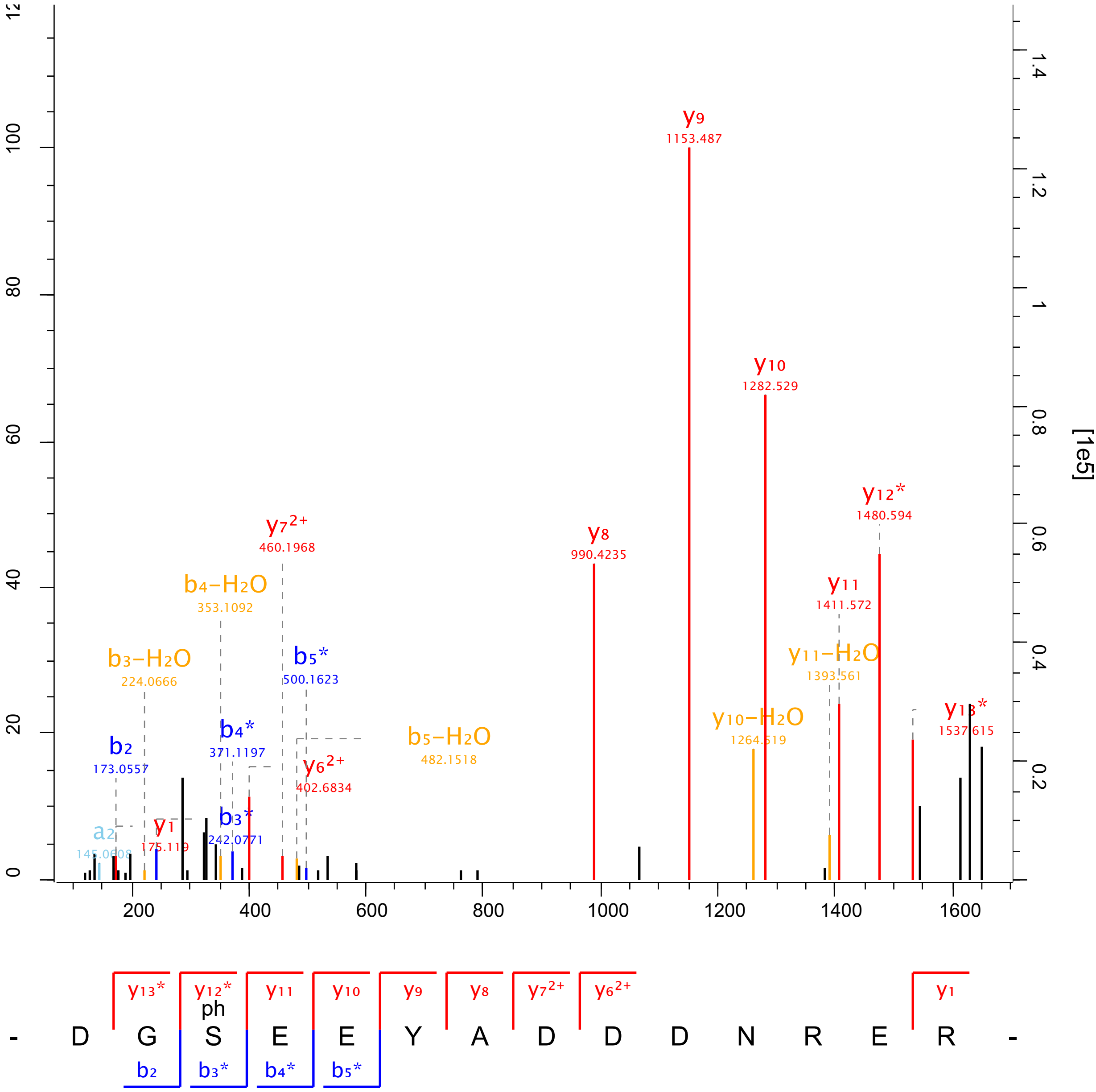

| Raw file | Scan | Method | Score | m/z |
|---|---|---|---|---|
| sys_15_1 | 6215 | FTMS; HCD | 110.12 | 644.79 |

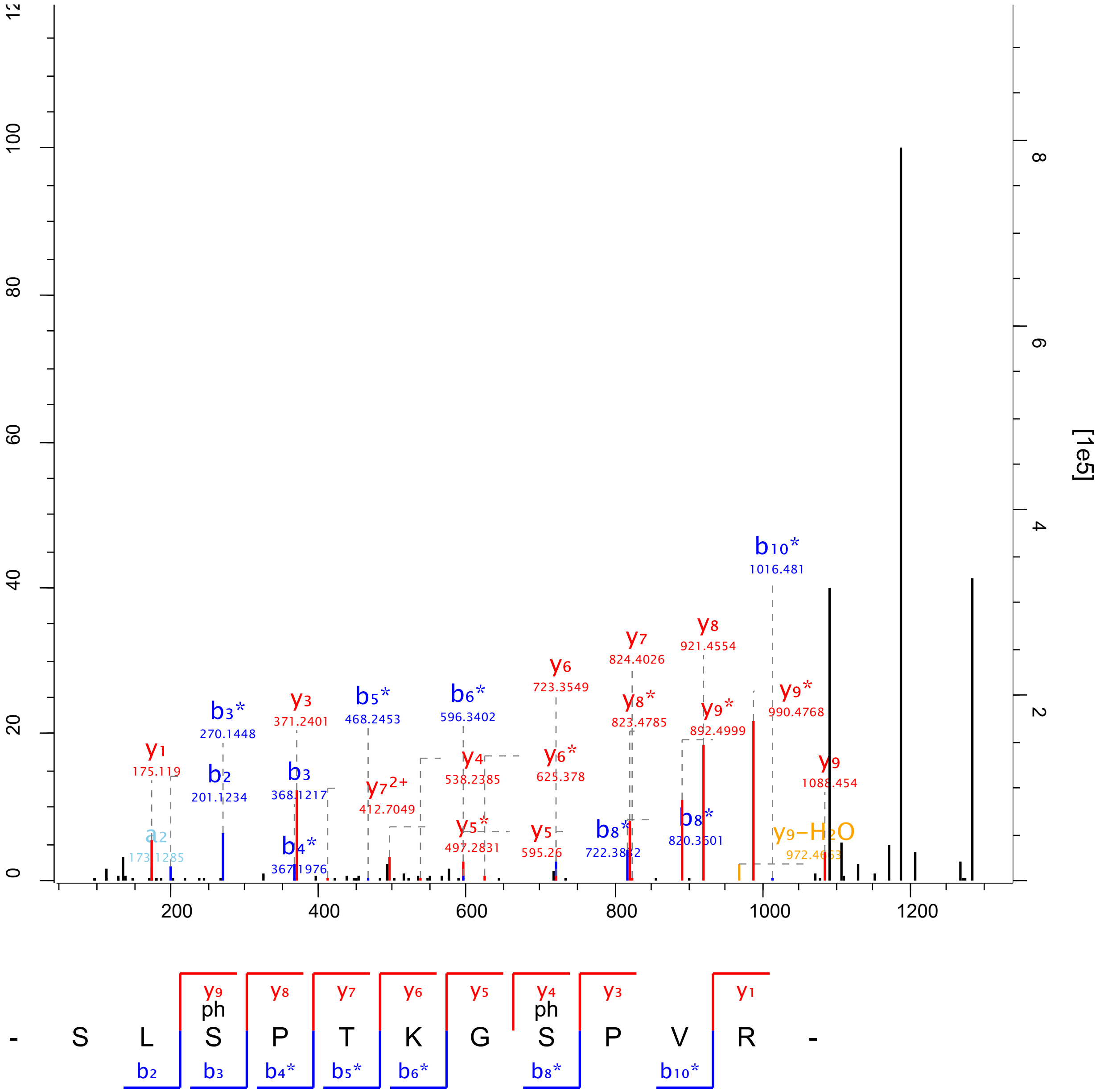

| Raw file | Scan | Method | Score | m/z |
|---|---|---|---|---|
| sys_15_1 | 6484 | FTMS; HCD | 217.67 | 644.29 |

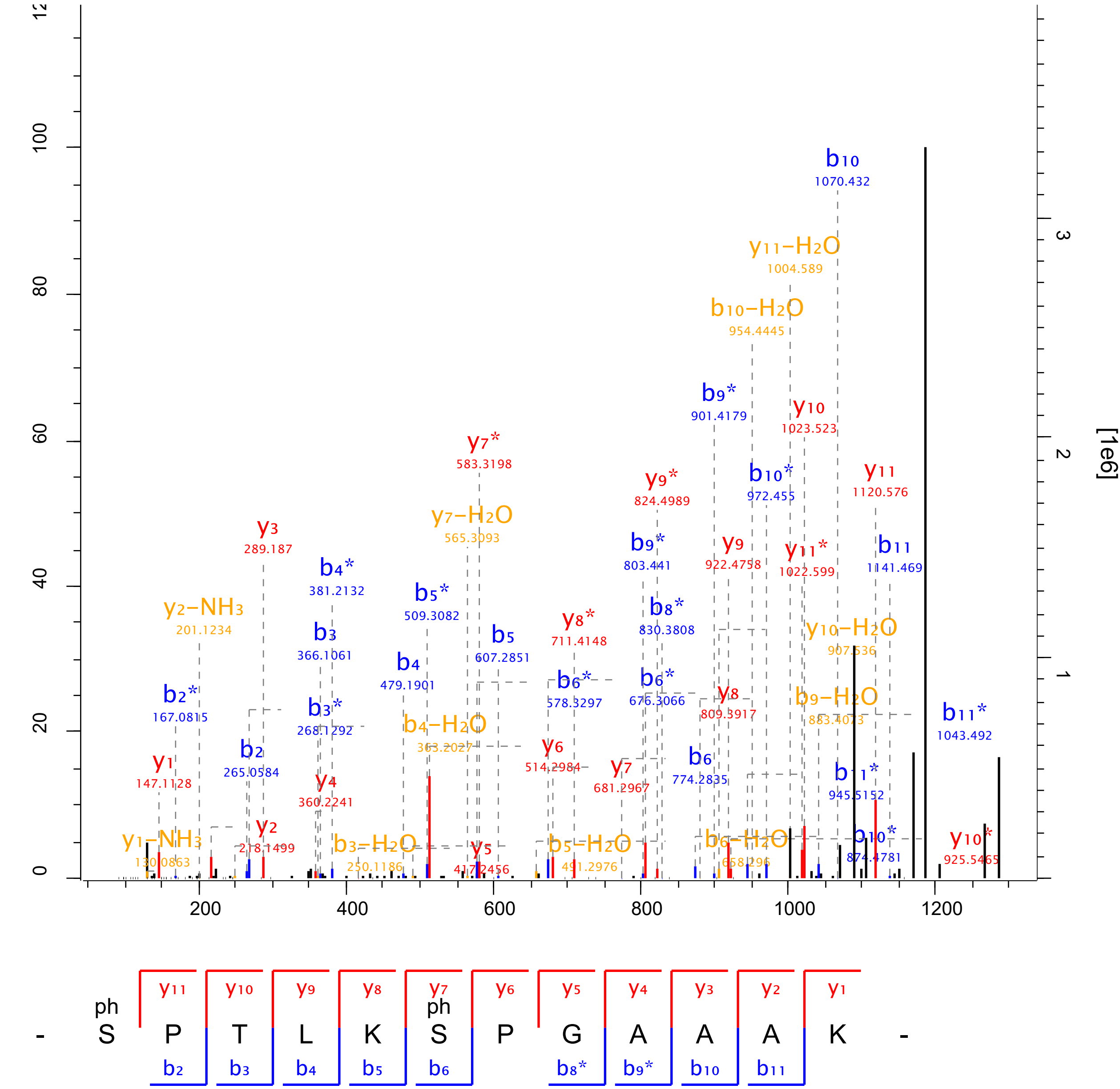

| Raw file | Scan | Method | Score | m/z |
| --- | --- | --- | --- | --- |
| sys_15_1 | 6488 | FTMS; HCD | 121.03 | 695.75 |

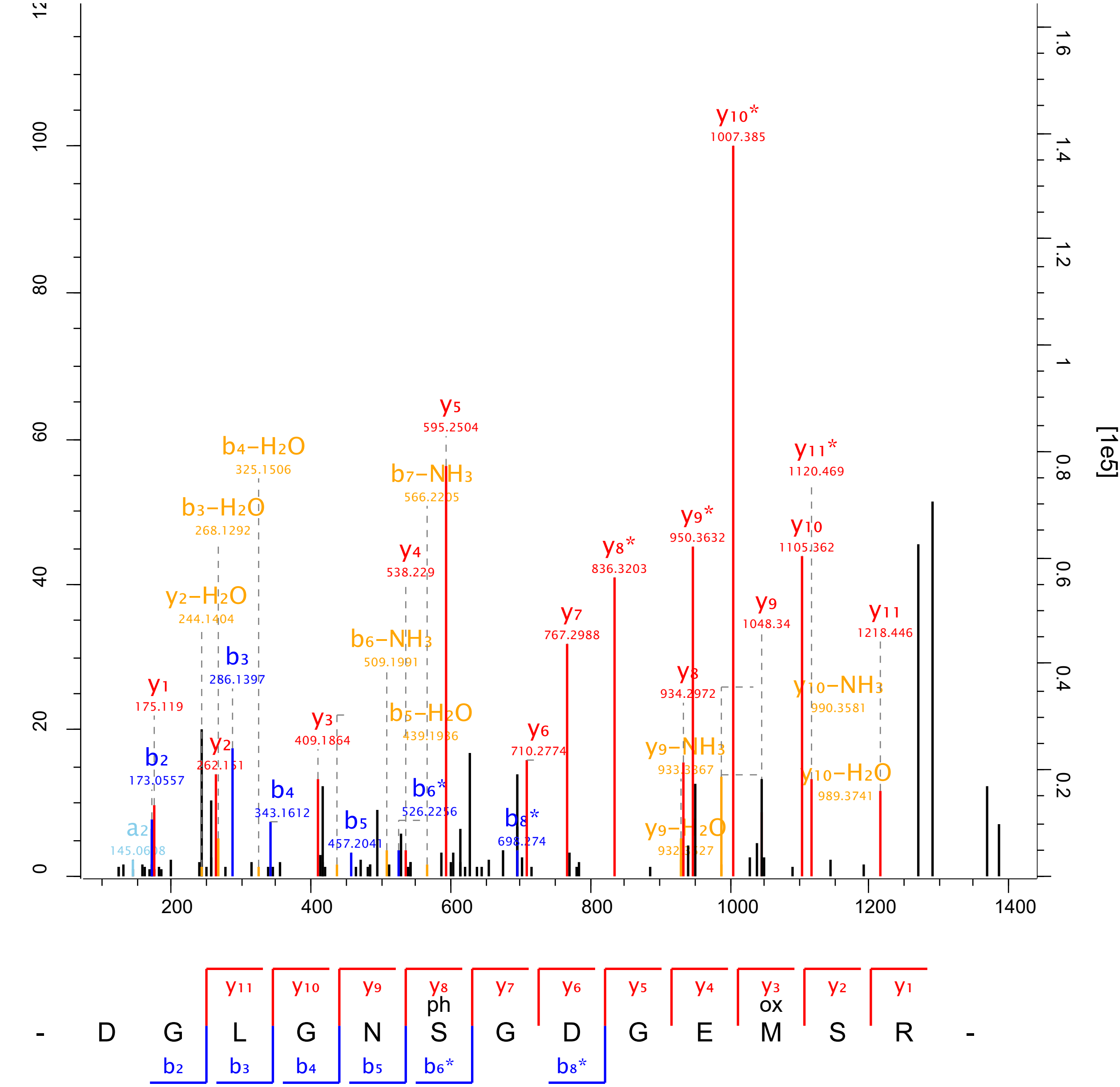

| Raw file | Scan | Method | Score | m/z |
| --- | --- | --- | --- | --- |
| sys_15_1 | 6498 | FTMS; HCD | 191.93 | 777.32 |

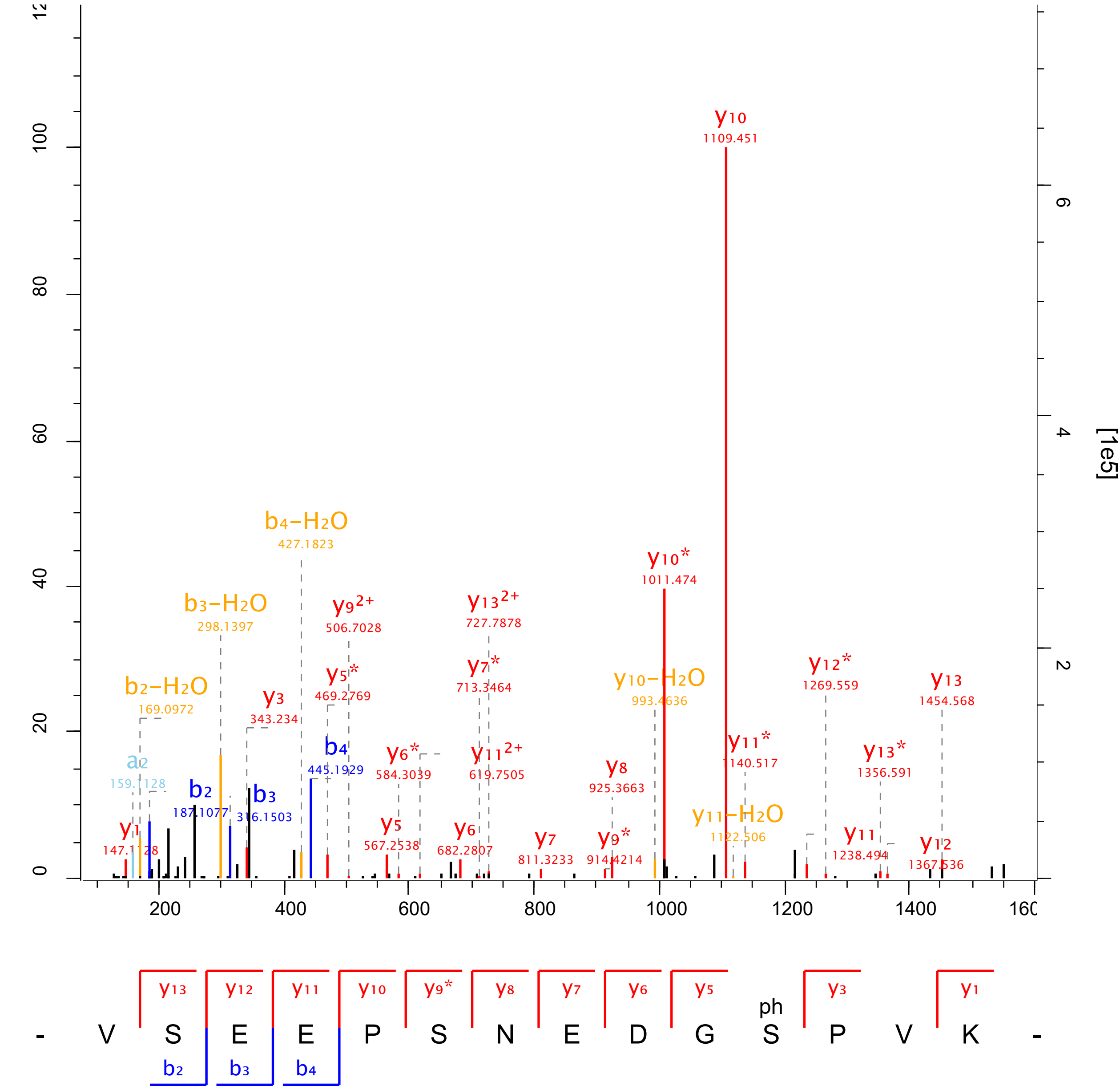

| Raw file | Scan | Method | Score | m/z |
|---|---|---|---|---|
| sys_15_1 | 6565 | FTMS; HCD | 65.54 | 445.2 |

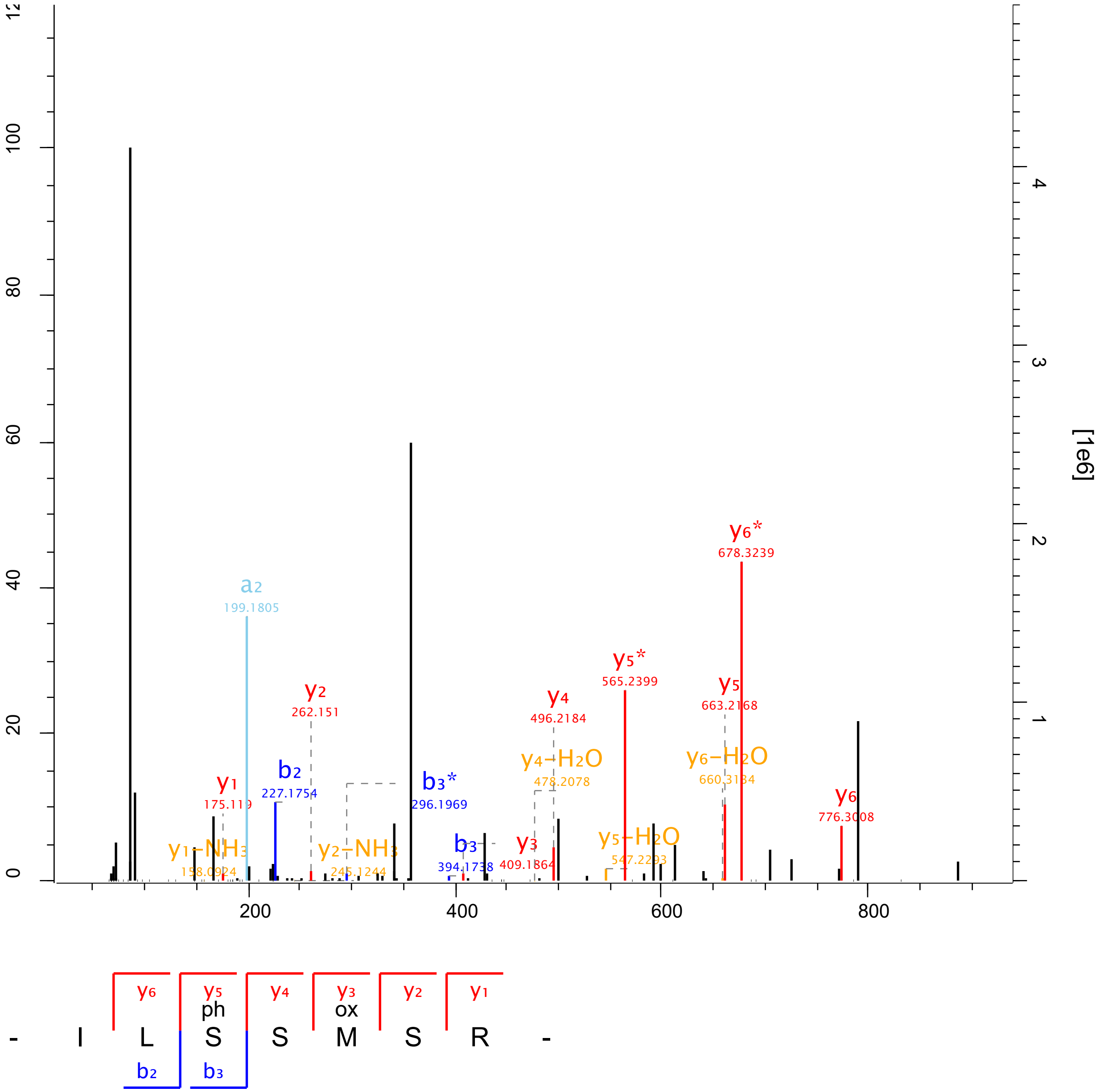

| Raw file | Scan | Method | Score | m/z |
| --- | --- | --- | --- | --- |
| sys_15_1 | 6585 | FTMS; HCD | 57.16 | 600.74 |

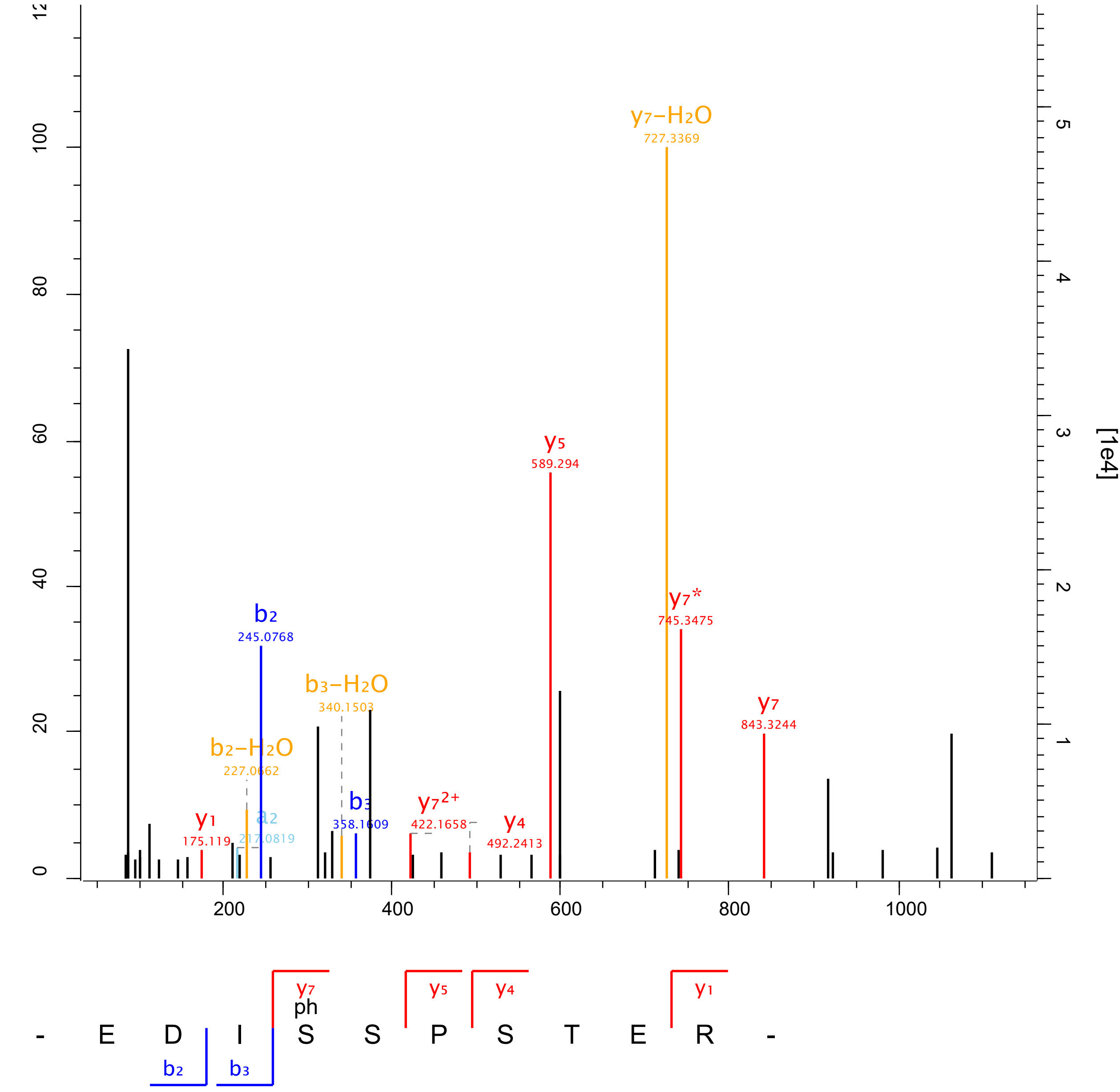

| Raw file | Scan | Method | Score | m/z |
| --- | --- | --- | --- | --- |
| sys_15_1 | 6603 | FTMS; HCD | 49.3 | 610.26 |

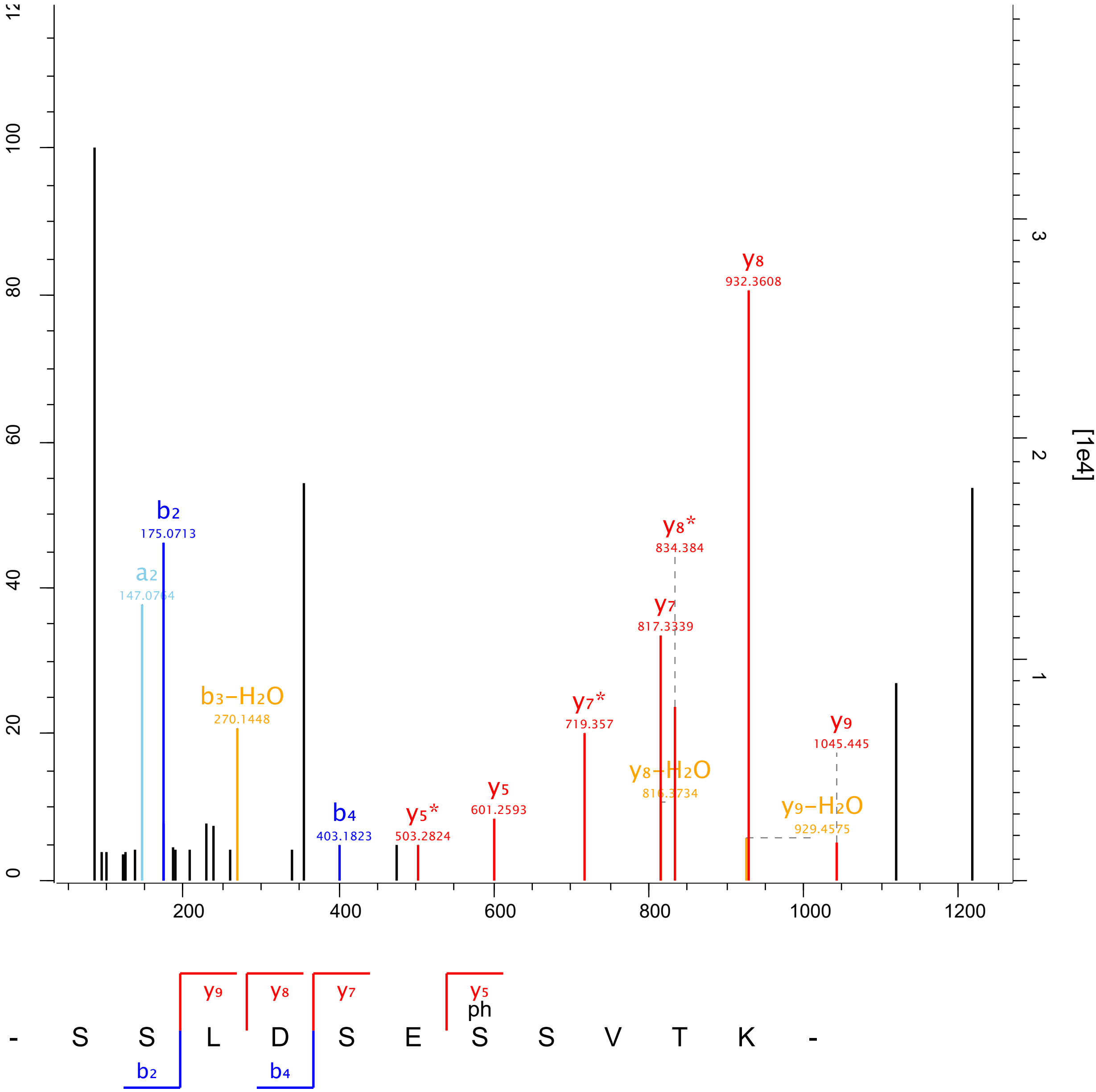

| Raw file | Scan | Method | Score | m/z |
|---|---|---|---|---|
| sys_15_1 | 6715 | FTMS; HCD | 169.54 | 571.23 |

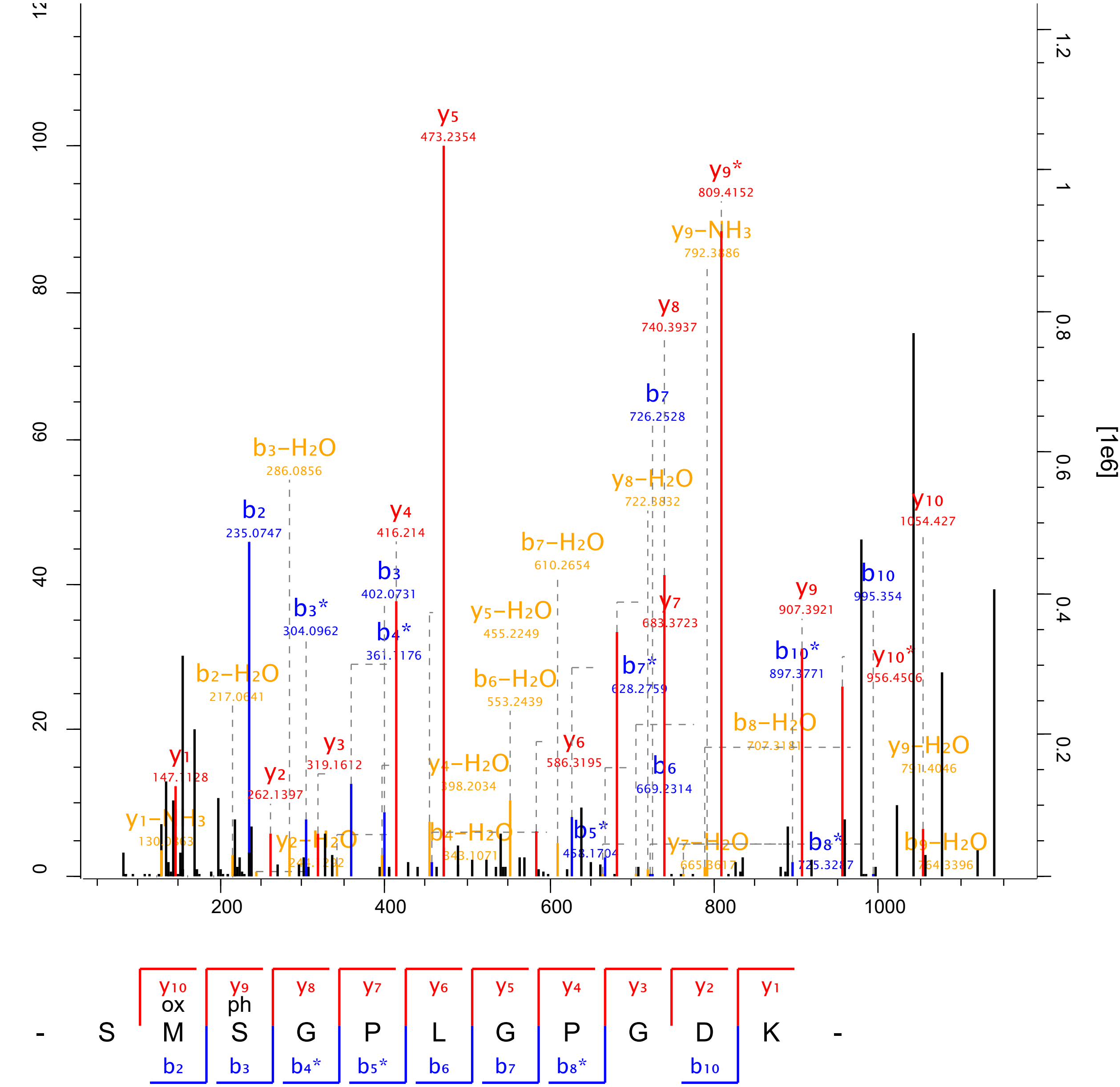

| Raw file | Scan | Method | Score | m/z |
|---|---|---|---|---|
| sys_15_1 | 6718 | FTMS; HCD | 182.02 | 675.76 |

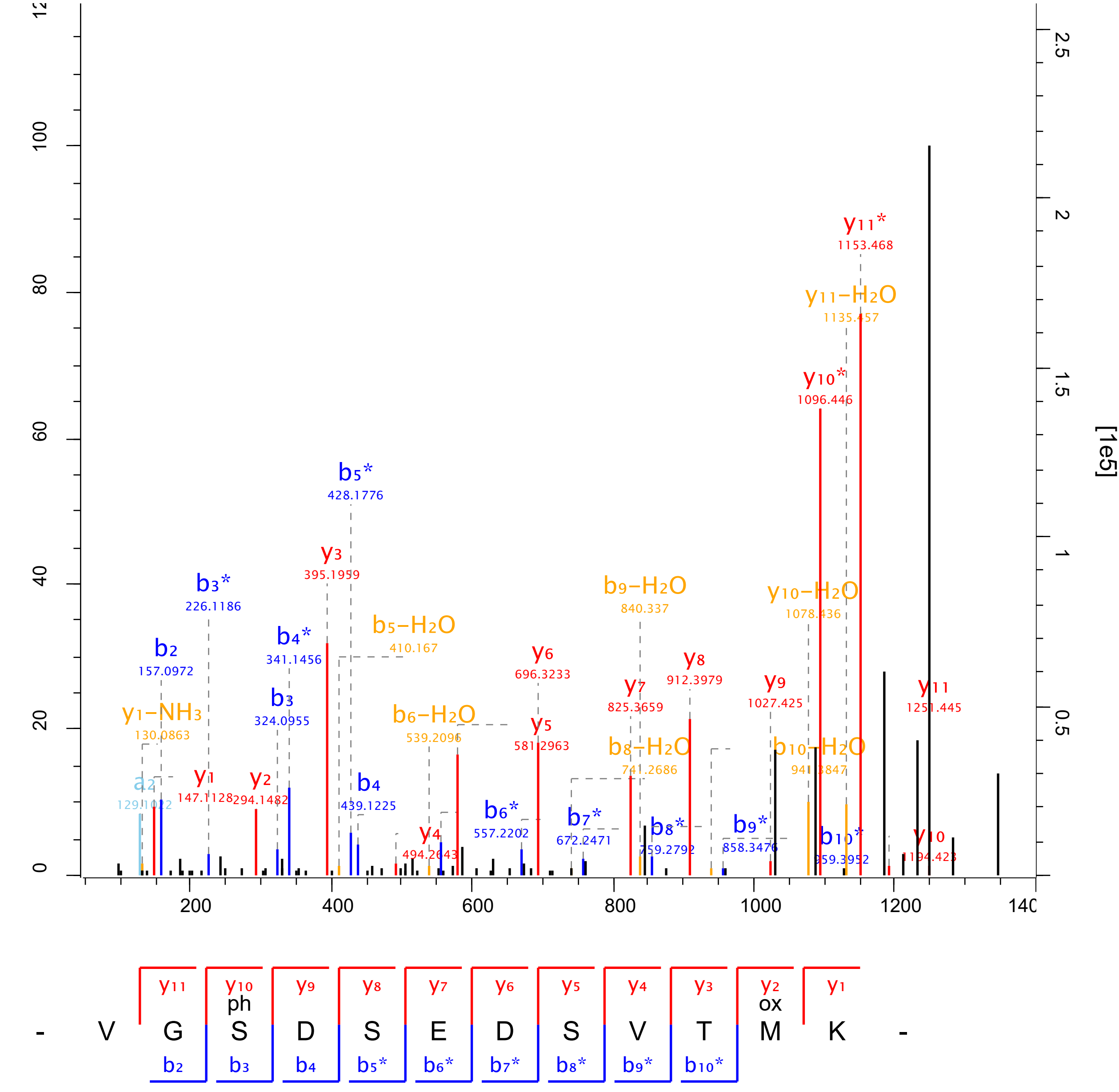

| Raw file | Scan | Method | Score | m/z |
|---|---|---|---|---|
| sys_15_1 | 6732 | FTMS; HCD | 210.98 | 762.35 |

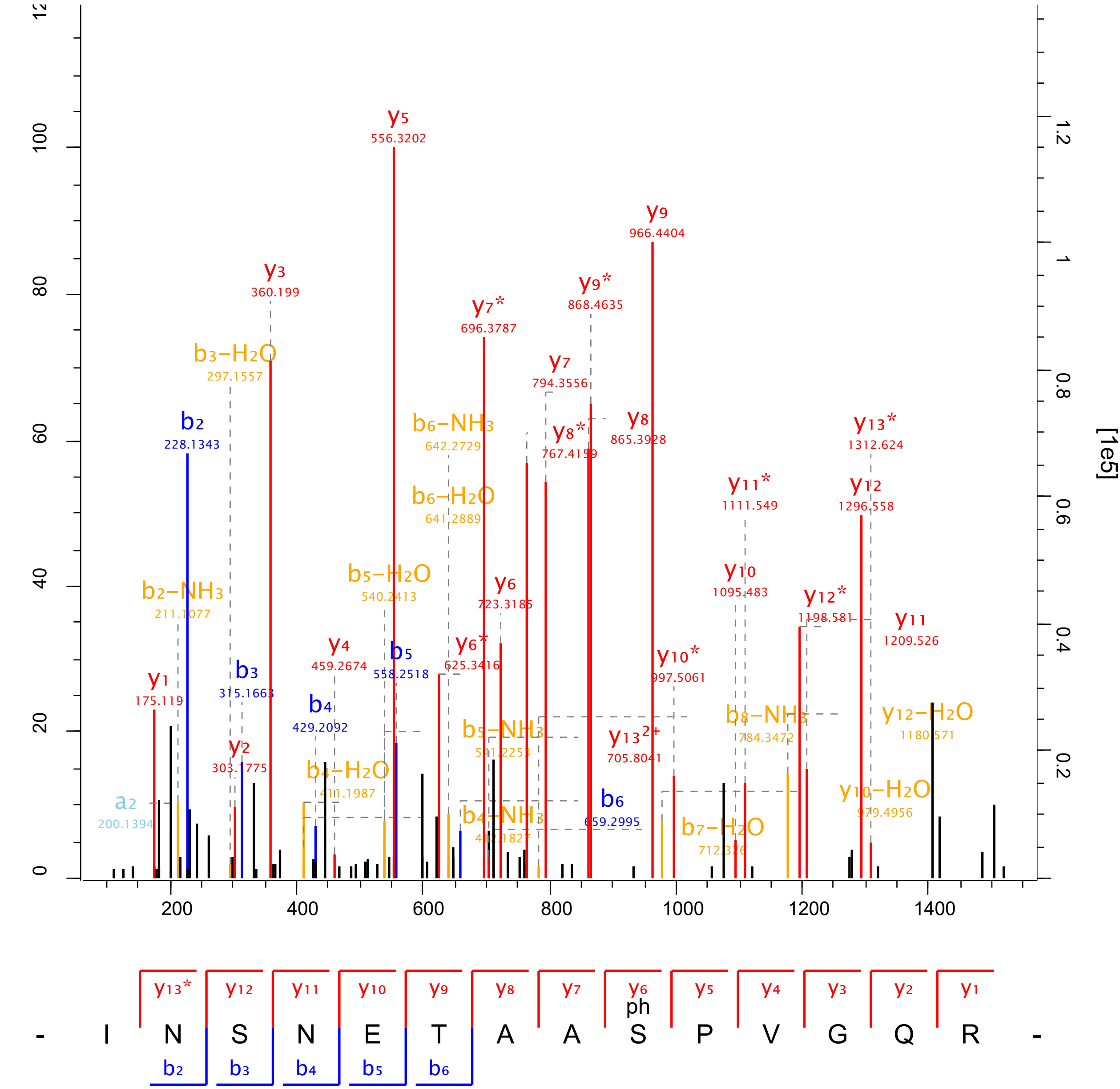

| Raw file | Scan | Method | Score | m/z |
|---|---|---|---|---|
| sys_15_1 | 6849 | FTMS; HCD | 69.03 | 638.72 |

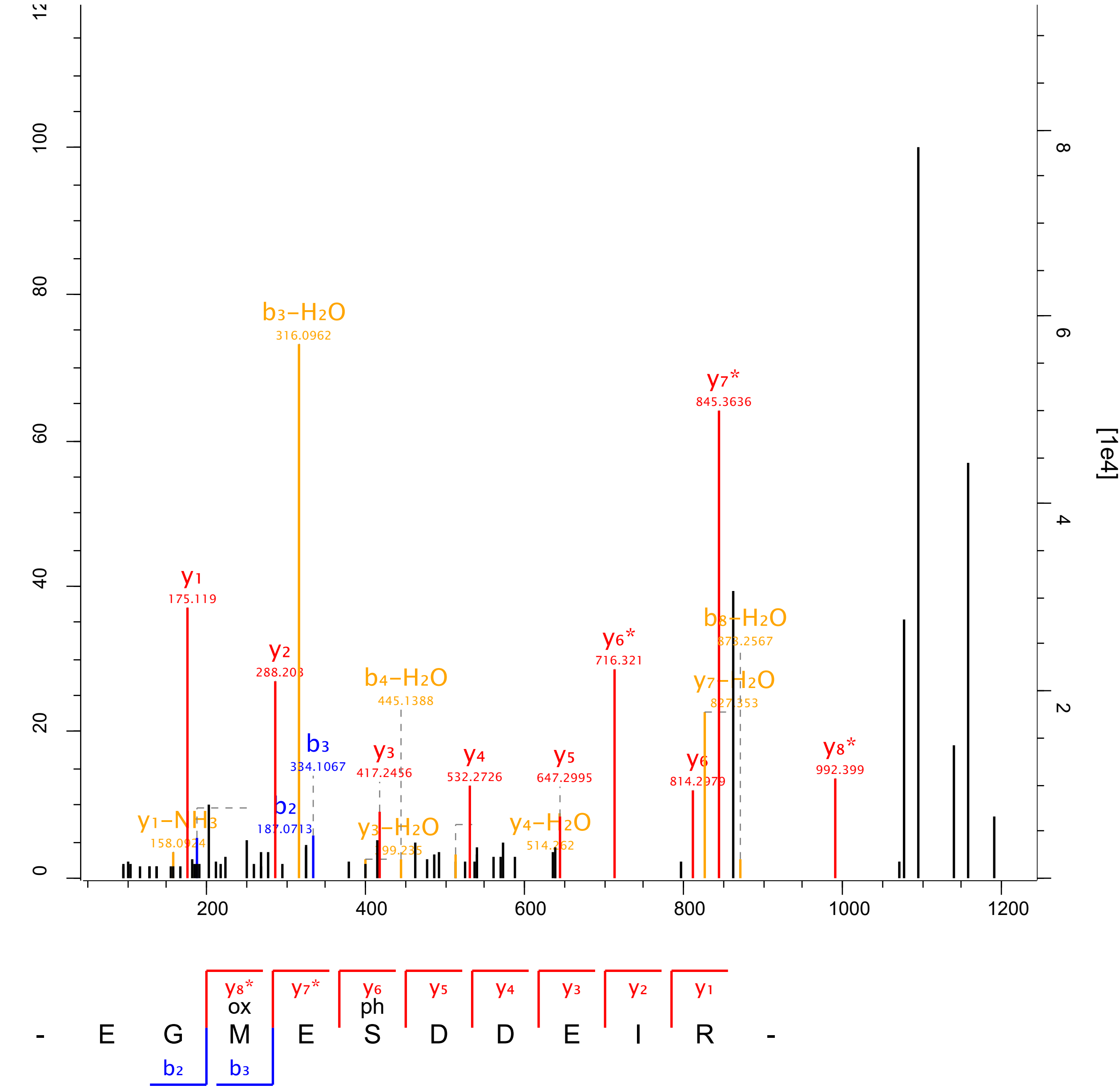

| Raw file | Scan | Method | Score | m/z |
|---|---|---|---|---|
| sys_15_1 | 6870 | FTMS; HCD | 41.62 | 623.25 |

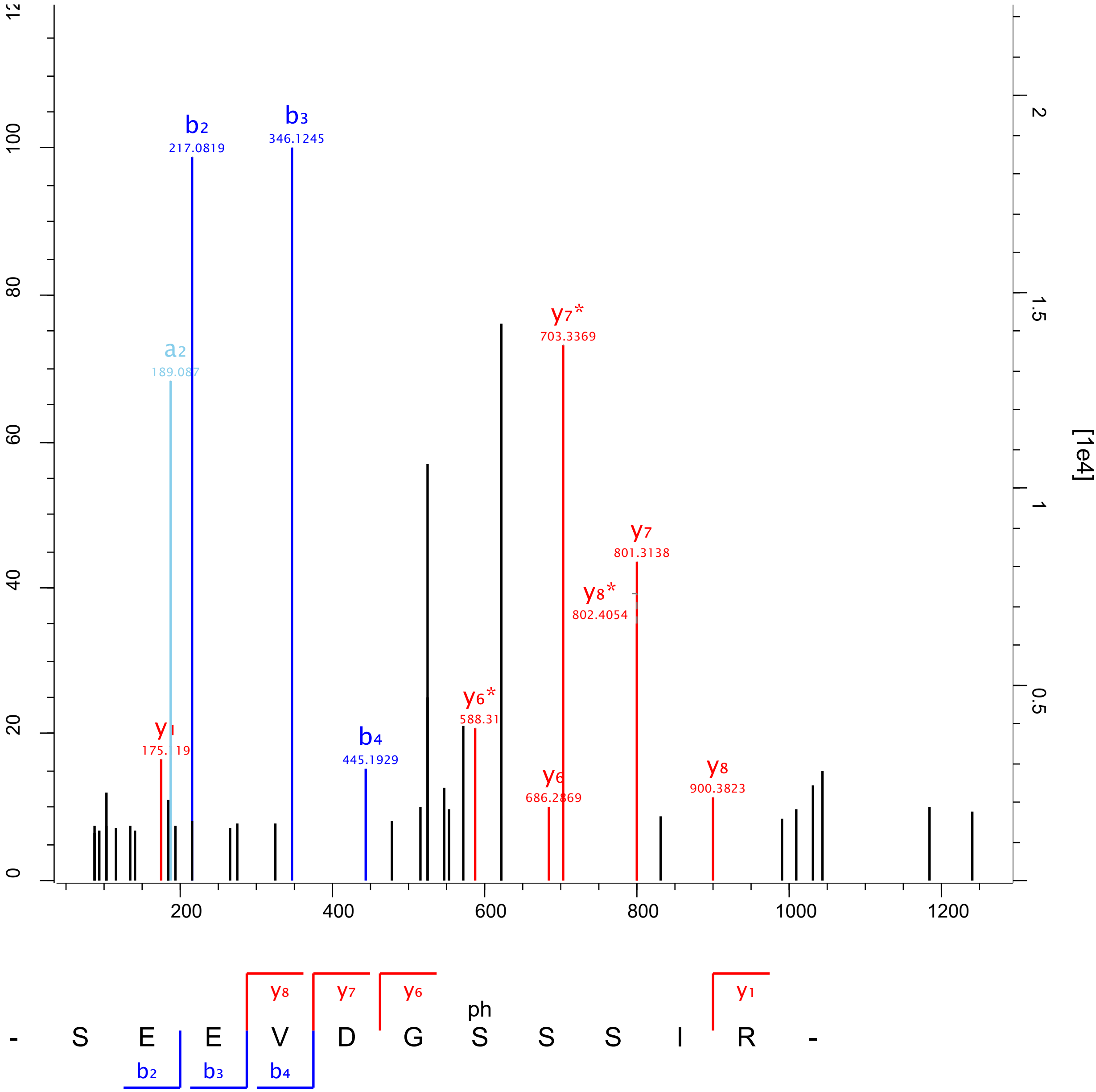

| Raw file | Scan | Method | Score | m/z |
|---|---|---|---|---|
| sys_15_1 | 7019 | FTMS; HCD | 73.63 | 525.2 |

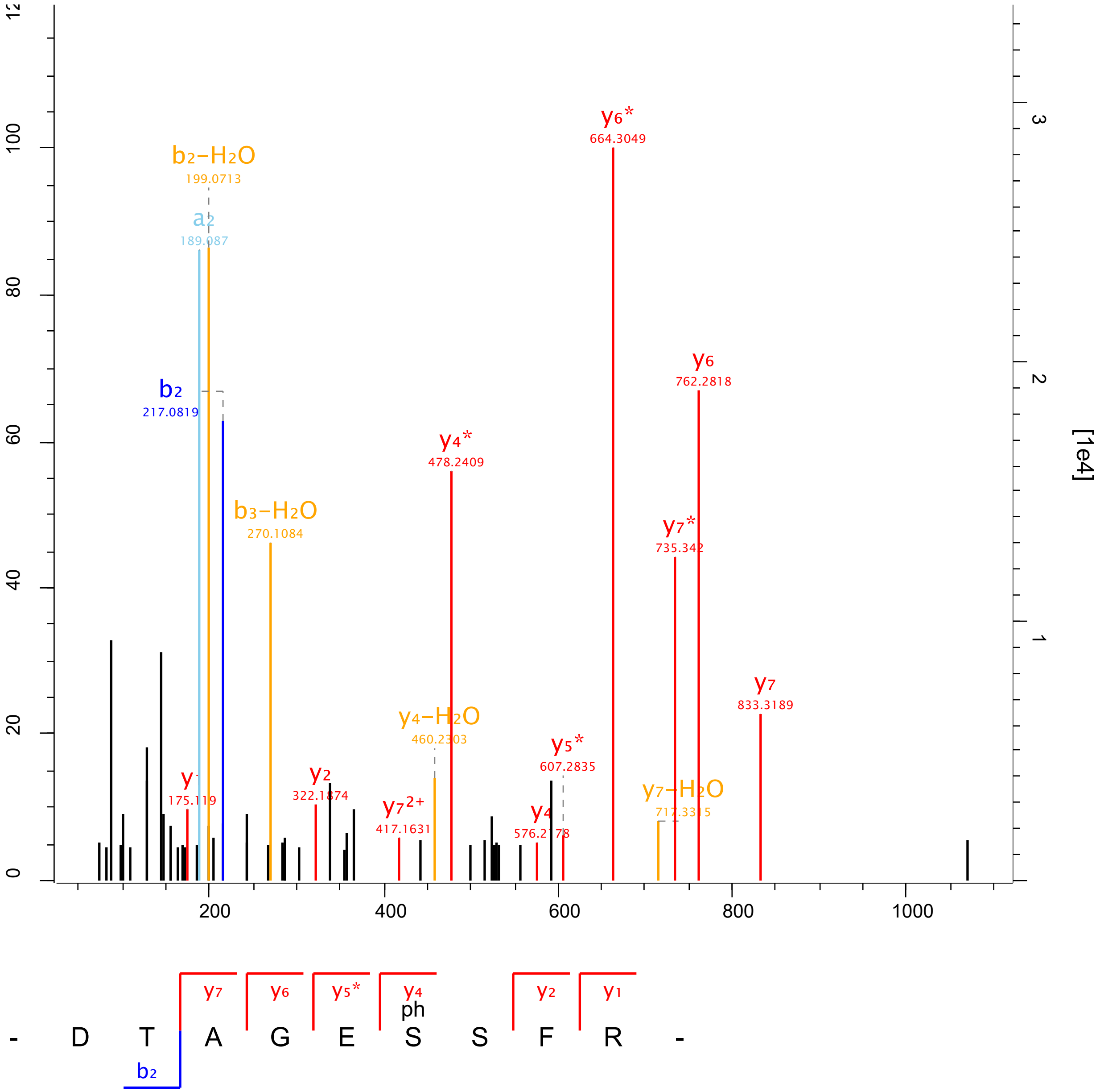

| Raw file | Scan | Method | Score | m/z |
|---|---|---|---|---|
| sys_15_1 | 7060 | FTMS; HCD | 41.62 | 793.86 |

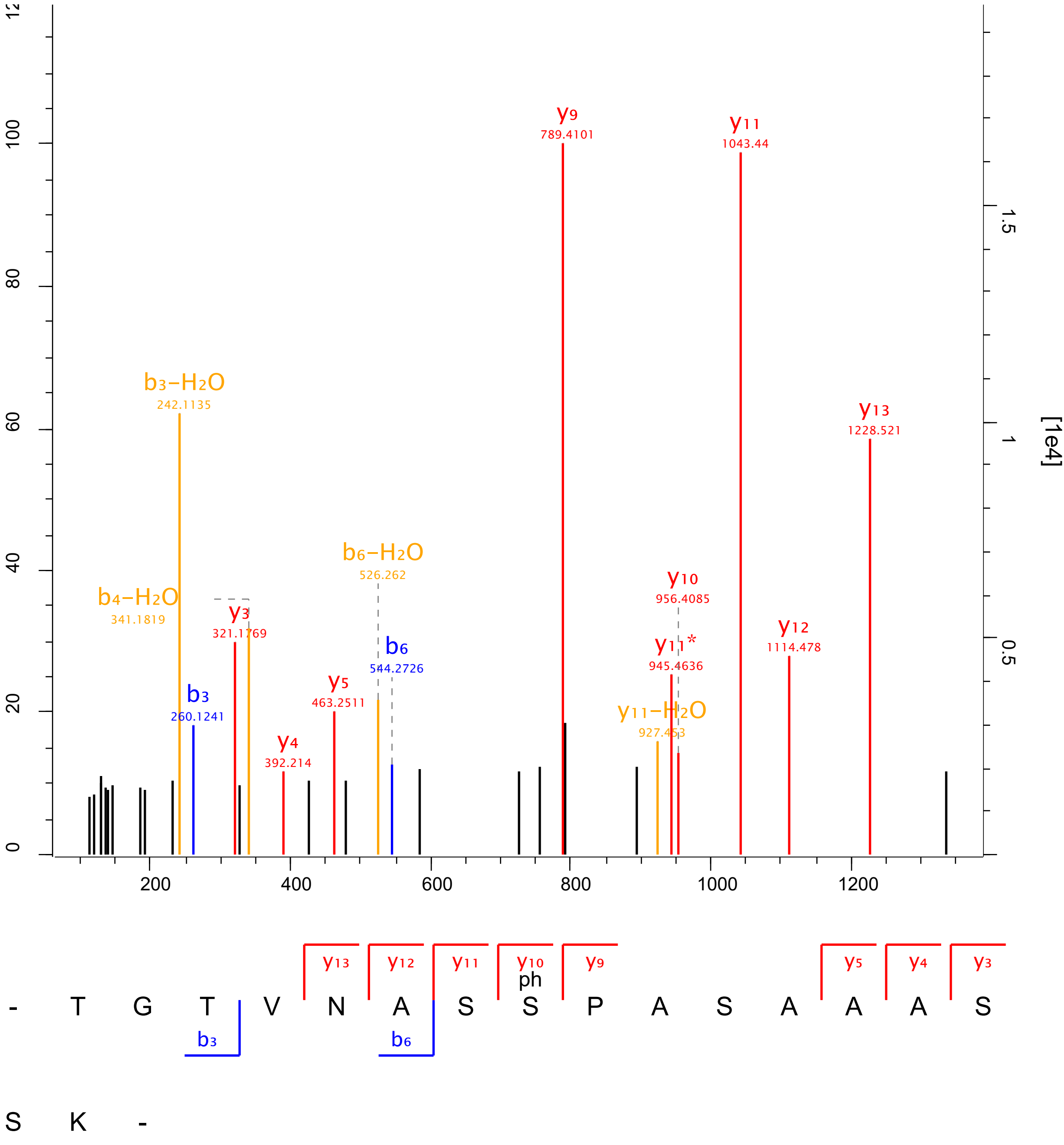

| Raw file | Scan | Method | Score | m/z |
|---|---|---|---|---|
| sys_15_1 | 7062 | FTMS; HCD | 43.76 | 688.27 |

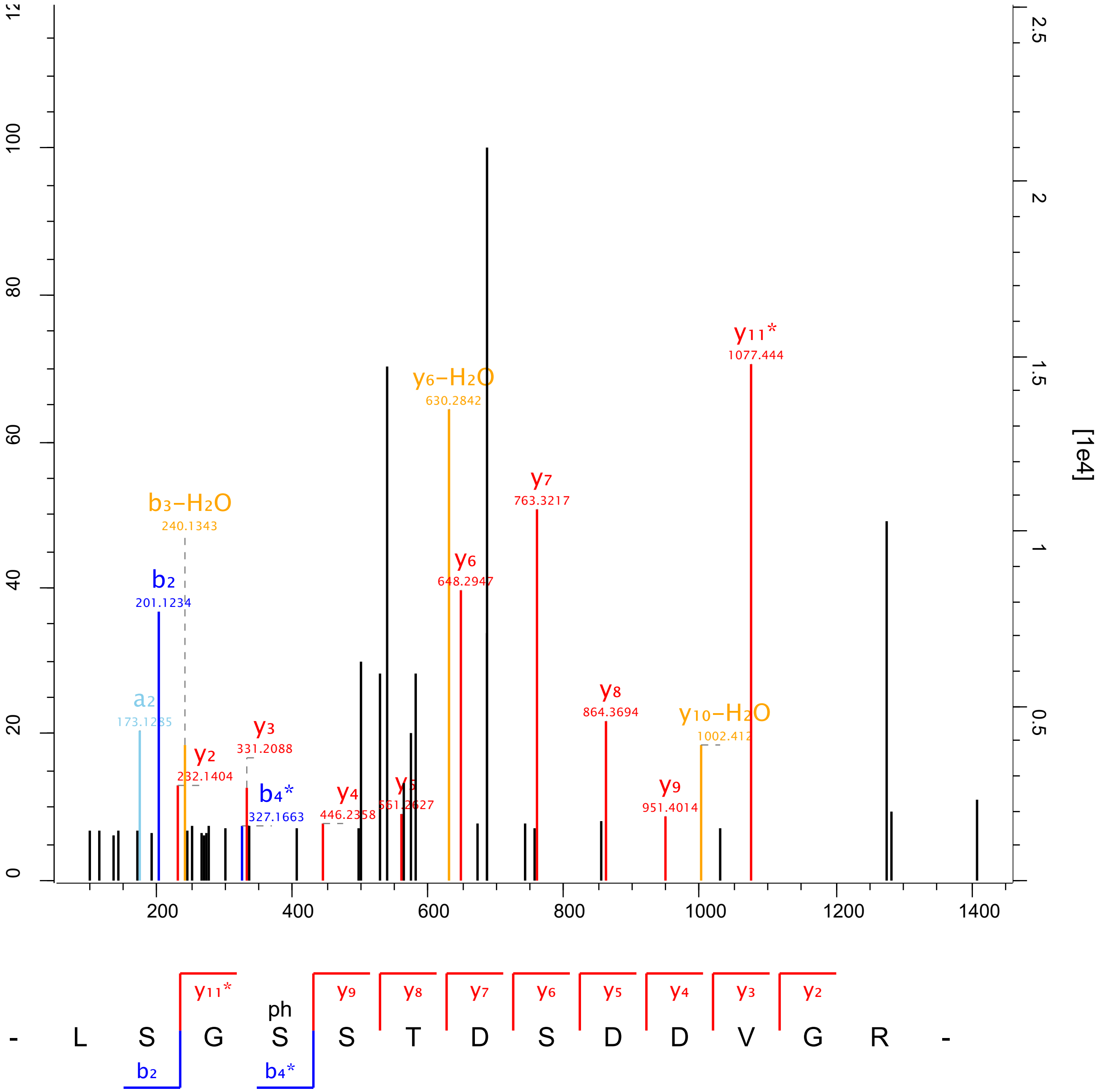

| Raw file | Scan | Method | Score | m/z |
|---|---|---|---|---|
| sys_15_1 | 7068 | FTMS; HCD | 162.31 | 709.25 |

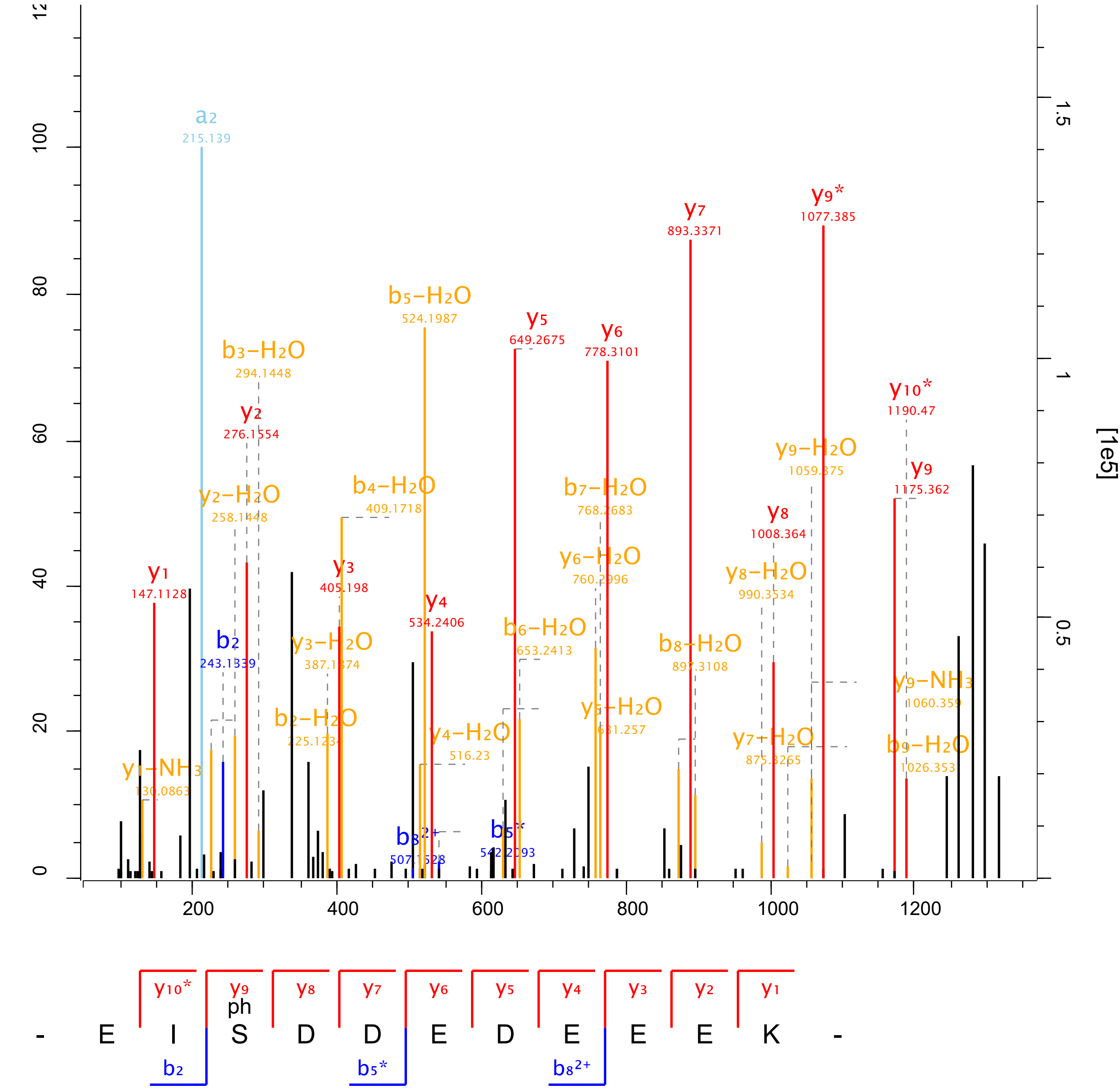

| Raw file | Scan | Method | Score | m/z |
| --- | --- | --- | --- | --- |
| sys_15_1 | 7132 | FTMS; HCD | 87.18 | 499.22 |

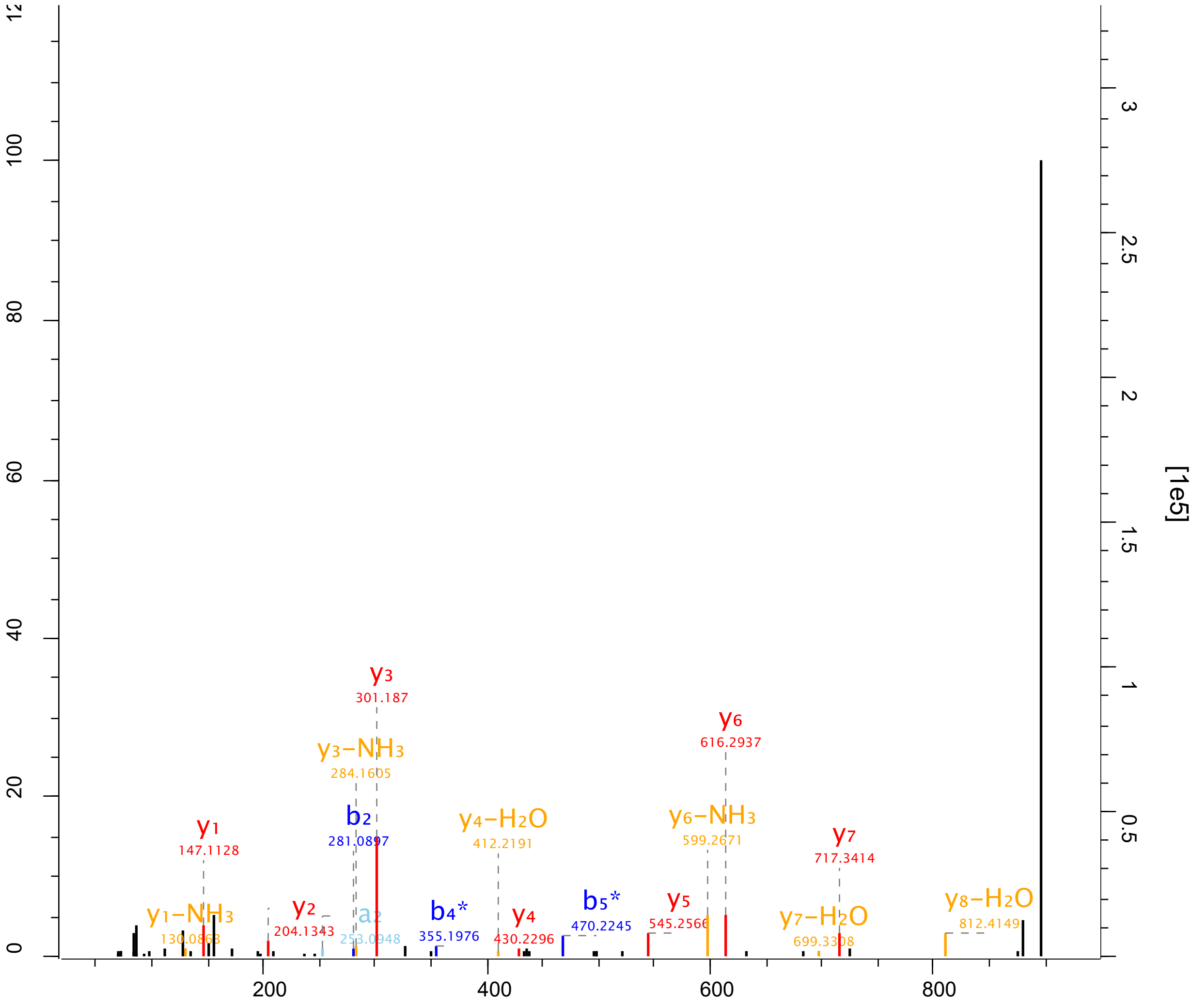


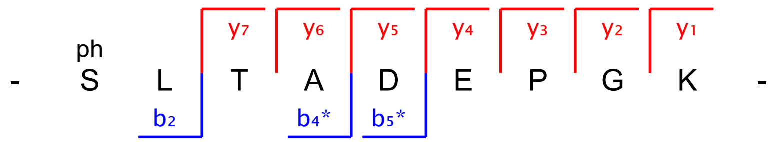

| Raw file | Scan | Method | Score | m/z |
|---|---|---|---|---|
| sys_15_1 | 7194 | FTMS; HCD | 72.29 | 550.25 |

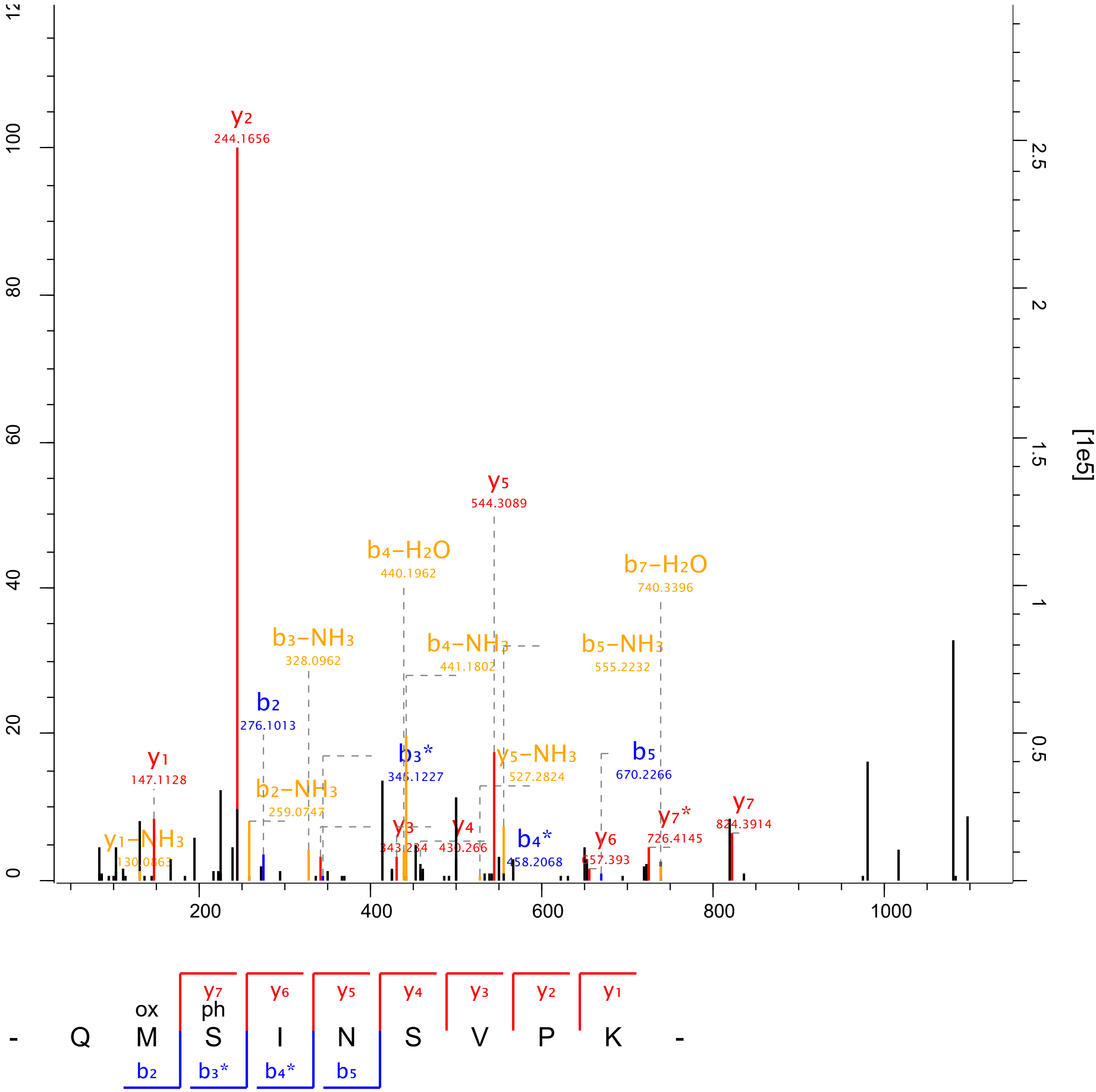

| Raw file | Scan | Method | Score | m/z |
|---|---|---|---|---|
| sys_15_1 | 7232 | FTMS; HCD | 72.89 | 591.71 |

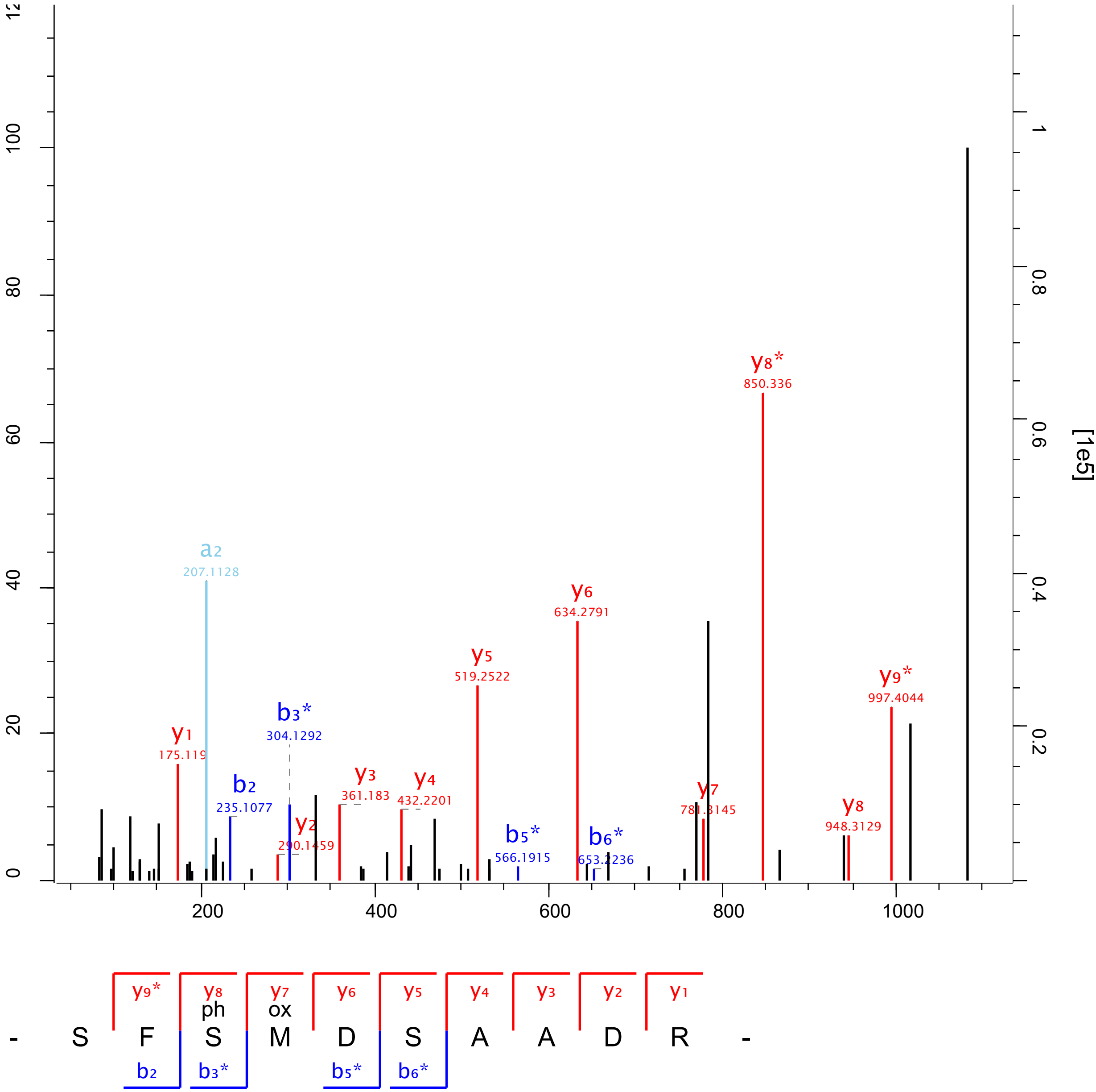

| Raw file | Scan | Method | Score | m/z |
|---|---|---|---|---|
| sys_15_1 | 7368 | FTMS; HCD | 85.91 | 643.74 |

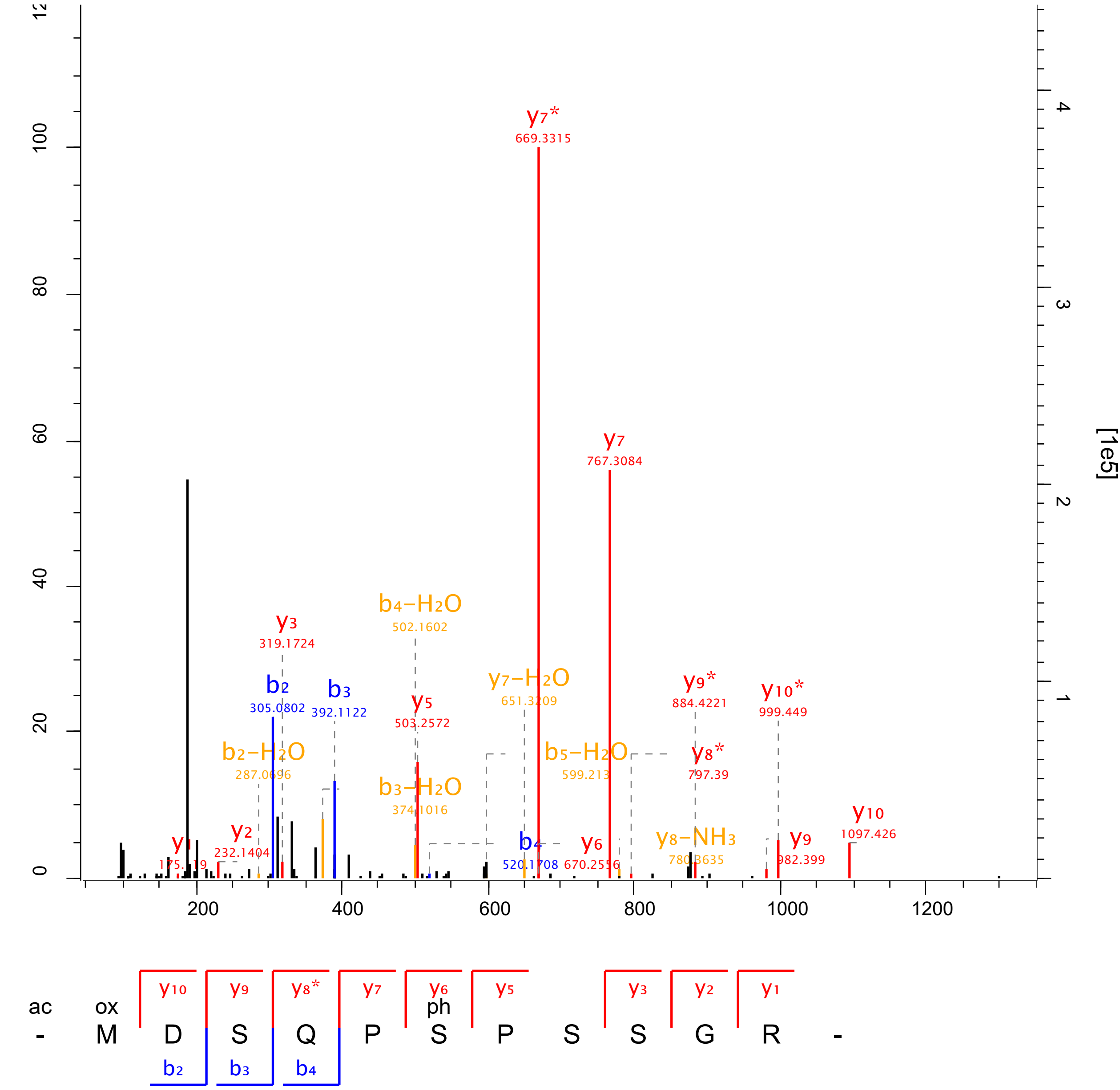

| Raw file | Scan | Method | Score | m/z |
|---|---|---|---|---|
| sys_15_1 | 7398 | FTMS; HCD | 101.75 | 690.28 |

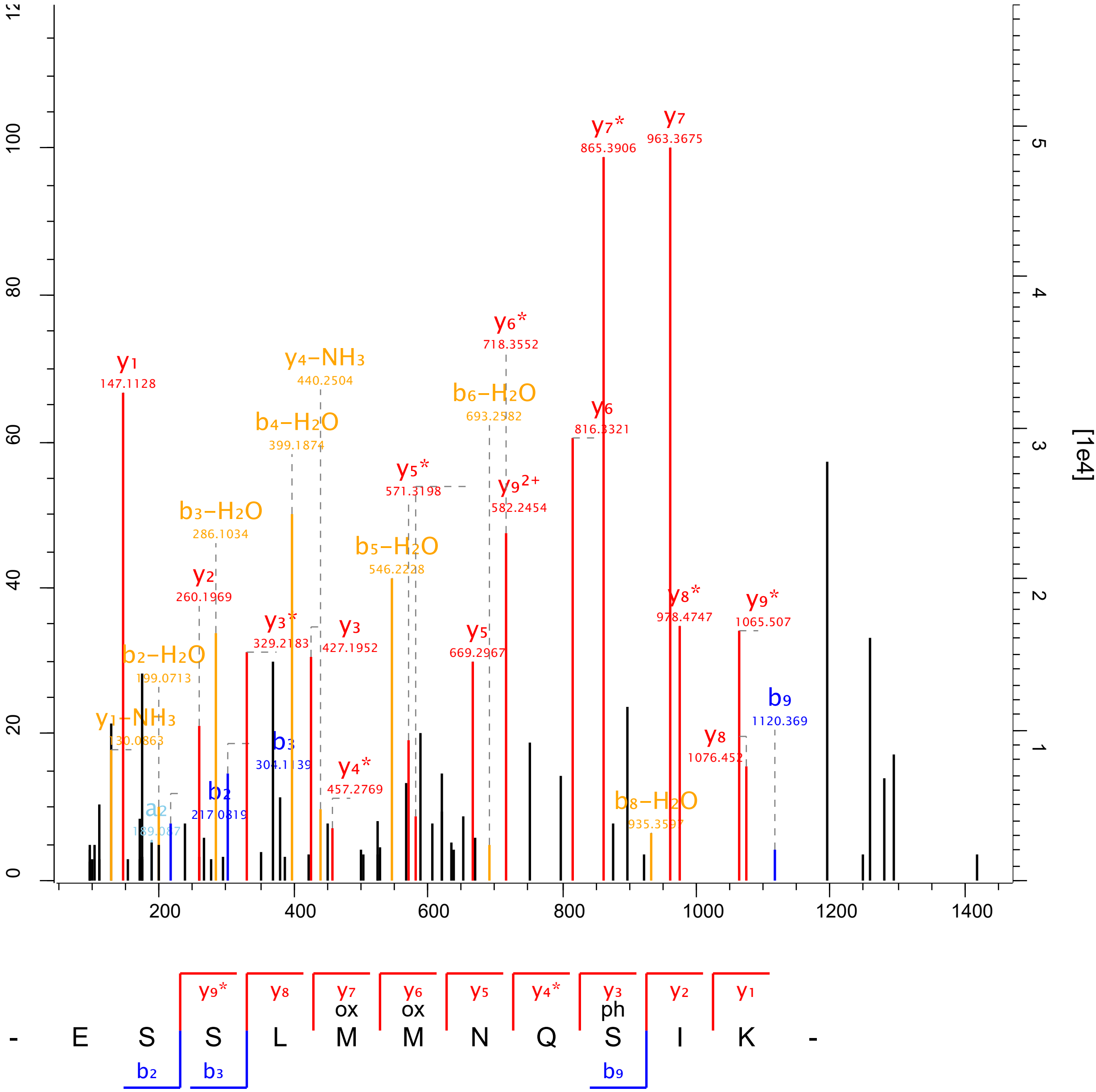

| Raw file | Scan | Method | Score | m/z |
| --- | --- | --- | --- | --- |
| sys_15_1 | 7409 | FTMS; HCD | 68.48 | 756.32 |

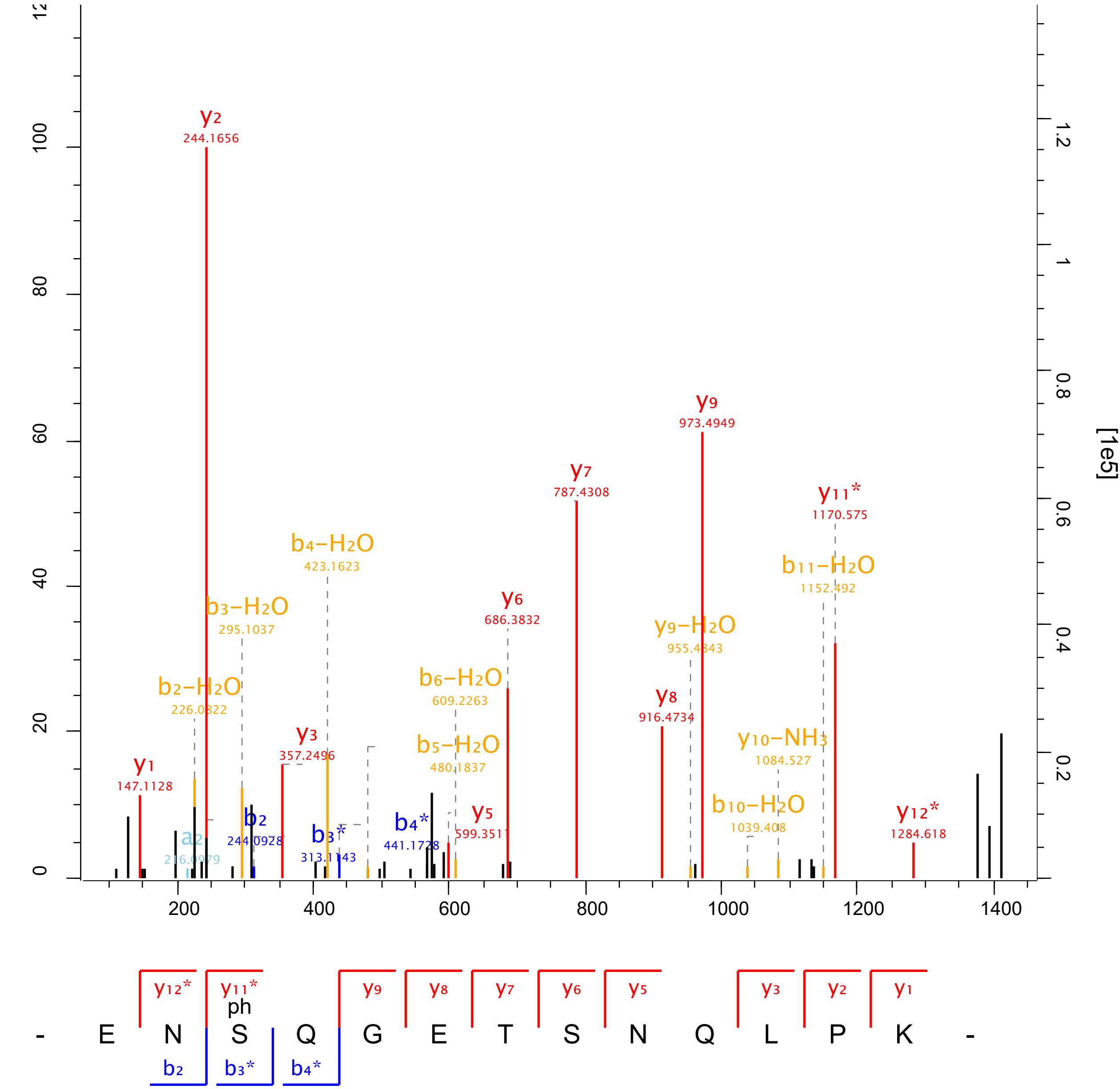

| Raw file | Scan | Method | Score | m/z |
|---|---|---|---|---|
| sys_15_1 | 7423 | FTMS; HCD | 99.06 | 658.29 |

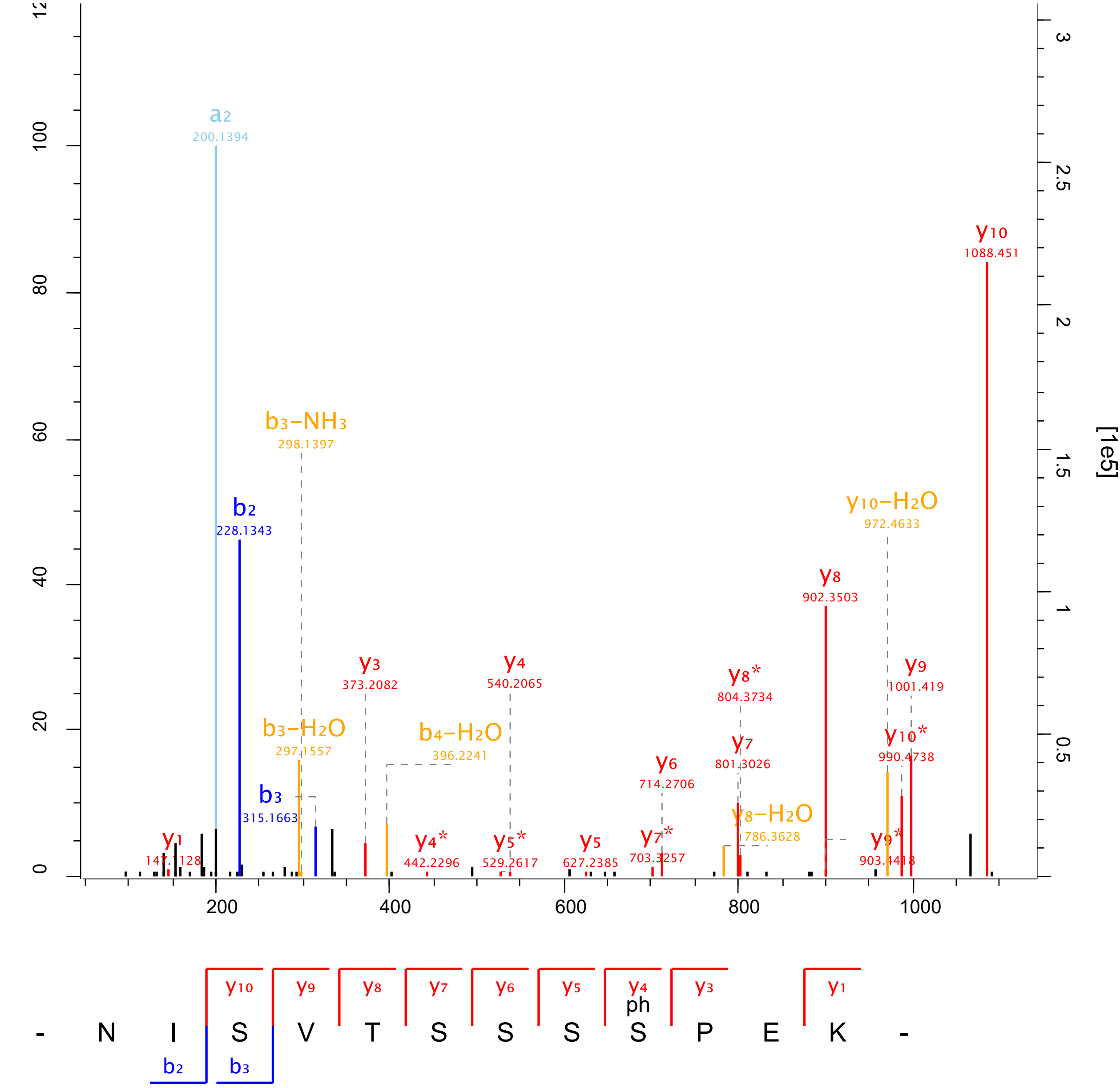

| Raw file | Scan | Method | Score | m/z |
|---|---|---|---|---|
| sys_15_1 | 7559 | FTMS; HCD | 61.59 | 452.2 |

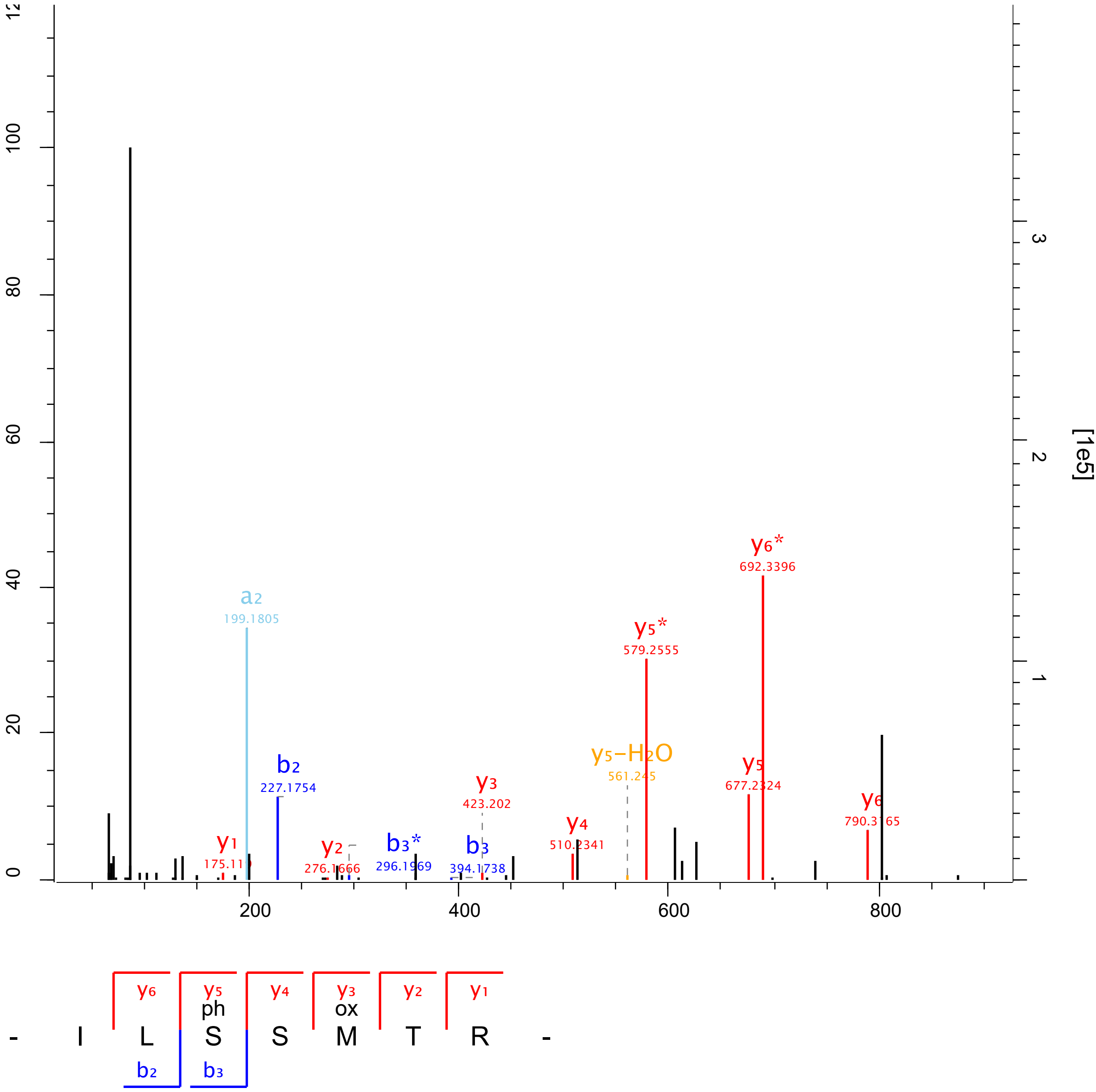

| Raw file | Scan | Method | Score | m/z |
|---|---|---|---|---|
| sys_15_1 | 7591 | FTMS; HCD | 51.76 | 768.77 |

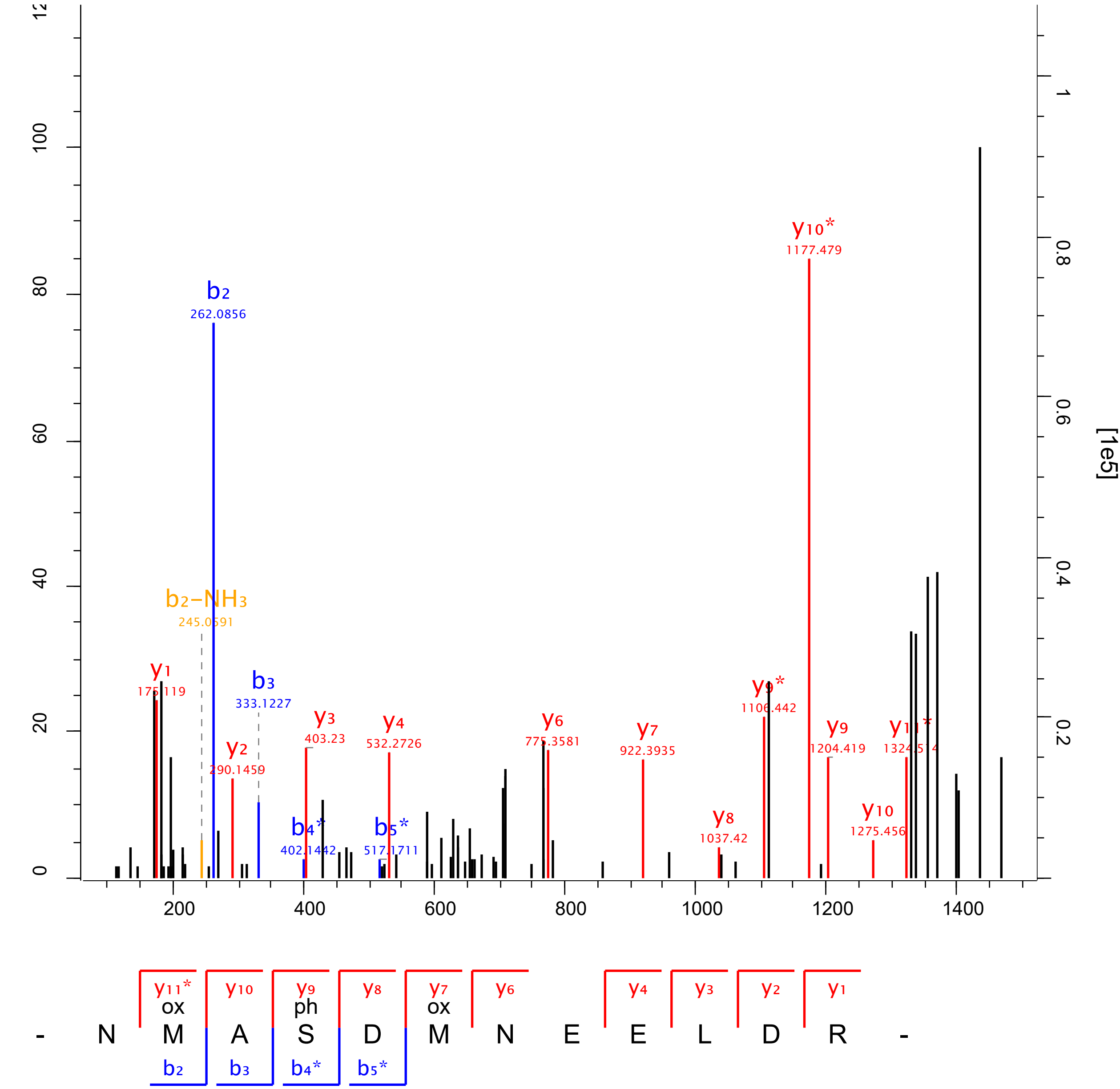

| Raw file | Scan | Method | Score | m/z |
|---|---|---|---|---|
| sys_15_1 | 7627 | FTMS; HCD | 67.14 | 673.74 |

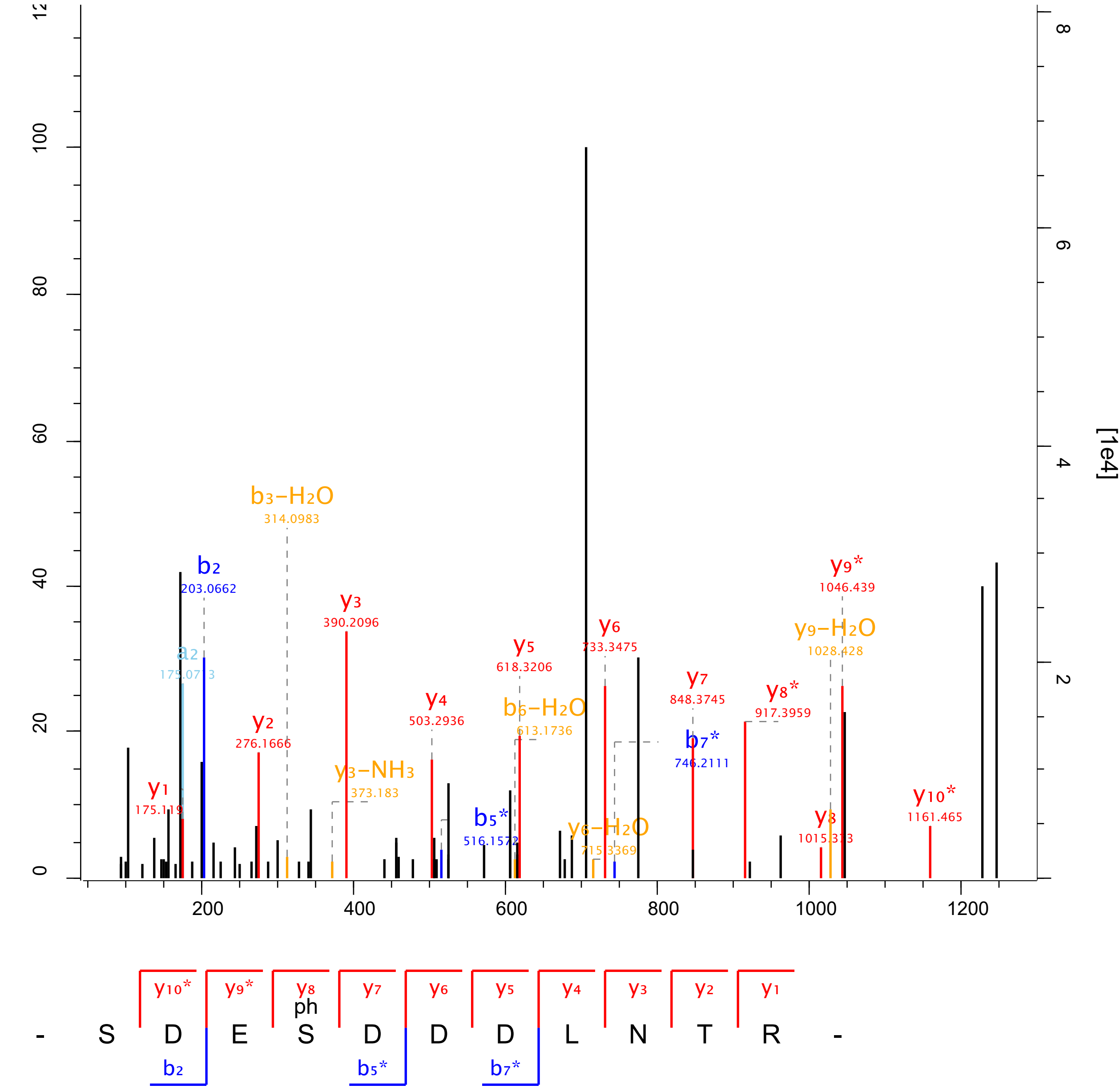

| Raw file | Scan | Method | Score | m/z |
|---|---|---|---|---|
| sys_15_1 | 7720 | FTMS; HCD | 139.86 | 597.26 |

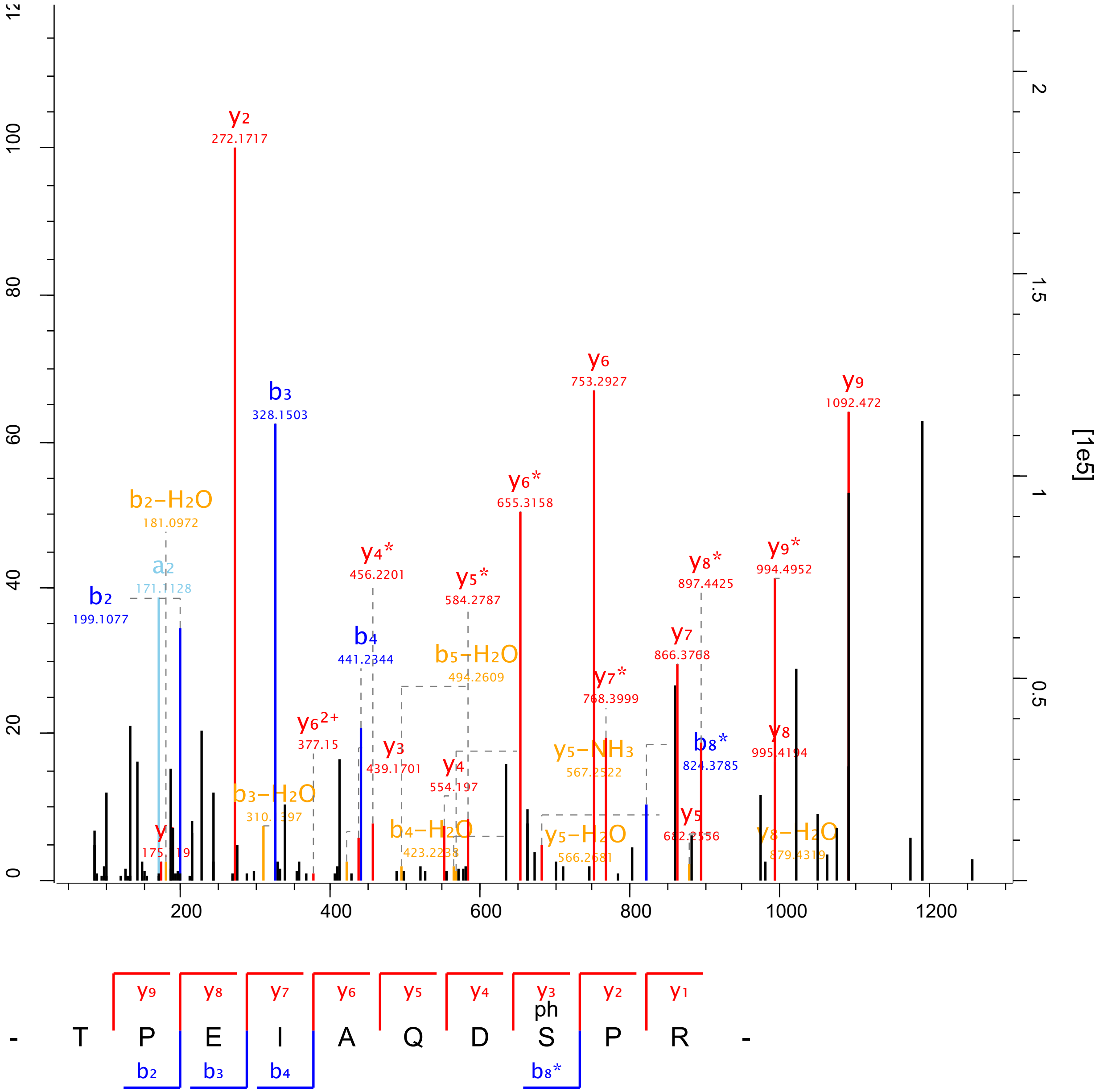

| Raw file | Scan | Method | Score | m/z |
|---|---|---|---|---|
| sys_15_1 | 7756 | FTMS; HCD | 109.72 | 678.8 |

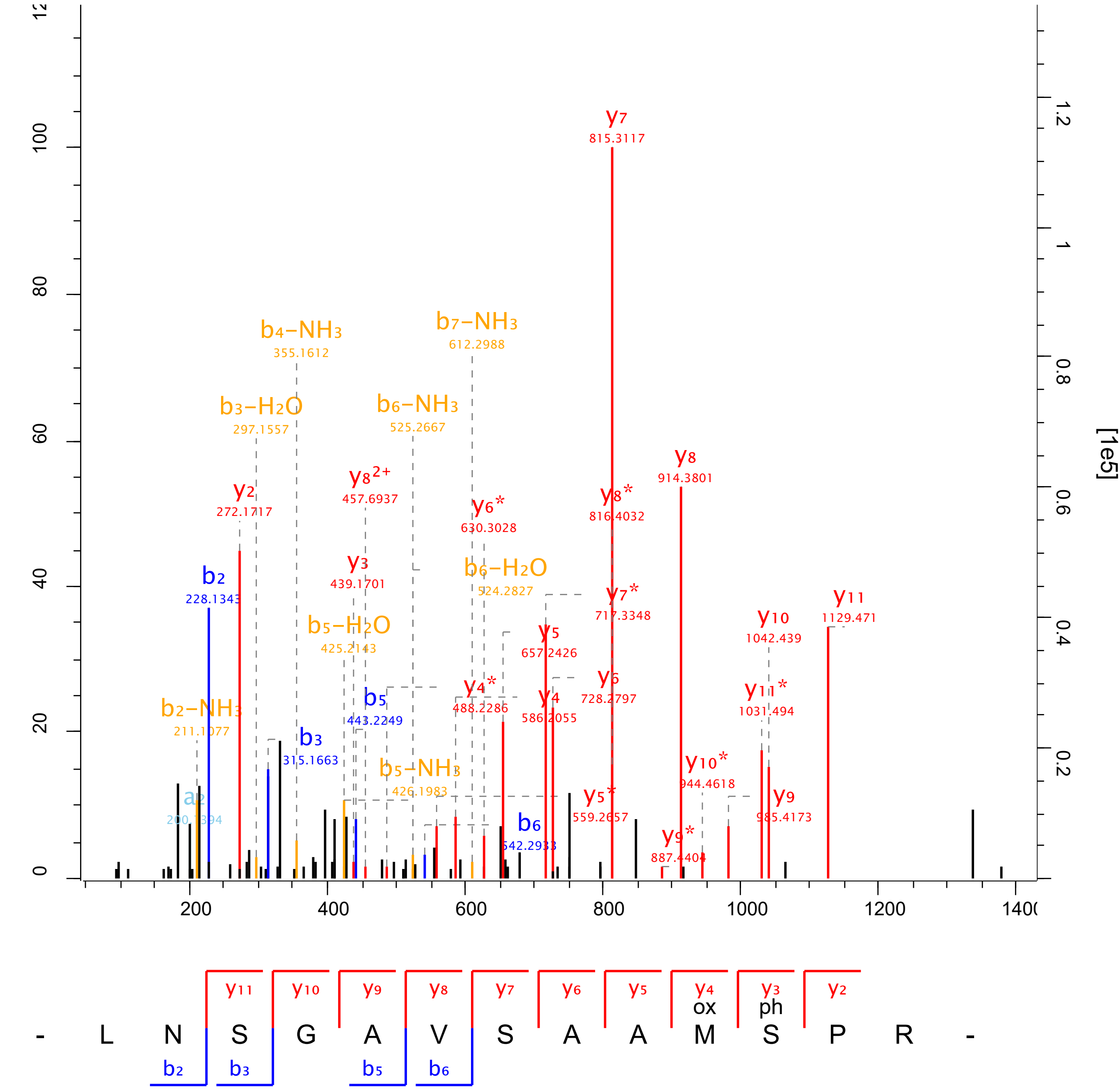

| Raw file | Scan | Method | Score | m/z |
|---|---|---|---|---|
| sys_15_1 | 7768 | FTMS; HCD | 118.31 | 530.74 |

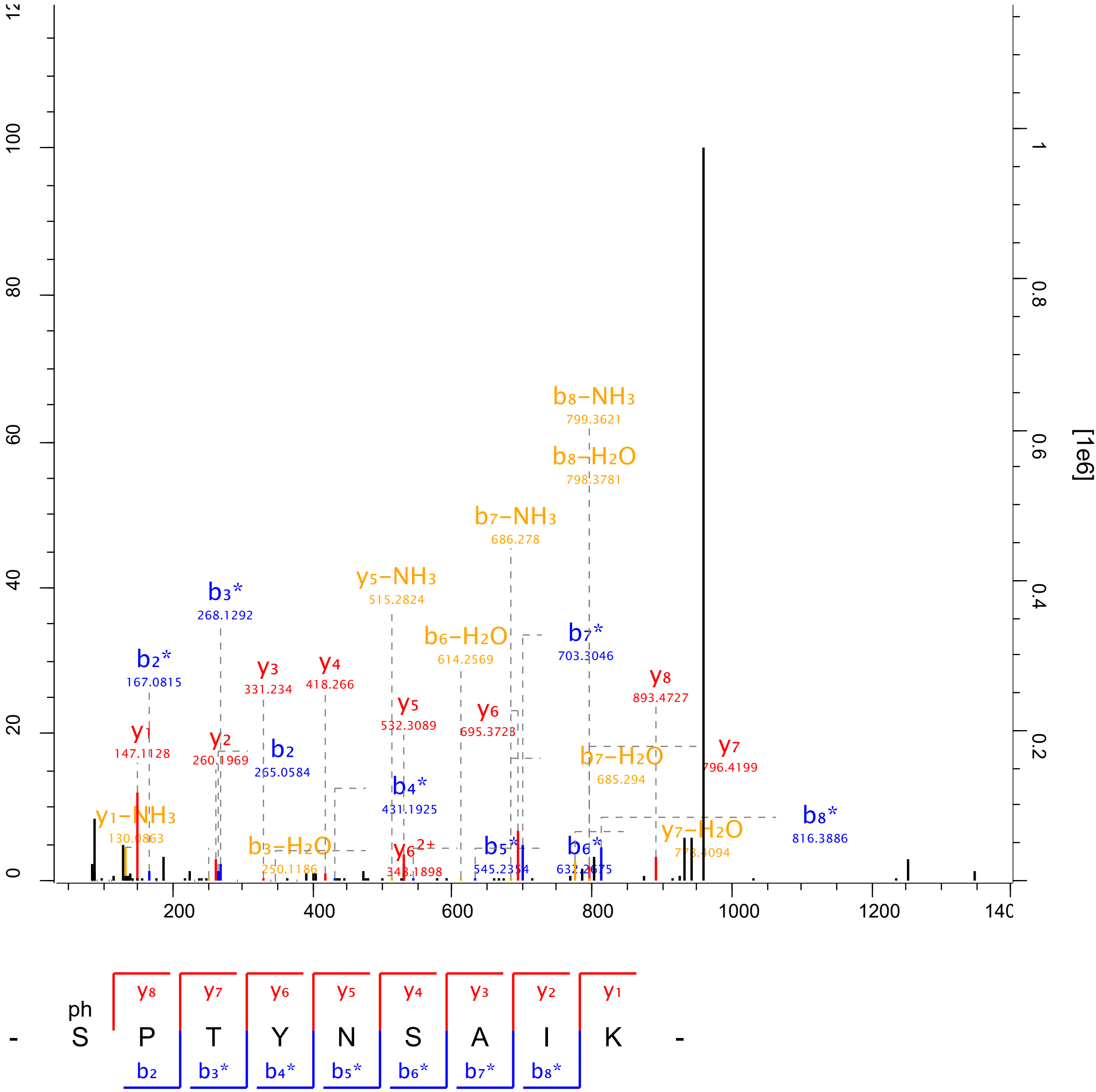

| Raw file | Scan | Method | Score | m/z |
|---|---|---|---|---|
| sys_15_1 | 7846 | FTMS; HCD | 67.14 | 680.76 |

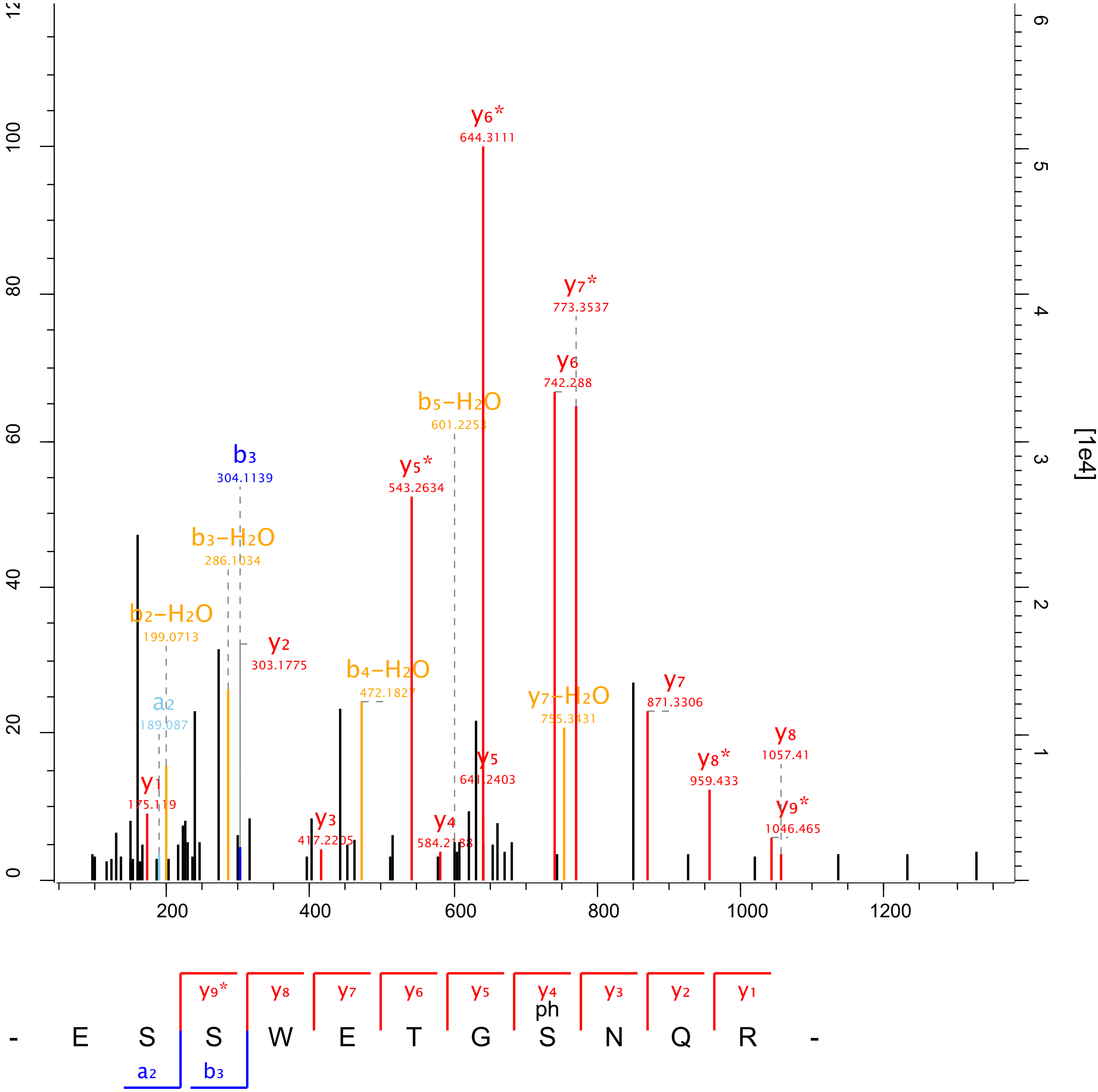

| Raw file | Scan | Method | Score | m/z |
|---|---|---|---|---|
| sys_15_1 | 7861 | FTMS; HCD | 47.6 | 528.23 |

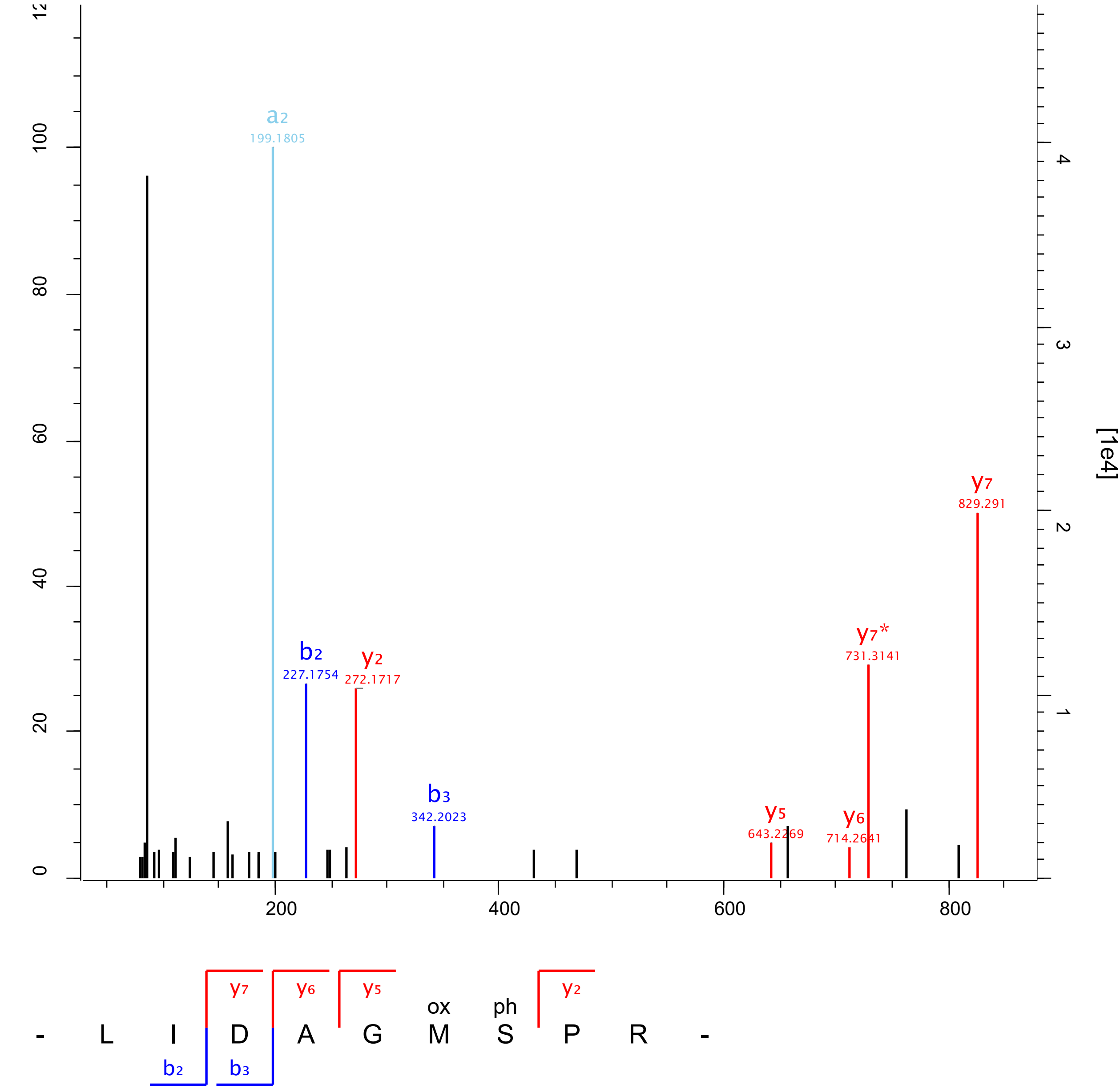

| Raw file | Scan | Method | Score | m/z |
| --- | --- | --- | --- | --- |
| sys_15_1 | 7873 | FTMS; HCD | 70.89 | 686.77 |

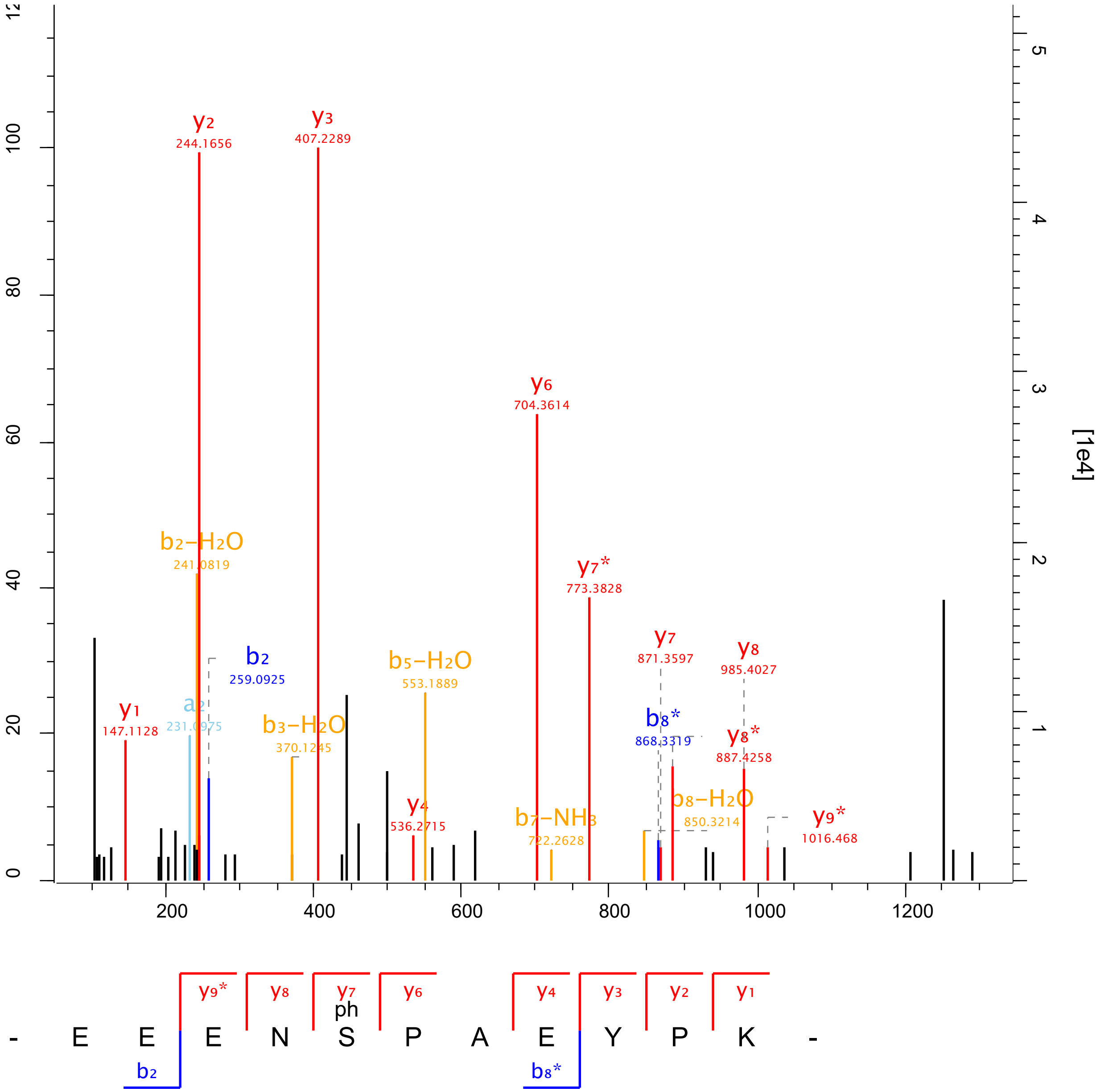

| Raw file | Scan | Method | Score | m/z |
|---|---|---|---|---|
| sys_15_1 | 7904 | FTMS; HCD | 88.02 | 603.78 |

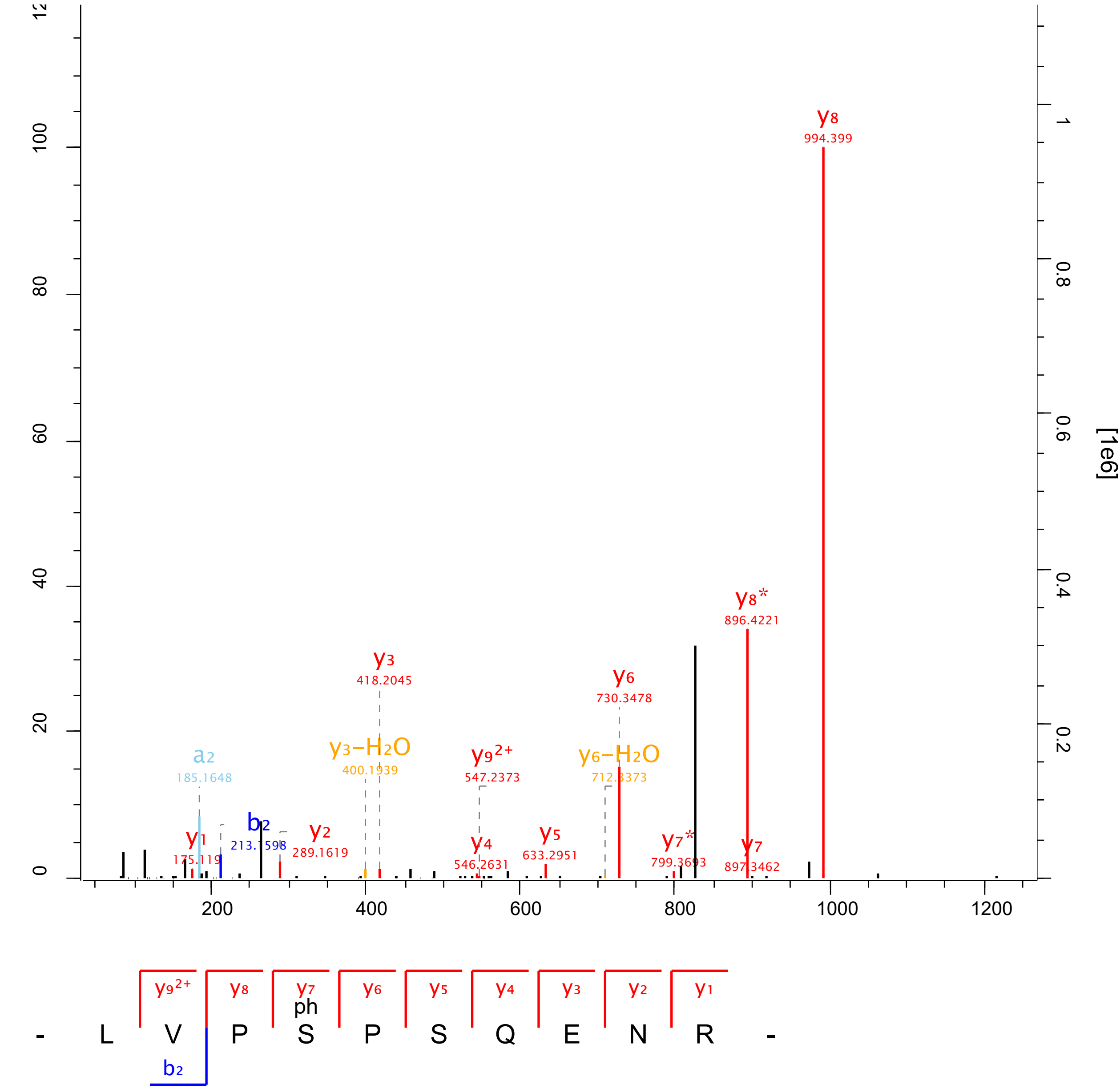

| Raw file | Scan | Method | Score | m/z |
|---|---|---|---|---|
| sys_15_1 | 7980 | FTMS; HCD | 179.14 | 783.79 |

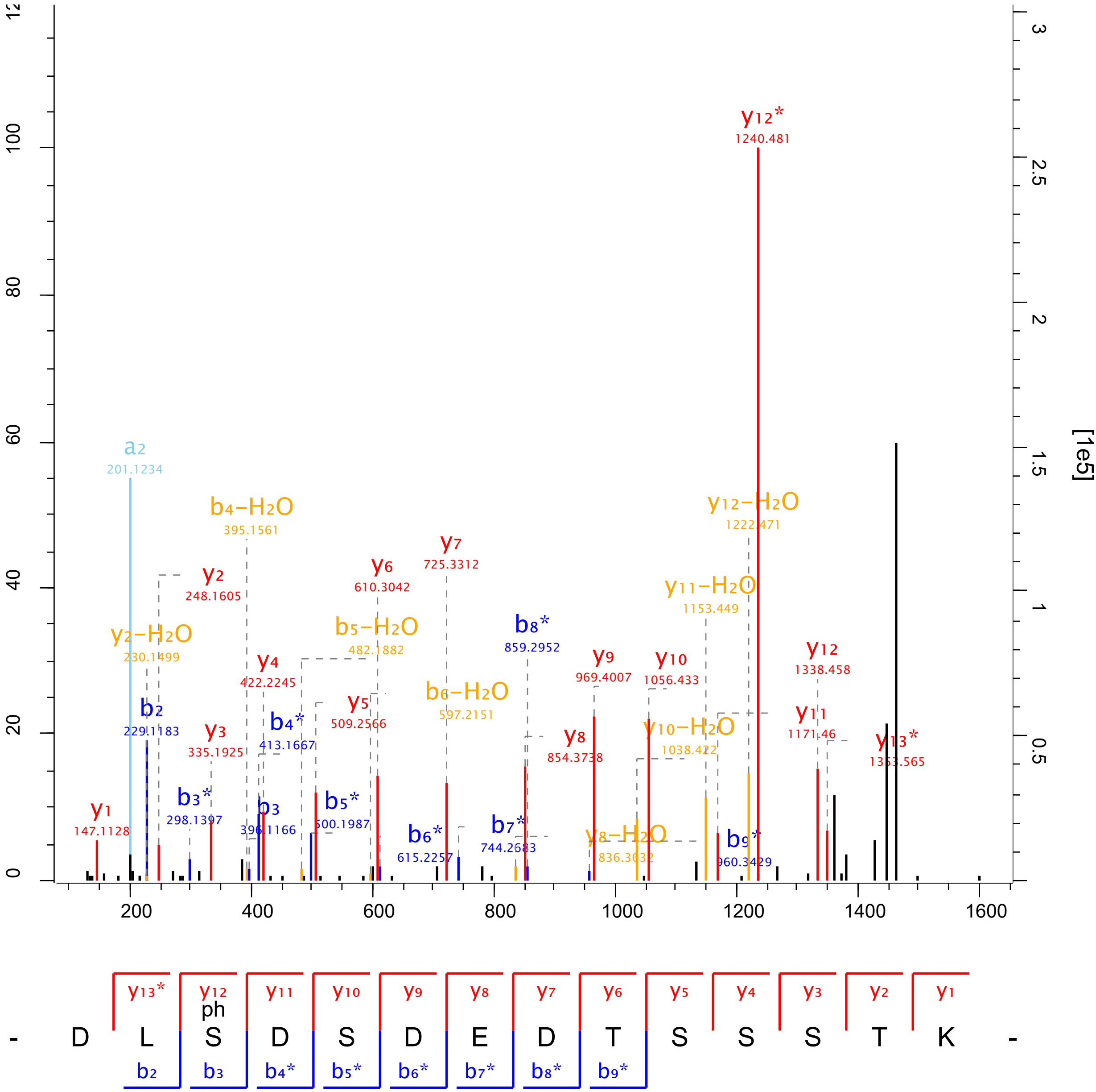

| Raw file | Scan | Method | Score | m/z |
| --- | --- | --- | --- | --- |
| sys_15_1 | 8139 | FTMS; HCD | 84.76 | 636.26 |

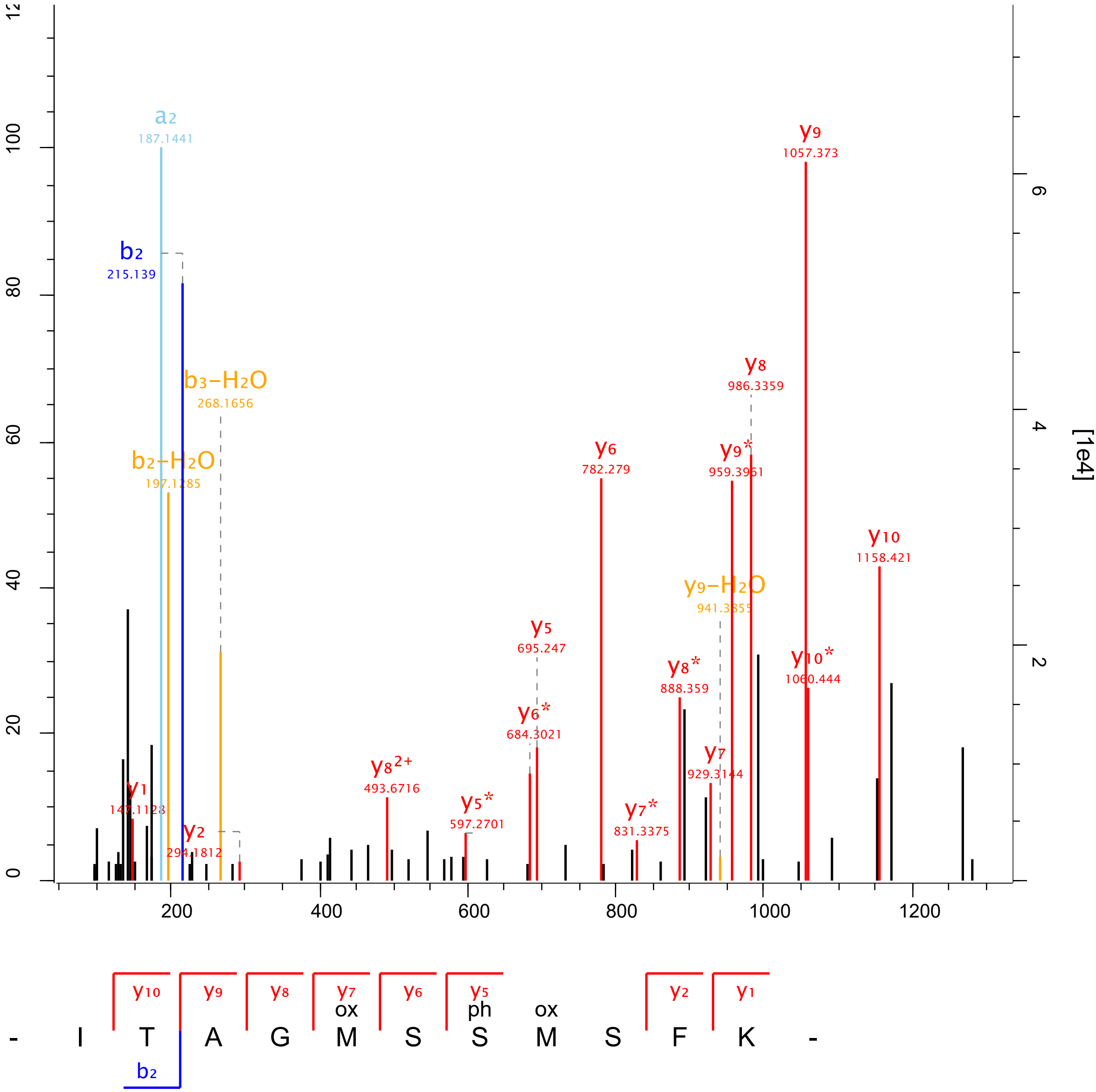

| Raw file | Scan | Method | Score | m/z |
|---|---|---|---|---|
| sys_15_1 | 8143 | FTMS; HCD | 65.04 | 659.75 |

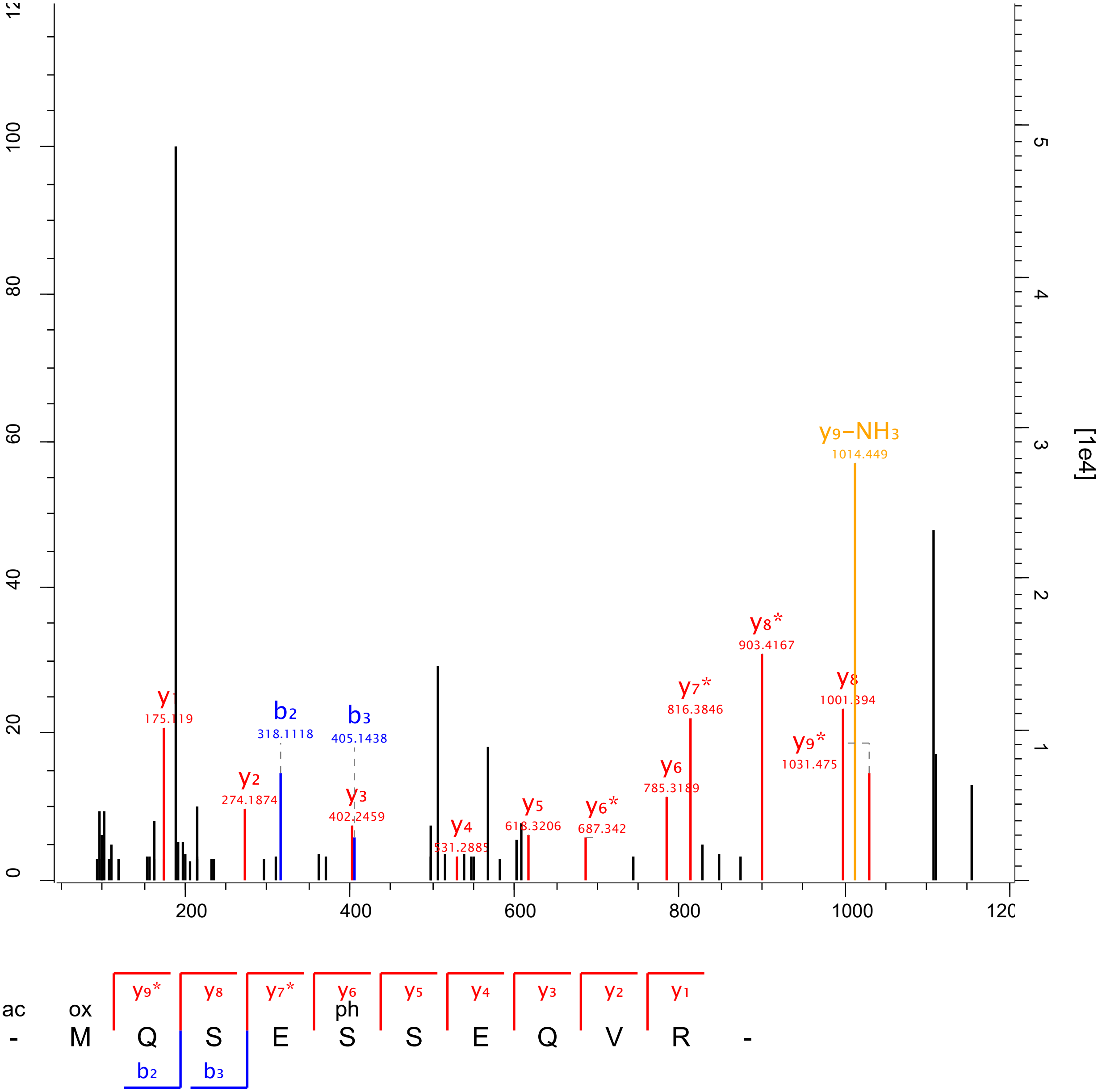

| Raw file | Scan | Method | Score | m/z |
|---|---|---|---|---|
| sys_15_1 | 8169 | FTMS; HCD | 198.04 | 735.8 |

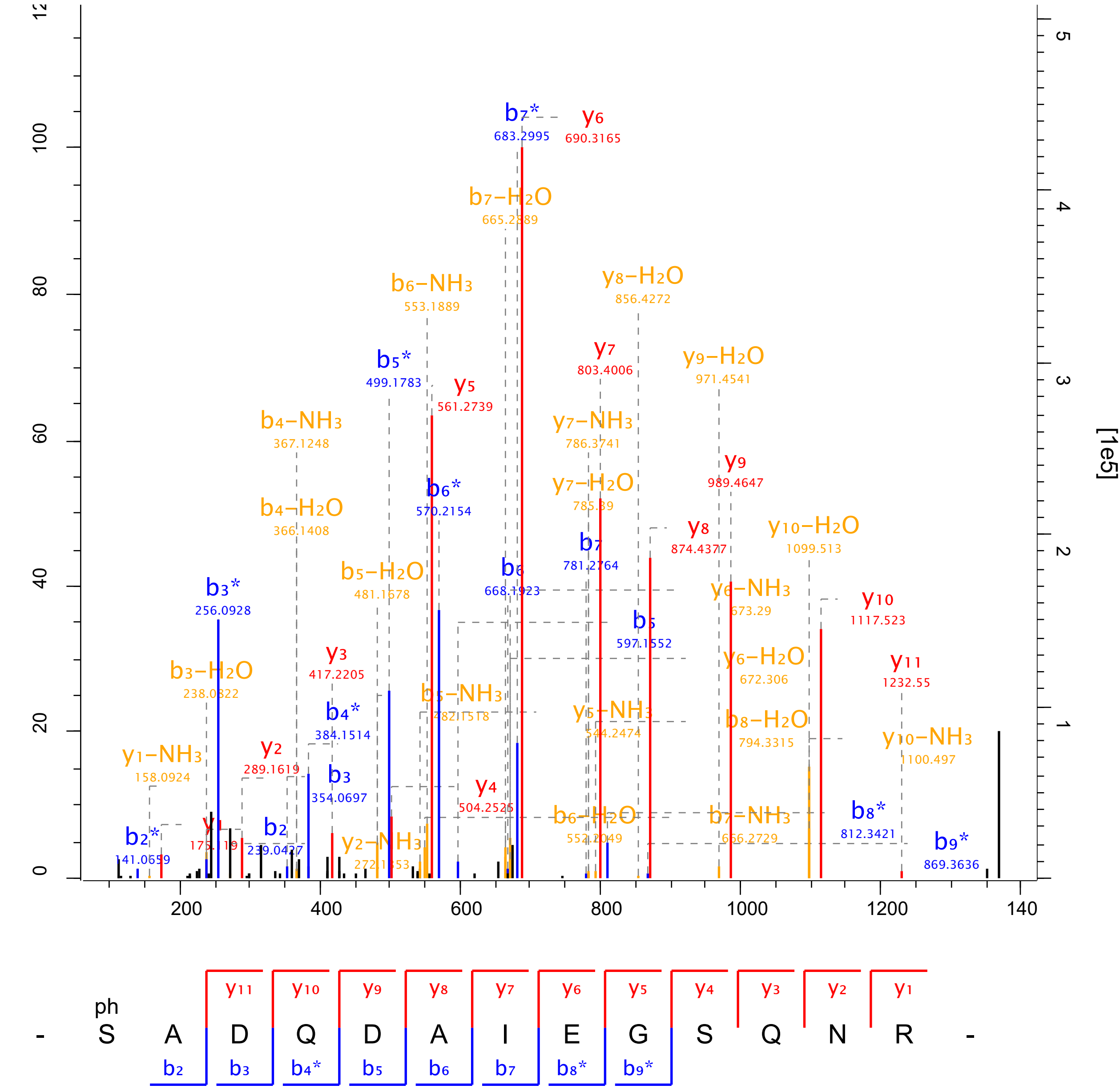

| Raw file | Scan | Method | Score | m/z |
| --- | --- | --- | --- | --- |
| sys_15_1 | 8189 | FTMS; HCD | 51.29 | 541.68 |

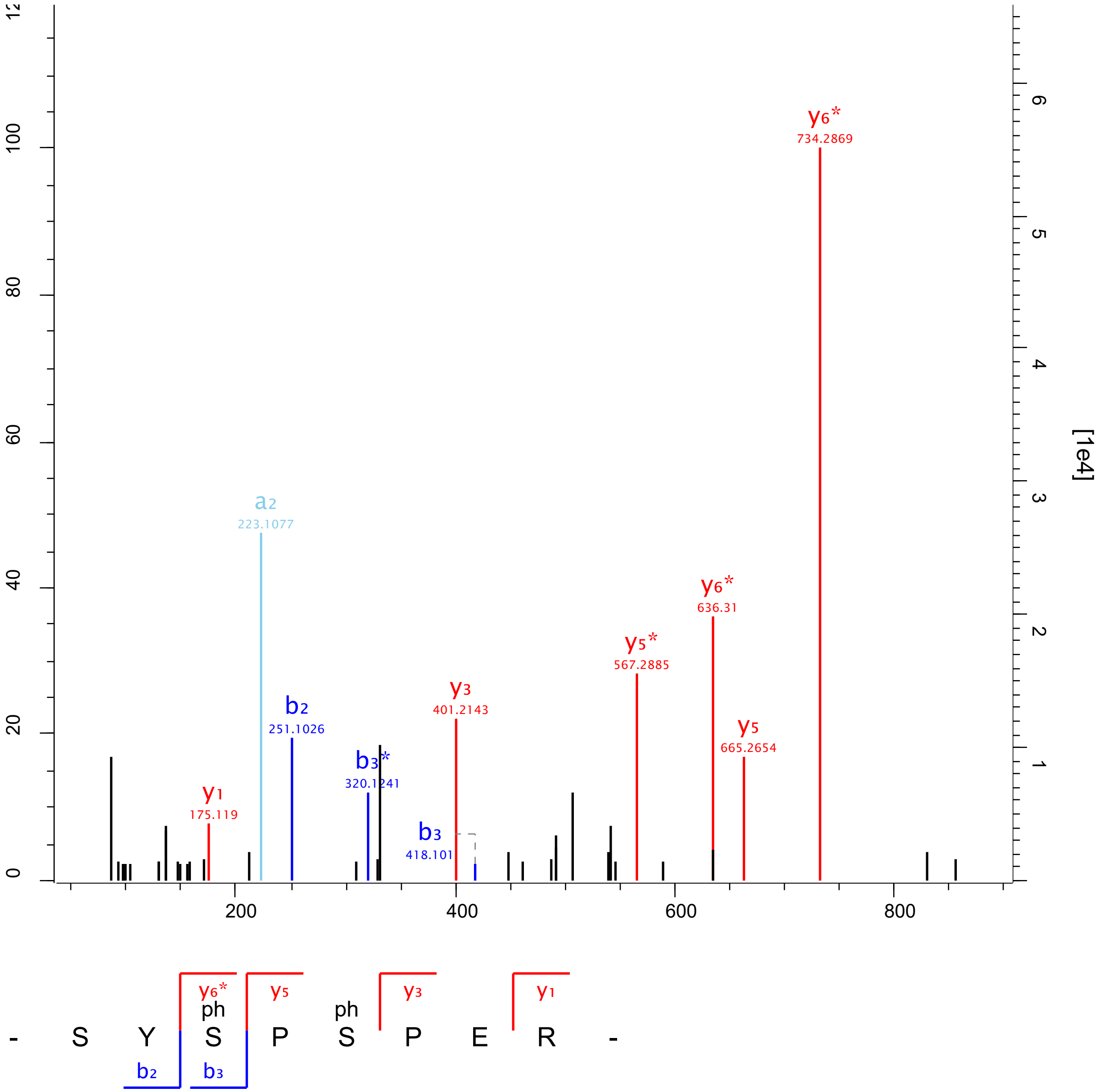

| Raw file | Scan | Method | Score | m/z |
| --- | --- | --- | --- | --- |
| sys_15_1 | 8202 | FTMS; HCD | 125.82 | 587.75 |

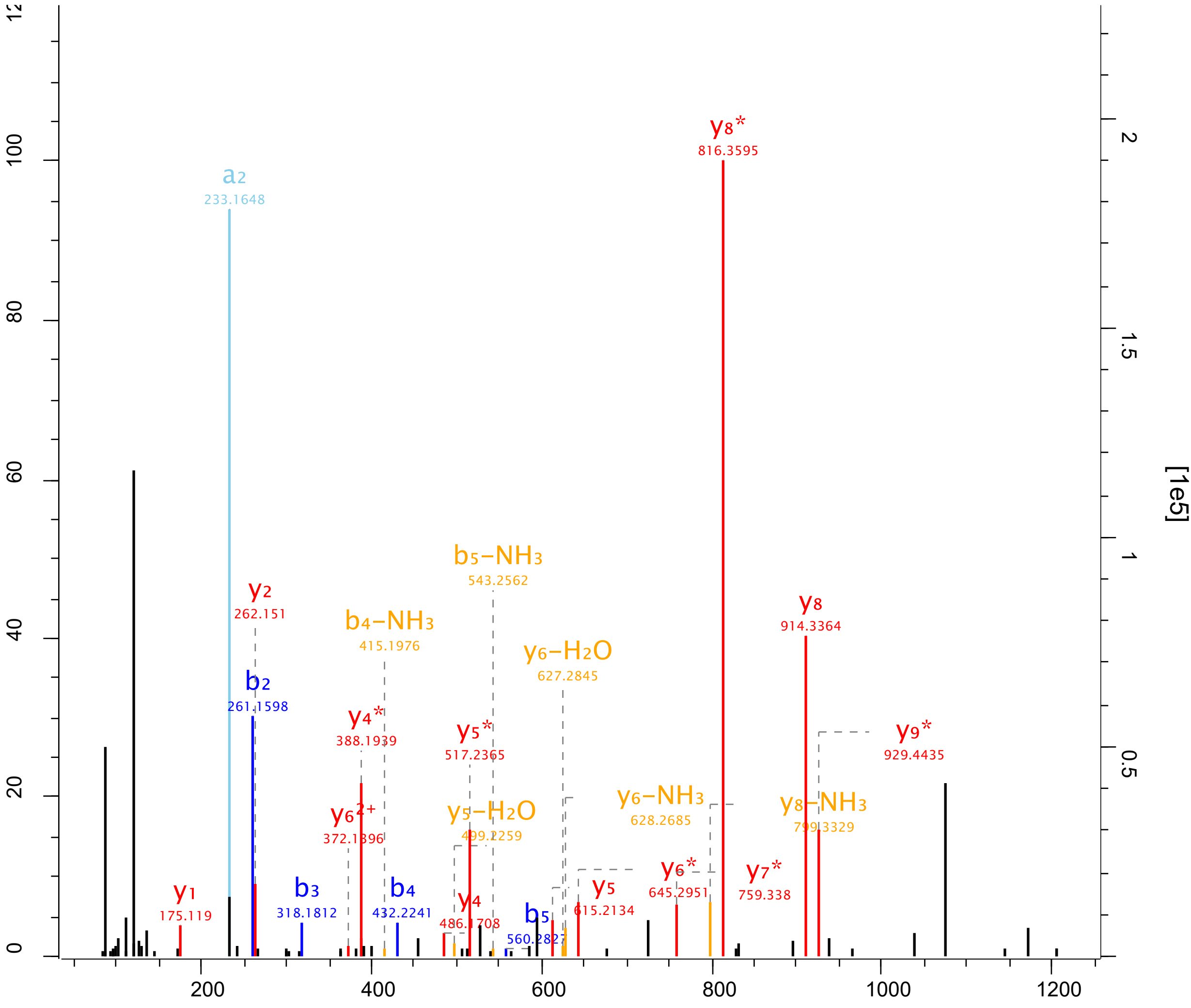


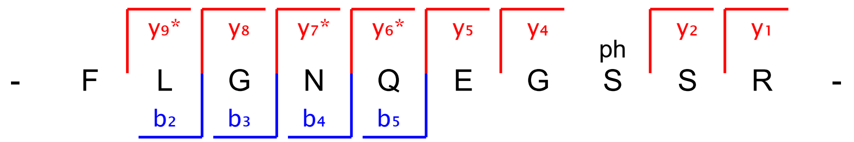

| Raw file | Scan | Method | Score | m/z |
|---|---|---|---|---|
| sys_15_1 | 8239 | FTMS; HCD | 54.09 | 643.58 |

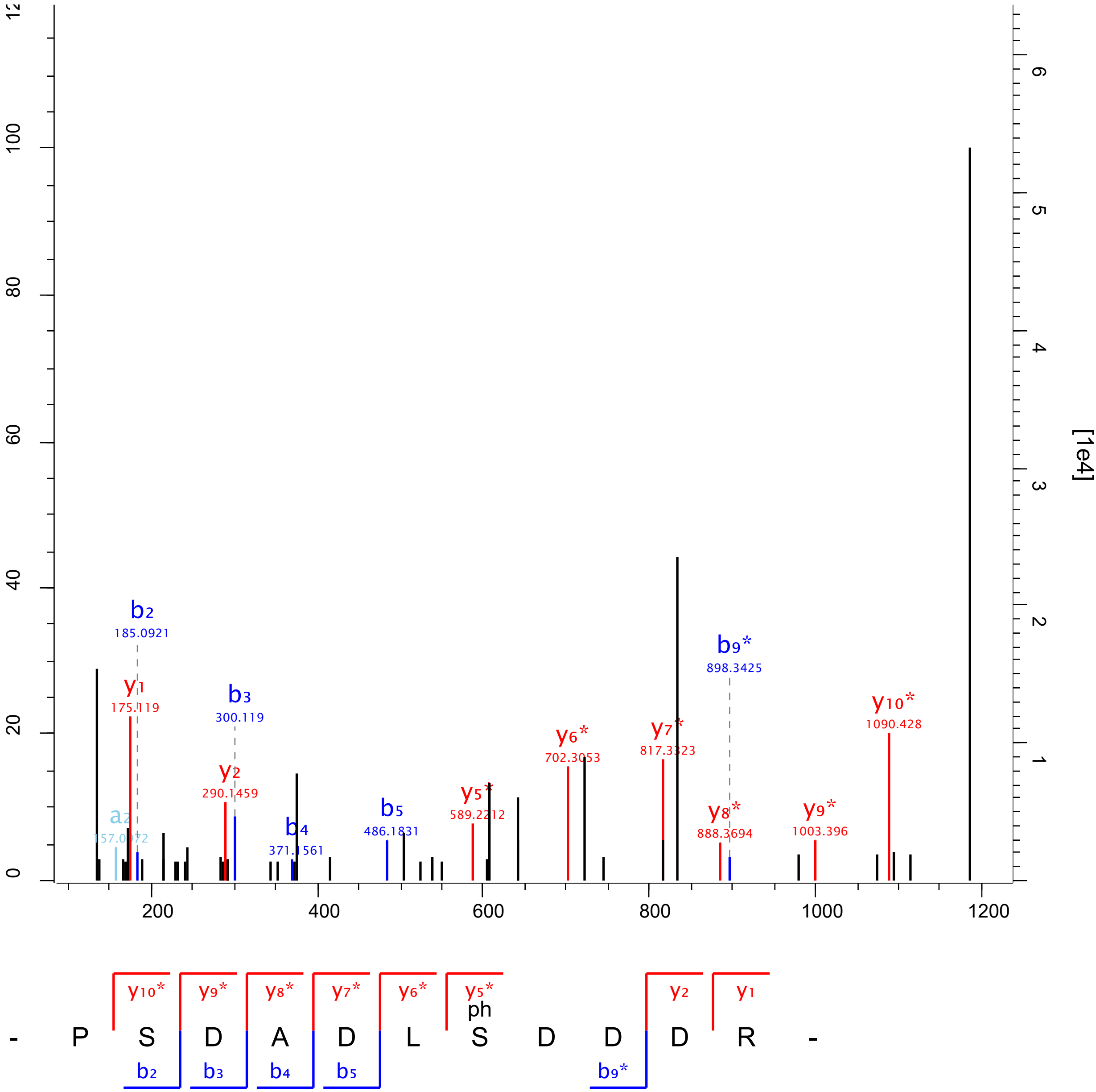

| Raw file | Scan | Method | Score | m/z |
|---|---|---|---|---|
| sys_15_1 | 8287 | FTMS; HCD | 161.2 | 869.35 |

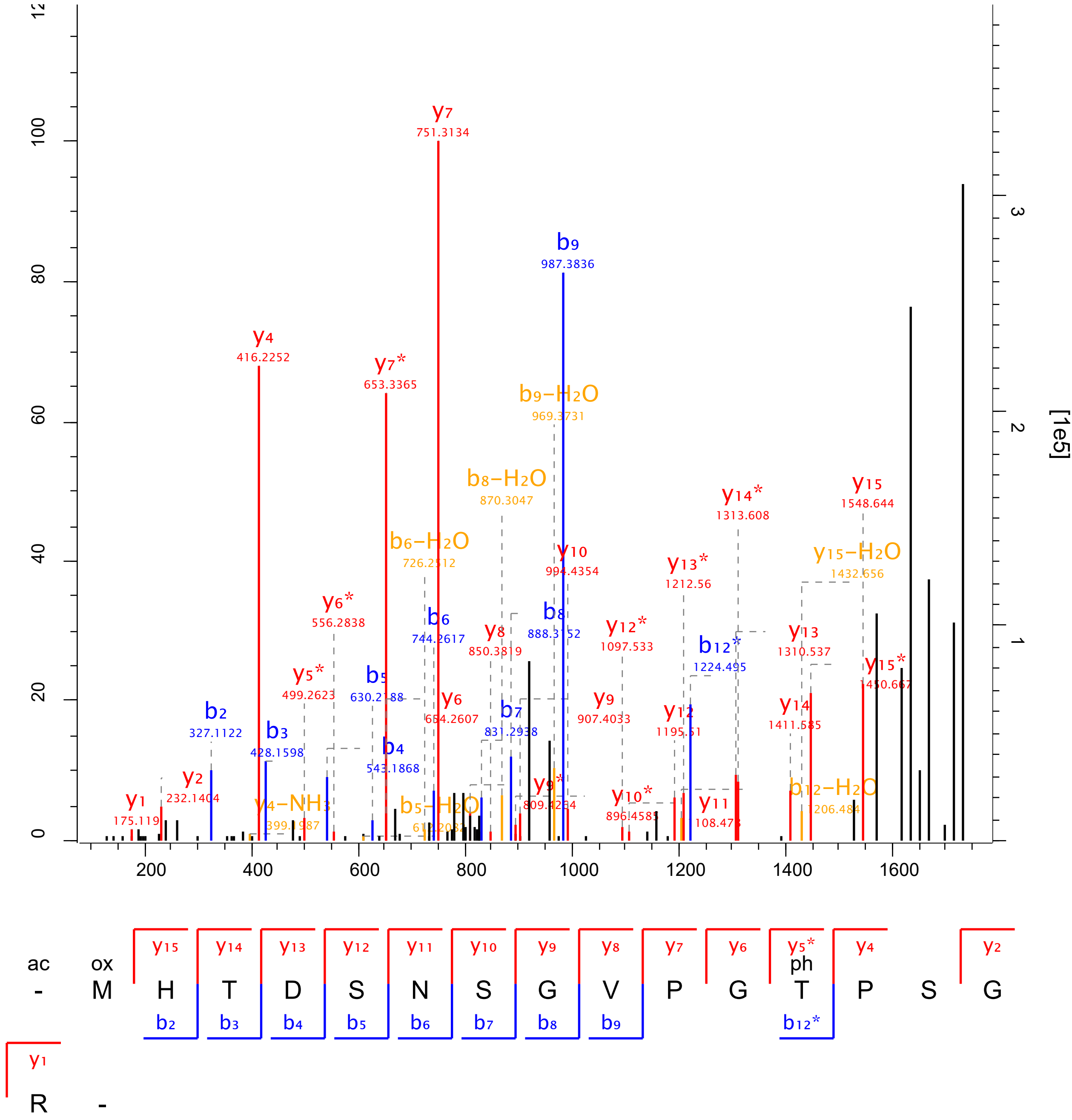

| Raw file | Scan | Method | Score | m/z |
|---|---|---|---|---|
| sys_15_1 | 8369 | FTMS; HCD | 191.47 | 655.77 |

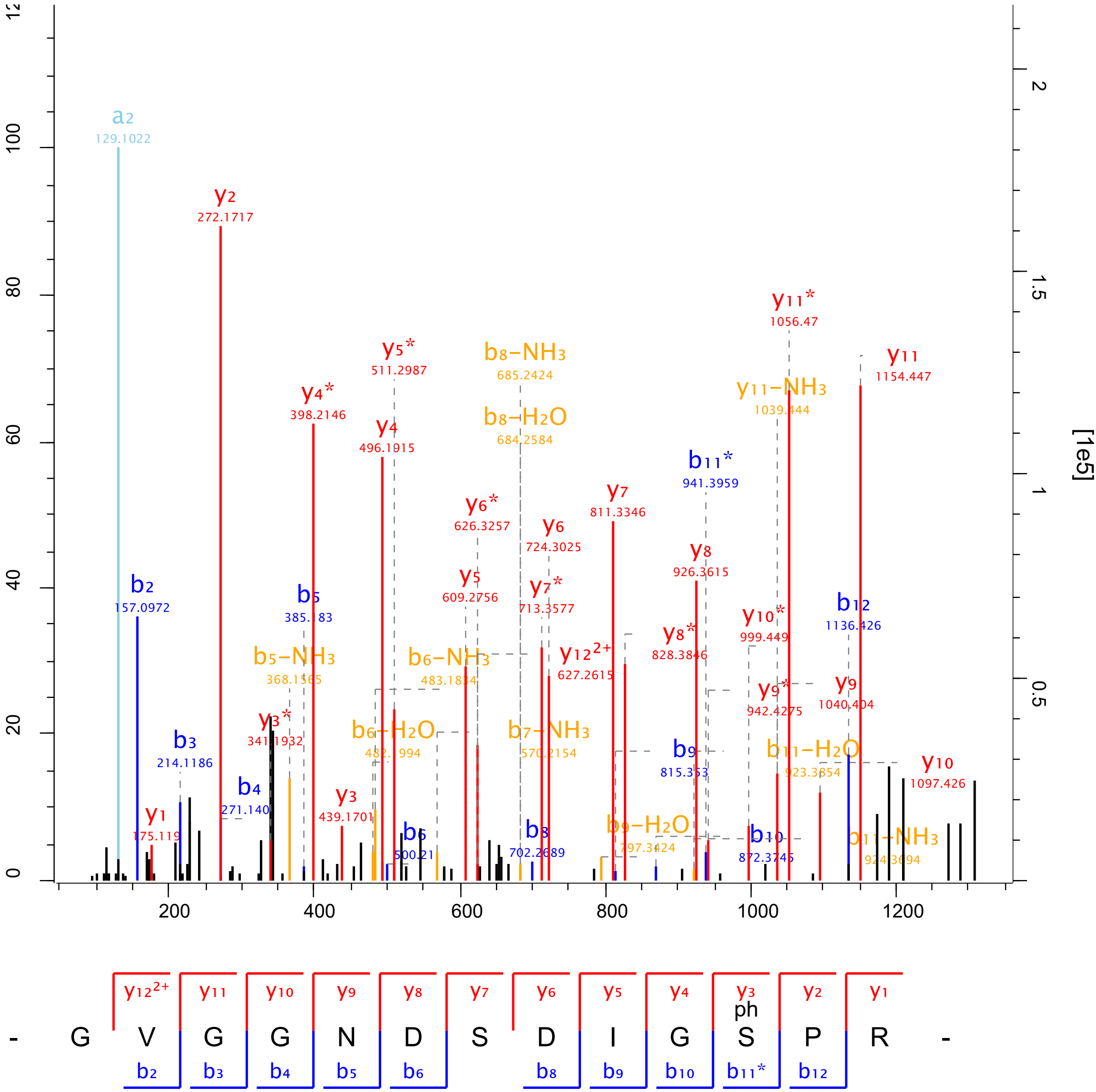

| Raw file | Scan | Method | Score | m/z |
|---|---|---|---|---|
| sys_15_1 | 8523 | FTMS; HCD | 44.85 | 559.23 |

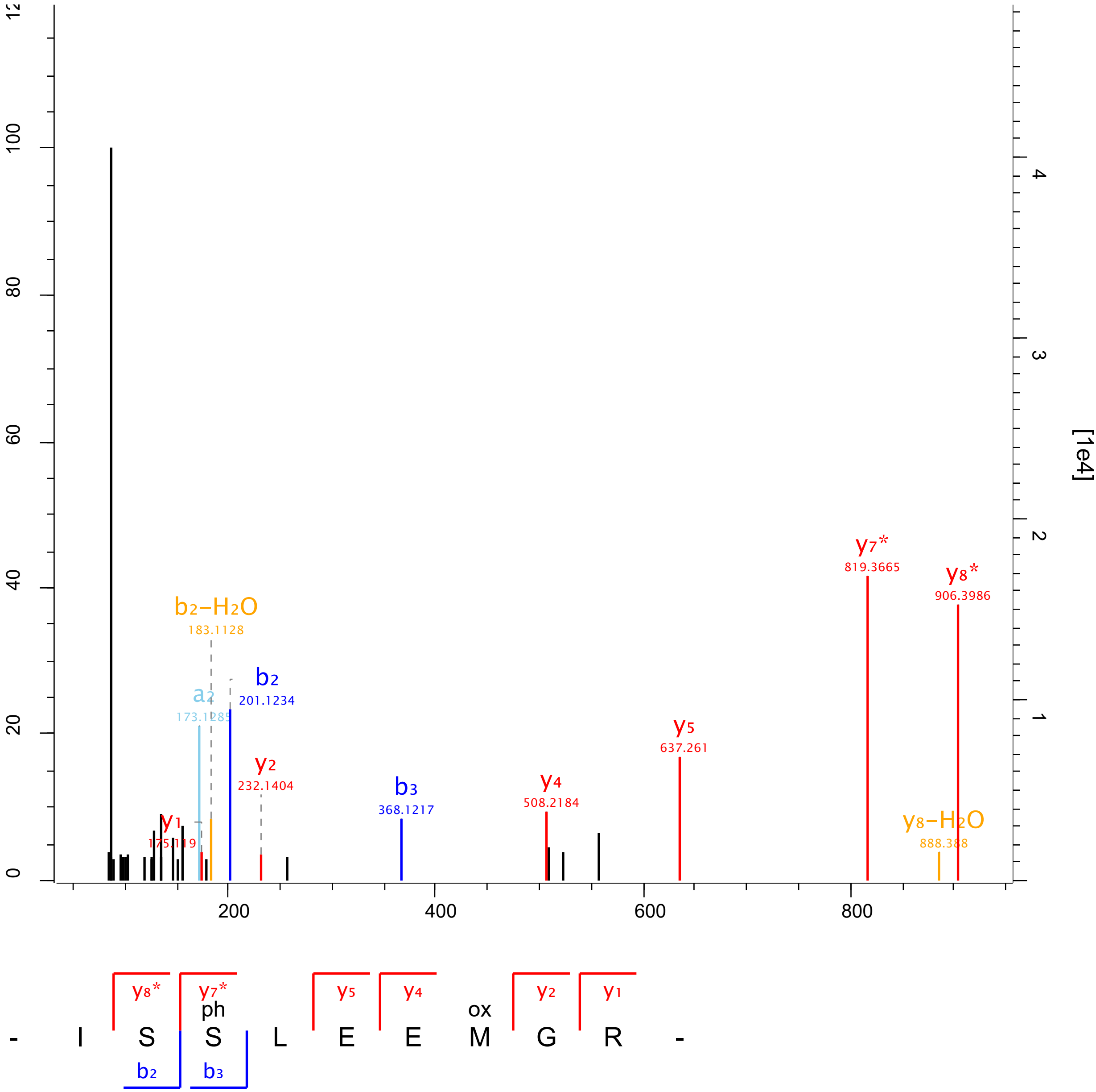

| Raw file | Scan | Method | Score | m/z |
|---|---|---|---|---|
| sys_15_1 | 8632 | FTMS; HCD | 210.91 | 622.28 |

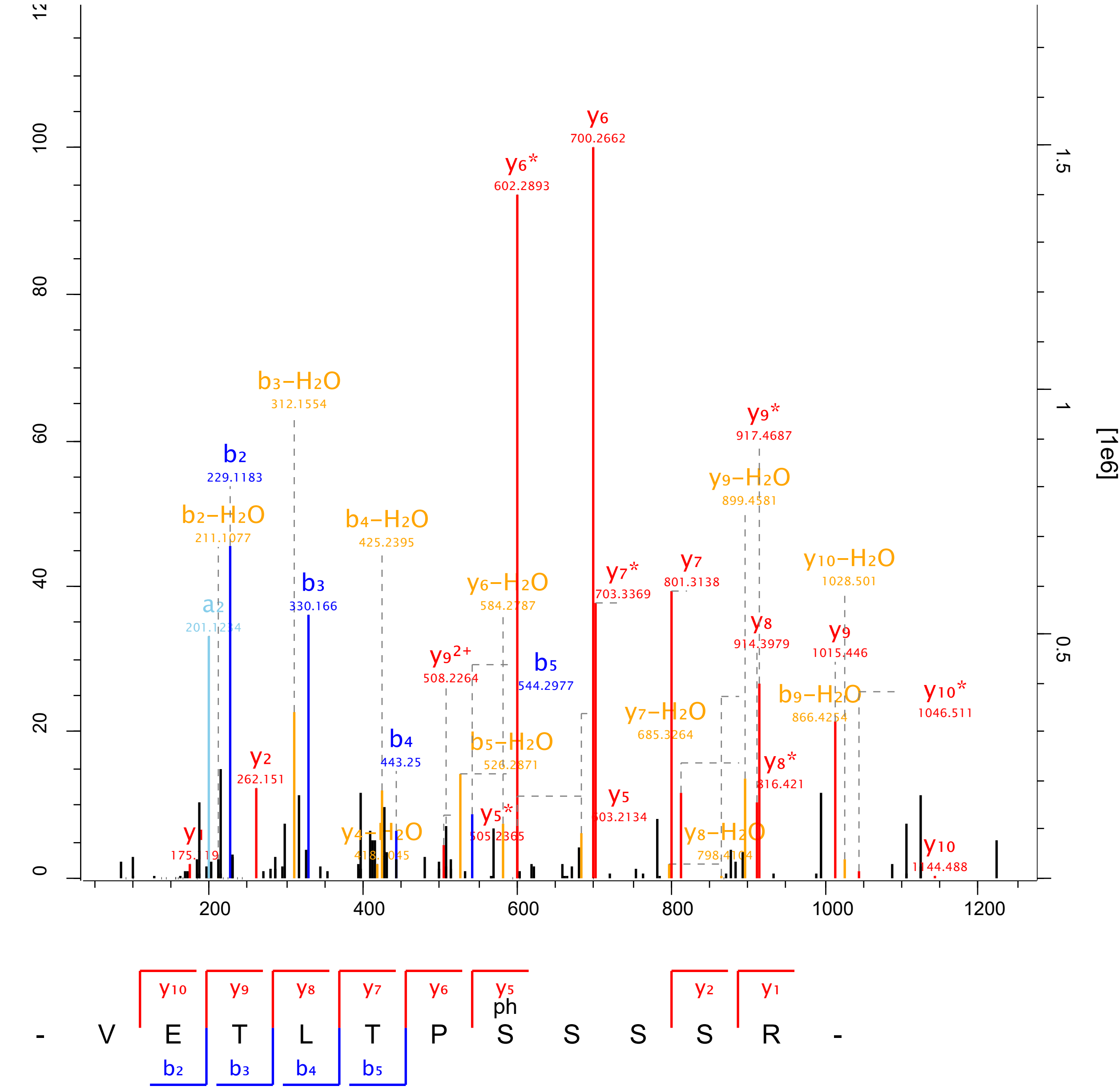

| Raw file | Scan | Method | Score | m/z |
| --- | --- | --- | --- | --- |
| sys_15_1 | 8737 | FTMS; HCD | 63.68 | 649.31 |

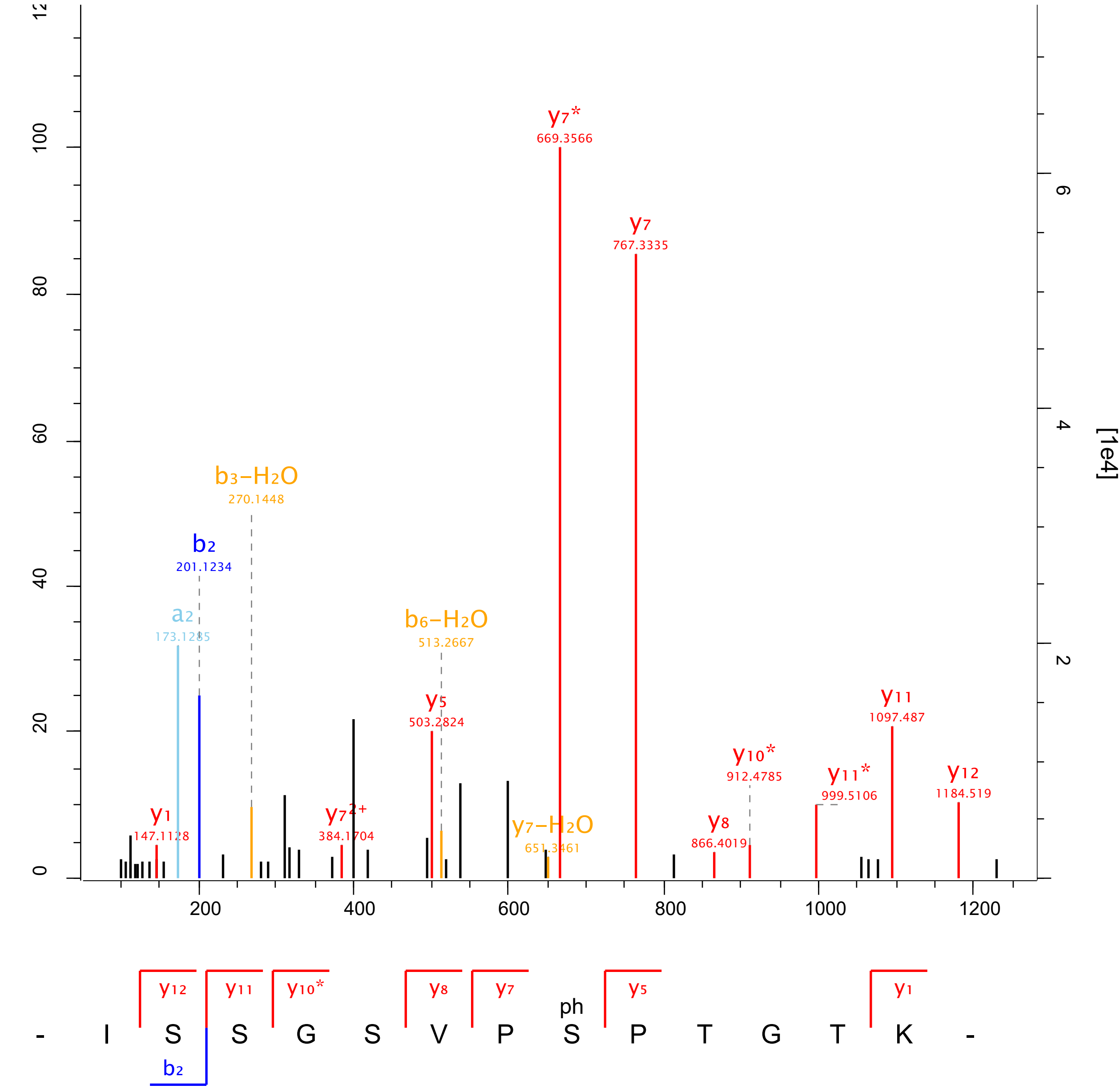

| Raw file | Scan | Method | Score | m/z |
| --- | --- | --- | --- | --- |
| sys_15_1 | 8780 | FTMS; HCD | 103.9 | 880.86 |

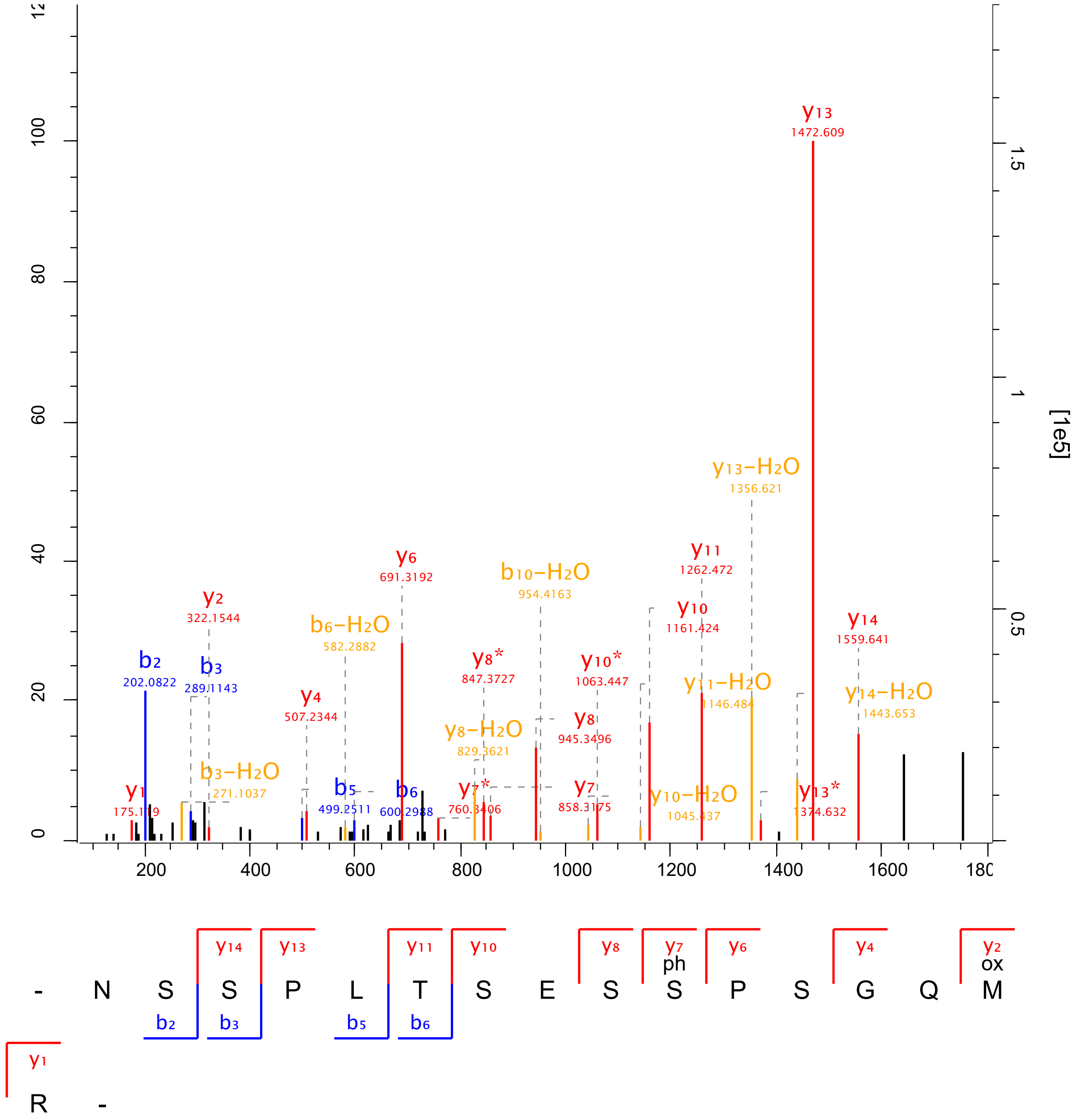

| Raw file | Scan | Method | Score | m/z |
| --- | --- | --- | --- | --- |
| sys_15_1 | 8826 | FTMS; HCD | 43.76 | 574.24 |

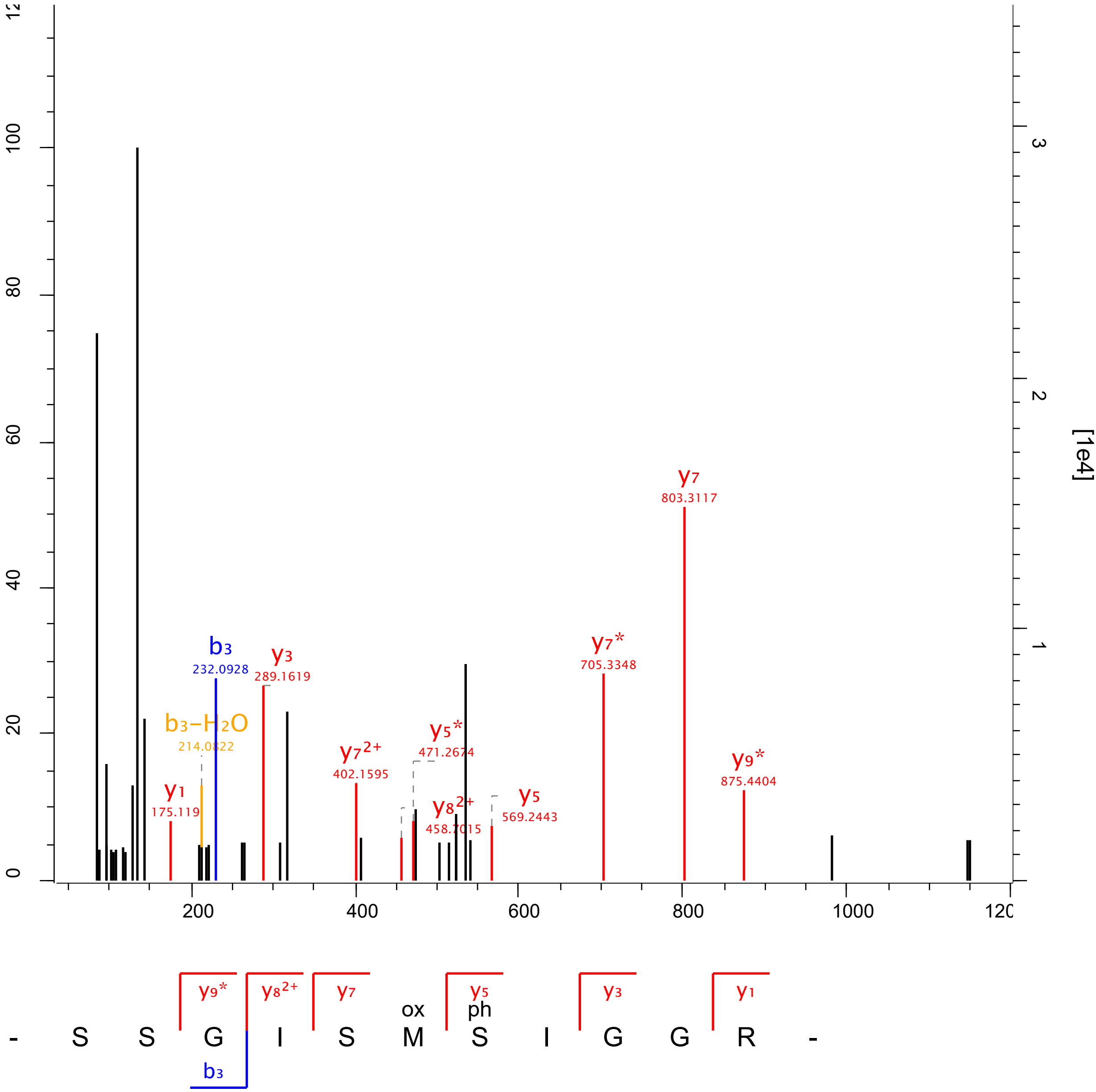

| Raw file | Scan | Method | Score | m/z |
|---|---|---|---|---|
| sys_15_1 | 8840 | FTMS; HCD | 93.92 | 638.59 |

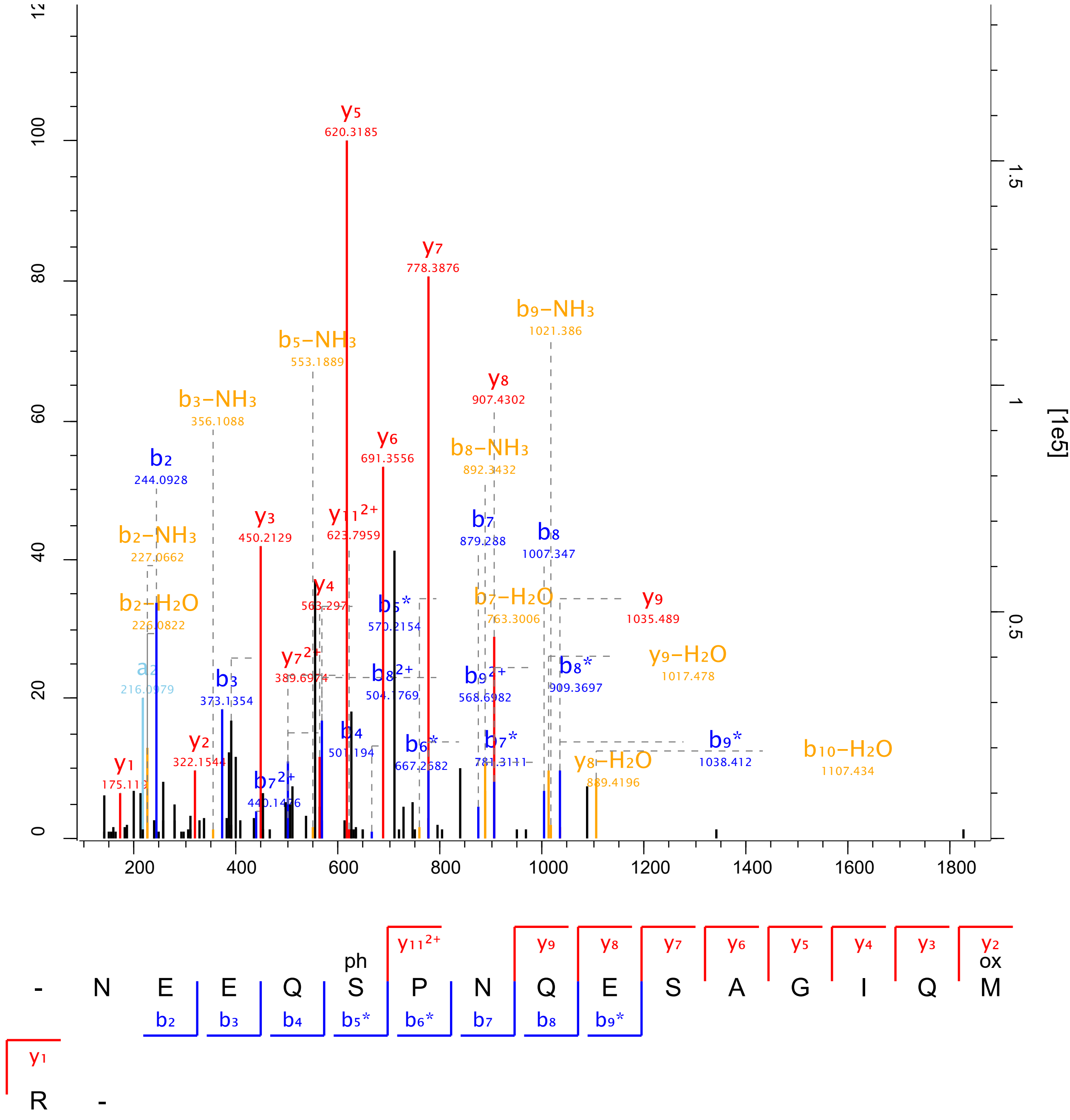

| Raw file | Scan | Method | Score | m/z |
|---|---|---|---|---|
| sys_15_1 | 8973 | FTMS; HCD | 61.26 | 758.82 |

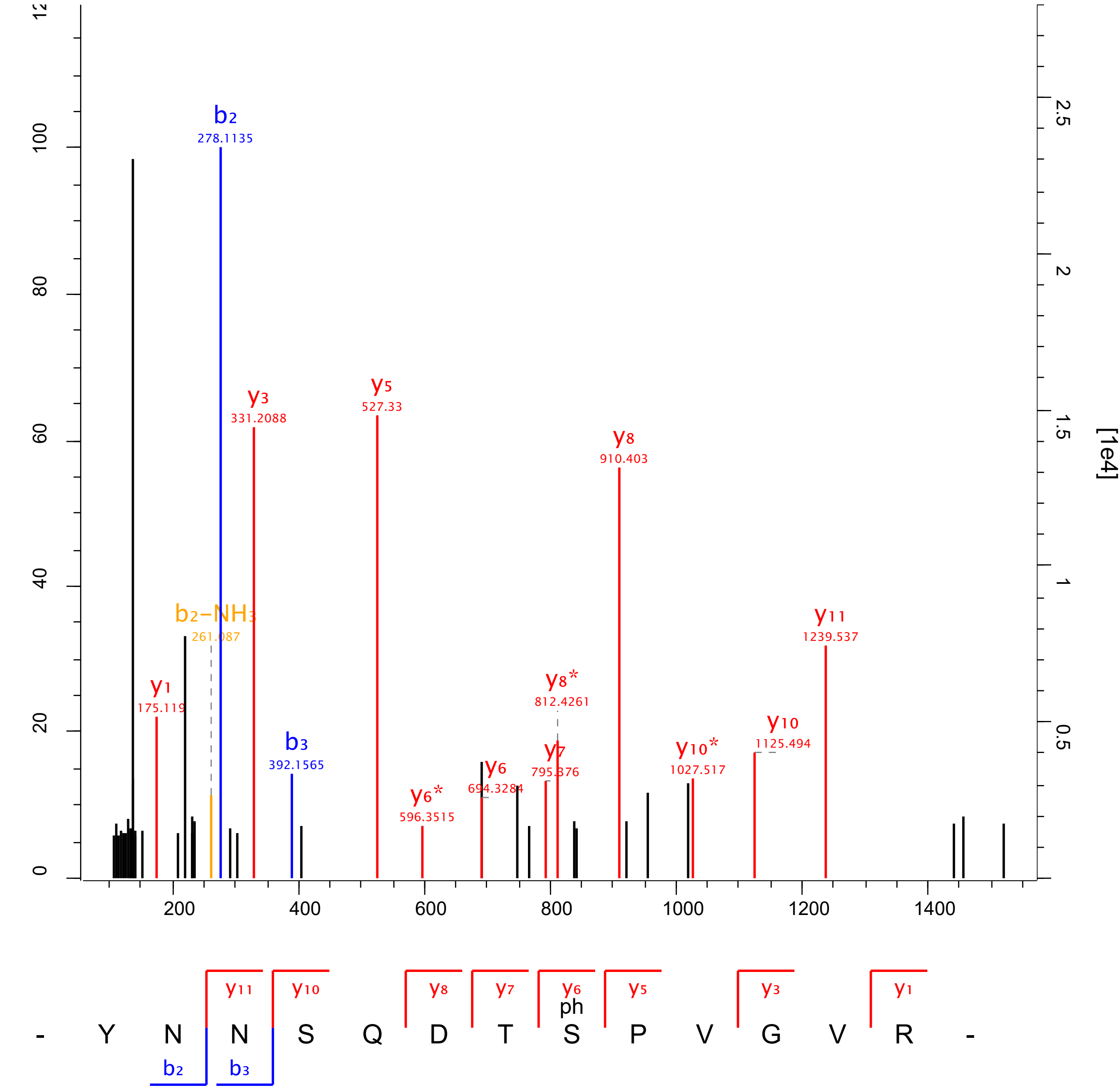

| Raw file | Scan | Method | Score | m/z |
|---|---|---|---|---|
| sys_15_1 | 9007 | FTMS; HCD | 74.46 | 454.19 |

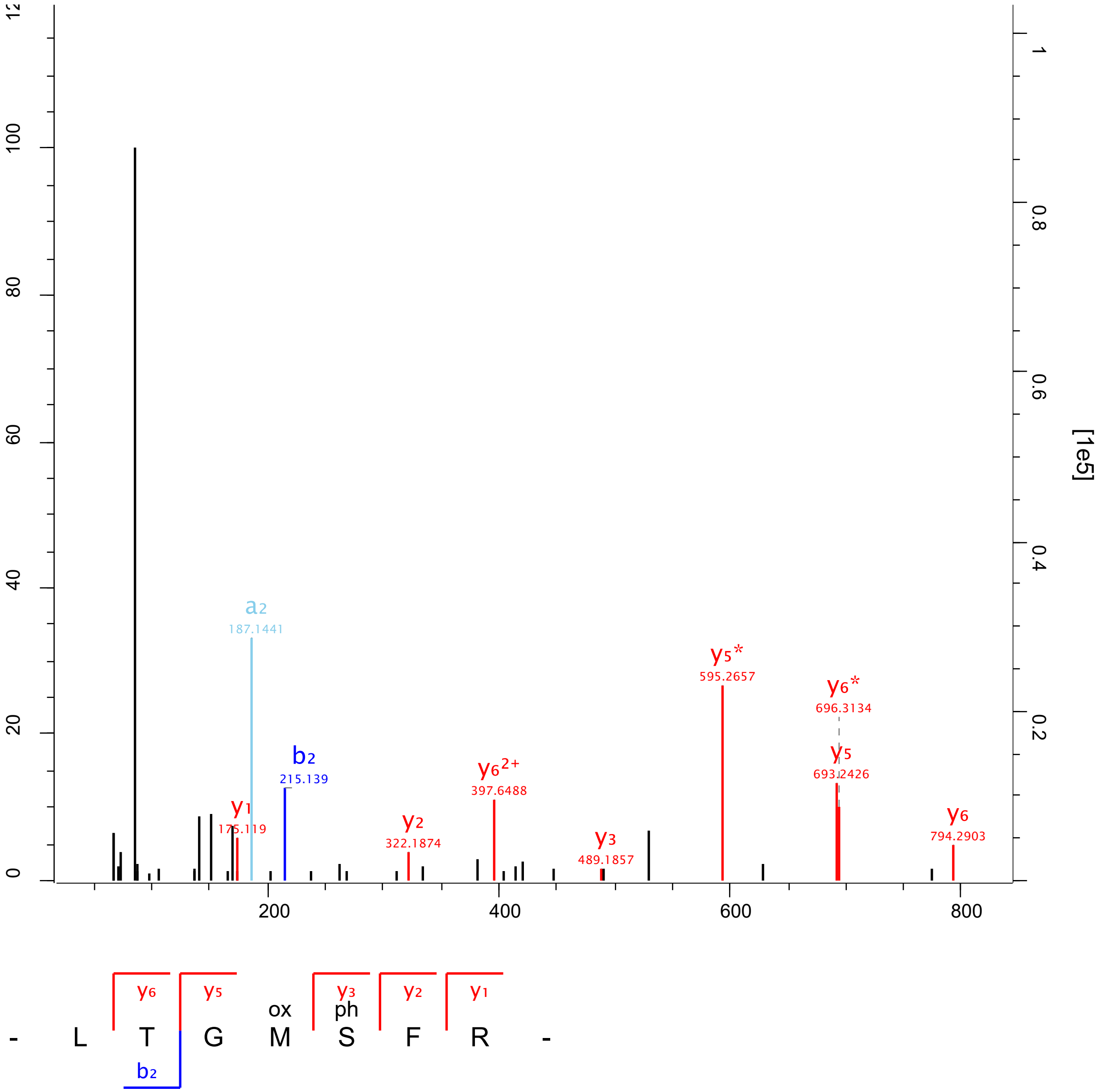

| Raw file | Scan | Method | Score | m/z |
|---|---|---|---|---|
| sys_15_1 | 9103 | FTMS; HCD | 81.68 | 524.23 |

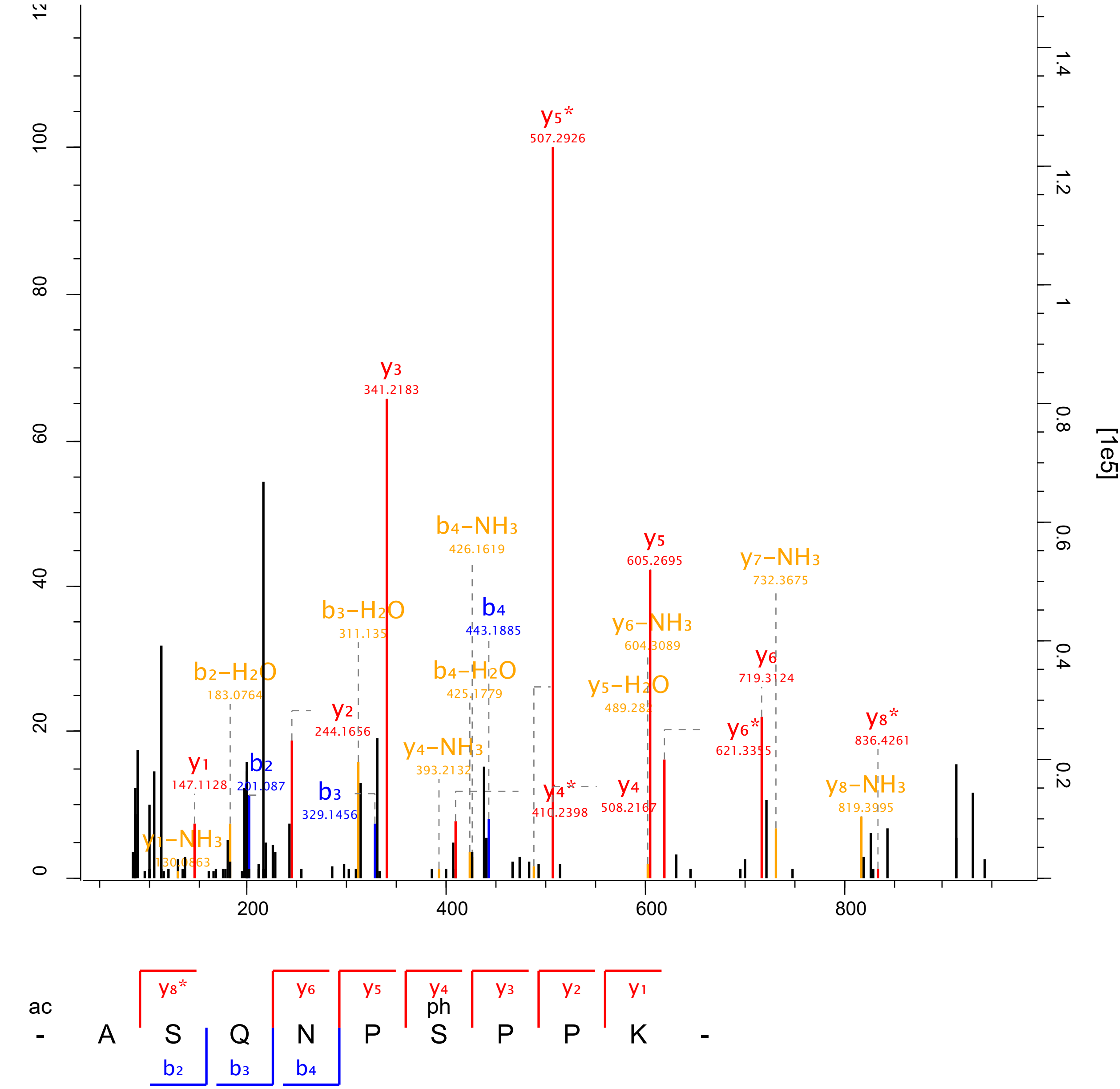

| Raw file | Scan | Method | Score | m/z |
|---|---|---|---|---|
| sys_15_1 | 9115 | FTMS; HCD | 197.41 | 888.84 |

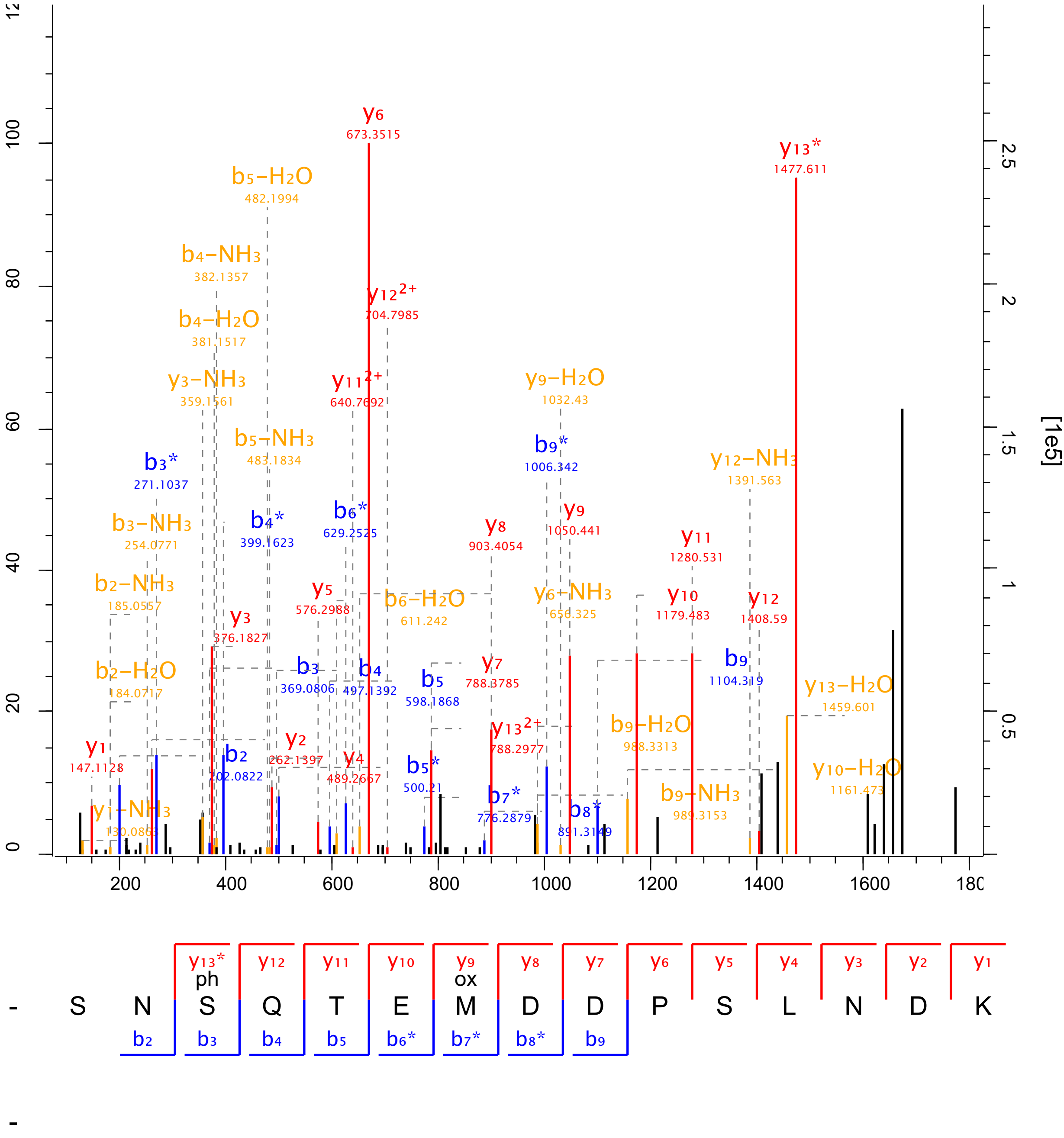

| Raw file | Scan | Method | Score | m/z |
| --- | --- | --- | --- | --- |
| sys_15_1 | 9140 | FTMS; HCD | 42.34 | 452.72 |

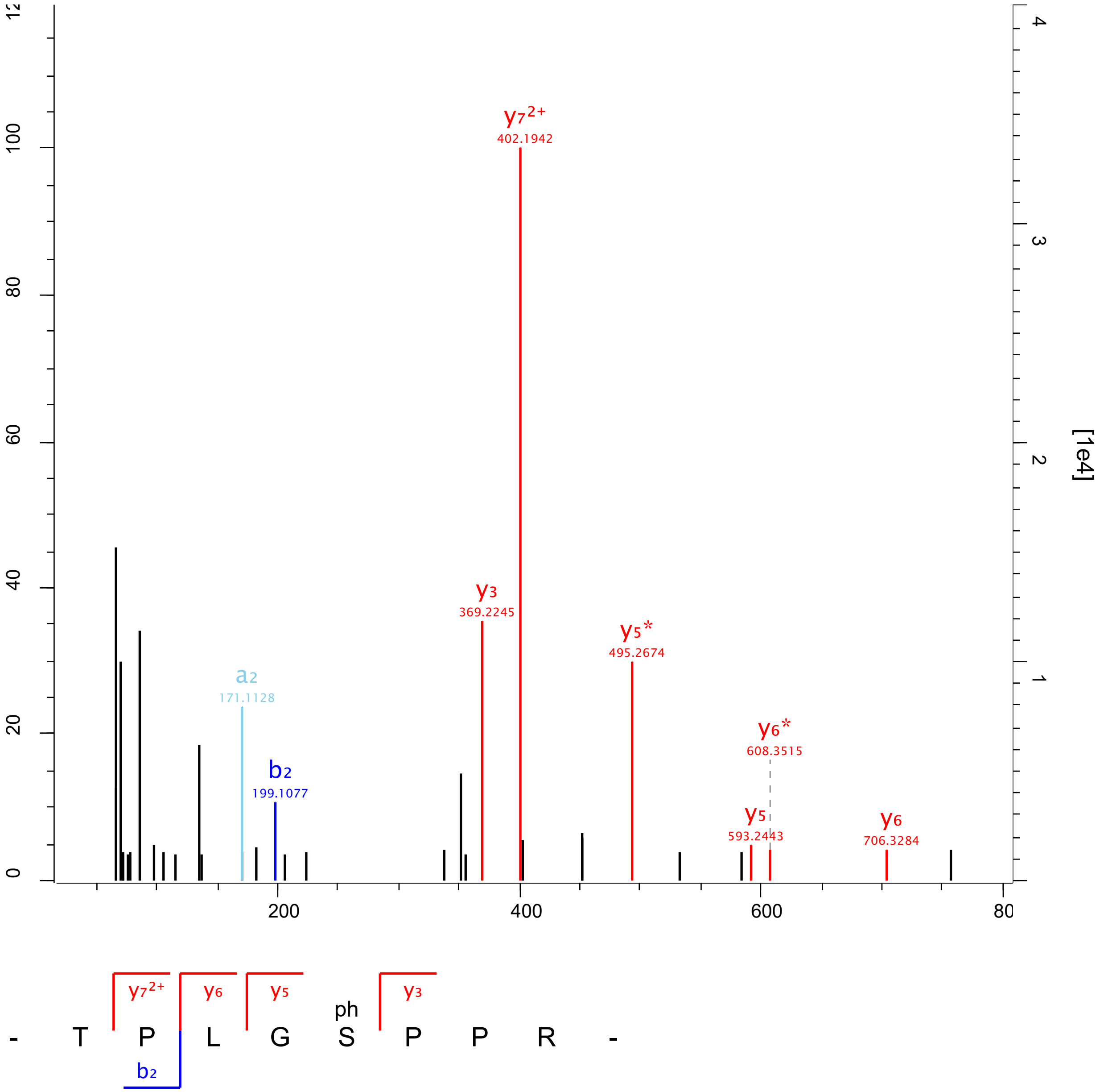

| Raw file | Scan | Method | Score | m/z |
|---|---|---|---|---|
| sys_15_1 | 9177 | FTMS; HCD | 174.59 | 734.78 |

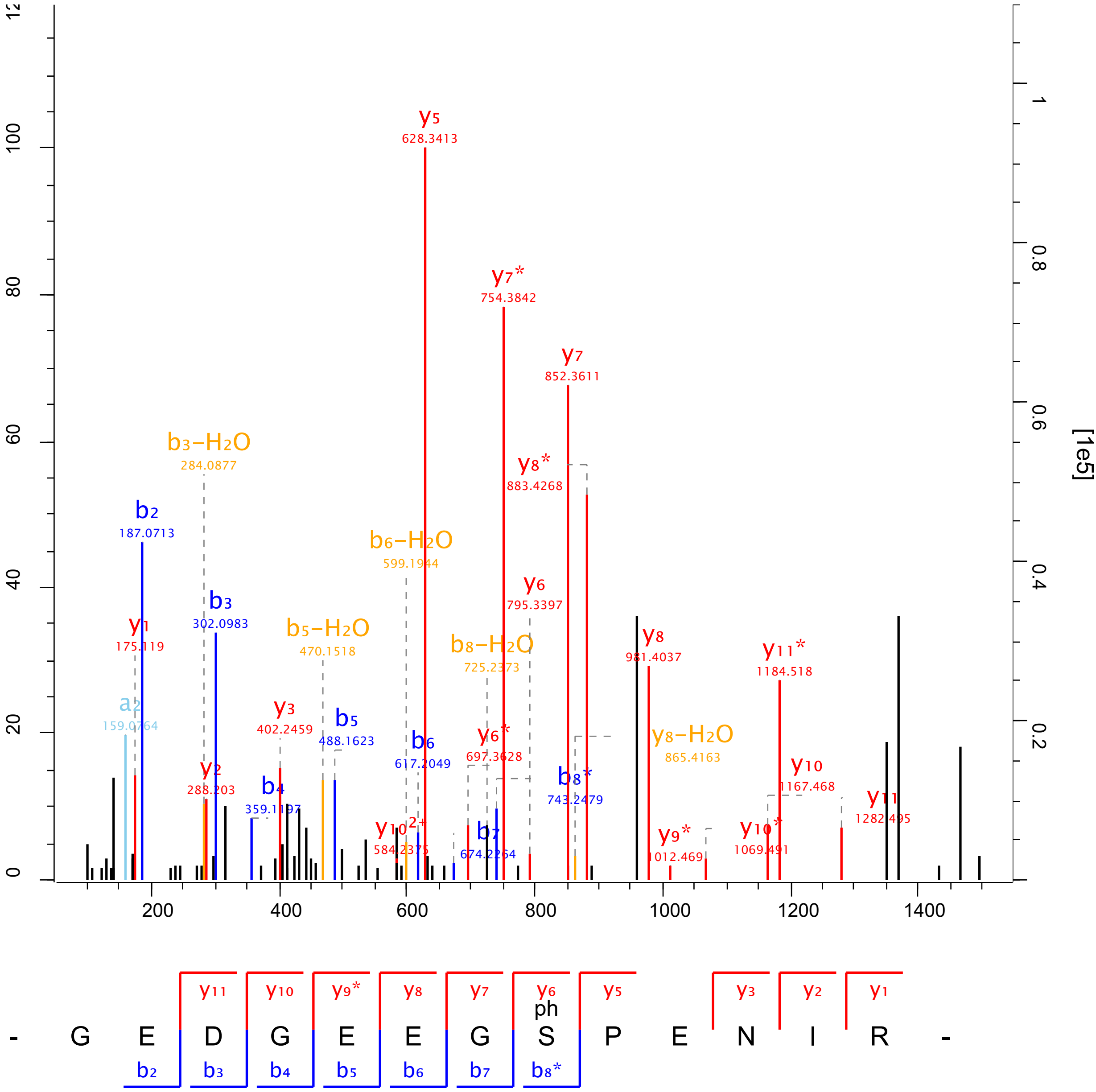

| Raw file | Scan | Method | Score | m/z |
|---|---|---|---|---|
| sys_15_1 | 9212 | FTMS; HCD | 155.57 | 663.79 |

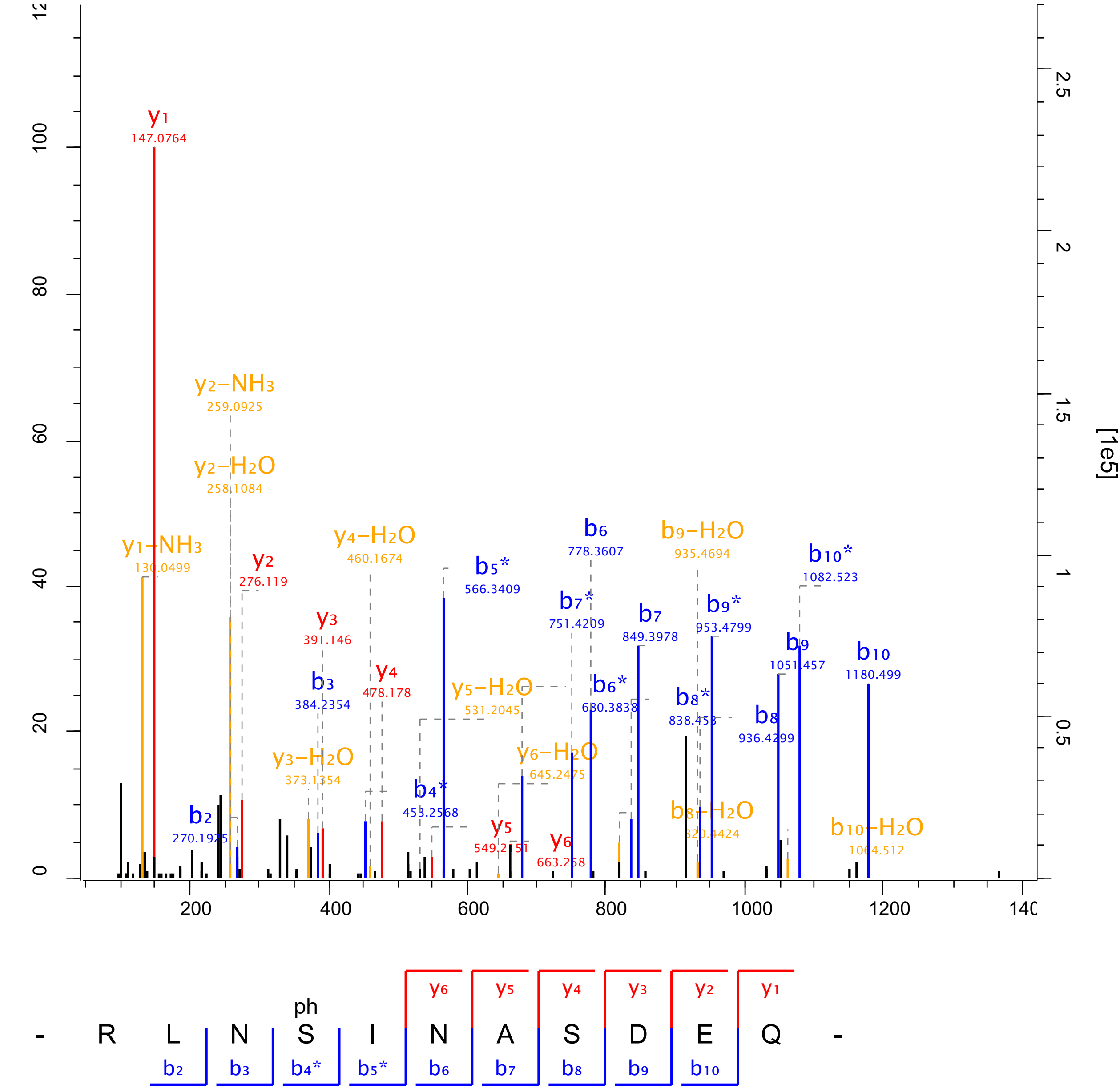

| Raw file | Scan | Method | Score | m/z |
| --- | --- | --- | --- | --- |
| sys_15_1 | 9379 | FTMS; HCD | 160.22 | 563.23 |

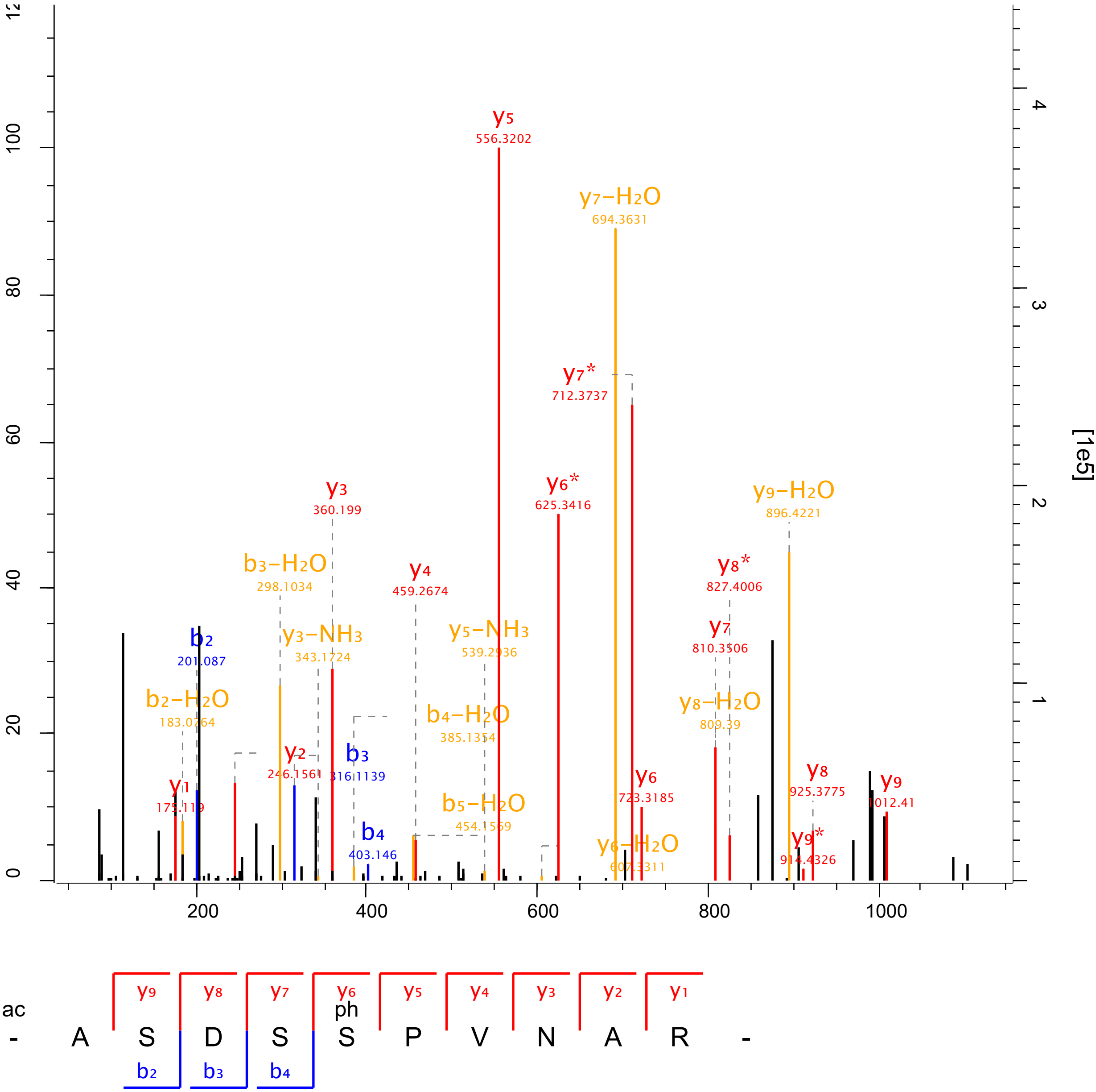

| Raw file | Scan | Method | Score | m/z |
|---|---|---|---|---|
| sys_15_1 | 9389 | FTMS; HCD | 148.52 | 630.77 |

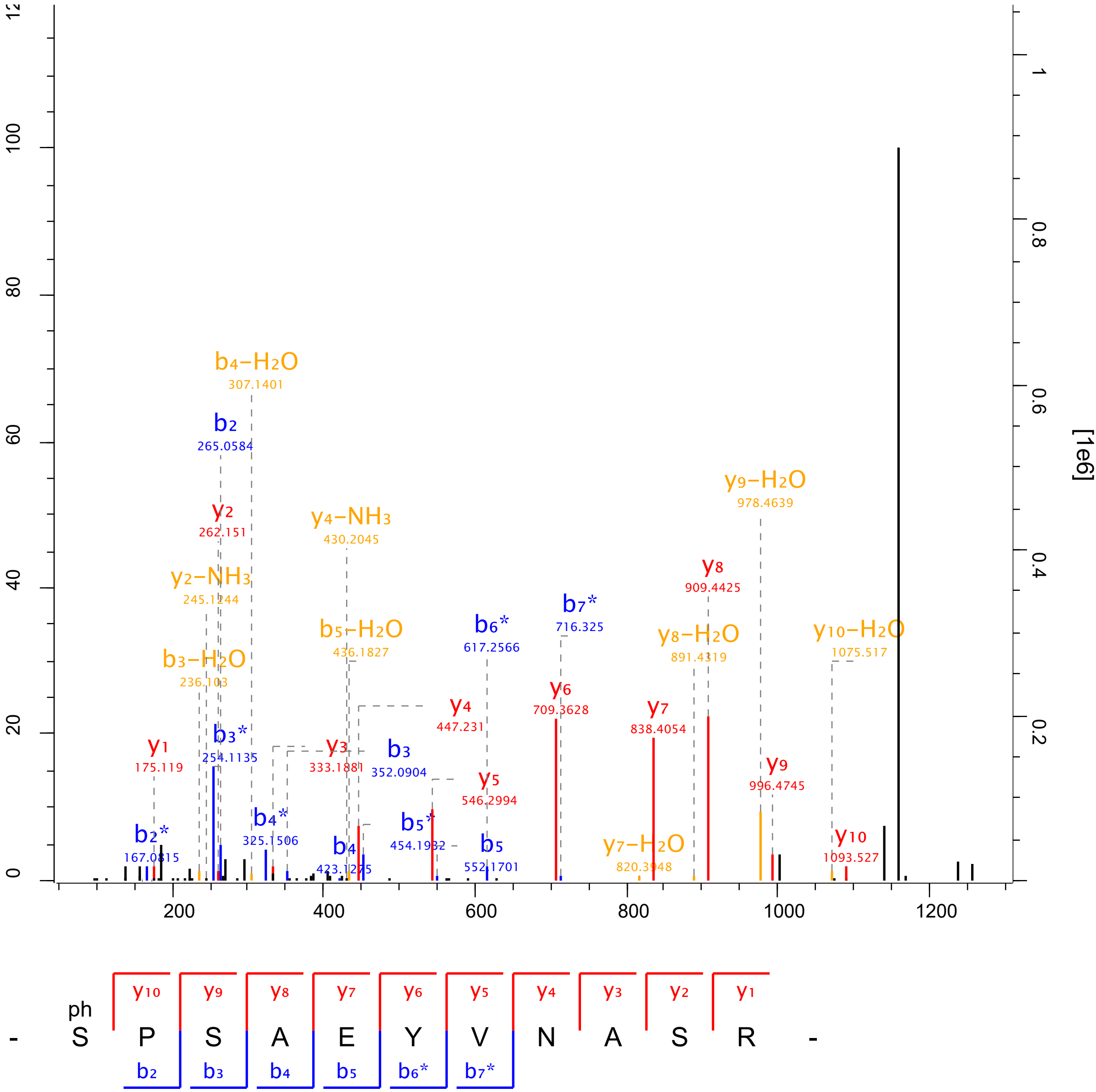

| Raw file | Scan | Method | Score | m/z |
| --- | --- | --- | --- | --- |
| sys_15_1 | 9410 | FTMS; HCD | 125.97 | 733.24 |

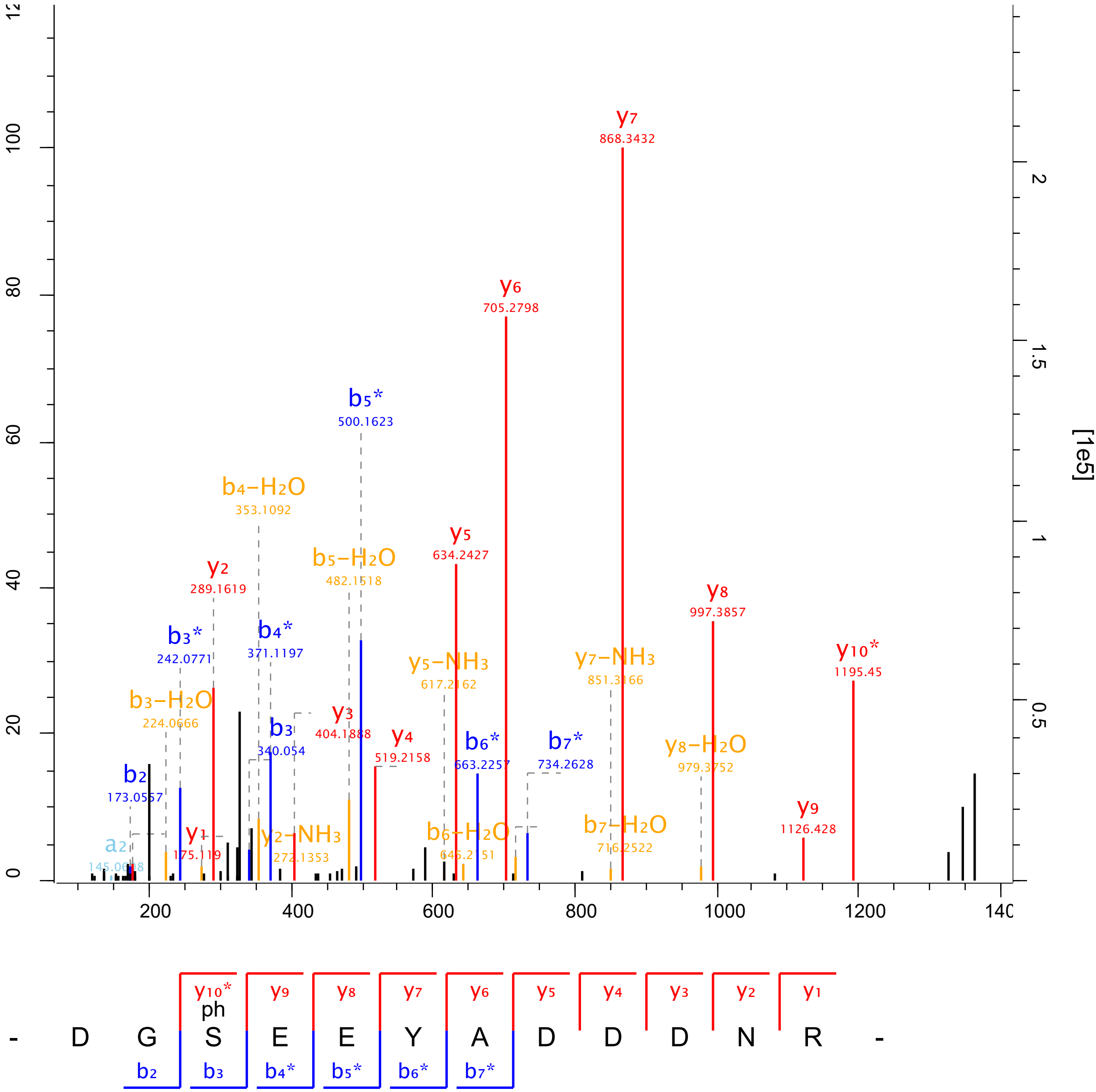

| Raw file | Scan | Method | Score | m/z |
| --- | --- | --- | --- | --- |
| sys_15_1 | 9413 | FTMS; HCD | 121.82 | 652.77 |

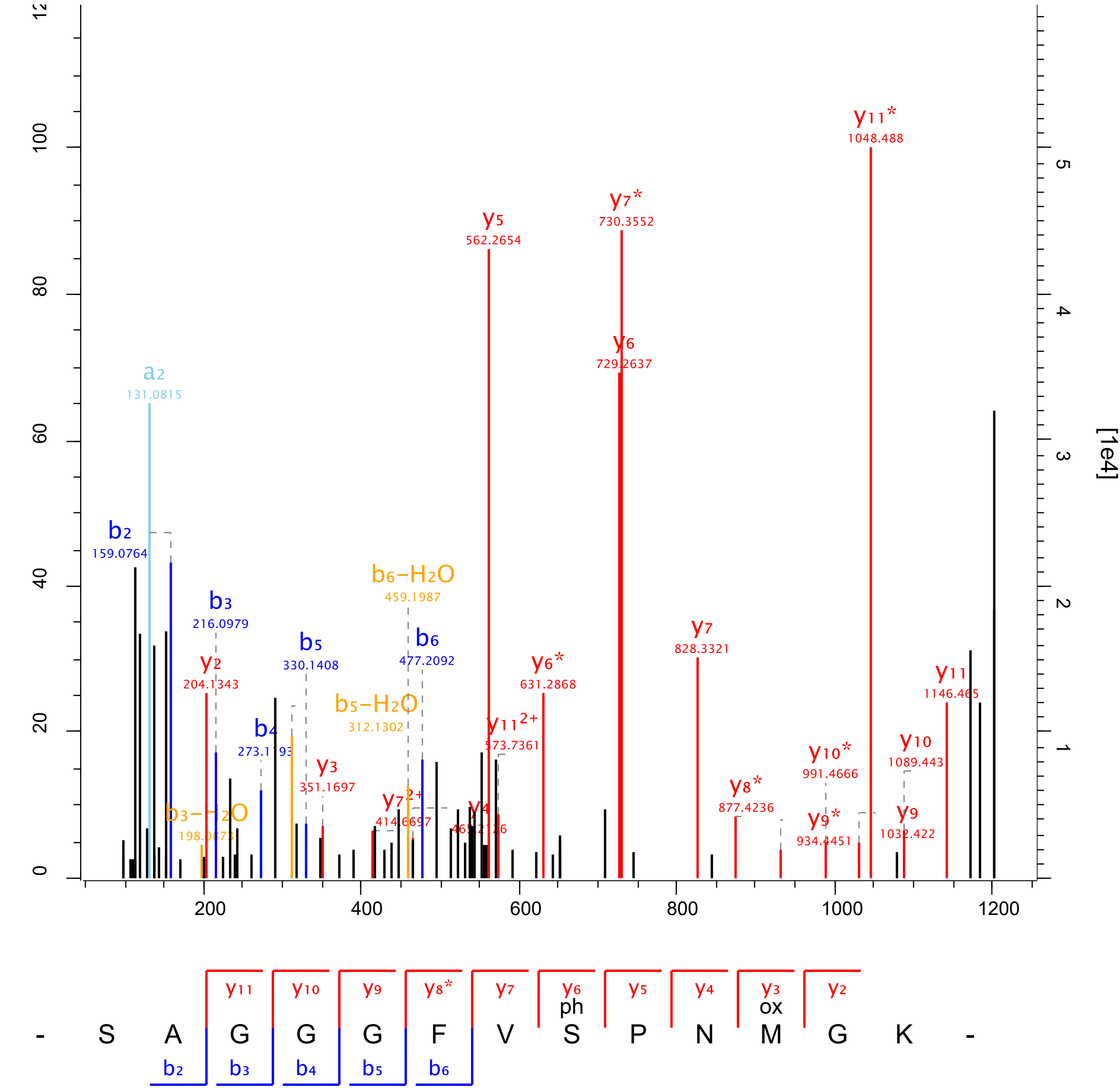

| Raw file | Scan | Method | Score | m/z |
|---|---|---|---|---|
| sys_15_1 | 9477 | FTMS; HCD | 174.18 | 777.79 |

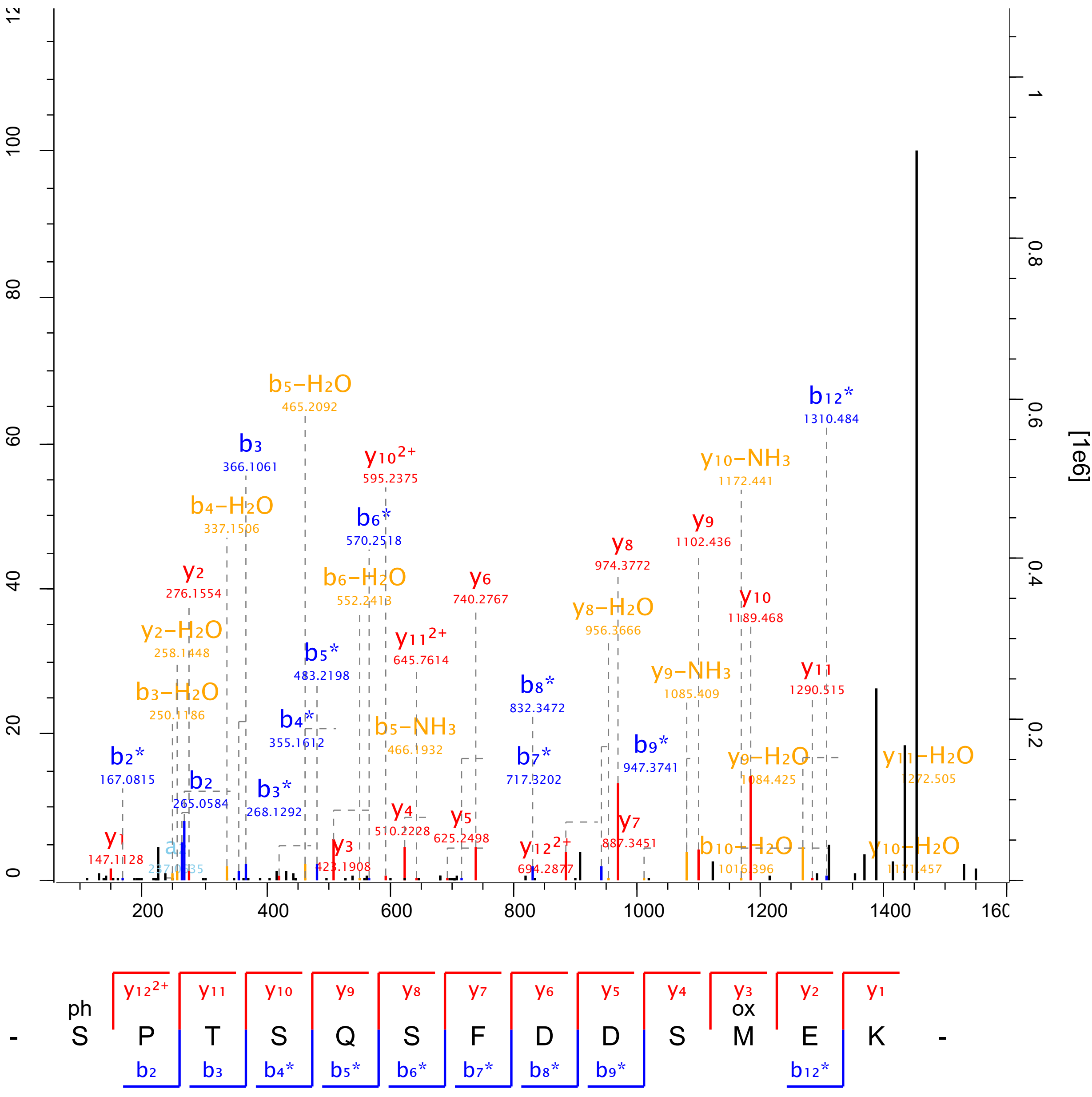

| Raw file | Scan | Method | Score | m/z |
| --- | --- | --- | --- | --- |
| sys_15_1 | 9494 | FTMS; HCD | 253.73 | 763.85 |

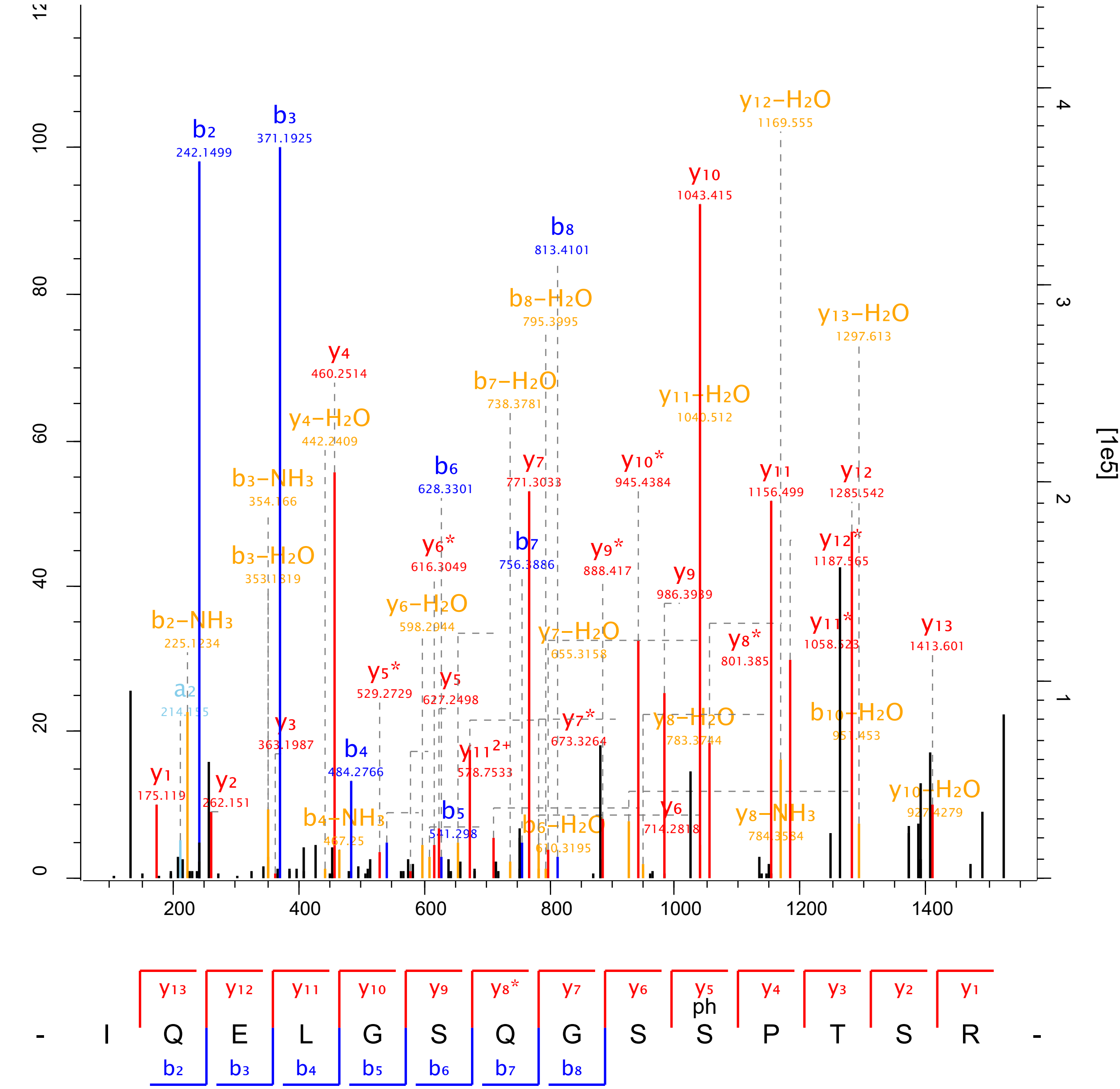

| Raw file | Scan | Method | Score | m/z |
| --- | --- | --- | --- | --- |
| sys_15_1 | 9500 | FTMS; HCD | 81.63 | 602.78 |

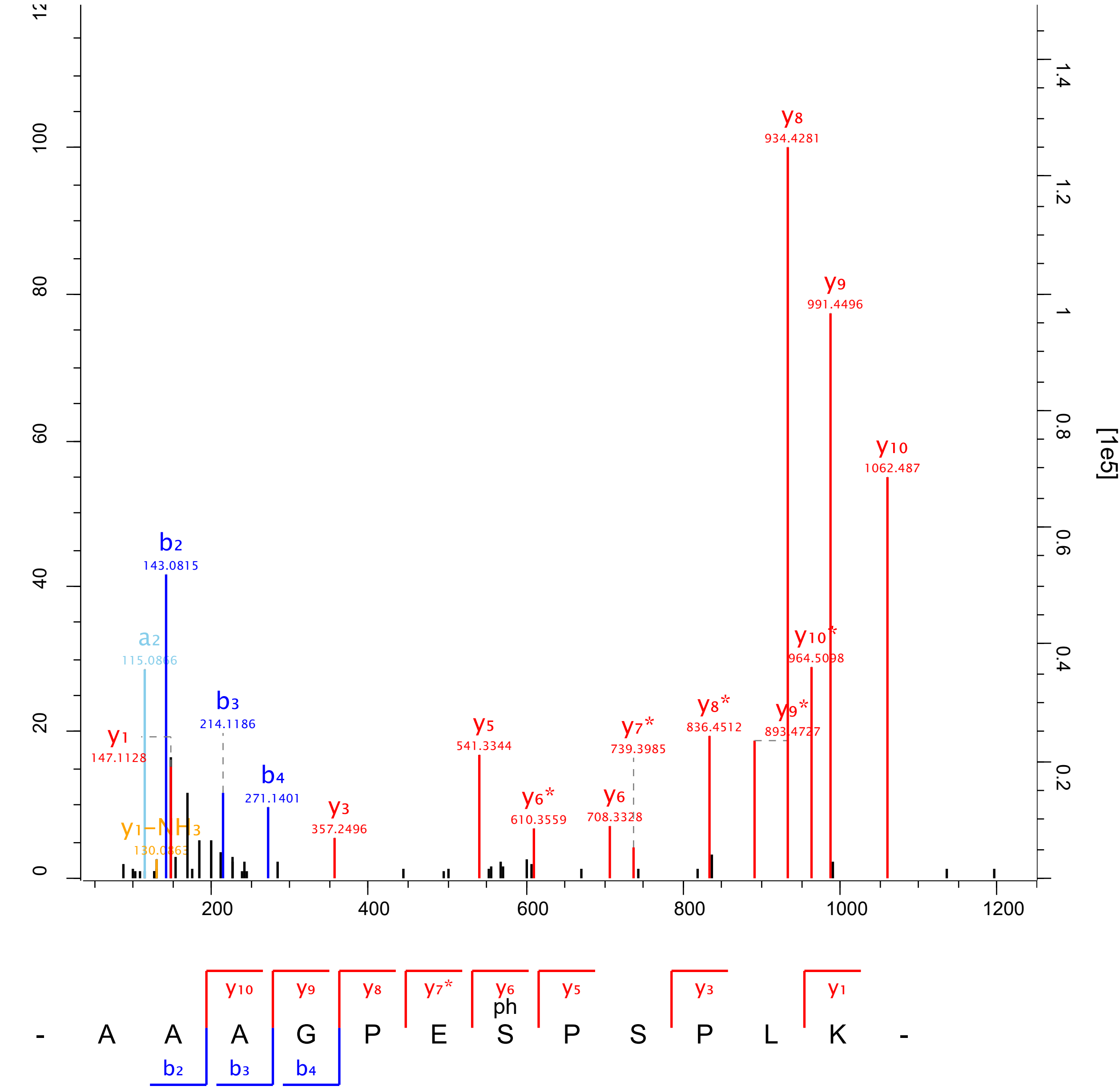

| Raw file | Scan | Method | Score | m/z |
| --- | --- | --- | --- | --- |
| sys_15_1 | 9506 | FTMS; HCD | 60.94 | 818.3 |

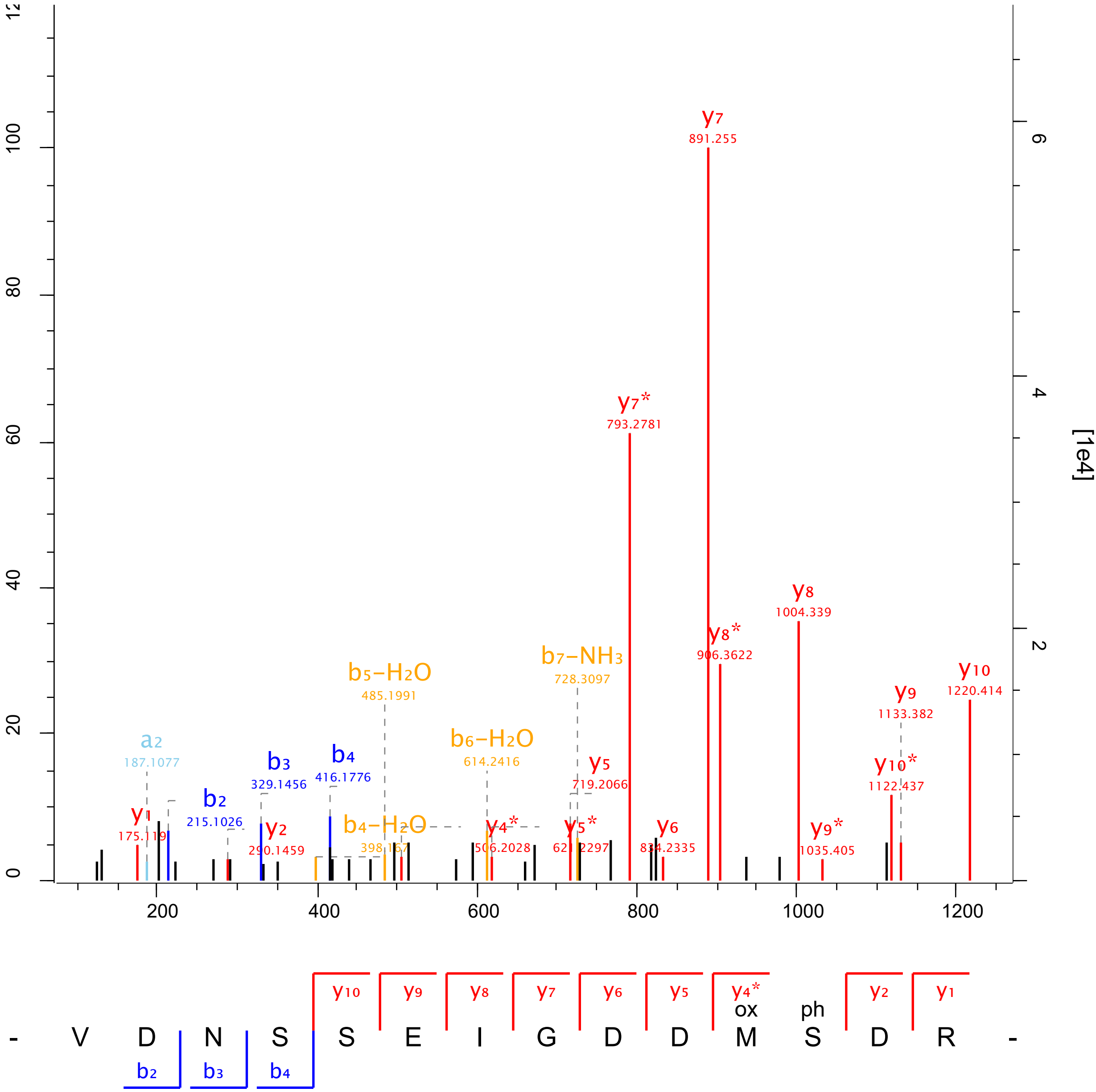

| Raw file | Scan | Method | Score | m/z |
|---|---|---|---|---|
| sys_15_1 | 9541 | FTMS; HCD | 66.25 | 695.3 |

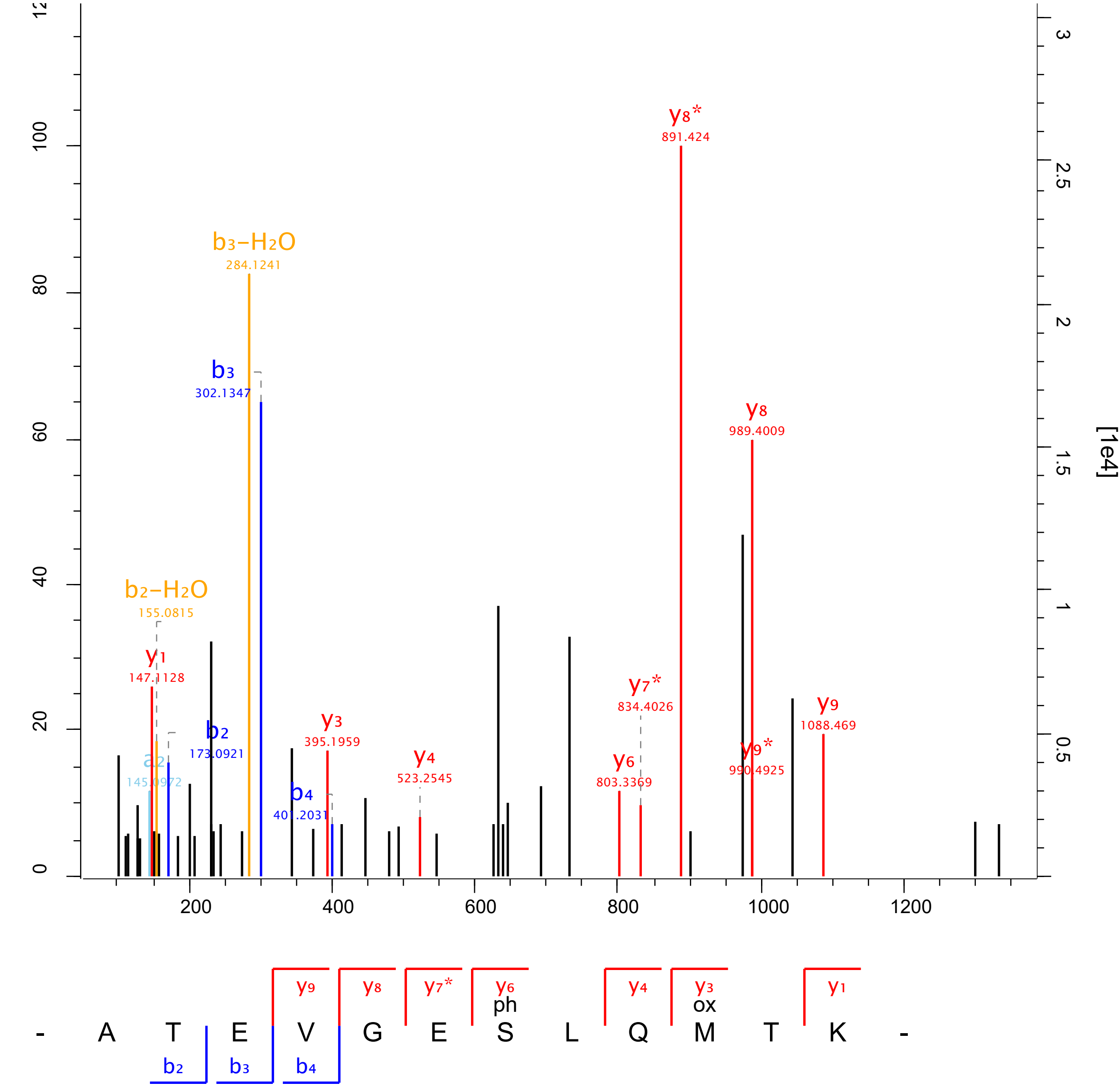

| Raw file | Scan | Method | Score | m/z |
|---|---|---|---|---|
| sys_15_1 | 9560 | FTMS; HCD | 94.2 | 716.31 |

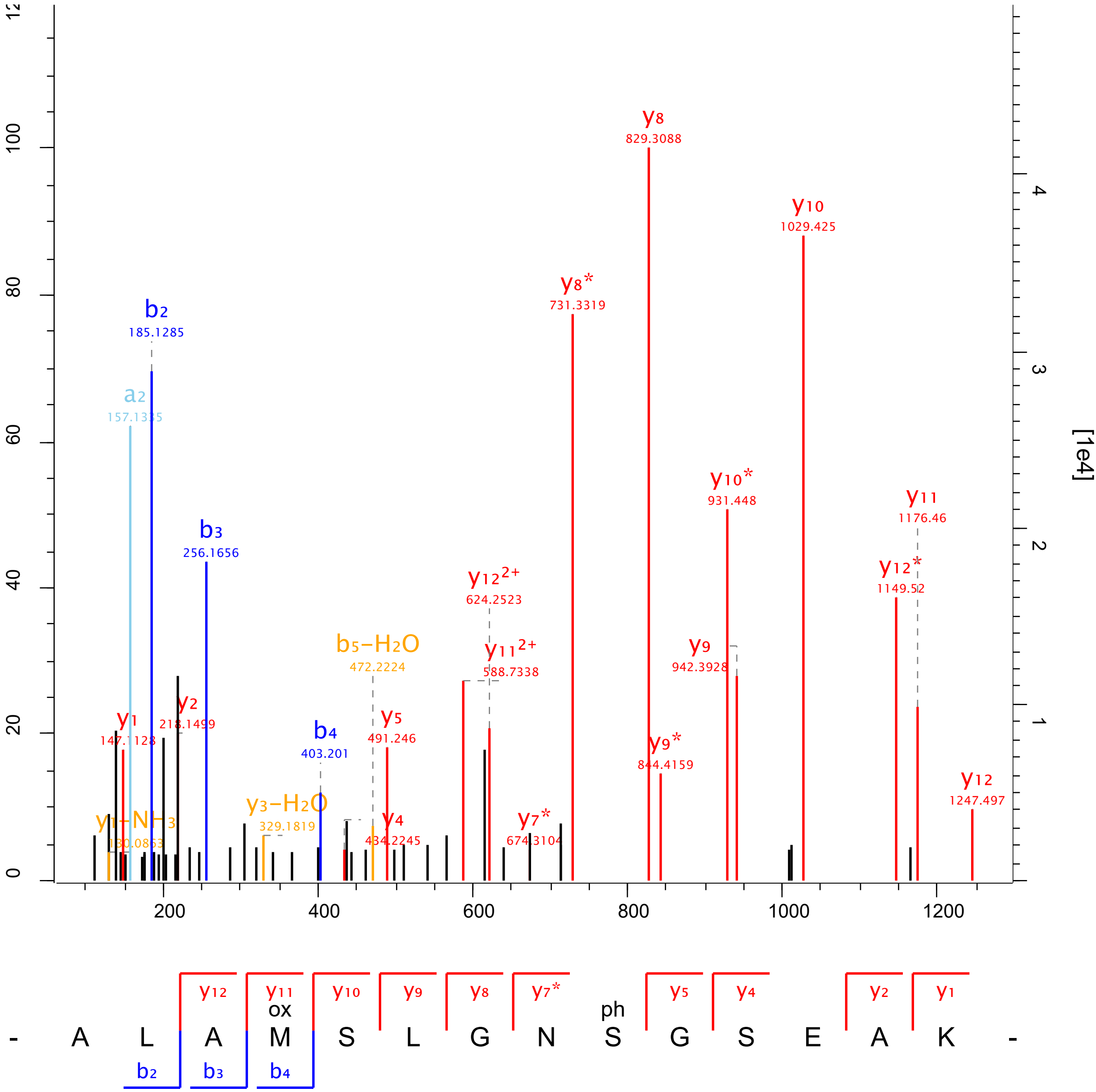

| Raw file | Scan | Method | Score | m/z |
| --- | --- | --- | --- | --- |
| sys_15_1 | 9581 | FTMS; HCD | 222.54 | 692.94 |

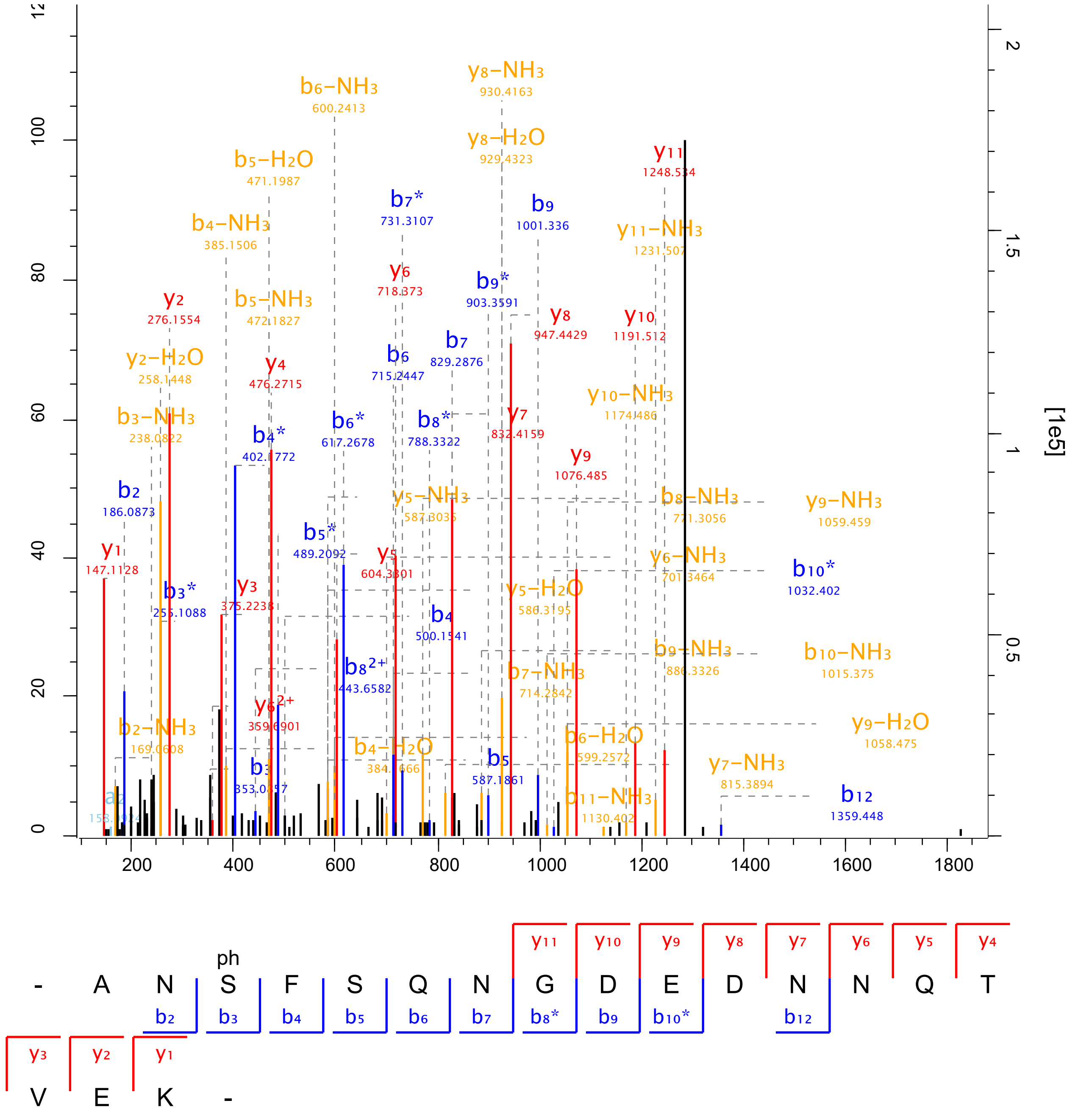

| Raw file | Scan | Method | Score | m/z |
| --- | --- | --- | --- | --- |
| sys_15_1 | 9626 | FTMS; HCD | 248.23 | 824.83 |

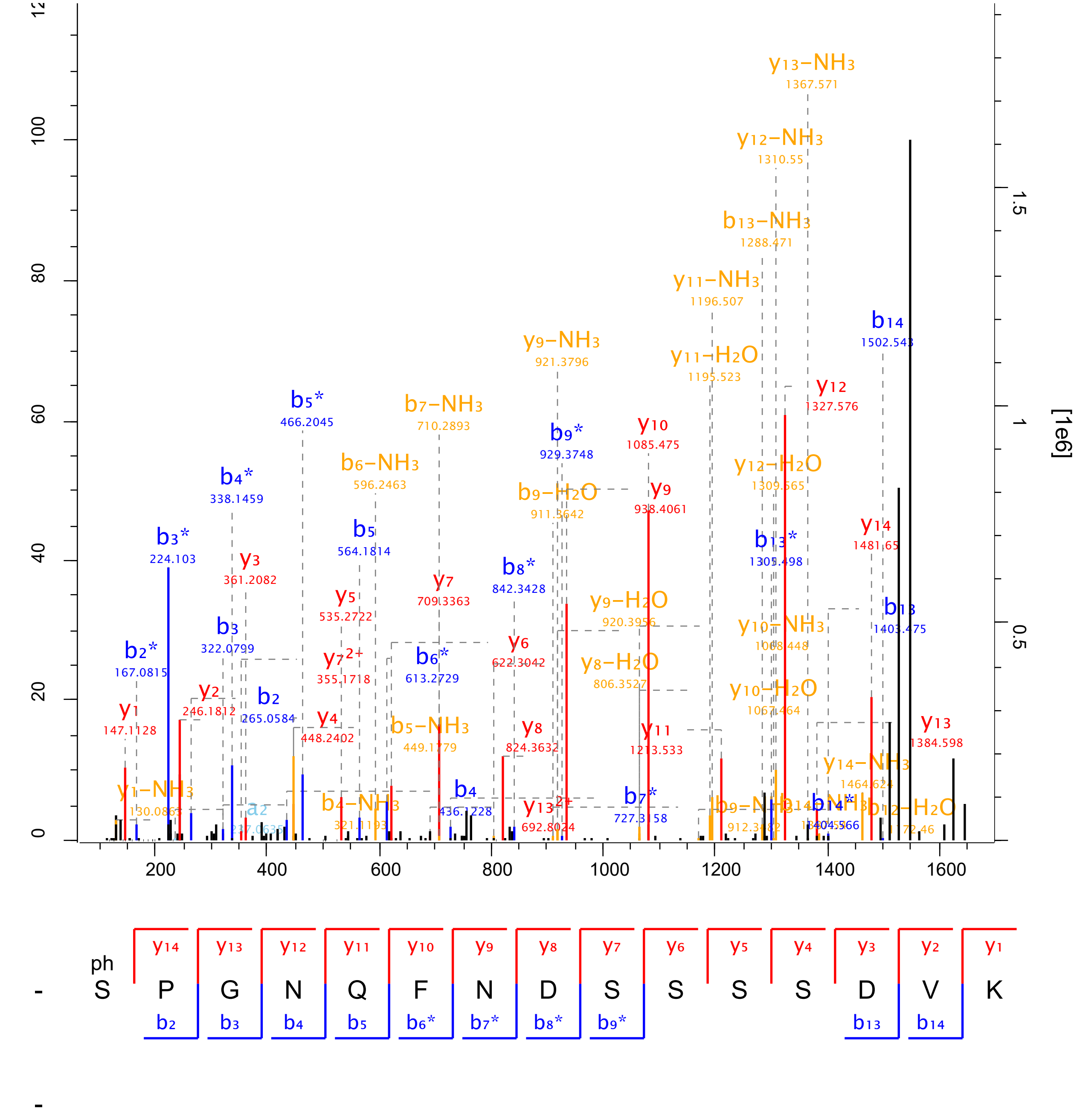

| Raw file | Scan | Method | Score | m/z |
|---|---|---|---|---|
| sys_15_1 | 9633 | FTMS; HCD | 70.09 | 514.72 |

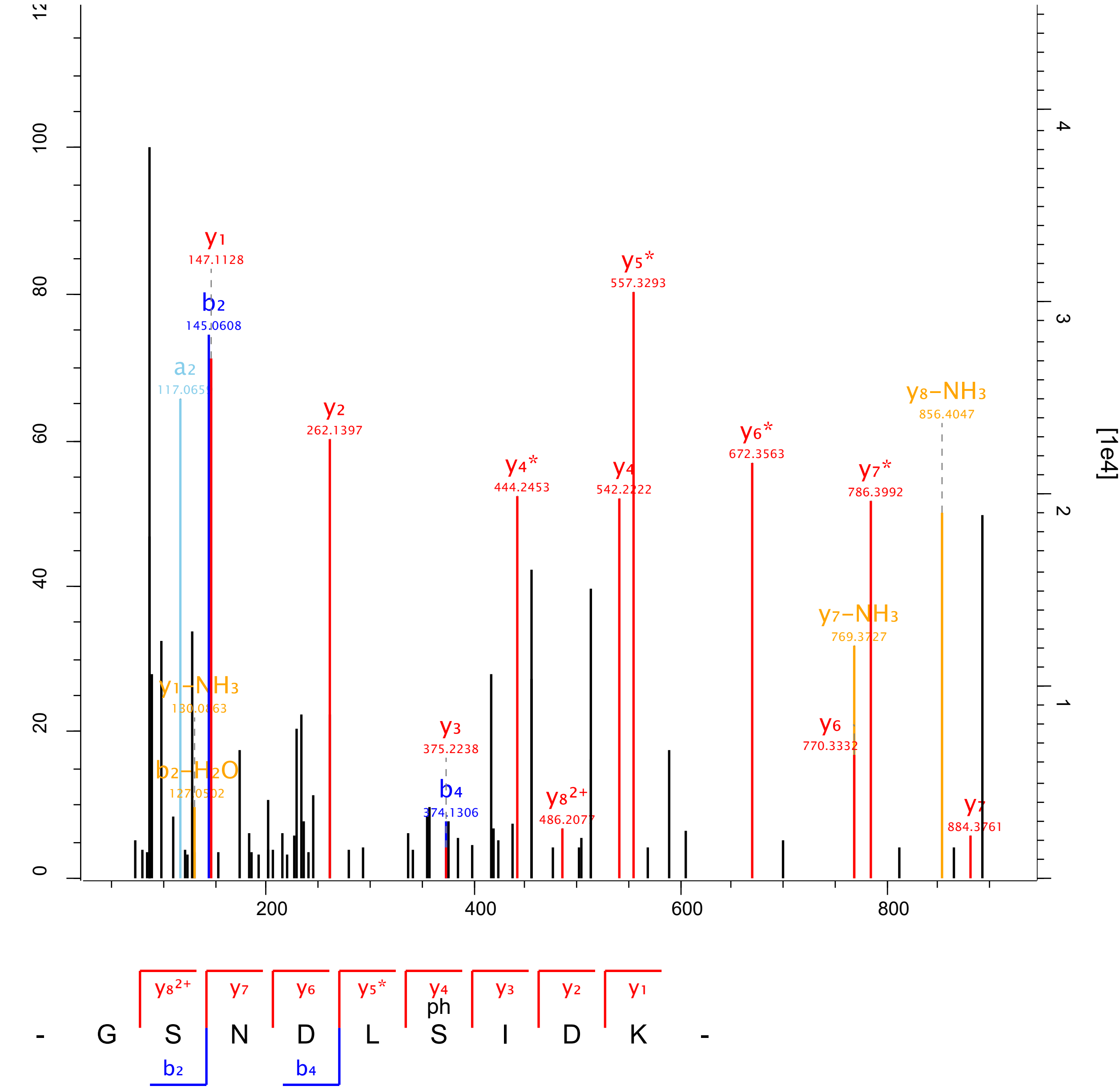

| Raw file | Scan | Method | Score | m/z |
|---|---|---|---|---|
| sys_15_1 | 9646 | FTMS; HCD | 138.55 | 621.28 |

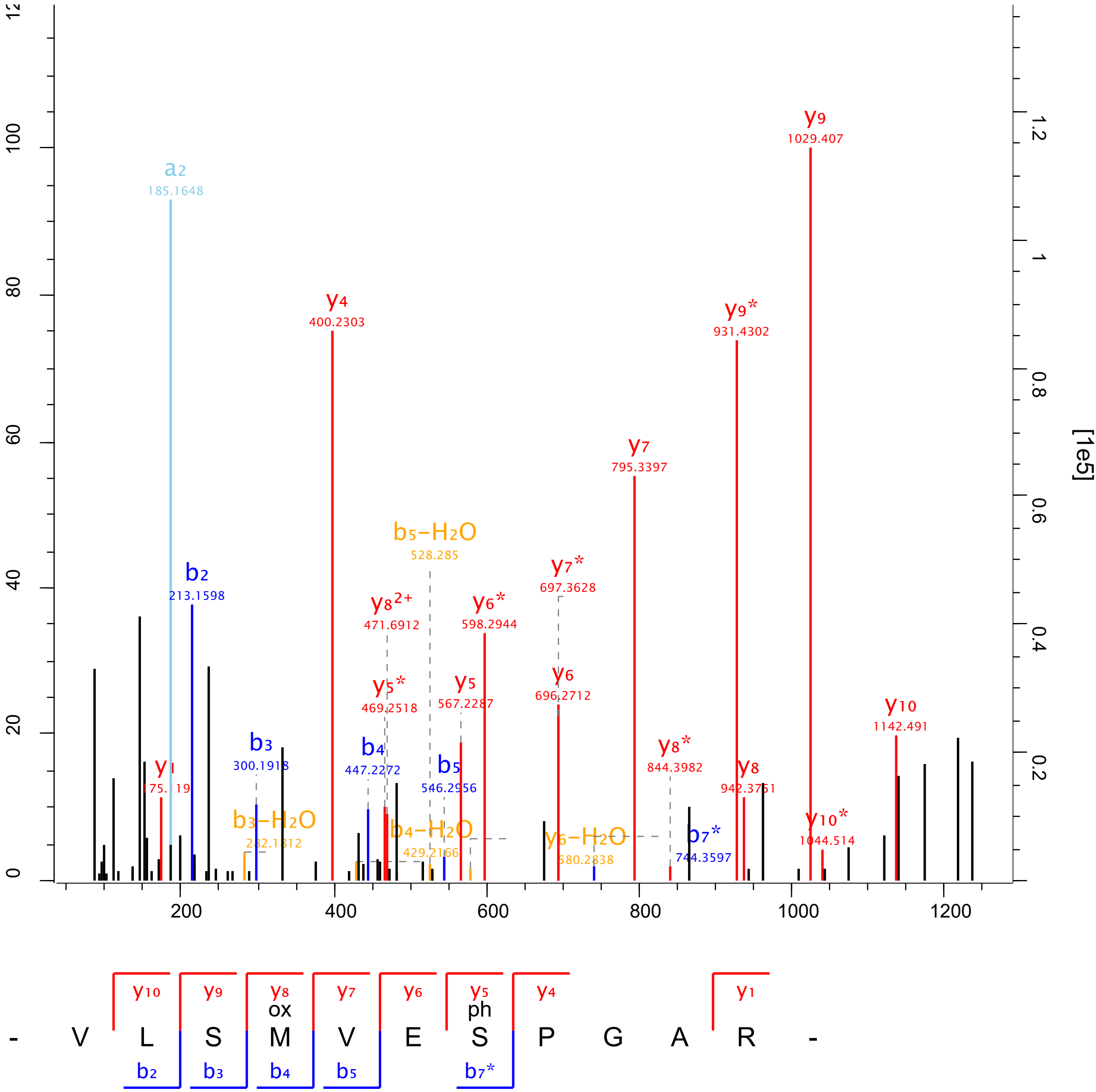

| Raw file | Scan | Method | Score | m/z |
|---|---|---|---|---|
| sys_15_1 | 9647 | FTMS; HCD | 73.93 | 586.75 |

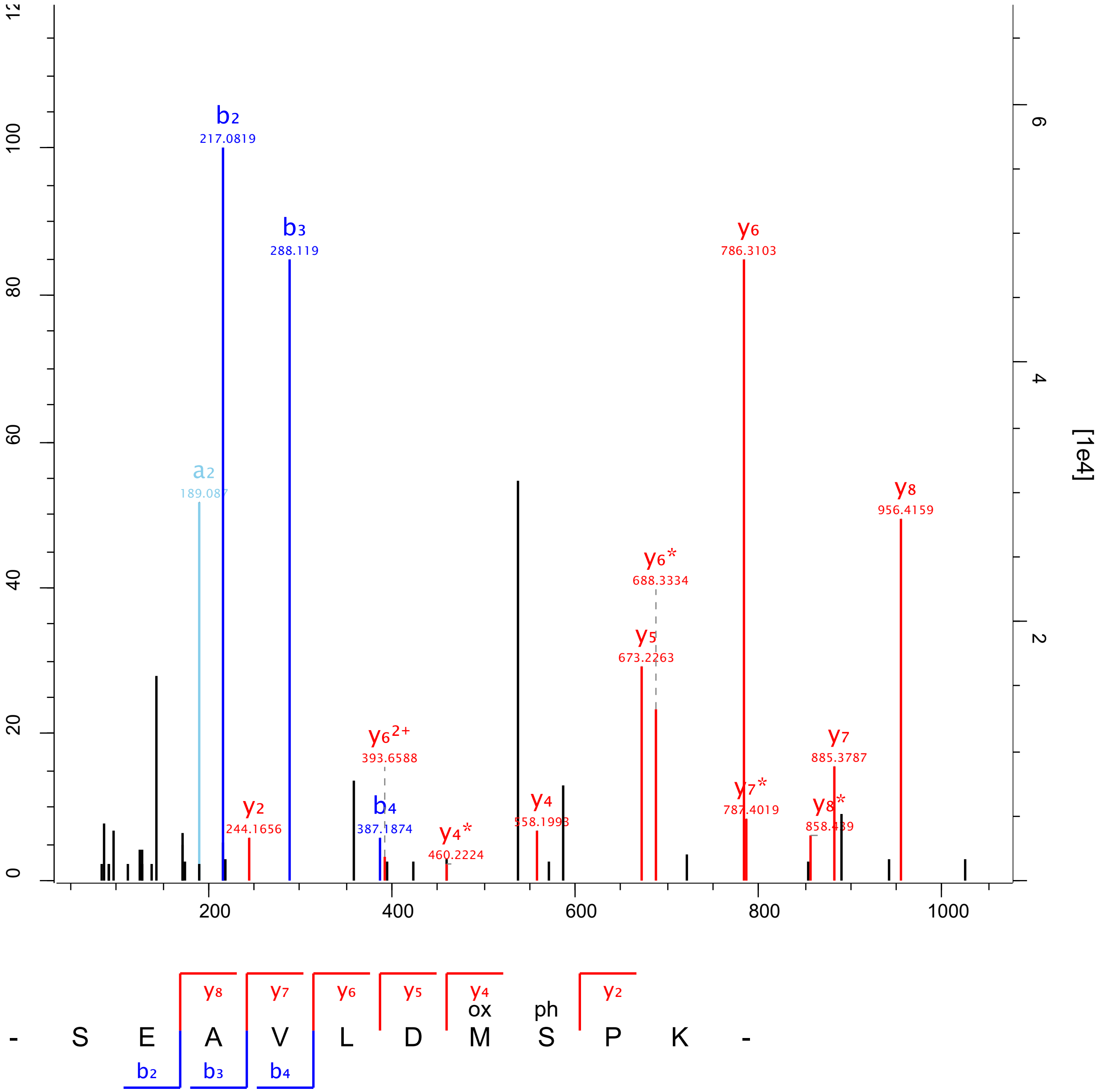

| Raw file | Scan | Method | Score | m/z |
| --- | --- | --- | --- | --- |
| sys_15_1 | 9650 | FTMS; HCD | 63.69 | 690.26 |

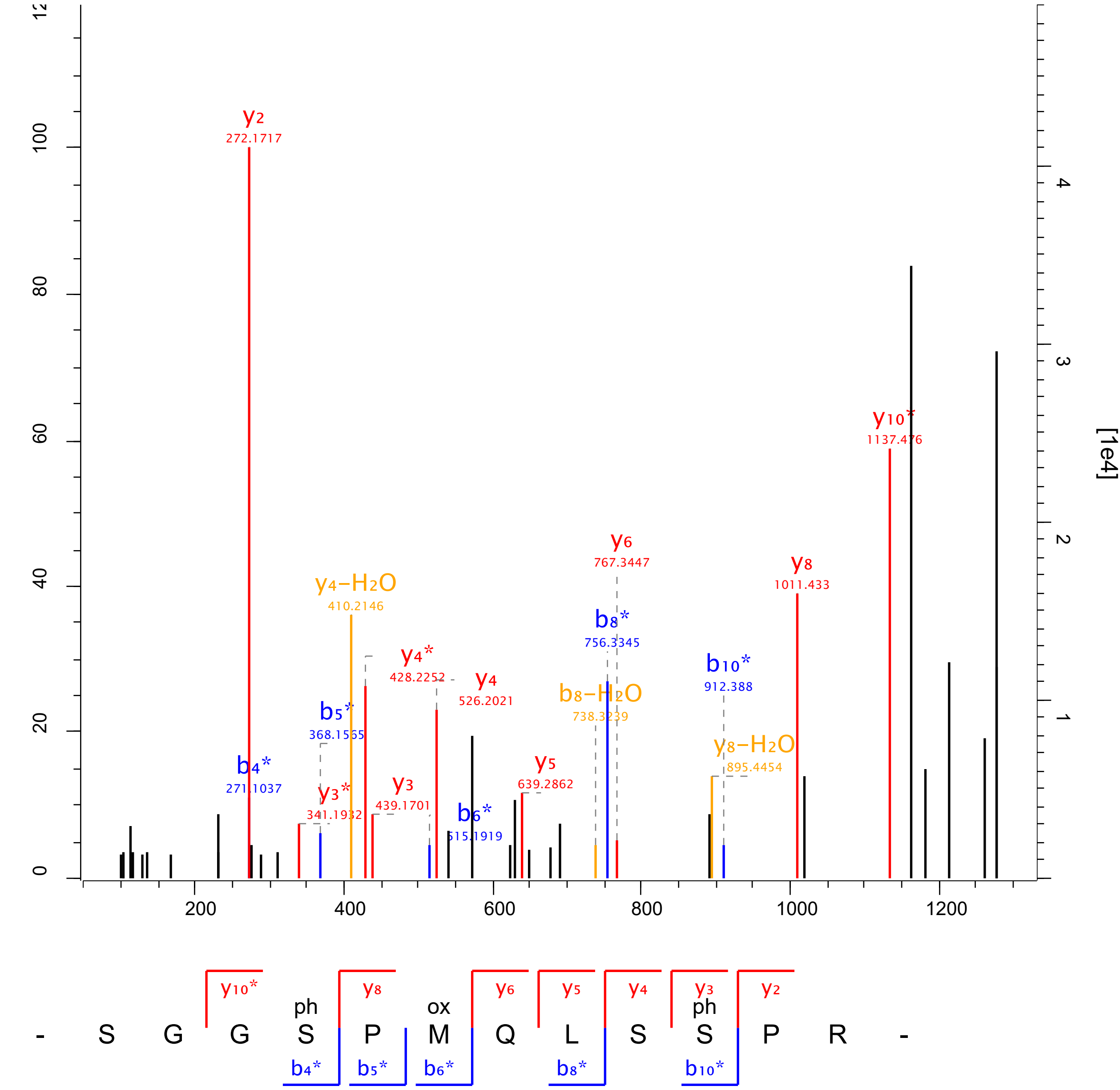

| Raw file | Scan | Method | Score | m/z |
|---|---|---|---|---|
| sys_15_1 | 9755 | FTMS; HCD | 43.03 | 594.25 |

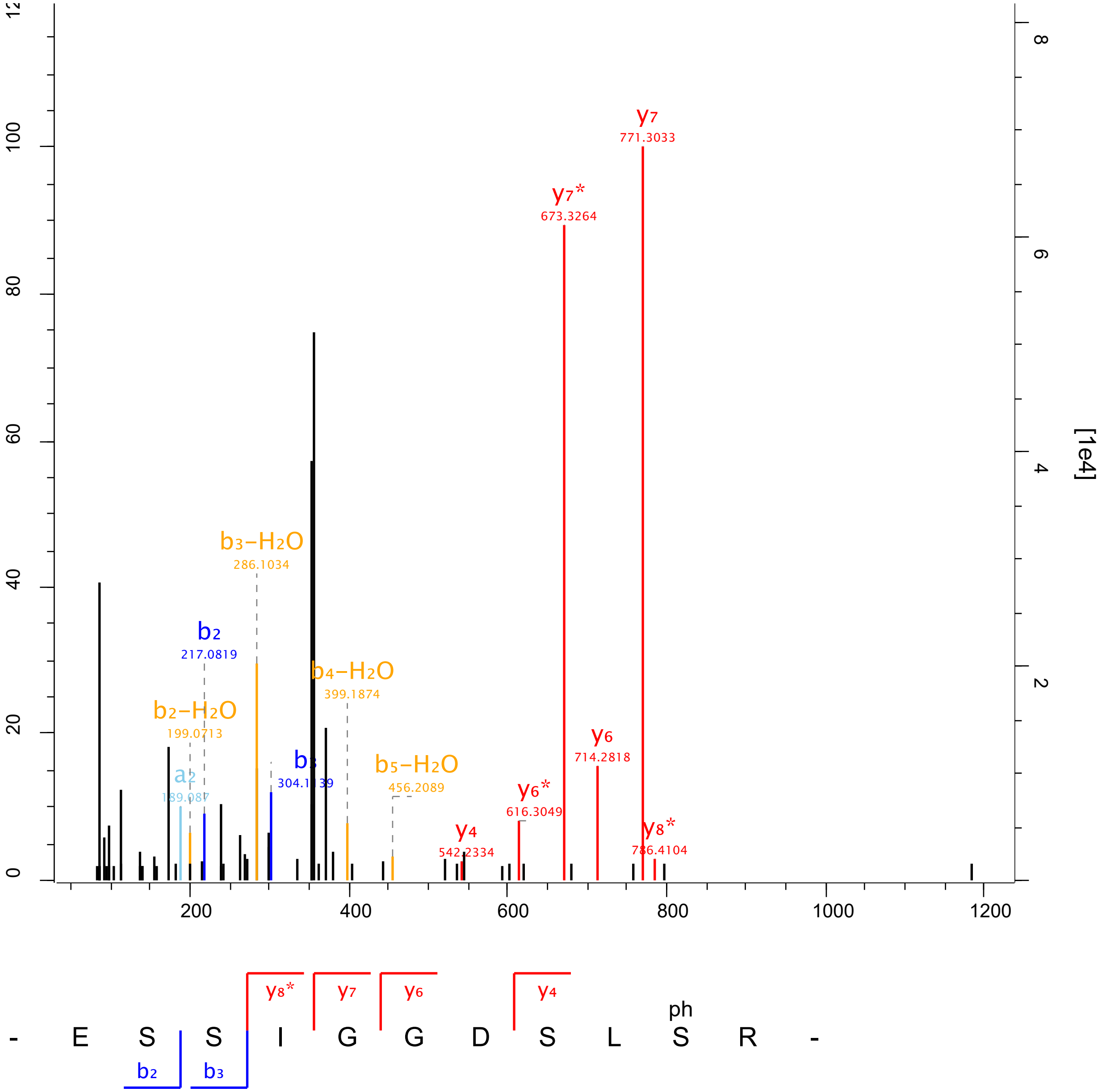

| Raw file | Scan | Method | Score | m/z |
|---|---|---|---|---|
| sys_15_1 | 9763 | FTMS; HCD | 92.38 | 484.87 |

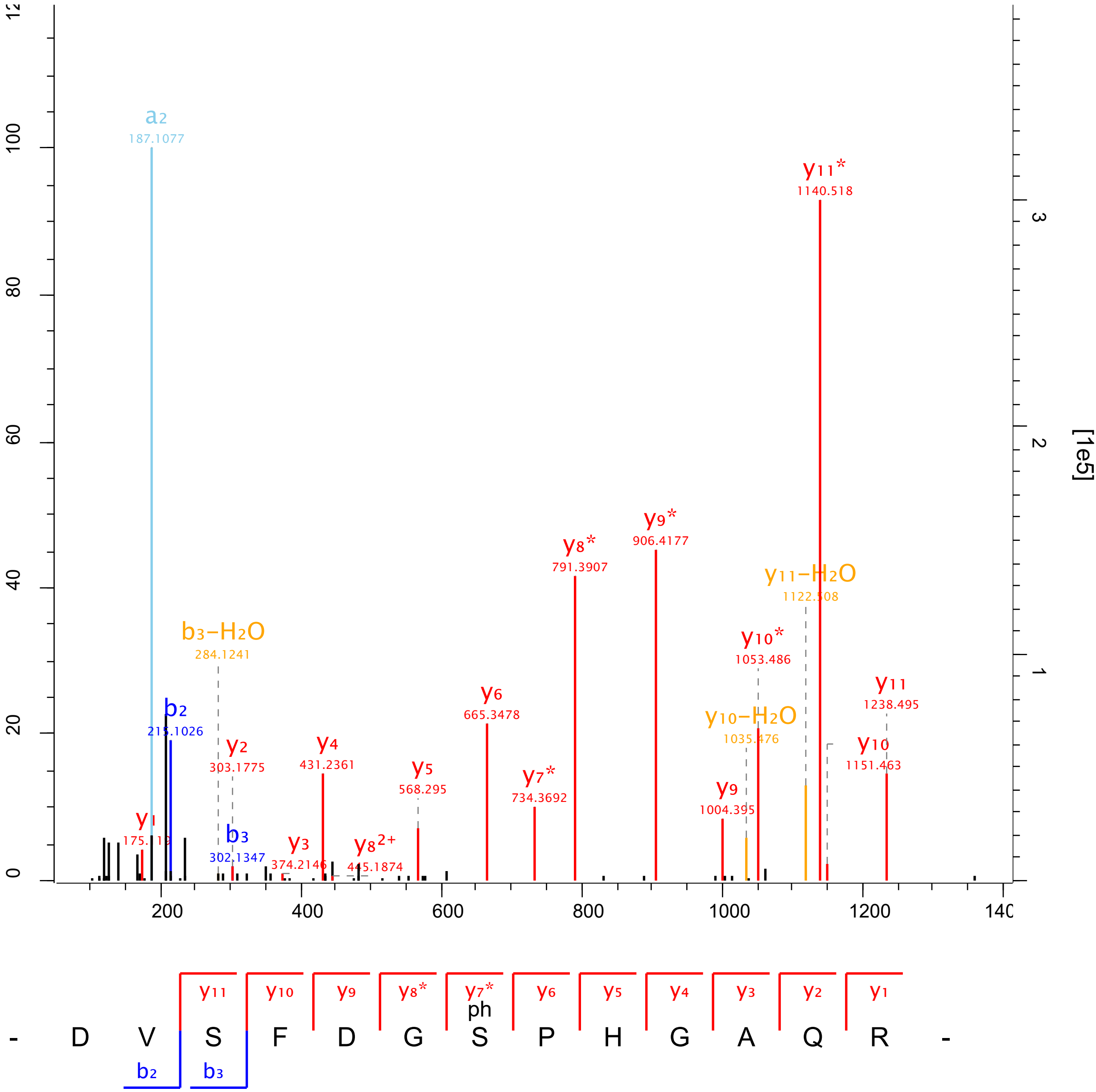

| Raw file | Scan | Method | Score | m/z |
|---|---|---|---|---|
| sys_15_1 | 9783 | FTMS; HCD | 52.17 | 693.3 |

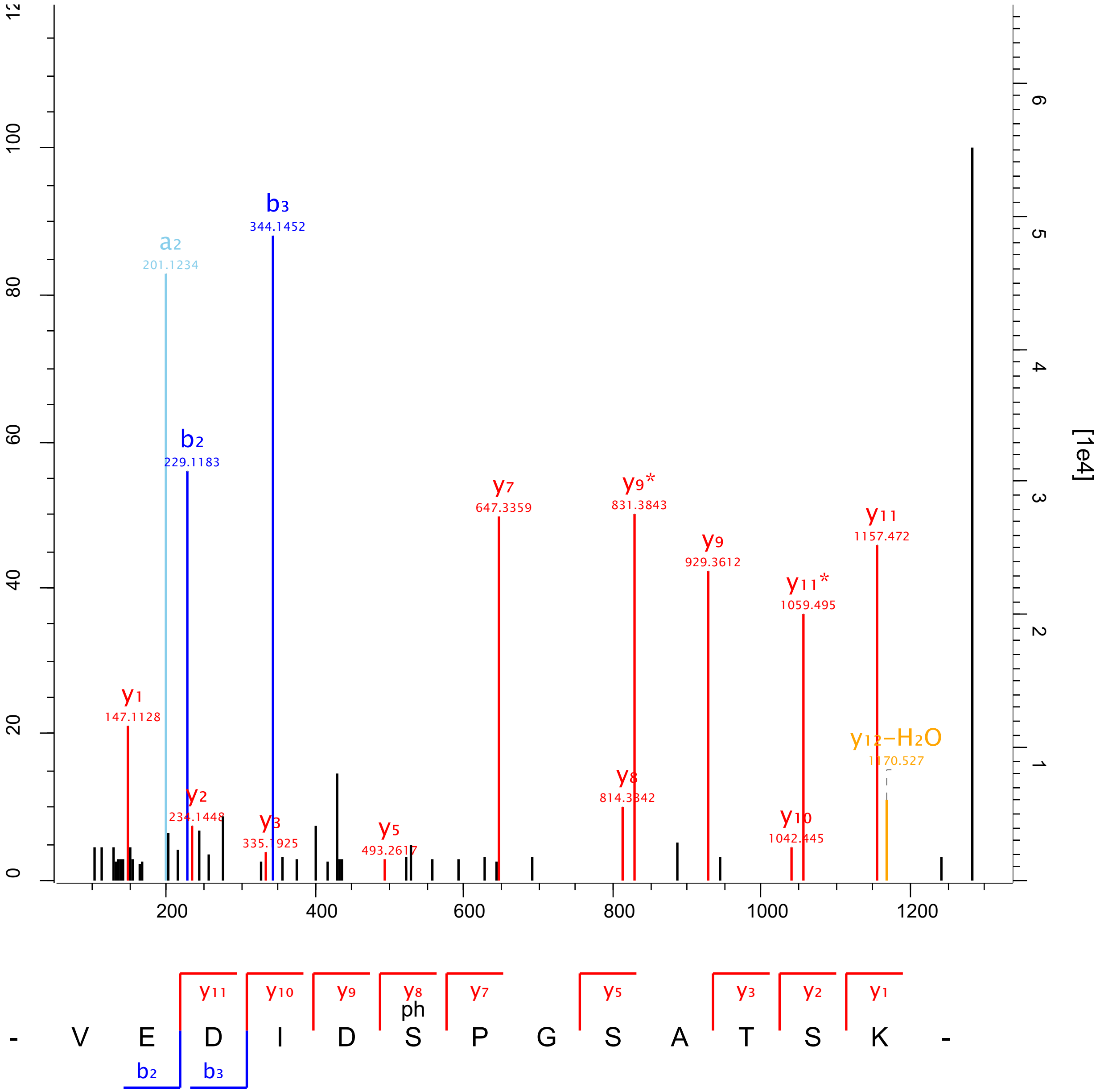

| Raw file | Scan | Method | Score | m/z |
|---|---|---|---|---|
| sys_15_1 | 9788 | FTMS; HCD | 86.21 | 624.6 |

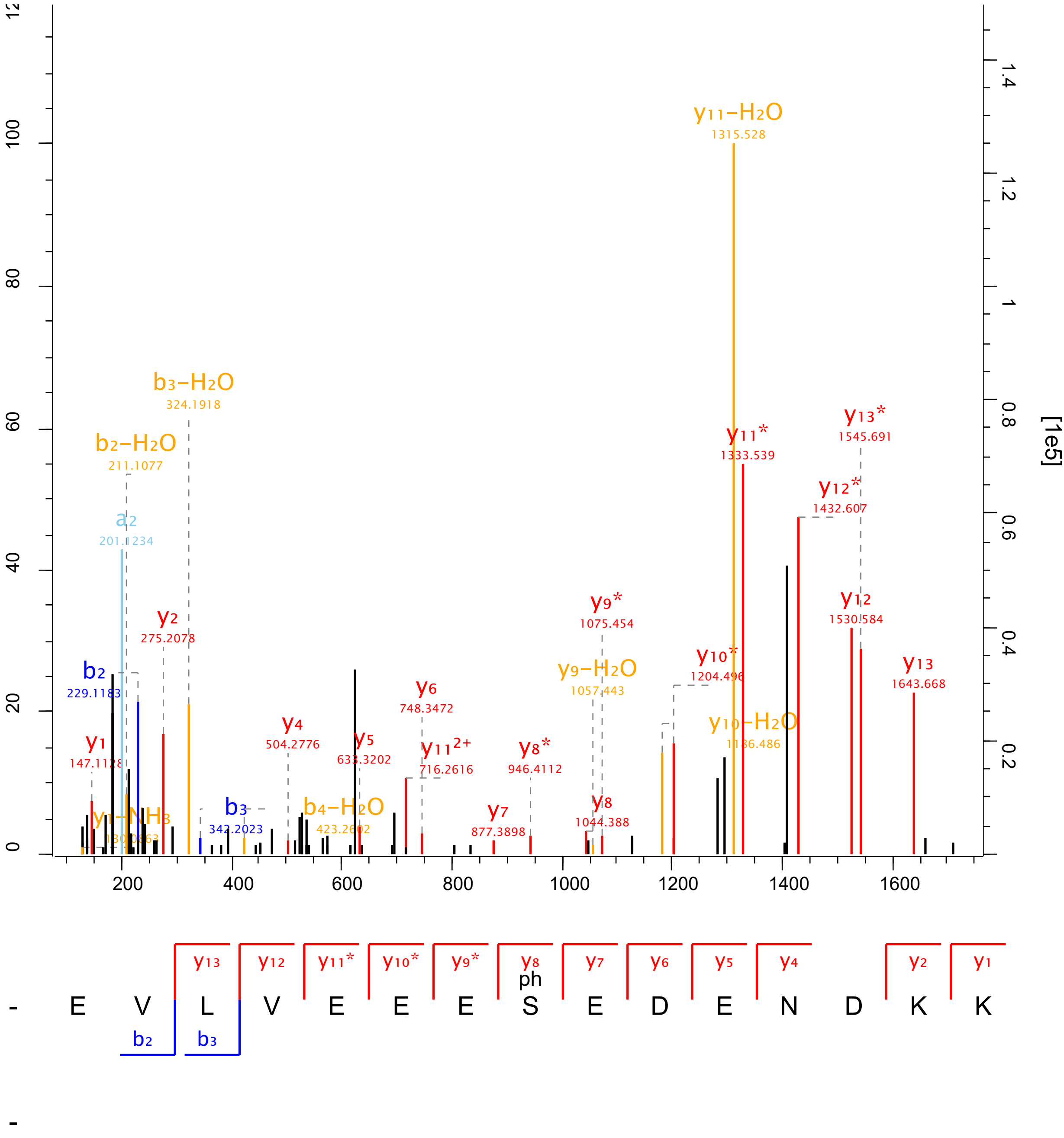

| Raw file | Scan | Method | Score | m/z |
|---|---|---|---|---|
| sys_15_1 | 9832 | FTMS; HCD | 50.04 | 644.79 |

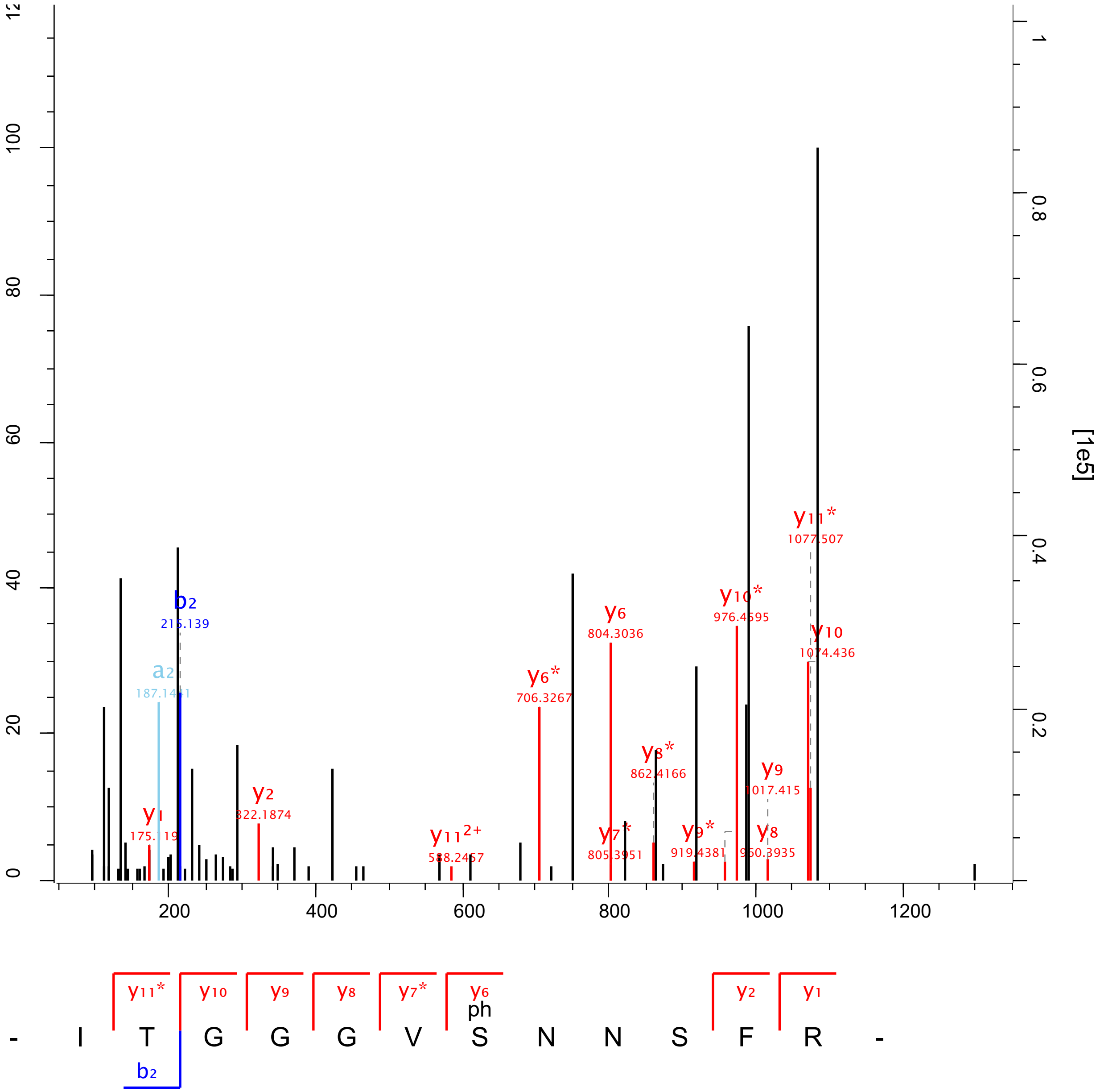

| Raw file | Scan | Method | Score | m/z |
| --- | --- | --- | --- | --- |
| sys_15_1 | 9901 | FTMS; HCD | 80.23 | 418.22 |

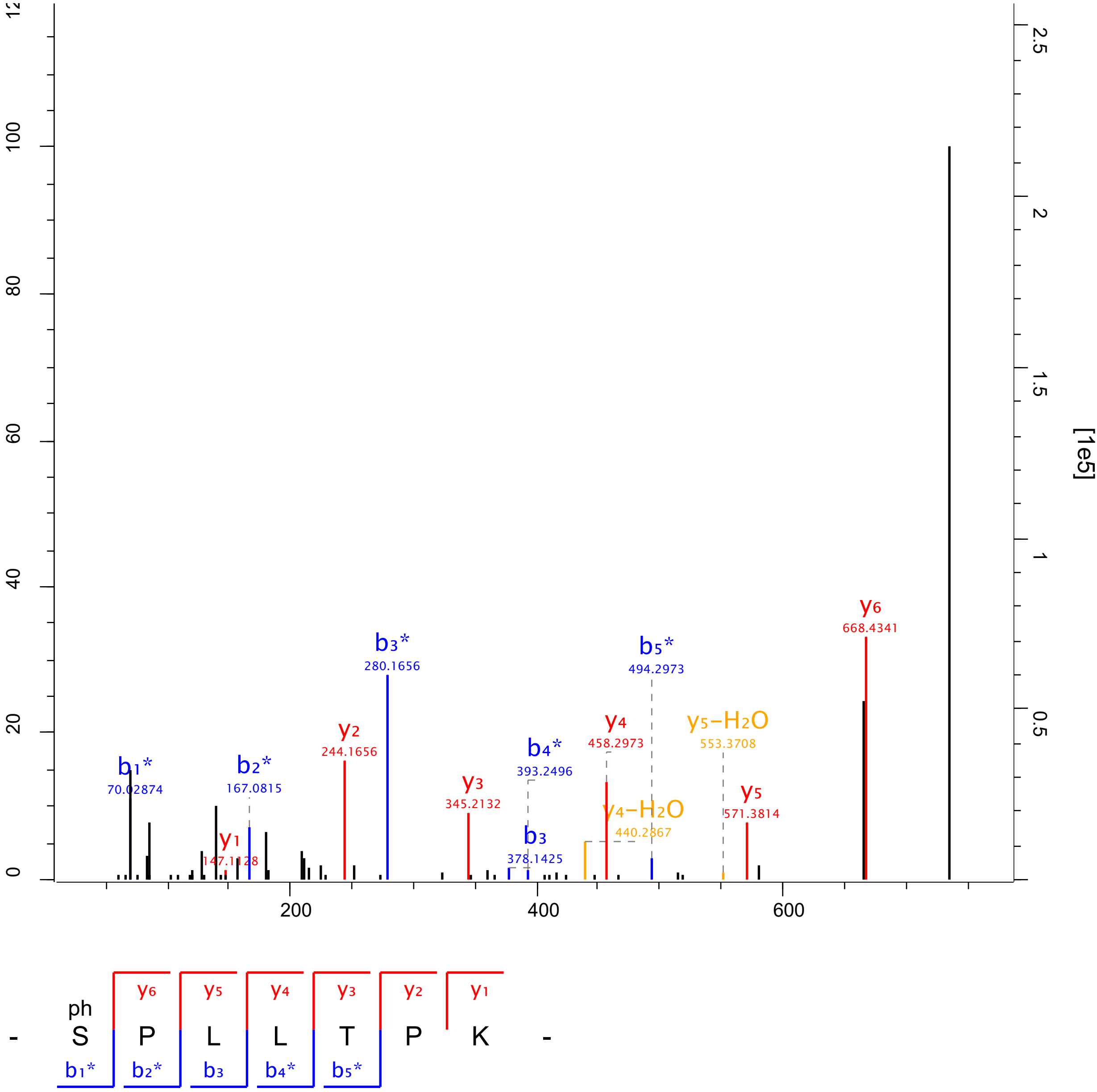

| Raw file | Scan | Method | Score | m/z |
|---|---|---|---|---|
| sys_15_1 | 9961 | FTMS; HCD | 96.49 | 635.26 |

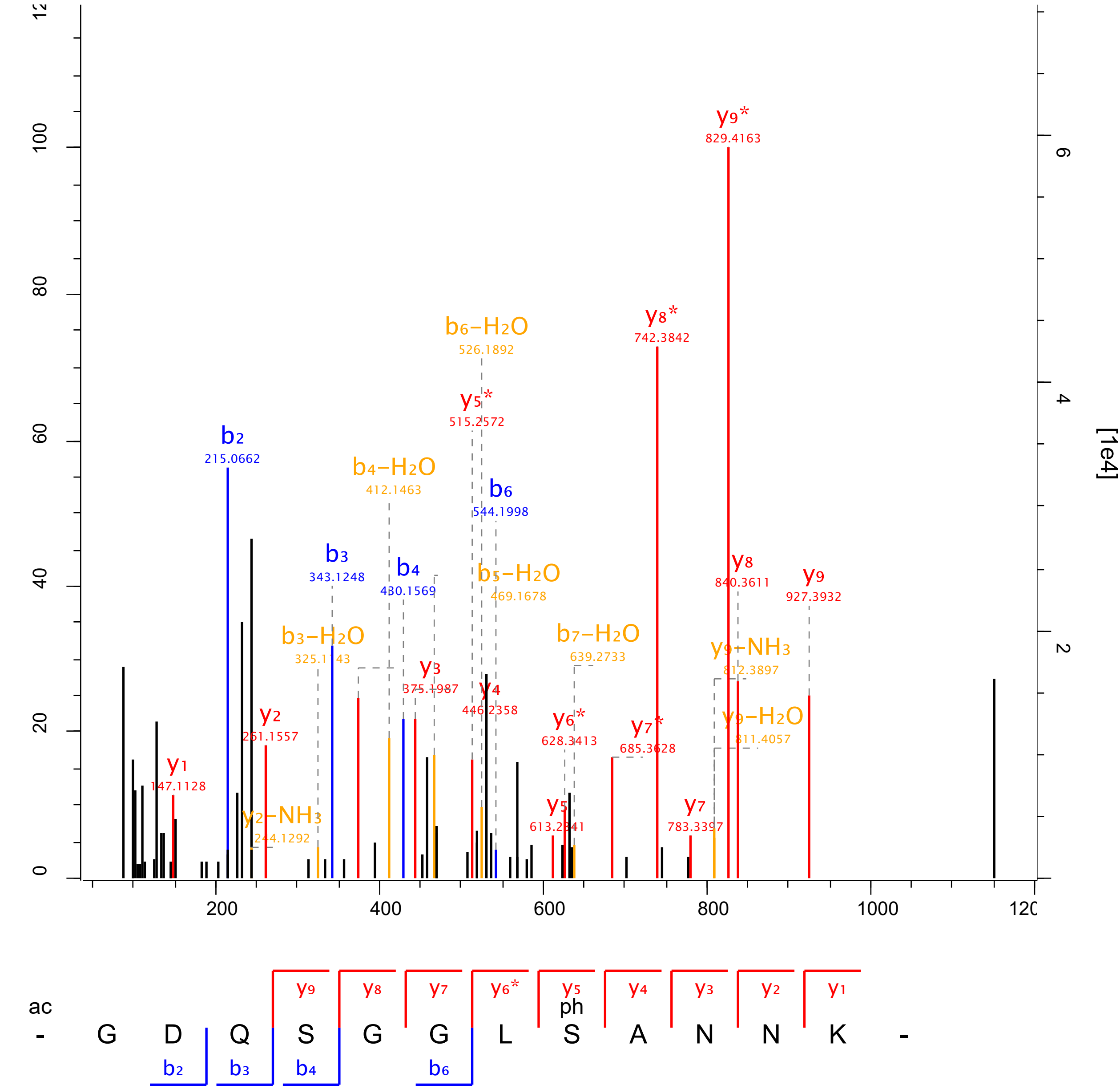

| Raw file | Scan | Method | Score | m/z |
| --- | --- | --- | --- | --- |
| sys_15_1 | 9998 | FTMS; HCD | 79.92 | 707.82 |

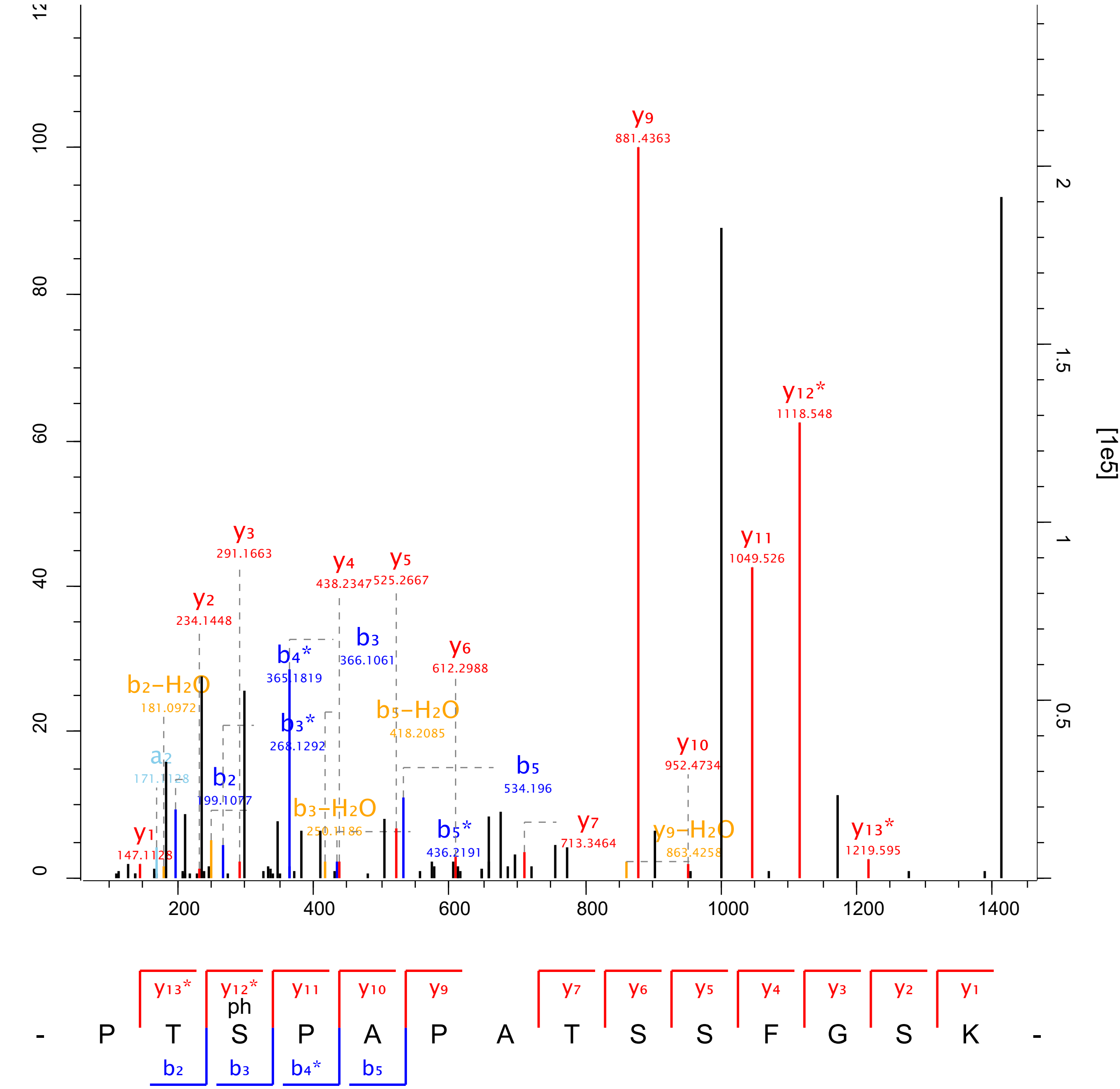

| Raw file | Scan | Method | Score | m/z |
|---|---|---|---|---|
| sys_15_2 | 30107 | FTMS; HCD | 185.25 | 798.87 |

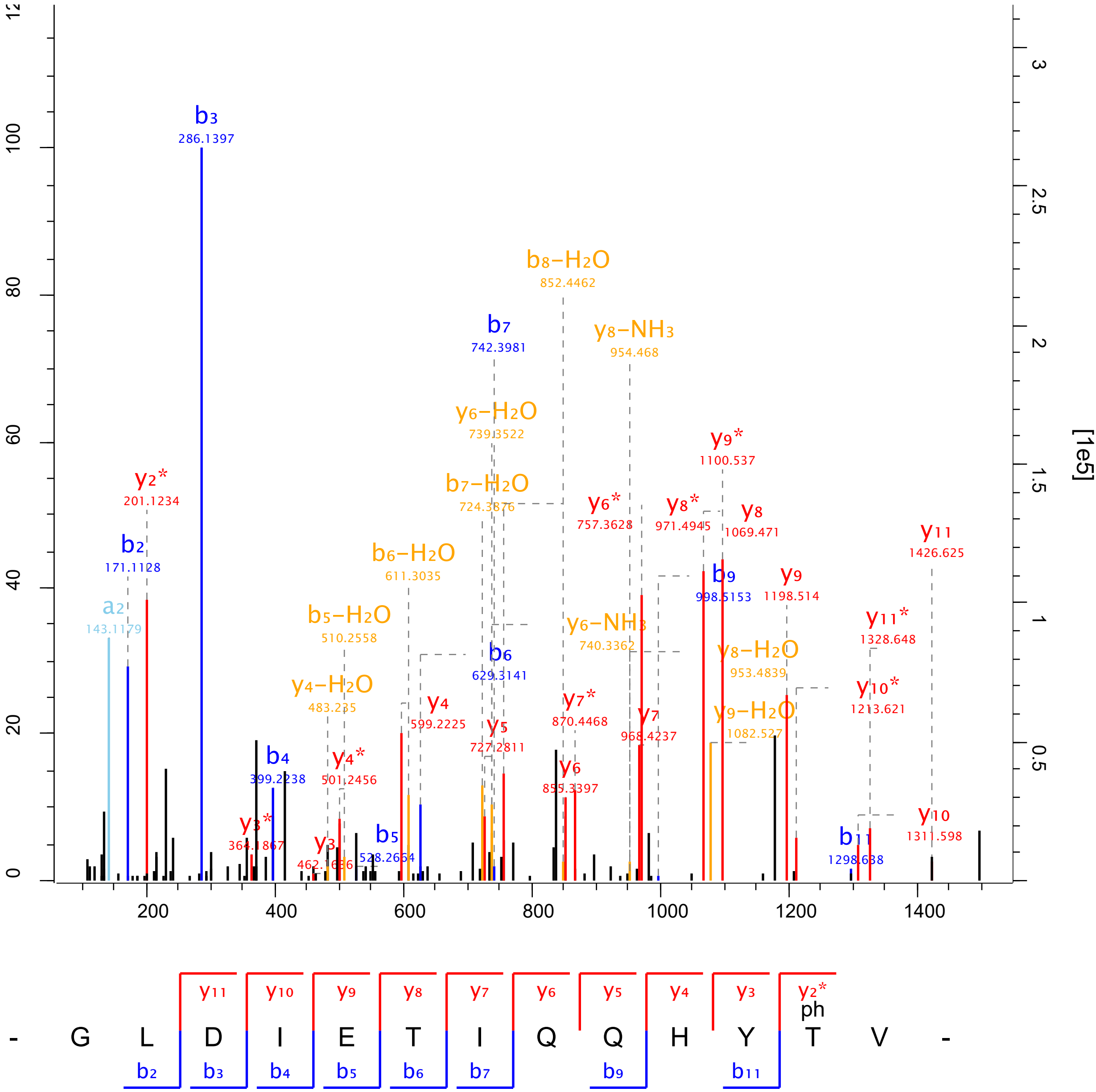

| Raw file | Scan | Method | Score | m/z |
|---|---|---|---|---|
| sys_15_2 | 30707 | FTMS; HCD | 109.53 | 832.7 |

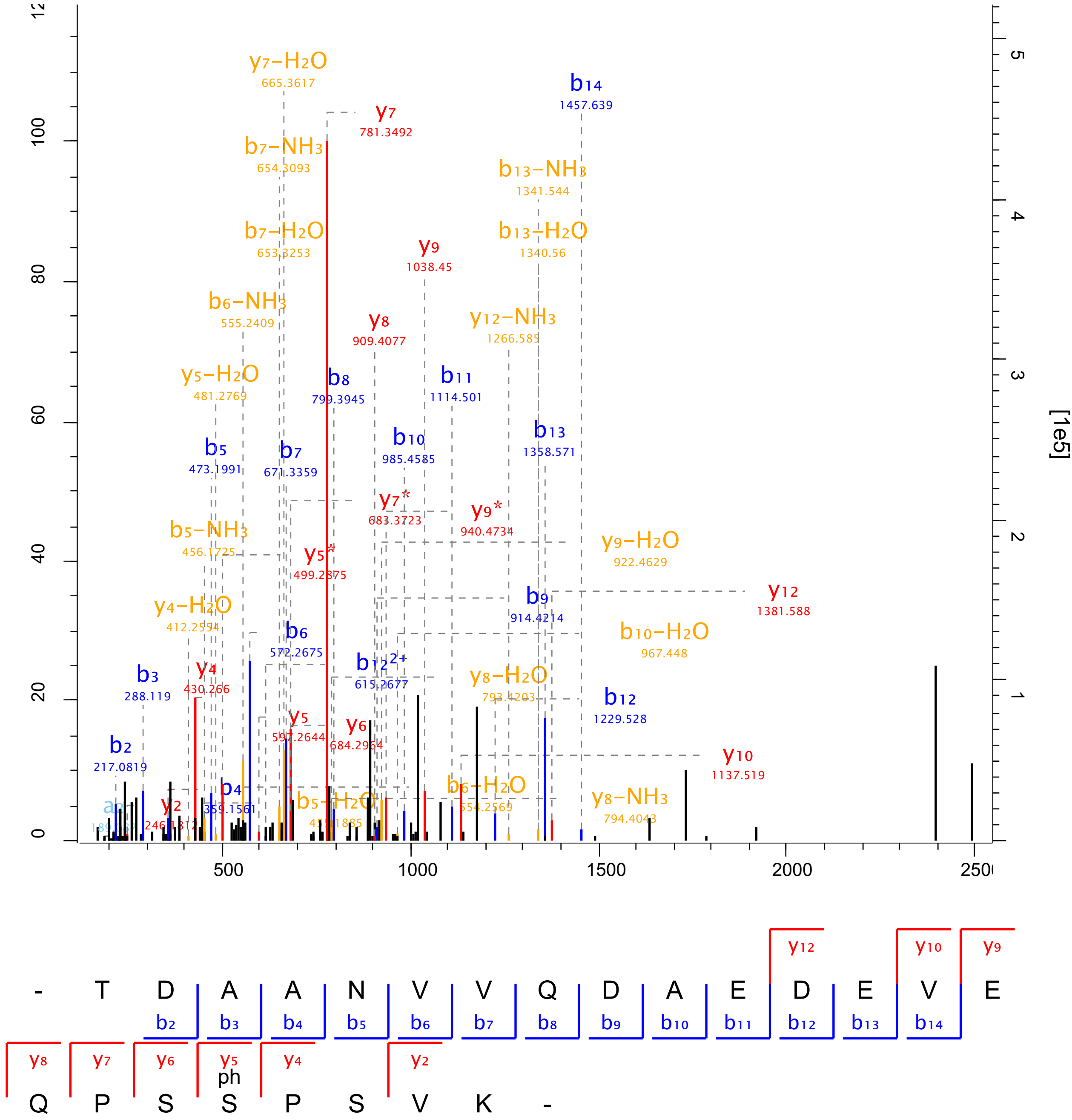

| Raw file | Scan | Method | Score | m/z |
|---|---|---|---|---|
| sys_15_2 | 33237 | FTMS; HCD | 54.47 | 1184.18 |

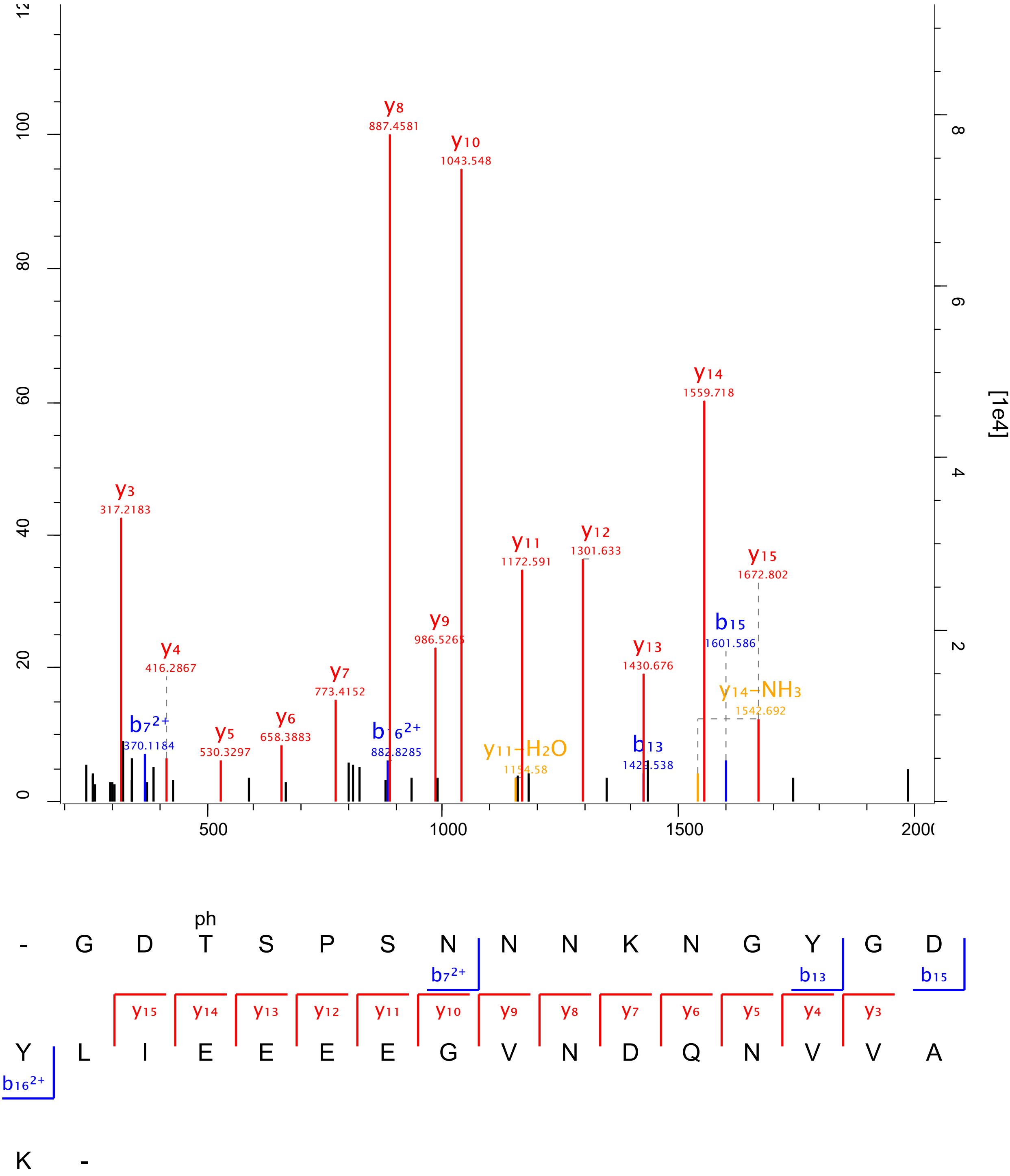

| Raw file | Scan | Method | Score | m/z |
|---|---|---|---|---|
| sys_15_3 | 10504 | FTMS; HCD | 174.59 | 692.3 |

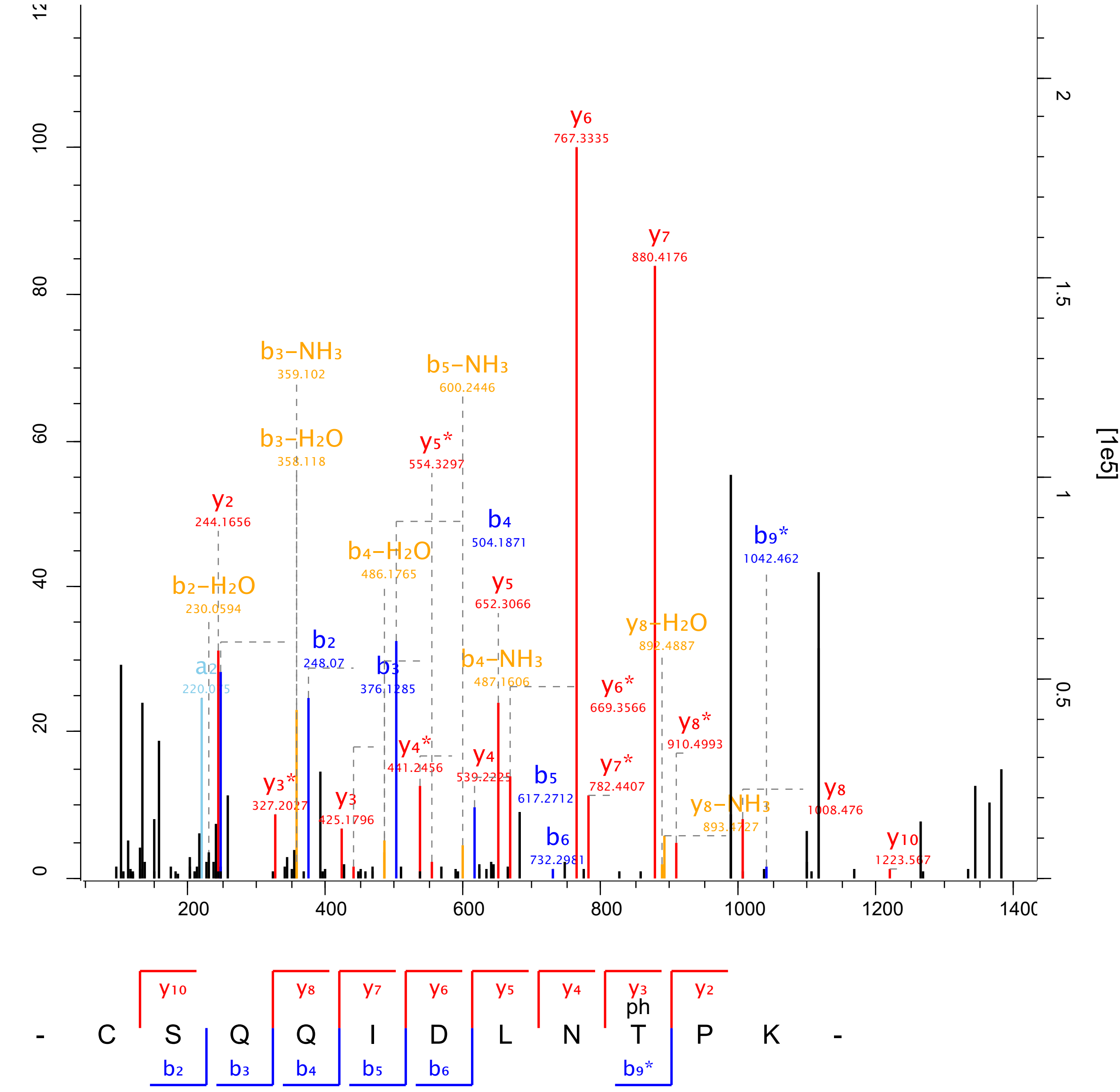

| Raw file | Scan | Method | Score | m/z |
| --- | --- | --- | --- | --- |
| sys_15_3 | 10529 | FTMS; HCD | 100.02 | 450.2 |

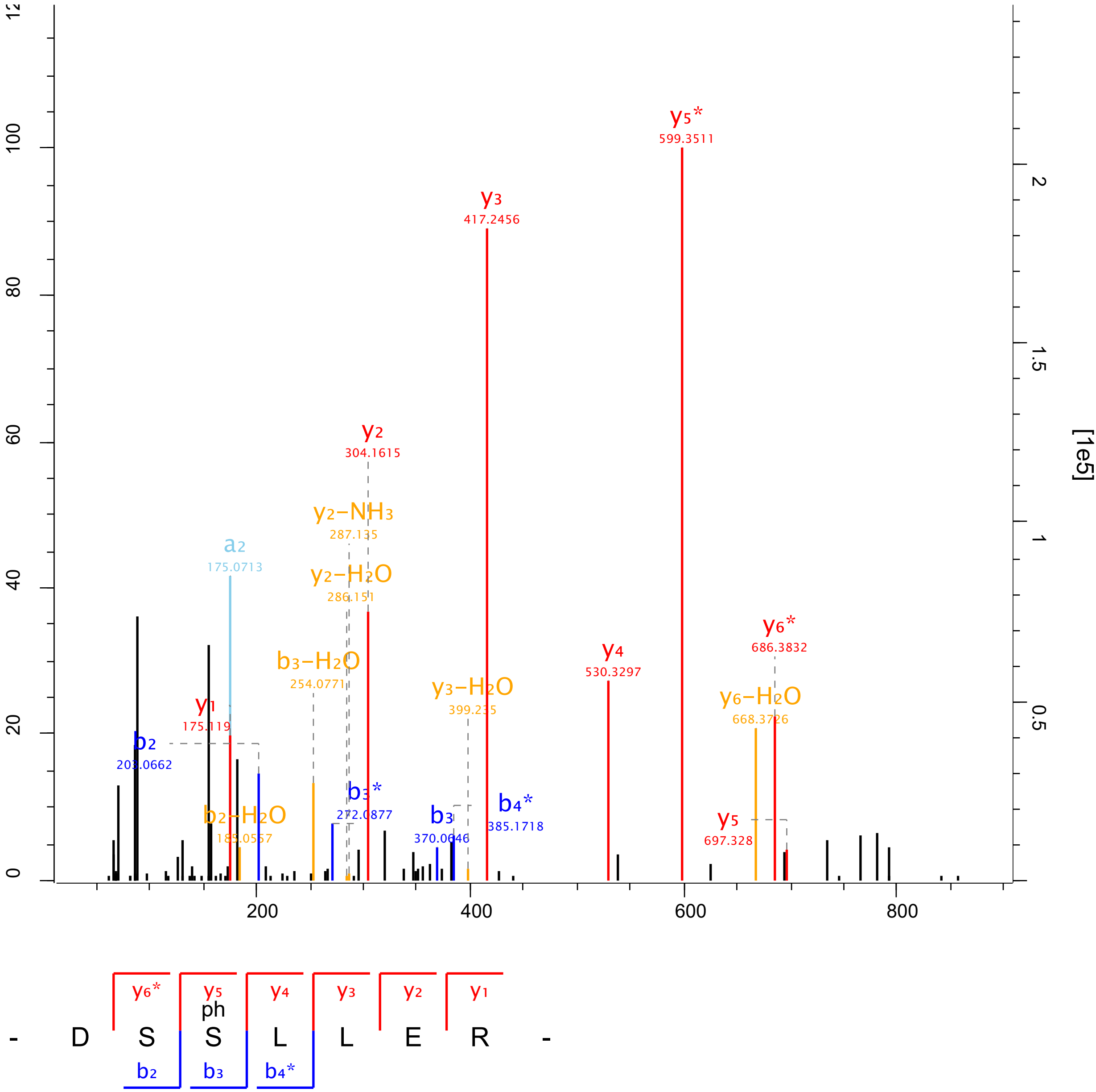

| Raw file | Scan | Method | Score | m/z |
|---|---|---|---|---|
| sys_15_3 | 10634 | FTMS; HCD | 119.74 | 492.24 |

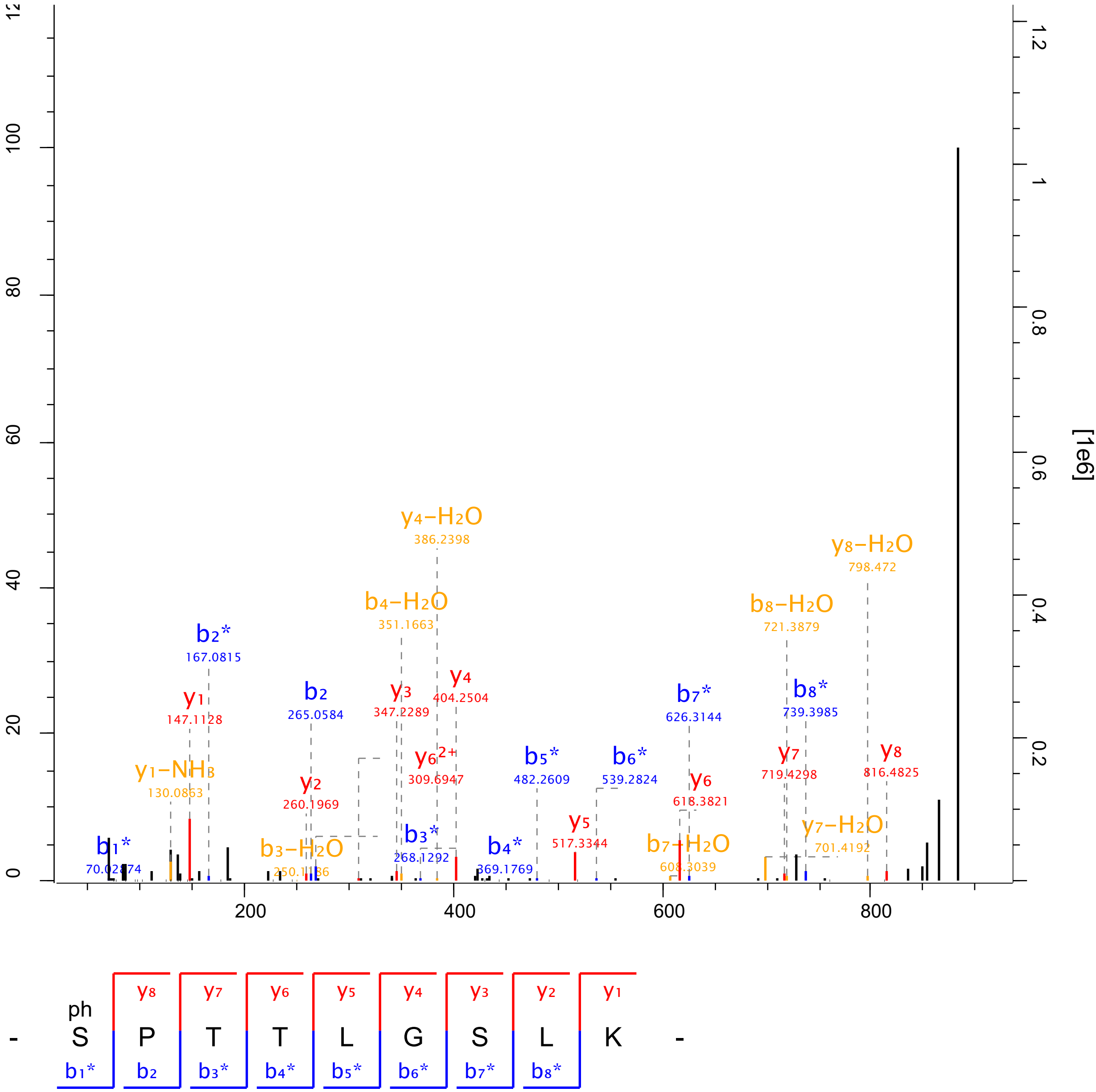

| Raw file | Scan | Method | Score | m/z |
| --- | --- | --- | --- | --- |
| sys_15_3 | 10648 | FTMS; HCD | 302.34 | 763.82 |

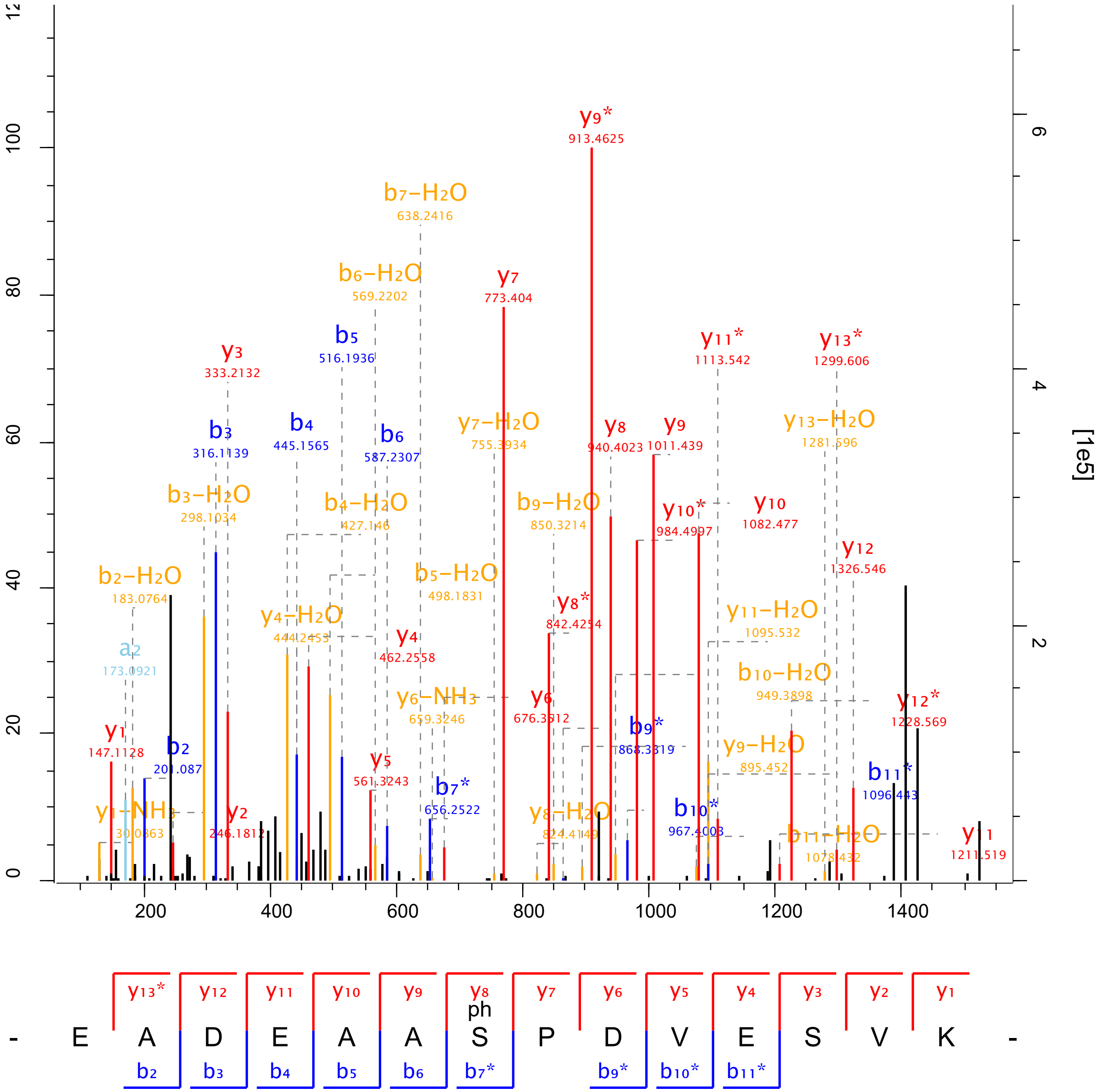

| Raw file | Scan | Method | Score | m/z |
|---|---|---|---|---|
| sys_15_3 | 11470 | FTMS; HCD | 60.55 | 570.75 |

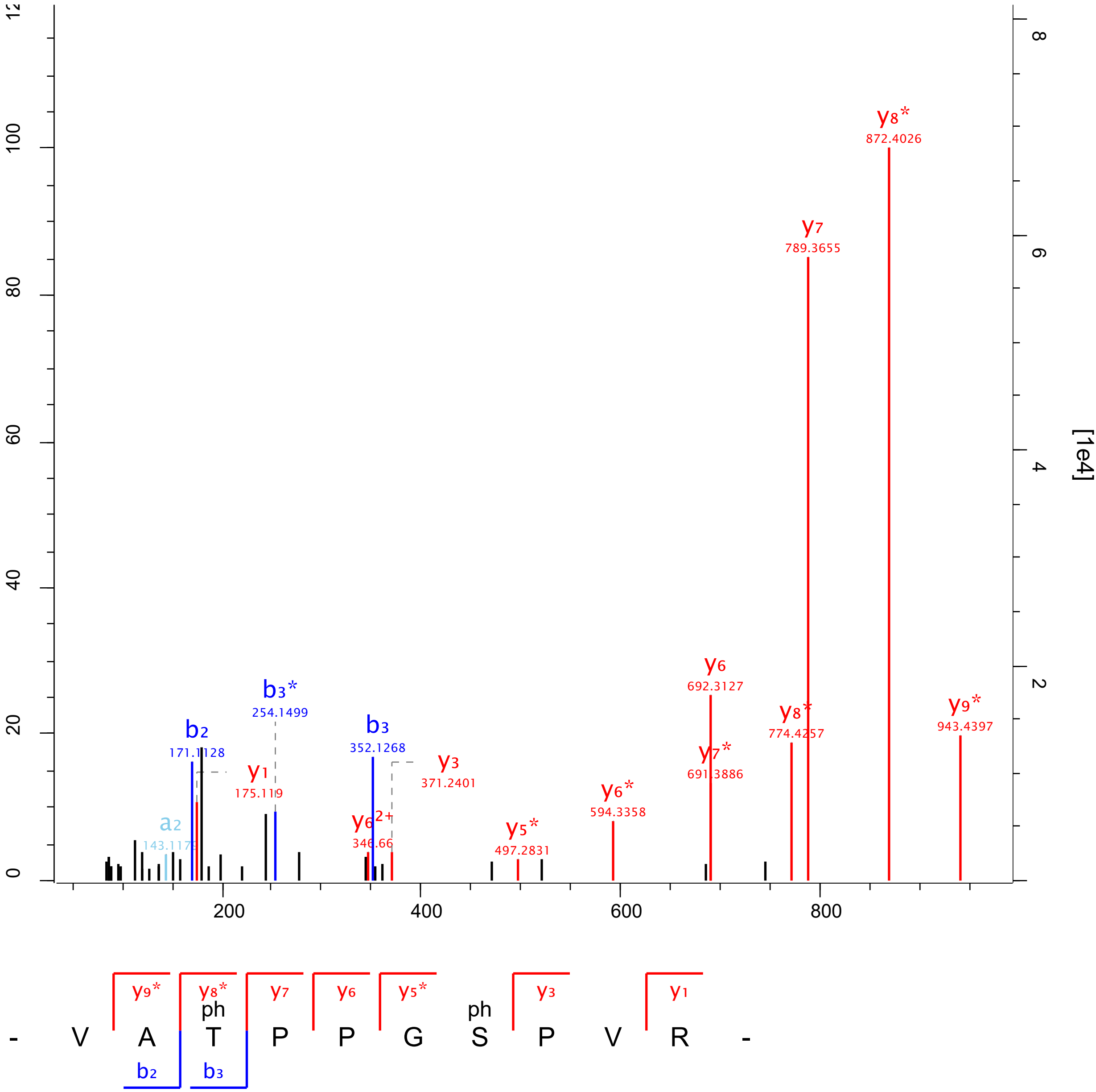

| Raw file | Scan | Method | Score | m/z |
|---|---|---|---|---|
| sys_15_3 | 11940 | FTMS; HCD | 48.4 | 598.8 |

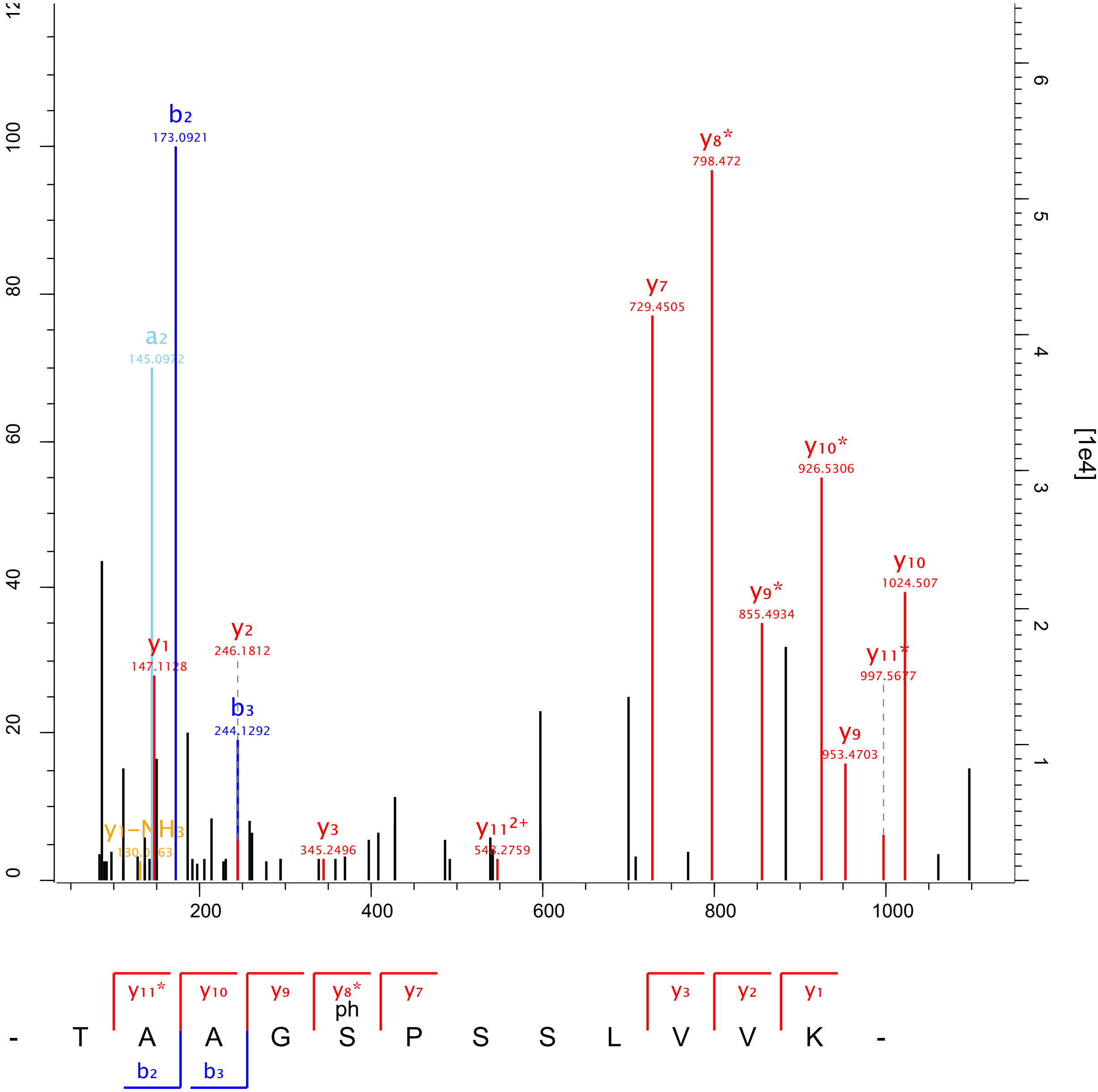

| Raw file | Scan | Method | Score | m/z |
| --- | --- | --- | --- | --- |
| sys_15_3 | 12055 | FTMS; HCD | 45.28 | 583.27 |

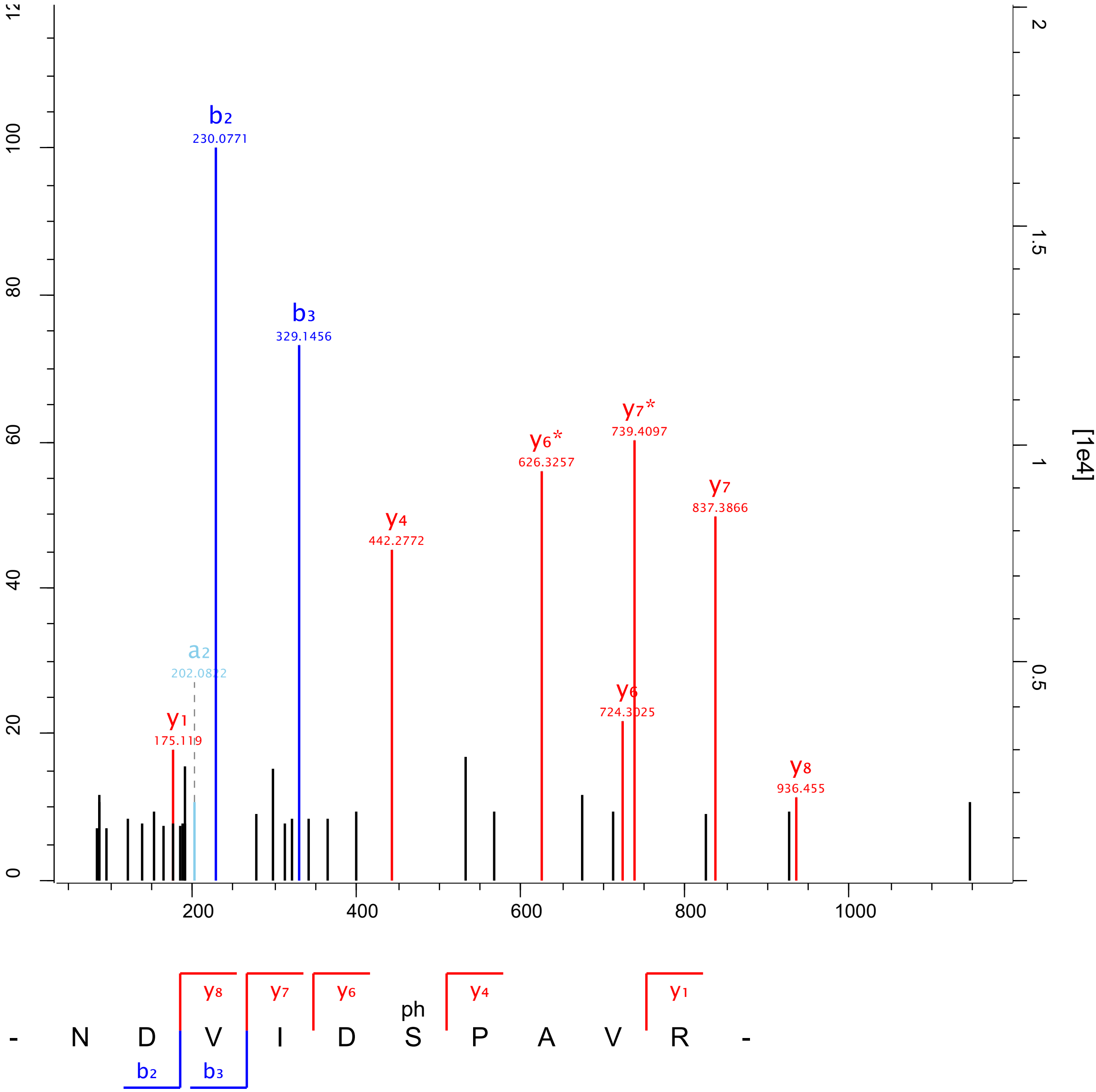

| Raw file | Scan | Method | Score | m/z |
|---|---|---|---|---|
| sys_15_3 | 12963 | FTMS; HCD | 48.95 | 568.21 |

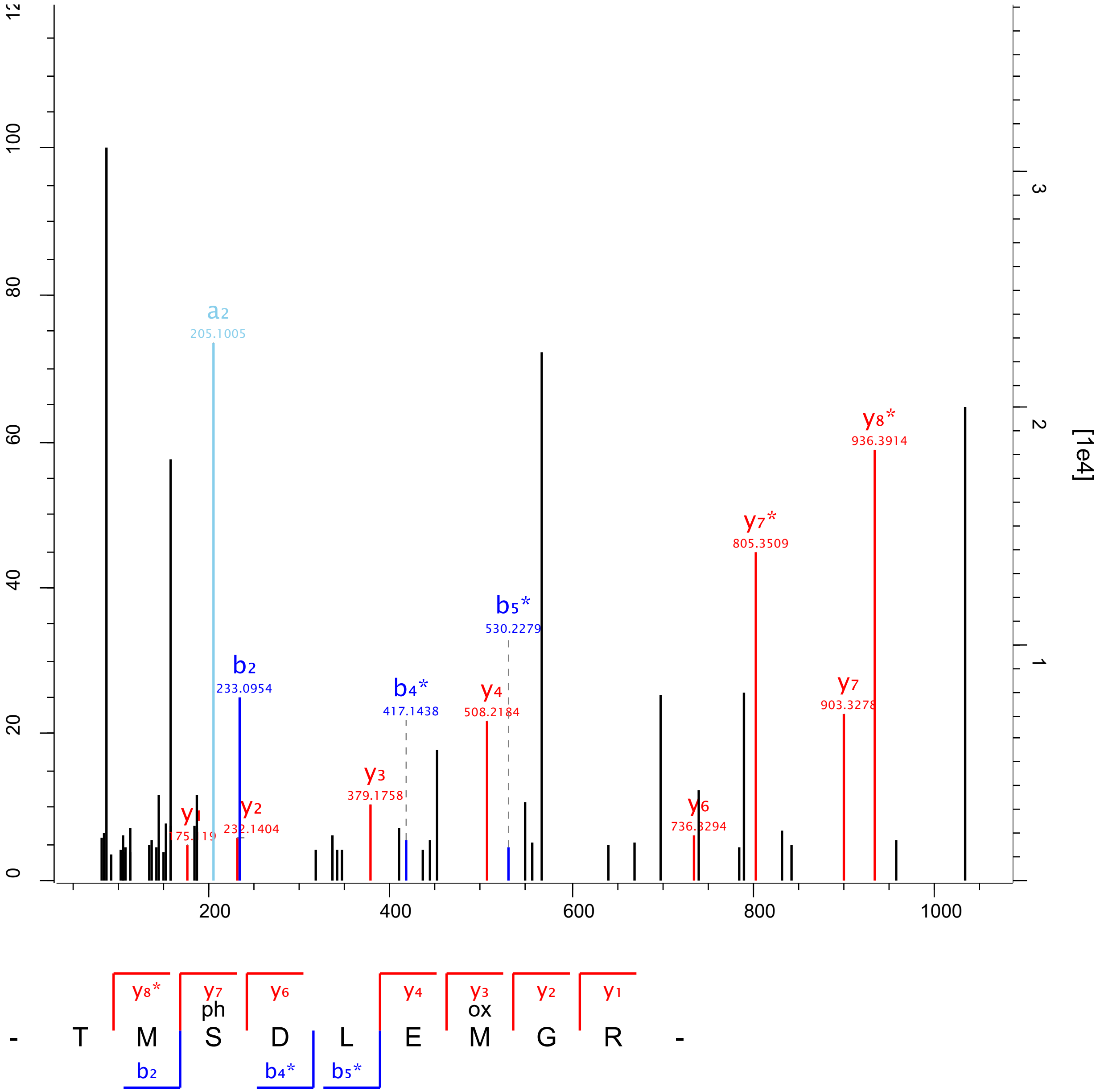

| Raw file | Scan | Method | Score | m/z |
| --- | --- | --- | --- | --- |
| sys_15_3 | 14088 | FTMS; HCD | 71.18 | 551.75 |

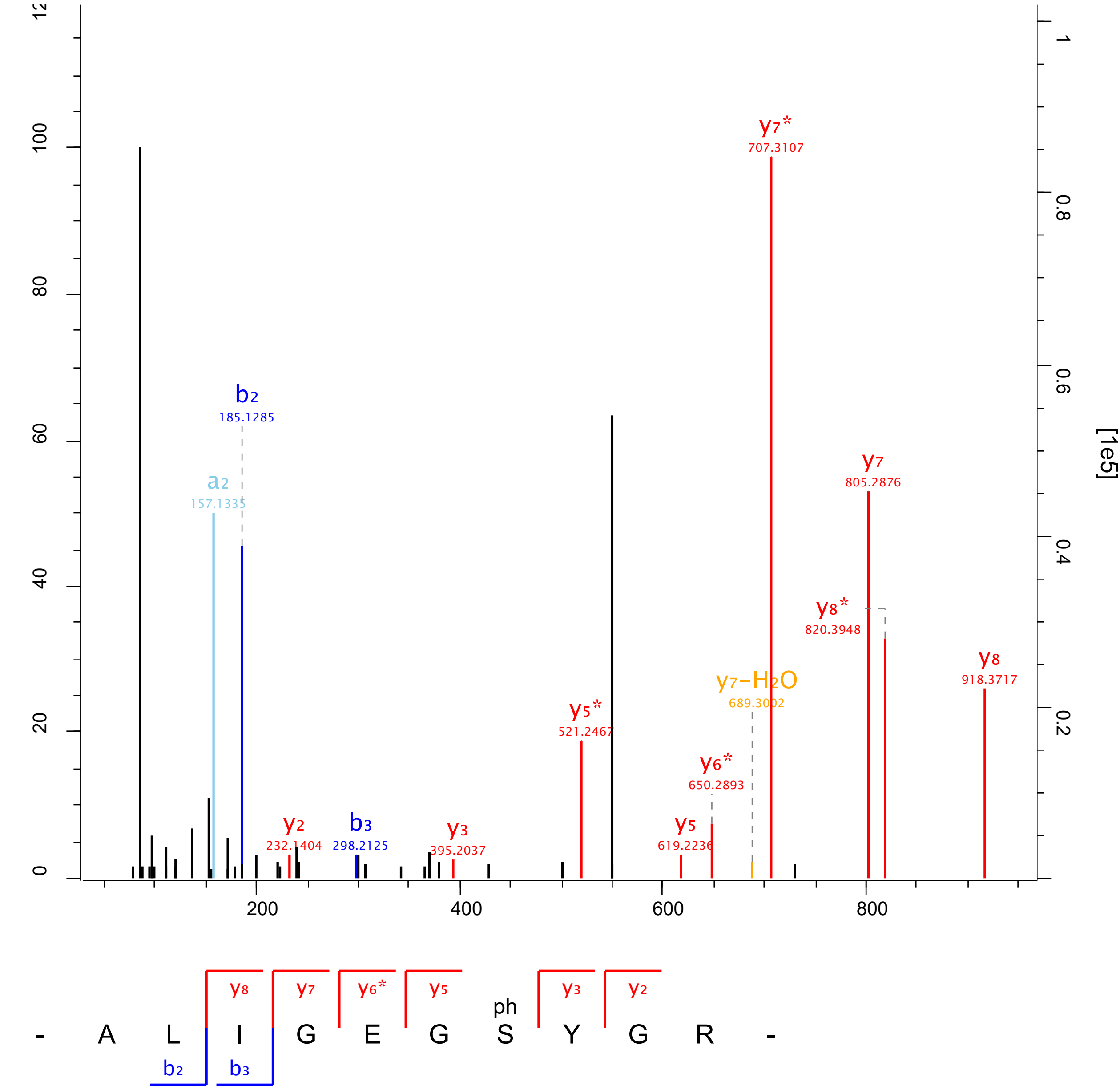

| Raw file | Scan | Method | Score | m/z |
|---|---|---|---|---|
| sys_15_3 | 14270 | FTMS; HCD | 42.98 | 521.22 |

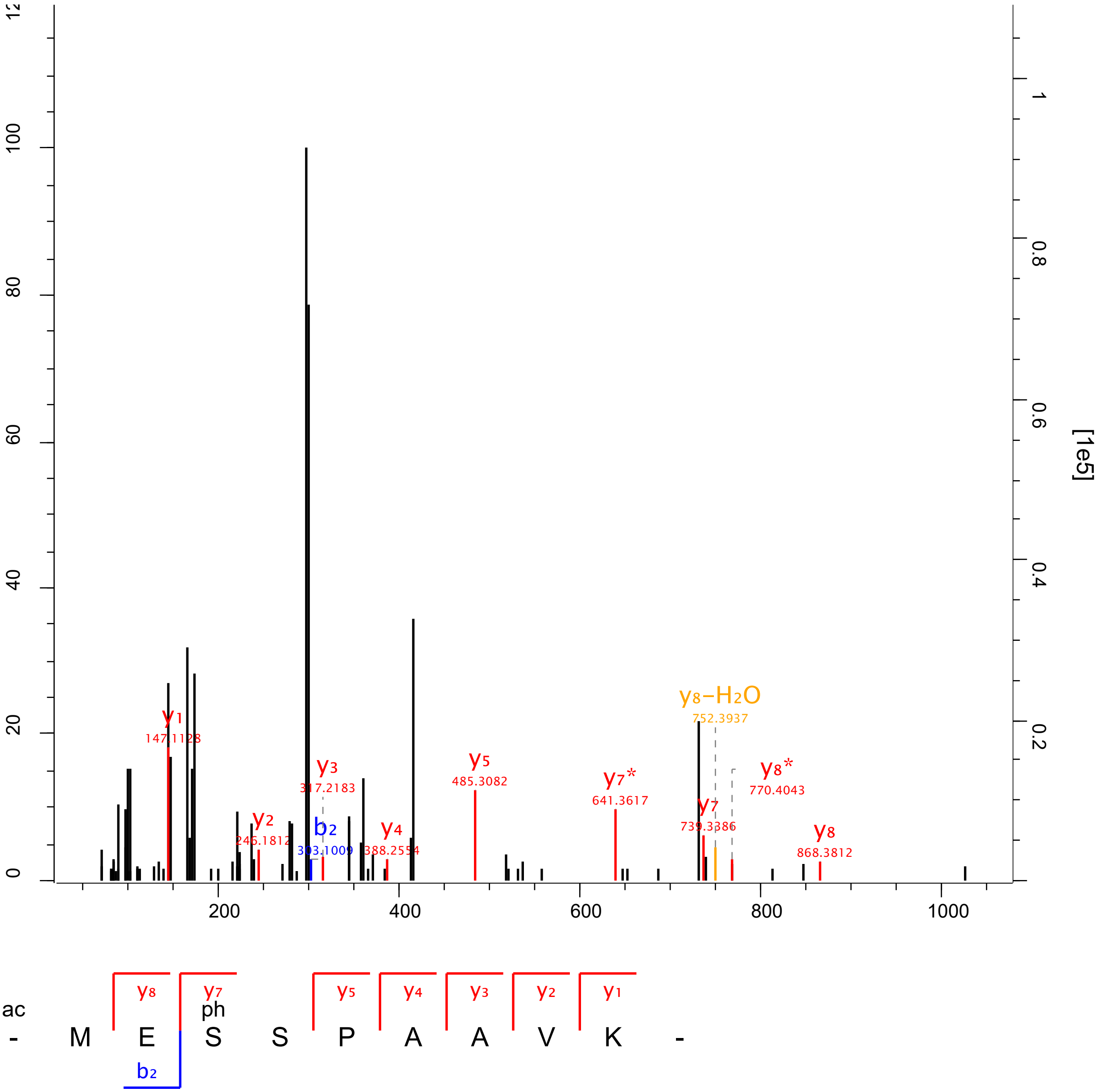

| Raw file | Scan | Method | Score | m/z |
|---|---|---|---|---|
| sys_15_3 | 14303 | FTMS; HCD | 119.62 | 437.2 |

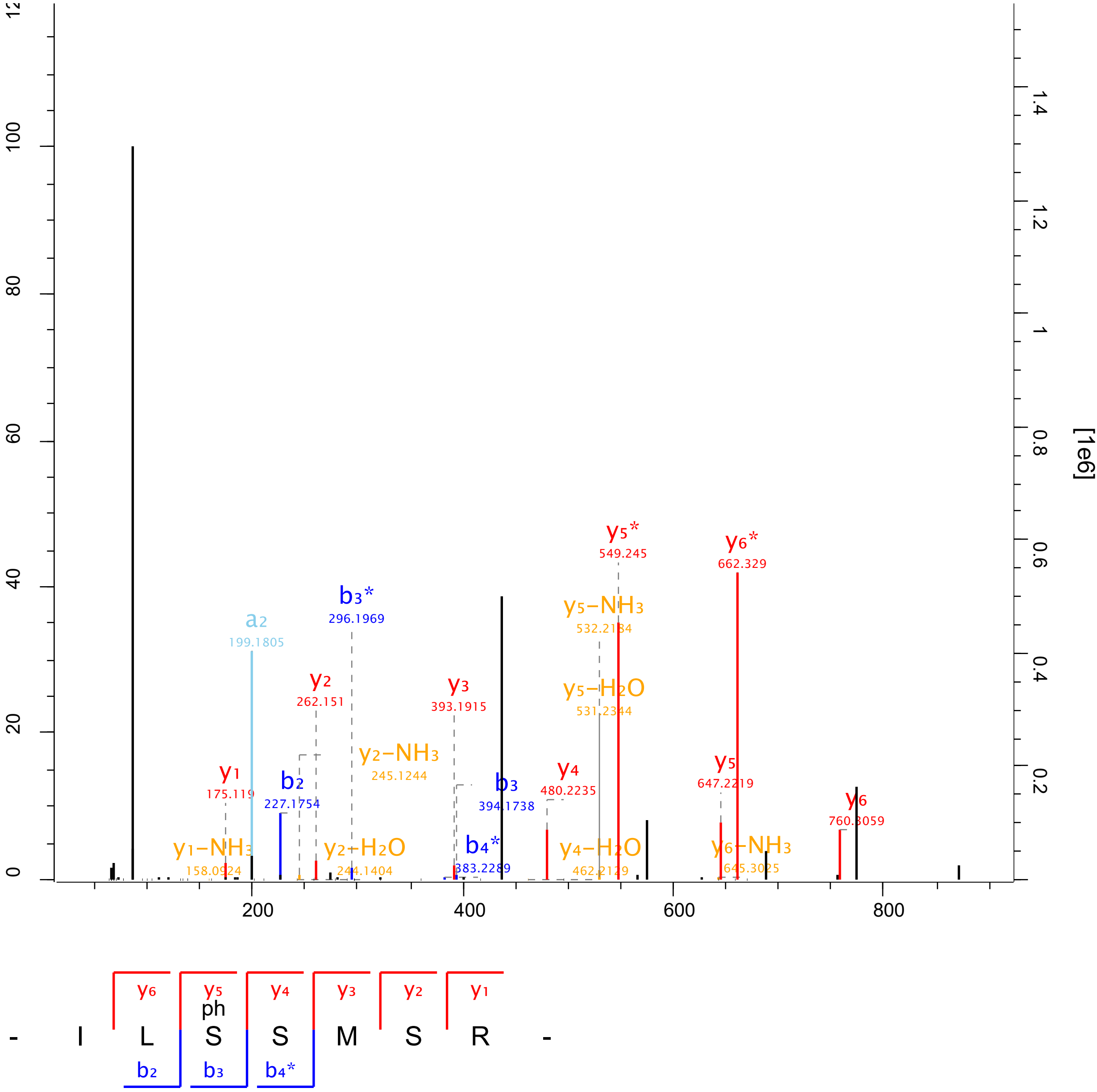

| Raw file | Scan | Method | Score | m/z |
|---|---|---|---|---|
| sys_15_3 | 15358 | FTMS; HCD | 69.72 | 542.78 |

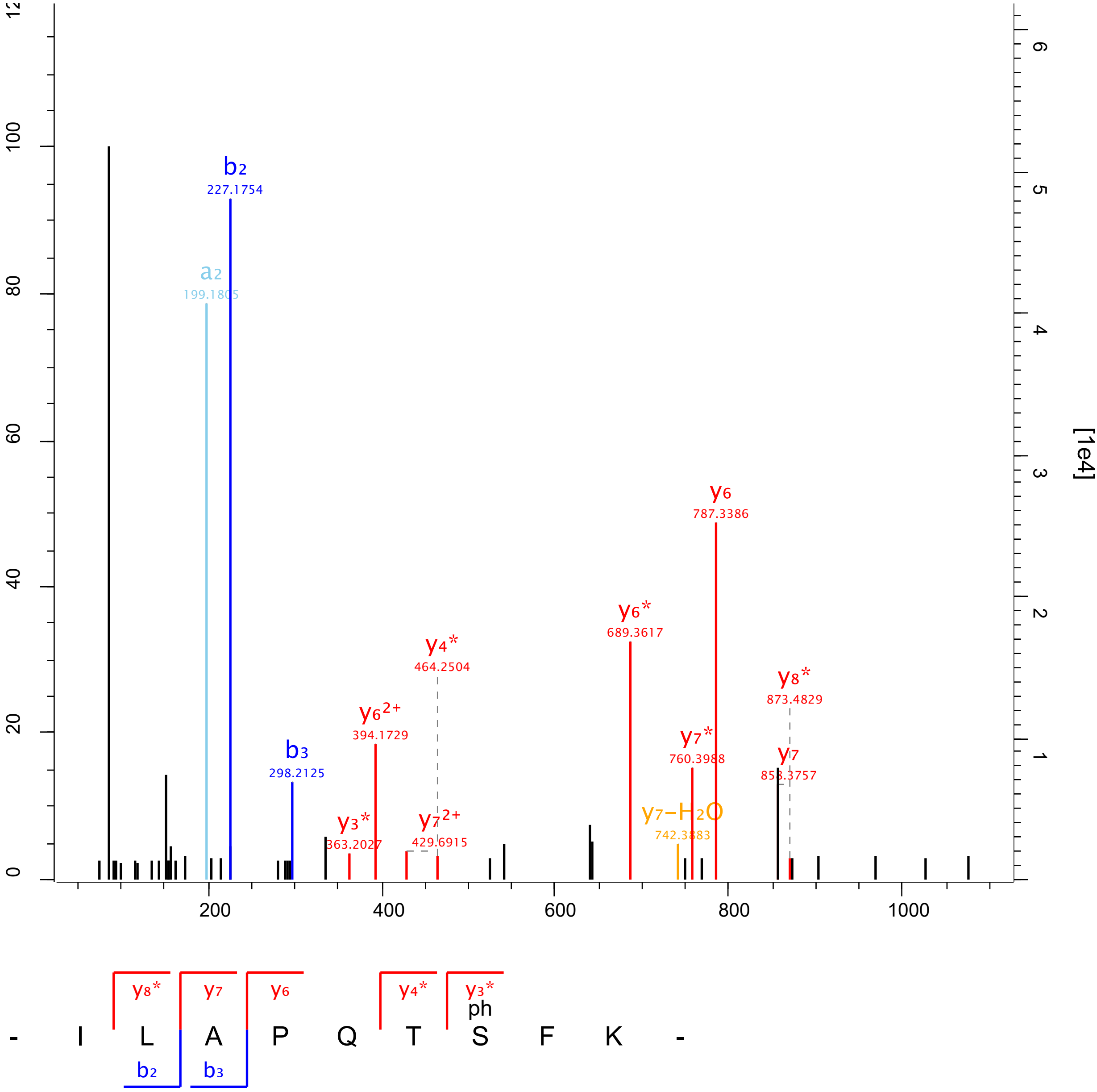

| Raw file | Scan | Method | Score | m/z |
|---|---|---|---|---|
| sys_15_3 | 17757 | FTMS; HCD | 201.15 | 938.36 |

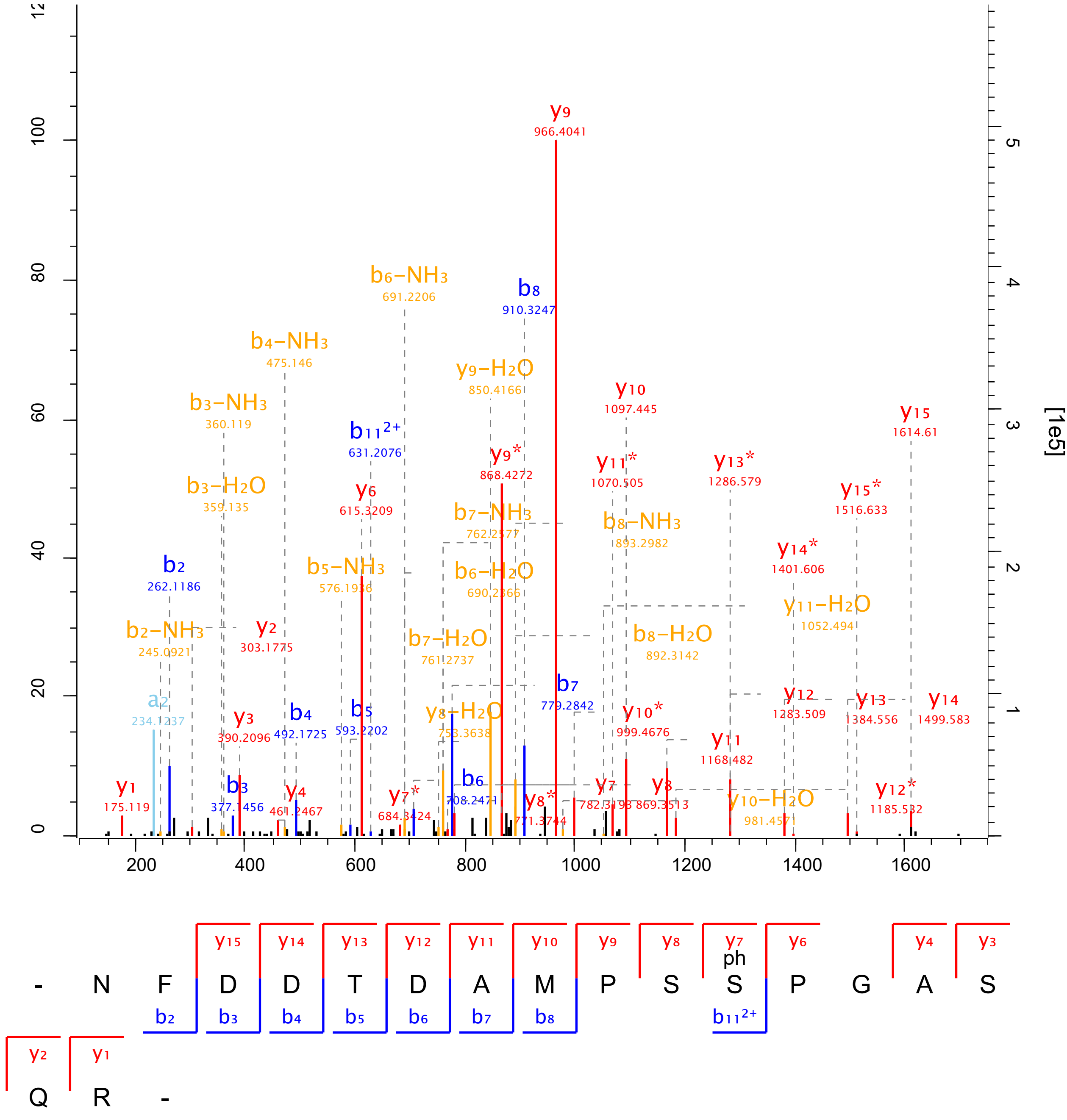

| Raw file | Scan | Method | Score | m/z |
|---|---|---|---|---|
| sys_15_3 | 17903 | FTMS; HCD | 45.61 | 569.27 |

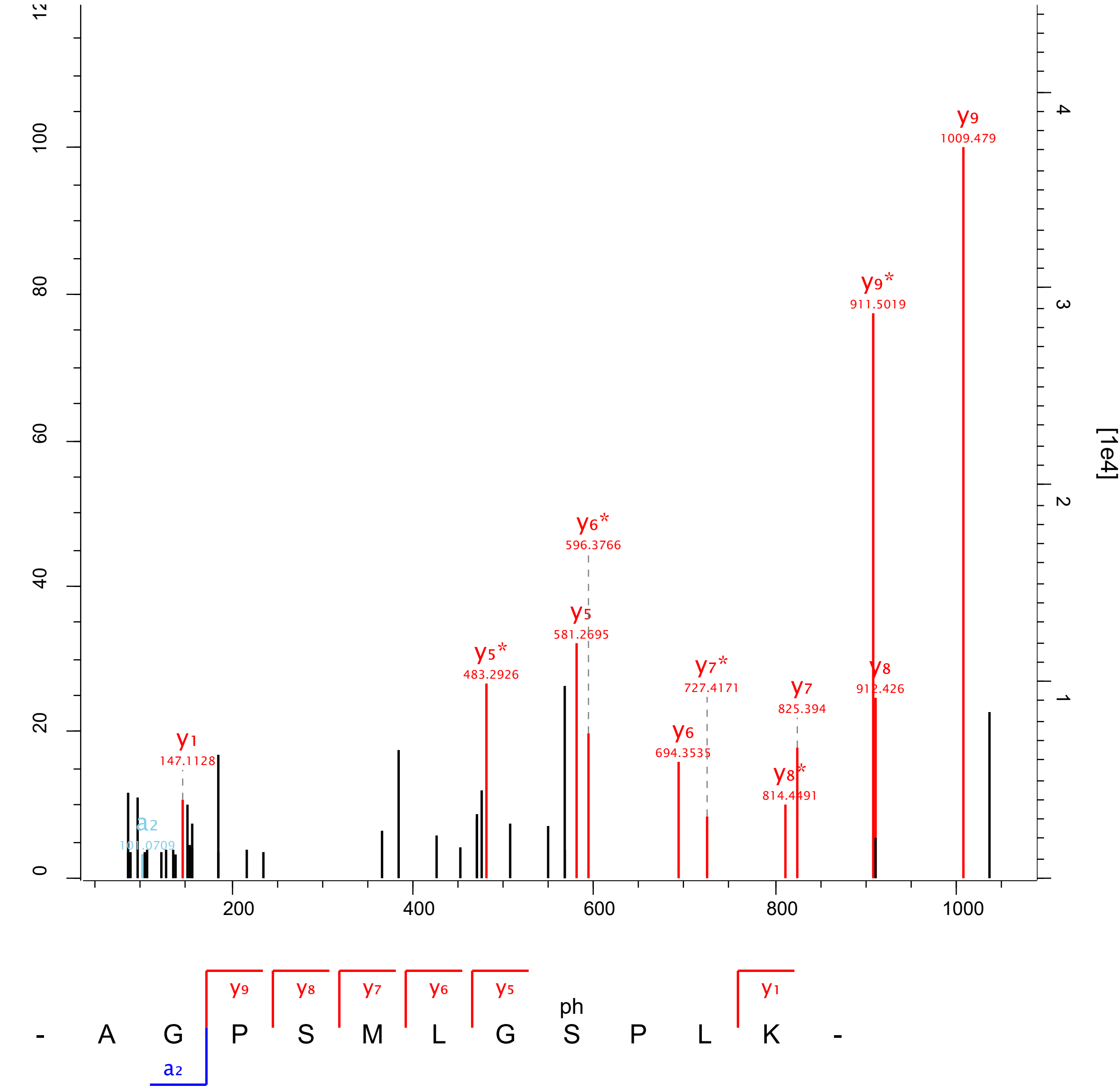

| Raw file | Scan | Method | Score | m/z |
|---|---|---|---|---|
| sys_15_3 | 19207 | FTMS; HCD | 87.43 | 712.83 |

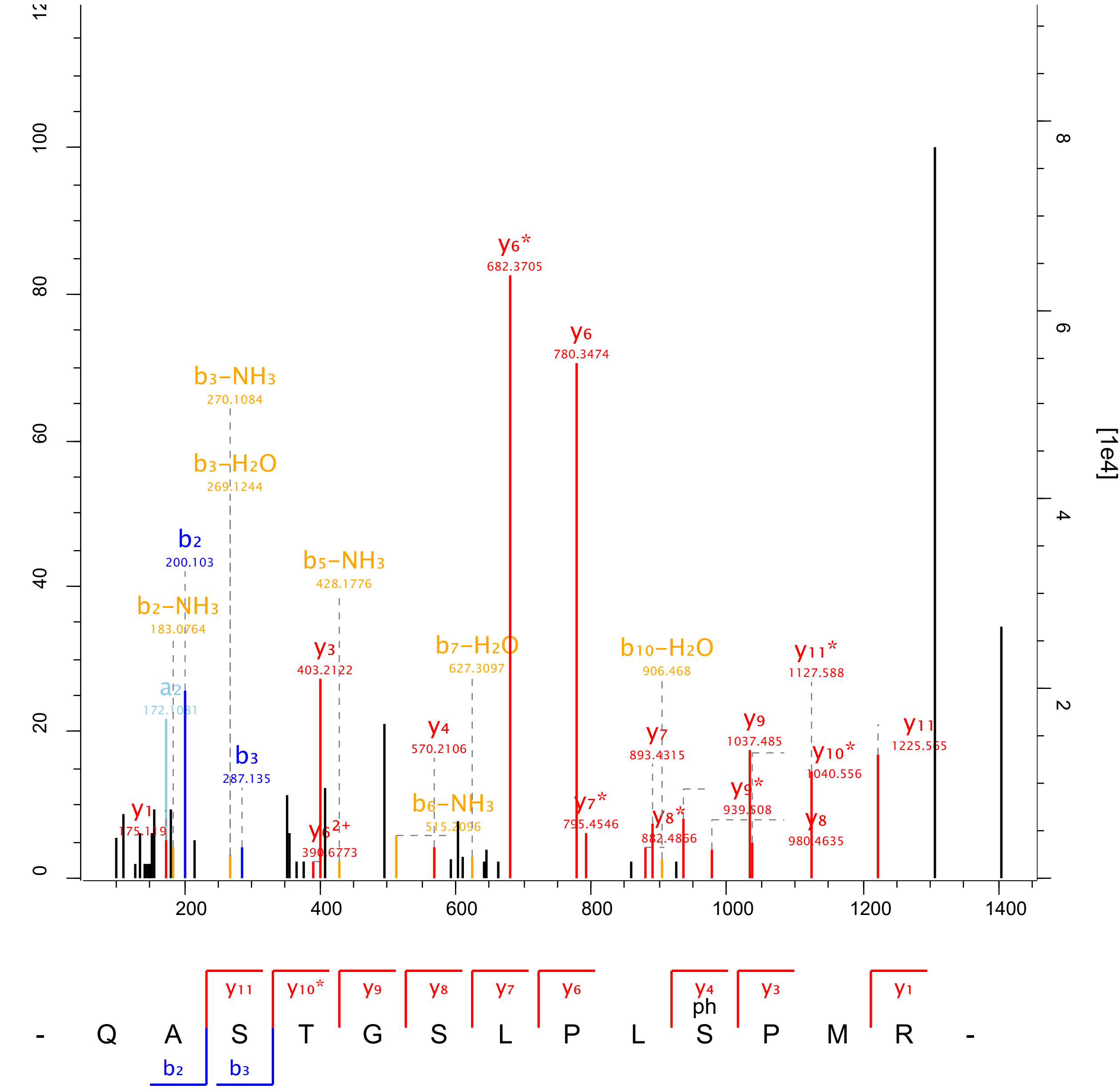

| Raw file | Scan | Method | Score | m/z |
|---|---|---|---|---|
| sys_15_3 | 21149 | FTMS; HCD | 75.59 | 671.27 |

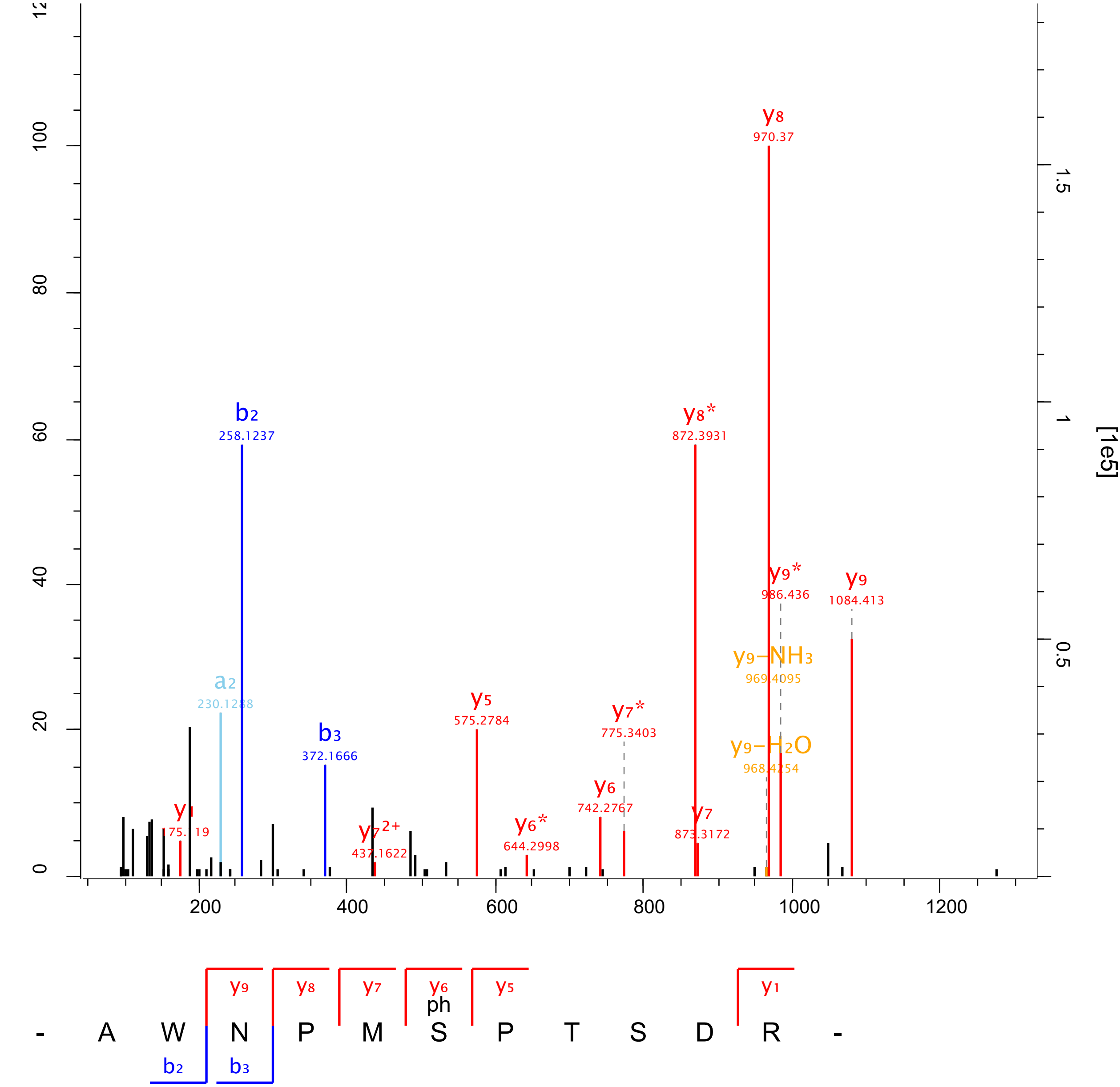

| Raw file | Scan | Method | Score | m/z |
|---|---|---|---|---|
| sys_15_3 | 35303 | FTMS; HCD | 81.81 | 969.39 |

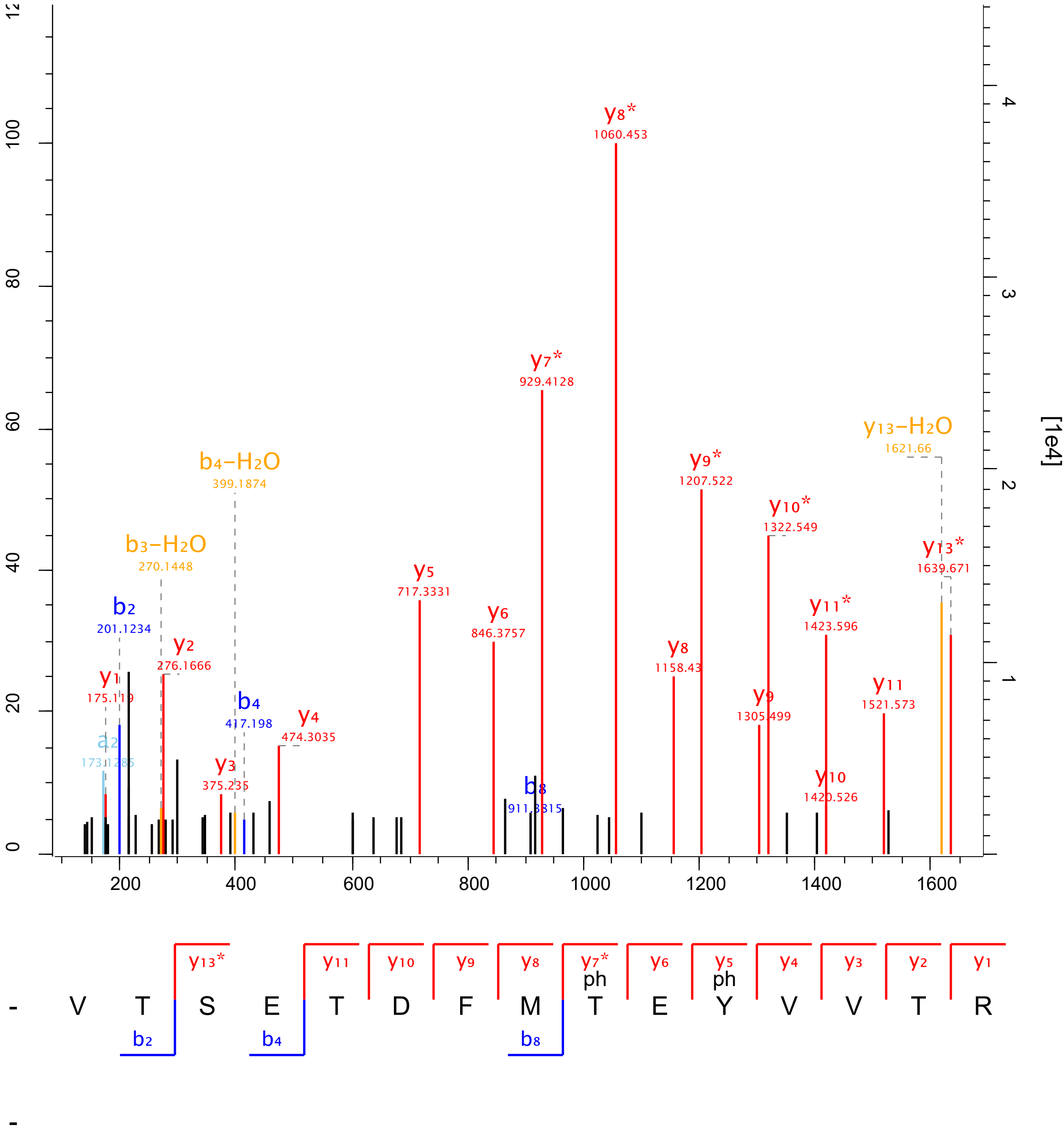

| Raw file | Scan | Method | Score | m/z |
|---|---|---|---|---|
| sys_15_3 | 5952 | FTMS; HCD | 62.34 | 529.74 |

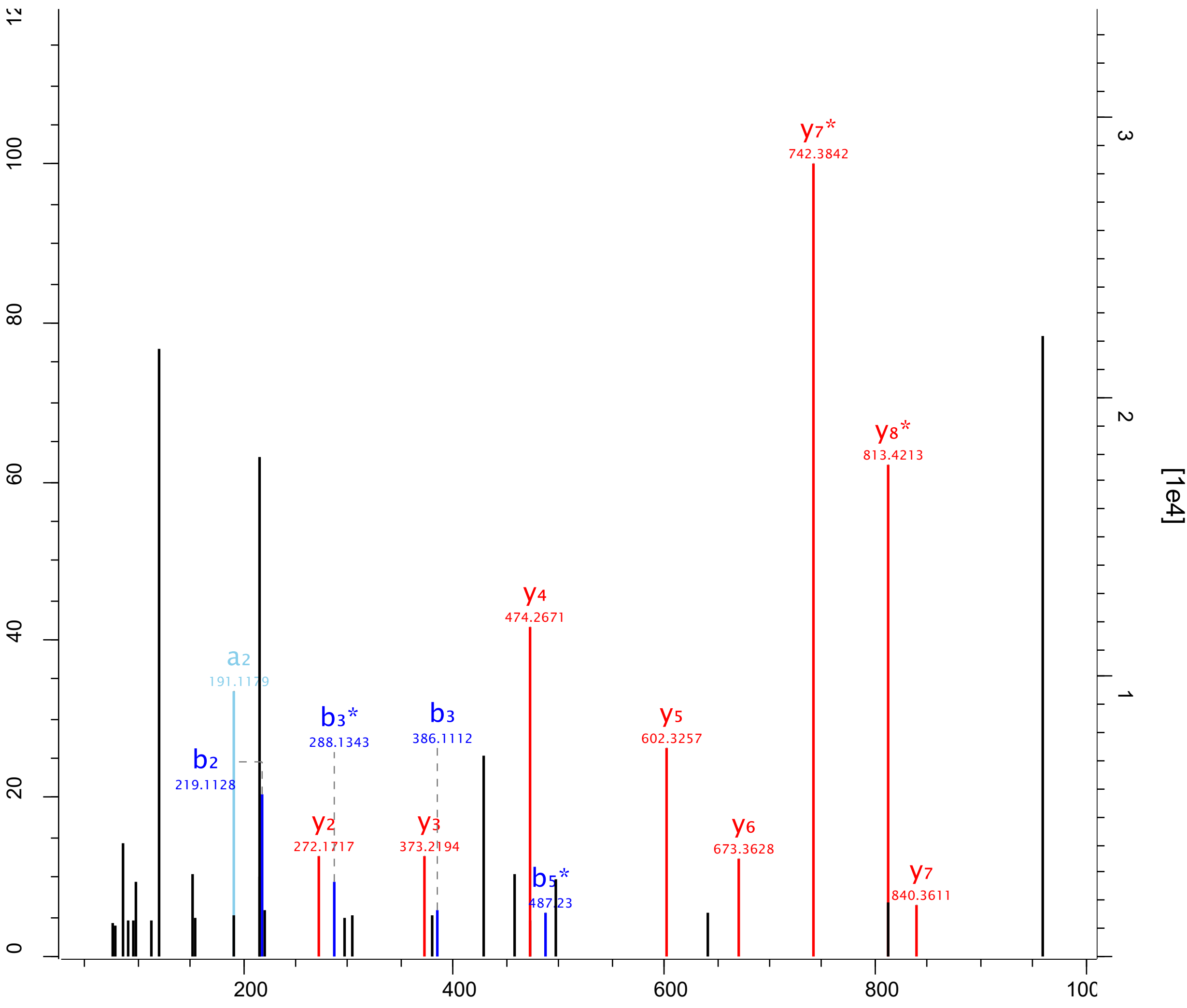


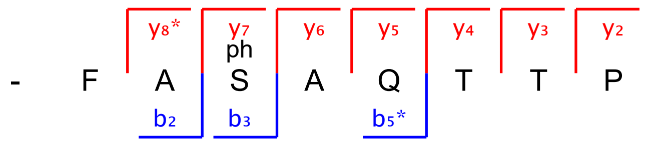

| Raw file | Scan | Method | Score | m/z |
|---|---|---|---|---|
| sys_15_3 | 7075 | FTMS; HCD | 63.69 | 515.23 |

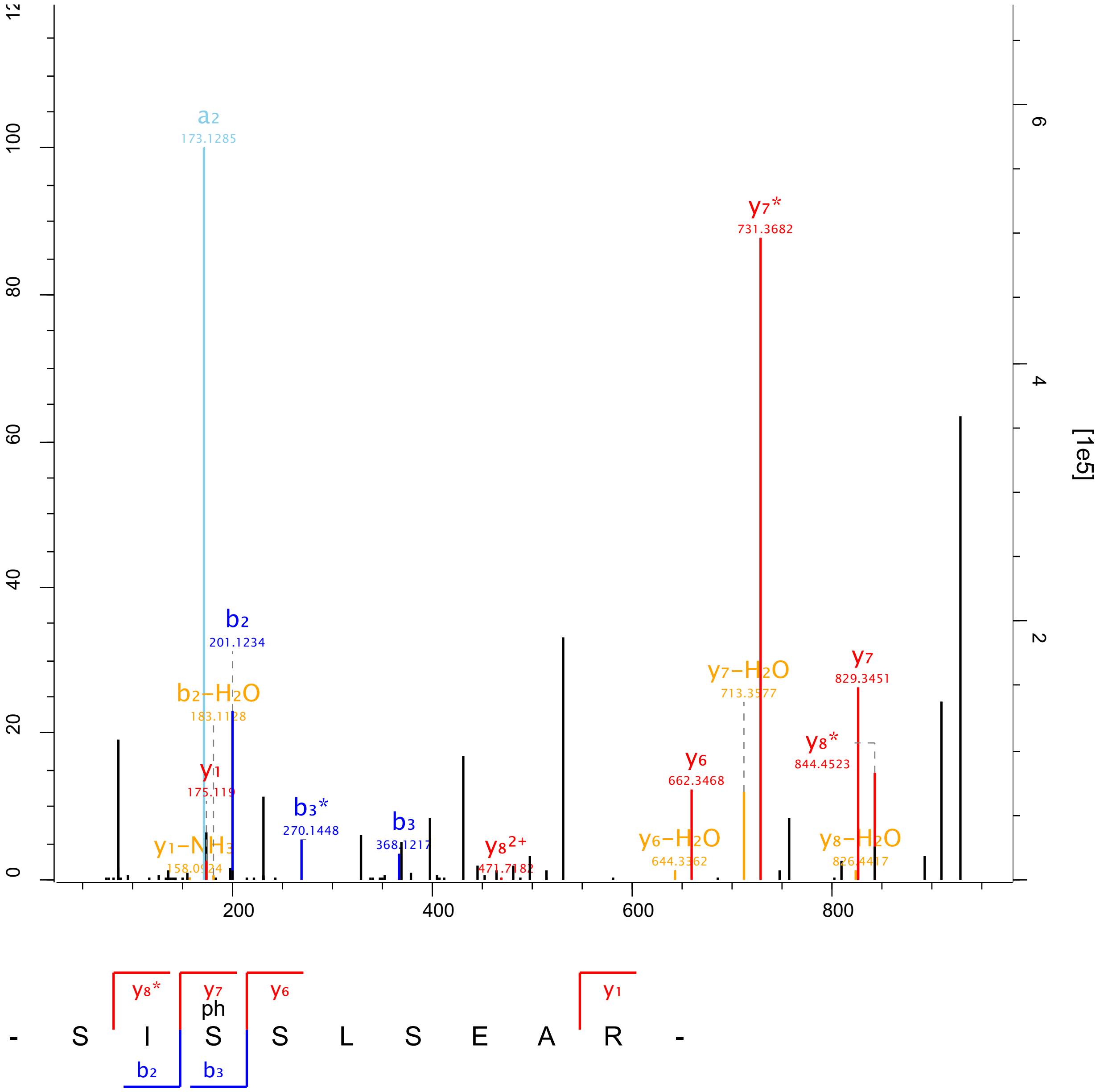

| Raw file | Scan | Method | Score | m/z |
|---|---|---|---|---|
| sys_15_3 | 8180 | FTMS; HCD | 93.1 | 525.21 |

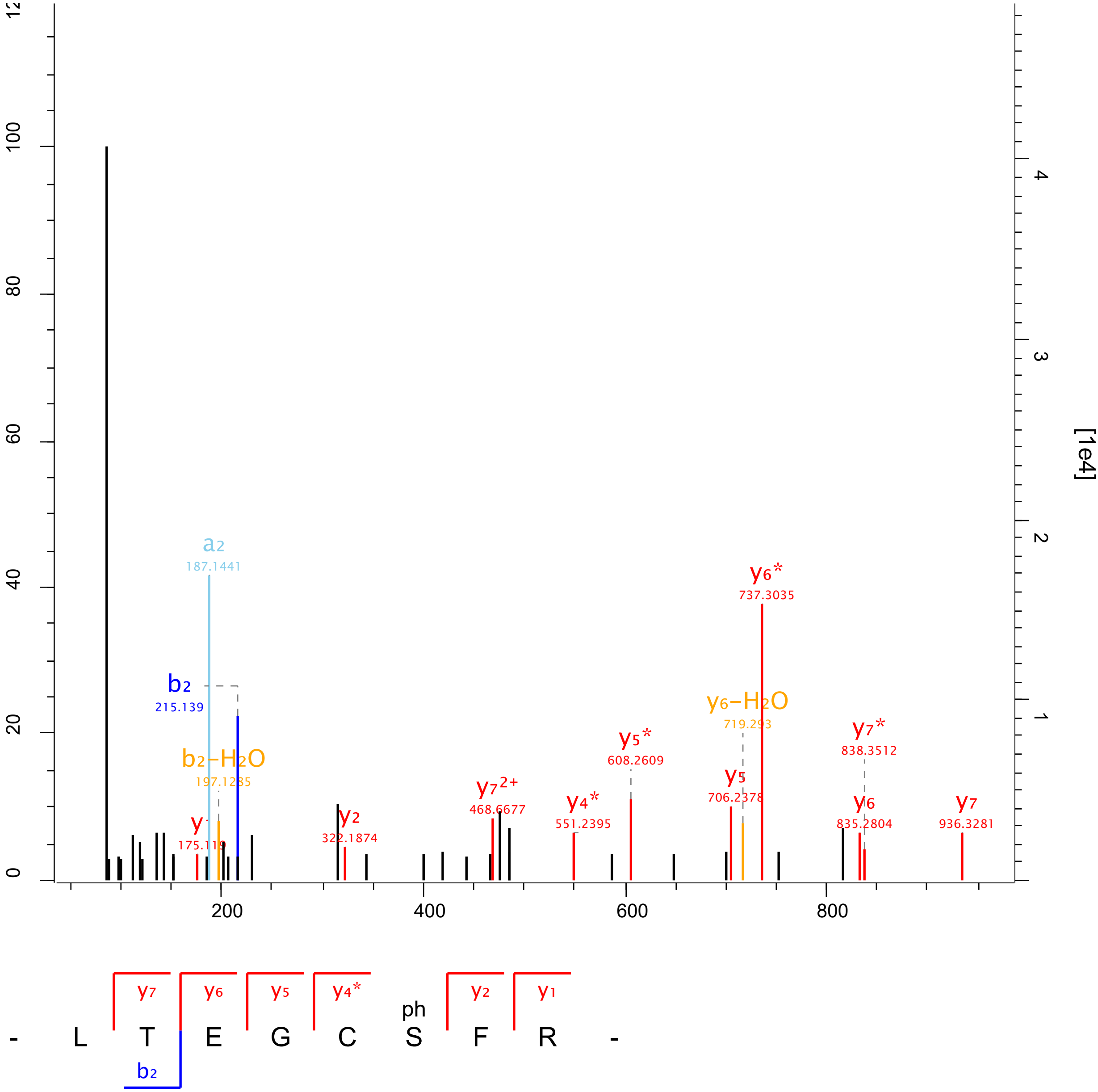

| Raw file | Scan | Method | Score | m/z |
|---|---|---|---|---|
| sys_15_3 | 8618 | FTMS; HCD | 99.82 | 486.23 |

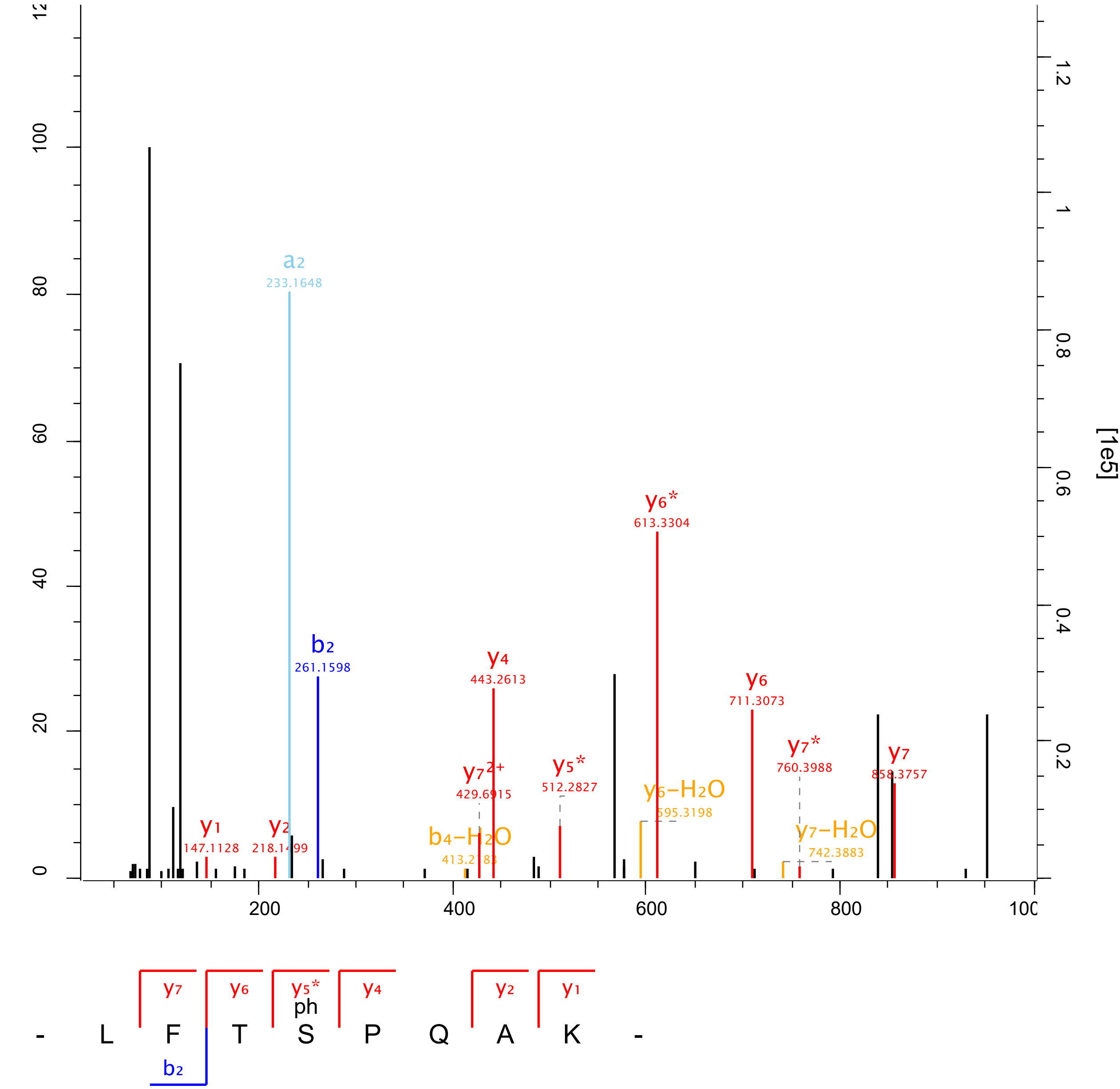

| Raw file | Scan | Method | Score | m/z |
| --- | --- | --- | --- | --- |
| sys_15_3 | 9628 | FTMS; HCD | 160.63 | 548.28 |

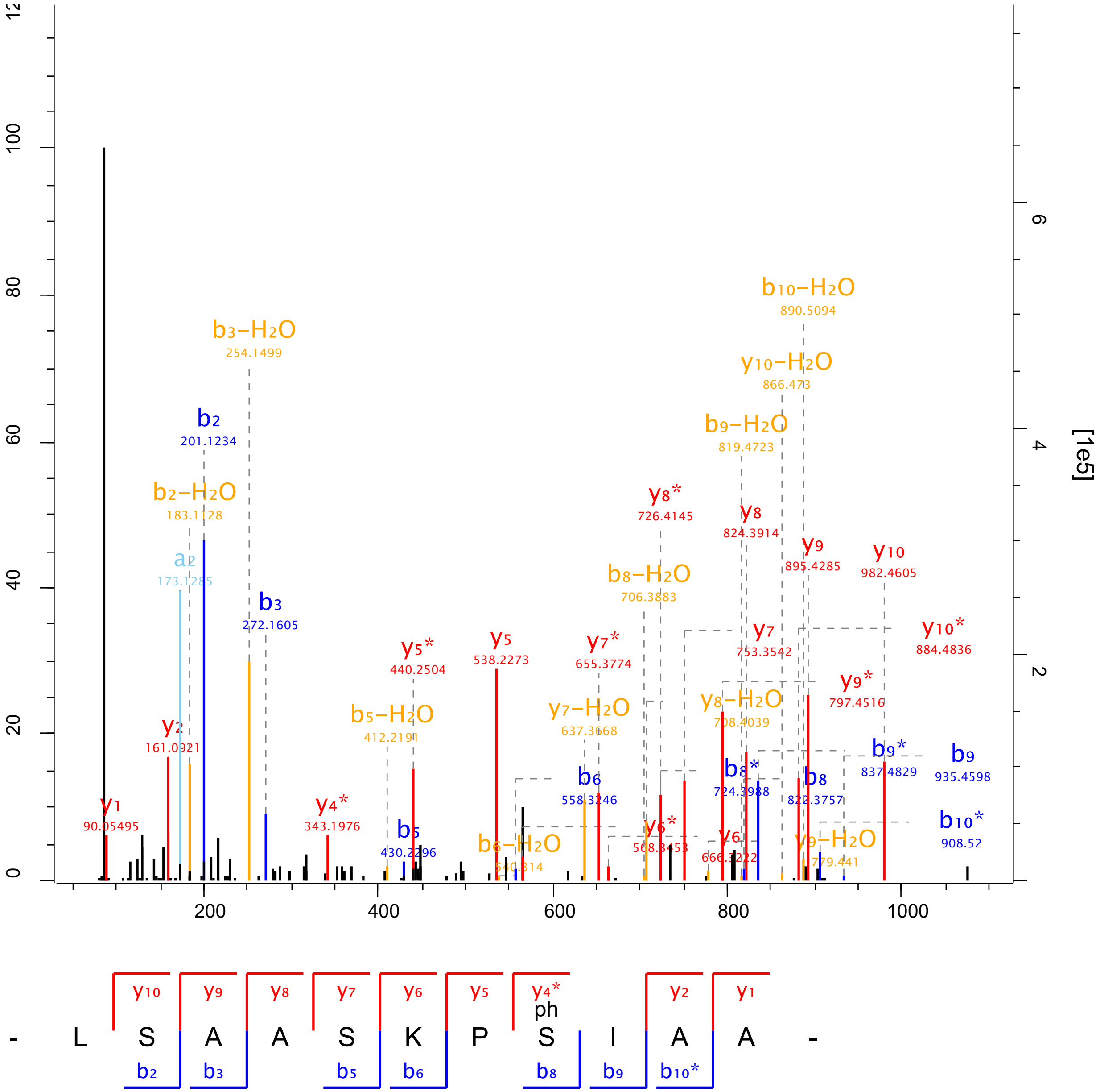

| Raw file | Scan | Method | Score | m/z |
|---|---|---|---|---|
| sys_15_3 | 9648 | FTMS; HCD | 69.03 | 524.21 |

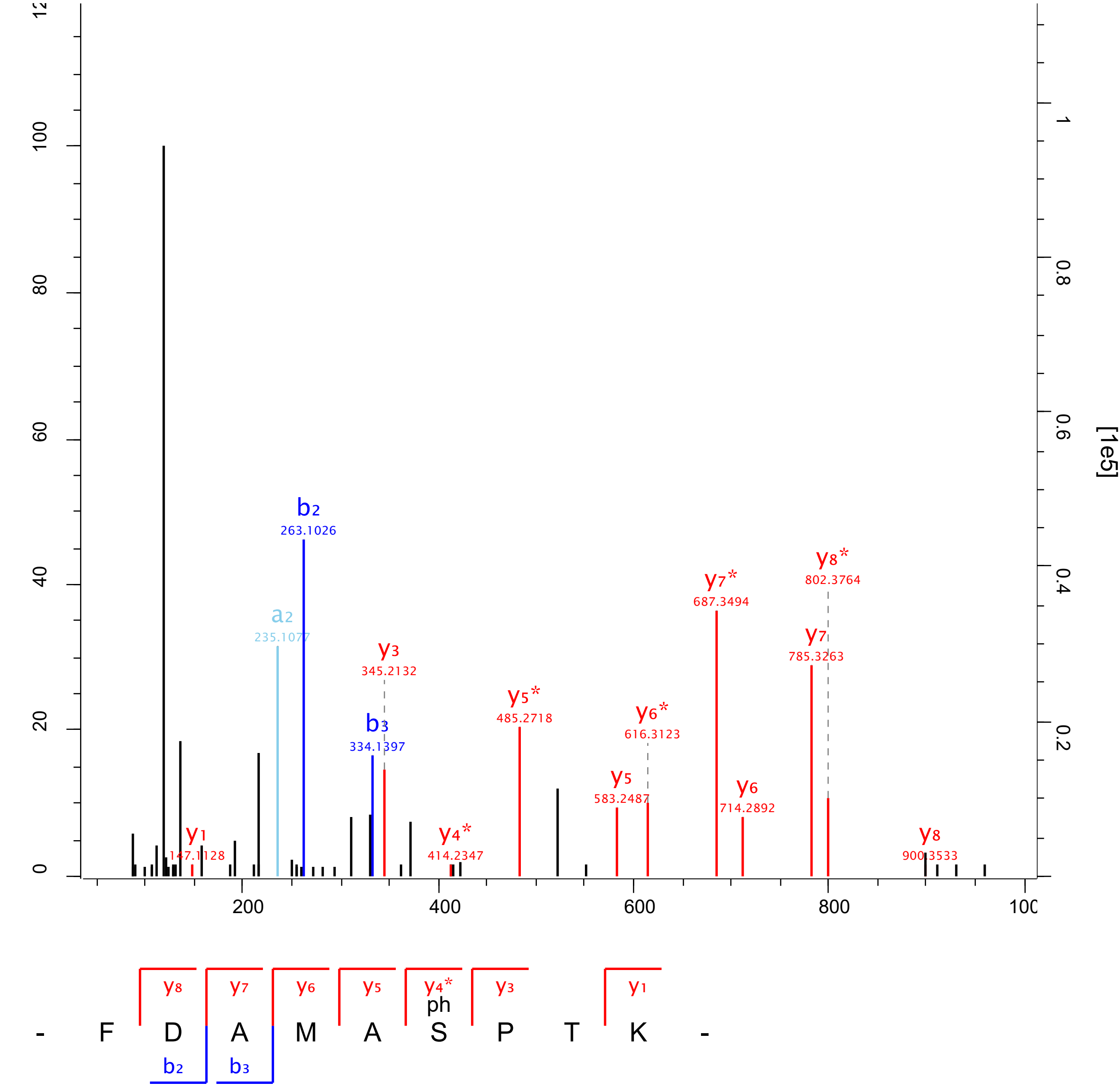

| Raw file | Scan | Method | Score | m/z |
|---|---|---|---|---|
| sys_30_1 | 14749 | FTMS; HCD | 120.63 | 785.82 |

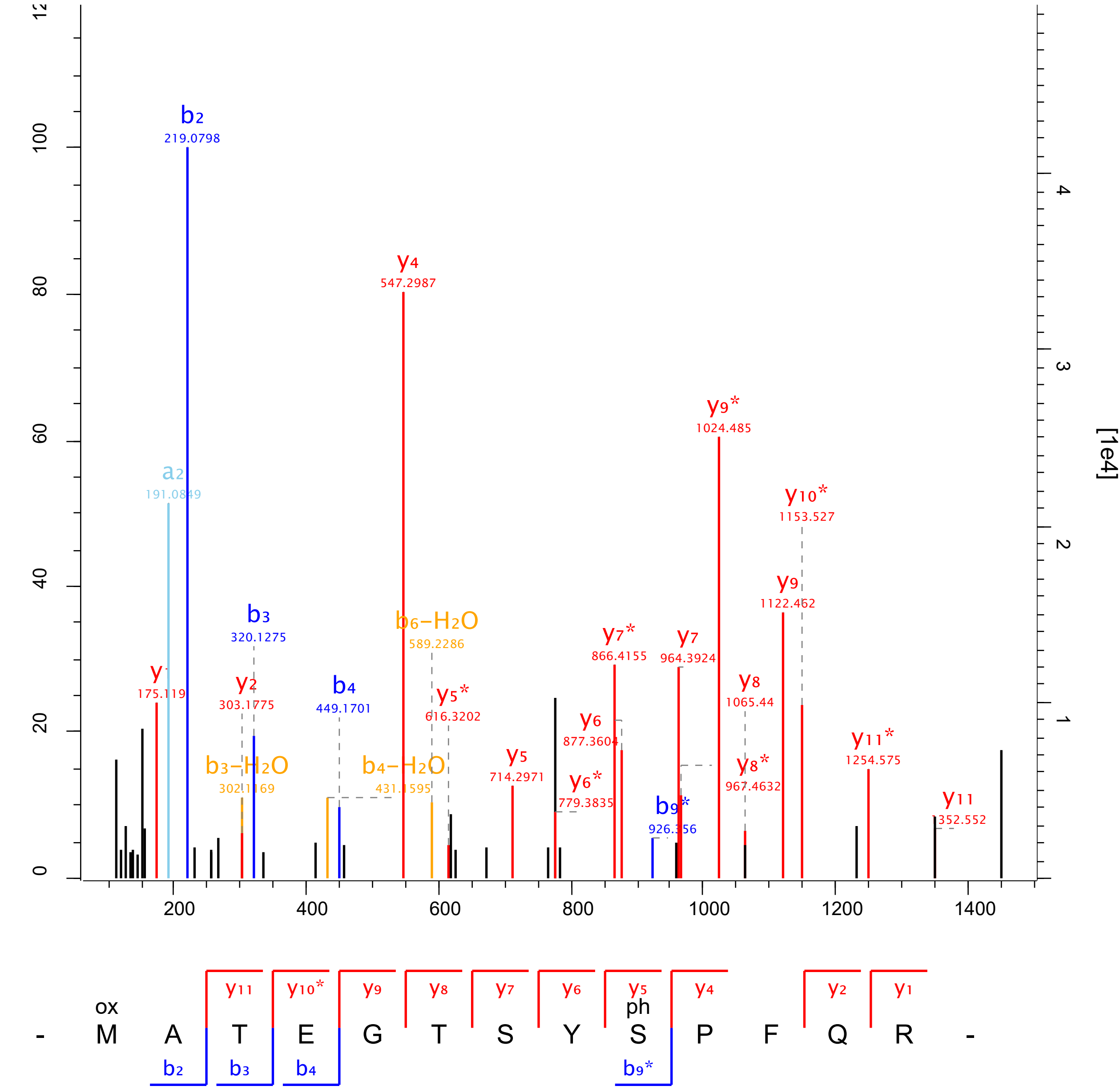

| Raw file | Scan | Method | Score | m/z |
|---|---|---|---|---|
| sys_30_1 | 16440 | FTMS; HCD | 139.5 | 749.81 |

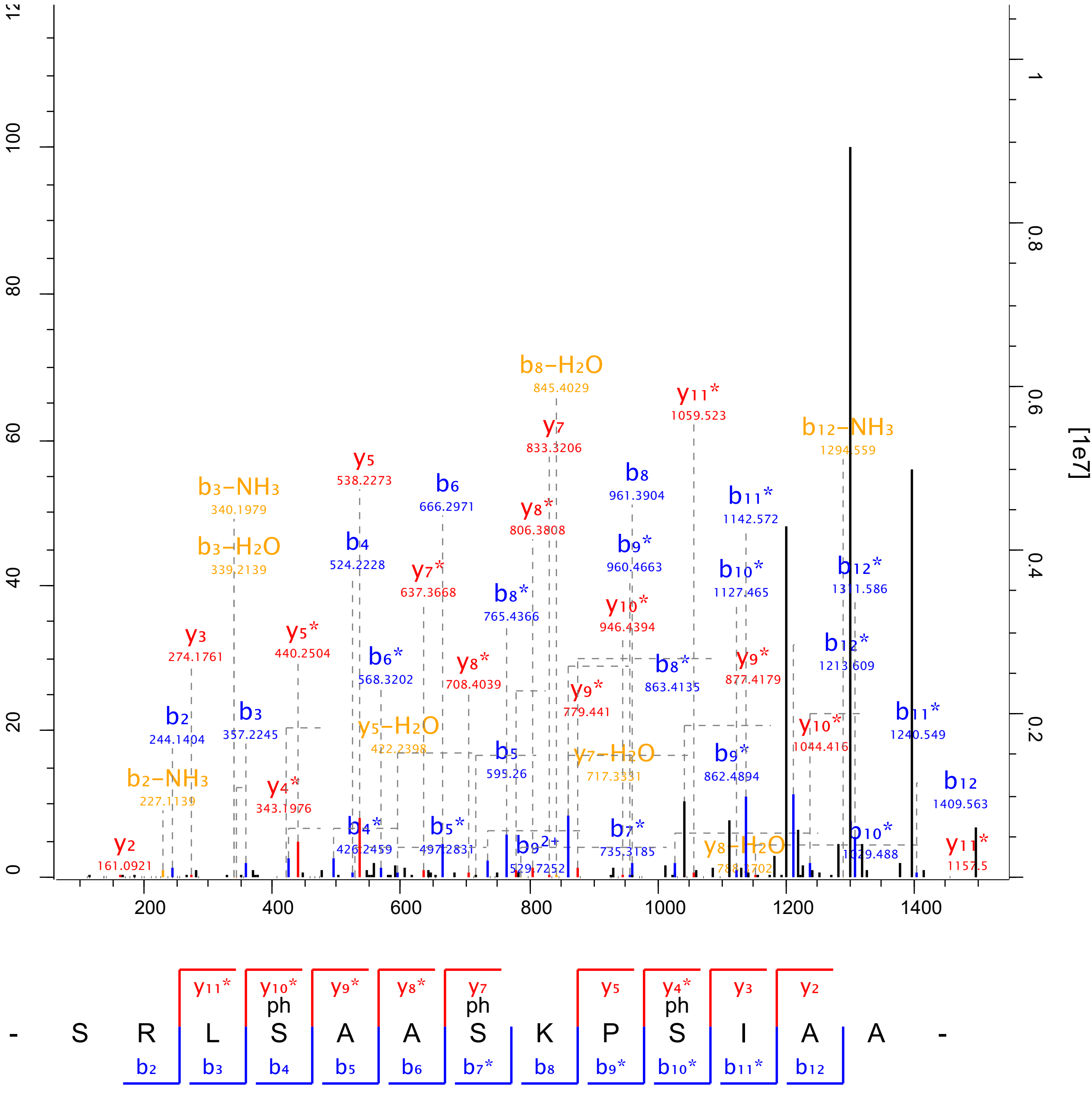

| Raw file | Scan | Method | Score | m/z |
| --- | --- | --- | --- | --- |
| sys_30_1 | 26002 | FTMS; HCD | 126.25 | 725.8 |

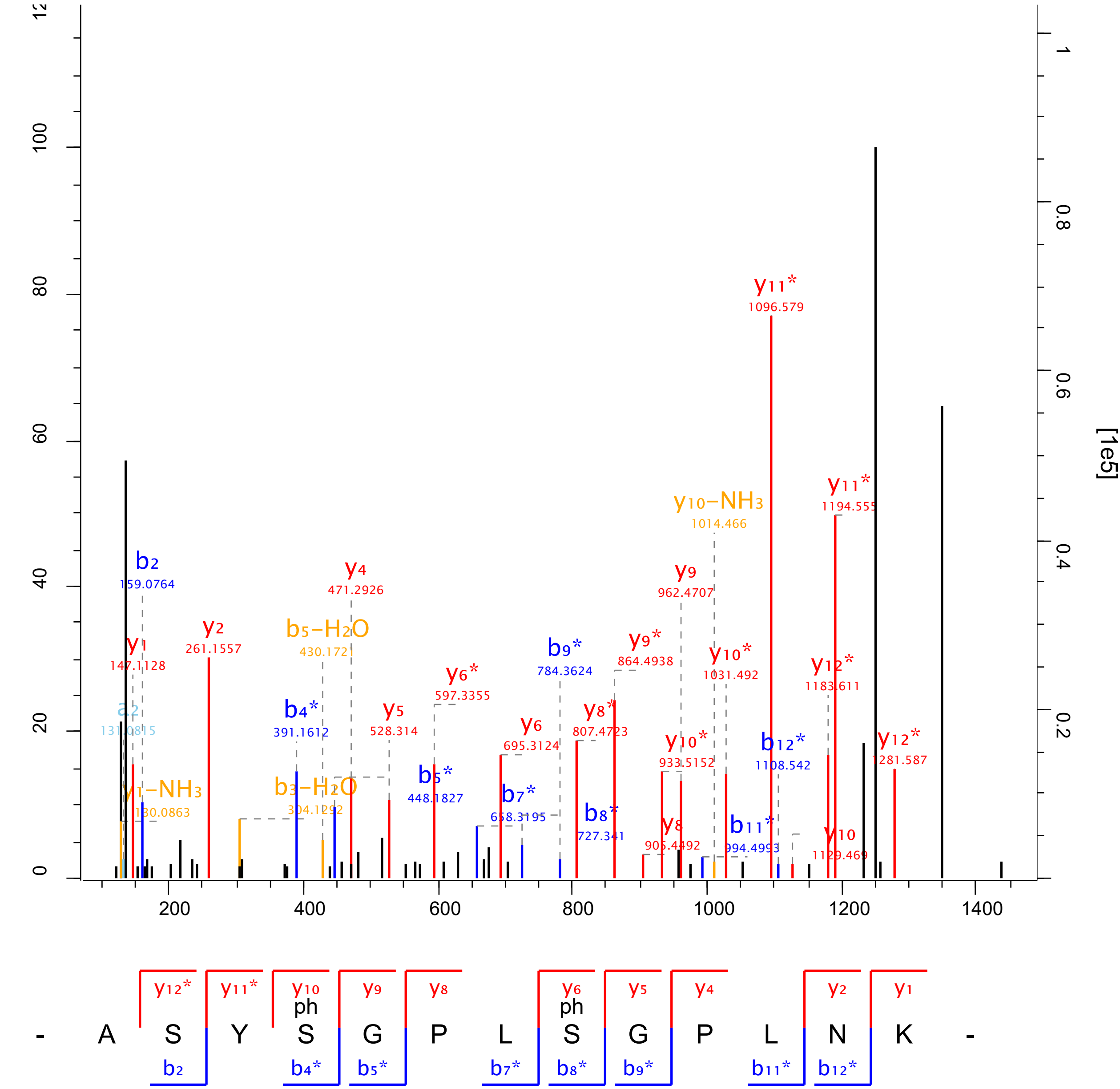

| Raw file | Scan | Method | Score | m/z |
| --- | --- | --- | --- | --- |
| sys_30_1 | 36281 | FTMS; HCD | 141.13 | 736.32 |

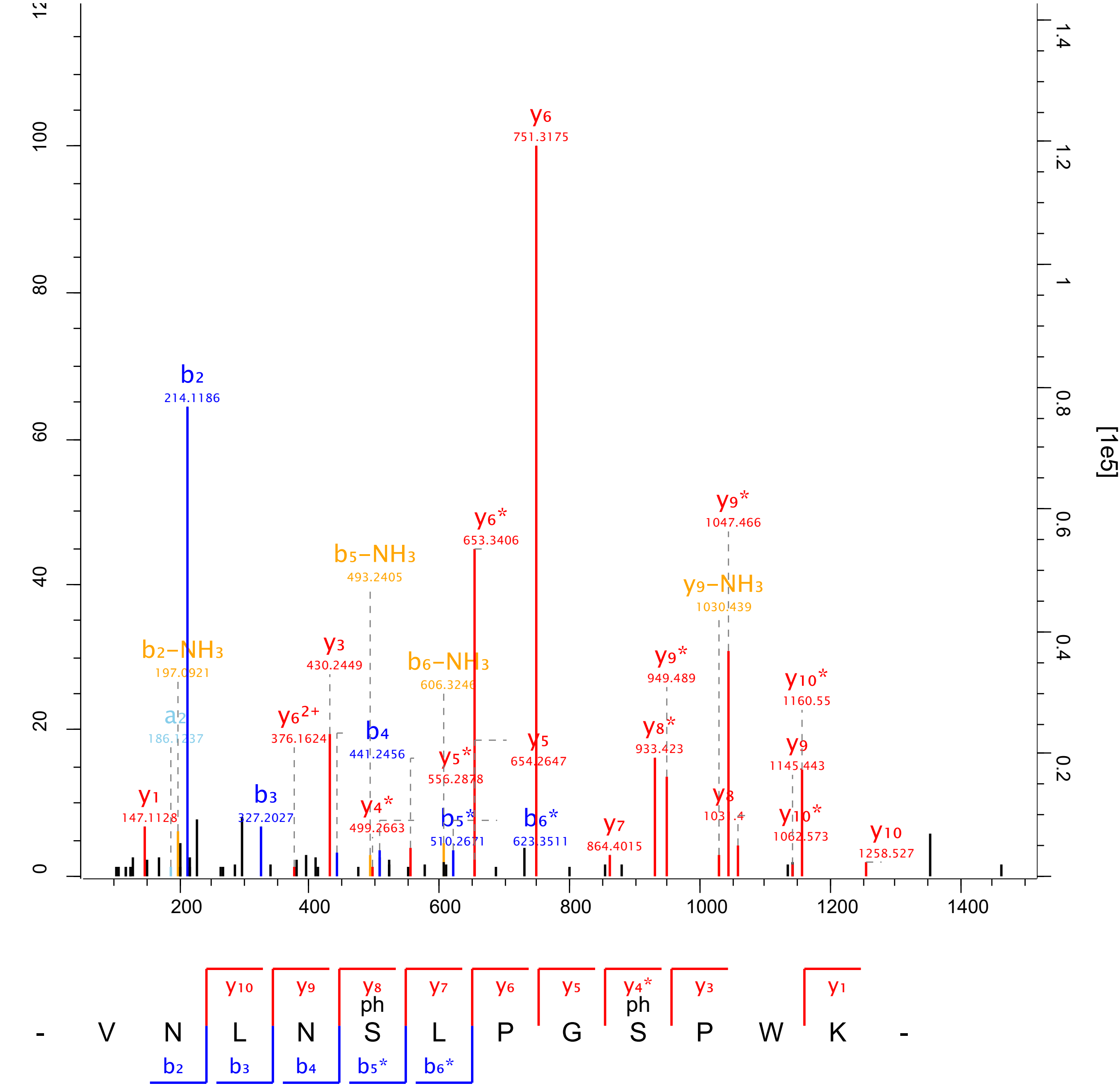

| Raw file | Scan | Method | Score | m/z |
|---|---|---|---|---|
| sys_30_1 | 8283 | FTMS; HCD | 114.97 | 450.7 |

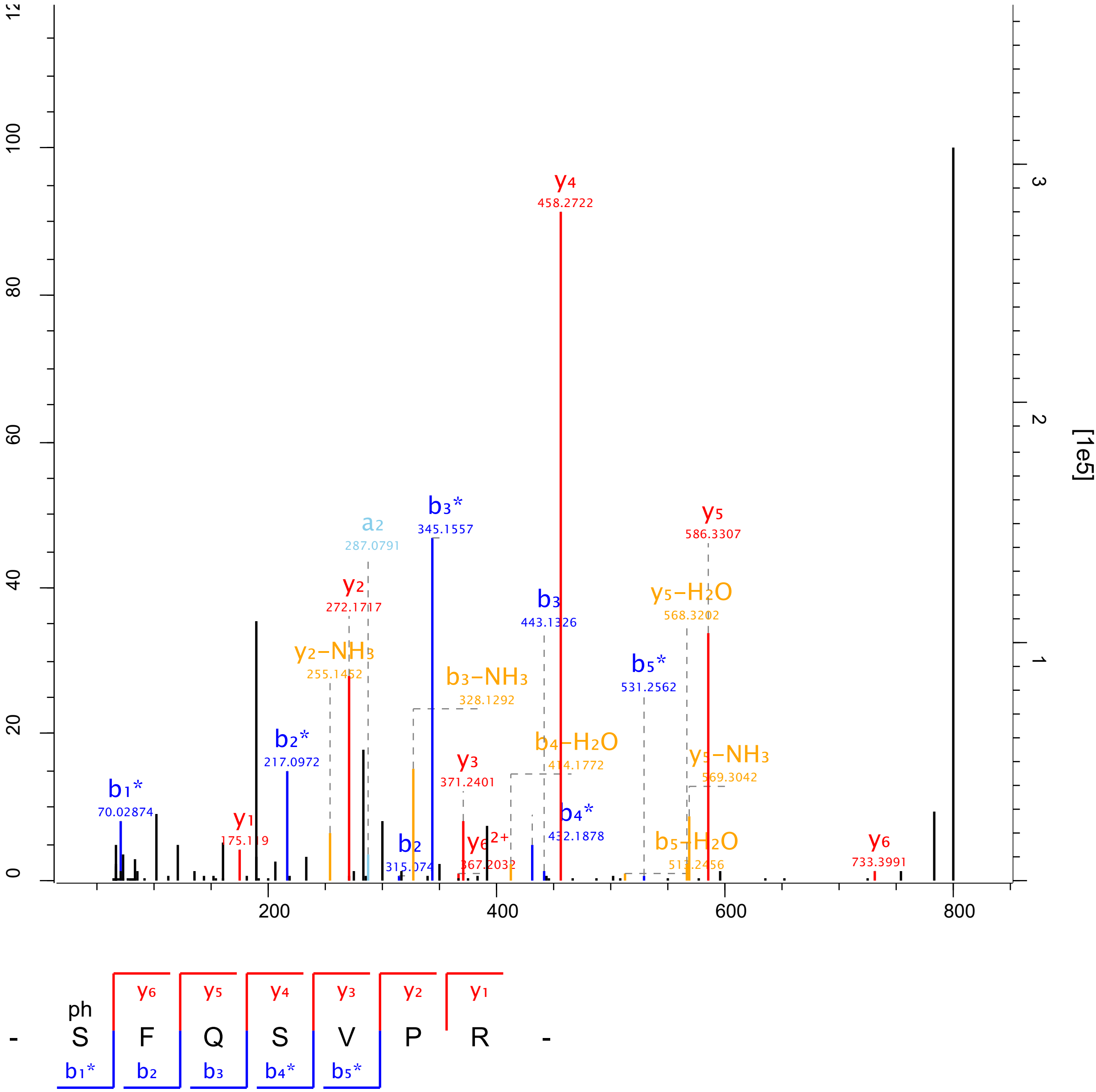

| Raw file | Scan | Method | Score | m/z |
| --- | --- | --- | --- | --- |
| sys_30_1 | 8449 | FTMS; HCD | 47.6 | 427.22 |

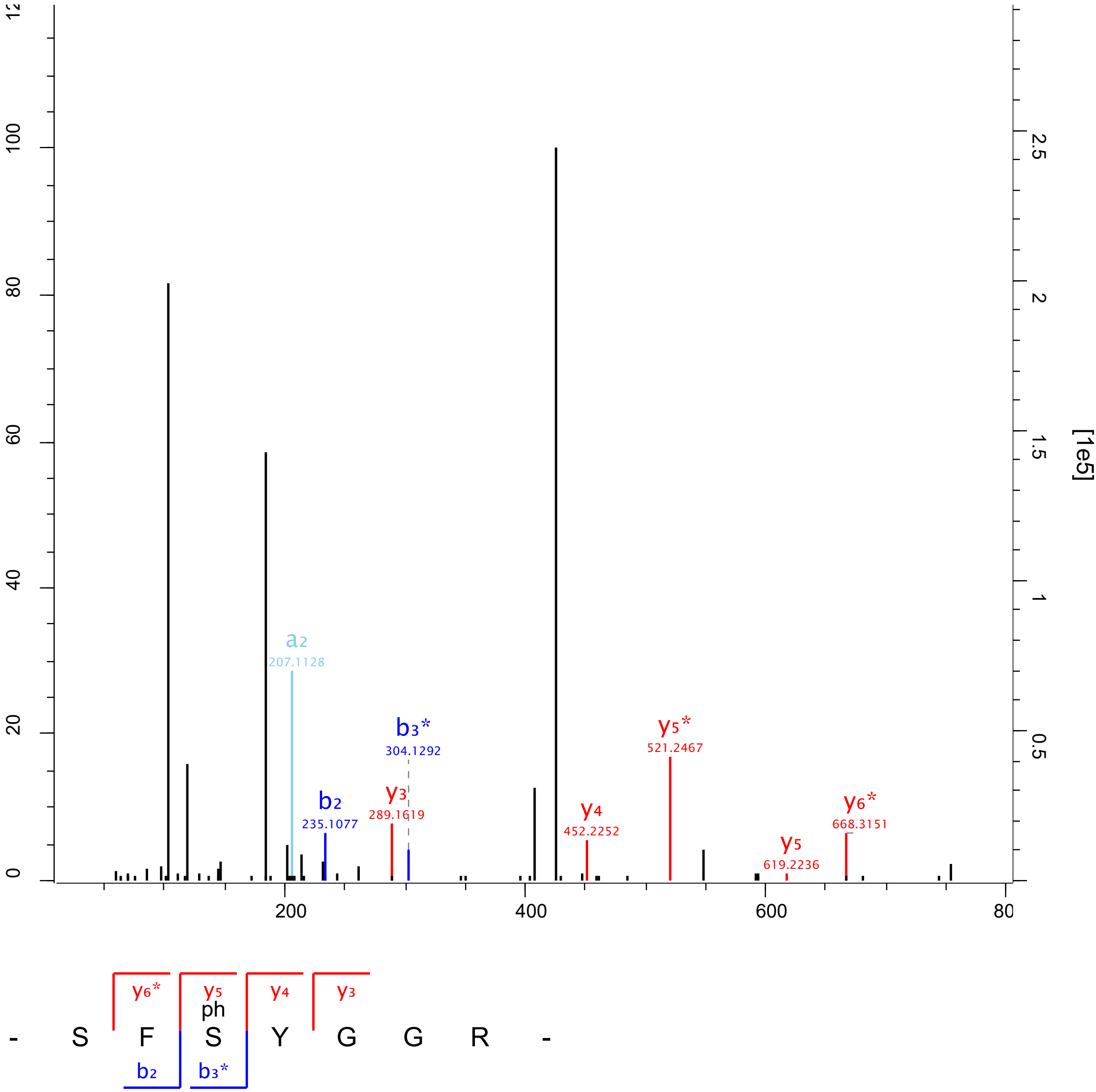

| Raw file | Scan | Method | Score | m/z |
| --- | --- | --- | --- | --- |
| sys_30_1 | 8770 | FTMS; HCD | 153.24 | 441.71 |

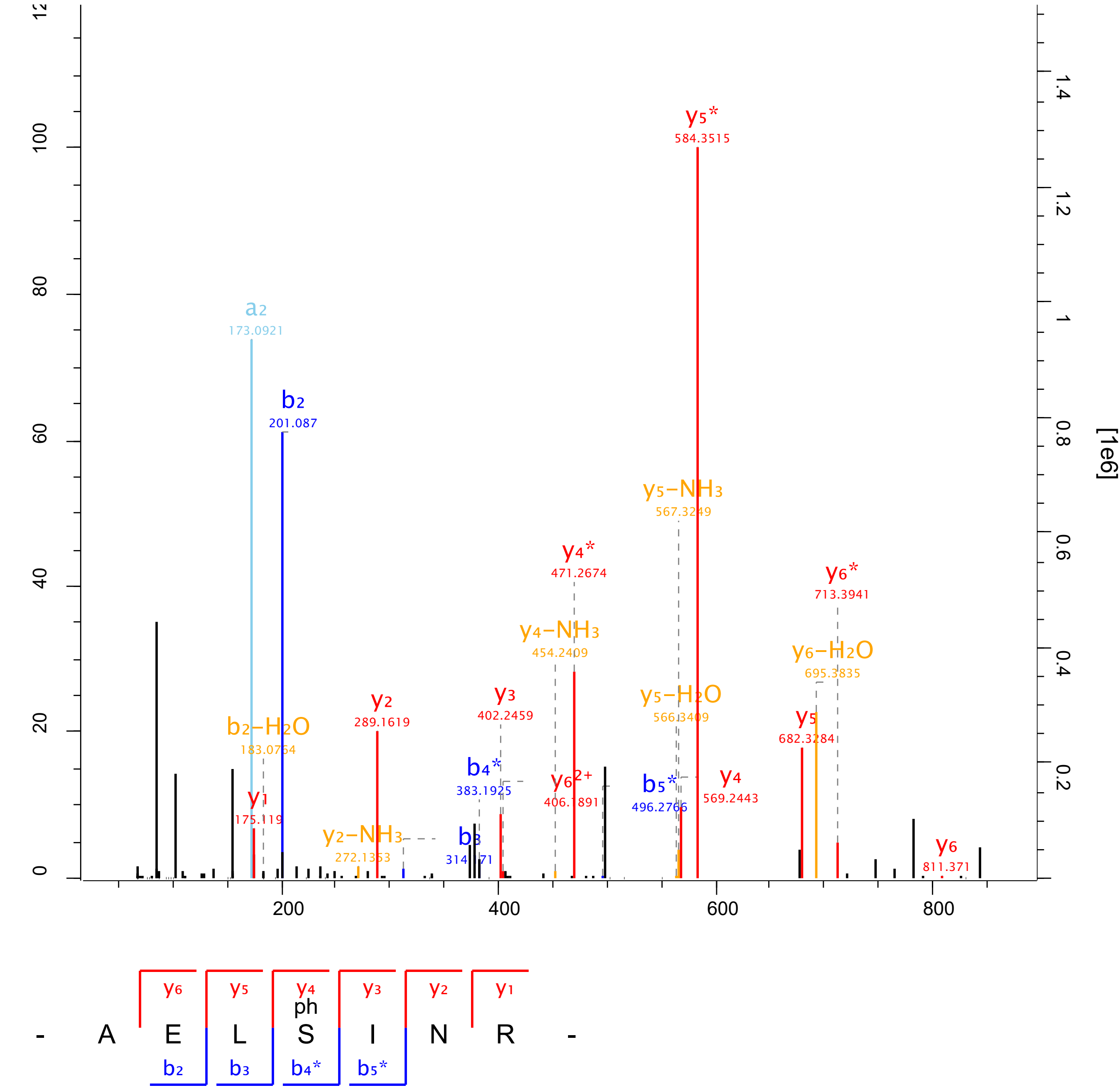

| Raw file | Scan | Method | Score | m/z |
|---|---|---|---|---|
| sys_30_1 | 8861 | FTMS; HCD | 75.82 | 525.7 |

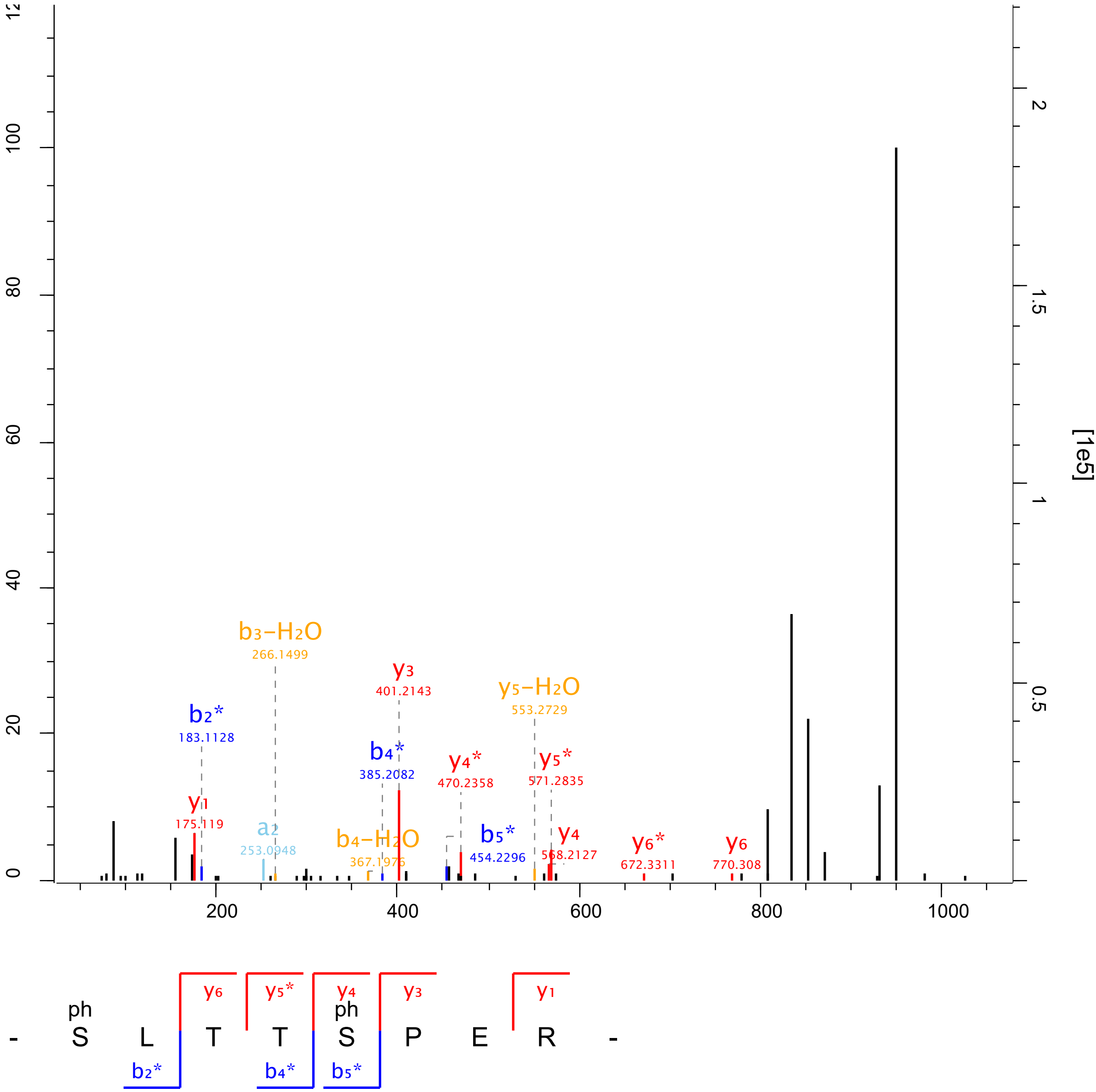

| Raw file | Scan | Method | Score | m/z |
|---|---|---|---|---|
| sys_30_1 | 9198 | FTMS; HCD | 153.54 | 534.74 |

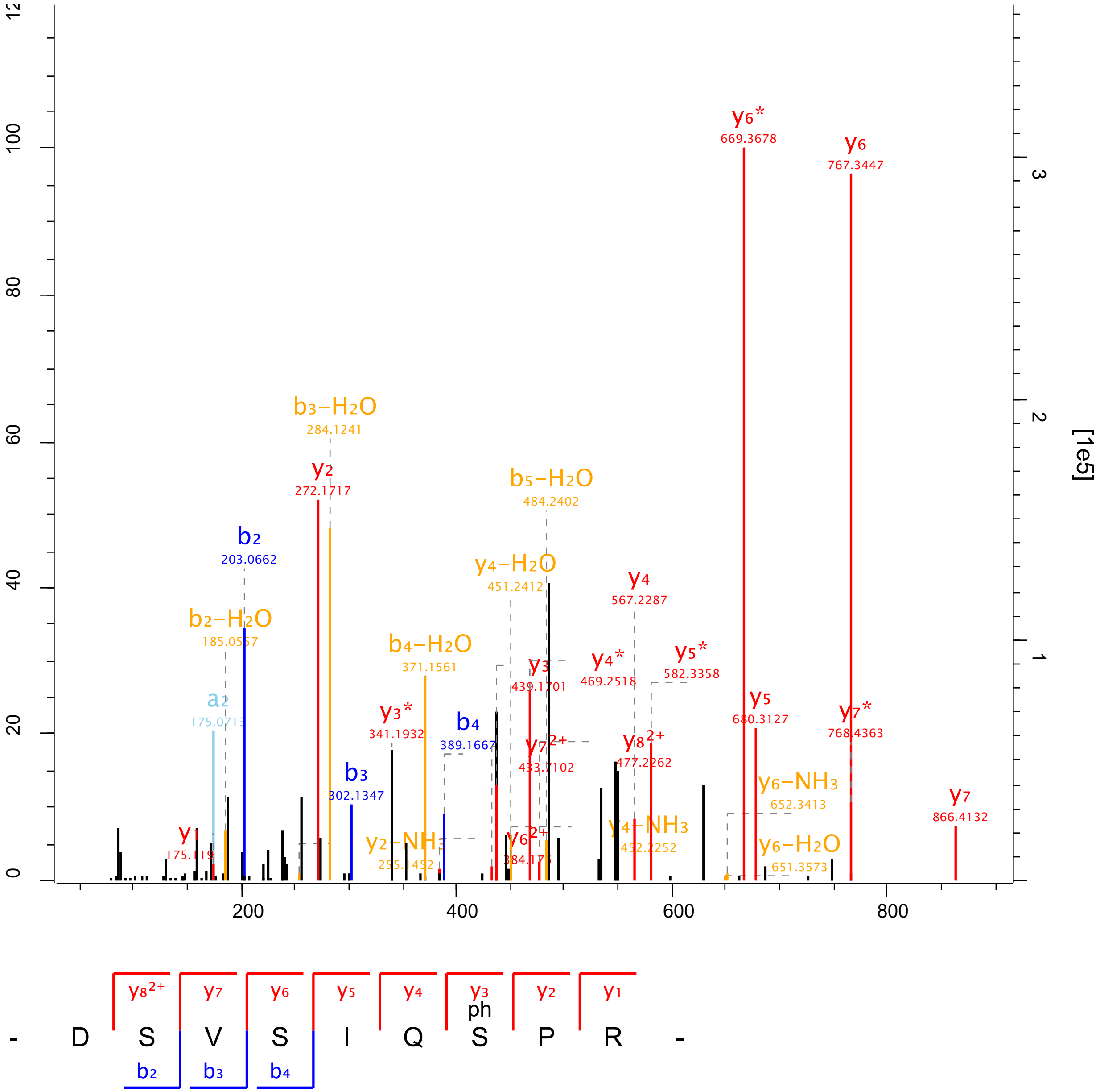

| Raw file | Scan | Method | Score | m/z |
|---|---|---|---|---|
| sys_30_2 | 10272 | FTMS; HCD | 218.01 | 602.27 |

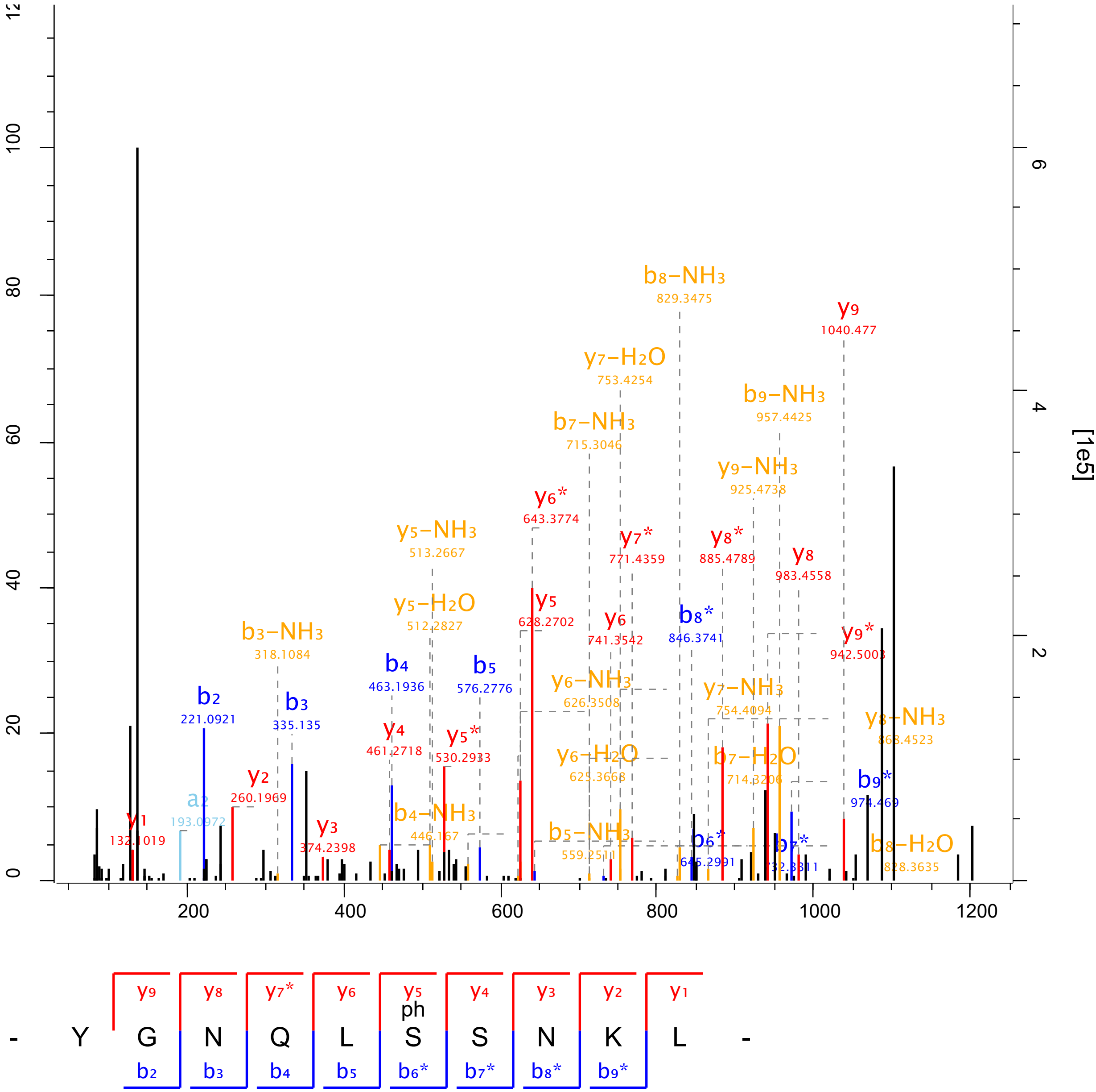

| Raw file | Scan | Method | Score | m/z |
|---|---|---|---|---|
| sys_30_2 | 20145 | FTMS; HCD | 102.55 | 459.19 |

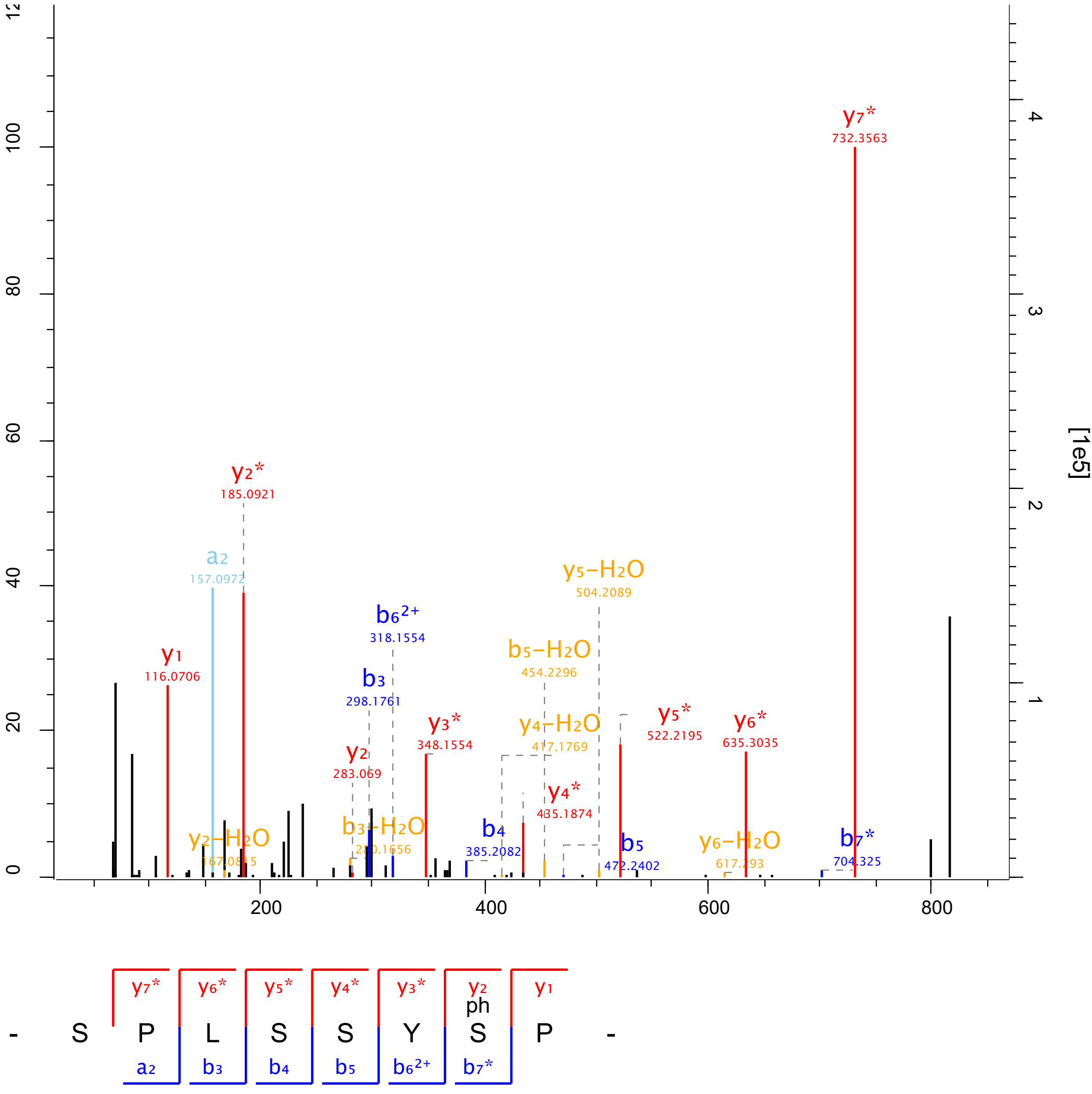

| Raw file | Scan | Method | Score | m/z |
|---|---|---|---|---|
| sys_30_2 | 3964 | FTMS; HCD | 81.92 | 512.74 |

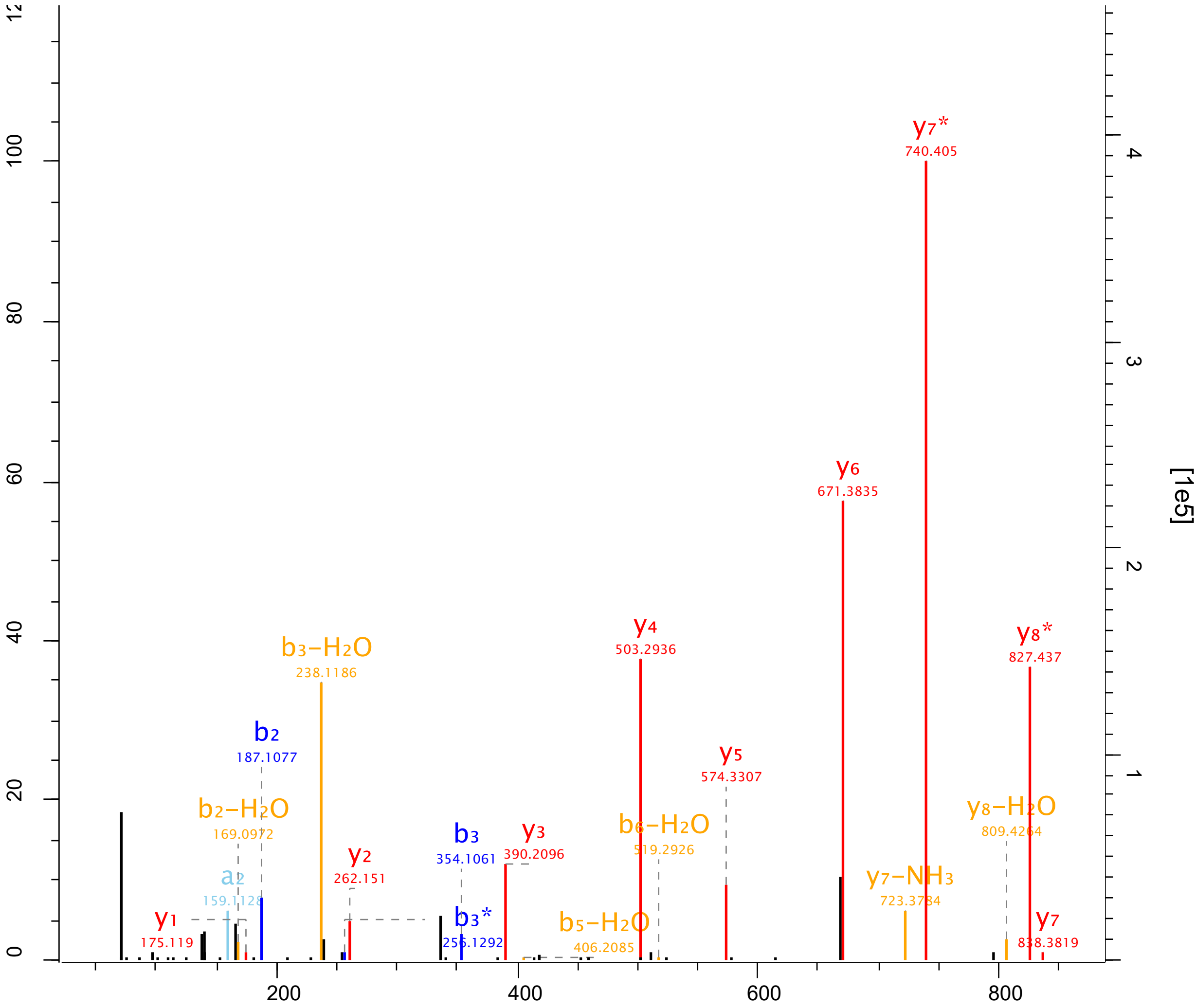


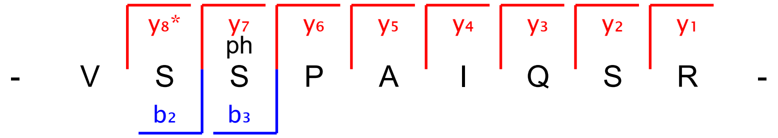

| Raw file | Scan | Method | Score | m/z |
|---|---|---|---|---|
| sys_30_2 | 5526 | FTMS; HCD | 66.27 | 407.2 |

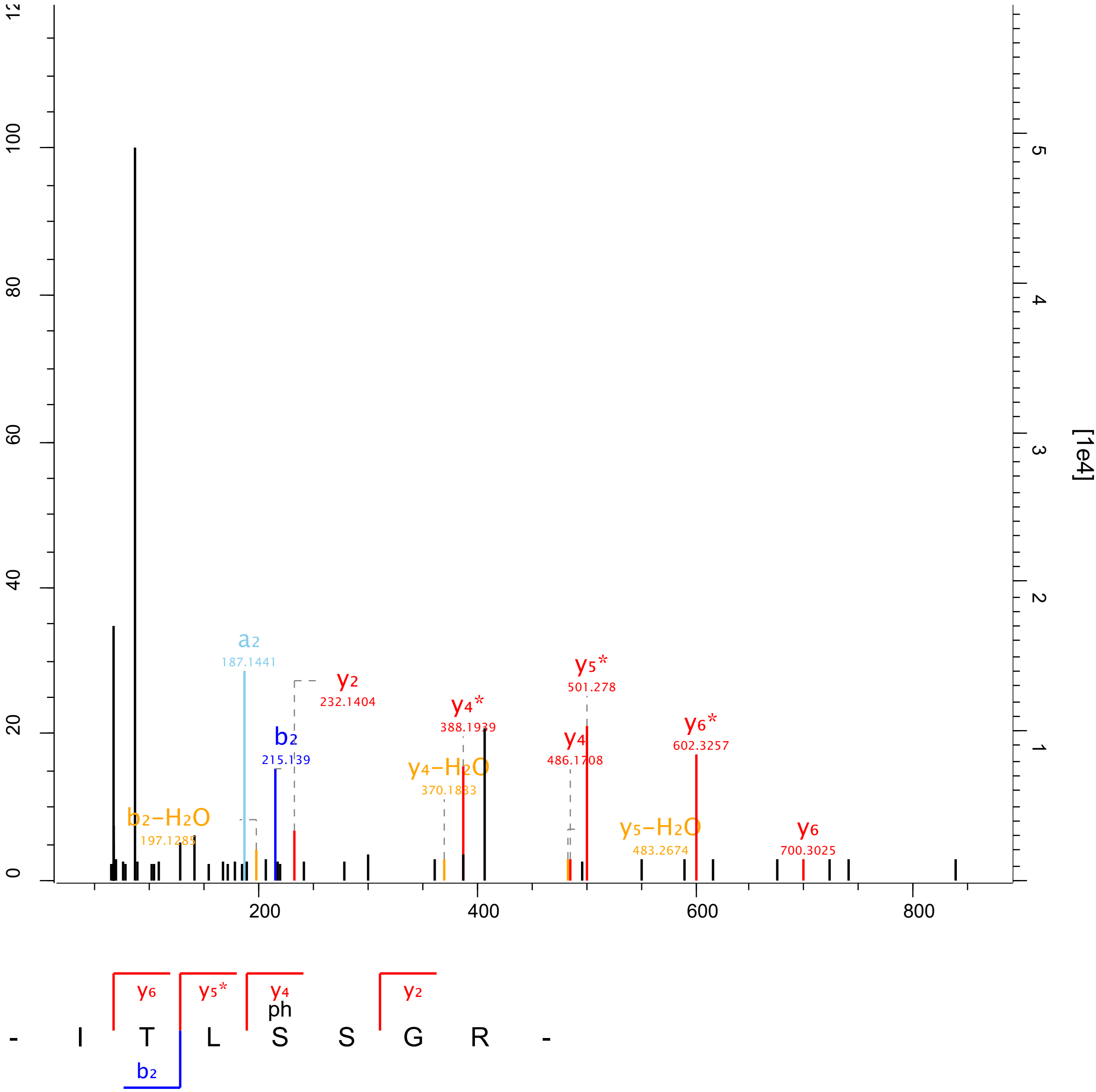

| Raw file | Scan | Method | Score | m/z |
|---|---|---|---|---|
| sys_30_2 | 6657 | FTMS; HCD | 119.75 | 441.22 |

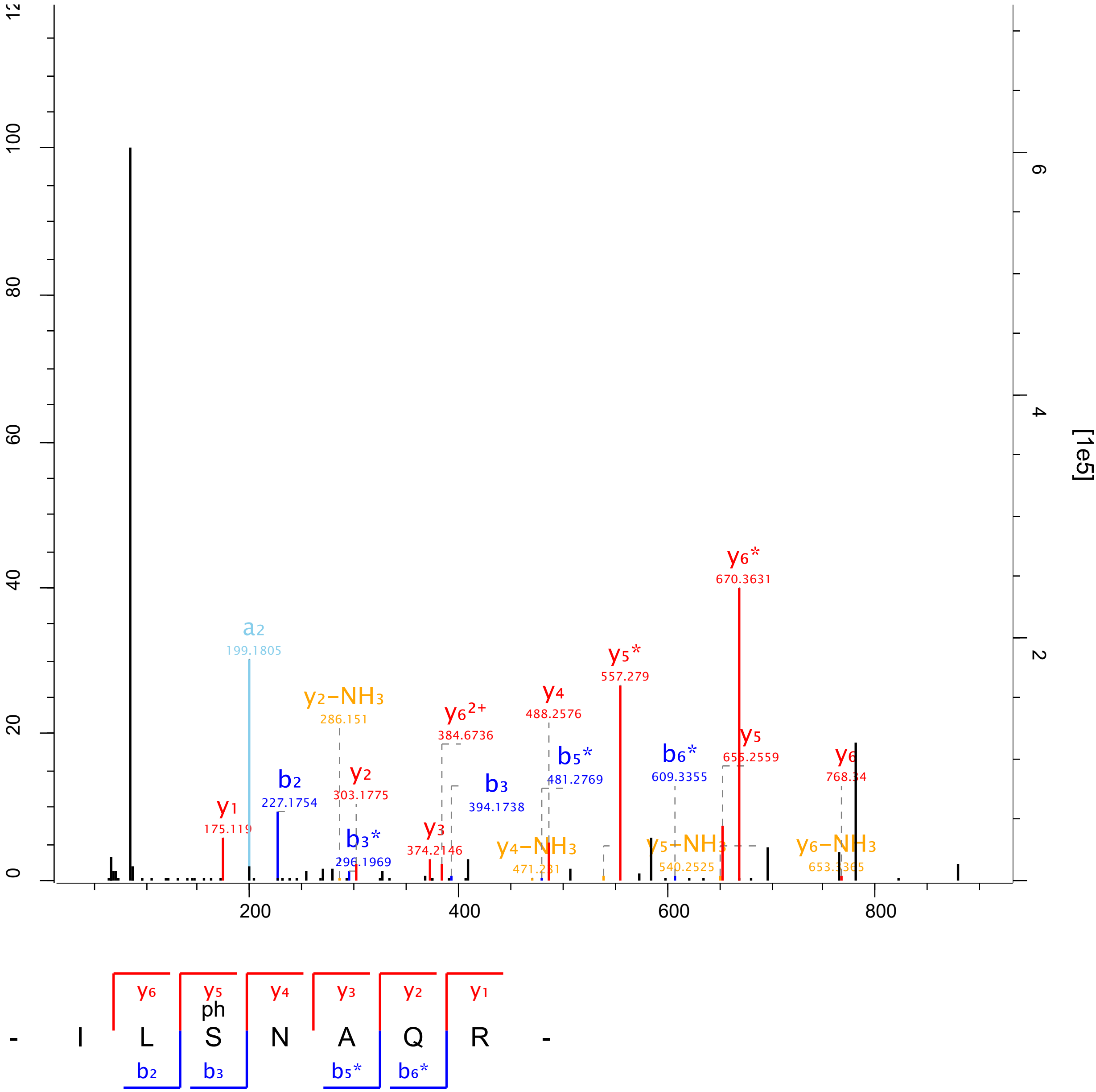

| Raw file | Scan | Method | Score | m/z |
|---|---|---|---|---|
| sys_30_2 | 8708 | FTMS; HCD | 152.23 | 501.72 |

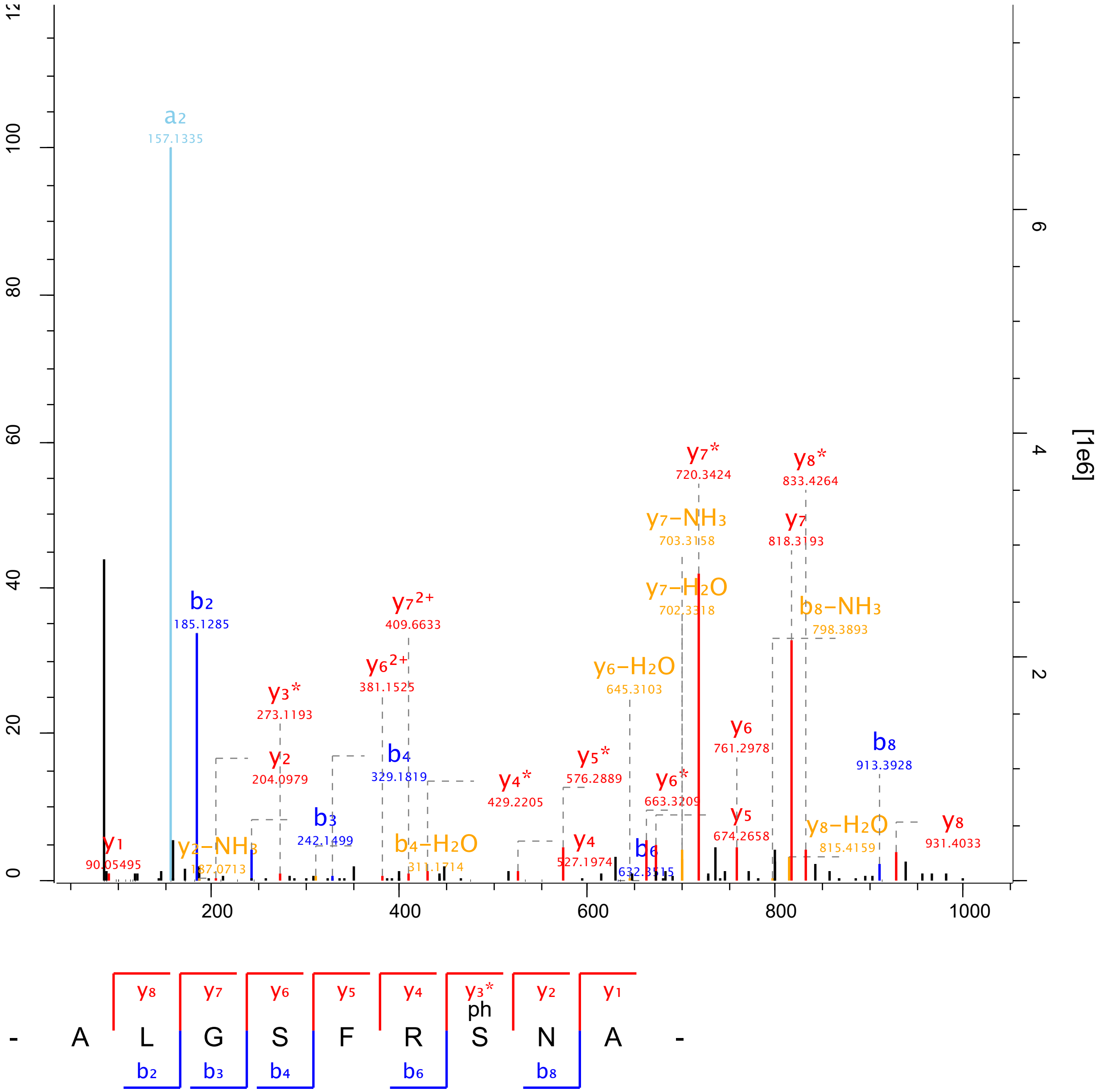

| Raw file | Scan | Method | Score | m/z |
|---|---|---|---|---|
| sys_30_3 | 10802 | FTMS; HCD | 150.13 | 709.83 |

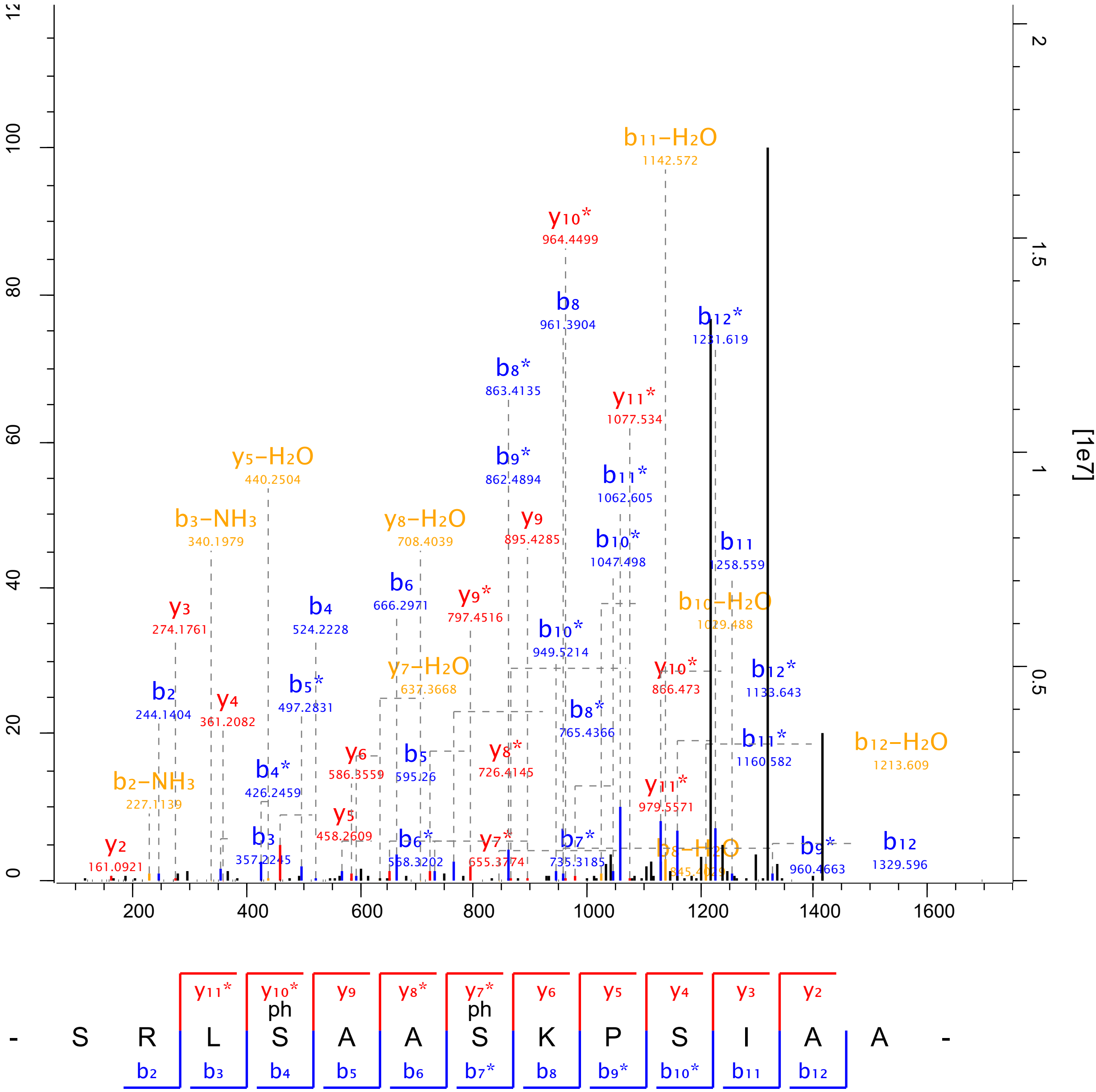

| Raw file | Scan | Method | Score | m/z |
|---|---|---|---|---|
| sys_30_3 | 13930 | FTMS; HCD | 57.53 | 598.77 |

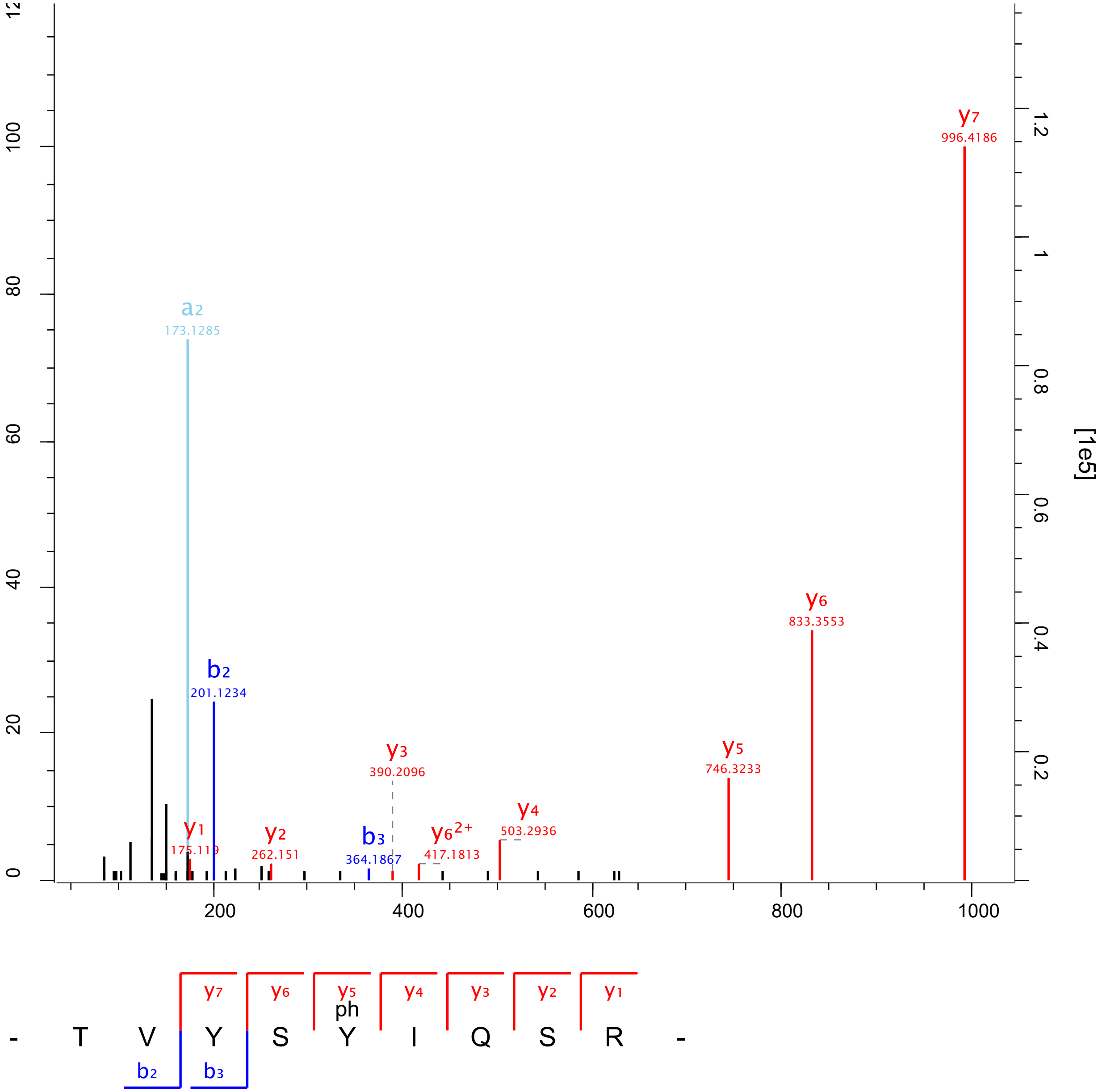

| Raw file | Scan | Method | Score | m/z |
|---|---|---|---|---|
| sys_30_3 | 19654 | FTMS; HCD | 47.55 | 722.82 |

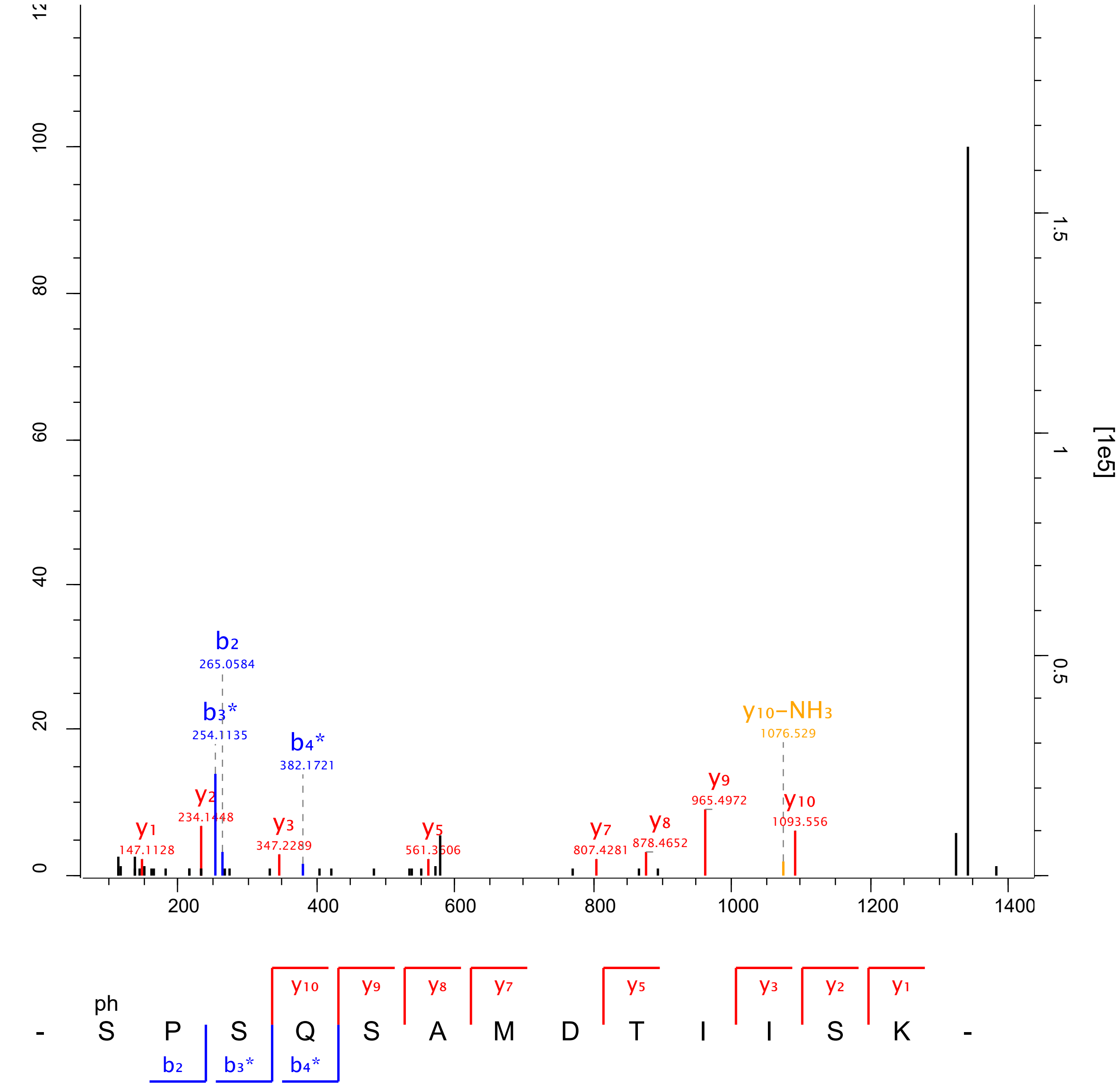

| Raw file | Scan | Method | Score | m/z |
| --- | --- | --- | --- | --- |
| sys_30_3 | 25568 | FTMS; HCD | 106.31 | 628.24 |

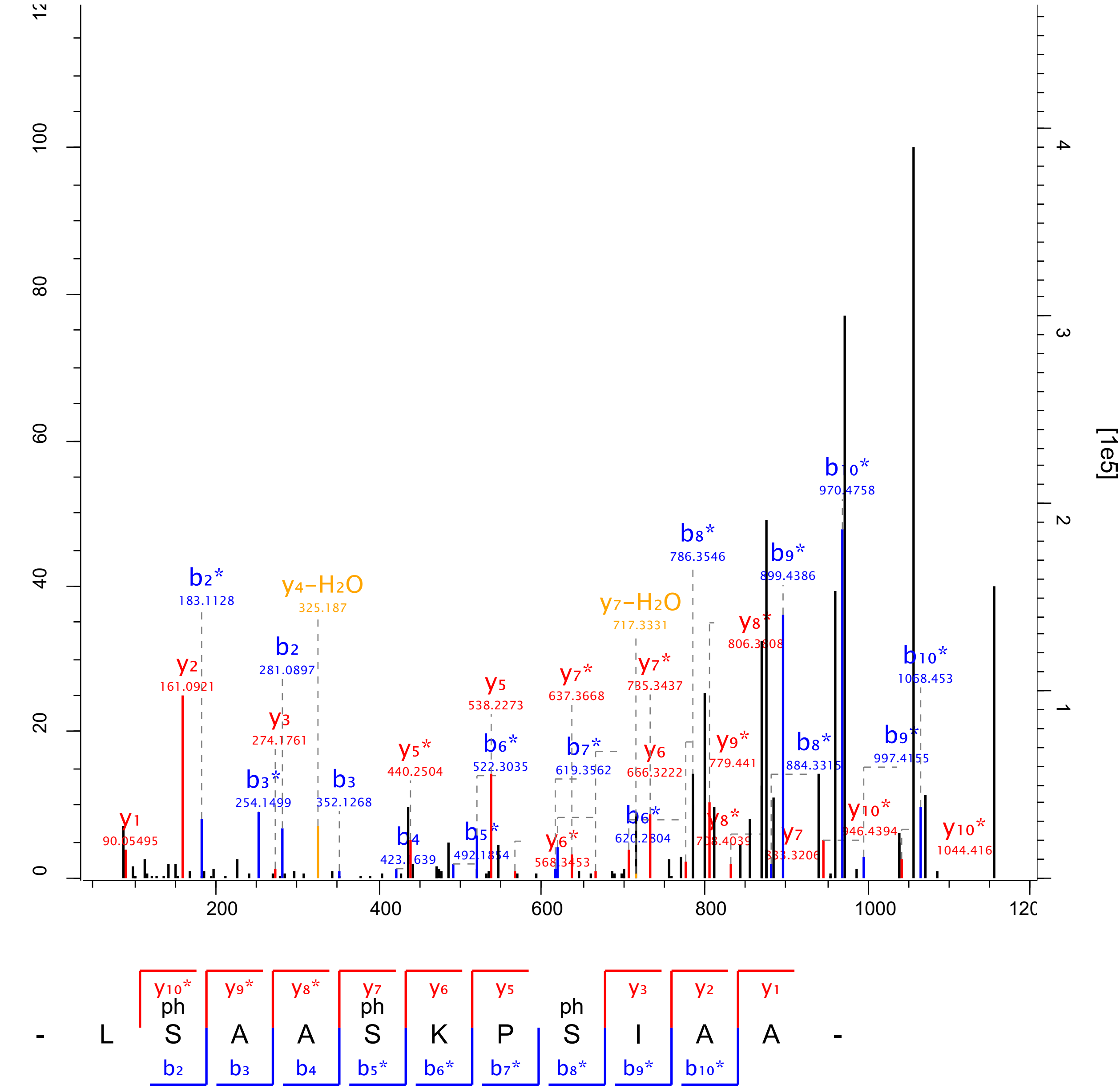

| Raw file | Scan | Method | Score | m/z |
|---|---|---|---|---|
| sys_30_3 | 3325 | FTMS; HCD | 66.57 | 422.17 |

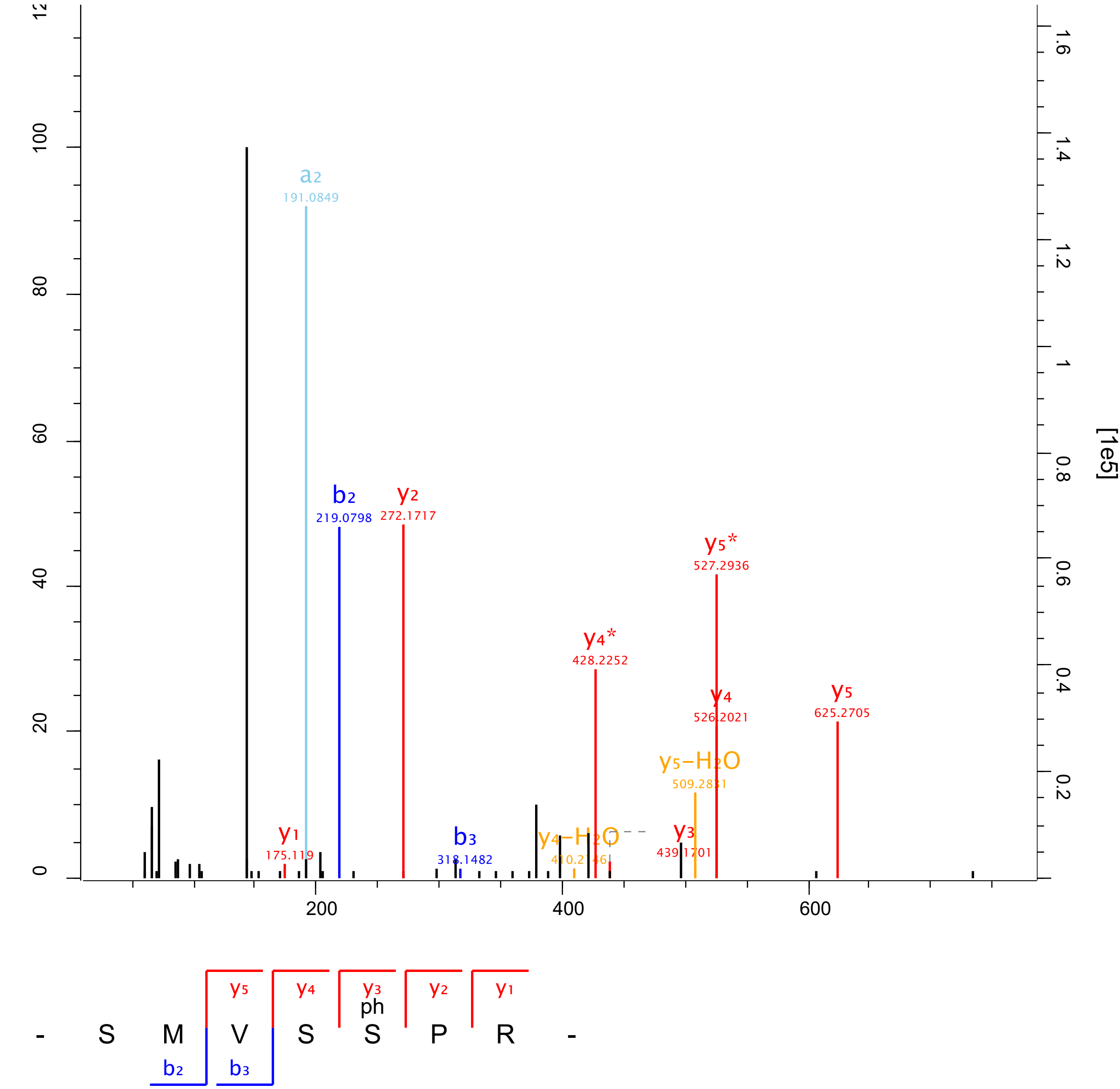

| Raw file | Scan | Method | Score | m/z |
| --- | --- | --- | --- | --- |
| sys_30_3 | 5822 | FTMS; HCD | 78.81 | 570.75 |

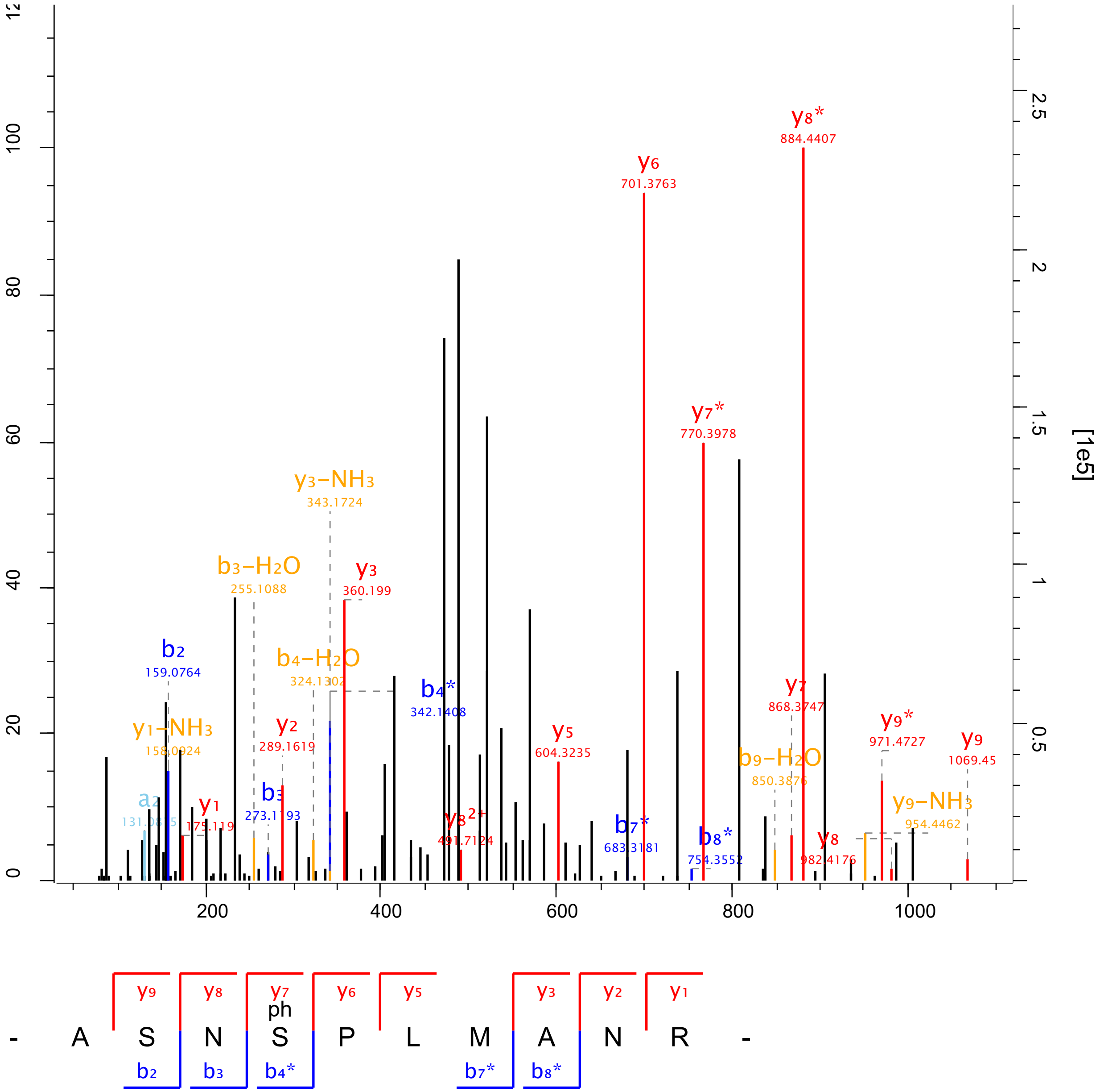

| Raw file | Scan | Method | Score | m/z |
|---|---|---|---|---|
| sys_30_3 | 8493 | FTMS; HCD | 139.86 | 480.22 |

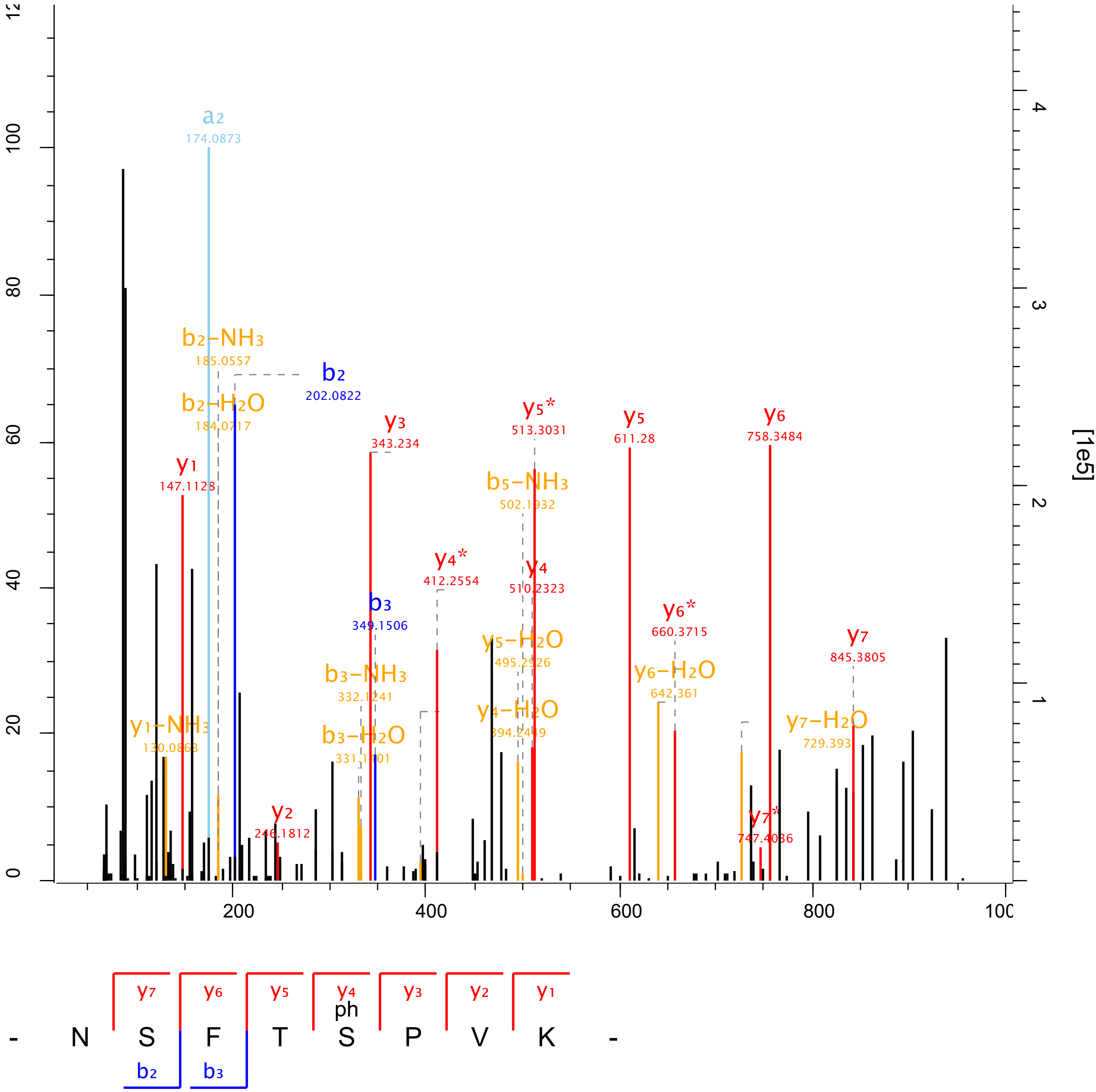

| Raw file | Scan | Method | Score | m/z |
|---|---|---|---|---|
| sys_30_3 | 9507 | FTMS; HCD | 140.31 | 621.28 |

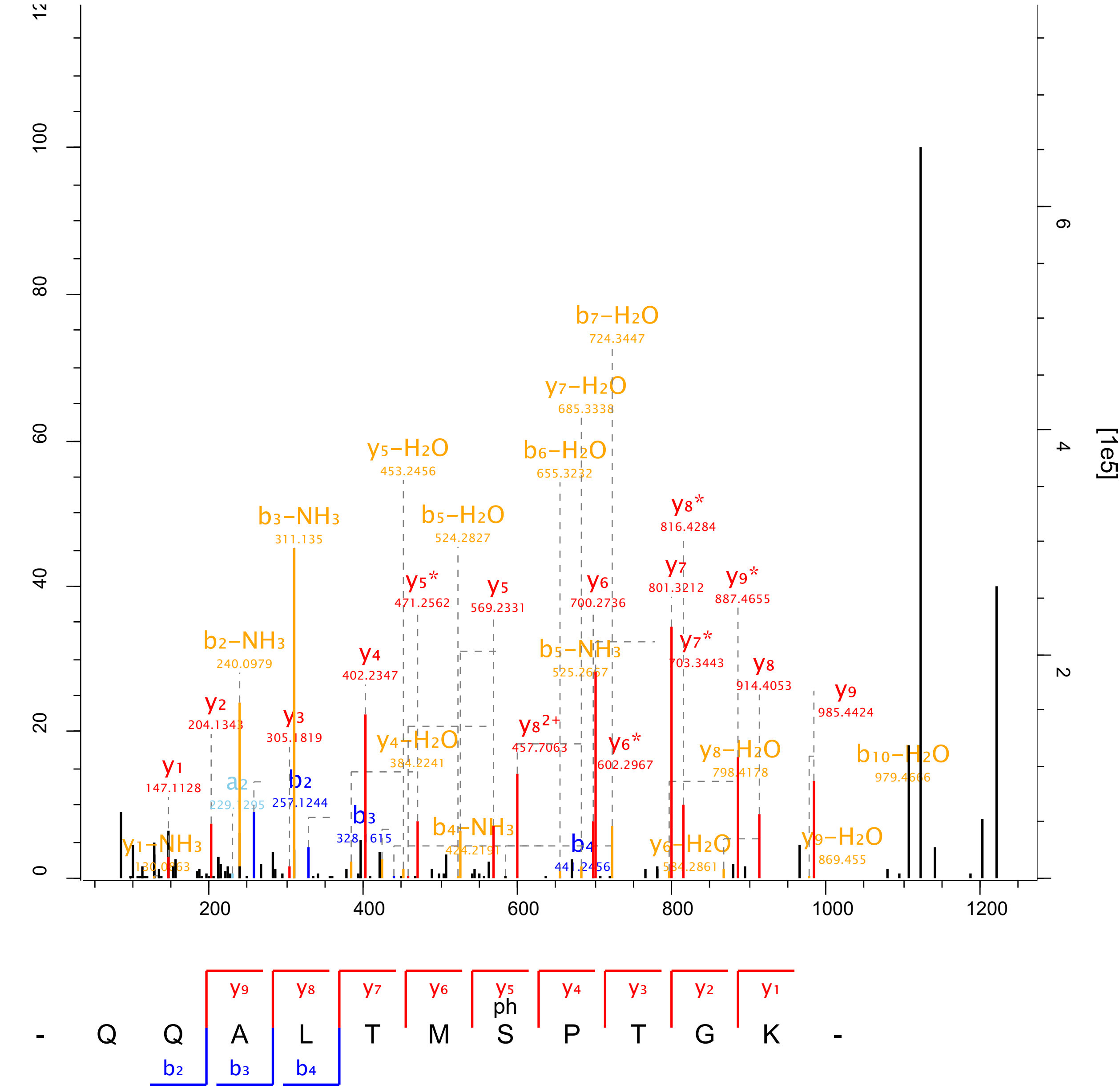

| Raw file | Scan | Method | Score | m/z |
|---|---|---|---|---|
| water_00_1 | 10060 | FTMS; HCD | 44.51 | 470.22 |

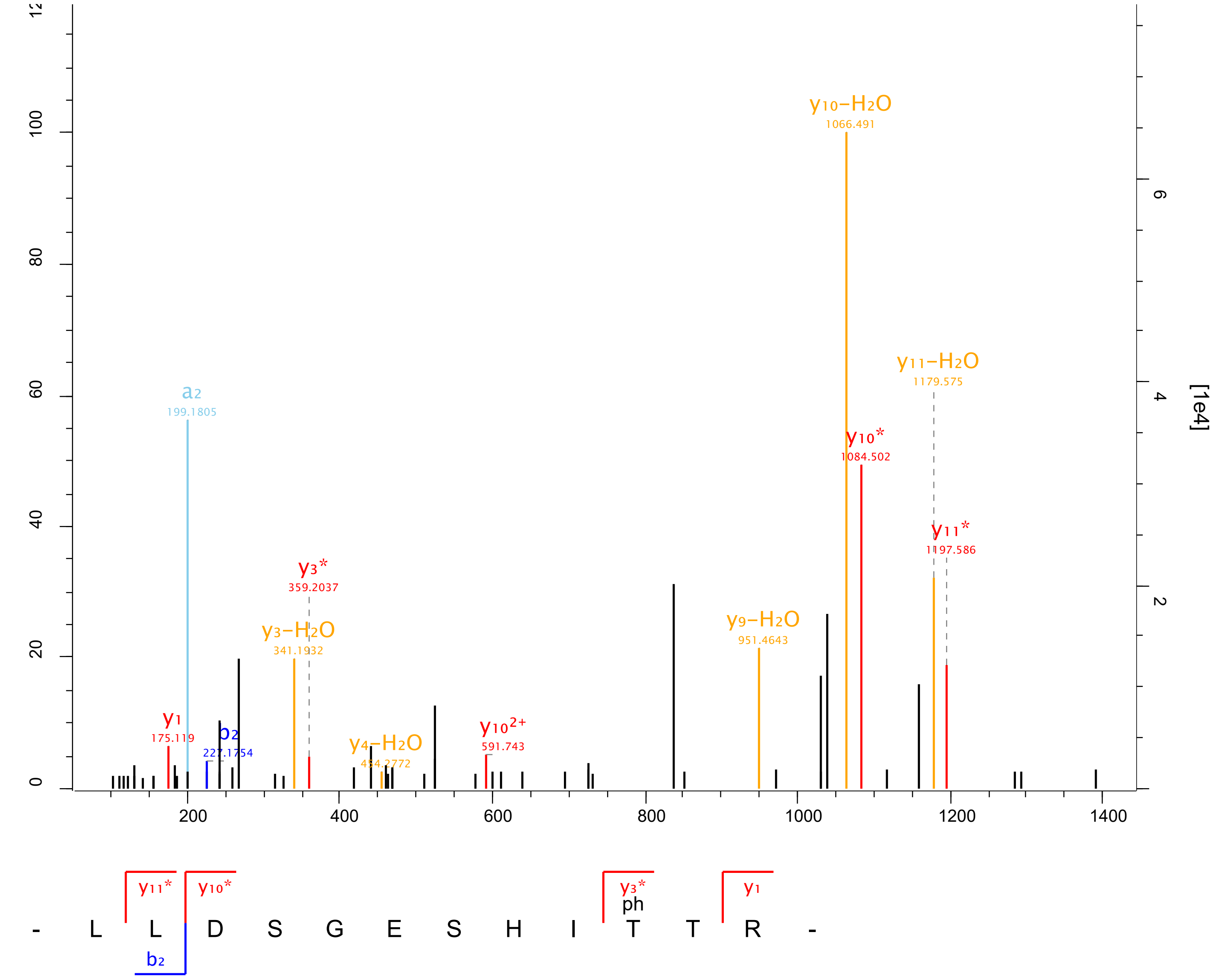

| Raw file | Scan | Method | Score | m/z |
|---|---|---|---|---|
| water_00_1 | 10428 | FTMS; HCD | 98.04 | 436.19 |

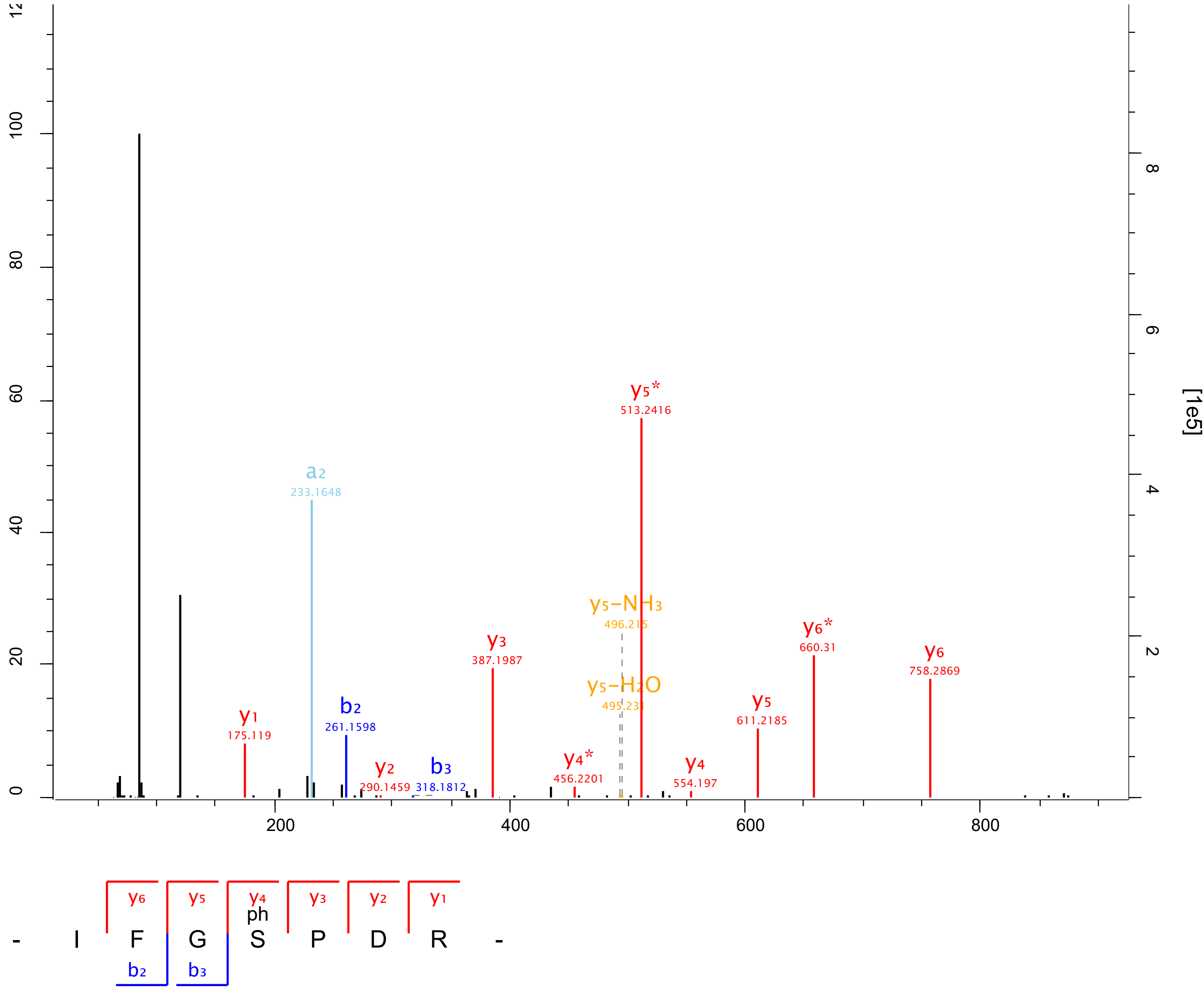

| Raw file | Scan | Method | Score | m/z |
| --- | --- | --- | --- | --- |
| water_00_1 | 10441 | FTMS; HCD | 72.9 | 556.24 |

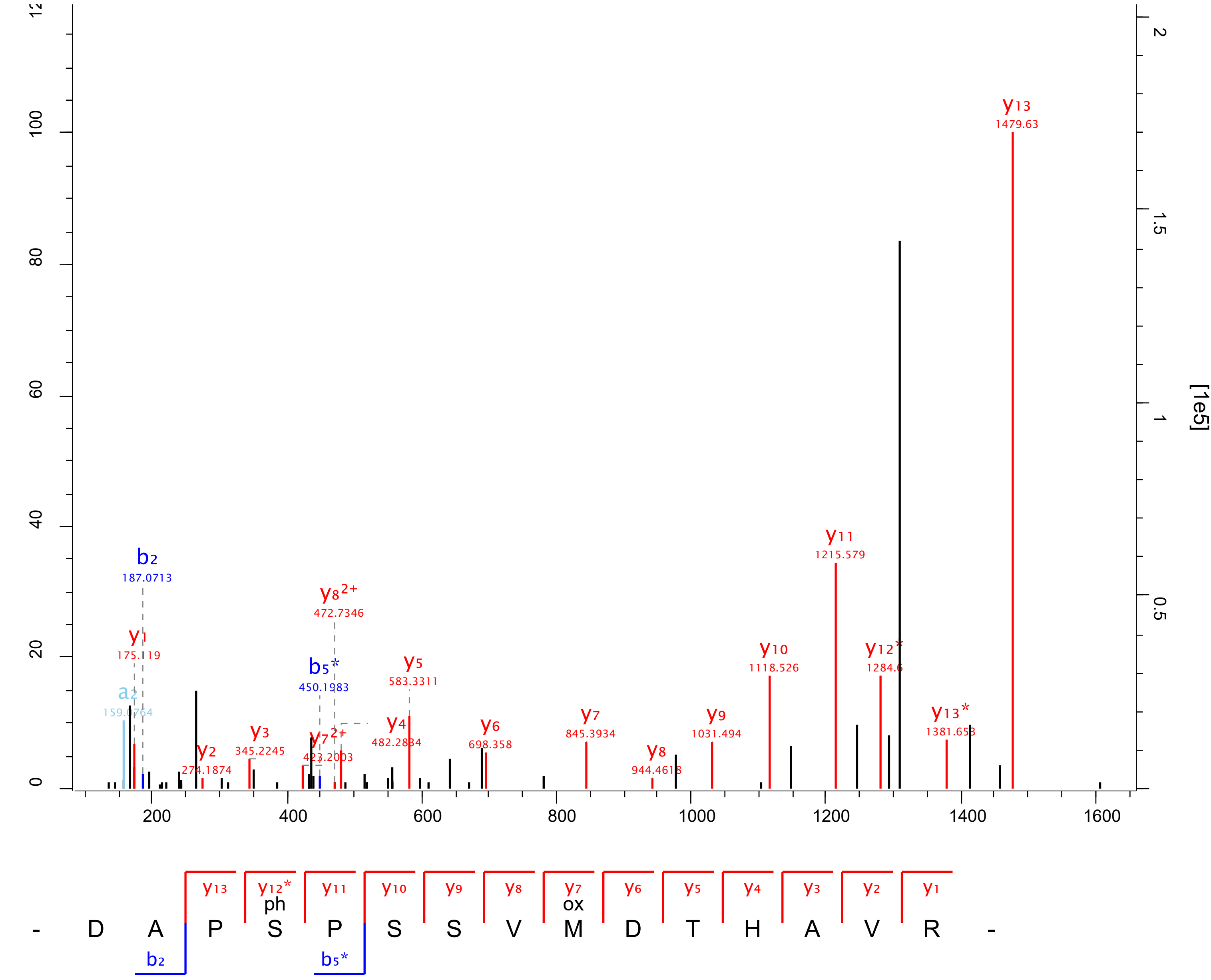

| Raw file | Scan | Method | Score | m/z |
|---|---|---|---|---|
| water_00_1 | 10625 | FTMS; HCD | 57.43 | 650.27 |

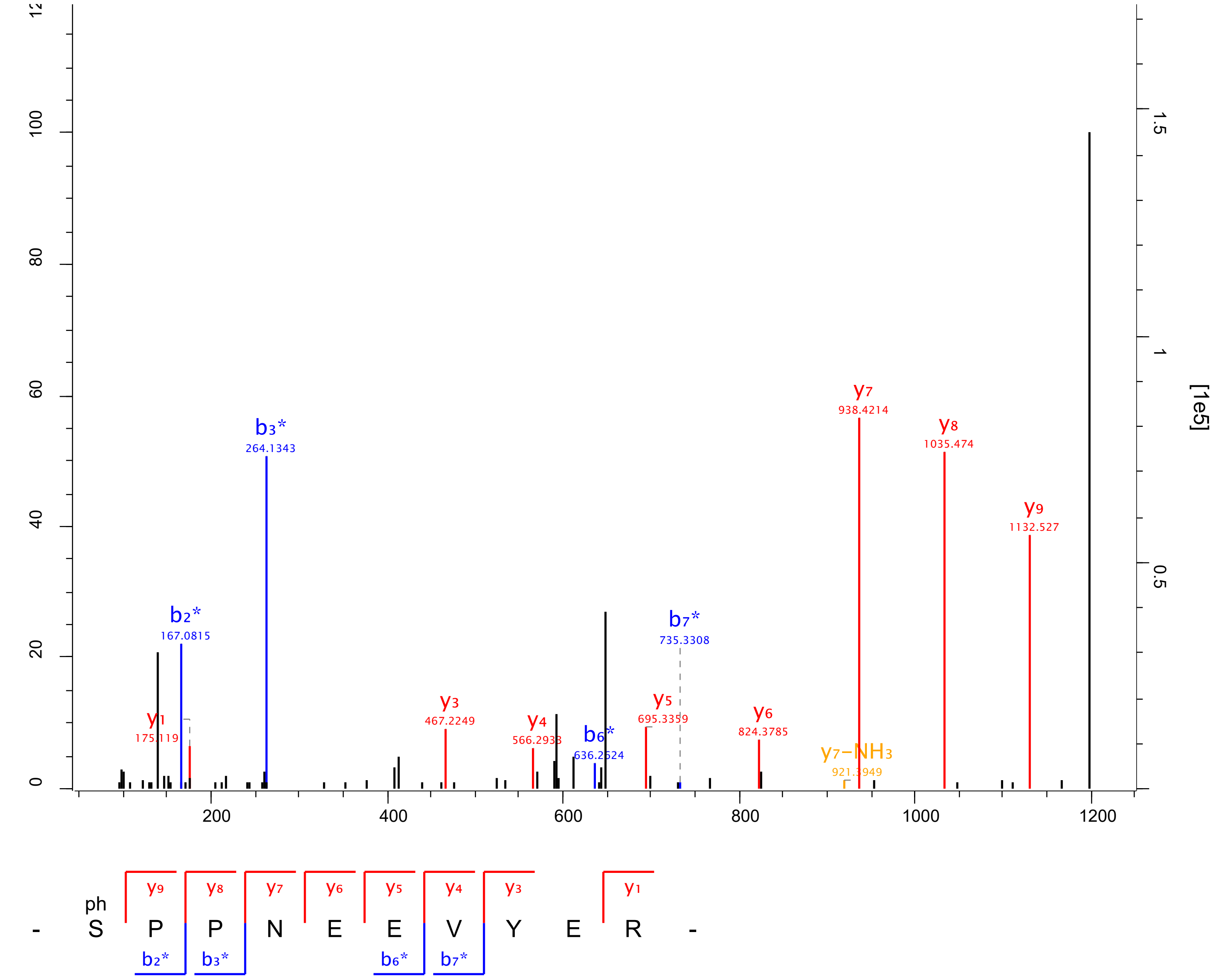

| Raw file | Scan | Method | Score | m/z |
|---|---|---|---|---|
| water_00_1 | 10688 | FTMS; HCD | 44.25 | 680.26 |

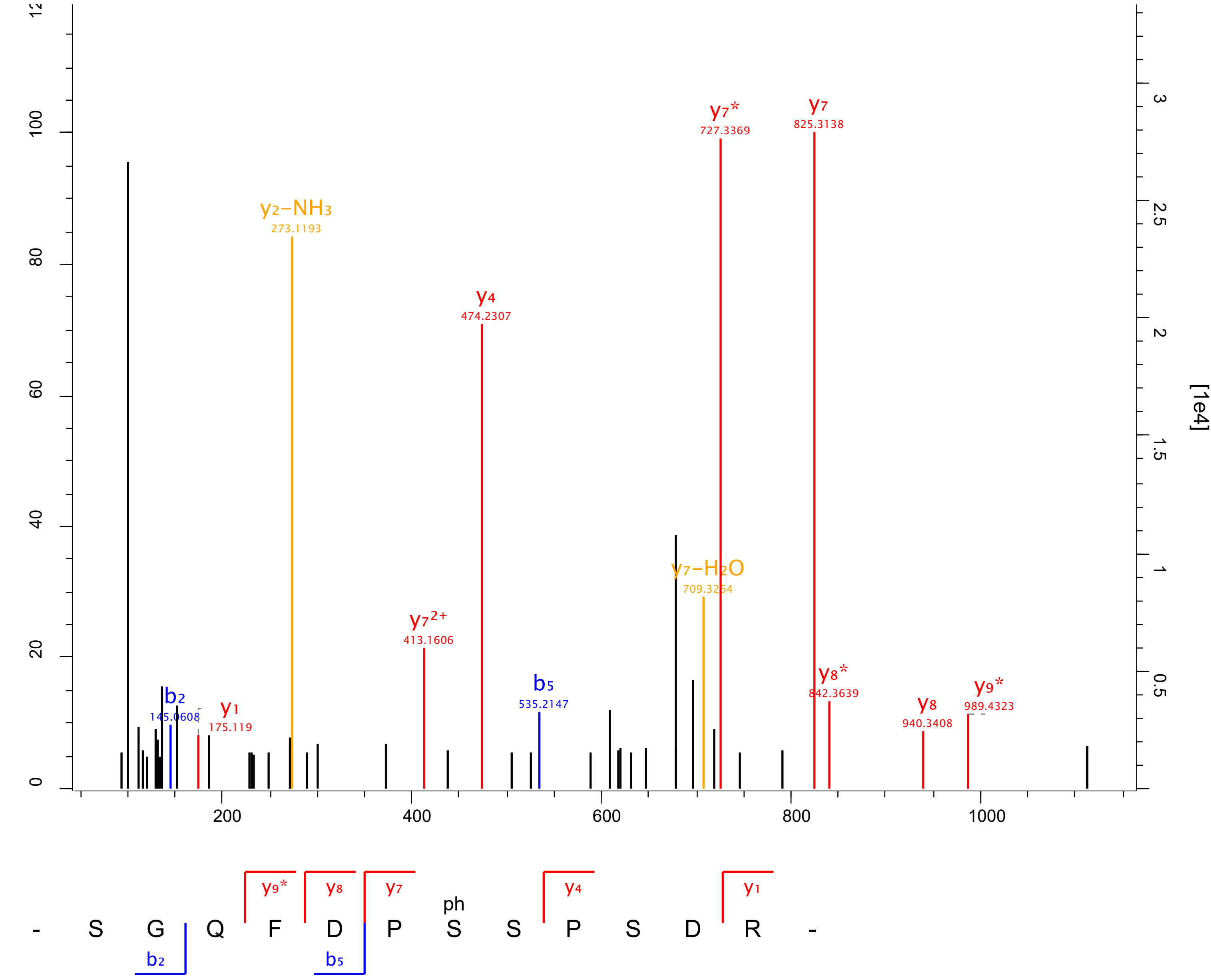

| Raw file | Scan | Method | Score | m/z |
|---|---|---|---|---|
| water_00_1 | 10799 | FTMS; HCD | 58.32 | 621.29 |

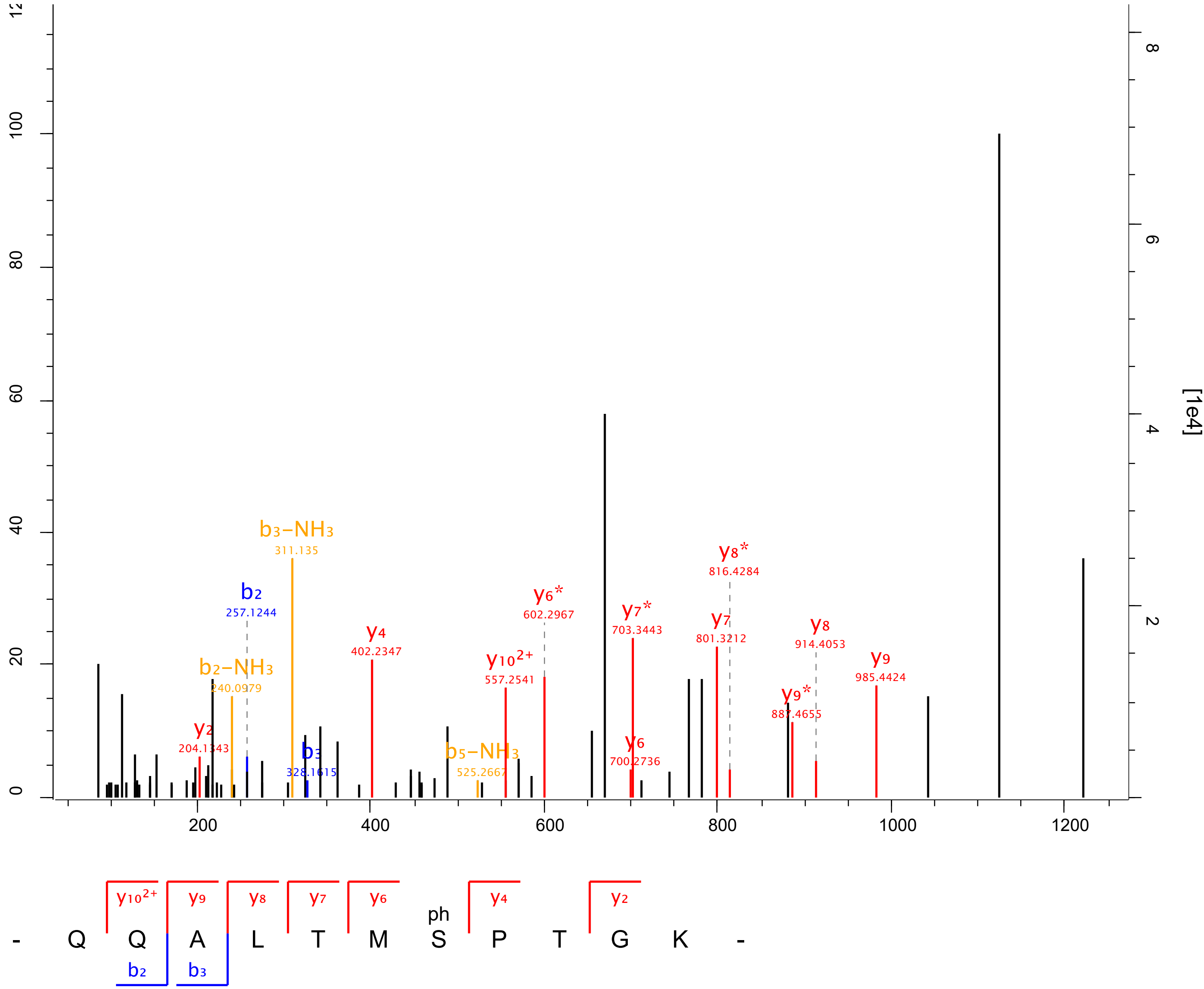

| Raw file | Scan | Method | Score | m/z |
|---|---|---|---|---|
| water_00_1 | 11014 | FTMS; HCD | 189.11 | 678.3 |

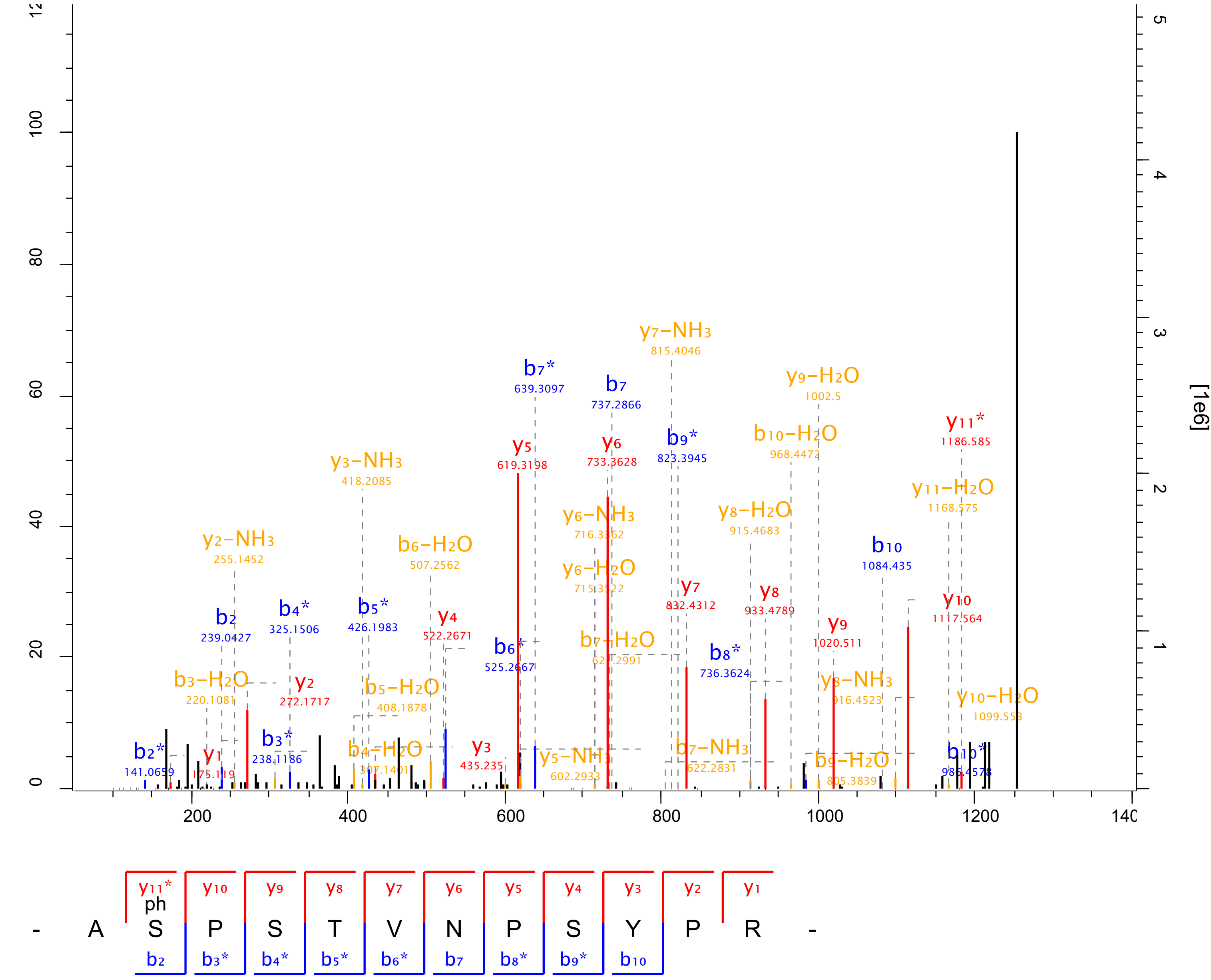

| Raw file | Scan | Method | Score | m/z |
|---|---|---|---|---|
| water_00_1 | 11138 | FTMS; HCD | 63.82 | 505.72 |

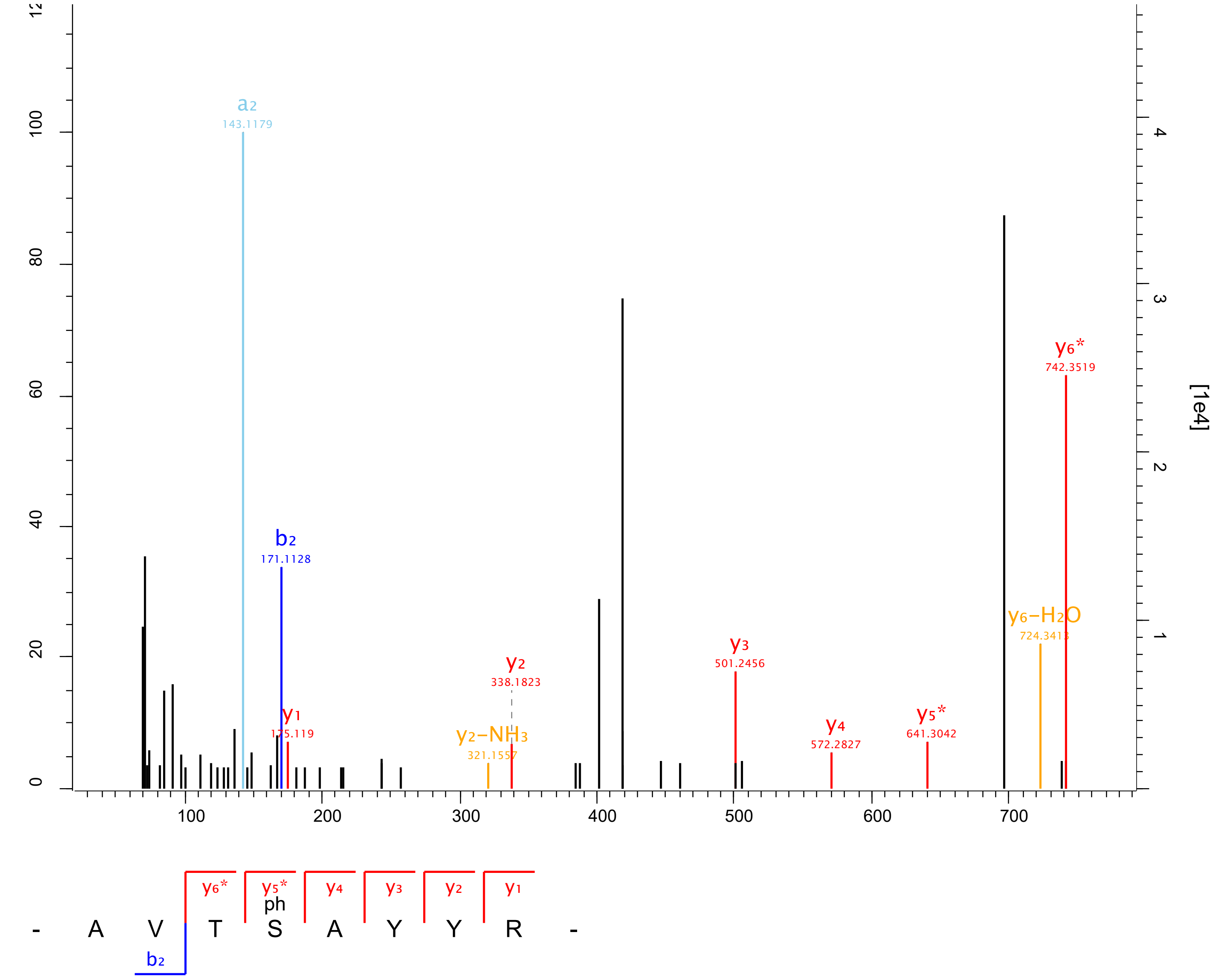

| Raw file | Scan | Method | Score | m/z |
|---|---|---|---|---|
| water_00_1 | 11212 | FTMS; HCD | 273.78 | 790.84 |

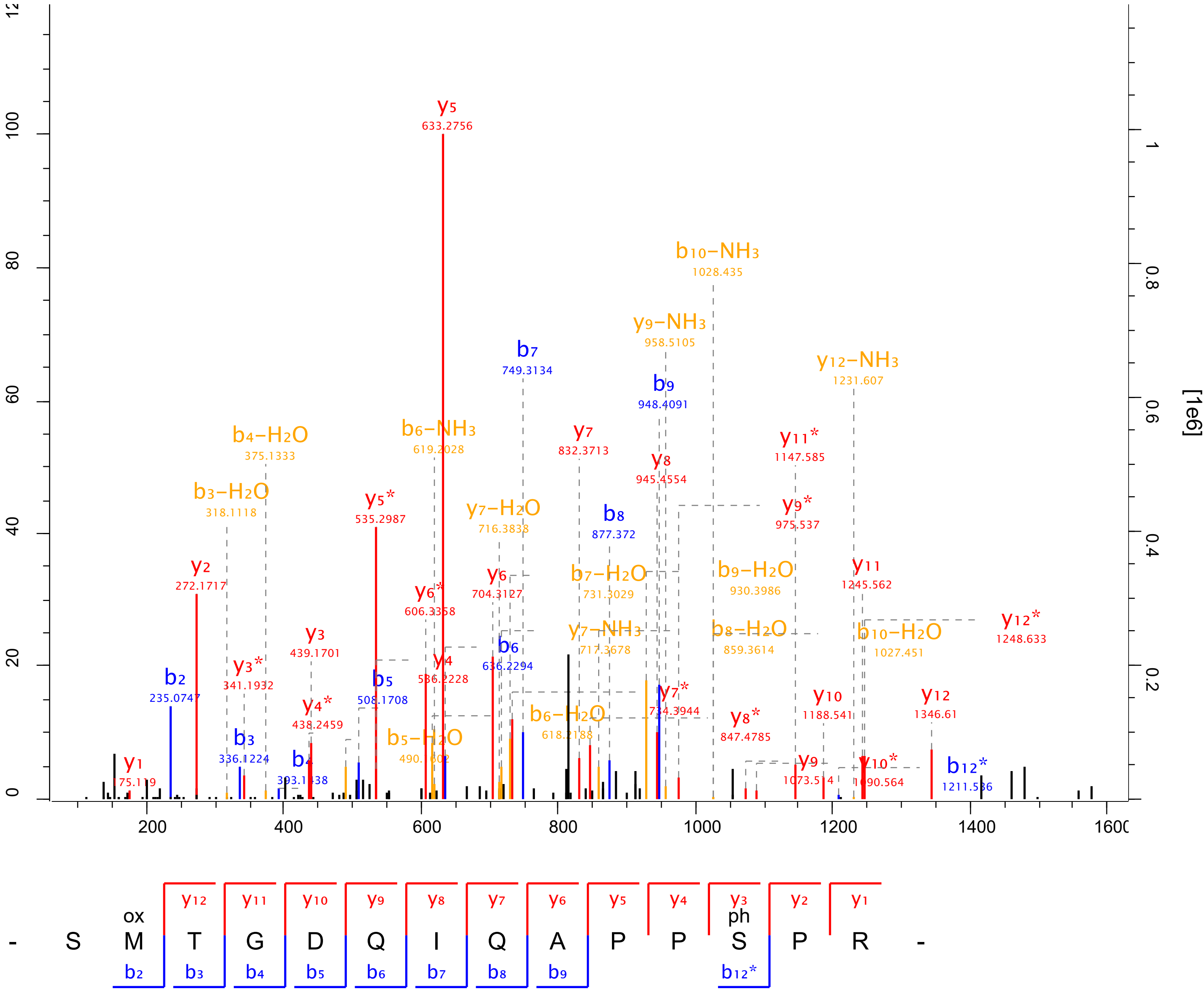

| Raw file | Scan | Method | Score | m/z |
|---|---|---|---|---|
| water_00_1 | 11360 | FTMS; HCD | 72.93 | 466.18 |

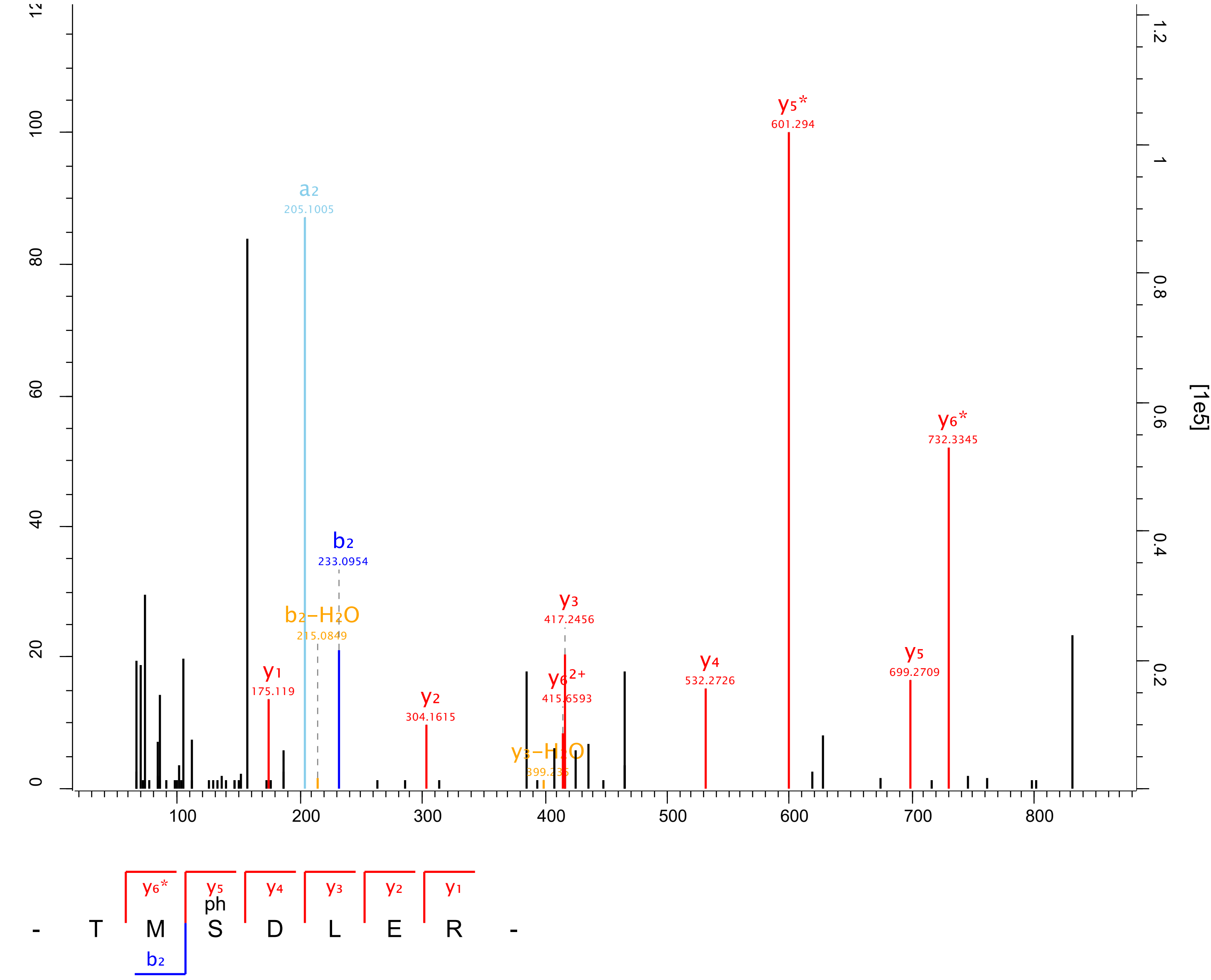

| Raw file | Scan | Method | Score | m/z |
|---|---|---|---|---|
| water_00_1 | 11454 | FTMS; HCD | 128.06 | 662.31 |

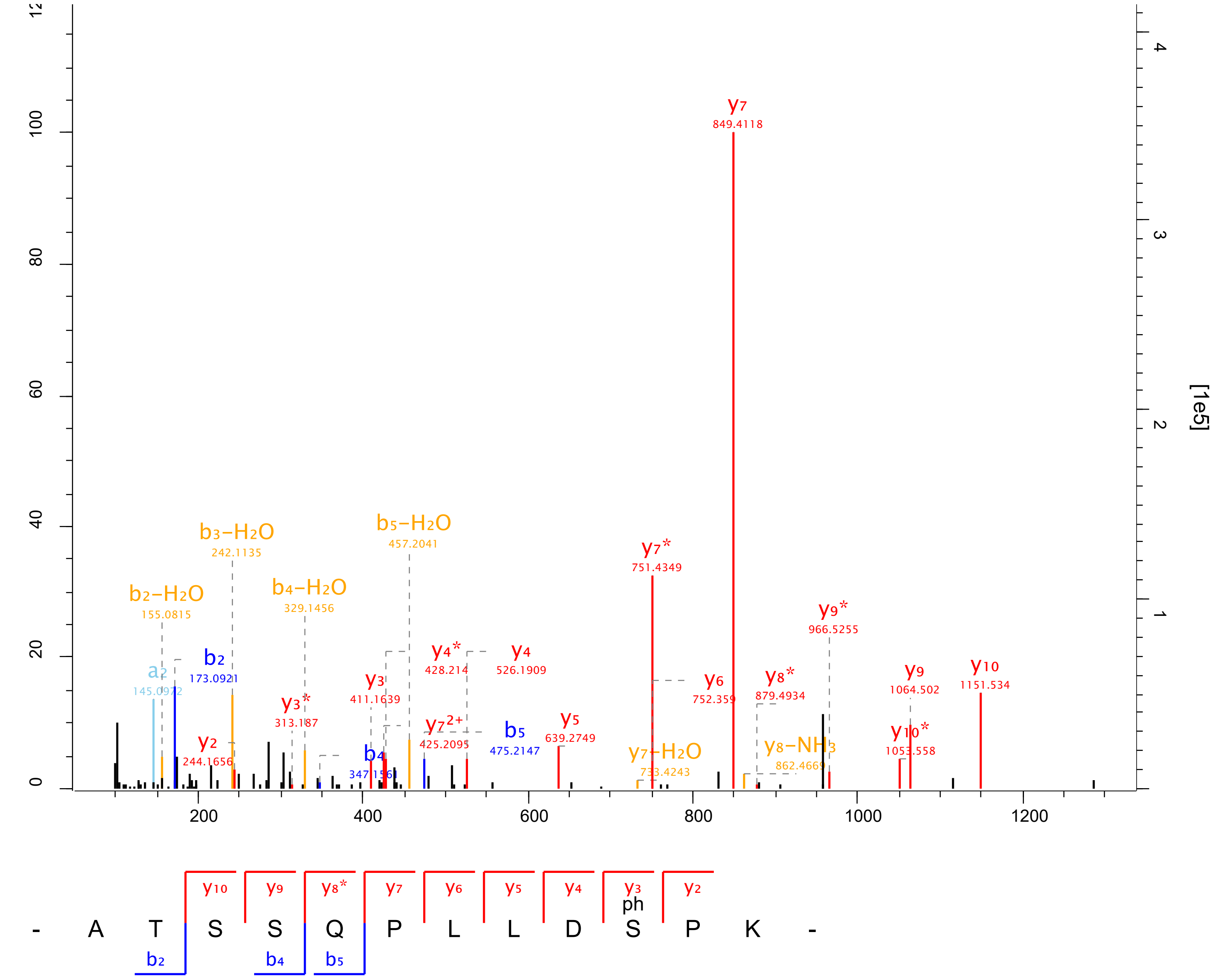

| Raw file | Scan | Method | Score | m/z |
|---|---|---|---|---|
| water_00_1 | 11710 | FTMS; HCD | 90.61 | 547.22 |

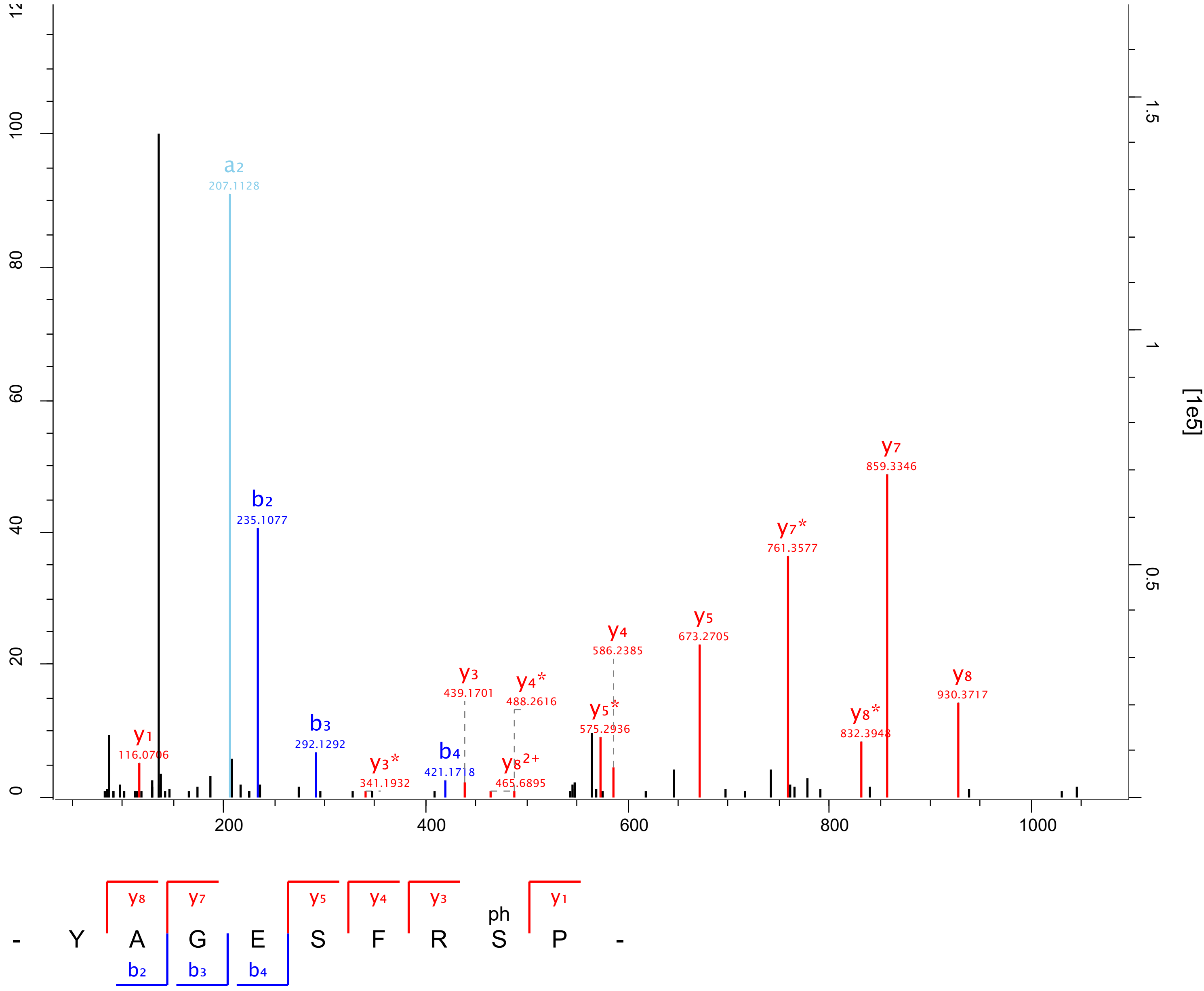

| Raw file | Scan | Method | Score | m/z |
|---|---|---|---|---|
| water_00_1 | 11895 | FTMS; HCD | 87.52 | 767.82 |

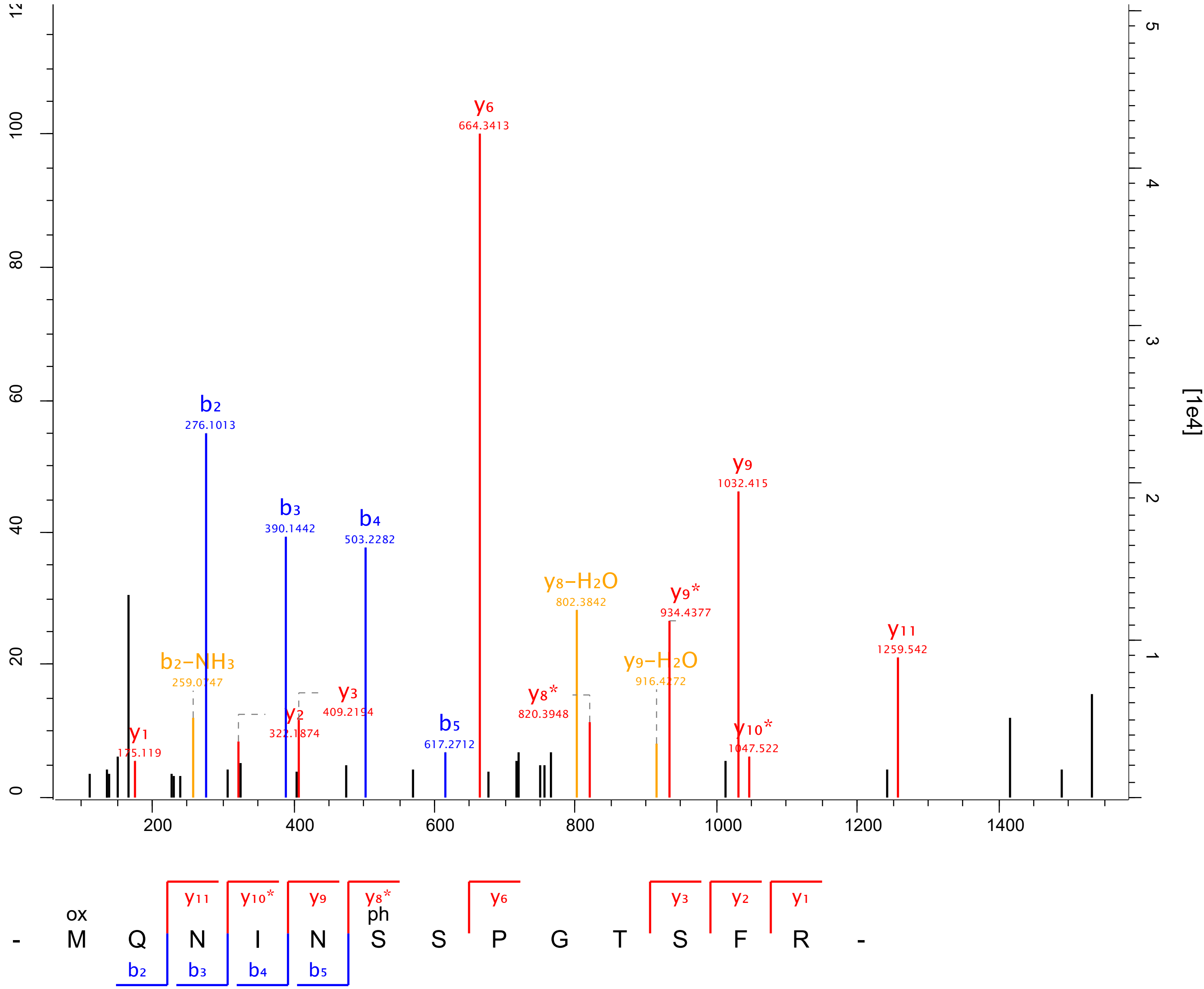

| Raw file | Scan | Method | Score | m/z |
|---|---|---|---|---|
| water_00_1 | 11941 | FTMS; HCD | 100.02 | 452.71 |

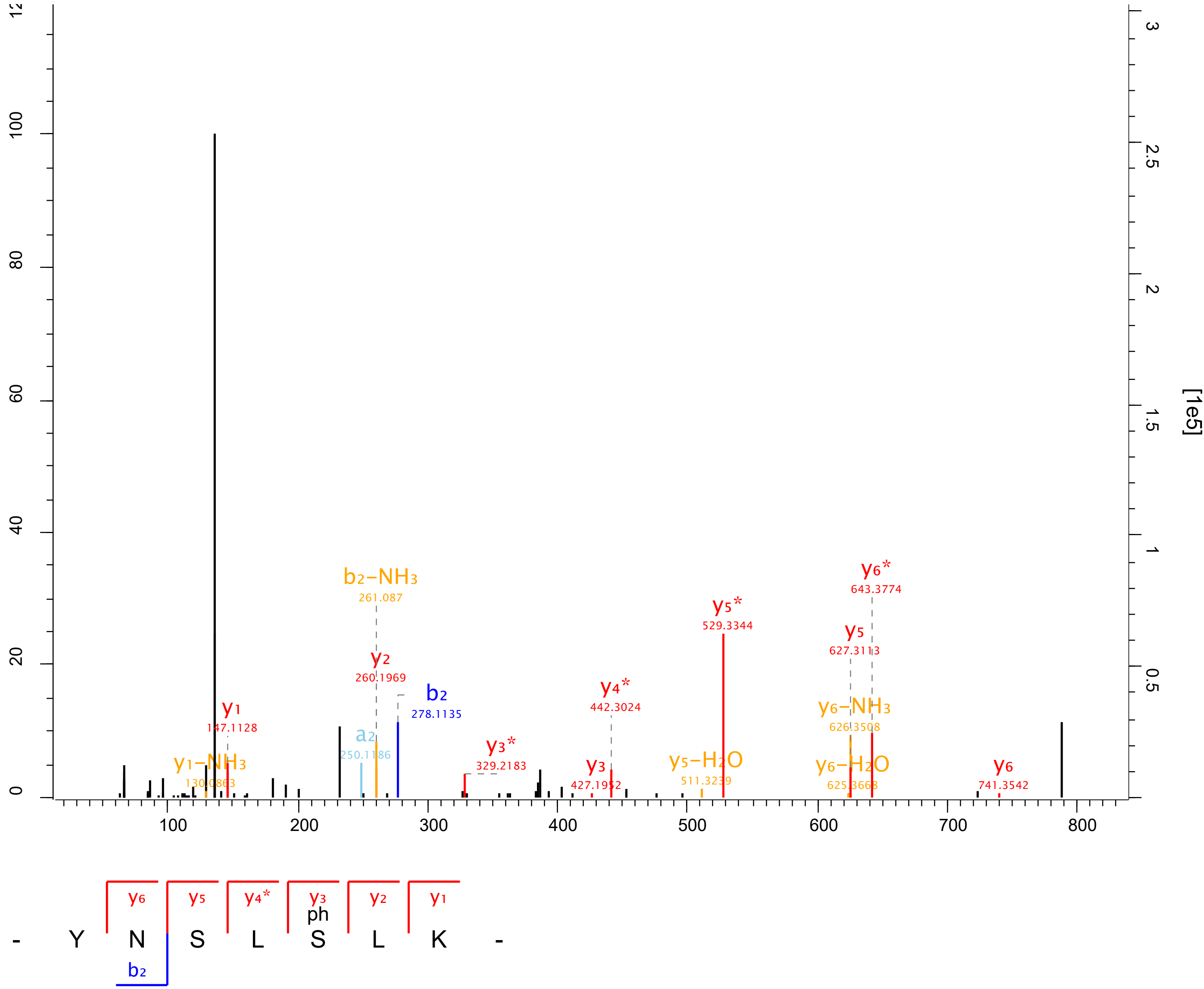

| Raw file | Scan | Method | Score | m/z |
| --- | --- | --- | --- | --- |
| water_00_1 | 11974 | FTMS; HCD | 106.33 | 692.3 |

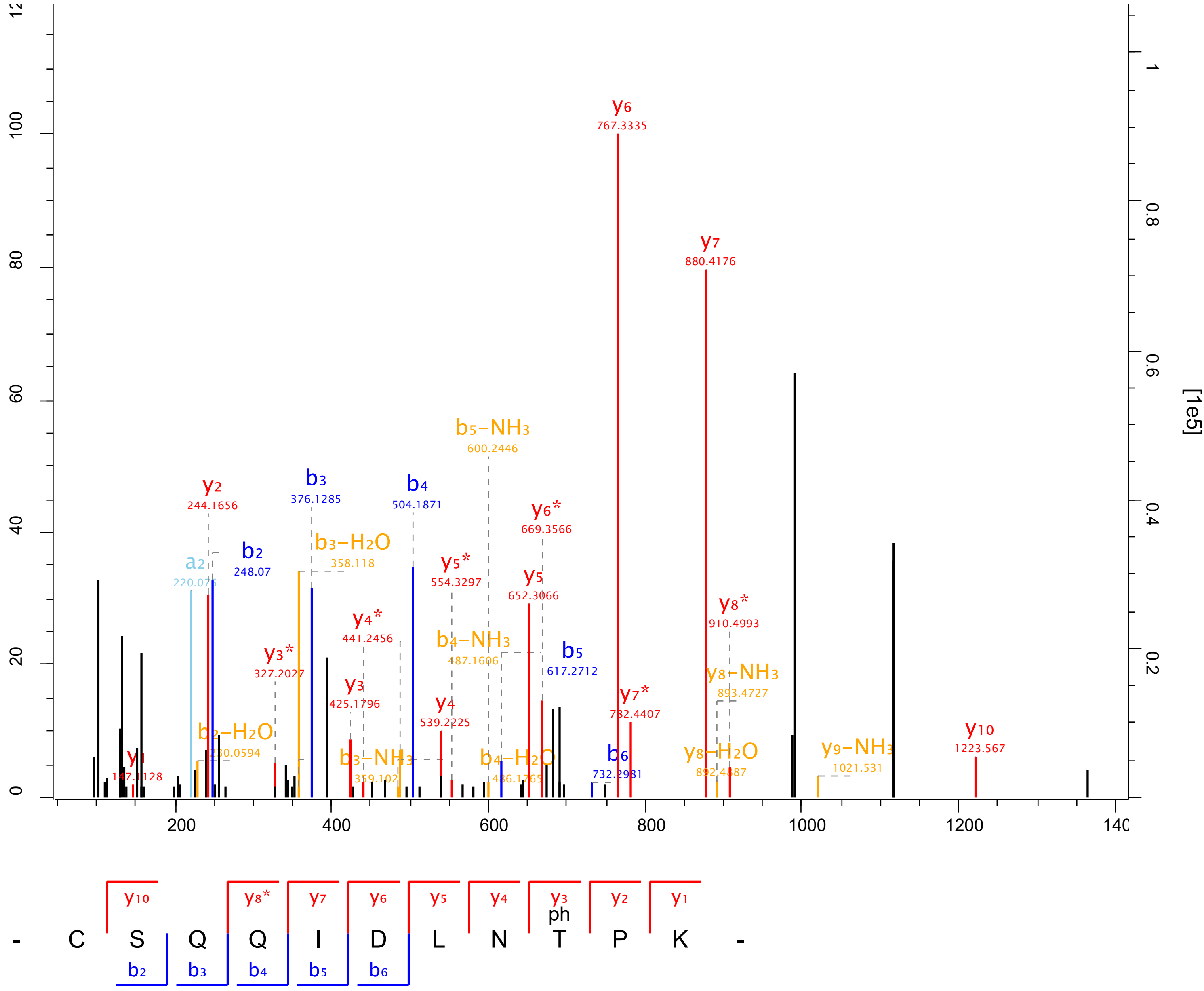

| Raw file | Scan | Method | Score | m/z |
|---|---|---|---|---|
| water_00_1 | 12077 | FTMS; HCD | 46.07 | 622.77 |

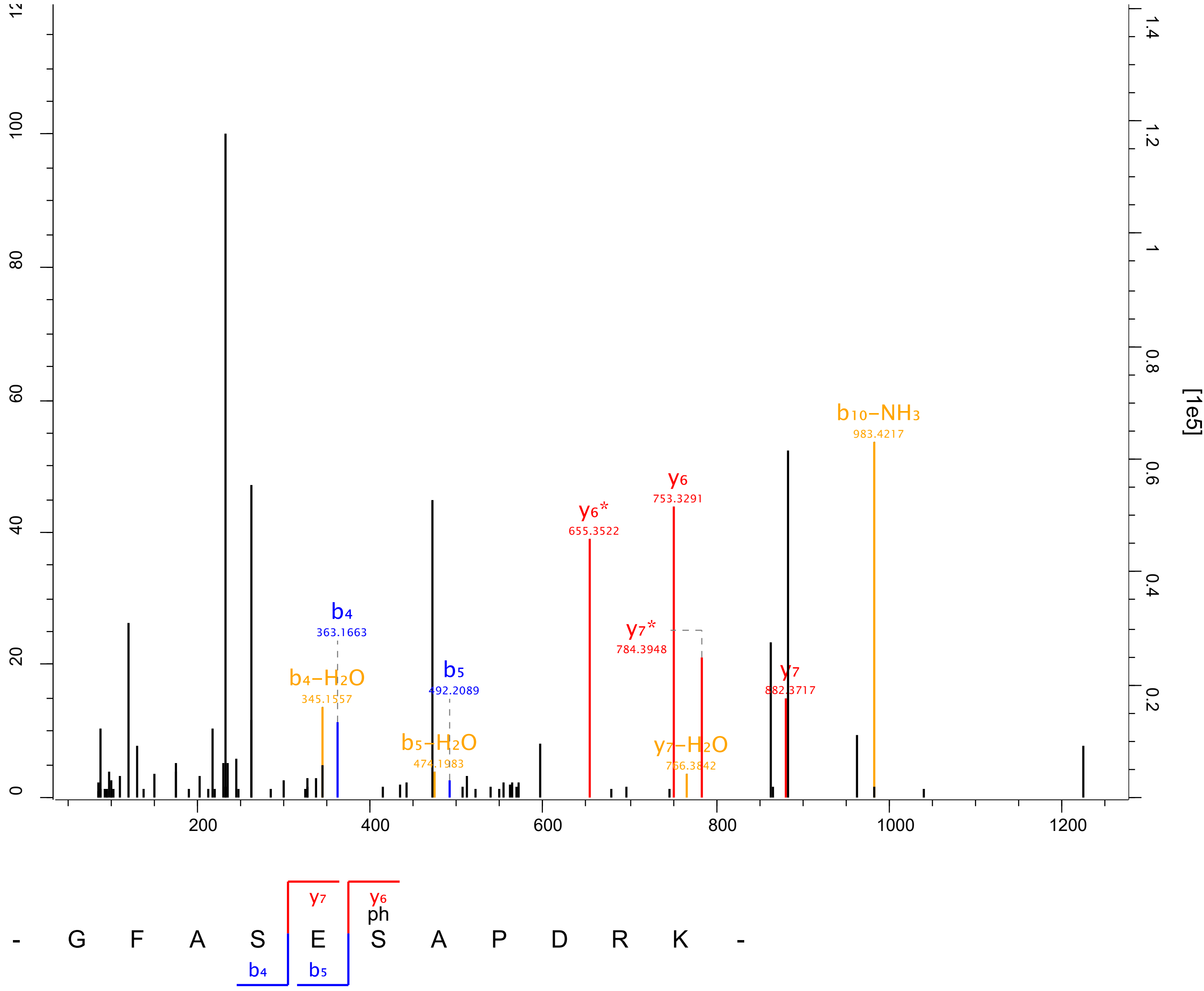

| Raw file | Scan | Method | Score | m/z |
| --- | --- | --- | --- | --- |
| water_00_1 | 12145 | FTMS; HCD | 42.42 | 532.72 |

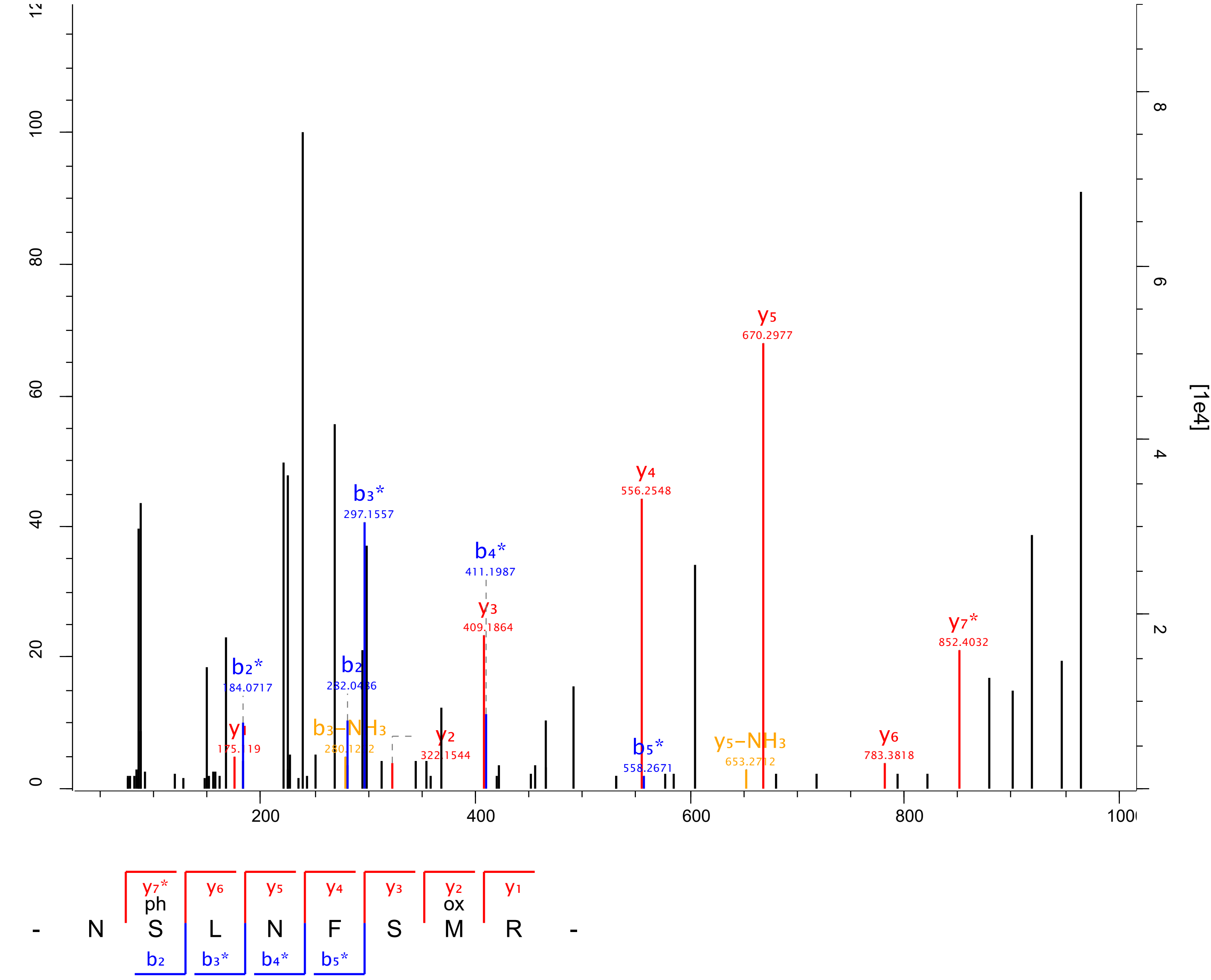

| Raw file | Scan | Method | Score | m/z |
| --- | --- | --- | --- | --- |
| water_00_1 | 12234 | FTMS; HCD | 119.41 | 671.63 |

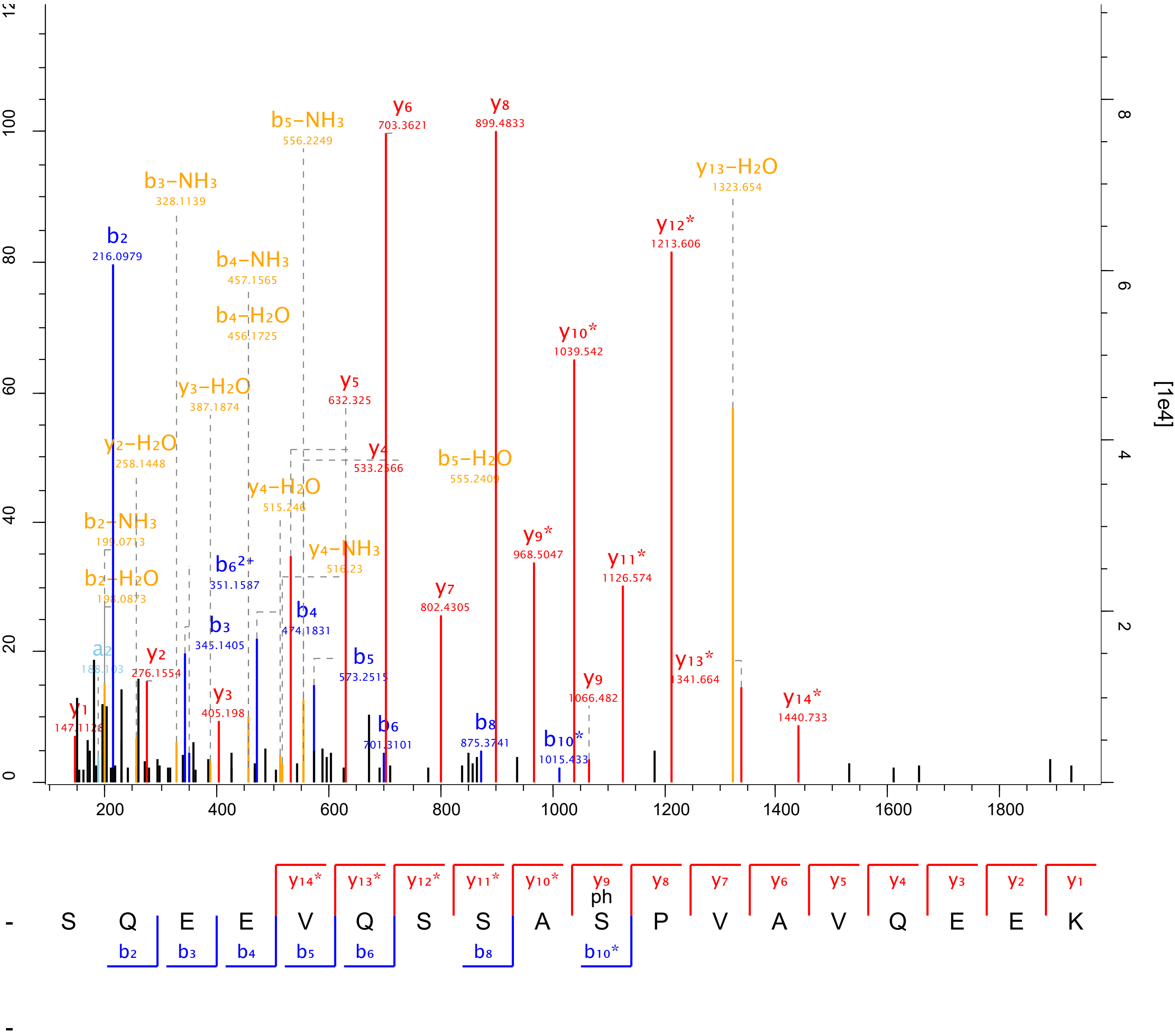

| Raw file | Scan | Method | Score | m/z |
|---|---|---|---|---|
| water_00_1 | 12460 | FTMS; HCD | 67.38 | 500.22 |

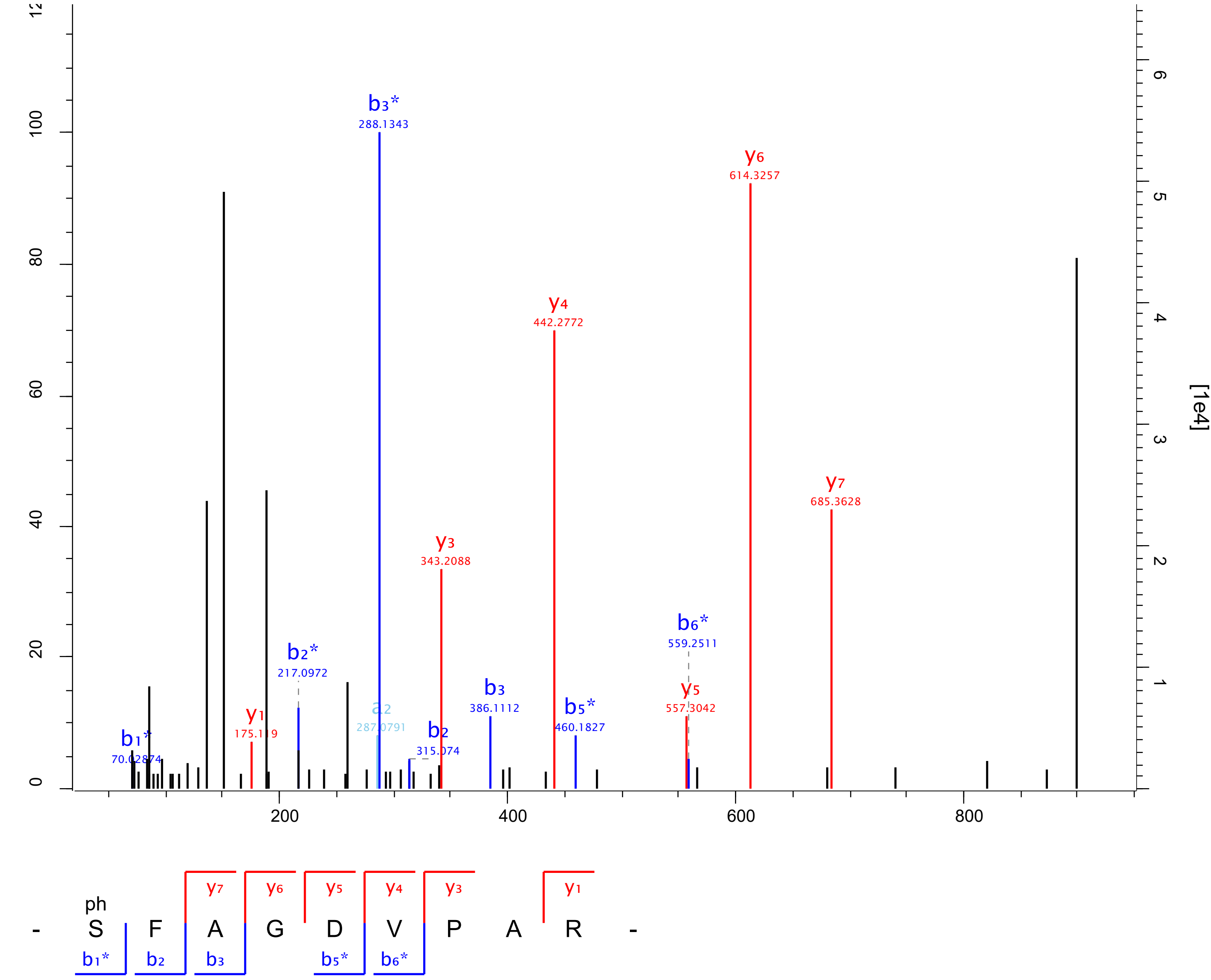

| Raw file | Scan | Method | Score | m/z |
|---|---|---|---|---|
| water_00_1 | 12468 | FTMS; HCD | 53.45 | 563.76 |

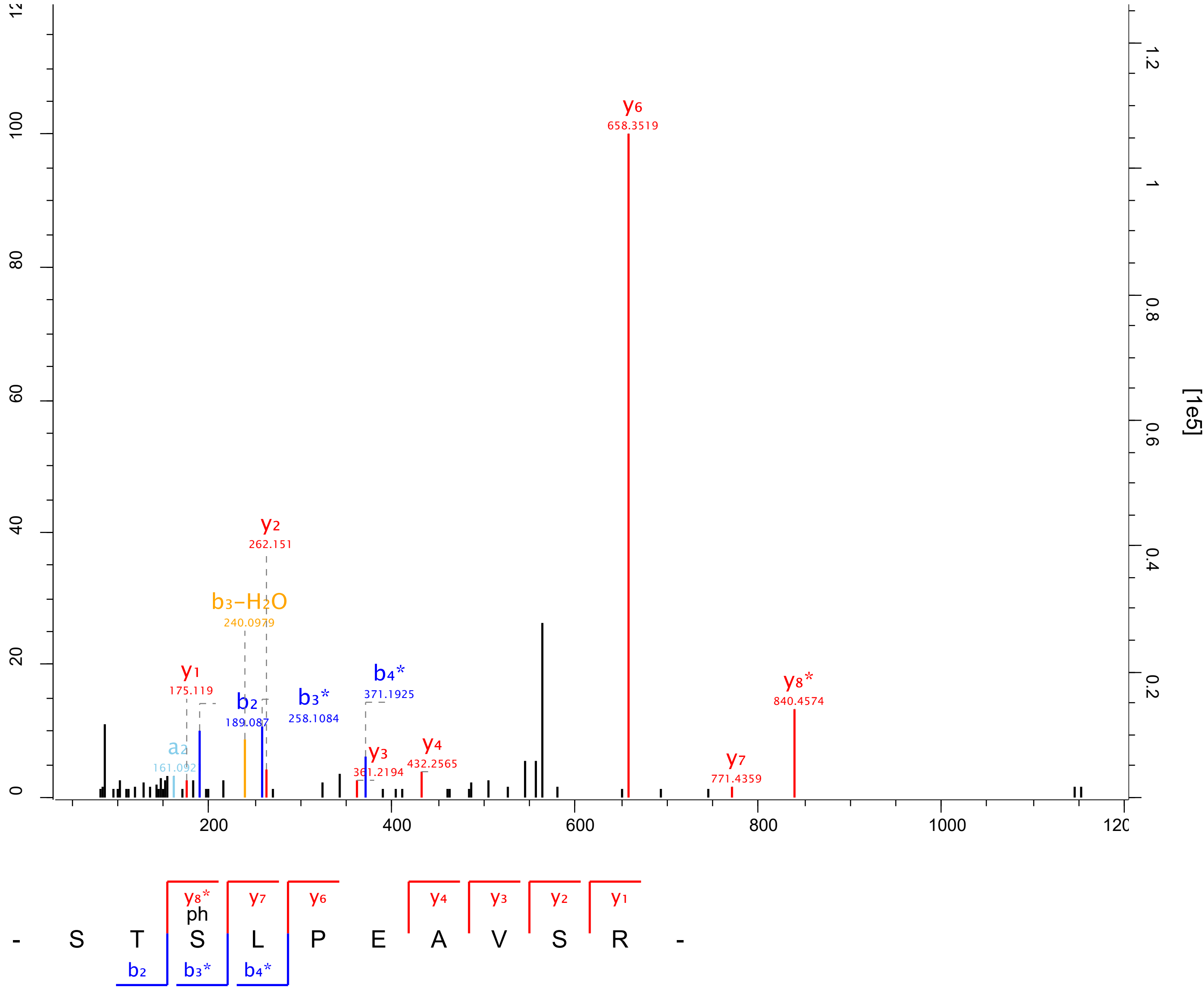

| Raw file | Scan | Method | Score | m/z |
|---|---|---|---|---|
| water_00_1 | 12519 | FTMS; HCD | 181.44 | 514.22 |

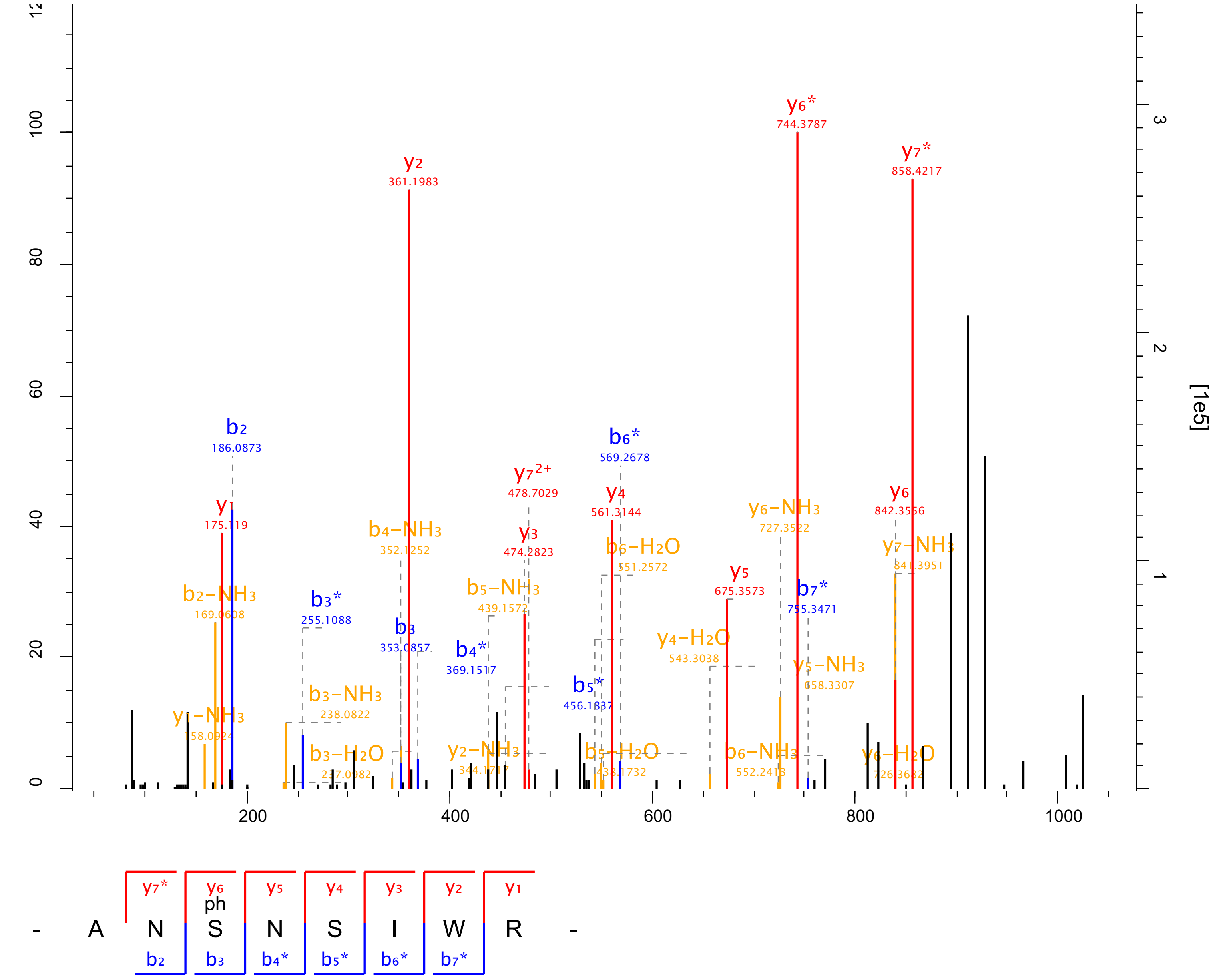

| Raw file | Scan | Method | Score | m/z |
|---|---|---|---|---|
| water_00_1 | 12540 | FTMS; HCD | 157.56 | 577.28 |

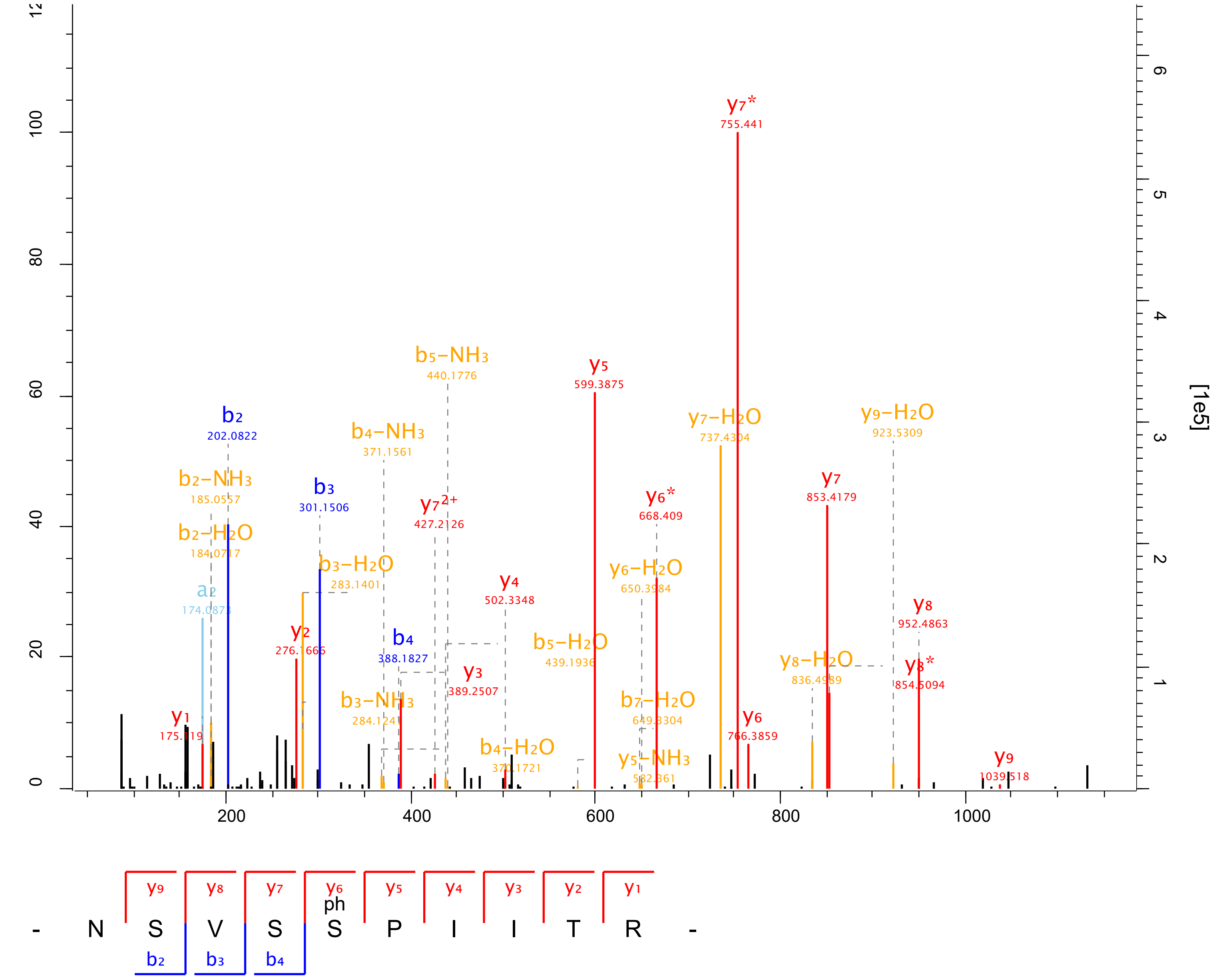

| Raw file | Scan | Method | Score | m/z |
|---|---|---|---|---|
| water_00_1 | 12822 | FTMS; HCD | 141.13 | 728.34 |

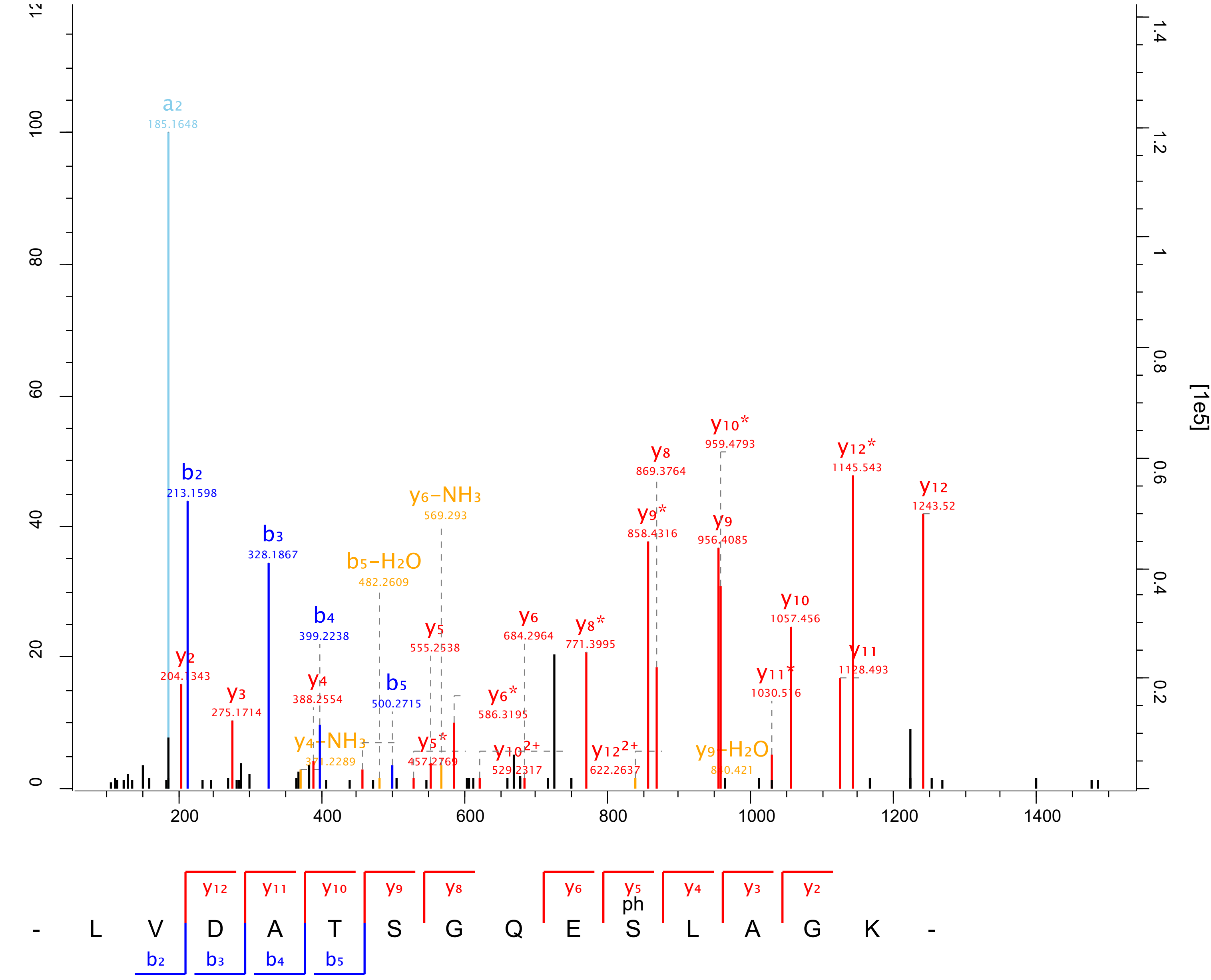

| Raw file | Scan | Method | Score | m/z |
| --- | --- | --- | --- | --- |
| water_00_1 | 12919 | FTMS; HCD | 43.76 | 620.8 |

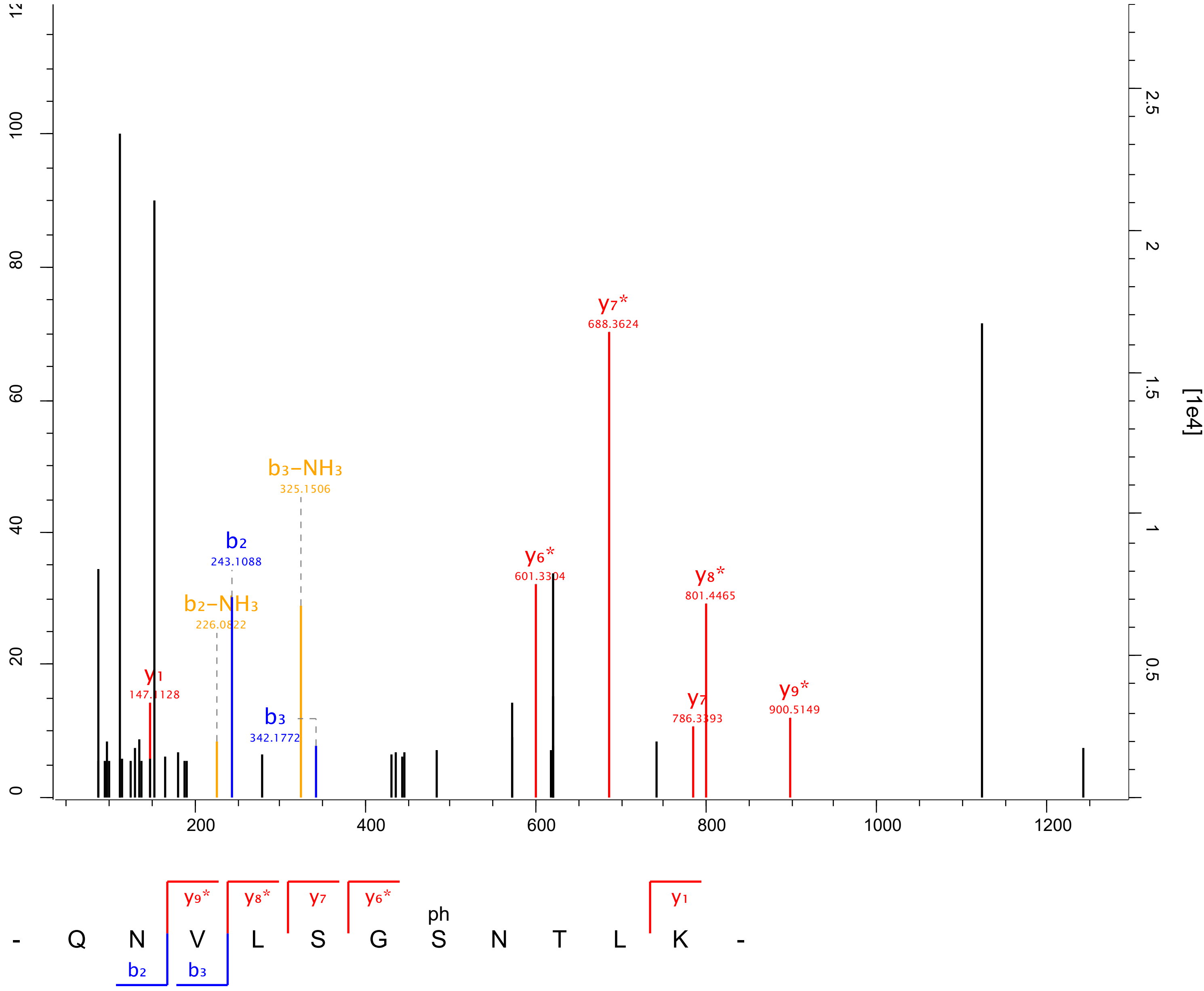

| Raw file | Scan | Method | Score | m/z |
| --- | --- | --- | --- | --- |
| water_00_1 | 13148 | FTMS; HCD | 90.05 | 521.73 |

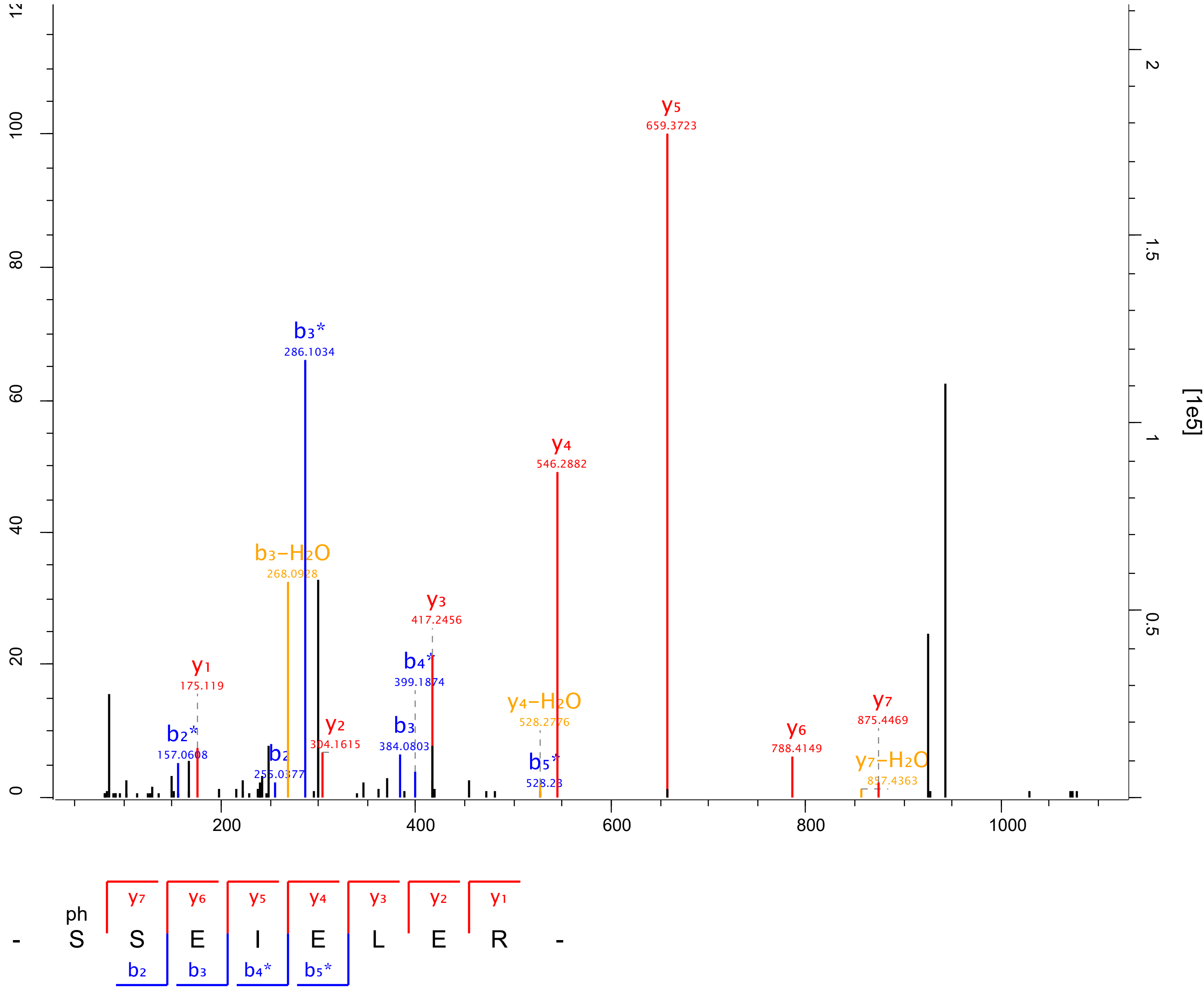

| Raw file | Scan | Method | Score | m/z |
| --- | --- | --- | --- | --- |
| water_00_1 | 13181 | FTMS; HCD | 144.3 | 745.33 |

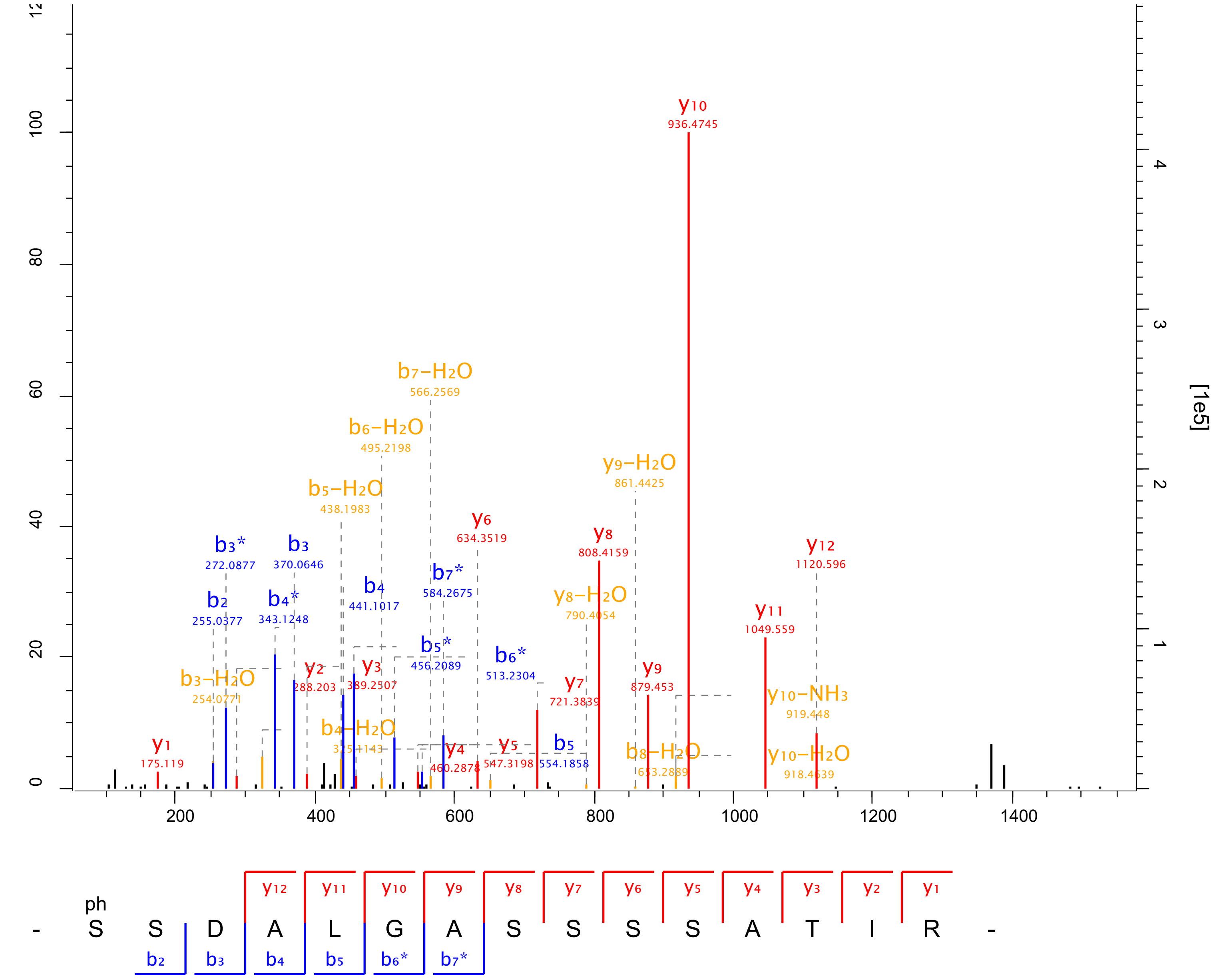

| Raw file | Scan | Method | Score | m/z |
| --- | --- | --- | --- | --- |
| water_00_1 | 13238 | FTMS; HCD | 46.16 | 677.95 |

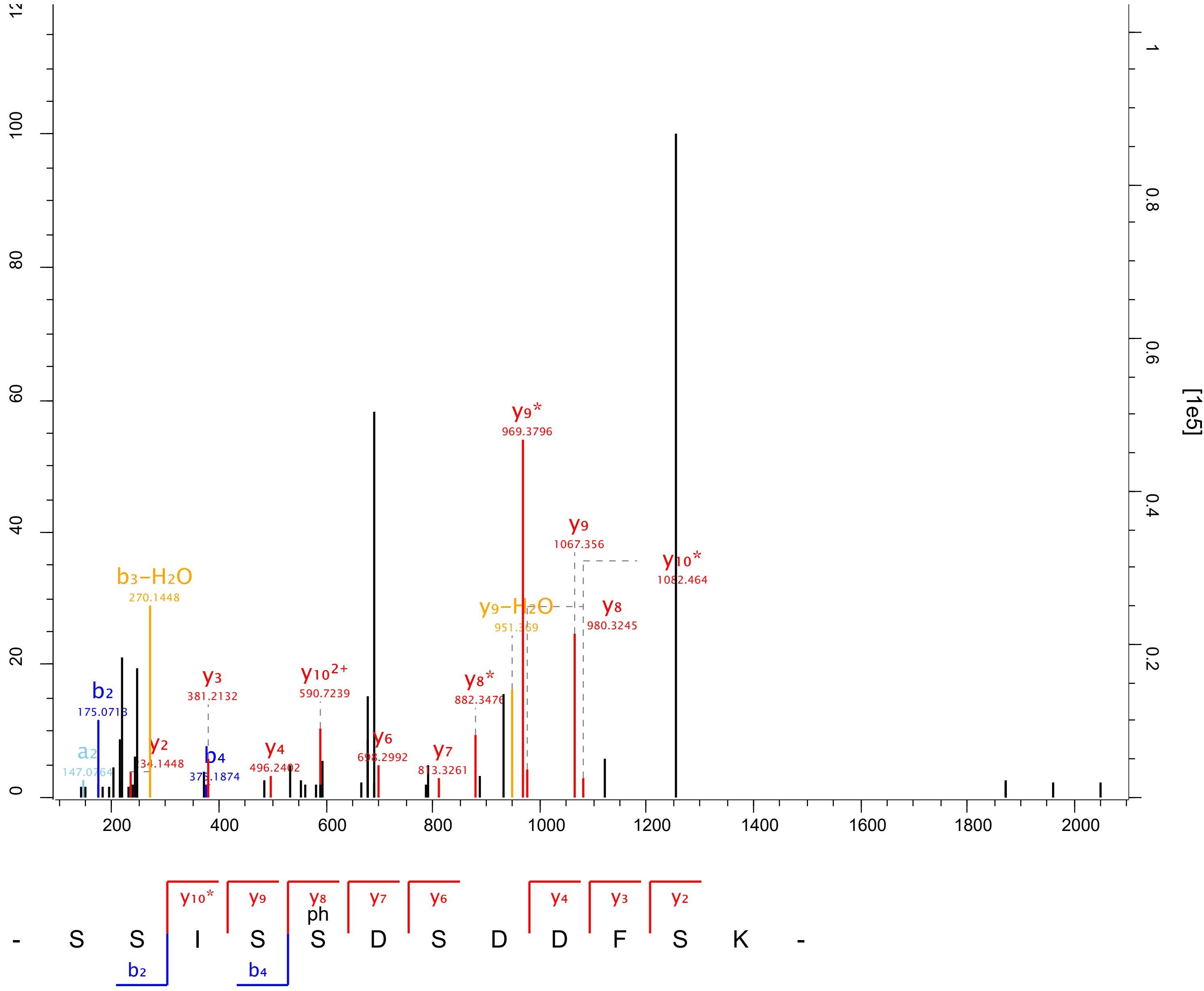

| Raw file | Scan | Method | Score | m/z |
|---|---|---|---|---|
| water_00_1 | 13380 | FTMS; HCD | 305.14 | 703.31 |

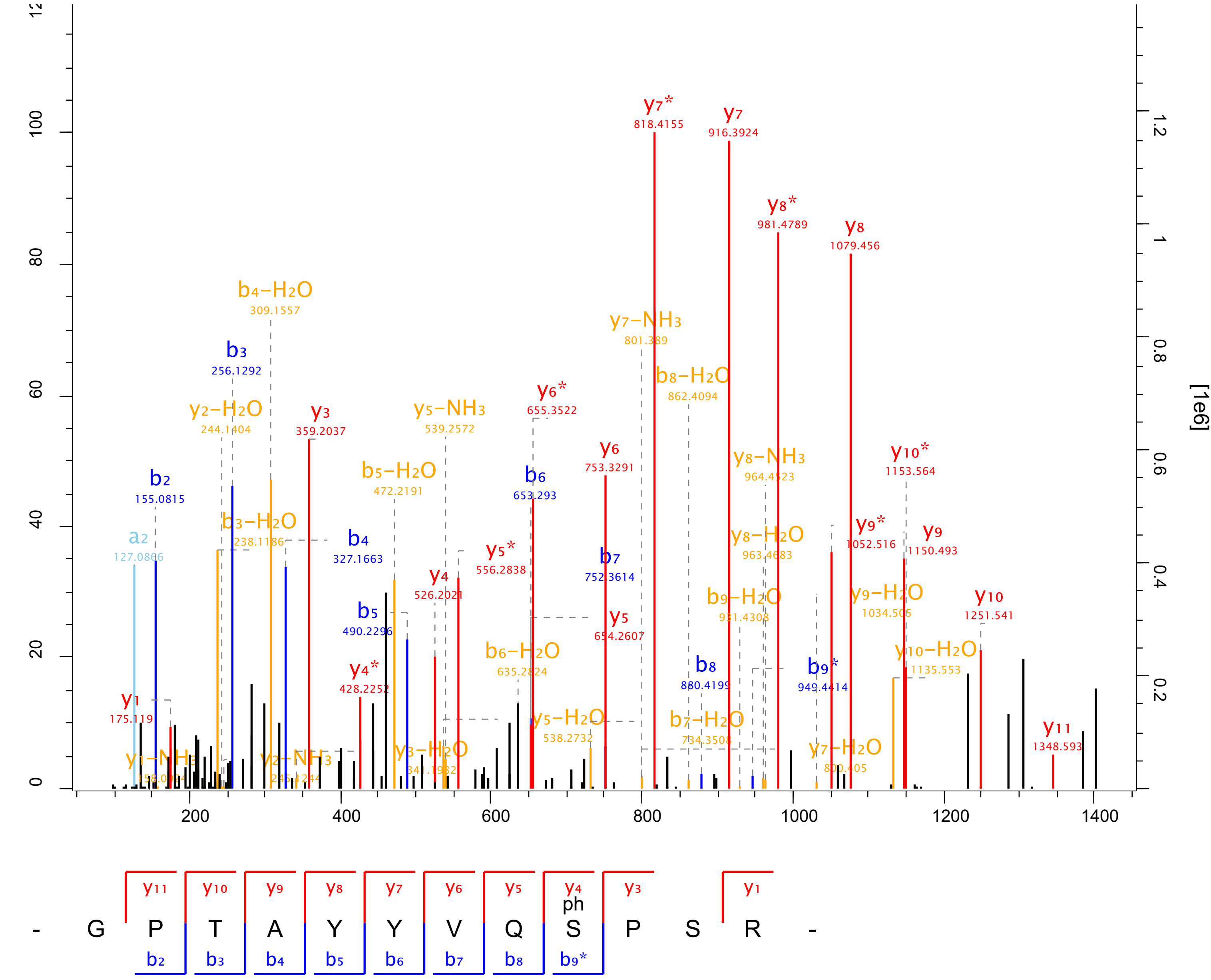

| Raw file | Scan | Method | Score | m/z |
|---|---|---|---|---|
| water_00_1 | 13427 | FTMS; HCD | 104.17 | 562.75 |

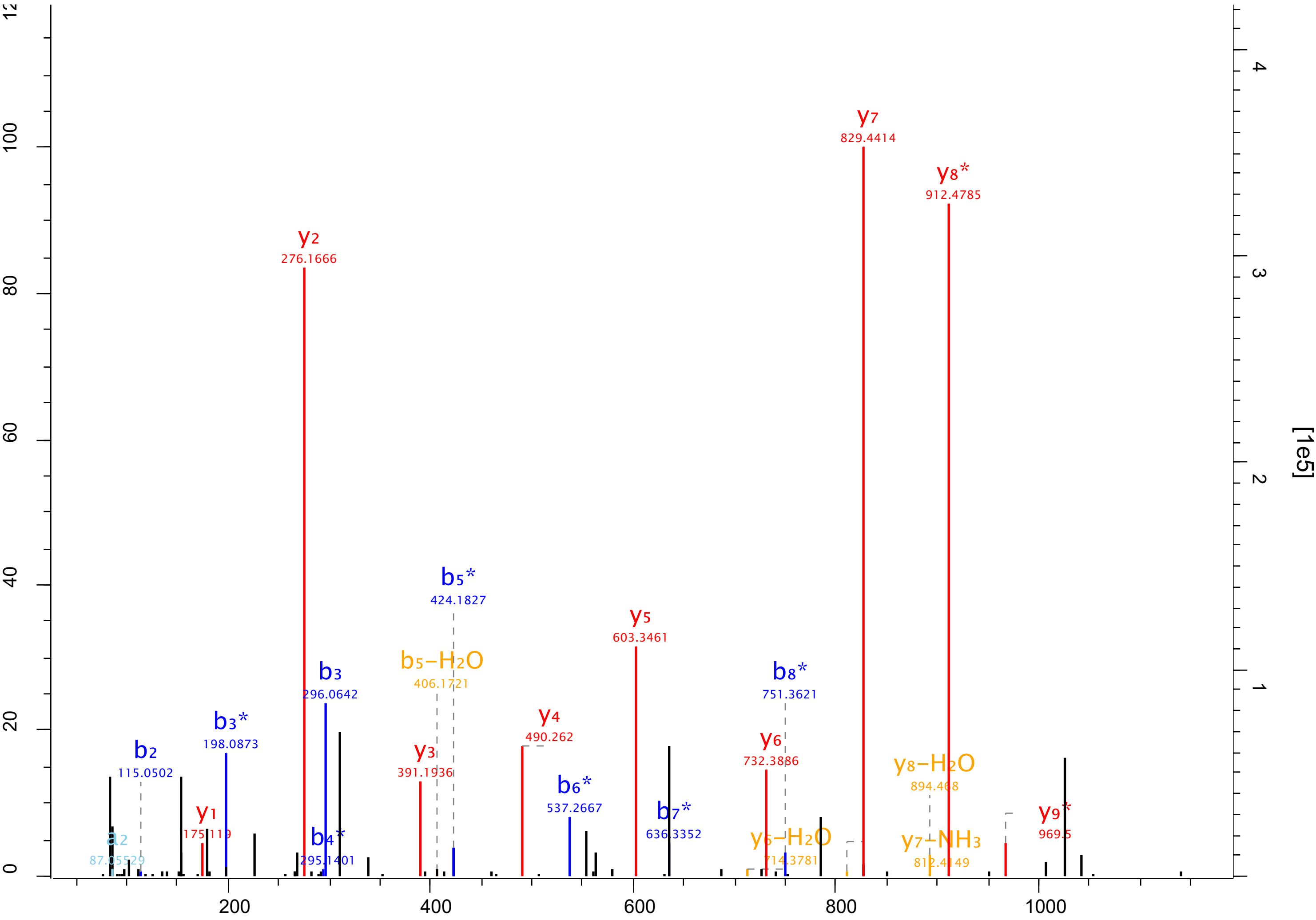


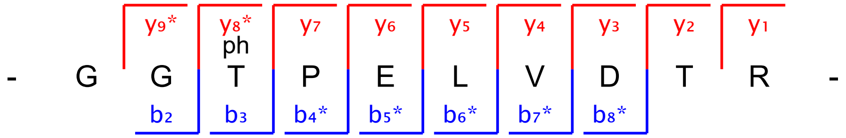

| Raw file | Scan | Method | Score | m/z |
| --- | --- | --- | --- | --- |
| water_00_1 | 13530 | FTMS; HCD | 88.28 | 828.87 |

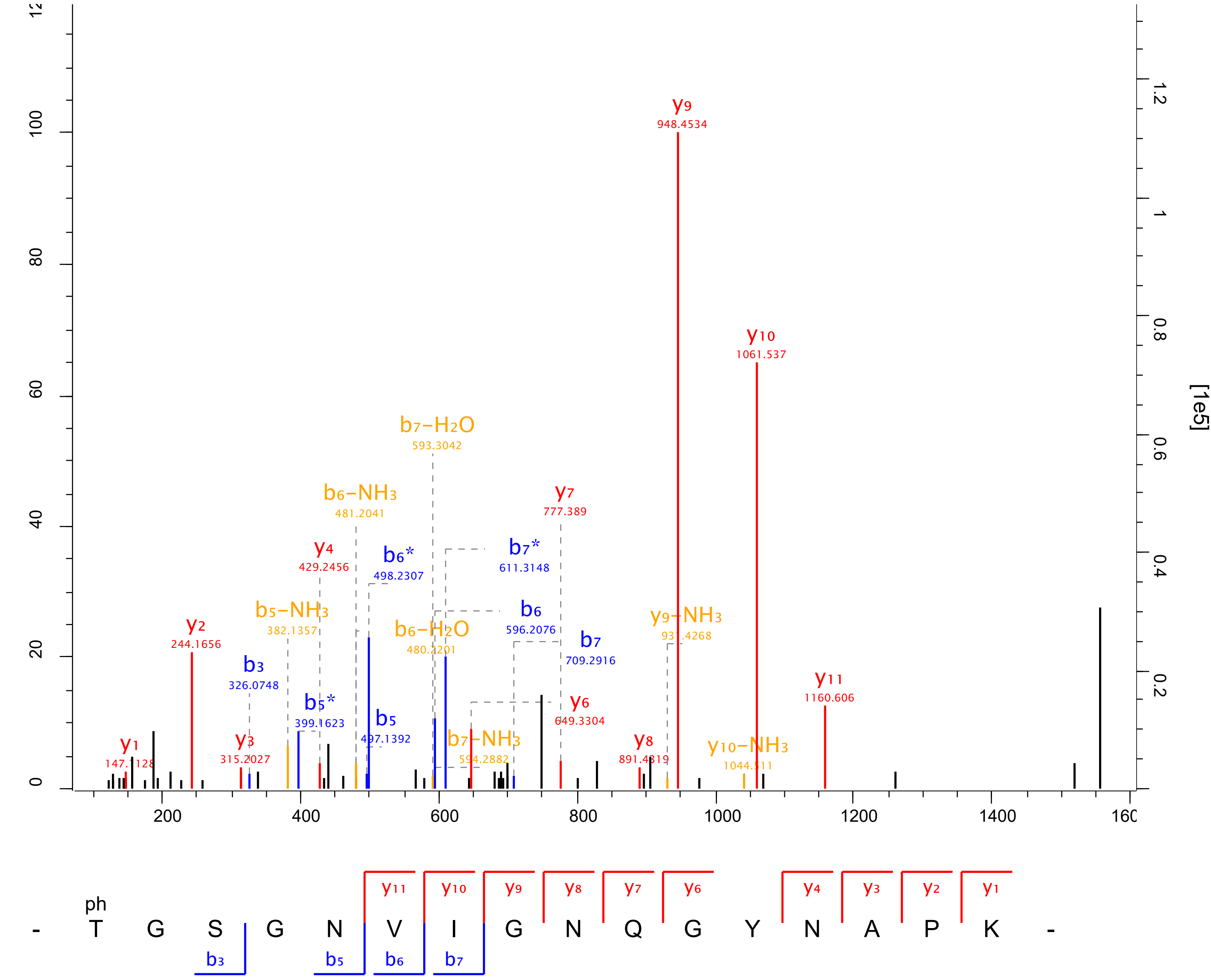

| Raw file | Scan | Method | Score | m/z |
|---|---|---|---|---|
| water_00_1 | 13664 | FTMS; HCD | 43.03 | 806.36 |

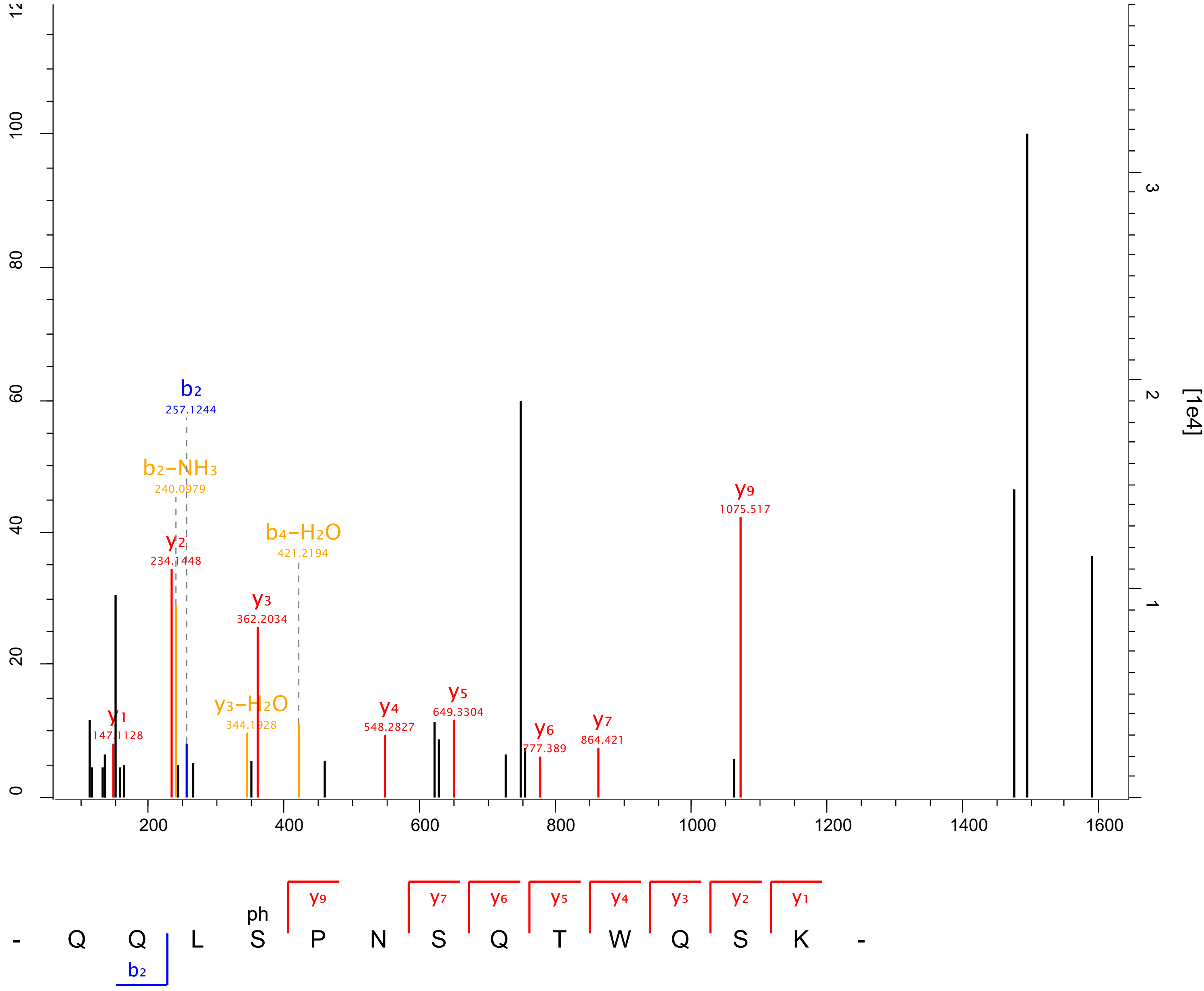

| Raw file | Scan | Method | Score | m/z |
|---|---|---|---|---|
| water_00_1 | 13681 | FTMS; HCD | 109.72 | 606.93 |

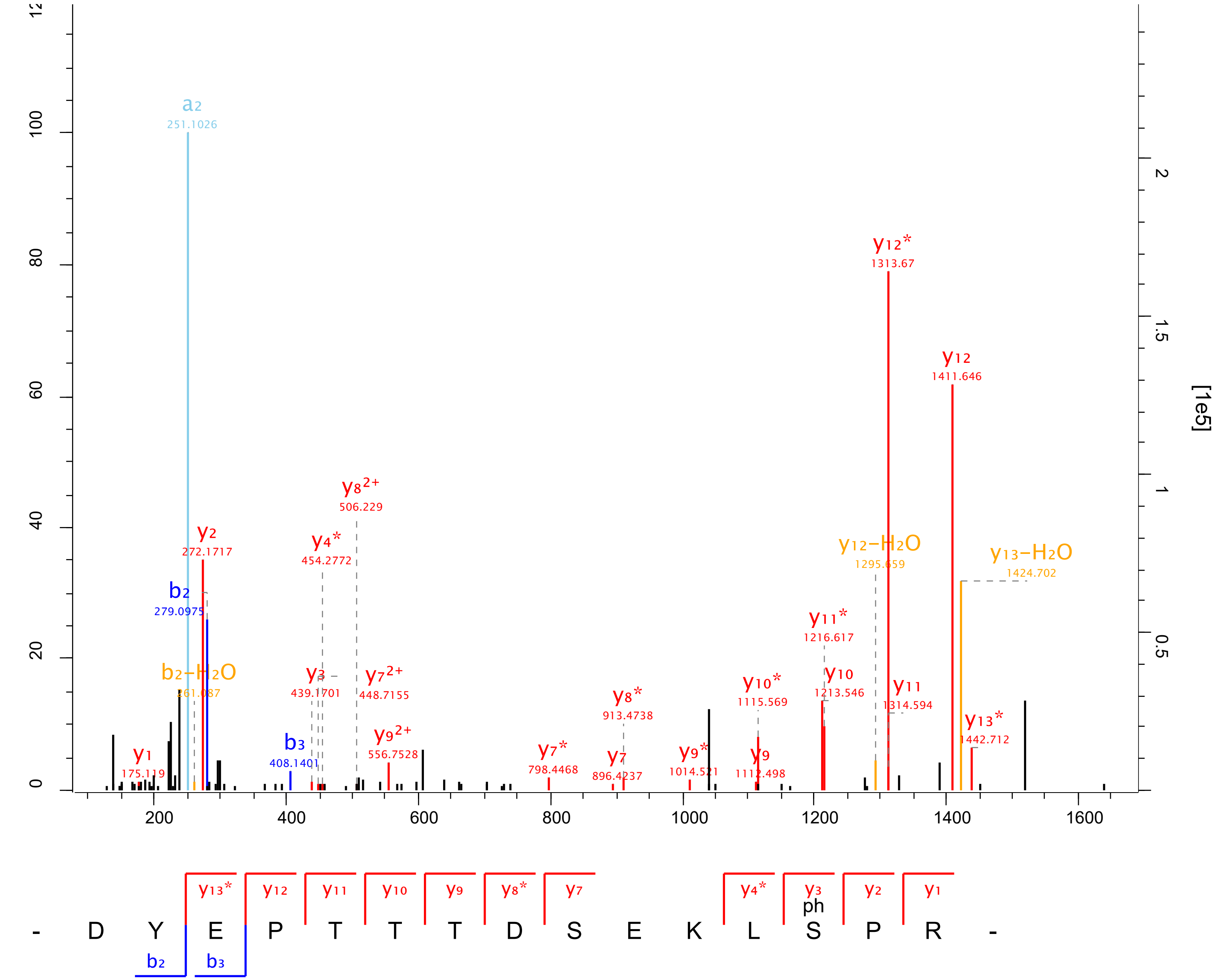

| Raw file | Scan | Method | Score | m/z |
| --- | --- | --- | --- | --- |
| water_00_1 | 13923 | FTMS; HCD | 94.09 | 679.27 |

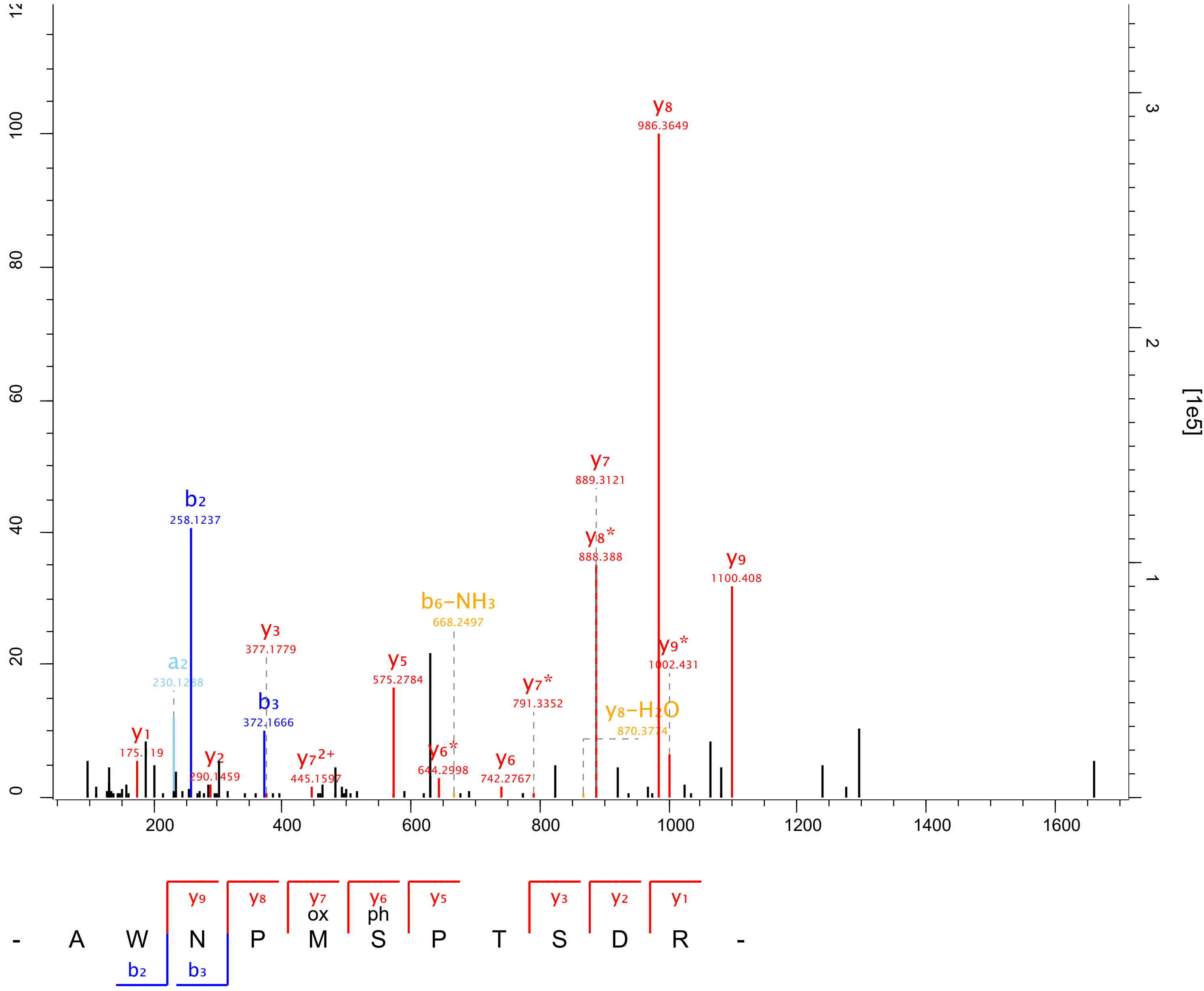

| Raw file | Scan | Method | Score | m/z |
| --- | --- | --- | --- | --- |
| water_00_1 | 13985 | FTMS; HCD | 354.55 | 998.36 |

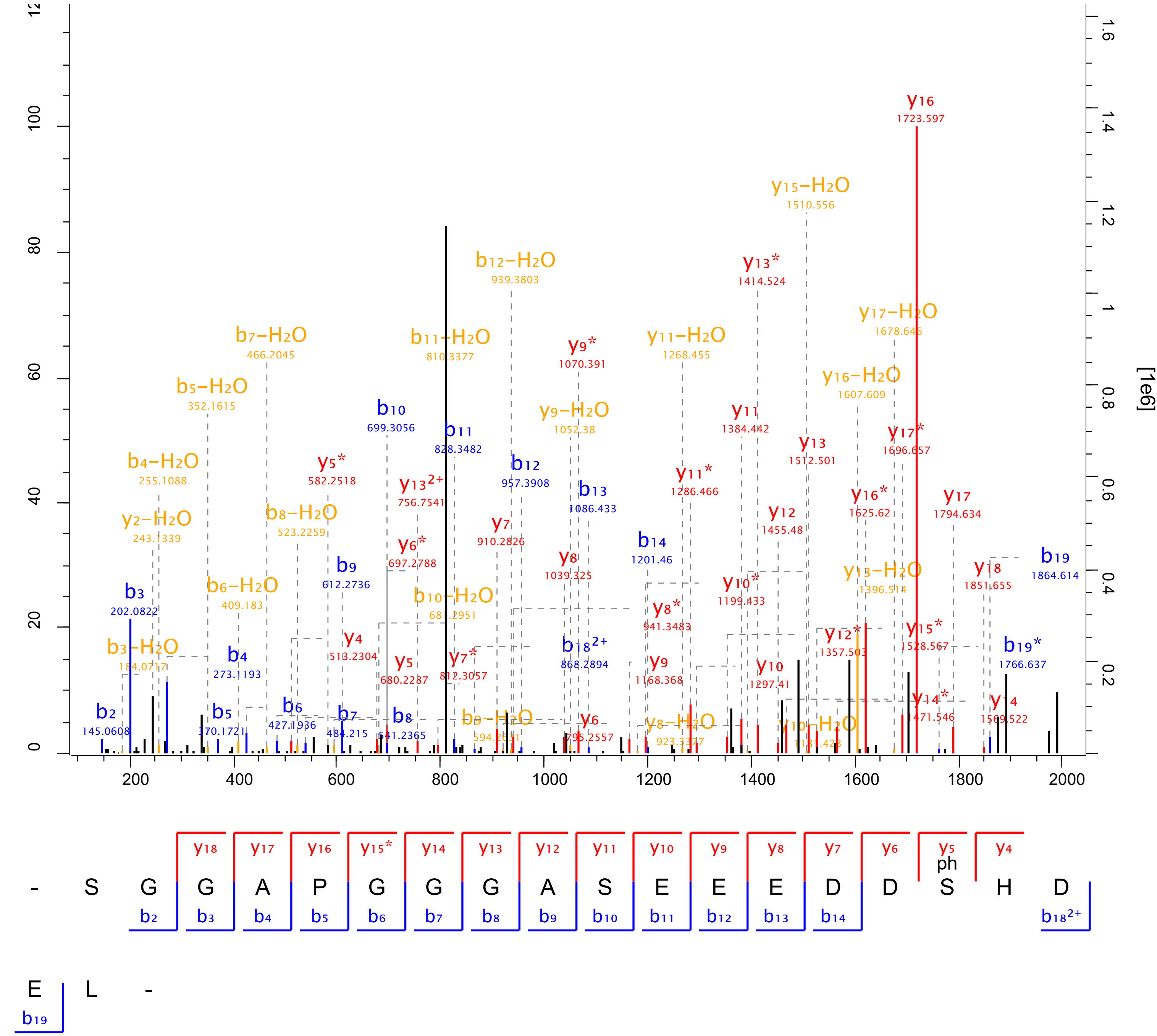

| Raw file | Scan | Method | Score | m/z |
|---|---|---|---|---|
| water_00_1 | 14219 | FTMS; HCD | 126.19 | 550.76 |

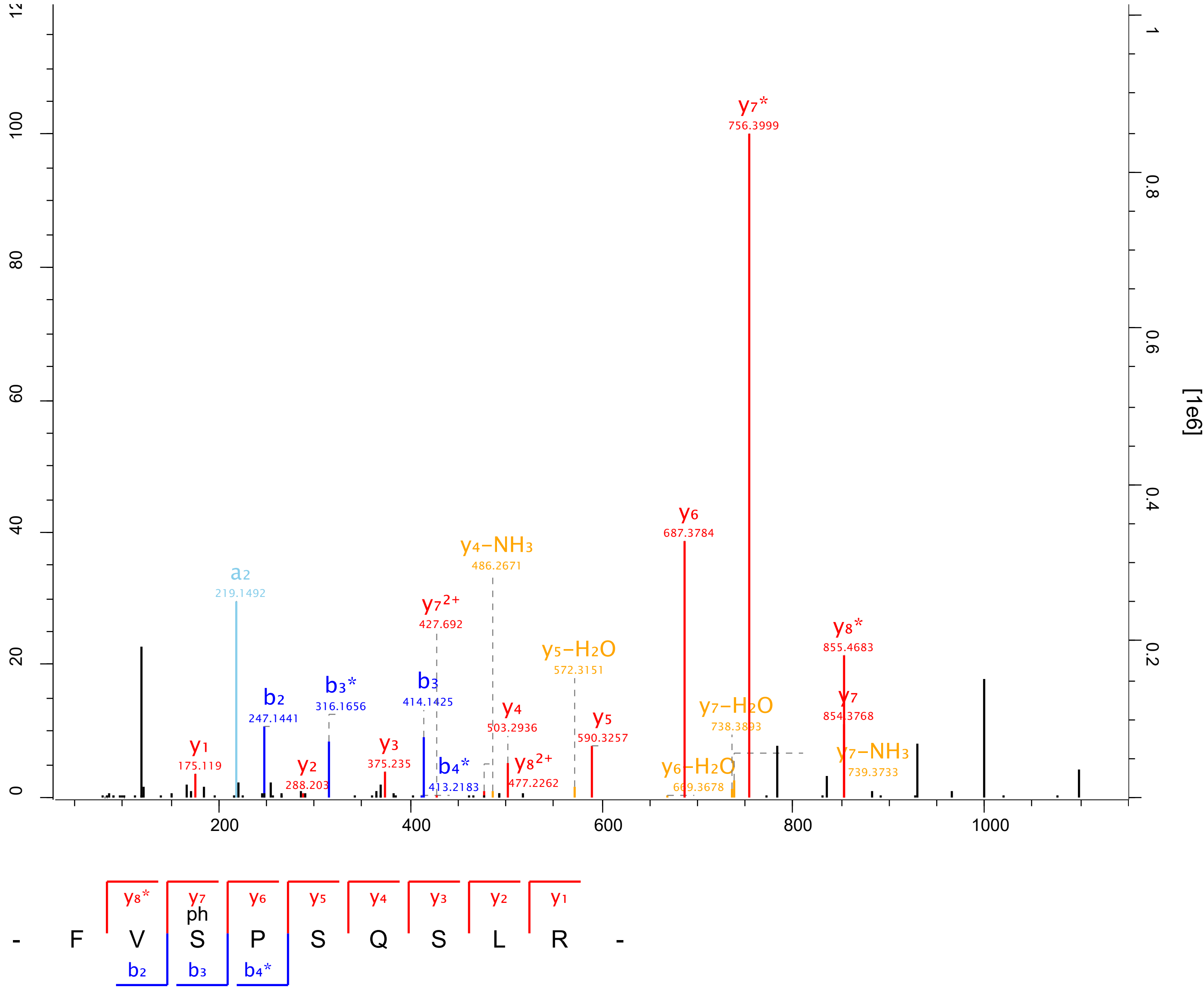

| Raw file | Scan | Method | Score | m/z |
| --- | --- | --- | --- | --- |
| water_00_1 | 14607 | FTMS; HCD | 43.68 | 593.75 |

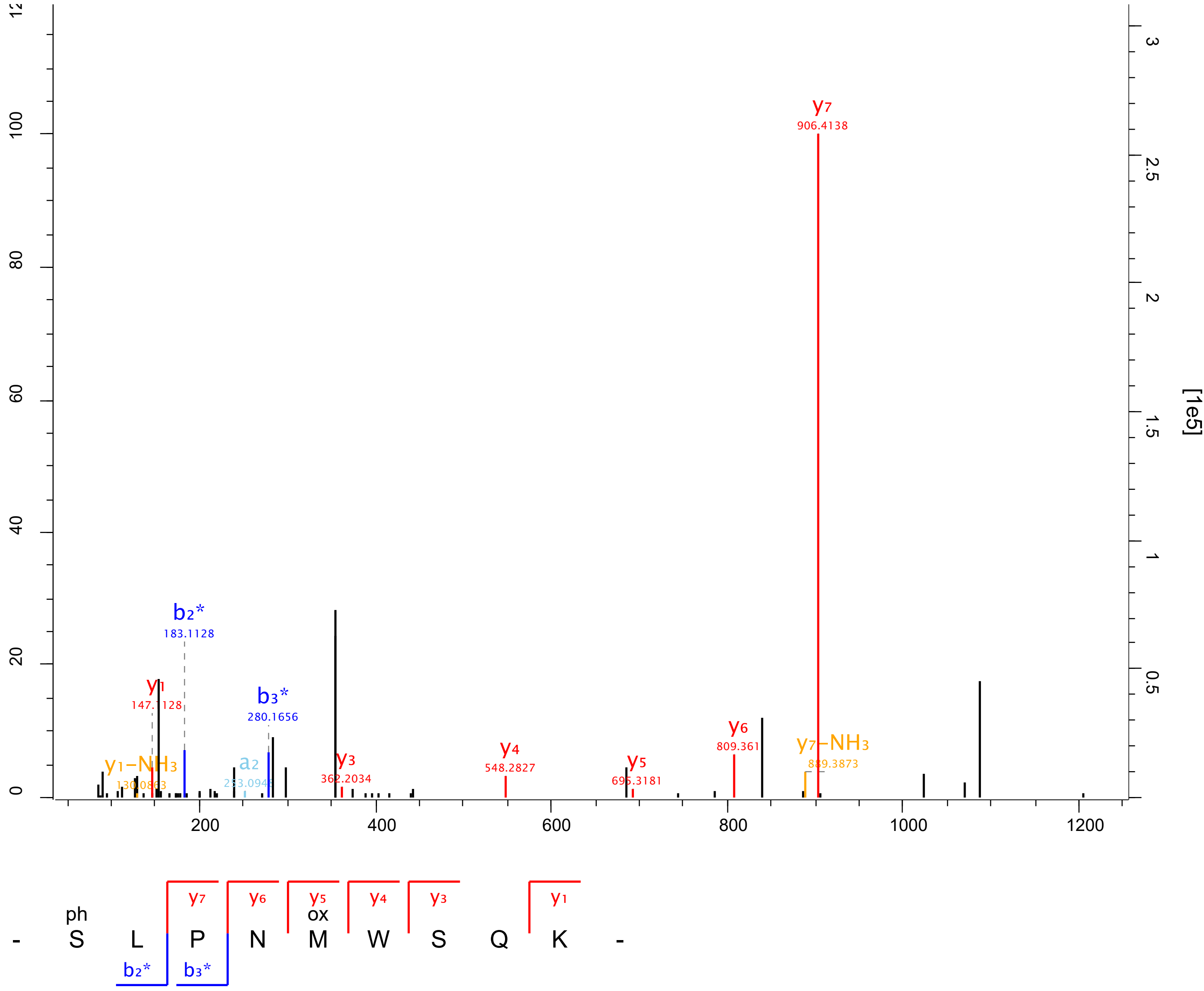

| Raw file | Scan | Method | Score | m/z |
| --- | --- | --- | --- | --- |
| water_00_1 | 14646 | FTMS; HCD | 161.47 | 908.41 |

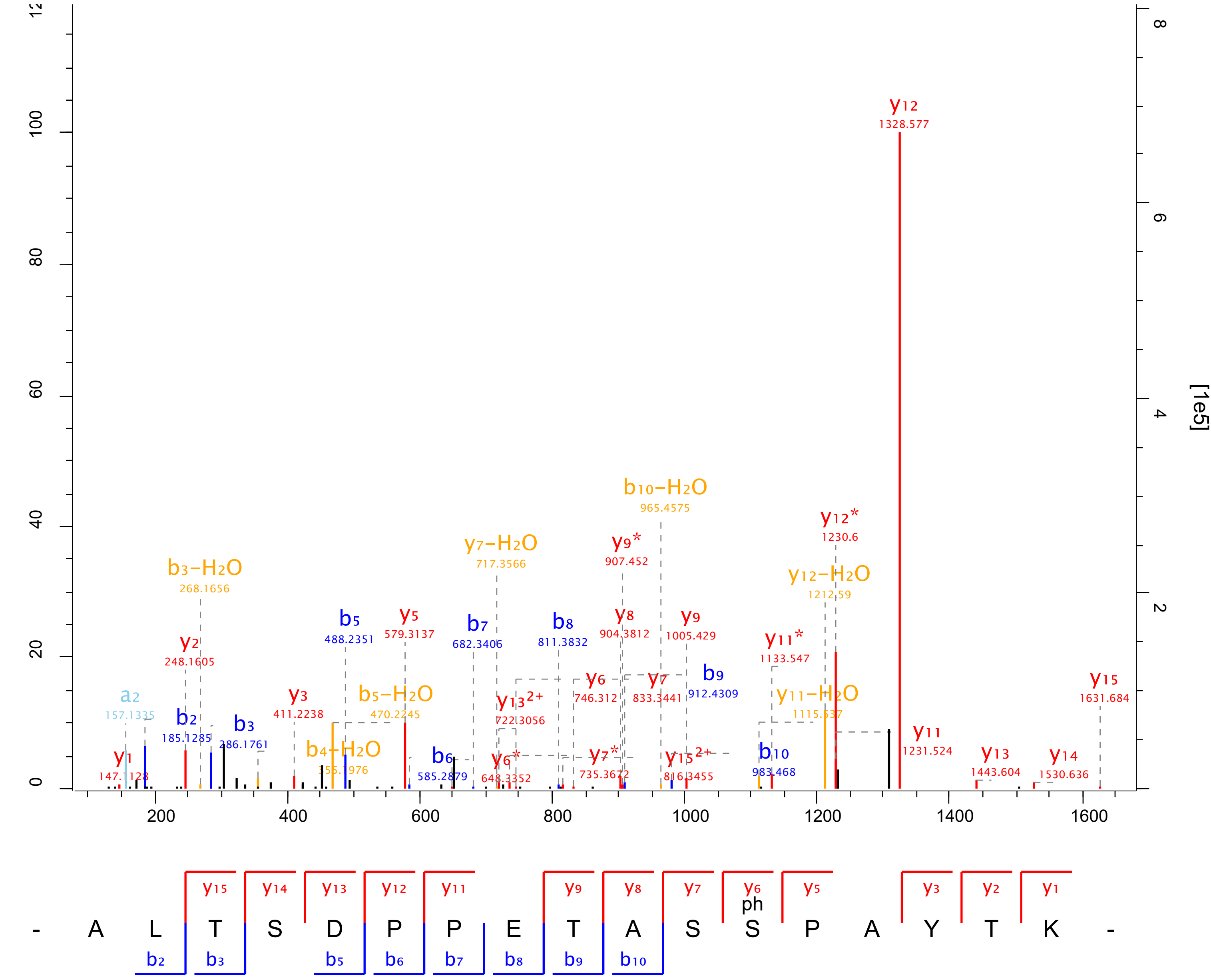

| Raw file | Scan | Method | Score | m/z |
|---|---|---|---|---|
| water_00_1 | 14946 | FTMS; HCD | 80.69 | 607.73 |

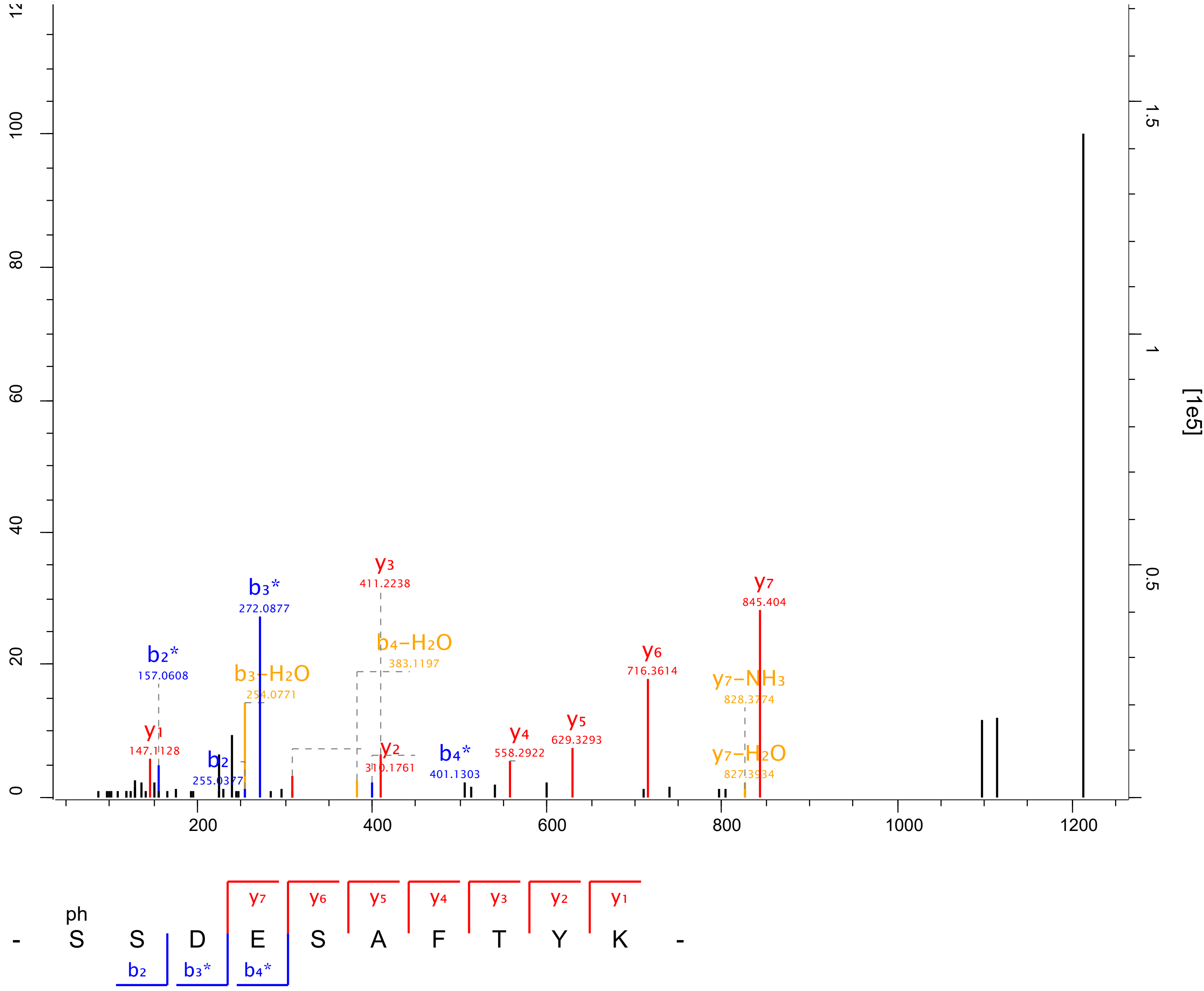

| Raw file | Scan | Method | Score | m/z |
| --- | --- | --- | --- | --- |
| water_00_1 | 15034 | FTMS; HCD | 63.69 | 802.32 |

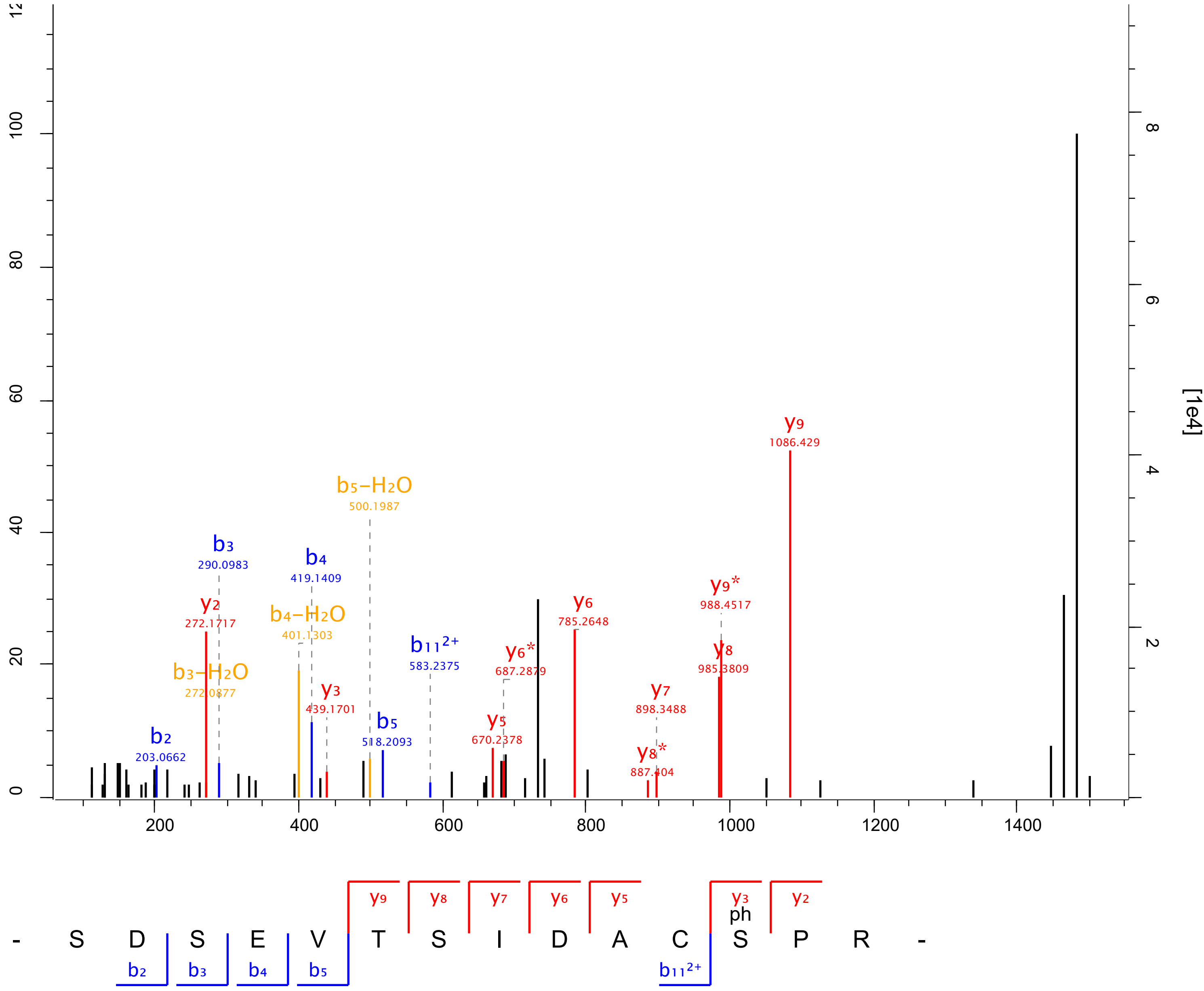

| Raw file | Scan | Method | Score | m/z |
| --- | --- | --- | --- | --- |
| water_00_1 | 15081 | FTMS; HCD | 80.69 | 512.73 |

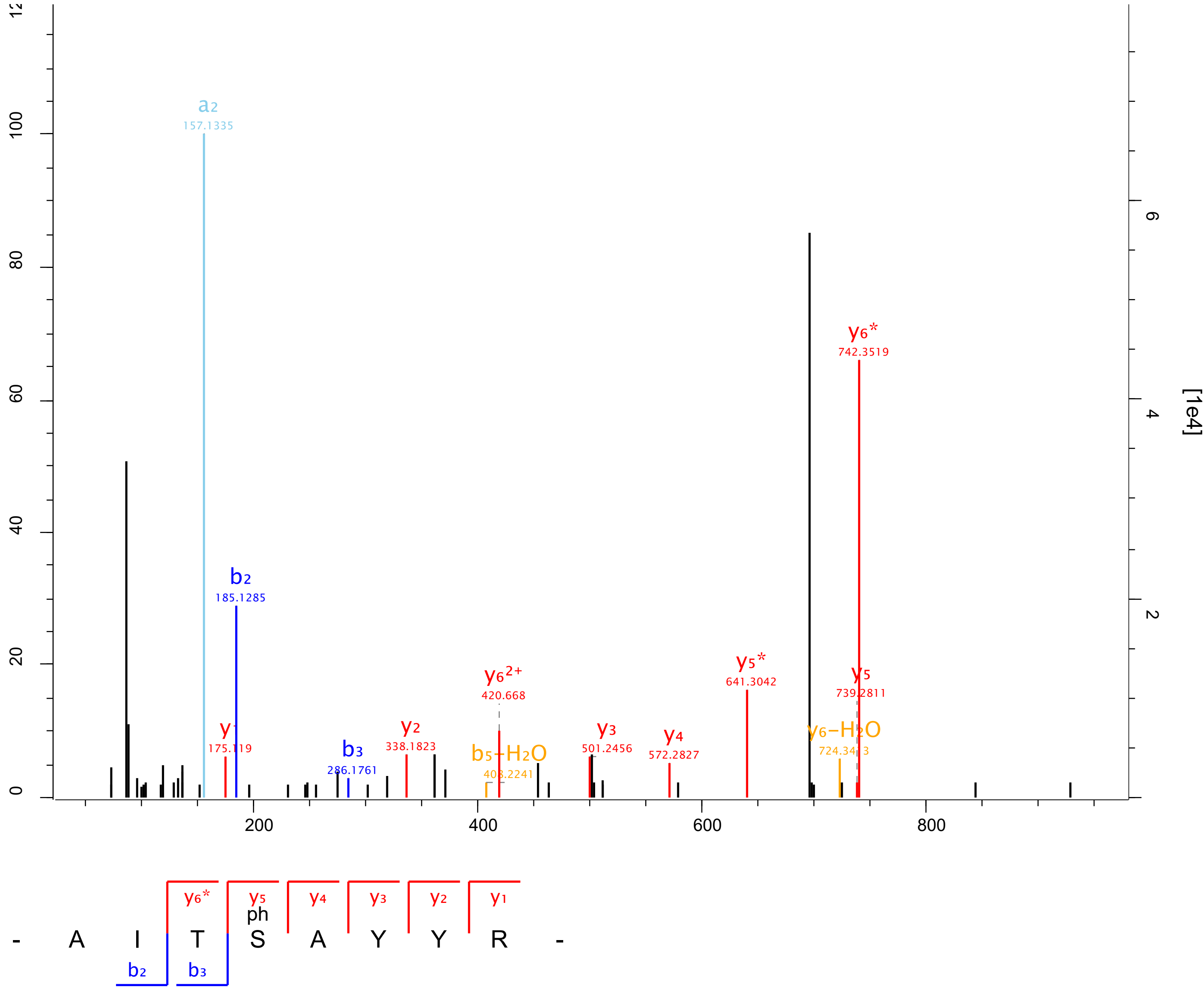

| Raw file | Scan | Method | Score | m/z |
|---|---|---|---|---|
| water_00_1 | 15639 | FTMS; HCD | 115.68 | 603.27 |

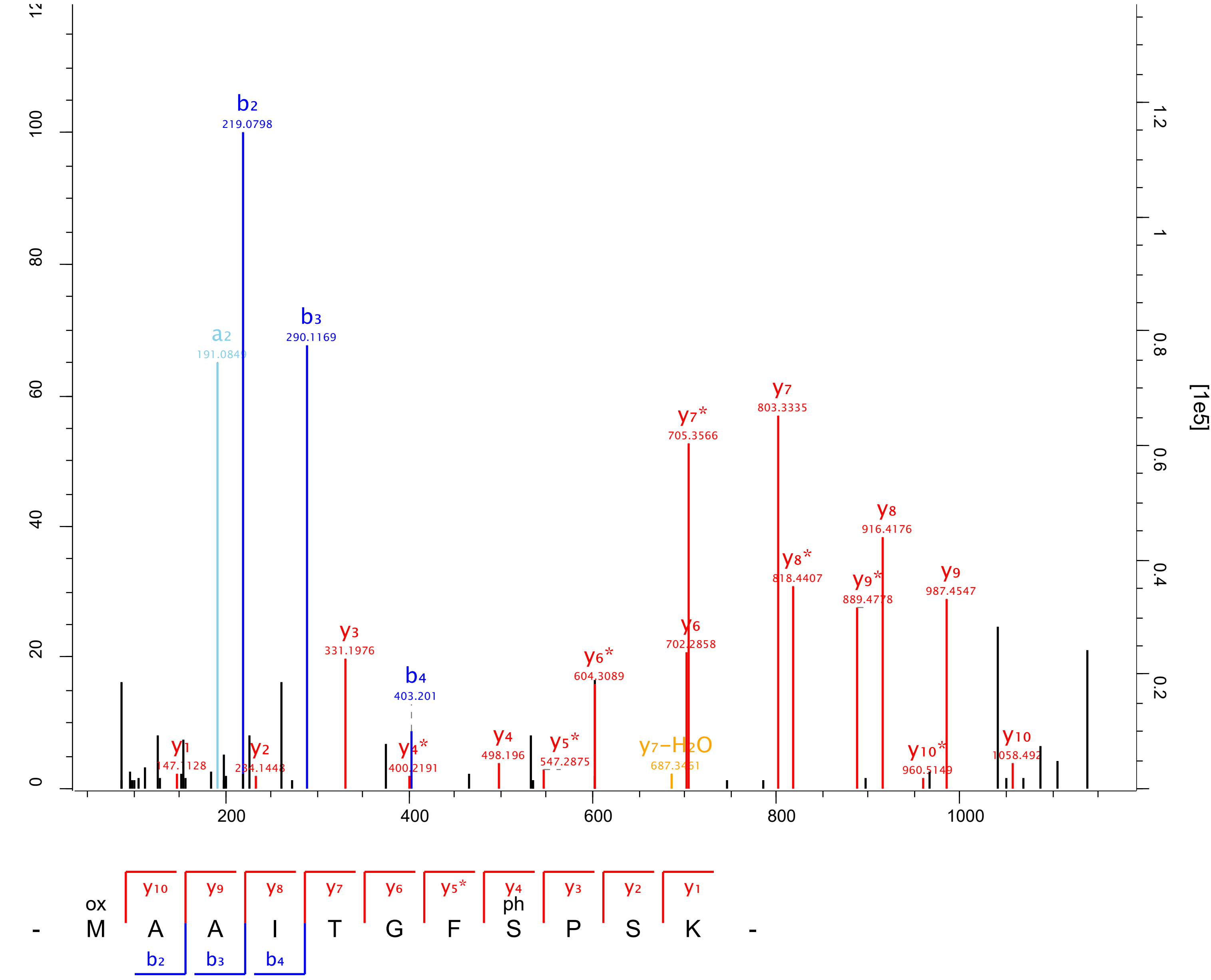

| Raw file | Scan | Method | Score | m/z |
|---|---|---|---|---|
| water_00_1 | 15750 | FTMS; HCD | 144.18 | 694.79 |

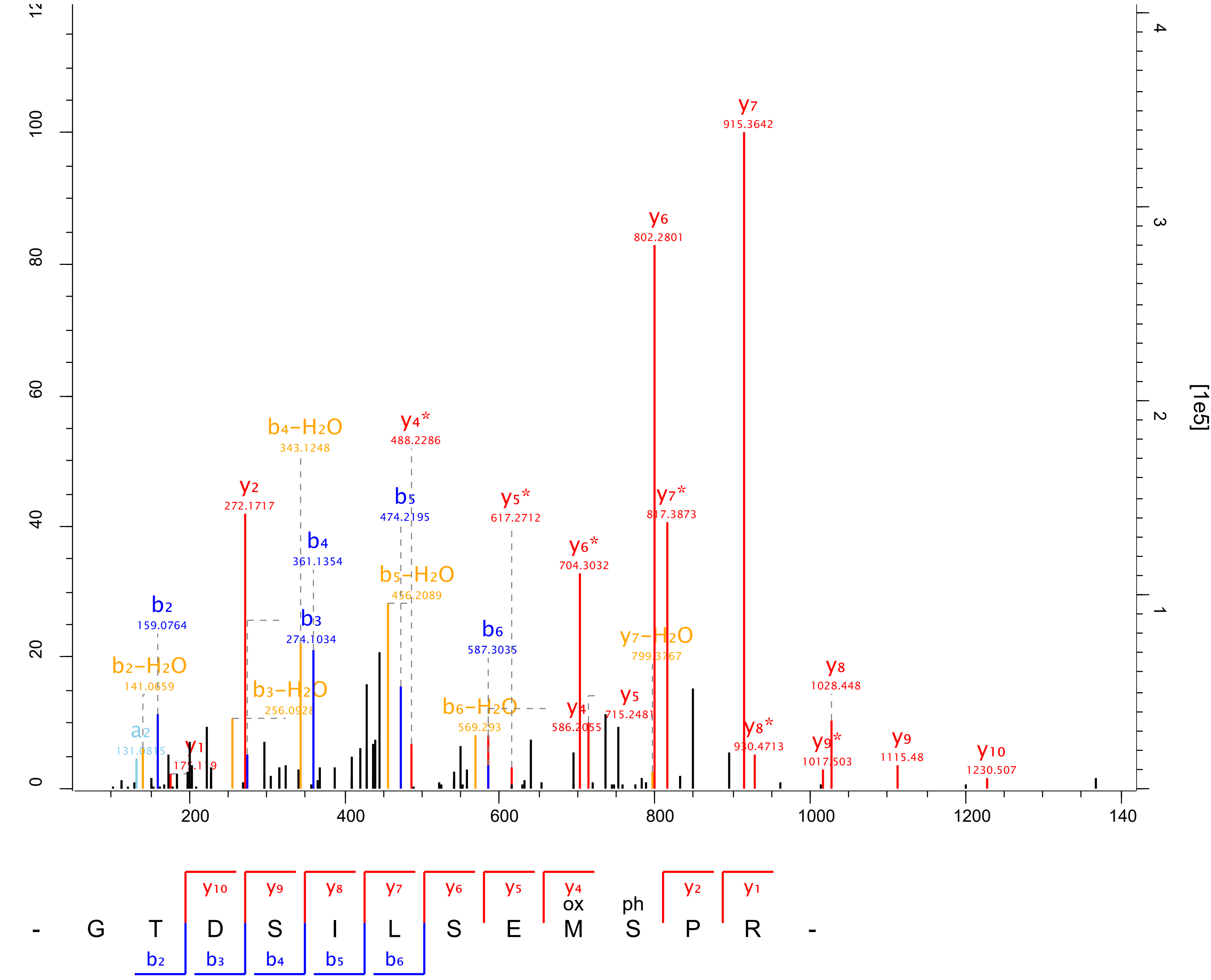

| Raw file | Scan | Method | Score | m/z |
| --- | --- | --- | --- | --- |
| water_00_1 | 15805 | FTMS; HCD | 79.61 | 591.26 |

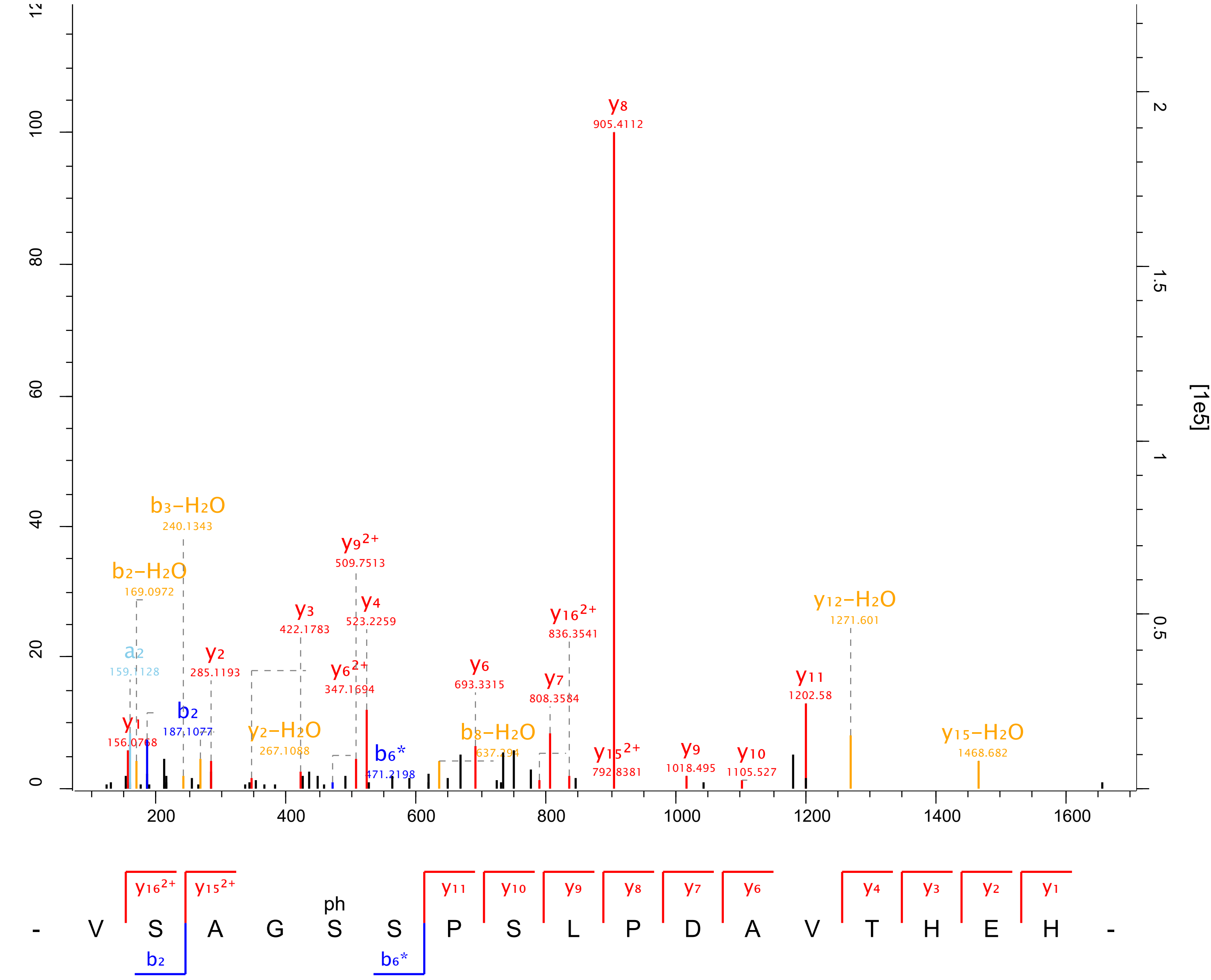

| Raw file | Scan | Method | Score | m/z |
| --- | --- | --- | --- | --- |
| water_00_1 | 15823 | FTMS; HCD | 99.17 | 500.75 |

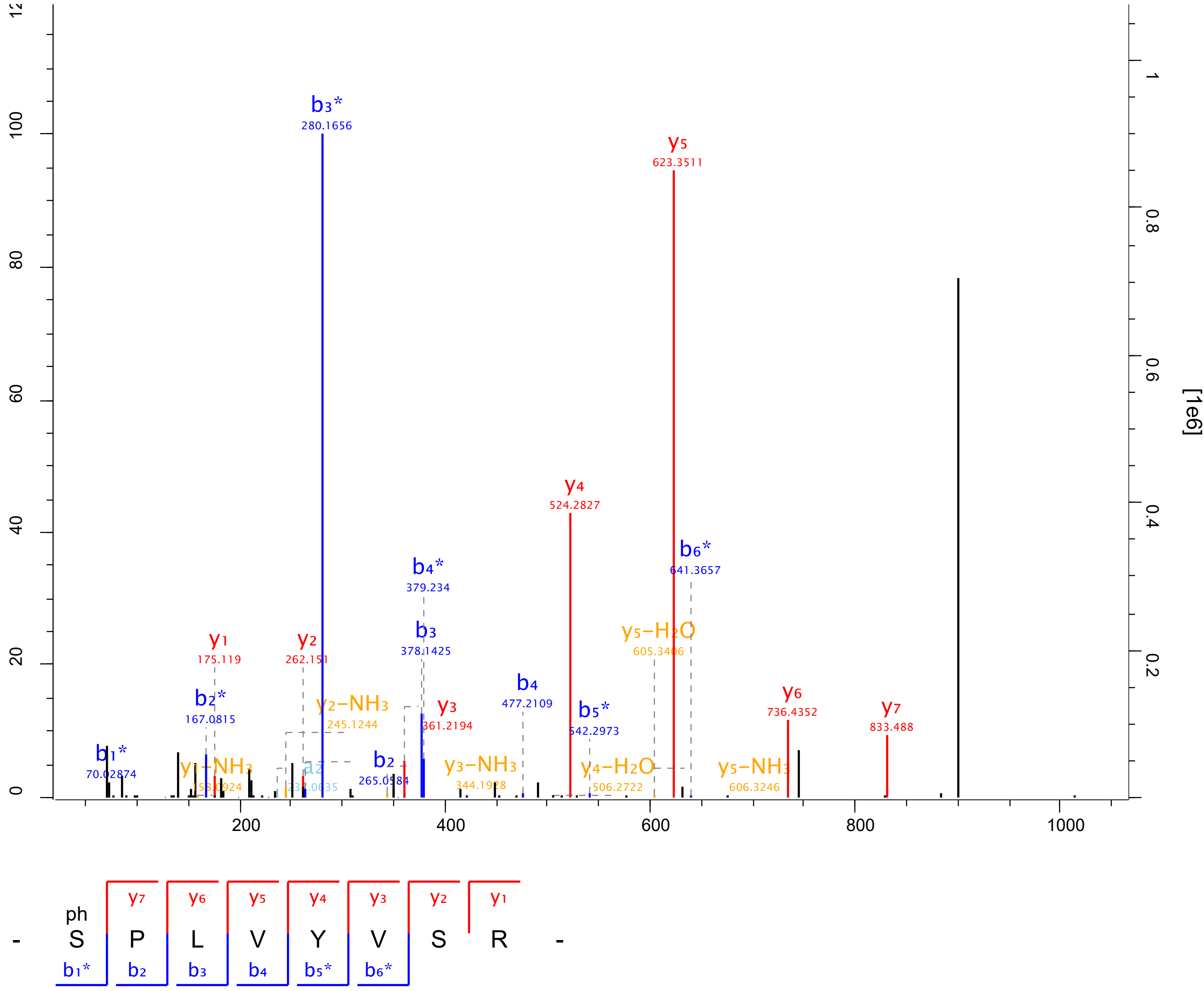

| Raw file | Scan | Method | Score | m/z |
| --- | --- | --- | --- | --- |
| water_00_1 | 15912 | FTMS; HCD | 67.23 | 952.89 |

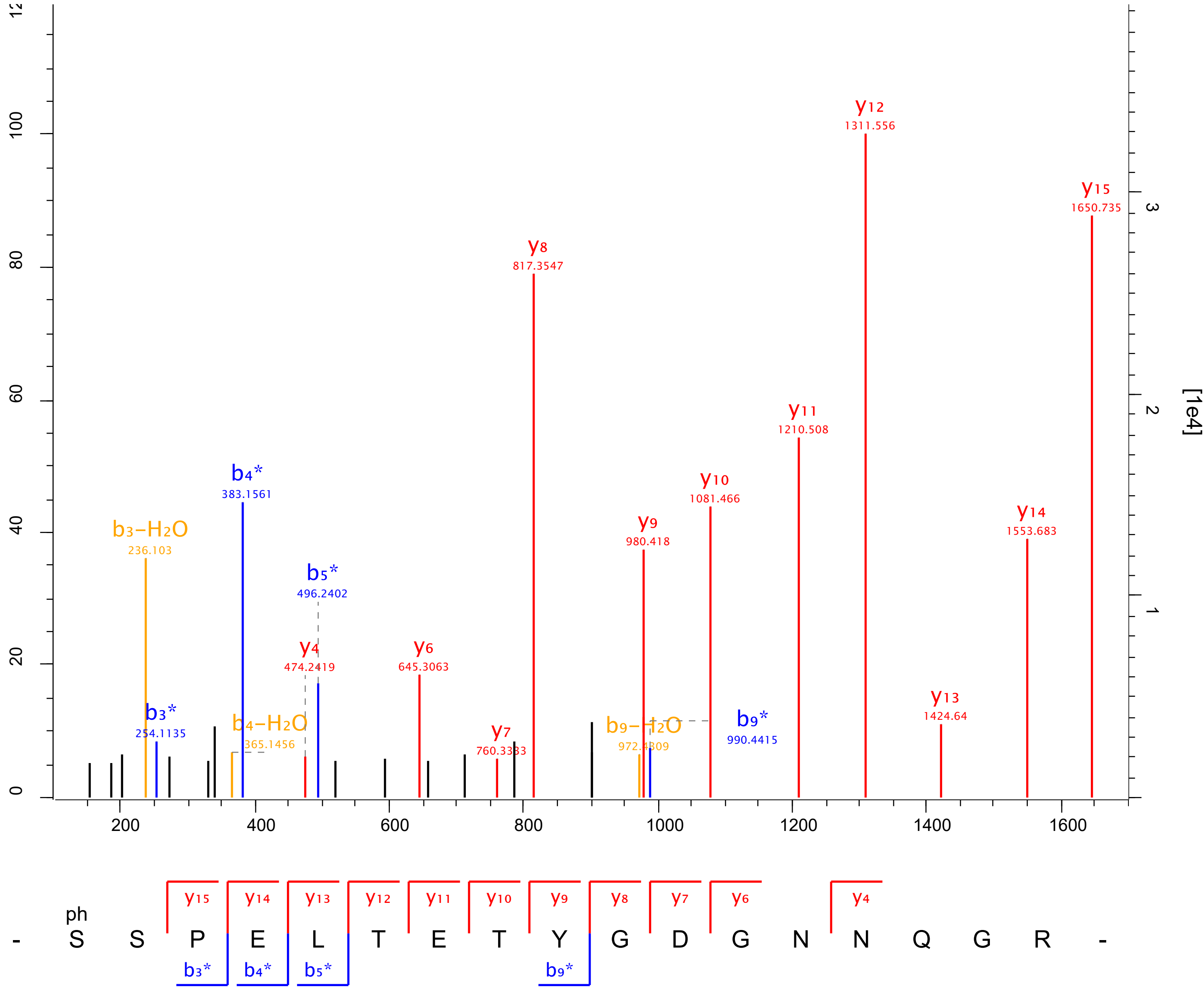

| Raw file | Scan | Method | Score | m/z |
|---|---|---|---|---|
| water_00_1 | 1592 | FTMS; HCD | 66.06 | 365.18 |

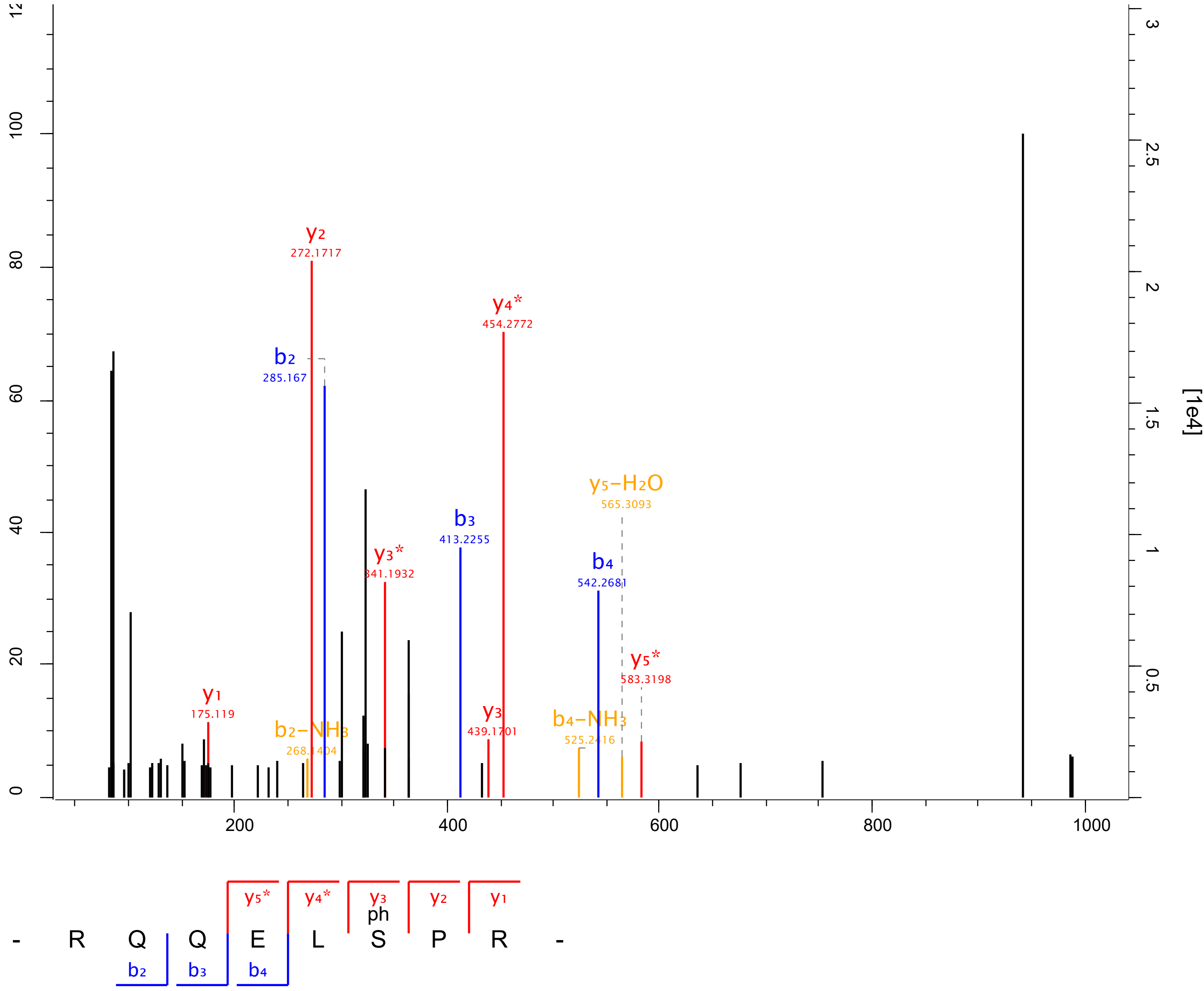

| Raw file | Scan | Method | Score | m/z |
| --- | --- | --- | --- | --- |
| water_00_1 | 15960 | FTMS; HCD | 71.08 | 503.77 |

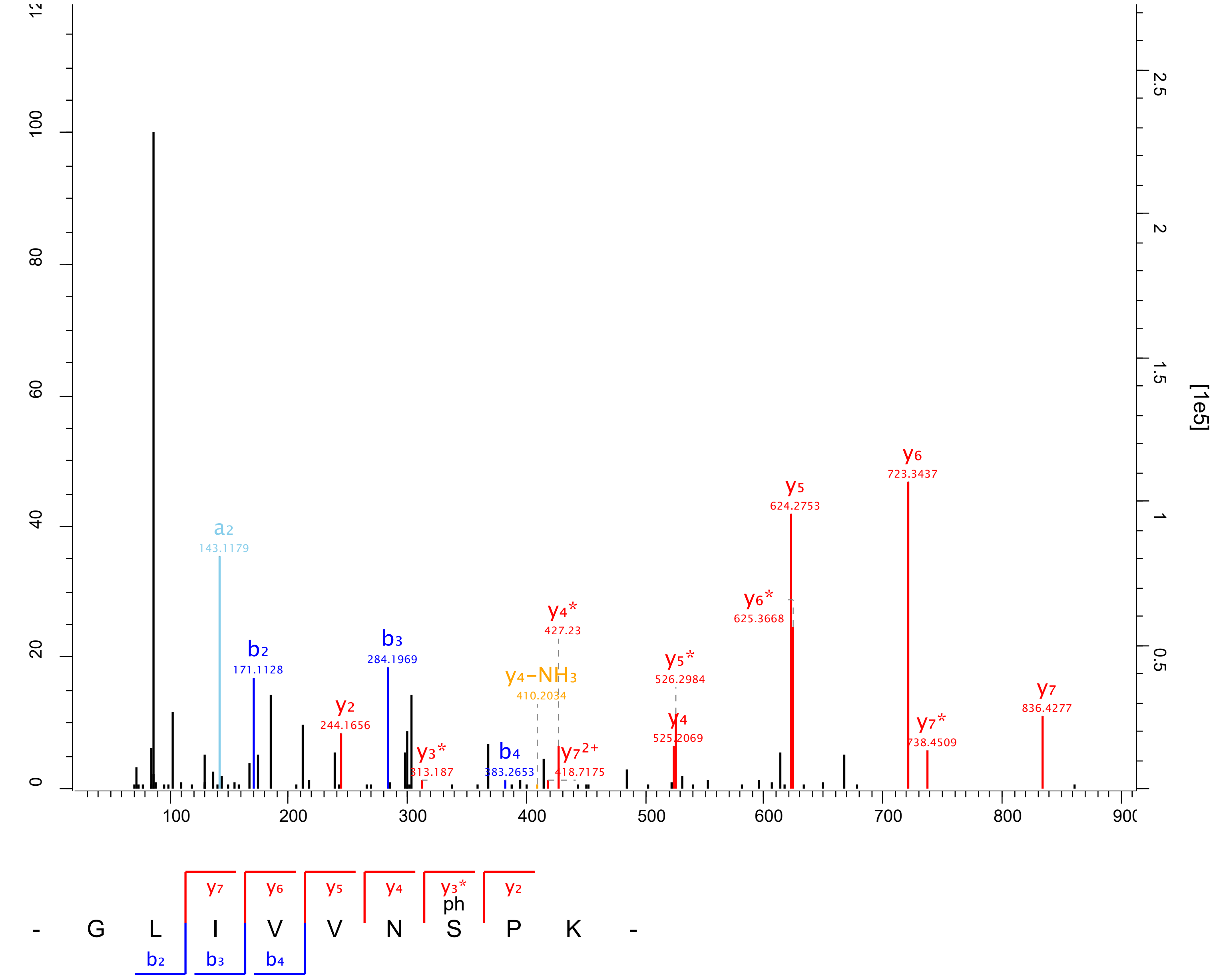

| Raw file | Scan | Method | Score | m/z |
|---|---|---|---|---|
| water_00_1 | 16159 | FTMS; HCD | 77.42 | 680.31 |

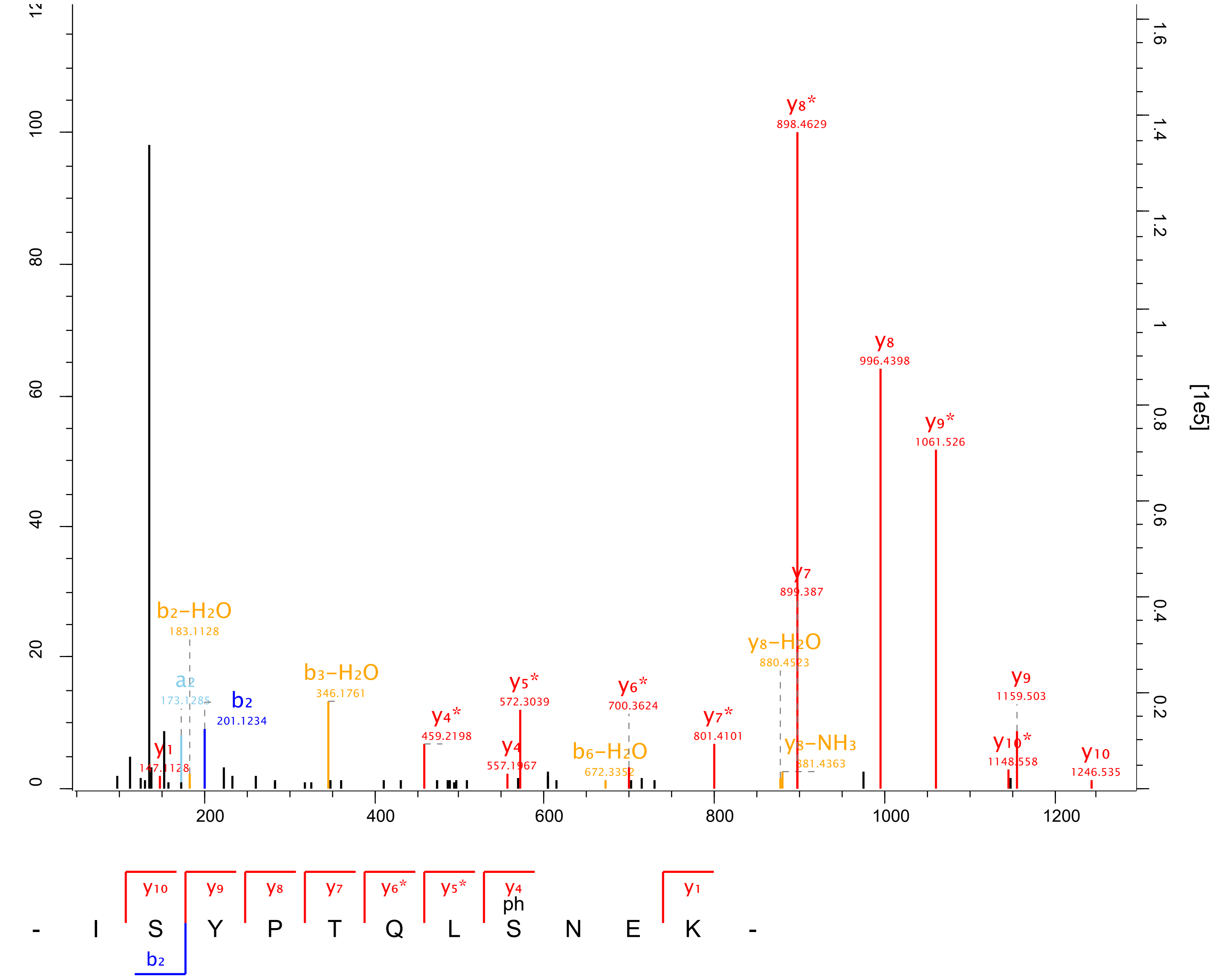

| Raw file | Scan | Method | Score | m/z |
|---|---|---|---|---|
| water_00_1 | 1630 | FTMS; HCD | 59.91 | 476.25 |

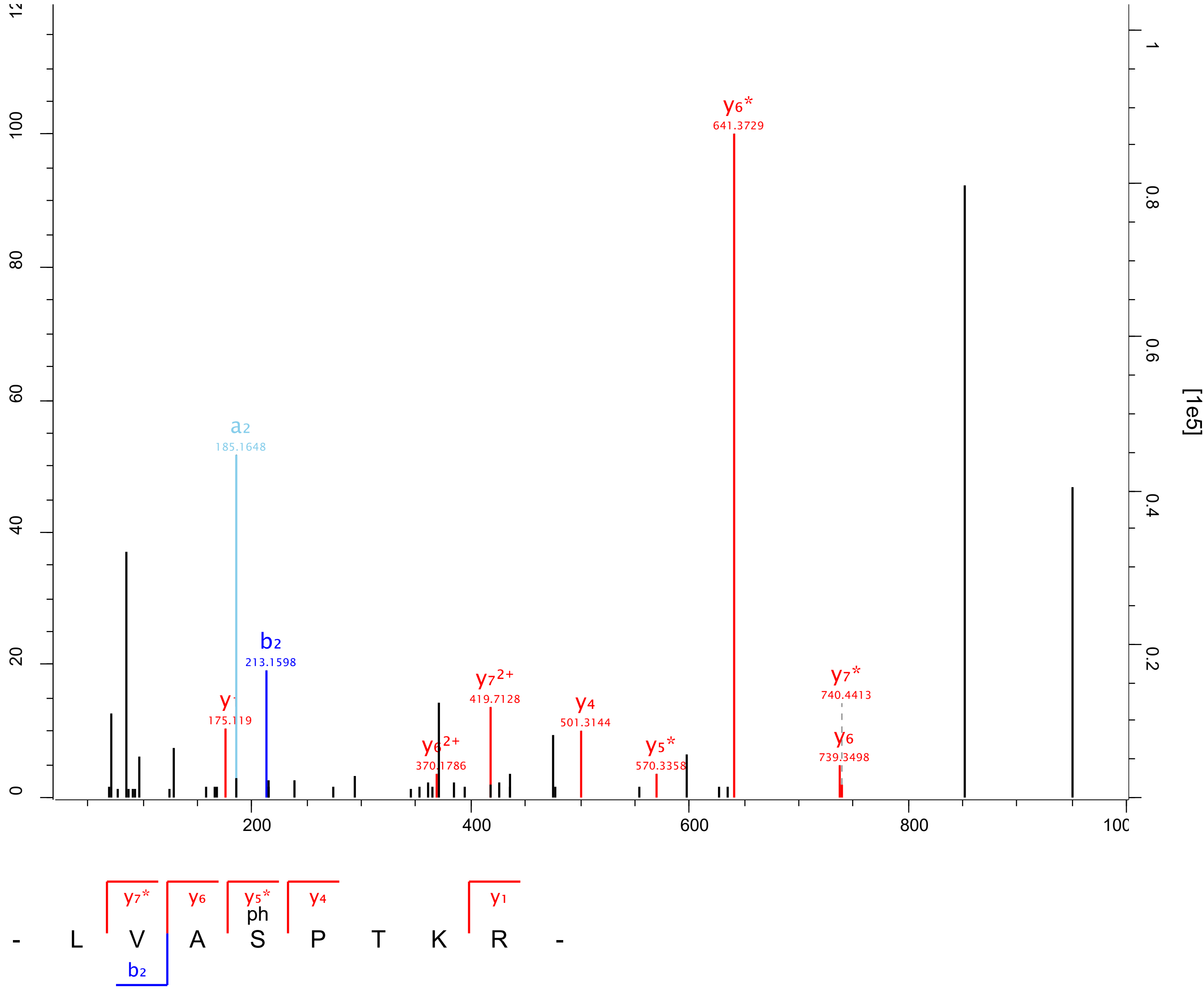

| Raw file | Scan | Method | Score | m/z |
|---|---|---|---|---|
| water_00_1 | 1667 | FTMS; HCD | 46.32 | 434.19 |

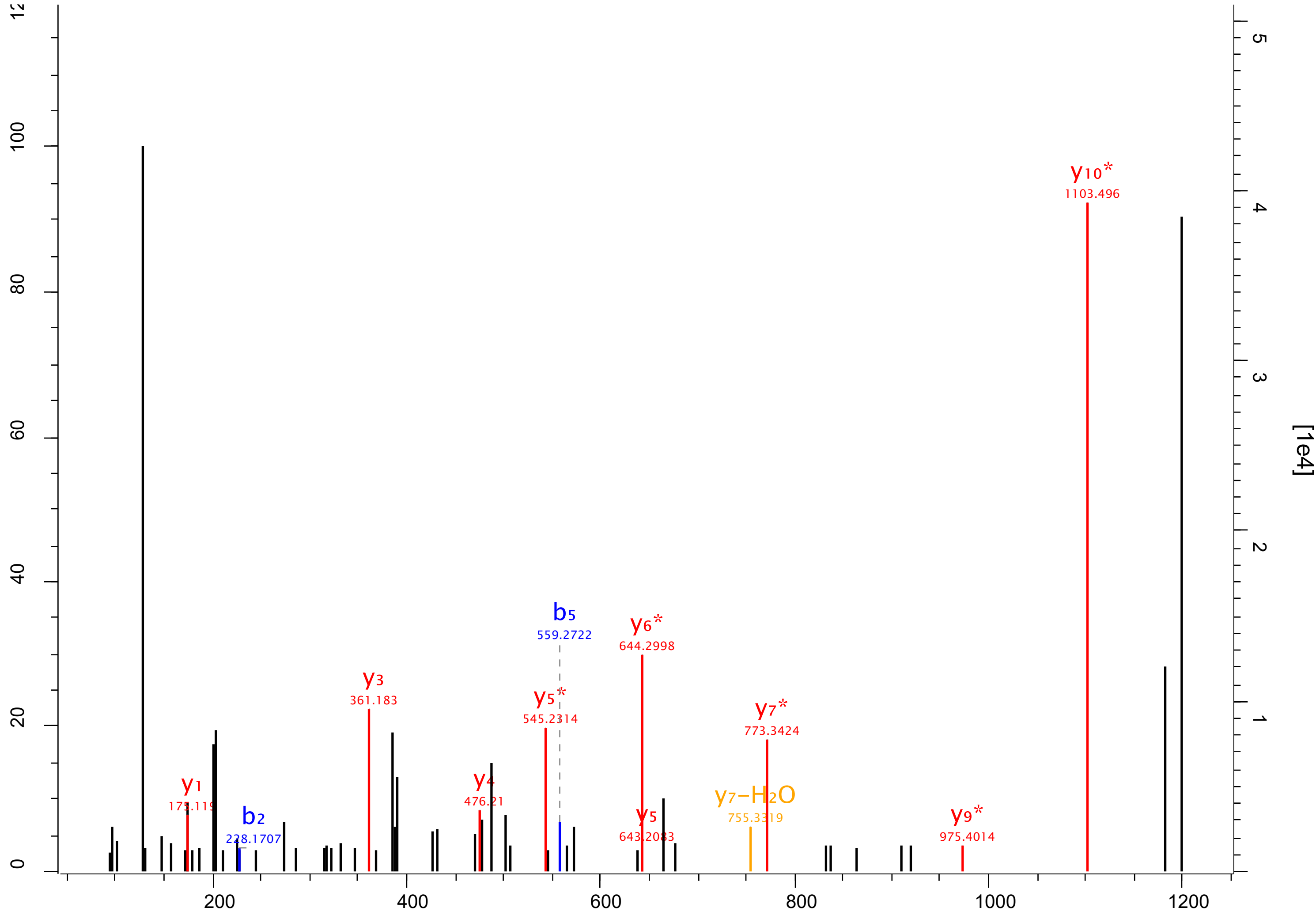


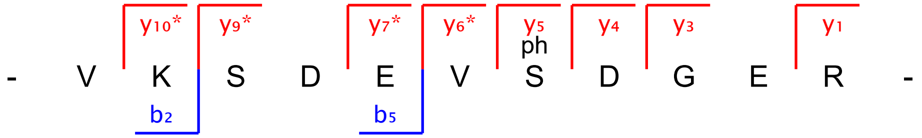

| Raw file | Scan | Method | Score | m/z |
| --- | --- | --- | --- | --- |
| water_00_1 | 16912 | FTMS; HCD | 46.32 | 692.28 |

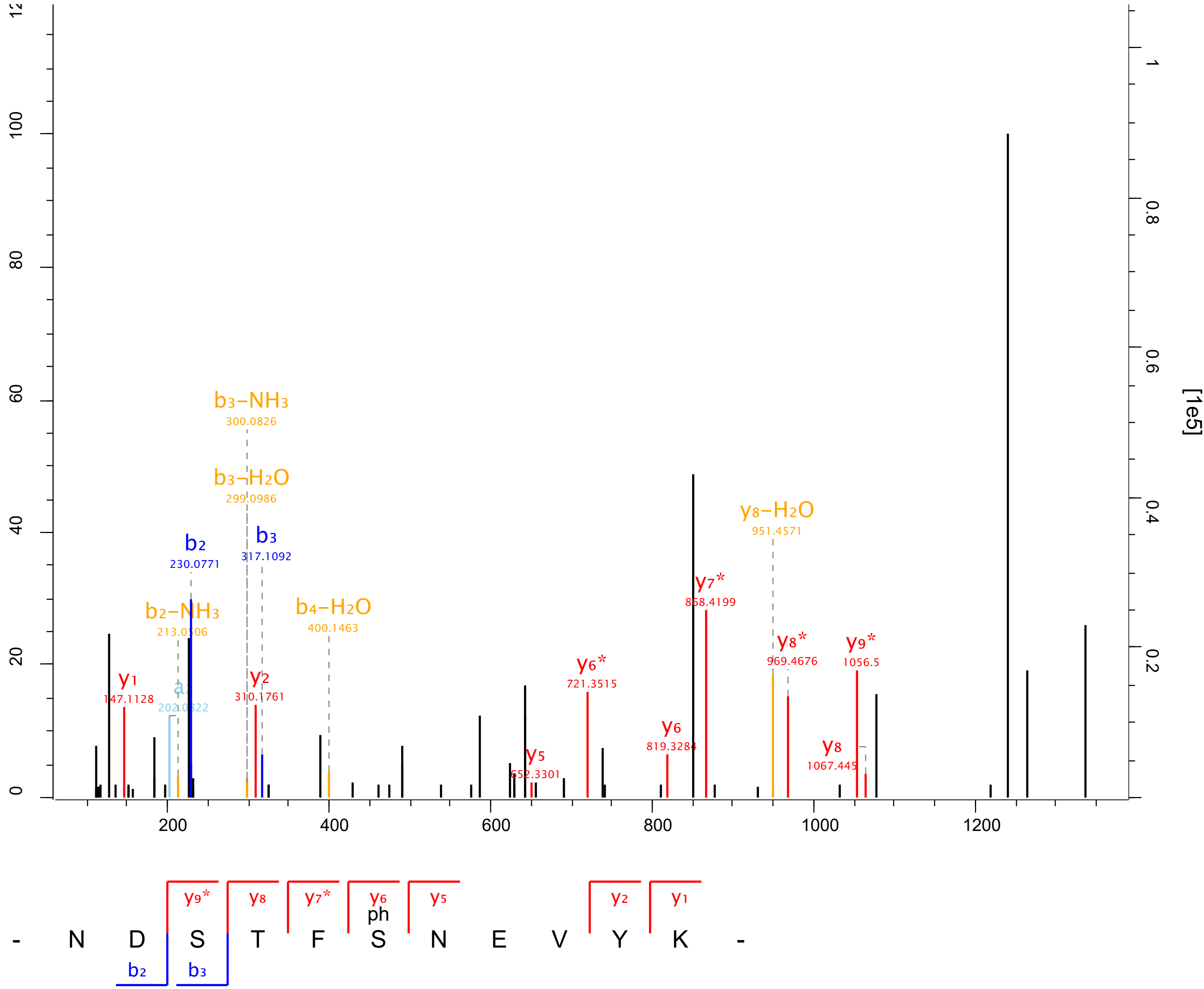

| Raw file | Scan | Method | Score | m/z |
| --- | --- | --- | --- | --- |
| water_00_1 | 1694 | FTMS; HCD | 65.84 | 442.2 |

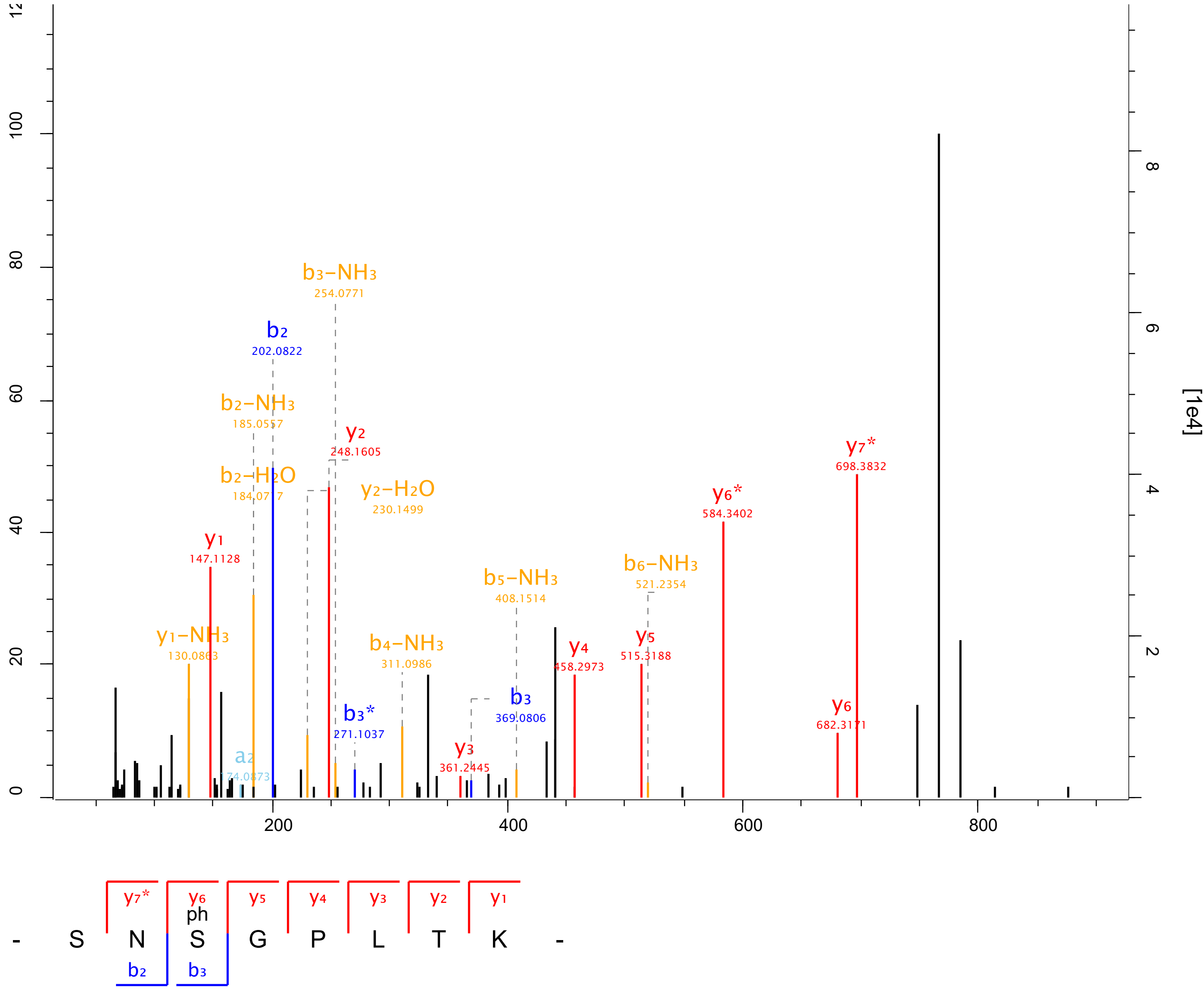

| Raw file | Scan | Method | Score | m/z |
|---|---|---|---|---|
| water_00_1 | 1697 | FTMS; HCD | 54.9 | 428.52 |

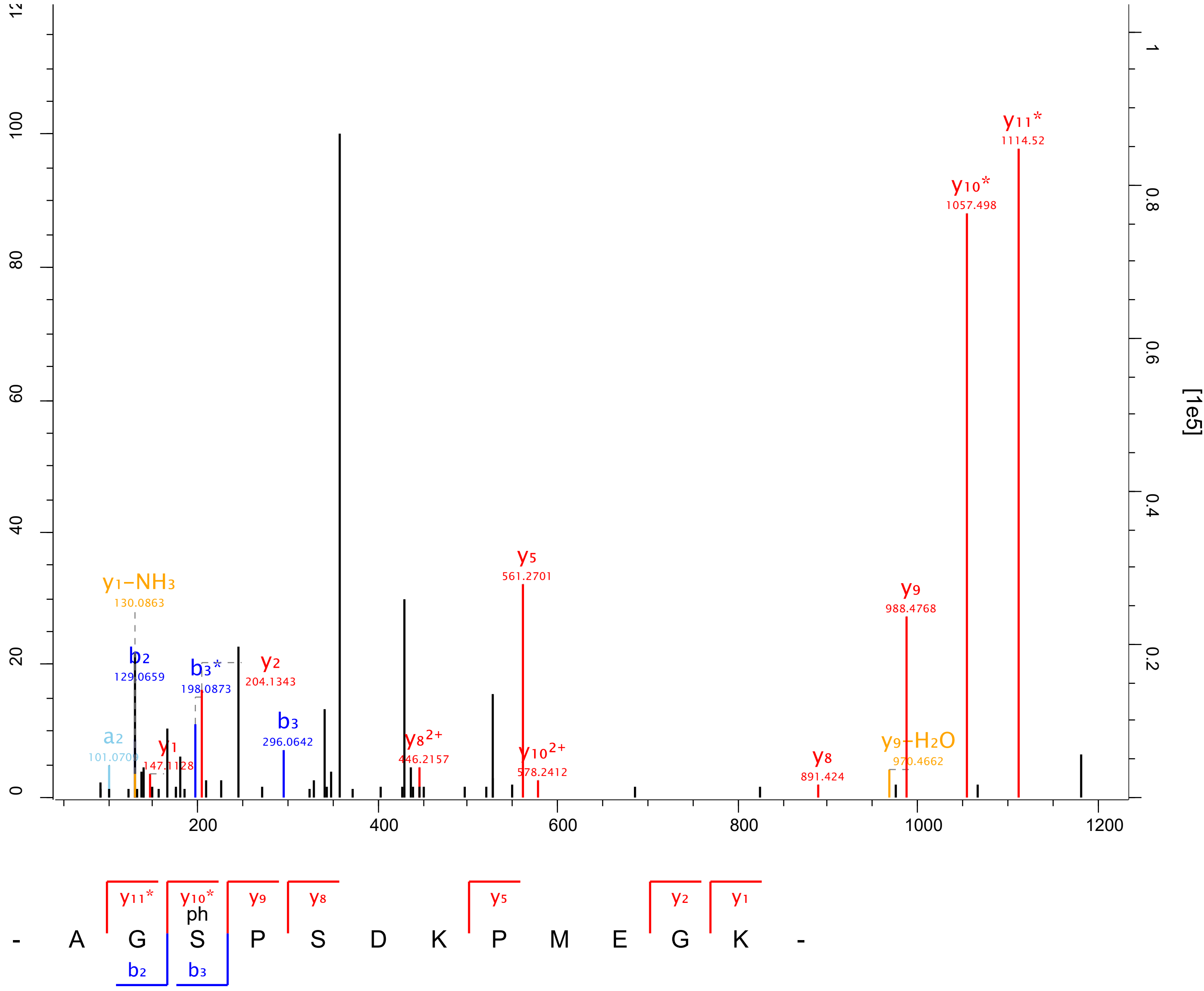

| Raw file | Scan | Method | Score | m/z |
|---|---|---|---|---|
| water_00_1 | 1702 | FTMS; HCD | 51.29 | 546.23 |

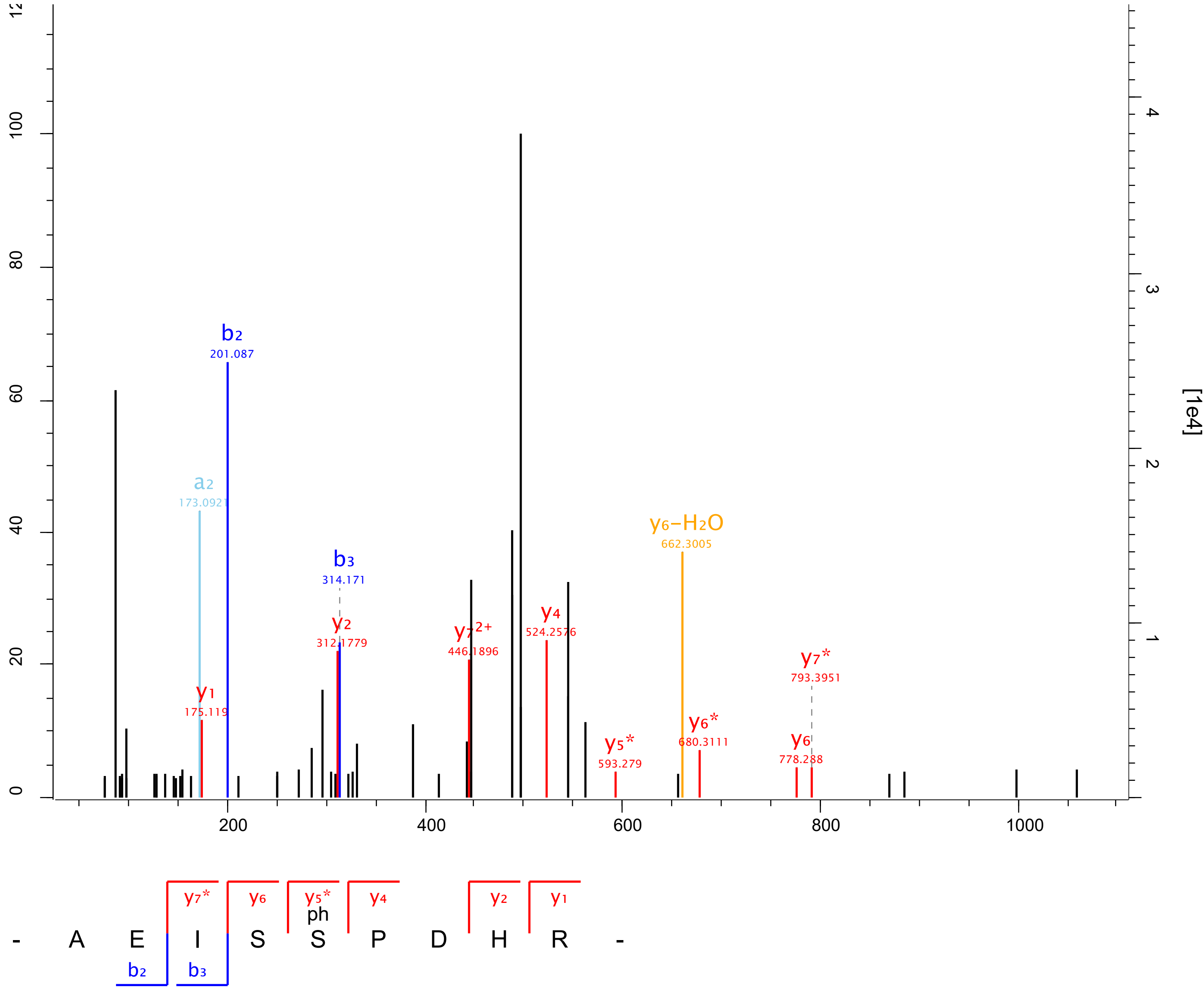

| Raw file | Scan | Method | Score | m/z |
|---|---|---|---|---|
| water_00_1 | 1707 | FTMS; HCD | 44.56 | 527.24 |

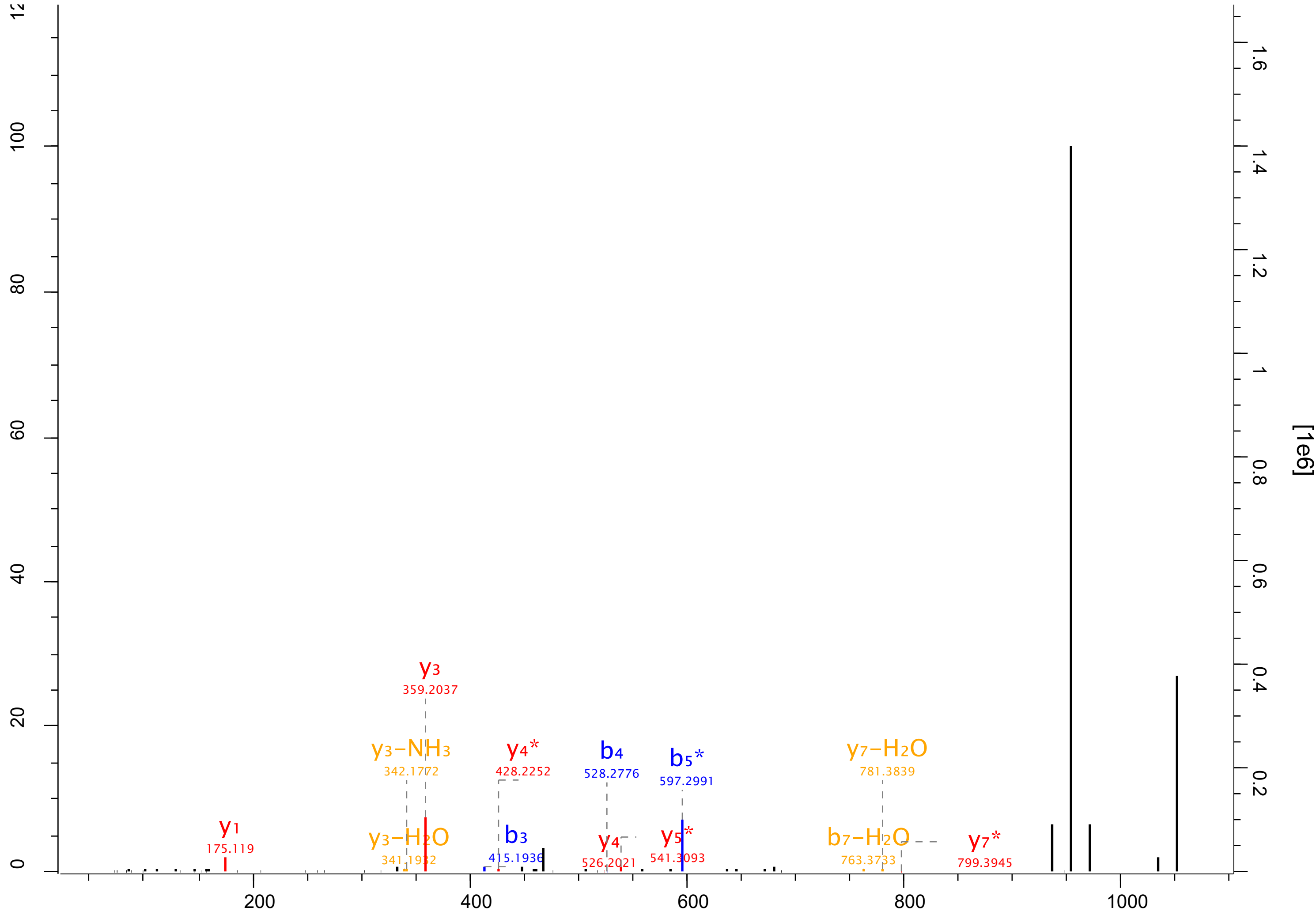


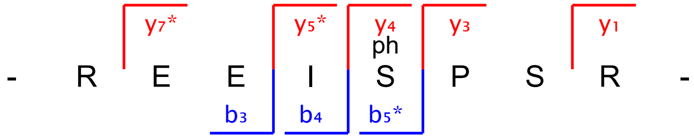

| Raw file | Scan | Method | Score | m/z |
| --- | --- | --- | --- | --- |
| water_00_1 | 17166 | FTMS; HCD | 102.64 | 690.3 |

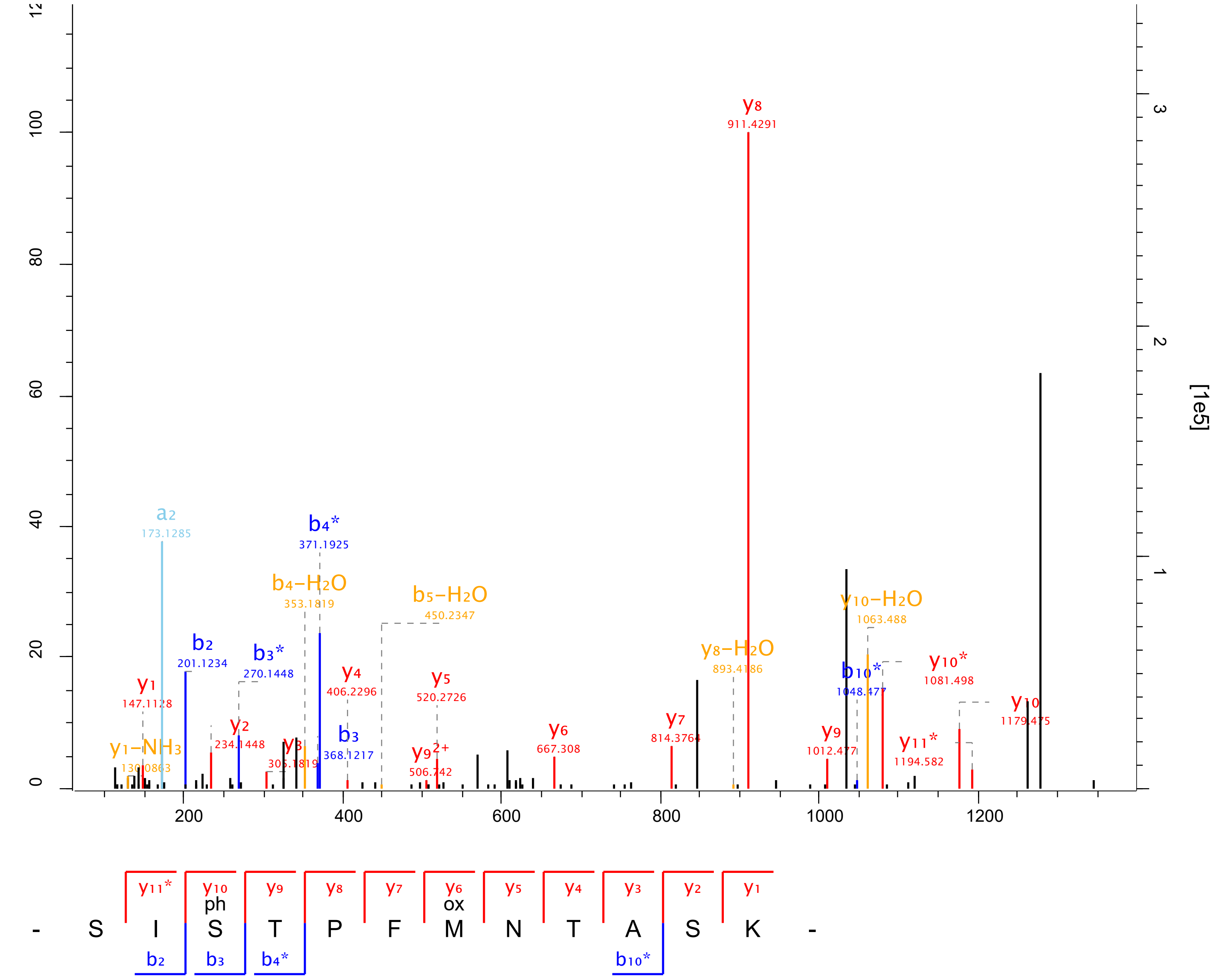

| Raw file | Scan | Method | Score | m/z |
|---|---|---|---|---|
| water_00_1 | 17182 | FTMS; HCD | 53.87 | 750.33 |

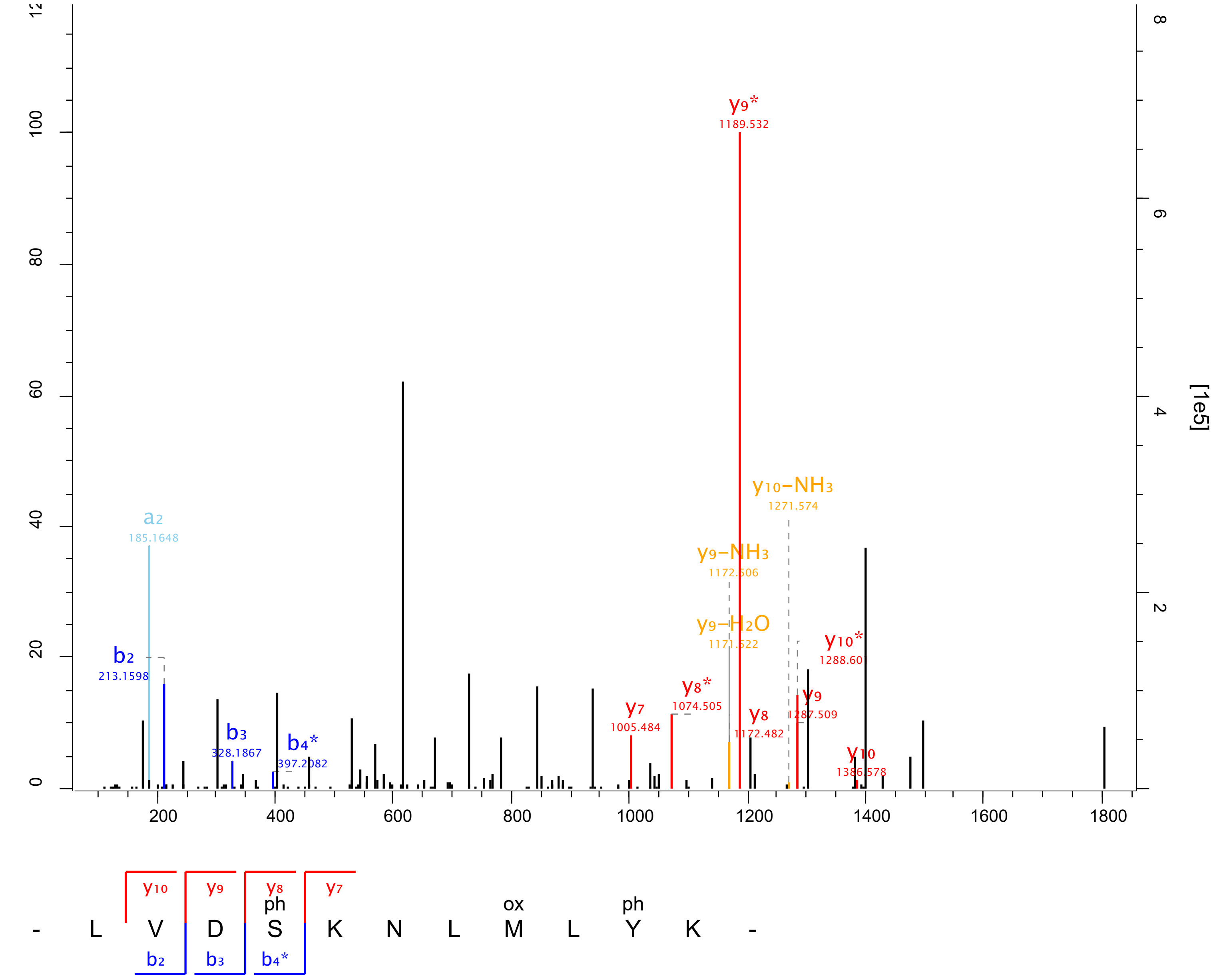

| Raw file | Scan | Method | Score | m/z |
|---|---|---|---|---|
| water_00_1 | 17248 | FTMS; HCD | 285.5 | 835.32 |

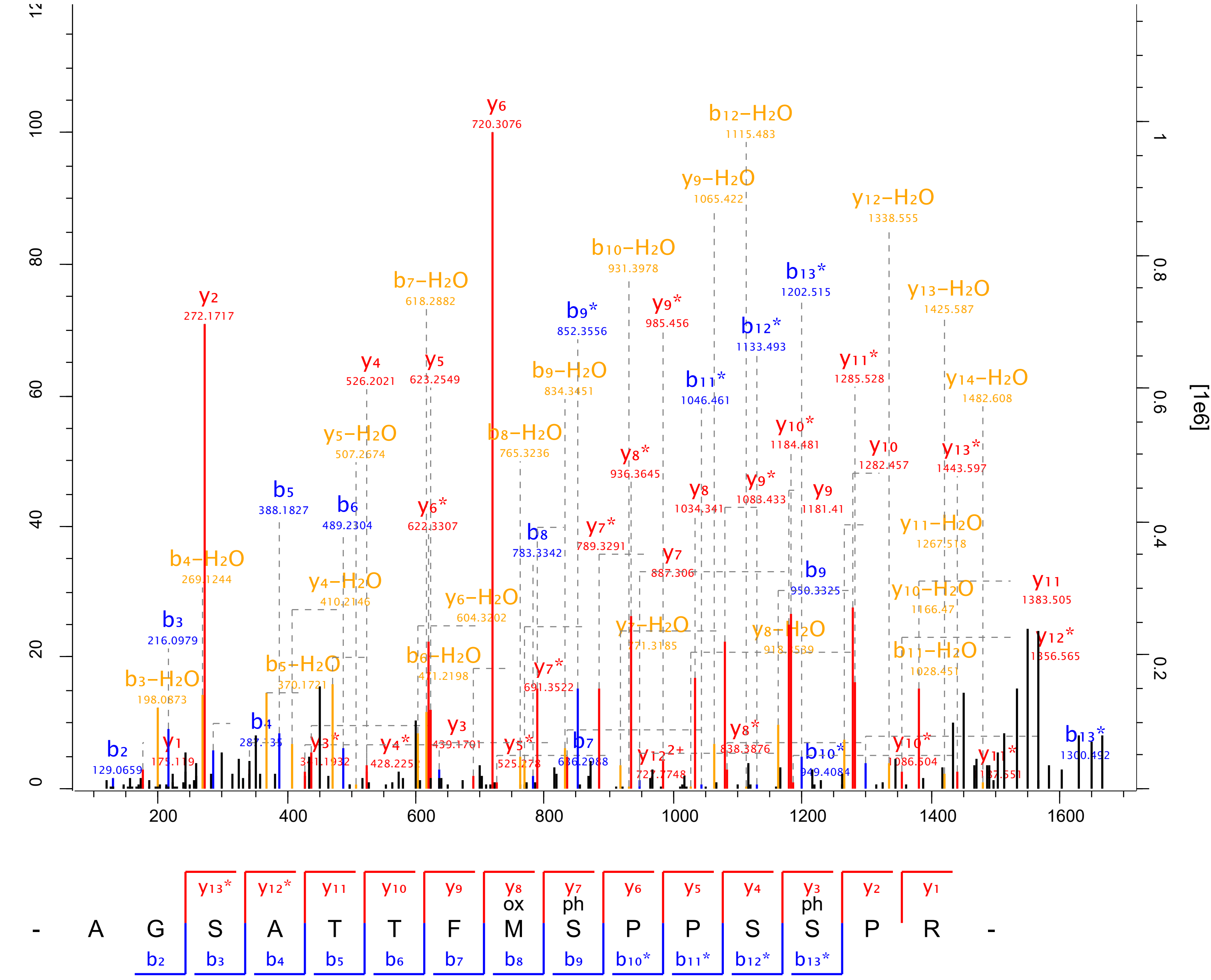

| Raw file | Scan | Method | Score | m/z |
|---|---|---|---|---|
| water_00_1 | 17265 | FTMS; HCD | 241.77 | 927.35 |

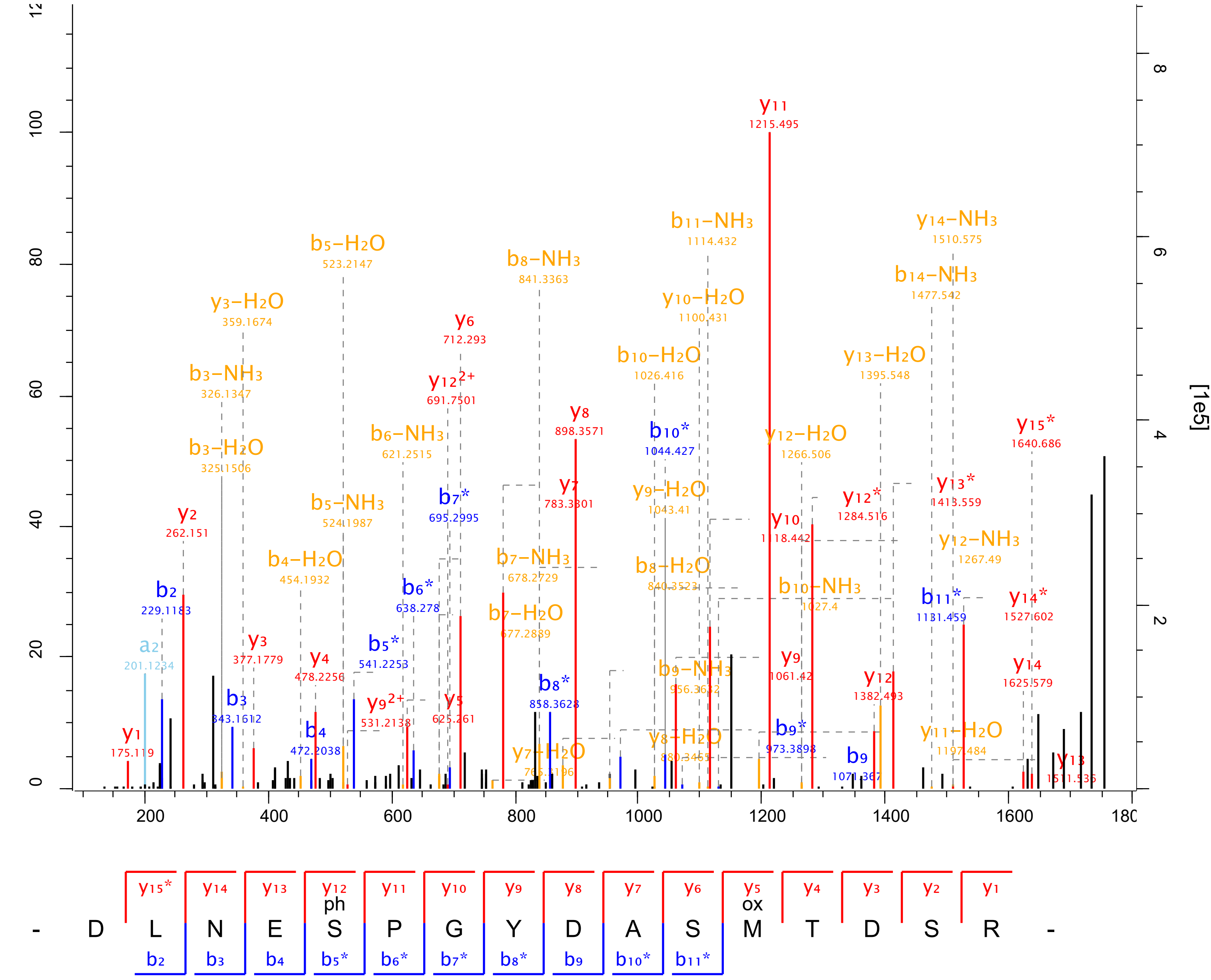

| Raw file | Scan | Method | Score | m/z |
|---|---|---|---|---|
| water_00_1 | 17277 | FTMS; HCD | 92.65 | 454.52 |

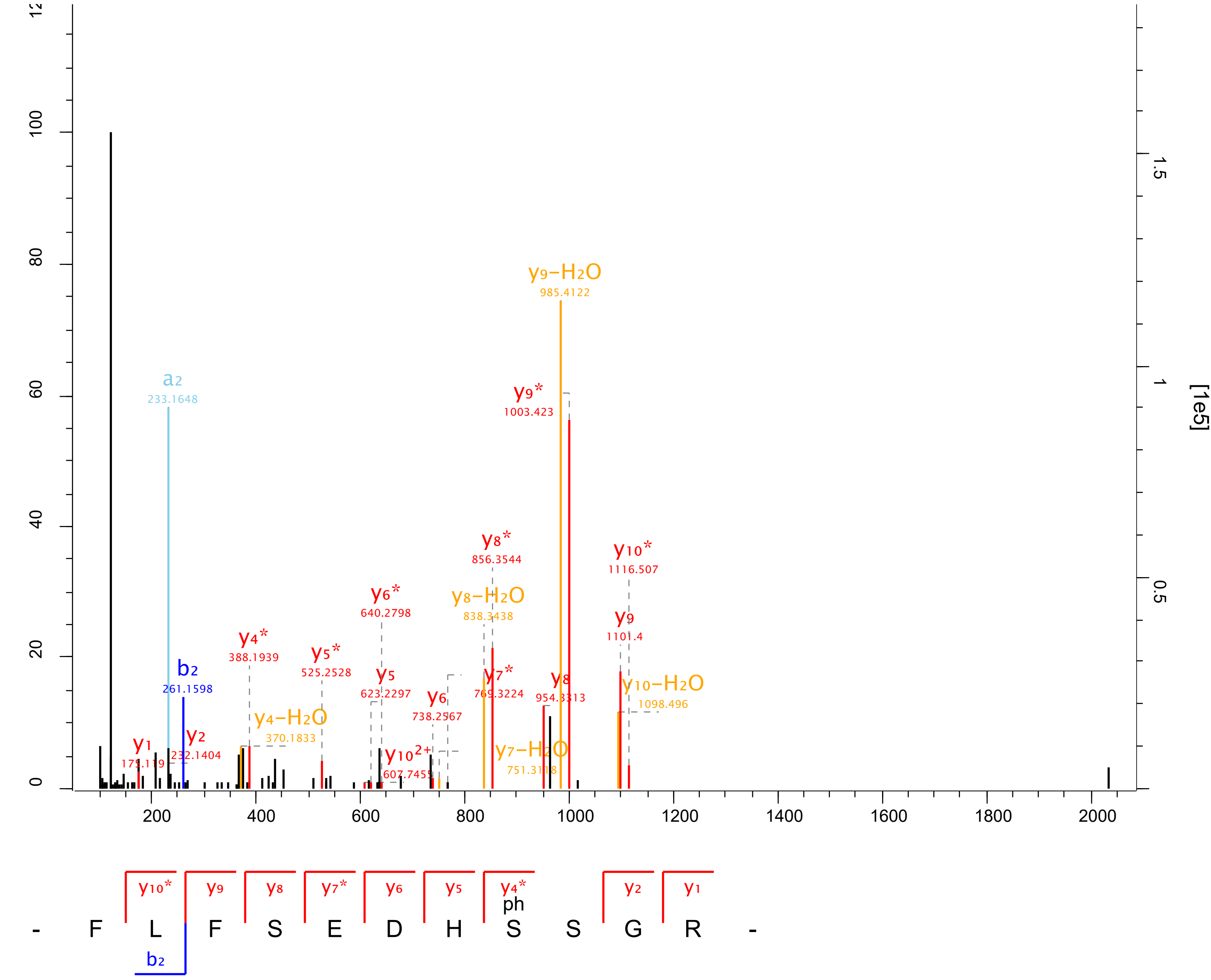

| Raw file | Scan | Method | Score | m/z |
|---|---|---|---|---|
| water_00_1 | 17587 | FTMS; HCD | 121.99 | 624.31 |

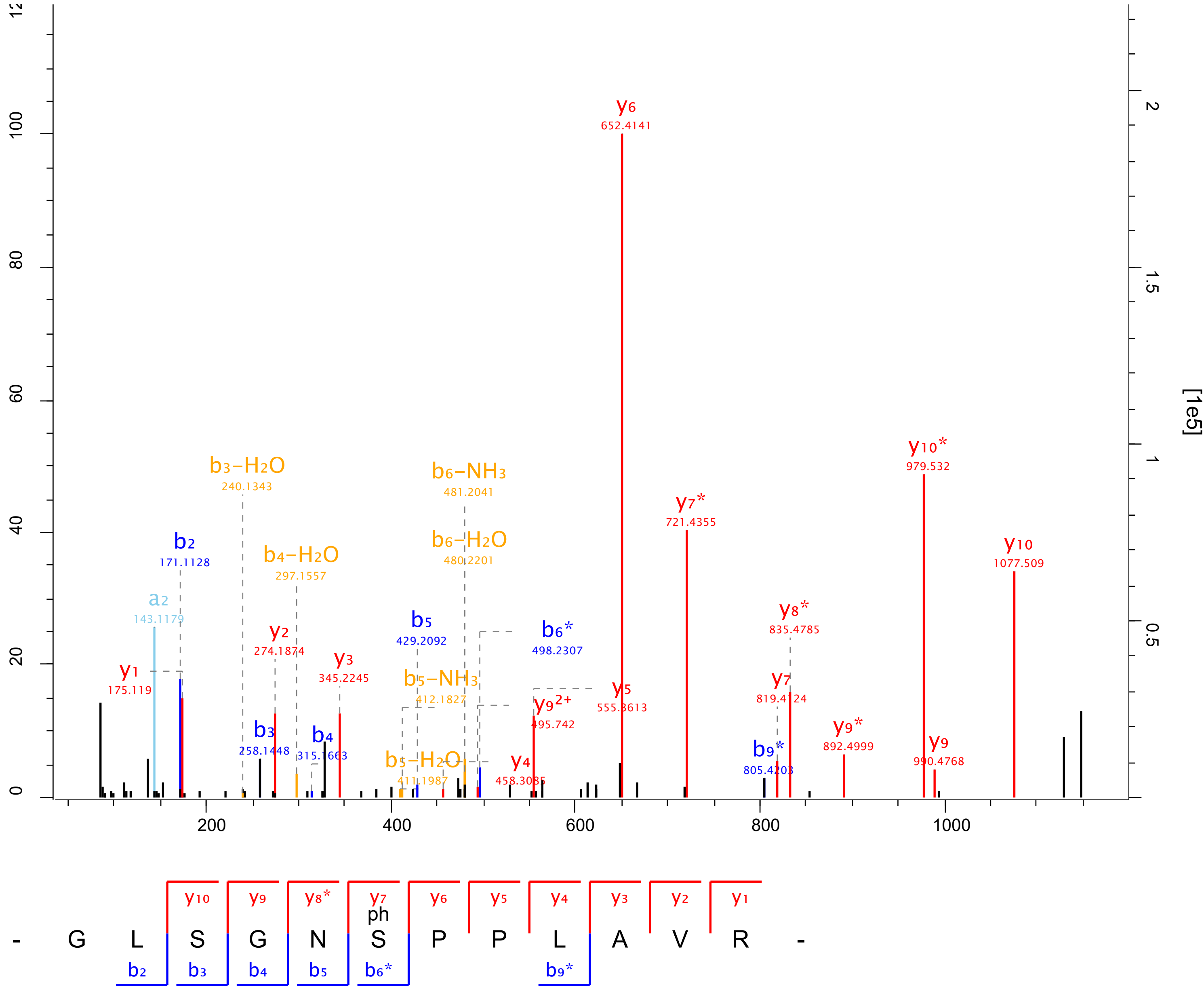

| Raw file | Scan | Method | Score | m/z |
| --- | --- | --- | --- | --- |
| water_00_1 | 17589 | FTMS; HCD | 53.36 | 556.93 |

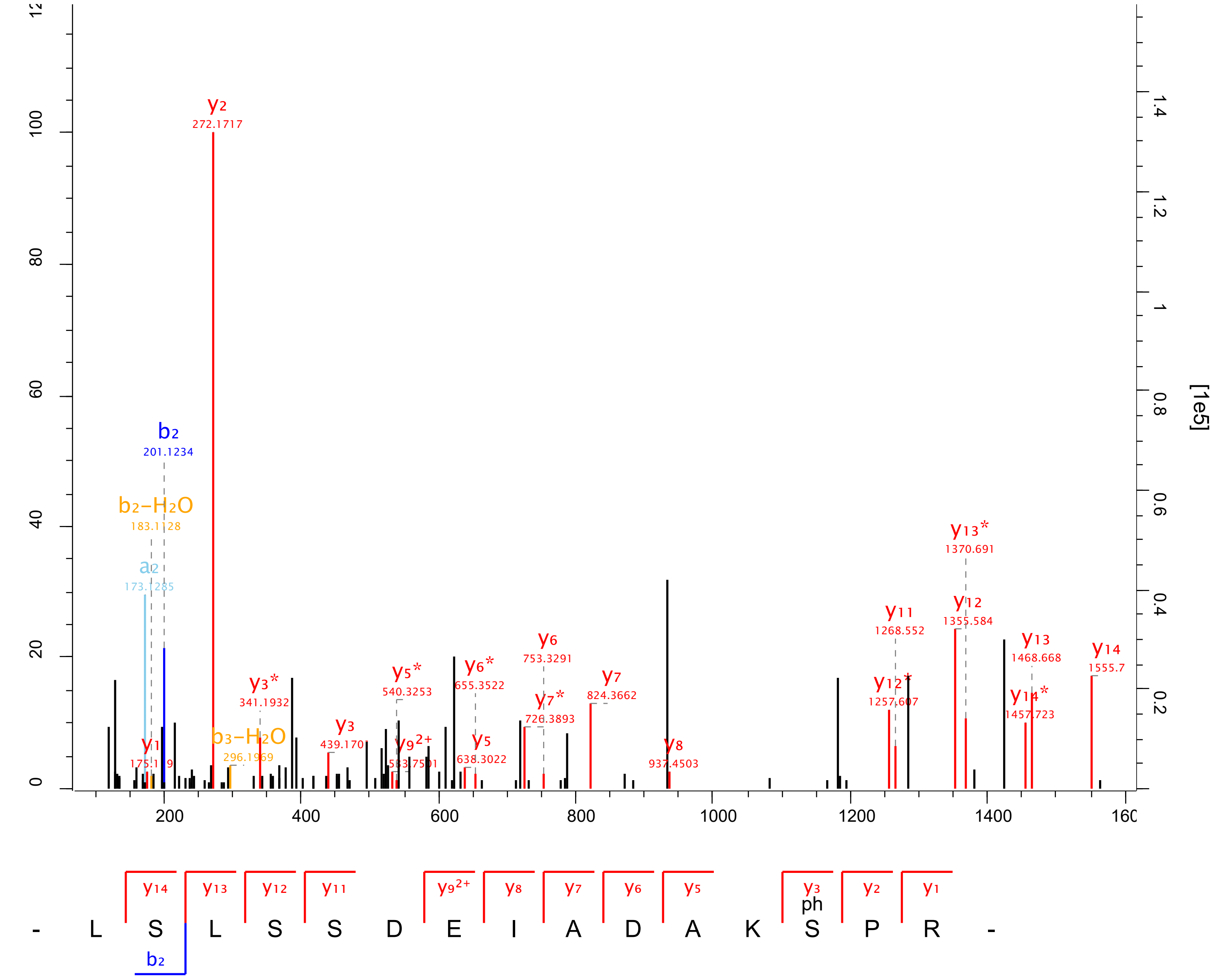

| Raw file | Scan | Method | Score | m/z |
|---|---|---|---|---|
| water_00_1 | 1762 | FTMS; HCD | 105.98 | 487.2 |

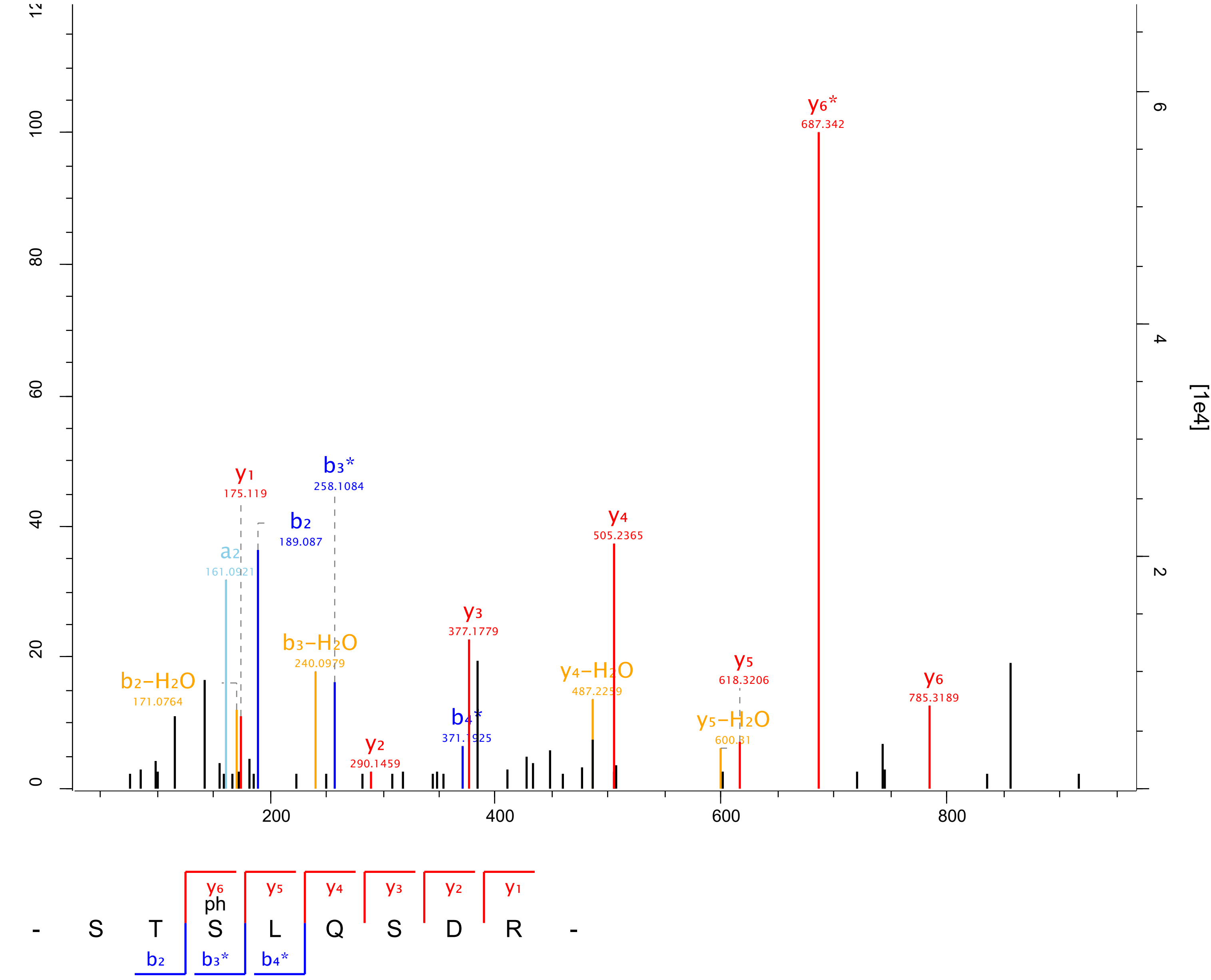

| Raw file | Scan | Method | Score | m/z |
| --- | --- | --- | --- | --- |
| water_00_1 | 17649 | FTMS; HCD | 152.88 | 668.77 |

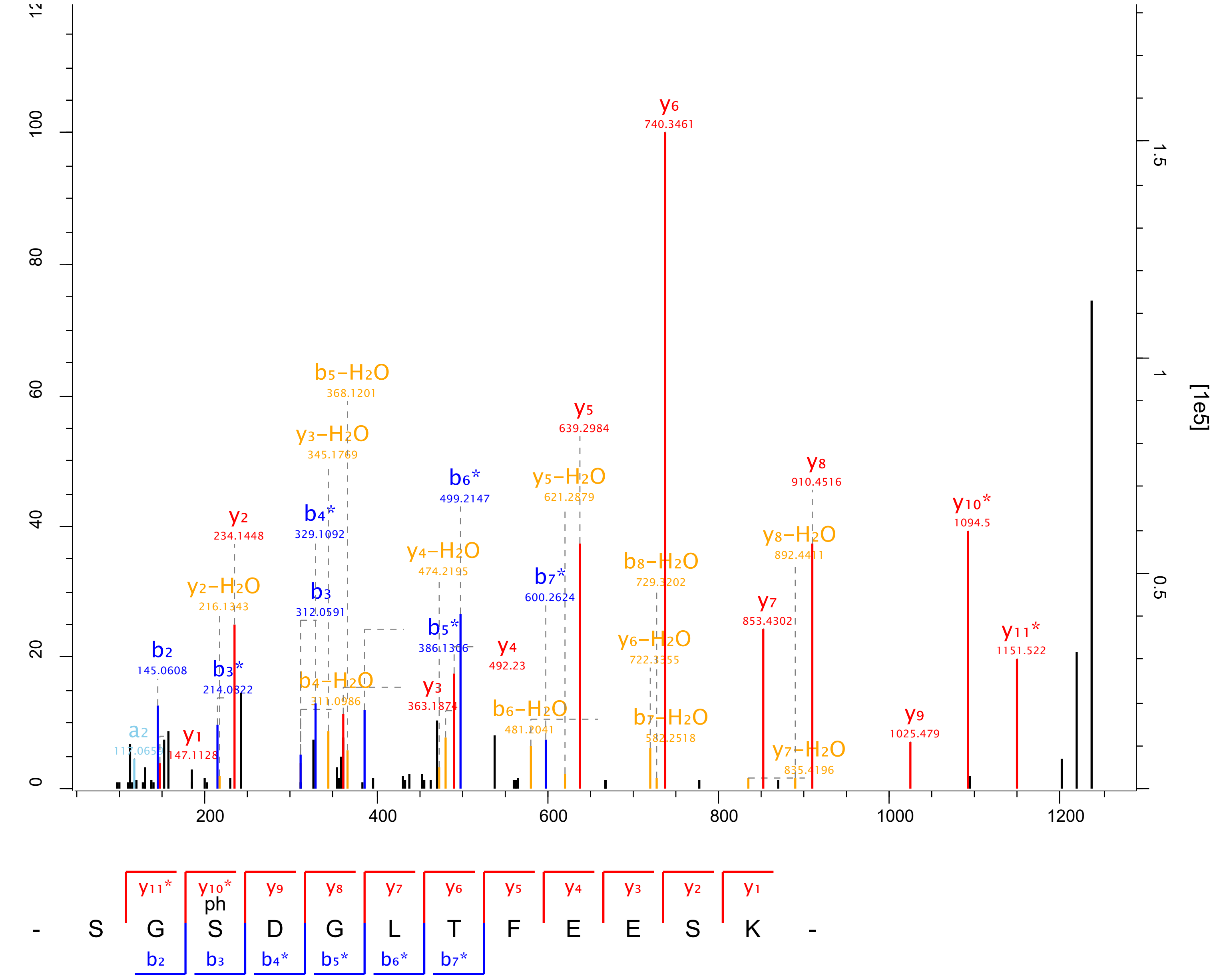

| Raw file | Scan | Method | Score | m/z |
|---|---|---|---|---|
| water_00_1 | 1767 | FTMS; HCD | 57.55 | 416.72 |

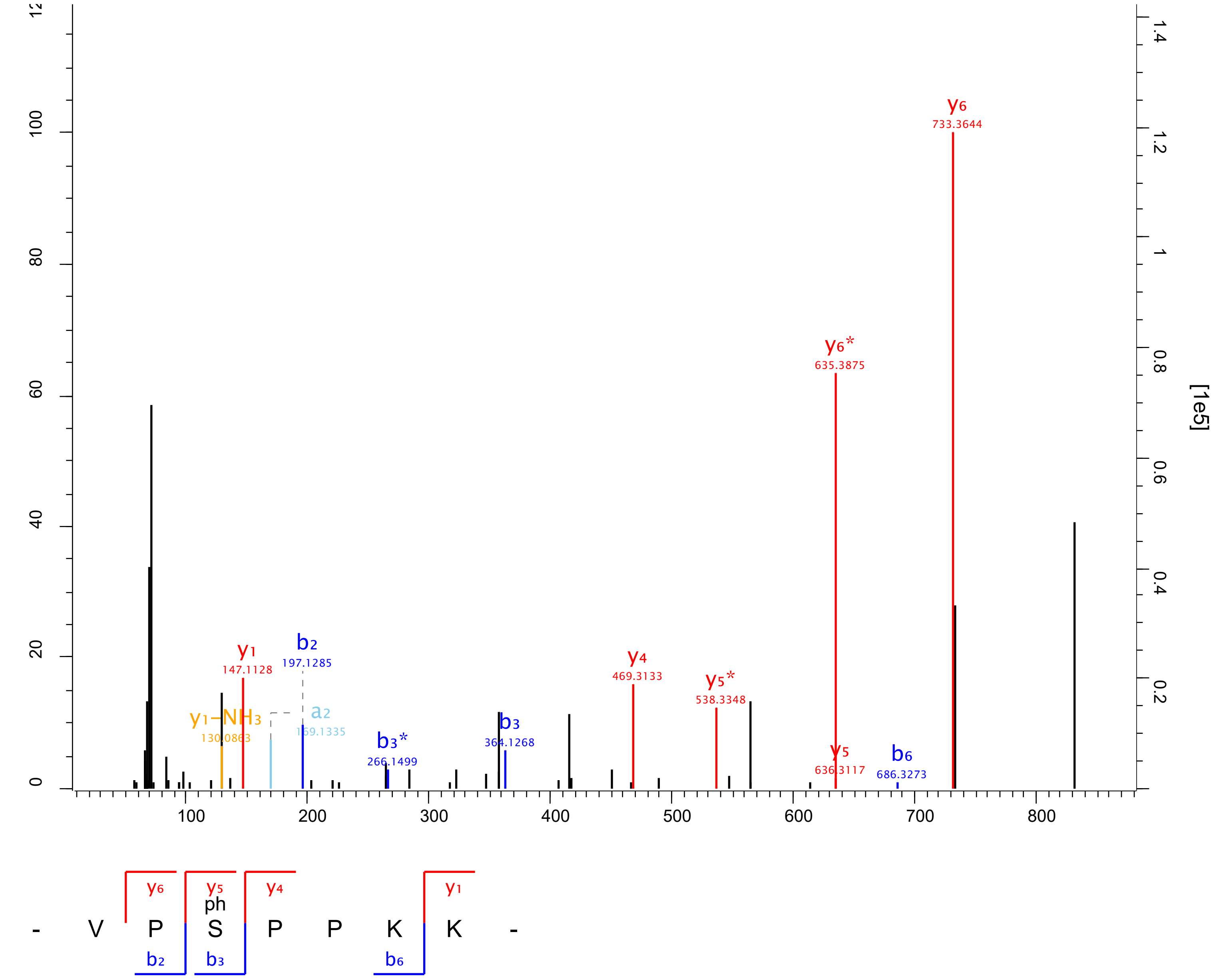

| Raw file | Scan | Method | Score | m/z |
| --- | --- | --- | --- | --- |
| water_00_1 | 17961 | FTMS; HCD | 121.13 | 626.81 |

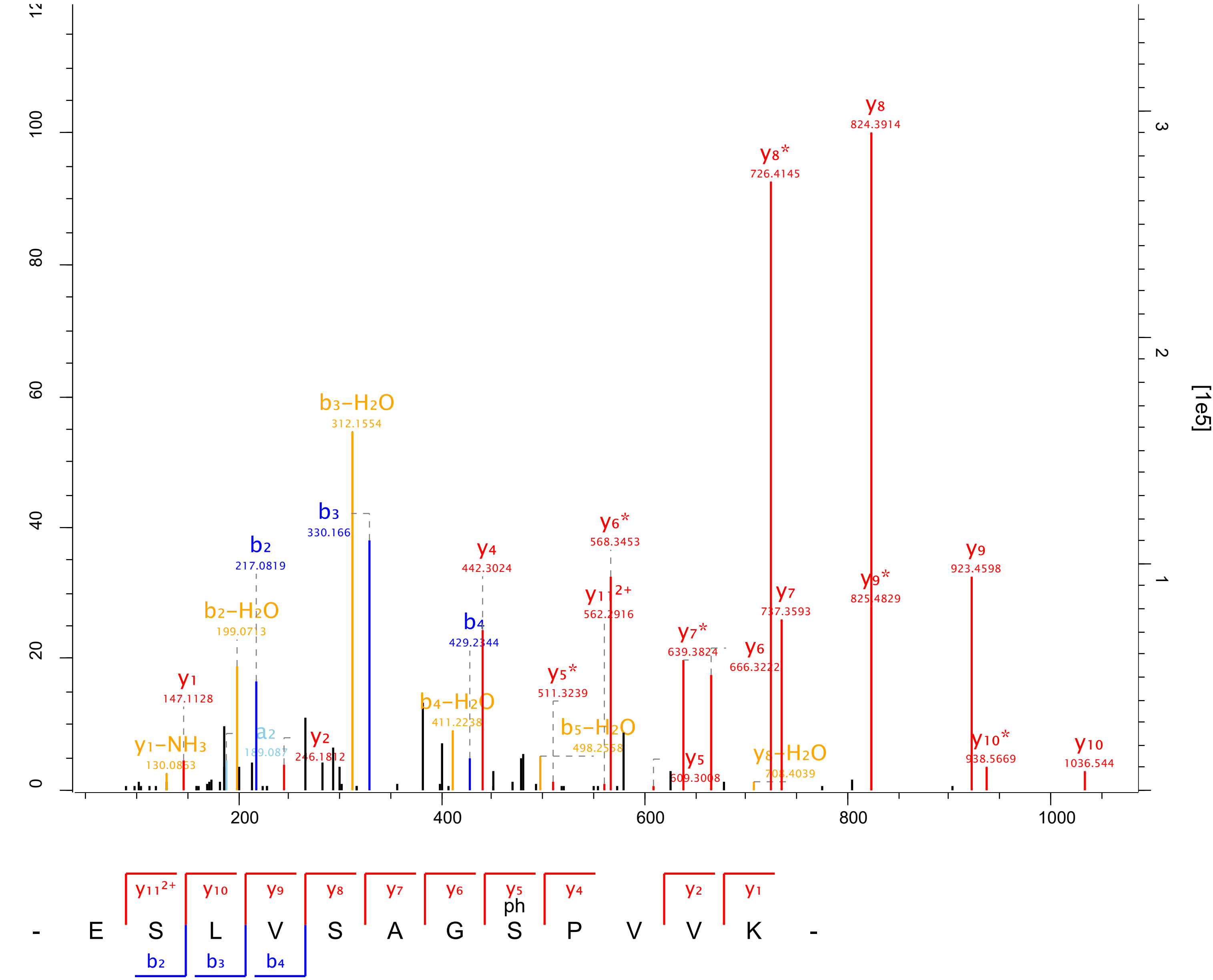

| Raw file | Scan | Method | Score | m/z |
|---|---|---|---|---|
| water_00_1 | 18143 | FTMS; HCD | 209.53 | 645.32 |

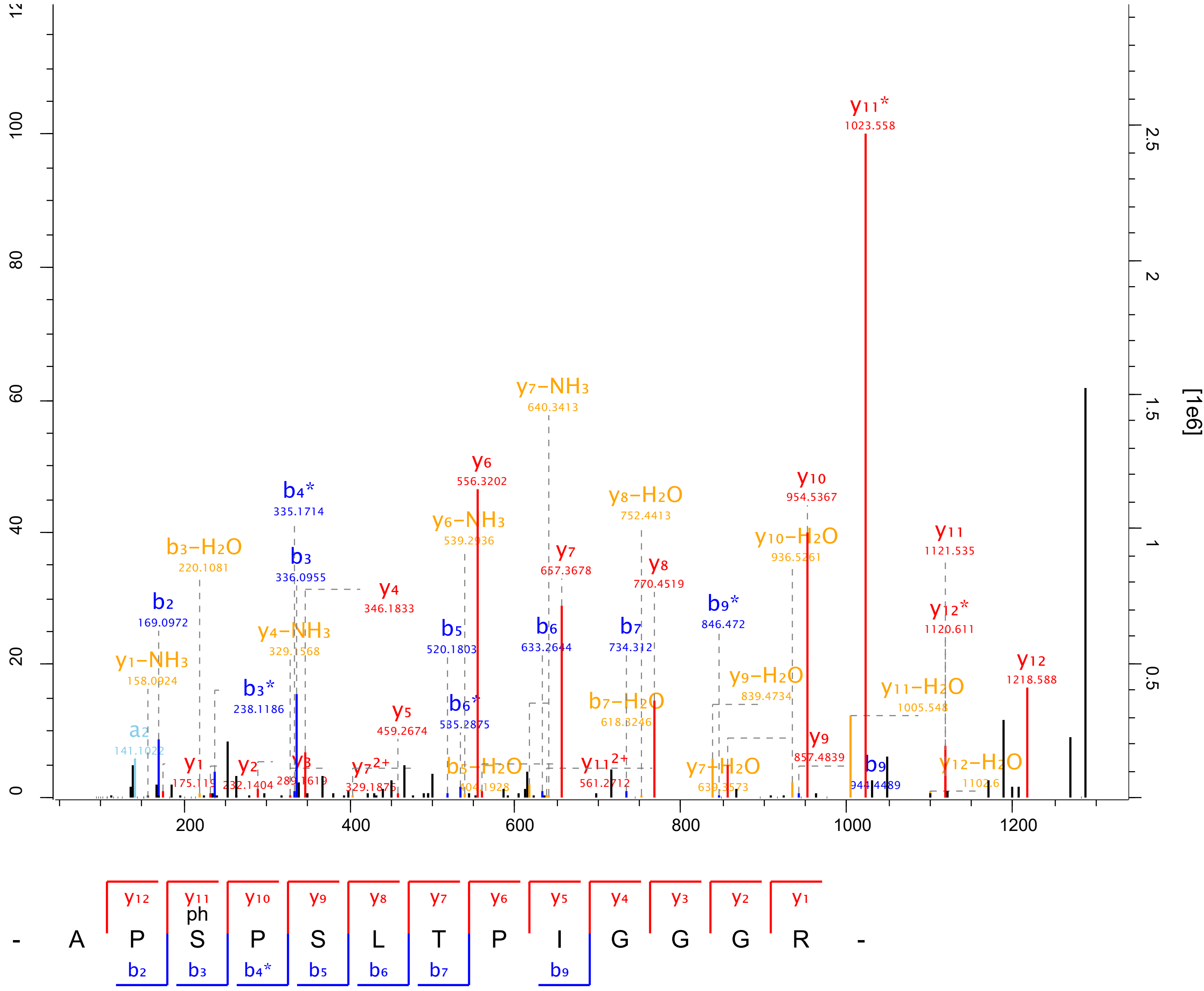

| Raw file | Scan | Method | Score | m/z |
| --- | --- | --- | --- | --- |
| water_00_1 | 18367 | FTMS; HCD | 137.69 | 838.85 |

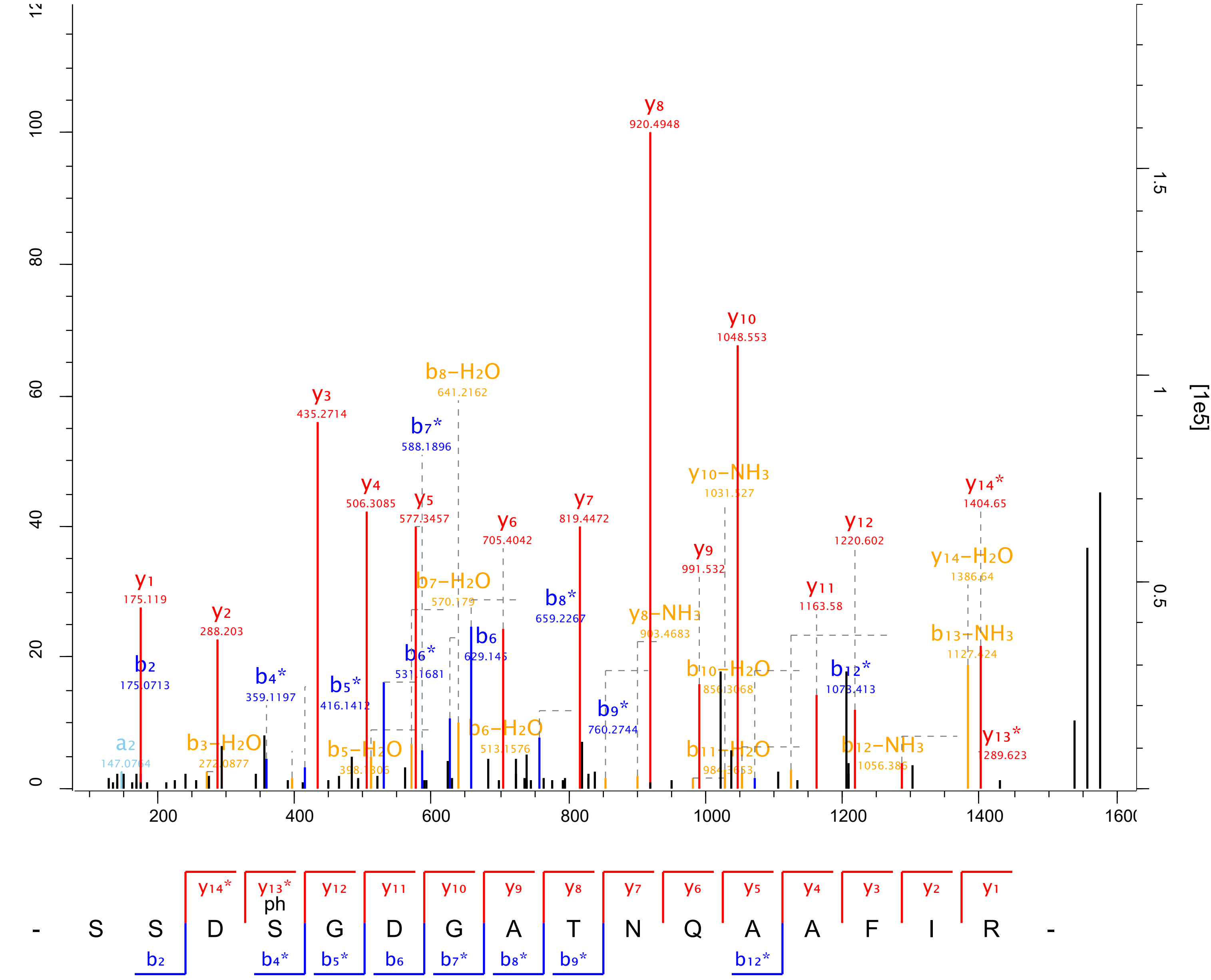

| Raw file | Scan | Method | Score | m/z |
|---|---|---|---|---|
| water_00_1 | 18418 | FTMS; HCD | 145.91 | 806.34 |

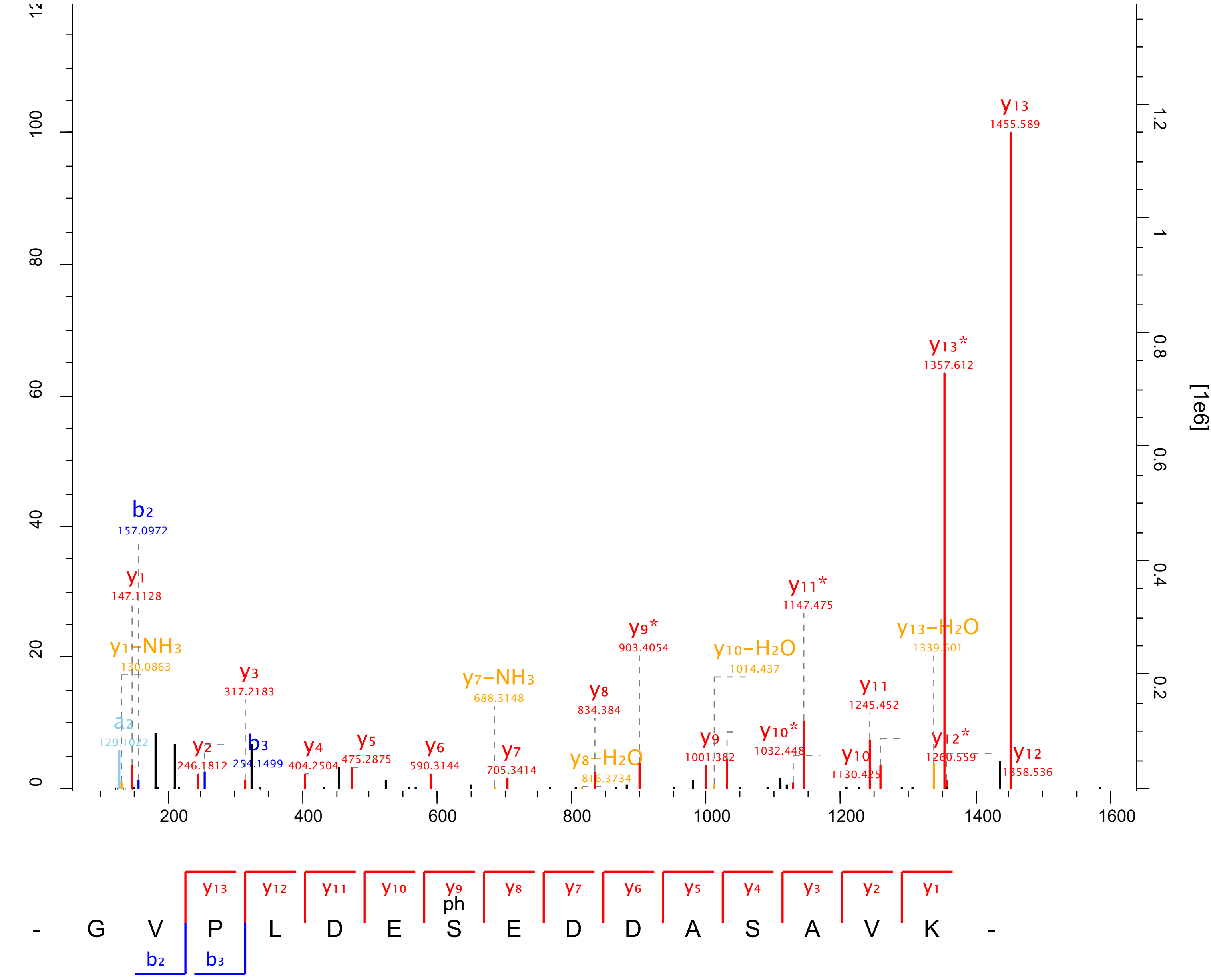

| Raw file | Scan | Method | Score | m/z |
|---|---|---|---|---|
| water_00_1 | 1855 | FTMS; HCD | 134.06 | 598.75 |

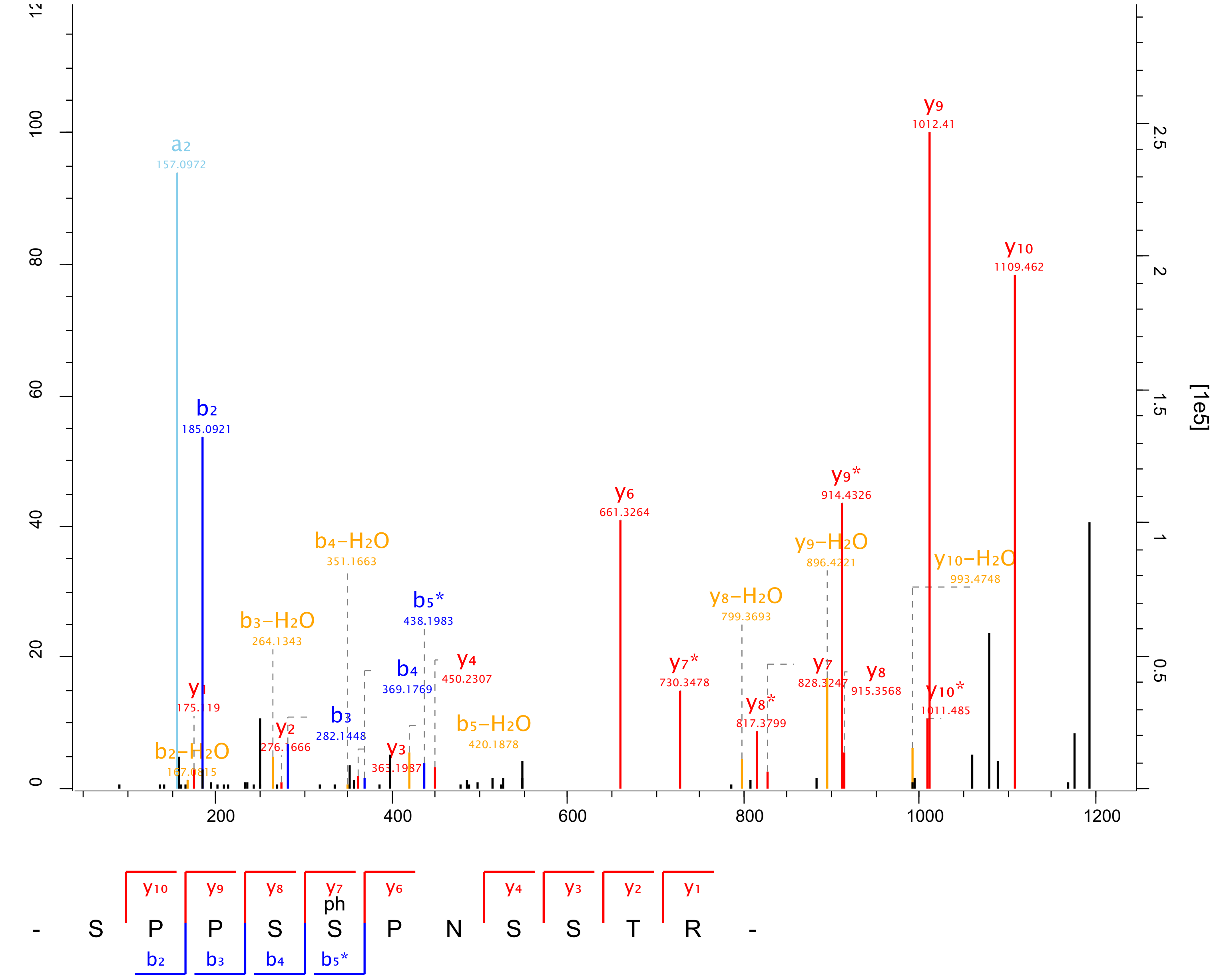

| Raw file | Scan | Method | Score | m/z |
| --- | --- | --- | --- | --- |
| water_00_1 | 19096 | FTMS; HCD | 186.62 | 1057.92 |

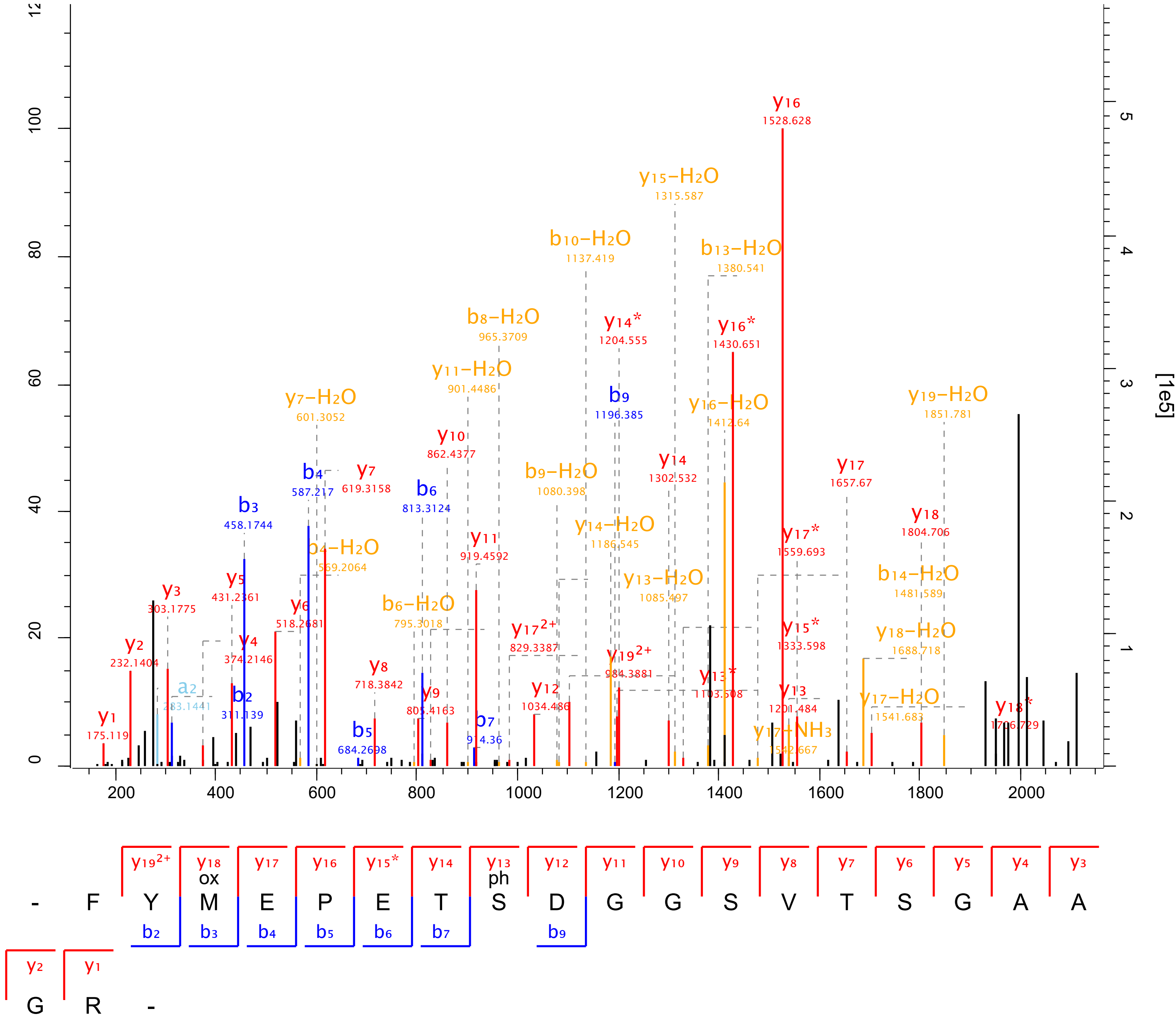

| Raw file | Scan | Method | Score | m/z |
|---|---|---|---|---|
| water_00_1 | 19112 | FTMS; HCD | 53.76 | 492.7 |

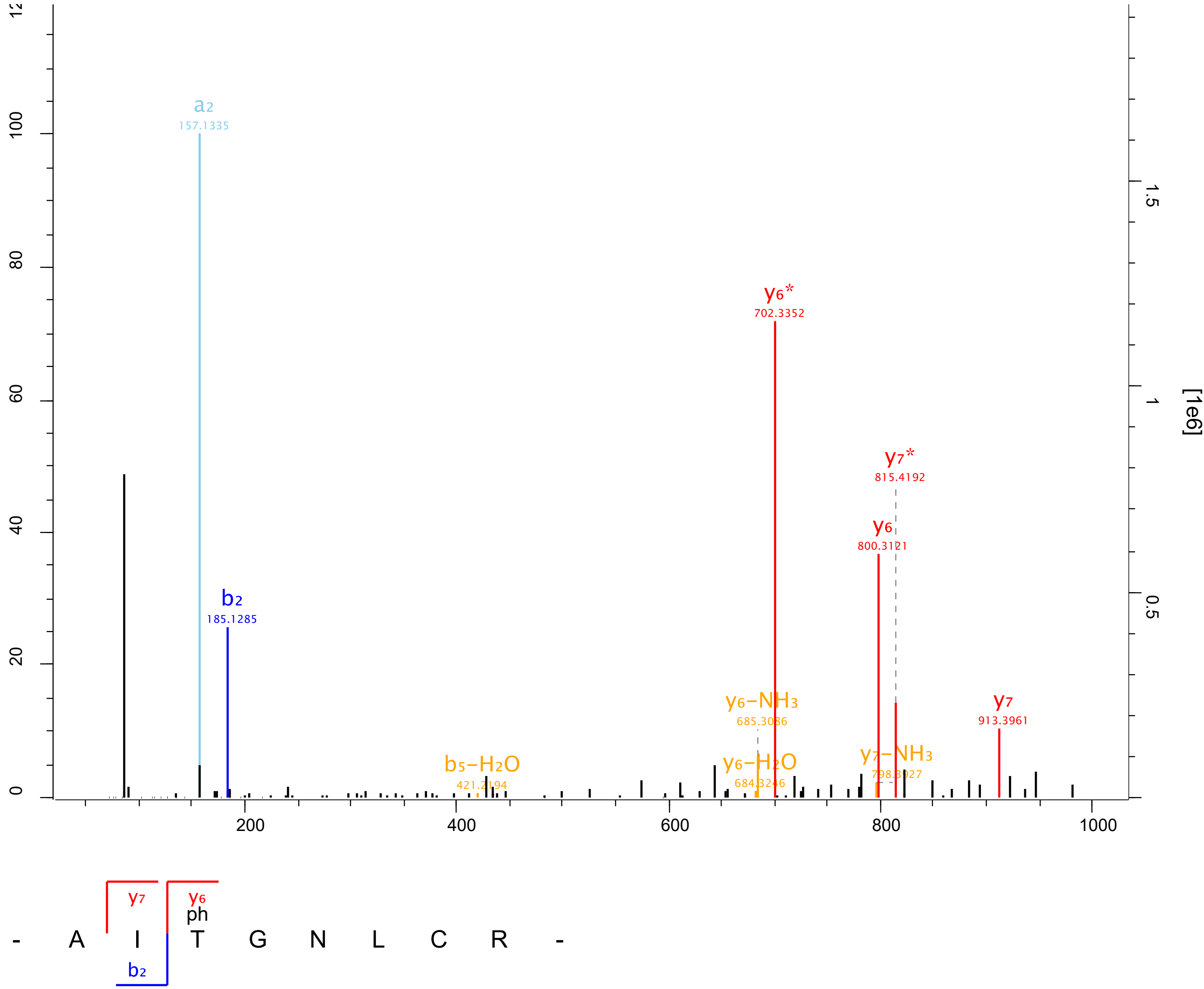

| Raw file | Scan | Method | Score | m/z |
| --- | --- | --- | --- | --- |
| water_00_1 | 19572 | FTMS; HCD | 127.01 | 694.63 |

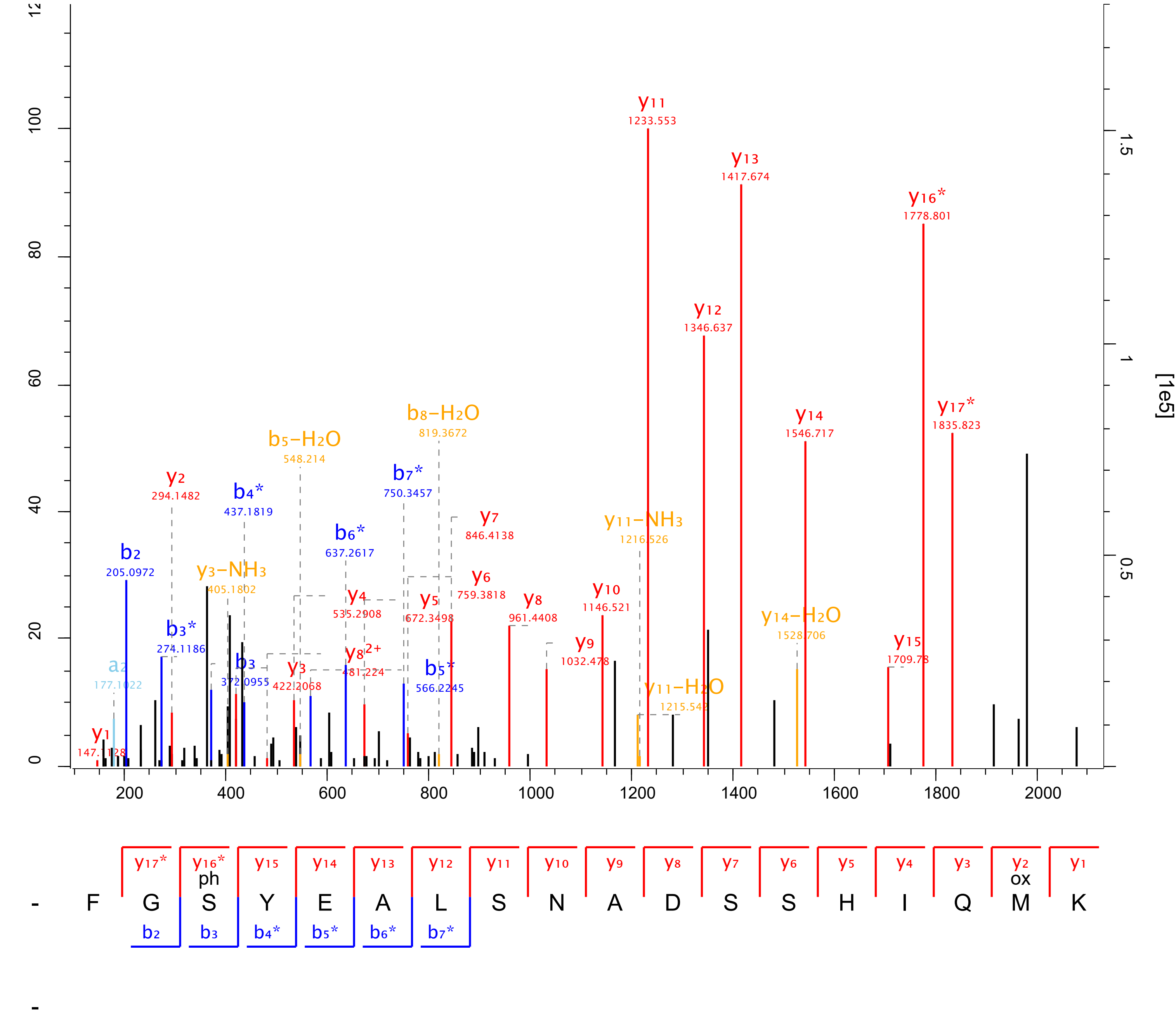

| Raw file | Scan | Method | Score | m/z |
|---|---|---|---|---|
| water_00_1 | 19999 | FTMS; HCD | 112.08 | 551.75 |

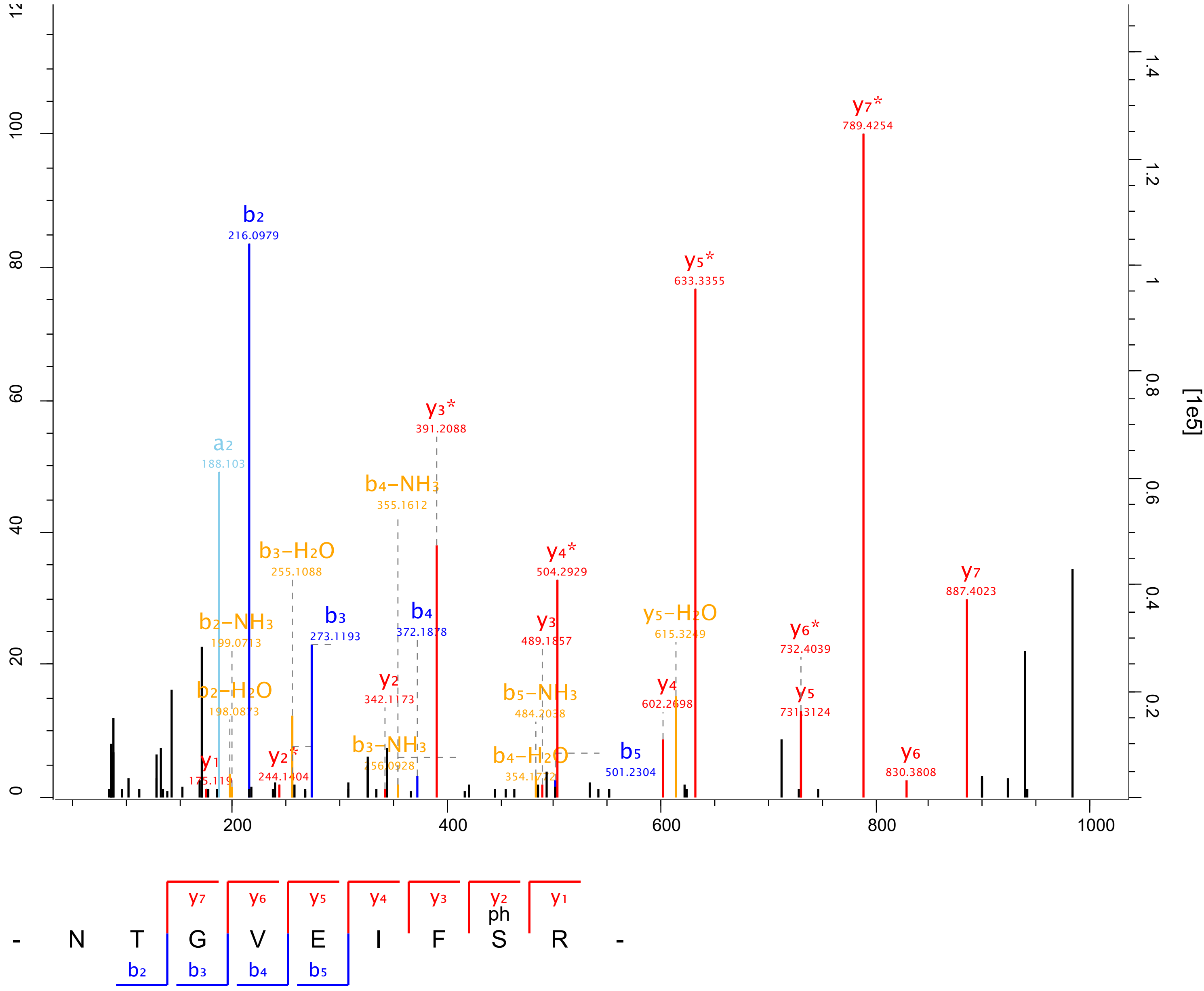

| Raw file | Scan | Method | Score | m/z |
| --- | --- | --- | --- | --- |
| water_00_1 | 20006 | FTMS; HCD | 58.17 | 577.78 |

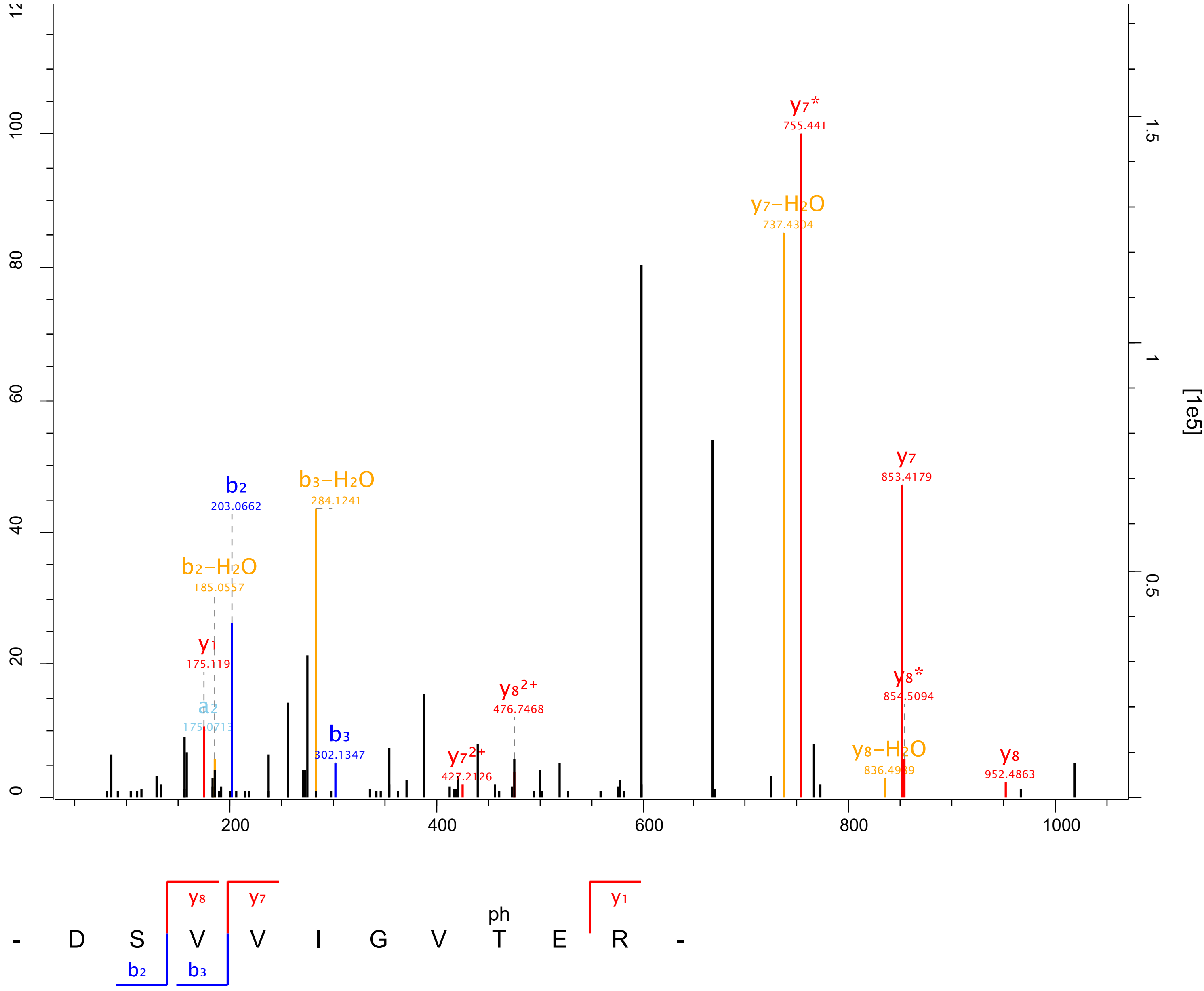

| Raw file | Scan | Method | Score | m/z |
| --- | --- | --- | --- | --- |
| water_00_1 | 20400 | FTMS; HCD | 100.02 | 851.38 |

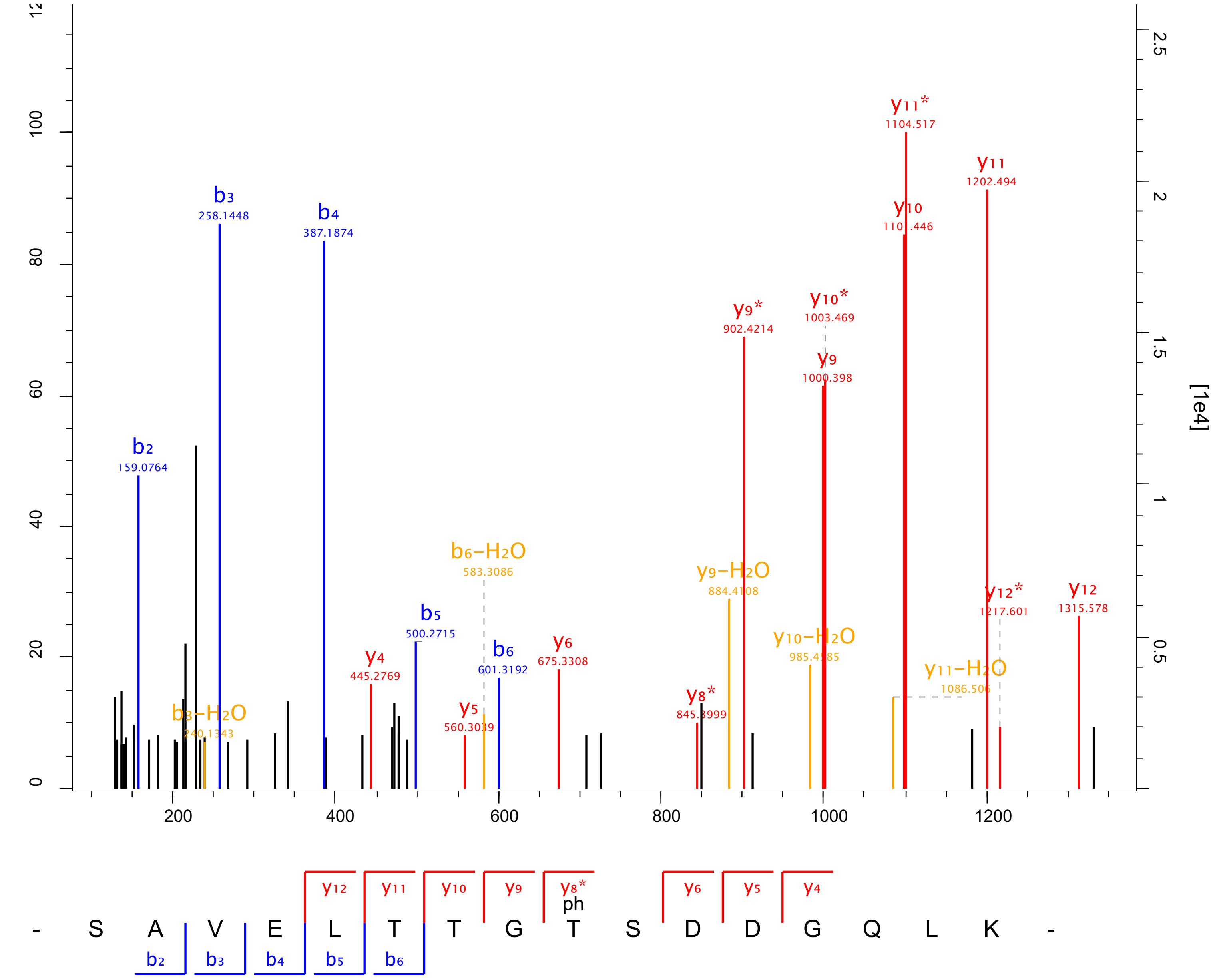

| Raw file | Scan | Method | Score | m/z |
|---|---|---|---|---|
| water_00_1 | 20500 | FTMS; HCD | 78.92 | 758.33 |

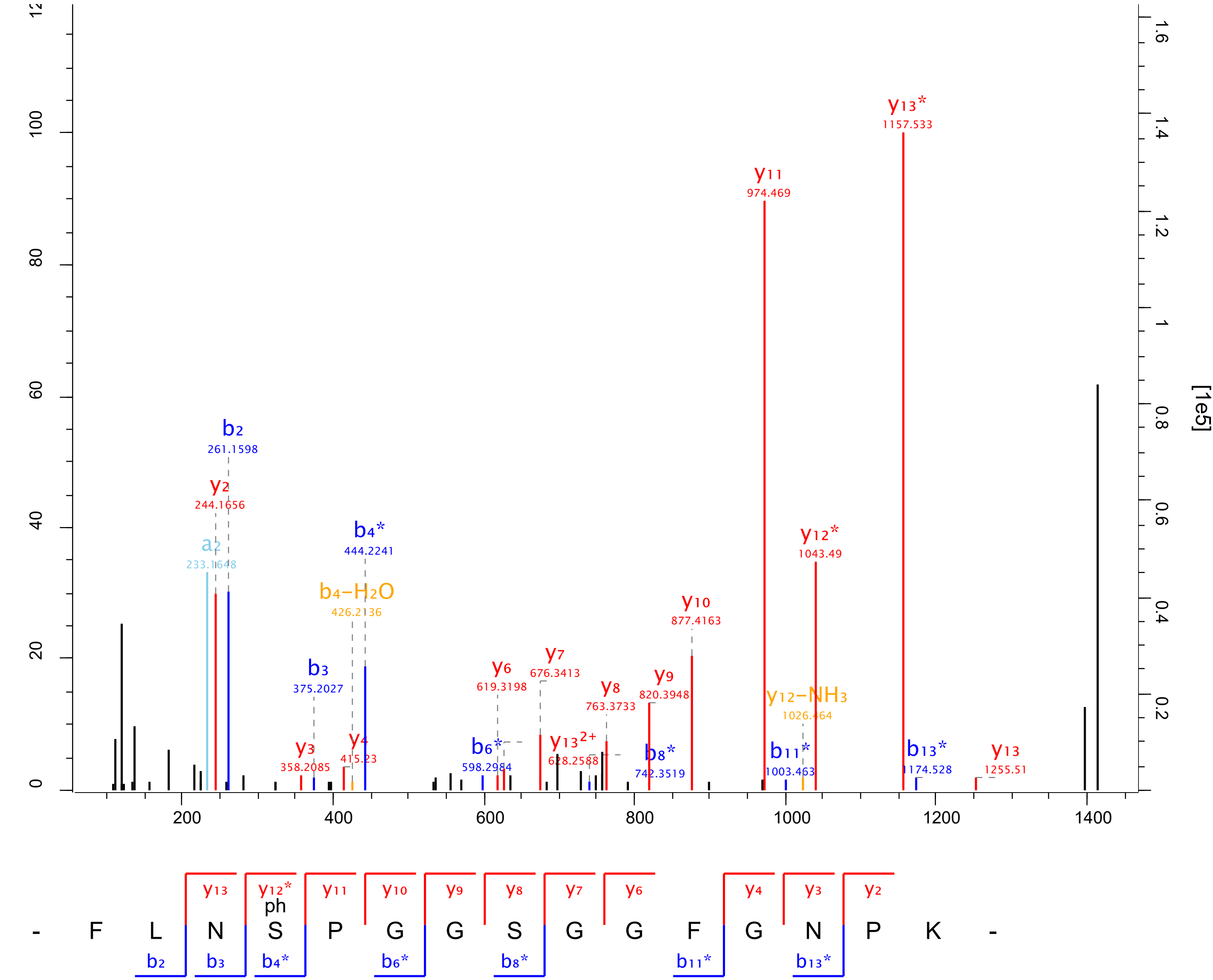

| Raw file | Scan | Method | Score | m/z |
|---|---|---|---|---|
| water_00_1 | 2055 | FTMS; HCD | 59.23 | 597.78 |

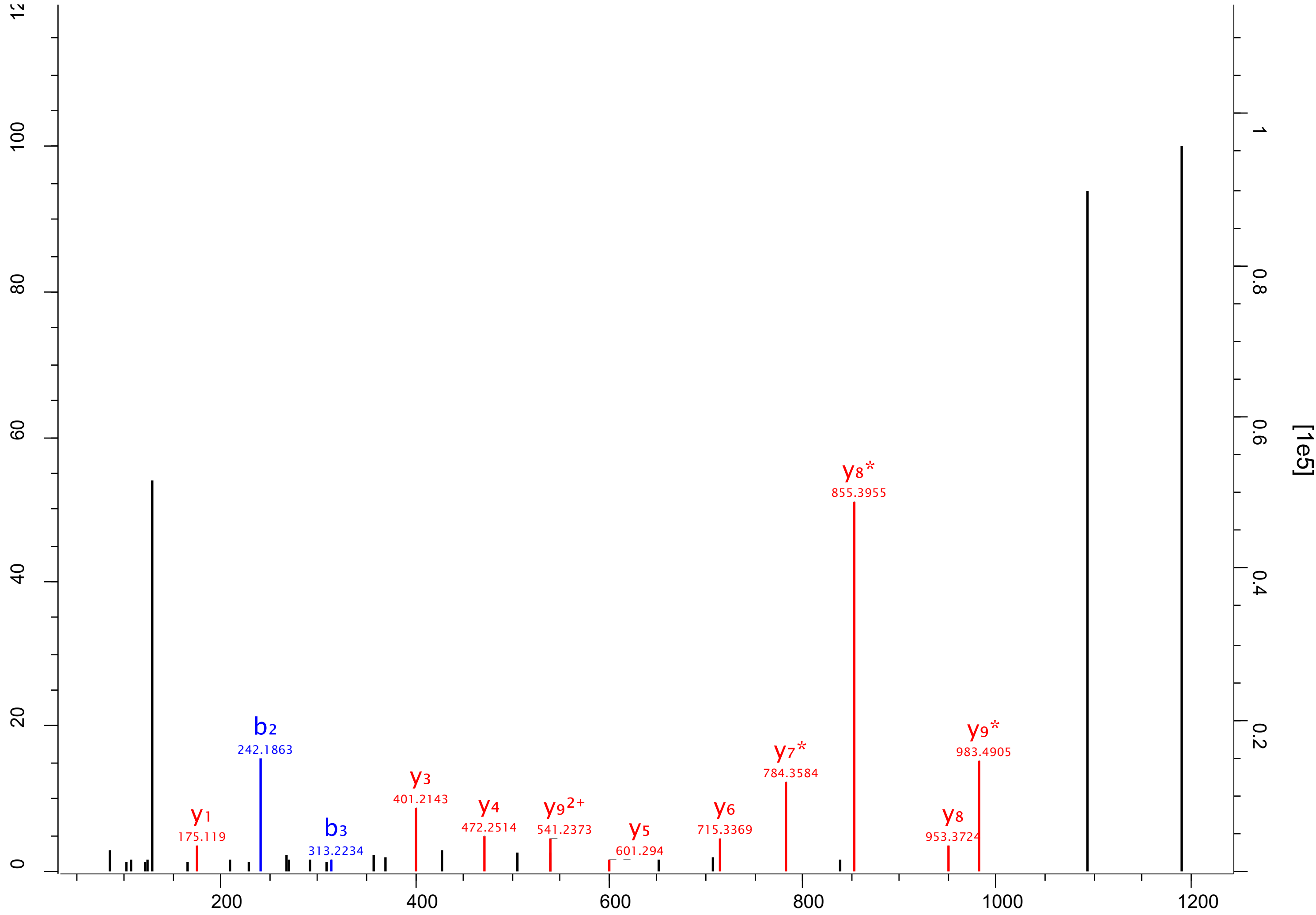


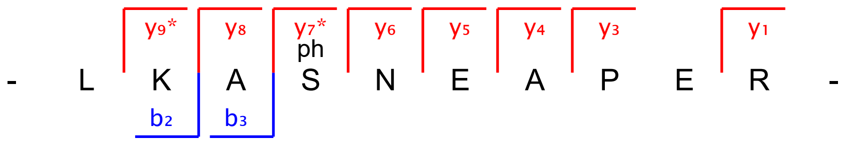

| Raw file | Scan | Method | Score | m/z |
|---|---|---|---|---|
| water_00_1 | 2061 | FTMS; HCD | 52.86 | 430.85 |

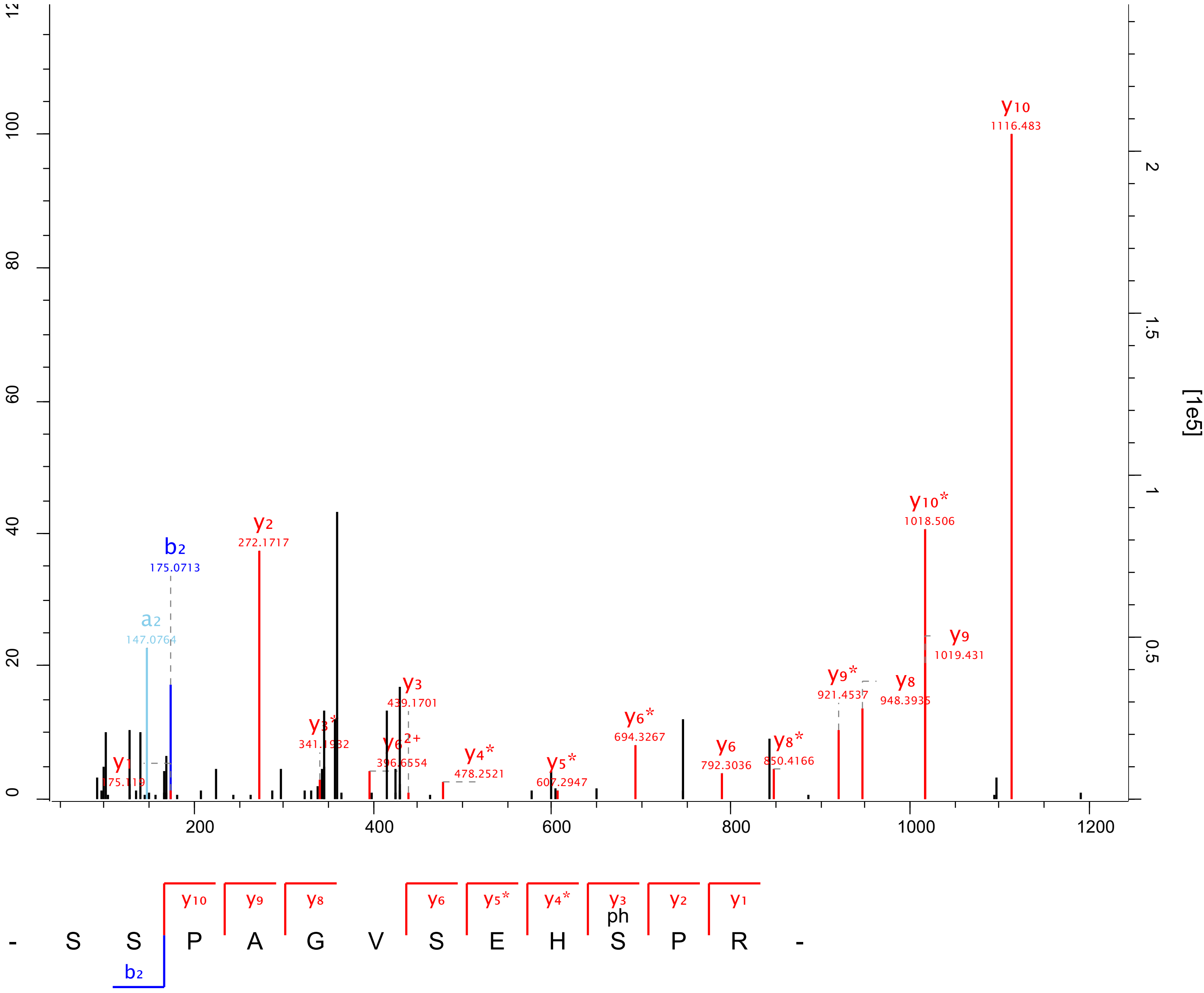

| Raw file | Scan | Method | Score | m/z |
|---|---|---|---|---|
| water_00_1 | 20639 | FTMS; HCD | 102.89 | 587.63 |

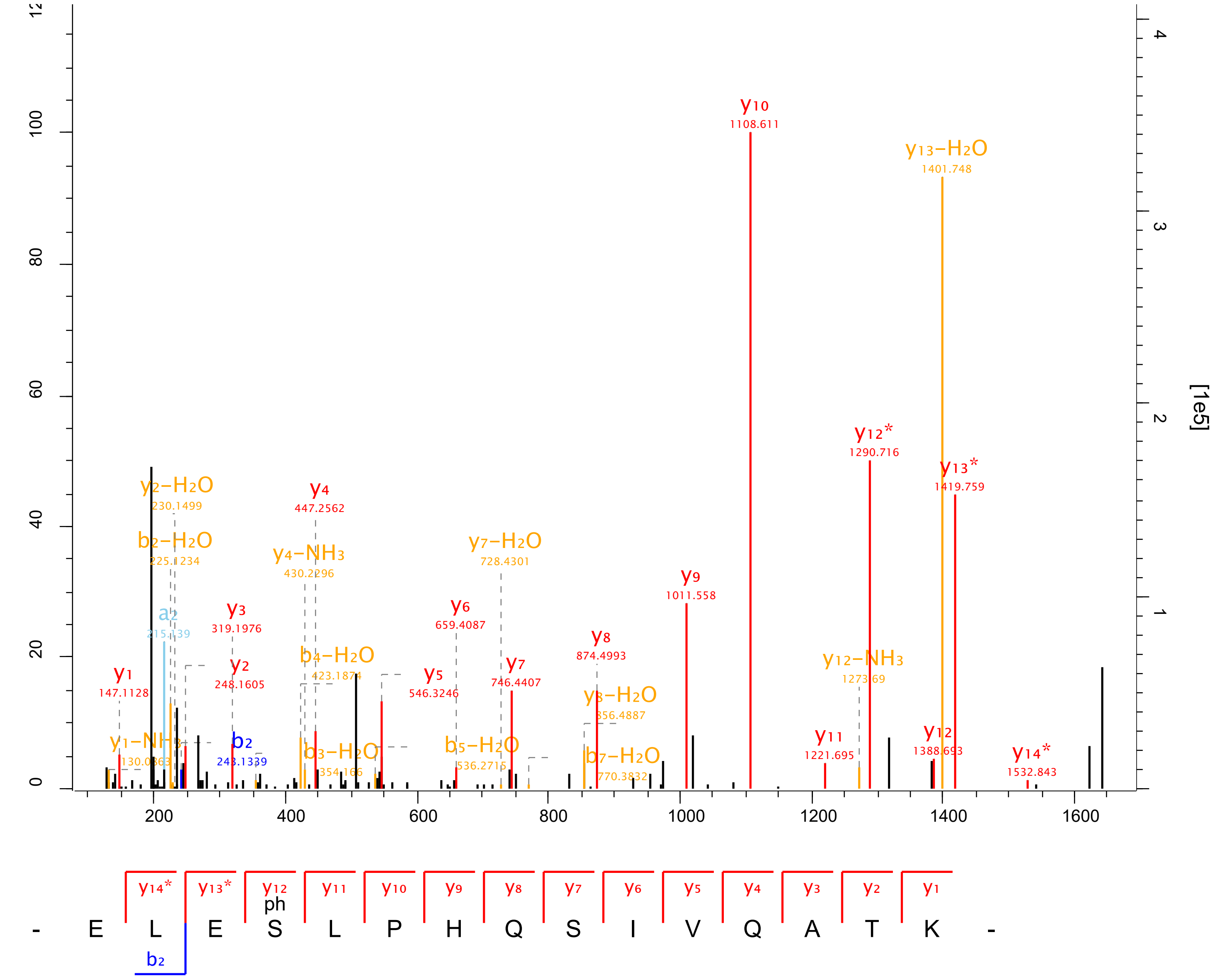

| Raw file | Scan | Method | Score | m/z |
|---|---|---|---|---|
| water_00_1 | 20965 | FTMS; HCD | 163.45 | 807.36 |

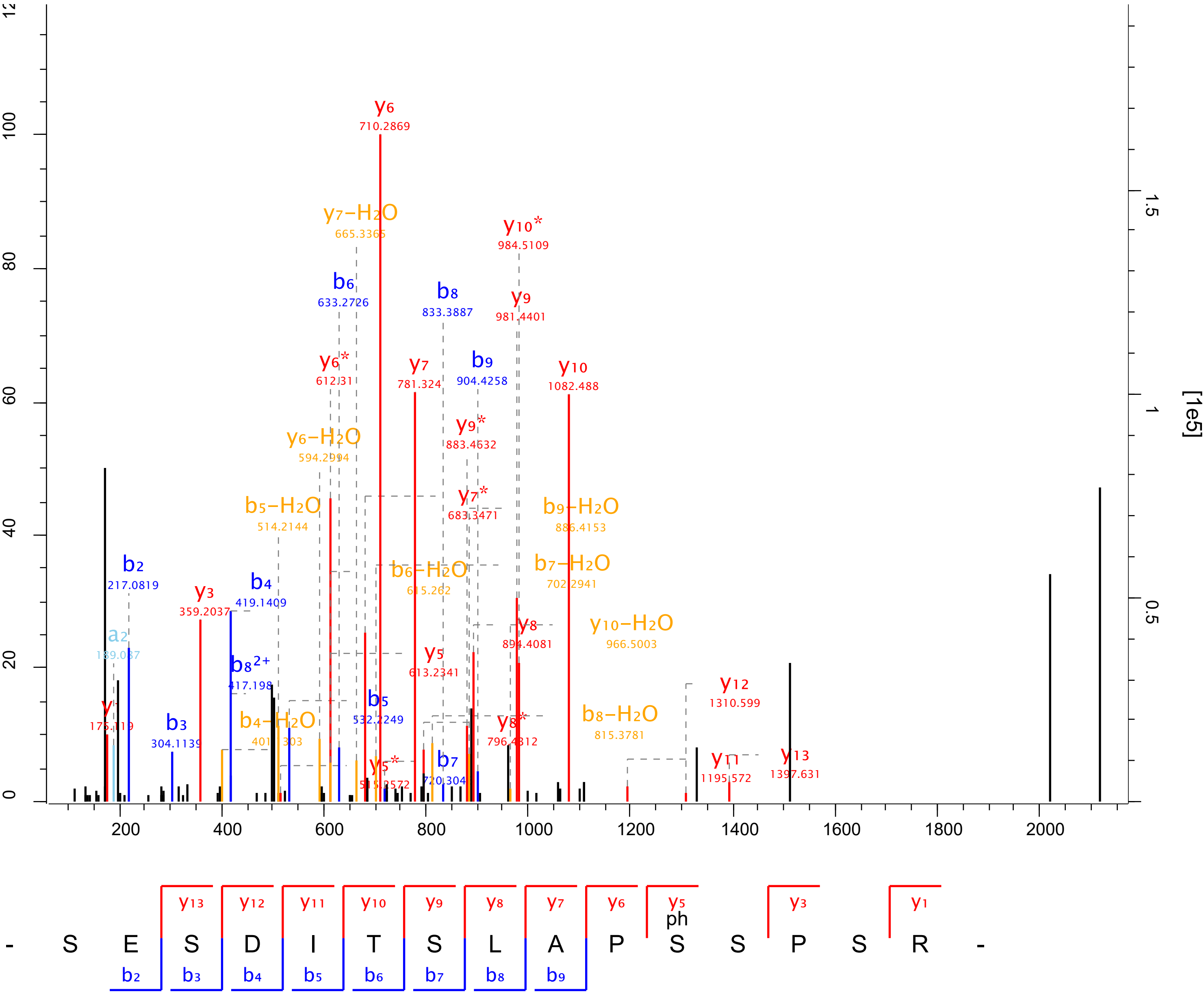

| Raw file | Scan | Method | Score | m/z |
|---|---|---|---|---|
| water_00_1 | 21162 | FTMS; HCD | 172.33 | 628.77 |

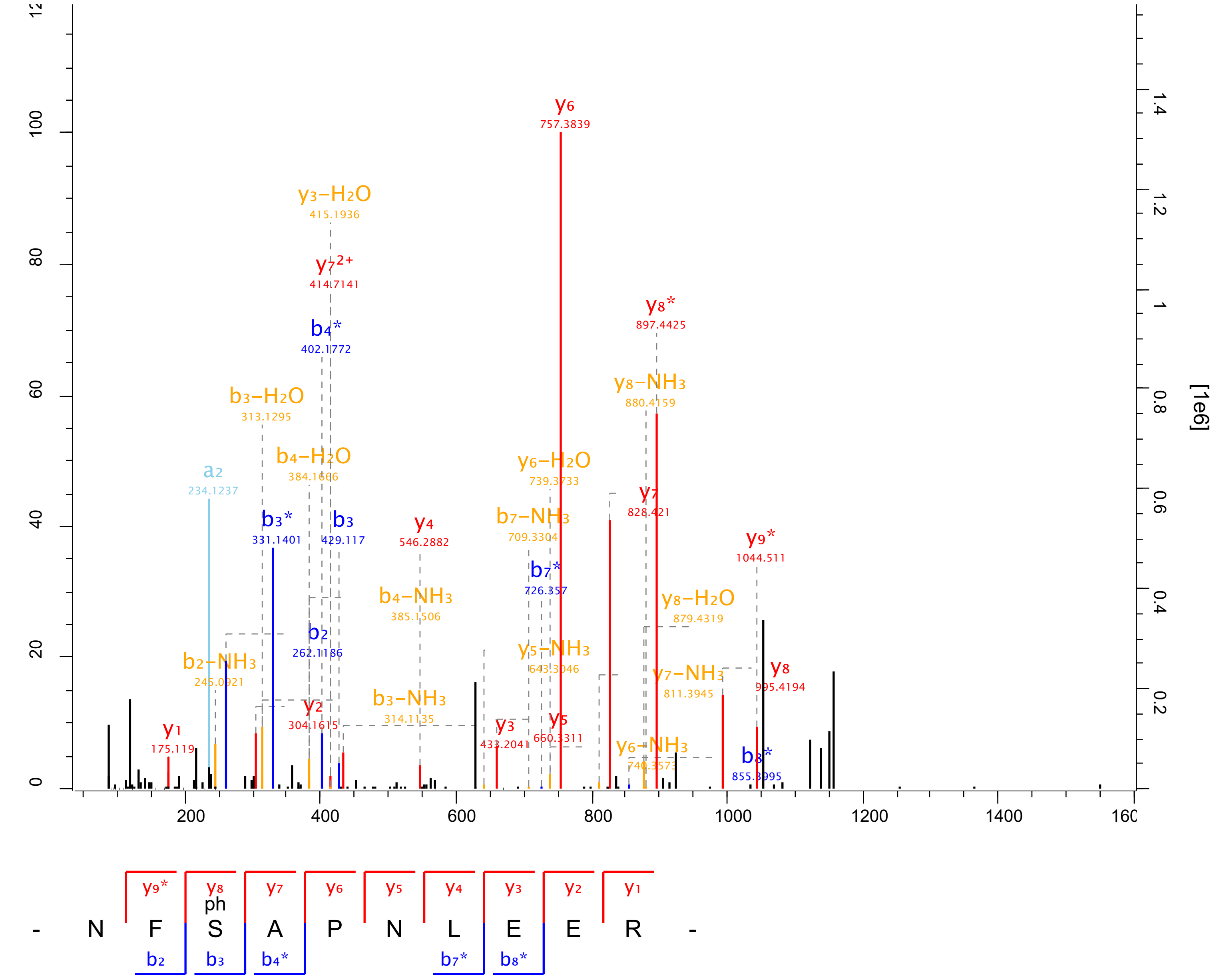

| Raw file | Scan | Method | Score | m/z |
|---|---|---|---|---|
| water_00_1 | 21200 | FTMS; HCD | 191.57 | 708.32 |

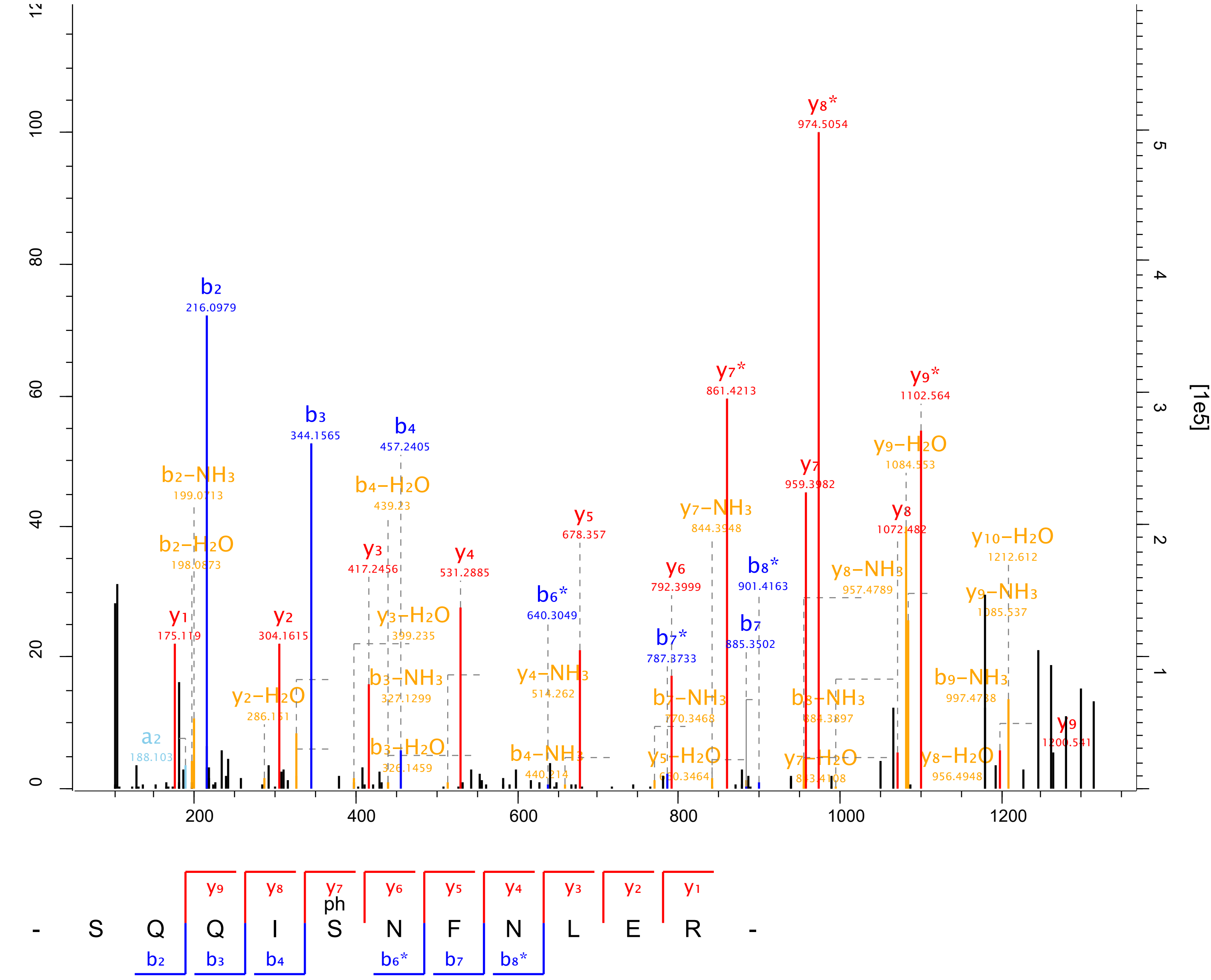

| Raw file | Scan | Method | Score | m/z |
|---|---|---|---|---|
| water_00_1 | 21536 | FTMS; HCD | 96.21 | 812.34 |

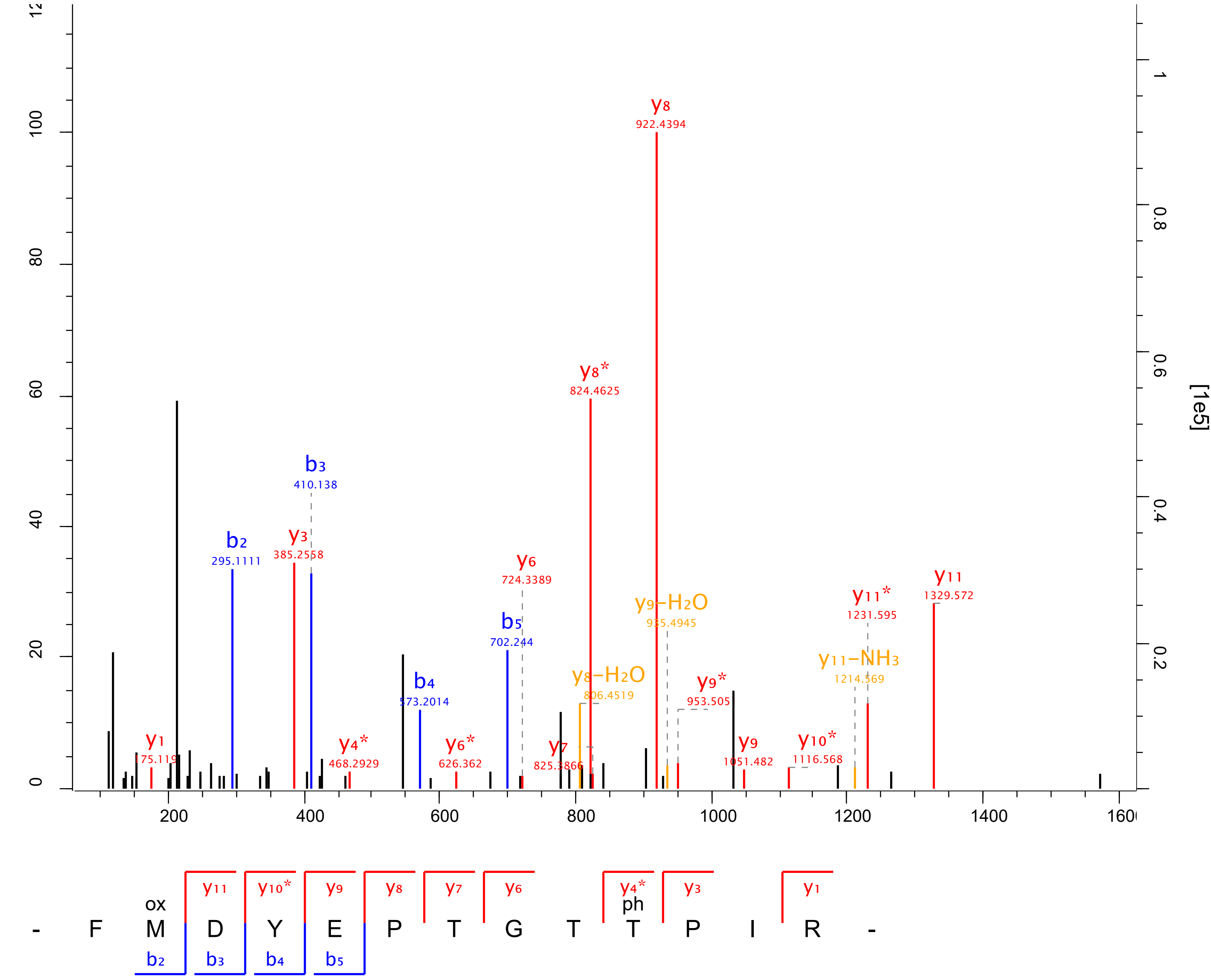

| Raw file | Scan | Method | Score | m/z |
|---|---|---|---|---|
| water_00_1 | 21574 | FTMS; HCD | 45.21 | 462.24 |

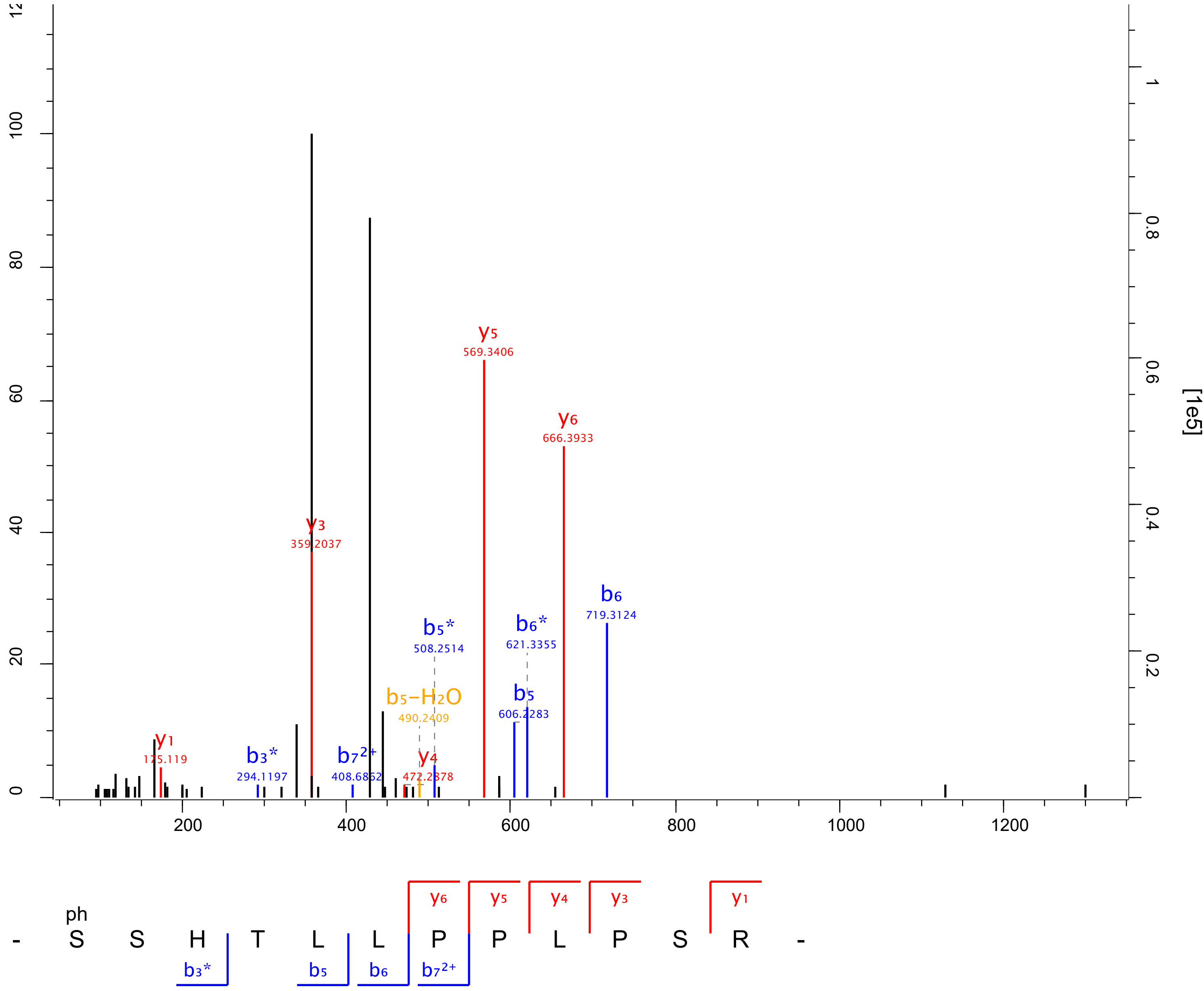

| Raw file | Scan | Method | Score | m/z |
| --- | --- | --- | --- | --- |
| water_00_1 | 21635 | FTMS; HCD | 93.4 | 709.3 |

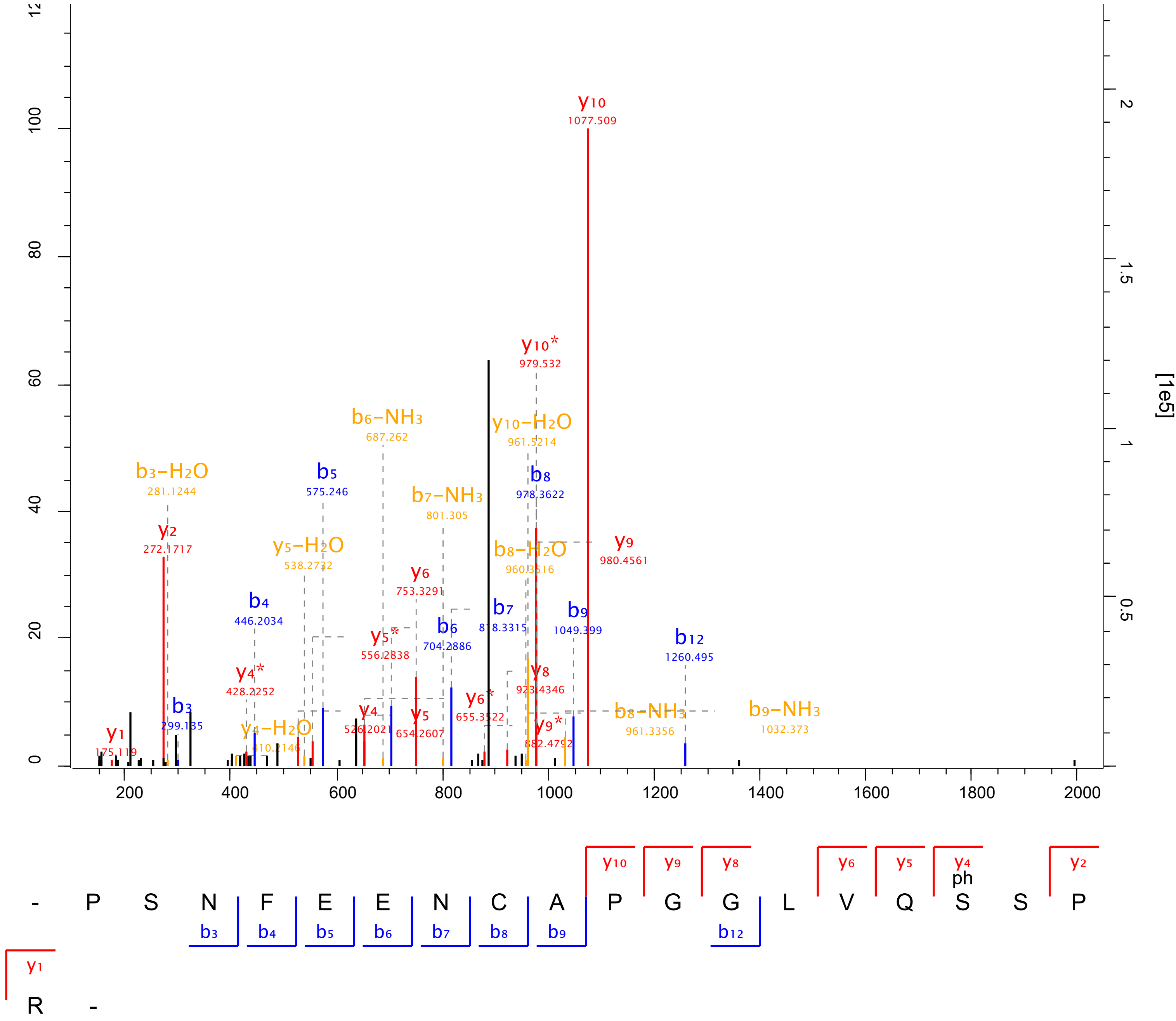

| Raw file | Scan | Method | Score | m/z |
| --- | --- | --- | --- | --- |
| water_00_1 | 21799 | FTMS; HCD | 68.48 | 685.82 |

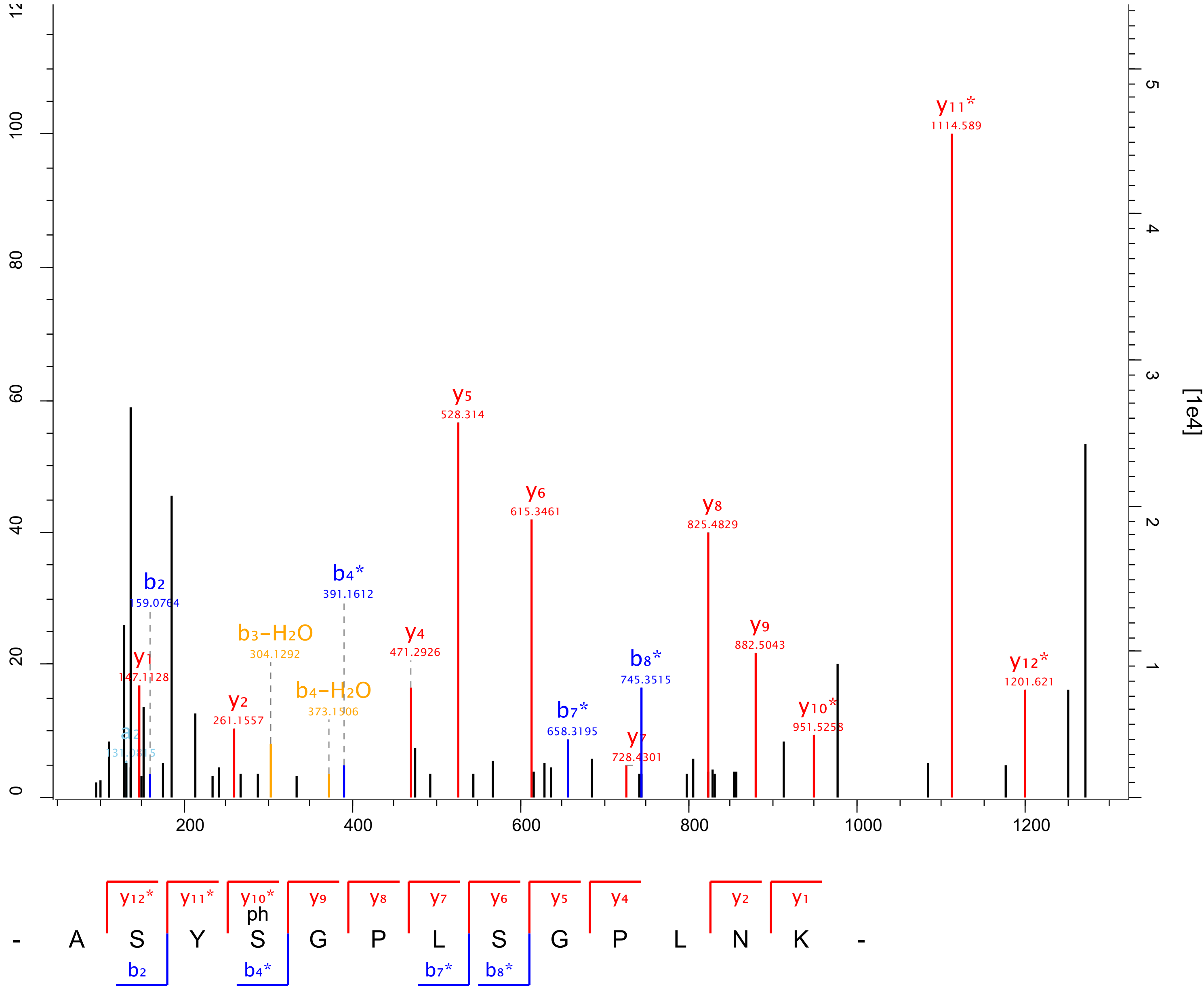

| Raw file | Scan | Method | Score | m/z |
|---|---|---|---|---|
| water_00_1 | 22573 | FTMS; HCD | 103.76 | 672.8 |

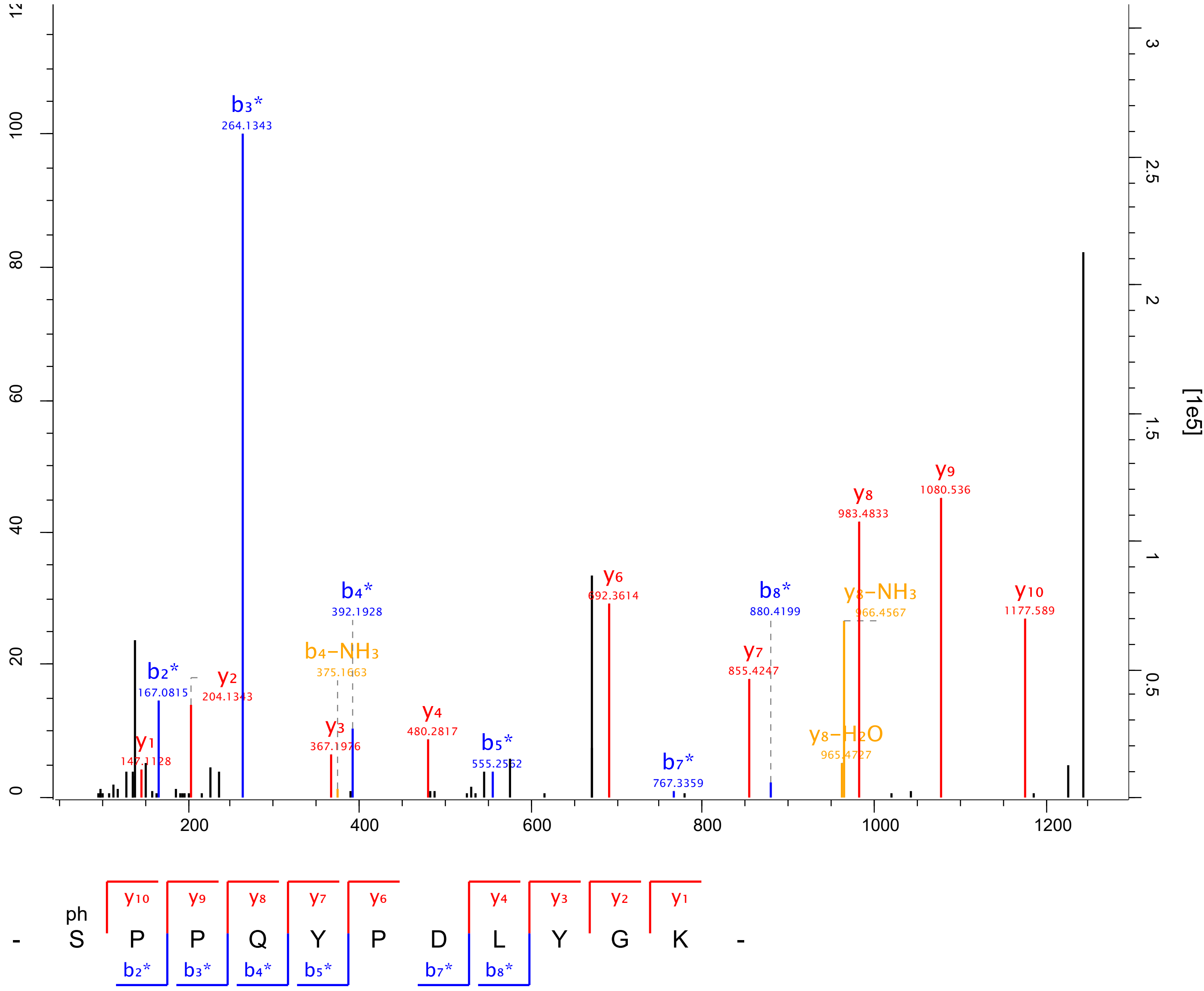

| Raw file | Scan | Method | Score | m/z |
| --- | --- | --- | --- | --- |
| water_00_1 | 22810 | FTMS; HCD | 182.57 | 691.79 |

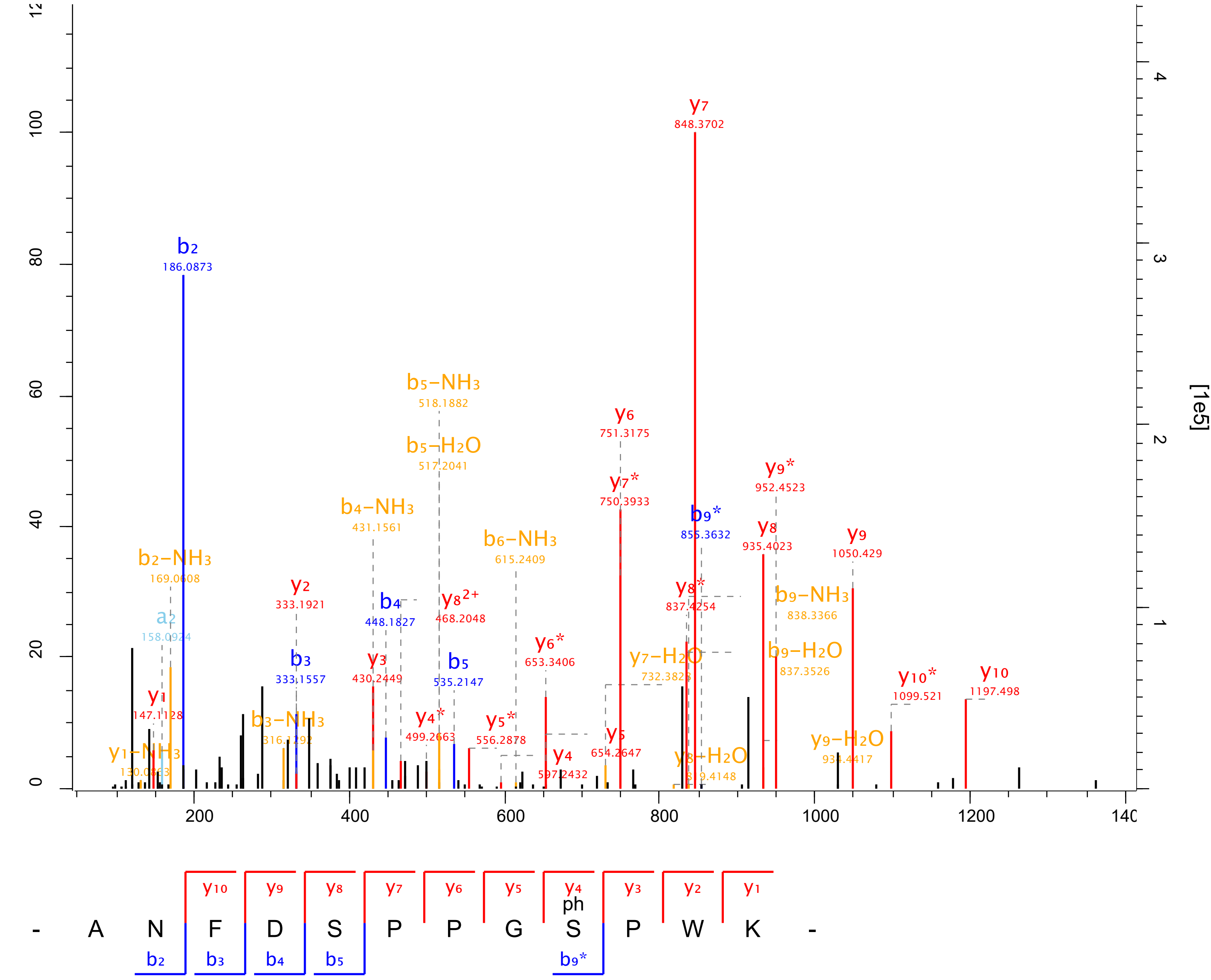

| Raw file | Scan | Method | Score | m/z |
|---|---|---|---|---|
| water_00_1 | 22868 | FTMS; HCD | 82.88 | 890.41 |

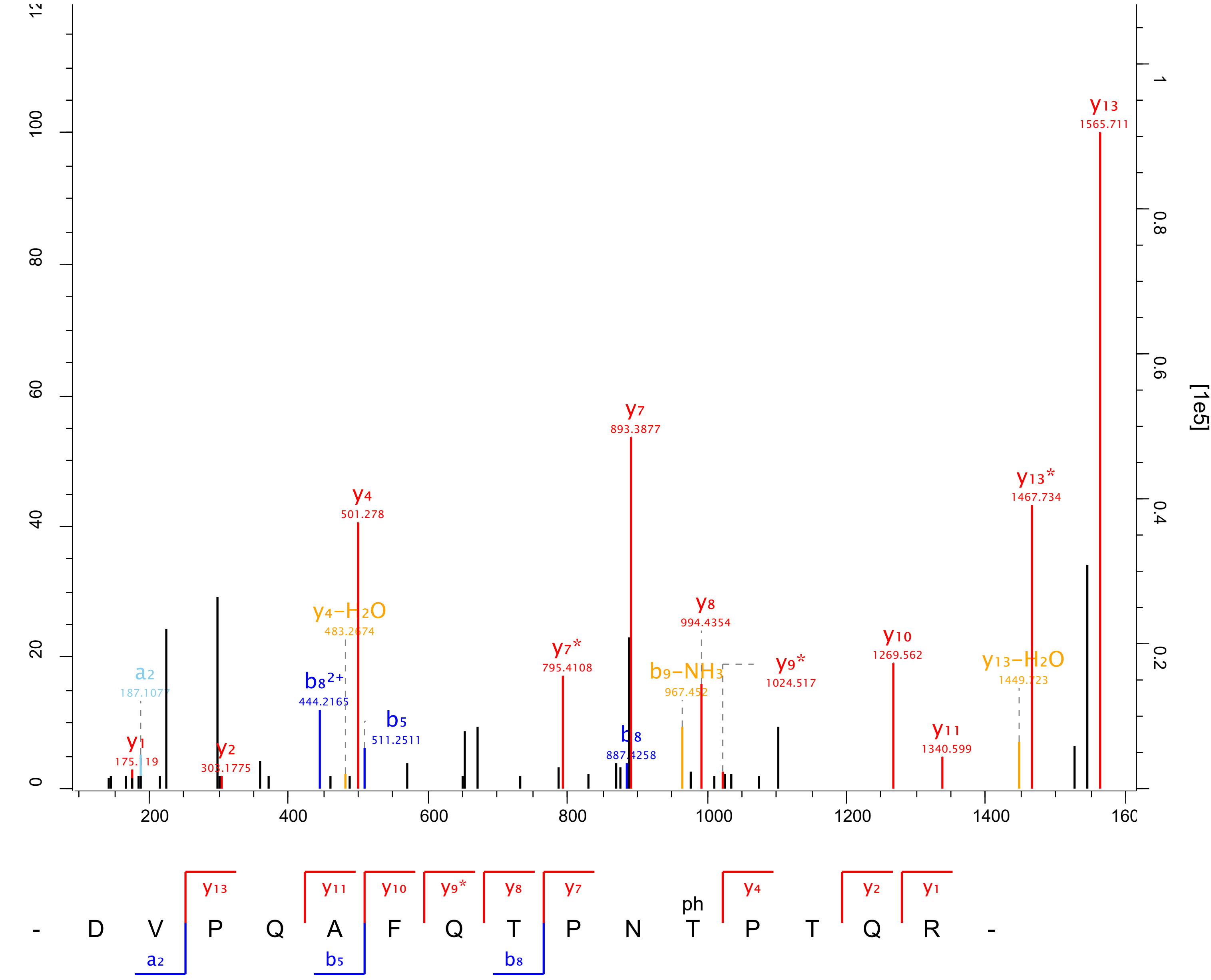

| Raw file | Scan | Method | Score | m/z |
| --- | --- | --- | --- | --- |
| water_00_1 | 2314 | FTMS; HCD | 51.28 | 448.2 |

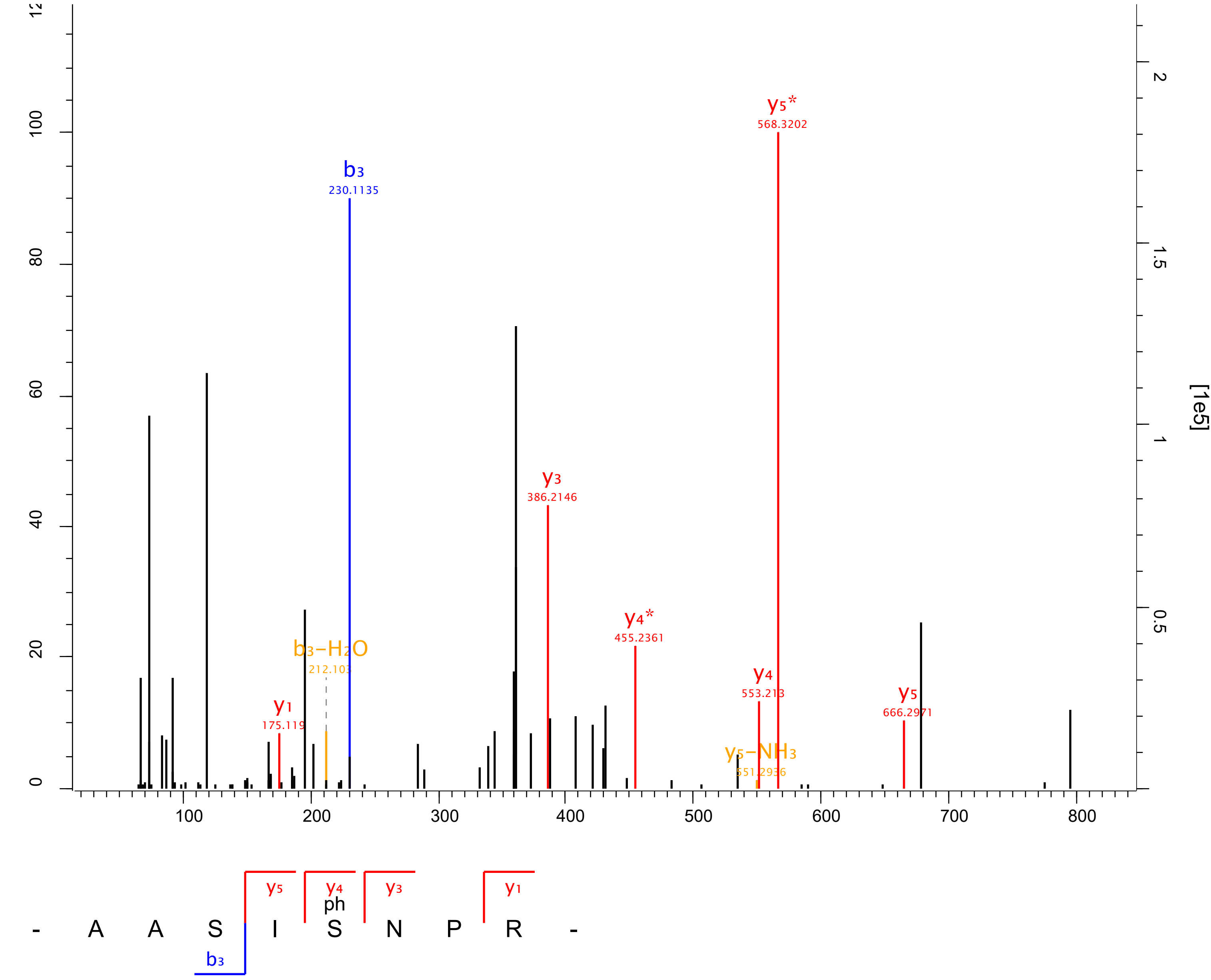

| Raw file | Scan | Method | Score | m/z |
| --- | --- | --- | --- | --- |
| water_00_1 | 23393 | FTMS; HCD | 81.32 | 832.02 |

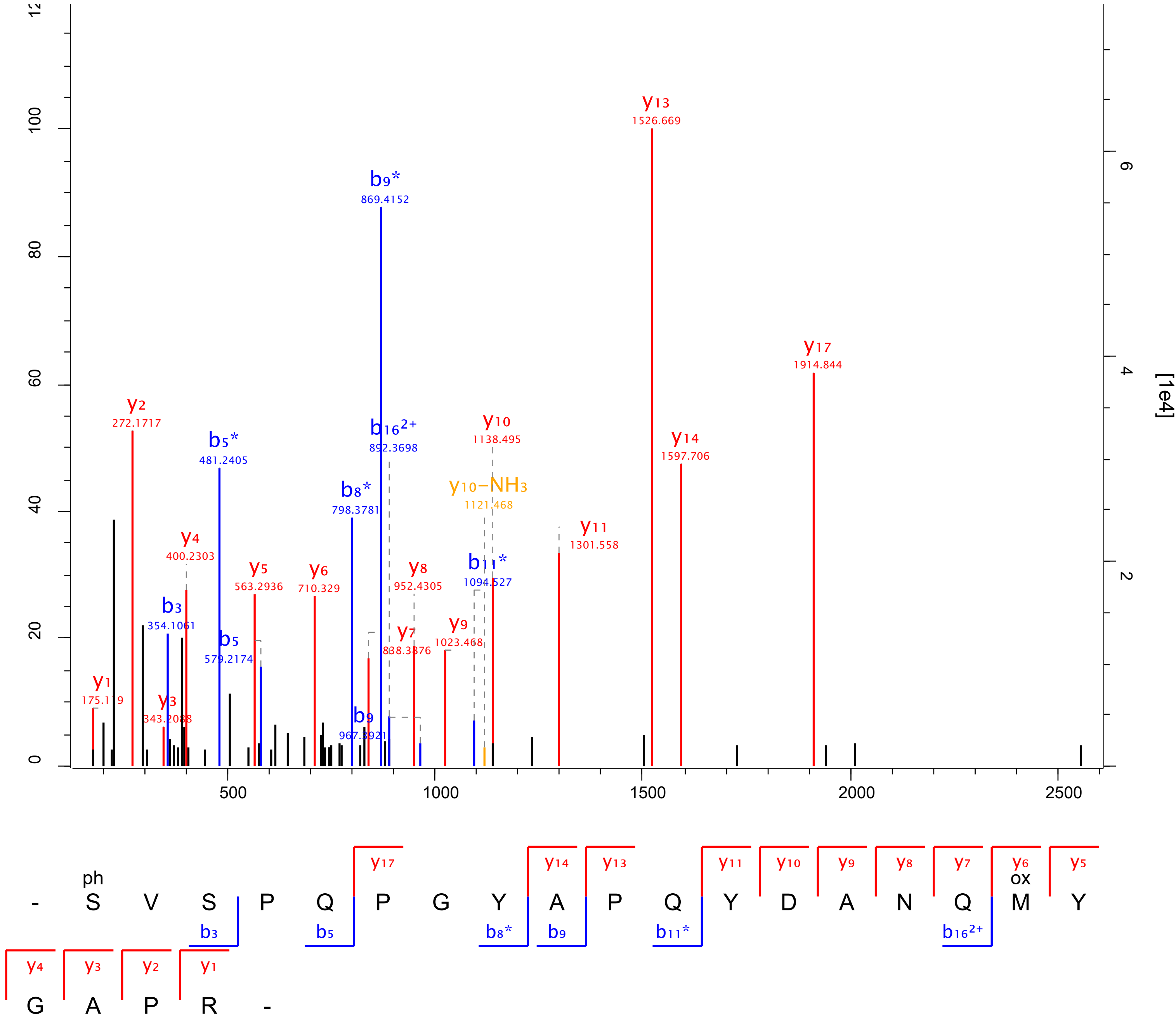

| Raw file | Scan | Method | Score | m/z |
|---|---|---|---|---|
| water_00_1 | 23504 | FTMS; HCD | 41.67 | 727 |

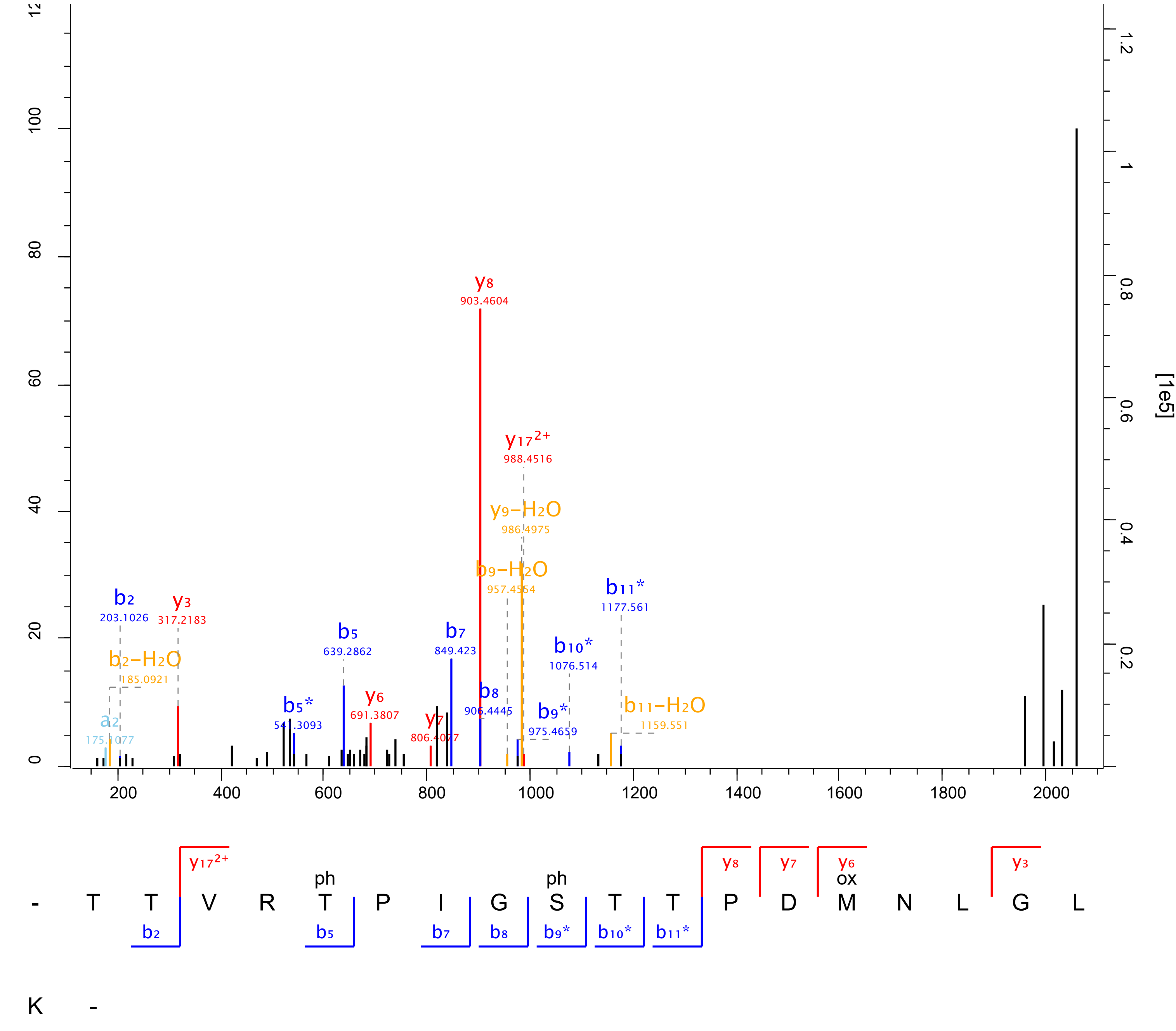

| Raw file | Scan | Method | Score | m/z |
|---|---|---|---|---|
| water_00_1 | 24034 | FTMS; HCD | 42.91 | 689.3 |

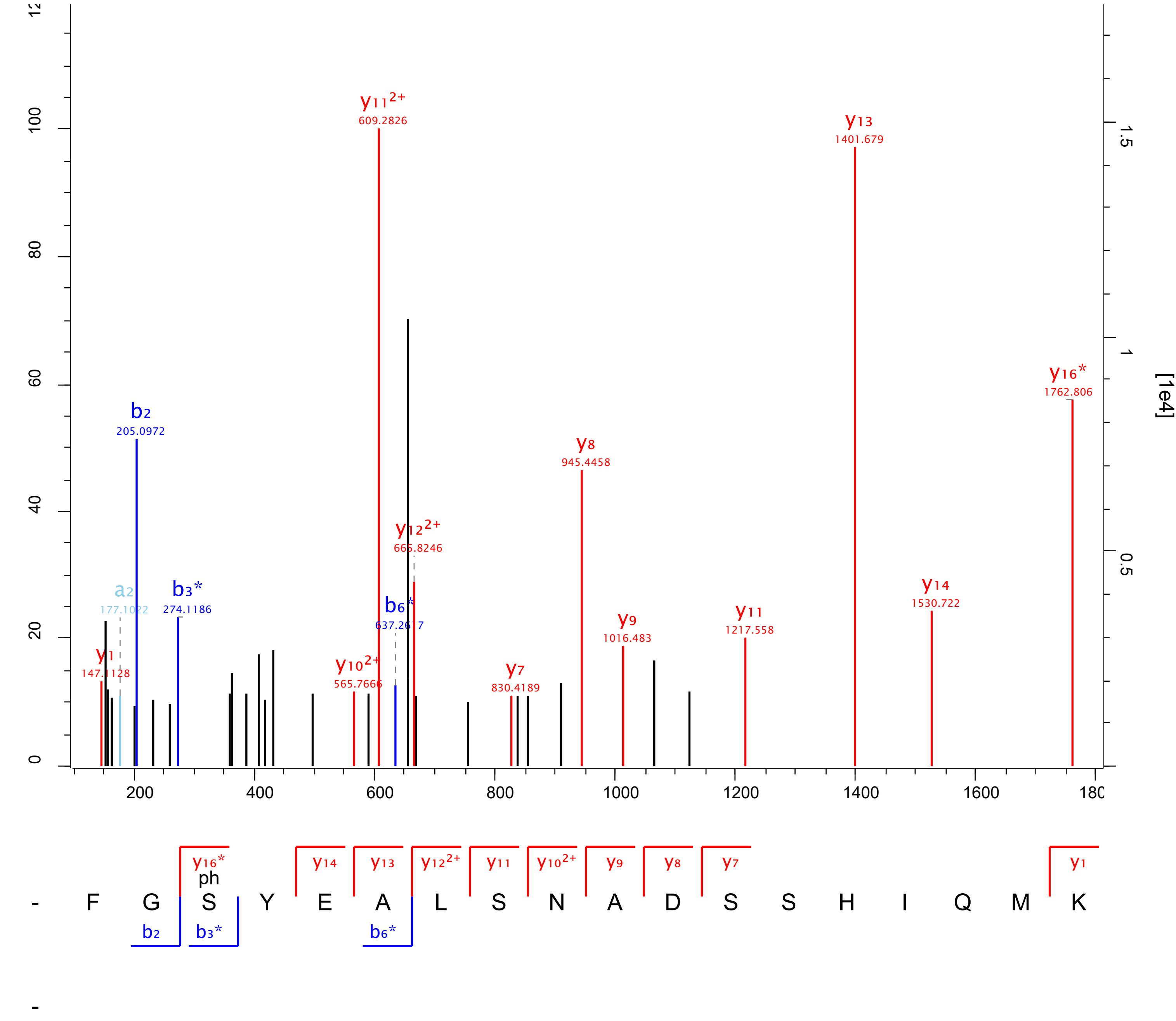

| Raw file | Scan | Method | Score | m/z |
|---|---|---|---|---|
| water_00_1 | 2430 | FTMS; HCD | 72.43 | 670.78 |

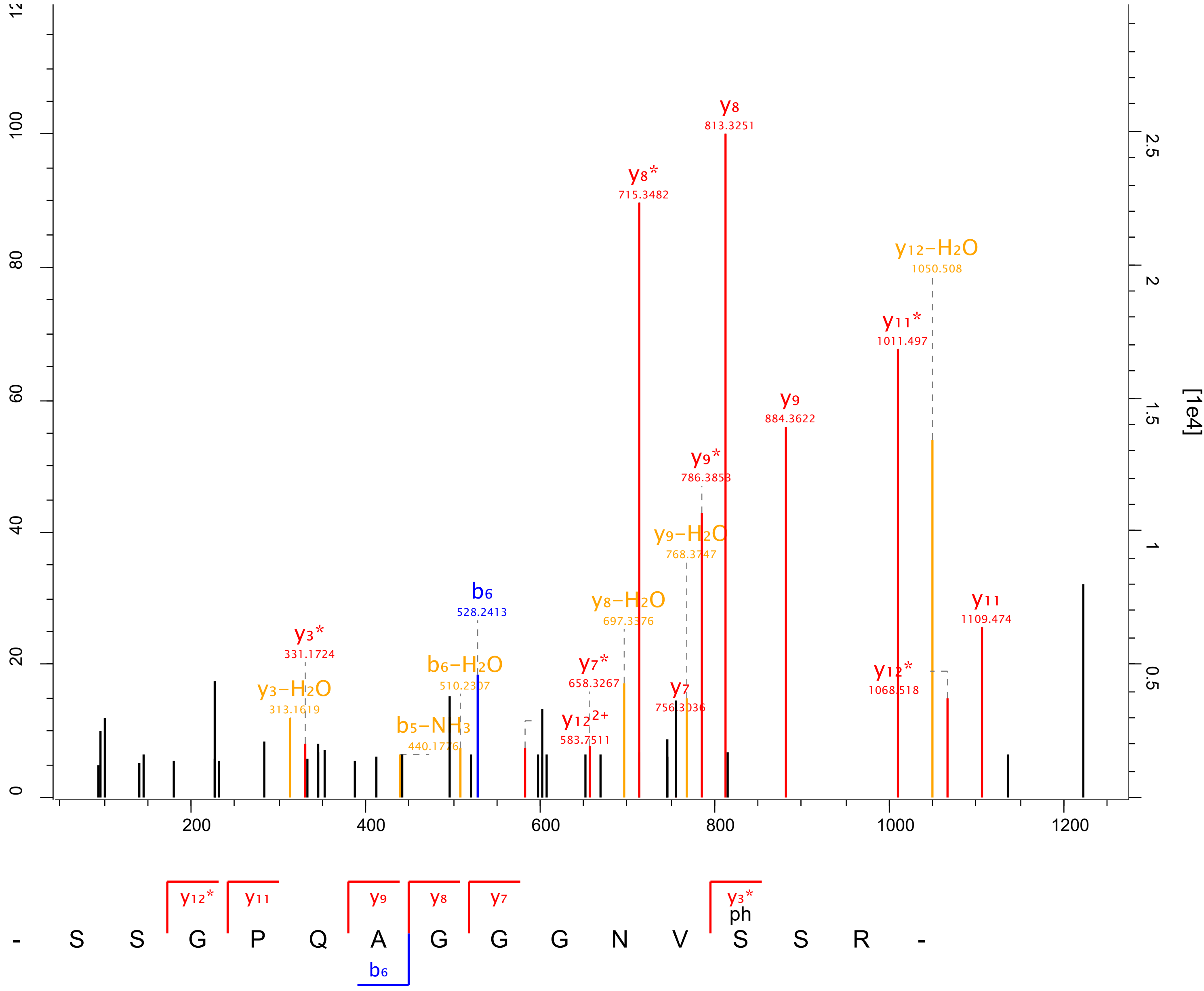

| Raw file | Scan | Method | Score | m/z |
|---|---|---|---|---|
| water_00_1 | 24754 | FTMS; HCD | 89.23 | 675.29 |

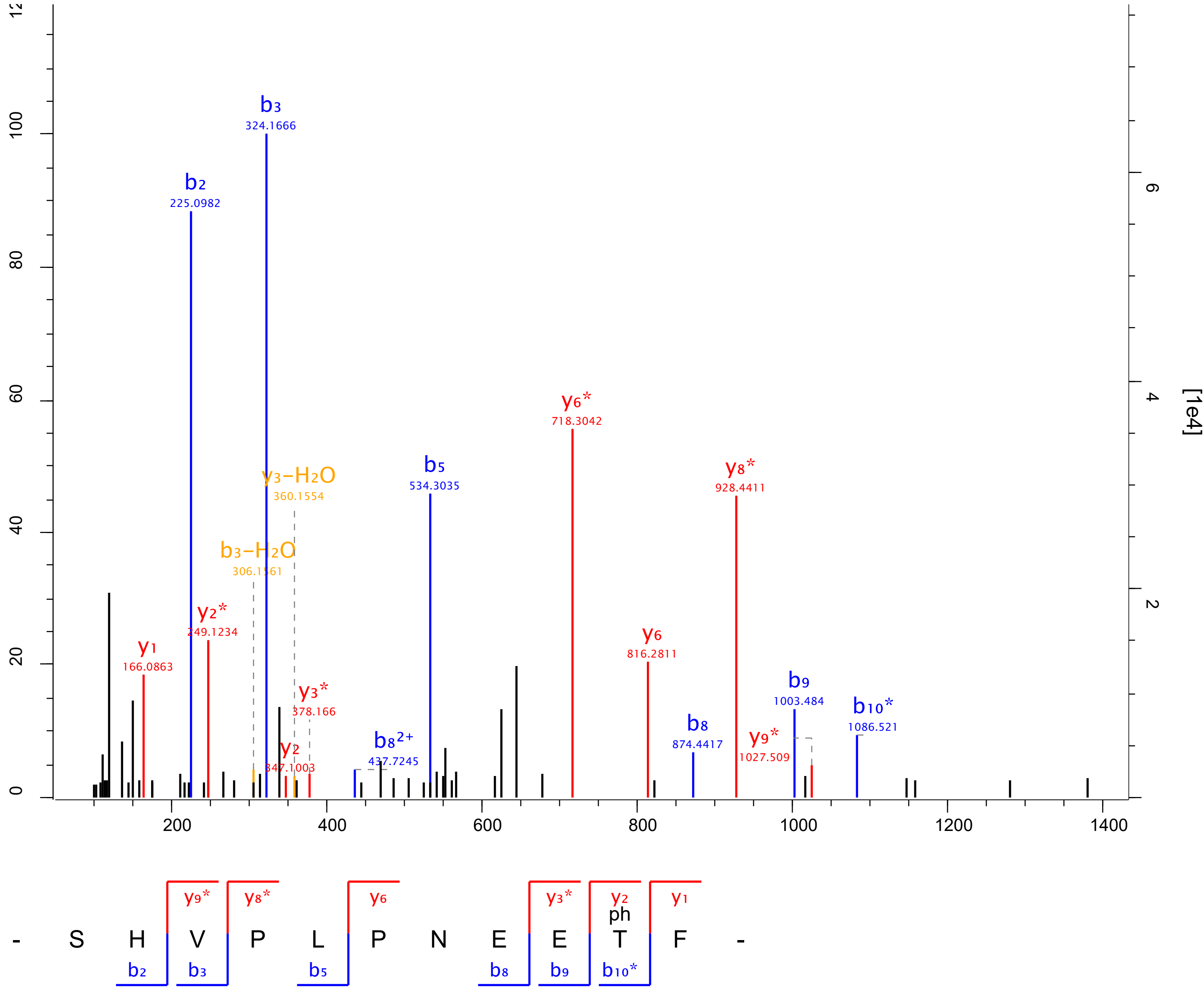

| Raw file | Scan | Method | Score | m/z |
| --- | --- | --- | --- | --- |
| water_00_1 | 24826 | FTMS; HCD | 231.37 | 718.86 |

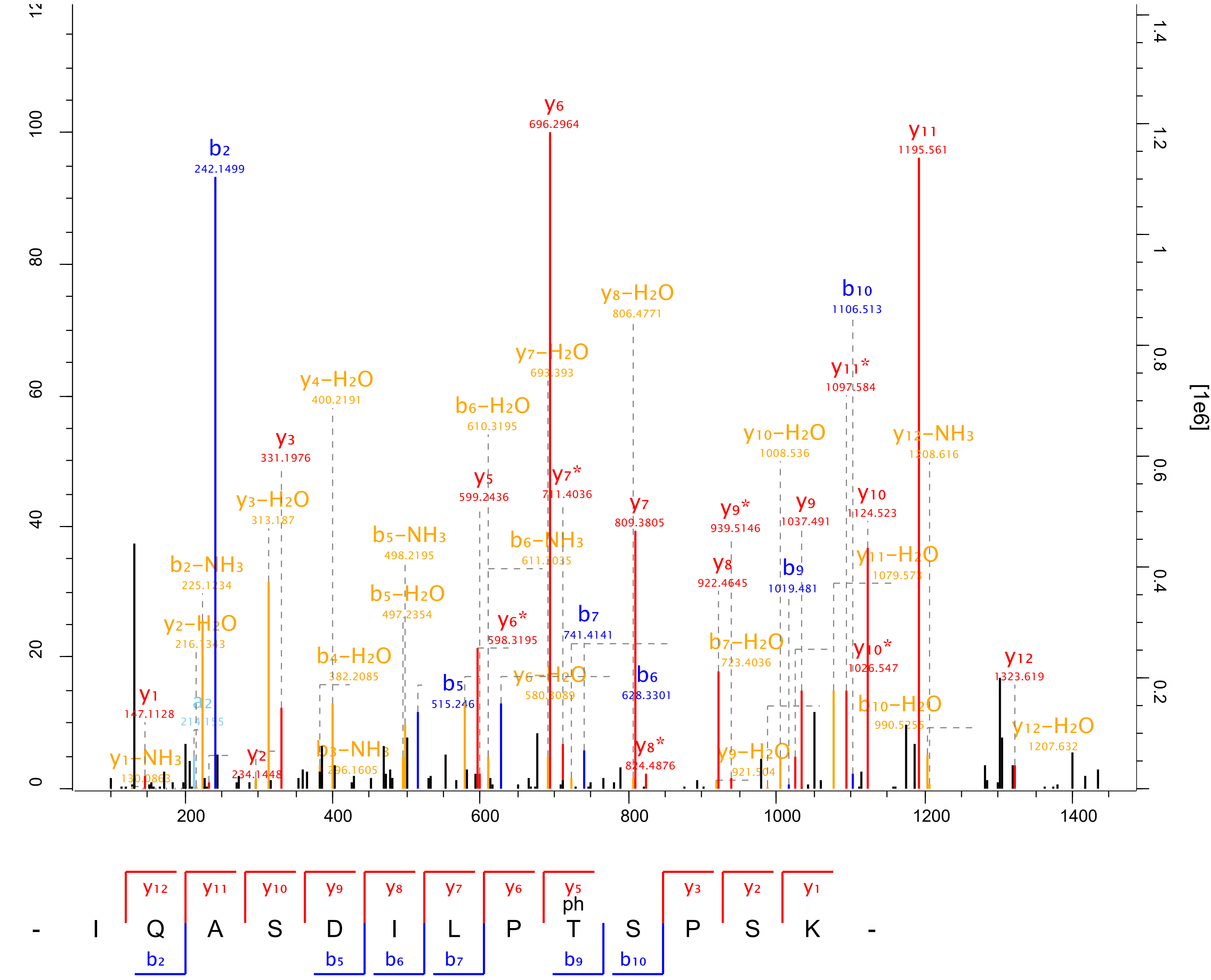

| Raw file | Scan | Method | Score | m/z |
|---|---|---|---|---|
| water_00_1 | 24887 | FTMS; HCD | 75.02 | 801.04 |

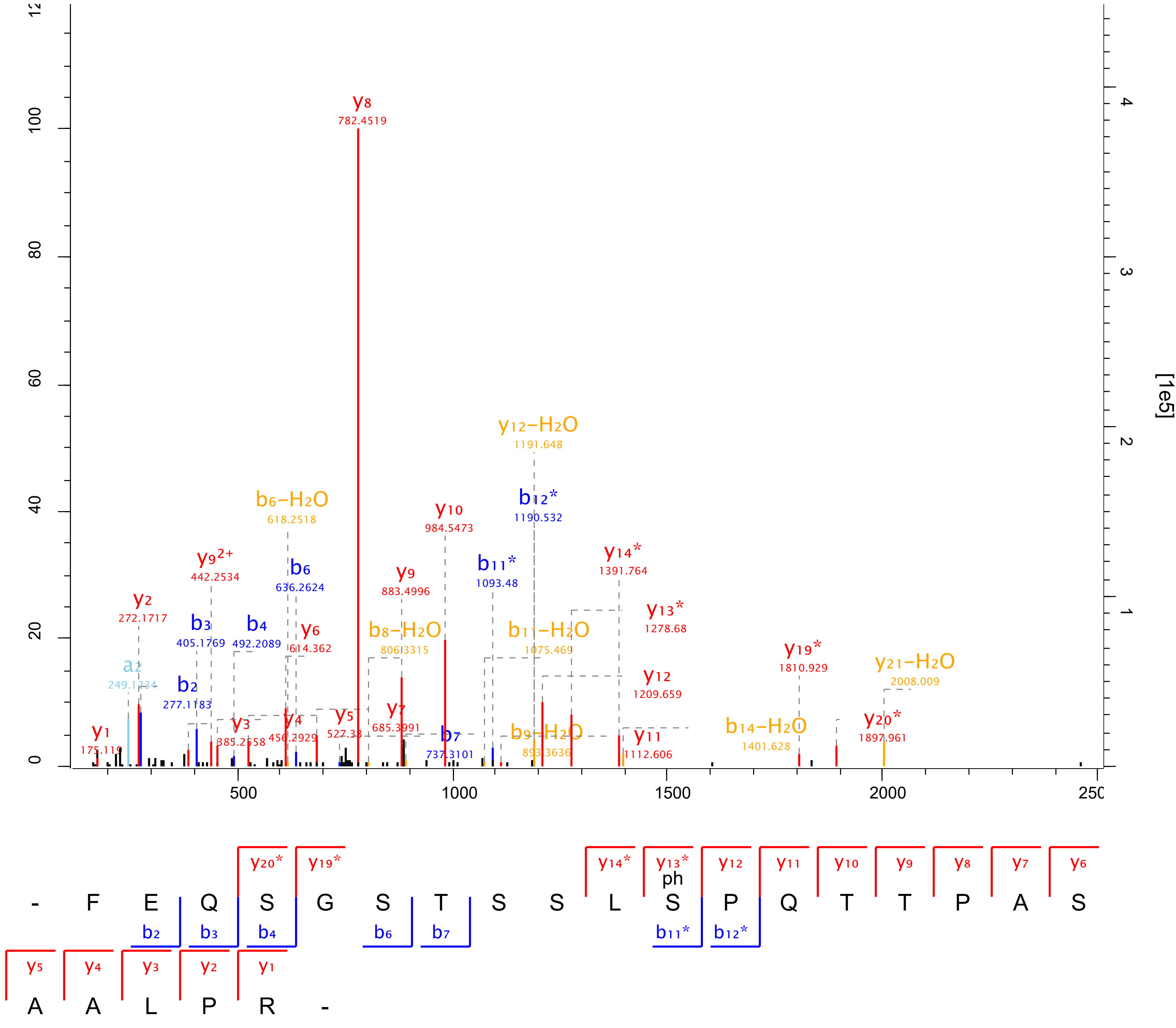

| Raw file | Scan | Method | Score | m/z |
|---|---|---|---|---|
| water_00_1 | 2499 | FTMS; HCD | 79.2 | 447.7 |

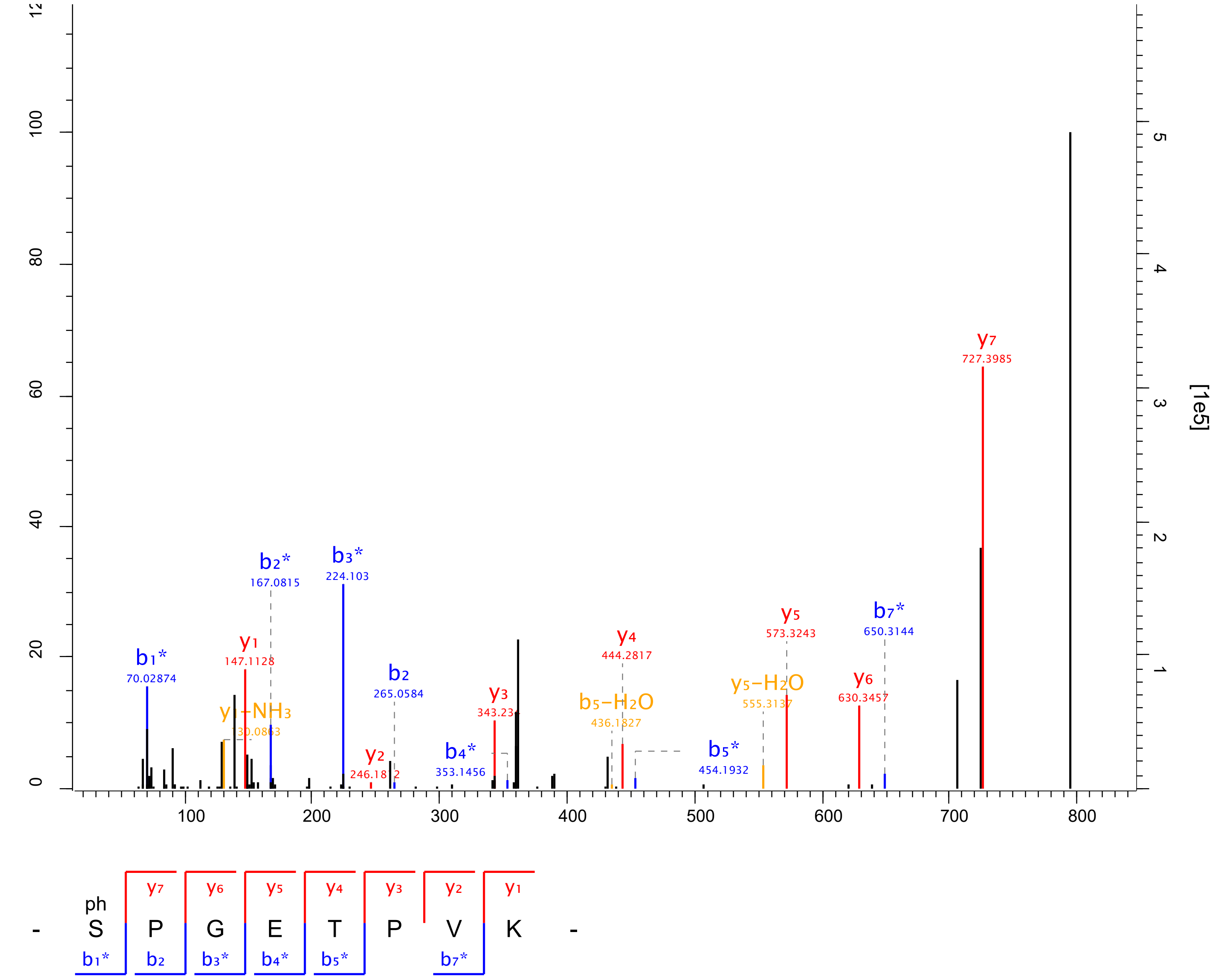

| Raw file | Scan | Method | Score | m/z |
| --- | --- | --- | --- | --- |
| water_00_1 | 25086 | FTMS; HCD | 52.9 | 806.39 |

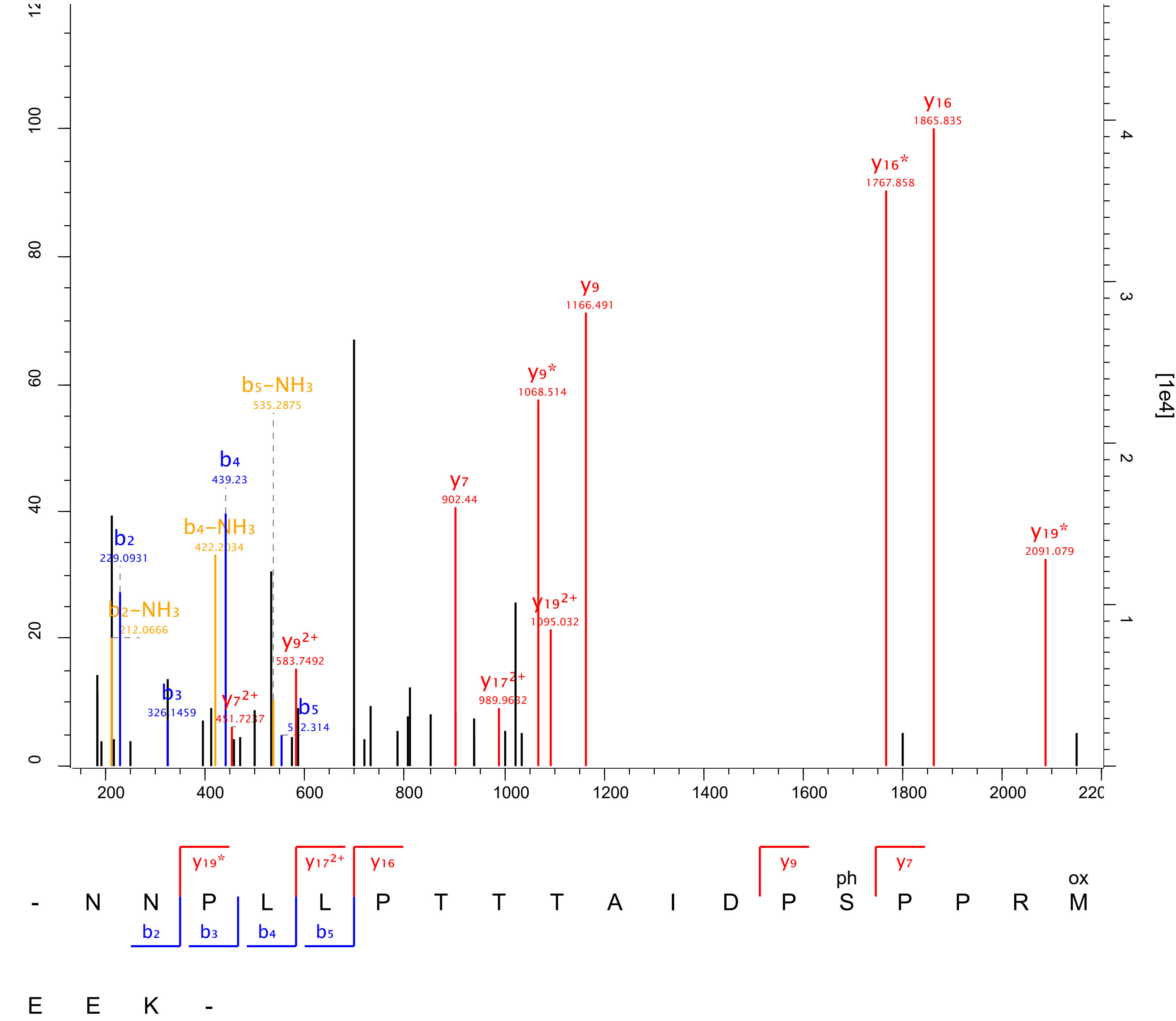

| Raw file | Scan | Method | Score | m/z |
|---|---|---|---|---|
| water_00_1 | 25165 | FTMS; HCD | 103.88 | 817.4 |

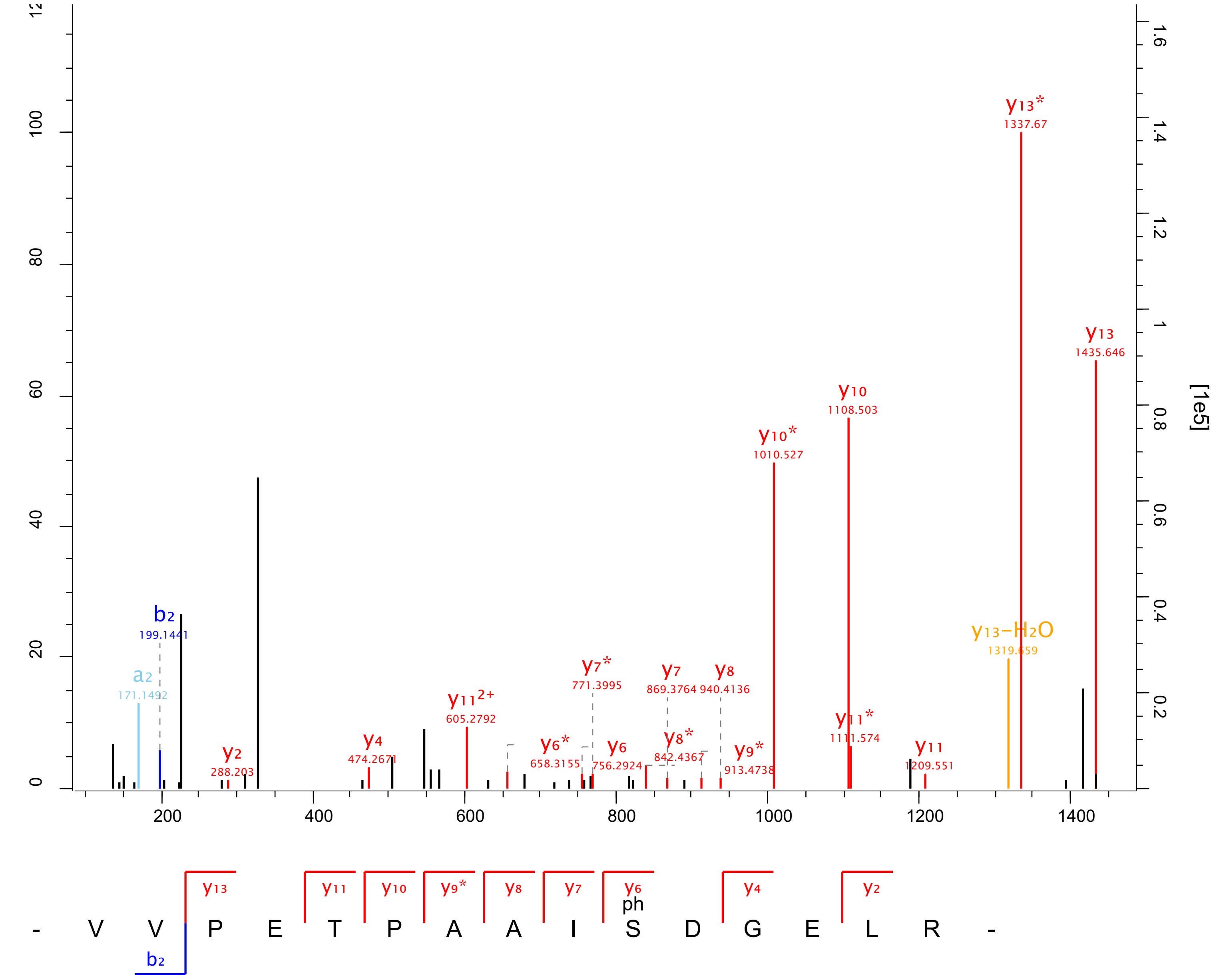

| Raw file | Scan | Method | Score | m/z |
|---|---|---|---|---|
| water_00_1 | 25658 | FTMS; HCD | 106.57 | 912.43 |

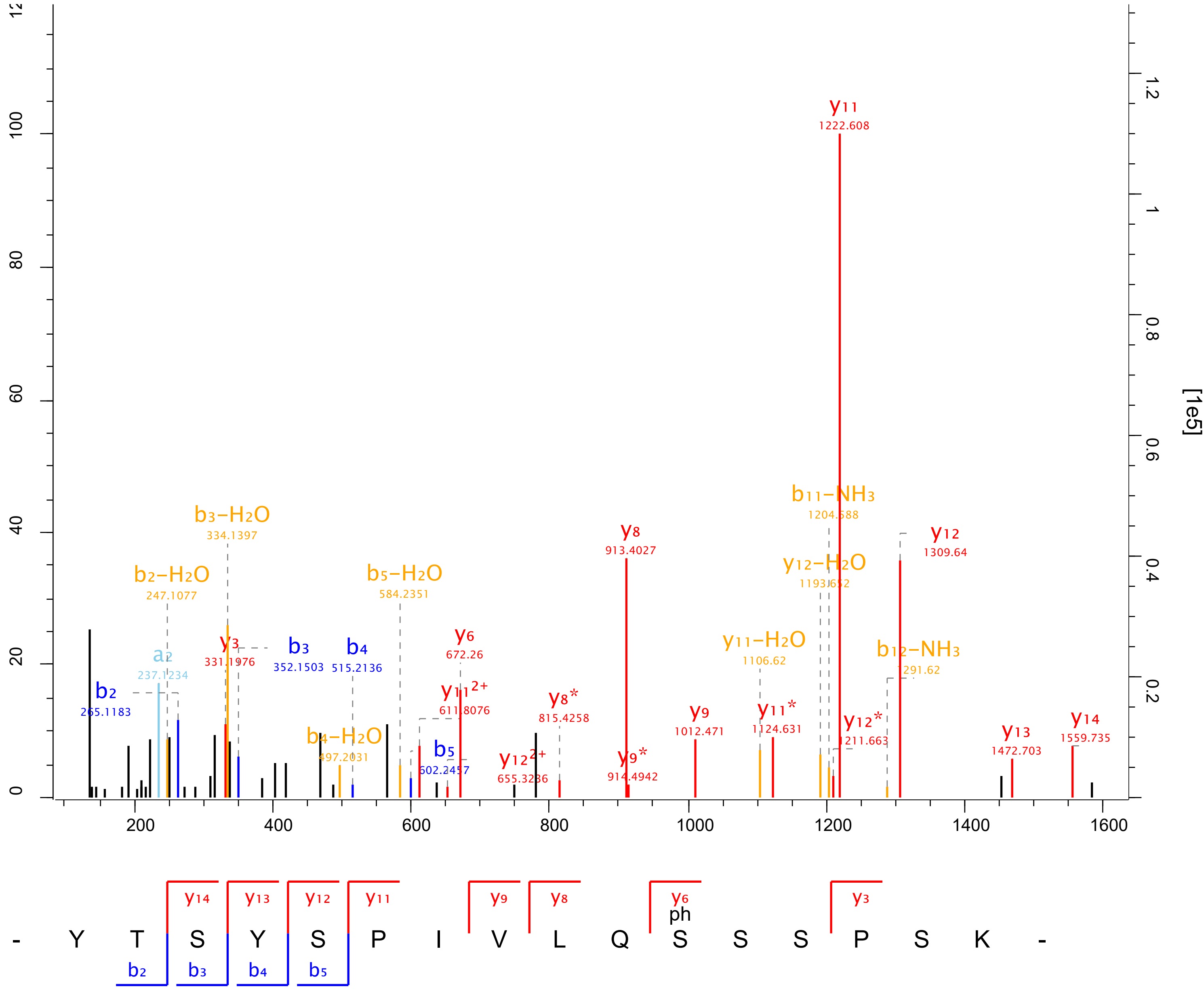

| Raw file | Scan | Method | Score | m/z |
|---|---|---|---|---|
| water_00_1 | 25995 | FTMS; HCD | 138.34 | 950.08 |

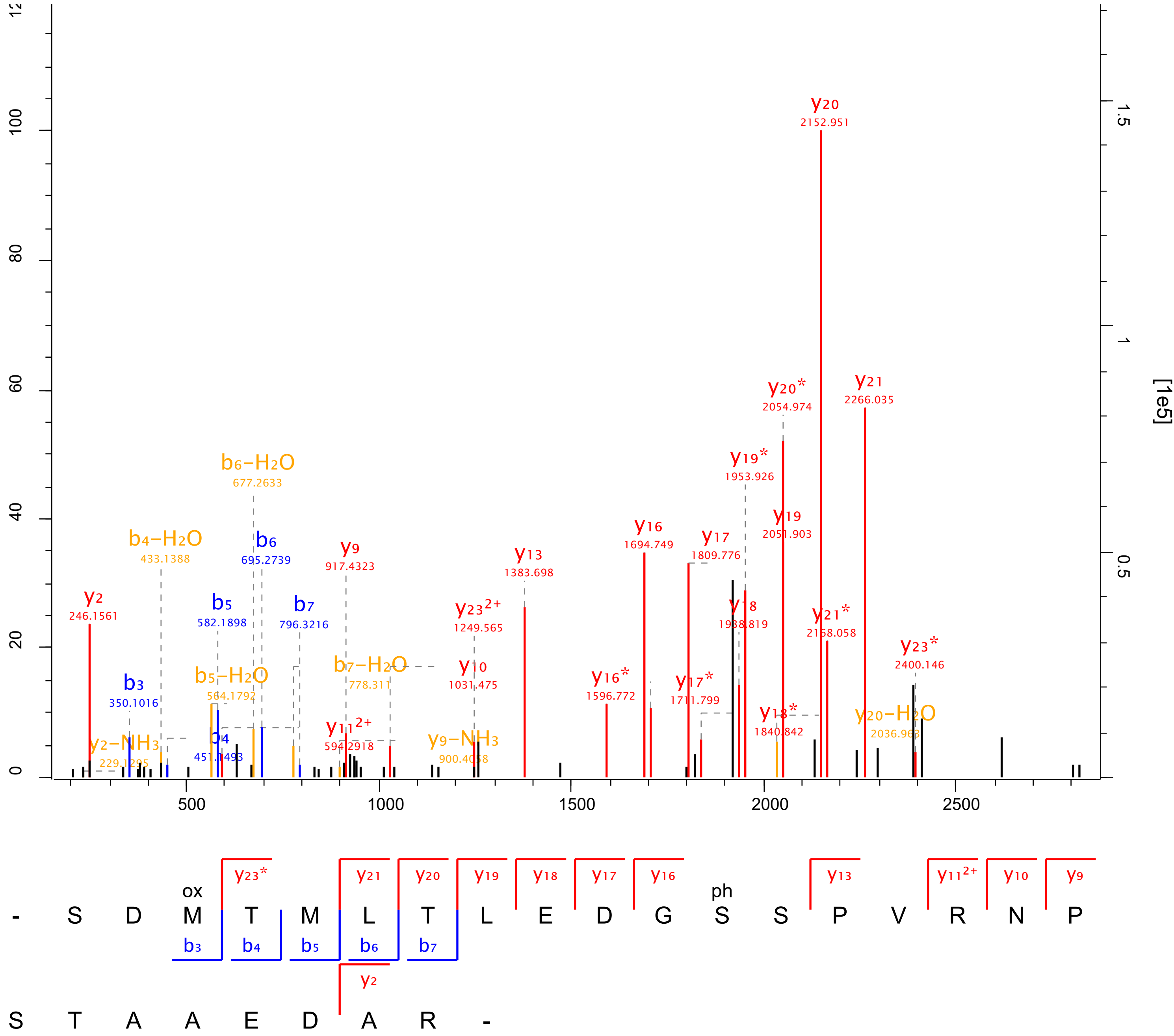

| Raw file | Scan | Method | Score | m/z |
| --- | --- | --- | --- | --- |
| water_00_1 | 26101 | FTMS; HCD | 129.77 | 633.31 |

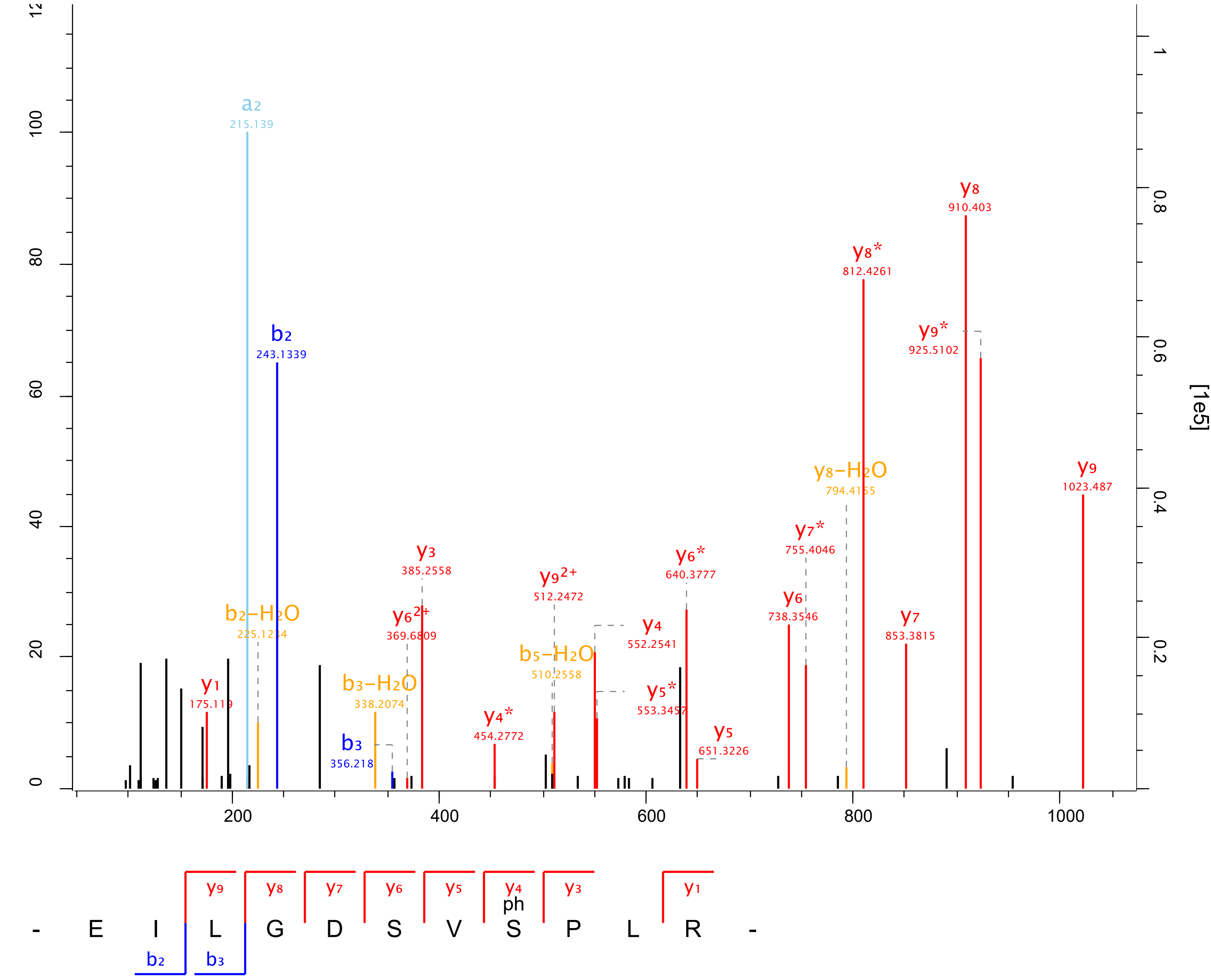

| Raw file | Scan | Method | Score | m/z |
| --- | --- | --- | --- | --- |
| water_00_1 | 26136 | FTMS; HCD | 121.64 | 444.71 |

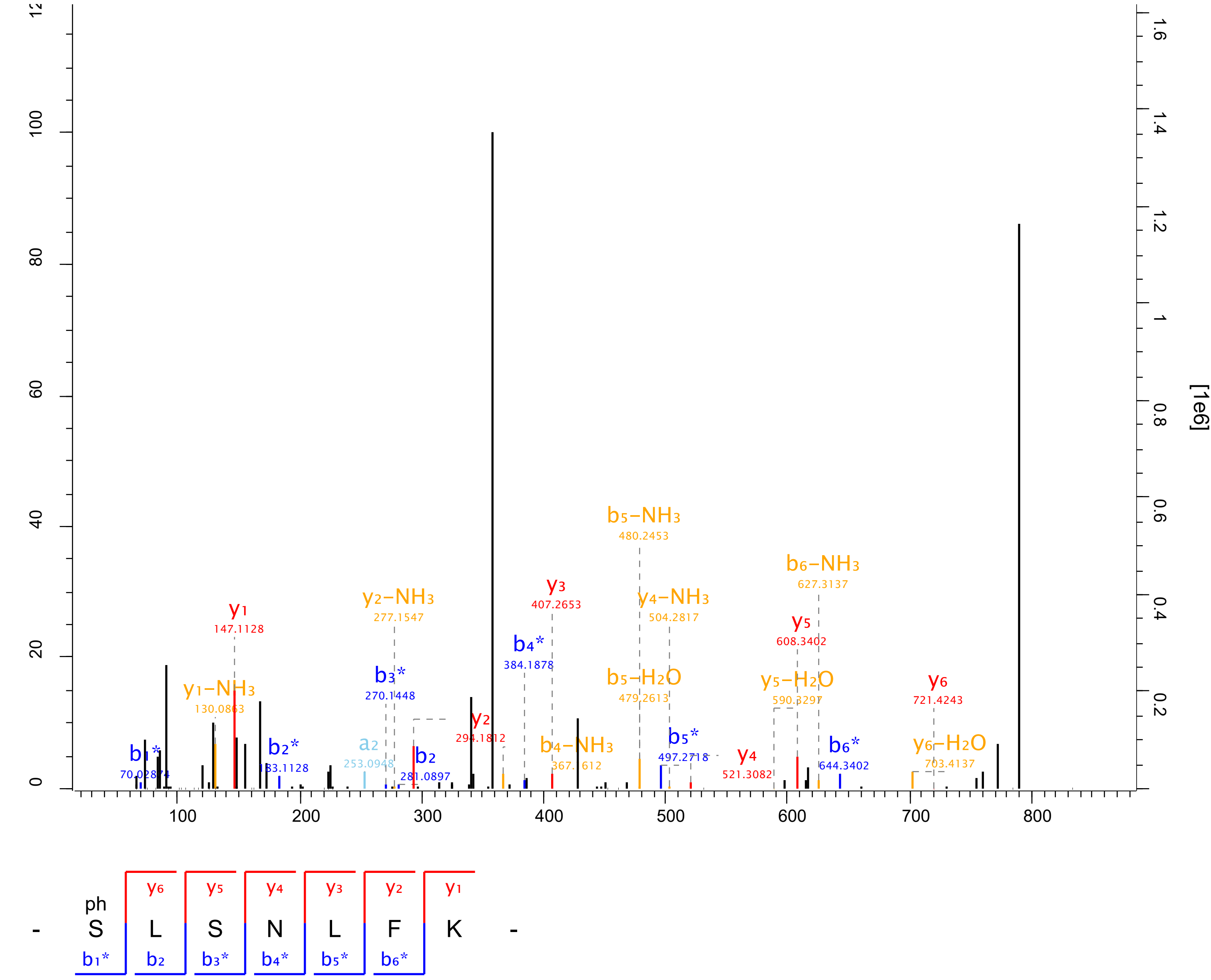

| Raw file | Scan | Method | Score | m/z |
| --- | --- | --- | --- | --- |
| water_00_1 | 26471 | FTMS; HCD | 203.32 | 722.8 |

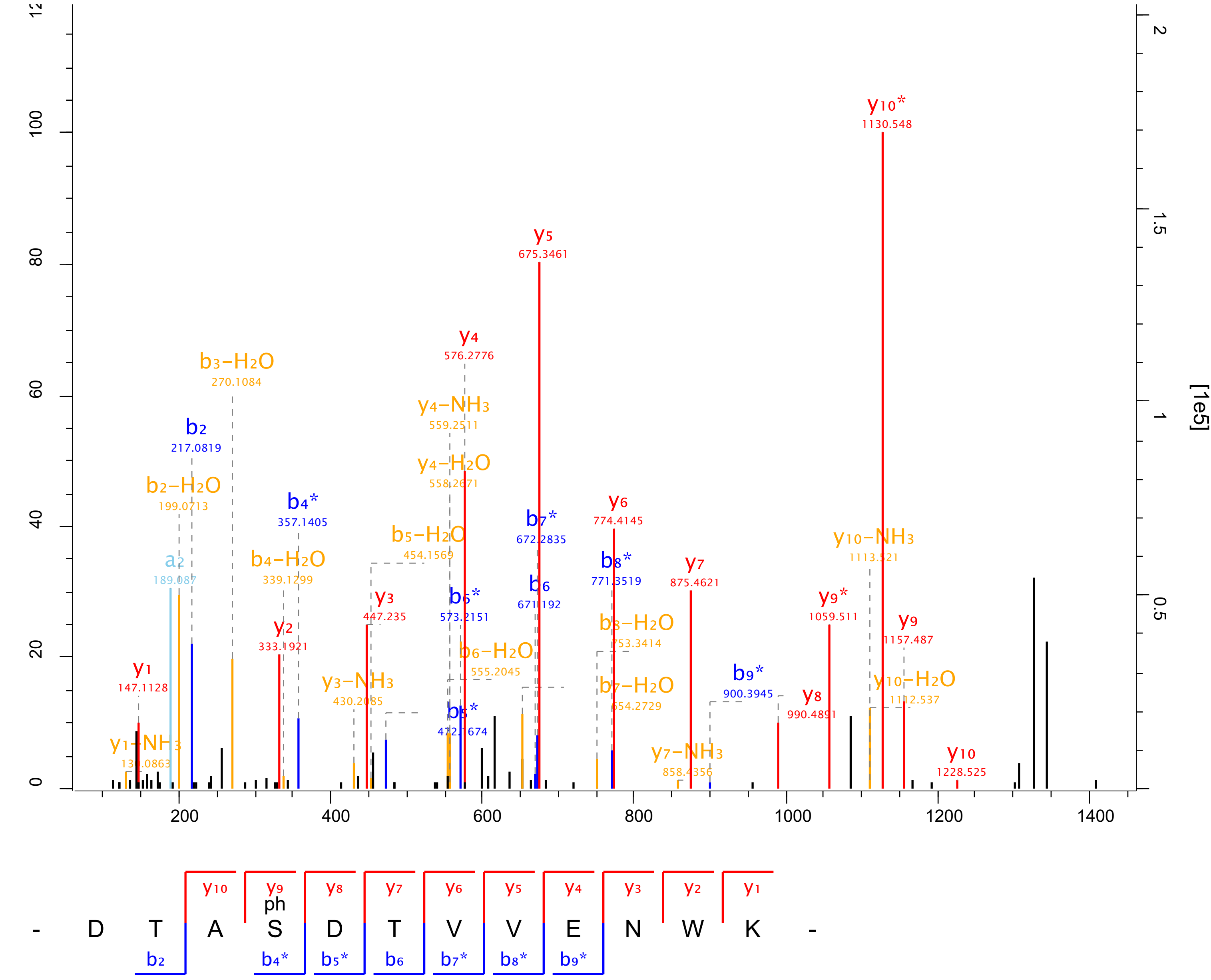

| Raw file | Scan | Method | Score | m/z |
|---|---|---|---|---|
| water_00_1 | 26473 | FTMS; HCD | 118.15 | 894.38 |

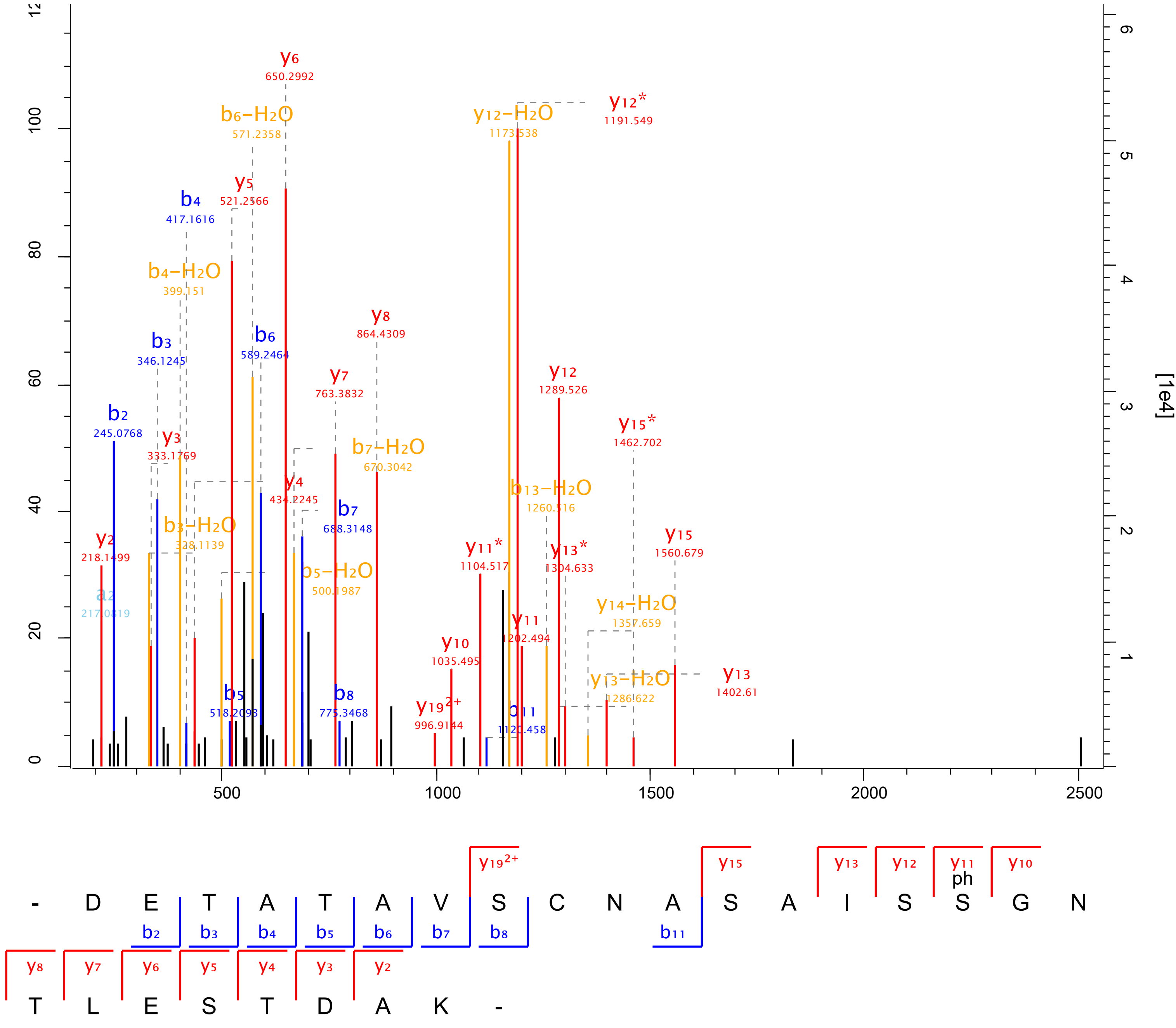

| Raw file | Scan | Method | Score | m/z |
|---|---|---|---|---|
| water_00_1 | 26535 | FTMS; HCD | 121.18 | 645.29 |

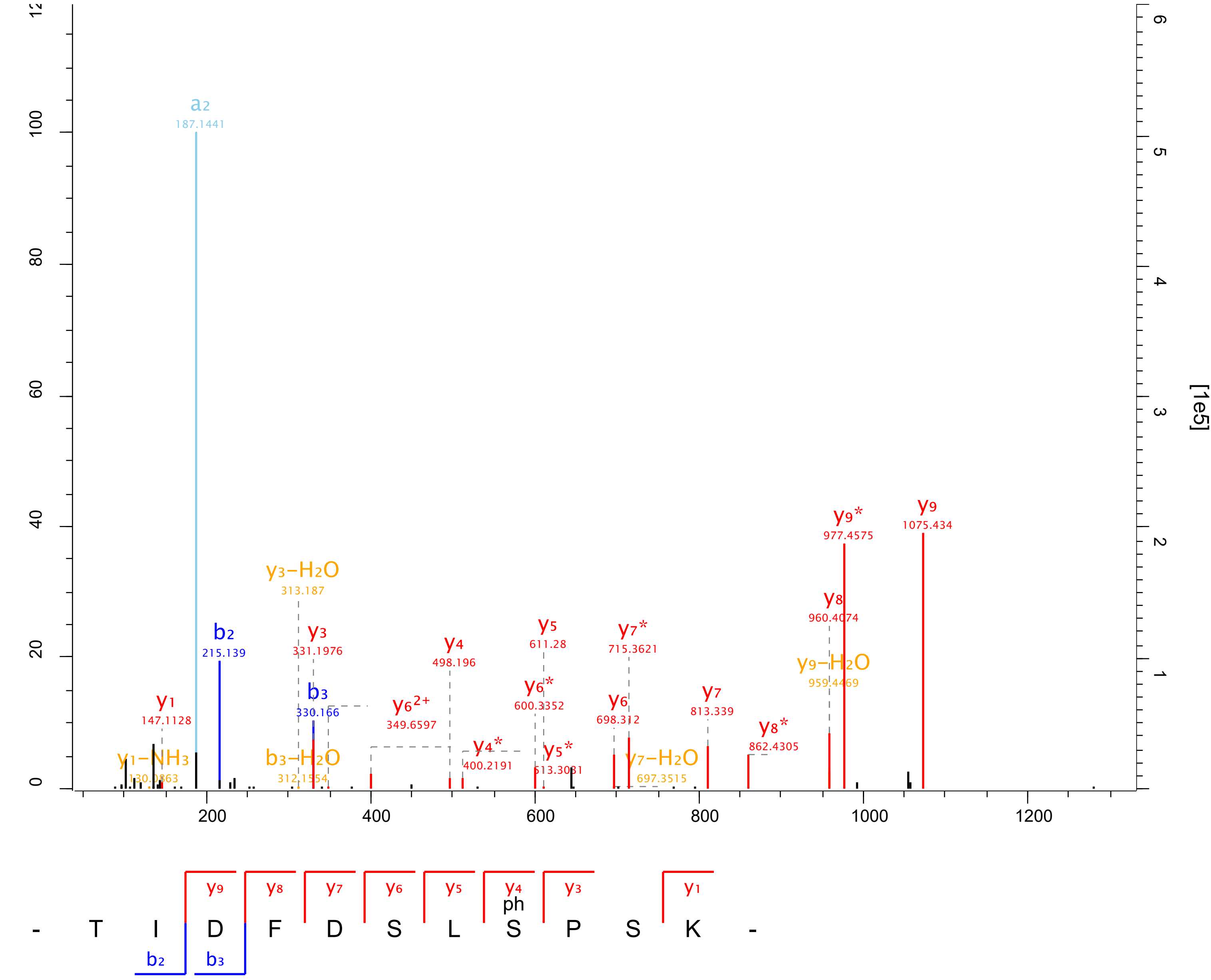

| Raw file | Scan | Method | Score | m/z |
|---|---|---|---|---|
| water_00_1 | 26674 | FTMS; HCD | 225.19 | 637.74 |

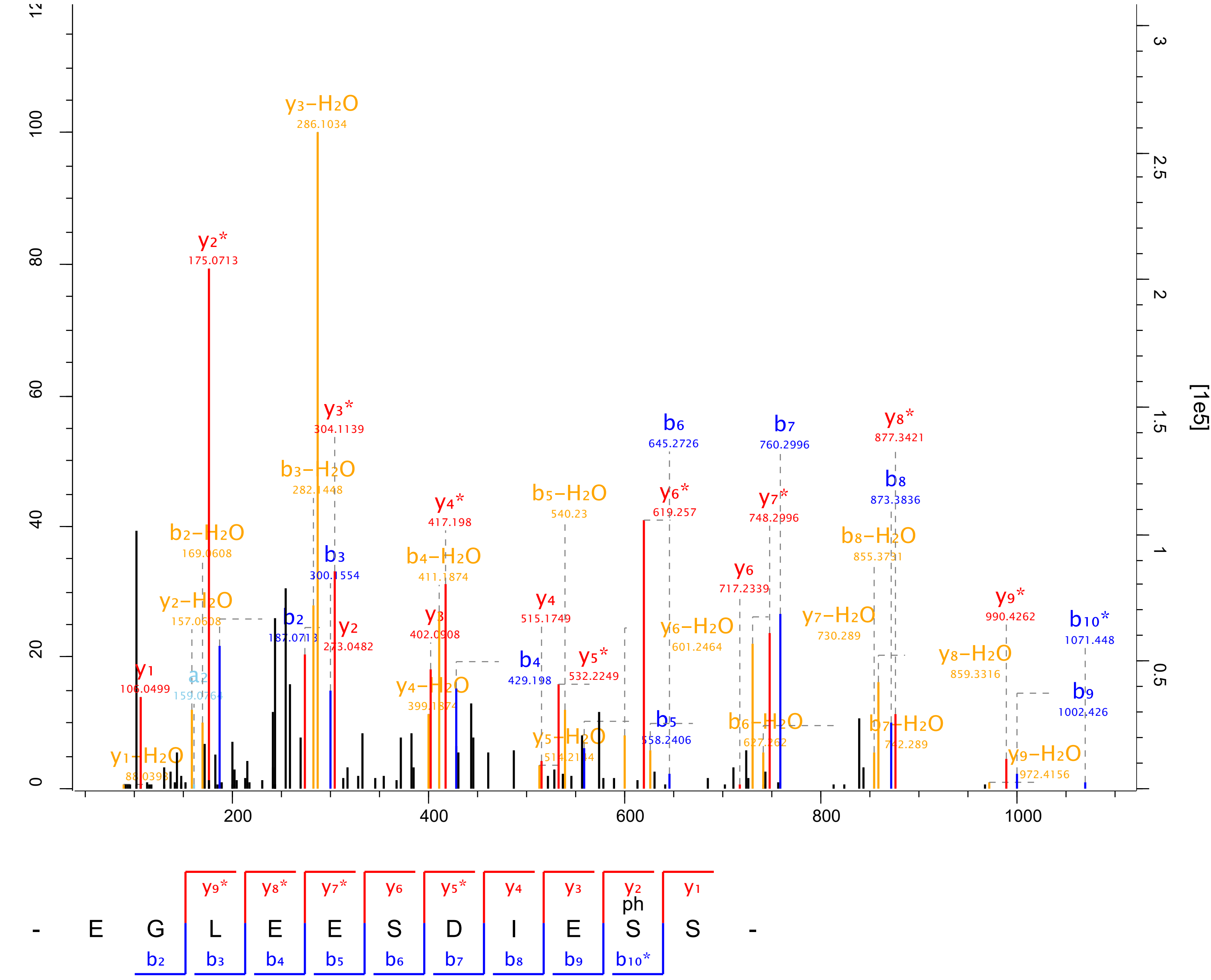

| Raw file | Scan | Method | Score | m/z |
|---|---|---|---|---|
| water_00_1 | 2670 | FTMS; HCD | 78.94 | 437.18 |

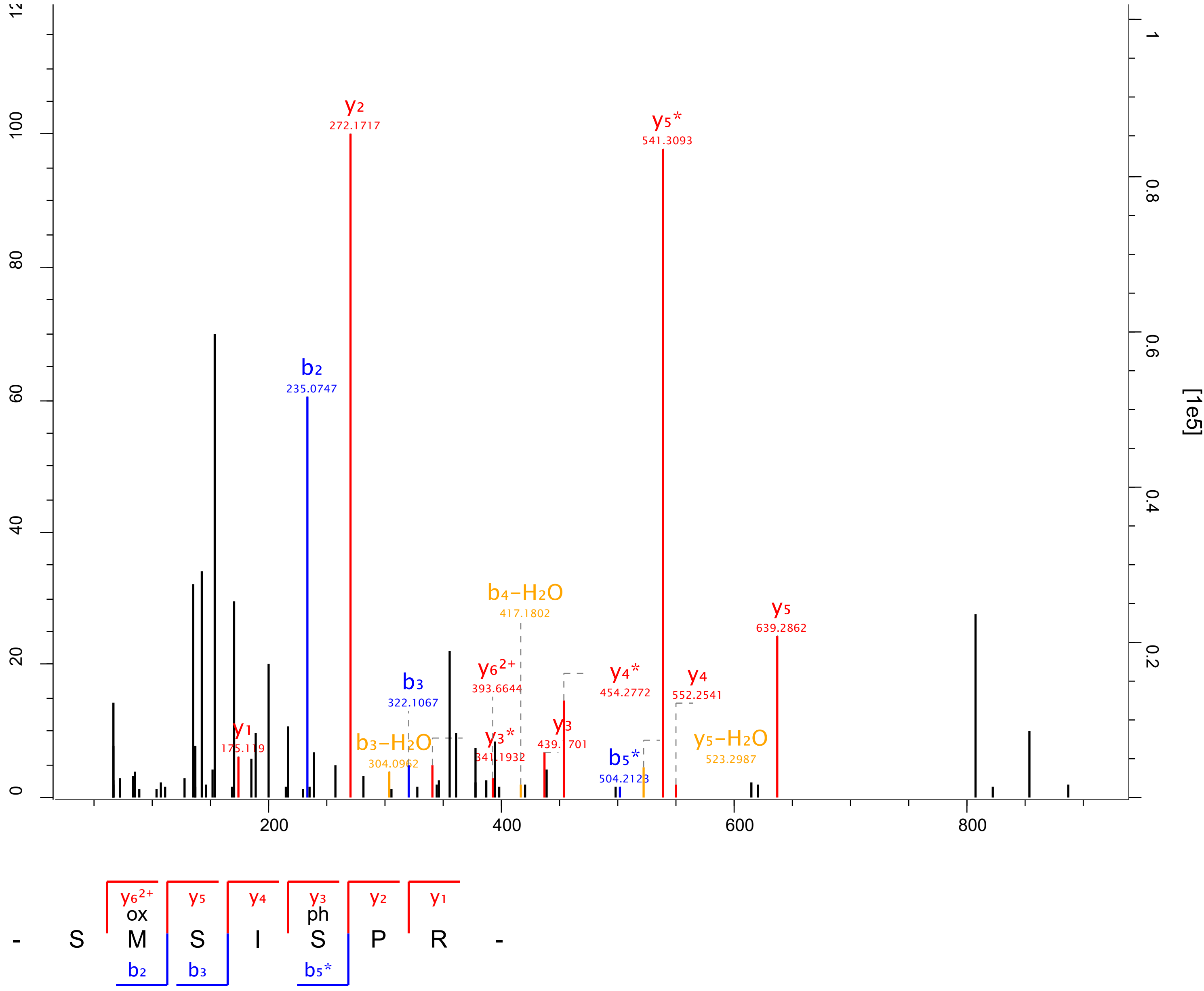

| Raw file | Scan | Method | Score | m/z |
|---|---|---|---|---|
| water_00_1 | 26791 | FTMS; HCD | 160.46 | 688.27 |

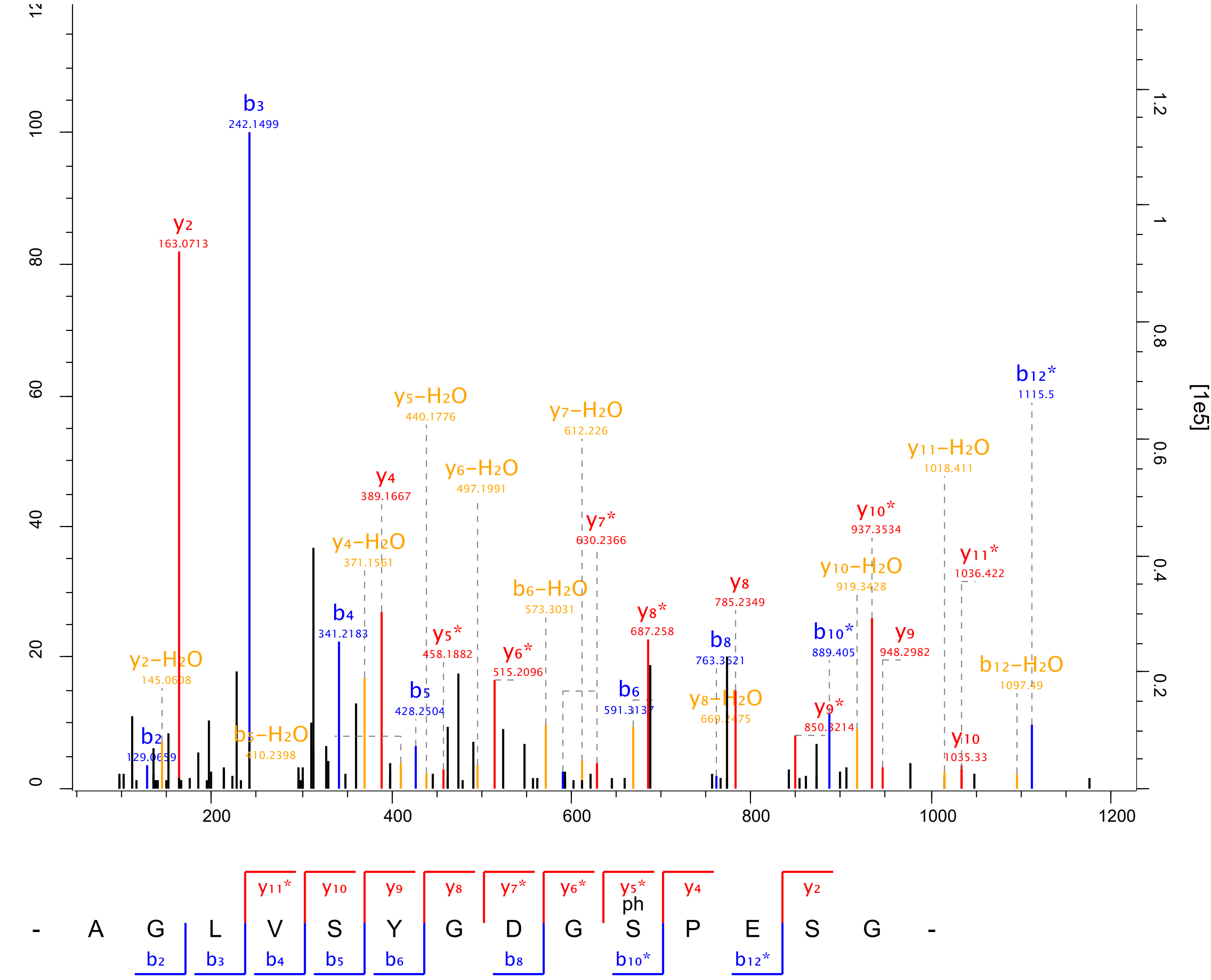

| Raw file | Scan | Method | Score | m/z |
|---|---|---|---|---|
| water_00_1 | 26825 | FTMS; HCD | 43.7 | 729.69 |

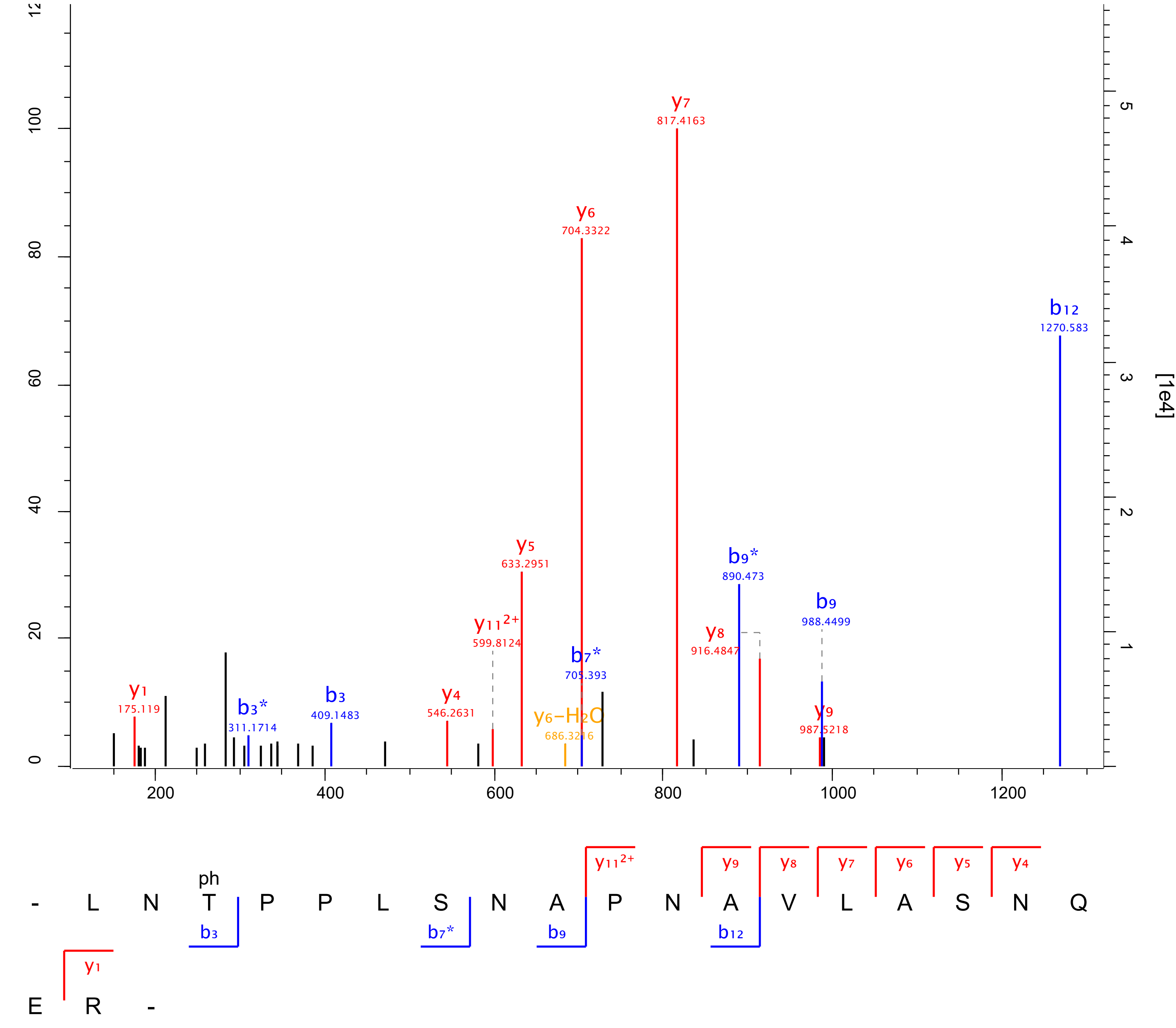

| Raw file | Scan | Method | Score | m/z |
|---|---|---|---|---|
| water_00_1 | 26837 | FTMS; HCD | 149.29 | 801.37 |

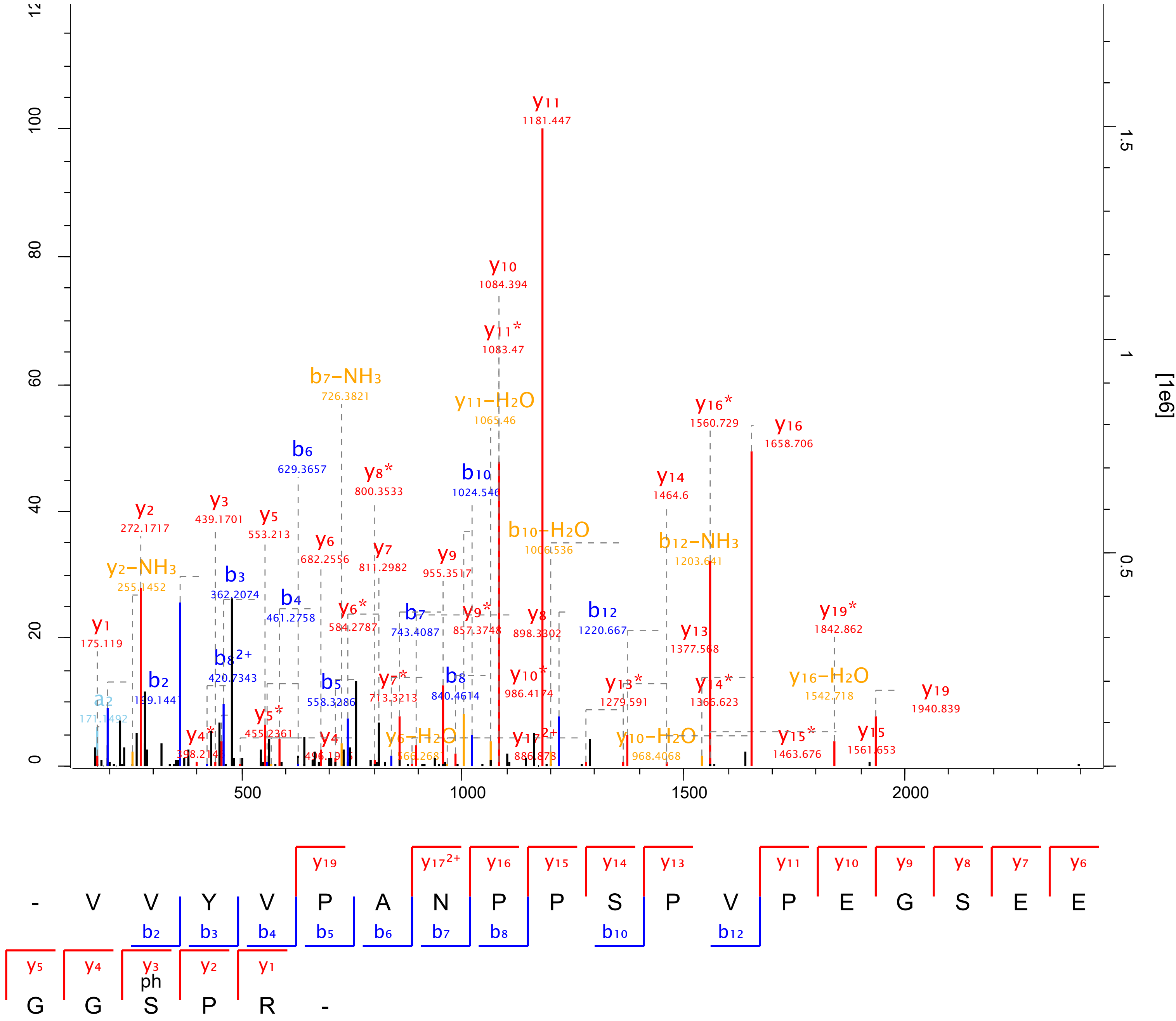

| Raw file | Scan | Method | Score | m/z |
|---|---|---|---|---|
| water_00_1 | 2690 | FTMS; HCD | 59.37 | 498.73 |

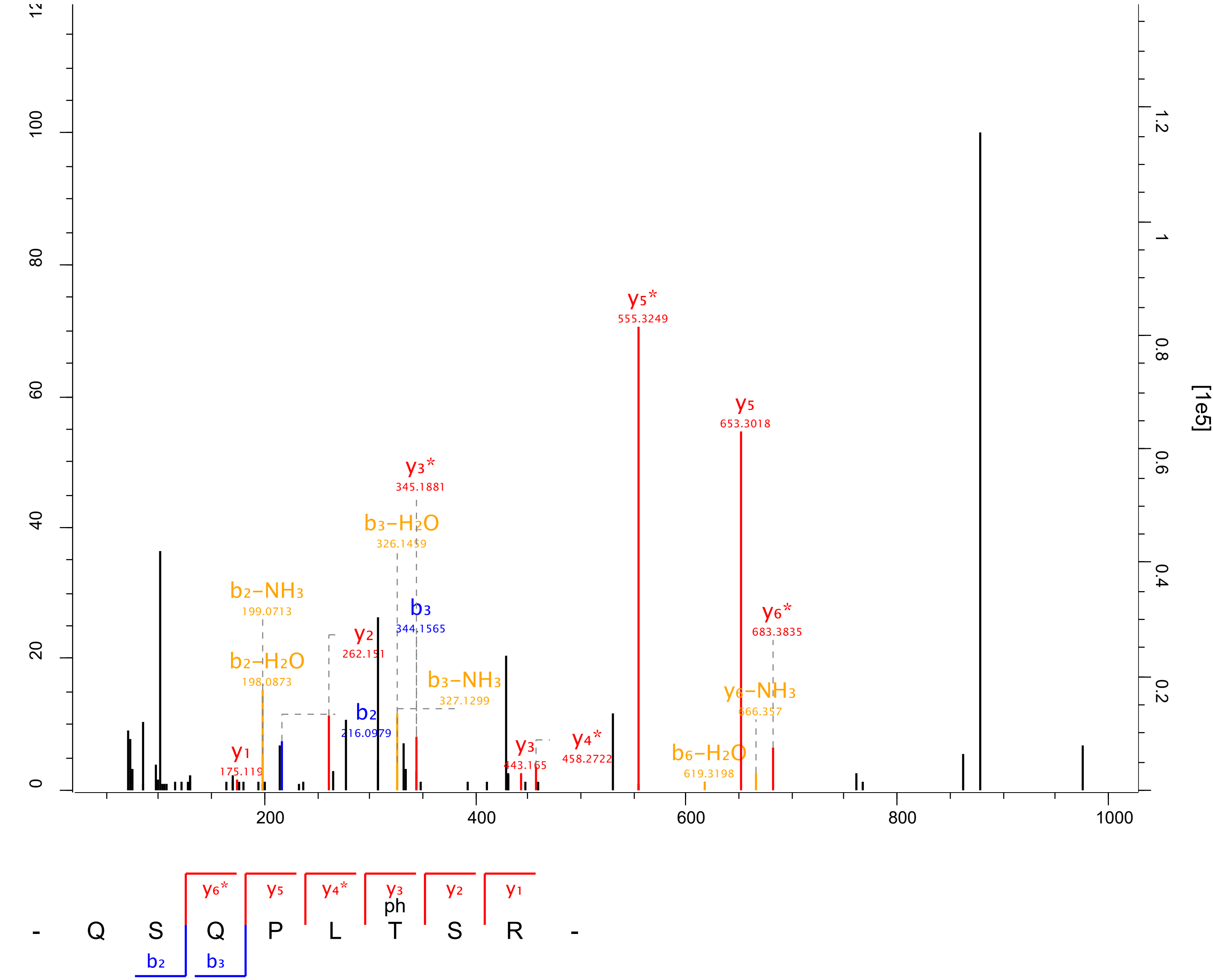

| Raw file | Scan | Method | Score | m/z |
|---|---|---|---|---|
| water_00_1 | 27479 | FTMS; HCD | 96.79 | 1089.01 |

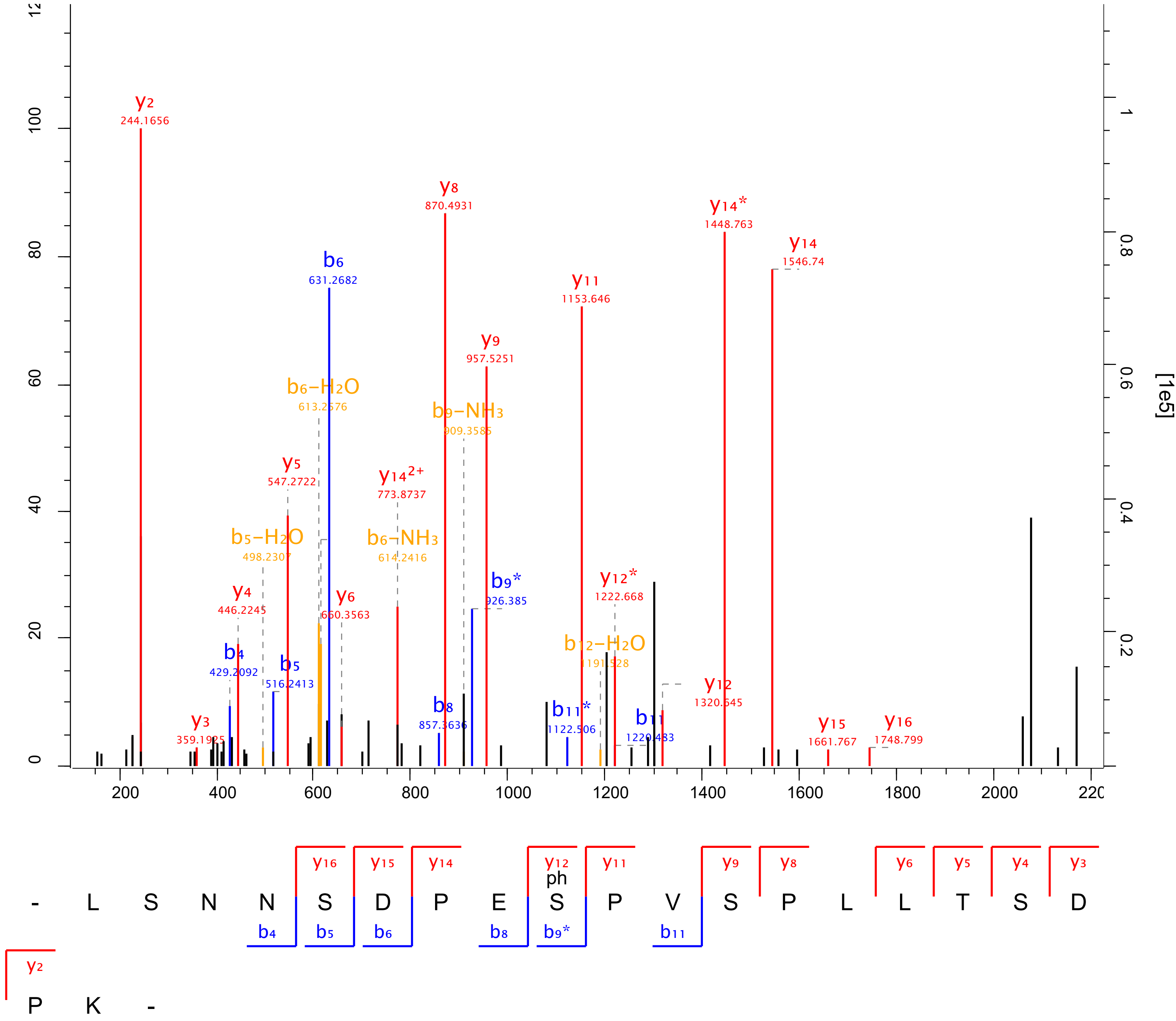

| Raw file | Scan | Method | Score | m/z |
| --- | --- | --- | --- | --- |
| water_00_1 | 2755 | FTMS; HCD | 41.24 | 586.75 |

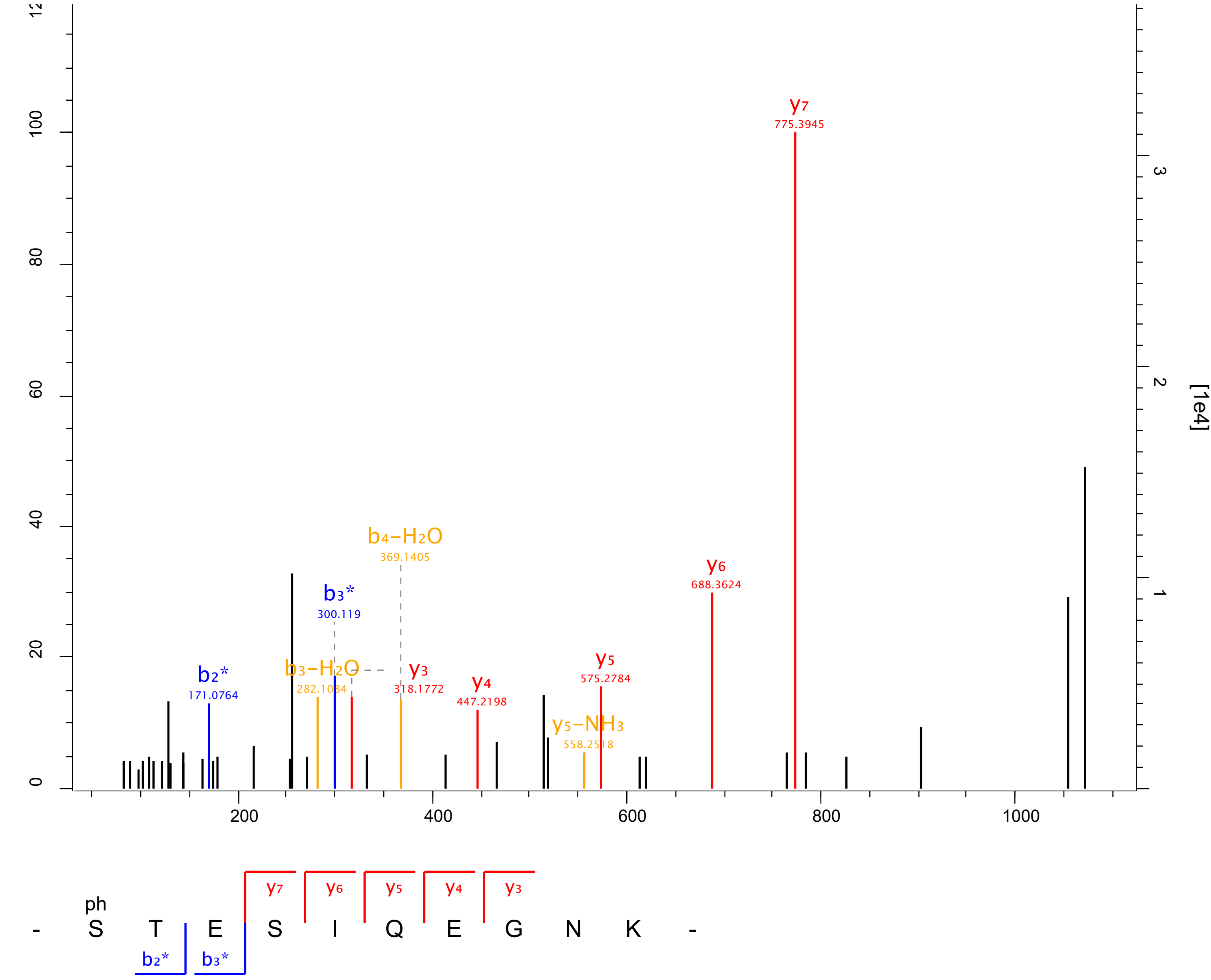

| Raw file | Scan | Method | Score | m/z |
| --- | --- | --- | --- | --- |
| water_00_1 | 27577 | FTMS; HCD | 57.29 | 895.82 |

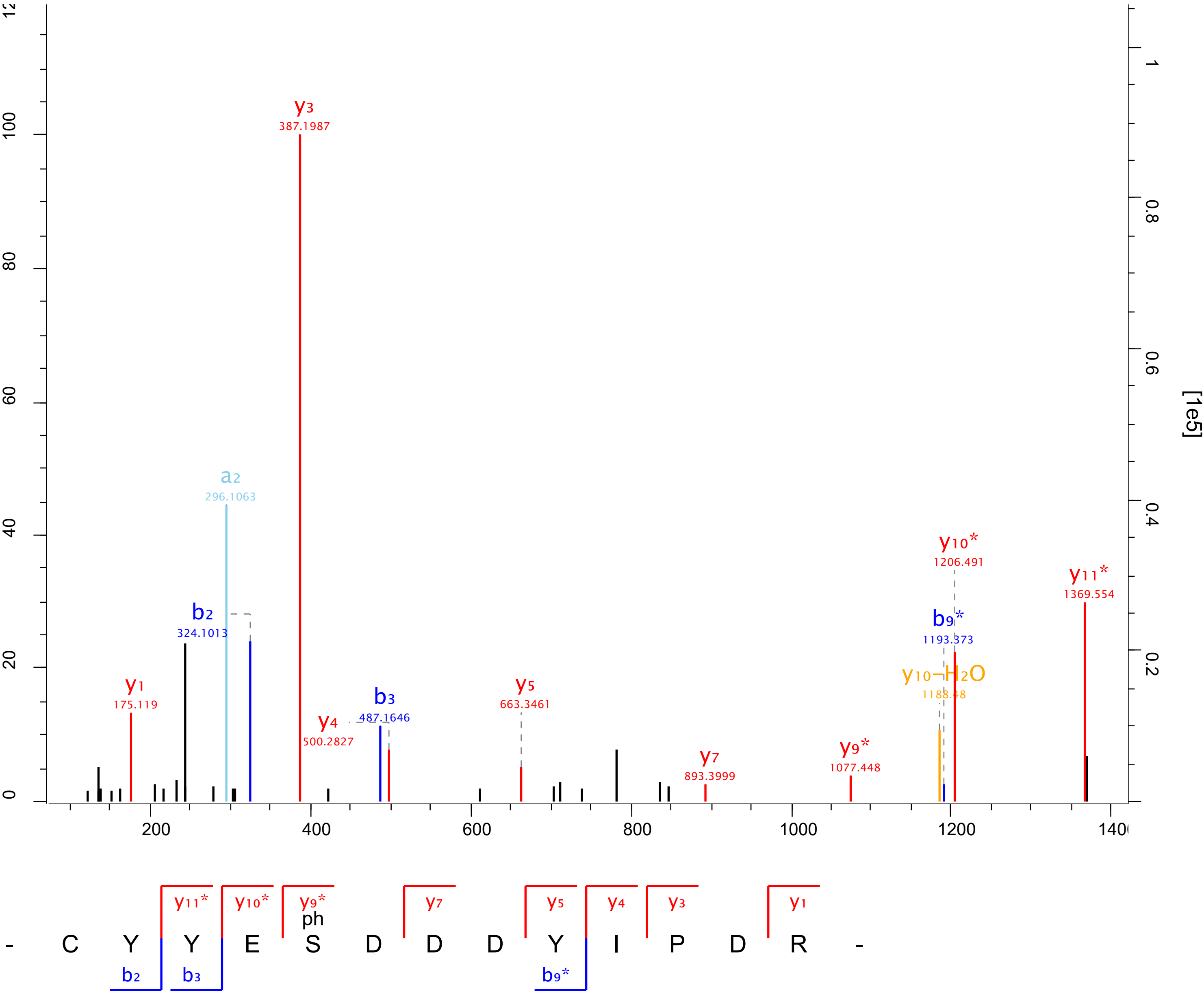

| Raw file | Scan | Method | Score | m/z |
|---|---|---|---|---|
| water_00_1 | 27912 | FTMS; HCD | 46.42 | 933.44 |

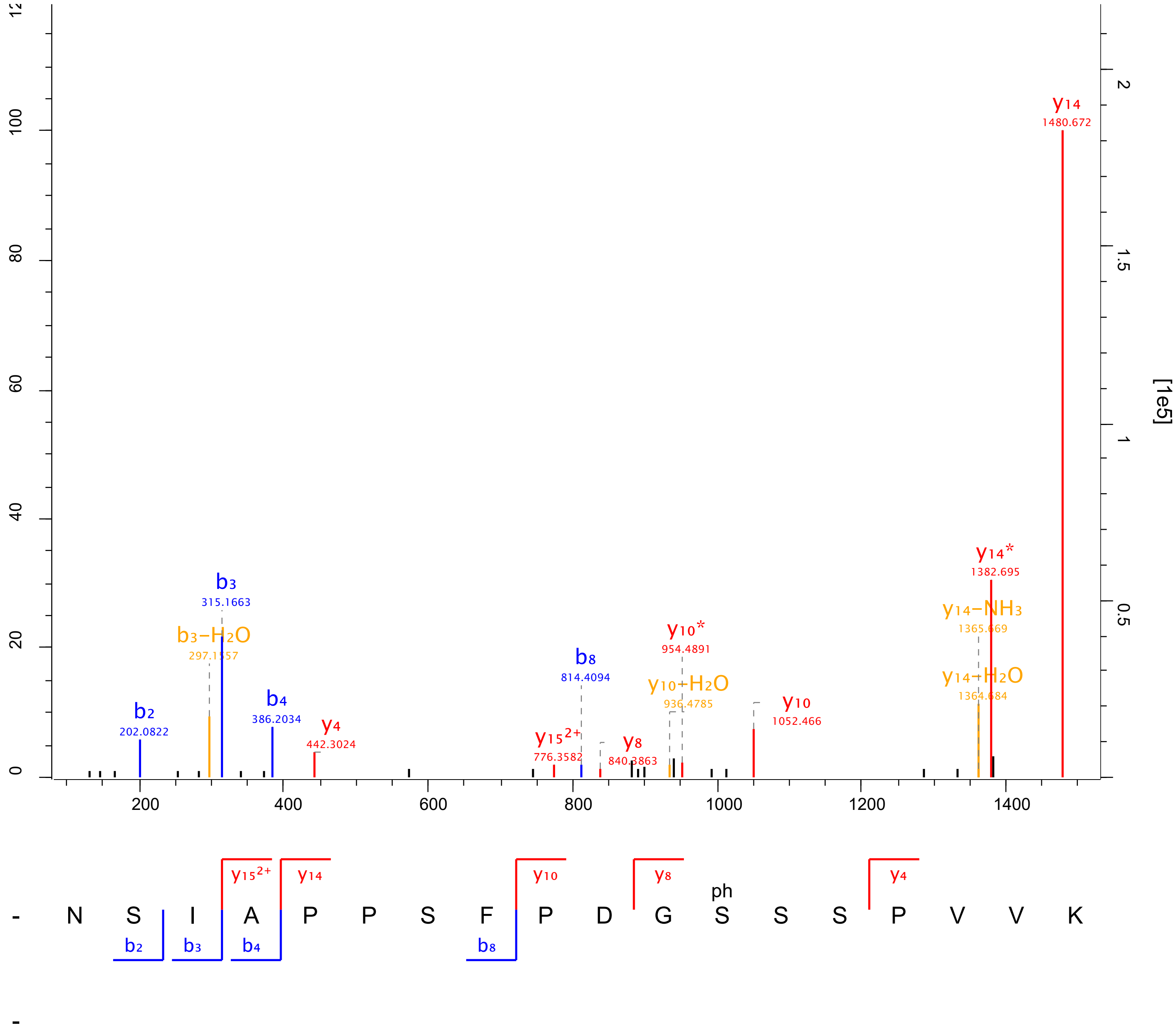

| Raw file | Scan | Method | Score | m/z |
|---|---|---|---|---|
| water_00_1 | 27940 | FTMS; HCD | 128.66 | 833.41 |

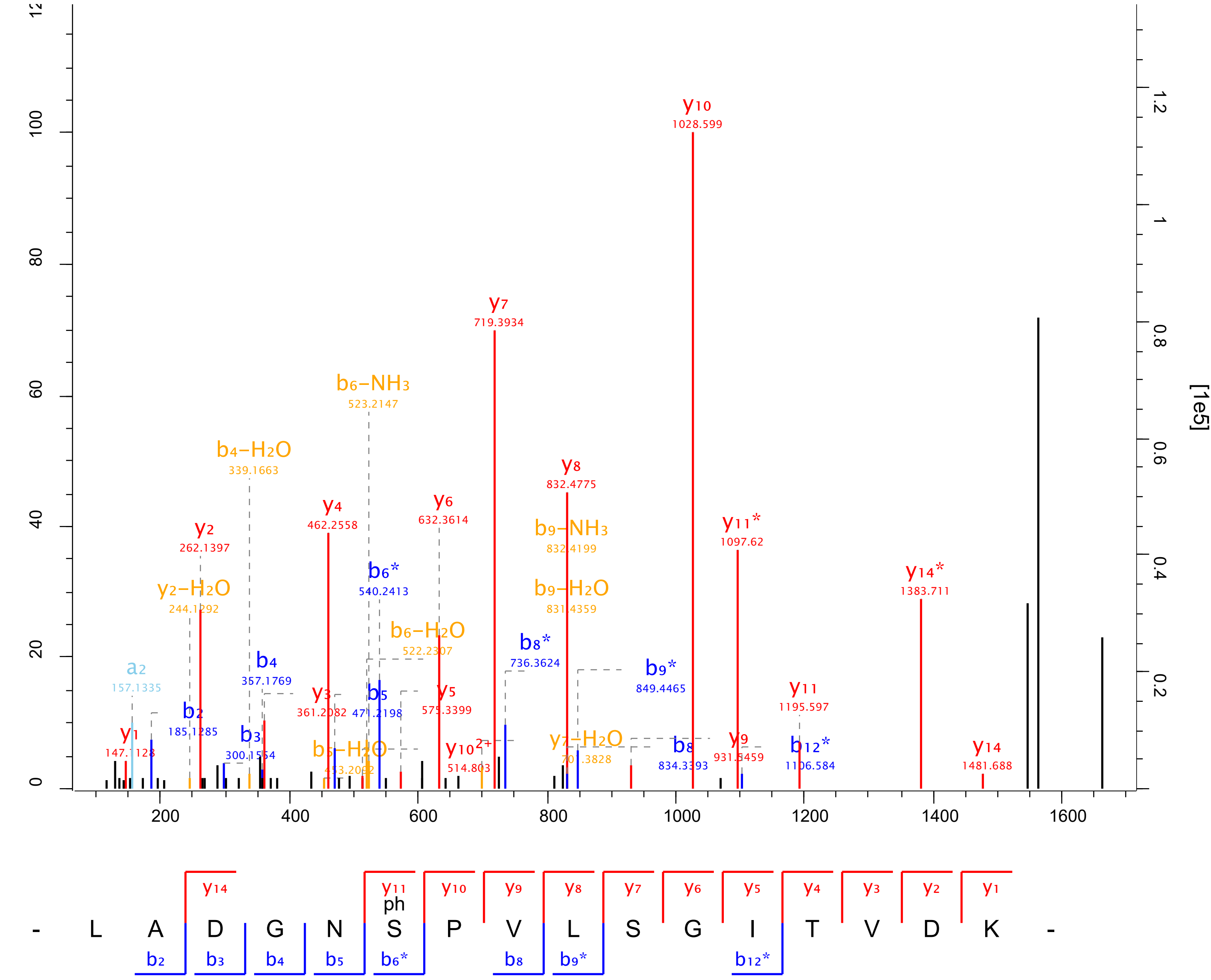

| Raw file | Scan | Method | Score | m/z |
|---|---|---|---|---|
| water_00_1 | 28227 | FTMS; HCD | 60.16 | 758.33 |

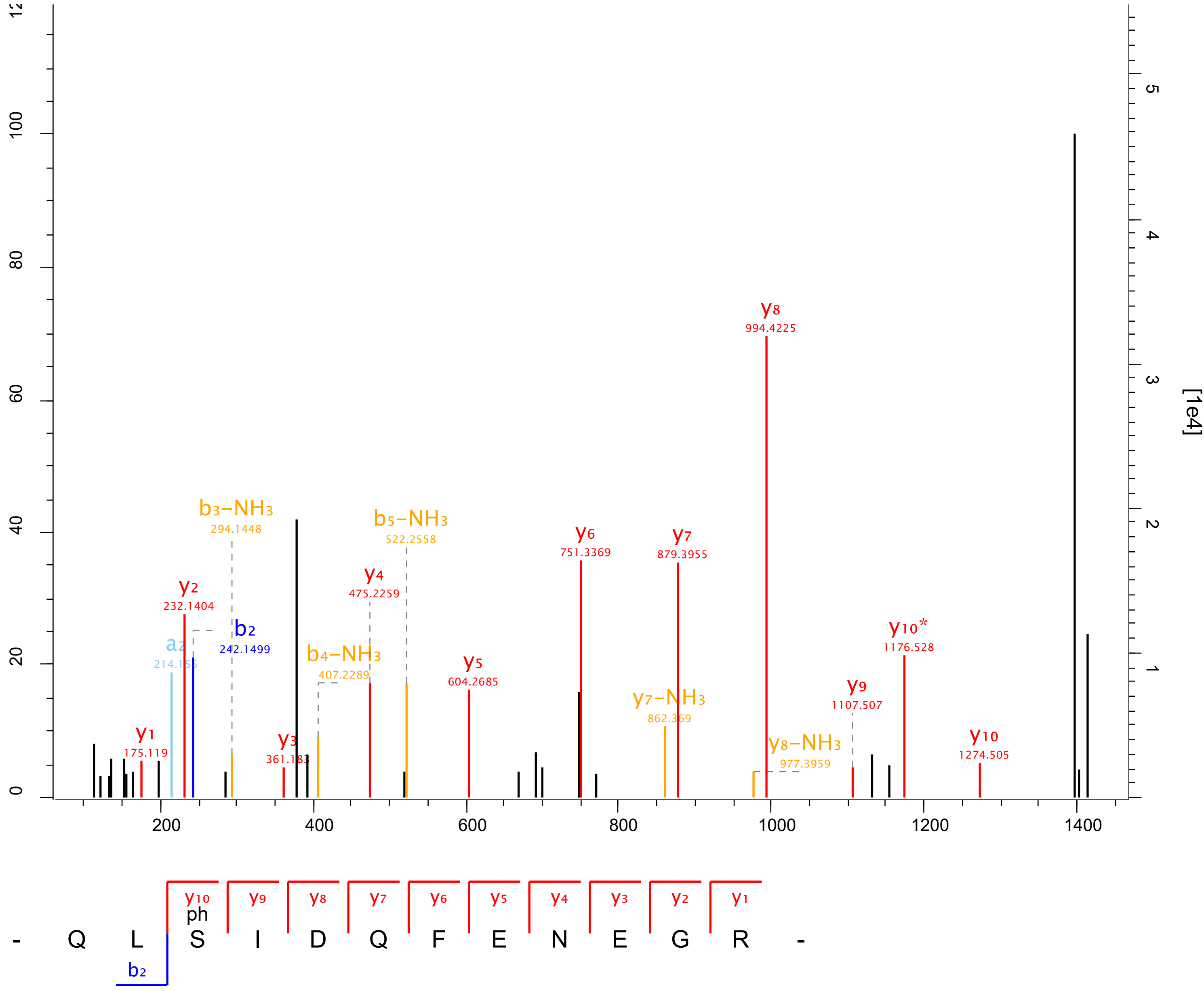

| Raw file | Scan | Method | Score | m/z |
| --- | --- | --- | --- | --- |
| water_00_1 | 2879 | FTMS; HCD | 93.91 | 517.87 |

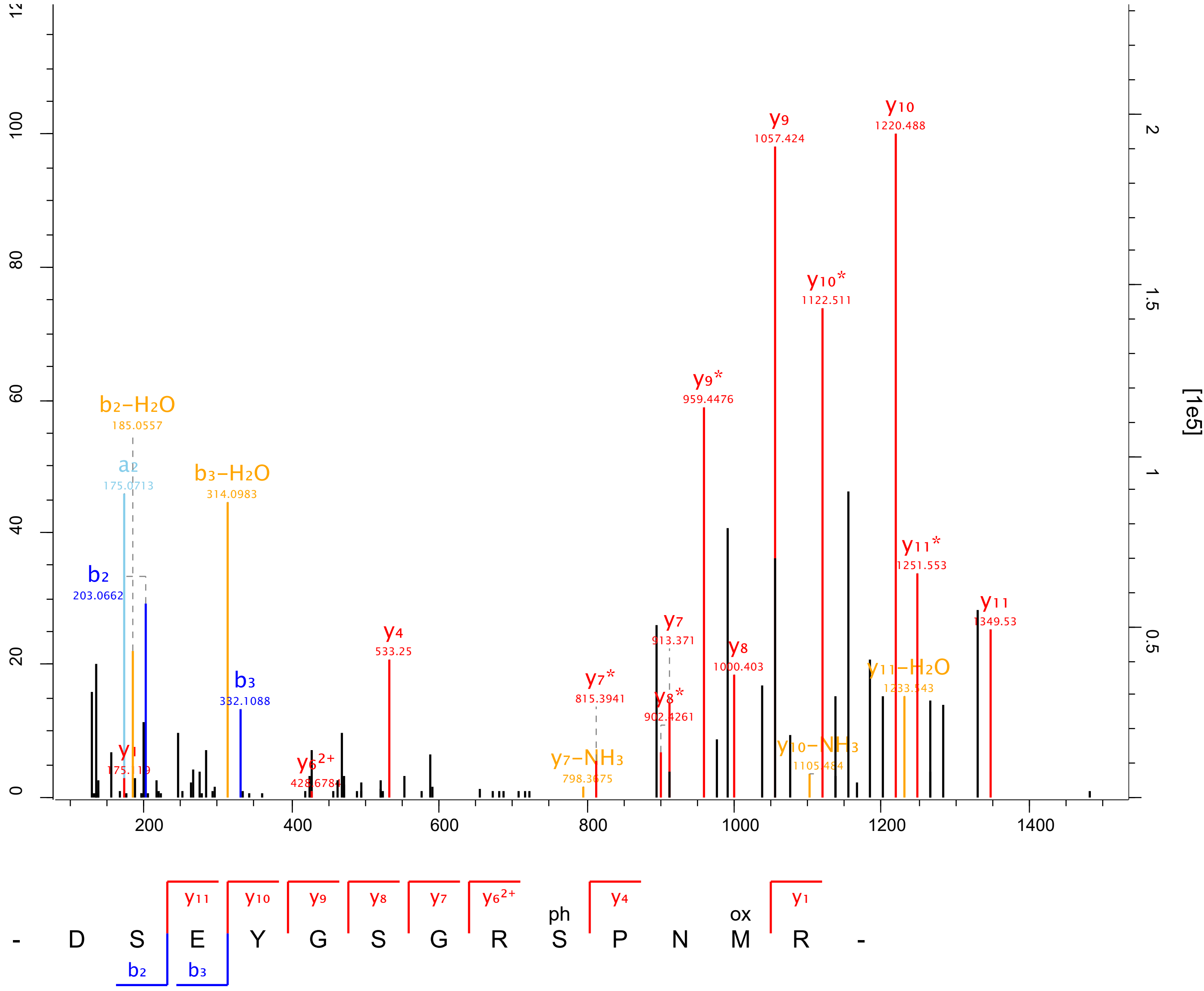

| Raw file | Scan | Method | Score | m/z |
|---|---|---|---|---|
| water_00_1 | 28822 | FTMS; HCD | 254.56 | 698.82 |

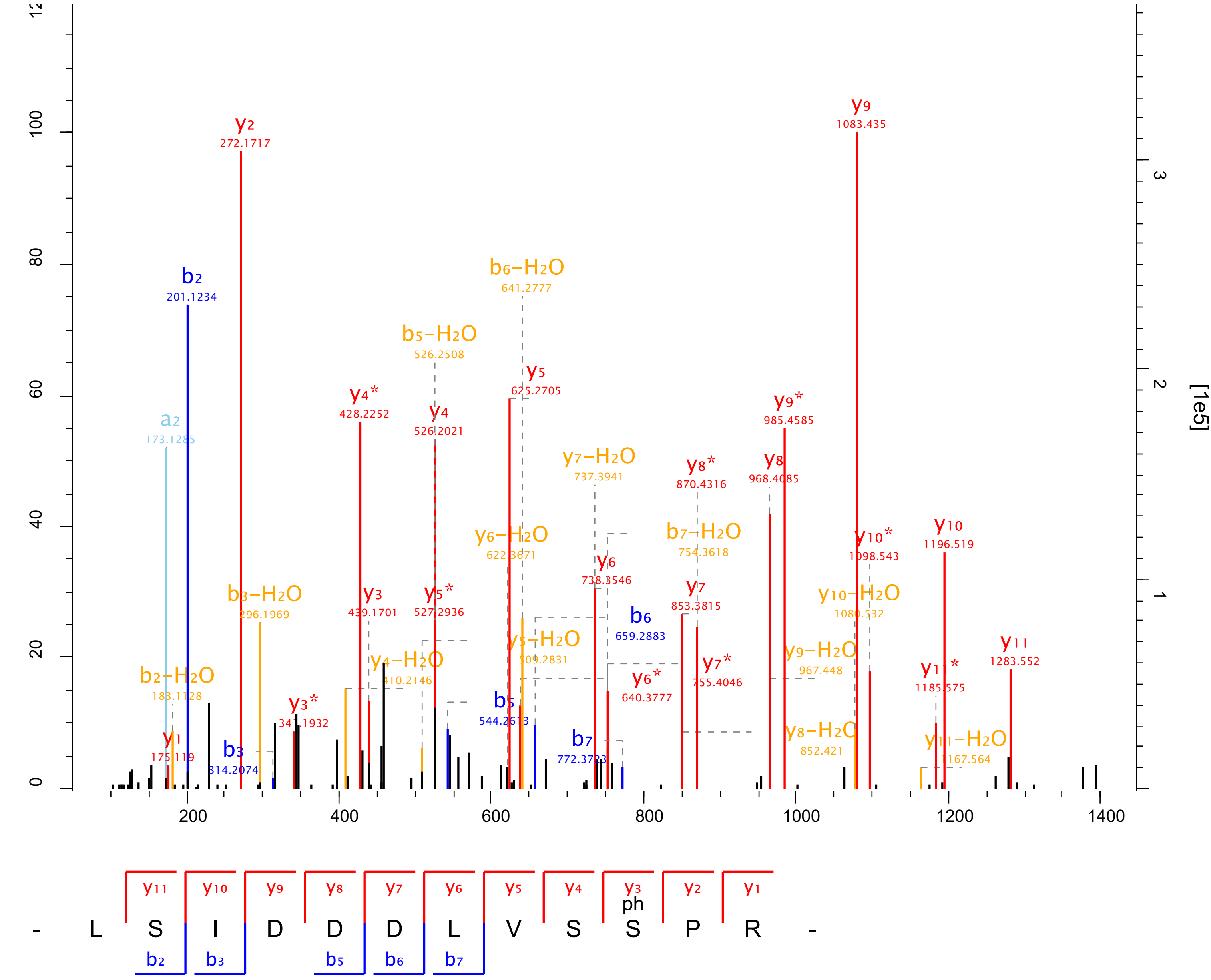

| Raw file | Scan | Method | Score | m/z |
|---|---|---|---|---|
| water_00_1 | 2900 | FTMS; HCD | 132.84 | 398.2 |

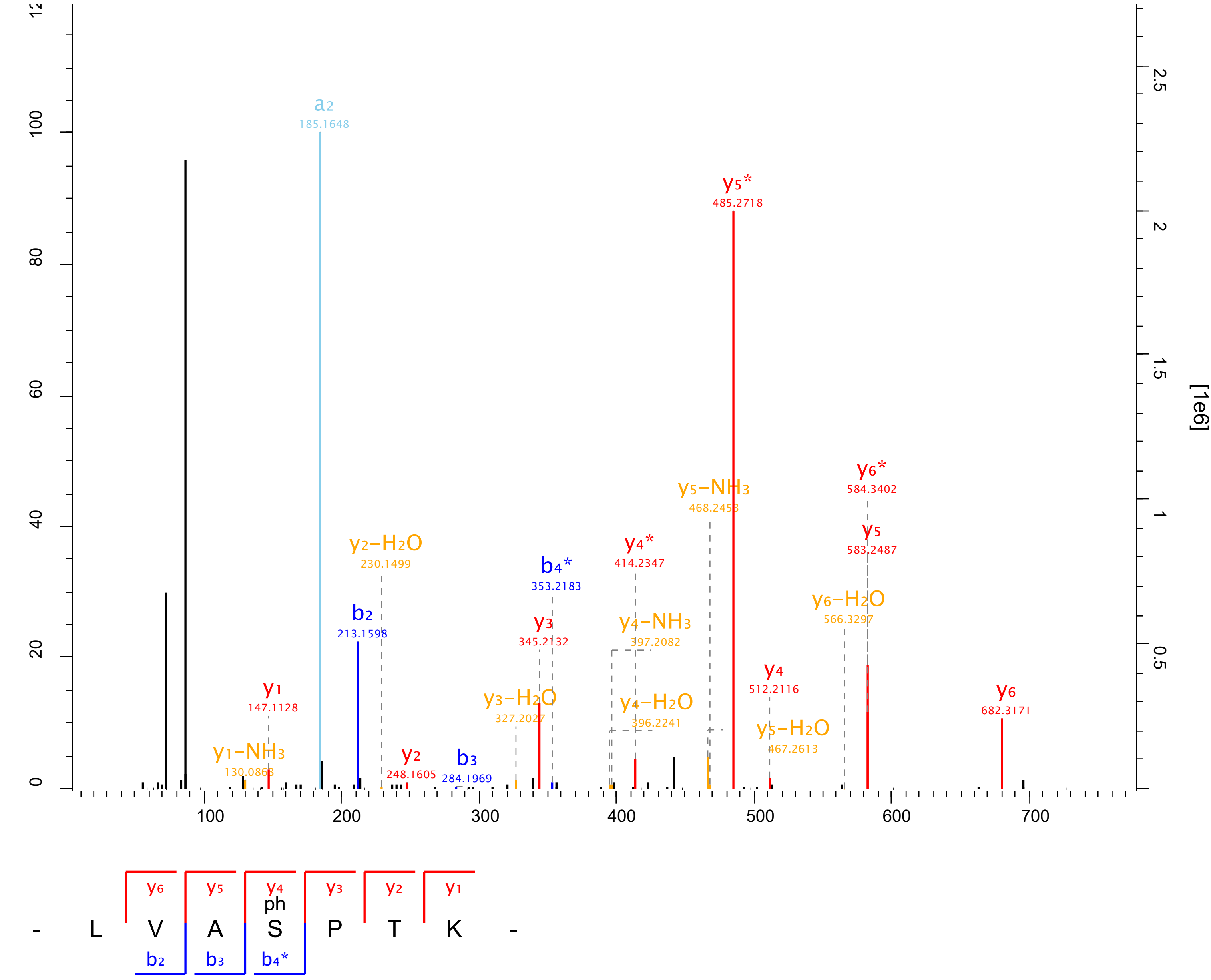

| Raw file | Scan | Method | Score | m/z |
|---|---|---|---|---|
| water_00_1 | 2911 | FTMS; HCD | 75.53 | 737.81 |

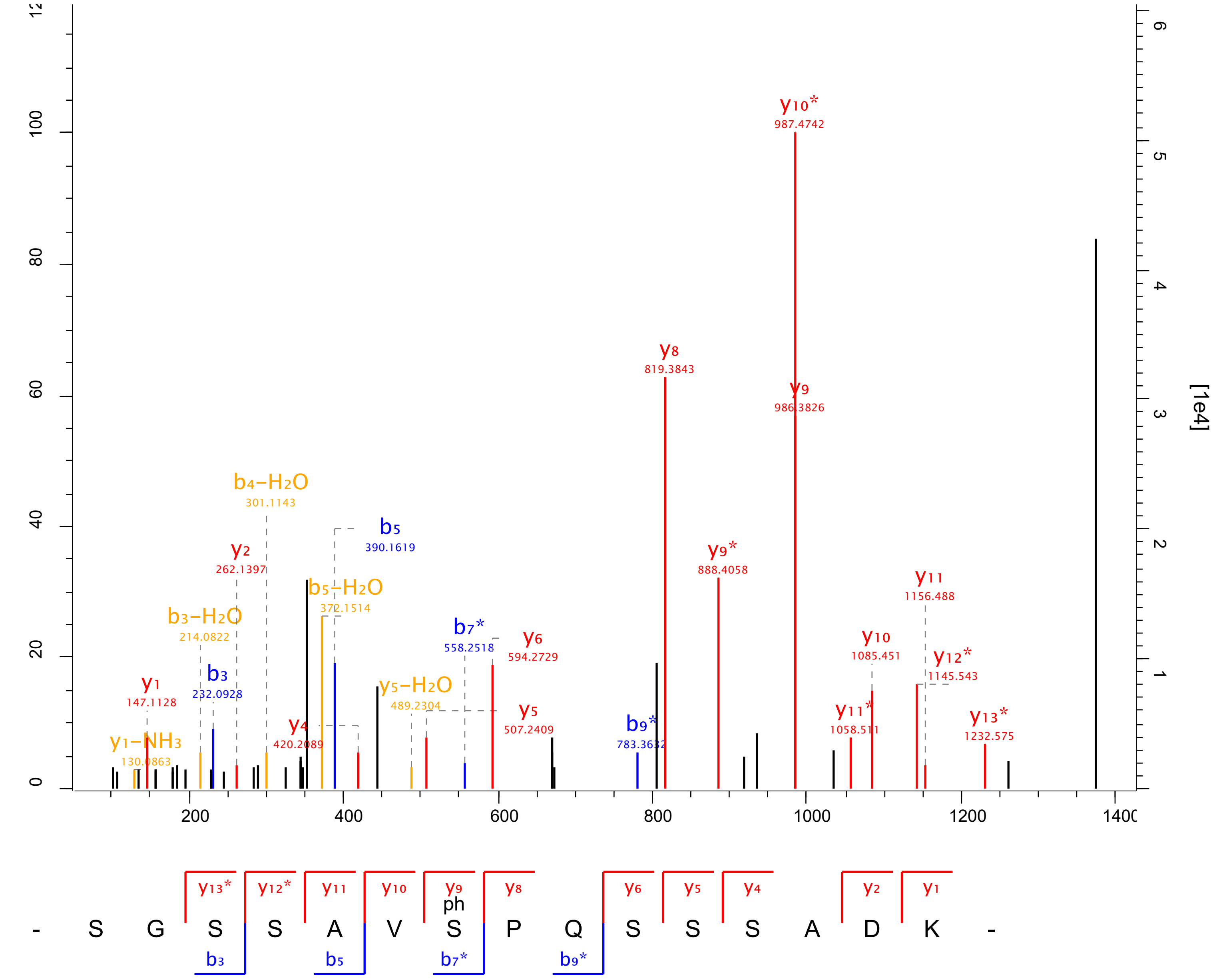

| Raw file | Scan | Method | Score | m/z |
|---|---|---|---|---|
| water_00_1 | 2936 | FTMS; HCD | 117 | 655.78 |

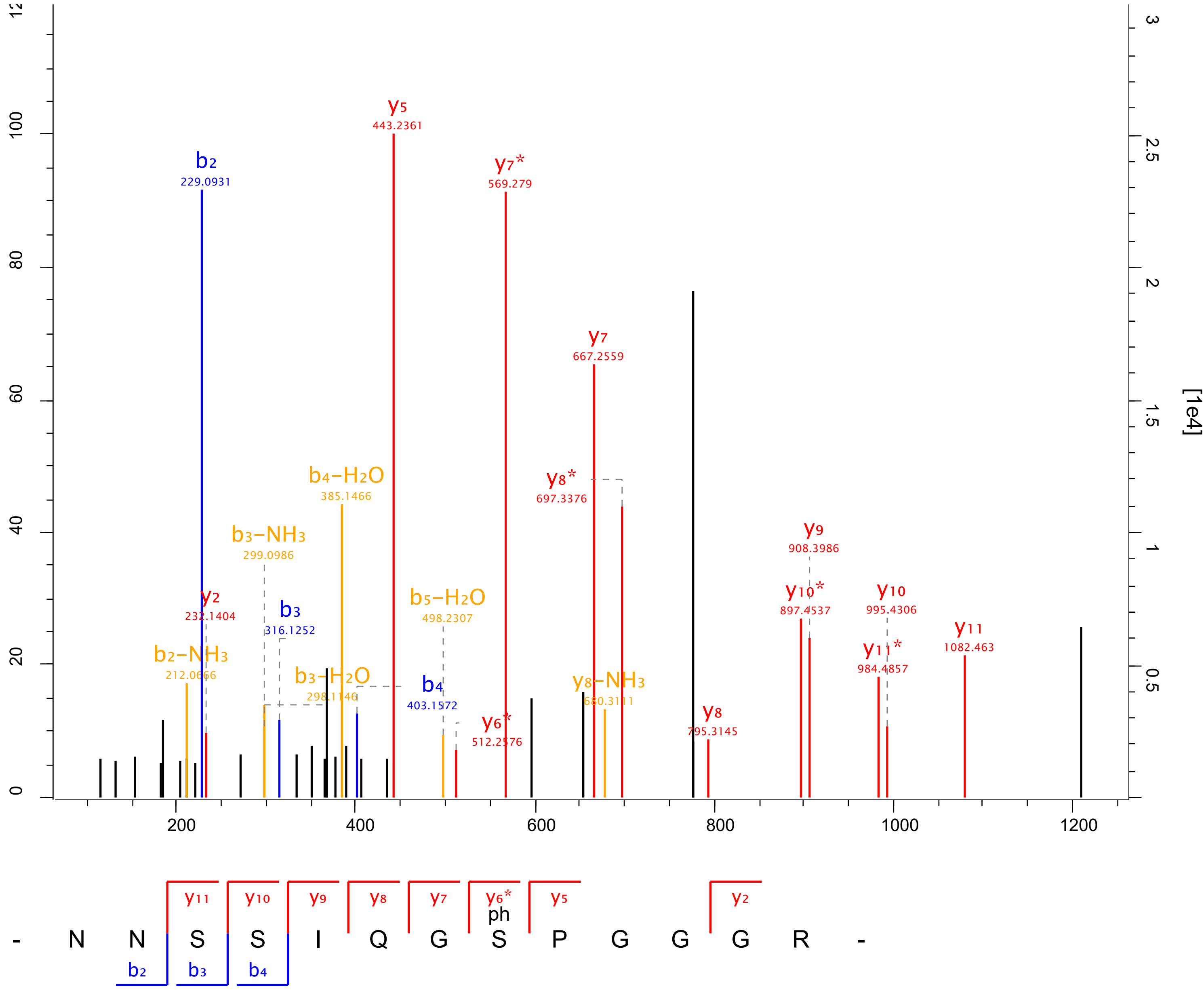

| Raw file | Scan | Method | Score | m/z |
| --- | --- | --- | --- | --- |
| water_00_1 | 2949 | FTMS; HCD | 92.87 | 508.23 |

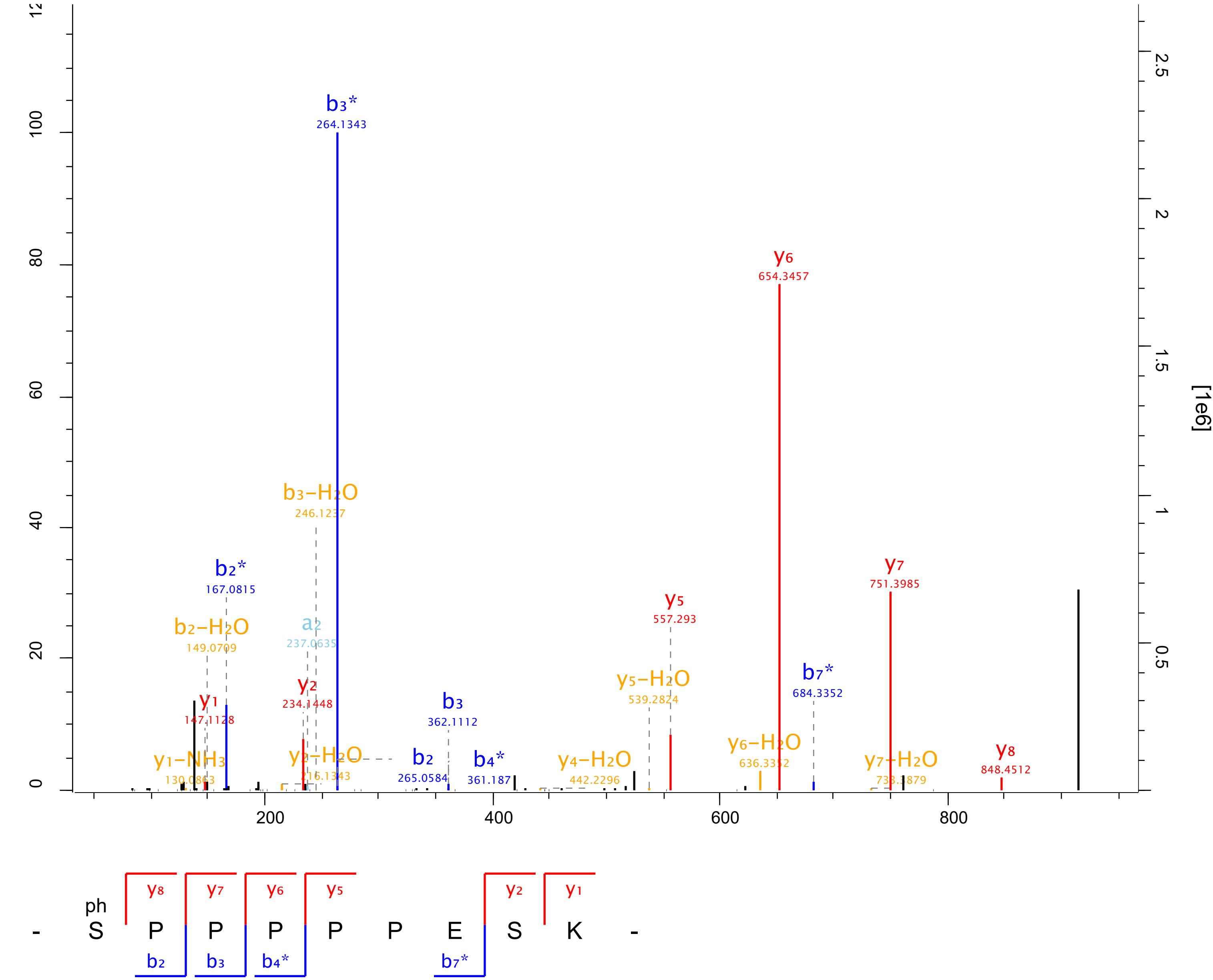

| Raw file | Scan | Method | Score | m/z |
|---|---|---|---|---|
| water_00_1 | 29508 | FTMS; HCD | 69.72 | 515.26 |

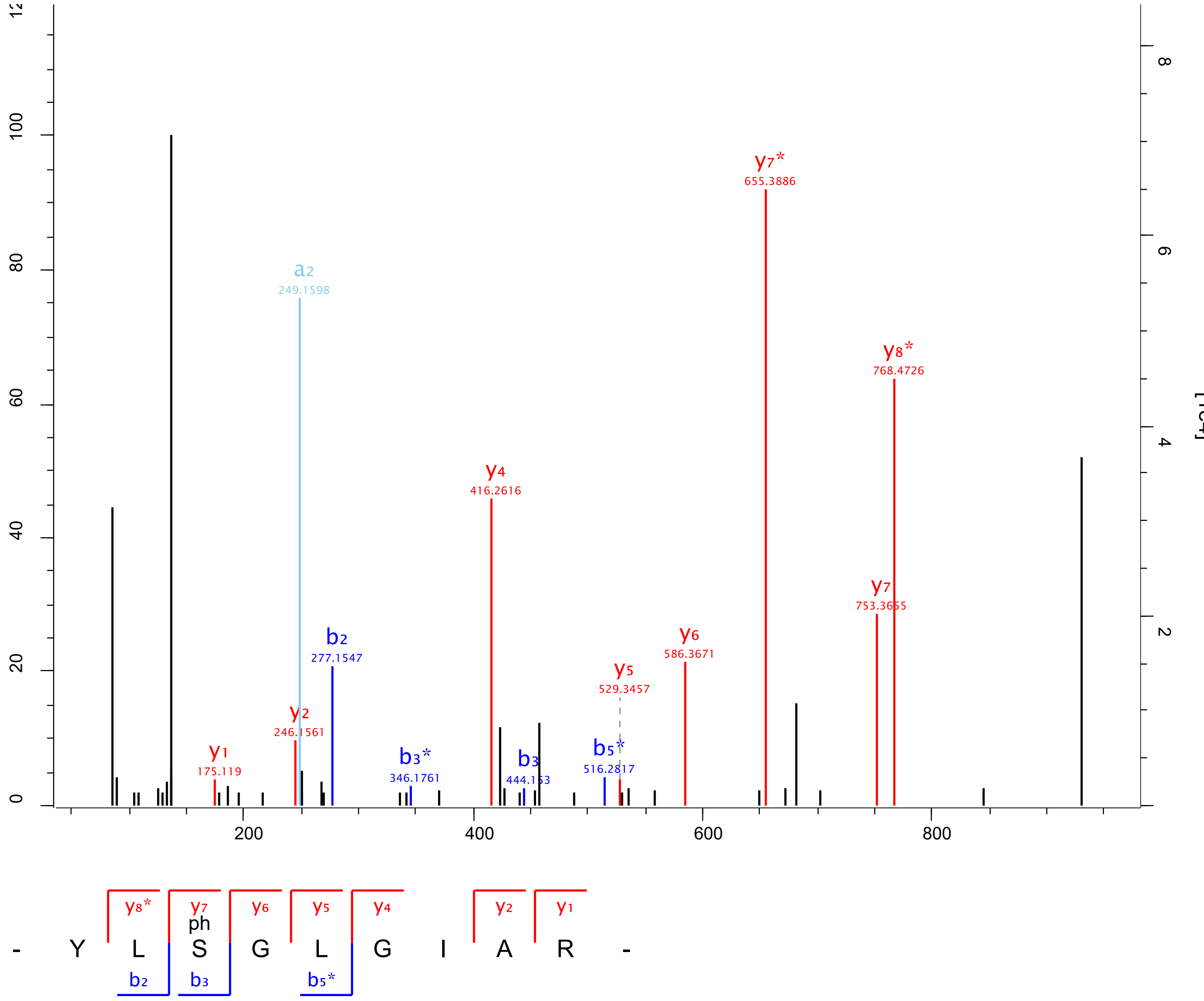

| Raw file | Scan | Method | Score | m/z |
| --- | --- | --- | --- | --- |
| water_00_1 | 29794 | FTMS; HCD | 135.91 | 846.4 |

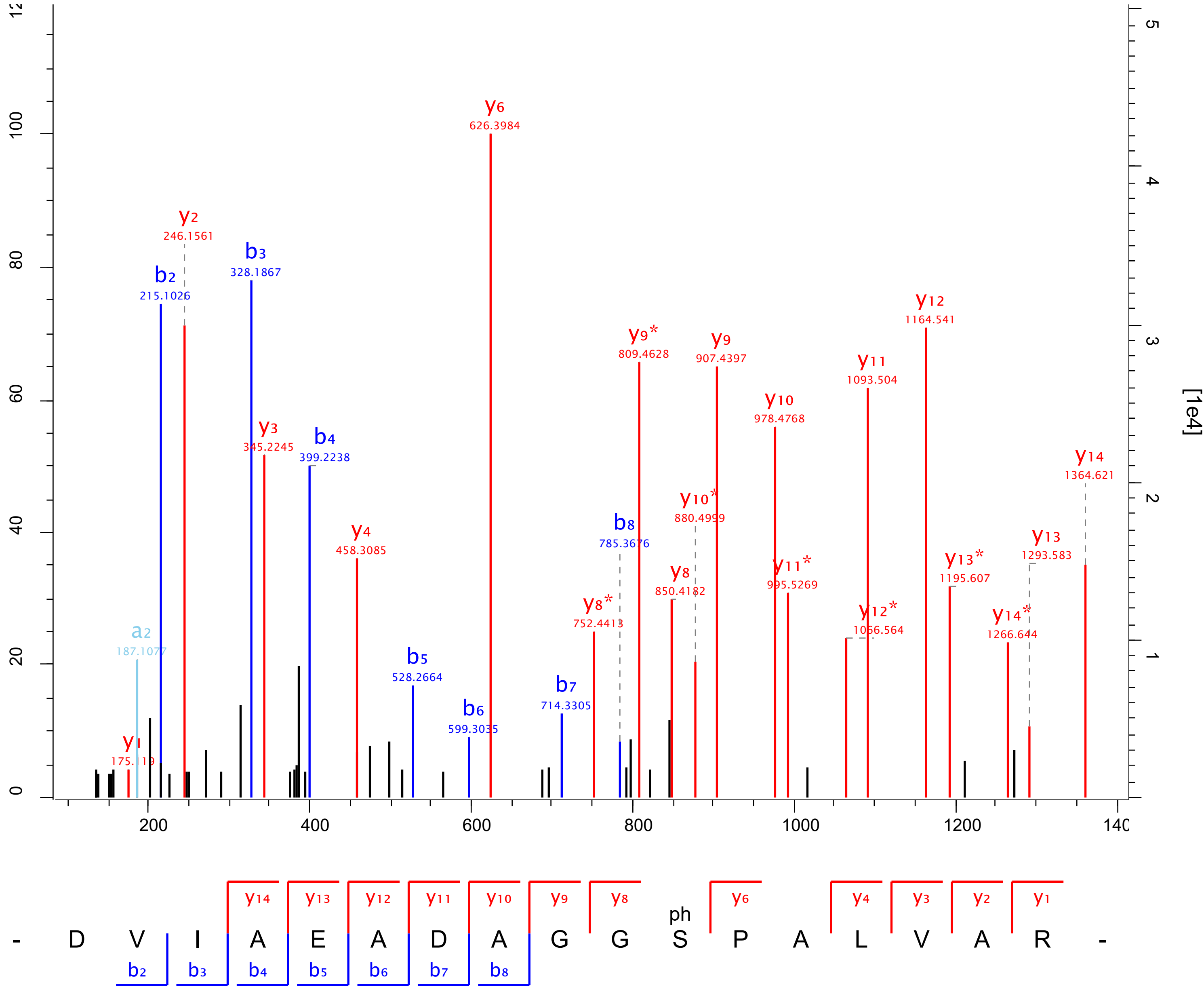

| Raw file | Scan | Method | Score | m/z |
|---|---|---|---|---|
| water_00_1 | 29859 | FTMS; HCD | 117.83 | 741.35 |

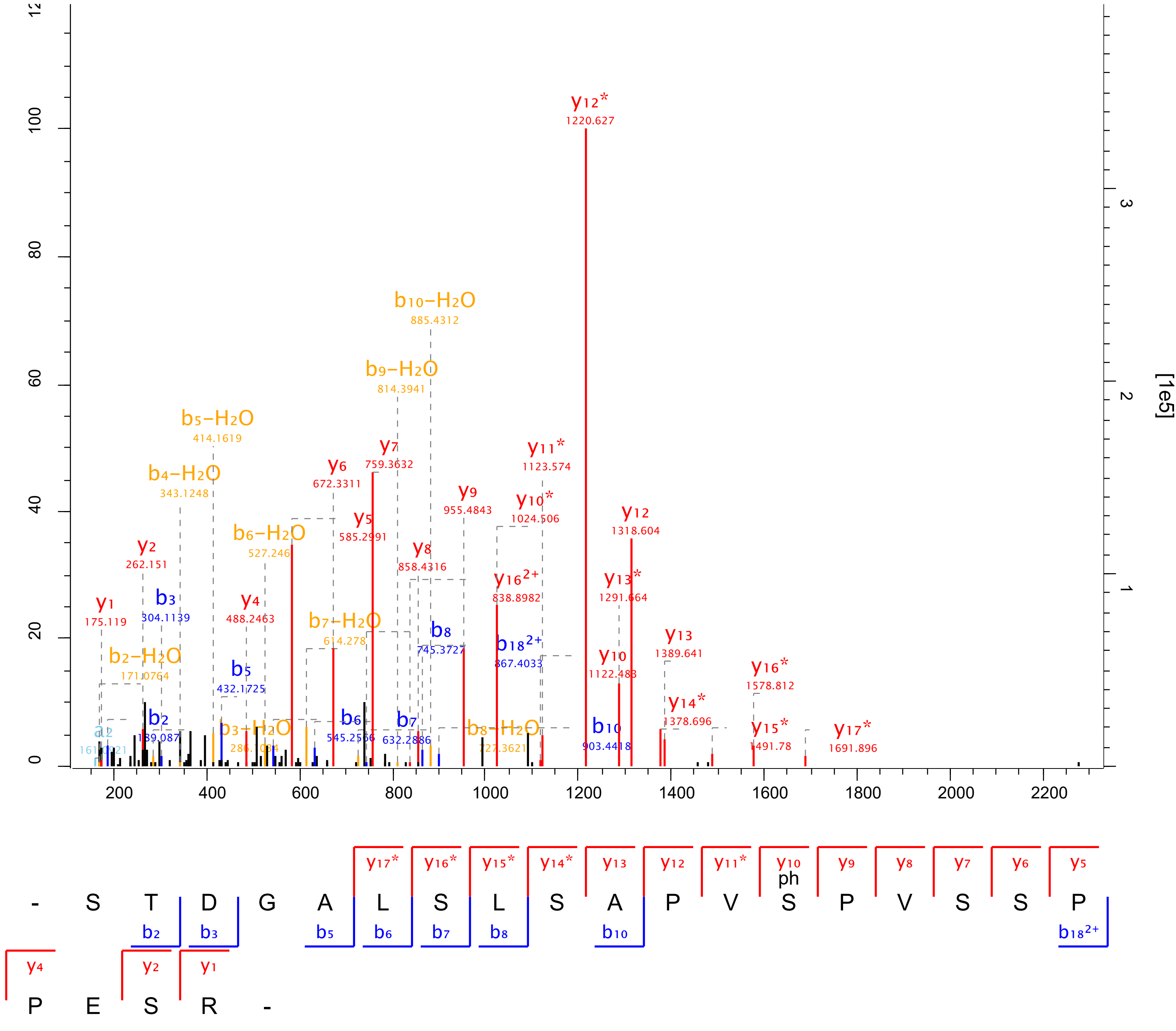

| Raw file | Scan | Method | Score | m/z |
|---|---|---|---|---|
| water_00_1 | 29863 | FTMS; HCD | 66.25 | 684.38 |

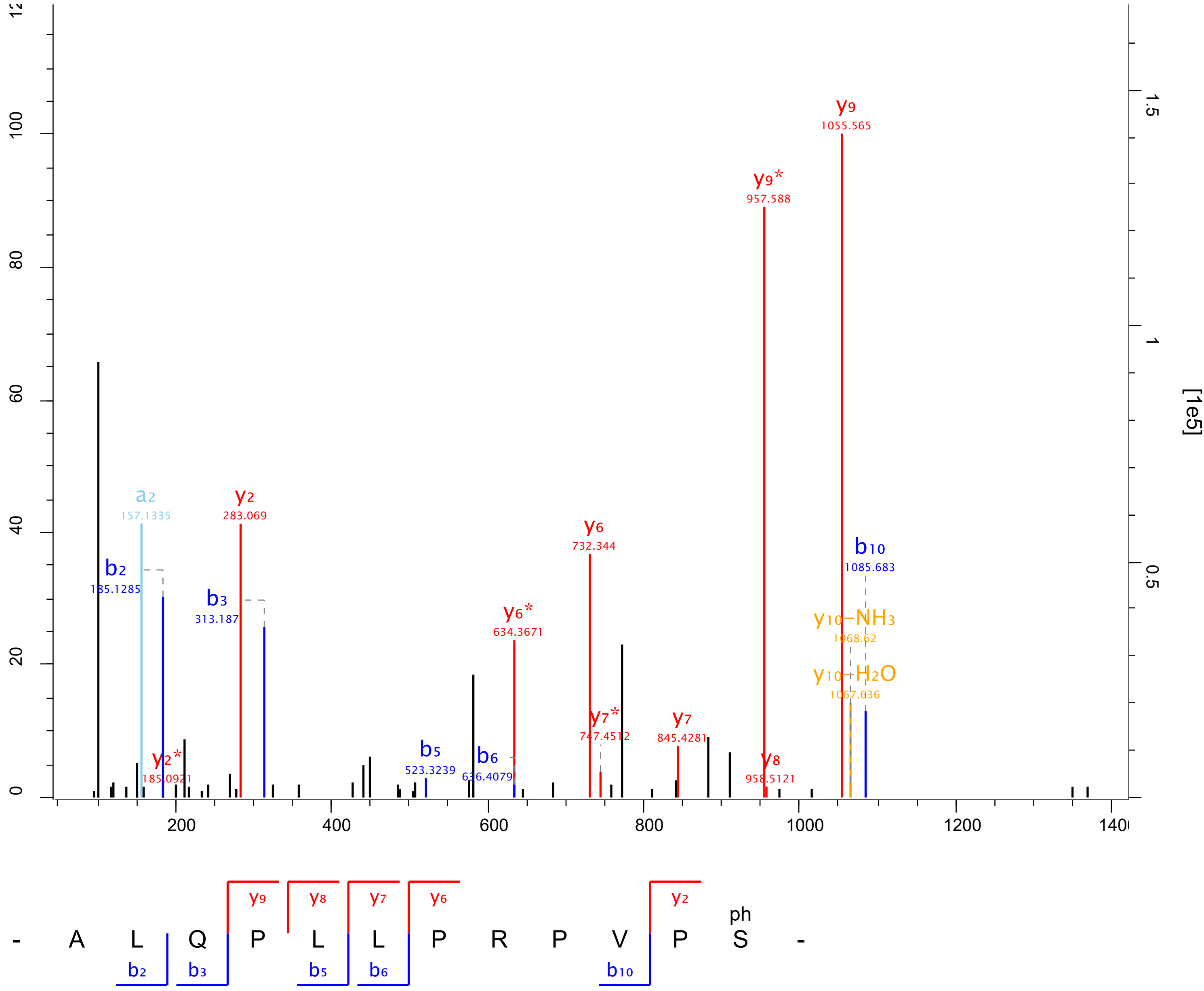

| Raw file | Scan | Method | Score | m/z |
| --- | --- | --- | --- | --- |
| water_00_1 | 29989 | FTMS; HCD | 46.56 | 1015.97 |

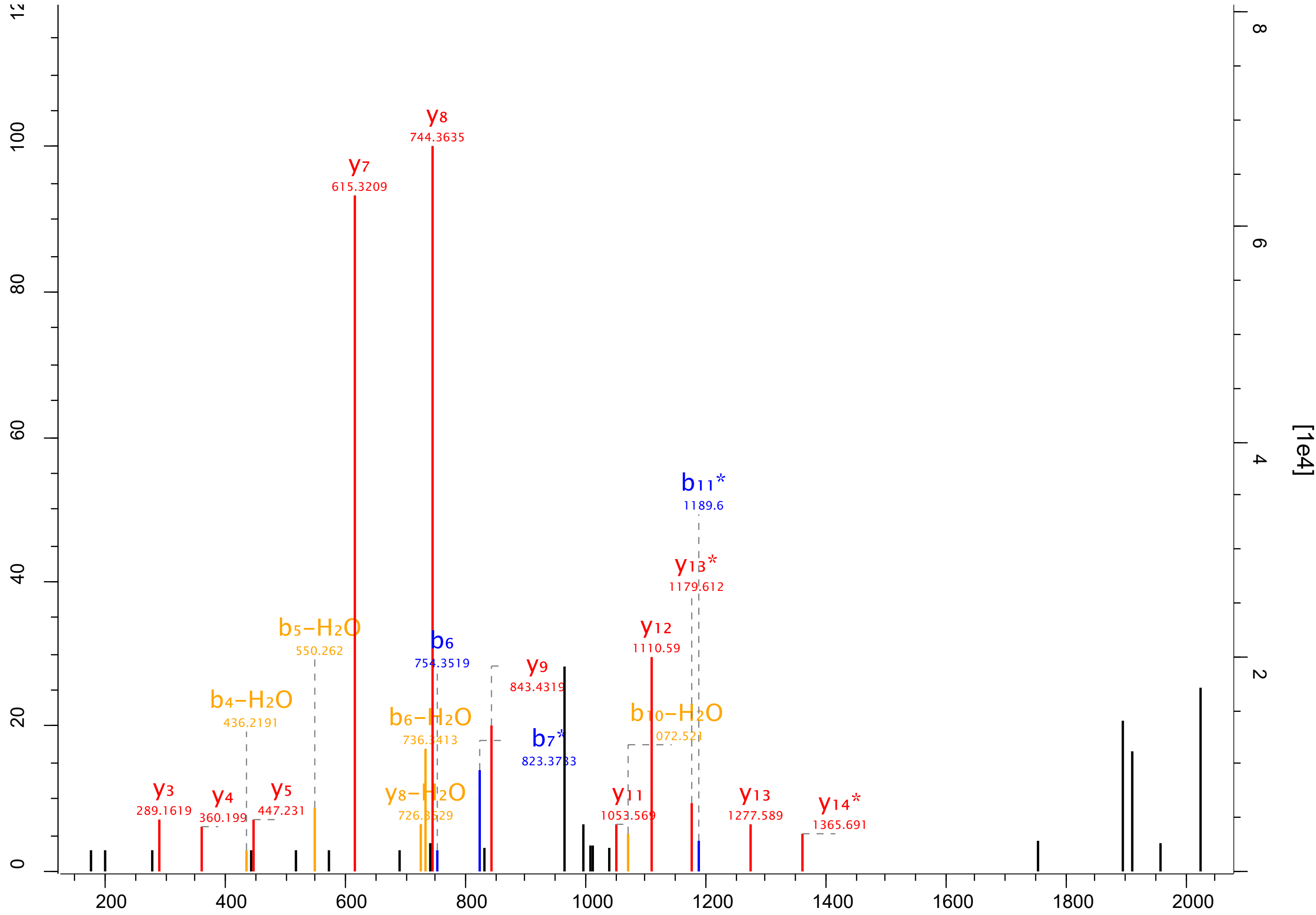


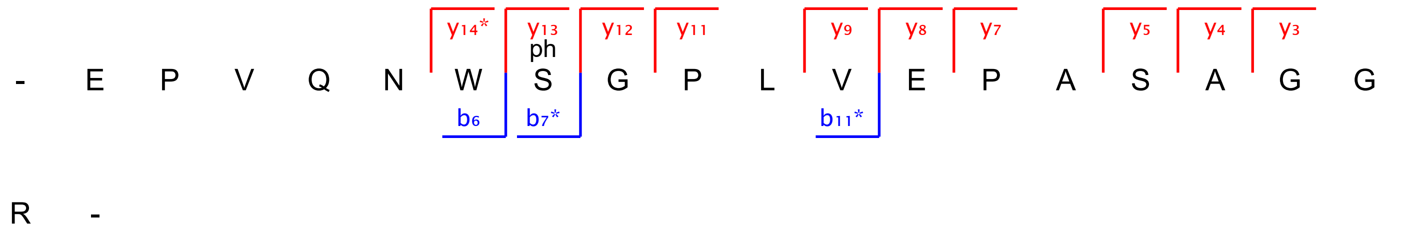

| Raw file | Scan | Method | Score | m/z |
| --- | --- | --- | --- | --- |
| water_00_1 | 2999 | FTMS; HCD | 52.58 | 709.27 |

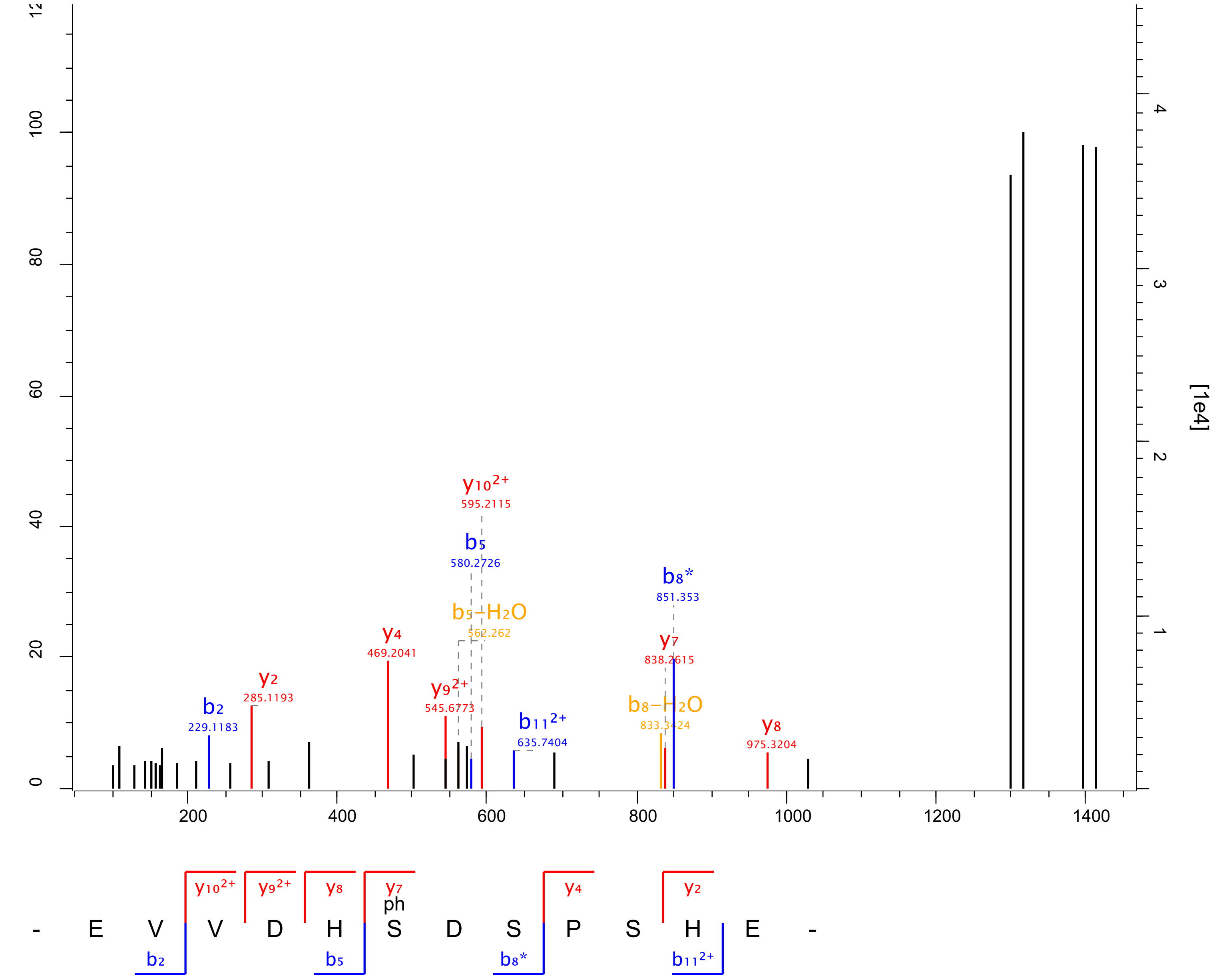

| Raw file | Scan | Method | Score | m/z |
|---|---|---|---|---|
| water_00_1 | 3041 | FTMS; HCD | 80.75 | 628.27 |

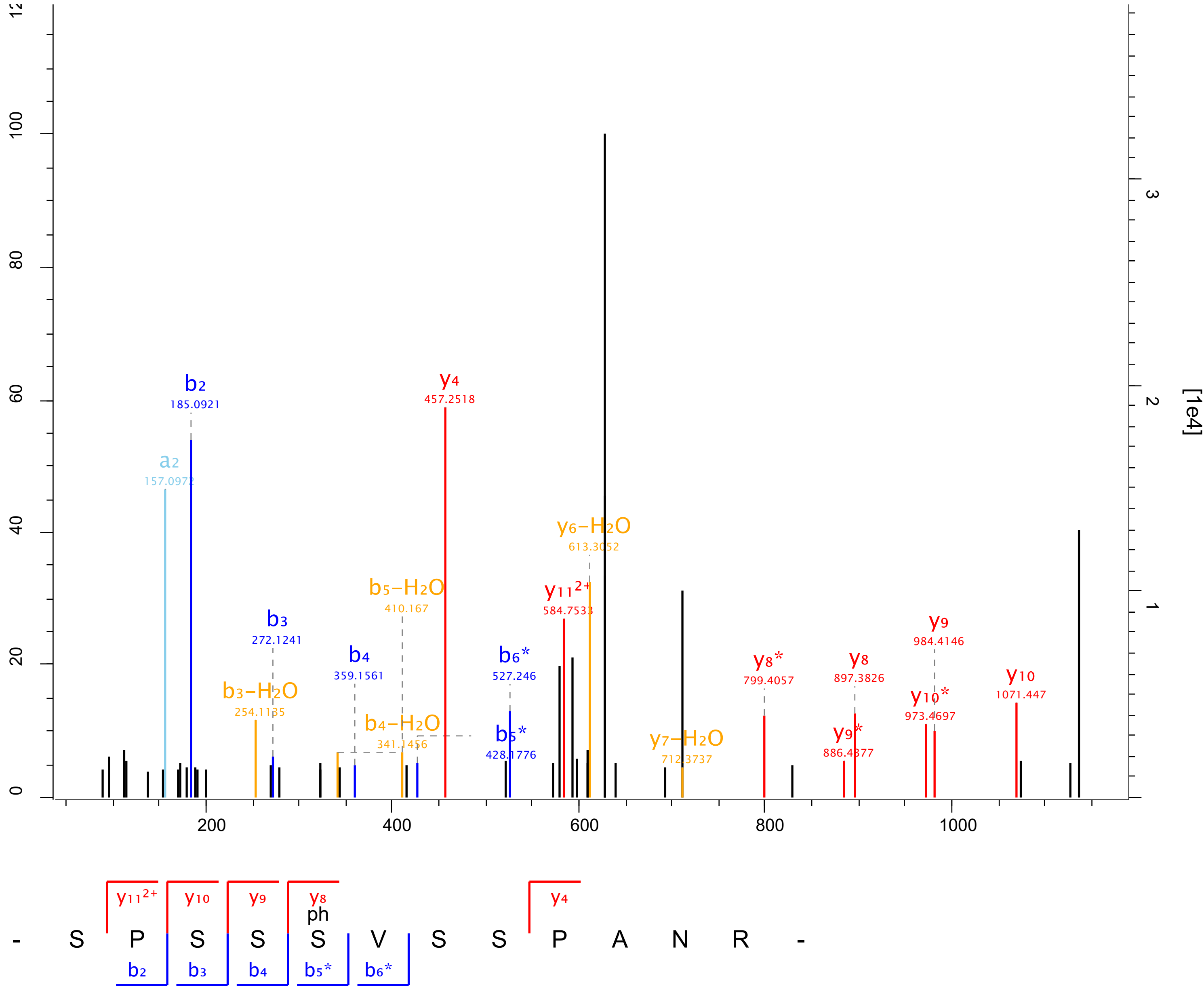

| Raw file | Scan | Method | Score | m/z |
|---|---|---|---|---|
| water_00_1 | 3078 | FTMS; HCD | 47.3 | 645.27 |

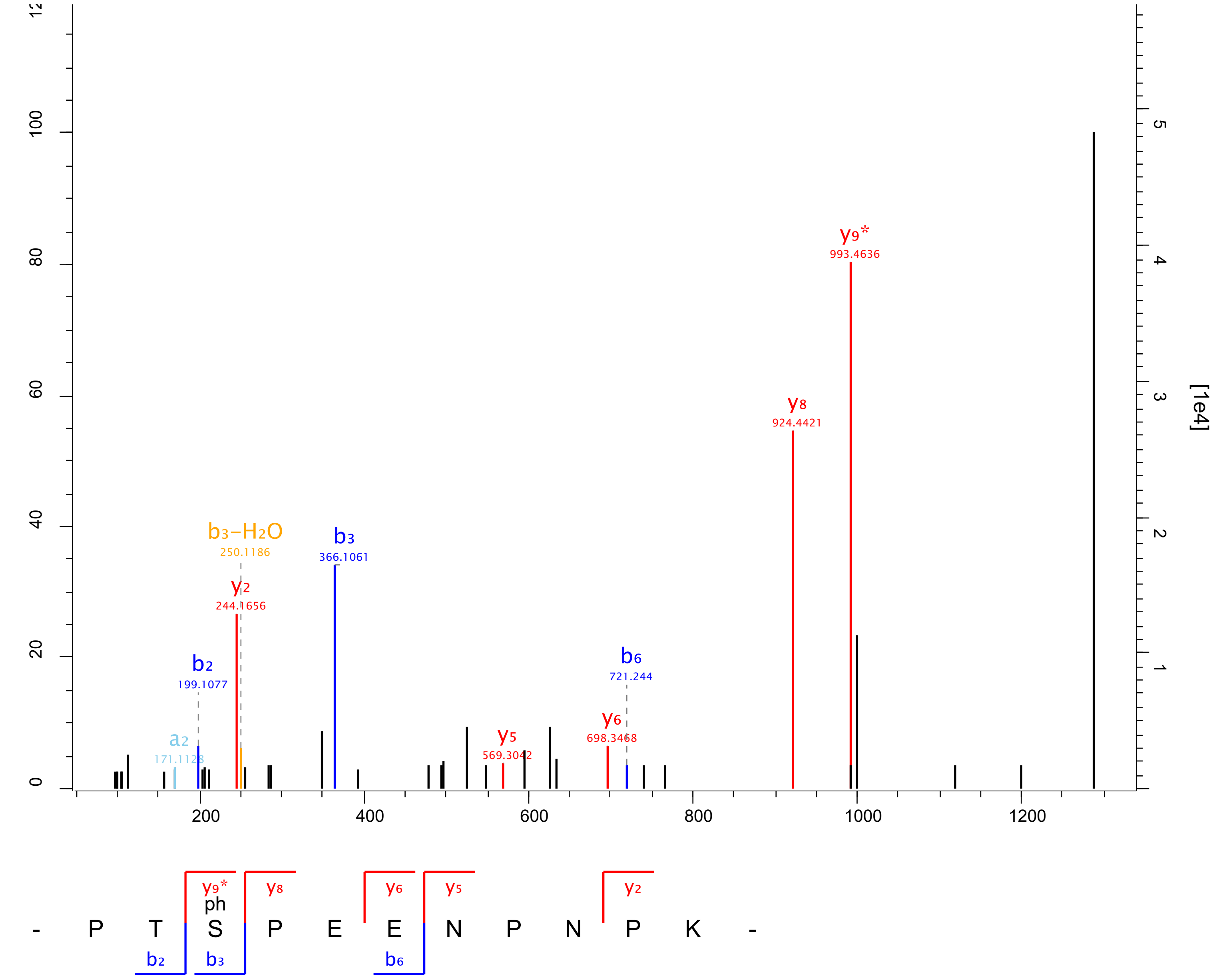

| Raw file | Scan | Method | Score | m/z |
|---|---|---|---|---|
| water_00_1 | 30940 | FTMS; HCD | 149.73 | 869.4 |

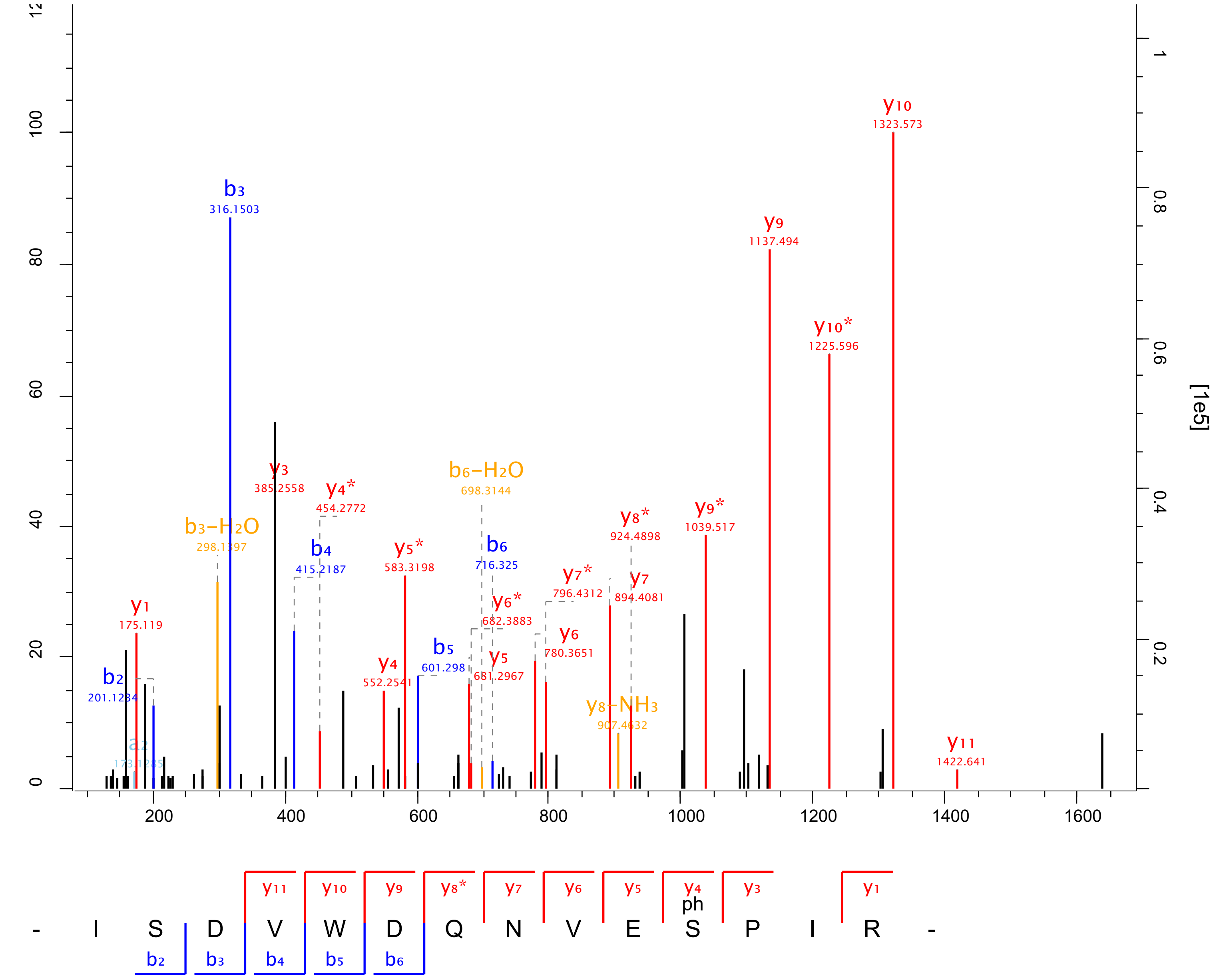

| Raw file | Scan | Method | Score | m/z |
|---|---|---|---|---|
| water_00_1 | 31047 | FTMS; HCD | 176.85 | 876.41 |

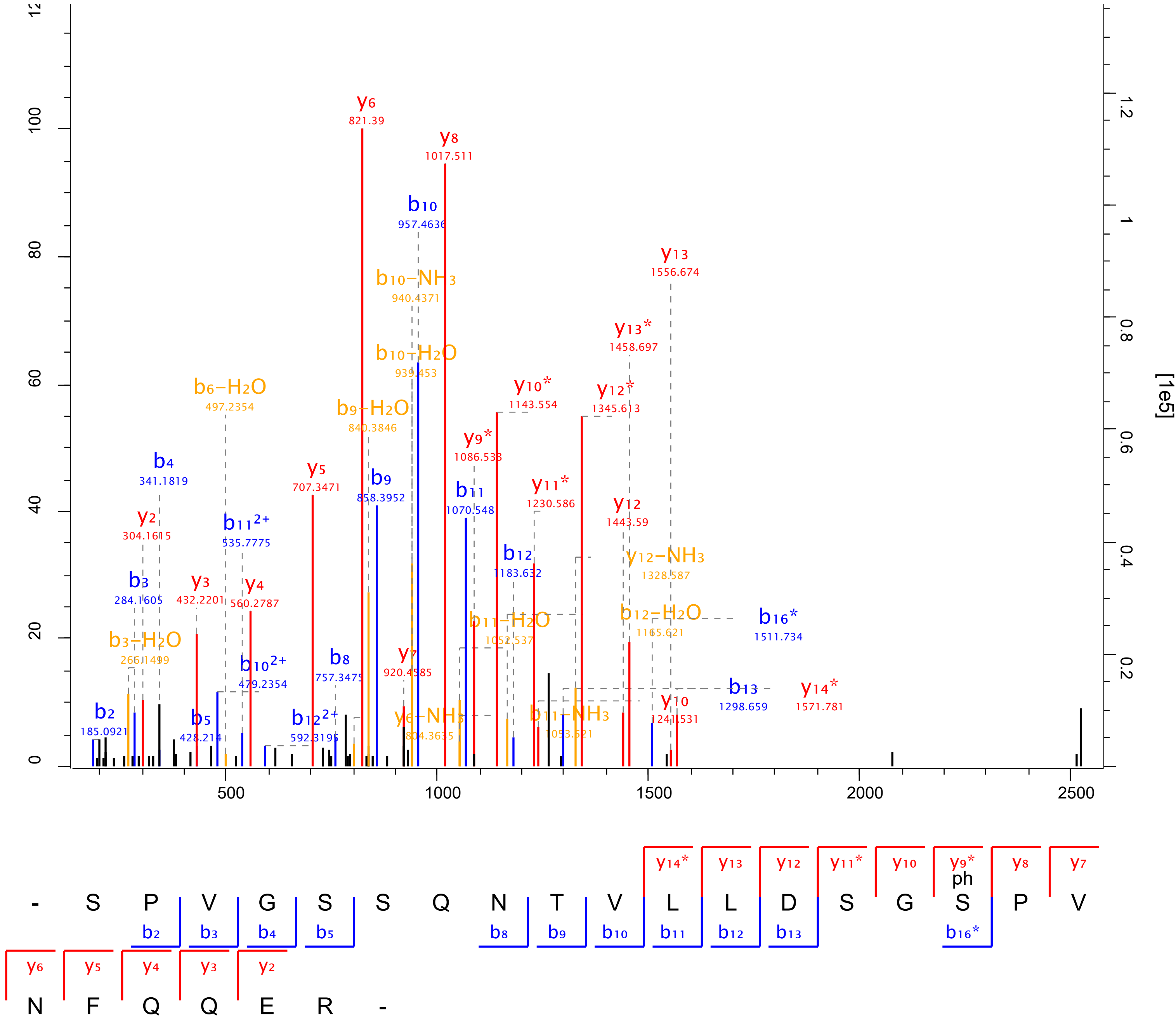

| Raw file | Scan | Method | Score | m/z |
| --- | --- | --- | --- | --- |
| water_00_1 | 3123 | FTMS; HCD | 82.85 | 593.76 |

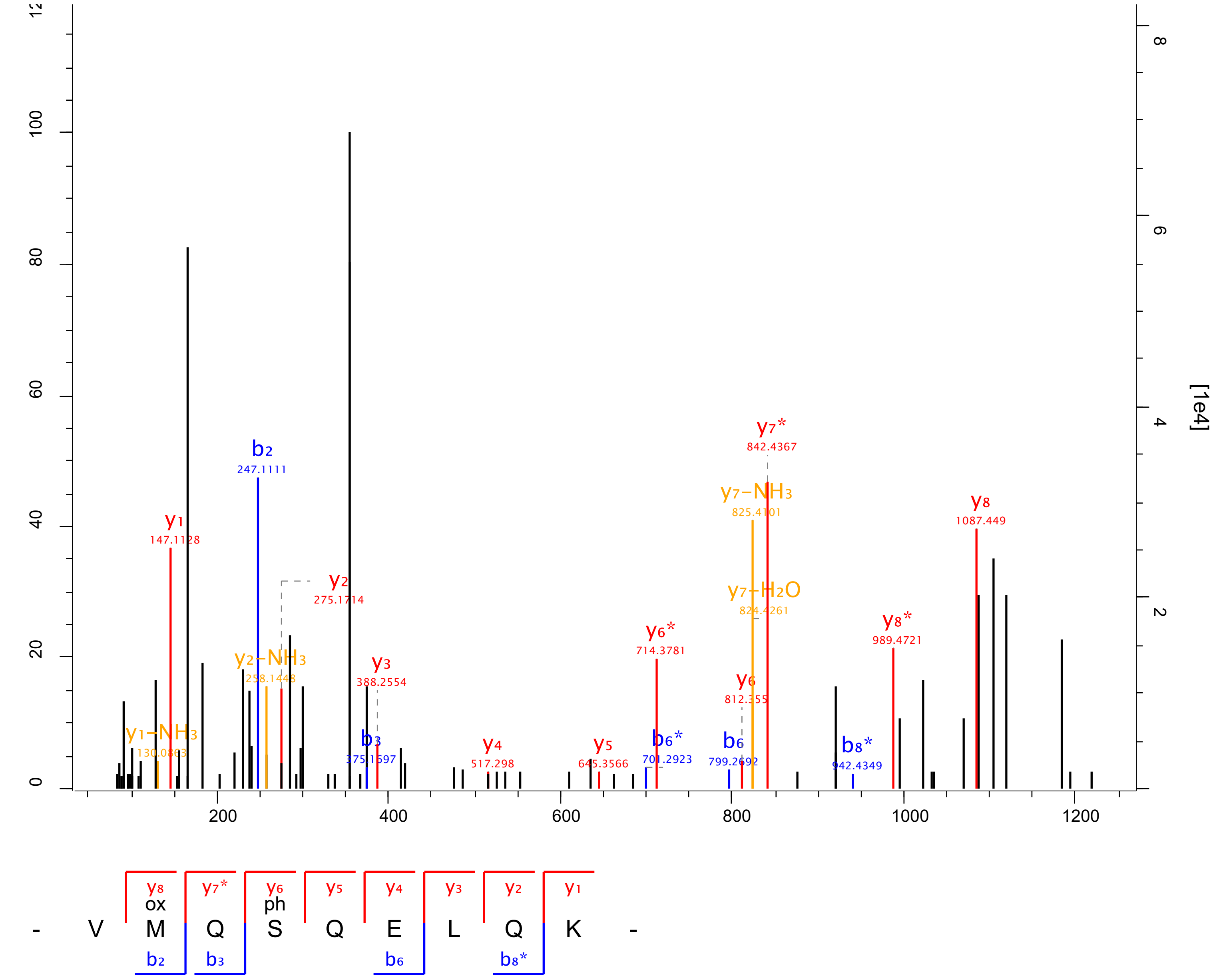

| Raw file | Scan | Method | Score | m/z |
| --- | --- | --- | --- | --- |
| water_00_1 | 3154 | FTMS; HCD | 134.04 | 632.9 |

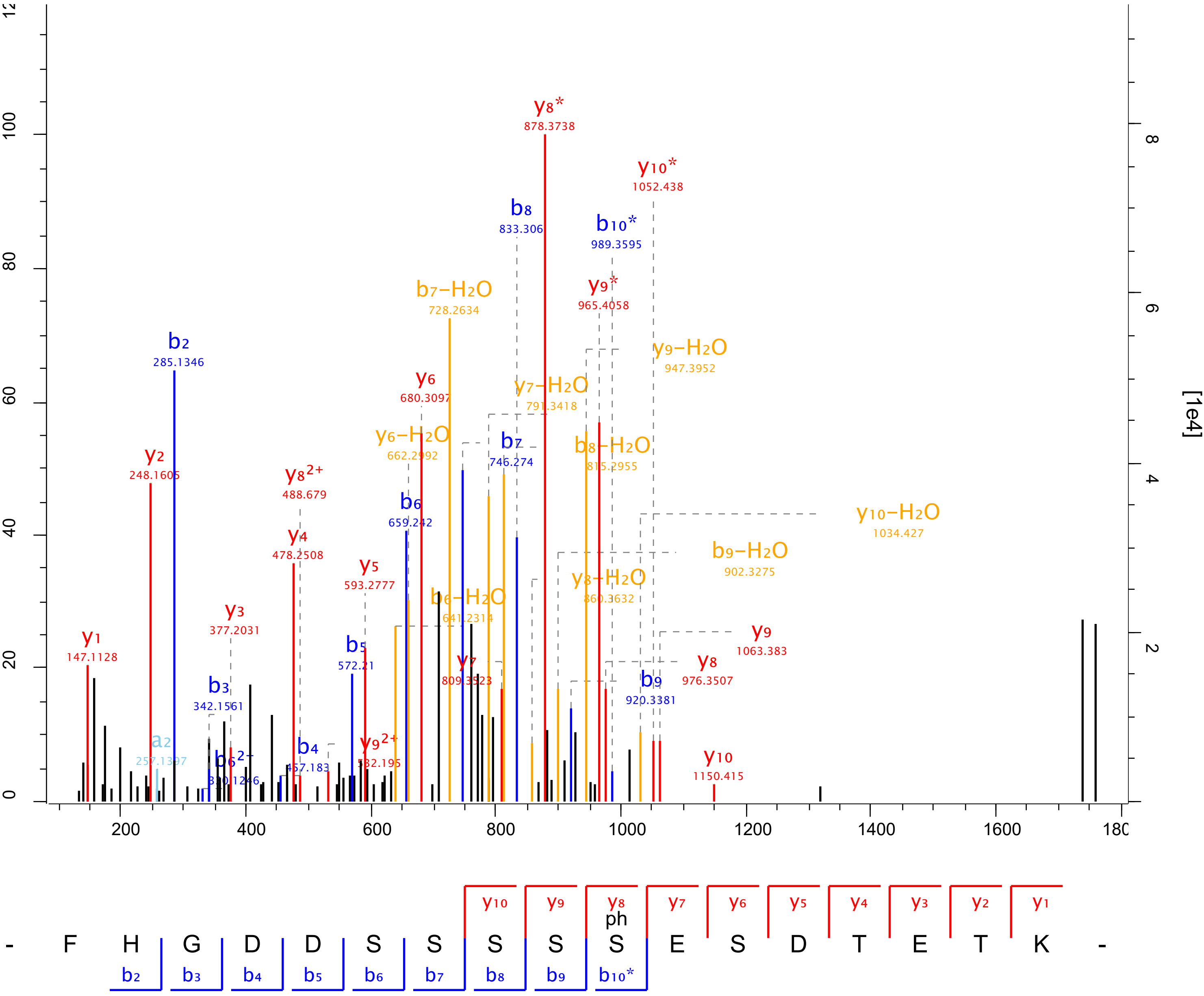

| Raw file | Scan | Method | Score | m/z |
|---|---|---|---|---|
| water_00_1 | 31767 | FTMS; HCD | 93.37 | 769.9 |

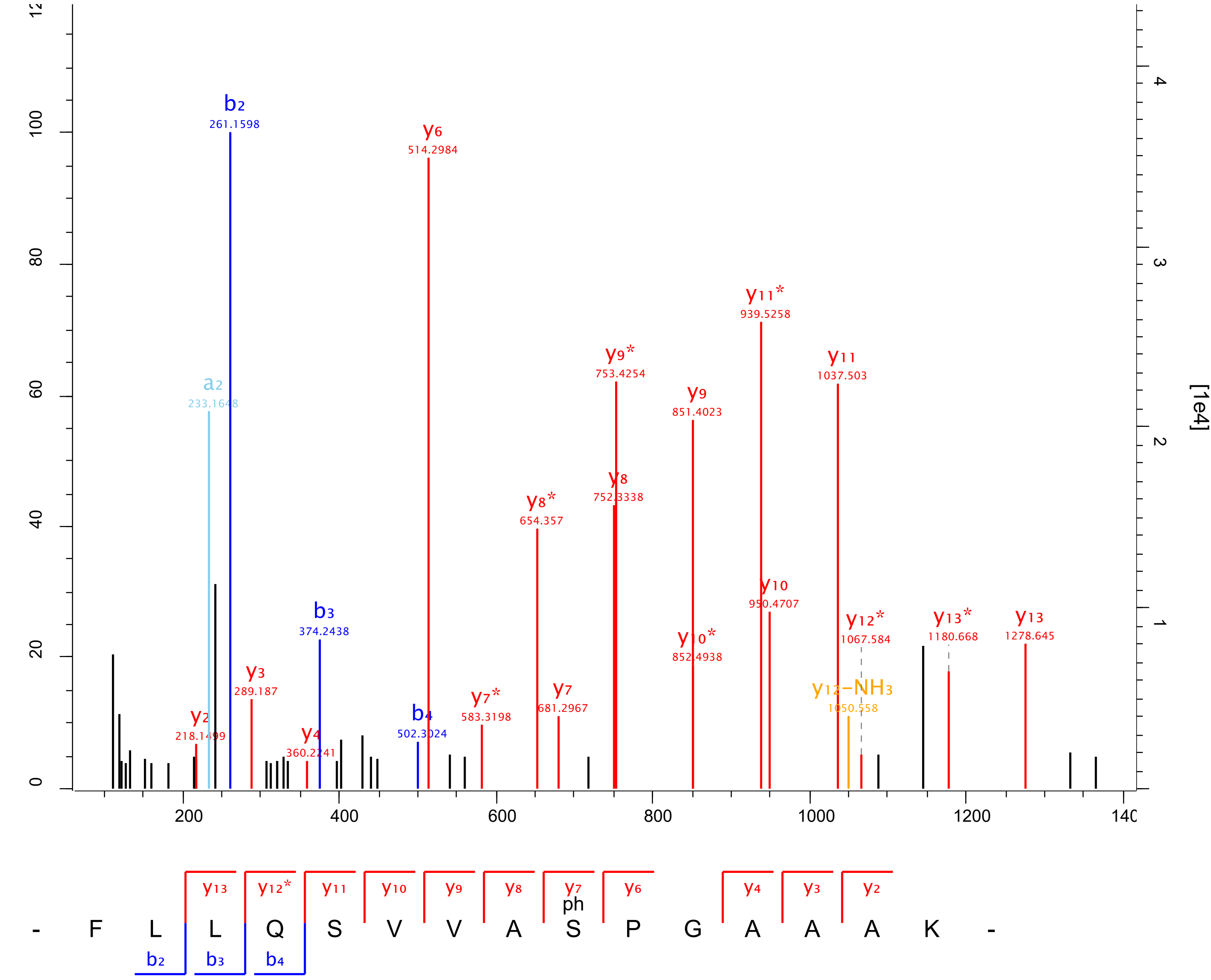

| Raw file | Scan | Method | Score | m/z |
| --- | --- | --- | --- | --- |
| water_00_1 | 3212 | FTMS; HCD | 56.51 | 670.3 |

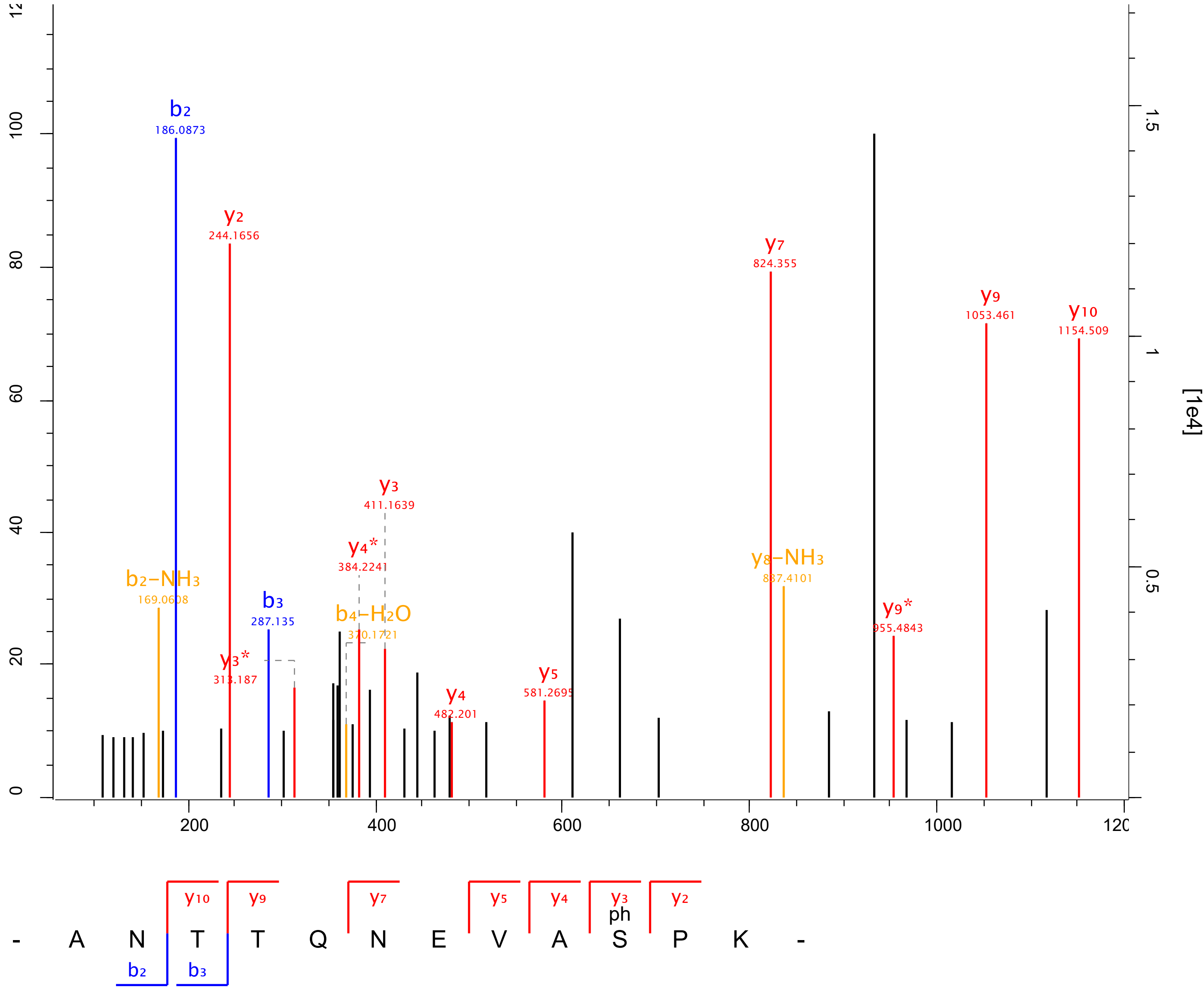

| Raw file | Scan | Method | Score | m/z |
|---|---|---|---|---|
| water_00_1 | 32454 | FTMS; HCD | 121.95 | 844.83 |

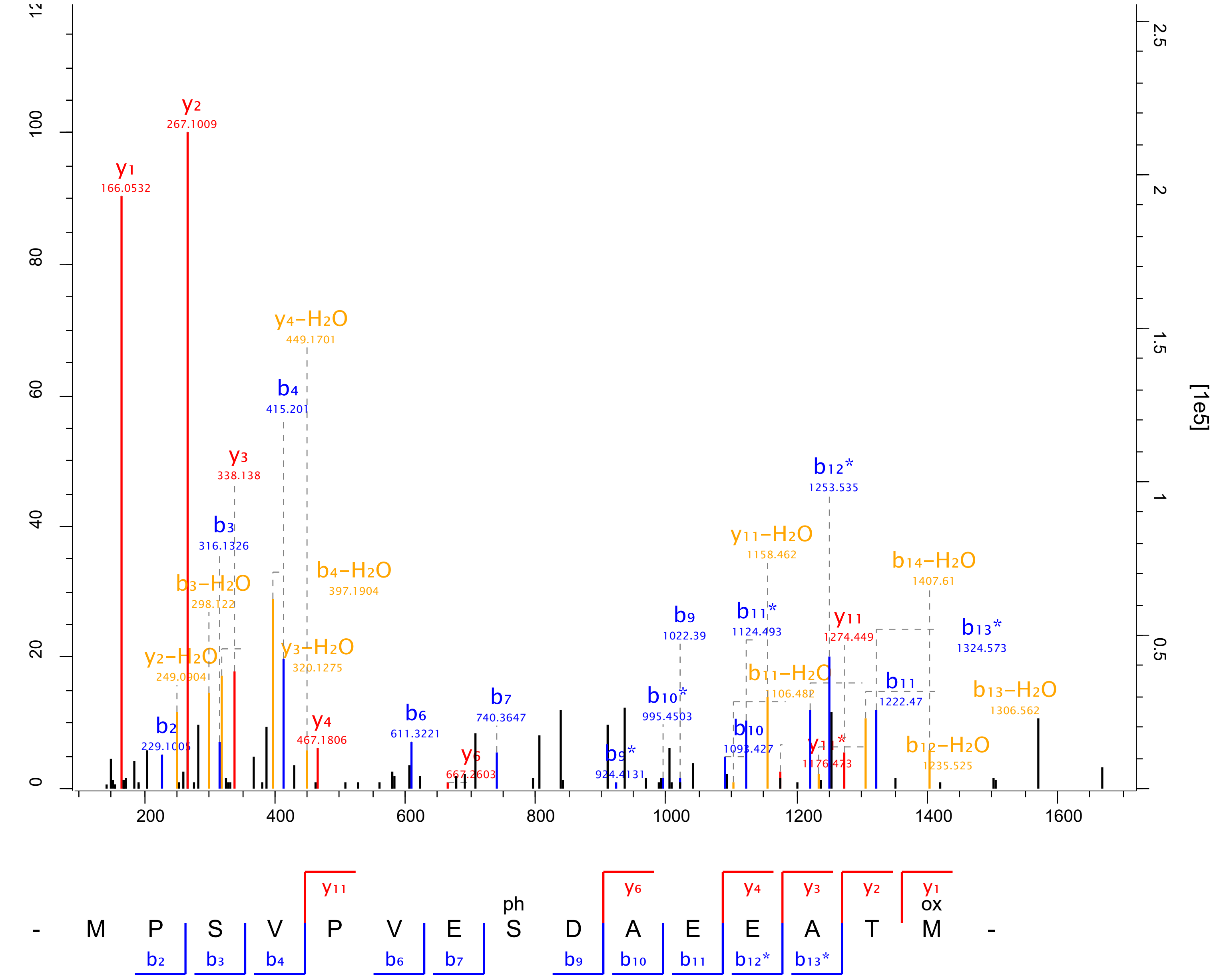

| Raw file | Scan | Method | Score | m/z |
|---|---|---|---|---|
| water_00_1 | 3291 | FTMS; HCD | 44.61 | 563.24 |

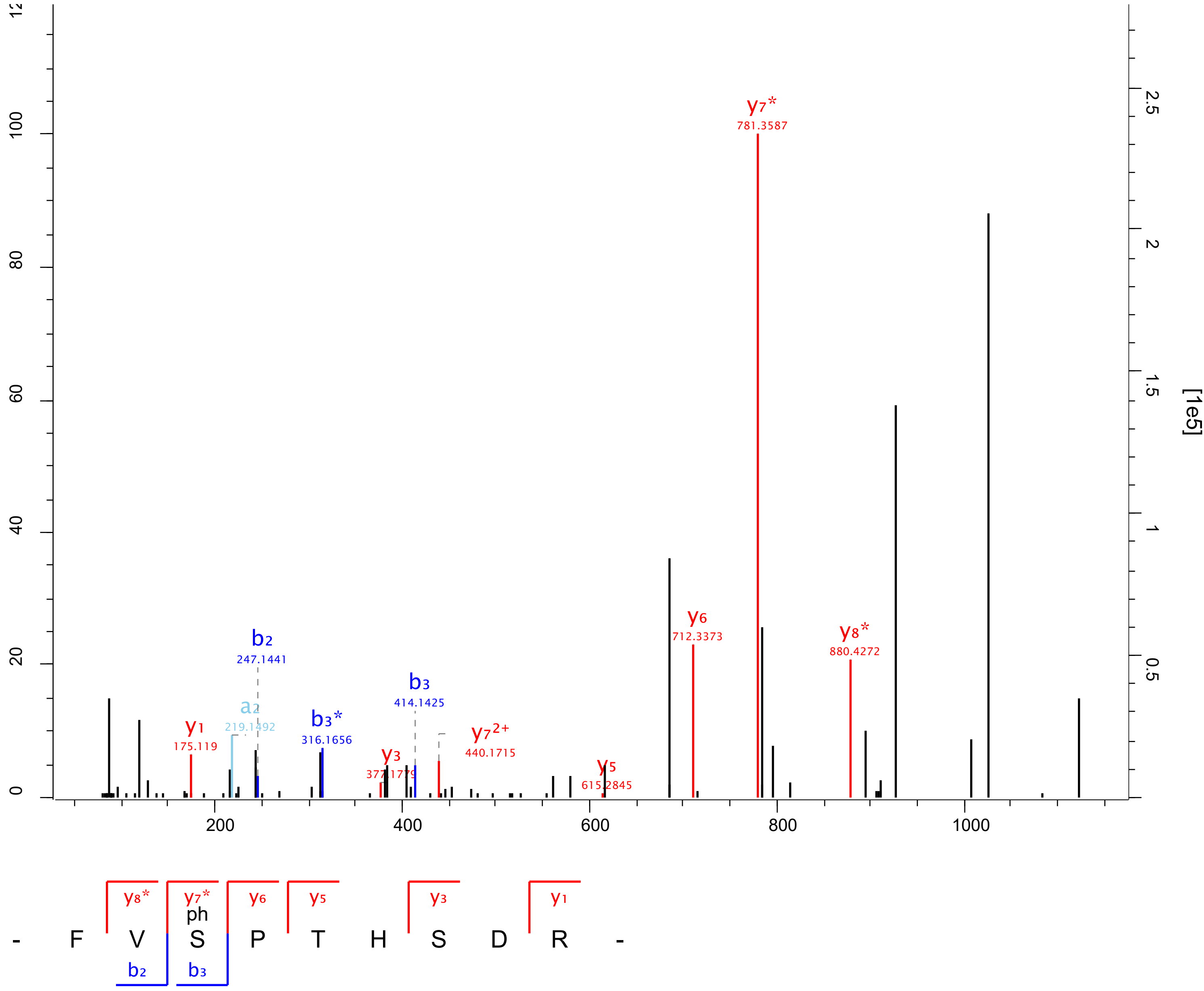

| Raw file | Scan | Method | Score | m/z |
| --- | --- | --- | --- | --- |
| water_00_1 | 33154 | FTMS; HCD | 51.52 | 806.72 |

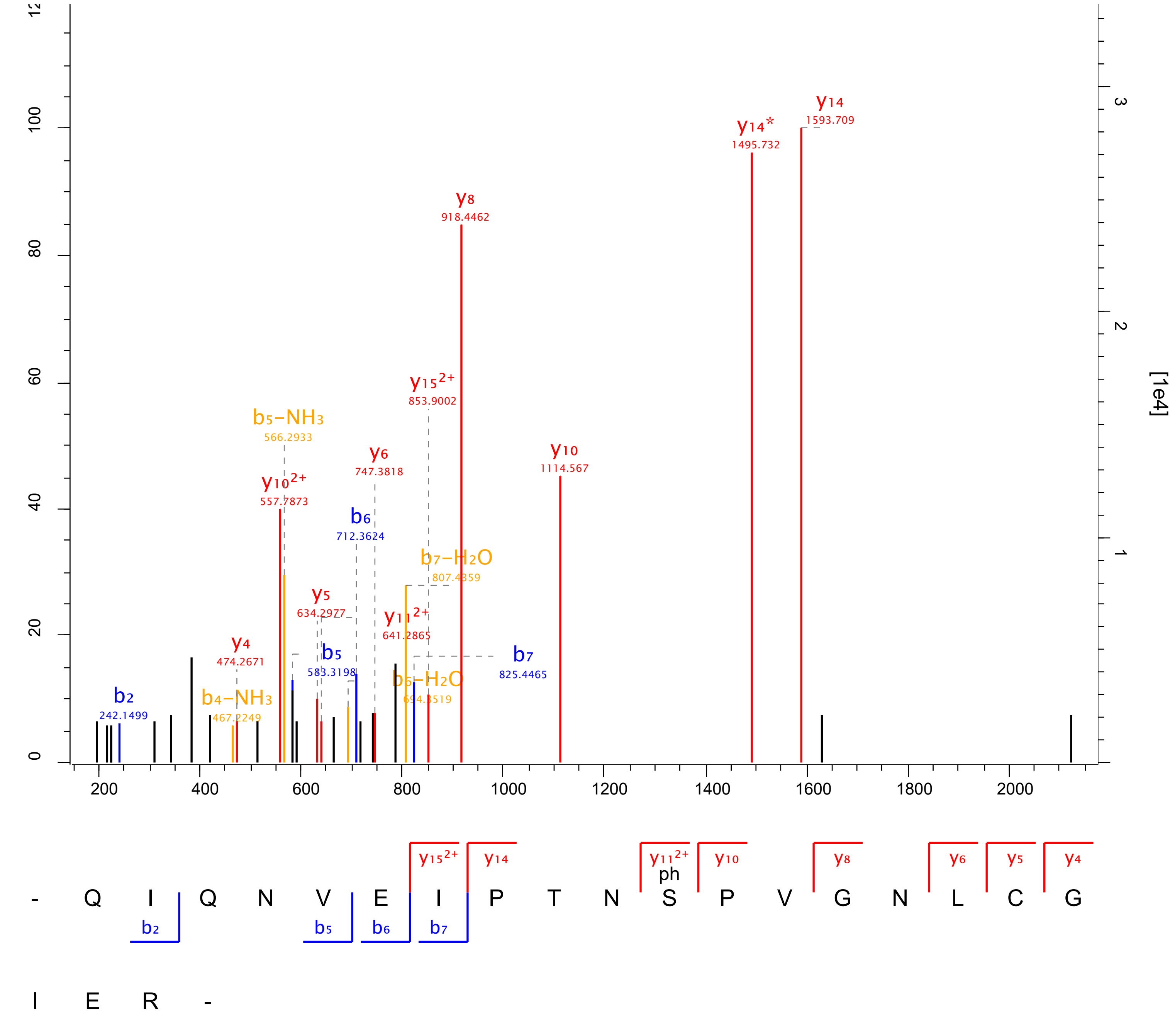

| Raw file | Scan | Method | Score | m/z |
| --- | --- | --- | --- | --- |
| water_00_1 | 33173 | FTMS; HCD | 47.84 | 535.18 |

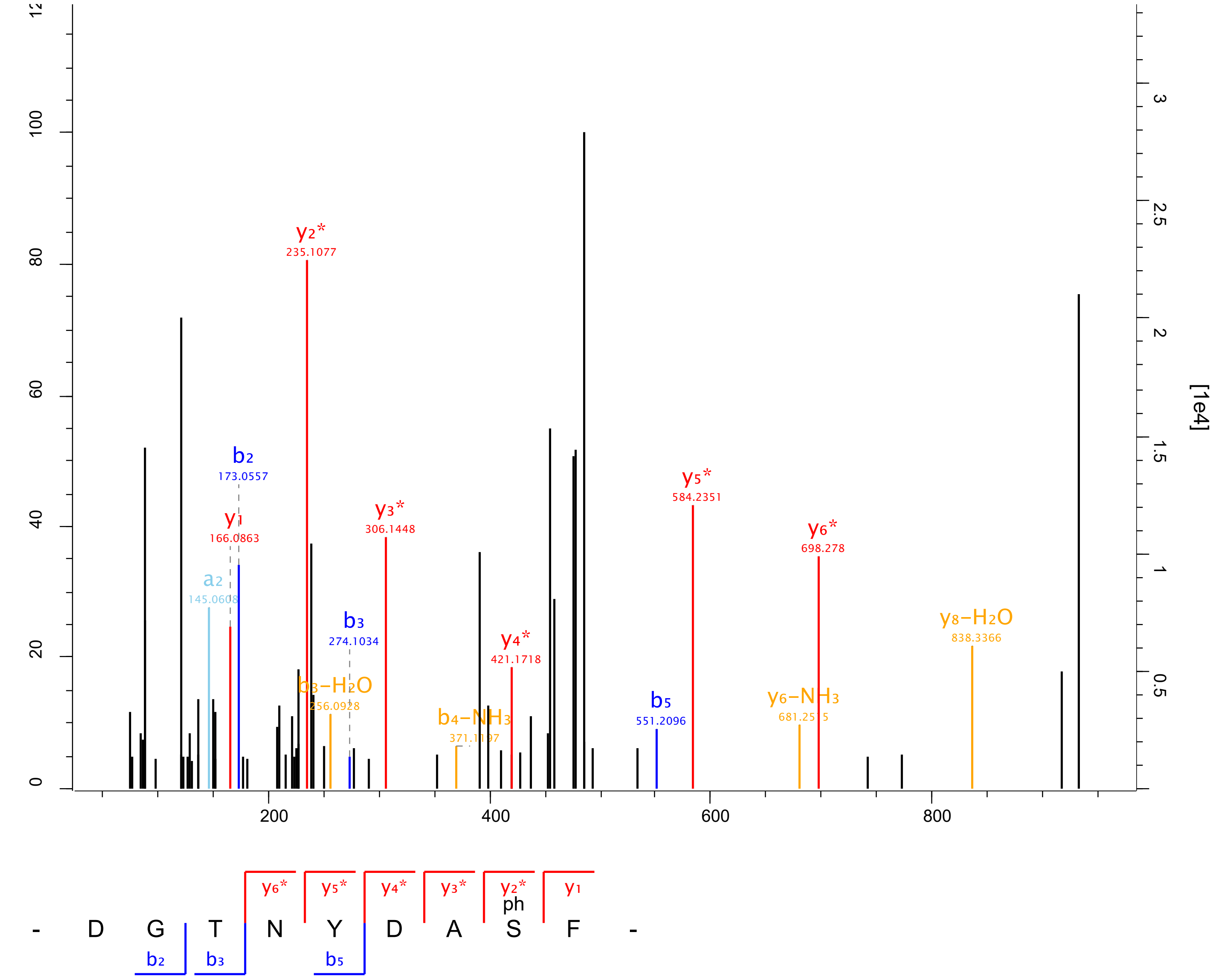

| Raw file | Scan | Method | Score | m/z |
|---|---|---|---|---|
| water_00_1 | 33712 | FTMS; HCD | 52.7 | 869.39 |

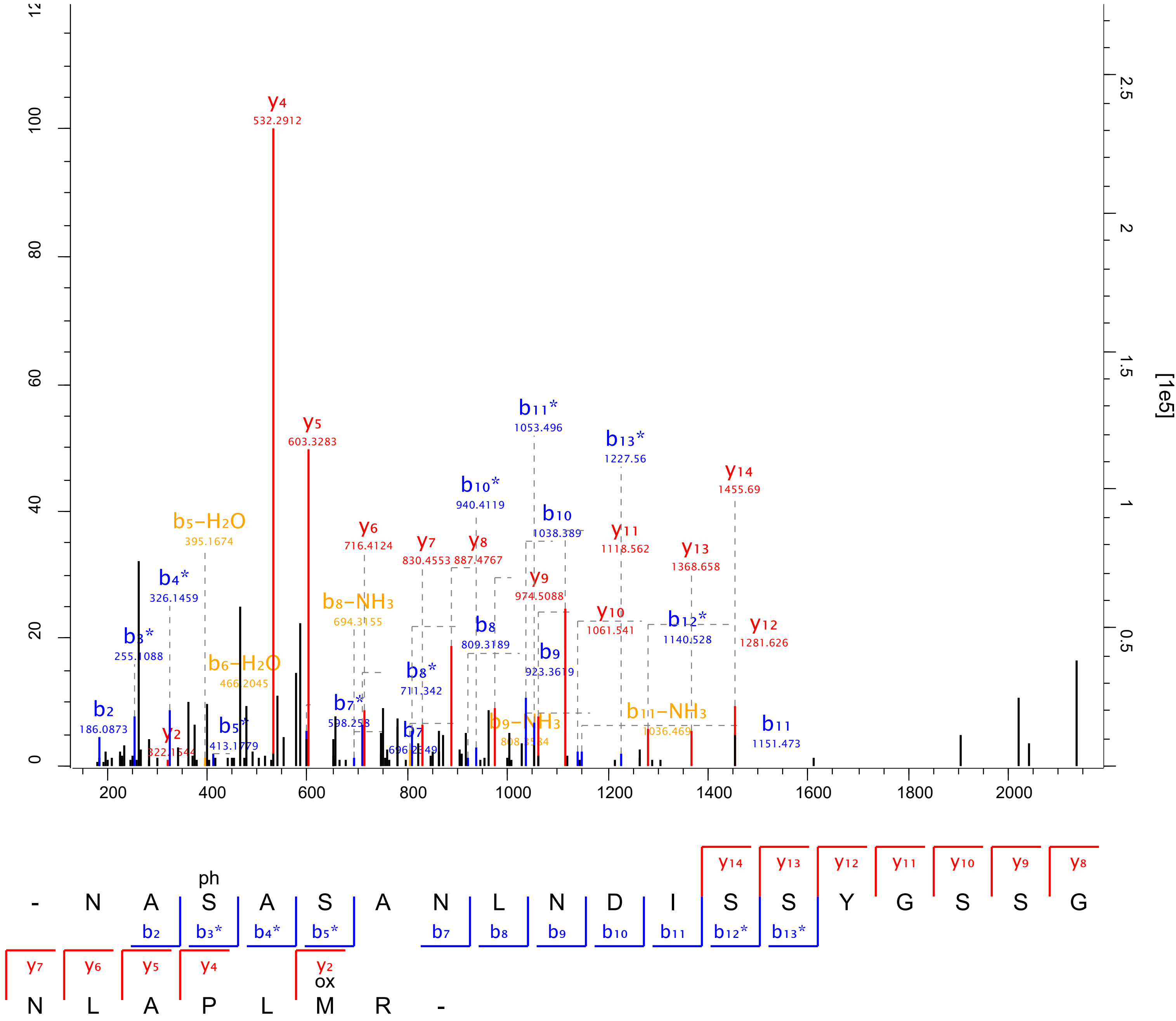

| Raw file | Scan | Method | Score | m/z |
|---|---|---|---|---|
| water_00_1 | 33803 | FTMS; HCD | 402.59 | 1070.96 |

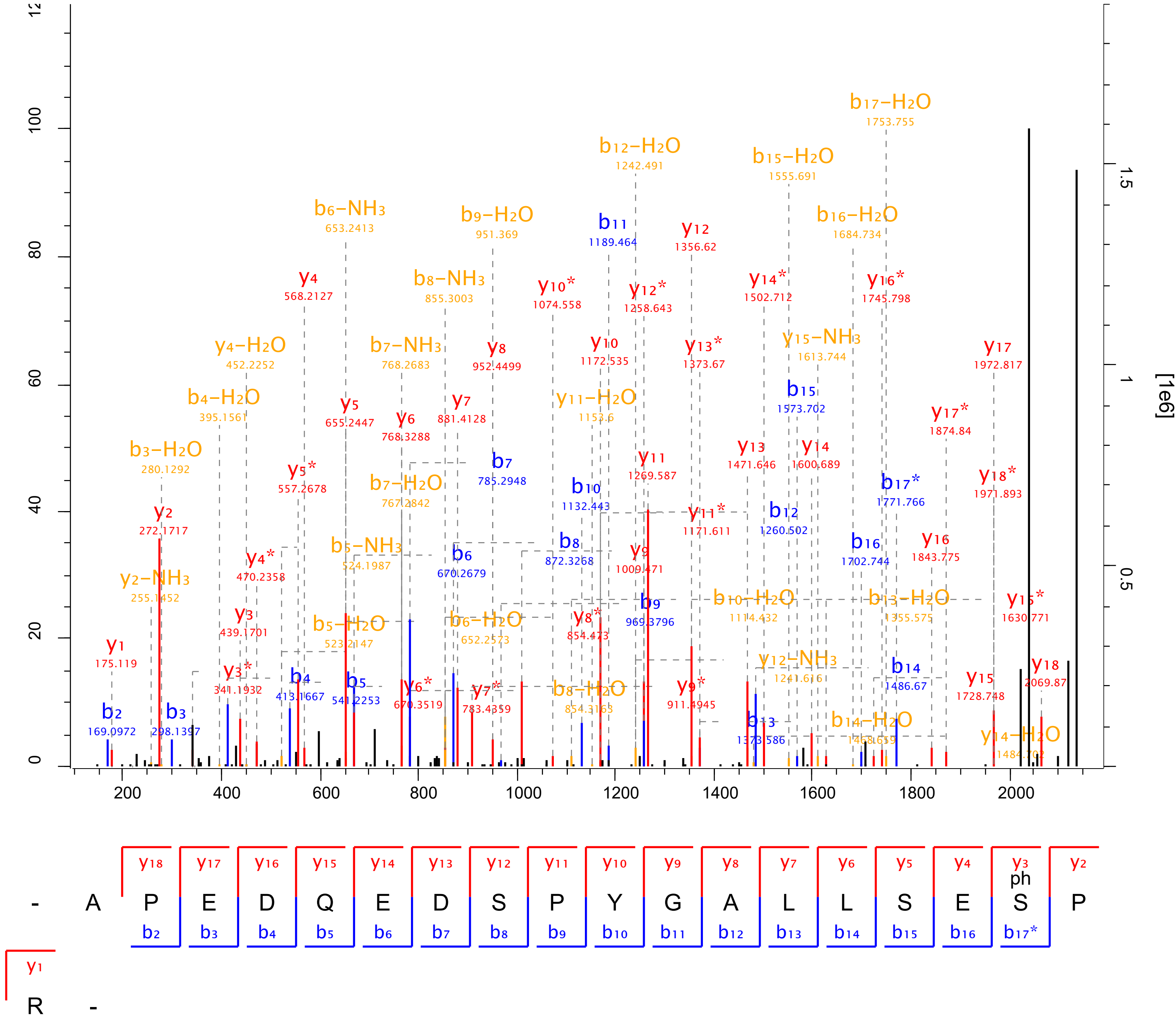

| Raw file | Scan | Method | Score | m/z |
| --- | --- | --- | --- | --- |
| water_00_1 | 33876 | FTMS; HCD | 78.95 | 815.87 |

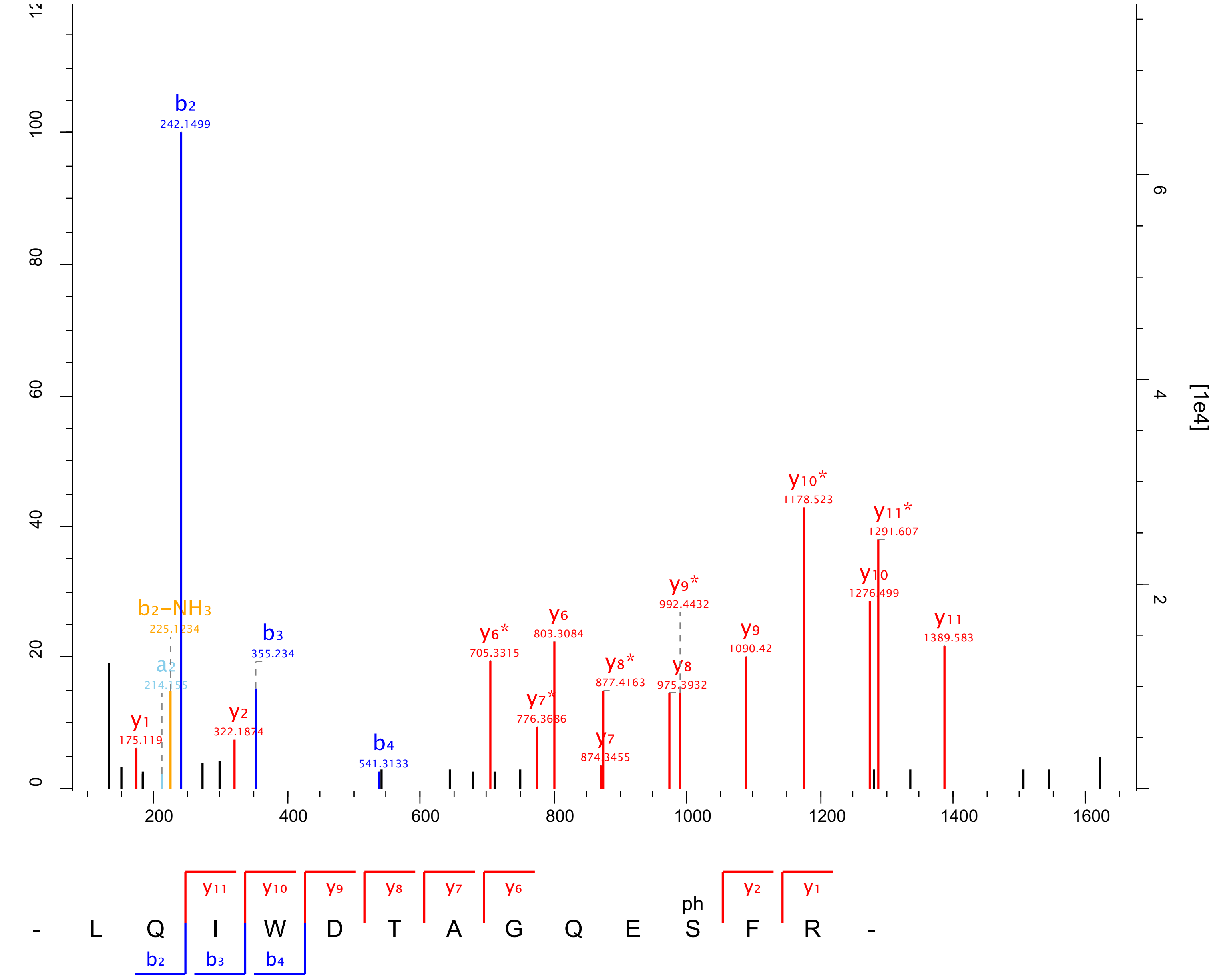

| Raw file | Scan | Method | Score | m/z |
|---|---|---|---|---|
| water_00_1 | 33973 | FTMS; HCD | 234.36 | 798.87 |

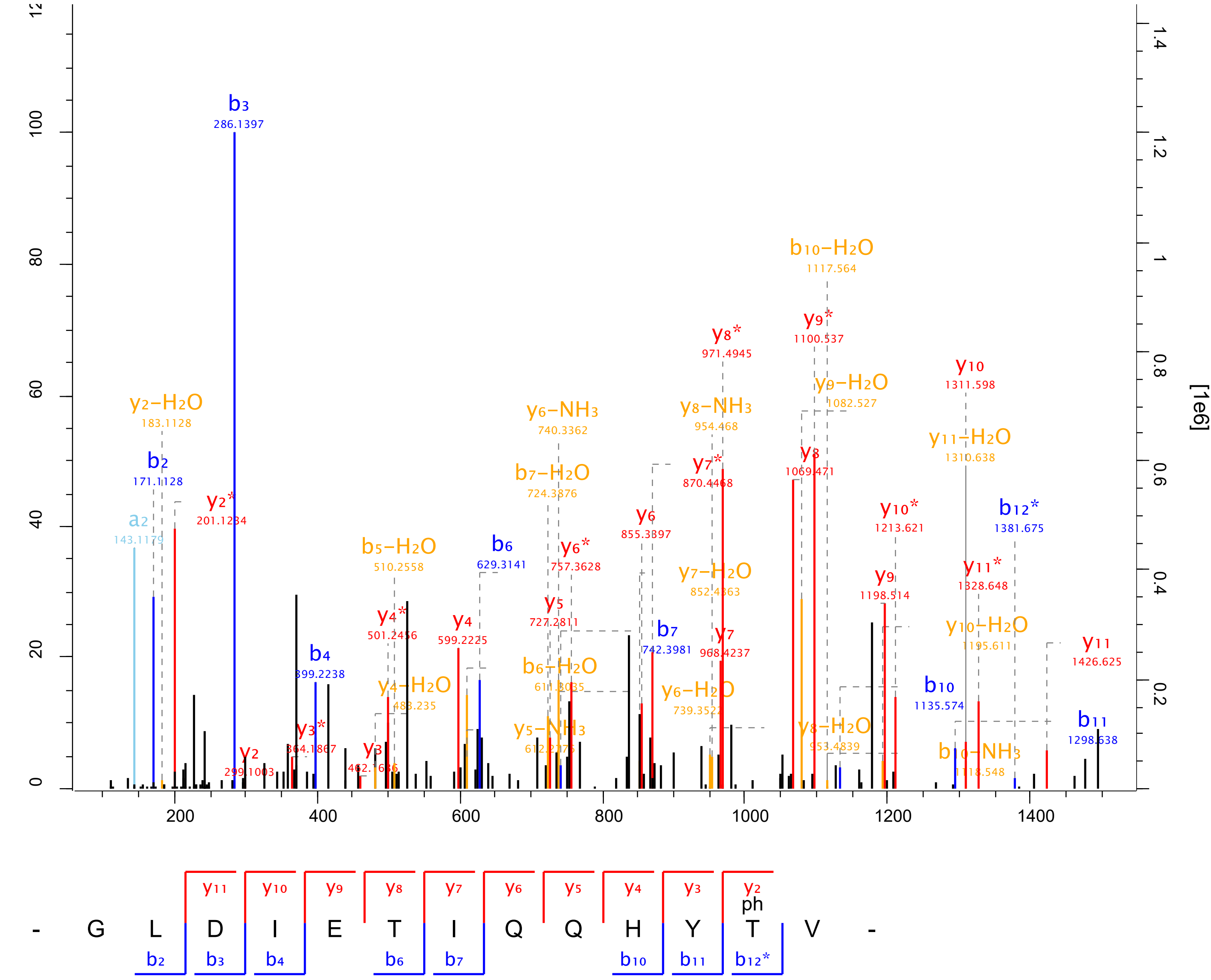

| Raw file | Scan | Method | Score | m/z |
|---|---|---|---|---|
| water_00_1 | 3412 | FTMS; HCD | 78.94 | 469.21 |

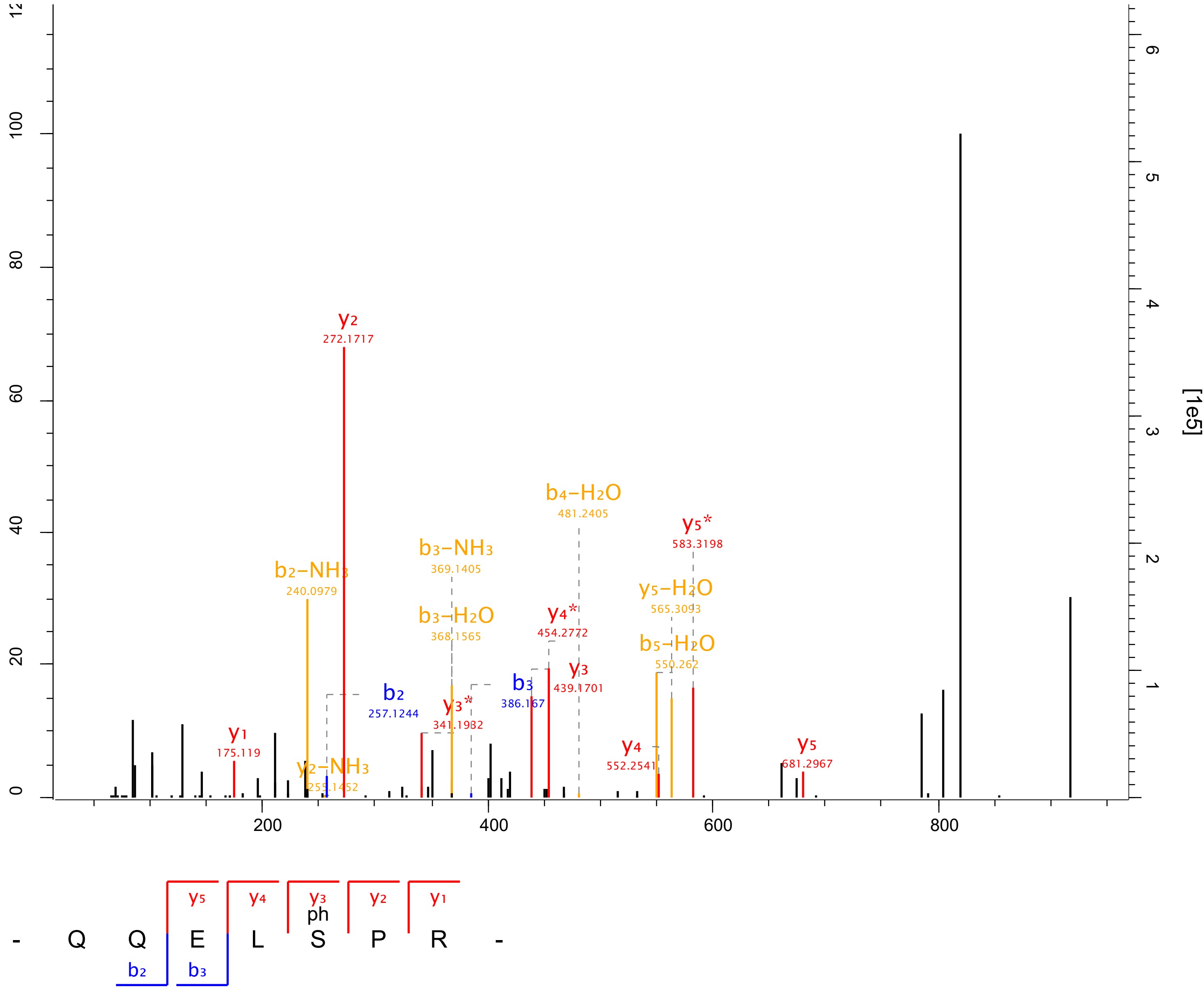

| Raw file | Scan | Method | Score | m/z |
|---|---|---|---|---|
| water_00_1 | 34314 | FTMS; HCD | 121.25 | 743.04 |

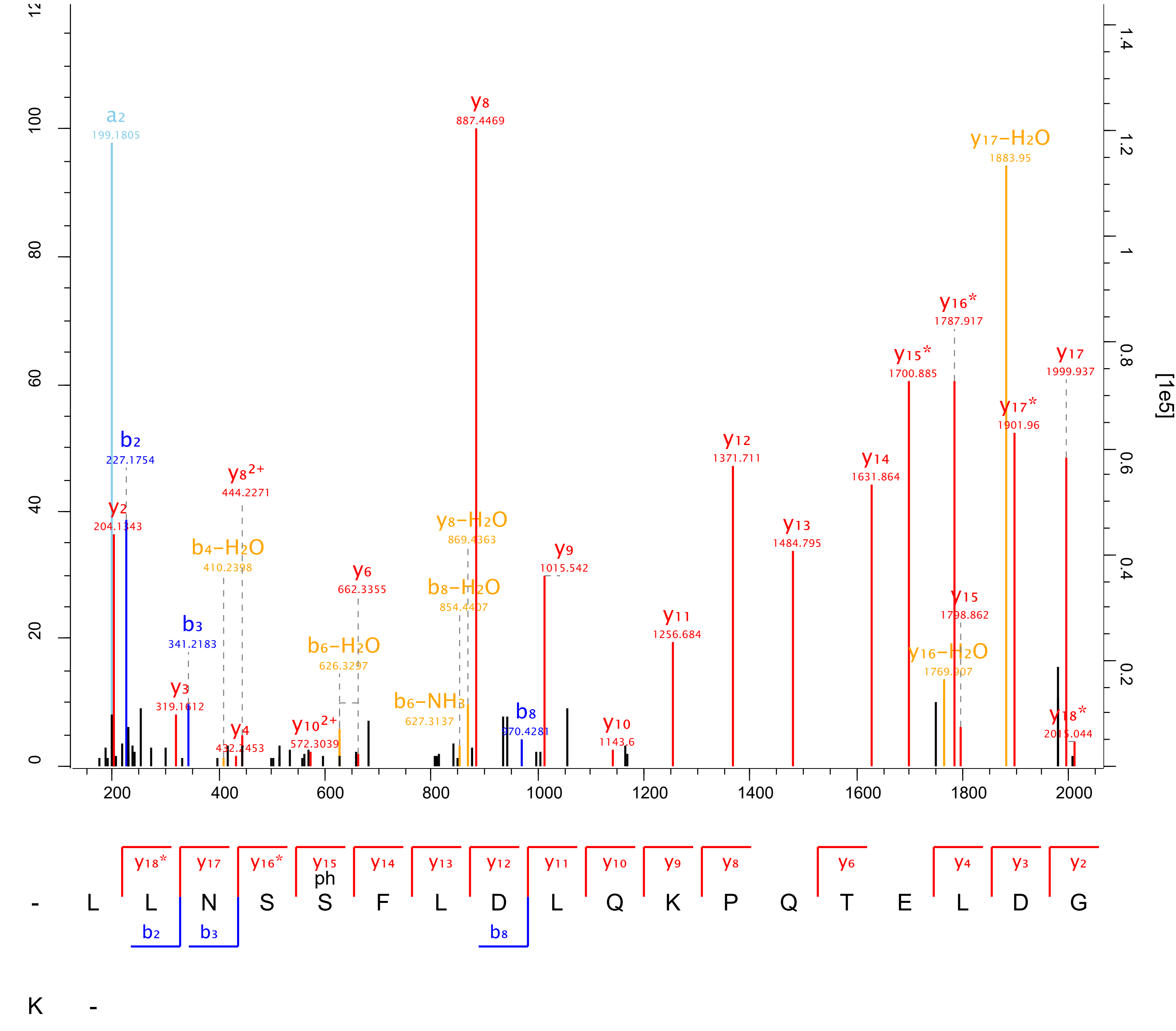

| Raw file | Scan | Method | Score | m/z |
|---|---|---|---|---|
| water_00_1 | 3439 | FTMS; HCD | 223.76 | 587.58 |

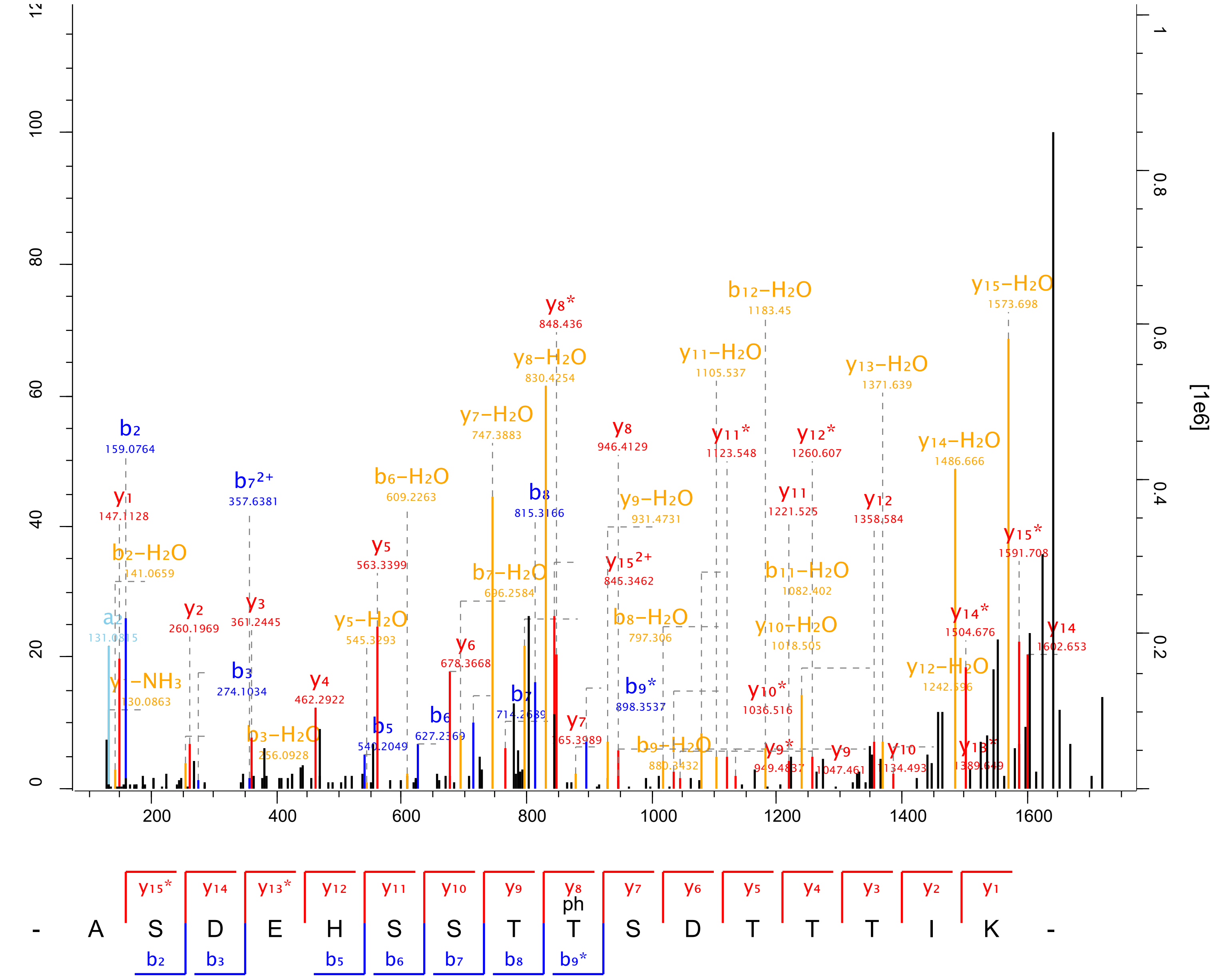

| Raw file | Scan | Method | Score | m/z |
| --- | --- | --- | --- | --- |
| water_00_1 | 34422 | FTMS; HCD | 66.29 | 904.42 |

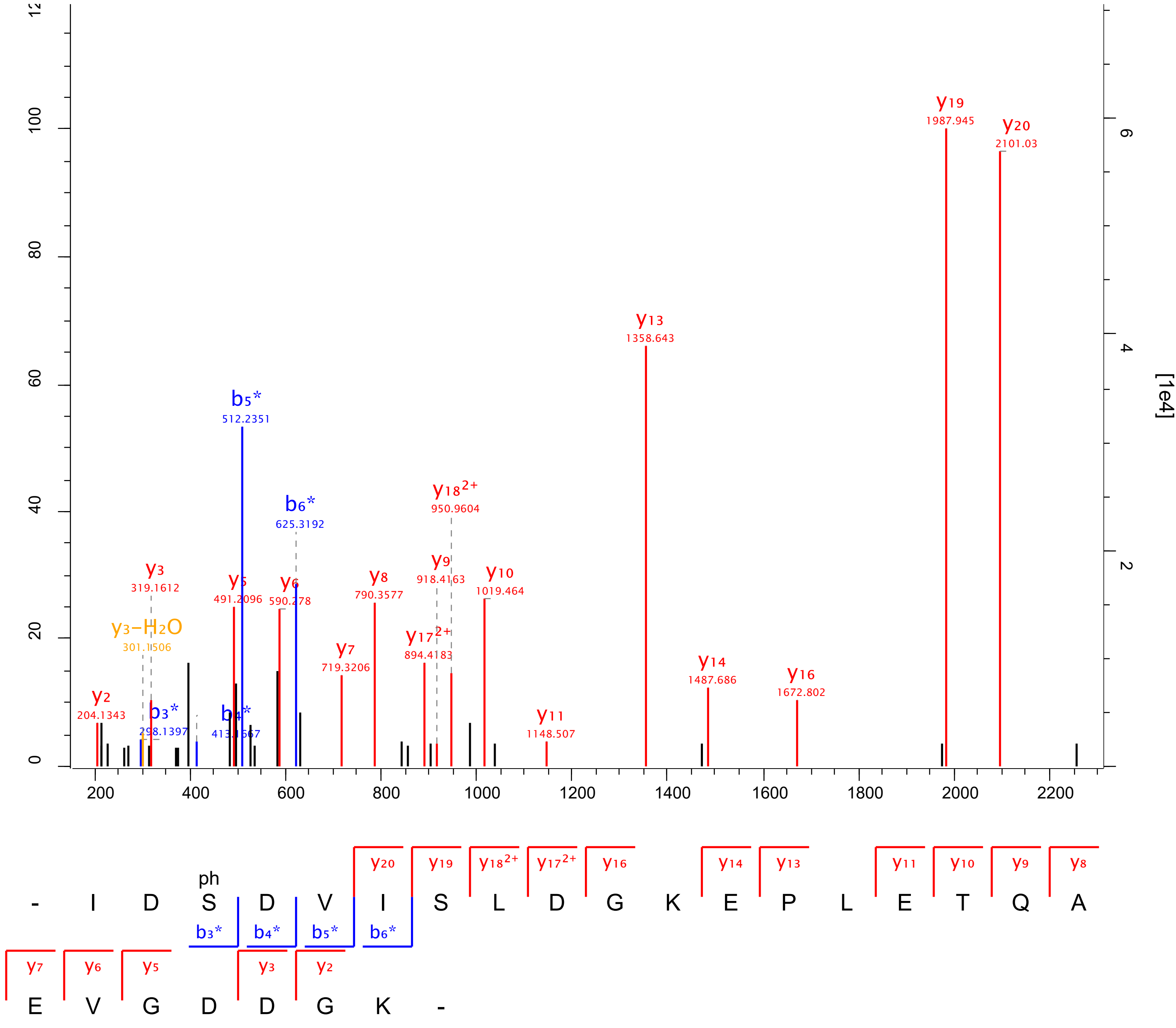

| Raw file | Scan | Method | Score | m/z |
|---|---|---|---|---|
| water_00_1 | 3524 | FTMS; HCD | 50.35 | 476.72 |

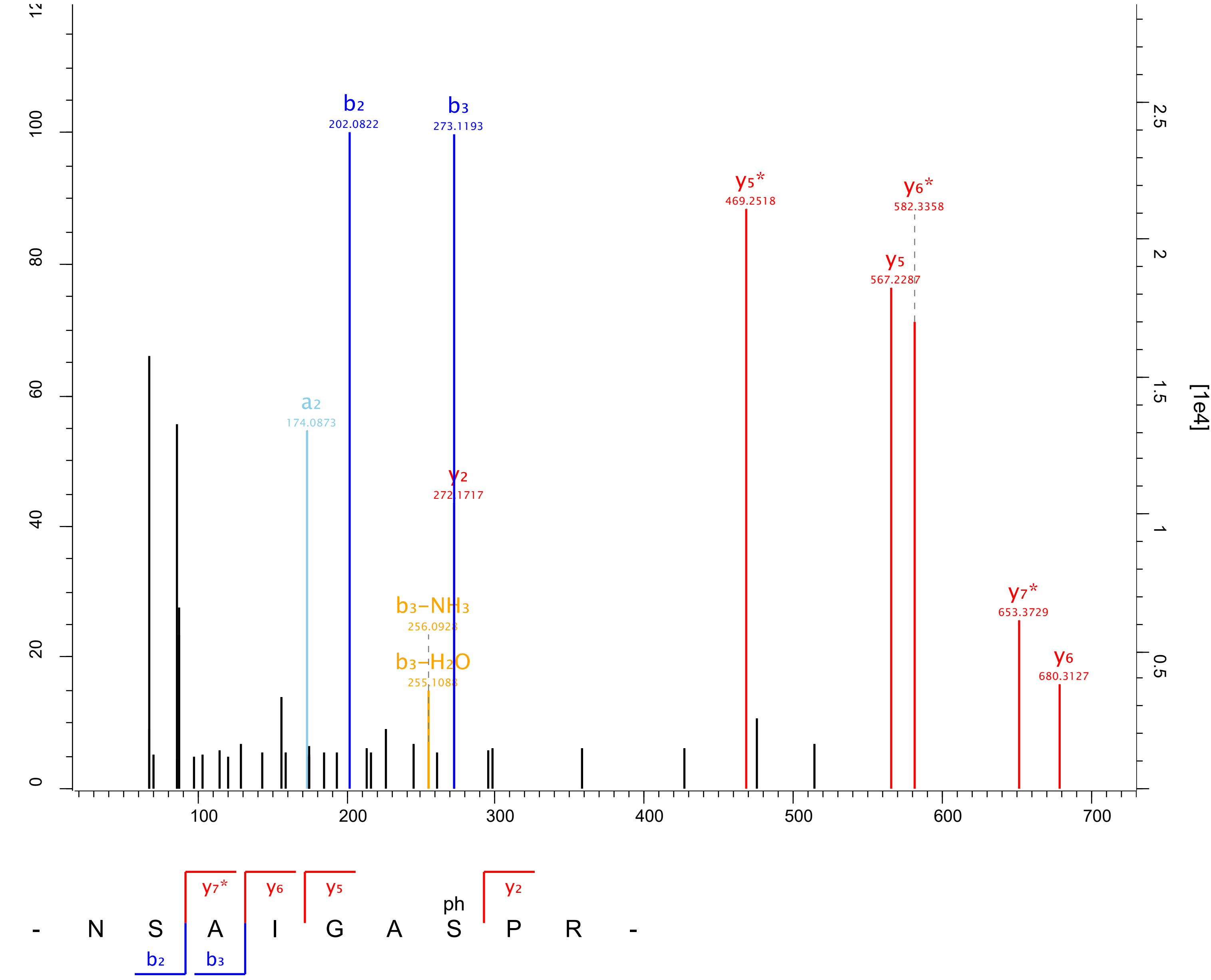

| Raw file | Scan | Method | Score | m/z |
| --- | --- | --- | --- | --- |
| water_00_1 | 3530 | FTMS; HCD | 64.22 | 583.75 |

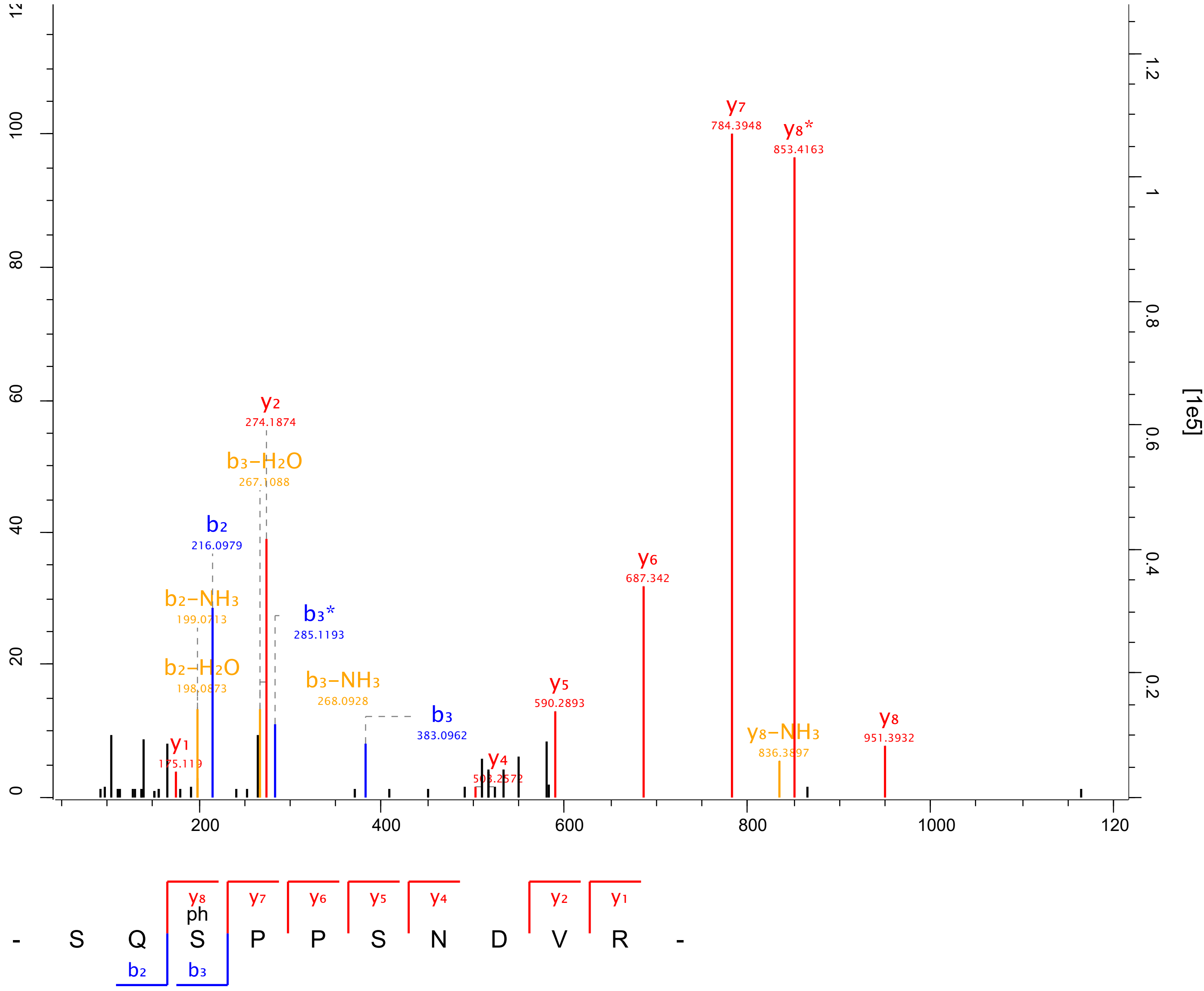

| Raw file | Scan | Method | Score | m/z |
|---|---|---|---|---|
| water_00_1 | 3578 | FTMS; HCD | 134.26 | 416.17 |

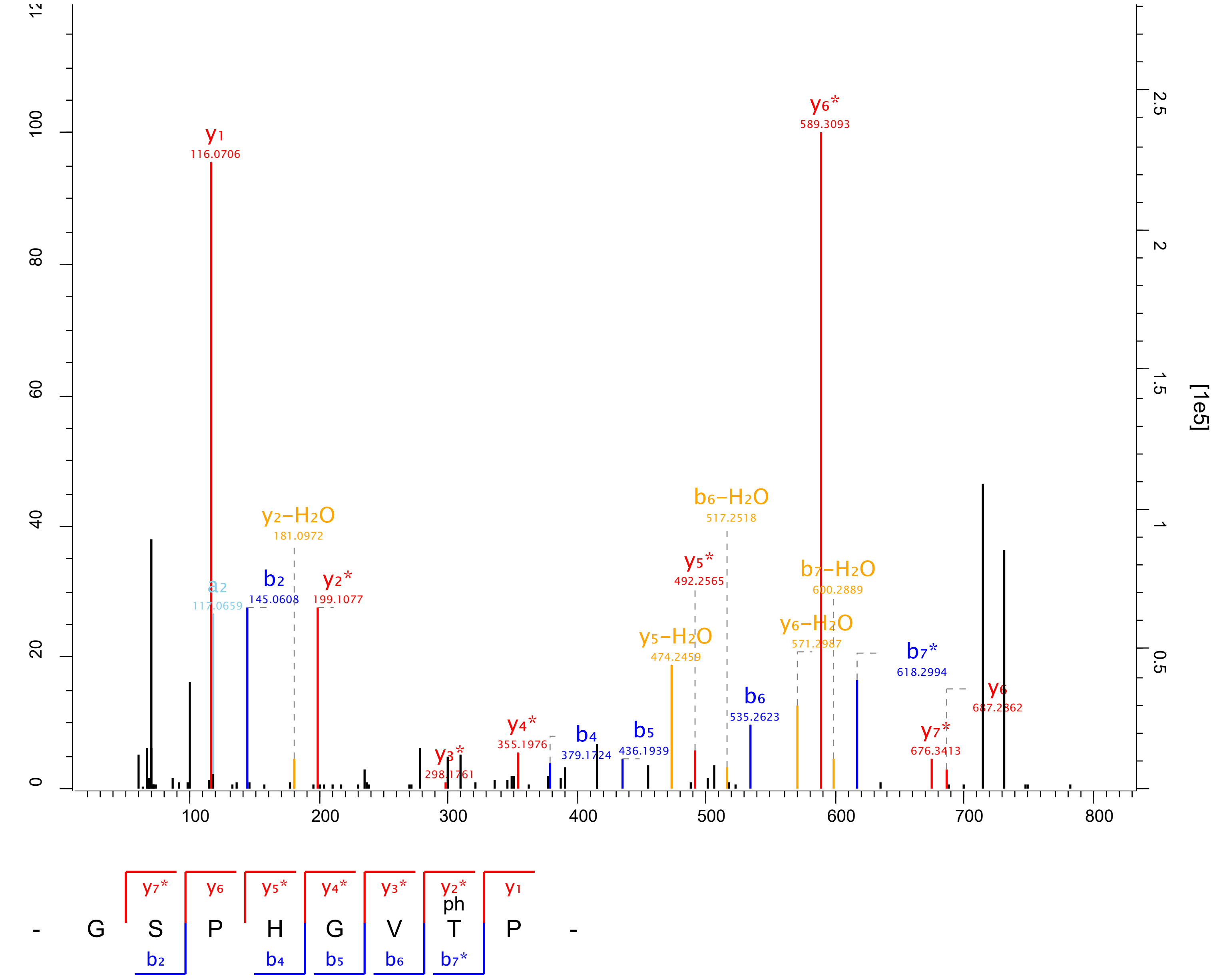

| Raw file | Scan | Method | Score | m/z |
|---|---|---|---|---|
| water_00_1 | 36060 | FTMS; HCD | 45.92 | 907.44 |

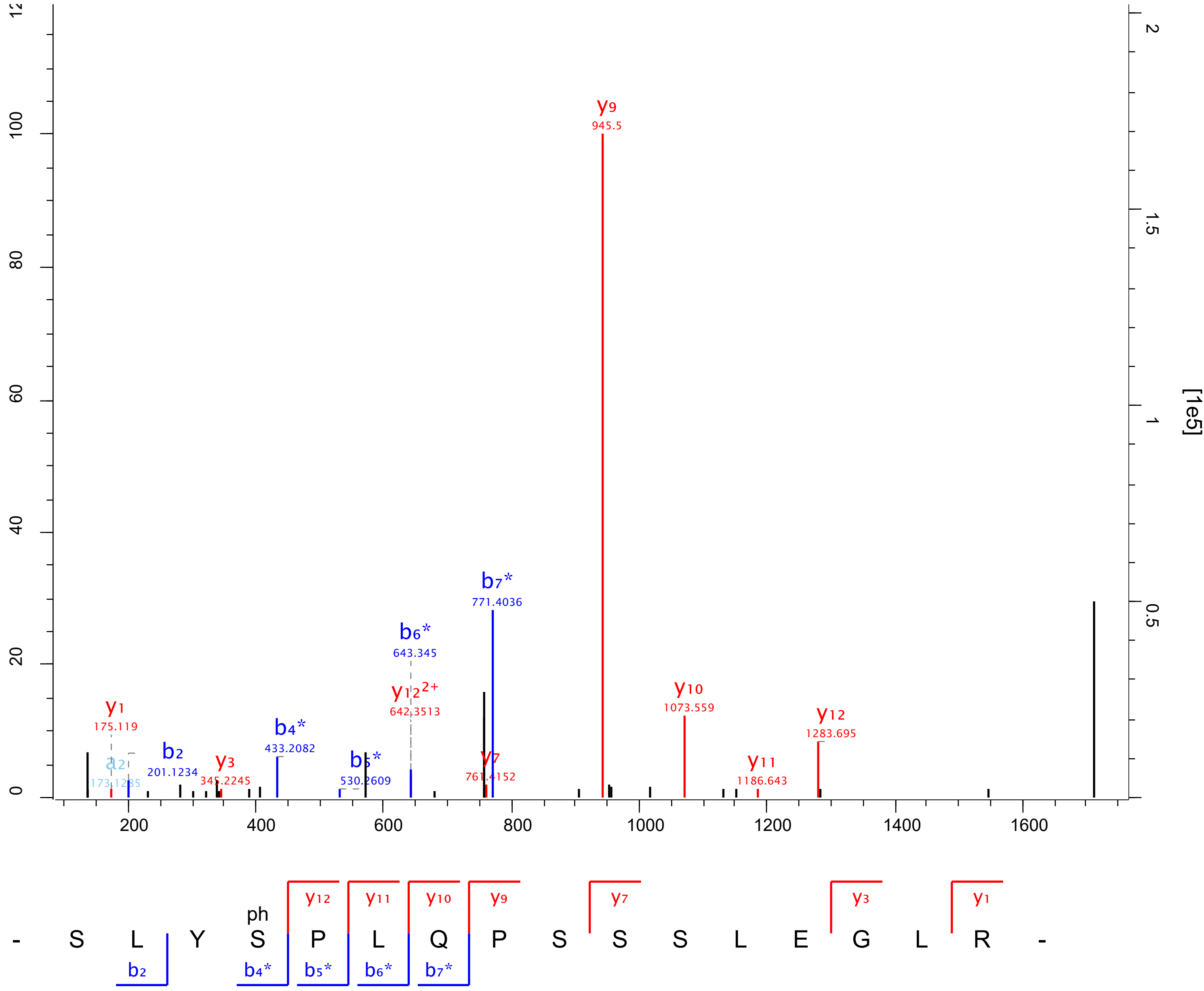

| Raw file | Scan | Method | Score | m/z |
| --- | --- | --- | --- | --- |
| water_00_1 | 36642 | FTMS; HCD | 57.53 | 550.7 |

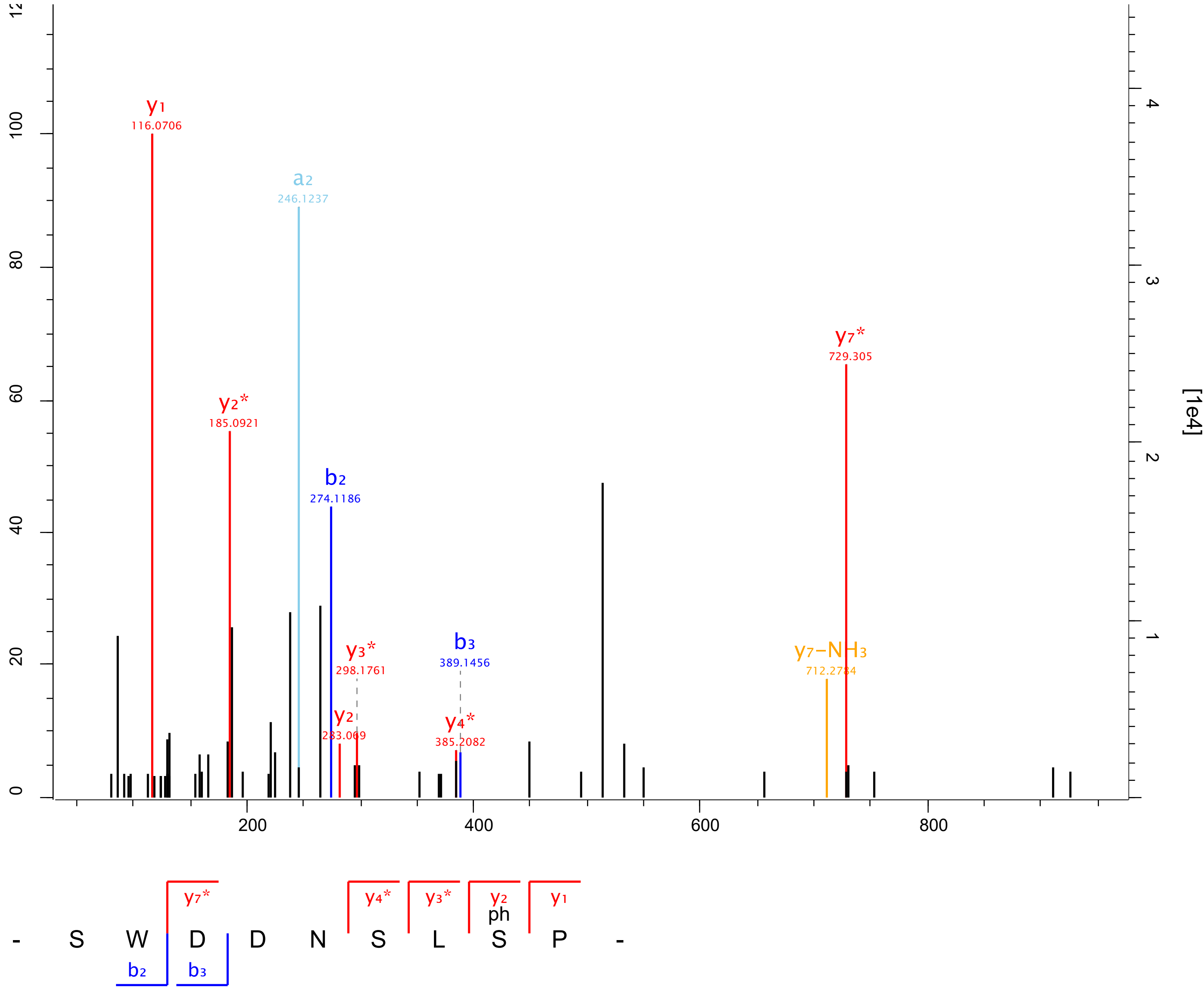

| Raw file | Scan | Method | Score | m/z |
|---|---|---|---|---|
| water_00_1 | 36869 | FTMS; HCD | 93.58 | 724.66 |

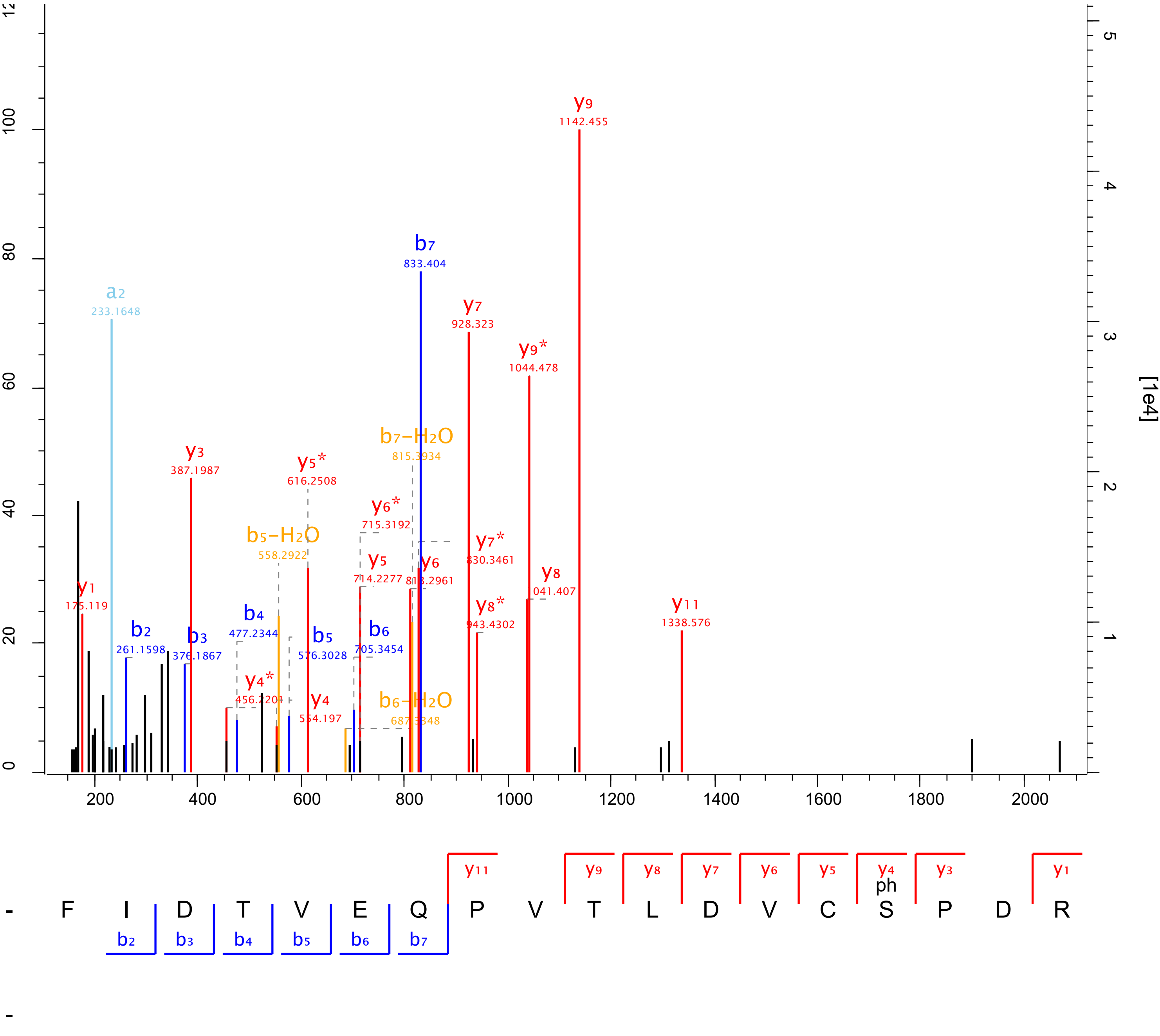

| Raw file | Scan | Method | Score | m/z |
| --- | --- | --- | --- | --- |
| water_00_1 | 3714 | FTMS; HCD | 101.5 | 605.92 |

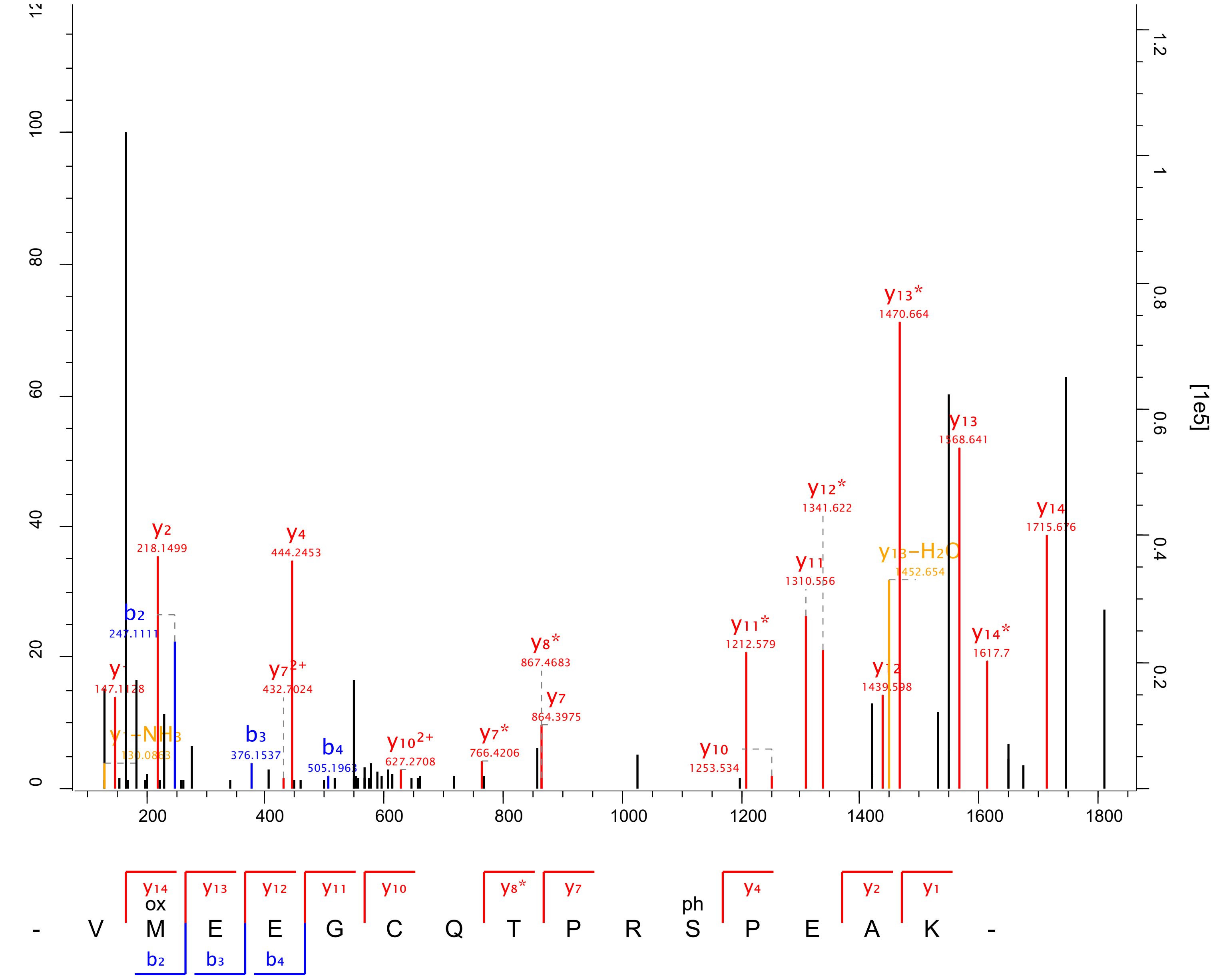

| Raw file | Scan | Method | Score | m/z |
|---|---|---|---|---|
| water_00_1 | 37166 | FTMS; HCD | 152.16 | 902.37 |

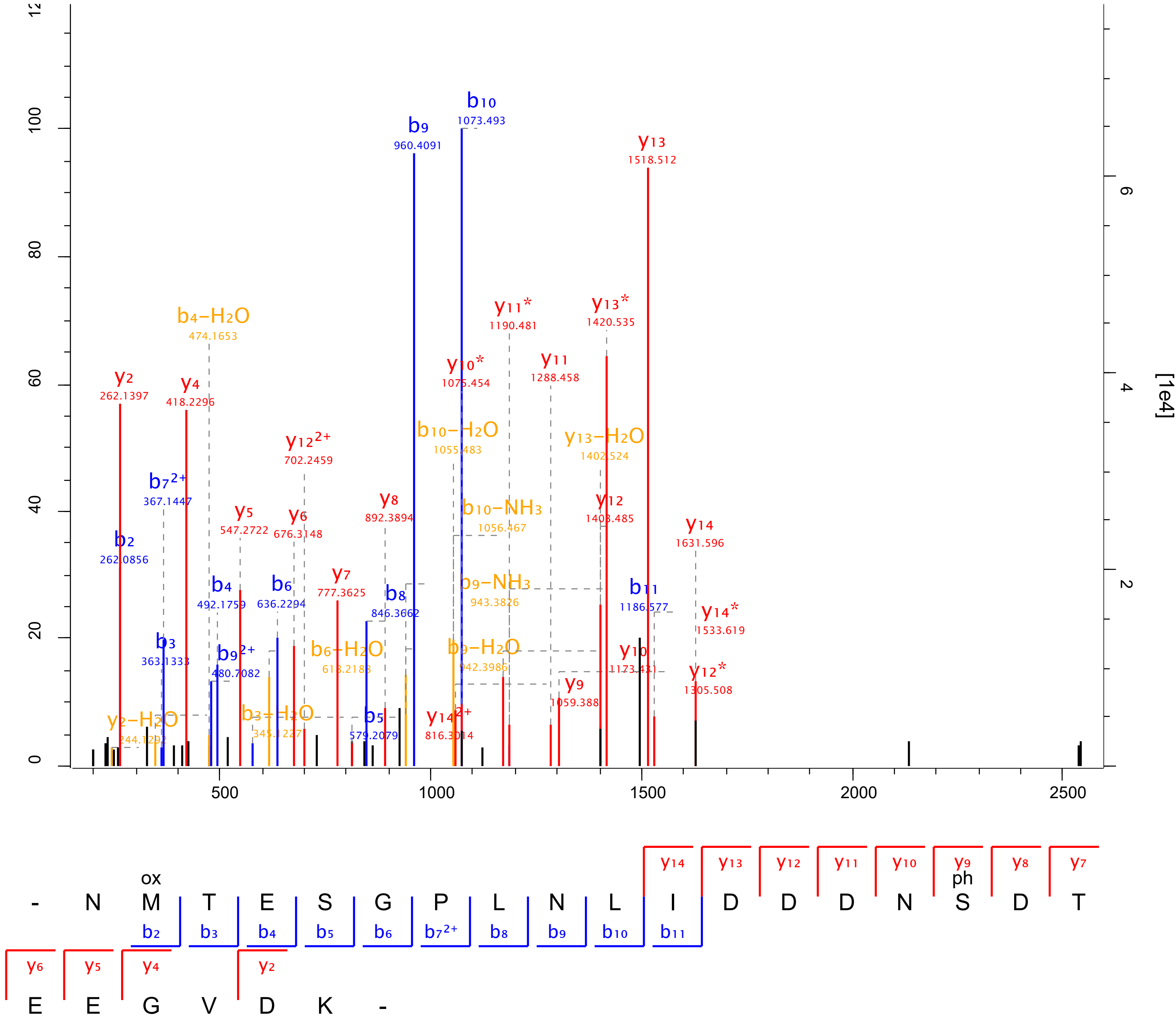

| Raw file | Scan | Method | Score | m/z |
| --- | --- | --- | --- | --- |
| water_00_1 | 3765 | FTMS; HCD | 77.89 | 526.74 |

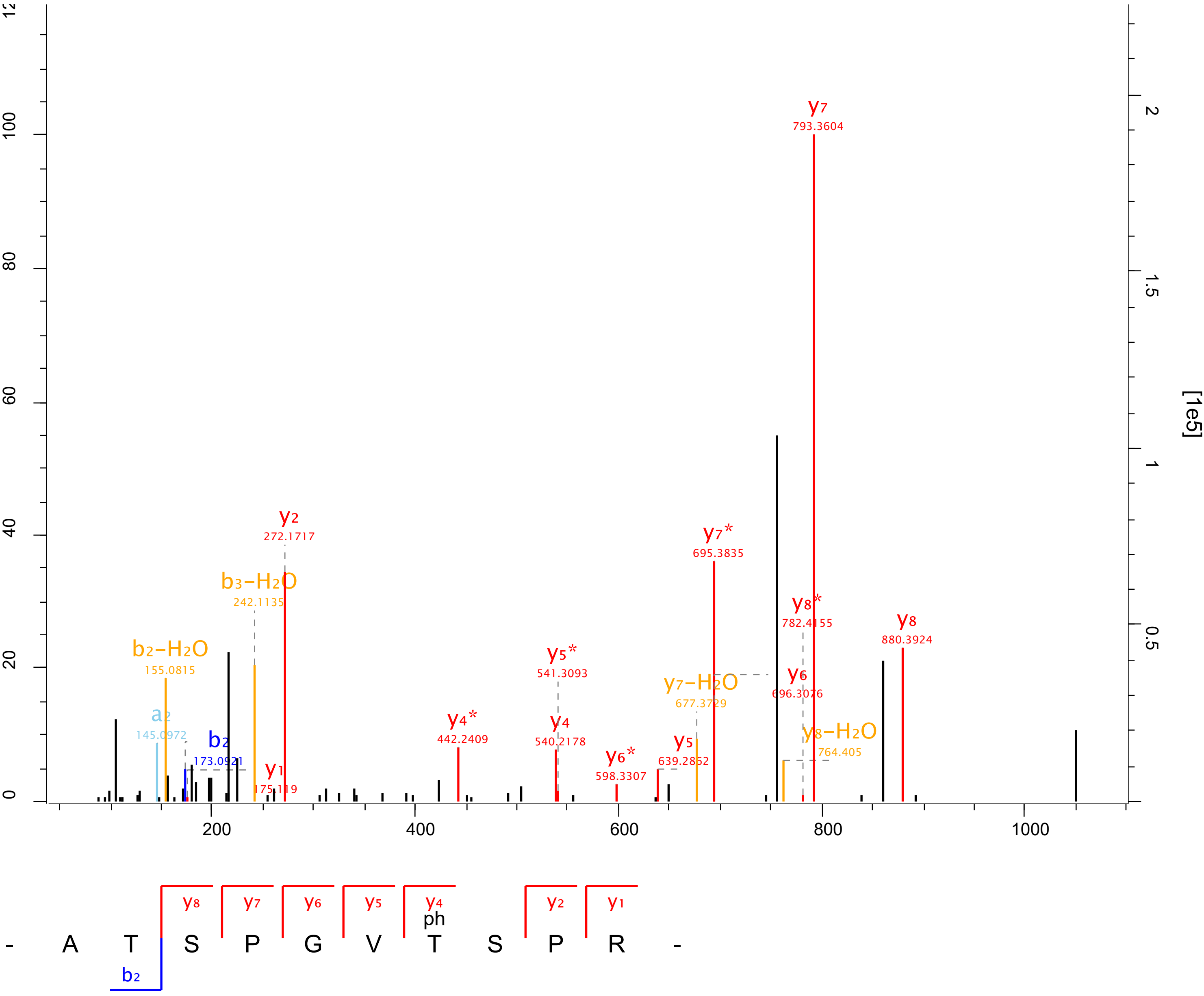

| Raw file | Scan | Method | Score | m/z |
| --- | --- | --- | --- | --- |
| water_00_1 | 37804 | FTMS; HCD | 71.79 | 773.7 |

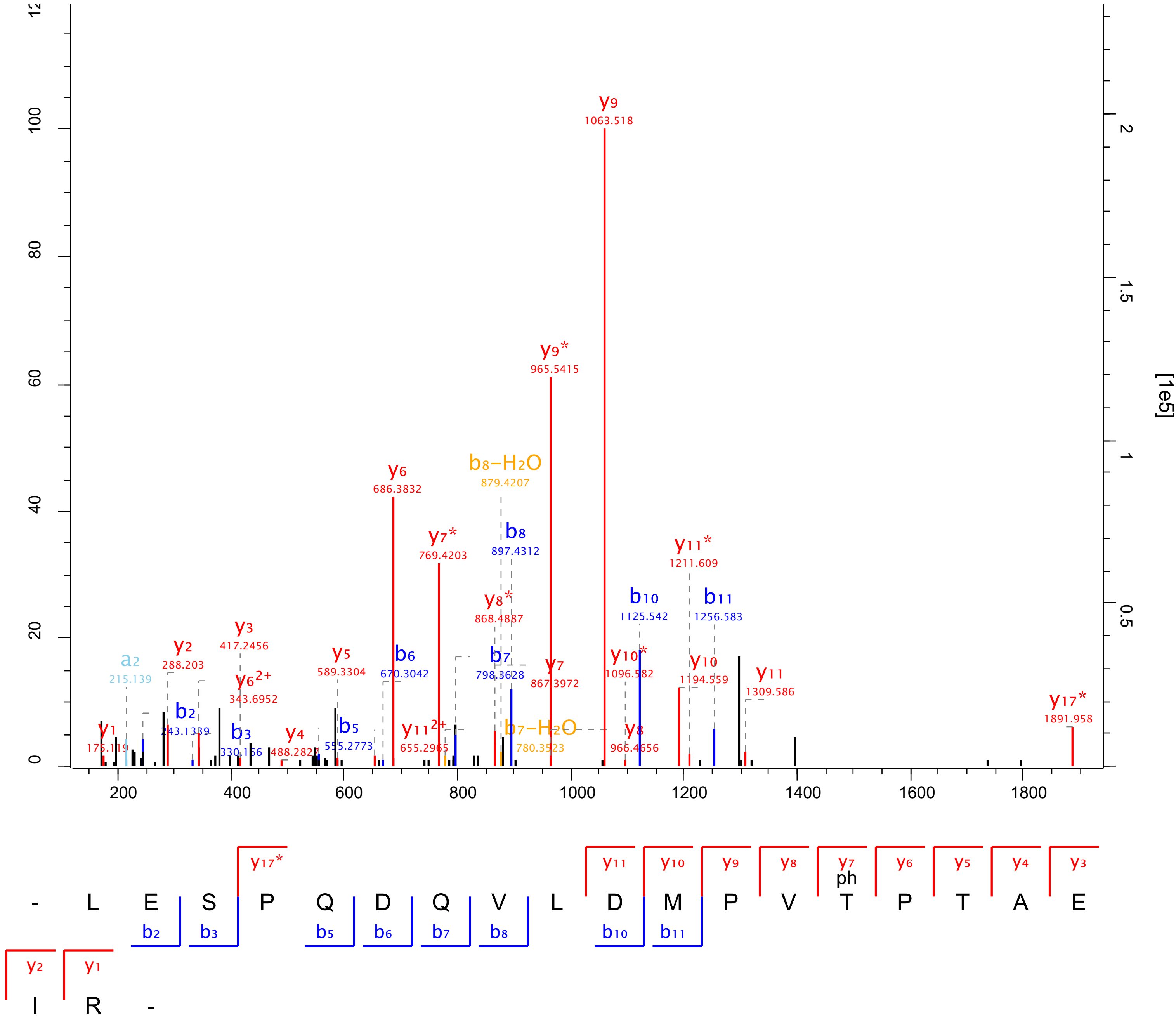

| Raw file | Scan | Method | Score | m/z |
|---|---|---|---|---|
| water_00_1 | 3831 | FTMS; HCD | 45.36 | 457.21 |

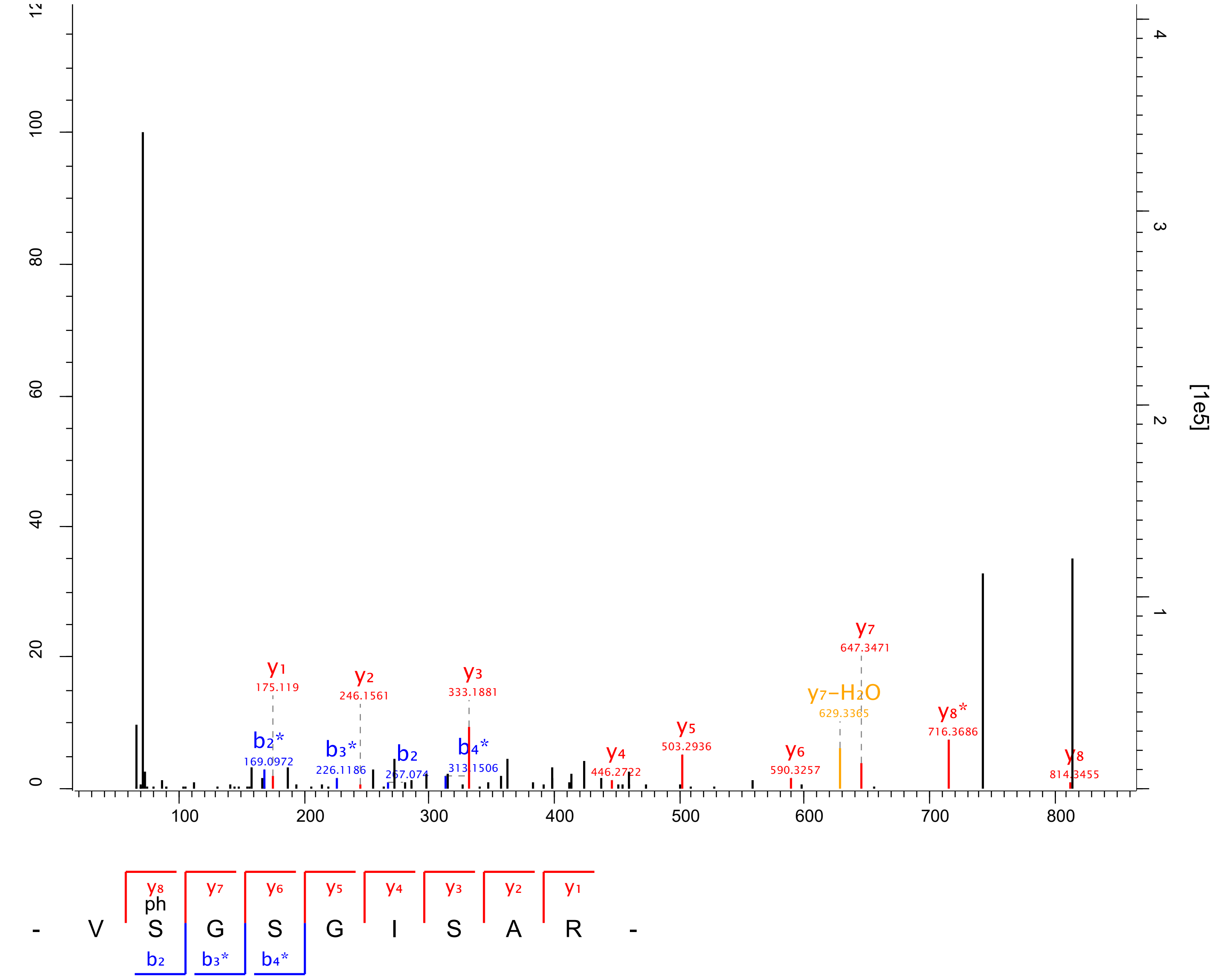

| Raw file | Scan | Method | Score | m/z |
| --- | --- | --- | --- | --- |
| water_00_1 | 39072 | FTMS; HCD | 70.89 | 828.36 |

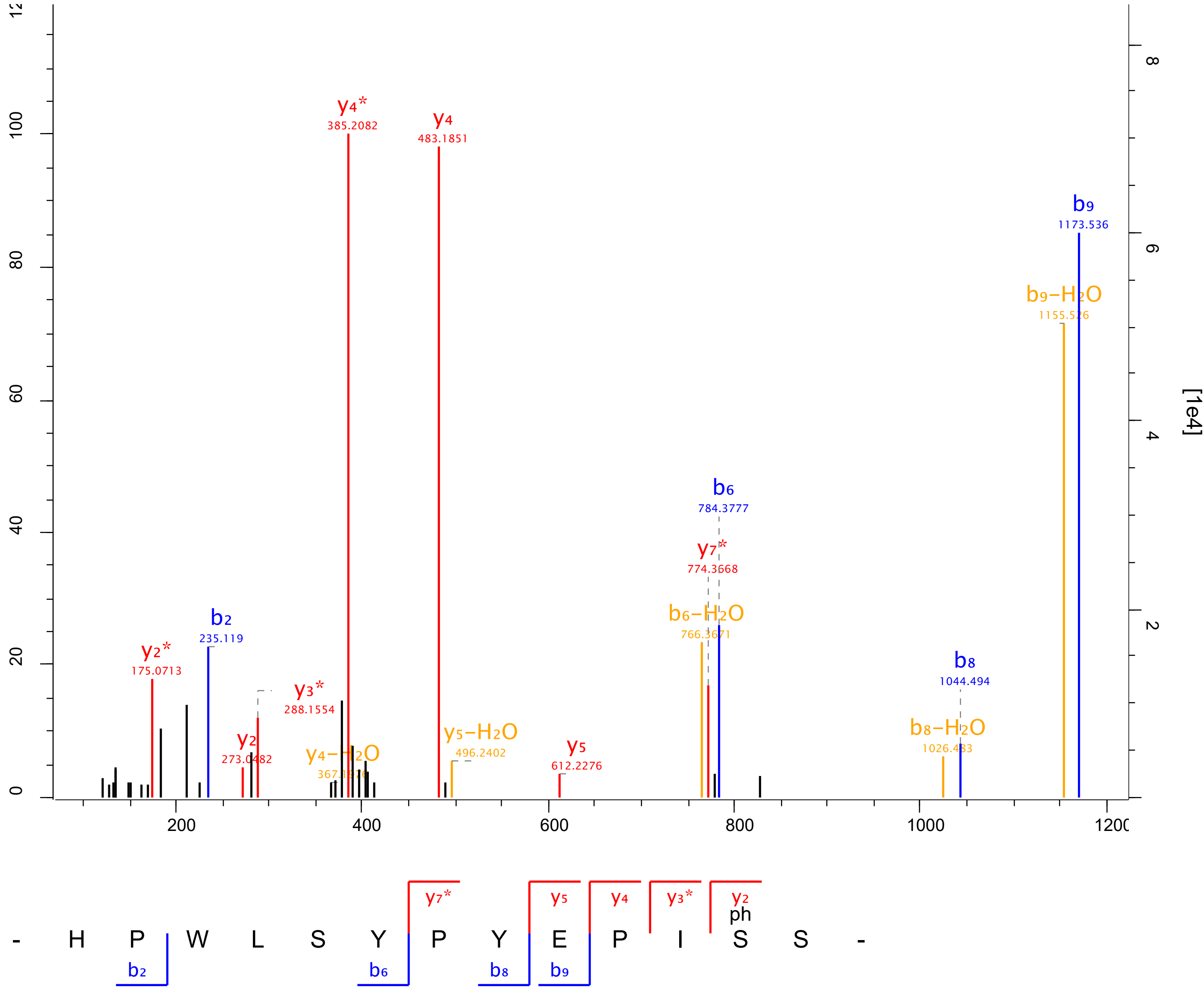

| Raw file | Scan | Method | Score | m/z |
|---|---|---|---|---|
| water_00_1 | 3928 | FTMS; HCD | 68.04 | 644.26 |

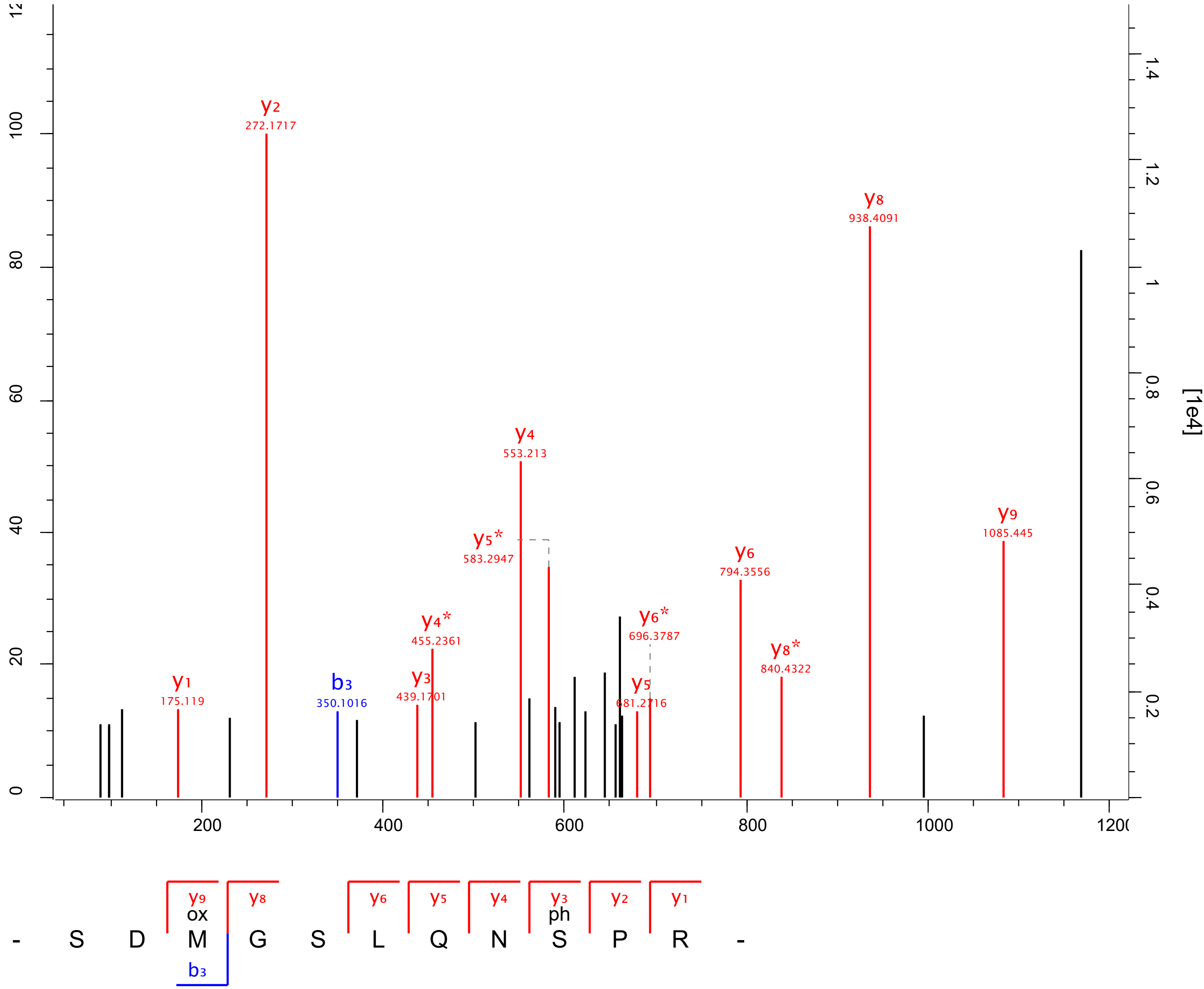

| Raw file | Scan | Method | Score | m/z |
|---|---|---|---|---|
| water_00_1 | 39521 | FTMS; HCD | 177.12 | 800.41 |

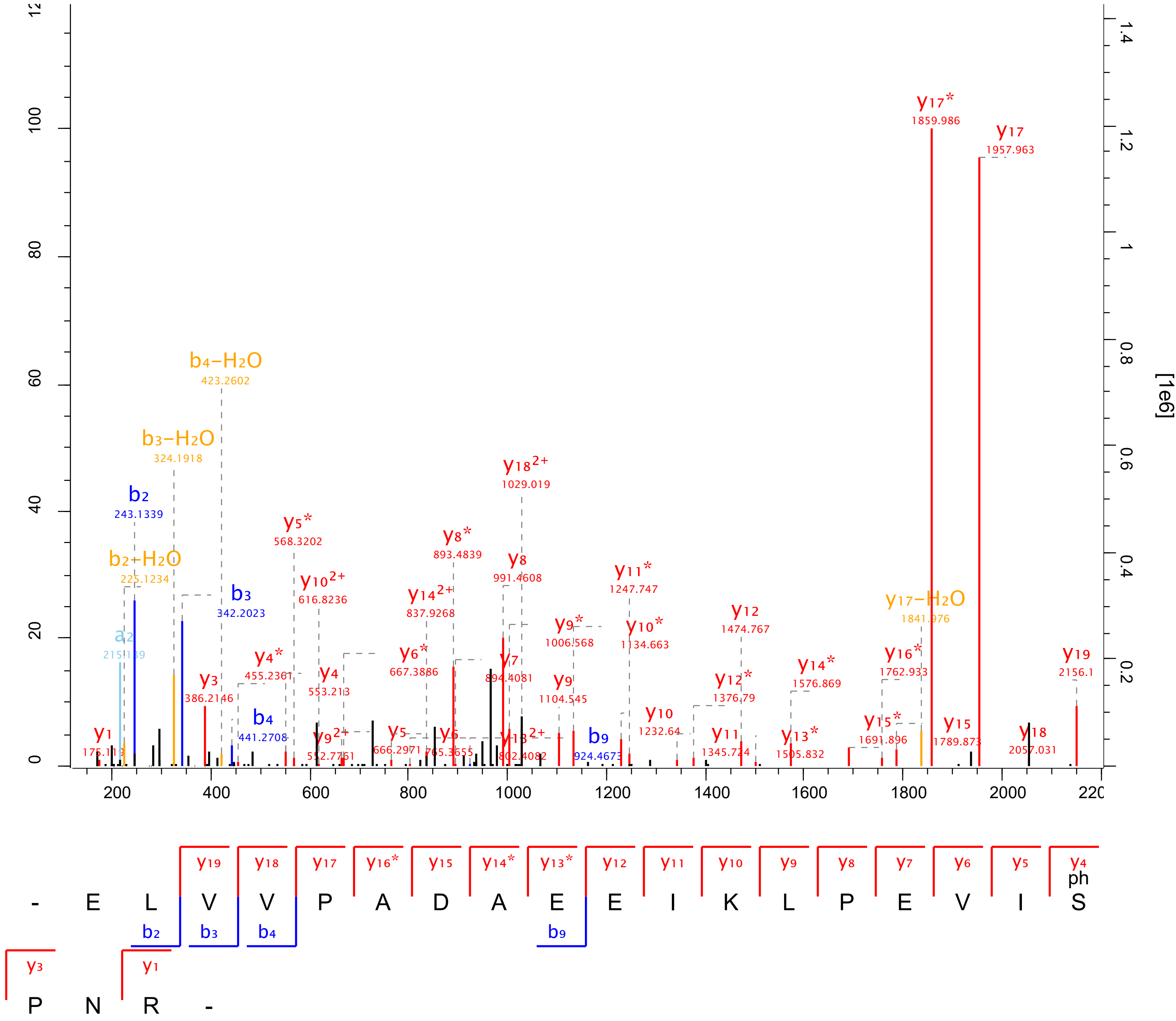

| Raw file | Scan | Method | Score | m/z |
|---|---|---|---|---|
| water_00_1 | 39540 | FTMS; HCD | 139.88 | 847.4 |

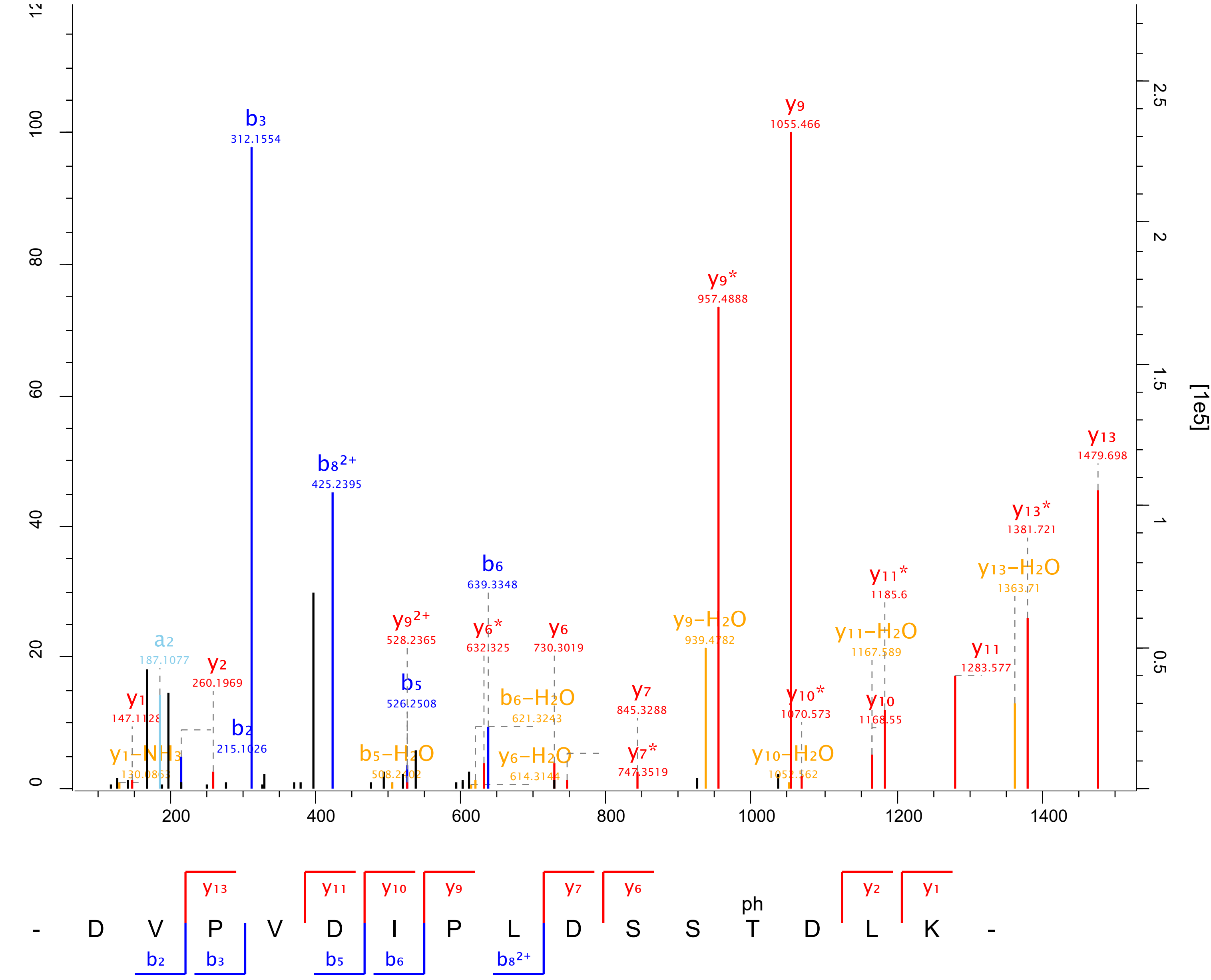

| Raw file | Scan | Method | Score | m/z |
|---|---|---|---|---|
| water_00_1 | 3999 | FTMS; HCD | 77.64 | 570.76 |

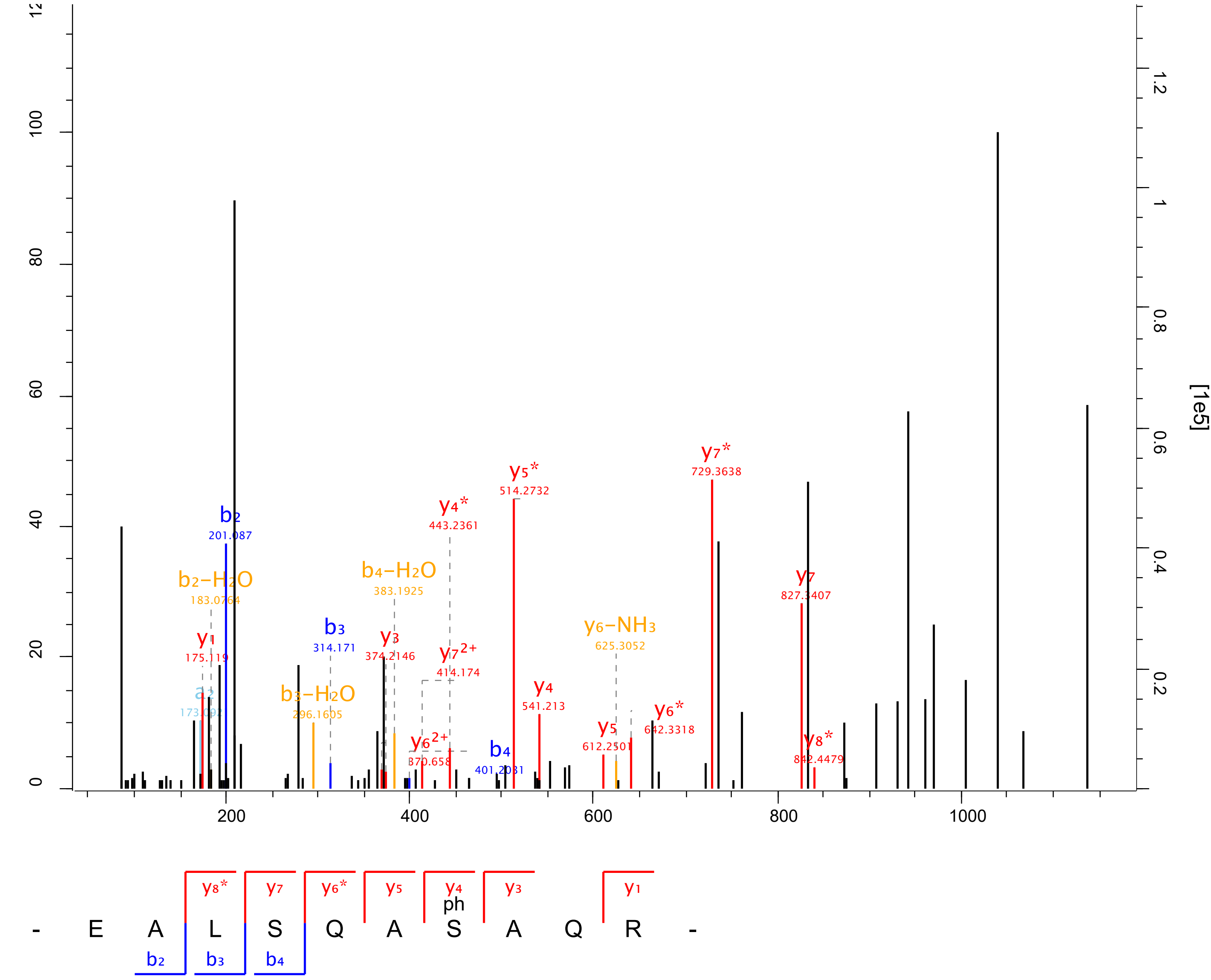

| Raw file | Scan | Method | Score | m/z |
|---|---|---|---|---|
| water_00_1 | 40190 | FTMS; HCD | 62.19 | 937.43 |

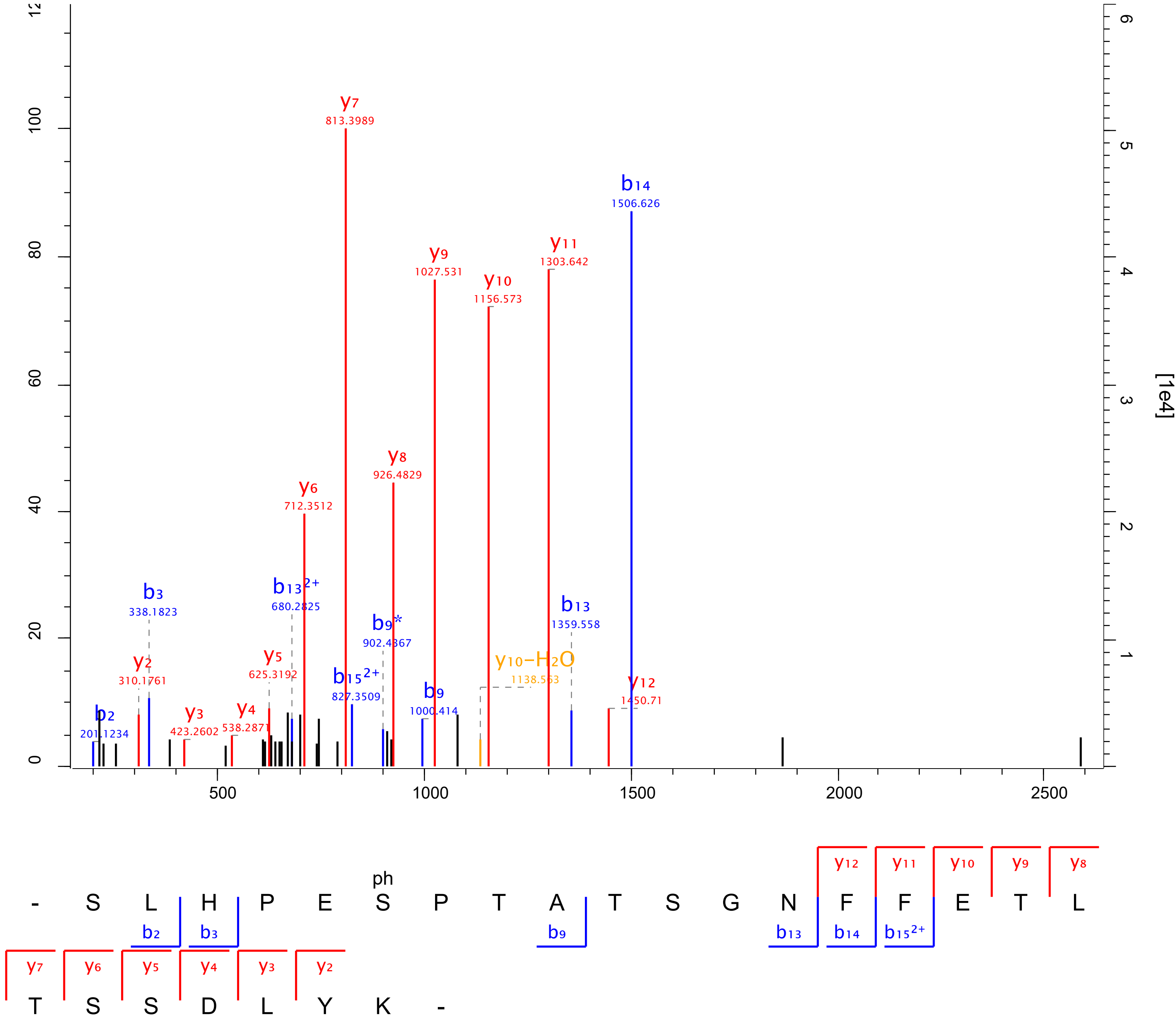

| Raw file | Scan | Method | Score | m/z |
|---|---|---|---|---|
| water_00_1 | 40577 | FTMS; HCD | 49.09 | 1038.82 |

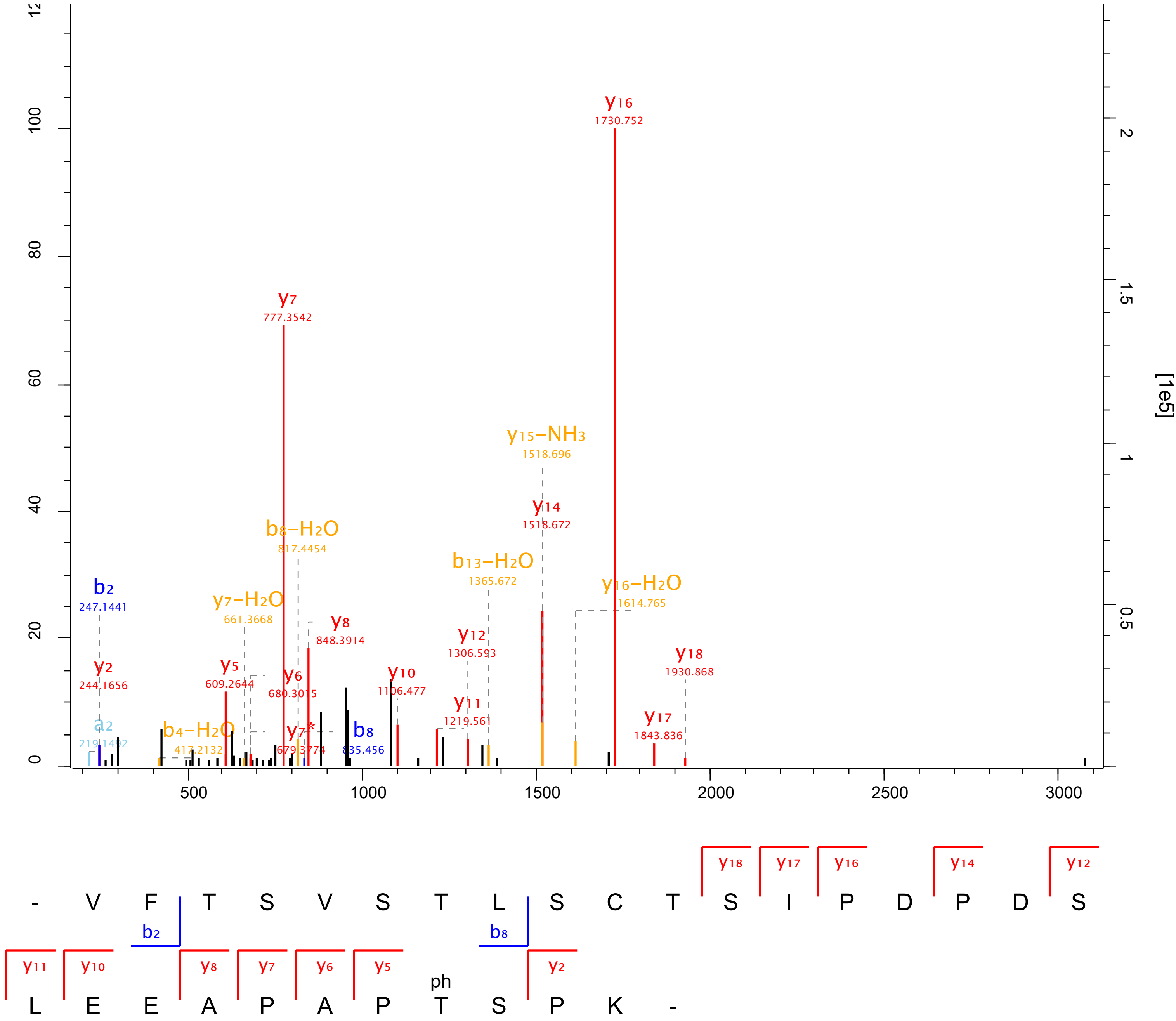

| Raw file | Scan | Method | Score | m/z |
| --- | --- | --- | --- | --- |
| water_00_1 | 4072 | FTMS; HCD | 90.39 | 636.76 |

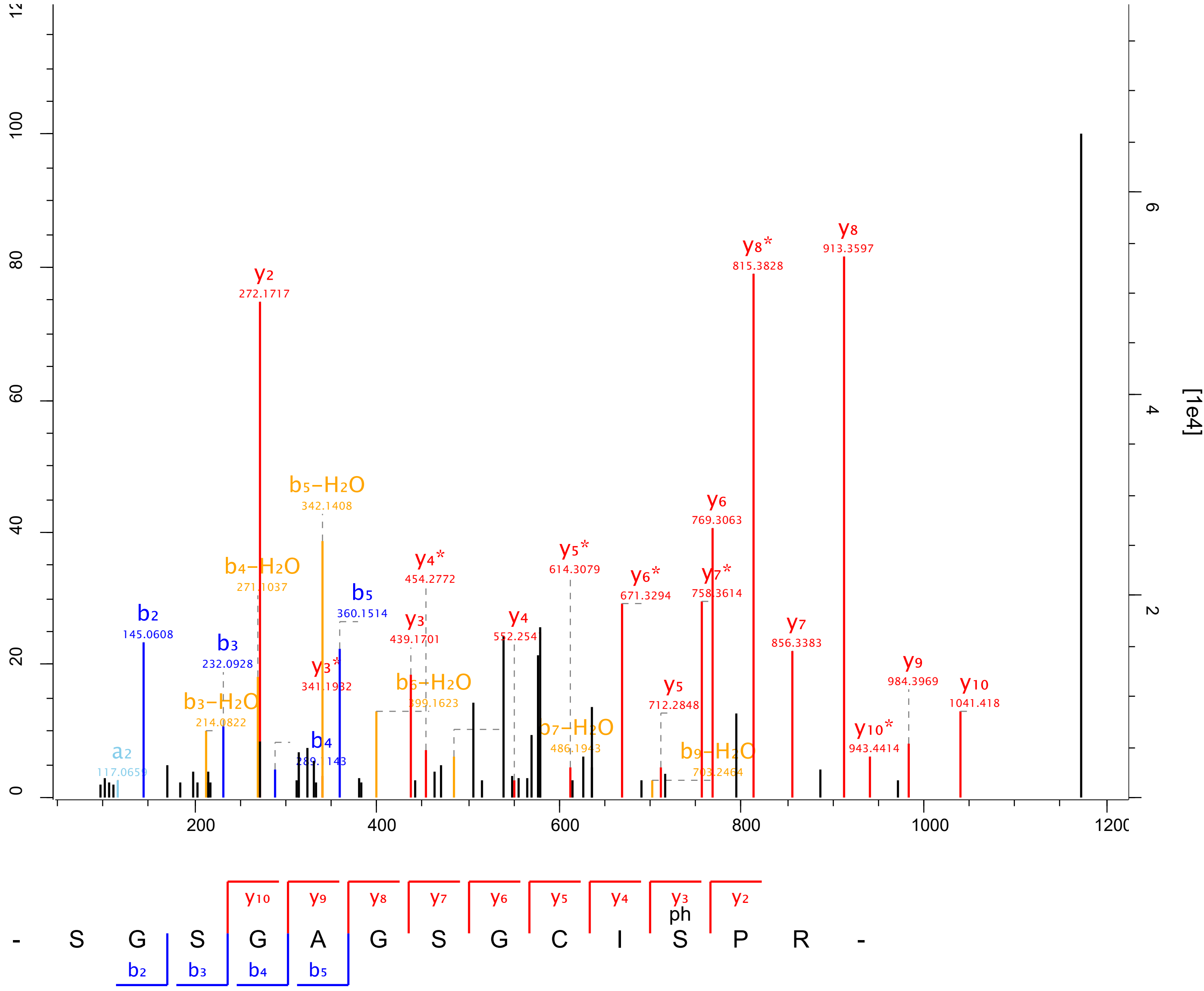

| Raw file | Scan | Method | Score | m/z |
|---|---|---|---|---|
| water_00_1 | 4098 | FTMS; HCD | 67.14 | 738.79 |

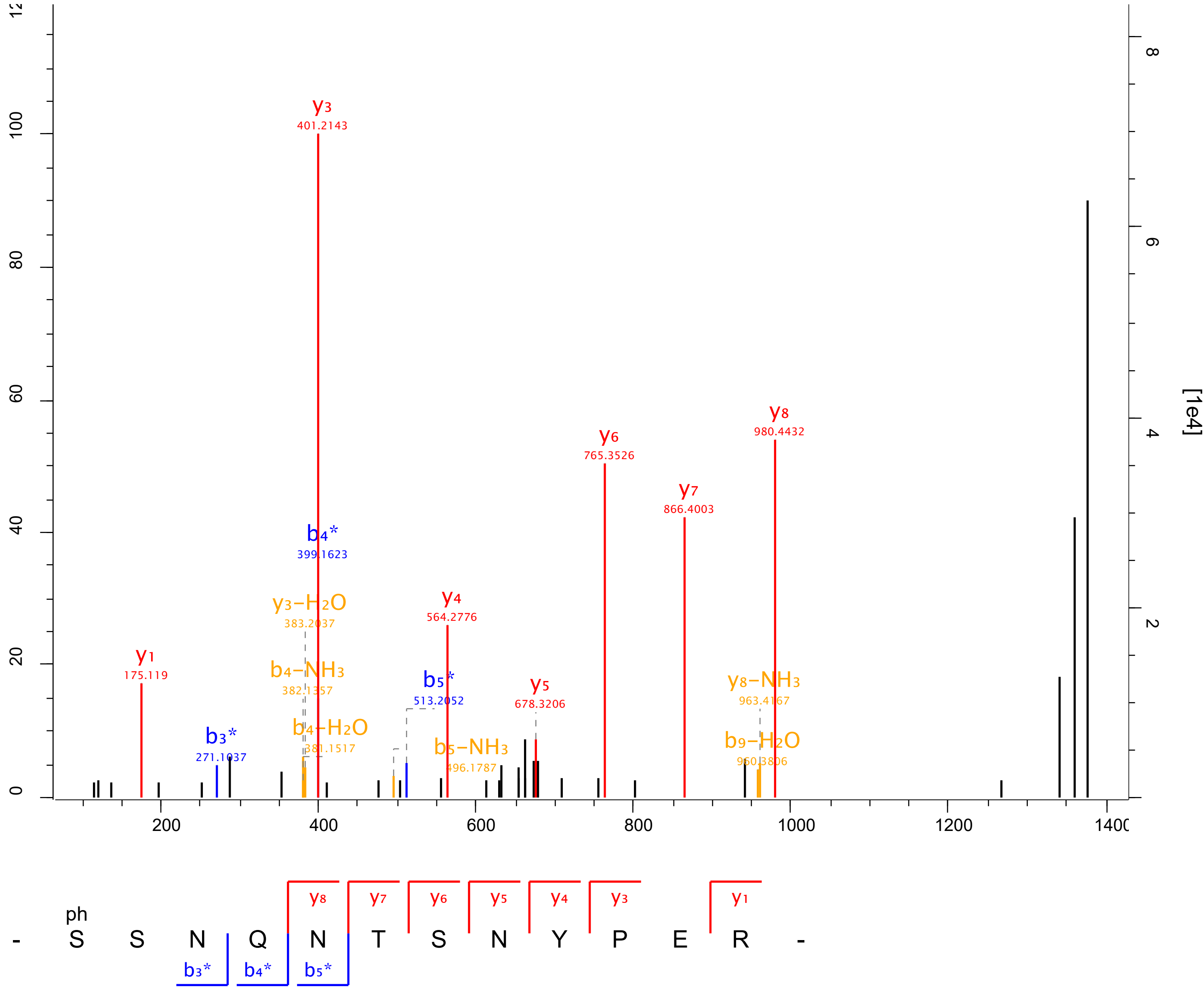

| Raw file | Scan | Method | Score | m/z |
|---|---|---|---|---|
| water_00_1 | 4123 | FTMS; HCD | 223.93 | 758.81 |

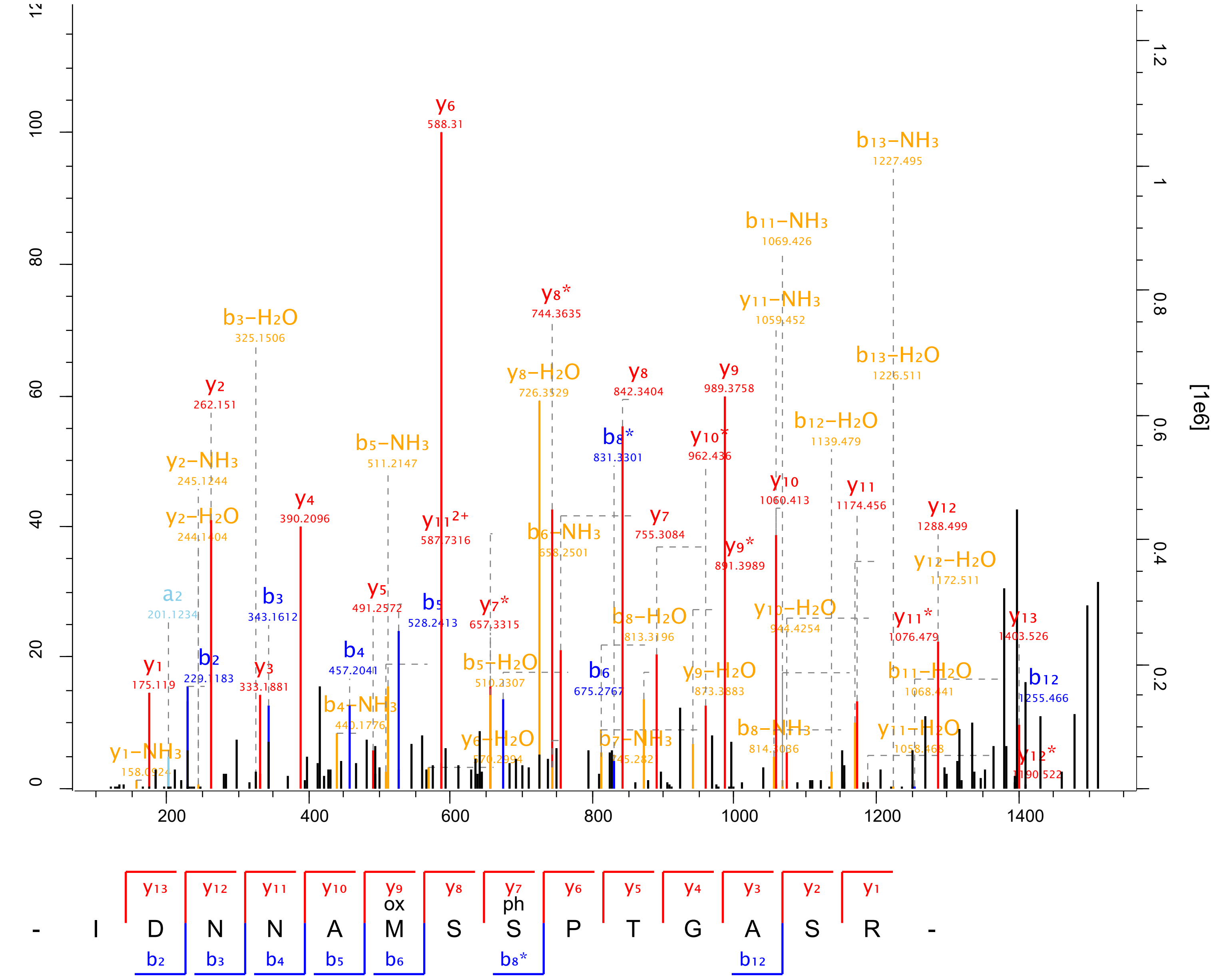

| Raw file | Scan | Method | Score | m/z |
|---|---|---|---|---|
| water_00_1 | 4127 | FTMS; HCD | 79.87 | 548.74 |

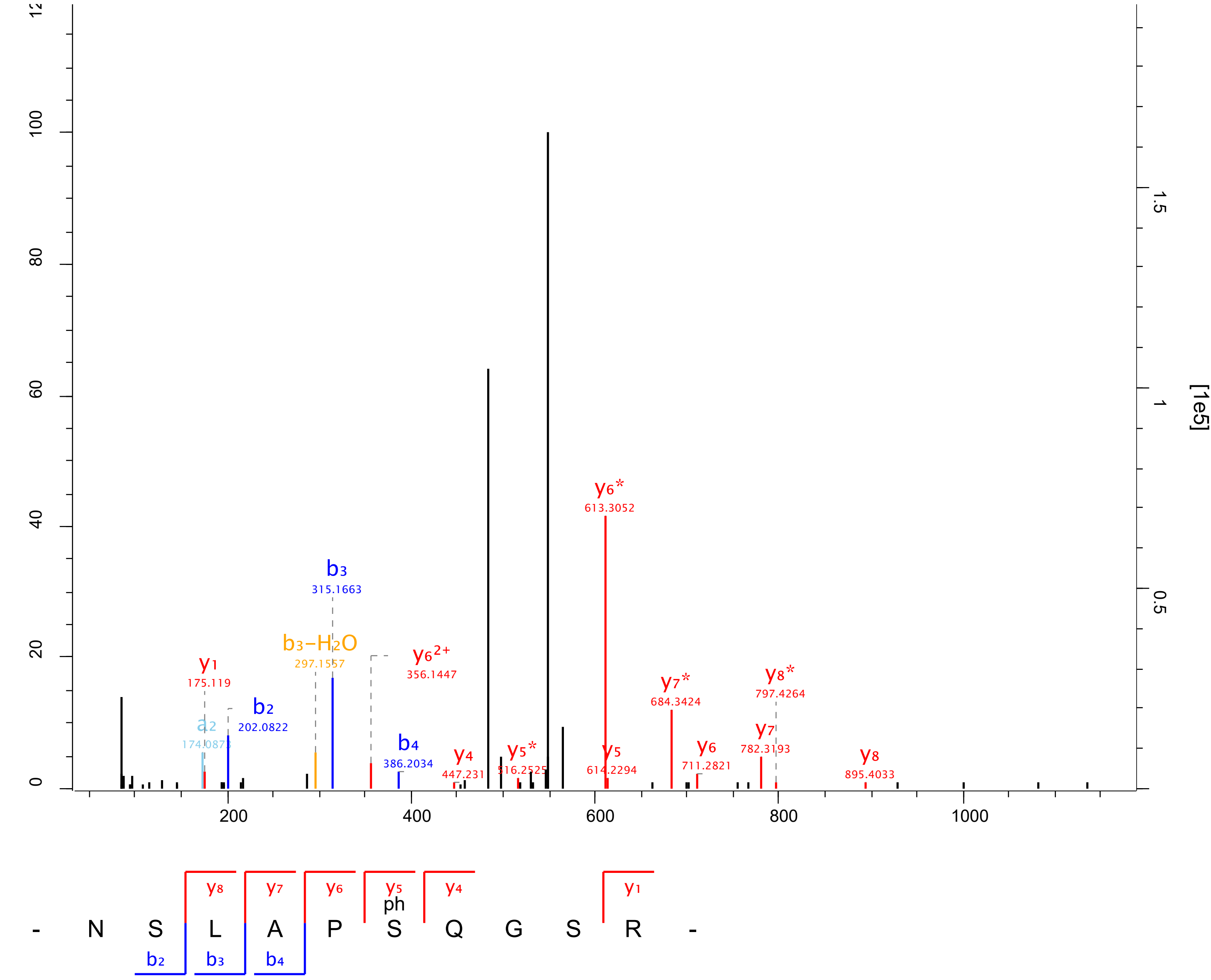

| Raw file | Scan | Method | Score | m/z |
|---|---|---|---|---|
| water_00_1 | 4156 | FTMS; HCD | 150.34 | 585.26 |

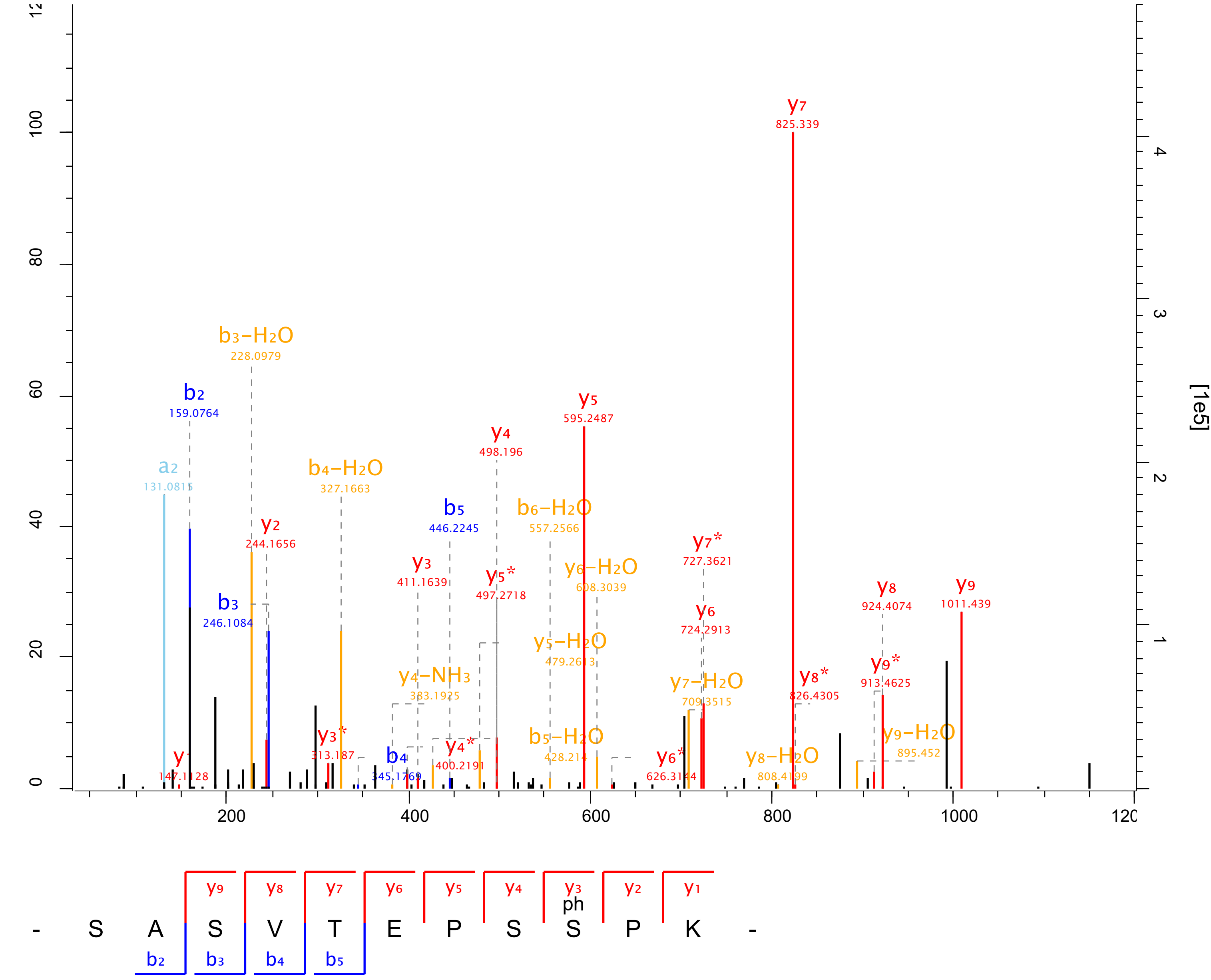

| Raw file | Scan | Method | Score | m/z |
|---|---|---|---|---|
| water_00_1 | 4244 | FTMS; HCD | 49.3 | 501.7 |

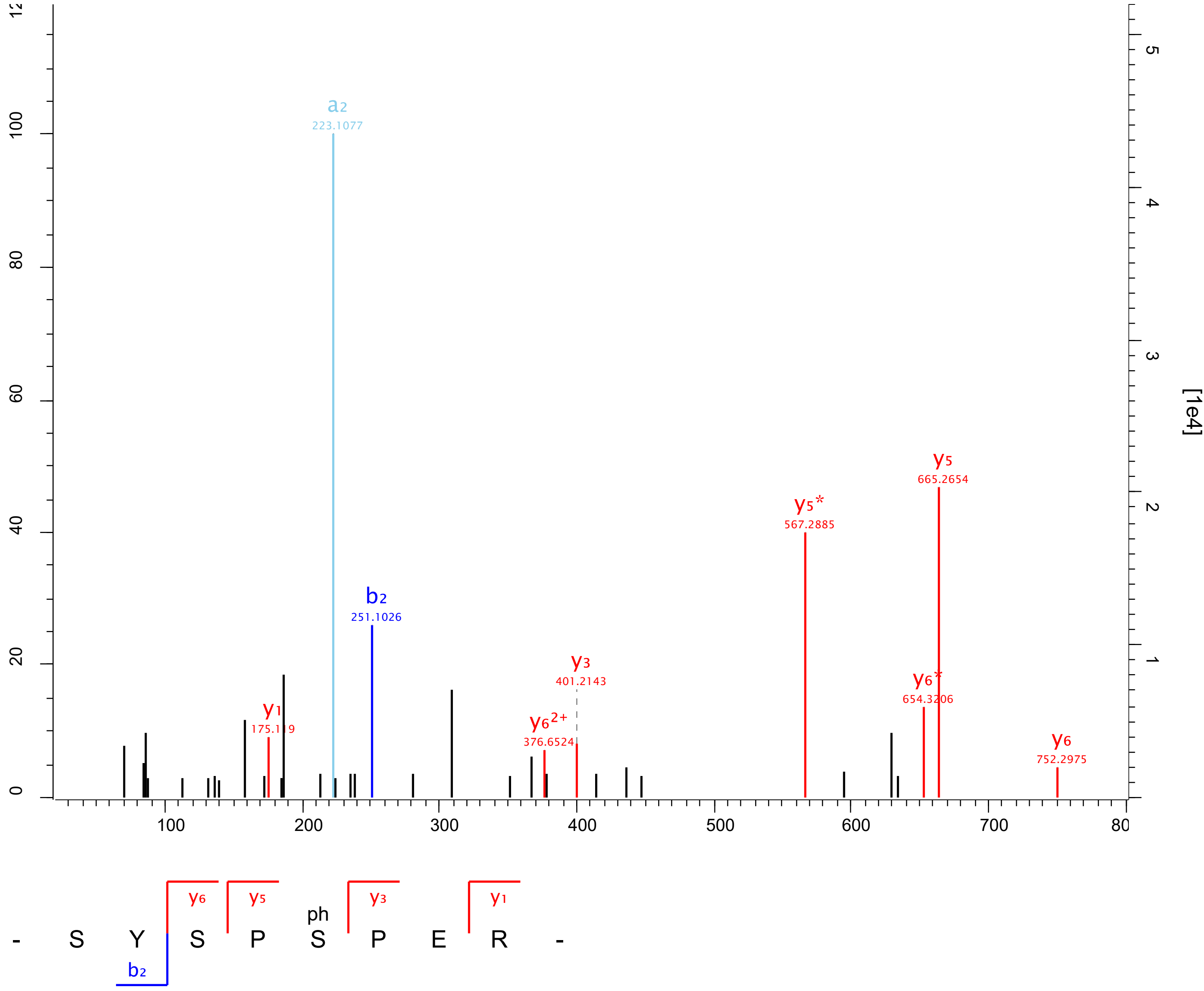

| Raw file | Scan | Method | Score | m/z |
|---|---|---|---|---|
| water_00_1 | 4246 | FTMS; HCD | 59.12 | 569.26 |

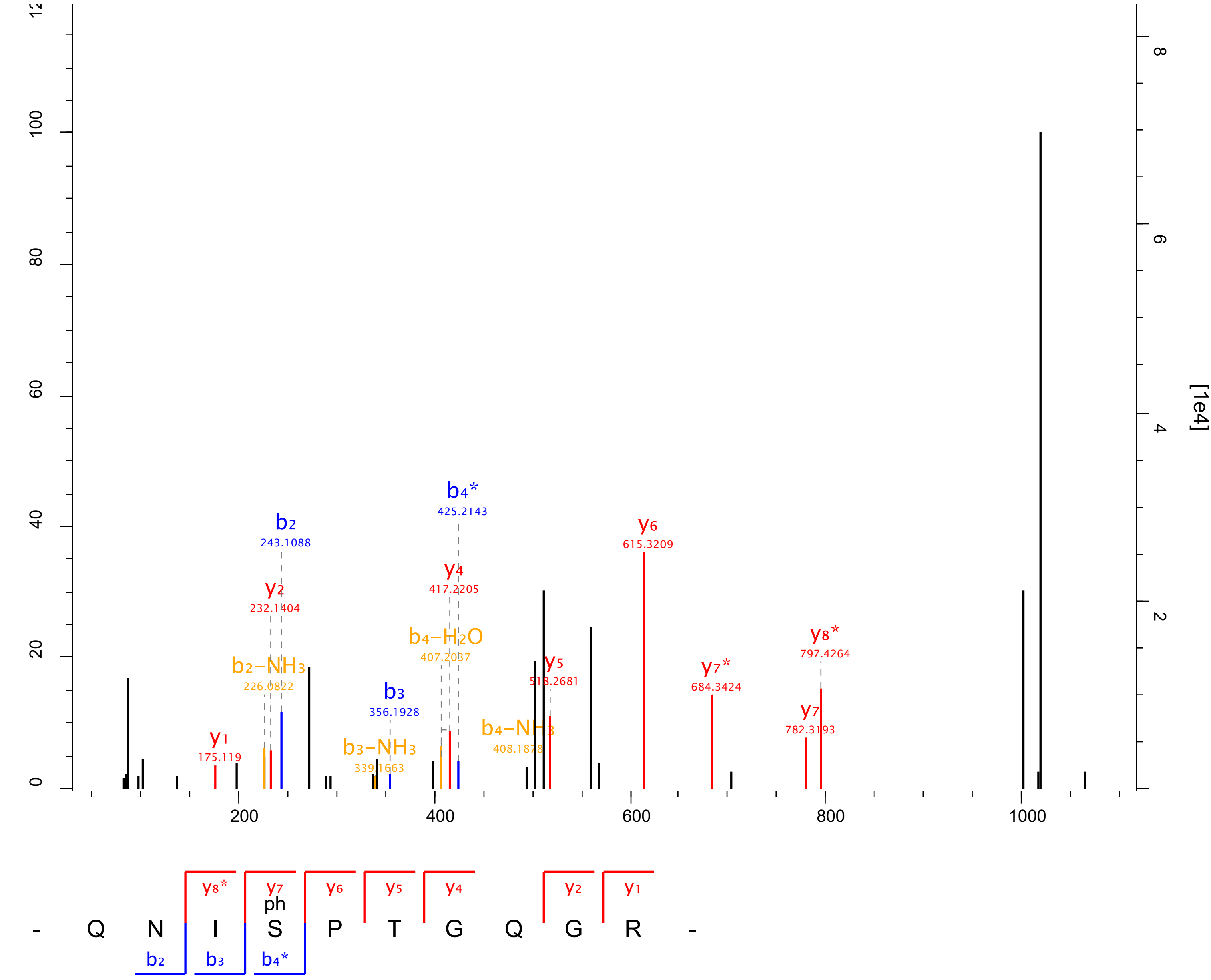

| Raw file | Scan | Method | Score | m/z |
|---|---|---|---|---|
| water_00_1 | 43060 | FTMS; HCD | 120.23 | 940.42 |

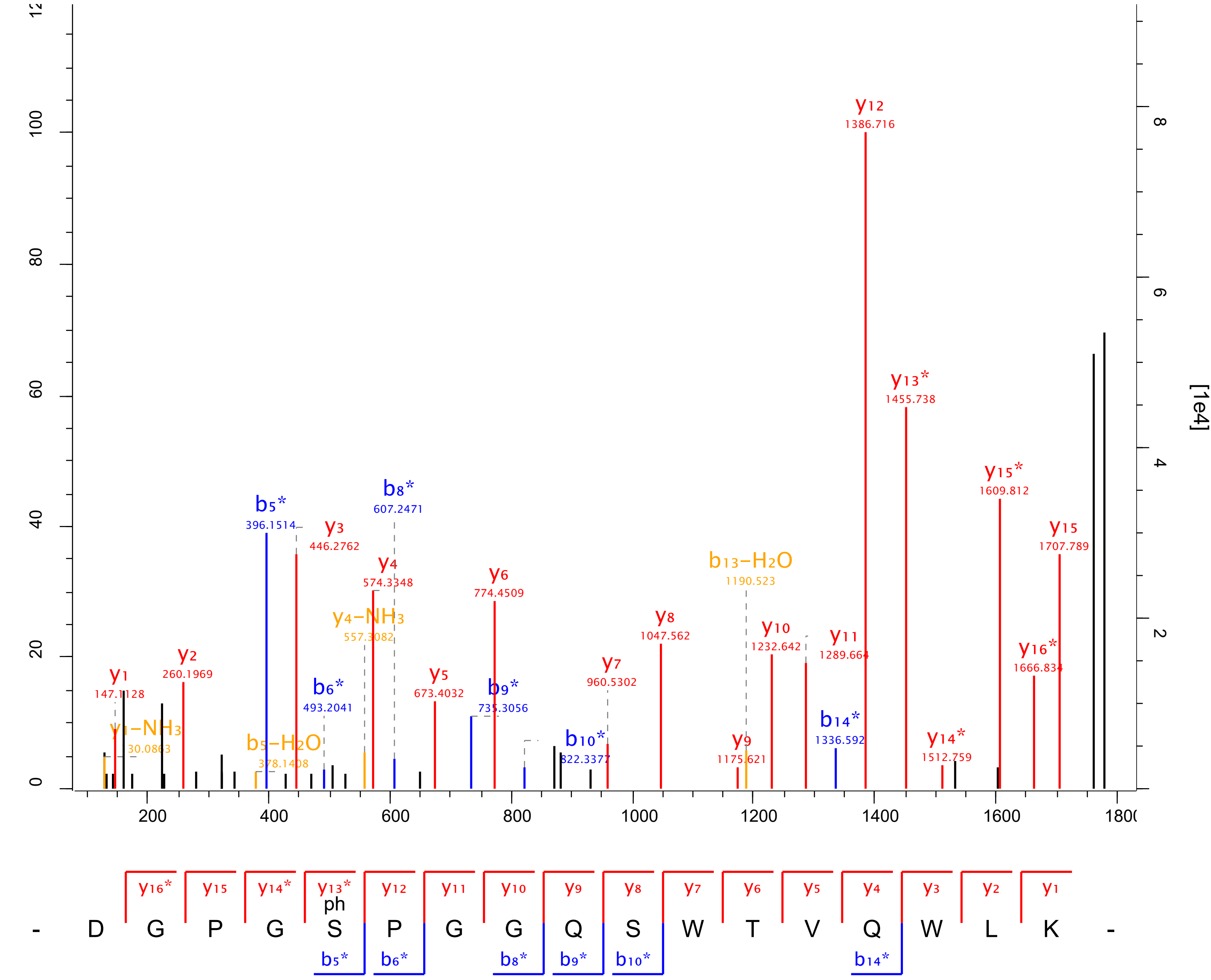

| Raw file | Scan | Method | Score | m/z |
| --- | --- | --- | --- | --- |
| water_00_1 | 4385 | FTMS; HCD | 97.81 | 485.72 |

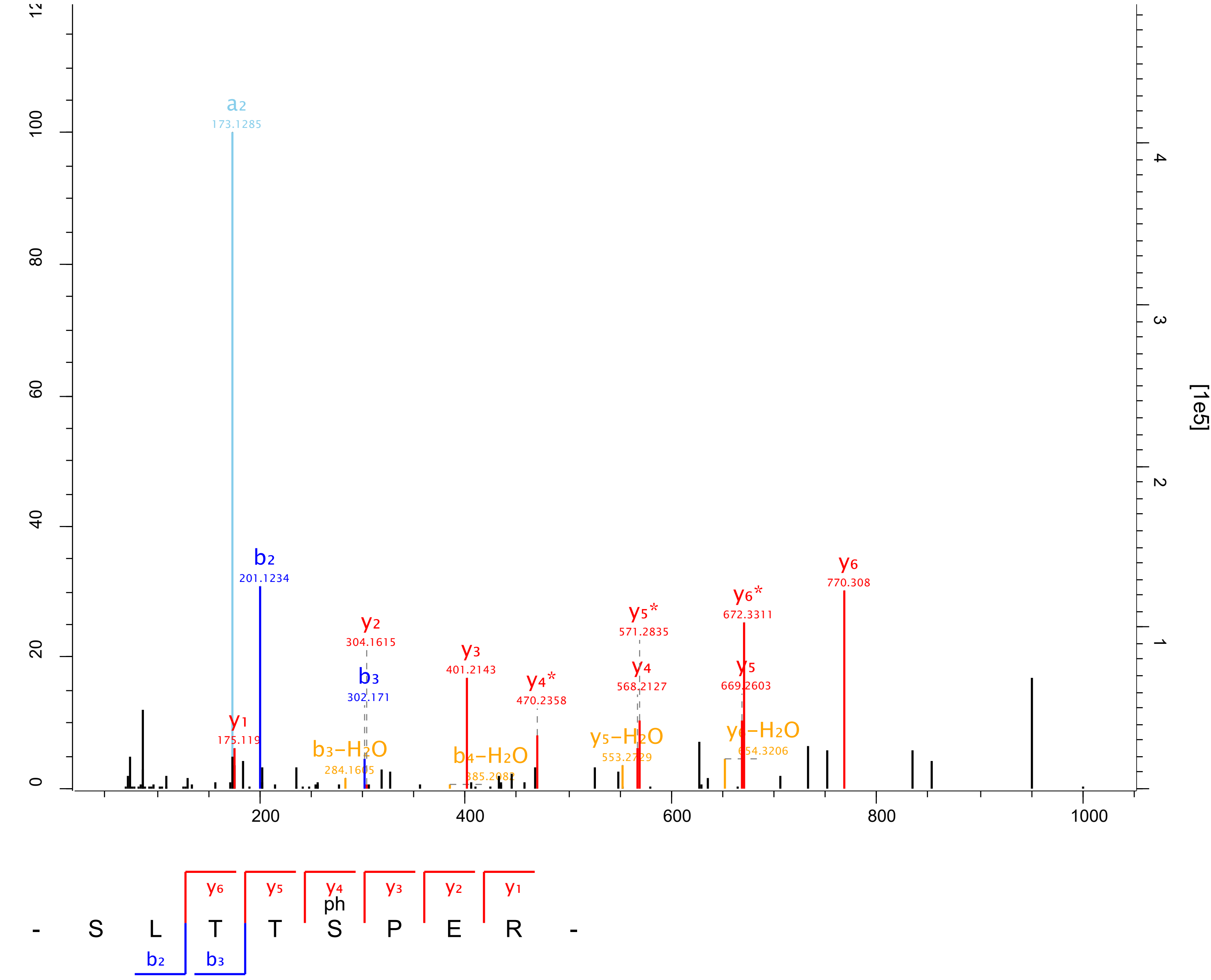

| Raw file | Scan | Method | Score | m/z |
|---|---|---|---|---|
| water_00_1 | 4394 | FTMS; HCD | 56.4 | 456.22 |

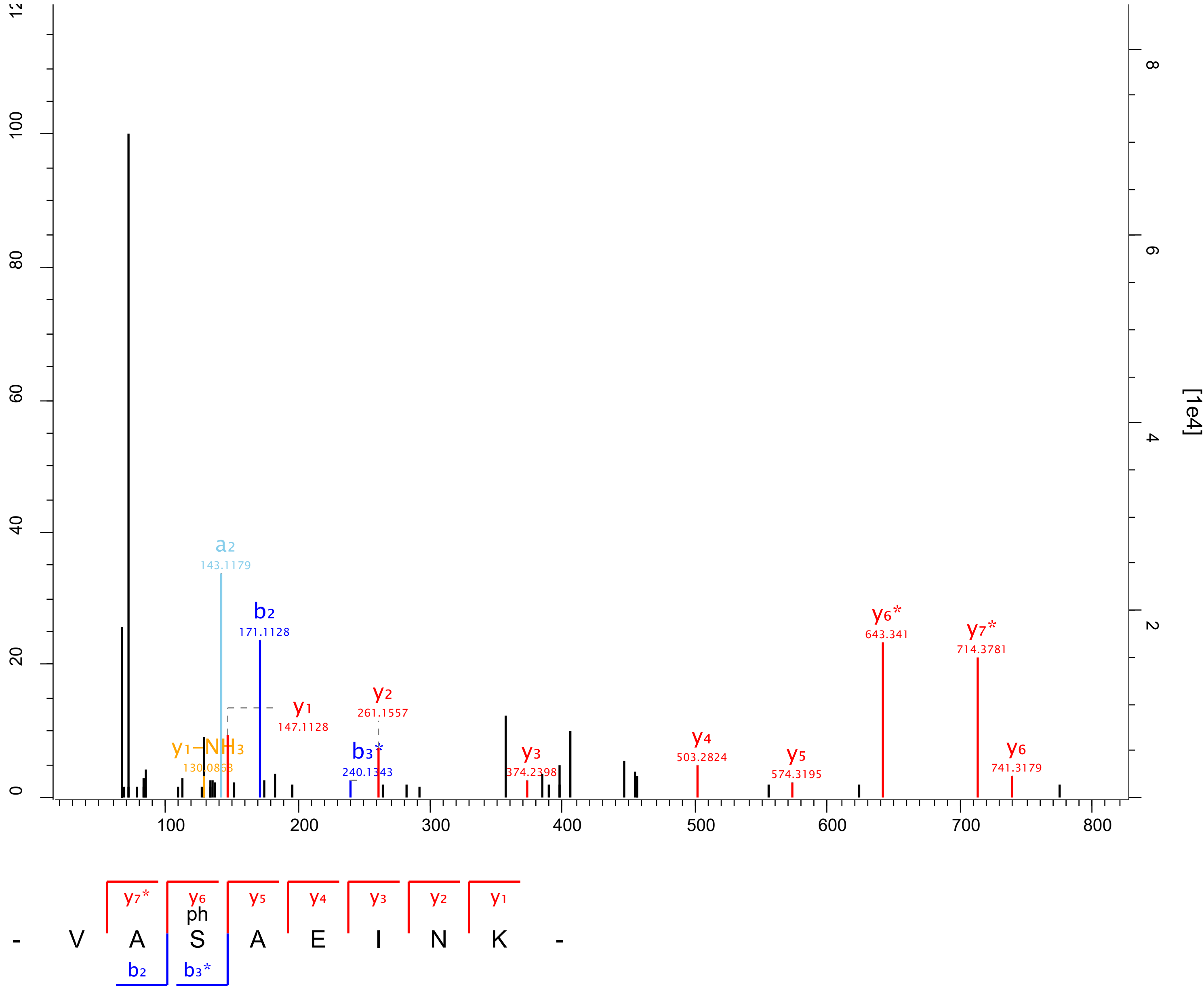

| Raw file | Scan | Method | Score | m/z |
|---|---|---|---|---|
| water_00_1 | 4419 | FTMS; HCD | 49.3 | 482.69 |

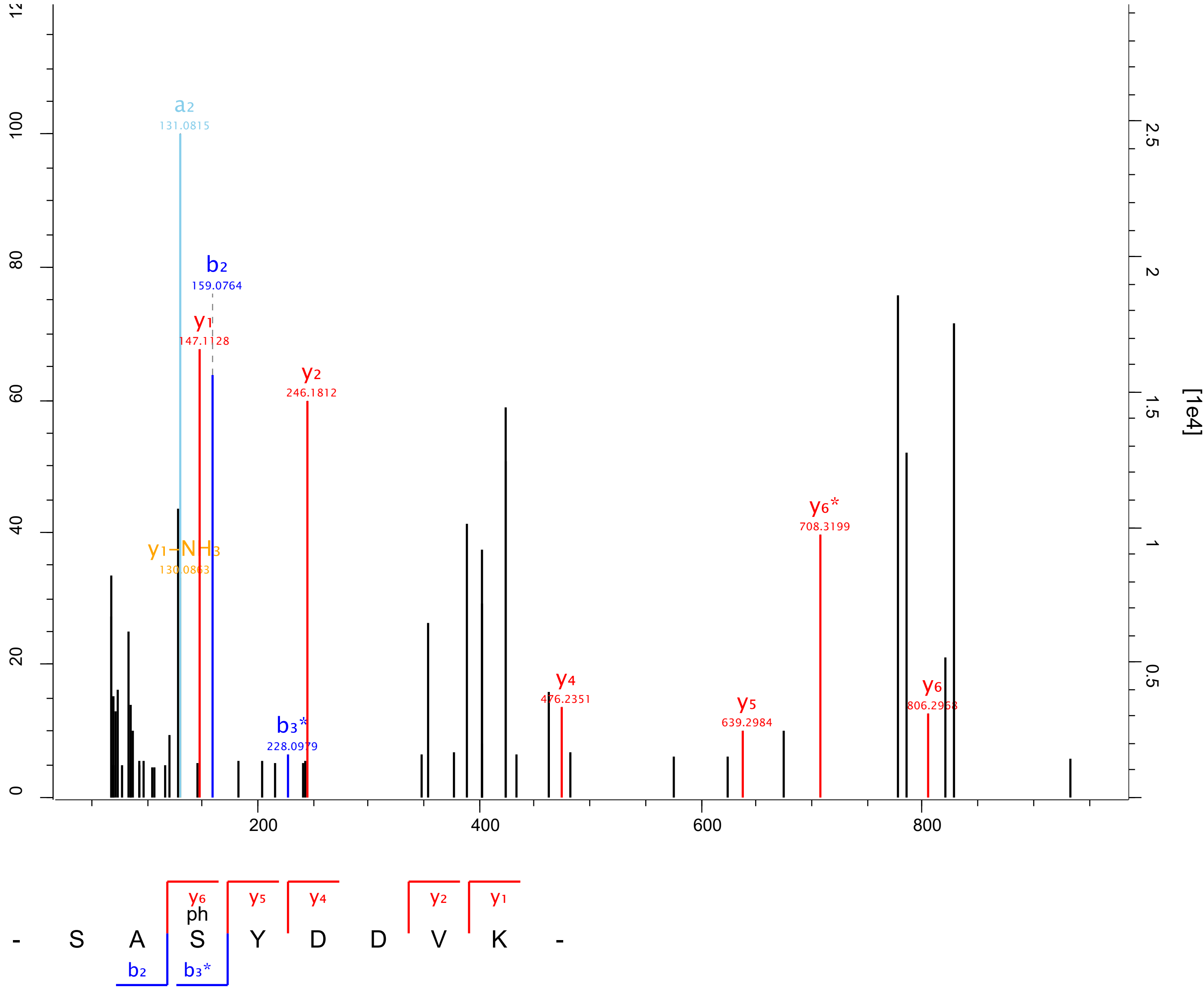

| Raw file | Scan | Method | Score | m/z |
|---|---|---|---|---|
| water_00_1 | 44529 | FTMS; HCD | 297.96 | 773.86 |

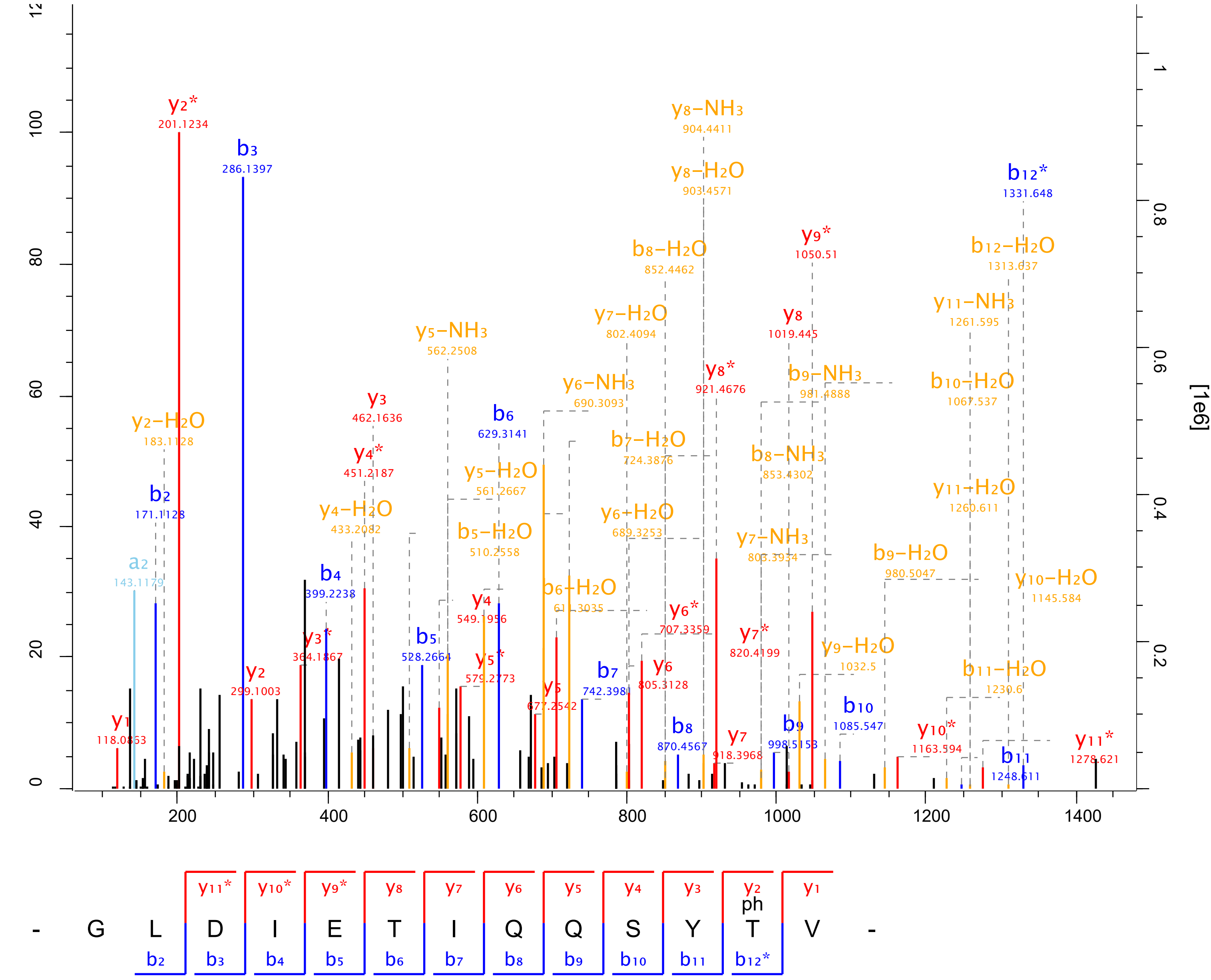

| Raw file | Scan | Method | Score | m/z |
| --- | --- | --- | --- | --- |
| water_00_1 | 44646 | FTMS; HCD | 113.91 | 885.41 |

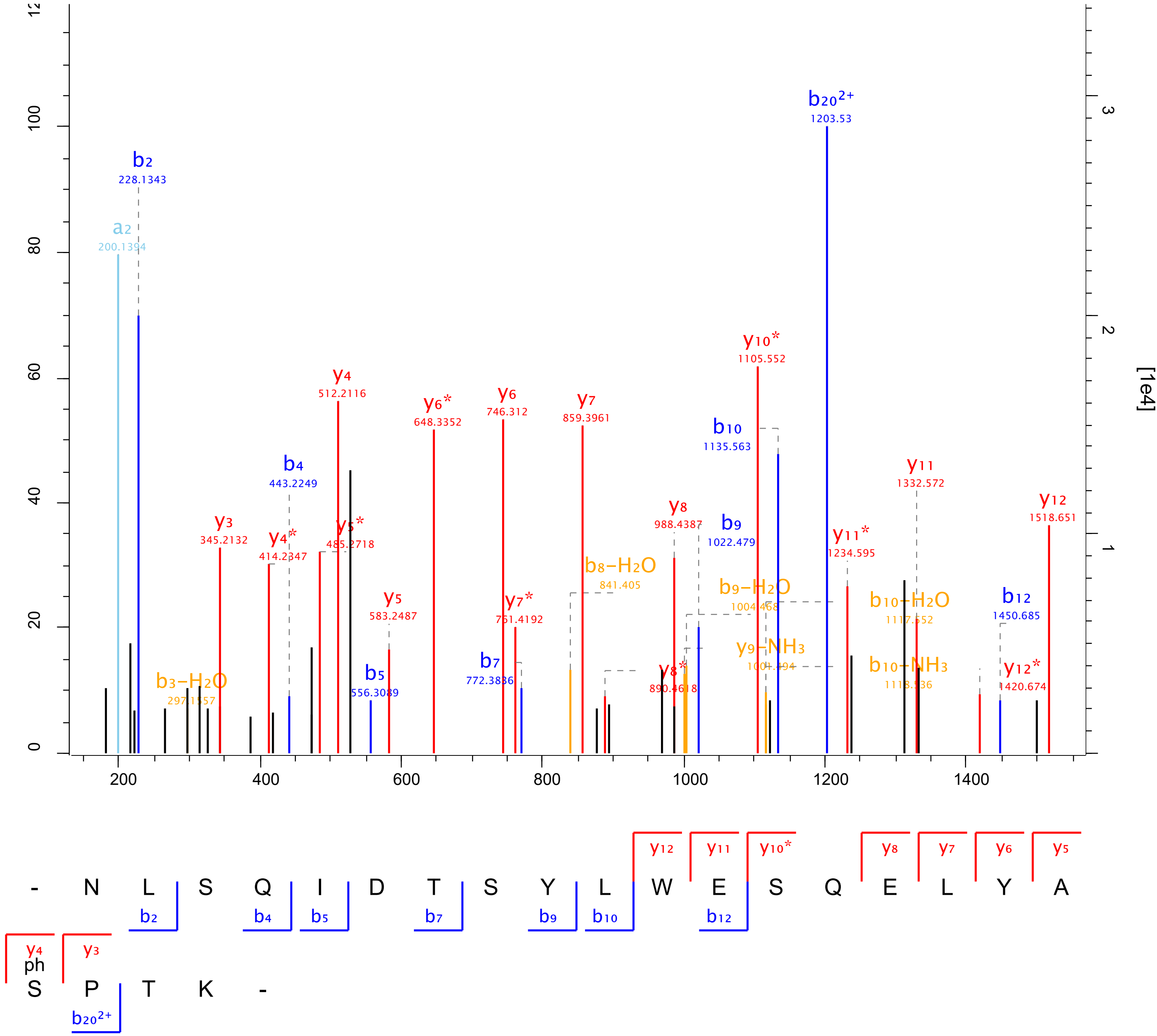

| Raw file | Scan | Method | Score | m/z |
|---|---|---|---|---|
| water_00_1 | 4477 | FTMS; HCD | 55.81 | 531.74 |

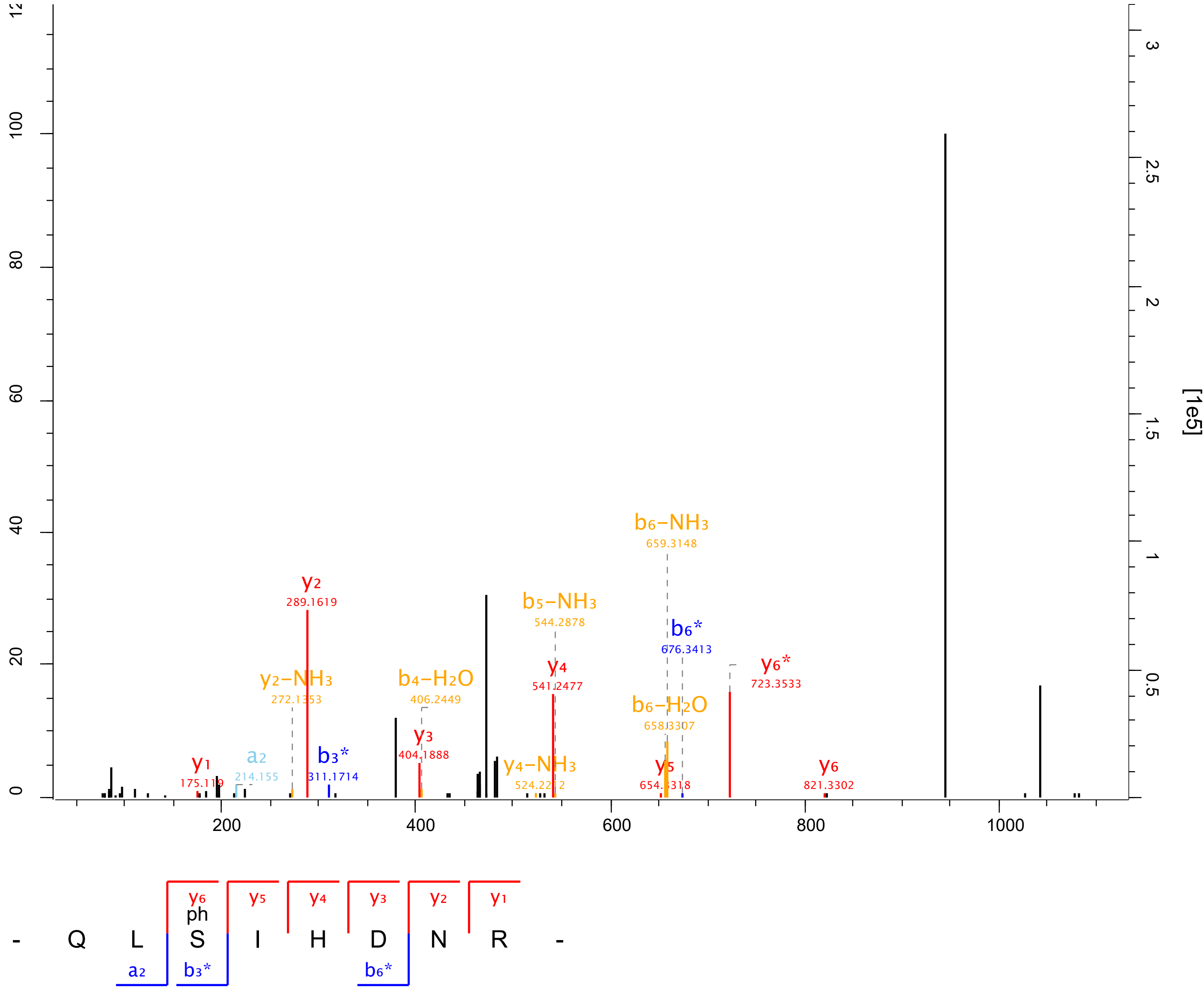

| Raw file | Scan | Method | Score | m/z |
| --- | --- | --- | --- | --- |
| water_00_1 | 45119 | FTMS; HCD | 102.6 | 848.09 |

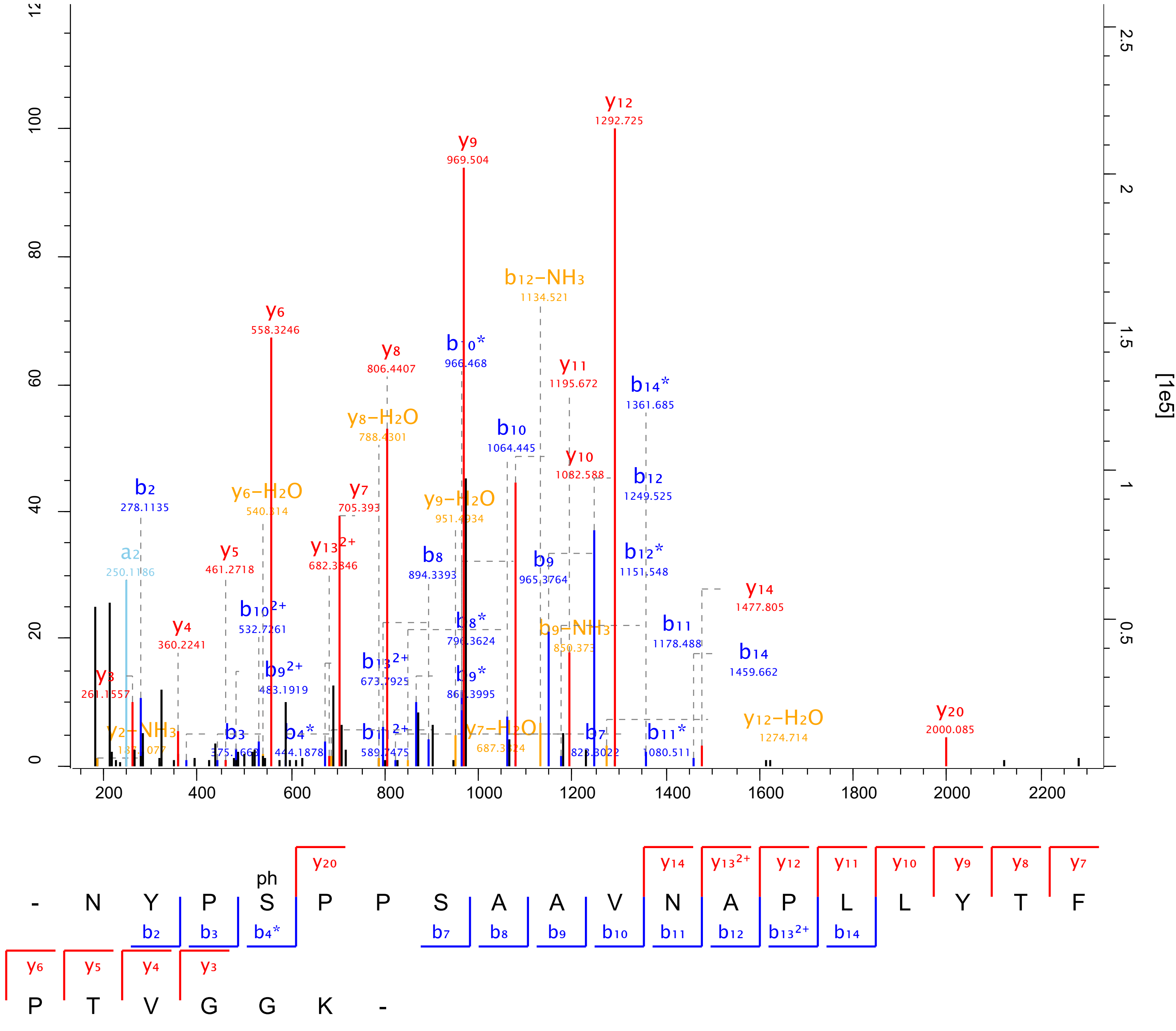

| Raw file | Scan | Method | Score | m/z |
|---|---|---|---|---|
| water_00_1 | 4517 | FTMS; HCD | 47.77 | 649.78 |

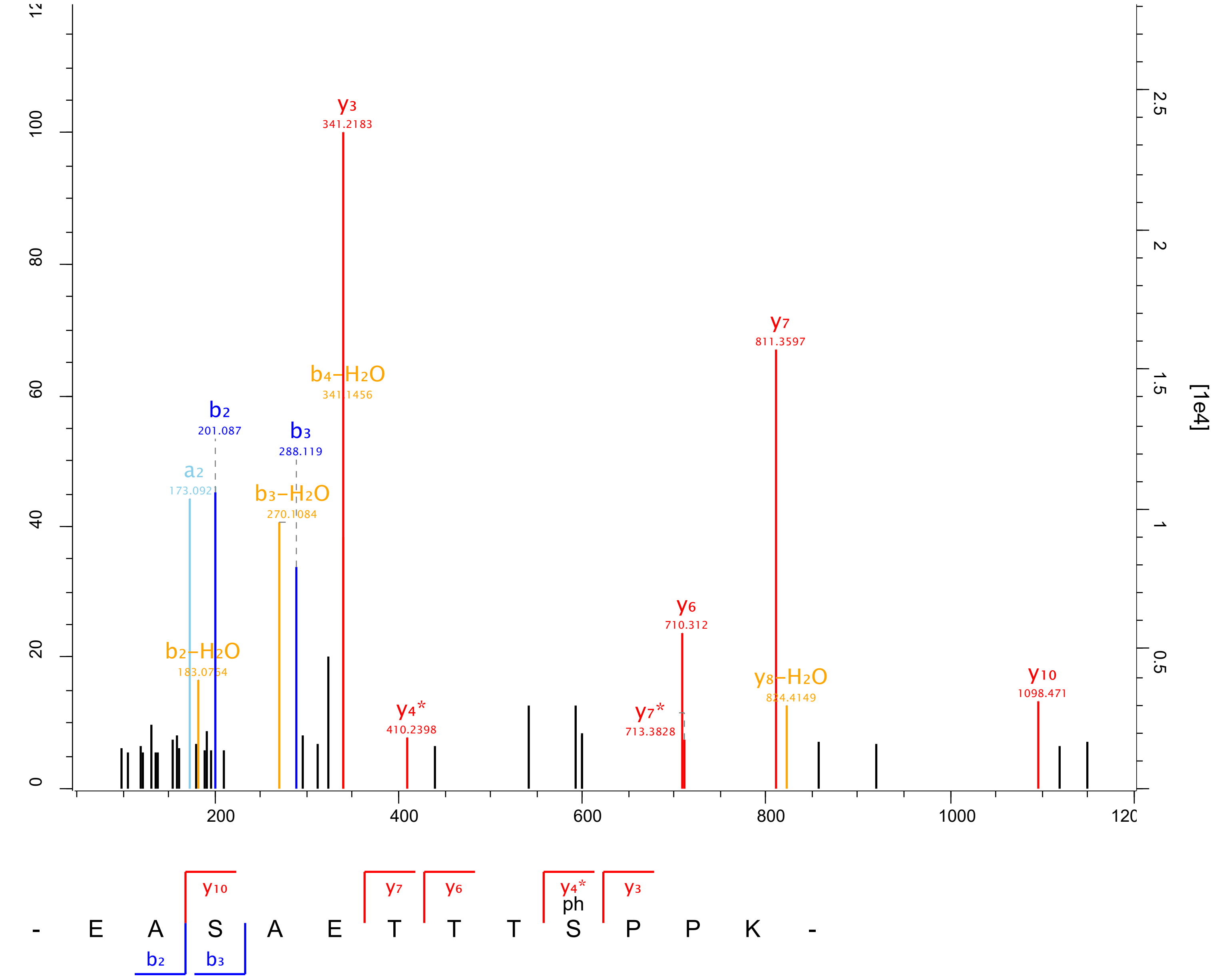

| Raw file | Scan | Method | Score | m/z |
|---|---|---|---|---|
| water_00_1 | 45314 | FTMS; HCD | 52.25 | 591.8 |

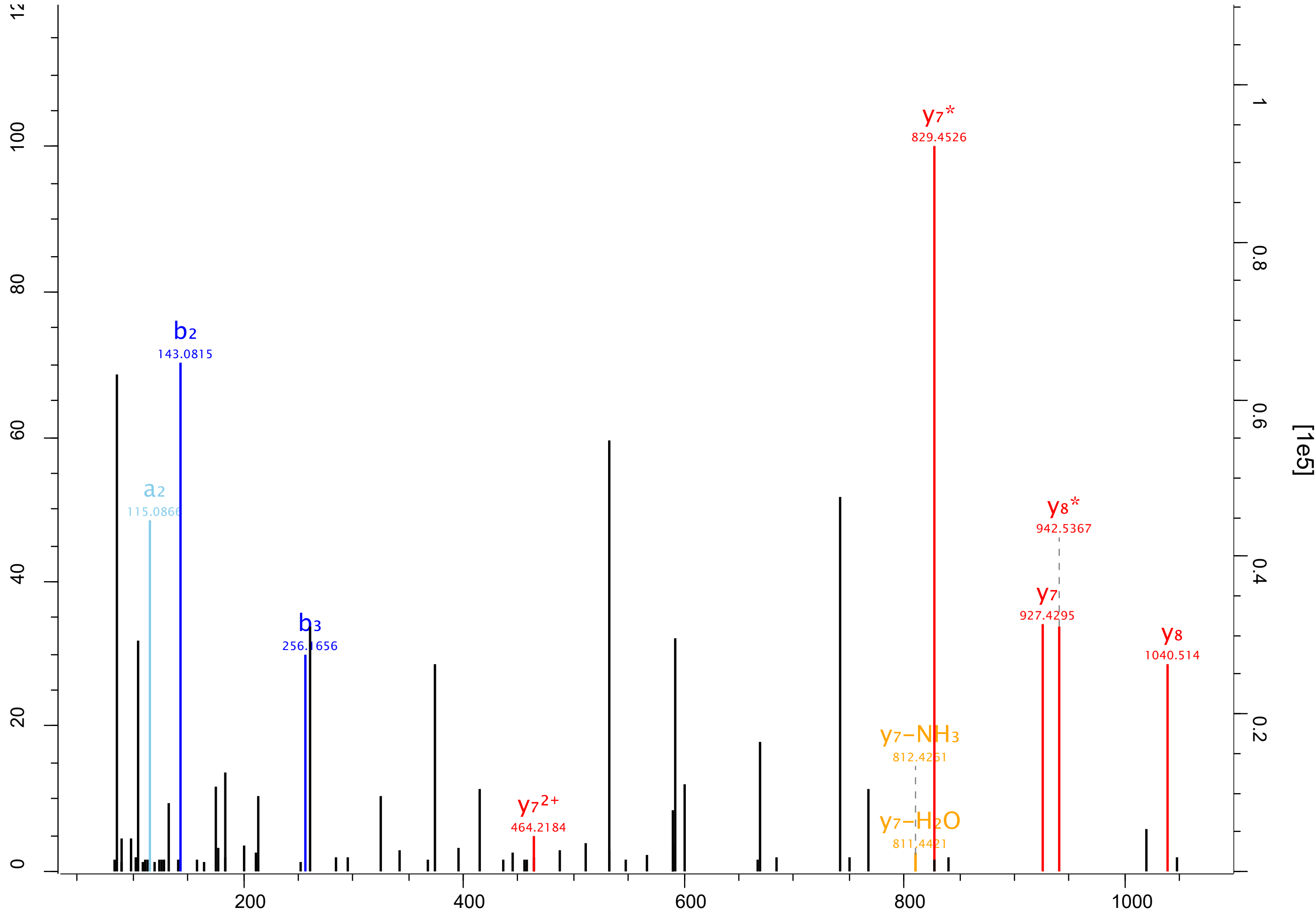


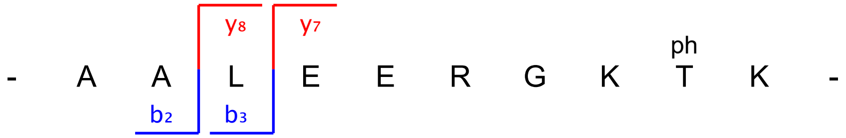

| Raw file | Scan | Method | Score | m/z |
|---|---|---|---|---|
| water_00_1 | 45460 | FTMS; HCD | 92.96 | 936.49 |

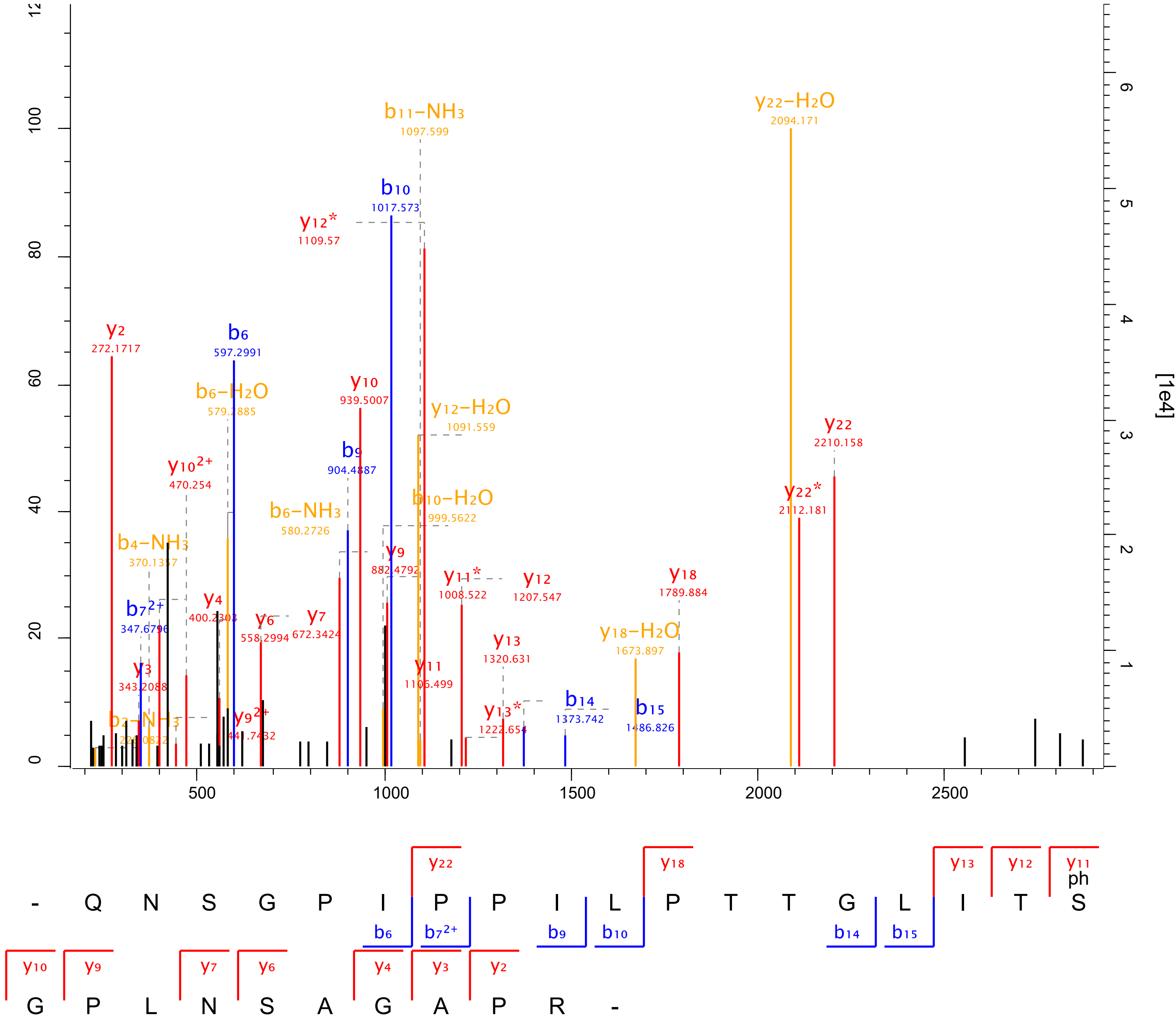

| Raw file | Scan | Method | Score | m/z |
|---|---|---|---|---|
| water_00_1 | 46719 | FTMS; HCD | 41.05 | 841.72 |

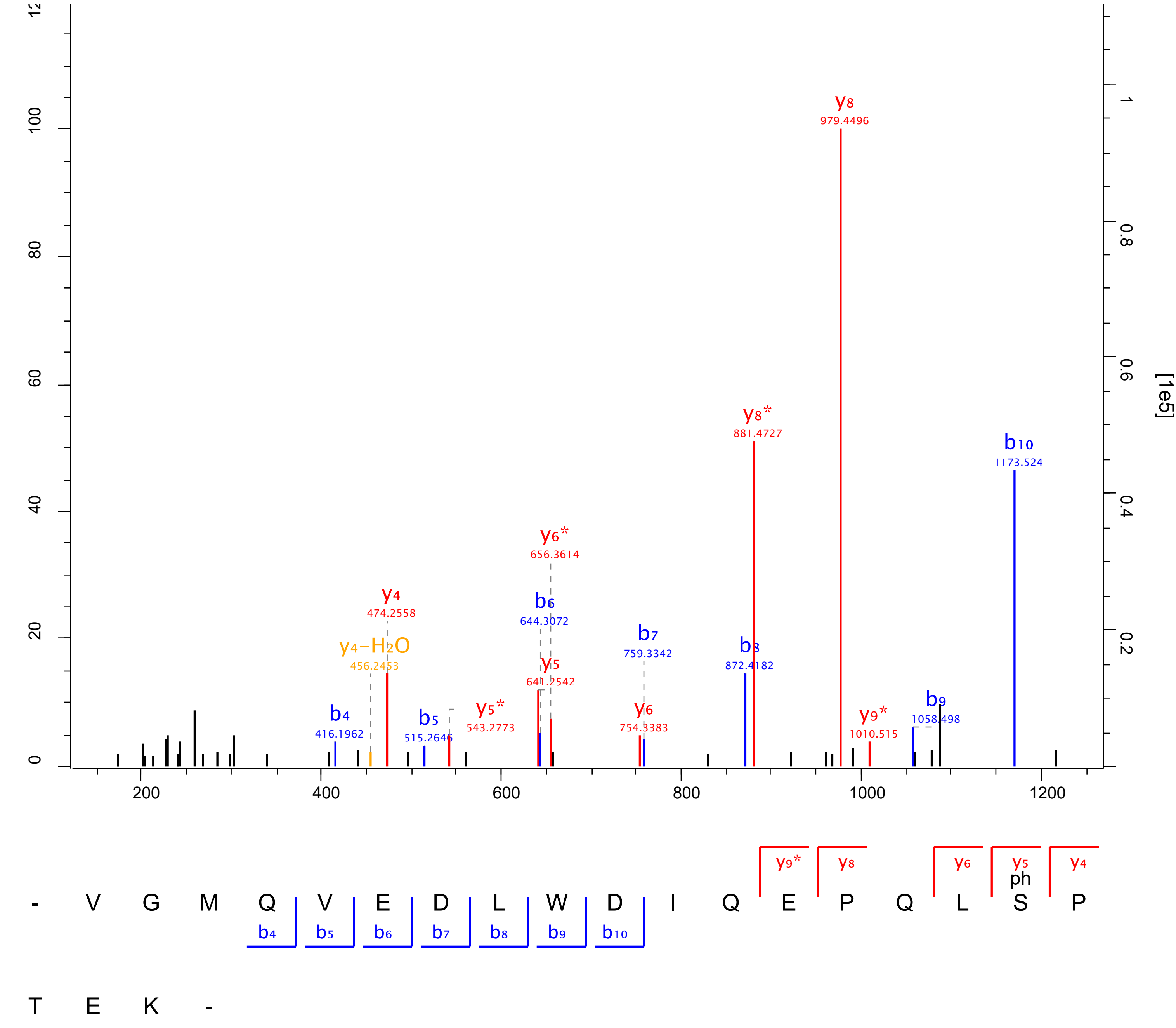

| Raw file | Scan | Method | Score | m/z |
| --- | --- | --- | --- | --- |
| water_00_1 | 47329 | FTMS; HCD | 50.3 | 733.35 |

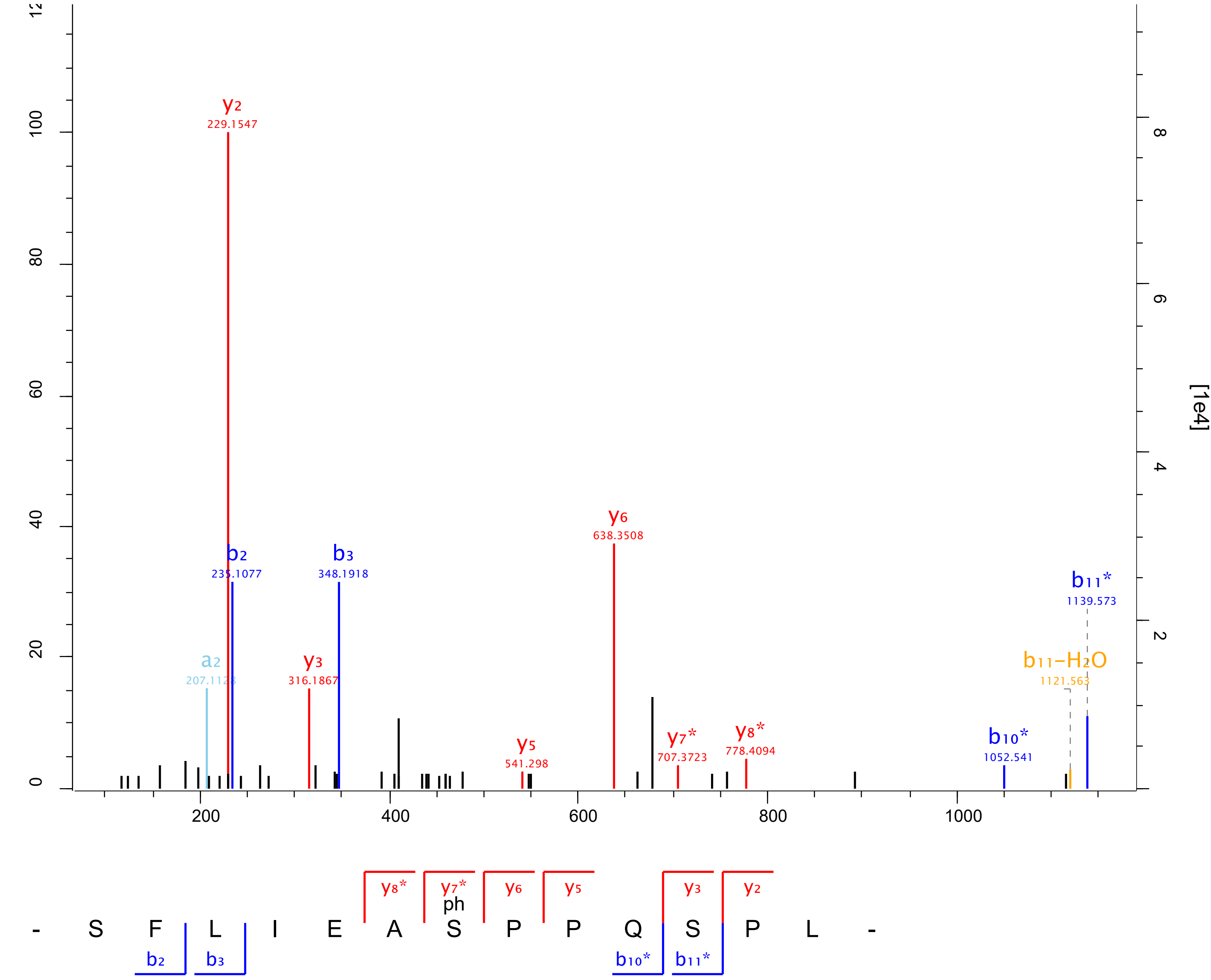

| Raw file | Scan | Method | Score | m/z |
| --- | --- | --- | --- | --- |
| water_00_1 | 48083 | FTMS; HCD | 154.34 | 910.43 |

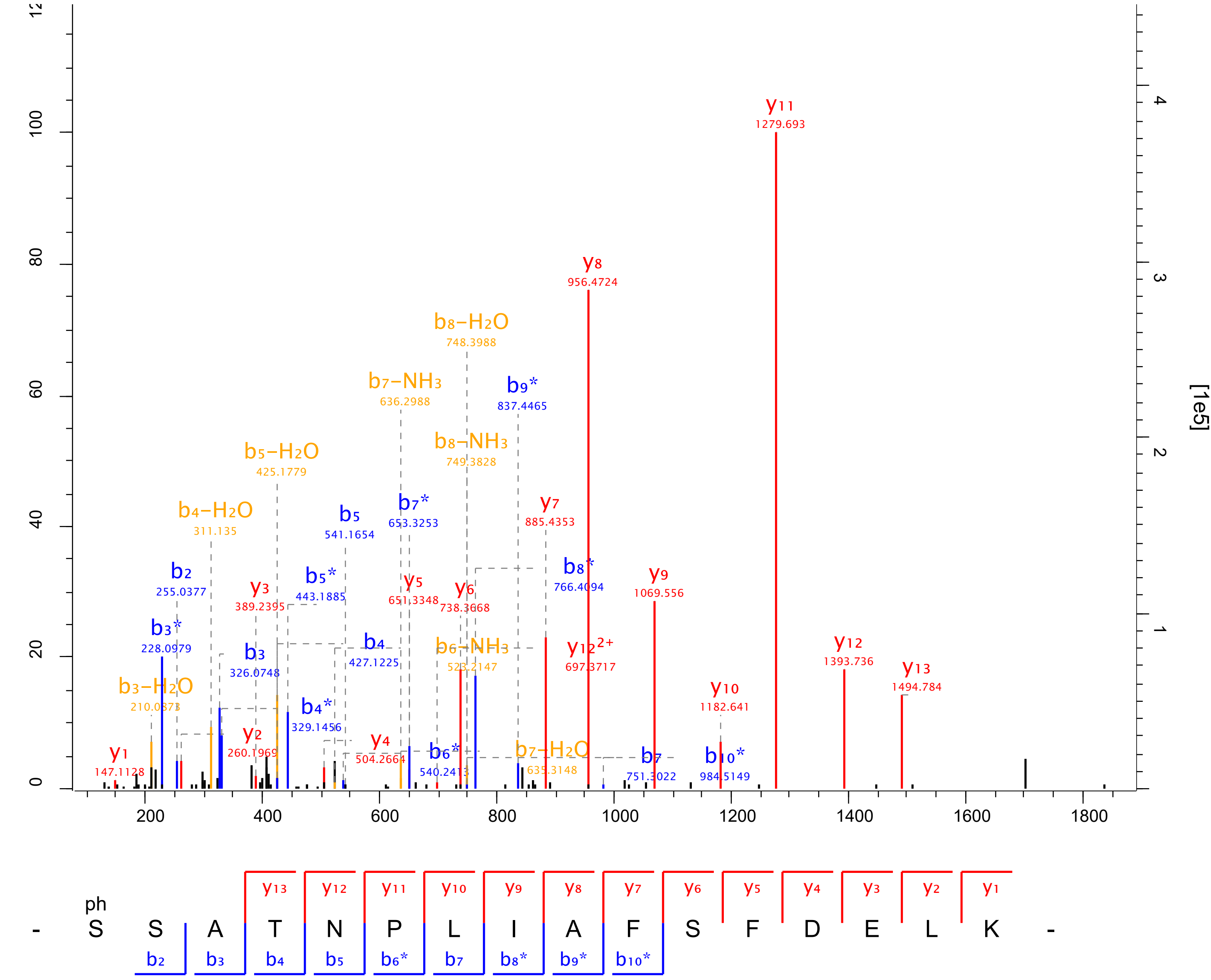

| Raw file | Scan | Method | Score | m/z |
|---|---|---|---|---|
| water_00_1 | 4813 | FTMS; HCD | 125.29 | 629.28 |

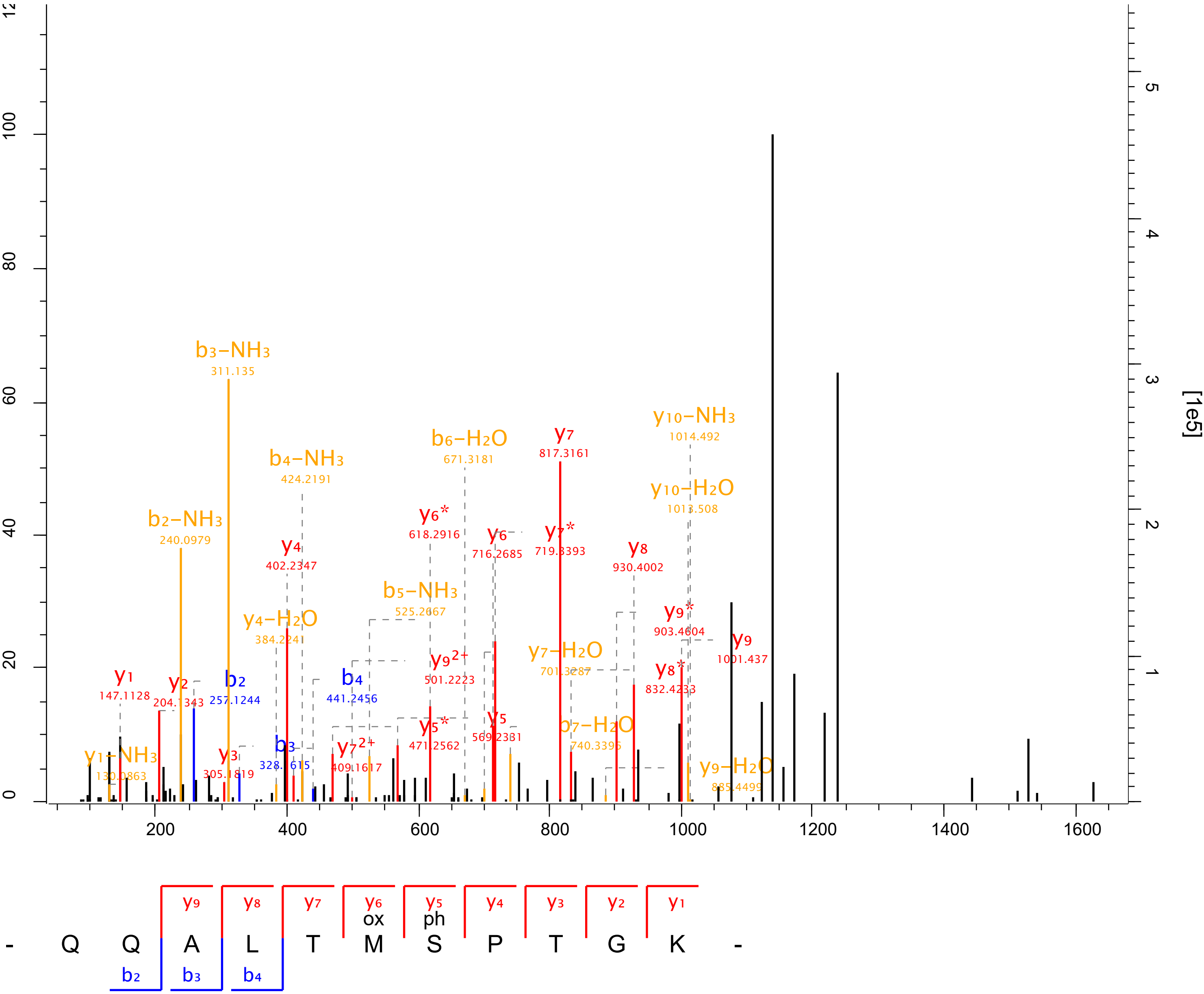

| Raw file | Scan | Method | Score | m/z |
|---|---|---|---|---|
| water_00_1 | 48244 | FTMS; HCD | 128.12 | 1008.79 |

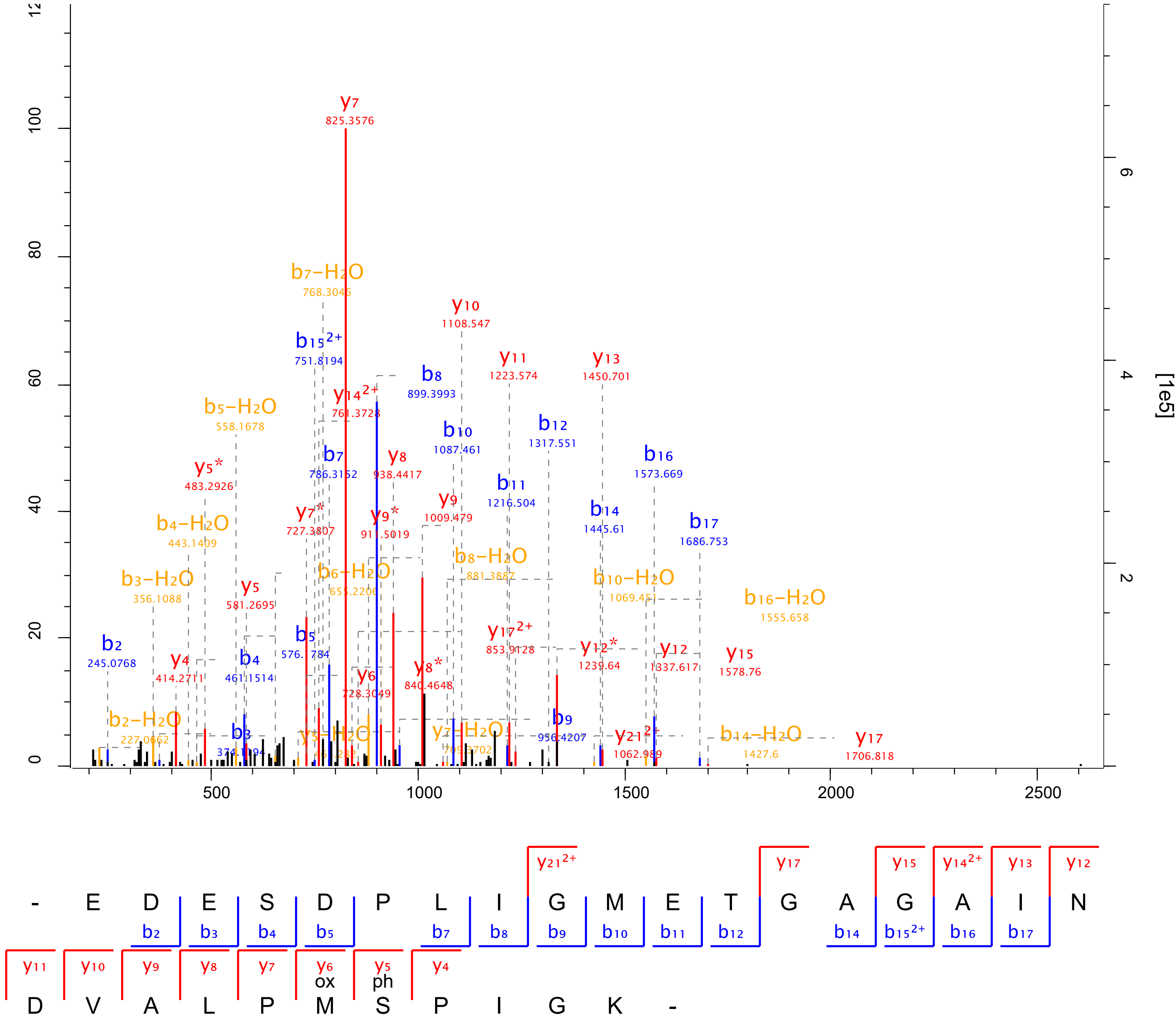

| Raw file | Scan | Method | Score | m/z |
| --- | --- | --- | --- | --- |
| water_00_1 | 4891 | FTMS; HCD | 93.78 | 632.58 |

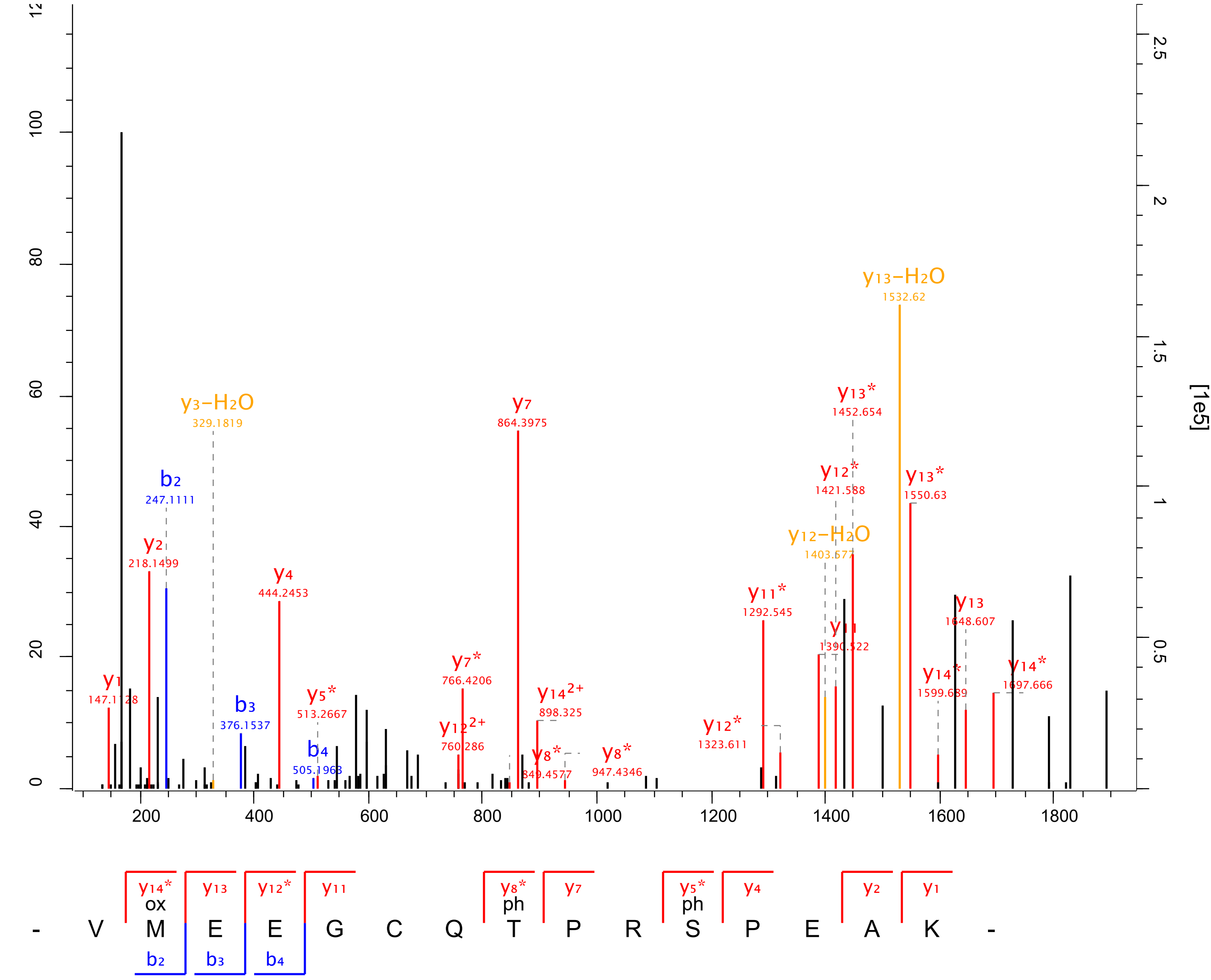

| Raw file | Scan | Method | Score | m/z |
| --- | --- | --- | --- | --- |
| water_00_1 | 4904 | FTMS; HCD | 51.28 | 461.72 |

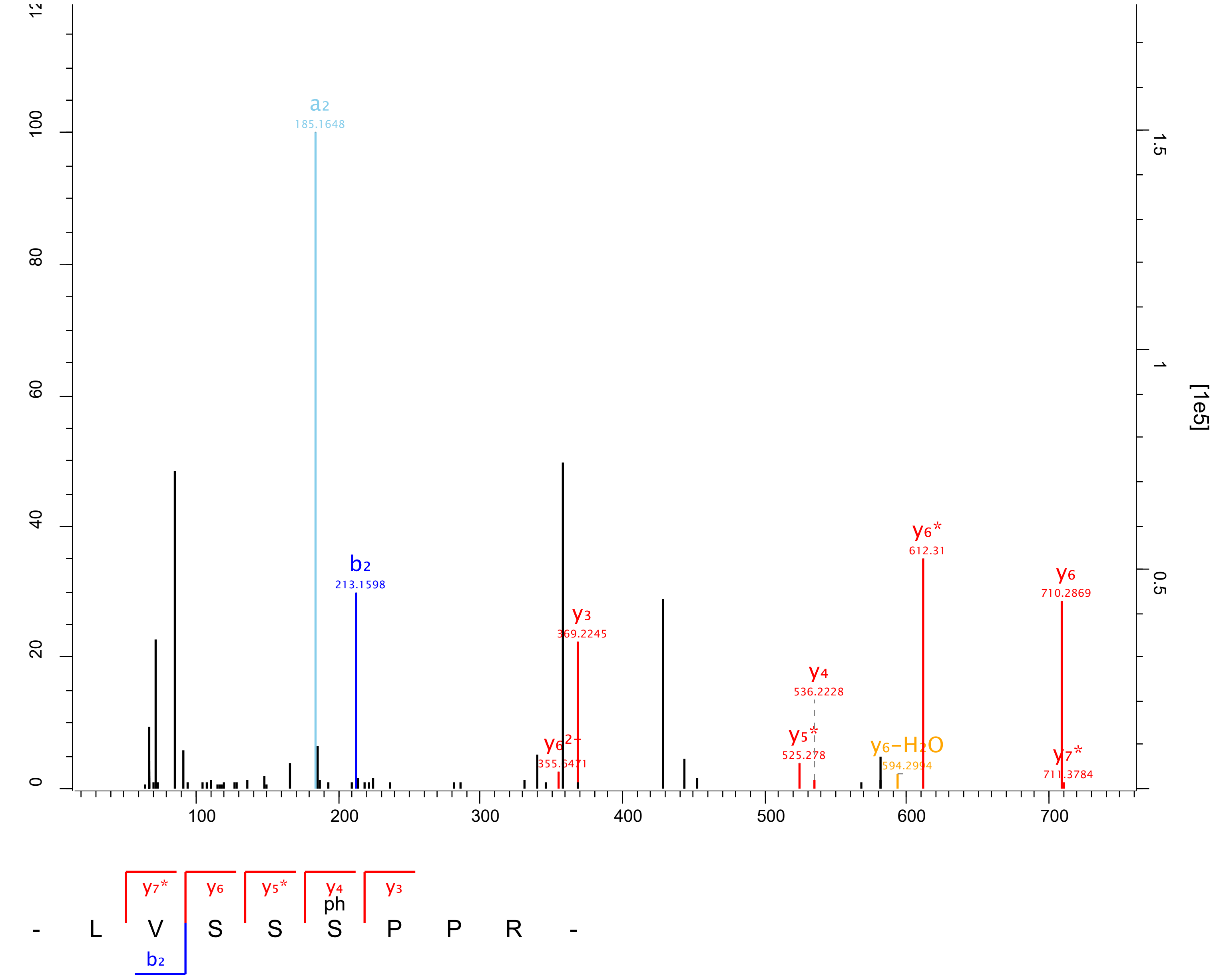

| Raw file | Scan | Method | Score | m/z |
| --- | --- | --- | --- | --- |
| water_00_1 | 5099 | FTMS; HCD | 64.82 | 489.7 |

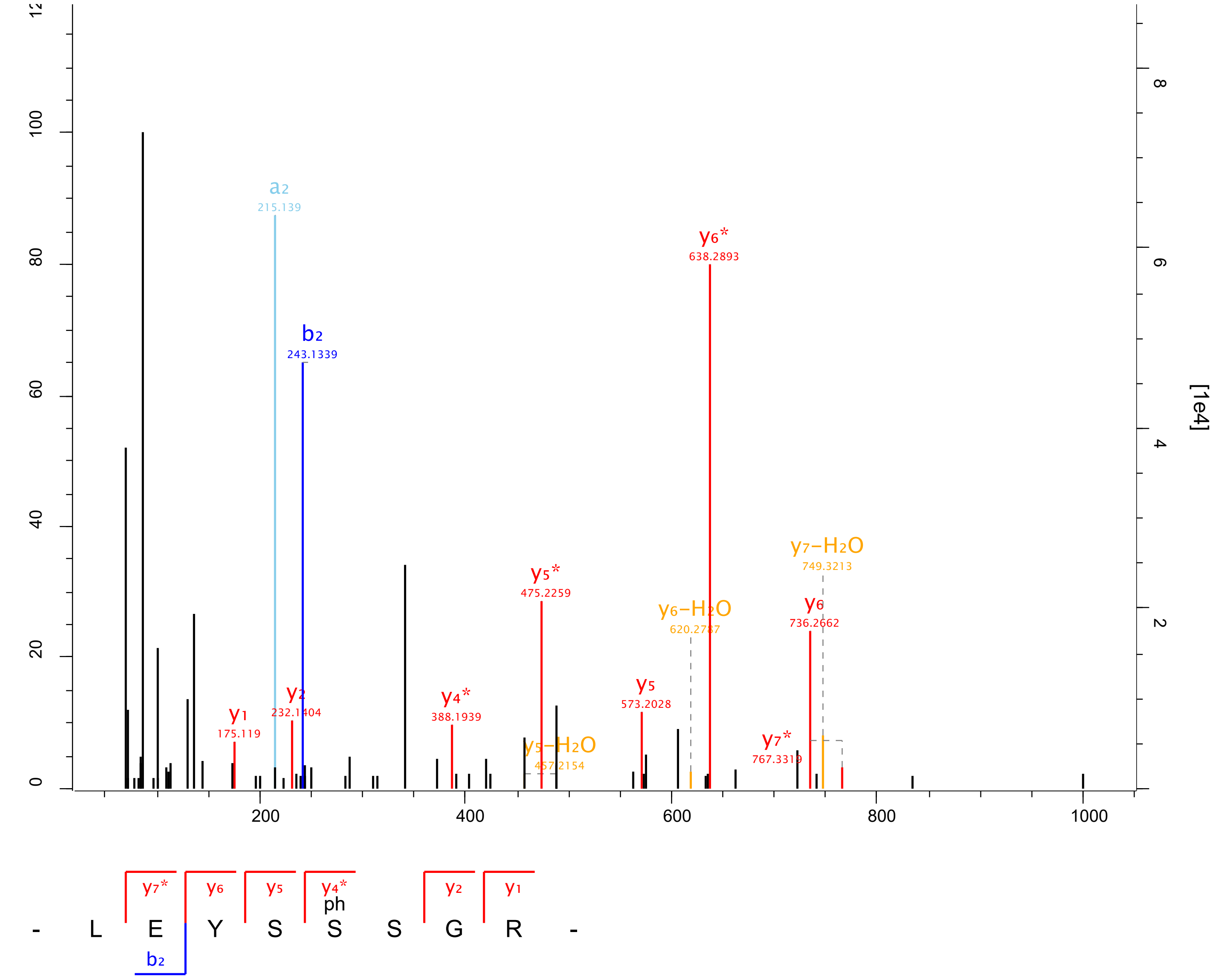

| Raw file | Scan | Method | Score | m/z |
| --- | --- | --- | --- | --- |
| water_00_1 | 5248 | FTMS; HCD | 89.51 | 541.24 |

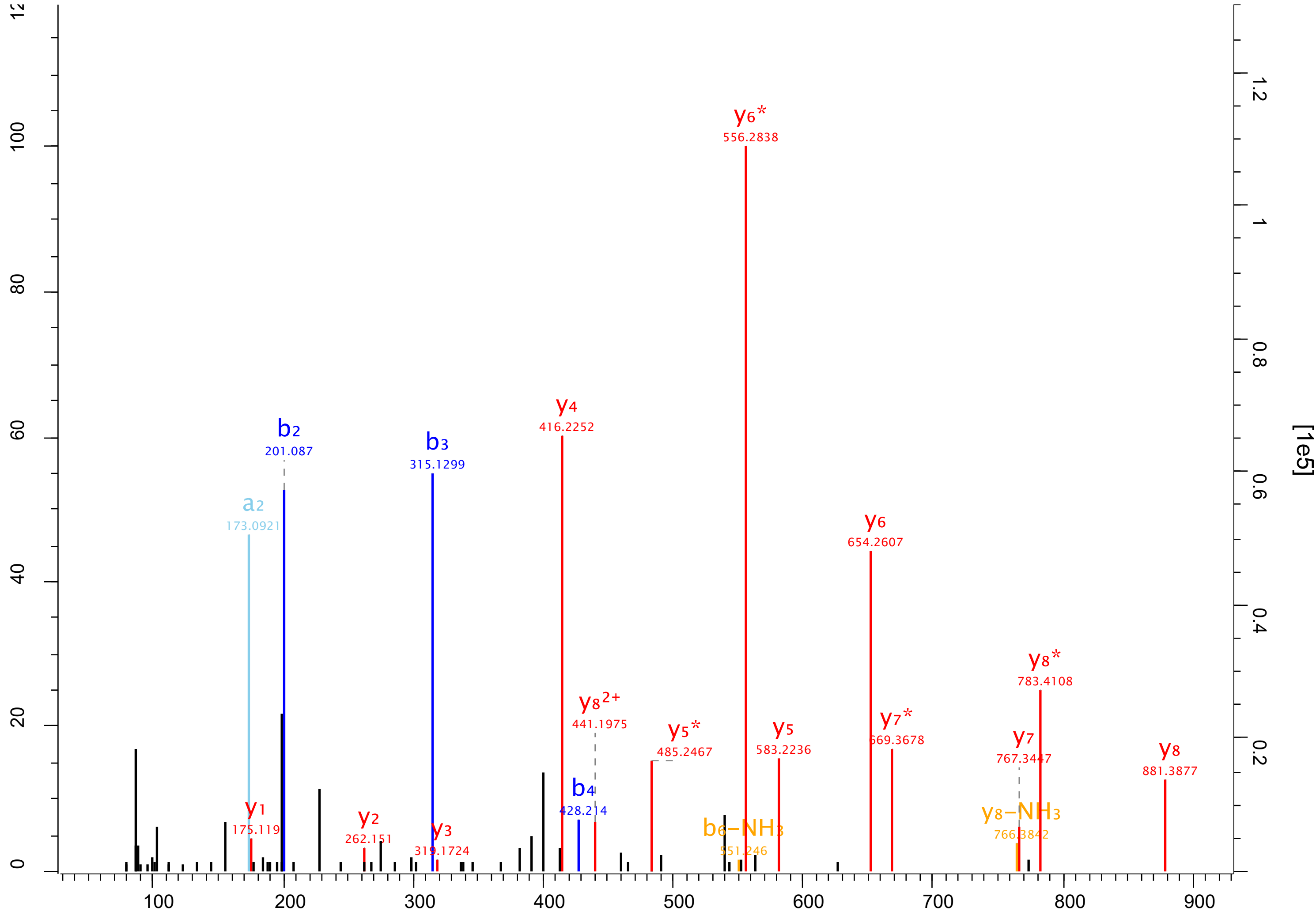


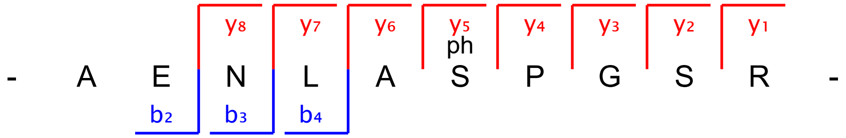

| Raw file | Scan | Method | Score | m/z |
|---|---|---|---|---|
| water_00_1 | 528 | FTMS; HCD | 61.16 | 523.24 |

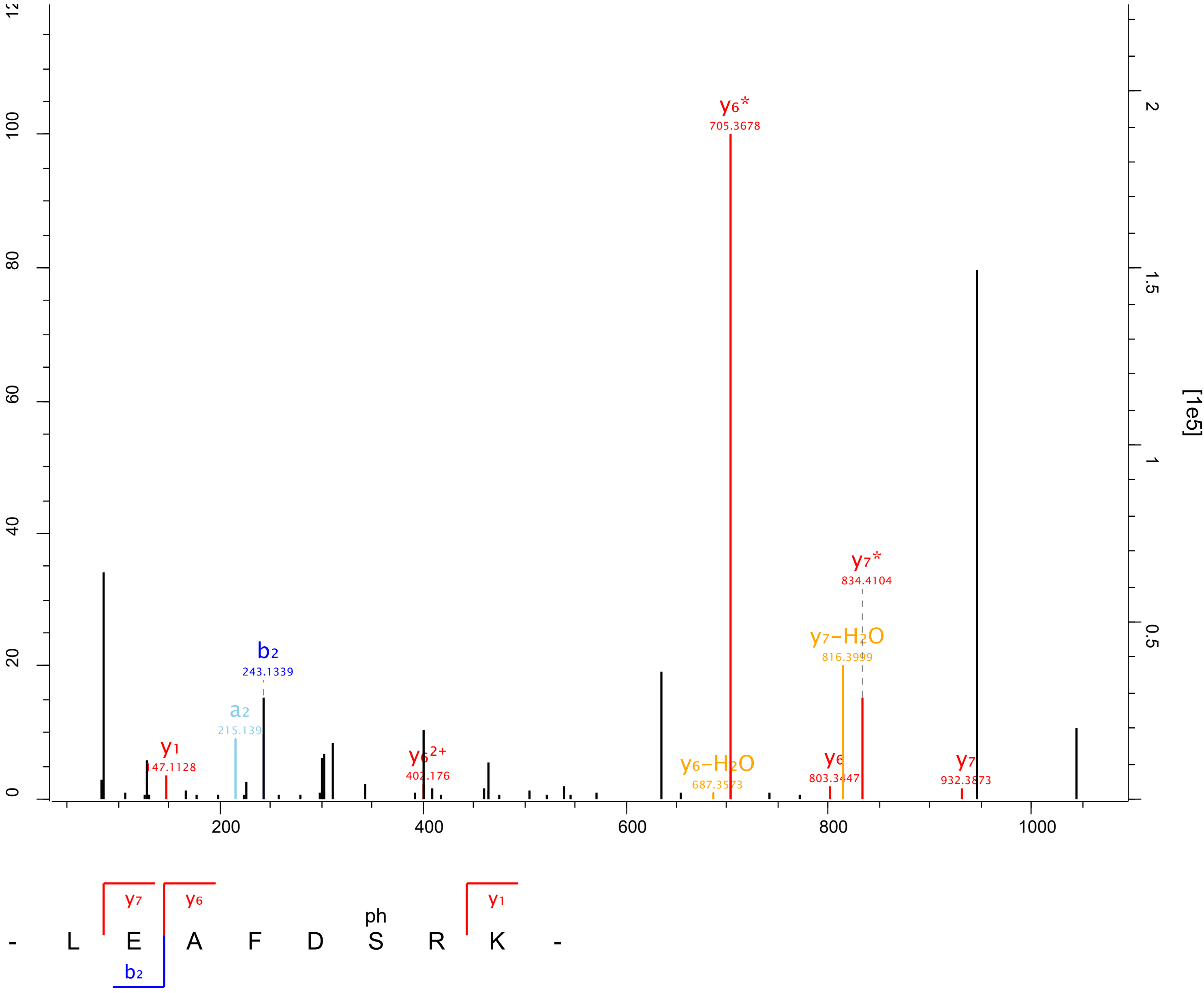

| Raw file | Scan | Method | Score | m/z |
|---|---|---|---|---|
| water_00_1 | 5288 | FTMS; HCD | 99.57 | 457.71 |

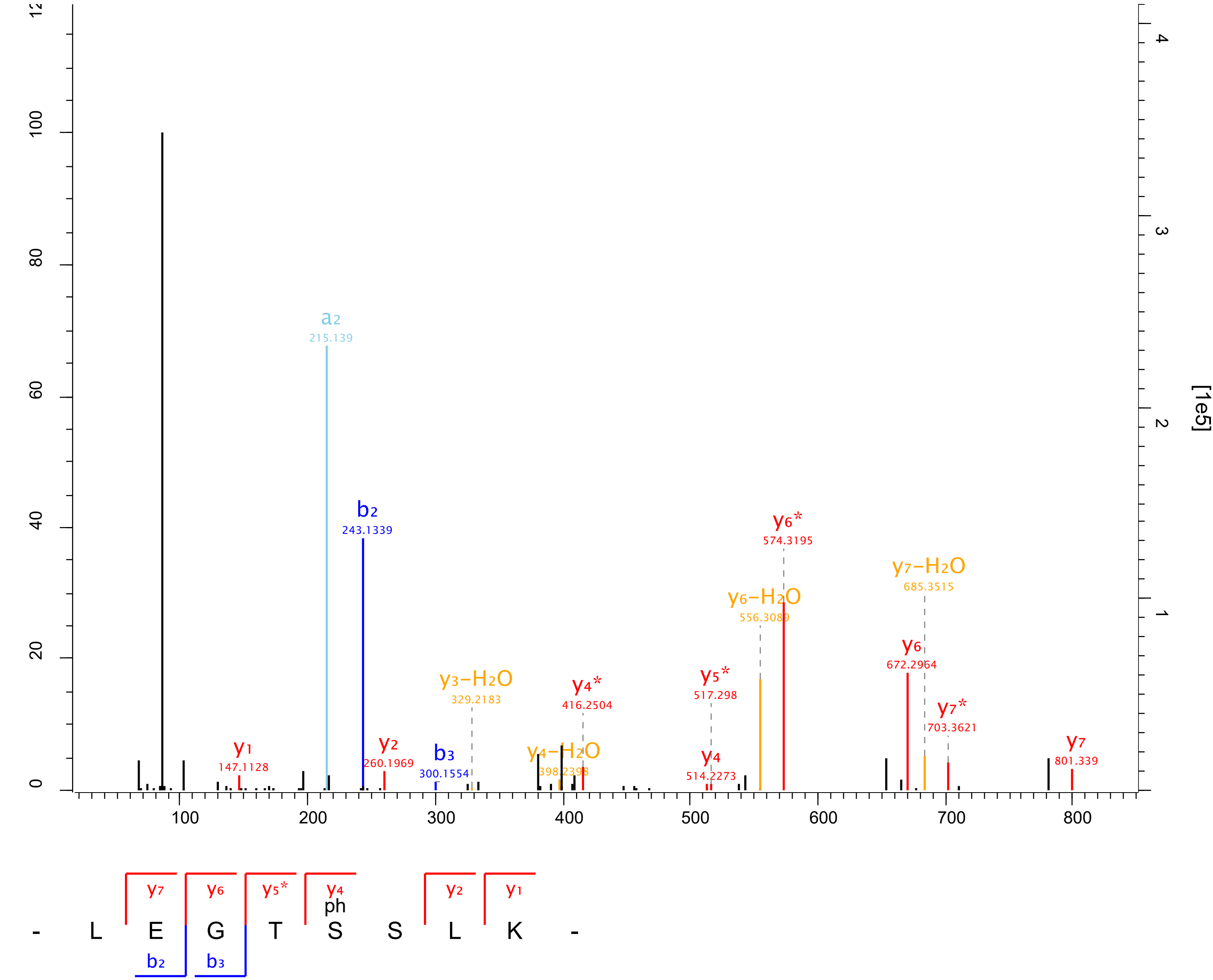

| Raw file | Scan | Method | Score | m/z |
|---|---|---|---|---|
| water_00_1 | 5298 | FTMS; HCD | 230.26 | 665.26 |

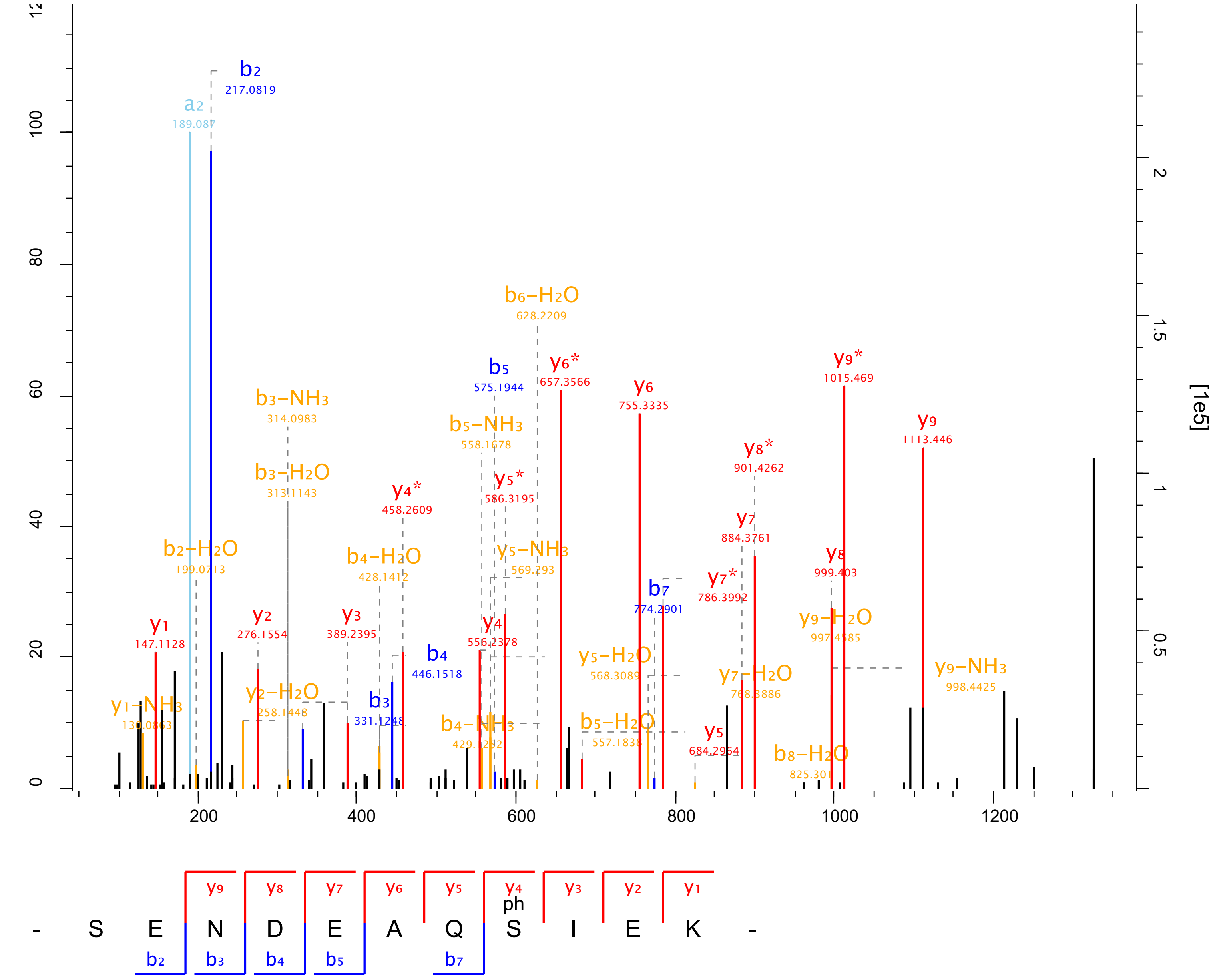

| Raw file | Scan | Method | Score | m/z |
|---|---|---|---|---|
| water_00_1 | 5304 | FTMS; HCD | 70.94 | 452.69 |

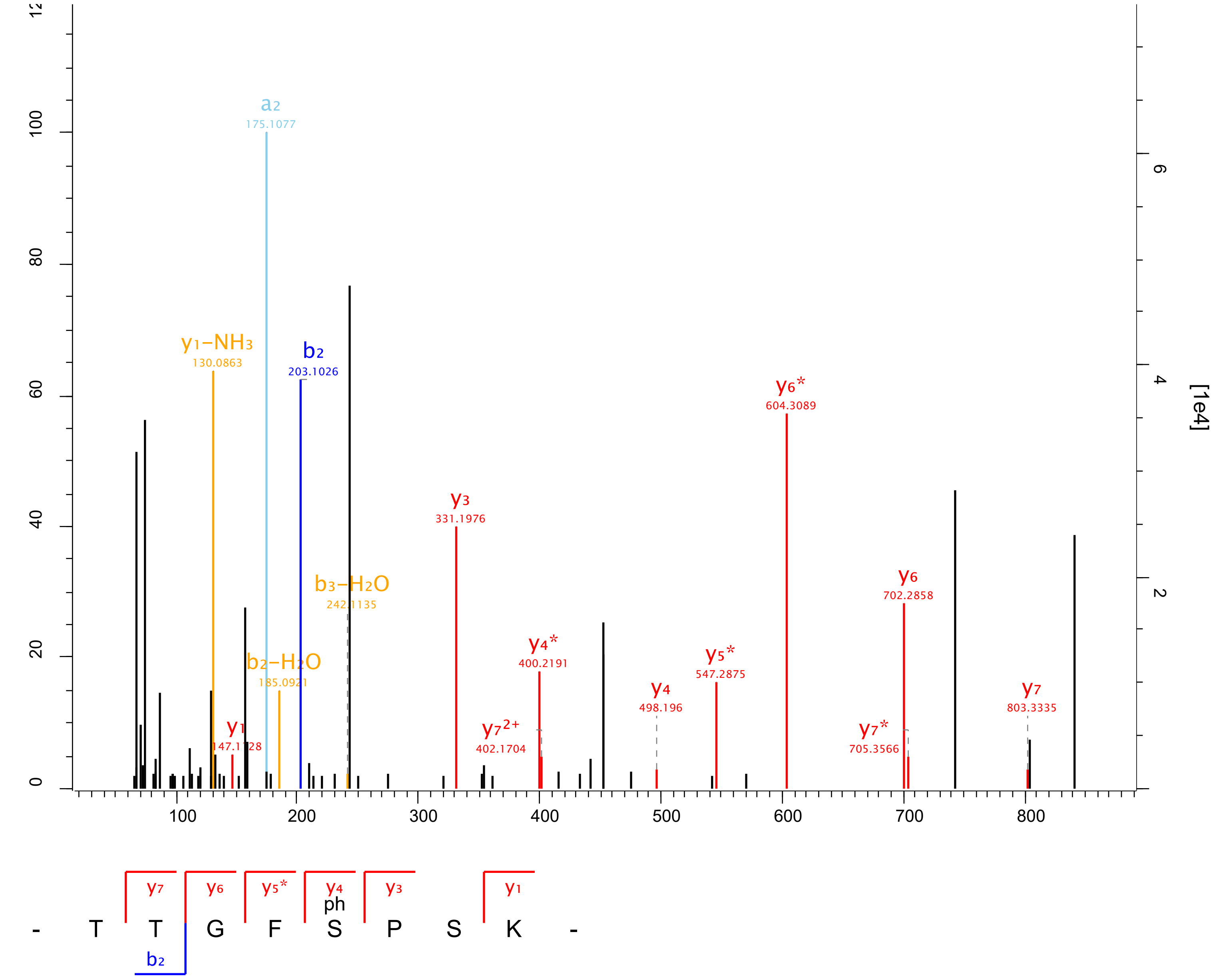

| Raw file | Scan | Method | Score | m/z |
|---|---|---|---|---|
| water_00_1 | 5425 | FTMS; HCD | 93.78 | 829.32 |

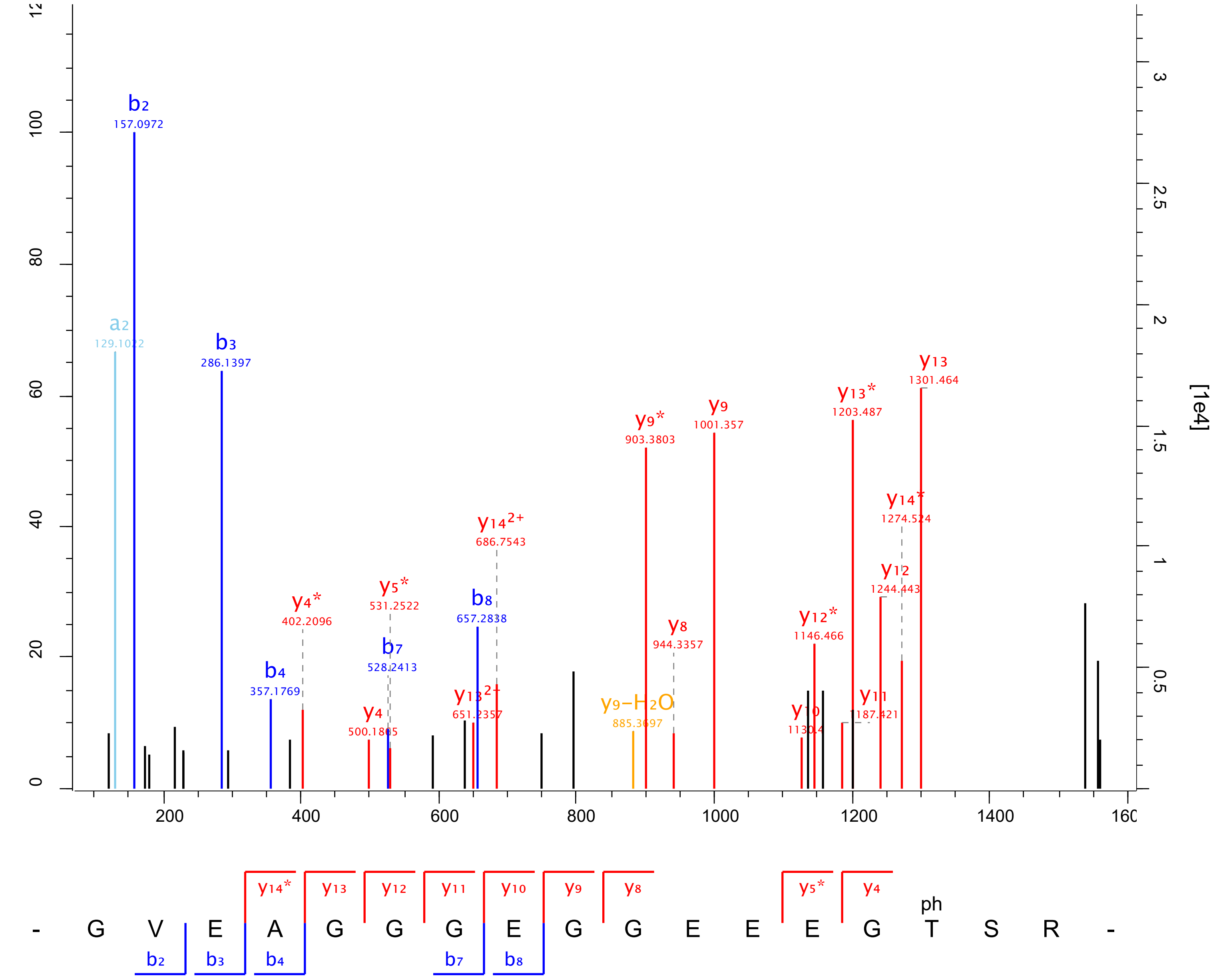

| Raw file | Scan | Method | Score | m/z |
| --- | --- | --- | --- | --- |
| water_00_1 | 5450 | FTMS; HCD | 224.59 | 769.81 |

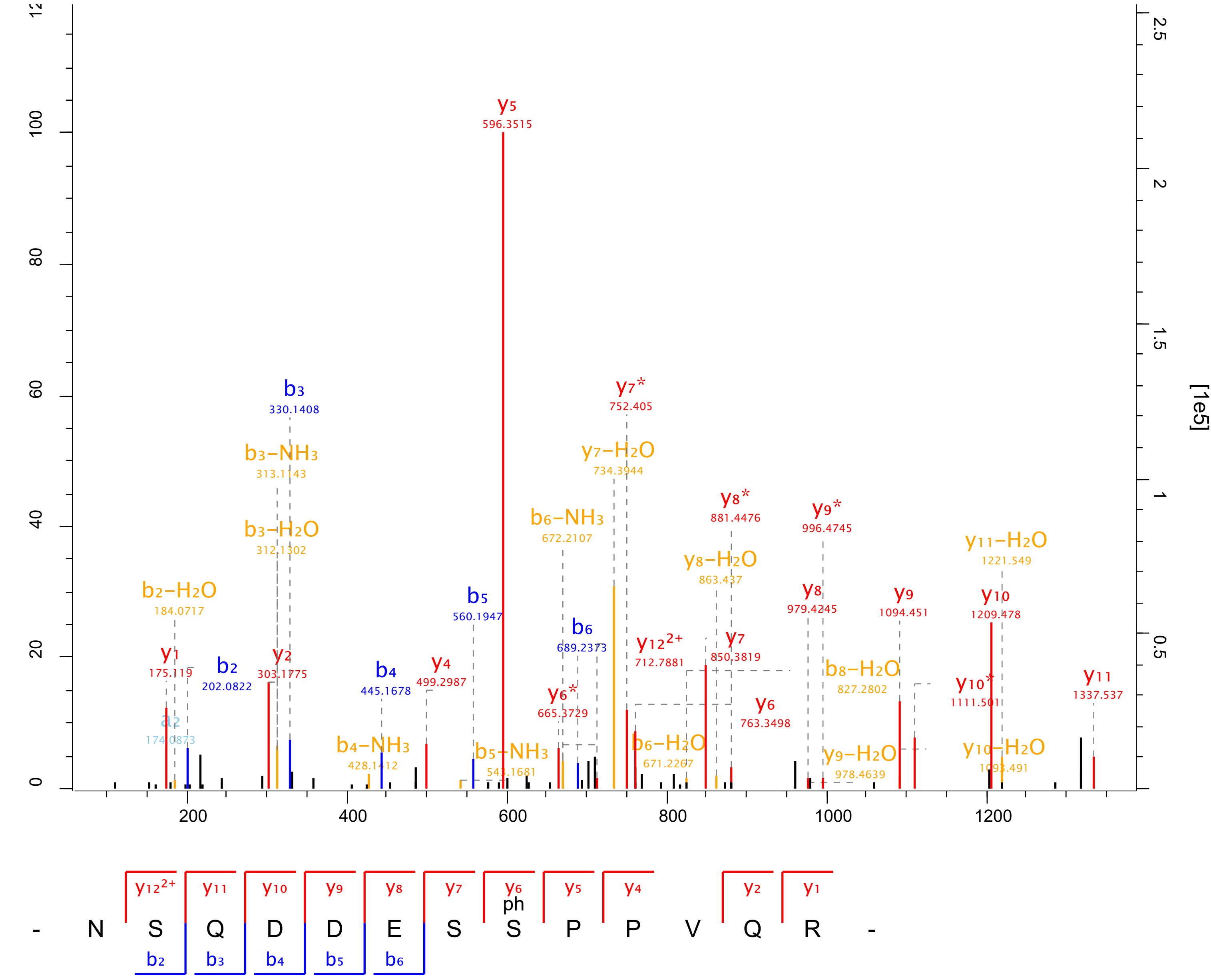

| Raw file | Scan | Method | Score | m/z |
|---|---|---|---|---|
| water_00_1 | 5515 | FTMS; HCD | 90.05 | 493.25 |

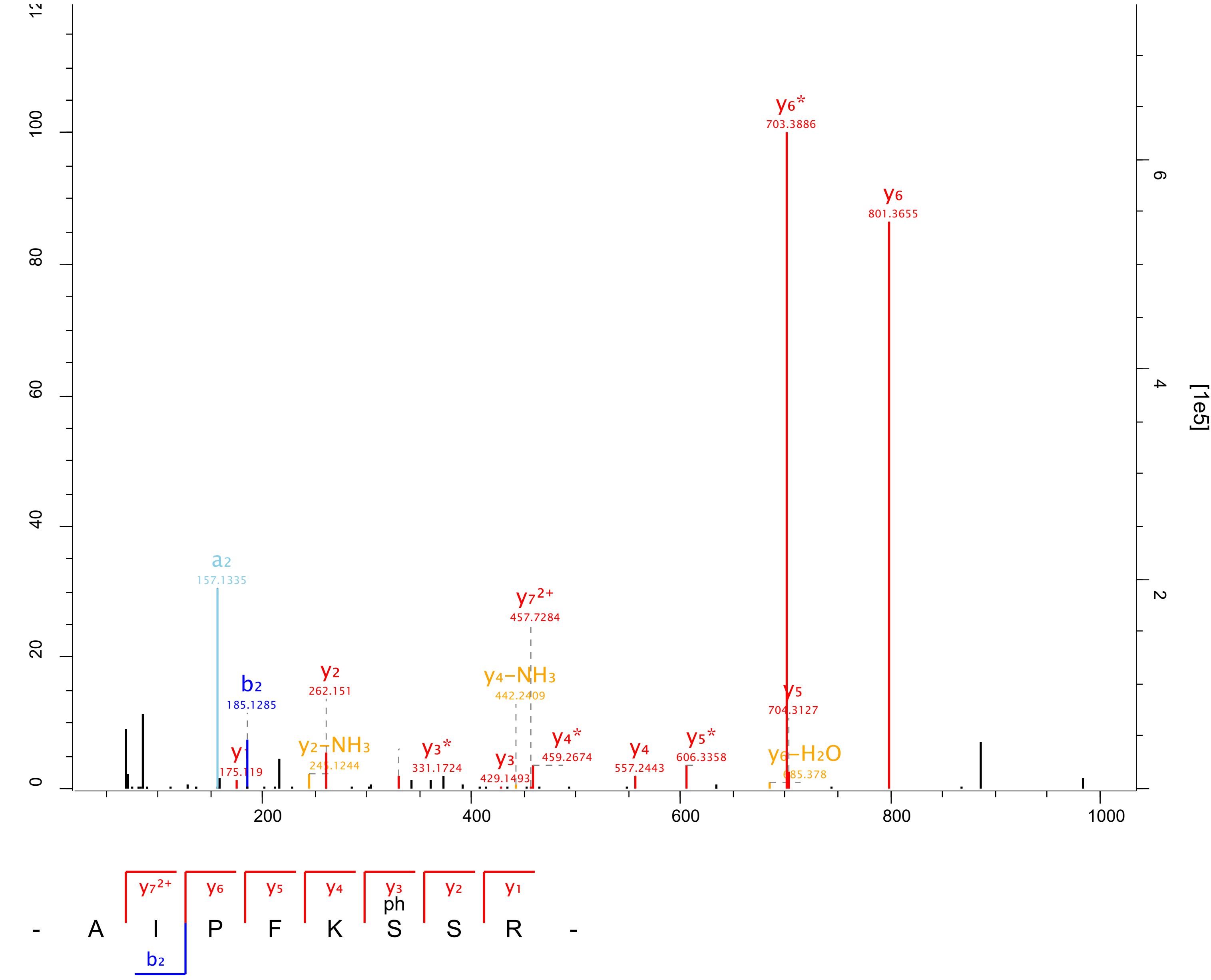

| Raw file | Scan | Method | Score | m/z |
| --- | --- | --- | --- | --- |
| water_00_1 | 5582 | FTMS; HCD | 158.58 | 684.3 |

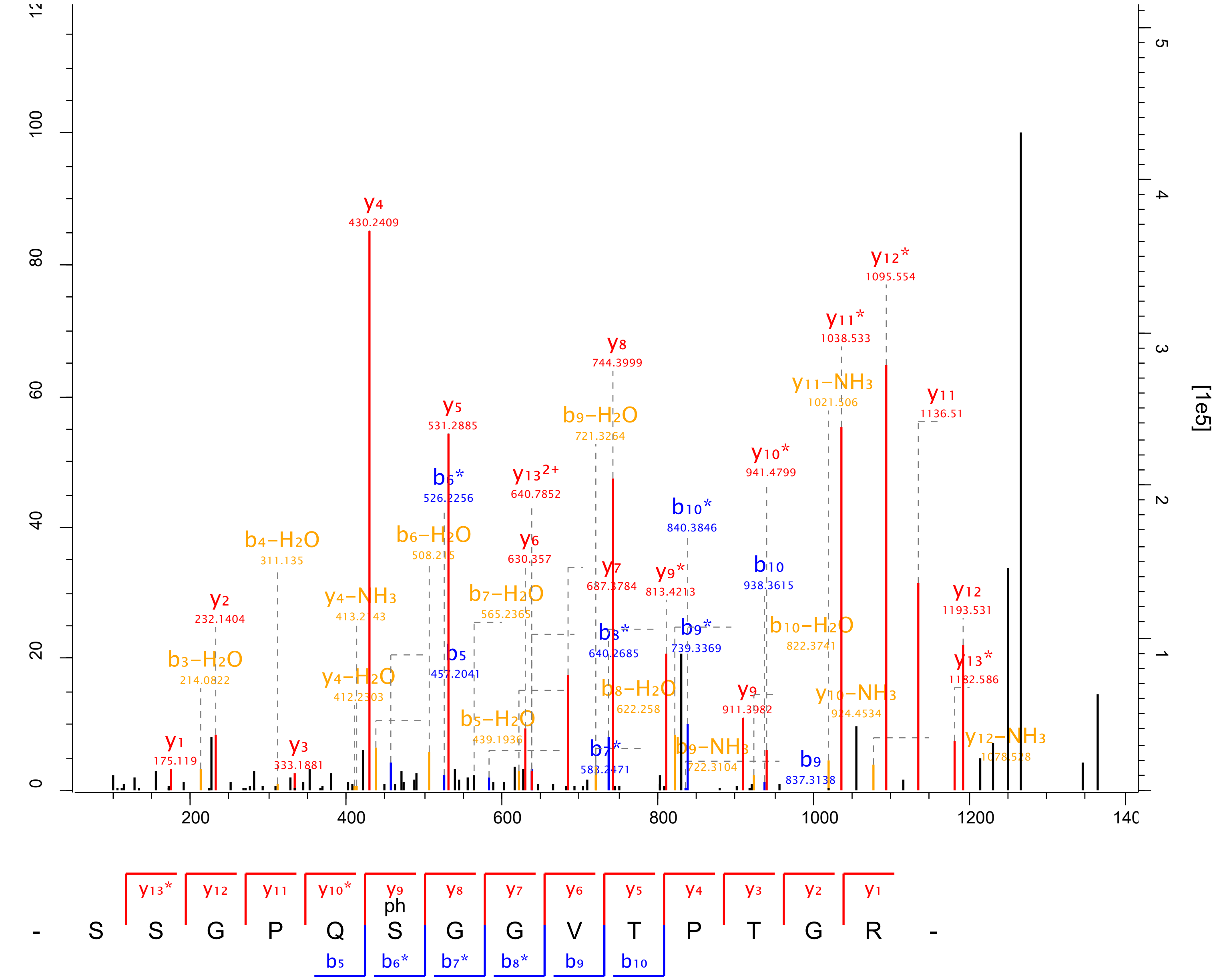

| Raw file | Scan | Method | Score | m/z |
| --- | --- | --- | --- | --- |
| water_00_1 | 5607 | FTMS; HCD | 60.64 | 459.7 |

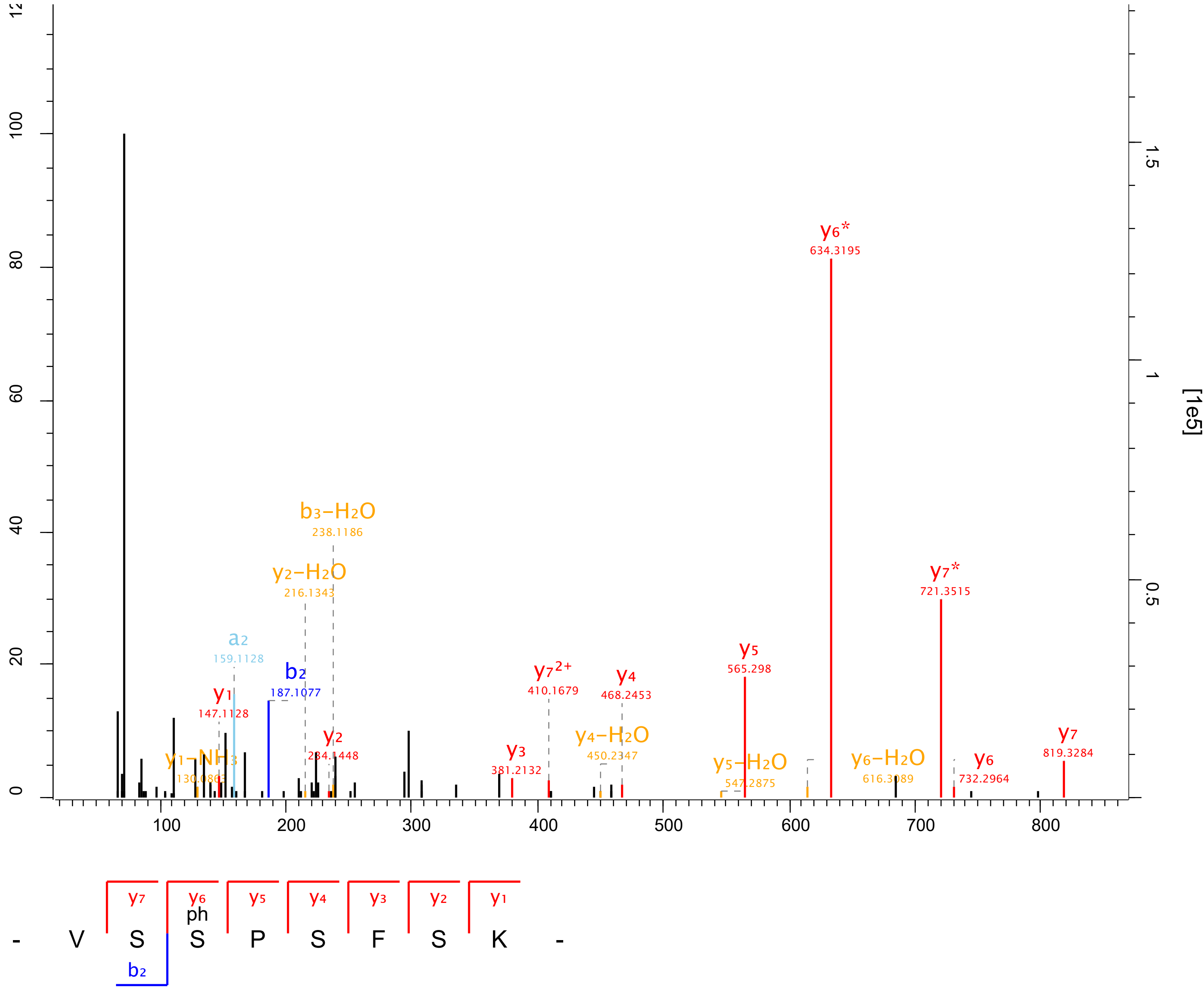

| Raw file | Scan | Method | Score | m/z |
|---|---|---|---|---|
| water_00_1 | 5610 | FTMS; HCD | 81.1 | 436.67 |

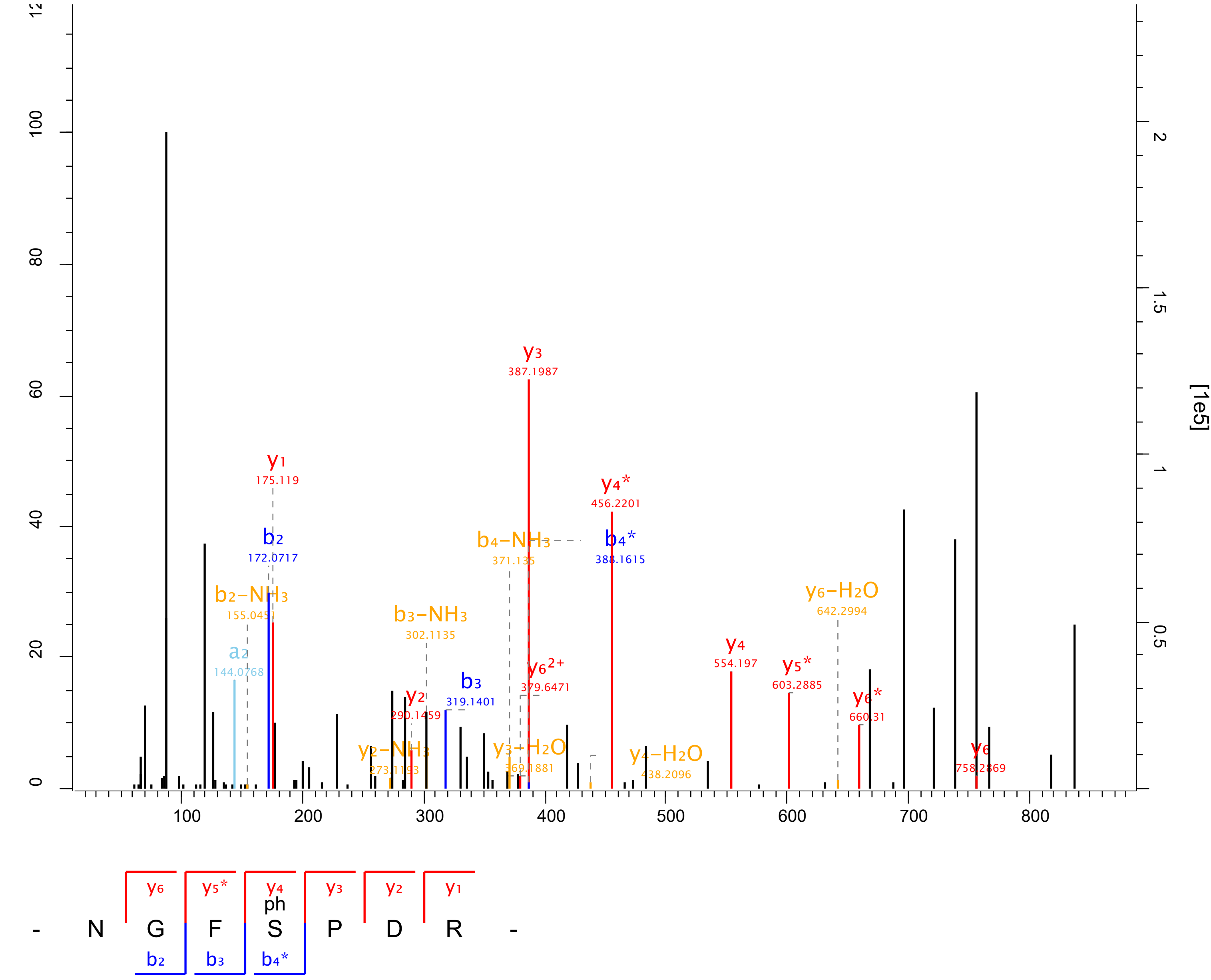

| Raw file | Scan | Method | Score | m/z |
|---|---|---|---|---|
| water_00_1 | 5649 | FTMS; HCD | 59.71 | 618.77 |

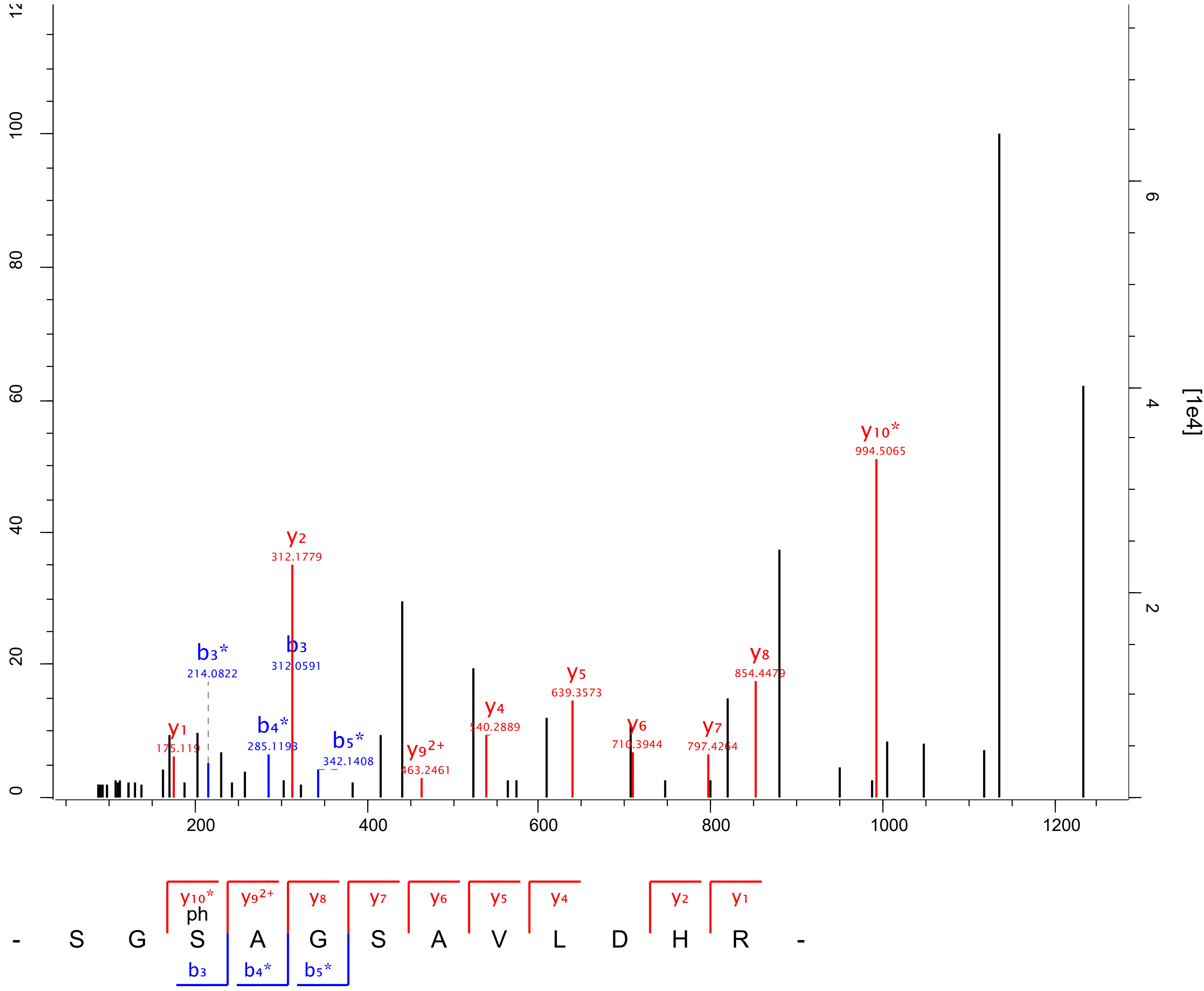

| Raw file | Scan | Method | Score | m/z |
| --- | --- | --- | --- | --- |
| water_00_1 | 5896 | FTMS; HCD | 131.4 | 493.56 |

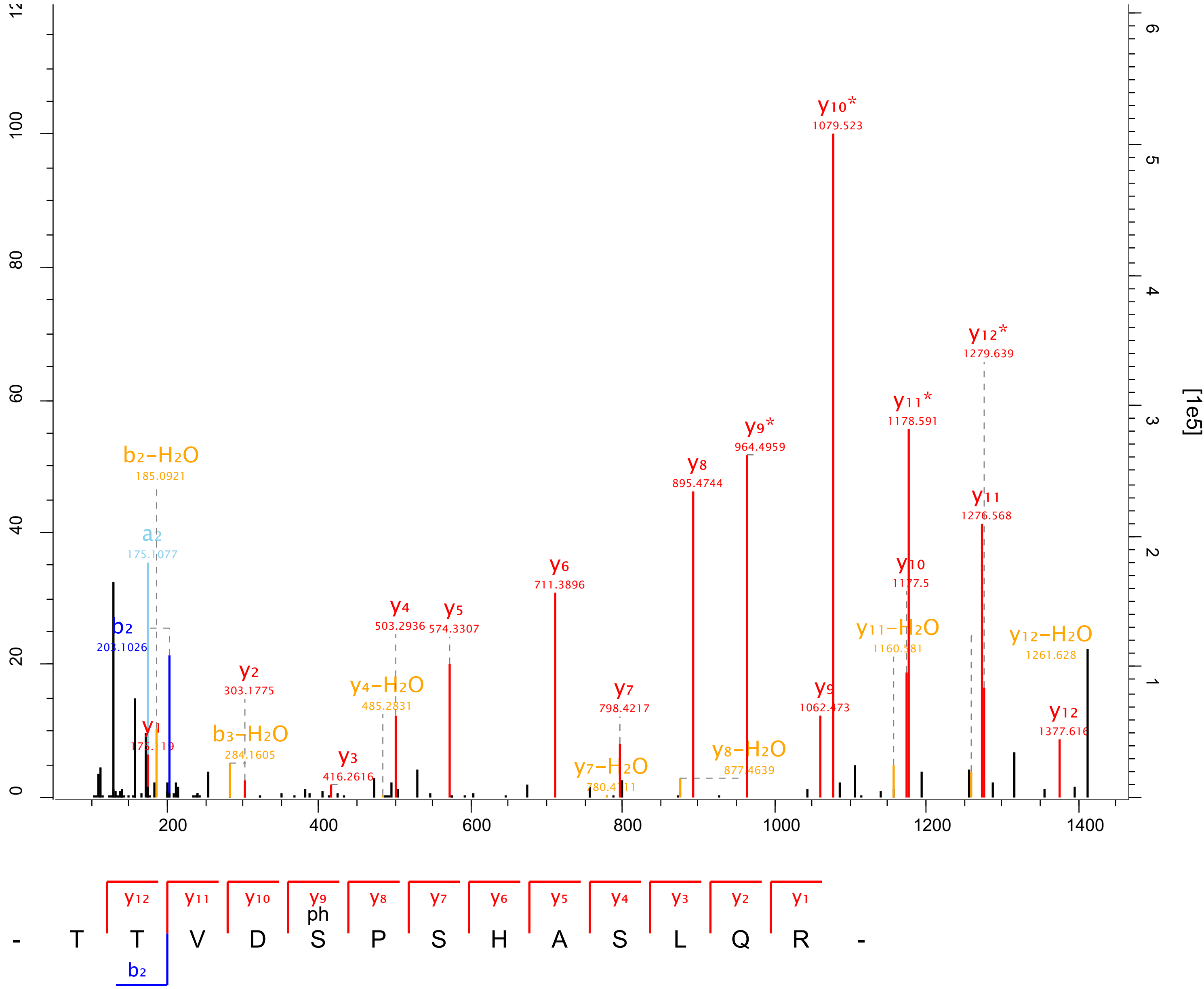

| Raw file | Scan | Method | Score | m/z |
| --- | --- | --- | --- | --- |
| water_00_1 | 5897 | FTMS; HCD | 81.57 | 532.21 |

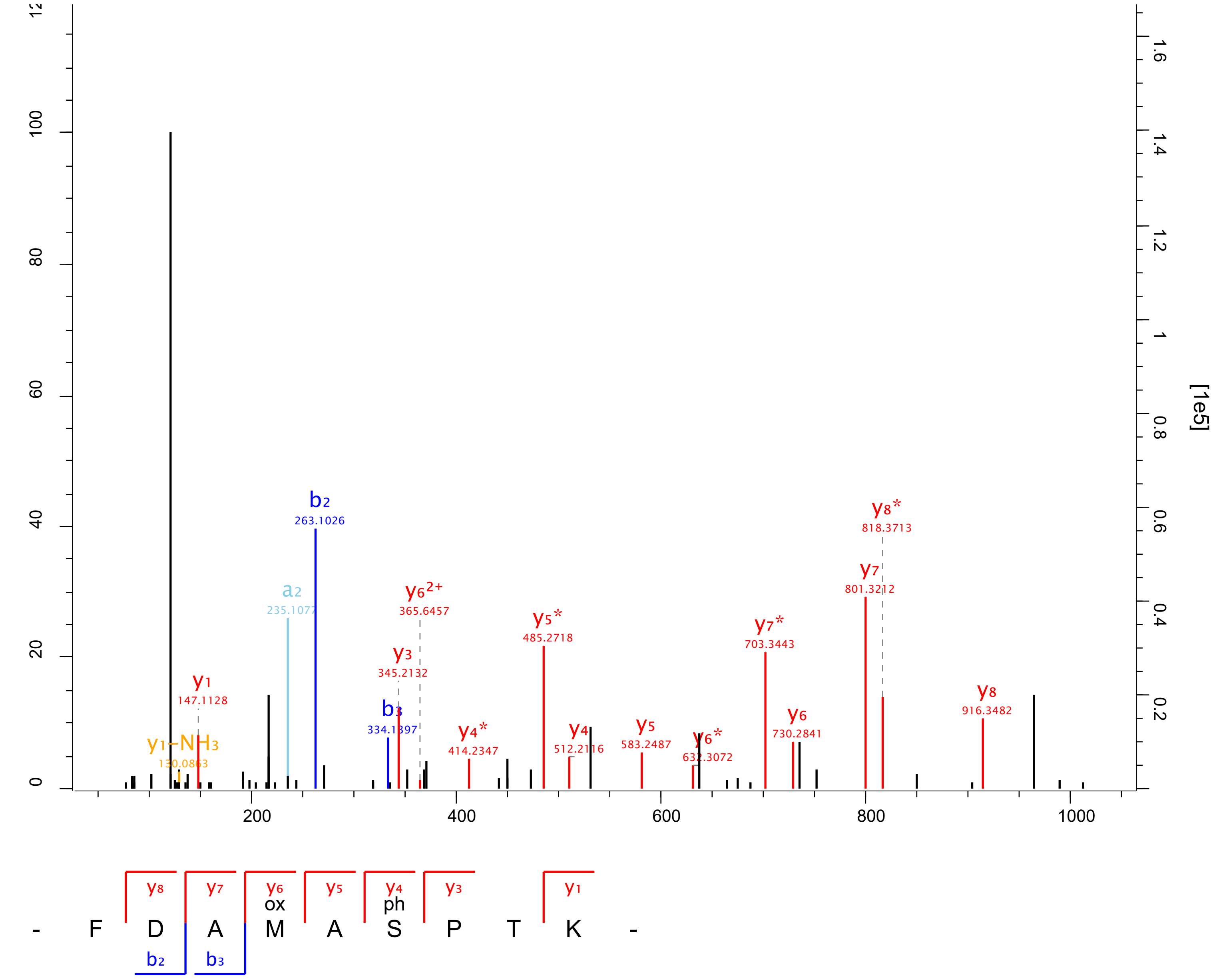

| Raw file | Scan | Method | Score | m/z |
| --- | --- | --- | --- | --- |
| water_00_1 | 6137 | FTMS; HCD | 109.79 | 515.88 |

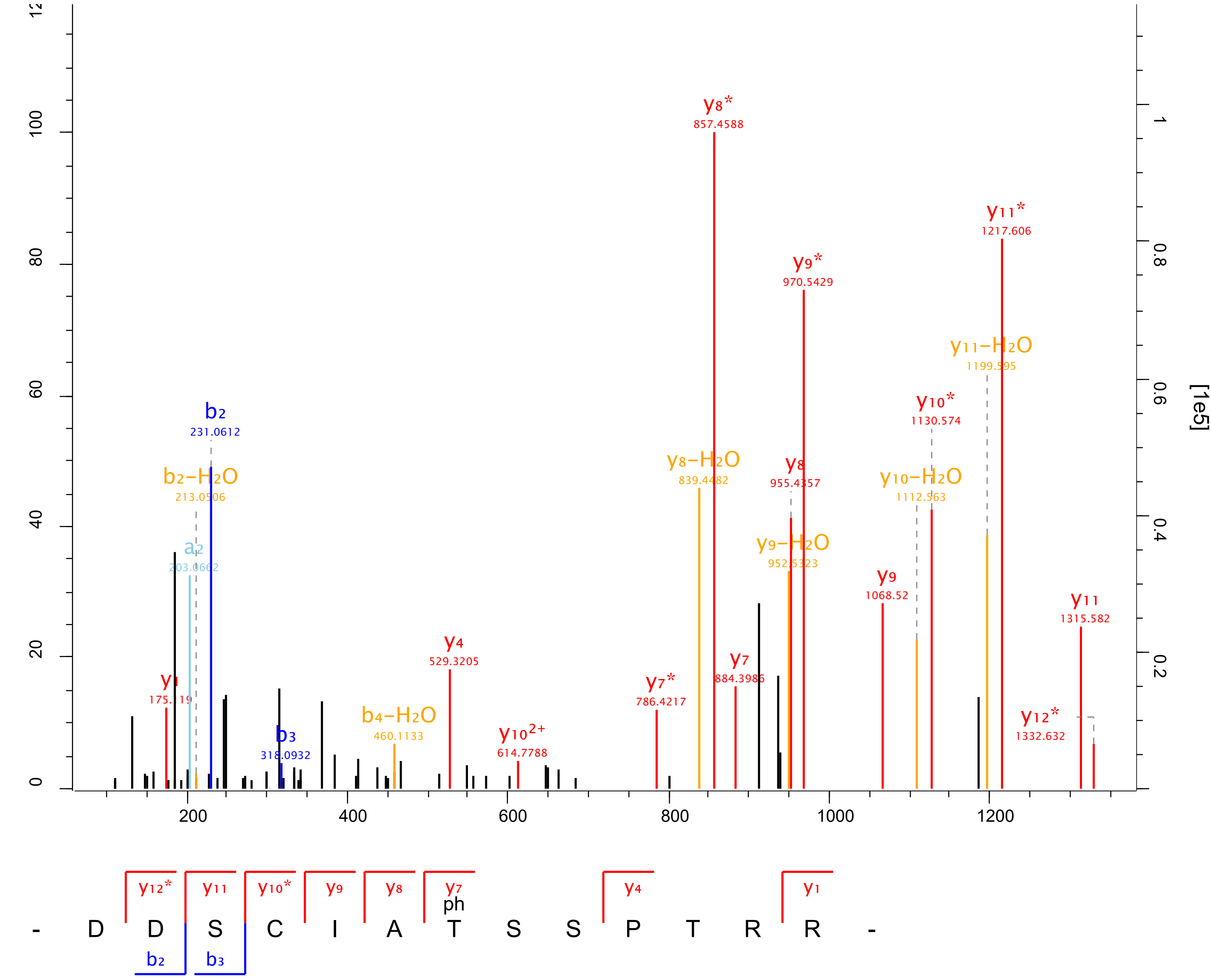

| Raw file | Scan | Method | Score | m/z |
|---|---|---|---|---|
| water_00_1 | 6254 | FTMS; HCD | 95.61 | 434.7 |

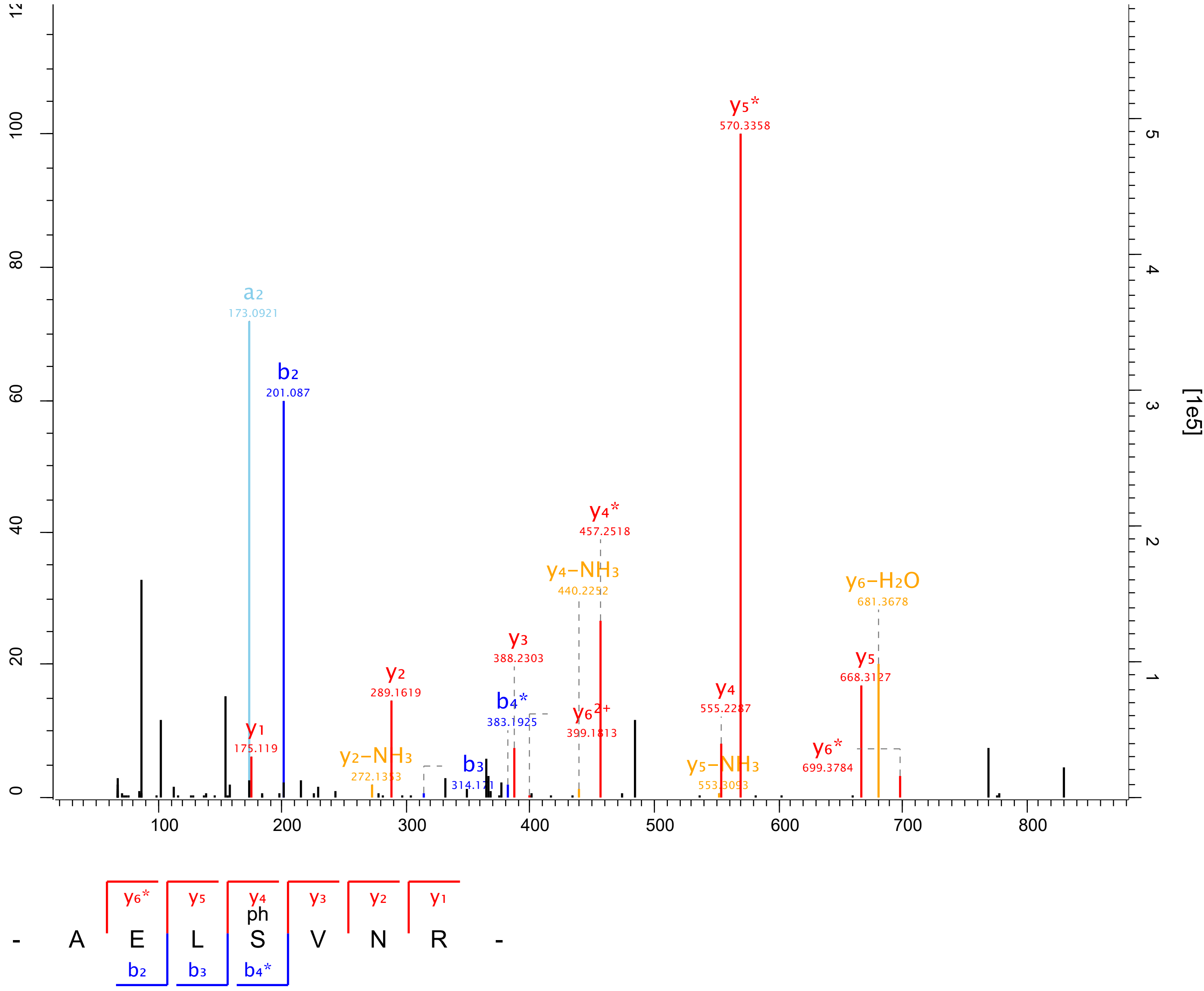

| Raw file | Scan | Method | Score | m/z |
|---|---|---|---|---|
| water_00_1 | 6335 | FTMS; HCD | 46.35 | 620.22 |

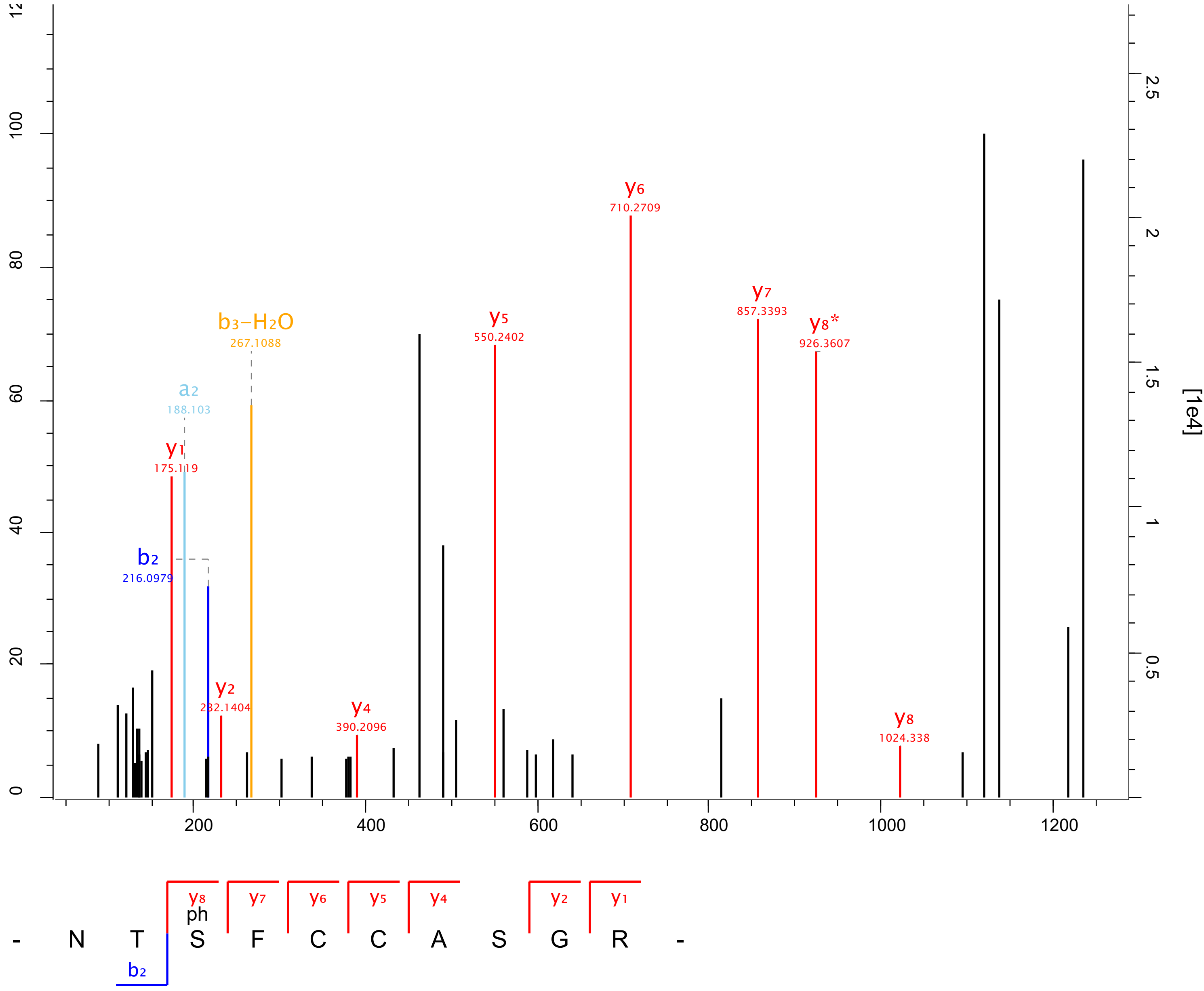

| Raw file | Scan | Method | Score | m/z |
|---|---|---|---|---|
| water_00_1 | 6430 | FTMS; HCD | 84.89 | 538.72 |

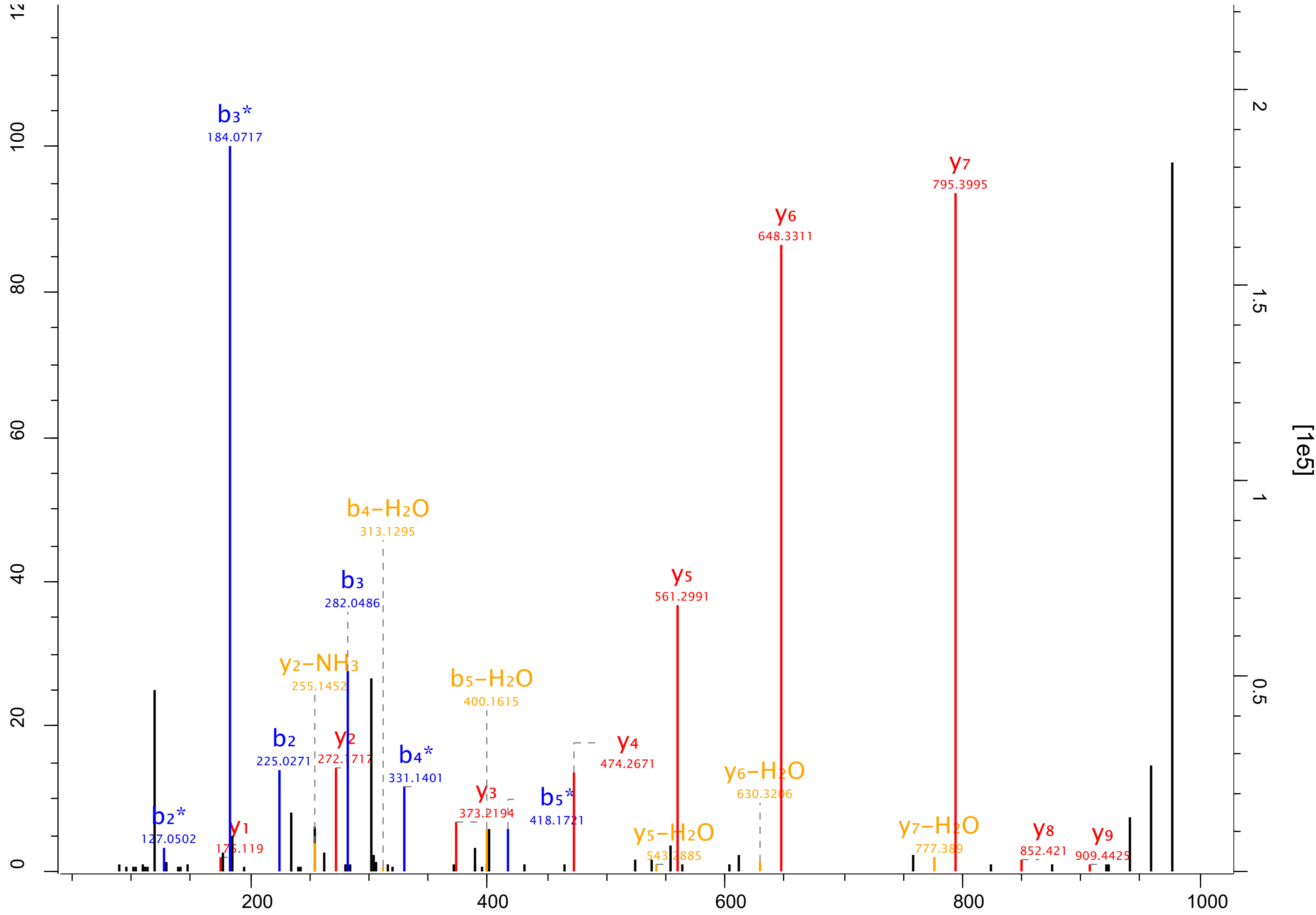


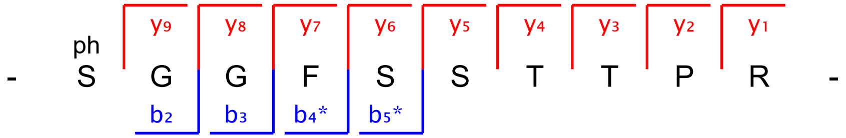

| Raw file | Scan | Method | Score | m/z |
|---|---|---|---|---|
| water_00_1 | 6483 | FTMS; HCD | 114.64 | 675.76 |

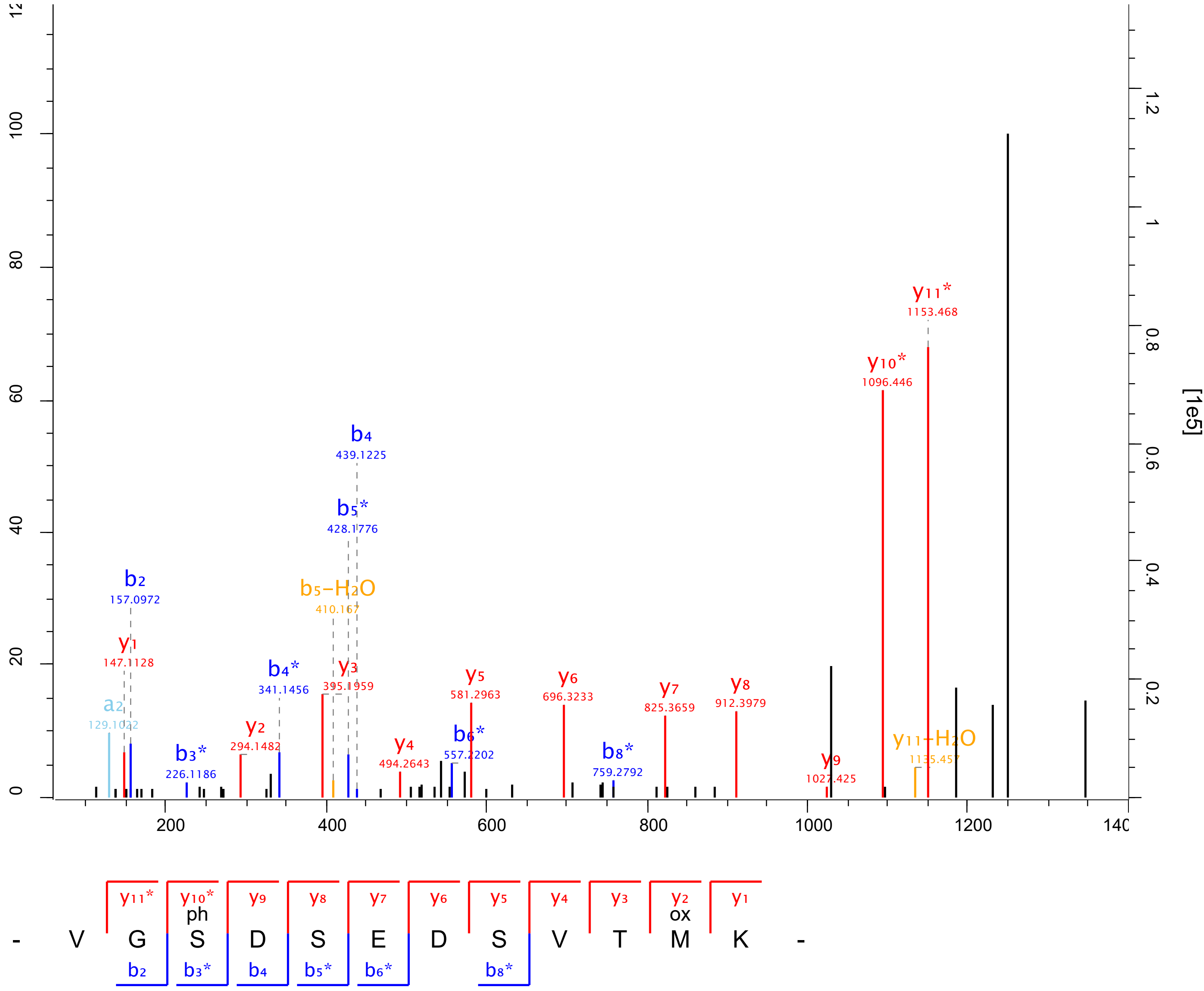

| Raw file | Scan | Method | Score | m/z |
|---|---|---|---|---|
| water_00_1 | 6635 | FTMS; HCD | 77.74 | 476.71 |

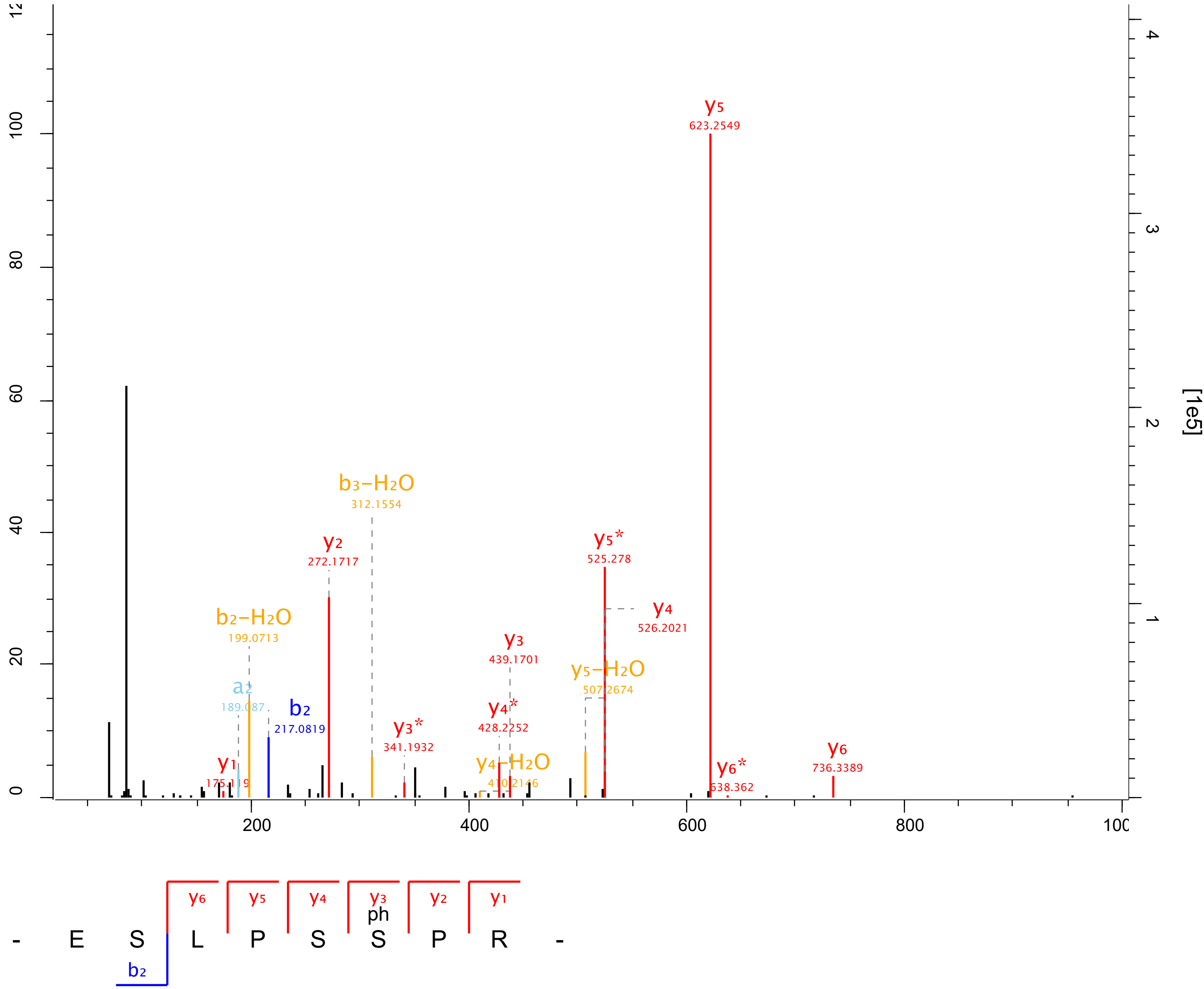

| Raw file | Scan | Method | Score | m/z |
|---|---|---|---|---|
| water_00_1 | 6712 | FTMS; HCD | 103.44 | 541.73 |

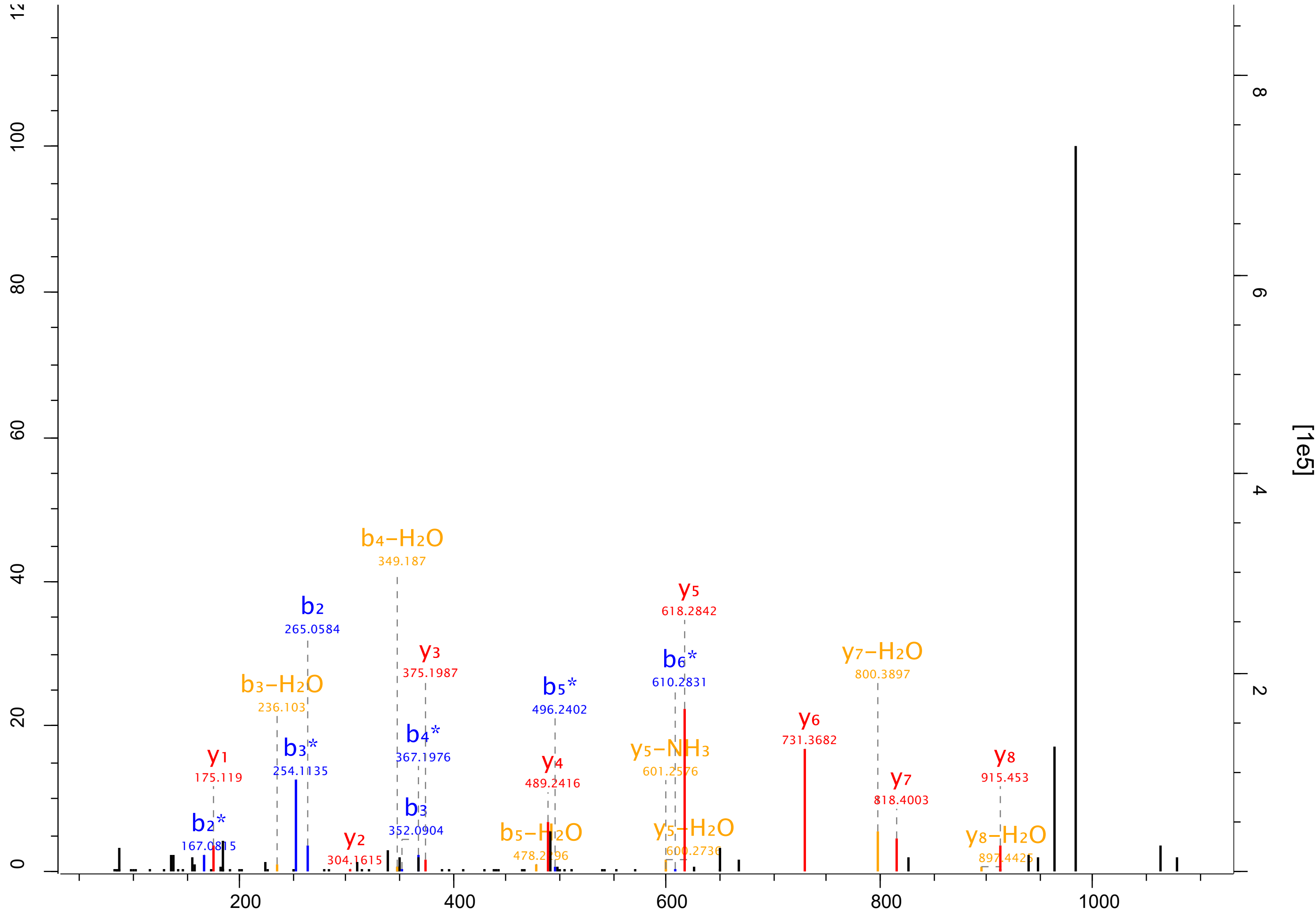


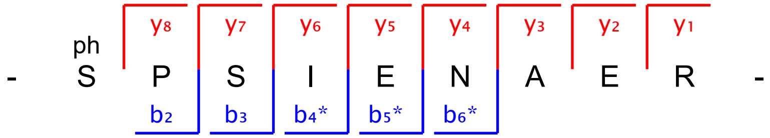

| Raw file | Scan | Method | Score | m/z |
|---|---|---|---|---|
| water_00_1 | 7027 | FTMS; HCD | 73.5 | 477.16 |

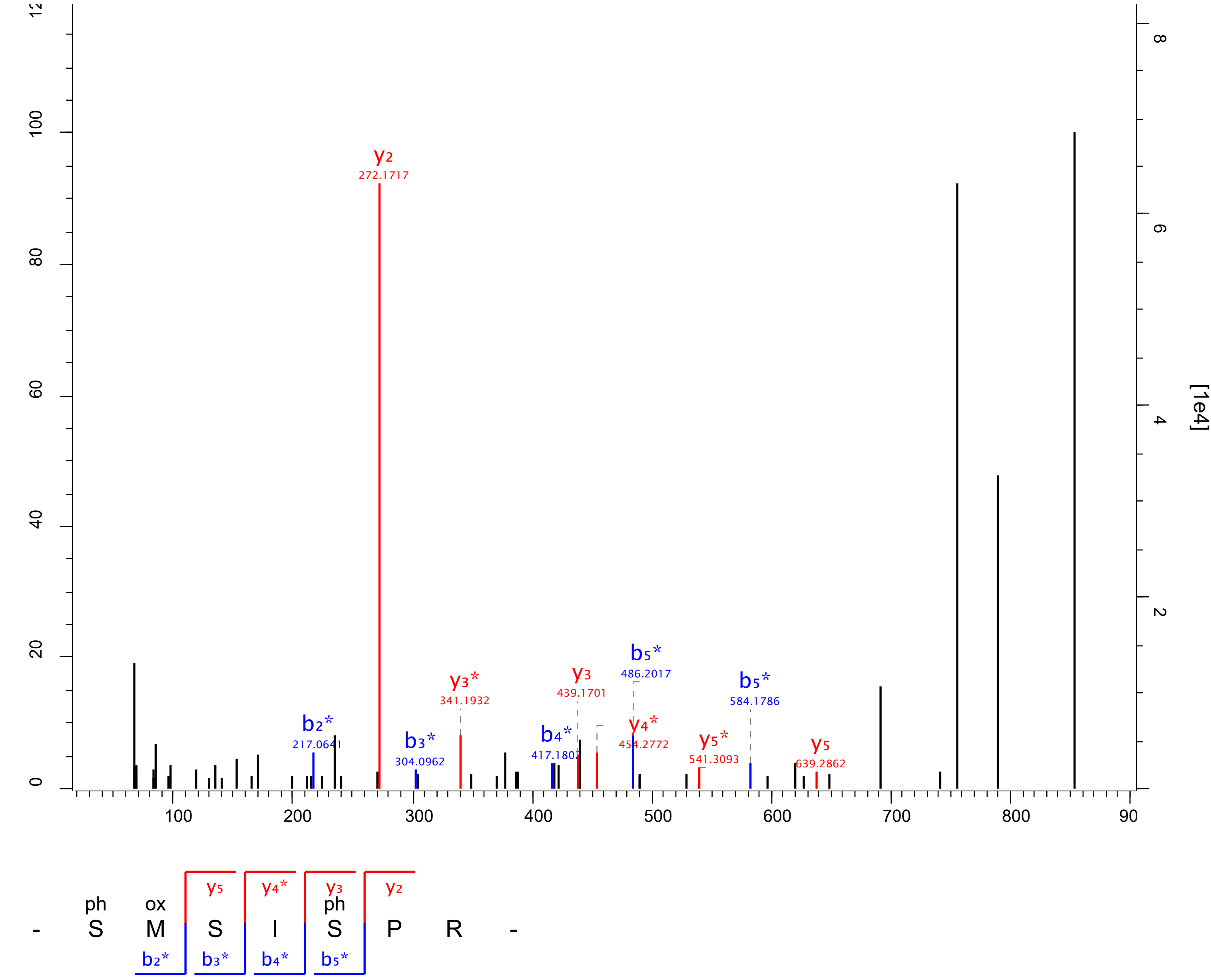

| Raw file | Scan | Method | Score | m/z |
| --- | --- | --- | --- | --- |
| water_00_1 | 7183 | FTMS; HCD | 93.77 | 540.76 |

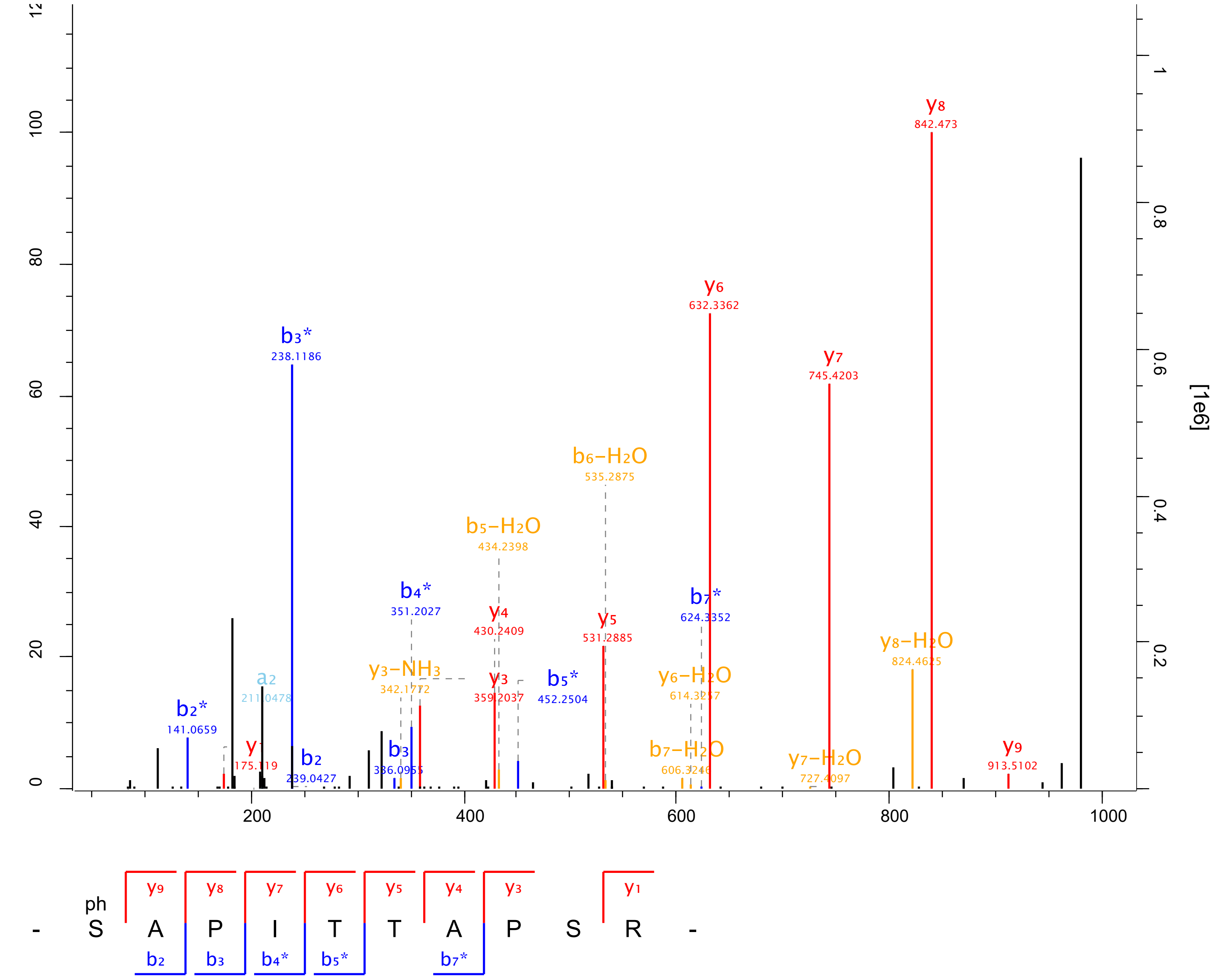

| Raw file | Scan | Method | Score | m/z |
|---|---|---|---|---|
| water_00_1 | 7261 | FTMS; HCD | 51.13 | 429.85 |

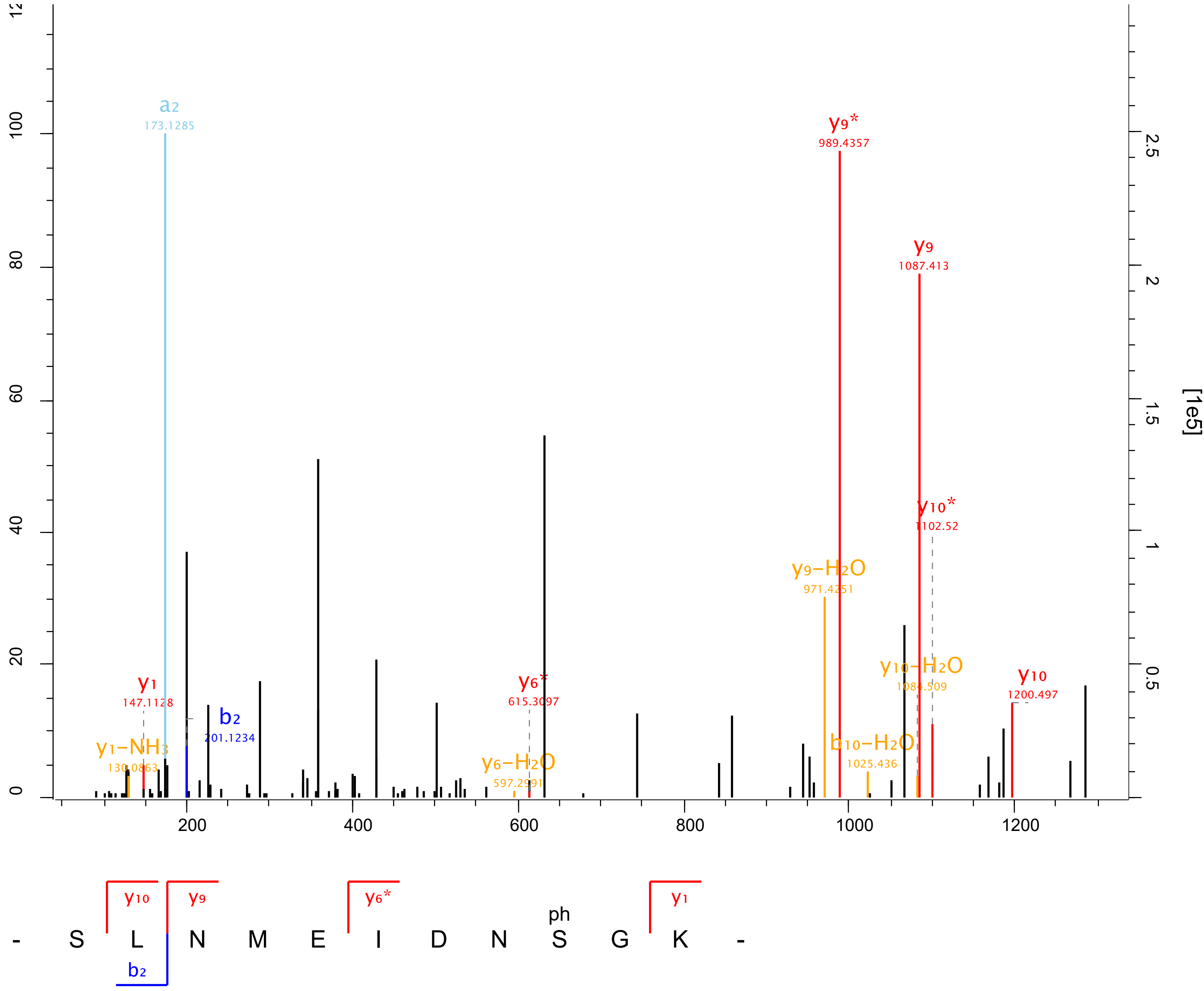

| Raw file | Scan | Method | Score | m/z |
|---|---|---|---|---|
| water_00_1 | 7425 | FTMS; HCD | 75.76 | 649.78 |

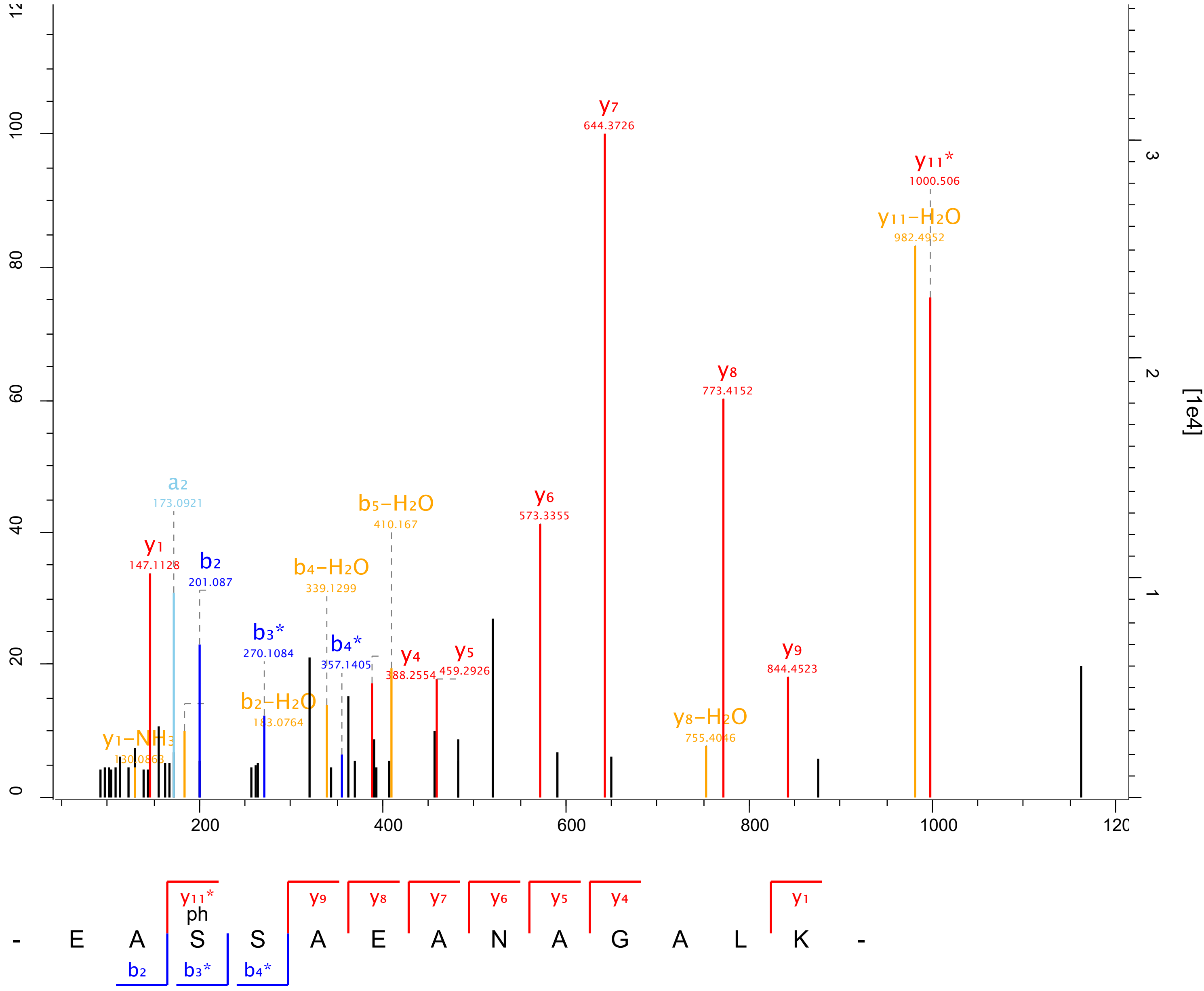

| Raw file | Scan | Method | Score | m/z |
|---|---|---|---|---|
| water_00_1 | 7438 | FTMS; HCD | 117.3 | 678.8 |

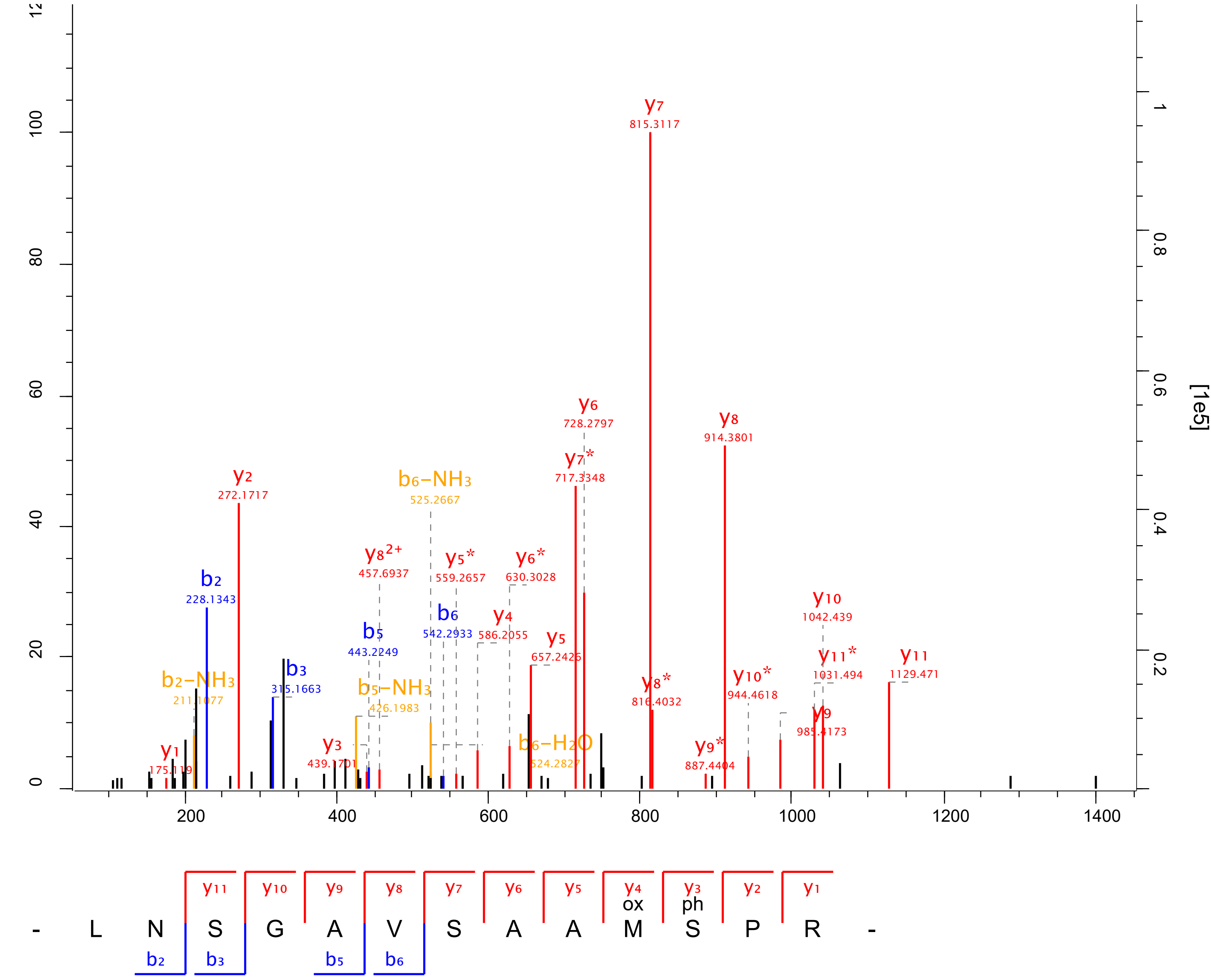

| Raw file | Scan | Method | Score | m/z |
| --- | --- | --- | --- | --- |
| water_00_1 | 7455 | FTMS; HCD | 69.92 | 680.76 |

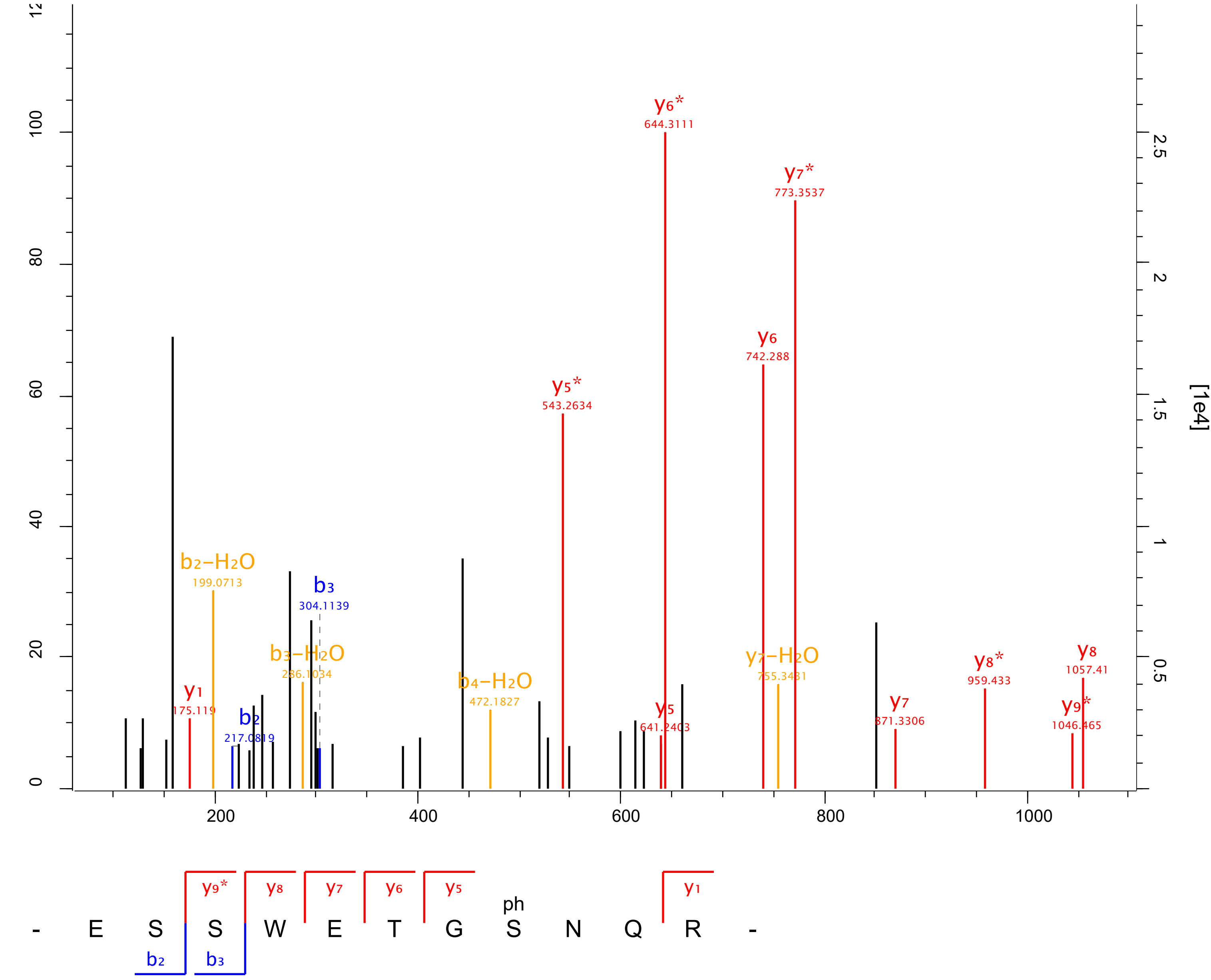

| Raw file | Scan | Method | Score | m/z |
|---|---|---|---|---|
| water_00_1 | 7560 | FTMS; HCD | 52.53 | 652.27 |

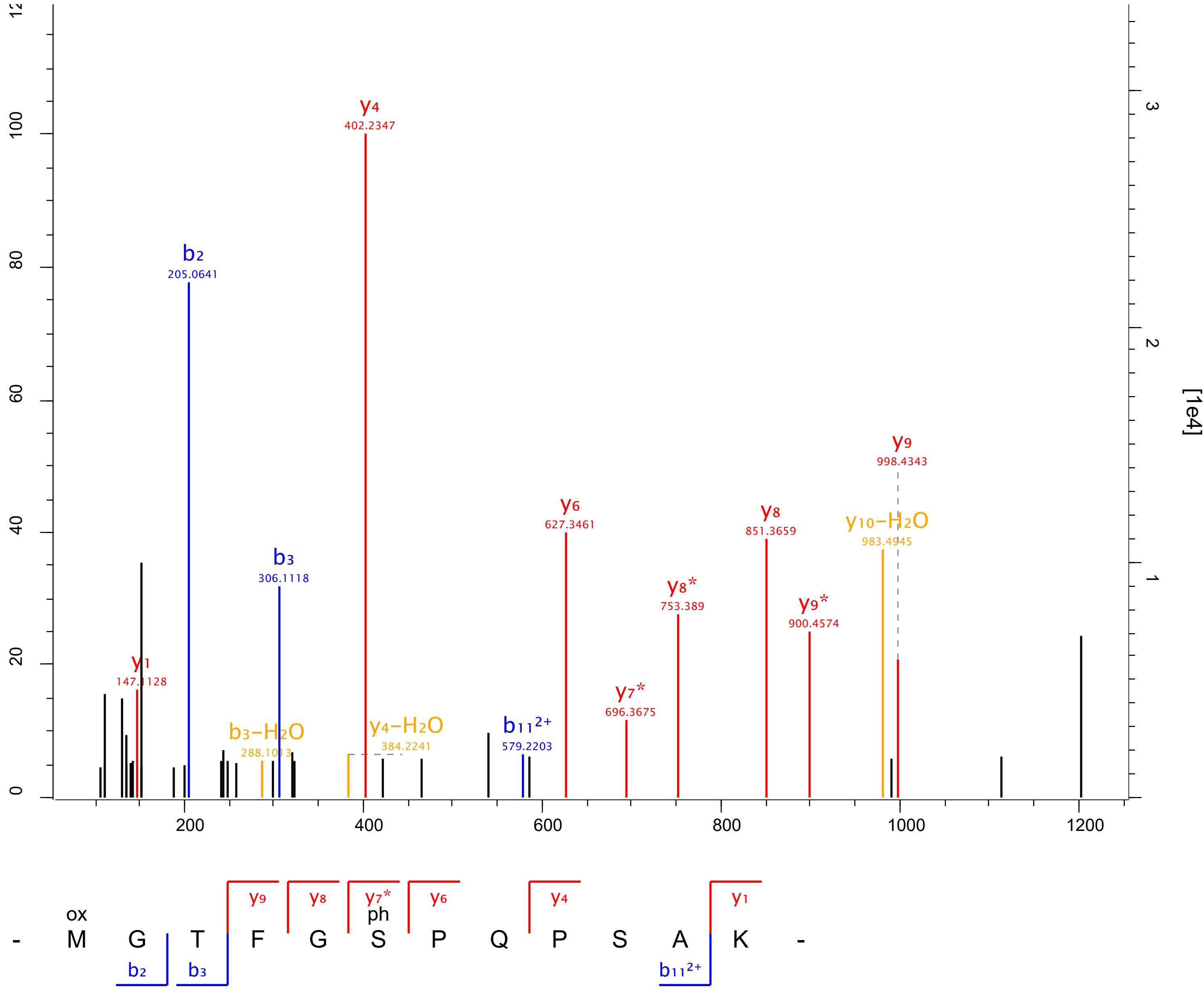

| Raw file | Scan | Method | Score | m/z |
| --- | --- | --- | --- | --- |
| water_00_1 | 7822 | FTMS; HCD | 266.28 | 826.33 |

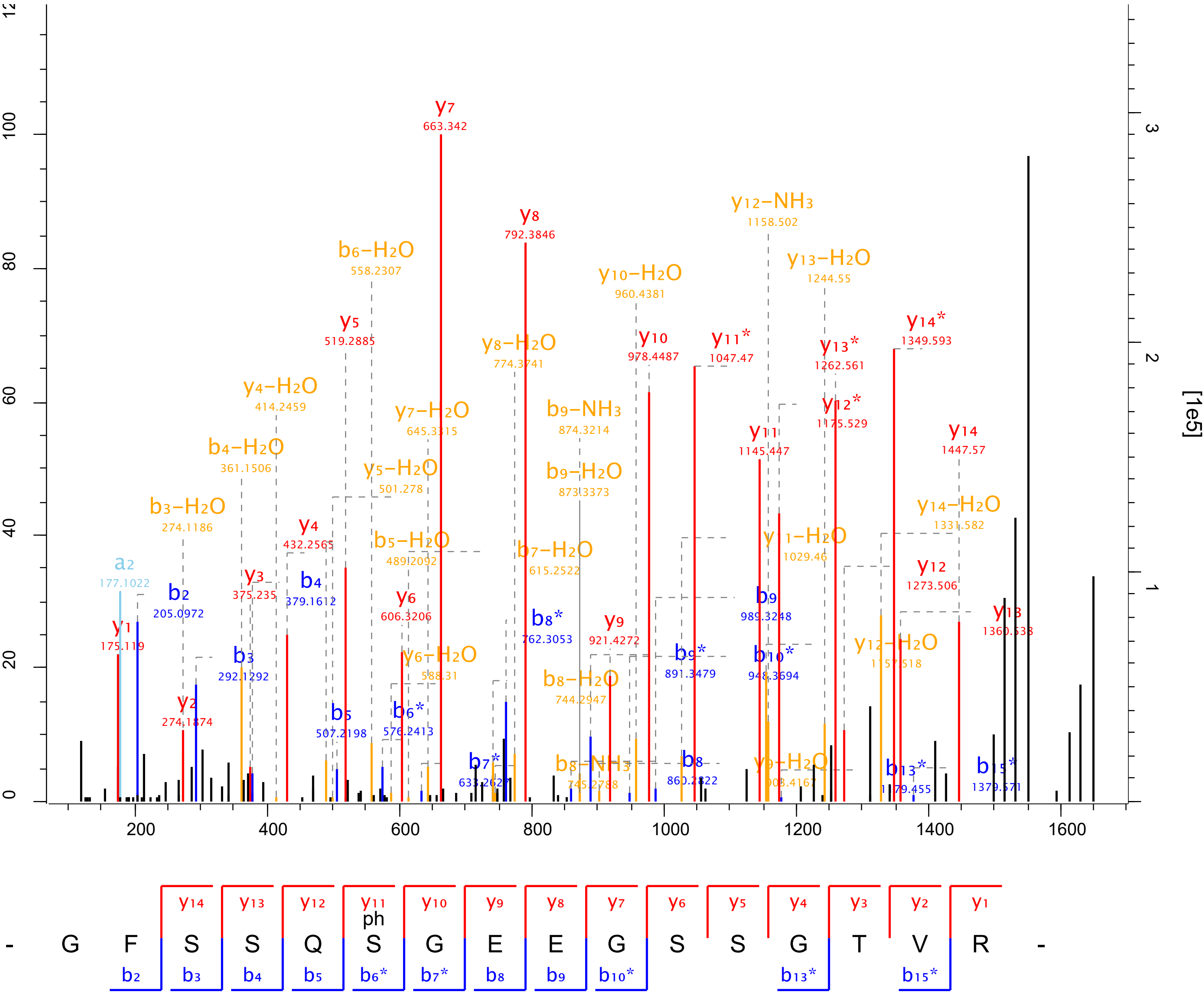

| Raw file | Scan | Method | Score | m/z |
| --- | --- | --- | --- | --- |
| water_00_1 | 7878 | FTMS; HCD | 85.81 | 451.2 |

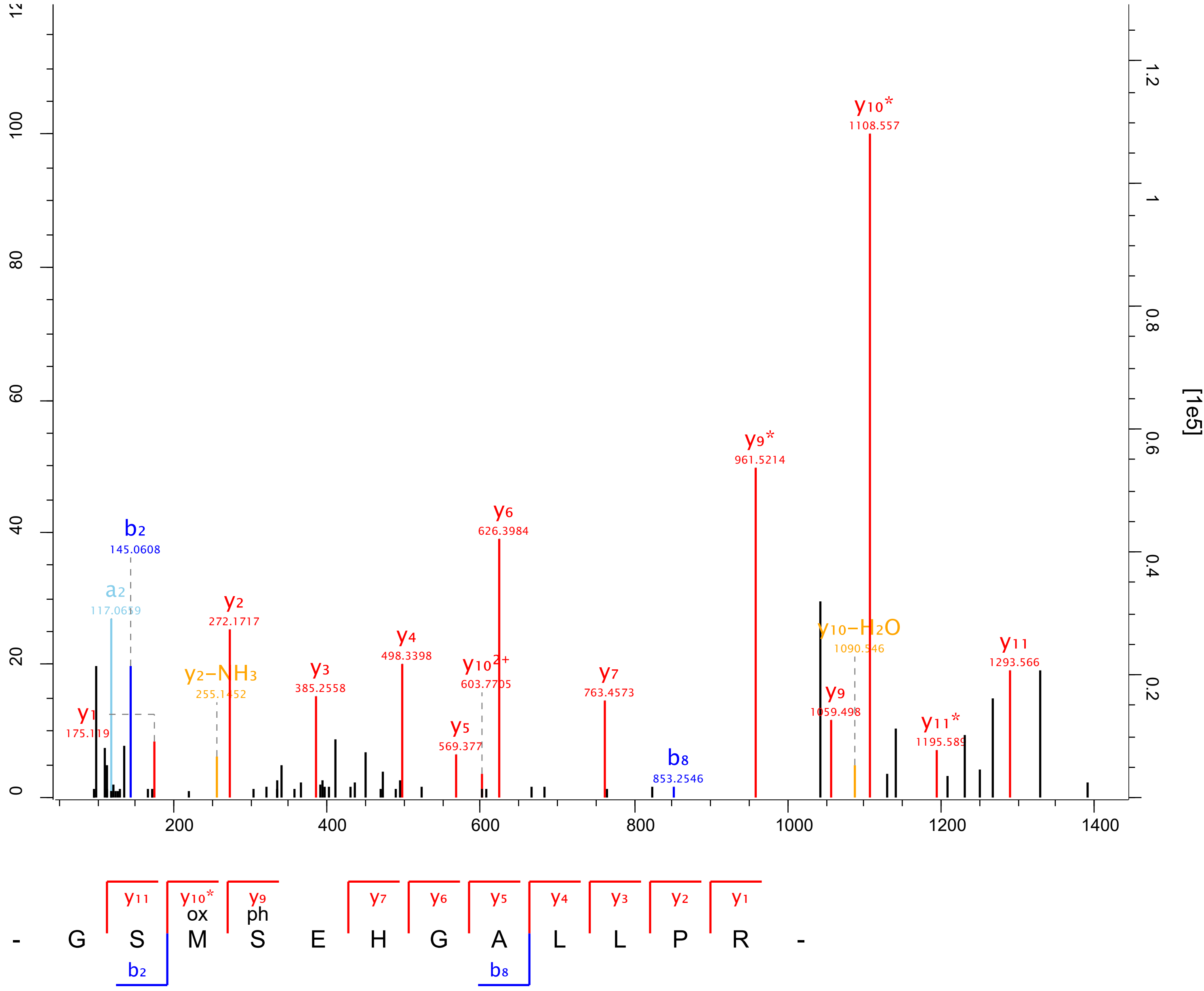

| Raw file | Scan | Method | Score | m/z |
|---|---|---|---|---|
| water_00_1 | 7963 | FTMS; HCD | 74.14 | 678.77 |

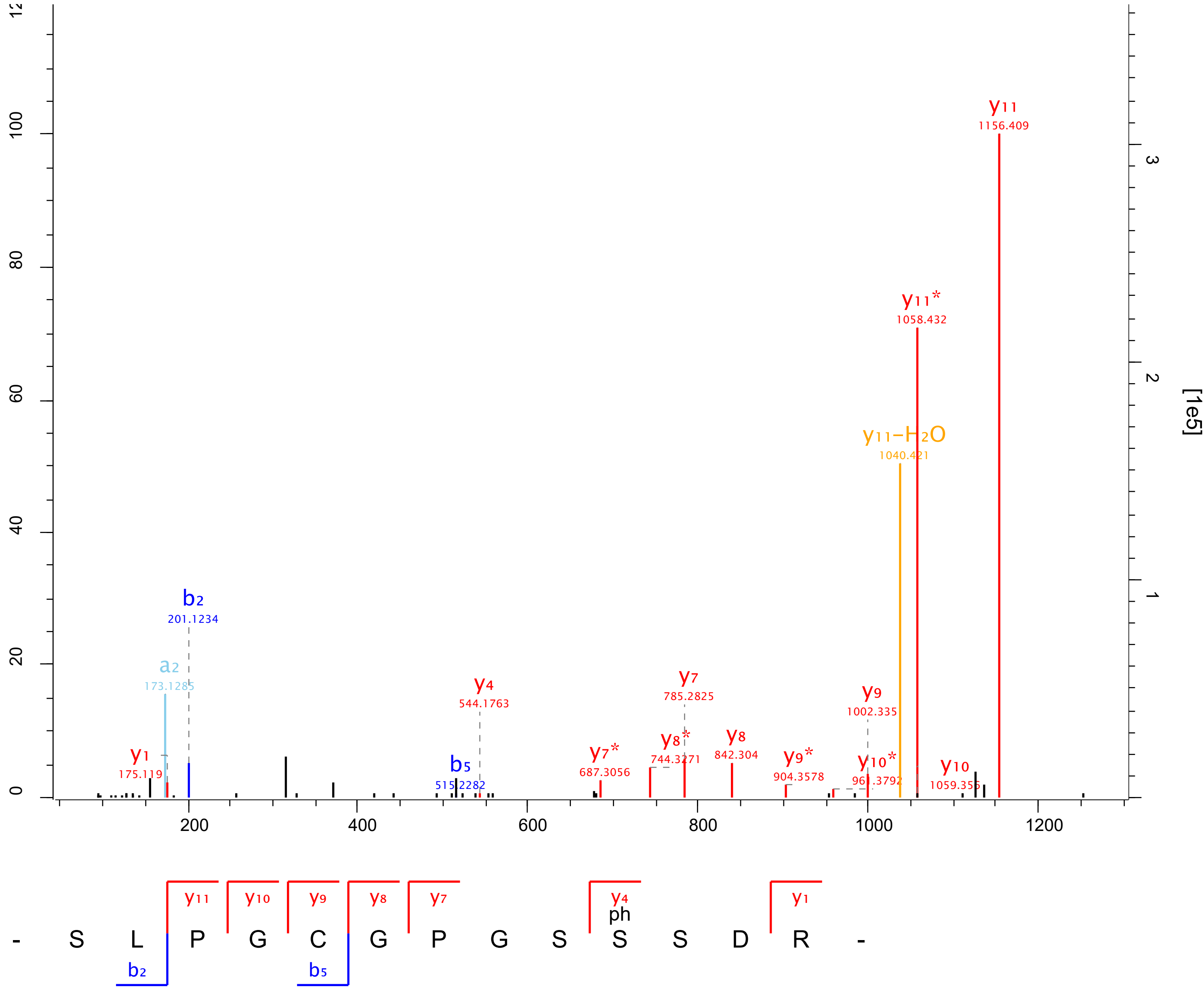

| Raw file | Scan | Method | Score | m/z |
|---|---|---|---|---|
| water_00_1 | 7969 | FTMS; HCD | 91.55 | 515.22 |

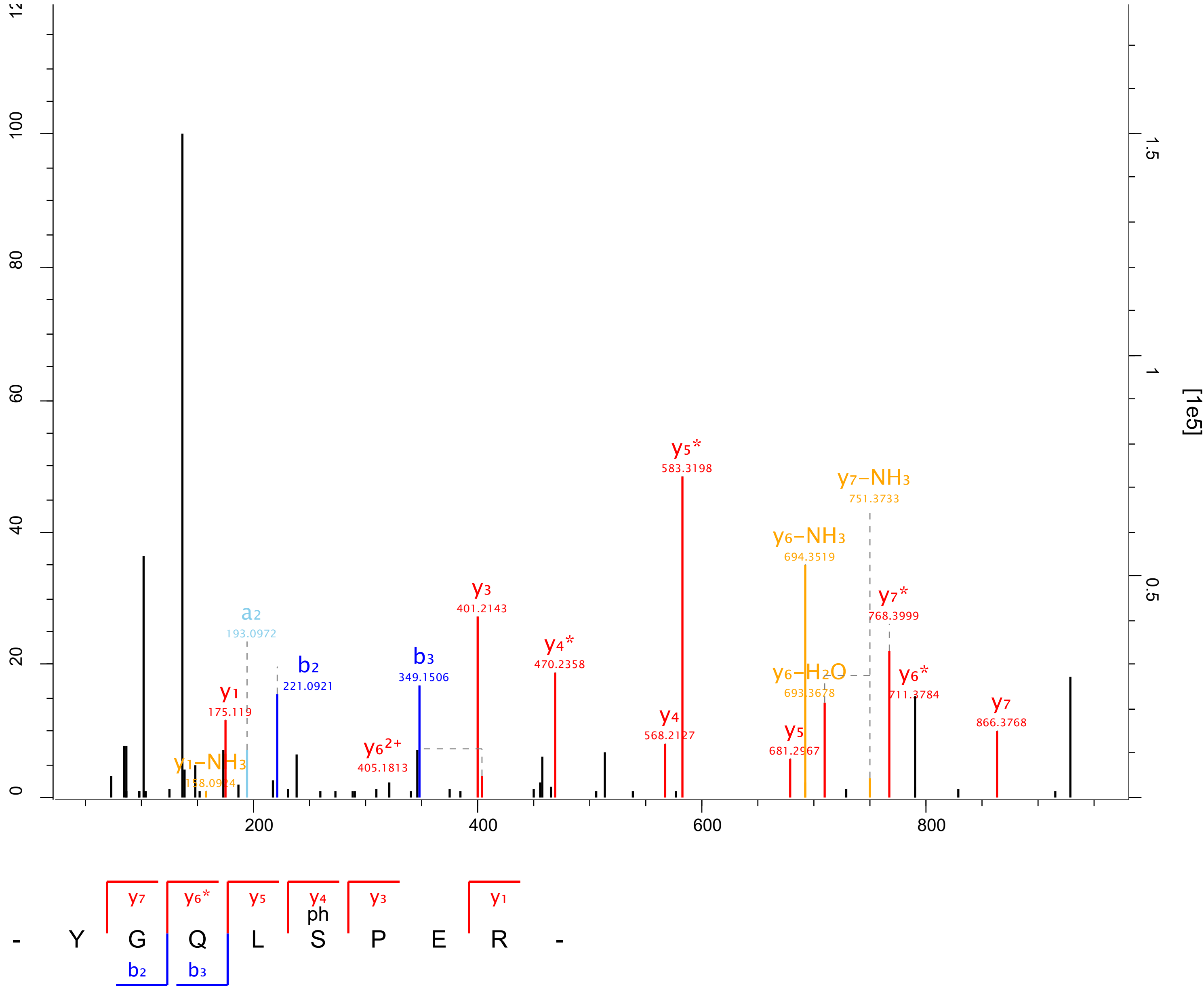

| Raw file | Scan | Method | Score | m/z |
|---|---|---|---|---|
| water_00_1 | 8044 | FTMS; HCD | 145.46 | 719.81 |

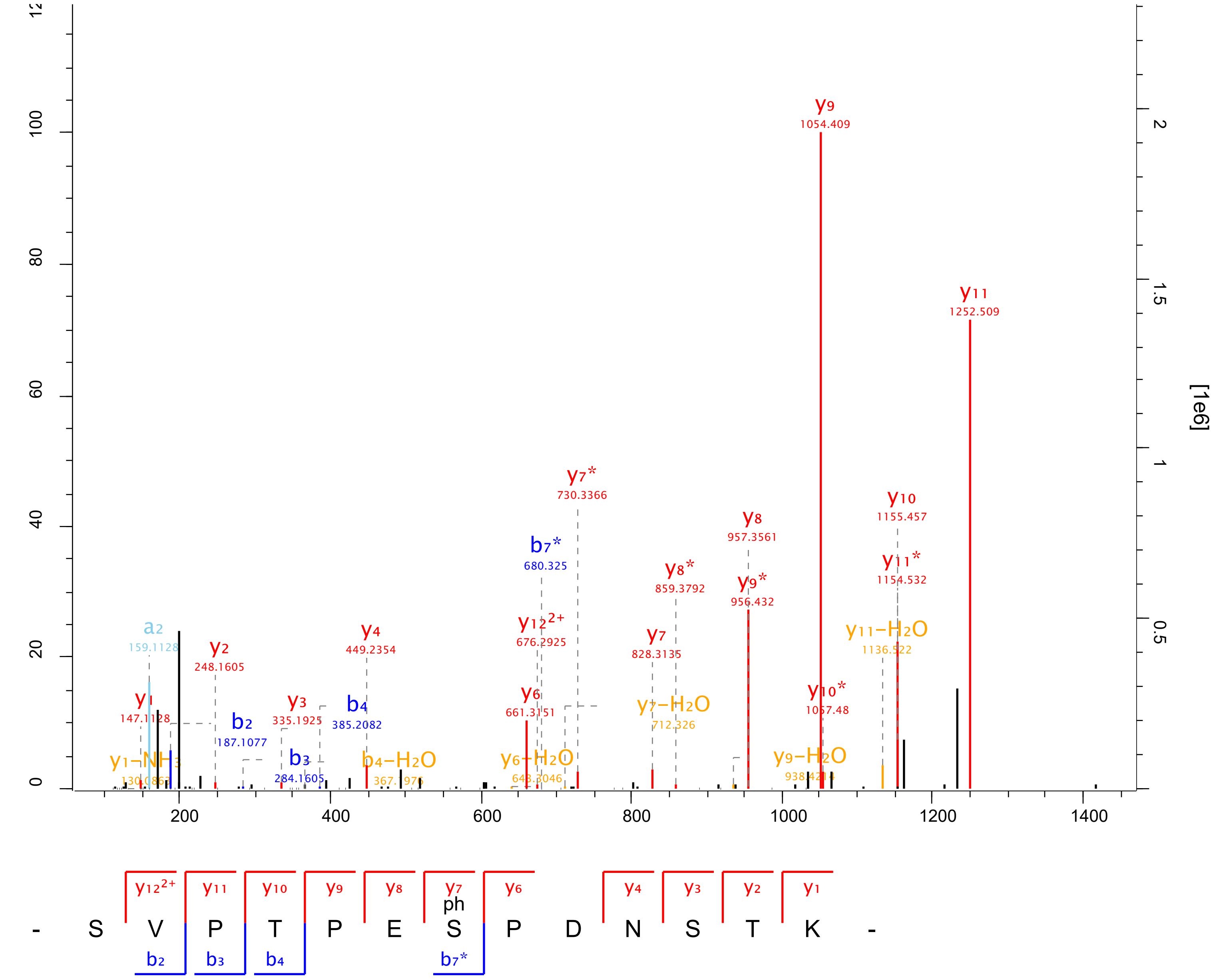

| Raw file | Scan | Method | Score | m/z |
| --- | --- | --- | --- | --- |
| water_00_1 | 8189 | FTMS; HCD | 59.73 | 553.58 |

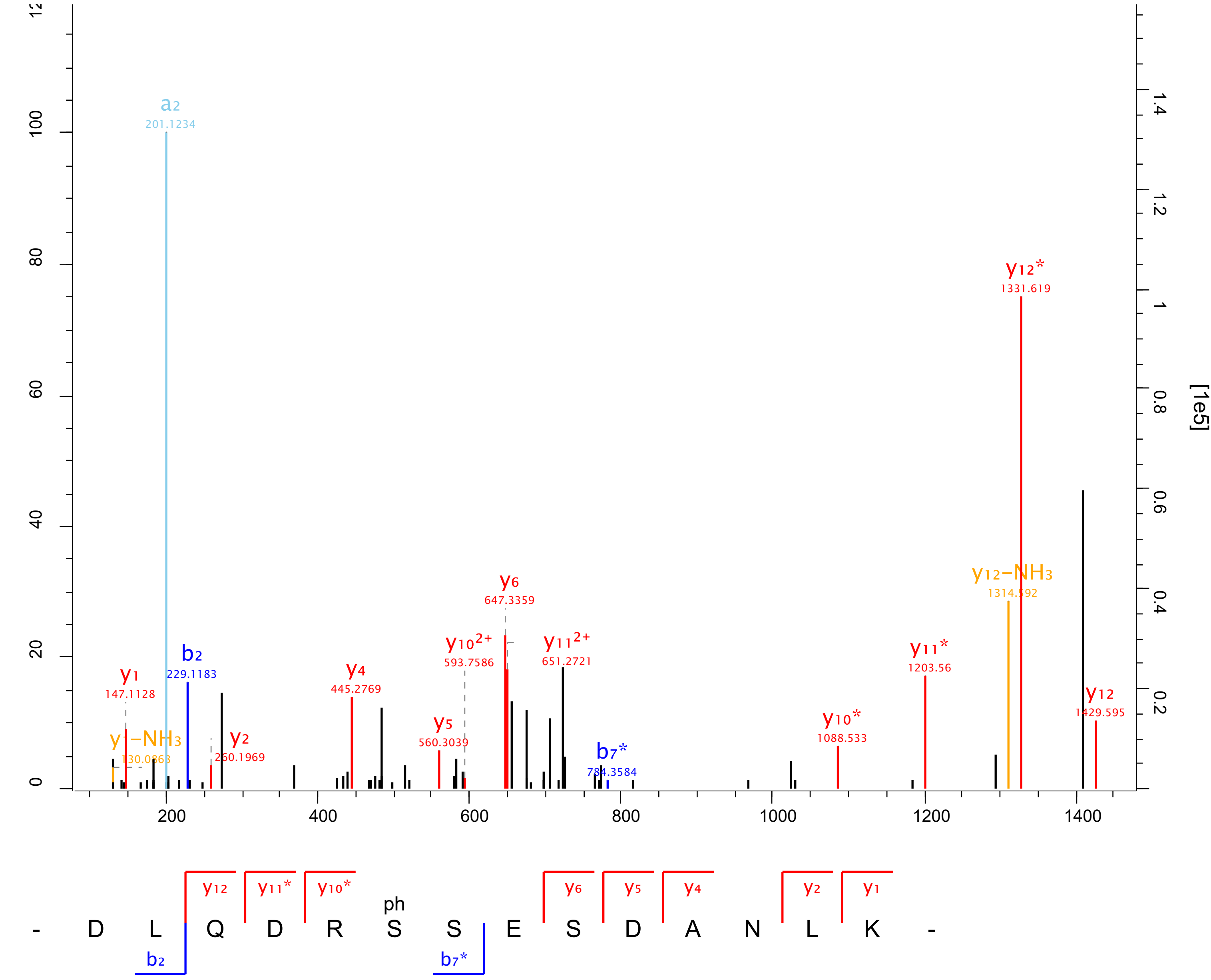

| Raw file | Scan | Method | Score | m/z |
| --- | --- | --- | --- | --- |
| water_00_1 | 8195 | FTMS; HCD | 51.29 | 559.23 |

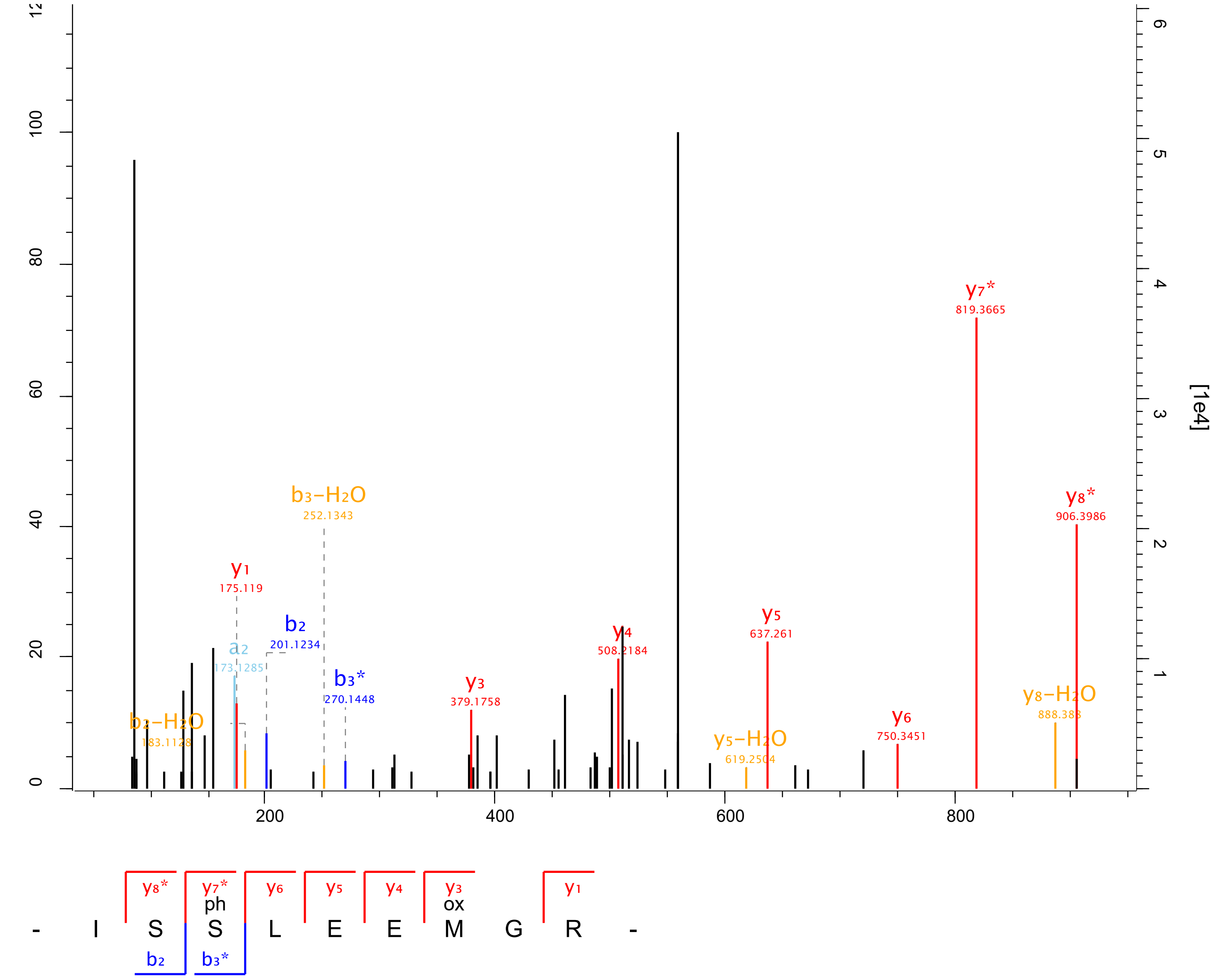

| Raw file | Scan | Method | Score | m/z |
|---|---|---|---|---|
| water_00_1 | 8371 | FTMS; HCD | 124.08 | 567.93 |

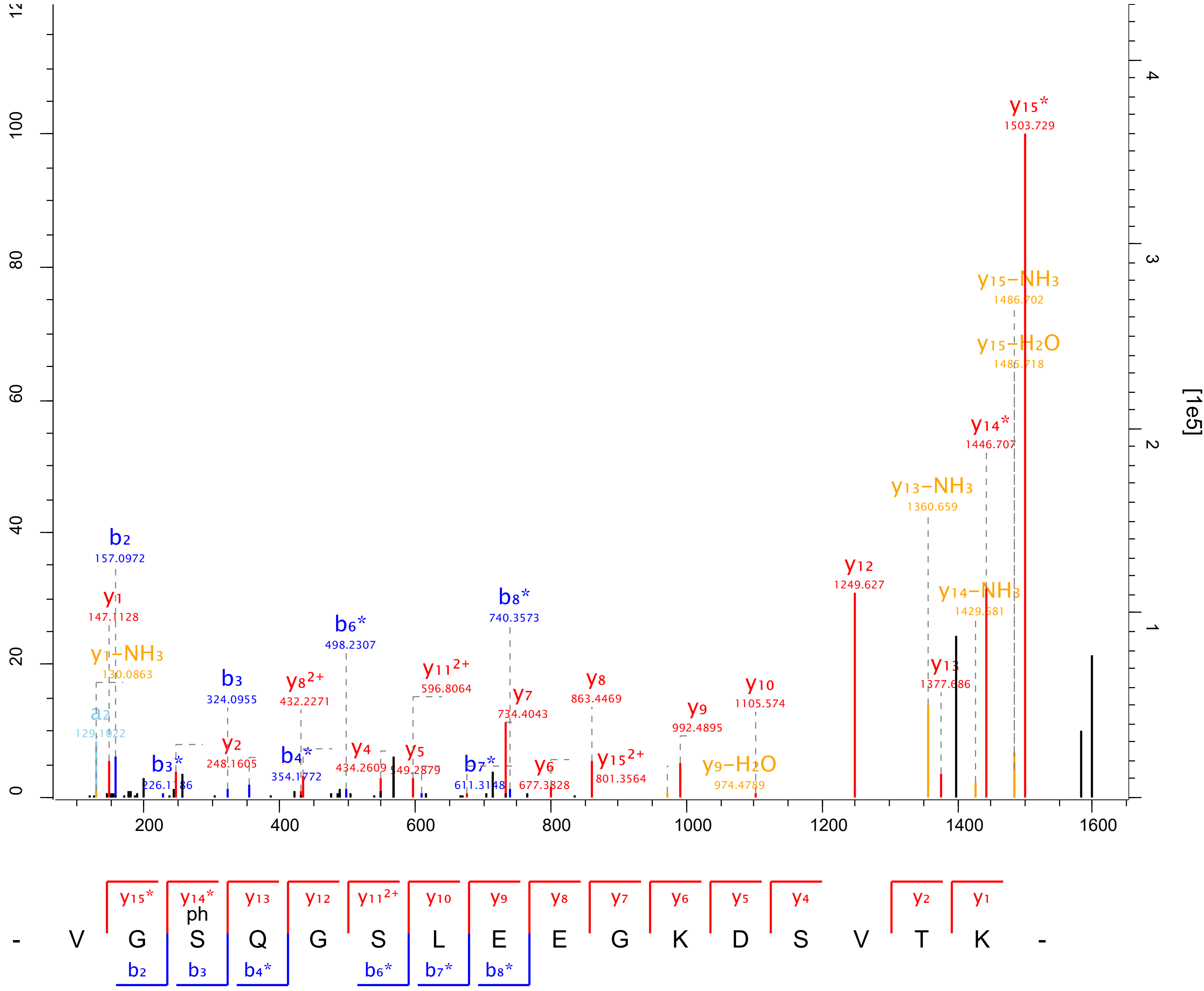

| Raw file | Scan | Method | Score | m/z |
|---|---|---|---|---|
| water_00_1 | 8424 | FTMS; HCD | 166.86 | 474.2 |

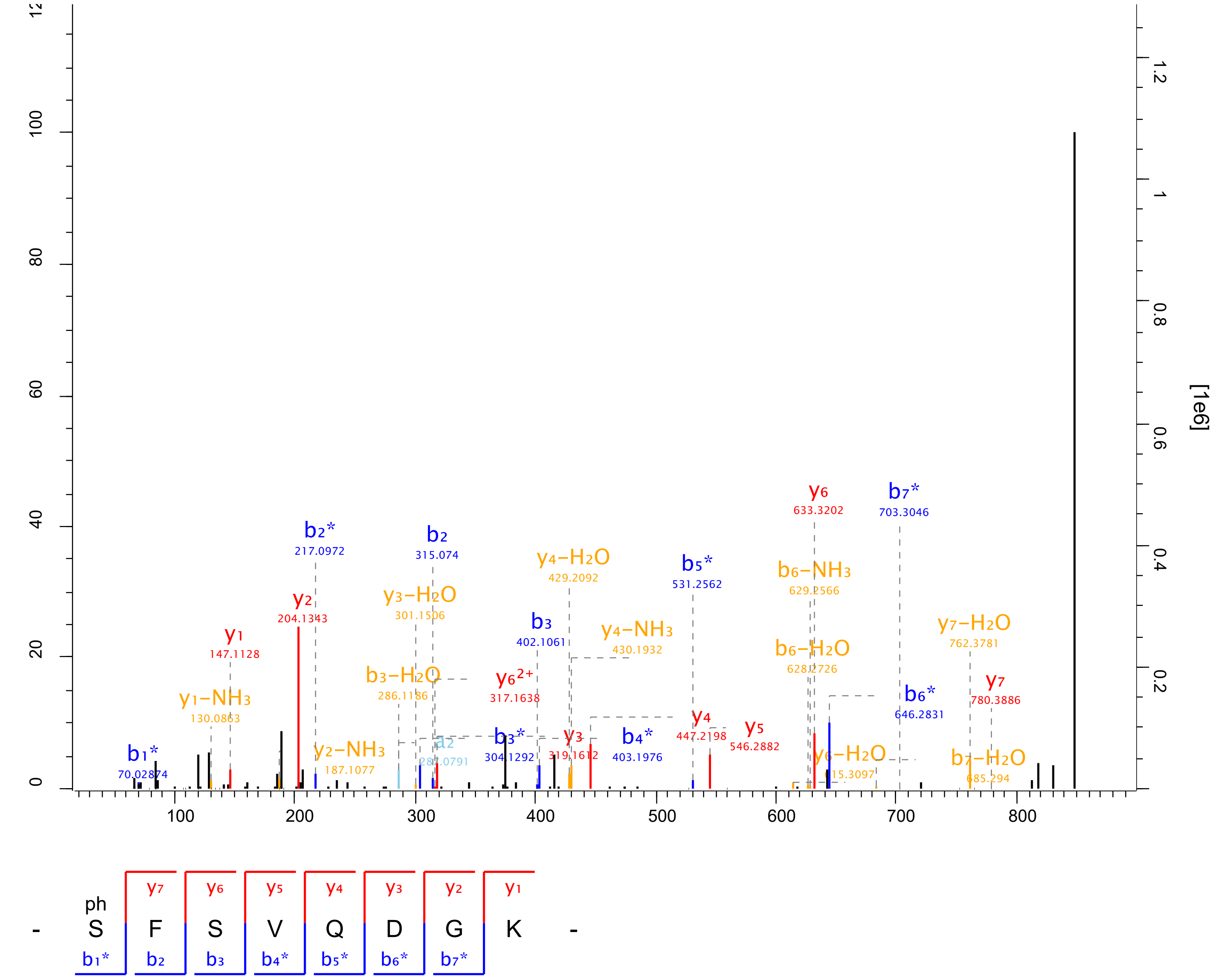

| Raw file | Scan | Method | Score | m/z |
|---|---|---|---|---|
| water_00_1 | 8500 | FTMS; HCD | 94.63 | 638.93 |

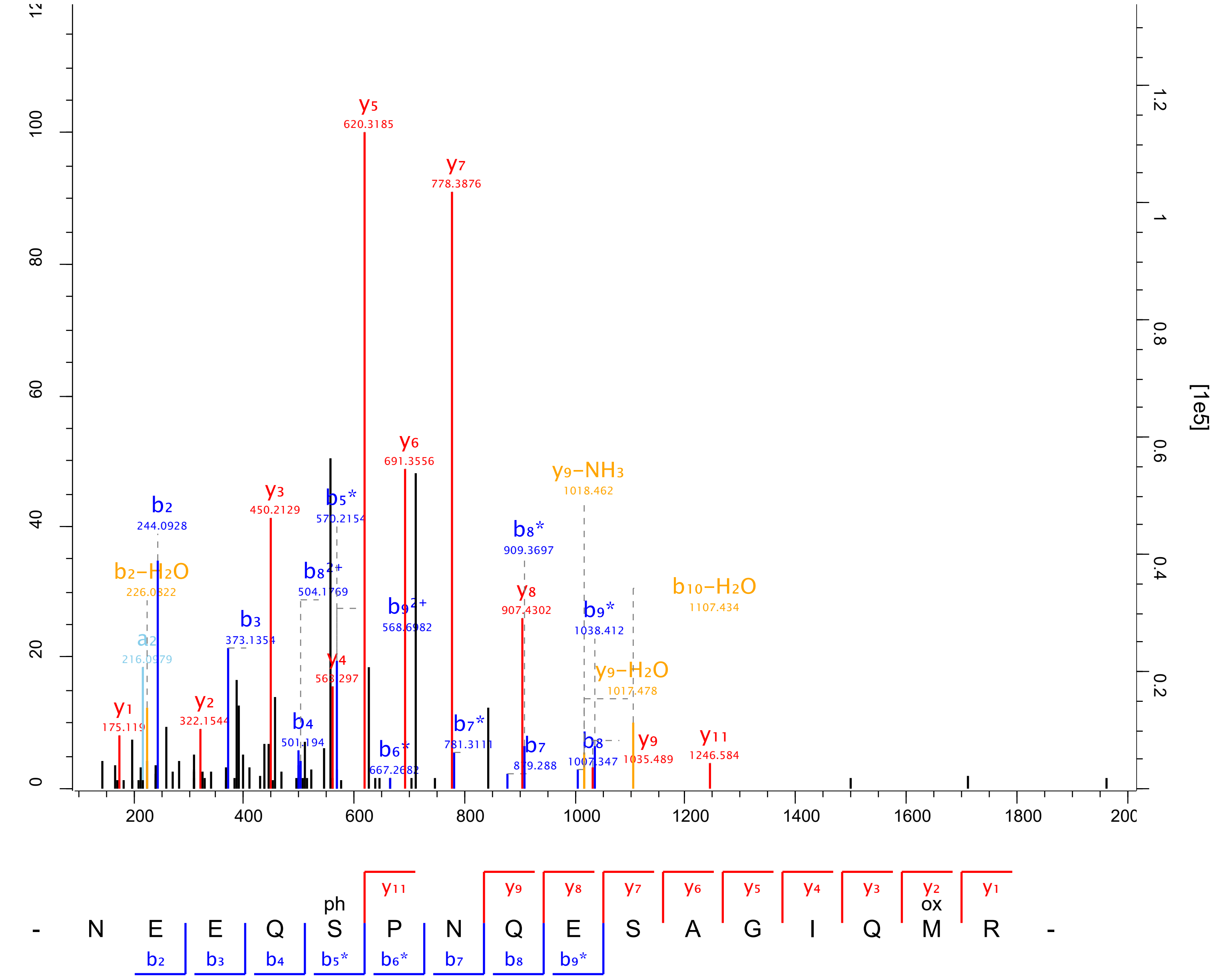

| Raw file | Scan | Method | Score | m/z |
|---|---|---|---|---|
| water_00_1 | 8865 | FTMS; HCD | 62.47 | 616.79 |

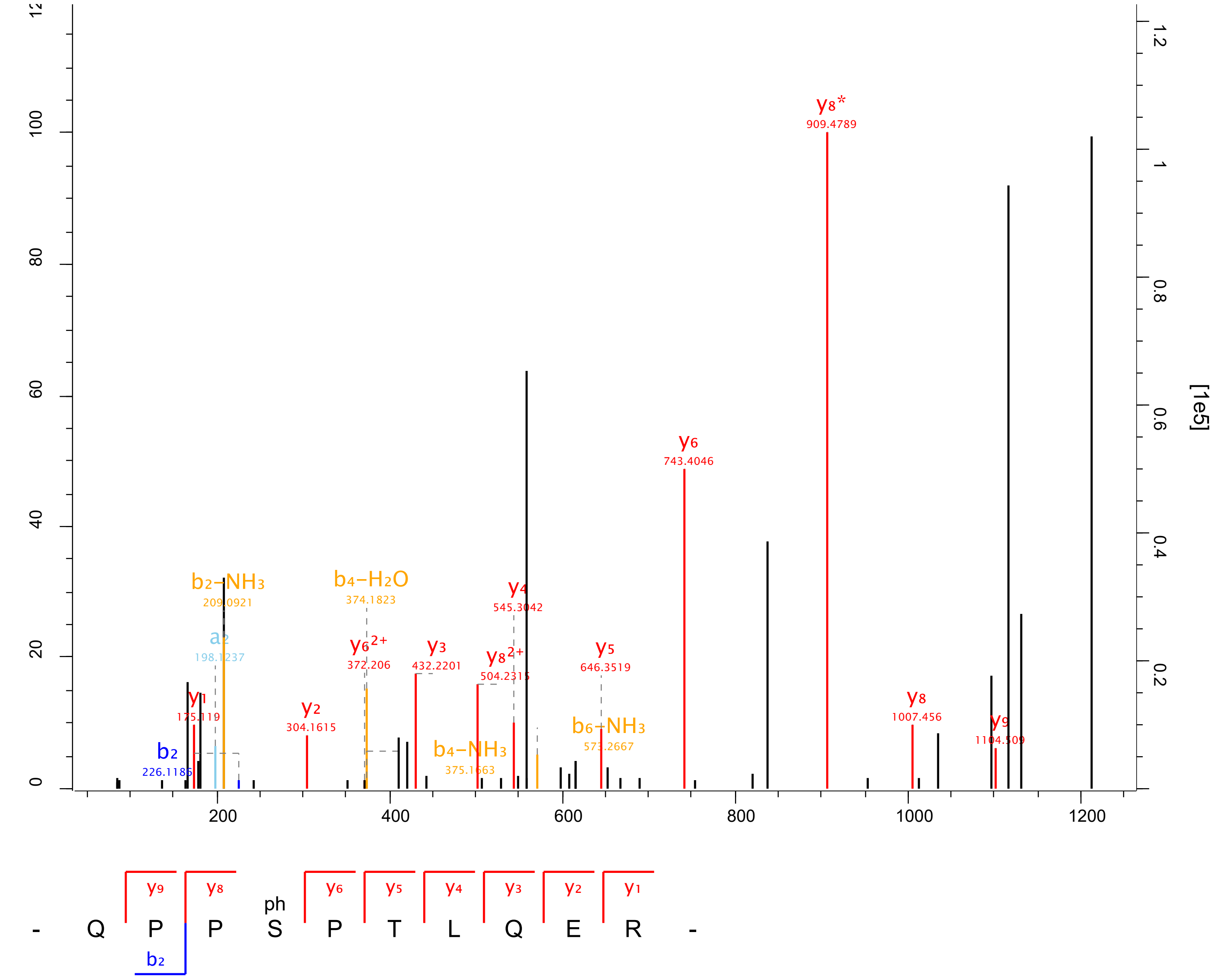

| Raw file | Scan | Method | Score | m/z |
| --- | --- | --- | --- | --- |
| water_00_1 | 8950 | FTMS; HCD | 126.31 | 450.7 |

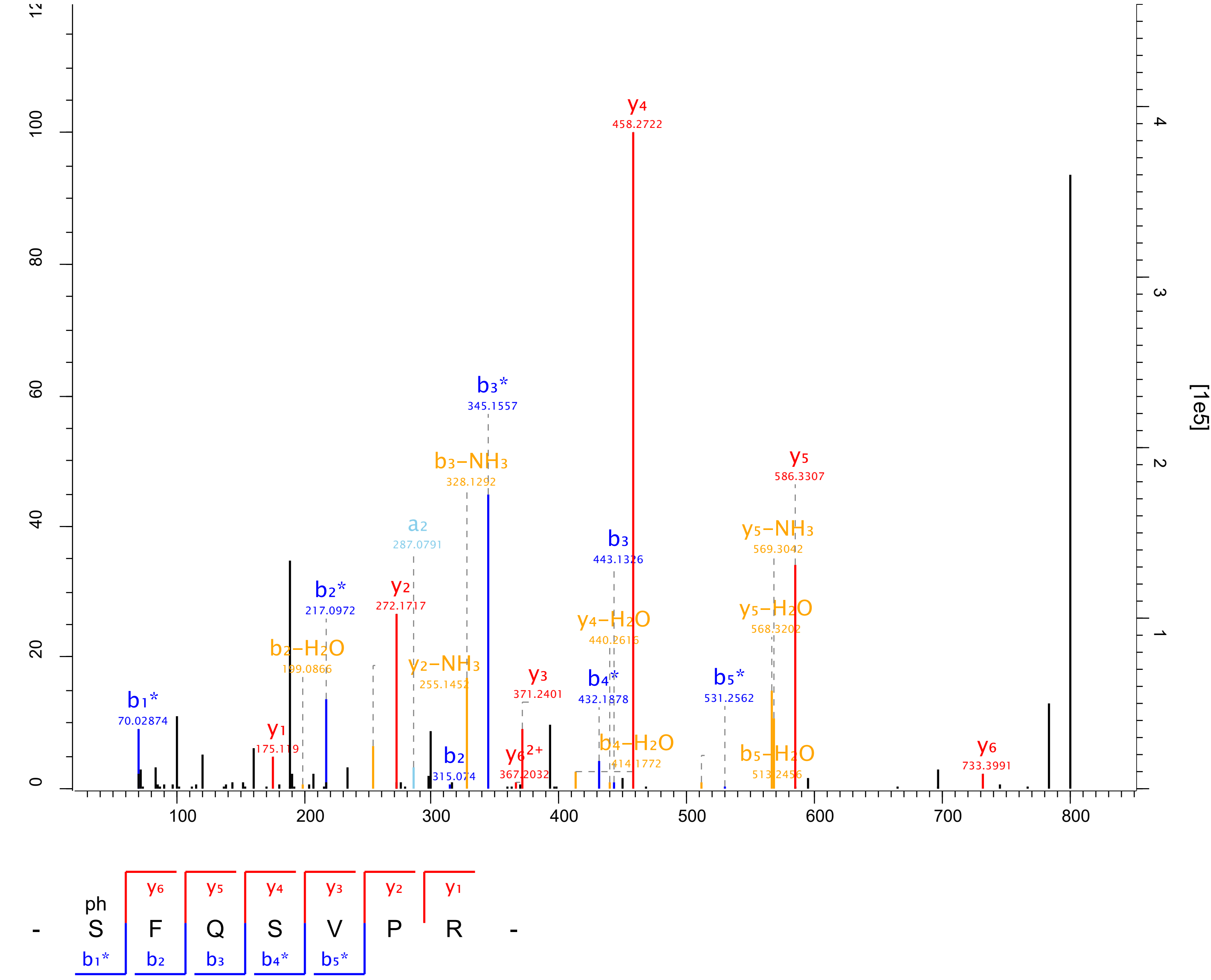

| Raw file | Scan | Method | Score | m/z |
| --- | --- | --- | --- | --- |
| water_00_1 | 9149 | FTMS; HCD | 56.57 | 462.53 |

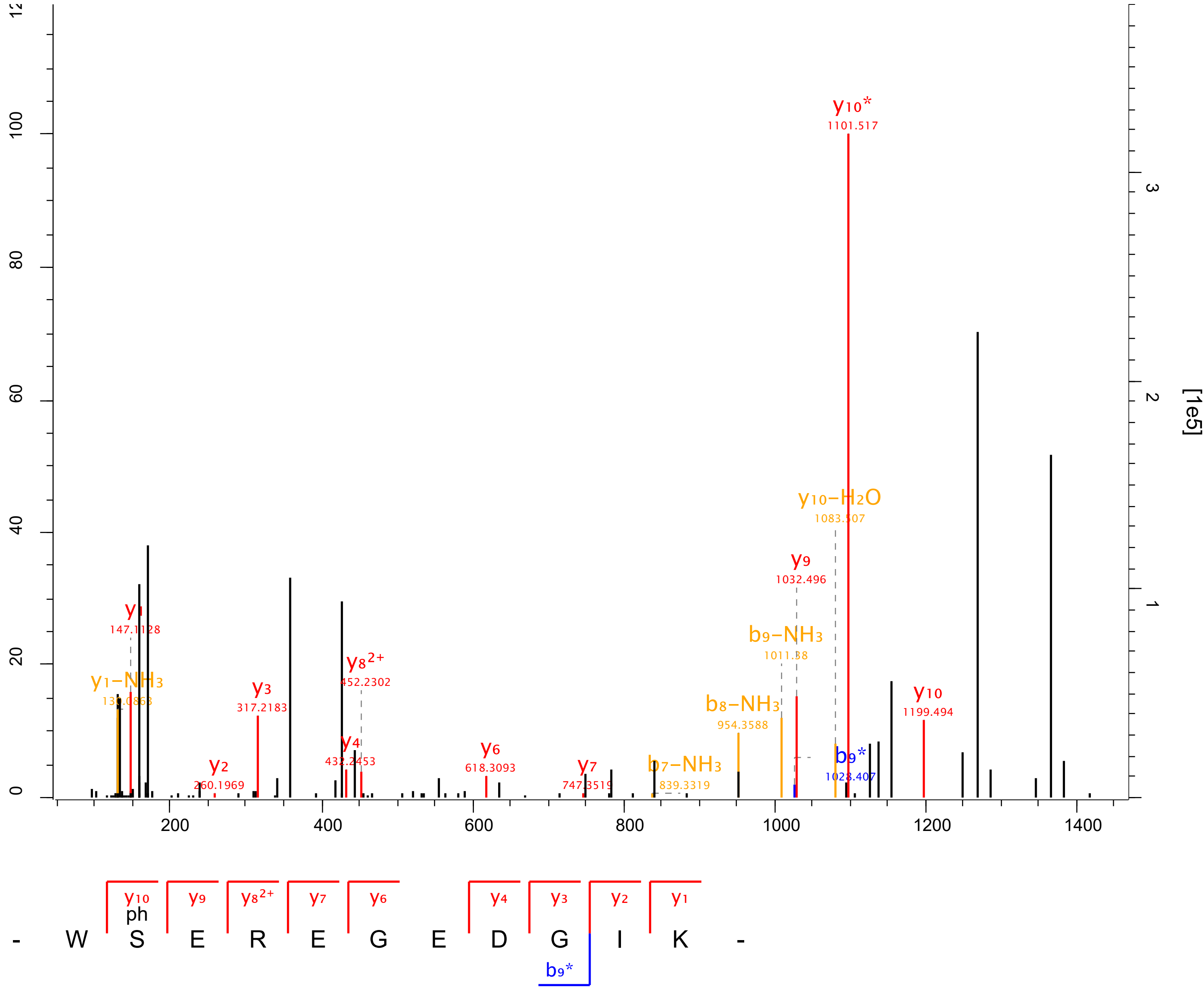

| Raw file | Scan | Method | Score | m/z |
|---|---|---|---|---|
| water_00_1 | 9158 | FTMS; HCD | 77.78 | 626.76 |

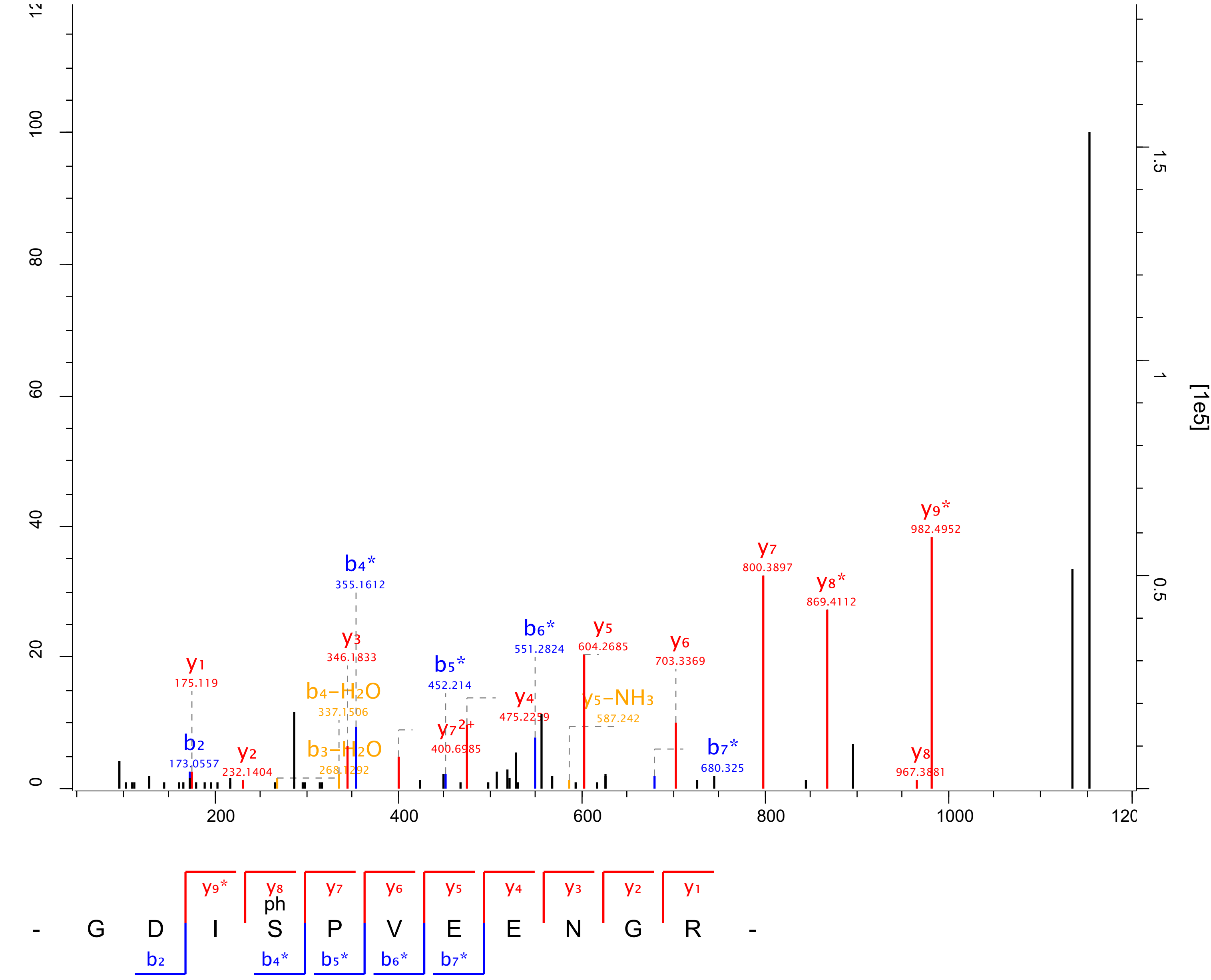

| Raw file | Scan | Method | Score | m/z |
|---|---|---|---|---|
| water_00_1 | 9188 | FTMS; HCD | 247.28 | 824.83 |

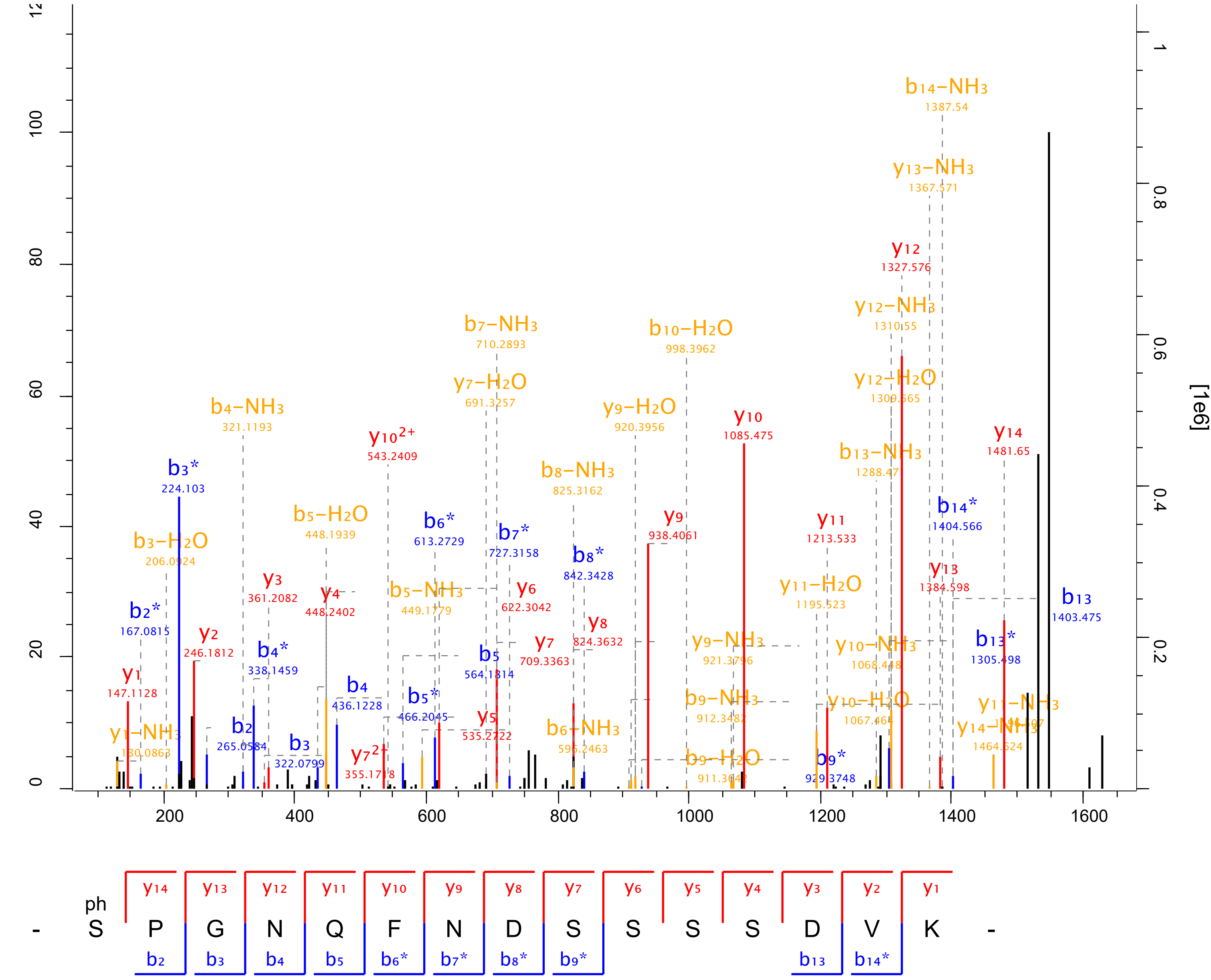

| Raw file | Scan | Method | Score | m/z |
|---|---|---|---|---|
| water_00_1 | 9281 | FTMS; HCD | 72.2 | 515.72 |

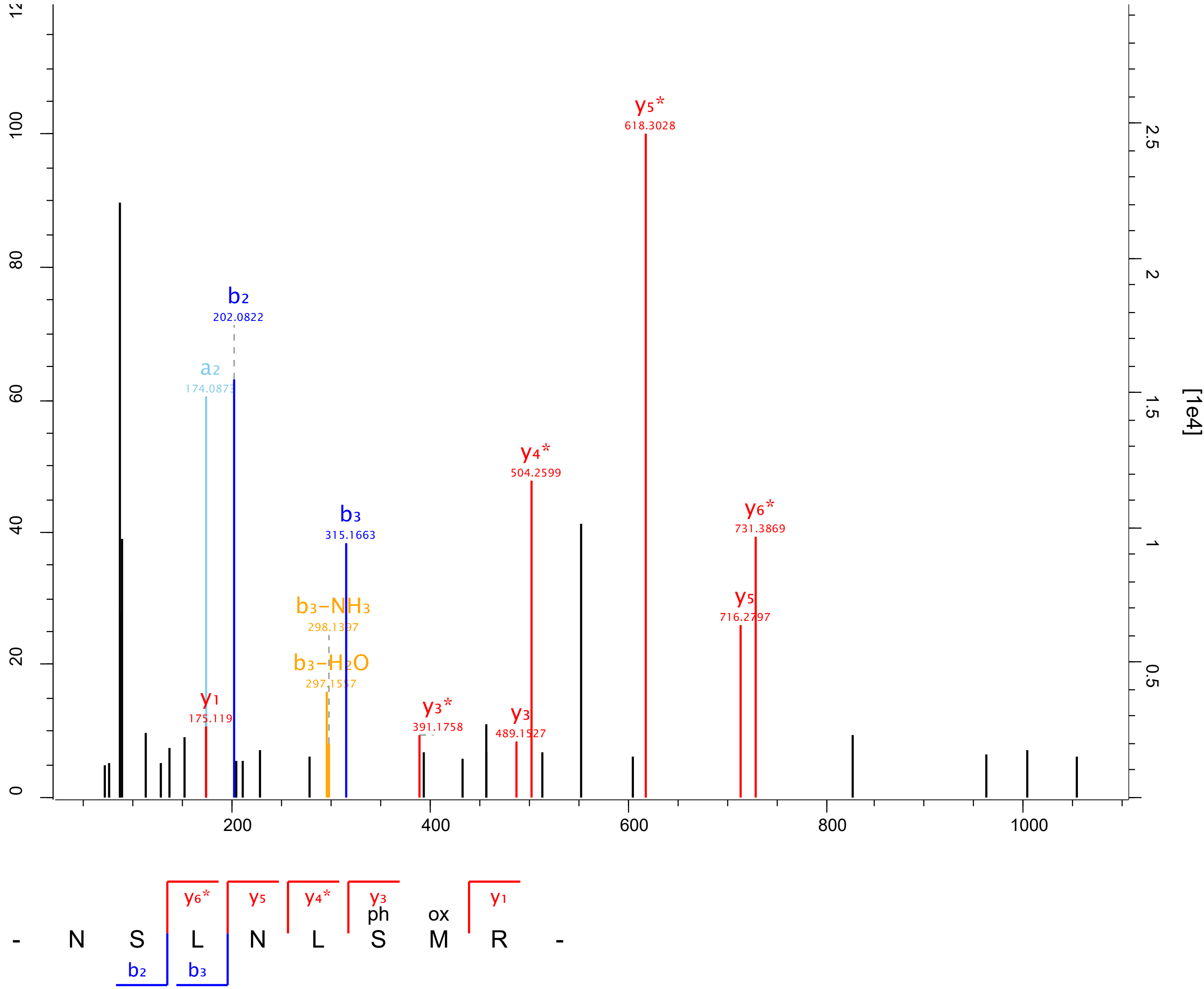

| Raw file | Scan | Method | Score | m/z |
|---|---|---|---|---|
| water_00_1 | 9300 | FTMS; HCD | 97.69 | 581.28 |

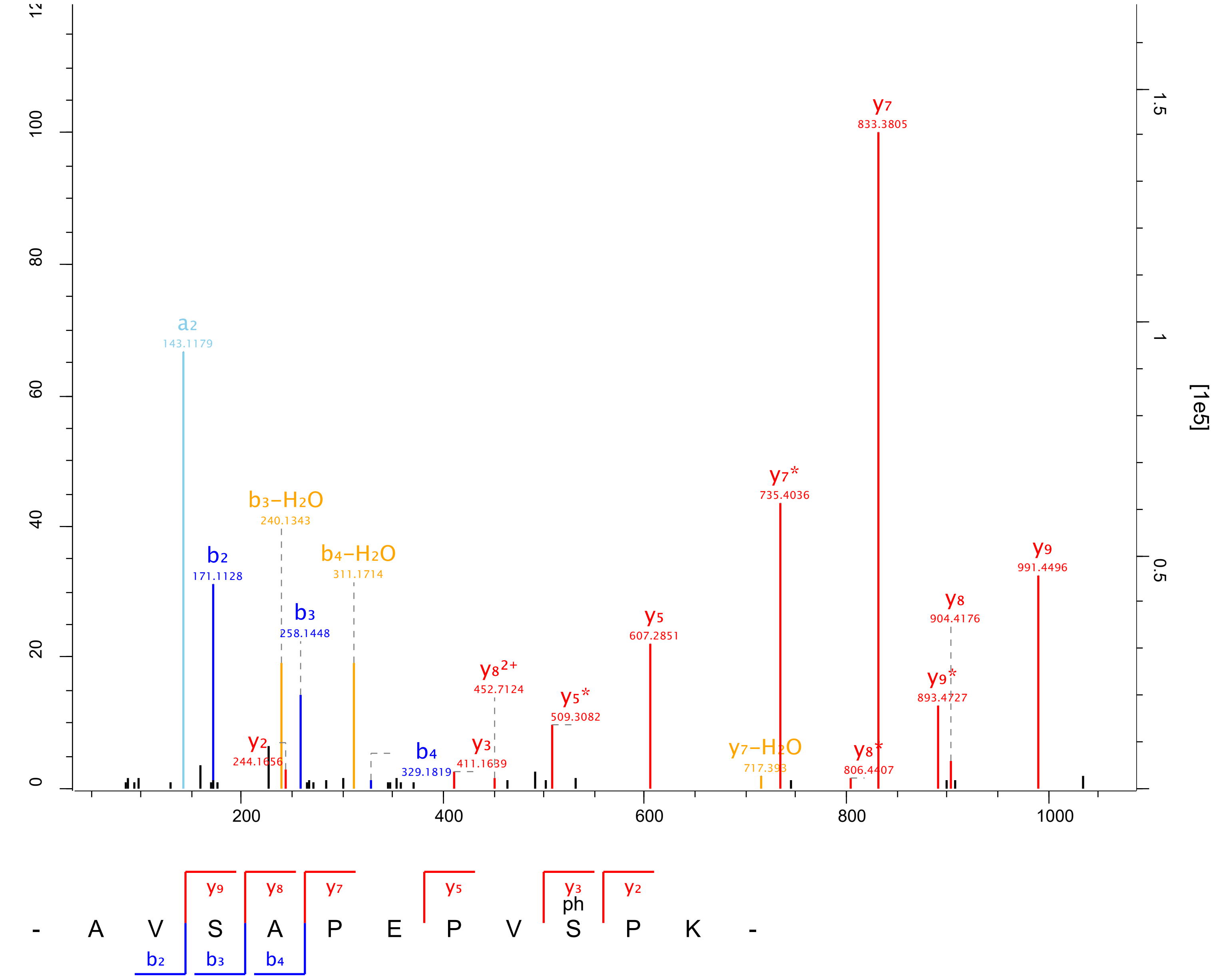

| Raw file | Scan | Method | Score | m/z |
|---|---|---|---|---|
| water_00_1 | 9321 | FTMS; HCD | 145.61 | 561.27 |

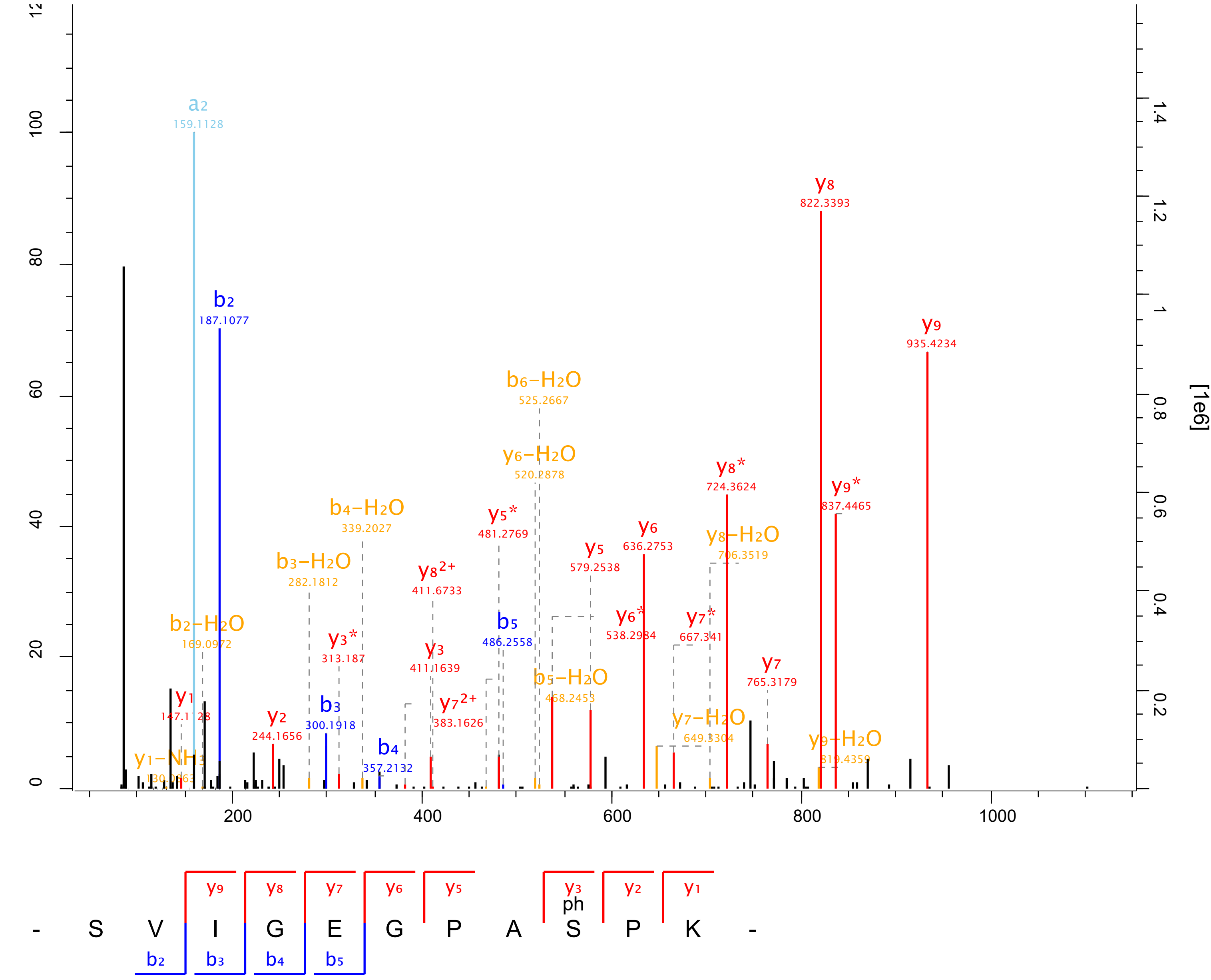

| Raw file | Scan | Method | Score | m/z |
| --- | --- | --- | --- | --- |
| water_00_1 | 9430 | FTMS; HCD | 159.58 | 578.74 |

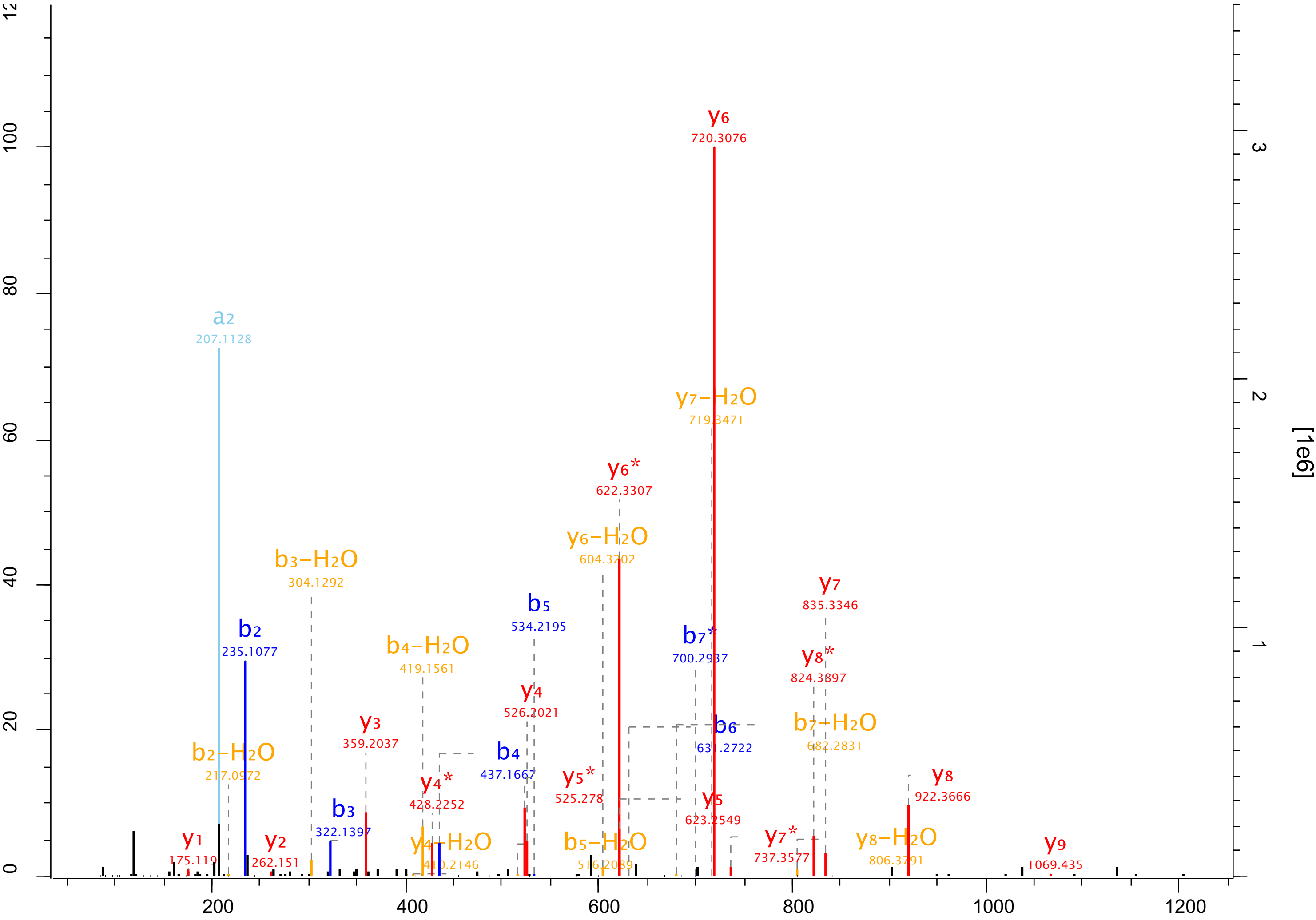


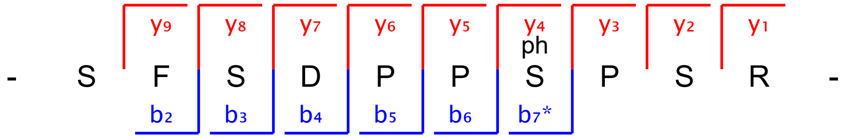

| Raw file | Scan | Method | Score | m/z |
|---|---|---|---|---|
| water_00_1 | 9593 | FTMS; HCD | 130.01 | 480.22 |

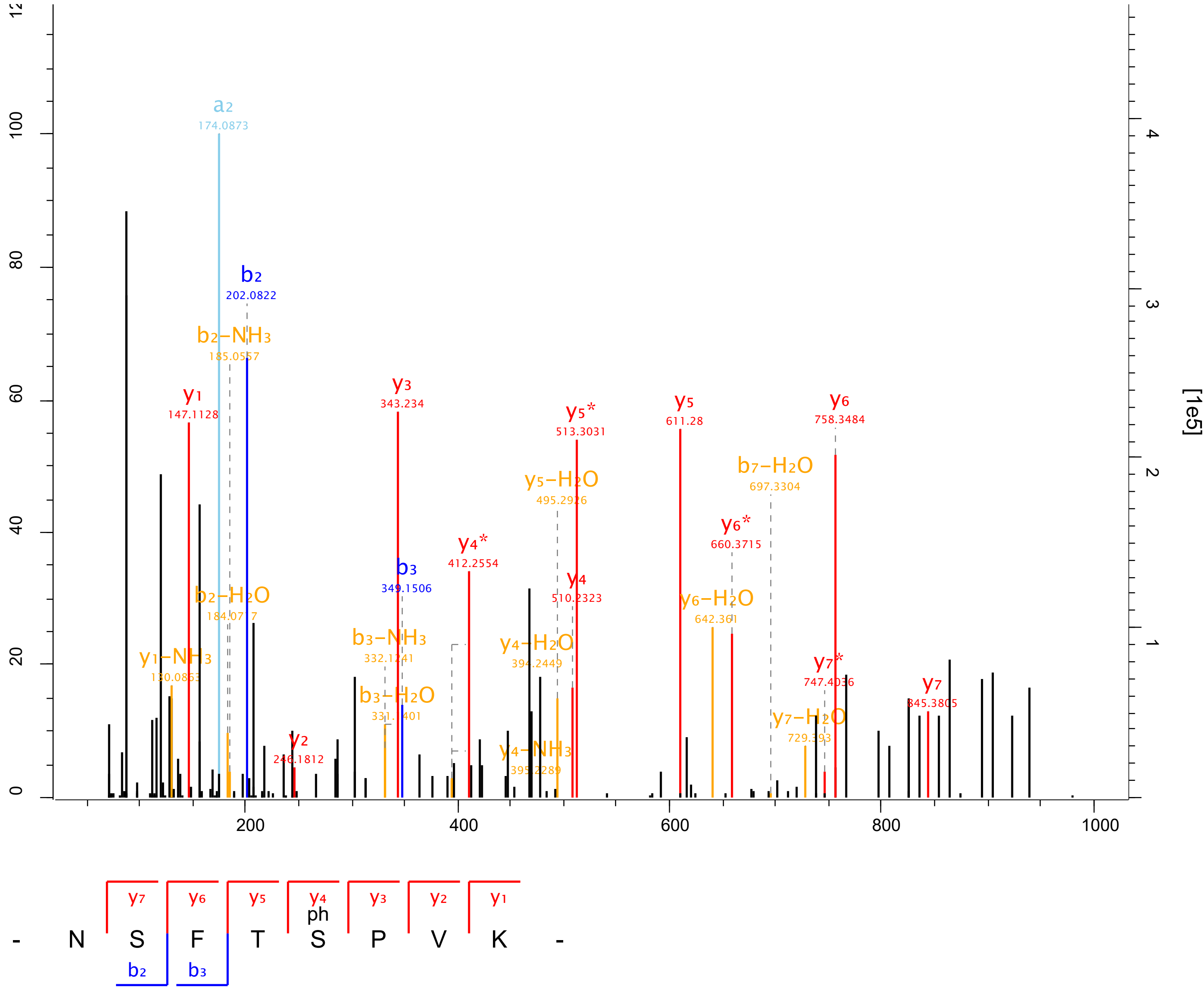

| Raw file | Scan | Method | Score | m/z |
|---|---|---|---|---|
| water_00_1 | 9608 | FTMS; HCD | 90.83 | 517.71 |

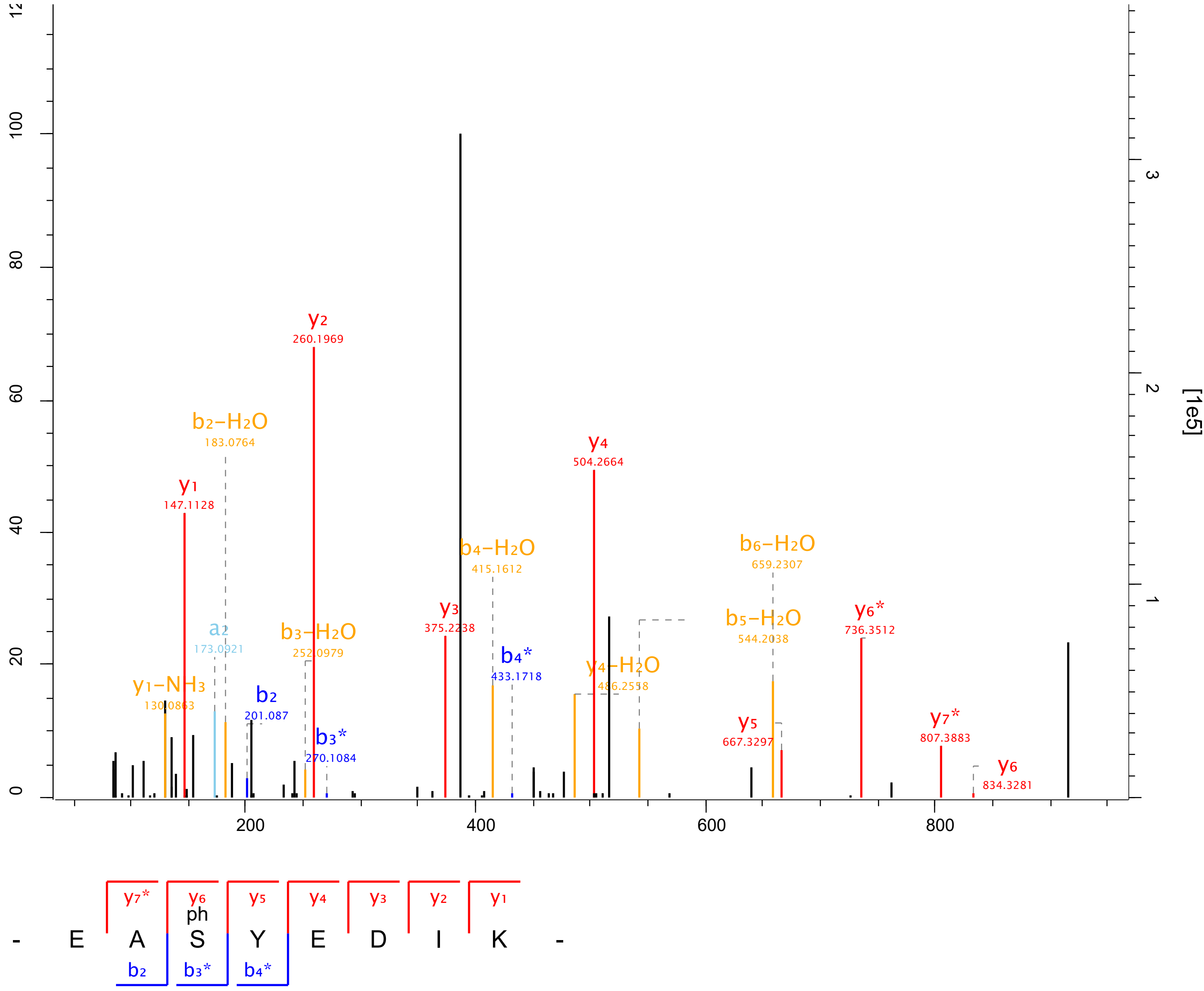

| Raw file | Scan | Method | Score | m/z |
|---|---|---|---|---|
| water_00_1 | 9803 | FTMS; HCD | 132.08 | 695.27 |

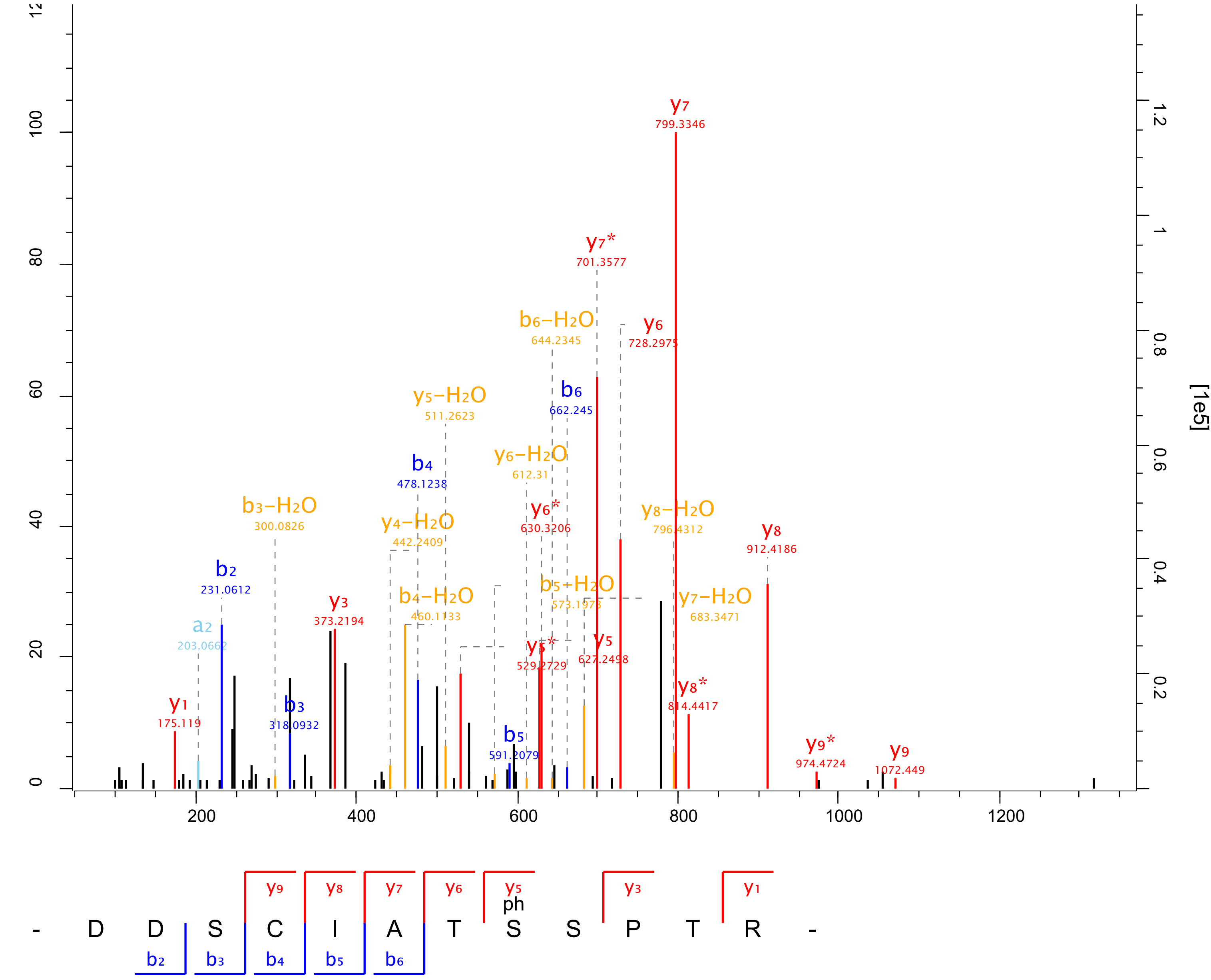

| Raw file | Scan | Method | Score | m/z |
|---|---|---|---|---|
| water_00_1 | 9812 | FTMS; HCD | 168.61 | 592.27 |

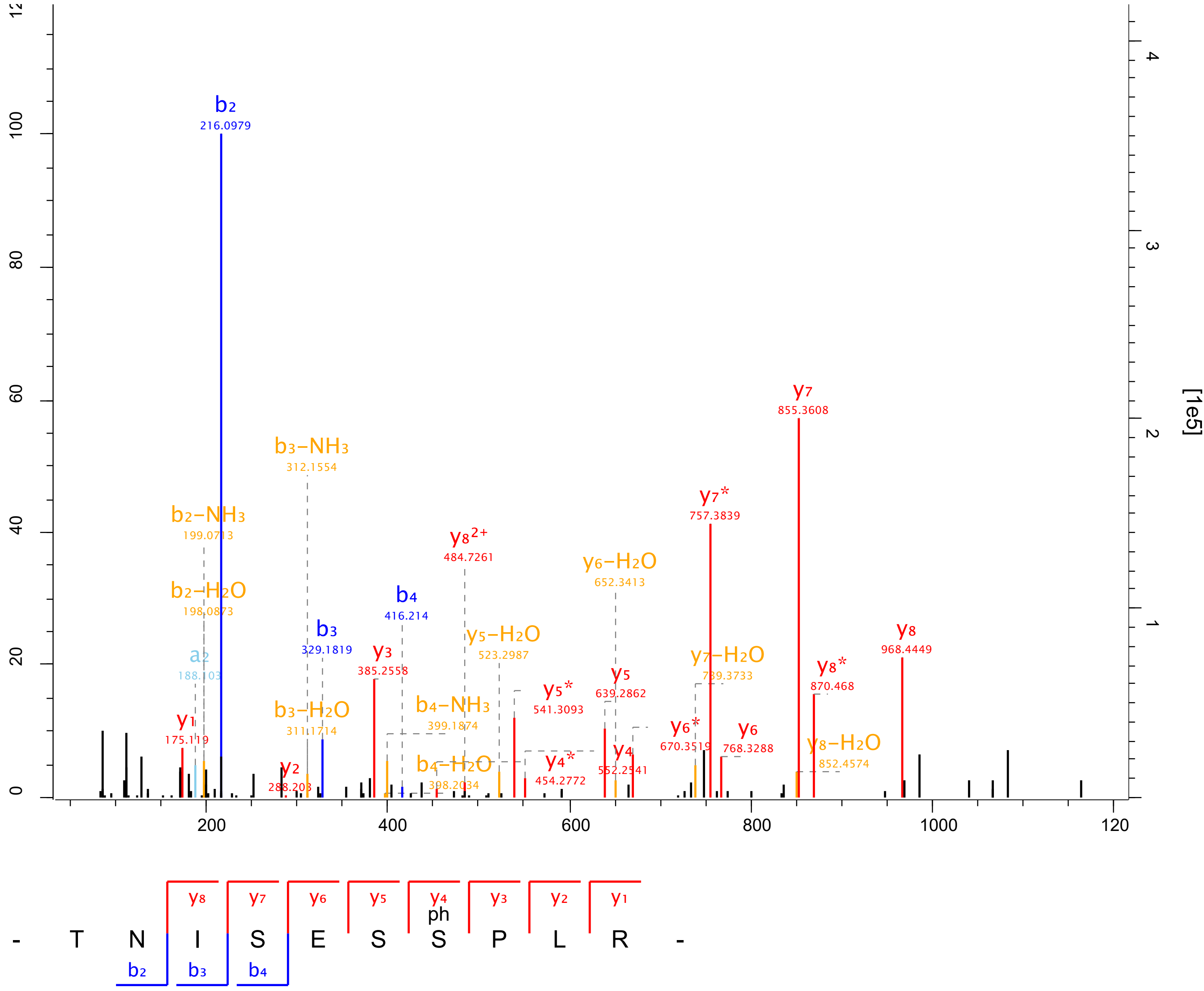

| Raw file | Scan | Method | Score | m/z |
|---|---|---|---|---|
| water_00_1 | 9875 | FTMS; HCD | 342.97 | 759.29 |

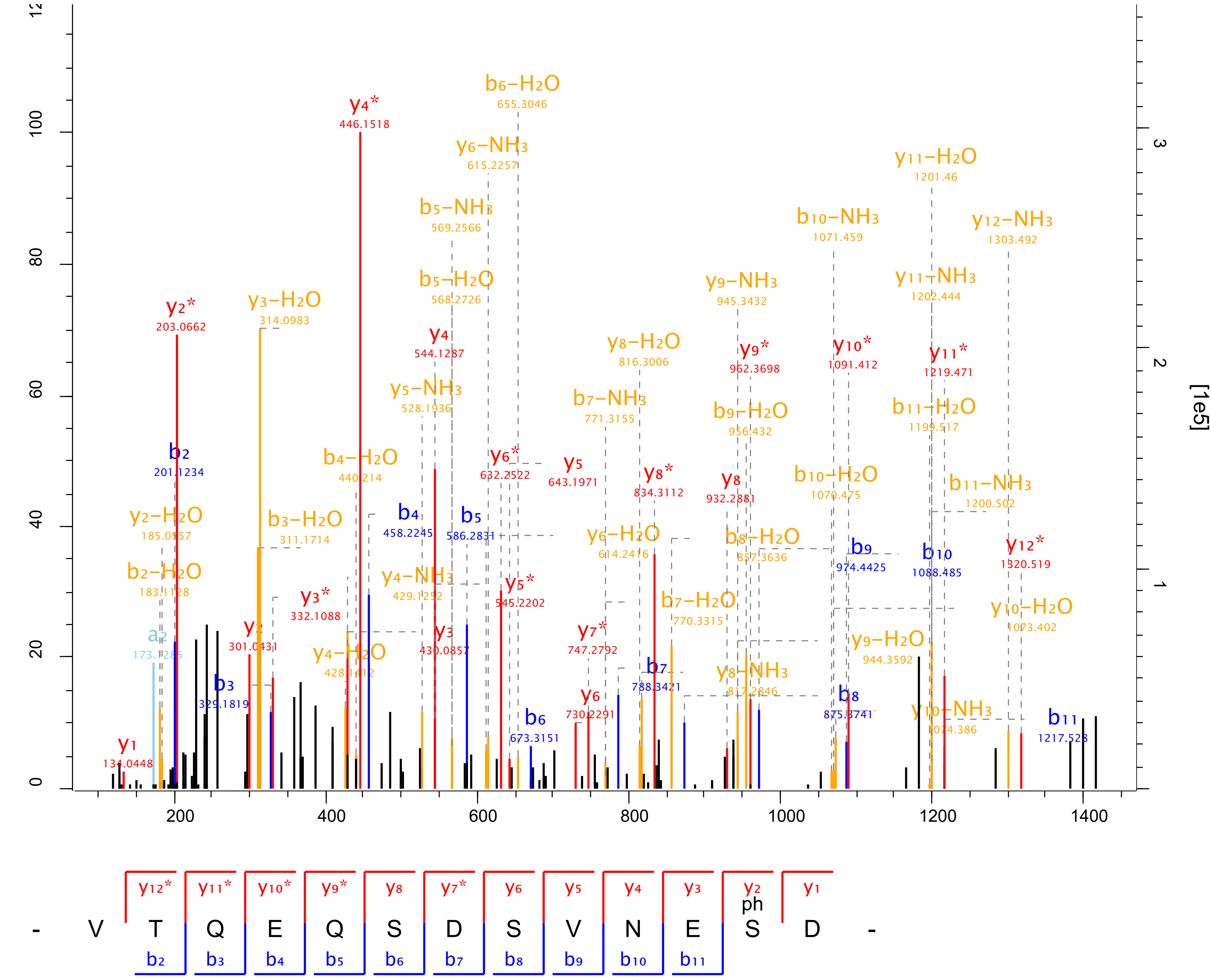

| Raw file | Scan | Method | Score | m/z |
|---|---|---|---|---|
| water_00_2 | 15732 | FTMS; HCD | 57.59 | 433.23 |

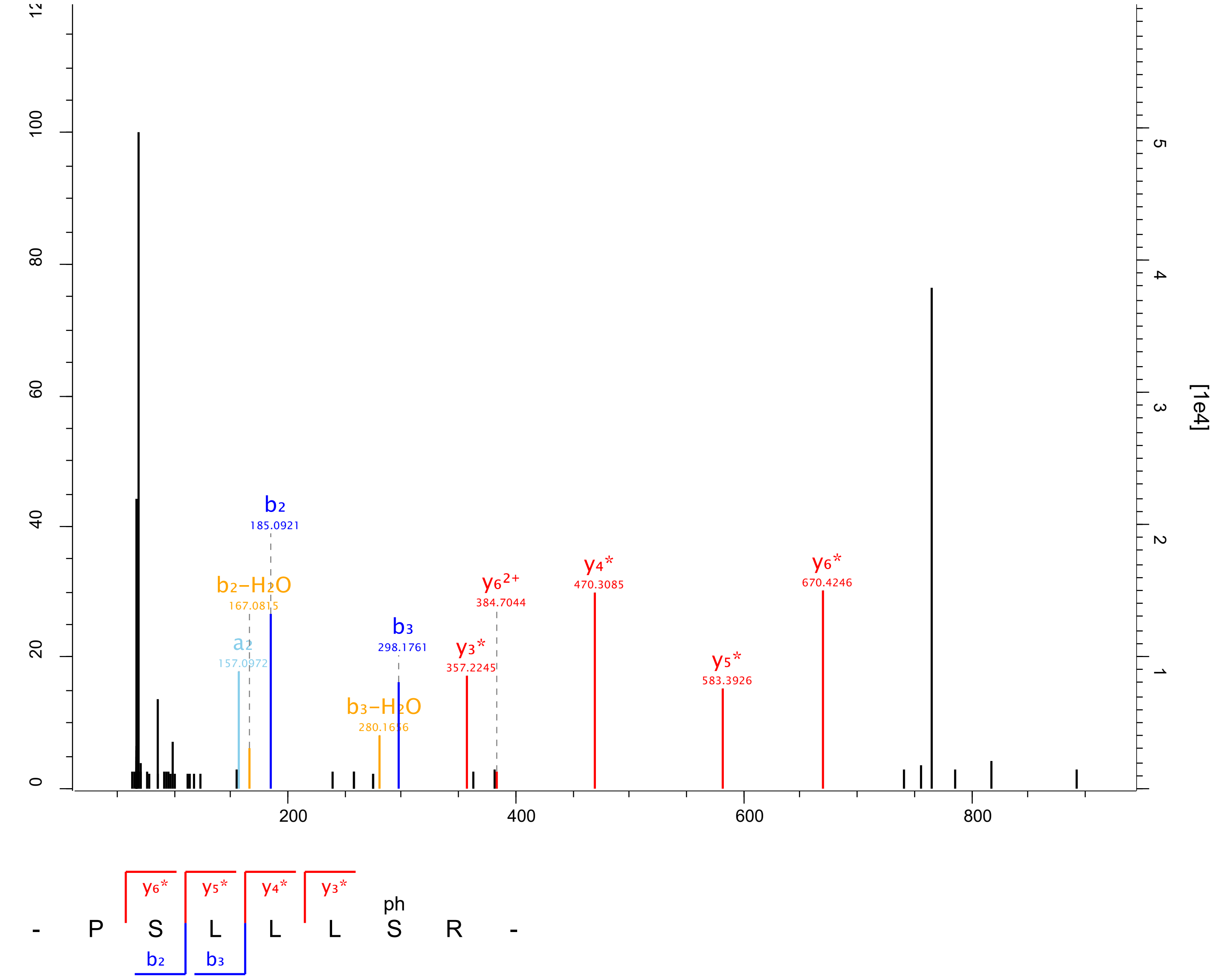

| Raw file | Scan | Method | Score | m/z |
|---|---|---|---|---|
| water_00_2 | 3639 | FTMS; HCD | 145.6 | 586.27 |

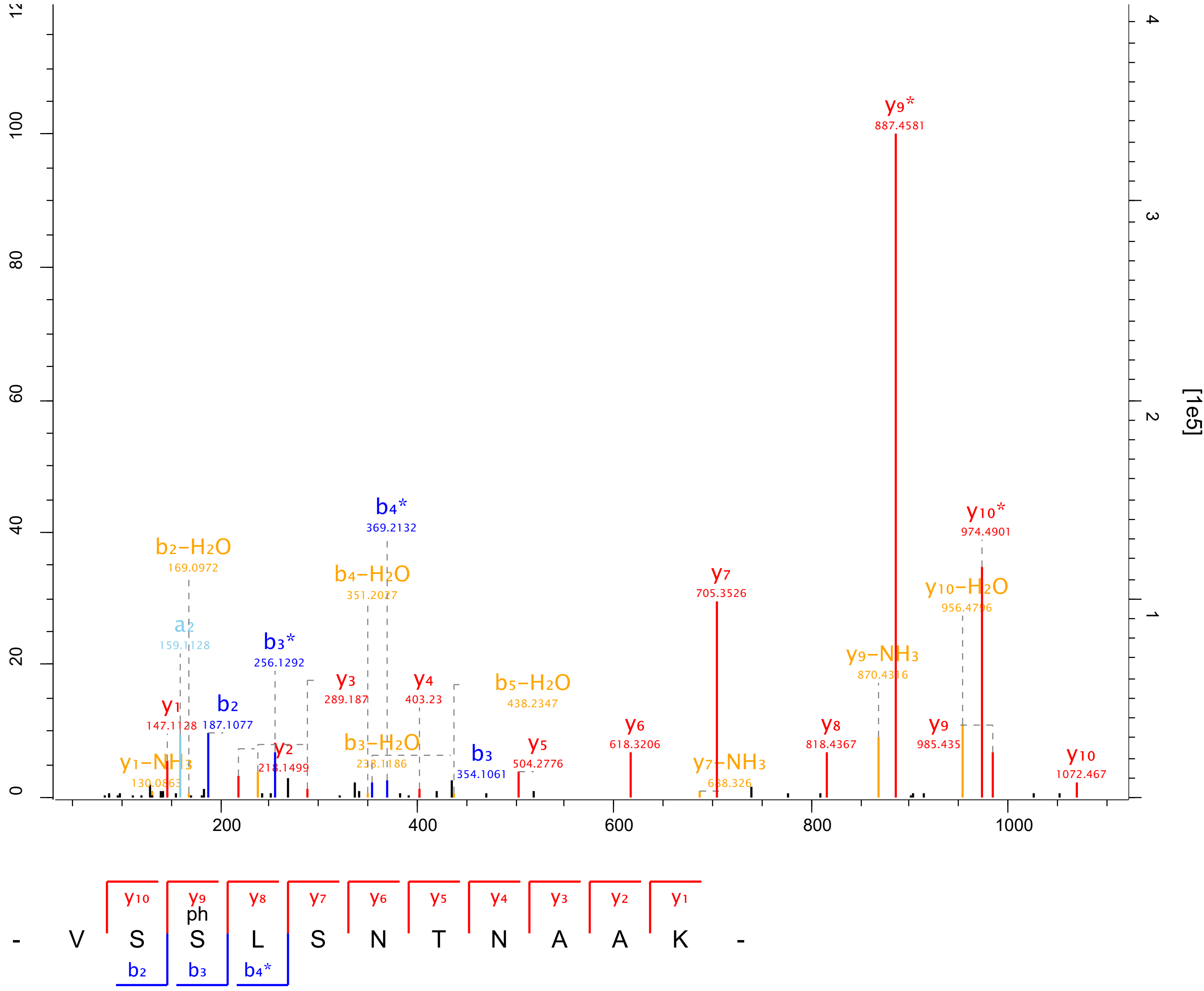

| Raw file | Scan | Method | Score | m/z |
|---|---|---|---|---|
| water_00_2 | 4665 | FTMS; HCD | 176.94 | 634.26 |

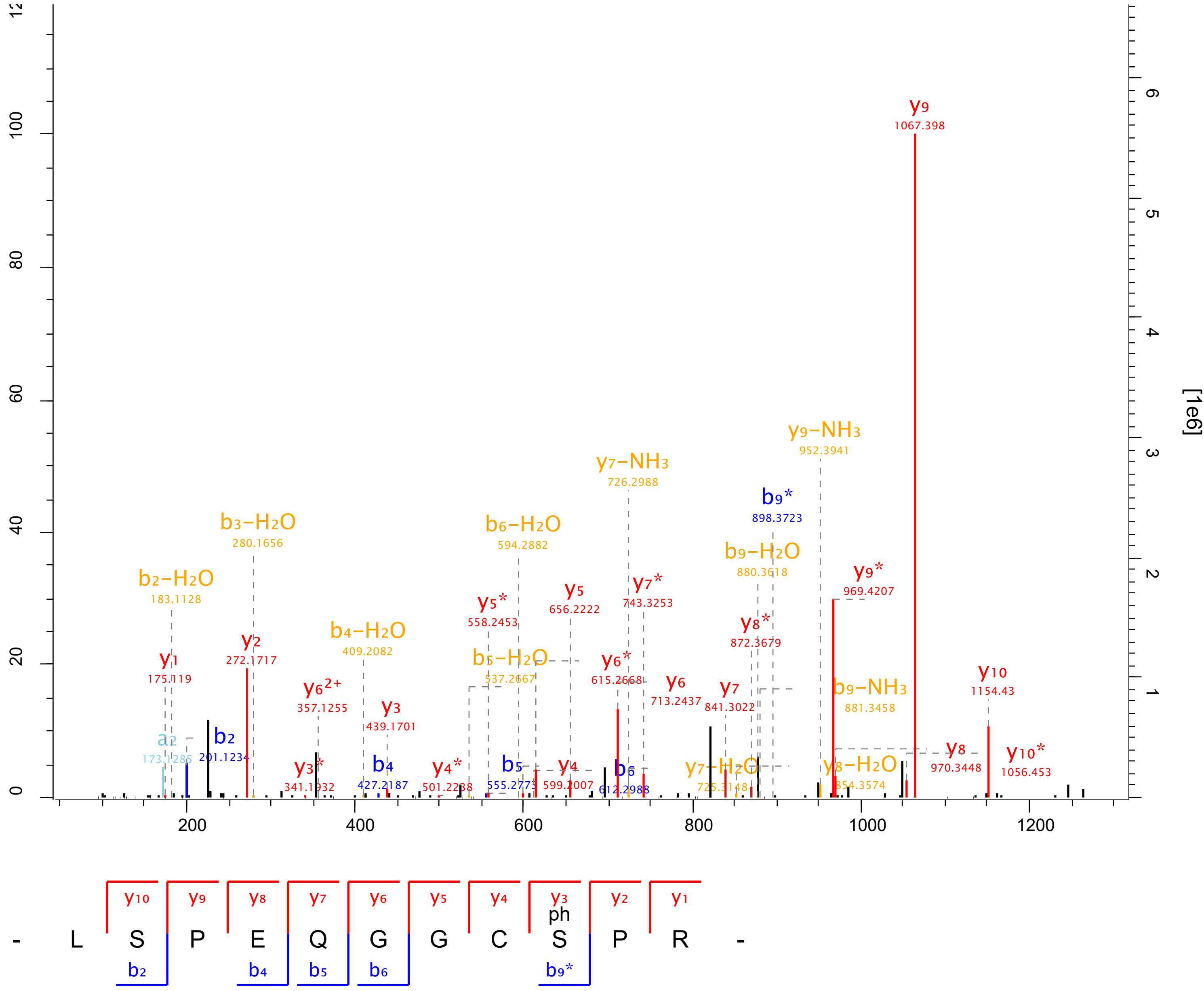

| Raw file | Scan | Method | Score | m/z |
|---|---|---|---|---|
| water_00_2 | 9067 | FTMS; HCD | 188.96 | 530.25 |

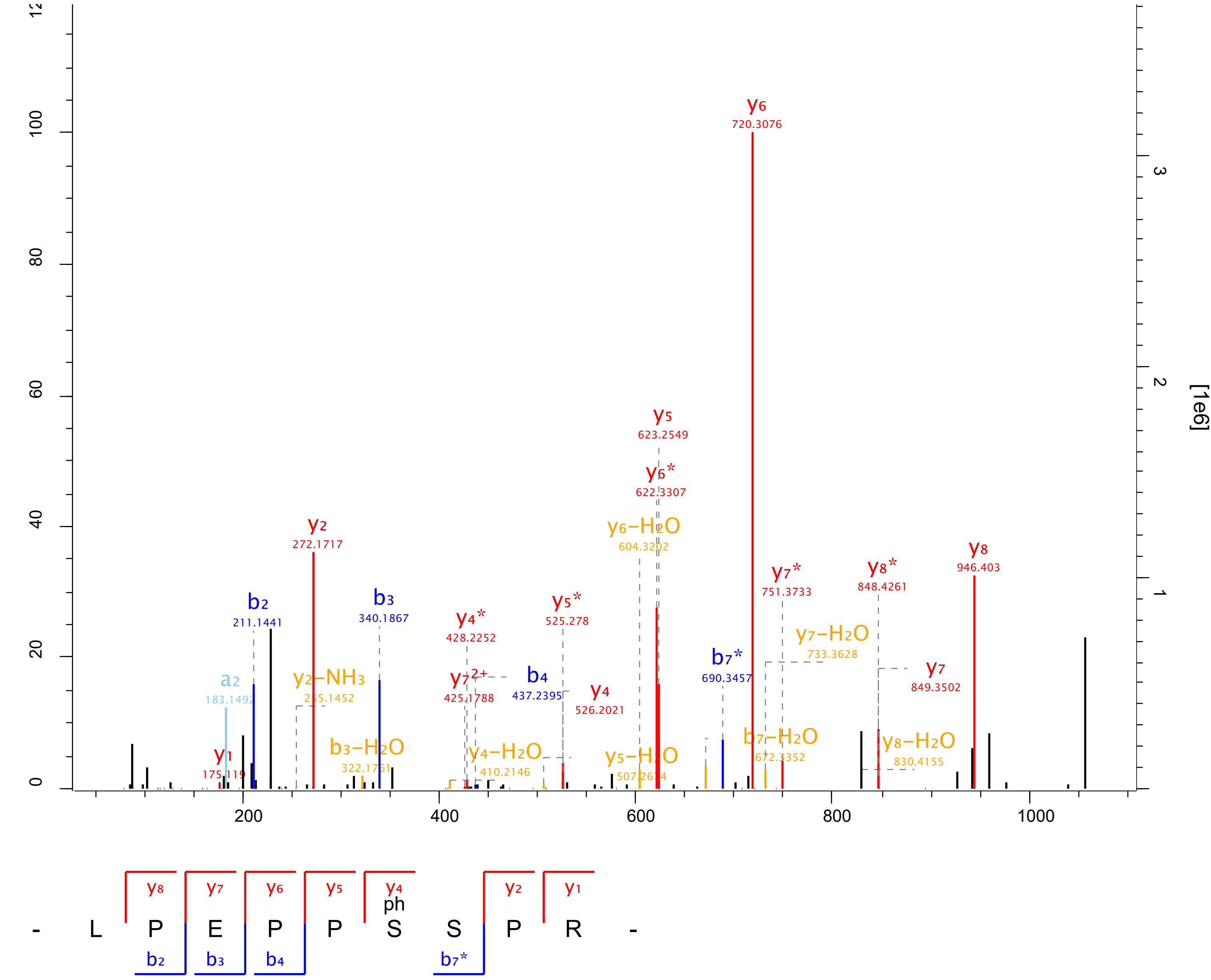

| Raw file | Scan | Method | Score | m/z |
|---|---|---|---|---|
| water_00_3short | 12201 | FTMS; HCD | 137.62 | 584.22 |

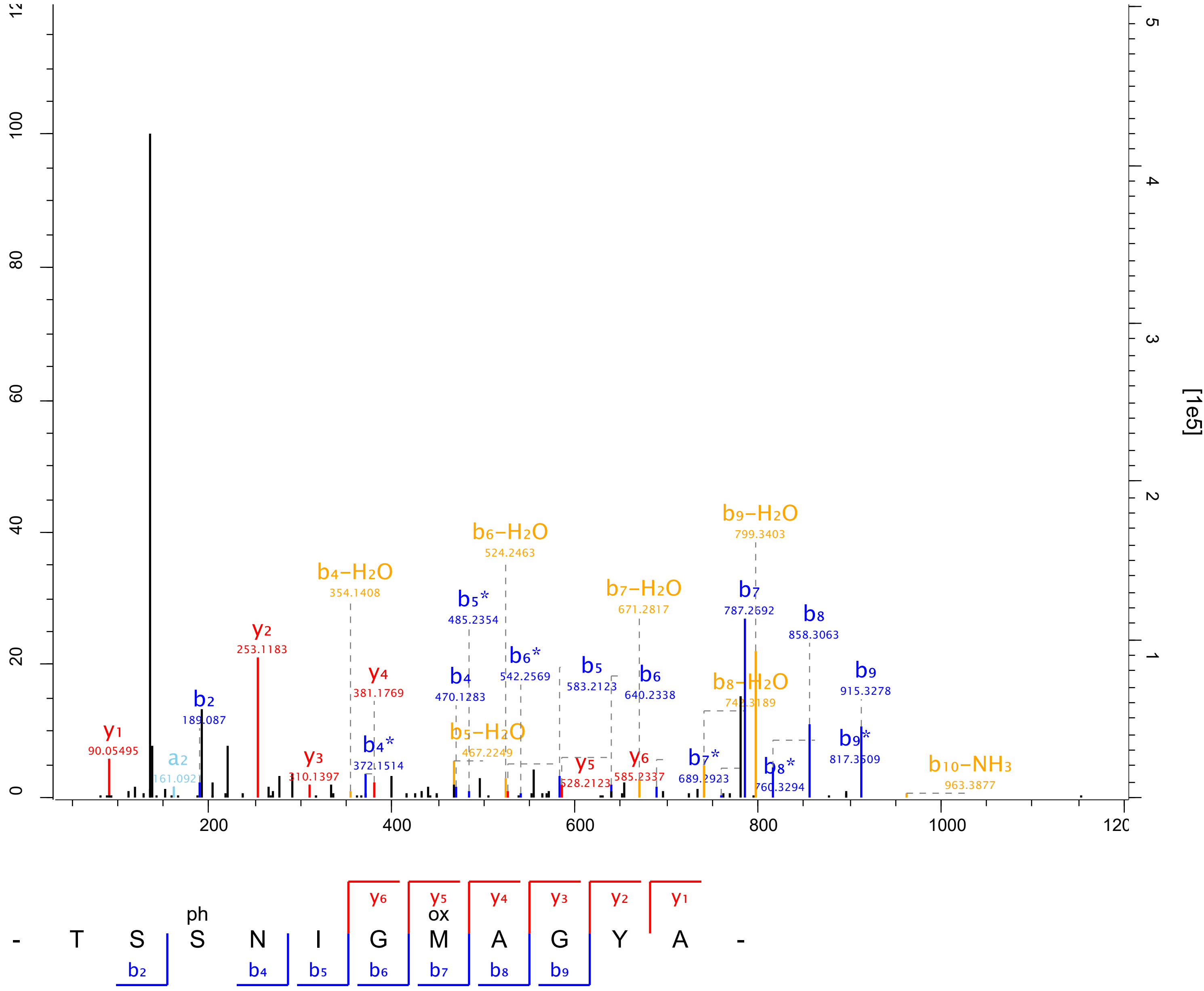

| Raw file | Scan | Method | Score | m/z |
|---|---|---|---|---|
| water_00_3short | 5770 | FTMS; HCD | 118.5 | 412.22 |

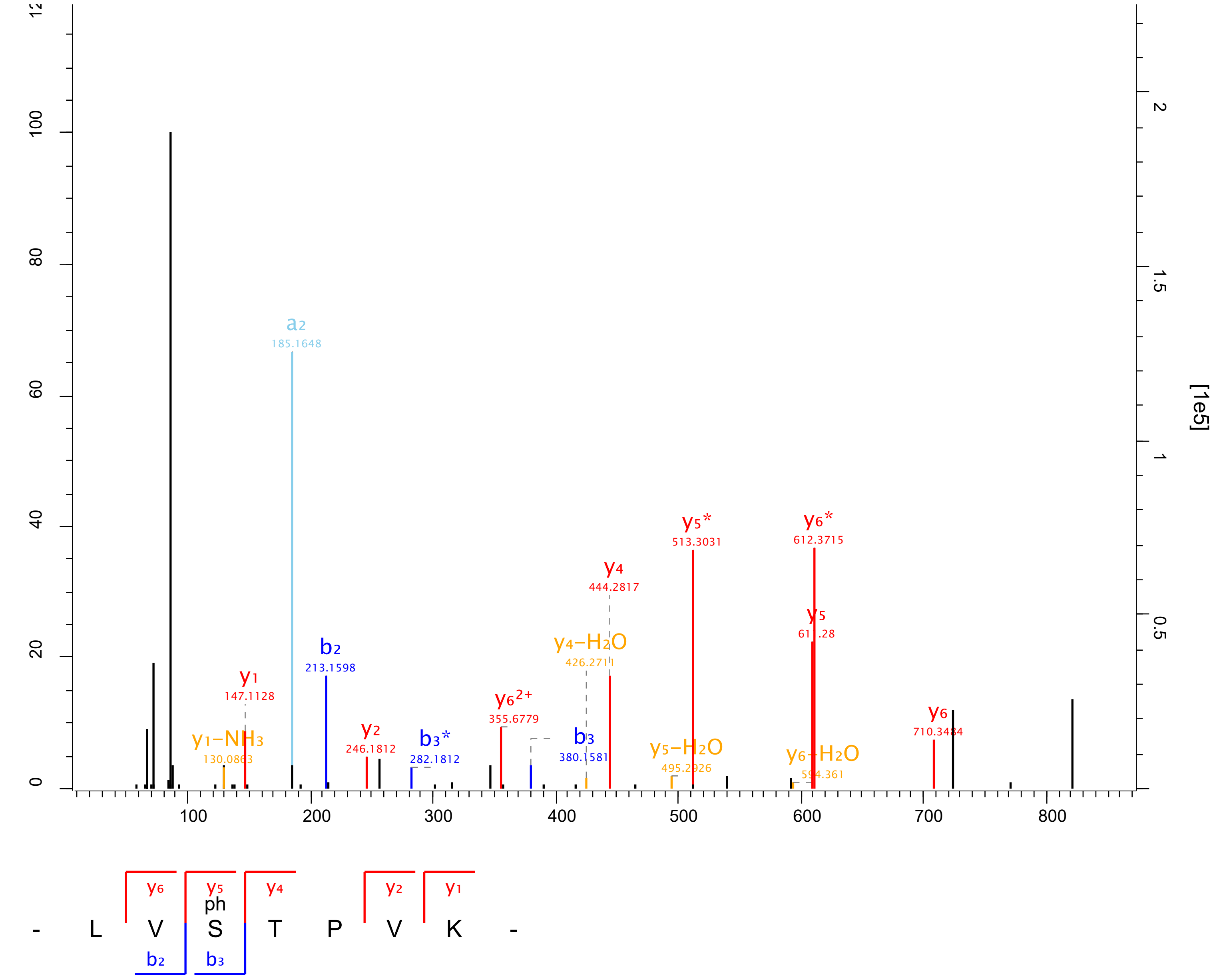

| Raw file | Scan | Method | Score | m/z |
|---|---|---|---|---|
| water_02_1 | 10033 | FTMS; HCD | 69.98 | 768.26 |

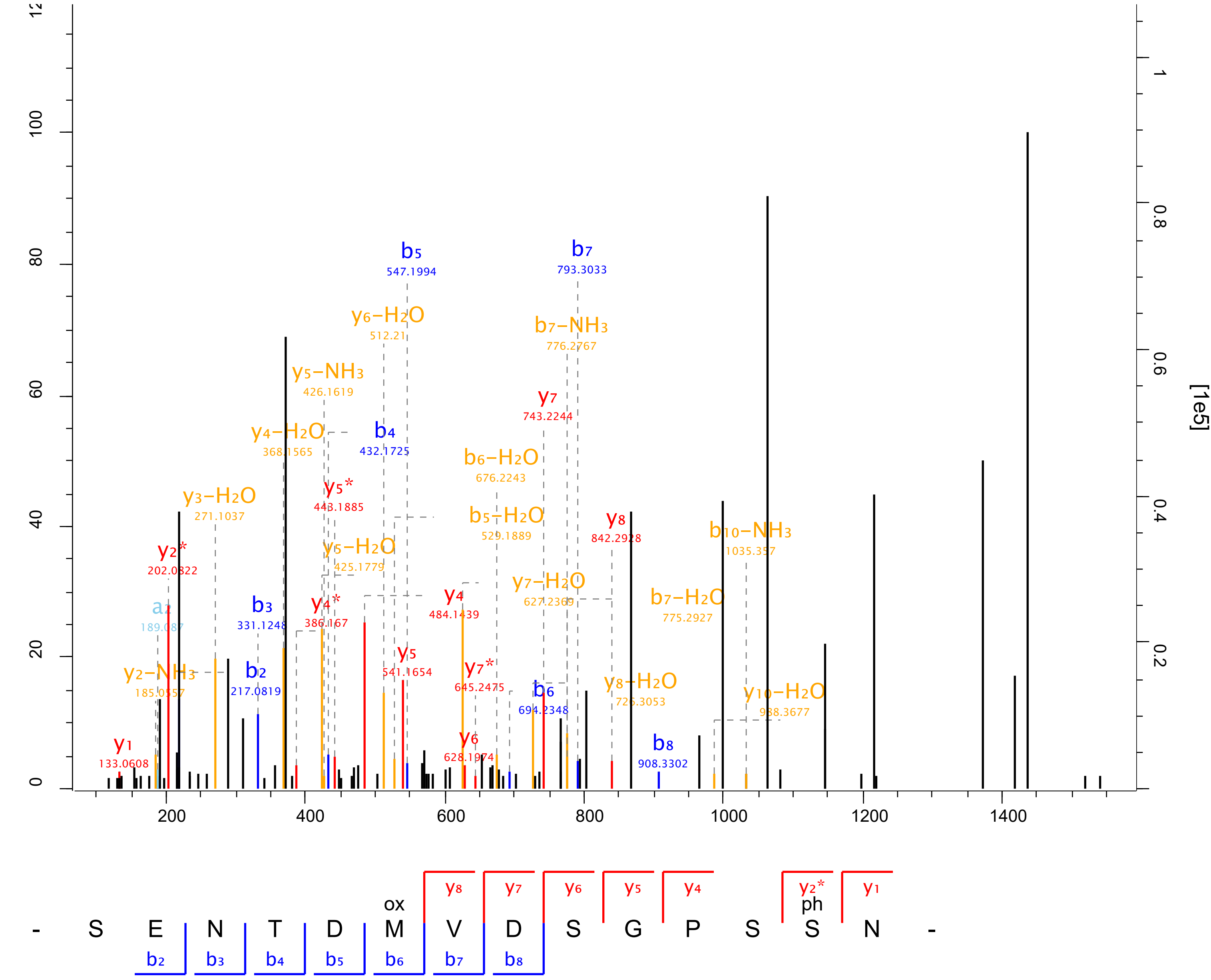

| Raw file | Scan | Method | Score | m/z |
| --- | --- | --- | --- | --- |
| water_02_1 | 10044 | FTMS; HCD | 124.68 | 794.31 |

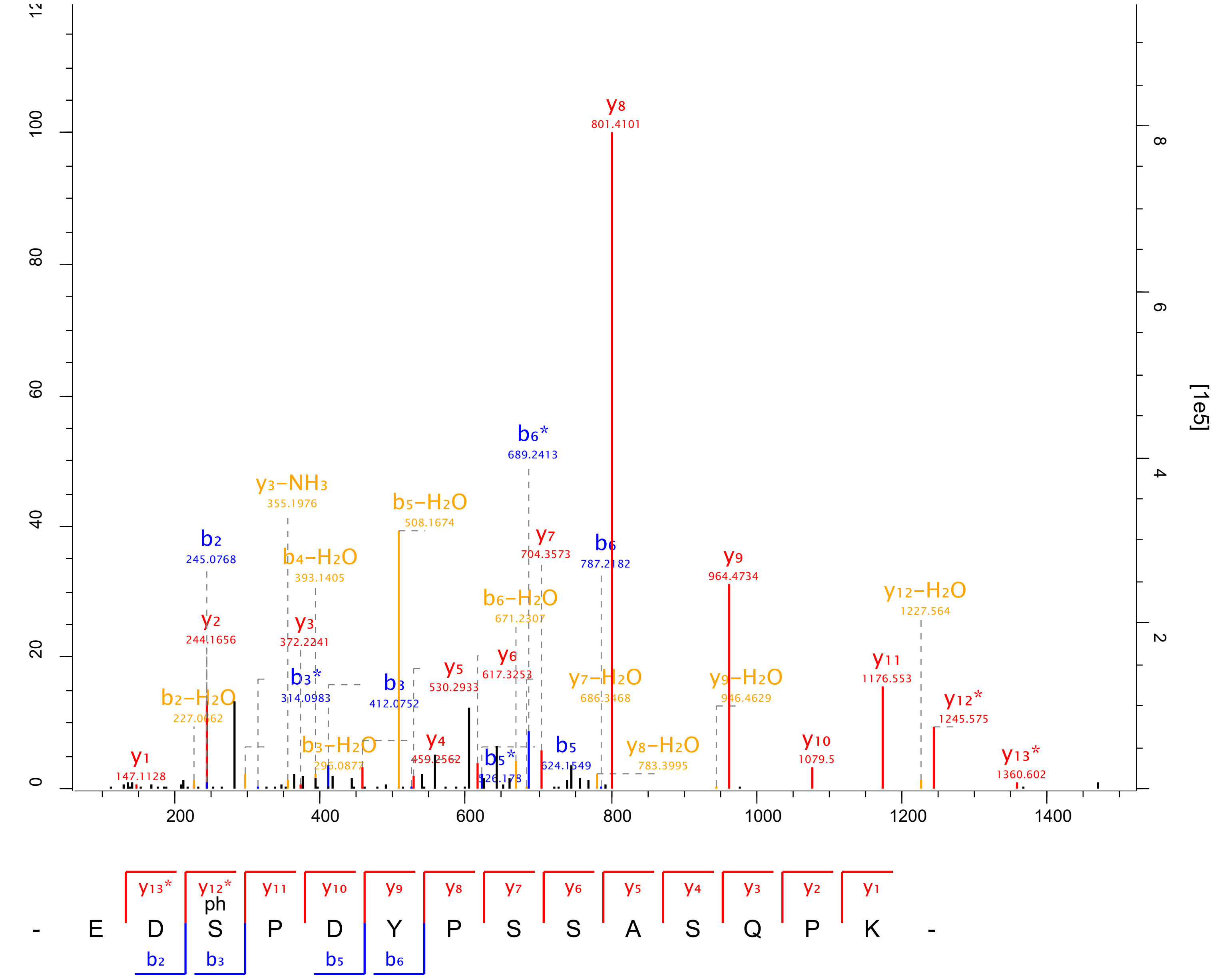

| Raw file | Scan | Method | Score | m/z |
| --- | --- | --- | --- | --- |
| water_02_1 | 10050 | FTMS; HCD | 168.61 | 570.77 |

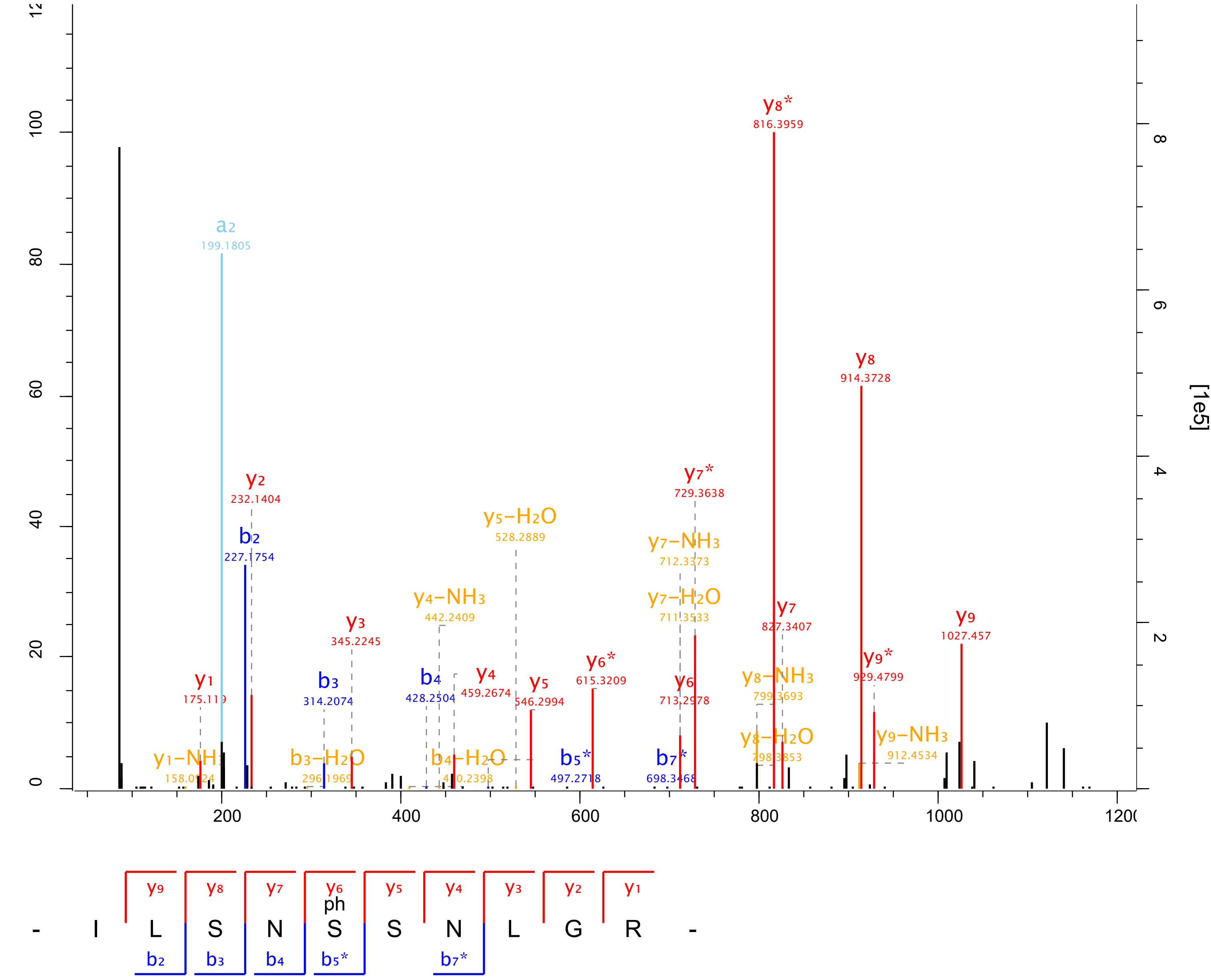

| Raw file | Scan | Method | Score | m/z |
|---|---|---|---|---|
| water_02_1 | 10105 | FTMS; HCD | 74.79 | 580.59 |

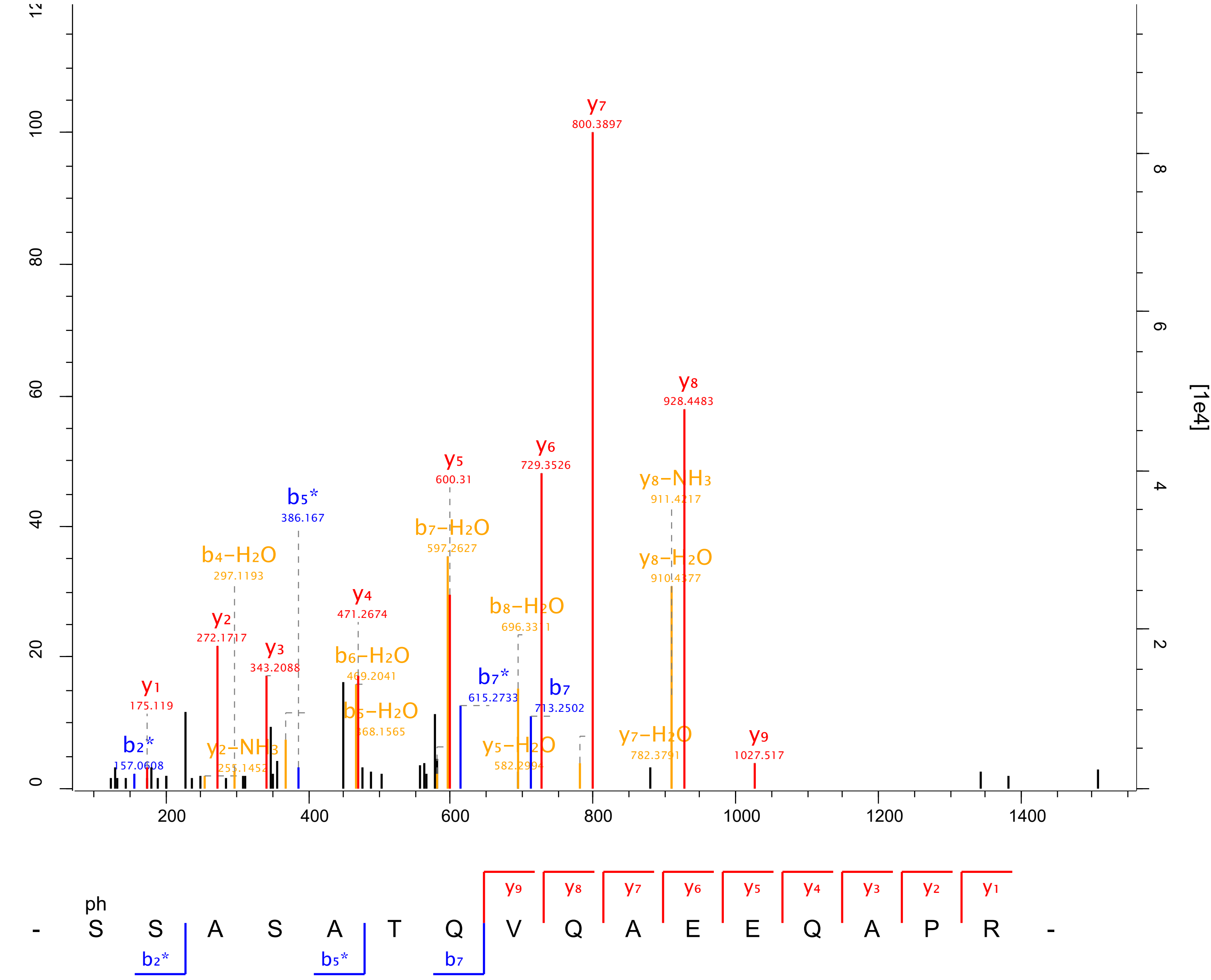

| Raw file | Scan | Method | Score | m/z |
| --- | --- | --- | --- | --- |
| water_02_1 | 10121 | FTMS; HCD | 146.5 | 501.72 |

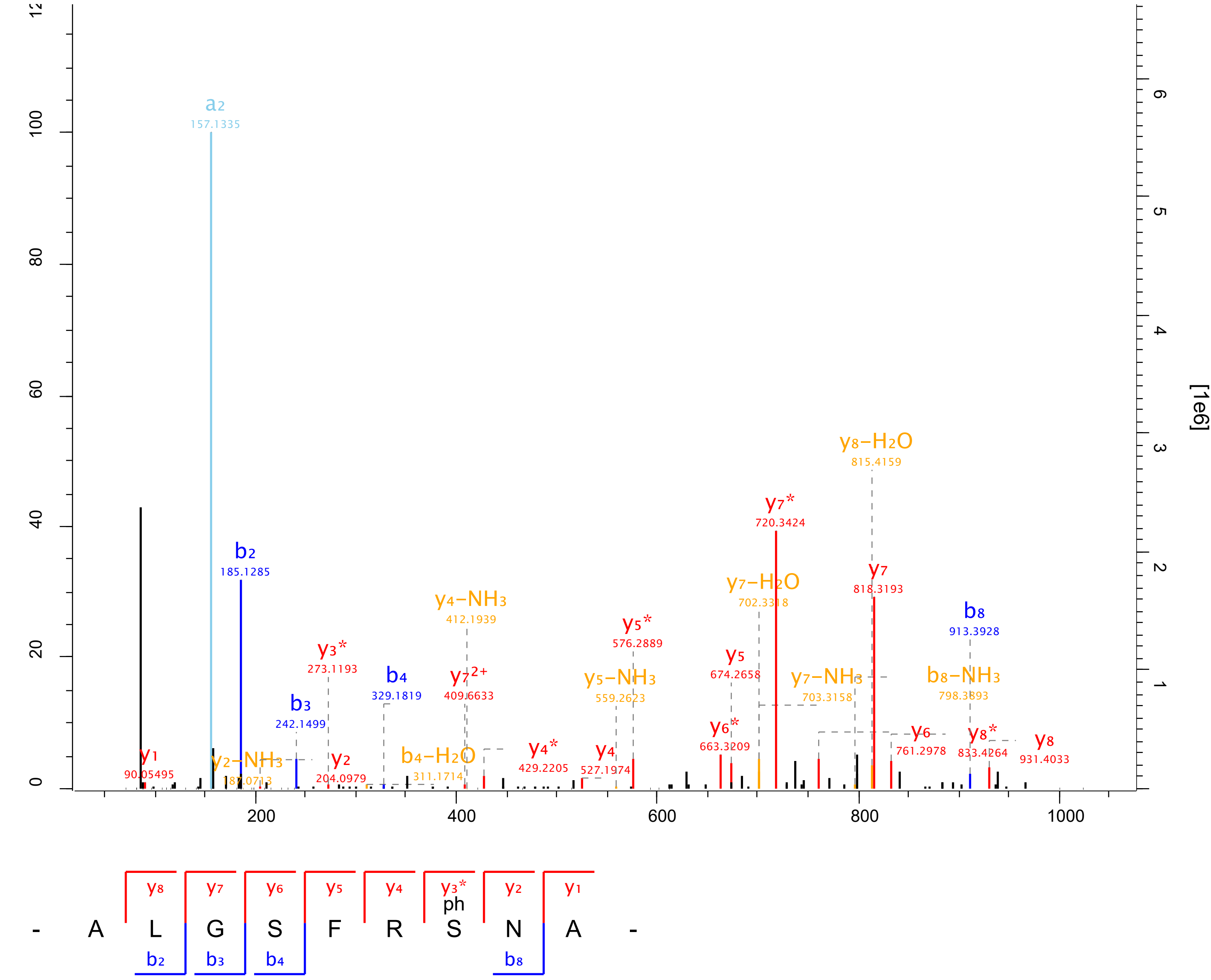

| Raw file | Scan | Method | Score | m/z |
| --- | --- | --- | --- | --- |
| water_02_1 | 10243 | FTMS; HCD | 200.88 | 706.31 |

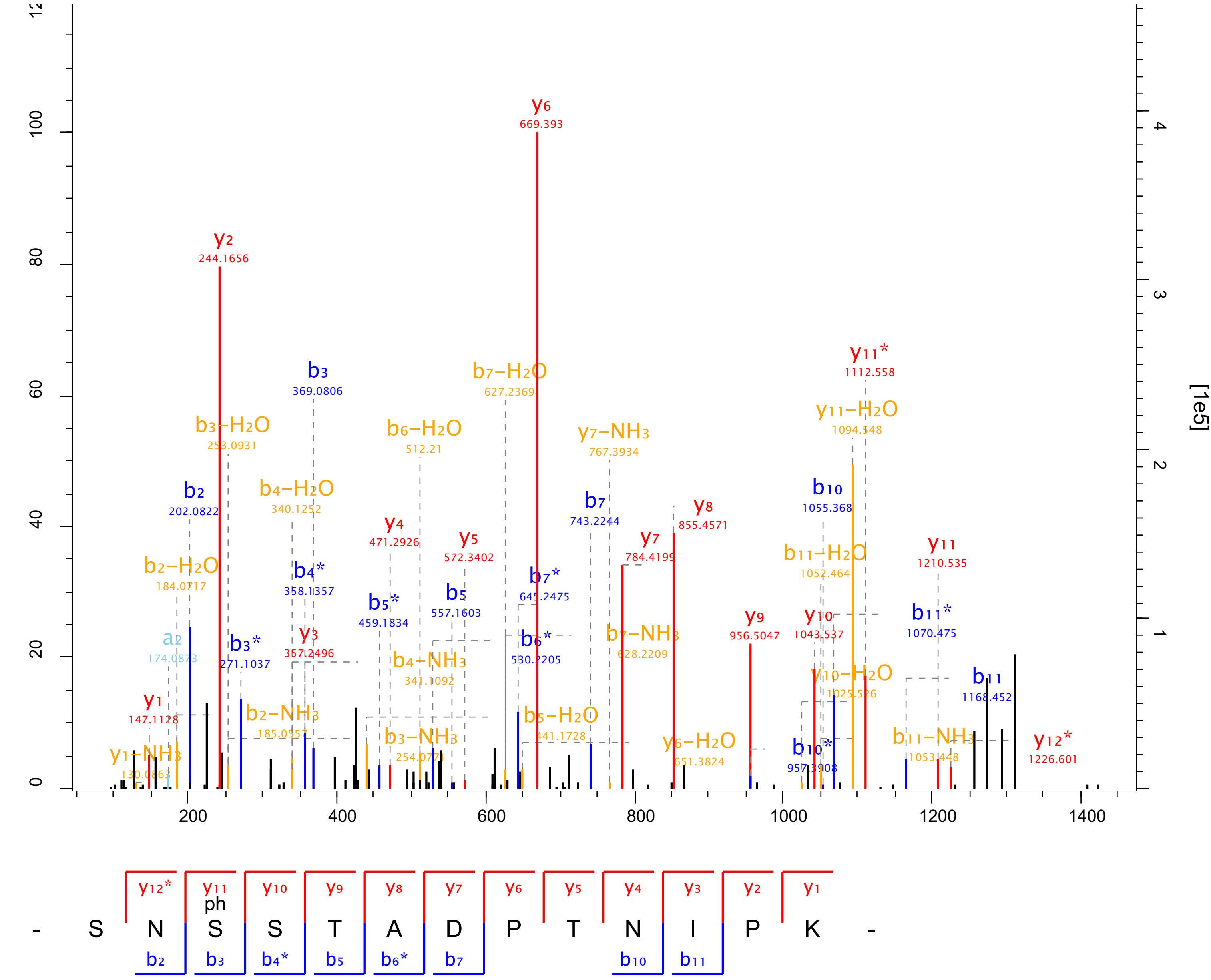

| Raw file | Scan | Method | Score | m/z |
| --- | --- | --- | --- | --- |
| water_02_1 | 10329 | FTMS; HCD | 64.8 | 618.79 |

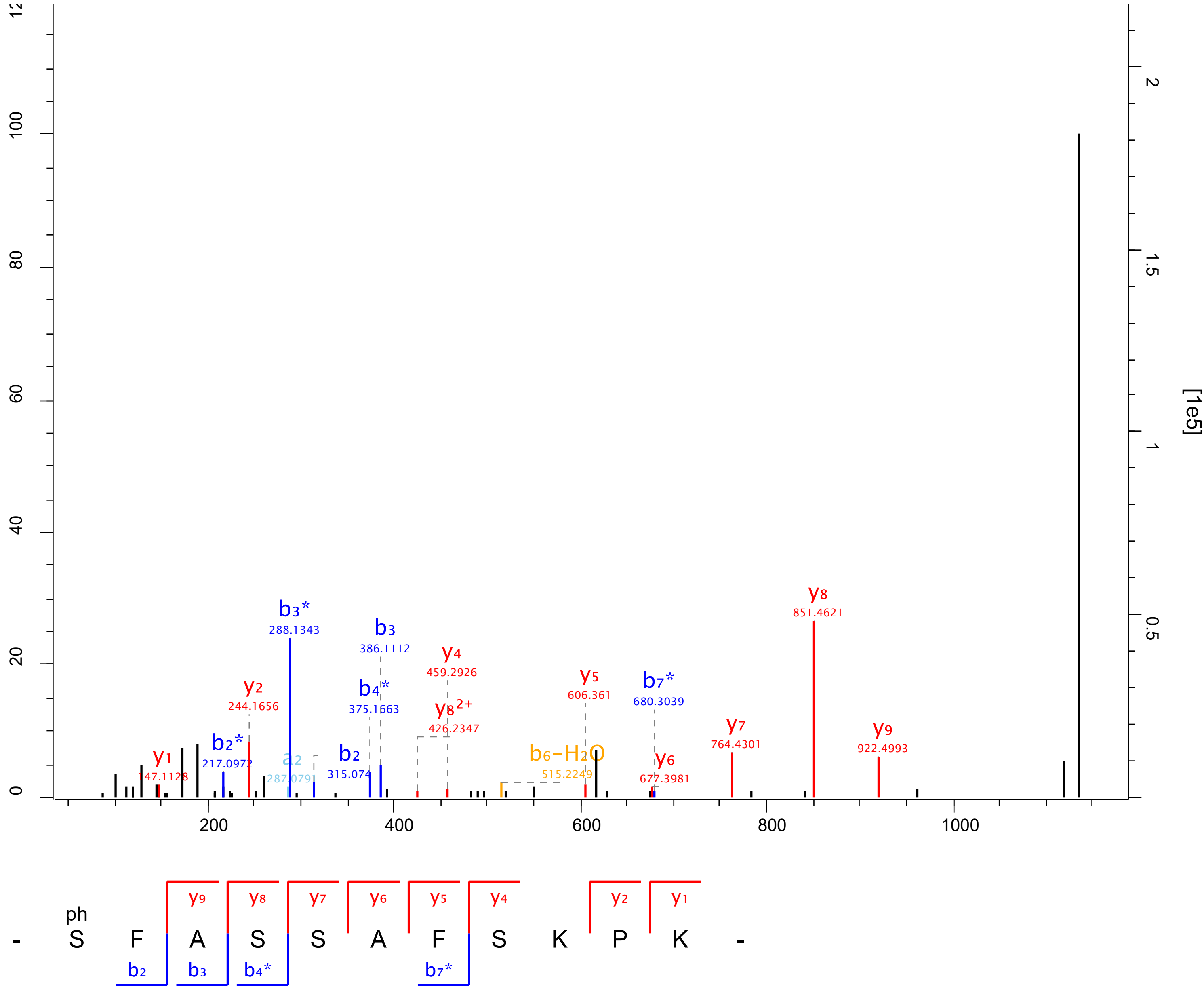

| Raw file | Scan | Method | Score | m/z |
|---|---|---|---|---|
| water_02_1 | 10395 | FTMS; HCD | 41.24 | 487.21 |

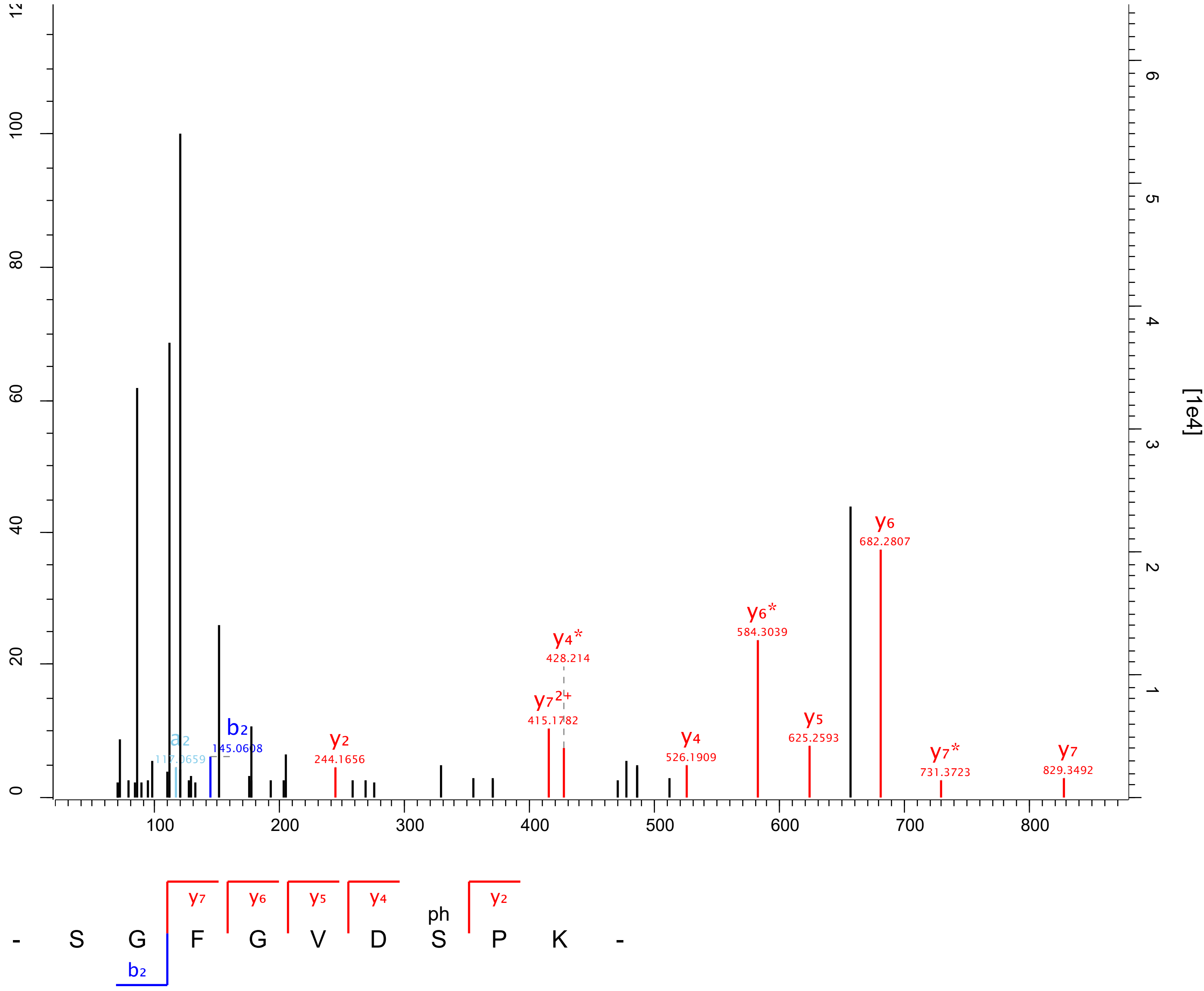

| Raw file | Scan | Method | Score | m/z |
|---|---|---|---|---|
| water_02_1 | 10399 | FTMS; HCD | 75.76 | 524.87 |

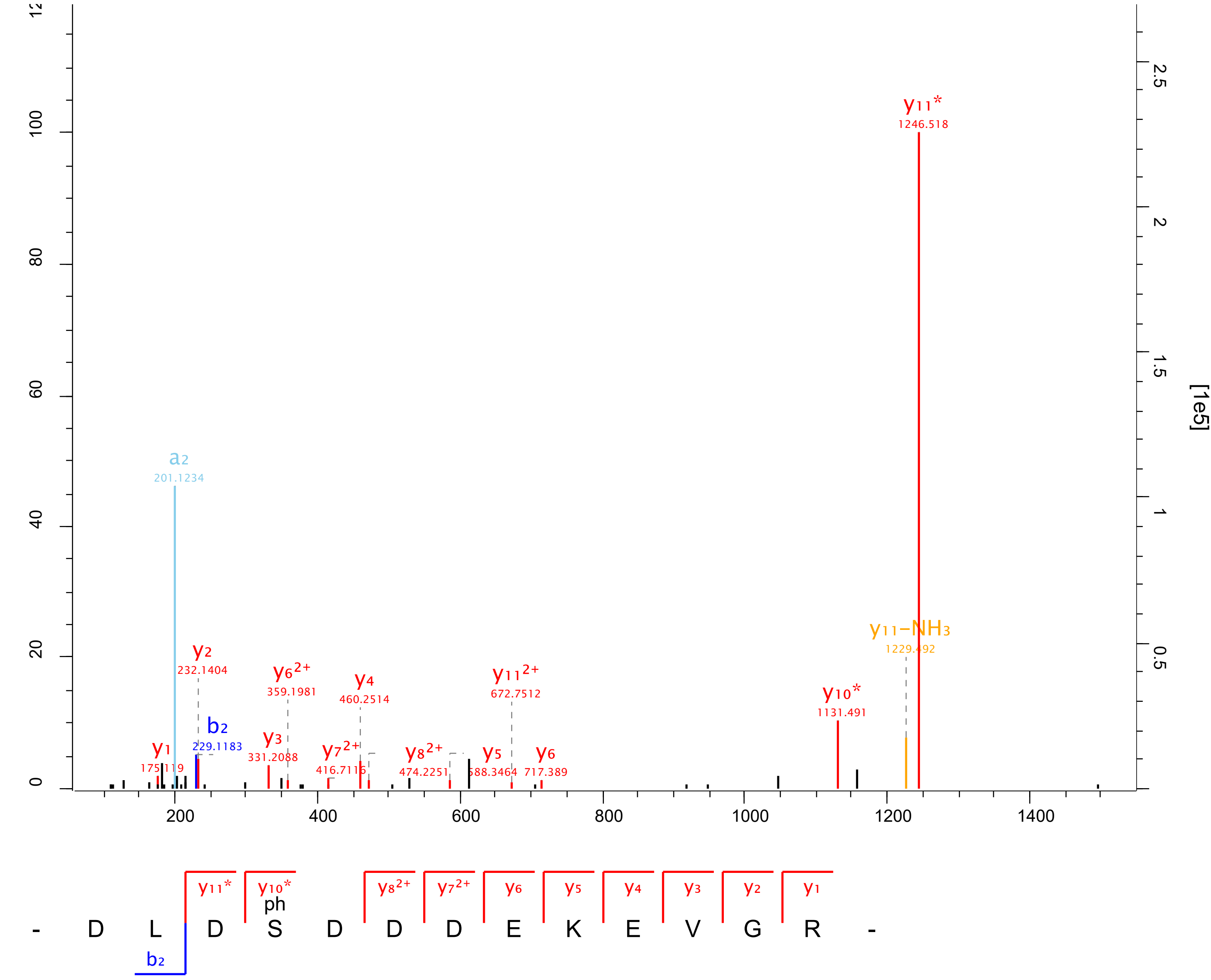

| Raw file | Scan | Method | Score | m/z |
| --- | --- | --- | --- | --- |
| water_02_1 | 10412 | FTMS; HCD | 47.71 | 616.78 |

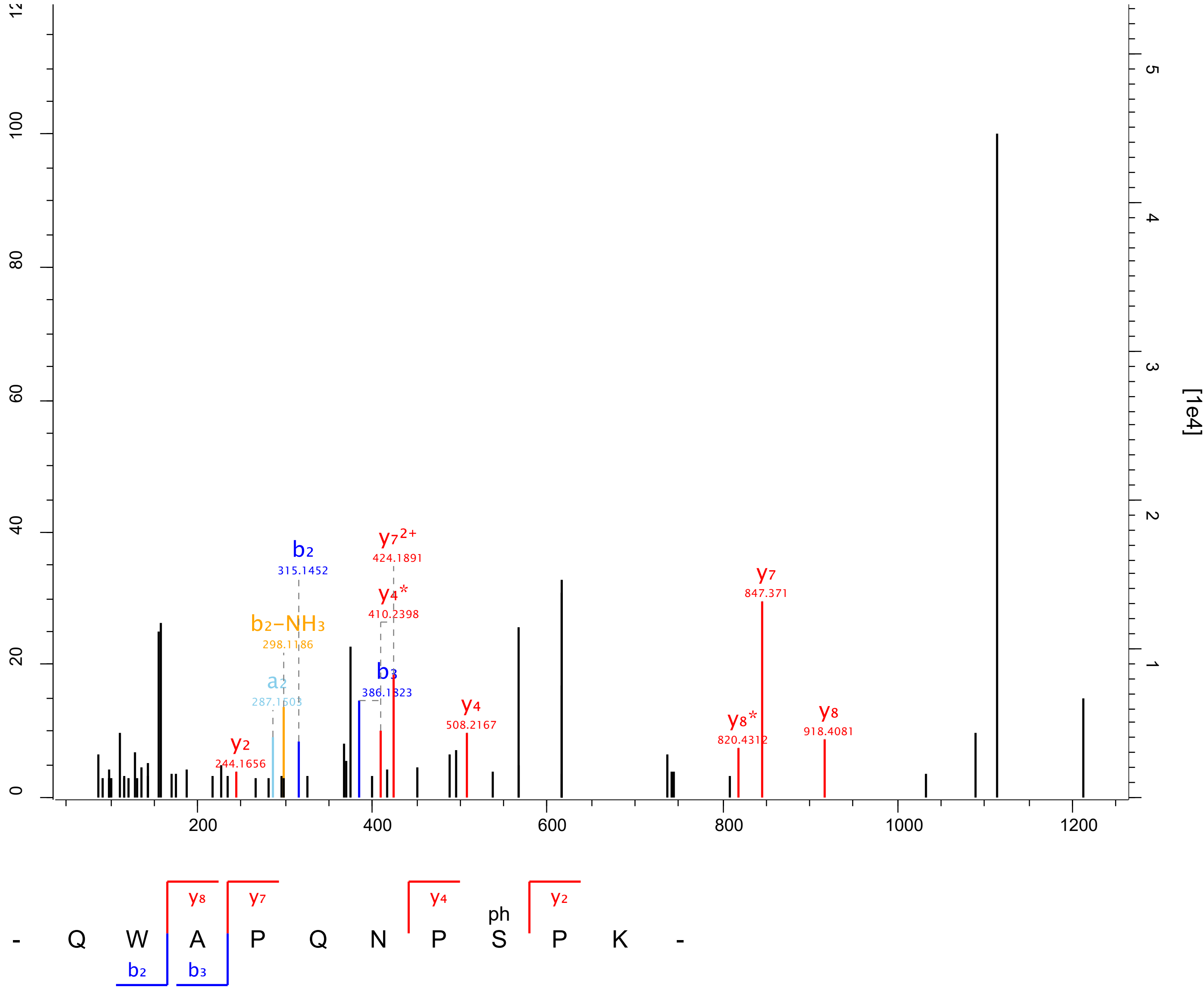

| Raw file | Scan | Method | Score | m/z |
|---|---|---|---|---|
| water_02_1 | 10443 | FTMS; HCD | 68.48 | 678.8 |

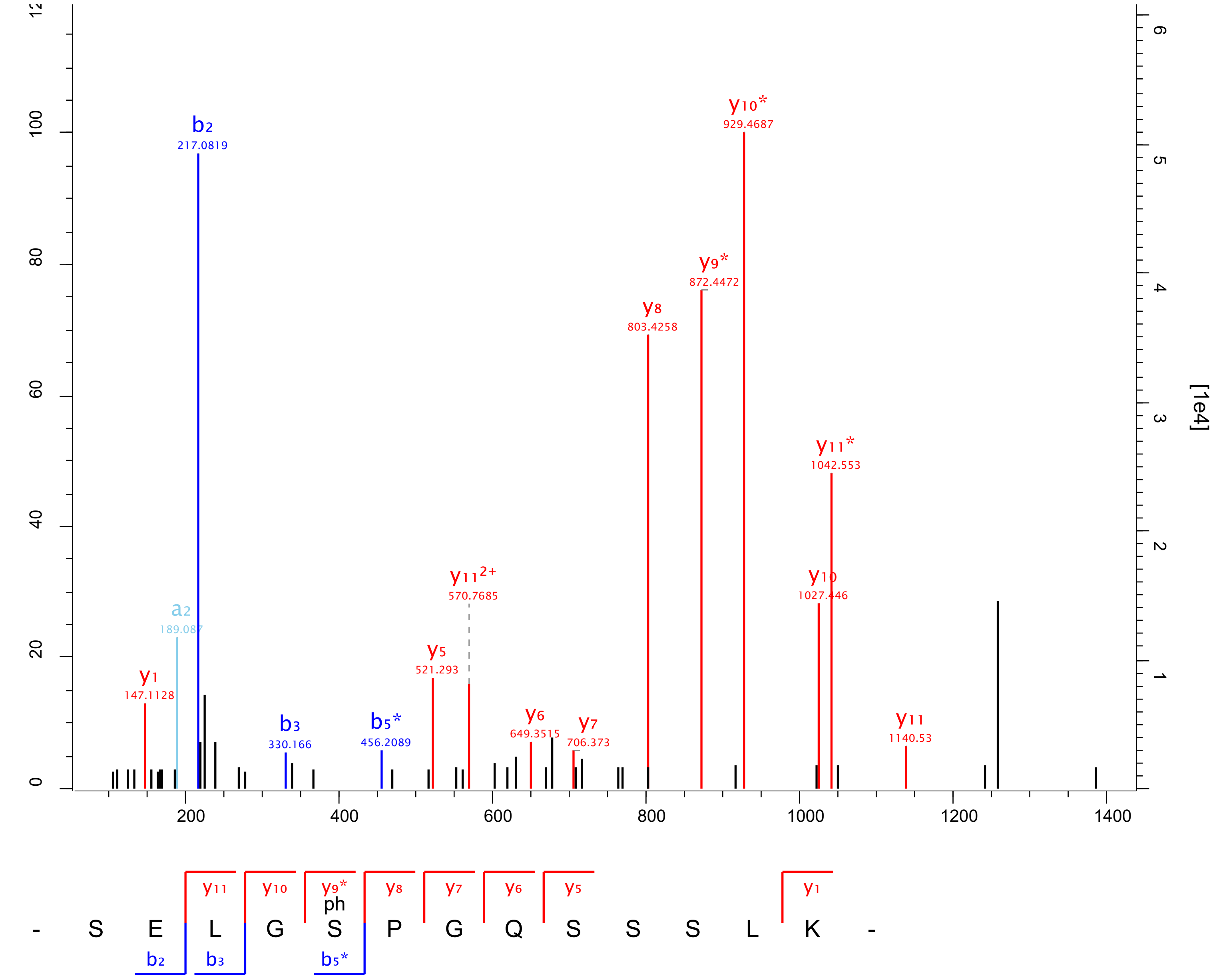

| Raw file | Scan | Method | Score | m/z |
|---|---|---|---|---|
| water_02_1 | 10469 | FTMS; HCD | 94.02 | 544.25 |

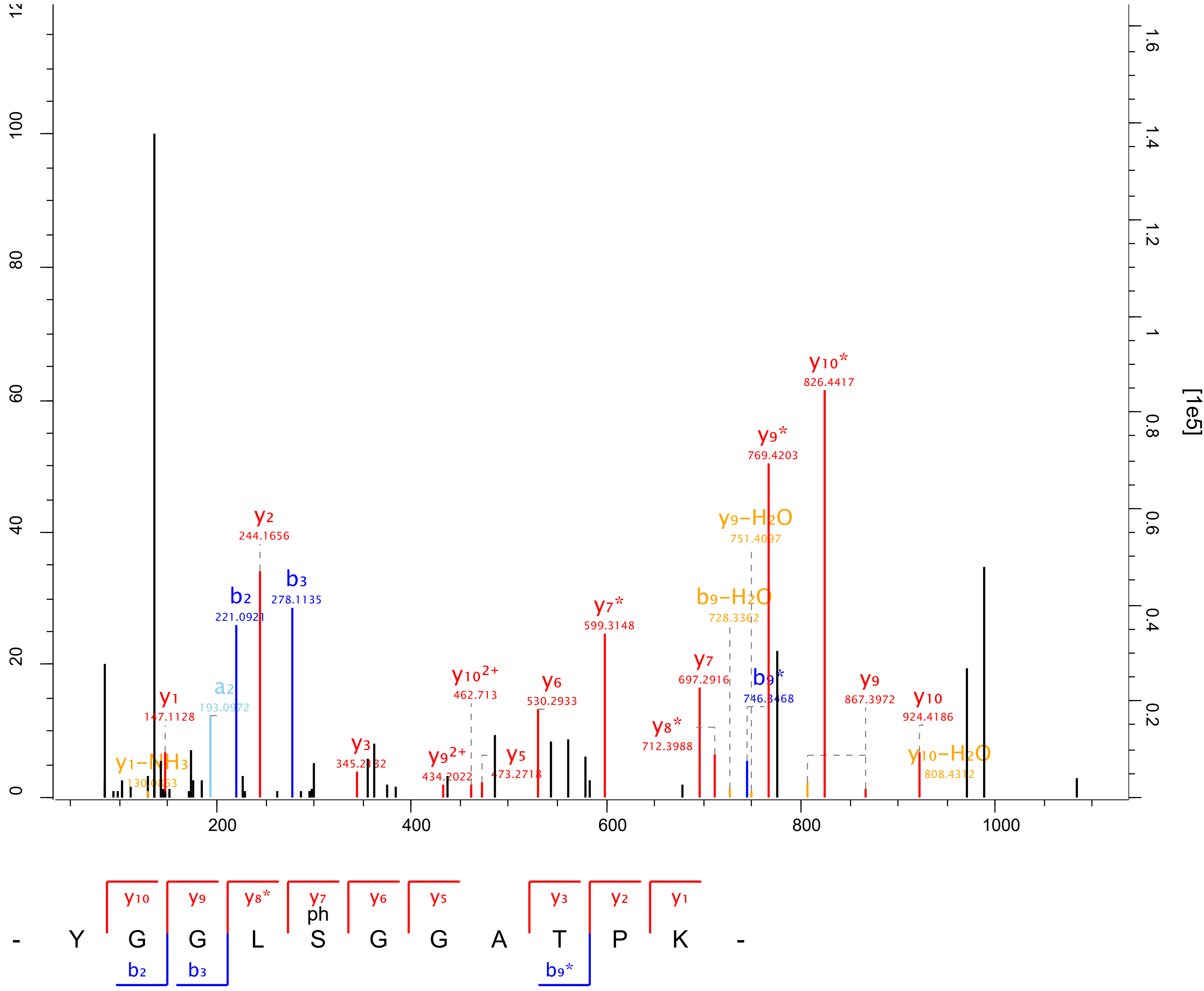

| Raw file | Scan | Method | Score | m/z |
|---|---|---|---|---|
| water_02_1 | 10500 | FTMS; HCD | 49.33 | 677.77 |

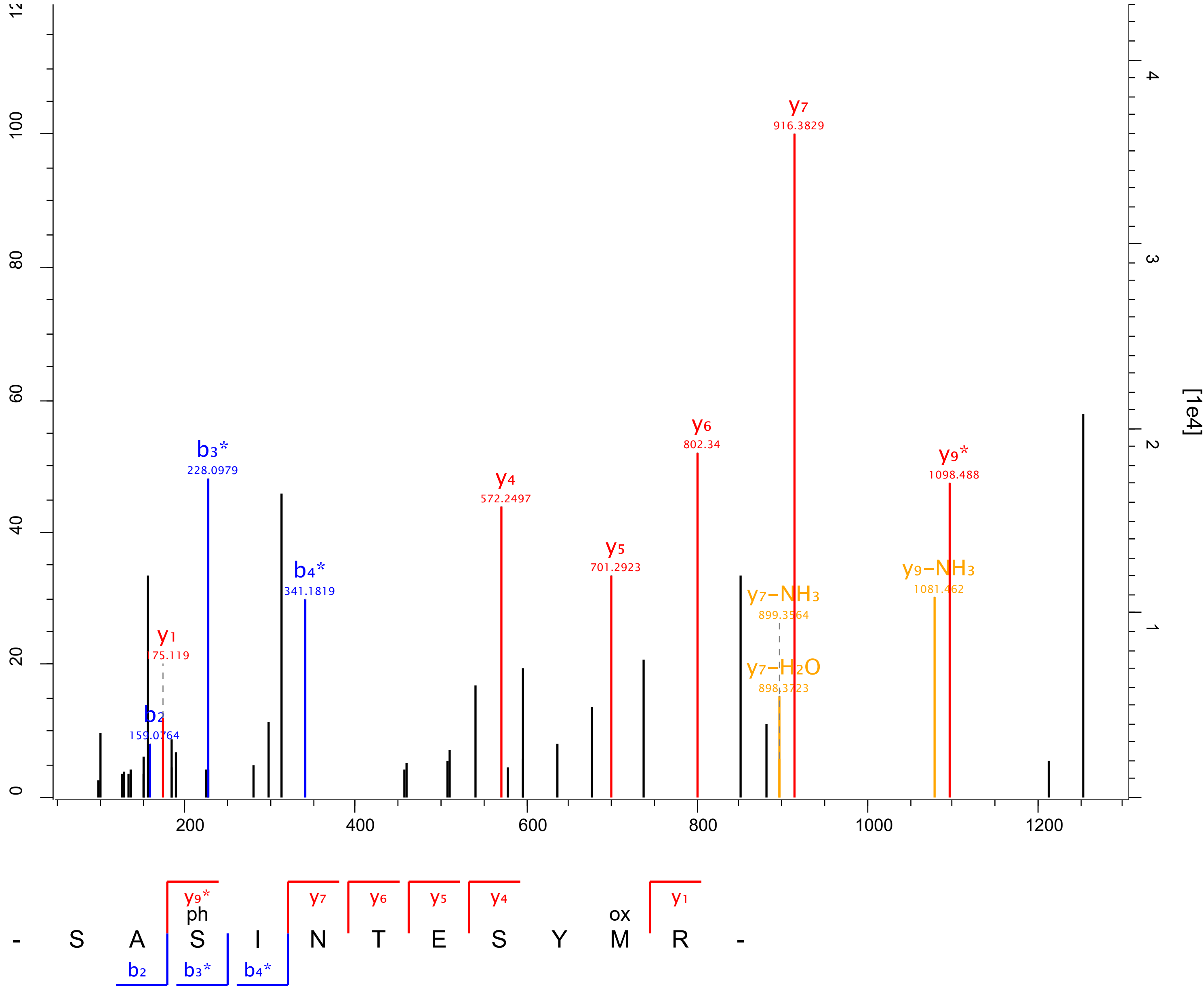

| Raw file | Scan | Method | Score | m/z |
|---|---|---|---|---|
| water_02_1 | 10502 | FTMS; HCD | 124.08 | 819.86 |

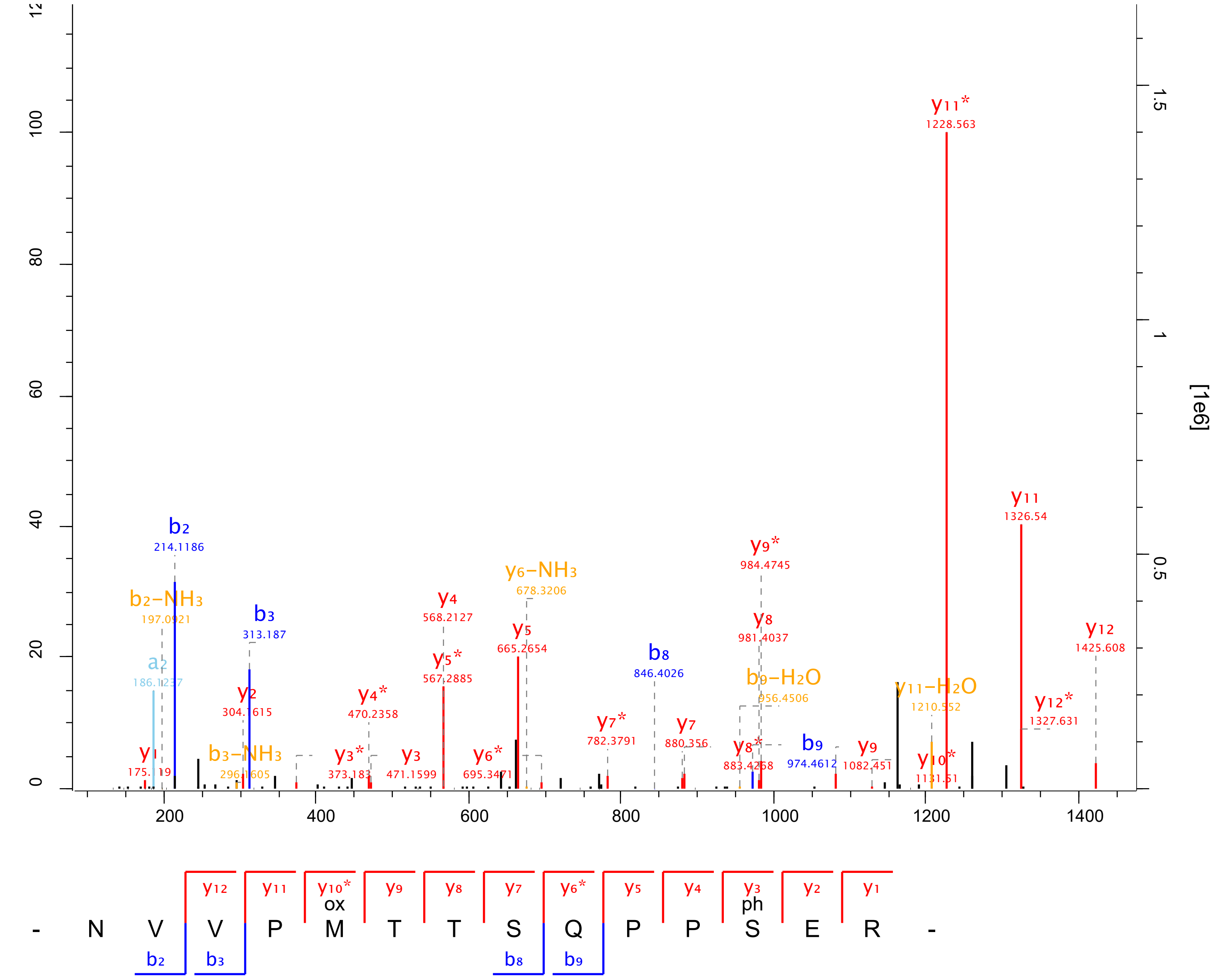

| Raw file | Scan | Method | Score | m/z |
|---|---|---|---|---|
| water_02_1 | 10534 | FTMS; HCD | 88.28 | 608.24 |

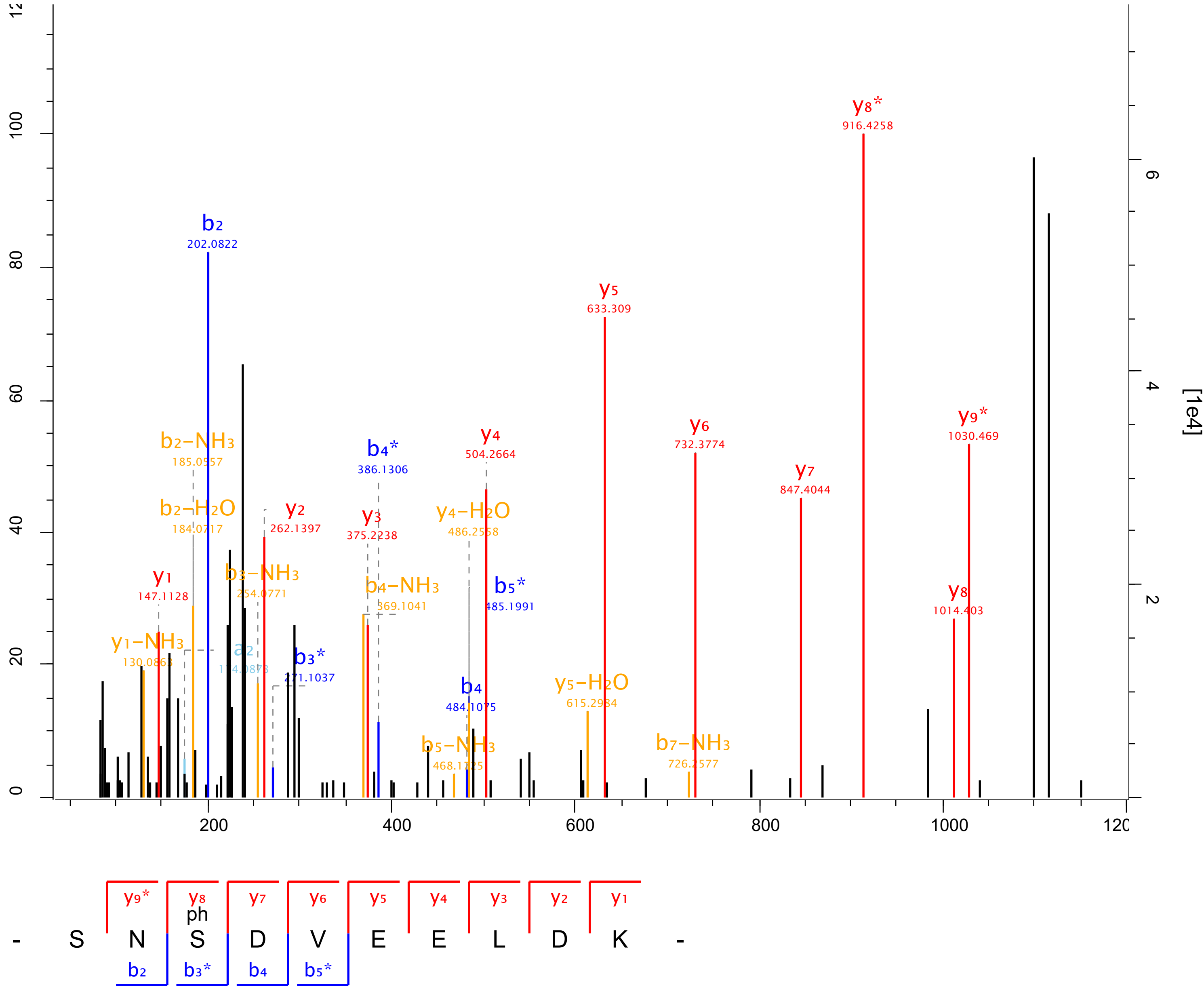

| Raw file | Scan | Method | Score | m/z |
|---|---|---|---|---|
| water_02_1 | 10579 | FTMS; HCD | 58.7 | 564.76 |

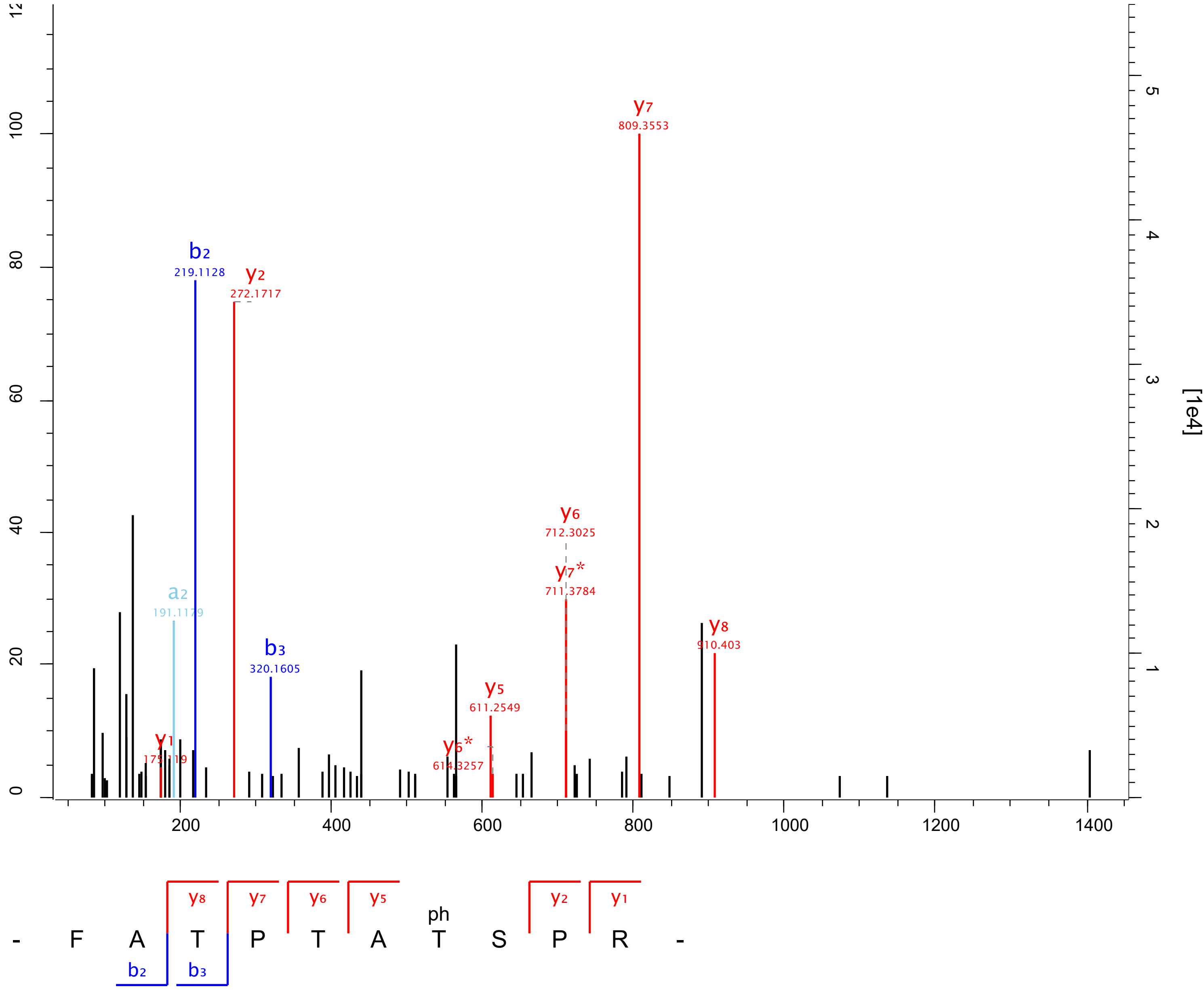

| Raw file | Scan | Method | Score | m/z |
|---|---|---|---|---|
| water_02_1 | 10608 | FTMS; HCD | 96.14 | 743.83 |

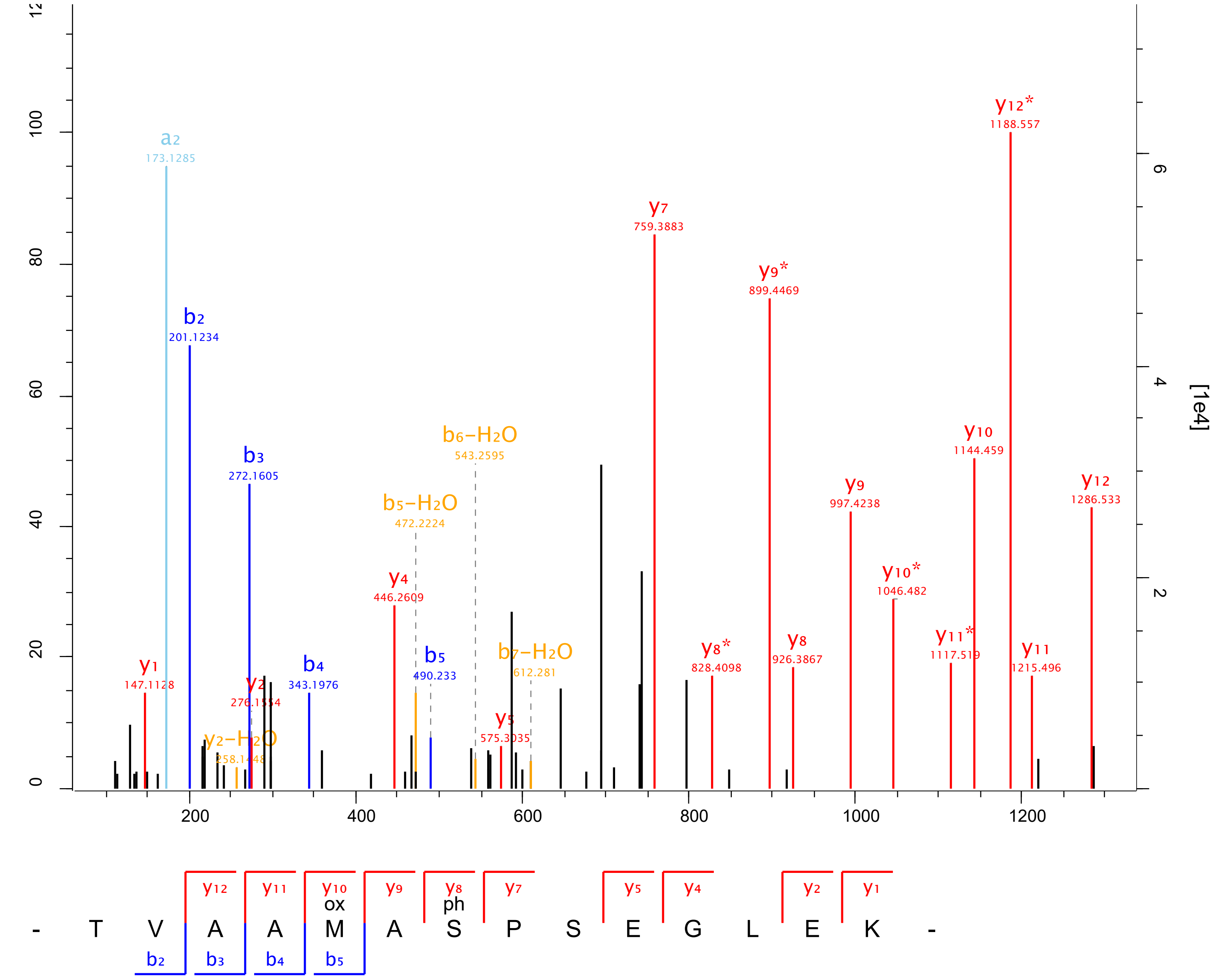

| Raw file | Scan | Method | Score | m/z |
| --- | --- | --- | --- | --- |
| water_02_1 | 10656 | FTMS; HCD | 116.71 | 577.77 |

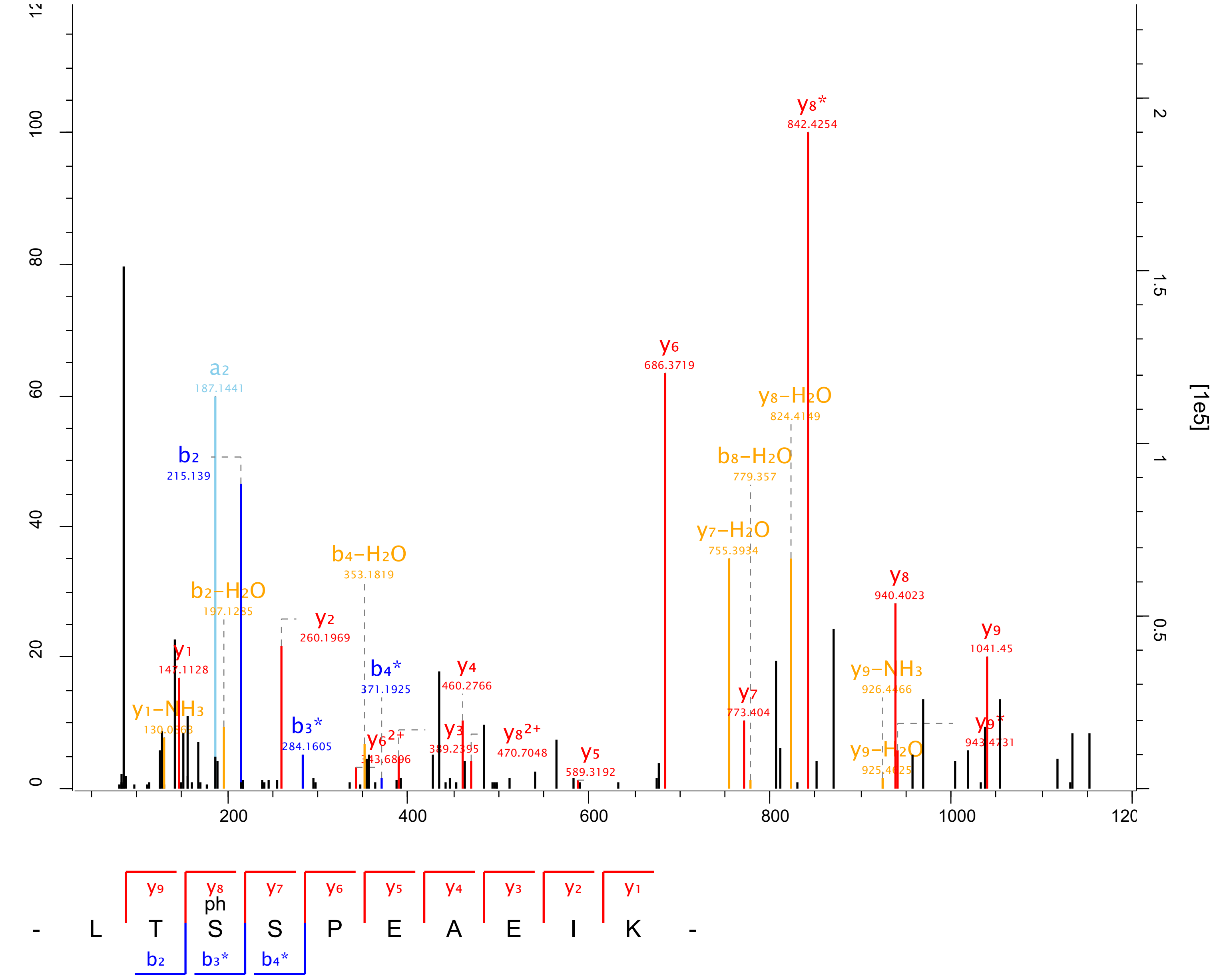

| Raw file | Scan | Method | Score | m/z |
|---|---|---|---|---|
| water_02_1 | 10670 | FTMS; HCD | 96.17 | 503.73 |

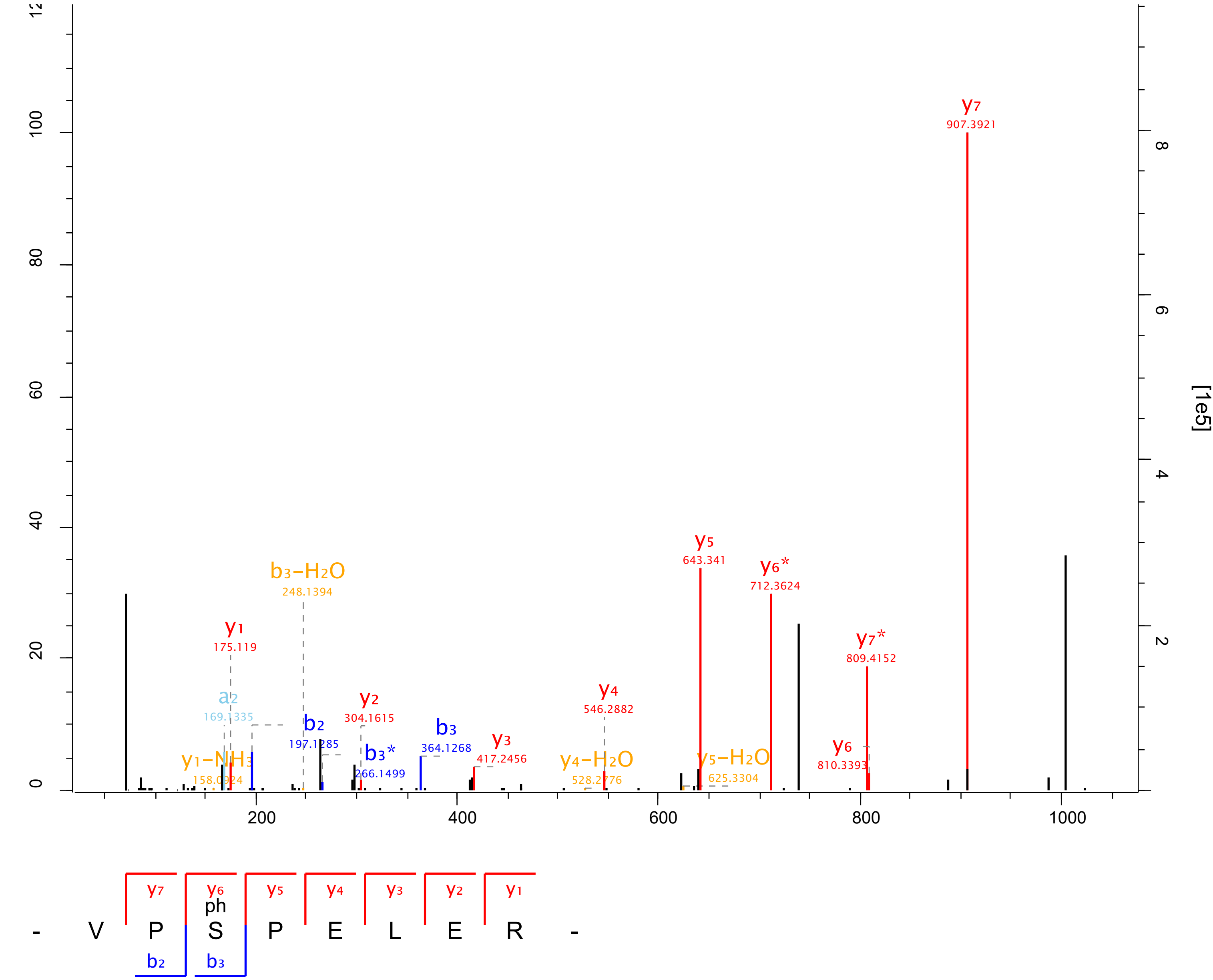

| Raw file | Scan | Method | Score | m/z |
| --- | --- | --- | --- | --- |
| water_02_1 | 10750 | FTMS; HCD | 169.51 | 671.58 |

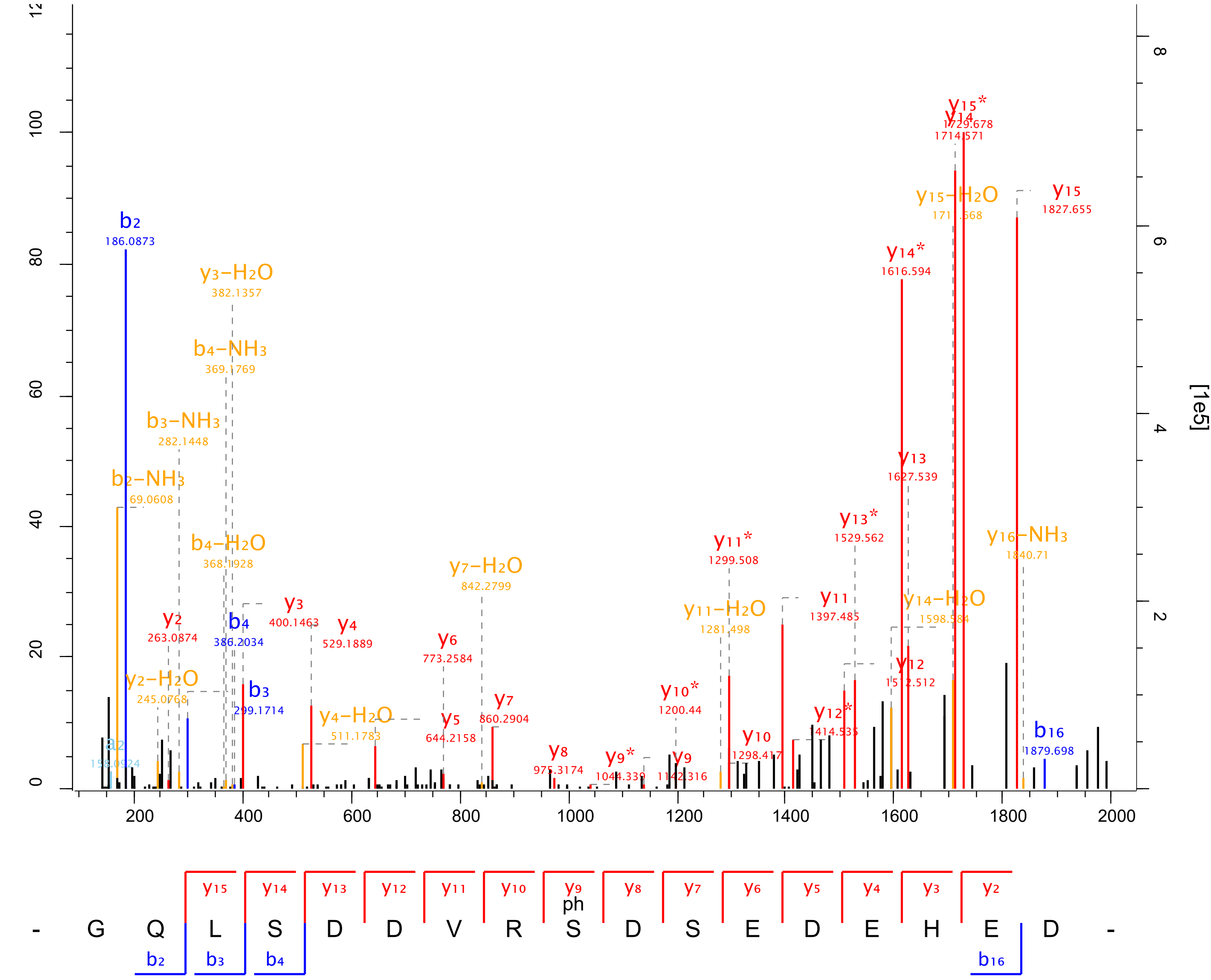

| Raw file | Scan | Method | Score | m/z |
|---|---|---|---|---|
| water_02_1 | 10770 | FTMS; HCD | 105.65 | 588.79 |

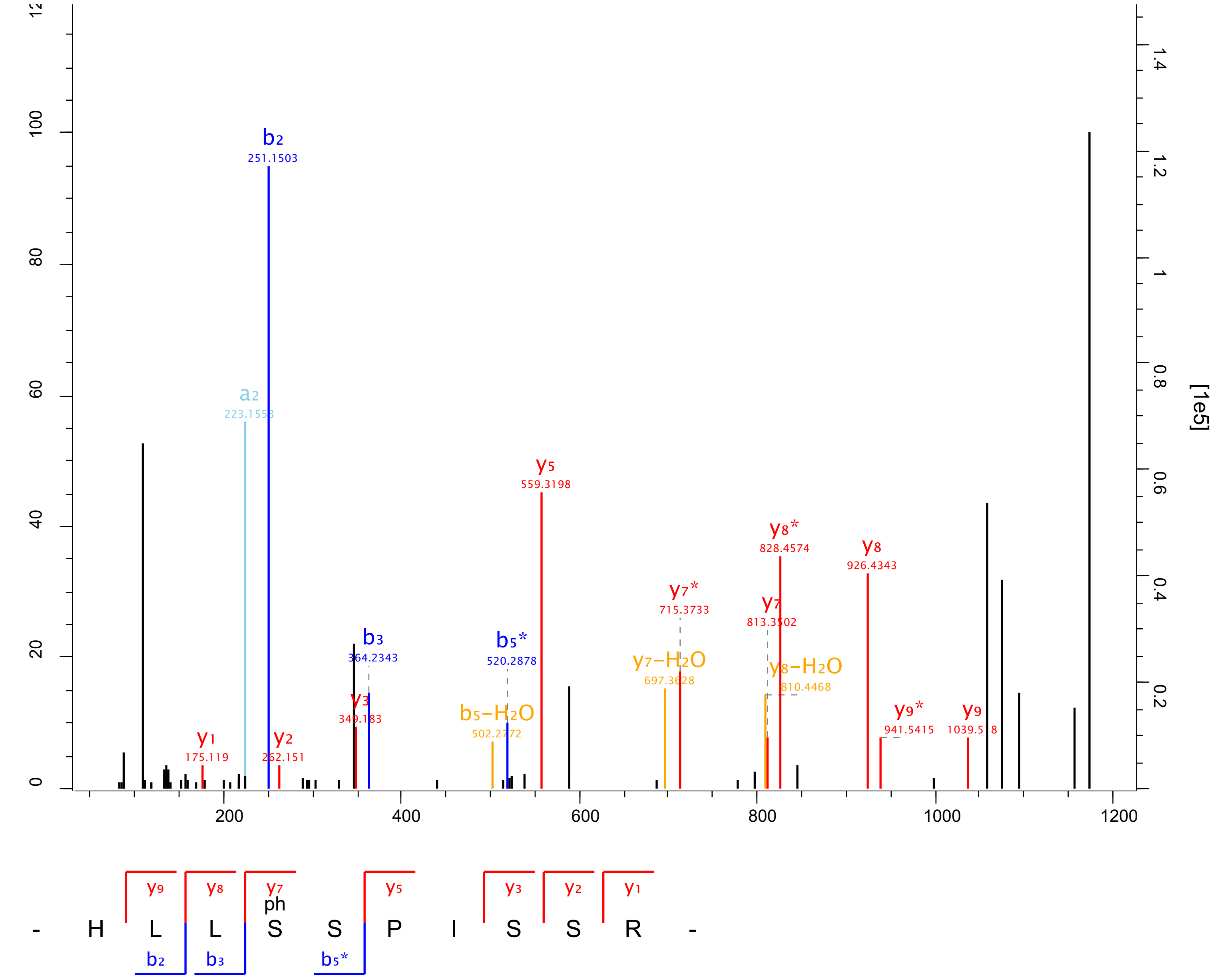

| Raw file | Scan | Method | Score | m/z |
| --- | --- | --- | --- | --- |
| water_02_1 | 10797 | FTMS; HCD | 76.23 | 495.75 |

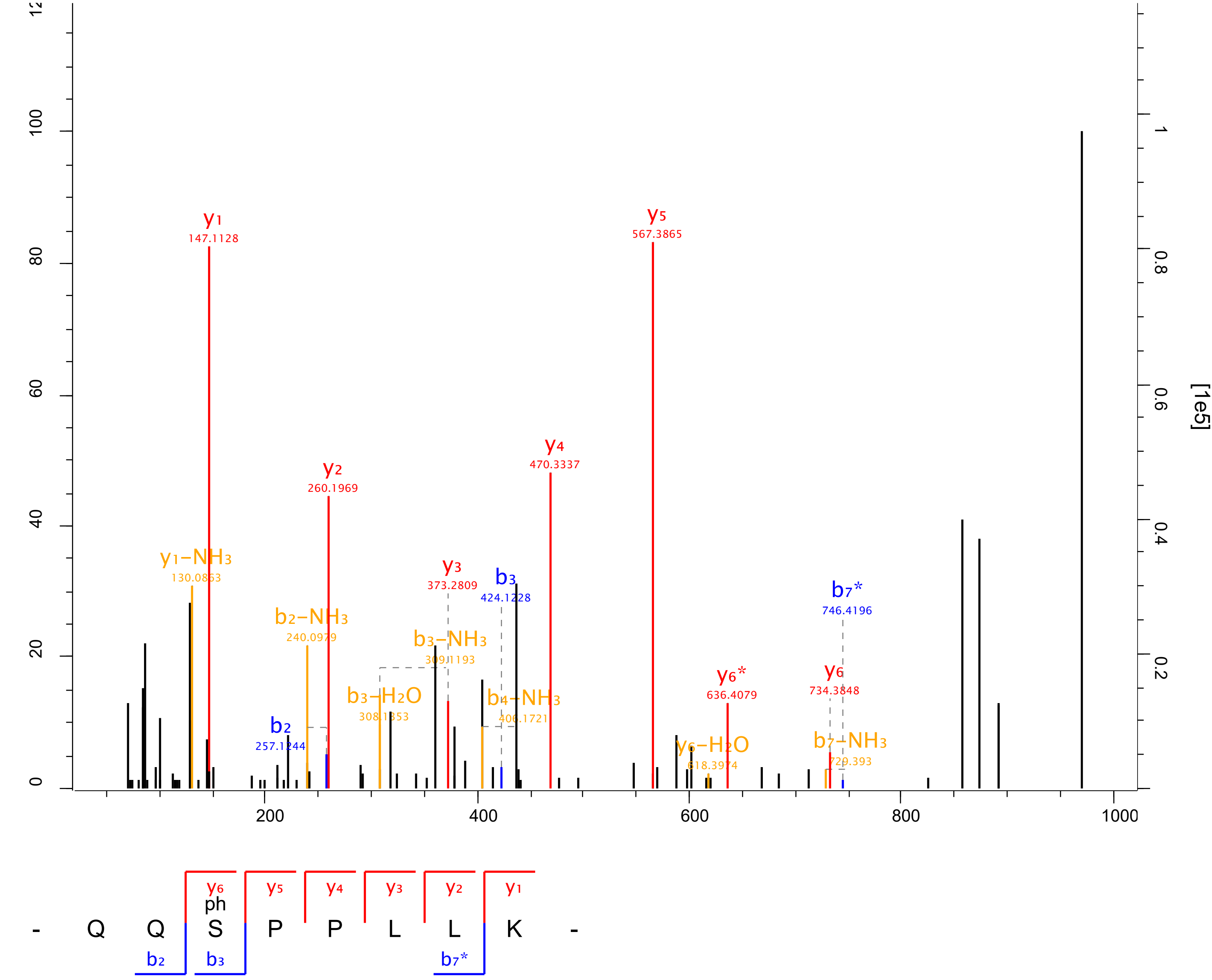

| Raw file | Scan | Method | Score | m/z |
| --- | --- | --- | --- | --- |
| water_02_1 | 10870 | FTMS; HCD | 99.75 | 562.27 |

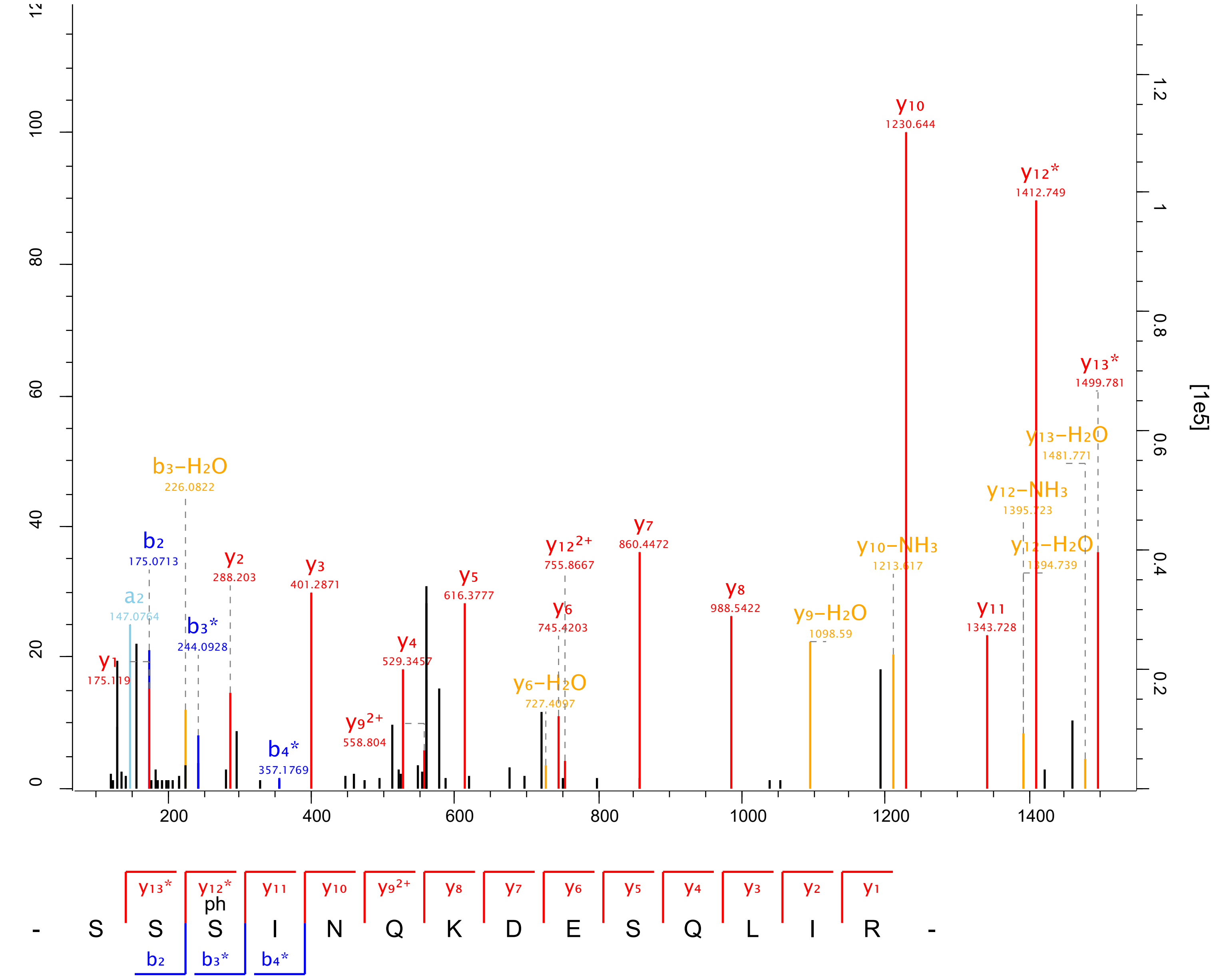

| Raw file | Scan | Method | Score | m/z |
|---|---|---|---|---|
| water_02_1 | 10919 | FTMS; HCD | 73.17 | 550.56 |

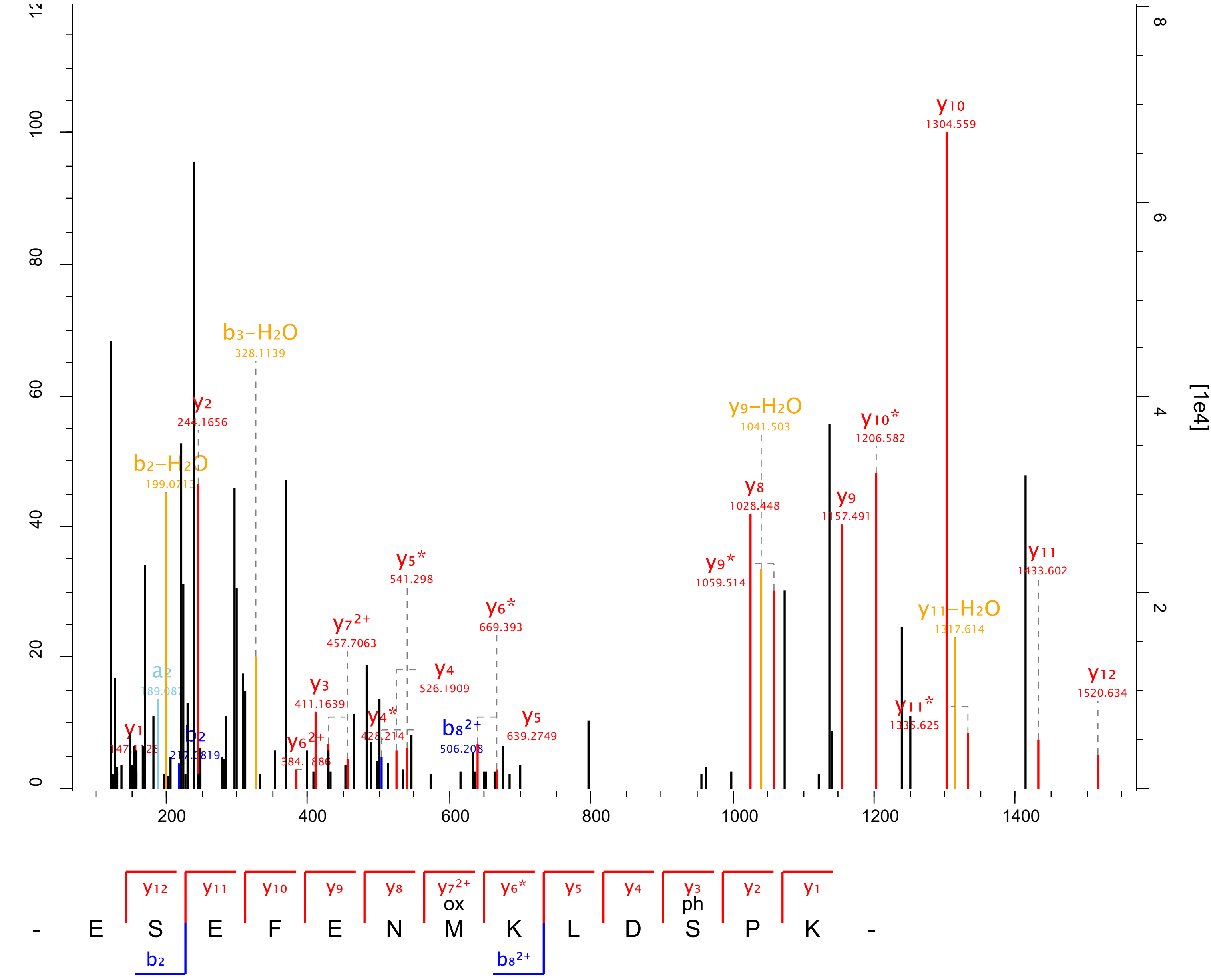

| Raw file | Scan | Method | Score | m/z |
| --- | --- | --- | --- | --- |
| water_02_1 | 10961 | FTMS; HCD | 43.9 | 499.75 |

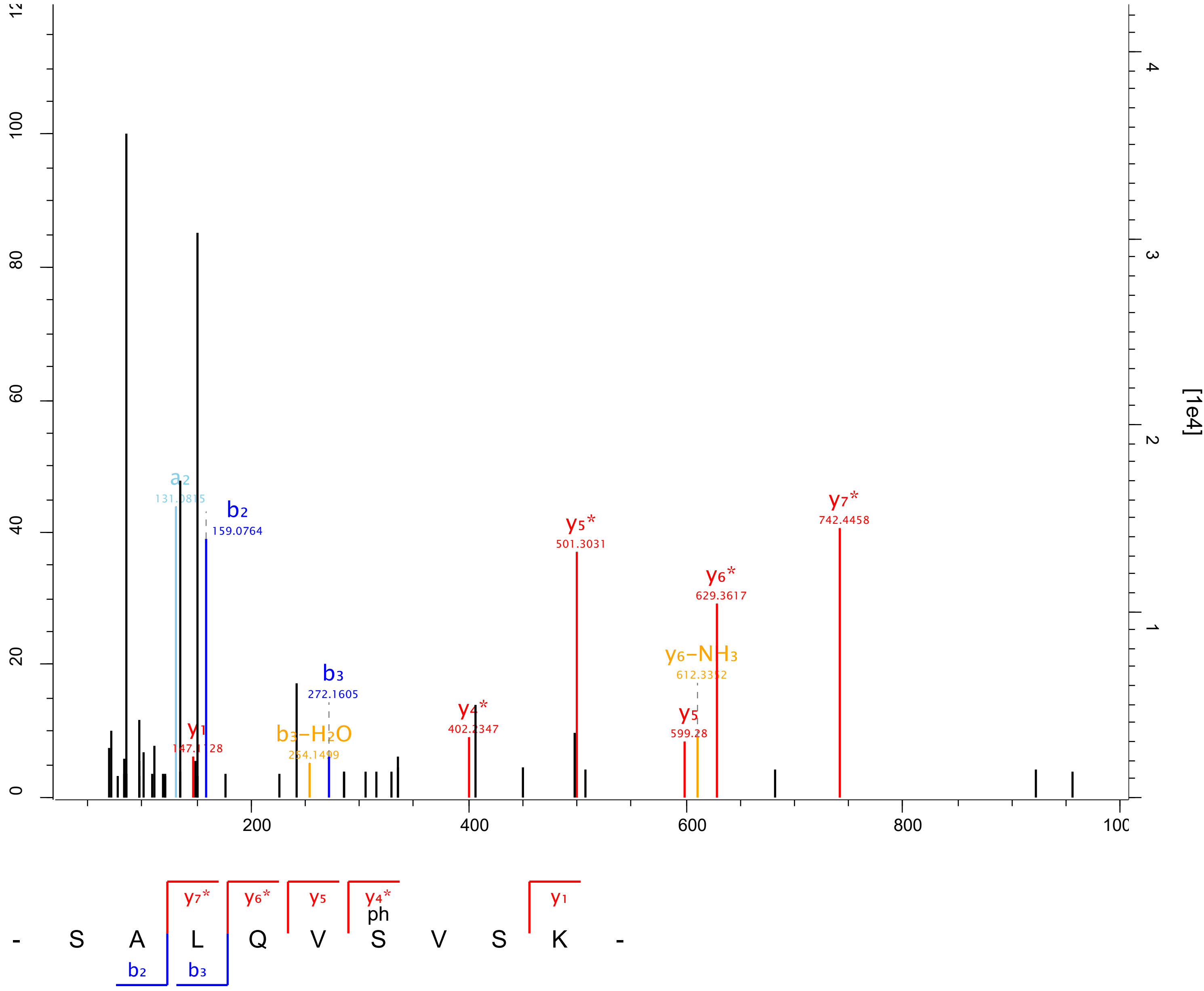

| Raw file | Scan | Method | Score | m/z |
|---|---|---|---|---|
| water_02_1 | 10988 | FTMS; HCD | 43.5 | 575.92 |

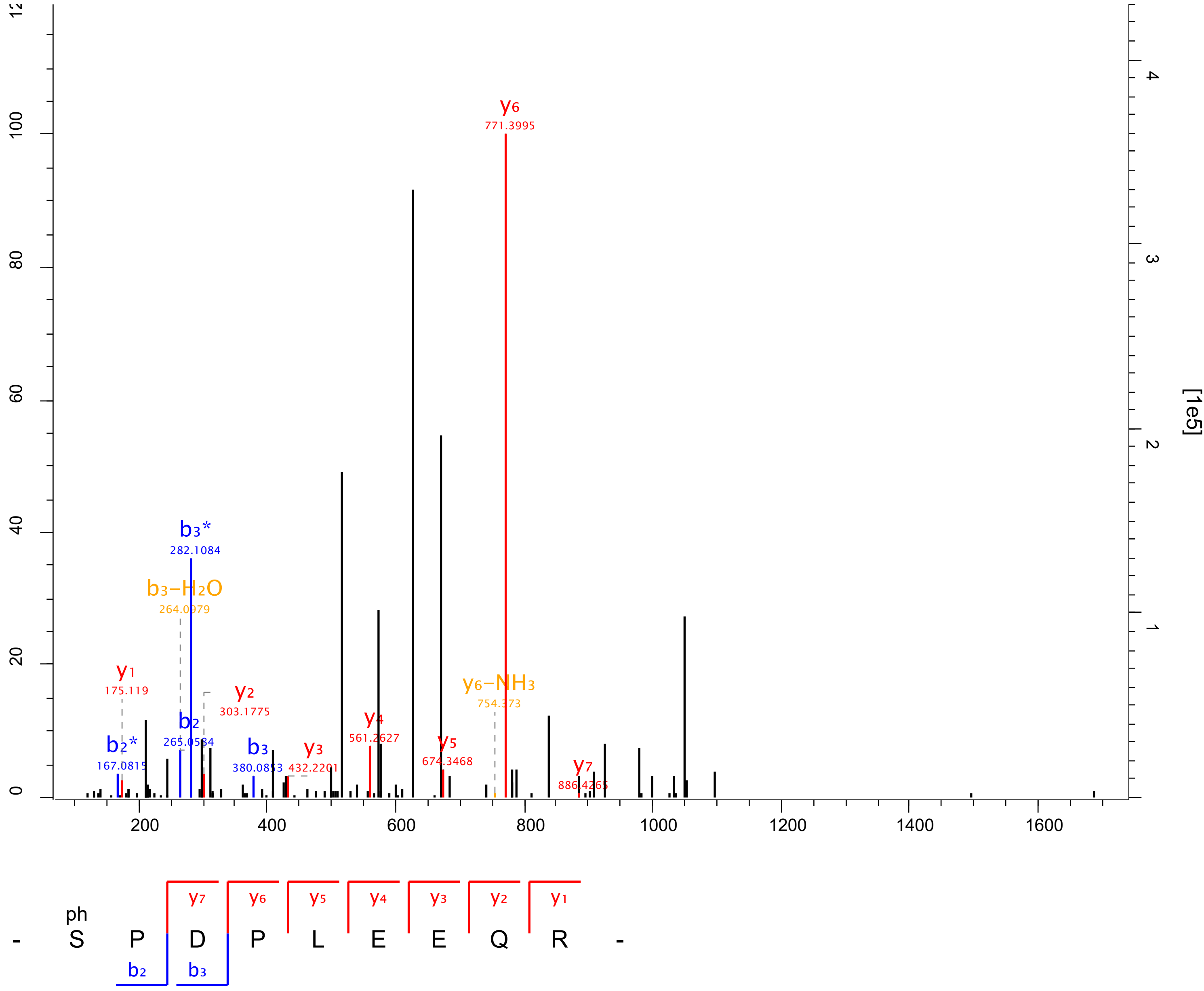

| Raw file | Scan | Method | Score | m/z |
|---|---|---|---|---|
| water_02_1 | 11029 | FTMS; HCD | 58.51 | 544.24 |

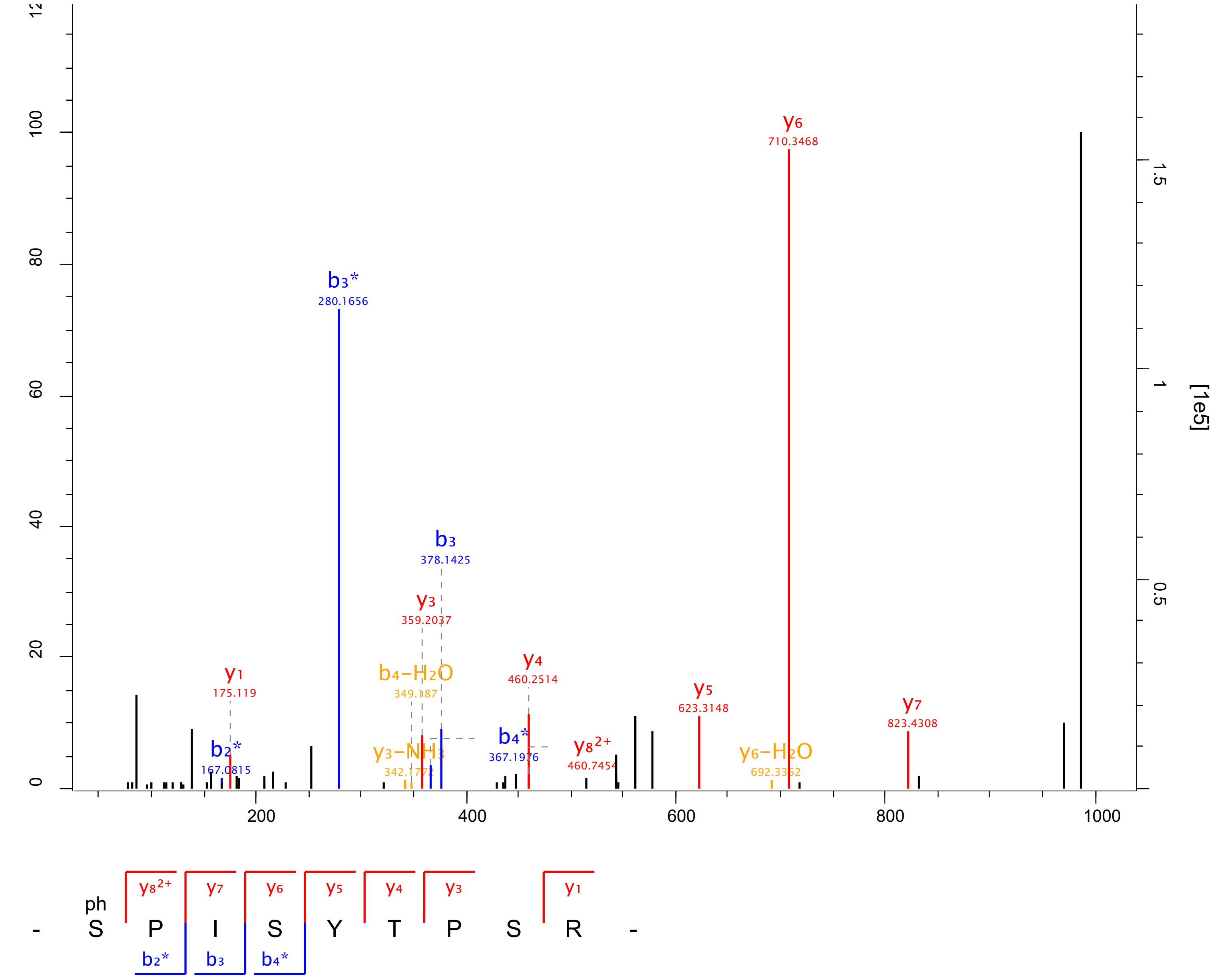

| Raw file | Scan | Method | Score | m/z |
|---|---|---|---|---|
| water_02_1 | 11053 | FTMS; HCD | 90.23 | 621.27 |

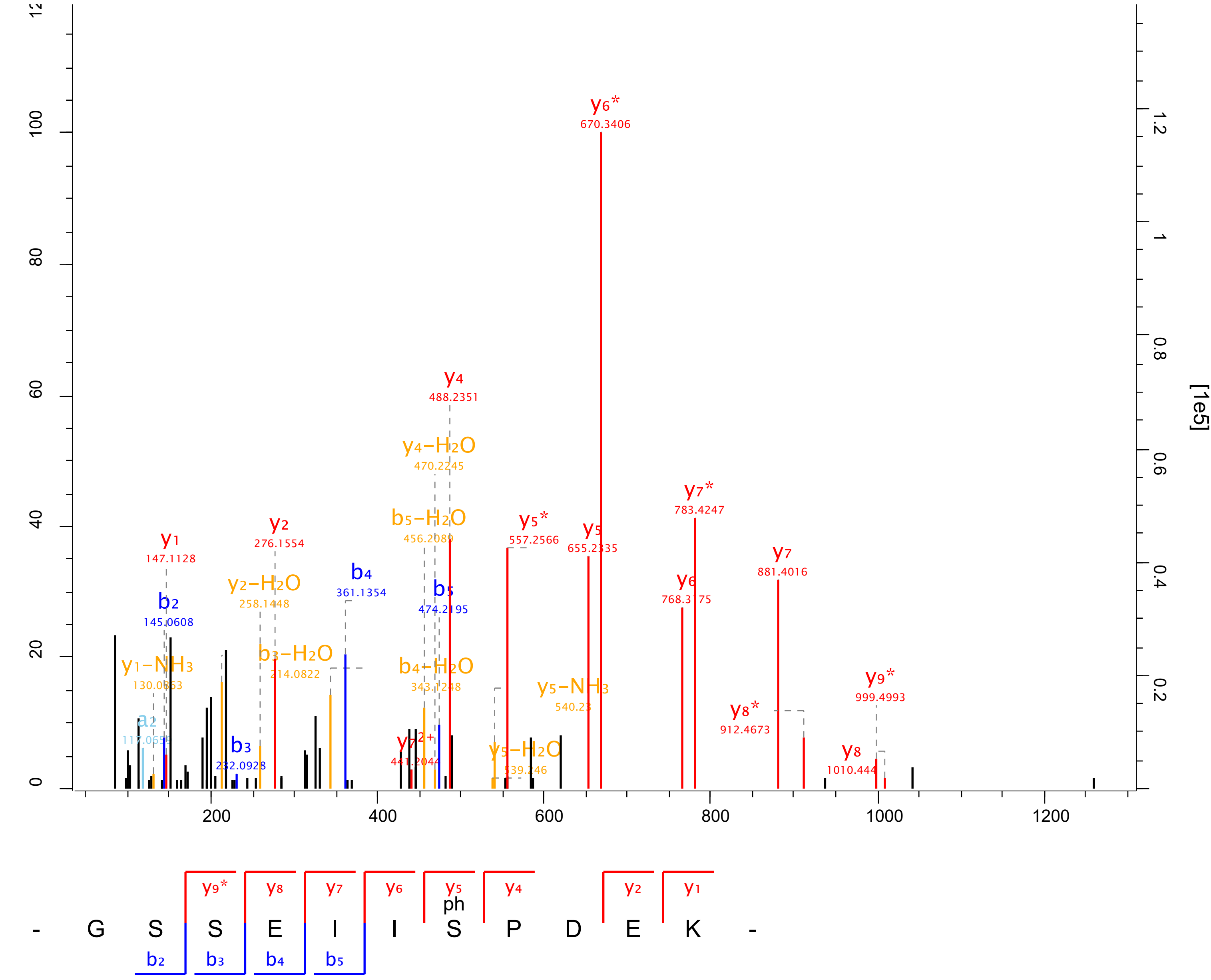

| Raw file | Scan | Method | Score | m/z |
| --- | --- | --- | --- | --- |
| water_02_1 | 11099 | FTMS; HCD | 63.43 | 437.71 |

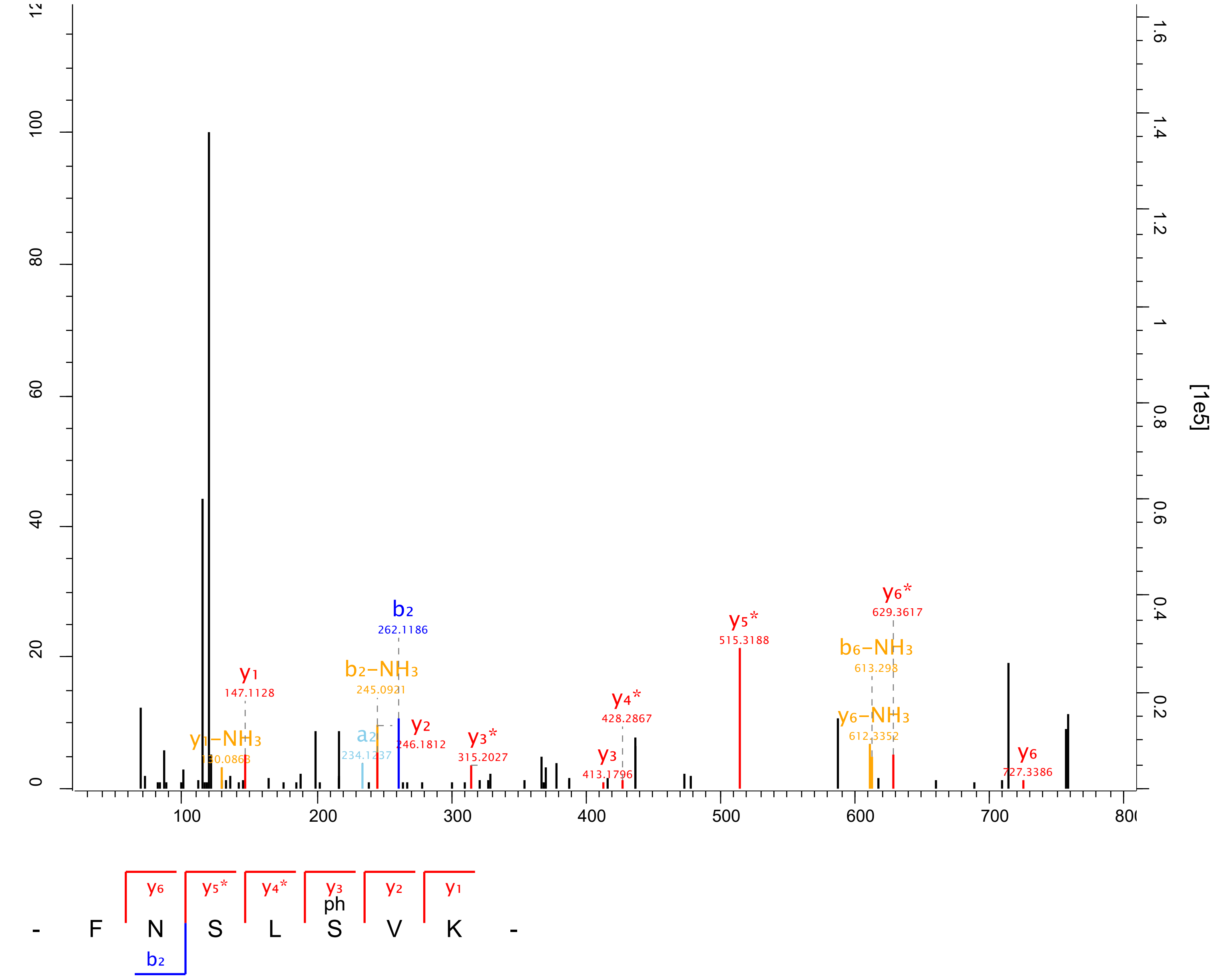

| Raw file | Scan | Method | Score | m/z |
| --- | --- | --- | --- | --- |
| water_02_1 | 11214 | FTMS; HCD | 71.08 | 699.33 |

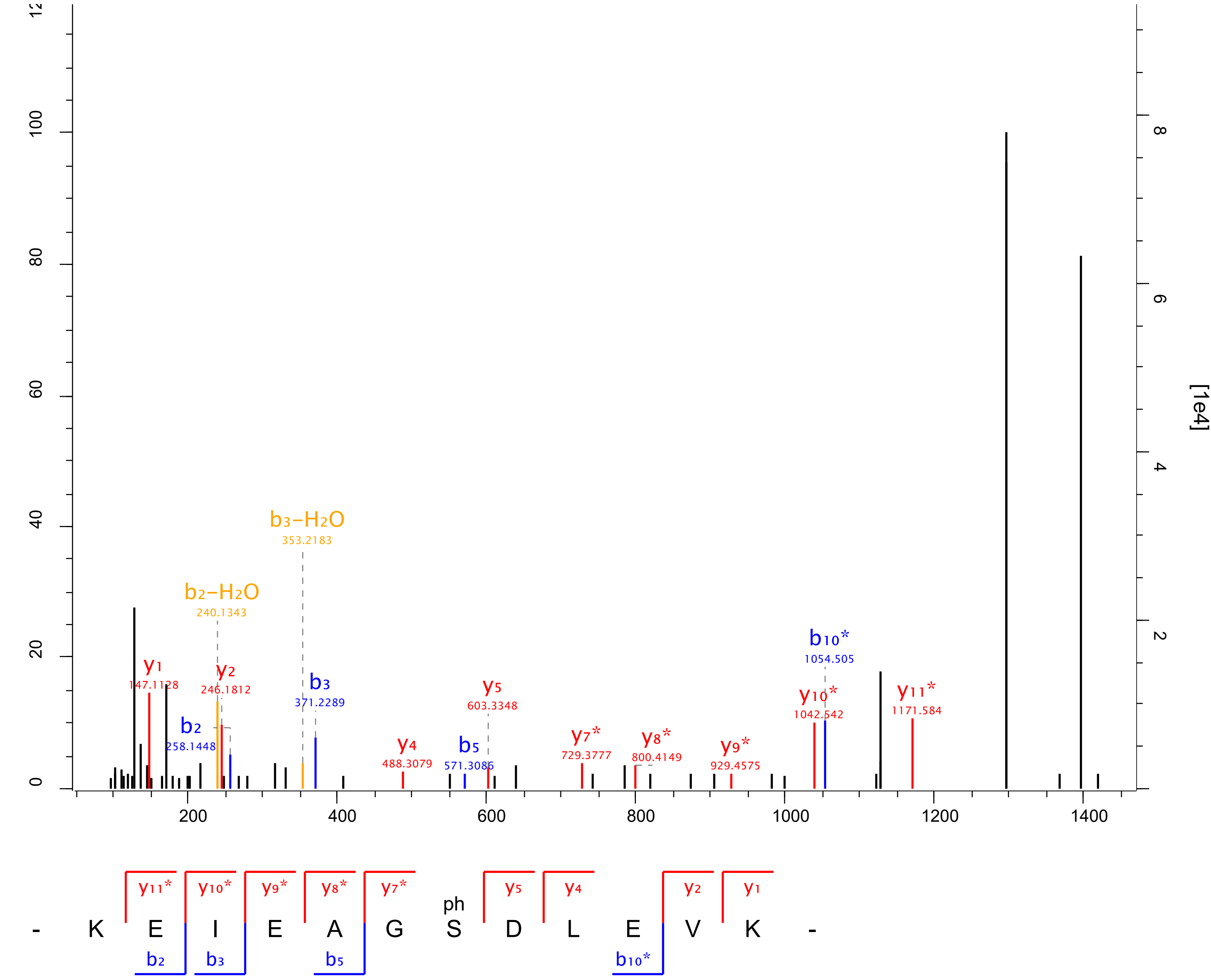

| Raw file | Scan | Method | Score | m/z |
|---|---|---|---|---|
| water_02_1 | 11264 | FTMS; HCD | 88.57 | 739.3 |

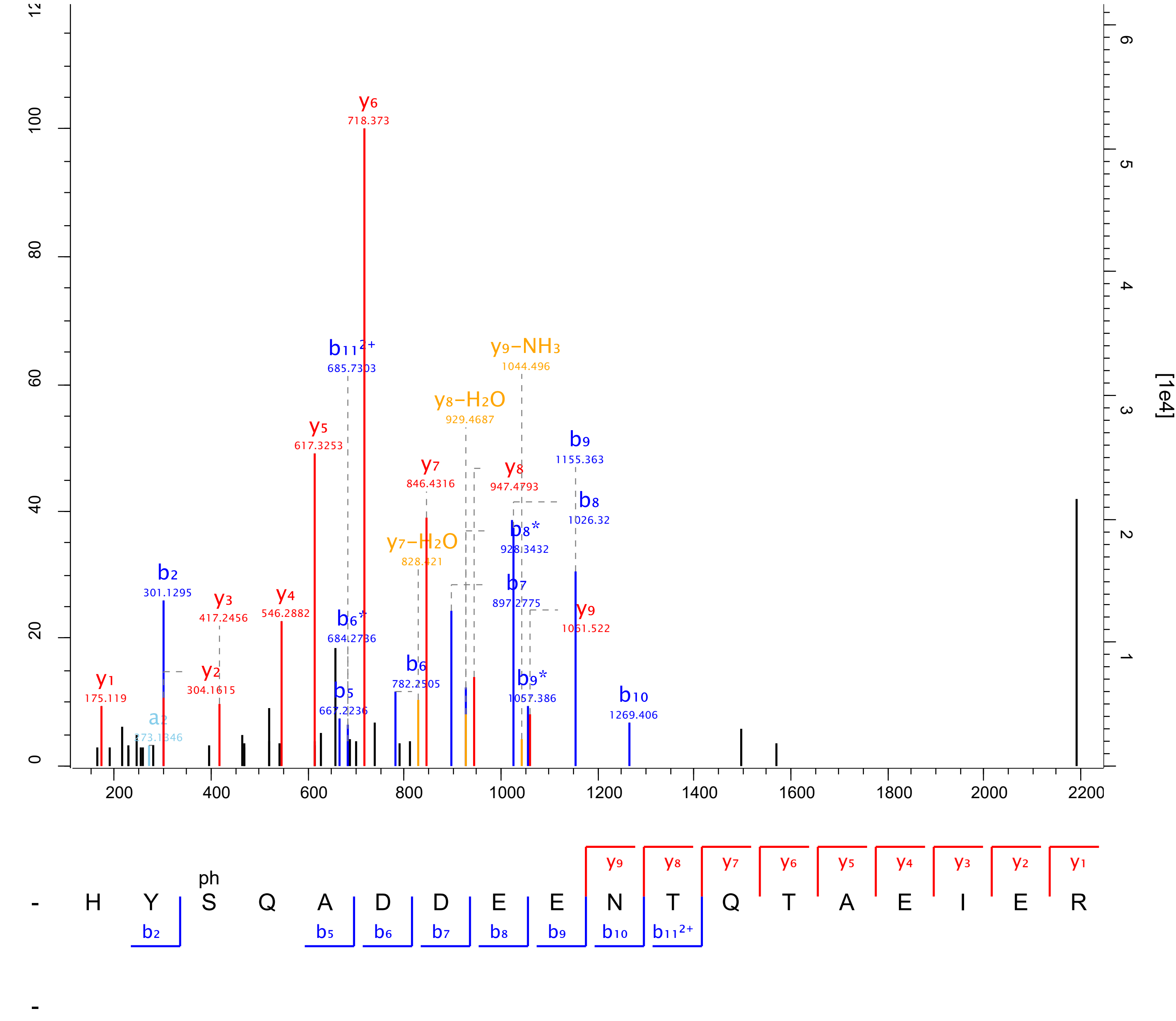

| Raw file | Scan | Method | Score | m/z |
|---|---|---|---|---|
| water_02_1 | 11331 | FTMS; HCD | 51.73 | 625.3 |

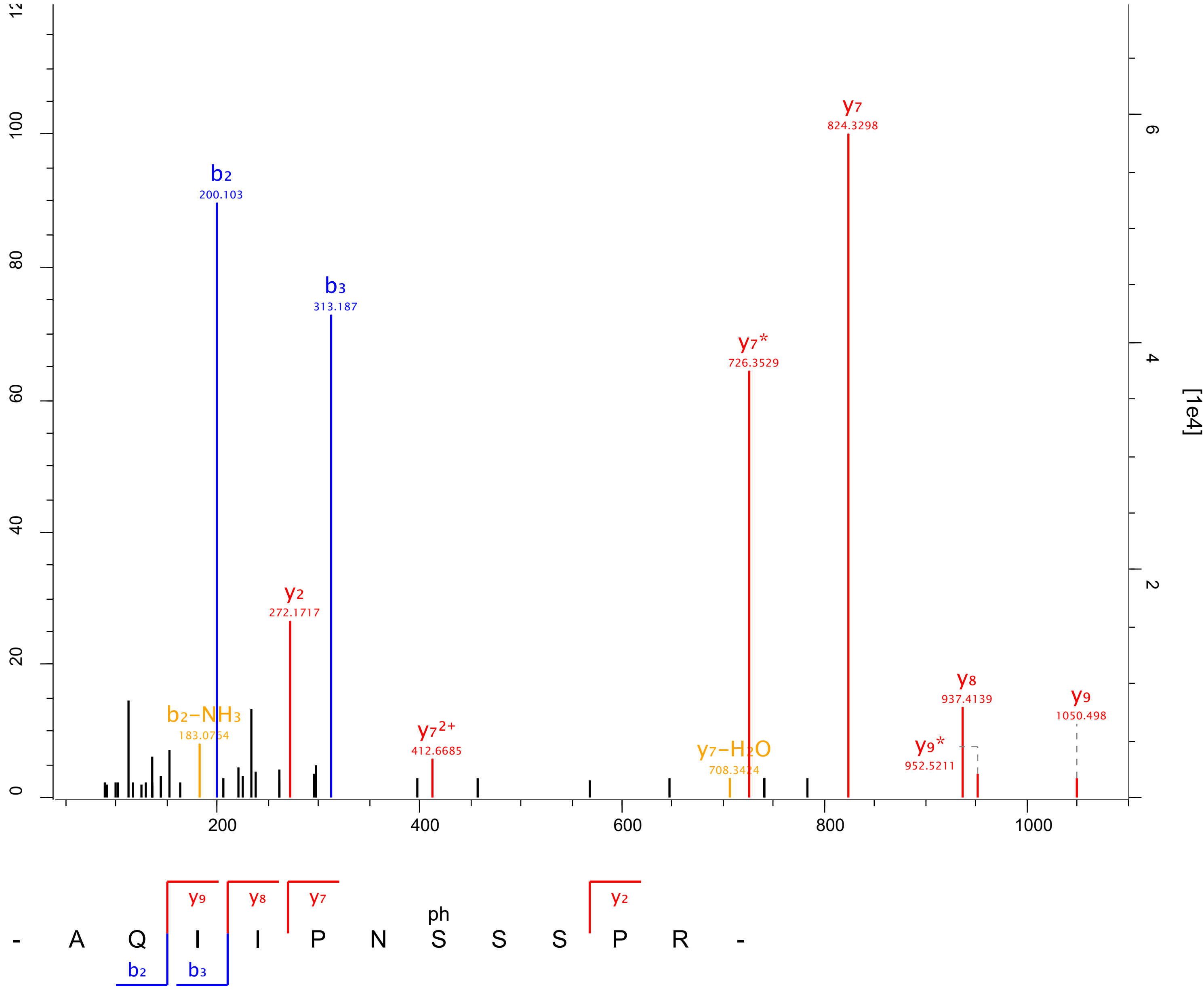

| Raw file | Scan | Method | Score | m/z |
|---|---|---|---|---|
| water_02_1 | 11344 | FTMS; HCD | 52.19 | 540.94 |

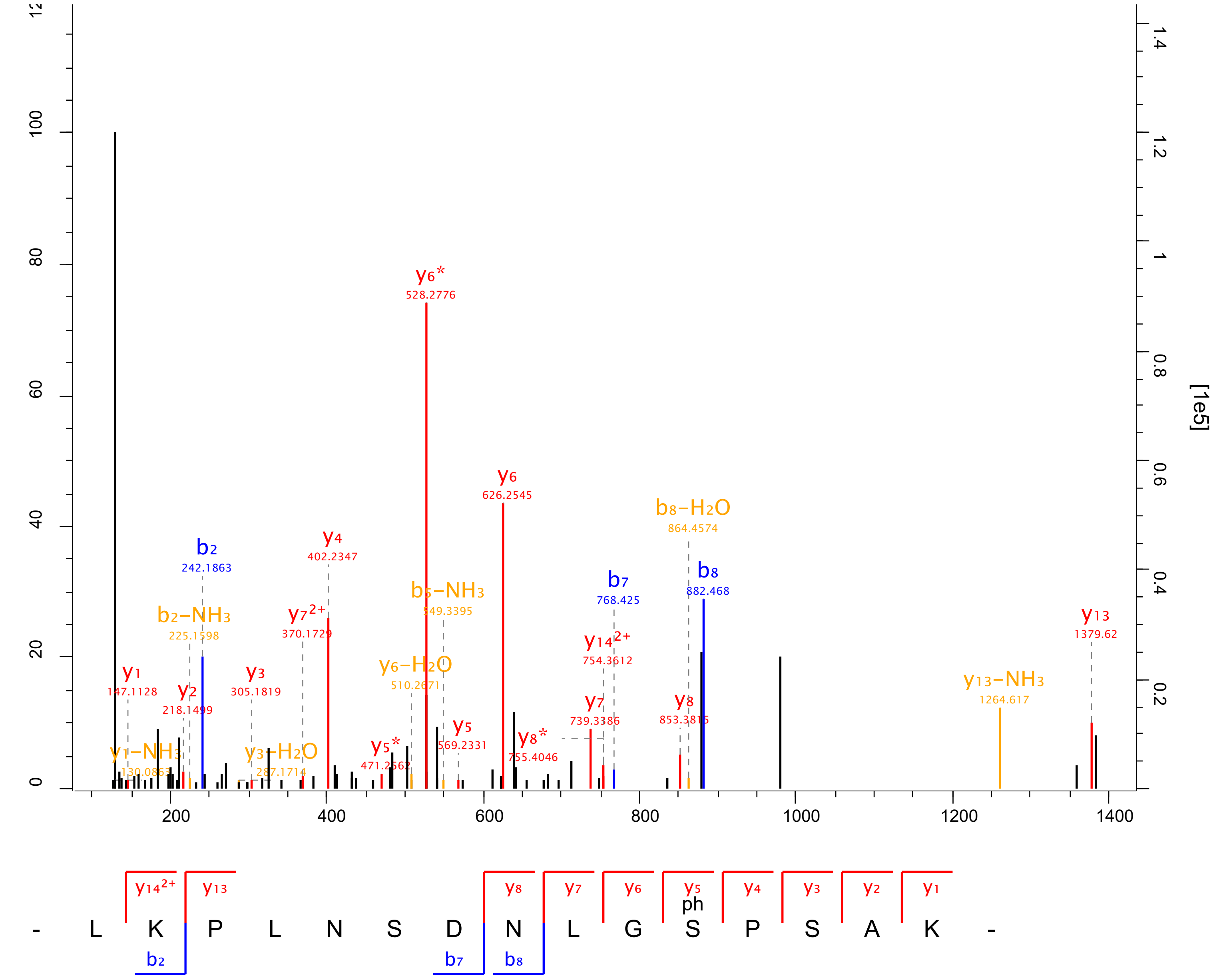

| Raw file | Scan | Method | Score | m/z |
|---|---|---|---|---|
| water_02_1 | 11507 | FTMS; HCD | 70.2 | 546.59 |

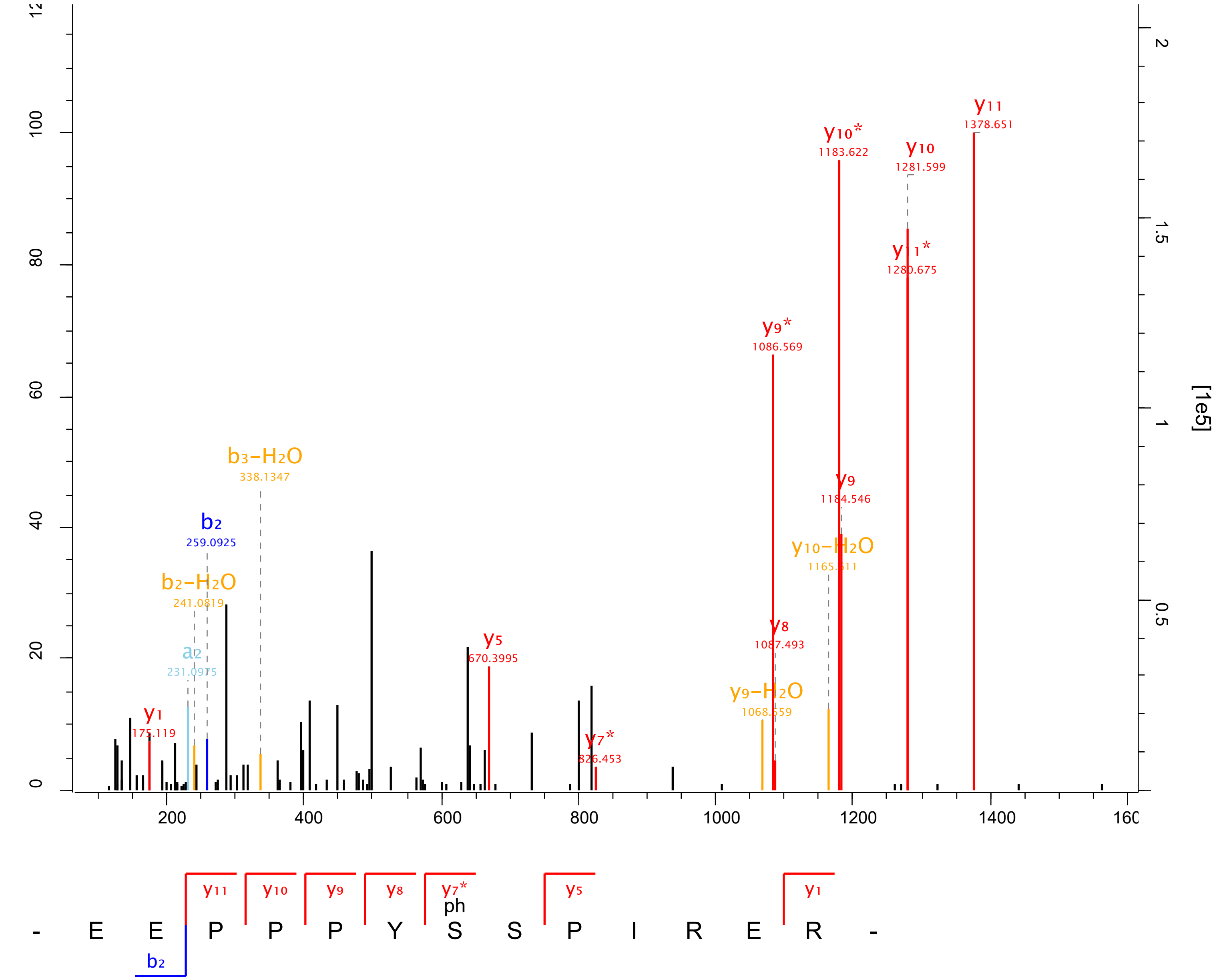

| Raw file | Scan | Method | Score | m/z |
|---|---|---|---|---|
| water_02_1 | 11595 | FTMS; HCD | 98.42 | 713.8 |

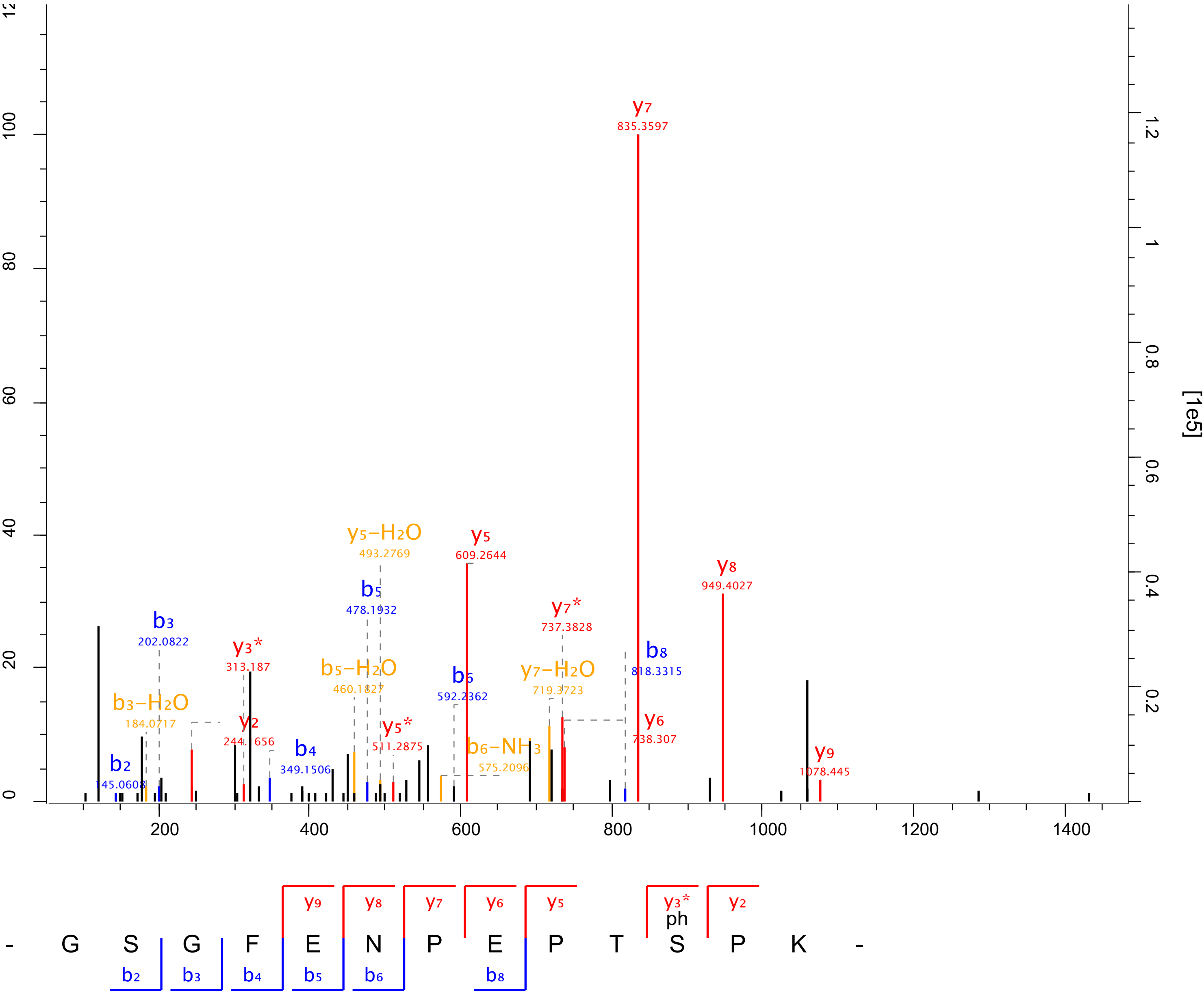

| Raw file | Scan | Method | Score | m/z |
| --- | --- | --- | --- | --- |
| water_02_1 | 11616 | FTMS; HCD | 206.64 | 770.33 |

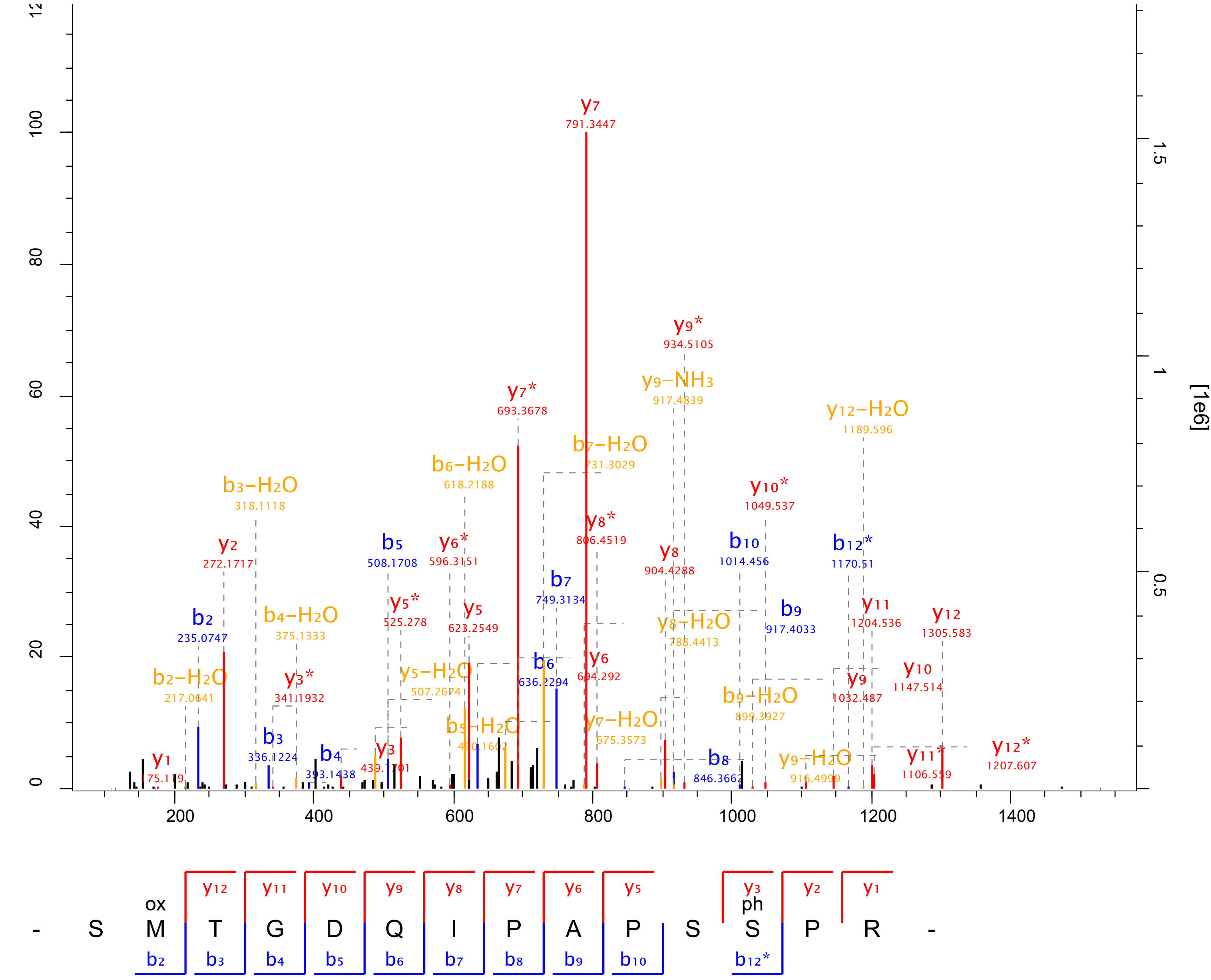

| Raw file | Scan | Method | Score | m/z |
| --- | --- | --- | --- | --- |
| water_02_1 | 11625 | FTMS; HCD | 45.43 | 511.26 |

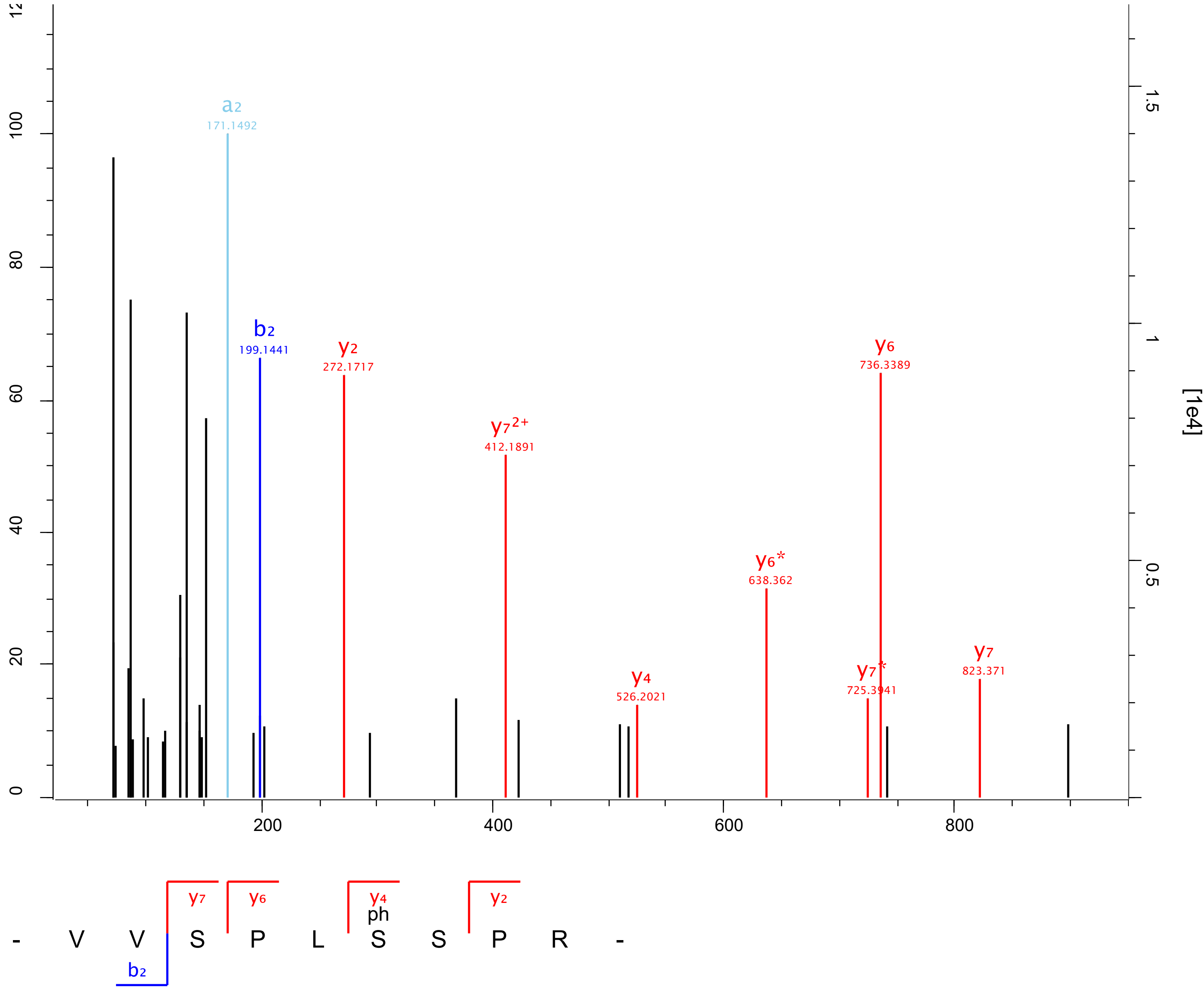

| Raw file | Scan | Method | Score | m/z |
|---|---|---|---|---|
| water_02_1 | 11629 | FTMS; HCD | 83.05 | 546.22 |

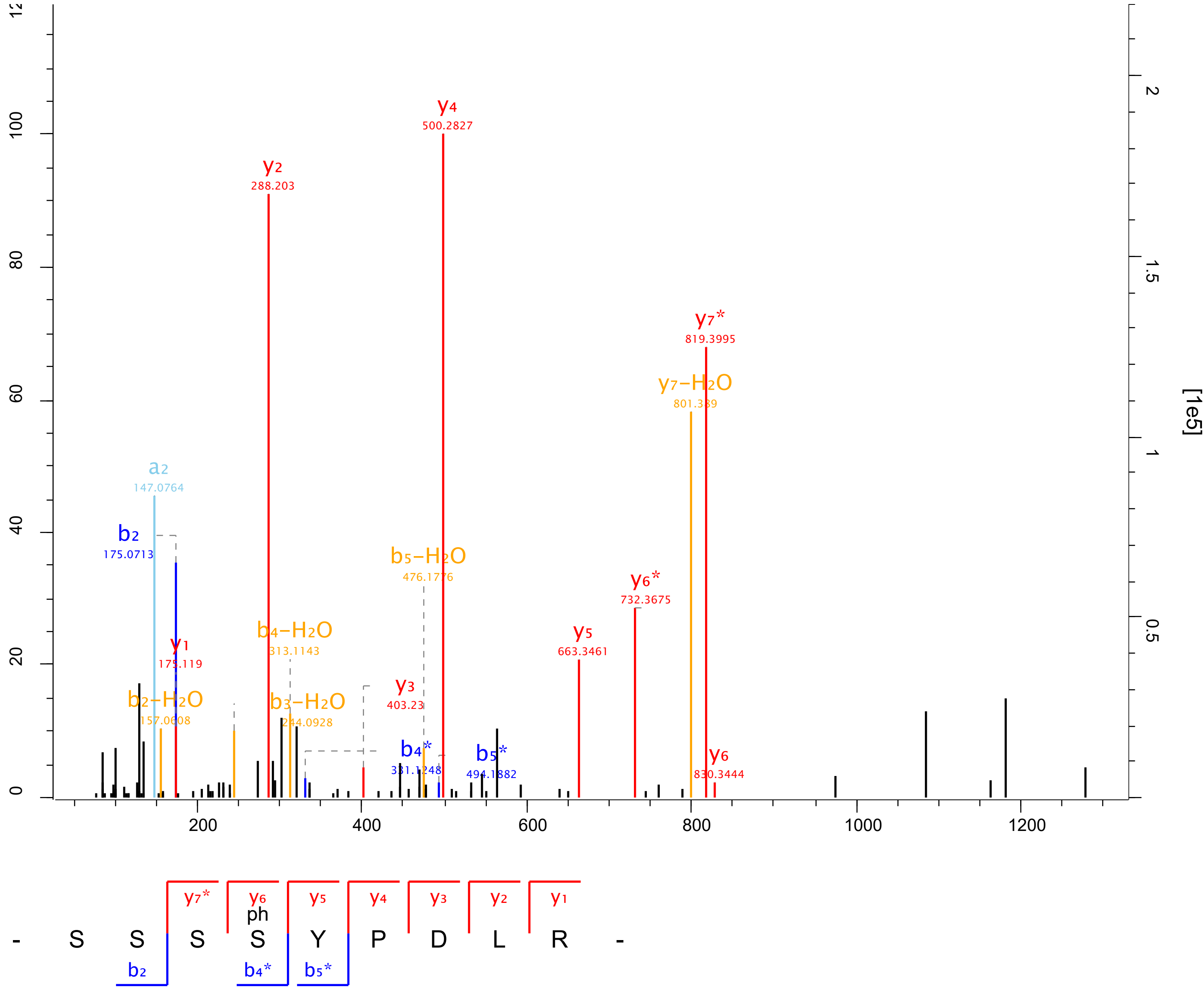

| Raw file | Scan | Method | Score | m/z |
|---|---|---|---|---|
| water_02_1 | 11682 | FTMS; HCD | 72.33 | 441.21 |

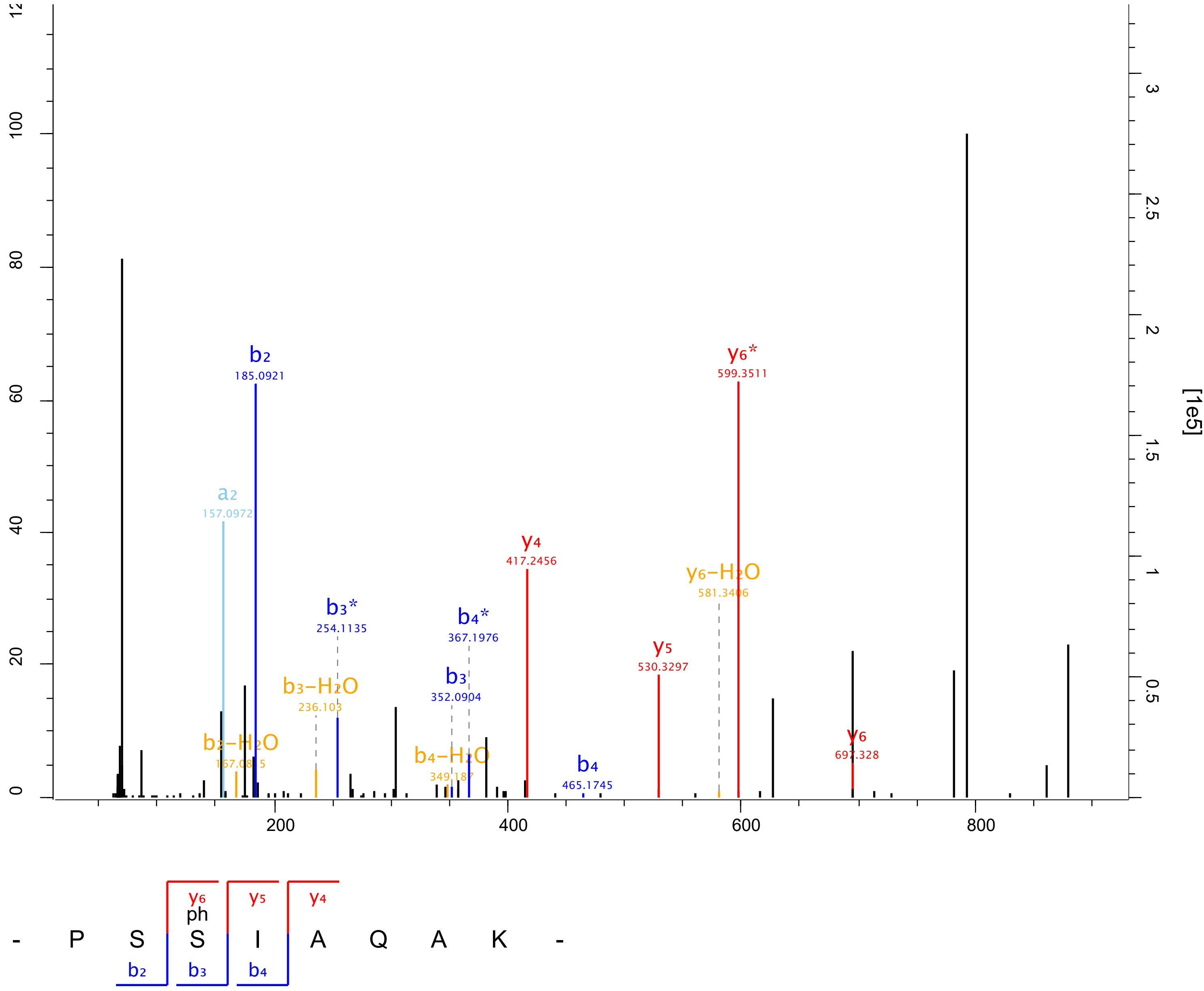

| Raw file | Scan | Method | Score | m/z |
|---|---|---|---|---|
| water_02_1 | 11695 | FTMS; HCD | 157.99 | 436.7 |

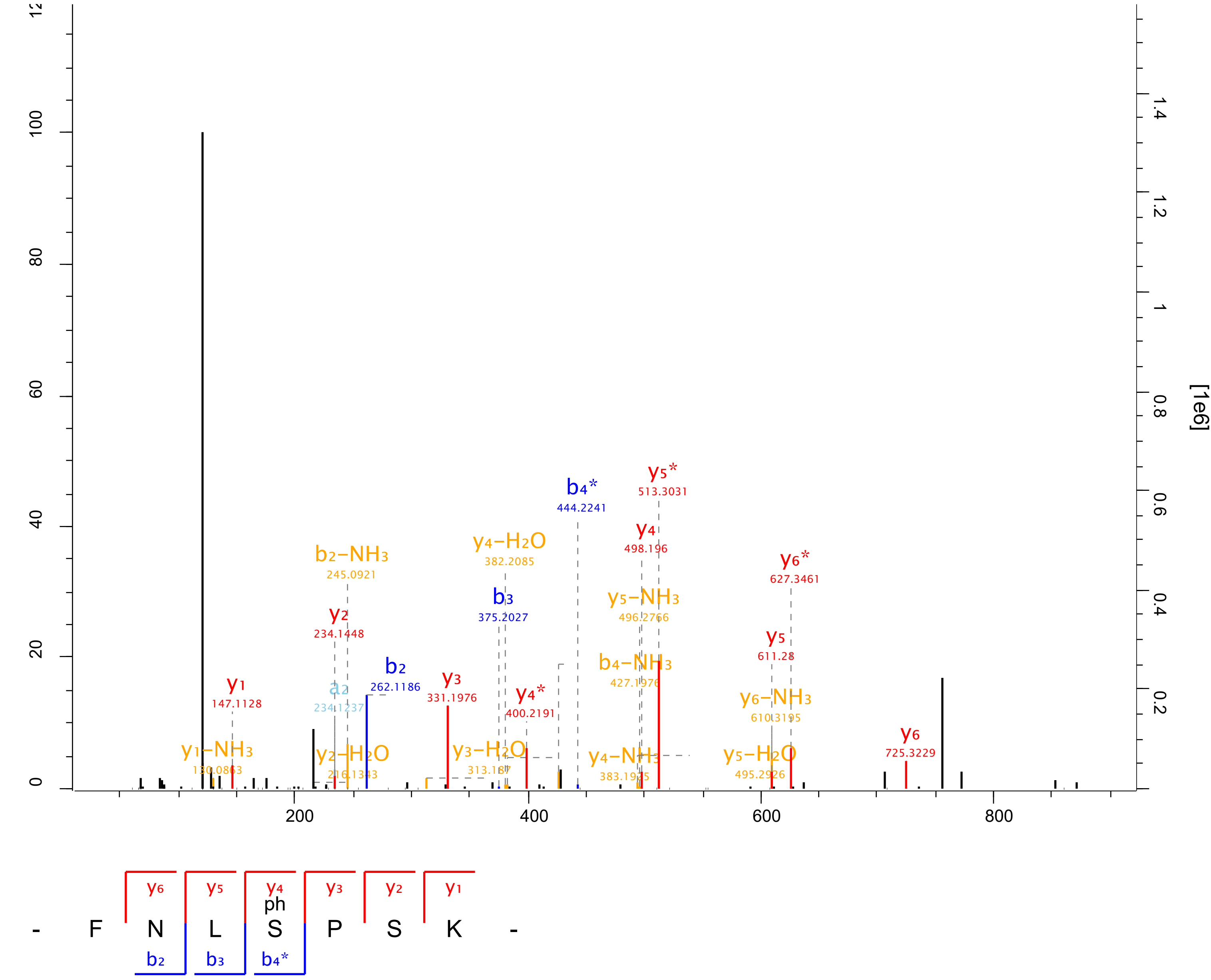

| Raw file | Scan | Method | Score | m/z |
|---|---|---|---|---|
| water_02_1 | 11700 | FTMS; HCD | 241.11 | 795.34 |

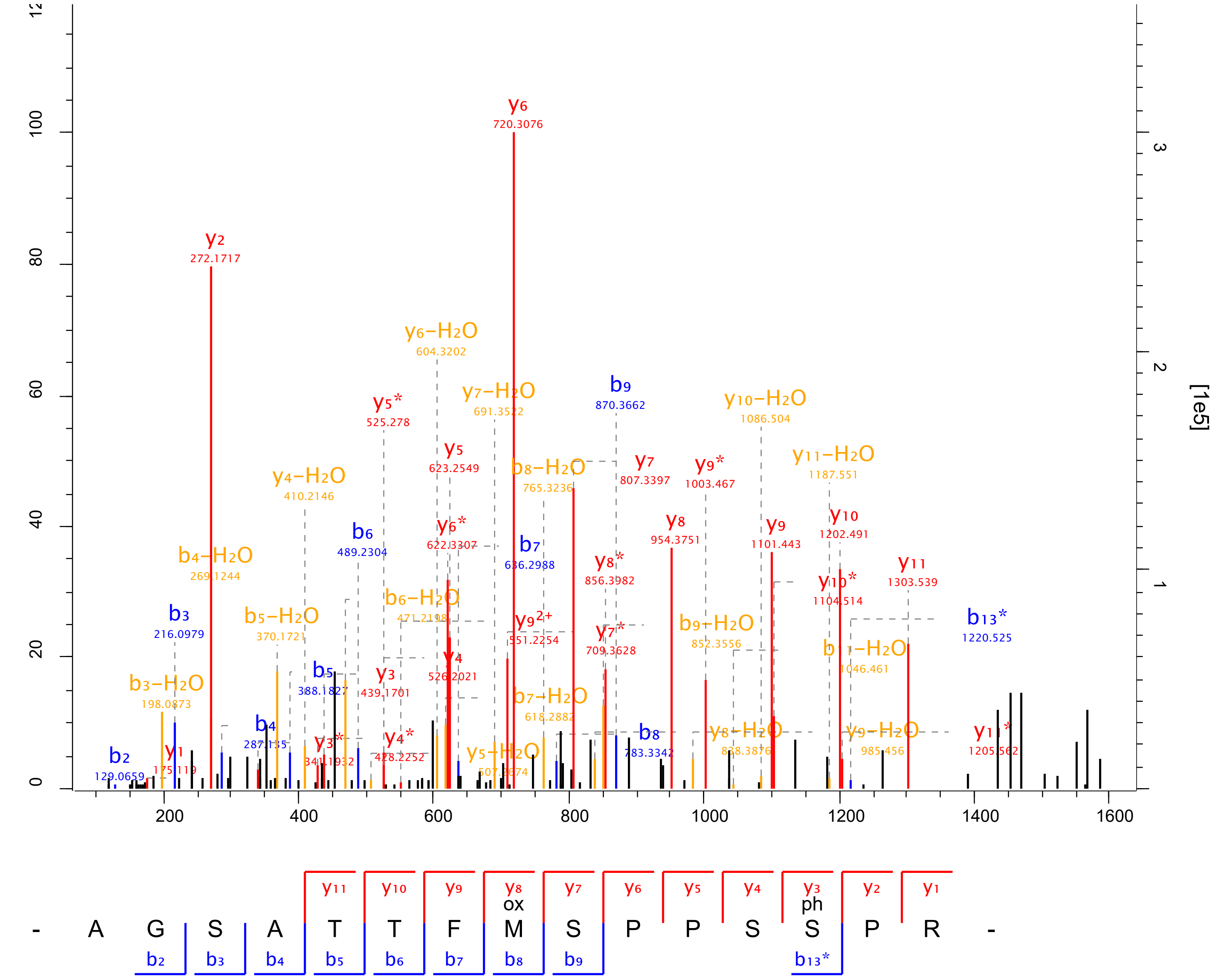

| Raw file | Scan | Method | Score | m/z |
|---|---|---|---|---|
| water_02_1 | 11720 | FTMS; HCD | 72.43 | 448.72 |

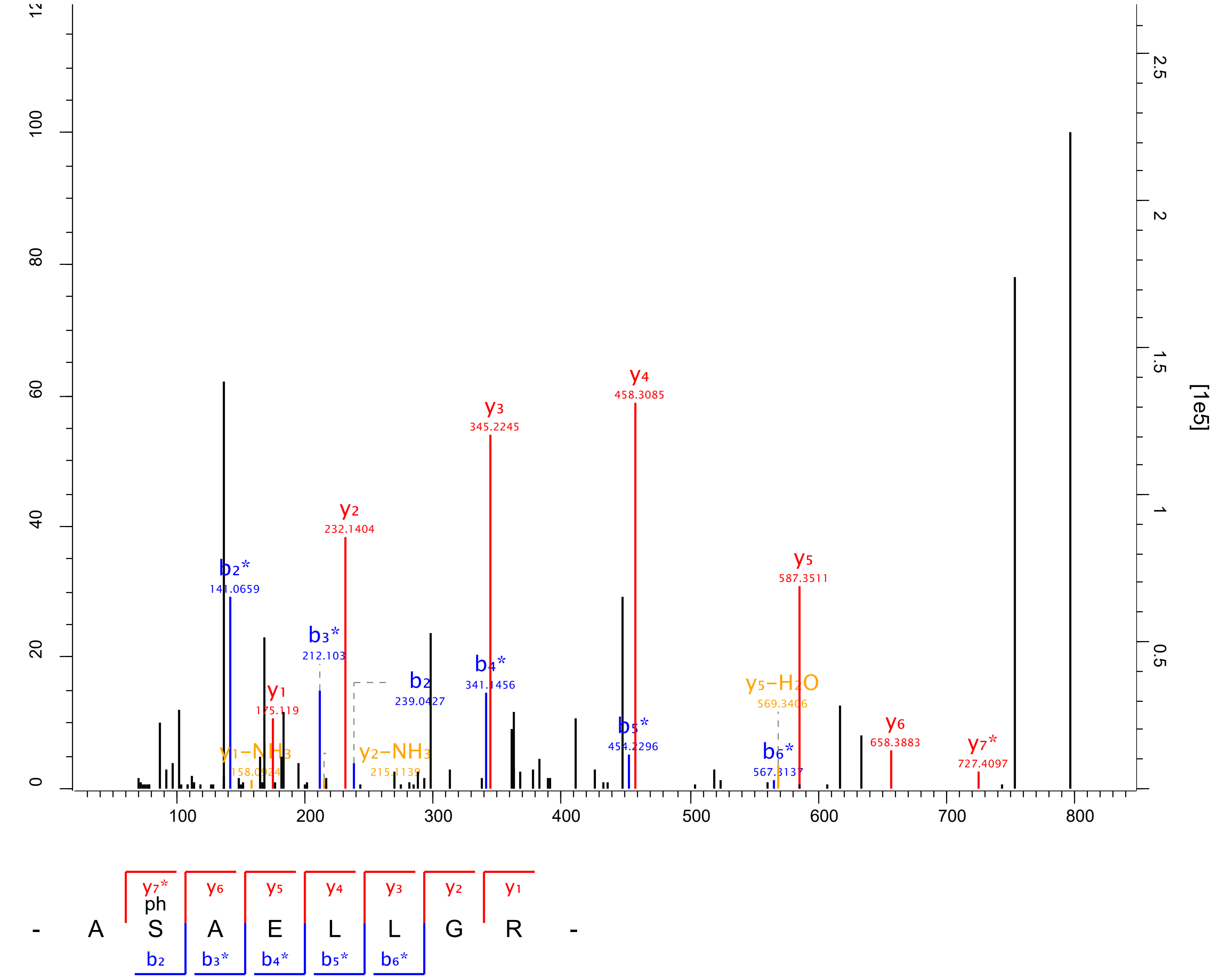

| Raw file | Scan | Method | Score | m/z |
|---|---|---|---|---|
| water_02_1 | 11726 | FTMS; HCD | 87.06 | 825.29 |

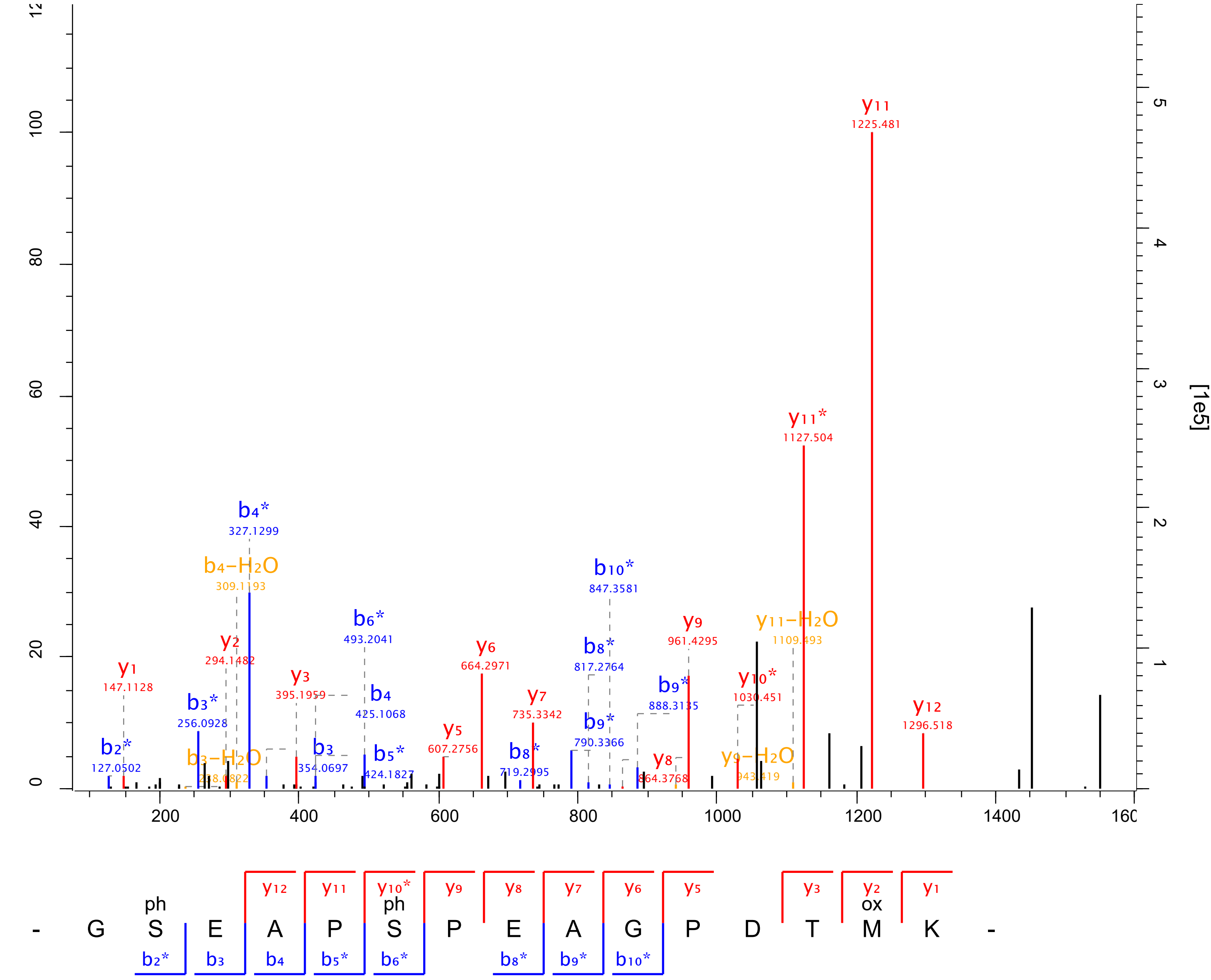

| Raw file | Scan | Method | Score | m/z |
|---|---|---|---|---|
| water_02_1 | 11739 | FTMS; HCD | 53.3 | 698.82 |

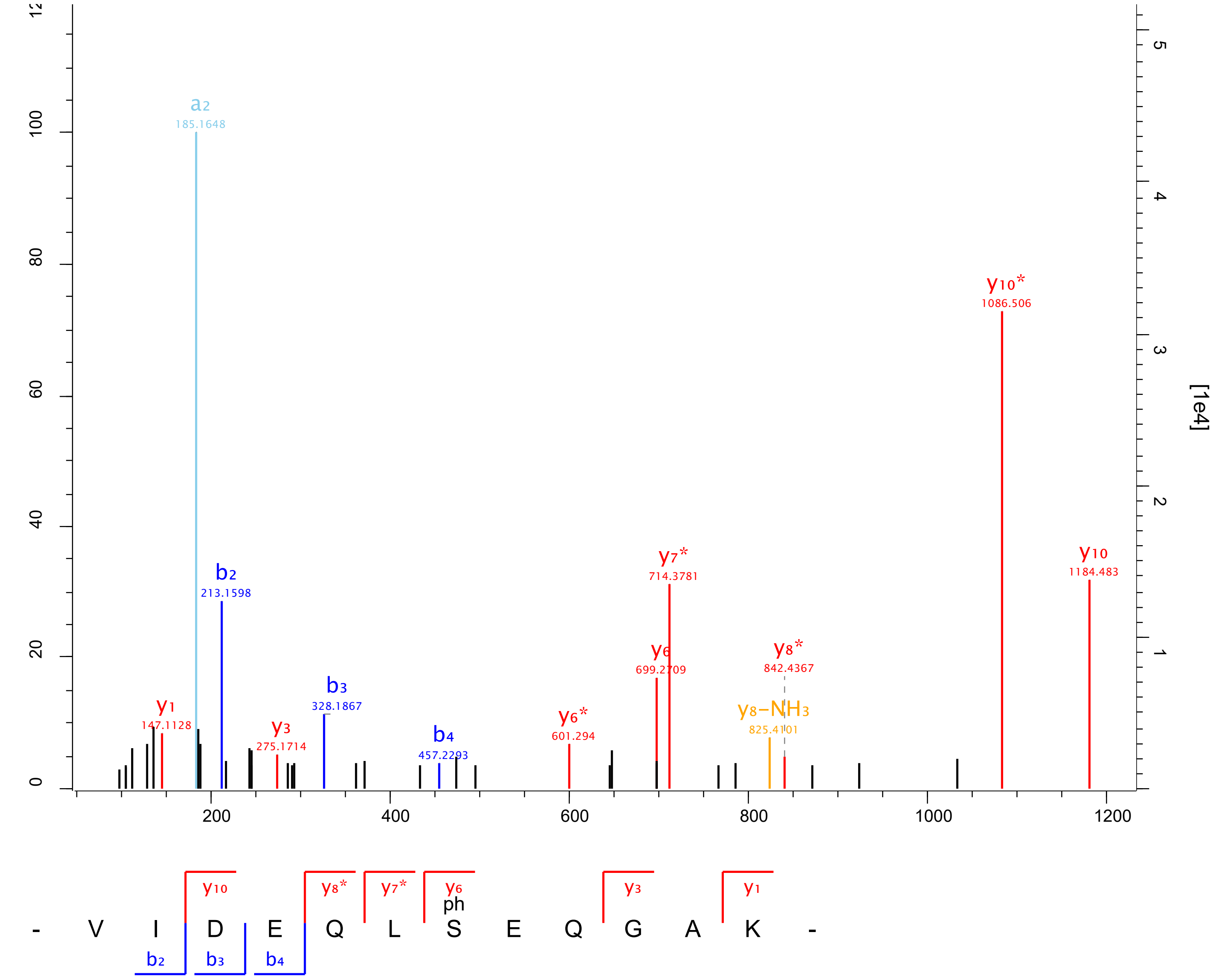

| Raw file | Scan | Method | Score | m/z |
|---|---|---|---|---|
| water_02_1 | 11768 | FTMS; HCD | 164.93 | 693.81 |

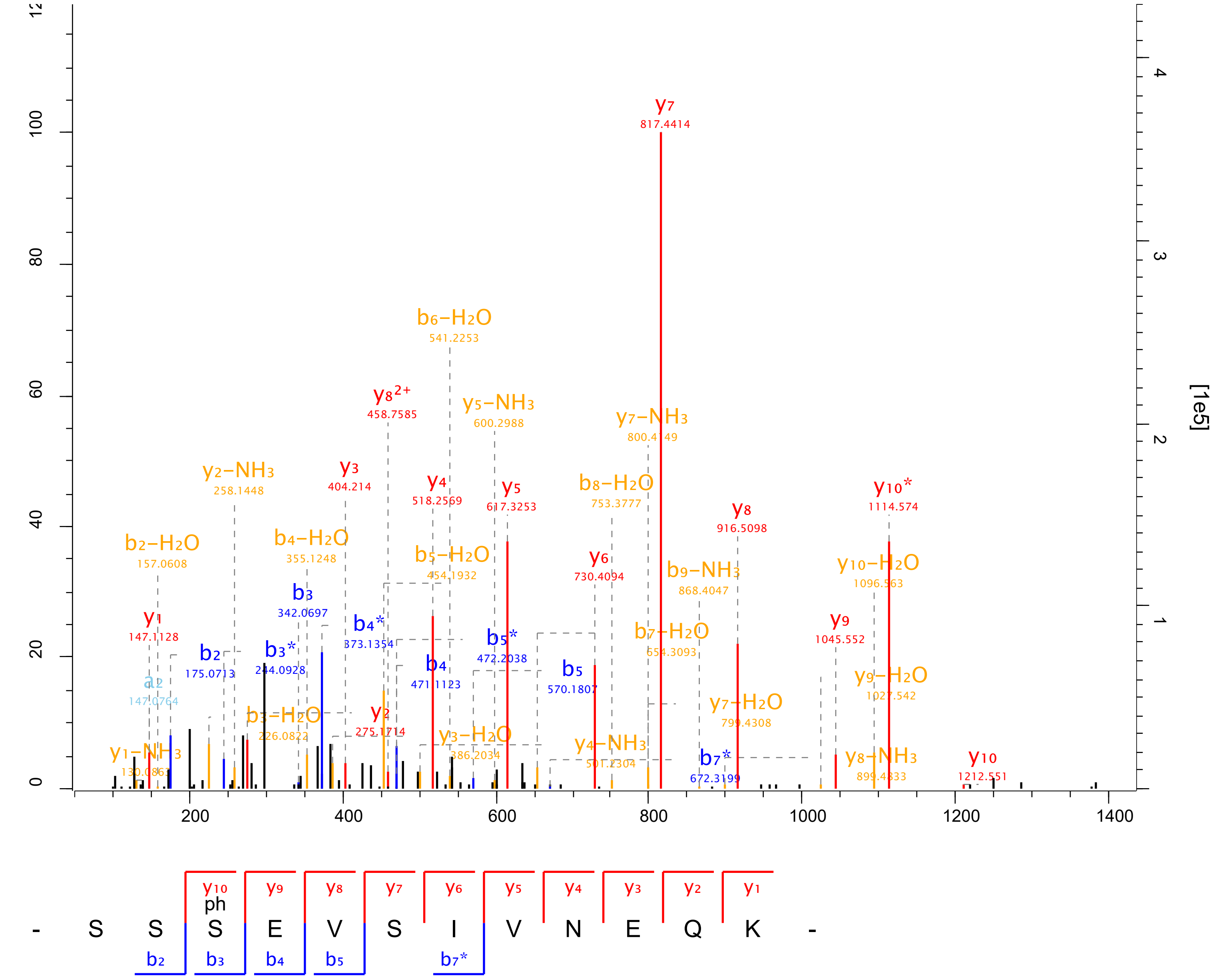

| Raw file | Scan | Method | Score | m/z |
| --- | --- | --- | --- | --- |
| water_02_1 | 11796 | FTMS; HCD | 132.48 | 690.65 |

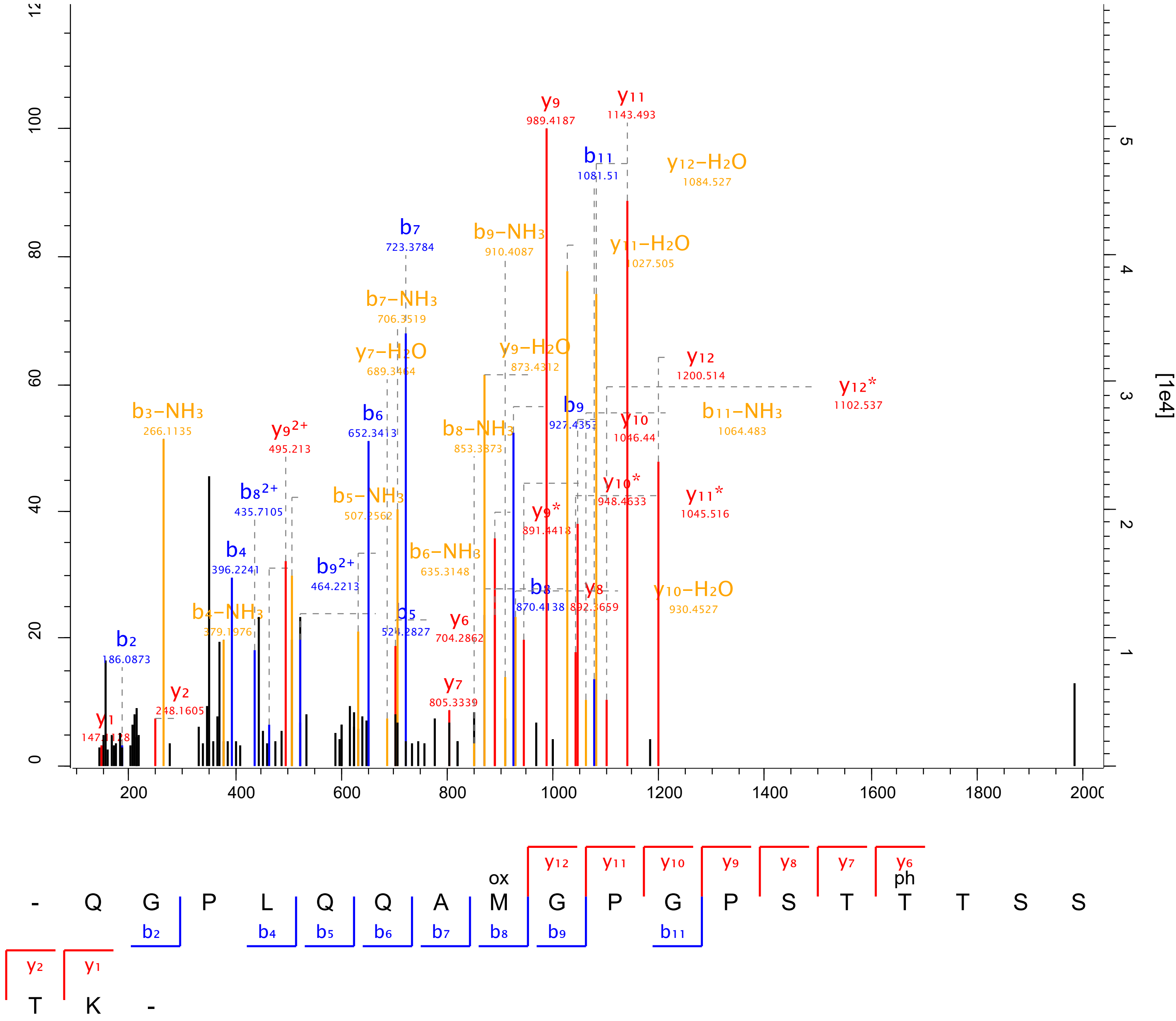

| Raw file | Scan | Method | Score | m/z |
|---|---|---|---|---|
| water_02_1 | 11816 | FTMS; HCD | 42.03 | 836.4 |

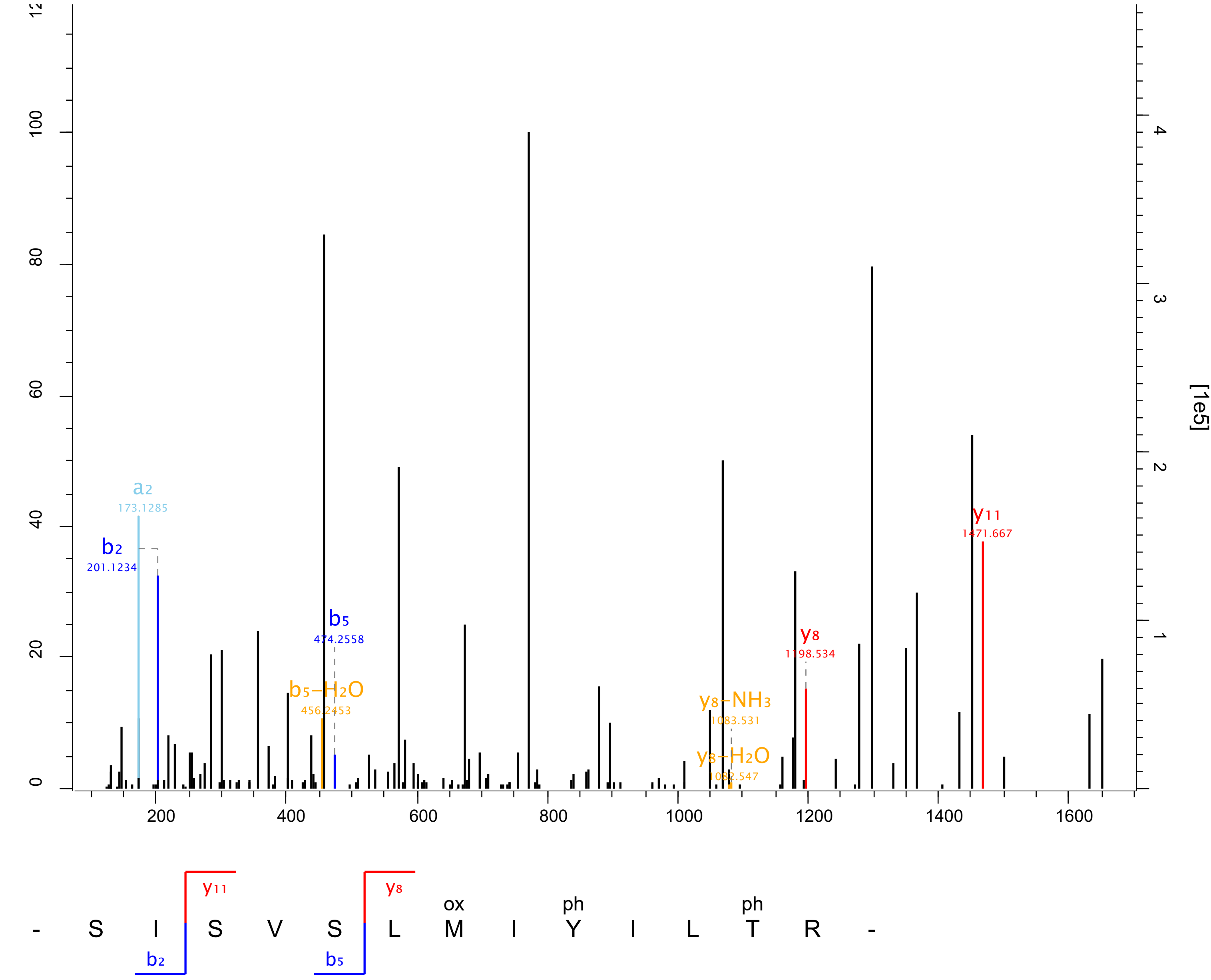

| Raw file | Scan | Method | Score | m/z |
|---|---|---|---|---|
| water_02_1 | 11874 | FTMS; HCD | 117.38 | 841.87 |

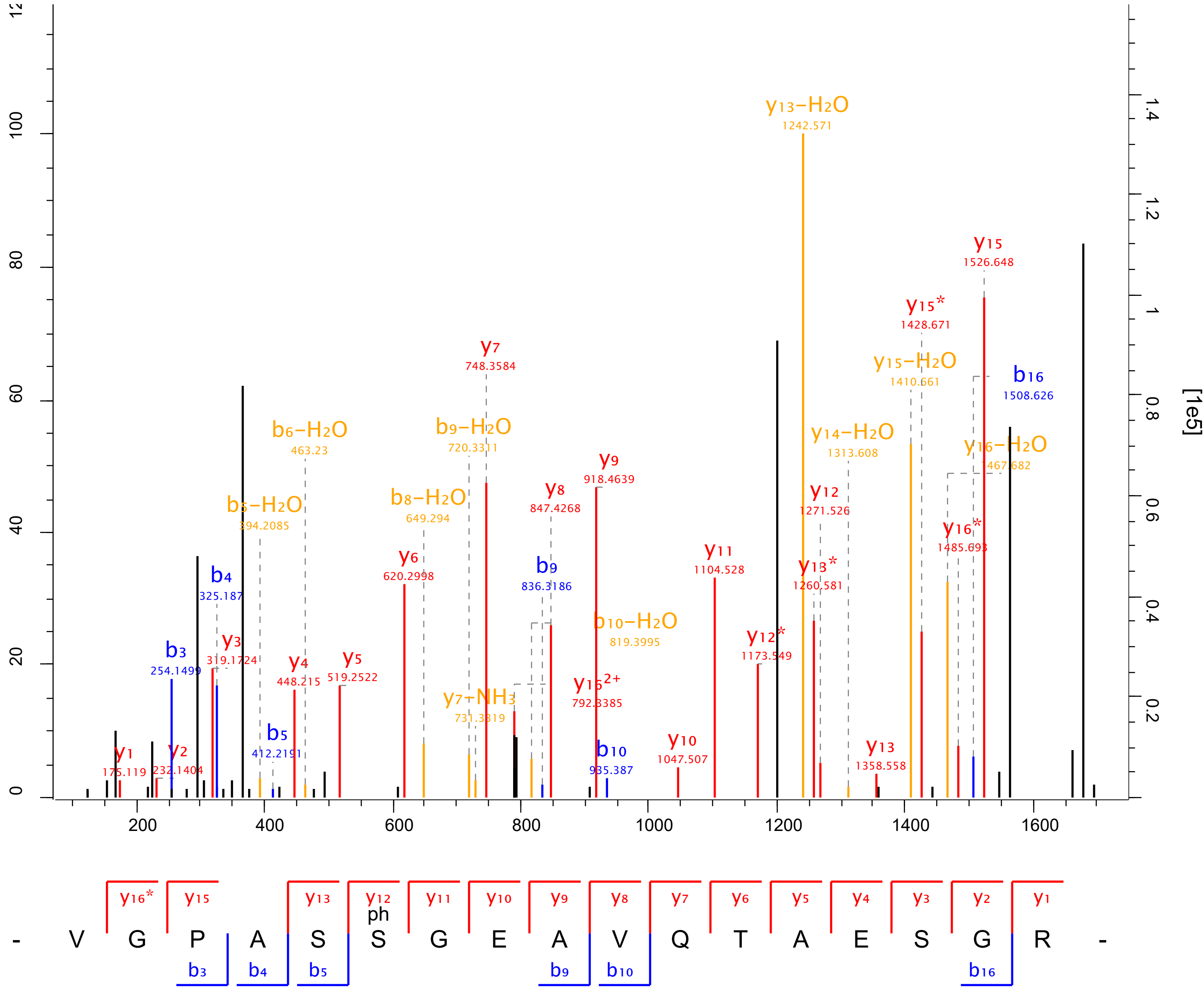

| Raw file | Scan | Method | Score | m/z |
|---|---|---|---|---|
| water_02_1 | 11912 | FTMS; HCD | 62.47 | 491.73 |

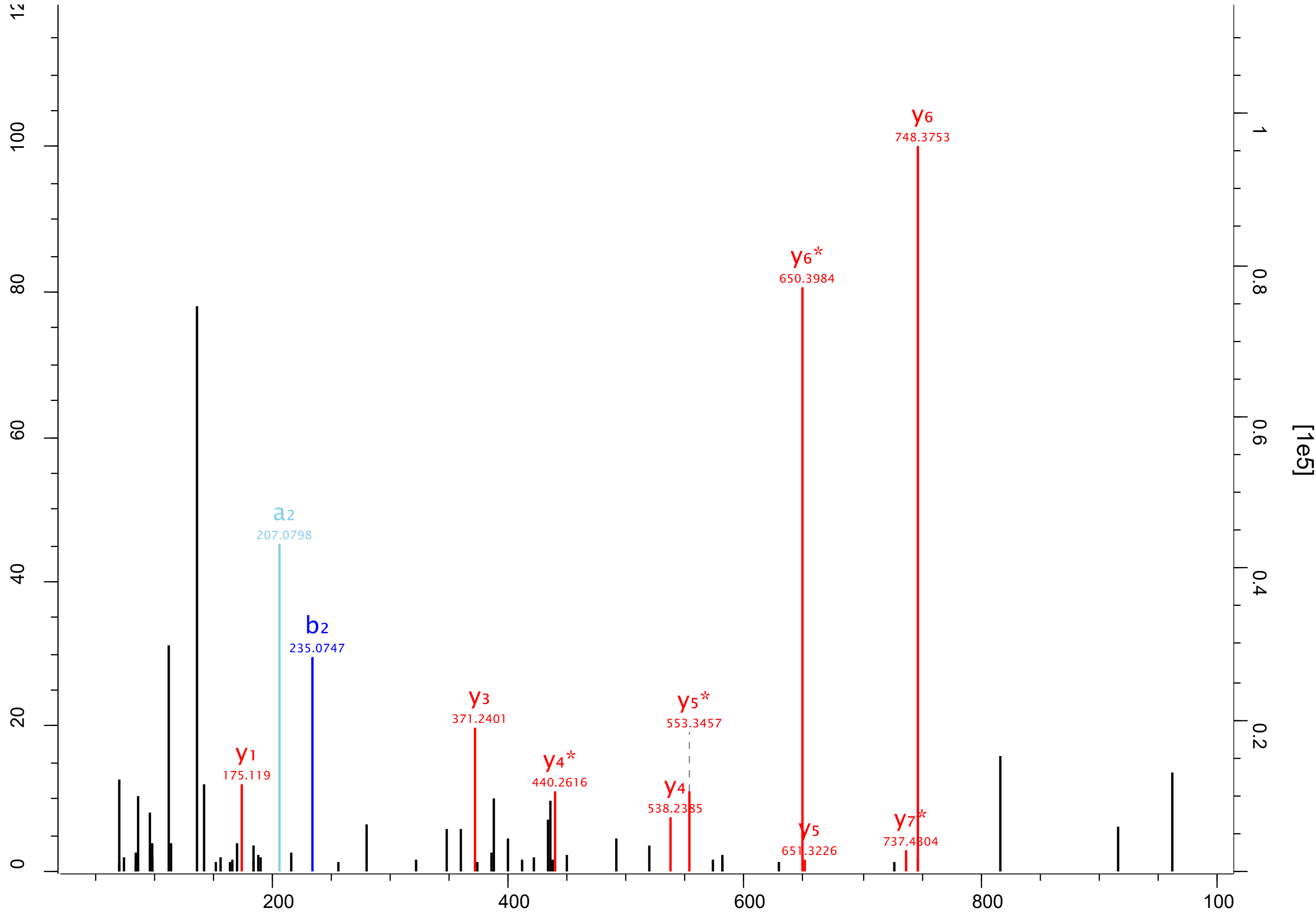


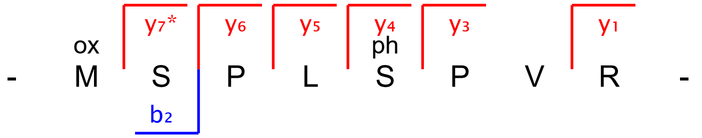

| Raw file | Scan | Method | Score | m/z |
| --- | --- | --- | --- | --- |
| water_02_1 | 11971 | FTMS; HCD | 41.62 | 656.3 |

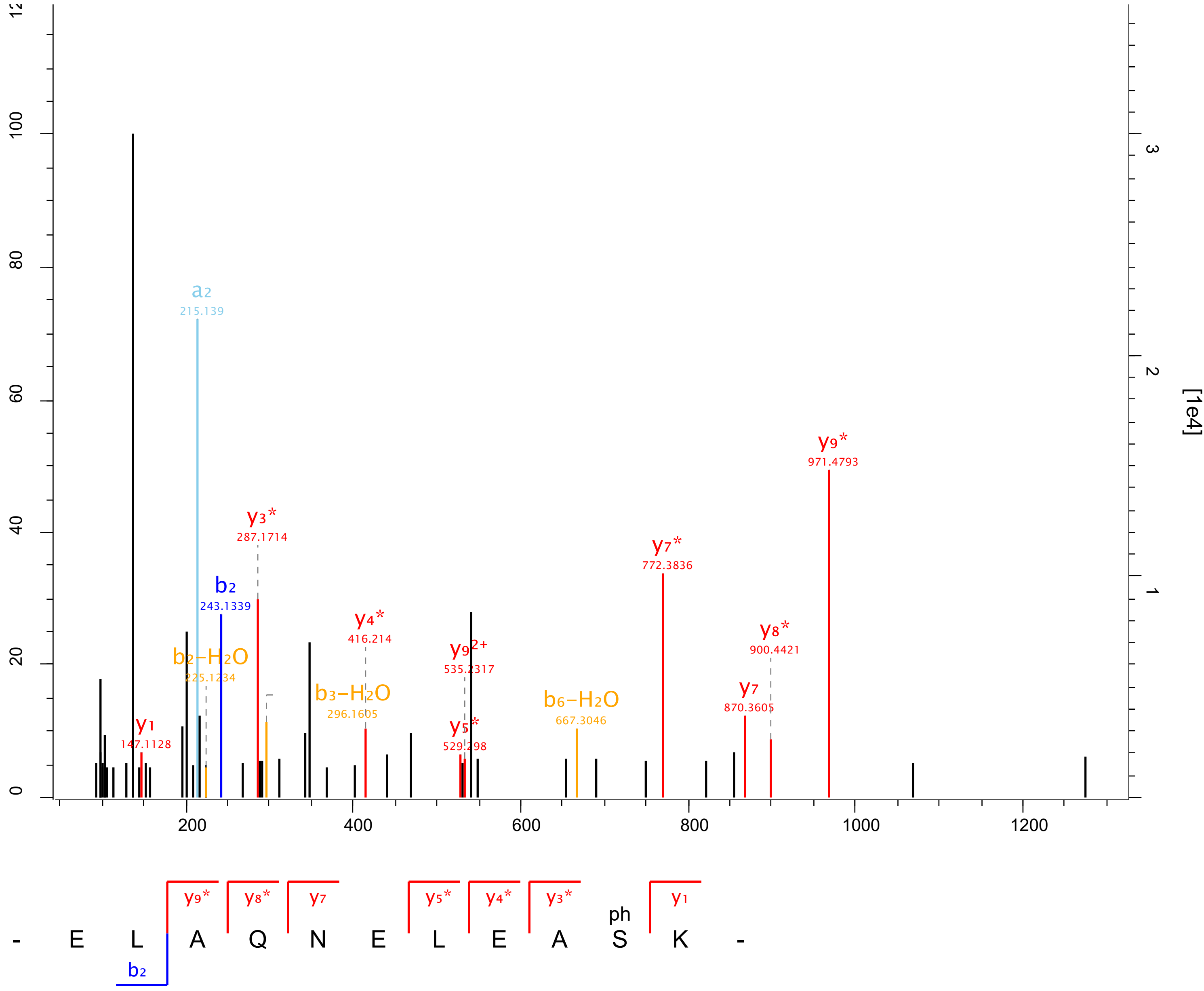

| Raw file | Scan | Method | Score | m/z |
| --- | --- | --- | --- | --- |
| water_02_1 | 11981 | FTMS; HCD | 100.22 | 595.75 |

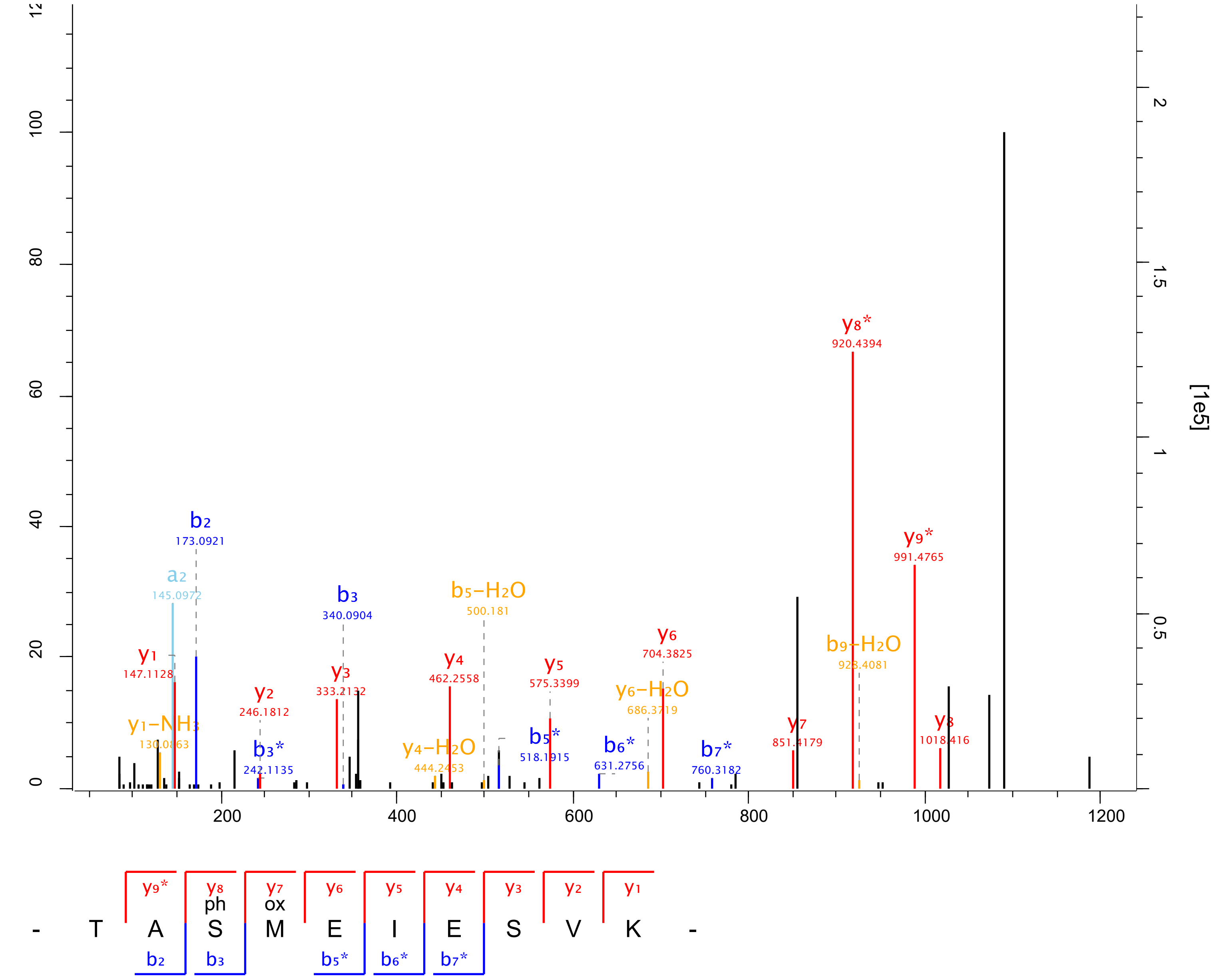

| Raw file | Scan | Method | Score | m/z |
| --- | --- | --- | --- | --- |
| water_02_1 | 12026 | FTMS; HCD | 155.26 | 456.73 |

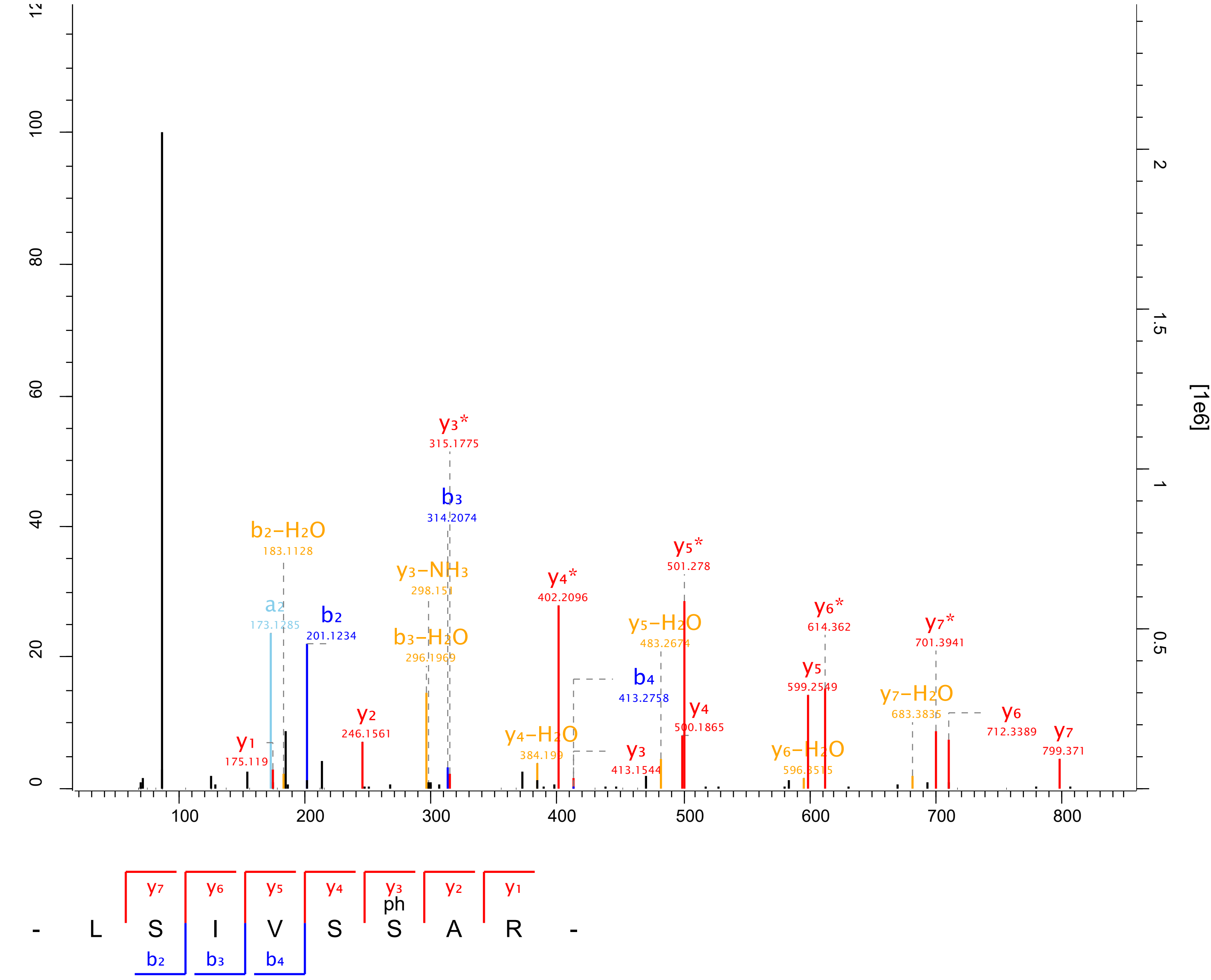

| Raw file | Scan | Method | Score | m/z |
|---|---|---|---|---|
| water_02_1 | 12080 | FTMS; HCD | 50.9 | 632.81 |

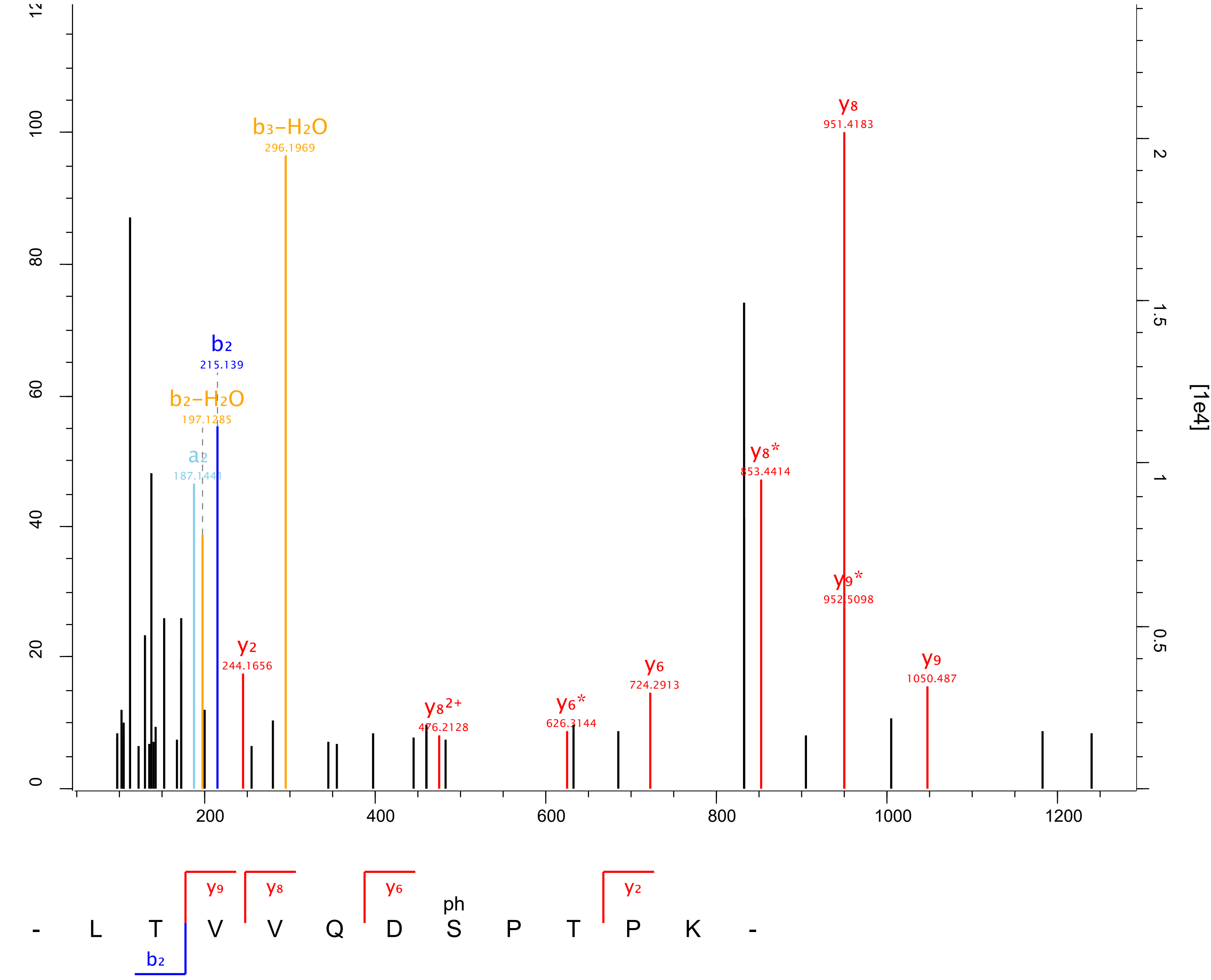

| Raw file | Scan | Method | Score | m/z |
|---|---|---|---|---|
| water_02_1 | 12180 | FTMS; HCD | 78.56 | 424.2 |

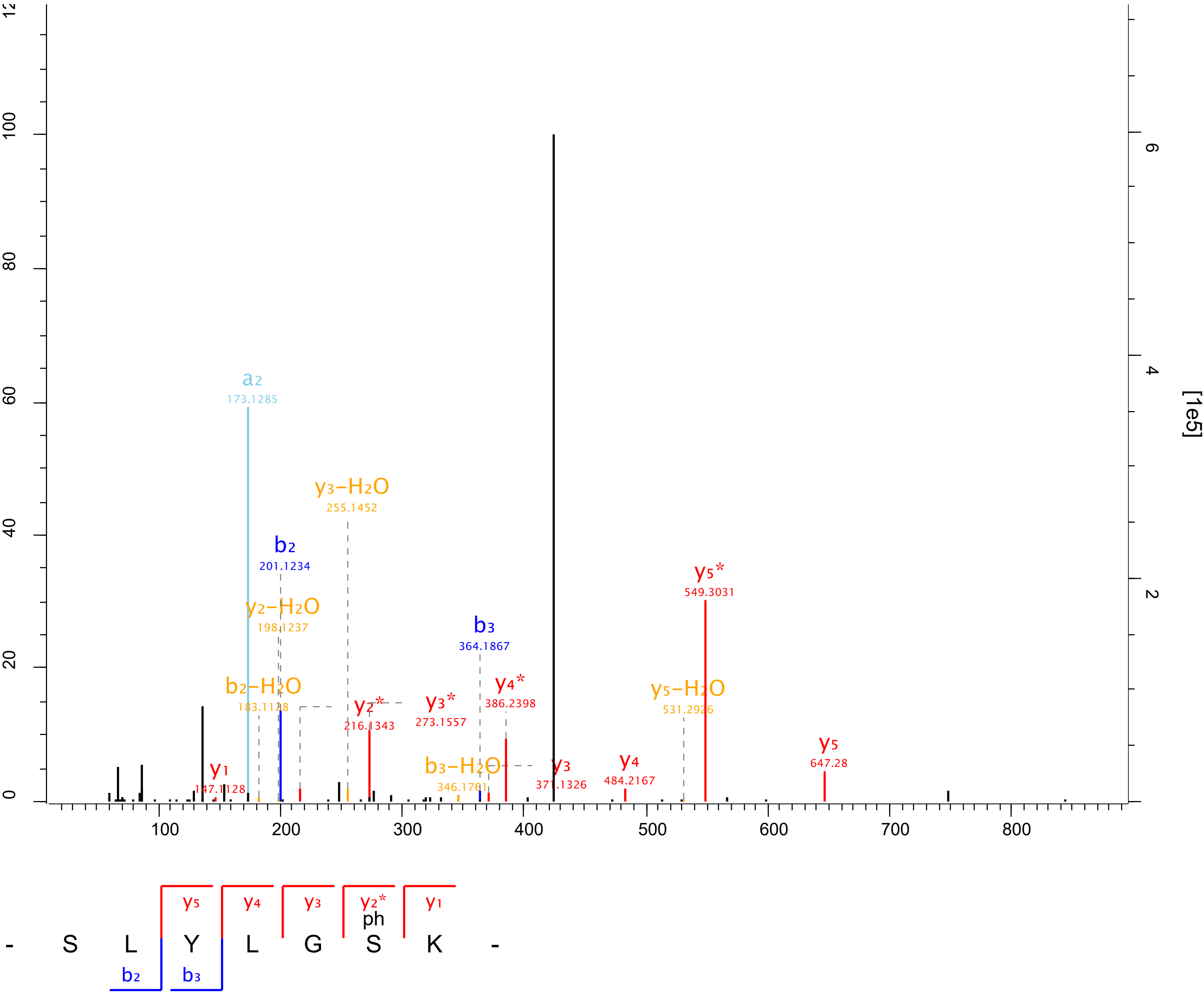

| Raw file | Scan | Method | Score | m/z |
|---|---|---|---|---|
| water_02_1 | 12276 | FTMS; HCD | 69.26 | 800.84 |

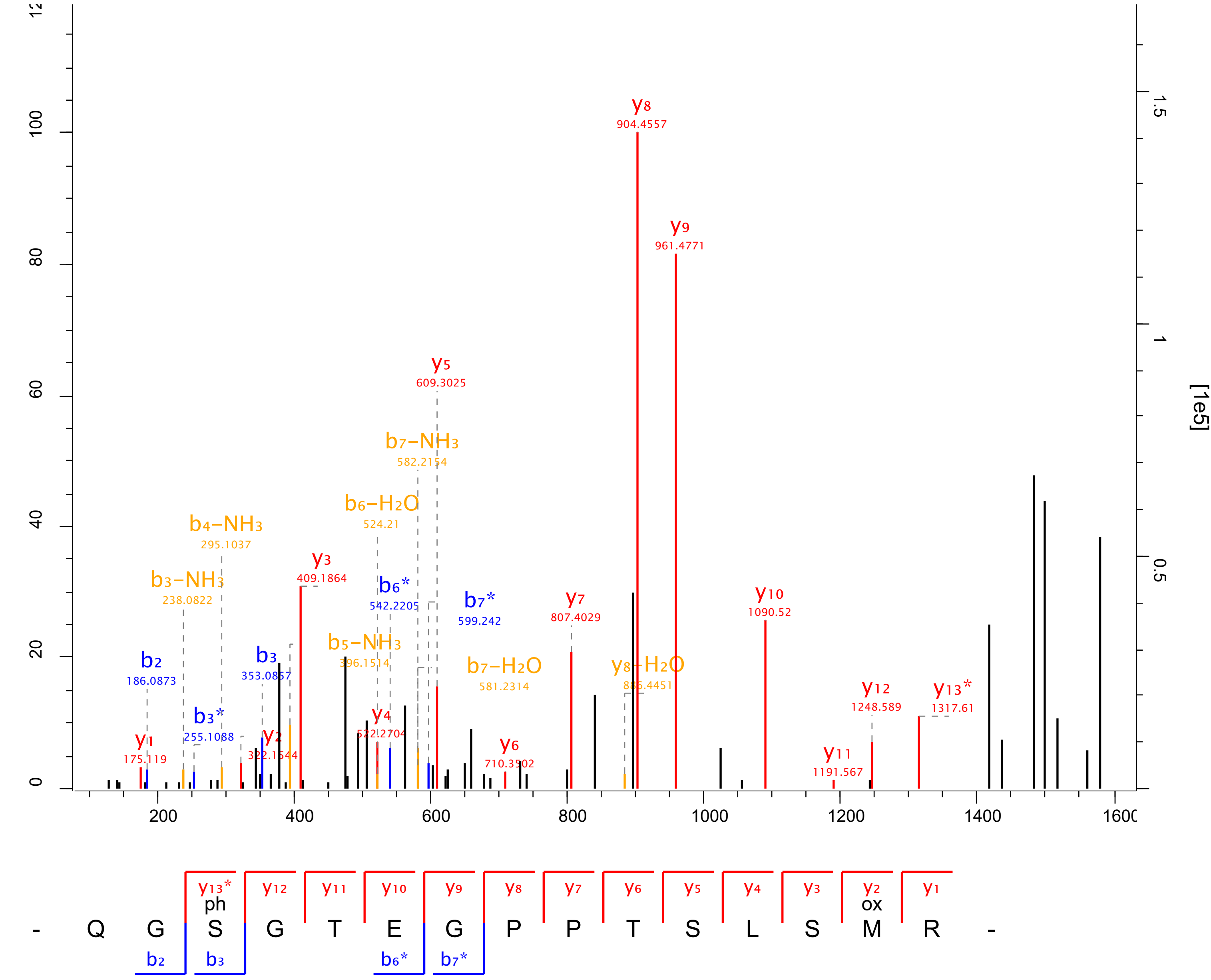

| Raw file | Scan | Method | Score | m/z |
|---|---|---|---|---|
| water_02_1 | 12305 | FTMS; HCD | 119.21 | 619.71 |

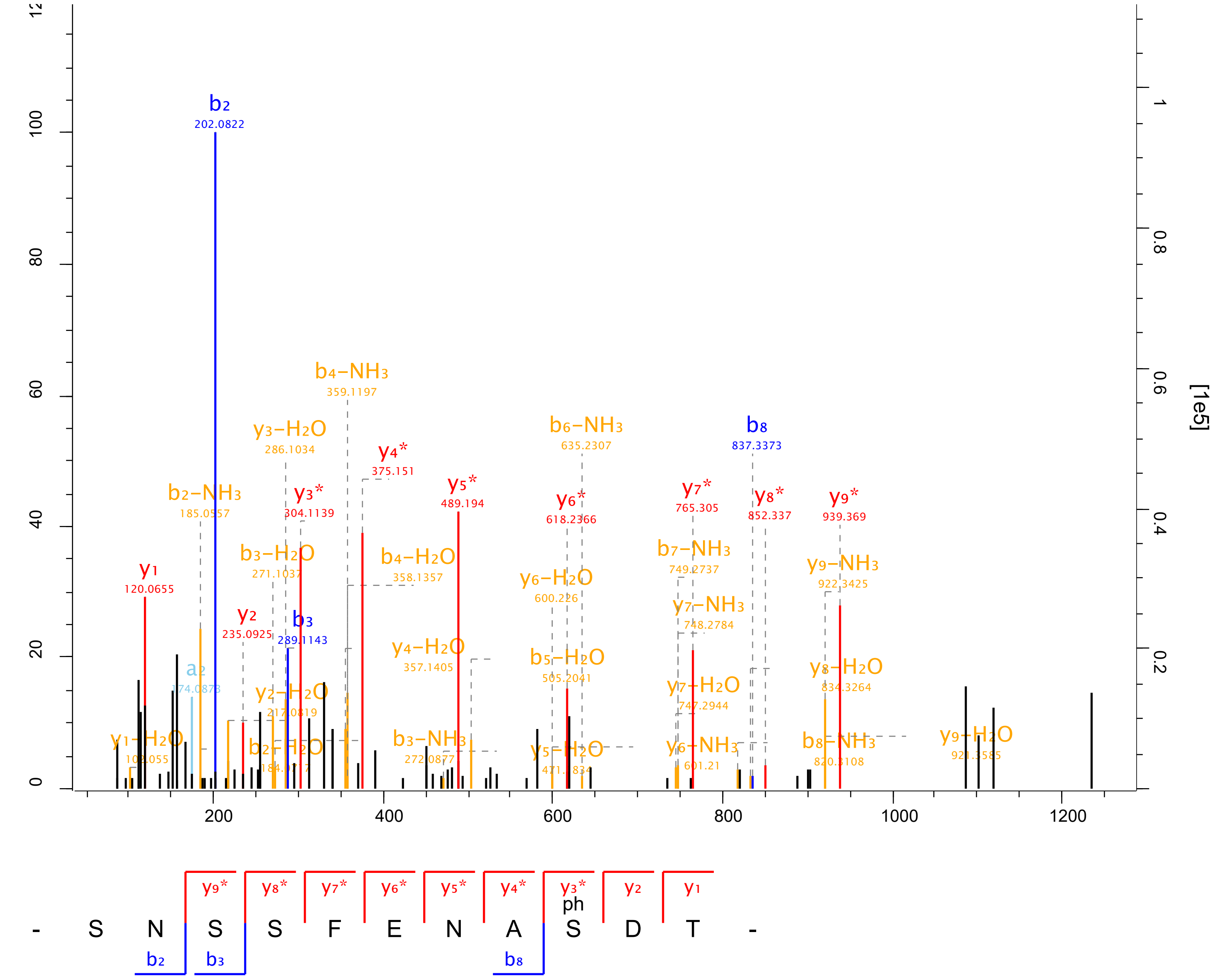

| Raw file | Scan | Method | Score | m/z |
|---|---|---|---|---|
| water_02_1 | 12346 | FTMS; HCD | 111.52 | 492.24 |

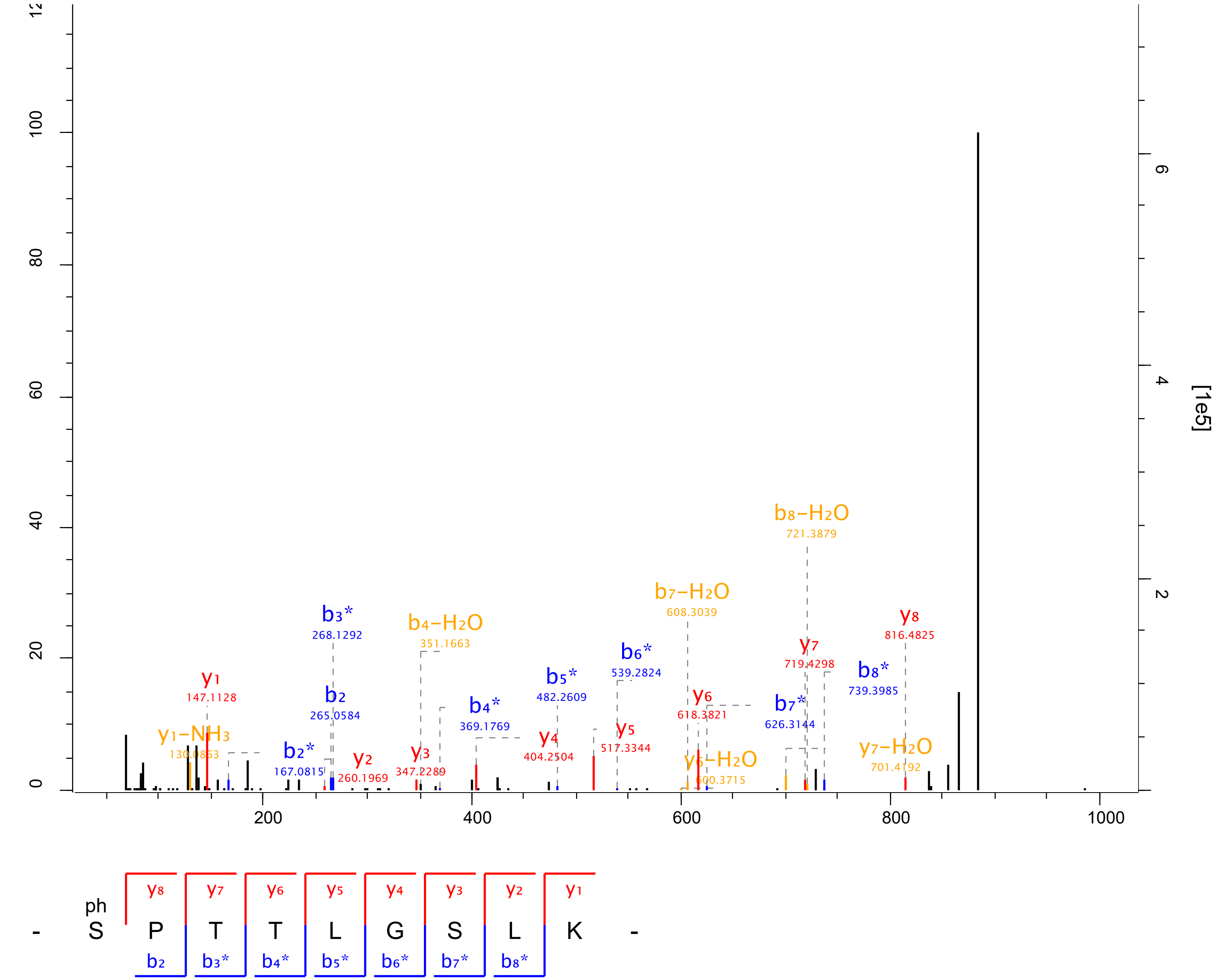

| Raw file | Scan | Method | Score | m/z |
|---|---|---|---|---|
| water_02_1 | 12463 | FTMS; HCD | 52.87 | 718.79 |

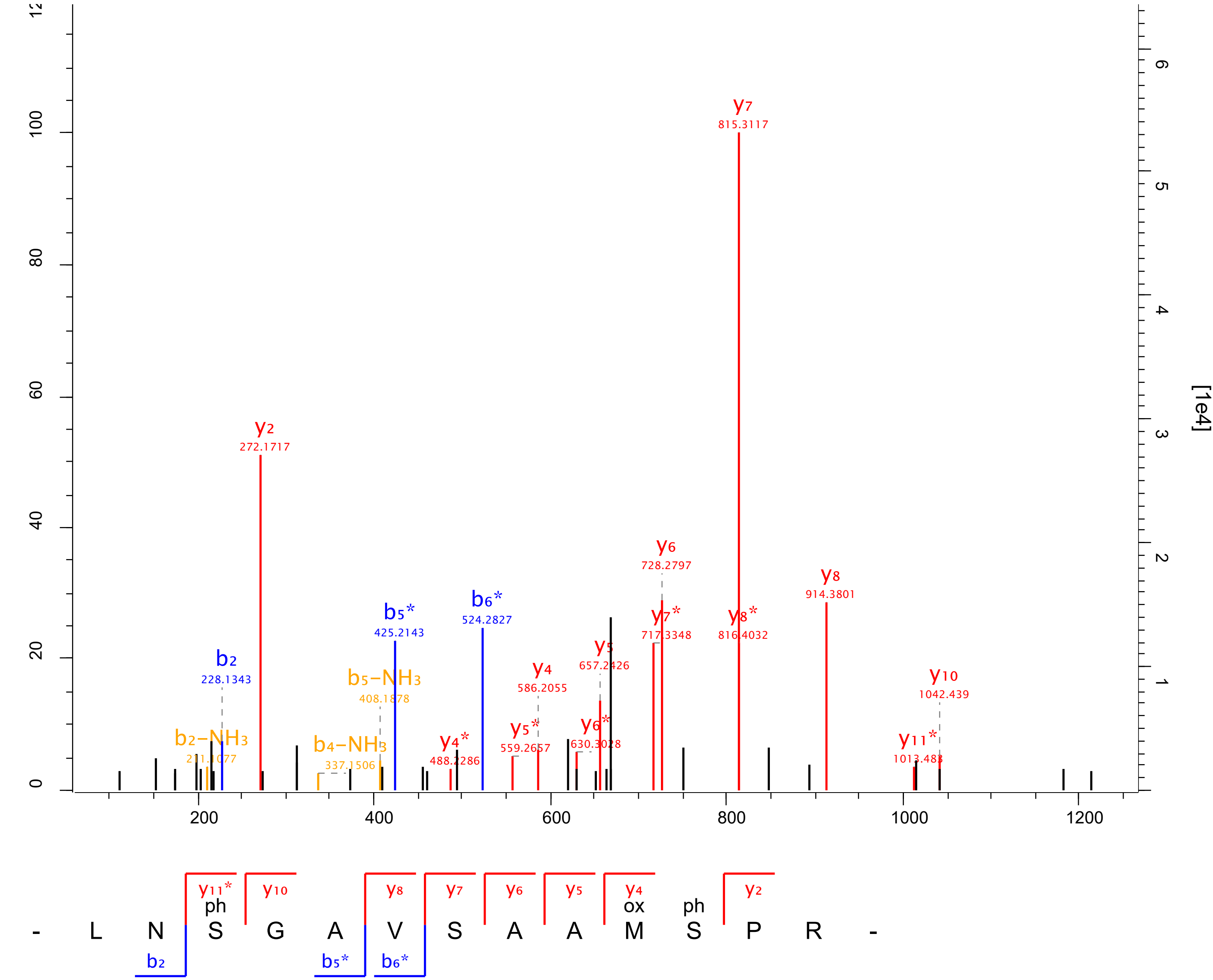

| Raw file | Scan | Method | Score | m/z |
| --- | --- | --- | --- | --- |
| water_02_1 | 12526 | FTMS; HCD | 62.2 | 823.85 |

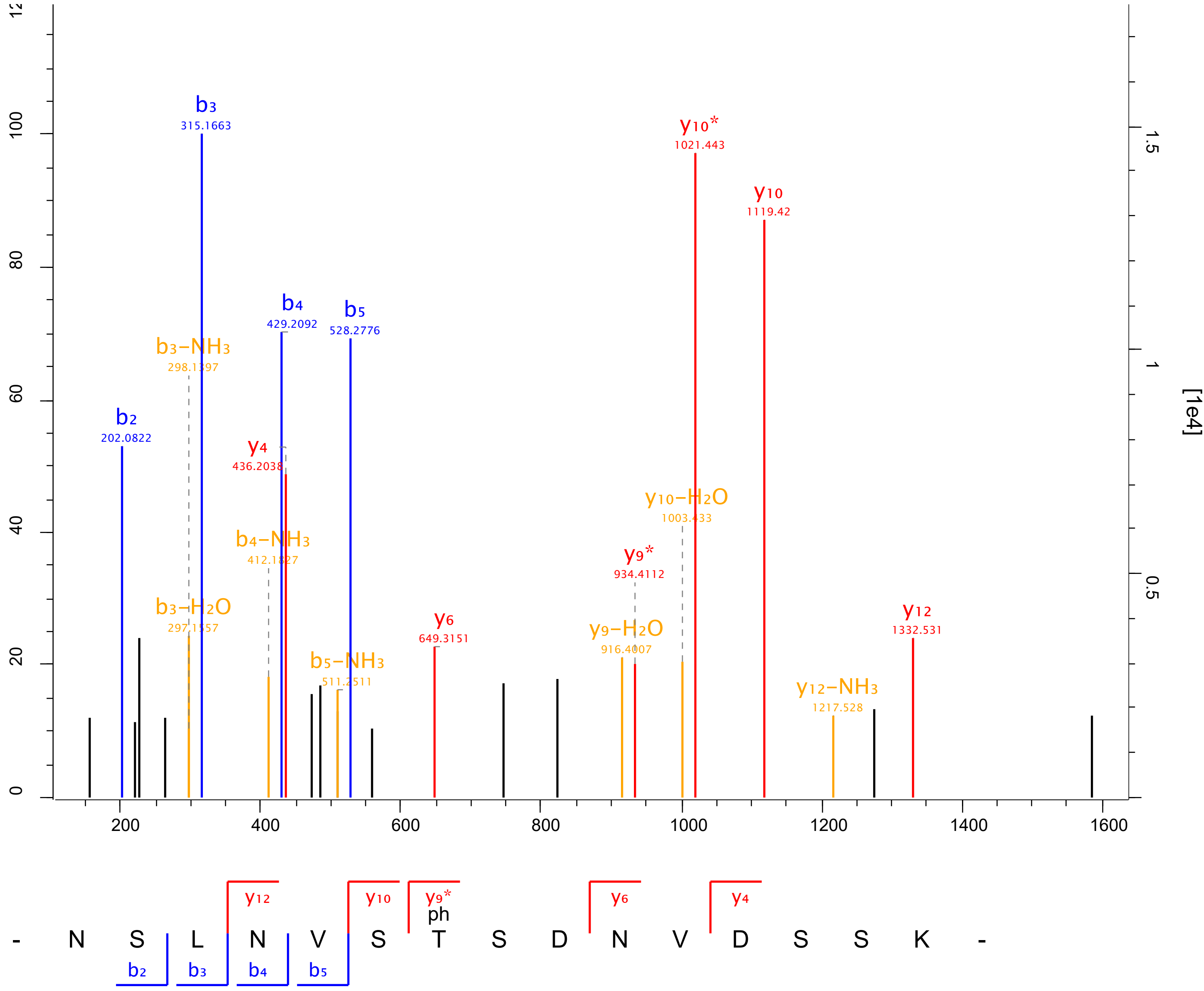

| Raw file | Scan | Method | Score | m/z |
|---|---|---|---|---|
| water_02_1 | 12655 | FTMS; HCD | 47.68 | 630.59 |

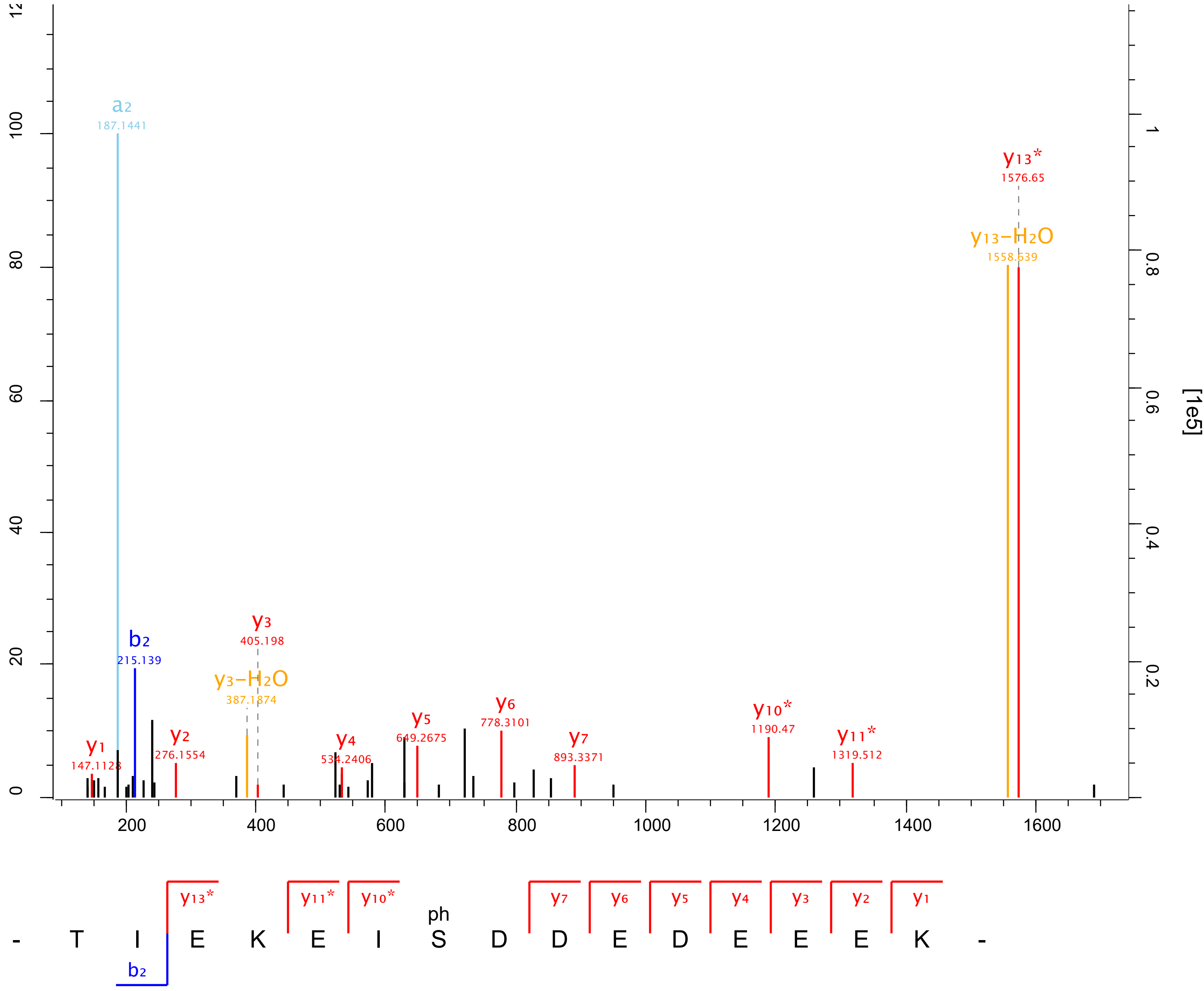

| Raw file | Scan | Method | Score | m/z |
| --- | --- | --- | --- | --- |
| water_02_1 | 12706 | FTMS; HCD | 91.59 | 855.89 |

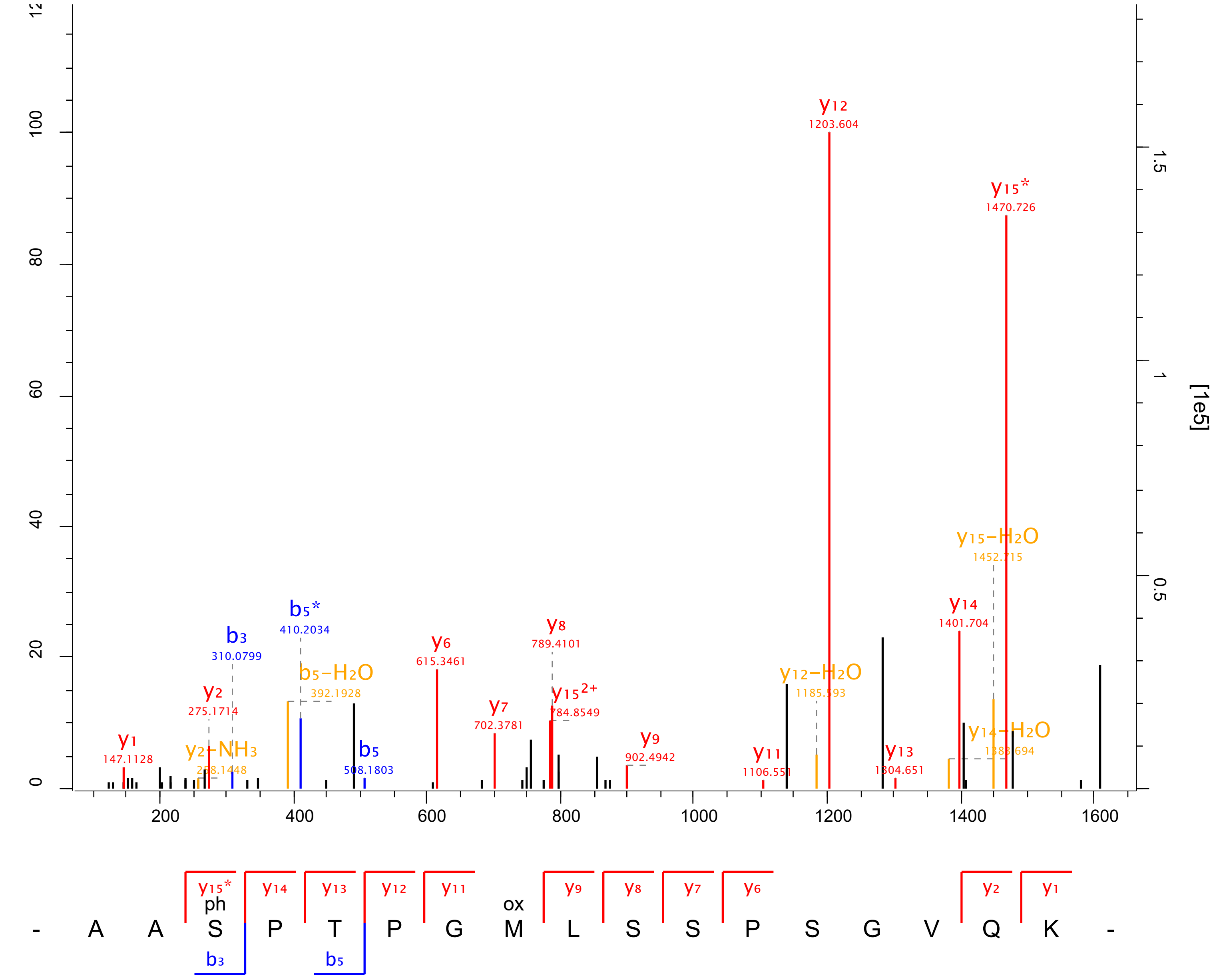

| Raw file | Scan | Method | Score | m/z |
|---|---|---|---|---|
| water_02_1 | 12905 | FTMS; HCD | 89.48 | 657.3 |

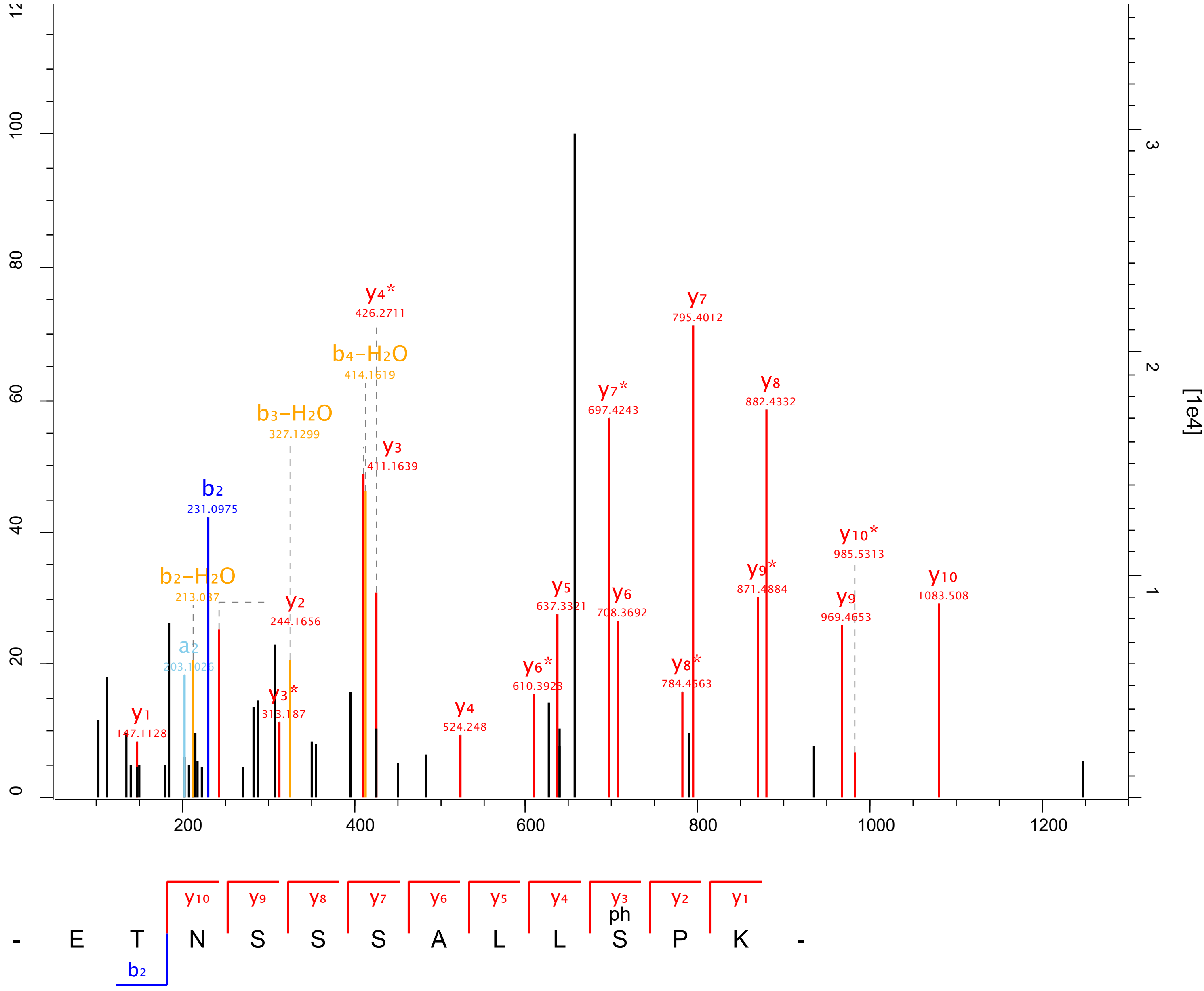

| Raw file | Scan | Method | Score | m/z |
|---|---|---|---|---|
| water_02_1 | 12911 | FTMS; HCD | 60.26 | 528.24 |

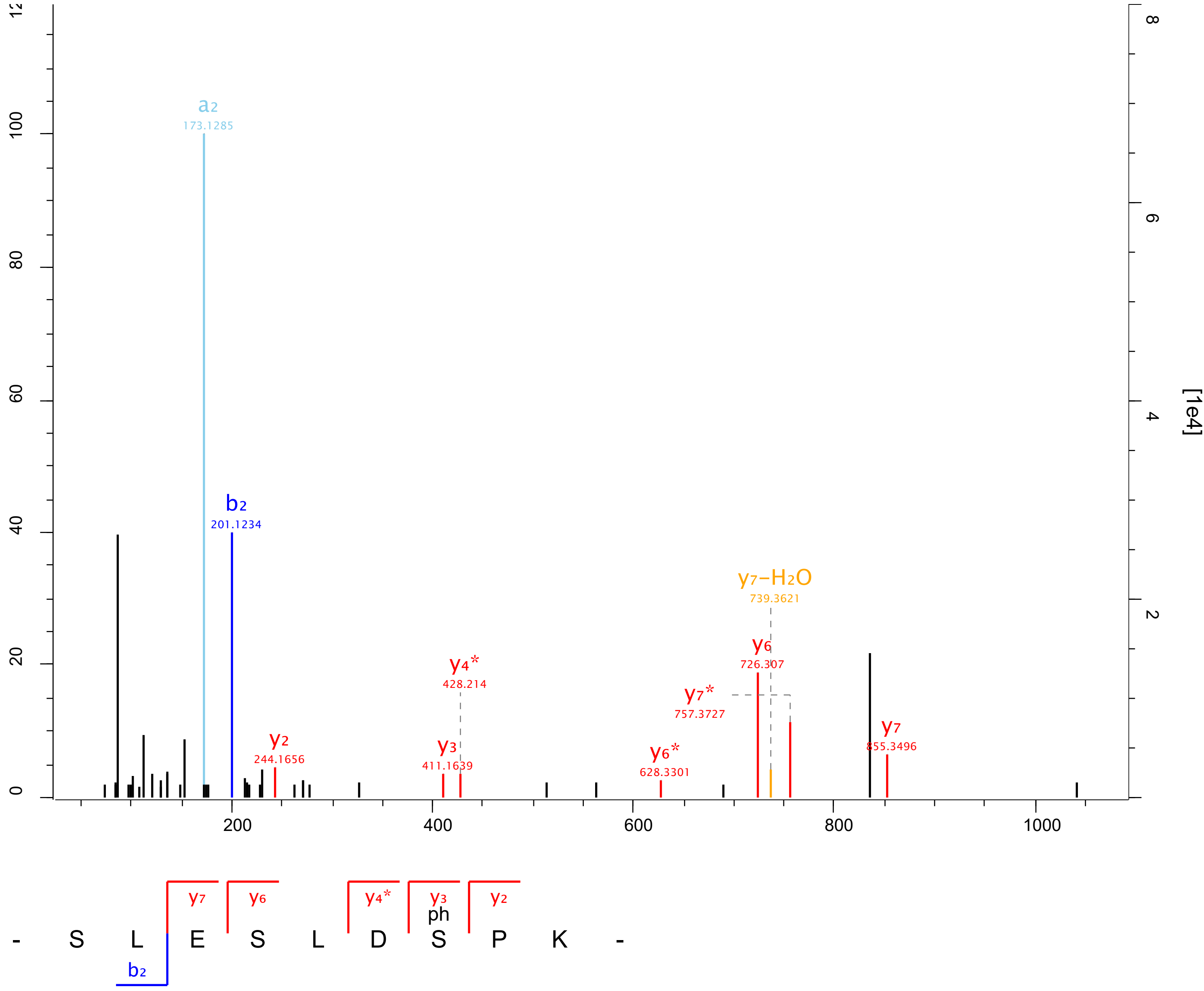

| Raw file | Scan | Method | Score | m/z |
| --- | --- | --- | --- | --- |
| water_02_1 | 12948 | FTMS; HCD | 67.38 | 573.73 |

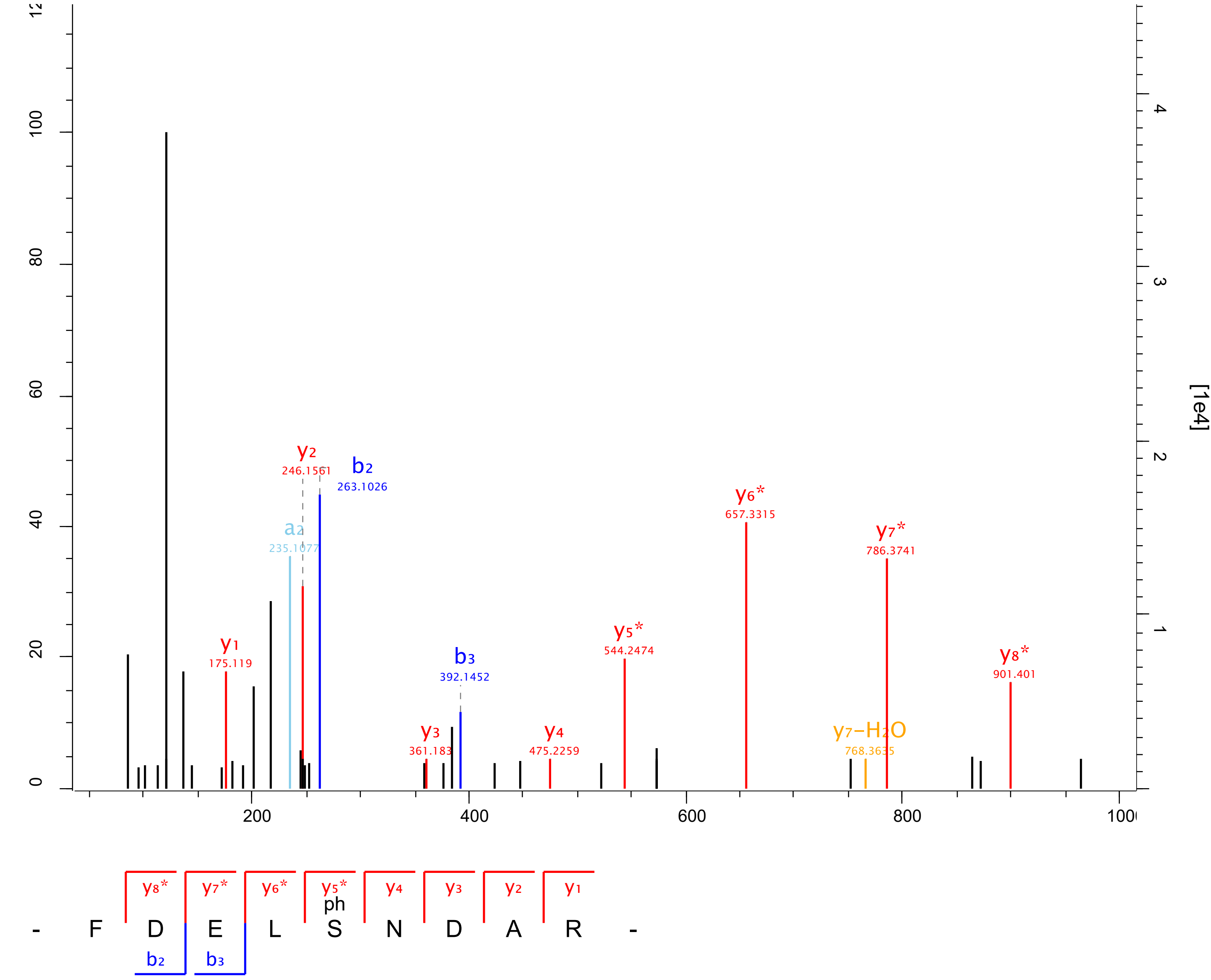

| Raw file | Scan | Method | Score | m/z |
|---|---|---|---|---|
| water_02_1 | 13018 | FTMS; HCD | 135.65 | 590.21 |

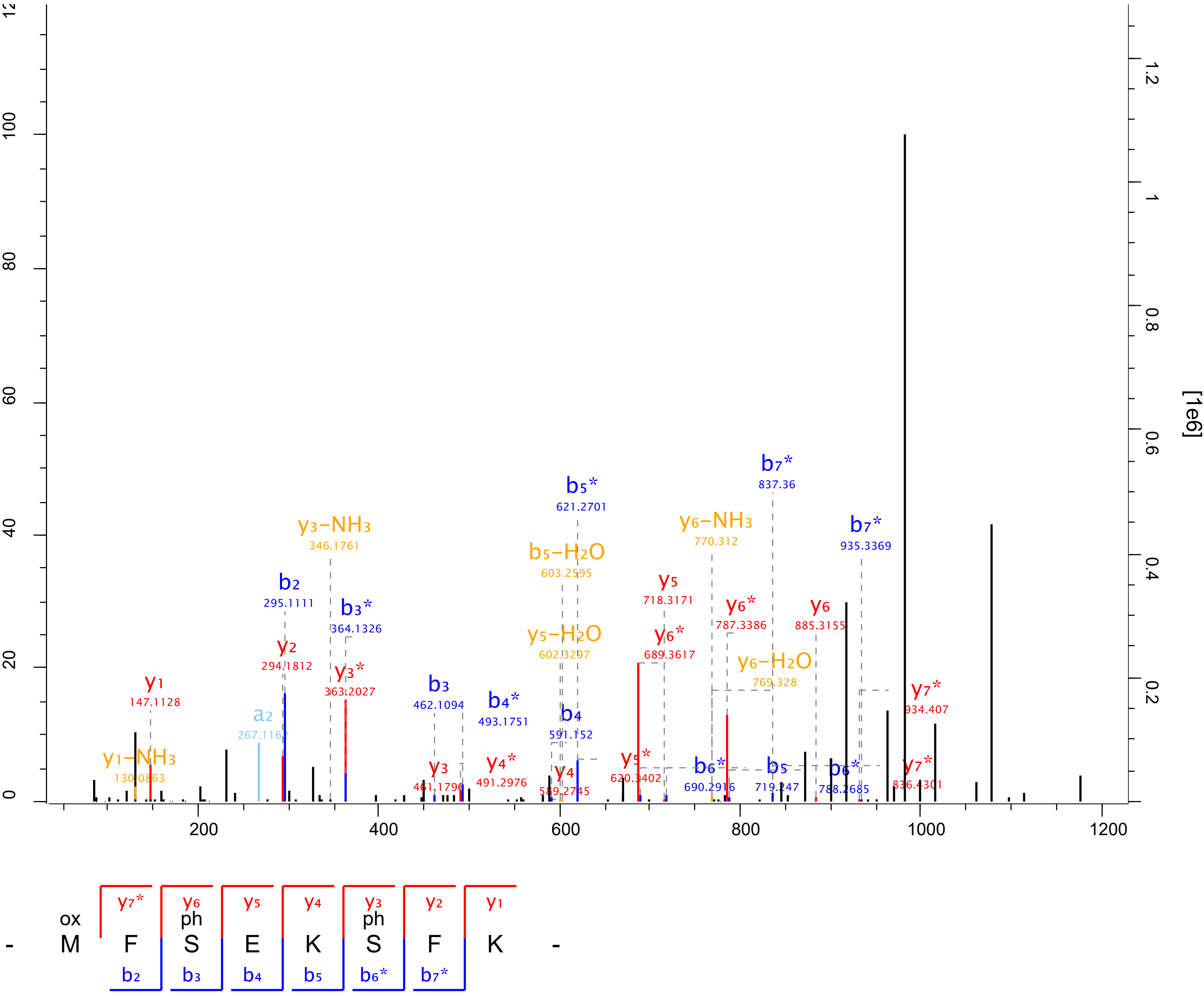

| Raw file | Scan | Method | Score | m/z |
| --- | --- | --- | --- | --- |
| water_02_1 | 13031 | FTMS; HCD | 52.25 | 545.23 |

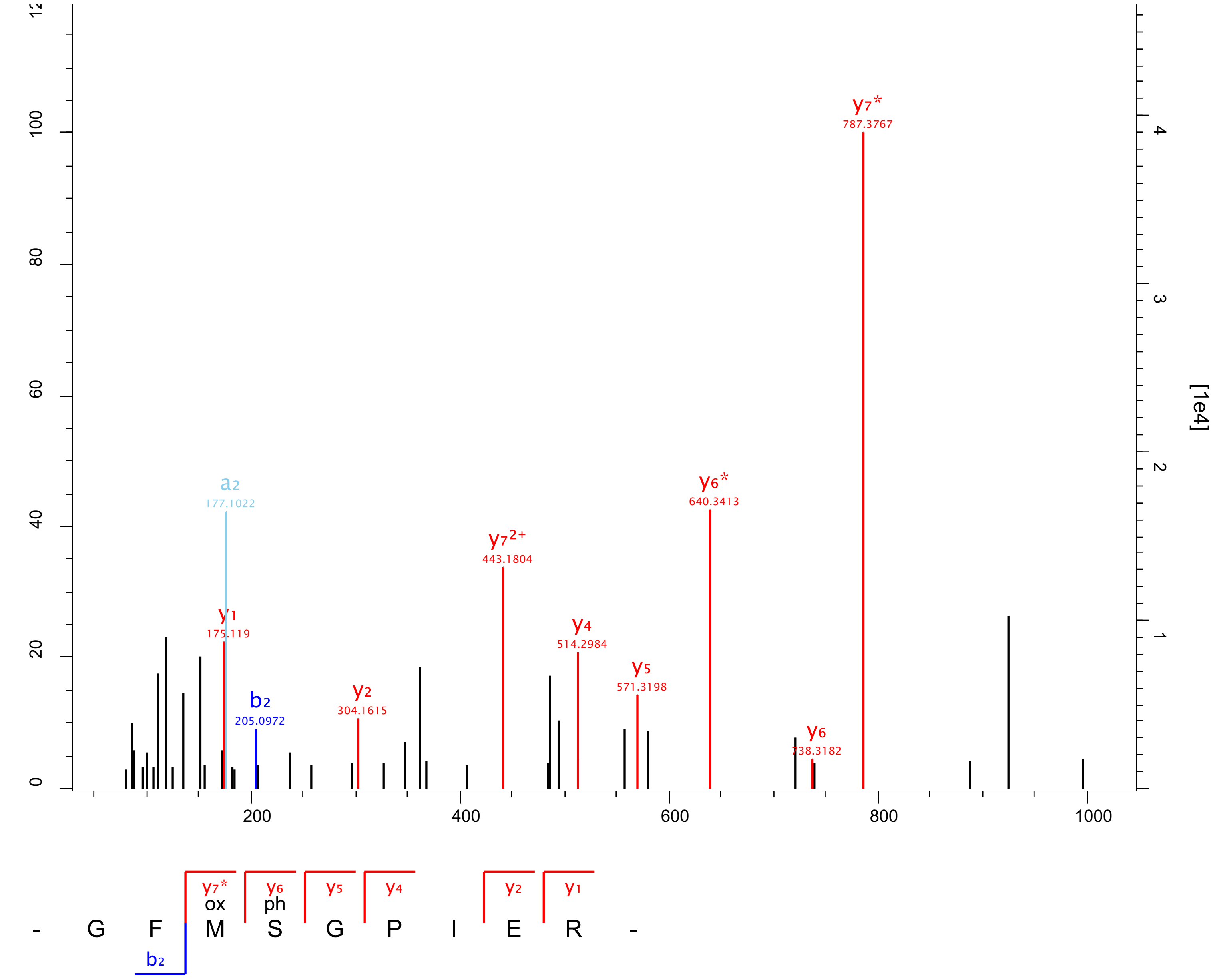

| Raw file | Scan | Method | Score | m/z |
| --- | --- | --- | --- | --- |
| water_02_1 | 13034 | FTMS; HCD | 104.76 | 539.27 |

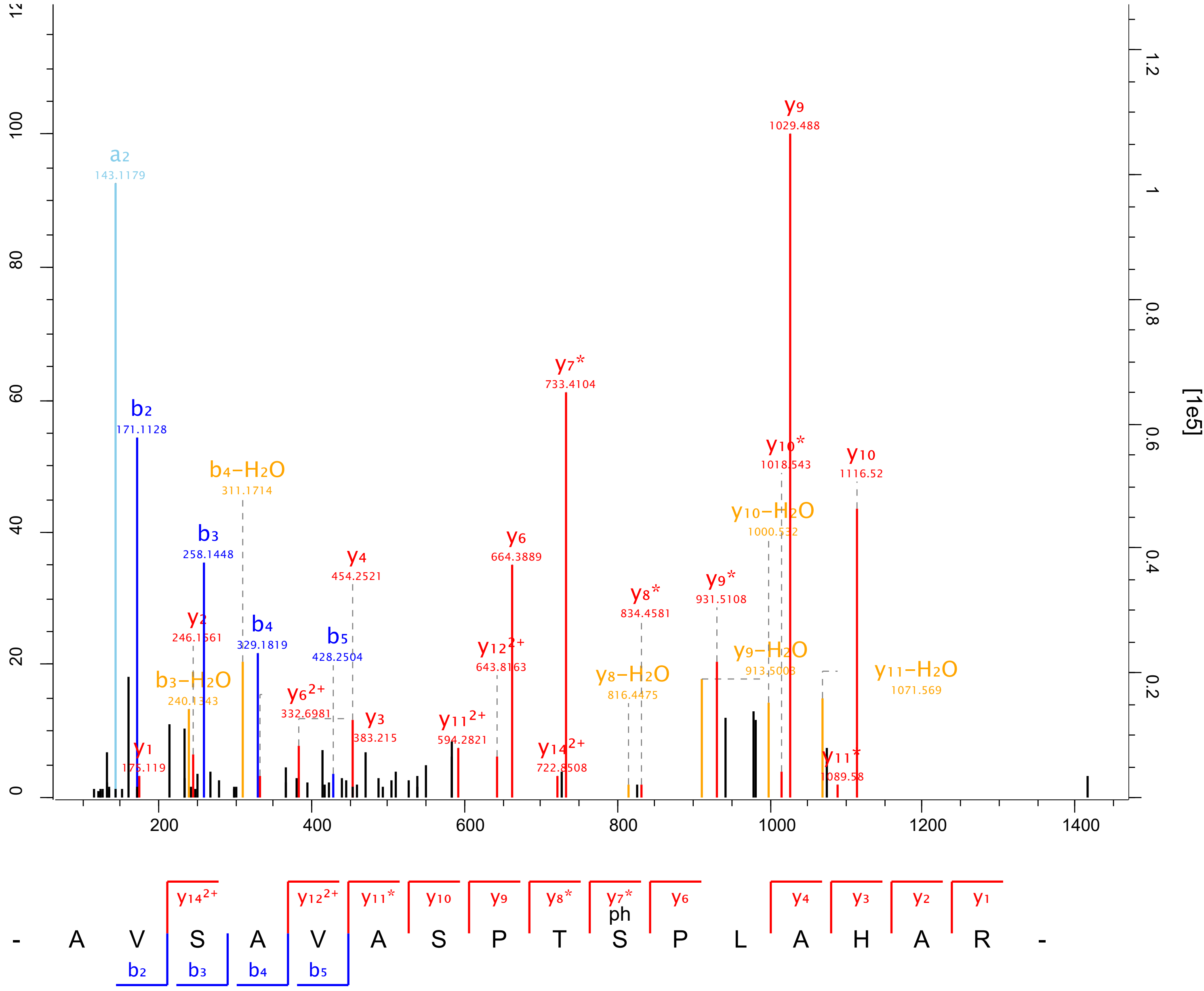

| Raw file | Scan | Method | Score | m/z |
|---|---|---|---|---|
| water_02_1 | 13088 | FTMS; HCD | 72.9 | 645.78 |

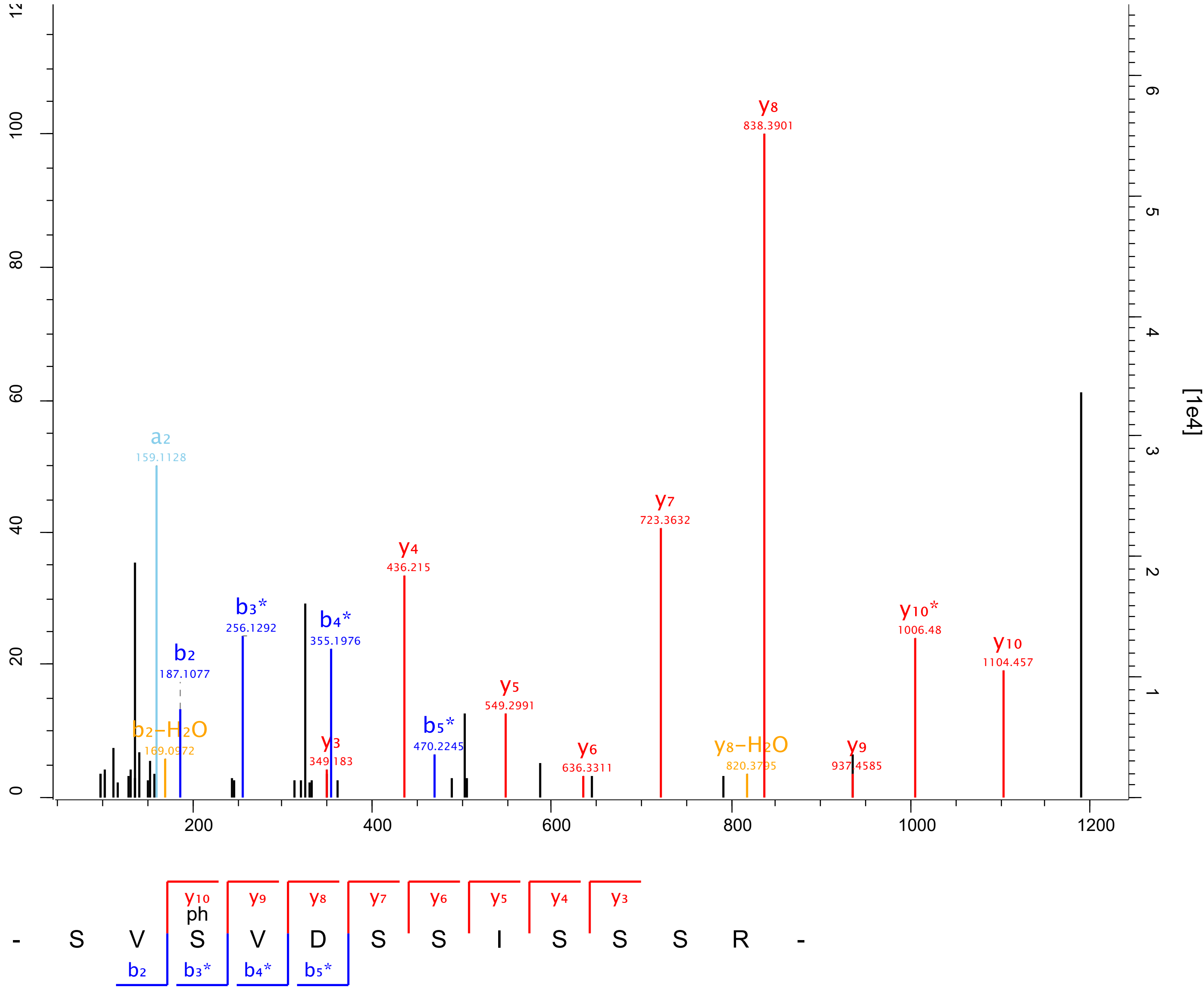

| Raw file | Scan | Method | Score | m/z |
|---|---|---|---|---|
| water_02_1 | 13221 | FTMS; HCD | 69.98 | 758.83 |

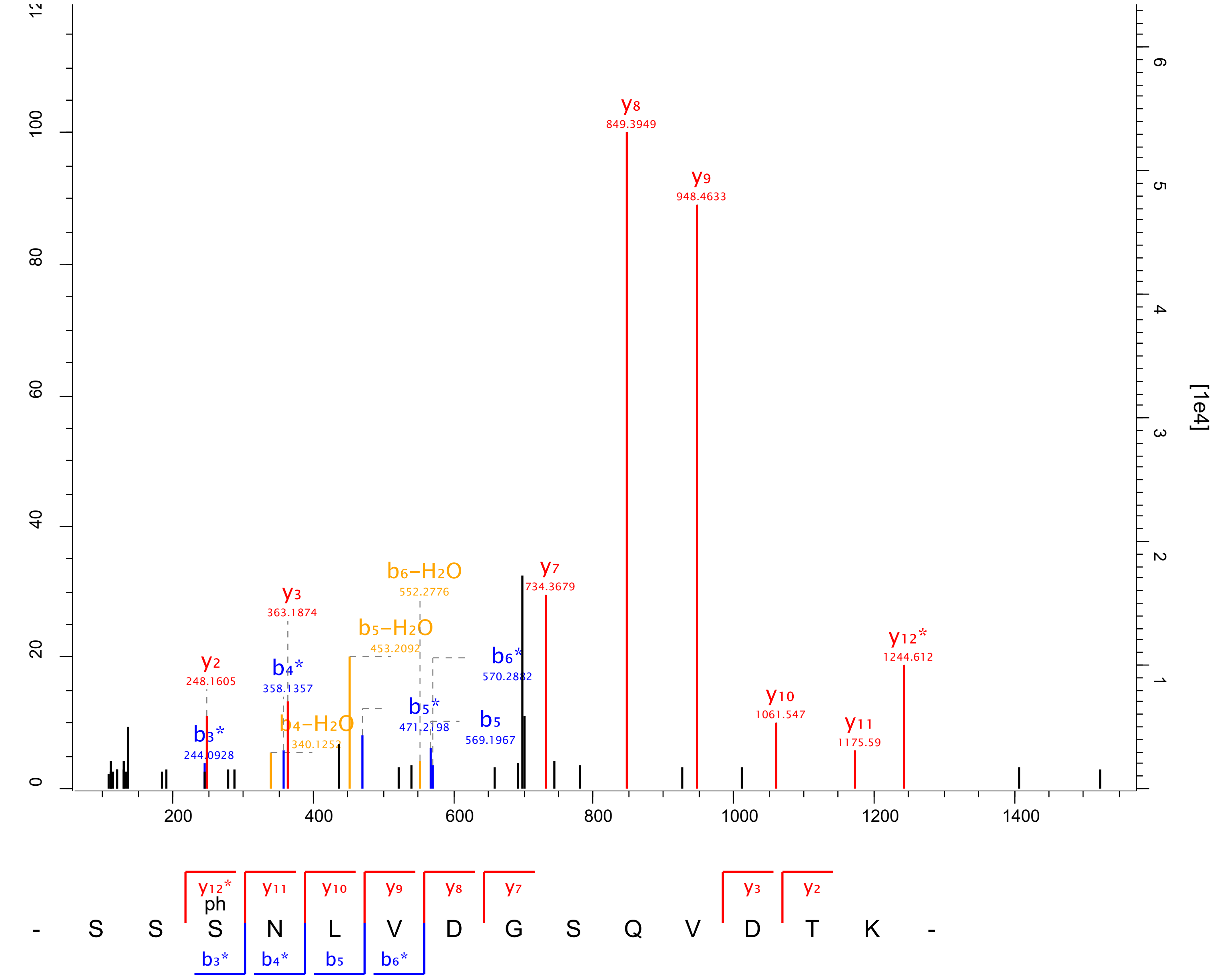

| Raw file | Scan | Method | Score | m/z |
|---|---|---|---|---|
| water_02_1 | 13247 | FTMS; HCD | 49.45 | 573.78 |

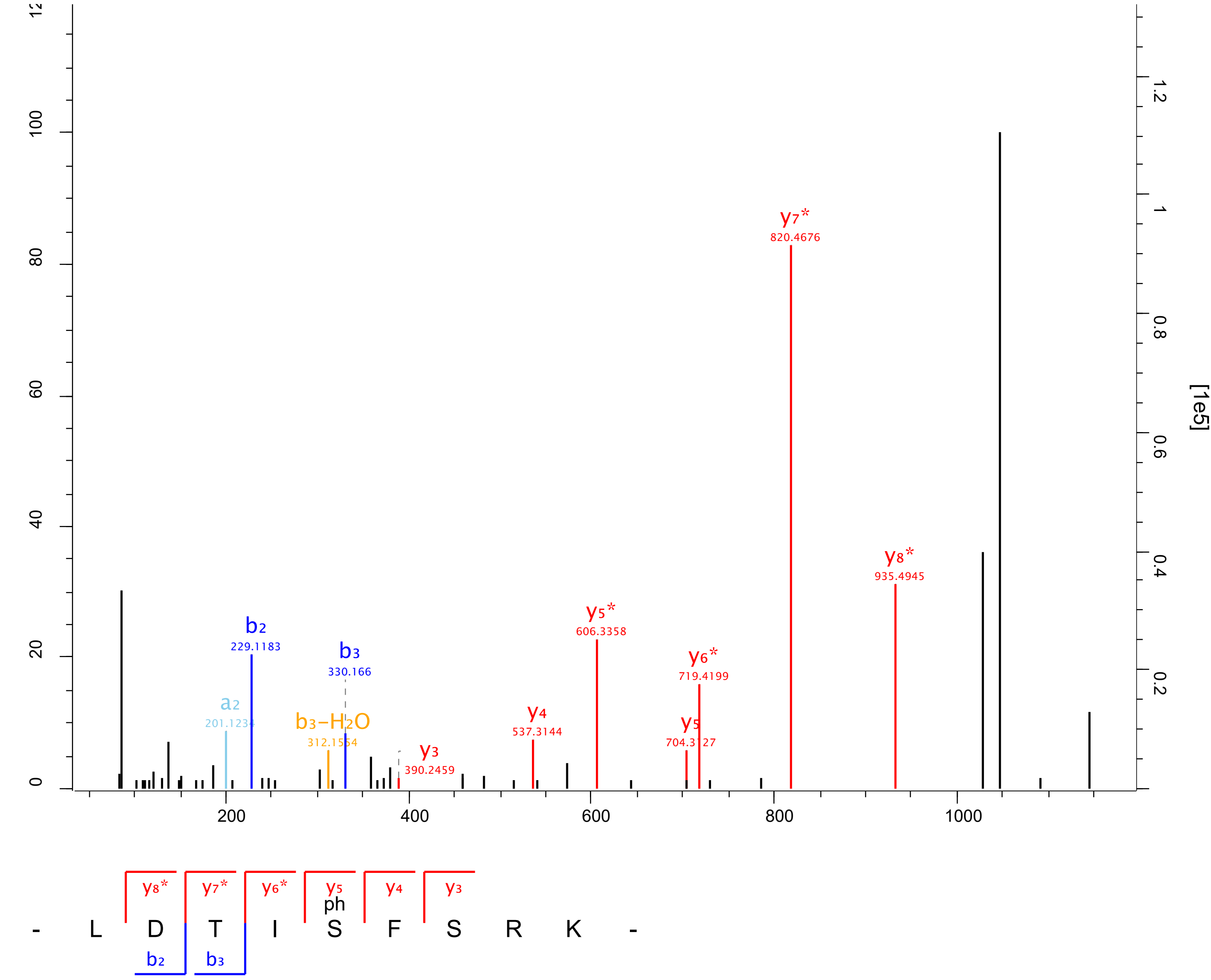

| Raw file | Scan | Method | Score | m/z |
| --- | --- | --- | --- | --- |
| water_02_1 | 13266 | FTMS; HCD | 172.82 | 693.31 |

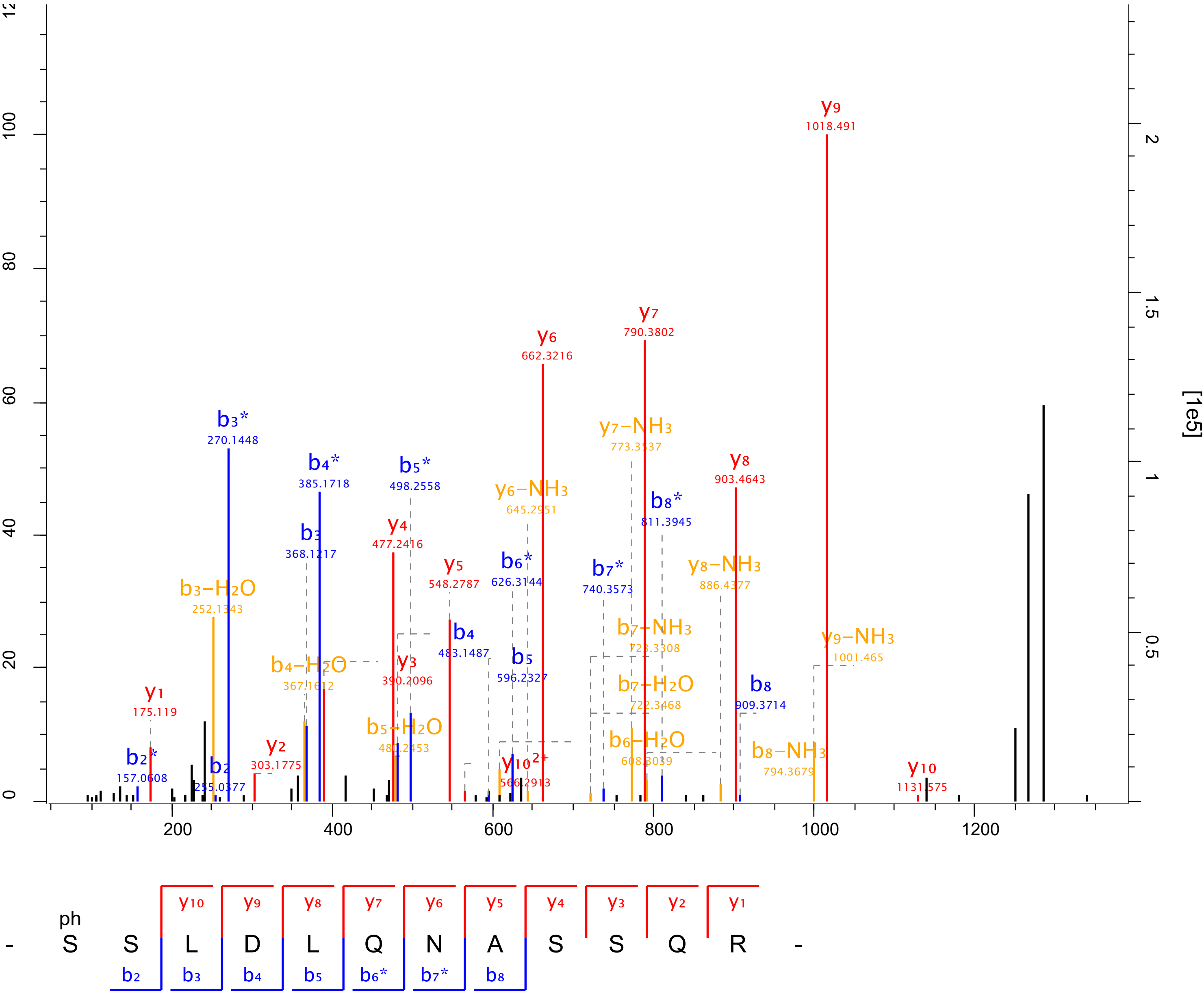

| Raw file | Scan | Method | Score | m/z |
| --- | --- | --- | --- | --- |
| water_02_1 | 13345 | FTMS; HCD | 50.83 | 736.36 |

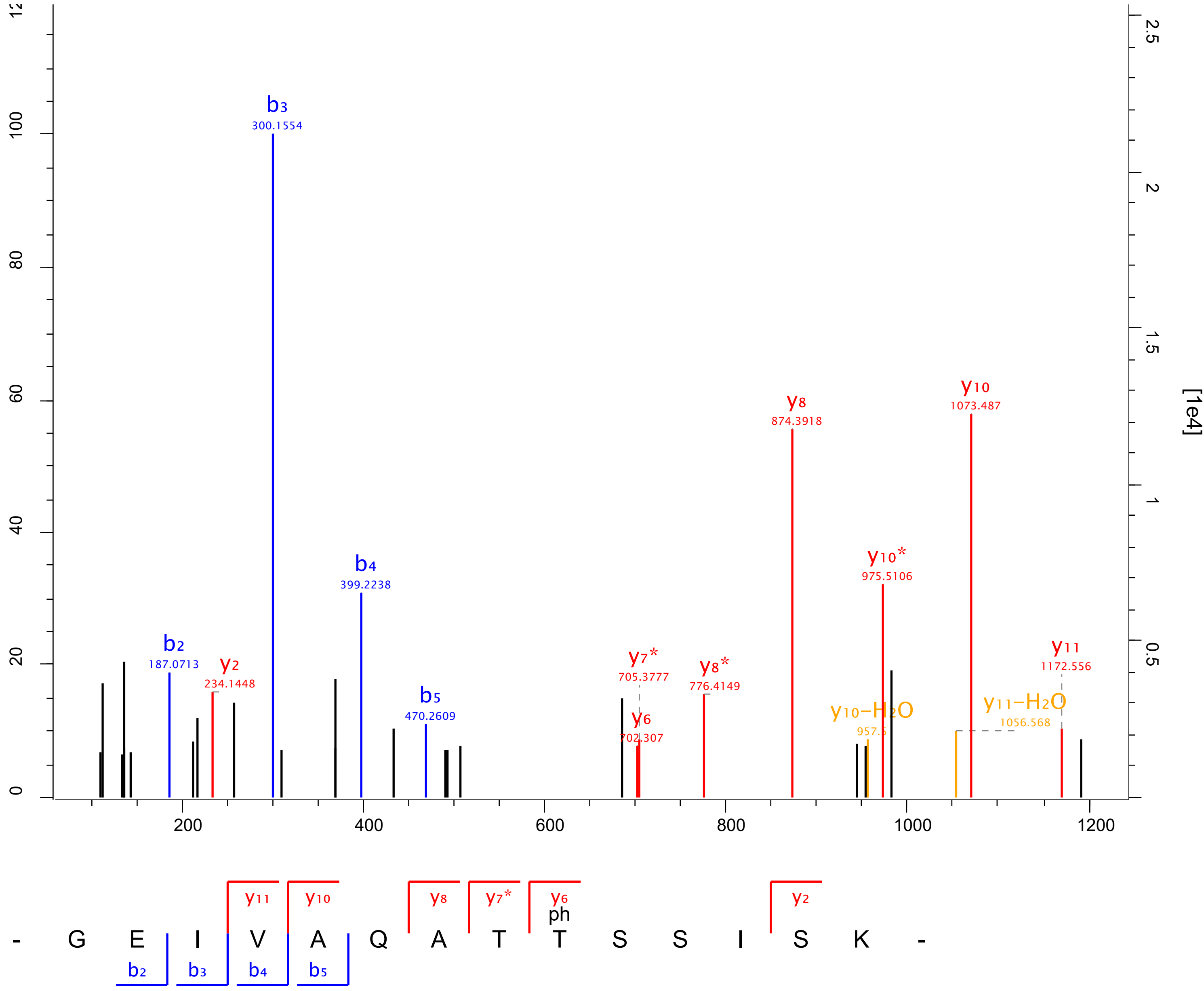

| Raw file | Scan | Method | Score | m/z |
| --- | --- | --- | --- | --- |
| water_02_1 | 13397 | FTMS; HCD | 329.19 | 946.36 |

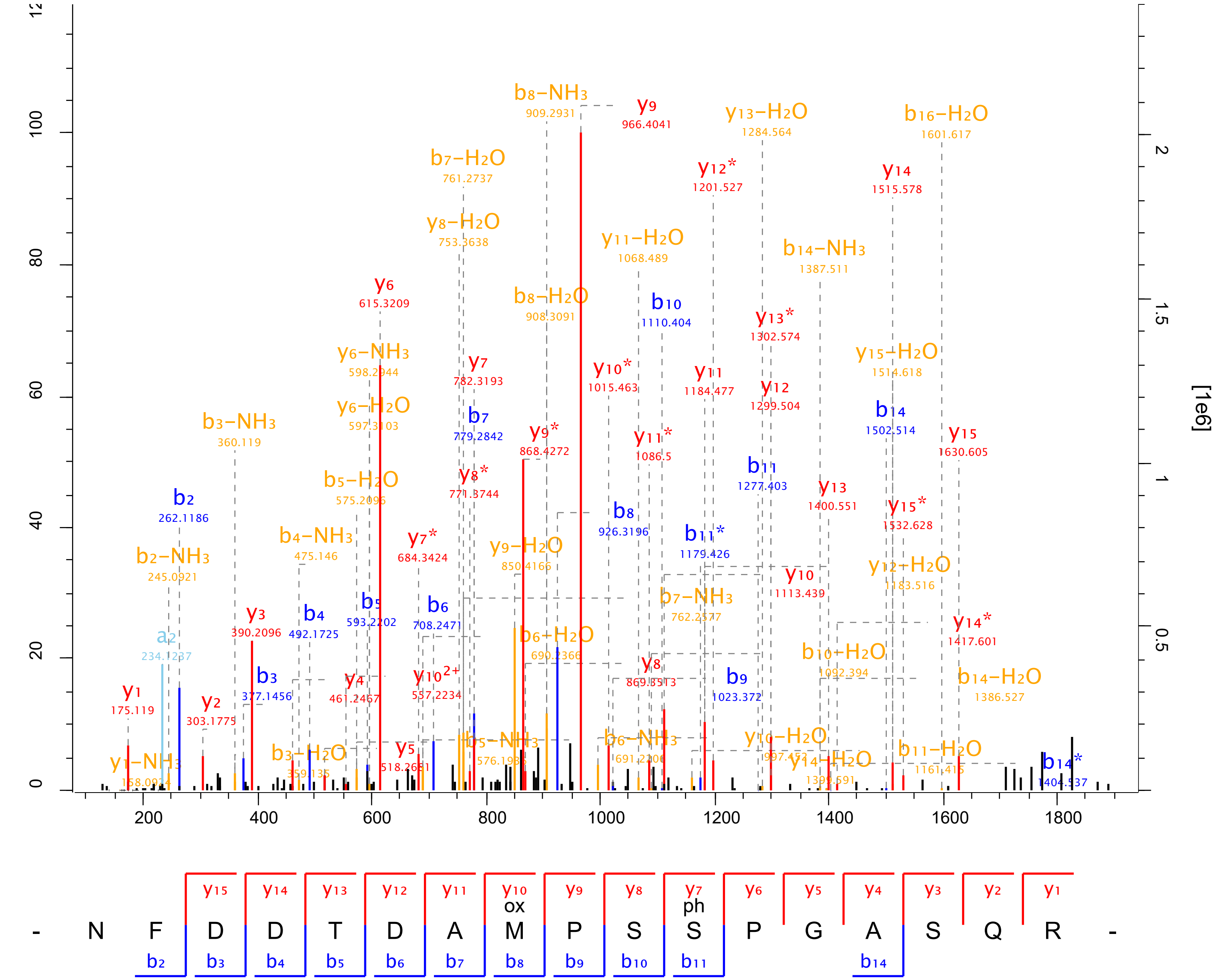

| Raw file | Scan | Method | Score | m/z |
| --- | --- | --- | --- | --- |
| water_02_1 | 13400 | FTMS; HCD | 57.29 | 723.82 |

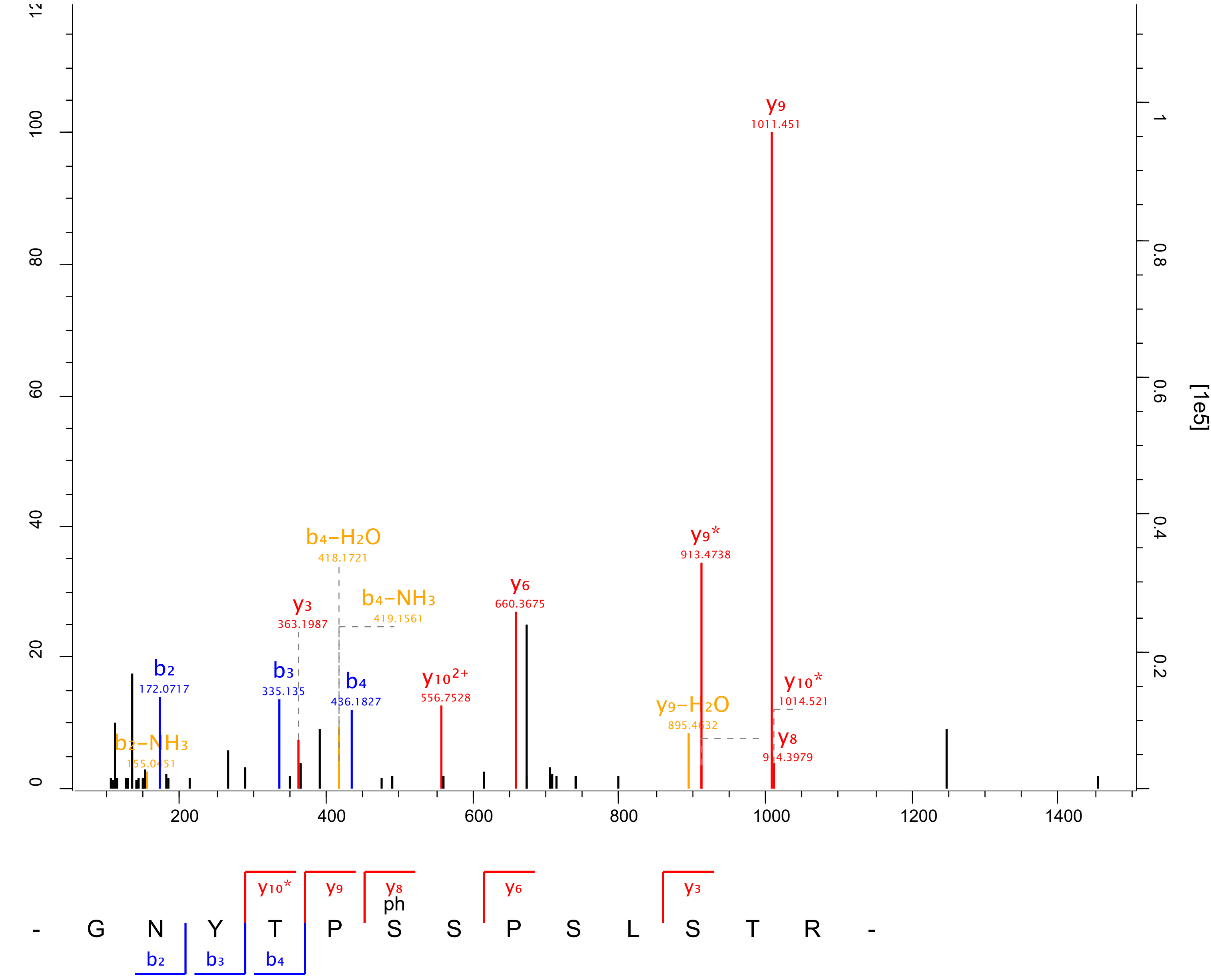

| Raw file | Scan | Method | Score | m/z |
| --- | --- | --- | --- | --- |
| water_02_1 | 13411 | FTMS; HCD | 135.02 | 662.24 |

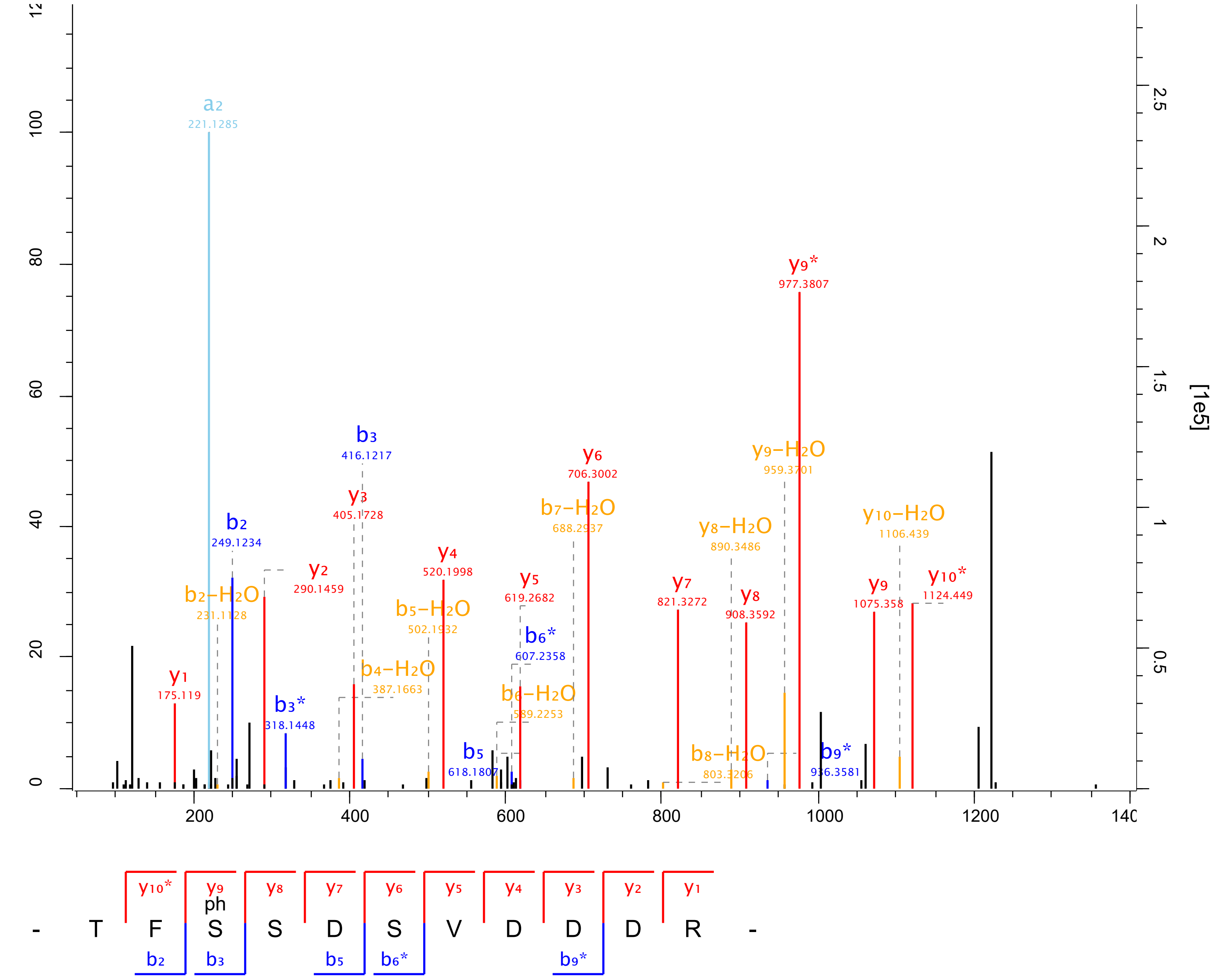

| Raw file | Scan | Method | Score | m/z |
|---|---|---|---|---|
| water_02_1 | 13474 | FTMS; HCD | 99.28 | 498.75 |

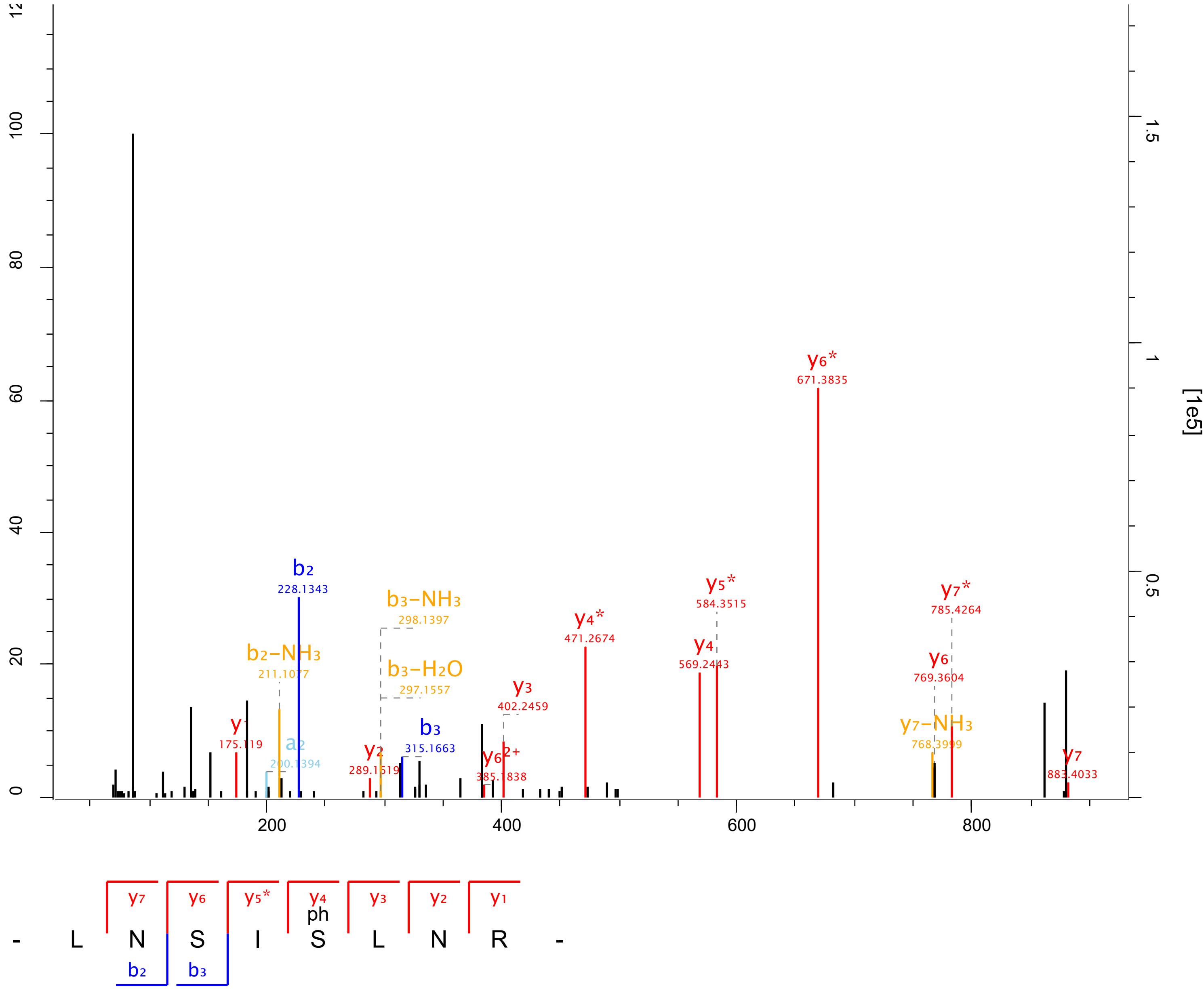

| Raw file | Scan | Method | Score | m/z |
|---|---|---|---|---|
| water_02_1 | 13495 | FTMS; HCD | 100.97 | 803.33 |

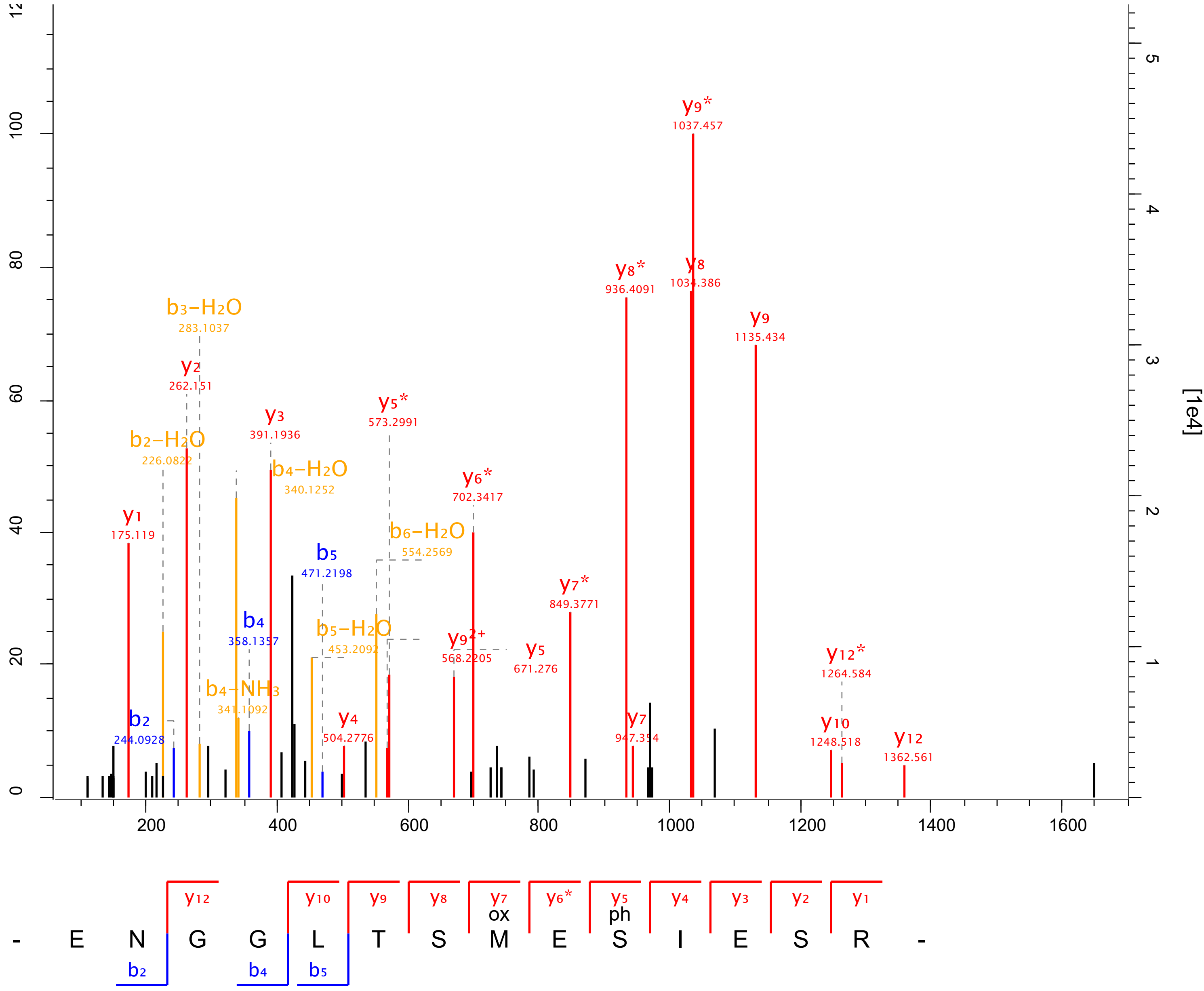

| Raw file | Scan | Method | Score | m/z |
|---|---|---|---|---|
| water_02_1 | 13561 | FTMS; HCD | 42.98 | 559.69 |

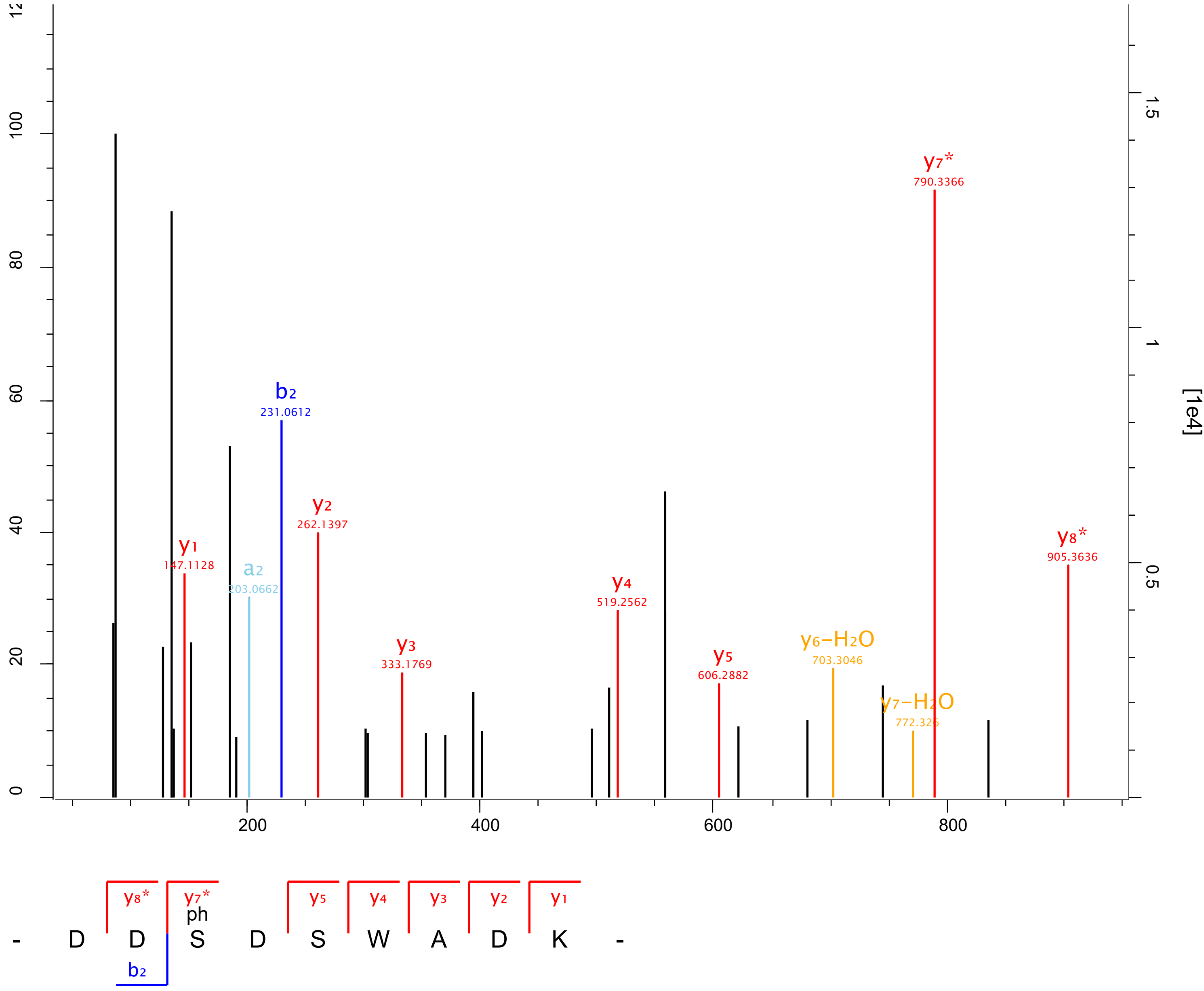

| Raw file | Scan | Method | Score | m/z |
|---|---|---|---|---|
| water_02_1 | 13659 | FTMS; HCD | 54 | 787.35 |

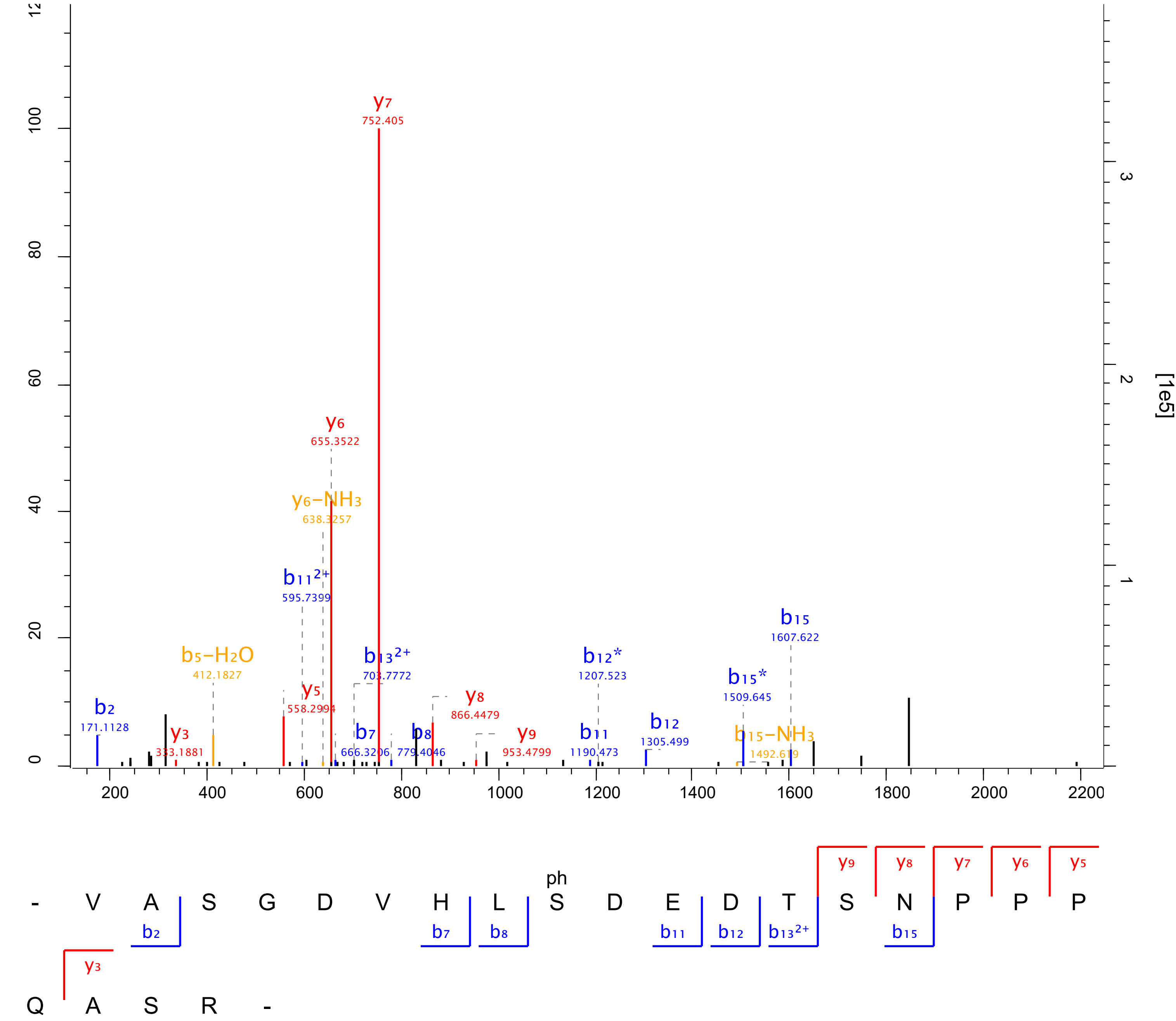

| Raw file | Scan | Method | Score | m/z |
| --- | --- | --- | --- | --- |
| water_02_1 | 13733 | FTMS; HCD | 243.93 | 891.87 |

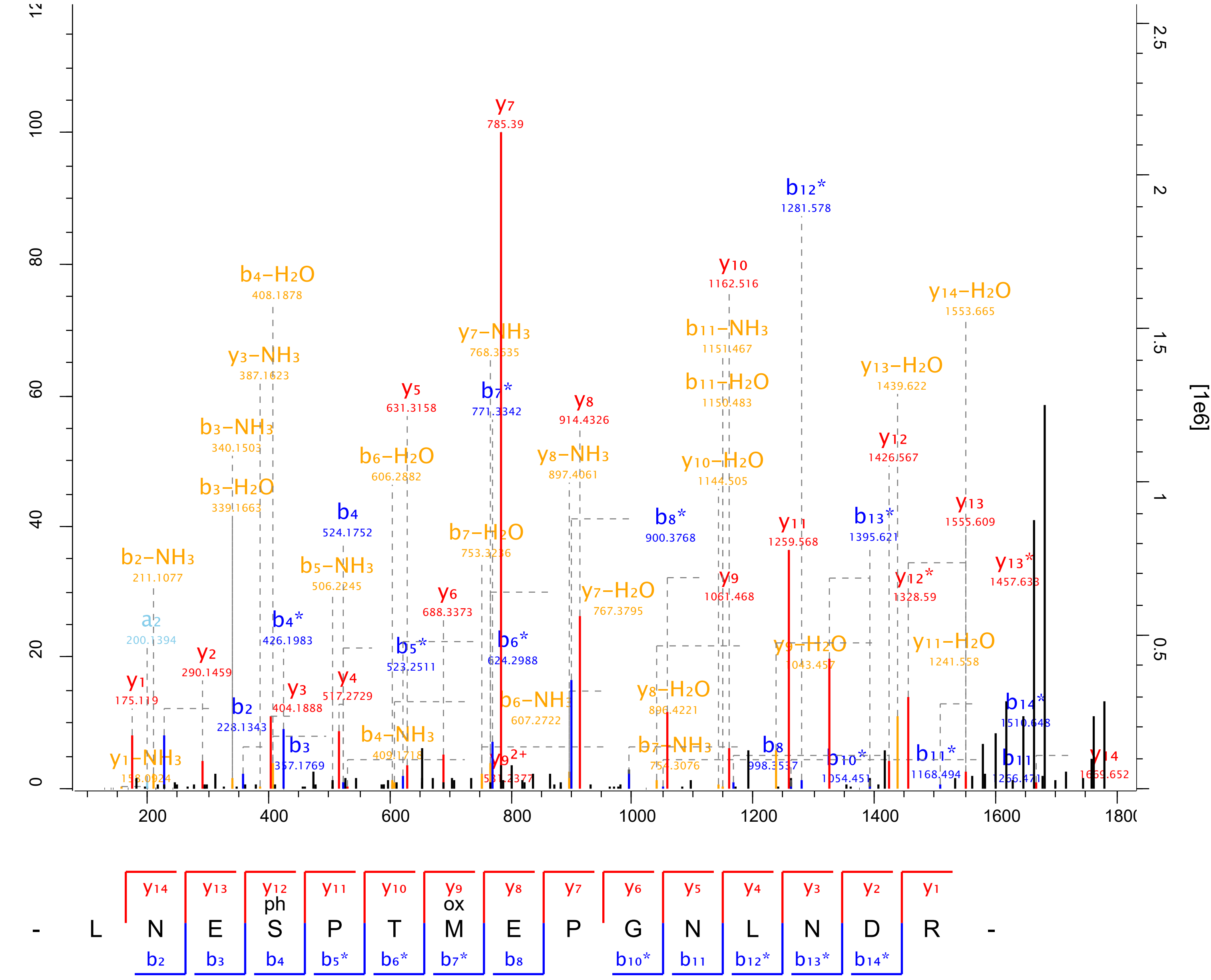

| Raw file | Scan | Method | Score | m/z |
|---|---|---|---|---|
| water_02_1 | 13851 | FTMS; HCD | 61.17 | 731.83 |

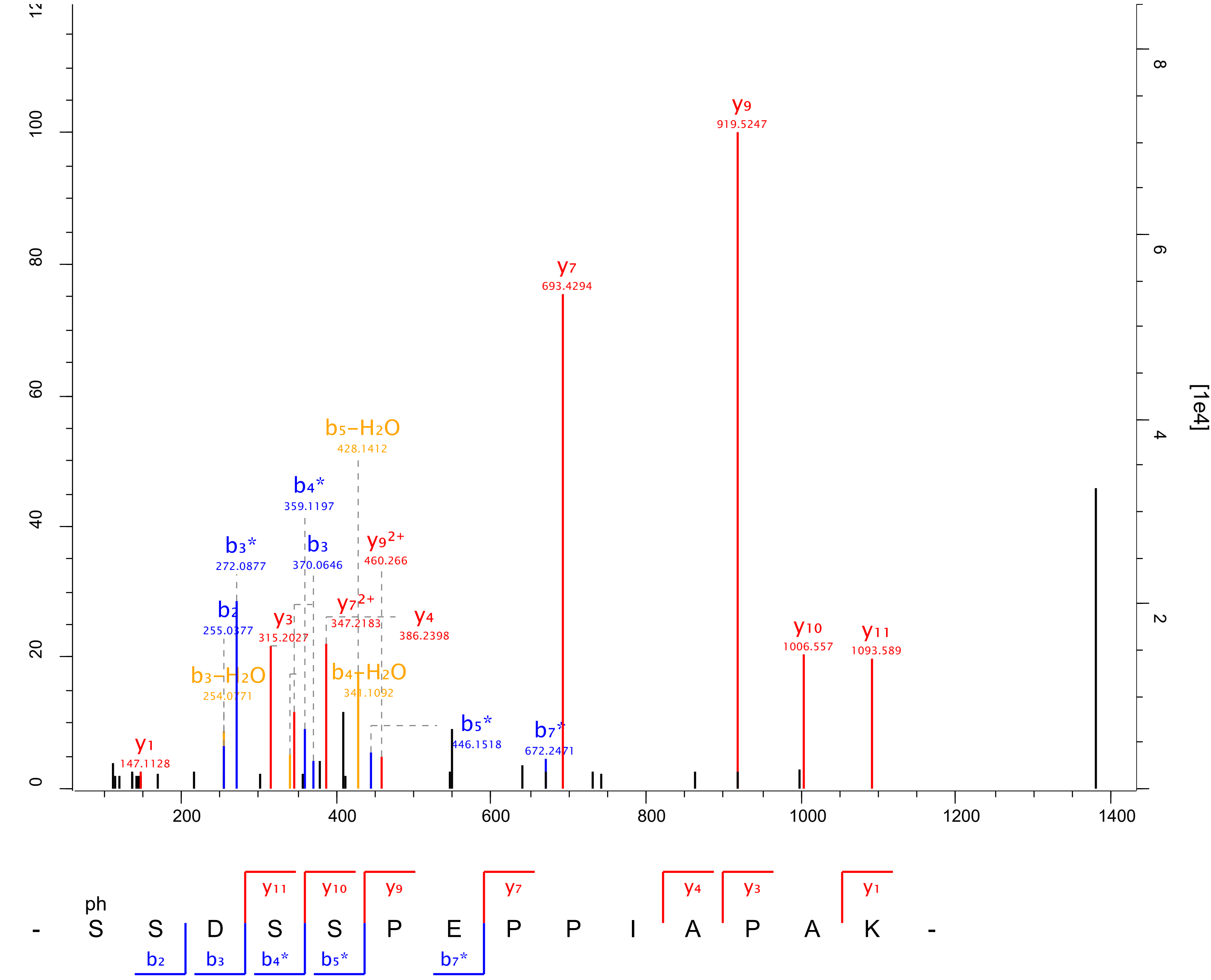

| Raw file | Scan | Method | Score | m/z |
|---|---|---|---|---|
| water_02_1 | 13872 | FTMS; HCD | 129.5 | 726.62 |

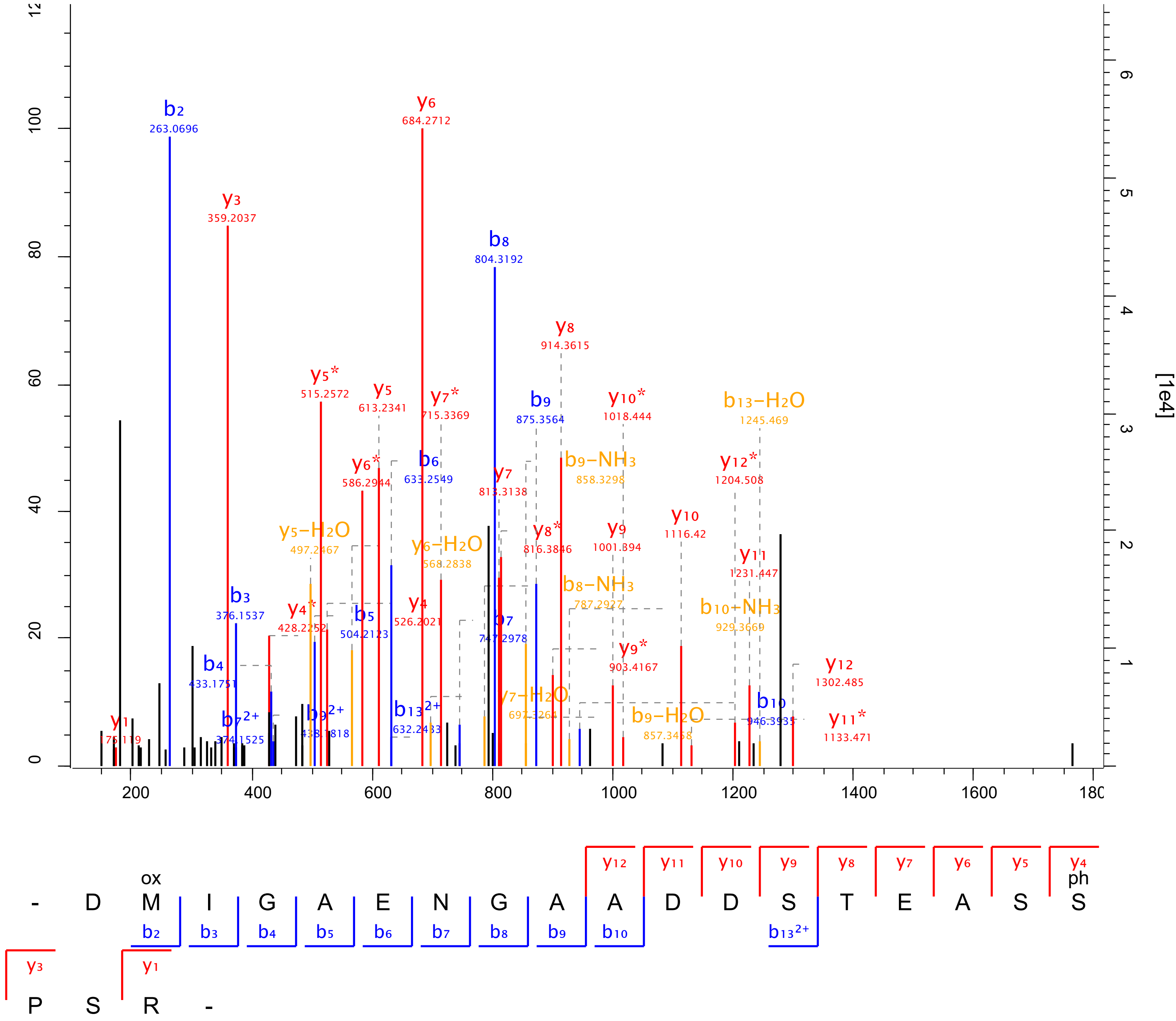

| Raw file | Scan | Method | Score | m/z |
| --- | --- | --- | --- | --- |
| water_02_1 | 13895 | FTMS; HCD | 333.46 | 858.82 |

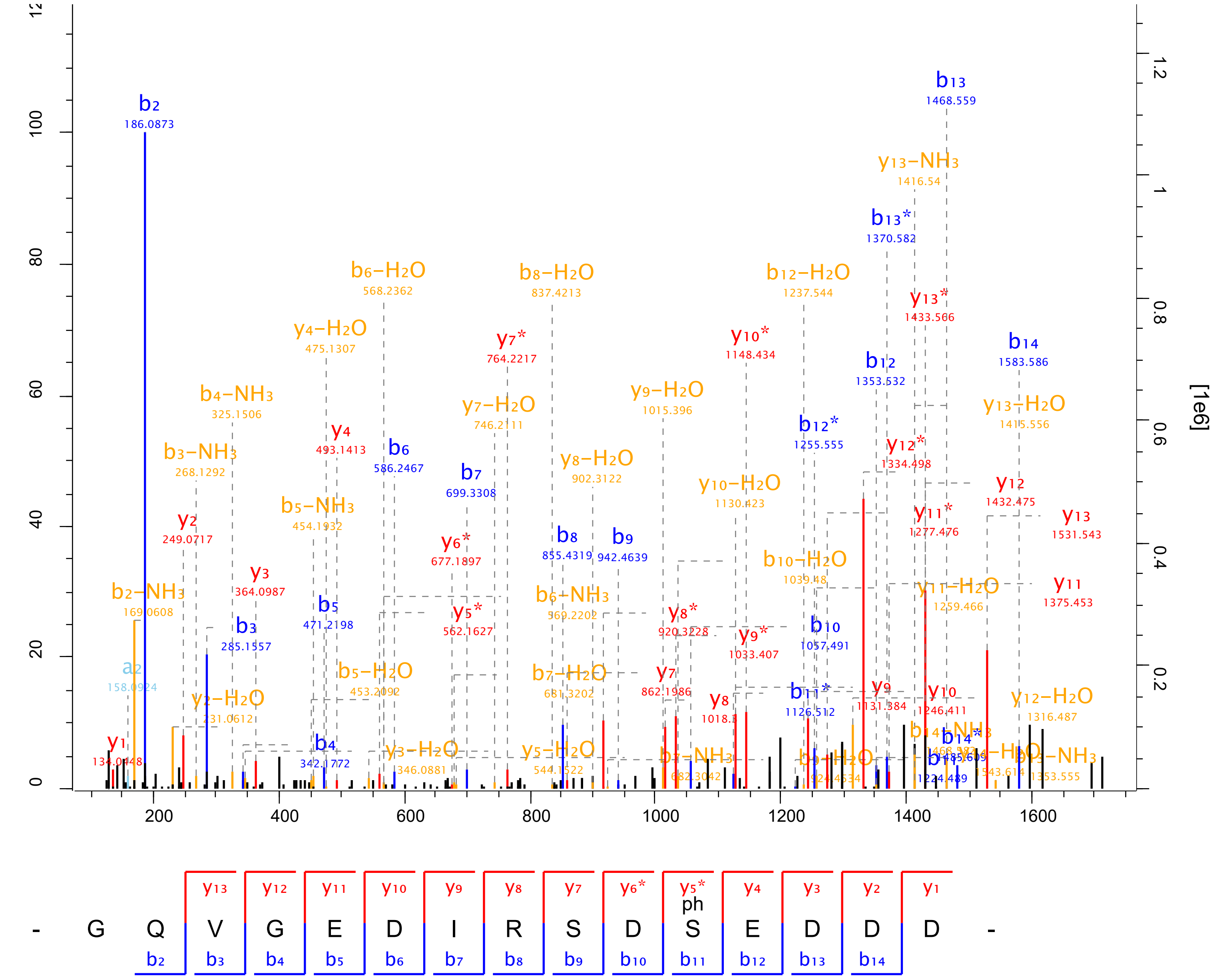

| Raw file | Scan | Method | Score | m/z |
| --- | --- | --- | --- | --- |
| water_02_1 | 13985 | FTMS; HCD | 42.31 | 964.91 |

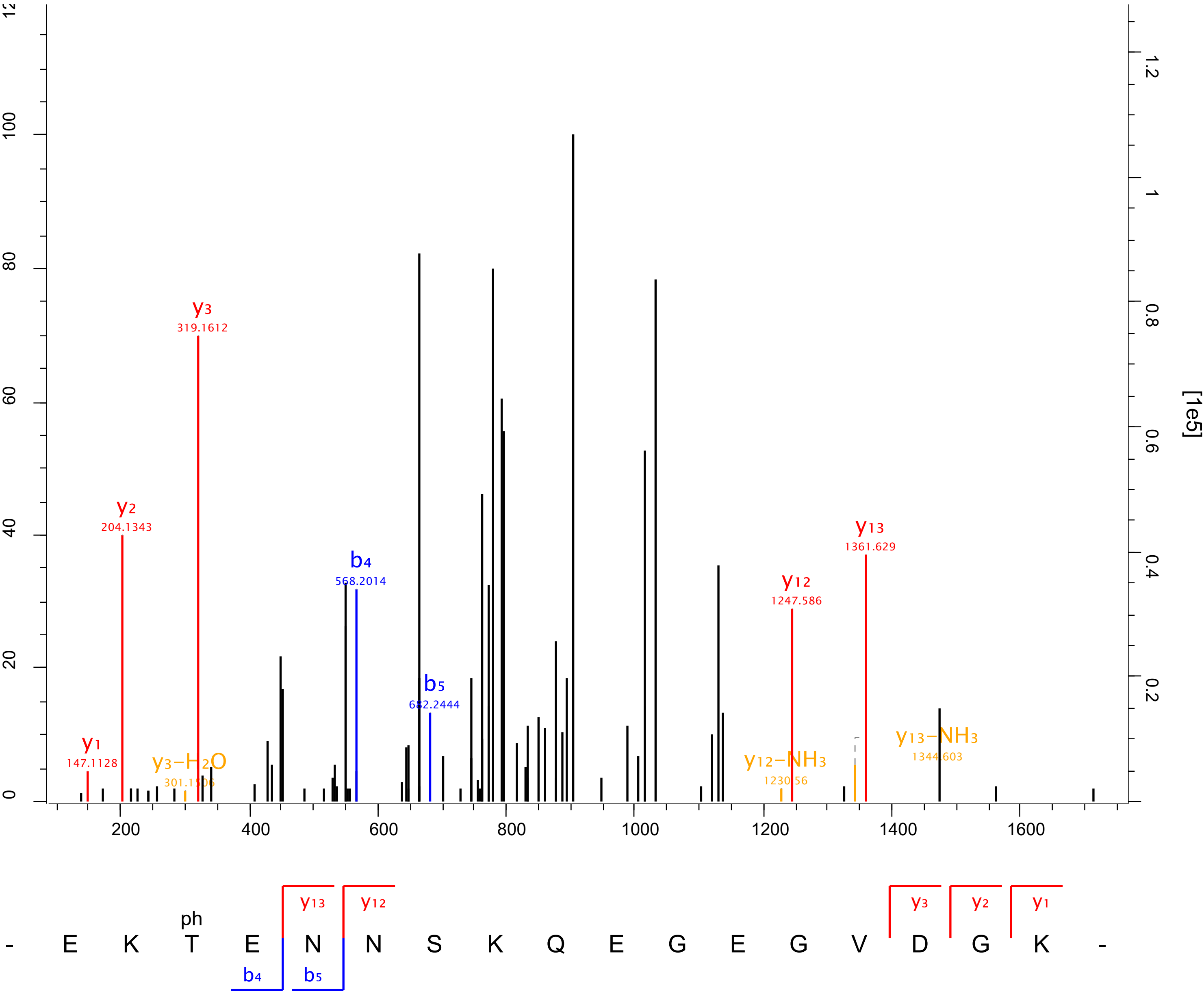

| Raw file | Scan | Method | Score | m/z |
| --- | --- | --- | --- | --- |
| water_02_1 | 14018 | FTMS; HCD | 61.26 | 476.93 |

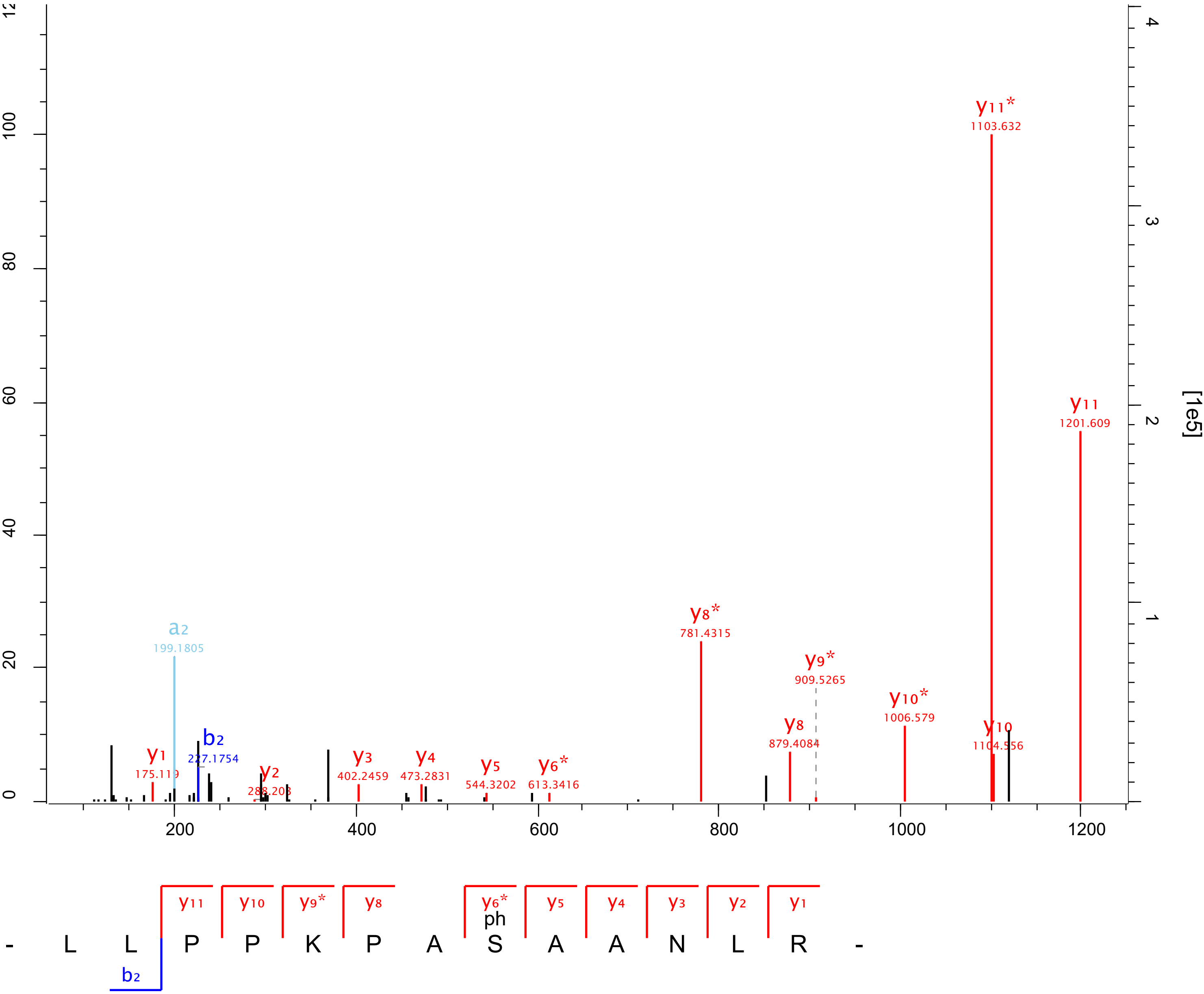

| Raw file | Scan | Method | Score | m/z |
| --- | --- | --- | --- | --- |
| water_02_1 | 14115 | FTMS; HCD | 99.94 | 637.93 |

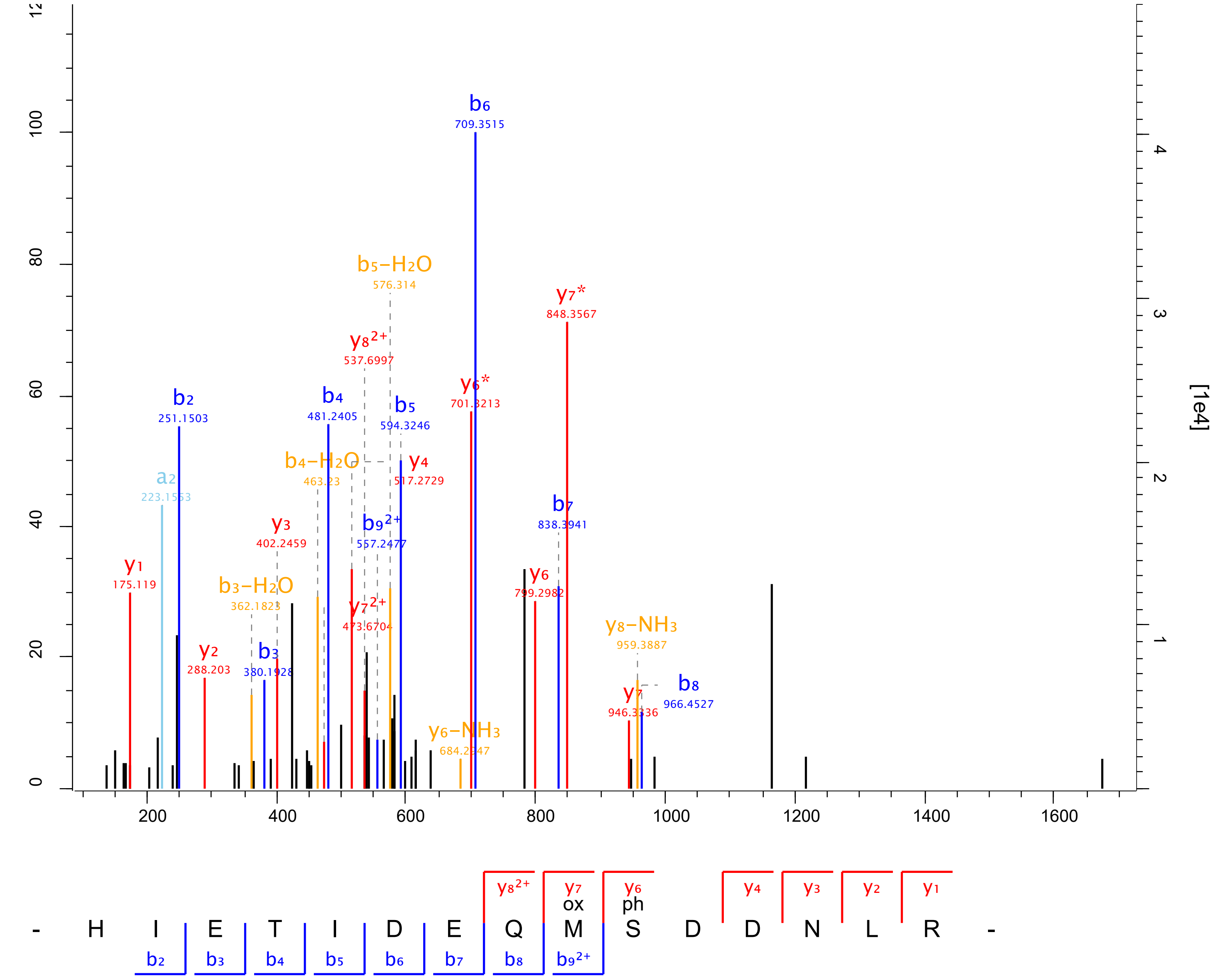

| Raw file | Scan | Method | Score | m/z |
| --- | --- | --- | --- | --- |
| water_02_1 | 14182 | FTMS; HCD | 94.26 | 840.88 |

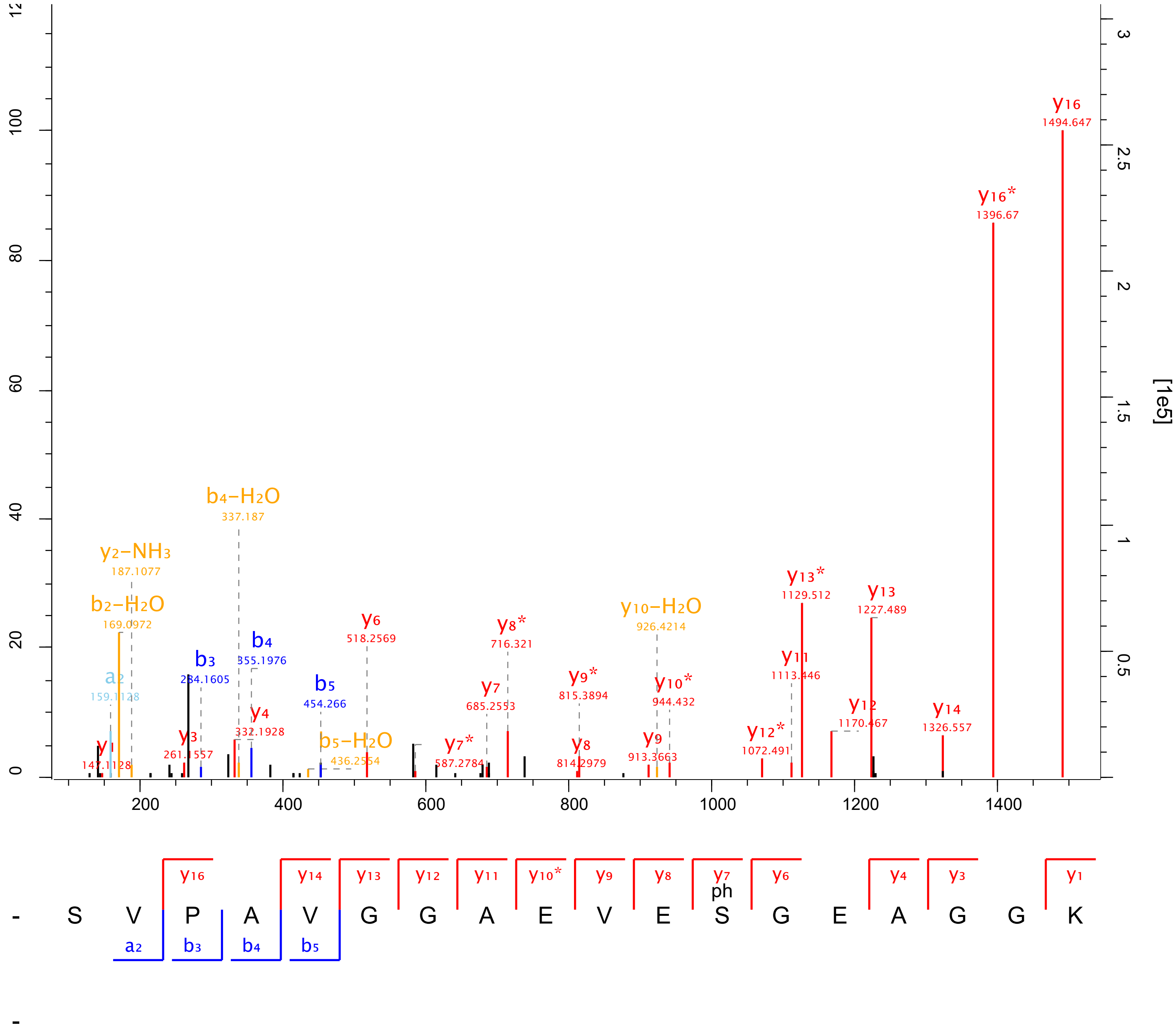

| Raw file | Scan | Method | Score | m/z |
|---|---|---|---|---|
| water_02_1 | 14330 | FTMS; HCD | 85.34 | 712.07 |

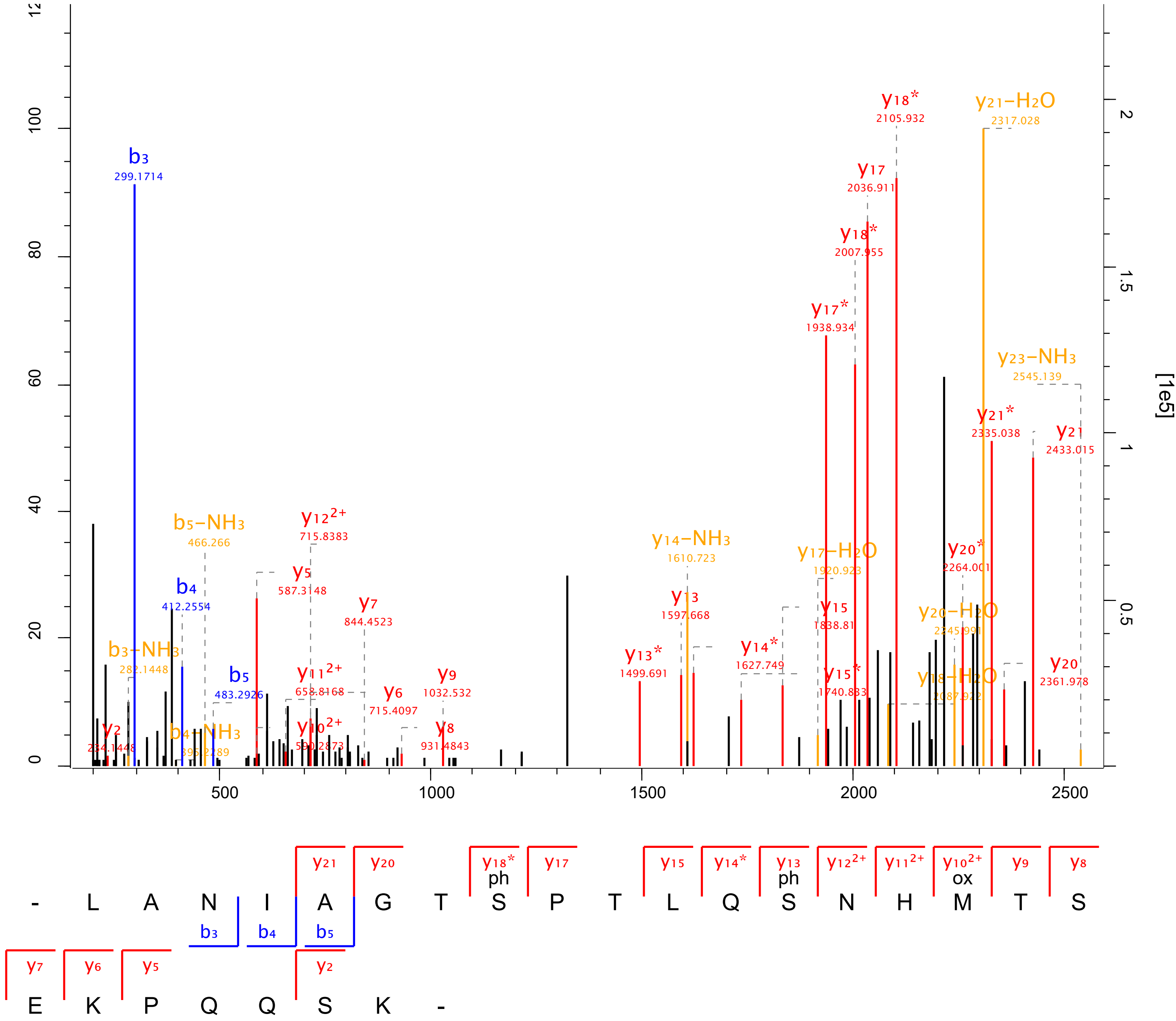

| Raw file | Scan | Method | Score | m/z |
|---|---|---|---|---|
| water_02_1 | 14379 | FTMS; HCD | 88.18 | 892.37 |

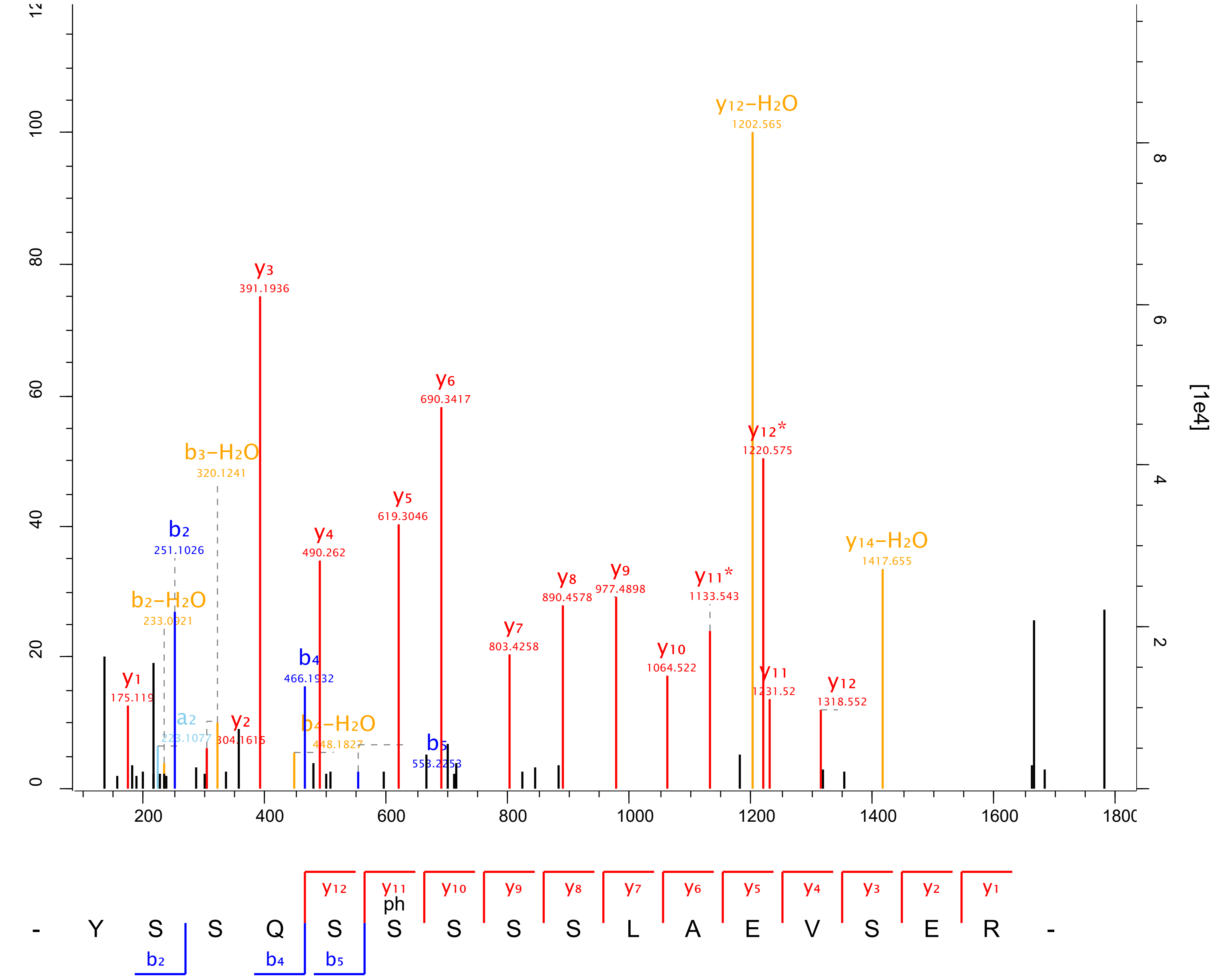

| Raw file | Scan | Method | Score | m/z |
|---|---|---|---|---|
| water_02_1 | 14523 | FTMS; HCD | 138.31 | 606.26 |

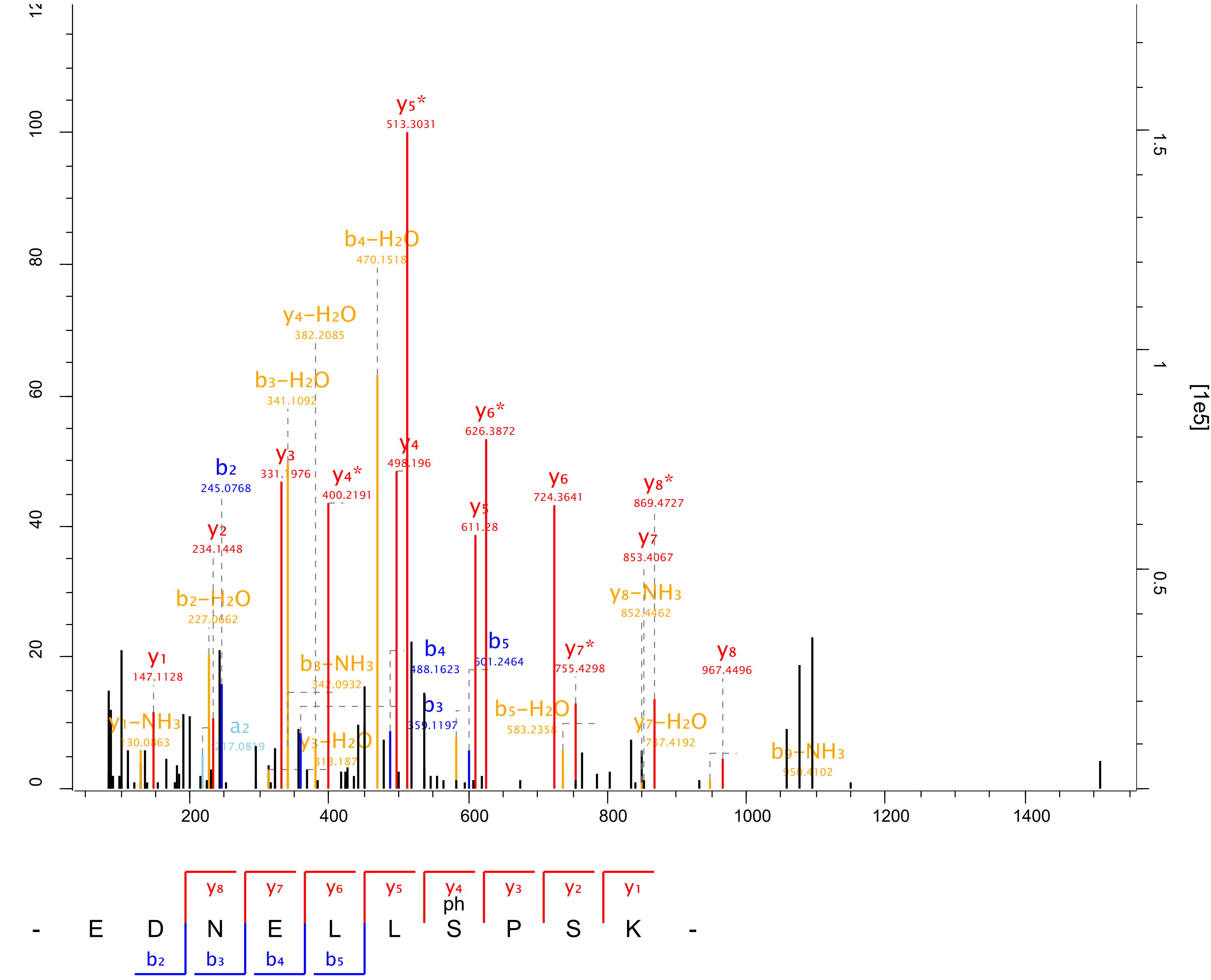

| Raw file | Scan | Method | Score | m/z |
|---|---|---|---|---|
| water_02_1 | 14545 | FTMS; HCD | 67.02 | 564.27 |

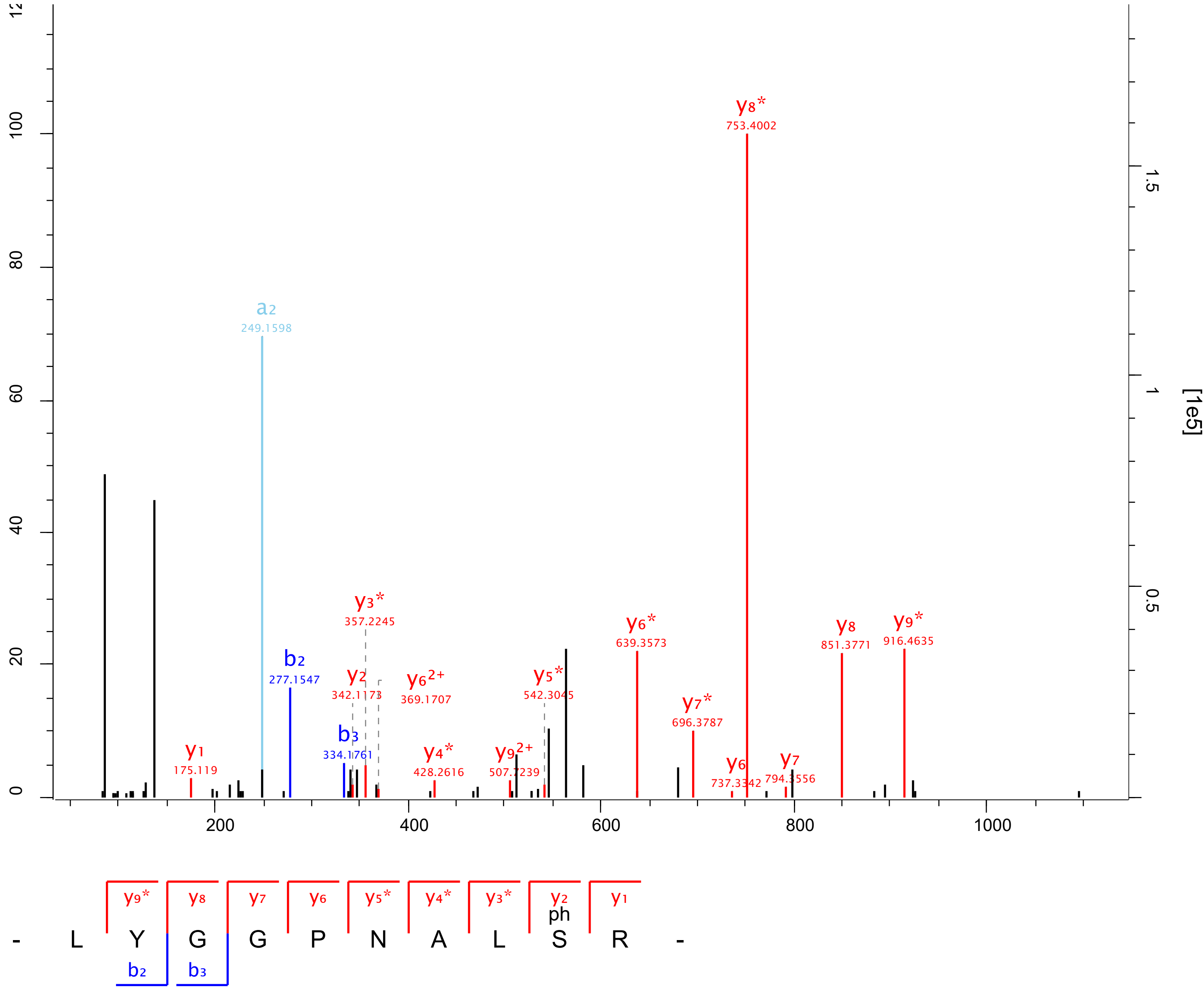

| Raw file | Scan | Method | Score | m/z |
|---|---|---|---|---|
| water_02_1 | 14588 | FTMS; HCD | 104.18 | 693.98 |

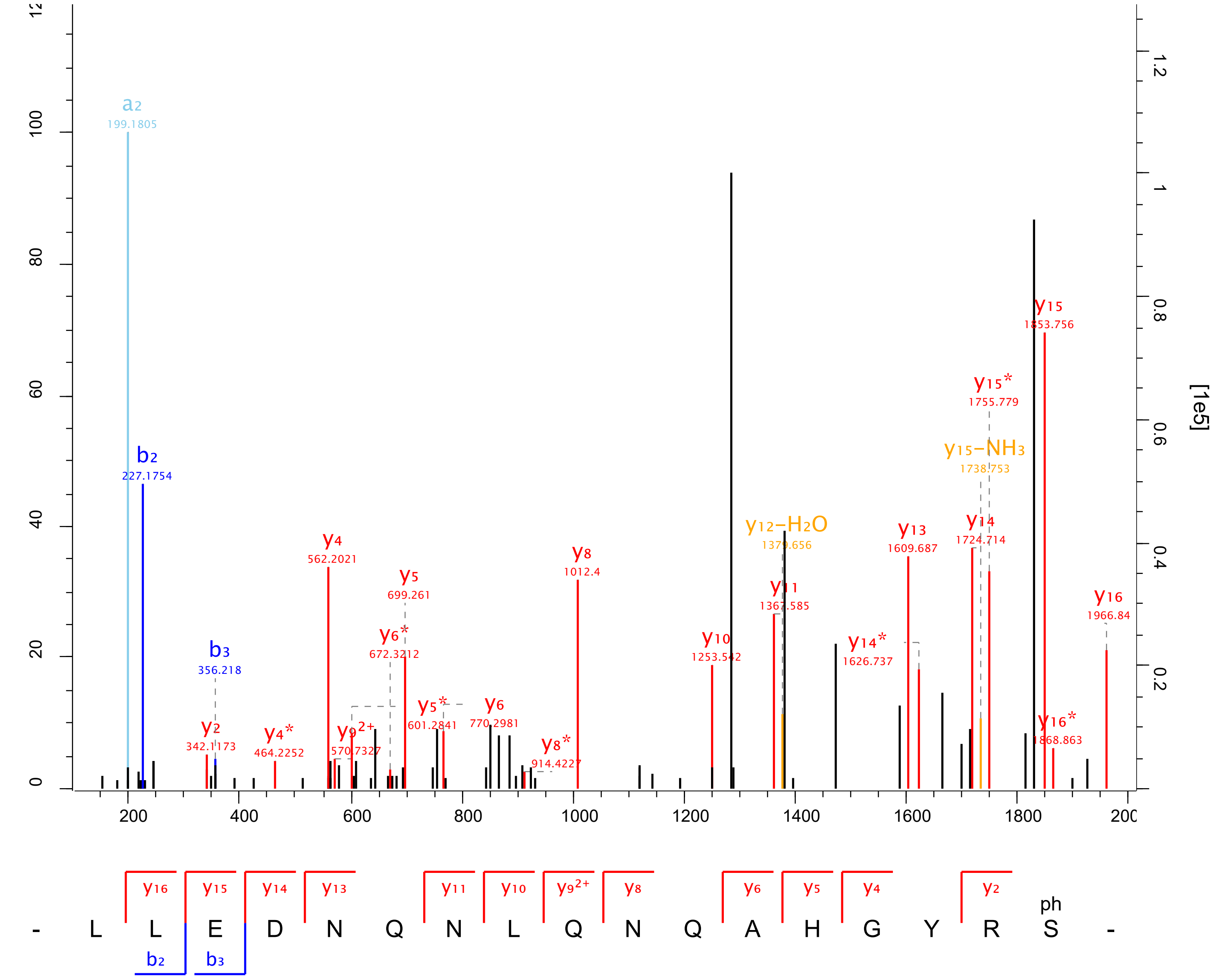

| Raw file | Scan | Method | Score | m/z |
|---|---|---|---|---|
| water_02_1 | 14619 | FTMS; HCD | 47.09 | 604.25 |

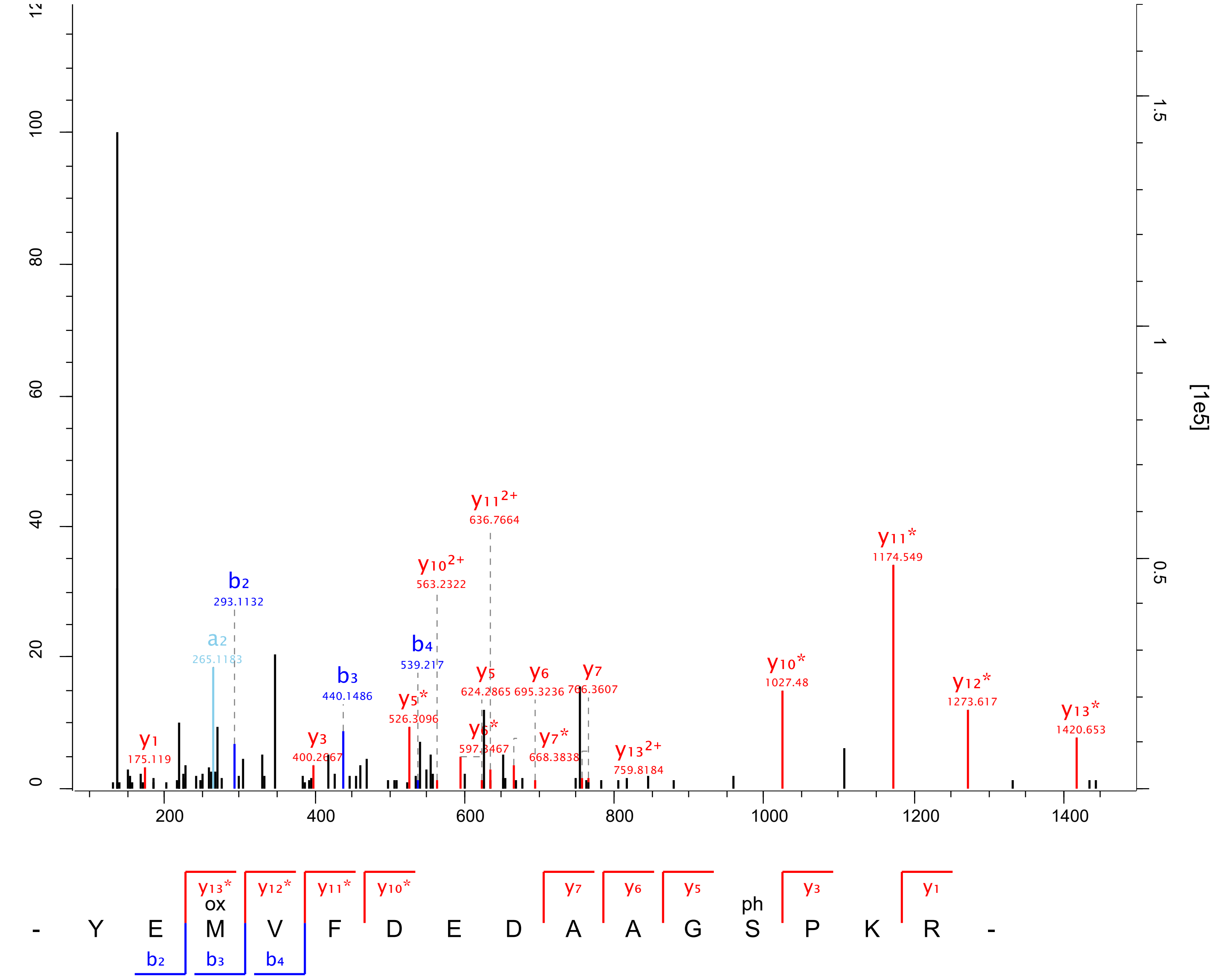

| Raw file | Scan | Method | Score | m/z |
|---|---|---|---|---|
| water_02_1 | 14629 | FTMS; HCD | 61.11 | 541.75 |

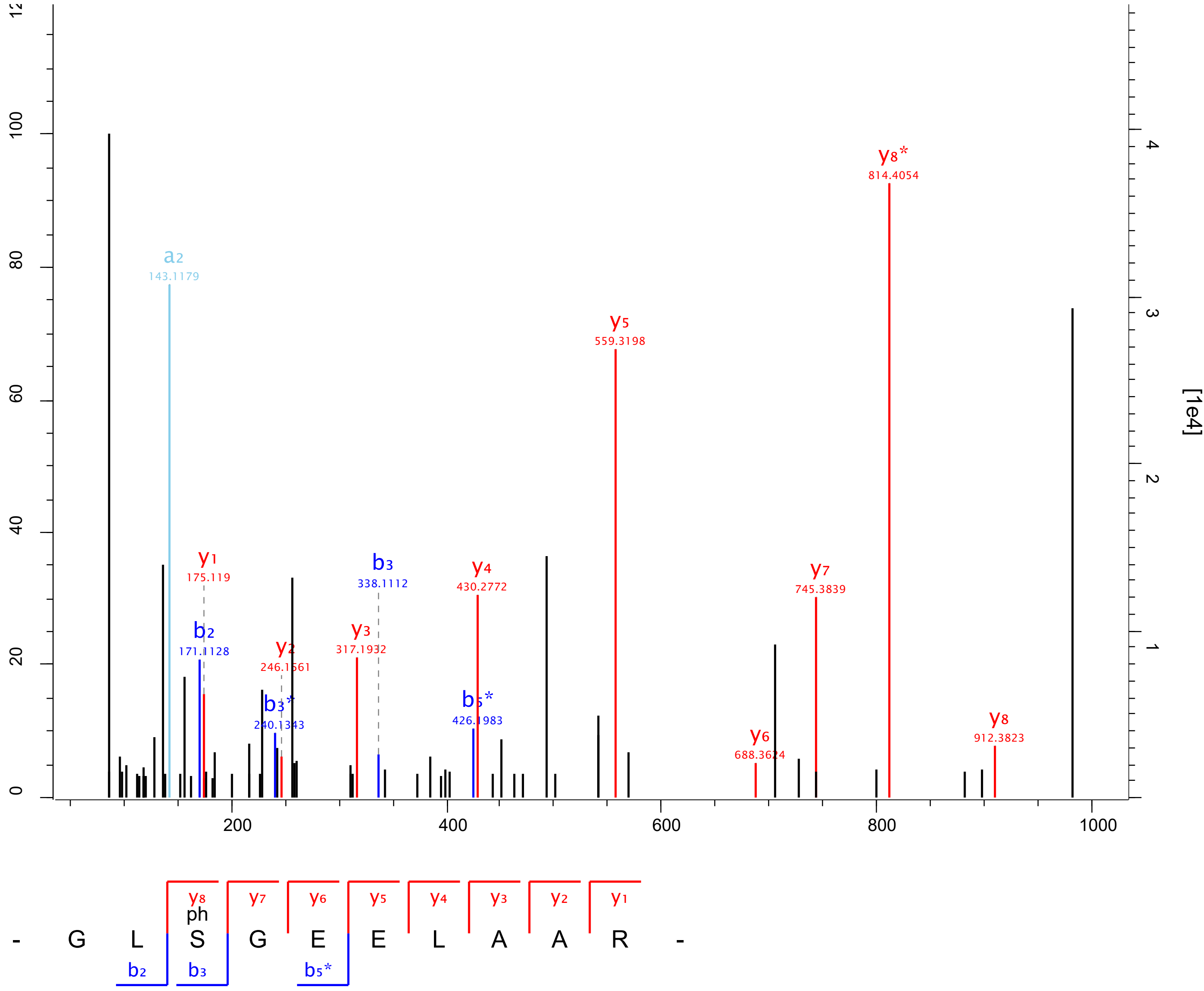

| Raw file | Scan | Method | Score | m/z |
|---|---|---|---|---|
| water_02_1 | 14662 | FTMS; HCD | 69.77 | 623.61 |

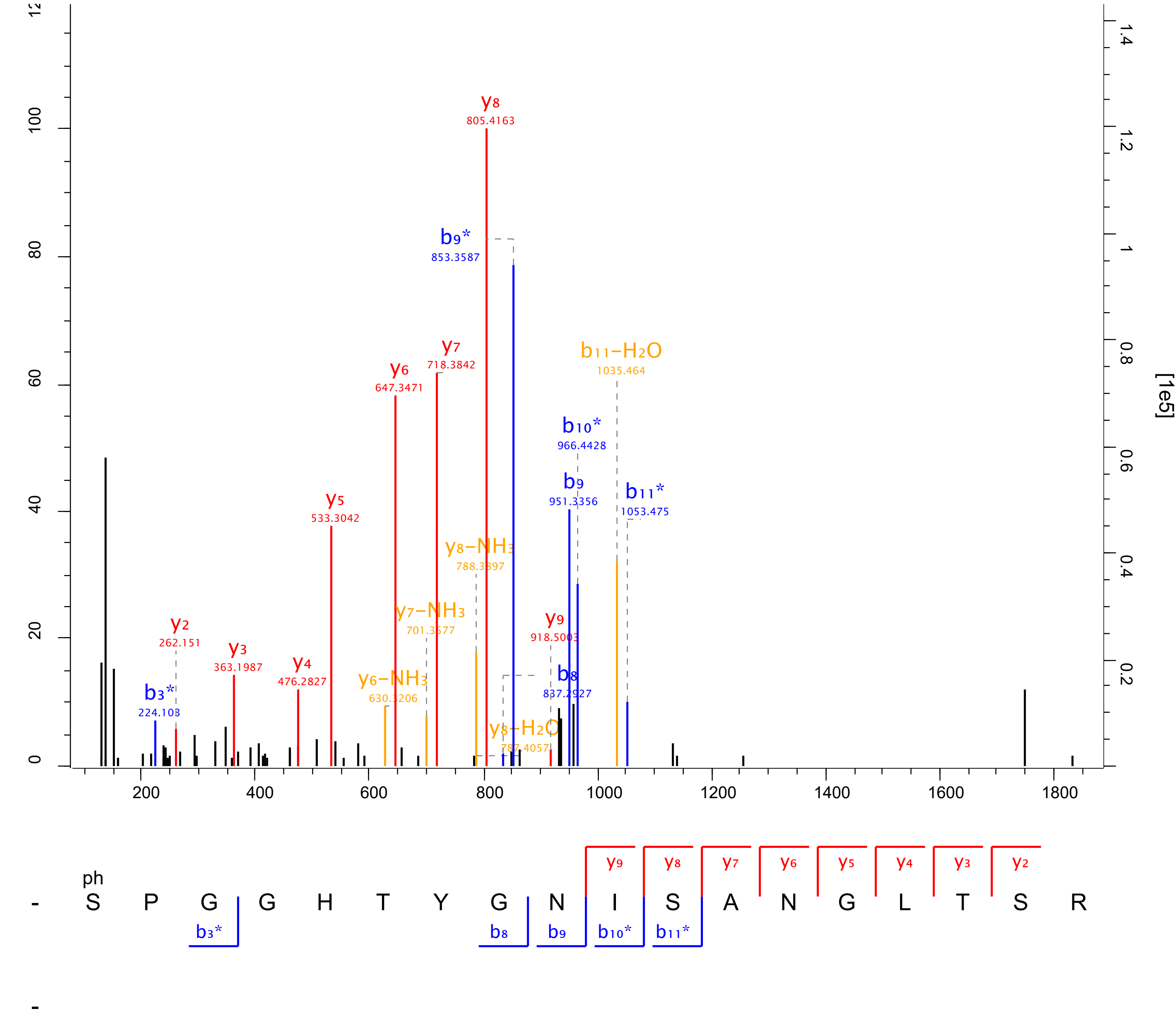

| Raw file | Scan | Method | Score | m/z |
| --- | --- | --- | --- | --- |
| water_02_1 | 14733 | FTMS; HCD | 121.44 | 630.25 |

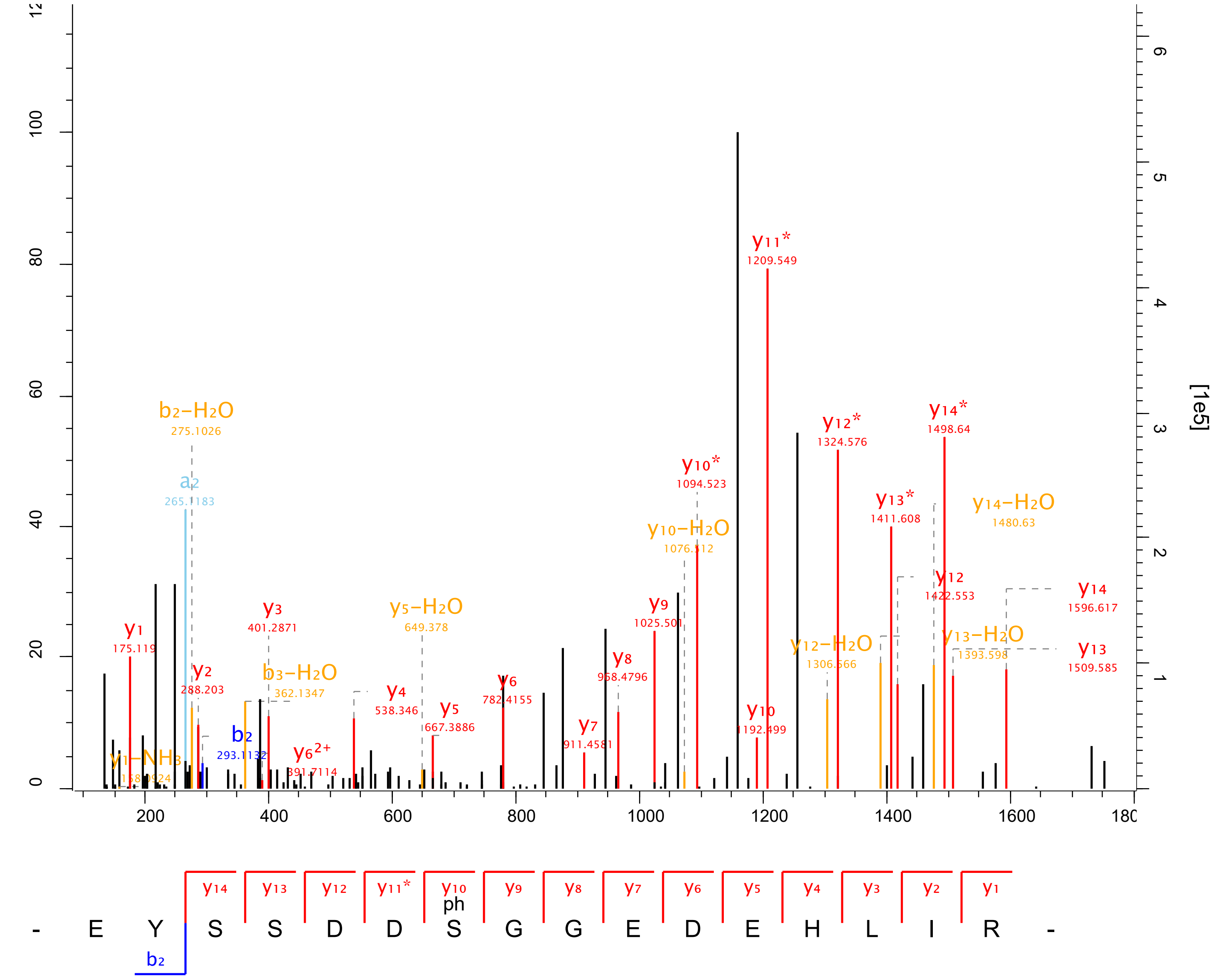

| Raw file | Scan | Method | Score | m/z |
| --- | --- | --- | --- | --- |
| water_02_1 | 14983 | FTMS; HCD | 44.31 | 587.92 |

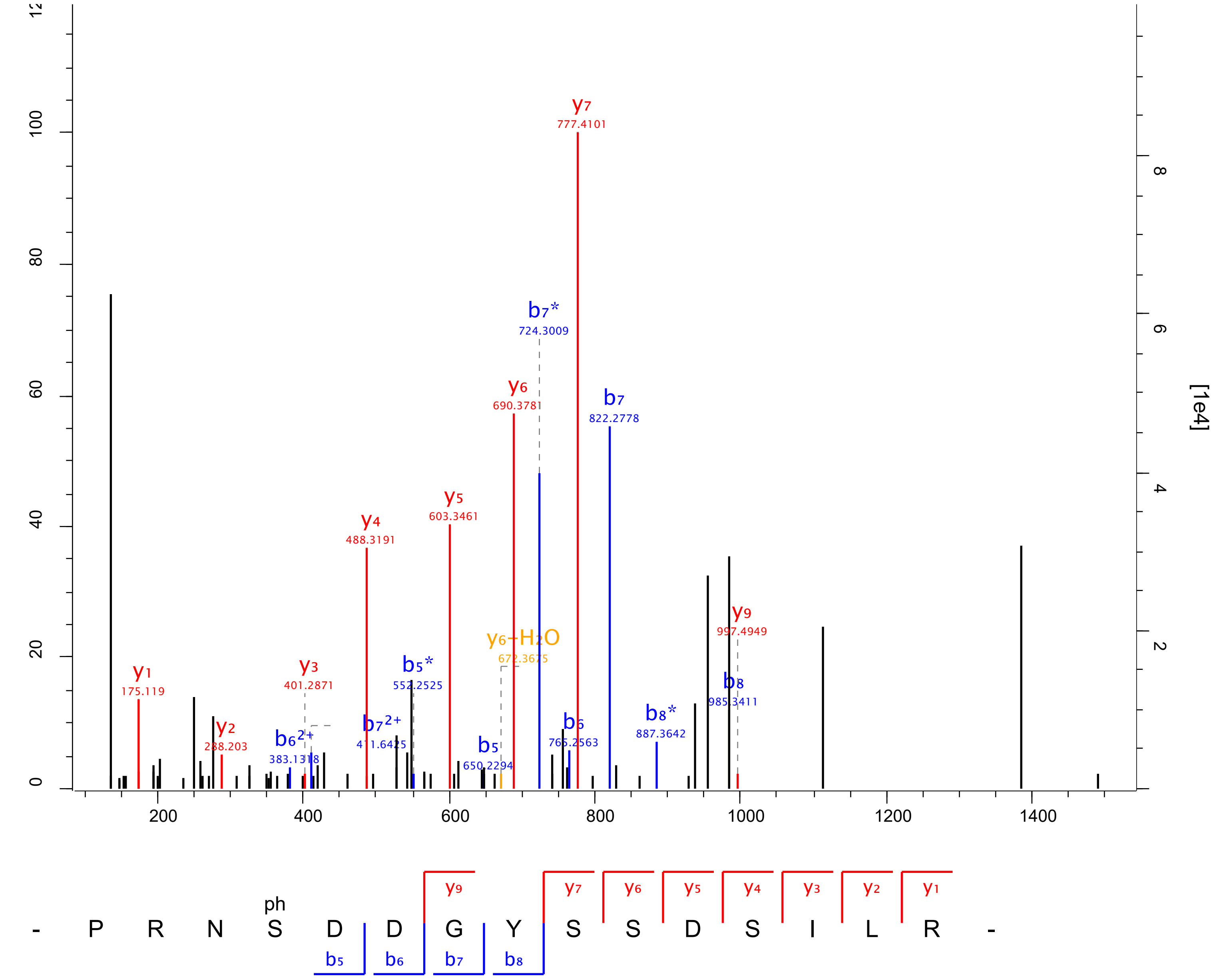

| Raw file | Scan | Method | Score | m/z |
| --- | --- | --- | --- | --- |
| water_02_1 | 15018 | FTMS; HCD | 81.88 | 661.6 |

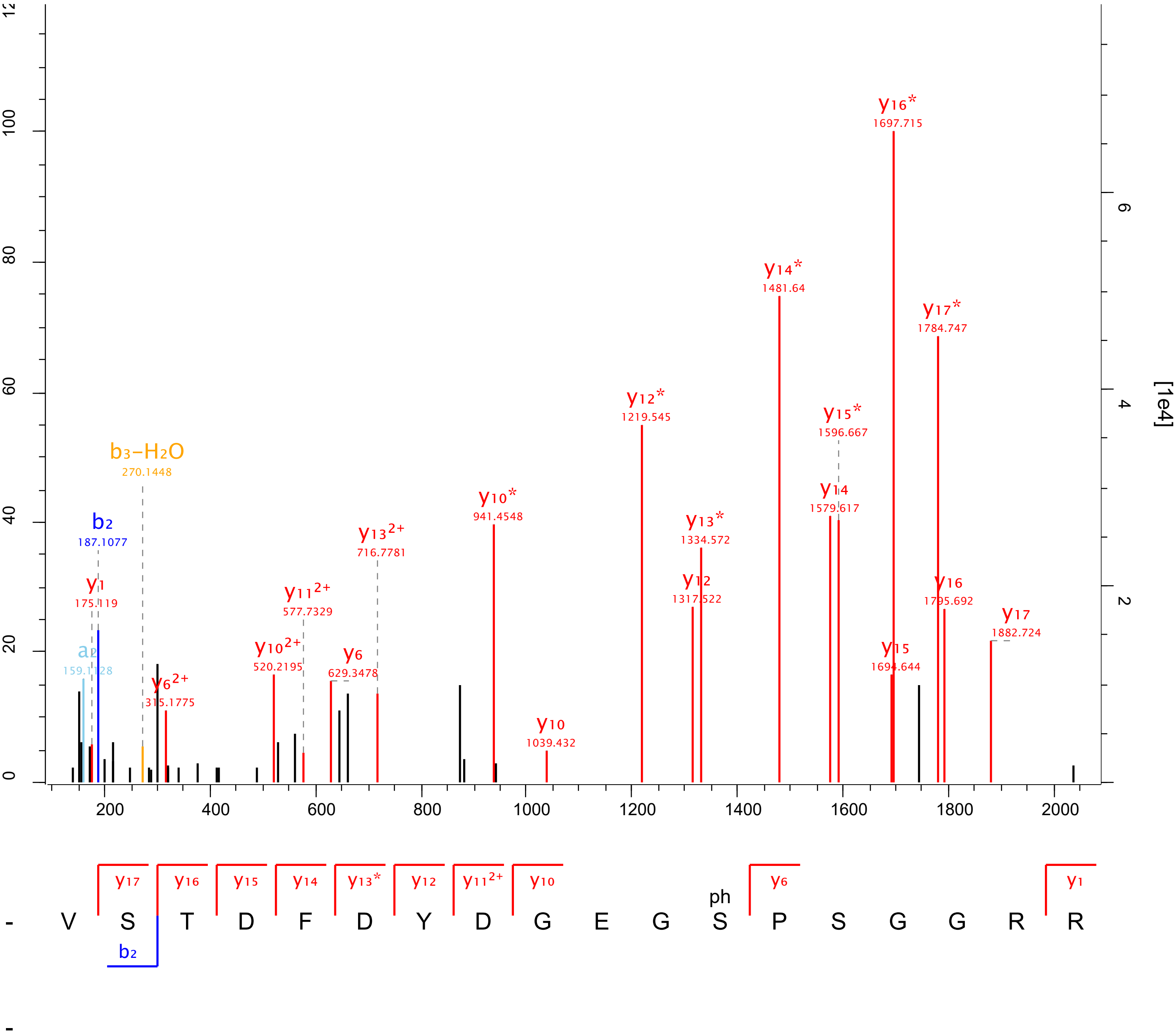

| Raw file | Scan | Method | Score | m/z |
|---|---|---|---|---|
| water_02_1 | 15022 | FTMS; HCD | 221.46 | 760.82 |

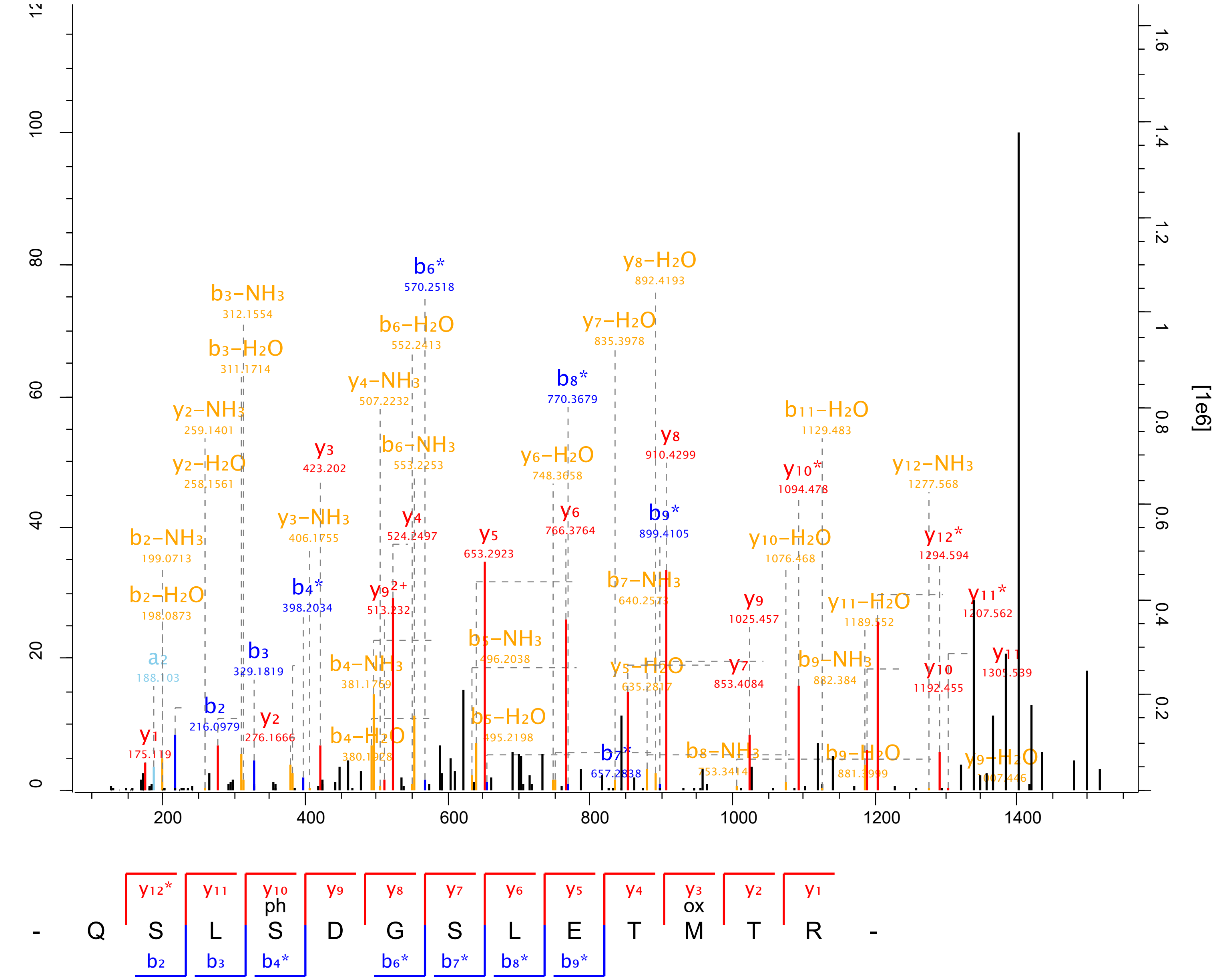

| Raw file | Scan | Method | Score | m/z |
| --- | --- | --- | --- | --- |
| water_02_1 | 15037 | FTMS; HCD | 60.49 | 664.28 |

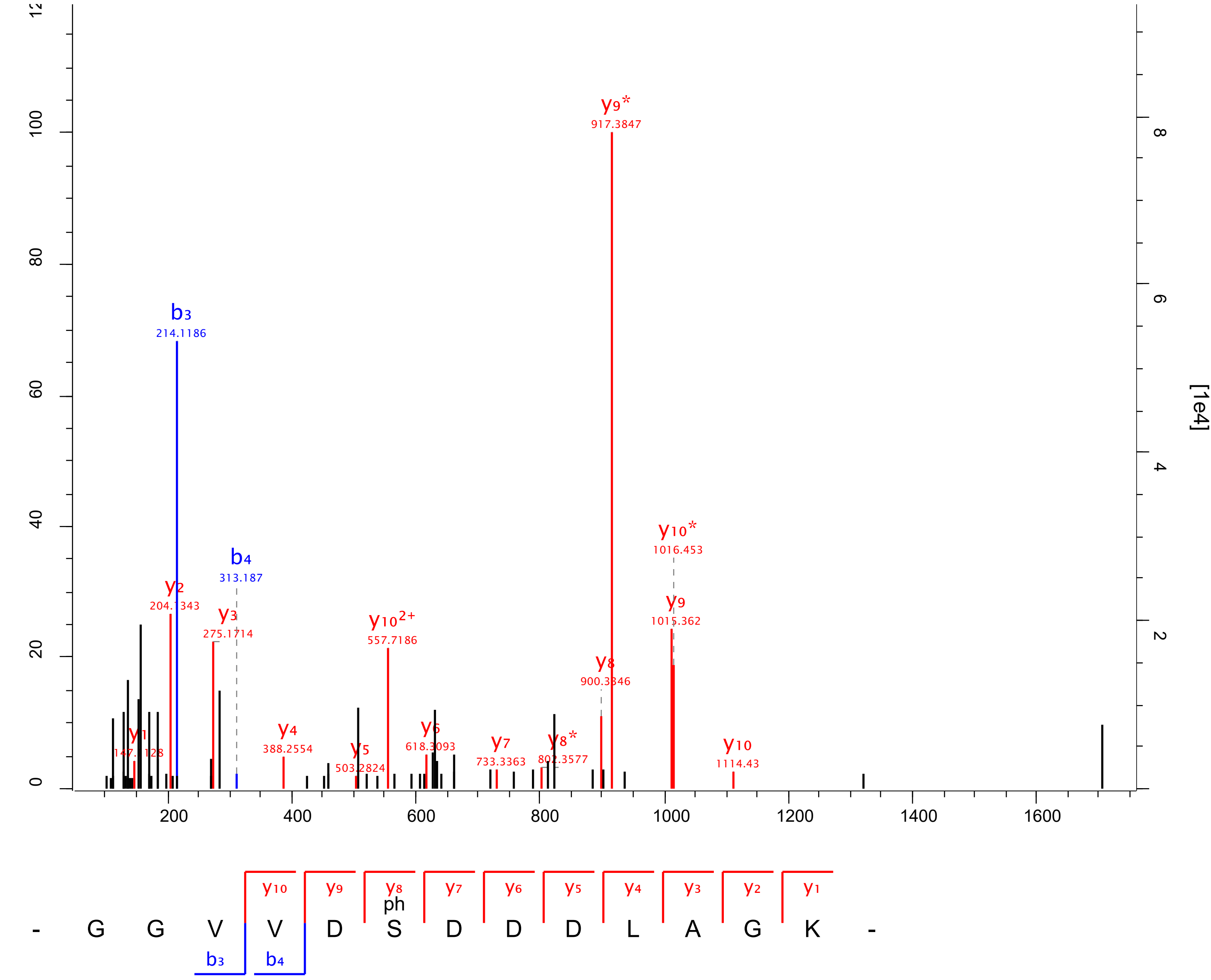

| Raw file | Scan | Method | Score | m/z |
|---|---|---|---|---|
| water_02_1 | 15062 | FTMS; HCD | 289.35 | 977.91 |

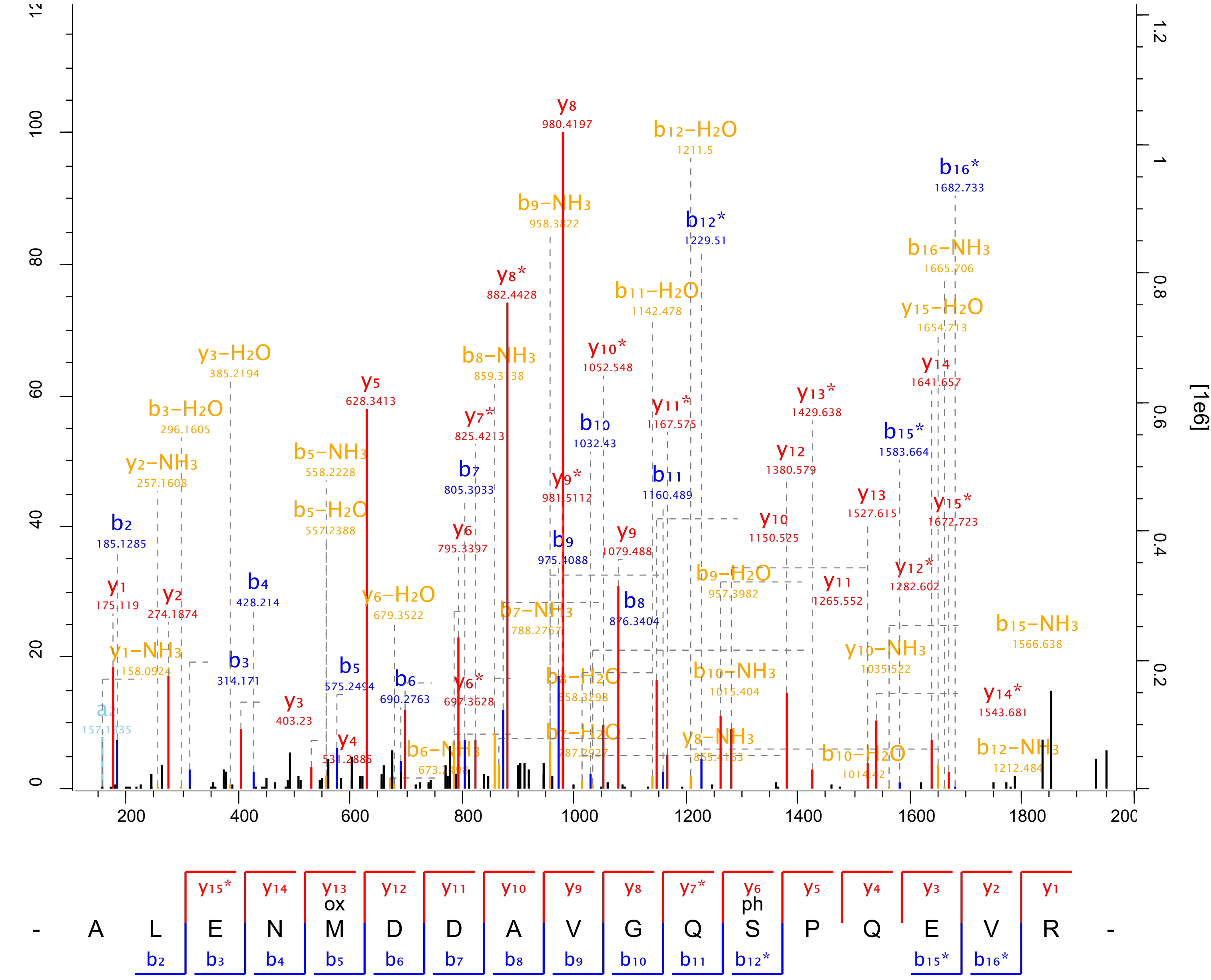

| Raw file | Scan | Method | Score | m/z |
|---|---|---|---|---|
| water_02_1 | 1510 | FTMS; HCD | 118.37 | 622.9 |

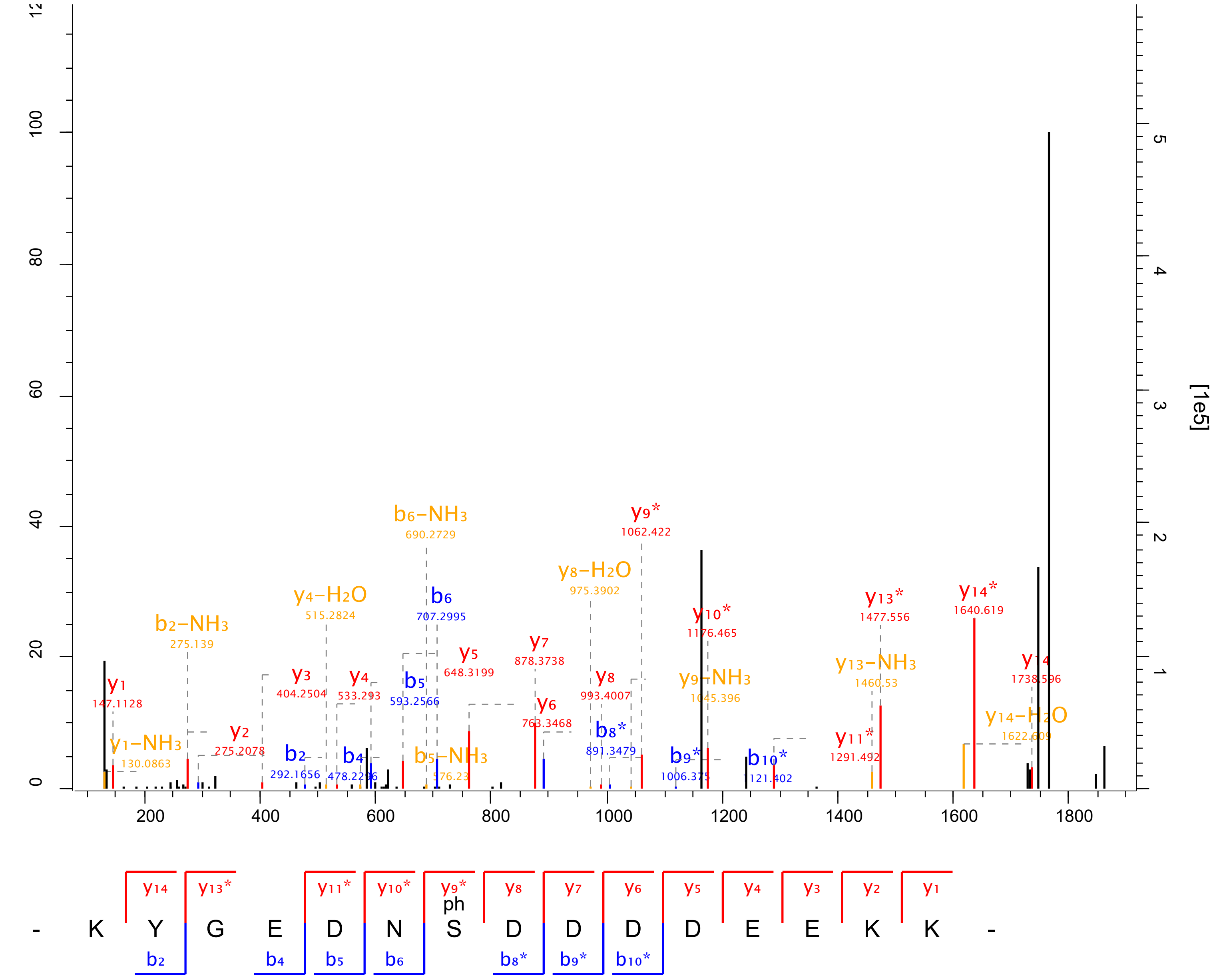

| Raw file | Scan | Method | Score | m/z |
|---|---|---|---|---|
| water_02_1 | 1522 | FTMS; HCD | 56.4 | 491.21 |

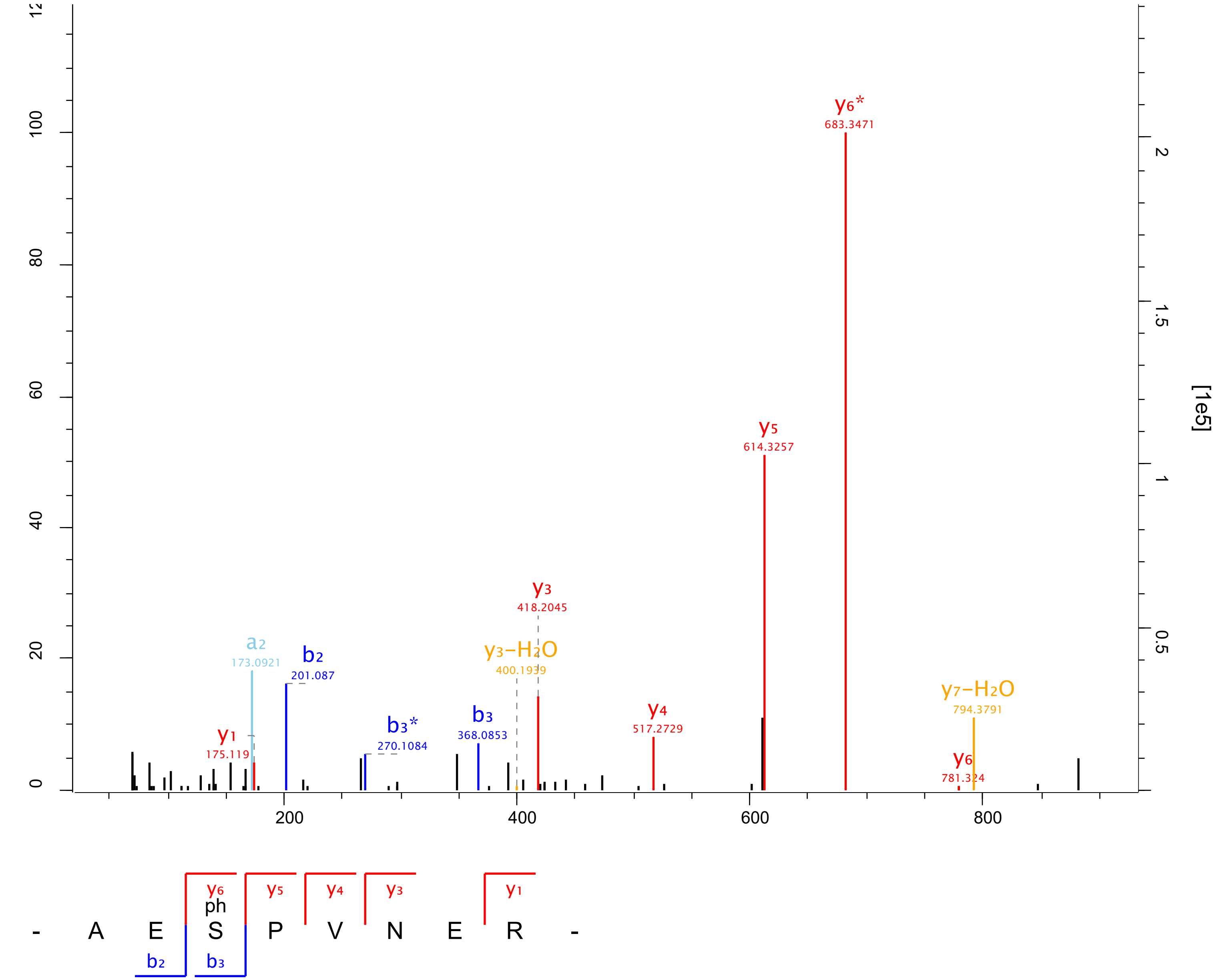

| Raw file | Scan | Method | Score | m/z |
|---|---|---|---|---|
| water_02_1 | 15251 | FTMS; HCD | 151.87 | 584.95 |

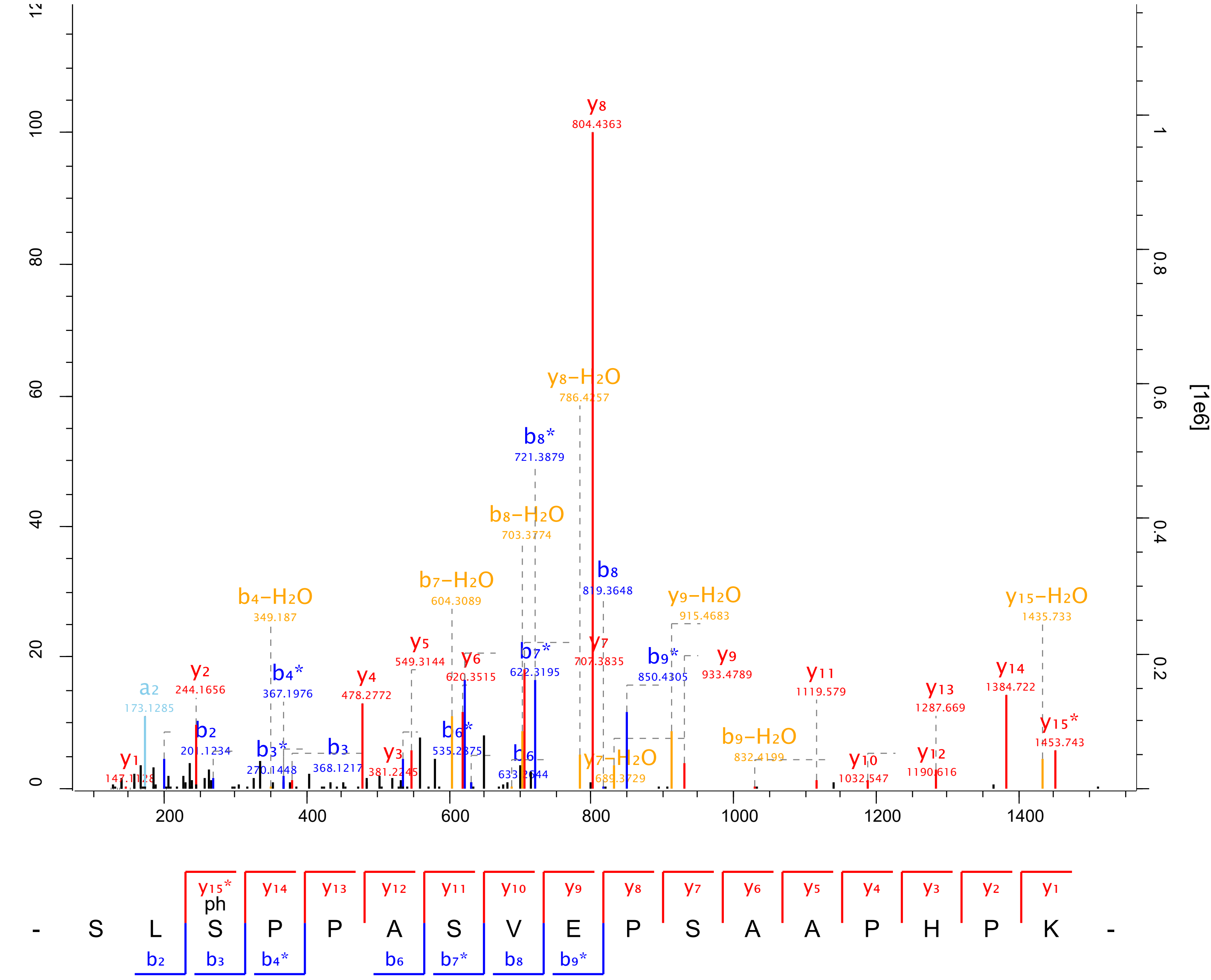

| Raw file | Scan | Method | Score | m/z |
| --- | --- | --- | --- | --- |
| water_02_1 | 15286 | FTMS; HCD | 99.34 | 842.33 |

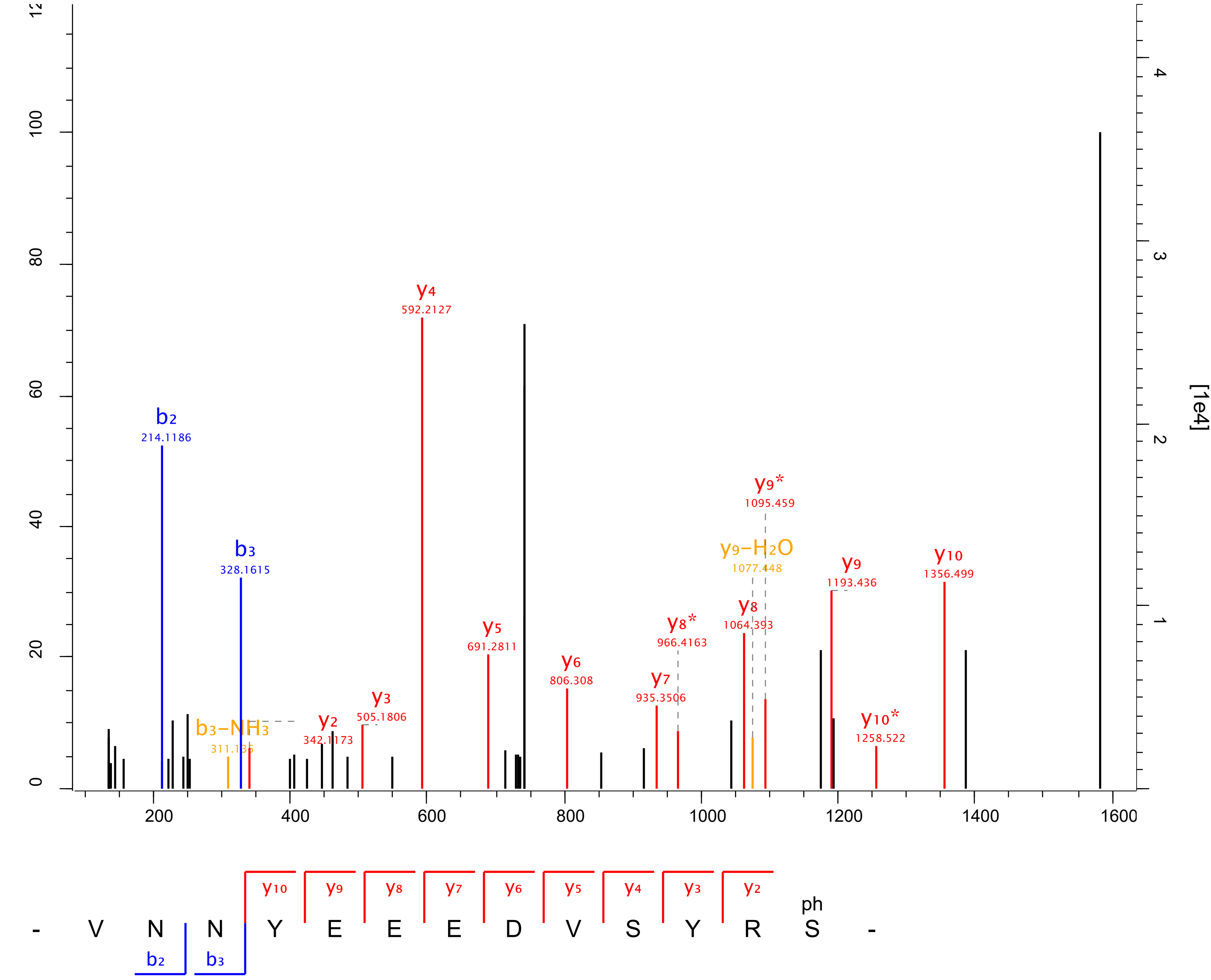

| Raw file | Scan | Method | Score | m/z |
| --- | --- | --- | --- | --- |
| water_02_1 | 15305 | FTMS; HCD | 79.39 | 741.31 |

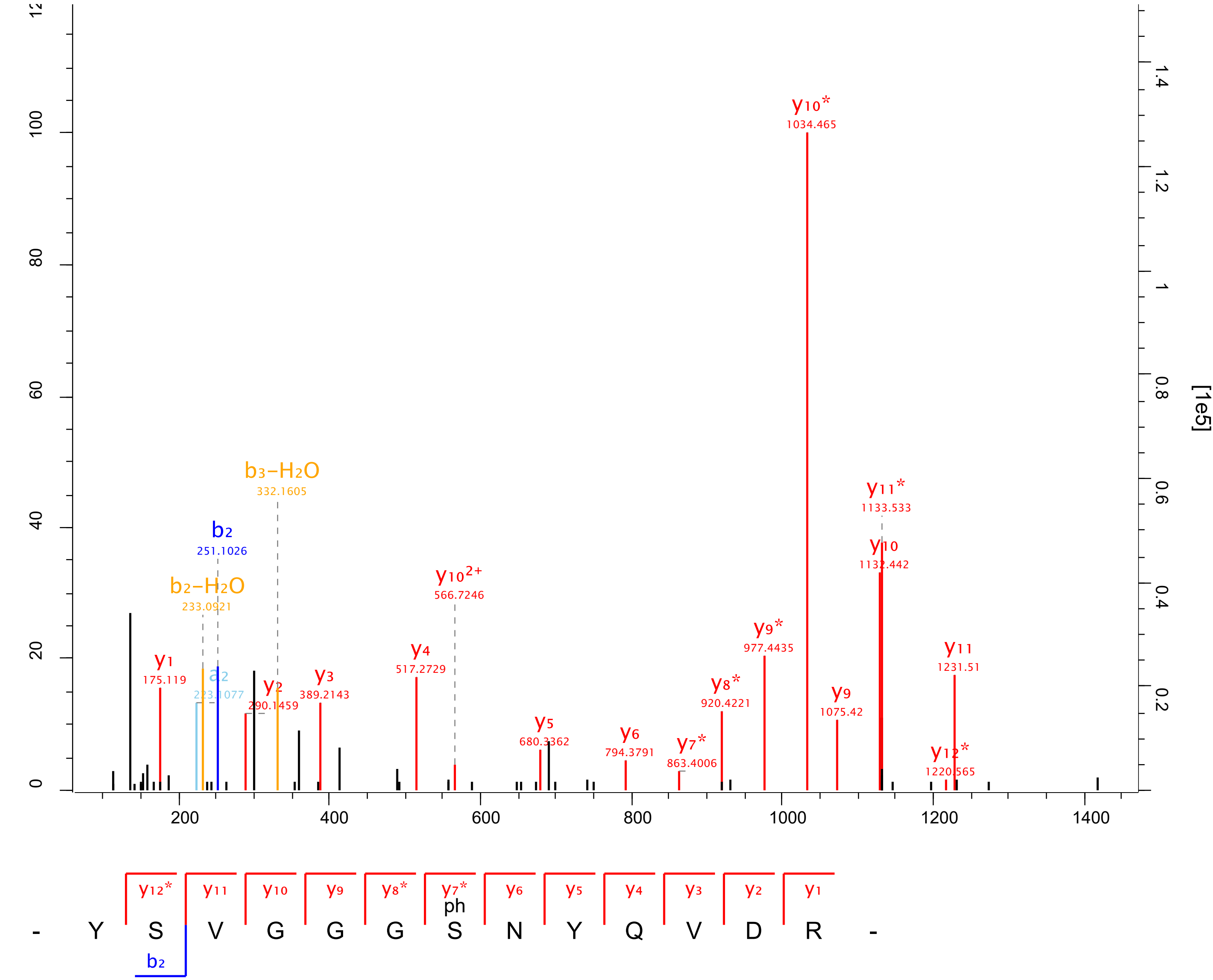

| Raw file | Scan | Method | Score | m/z |
| --- | --- | --- | --- | --- |
| water_02_1 | 1534 | FTMS; HCD | 56.82 | 540.55 |

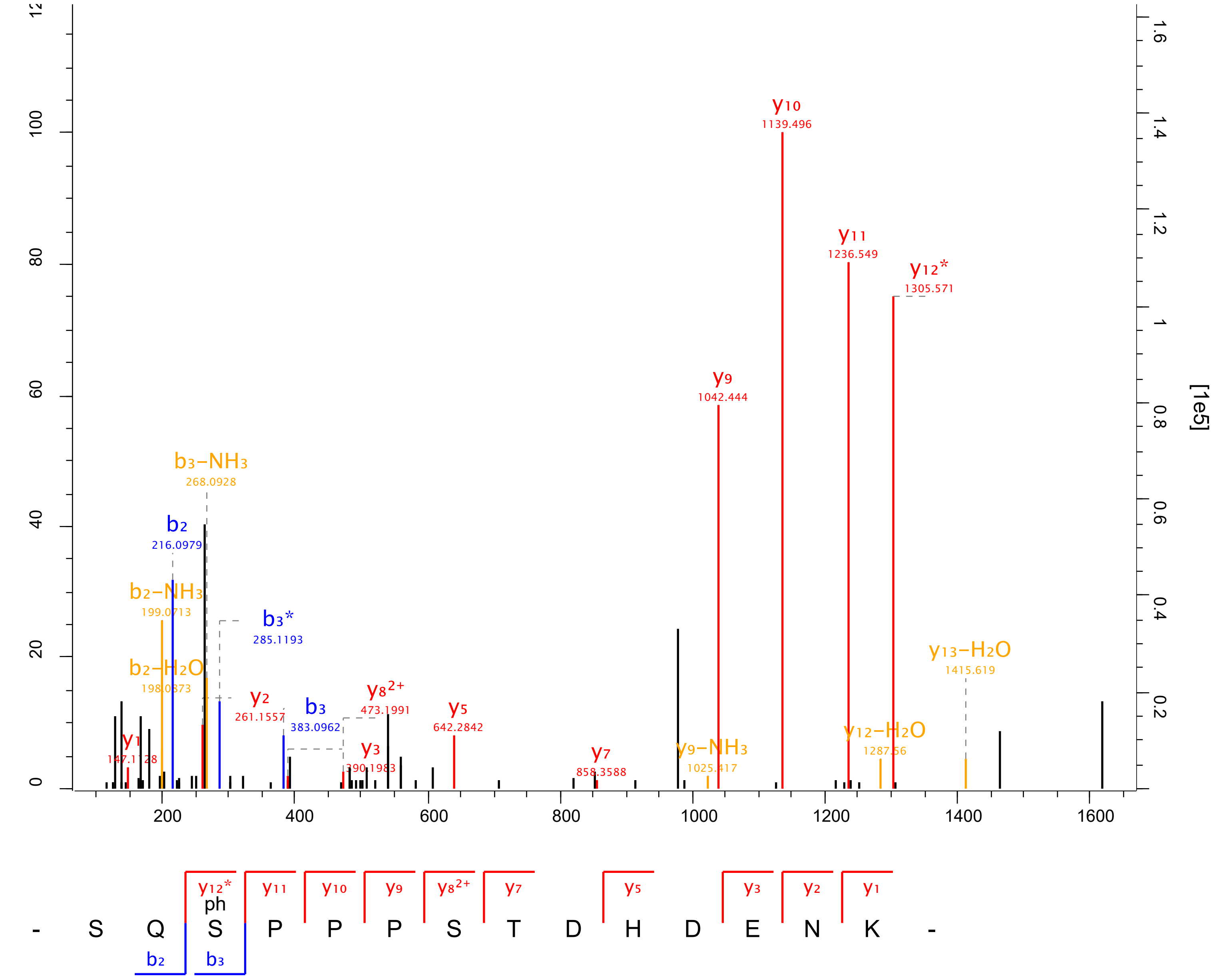

| Raw file | Scan | Method | Score | m/z |
|---|---|---|---|---|
| water_02_1 | 15343 | FTMS; HCD | 124.69 | 624.93 |

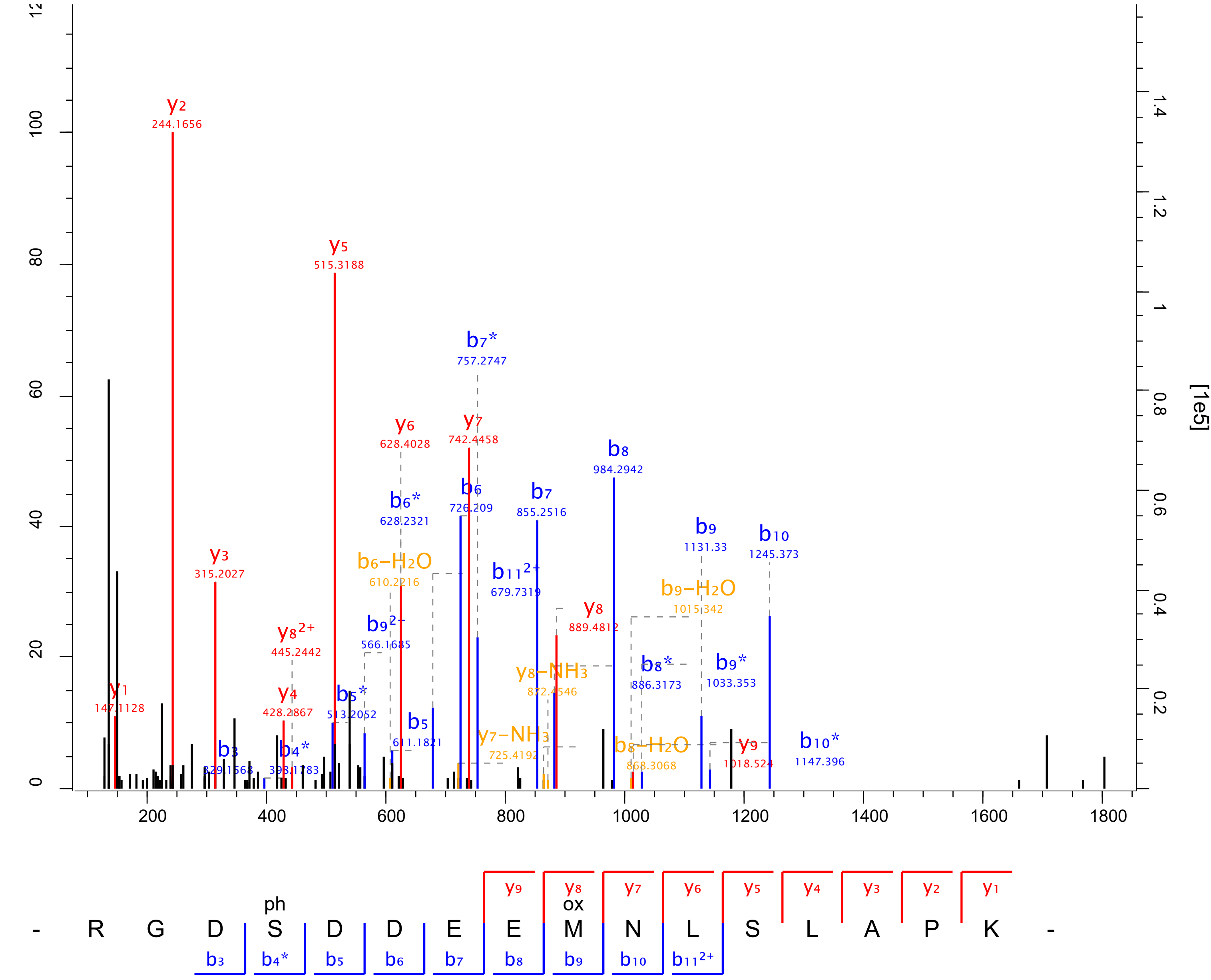

| Raw file | Scan | Method | Score | m/z |
|---|---|---|---|---|
| water_02_1 | 1545 | FTMS; HCD | 74.96 | 536.22 |

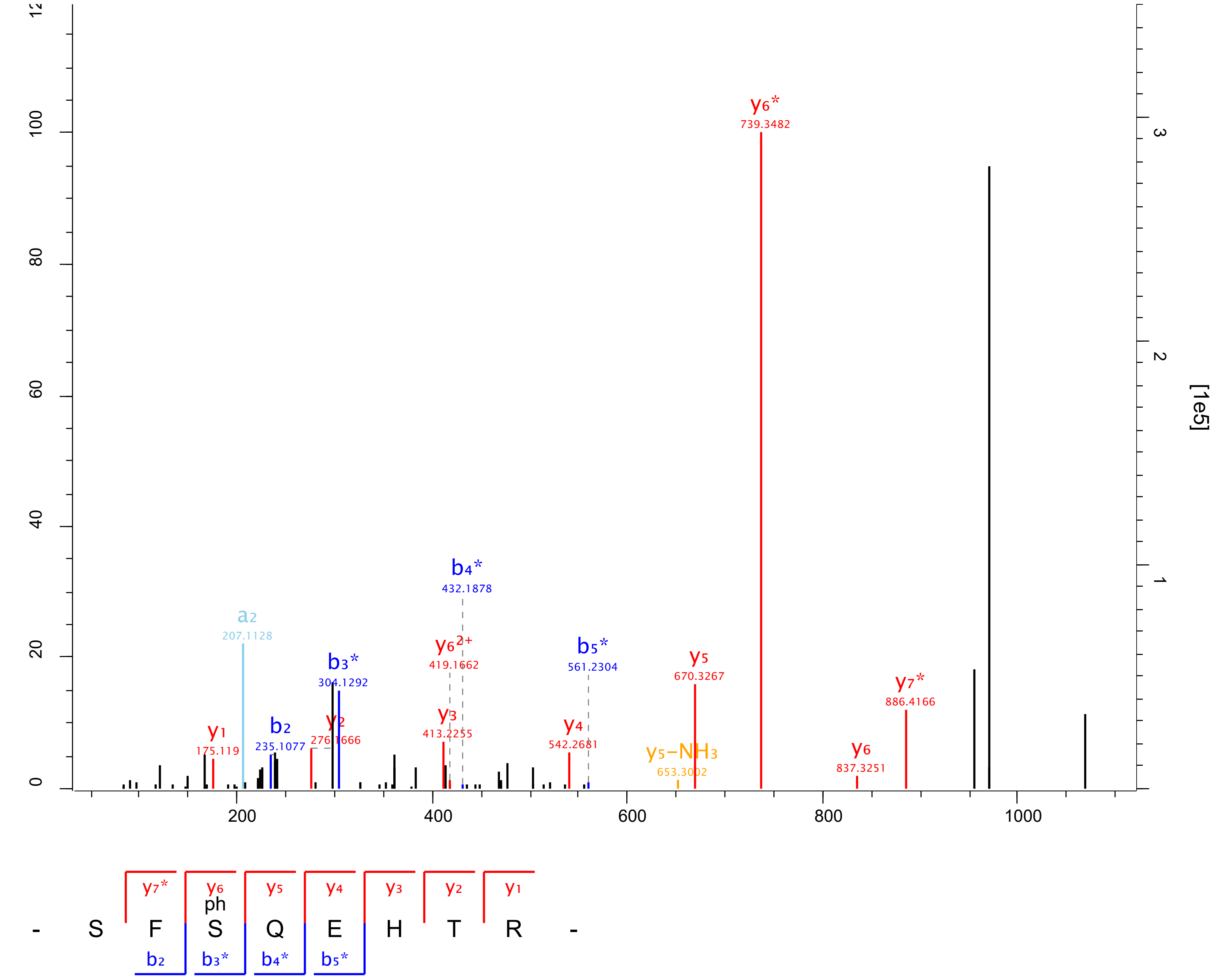

| Raw file | Scan | Method | Score | m/z |
|---|---|---|---|---|
| water_02_1 | 15491 | FTMS; HCD | 80.36 | 707.32 |

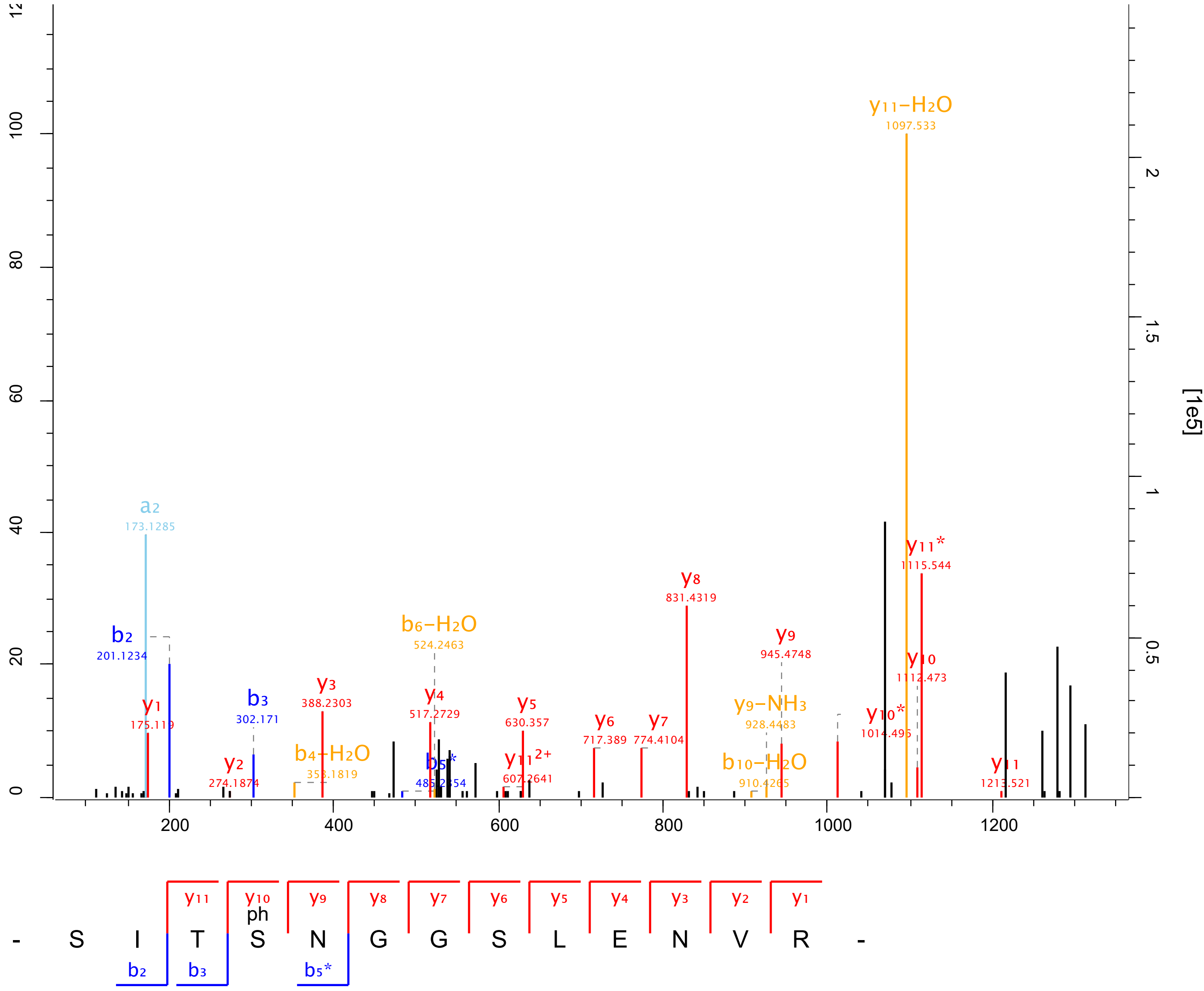

| Raw file | Scan | Method | Score | m/z |
| --- | --- | --- | --- | --- |
| water_02_1 | 1550 | FTMS; HCD | 92.77 | 462.23 |

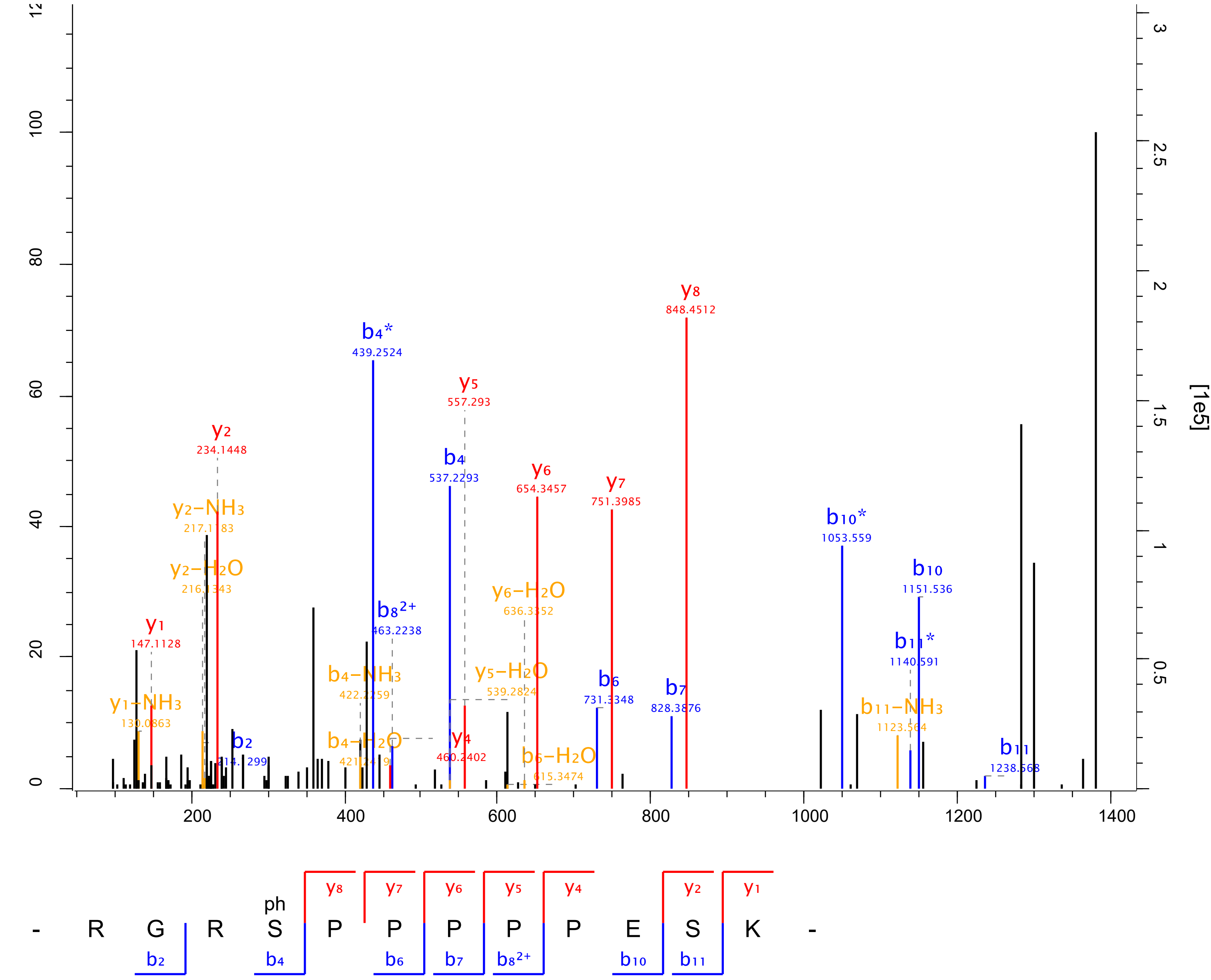

| Raw file | Scan | Method | Score | m/z |
| --- | --- | --- | --- | --- |
| water_02_1 | 15598 | FTMS; HCD | 70.37 | 488.91 |

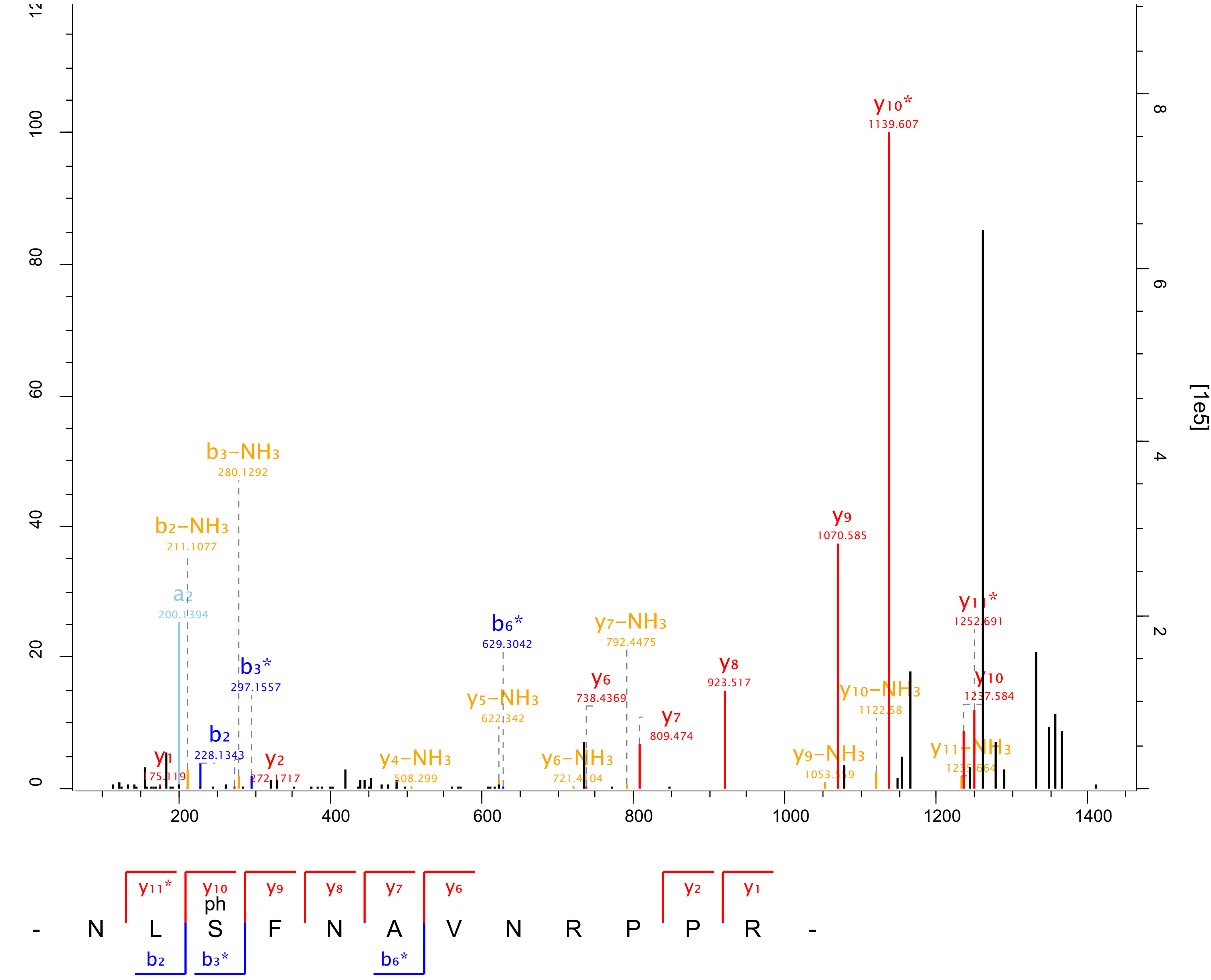

| Raw file | Scan | Method | Score | m/z |
| --- | --- | --- | --- | --- |
| water_02_1 | 15633 | FTMS; HCD | 124.84 | 507.59 |

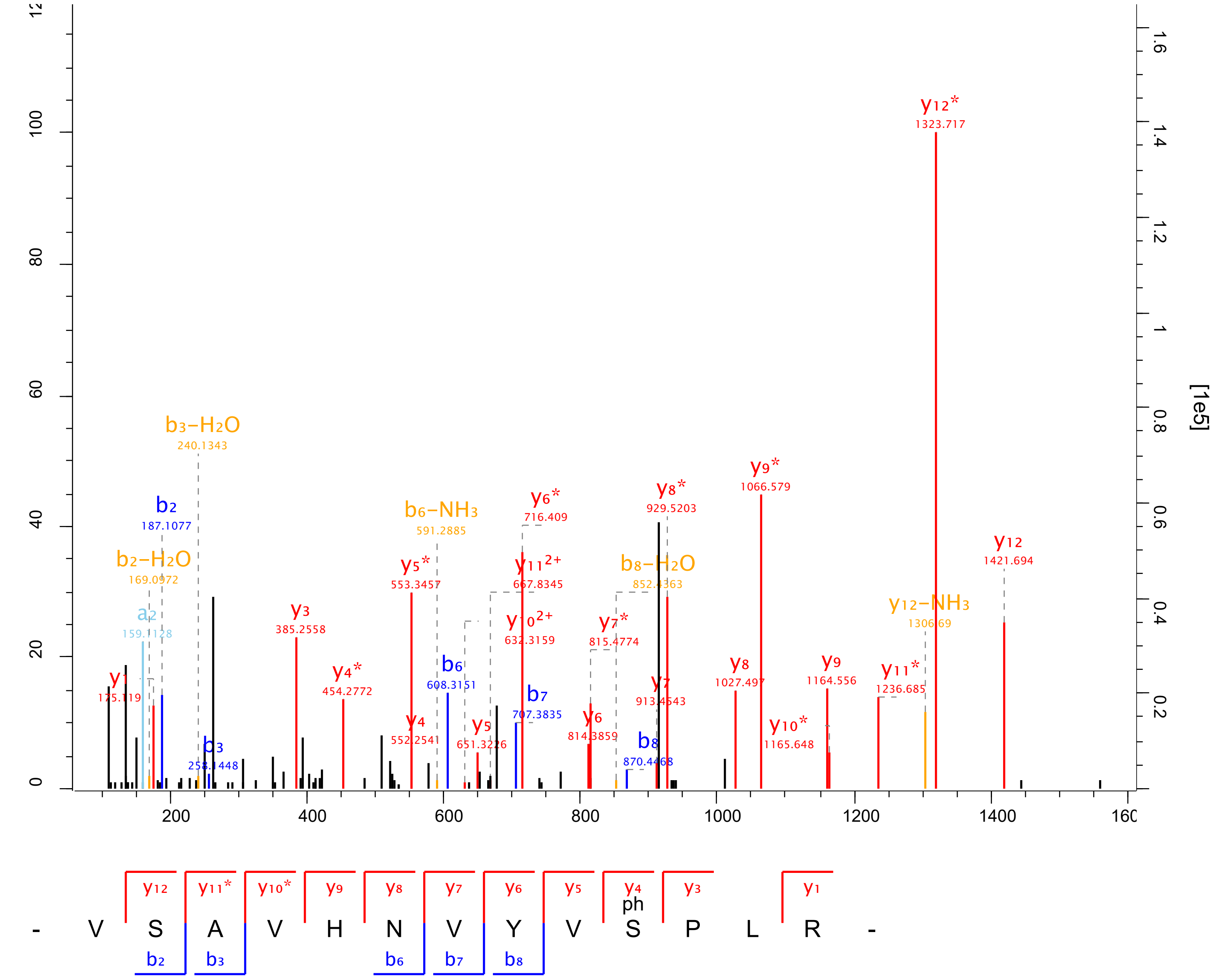

| Raw file | Scan | Method | Score | m/z |
| --- | --- | --- | --- | --- |
| water_02_1 | 15650 | FTMS; HCD | 73.83 | 416.55 |

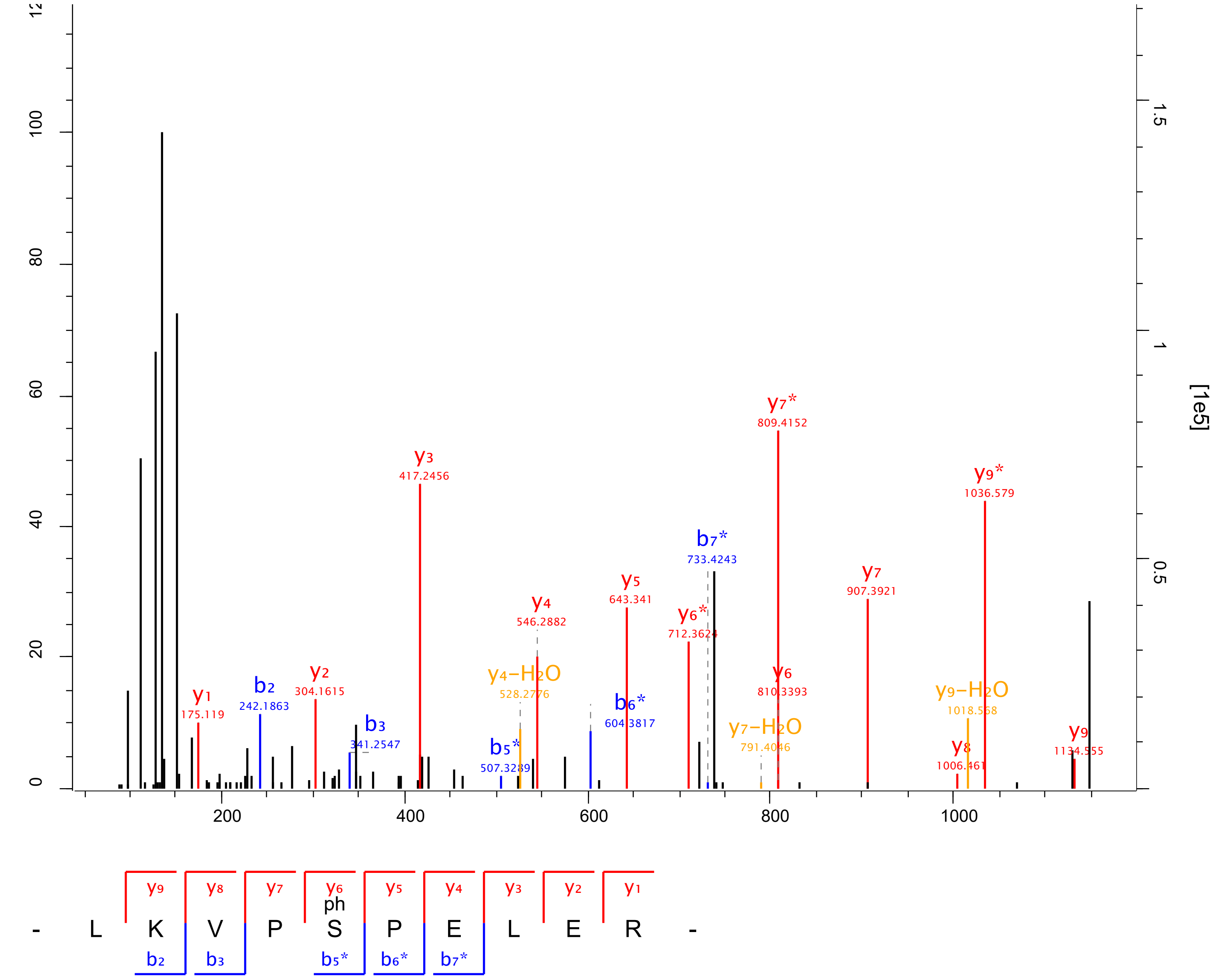

| Raw file | Scan | Method | Score | m/z |
| --- | --- | --- | --- | --- |
| water_02_1 | 15697 | FTMS; HCD | 143.54 | 780.67 |

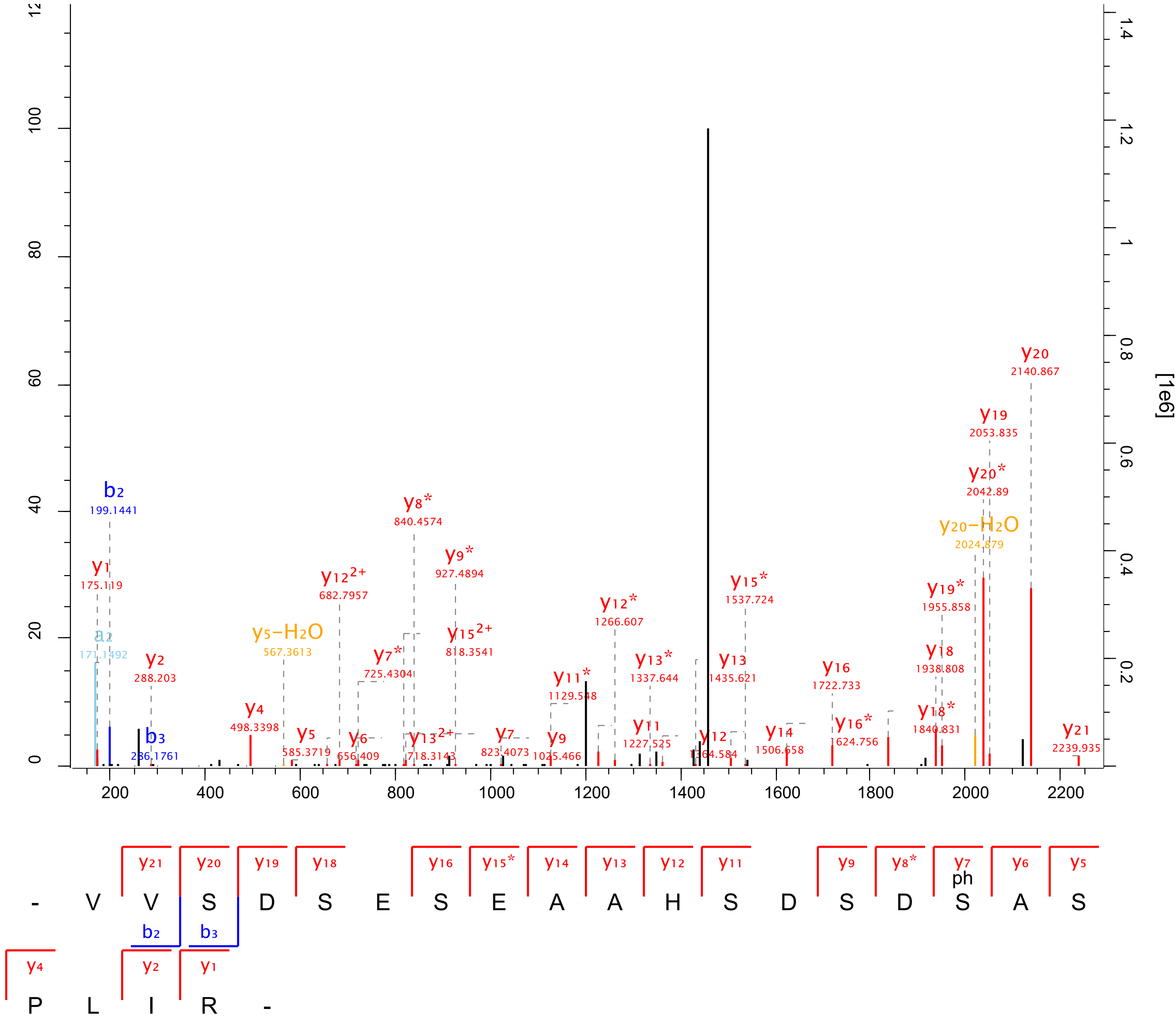

| Raw file | Scan | Method | Score | m/z |
| --- | --- | --- | --- | --- |
| water_02_1 | 1573 | FTMS; HCD | 146.19 | 611.58 |

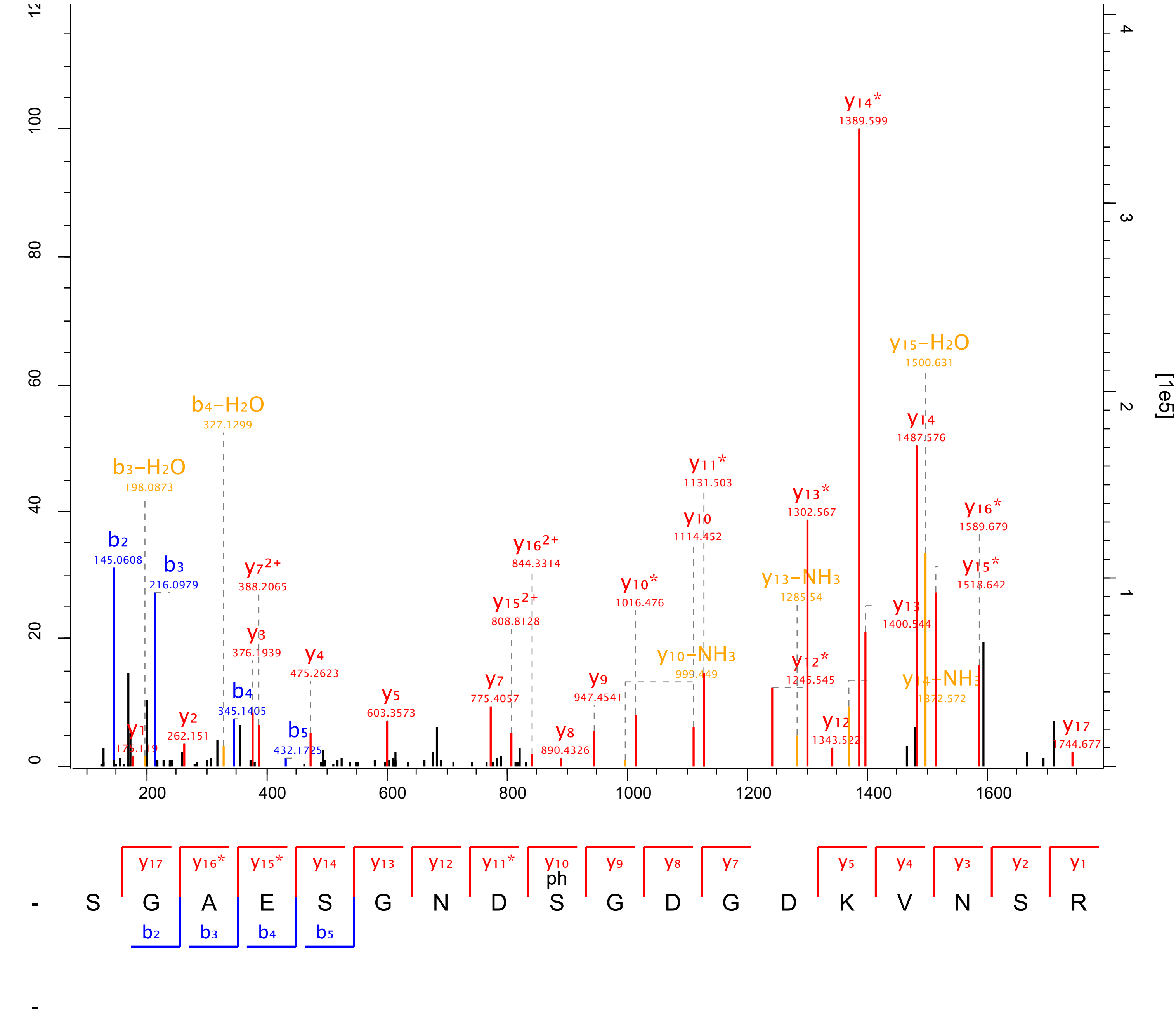

| Raw file | Scan | Method | Score | m/z |
|---|---|---|---|---|
| water_02_1 | 15810 | FTMS; HCD | 176.44 | 889.32 |

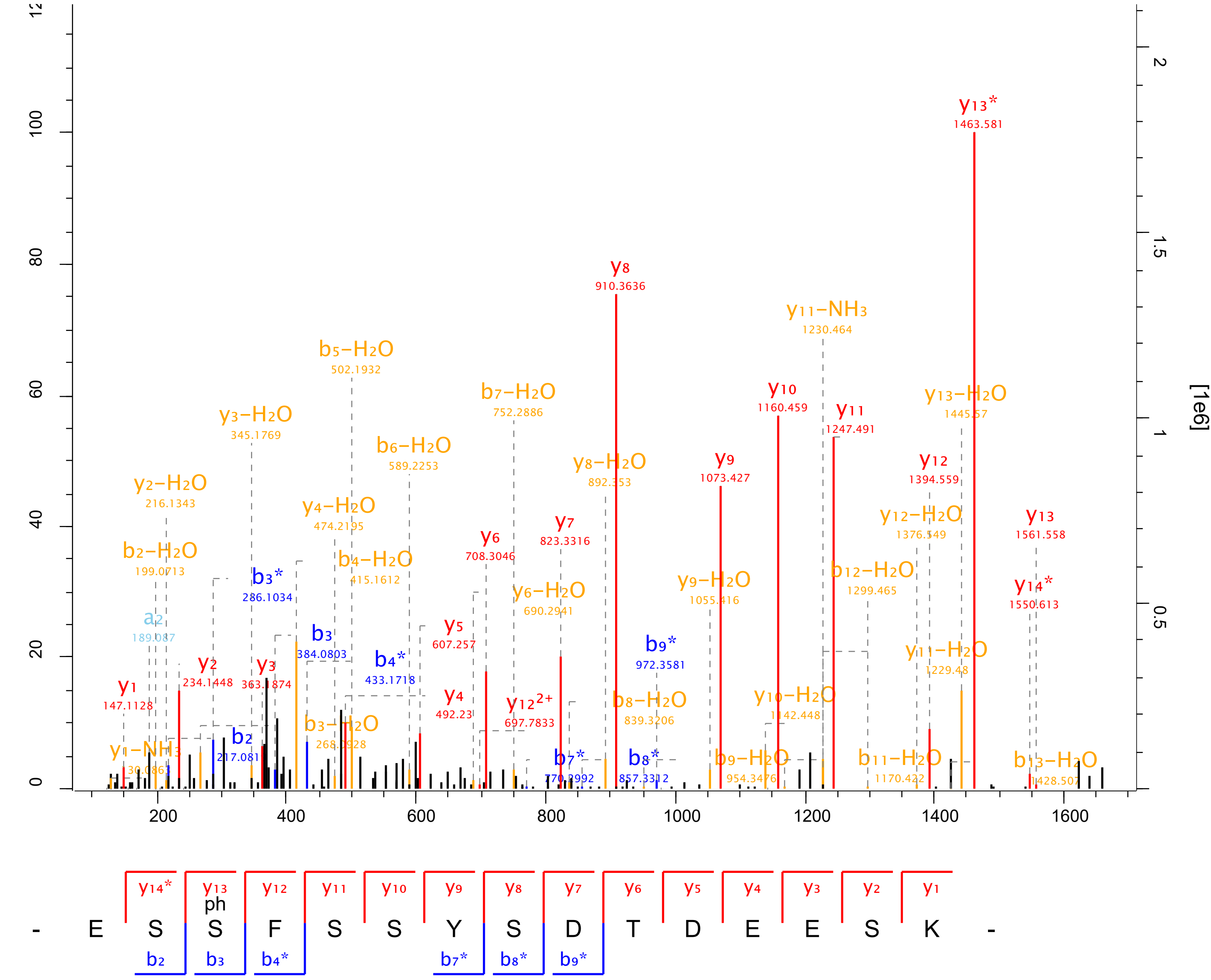

| Raw file | Scan | Method | Score | m/z |
|---|---|---|---|---|
| water_02_1 | 1582 | FTMS; HCD | 54.02 | 554.76 |

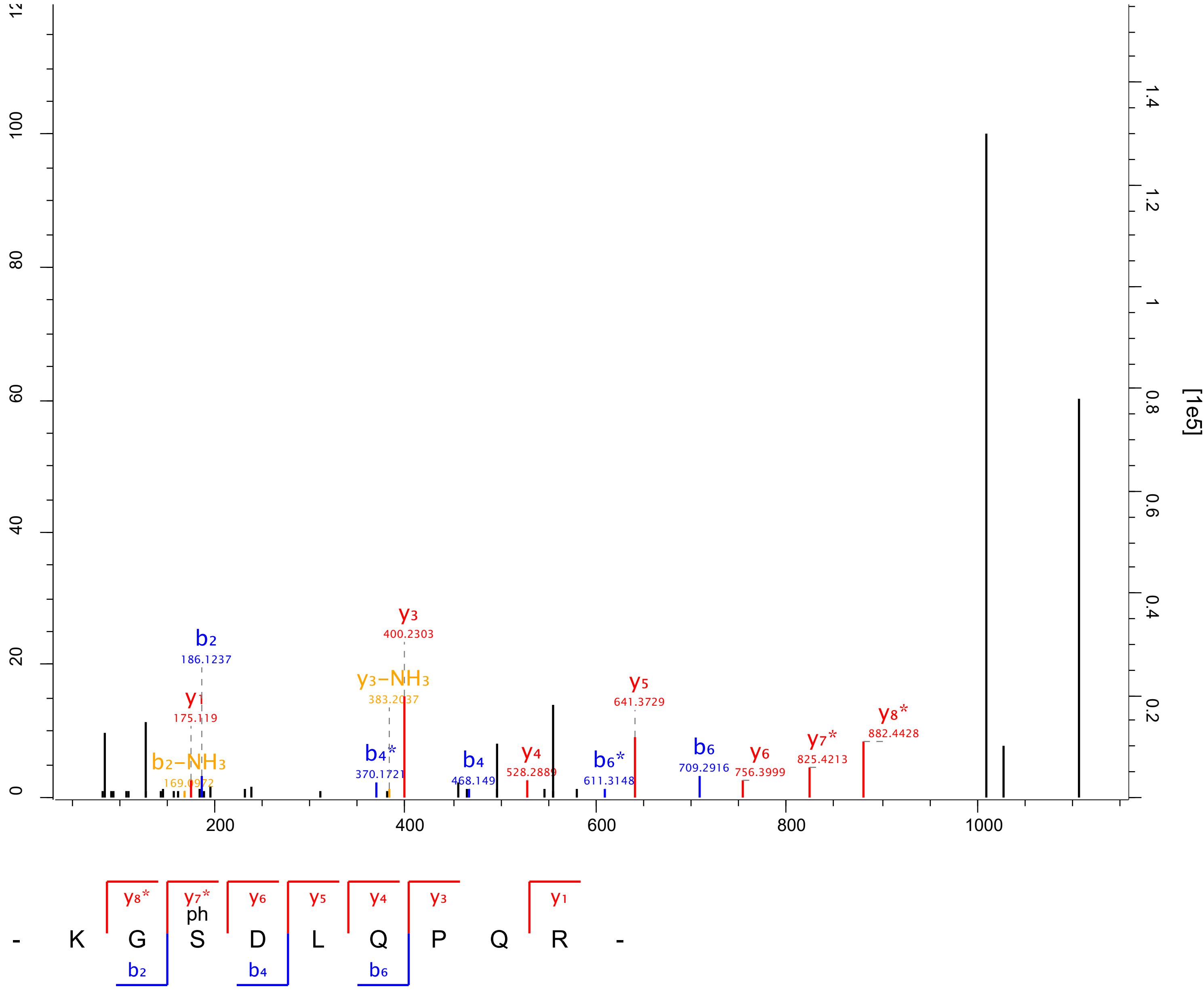

| Raw file | Scan | Method | Score | m/z |
| --- | --- | --- | --- | --- |
| water_02_1 | 15846 | FTMS; HCD | 67.23 | 873.36 |

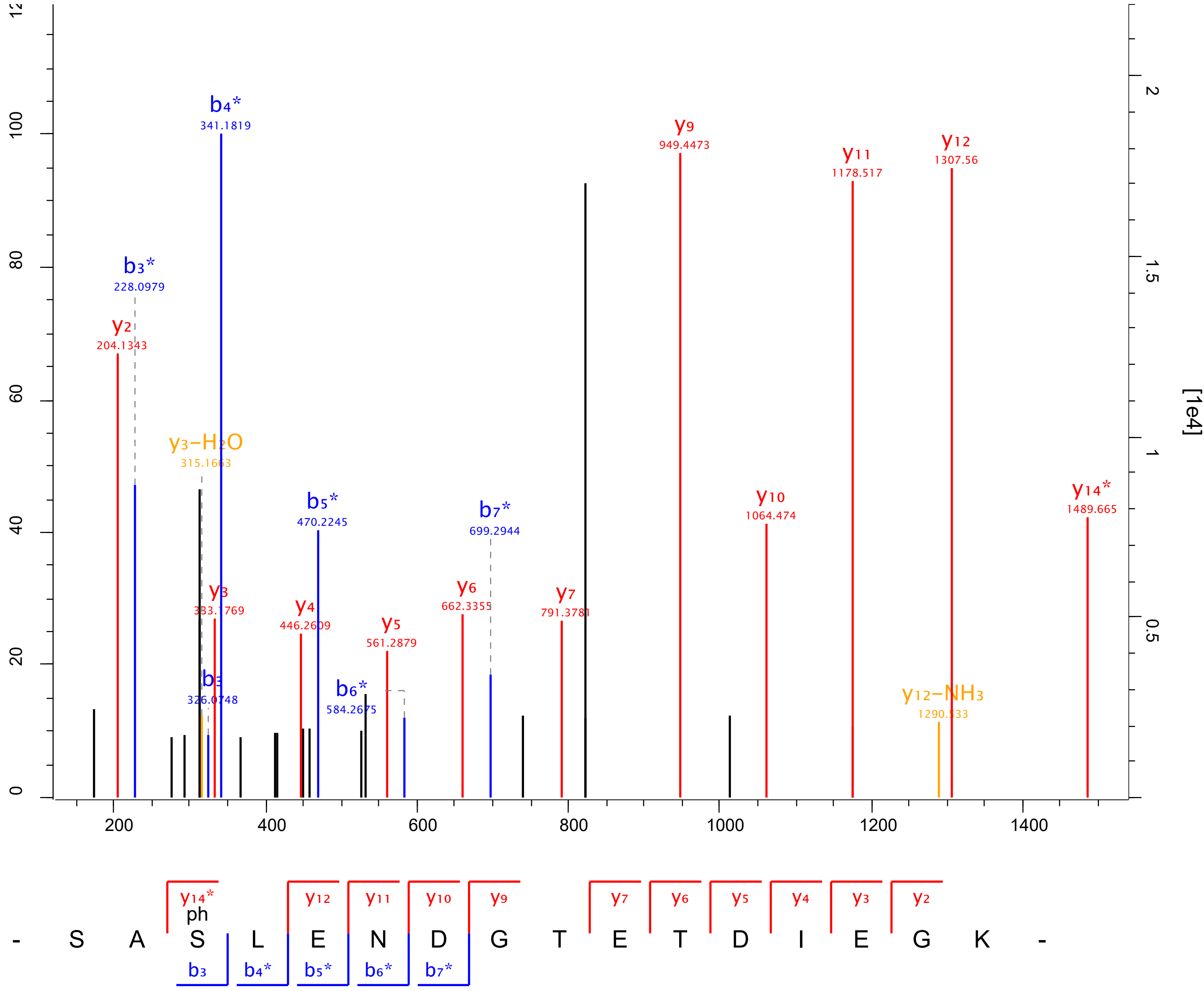

| Raw file | Scan | Method | Score | m/z |
|---|---|---|---|---|
| water_02_1 | 15865 | FTMS; HCD | 115.63 | 851.35 |

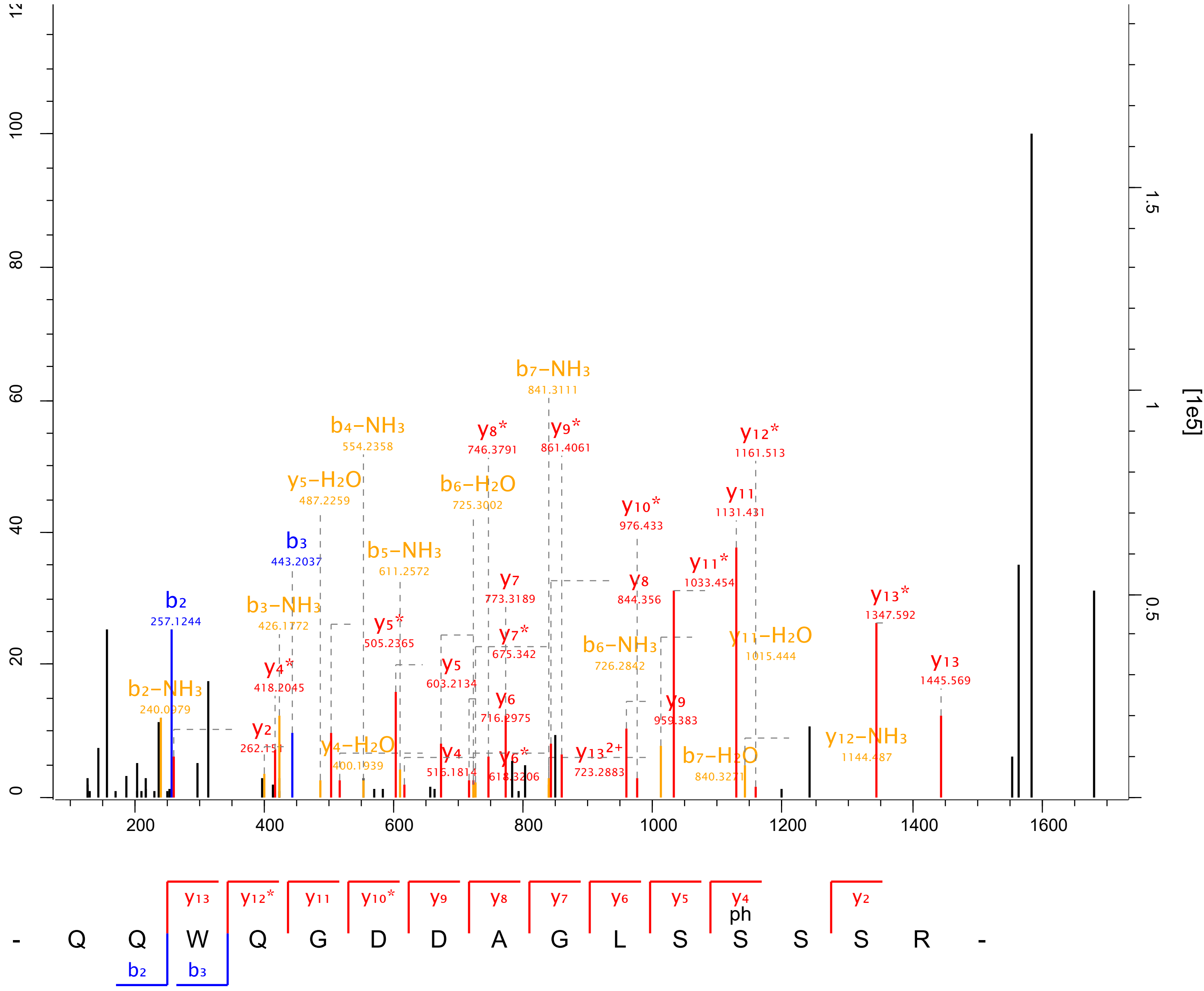

| Raw file | Scan | Method | Score | m/z |
| --- | --- | --- | --- | --- |
| water_02_1 | 1601 | FTMS; HCD | 57.35 | 457.71 |

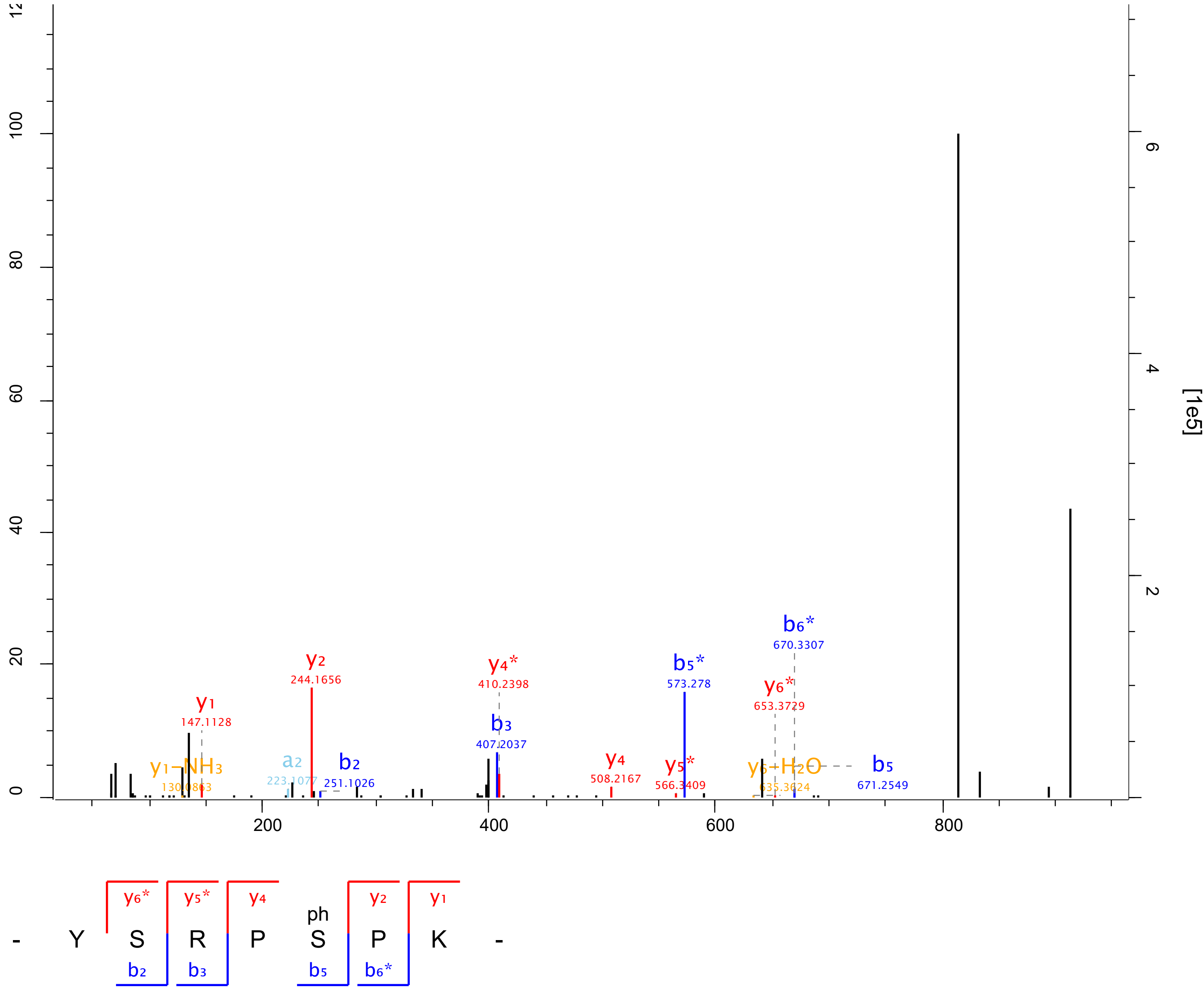

| Raw file | Scan | Method | Score | m/z |
| --- | --- | --- | --- | --- |
| water_02_1 | 1605 | FTMS; HCD | 41.16 | 348.49 |

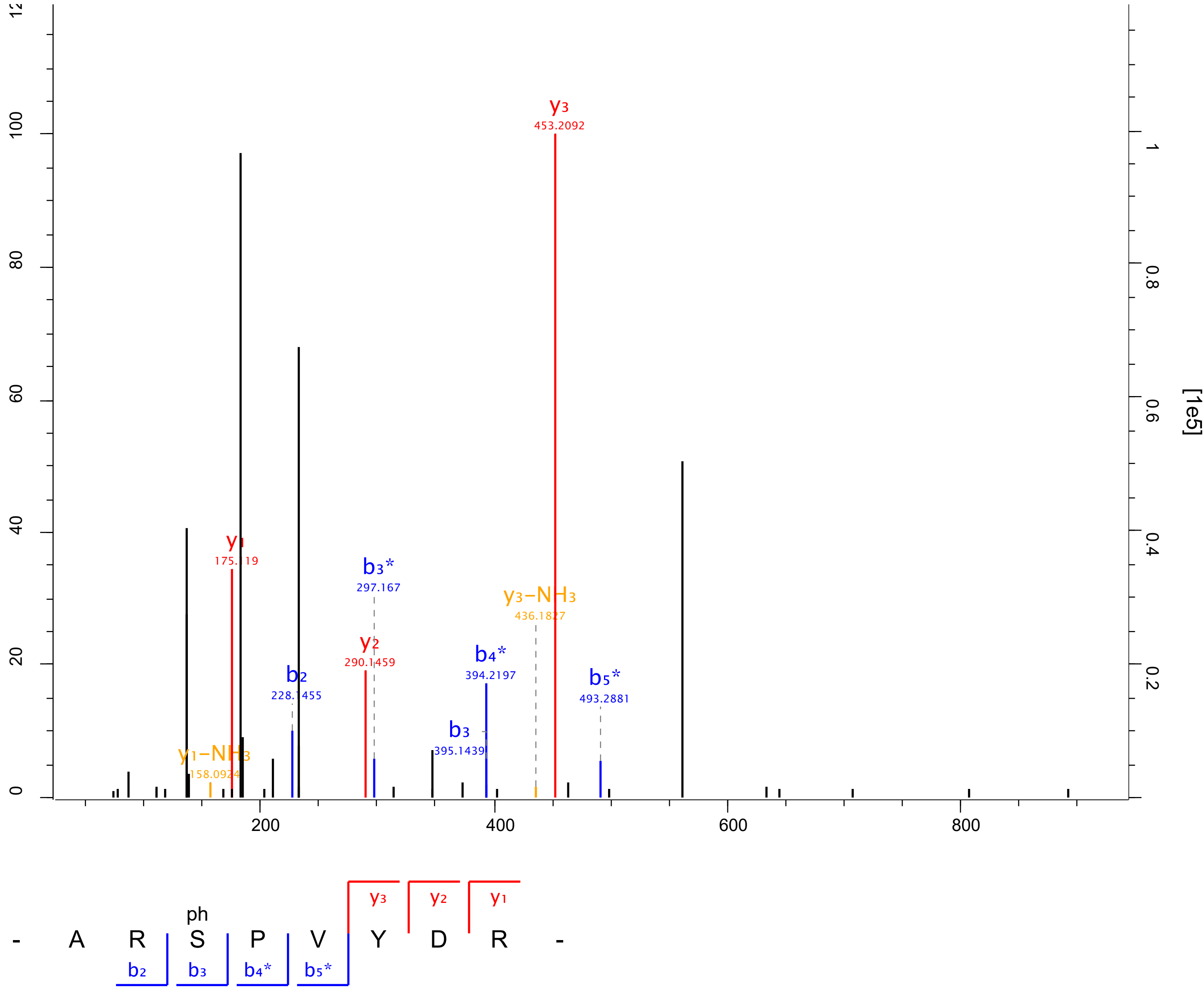

| Raw file | Scan | Method | Score | m/z |
|---|---|---|---|---|
| water_02_1 | 16067 | FTMS; HCD | 117.23 | 859.85 |

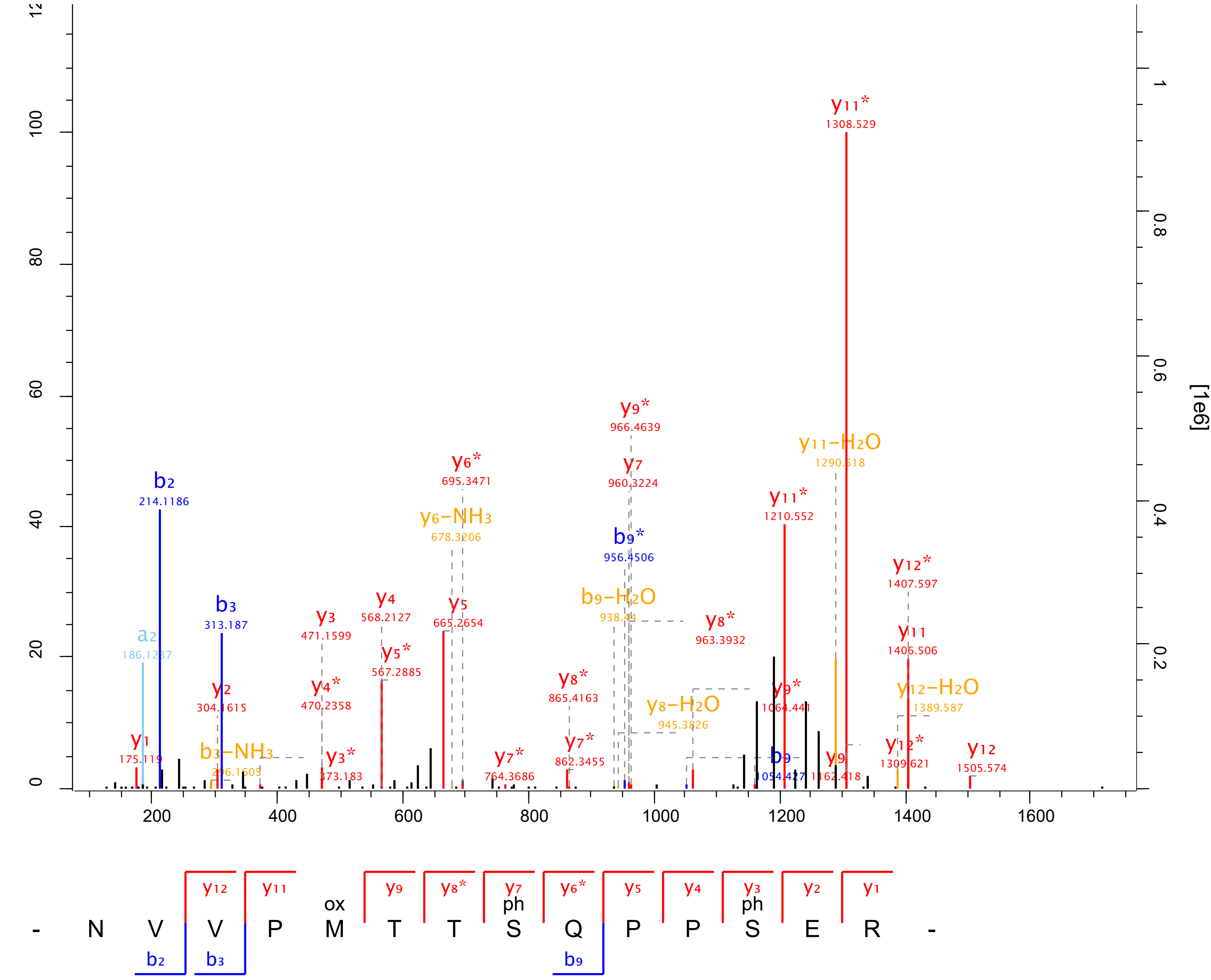

| Raw file | Scan | Method | Score | m/z |
|---|---|---|---|---|
| water_02_1 | 16076 | FTMS; HCD | 100.88 | 482.73 |

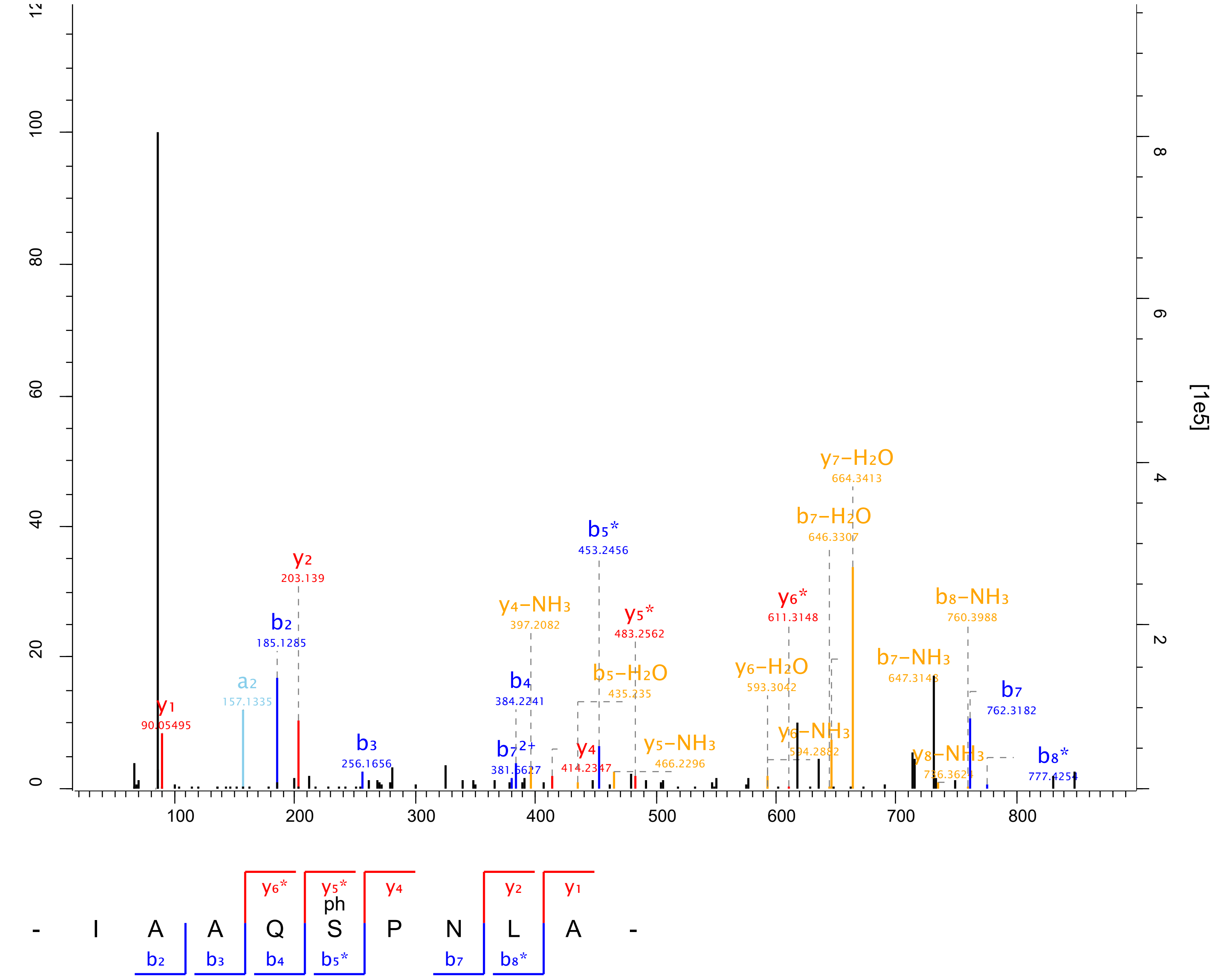

| Raw file | Scan | Method | Score | m/z |
|---|---|---|---|---|
| water_02_1 | 16159 | FTMS; HCD | 66.25 | 512.25 |

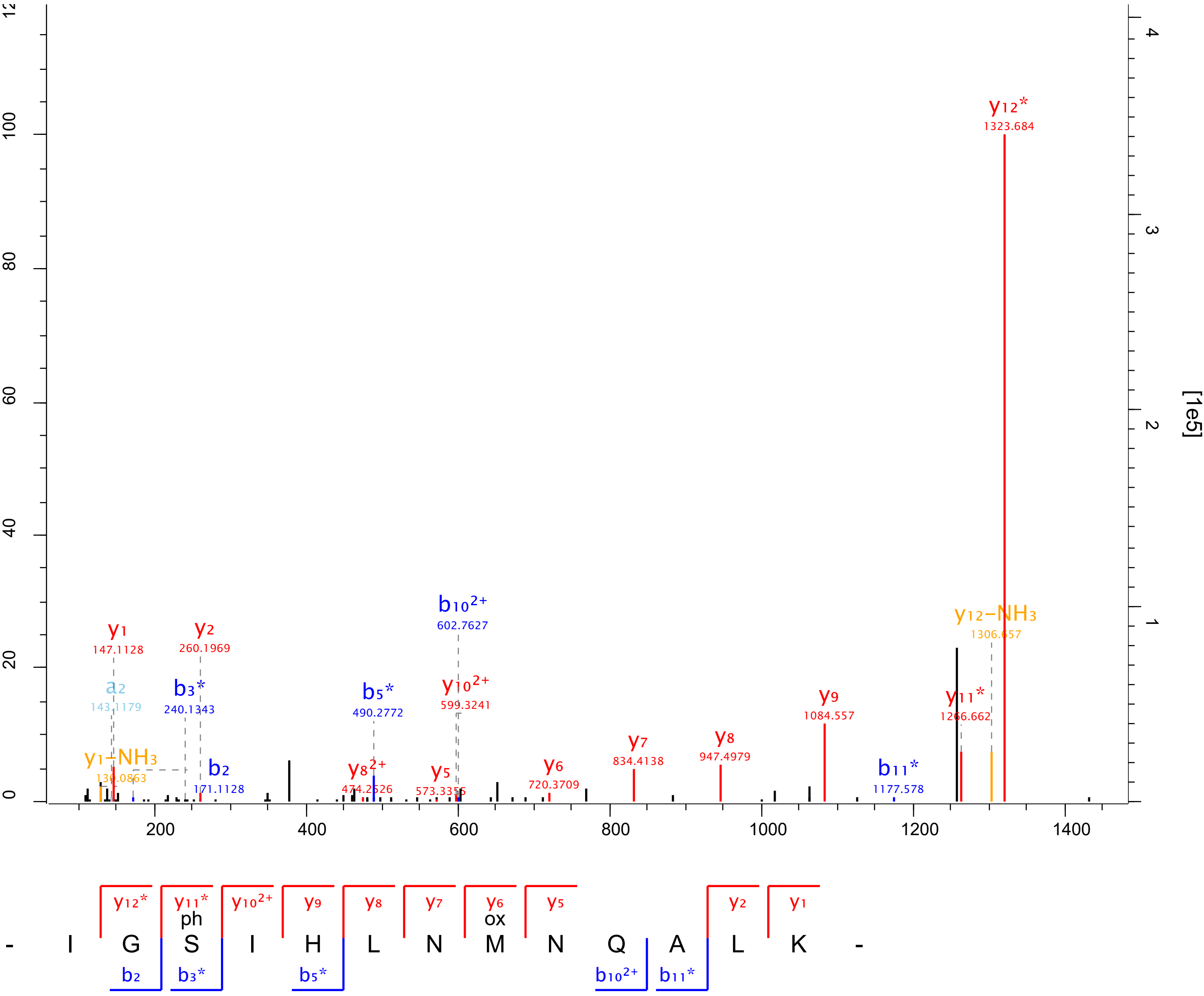

| Raw file | Scan | Method | Score | m/z |
| --- | --- | --- | --- | --- |
| water_02_1 | 16160 | FTMS; HCD | 72.99 | 843.02 |

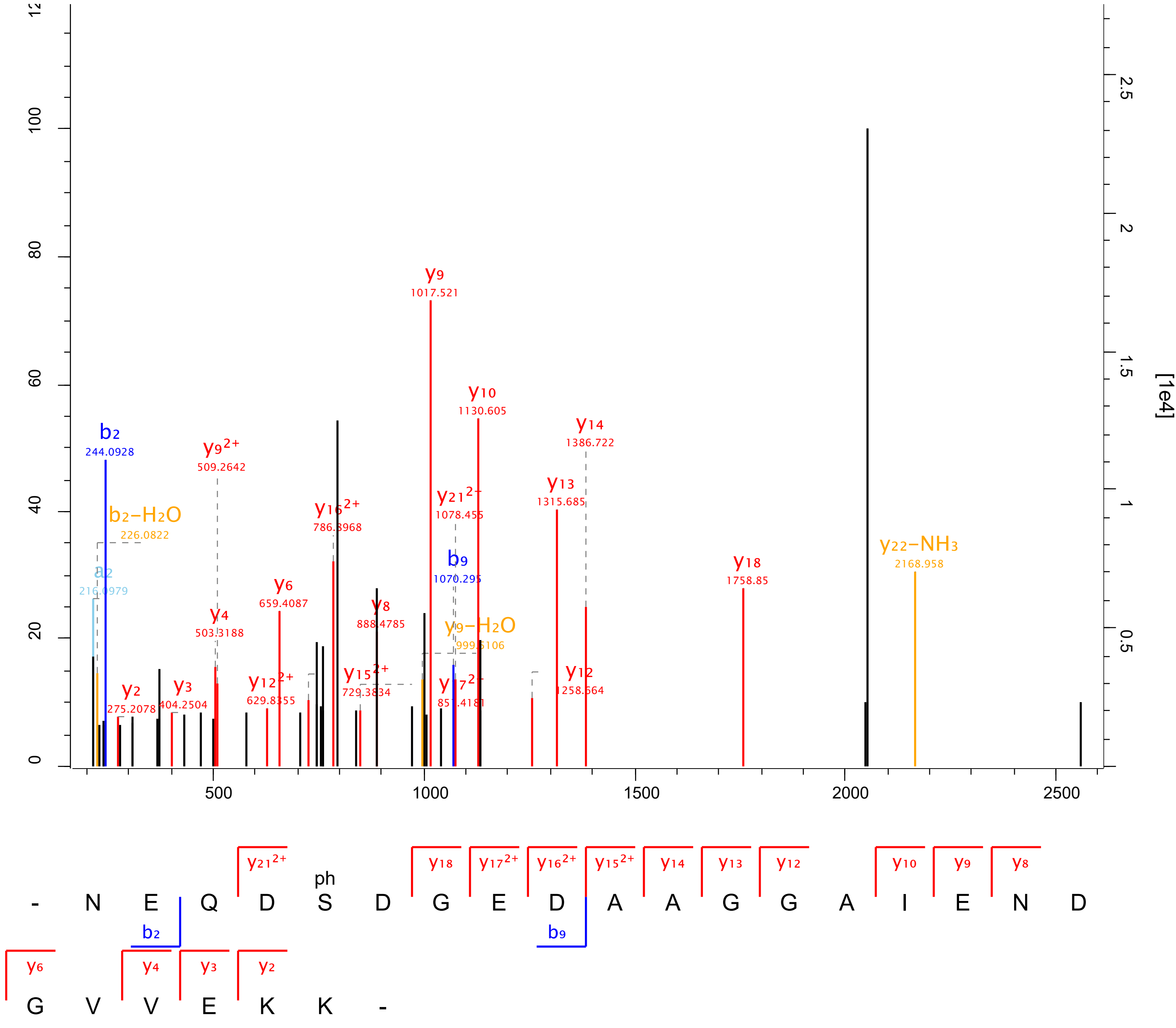

| Raw file | Scan | Method | Score | m/z |
|---|---|---|---|---|
| water_02_1 | 16294 | FTMS; HCD | 94.63 | 856.4 |

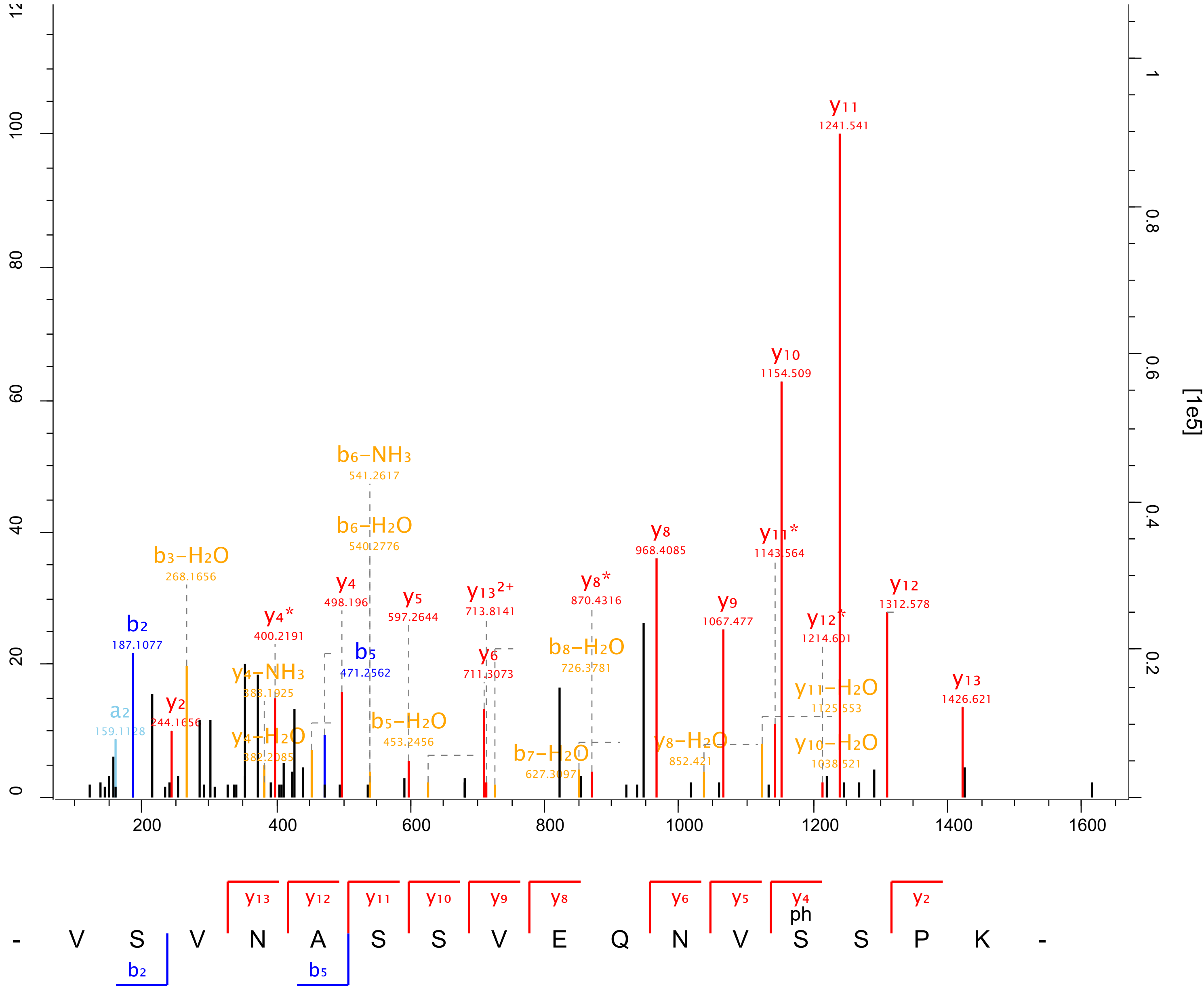

| Raw file | Scan | Method | Score | m/z |
| --- | --- | --- | --- | --- |
| water_02_1 | 16458 | FTMS; HCD | 157.5 | 503.86 |

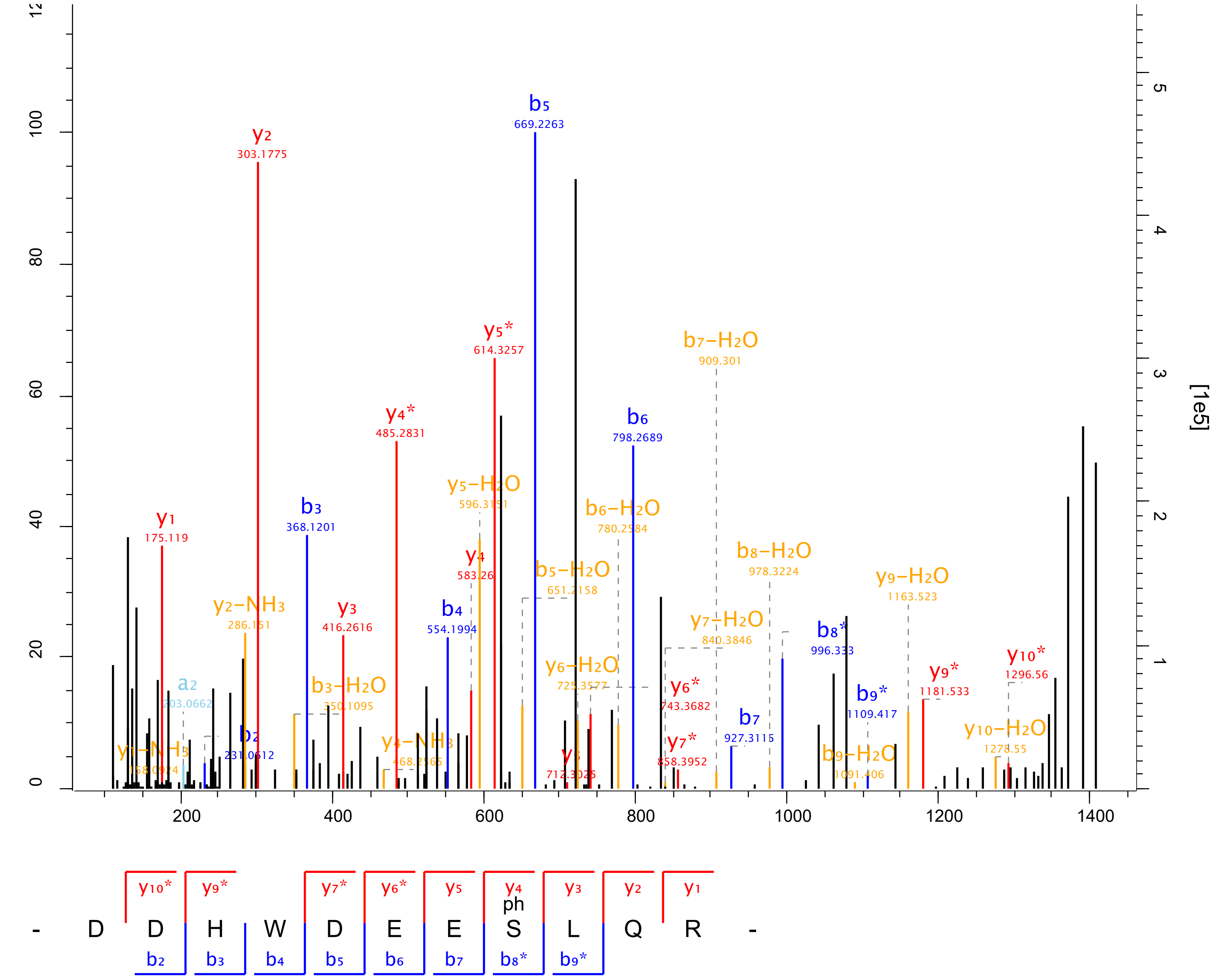

| Raw file | Scan | Method | Score | m/z |
| --- | --- | --- | --- | --- |
| water_02_1 | 16486 | FTMS; HCD | 165.3 | 794.35 |

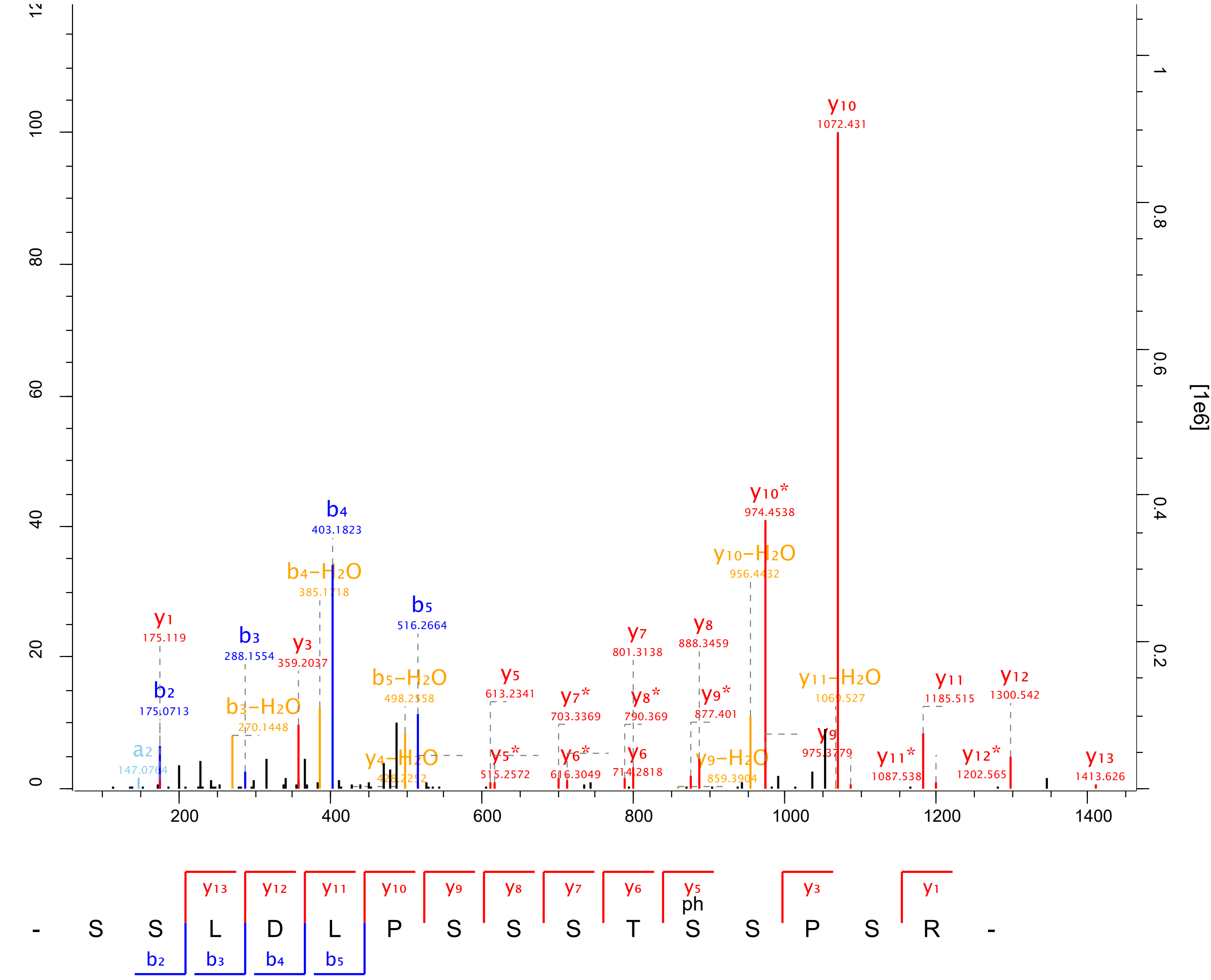

| Raw file | Scan | Method | Score | m/z |
|---|---|---|---|---|
| water_02_1 | 16524 | FTMS; HCD | 68.19 | 688.97 |

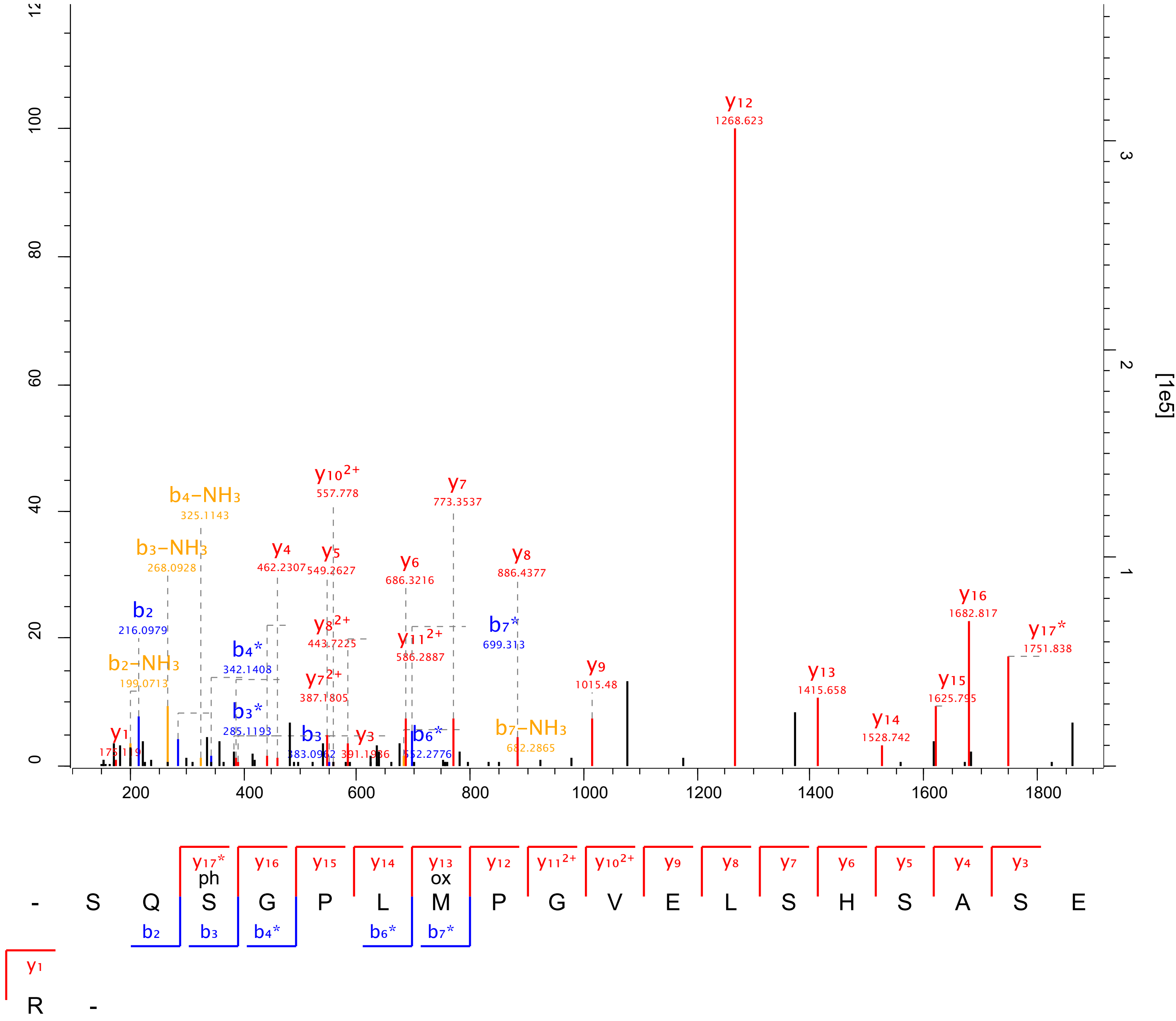

| Raw file | Scan | Method | Score | m/z |
|---|---|---|---|---|
| water_02_1 | 16559 | FTMS; HCD | 129.29 | 574.26 |

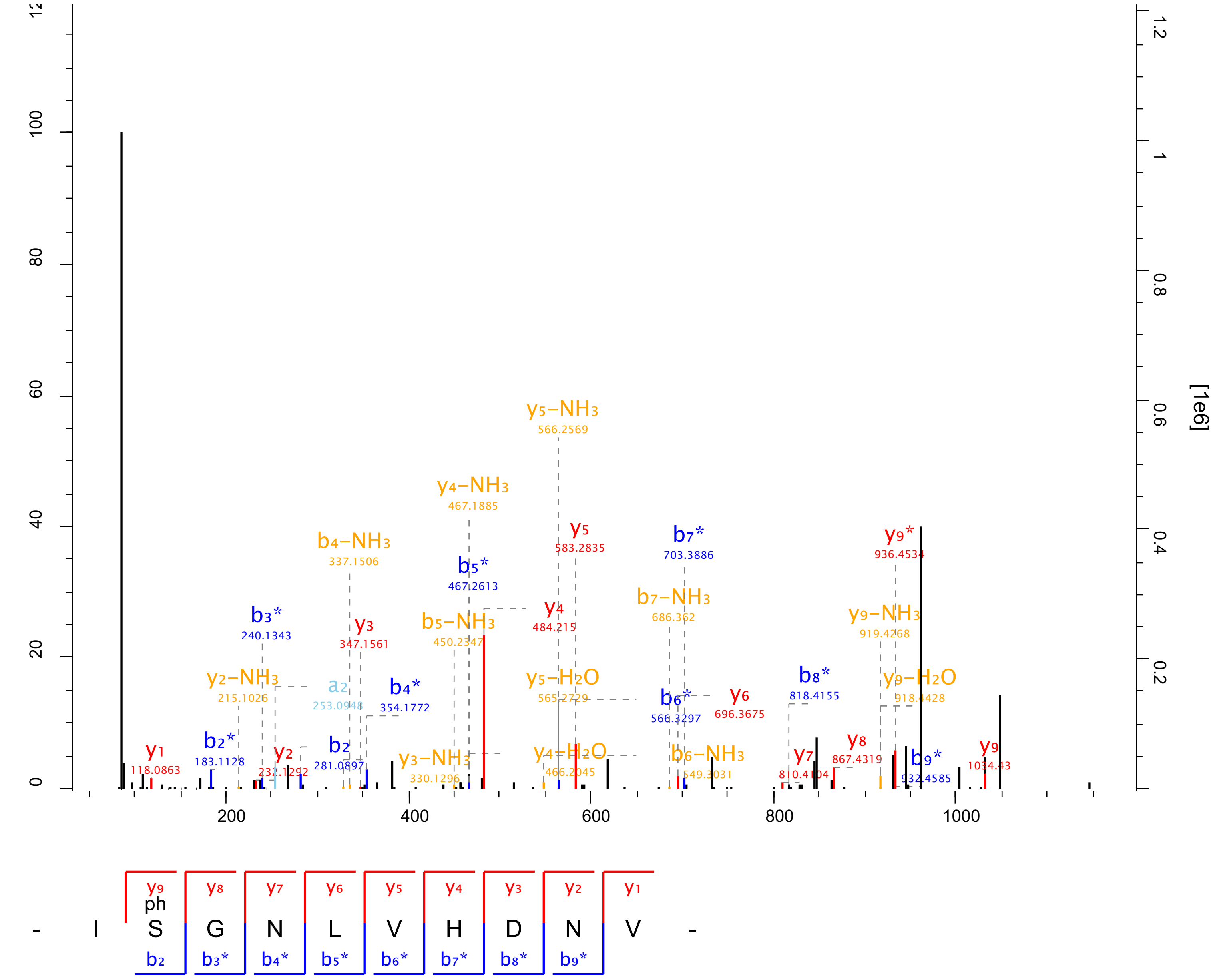

| Raw file | Scan | Method | Score | m/z |
| --- | --- | --- | --- | --- |
| water_02_1 | 16784 | FTMS; HCD | 239.43 | 865.85 |

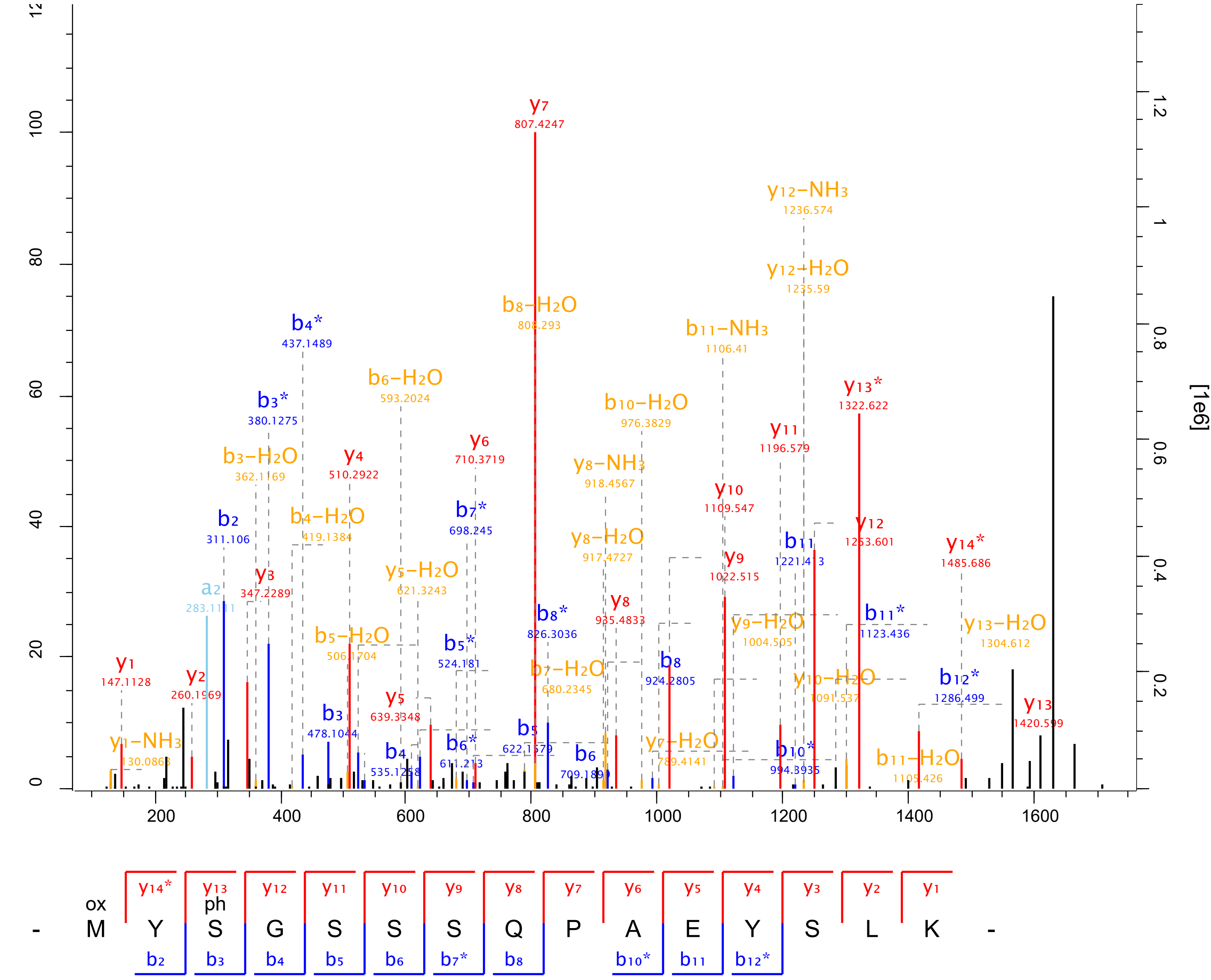

| Raw file | Scan | Method | Score | m/z |
| --- | --- | --- | --- | --- |
| water_02_1 | 16874 | FTMS; HCD | 94.2 | 848.86 |

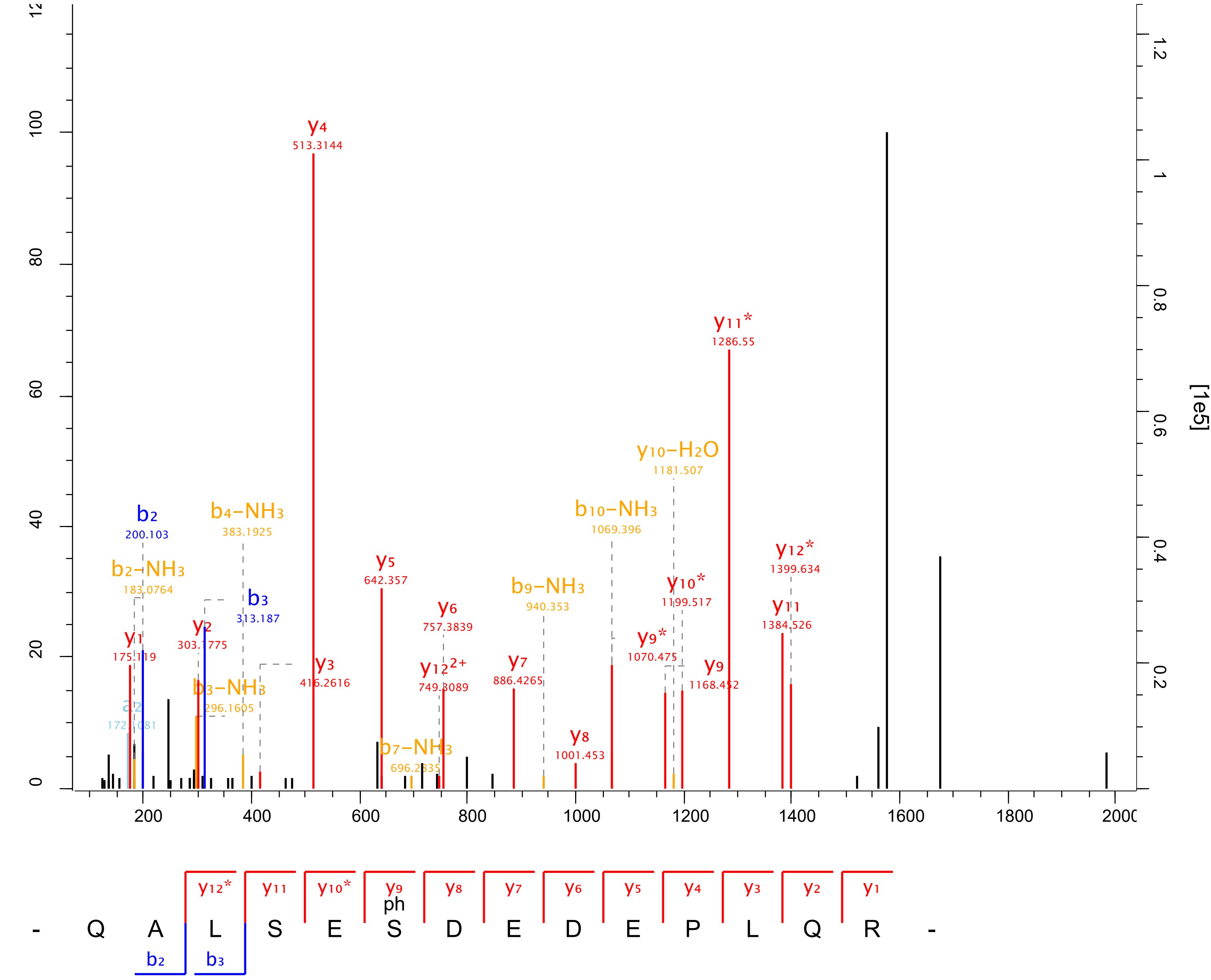

| Raw file | Scan | Method | Score | m/z |
|---|---|---|---|---|
| water_02_1 | 16906 | FTMS; HCD | 98.54 | 883.87 |

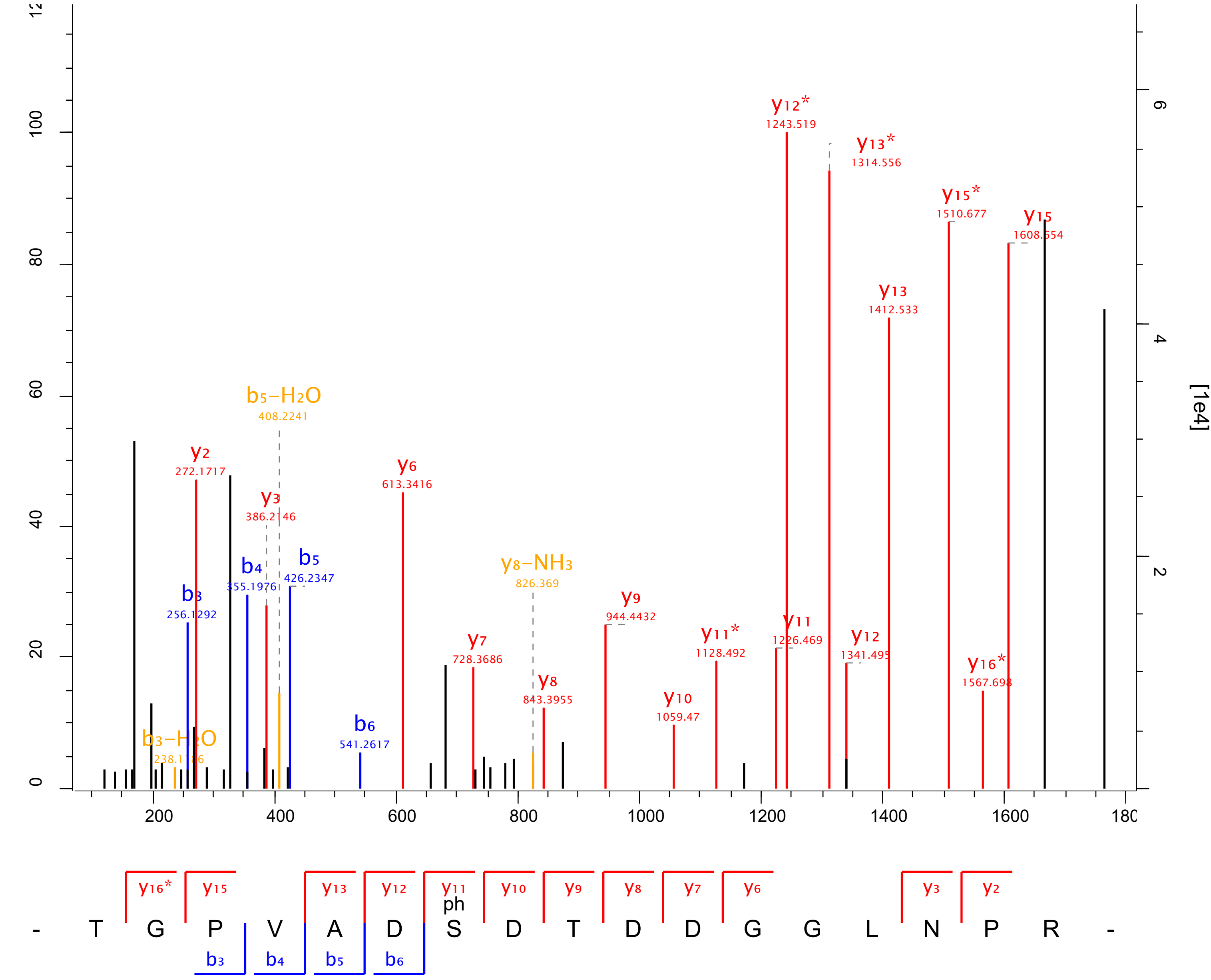

| Raw file | Scan | Method | Score | m/z |
|---|---|---|---|---|
| water_02_1 | 16927 | FTMS; HCD | 239.87 | 721.63 |

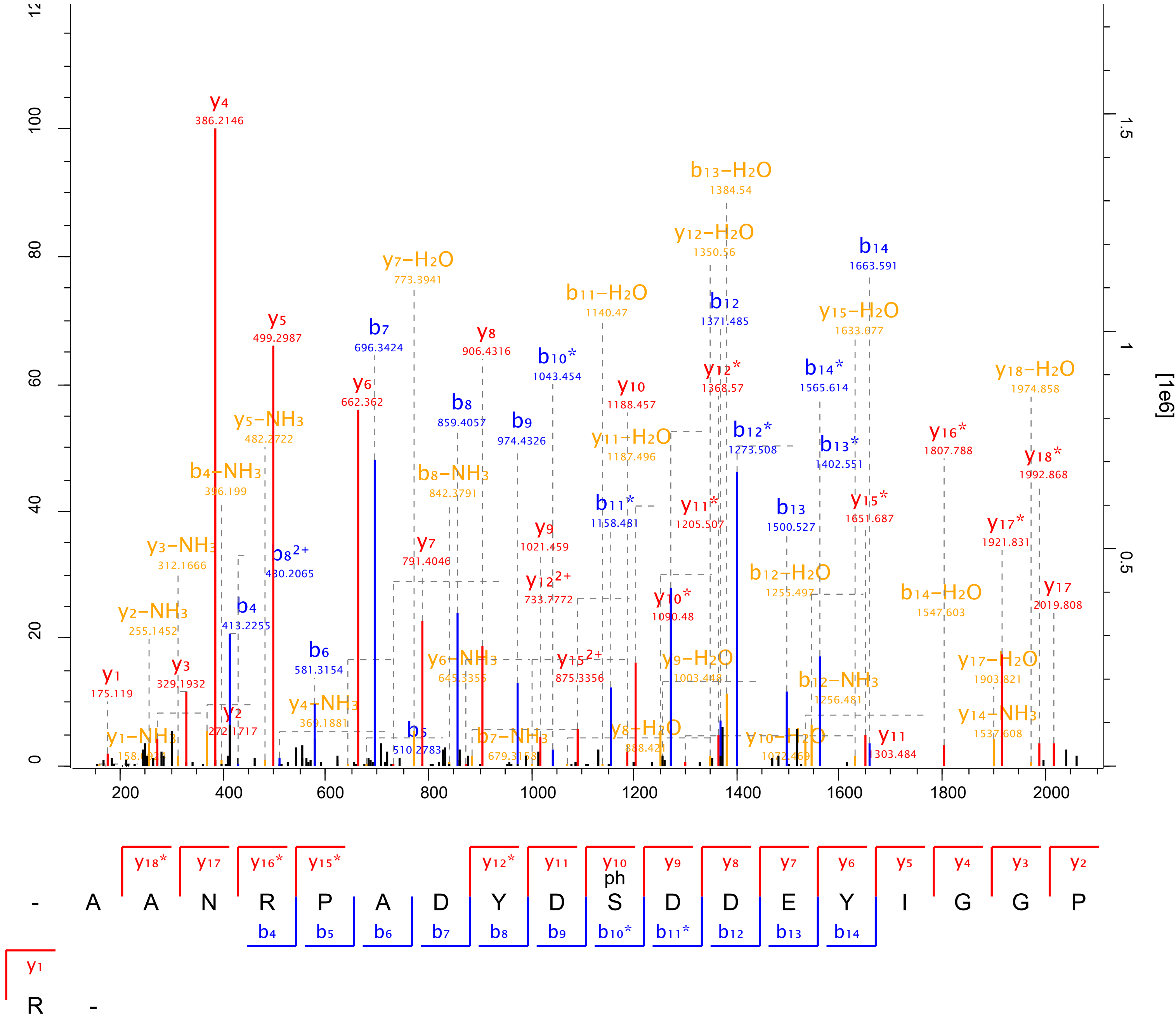

| Raw file | Scan | Method | Score | m/z |
|---|---|---|---|---|
| water_02_1 | 16958 | FTMS; HCD | 155.06 | 607.78 |

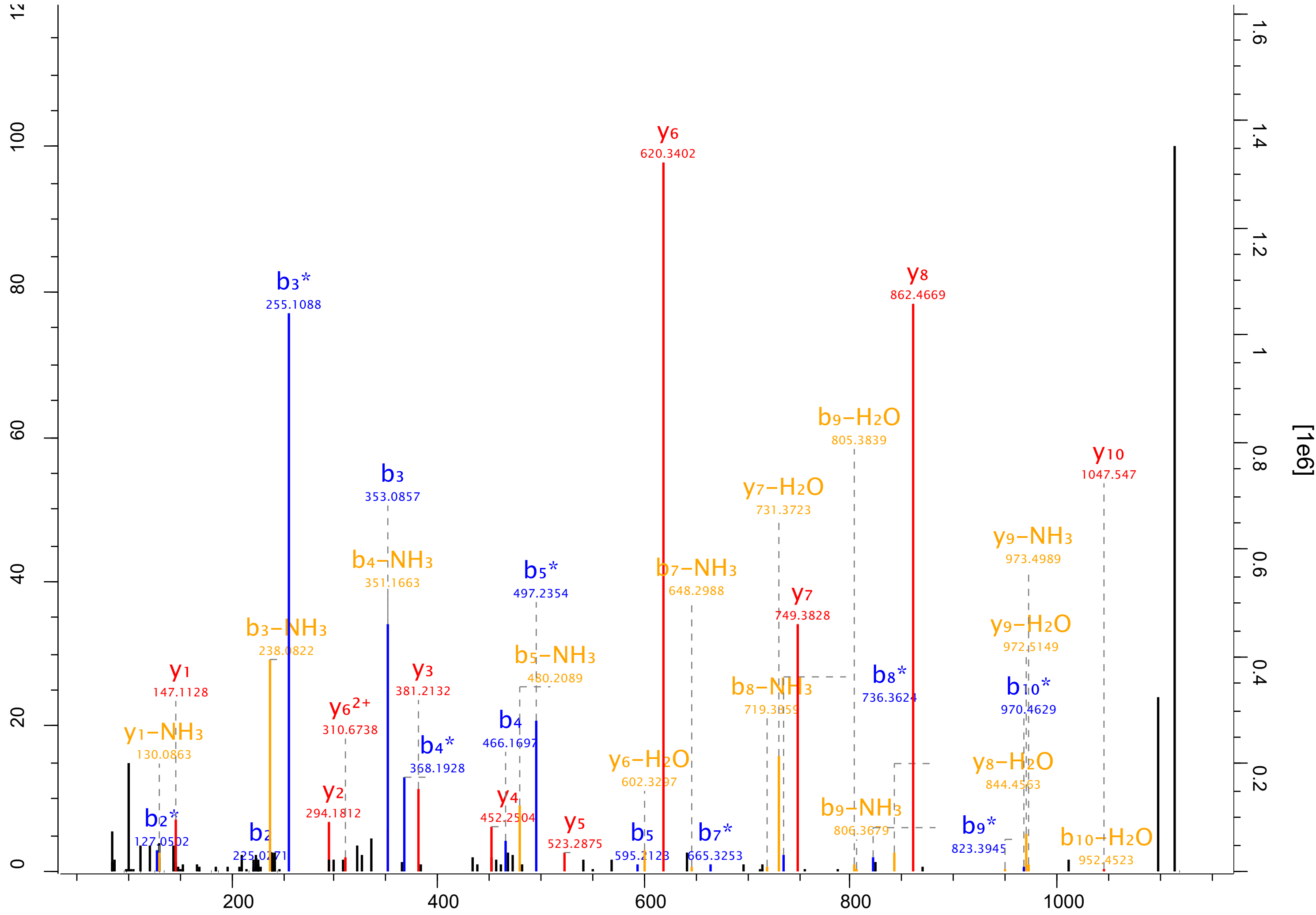


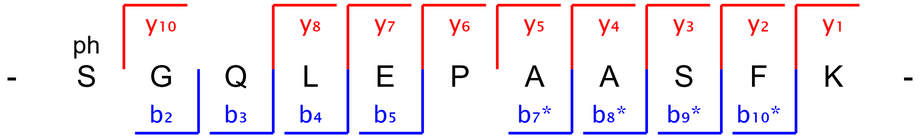

| Raw file | Scan | Method | Score | m/z |
|---|---|---|---|---|
| water_02_1 | 1696 | FTMS; HCD | 77.18 | 614.74 |

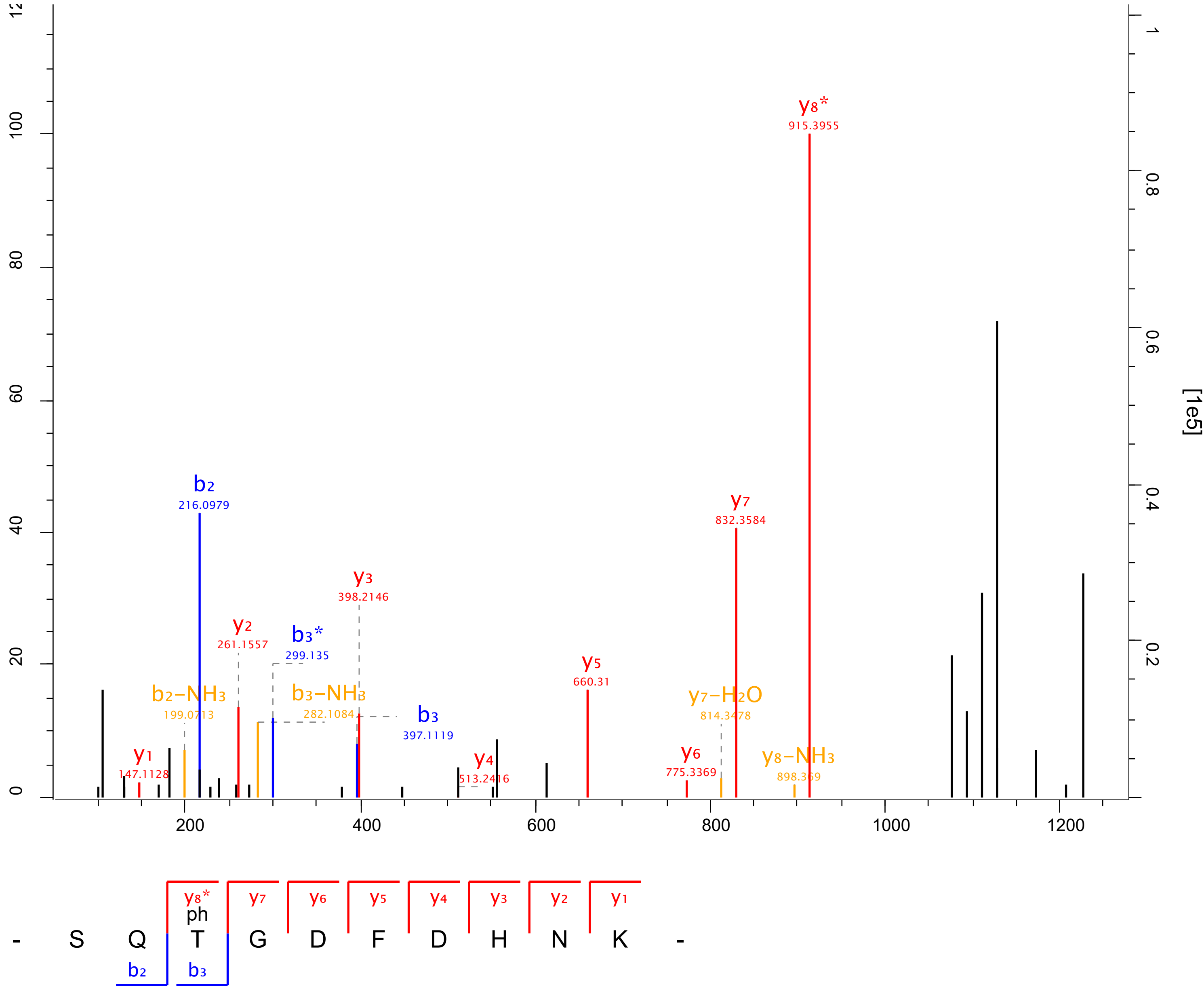

| Raw file | Scan | Method | Score | m/z |
|---|---|---|---|---|
| water_02_1 | 17010 | FTMS; HCD | 81.02 | 507.72 |

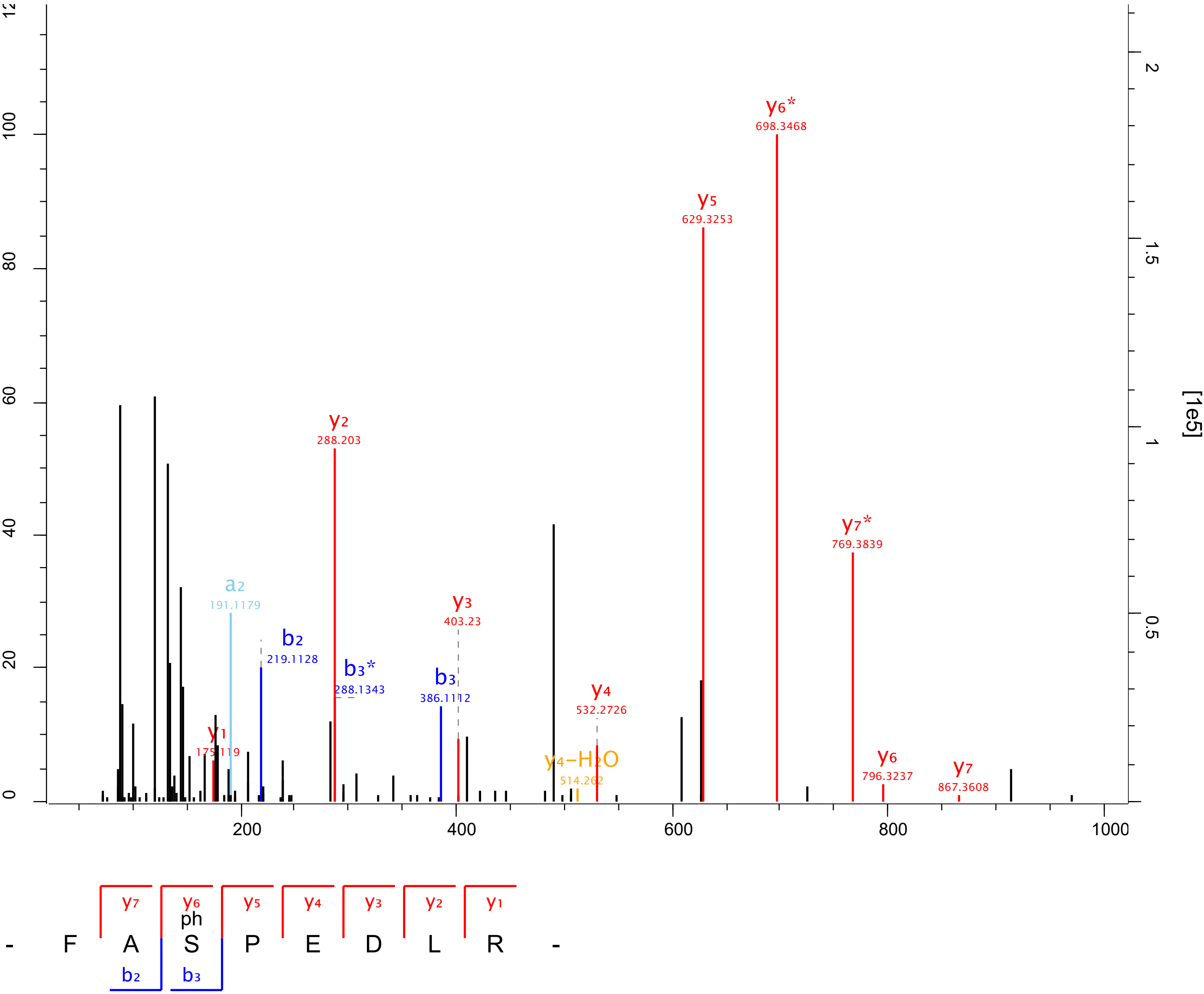

| Raw file | Scan | Method | Score | m/z |
|---|---|---|---|---|
| water_02_1 | 17021 | FTMS; HCD | 63.7 | 811.68 |

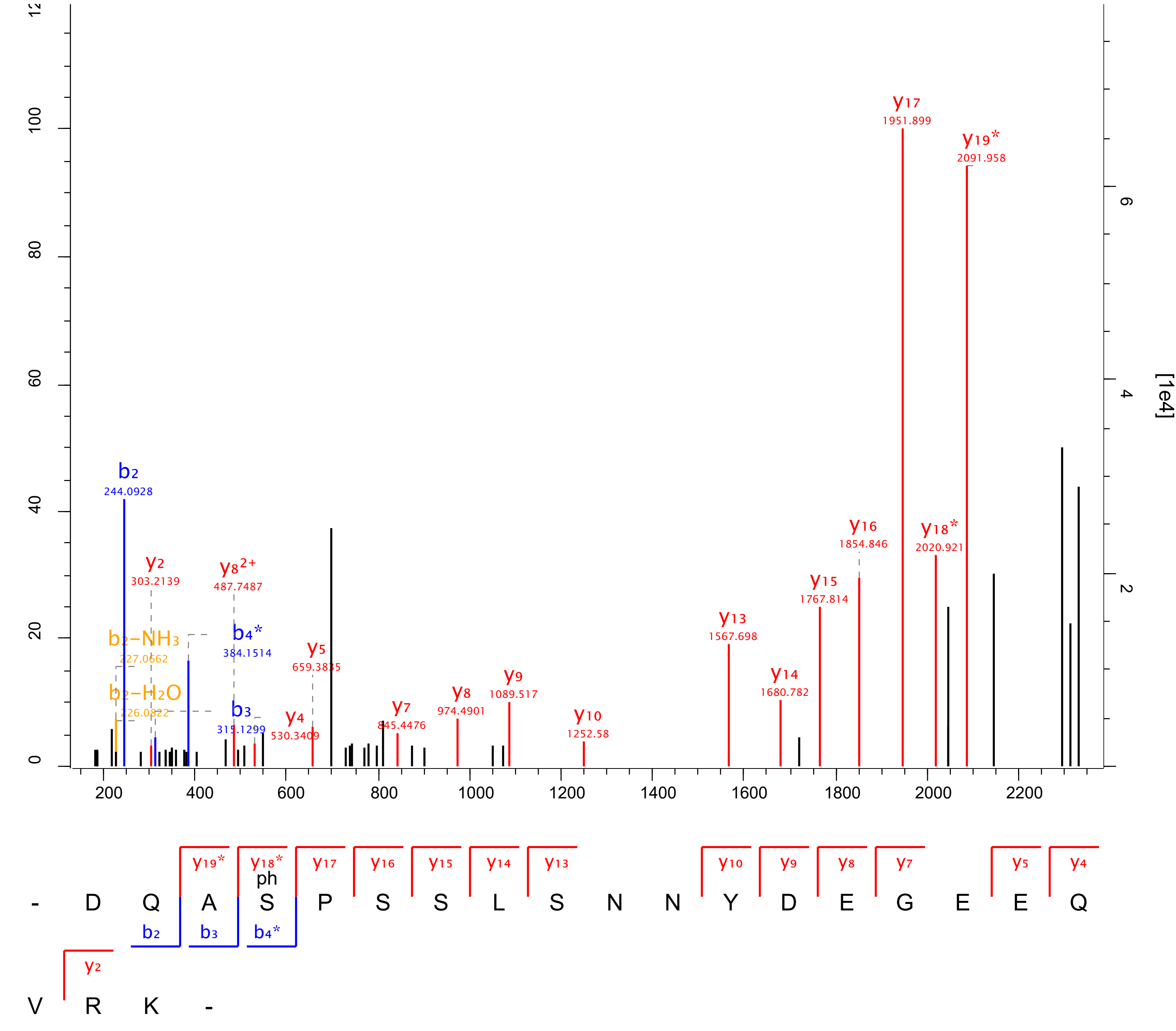

| Raw file | Scan | Method | Score | m/z |
| --- | --- | --- | --- | --- |
| water_02_1 | 17078 | FTMS; HCD | 153.56 | 726.29 |

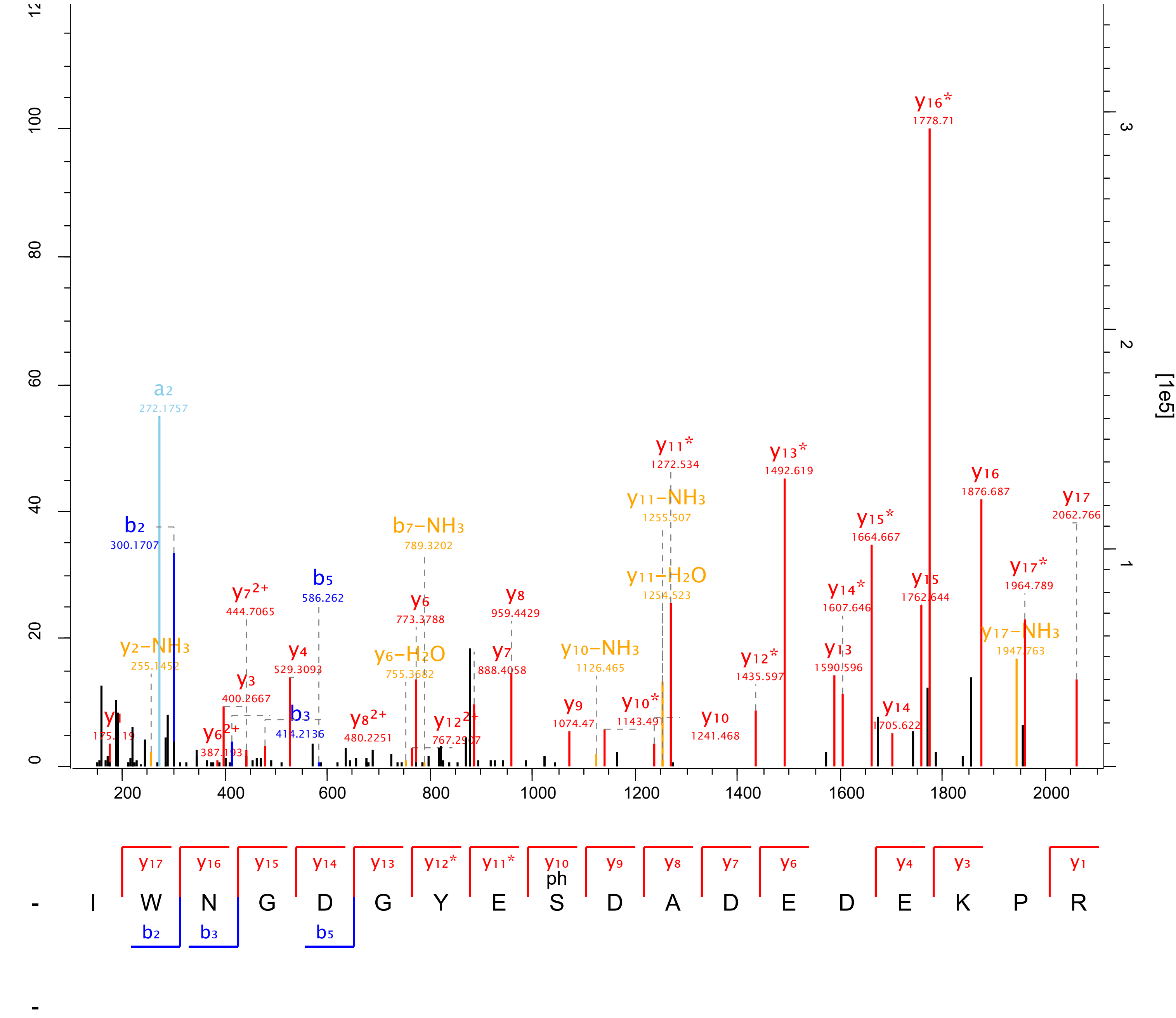

| Raw file | Scan | Method | Score | m/z |
| --- | --- | --- | --- | --- |
| water_02_1 | 17097 | FTMS; HCD | 159.06 | 591.6 |

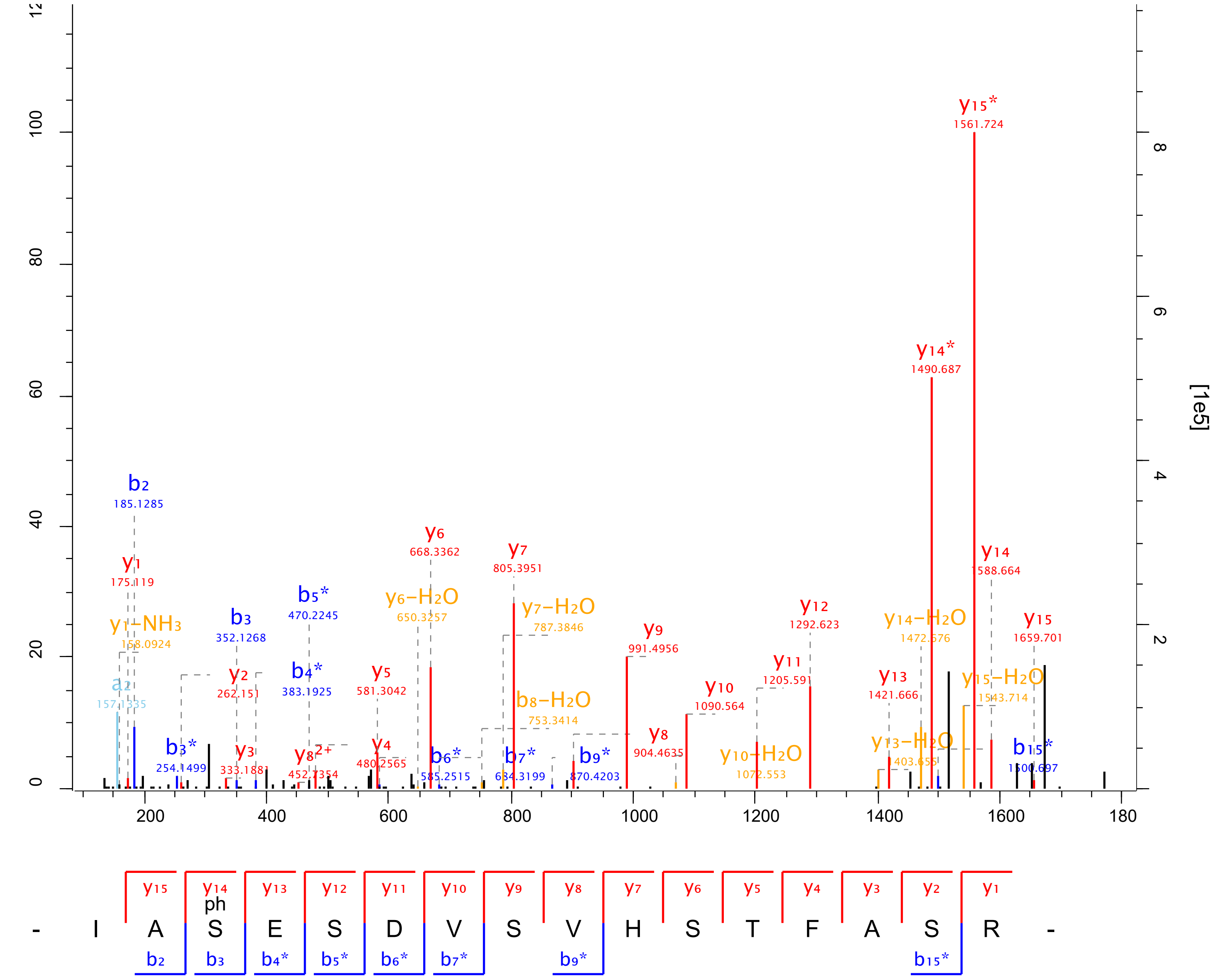

| Raw file | Scan | Method | Score | m/z |
| --- | --- | --- | --- | --- |
| water_02_1 | 1721 | FTMS; HCD | 106.6 | 711.79 |

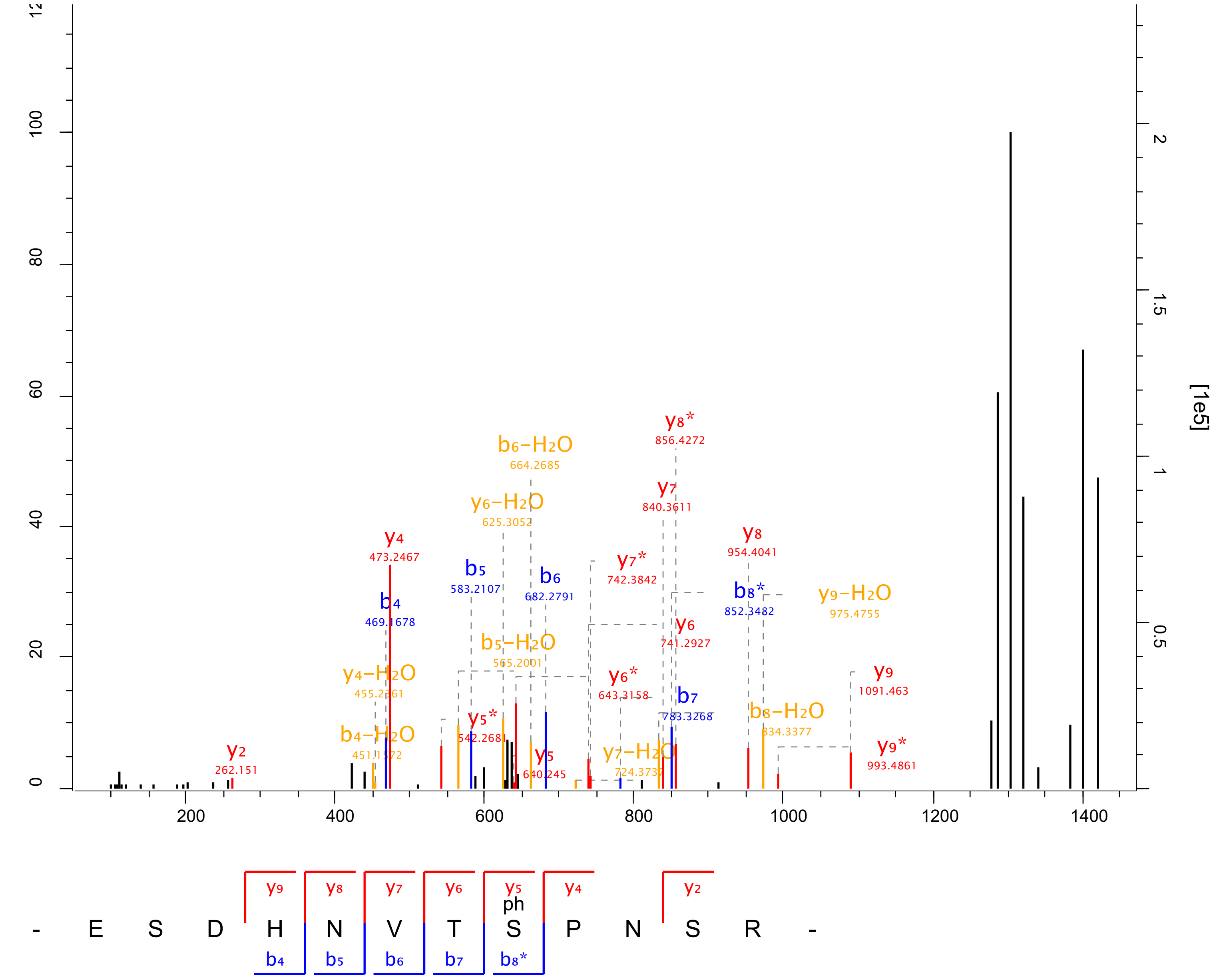

| Raw file | Scan | Method | Score | m/z |
| --- | --- | --- | --- | --- |
| water_02_1 | 1724 | FTMS; HCD | 52.52 | 679.78 |

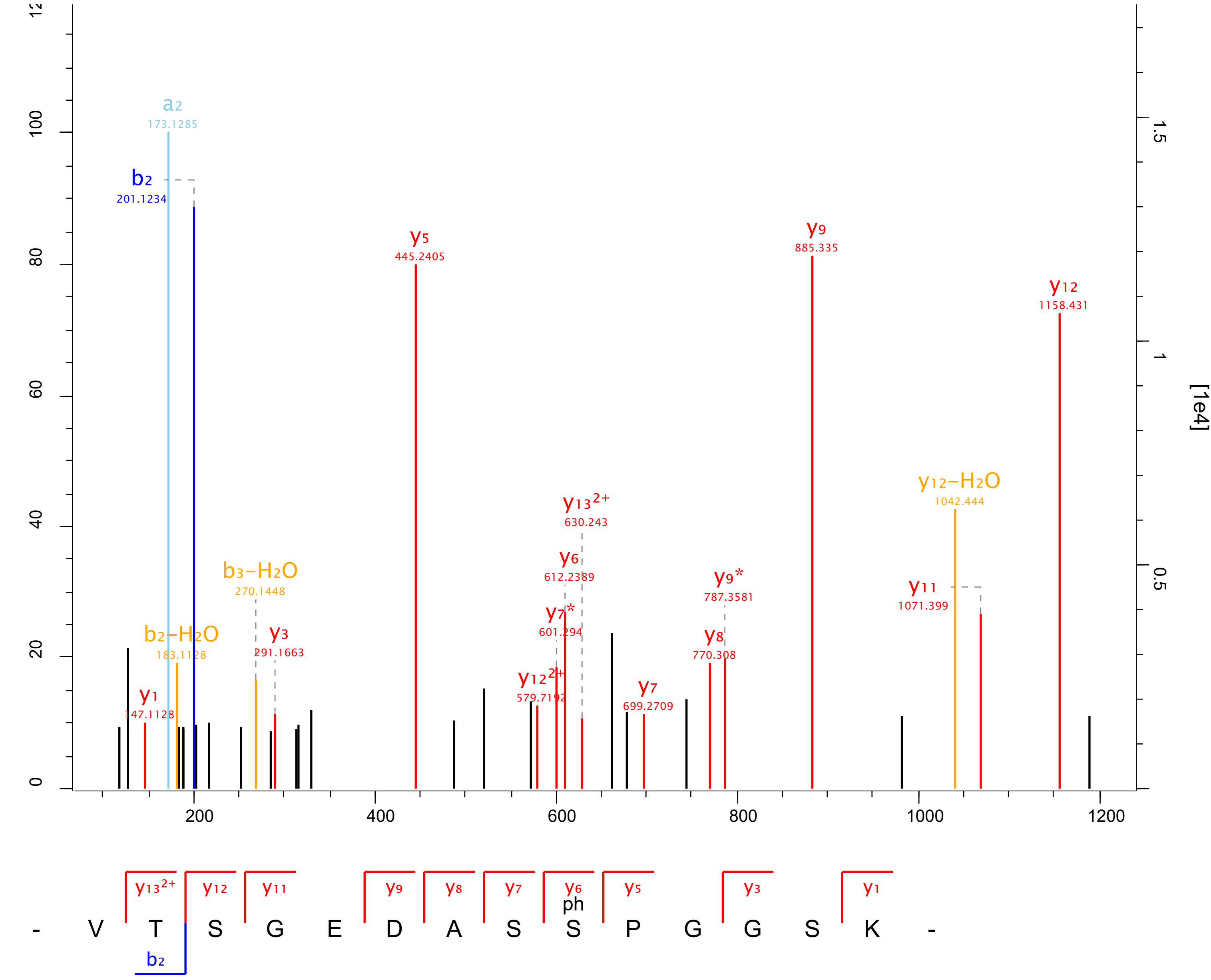

| Raw file | Scan | Method | Score | m/z |
|---|---|---|---|---|
| water_02_1 | 17273 | FTMS; HCD | 305.11 | 923.42 |

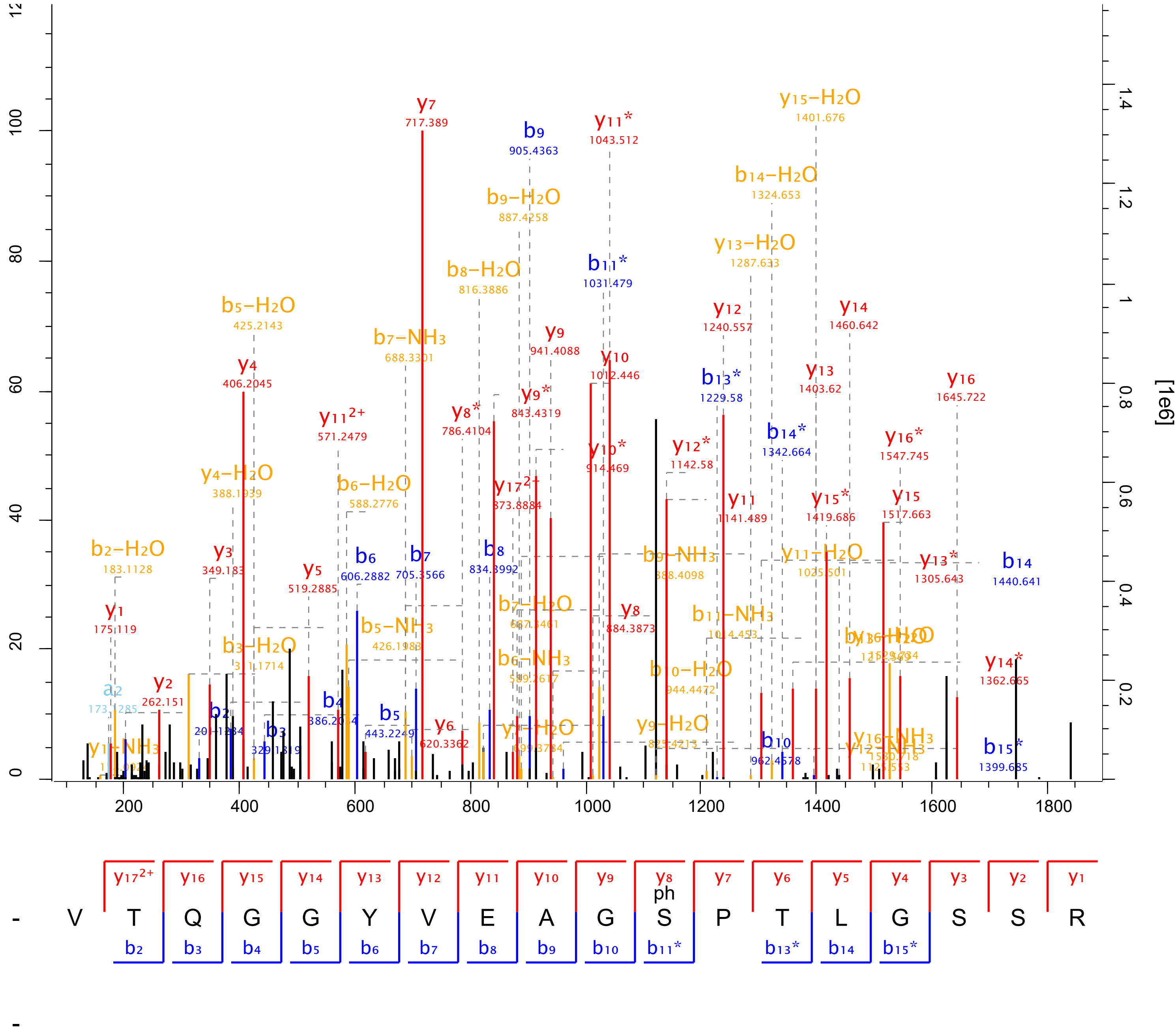

| Raw file | Scan | Method | Score | m/z |
| --- | --- | --- | --- | --- |
| water_02_1 | 17312 | FTMS; HCD | 142.32 | 795.01 |

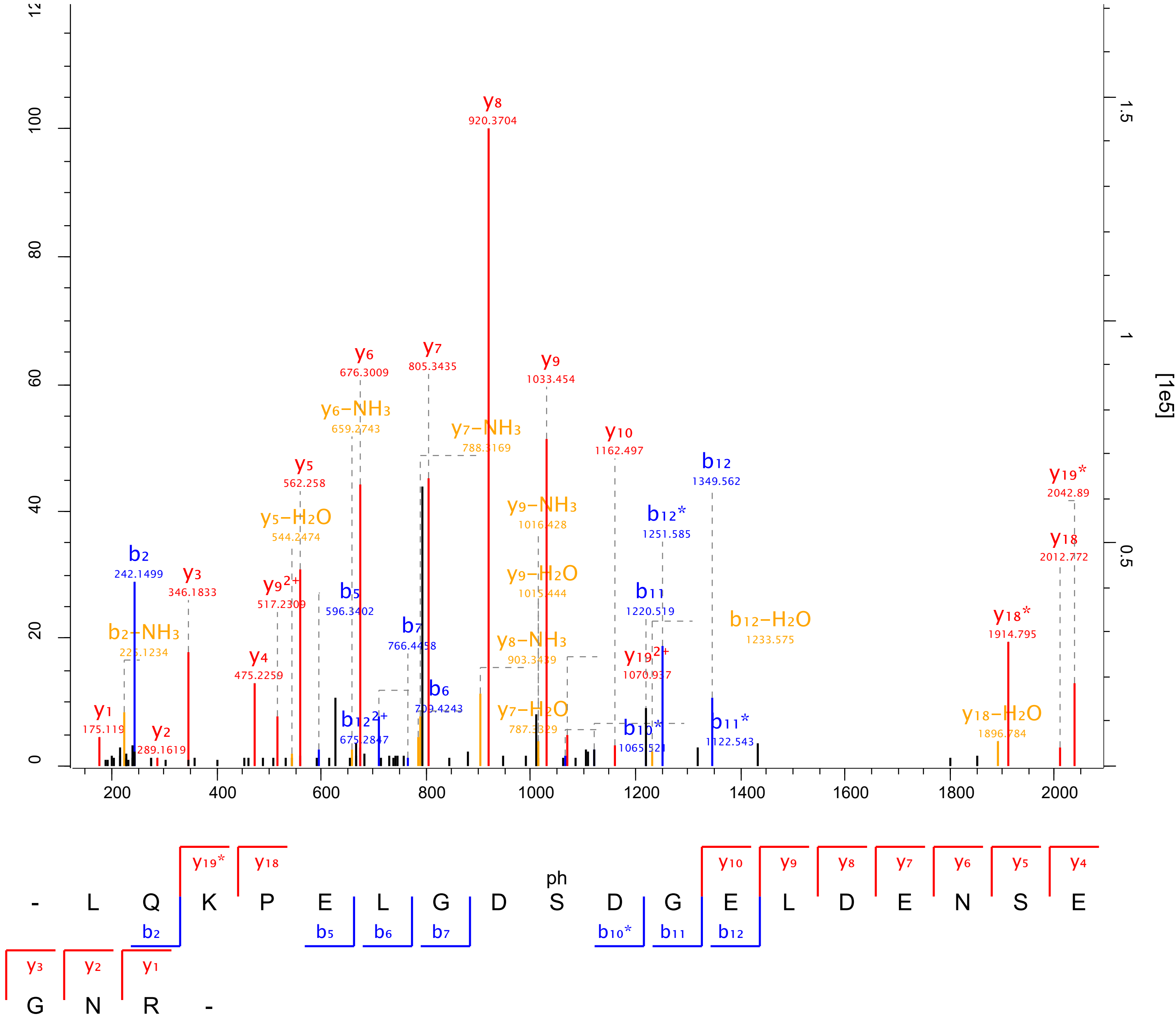

| Raw file | Scan | Method | Score | m/z |
| --- | --- | --- | --- | --- |
| water_02_1 | 17315 | FTMS; HCD | 49.3 | 563.78 |

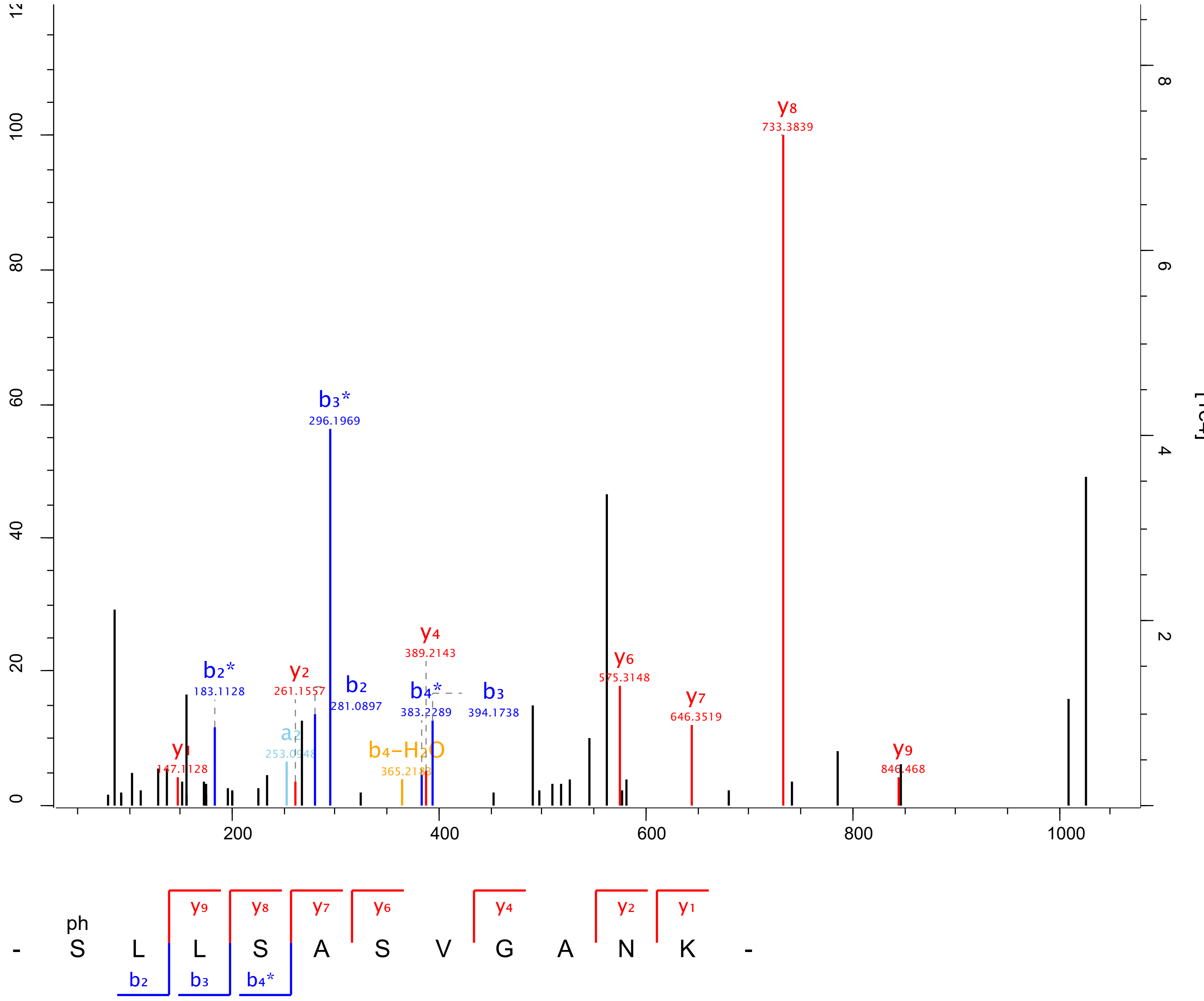

| Raw file | Scan | Method | Score | m/z |
| --- | --- | --- | --- | --- |
| water_02_1 | 1732 | FTMS; HCD | 47.6 | 437.18 |

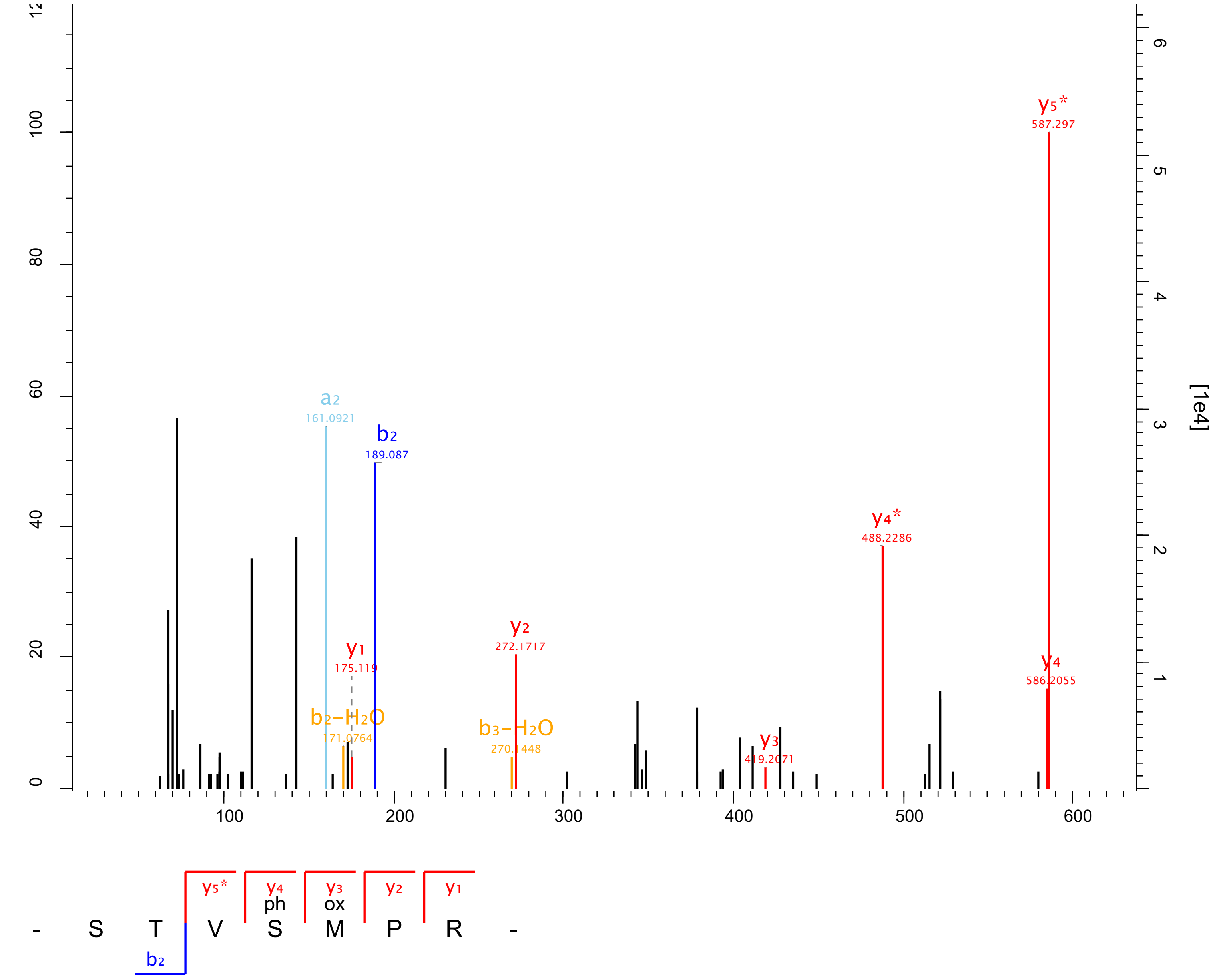

| Raw file | Scan | Method | Score | m/z |
| --- | --- | --- | --- | --- |
| water_02_1 | 1737 | FTMS; HCD | 59.73 | 721.8 |

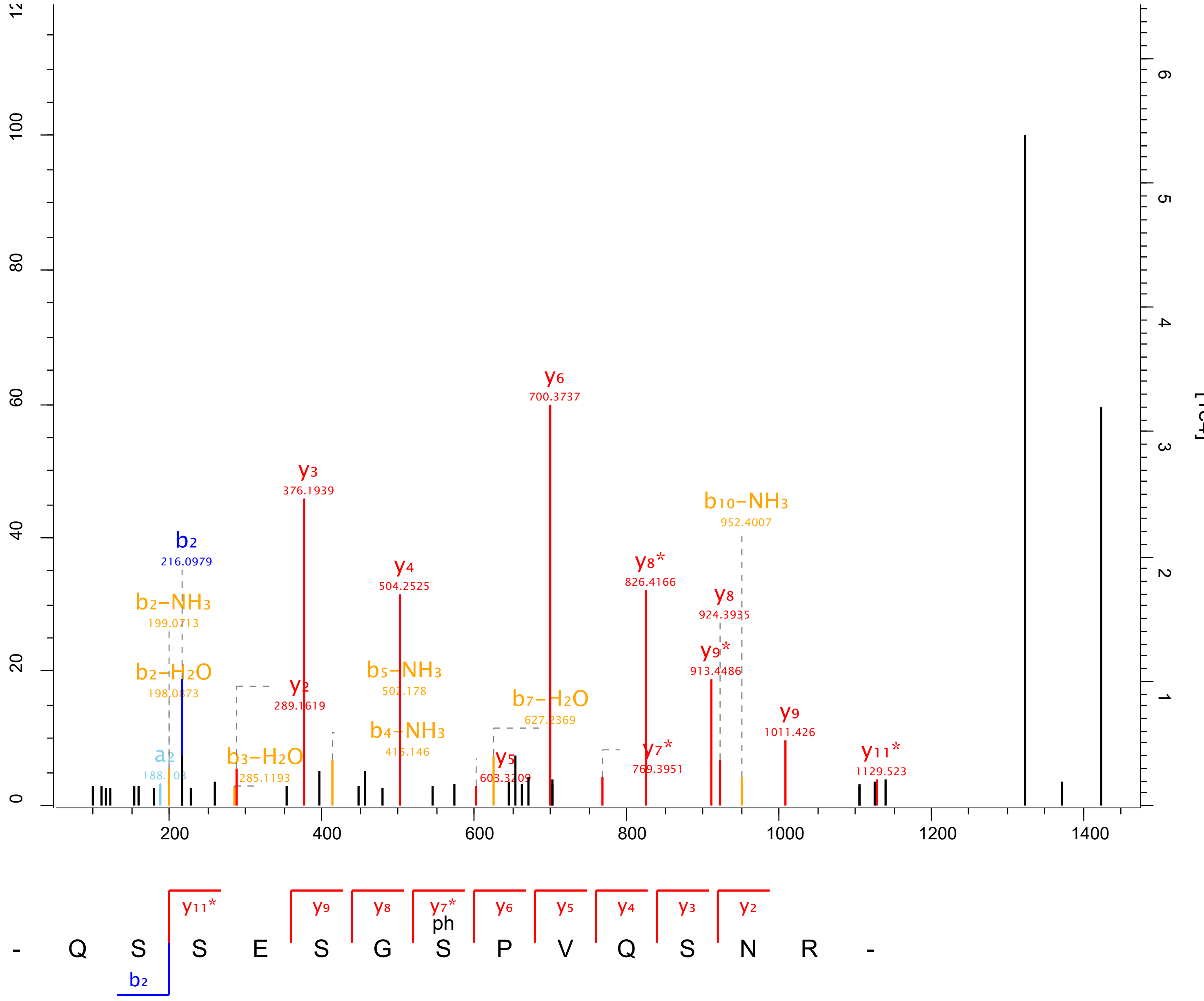

| Raw file | Scan | Method | Score | m/z |
|---|---|---|---|---|
| water_02_1 | 17499 | FTMS; HCD | 95.62 | 888.9 |

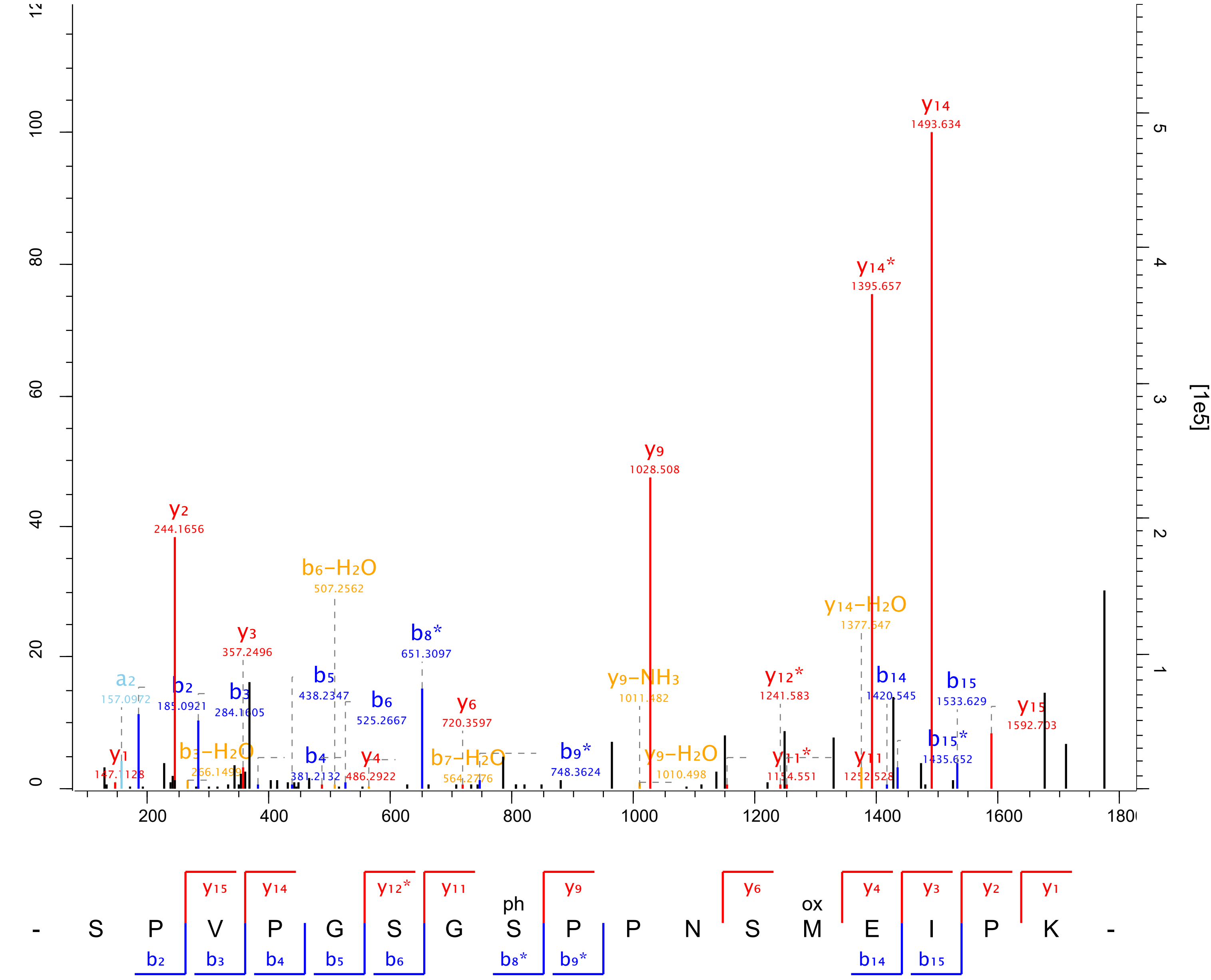

| Raw file | Scan | Method | Score | m/z |
| --- | --- | --- | --- | --- |
| water_02_1 | 17571 | FTMS; HCD | 40.43 | 918.86 |

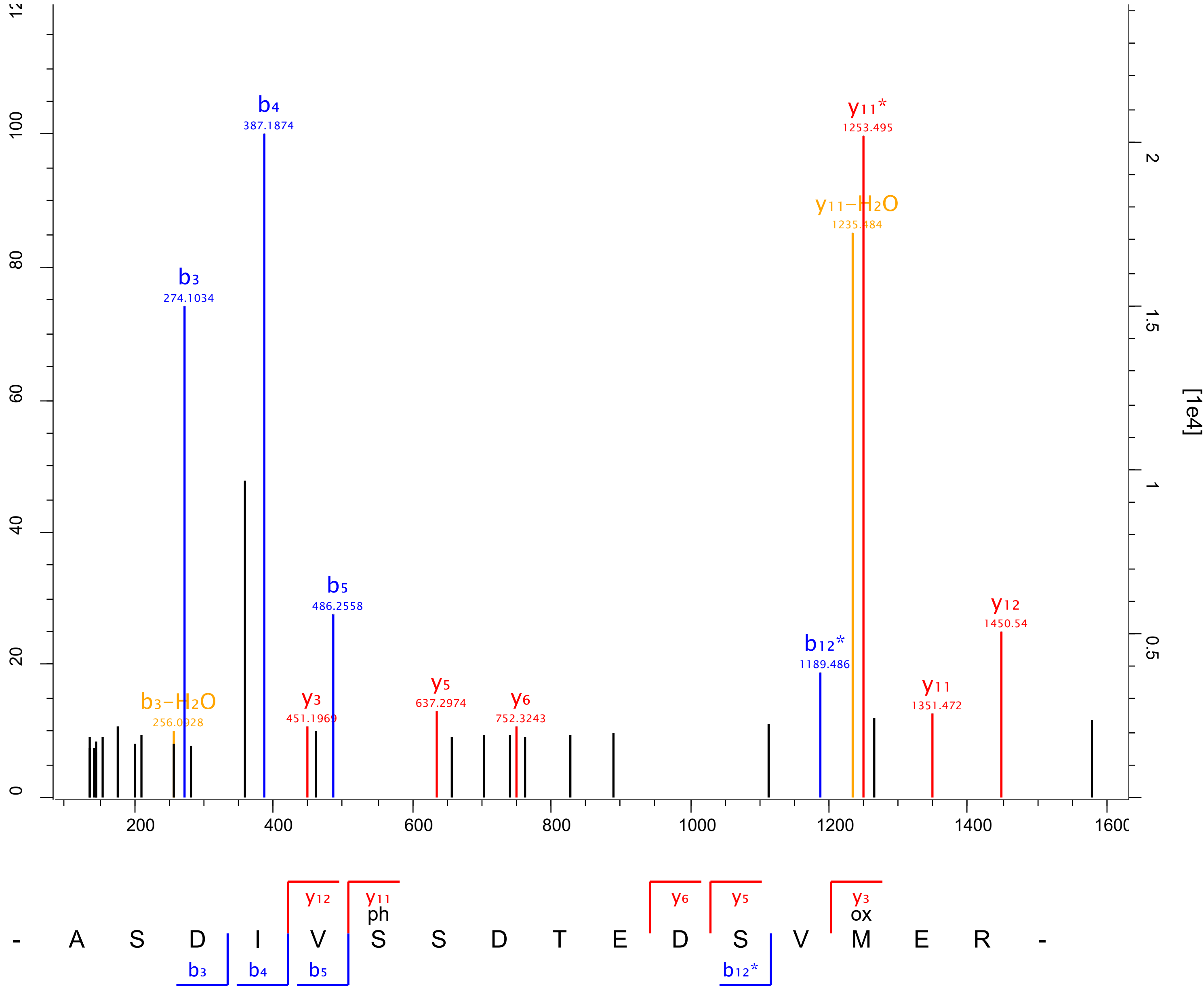

| Raw file | Scan | Method | Score | m/z |
|---|---|---|---|---|
| water_02_1 | 17588 | FTMS; HCD | 131.72 | 528.26 |

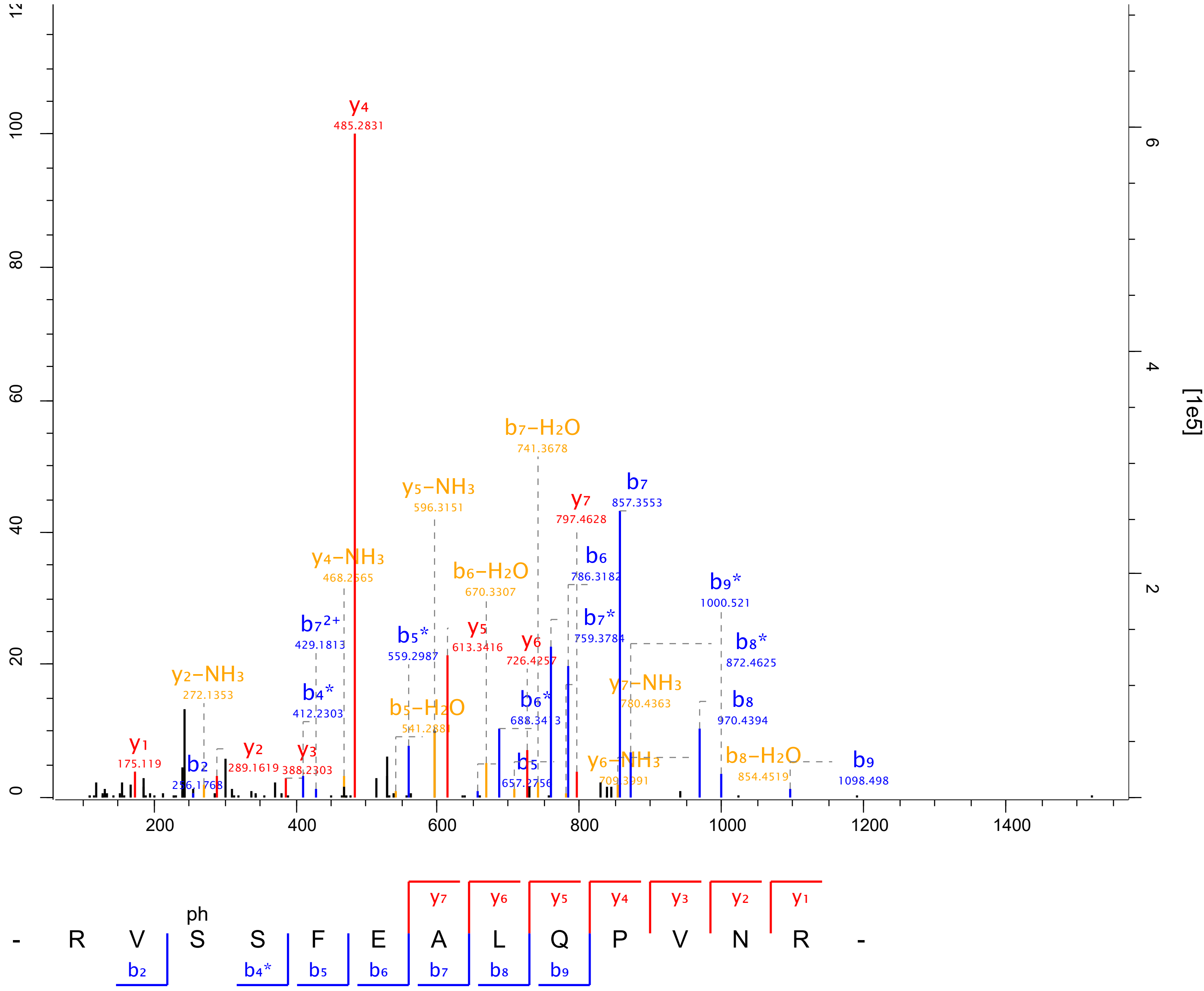

| Raw file | Scan | Method | Score | m/z |
|---|---|---|---|---|
| water_02_1 | 17634 | FTMS; HCD | 77.73 | 905.05 |

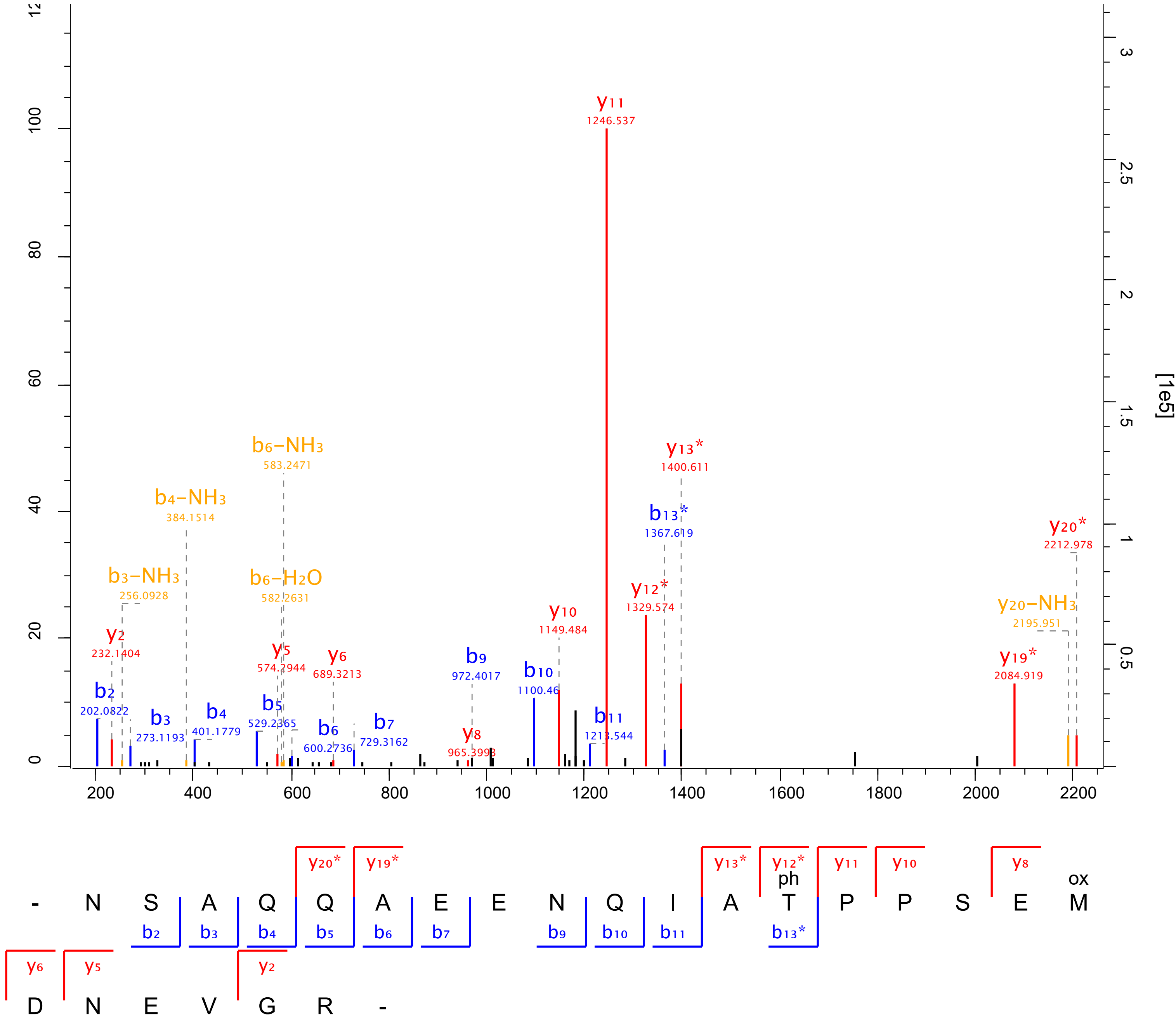

| Raw file | Scan | Method | Score | m/z |
| --- | --- | --- | --- | --- |
| water_02_1 | 17673 | FTMS; HCD | 40.85 | 858.04 |

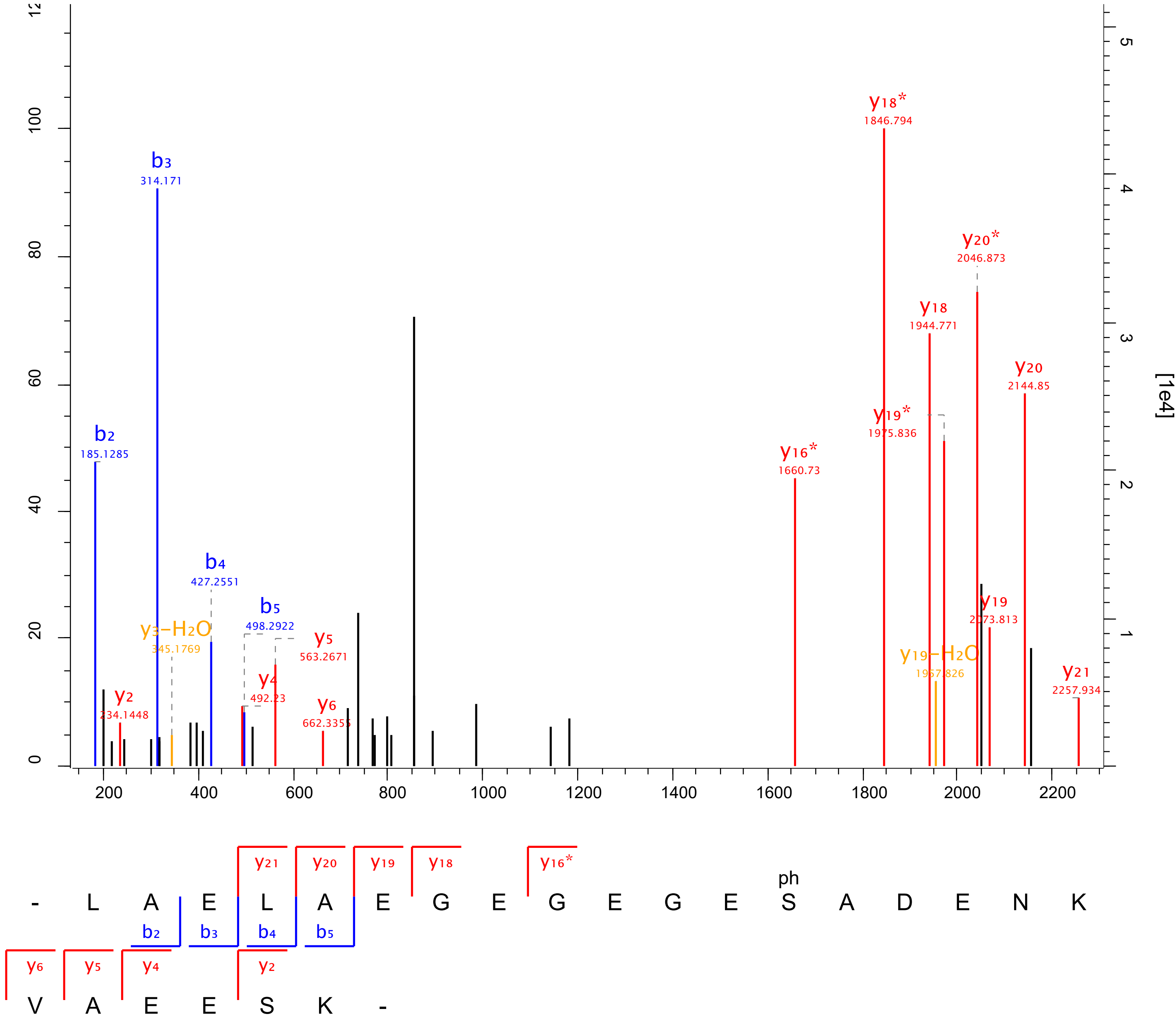

| Raw file | Scan | Method | Score | m/z |
|---|---|---|---|---|
| water_02_1 | 17674 | FTMS; HCD | 54.28 | 758.86 |

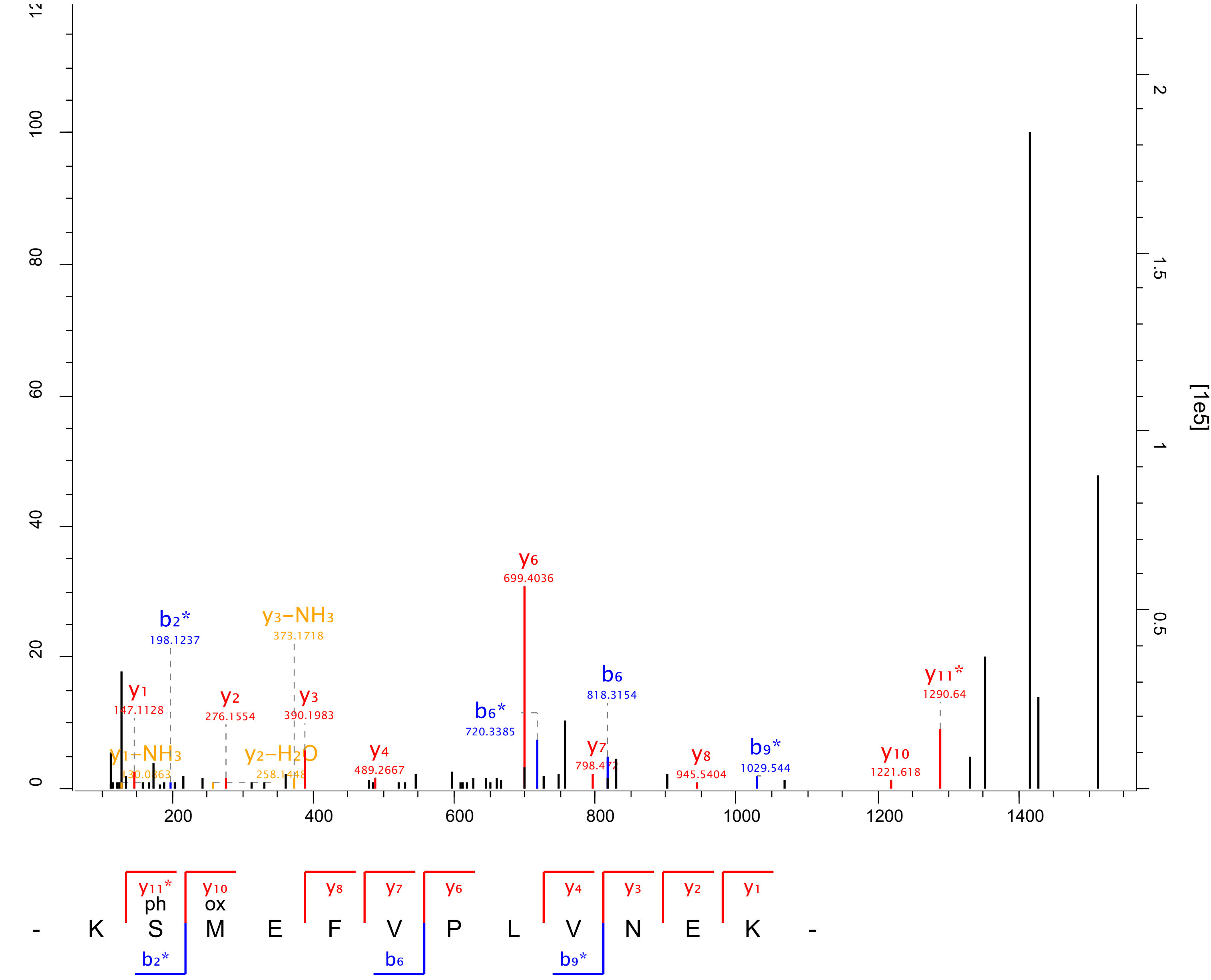

| Raw file | Scan | Method | Score | m/z |
|---|---|---|---|---|
| water_02_1 | 17713 | FTMS; HCD | 114.63 | 511.91 |

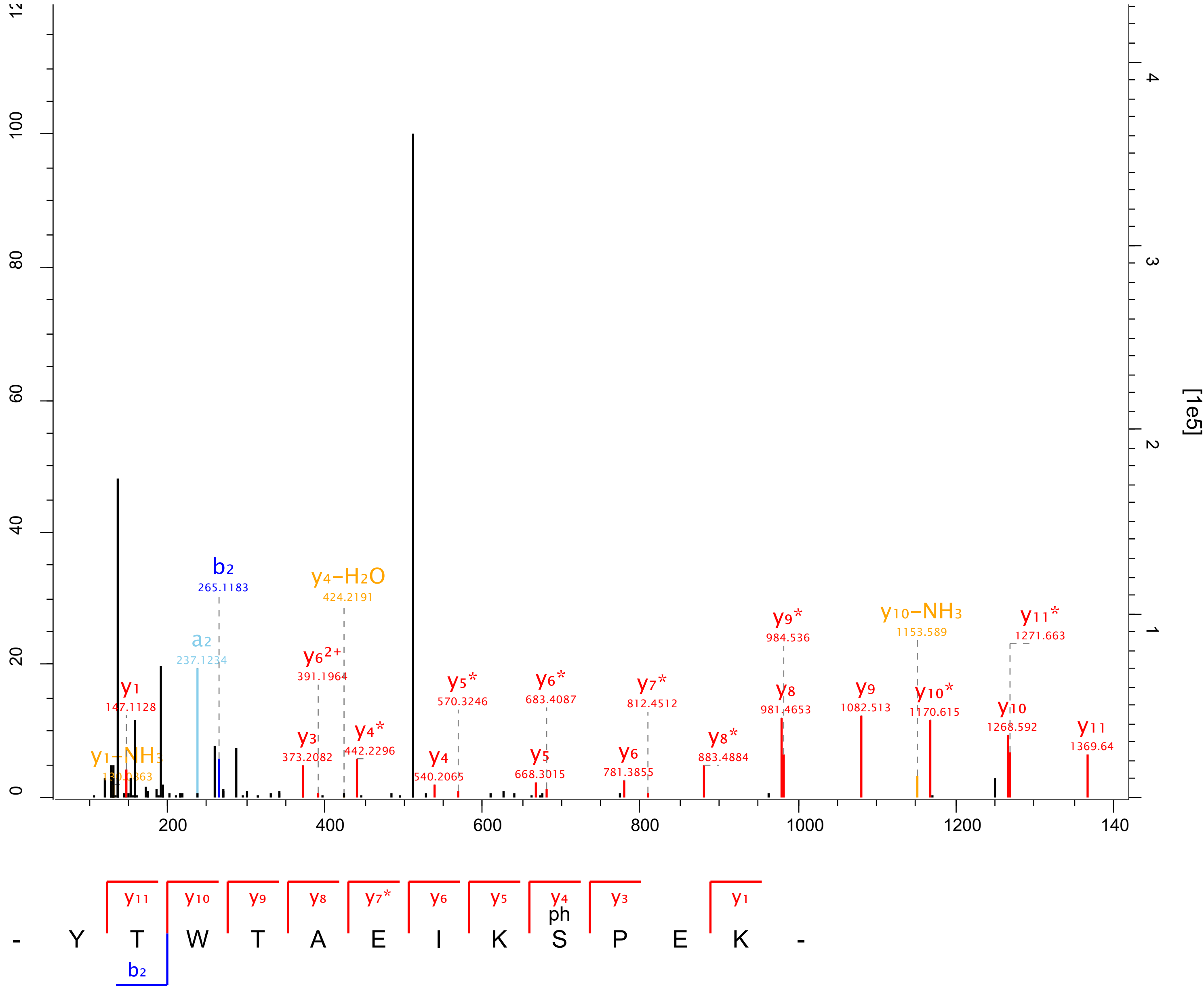

| Raw file | Scan | Method | Score | m/z |
|---|---|---|---|---|
| water_02_1 | 17728 | FTMS; HCD | 61.34 | 542.78 |

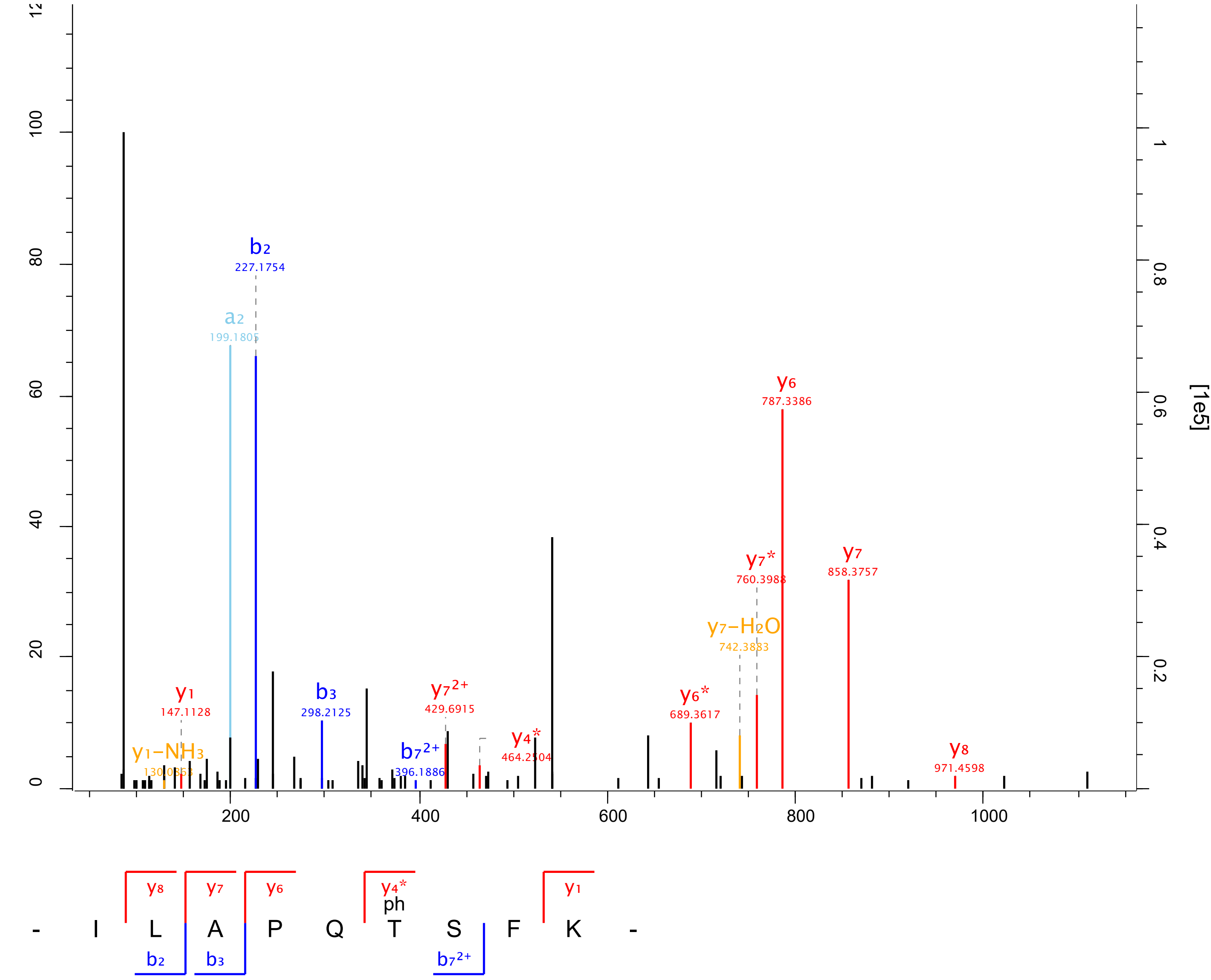

| Raw file | Scan | Method | Score | m/z |
| --- | --- | --- | --- | --- |
| water_02_1 | 17784 | FTMS; HCD | 69.77 | 950.42 |

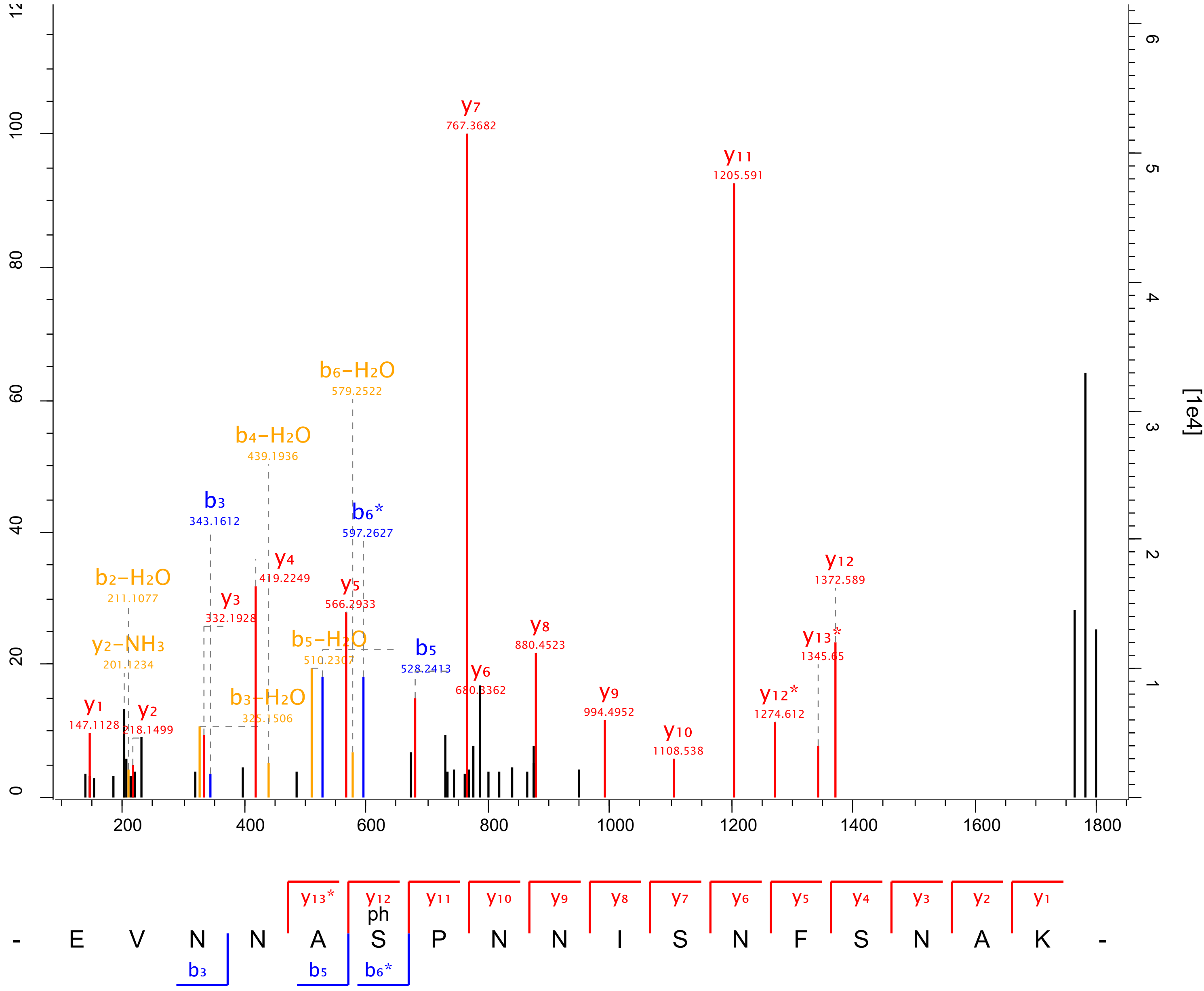

| Raw file | Scan | Method | Score | m/z |
| --- | --- | --- | --- | --- |
| water_02_1 | 17821 | FTMS; HCD | 59.12 | 552.78 |

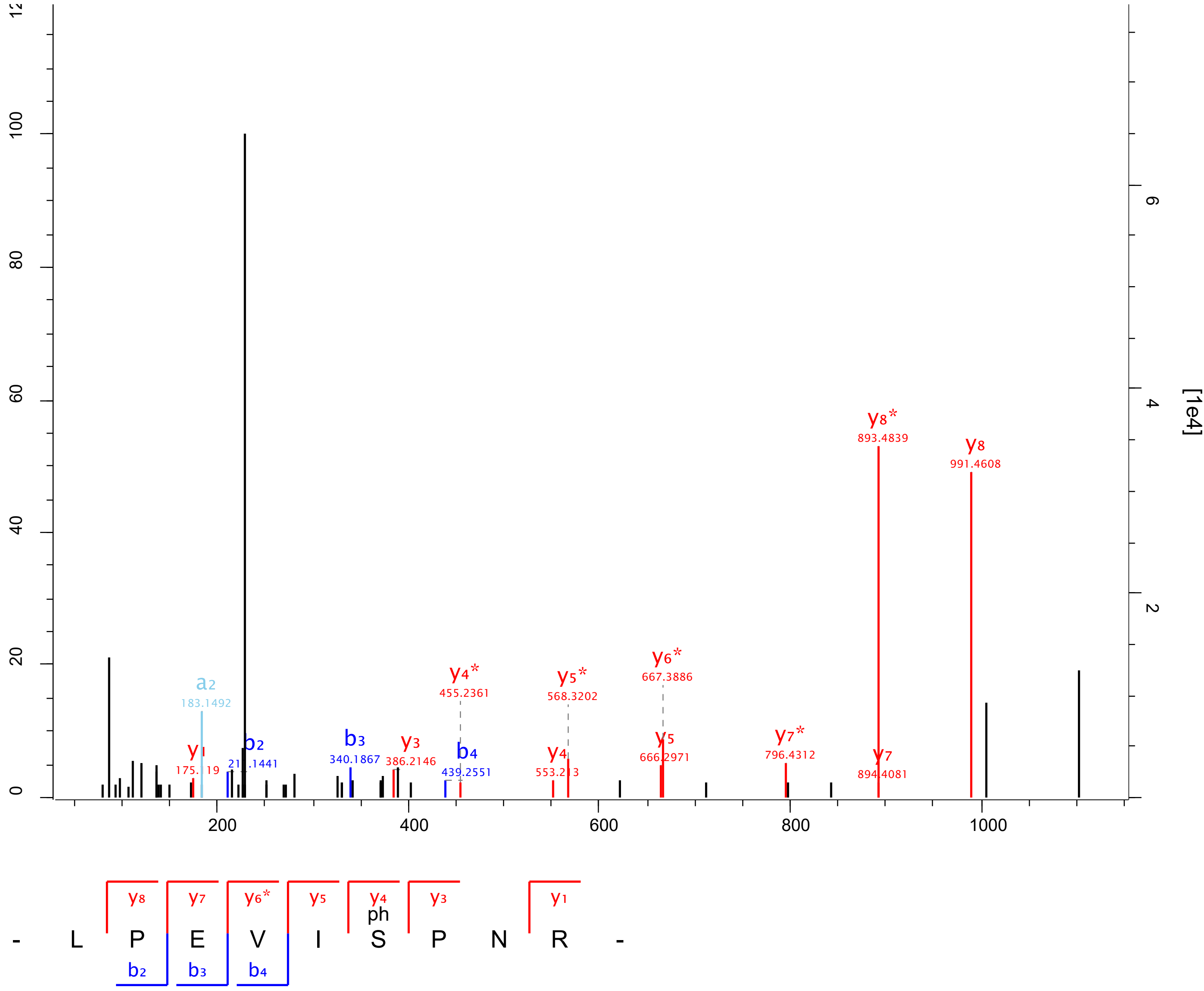

| Raw file | Scan | Method | Score | m/z |
|---|---|---|---|---|
| water_02_1 | 17943 | FTMS; HCD | 199.01 | 579.94 |

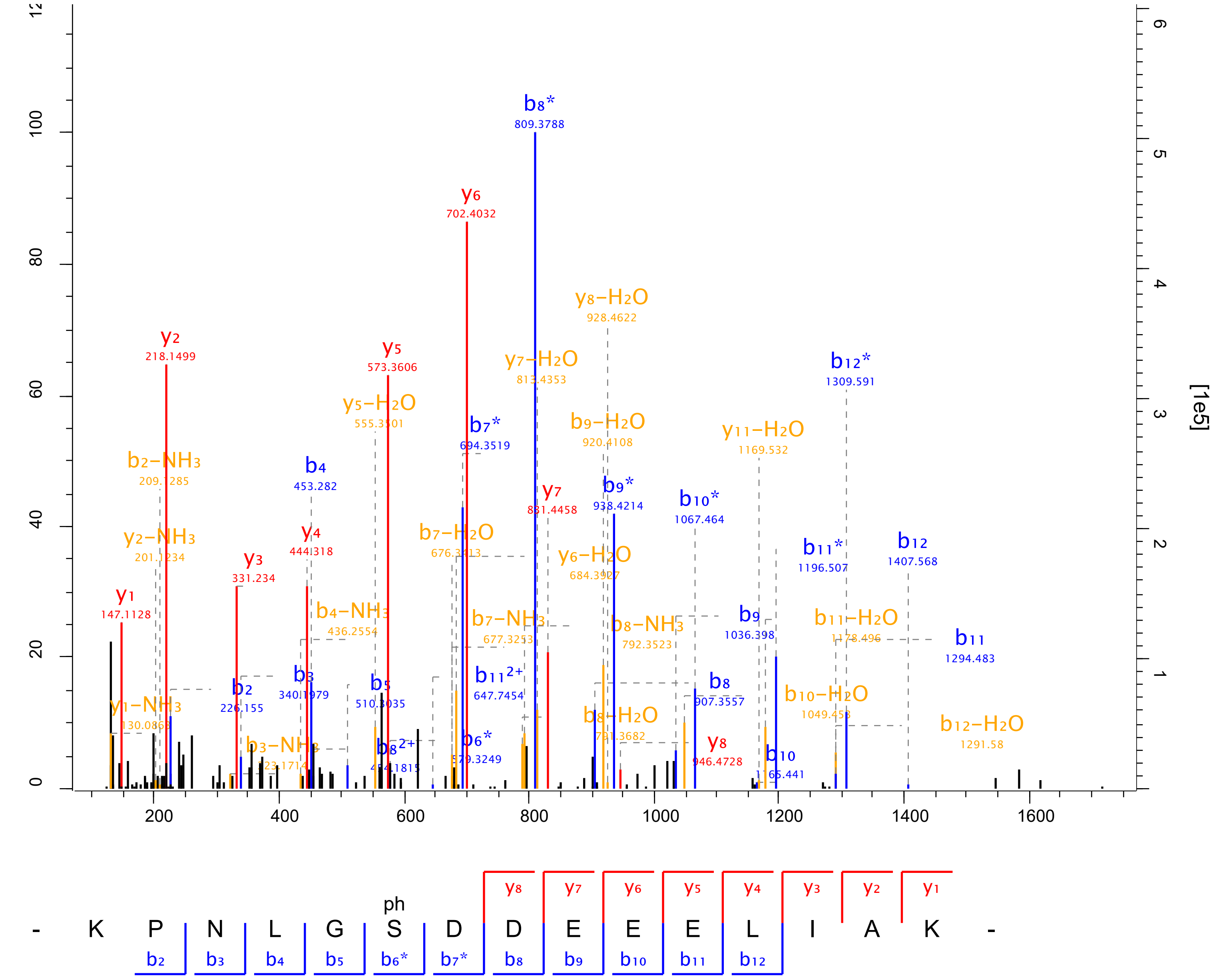

| Raw file | Scan | Method | Score | m/z |
|---|---|---|---|---|
| water_02_1 | 17979 | FTMS; HCD | 135.3 | 670.29 |

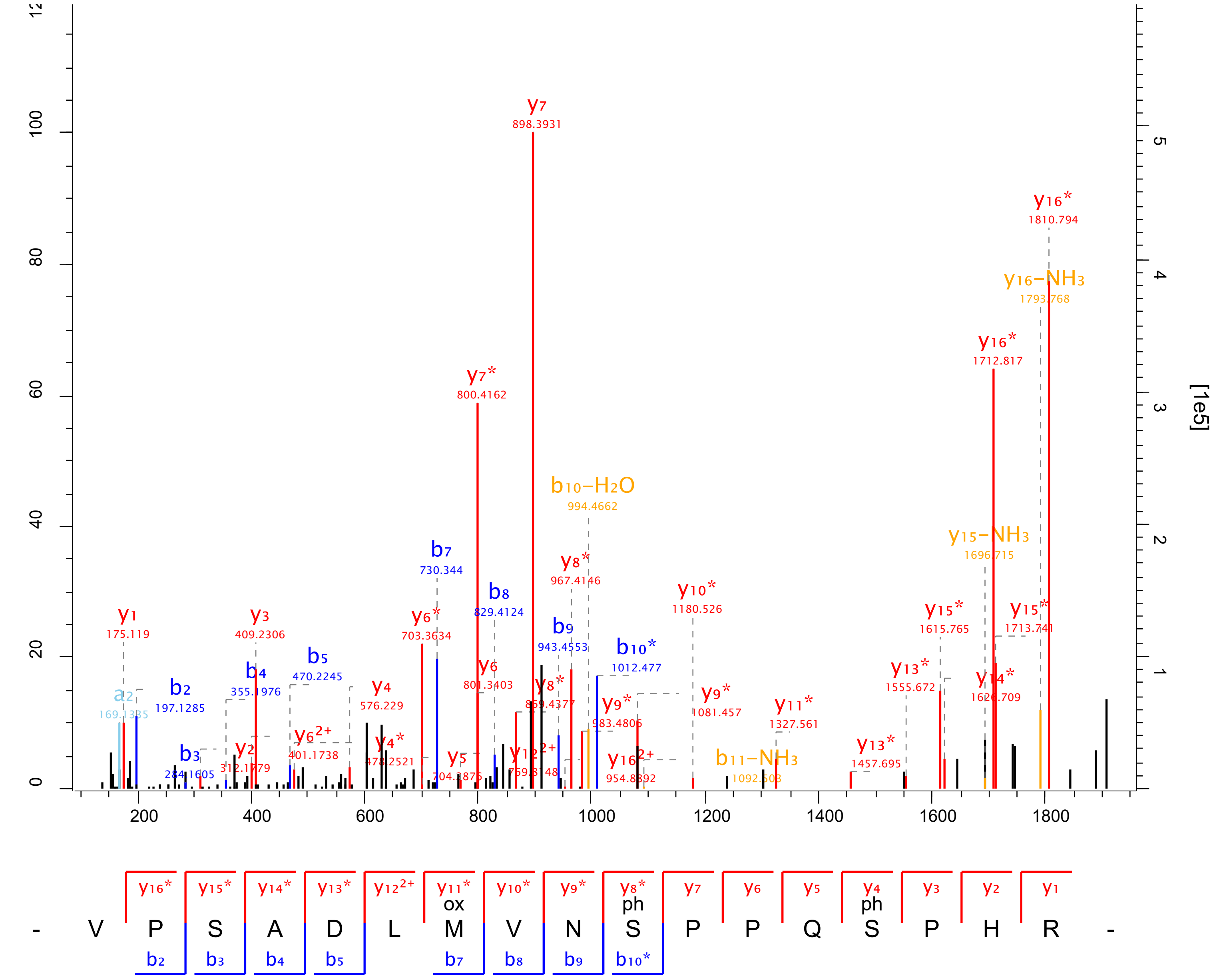

| Raw file | Scan | Method | Score | m/z |
|---|---|---|---|---|
| water_02_1 | 18004 | FTMS; HCD | 82.53 | 708.28 |

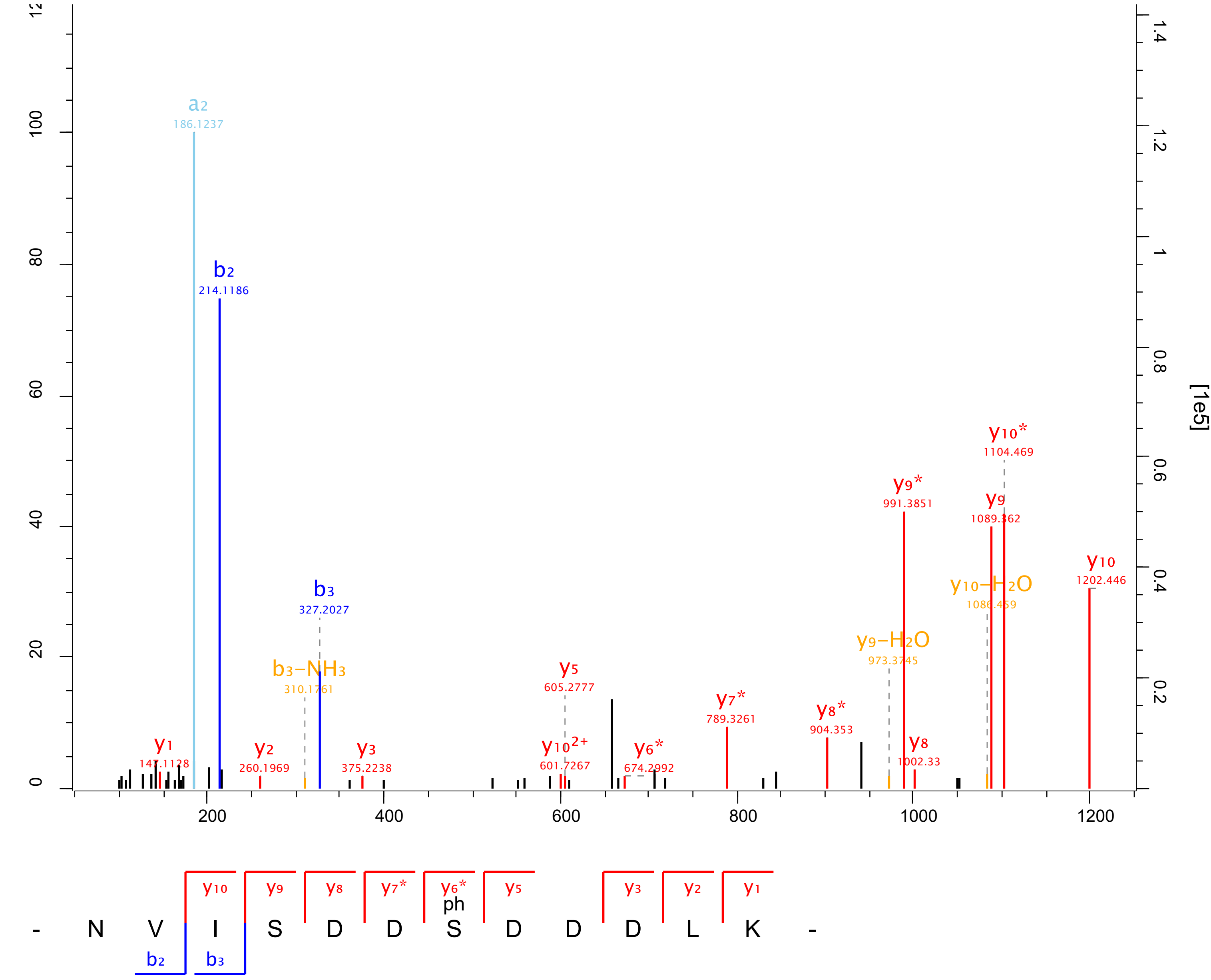

| Raw file | Scan | Method | Score | m/z |
|---|---|---|---|---|
| water_02_1 | 18040 | FTMS; HCD | 48.9 | 785.31 |

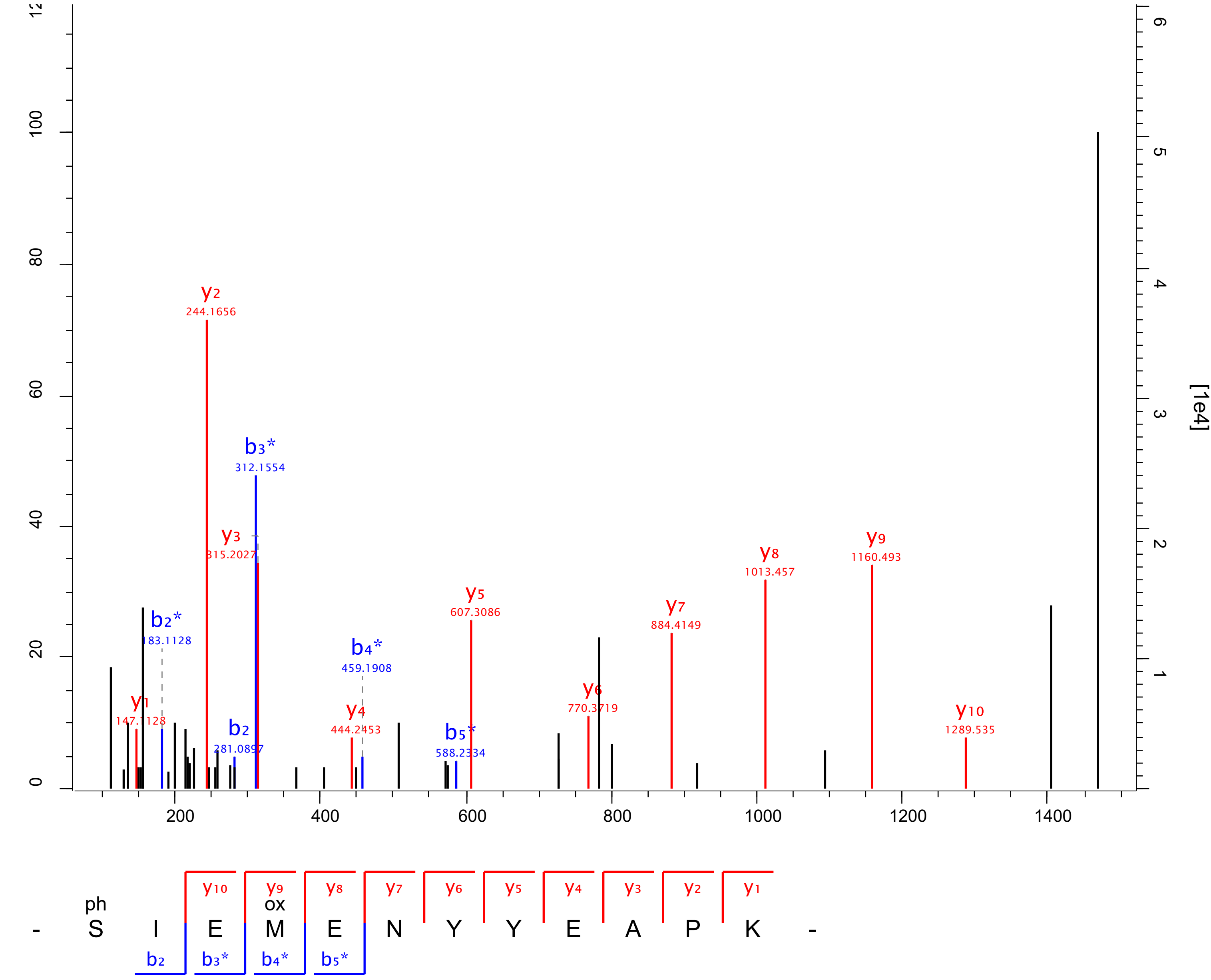

| Raw file | Scan | Method | Score | m/z |
|---|---|---|---|---|
| water_02_1 | 18049 | FTMS; HCD | 52.97 | 776.34 |

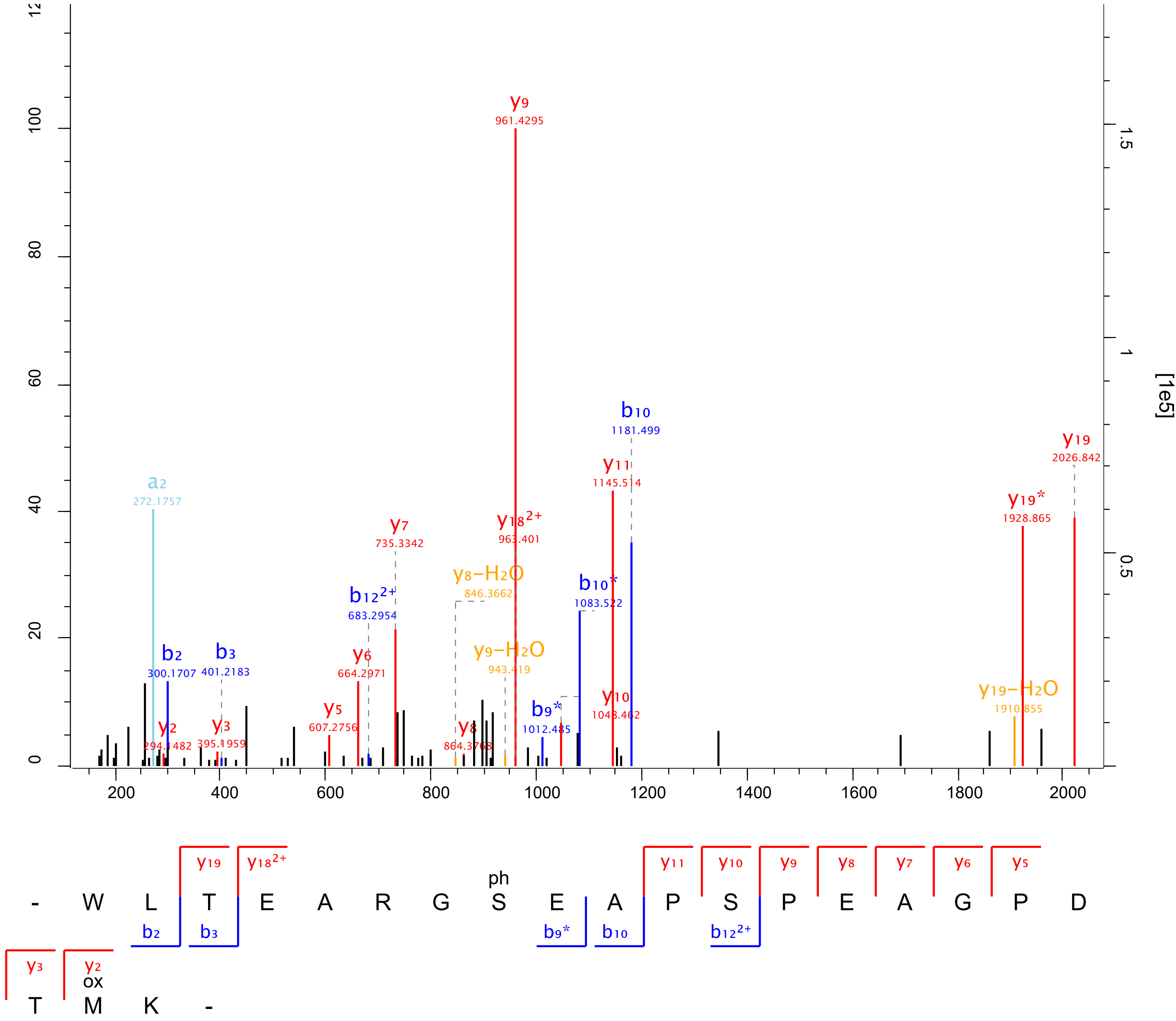

| Raw file | Scan | Method | Score | m/z |
|---|---|---|---|---|
| water_02_1 | 18092 | FTMS; HCD | 81 | 519.2 |

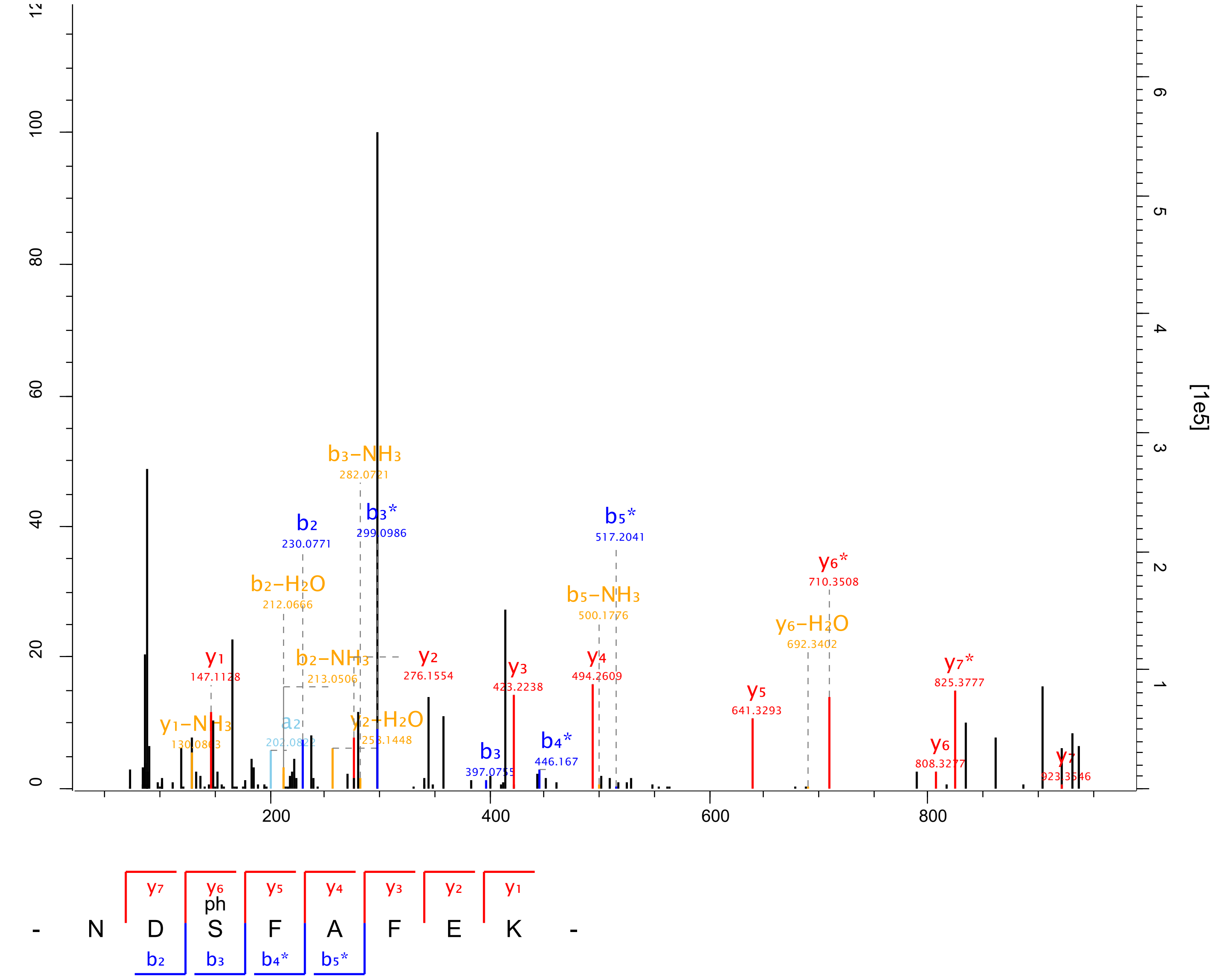

| Raw file | Scan | Method | Score | m/z |
| --- | --- | --- | --- | --- |
| water_02_1 | 18122 | FTMS; HCD | 196.78 | 895.88 |

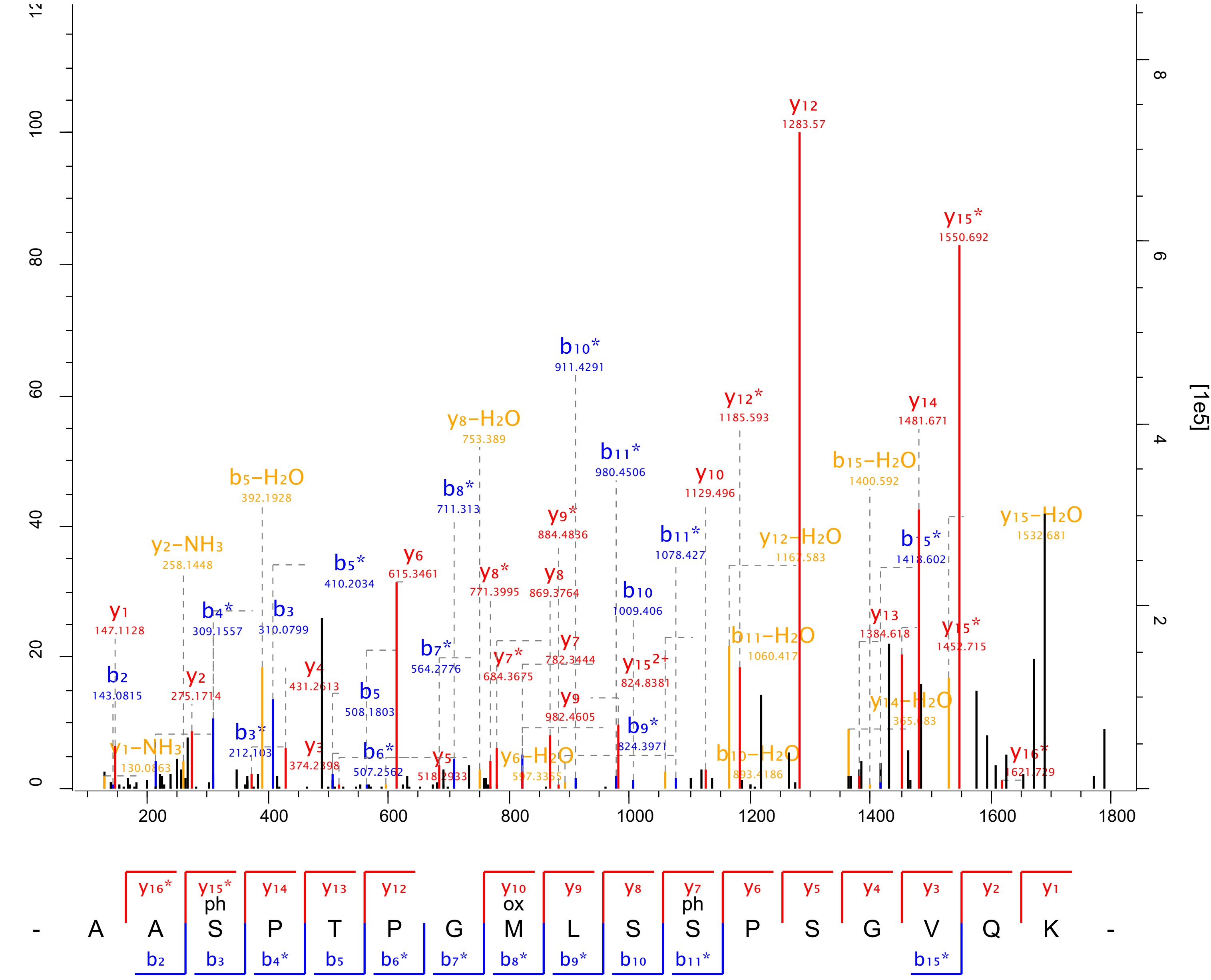

| Raw file | Scan | Method | Score | m/z |
| --- | --- | --- | --- | --- |
| water_02_1 | 18136 | FTMS; HCD | 76.01 | 667.29 |

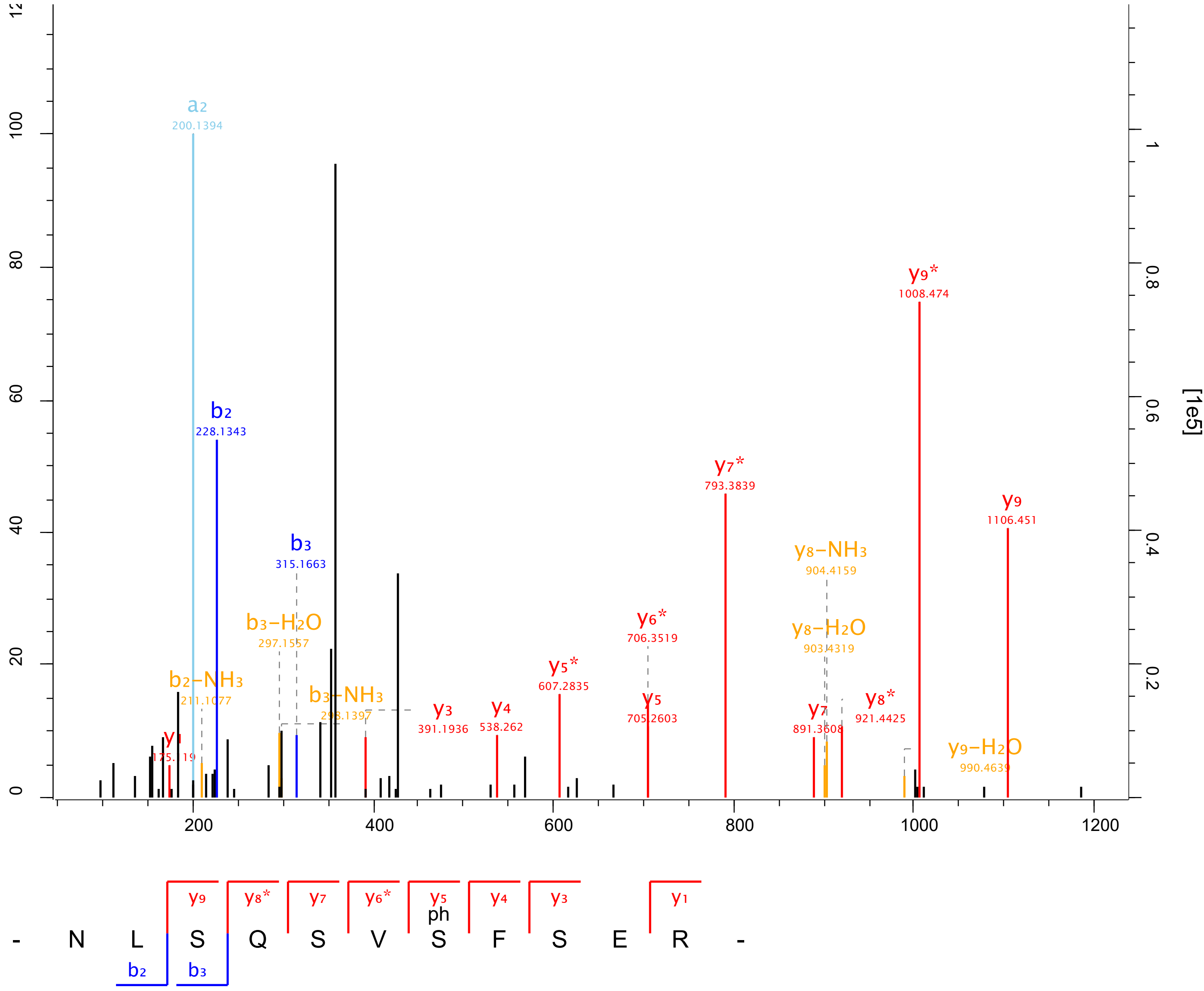

| Raw file | Scan | Method | Score | m/z |
|---|---|---|---|---|
| water_02_1 | 18234 | FTMS; HCD | 187.97 | 652.27 |

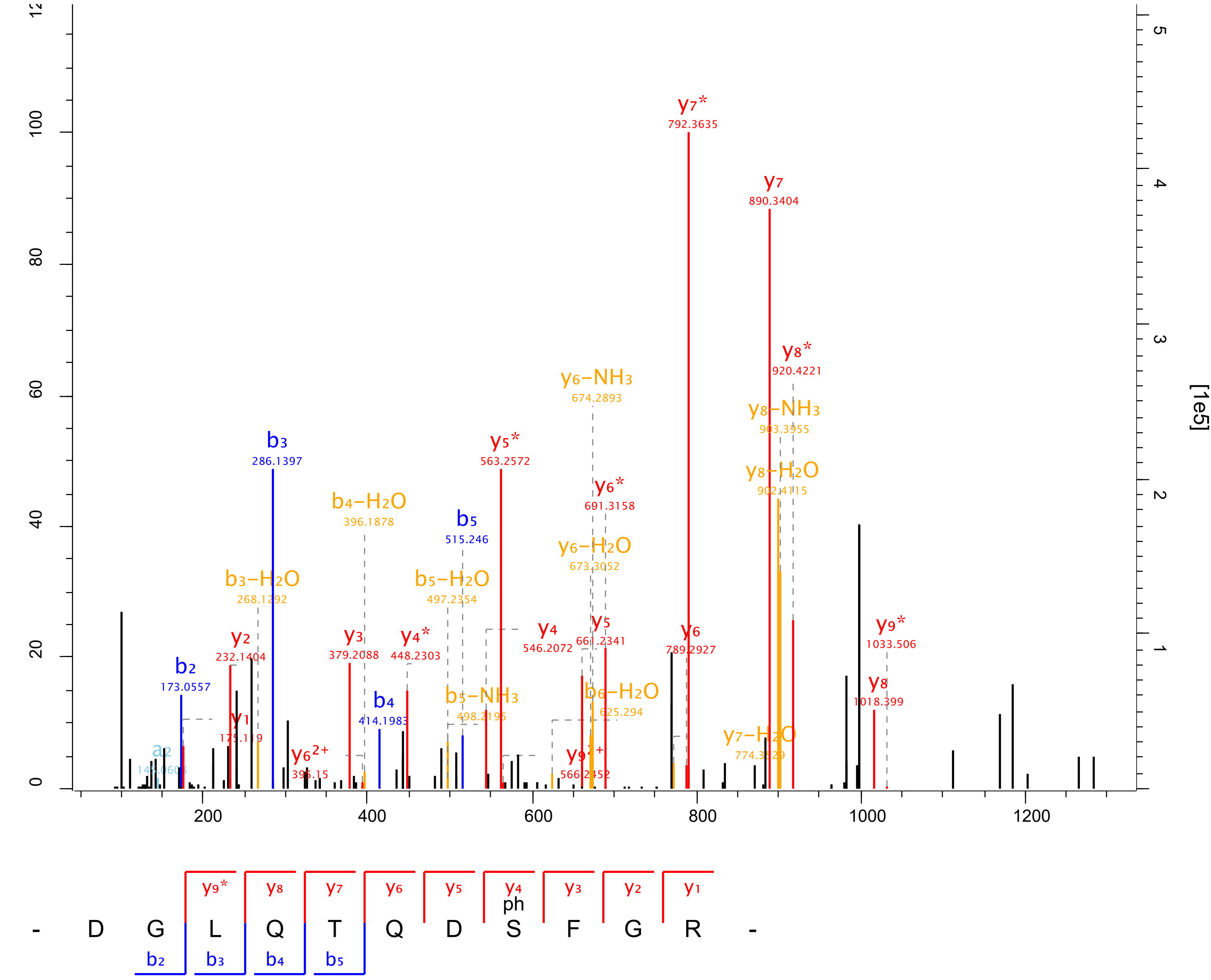

| Raw file | Scan | Method | Score | m/z |
| --- | --- | --- | --- | --- |
| water_02_1 | 1830 | FTMS; HCD | 52.81 | 835.81 |

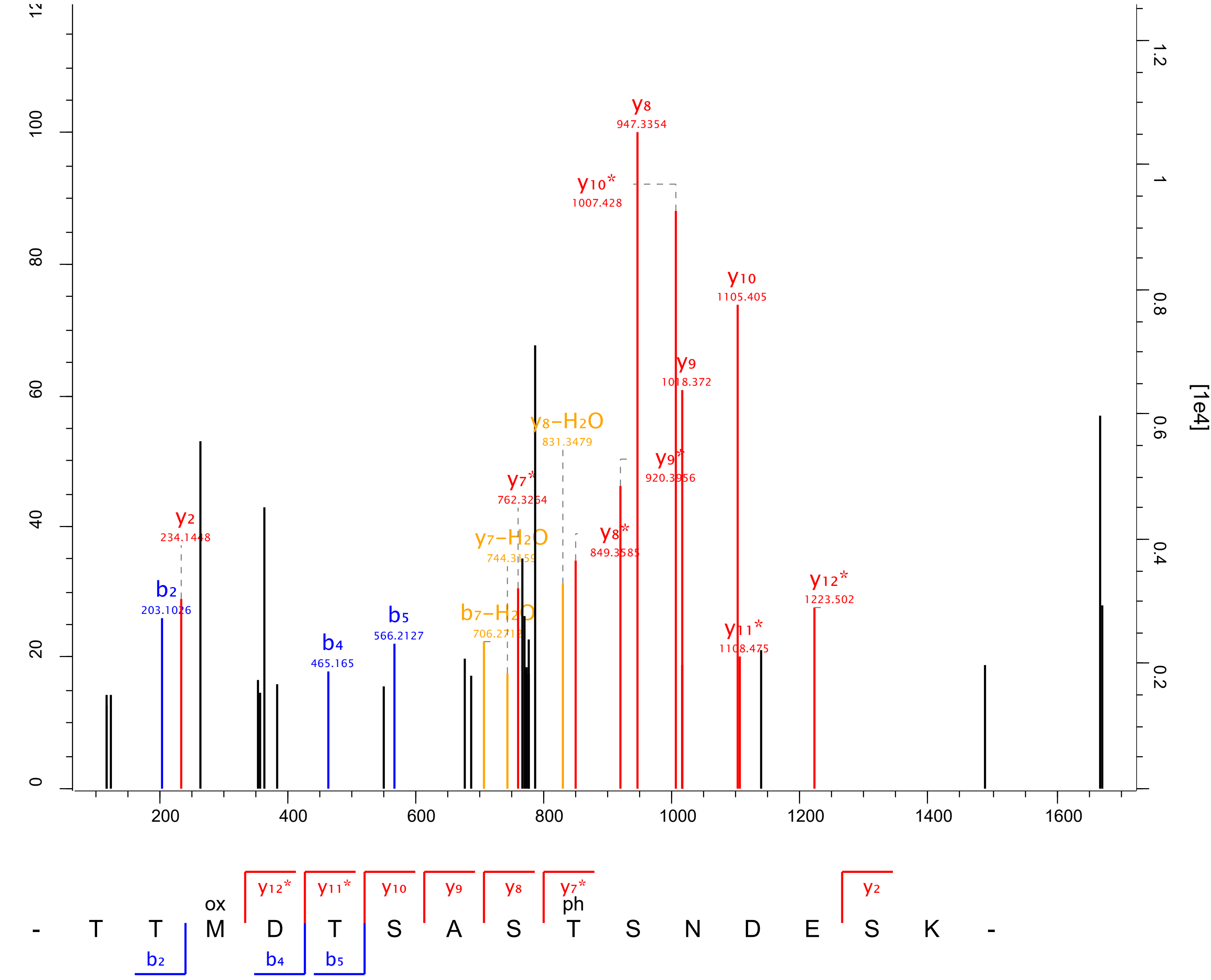

| Raw file | Scan | Method | Score | m/z |
| --- | --- | --- | --- | --- |
| water_02_1 | 18315 | FTMS; HCD | 58.98 | 832.83 |

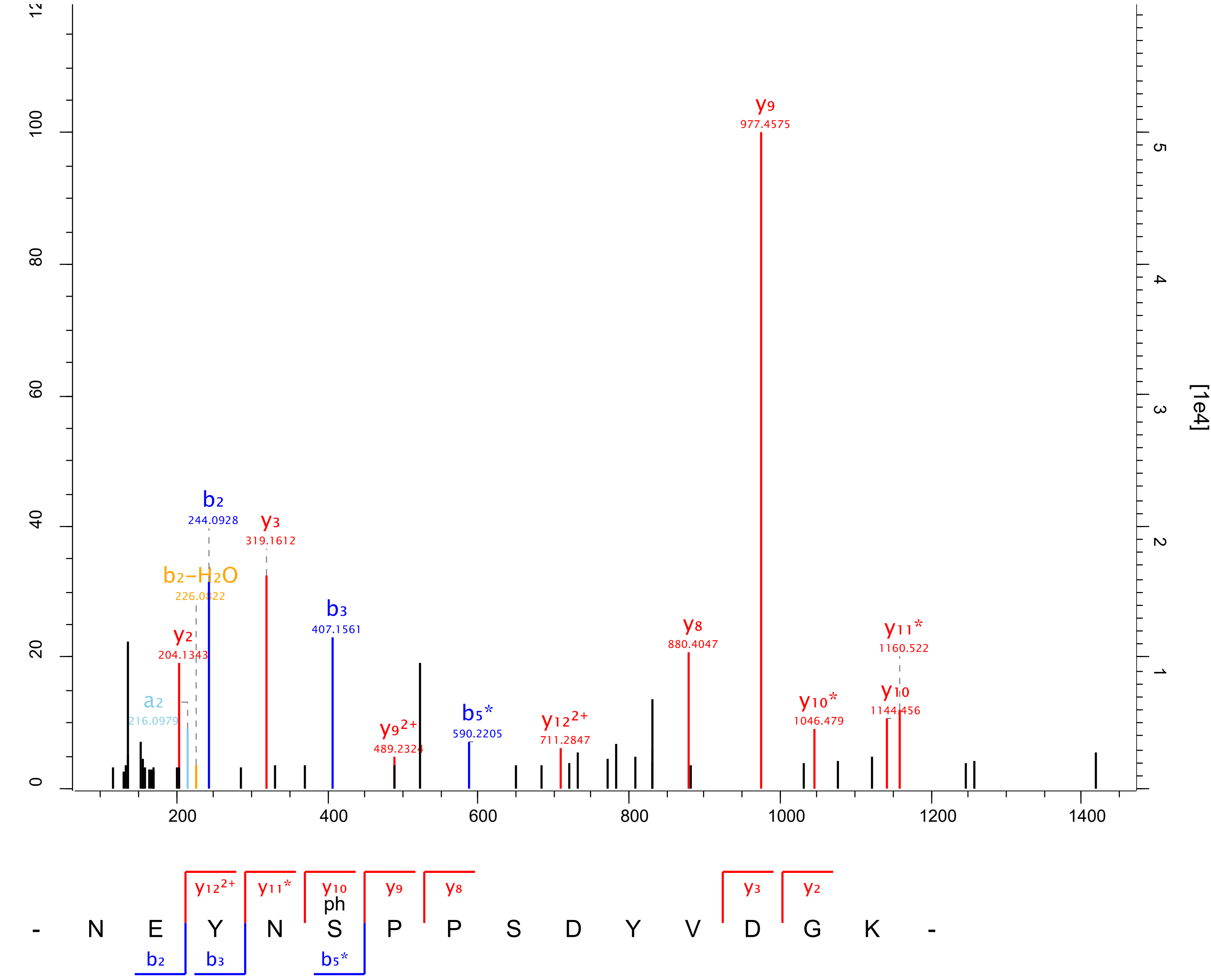

| Raw file | Scan | Method | Score | m/z |
|---|---|---|---|---|
| water_02_1 | 18381 | FTMS; HCD | 55.01 | 503.91 |

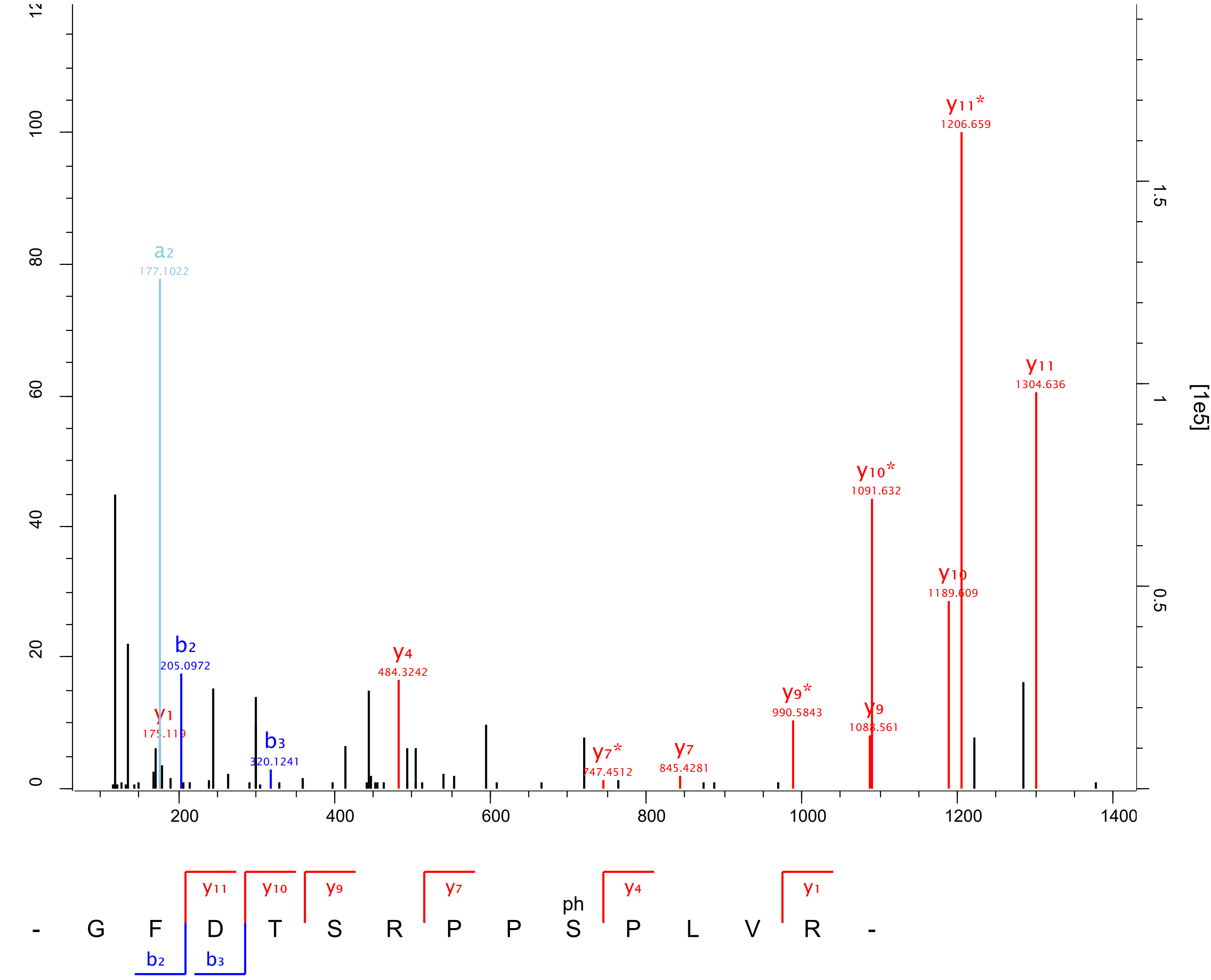

| Raw file | Scan | Method | Score | m/z |
|---|---|---|---|---|
| water_02_1 | 18404 | FTMS; HCD | 159.1 | 914.88 |

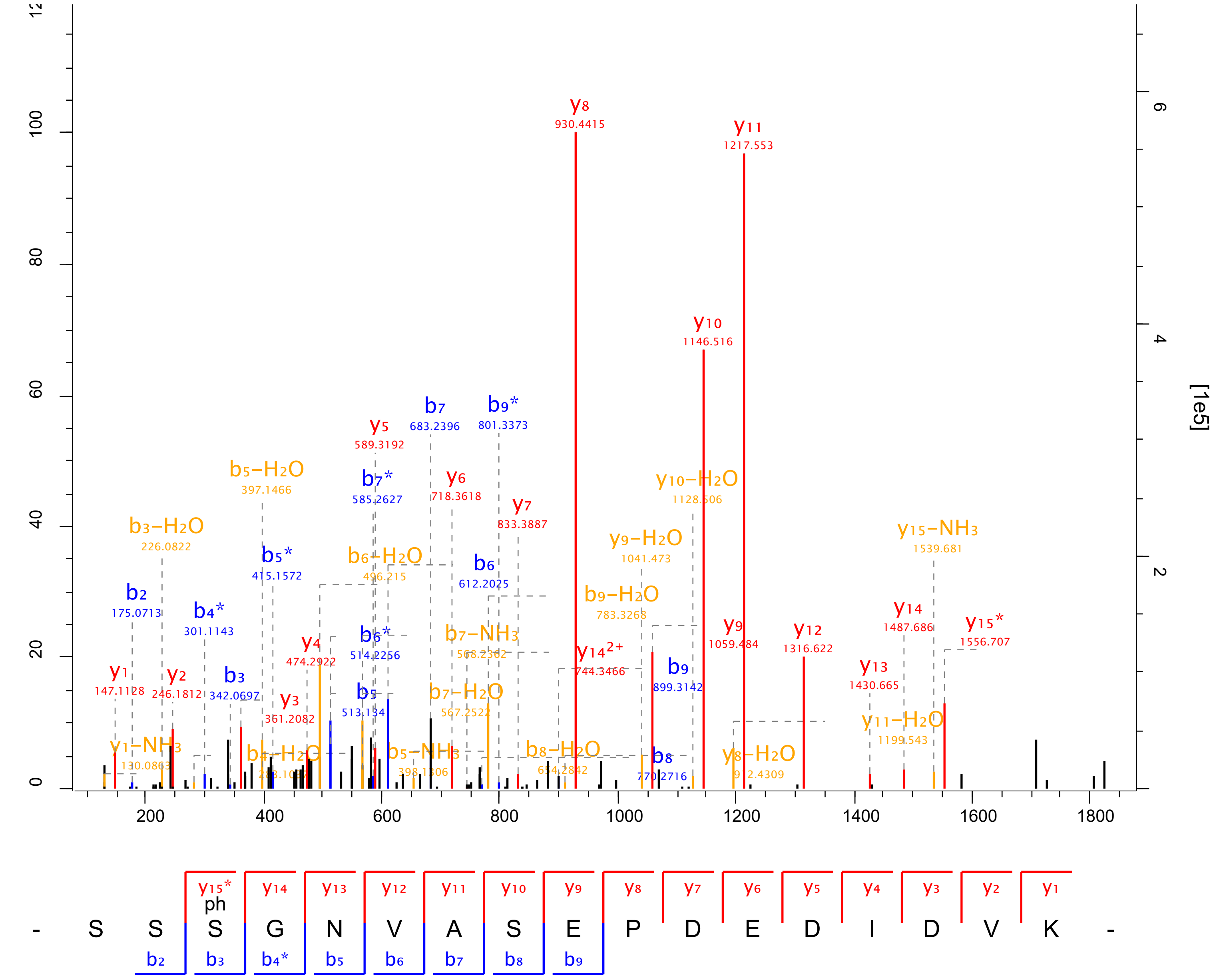

| Raw file | Scan | Method | Score | m/z |
|---|---|---|---|---|
| water_02_1 | 18446 | FTMS; HCD | 55.13 | 668.32 |

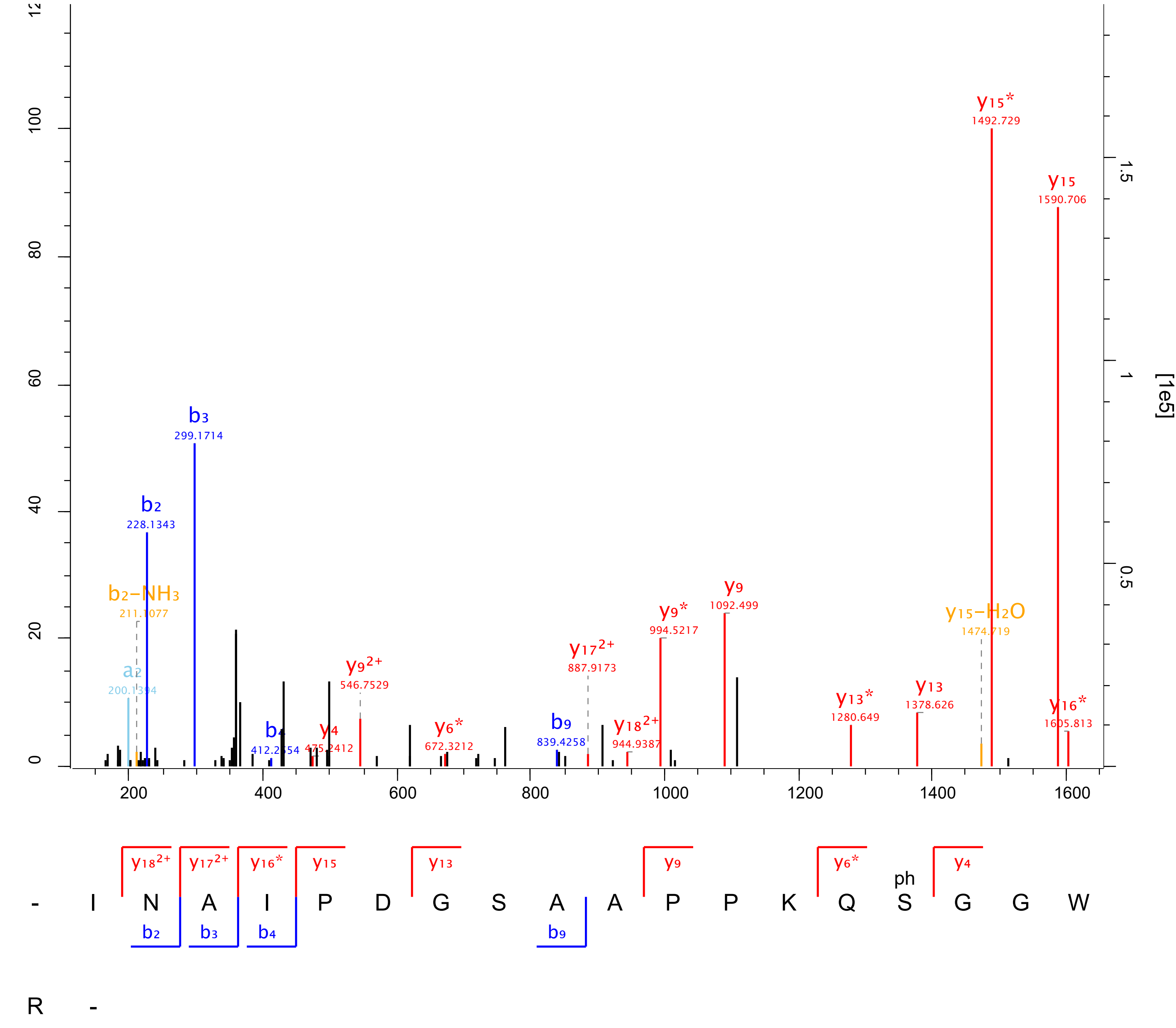

| Raw file | Scan | Method | Score | m/z |
|---|---|---|---|---|
| water_02_1 | 1857 | FTMS; HCD | 118.21 | 745.31 |

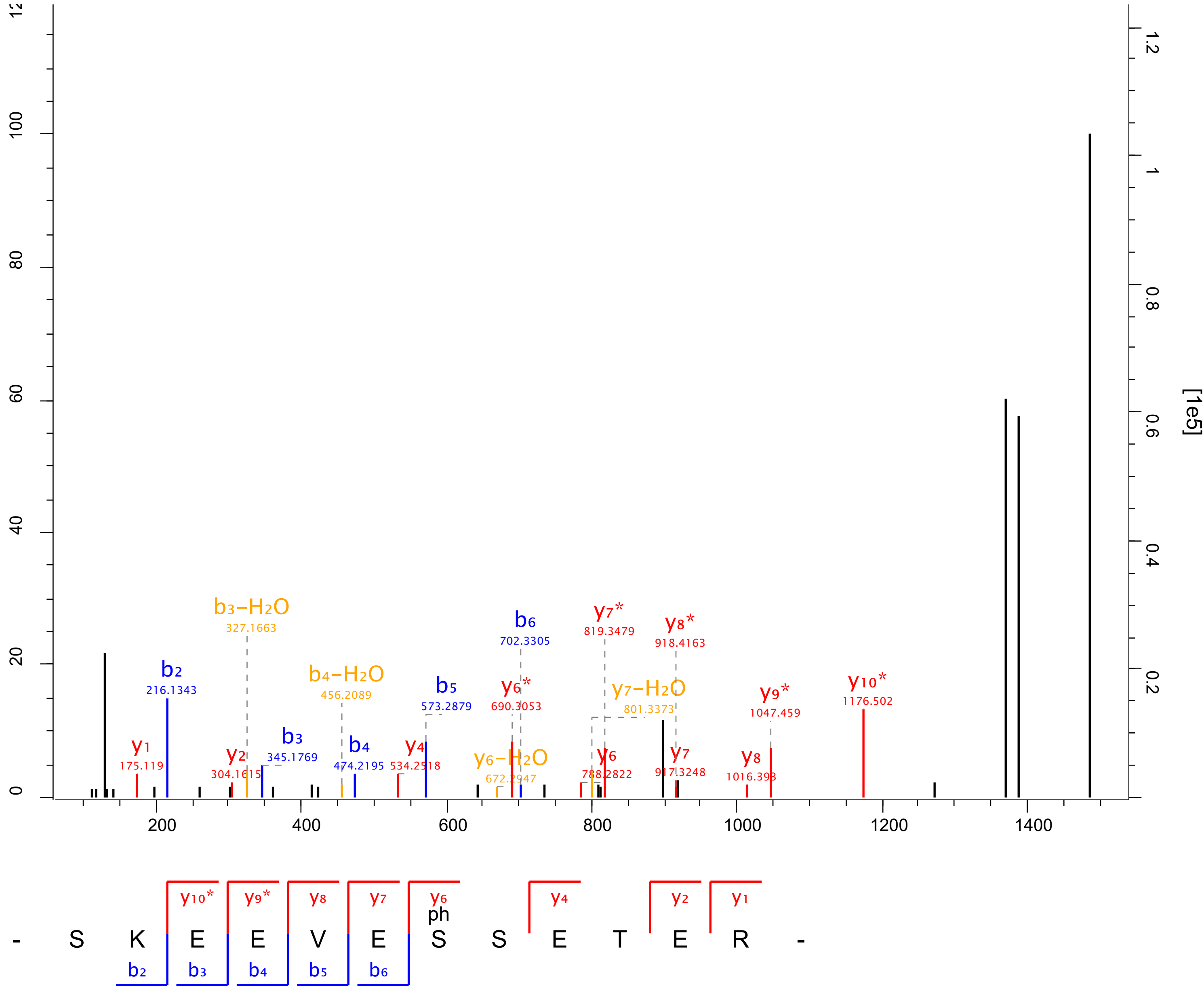

| Raw file | Scan | Method | Score | m/z |
| --- | --- | --- | --- | --- |
| water_02_1 | 18570 | FTMS; HCD | 105.03 | 560.59 |

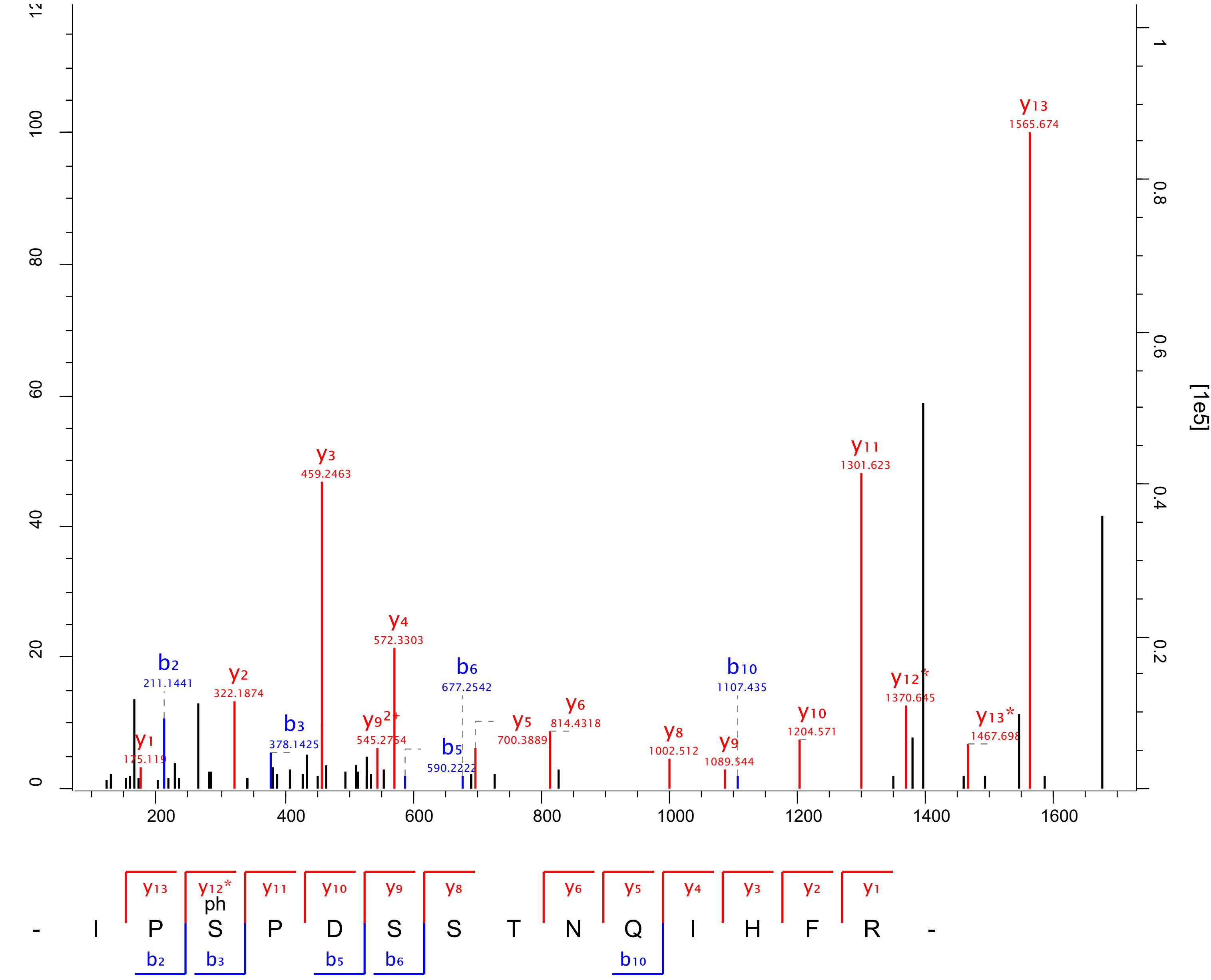

| Raw file | Scan | Method | Score | m/z |
|---|---|---|---|---|
| water_02_1 | 18759 | FTMS; HCD | 98.06 | 592.28 |

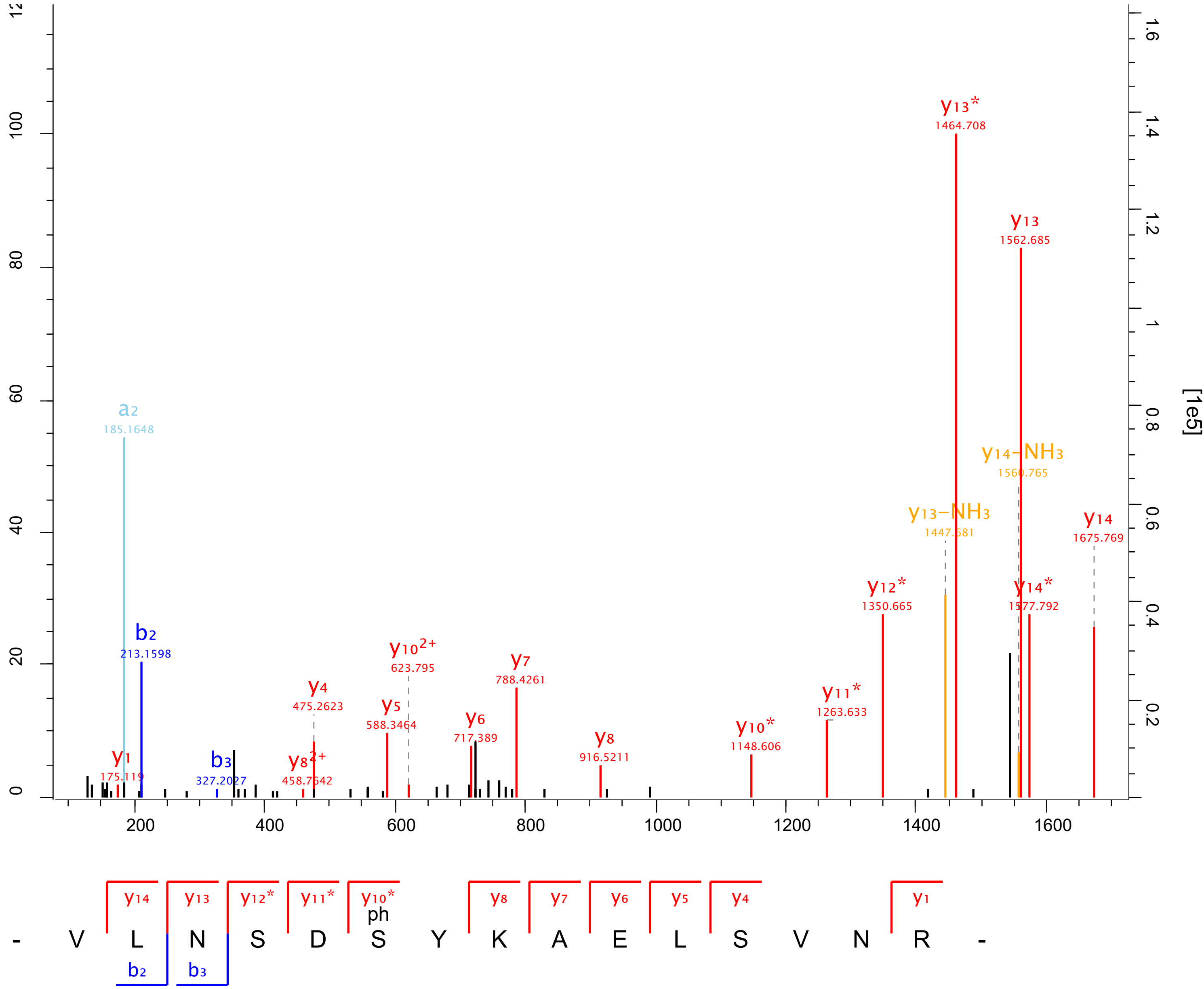

| Raw file | Scan | Method | Score | m/z |
| --- | --- | --- | --- | --- |
| water_02_1 | 18826 | FTMS; HCD | 129.46 | 728.84 |

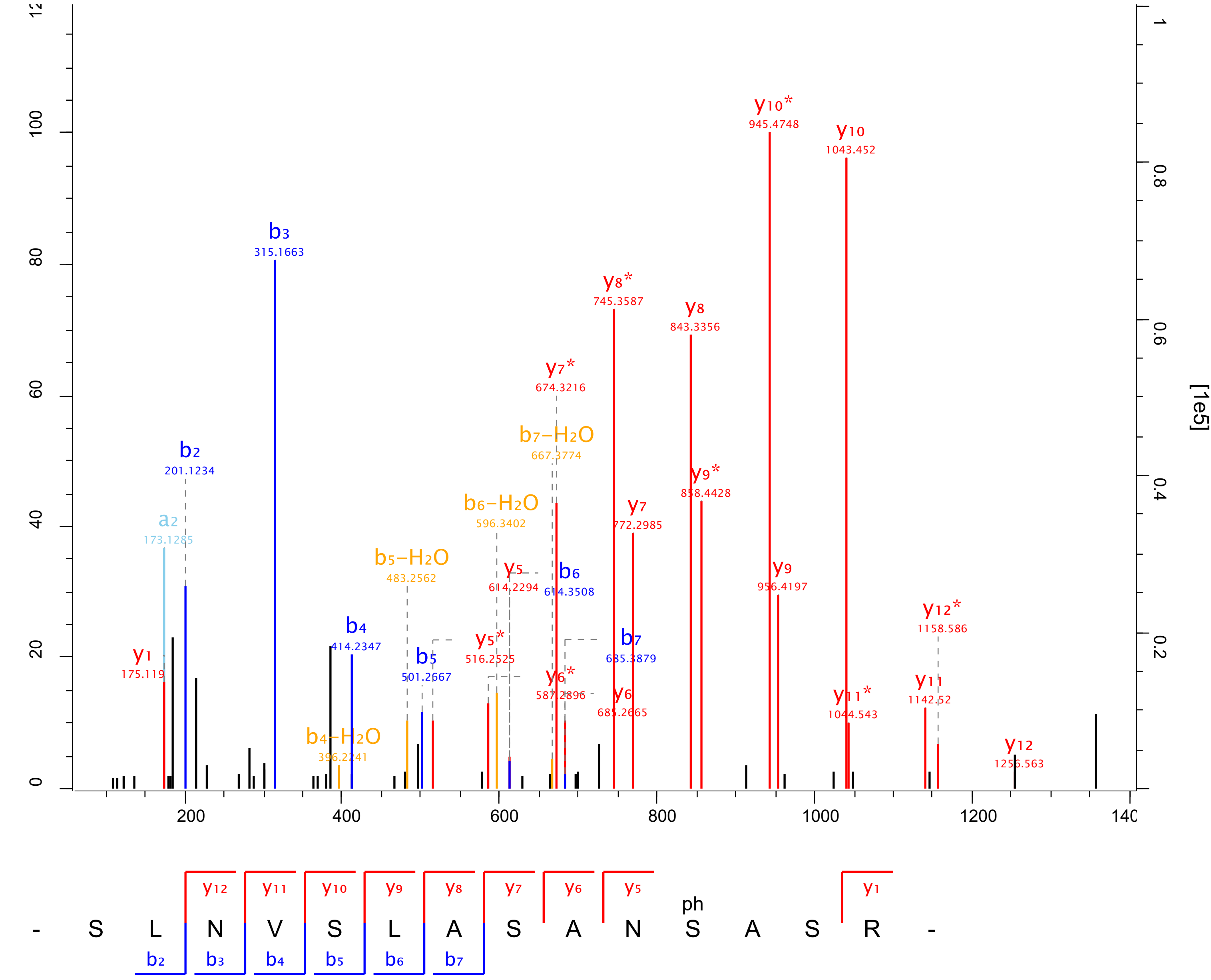

| Raw file | Scan | Method | Score | m/z |
|---|---|---|---|---|
| water_02_1 | 18868 | FTMS; HCD | 135.62 | 740.29 |

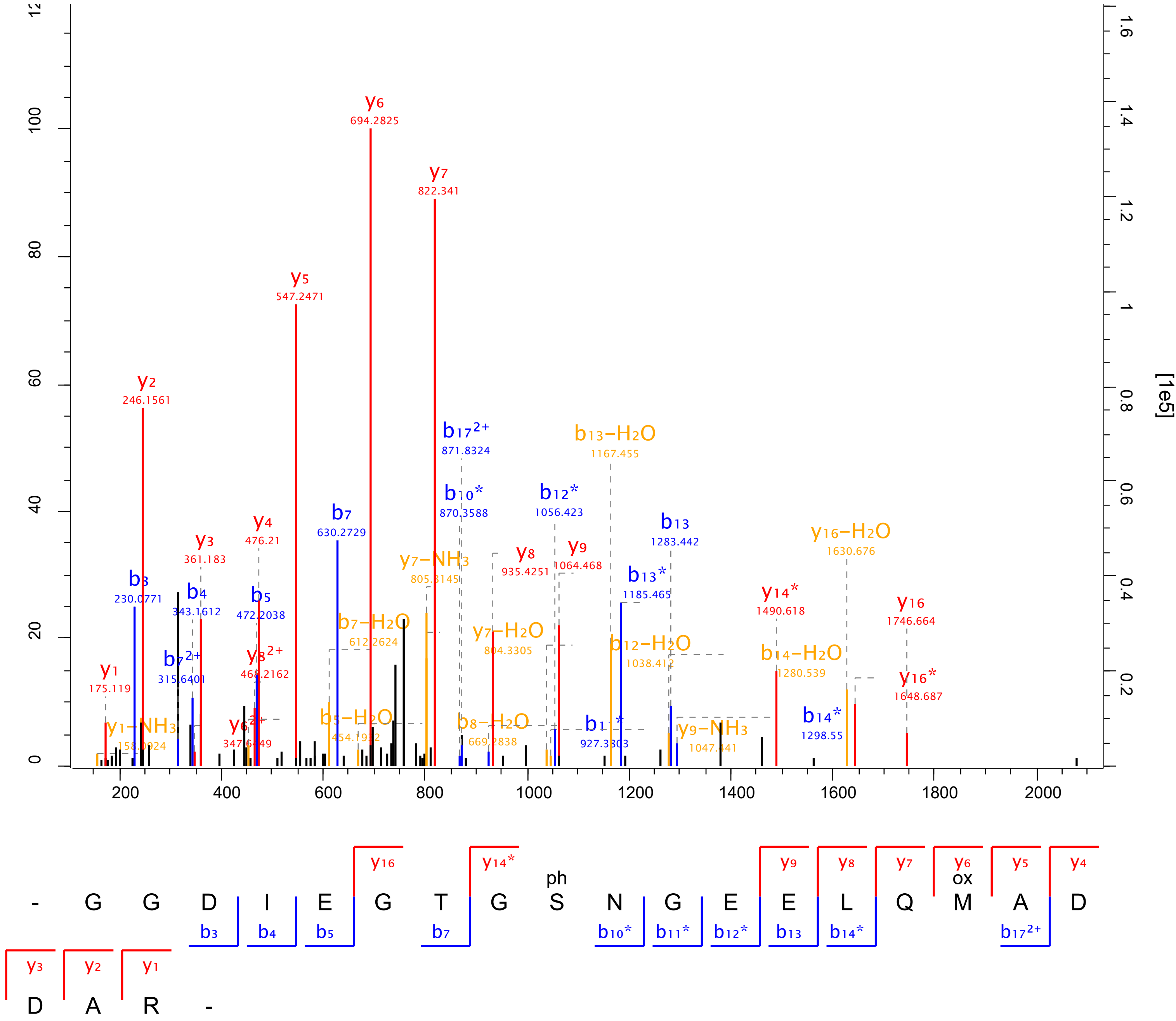

| Raw file | Scan | Method | Score | m/z |
|---|---|---|---|---|
| water_02_1 | 18915 | FTMS; HCD | 82.26 | 515.22 |

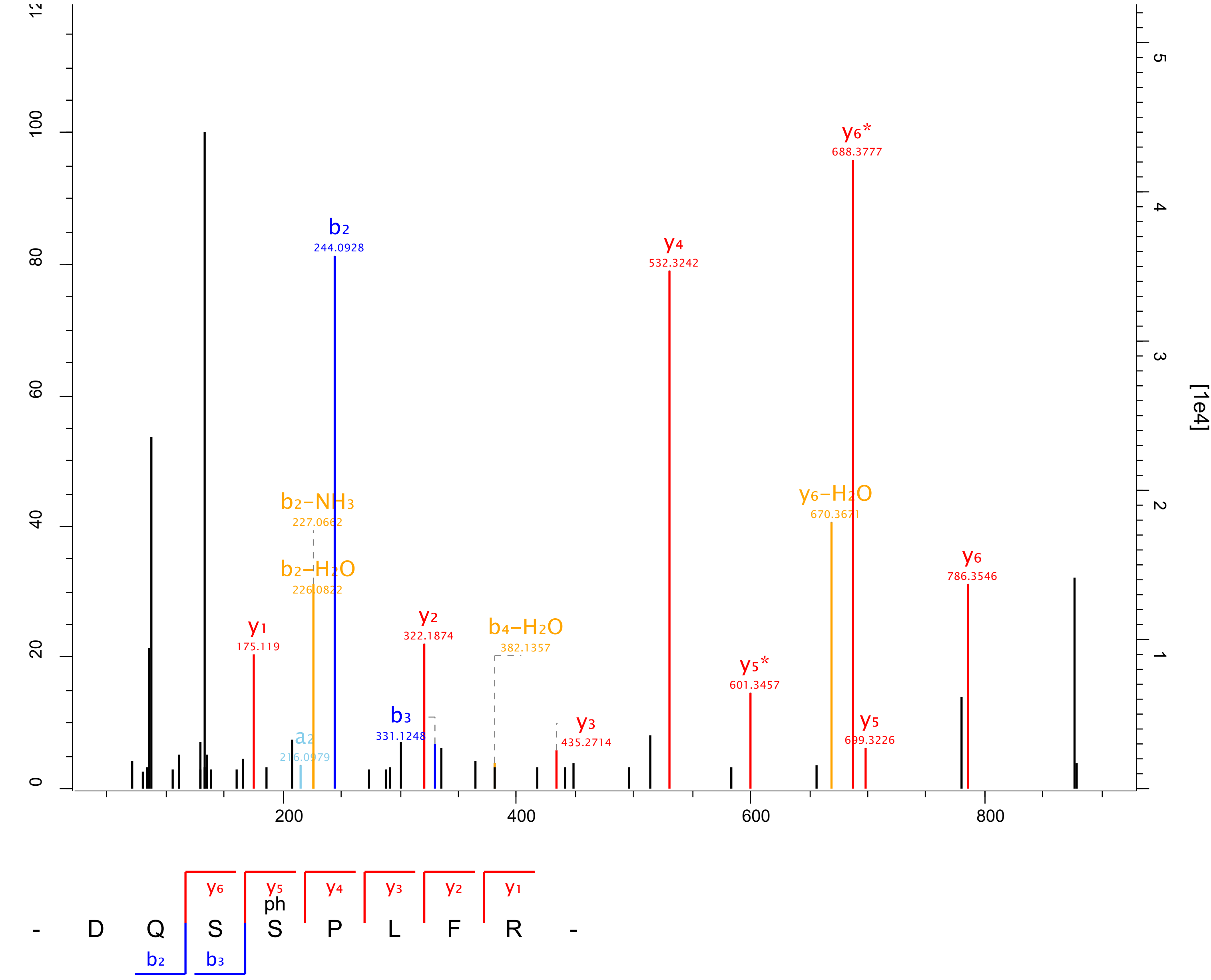

| Raw file | Scan | Method | Score | m/z |
| --- | --- | --- | --- | --- |
| water_02_1 | 18970 | FTMS; HCD | 141.37 | 572.76 |

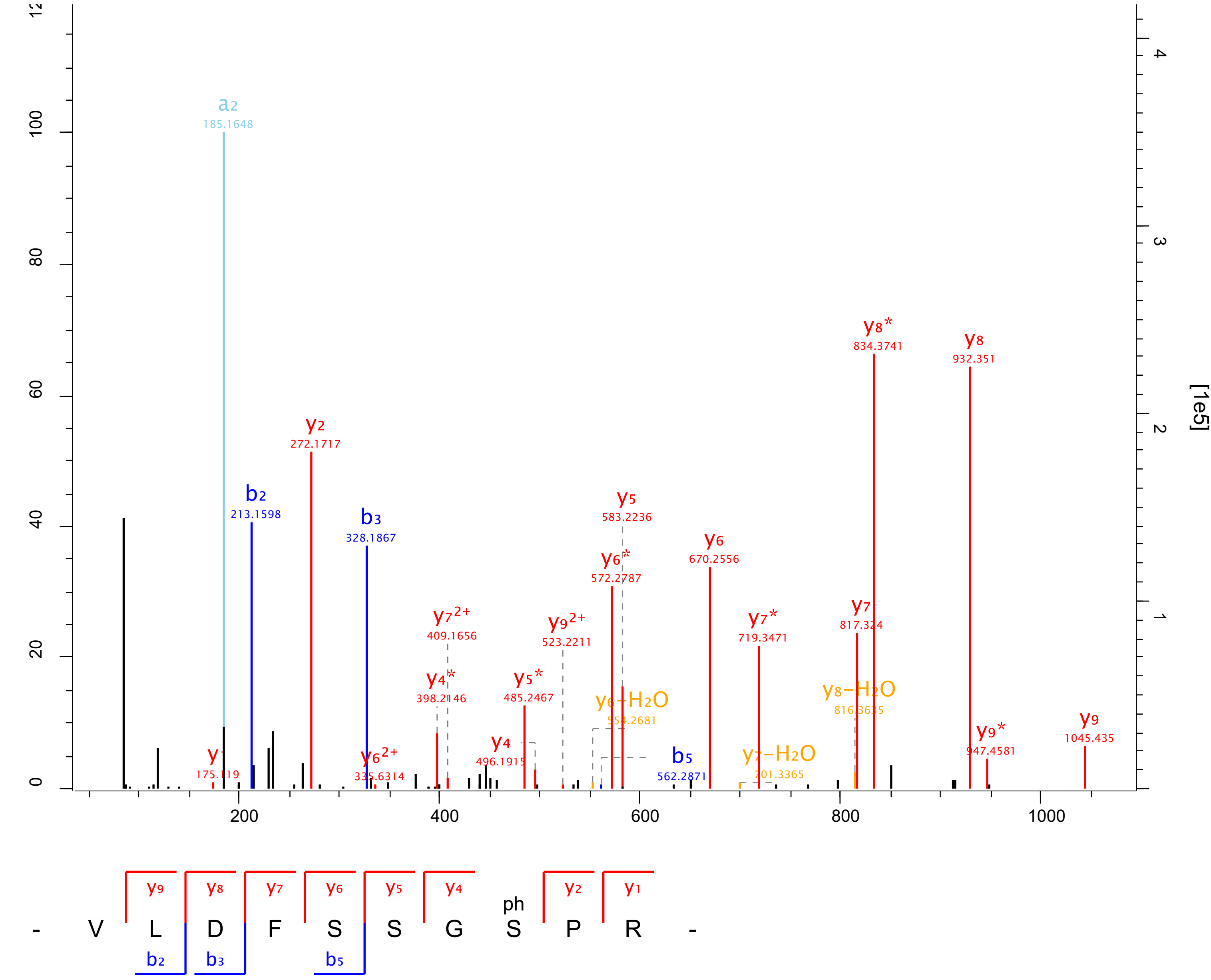

| Raw file | Scan | Method | Score | m/z |
| --- | --- | --- | --- | --- |
| water_02_1 | 19017 | FTMS; HCD | 169.8 | 780.97 |

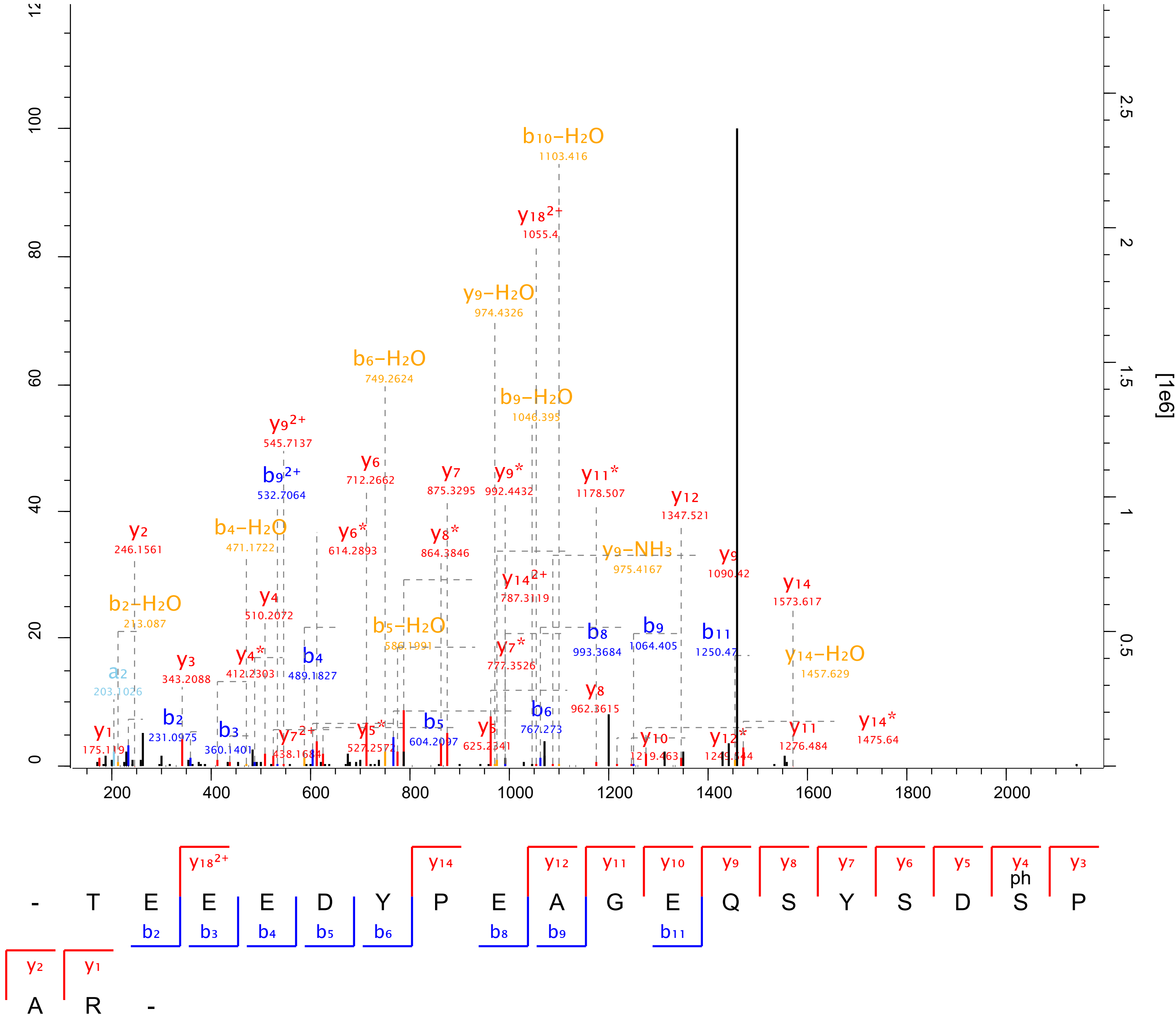

| Raw file | Scan | Method | Score | m/z |
|---|---|---|---|---|
| water_02_1 | 19028 | FTMS; HCD | 142.41 | 959.38 |

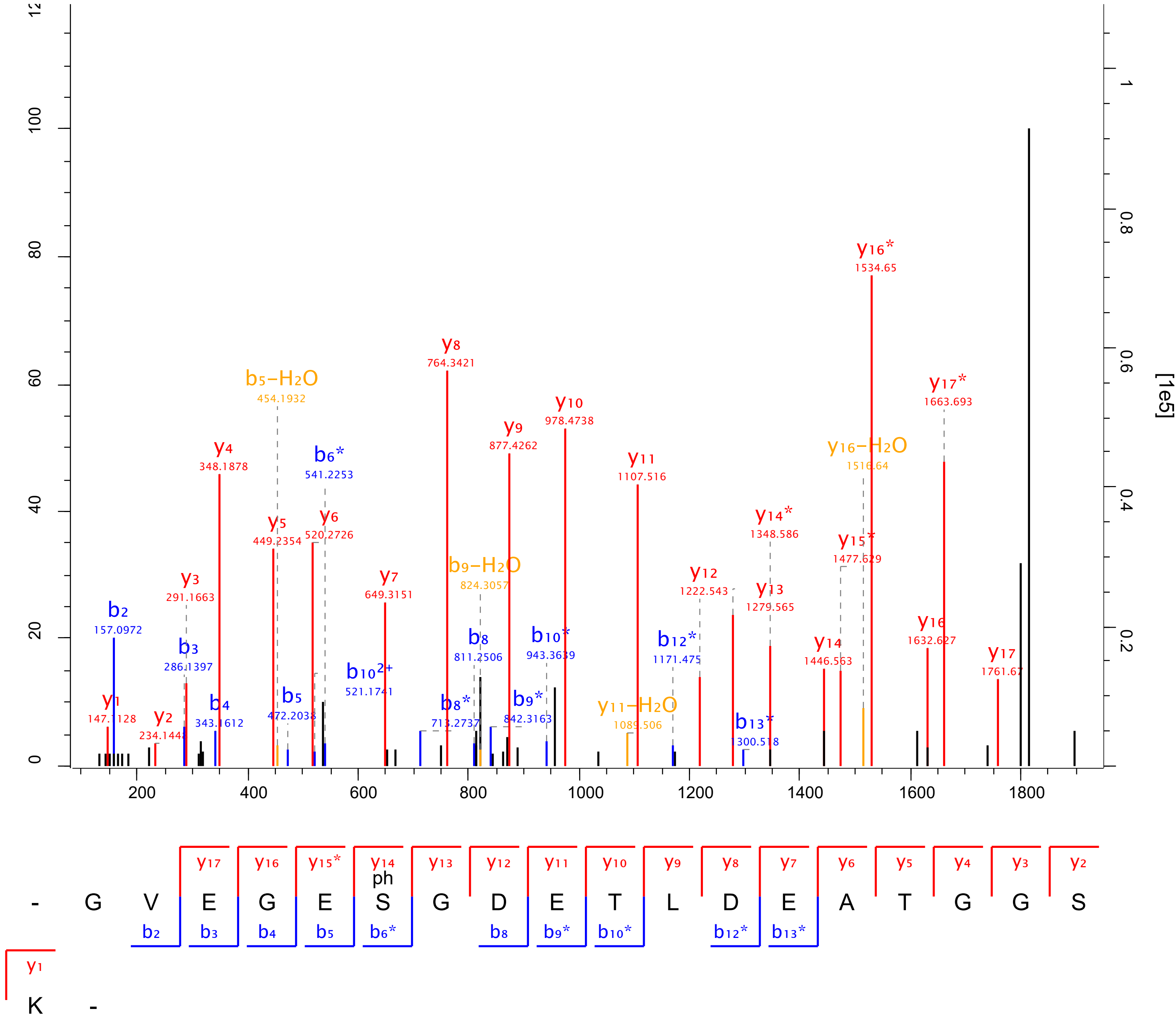

| Raw file | Scan | Method | Score | m/z |
|---|---|---|---|---|
| water_02_1 | 19039 | FTMS; HCD | 69.19 | 549.57 |

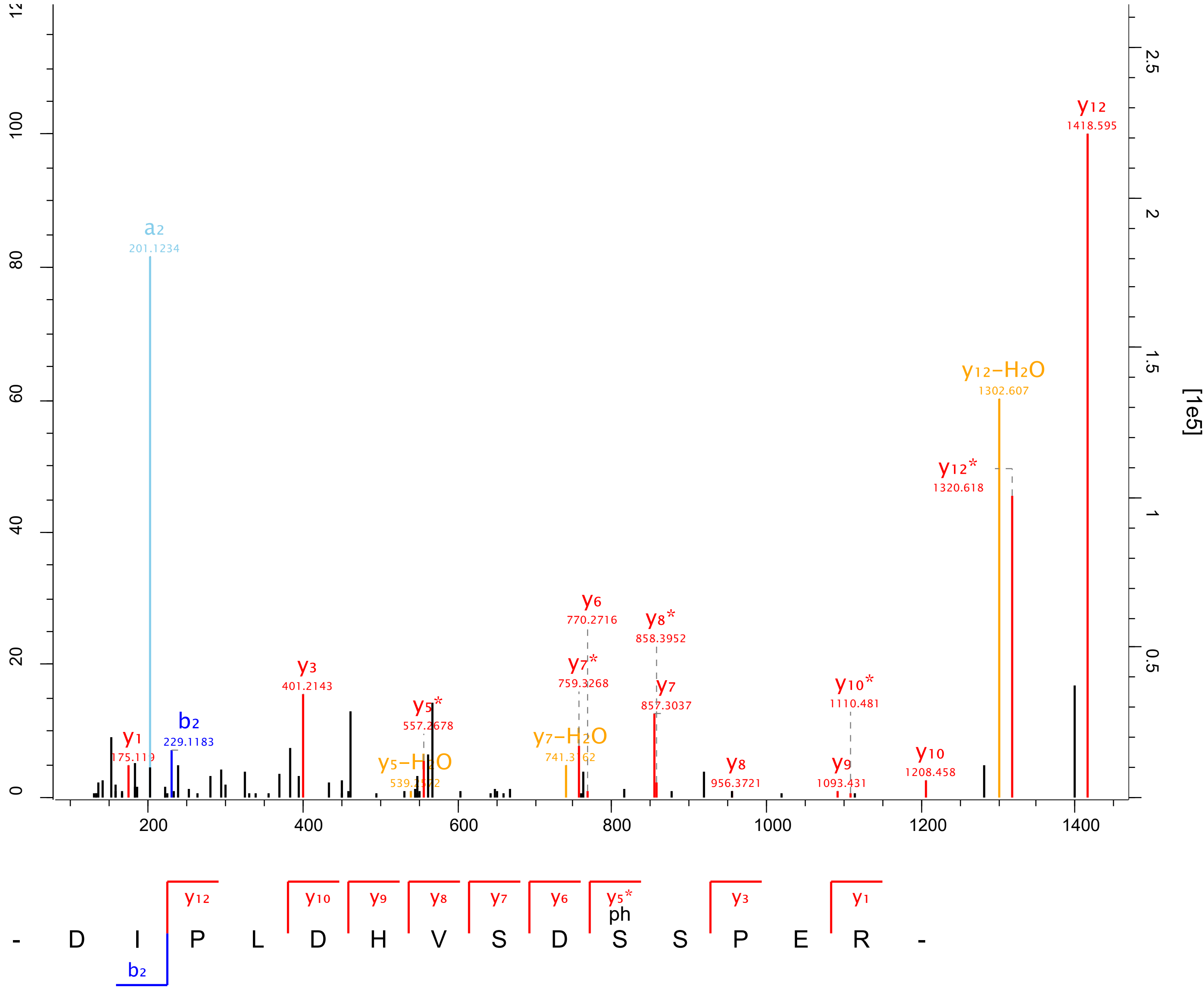

| Raw file | Scan | Method | Score | m/z |
| --- | --- | --- | --- | --- |
| water_02_1 | 19170 | FTMS; HCD | 85.06 | 739.31 |

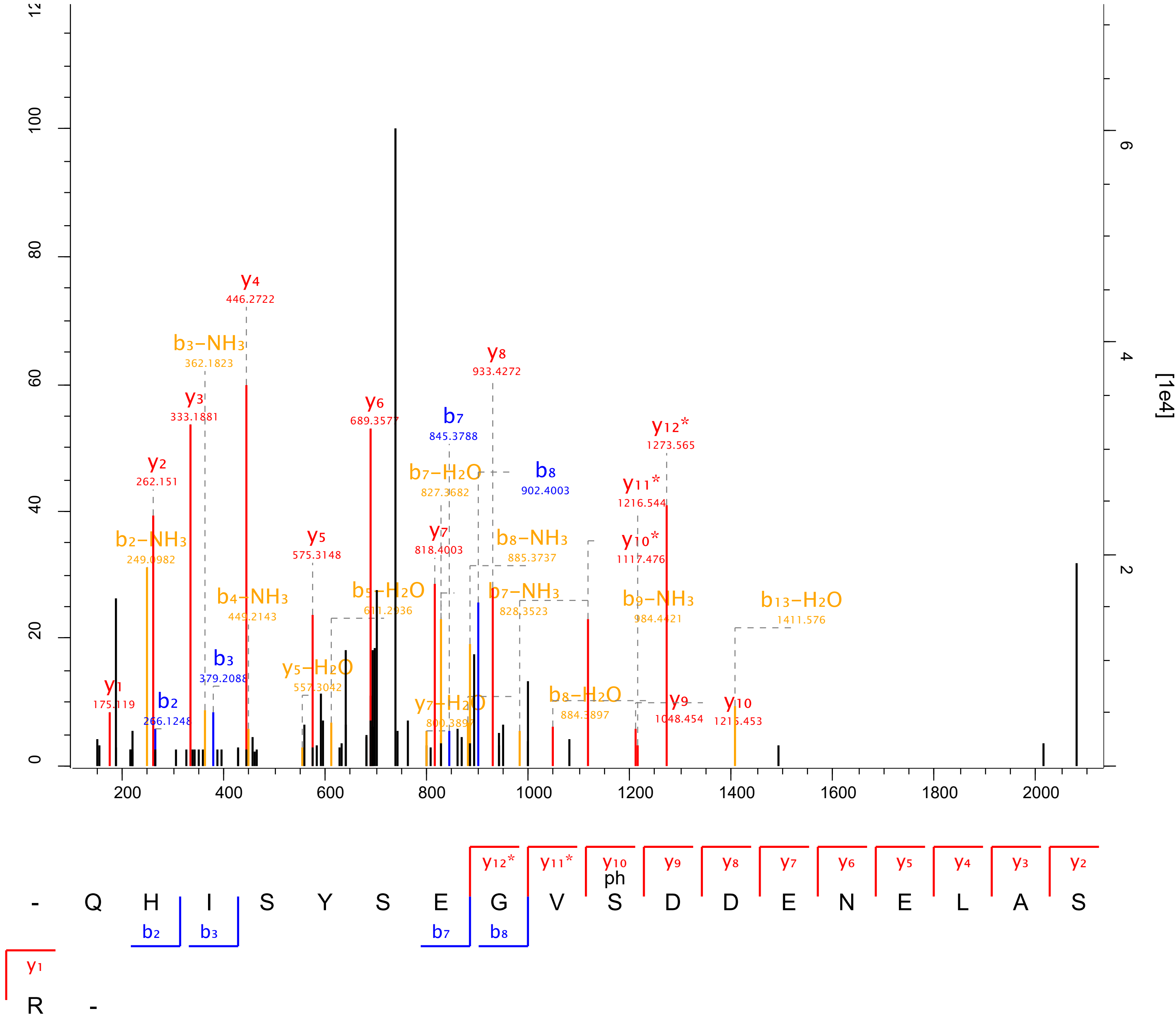

| Raw file | Scan | Method | Score | m/z |
|---|---|---|---|---|
| water_02_1 | 19187 | FTMS; HCD | 82.89 | 782.01 |

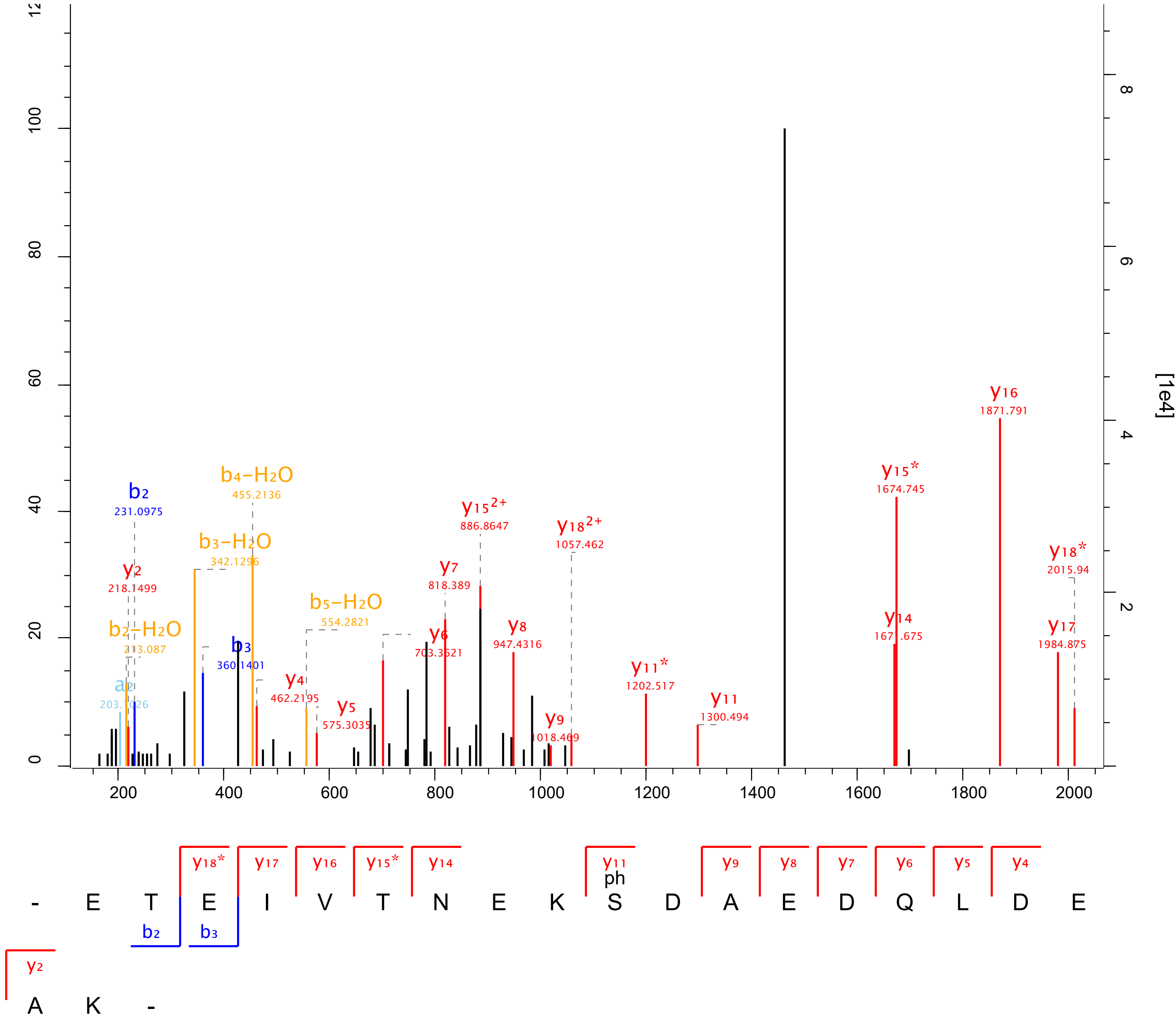

| Raw file | Scan | Method | Score | m/z |
| --- | --- | --- | --- | --- |
| water_02_1 | 19397 | FTMS; HCD | 44.25 | 716.35 |

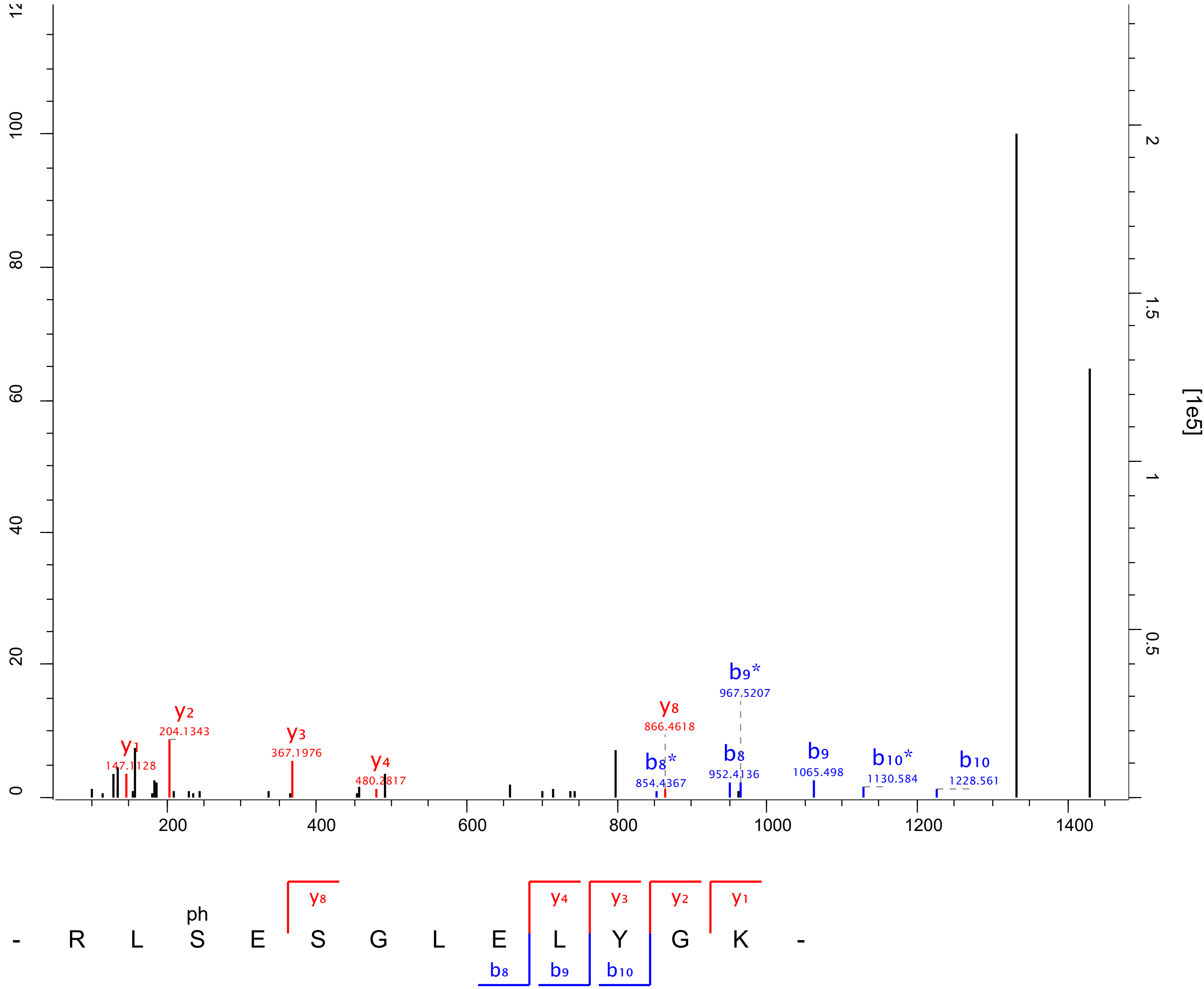

| Raw file | Scan | Method | Score | m/z |
| --- | --- | --- | --- | --- |
| water_02_1 | 19423 | FTMS; HCD | 65.85 | 662.64 |

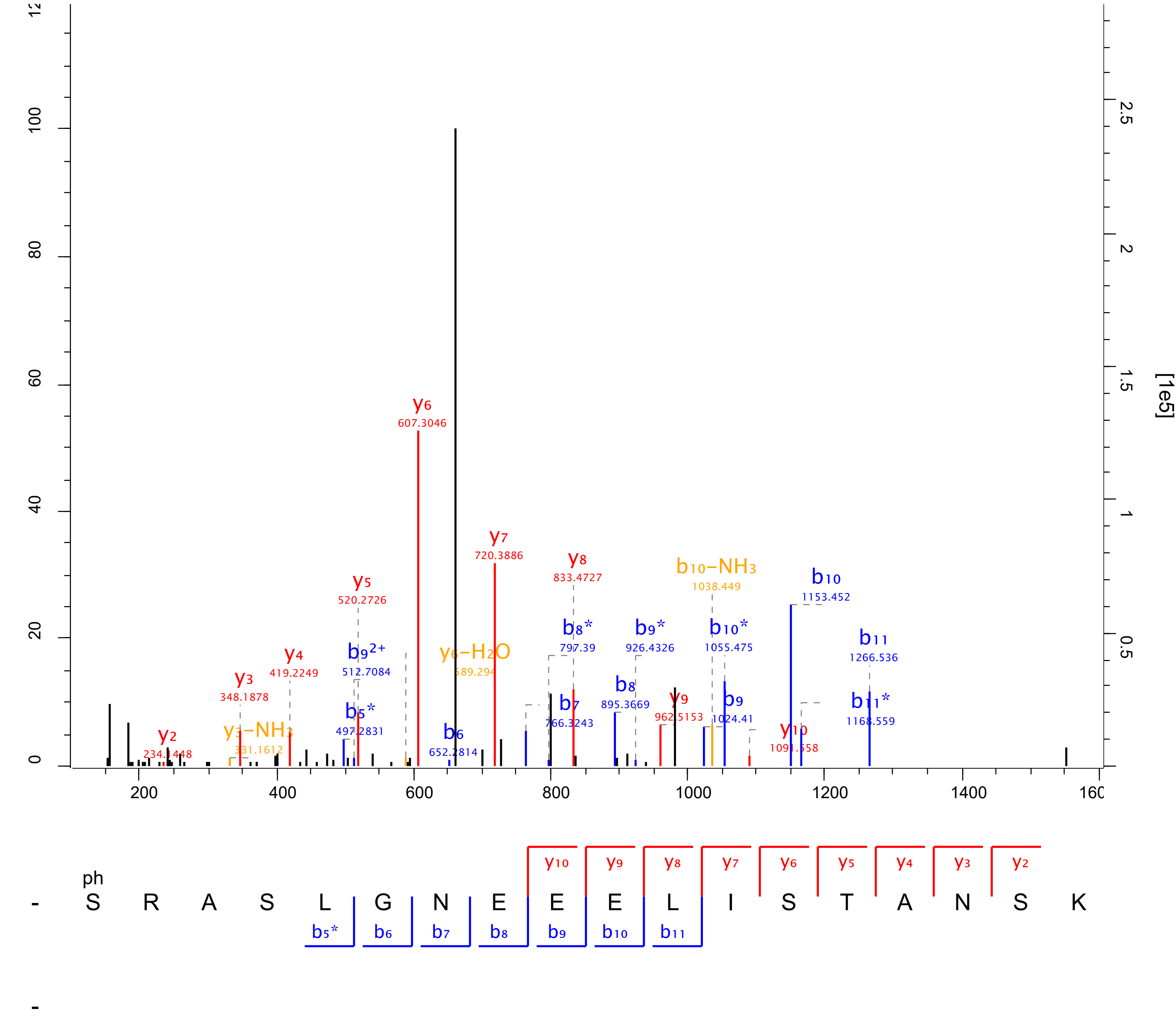

| Raw file | Scan | Method | Score | m/z |
| --- | --- | --- | --- | --- |
| water_02_1 | 19430 | FTMS; HCD | 76.07 | 439.86 |

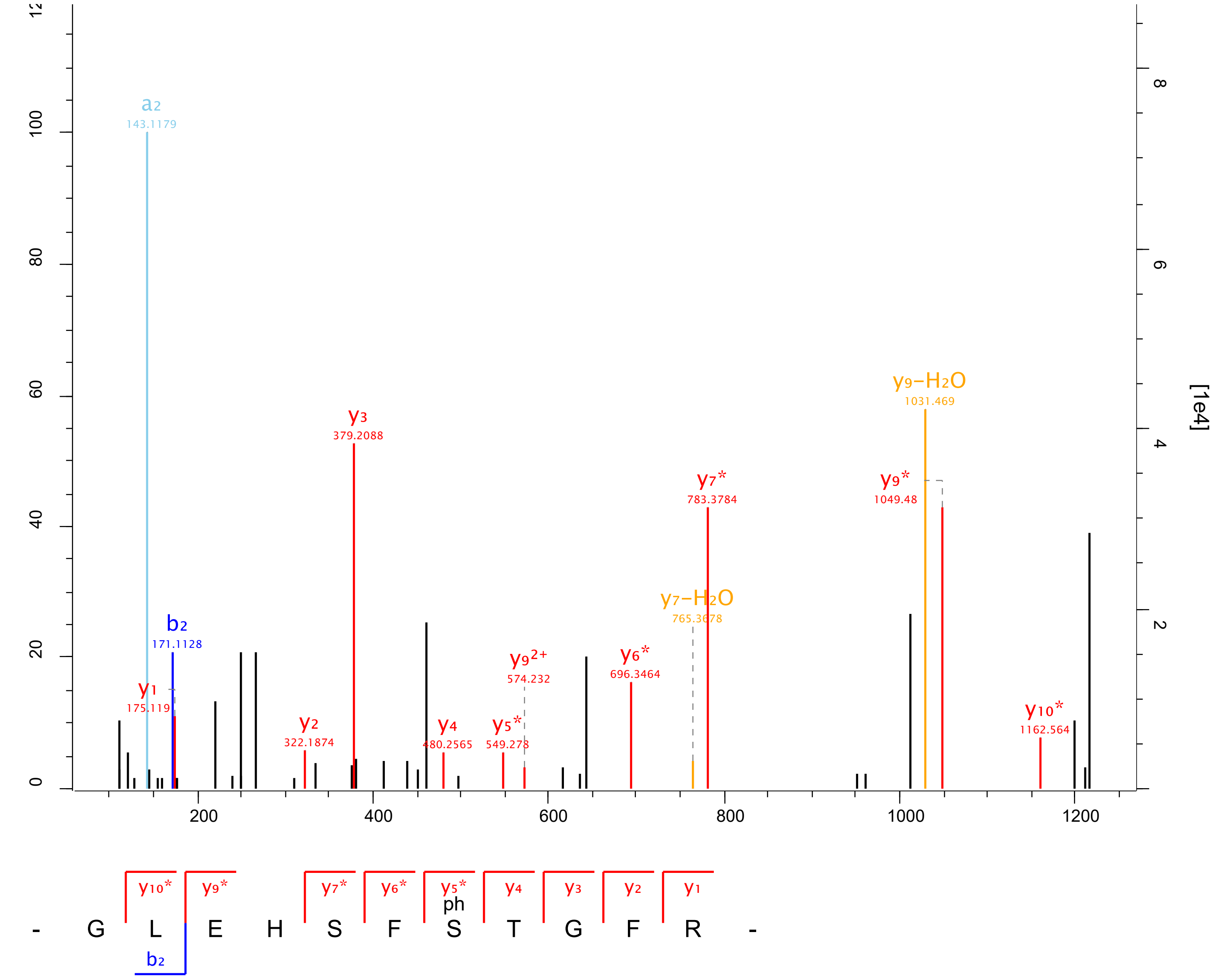

| Raw file | Scan | Method | Score | m/z |
| --- | --- | --- | --- | --- |
| water_02_1 | 19444 | FTMS; HCD | 68.38 | 492.71 |

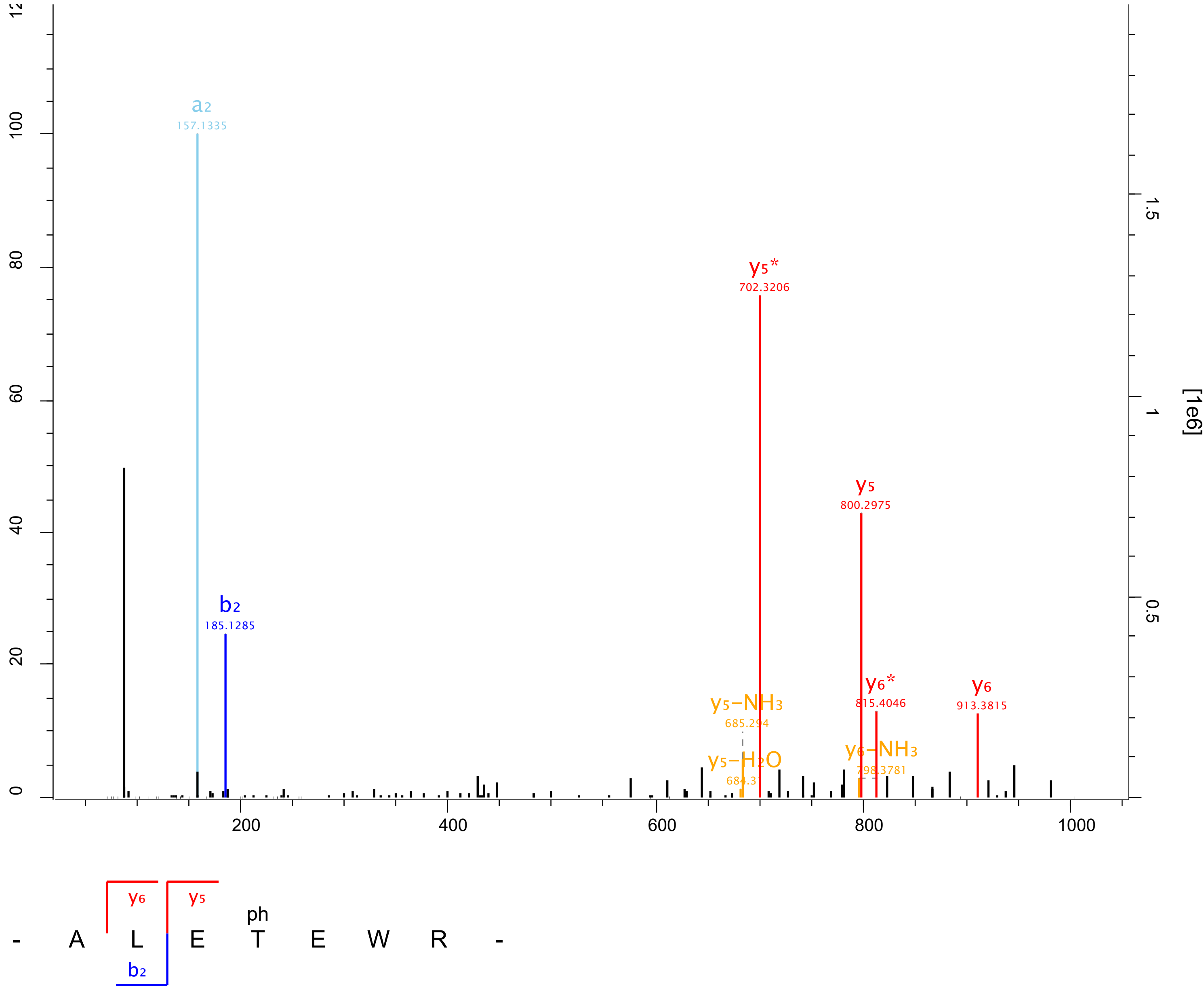

| Raw file | Scan | Method | Score | m/z |
| --- | --- | --- | --- | --- |
| water_02_1 | 19550 | FTMS; HCD | 139.86 | 657.79 |

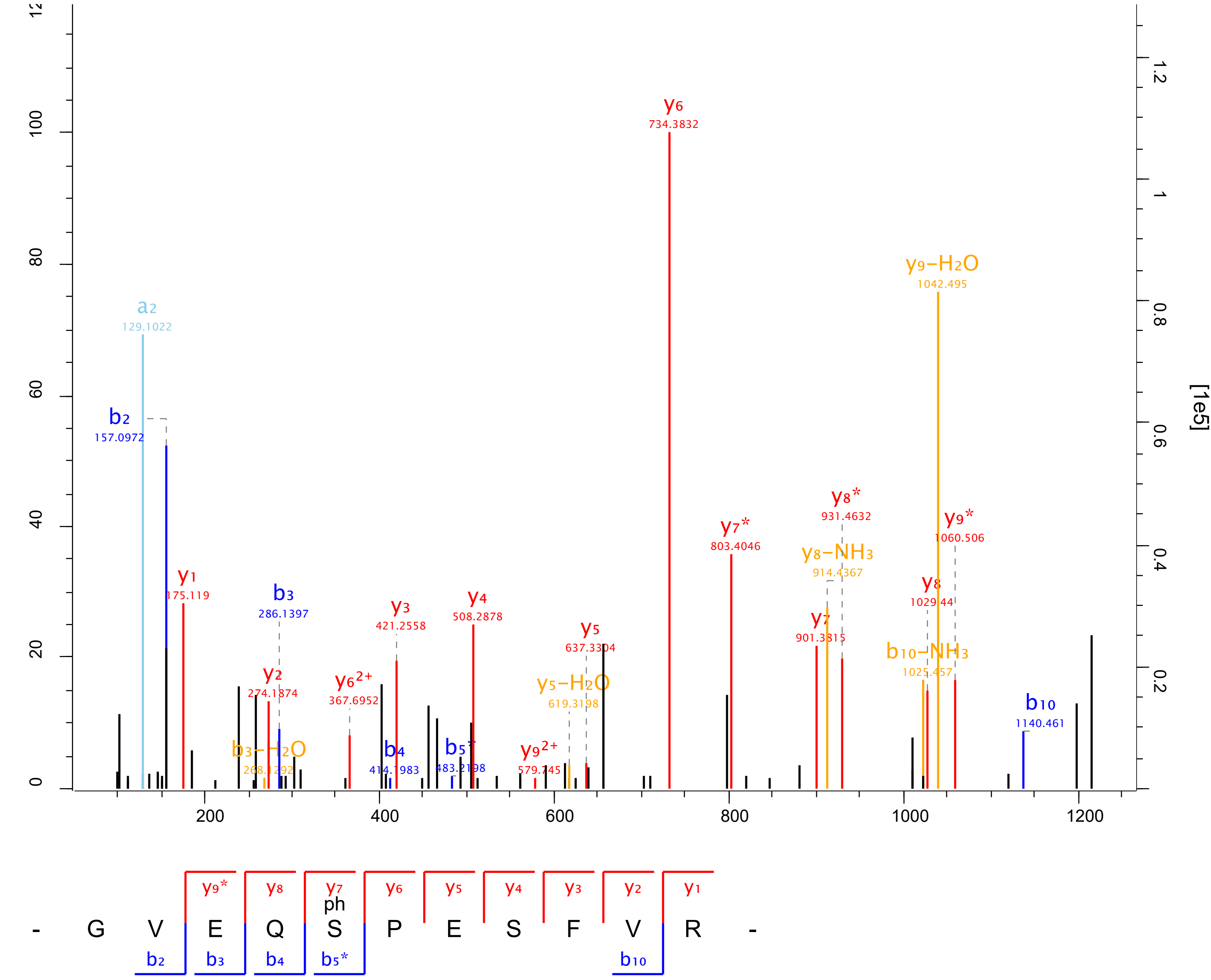

| Raw file | Scan | Method | Score | m/z |
| --- | --- | --- | --- | --- |
| water_02_1 | 19553 | FTMS; HCD | 64.57 | 451.24 |

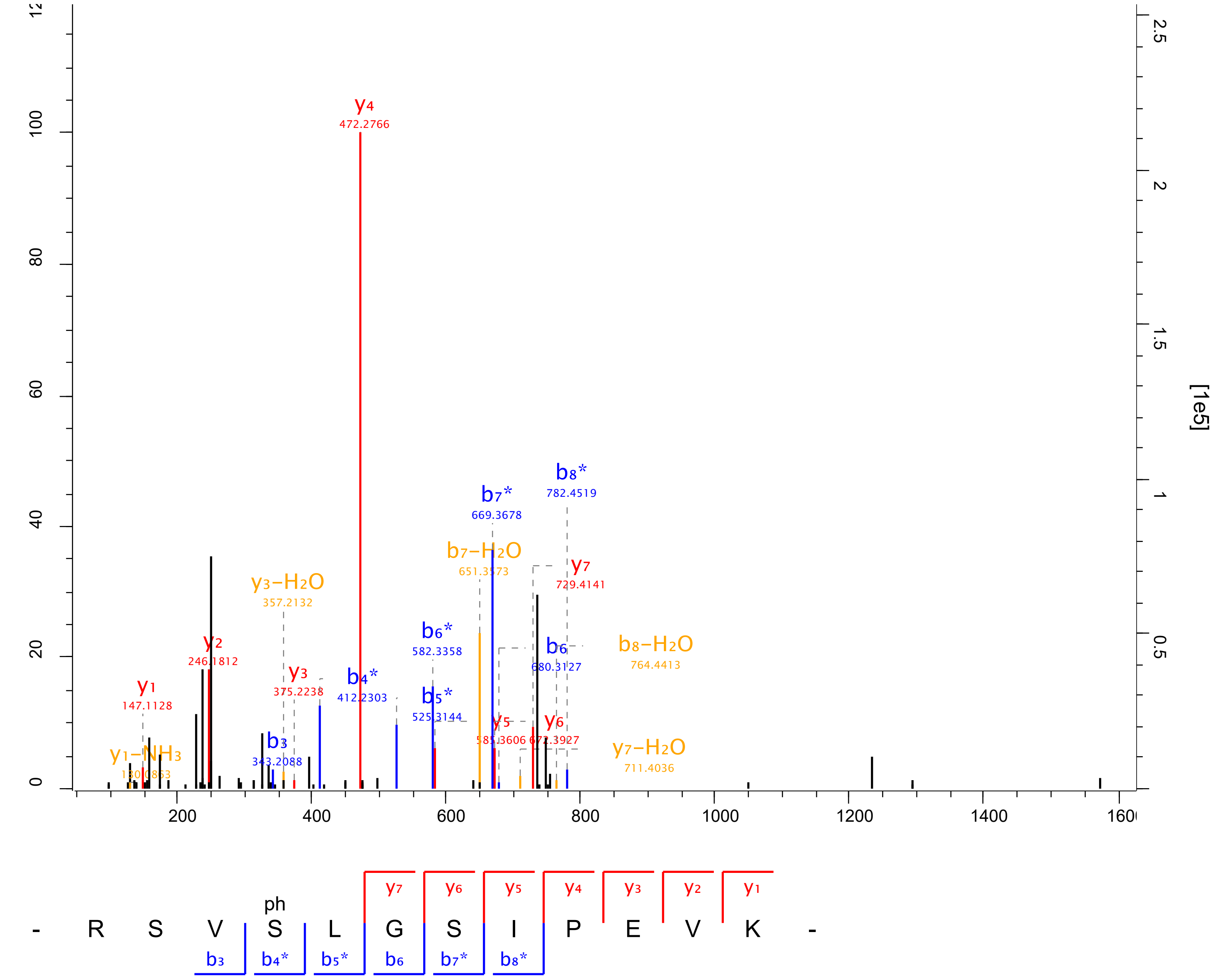

| Raw file | Scan | Method | Score | m/z |
|---|---|---|---|---|
| water_02_1 | 1960 | FTMS; HCD | 68.68 | 535.27 |

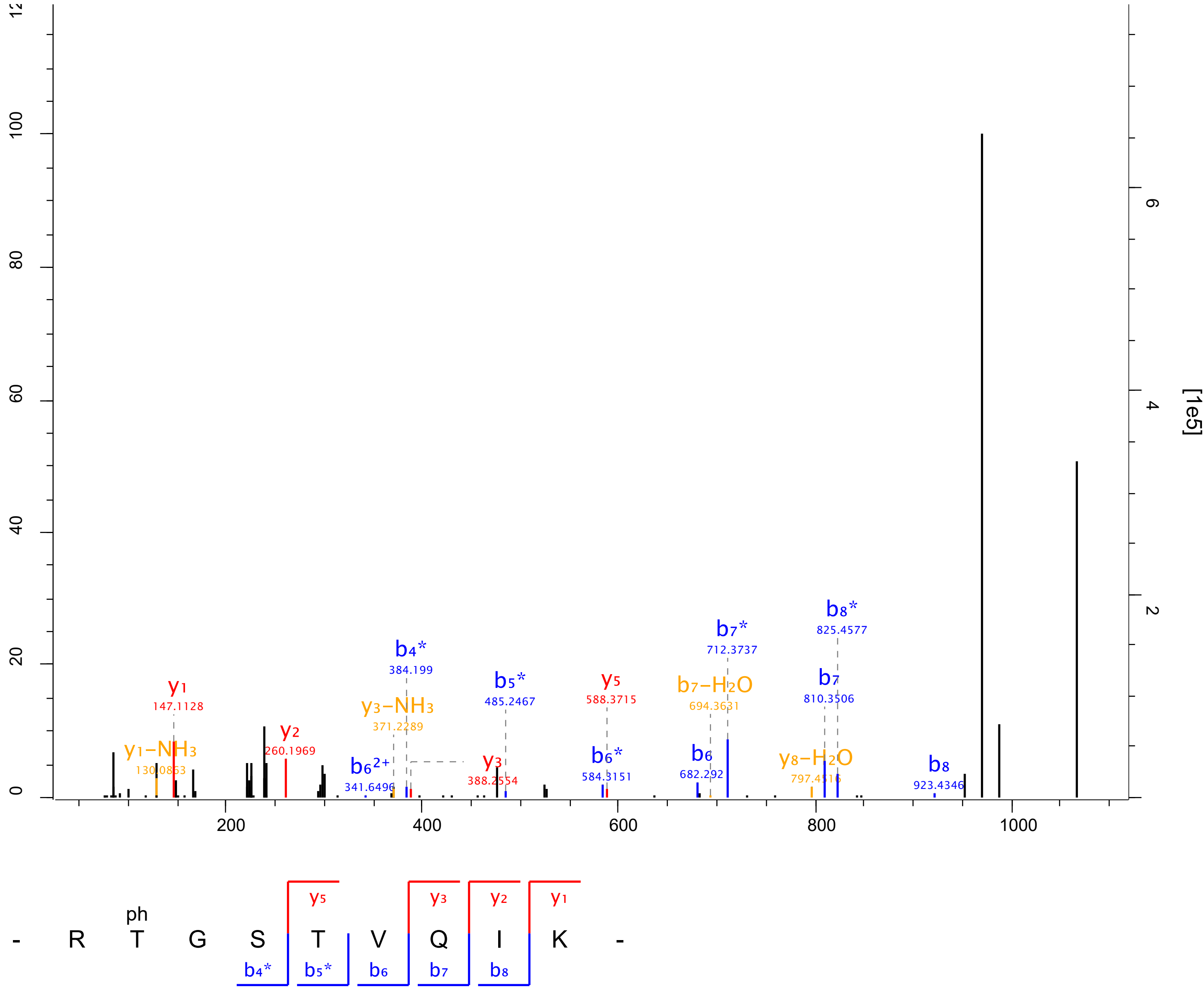

| Raw file | Scan | Method | Score | m/z |
|---|---|---|---|---|
| water_02_1 | 19649 | FTMS; HCD | 108.5 | 946.42 |

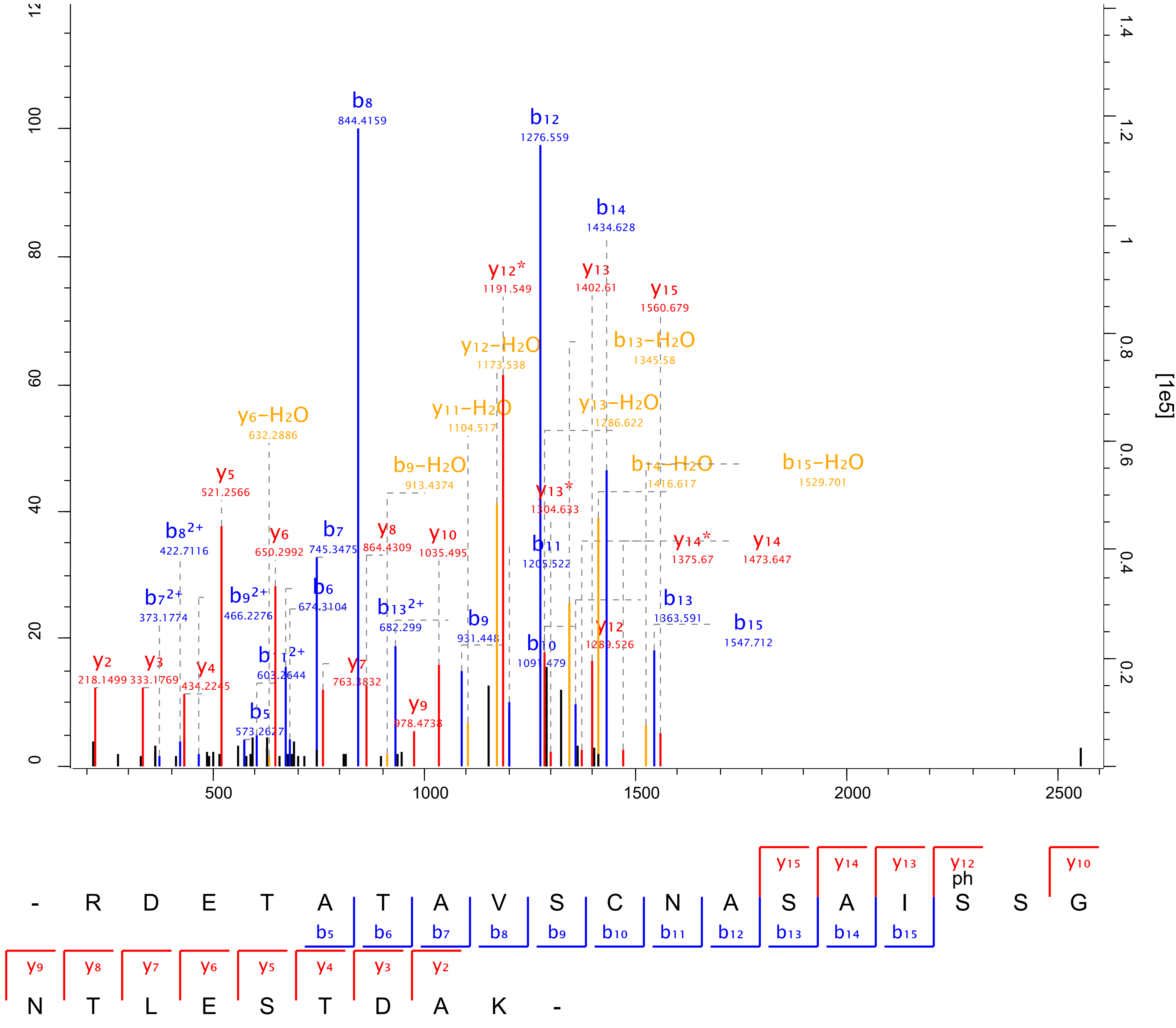

| Raw file | Scan | Method | Score | m/z |
|---|---|---|---|---|
| water_02_1 | 19723 | FTMS; HCD | 54.28 | 907.88 |

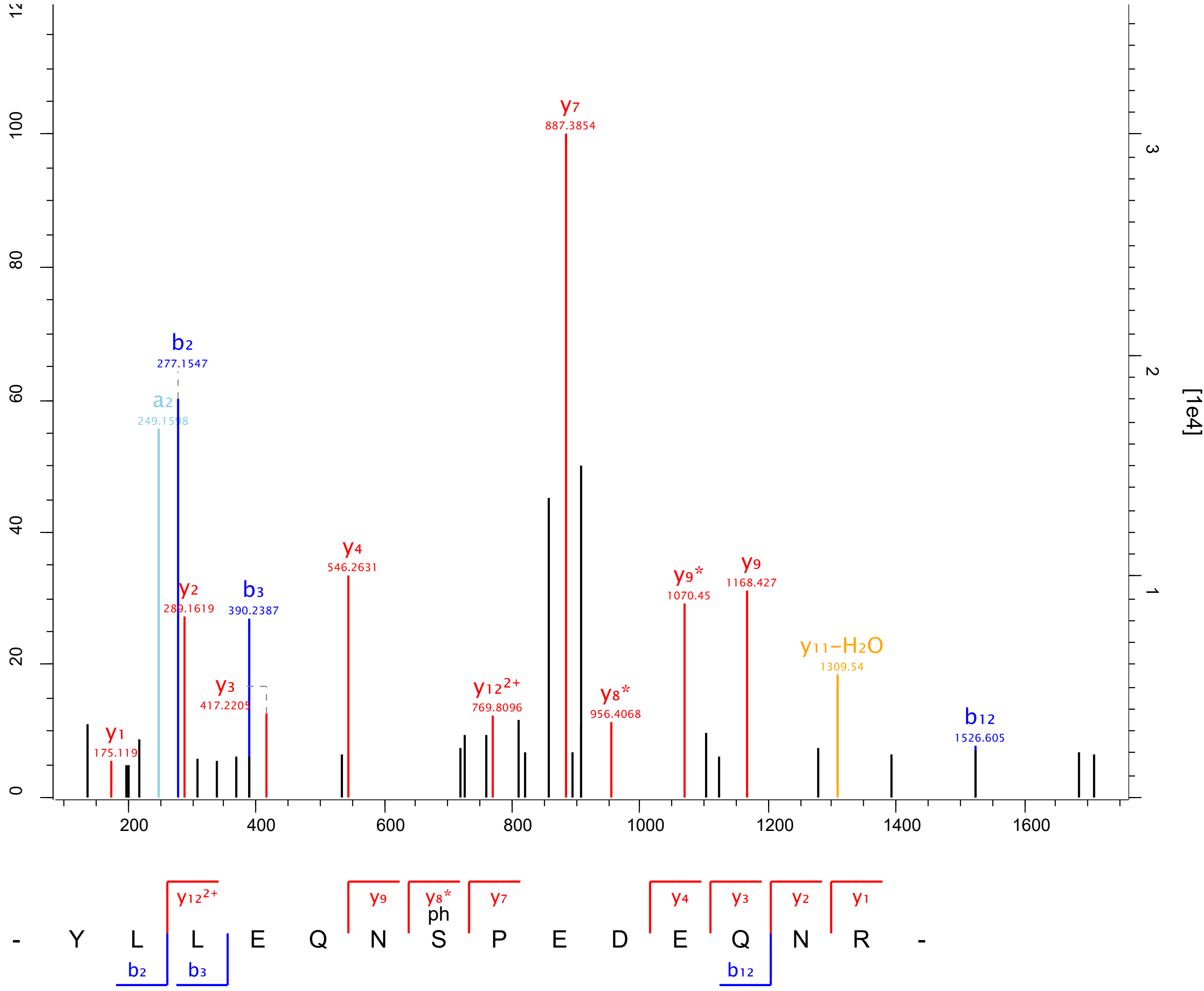

| Raw file | Scan | Method | Score | m/z |
|---|---|---|---|---|
| water_02_1 | 1974 | FTMS; HCD | 50.09 | 572.22 |

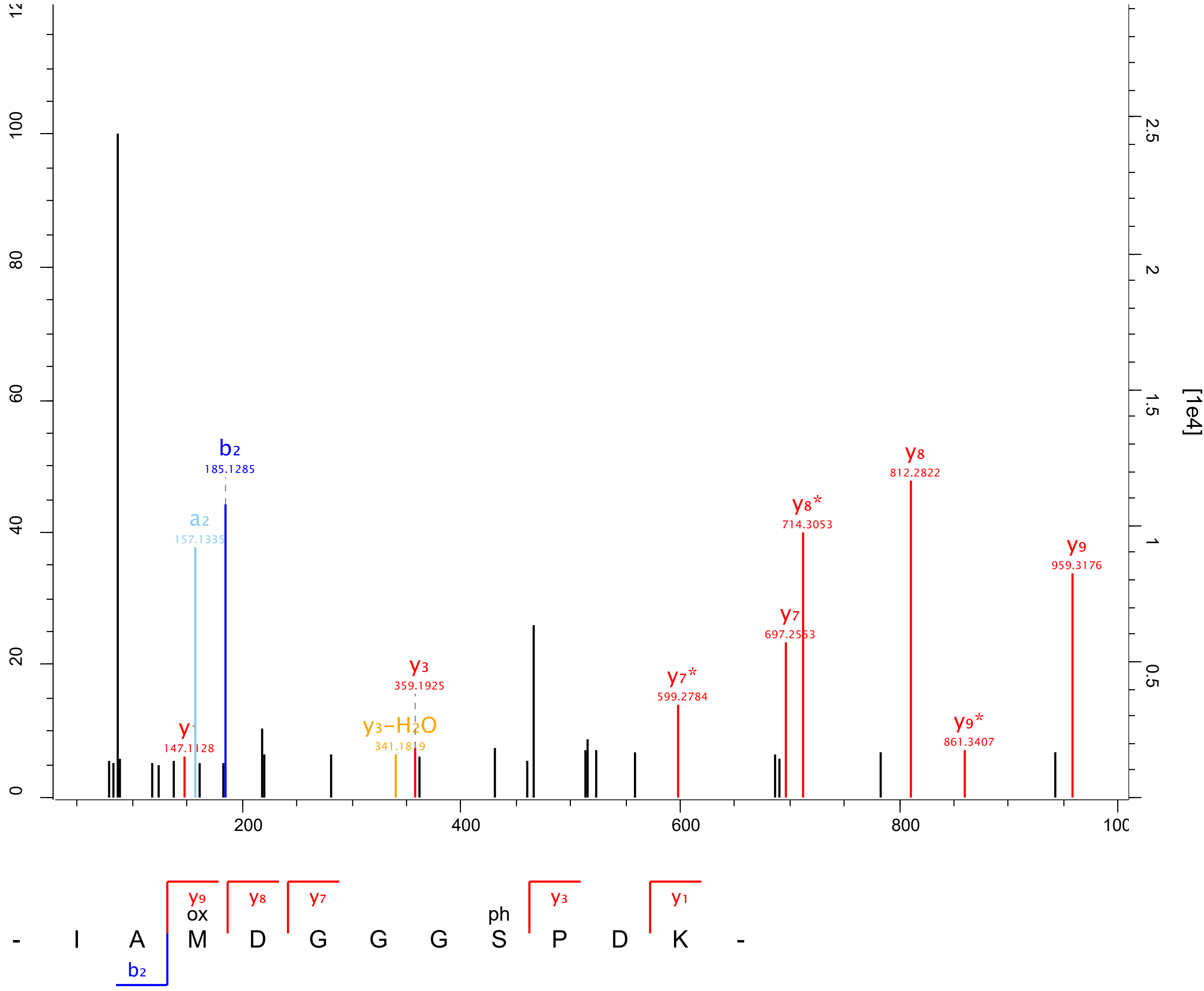

| Raw file | Scan | Method | Score | m/z |
|---|---|---|---|---|
| water_02_1 | 19746 | FTMS; HCD | 158.67 | 889.72 |

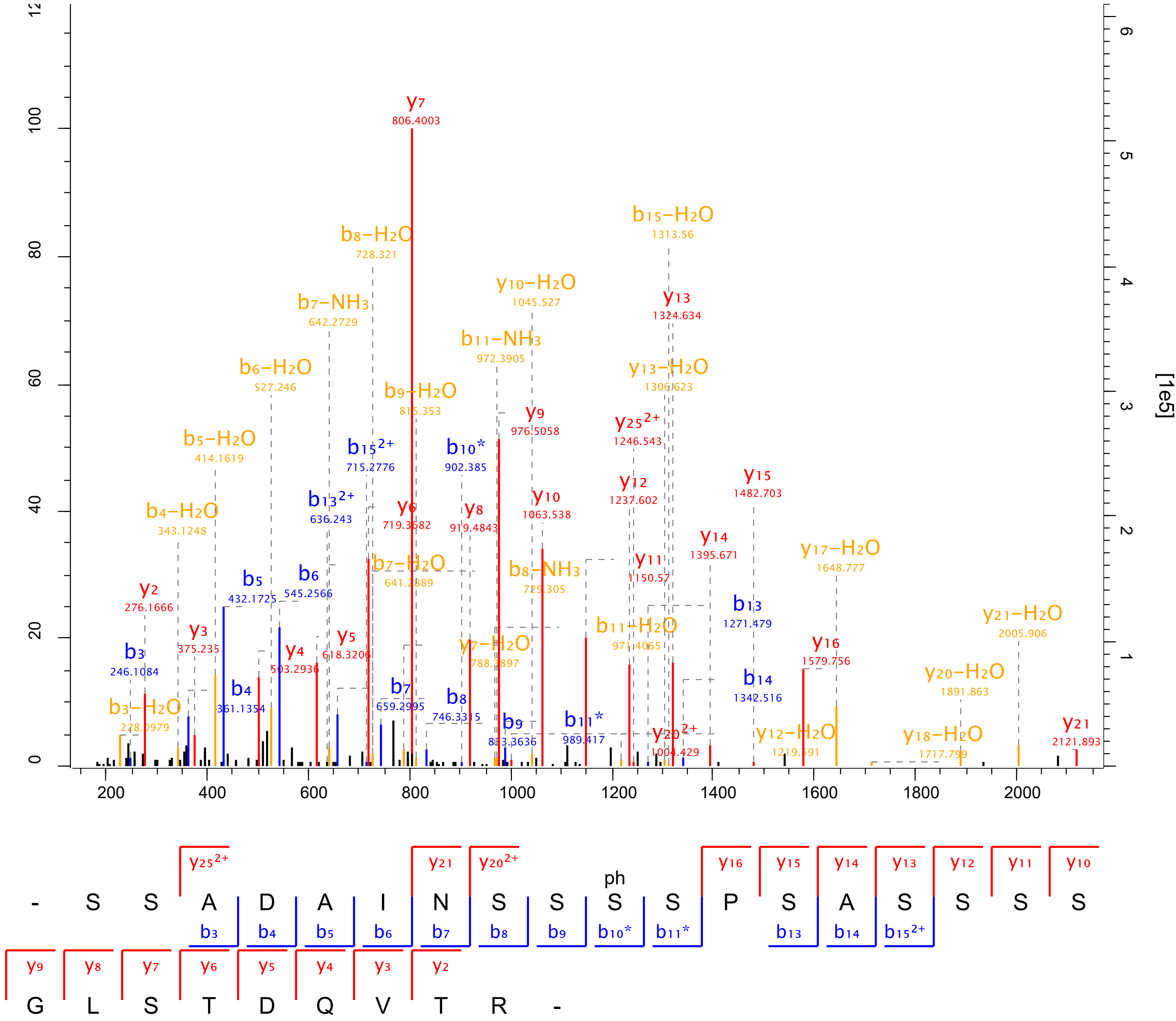

| Raw file | Scan | Method | Score | m/z |
| --- | --- | --- | --- | --- |
| water_02_1 | 1996 | FTMS; HCD | 42.03 | 368.15 |

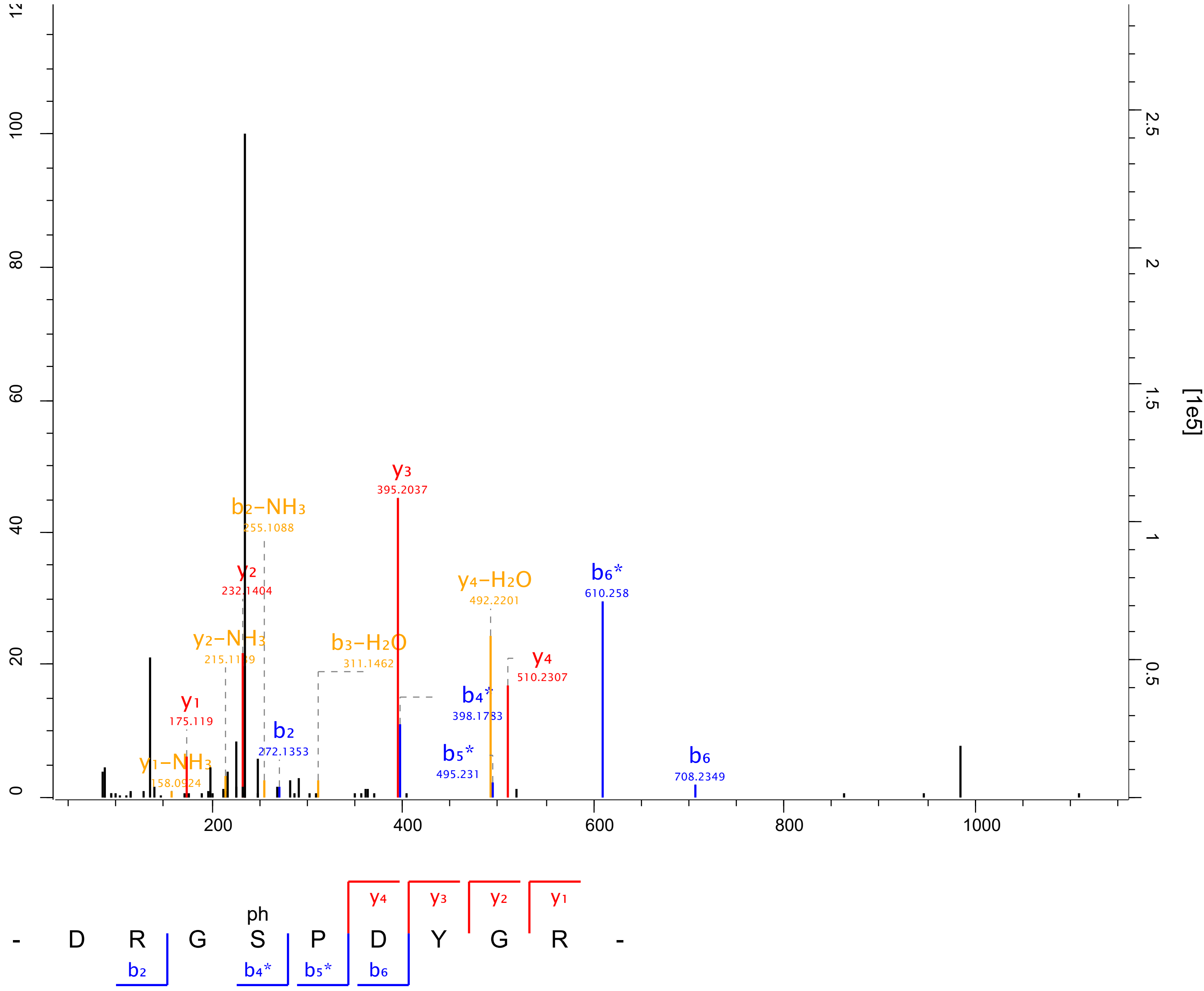

| Raw file | Scan | Method | Score | m/z |
|---|---|---|---|---|
| water_02_1 | 19966 | FTMS; HCD | 169.85 | 862.86 |

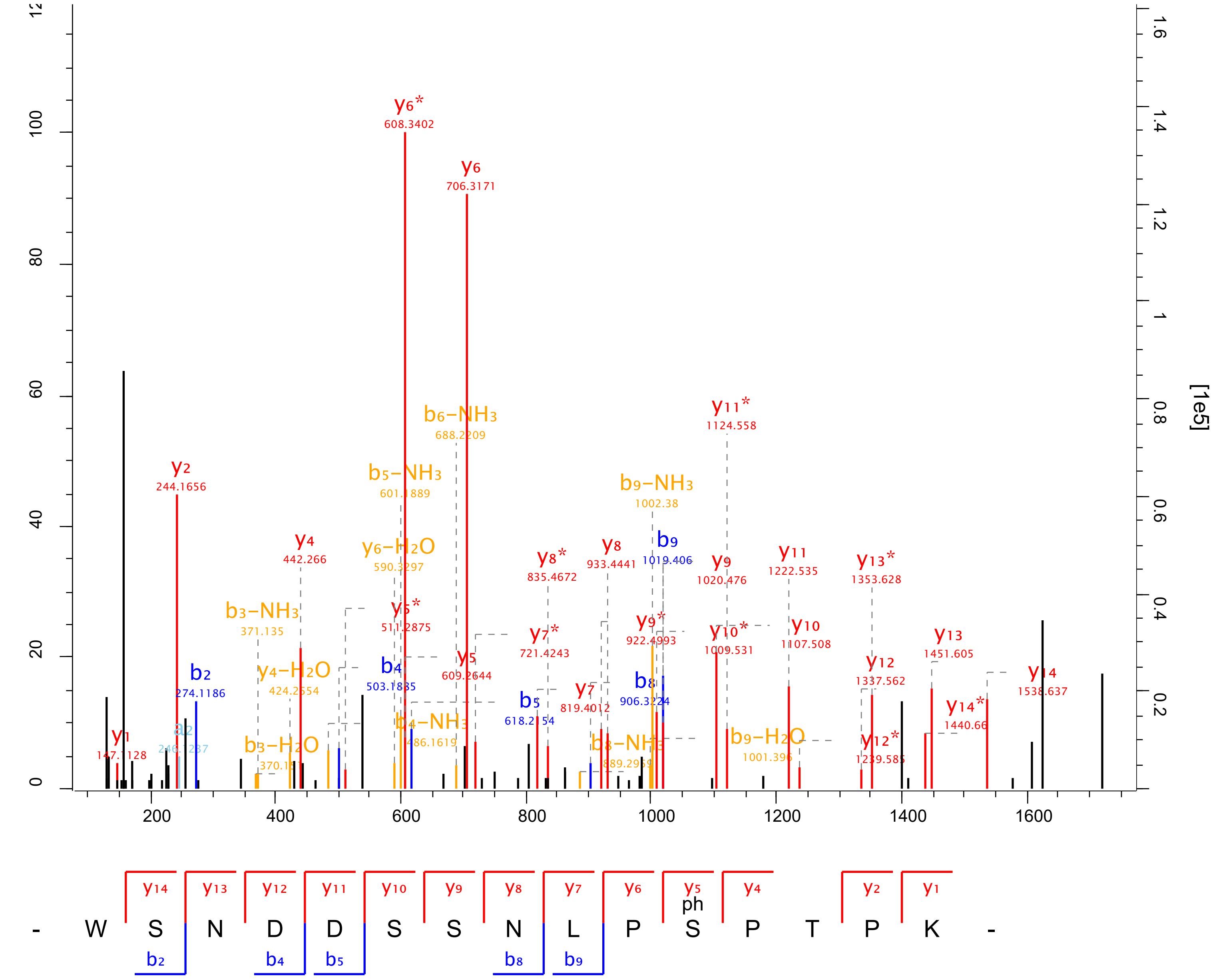

| Raw file | Scan | Method | Score | m/z |
| --- | --- | --- | --- | --- |
| water_02_1 | 19977 | FTMS; HCD | 189.11 | 632.25 |

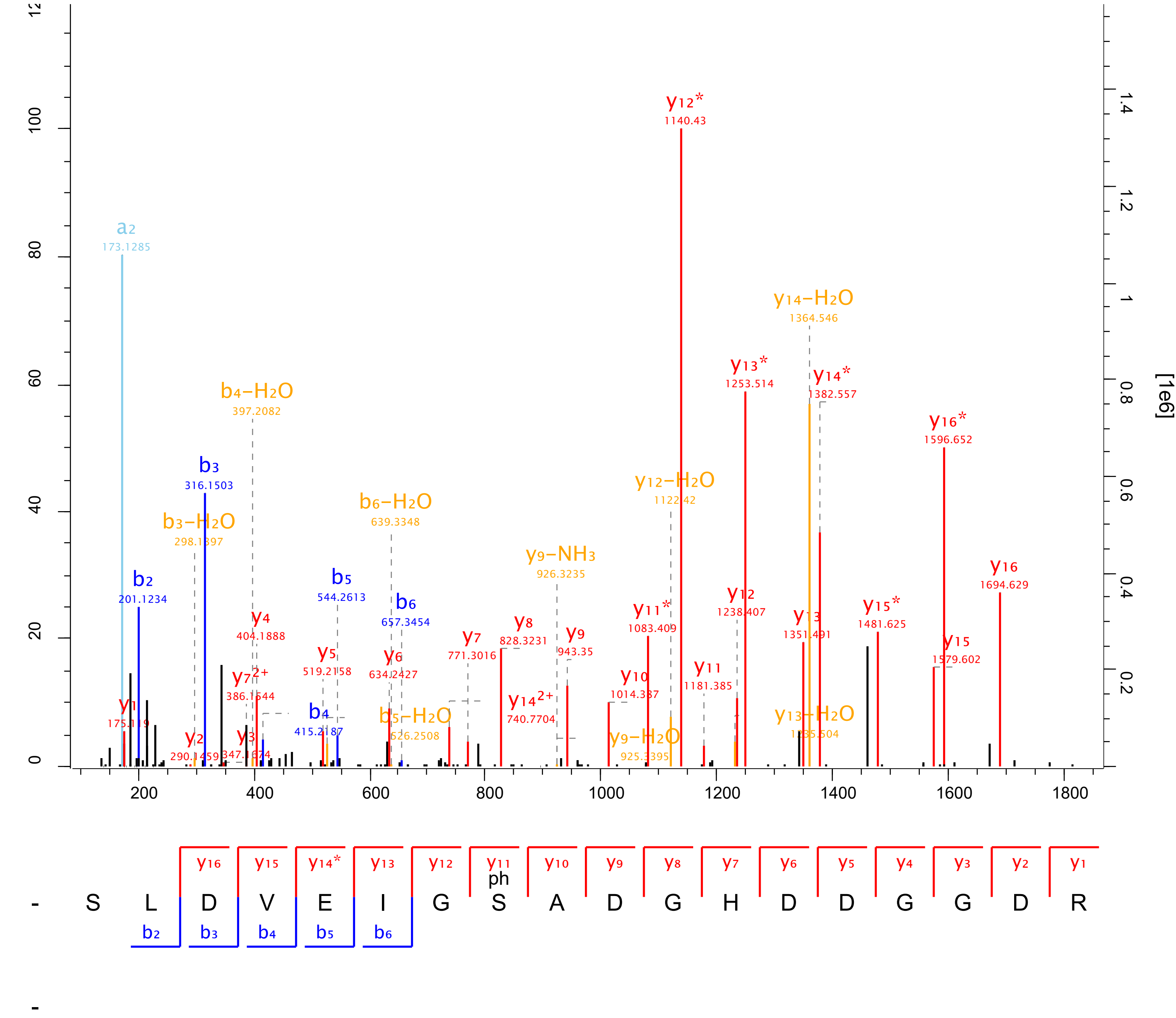

| Raw file | Scan | Method | Score | m/z |
|---|---|---|---|---|
| water_02_1 | 20005 | FTMS; HCD | 53.38 | 602.77 |

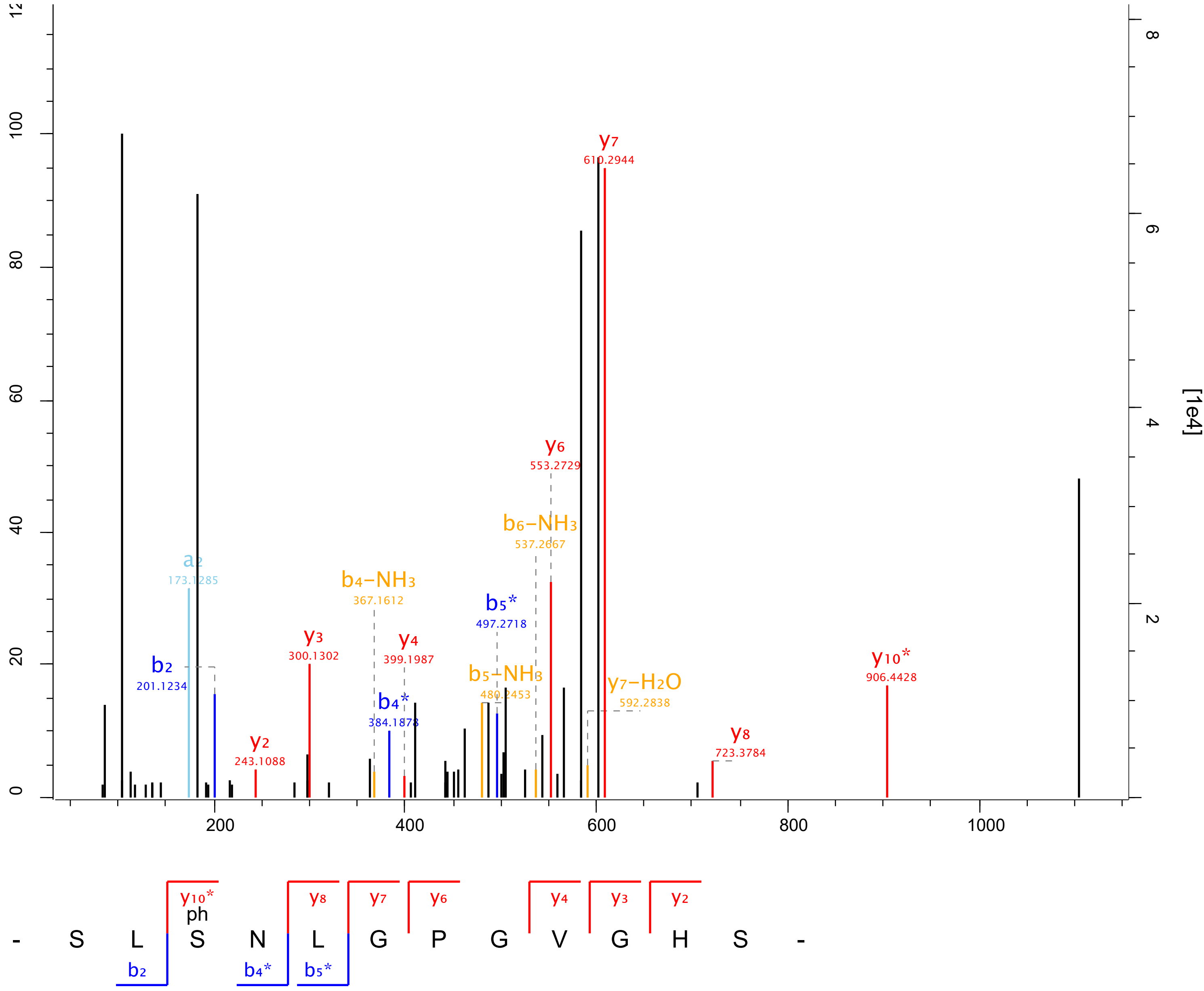

| Raw file | Scan | Method | Score | m/z |
|---|---|---|---|---|
| water_02_1 | 20023 | FTMS; HCD | 96.6 | 580.74 |

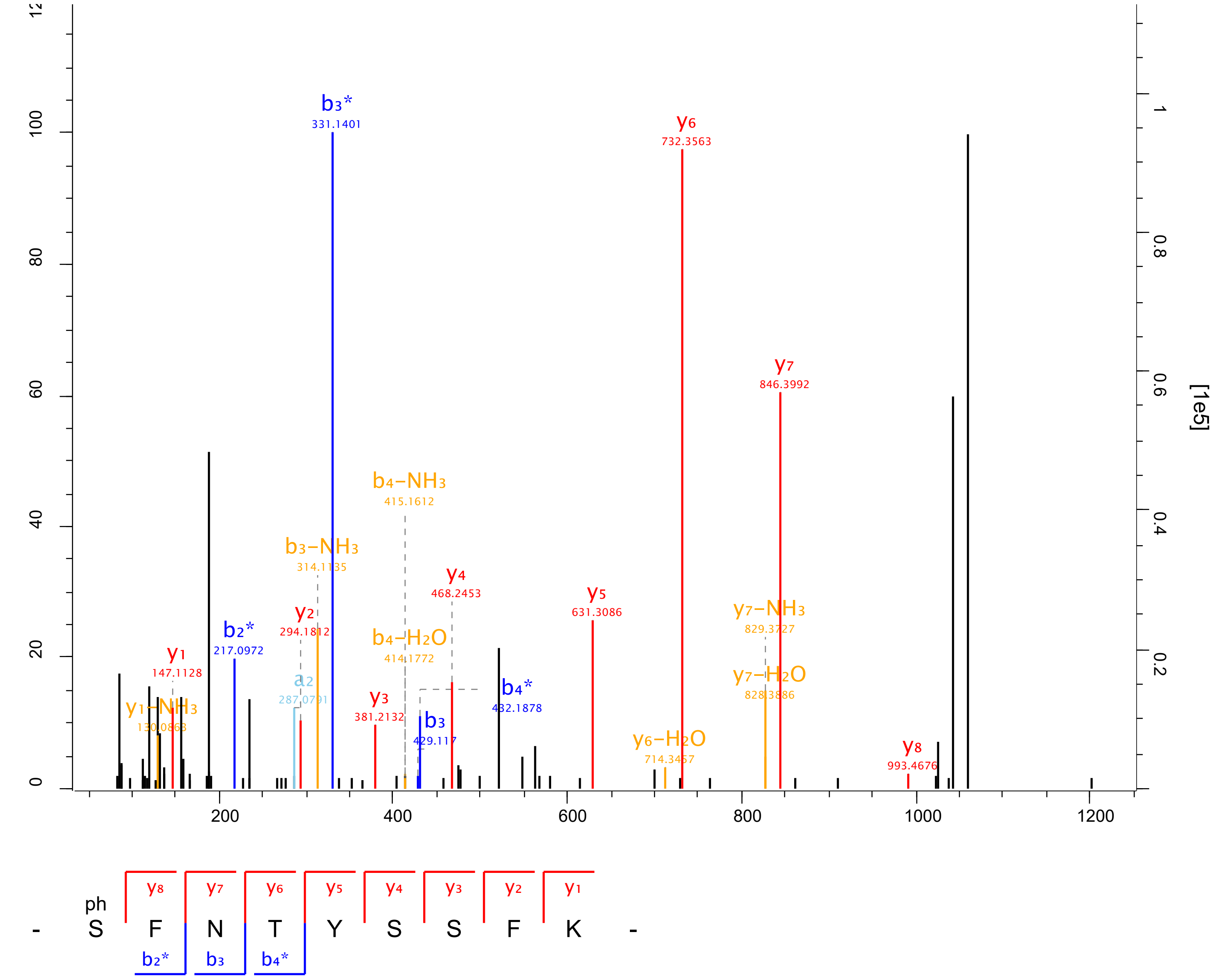

| Raw file | Scan | Method | Score | m/z |
|---|---|---|---|---|
| water_02_1 | 20135 | FTMS; HCD | 96.39 | 883.73 |

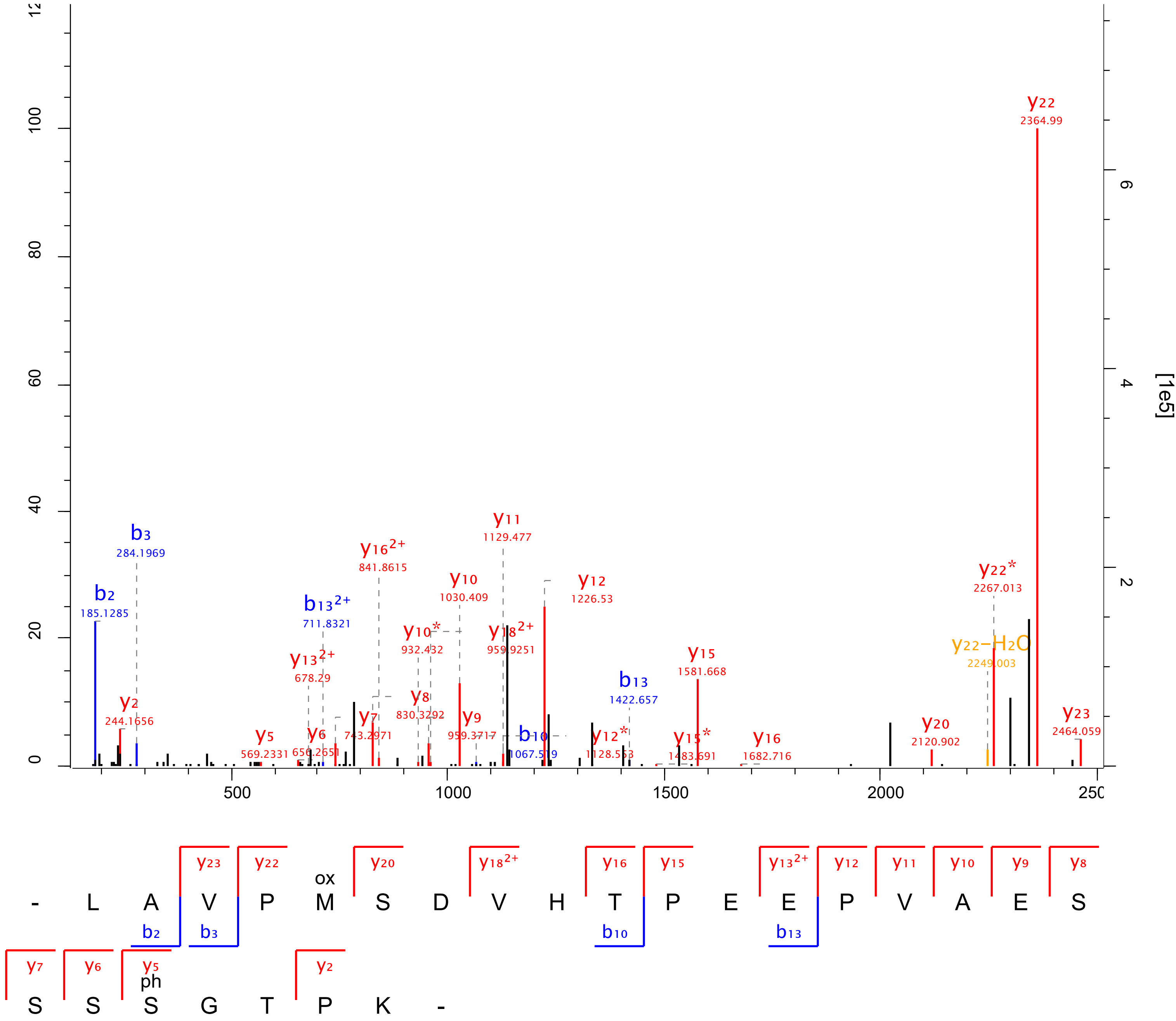

| Raw file | Scan | Method | Score | m/z |
|---|---|---|---|---|
| water_02_1 | 2028 | FTMS; HCD | 59.2 | 566.74 |

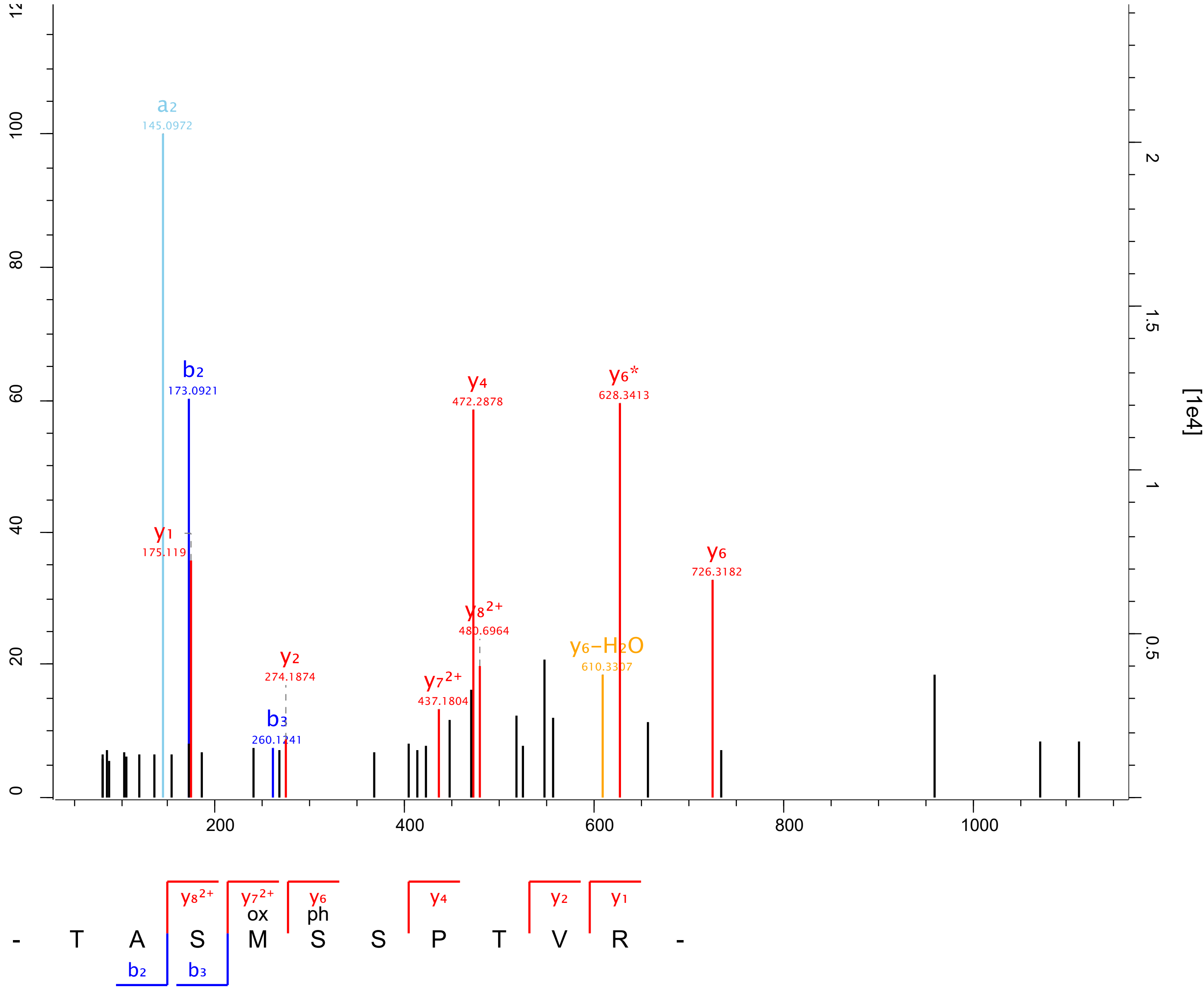

| Raw file | Scan | Method | Score | m/z |
| --- | --- | --- | --- | --- |
| water_02_1 | 2034 | FTMS; HCD | 40.48 | 405.5 |

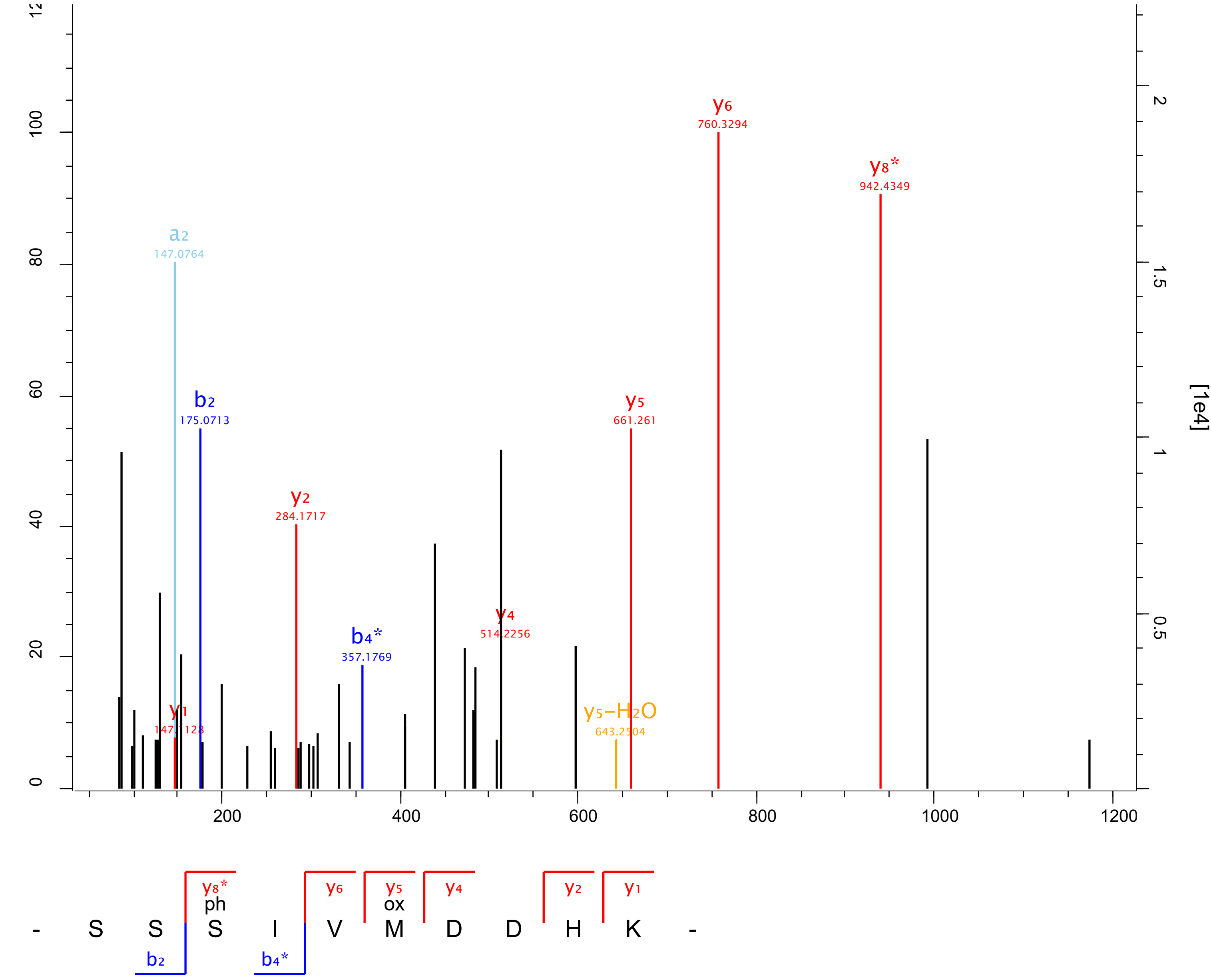

| Raw file | Scan | Method | Score | m/z |
|---|---|---|---|---|
| water_02_1 | 20394 | FTMS; HCD | 54 | 847.37 |

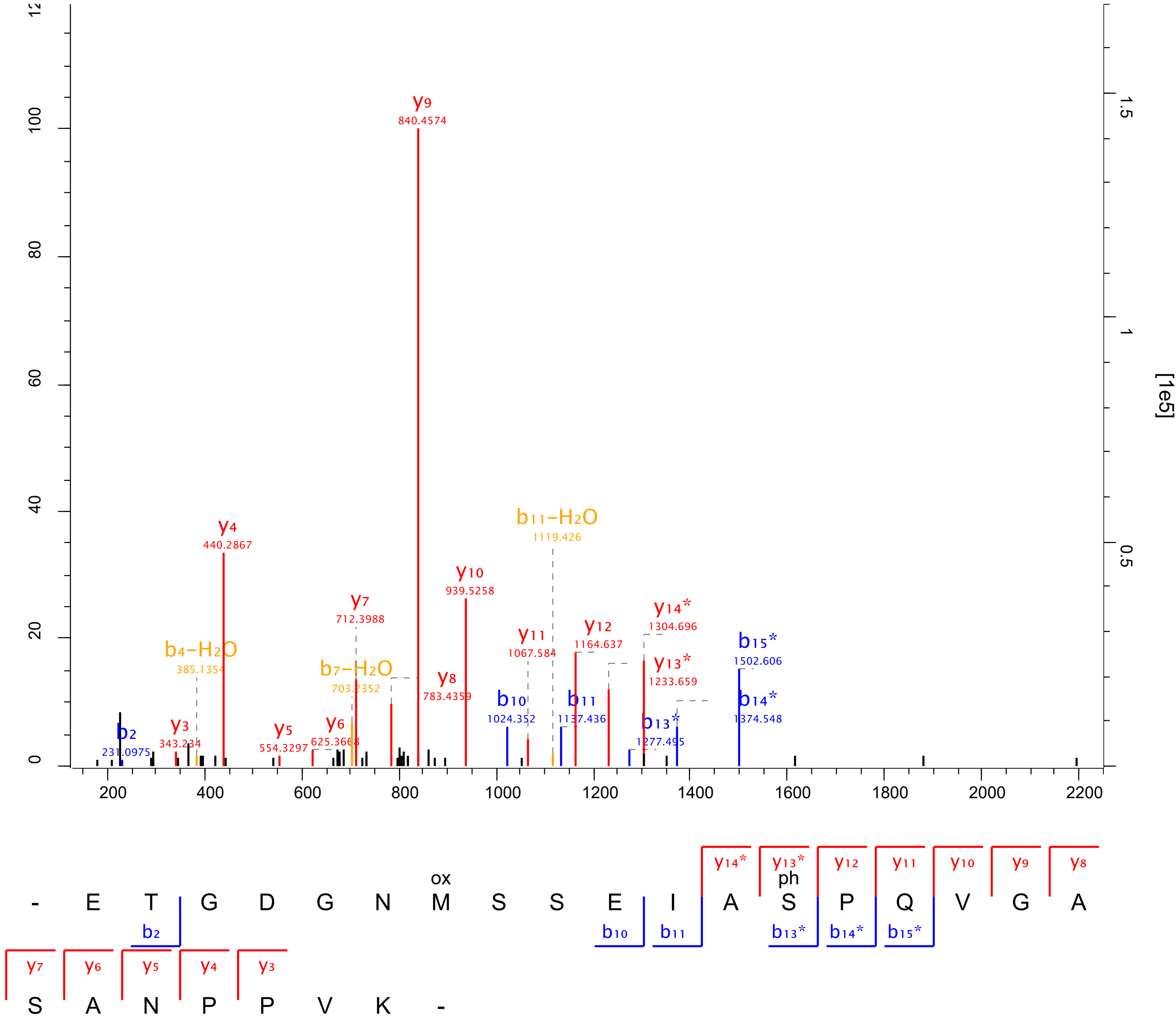

| Raw file | Scan | Method | Score | m/z |
|---|---|---|---|---|
| water_02_1 | 20406 | FTMS; HCD | 197.38 | 787.81 |

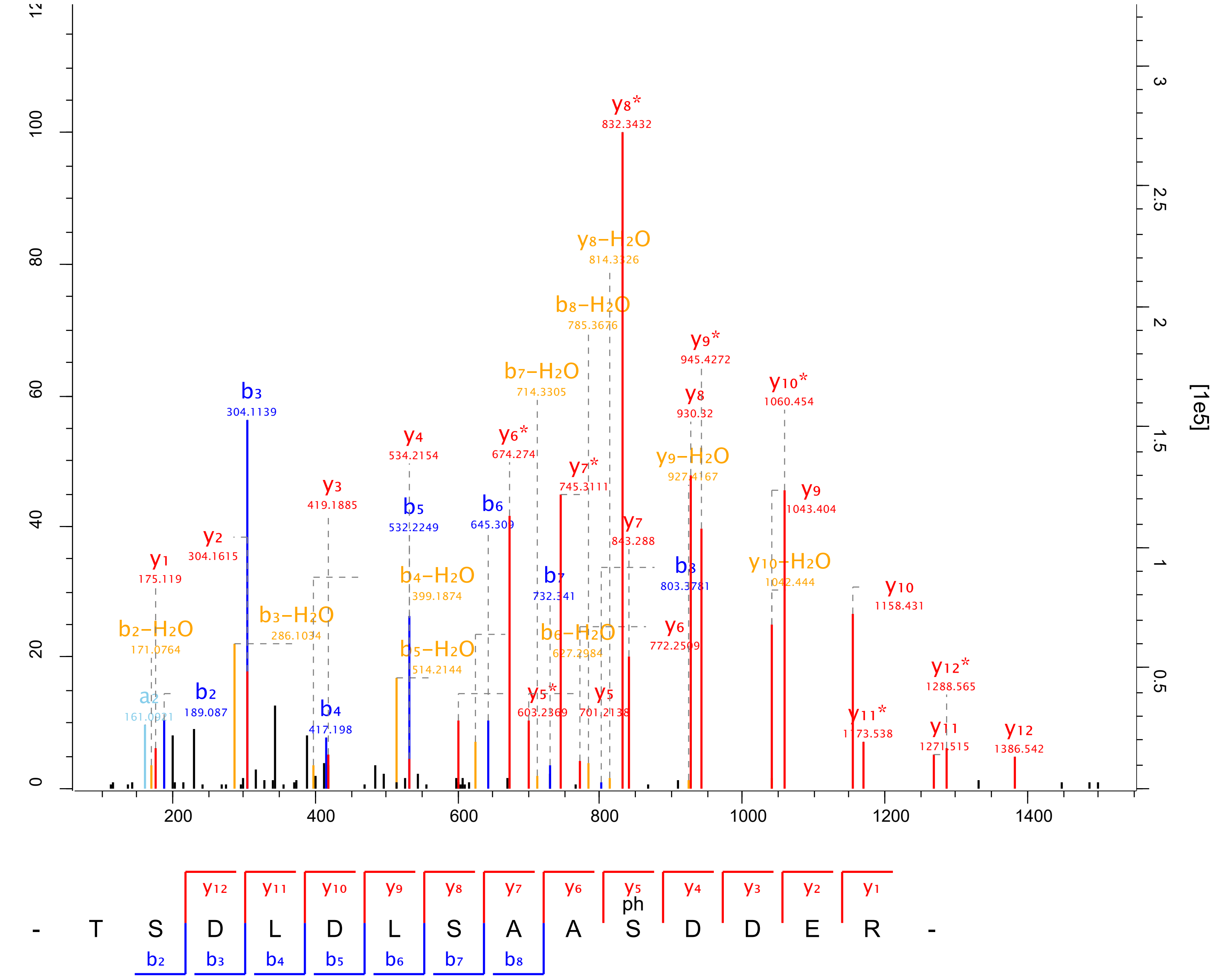

| Raw file | Scan | Method | Score | m/z |
|---|---|---|---|---|
| water_02_1 | 20424 | FTMS; HCD | 132.51 | 871.91 |

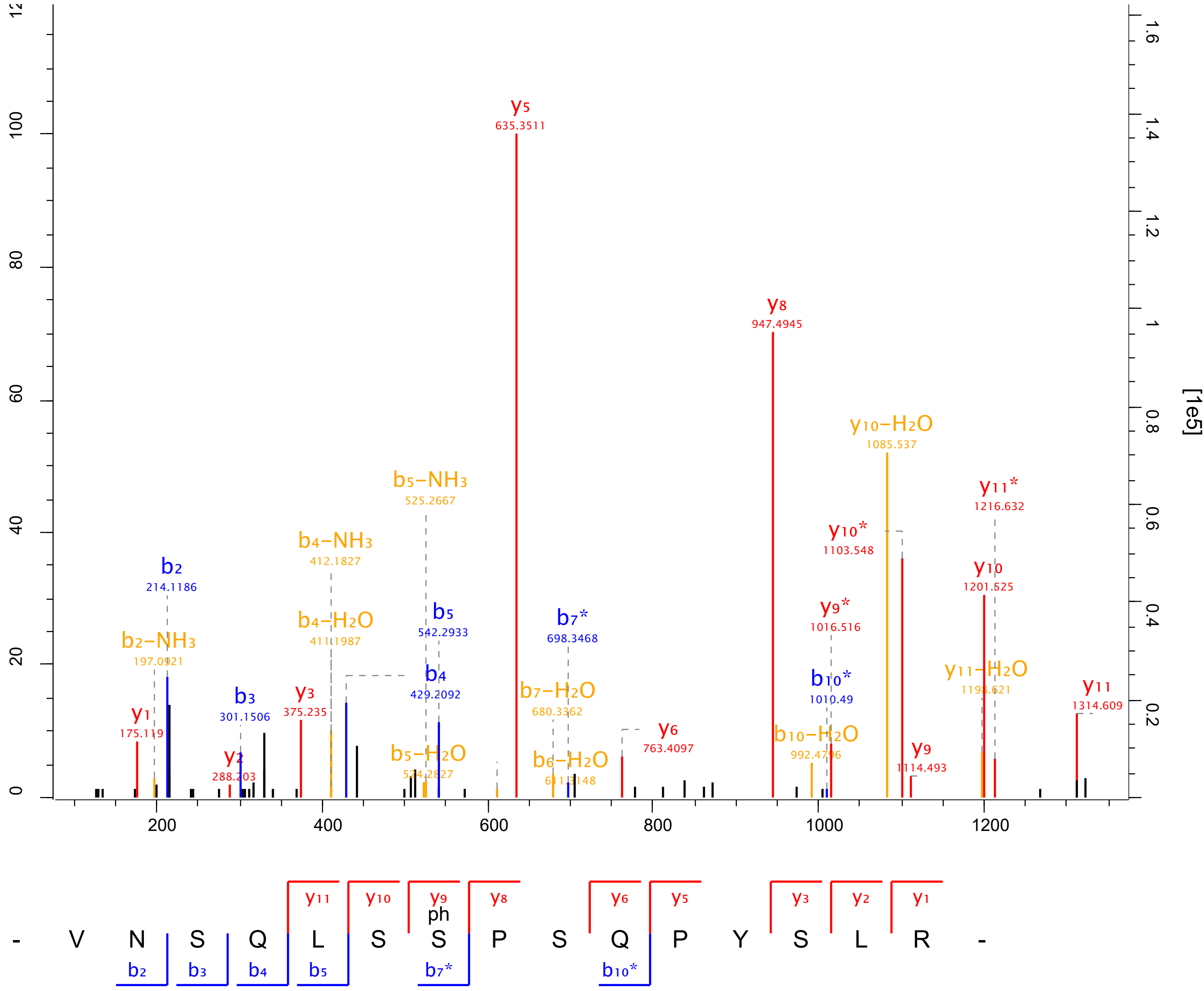

| Raw file | Scan | Method | Score | m/z |
|---|---|---|---|---|
| water_02_1 | 20578 | FTMS; HCD | 170.38 | 805.36 |

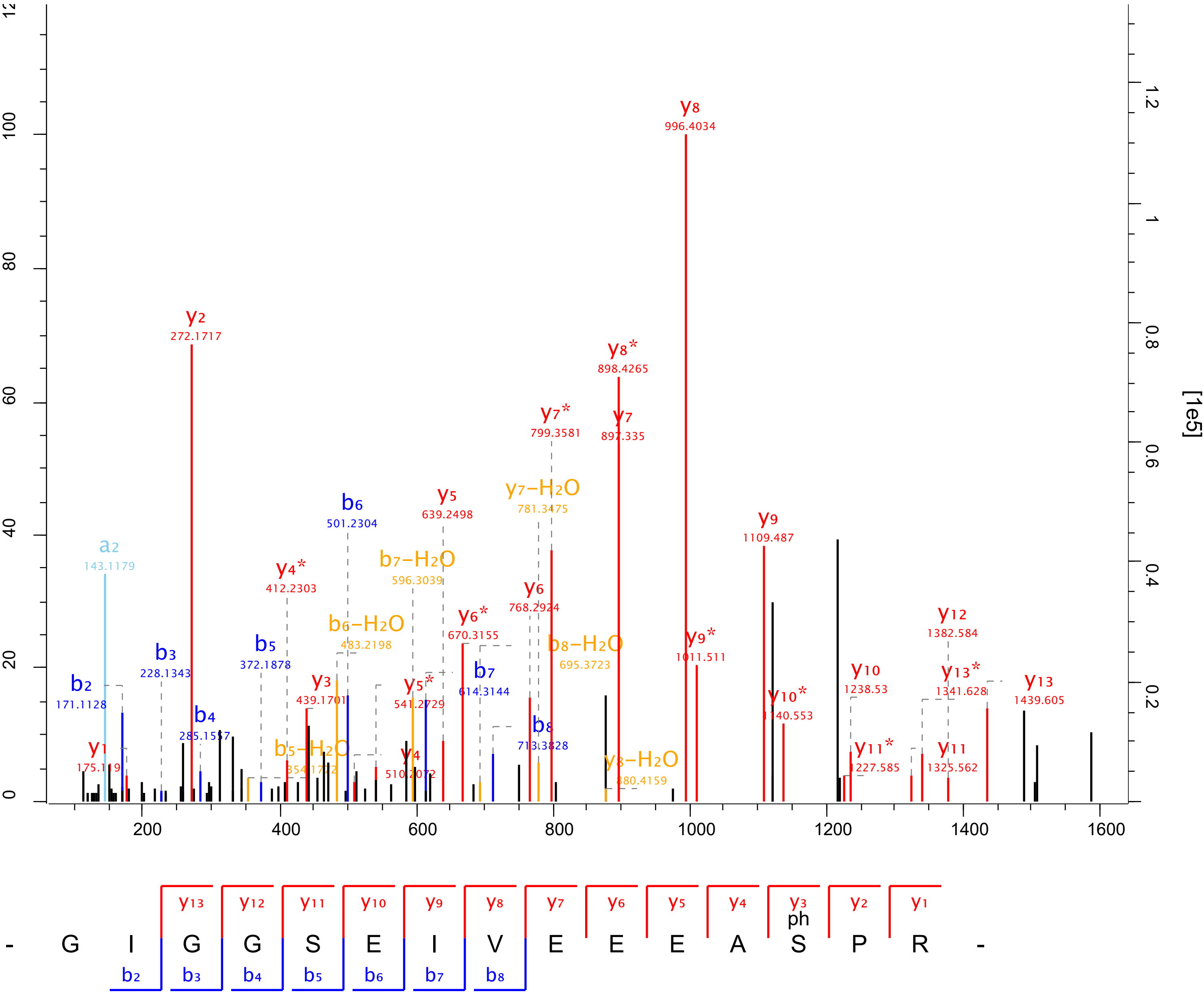

| Raw file | Scan | Method | Score | m/z |
| --- | --- | --- | --- | --- |
| water_02_1 | 20650 | FTMS; HCD | 57.53 | 638.29 |

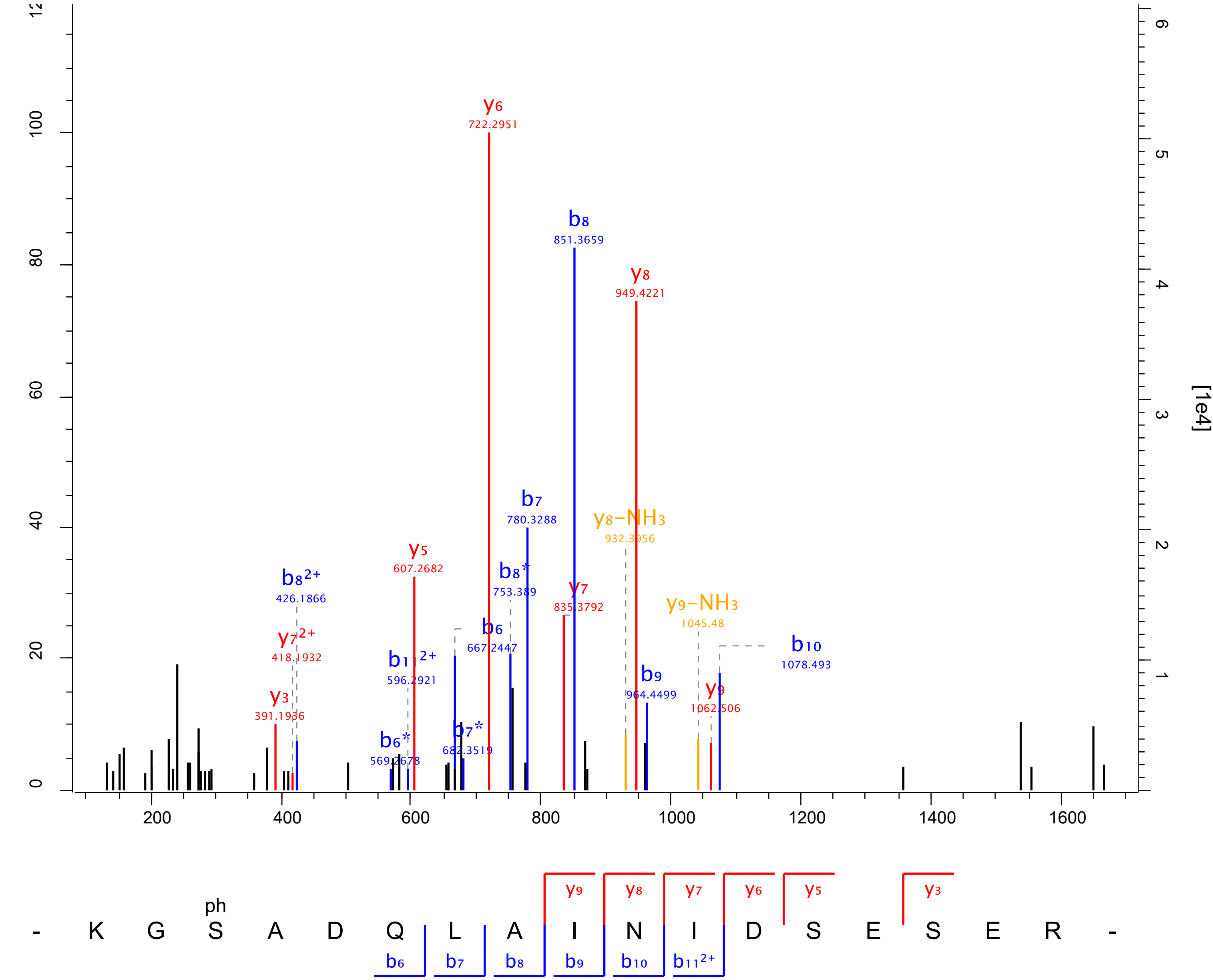

| Raw file | Scan | Method | Score | m/z |
|---|---|---|---|---|
| water_02_1 | 20669 | FTMS; HCD | 78.52 | 433.23 |

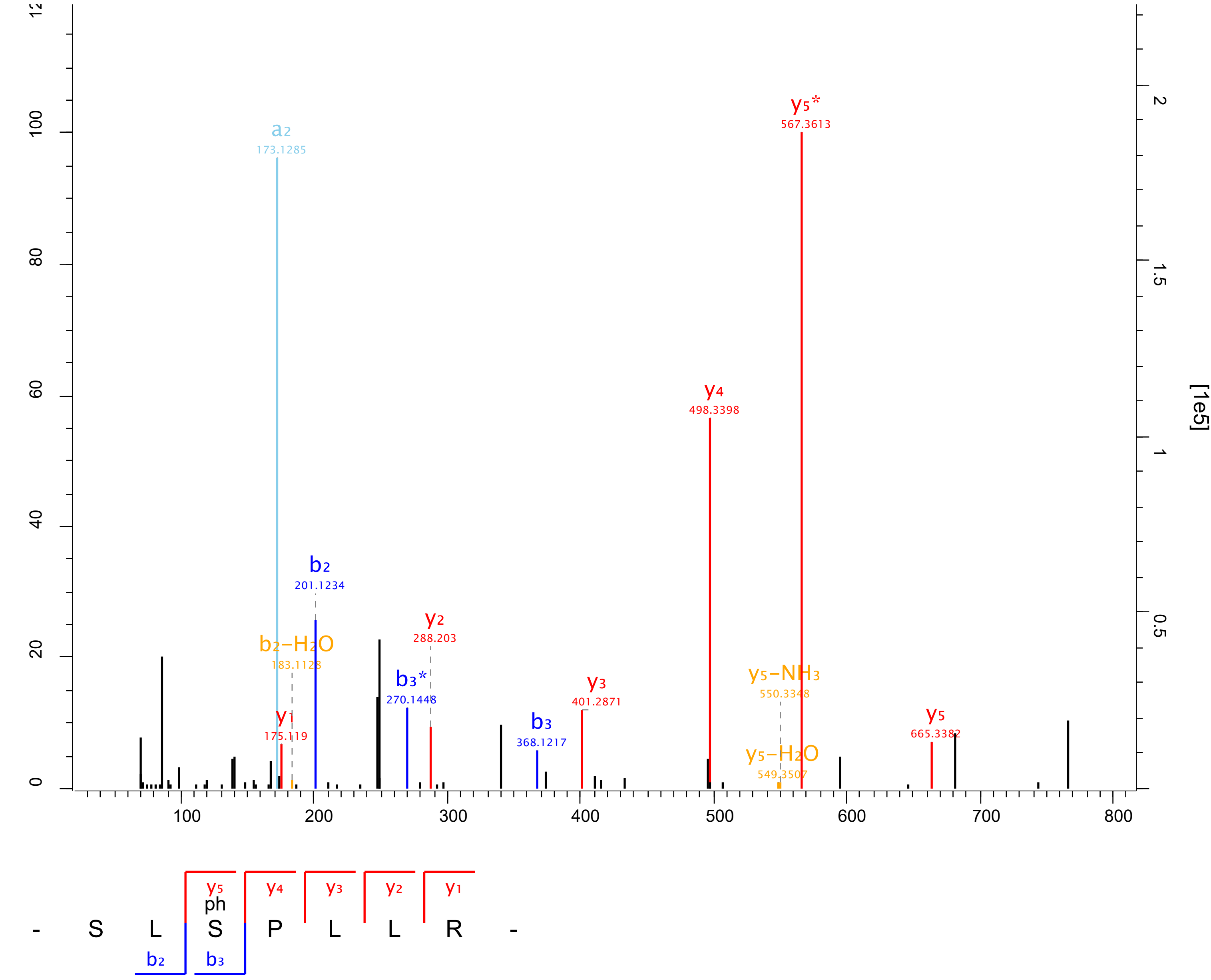

| Raw file | Scan | Method | Score | m/z |
|---|---|---|---|---|
| water_02_1 | 20876 | FTMS; HCD | 158.86 | 599.28 |

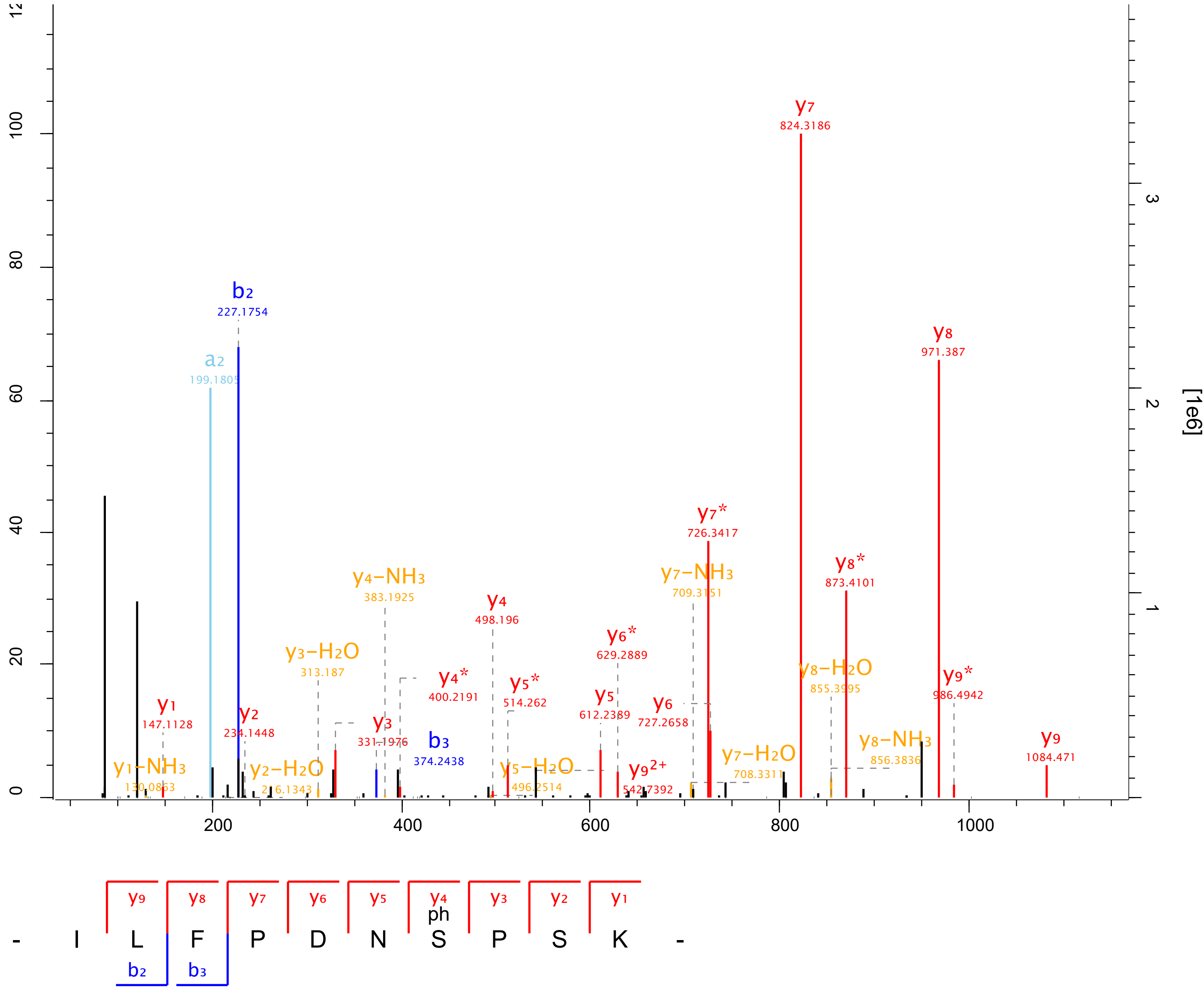

| Raw file | Scan | Method | Score | m/z |
| --- | --- | --- | --- | --- |
| water_02_1 | 20981 | FTMS; HCD | 68 | 533.77 |

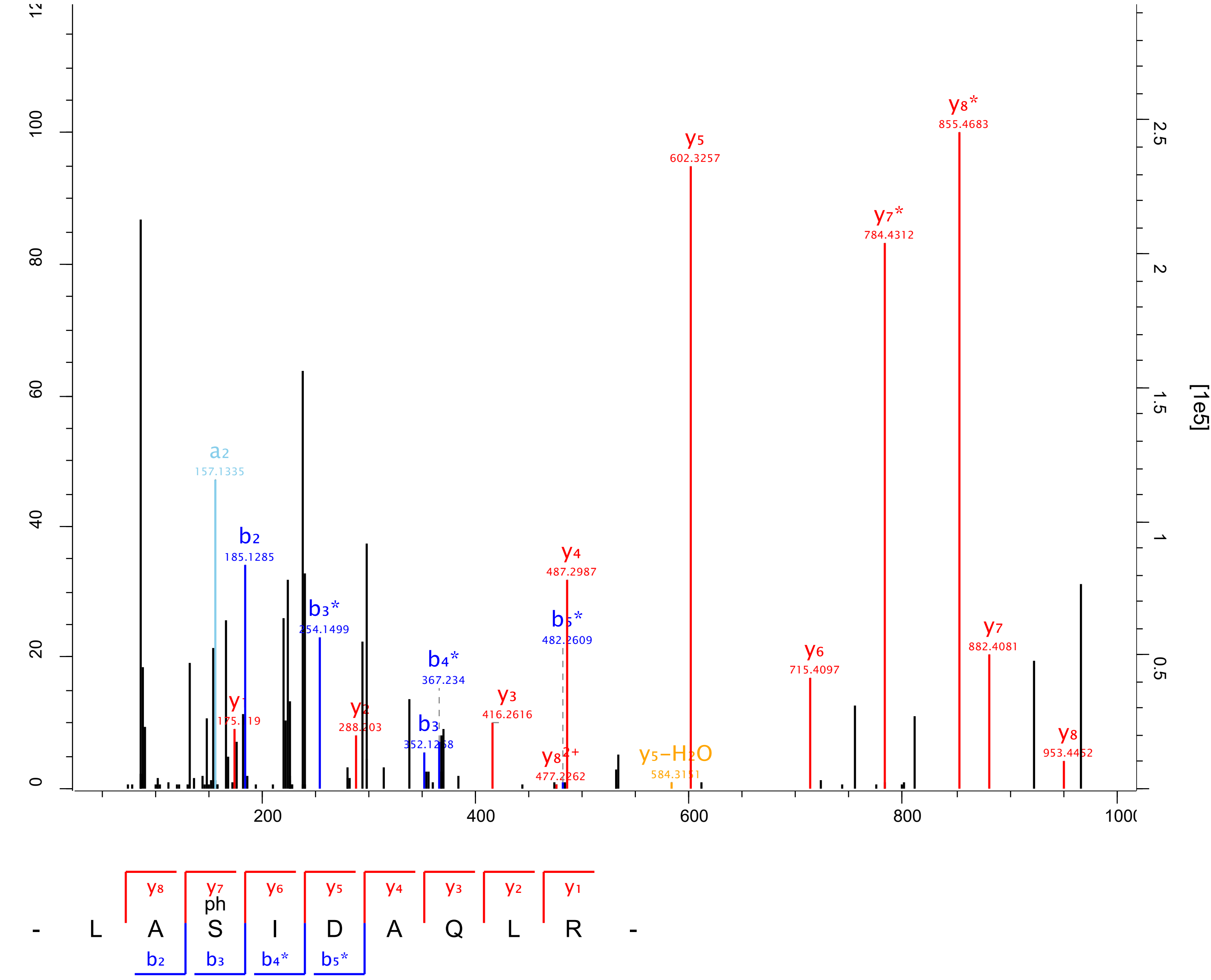

| Raw file | Scan | Method | Score | m/z |
| --- | --- | --- | --- | --- |
| water_02_1 | 20998 | FTMS; HCD | 212.35 | 696.31 |

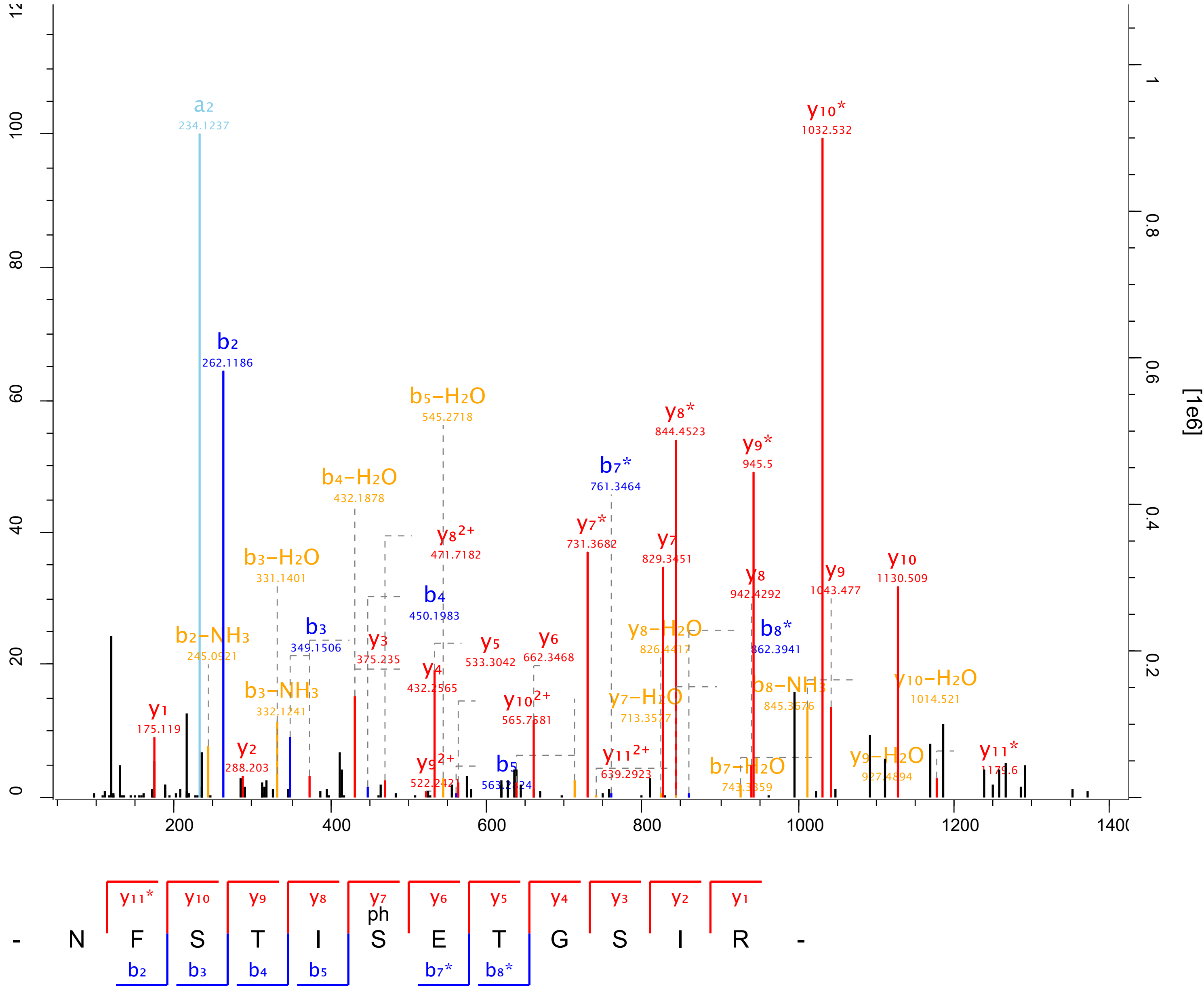

| Raw file | Scan | Method | Score | m/z |
| --- | --- | --- | --- | --- |
| water_02_1 | 21046 | FTMS; HCD | 126.96 | 680.31 |

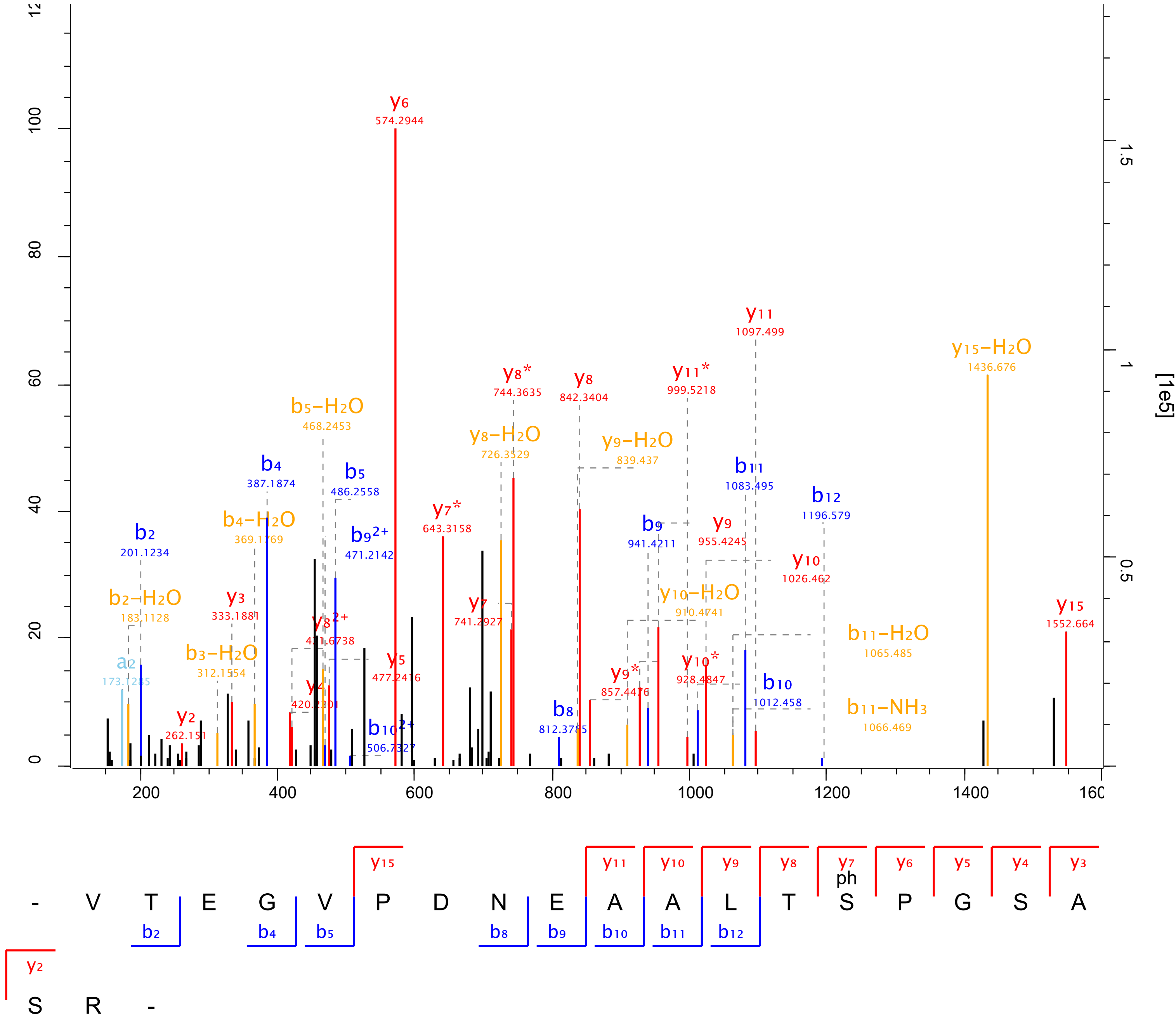

| Raw file | Scan | Method | Score | m/z |
|---|---|---|---|---|
| water_02_1 | 21056 | FTMS; HCD | 44.64 | 843.31 |

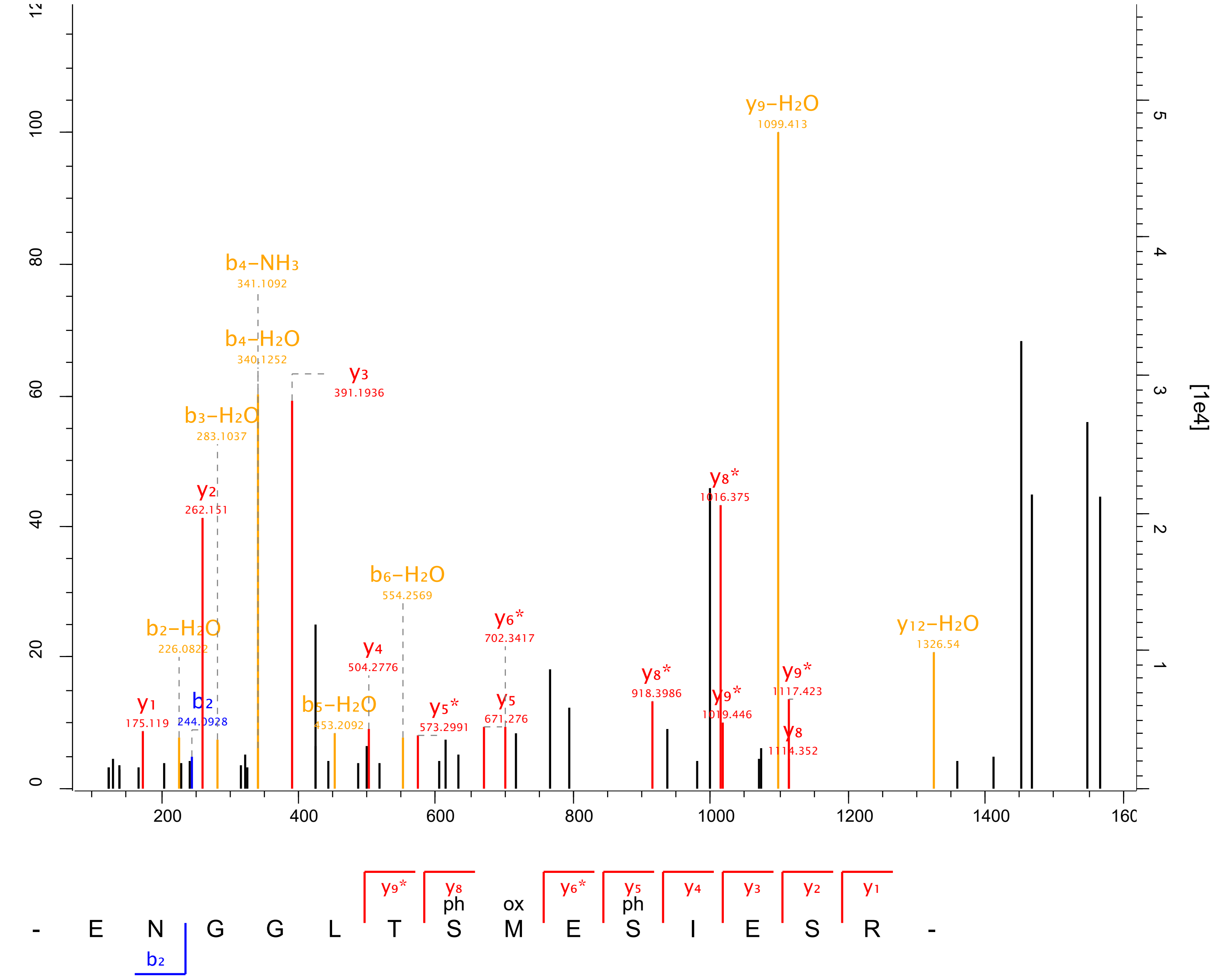

| Raw file | Scan | Method | Score | m/z |
| --- | --- | --- | --- | --- |
| water_02_1 | 21060 | FTMS; HCD | 152.06 | 939.44 |

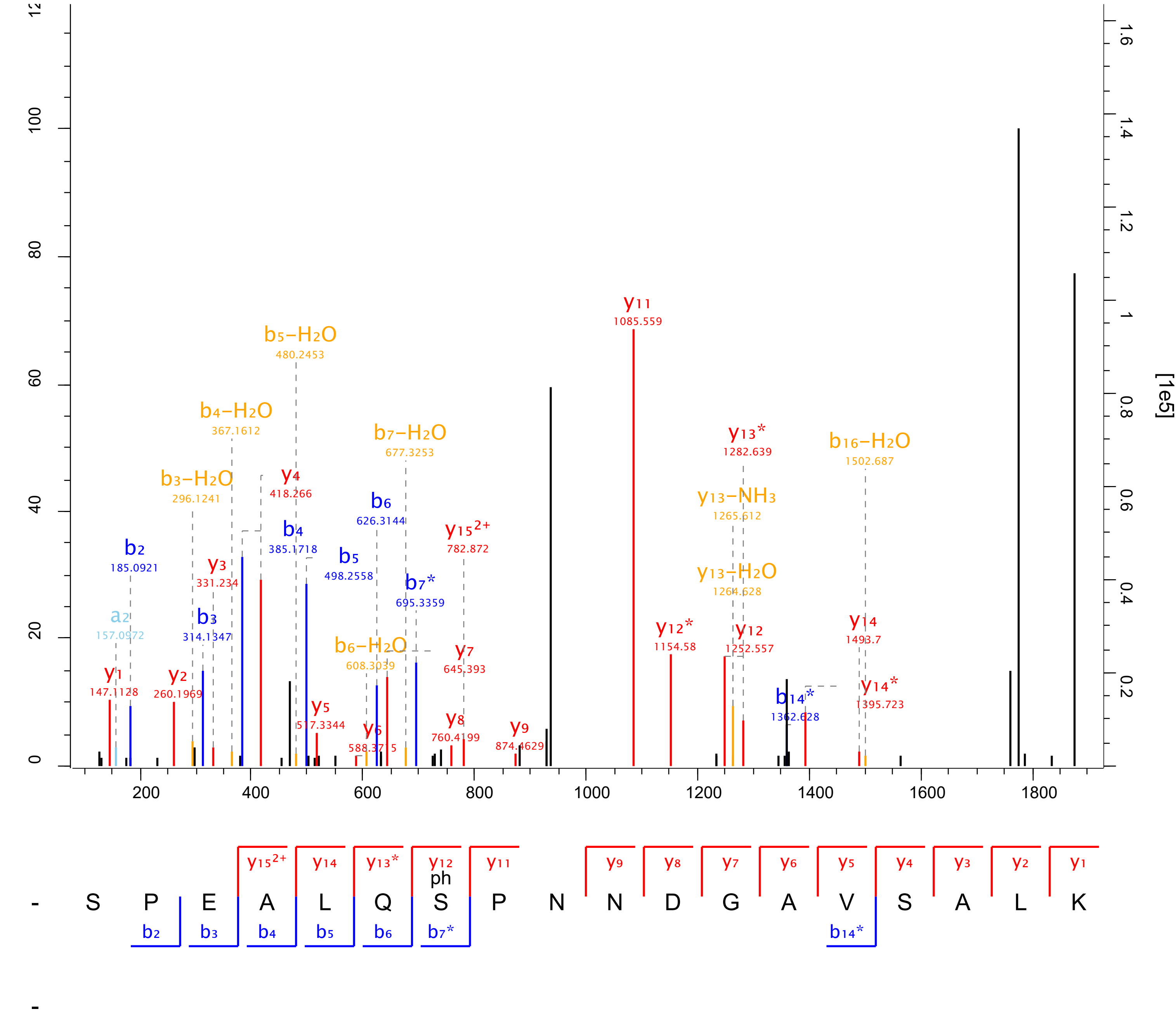

| Raw file | Scan | Method | Score | m/z |
|---|---|---|---|---|
| water_02_1 | 21065 | FTMS; HCD | 65.37 | 693.82 |

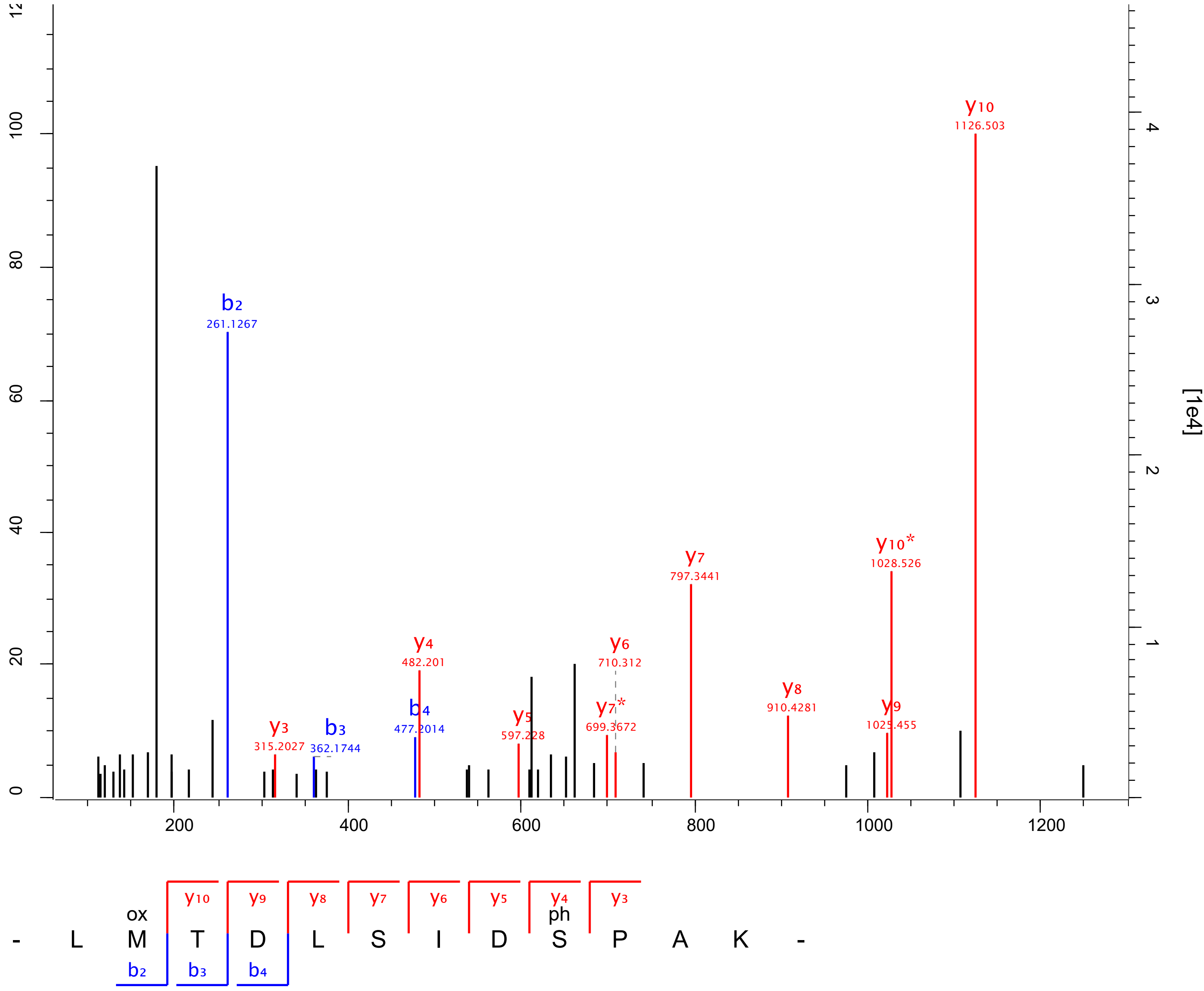

| Raw file | Scan | Method | Score | m/z |
|---|---|---|---|---|
| water_02_1 | 21094 | FTMS; HCD | 50.35 | 587.63 |

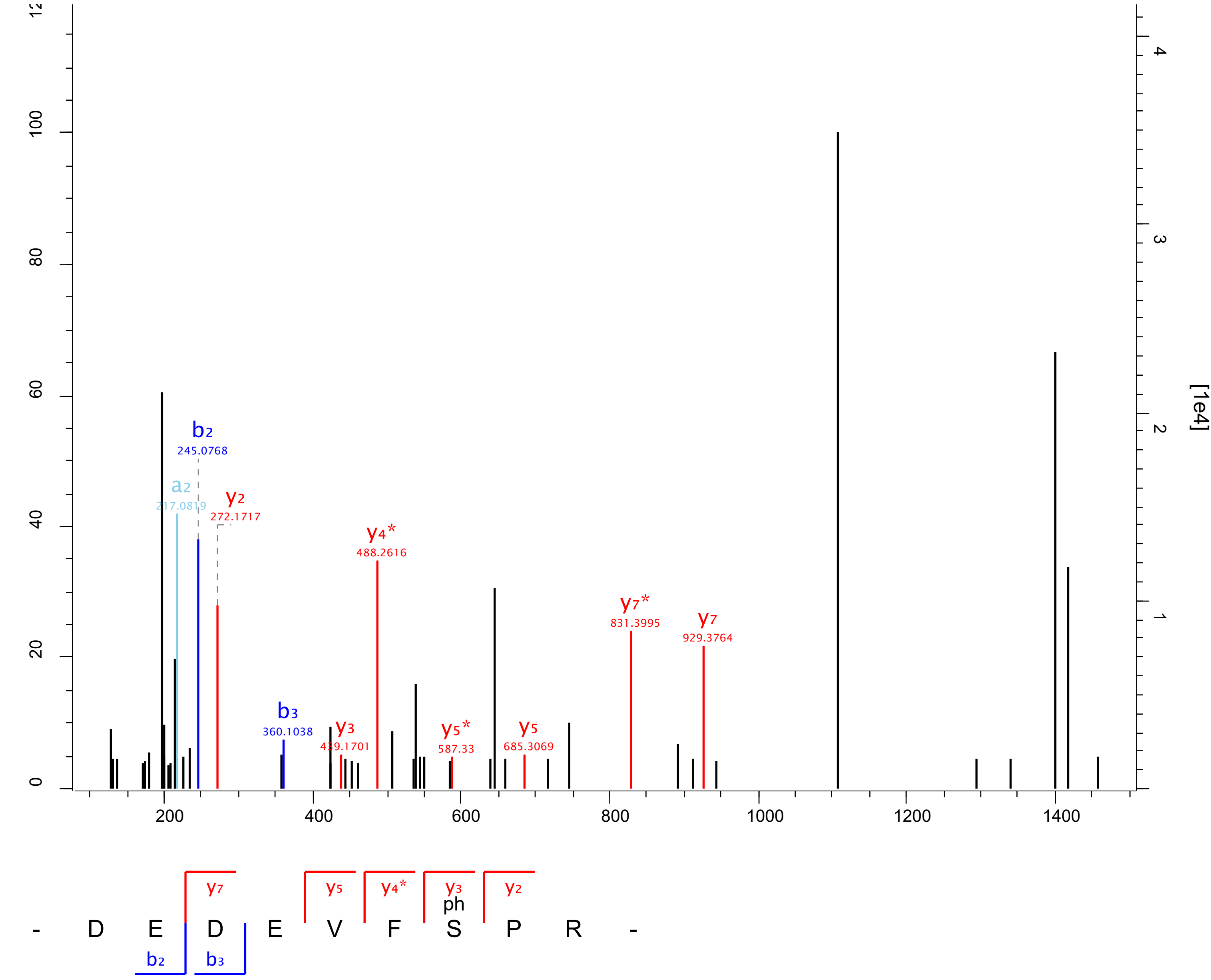

| Raw file | Scan | Method | Score | m/z |
|---|---|---|---|---|
| water_02_1 | 21099 | FTMS; HCD | 51.29 | 620.89 |

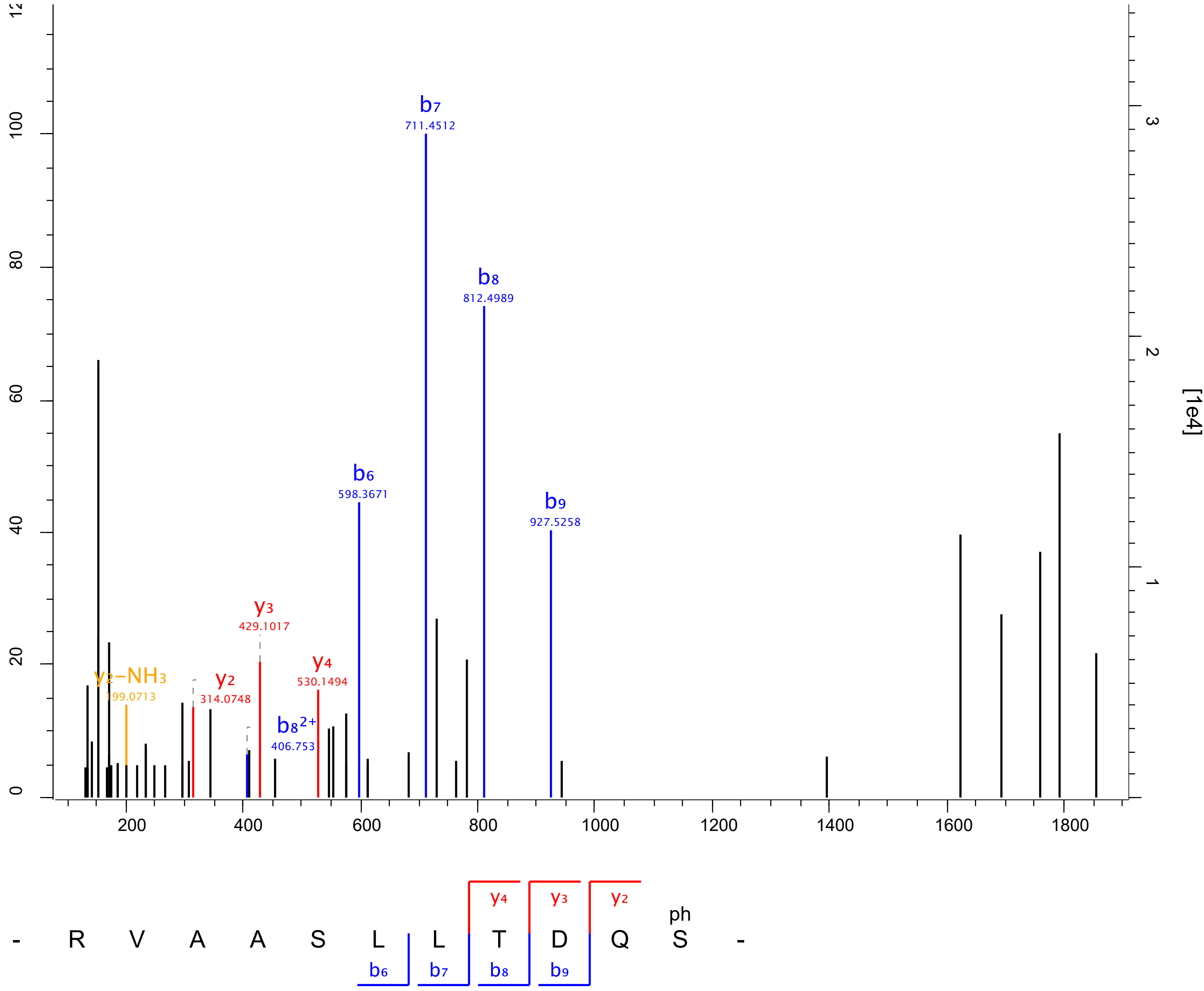

| Raw file | Scan | Method | Score | m/z |
|---|---|---|---|---|
| water_02_1 | 21203 | FTMS; HCD | 171.26 | 744.83 |

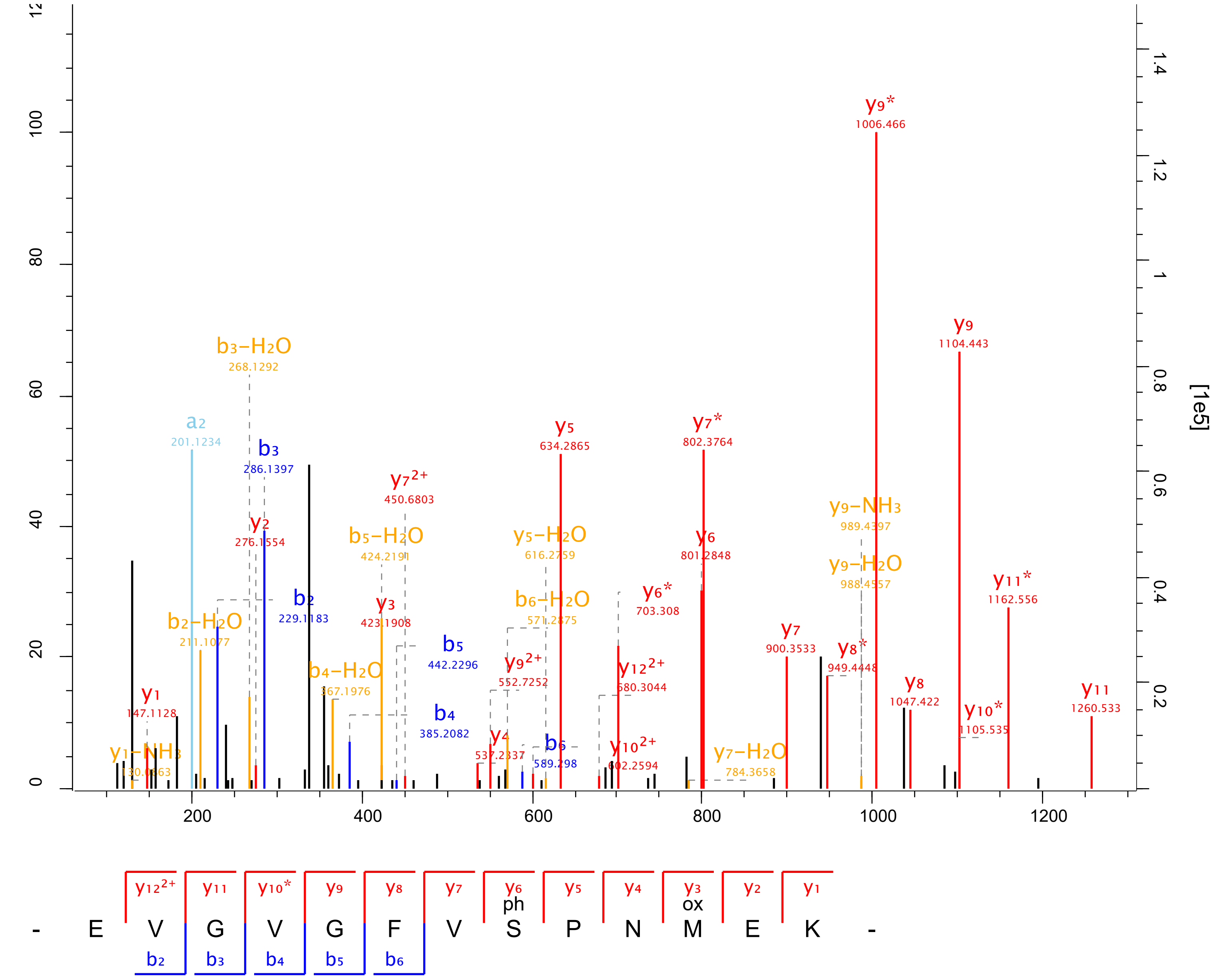

| Raw file | Scan | Method | Score | m/z |
| --- | --- | --- | --- | --- |
| water_02_1 | 21353 | FTMS; HCD | 52.81 | 596.96 |

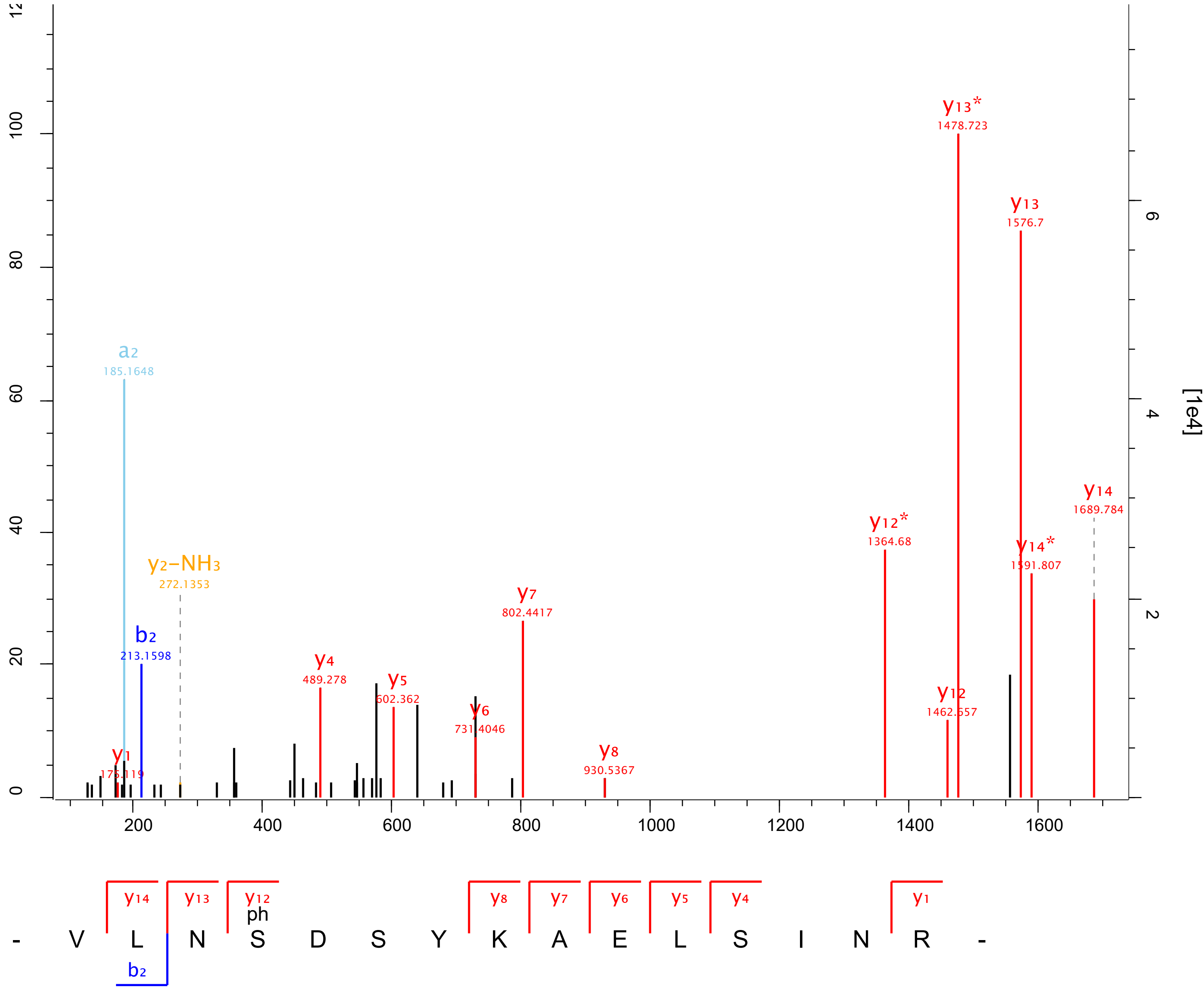

| Raw file | Scan | Method | Score | m/z |
| --- | --- | --- | --- | --- |
| water_02_1 | 21374 | FTMS; HCD | 79.65 | 599.6 |

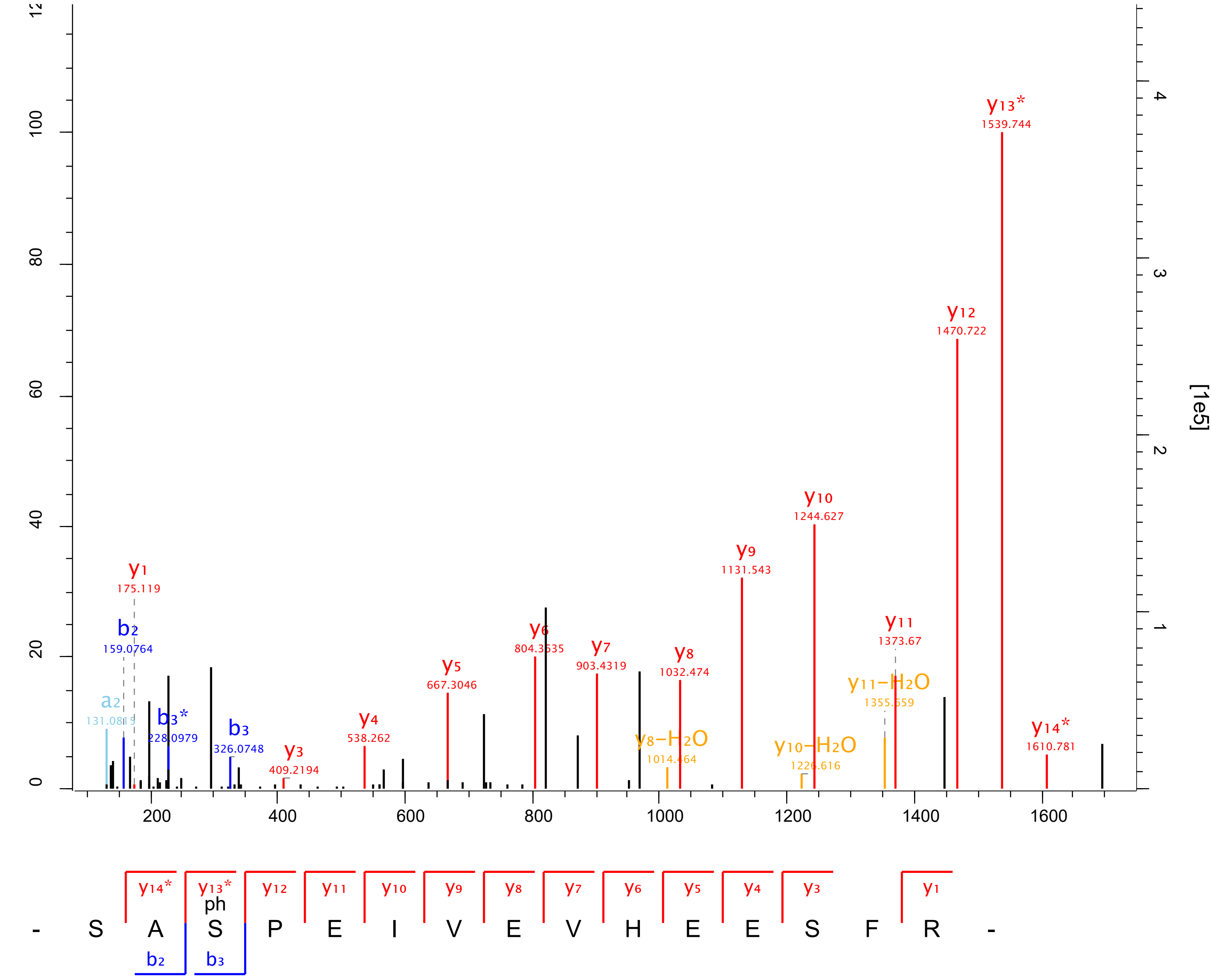

| Raw file | Scan | Method | Score | m/z |
|---|---|---|---|---|
| water_02_1 | 21425 | FTMS; HCD | 69.92 | 649.82 |

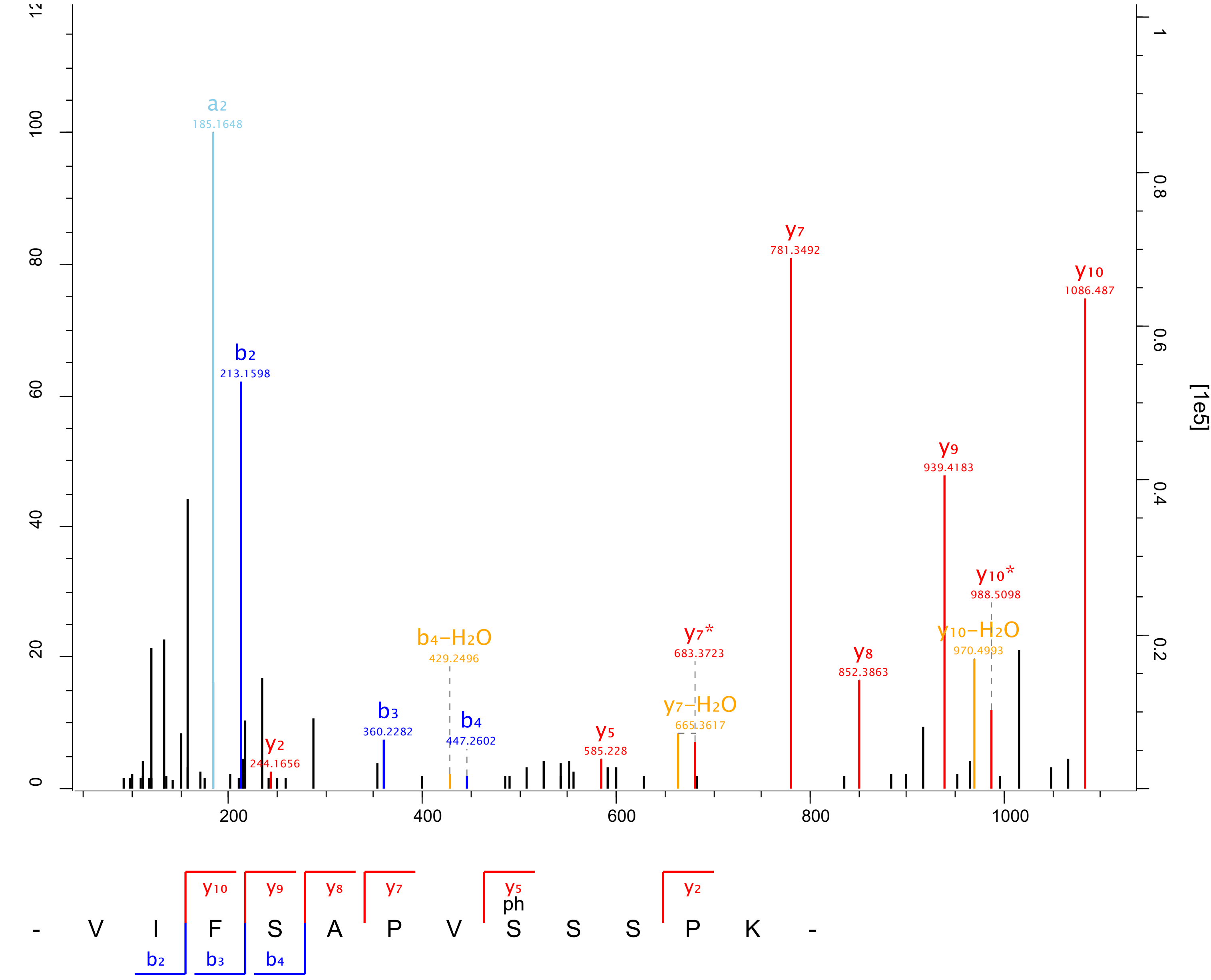

| Raw file | Scan | Method | Score | m/z |
| --- | --- | --- | --- | --- |
| water_02_1 | 21454 | FTMS; HCD | 54.76 | 853.86 |

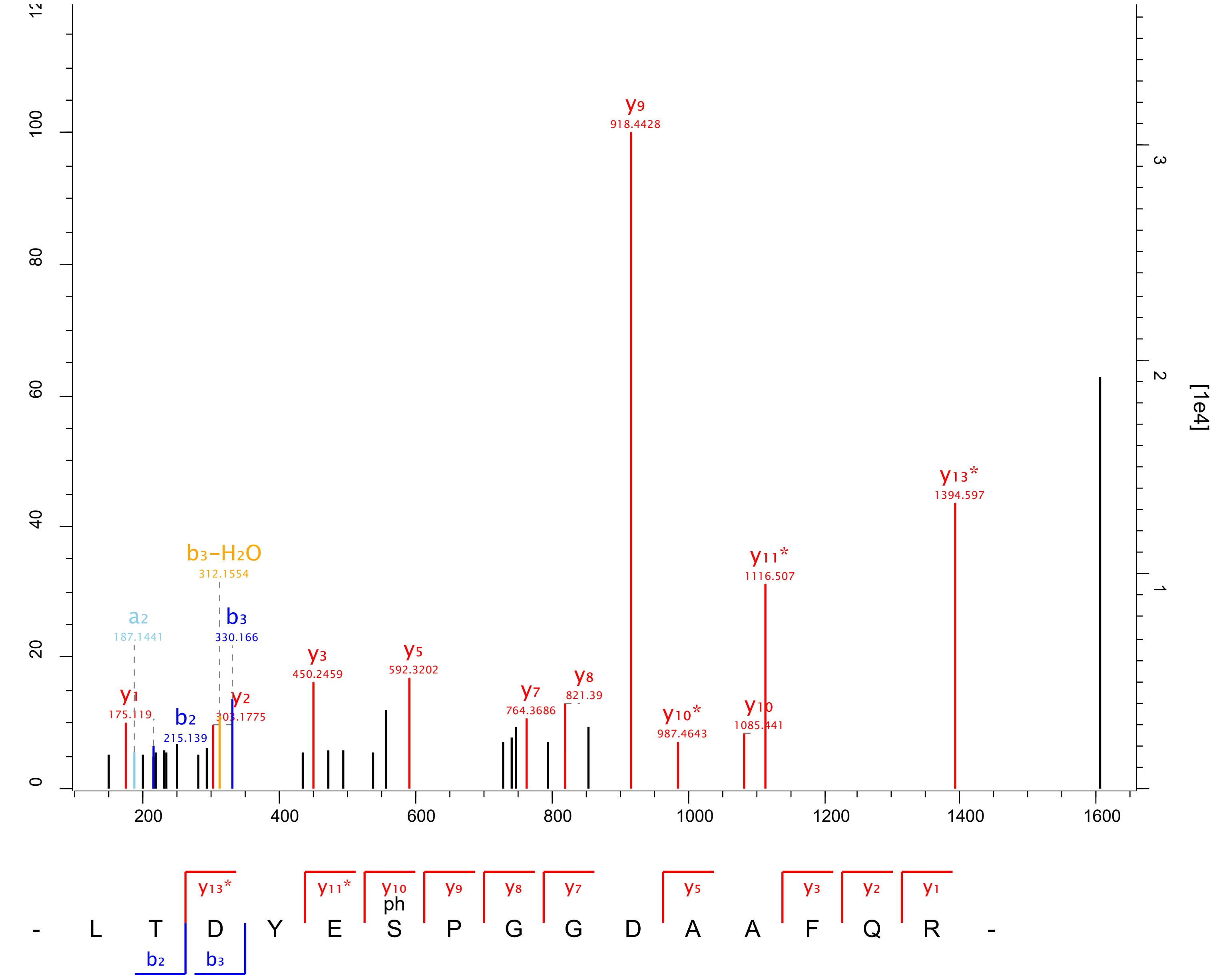

| Raw file | Scan | Method | Score | m/z |
| --- | --- | --- | --- | --- |
| water_02_1 | 21470 | FTMS; HCD | 46.88 | 609.77 |

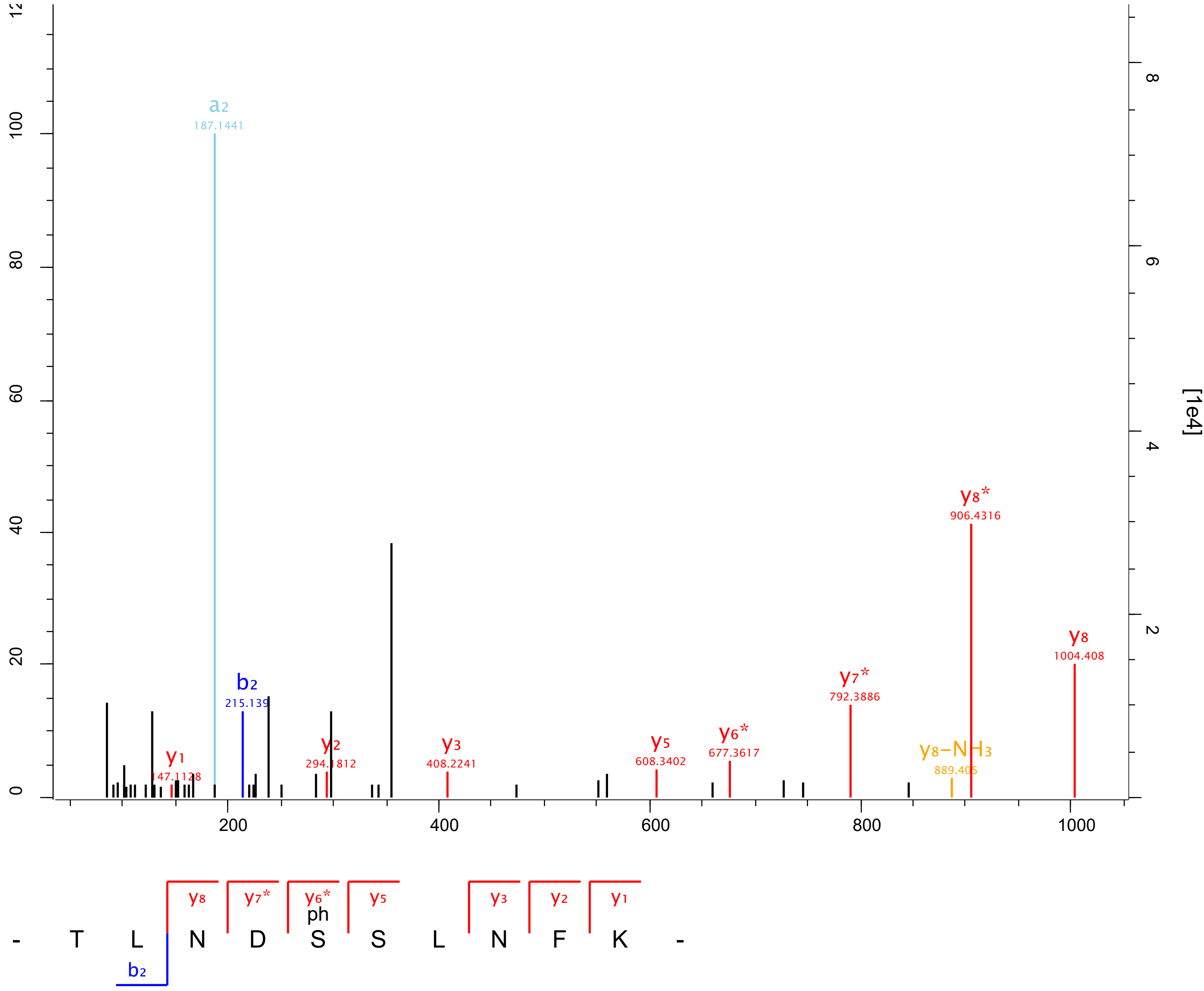

| Raw file | Scan | Method | Score | m/z |
| --- | --- | --- | --- | --- |
| water_02_1 | 2153 | FTMS; HCD | 106.26 | 543.72 |

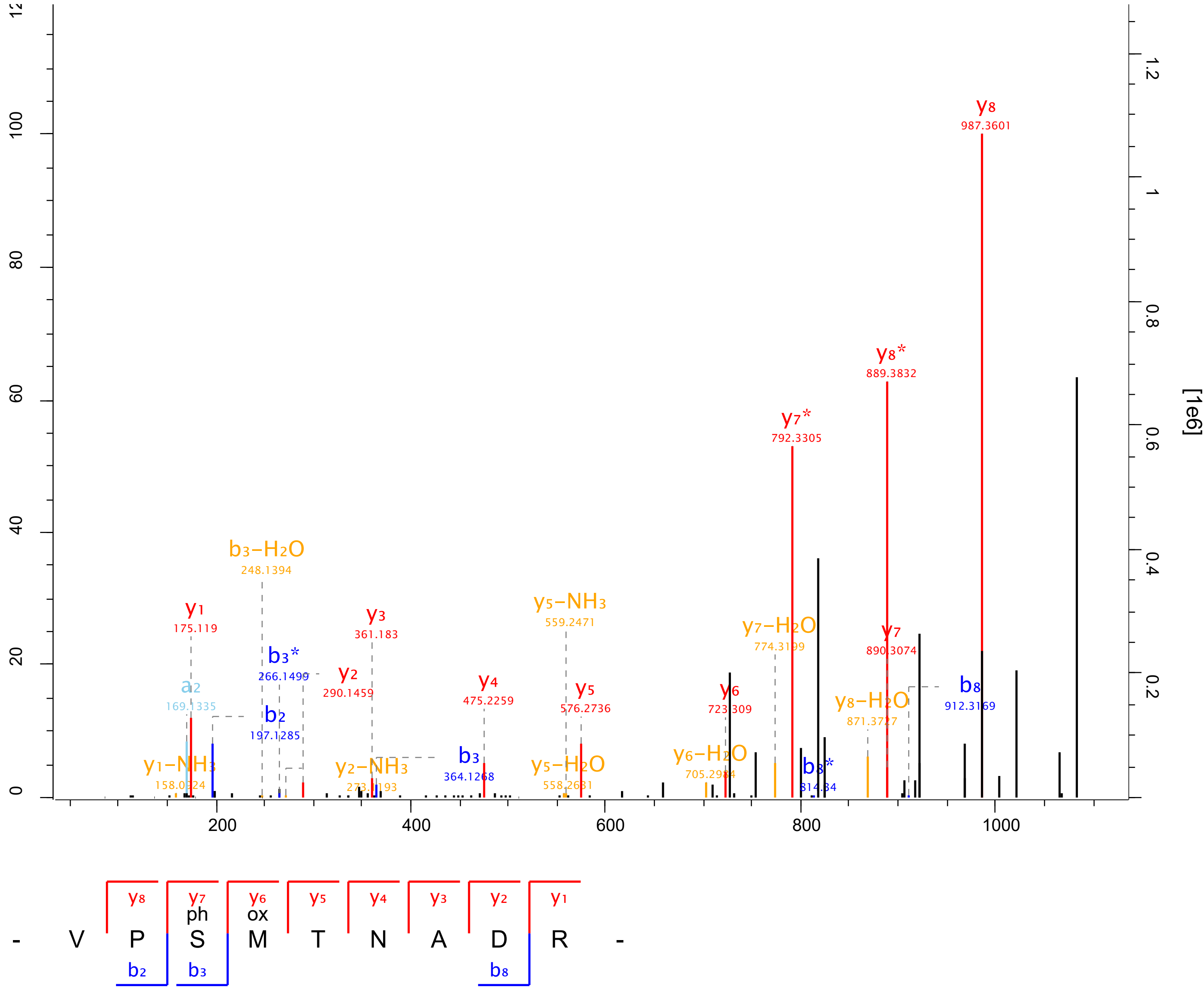

| Raw file | Scan | Method | Score | m/z |
|---|---|---|---|---|
| water_02_1 | 21632 | FTMS; HCD | 132.64 | 501.61 |

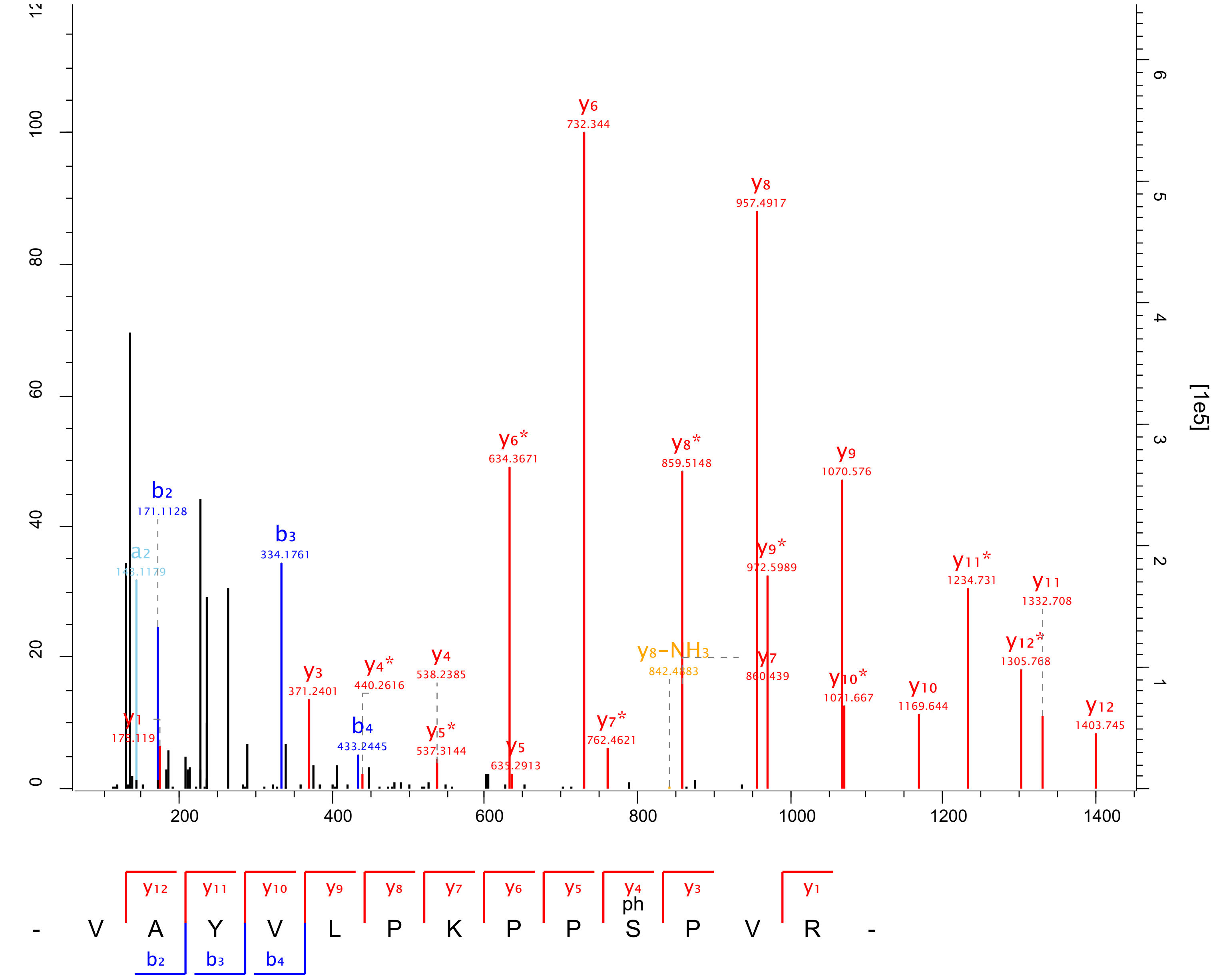

| Raw file | Scan | Method | Score | m/z |
|---|---|---|---|---|
| water_02_1 | 21671 | FTMS; HCD | 175.79 | 1049.47 |

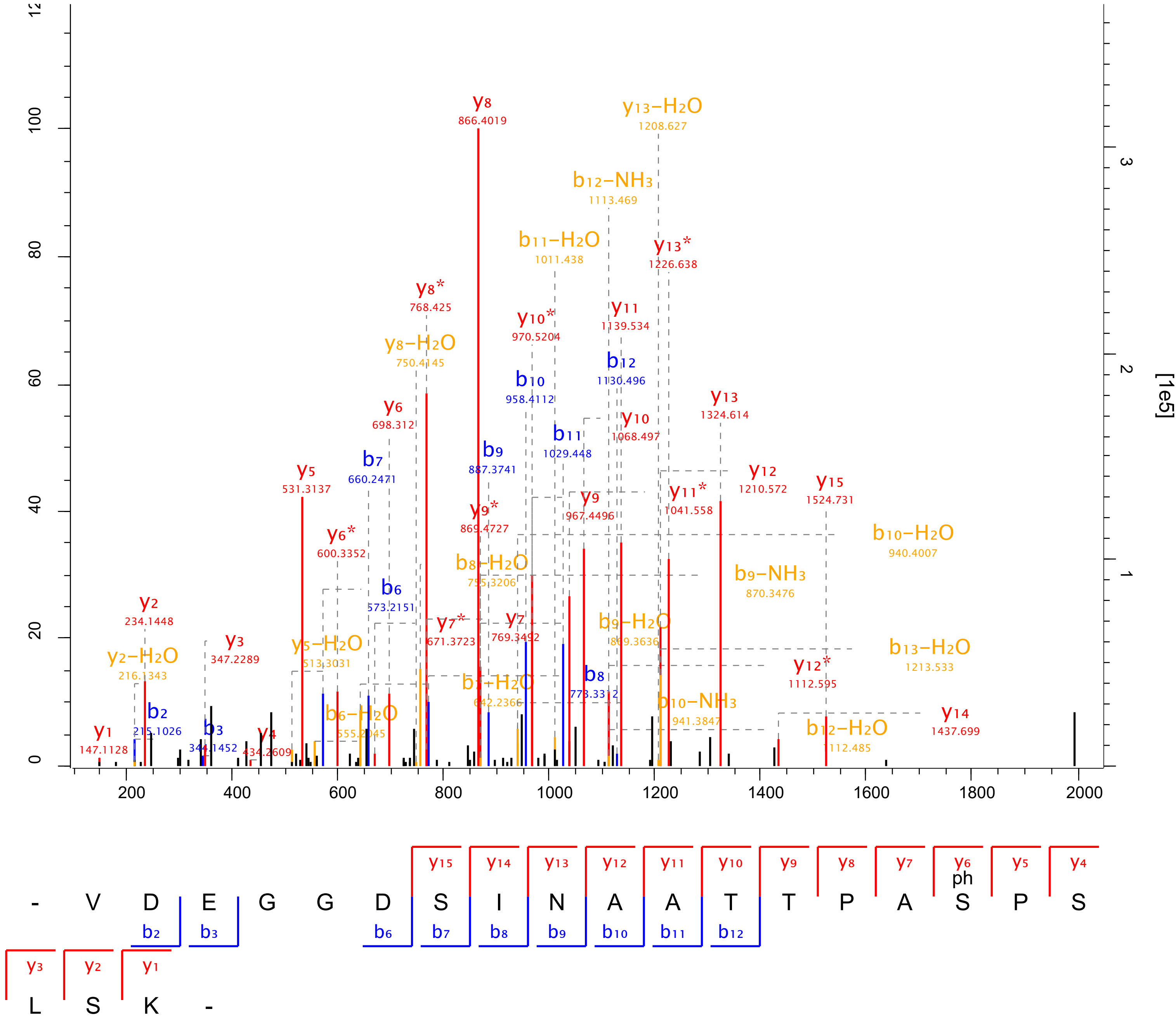

| Raw file | Scan | Method | Score | m/z |
| --- | --- | --- | --- | --- |
| water_02_1 | 21678 | FTMS; HCD | 72.9 | 673.78 |

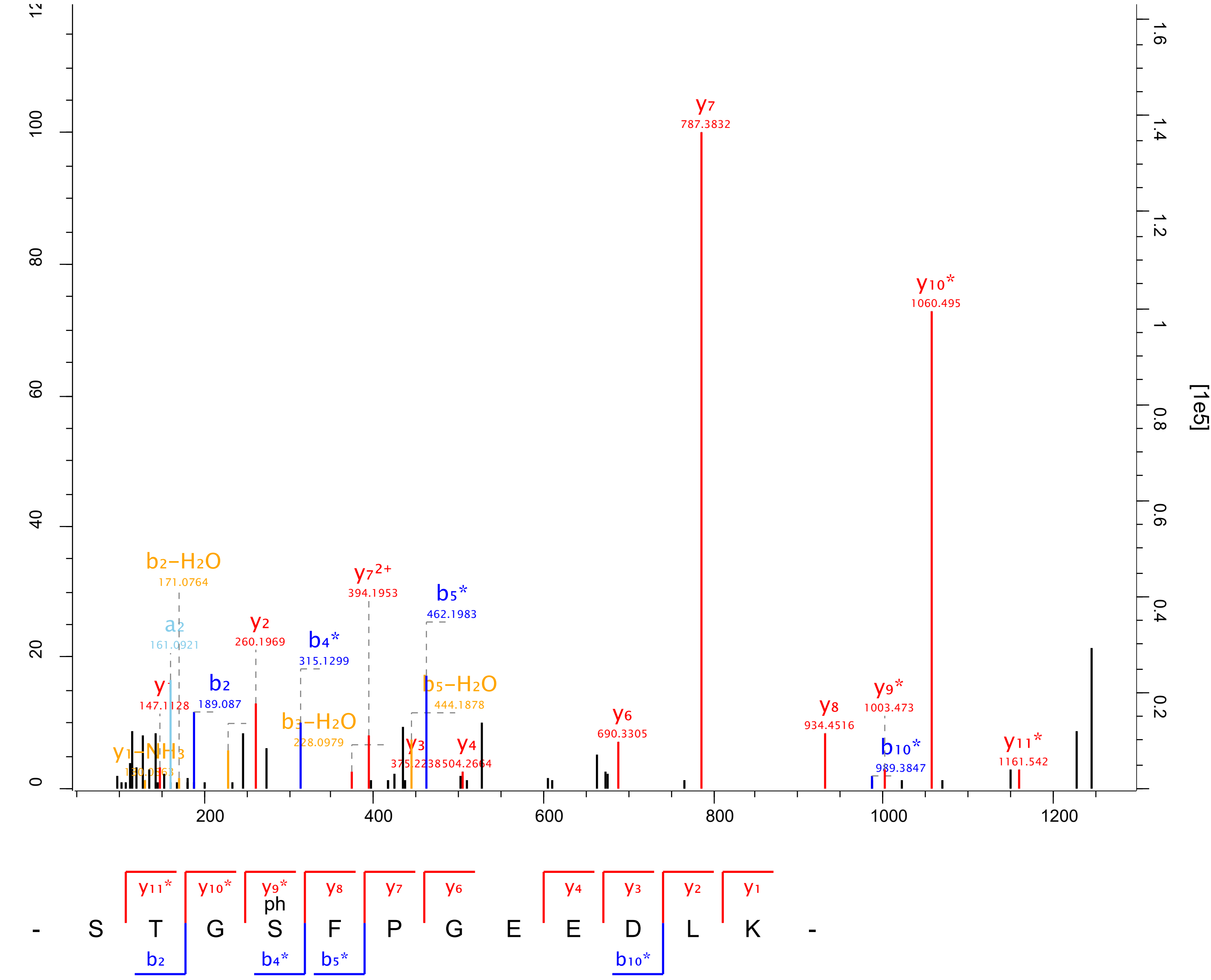

| Raw file | Scan | Method | Score | m/z |
|---|---|---|---|---|
| water_02_1 | 21702 | FTMS; HCD | 137.61 | 827.82 |

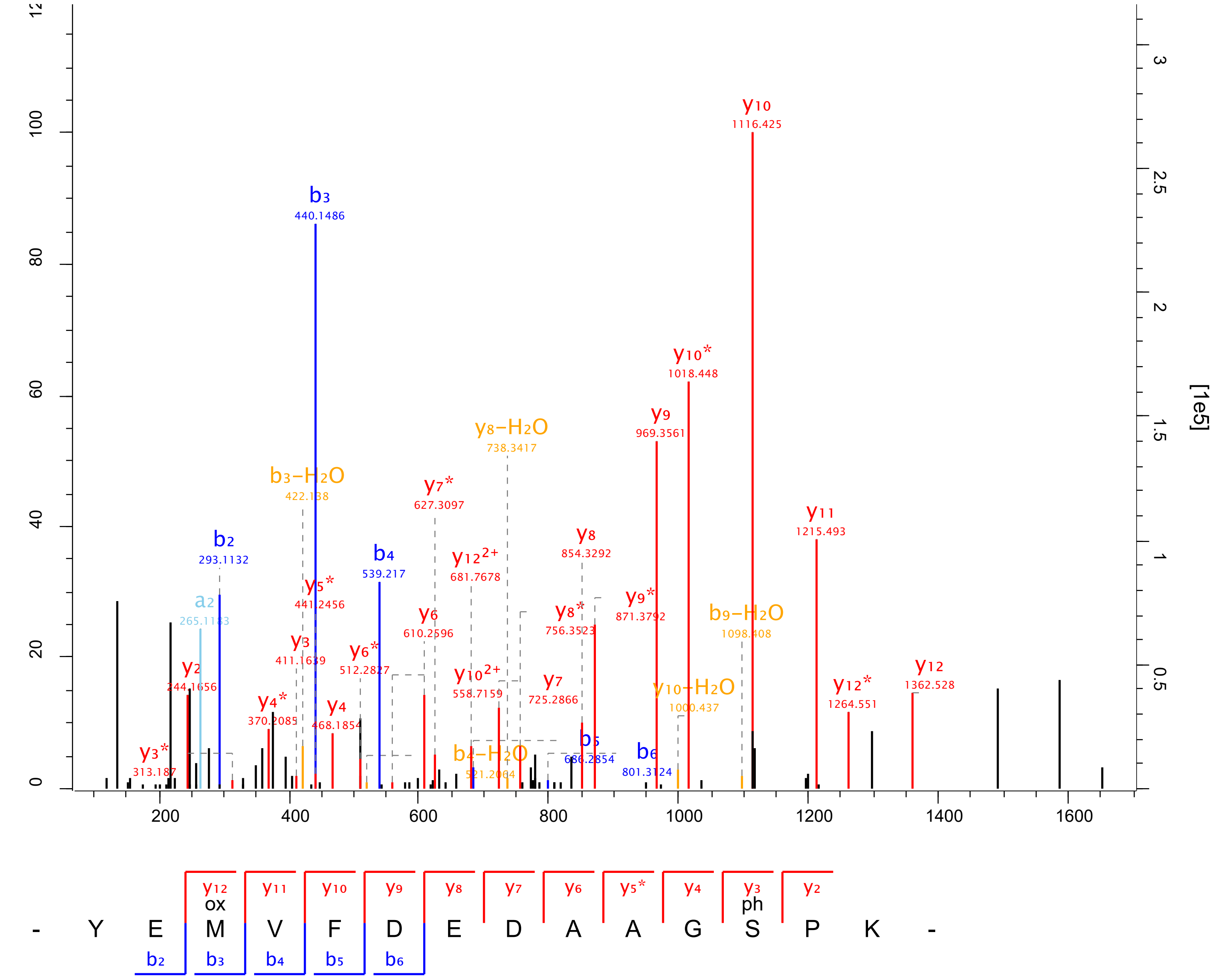

| Raw file | Scan | Method | Score | m/z |
|---|---|---|---|---|
| water_02_1 | 21723 | FTMS; HCD | 94.63 | 787.89 |

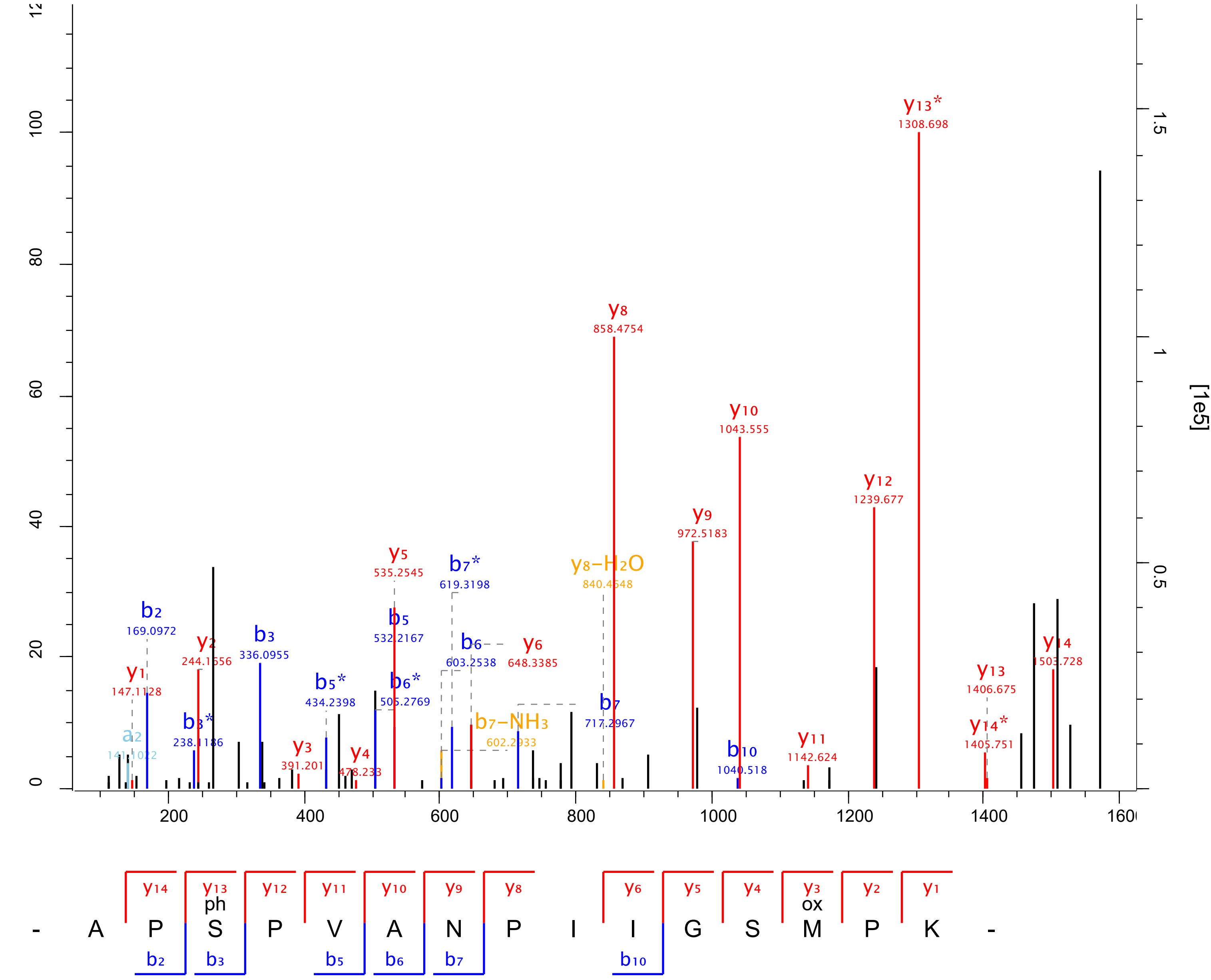

| Raw file | Scan | Method | Score | m/z |
|---|---|---|---|---|
| water_02_1 | 21840 | FTMS; HCD | 130.83 | 627.29 |

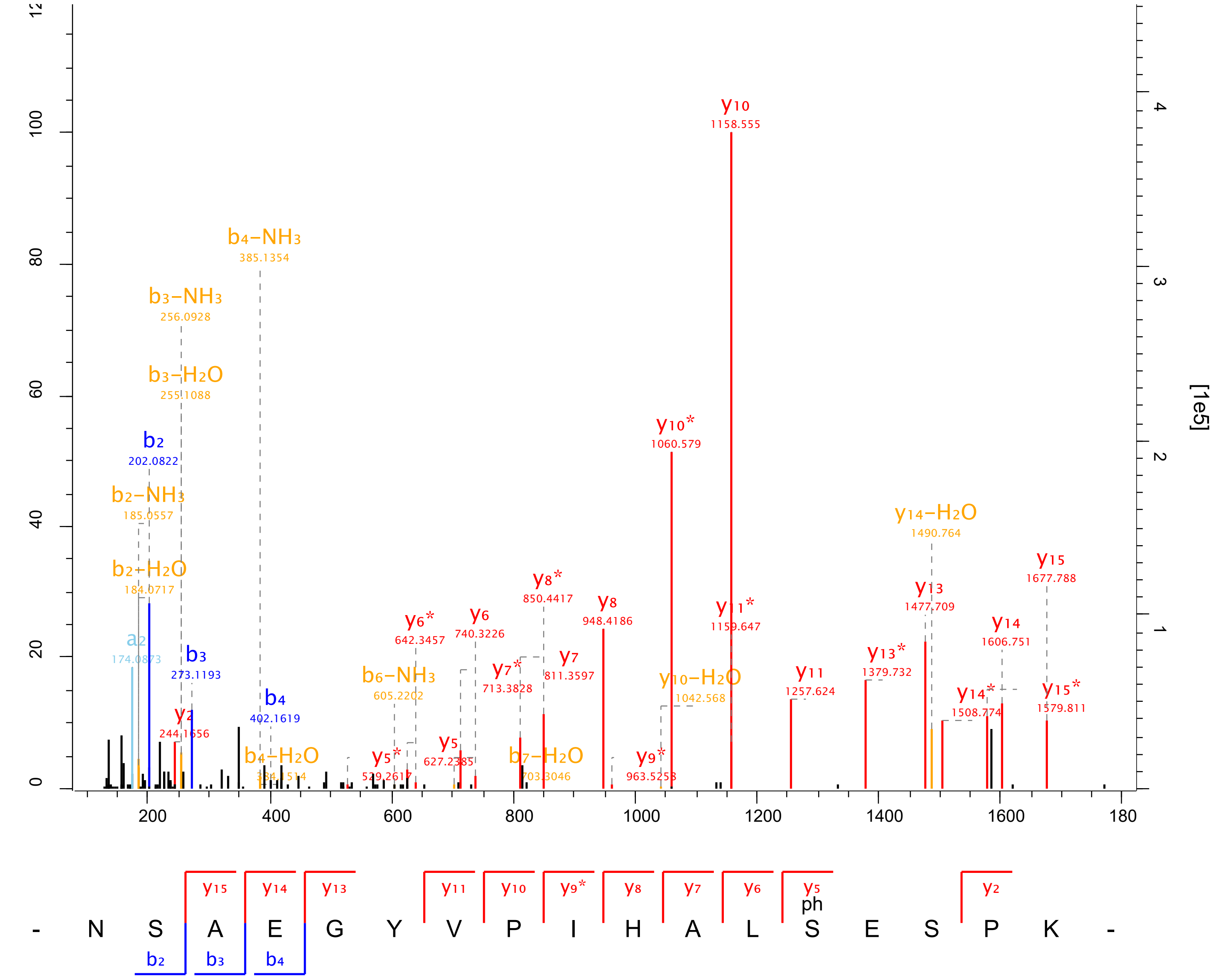

| Raw file | Scan | Method | Score | m/z |
| --- | --- | --- | --- | --- |
| water_02_1 | 21864 | FTMS; HCD | 118.54 | 795.67 |

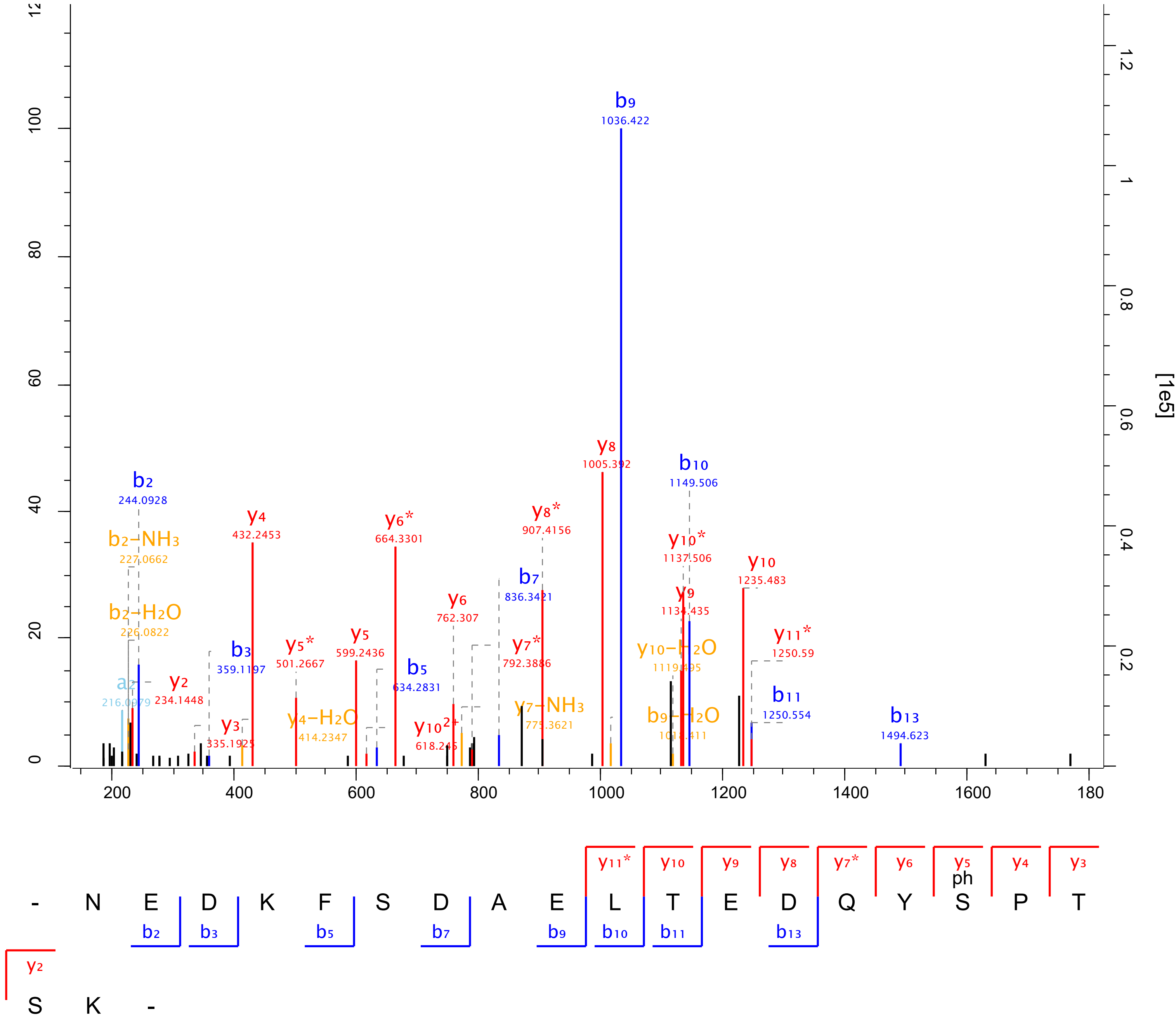

| Raw file | Scan | Method | Score | m/z |
| --- | --- | --- | --- | --- |
| water_02_1 | 21870 | FTMS; HCD | 70.89 | 436.91 |

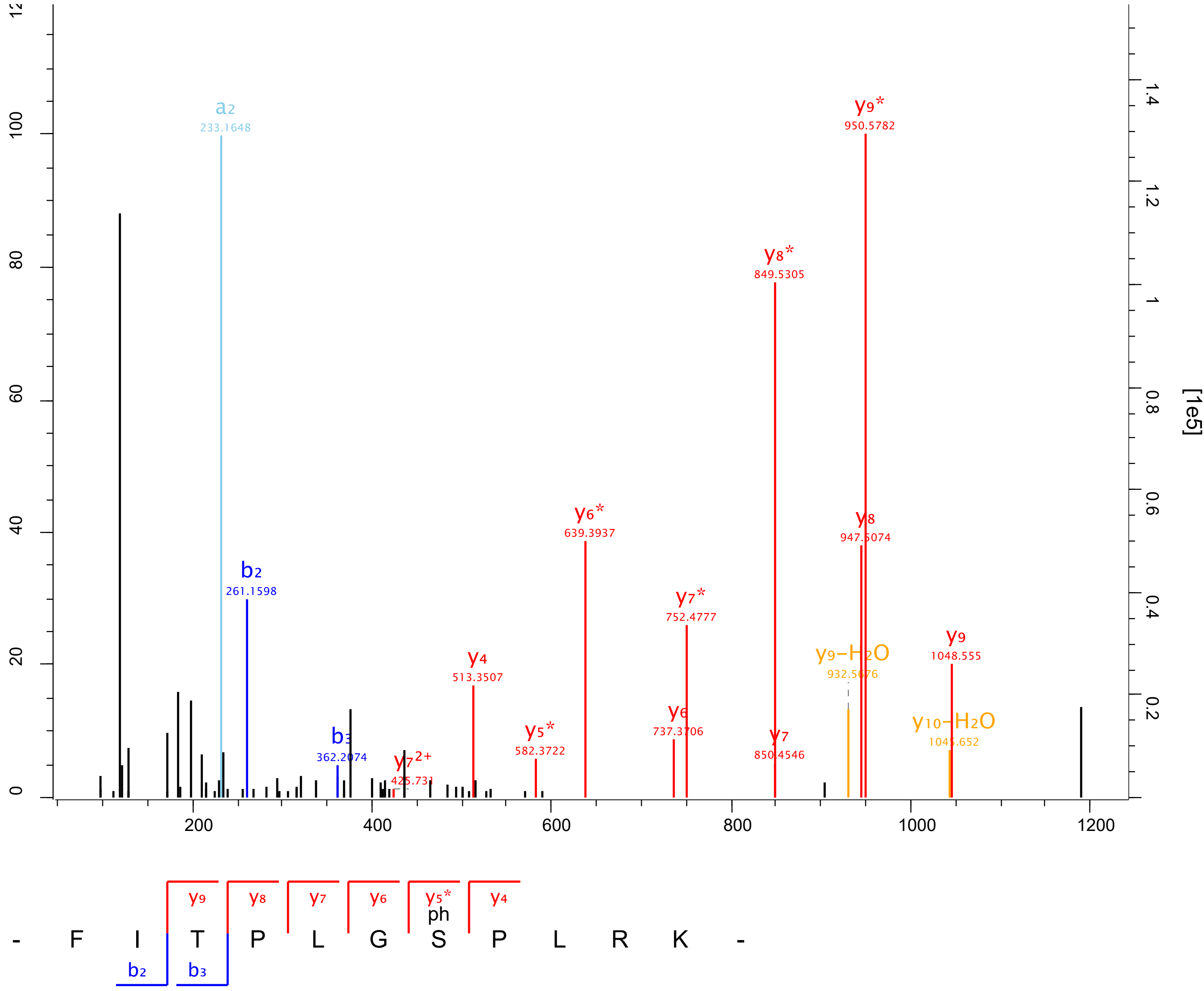

| Raw file | Scan | Method | Score | m/z |
|---|---|---|---|---|
| water_02_1 | 21976 | FTMS; HCD | 45.61 | 646.24 |

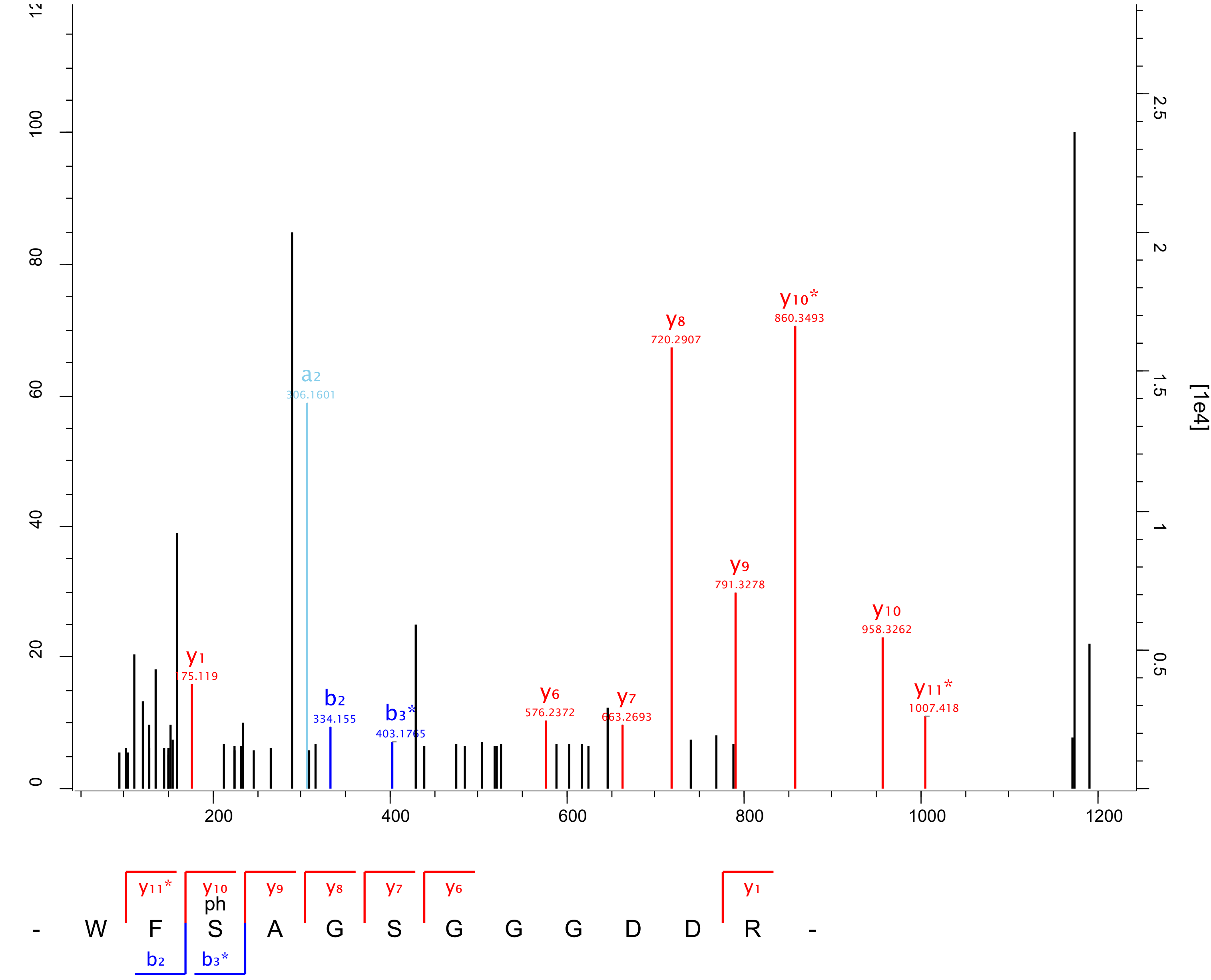

| Raw file | Scan | Method | Score | m/z |
| --- | --- | --- | --- | --- |
| water_02_1 | 21979 | FTMS; HCD | 97.45 | 639.27 |

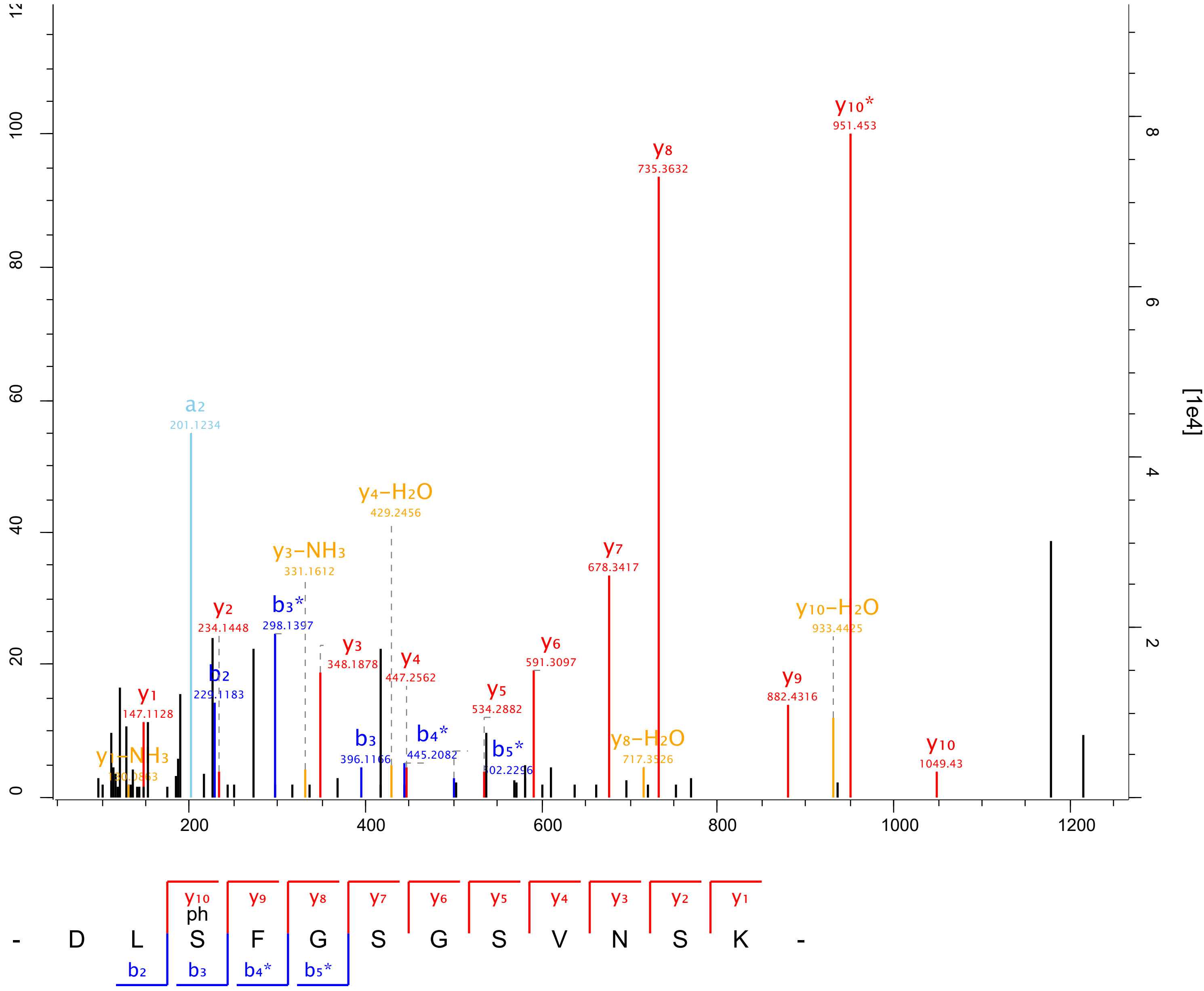

| Raw file | Scan | Method | Score | m/z |
|---|---|---|---|---|
| water_02_1 | 22086 | FTMS; HCD | 130.77 | 525.74 |

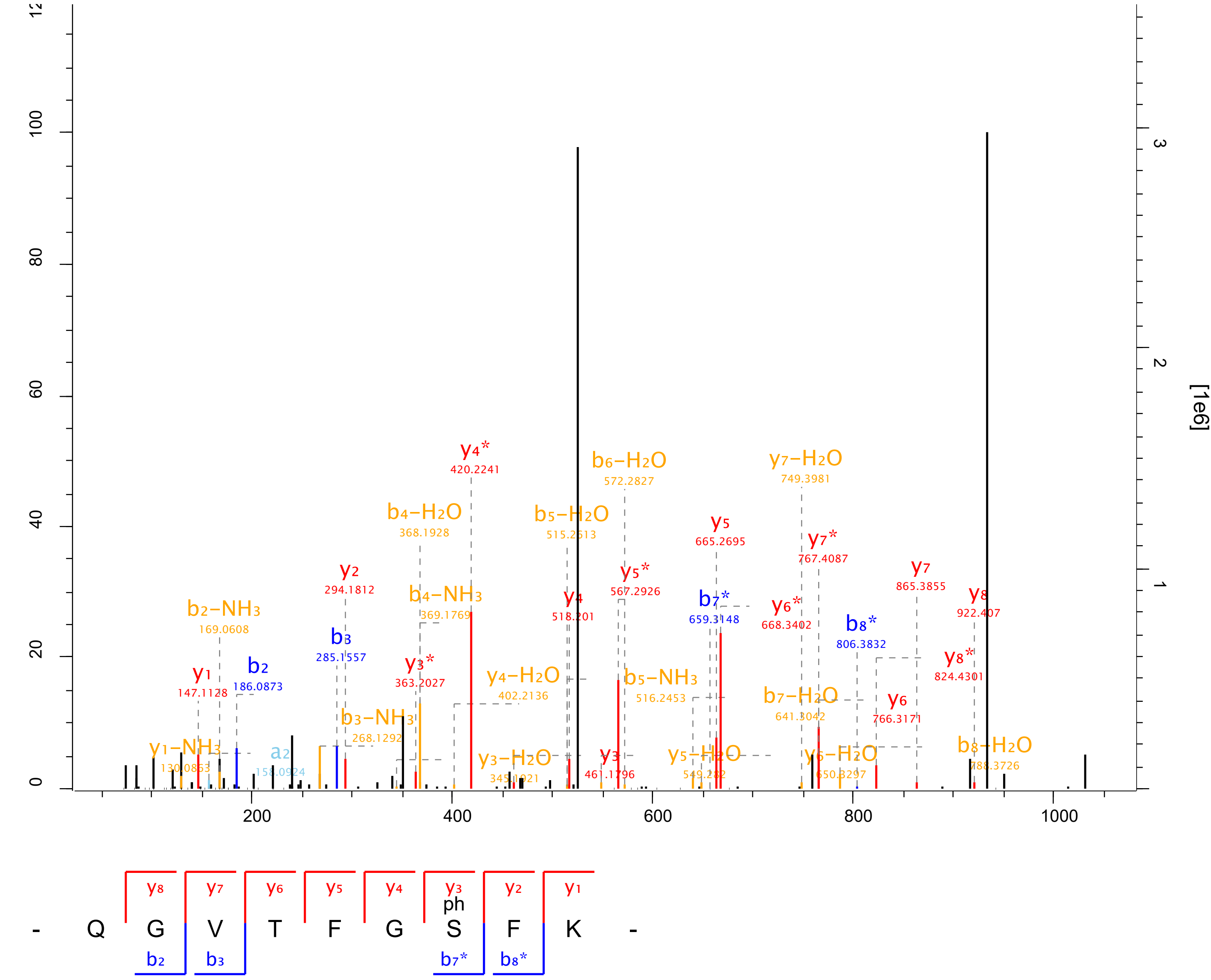

| Raw file | Scan | Method | Score | m/z |
|---|---|---|---|---|
| water_02_1 | 2210 | FTMS; HCD | 51.45 | 472.74 |

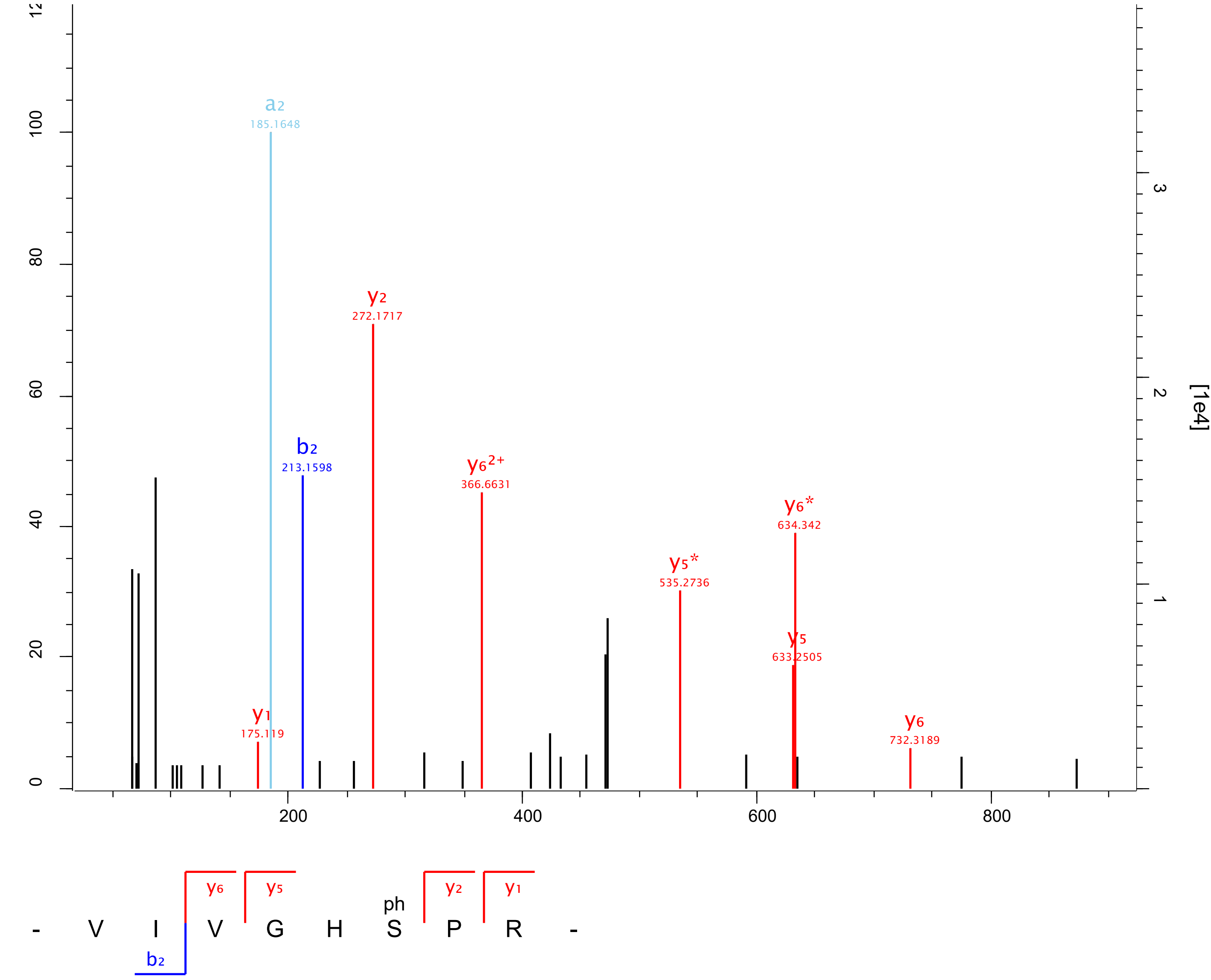

| Raw file | Scan | Method | Score | m/z |
| --- | --- | --- | --- | --- |
| water_02_1 | 22110 | FTMS; HCD | 72.99 | 735.98 |

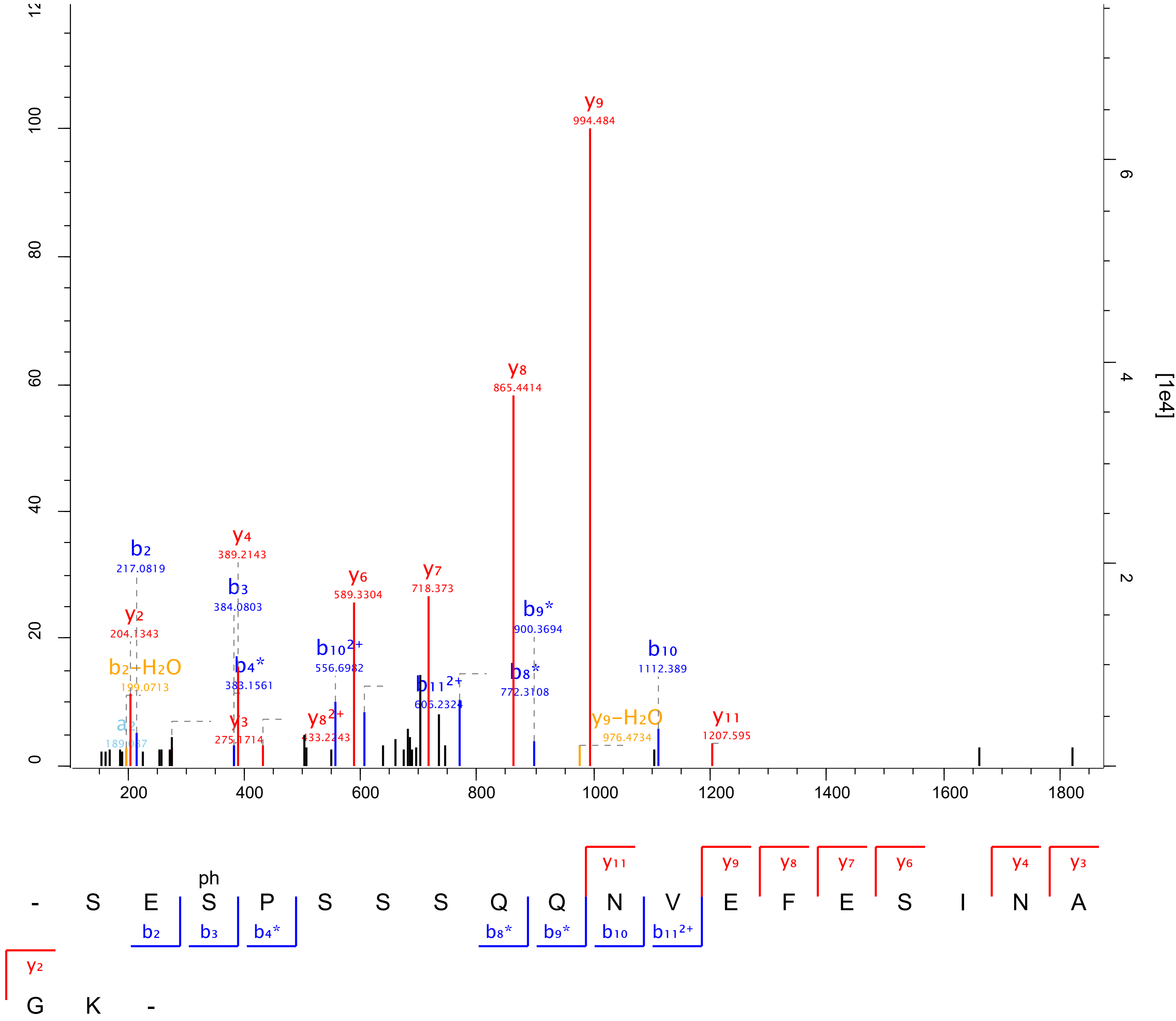

| Raw file | Scan | Method | Score | m/z |
|---|---|---|---|---|
| water_02_1 | 22112 | FTMS; HCD | 162.59 | 754.8 |

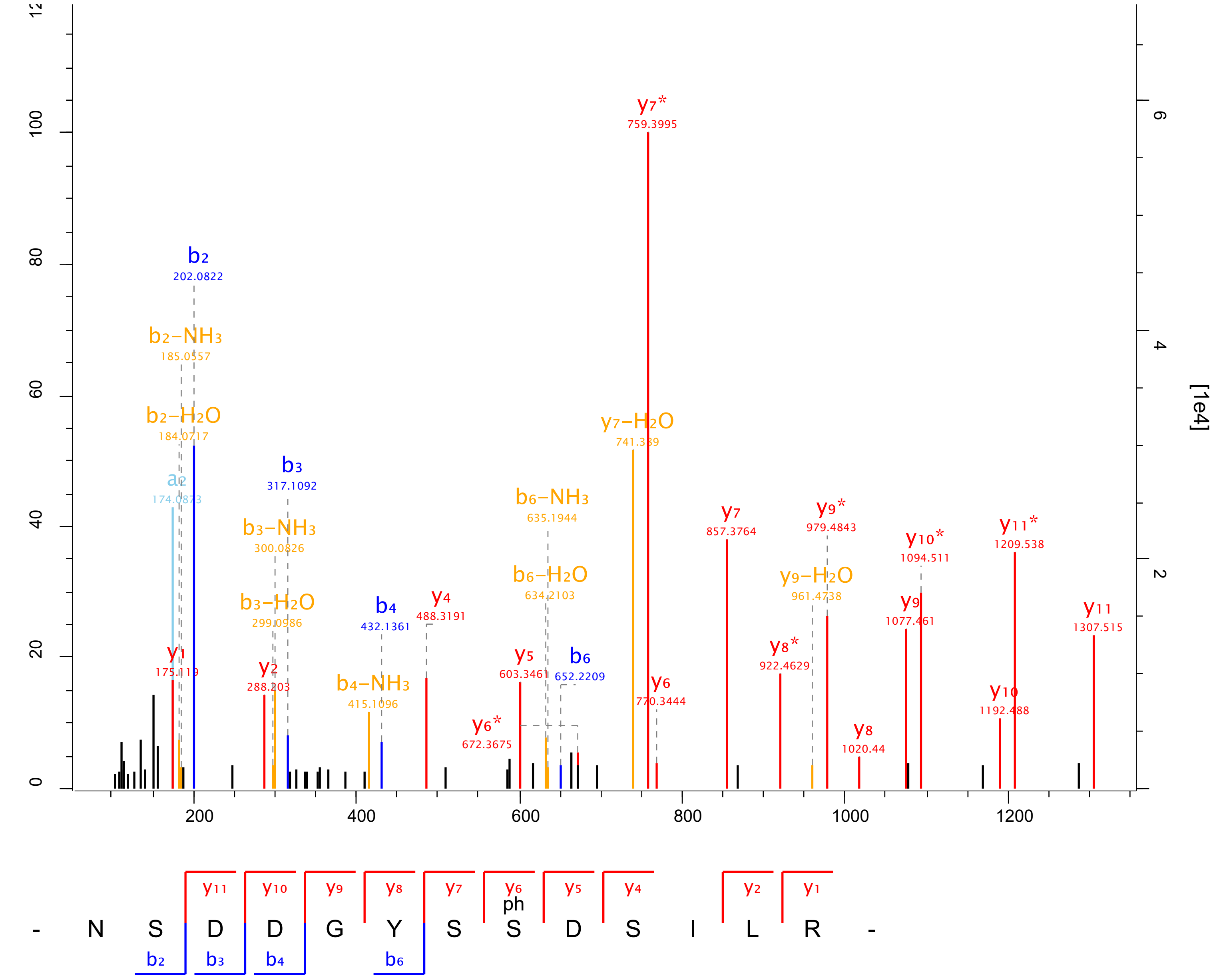

| Raw file | Scan | Method | Score | m/z |
| --- | --- | --- | --- | --- |
| water_02_1 | 22209 | FTMS; HCD | 344.39 | 913.35 |

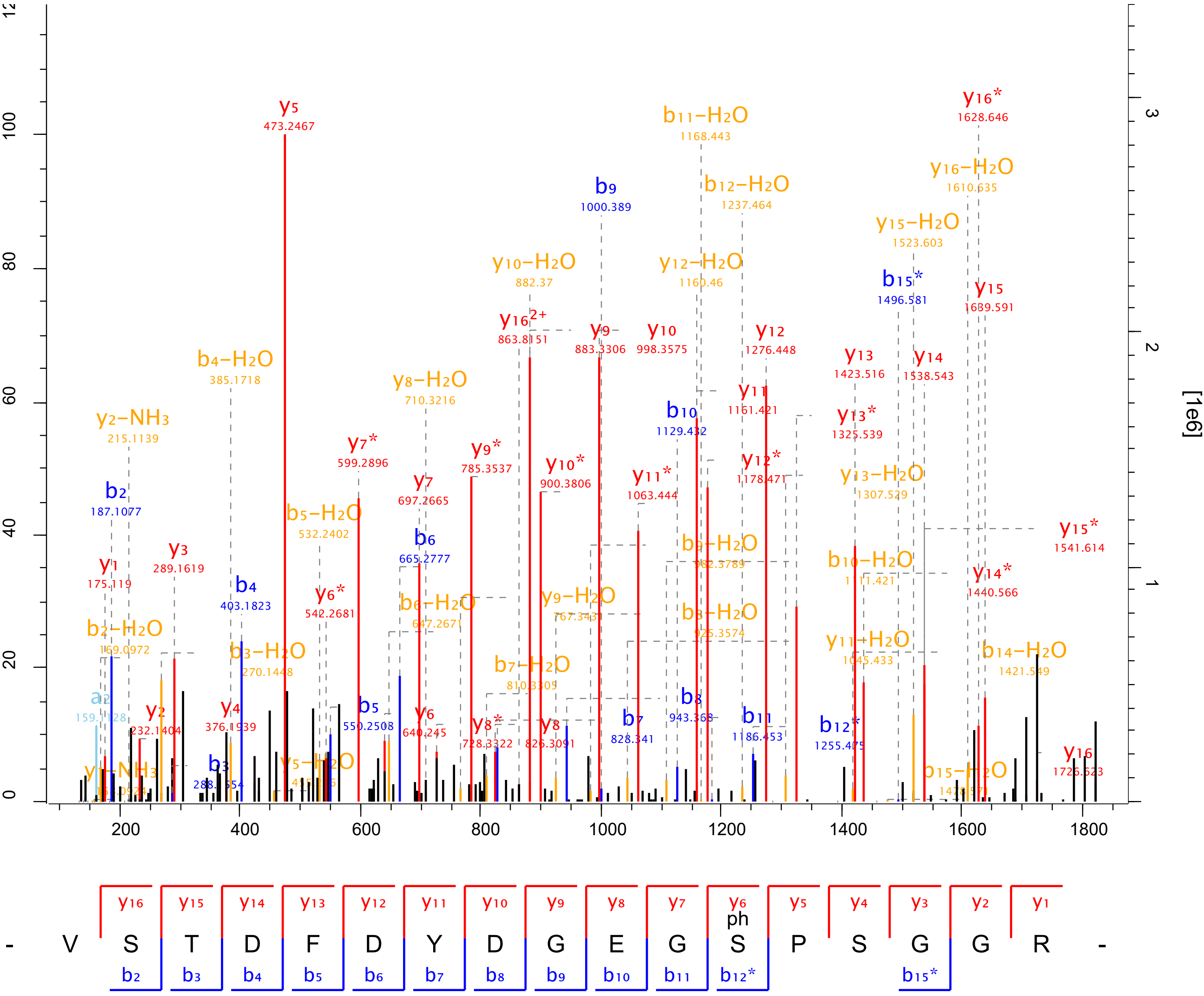

| Raw file | Scan | Method | Score | m/z |
| --- | --- | --- | --- | --- |
| water_02_1 | 22346 | FTMS; HCD | 133.89 | 509.73 |

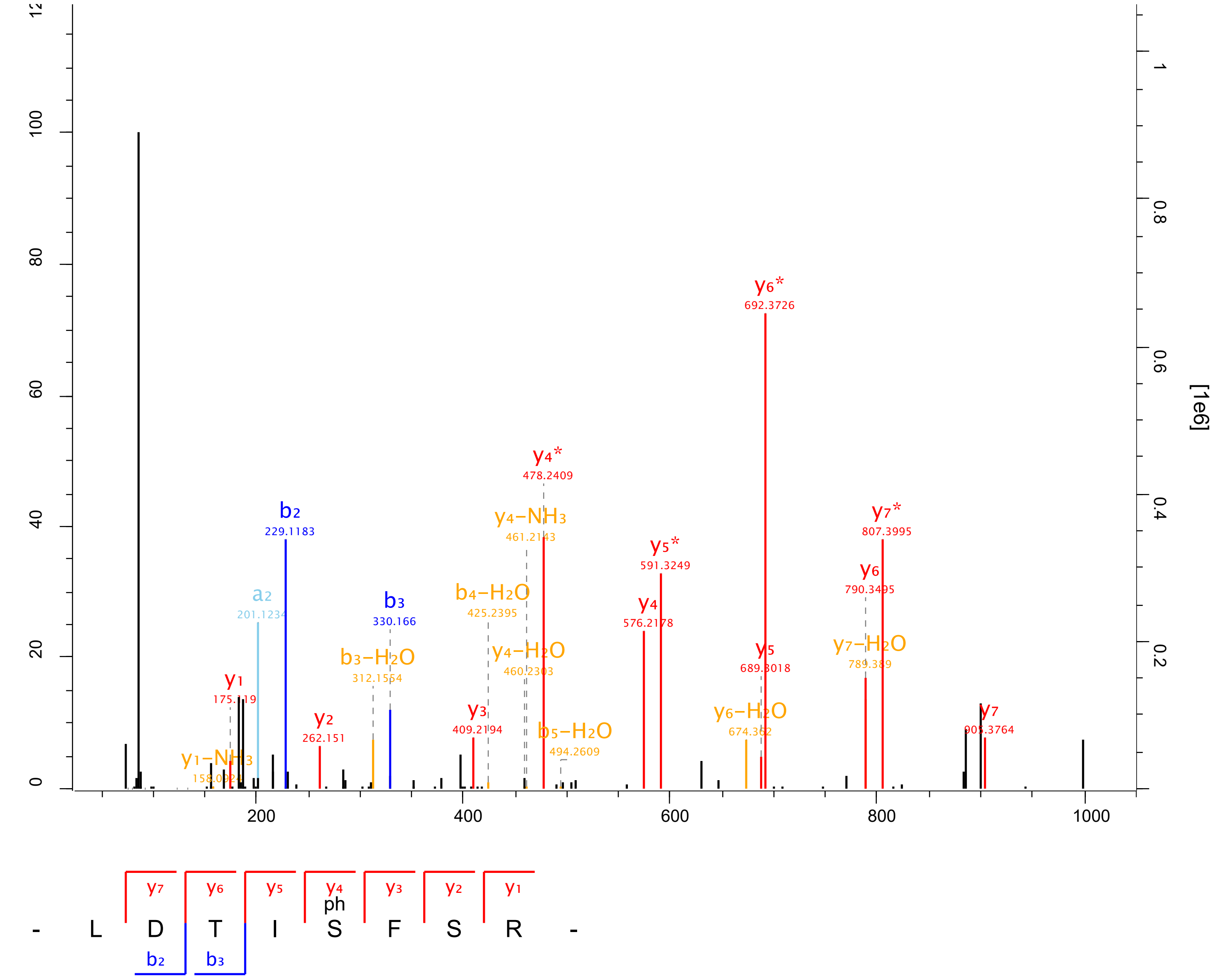

| Raw file | Scan | Method | Score | m/z |
|---|---|---|---|---|
| water_02_1 | 22368 | FTMS; HCD | 127.83 | 775.38 |

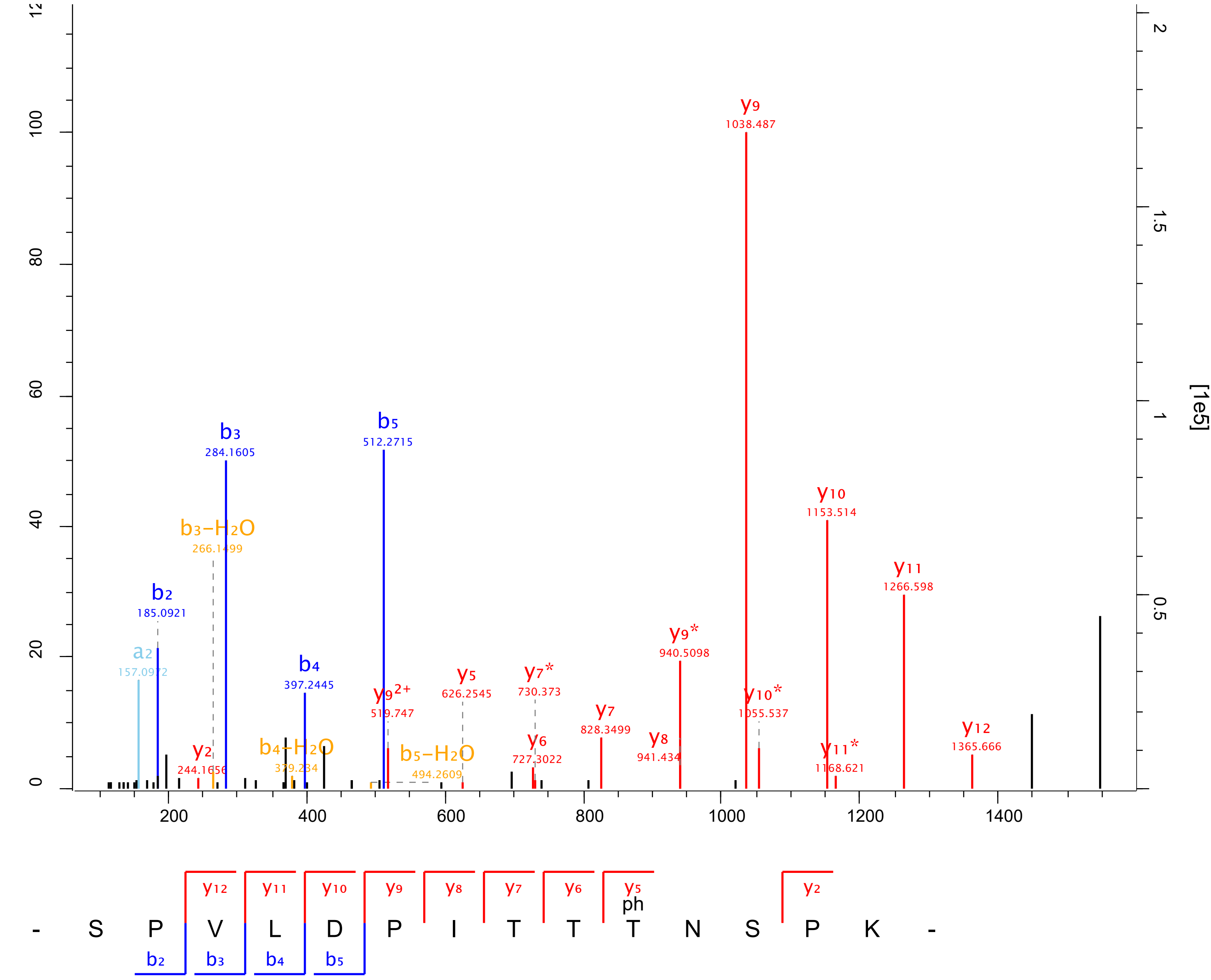

| Raw file | Scan | Method | Score | m/z |
| --- | --- | --- | --- | --- |
| water_02_1 | 22476 | FTMS; HCD | 63.87 | 676.28 |

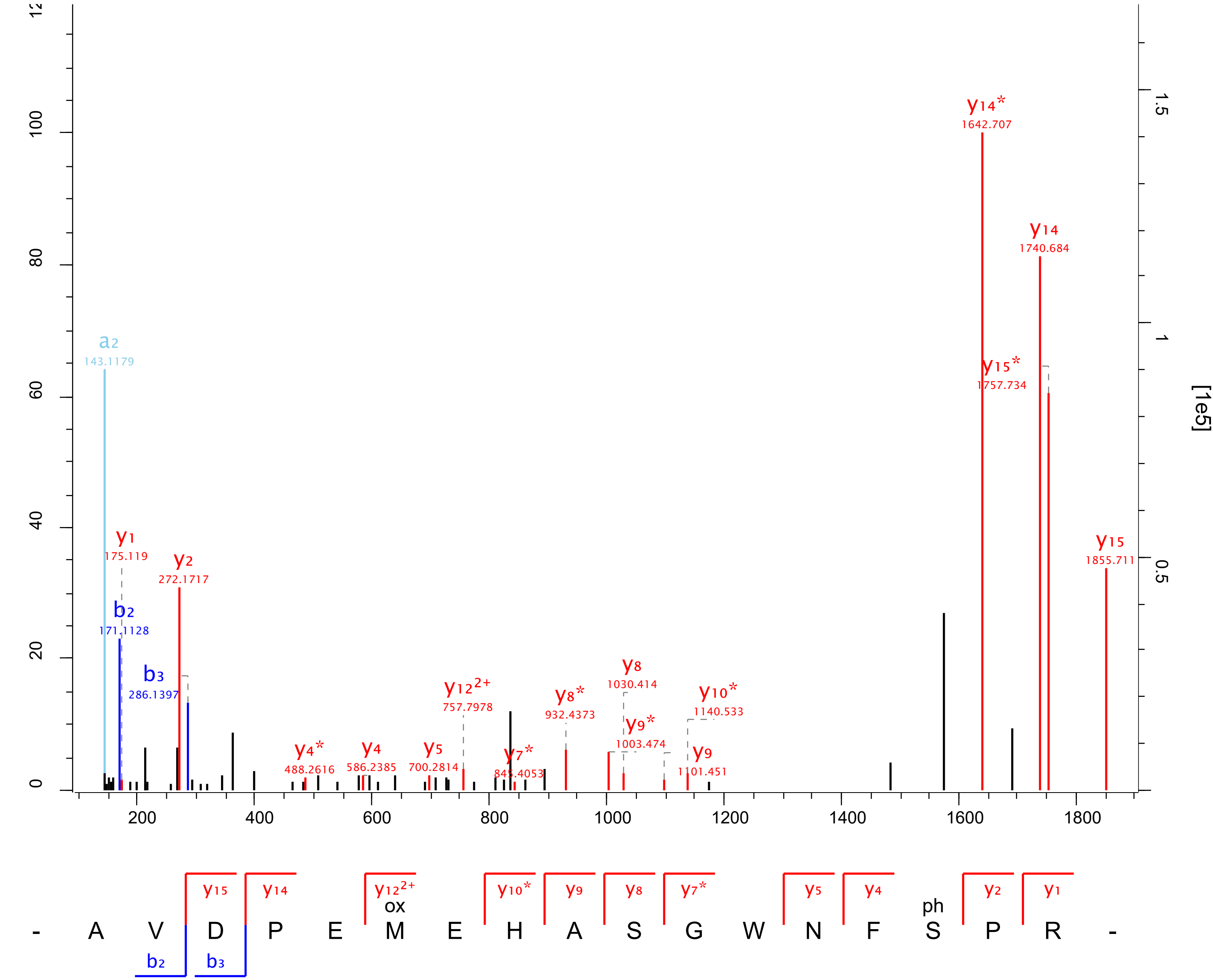

| Raw file | Scan | Method | Score | m/z |
|---|---|---|---|---|
| water_02_1 | 22489 | FTMS; HCD | 101.92 | 827.89 |

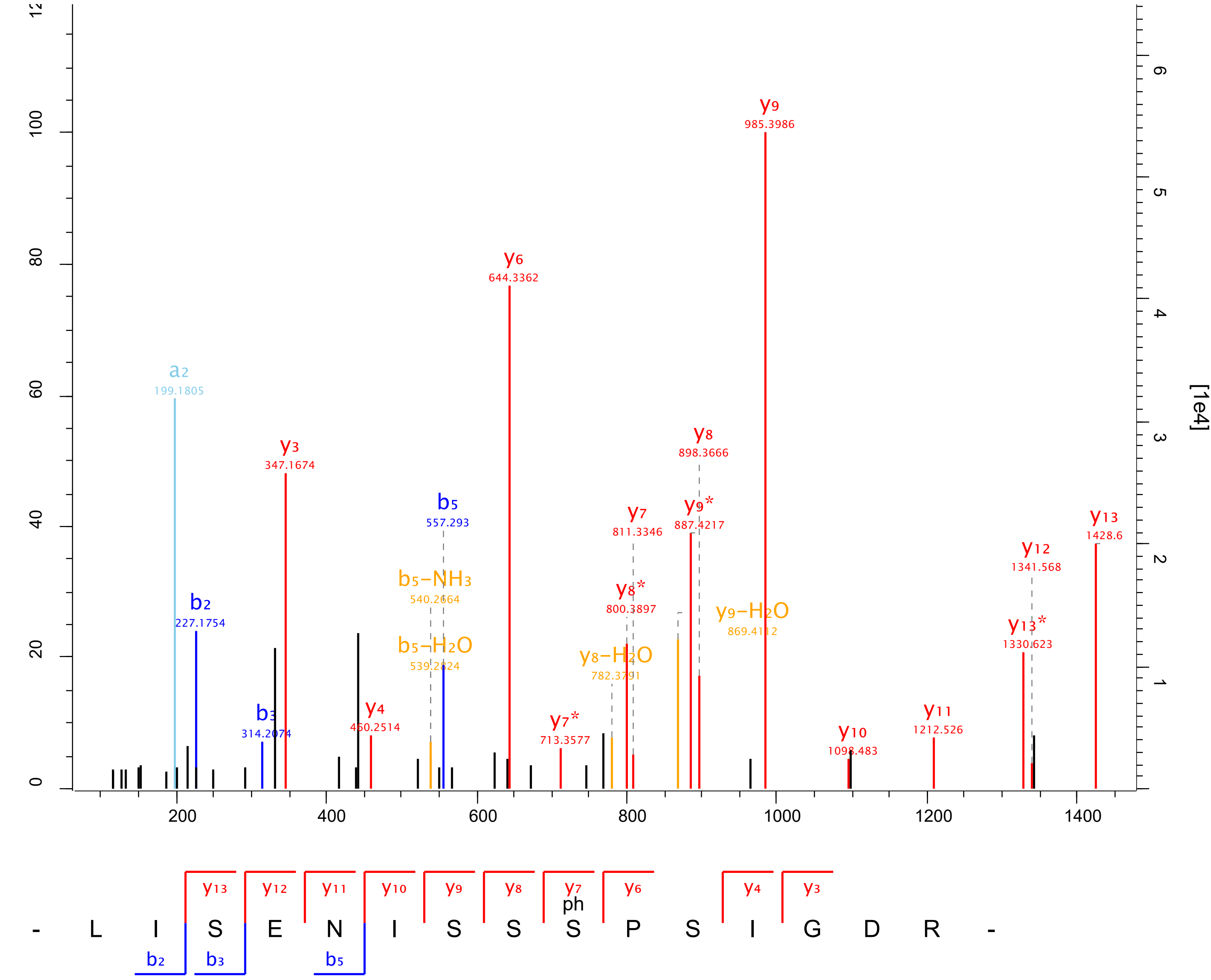

| Raw file | Scan | Method | Score | m/z |
|---|---|---|---|---|
| water_02_1 | 22543 | FTMS; HCD | 131.56 | 768.32 |

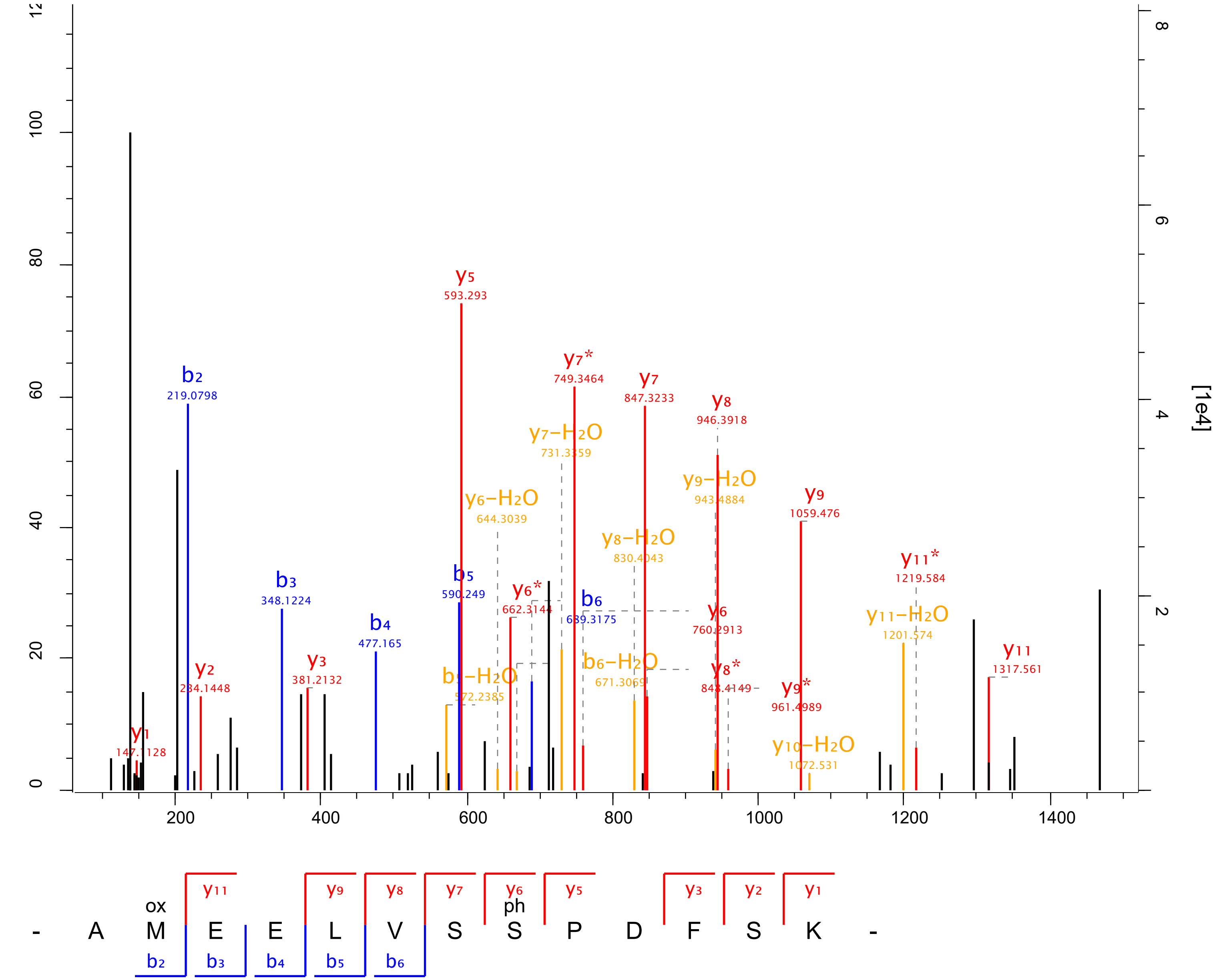

| Raw file | Scan | Method | Score | m/z |
|---|---|---|---|---|
| water_02_1 | 22553 | FTMS; HCD | 188.59 | 684.79 |

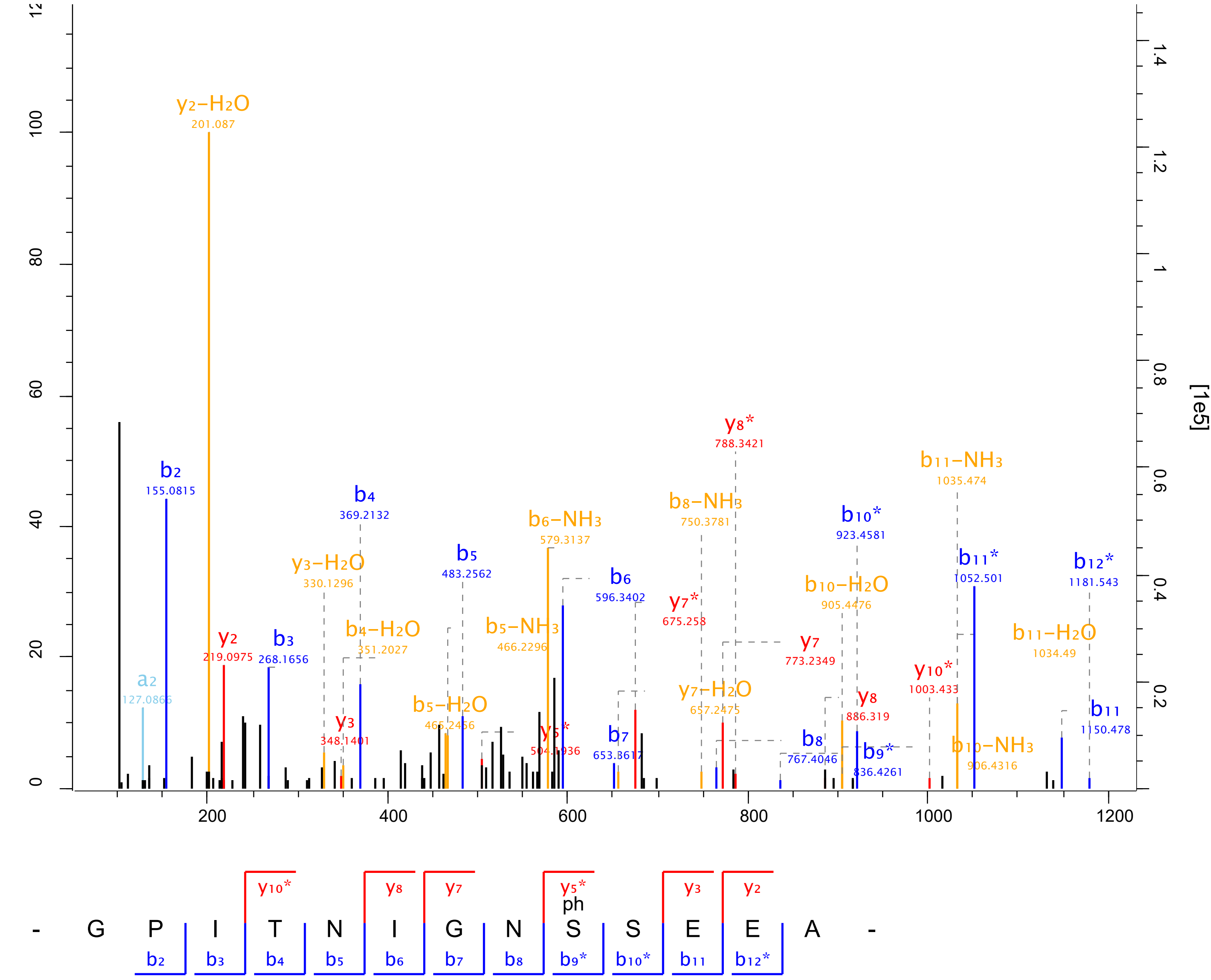

| Raw file | Scan | Method | Score | m/z |
|---|---|---|---|---|
| water_02_1 | 22656 | FTMS; HCD | 59.64 | 803 |

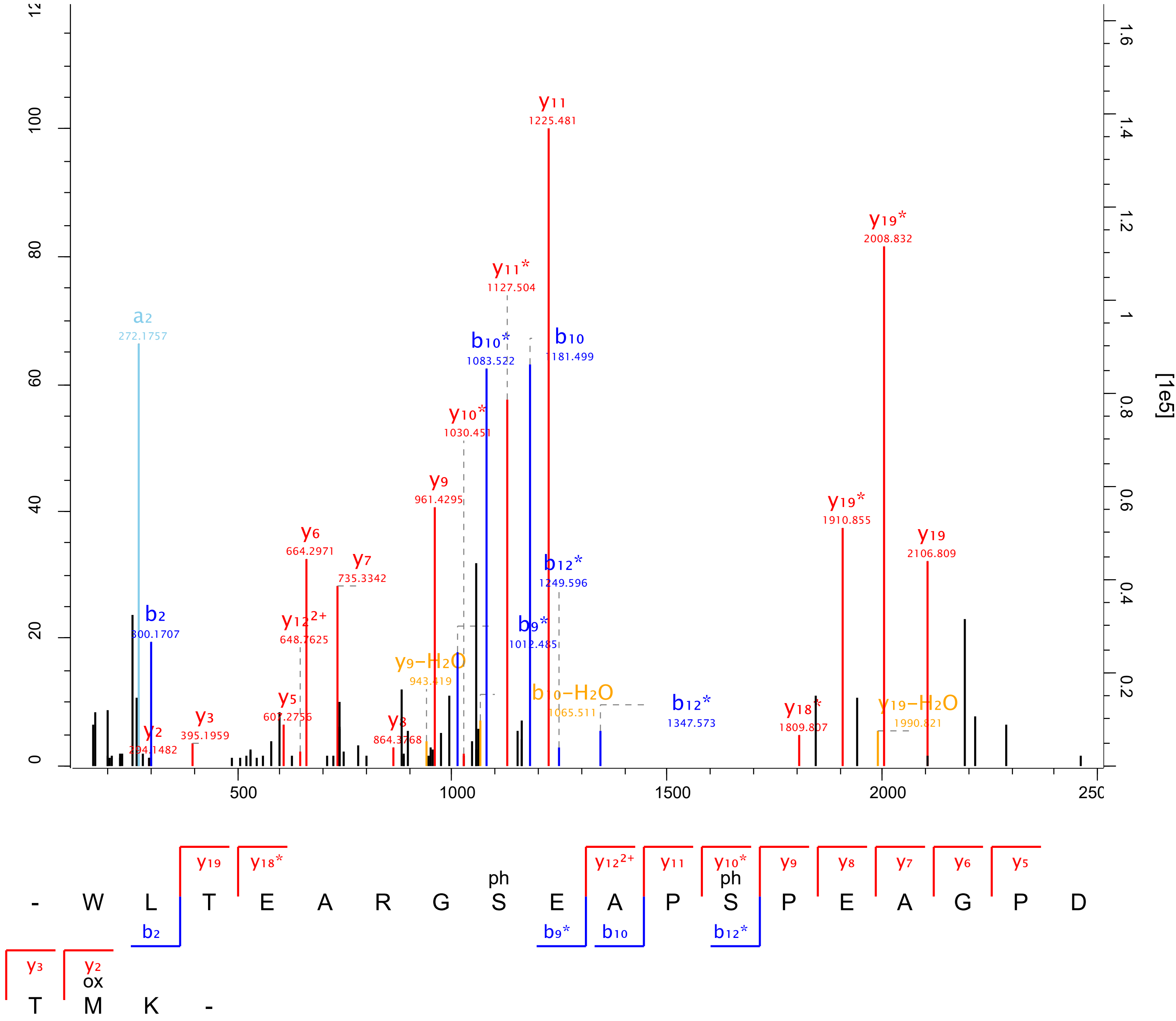

| Raw file | Scan | Method | Score | m/z |
|---|---|---|---|---|
| water_02_1 | 22660 | FTMS; HCD | 74.03 | 999.93 |

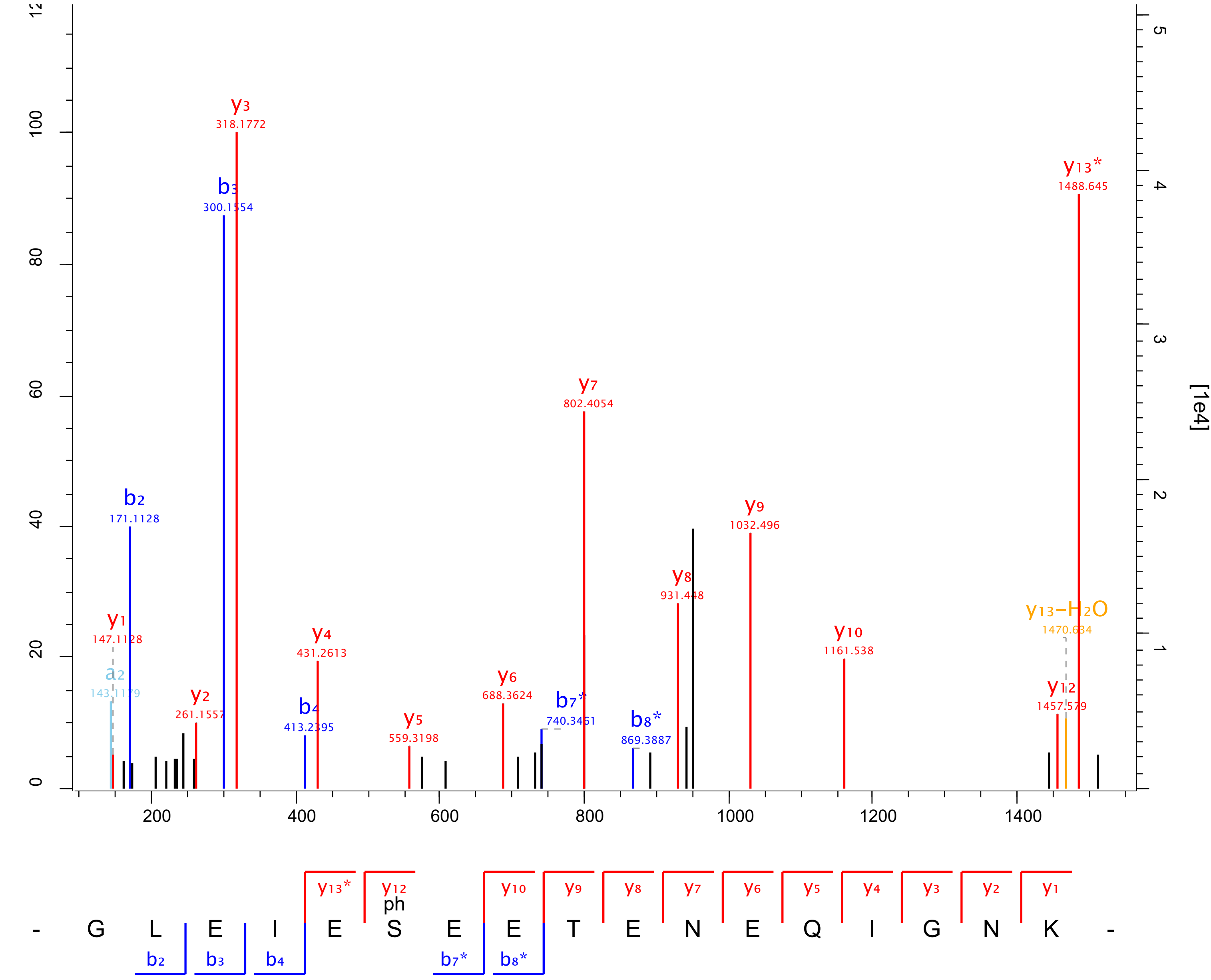

| Raw file | Scan | Method | Score | m/z |
| --- | --- | --- | --- | --- |
| water_02_1 | 22686 | FTMS; HCD | 96.03 | 532.73 |

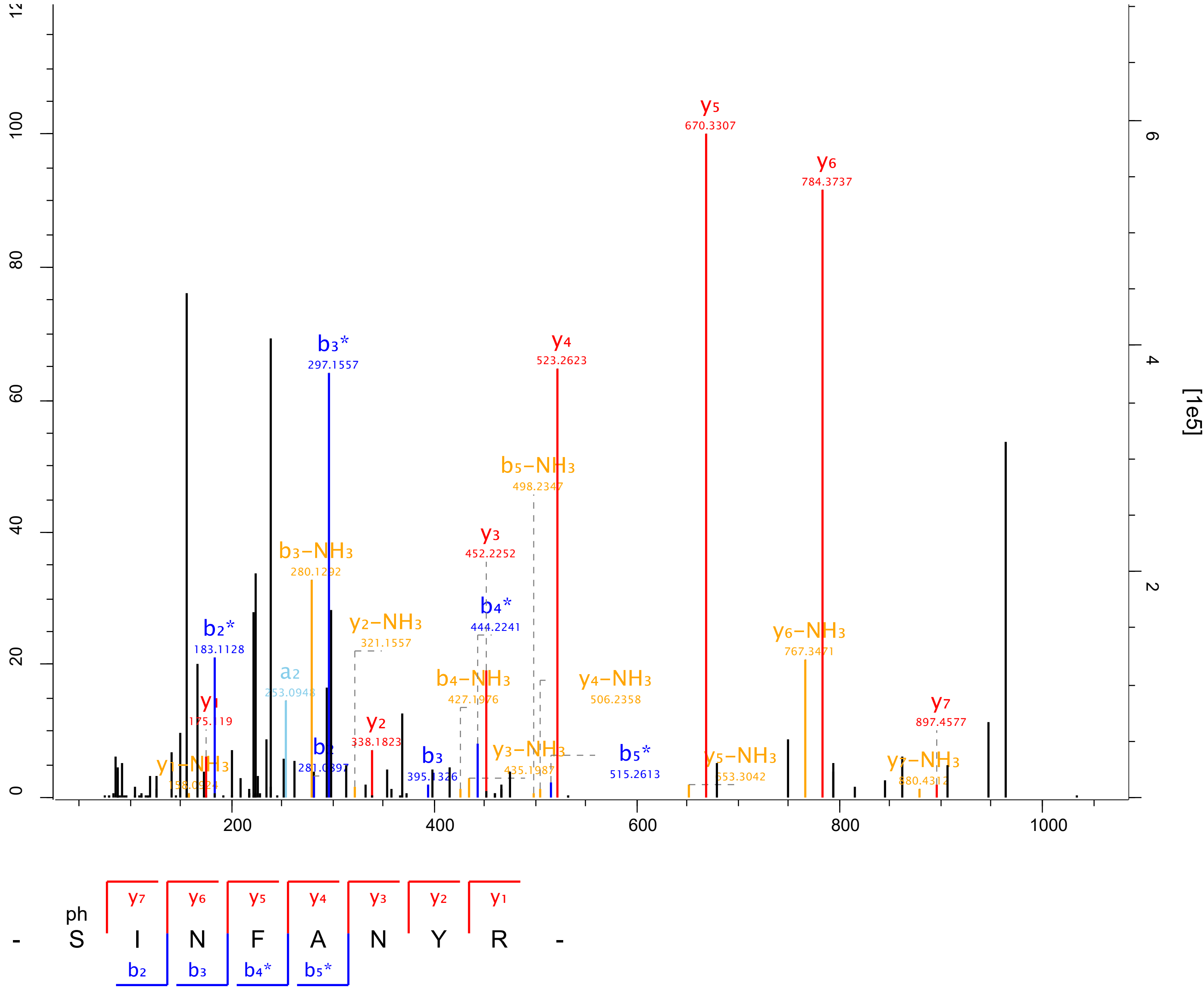

| Raw file | Scan | Method | Score | m/z |
|---|---|---|---|---|
| water_02_1 | 22794 | FTMS; HCD | 132.97 | 559.27 |

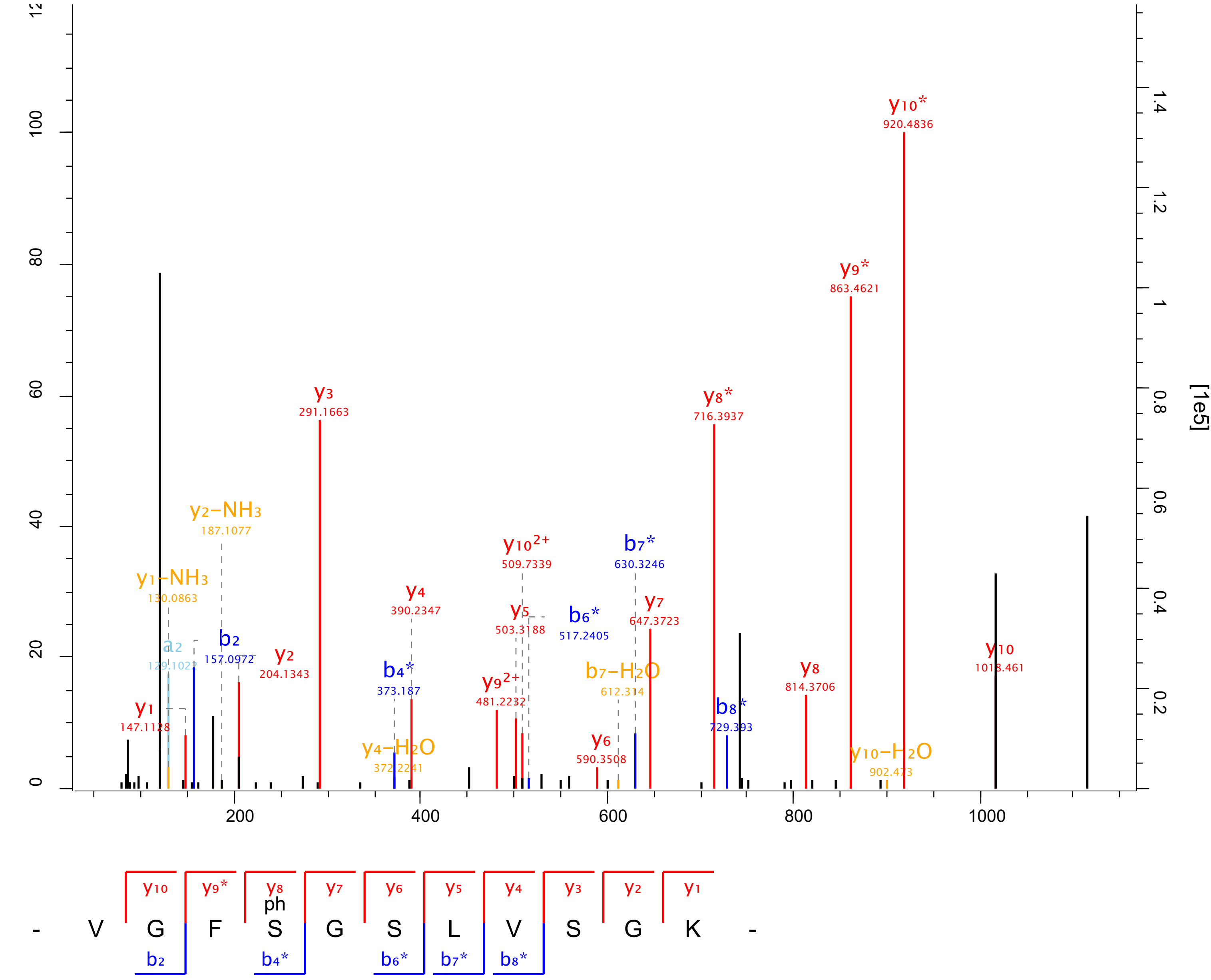

| Raw file | Scan | Method | Score | m/z |
| --- | --- | --- | --- | --- |
| water_02_1 | 2290 | FTMS; HCD | 65.18 | 537.19 |

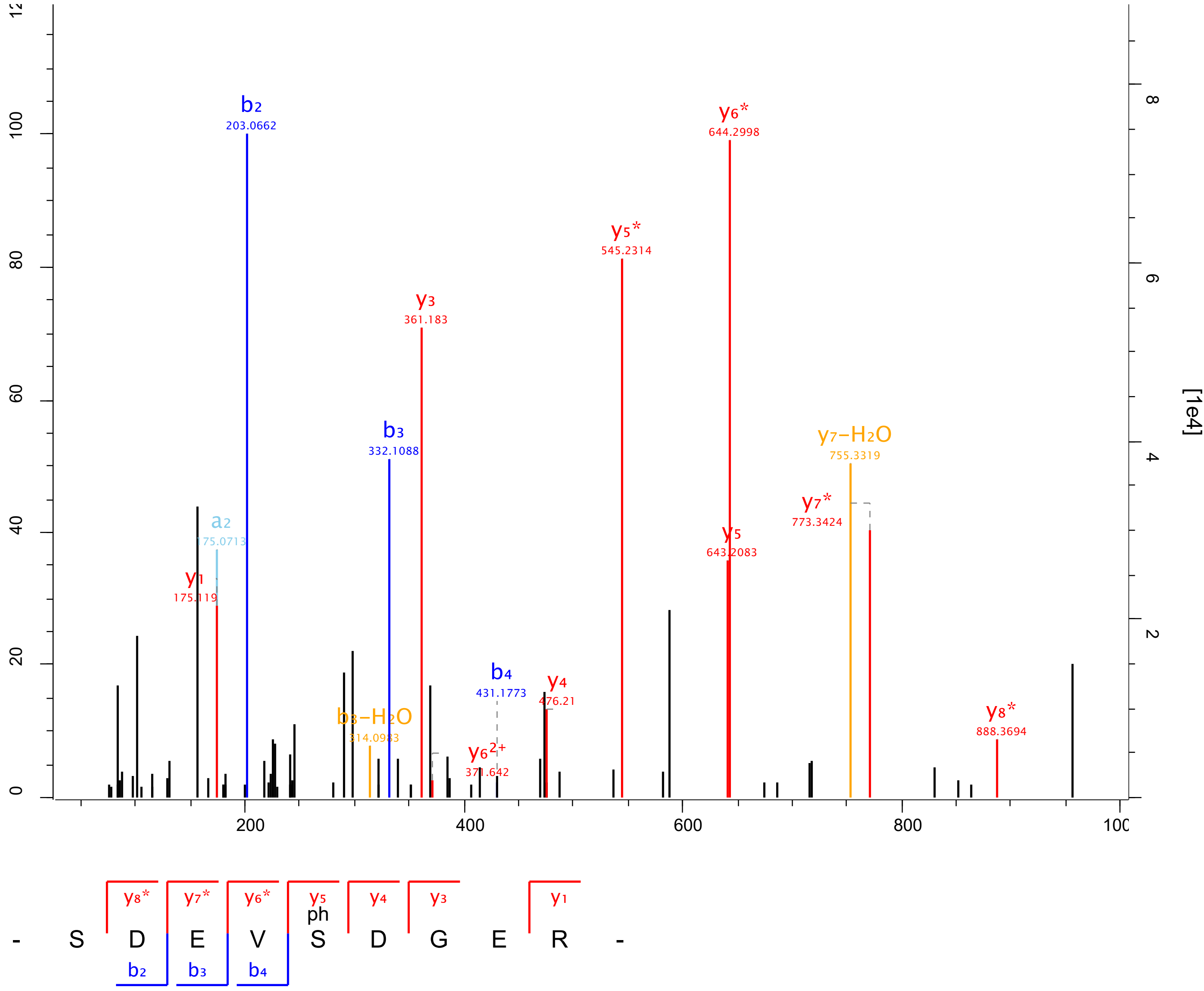

| Raw file | Scan | Method | Score | m/z |
|---|---|---|---|---|
| water_02_1 | 22989 | FTMS; HCD | 256.29 | 1199.98 |

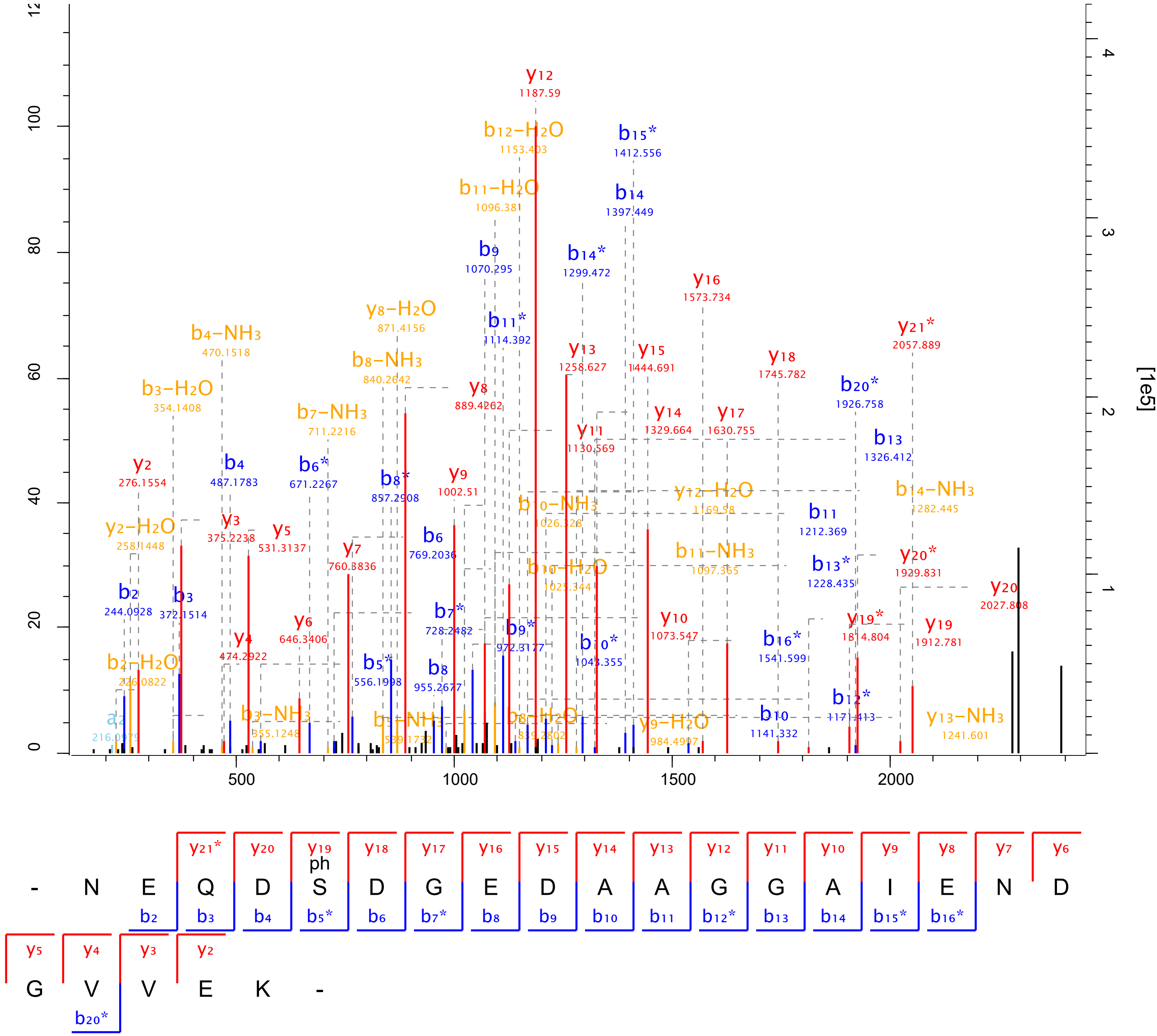

| Raw file | Scan | Method | Score | m/z |
|---|---|---|---|---|
| water_02_1 | 23040 | FTMS; HCD | 114.37 | 1078 |

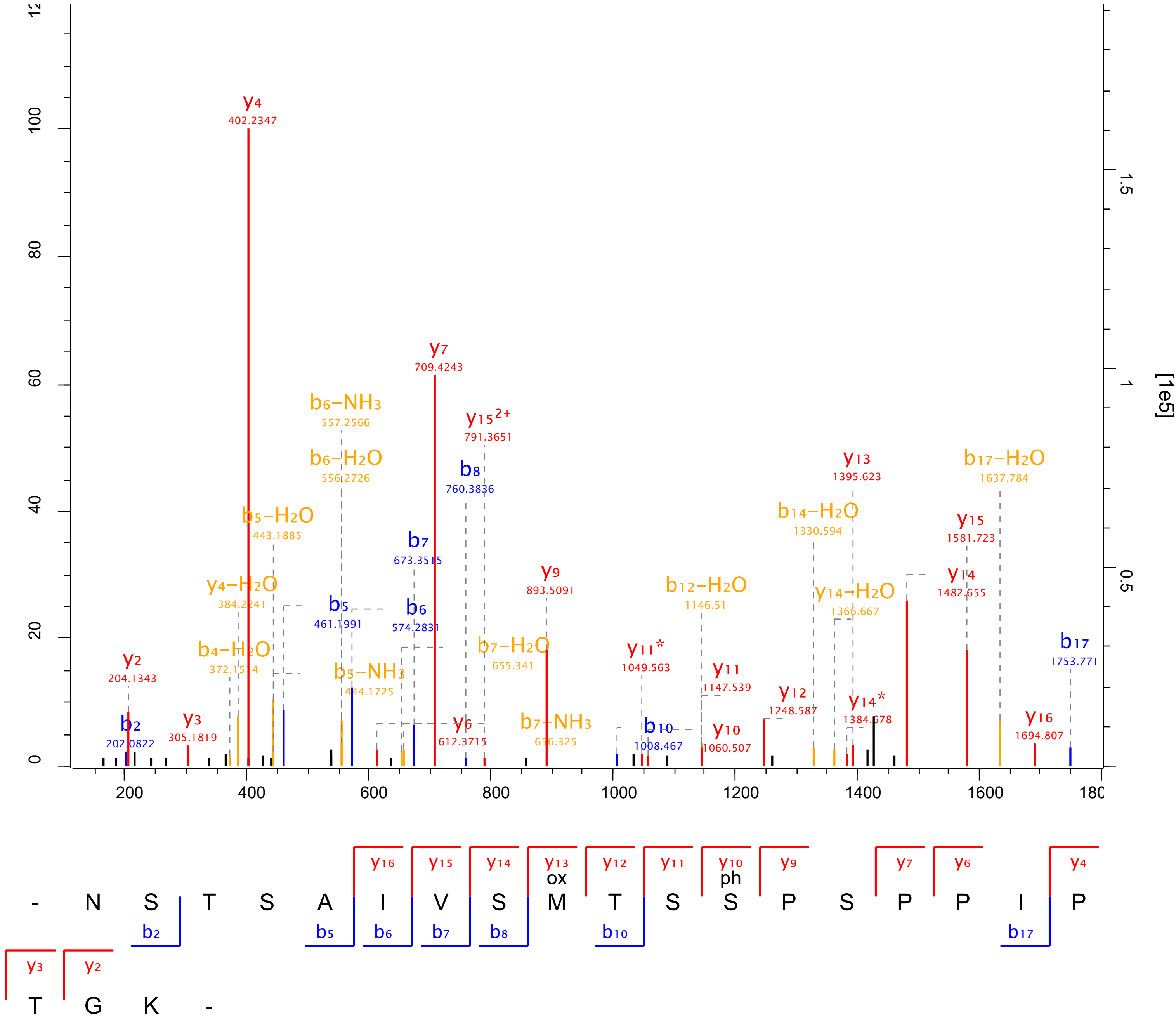

| Raw file | Scan | Method | Score | m/z |
|---|---|---|---|---|
| water_02_1 | 2310 | FTMS; HCD | 92.25 | 582.73 |

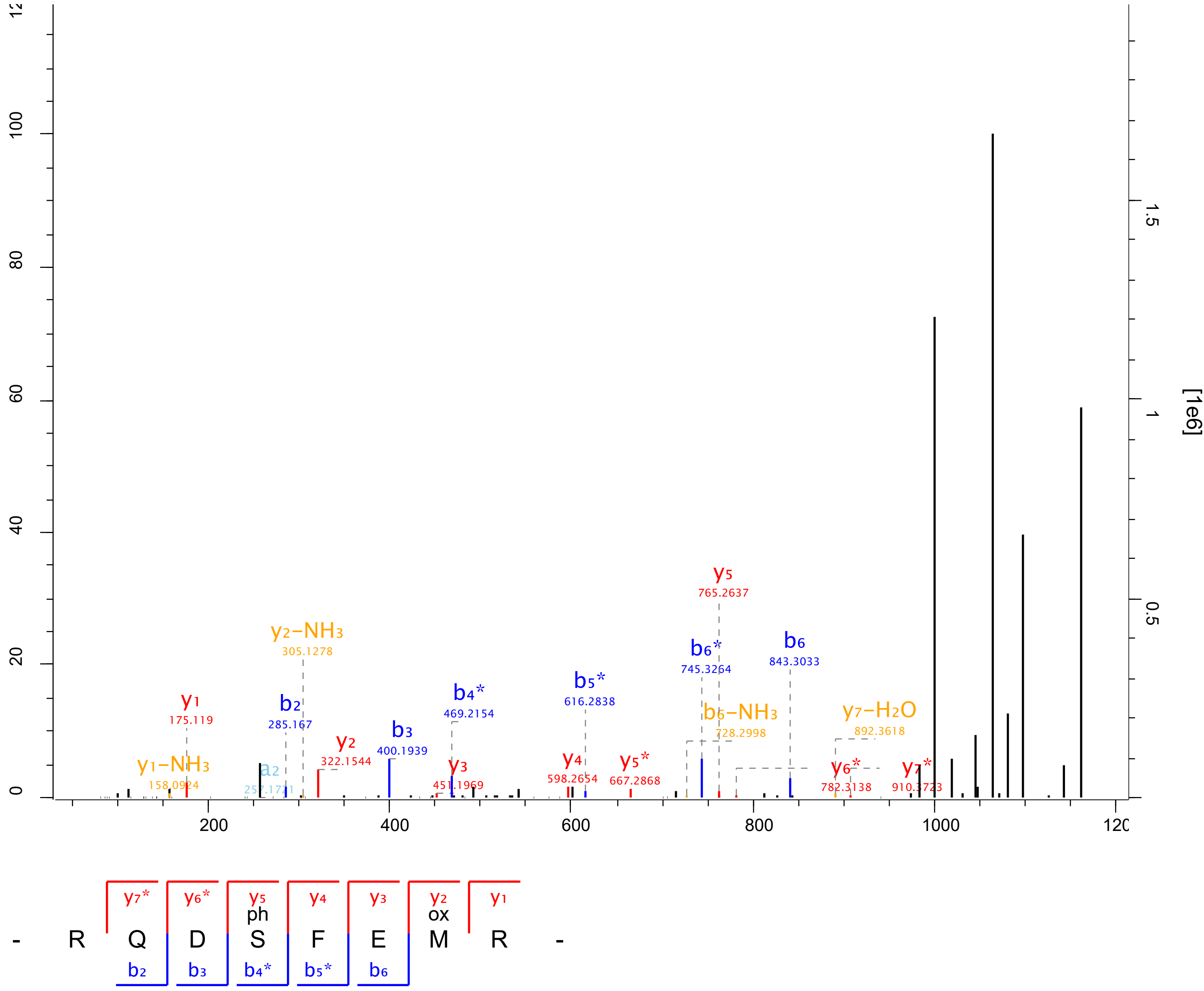

| Raw file | Scan | Method | Score | m/z |
| --- | --- | --- | --- | --- |
| water_02_1 | 23148 | FTMS; HCD | 114.96 | 762.36 |

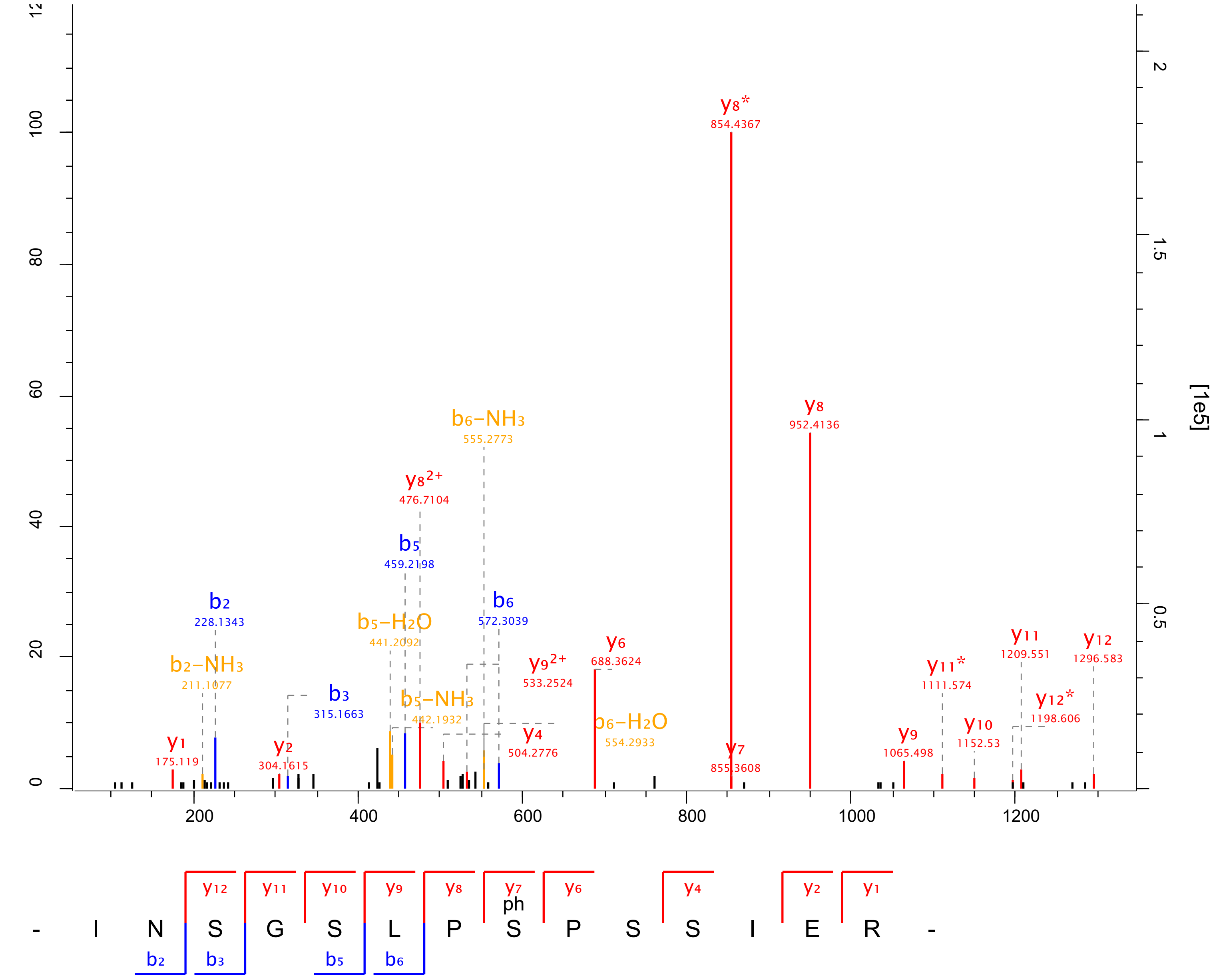

| Raw file | Scan | Method | Score | m/z |
|---|---|---|---|---|
| water_02_1 | 2315 | FTMS; HCD | 64.22 | 683.8 |

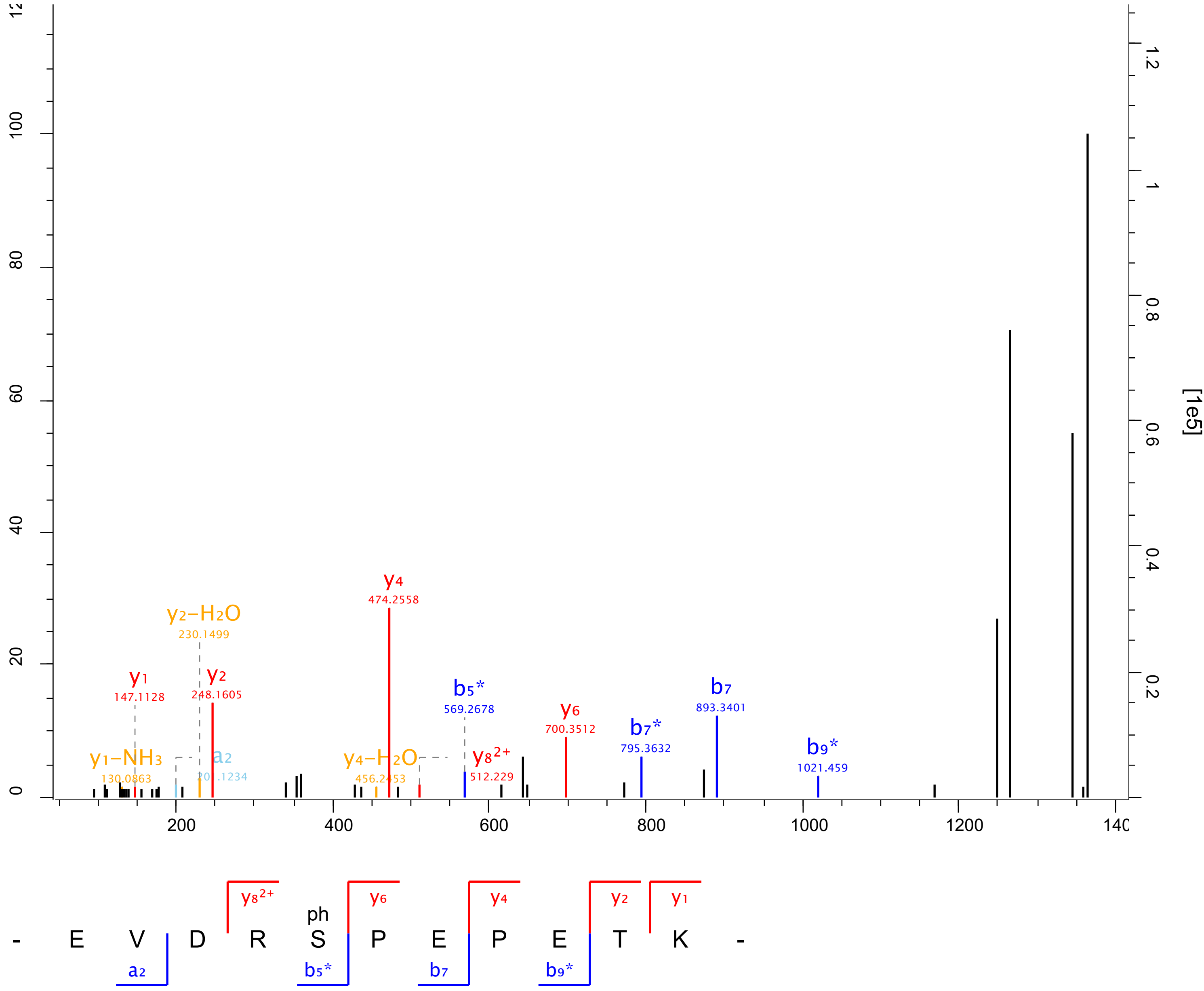

| Raw file | Scan | Method | Score | m/z |
| --- | --- | --- | --- | --- |
| water_02_1 | 23255 | FTMS; HCD | 46.88 | 572.76 |

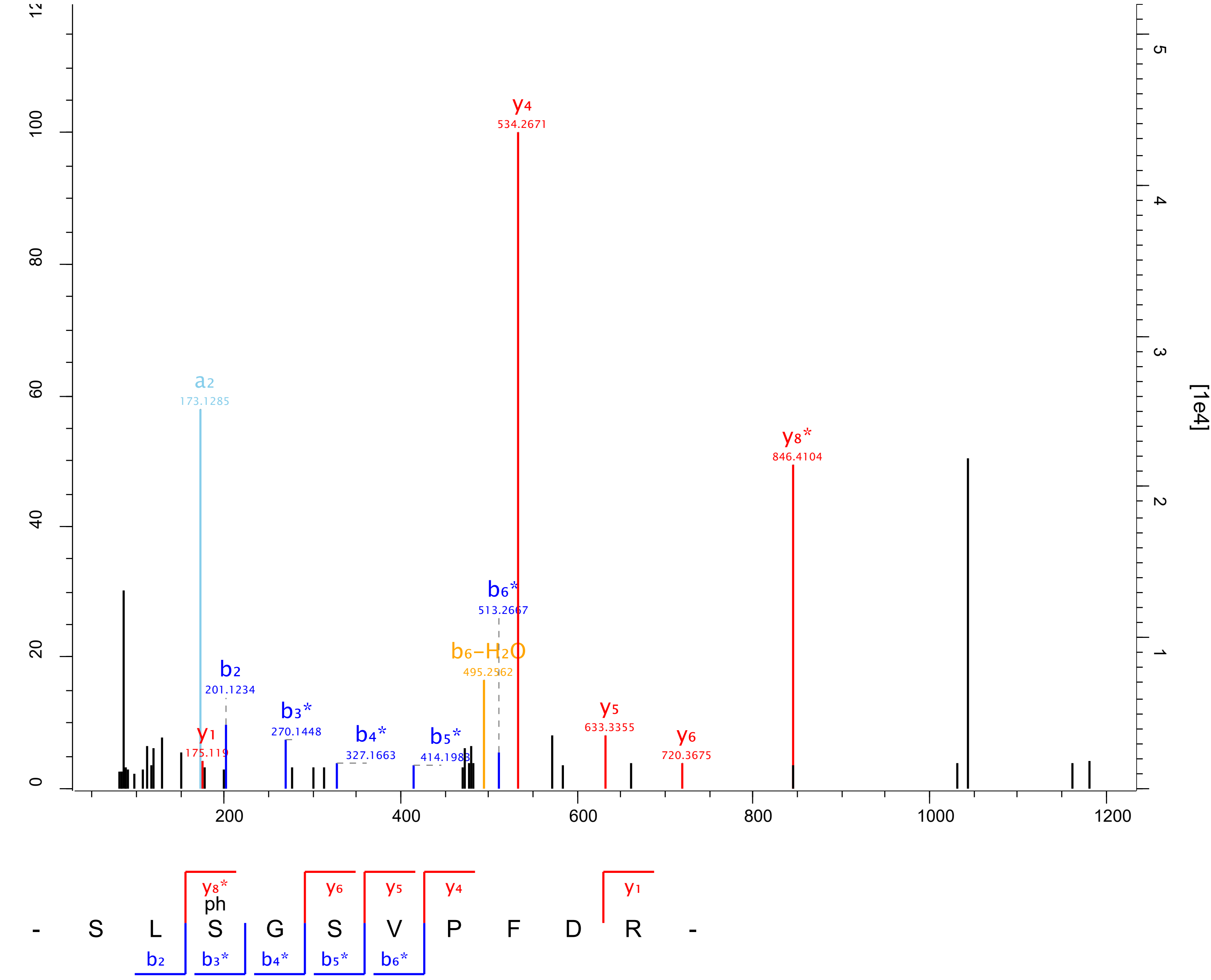

| Raw file | Scan | Method | Score | m/z |
| --- | --- | --- | --- | --- |
| water_02_1 | 23266 | FTMS; HCD | 72.34 | 498.55 |

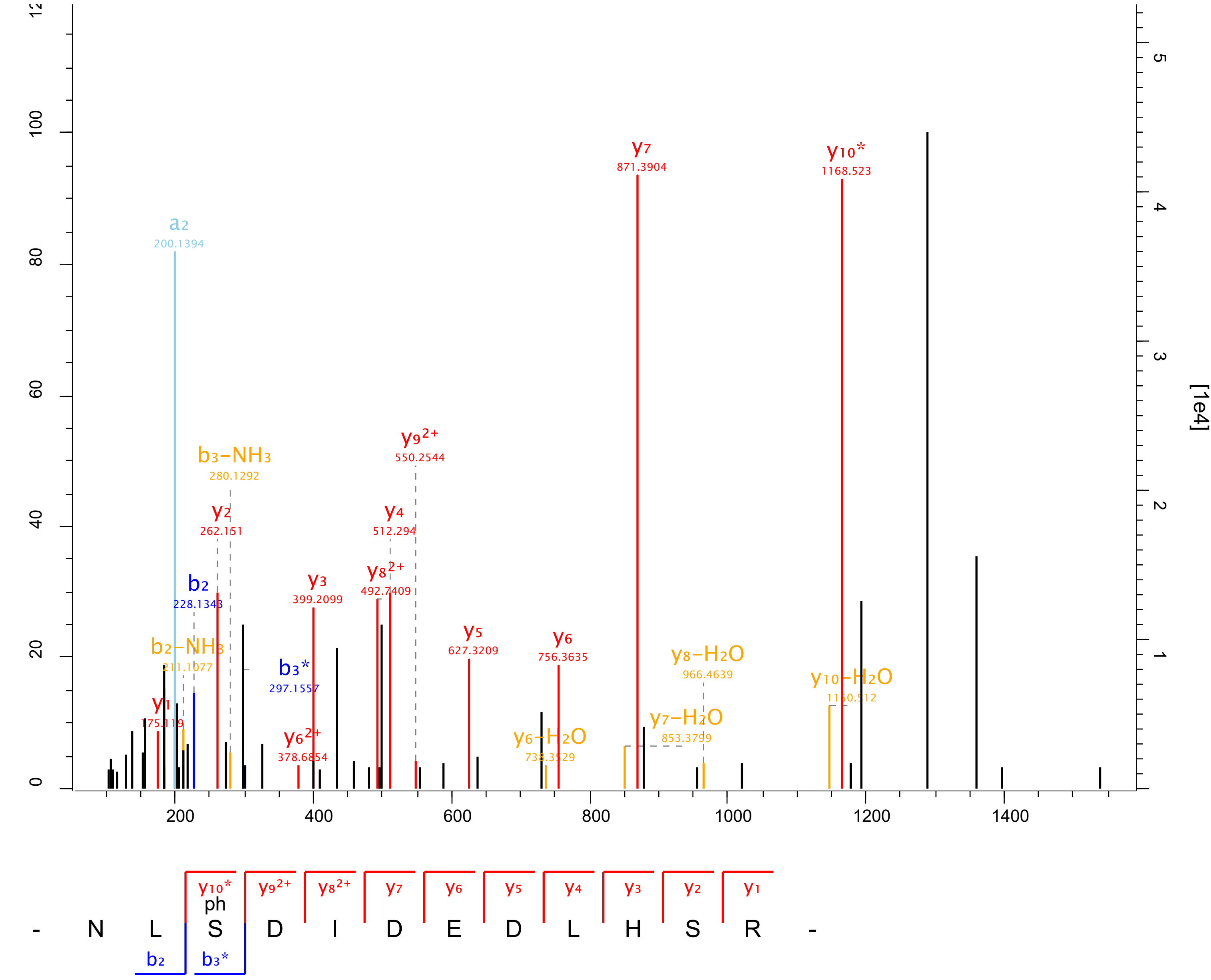

| Raw file | Scan | Method | Score | m/z |
| --- | --- | --- | --- | --- |
| water_02_1 | 2335 | FTMS; HCD | 85.55 | 555.7 |

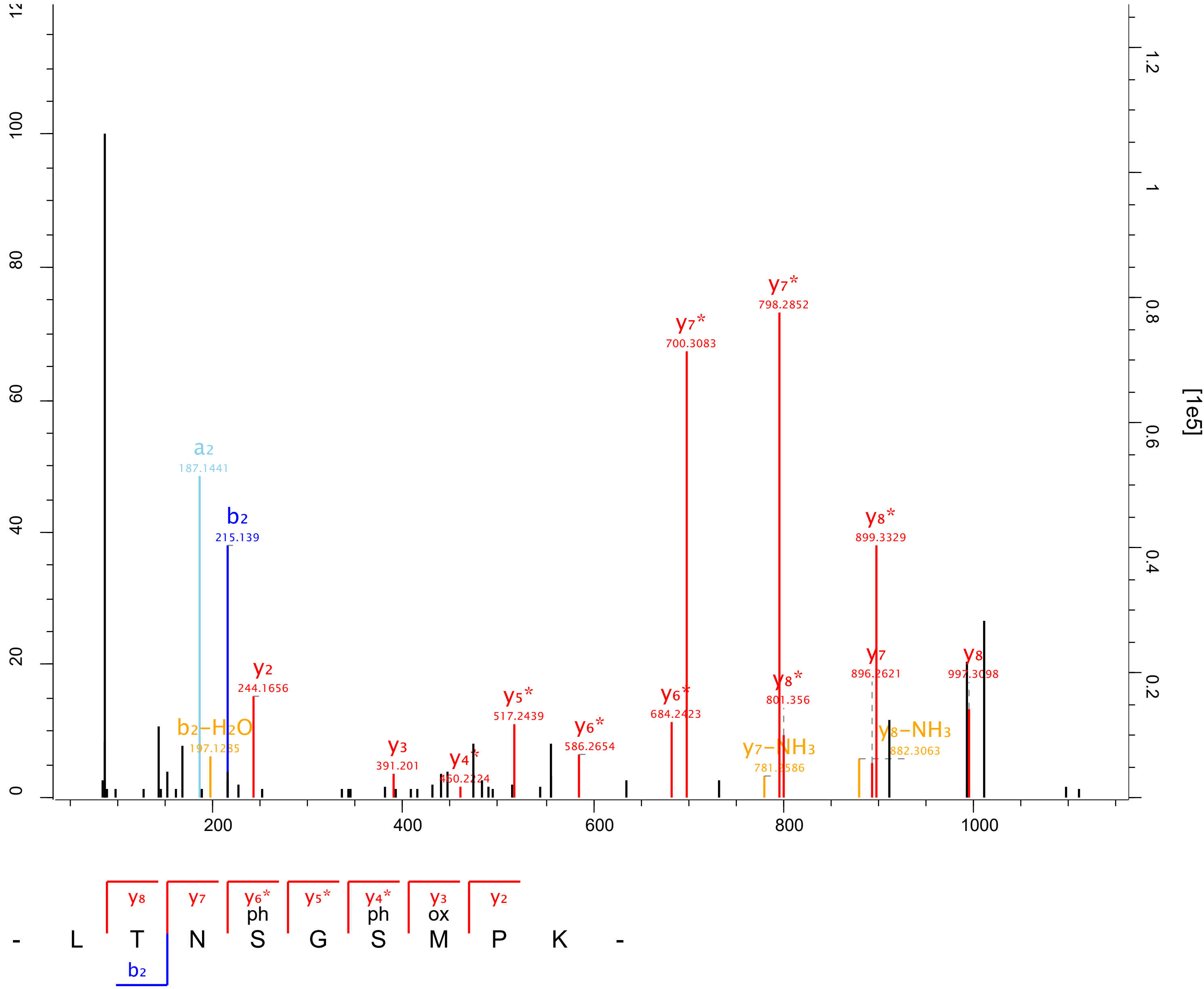

| Raw file | Scan | Method | Score | m/z |
| --- | --- | --- | --- | --- |
| water_02_1 | 23496 | FTMS; HCD | 59.73 | 766.32 |

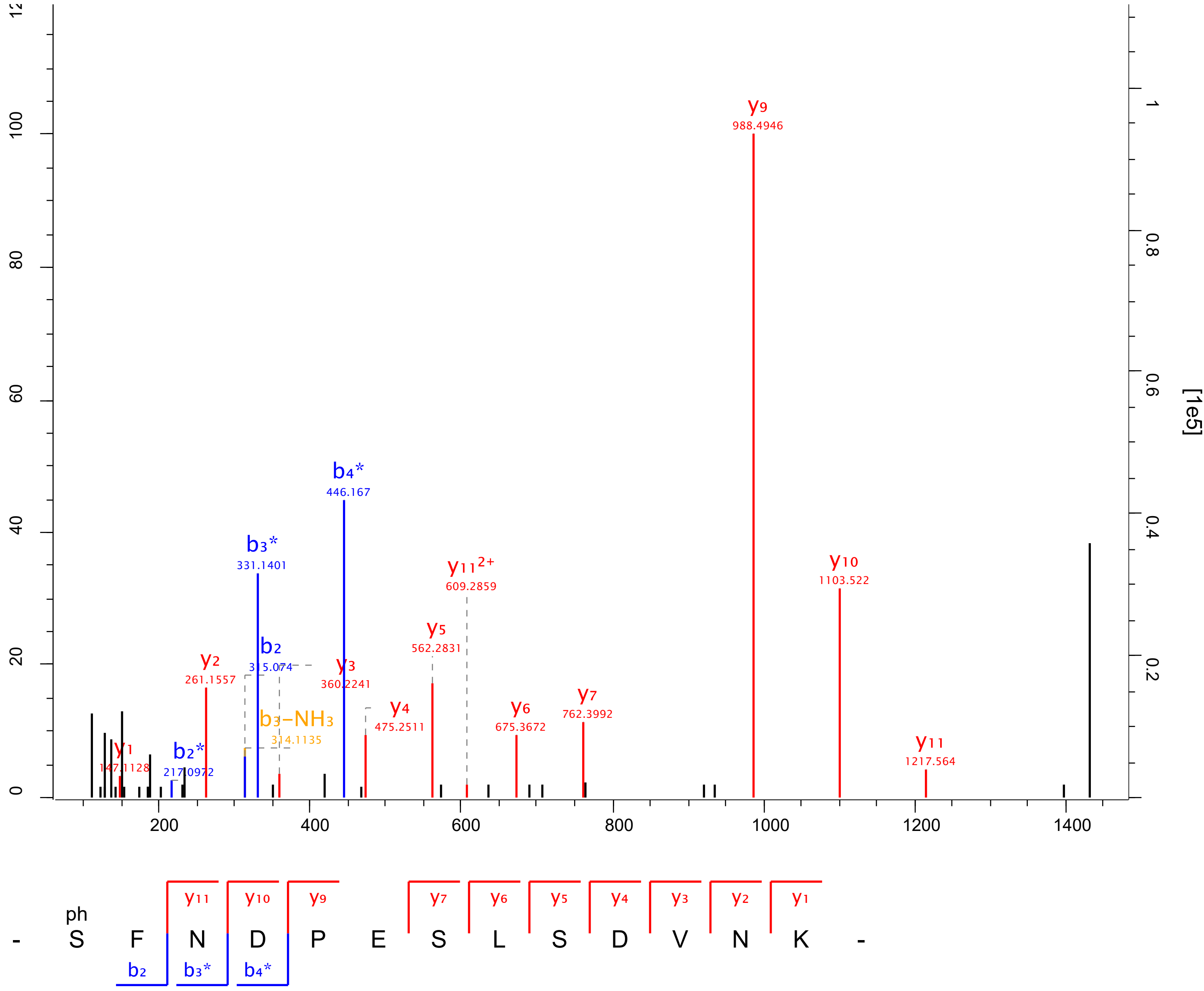

| Raw file | Scan | Method | Score | m/z |
| --- | --- | --- | --- | --- |
| water_02_1 | 23634 | FTMS; HCD | 80.63 | 740.27 |

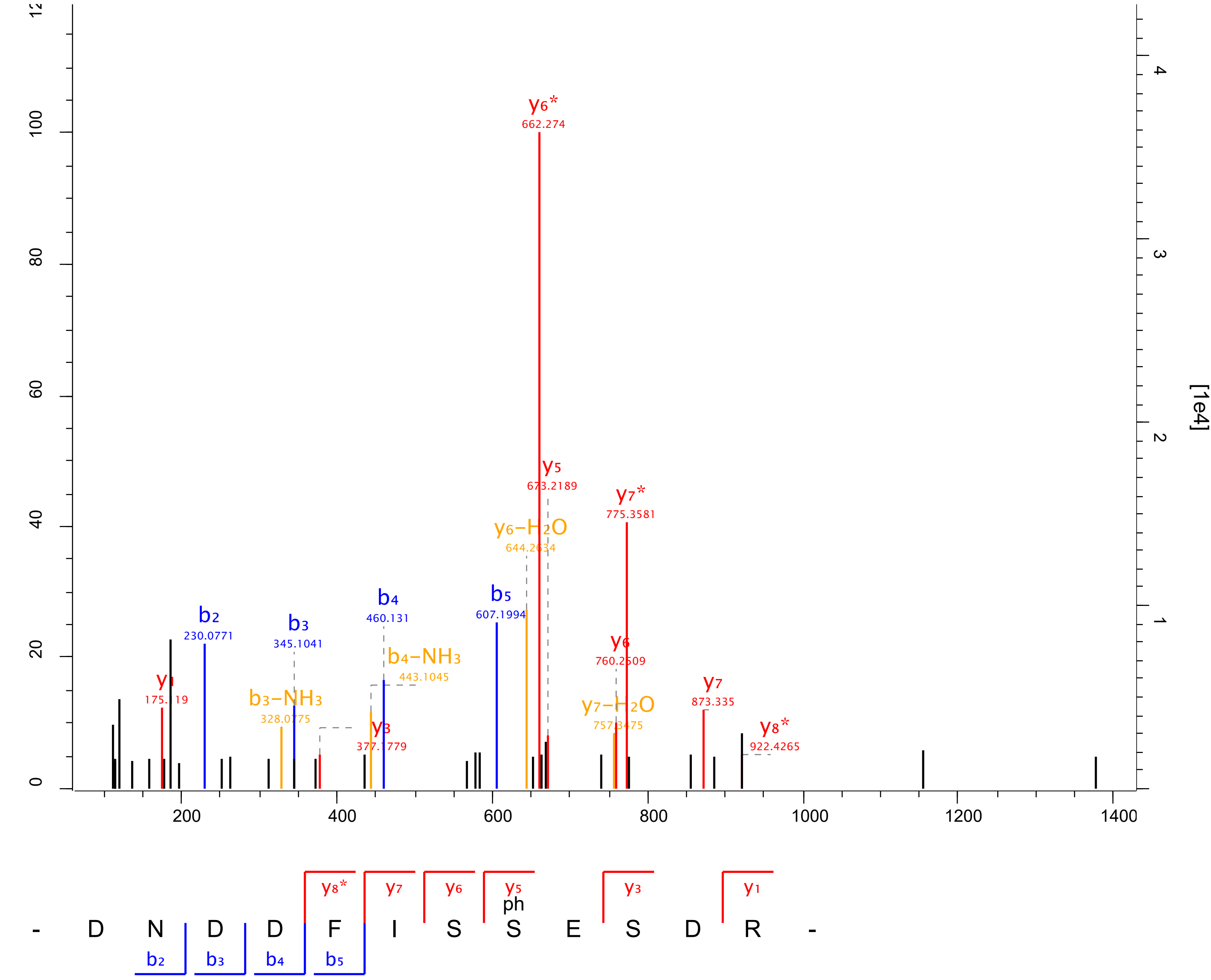

| Raw file | Scan | Method | Score | m/z |
|---|---|---|---|---|
| water_02_1 | 23686 | FTMS; HCD | 64.45 | 794.83 |

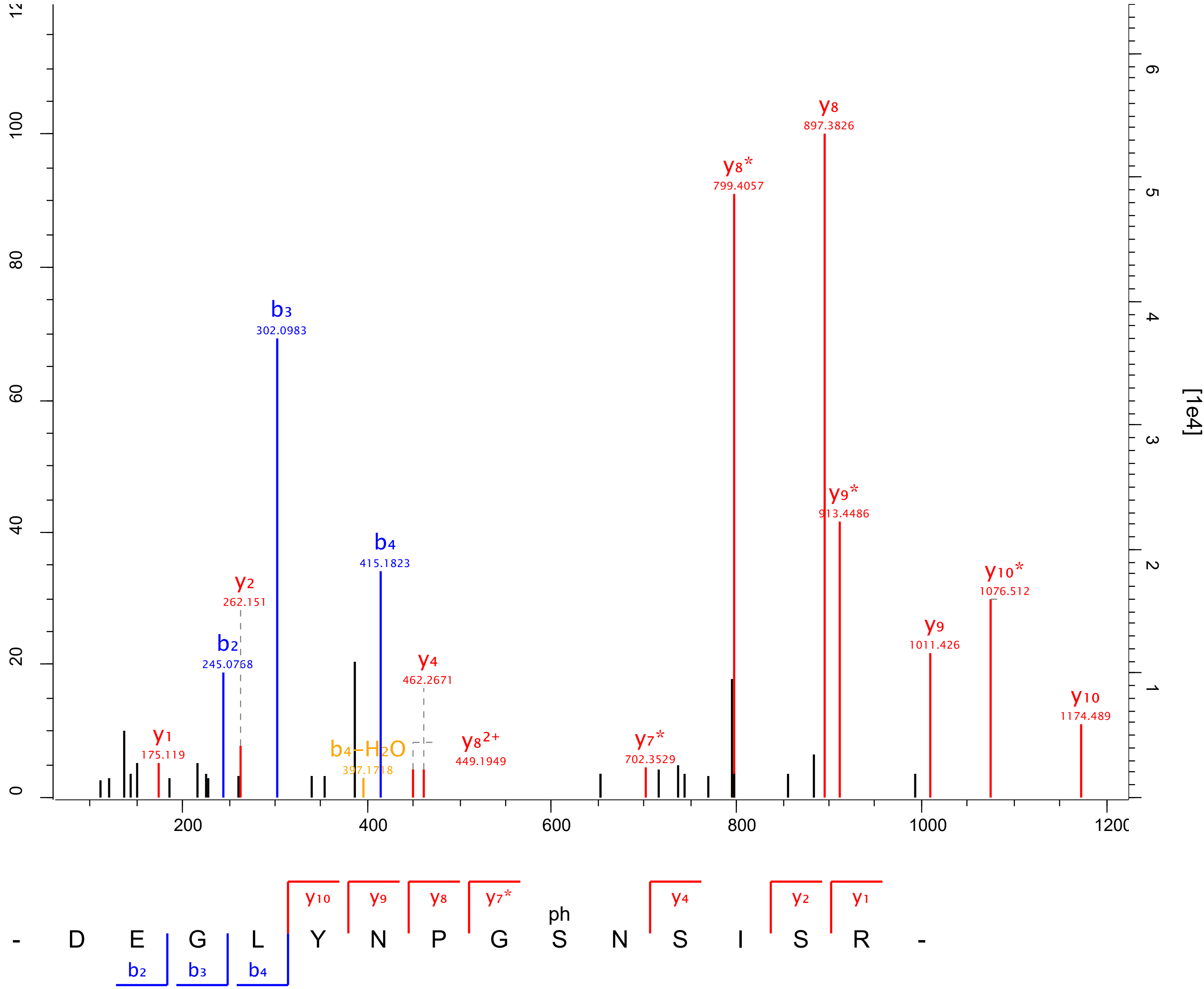

| Raw file | Scan | Method | Score | m/z |
| --- | --- | --- | --- | --- |
| water_02_1 | 24055 | FTMS; HCD | 75.55 | 875.39 |

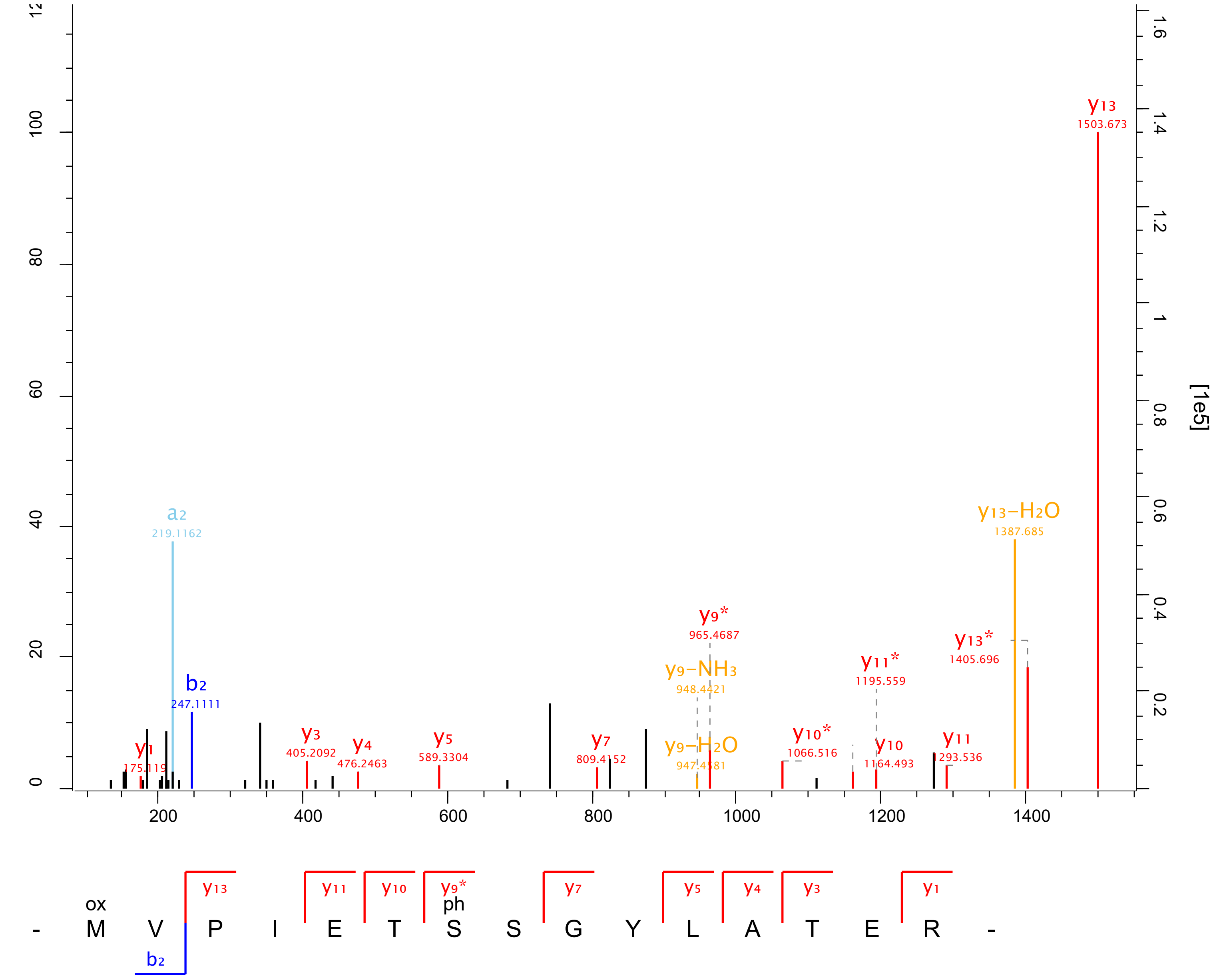

| Raw file | Scan | Method | Score | m/z |
|---|---|---|---|---|
| water_02_1 | 24057 | FTMS; HCD | 179.14 | 749.36 |

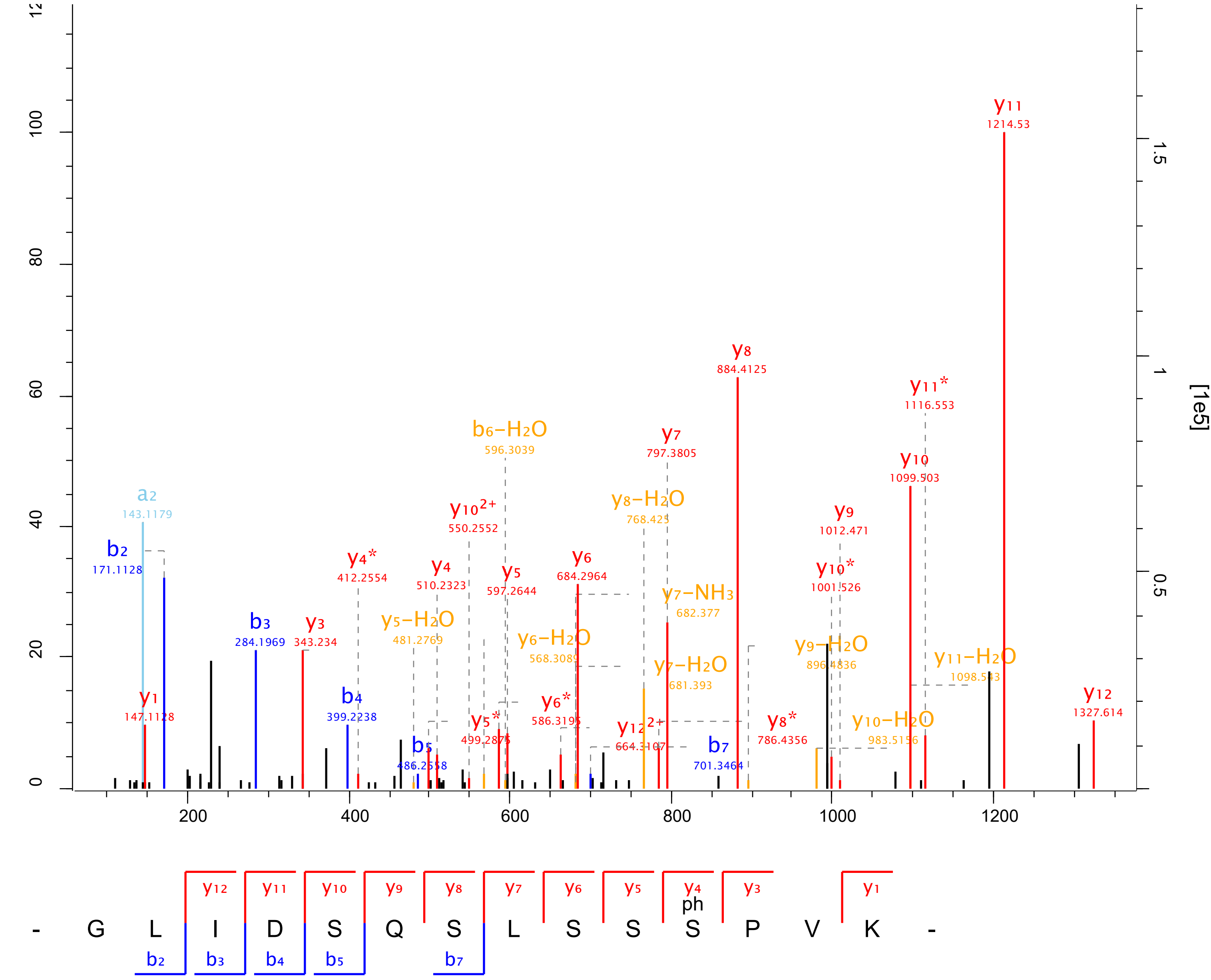

| Raw file | Scan | Method | Score | m/z |
| --- | --- | --- | --- | --- |
| water_02_1 | 24089 | FTMS; HCD | 78.67 | 550.61 |

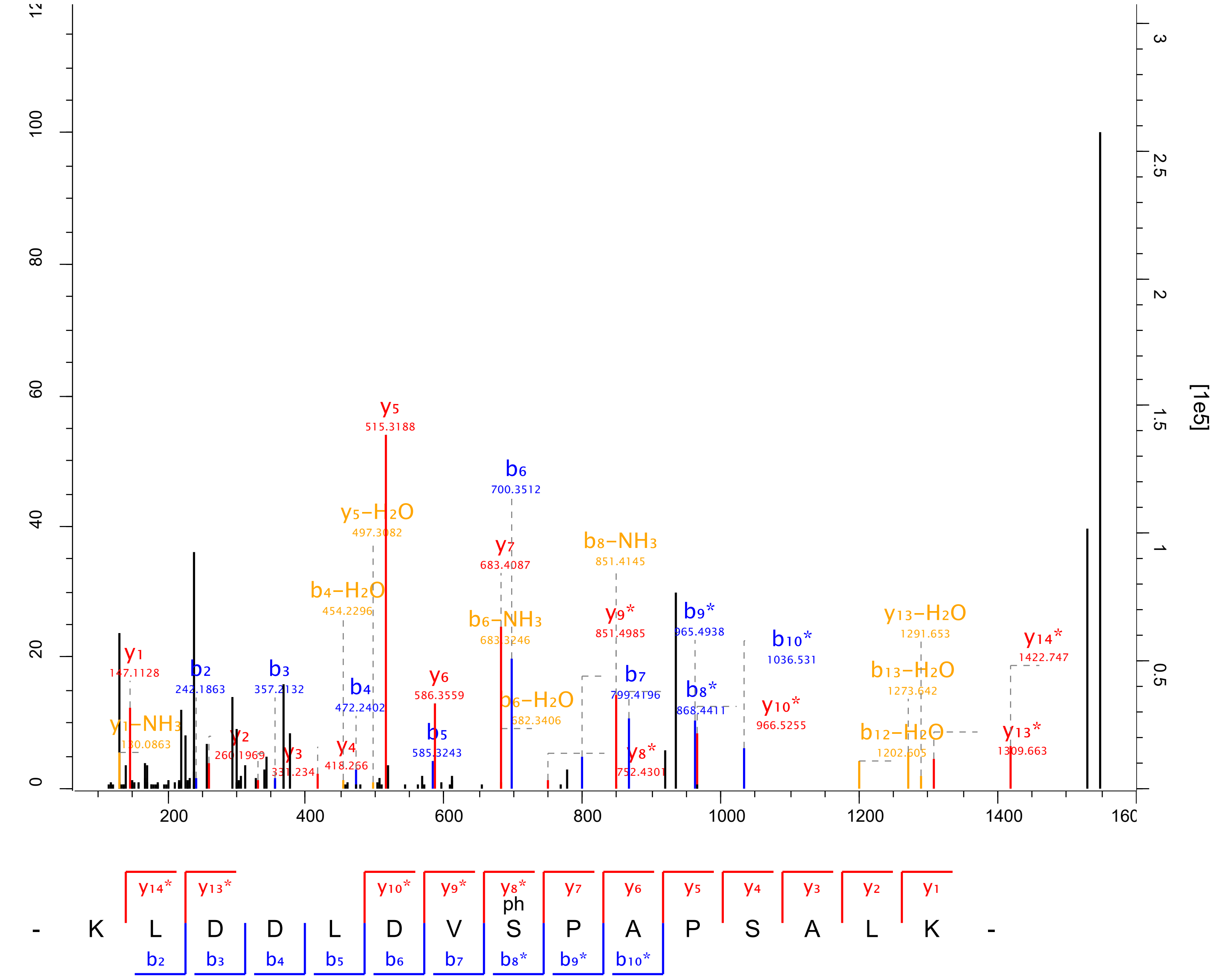

| Raw file | Scan | Method | Score | m/z |
| --- | --- | --- | --- | --- |
| water_02_1 | 24285 | FTMS; HCD | 45.04 | 817.06 |

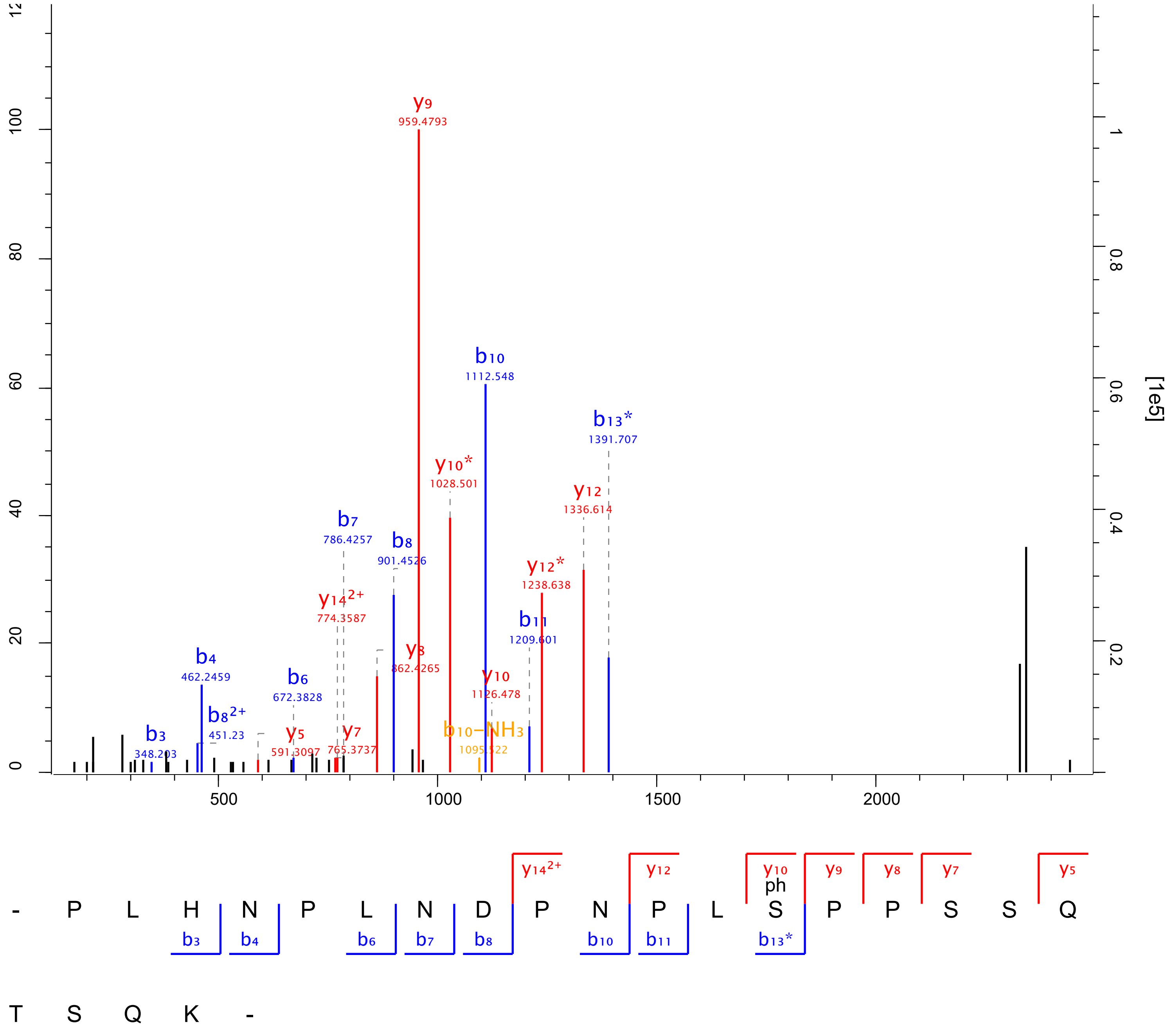

| Raw file | Scan | Method | Score | m/z |
|---|---|---|---|---|
| water_02_1 | 24412 | FTMS; HCD | 131.25 | 762.36 |

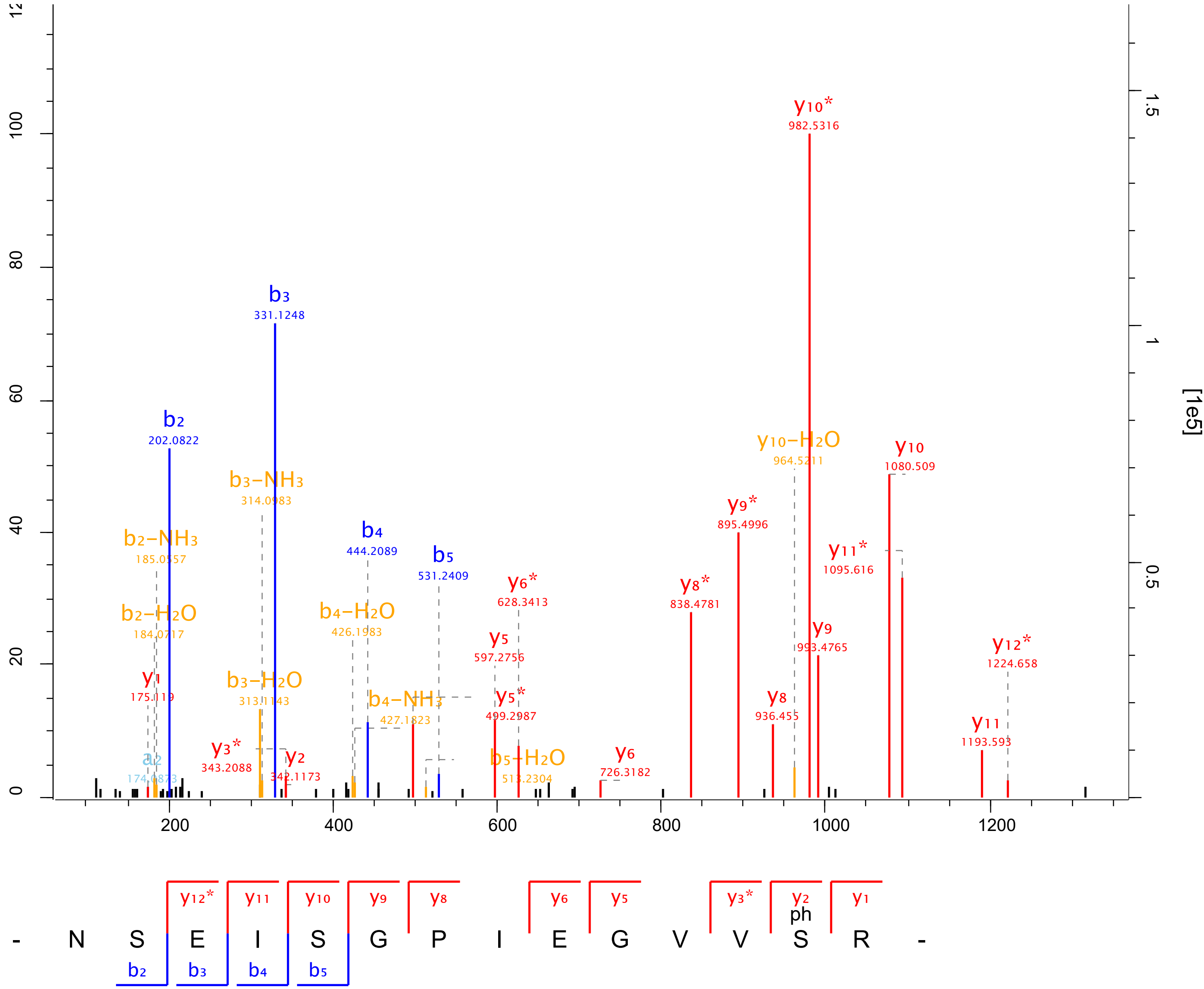

| Raw file | Scan | Method | Score | m/z |
|---|---|---|---|---|
| water_02_1 | 24492 | FTMS; HCD | 250.79 | 949.39 |

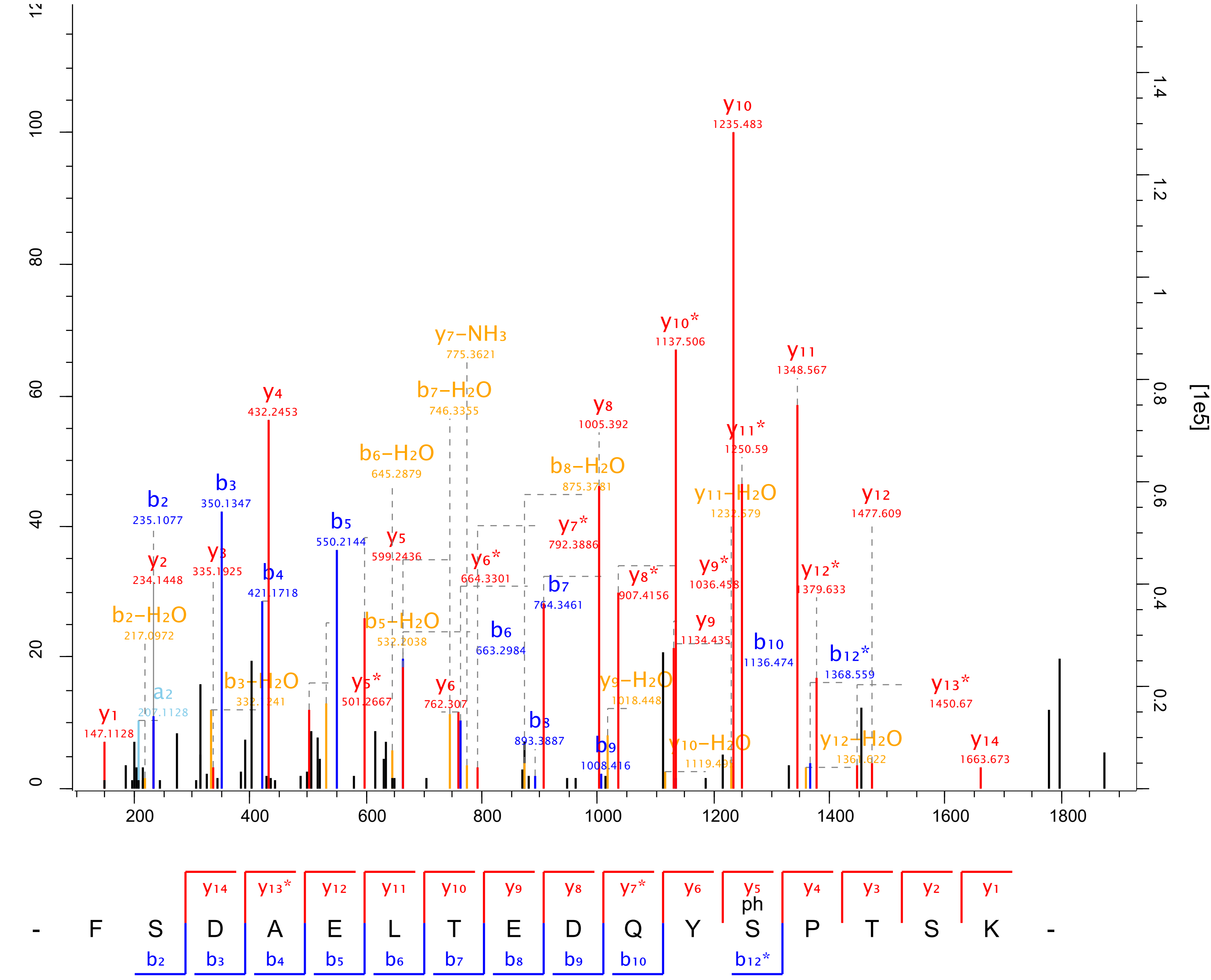

| Raw file | Scan | Method | Score | m/z |
| --- | --- | --- | --- | --- |
| water_02_1 | 24577 | FTMS; HCD | 181.81 | 1027.12 |

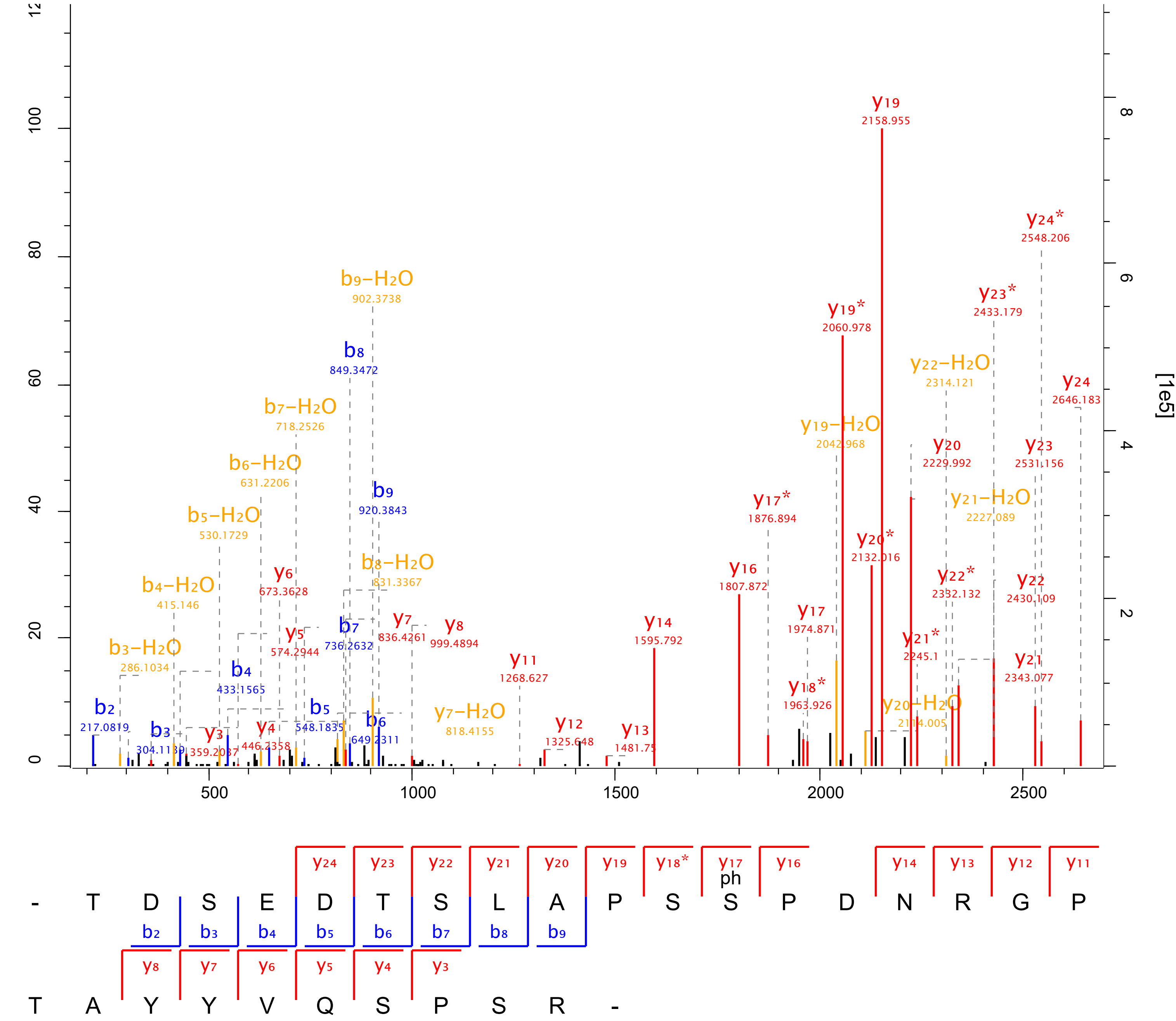

| Raw file | Scan | Method | Score | m/z |
|---|---|---|---|---|
| water_02_1 | 24598 | FTMS; HCD | 52.18 | 892.34 |

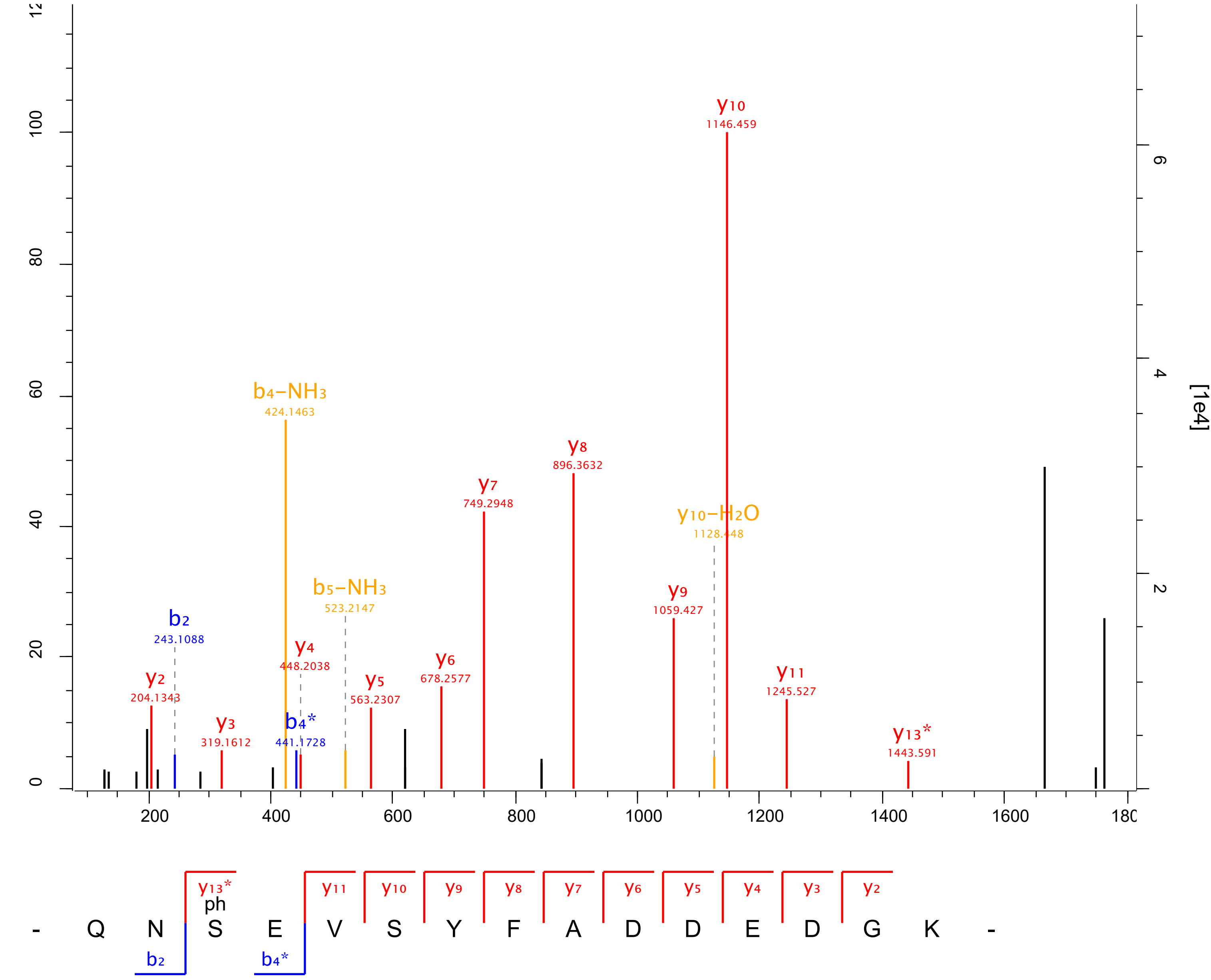

| Raw file | Scan | Method | Score | m/z |
|---|---|---|---|---|
| water_02_1 | 24652 | FTMS; HCD | 117.3 | 814.85 |

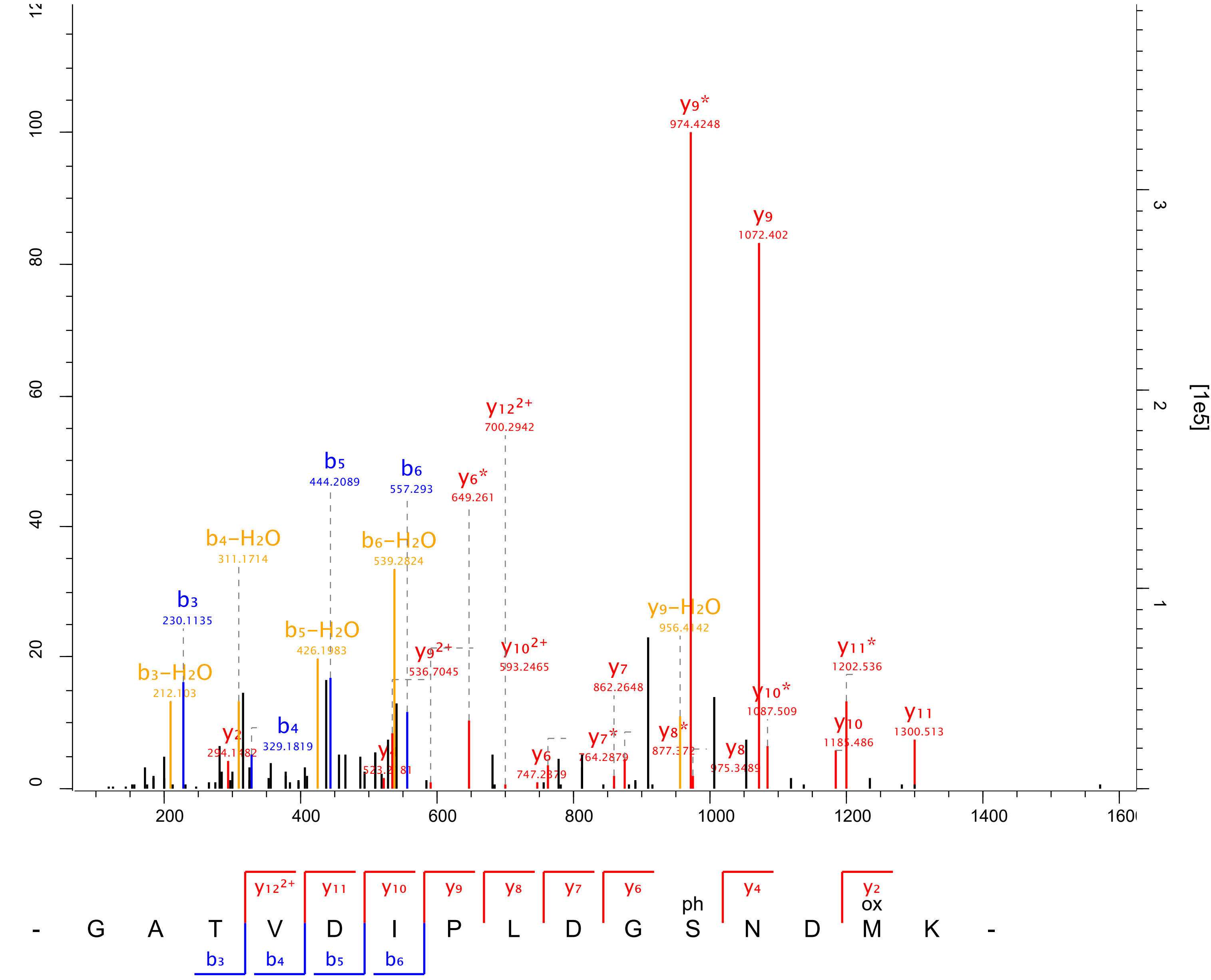

| Raw file | Scan | Method | Score | m/z |
|---|---|---|---|---|
| water_02_1 | 24660 | FTMS; HCD | 56.51 | 805.35 |

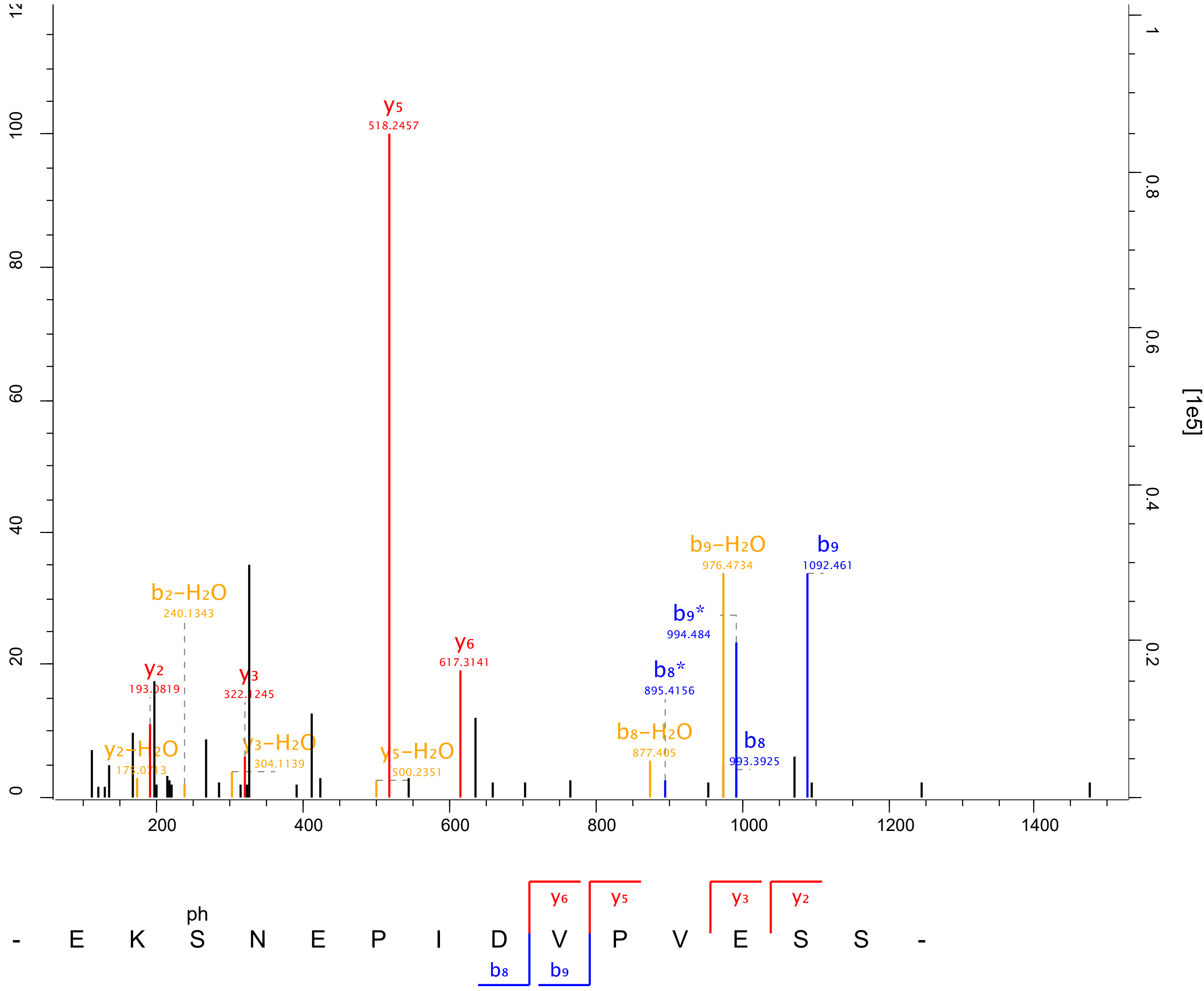

| Raw file | Scan | Method | Score | m/z |
| --- | --- | --- | --- | --- |
| water_02_1 | 24704 | FTMS; HCD | 189.51 | 889.84 |

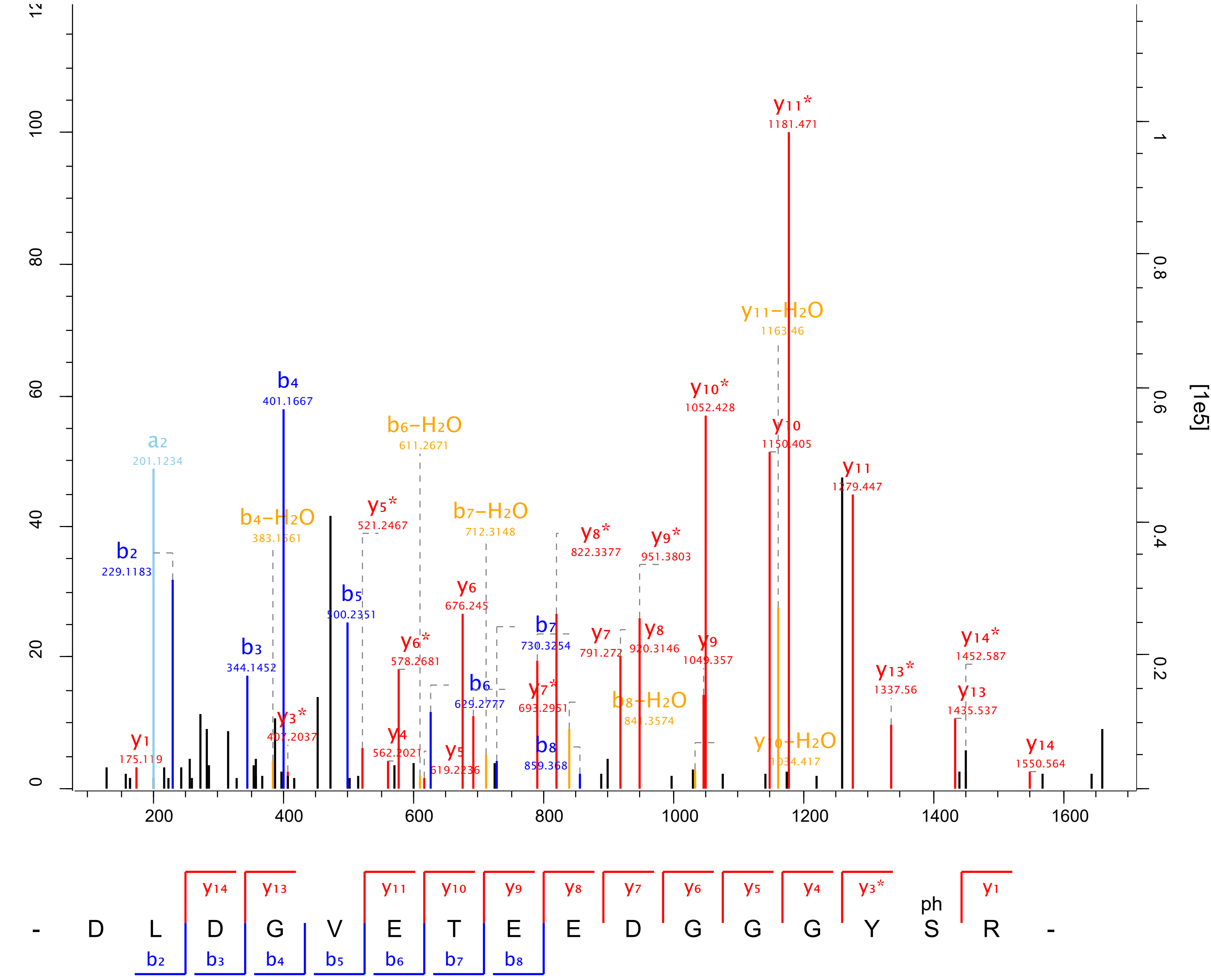

| Raw file | Scan | Method | Score | m/z |
| --- | --- | --- | --- | --- |
| water_02_1 | 24722 | FTMS; HCD | 59.37 | 824.37 |

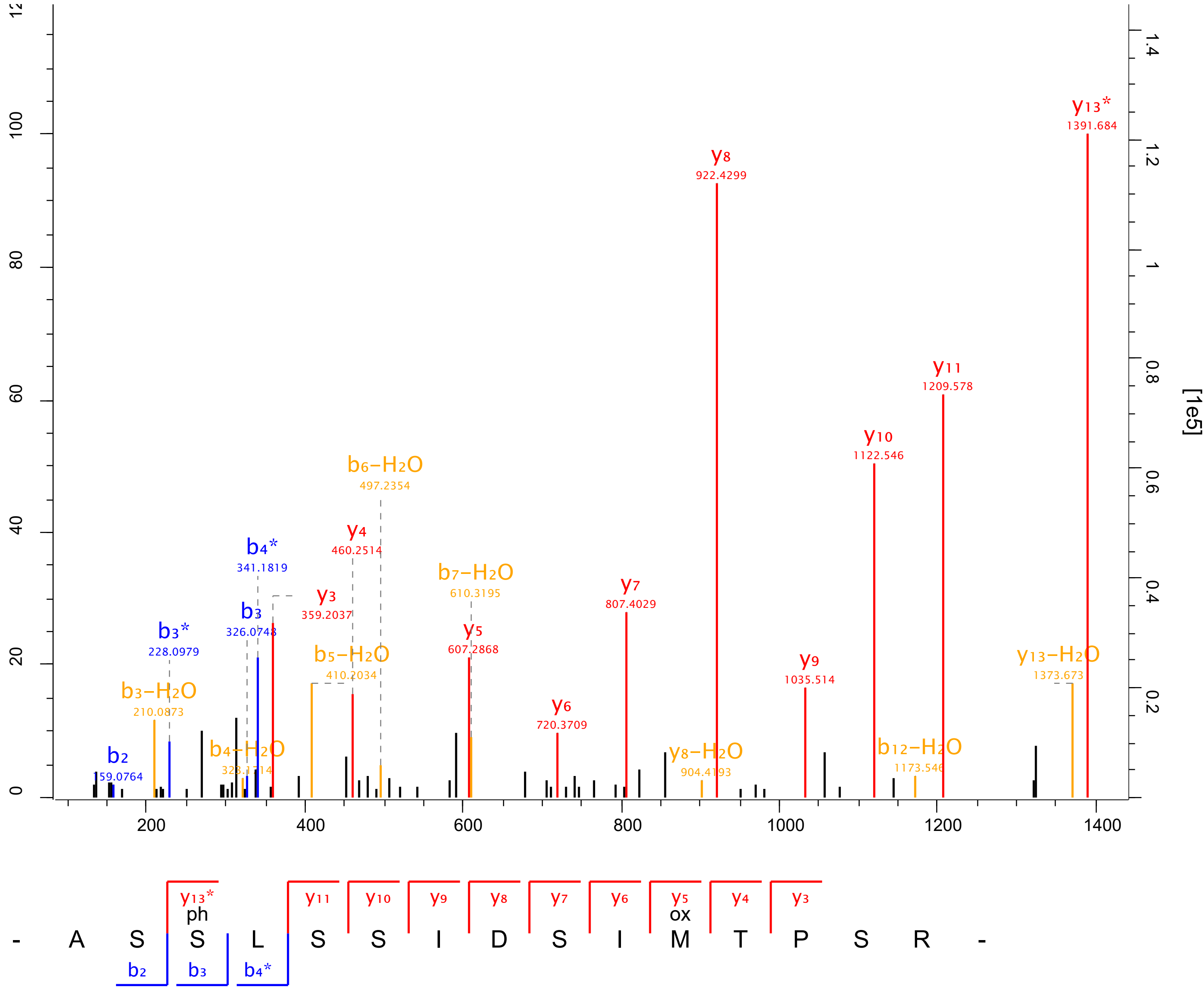

| Raw file | Scan | Method | Score | m/z |
| --- | --- | --- | --- | --- |
| water_02_1 | 24735 | FTMS; HCD | 91.59 | 773.65 |

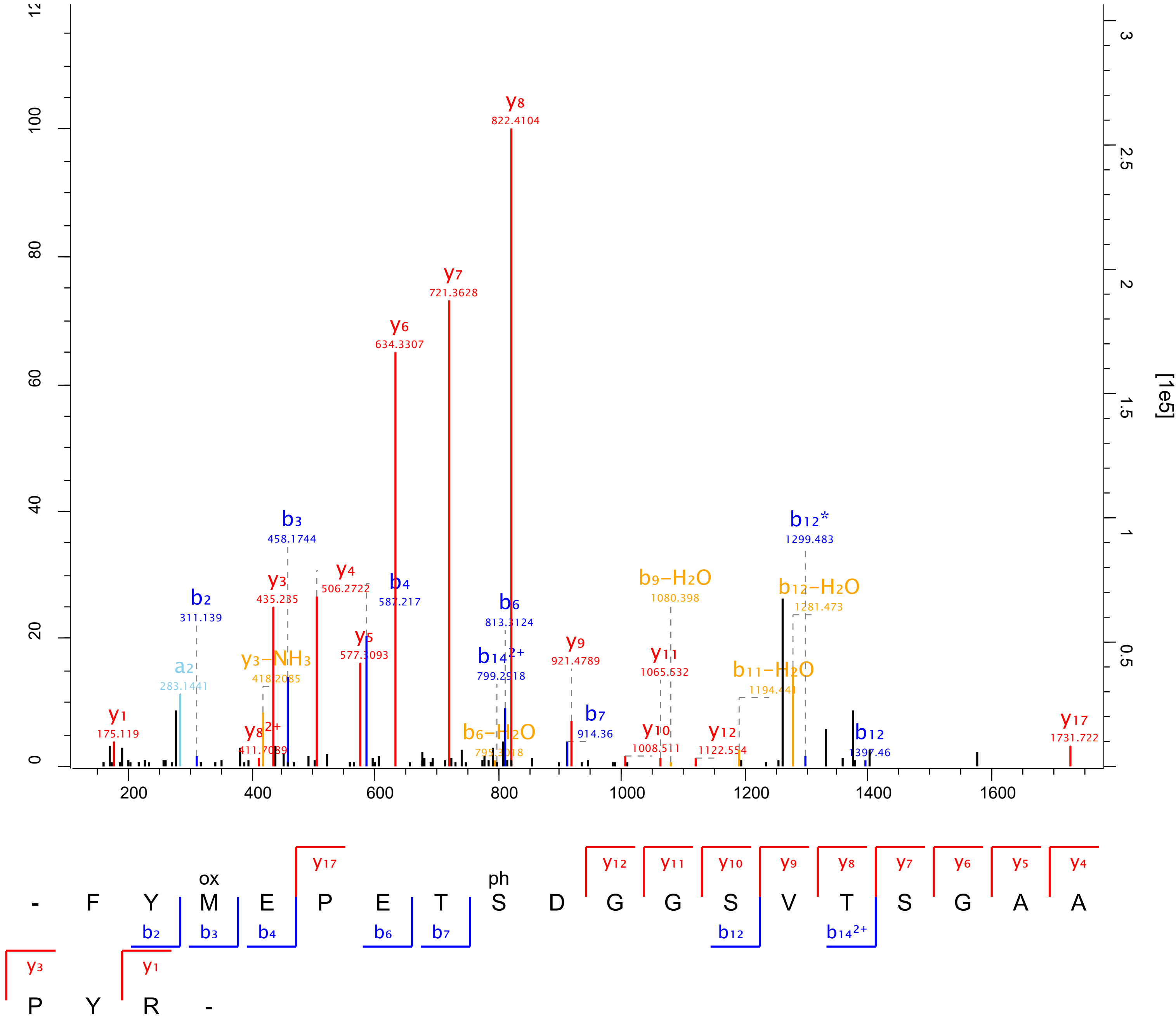

| Raw file | Scan | Method | Score | m/z |
| --- | --- | --- | --- | --- |
| water_02_1 | 24755 | FTMS; HCD | 50.9 | 598.3 |

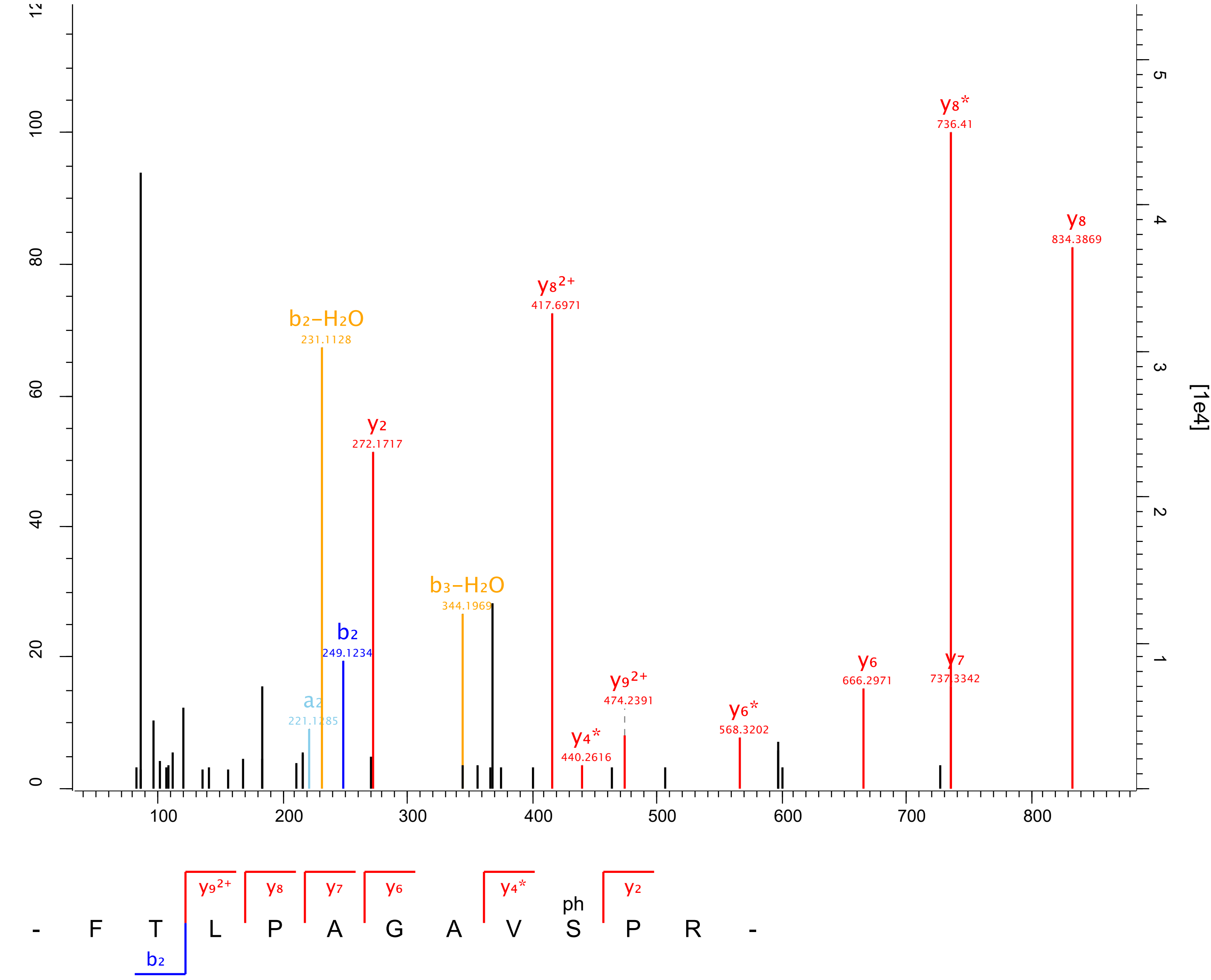

| Raw file | Scan | Method | Score | m/z |
| --- | --- | --- | --- | --- |
| water_02_1 | 24839 | FTMS; HCD | 120.6 | 555.78 |

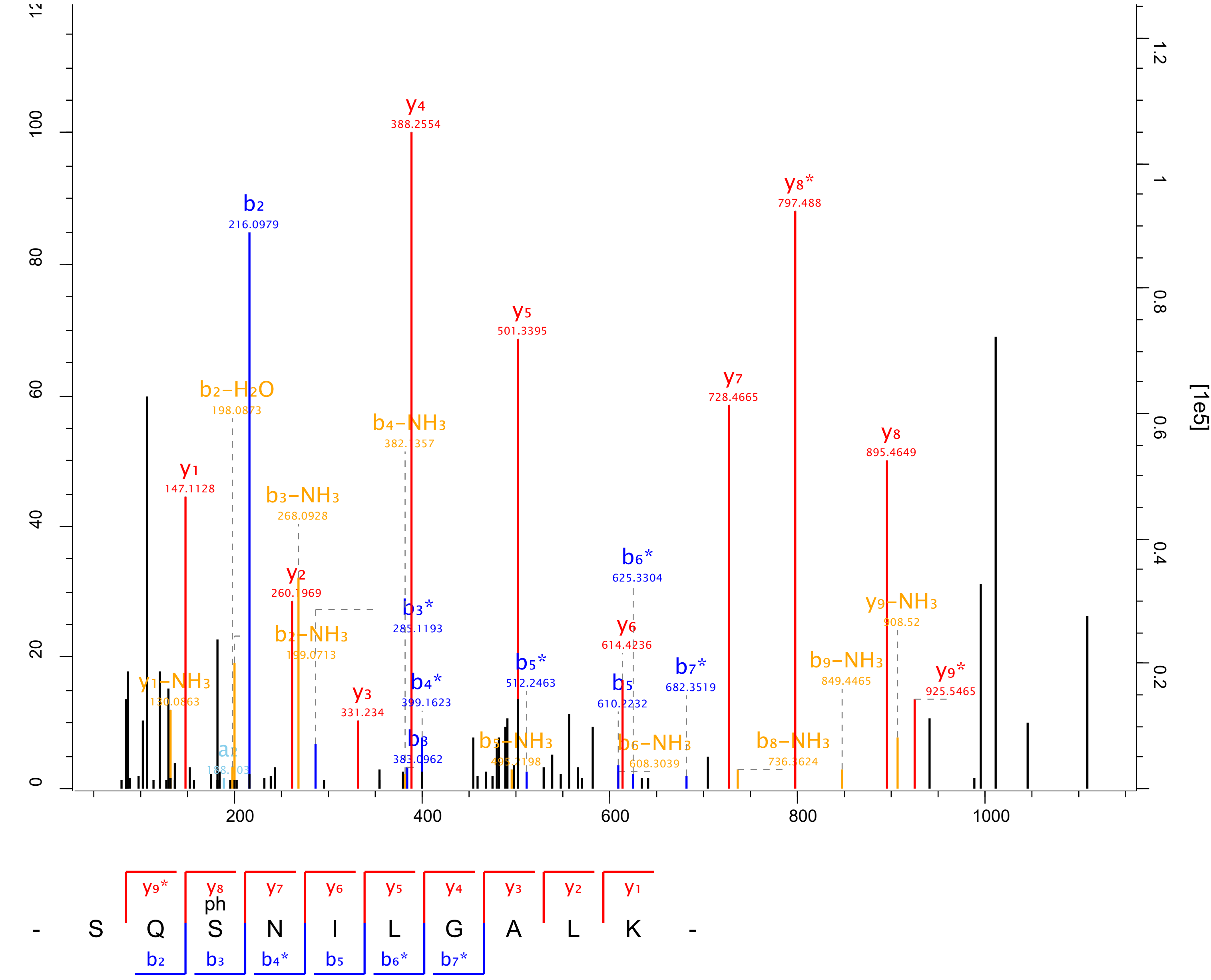

| Raw file | Scan | Method | Score | m/z |
| --- | --- | --- | --- | --- |
| water_02_1 | 24904 | FTMS; HCD | 170.11 | 832.36 |

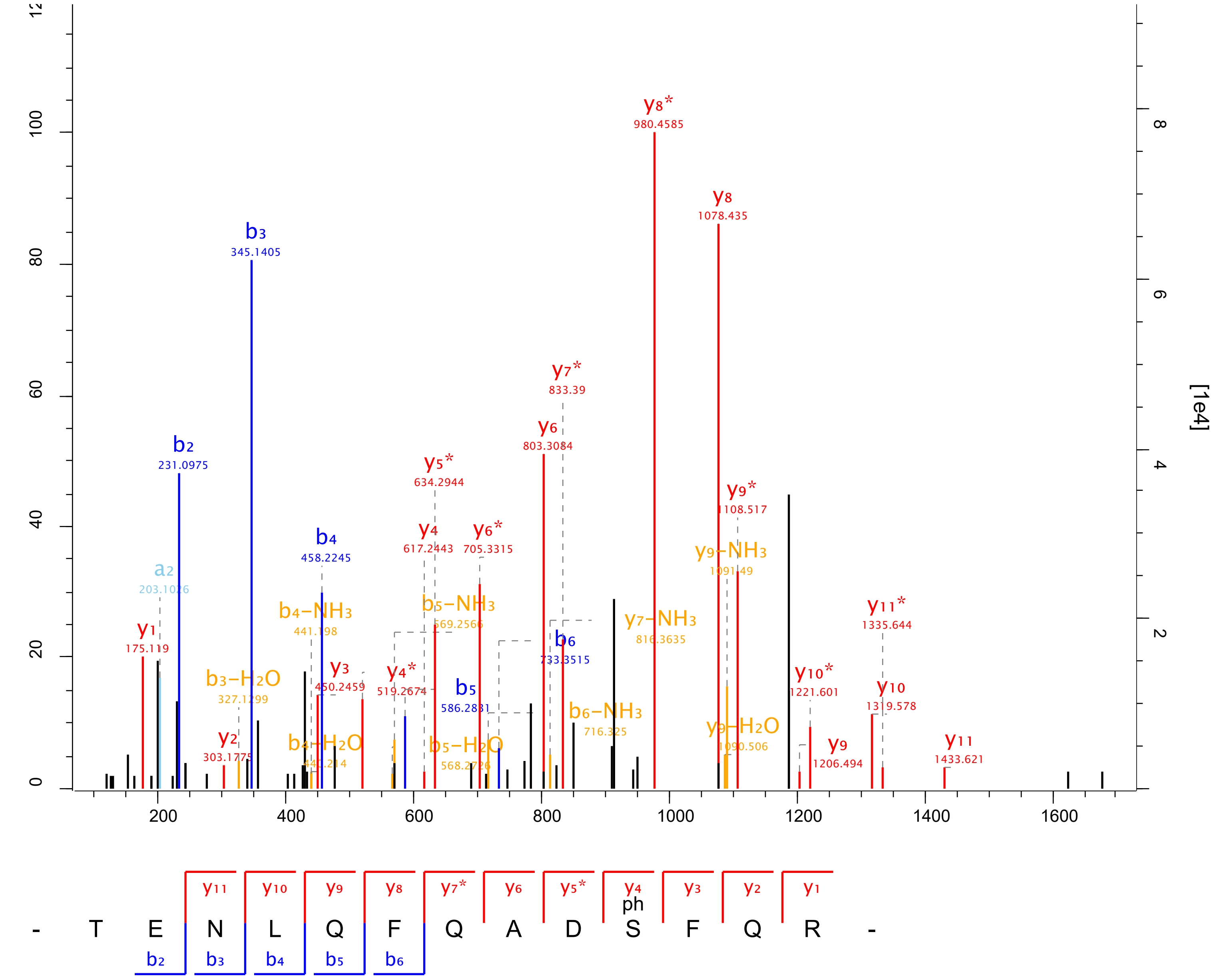

| Raw file | Scan | Method | Score | m/z |
|---|---|---|---|---|
| water_02_1 | 24958 | FTMS; HCD | 51.47 | 836.7 |

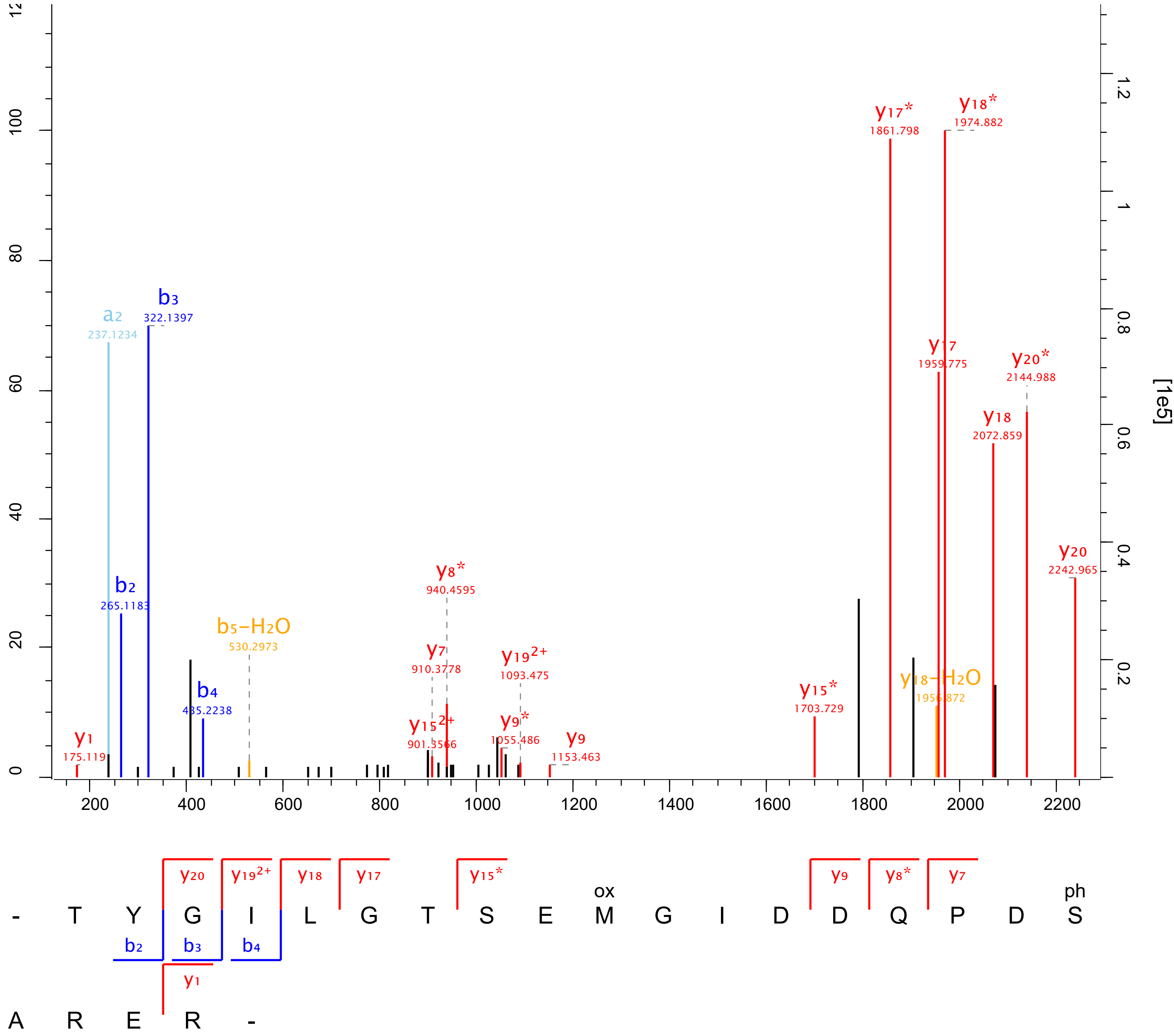

| Raw file | Scan | Method | Score | m/z |
|---|---|---|---|---|
| water_02_1 | 24967 | FTMS; HCD | 65.72 | 526.2 |

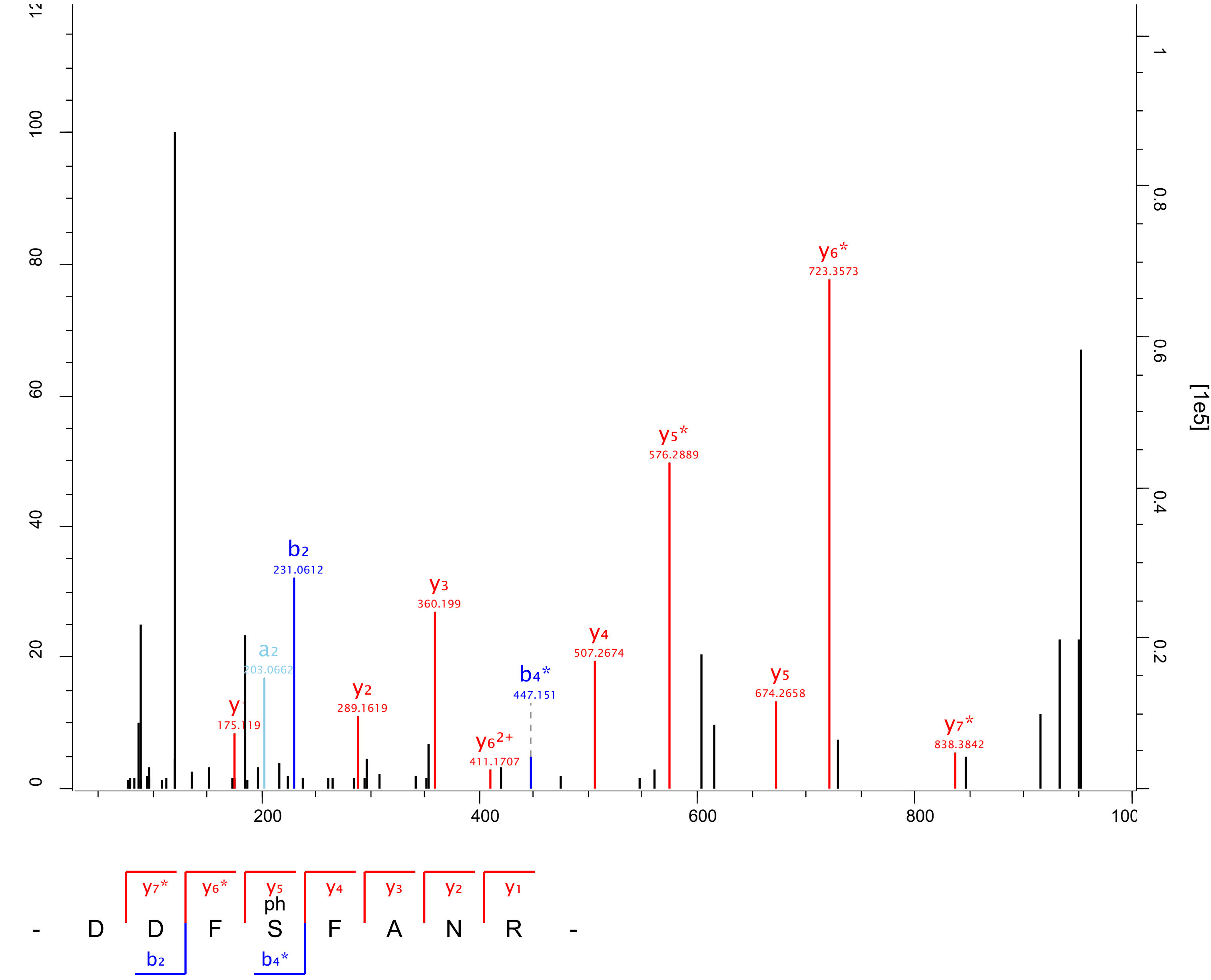

| Raw file | Scan | Method | Score | m/z |
| --- | --- | --- | --- | --- |
| water_02_1 | 24992 | FTMS; HCD | 82.66 | 765.81 |

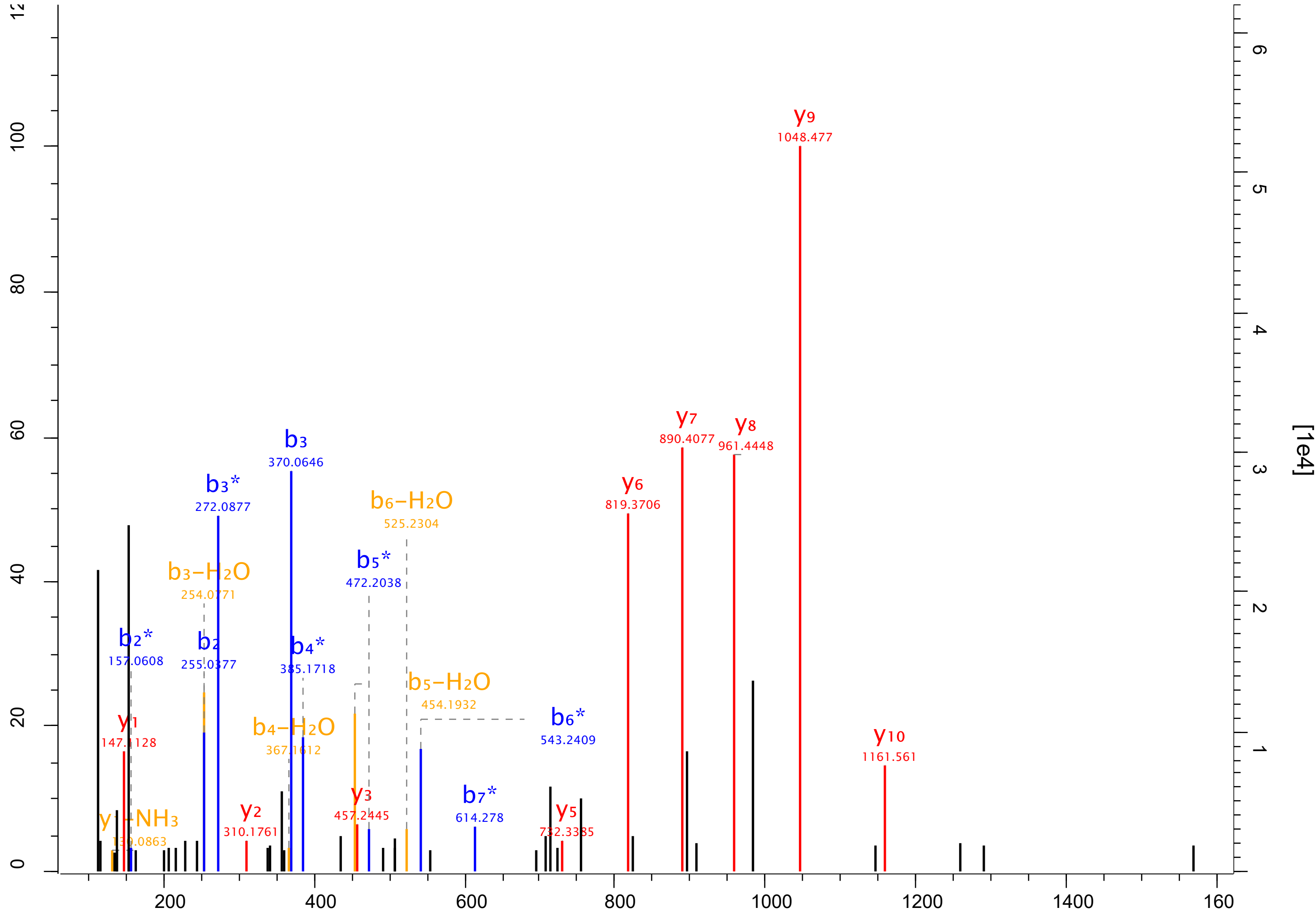


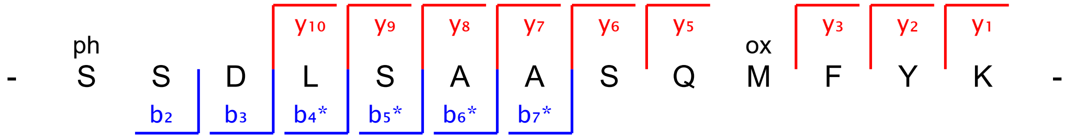

| Raw file | Scan | Method | Score | m/z |
|---|---|---|---|---|
| water_02_1 | 25097 | FTMS; HCD | 212.76 | 782.87 |

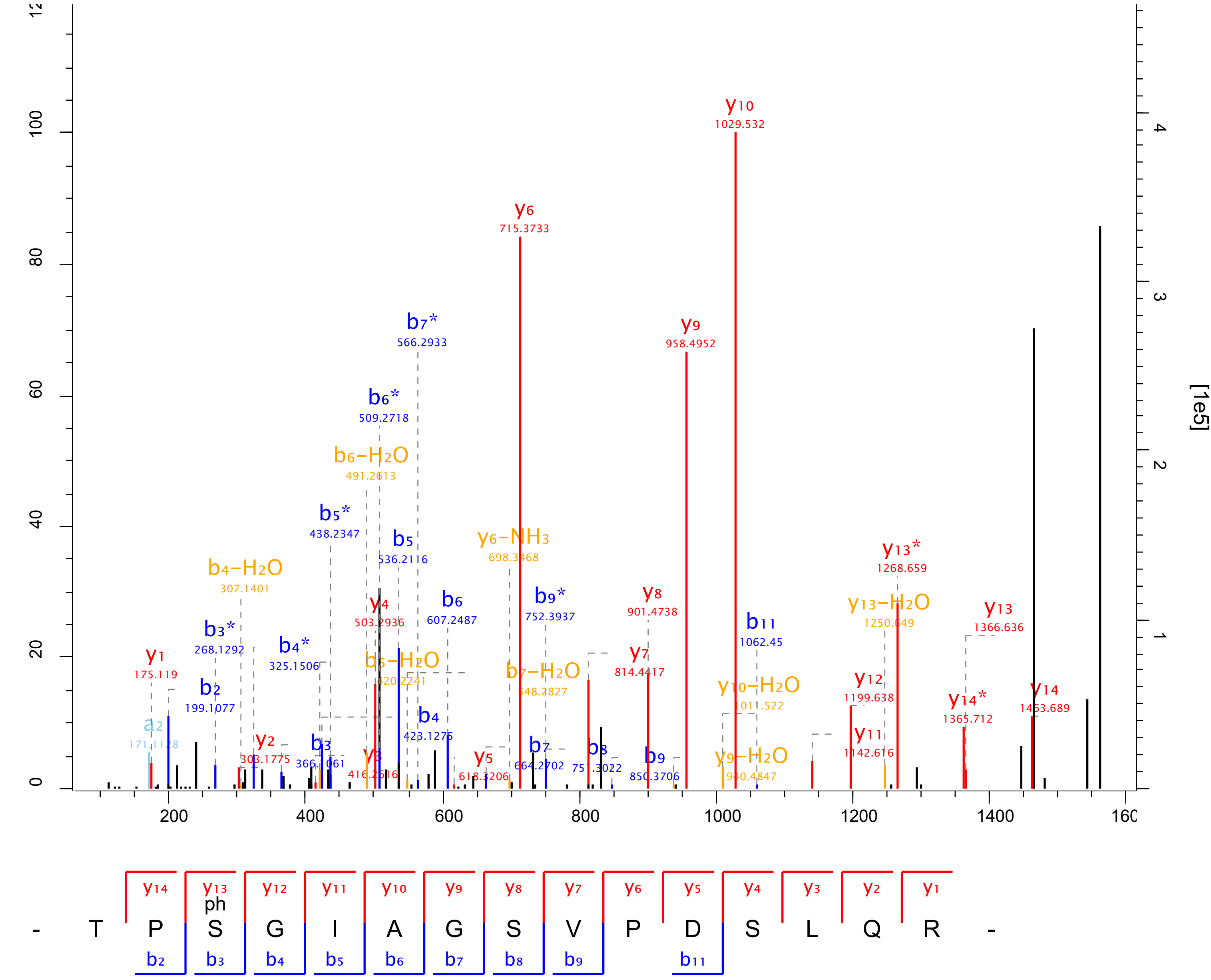

| Raw file | Scan | Method | Score | m/z |
| --- | --- | --- | --- | --- |
| water_02_1 | 25217 | FTMS; HCD | 42.84 | 550.77 |

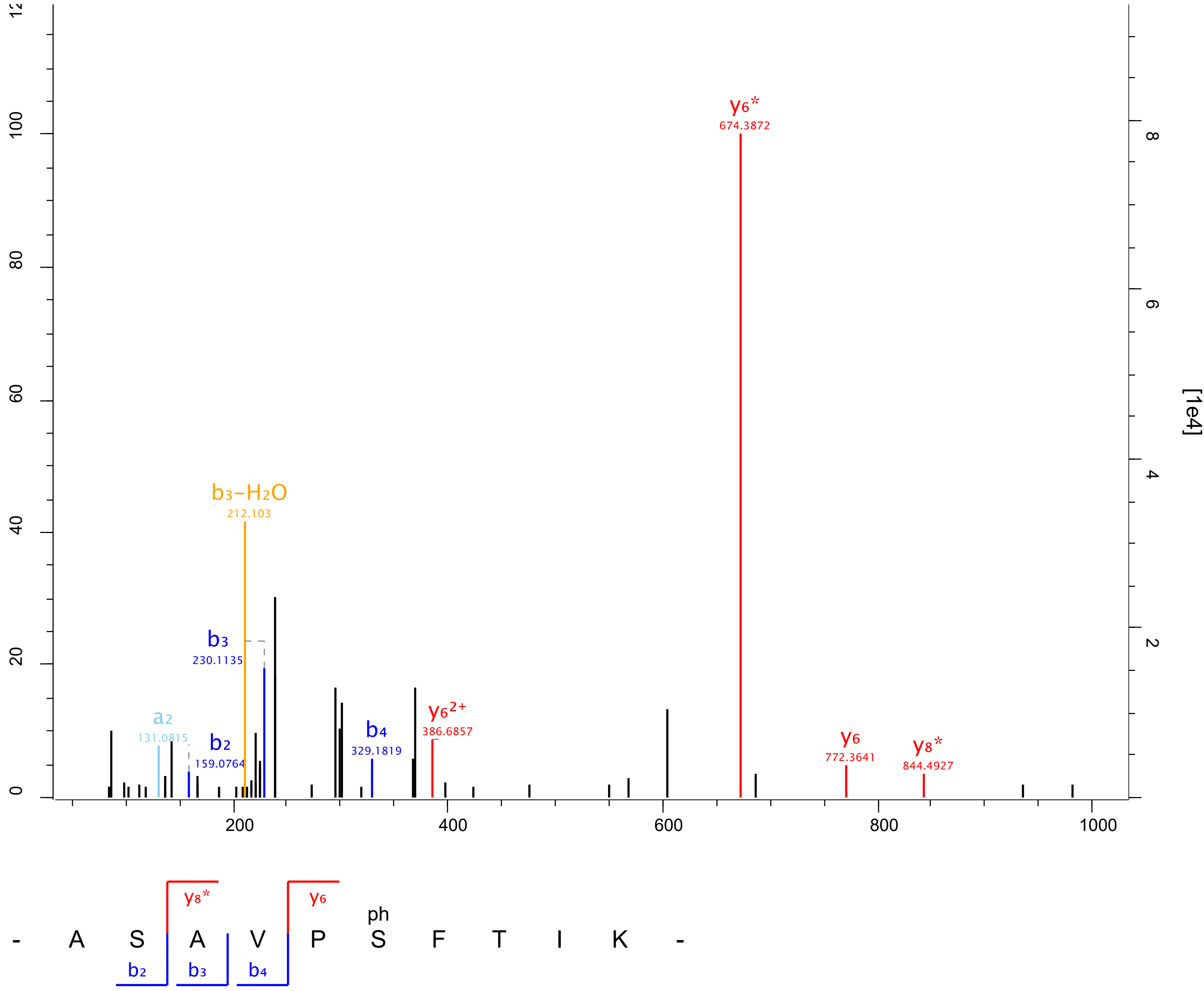

| Raw file | Scan | Method | Score | m/z |
|---|---|---|---|---|
| water_02_1 | 25231 | FTMS; HCD | 48.57 | 514.24 |

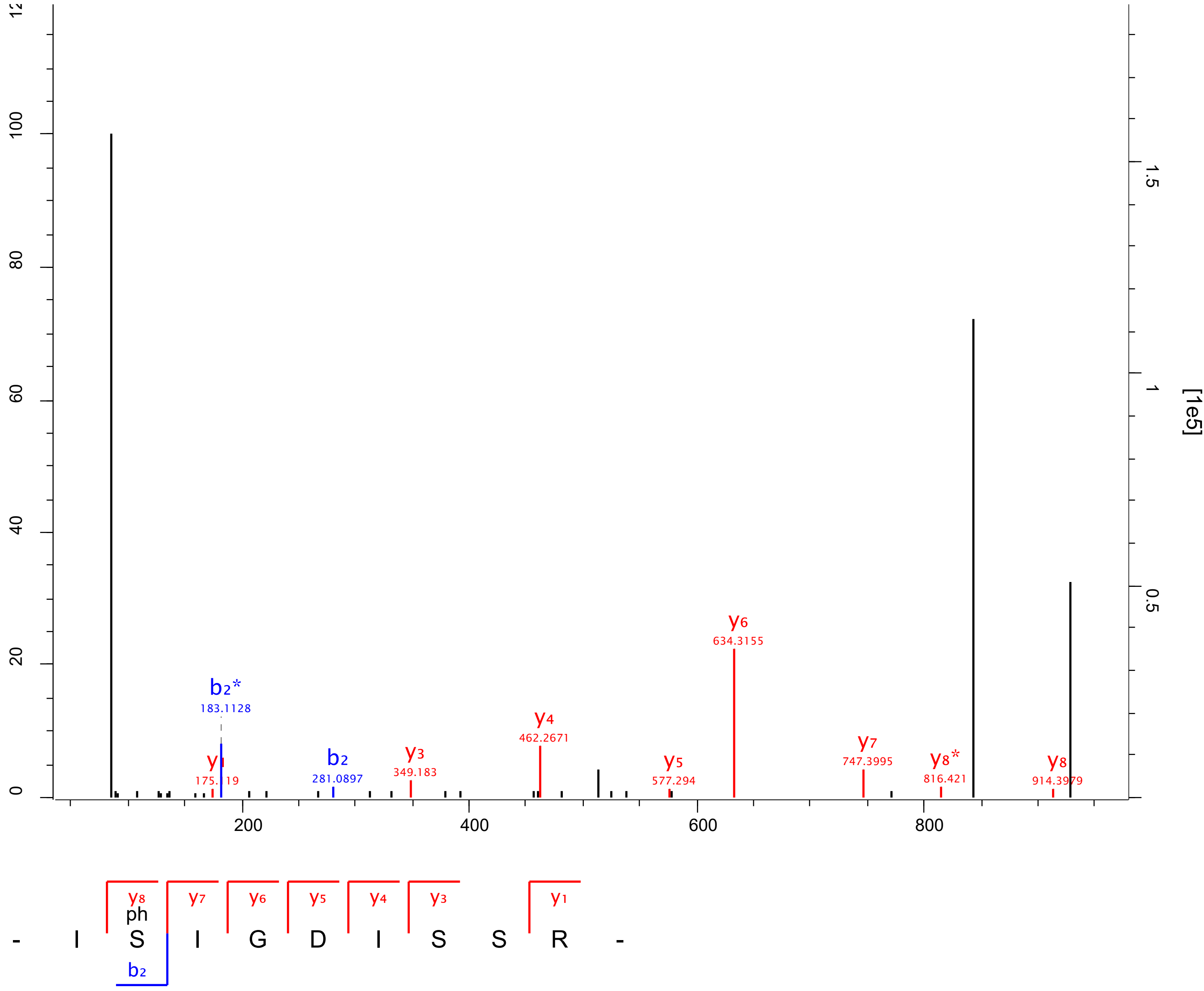

| Raw file | Scan | Method | Score | m/z |
|---|---|---|---|---|
| water_02_1 | 25267 | FTMS; HCD | 148.36 | 825.85 |

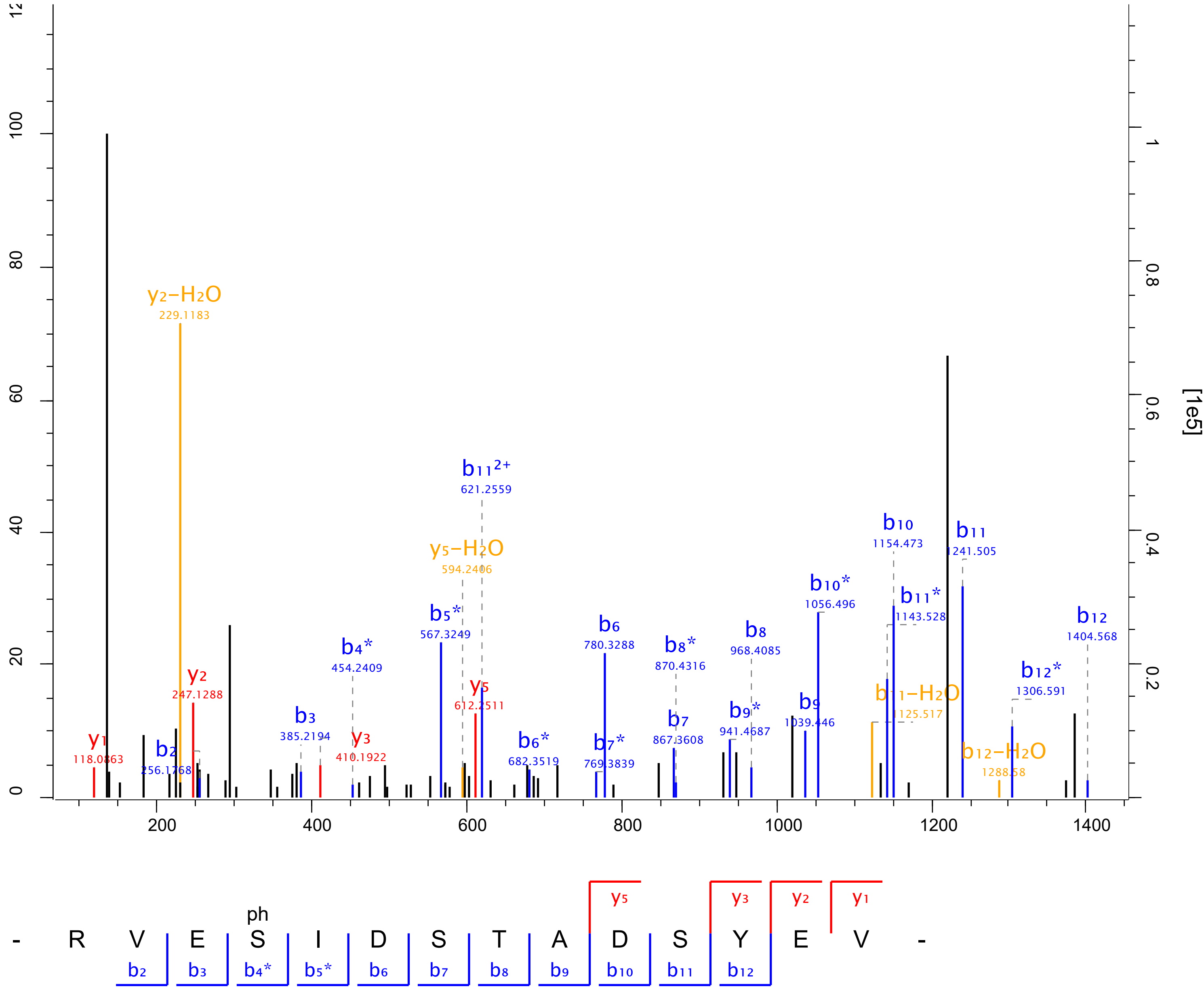

| Raw file | Scan | Method | Score | m/z |
| --- | --- | --- | --- | --- |
| water_02_1 | 25291 | FTMS; HCD | 68.02 | 478.71 |

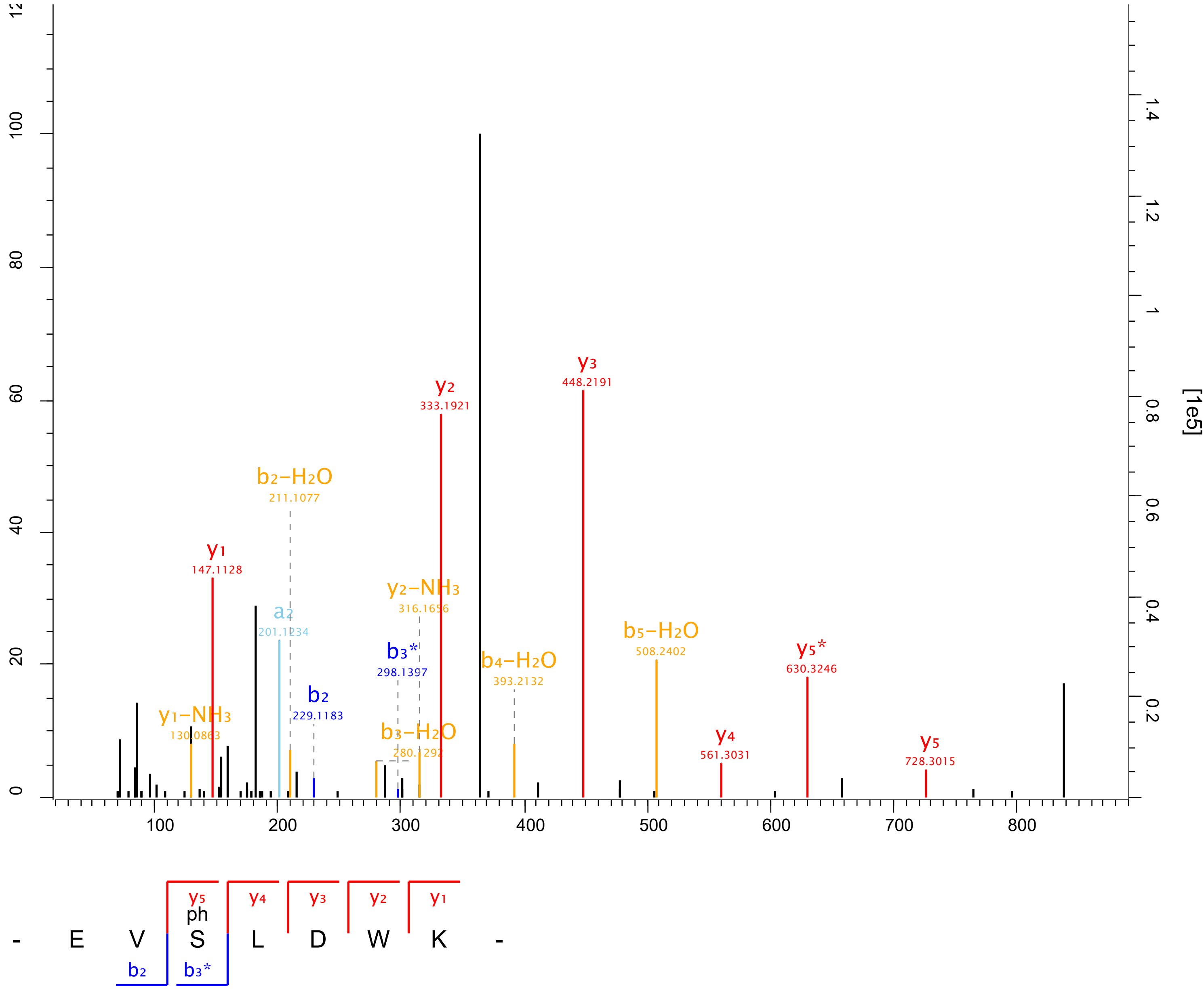

| Raw file | Scan | Method | Score | m/z |
| --- | --- | --- | --- | --- |
| water_02_1 | 2538 | FTMS; HCD | 43.03 | 658.23 |

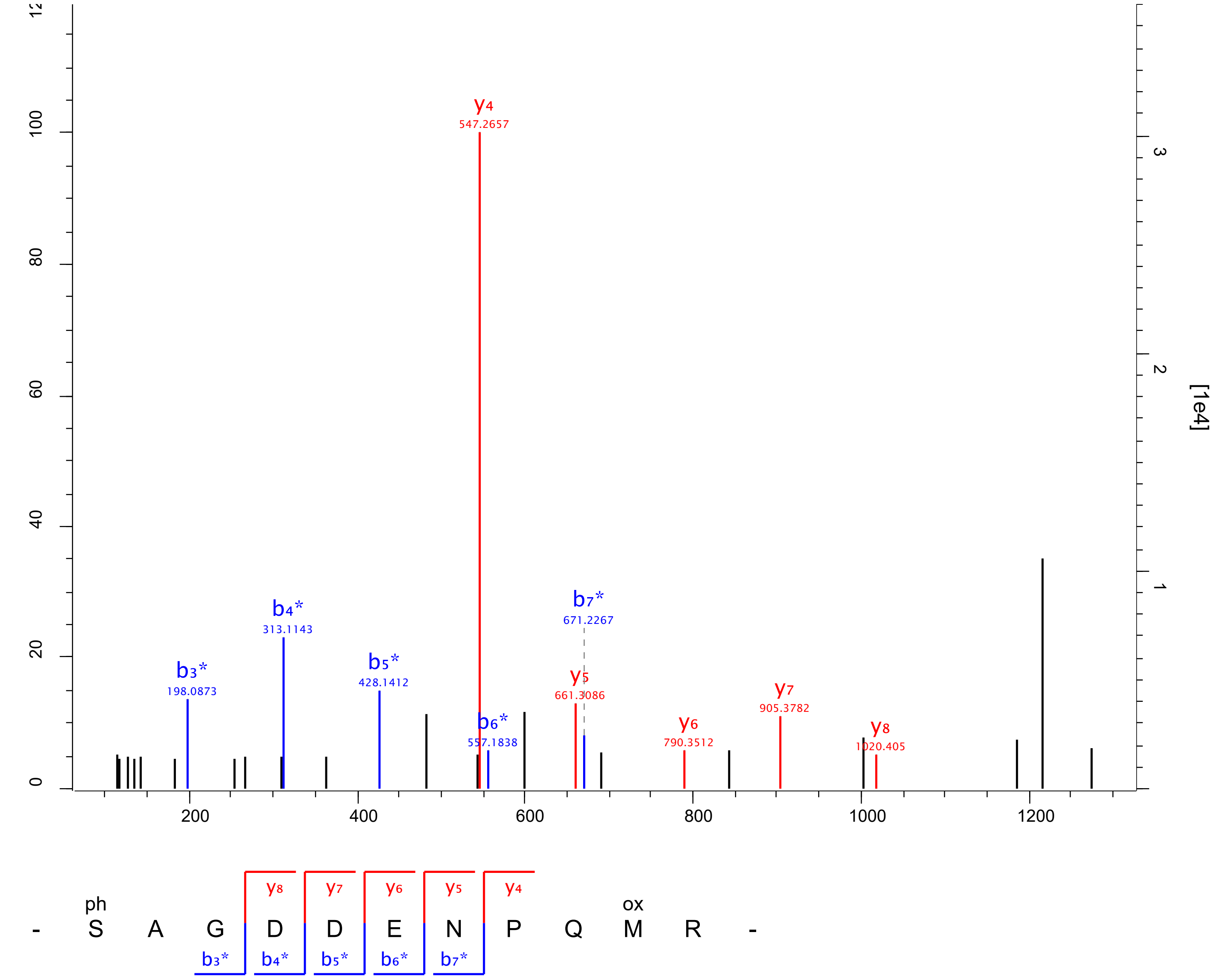

| Raw file | Scan | Method | Score | m/z |
| --- | --- | --- | --- | --- |
| water_02_1 | 25519 | FTMS; HCD | 122.18 | 623.63 |

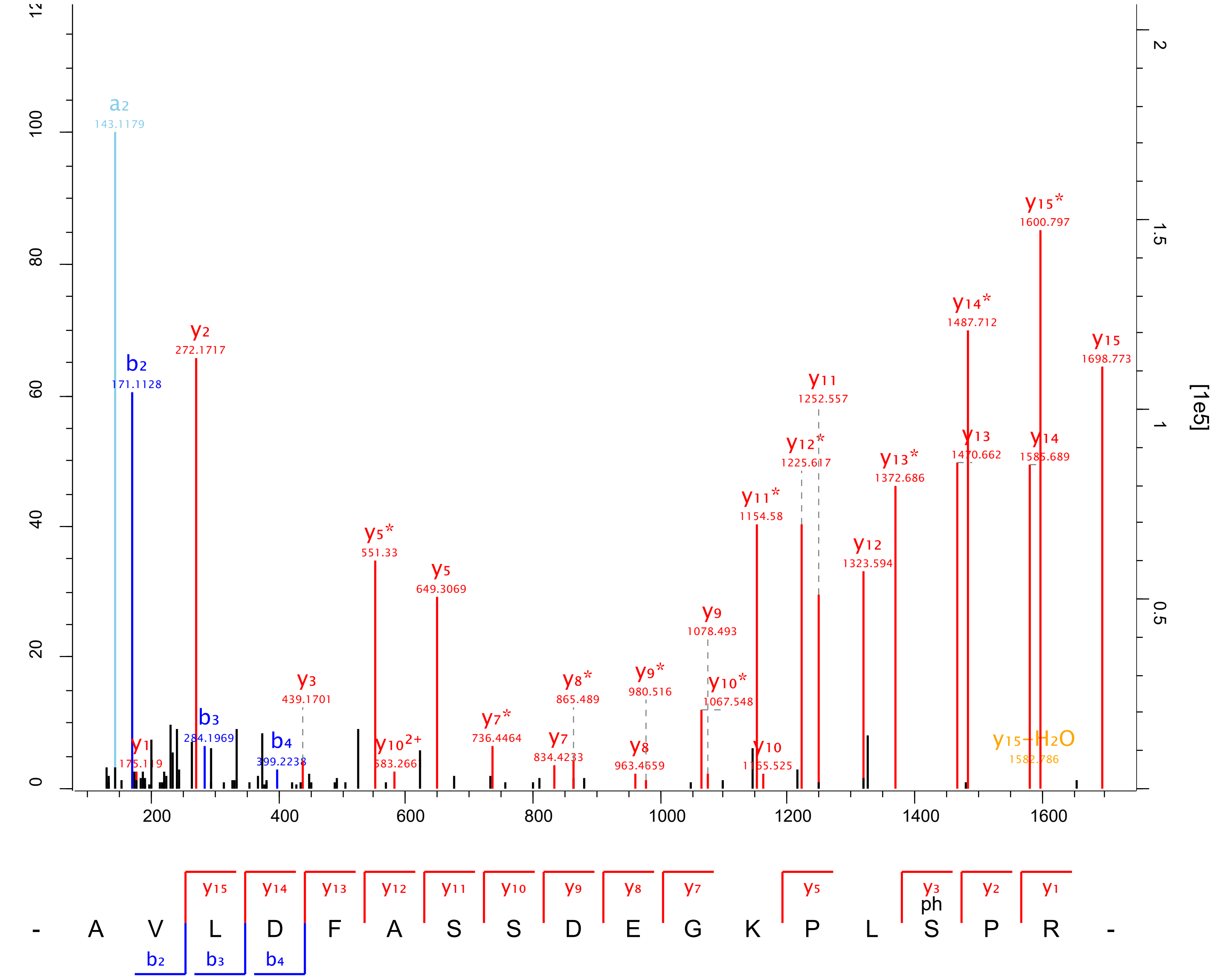

| Raw file | Scan | Method | Score | m/z |
| --- | --- | --- | --- | --- |
| water_02_1 | 25545 | FTMS; HCD | 113.29 | 1039.82 |

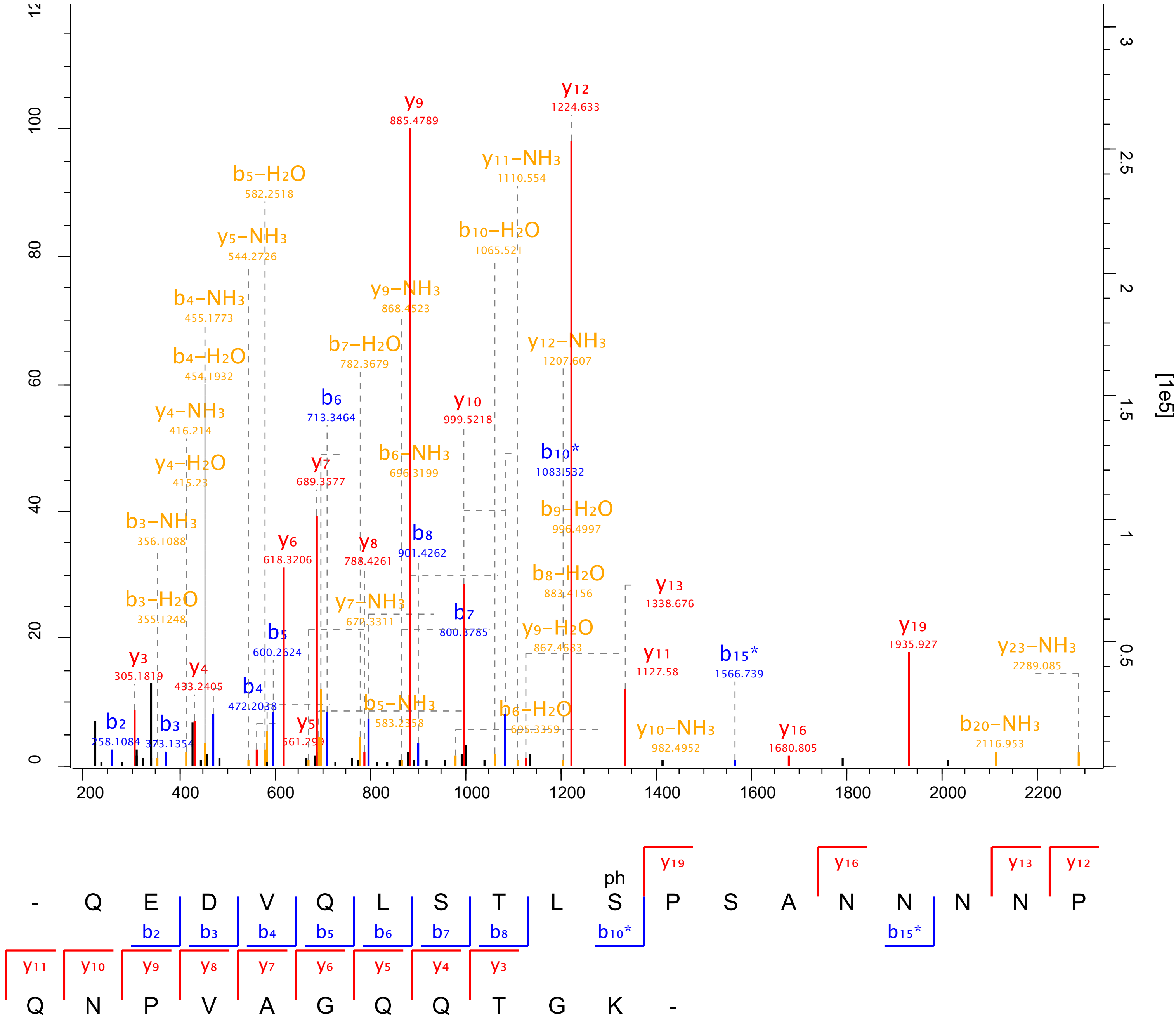

| Raw file | Scan | Method | Score | m/z |
|---|---|---|---|---|
| water_02_1 | 25573 | FTMS; HCD | 153.58 | 657.29 |

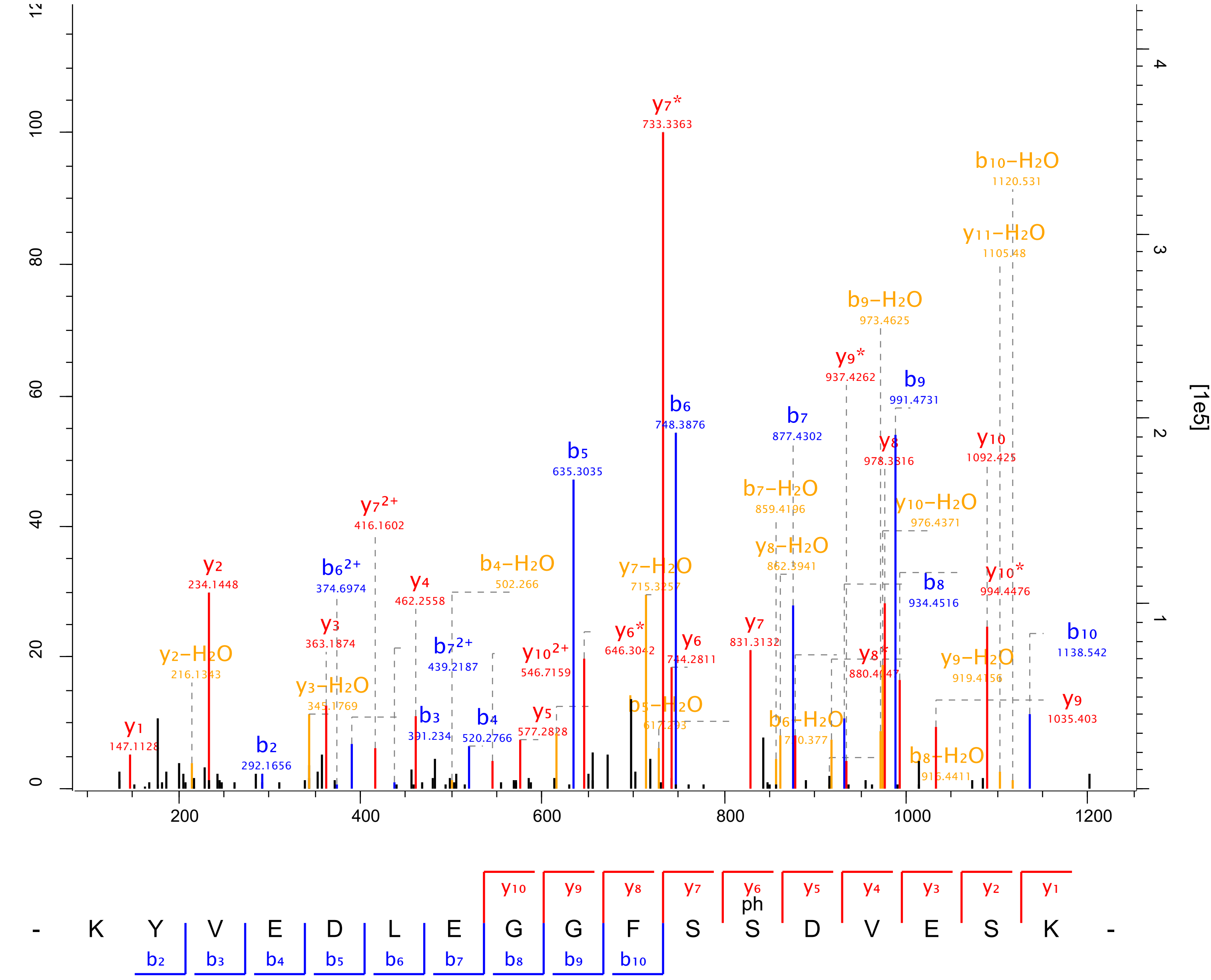

| Raw file | Scan | Method | Score | m/z |
| --- | --- | --- | --- | --- |
| water_02_1 | 25661 | FTMS; HCD | 182.76 | 951.87 |

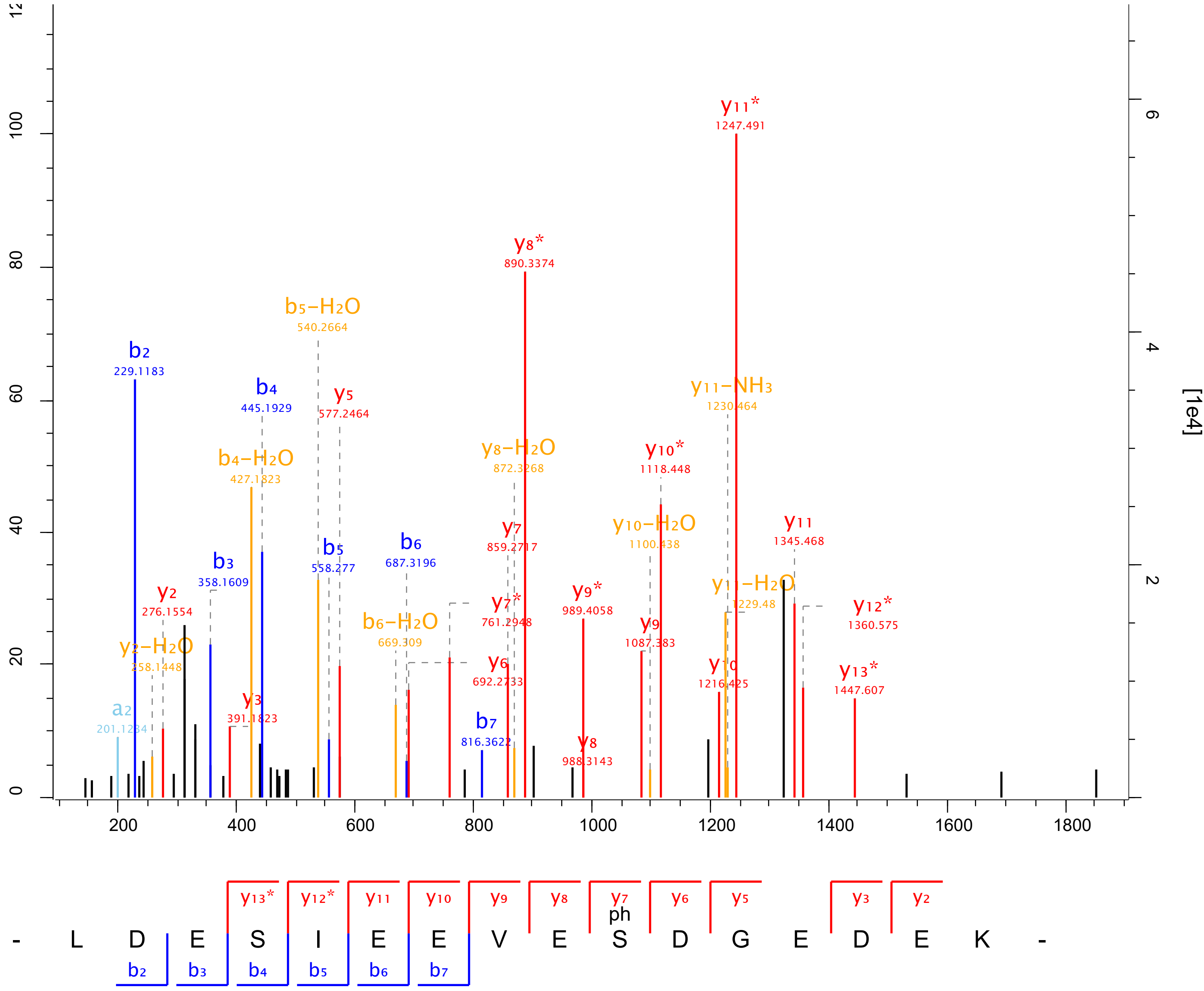

| Raw file | Scan | Method | Score | m/z |
| --- | --- | --- | --- | --- |
| water_02_1 | 25711 | FTMS; HCD | 180.47 | 590.75 |

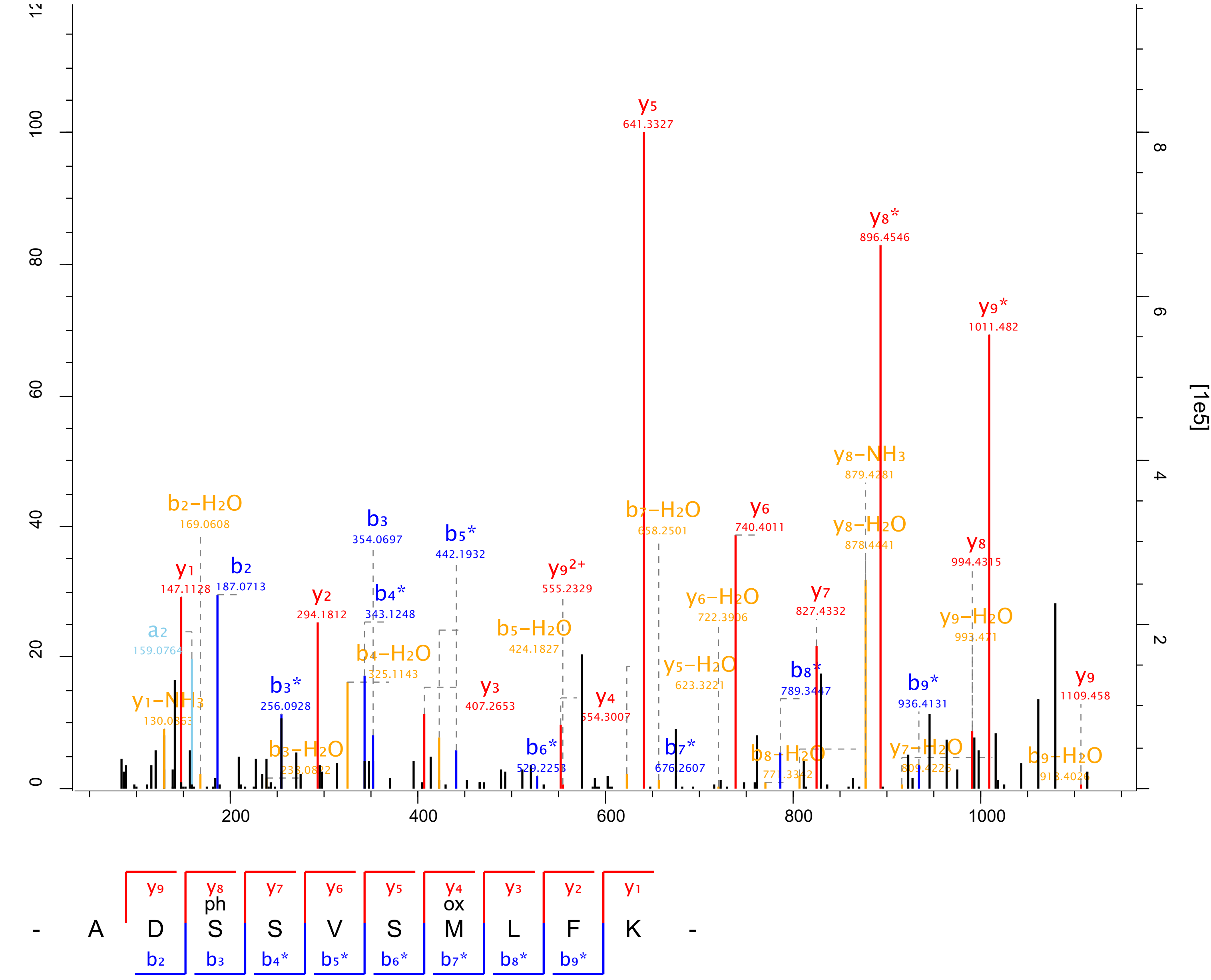

| Raw file | Scan | Method | Score | m/z |
|---|---|---|---|---|
| water_02_1 | 25717 | FTMS; HCD | 60.95 | 546.94 |

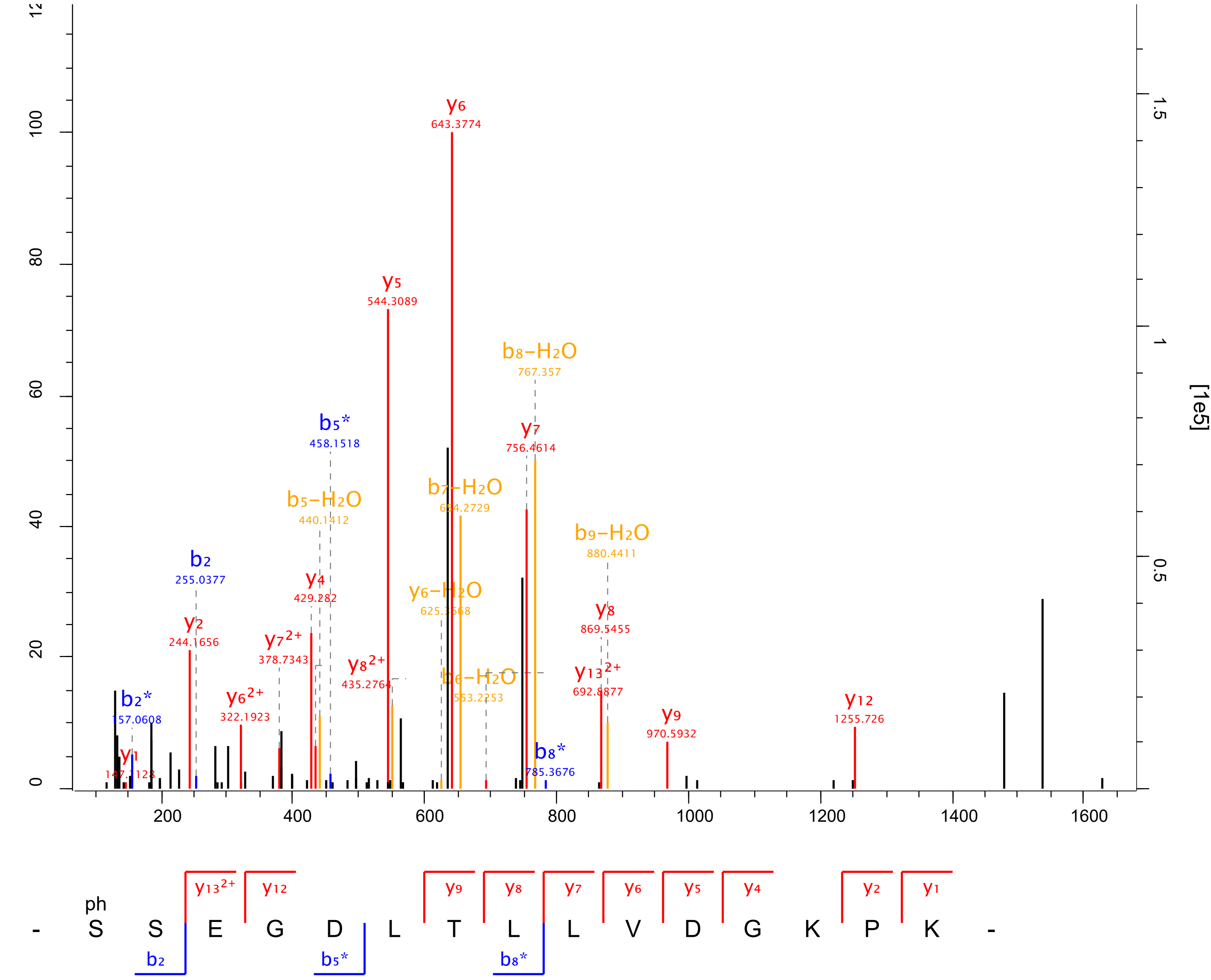

| Raw file | Scan | Method | Score | m/z |
|---|---|---|---|---|
| water_02_1 | 2574 | FTMS; HCD | 142.45 | 599.76 |

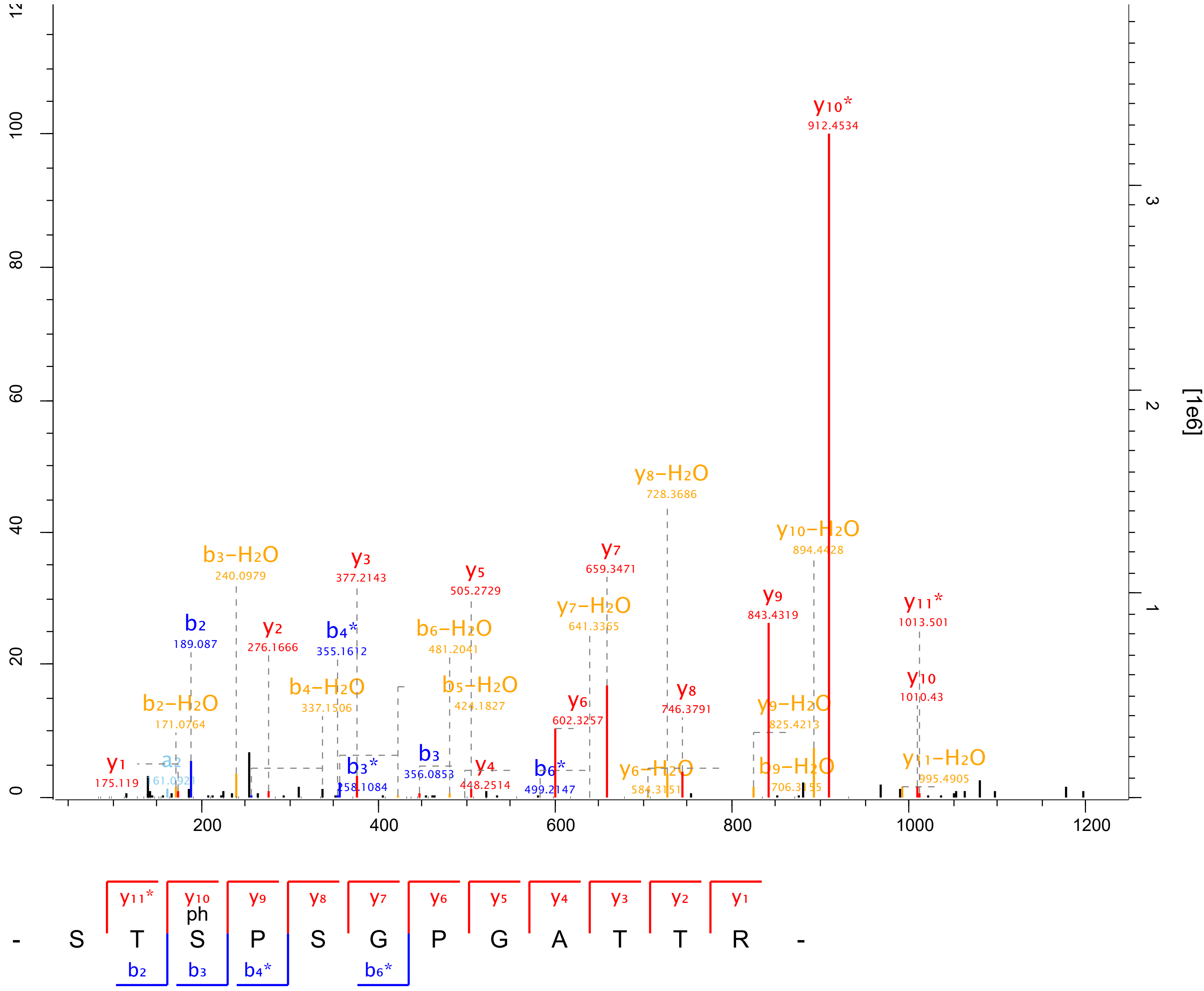

| Raw file | Scan | Method | Score | m/z |
| --- | --- | --- | --- | --- |
| water_02_1 | 2575 | FTMS; HCD | 62.34 | 395.86 |

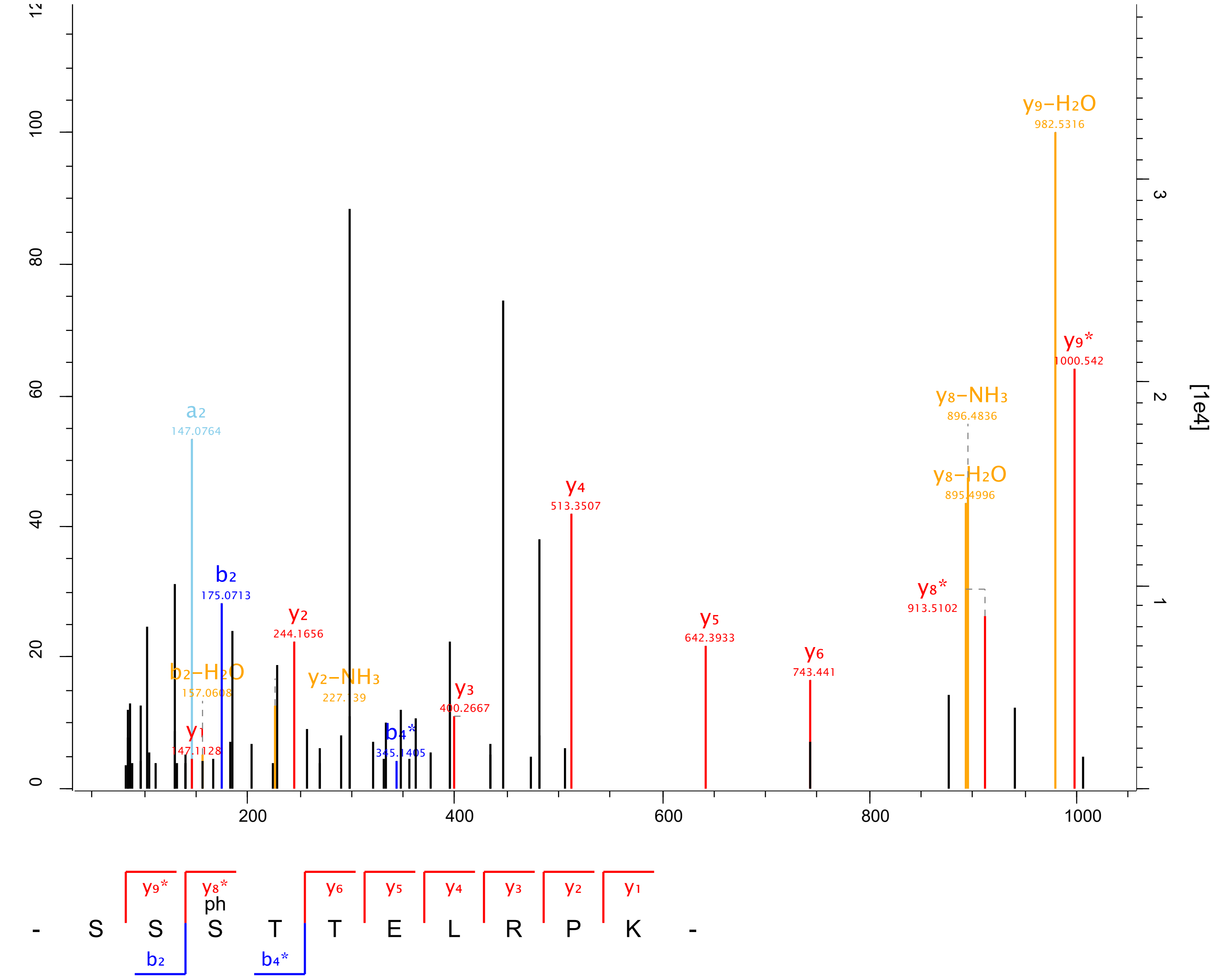

| Raw file | Scan | Method | Score | m/z |
| --- | --- | --- | --- | --- |
| water_02_1 | 25798 | FTMS; HCD | 123.02 | 794.37 |

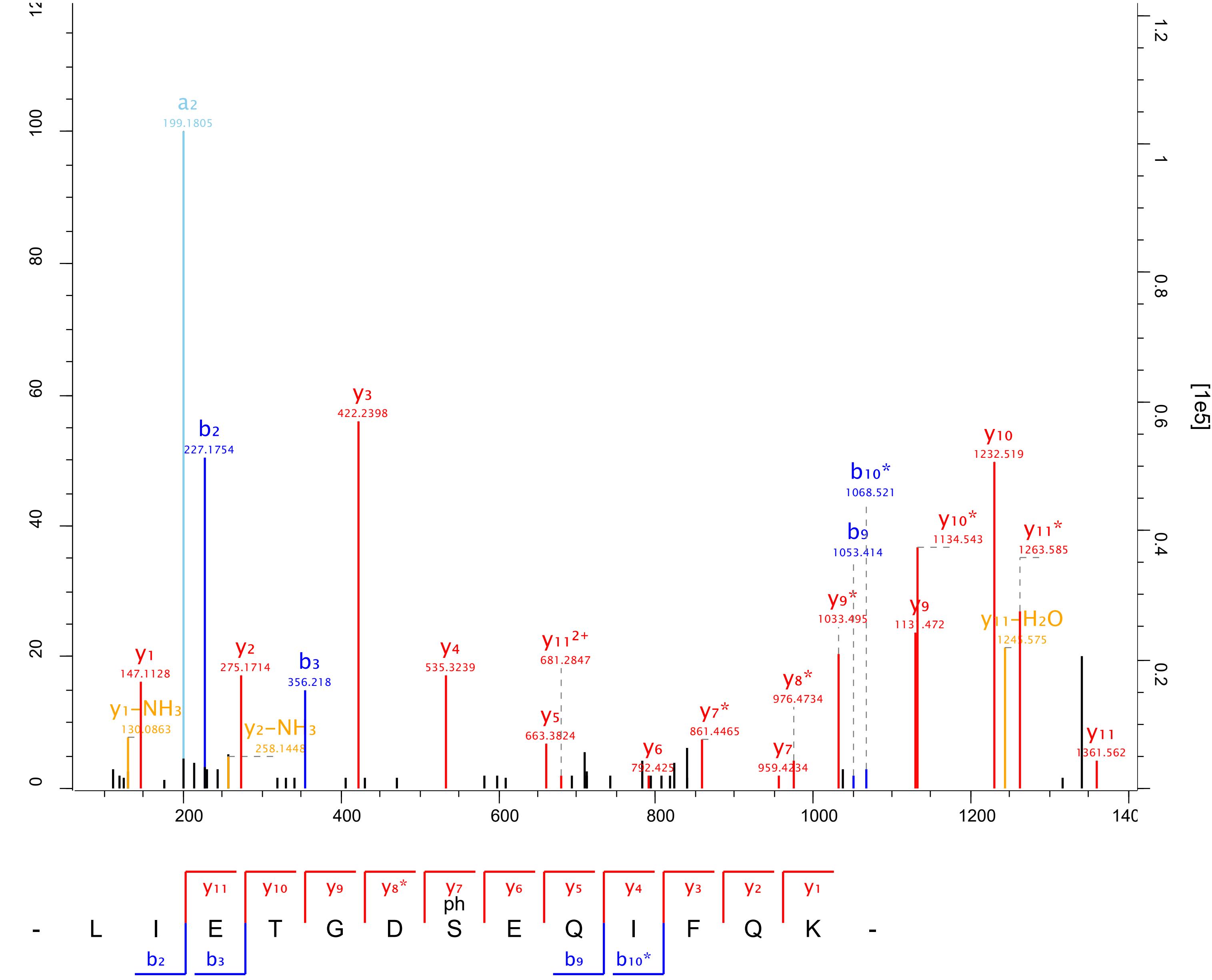

| Raw file | Scan | Method | Score | m/z |
| --- | --- | --- | --- | --- |
| water_02_1 | 25843 | FTMS; HCD | 105.72 | 722.96 |

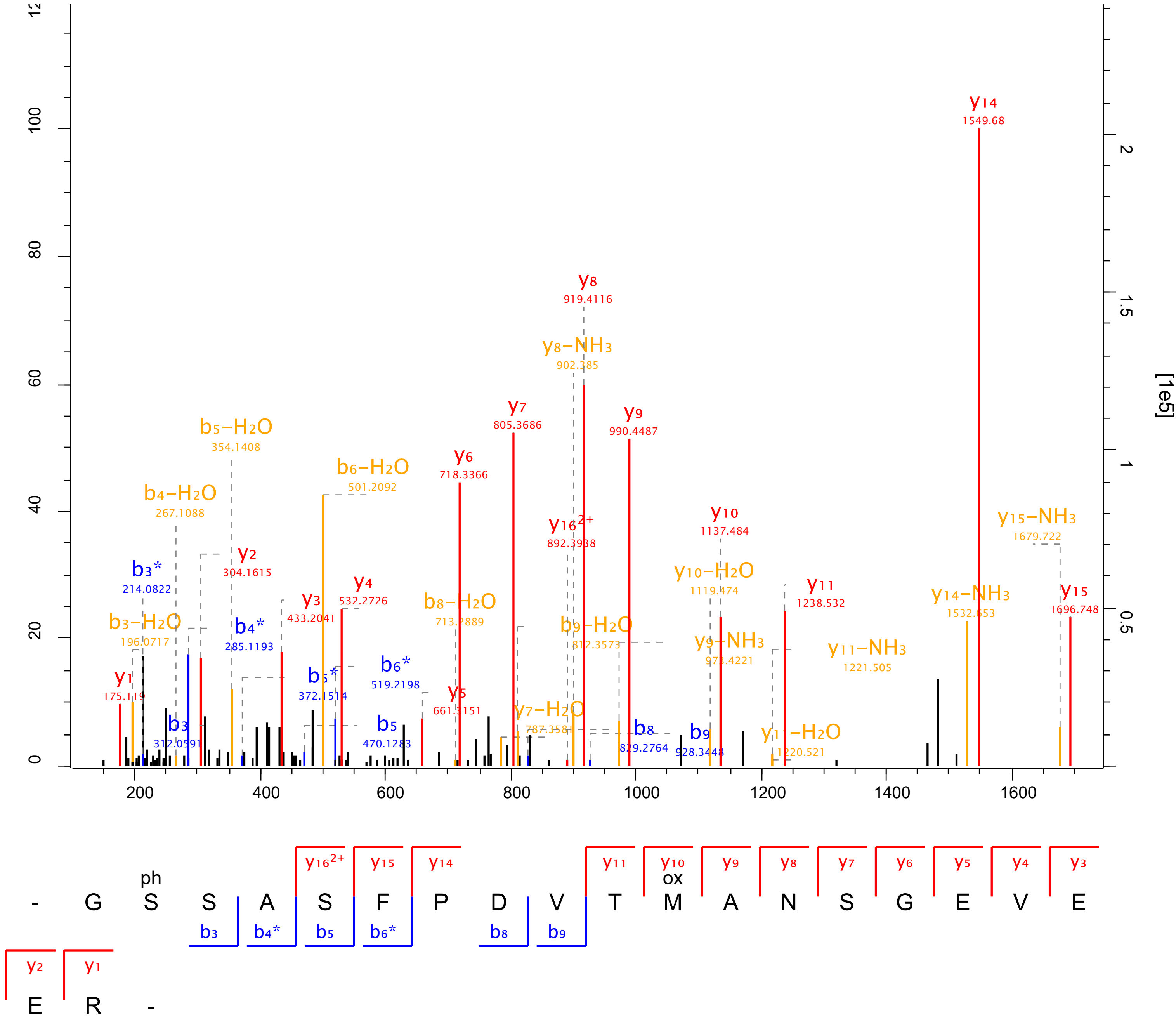

| Raw file | Scan | Method | Score | m/z |
|---|---|---|---|---|
| water_02_1 | 25963 | FTMS; HCD | 61.86 | 640.63 |

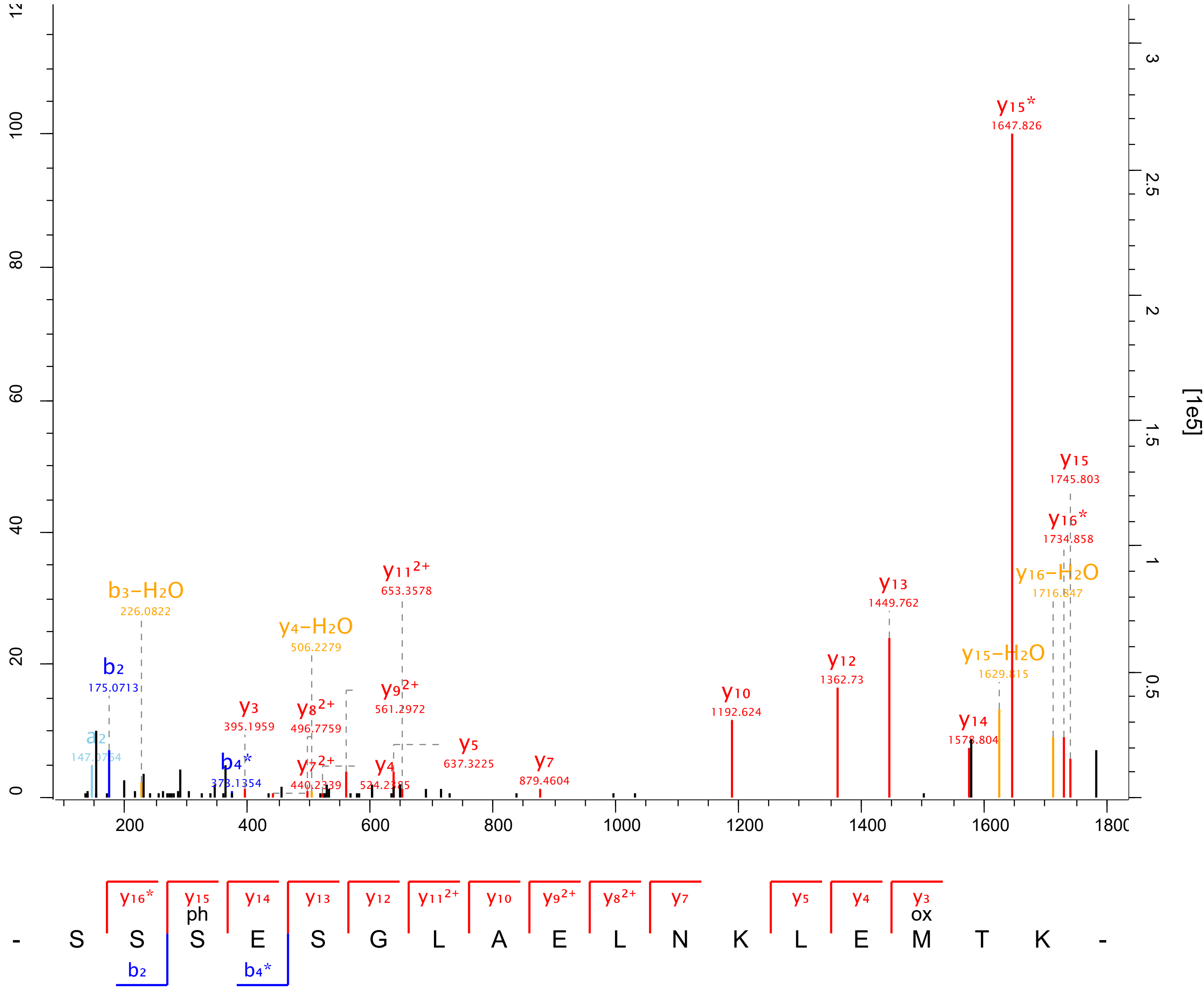

| Raw file | Scan | Method | Score | m/z |
| --- | --- | --- | --- | --- |
| water_02_1 | 25992 | FTMS; HCD | 59.71 | 847.39 |

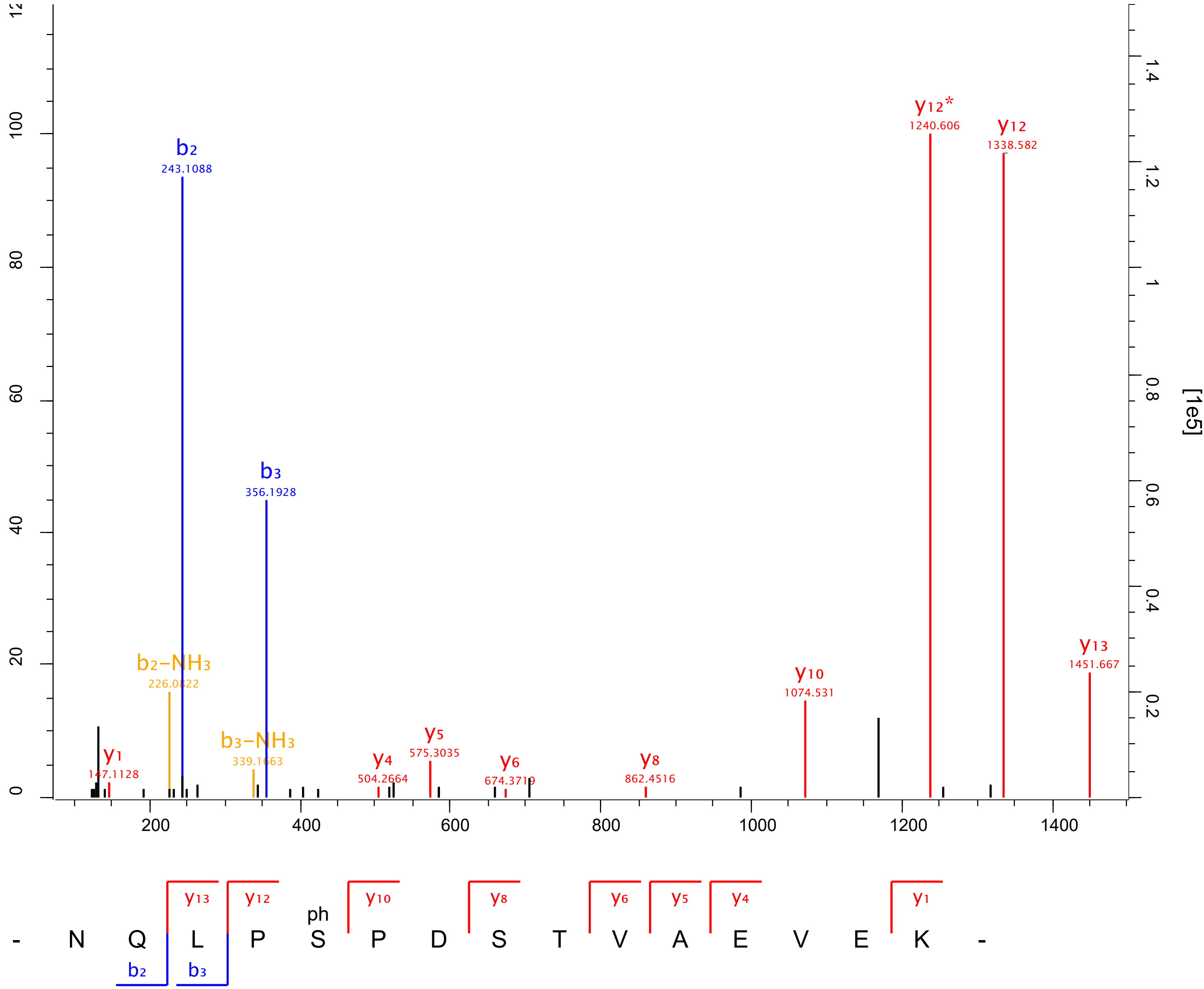

| Raw file | Scan | Method | Score | m/z |
|---|---|---|---|---|
| water_02_1 | 2600 | FTMS; HCD | 43.31 | 336.84 |

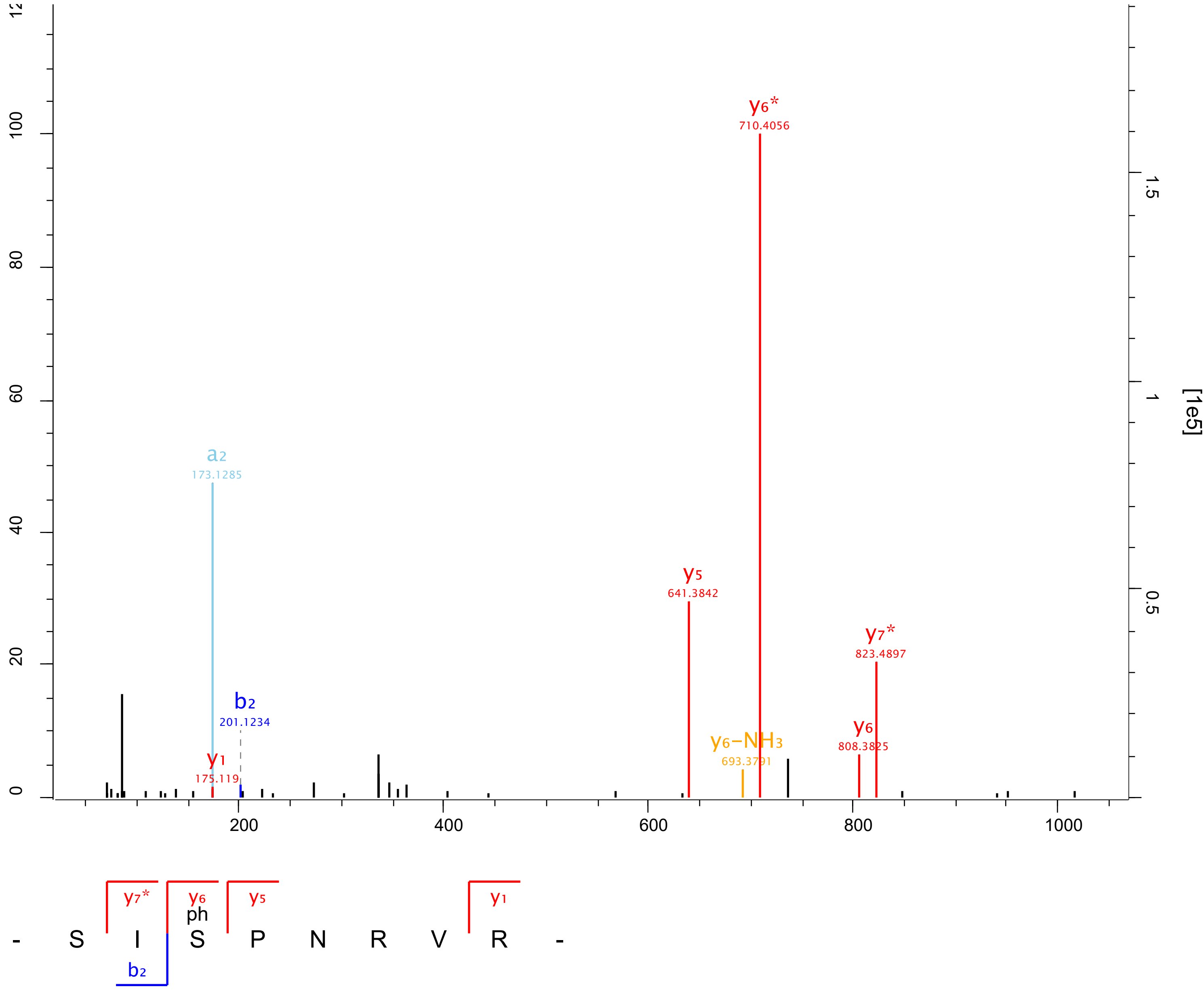

| Raw file | Scan | Method | Score | m/z |
|---|---|---|---|---|
| water_02_1 | 26019 | FTMS; HCD | 92.41 | 867.02 |

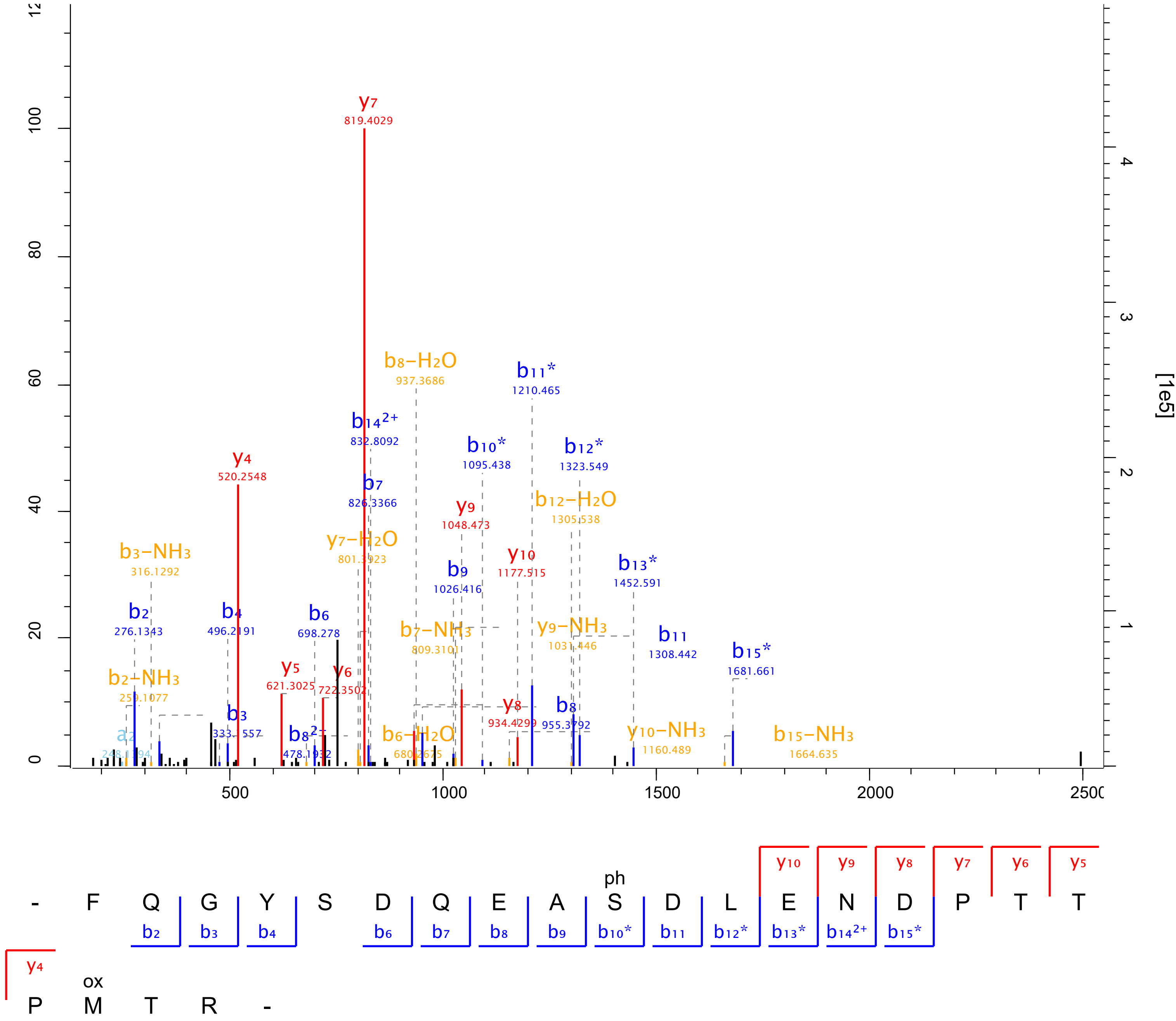

| Raw file | Scan | Method | Score | m/z |
|---|---|---|---|---|
| water_02_1 | 26040 | FTMS; HCD | 60.76 | 871.87 |

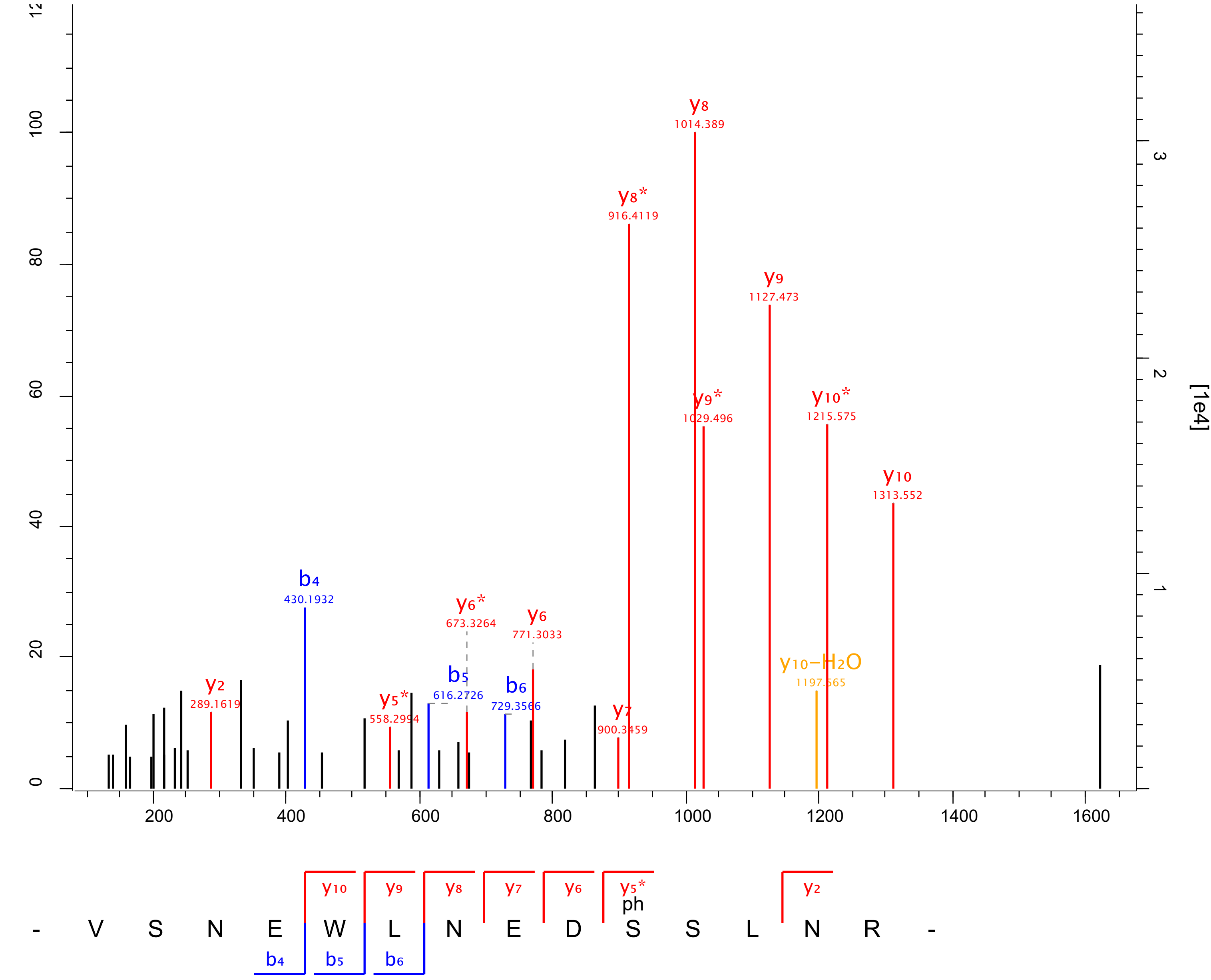

| Raw file | Scan | Method | Score | m/z |
| --- | --- | --- | --- | --- |
| water_02_1 | 26065 | FTMS; HCD | 75.54 | 761.03 |

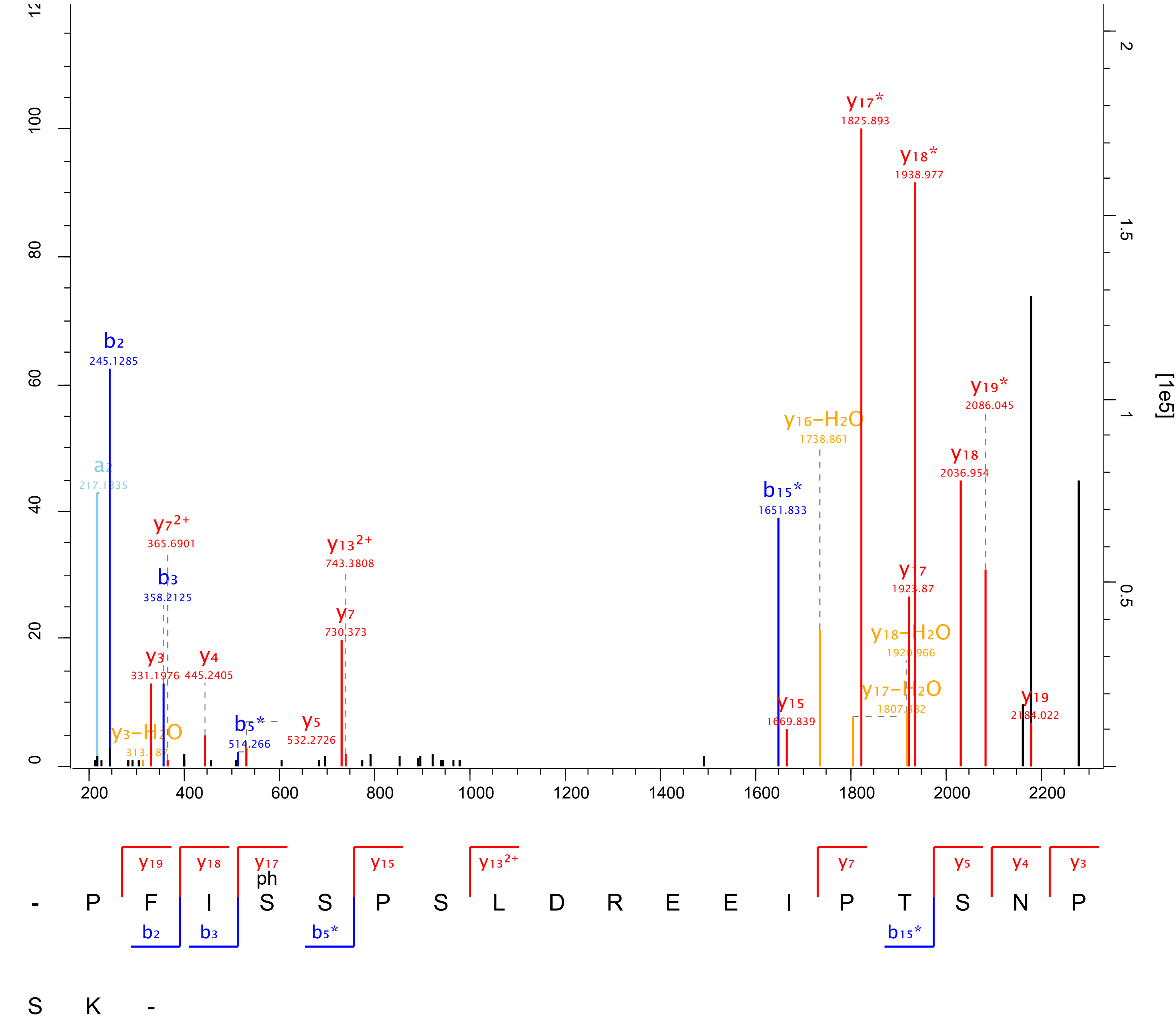

| Raw file | Scan | Method | Score | m/z |
| --- | --- | --- | --- | --- |
| water_02_1 | 26141 | FTMS; HCD | 125.48 | 874.14 |

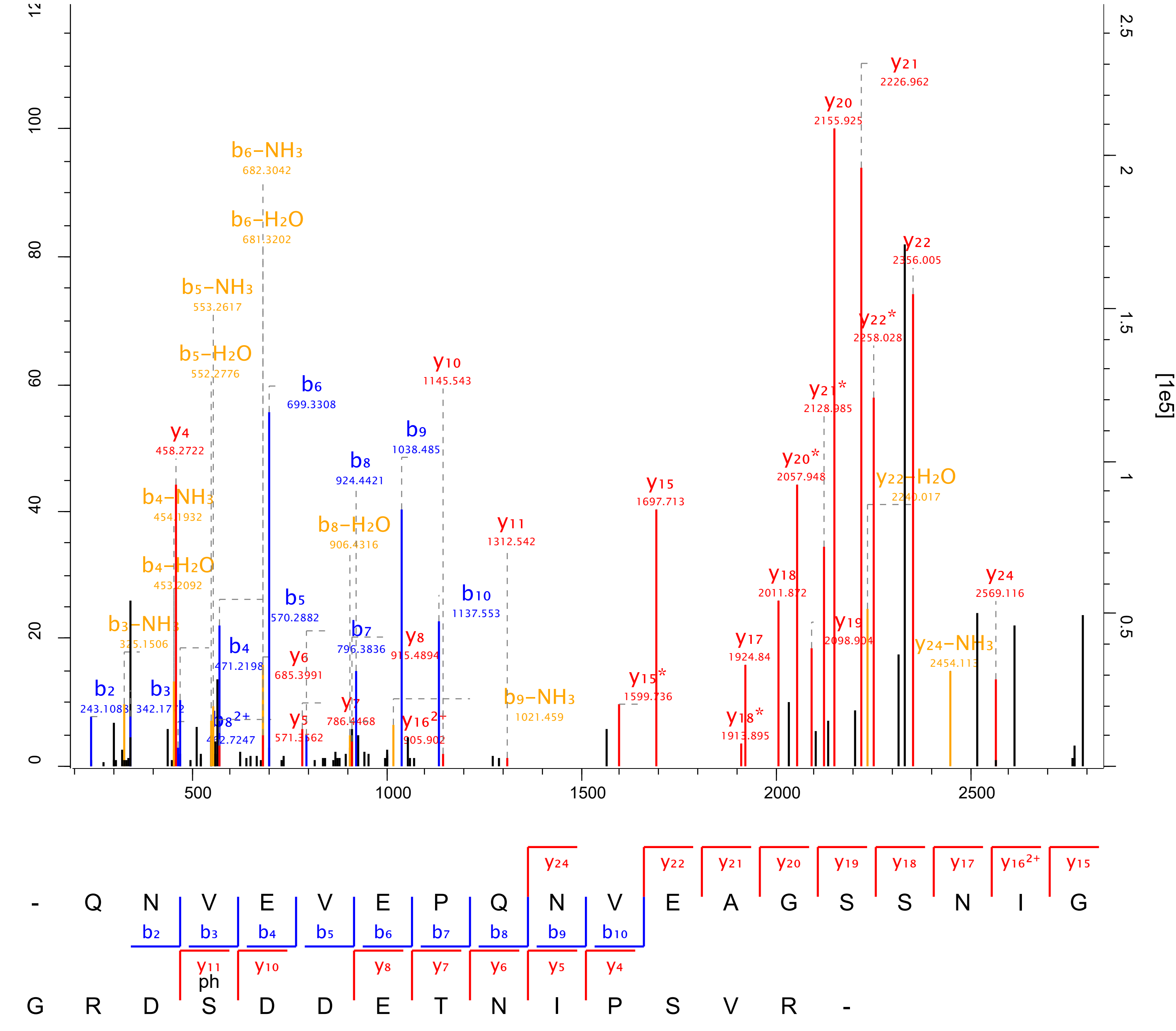

| Raw file | Scan | Method | Score | m/z |
|---|---|---|---|---|
| water_02_1 | 26253 | FTMS; HCD | 136.39 | 792.34 |

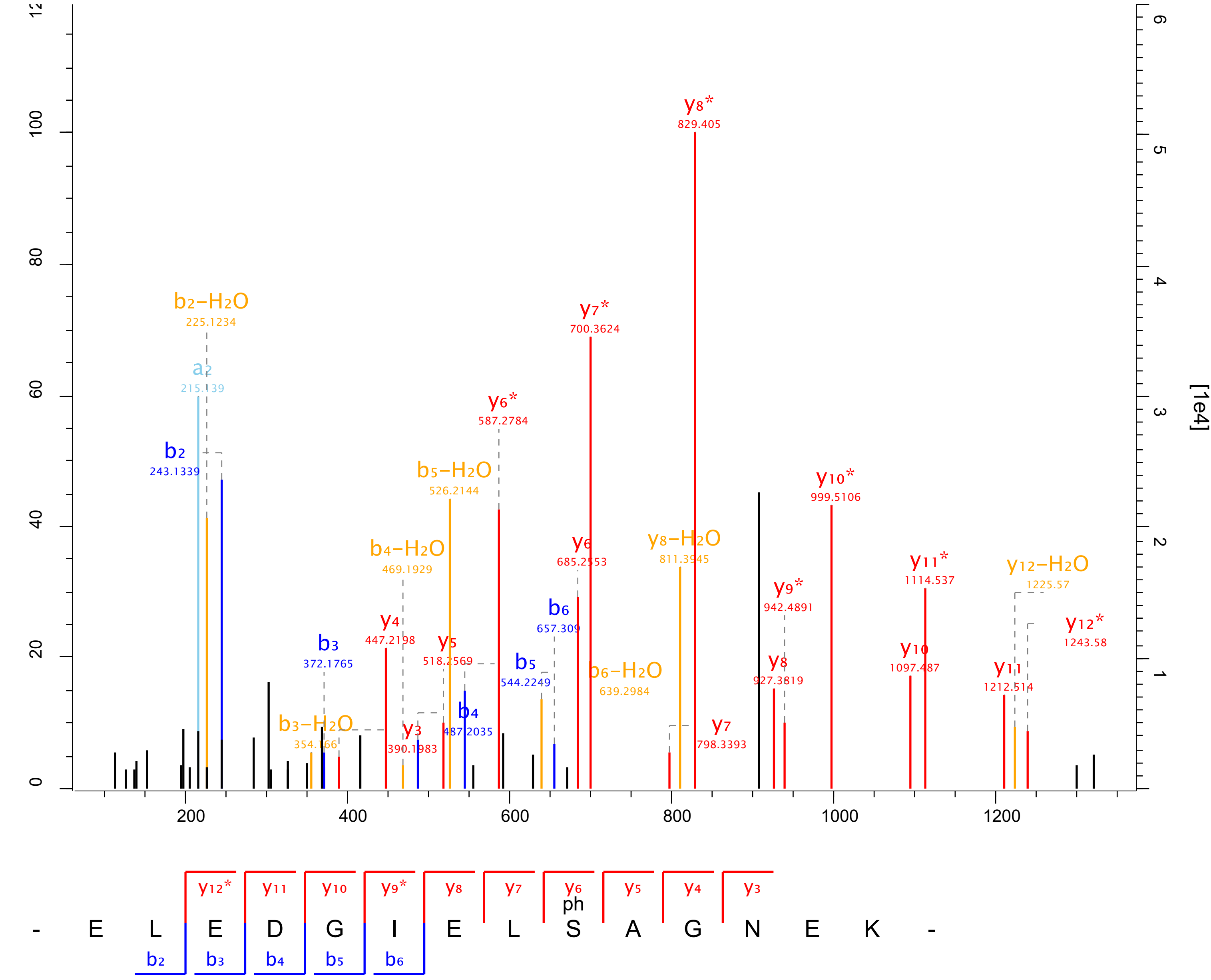

| Raw file | Scan | Method | Score | m/z |
| --- | --- | --- | --- | --- |
| water_02_1 | 2645 | FTMS; HCD | 47.3 | 502.69 |

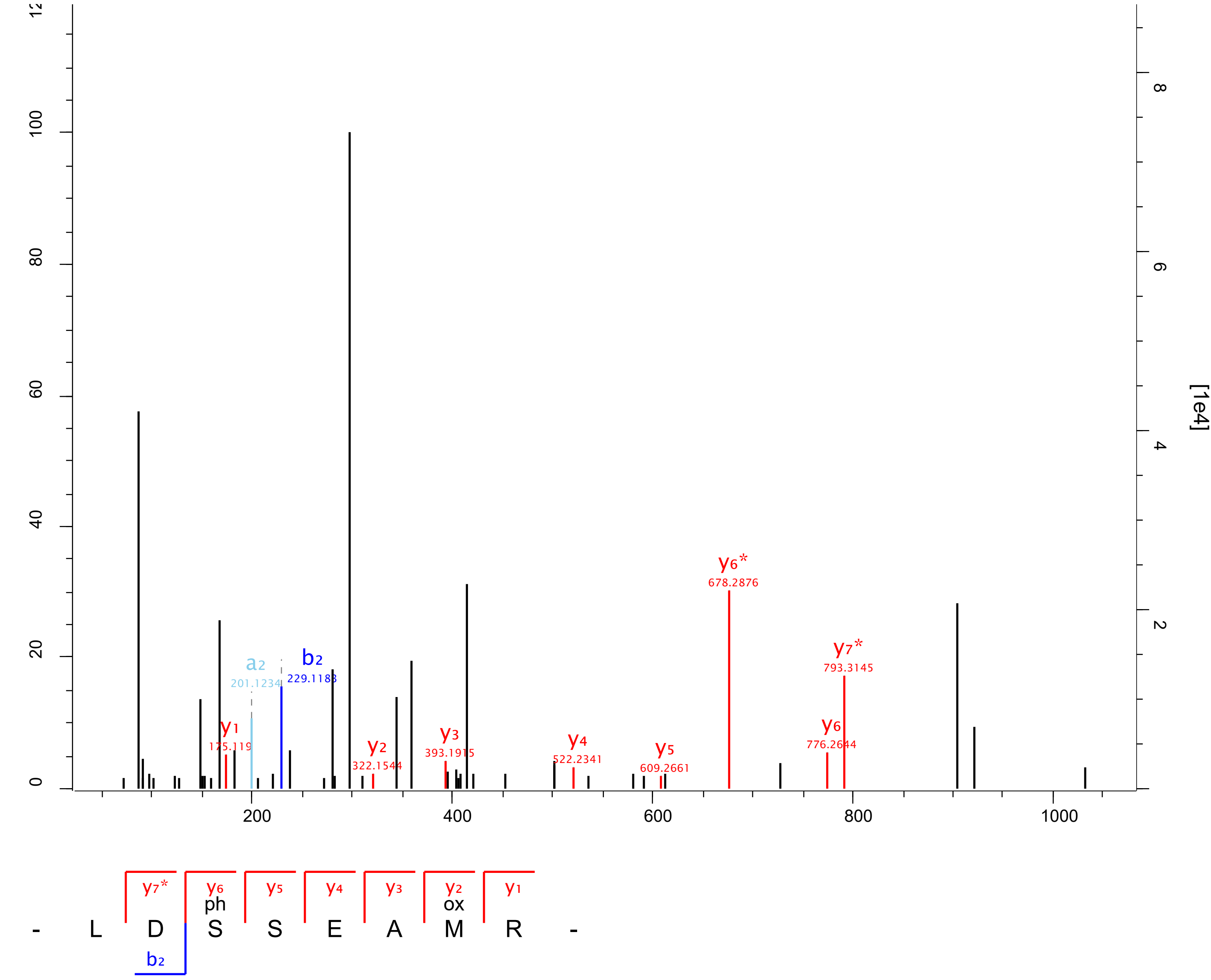

| Raw file | Scan | Method | Score | m/z |
|---|---|---|---|---|
| water_02_1 | 26536 | FTMS; HCD | 231.44 | 784.68 |

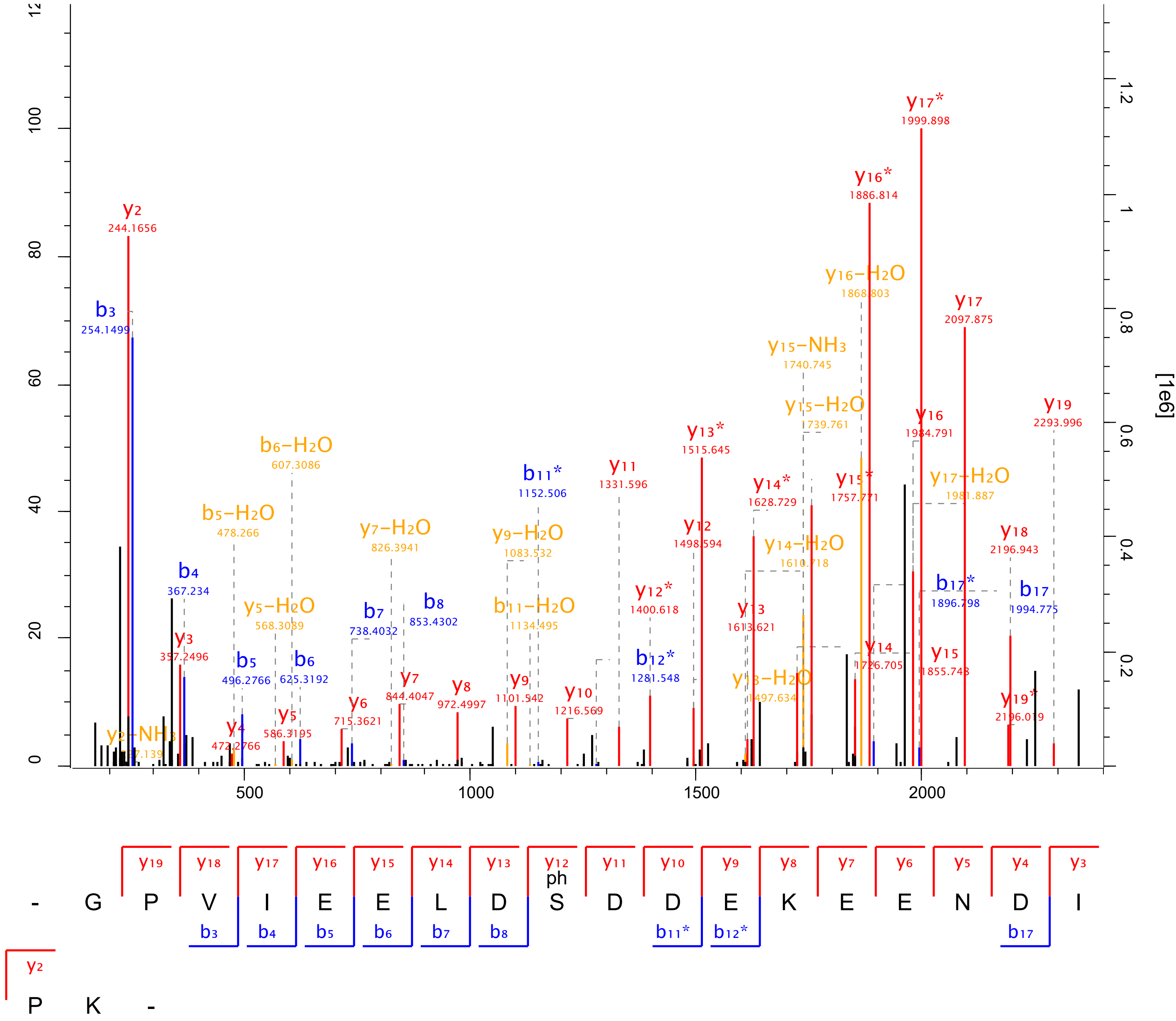

| Raw file | Scan | Method | Score | m/z |
| --- | --- | --- | --- | --- |
| water_02_1 | 26547 | FTMS; HCD | 88.24 | 626.28 |

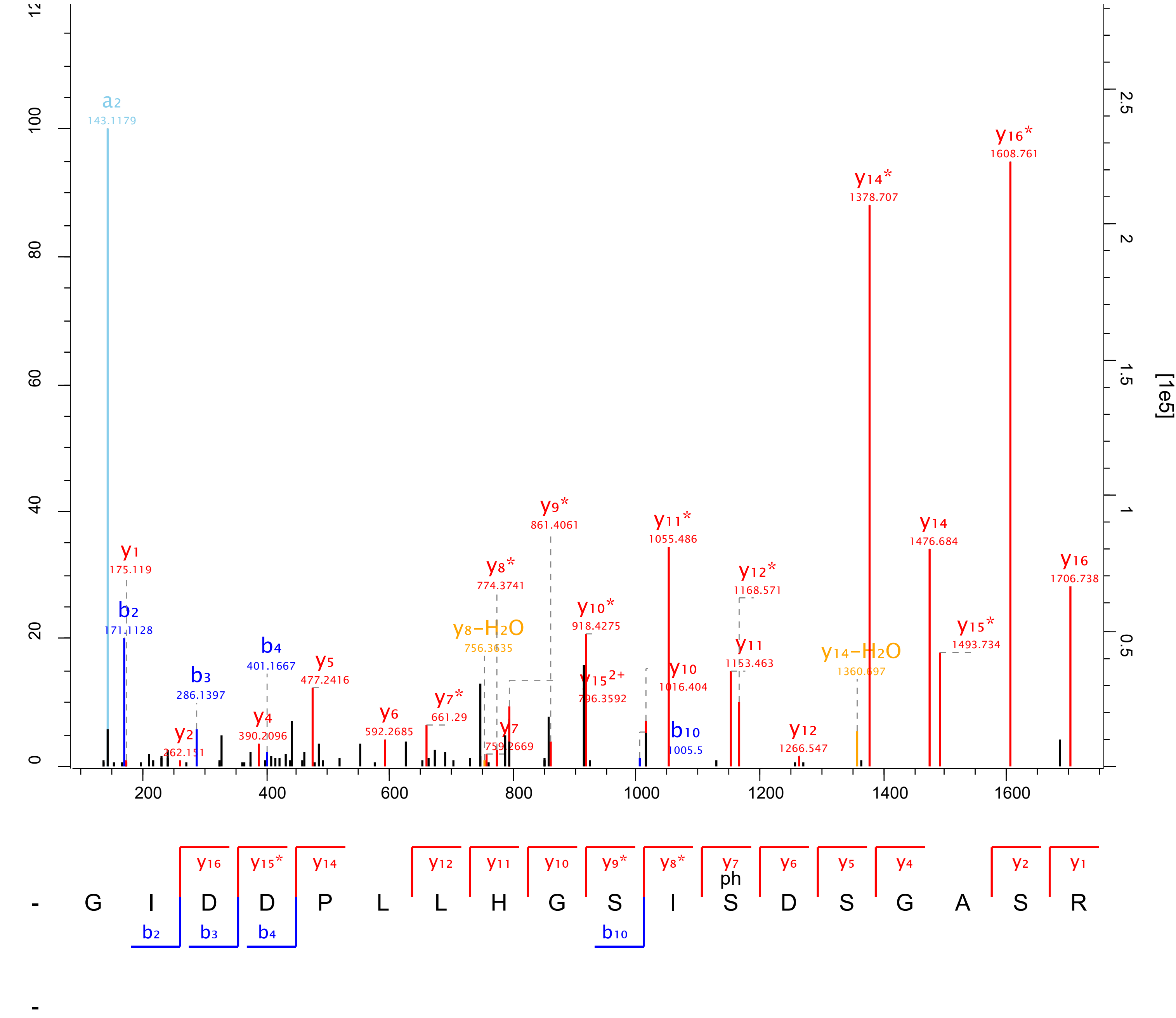

| Raw file | Scan | Method | Score | m/z |
| --- | --- | --- | --- | --- |
| water_02_1 | 26681 | FTMS; HCD | 115.54 | 709.34 |

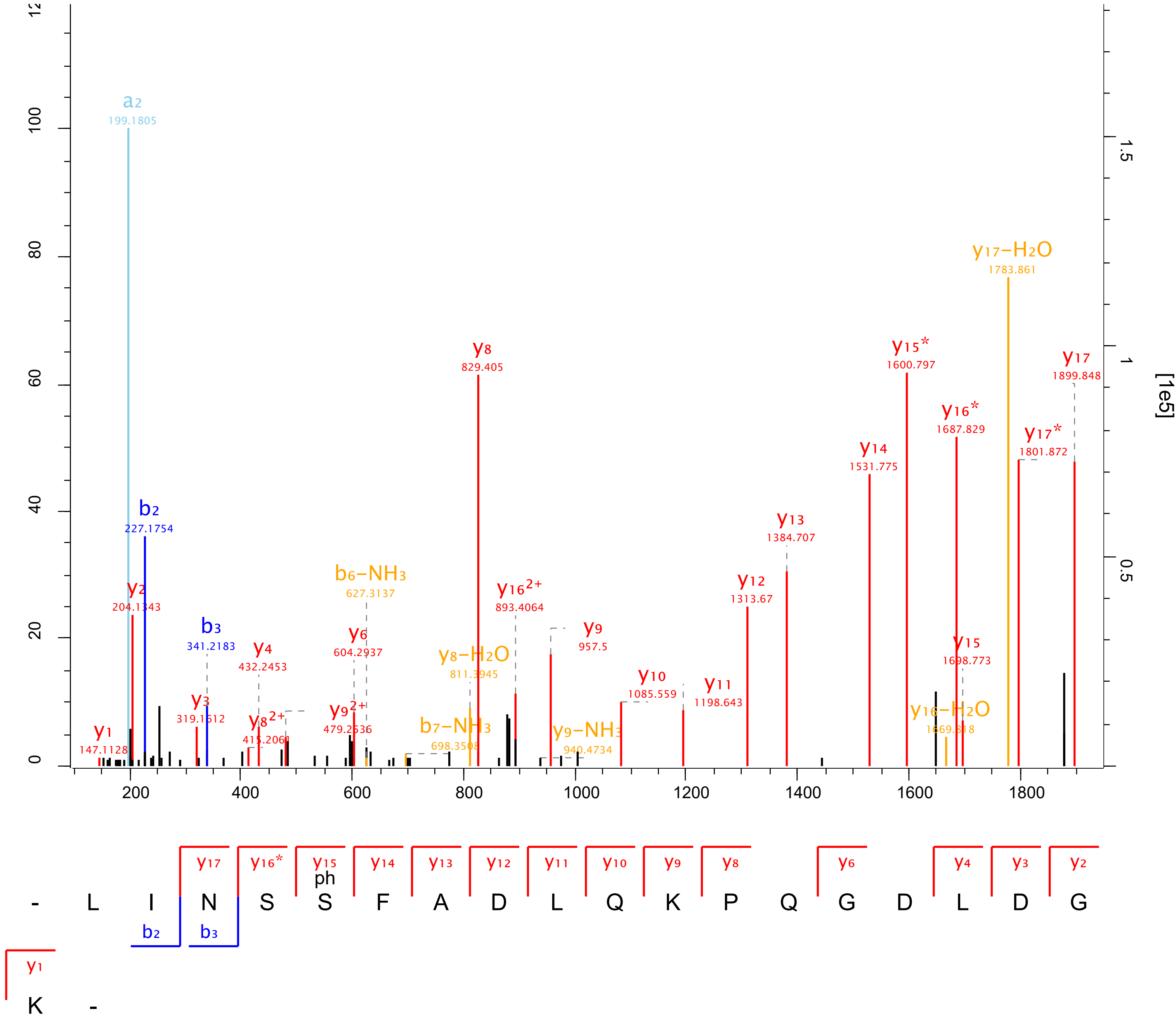

| Raw file | Scan | Method | Score | m/z |
|---|---|---|---|---|
| water_02_1 | 26772 | FTMS; HCD | 48.29 | 769.33 |

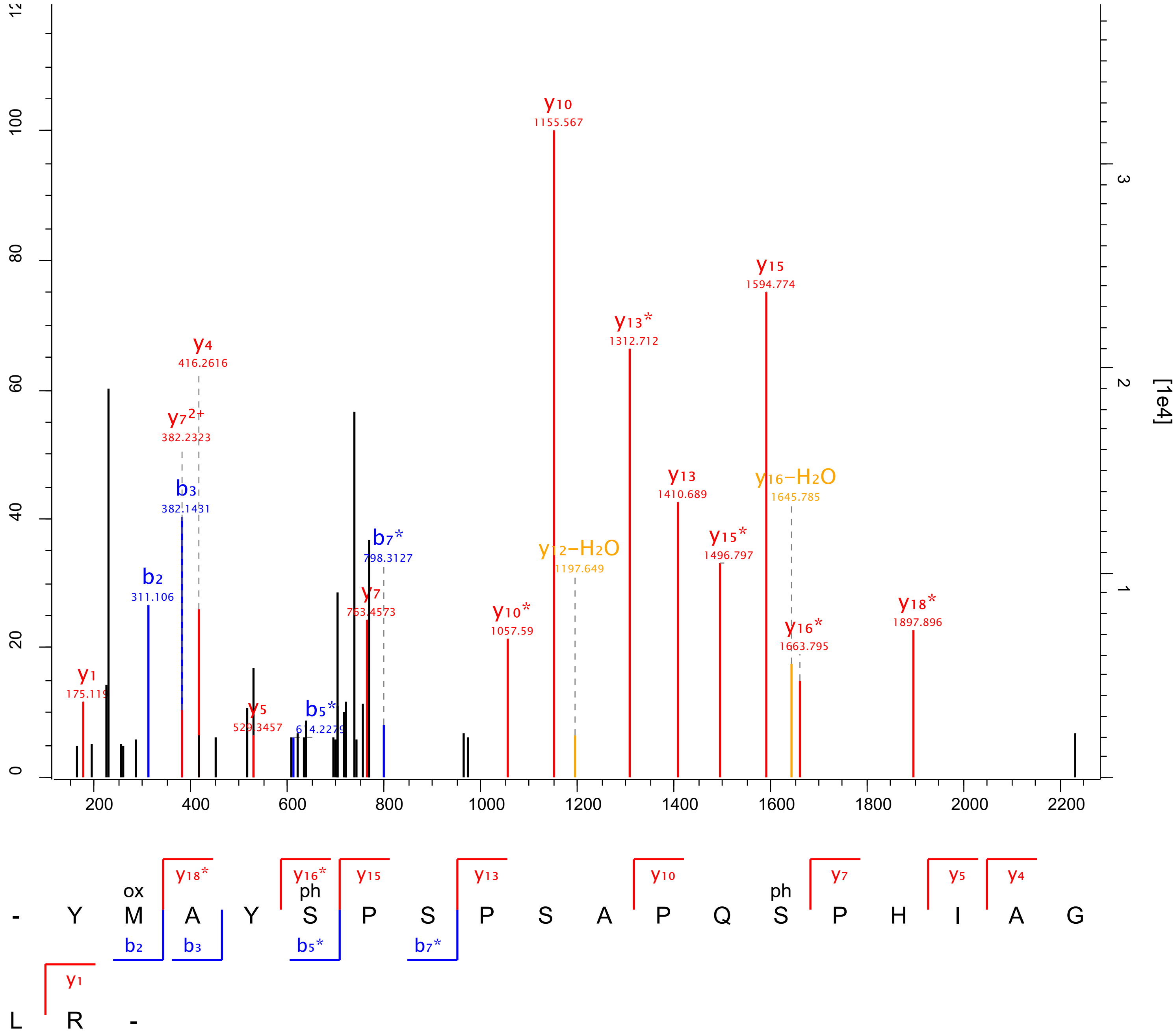

| Raw file | Scan | Method | Score | m/z |
| --- | --- | --- | --- | --- |
| water_02_1 | 26789 | FTMS; HCD | 69.33 | 741.32 |

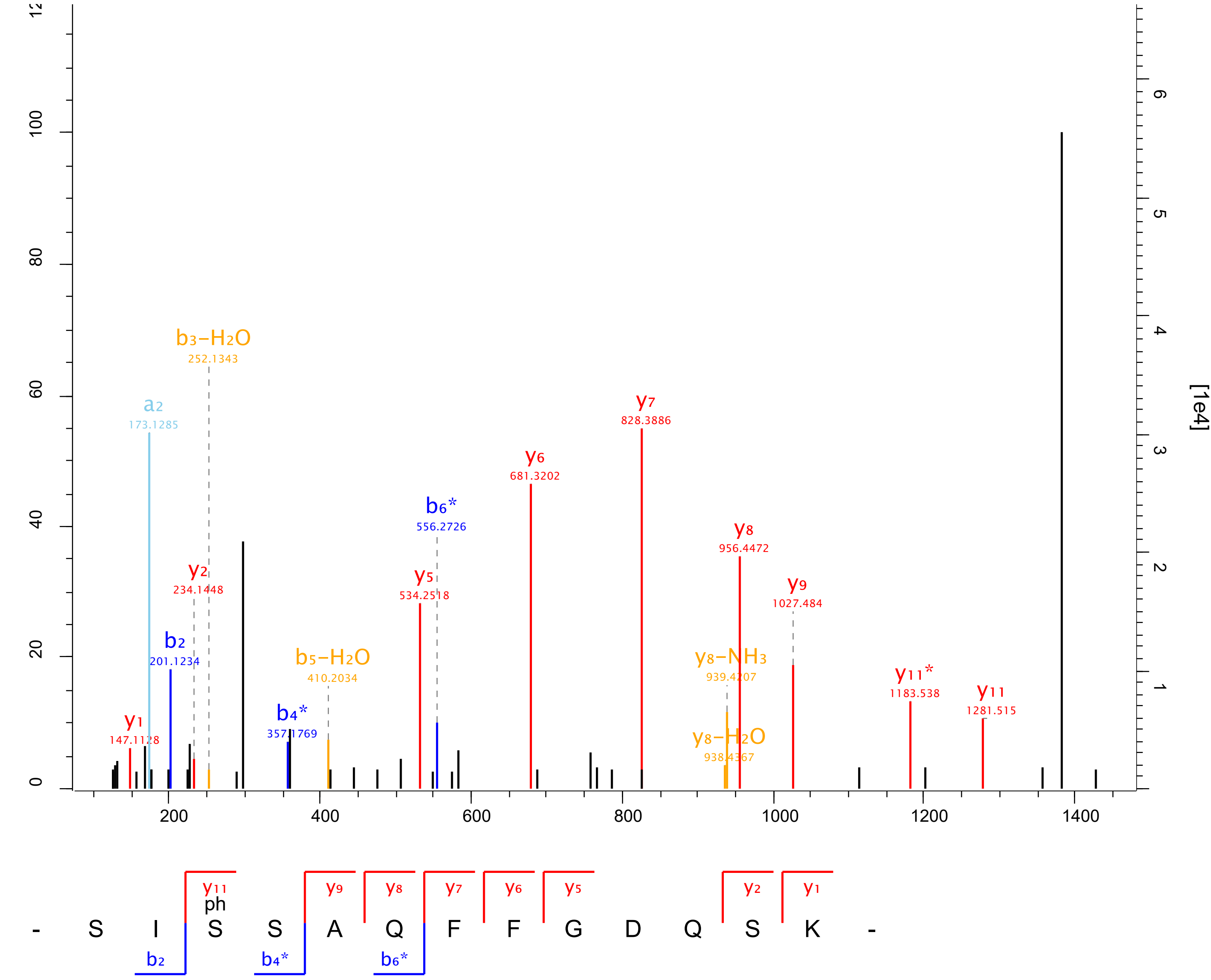

| Raw file | Scan | Method | Score | m/z |
|---|---|---|---|---|
| water_02_1 | 26890 | FTMS; HCD | 109 | 596.77 |

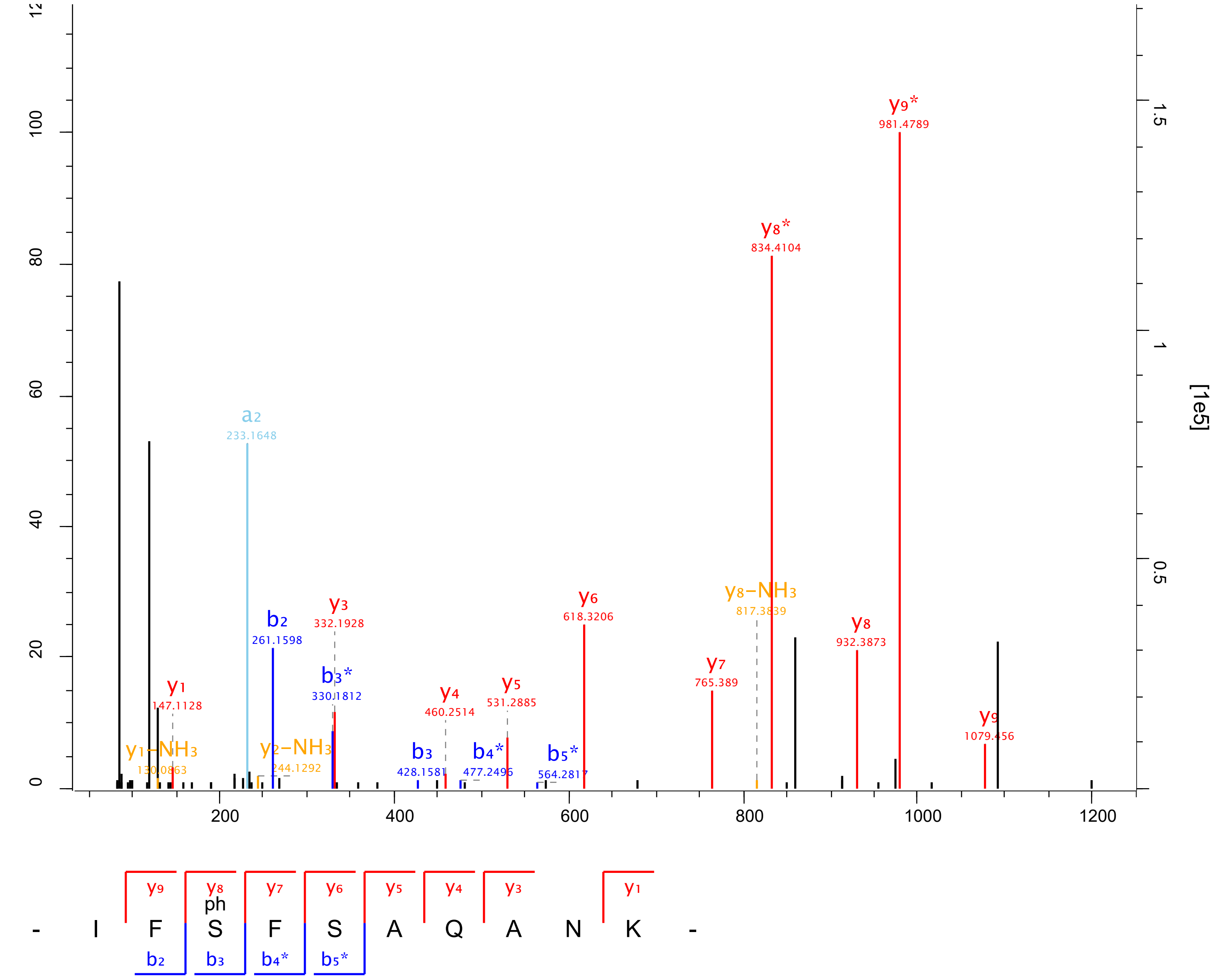

| Raw file | Scan | Method | Score | m/z |
|---|---|---|---|---|
| water_02_1 | 26961 | FTMS; HCD | 188.4 | 710.33 |

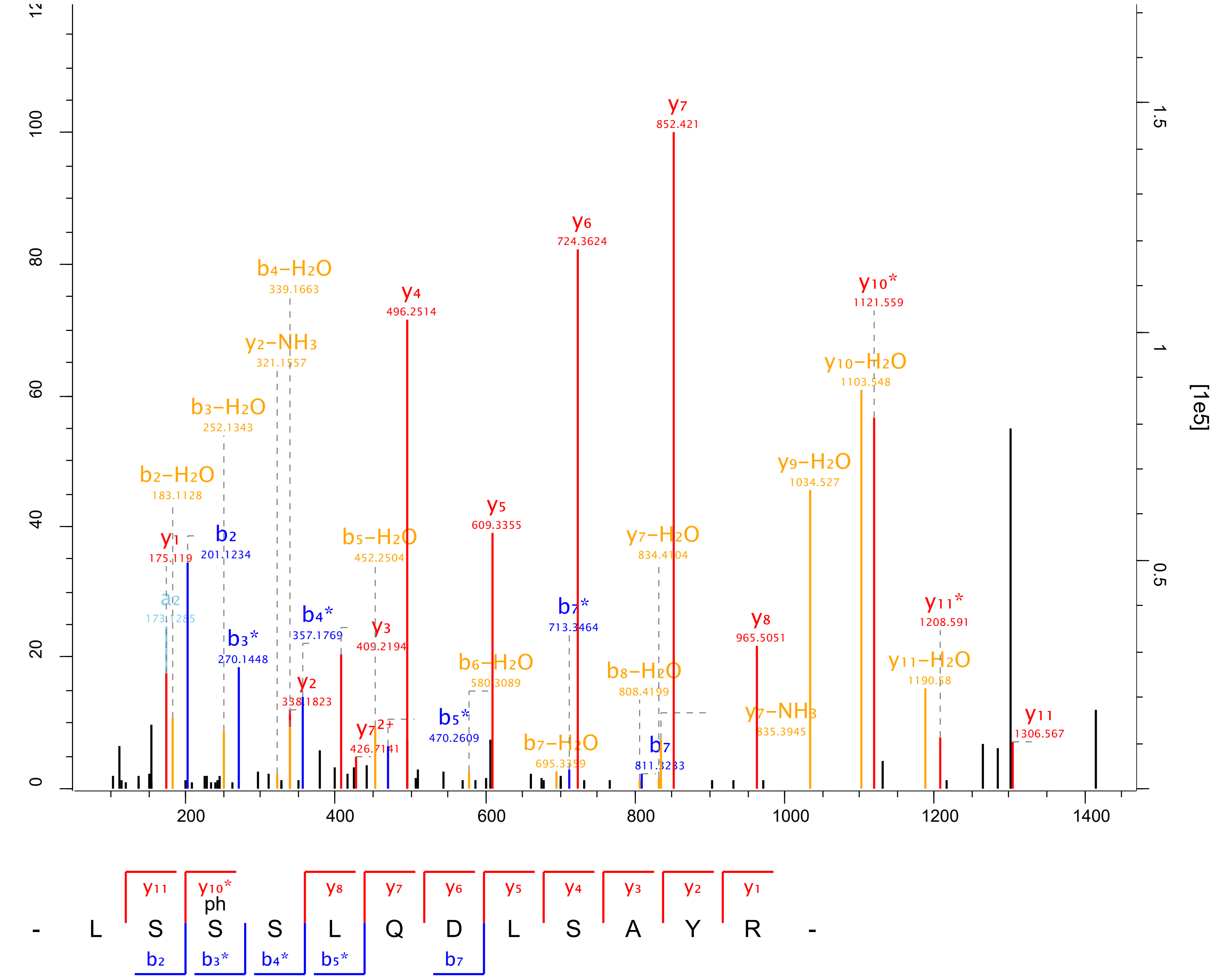

| Raw file | Scan | Method | Score | m/z |
| --- | --- | --- | --- | --- |
| water_02_1 | 27083 | FTMS; HCD | 69.98 | 579.93 |

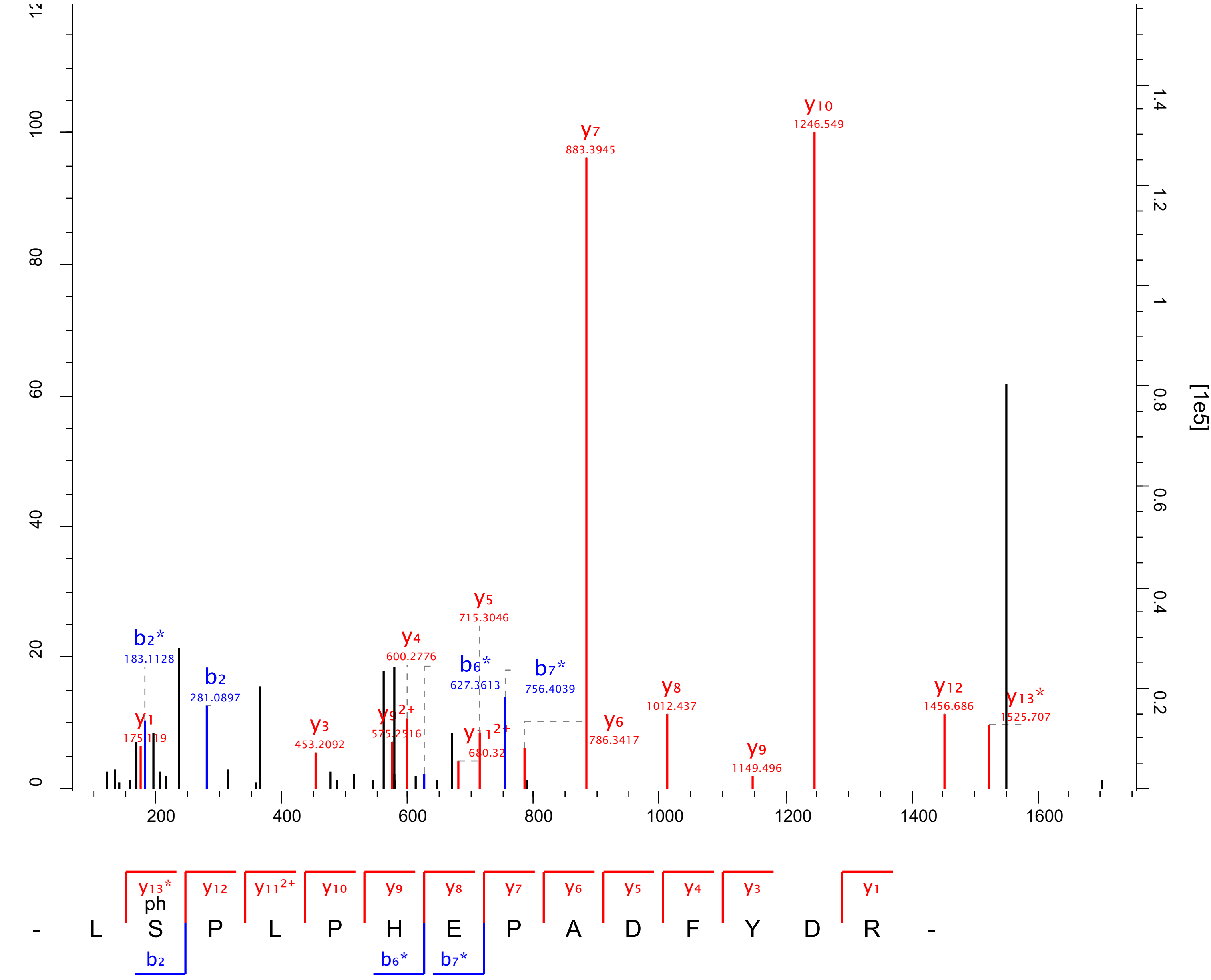

| Raw file | Scan | Method | Score | m/z |
| --- | --- | --- | --- | --- |
| water_02_1 | 27097 | FTMS; HCD | 184.34 | 961.91 |

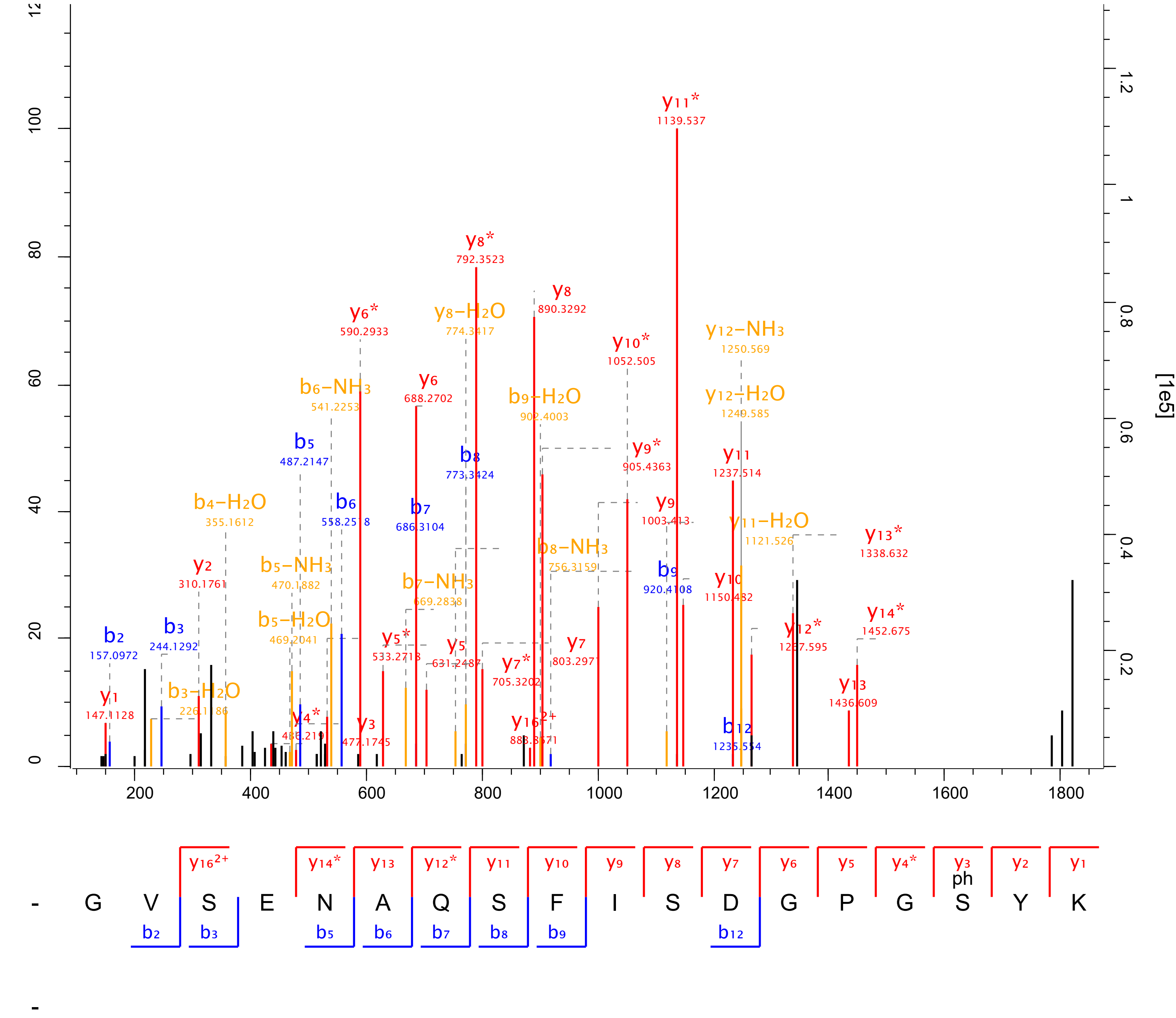

| Raw file | Scan | Method | Score | m/z |
|---|---|---|---|---|
| water_02_1 | 2711 | FTMS; HCD | 59.07 | 451.69 |

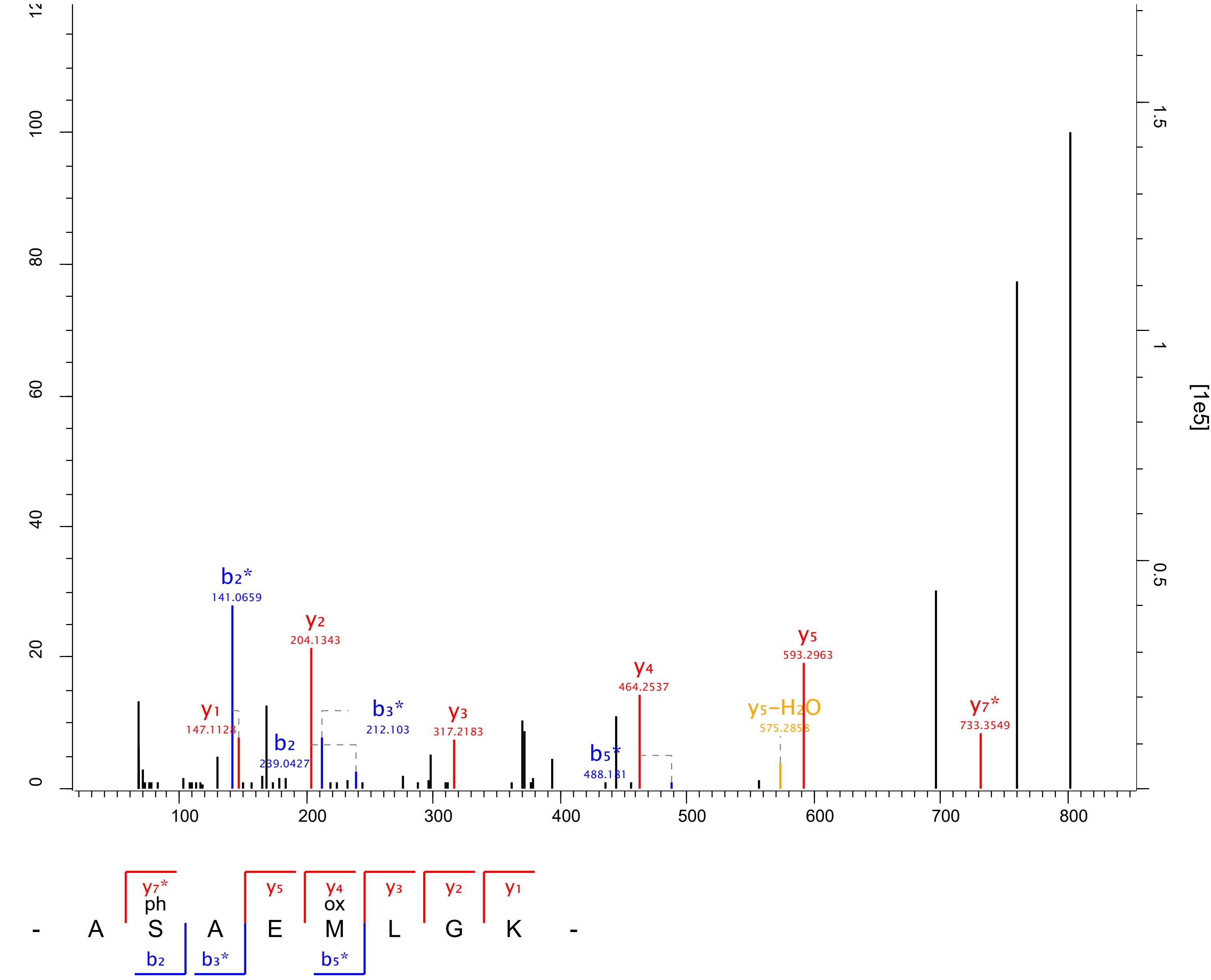

| Raw file | Scan | Method | Score | m/z |
|---|---|---|---|---|
| water_02_1 | 27382 | FTMS; HCD | 43.7 | 757.68 |

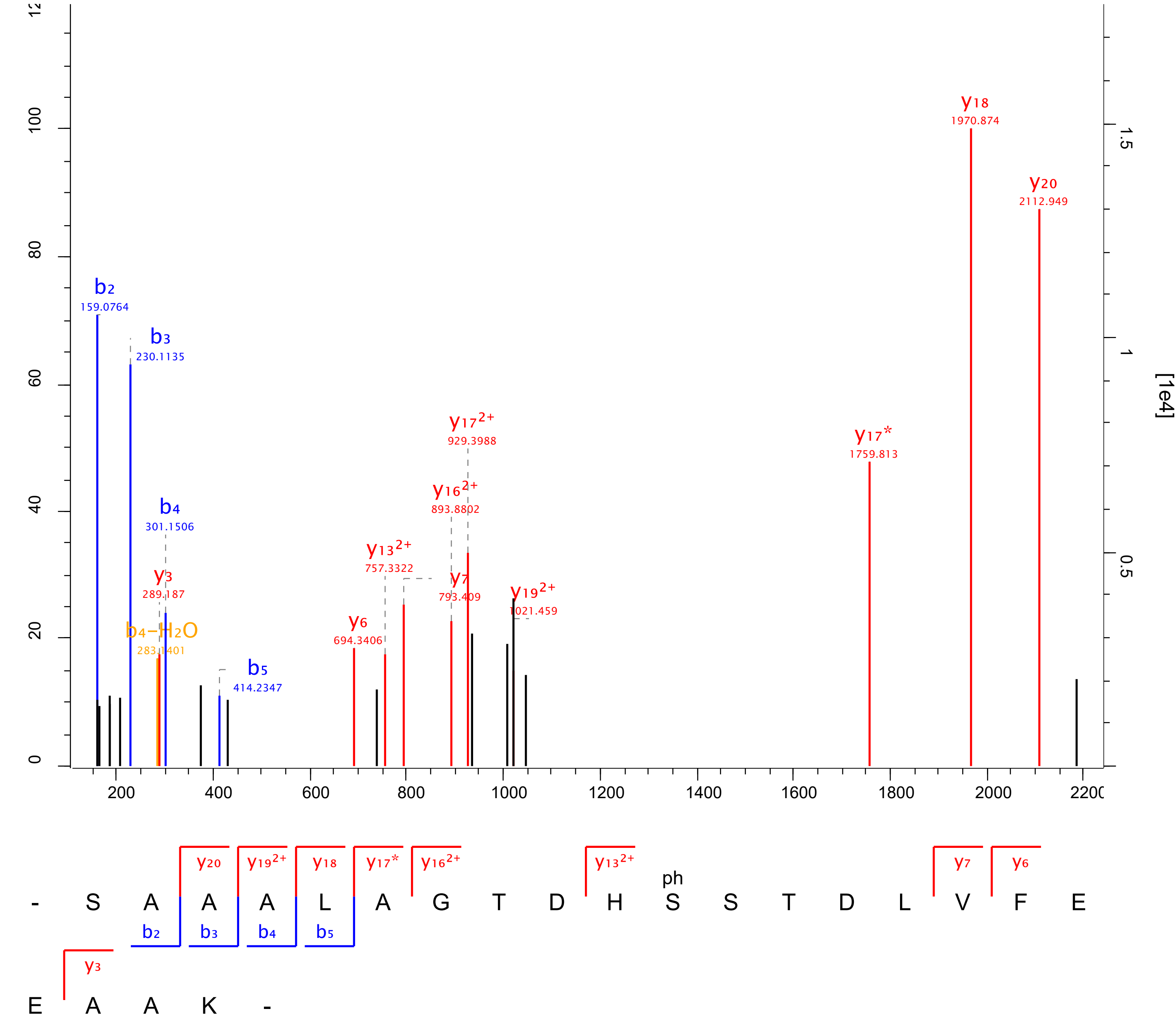

| Raw file | Scan | Method | Score | m/z |
|---|---|---|---|---|
| water_02_1 | 2752 | FTMS; HCD | 68.54 | 448.21 |

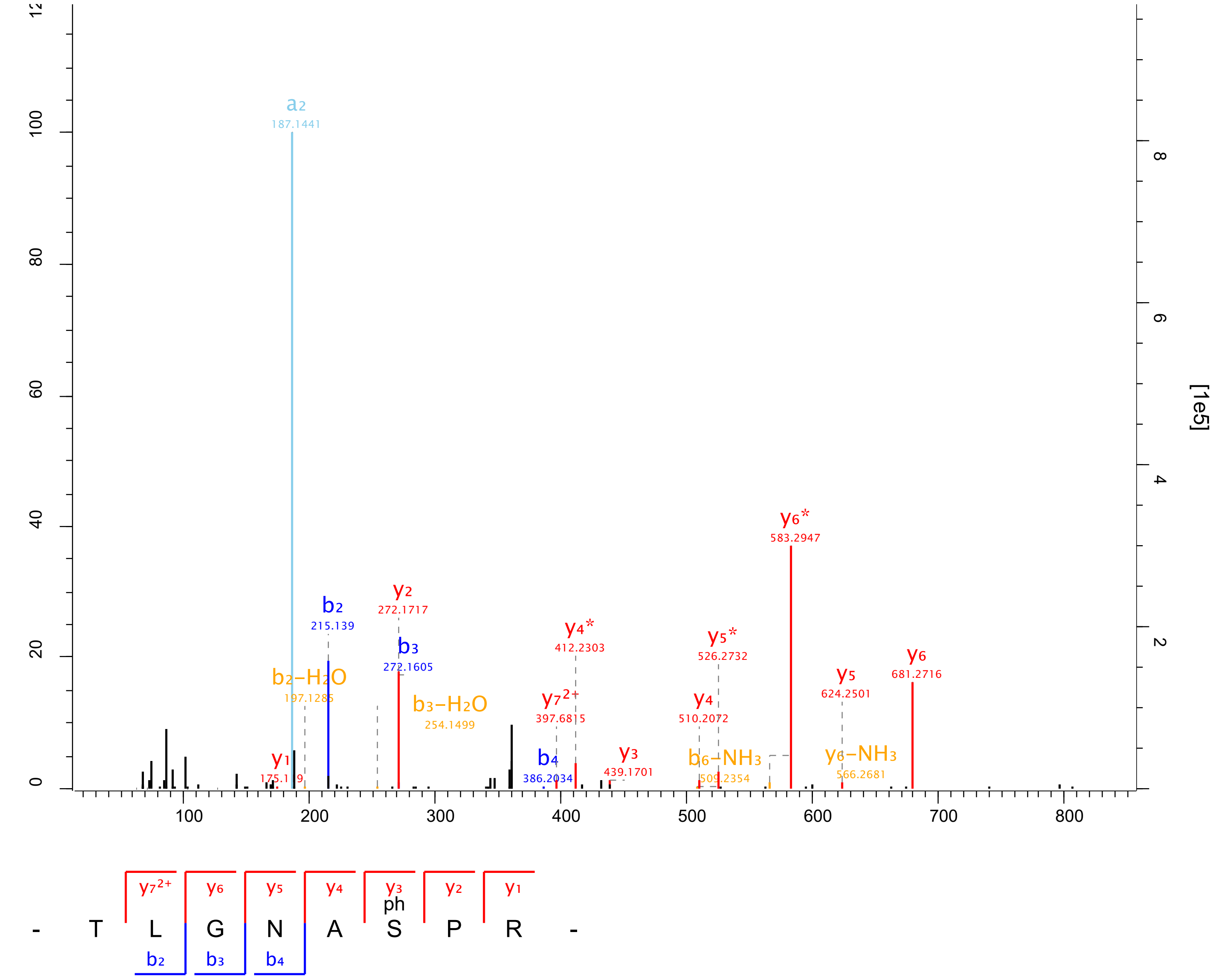

| Raw file | Scan | Method | Score | m/z |
|---|---|---|---|---|
| water_02_1 | 27551 | FTMS; HCD | 169.04 | 598.3 |

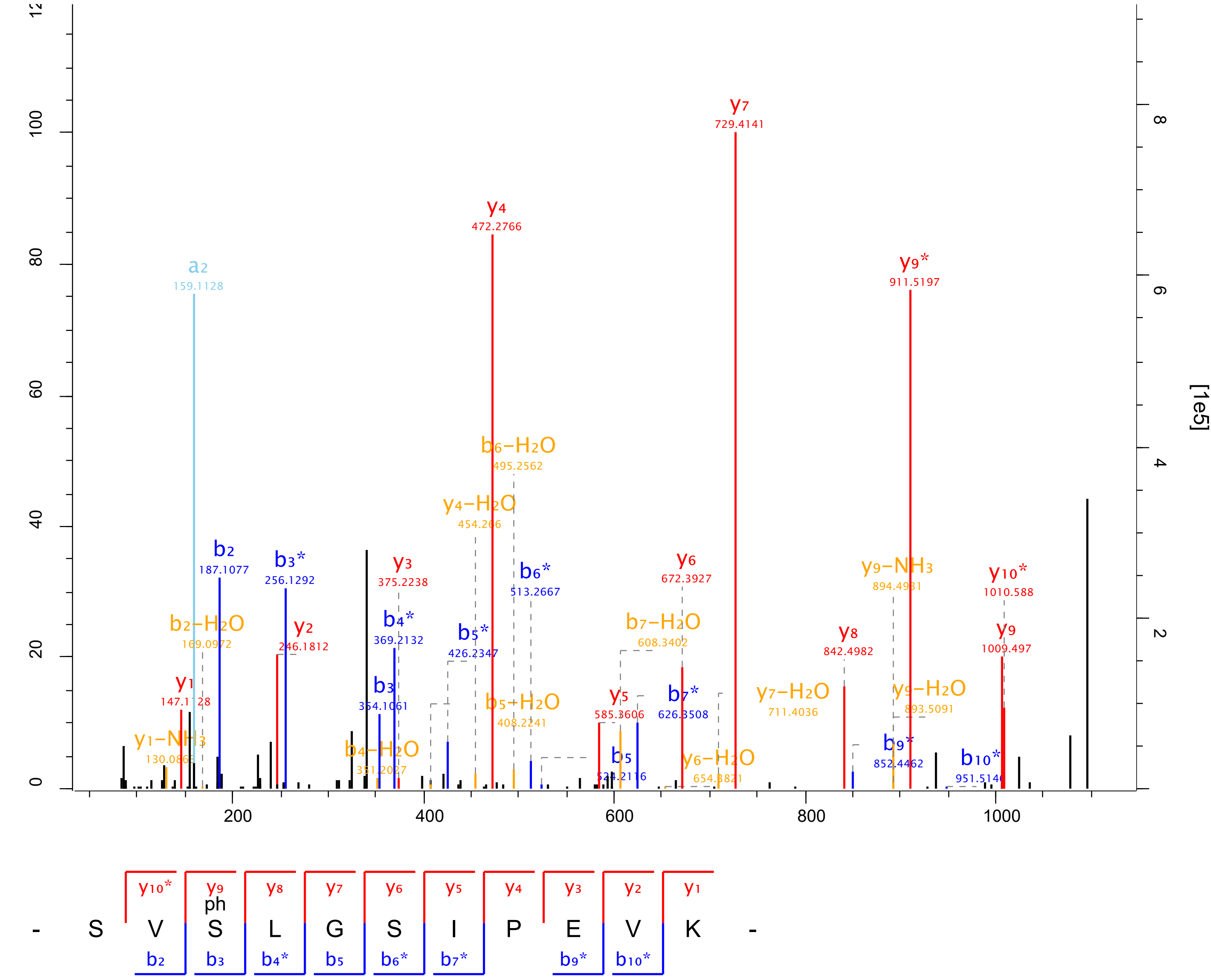

| Raw file | Scan | Method | Score | m/z |
|---|---|---|---|---|
| water_02_1 | 27570 | FTMS; HCD | 49.33 | 643.78 |

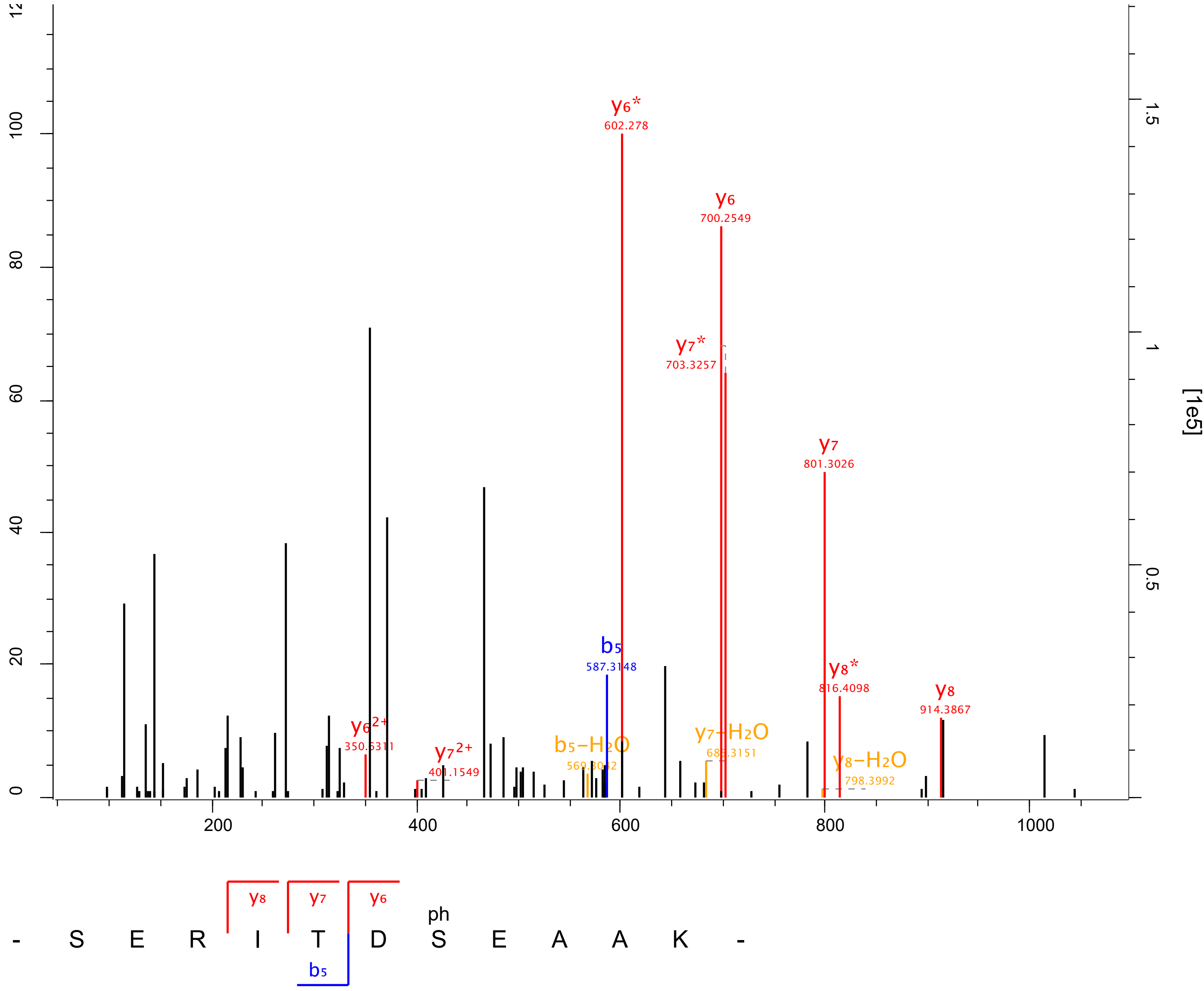

| Raw file | Scan | Method | Score | m/z |
|---|---|---|---|---|
| water_02_1 | 27618 | FTMS; HCD | 169.58 | 694.81 |

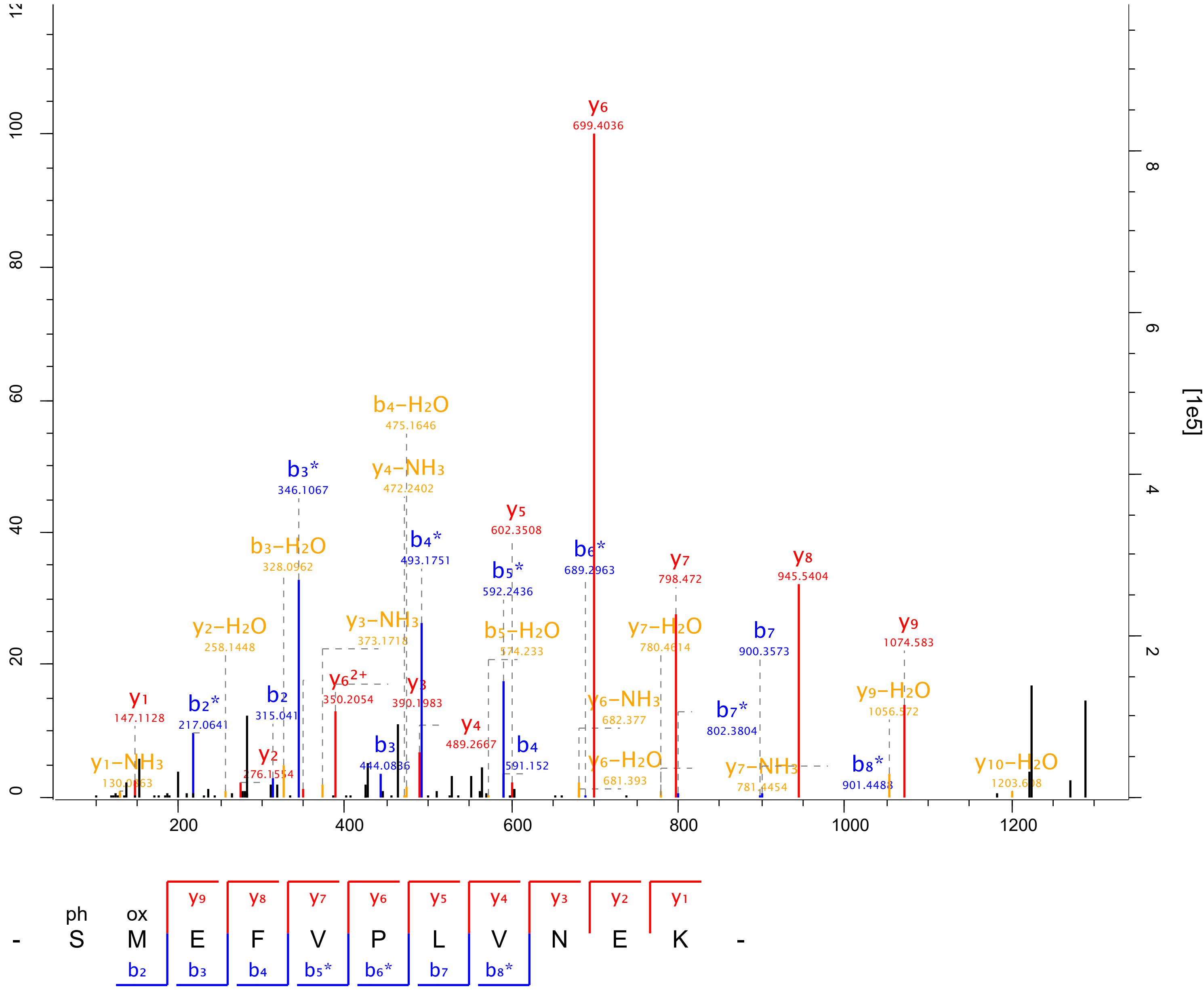

| Raw file | Scan | Method | Score | m/z |
|---|---|---|---|---|
| water_02_1 | 27701 | FTMS; HCD | 92.04 | 601.28 |

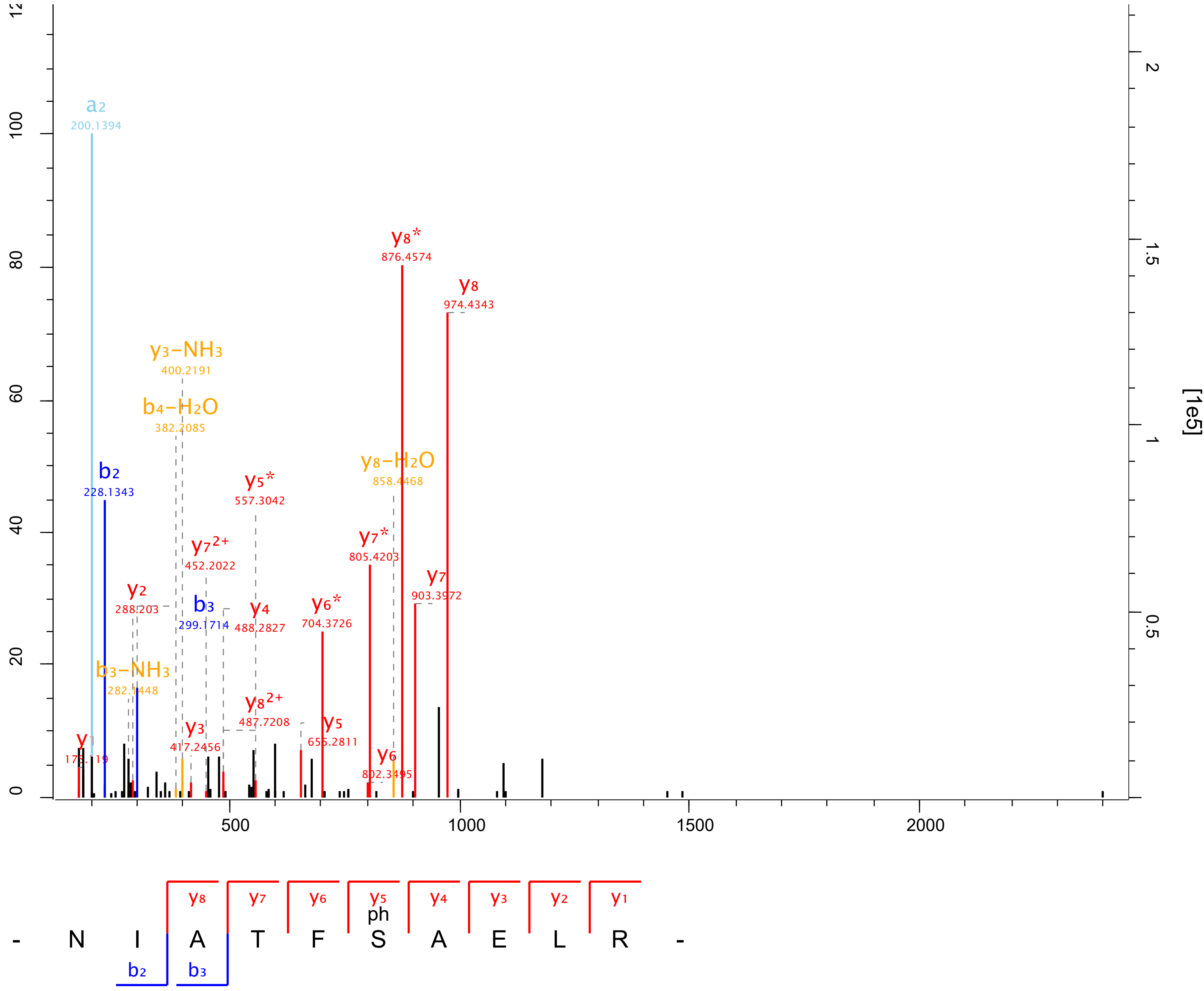

| Raw file | Scan | Method | Score | m/z |
|---|---|---|---|---|
| water_02_1 | 27764 | FTMS; HCD | 103.76 | 639.75 |

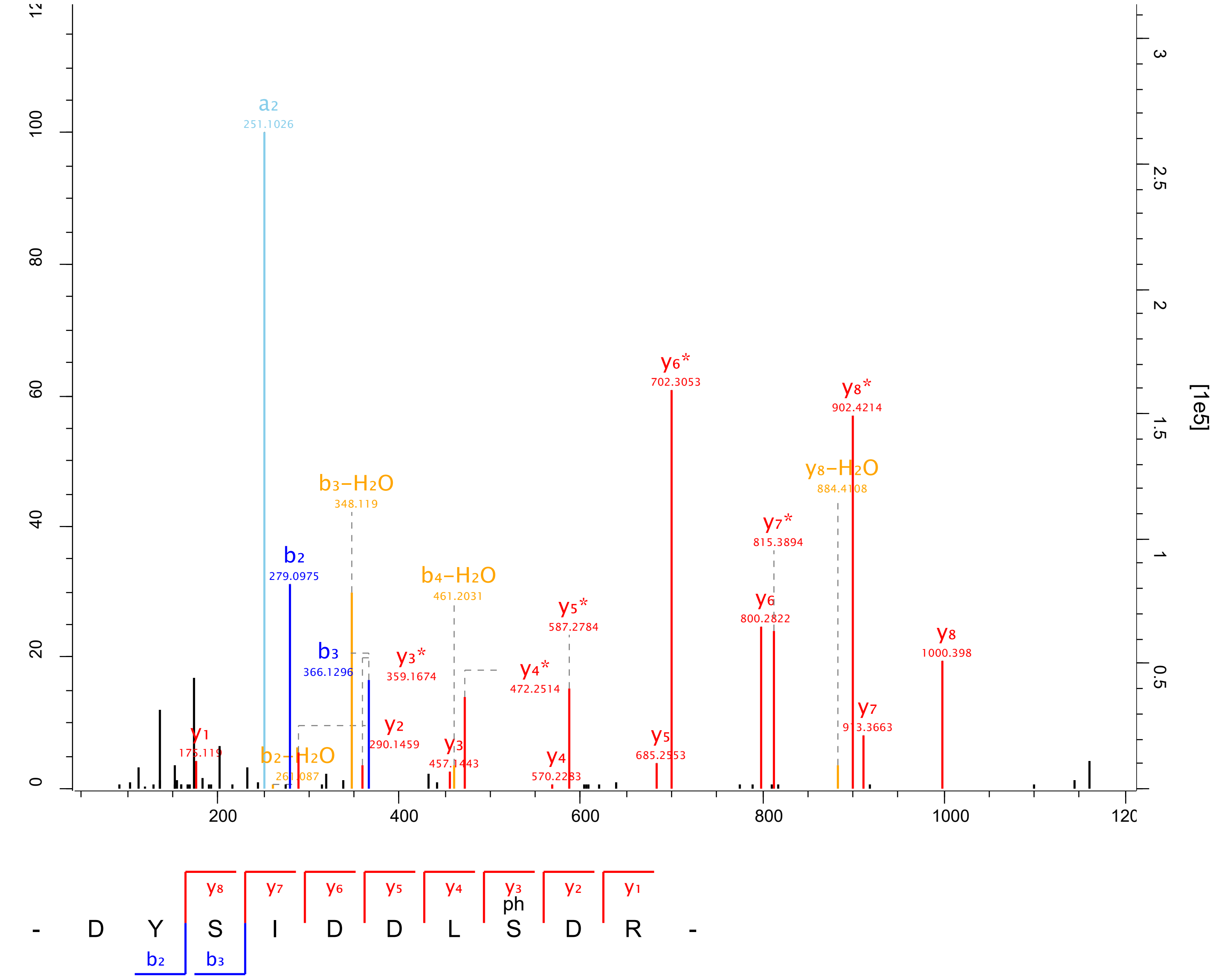

| Raw file | Scan | Method | Score | m/z |
|---|---|---|---|---|
| water_02_1 | 27781 | FTMS; HCD | 105.79 | 757.87 |

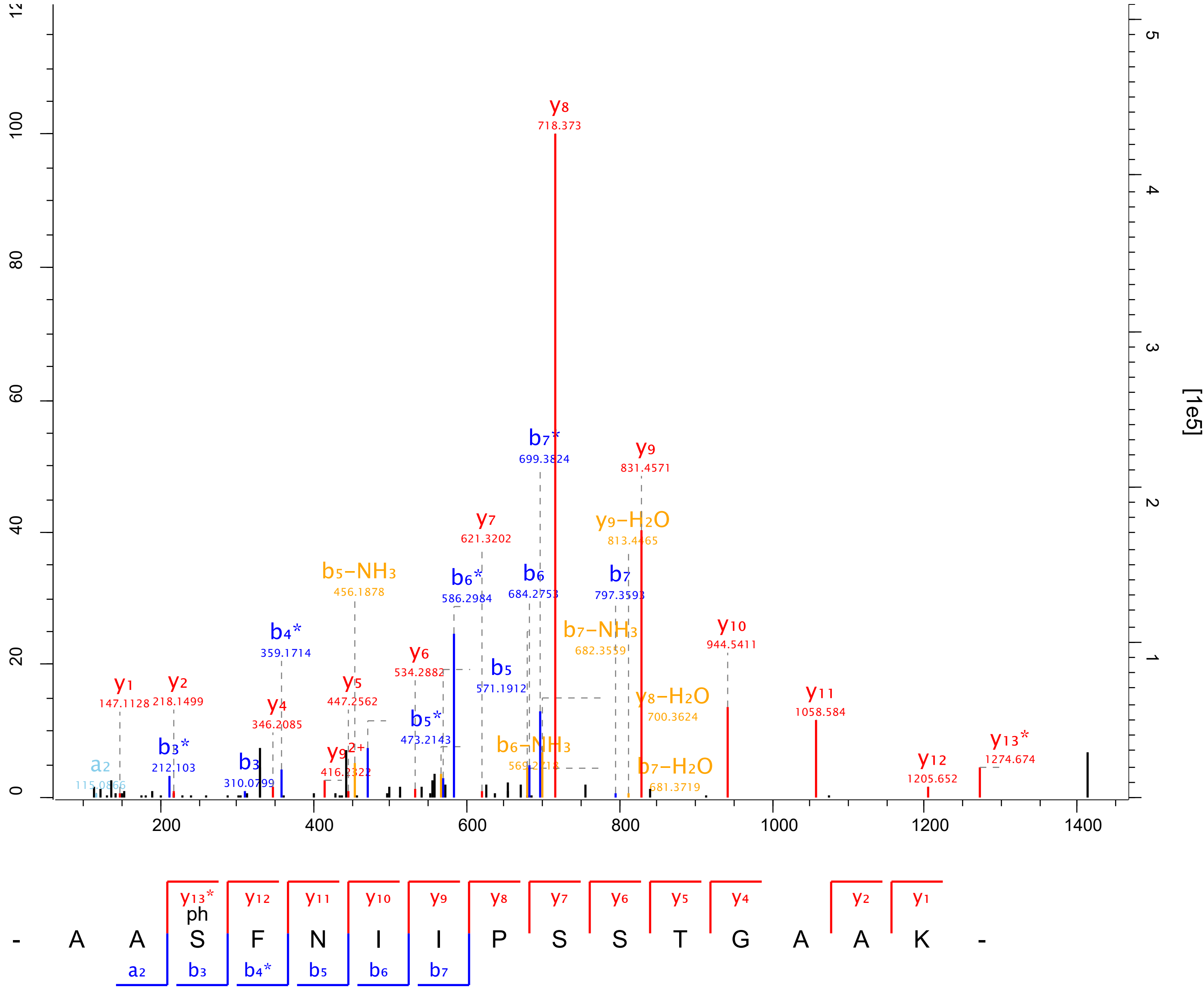

| Raw file | Scan | Method | Score | m/z |
| --- | --- | --- | --- | --- |
| water_02_1 | 2779 | FTMS; HCD | 45.43 | 506.71 |

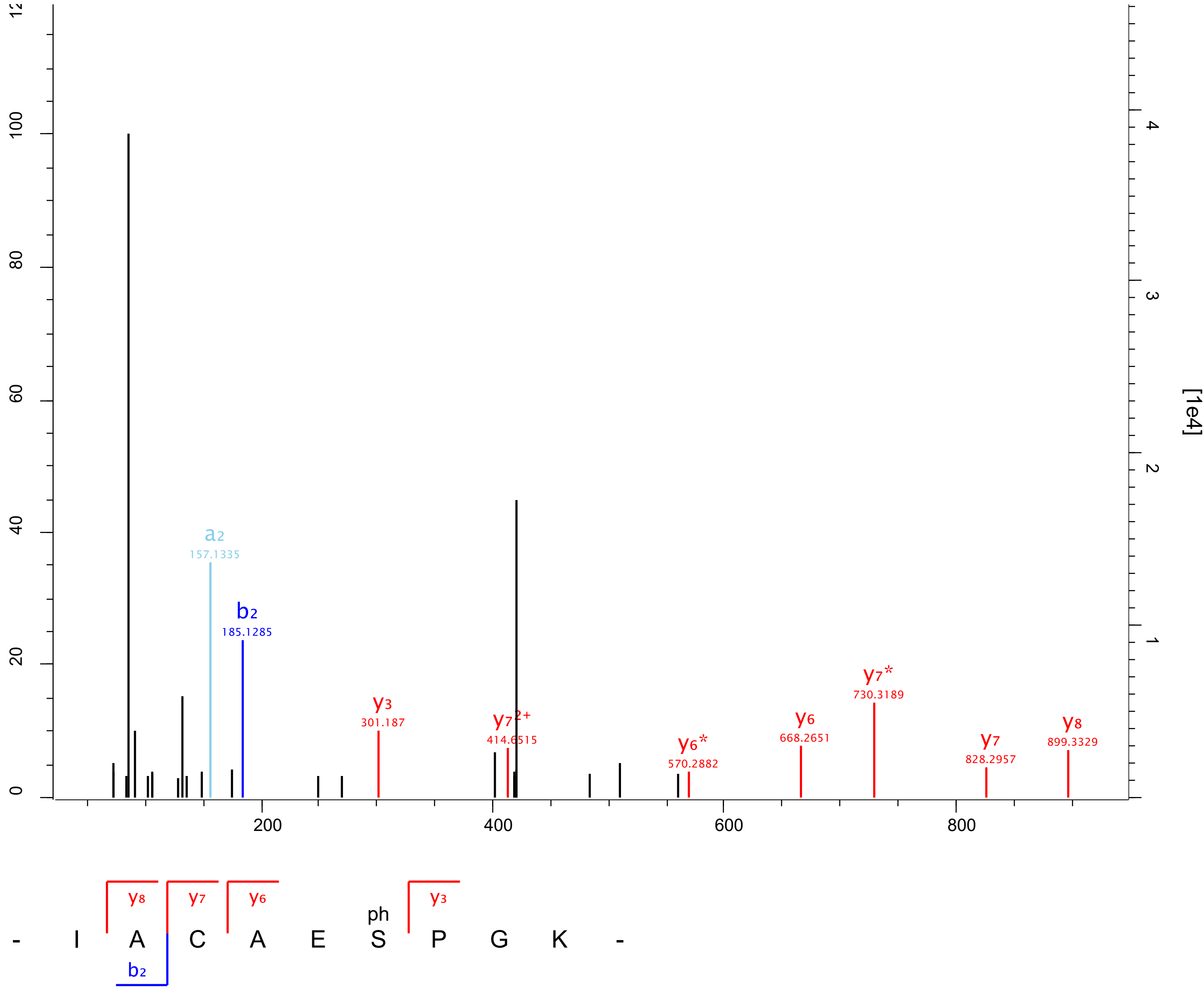

| Raw file | Scan | Method | Score | m/z |
| --- | --- | --- | --- | --- |
| water_02_1 | 27917 | FTMS; HCD | 174.04 | 658.33 |

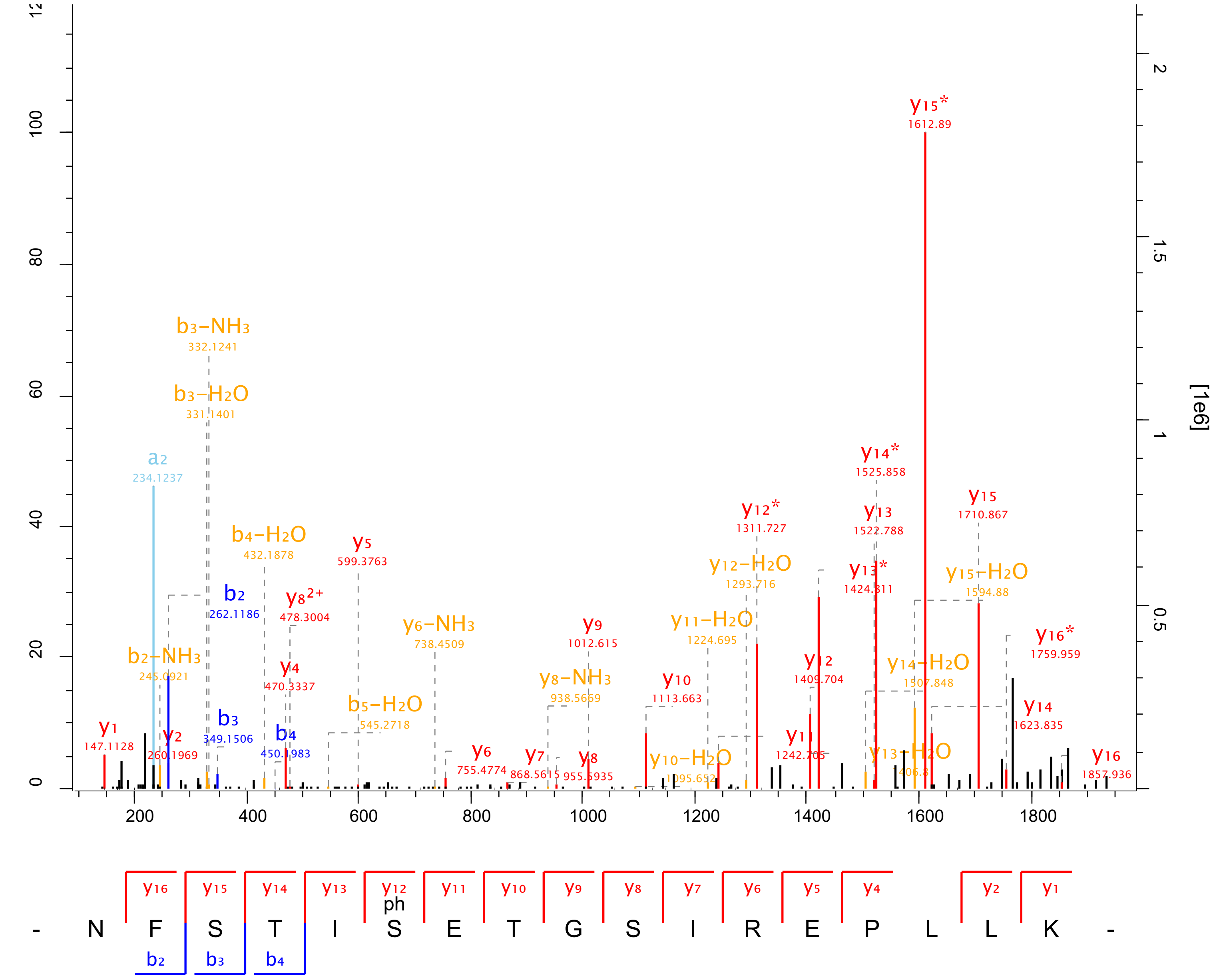

| Raw file | Scan | Method | Score | m/z |
| --- | --- | --- | --- | --- |
| water_02_1 | 2804 | FTMS; HCD | 98.56 | 616.57 |

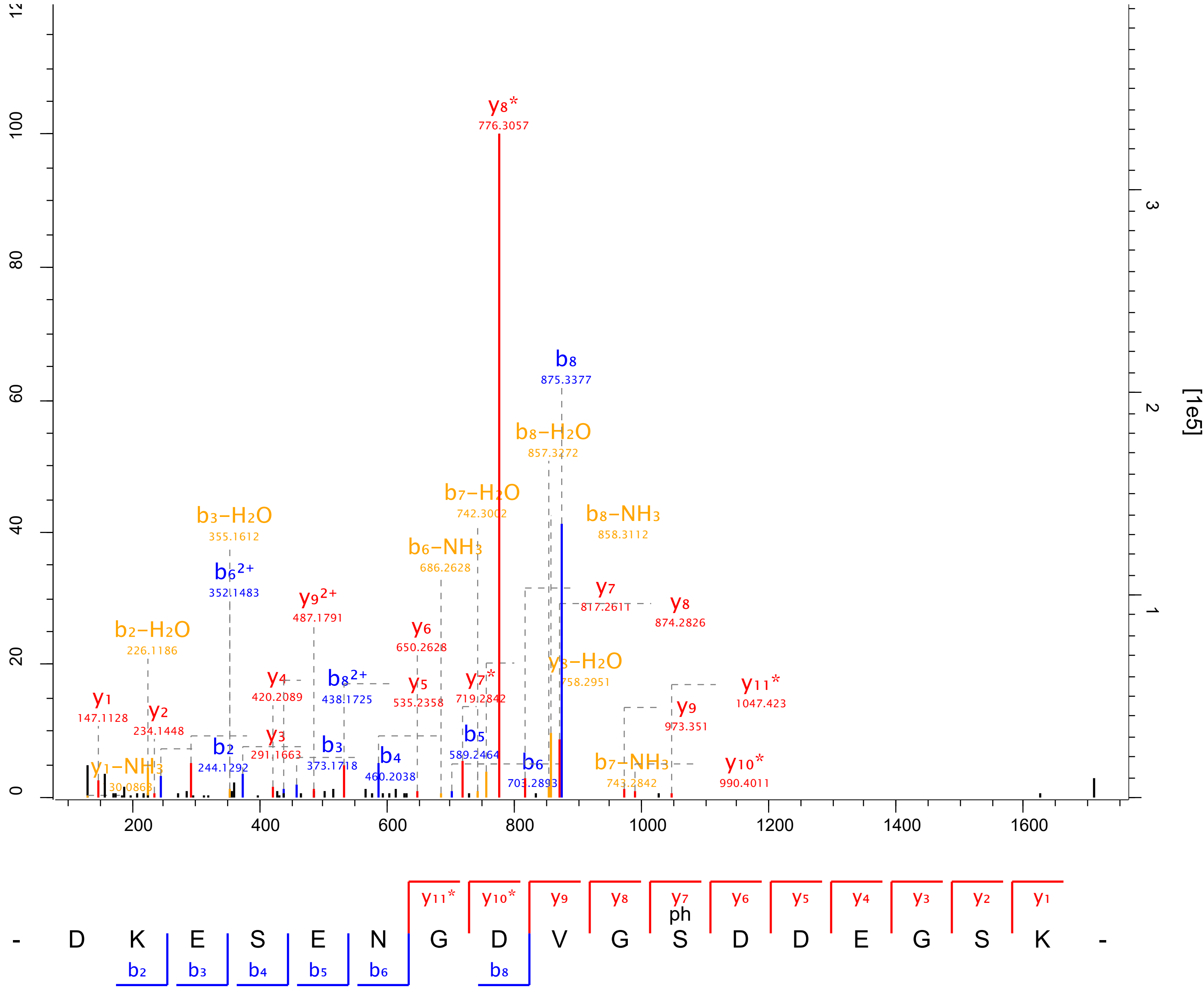

| Raw file | Scan | Method | Score | m/z |
|---|---|---|---|---|
| water_02_1 | 28160 | FTMS; HCD | 148.57 | 797.82 |

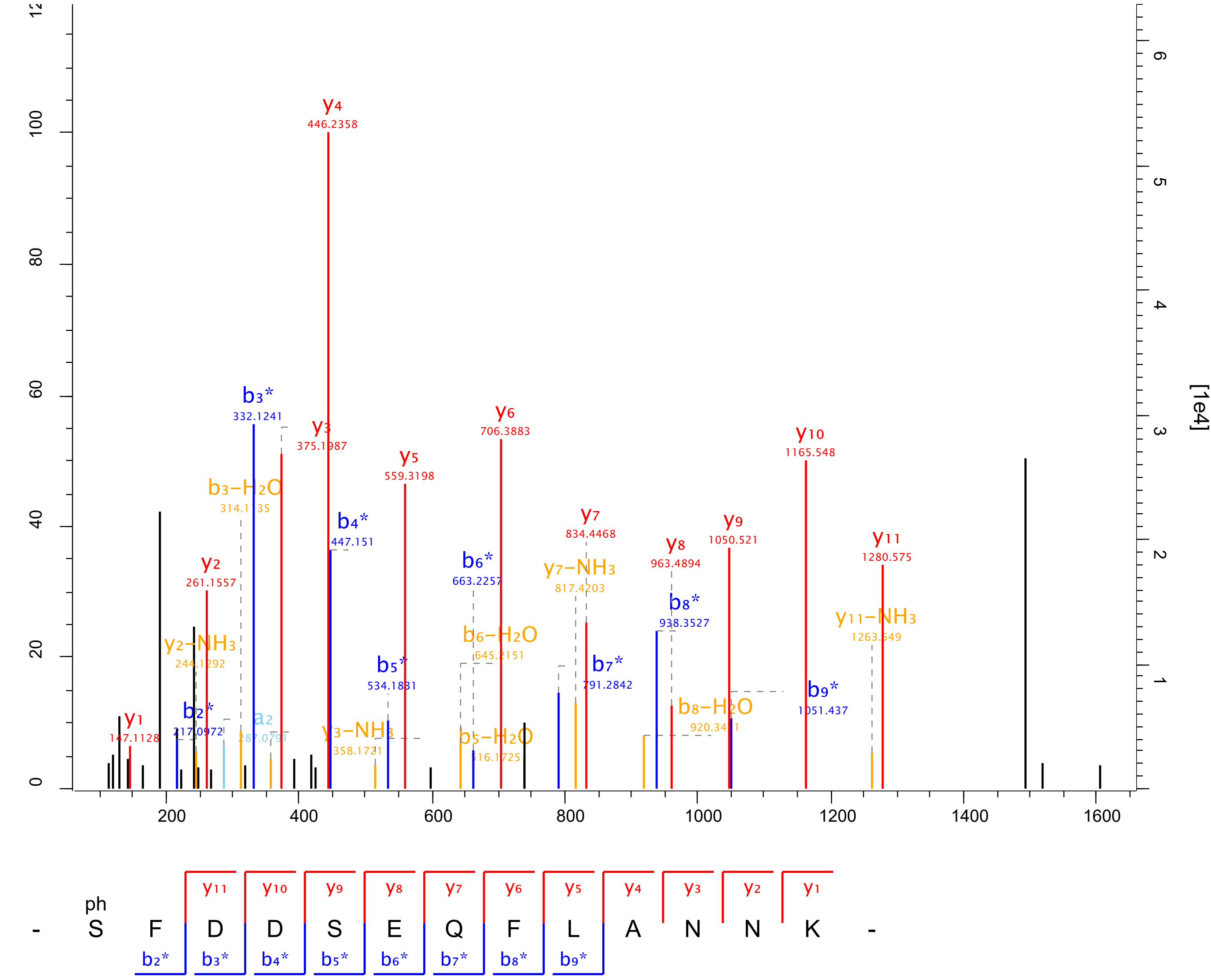

| Raw file | Scan | Method | Score | m/z |
|---|---|---|---|---|
| water_02_1 | 28170 | FTMS; HCD | 161.04 | 764.05 |

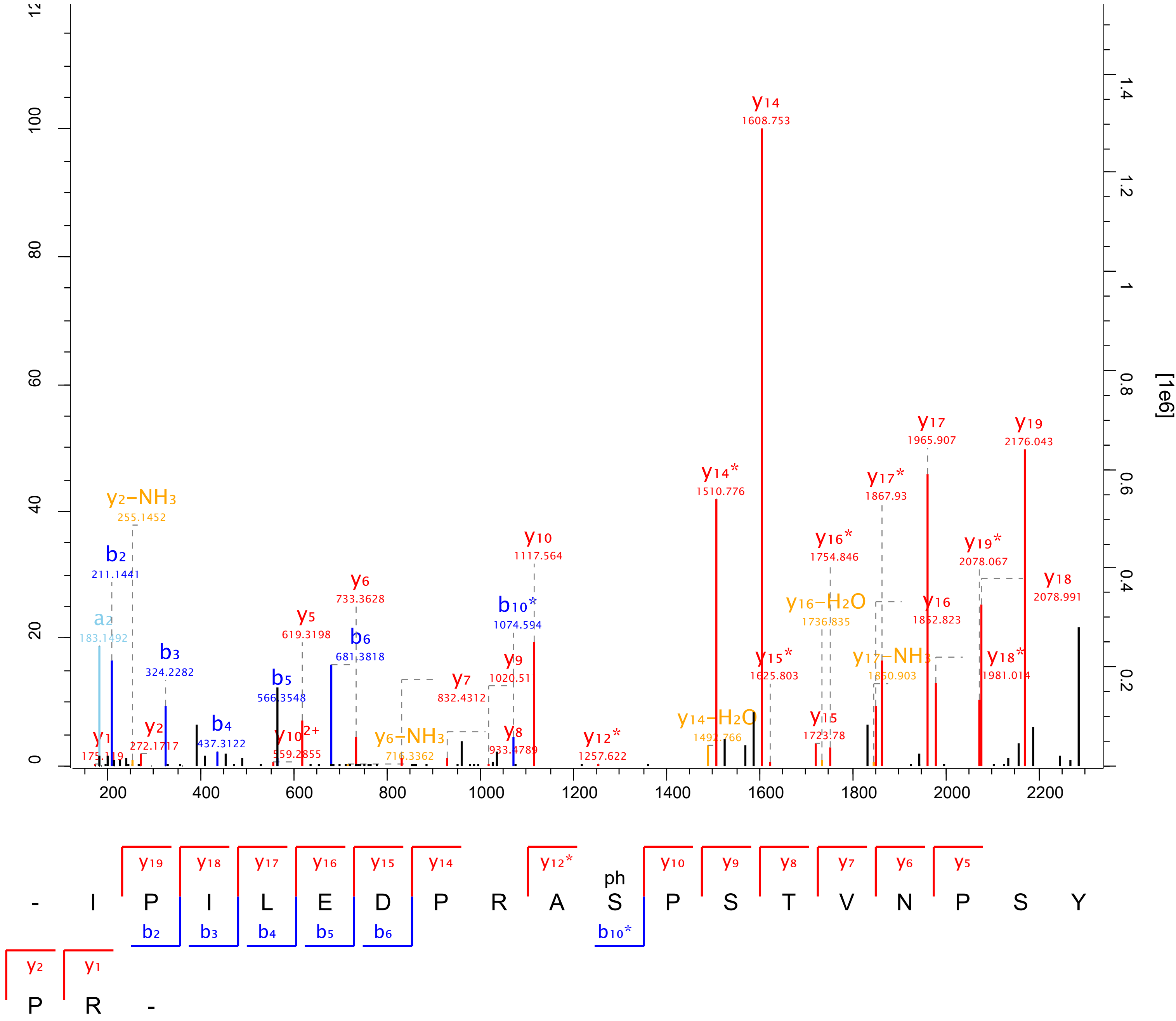

| Raw file | Scan | Method | Score | m/z |
|---|---|---|---|---|
| water_02_1 | 28202 | FTMS; HCD | 101.53 | 678.77 |

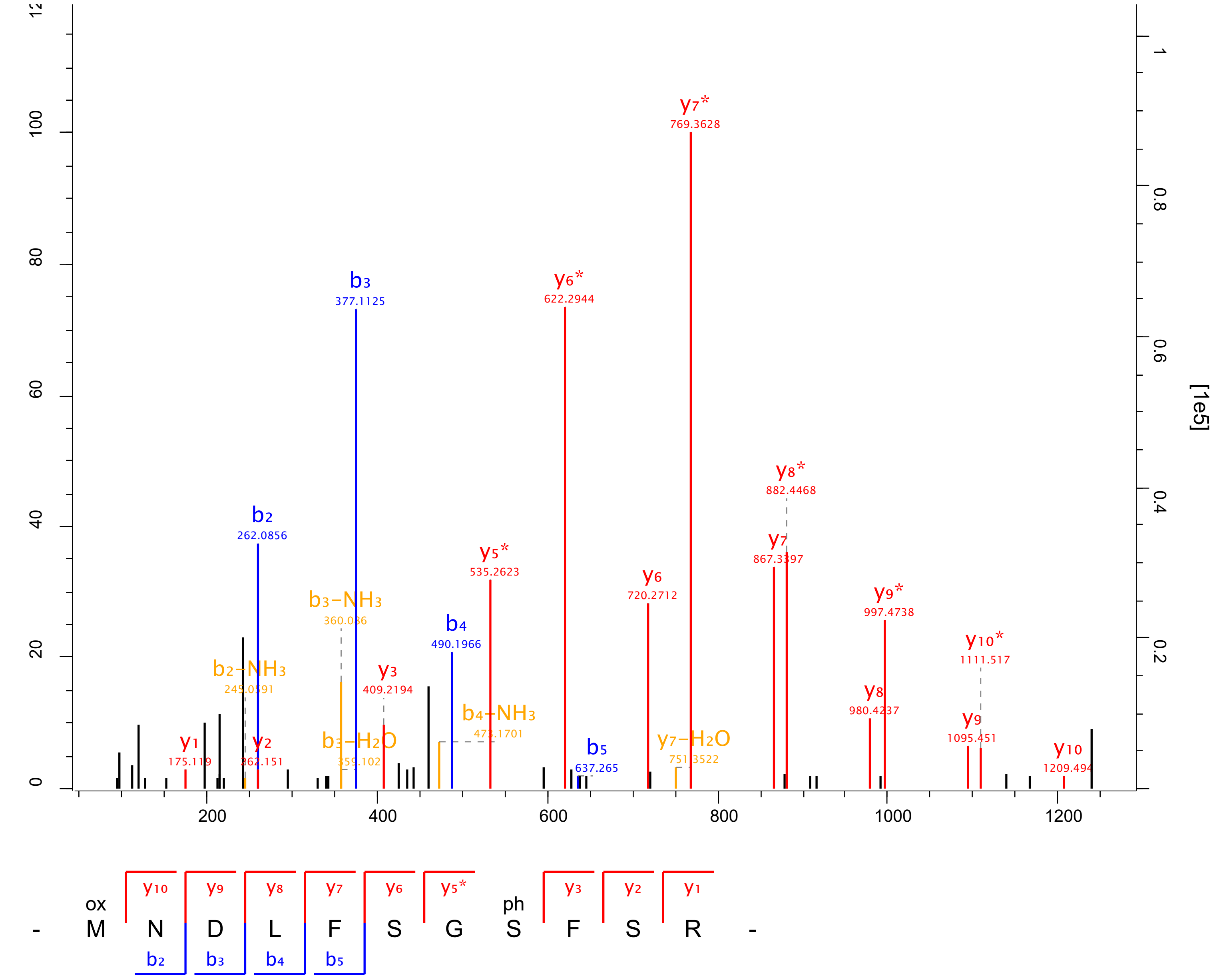

| Raw file | Scan | Method | Score | m/z |
|---|---|---|---|---|
| water_02_1 | 28408 | FTMS; HCD | 59.2 | 548.28 |

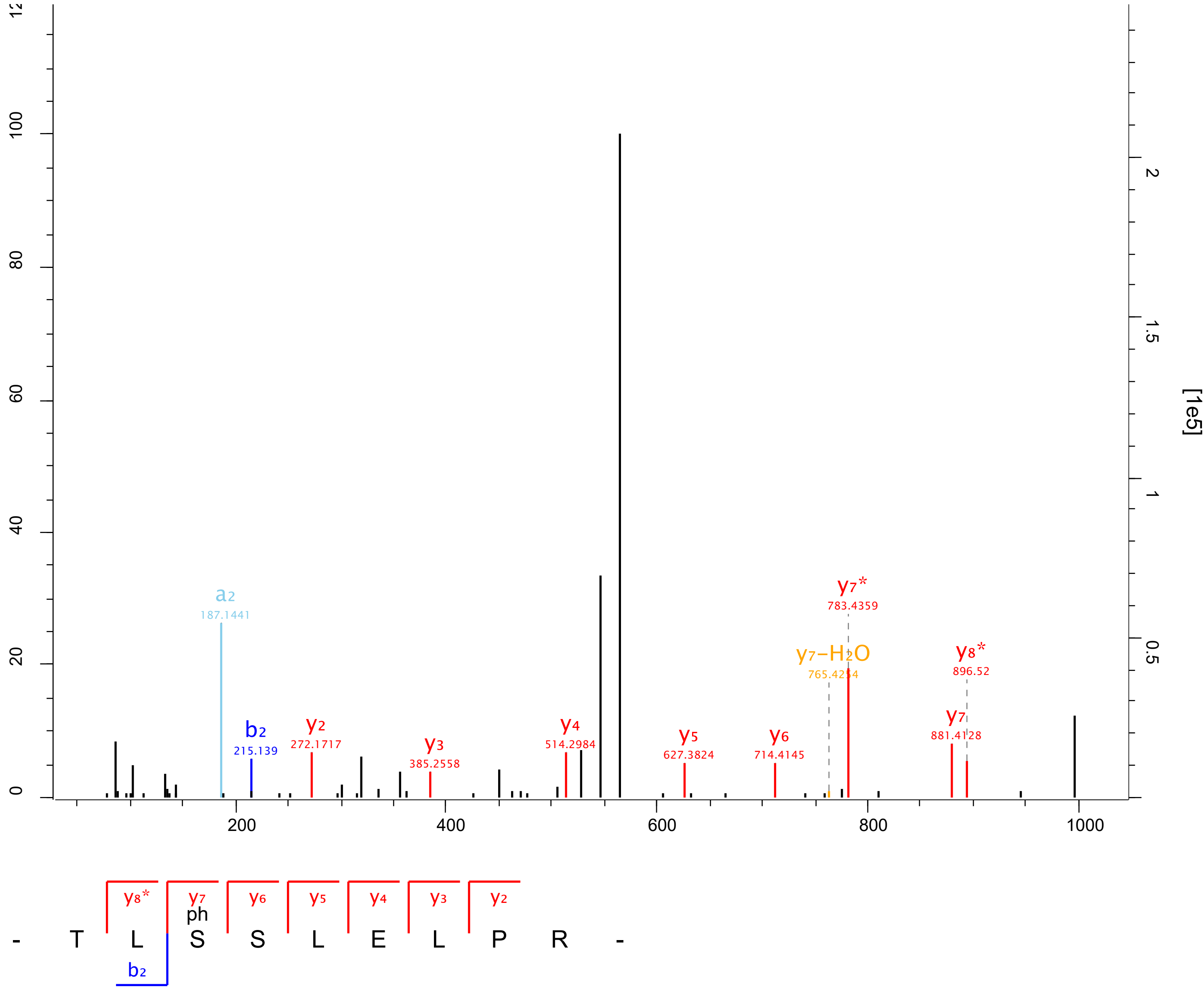

| Raw file | Scan | Method | Score | m/z |
| --- | --- | --- | --- | --- |
| water_02_1 | 2845 | FTMS; HCD | 89.51 | 590.27 |

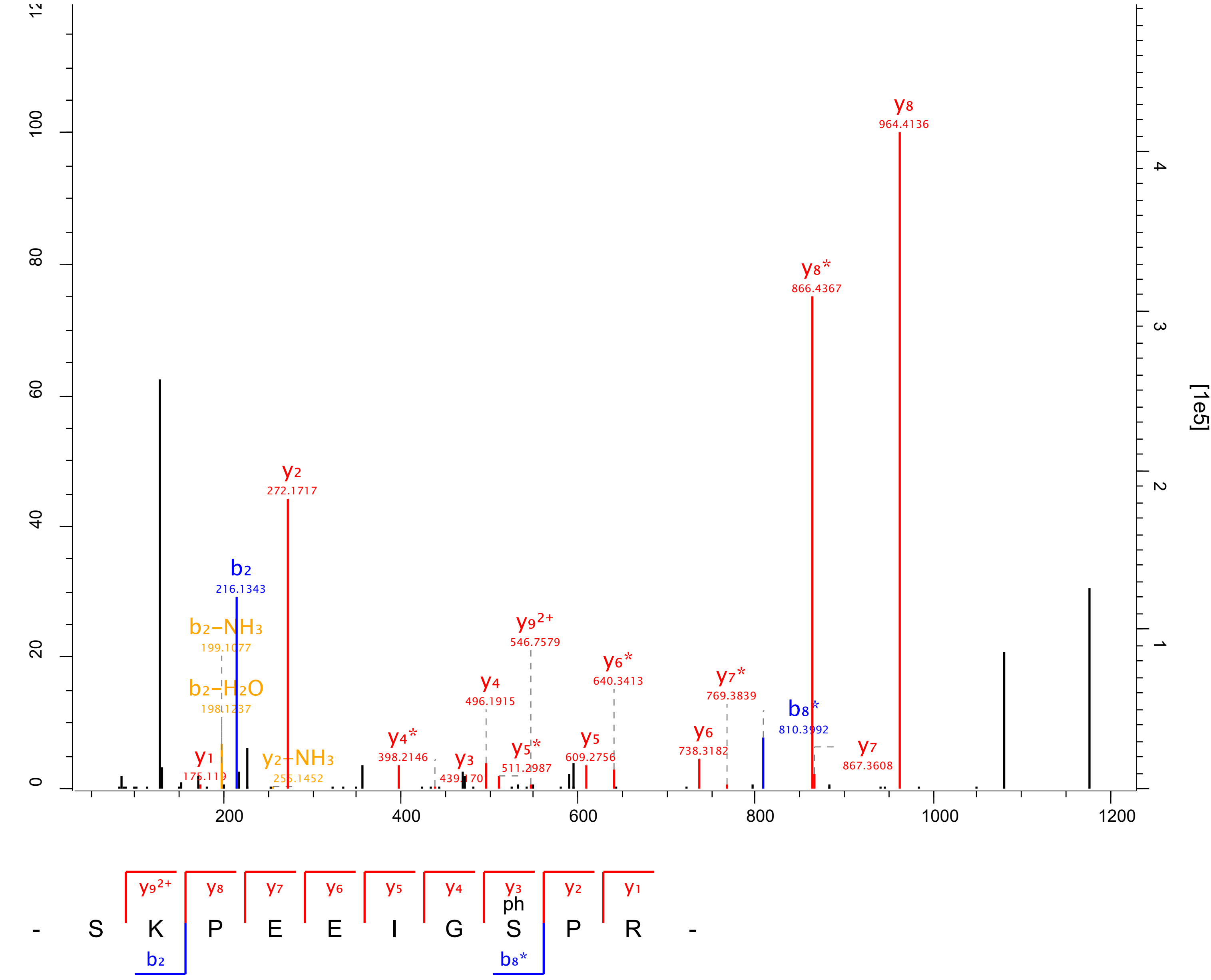

| Raw file | Scan | Method | Score | m/z |
|---|---|---|---|---|
| water_02_1 | 28452 | FTMS; HCD | 51.03 | 710.3 |

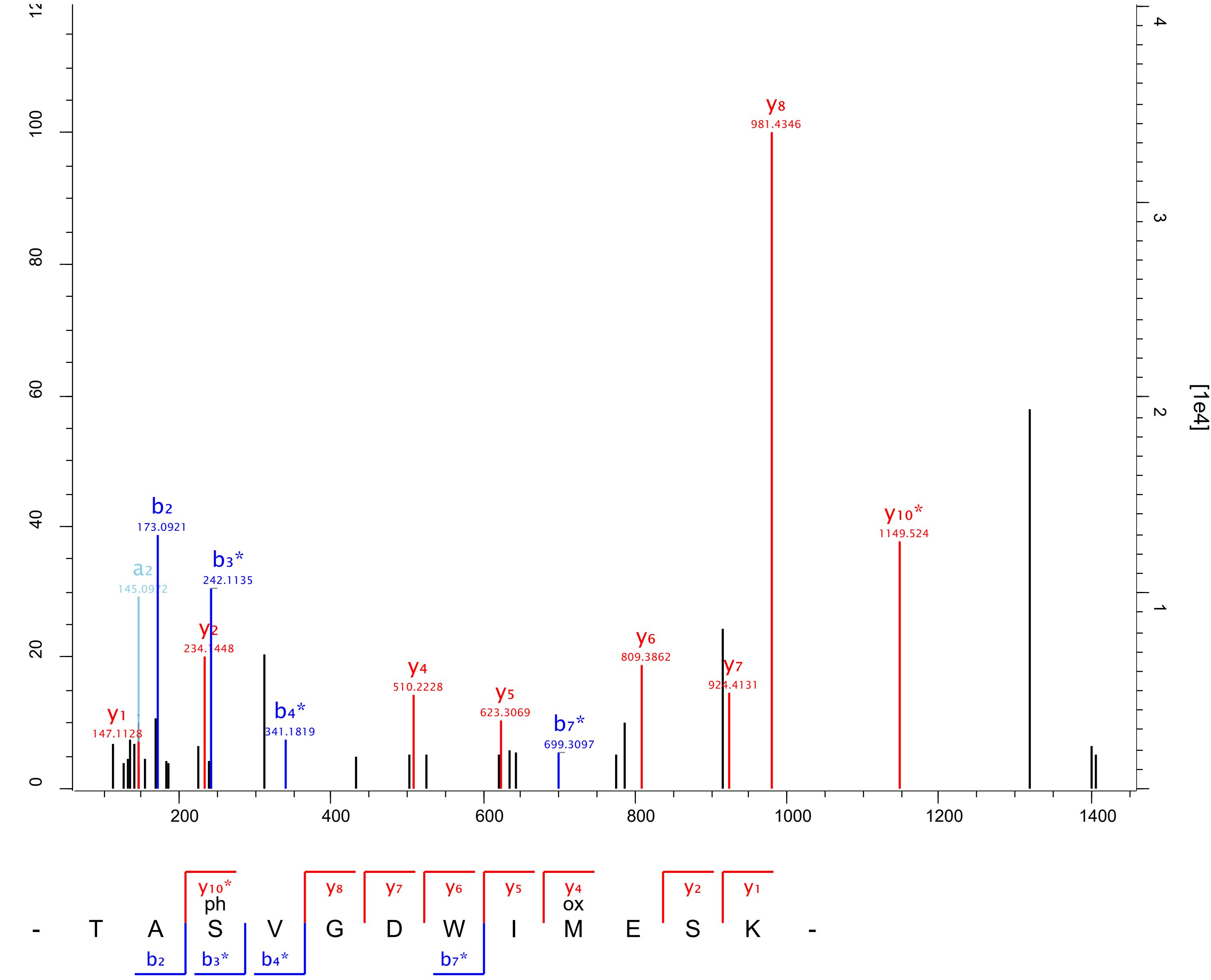

| Raw file | Scan | Method | Score | m/z |
| --- | --- | --- | --- | --- |
| water_02_1 | 28570 | FTMS; HCD | 180.16 | 921.42 |

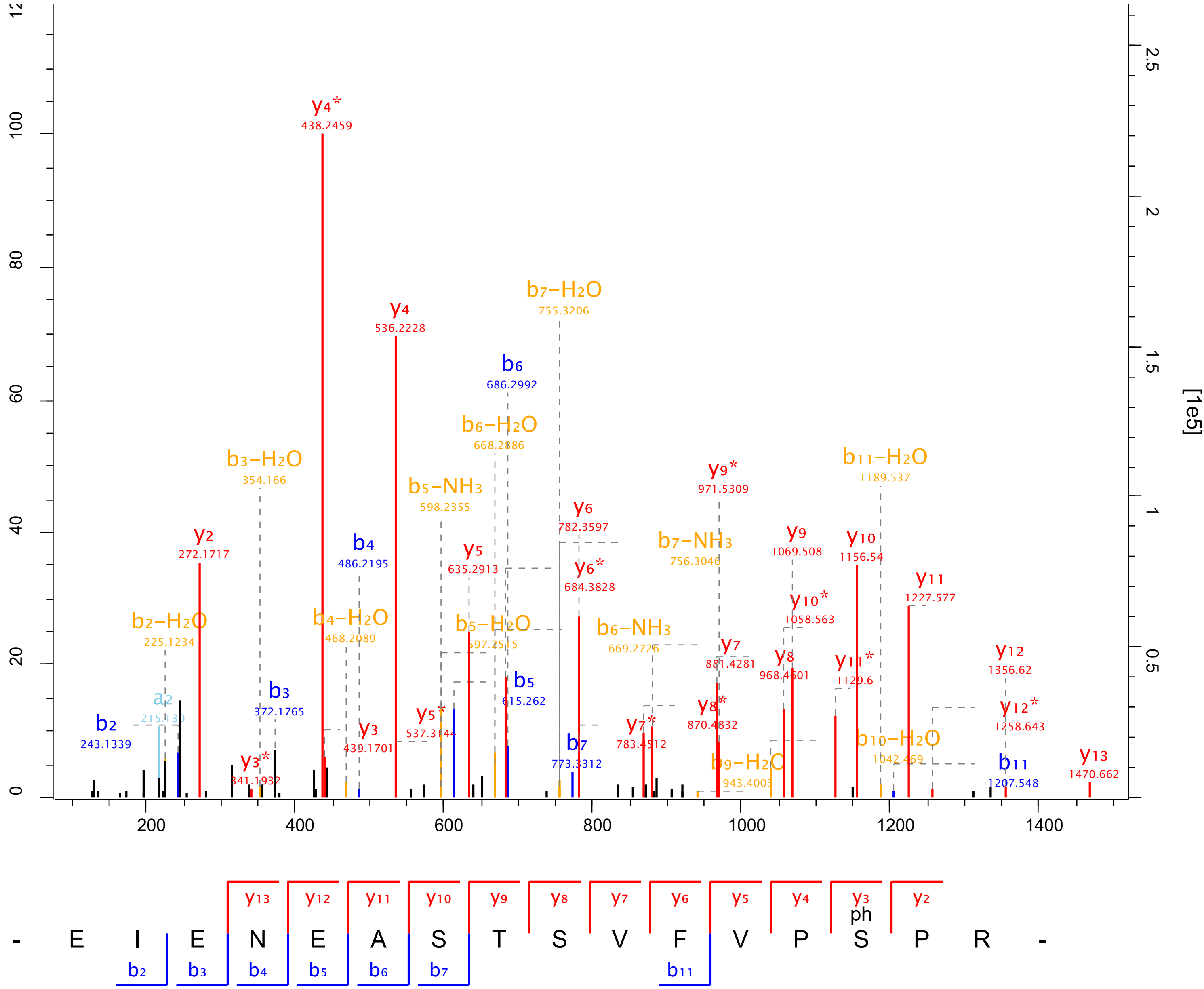

| Raw file | Scan | Method | Score | m/z |
|---|---|---|---|---|
| water_02_1 | 28615 | FTMS; HCD | 189.39 | 1079.98 |

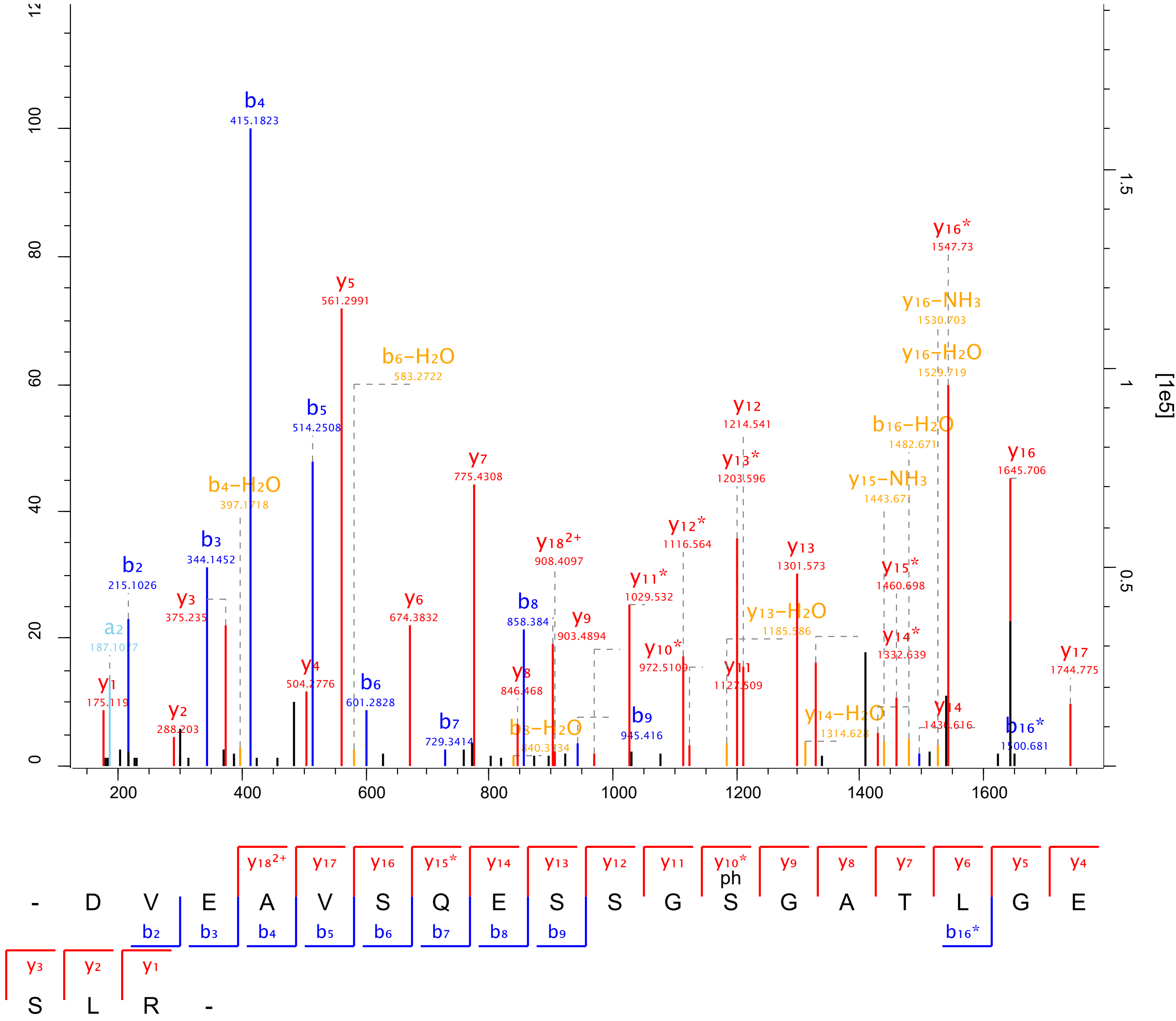

| Raw file | Scan | Method | Score | m/z |
| --- | --- | --- | --- | --- |
| water_02_1 | 28731 | FTMS; HCD | 75.56 | 616.98 |

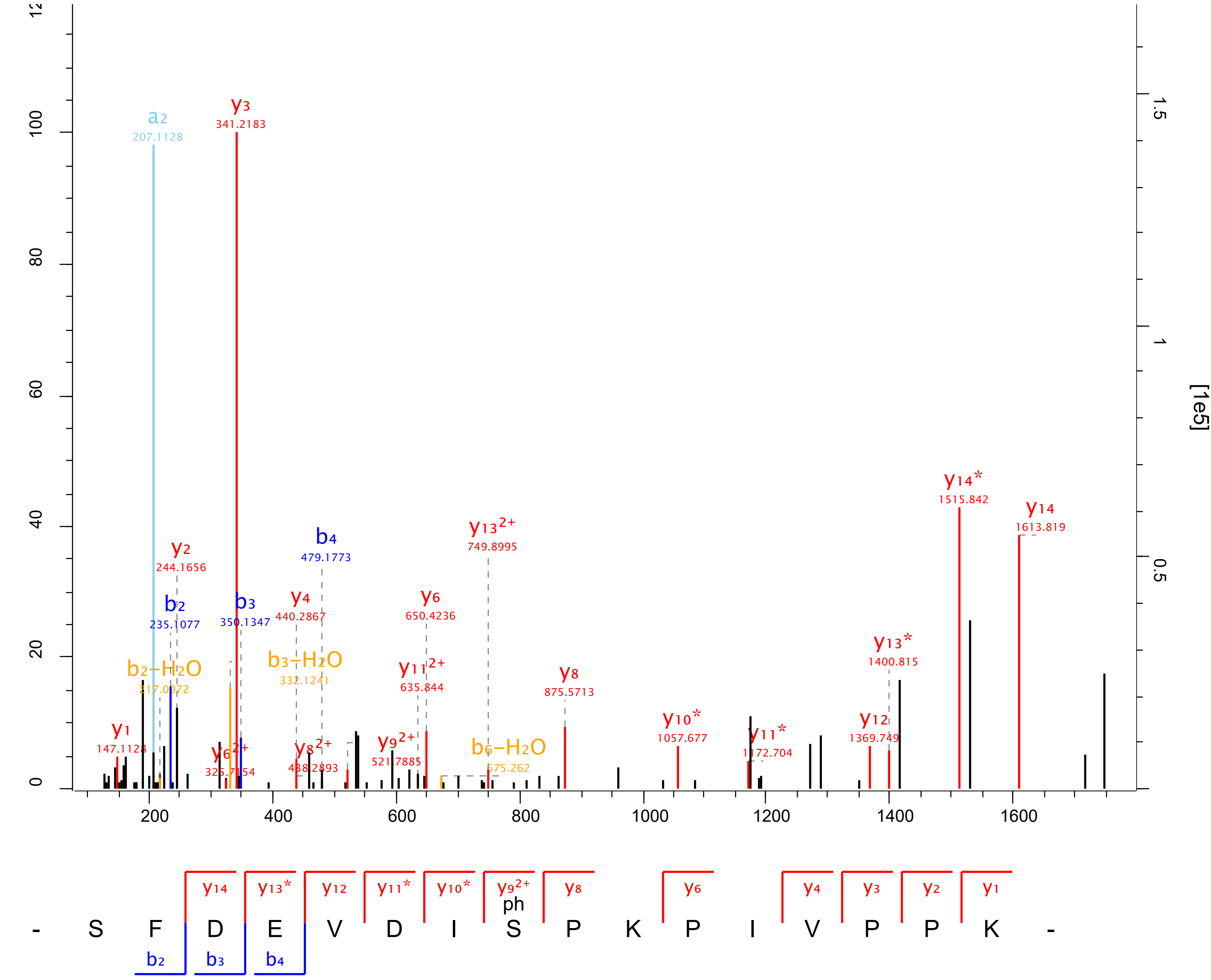

| Raw file | Scan | Method | Score | m/z |
|---|---|---|---|---|
| water_02_1 | 2882 | FTMS; HCD | 49.3 | 434.18 |

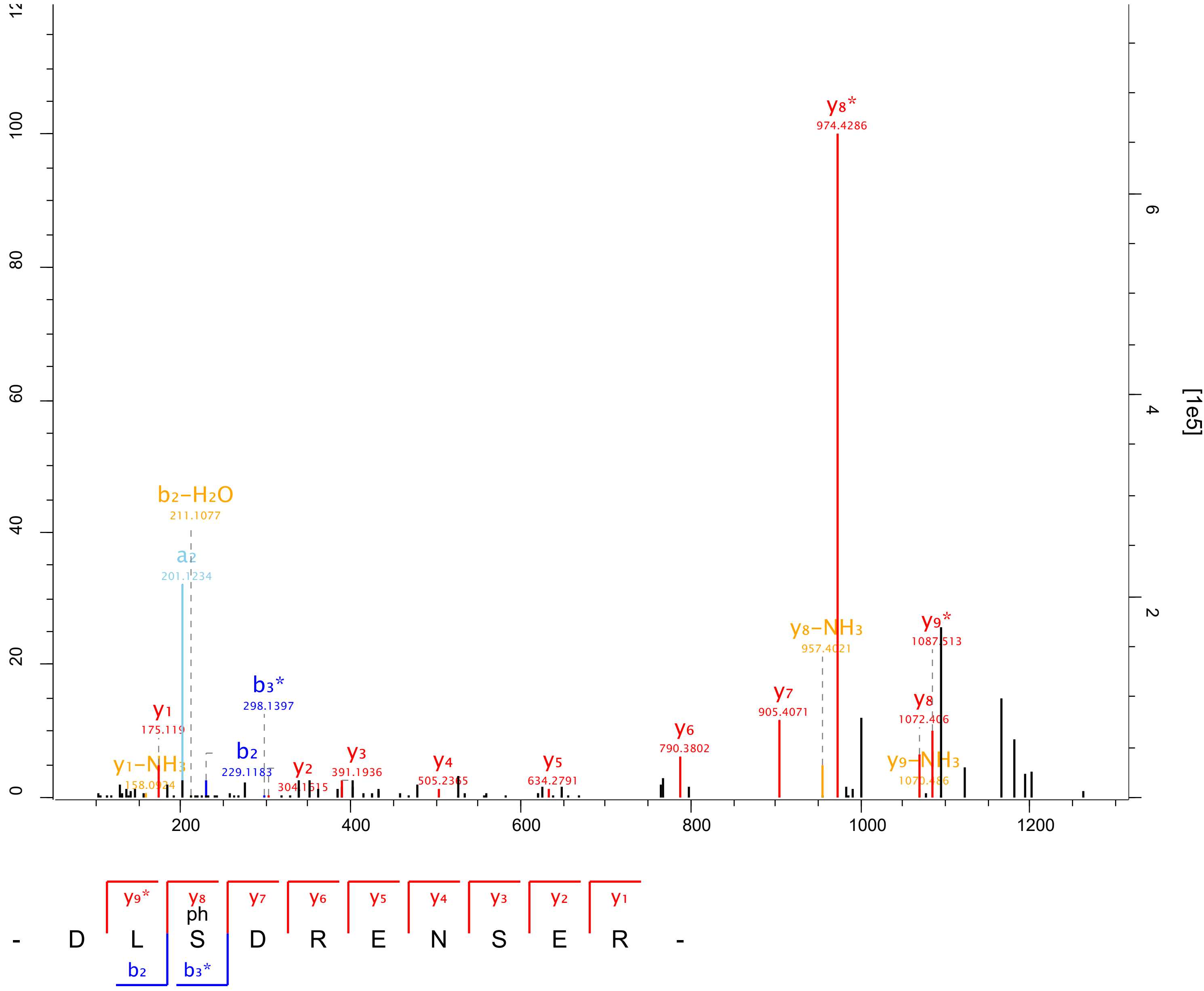

| Raw file | Scan | Method | Score | m/z |
|---|---|---|---|---|
| water_02_1 | 28848 | FTMS; HCD | 84.95 | 659.32 |

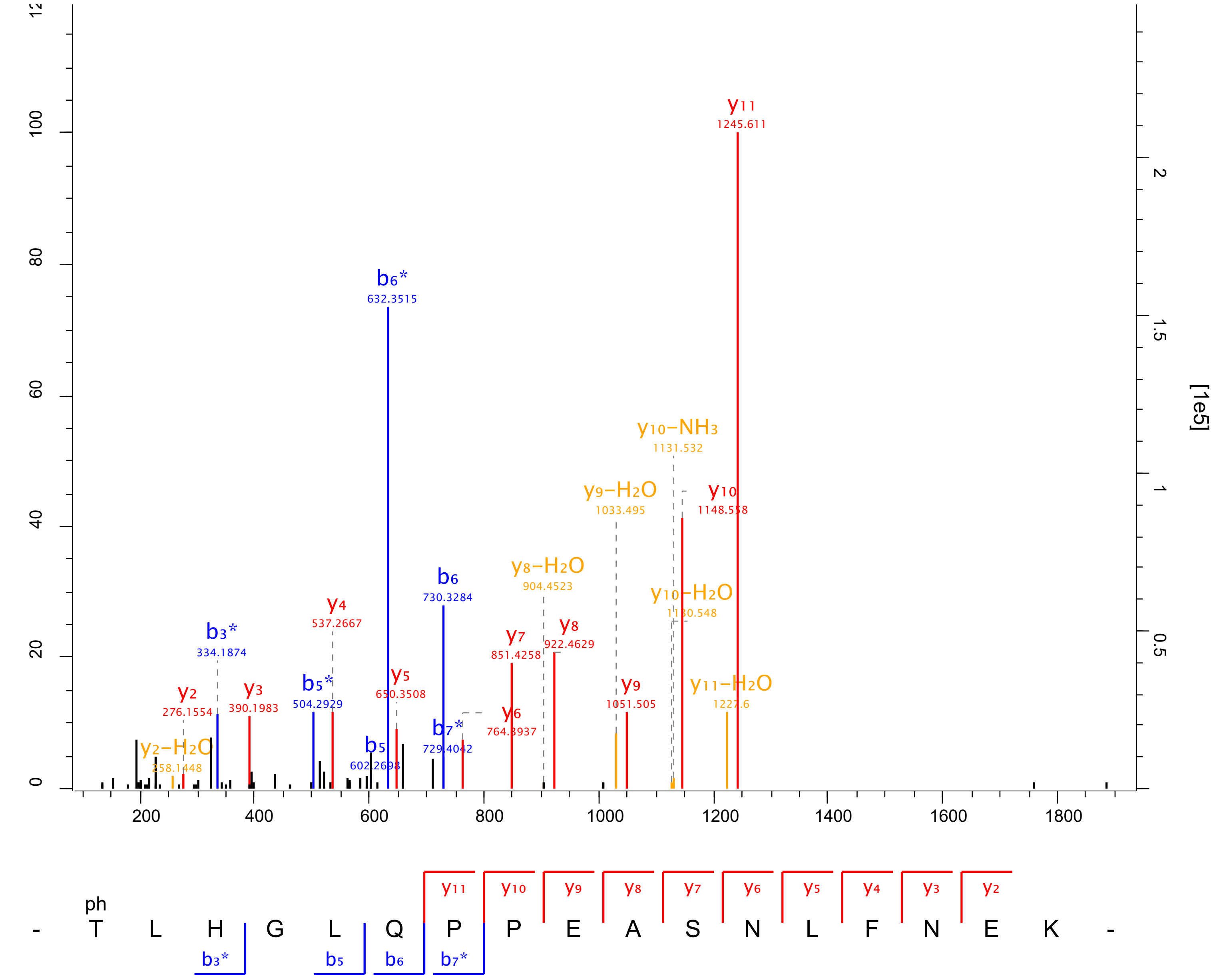

| Raw file | Scan | Method | Score | m/z |
| --- | --- | --- | --- | --- |
| water_02_1 | 28895 | FTMS; HCD | 118.51 | 728.69 |

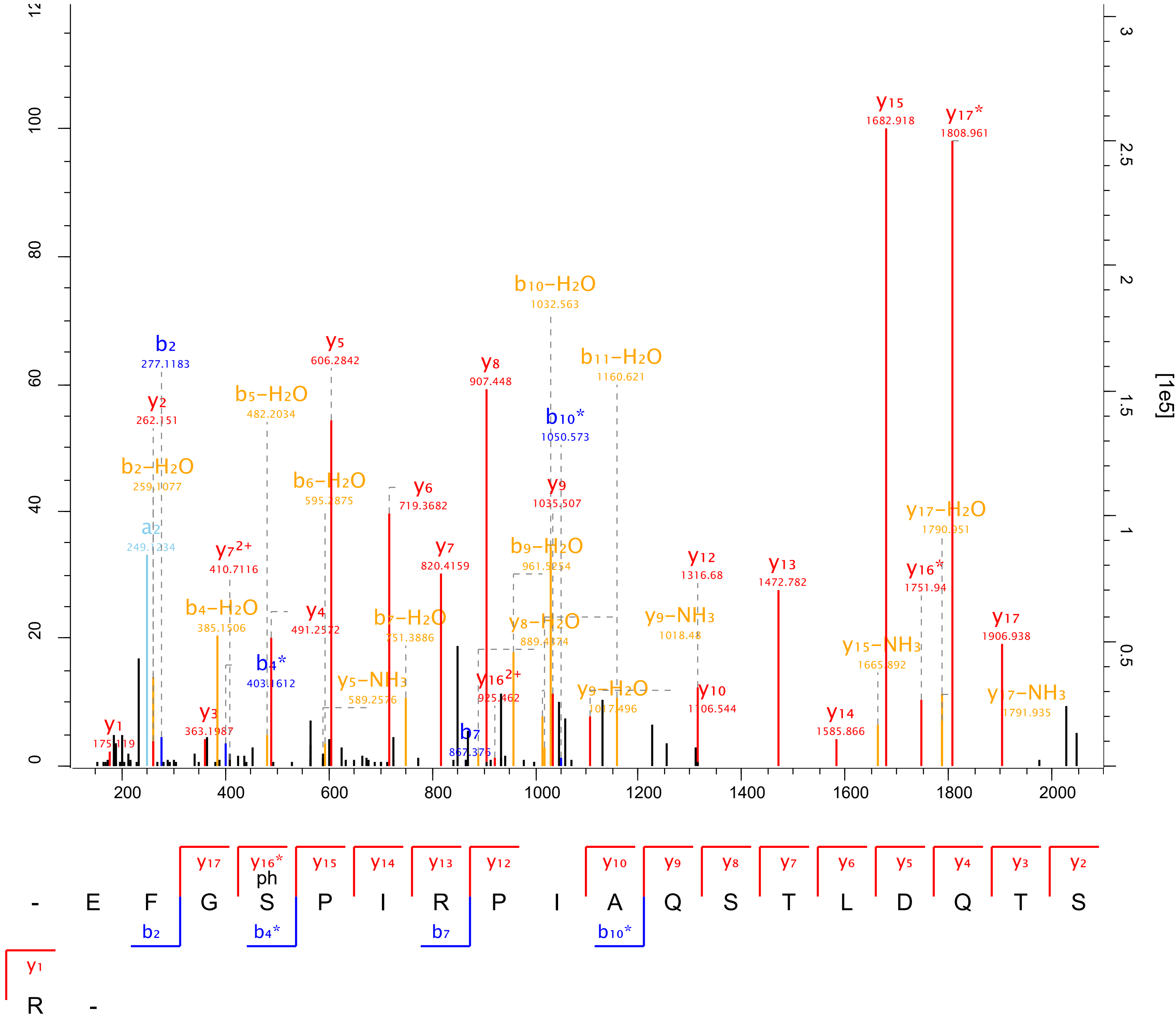

| Raw file | Scan | Method | Score | m/z |
|---|---|---|---|---|
| water_02_1 | 2896 | FTMS; HCD | 113.47 | 769.76 |

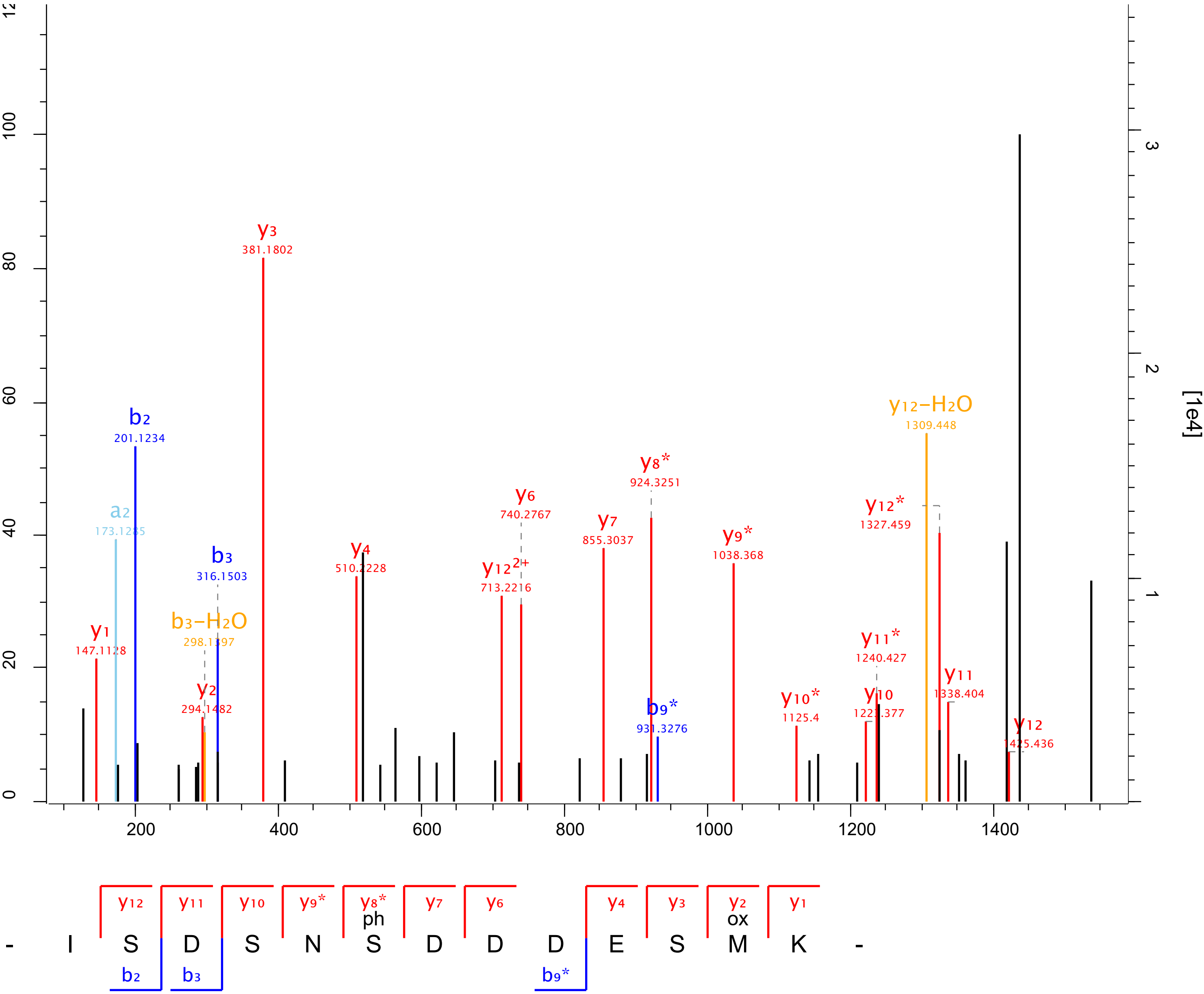

| Raw file | Scan | Method | Score | m/z |
|---|---|---|---|---|
| water_02_1 | 29037 | FTMS; HCD | 41.24 | 745.87 |

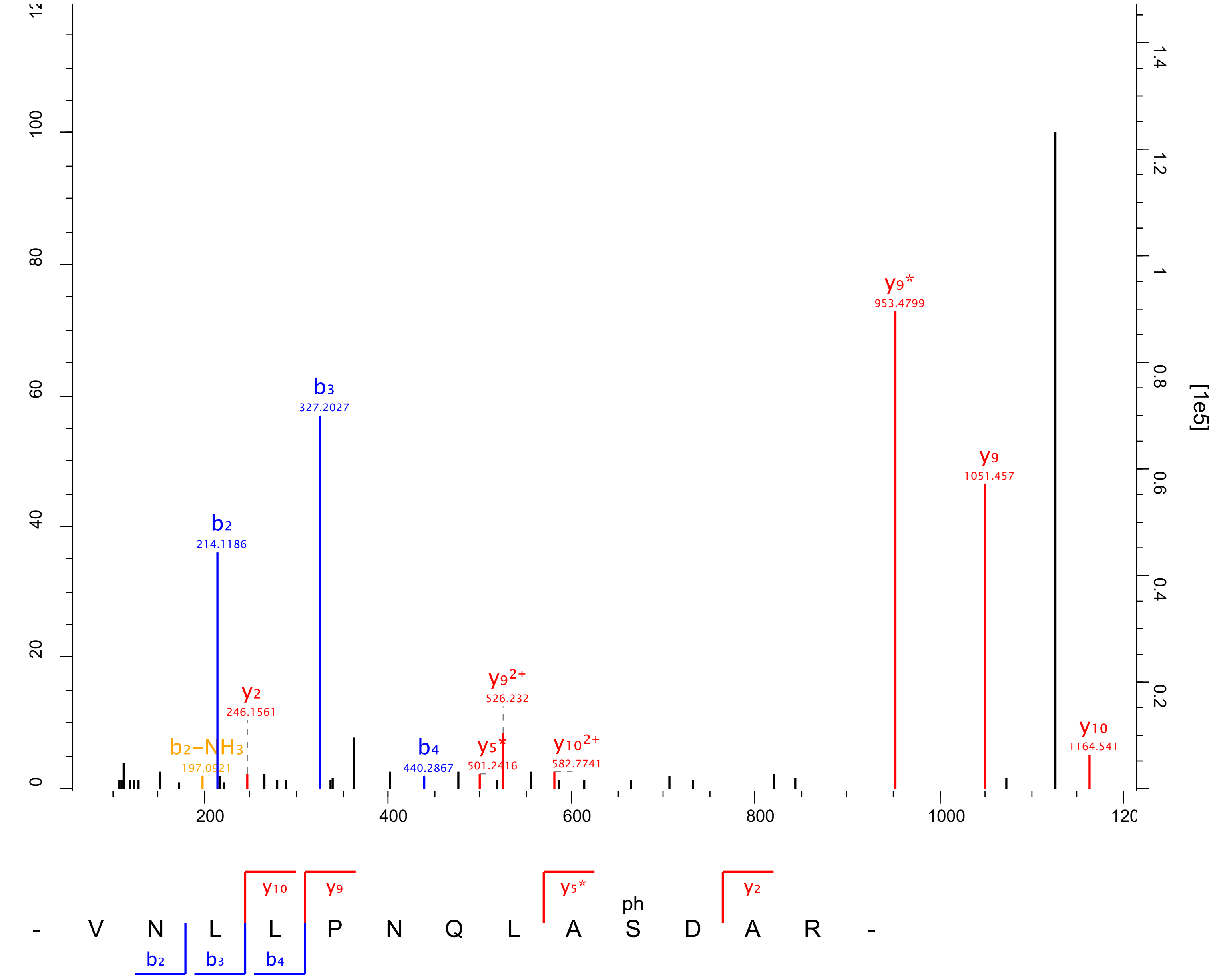

| Raw file | Scan | Method | Score | m/z |
|---|---|---|---|---|
| water_02_1 | 29203 | FTMS; HCD | 75.06 | 985.96 |

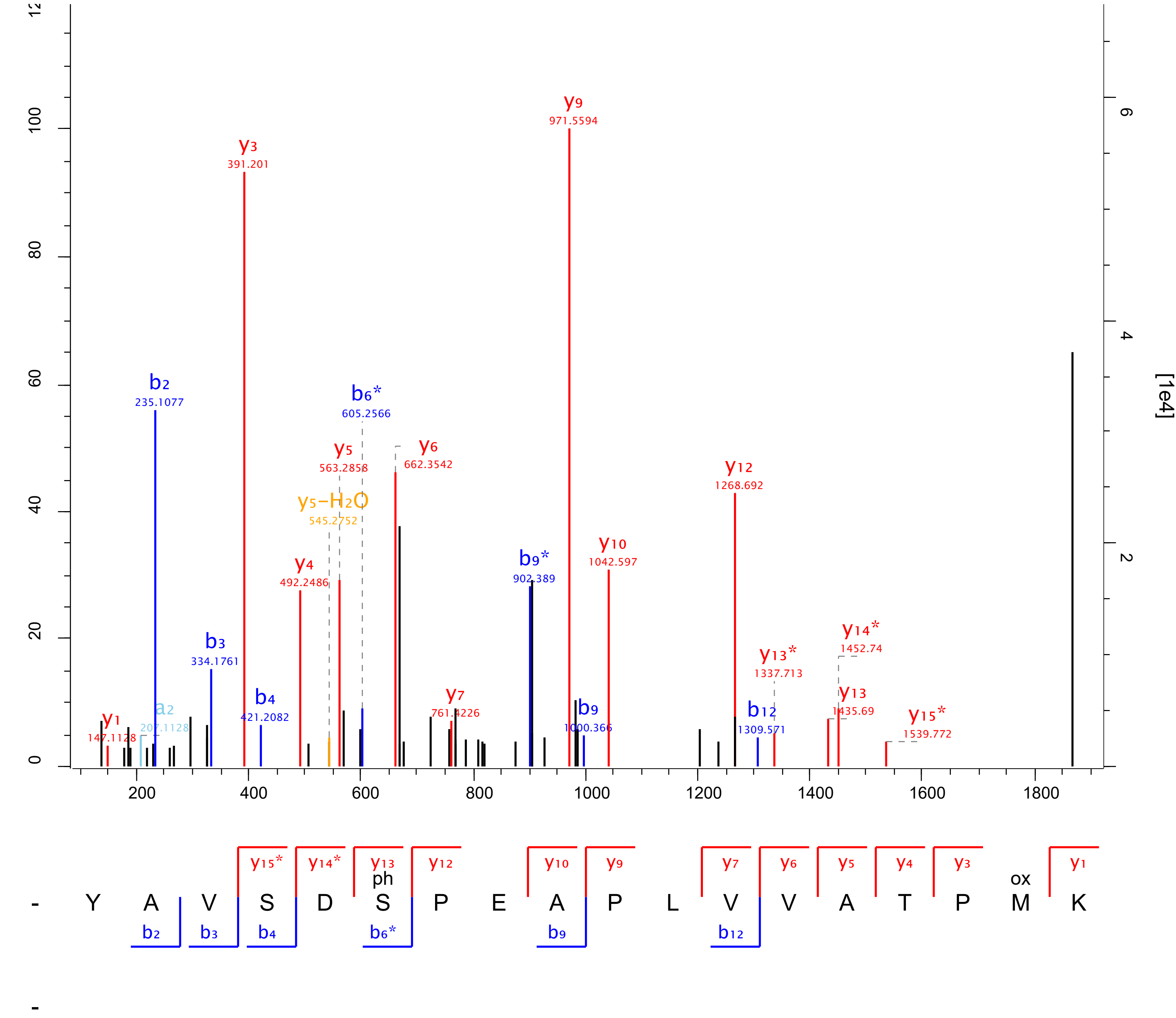

| Raw file | Scan | Method | Score | m/z |
|---|---|---|---|---|
| water_02_1 | 29225 | FTMS; HCD | 67.76 | 709.67 |

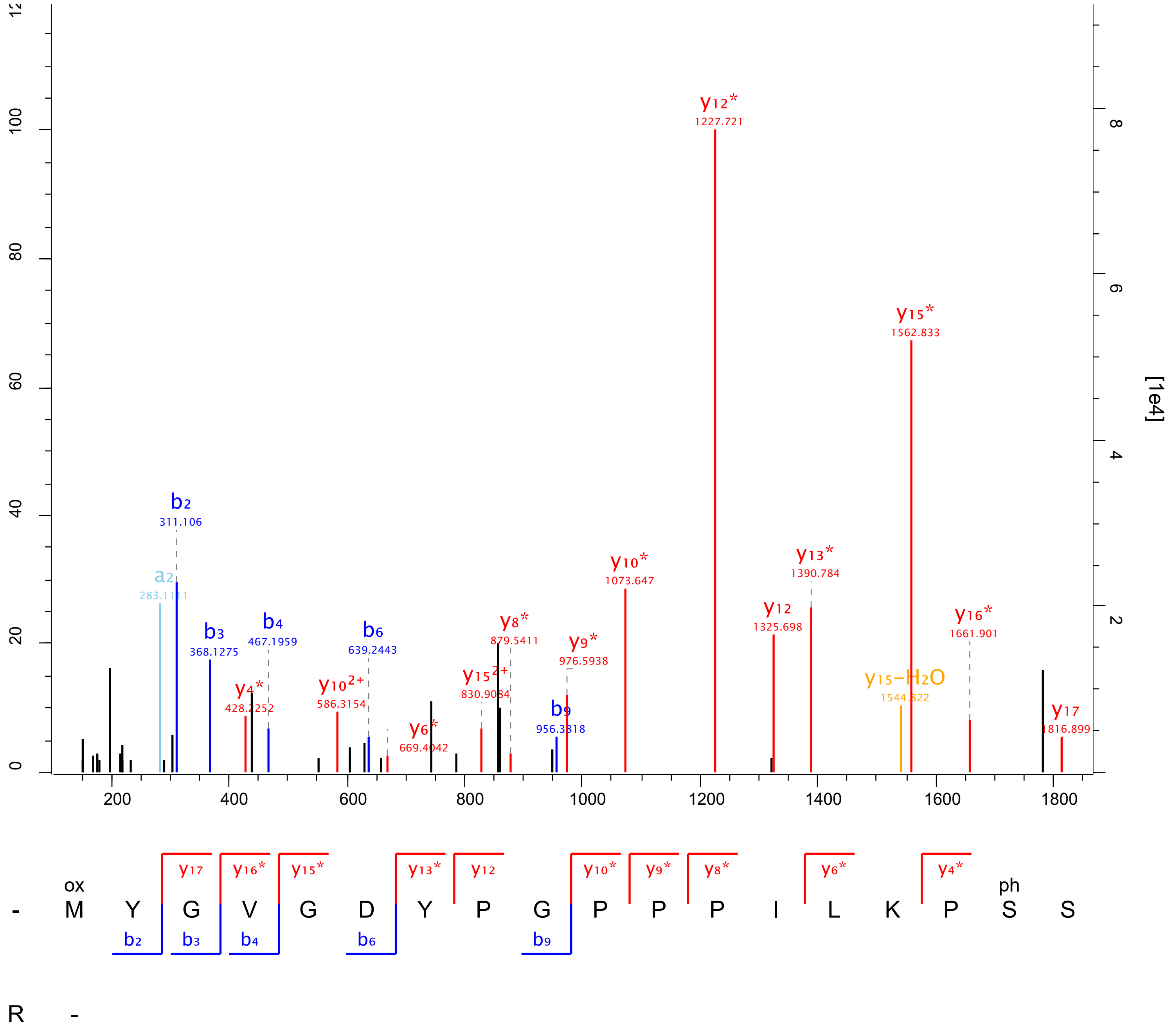

| Raw file | Scan | Method | Score | m/z |
|---|---|---|---|---|
| water_02_1 | 29245 | FTMS; HCD | 125.23 | 854.89 |

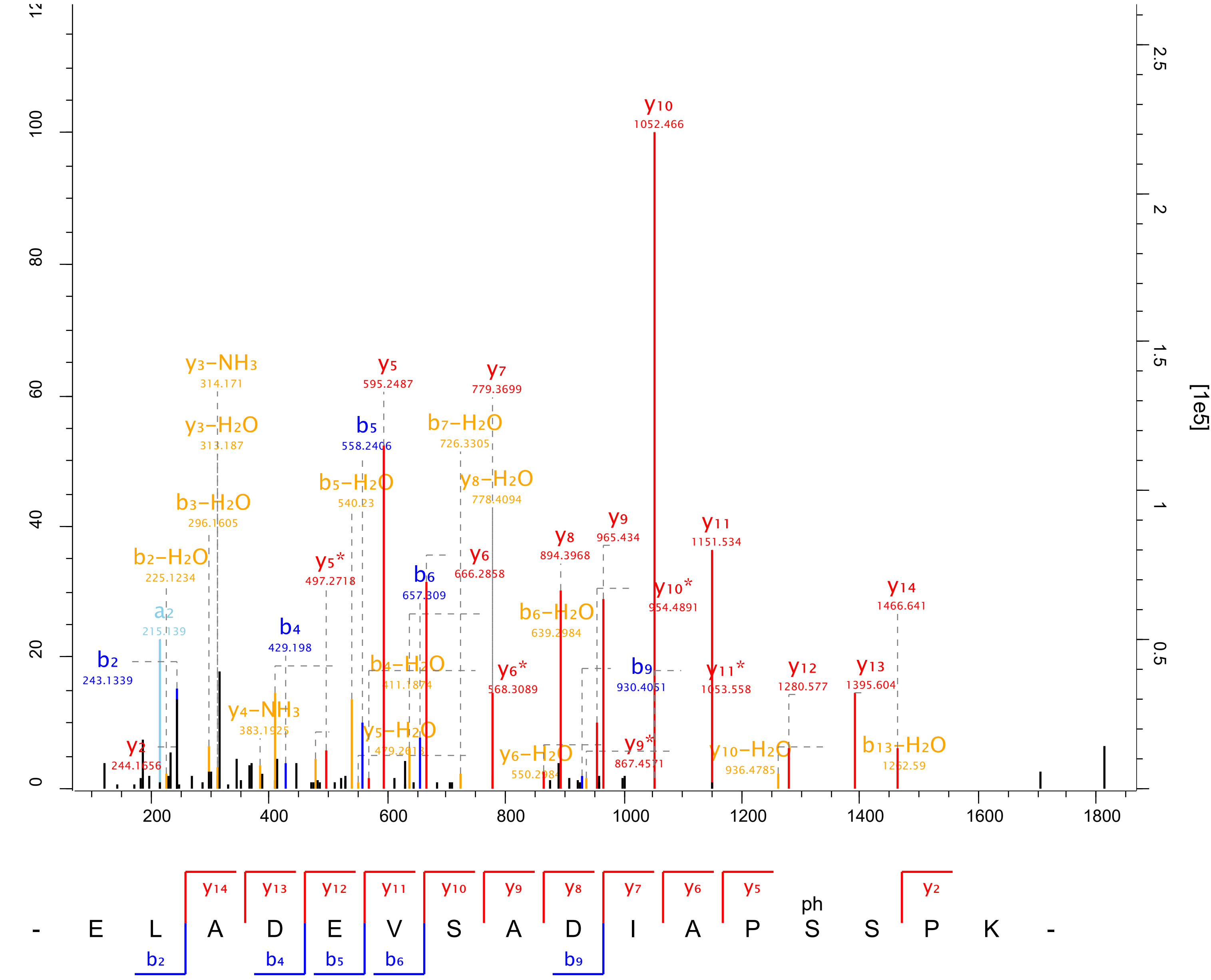

| Raw file | Scan | Method | Score | m/z |
|---|---|---|---|---|
| water_02_1 | 29313 | FTMS; HCD | 51.73 | 618.78 |

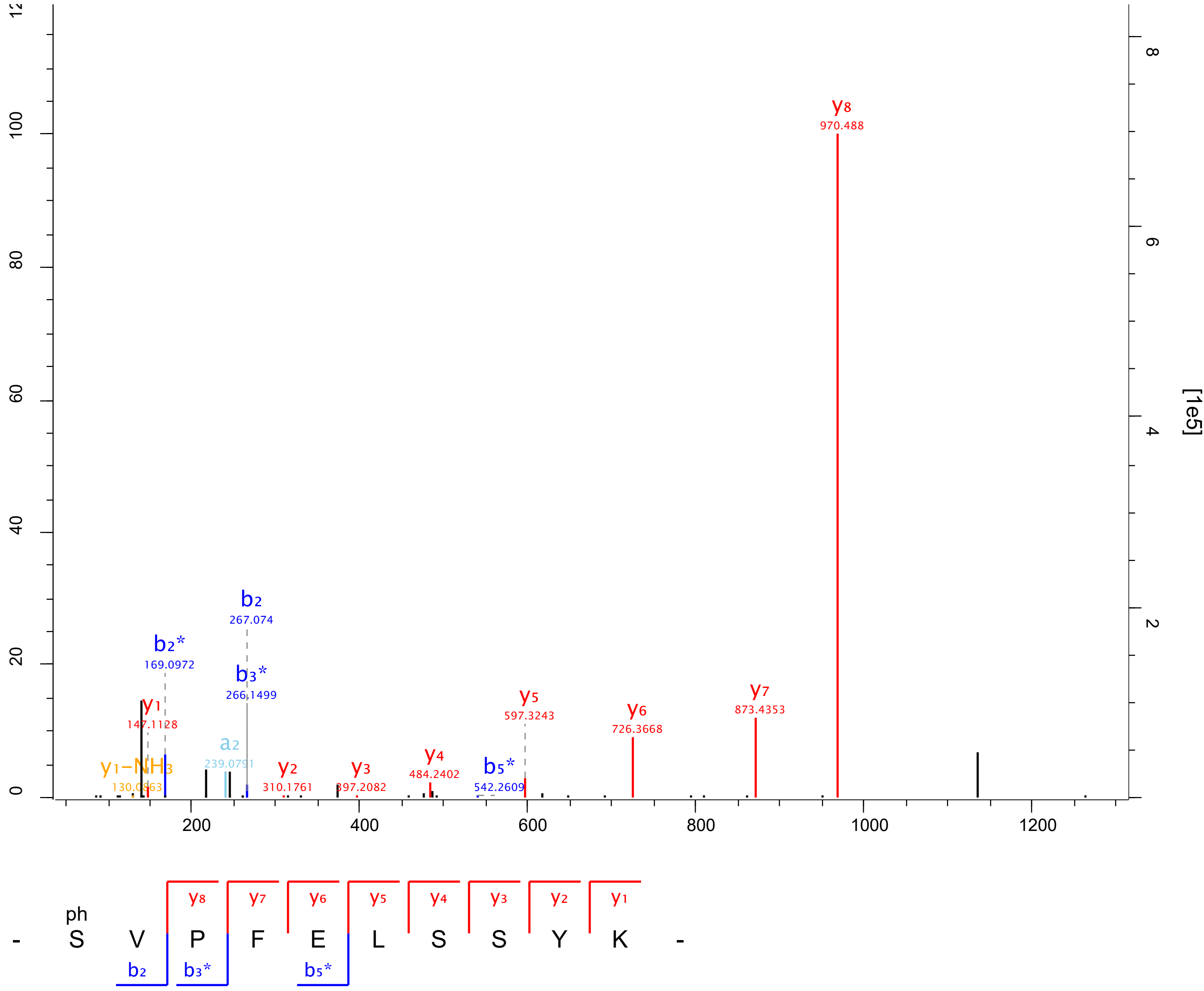

| Raw file | Scan | Method | Score | m/z |
|---|---|---|---|---|
| water_02_1 | 29394 | FTMS; HCD | 81.71 | 574.78 |

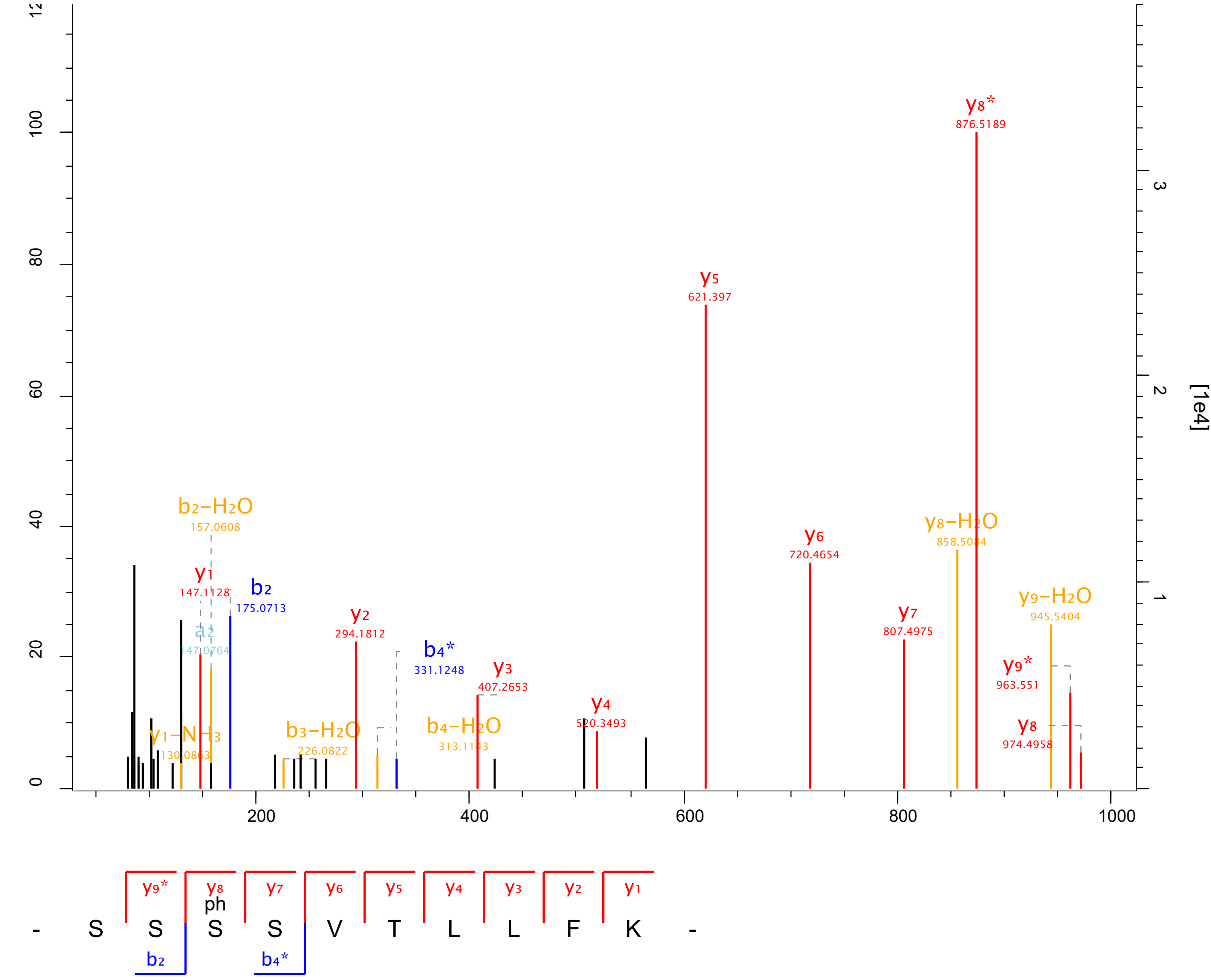

| Raw file | Scan | Method | Score | m/z |
|---|---|---|---|---|
| water_02_1 | 29601 | FTMS; HCD | 88.24 | 980.93 |

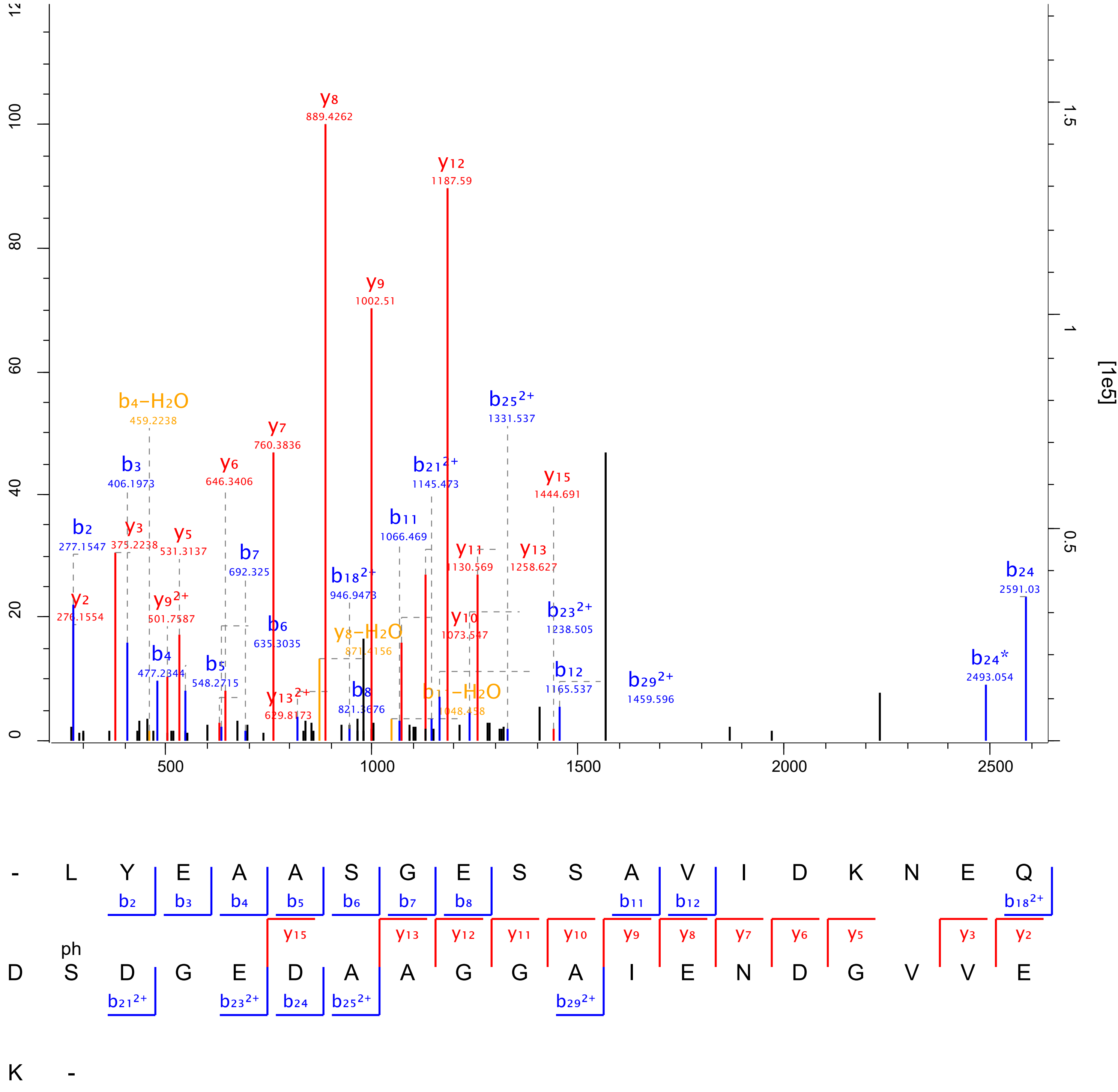

| Raw file | Scan | Method | Score | m/z |
| --- | --- | --- | --- | --- |
| water_02_1 | 2964 | FTMS; HCD | 176.98 | 596.26 |

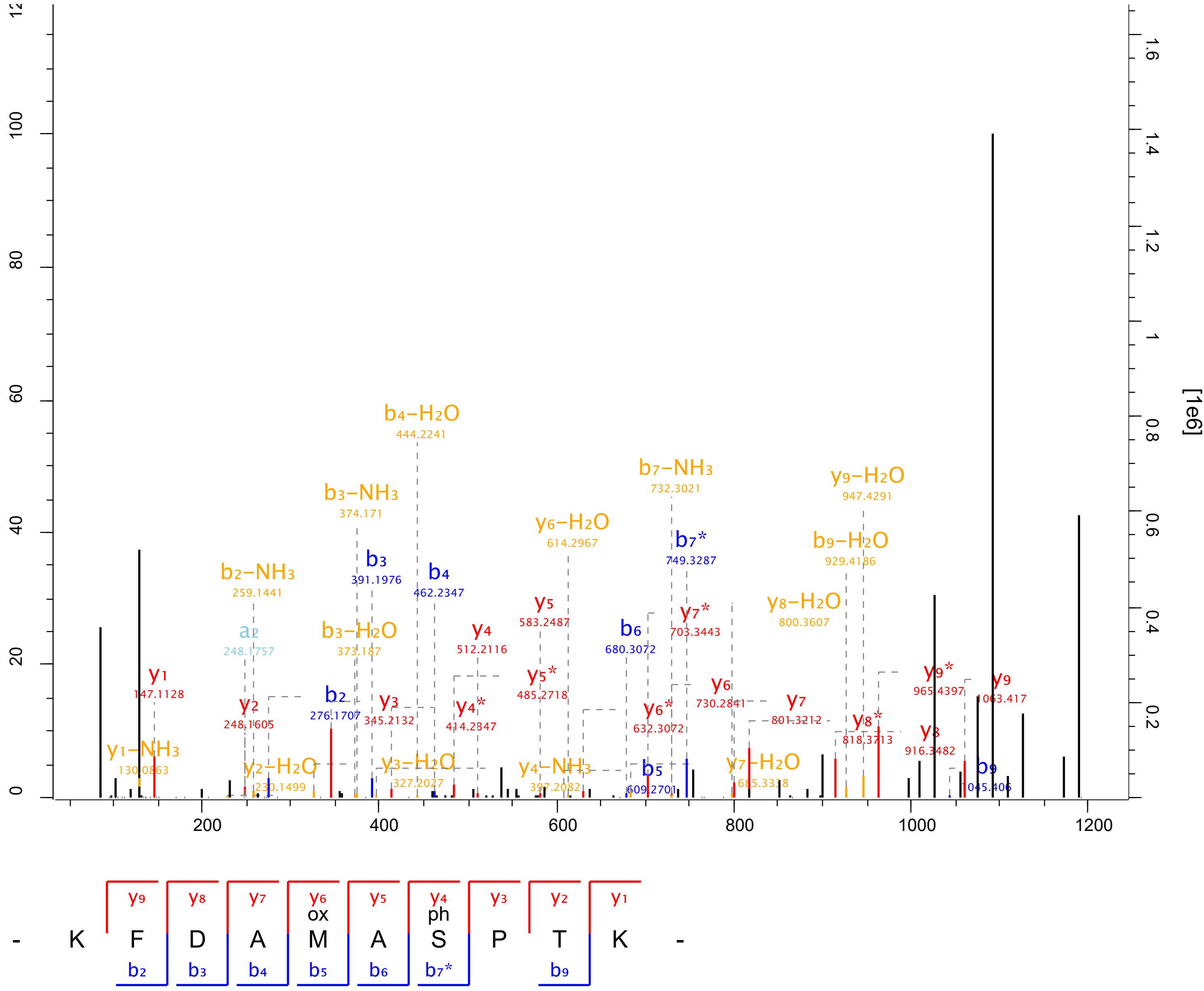

| Raw file | Scan | Method | Score | m/z |
|---|---|---|---|---|
| water_02_1 | 29707 | FTMS; HCD | 94.96 | 652.97 |

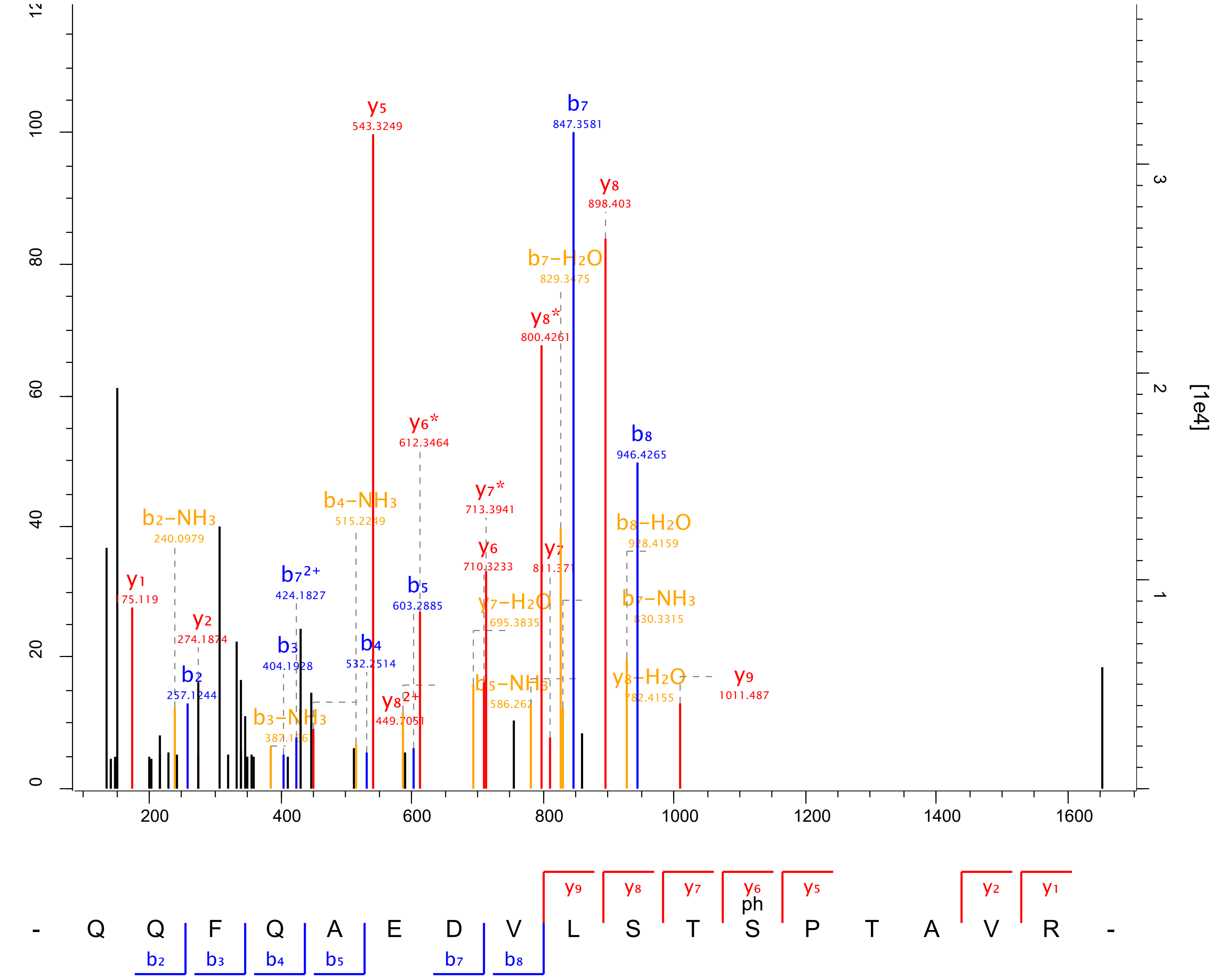

| Raw file | Scan | Method | Score | m/z |
| --- | --- | --- | --- | --- |
| water_02_1 | 29776 | FTMS; HCD | 89.48 | 755.87 |

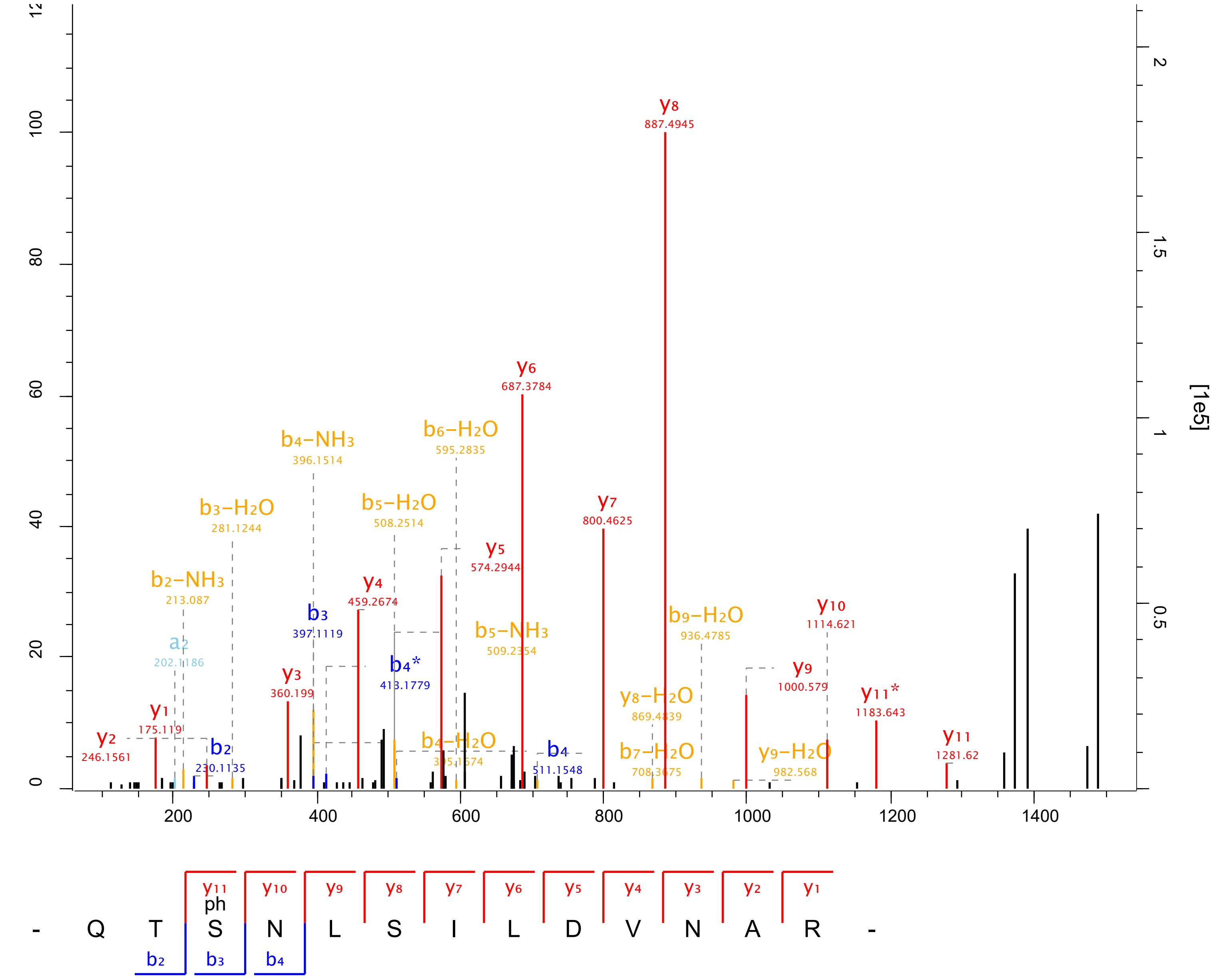

| Raw file | Scan | Method | Score | m/z |
|---|---|---|---|---|
| water_02_1 | 29789 | FTMS; HCD | 43.35 | 739.35 |

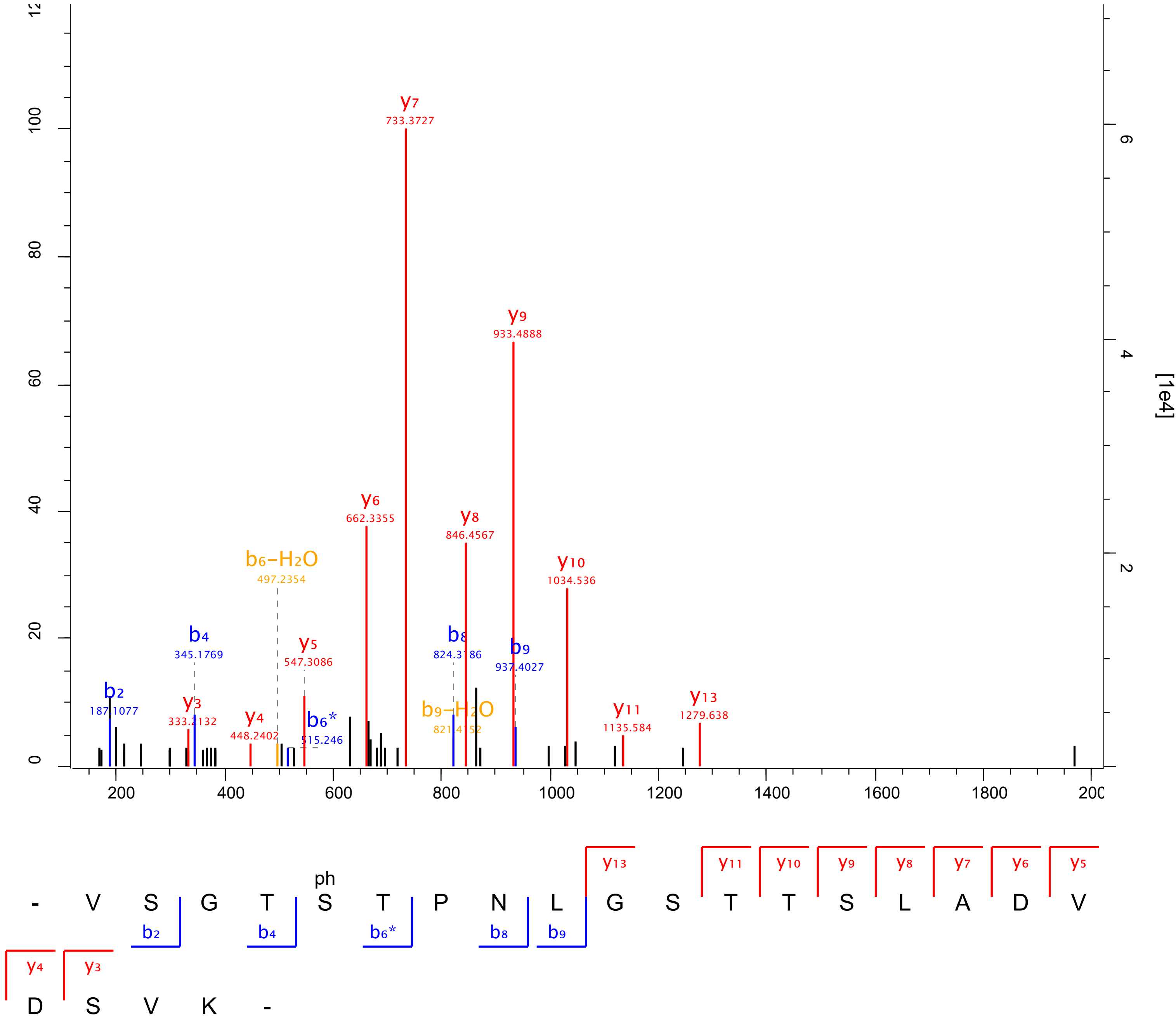

| Raw file | Scan | Method | Score | m/z |
|---|---|---|---|---|
| water_02_1 | 29795 | FTMS; HCD | 69.02 | 926.75 |

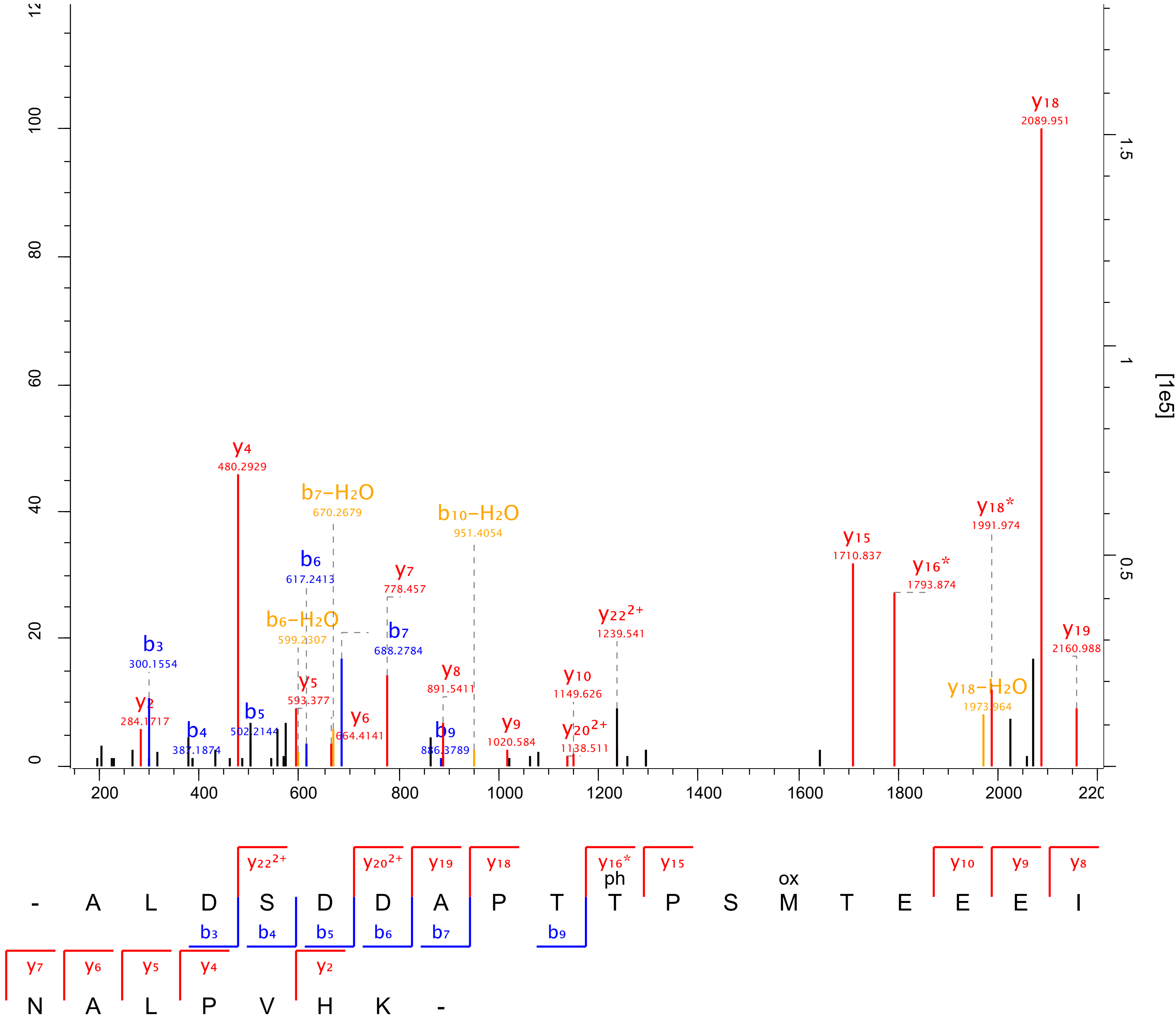

| Raw file | Scan | Method | Score | m/z |
|---|---|---|---|---|
| water_02_1 | 29843 | FTMS; HCD | 81 | 901.91 |

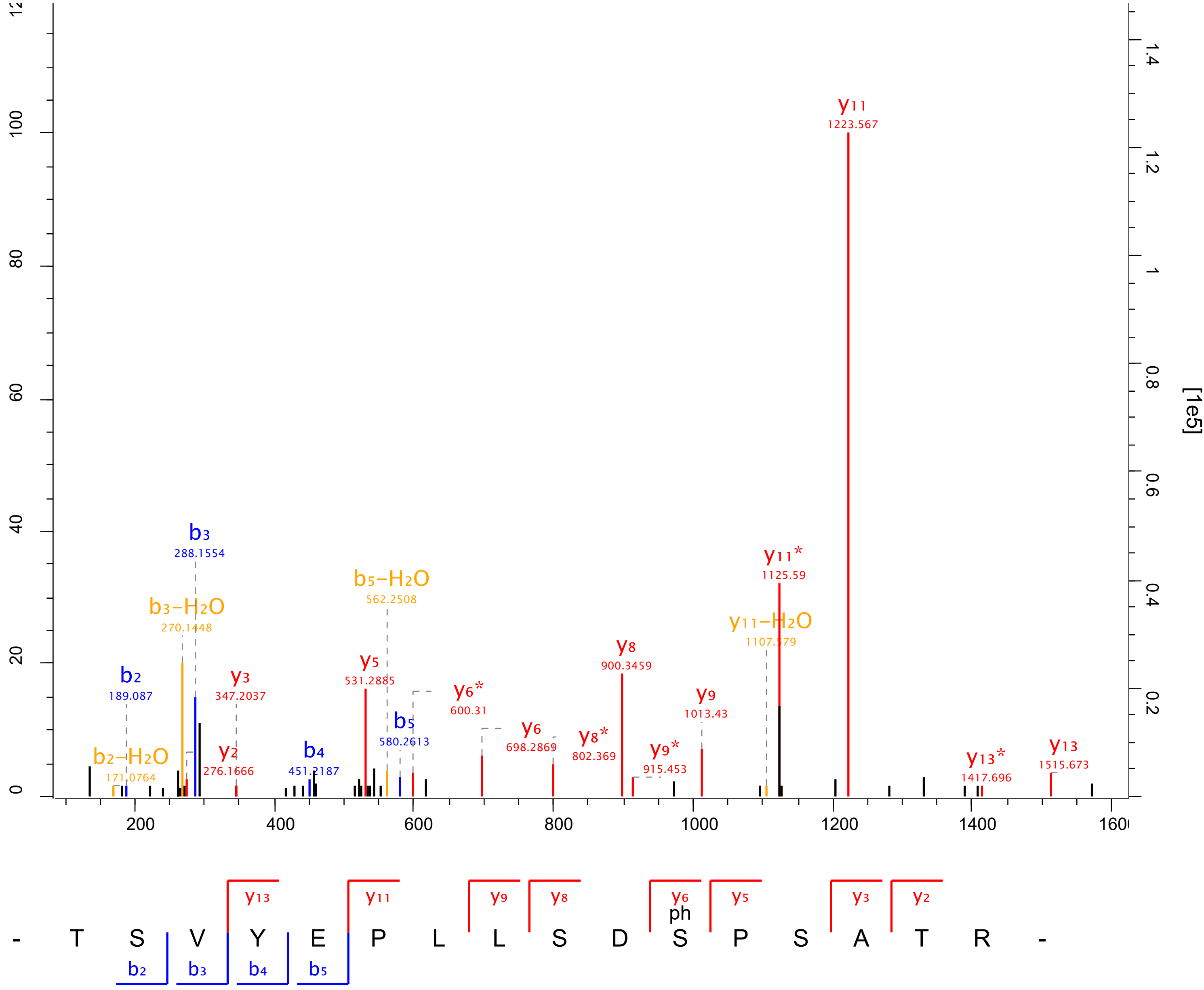

| Raw file | Scan | Method | Score | m/z |
| --- | --- | --- | --- | --- |
| water_02_1 | 30071 | FTMS; HCD | 142.45 | 773.33 |

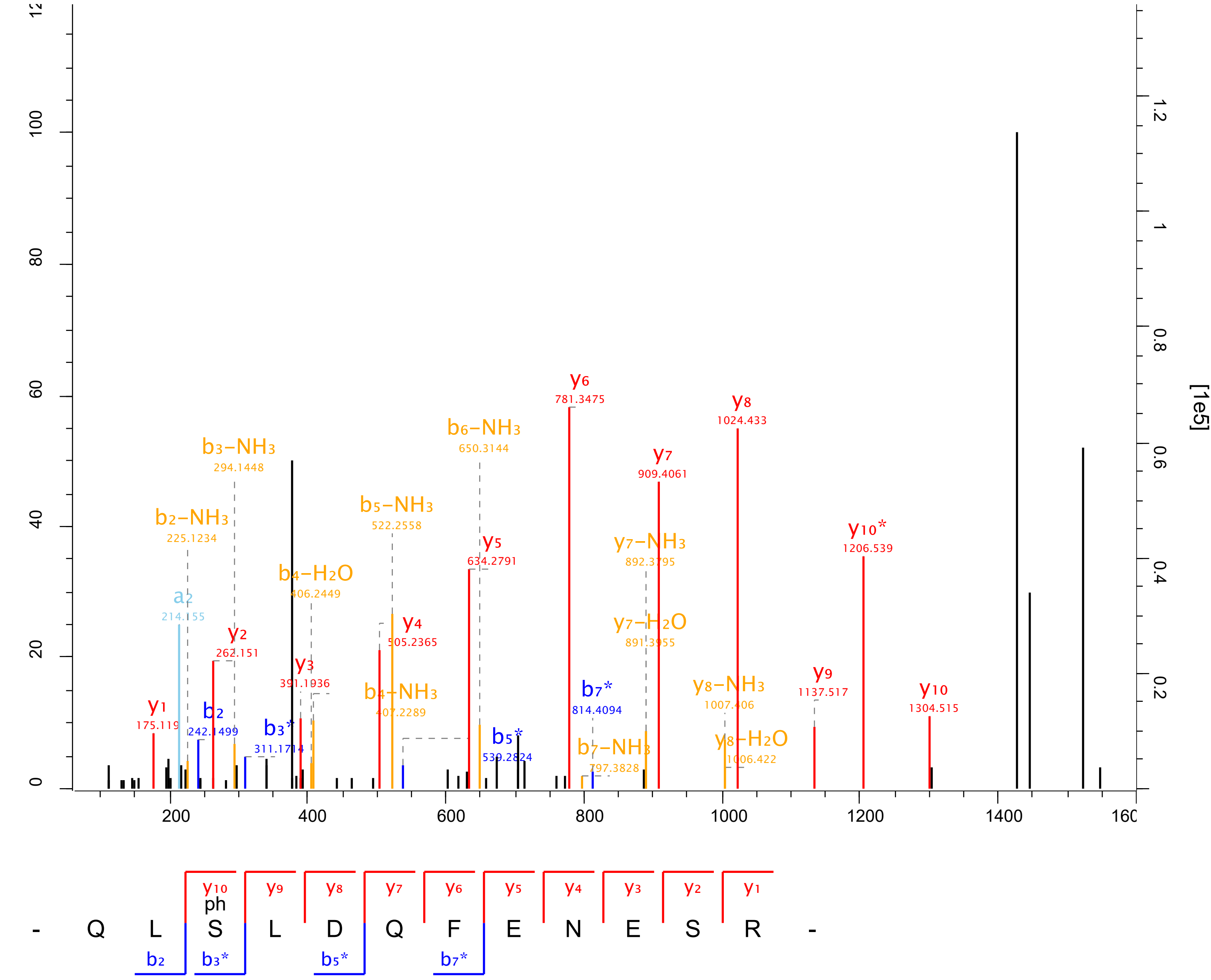

| Raw file | Scan | Method | Score | m/z |
| --- | --- | --- | --- | --- |
| water_02_1 | 30110 | FTMS; HCD | 65.18 | 673.32 |

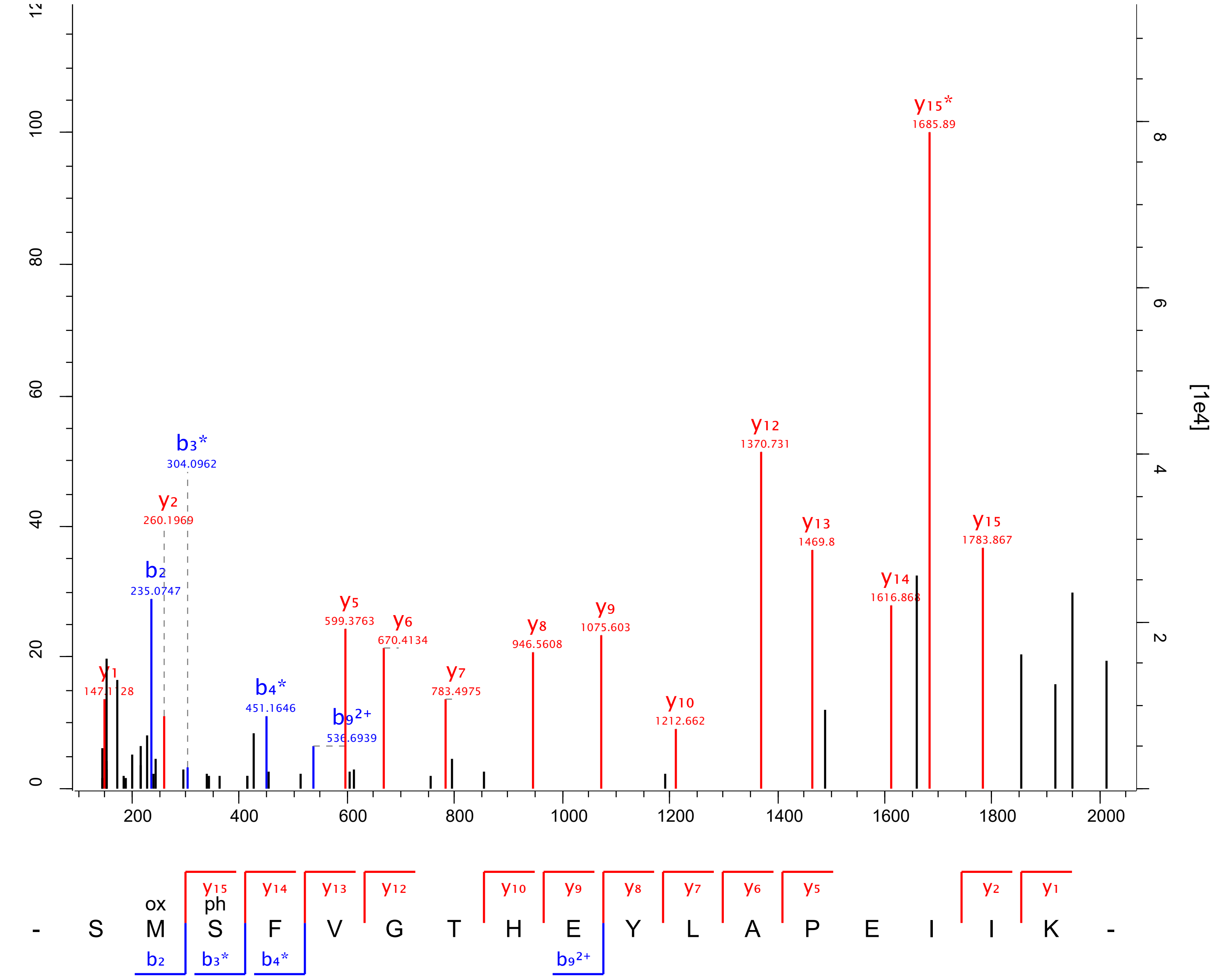

| Raw file | Scan | Method | Score | m/z |
|---|---|---|---|---|
| water_02_1 | 30172 | FTMS; HCD | 171.35 | 924.35 |

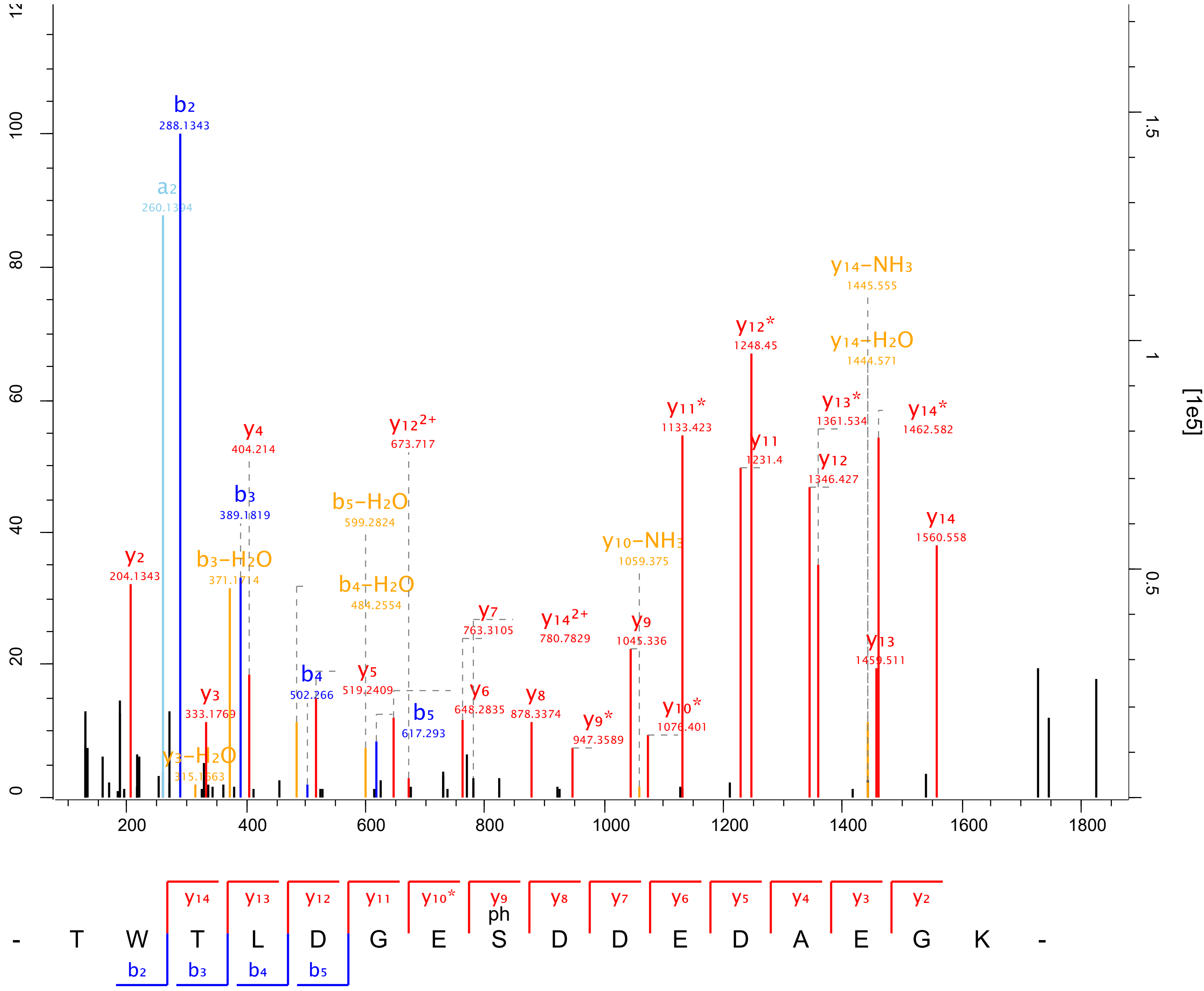

| Raw file | Scan | Method | Score | m/z |
|---|---|---|---|---|
| water_02_1 | 3025 | FTMS; HCD | 73.93 | 580.2 |

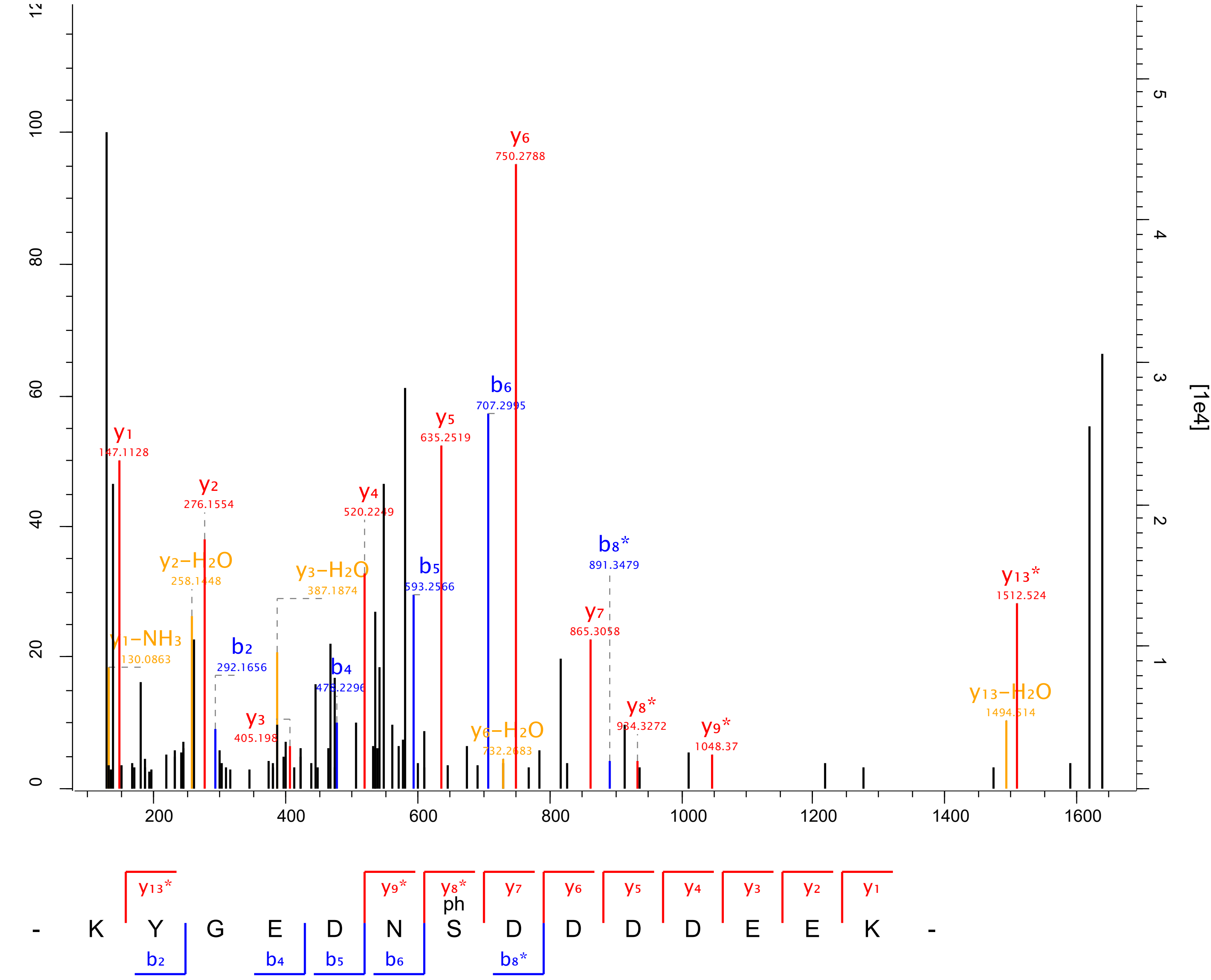

| Raw file | Scan | Method | Score | m/z |
|---|---|---|---|---|
| water_02_1 | 30312 | FTMS; HCD | 146.19 | 864.35 |

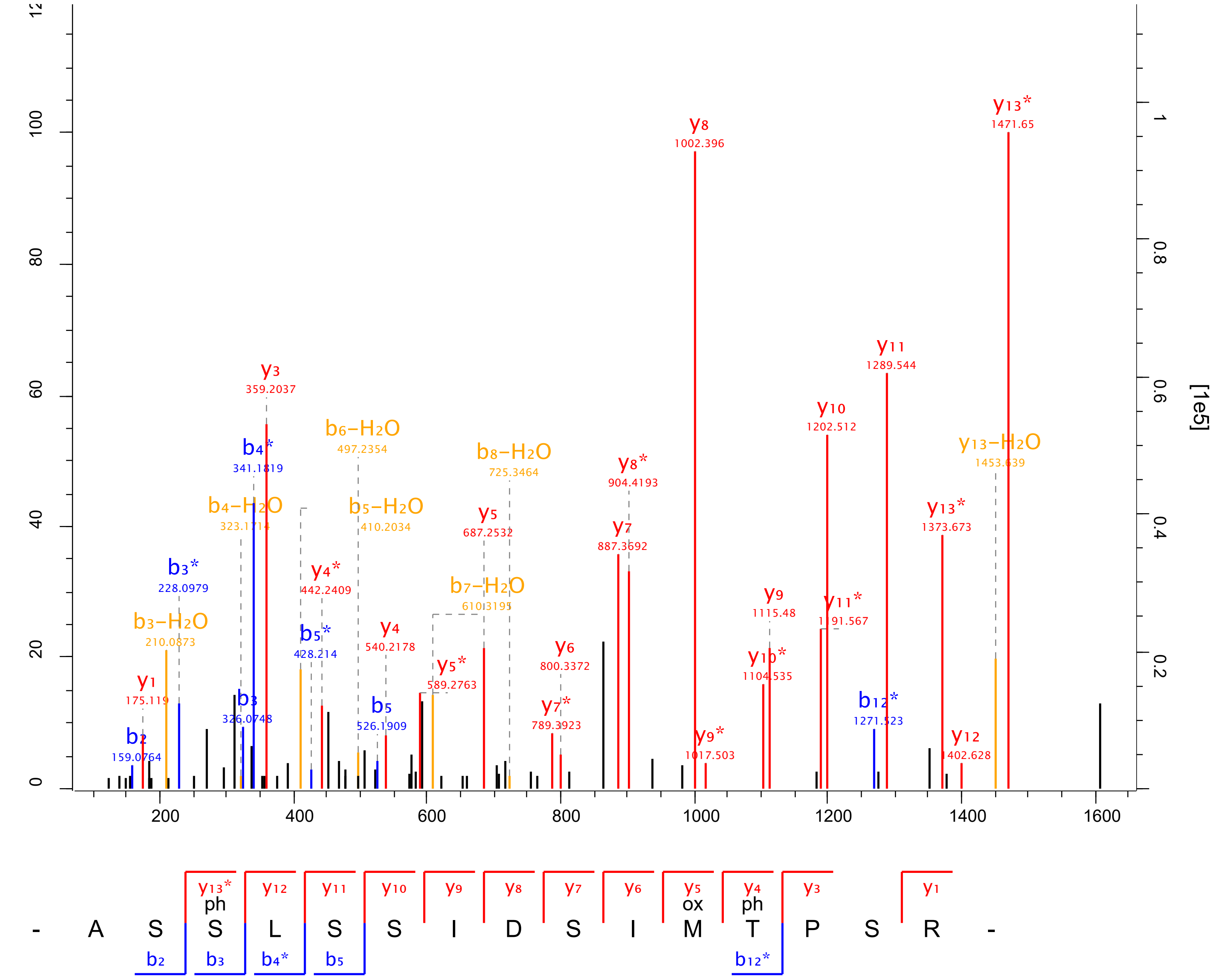

| Raw file | Scan | Method | Score | m/z |
| --- | --- | --- | --- | --- |
| water_02_1 | 3044 | FTMS; HCD | 110.32 | 679.8 |

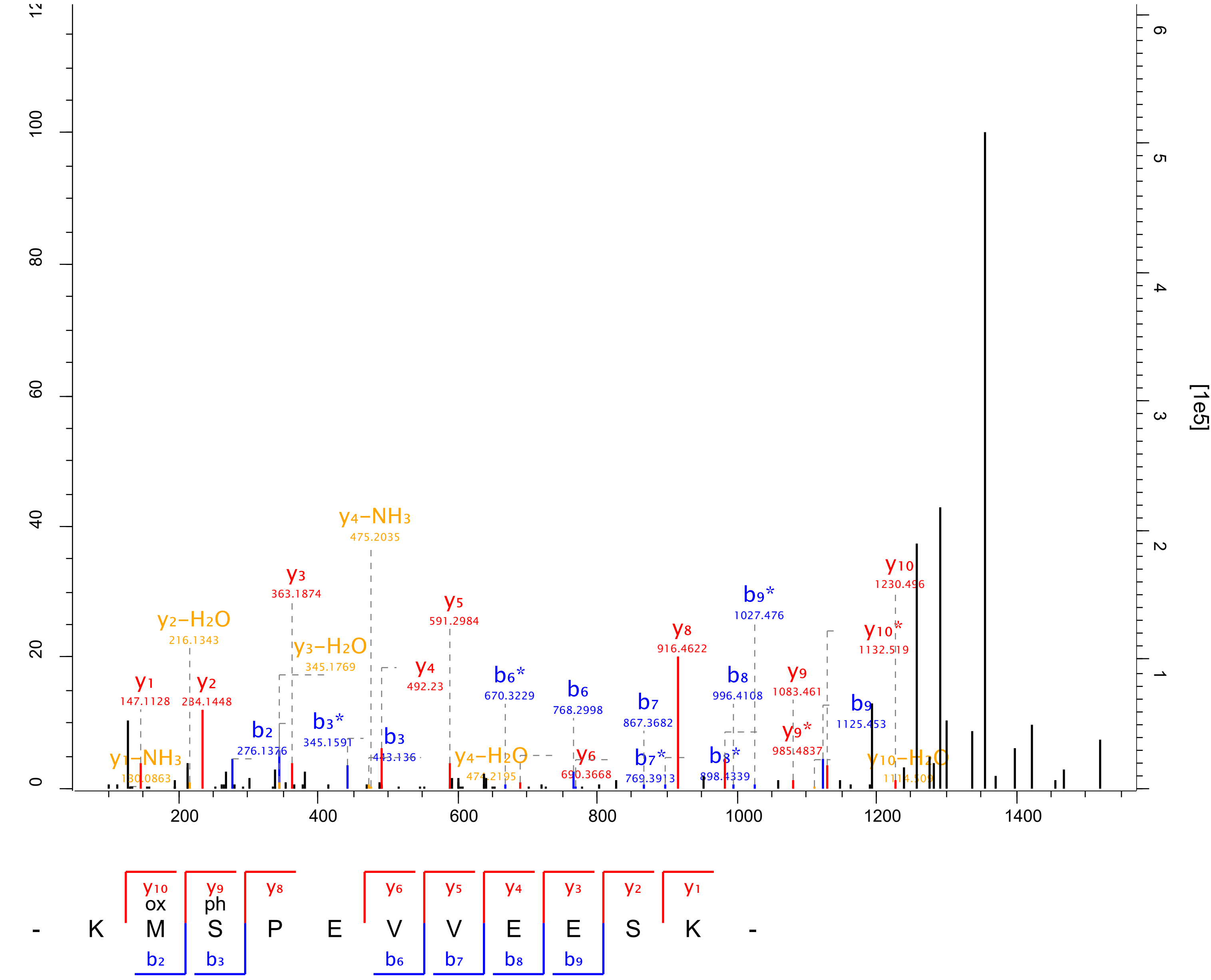

| Raw file | Scan | Method | Score | m/z |
| --- | --- | --- | --- | --- |
| water_02_1 | 30513 | FTMS; HCD | 47.77 | 803.42 |

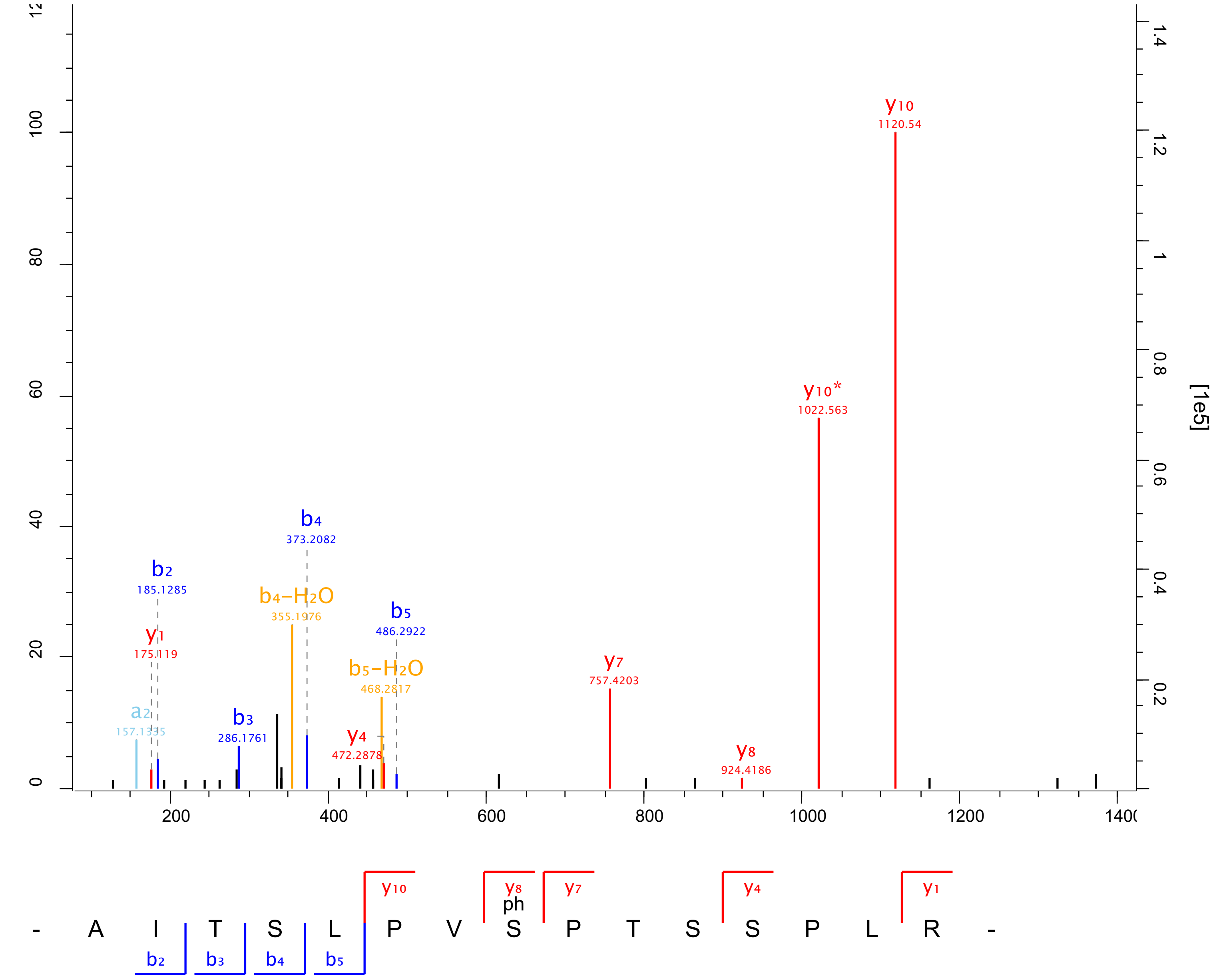

| Raw file | Scan | Method | Score | m/z |
|---|---|---|---|---|
| water_02_1 | 30639 | FTMS; HCD | 120.11 | 820.9 |

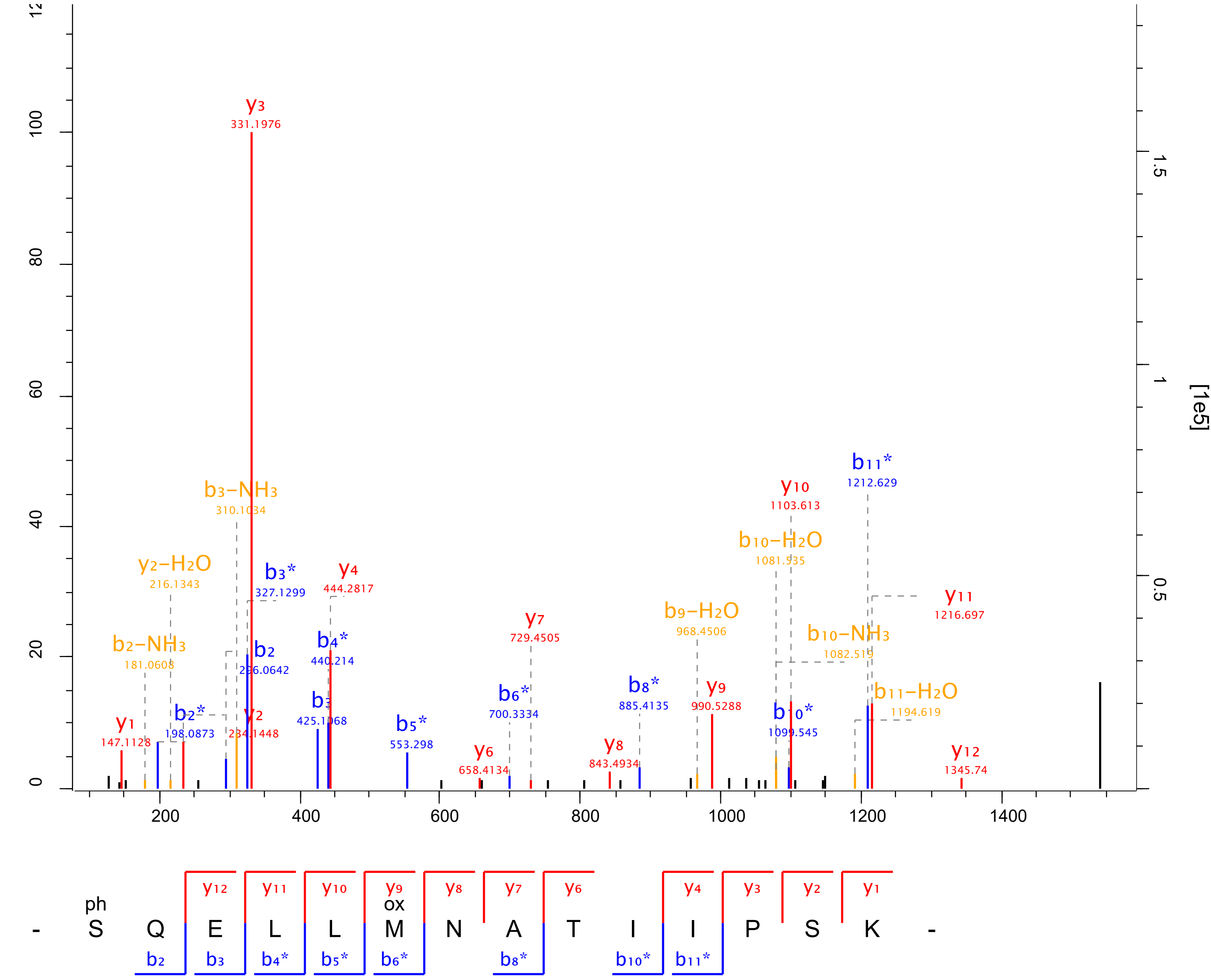

| Raw file | Scan | Method | Score | m/z |
| --- | --- | --- | --- | --- |
| water_02_1 | 30655 | FTMS; HCD | 82.45 | 813.86 |

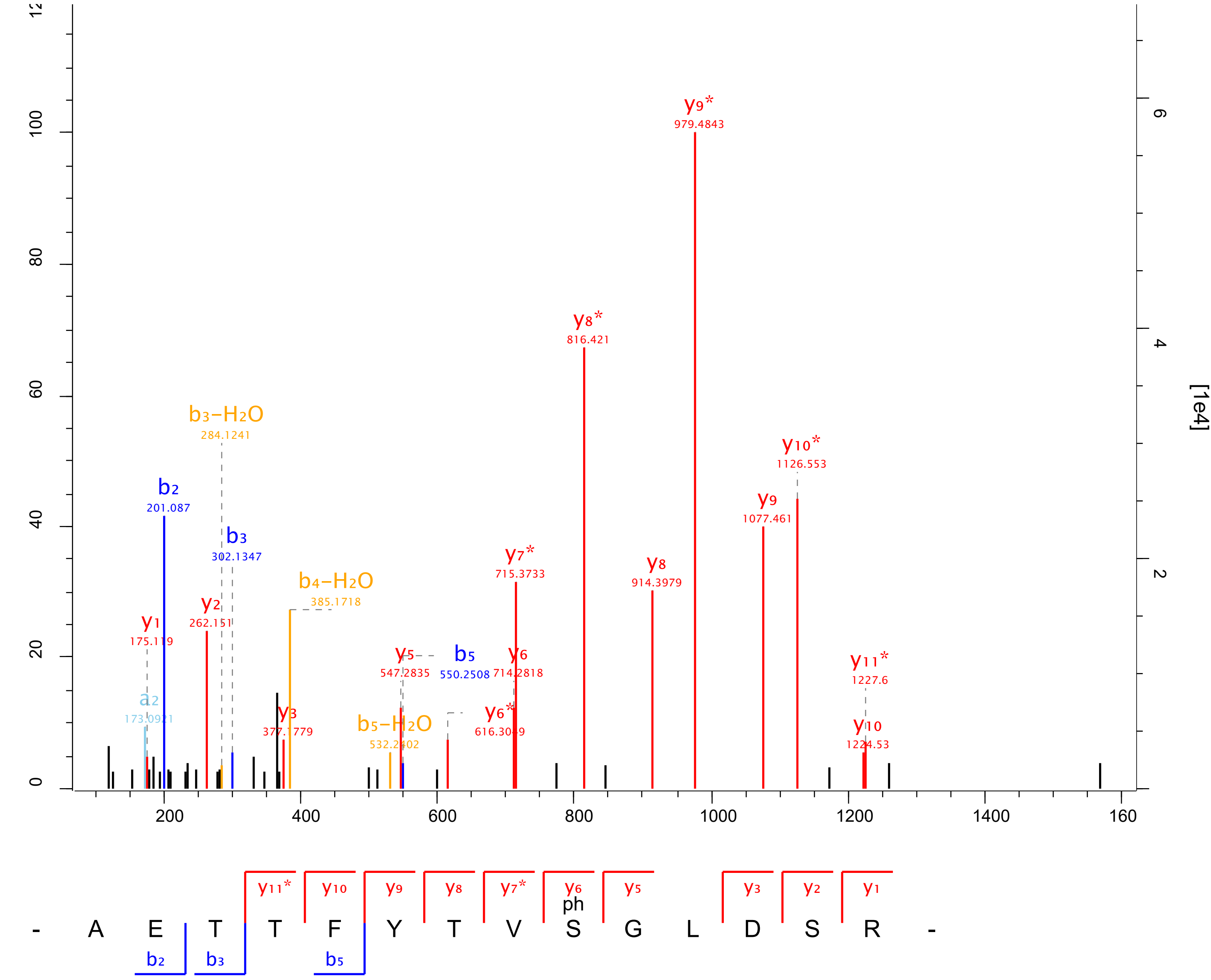

| Raw file | Scan | Method | Score | m/z |
|---|---|---|---|---|
| water_02_1 | 3076 | FTMS; HCD | 108.26 | 474.18 |

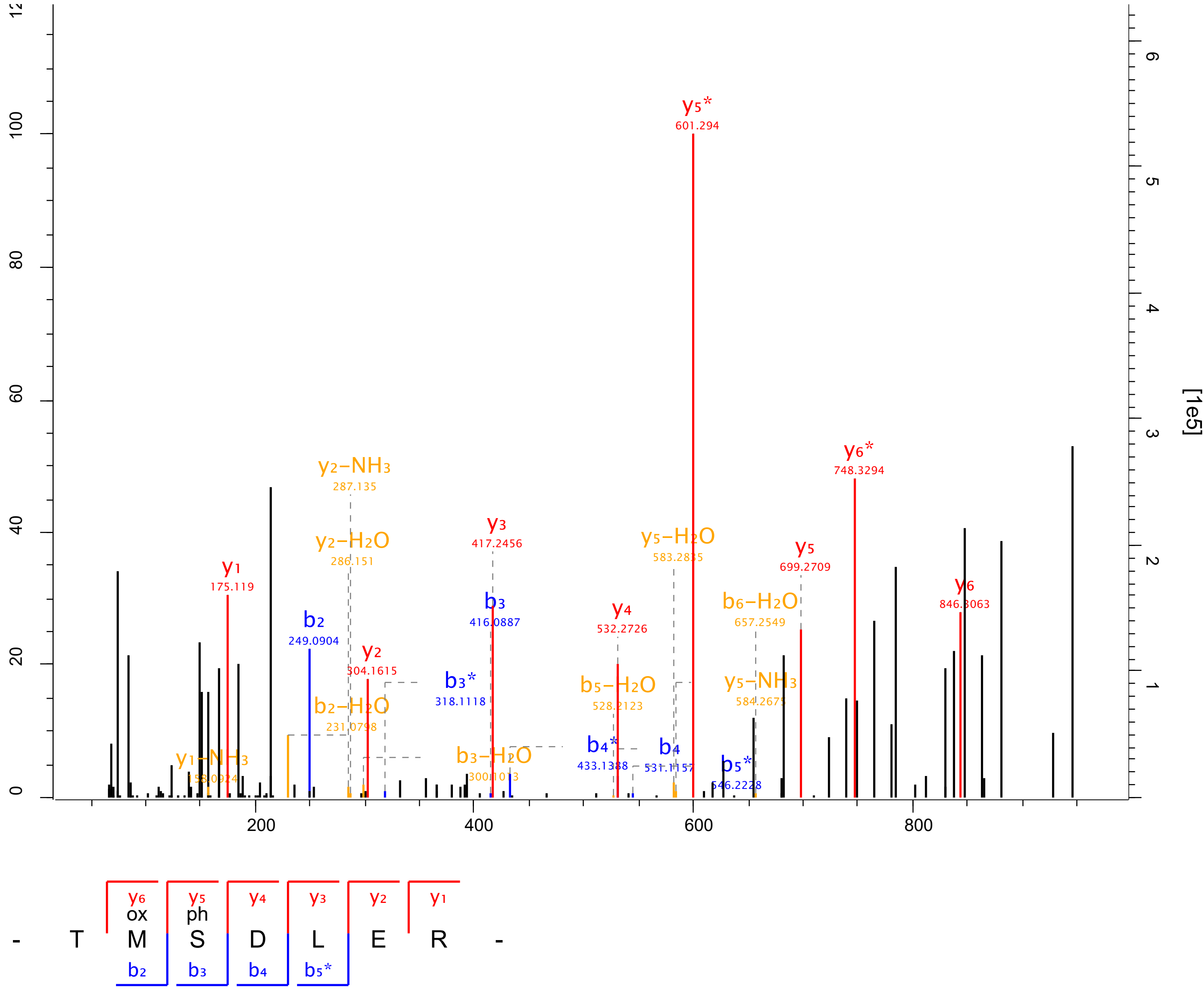

| Raw file | Scan | Method | Score | m/z |
|---|---|---|---|---|
| water_02_1 | 30781 | FTMS; HCD | 160.8 | 696.34 |

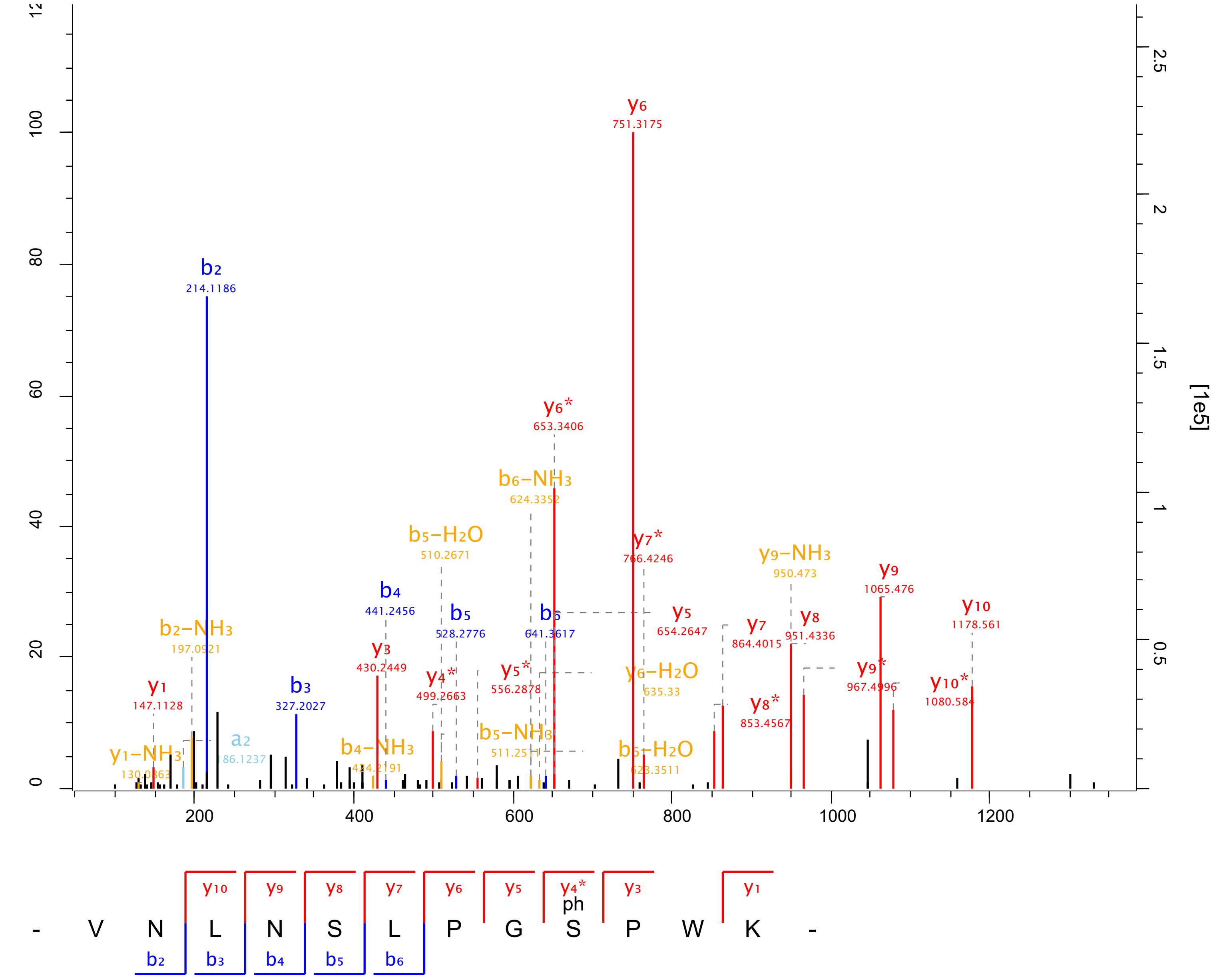

| Raw file | Scan | Method | Score | m/z |
|---|---|---|---|---|
| water_02_1 | 3079 | FTMS; HCD | 105.03 | 522.22 |

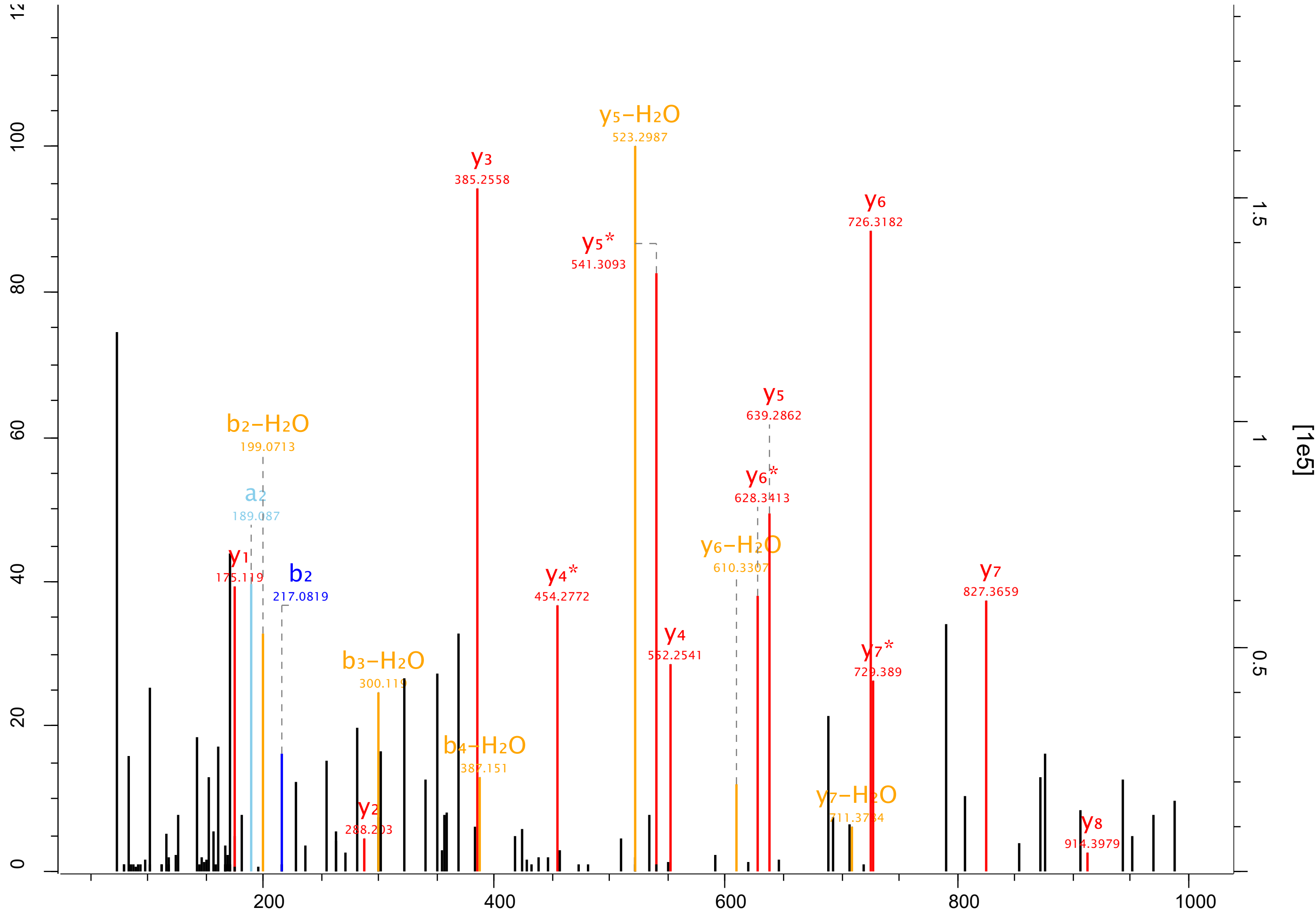


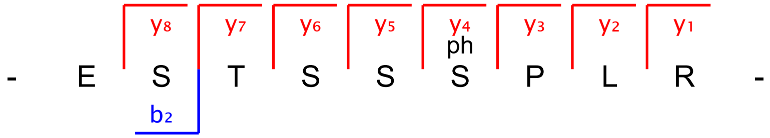

| Raw file | Scan | Method | Score | m/z |
|---|---|---|---|---|
| water_02_1 | 30908 | FTMS; HCD | 92.15 | 811.69 |

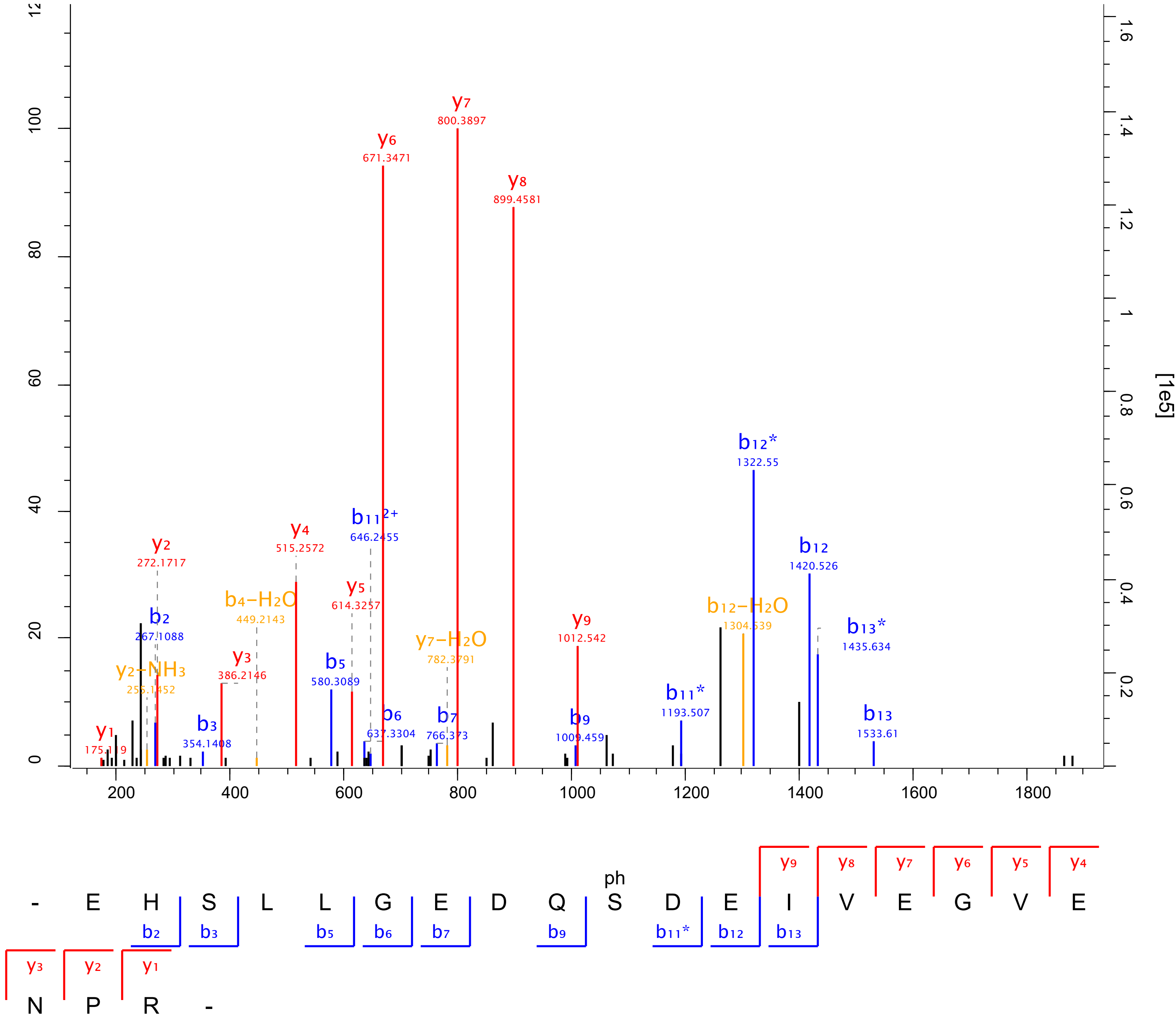

| Raw file | Scan | Method | Score | m/z |
|---|---|---|---|---|
| water_02_1 | 30935 | FTMS; HCD | 67.5 | 891.05 |

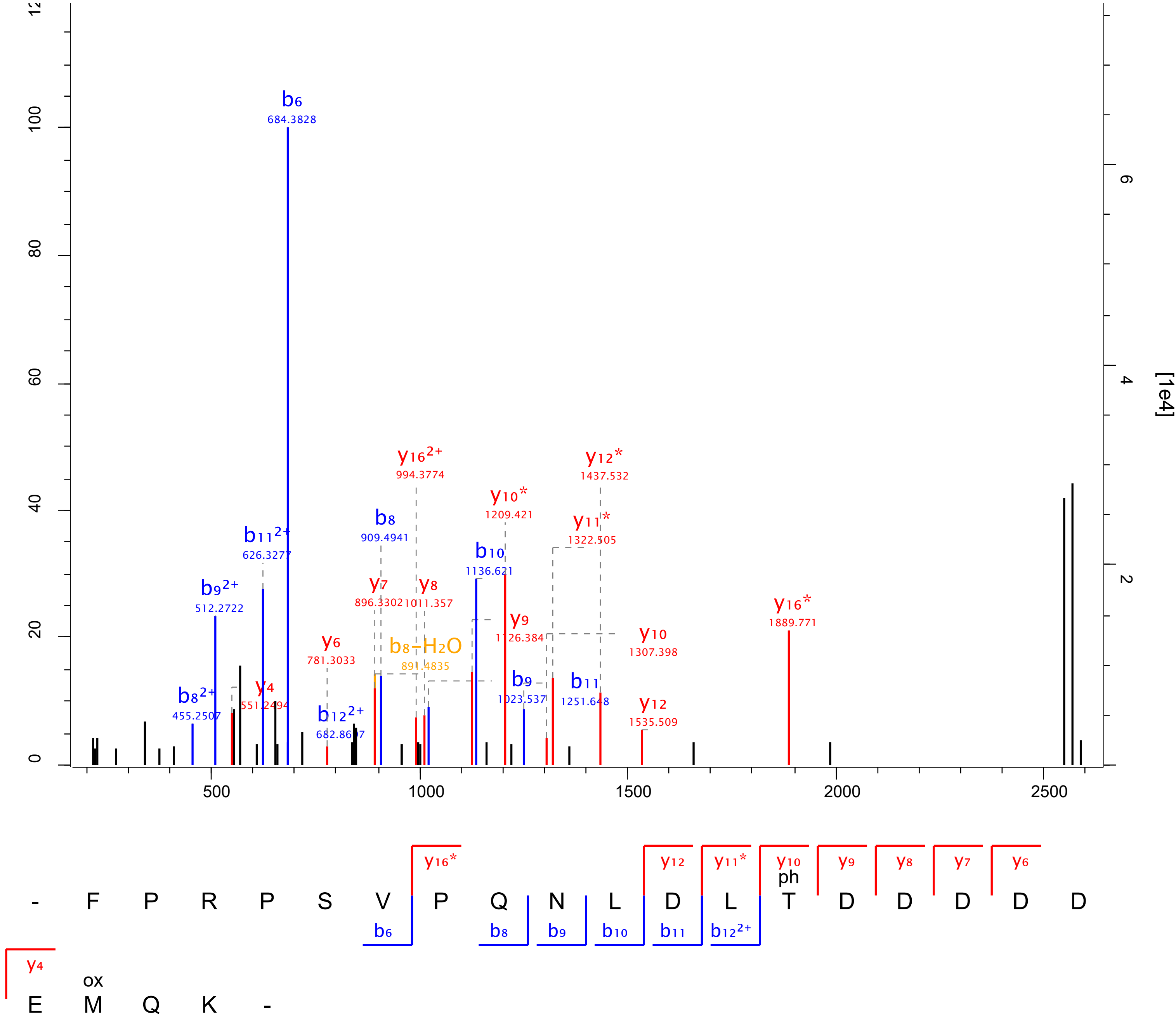

| Raw file | Scan | Method | Score | m/z |
| --- | --- | --- | --- | --- |
| water_02_1 | 3101 | FTMS; HCD | 48.57 | 436.85 |

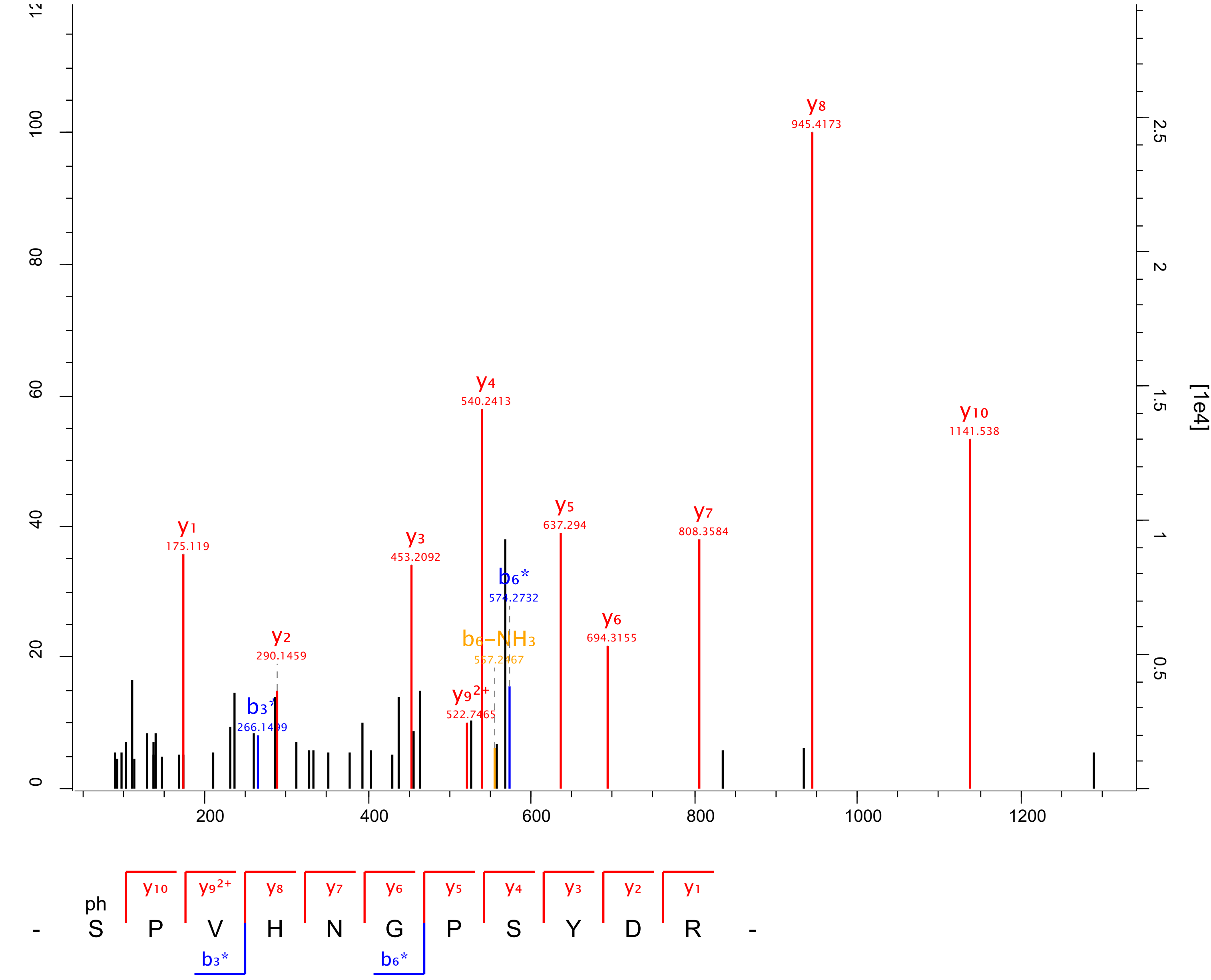

| Raw file | Scan | Method | Score | m/z |
|---|---|---|---|---|
| water_02_1 | 31178 | FTMS; HCD | 121.1 | 781.7 |

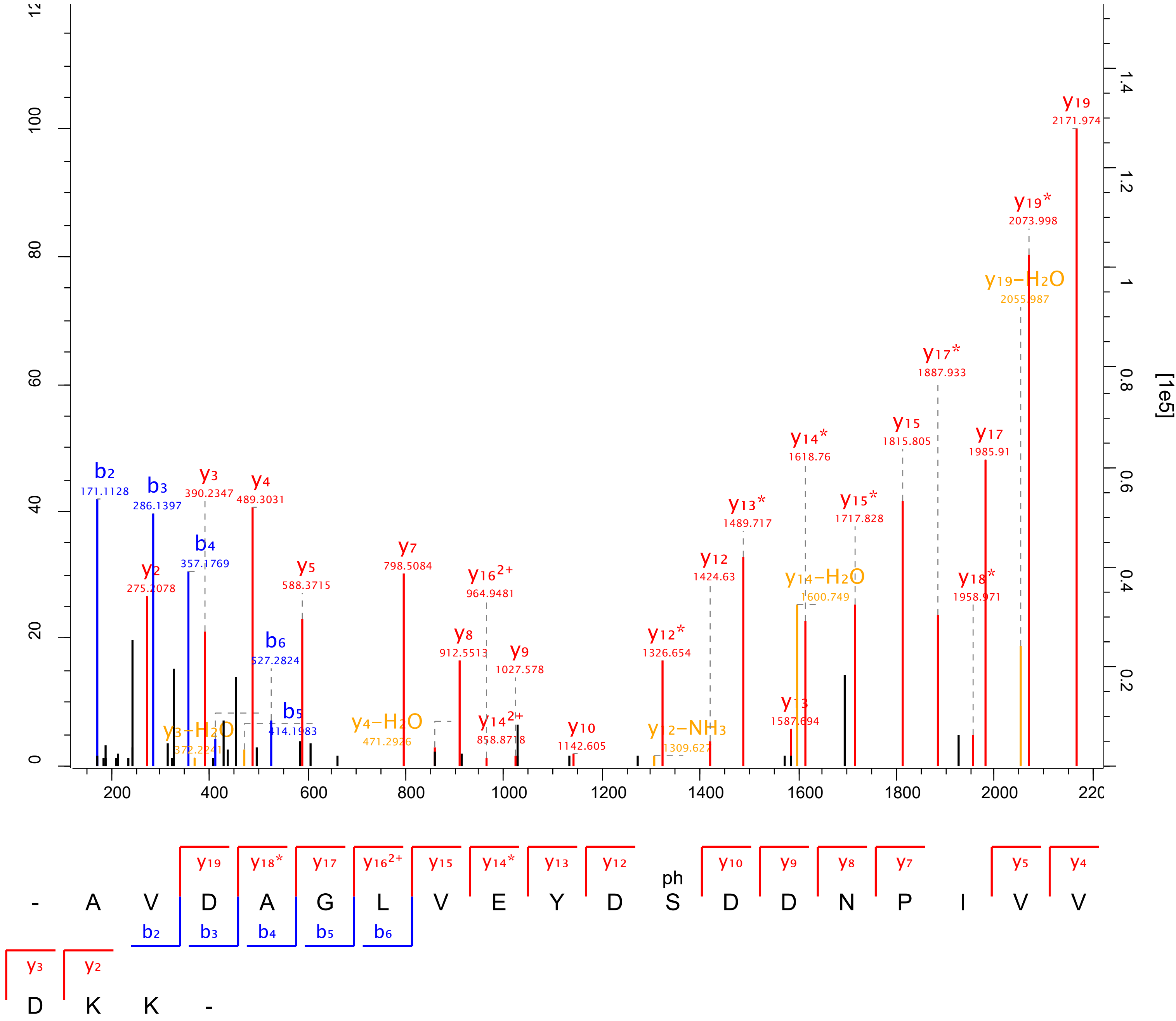

| Raw file | Scan | Method | Score | m/z |
| --- | --- | --- | --- | --- |
| water_02_1 | 3118 | FTMS; HCD | 83.18 | 571.74 |

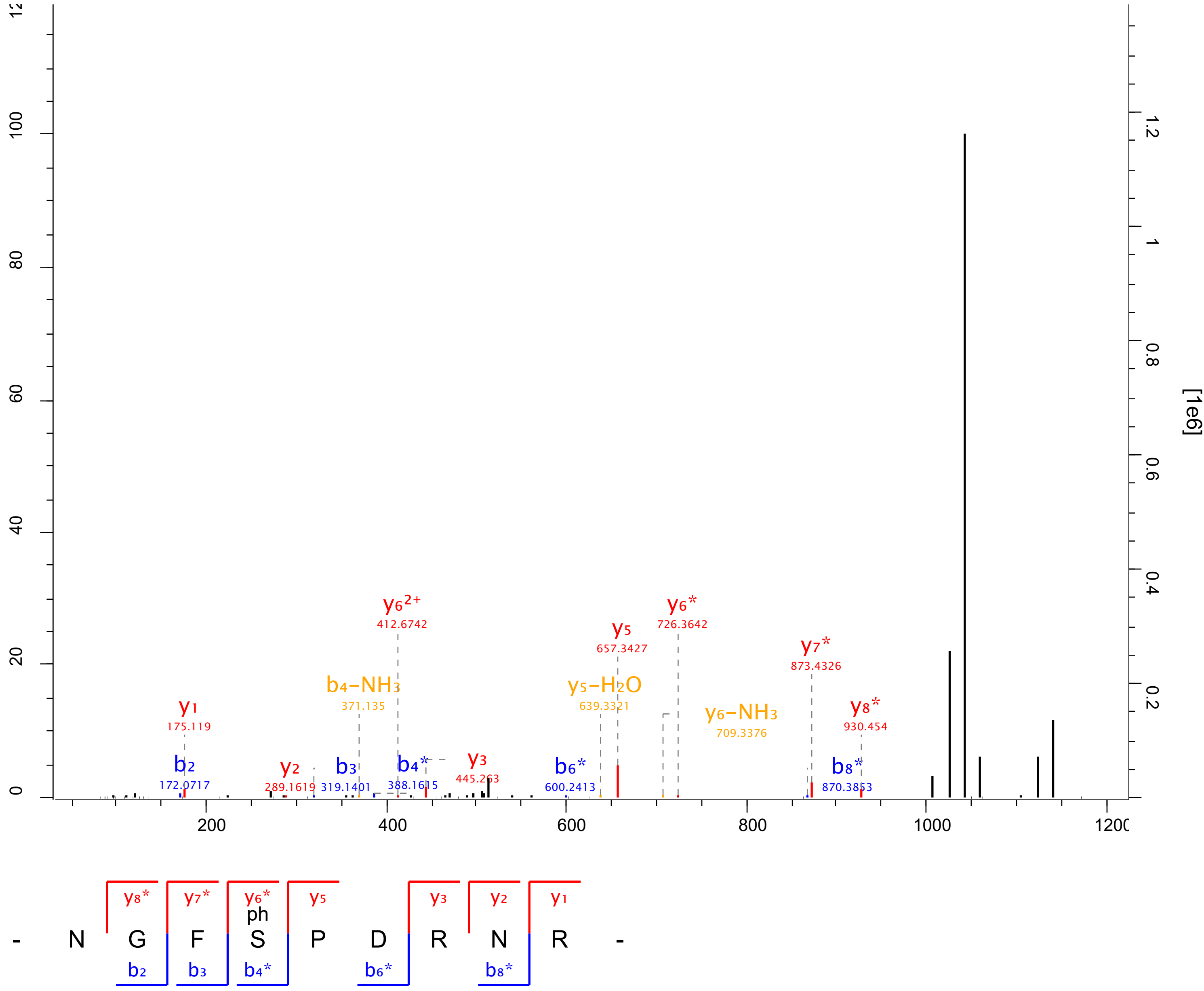

| Raw file | Scan | Method | Score | m/z |
| --- | --- | --- | --- | --- |
| water_02_1 | 31237 | FTMS; HCD | 118.31 | 750.87 |

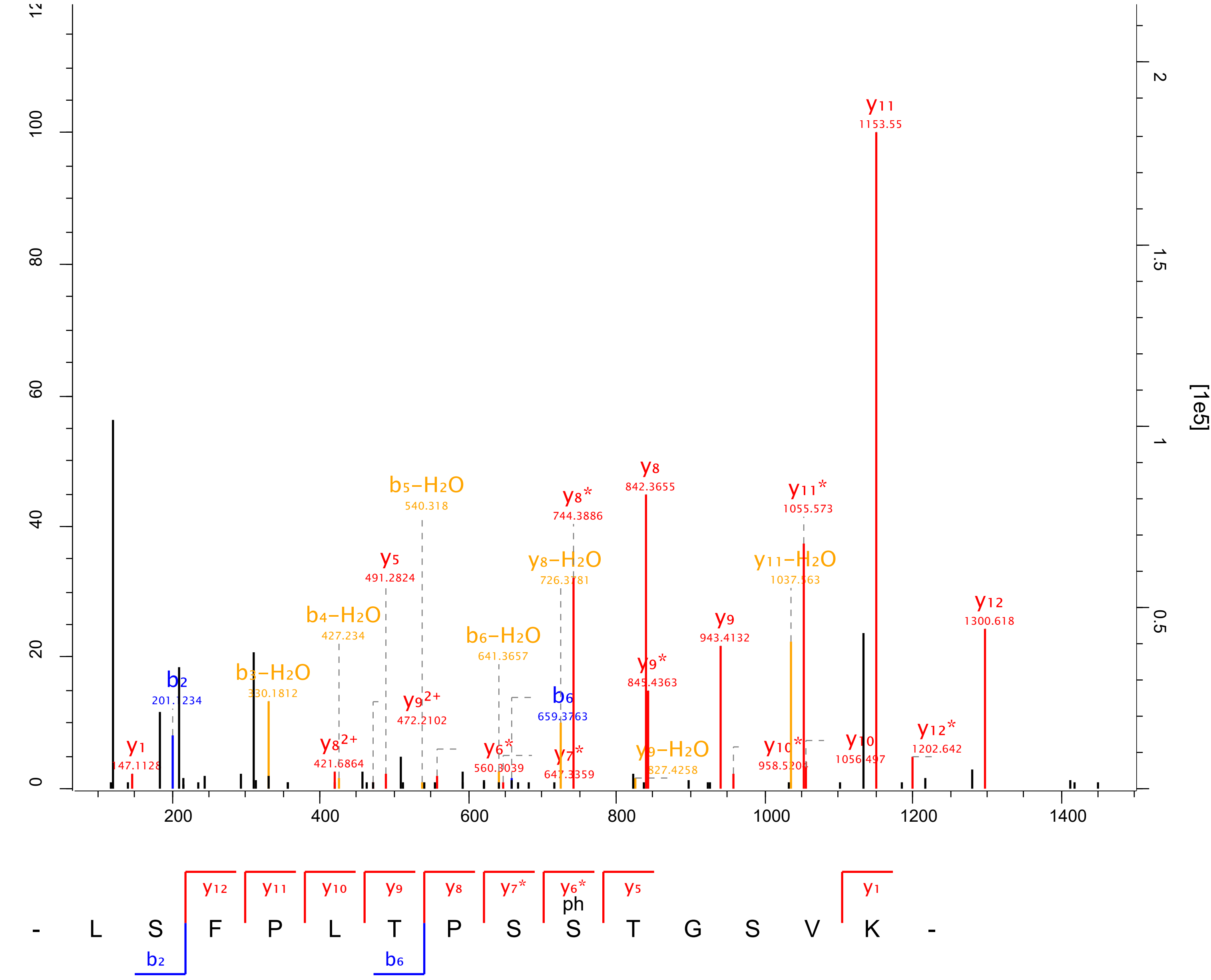

| Raw file | Scan | Method | Score | m/z |
| --- | --- | --- | --- | --- |
| water_02_1 | 31358 | FTMS; HCD | 143.19 | 827.04 |

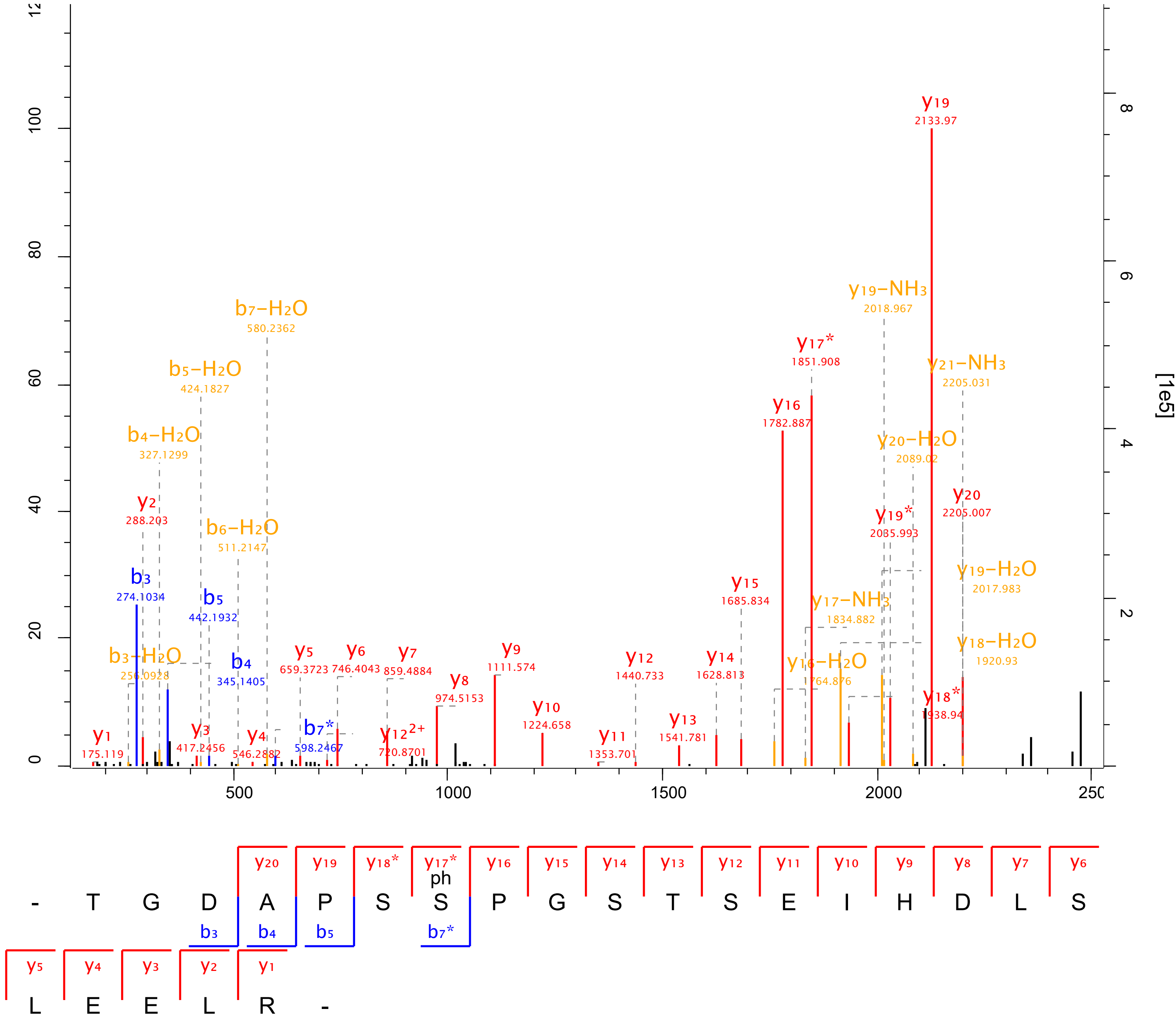

| Raw file | Scan | Method | Score | m/z |
| --- | --- | --- | --- | --- |
| water_02_1 | 31476 | FTMS; HCD | 41.62 | 916.41 |

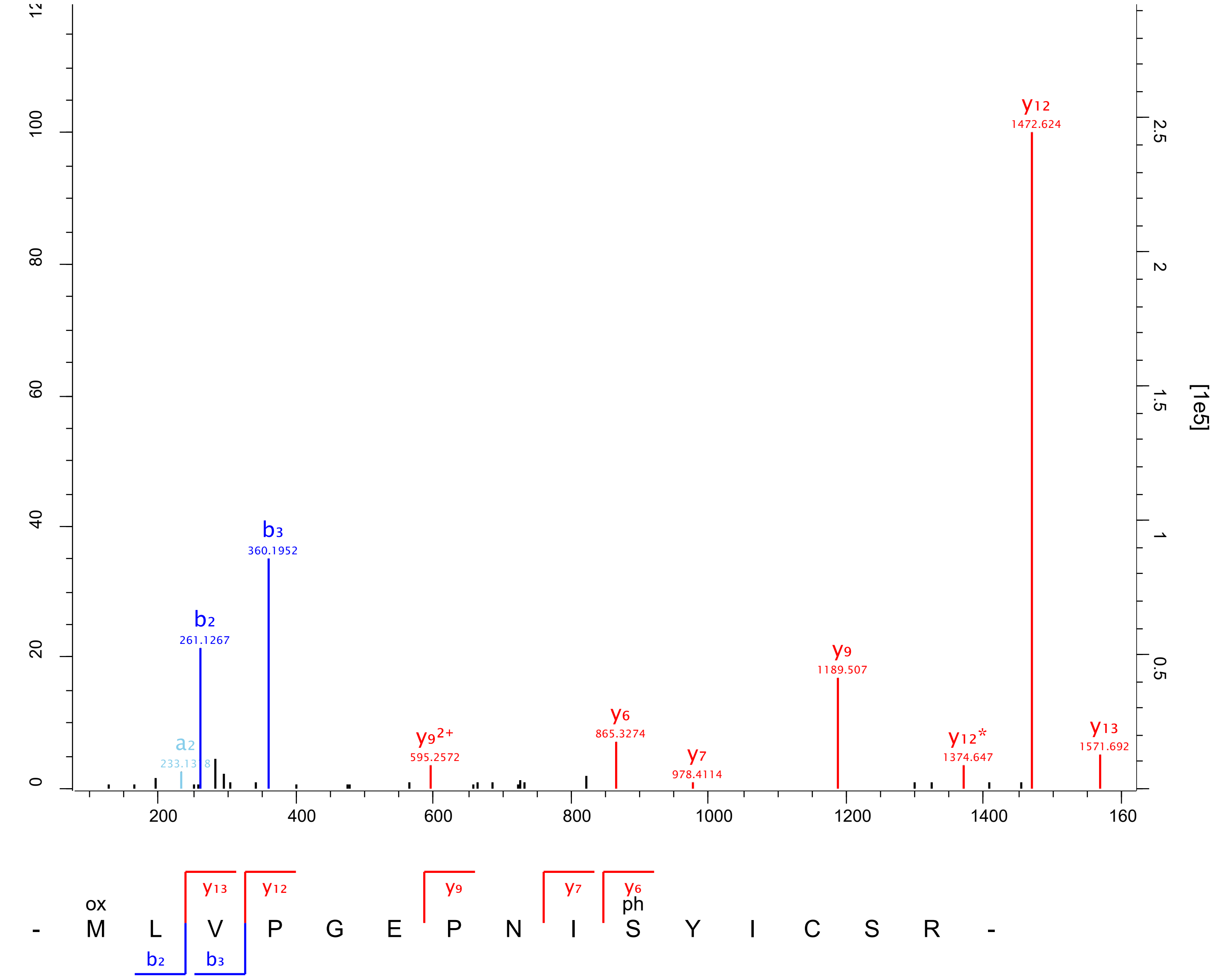

| Raw file | Scan | Method | Score | m/z |
|---|---|---|---|---|
| water_02_1 | 31613 | FTMS; HCD | 176.69 | 853.43 |

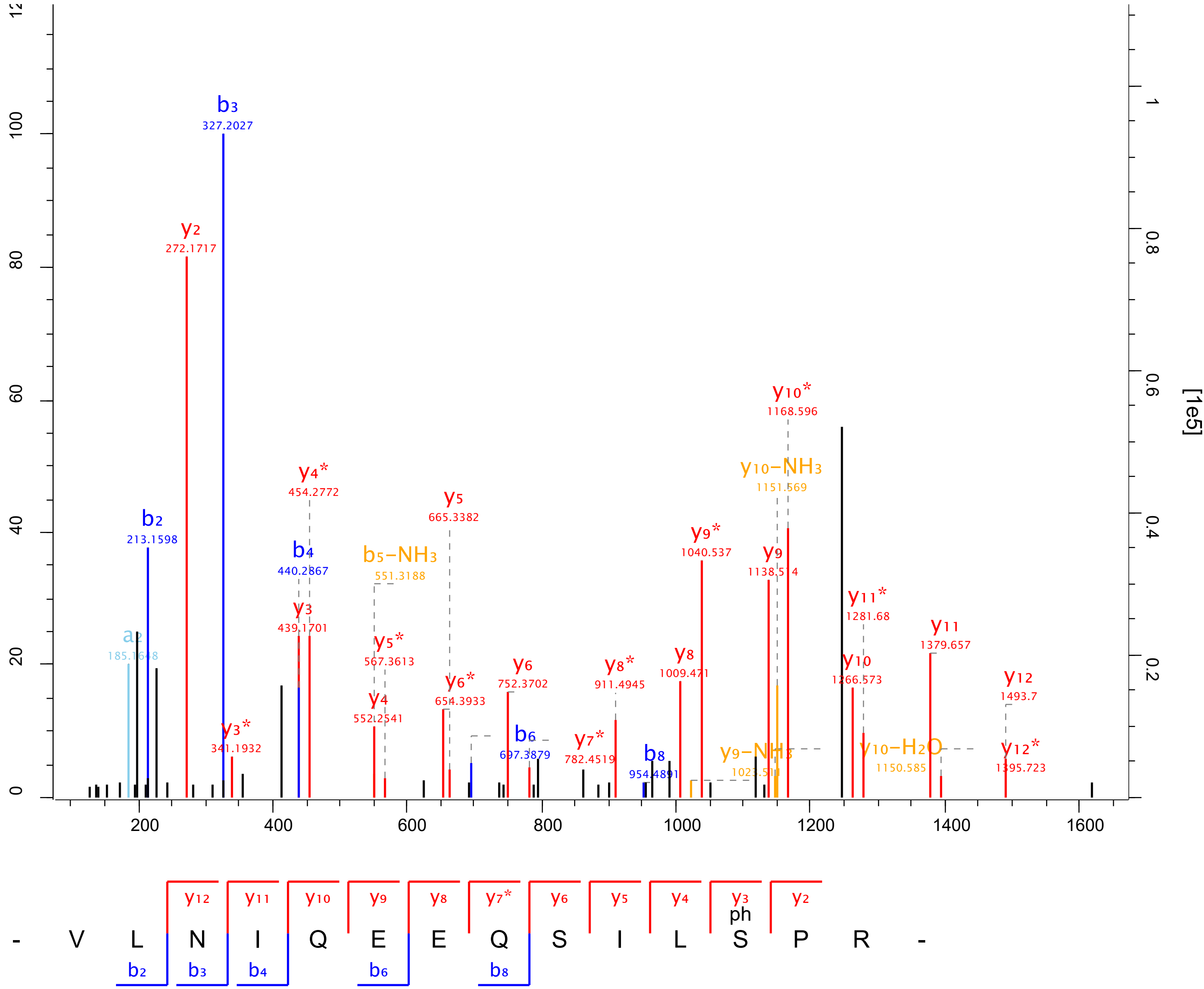

| Raw file | Scan | Method | Score | m/z |
| --- | --- | --- | --- | --- |
| water_02_1 | 31681 | FTMS; HCD | 74.48 | 729.84 |

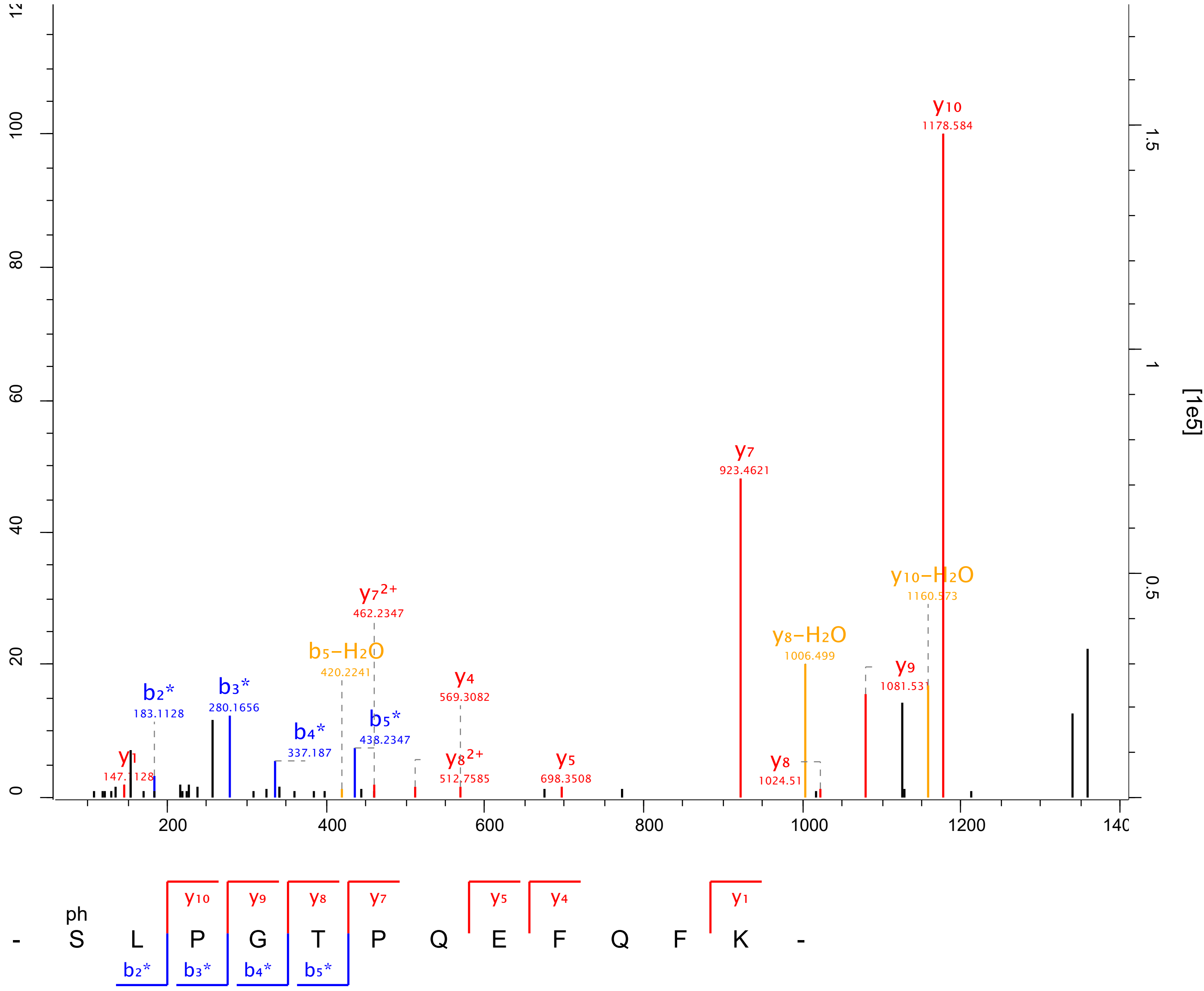

| Raw file | Scan | Method | Score | m/z |
| --- | --- | --- | --- | --- |
| water_02_1 | 31685 | FTMS; HCD | 134.25 | 1108.45 |

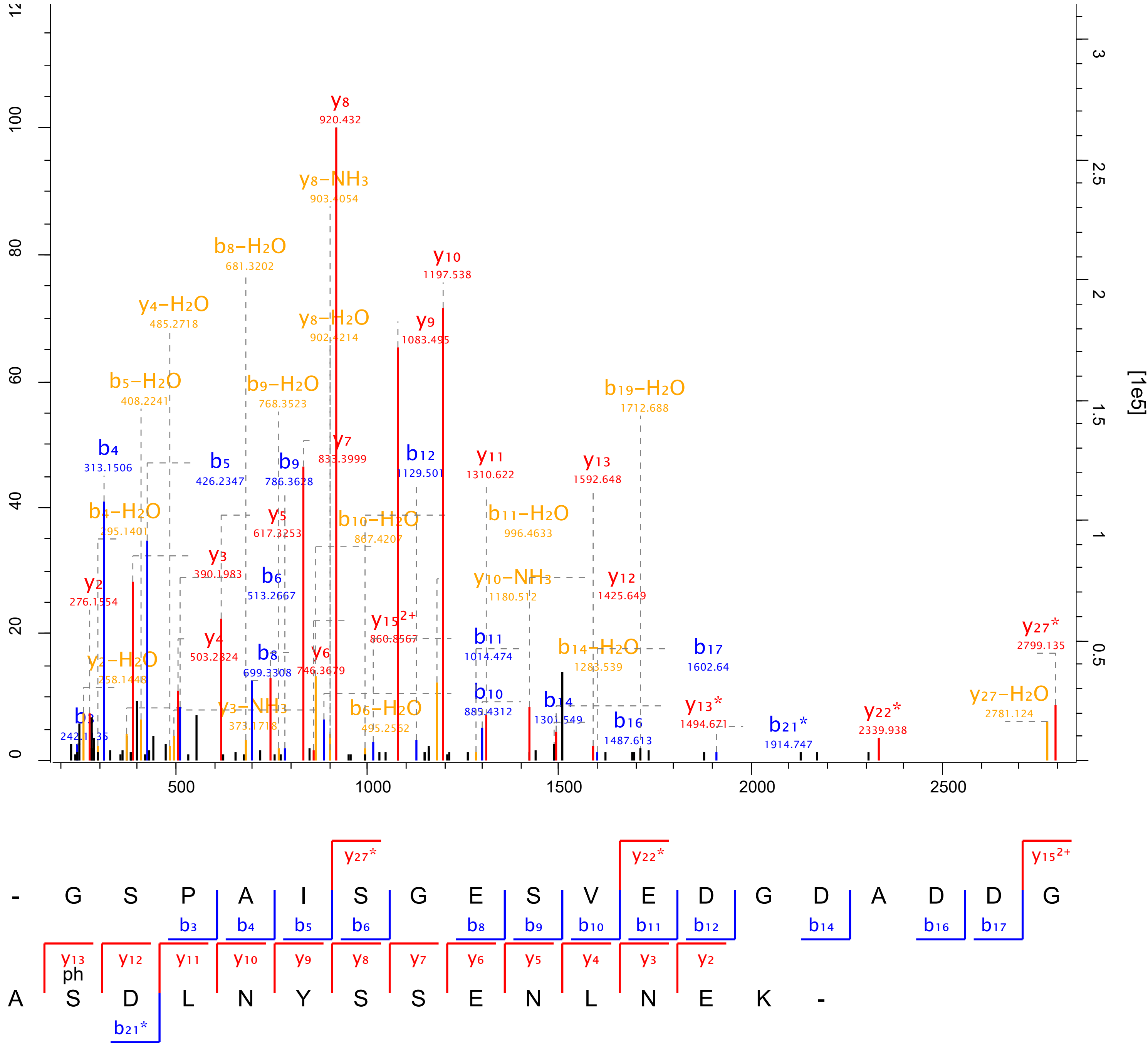

| Raw file | Scan | Method | Score | m/z |
|---|---|---|---|---|
| water_02_1 | 3185 | FTMS; HCD | 75.02 | 764.3 |

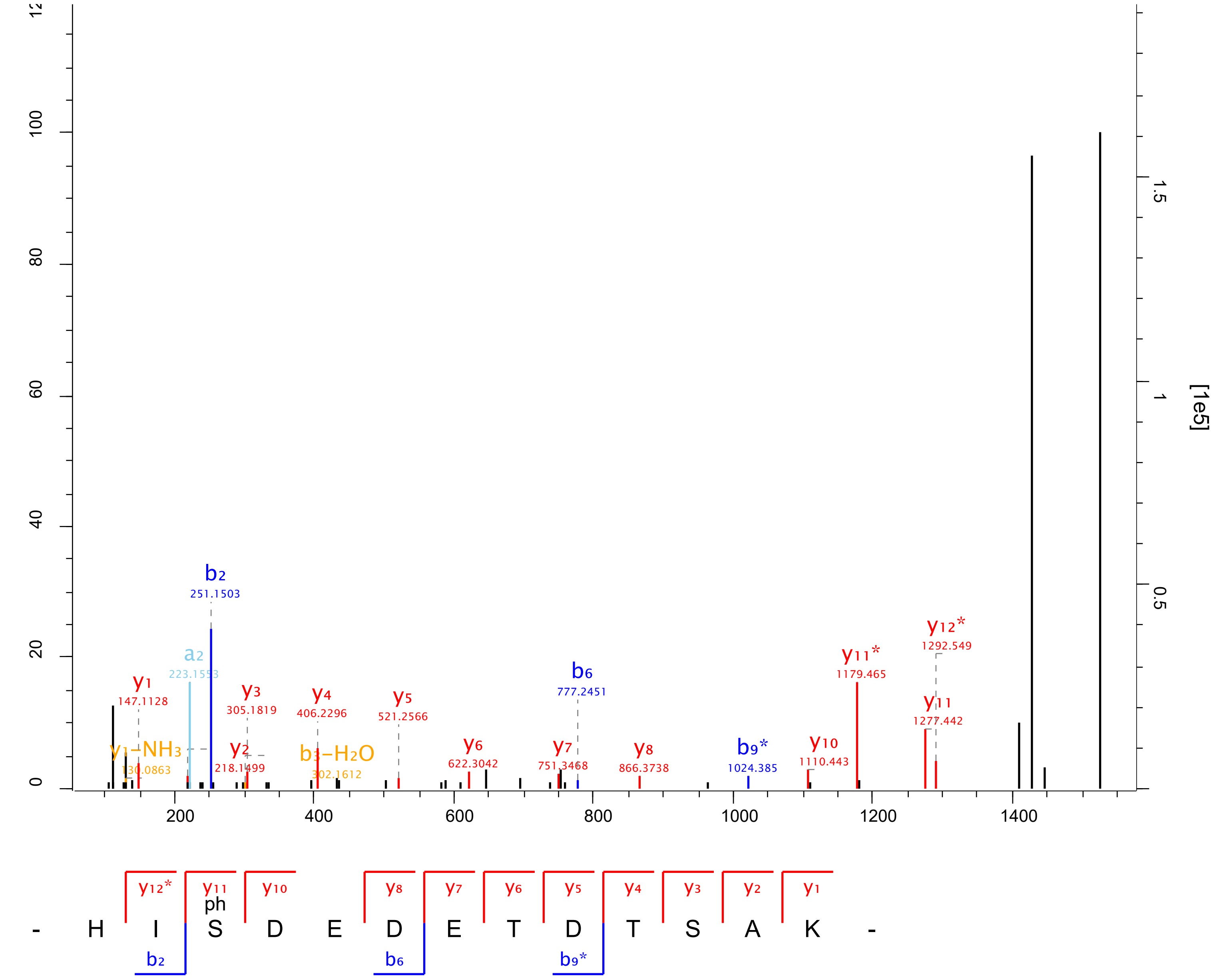

| Raw file | Scan | Method | Score | m/z |
| --- | --- | --- | --- | --- |
| water_02_1 | 31851 | FTMS; HCD | 103.75 | 755.33 |

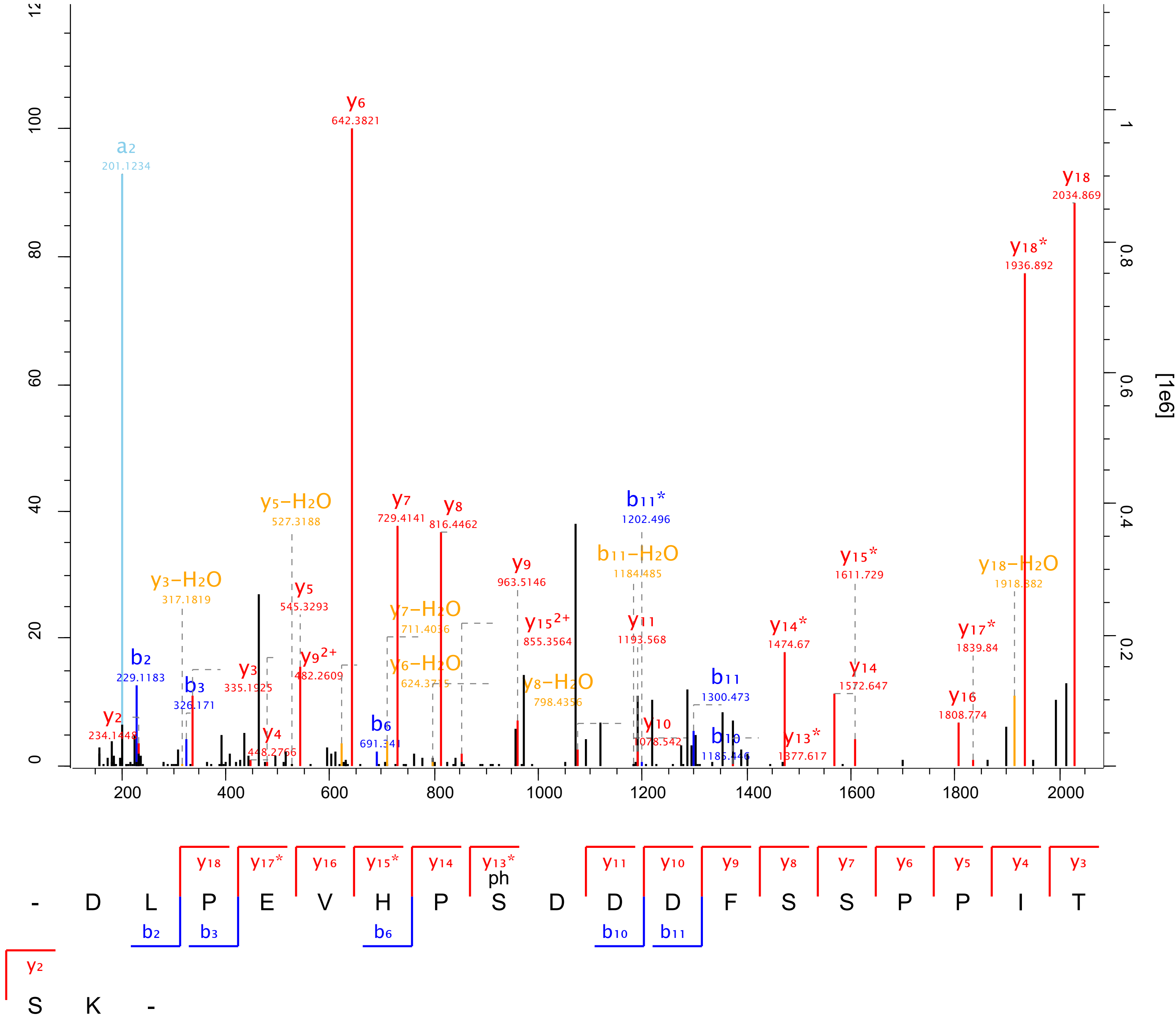

| Raw file | Scan | Method | Score | m/z |
| --- | --- | --- | --- | --- |
| water_02_1 | 31948 | FTMS; HCD | 83.24 | 1076.12 |

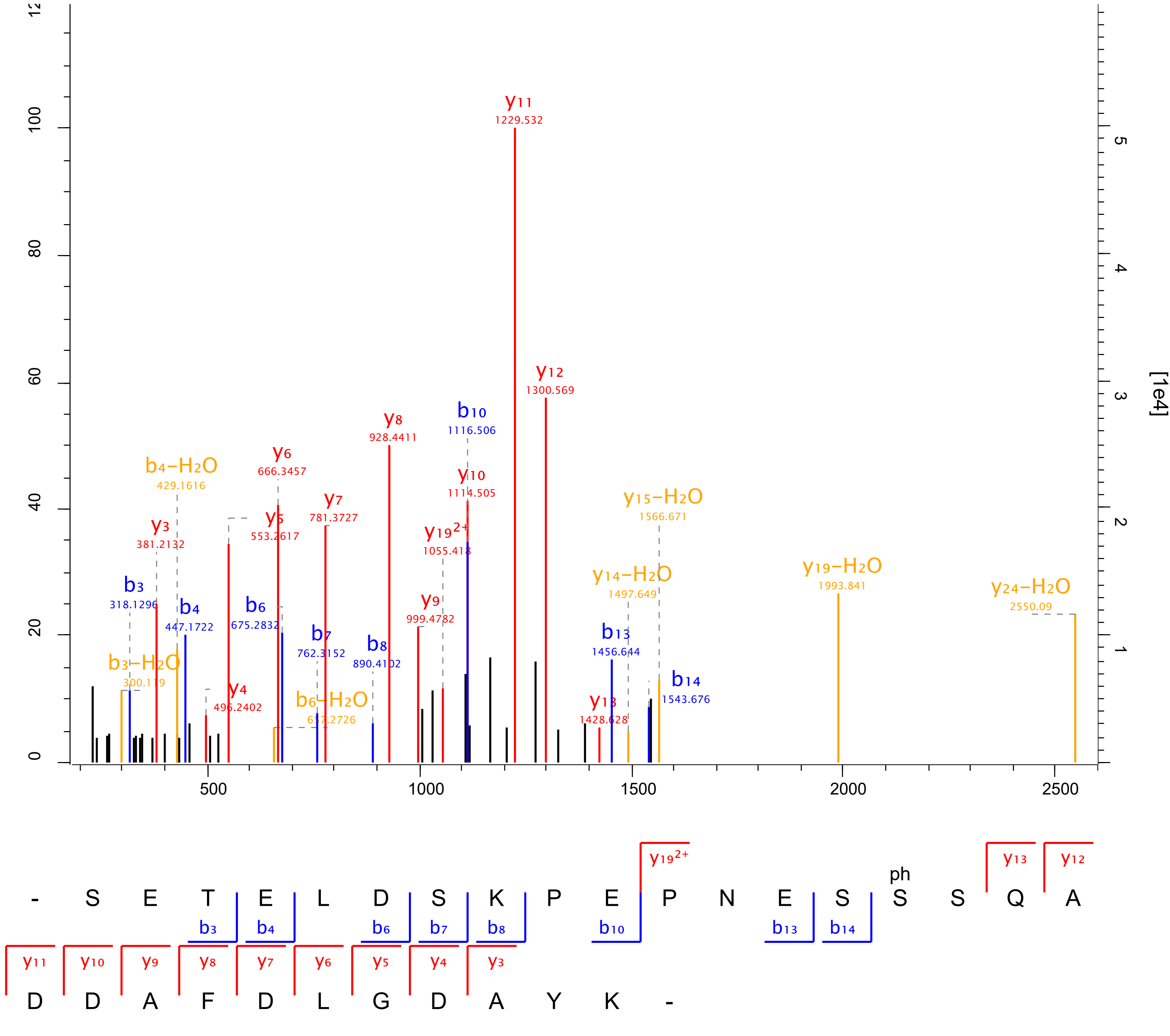

| Raw file | Scan | Method | Score | m/z |
| --- | --- | --- | --- | --- |
| water_02_1 | 3195 | FTMS; HCD | 111.64 | 736.32 |

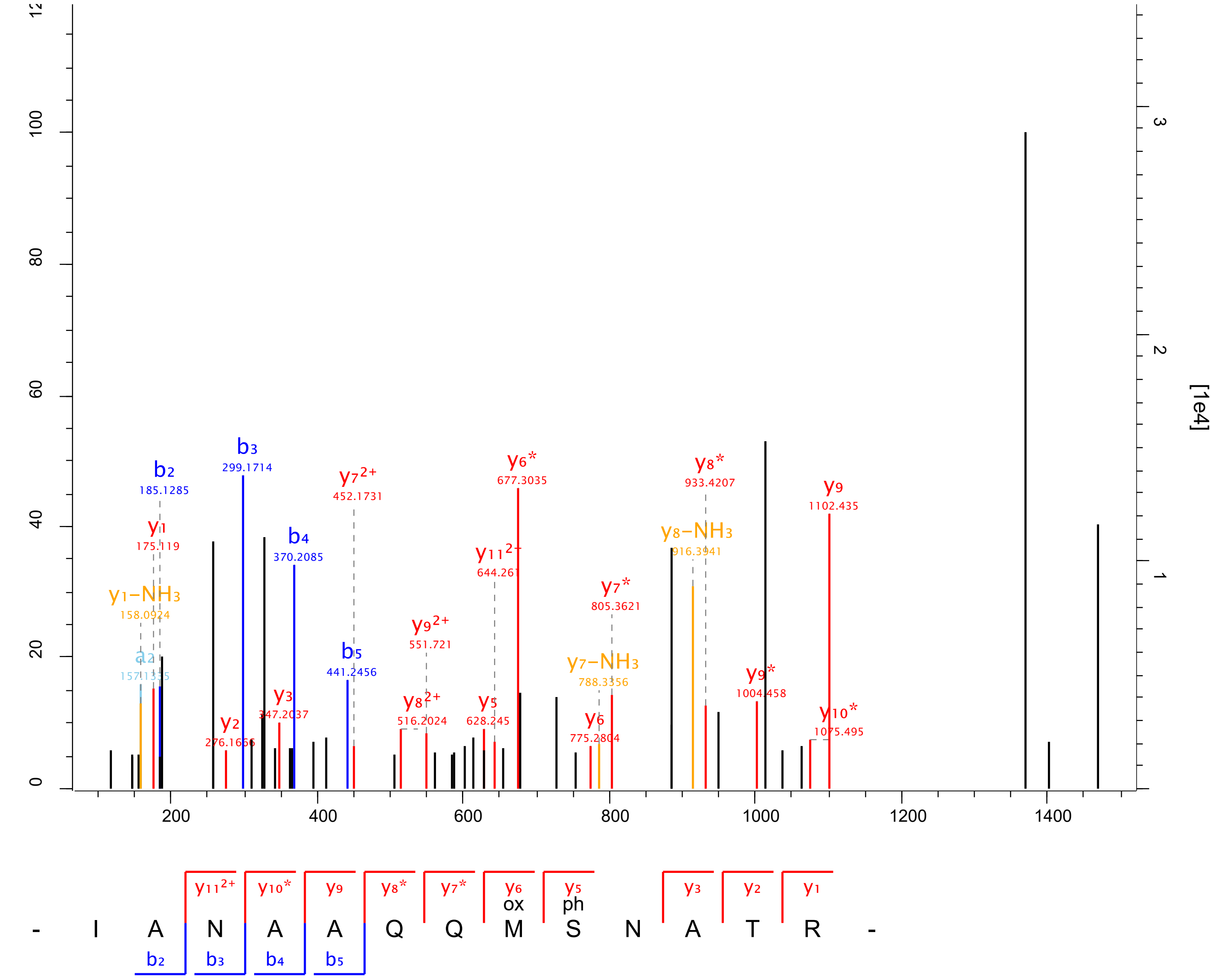

| Raw file | Scan | Method | Score | m/z |
|---|---|---|---|---|
| water_02_1 | 32064 | FTMS; HCD | 165.85 | 1023.42 |

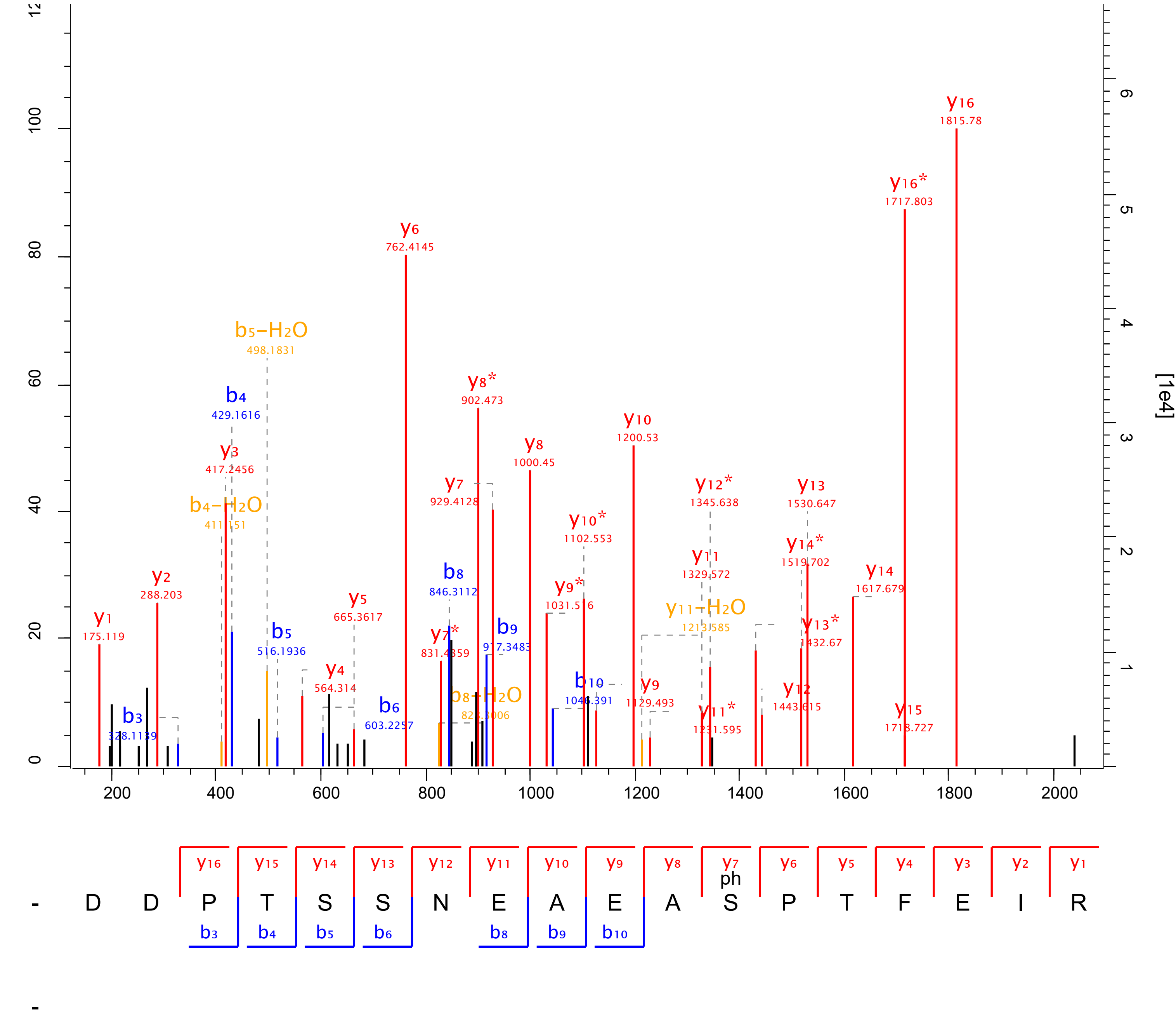

| Raw file | Scan | Method | Score | m/z |
| --- | --- | --- | --- | --- |
| water_02_1 | 32364 | FTMS; HCD | 40.53 | 976.09 |

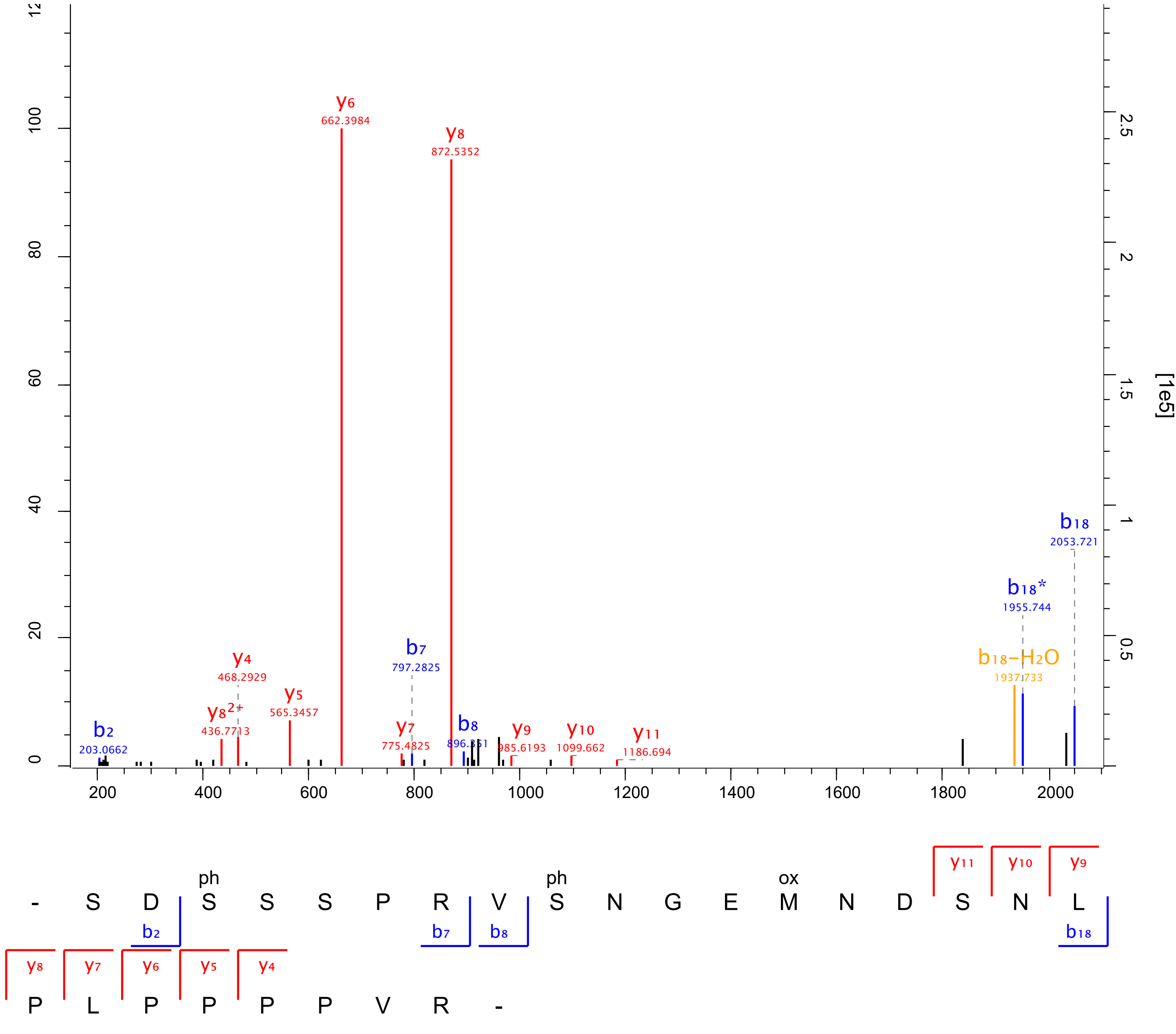

| Raw file | Scan | Method | Score | m/z |
| --- | --- | --- | --- | --- |
| water_02_1 | 32450 | FTMS; HCD | 44.16 | 875.03 |

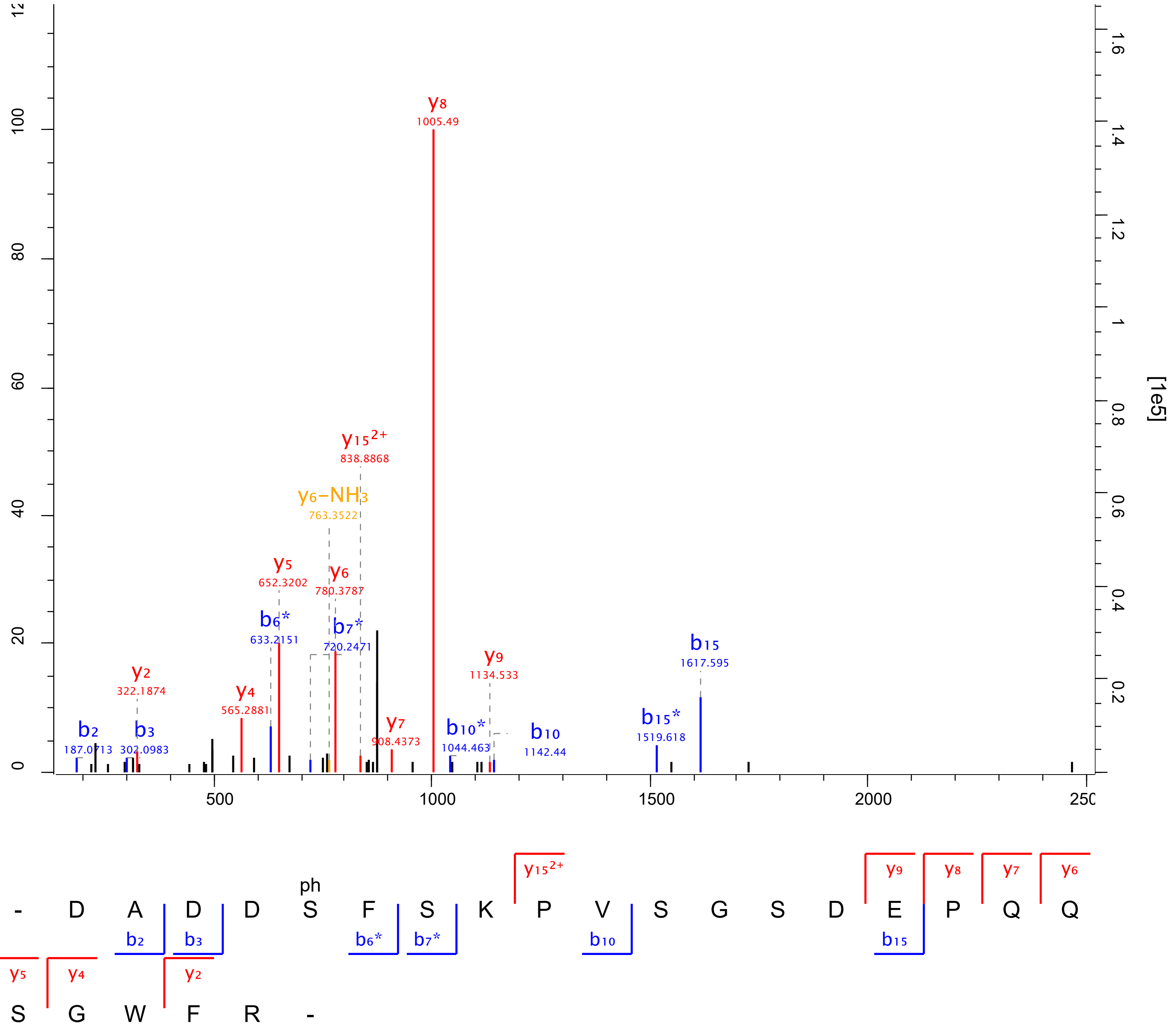

| Raw file | Scan | Method | Score | m/z |
| --- | --- | --- | --- | --- |
| water_02_1 | 32501 | FTMS; HCD | 279.16 | 774.35 |

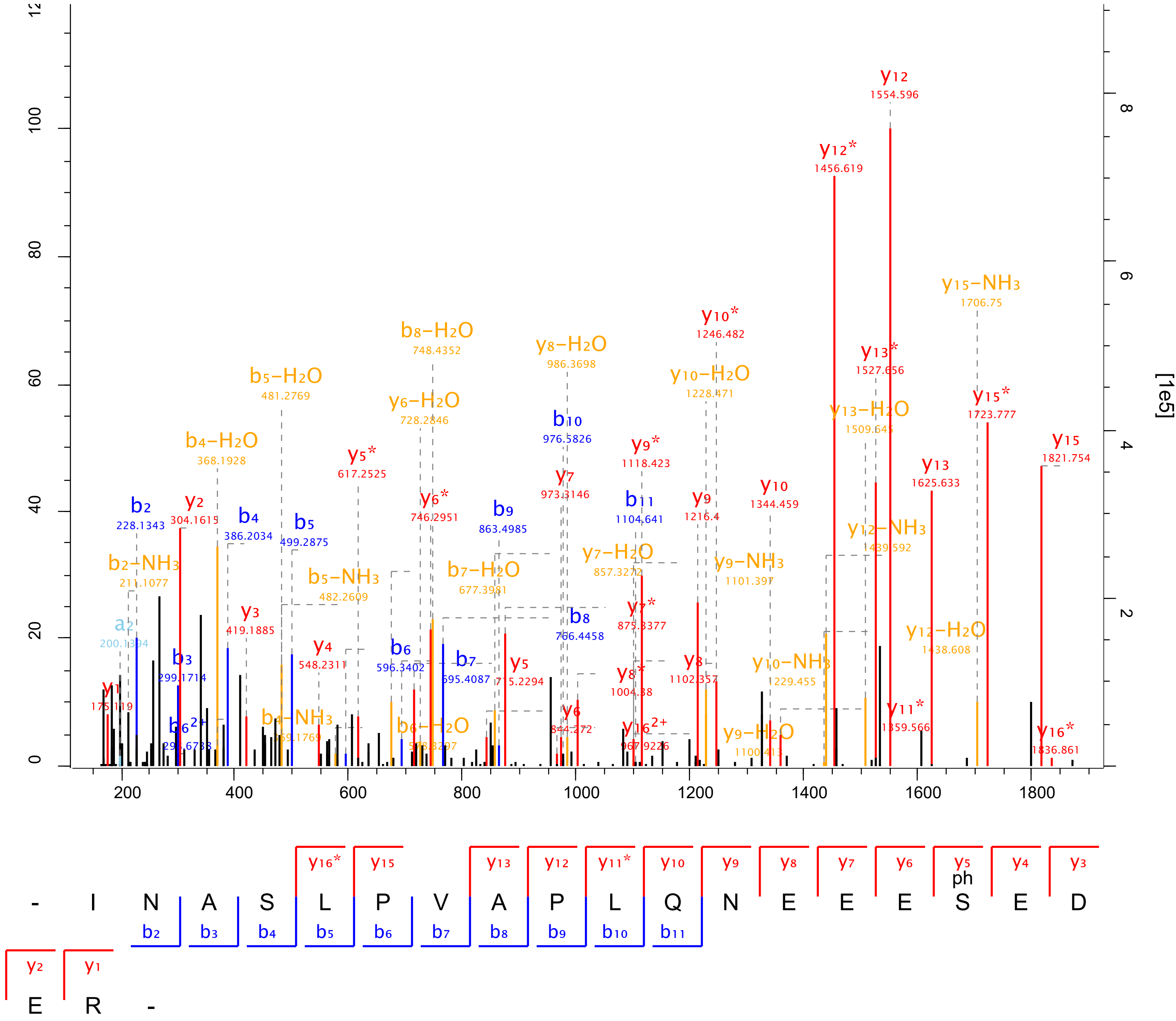

| Raw file | Scan | Method | Score | m/z |
|---|---|---|---|---|
| water_02_1 | 32760 | FTMS; HCD | 65.8 | 812.06 |

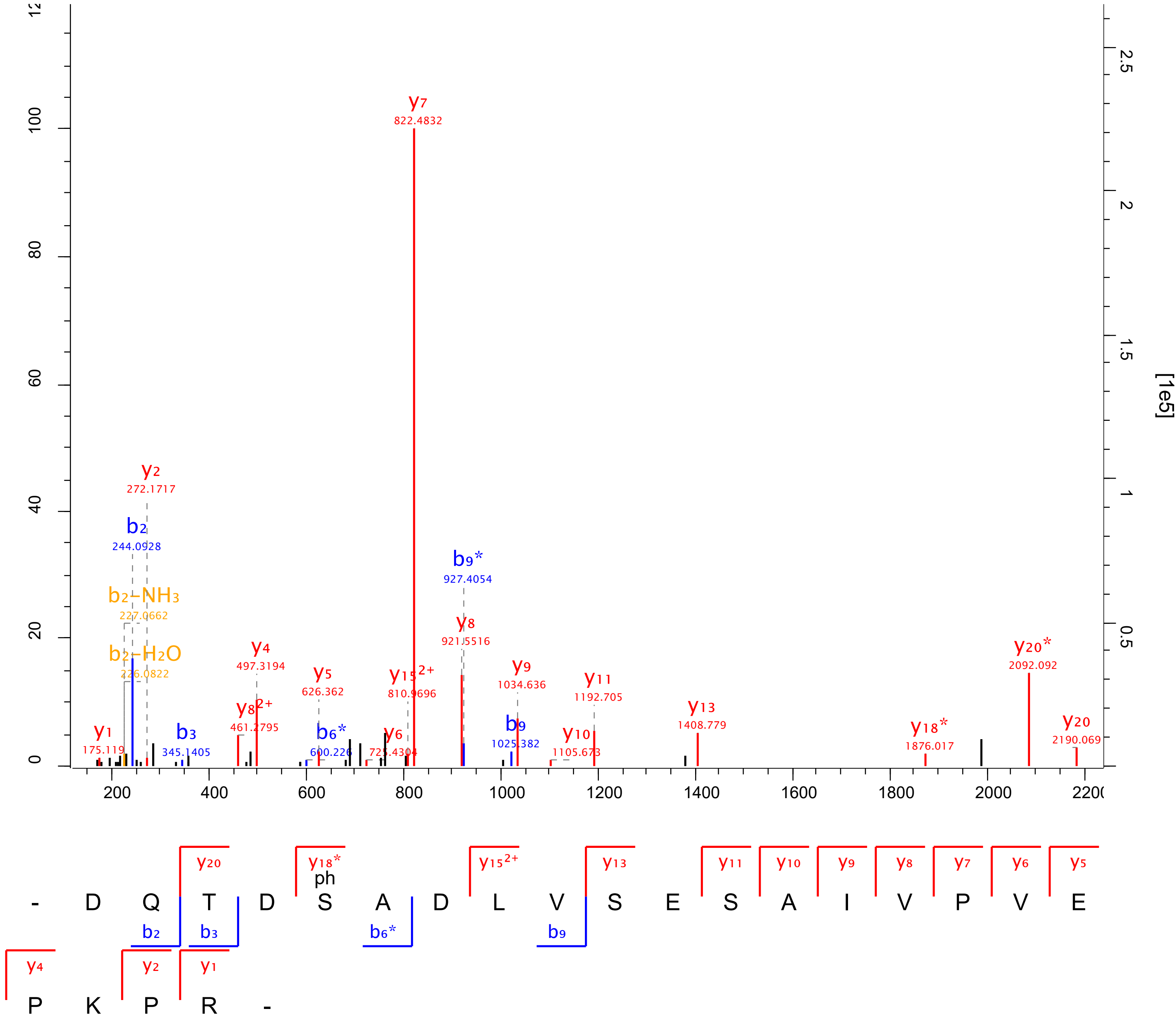

| Raw file | Scan | Method | Score | m/z |
| --- | --- | --- | --- | --- |
| water_02_1 | 32764 | FTMS; HCD | 106.99 | 838.01 |

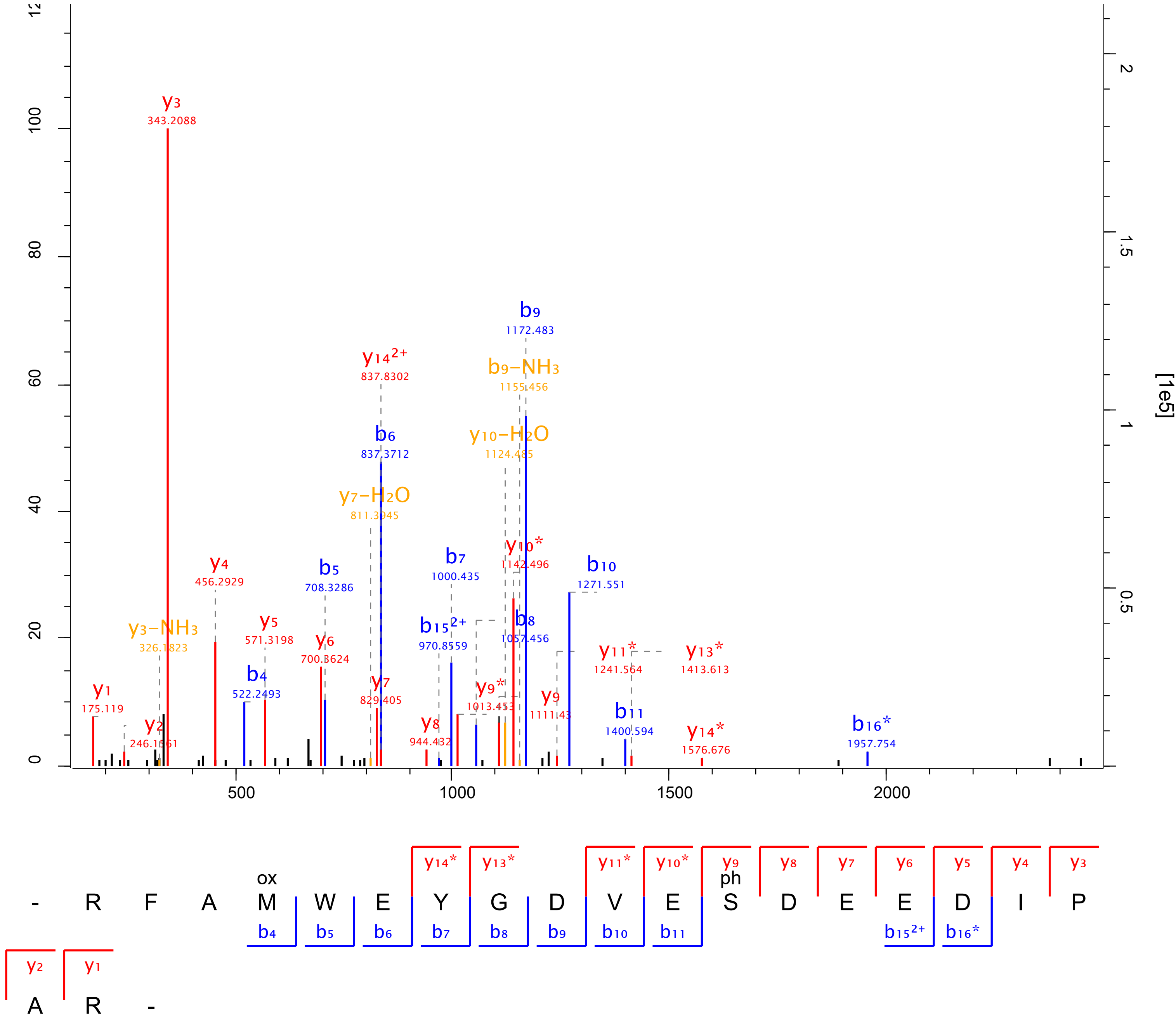

| Raw file | Scan | Method | Score | m/z |
| --- | --- | --- | --- | --- |
| water_02_1 | 32765 | FTMS; HCD | 94.28 | 865.36 |

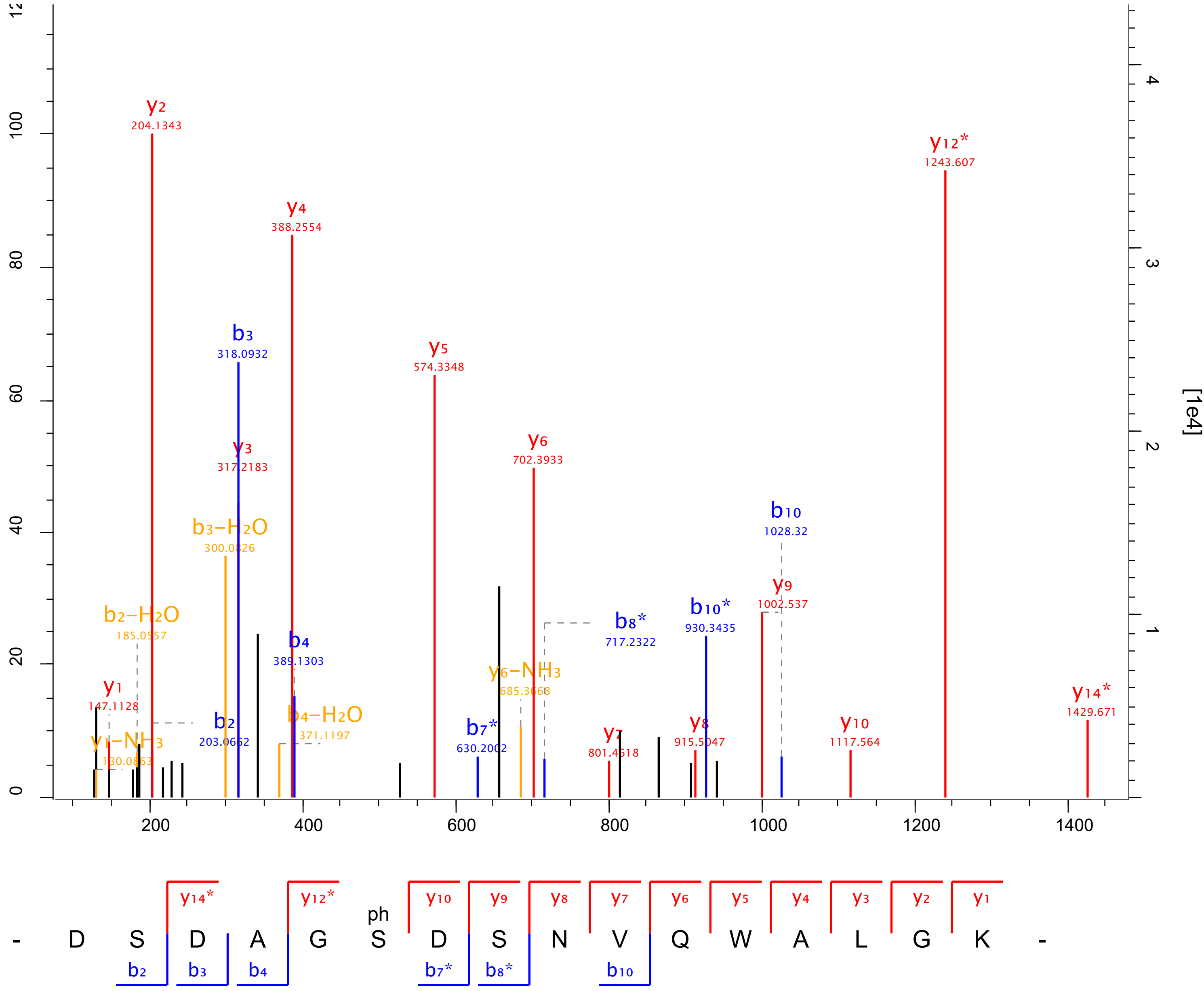

| Raw file | Scan | Method | Score | m/z |
| --- | --- | --- | --- | --- |
| water_02_1 | 32870 | FTMS; HCD | 189.02 | 779.37 |

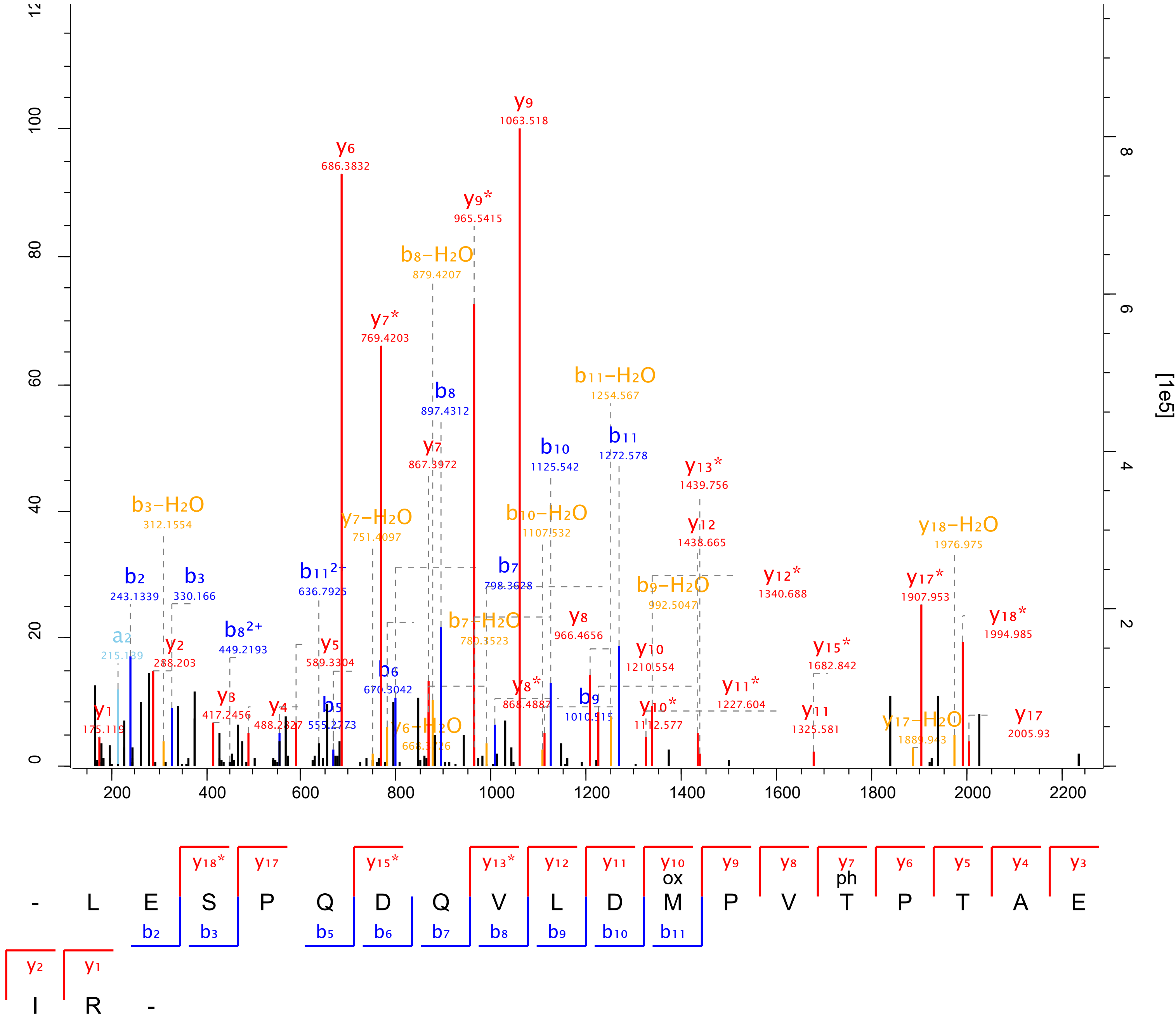

| Raw file | Scan | Method | Score | m/z |
| --- | --- | --- | --- | --- |
| water_02_1 | 3296 | FTMS; HCD | 69.38 | 431.87 |

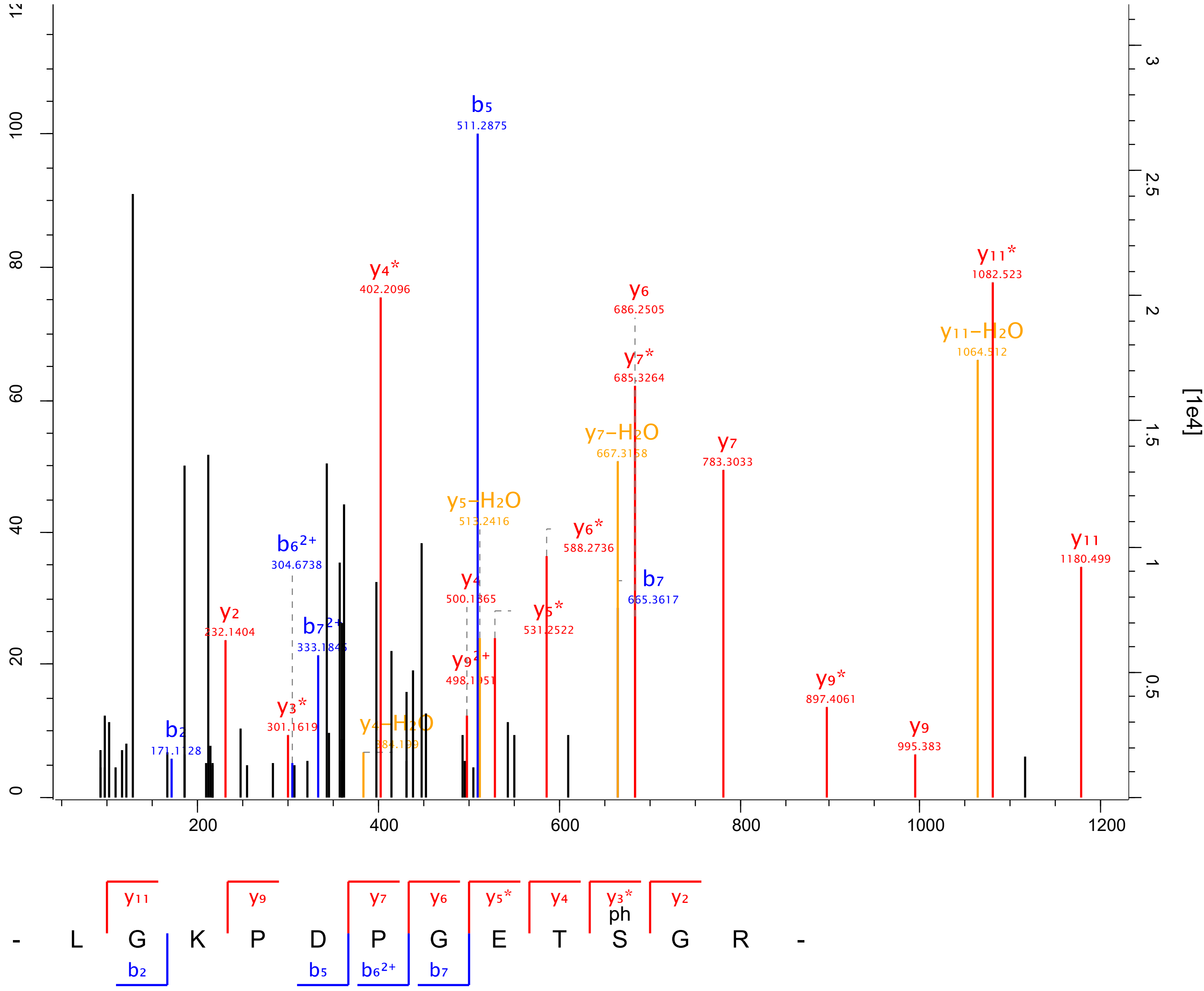

| Raw file | Scan | Method | Score | m/z |
| --- | --- | --- | --- | --- |
| water_02_1 | 32964 | FTMS; HCD | 197.28 | 696.7 |

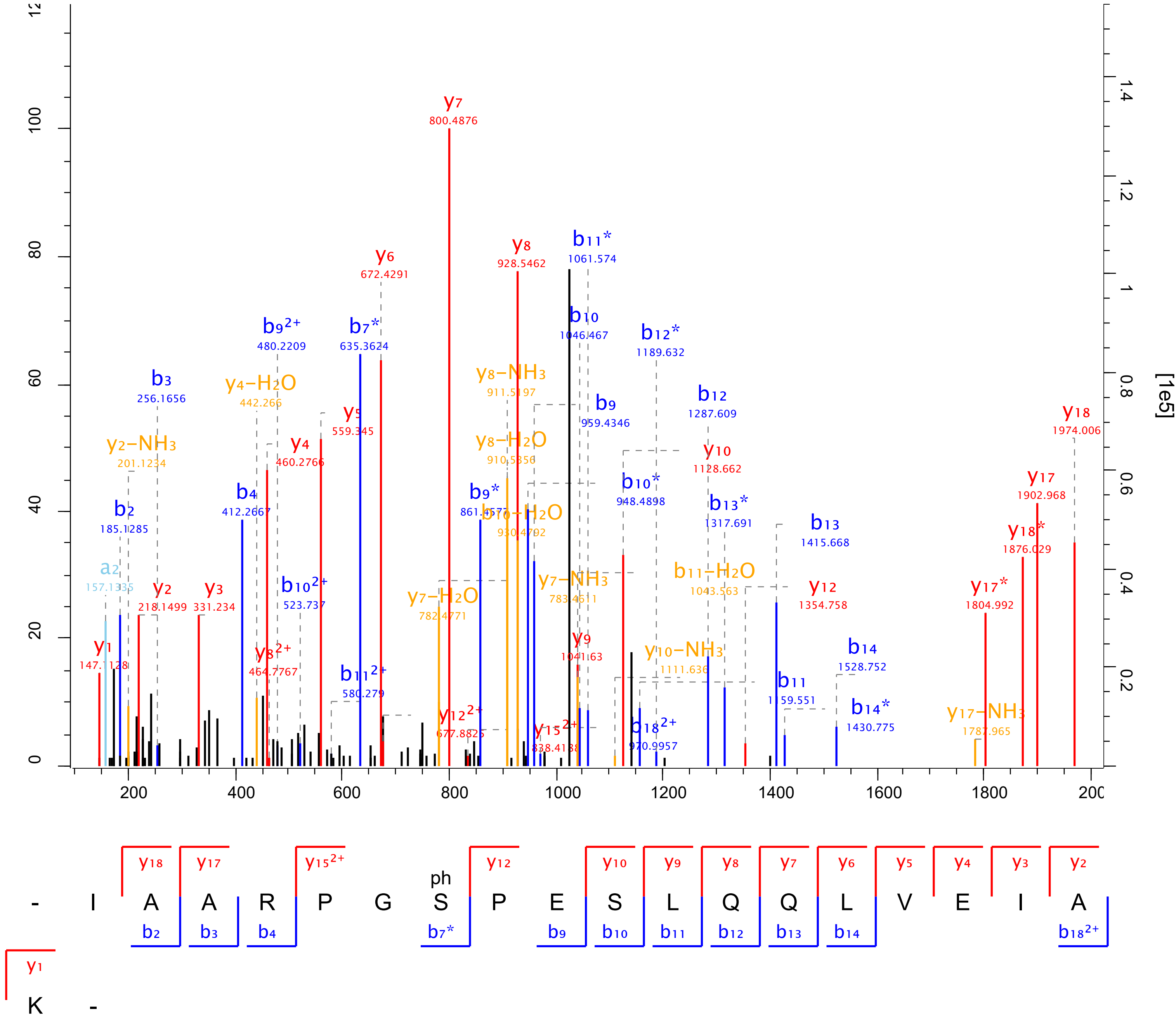

| Raw file | Scan | Method | Score | m/z |
|---|---|---|---|---|
| water_02_1 | 33129 | FTMS; HCD | 45.89 | 716.68 |

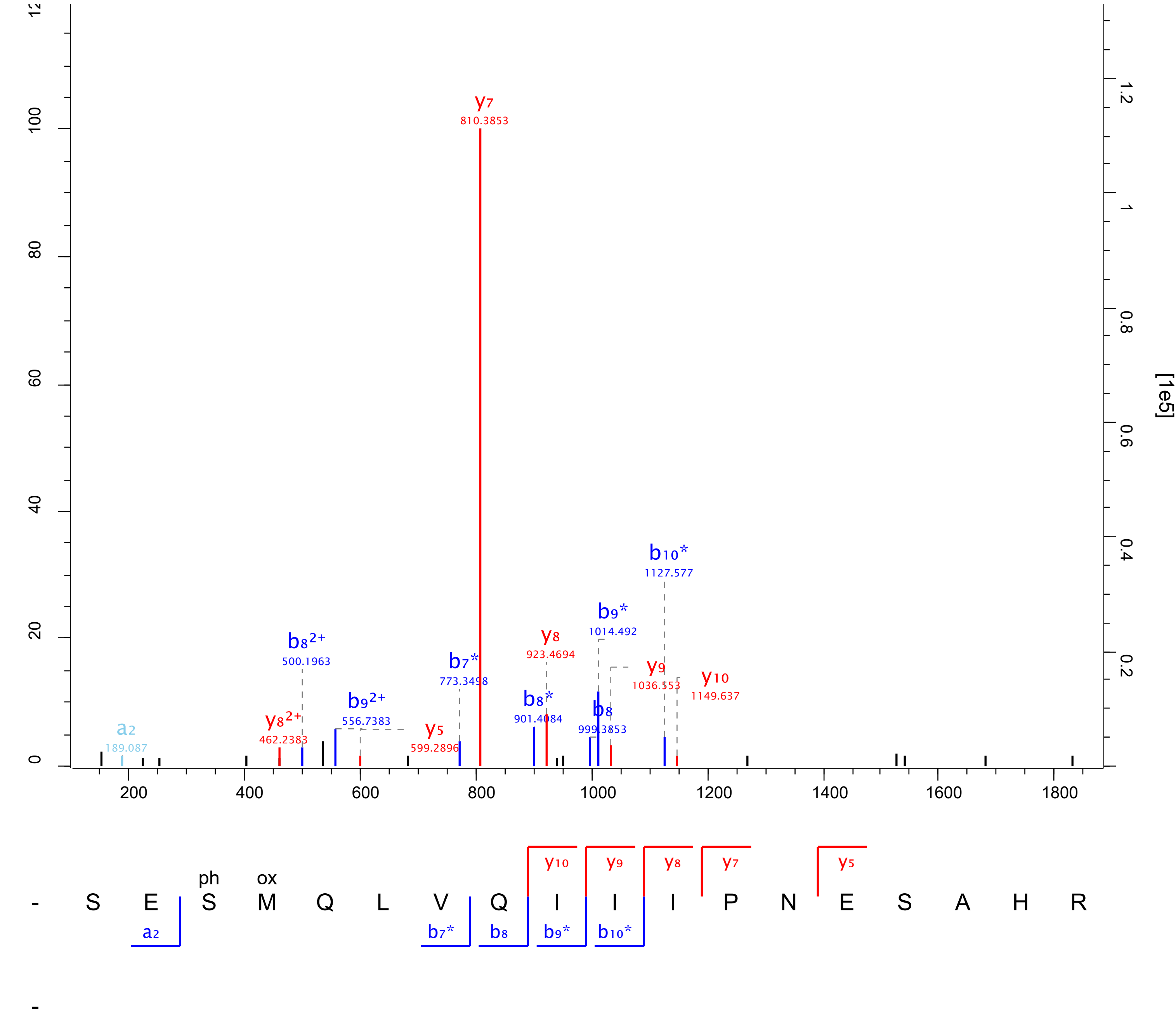

| Raw file | Scan | Method | Score | m/z |
|---|---|---|---|---|
| water_02_1 | 33142 | FTMS; HCD | 76.73 | 1088.83 |

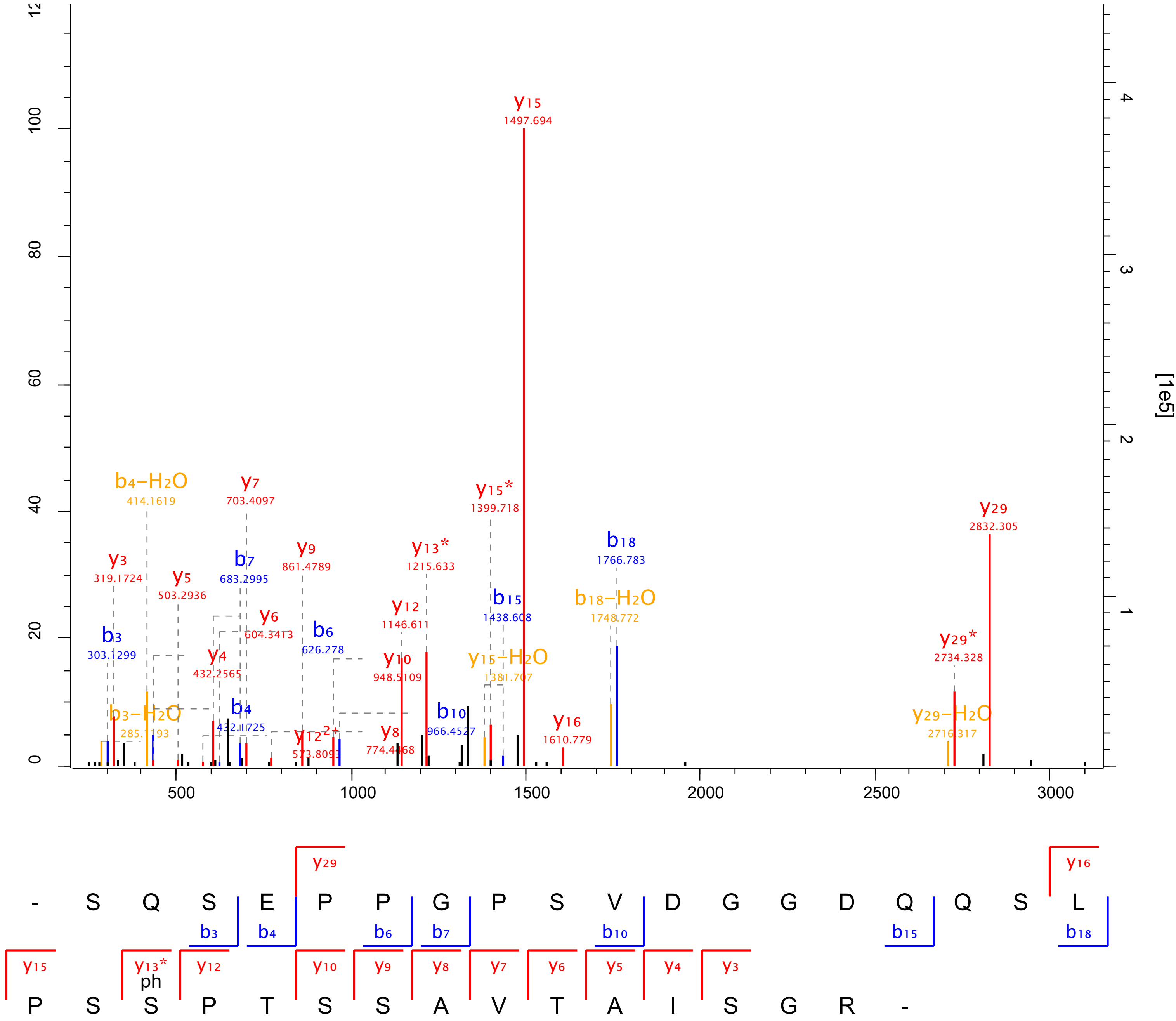

| Raw file | Scan | Method | Score | m/z |
|---|---|---|---|---|
| water_02_1 | 33282 | FTMS; HCD | 97.2 | 721.33 |

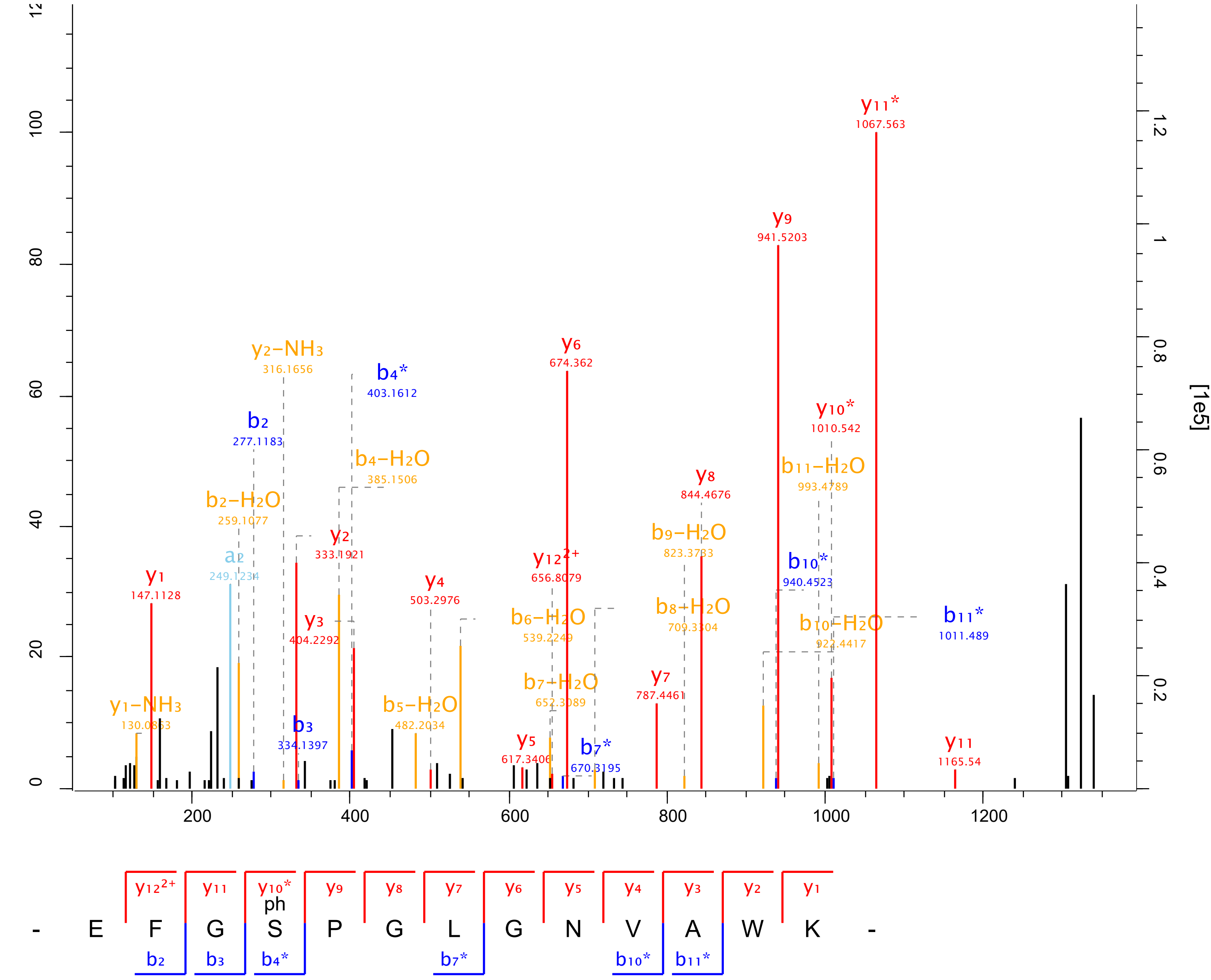

| Raw file | Scan | Method | Score | m/z |
| --- | --- | --- | --- | --- |
| water_02_1 | 33479 | FTMS; HCD | 71.08 | 804.36 |

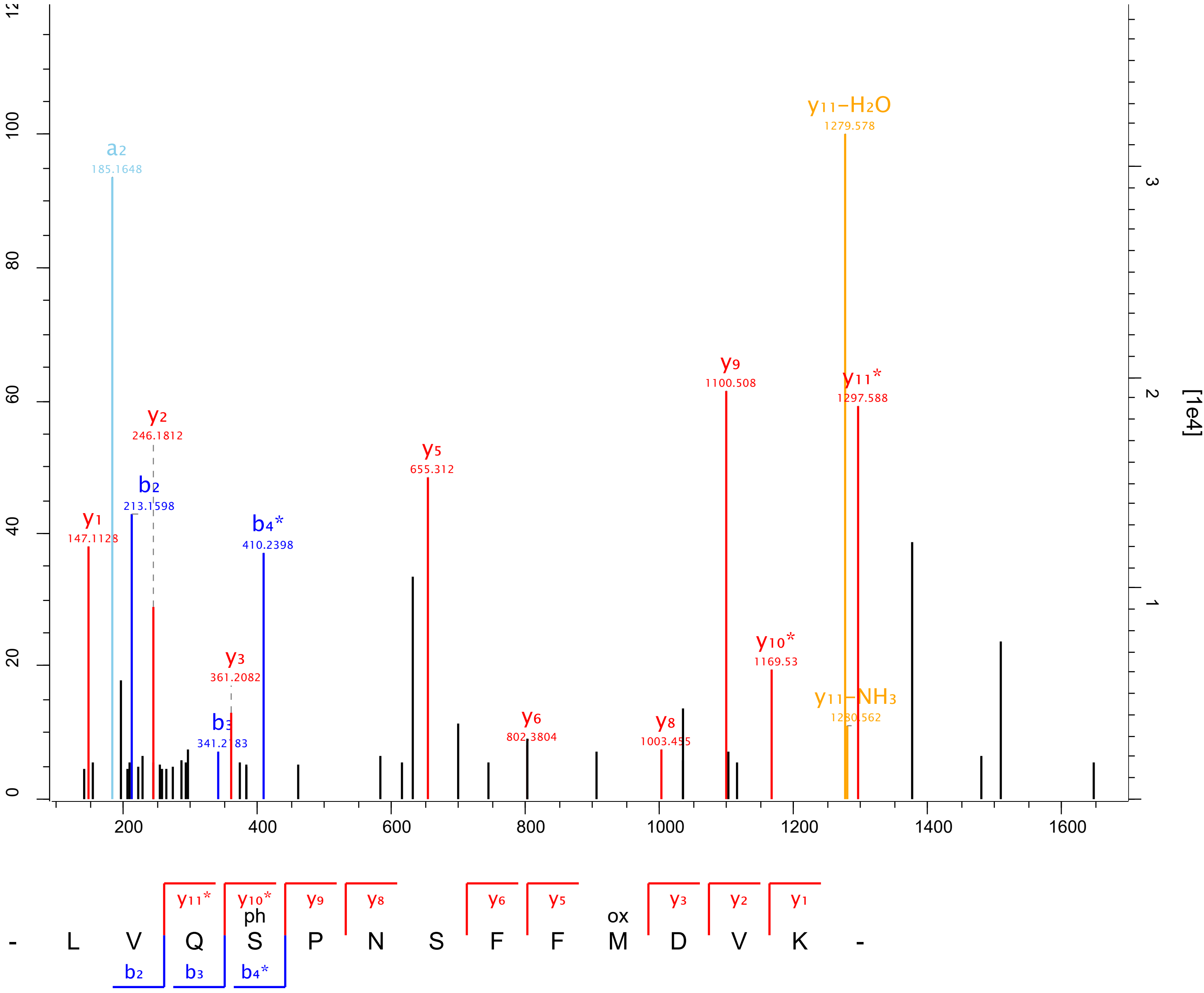

| Raw file | Scan | Method | Score | m/z |
| --- | --- | --- | --- | --- |
| water_02_1 | 33576 | FTMS; HCD | 75.44 | 890.35 |

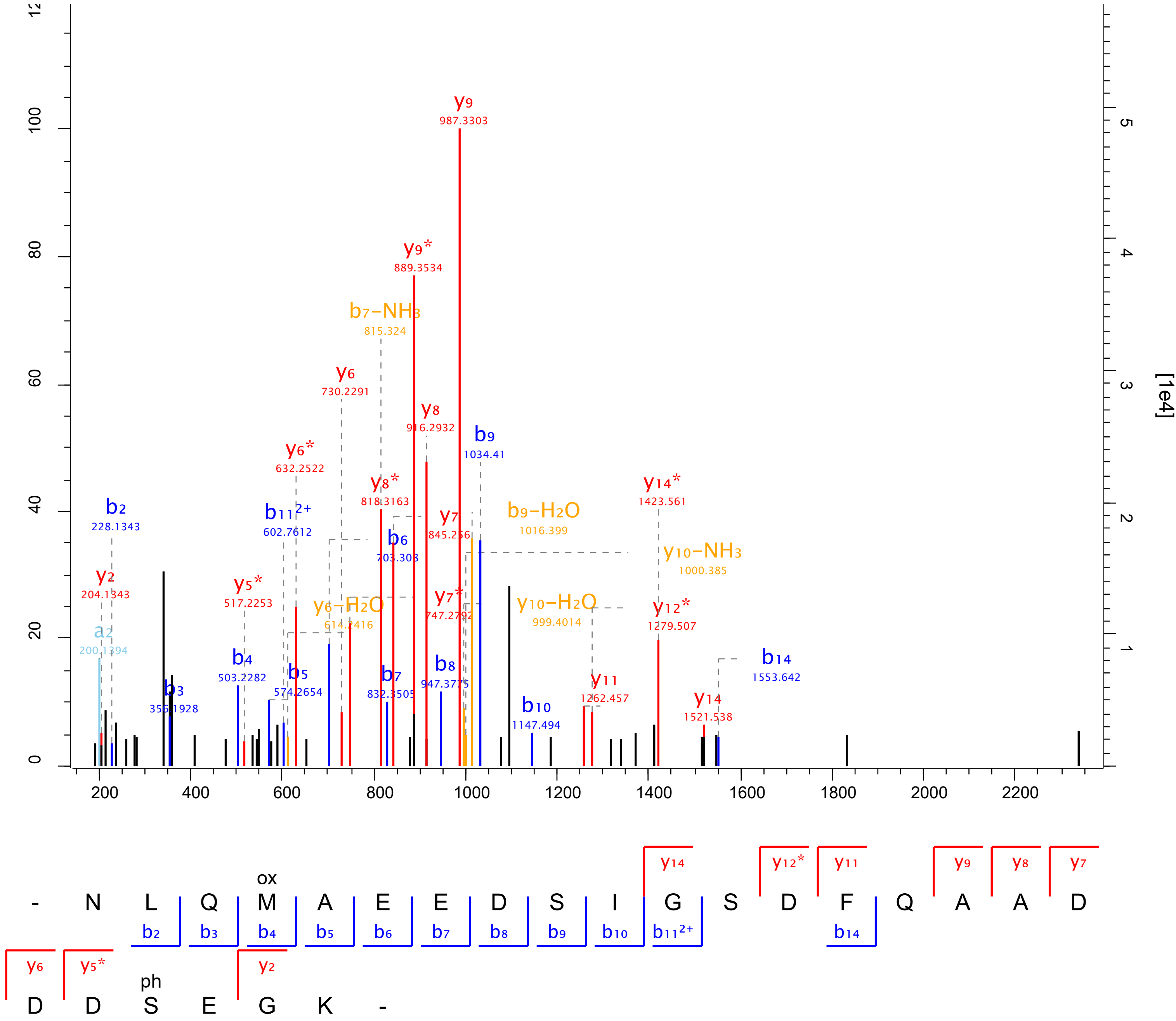

| Raw file | Scan | Method | Score | m/z |
|---|---|---|---|---|
| water_02_1 | 33689 | FTMS; HCD | 108.88 | 881.69 |

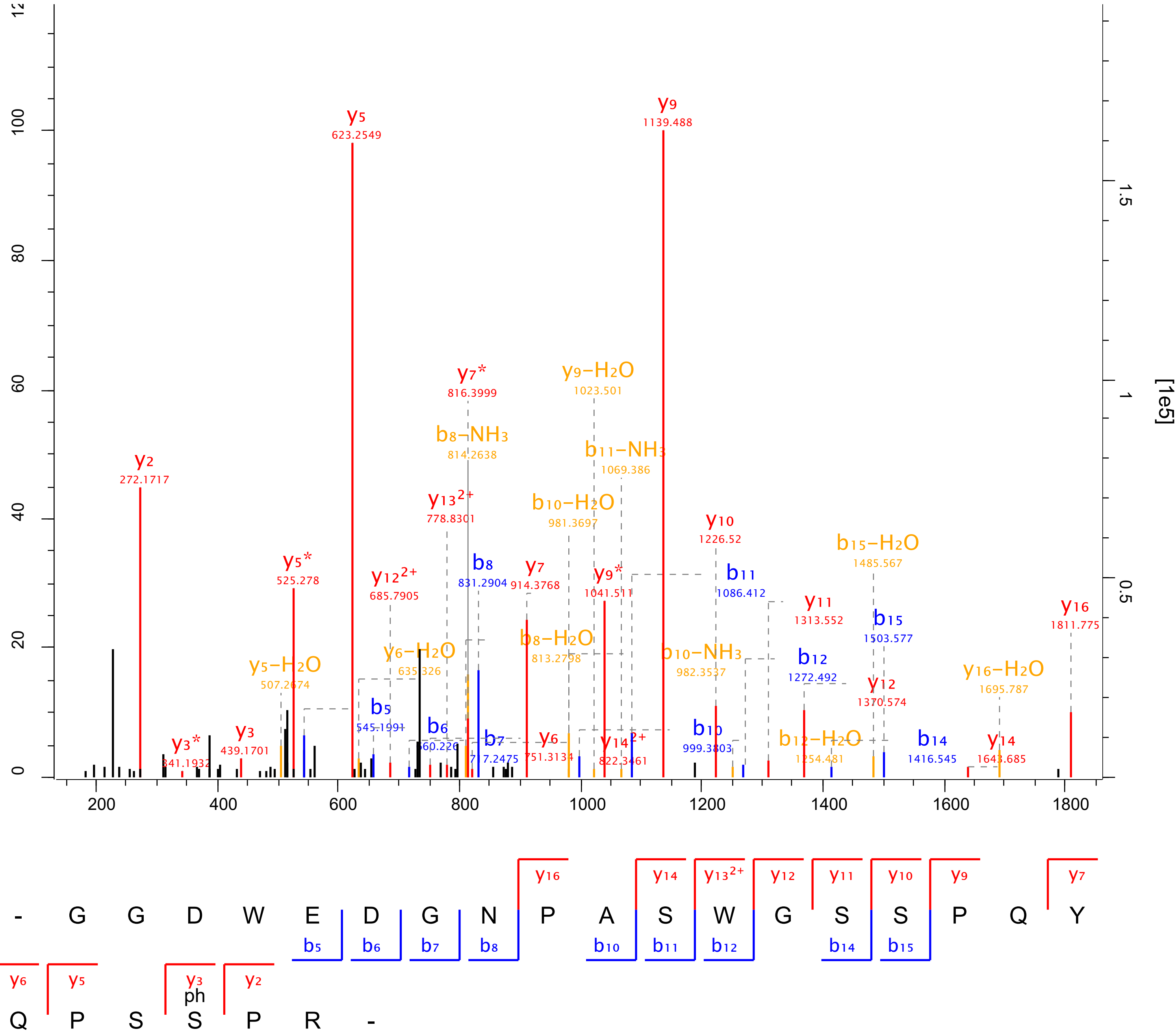

| Raw file | Scan | Method | Score | m/z |
| --- | --- | --- | --- | --- |
| water_02_1 | 33762 | FTMS; HCD | 146.6 | 1104.43 |

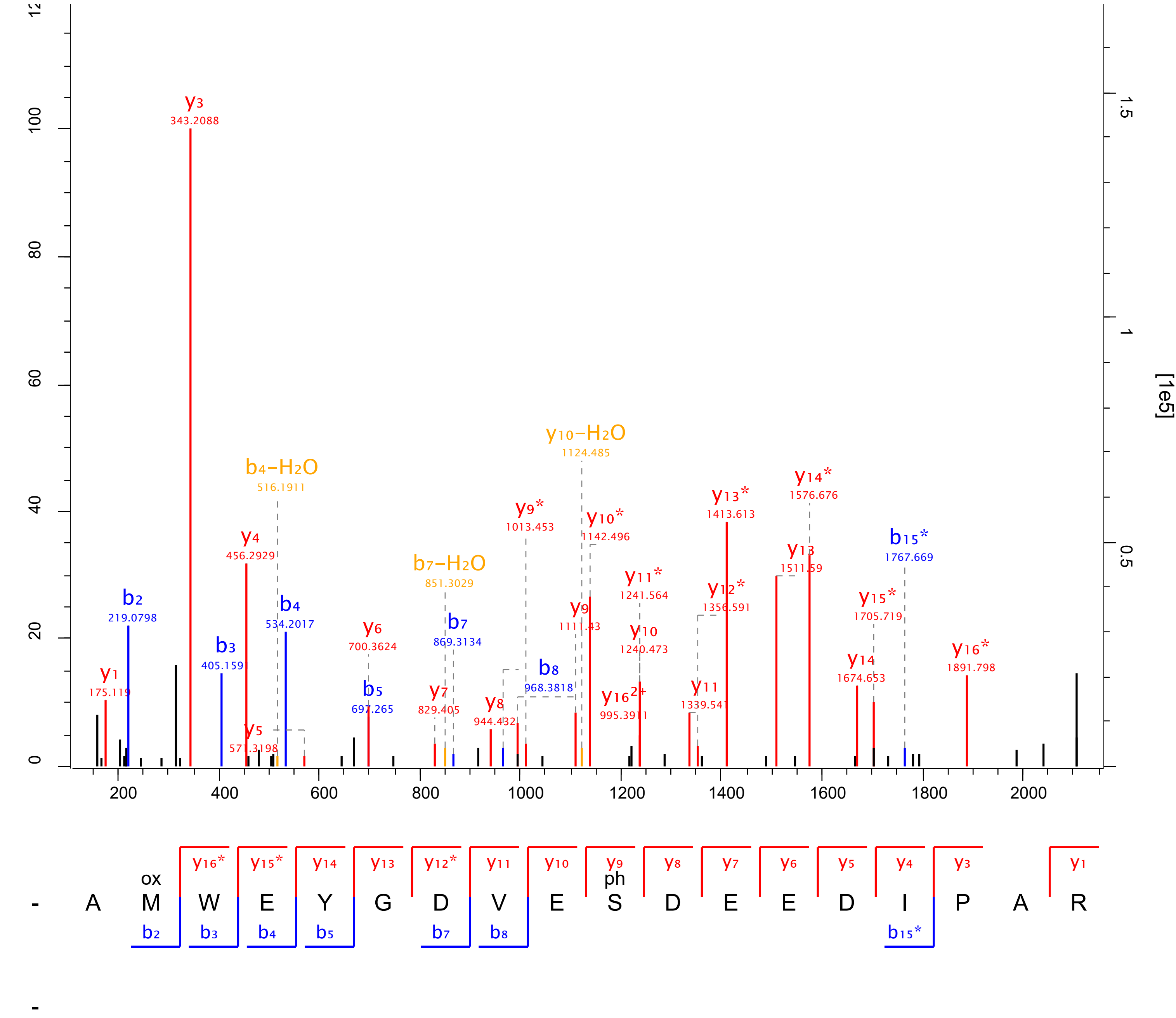

| Raw file | Scan | Method | Score | m/z |
|---|---|---|---|---|
| water_02_1 | 33892 | FTMS; HCD | 174.86 | 516.71 |

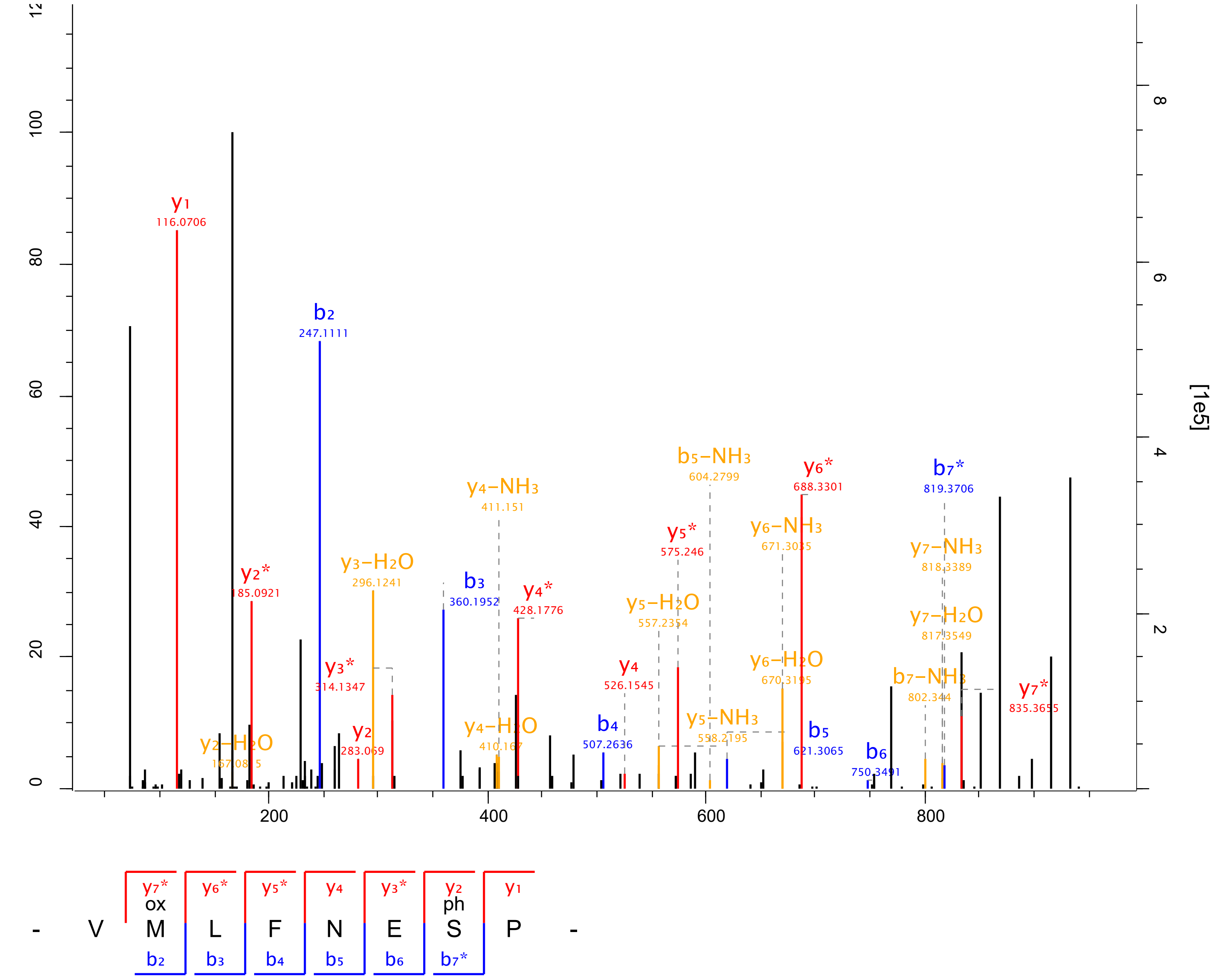

| Raw file | Scan | Method | Score | m/z |
| --- | --- | --- | --- | --- |
| water_02_1 | 34045 | FTMS; HCD | 49.66 | 740.96 |

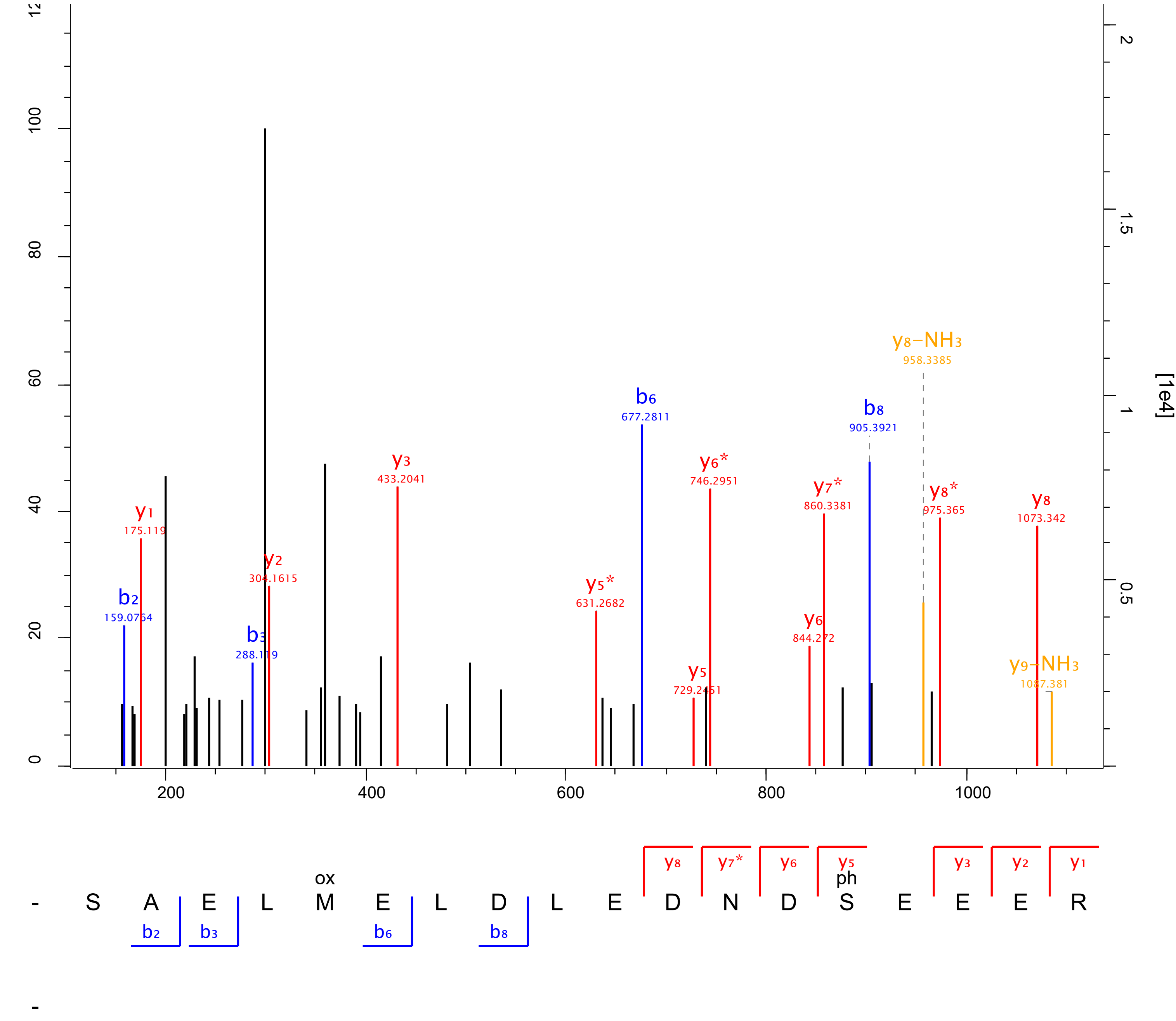

| Raw file | Scan | Method | Score | m/z |
| --- | --- | --- | --- | --- |
| water_02_1 | 34114 | FTMS; HCD | 99.04 | 787.35 |

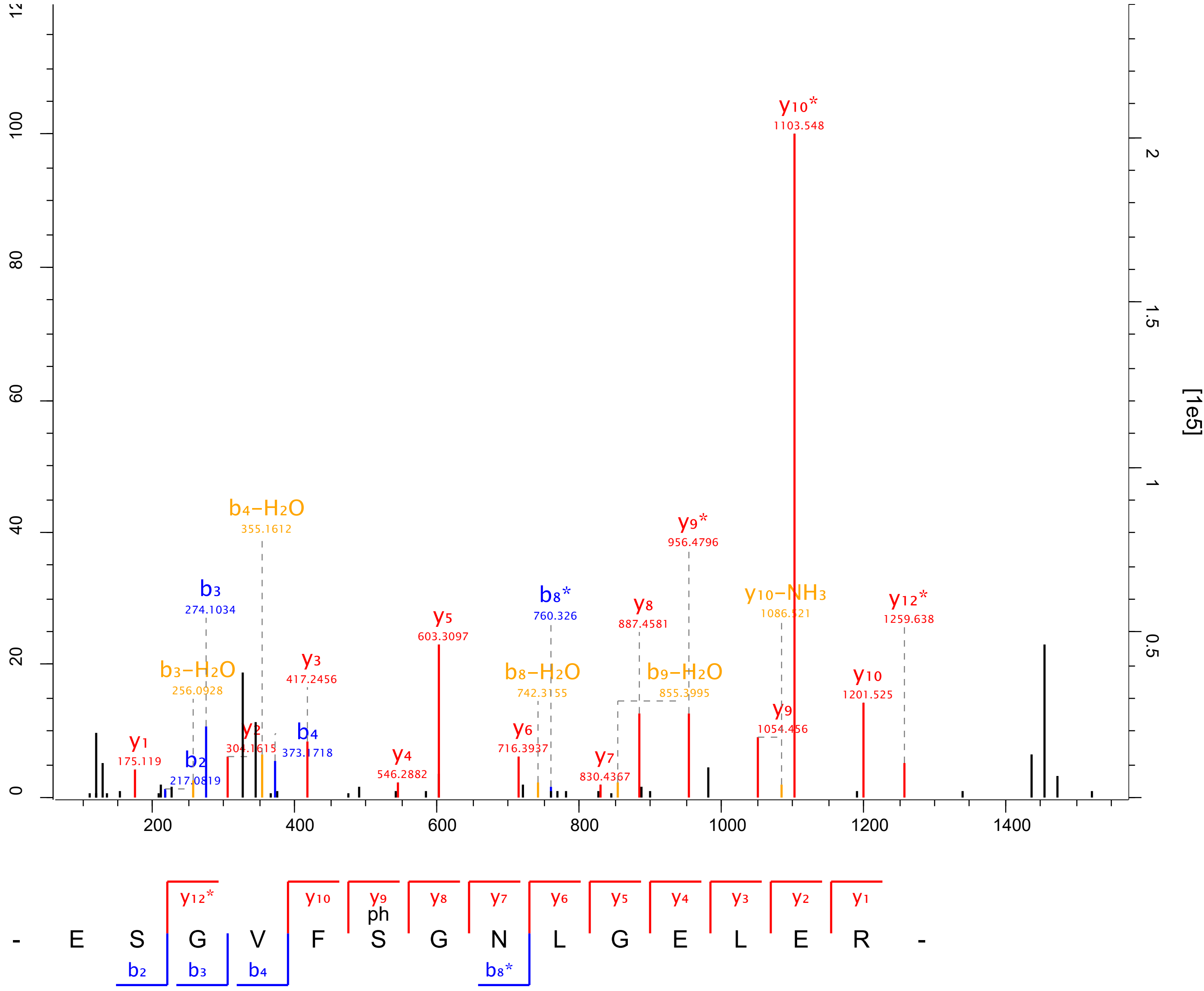

| Raw file | Scan | Method | Score | m/z |
|---|---|---|---|---|
| water_02_1 | 3414 | FTMS; HCD | 235.26 | 556.89 |

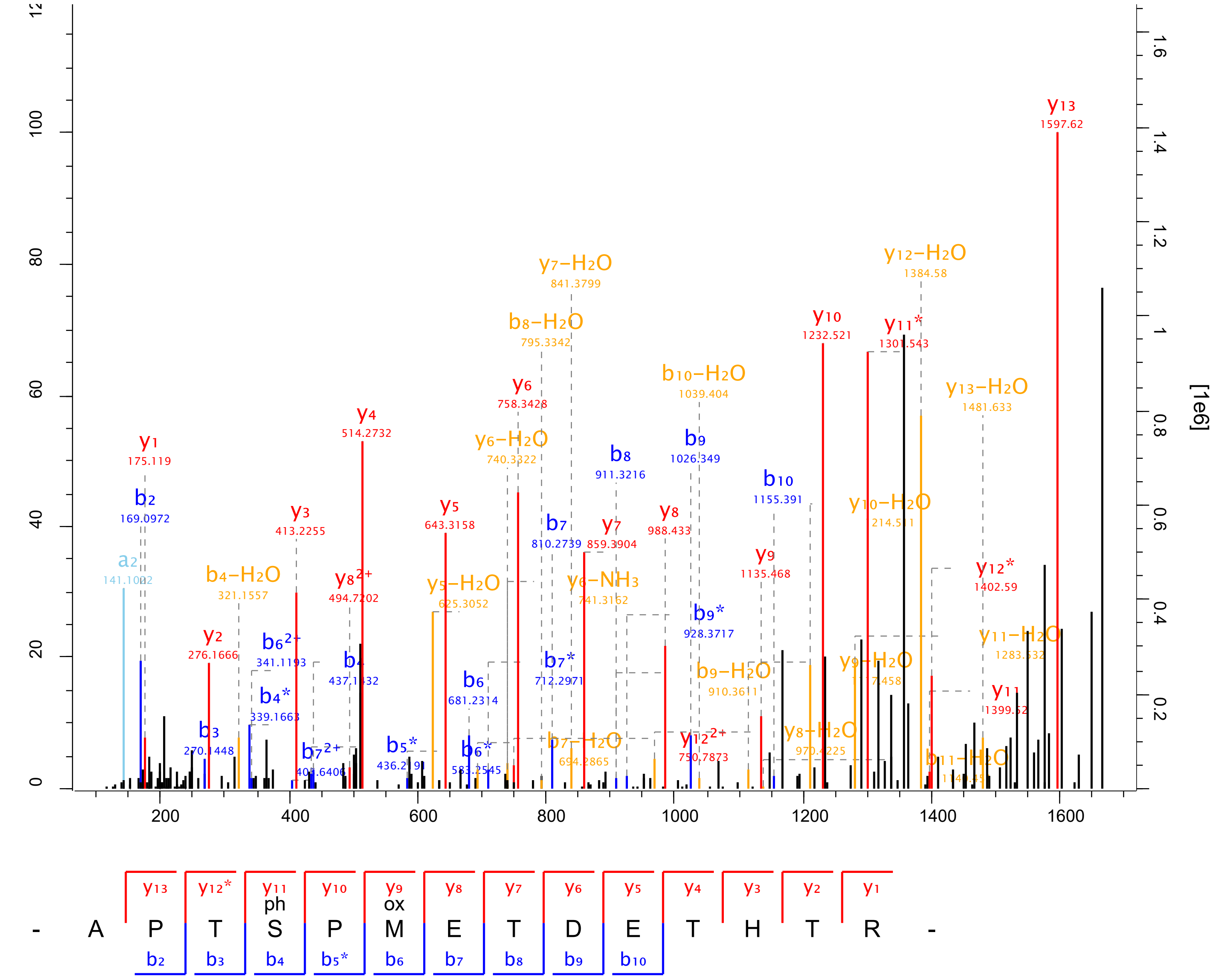

| Raw file | Scan | Method | Score | m/z |
| --- | --- | --- | --- | --- |
| water_02_1 | 3427 | FTMS; HCD | 98.9 | 554.25 |

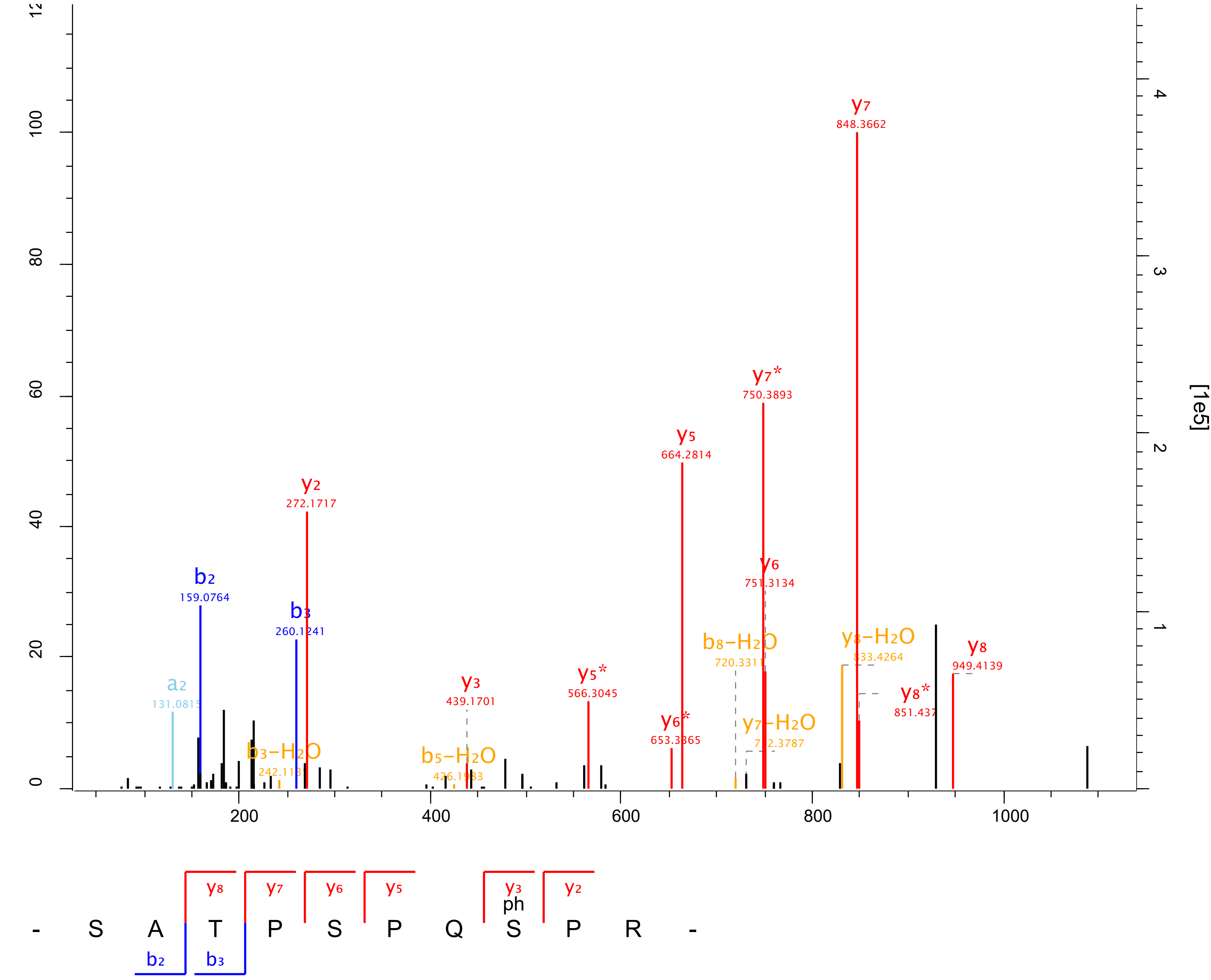

| Raw file | Scan | Method | Score | m/z |
|---|---|---|---|---|
| water_02_1 | 34408 | FTMS; HCD | 169.39 | 906.4 |

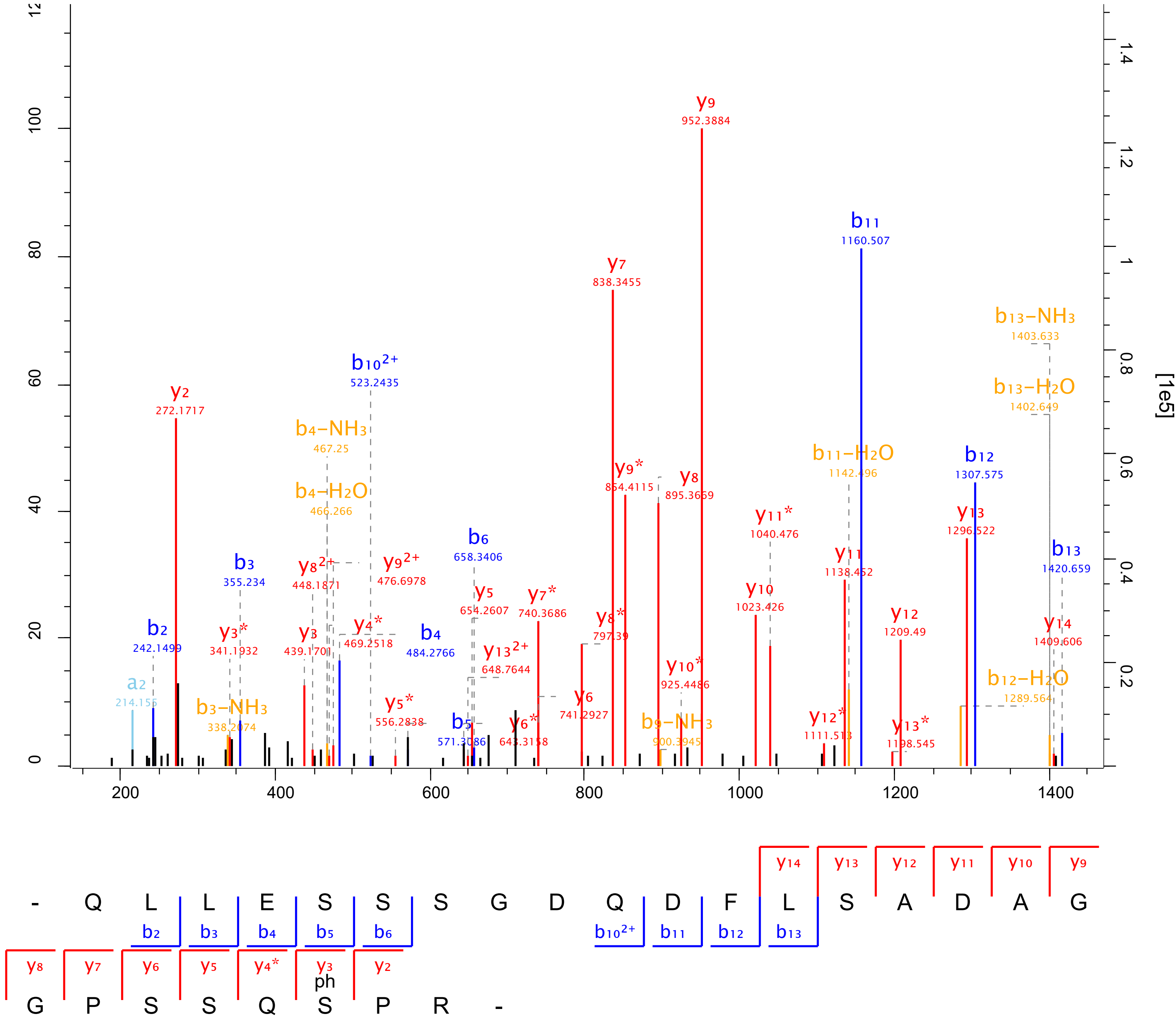

| Raw file | Scan | Method | Score | m/z |
|---|---|---|---|---|
| water_02_1 | 34451 | FTMS; HCD | 66.27 | 599.75 |

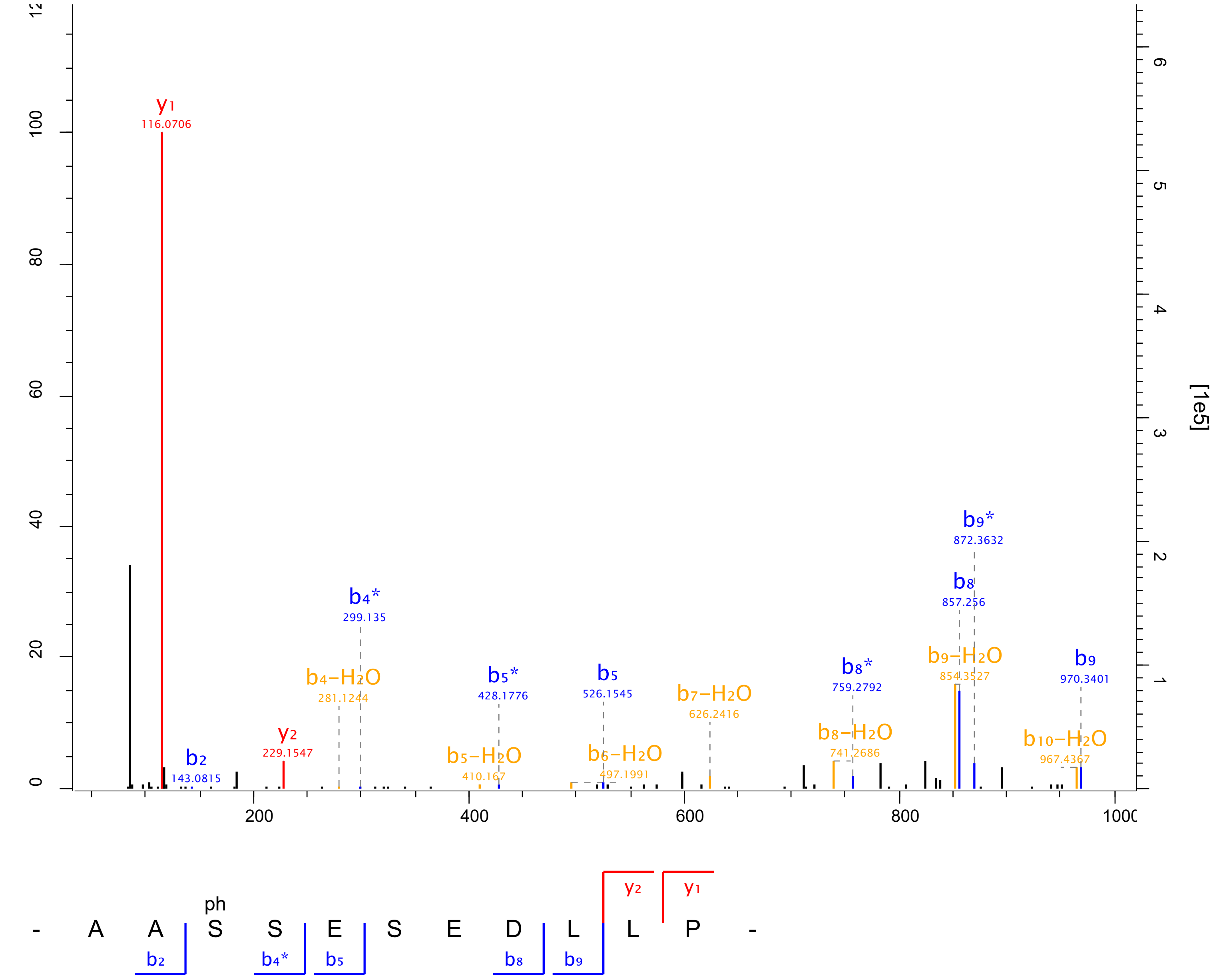

| Raw file | Scan | Method | Score | m/z |
| --- | --- | --- | --- | --- |
| water_02_1 | 3449 | FTMS; HCD | 71.85 | 851.83 |

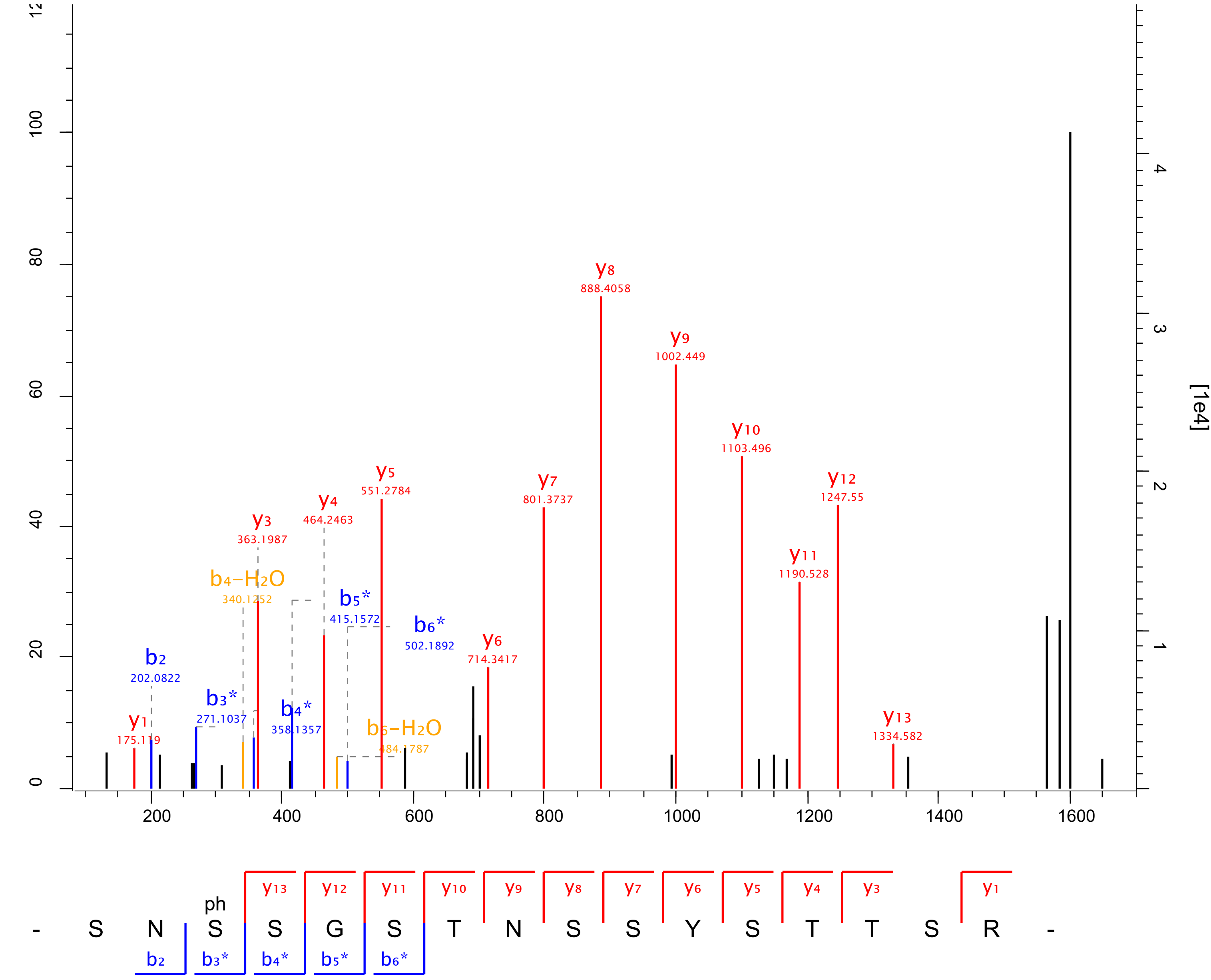

| Raw file | Scan | Method | Score | m/z |
|---|---|---|---|---|
| water_02_1 | 34558 | FTMS; HCD | 124.08 | 1136.47 |

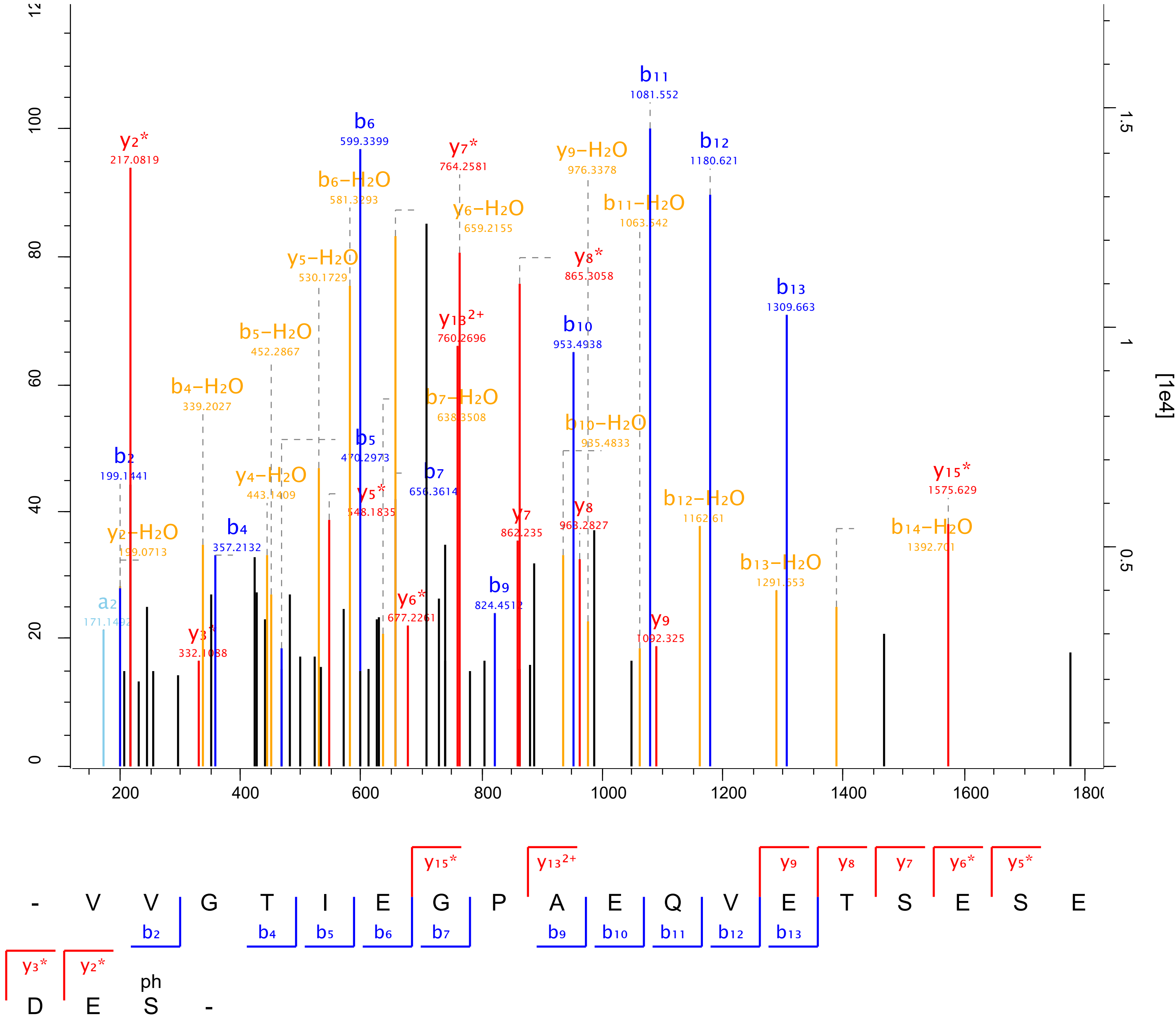

| Raw file | Scan | Method | Score | m/z |
|---|---|---|---|---|
| water_02_1 | 34602 | FTMS; HCD | 141.07 | 819.88 |

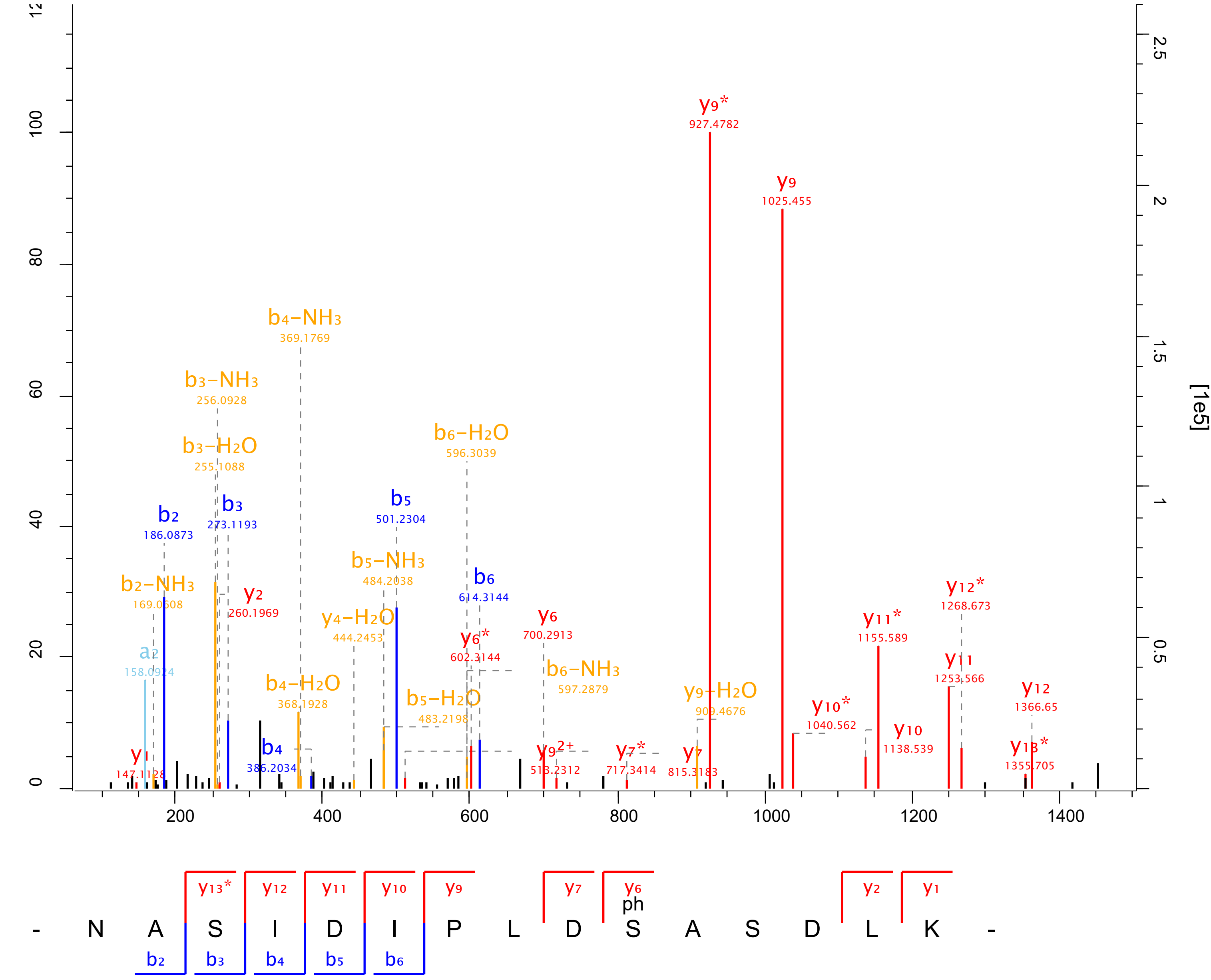

| Raw file | Scan | Method | Score | m/z |
|---|---|---|---|---|
| water_02_1 | 3463 | FTMS; HCD | 102.73 | 606.25 |

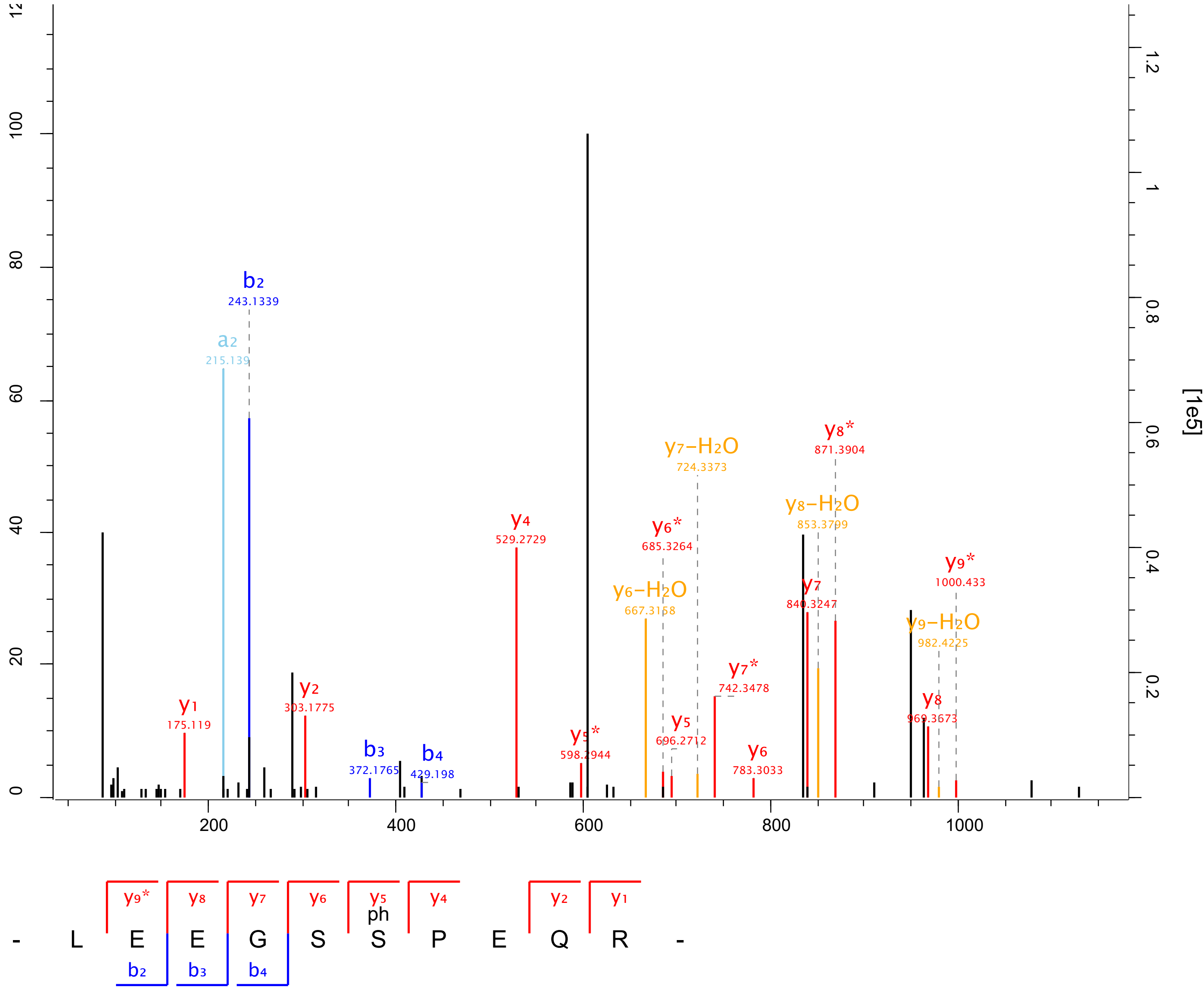

| Raw file | Scan | Method | Score | m/z |
| --- | --- | --- | --- | --- |
| water_02_1 | 34778 | FTMS; HCD | 72.55 | 1192.6 |

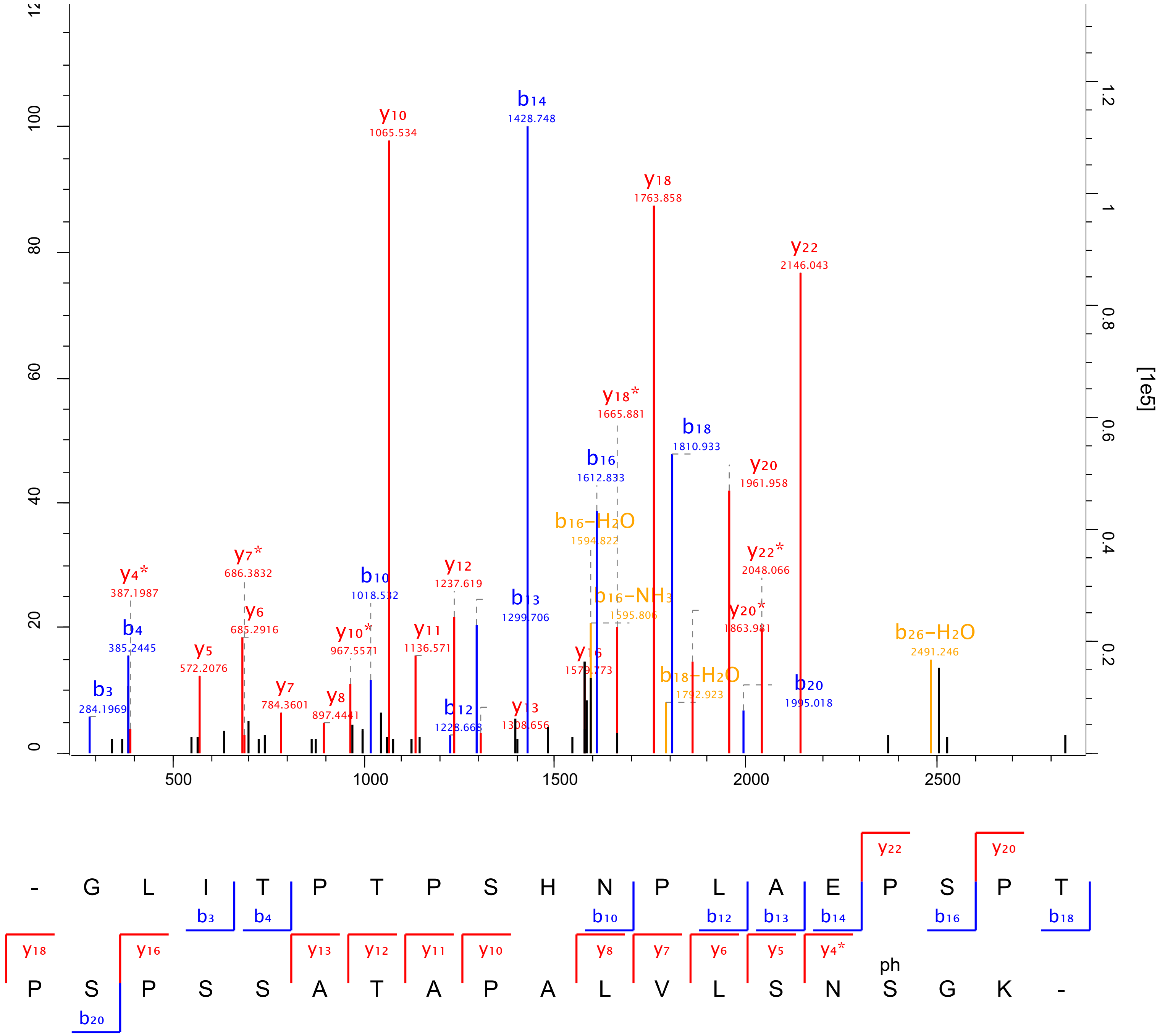

| Raw file | Scan | Method | Score | m/z |
|---|---|---|---|---|
| water_02_1 | 34873 | FTMS; HCD | 101.75 | 1108.43 |

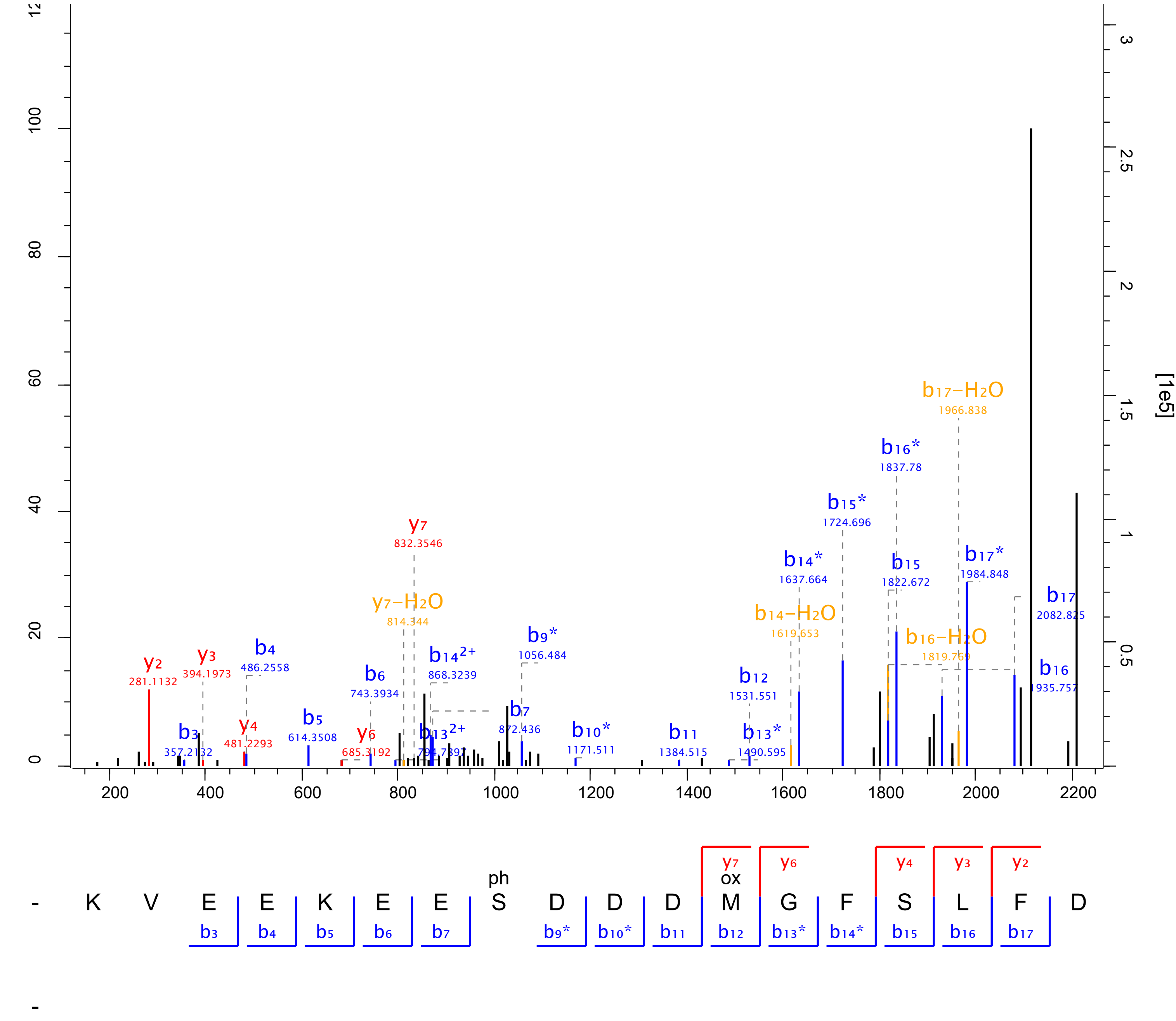

| Raw file | Scan | Method | Score | m/z |
|---|---|---|---|---|
| water_02_1 | 3500 | FTMS; HCD | 96.67 | 442.7 |

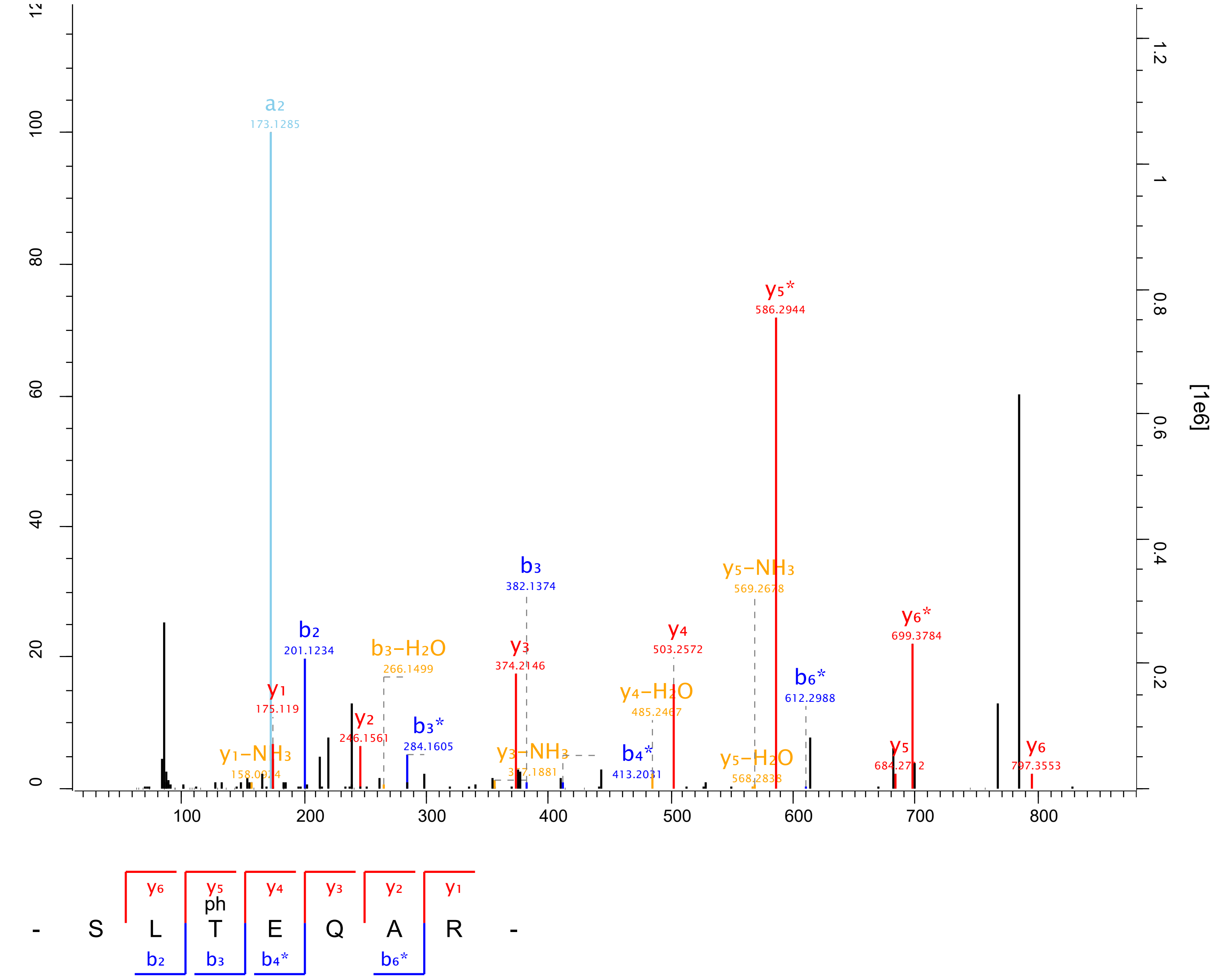

| Raw file | Scan | Method | Score | m/z |
|---|---|---|---|---|
| water_02_1 | 35049 | FTMS; HCD | 95.32 | 789.38 |

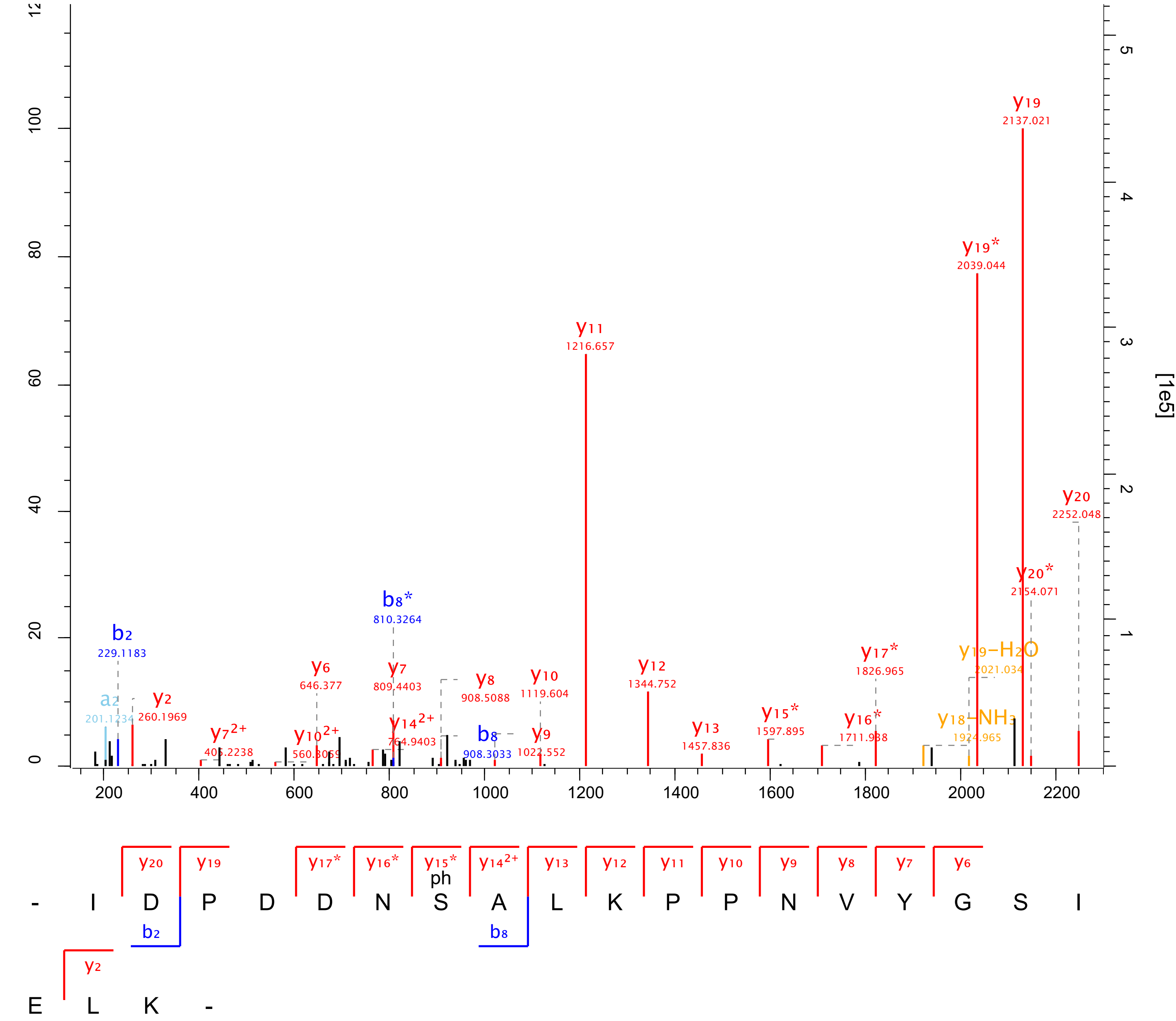

| Raw file | Scan | Method | Score | m/z |
| --- | --- | --- | --- | --- |
| water_02_1 | 35125 | FTMS; HCD | 166.03 | 650.79 |

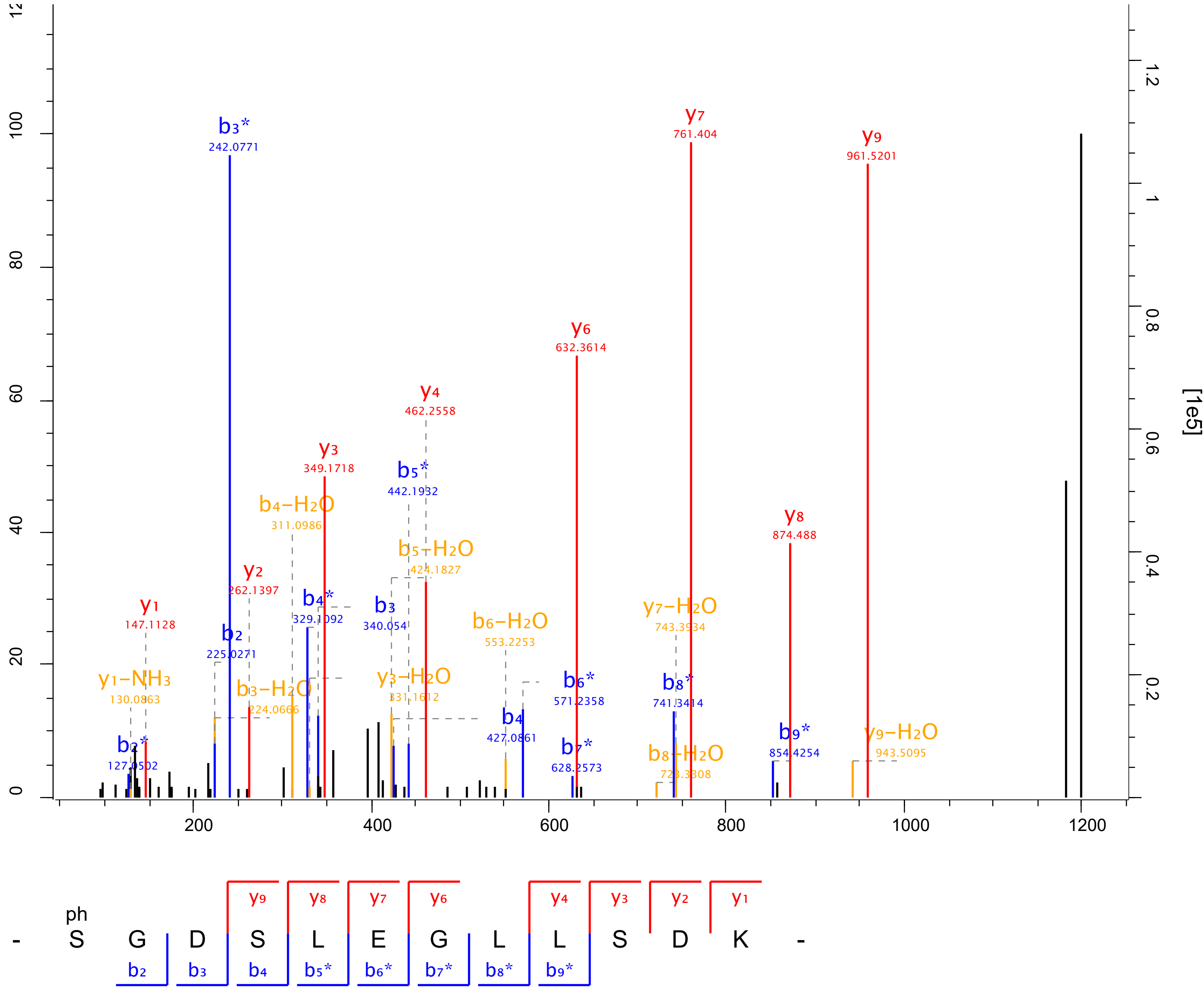

| Raw file | Scan | Method | Score | m/z |
|---|---|---|---|---|
| water_02_1 | 35135 | FTMS; HCD | 117.77 | 987.75 |

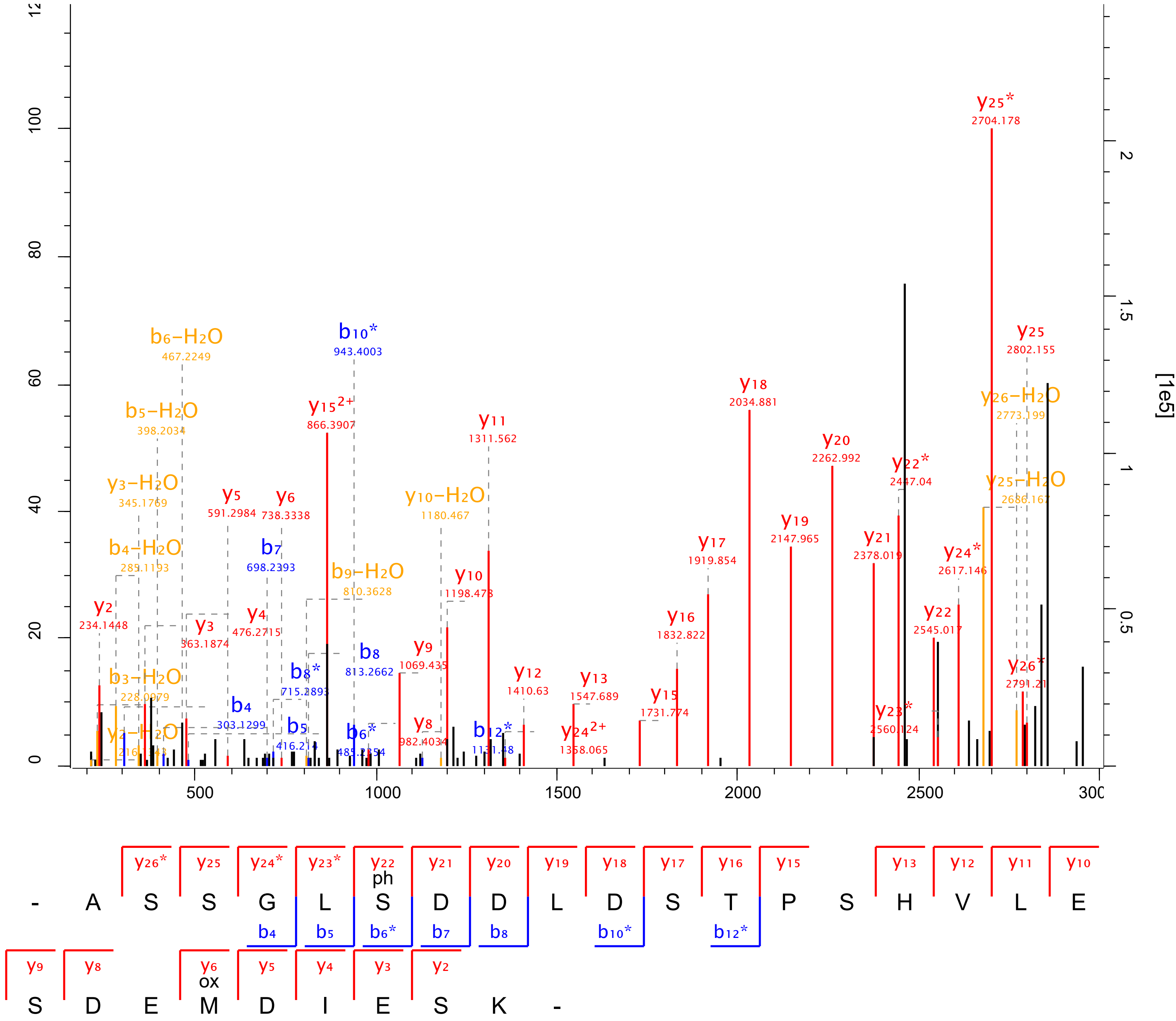

| Raw file | Scan | Method | Score | m/z |
|---|---|---|---|---|
| water_02_1 | 35193 | FTMS; HCD | 44.85 | 751 |

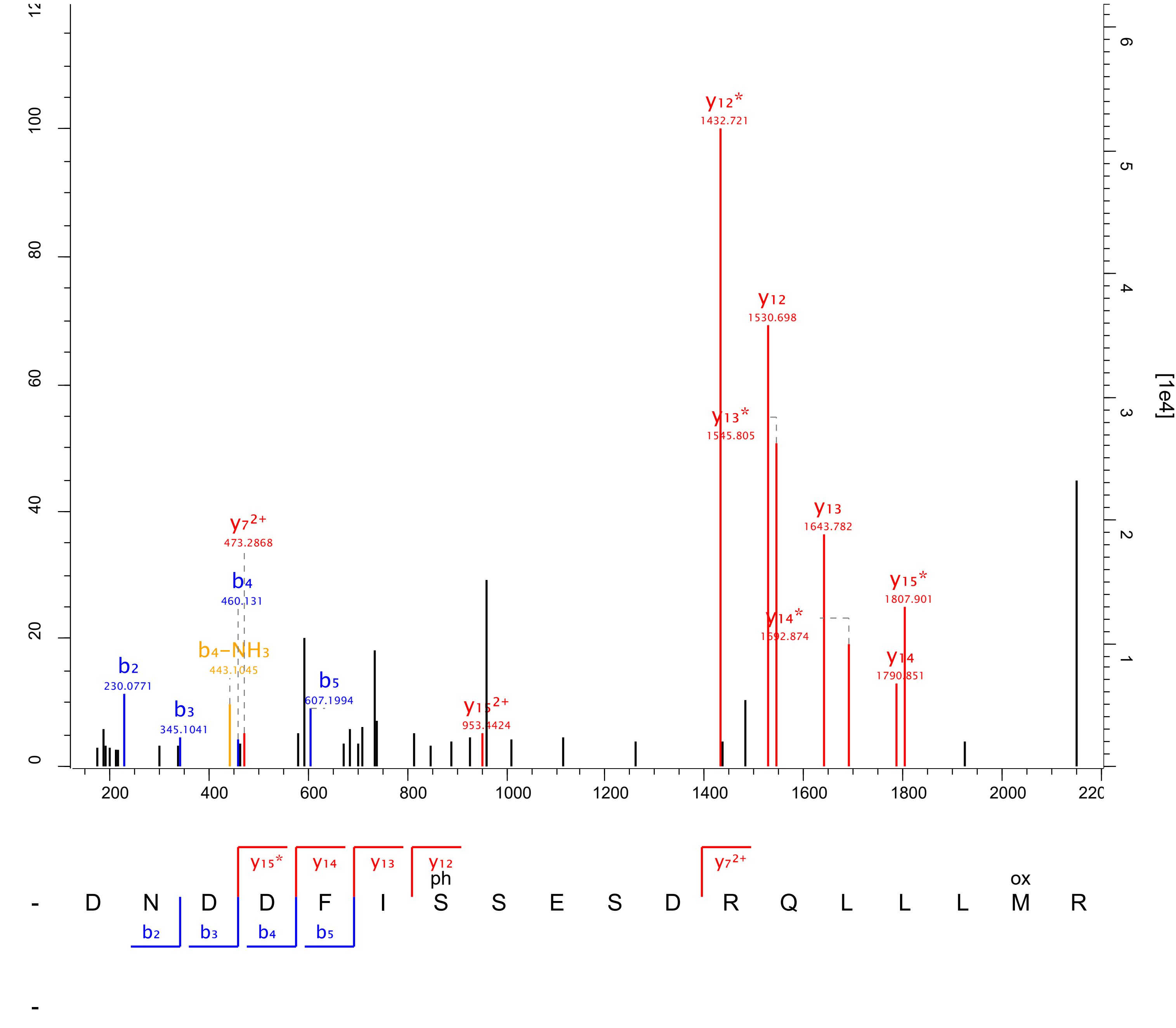

| Raw file | Scan | Method | Score | m/z |
|---|---|---|---|---|
| water_02_1 | 35198 | FTMS; HCD | 149.53 | 882.42 |

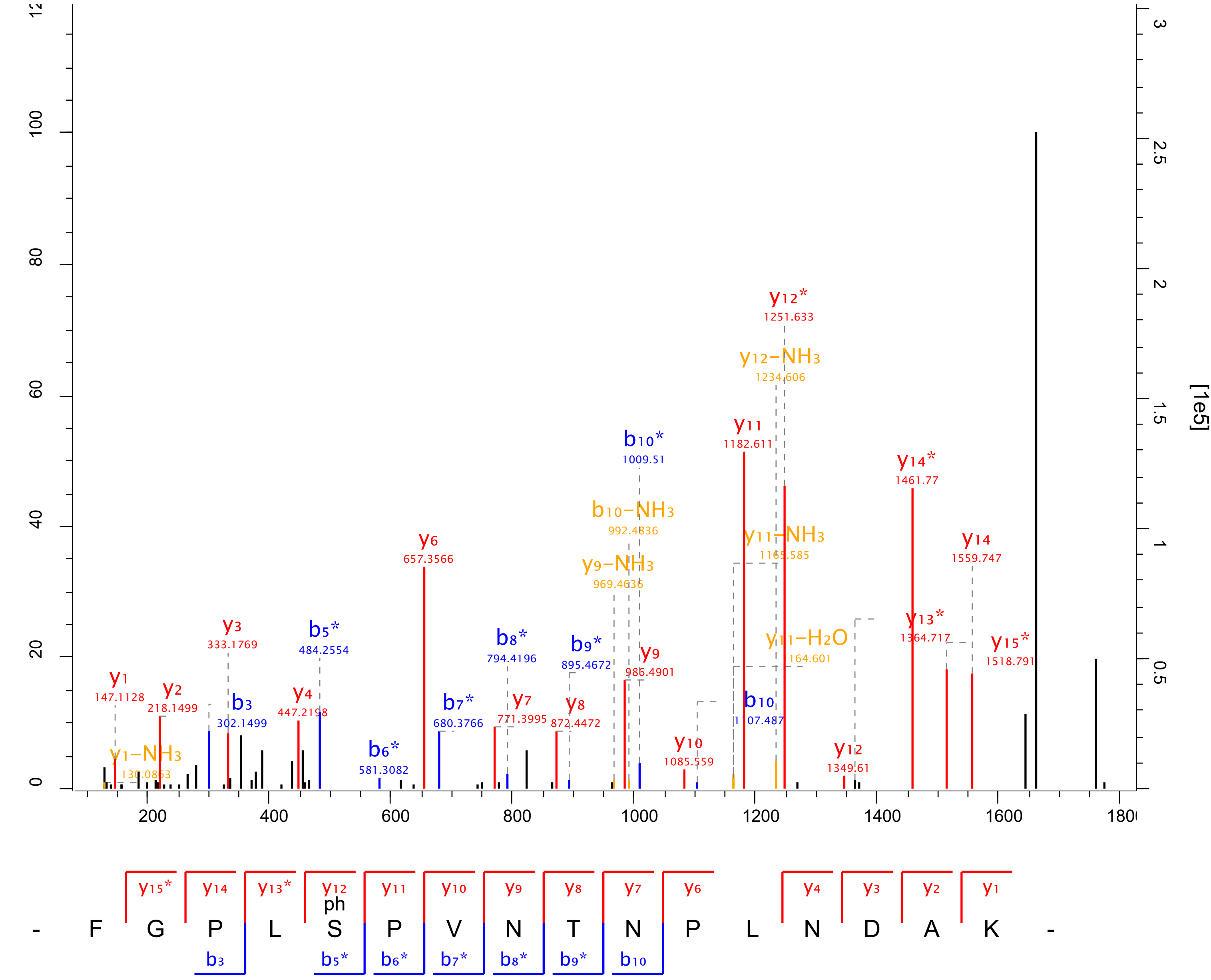

| Raw file | Scan | Method | Score | m/z |
| --- | --- | --- | --- | --- |
| water_02_1 | 3520 | FTMS; HCD | 42.63 | 577.23 |

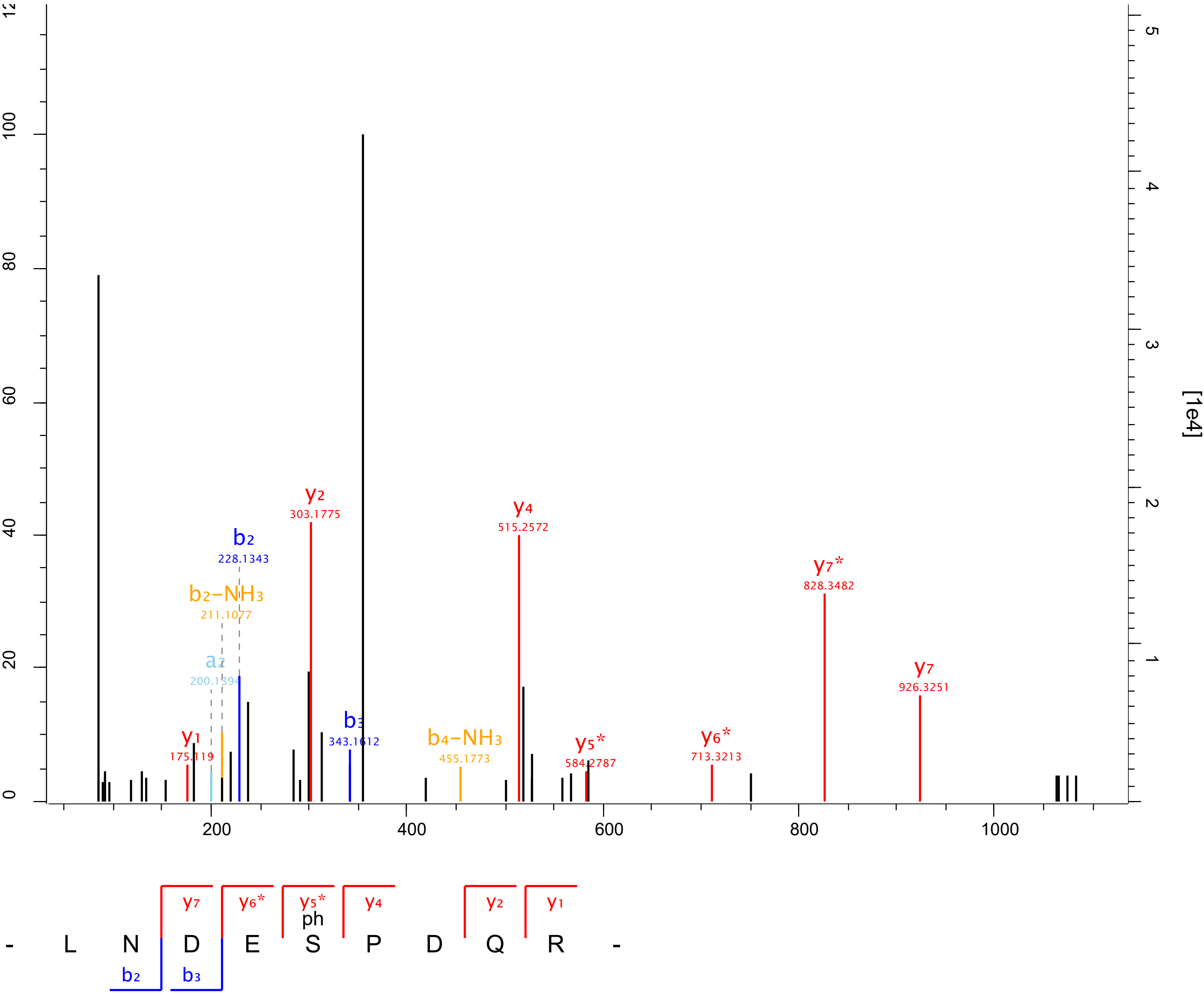

| Raw file | Scan | Method | Score | m/z |
| --- | --- | --- | --- | --- |
| water_02_1 | 35200 | FTMS; HCD | 43.7 | 1014.87 |

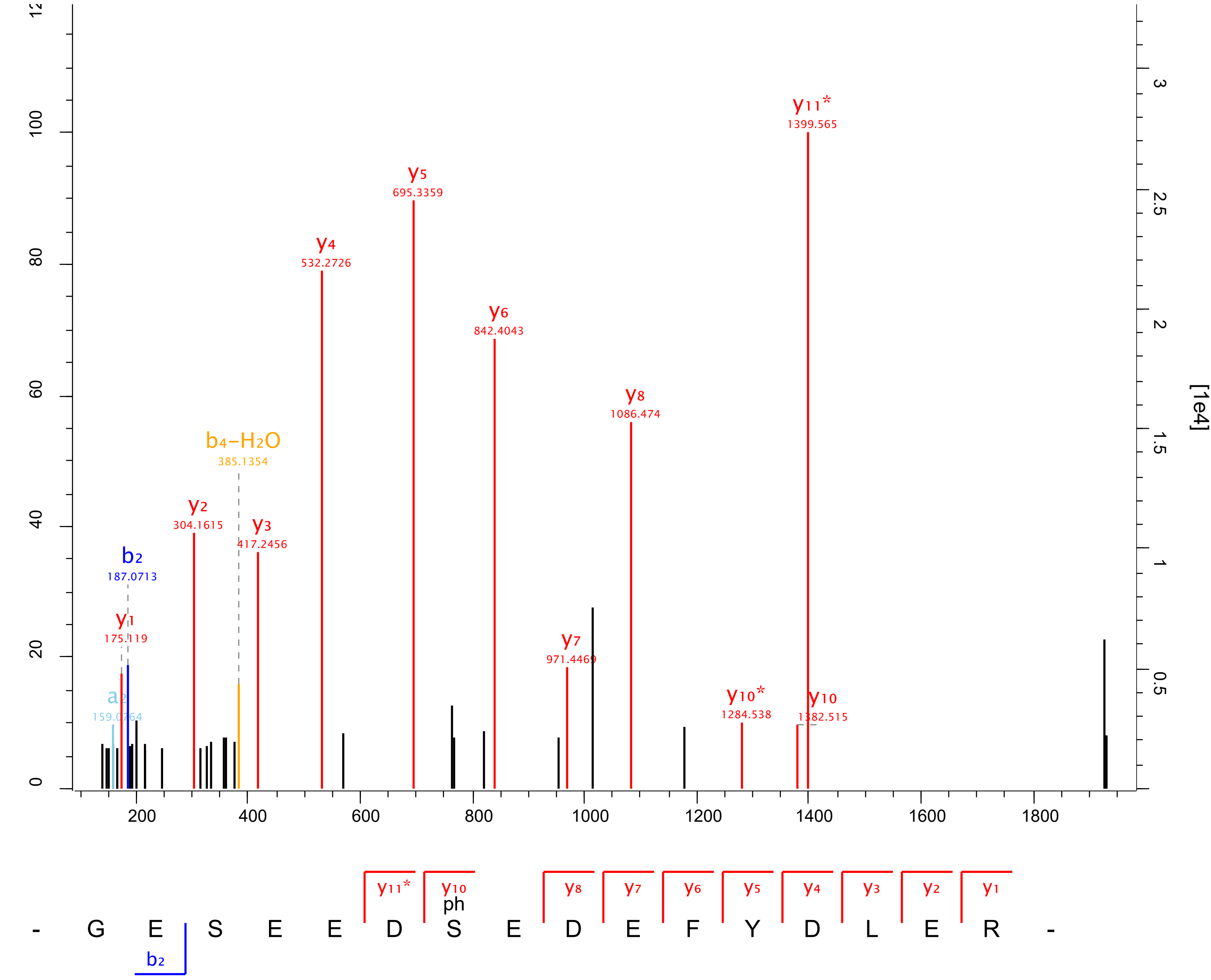

| Raw file | Scan | Method | Score | m/z |
|---|---|---|---|---|
| water_02_1 | 35231 | FTMS; HCD | 214.65 | 1010.92 |

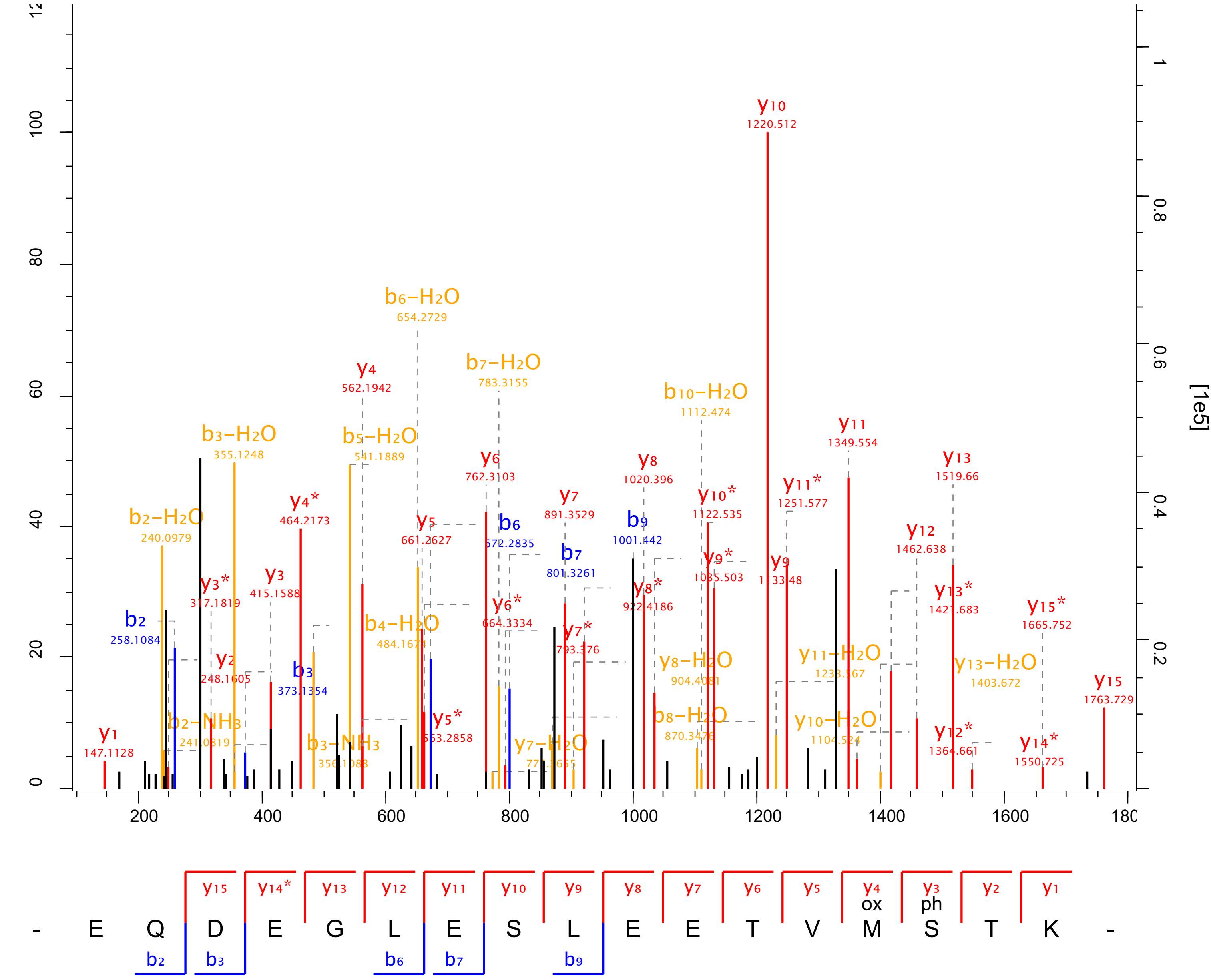

| Raw file | Scan | Method | Score | m/z |
|---|---|---|---|---|
| water_02_1 | 35308 | FTMS; HCD | 80.74 | 696.82 |

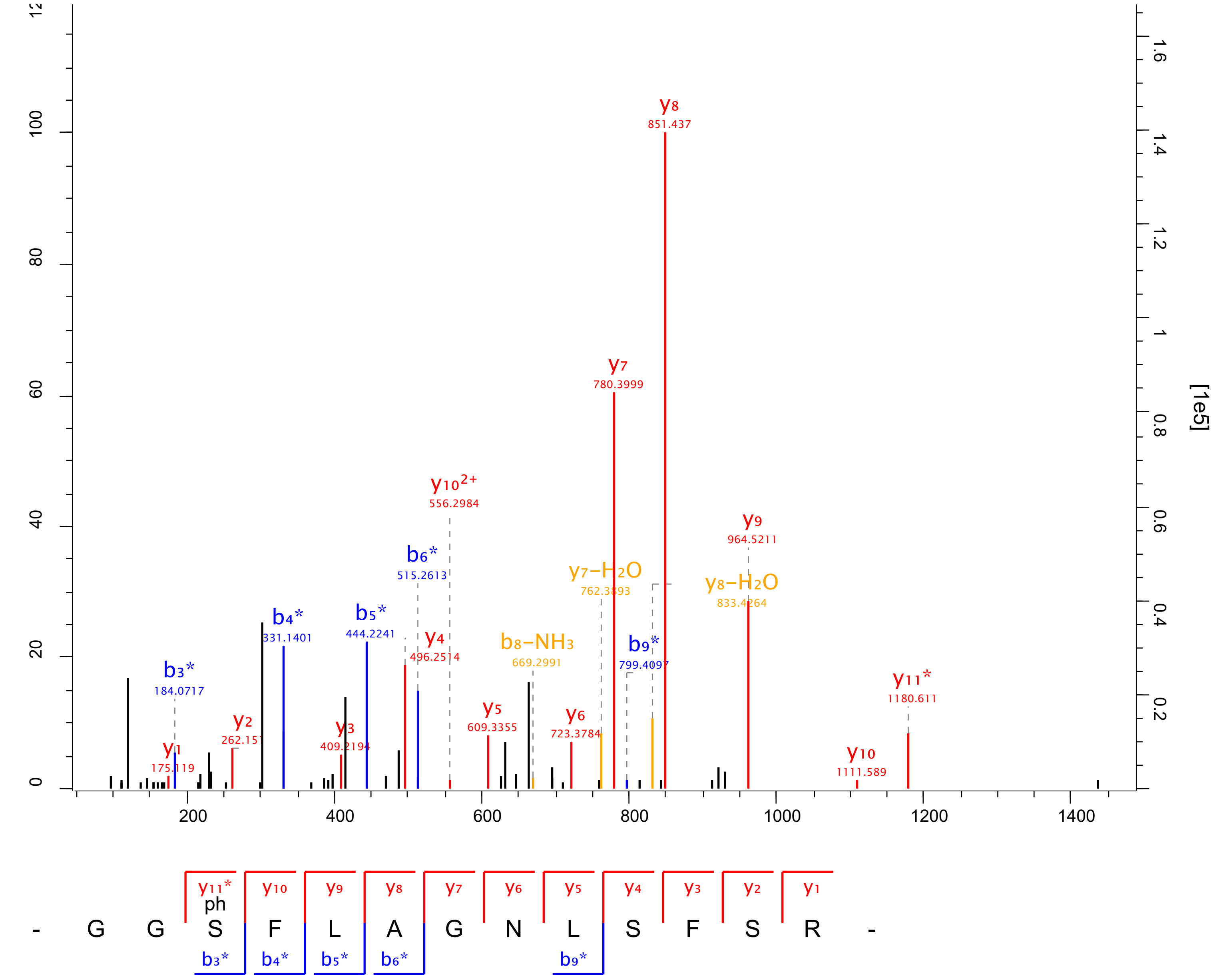

| Raw file | Scan | Method | Score | m/z |
|---|---|---|---|---|
| water_02_1 | 35326 | FTMS; HCD | 42.73 | 1086.11 |

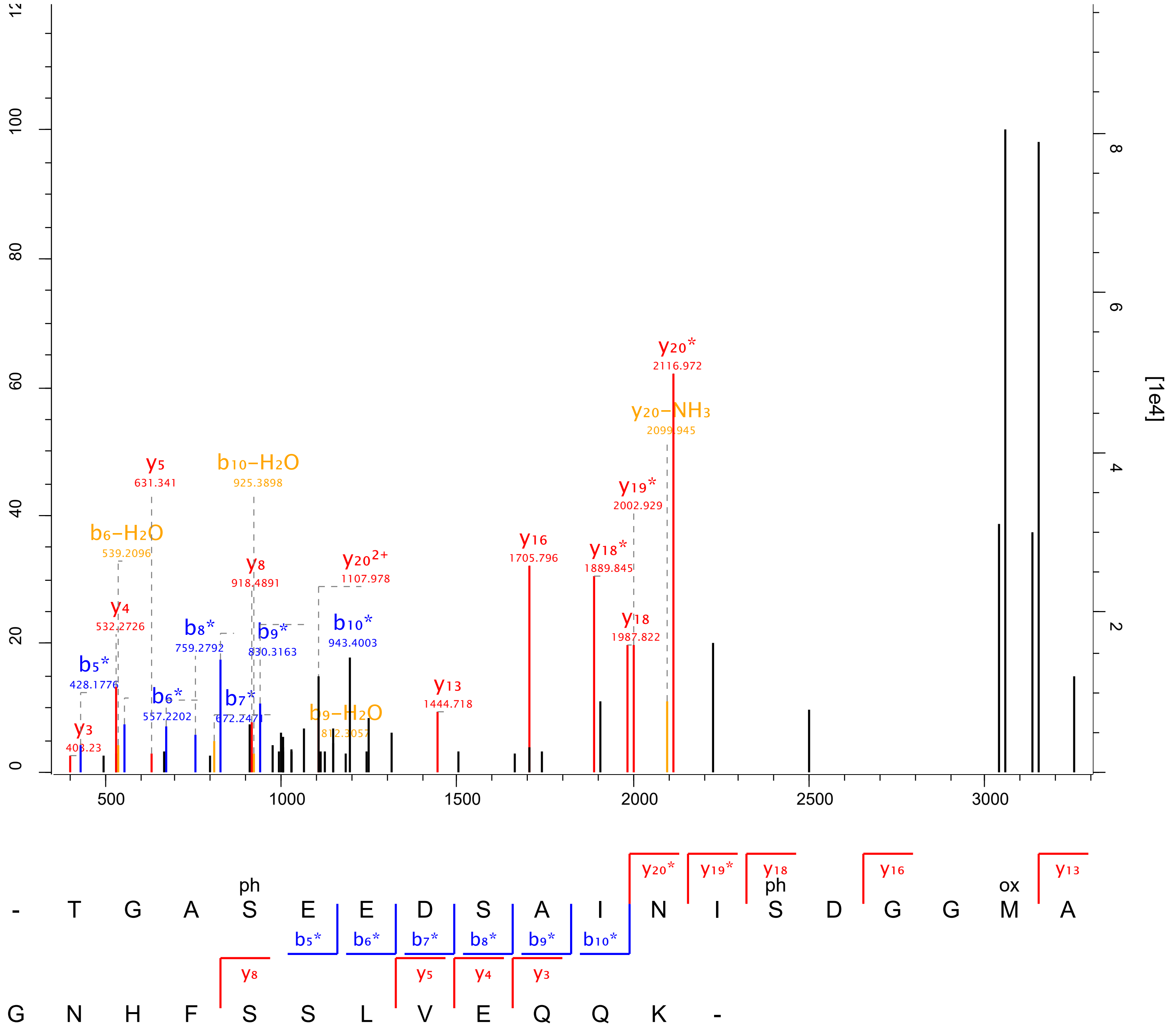

| Raw file | Scan | Method | Score | m/z |
|---|---|---|---|---|
| water_02_1 | 35402 | FTMS; HCD | 113.47 | 744.89 |

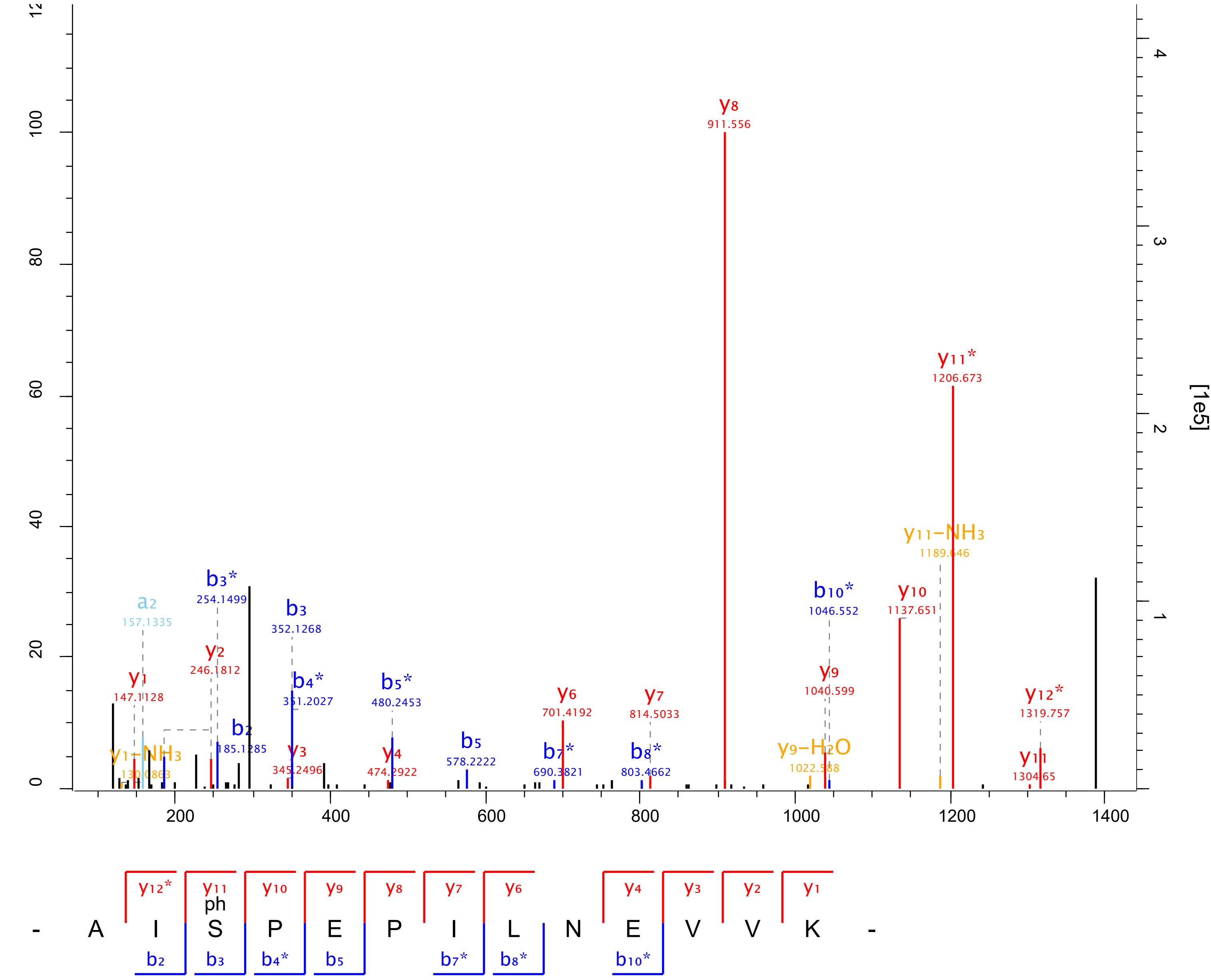

| Raw file | Scan | Method | Score | m/z |
| --- | --- | --- | --- | --- |
| water_02_1 | 35499 | FTMS; HCD | 96.12 | 904.42 |

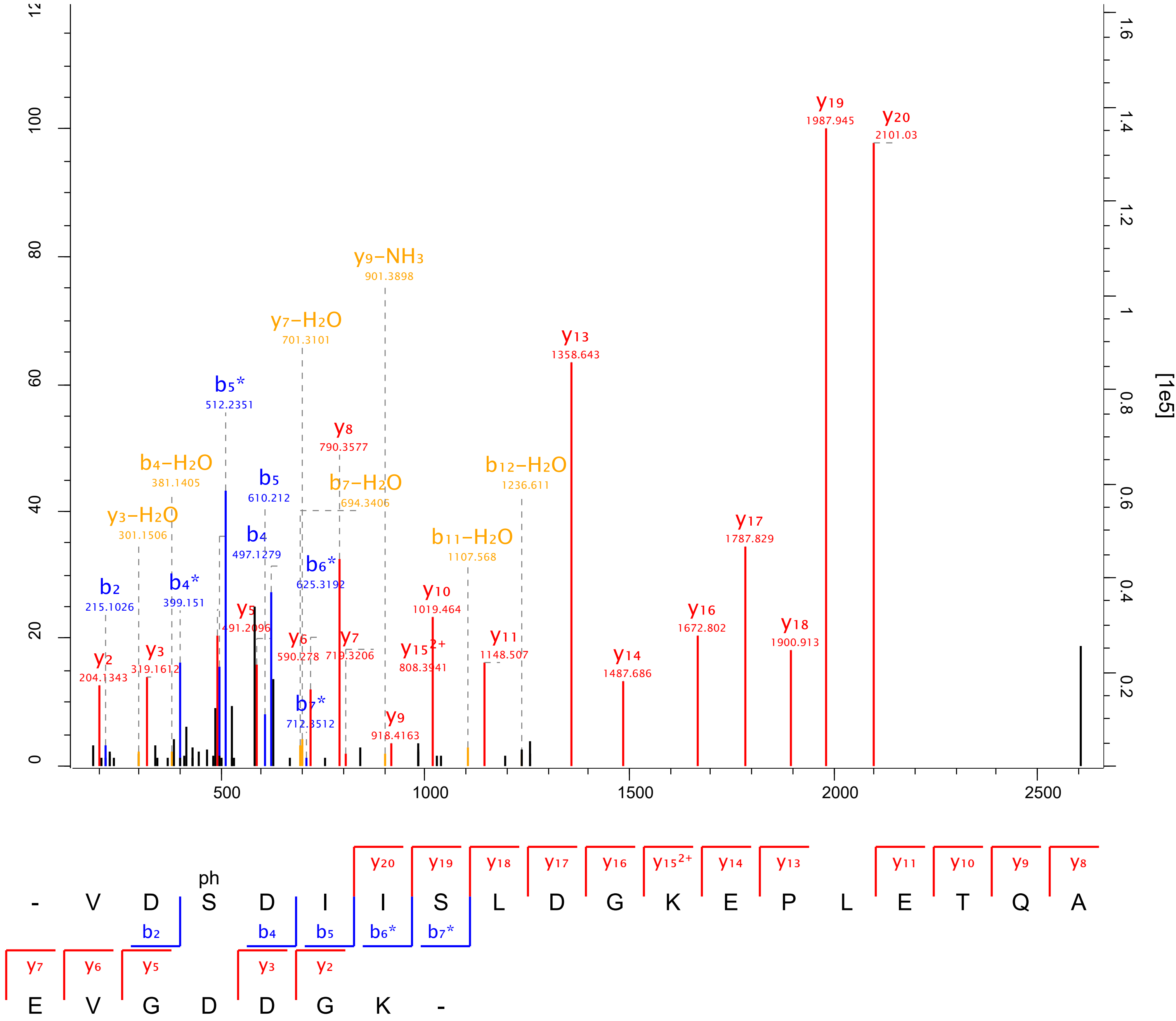

| Raw file | Scan | Method | Score | m/z |
|---|---|---|---|---|
| water_02_1 | 3551 | FTMS; HCD | 50.45 | 499.19 |

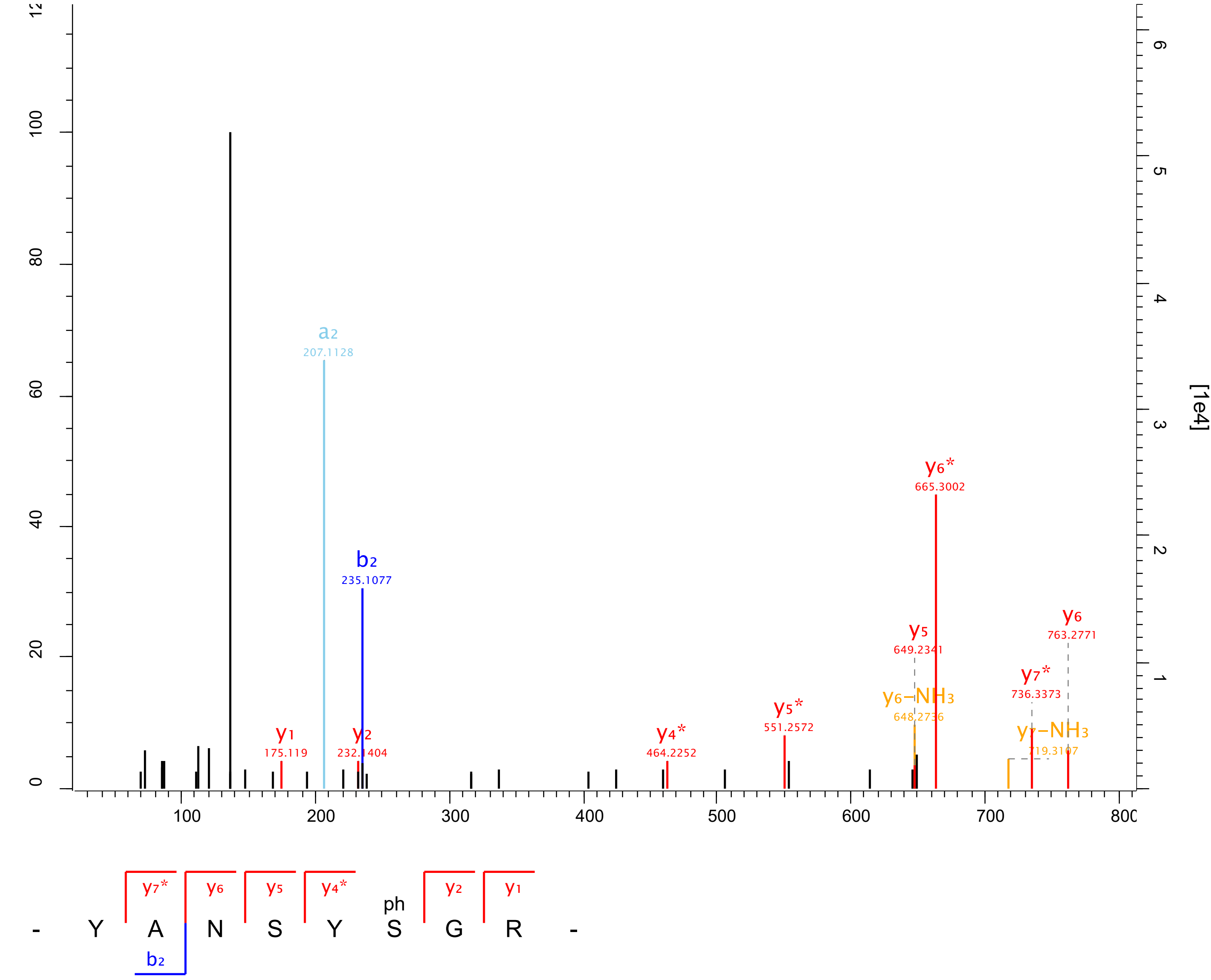

| Raw file | Scan | Method | Score | m/z |
| --- | --- | --- | --- | --- |
| water_02_1 | 35737 | FTMS; HCD | 116.09 | 923.86 |

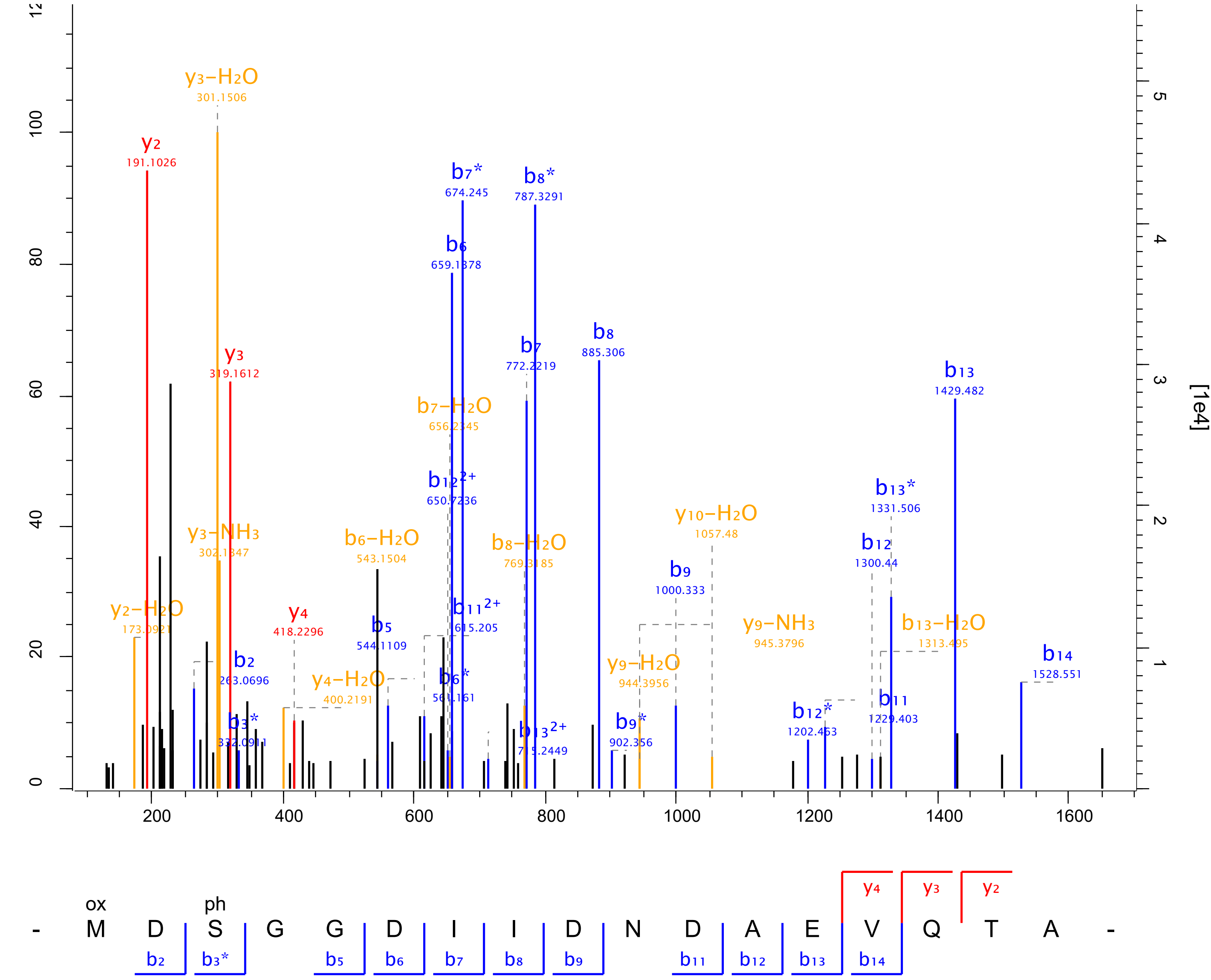

| Raw file | Scan | Method | Score | m/z |
| --- | --- | --- | --- | --- |
| water_02_1 | 35759 | FTMS; HCD | 44.79 | 663.95 |

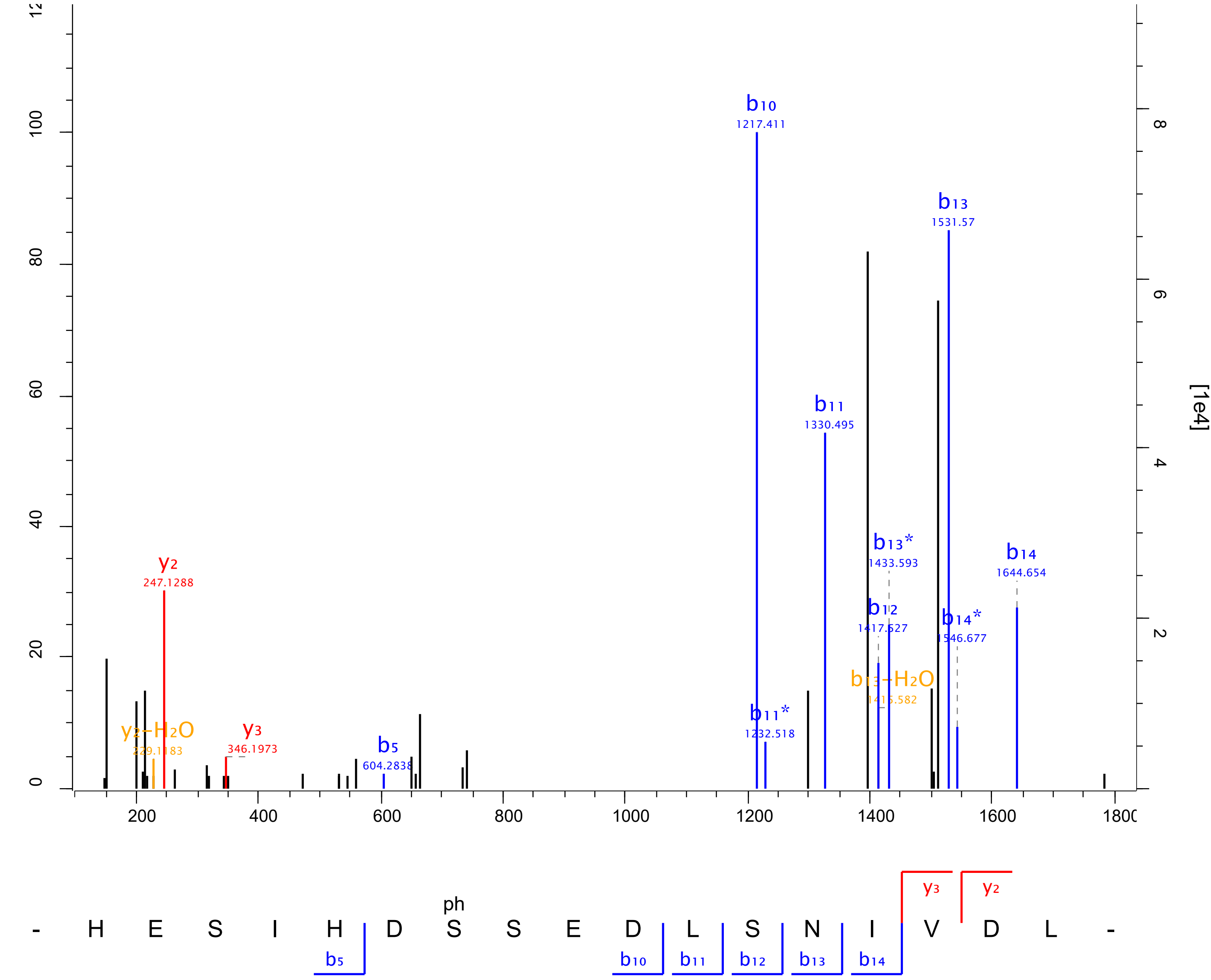

| Raw file | Scan | Method | Score | m/z |
| --- | --- | --- | --- | --- |
| water_02_1 | 3579 | FTMS; HCD | 94.58 | 485.7 |

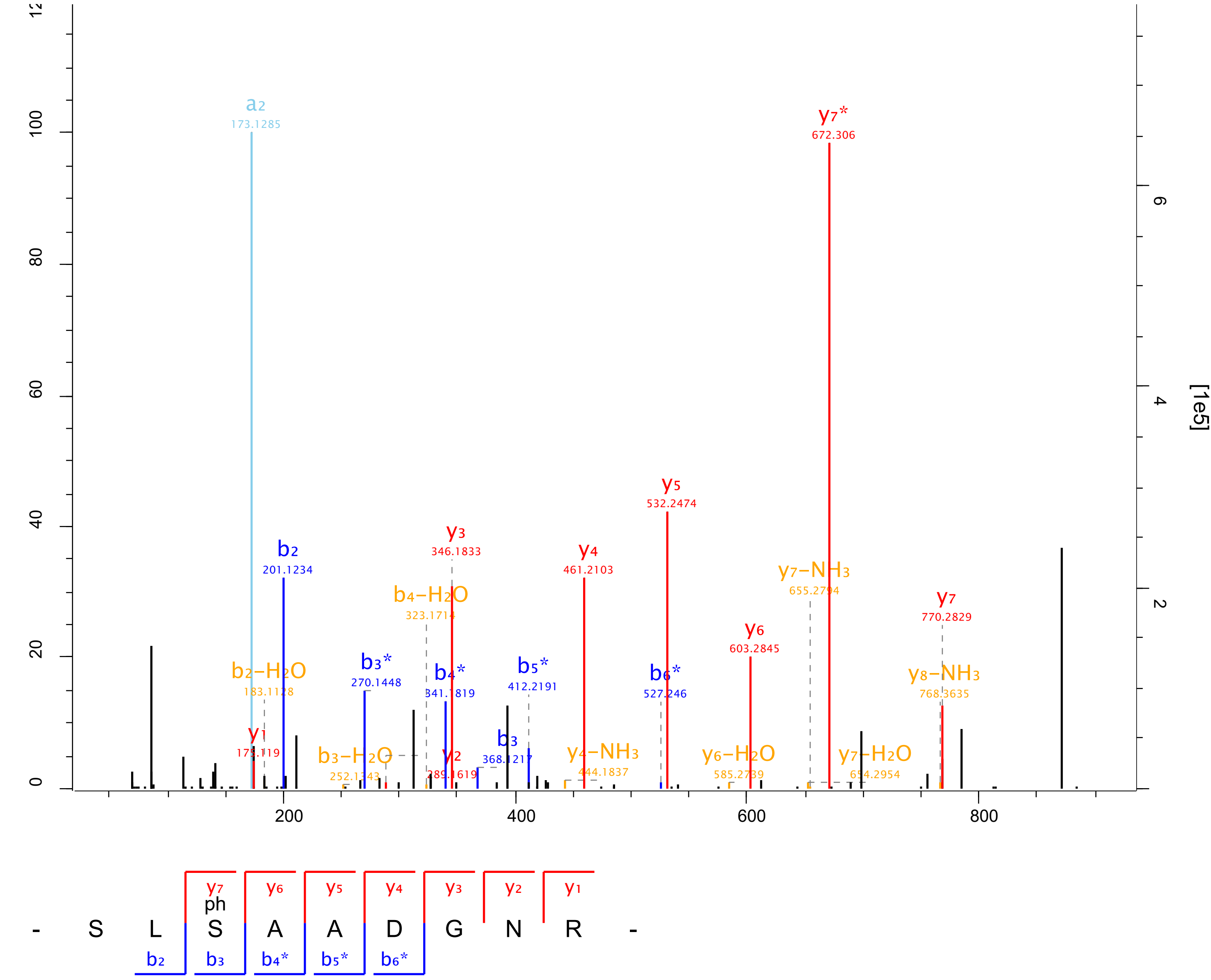

| Raw file | Scan | Method | Score | m/z |
|---|---|---|---|---|
| water_02_1 | 35873 | FTMS; HCD | 83.42 | 733.86 |

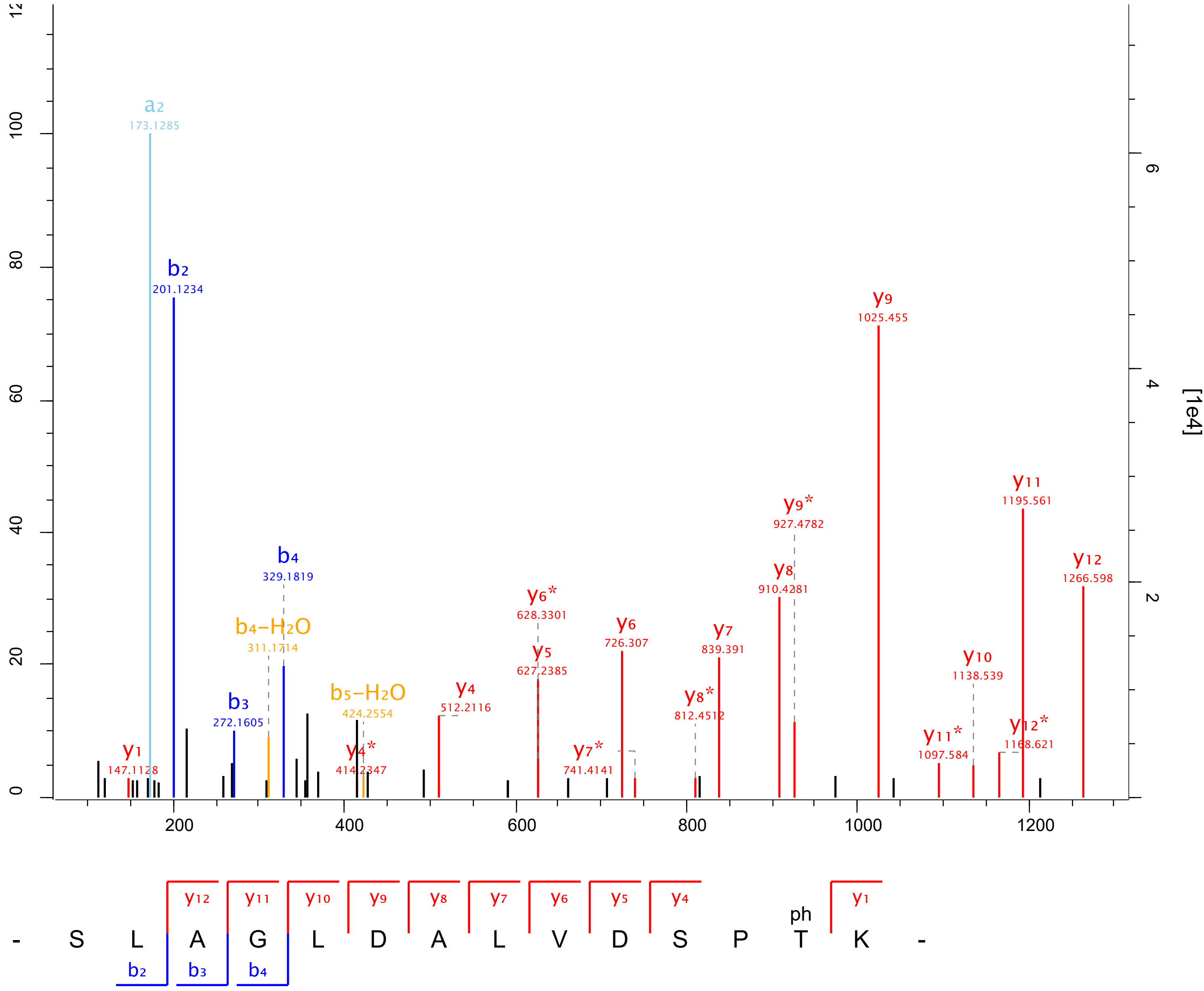

| Raw file | Scan | Method | Score | m/z |
|---|---|---|---|---|
| water_02_1 | 35921 | FTMS; HCD | 78.71 | 706.6 |

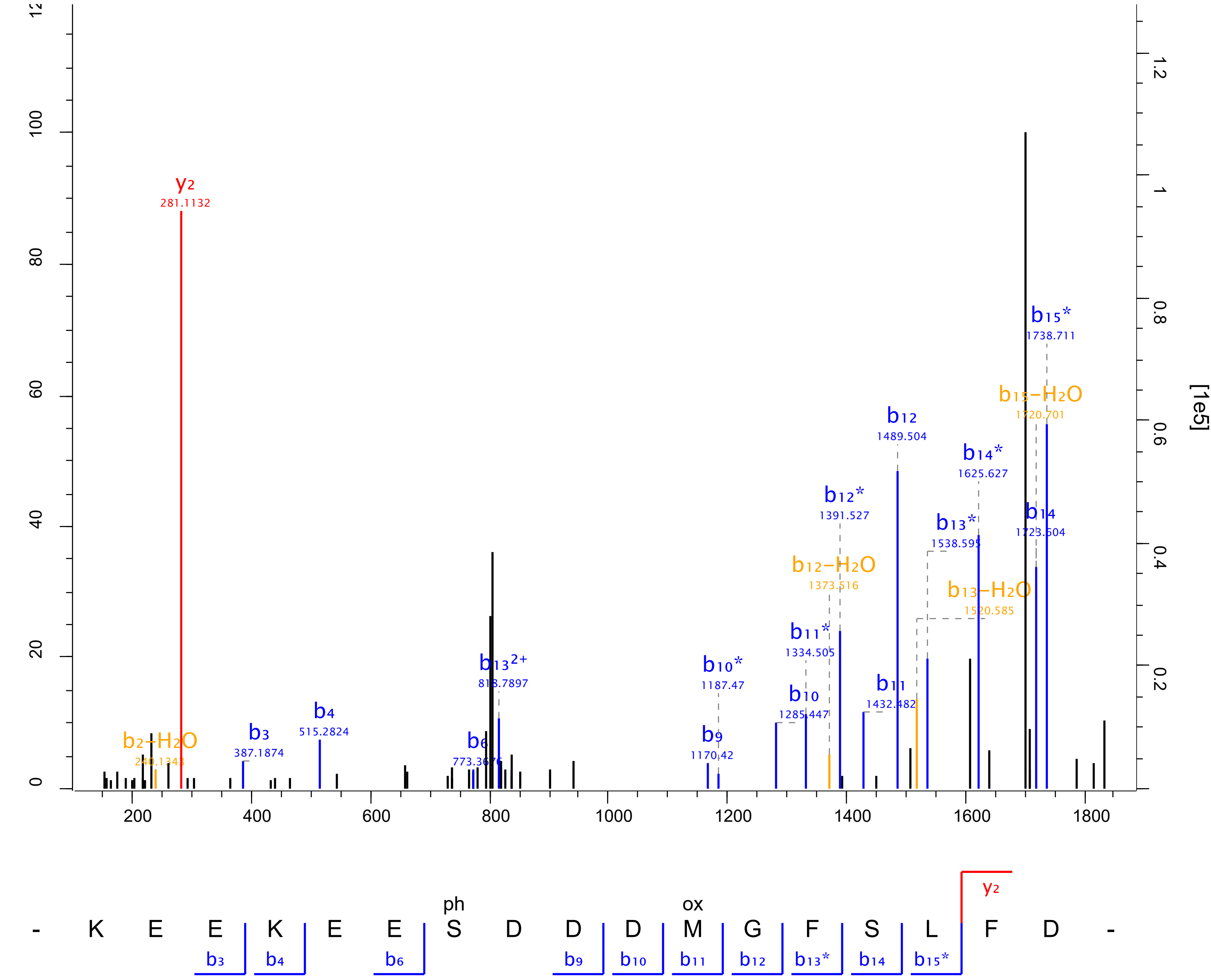

| Raw file | Scan | Method | Score | m/z |
| --- | --- | --- | --- | --- |
| water_02_1 | 36031 | FTMS; HCD | 104.38 | 750.31 |

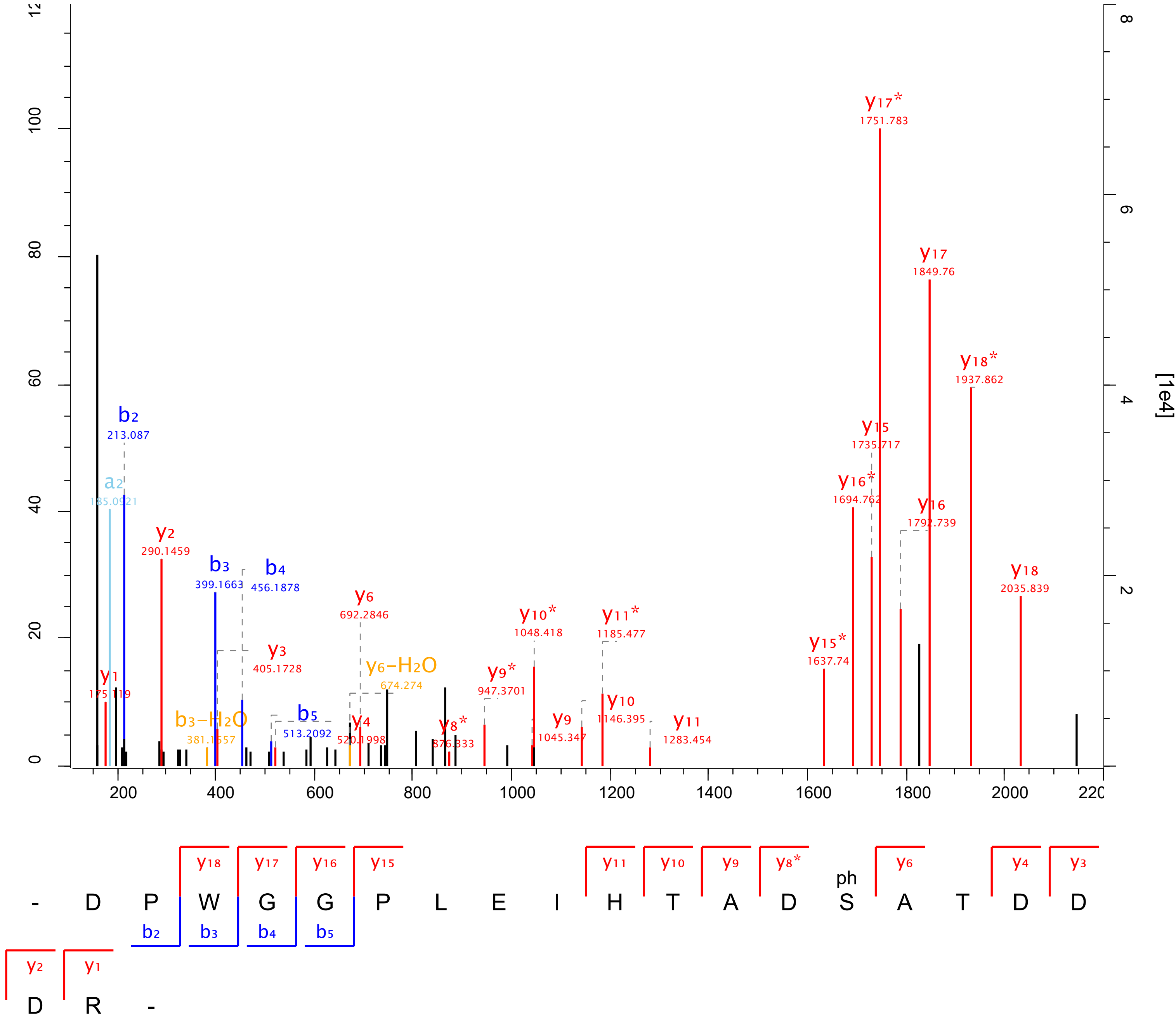

| Raw file | Scan | Method | Score | m/z |
| --- | --- | --- | --- | --- |
| water_02_1 | 36121 | FTMS; HCD | 46.15 | 956.78 |

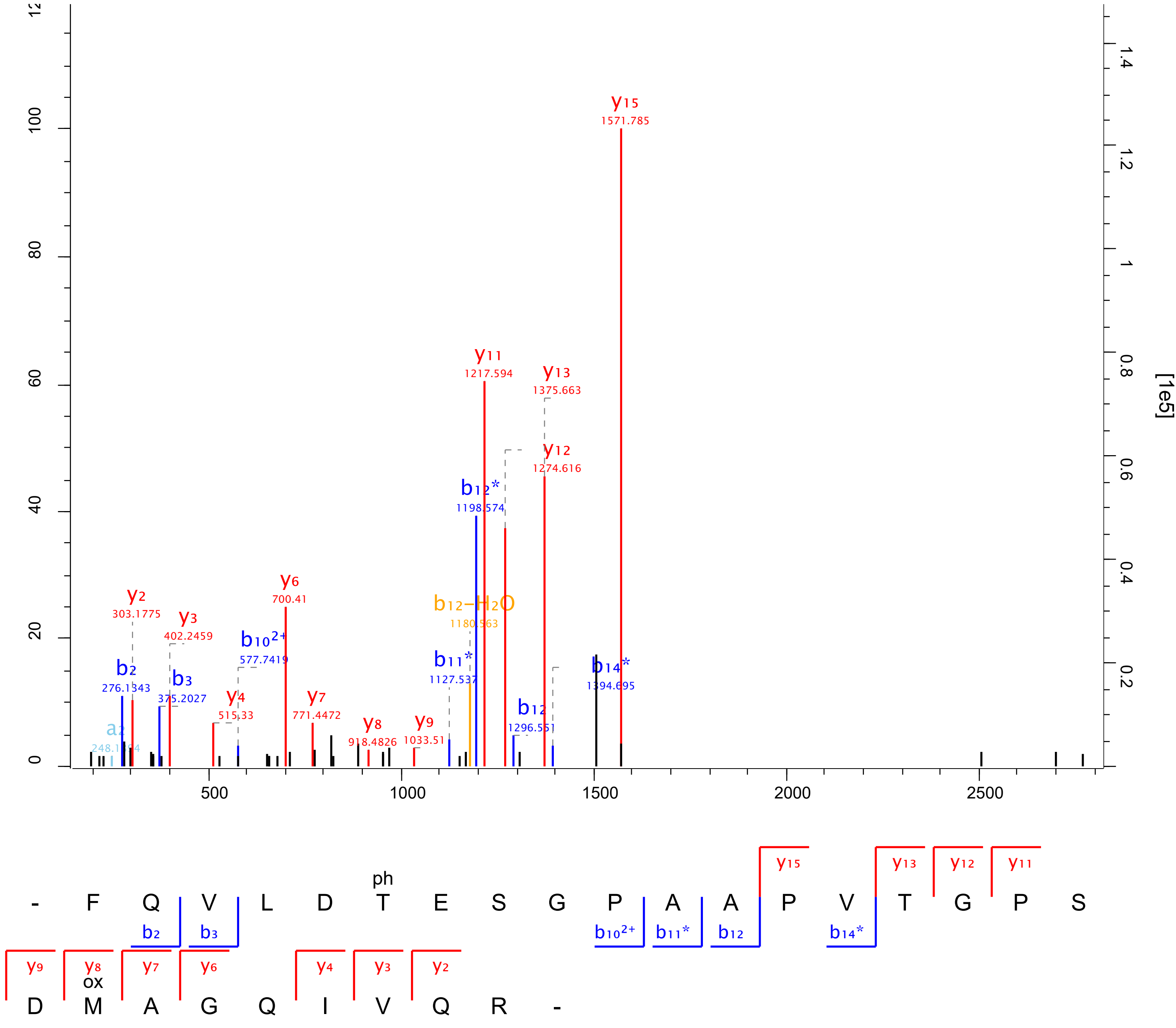

| Raw file | Scan | Method | Score | m/z |
| --- | --- | --- | --- | --- |
| water_02_1 | 36172 | FTMS; HCD | 52.25 | 651.29 |

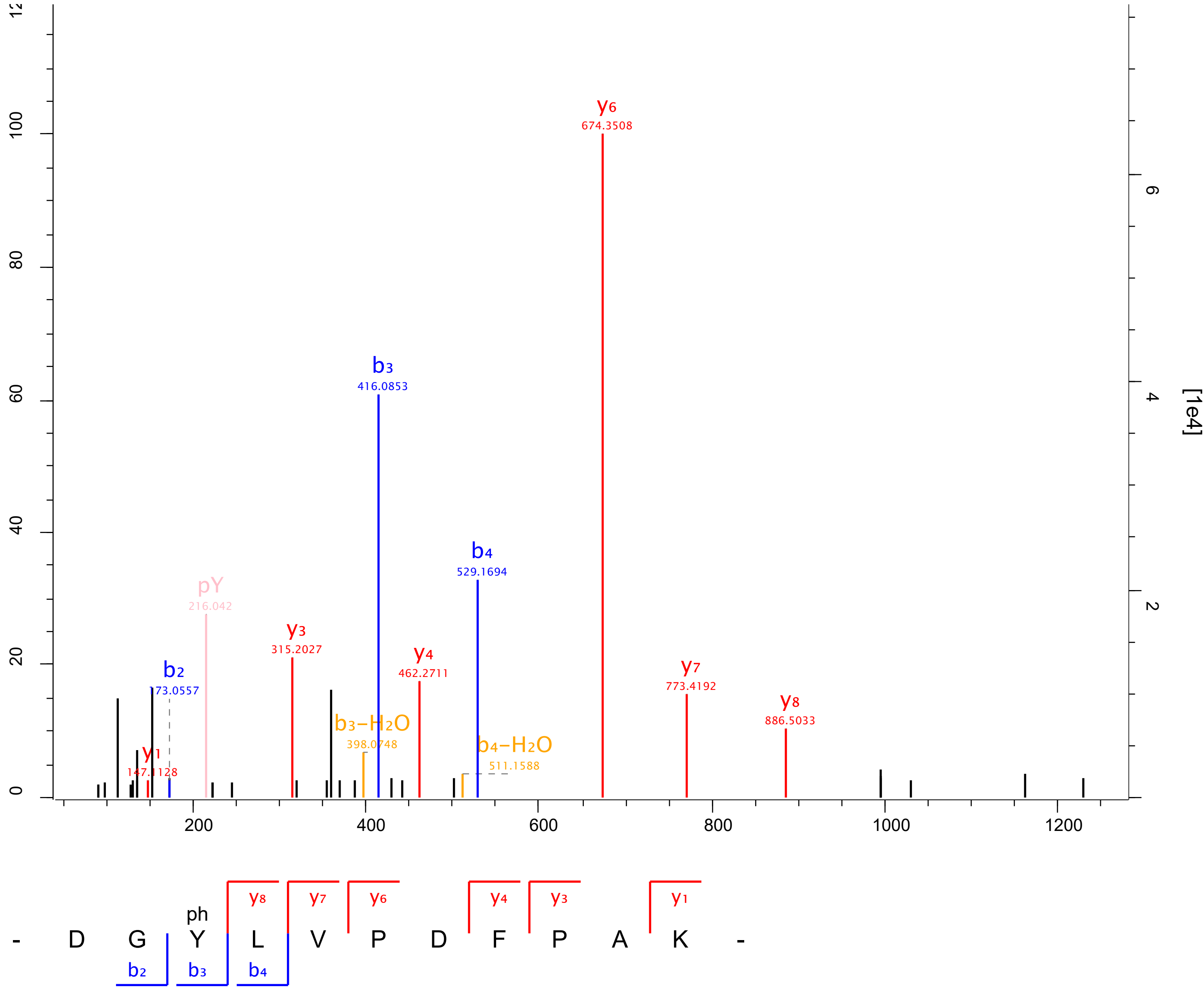

| Raw file | Scan | Method | Score | m/z |
| --- | --- | --- | --- | --- |
| water_02_1 | 36223 | FTMS; HCD | 65.23 | 807.38 |

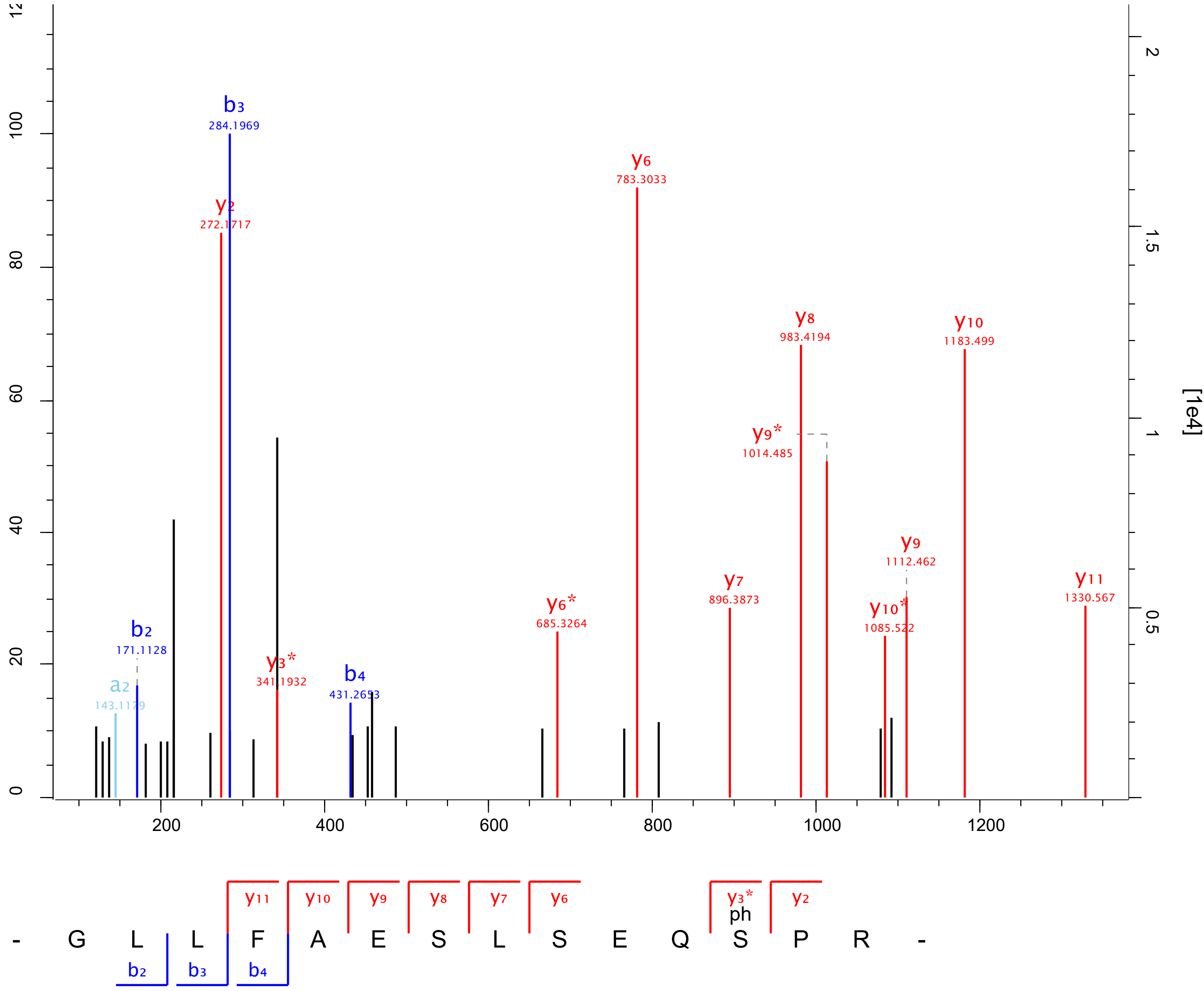

| Raw file | Scan | Method | Score | m/z |
| --- | --- | --- | --- | --- |
| water_02_1 | 36328 | FTMS; HCD | 110.26 | 1006.49 |

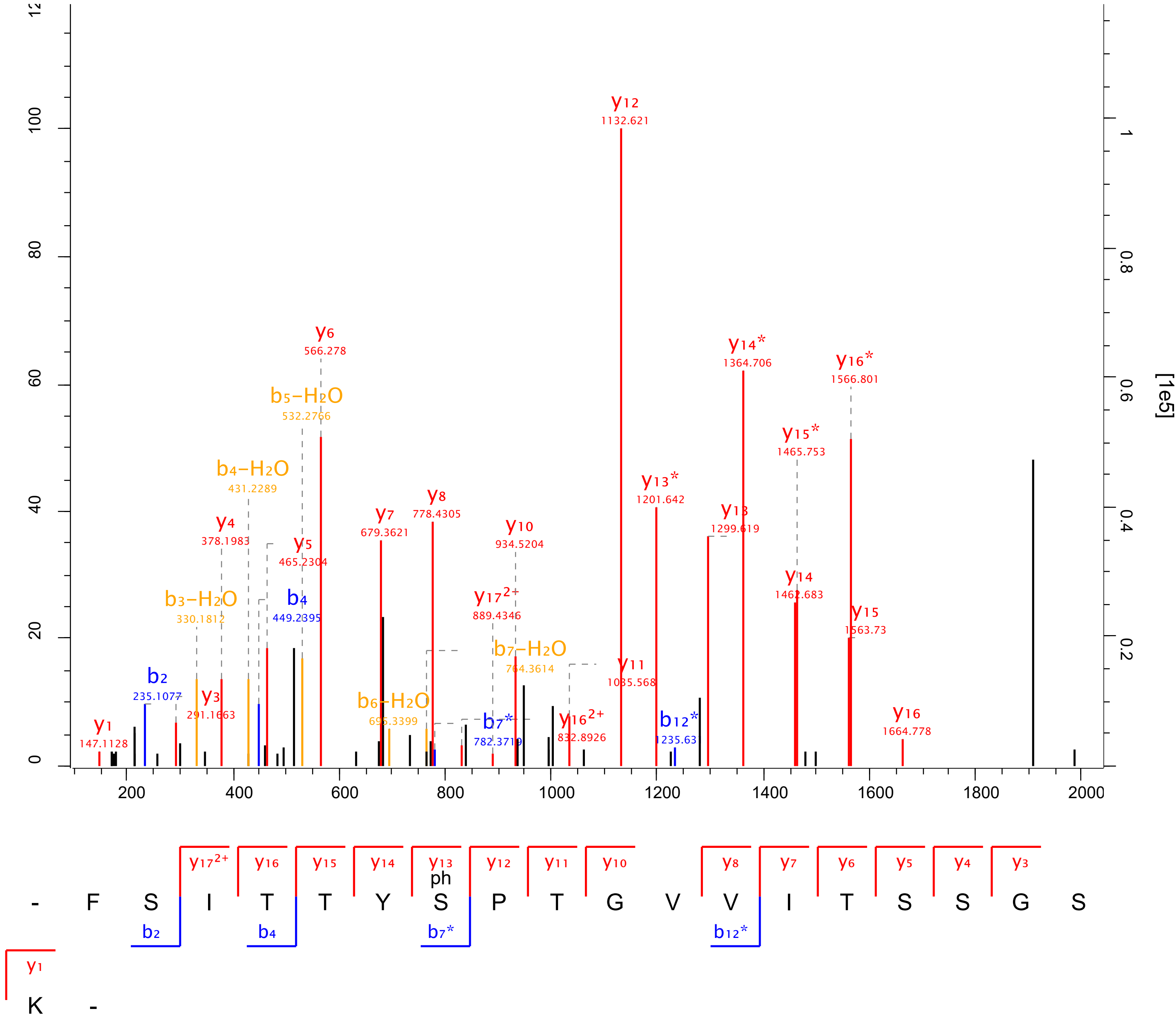

| Raw file | Scan | Method | Score | m/z |
| --- | --- | --- | --- | --- |
| water_02_1 | 36364 | FTMS; HCD | 146.15 | 812.36 |

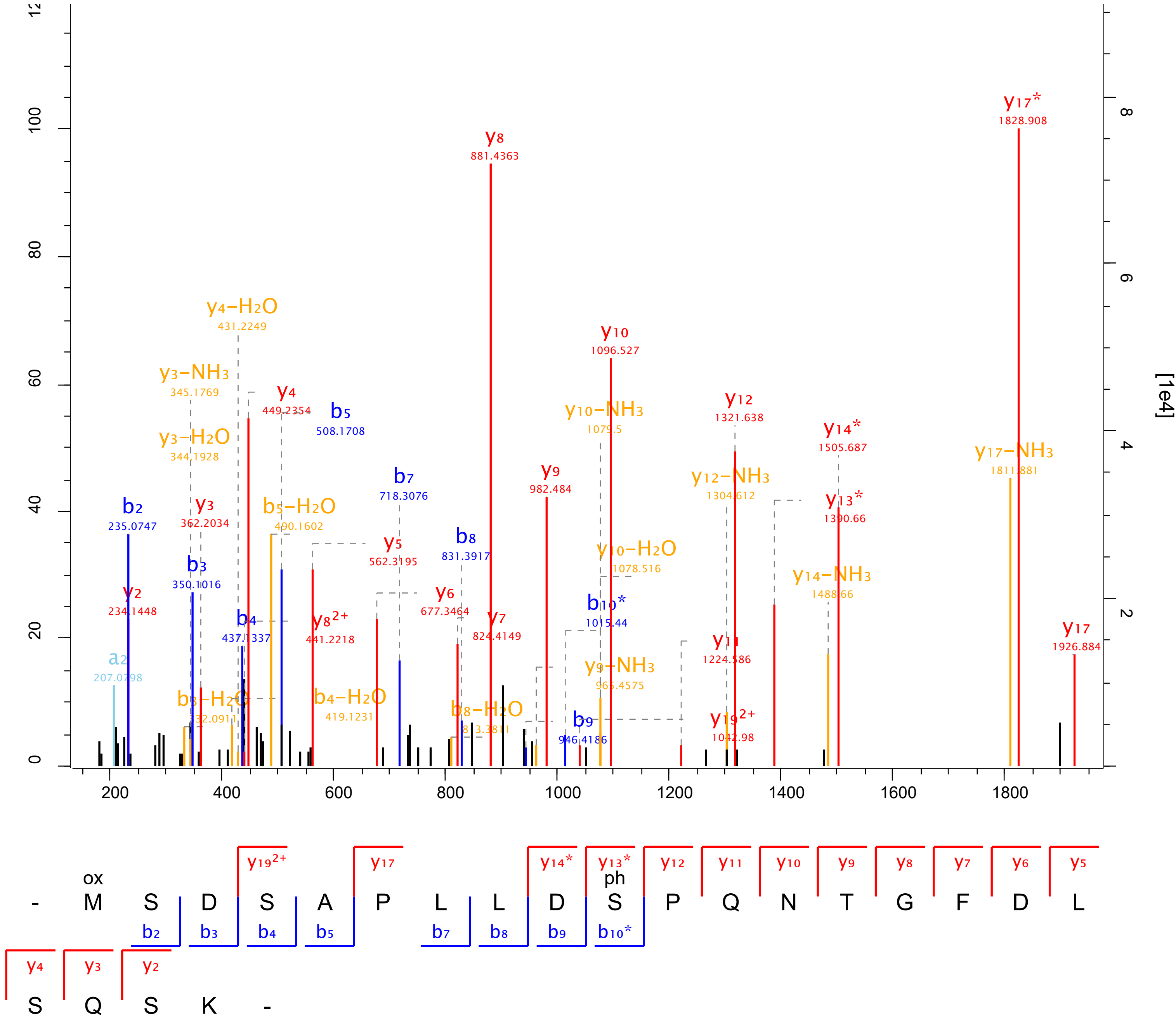

| Raw file | Scan | Method | Score | m/z |
|---|---|---|---|---|
| water_02_1 | 36457 | FTMS; HCD | 56.57 | 702.32 |

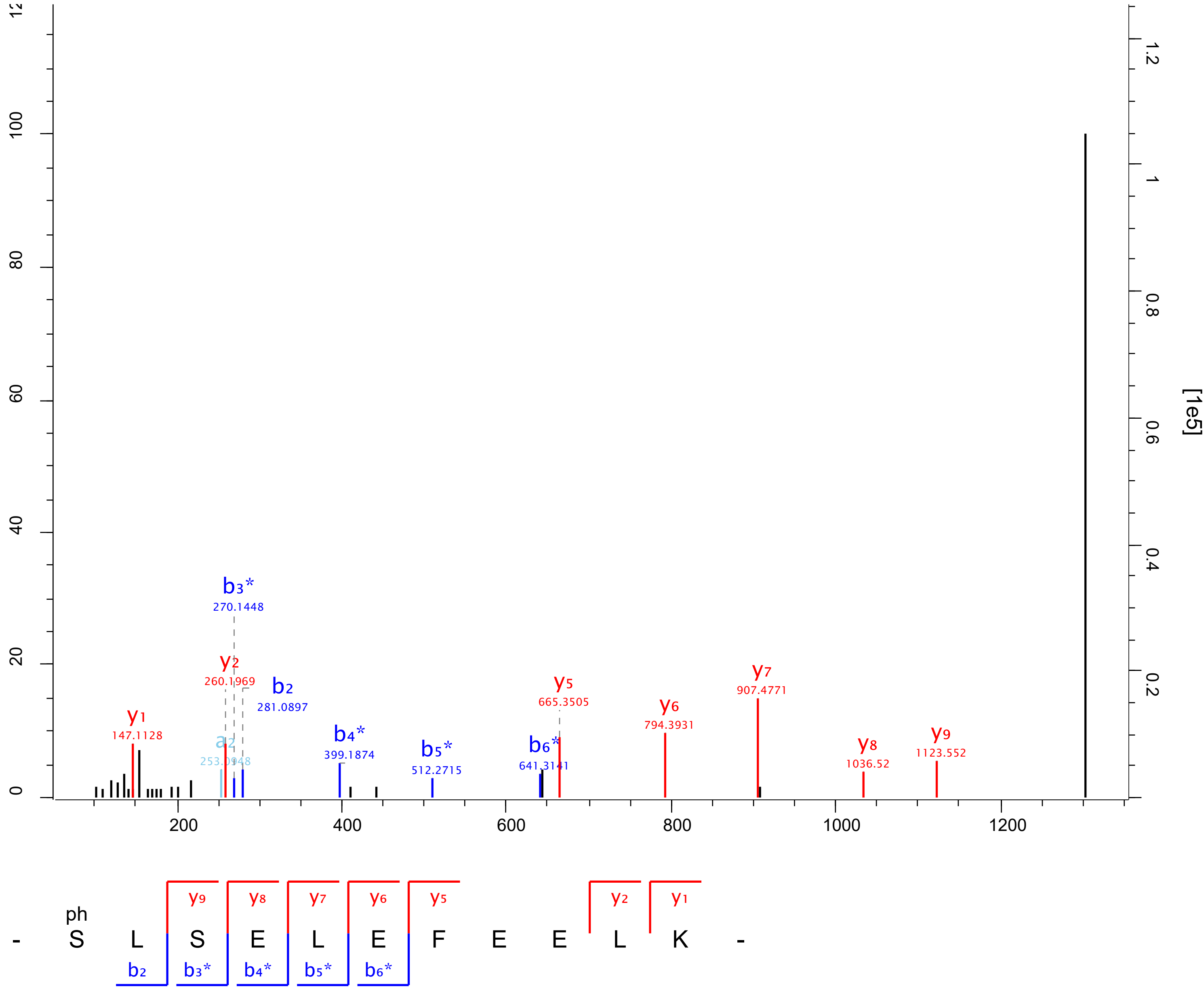

| Raw file | Scan | Method | Score | m/z |
|---|---|---|---|---|
| water_02_1 | 36475 | FTMS; HCD | 118.19 | 659.83 |

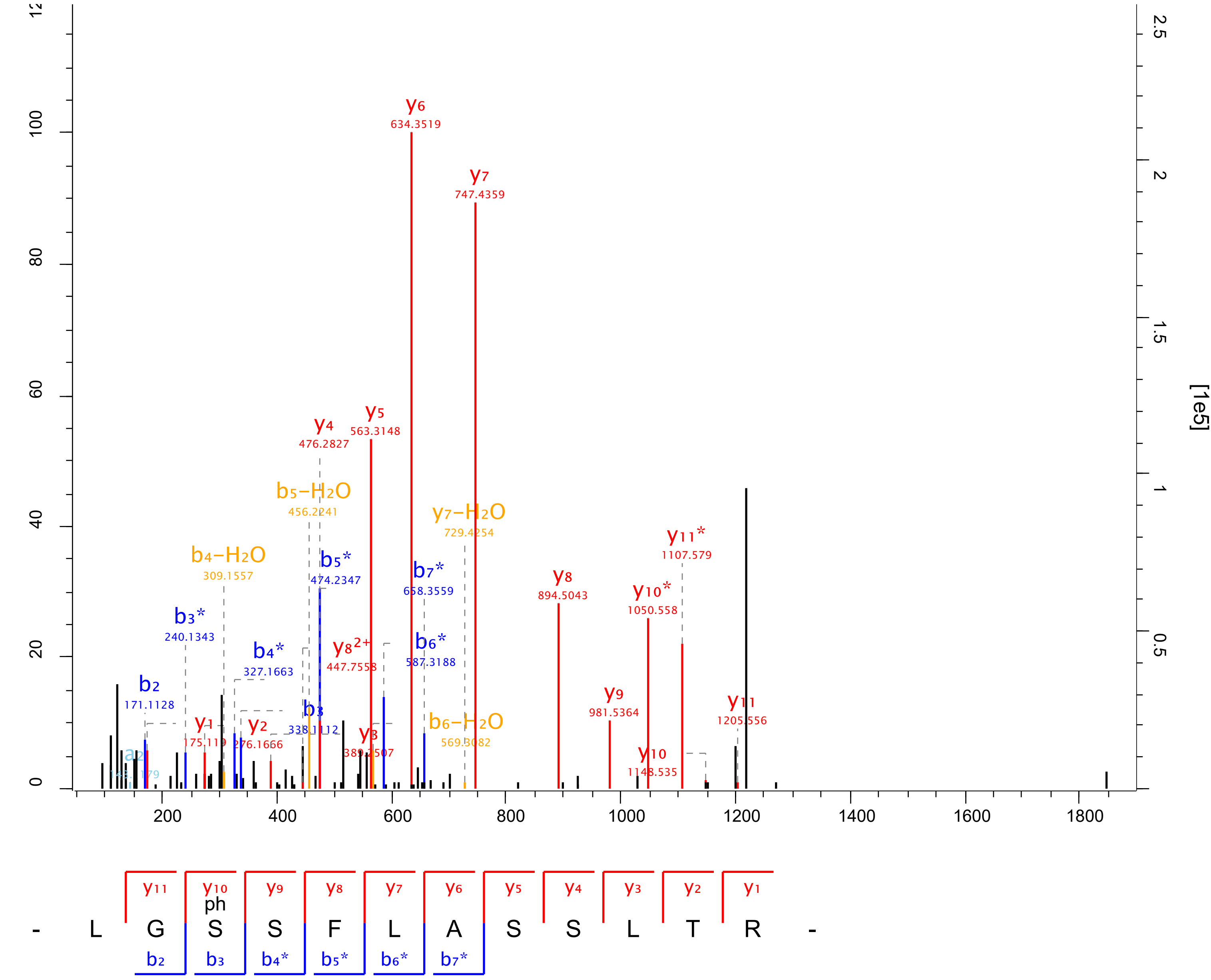

| Raw file | Scan | Method | Score | m/z |
|---|---|---|---|---|
| water_02_1 | 36484 | FTMS; HCD | 225.65 | 889.69 |

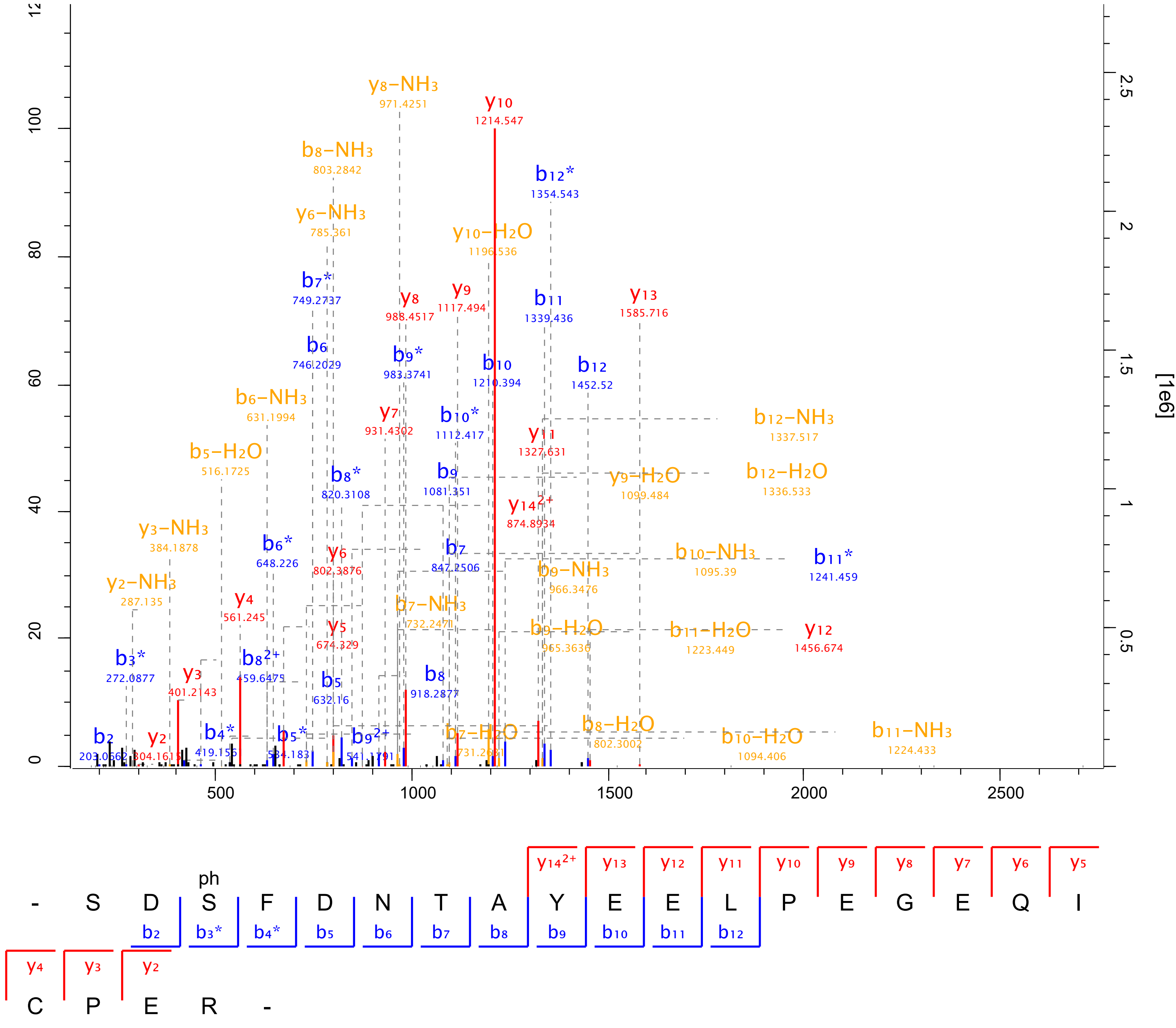

| Raw file | Scan | Method | Score | m/z |
| --- | --- | --- | --- | --- |
| water_02_1 | 36542 | FTMS; HCD | 71.42 | 860.12 |

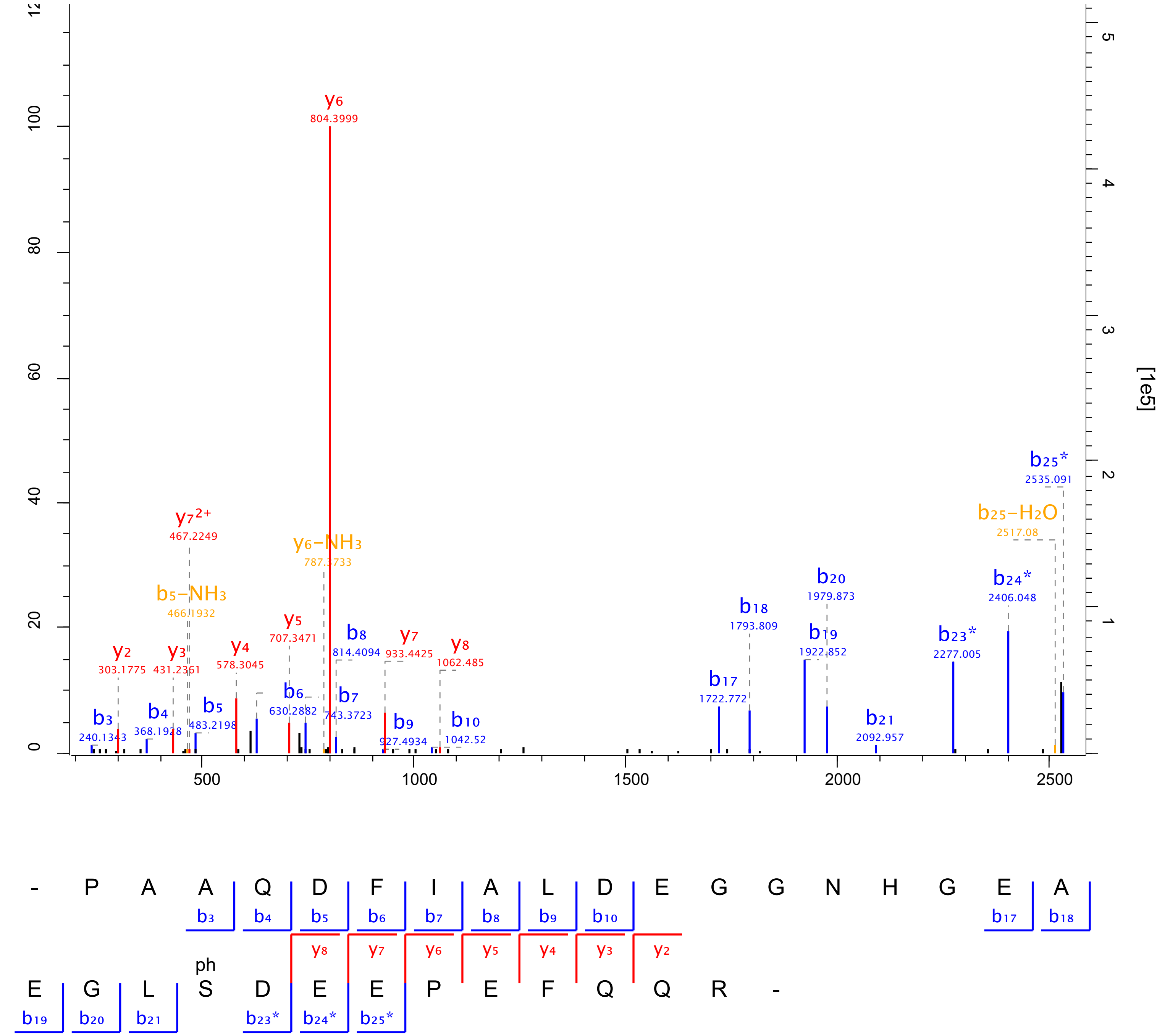

| Raw file | Scan | Method | Score | m/z |
|---|---|---|---|---|
| water_02_1 | 36549 | FTMS; HCD | 198.4 | 927.73 |

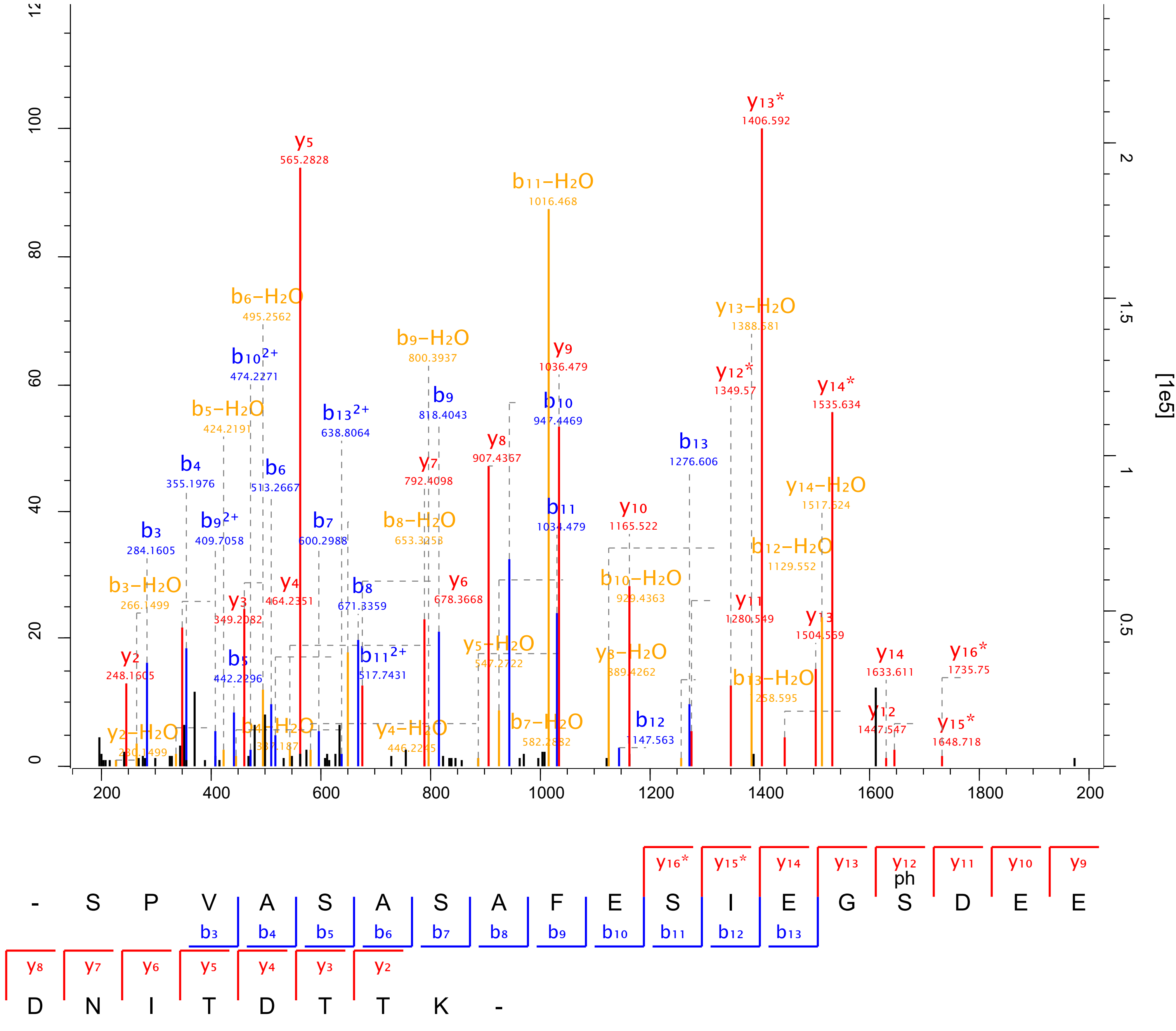

| Raw file | Scan | Method | Score | m/z |
|---|---|---|---|---|
| water_02_1 | 36559 | FTMS; HCD | 65.18 | 562.8 |

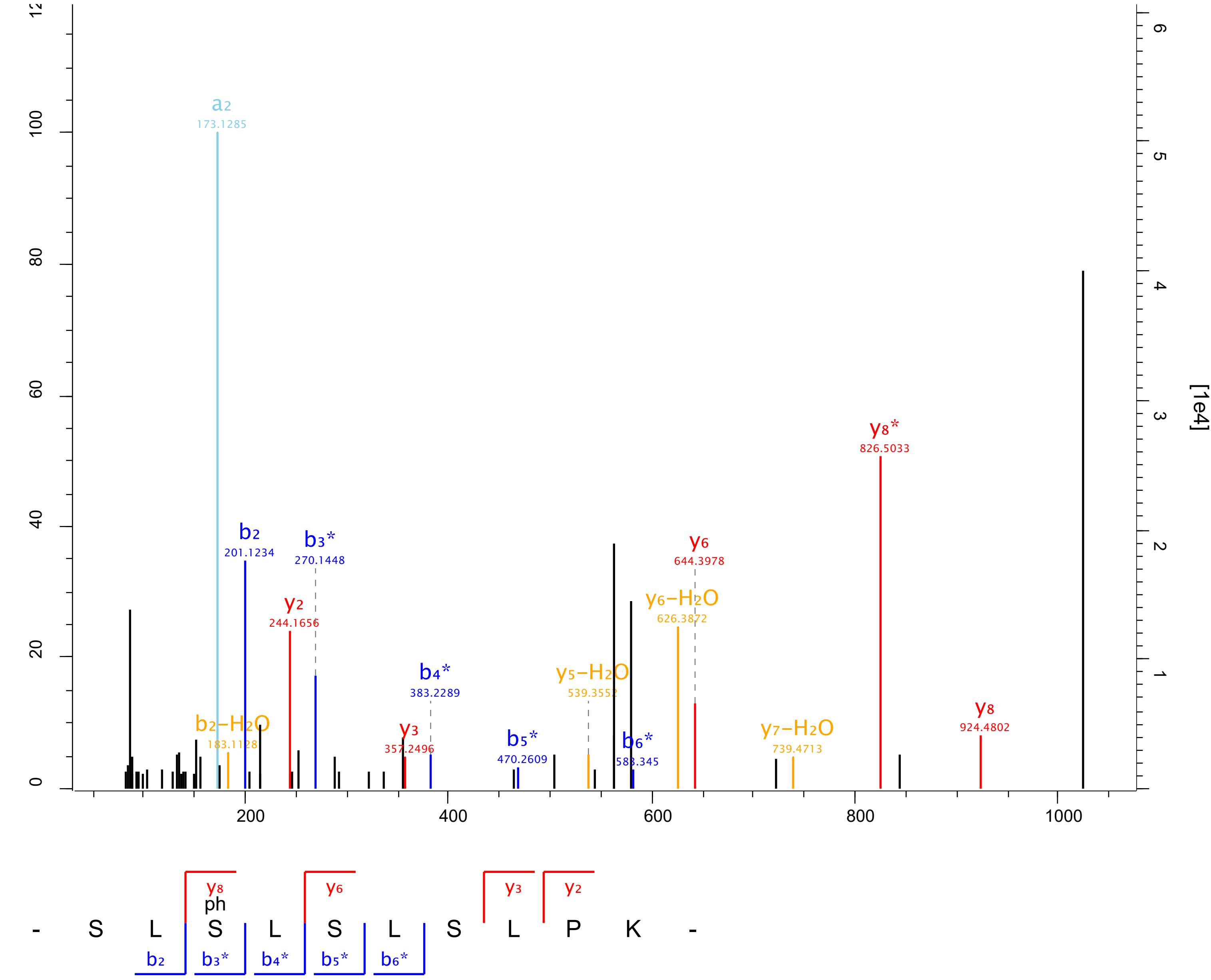

| Raw file | Scan | Method | Score | m/z |
|---|---|---|---|---|
| water_02_1 | 3659 | FTMS; HCD | 62.36 | 467.52 |

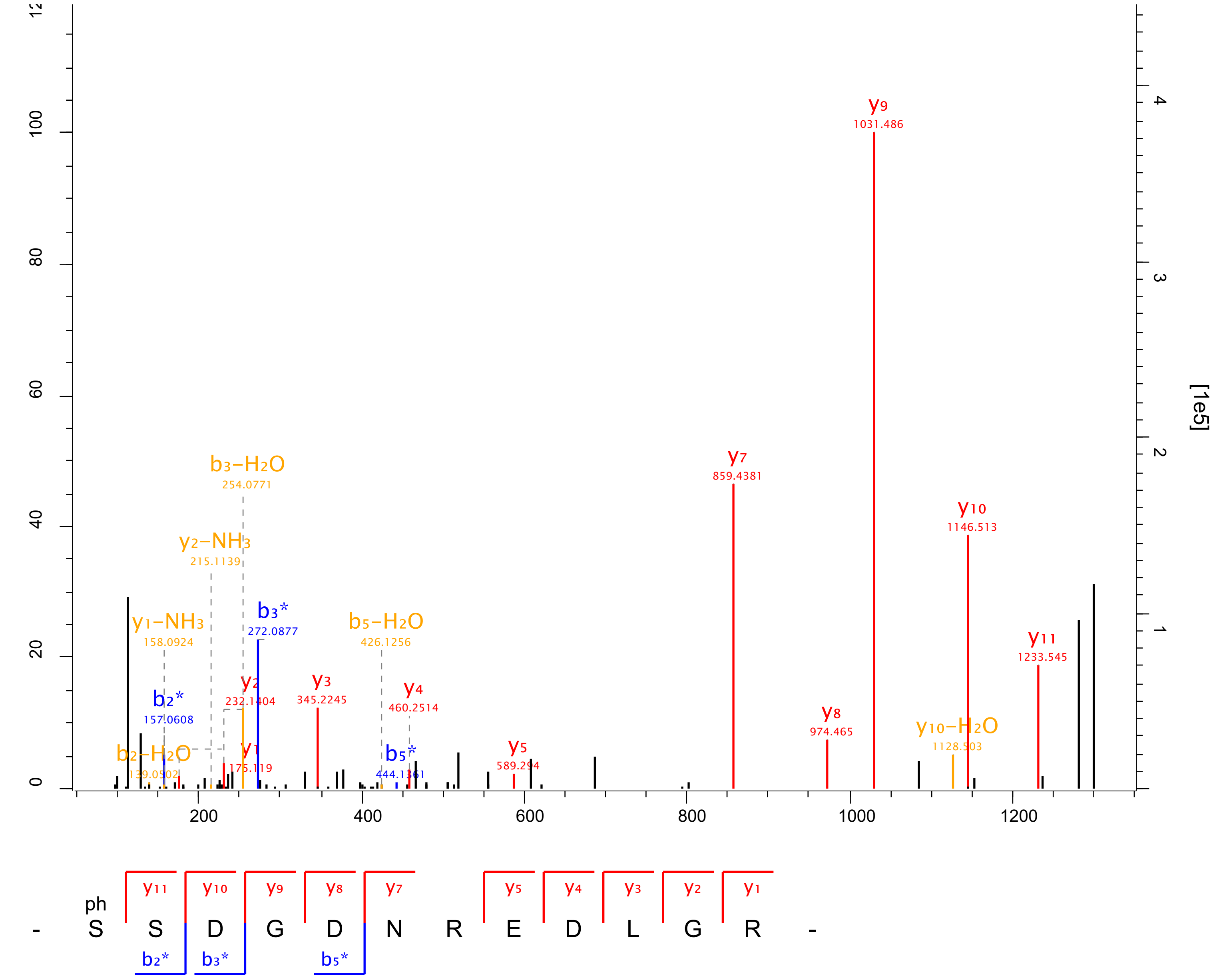

| Raw file | Scan | Method | Score | m/z |
|---|---|---|---|---|
| water_02_1 | 36680 | FTMS; HCD | 123.71 | 853.4 |

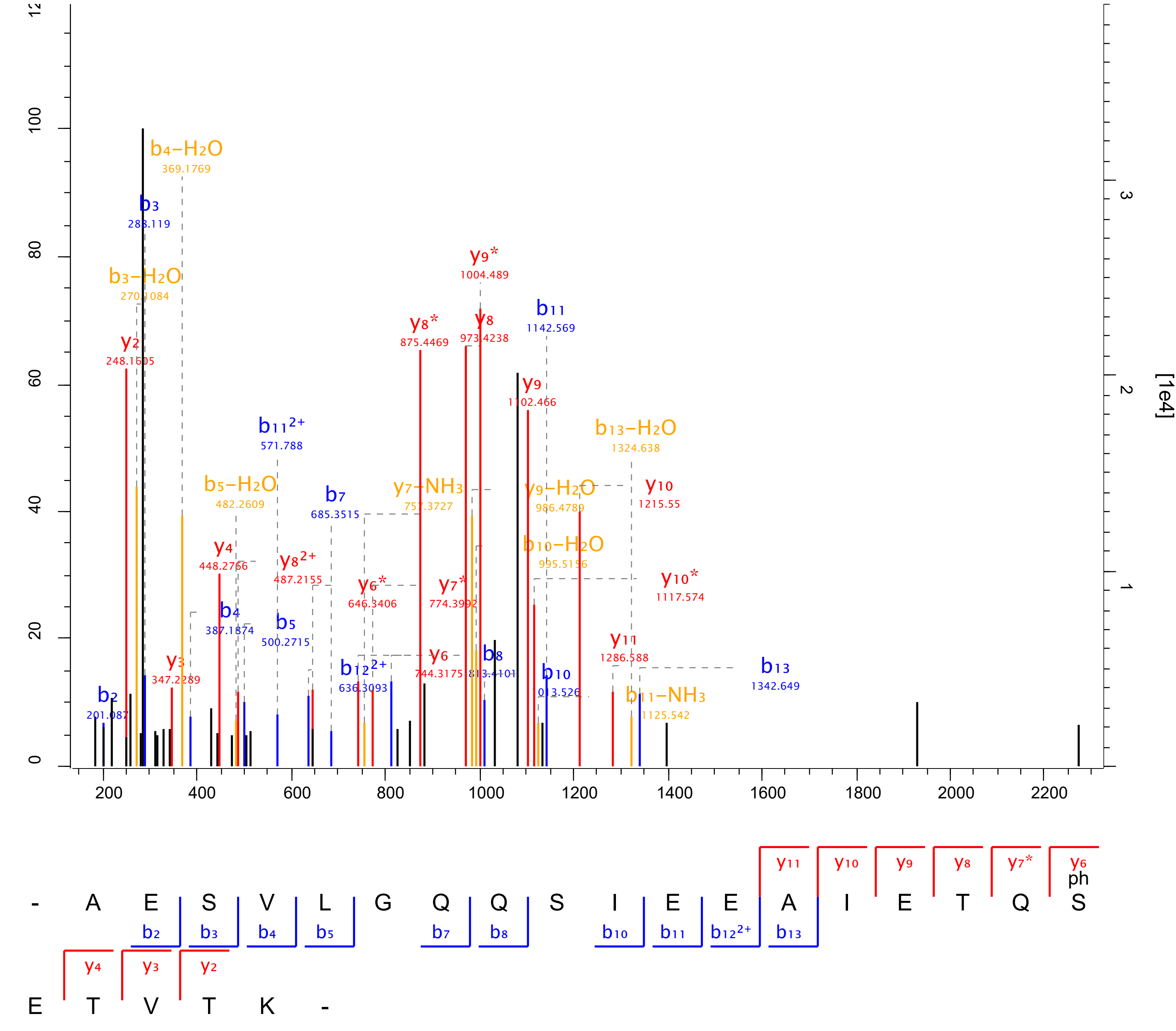

| Raw file | Scan | Method | Score | m/z |
|---|---|---|---|---|
| water_02_1 | 36754 | FTMS; HCD | 81.66 | 725.83 |

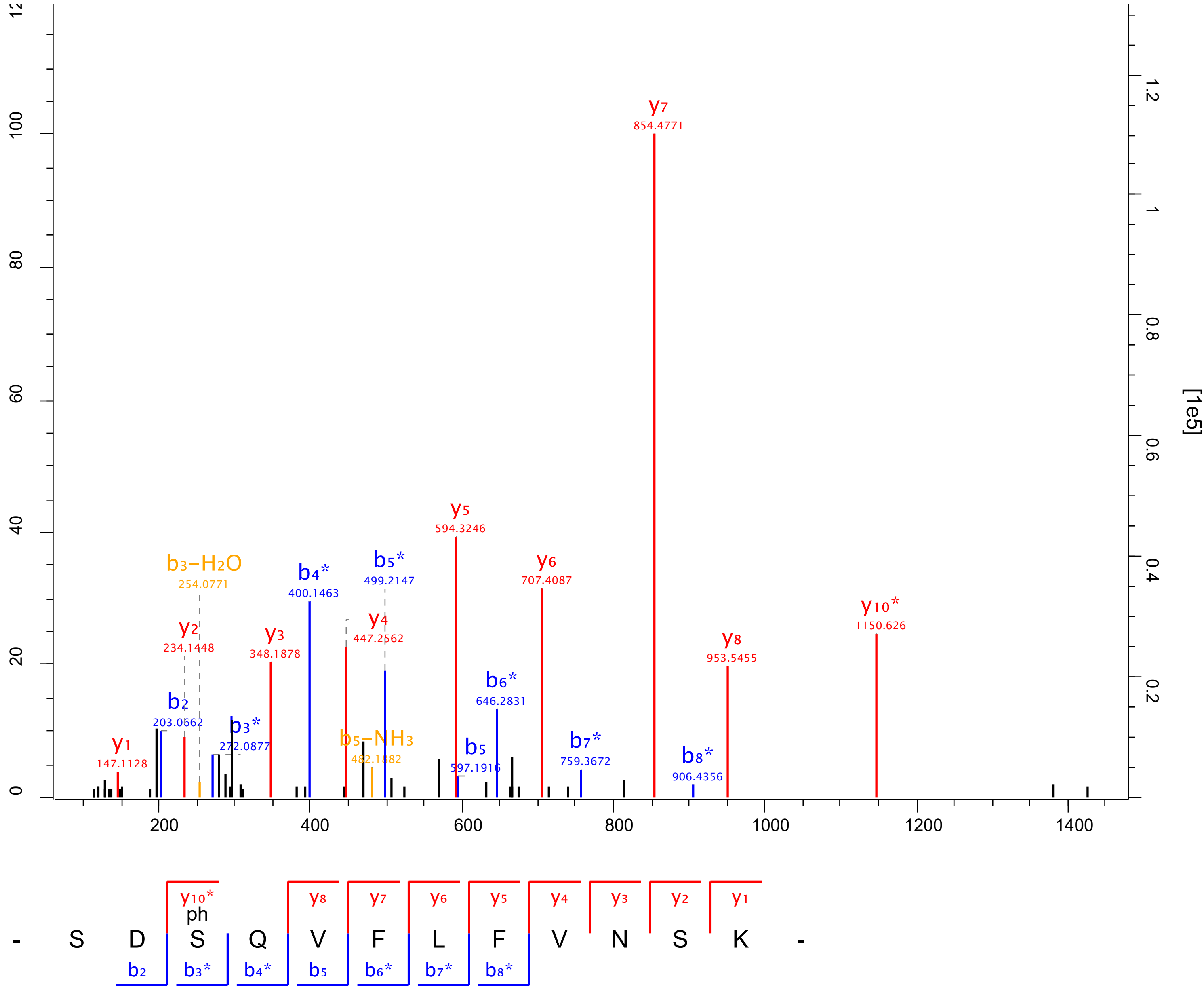

| Raw file | Scan | Method | Score | m/z |
| --- | --- | --- | --- | --- |
| water_02_1 | 36911 | FTMS; HCD | 105.72 | 986.5 |

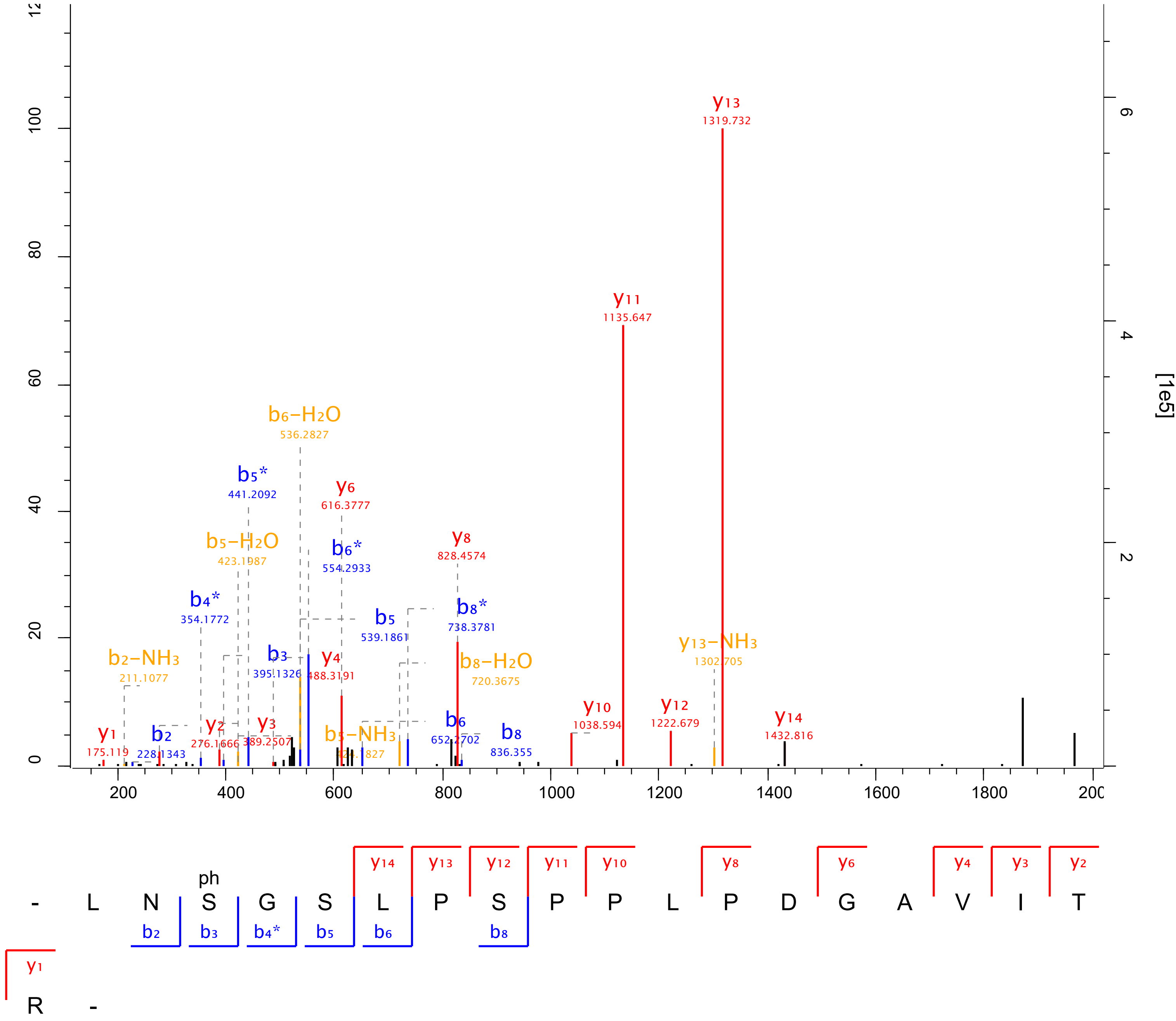

| Raw file | Scan | Method | Score | m/z |
| --- | --- | --- | --- | --- |
| water_02_1 | 3707 | FTMS; HCD | 65.37 | 594.76 |

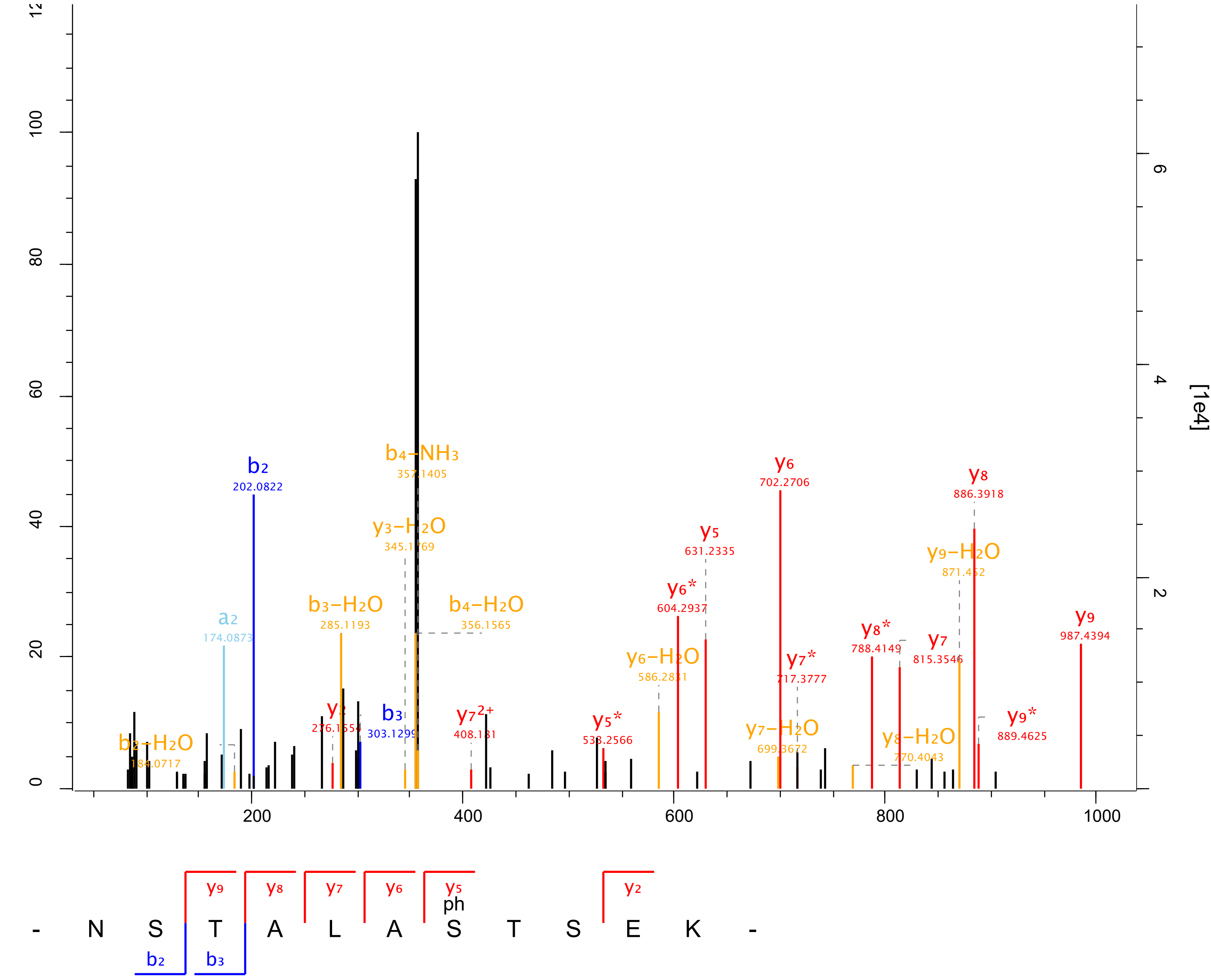

| Raw file | Scan | Method | Score | m/z |
|---|---|---|---|---|
| water_02_1 | 37116 | FTMS; HCD | 154.69 | 809.36 |

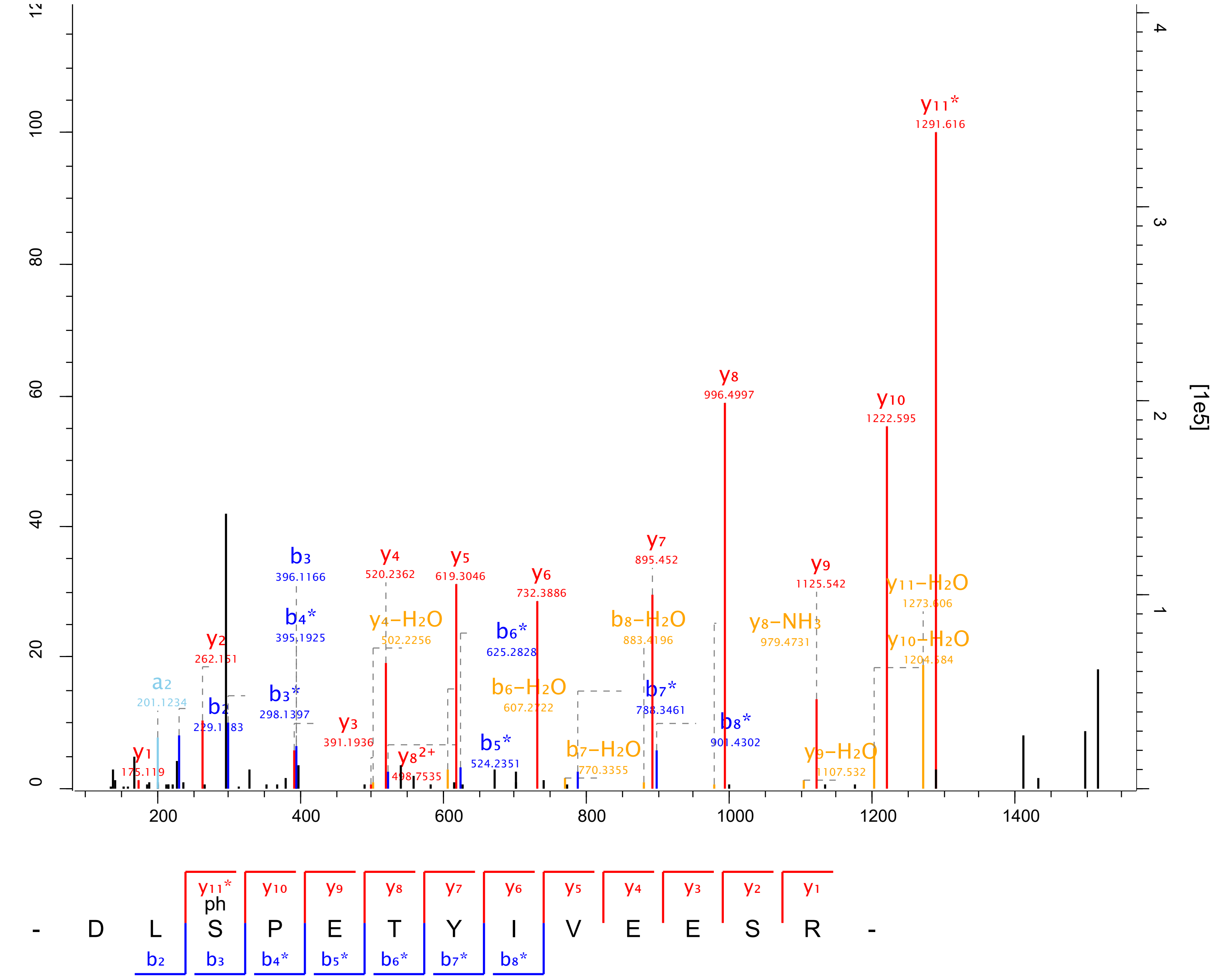

| Raw file | Scan | Method | Score | m/z |
|---|---|---|---|---|
| water_02_1 | 37176 | FTMS; HCD | 126.63 | 862.88 |

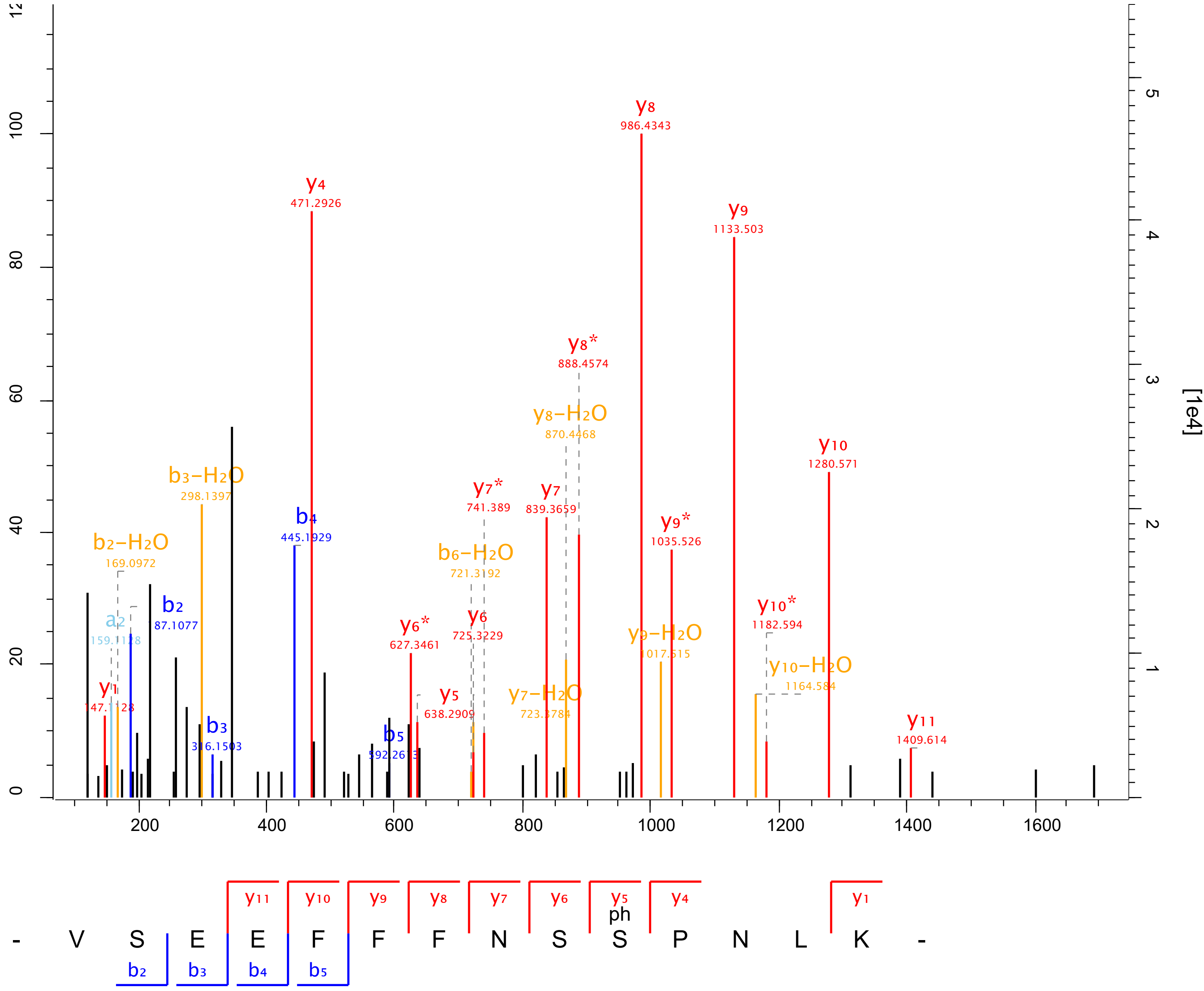

| Raw file | Scan | Method | Score | m/z |
| --- | --- | --- | --- | --- |
| water_02_1 | 3722 | FTMS; HCD | 48.57 | 548.74 |

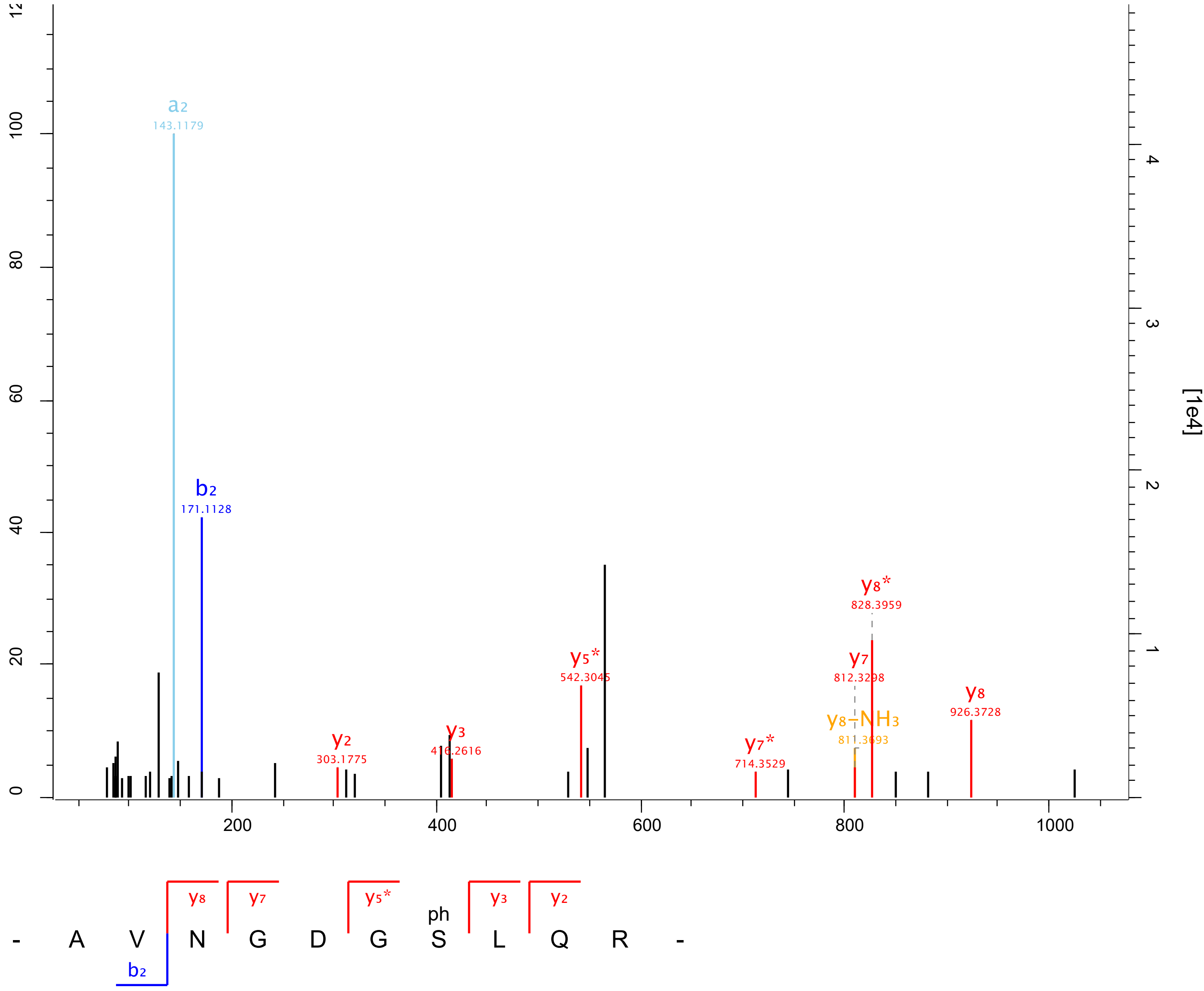

| Raw file | Scan | Method | Score | m/z |
|---|---|---|---|---|
| water_02_1 | 3729 | FTMS; HCD | 65.22 | 394.16 |

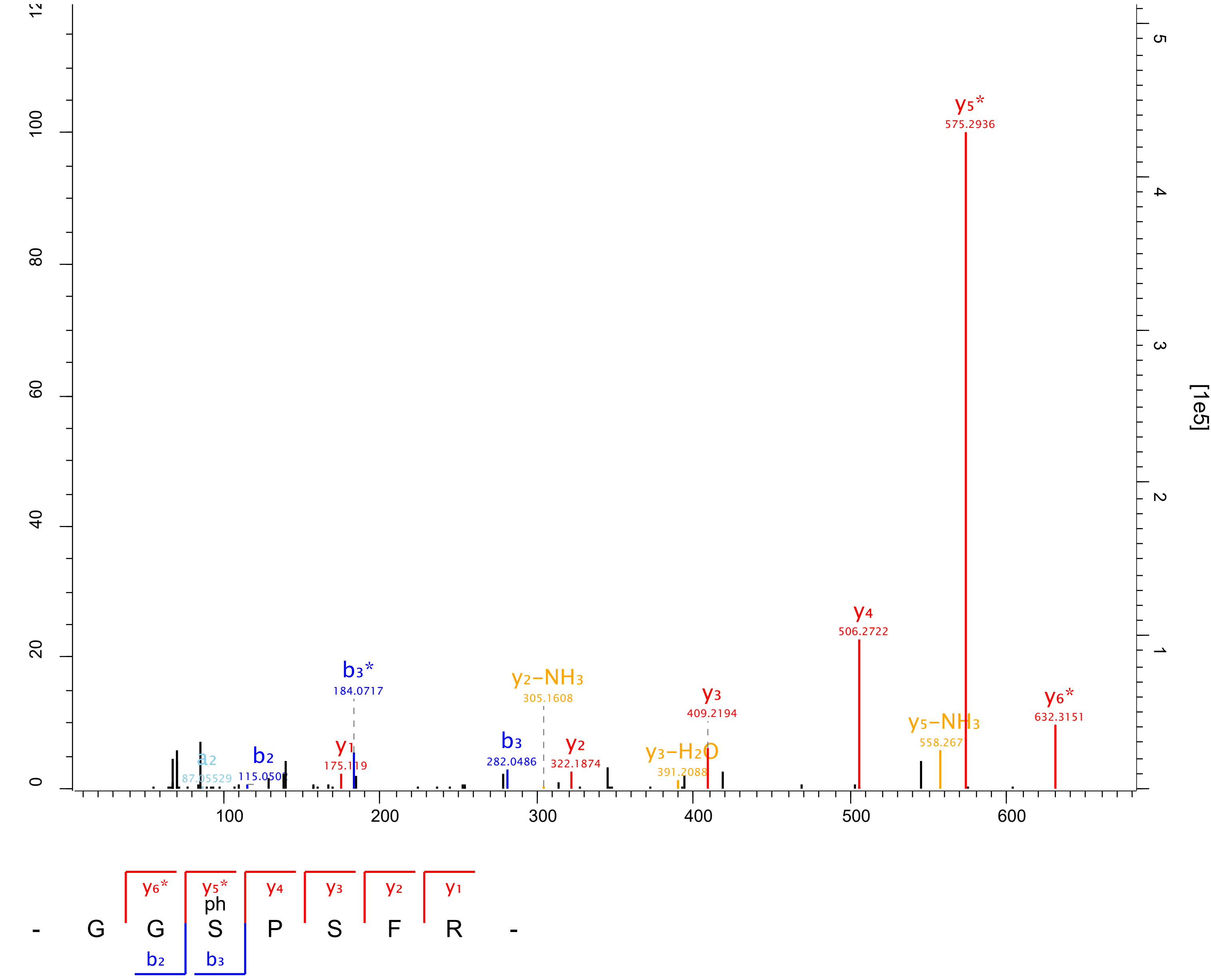

| Raw file | Scan | Method | Score | m/z |
|---|---|---|---|---|
| water_02_1 | 37292 | FTMS; HCD | 64.45 | 759.81 |

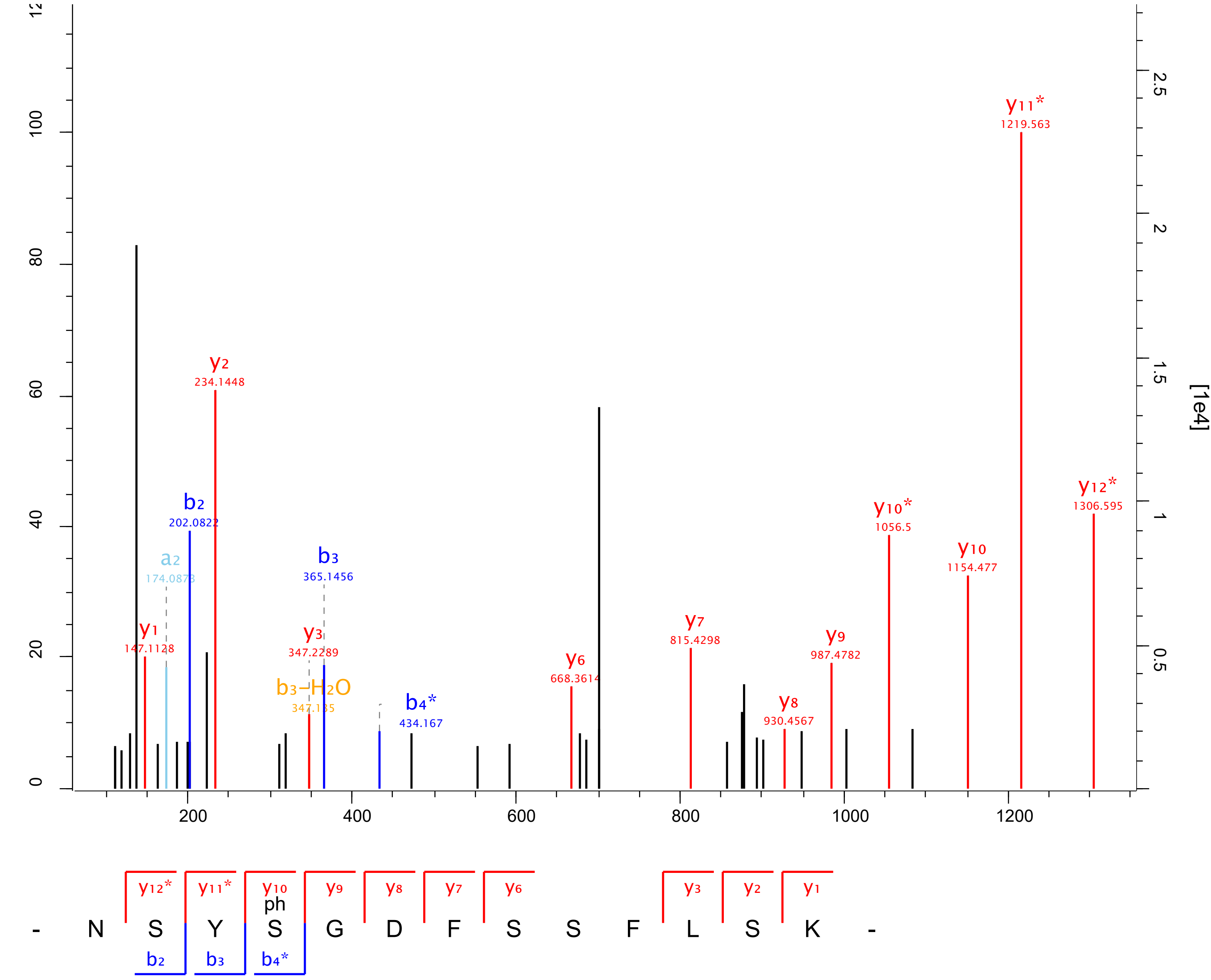

| Raw file | Scan | Method | Score | m/z |
| --- | --- | --- | --- | --- |
| water_02_1 | 37324 | FTMS; HCD | 95.2 | 797.89 |

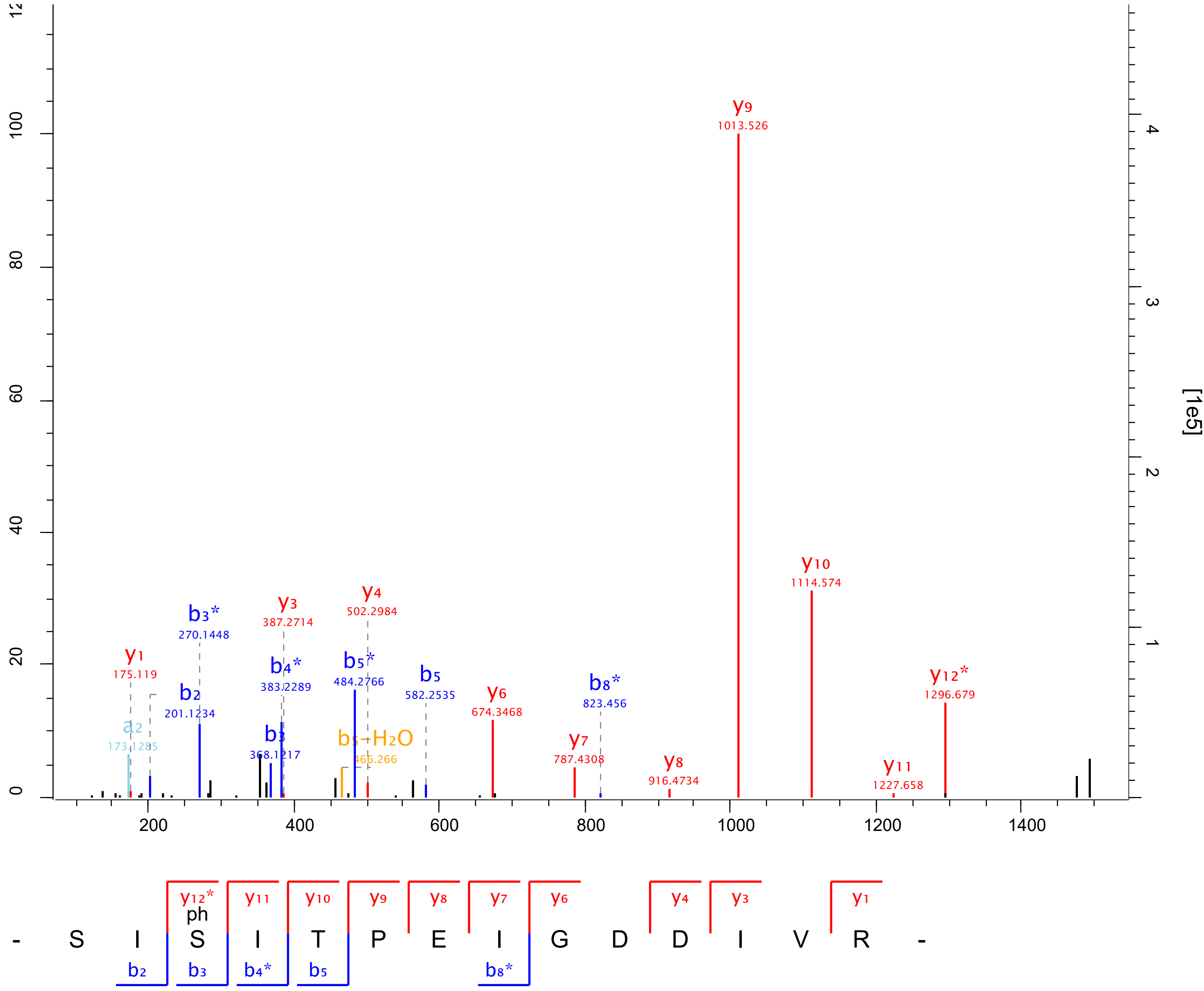

| Raw file | Scan | Method | Score | m/z |
|---|---|---|---|---|
| water_02_1 | 3734 | FTMS; HCD | 50.36 | 802.78 |

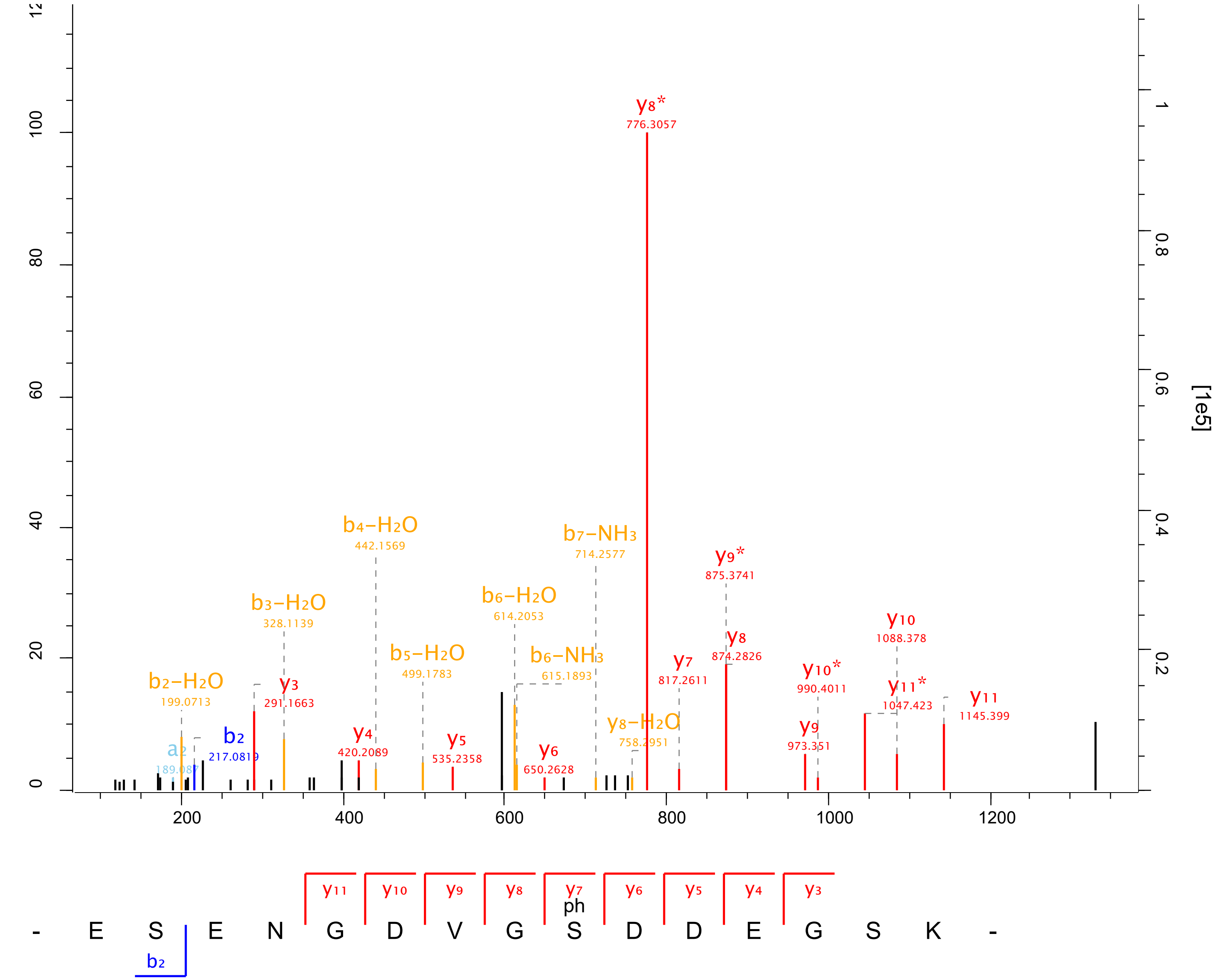

| Raw file | Scan | Method | Score | m/z |
| --- | --- | --- | --- | --- |
| water_02_1 | 37469 | FTMS; HCD | 171.05 | 814.03 |

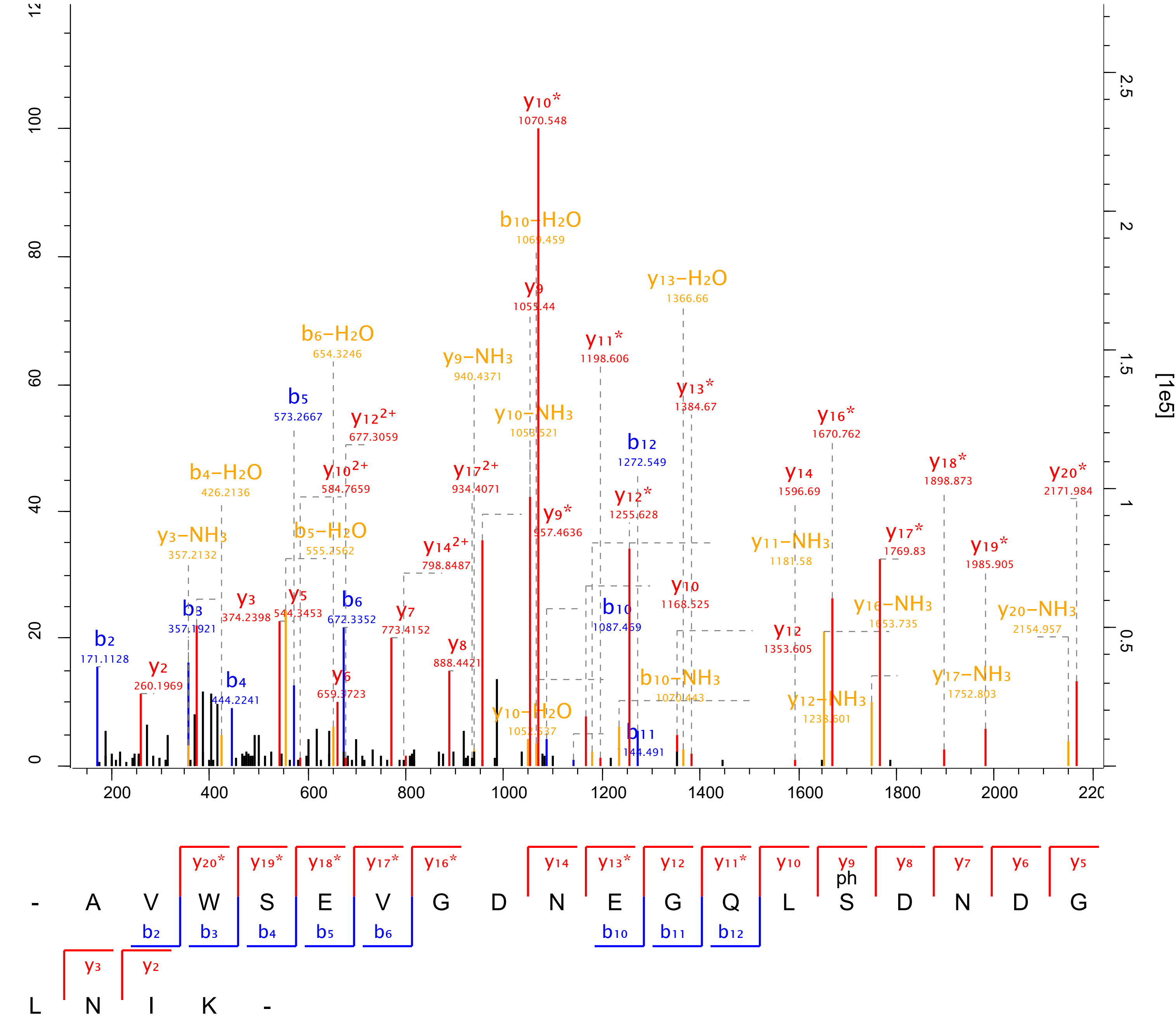

| Raw file | Scan | Method | Score | m/z |
| --- | --- | --- | --- | --- |
| water_02_1 | 37530 | FTMS; HCD | 81.88 | 958.92 |

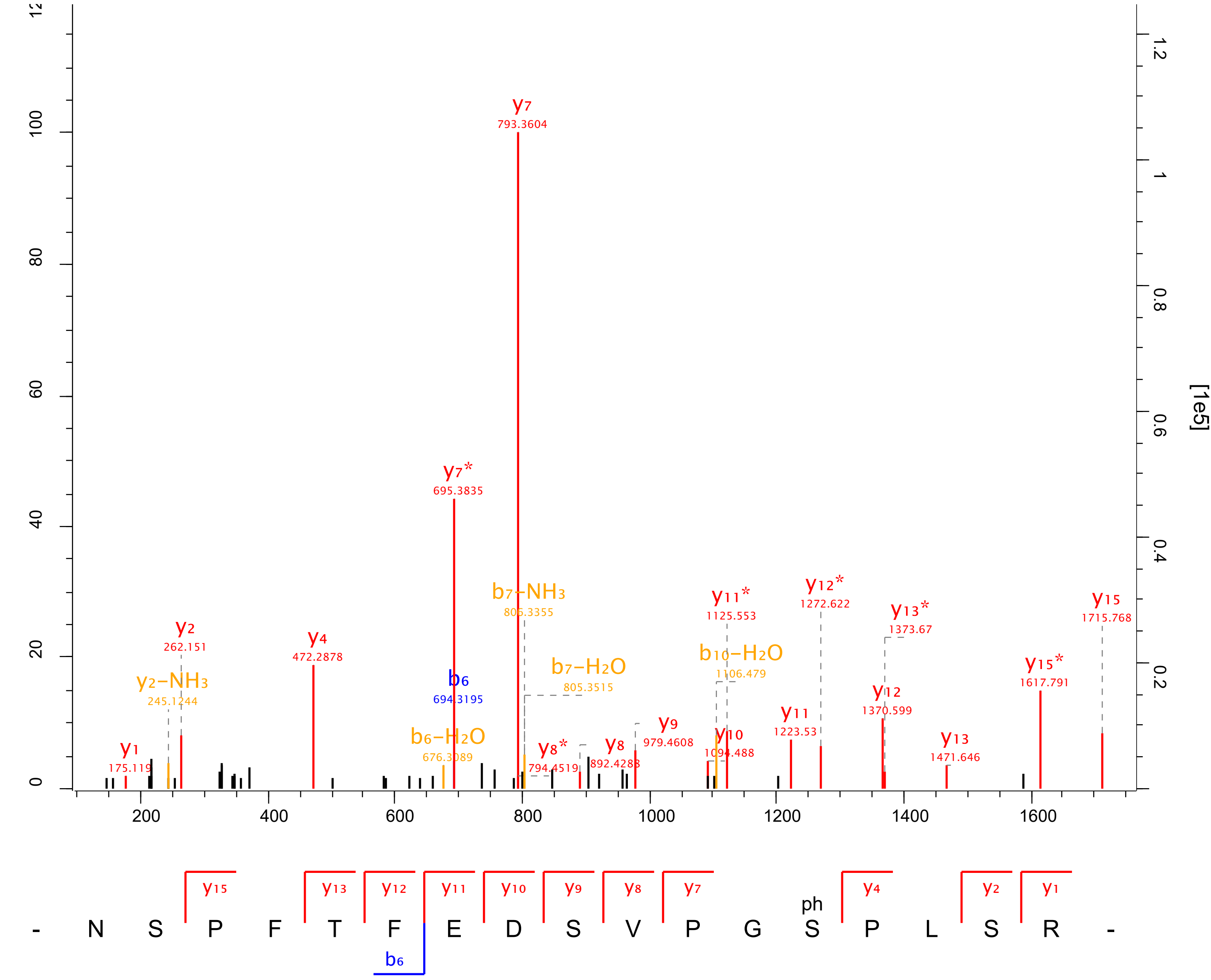

| Raw file | Scan | Method | Score | m/z |
|---|---|---|---|---|
| water_02_1 | 3762 | FTMS; HCD | 87.67 | 483.55 |

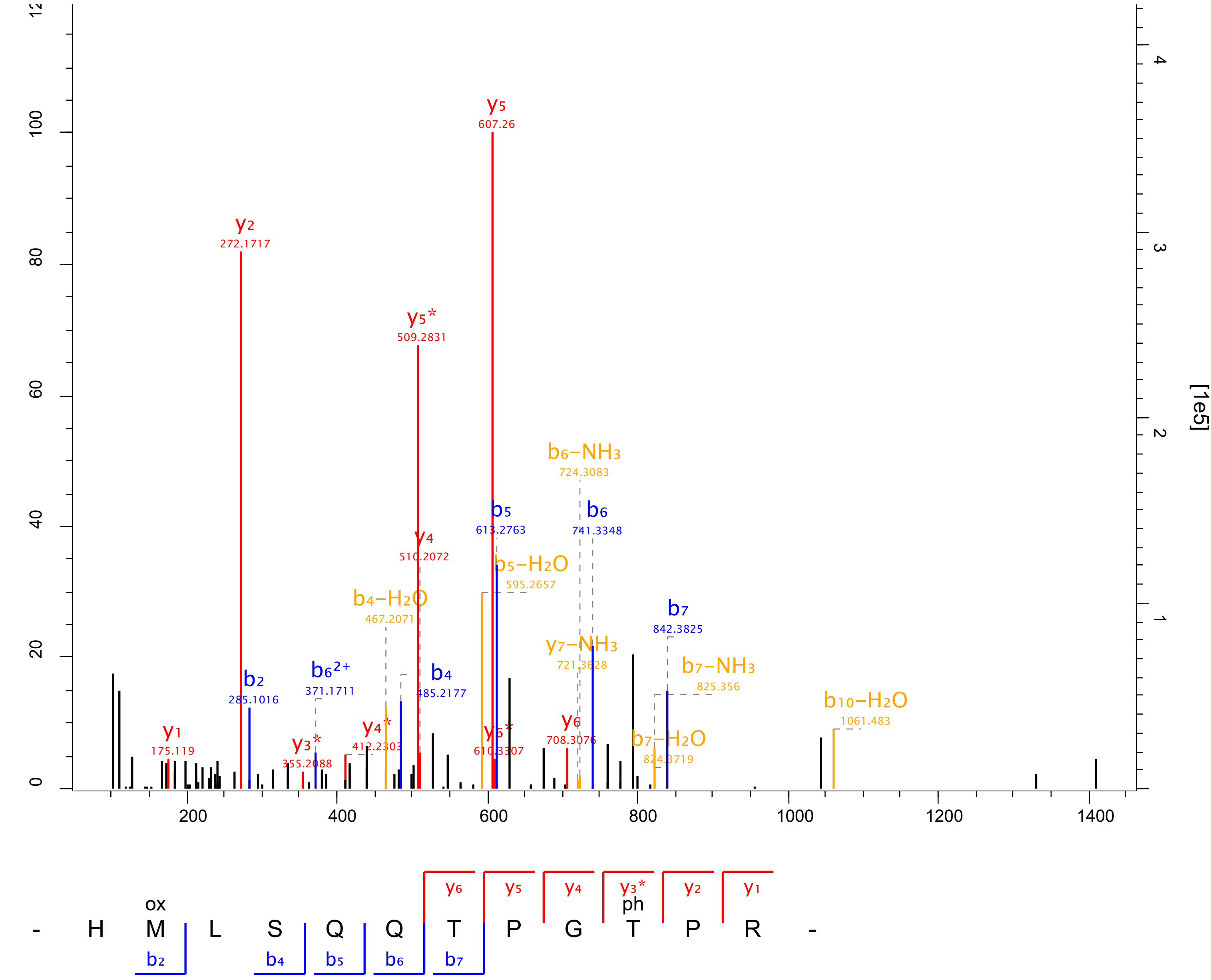

| Raw file | Scan | Method | Score | m/z |
| --- | --- | --- | --- | --- |
| water_02_1 | 37791 | FTMS; HCD | 42.32 | 1005.09 |

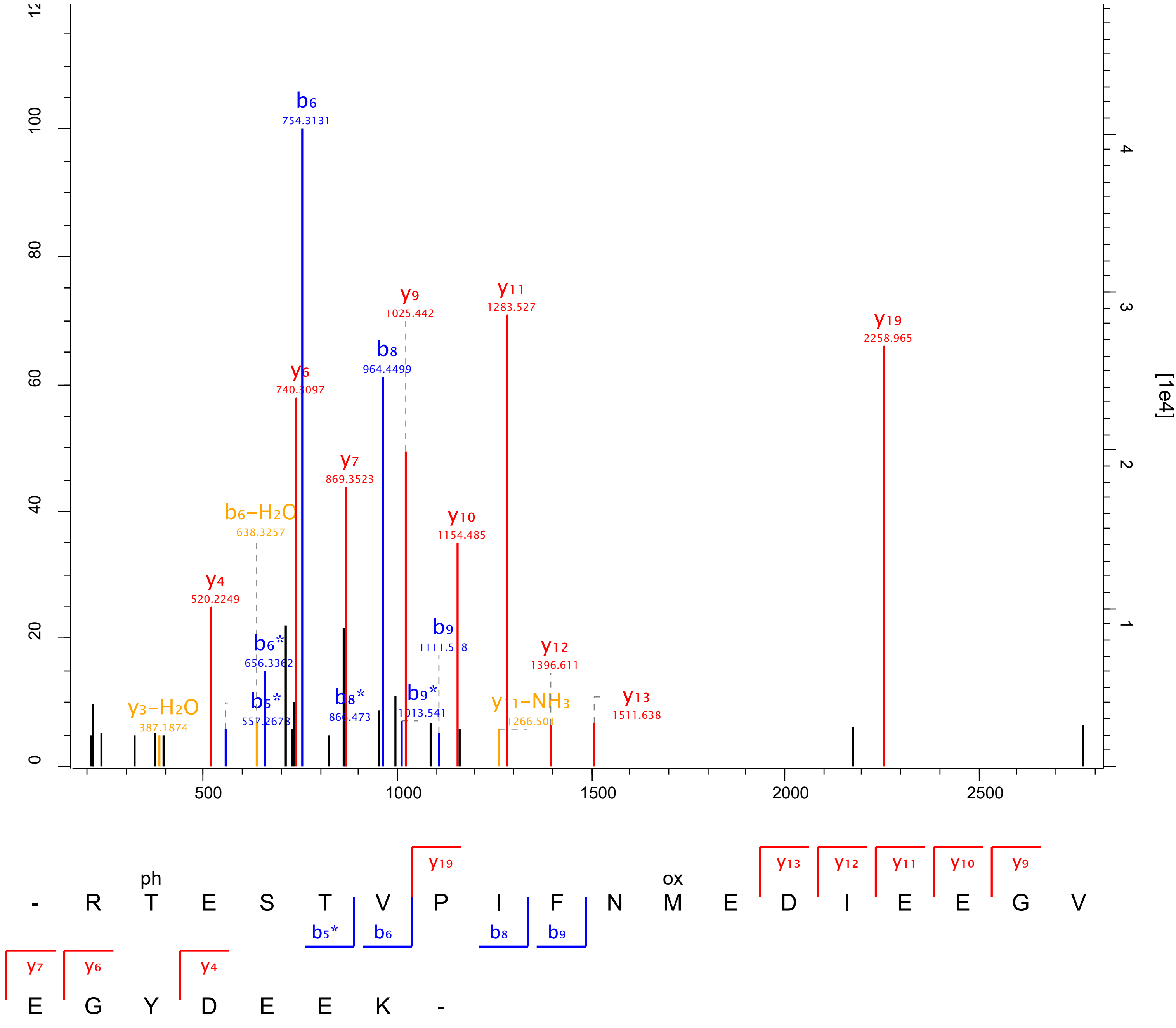

| Raw file | Scan | Method | Score | m/z |
|---|---|---|---|---|
| water_02_1 | 3795 | FTMS; HCD | 72.43 | 495.72 |

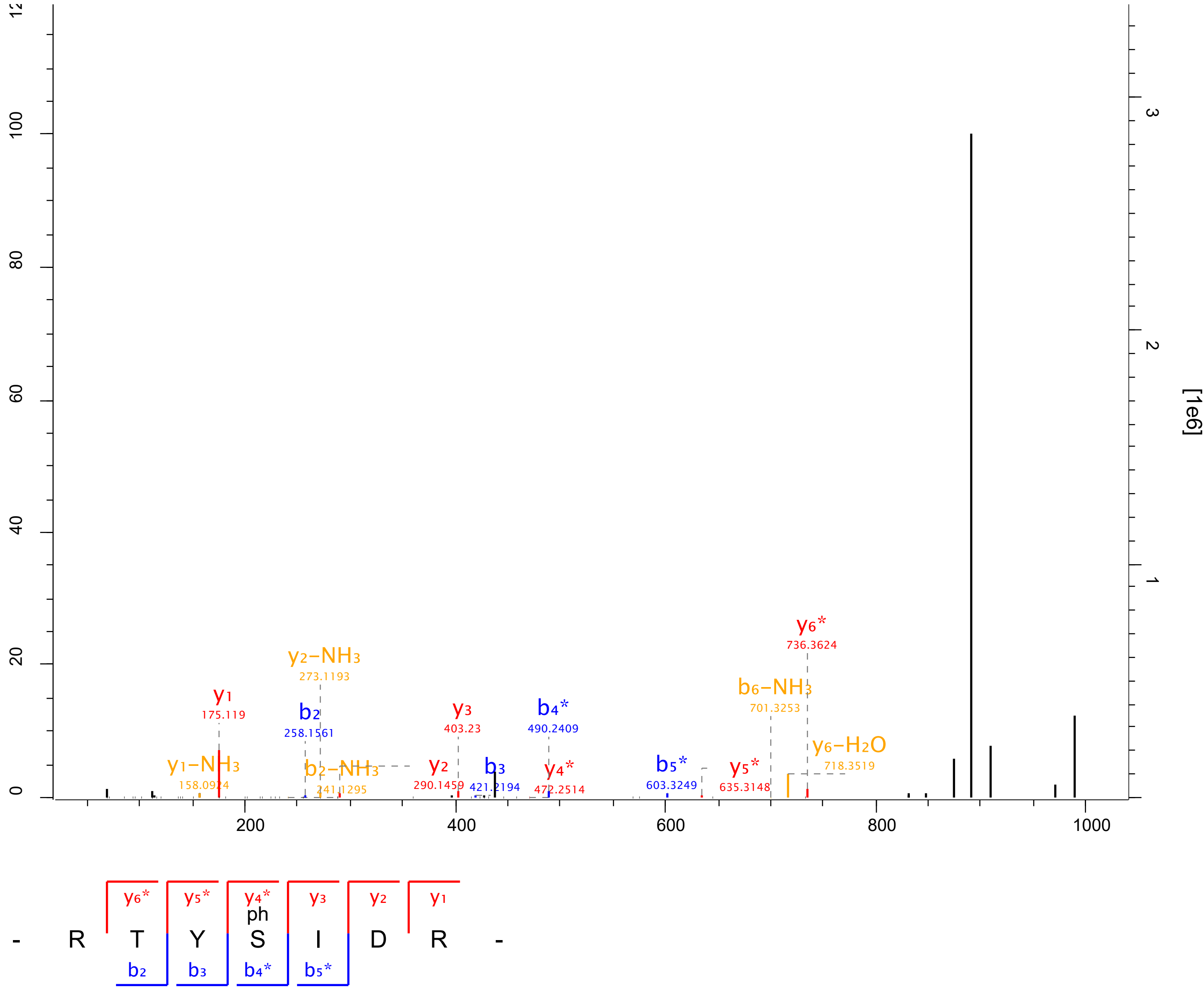

| Raw file | Scan | Method | Score | m/z |
|---|---|---|---|---|
| water_02_1 | 38026 | FTMS; HCD | 132.79 | 876.84 |

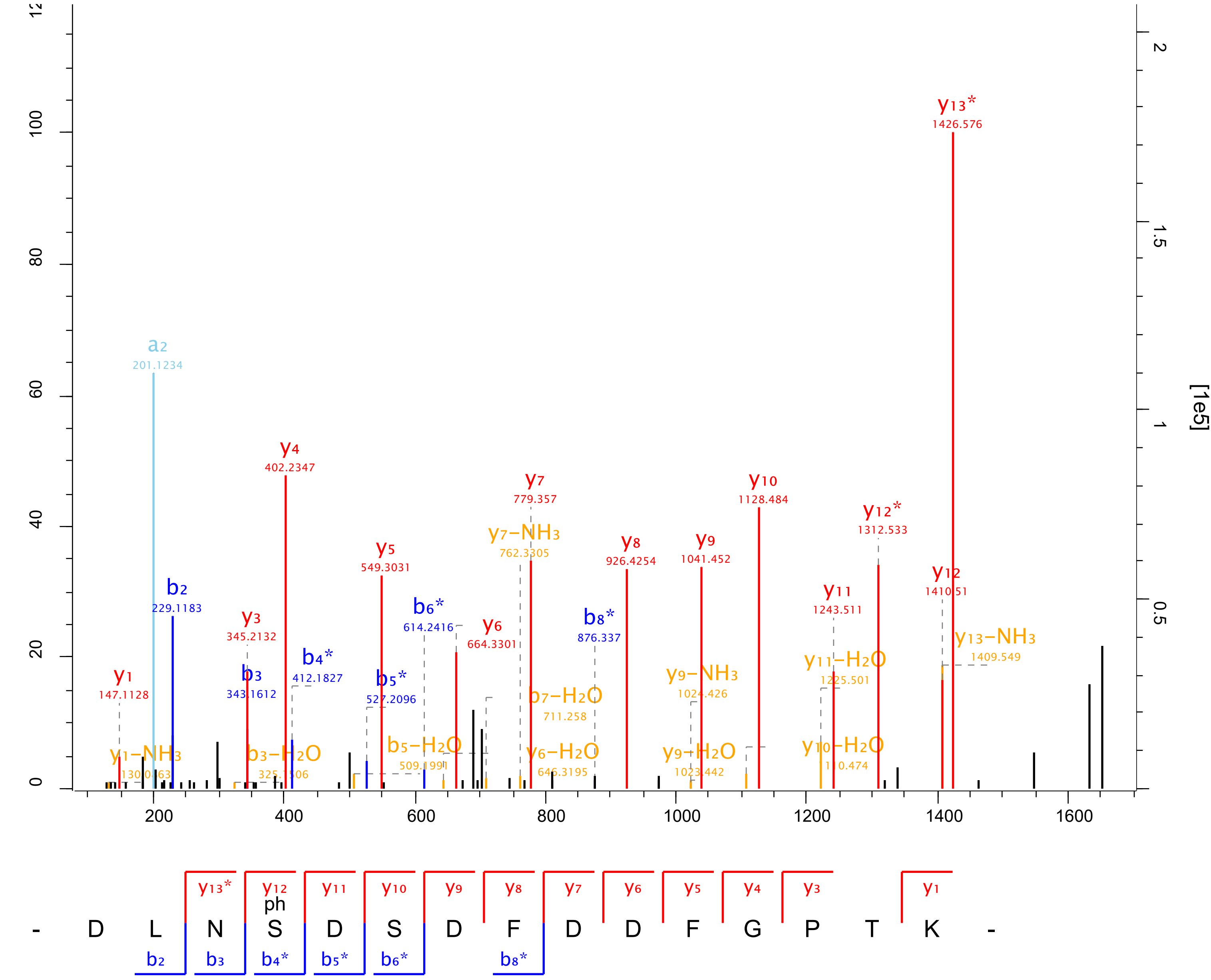

| Raw file | Scan | Method | Score | m/z |
|---|---|---|---|---|
| water_02_1 | 38035 | FTMS; HCD | 49.77 | 908.4 |

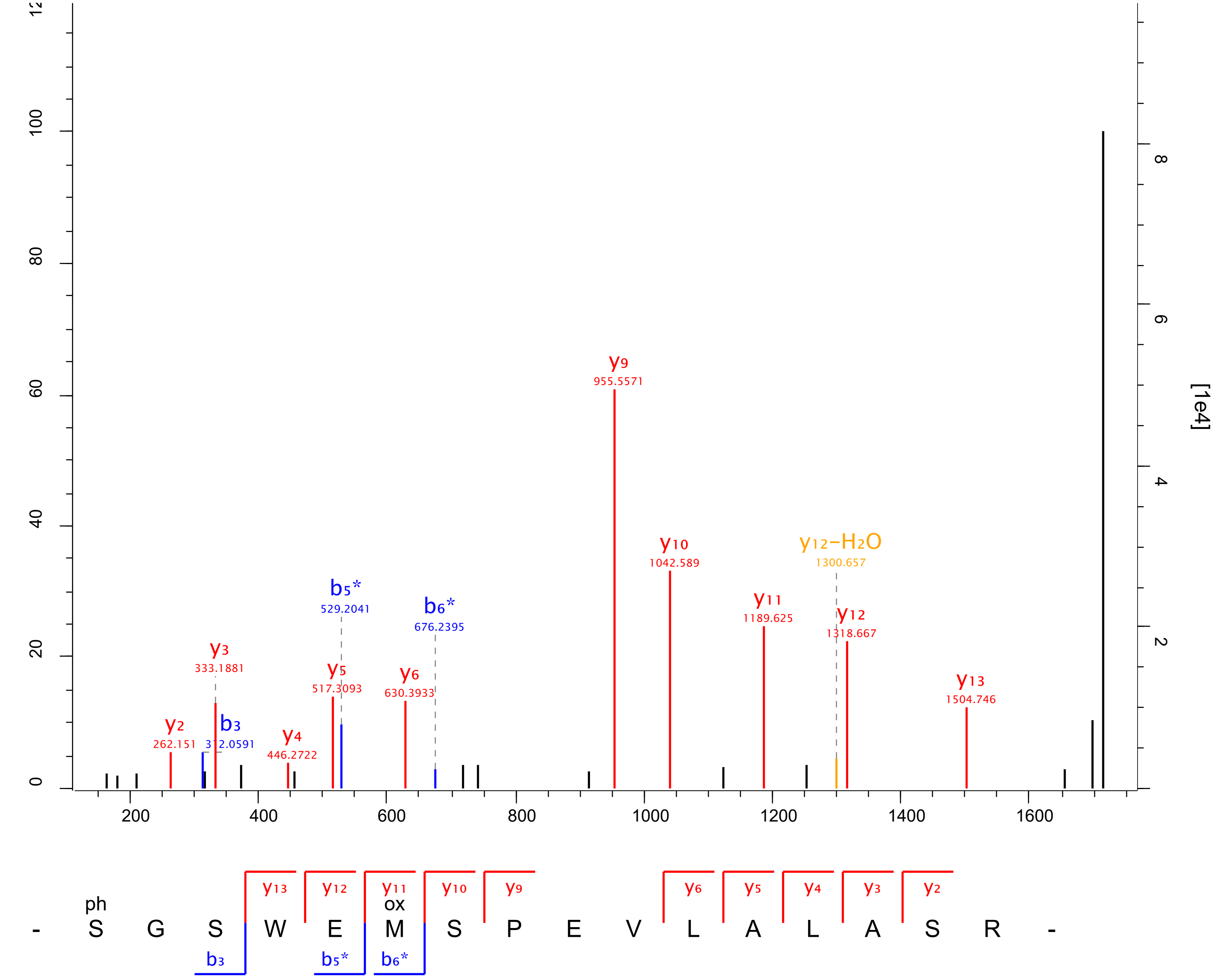

| Raw file | Scan | Method | Score | m/z |
|---|---|---|---|---|
| water_02_1 | 38073 | FTMS; HCD | 57.48 | 1020.47 |

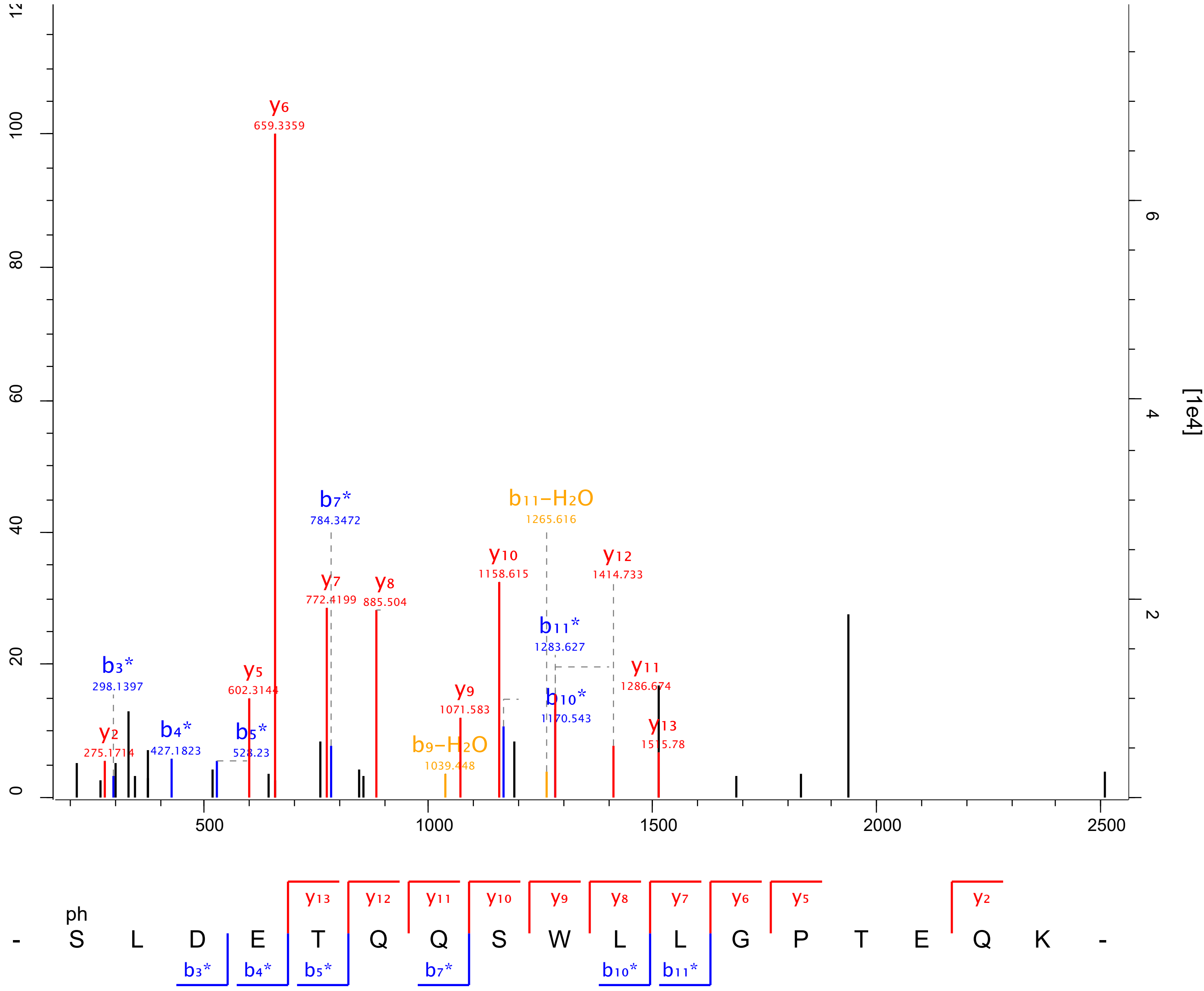

| Raw file | Scan | Method | Score | m/z |
| --- | --- | --- | --- | --- |
| water_02_1 | 38105 | FTMS; HCD | 78 | 983.77 |

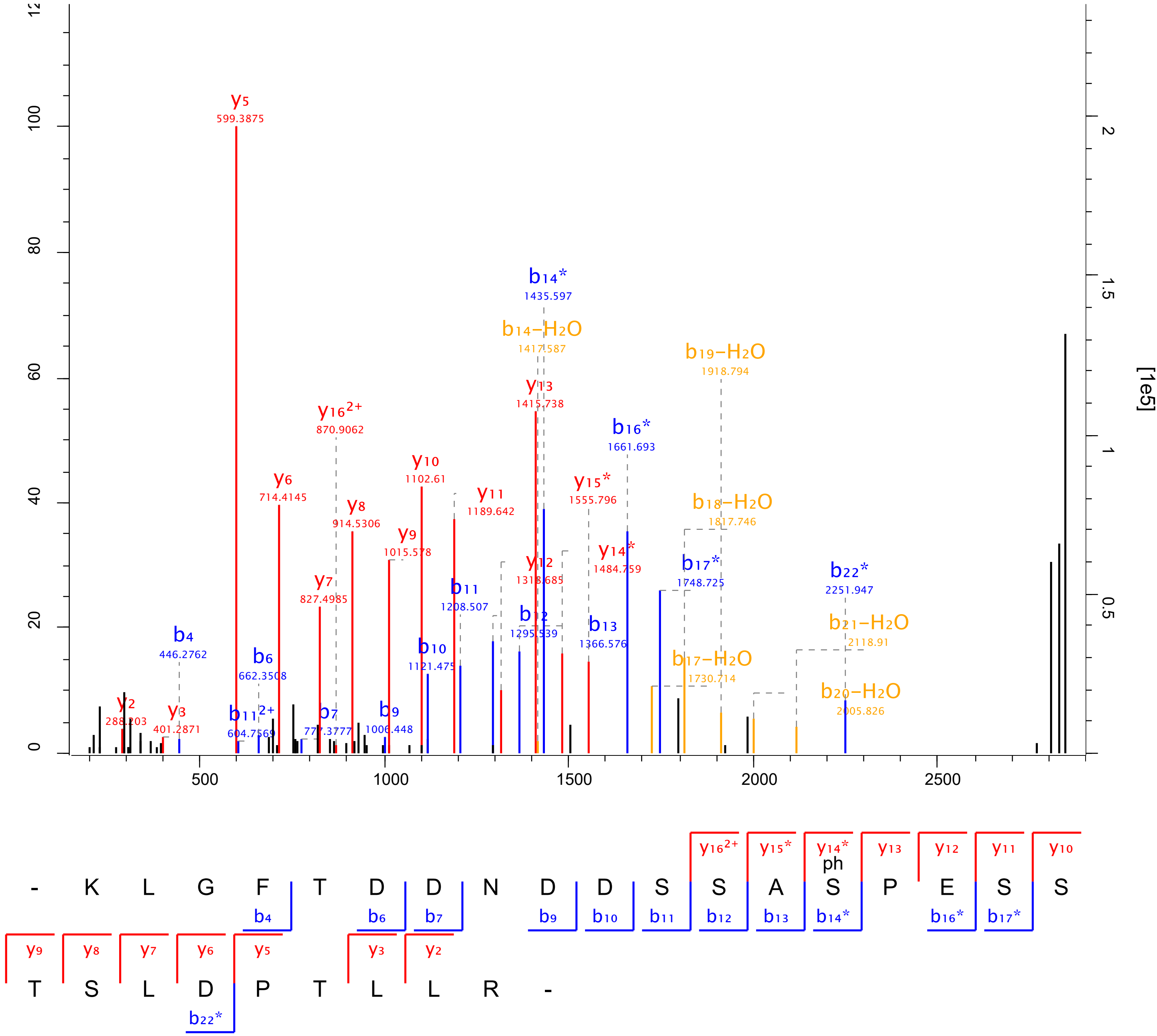

| Raw file | Scan | Method | Score | m/z |
|---|---|---|---|---|
| water_02_1 | 38180 | FTMS; HCD | 205.95 | 805.87 |

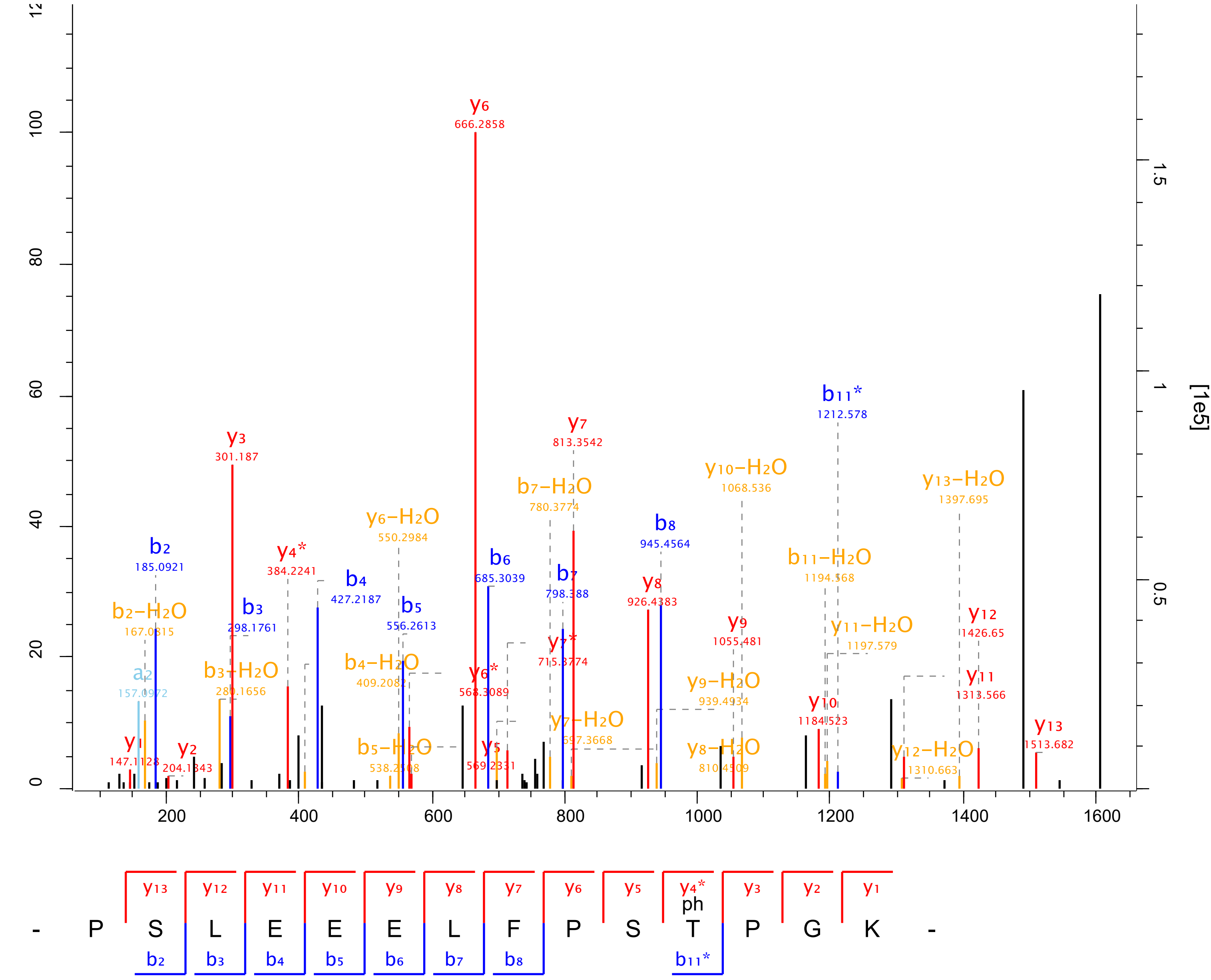

| Raw file | Scan | Method | Score | m/z |
| --- | --- | --- | --- | --- |
| water_02_1 | 38192 | FTMS; HCD | 138.9 | 887.4 |

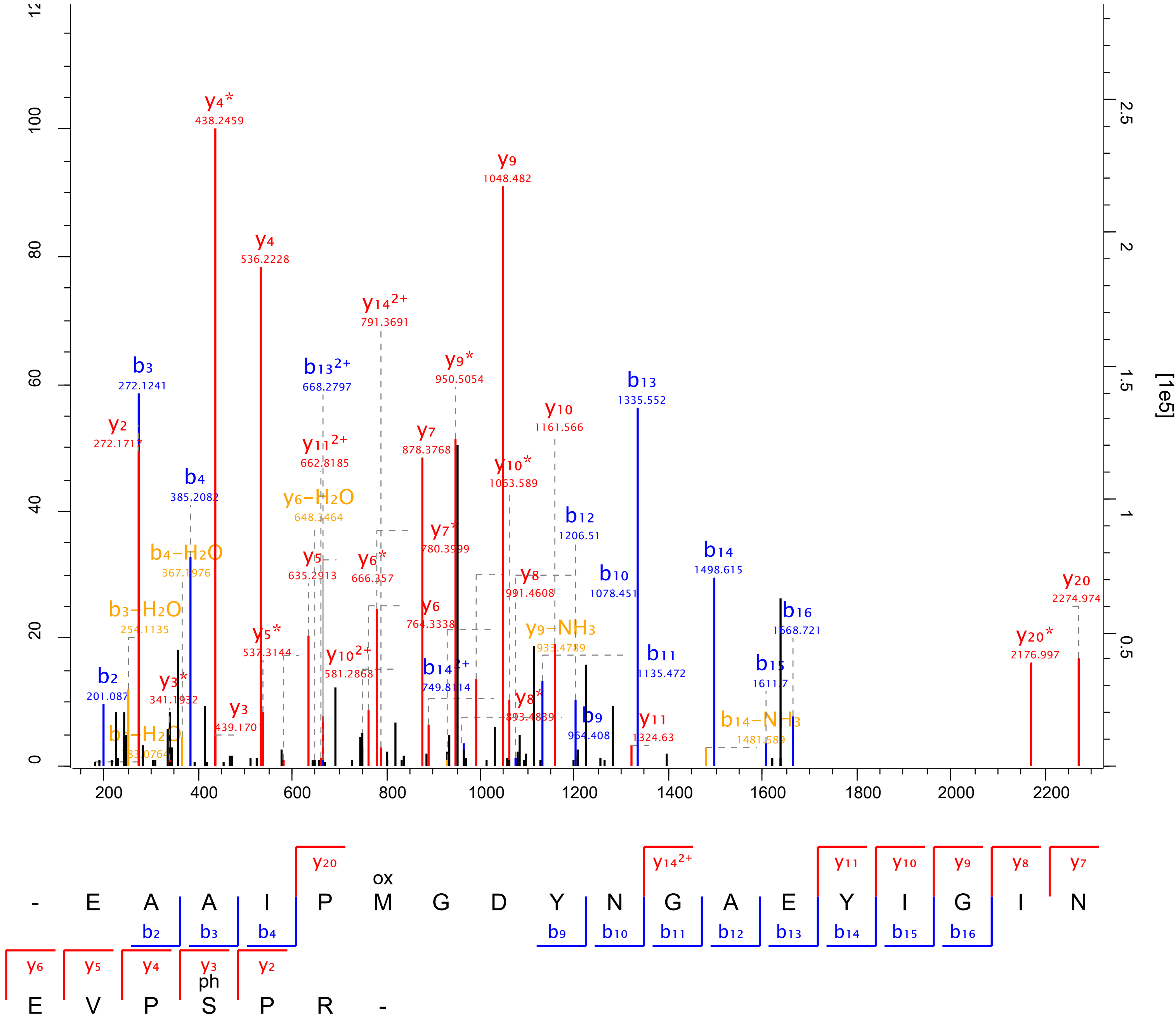

| Raw file | Scan | Method | Score | m/z |
|---|---|---|---|---|
| water_02_1 | 38508 | FTMS; HCD | 137.88 | 887.03 |

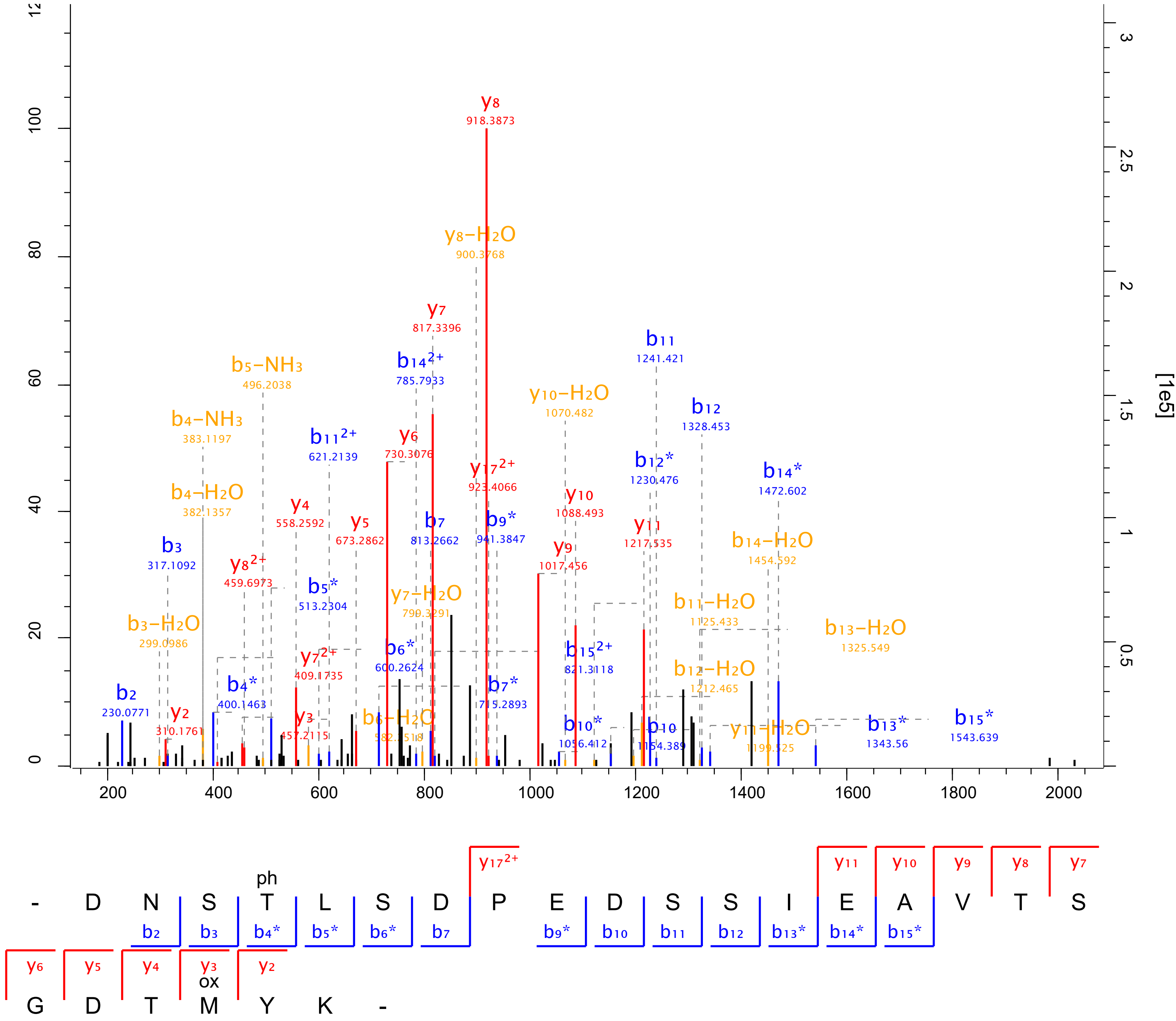

| Raw file | Scan | Method | Score | m/z |
|---|---|---|---|---|
| water_02_1 | 38604 | FTMS; HCD | 52.45 | 785.64 |

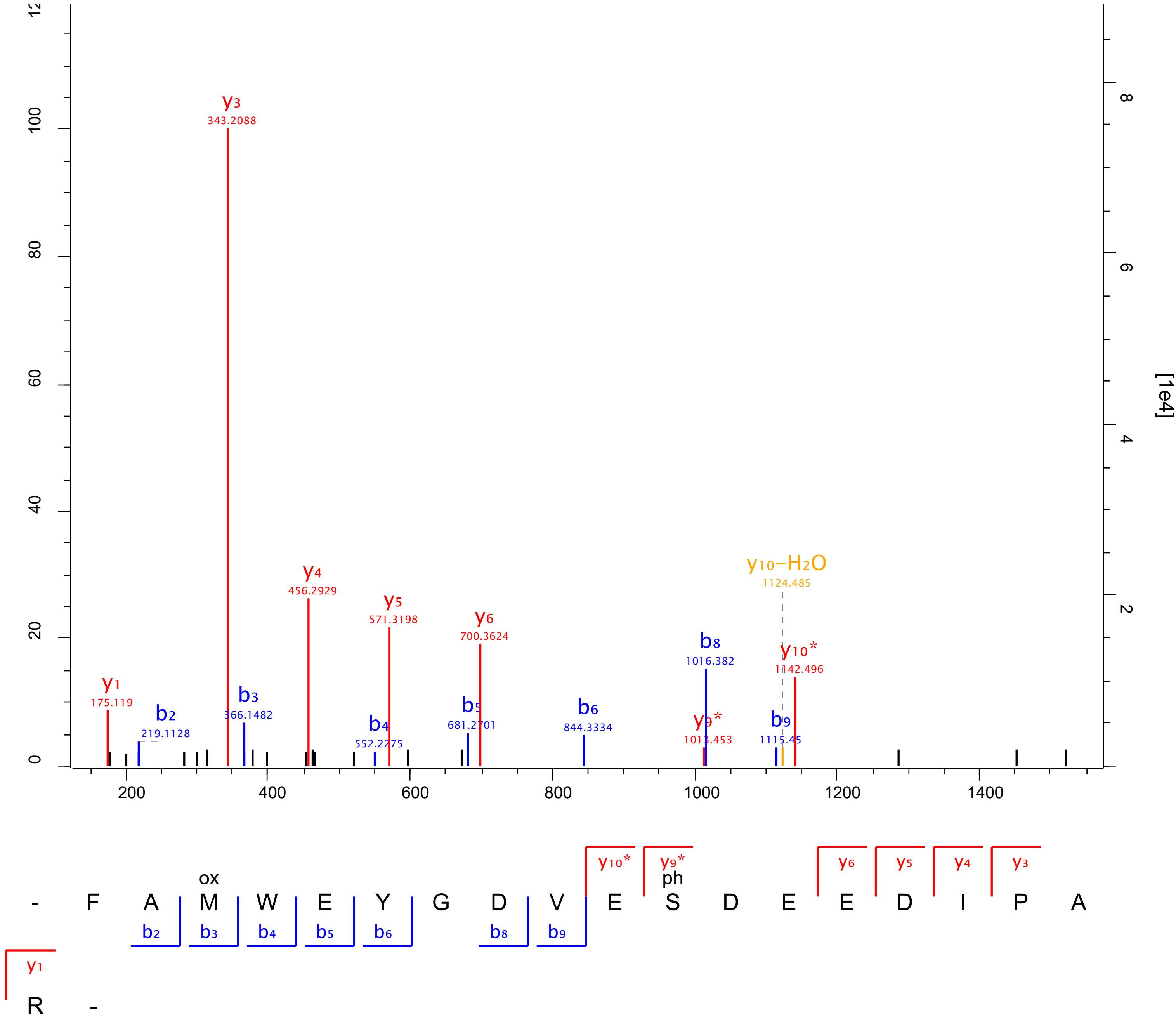

| Raw file | Scan | Method | Score | m/z |
| --- | --- | --- | --- | --- |
| water_02_1 | 38711 | FTMS; HCD | 144.74 | 1184.18 |

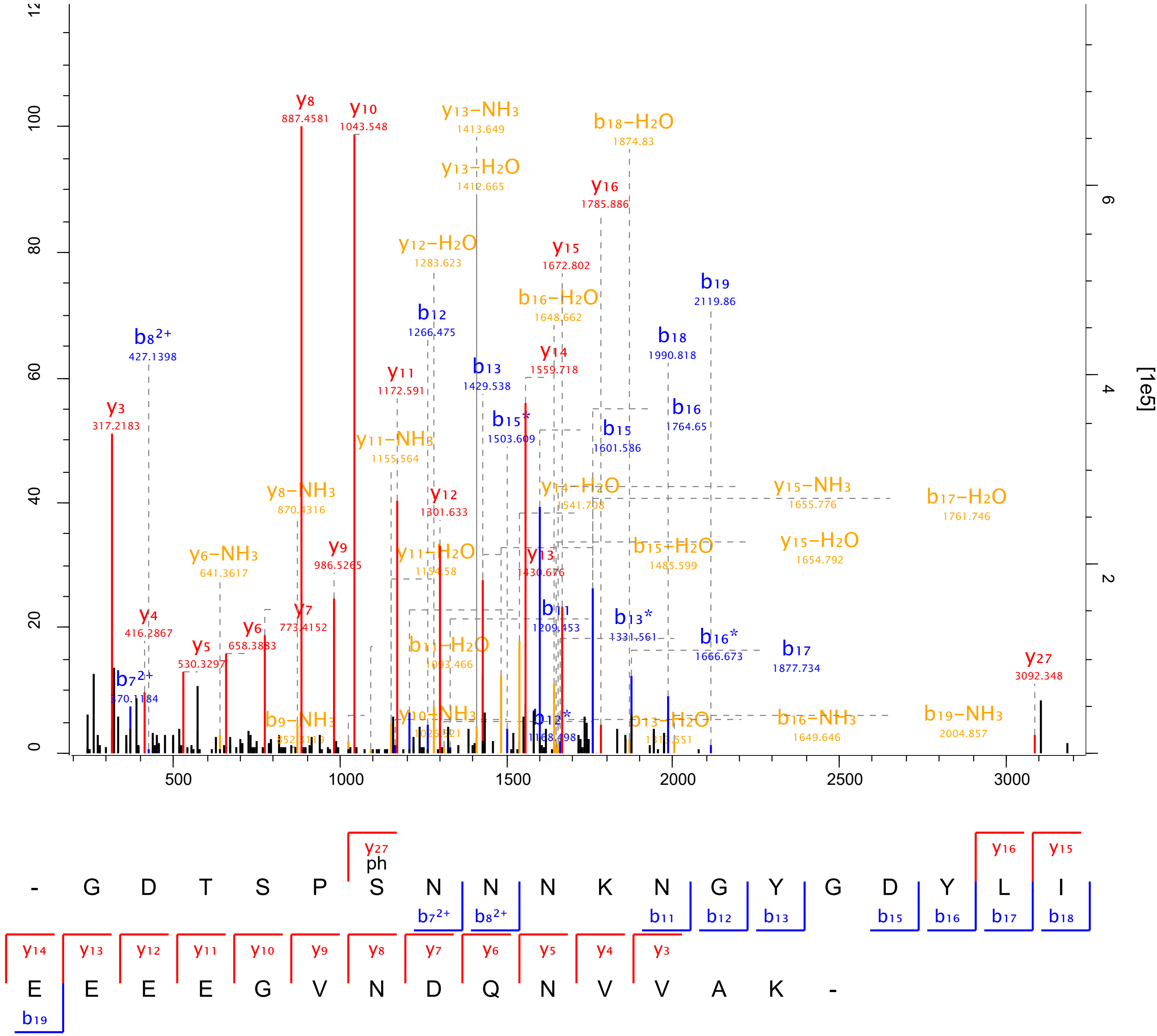

| Raw file | Scan | Method | Score | m/z |
|---|---|---|---|---|
| water_02_1 | 38913 | FTMS; HCD | 53.45 | 837.89 |

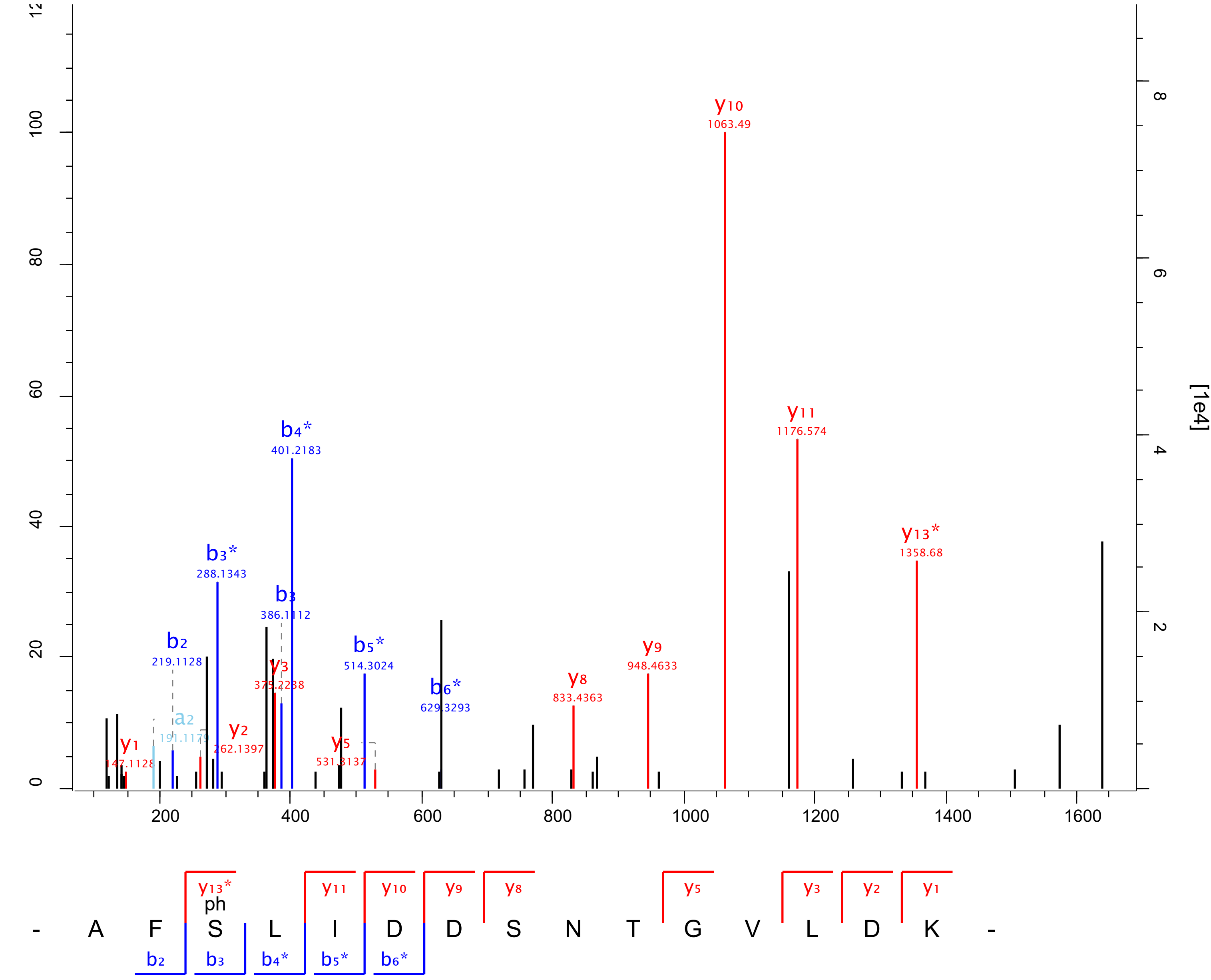

| Raw file | Scan | Method | Score | m/z |
|---|---|---|---|---|
| water_02_1 | 39063 | FTMS; HCD | 47.81 | 754.71 |

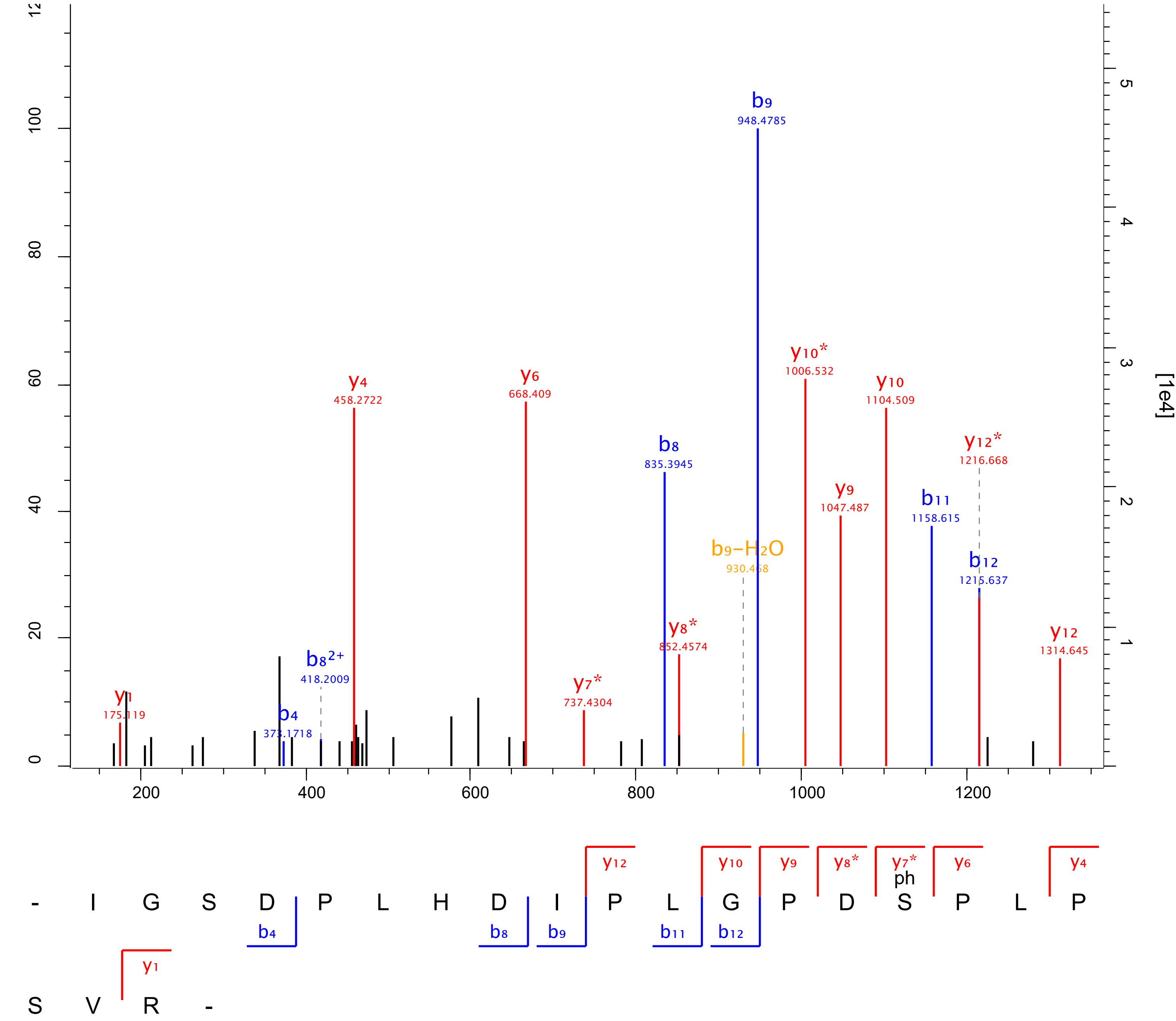

| Raw file | Scan | Method | Score | m/z |
|---|---|---|---|---|
| water_02_1 | 39125 | FTMS; HCD | 166.48 | 747.88 |

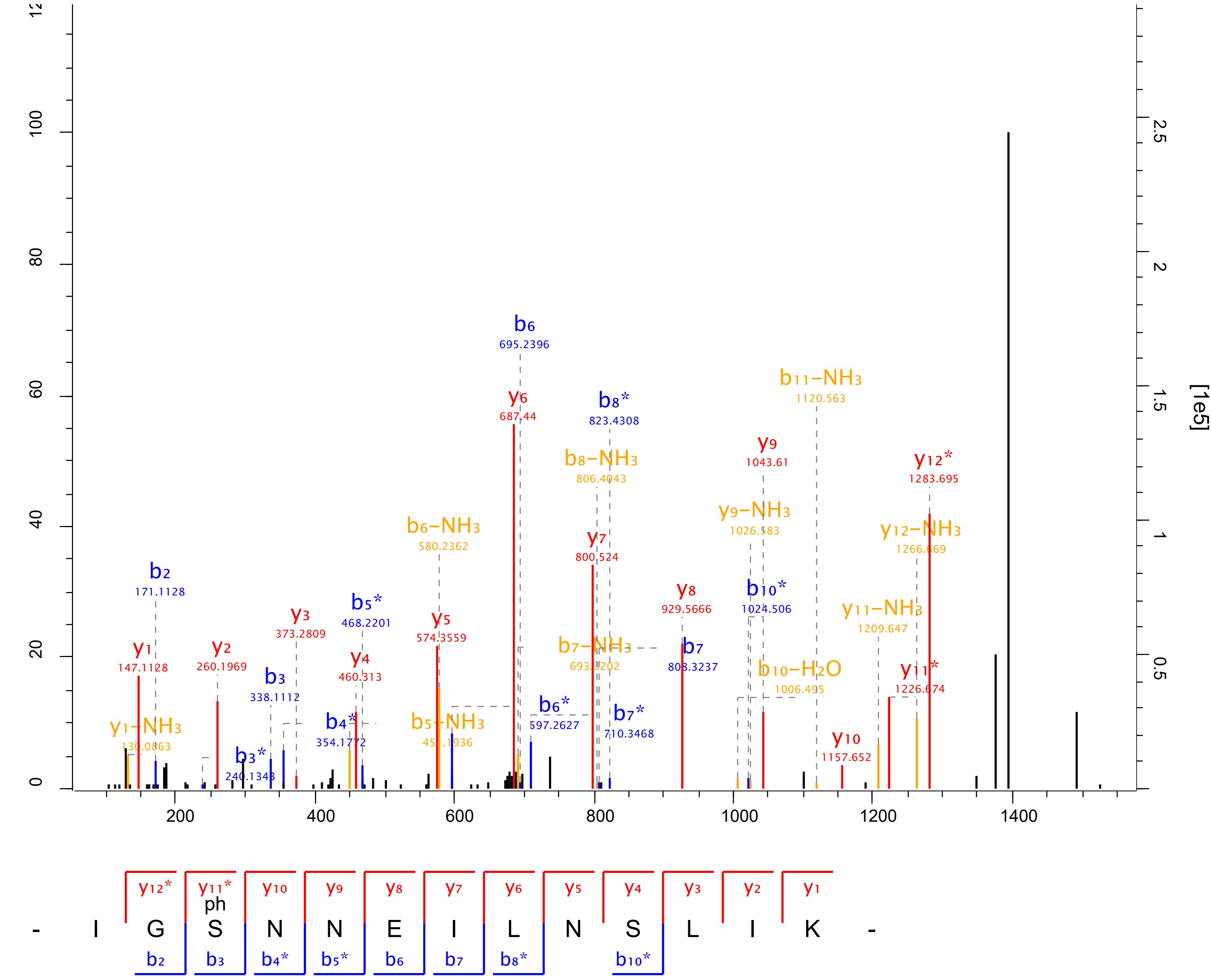

| Raw file | Scan | Method | Score | m/z |
|---|---|---|---|---|
| water_02_1 | 39164 | FTMS; HCD | 93.37 | 838.42 |

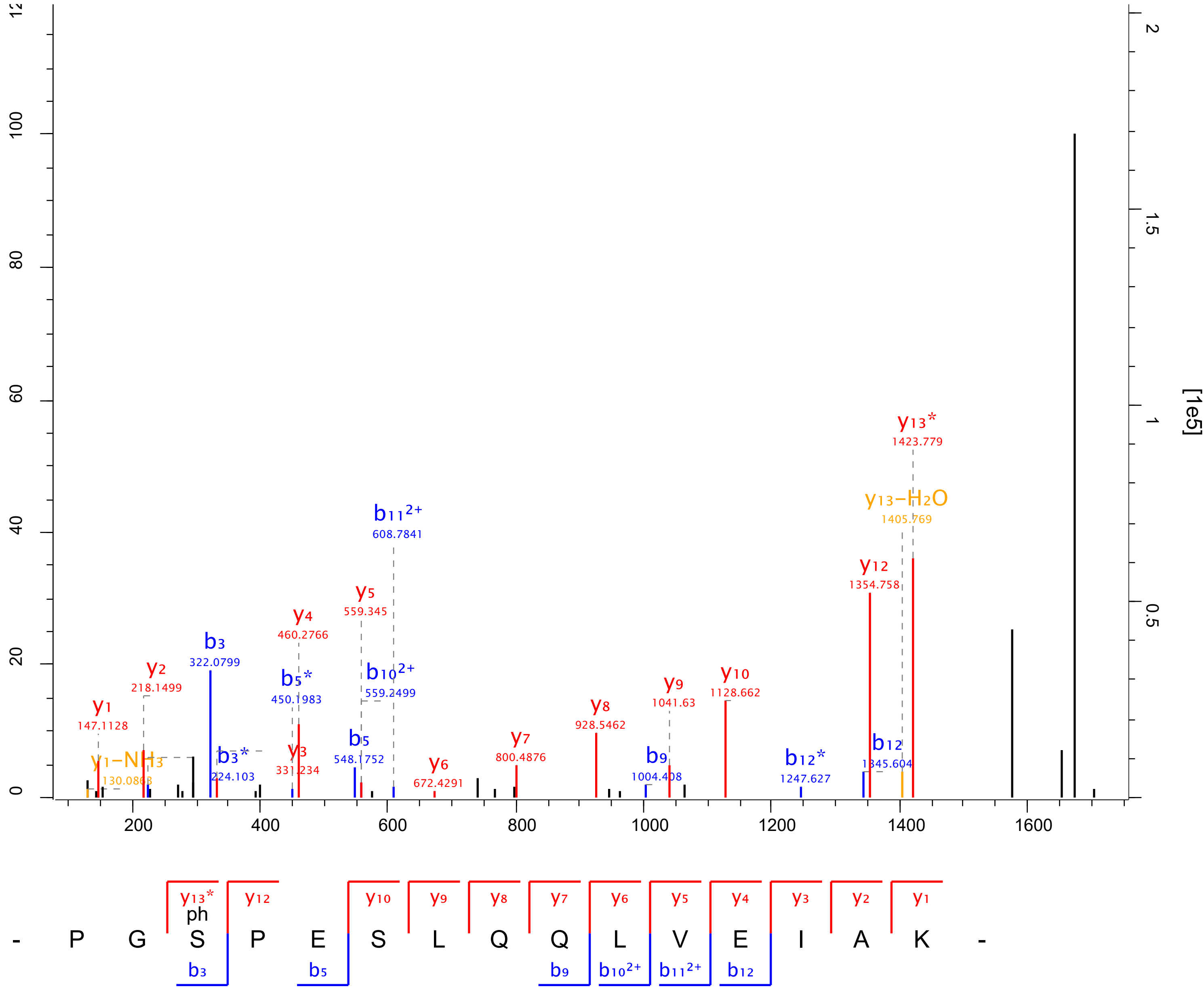

| Raw file | Scan | Method | Score | m/z |
|---|---|---|---|---|
| water_02_1 | 39288 | FTMS; HCD | 162 | 924.4 |

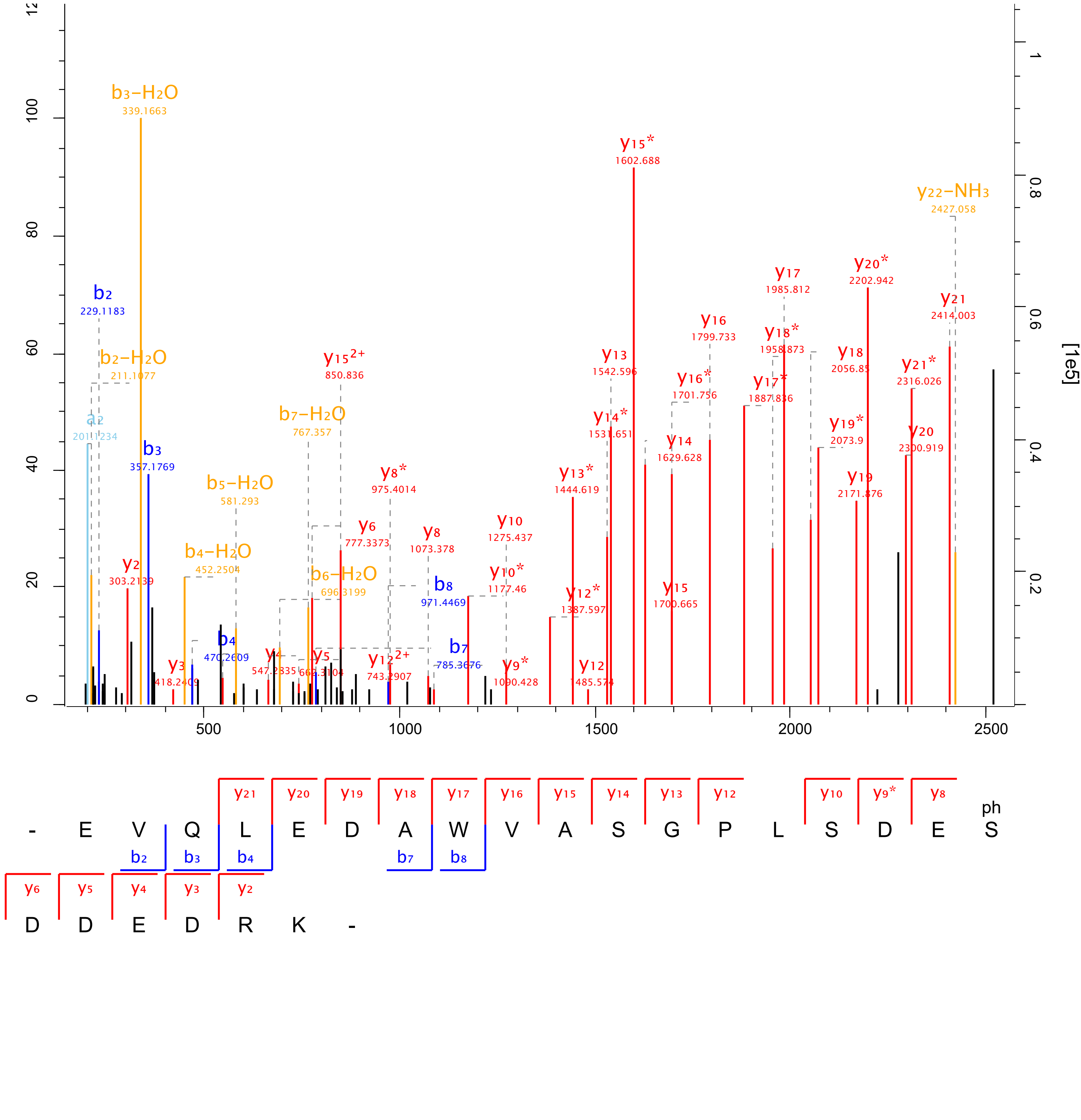

| Raw file | Scan | Method | Score | m/z |
|---|---|---|---|---|
| water_02_1 | 39302 | FTMS; HCD | 120.63 | 1108 |

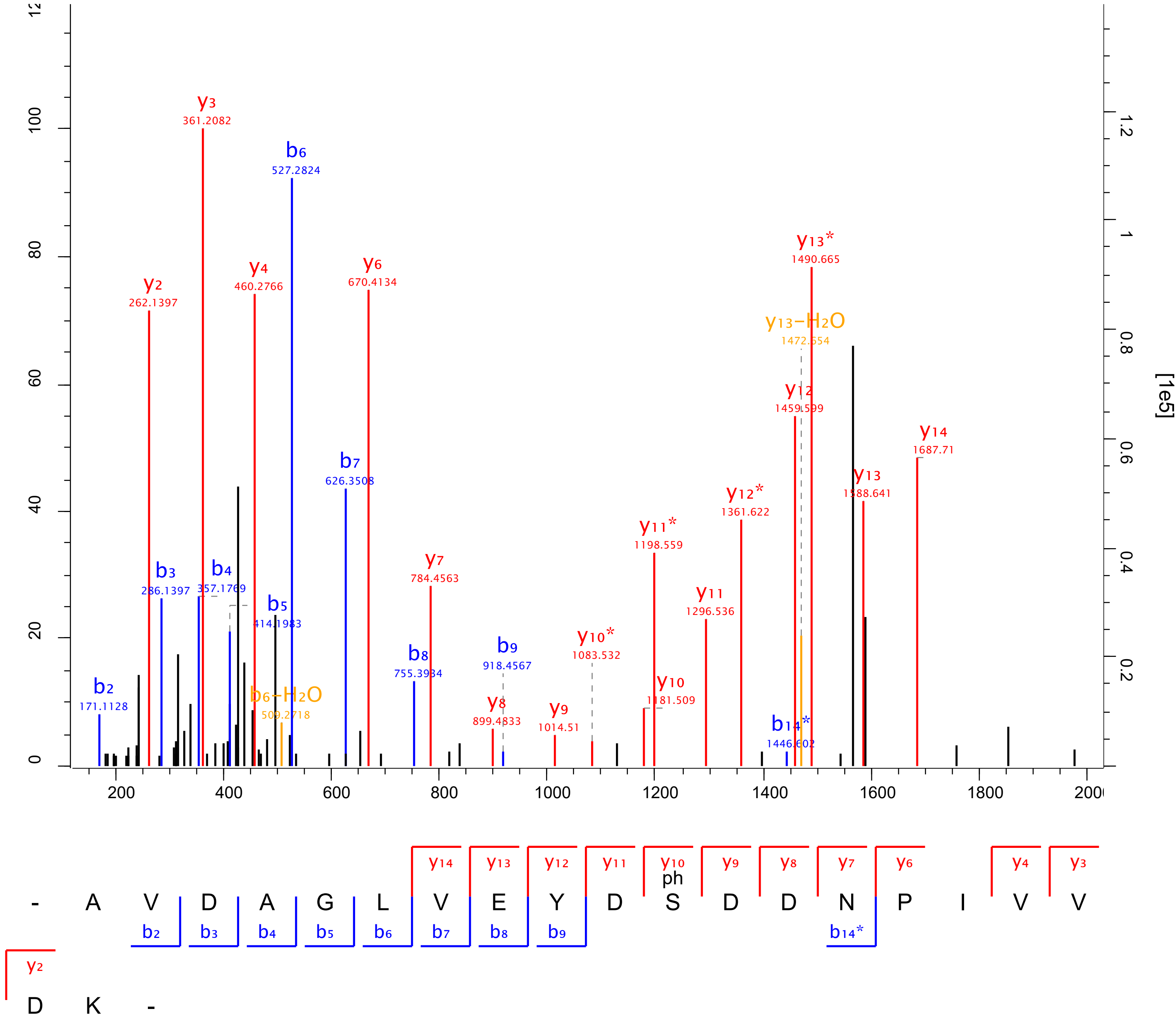

| Raw file | Scan | Method | Score | m/z |
| --- | --- | --- | --- | --- |
| water_02_1 | 39337 | FTMS; HCD | 136.43 | 919.92 |

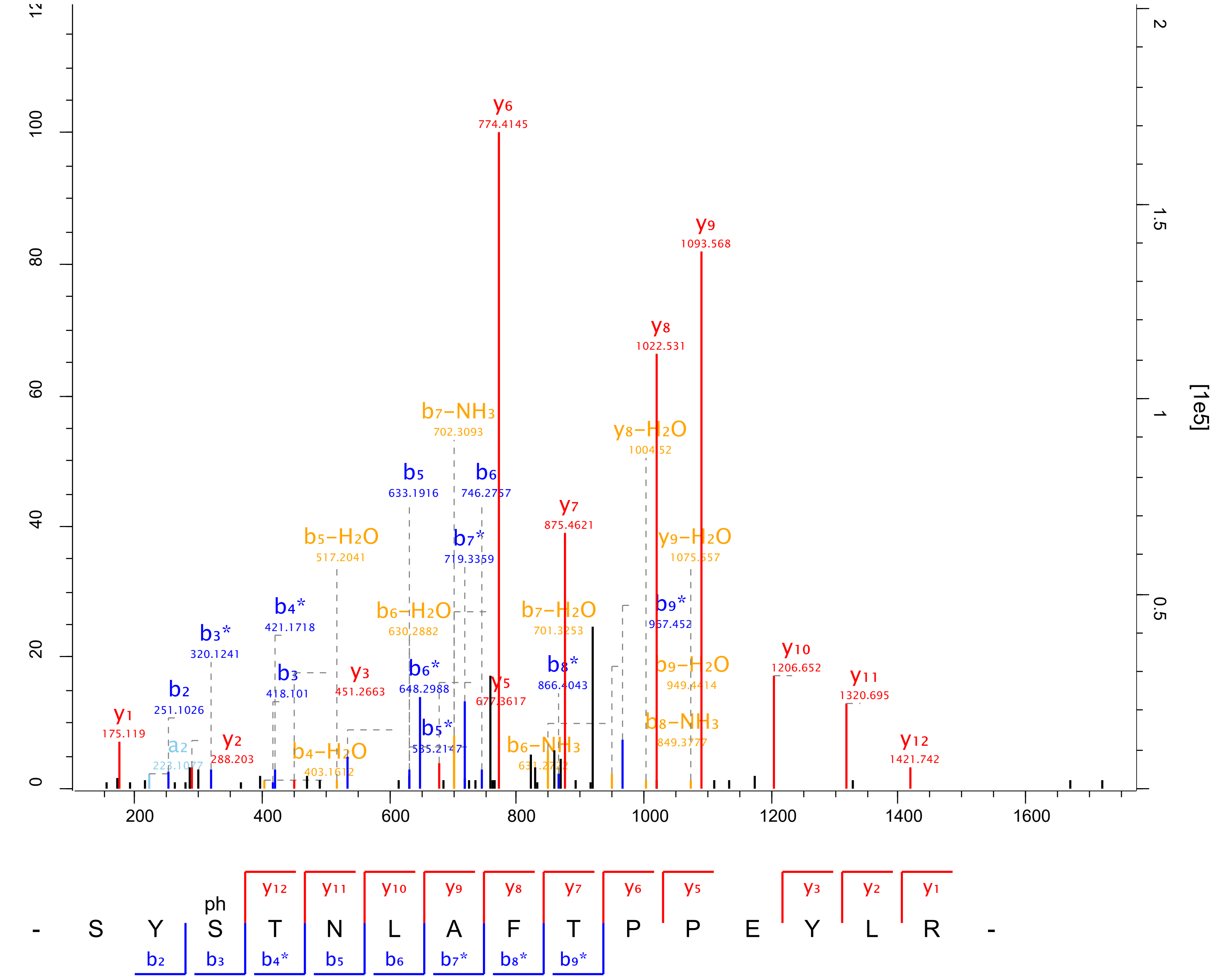

| Raw file | Scan | Method | Score | m/z |
|---|---|---|---|---|
| water_02_1 | 3946 | FTMS; HCD | 117.95 | 402.19 |

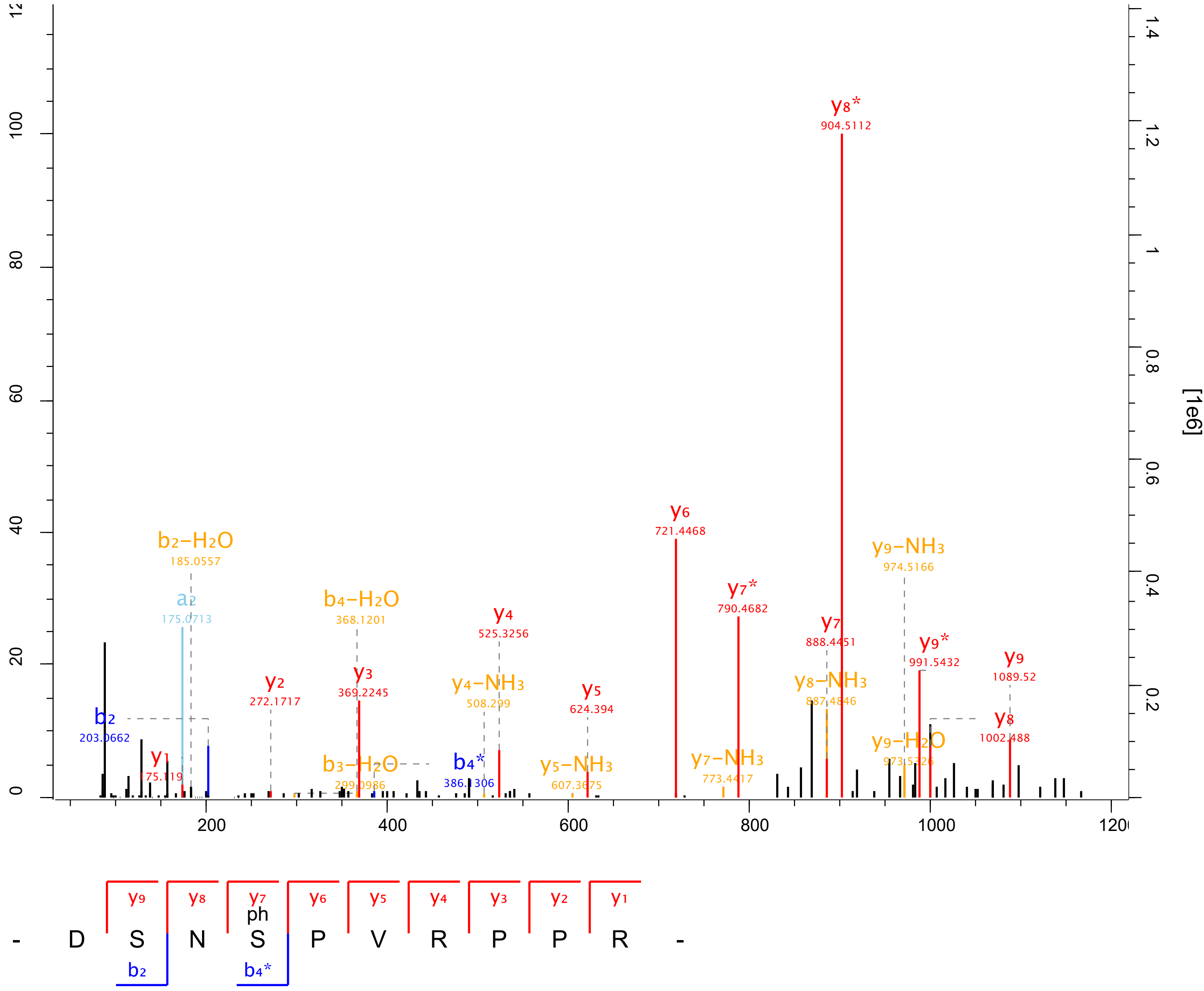

| Raw file | Scan | Method | Score | m/z |
|---|---|---|---|---|
| water_02_1 | 40088 | FTMS; HCD | 90.36 | 1064.45 |

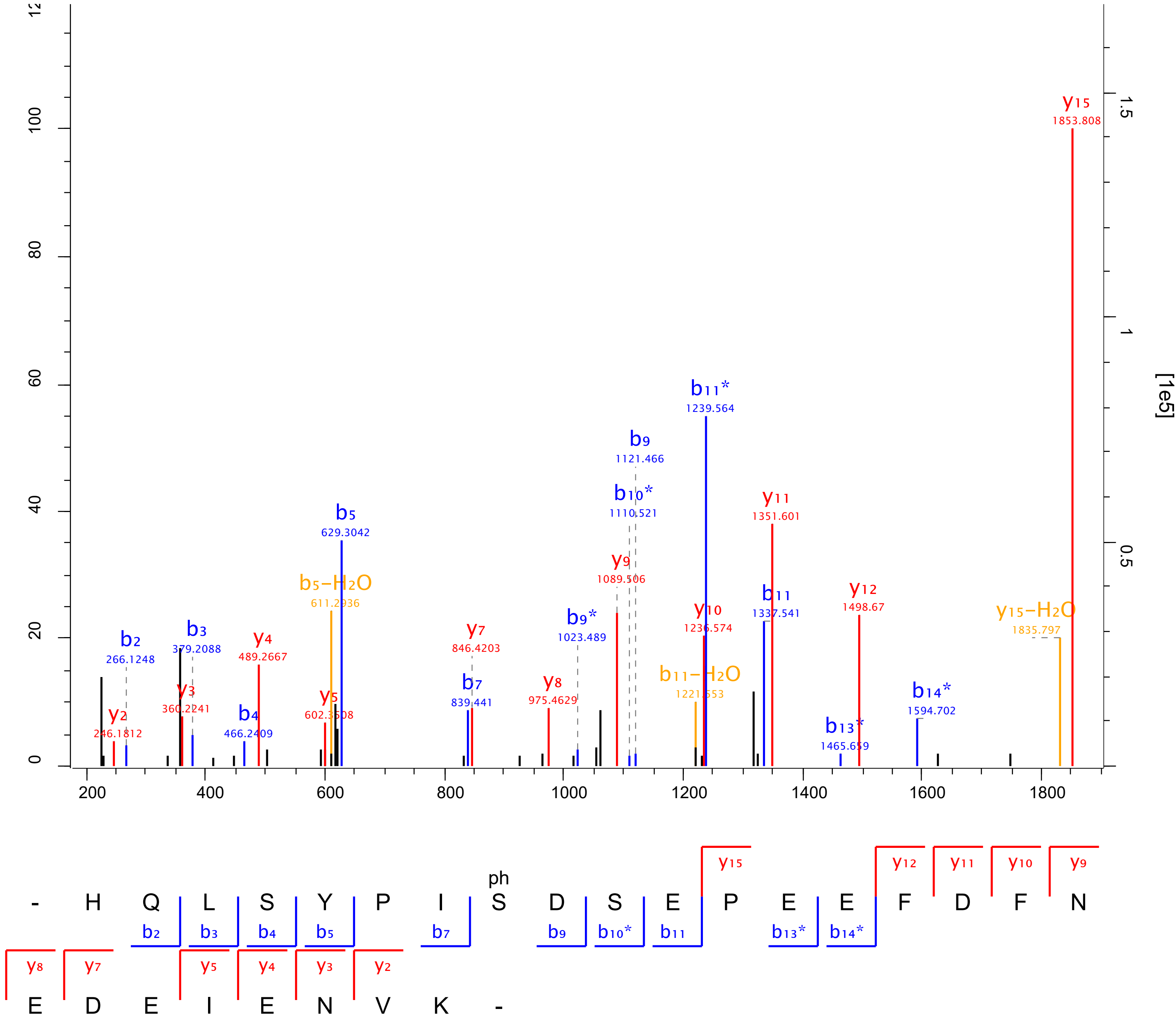

| Raw file | Scan | Method | Score | m/z |
|---|---|---|---|---|
| water_02_1 | 40141 | FTMS; HCD | 187.21 | 930.73 |

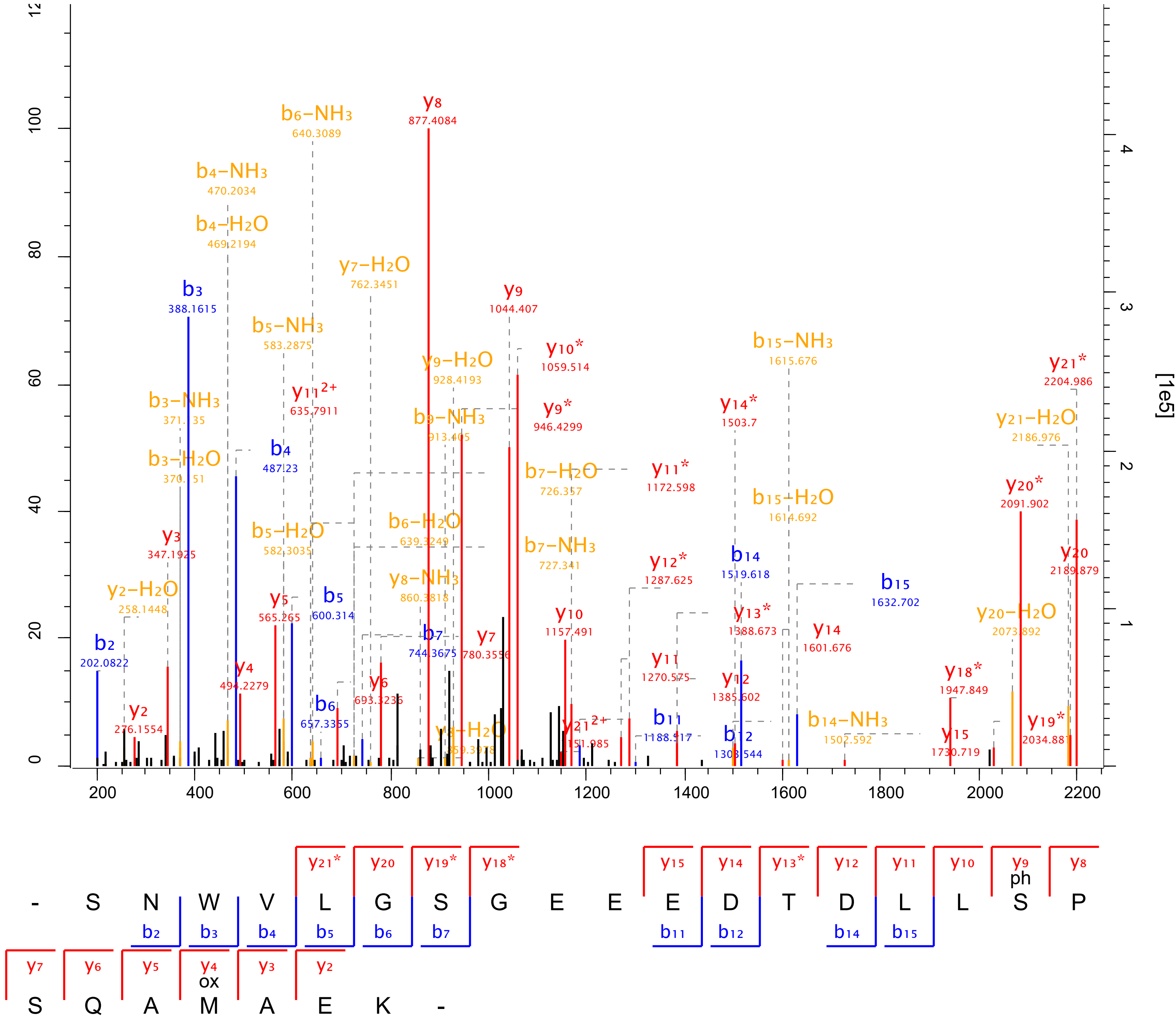

| Raw file | Scan | Method | Score | m/z |
|---|---|---|---|---|
| water_02_1 | 40186 | FTMS; HCD | 66.38 | 1012.76 |

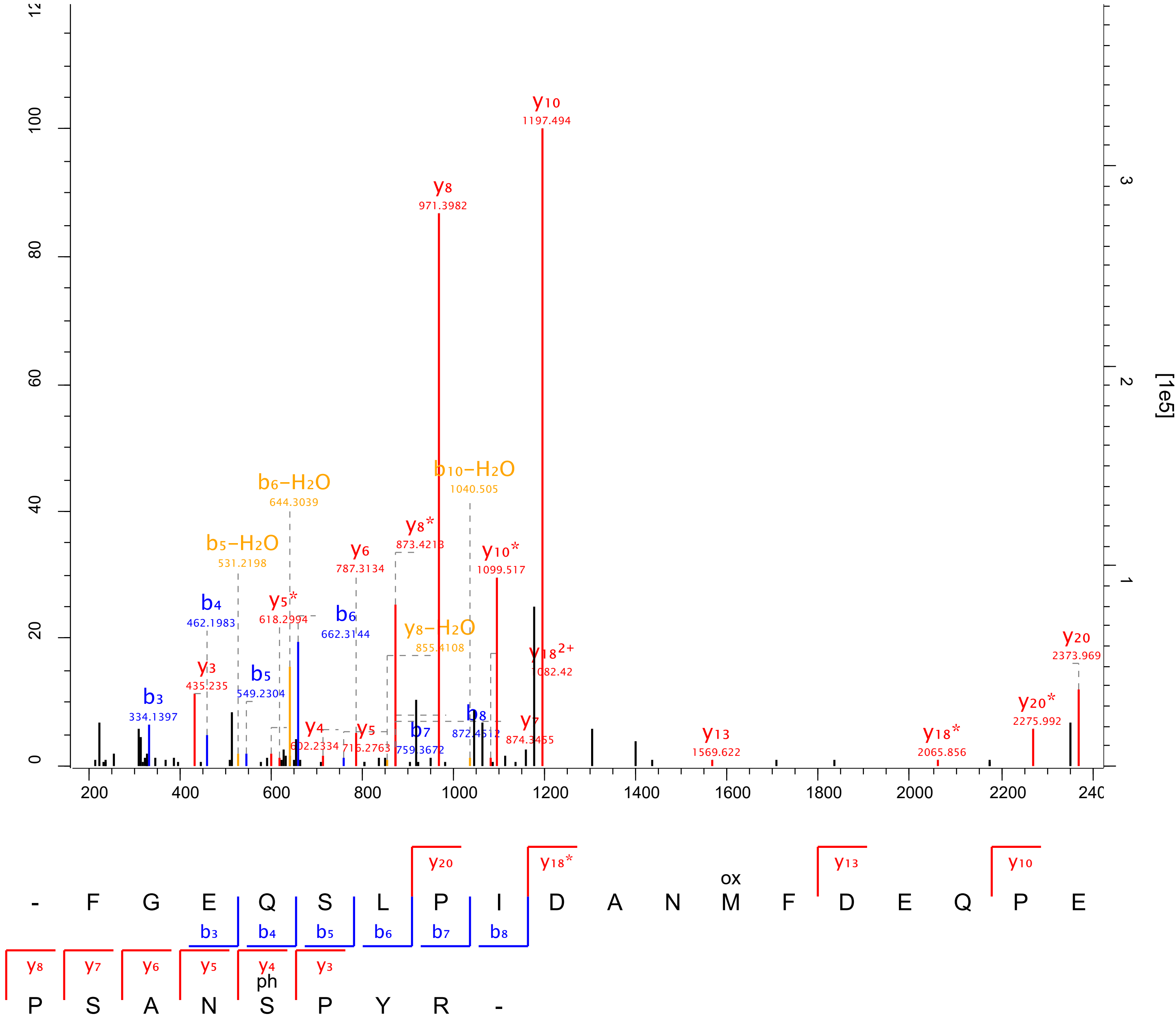

| Raw file | Scan | Method | Score | m/z |
|---|---|---|---|---|
| water_02_1 | 40369 | FTMS; HCD | 47.73 | 805.69 |

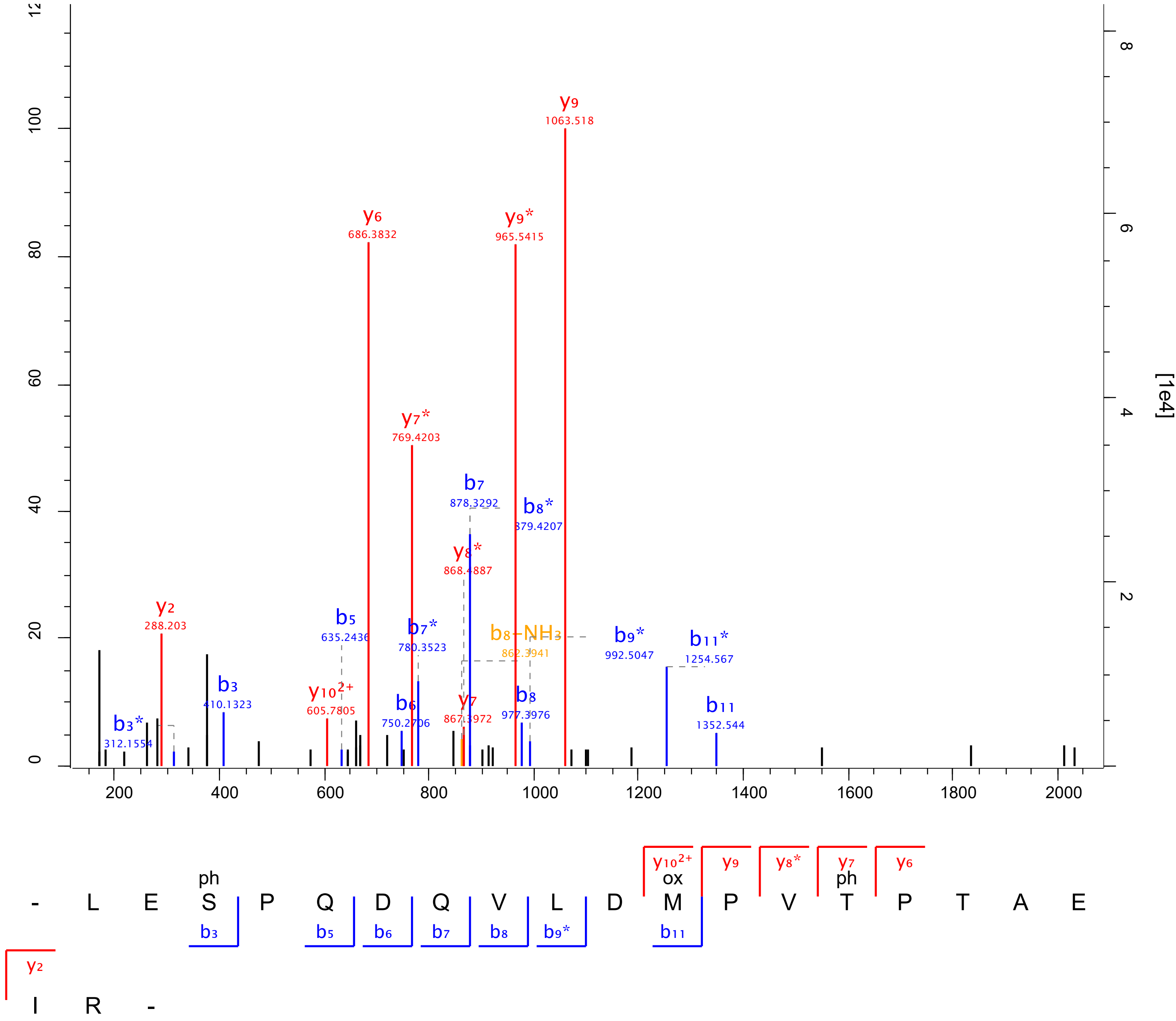

| Raw file | Scan | Method | Score | m/z |
| --- | --- | --- | --- | --- |
| water_02_1 | 4052 | FTMS; HCD | 137.78 | 486.7 |

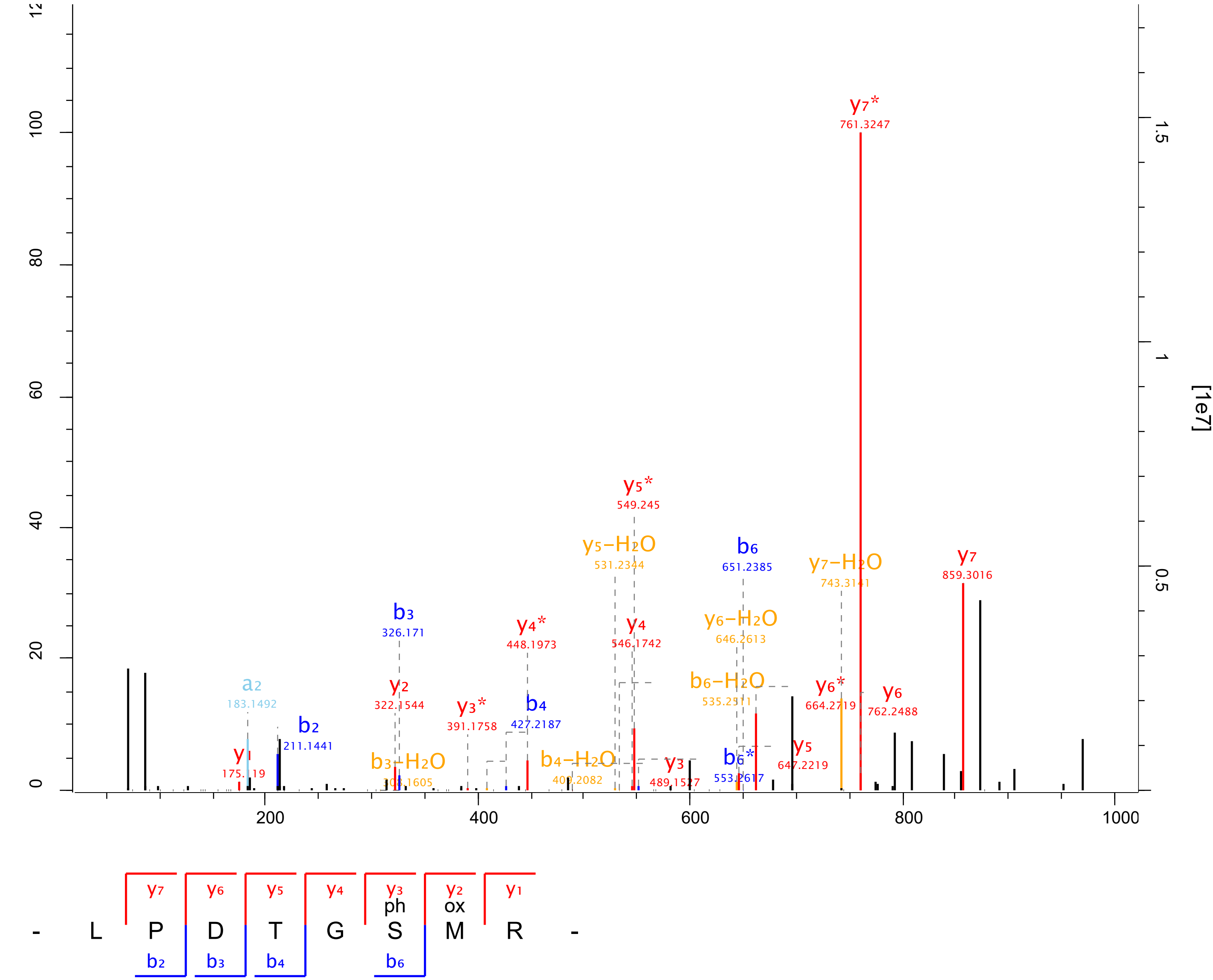

| Raw file | Scan | Method | Score | m/z |
|---|---|---|---|---|
| water_02_1 | 4053 | FTMS; HCD | 45.43 | 667.59 |

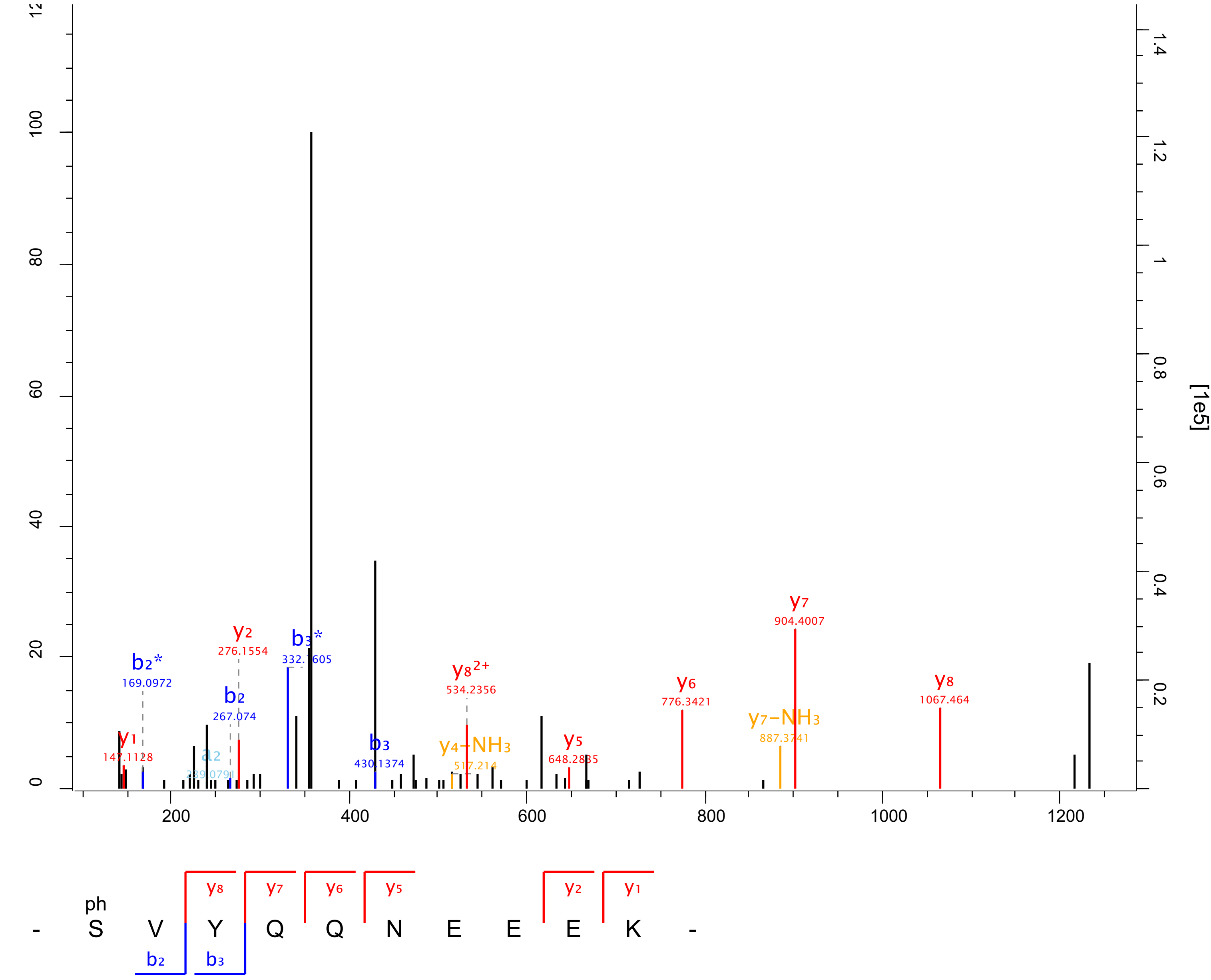

| Raw file | Scan | Method | Score | m/z |
|---|---|---|---|---|
| water_02_1 | 4058 | FTMS; HCD | 84.36 | 615.27 |

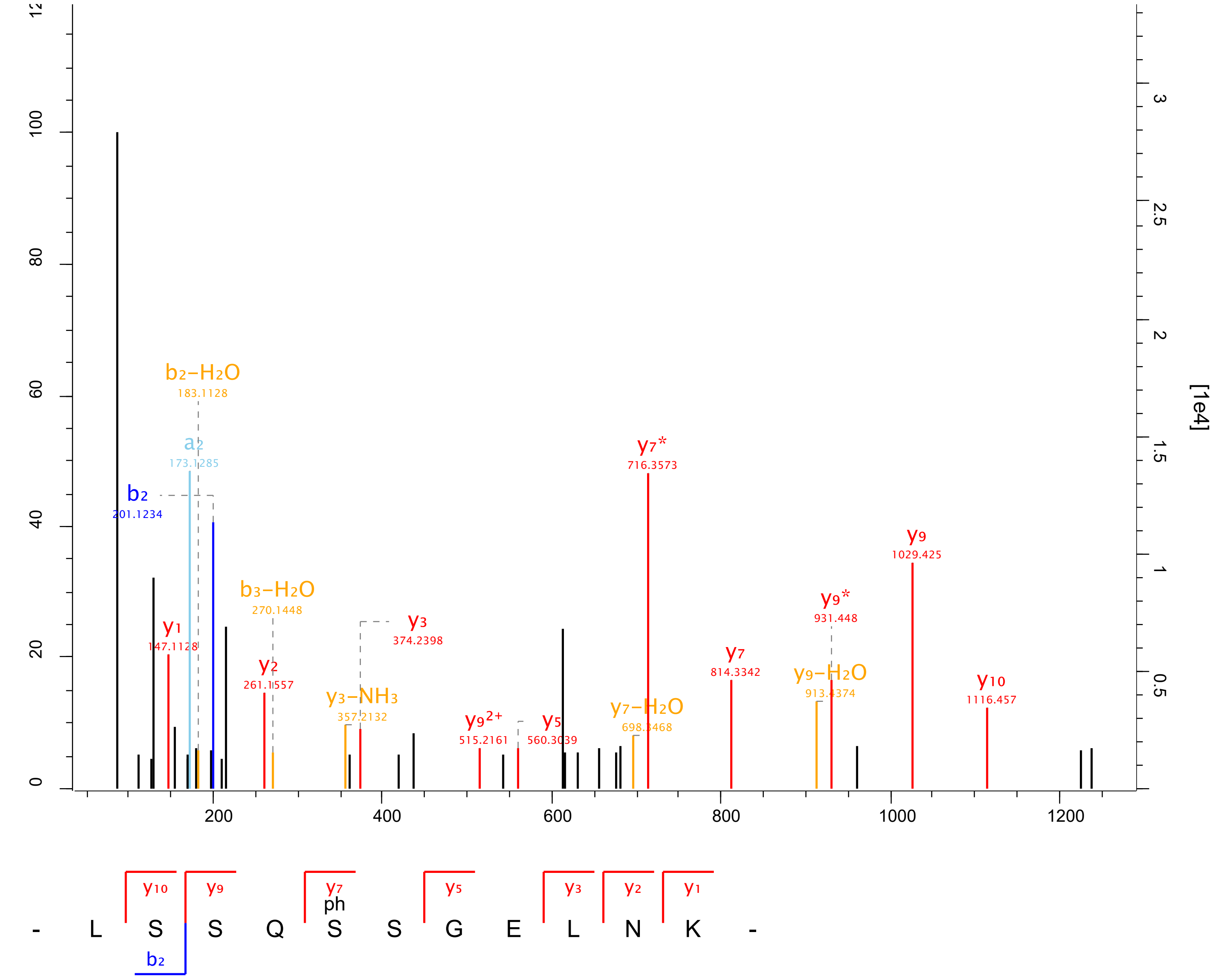

| Raw file | Scan | Method | Score | m/z |
| --- | --- | --- | --- | --- |
| water_02_1 | 40731 | FTMS; HCD | 41.45 | 802.02 |

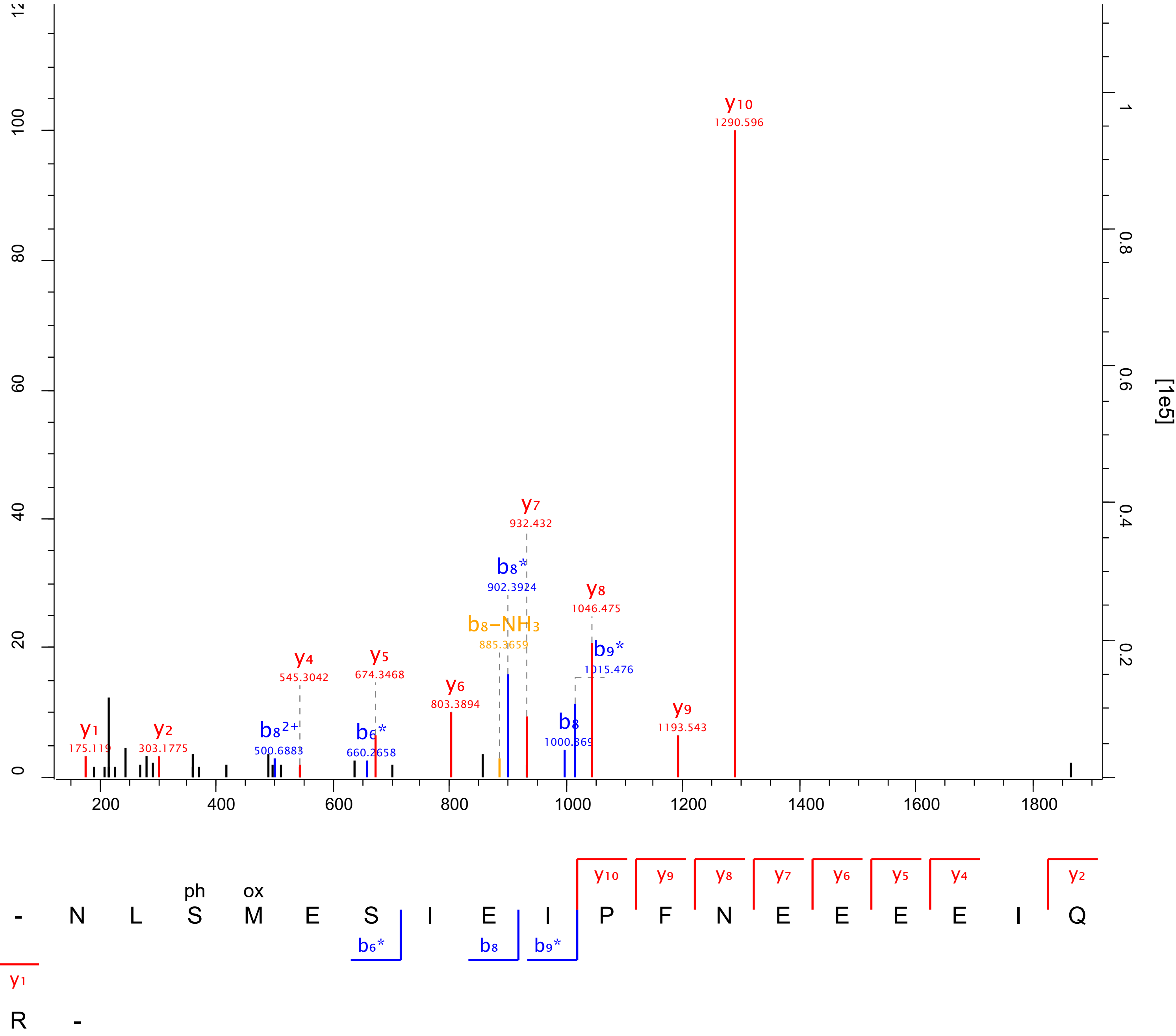

| Raw file | Scan | Method | Score | m/z |
| --- | --- | --- | --- | --- |
| water_02_1 | 4106 | FTMS; HCD | 43.81 | 504.7 |

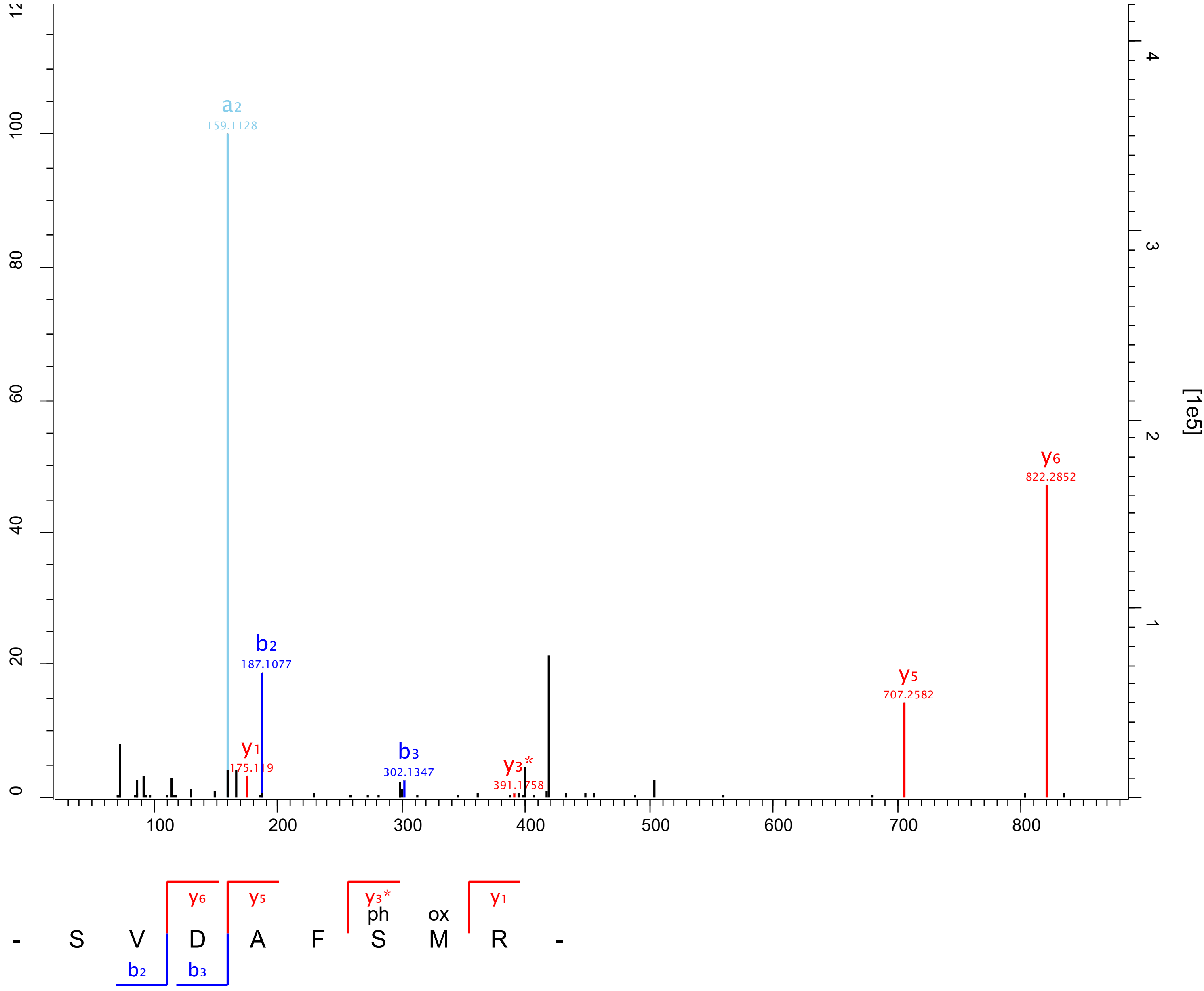

| Raw file | Scan | Method | Score | m/z |
|---|---|---|---|---|
| water_02_1 | 41102 | FTMS; HCD | 222.06 | 934.94 |

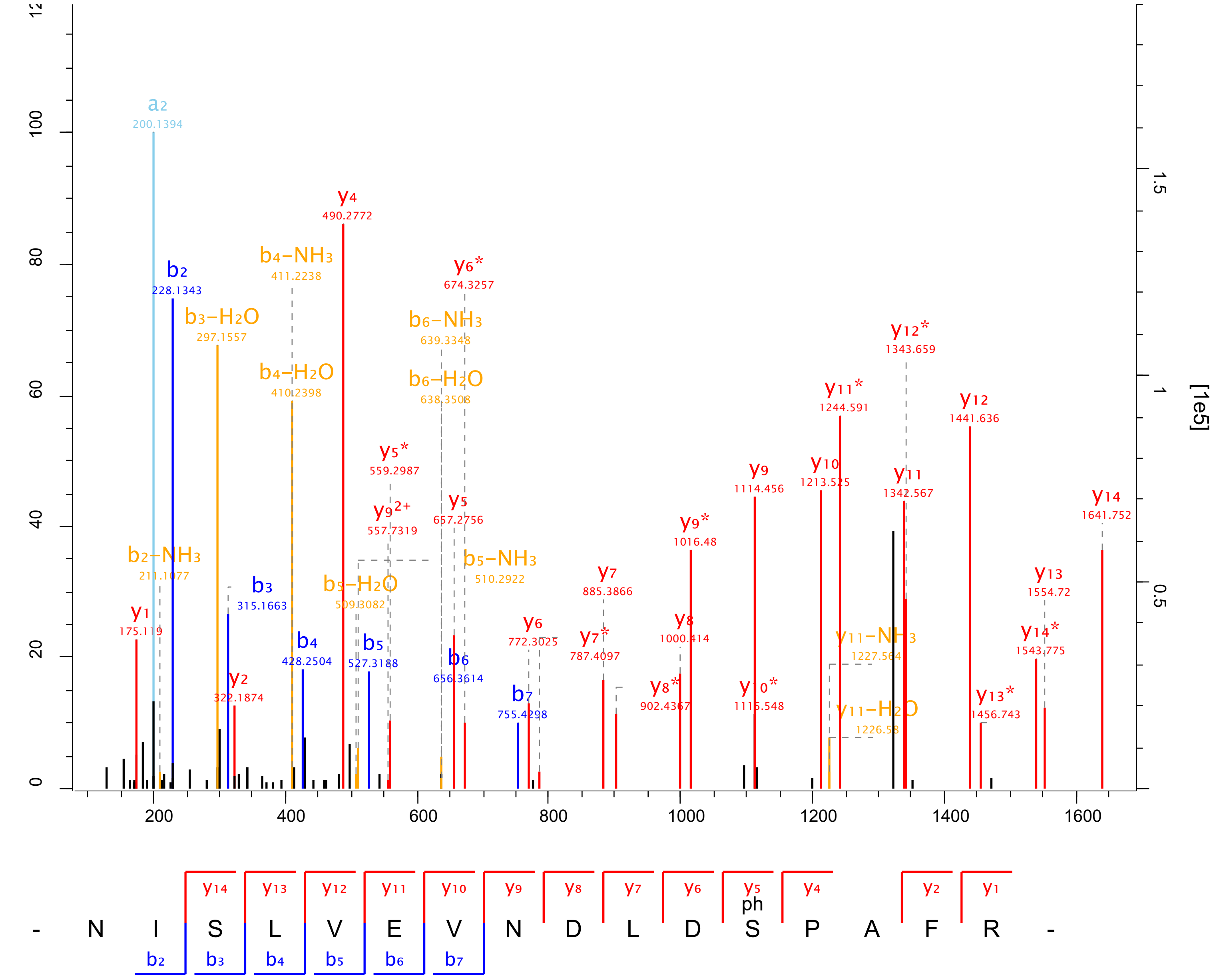

| Raw file | Scan | Method | Score | m/z |
| --- | --- | --- | --- | --- |
| water_02_1 | 41158 | FTMS; HCD | 161.34 | 1130.15 |

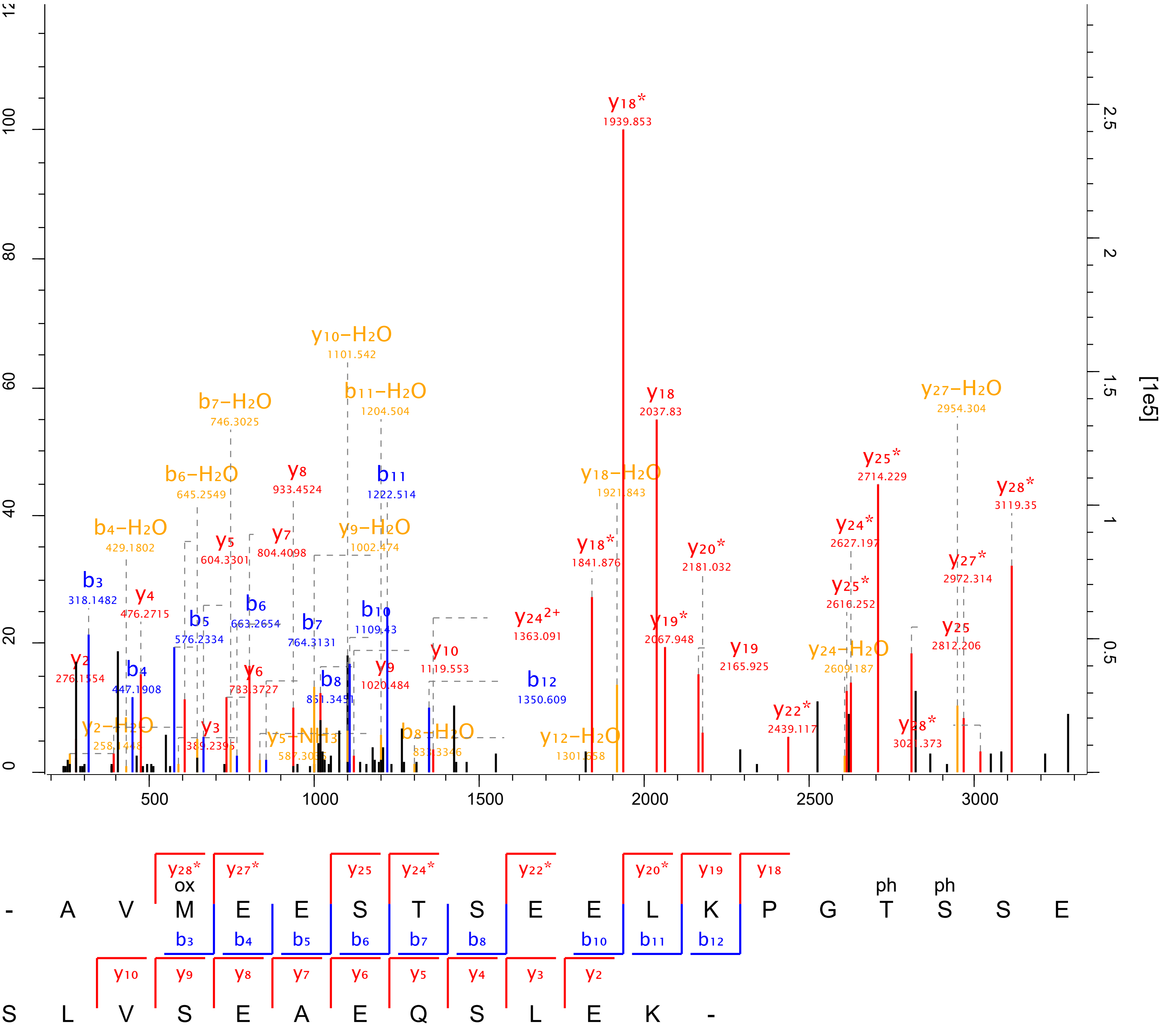

| Raw file | Scan | Method | Score | m/z |
|---|---|---|---|---|
| water_02_1 | 41339 | FTMS; HCD | 140.96 | 796.82 |

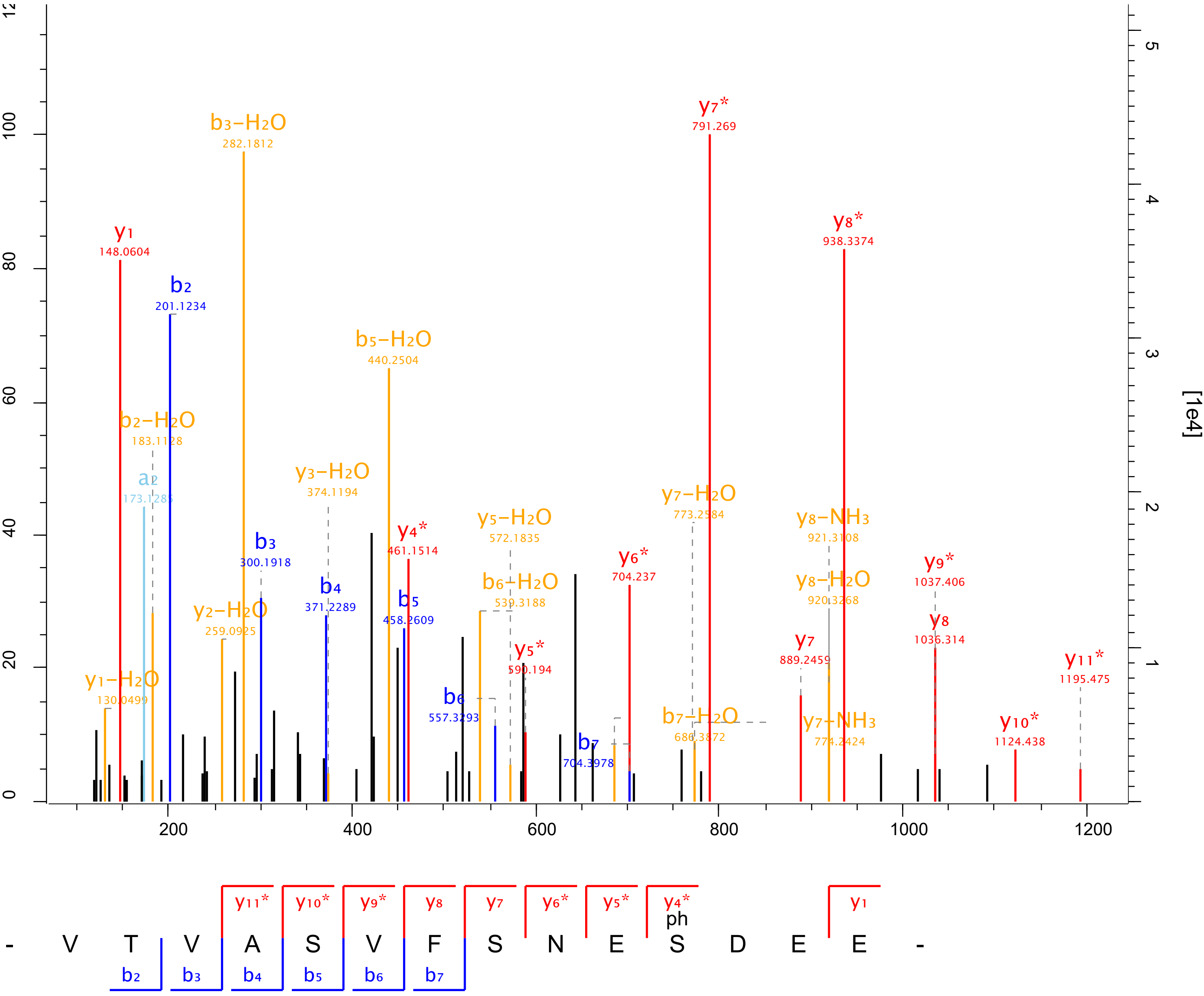

| Raw file | Scan | Method | Score | m/z |
| --- | --- | --- | --- | --- |
| water_02_1 | 41882 | FTMS; HCD | 108.5 | 978 |

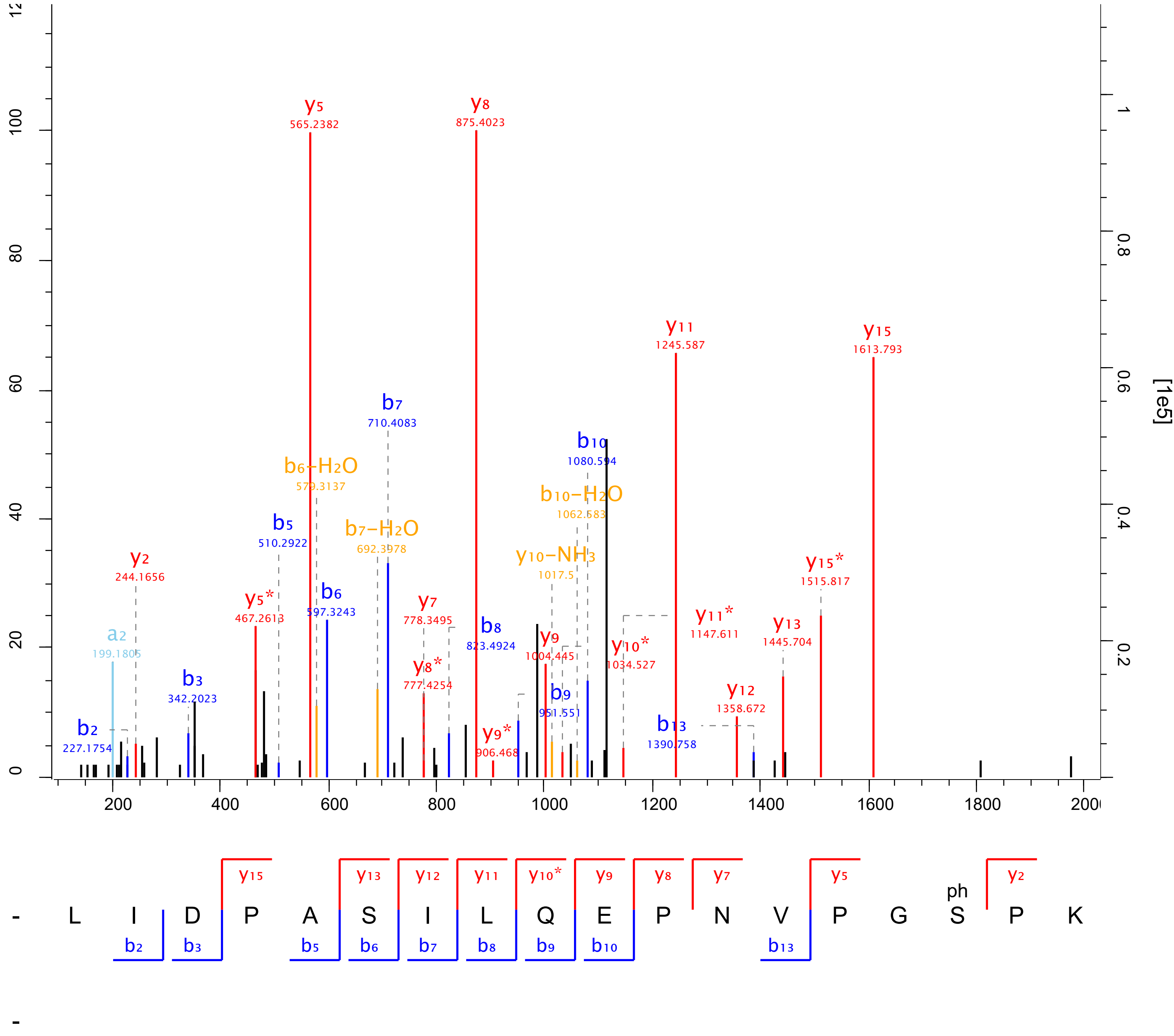

| Raw file | Scan | Method | Score | m/z |
| --- | --- | --- | --- | --- |
| water_02_1 | 41901 | FTMS; HCD | 61.48 | 897.39 |

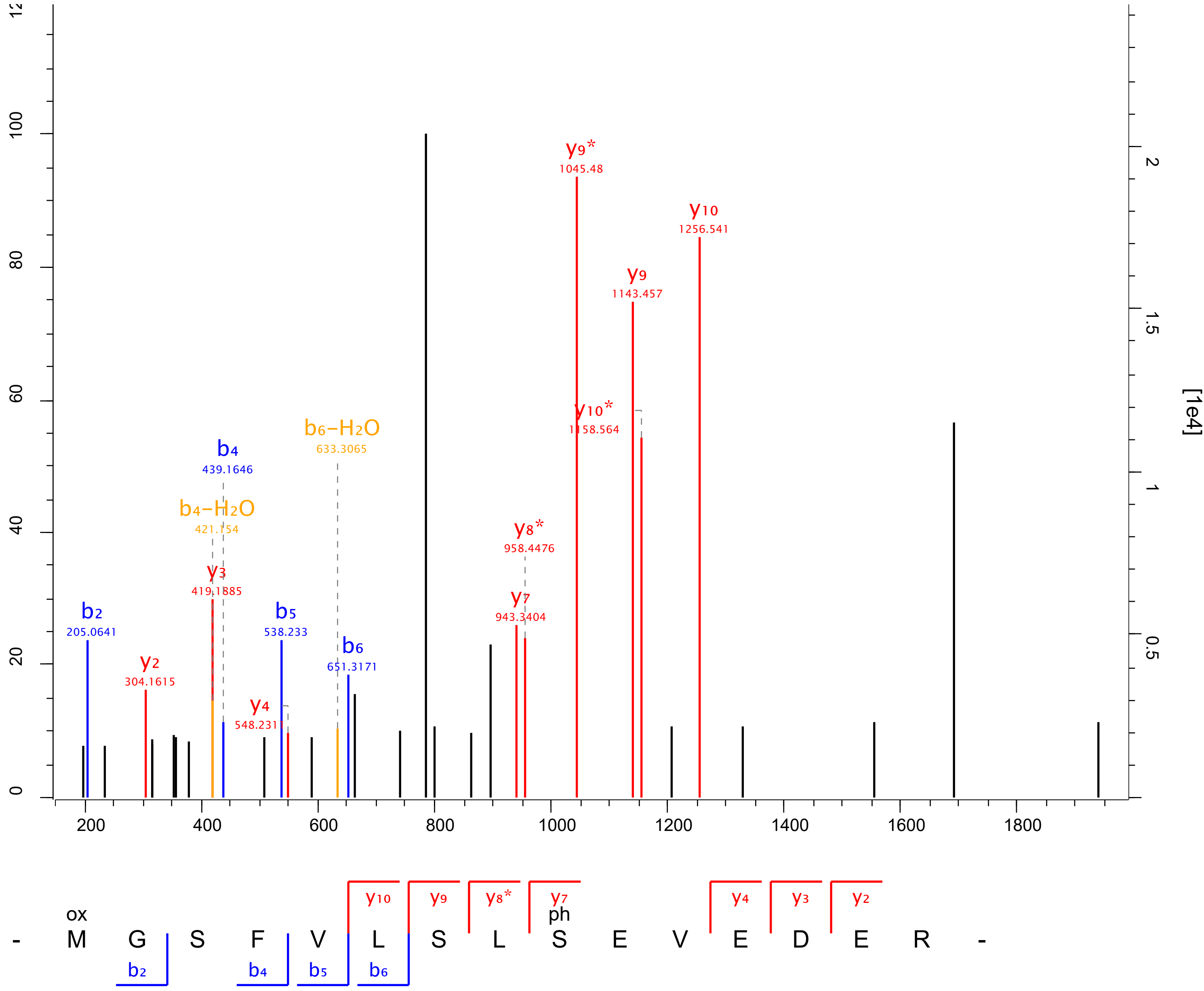

| Raw file | Scan | Method | Score | m/z |
|---|---|---|---|---|
| water_02_1 | 41946 | FTMS; HCD | 61.1 | 1058.45 |

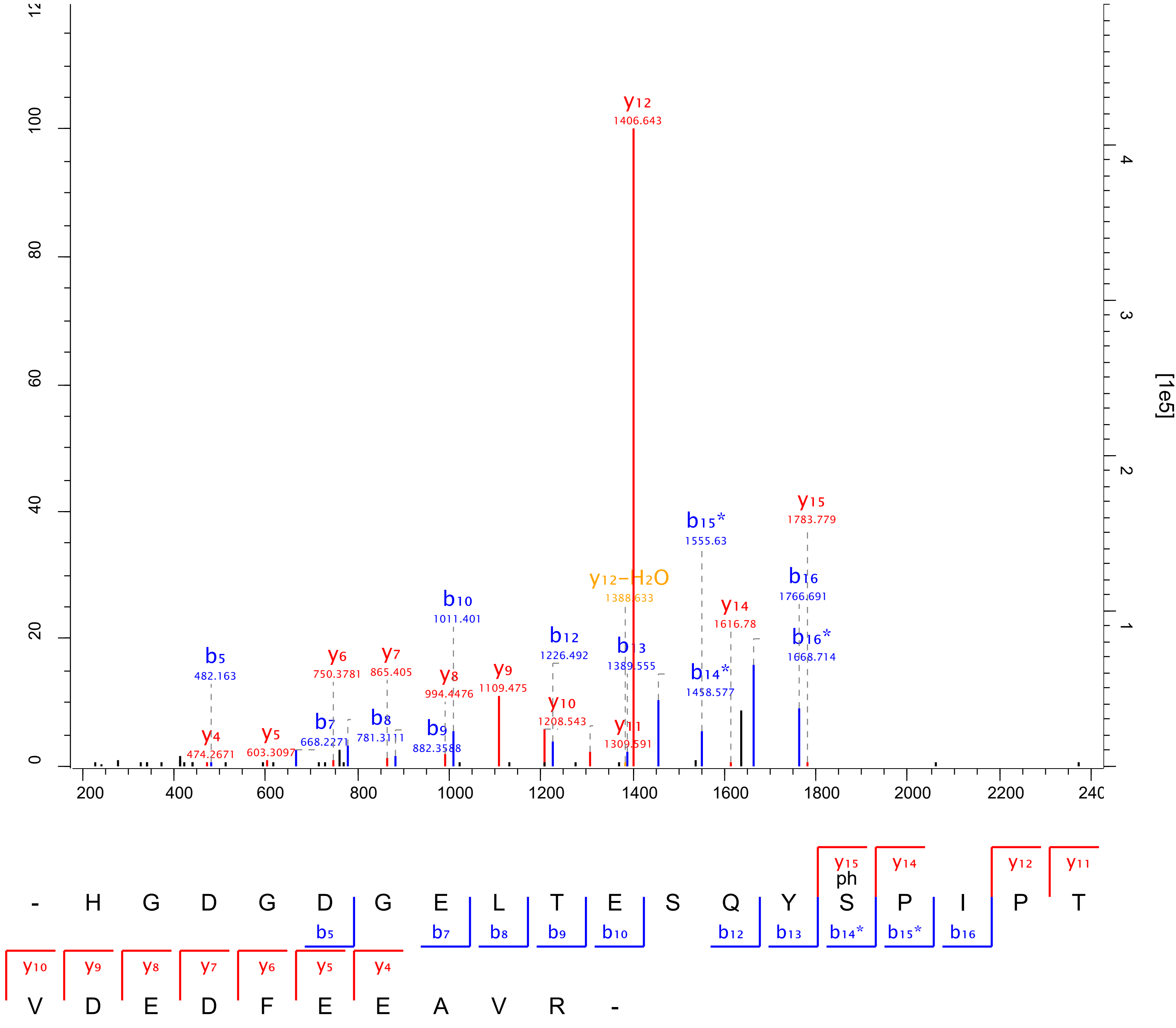

| Raw file | Scan | Method | Score | m/z |
| --- | --- | --- | --- | --- |
| water_02_1 | 42295 | FTMS; HCD | 102.69 | 861.36 |

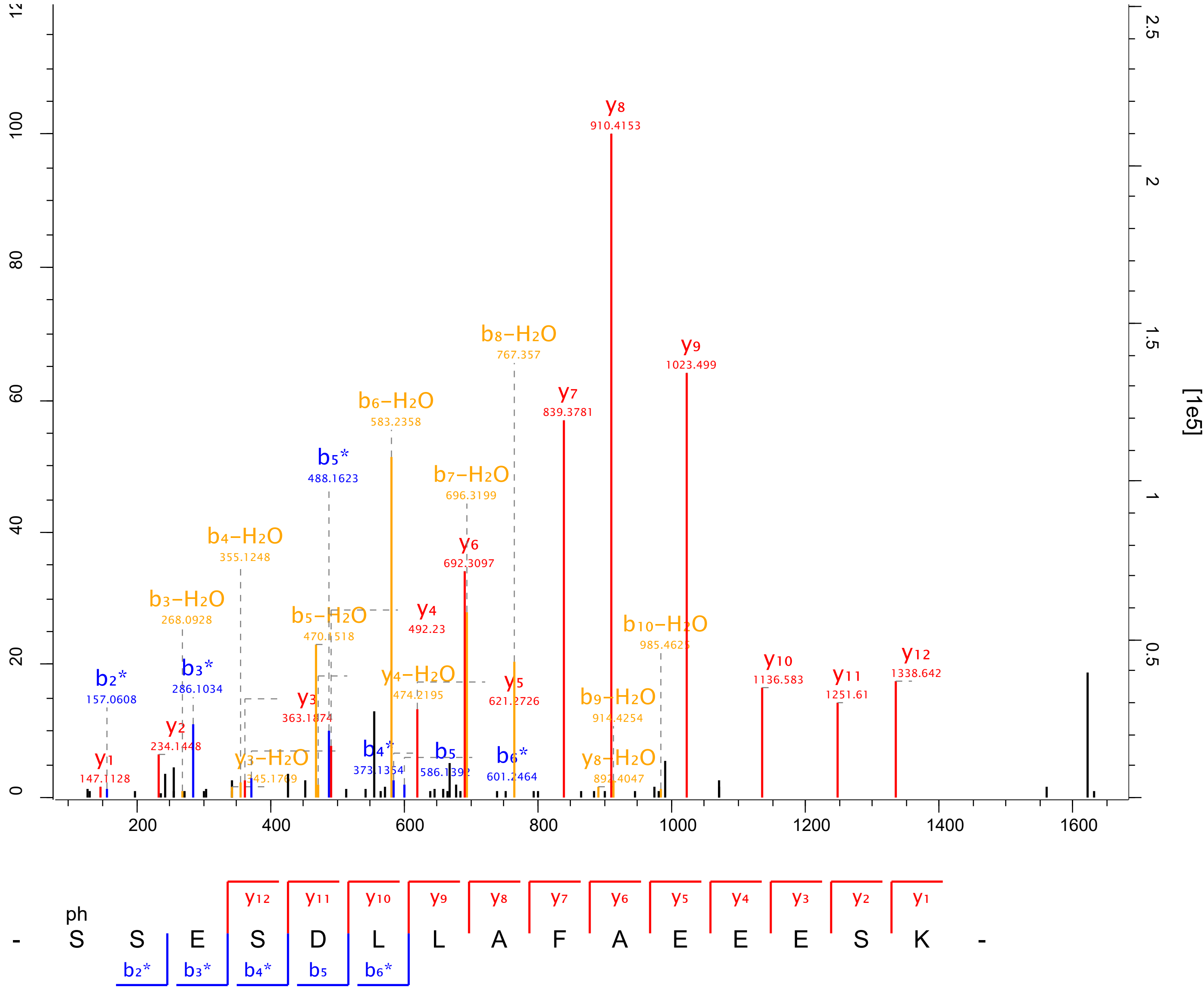

| Raw file | Scan | Method | Score | m/z |
|---|---|---|---|---|
| water_02_1 | 42319 | FTMS; HCD | 97.78 | 771.75 |

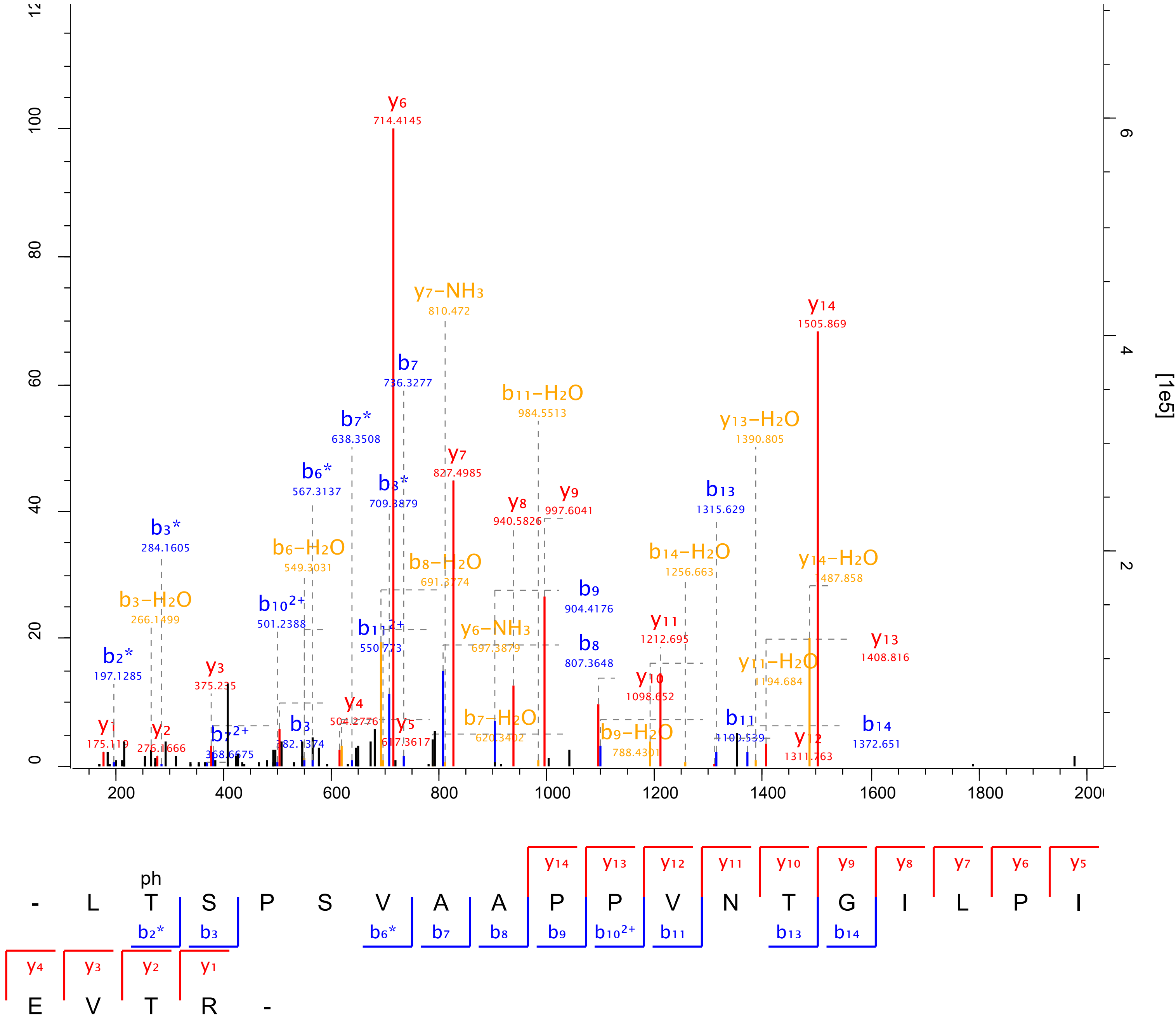

| Raw file | Scan | Method | Score | m/z |
|---|---|---|---|---|
| water_02_1 | 42530 | FTMS; HCD | 57.29 | 701.39 |

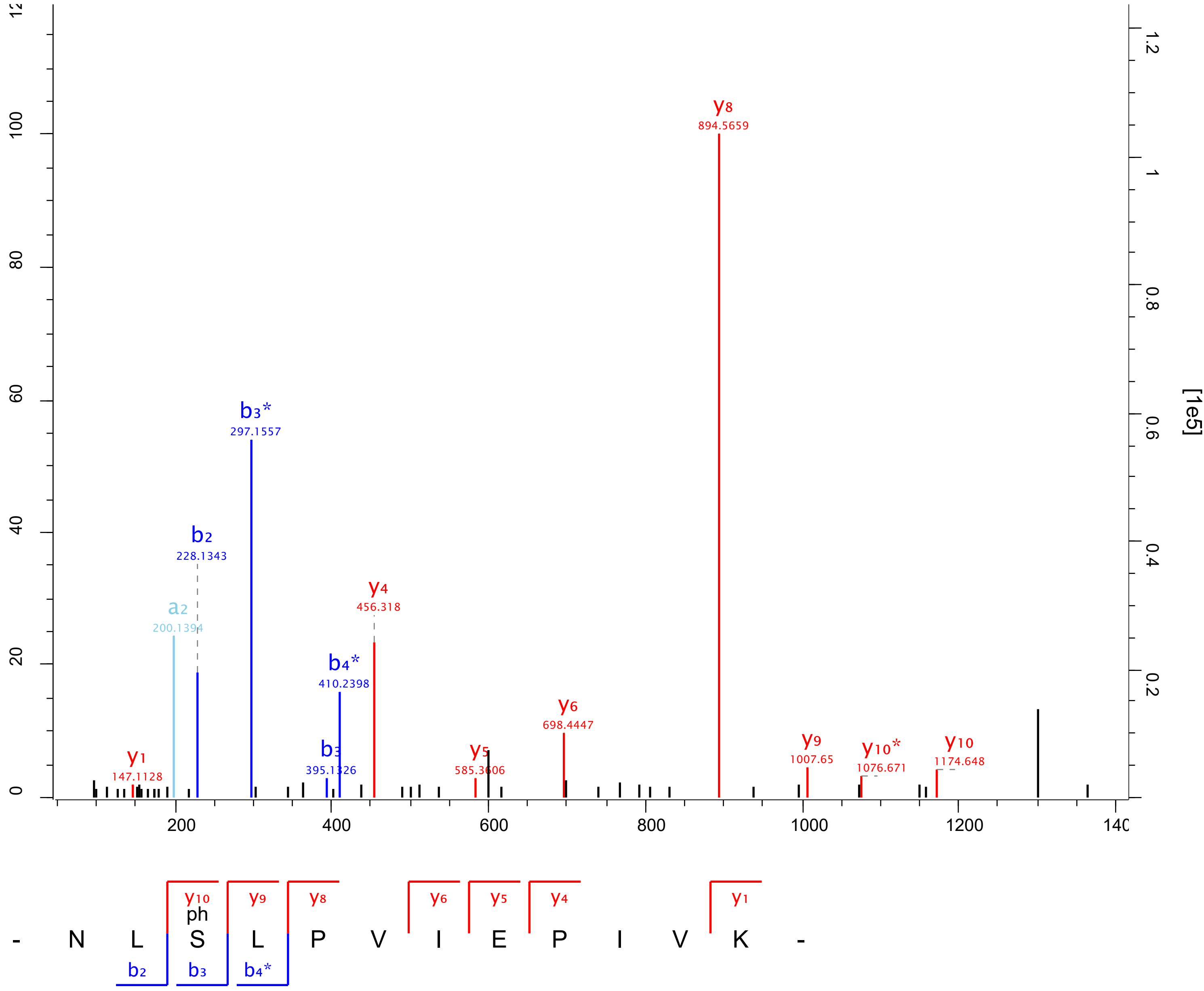

| Raw file | Scan | Method | Score | m/z |
| --- | --- | --- | --- | --- |
| water_02_1 | 43107 | FTMS; HCD | 62.66 | 579.27 |

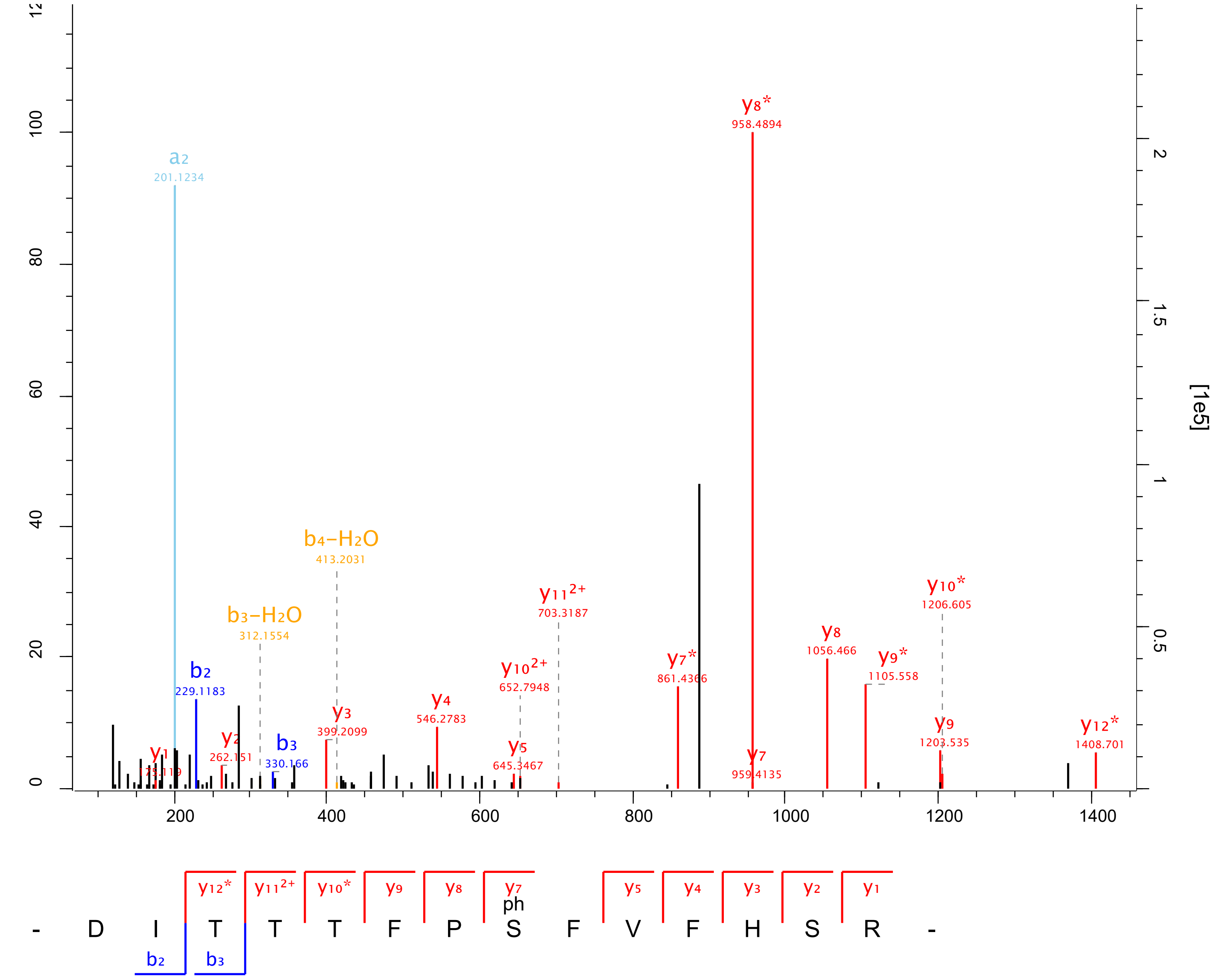

| Raw file | Scan | Method | Score | m/z |
|---|---|---|---|---|
| water_02_1 | 43205 | FTMS; HCD | 76.68 | 721.31 |

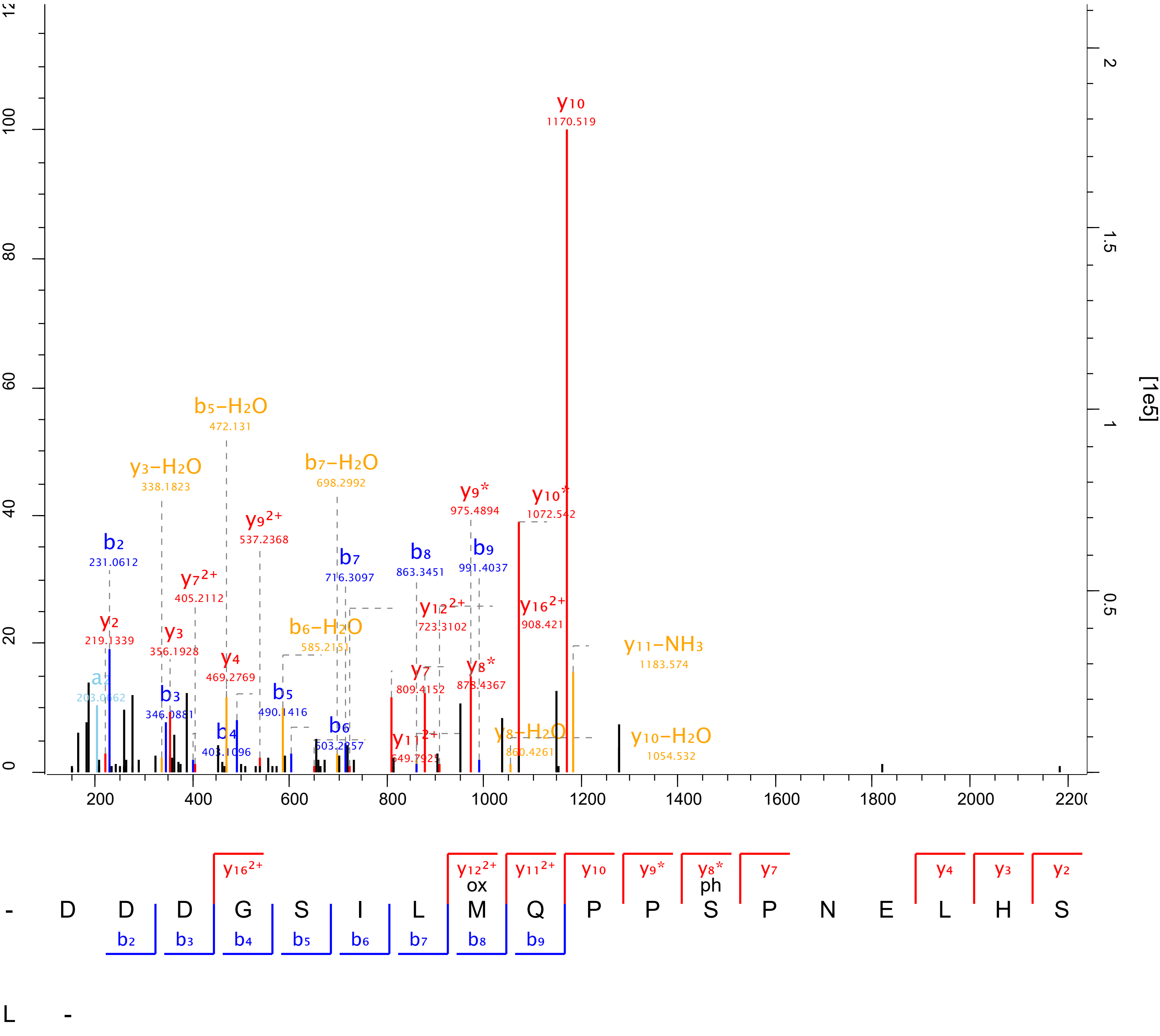

| Raw file | Scan | Method | Score | m/z |
|---|---|---|---|---|
| water_02_1 | 4332 | FTMS; HCD | 90.61 | 543.21 |

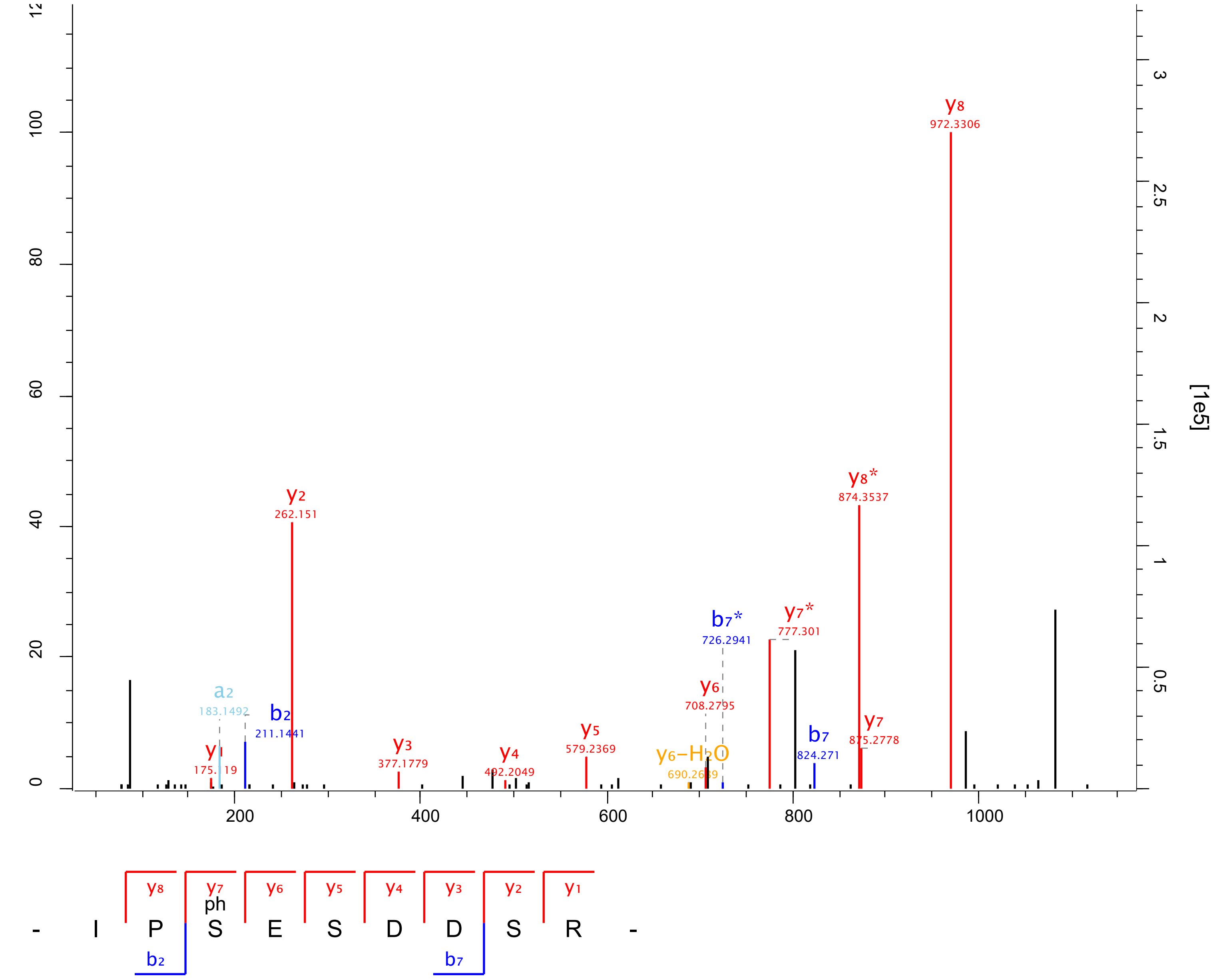

| Raw file | Scan | Method | Score | m/z |
|---|---|---|---|---|
| water_02_1 | 43372 | FTMS; HCD | 175.34 | 847.05 |

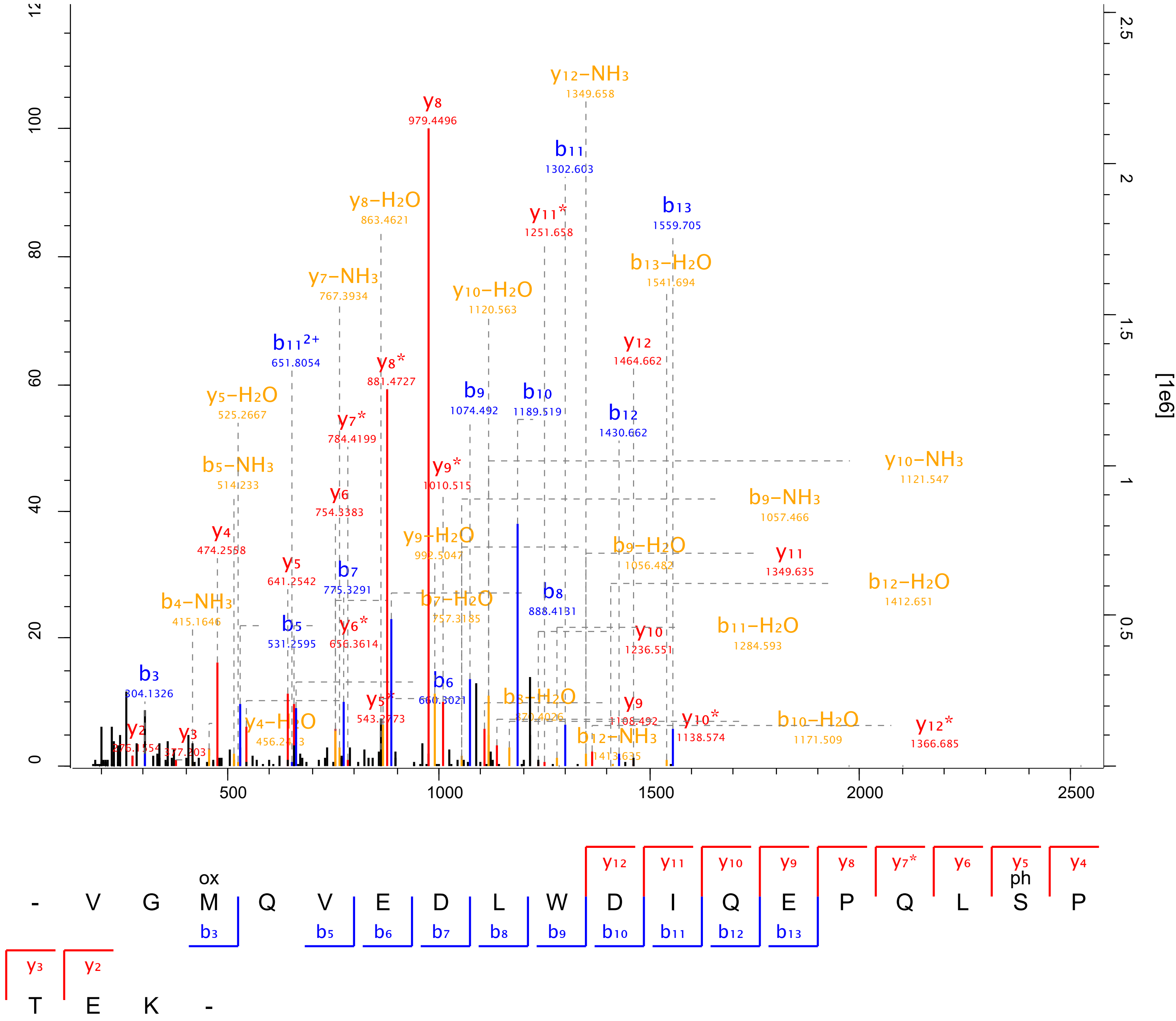

| Raw file | Scan | Method | Score | m/z |
|---|---|---|---|---|
| water_02_1 | 43659 | FTMS; HCD | 203.89 | 913.43 |

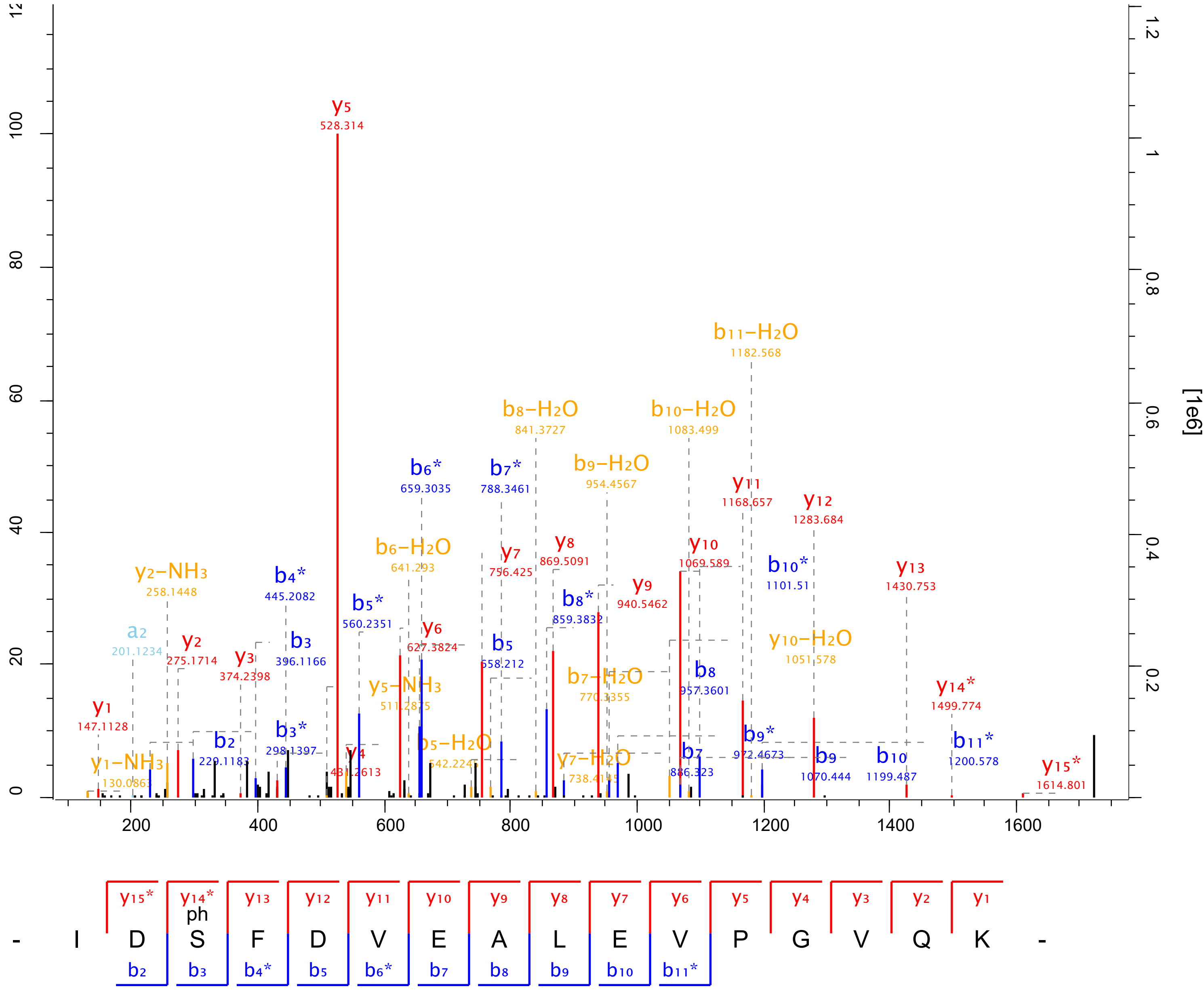

| Raw file | Scan | Method | Score | m/z |
|---|---|---|---|---|
| water_02_1 | 43879 | FTMS; HCD | 91.28 | 1223.62 |

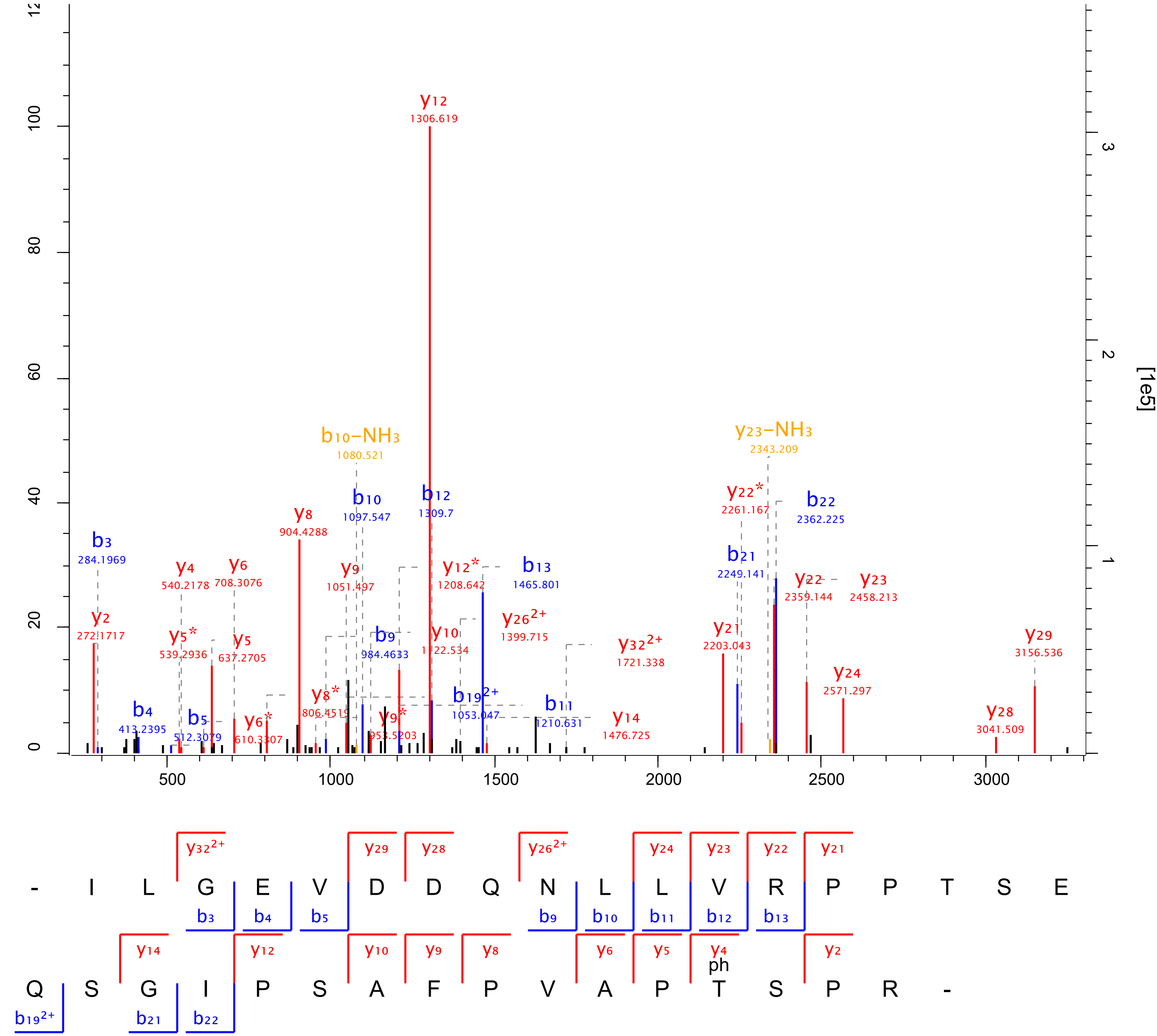

| Raw file | Scan | Method | Score | m/z |
|---|---|---|---|---|
| water_02_1 | 44036 | FTMS; HCD | 45.99 | 785.36 |

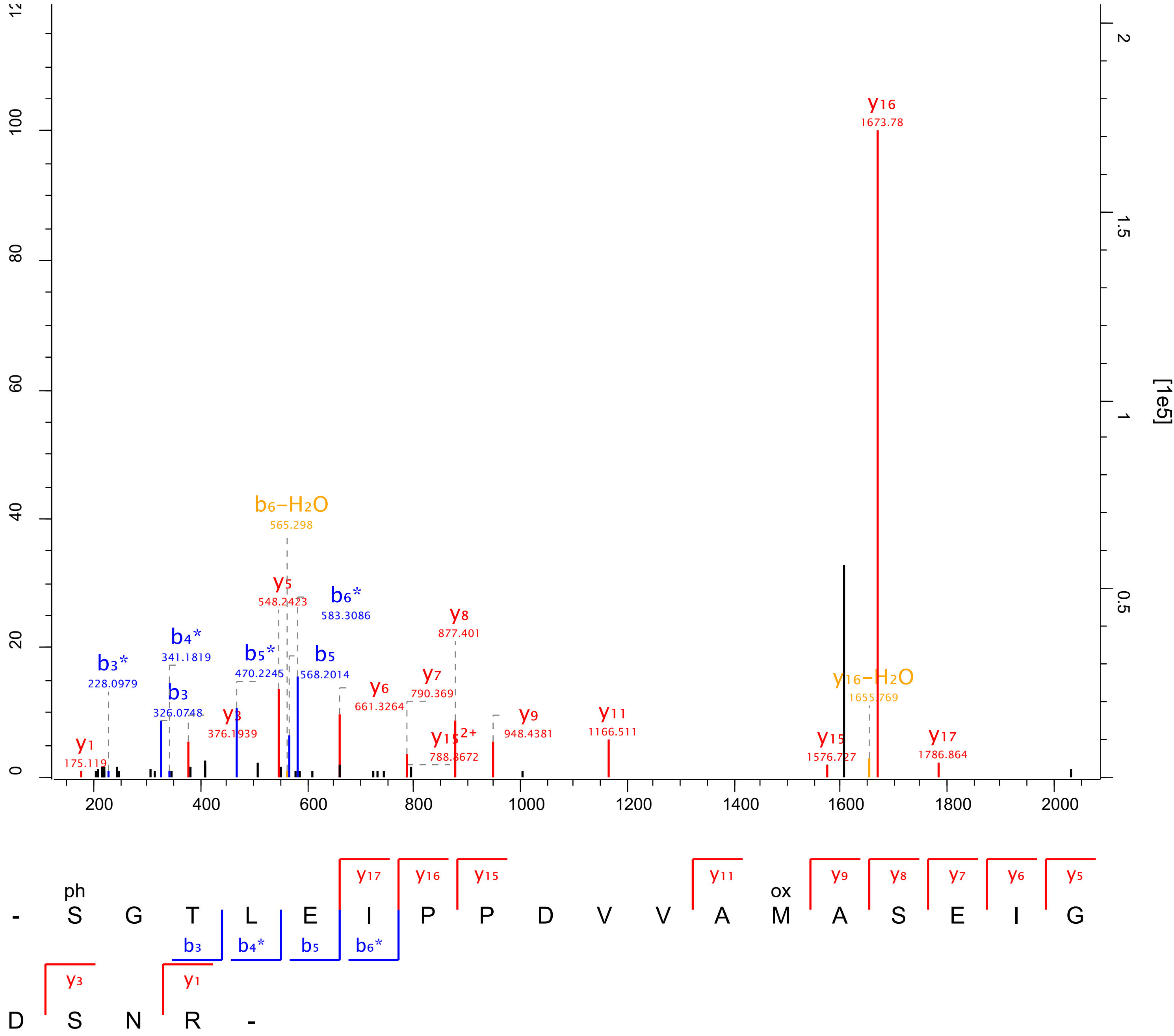

| Raw file | Scan | Method | Score | m/z |
|---|---|---|---|---|
| water_02_1 | 4406 | FTMS; HCD | 57.53 | 584.21 |

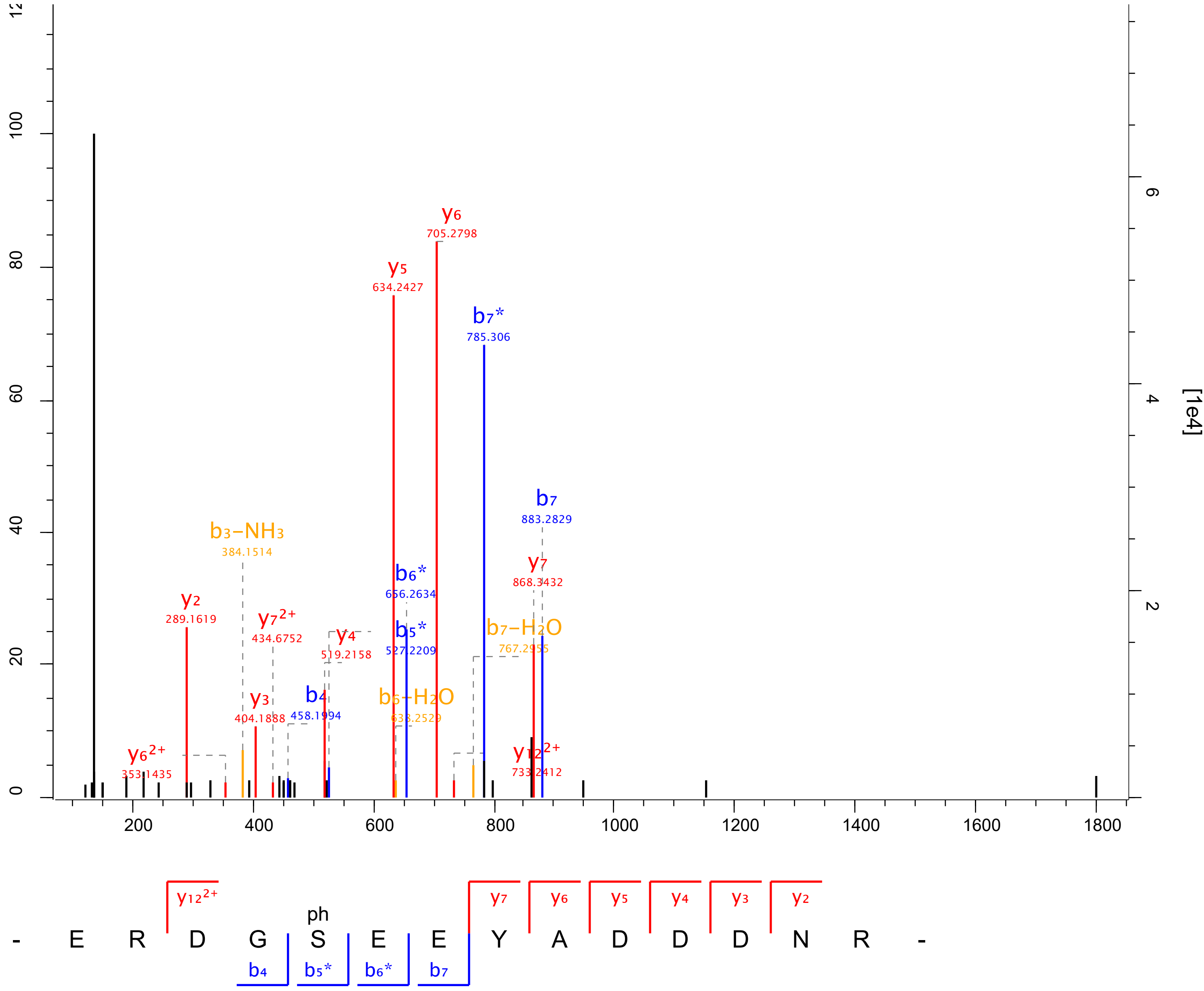

| Raw file | Scan | Method | Score | m/z |
|---|---|---|---|---|
| water_02_1 | 44128 | FTMS; HCD | 40.14 | 831.04 |

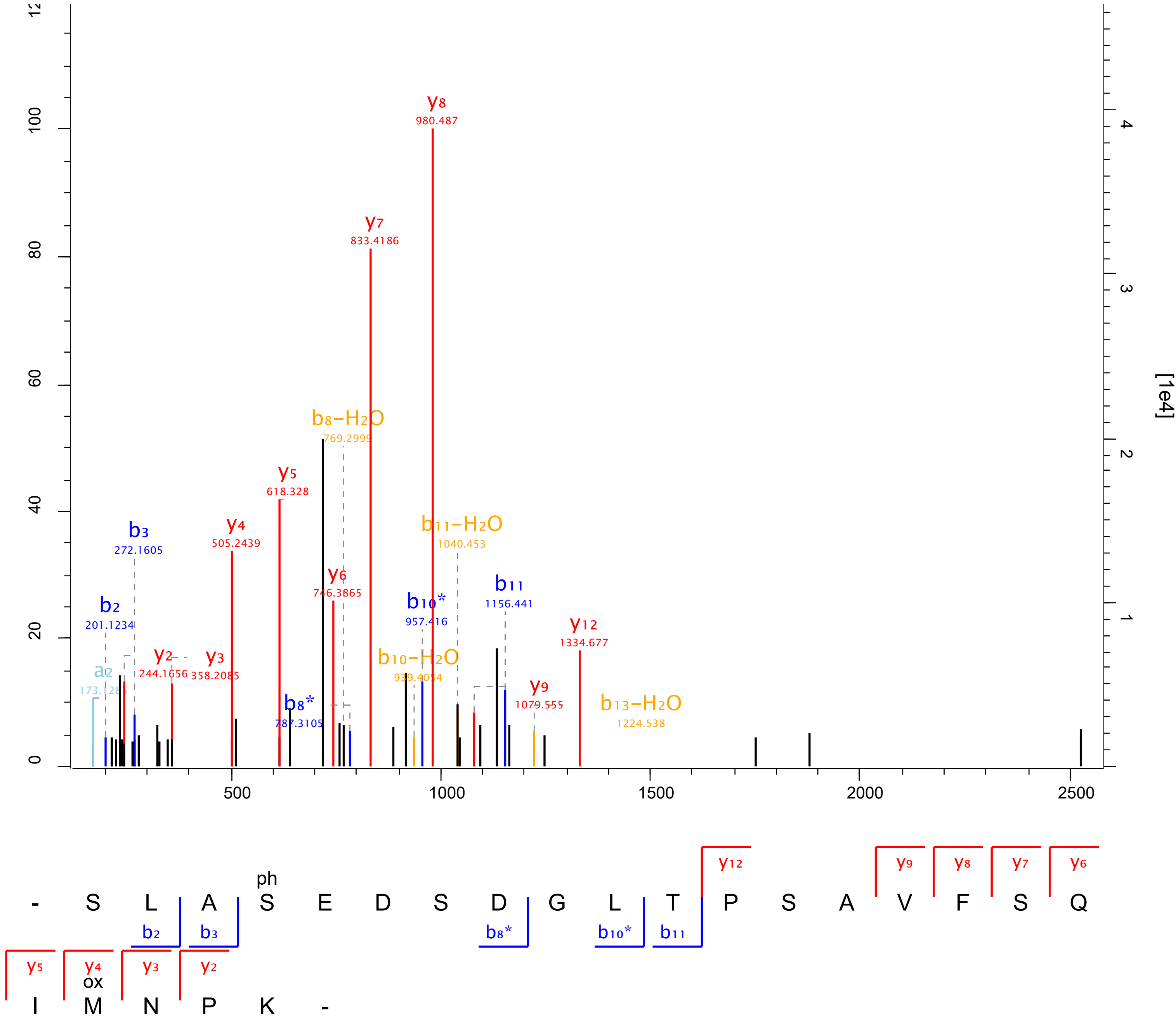

| Raw file | Scan | Method | Score | m/z |
| --- | --- | --- | --- | --- |
| water_02_1 | 4419 | FTMS; HCD | 107.03 | 506.73 |

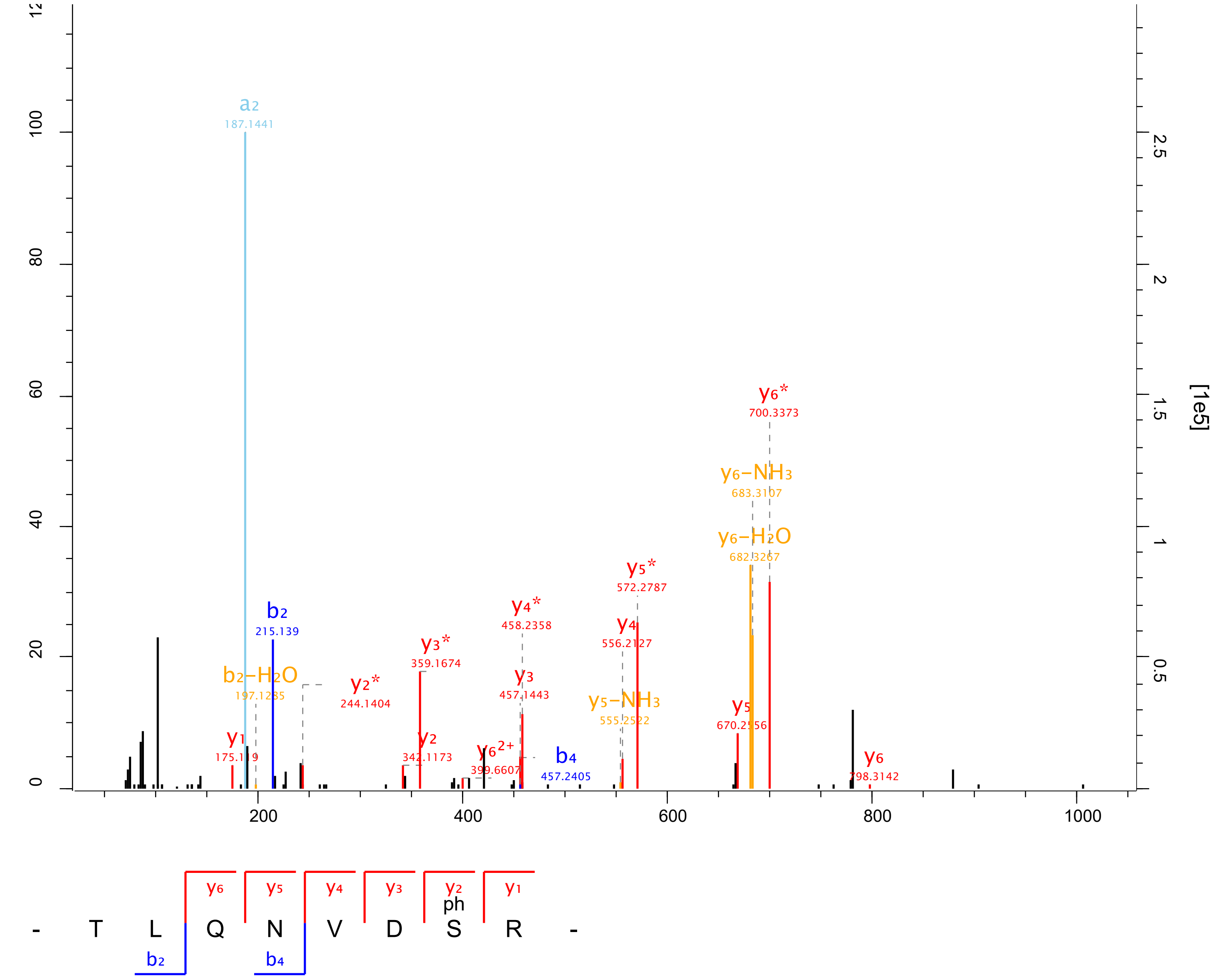

| Raw file | Scan | Method | Score | m/z |
|---|---|---|---|---|
| water_02_1 | 4420 | FTMS; HCD | 86.11 | 553.23 |

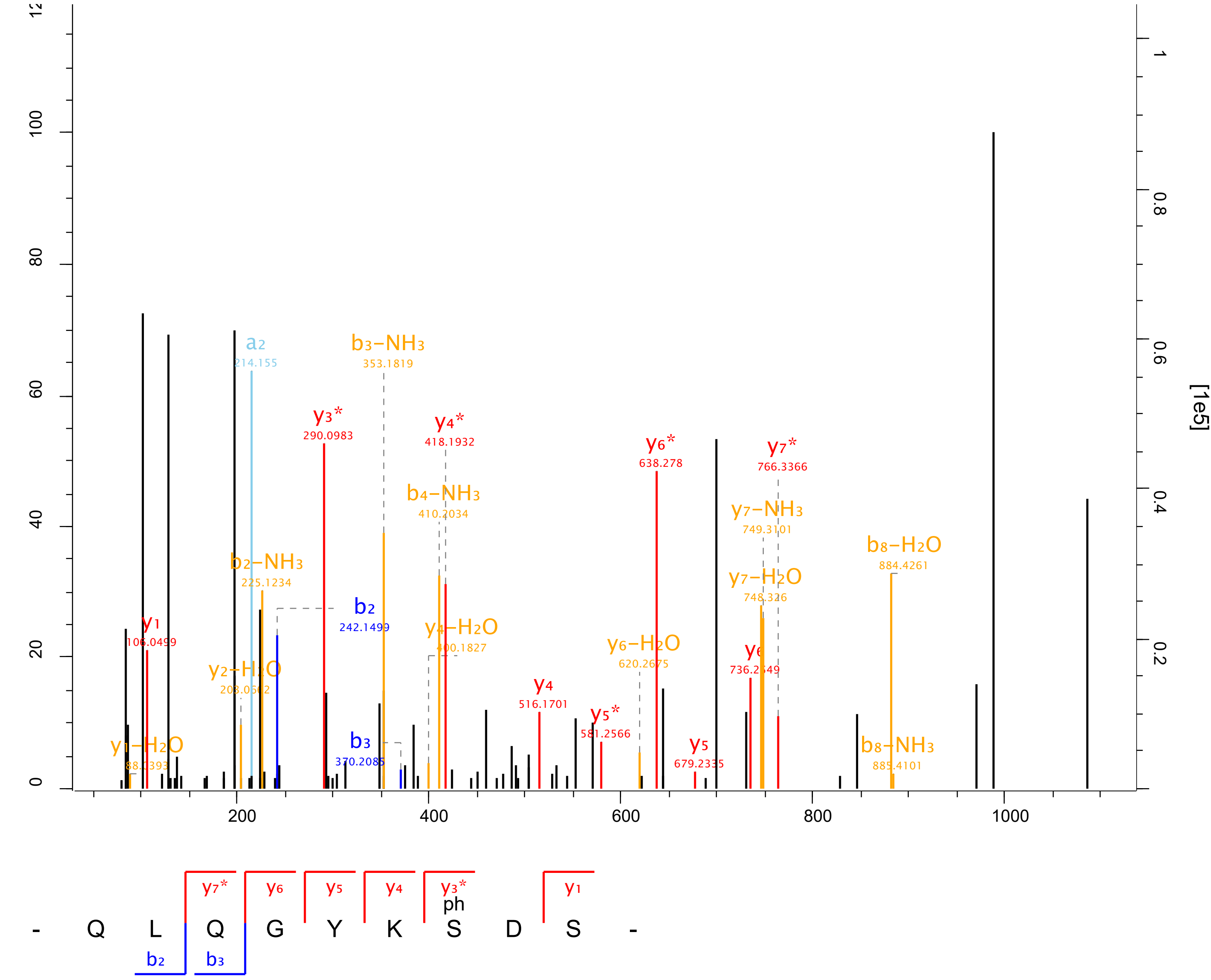

| Raw file | Scan | Method | Score | m/z |
|---|---|---|---|---|
| water_02_1 | 4440 | FTMS; HCD | 69.72 | 512.74 |

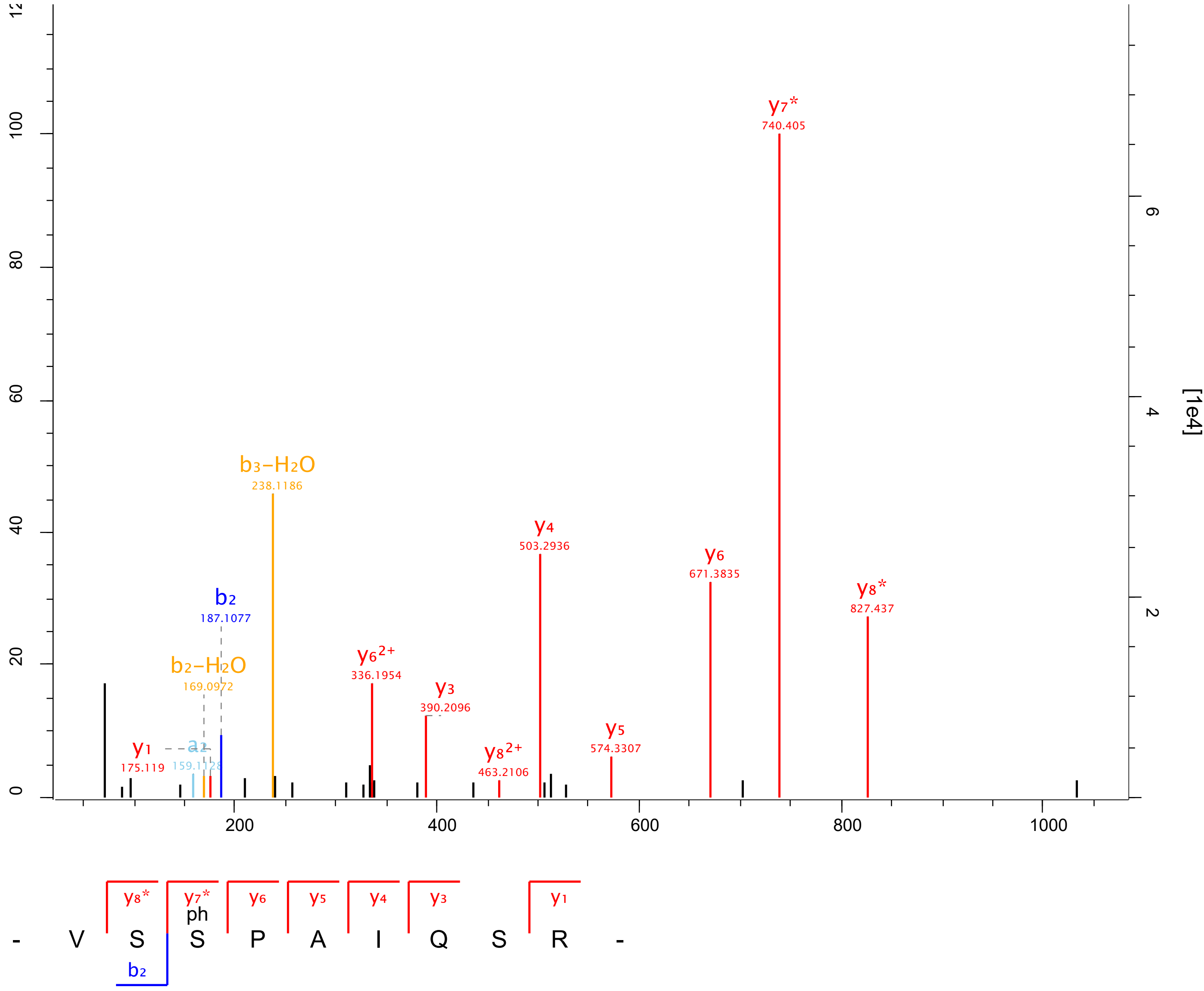

| Raw file | Scan | Method | Score | m/z |
|---|---|---|---|---|
| water_02_1 | 44435 | FTMS; HCD | 204.31 | 922.92 |

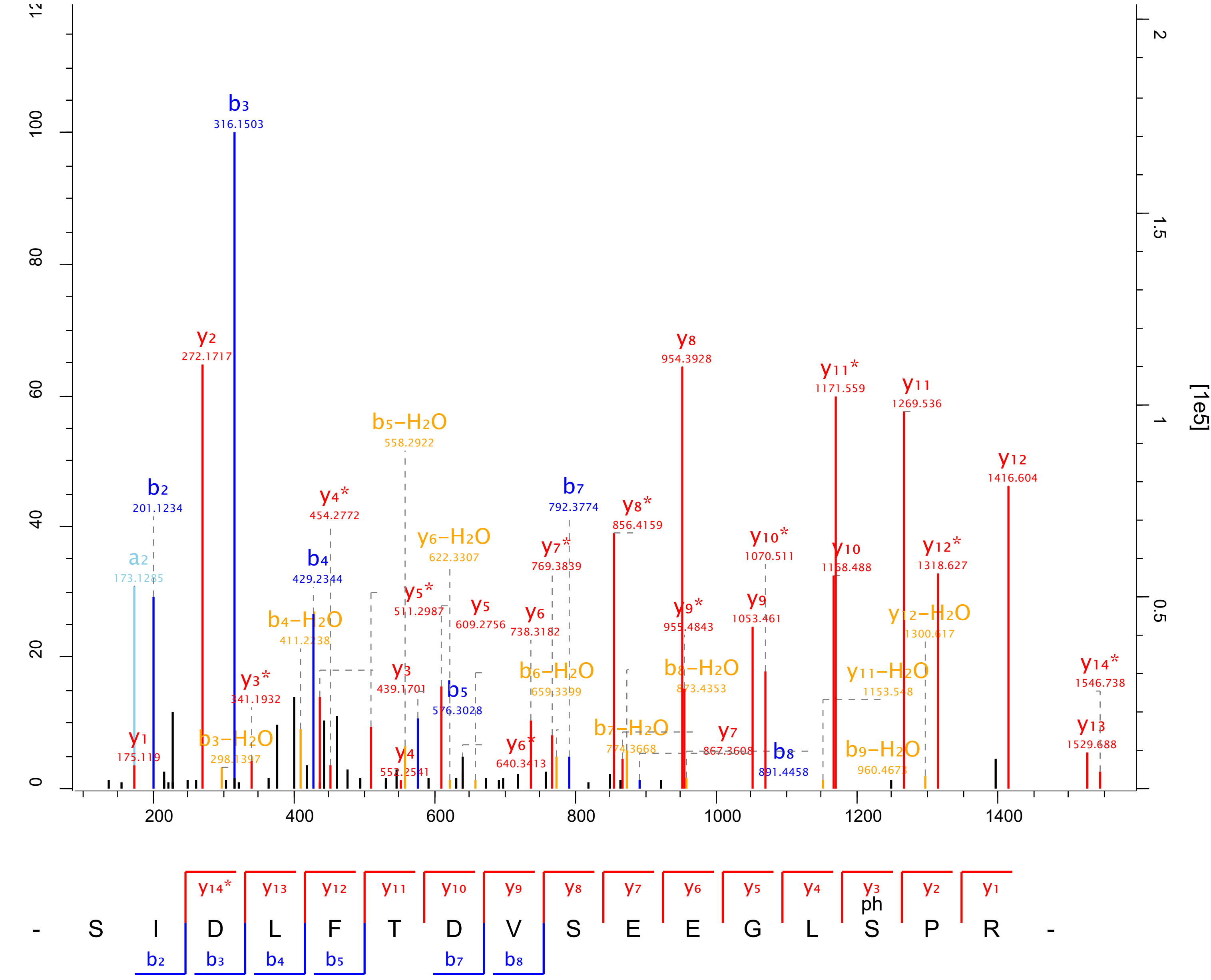

| Raw file | Scan | Method | Score | m/z |
|---|---|---|---|---|
| water_02_1 | 4453 | FTMS; HCD | 64.83 | 476.23 |

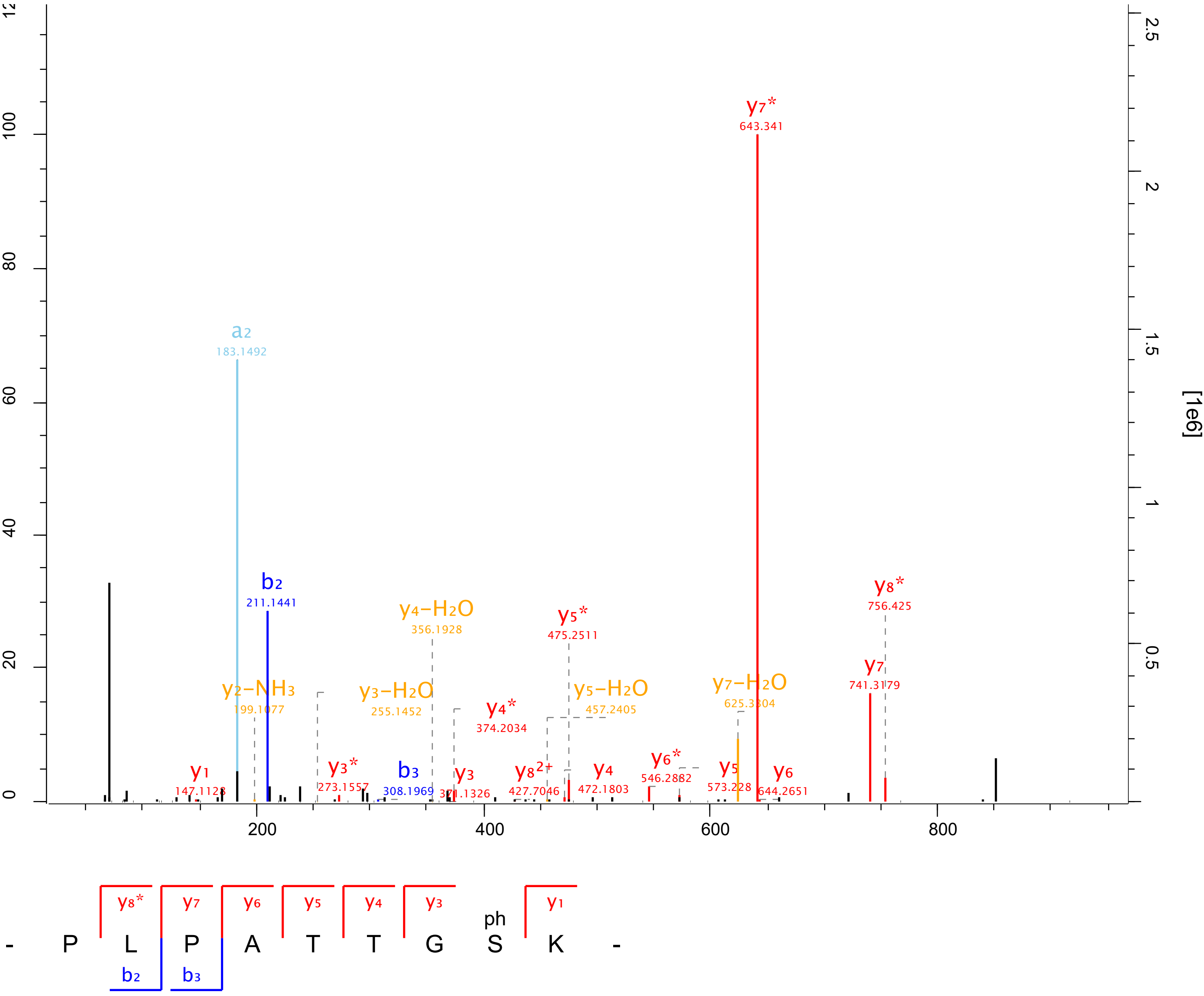

| Raw file | Scan | Method | Score | m/z |
|---|---|---|---|---|
| water_02_1 | 44564 | FTMS; HCD | 52.28 | 1030.48 |

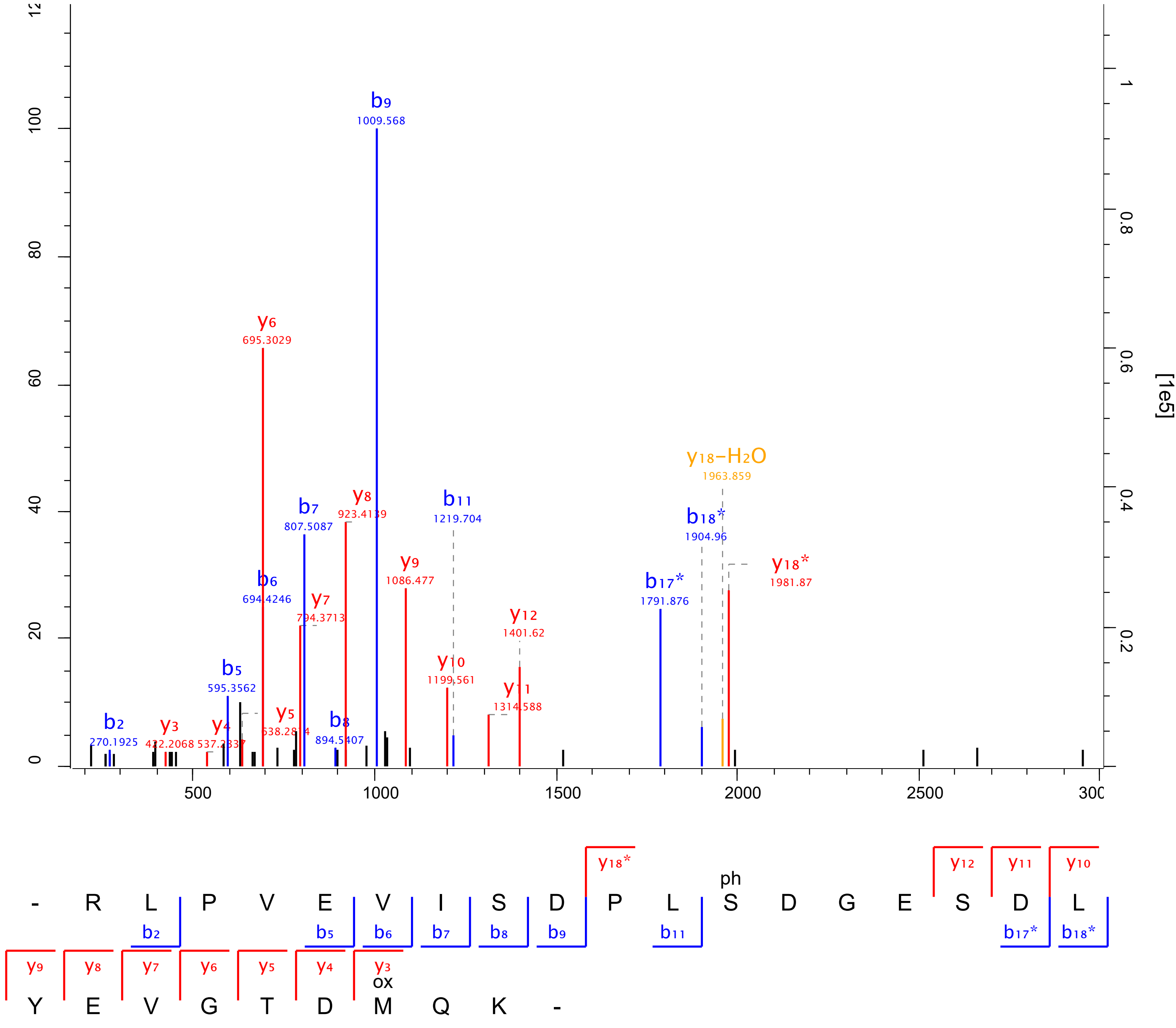

| Raw file | Scan | Method | Score | m/z |
| --- | --- | --- | --- | --- |
| water_02_1 | 44571 | FTMS; HCD | 45.35 | 829.4 |

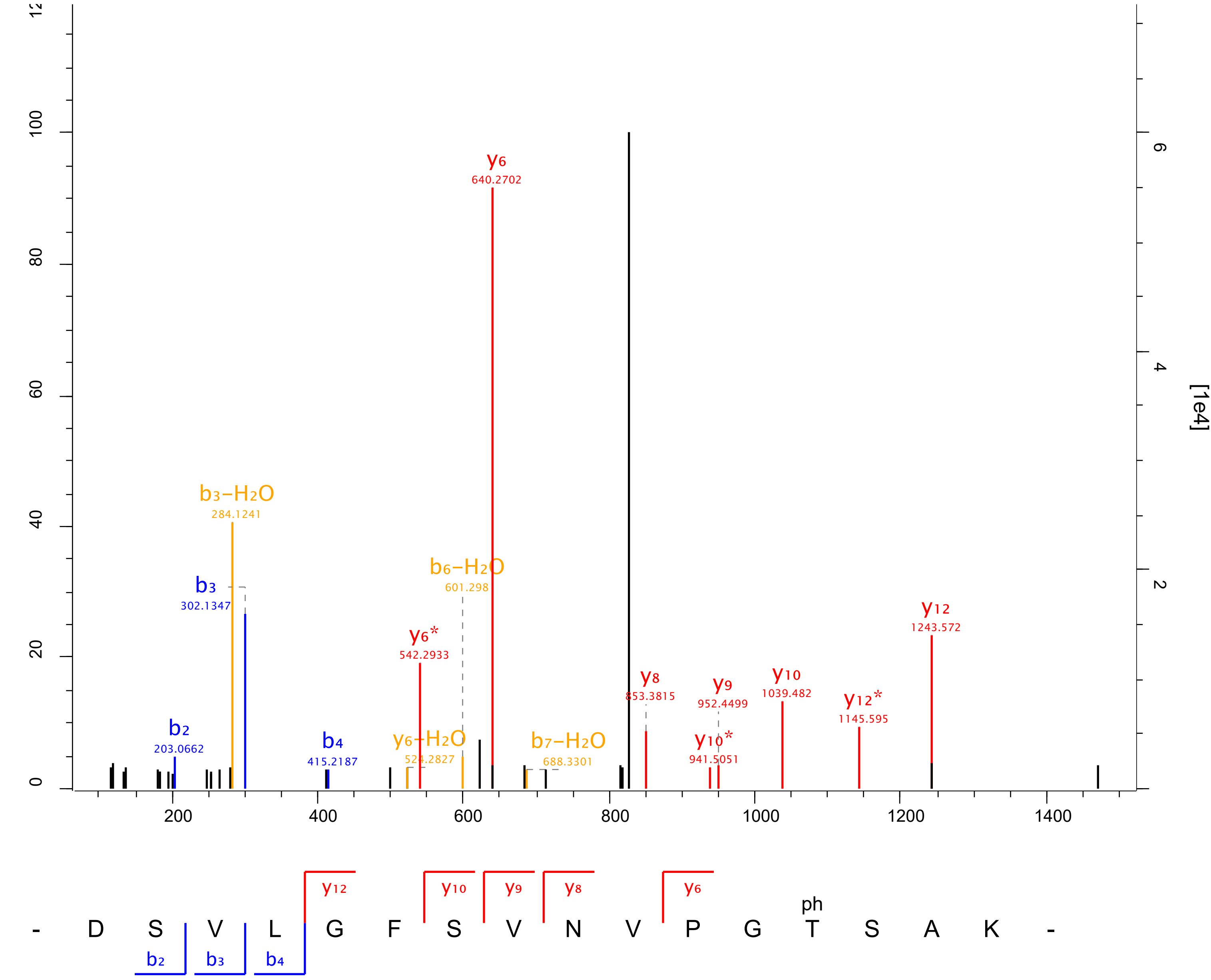

| Raw file | Scan | Method | Score | m/z |
|---|---|---|---|---|
| water_02_1 | 44622 | FTMS; HCD | 81.24 | 1023.76 |

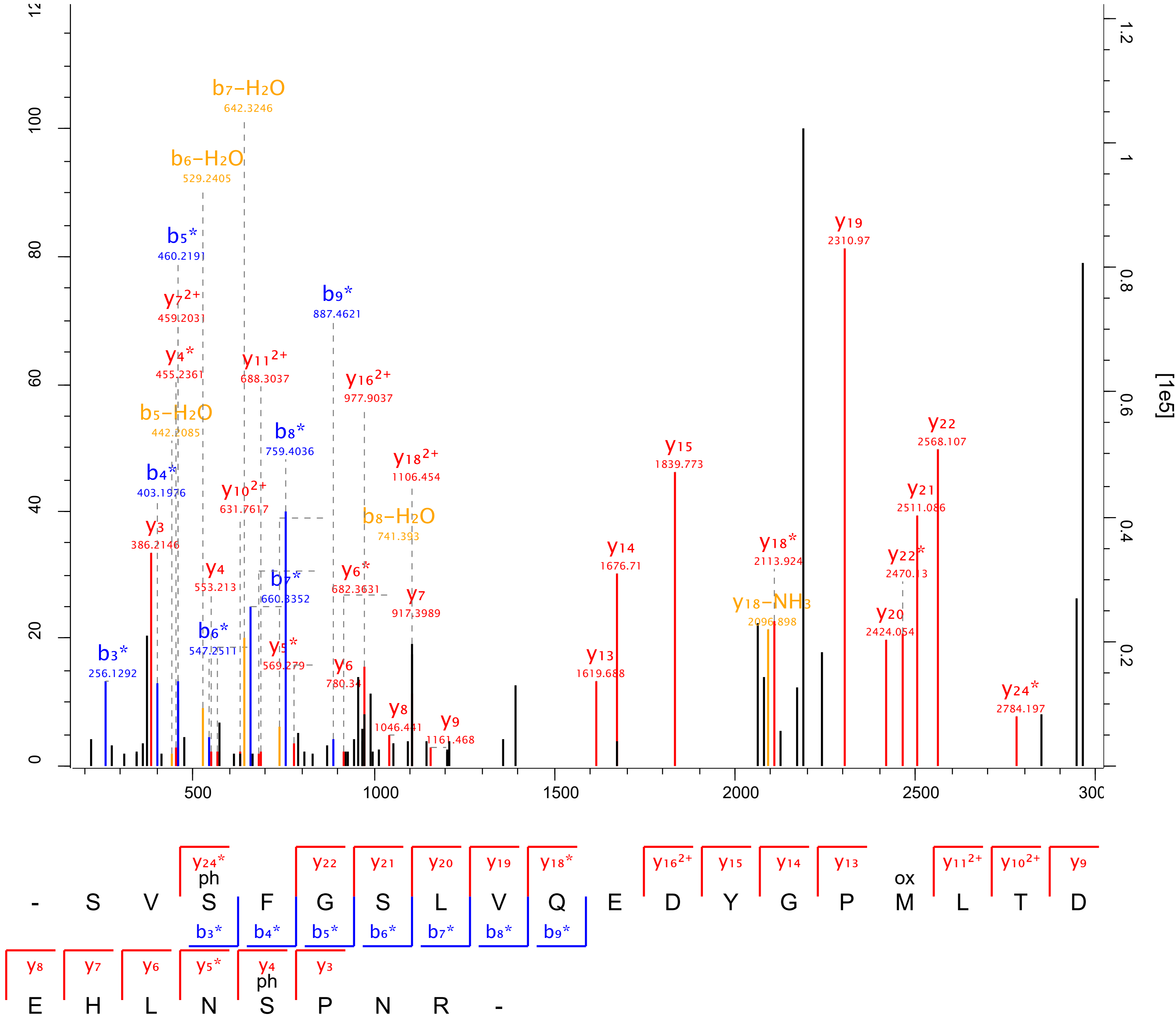

| Raw file | Scan | Method | Score | m/z |
| --- | --- | --- | --- | --- |
| water_02_1 | 4512 | FTMS; HCD | 79.61 | 594.22 |

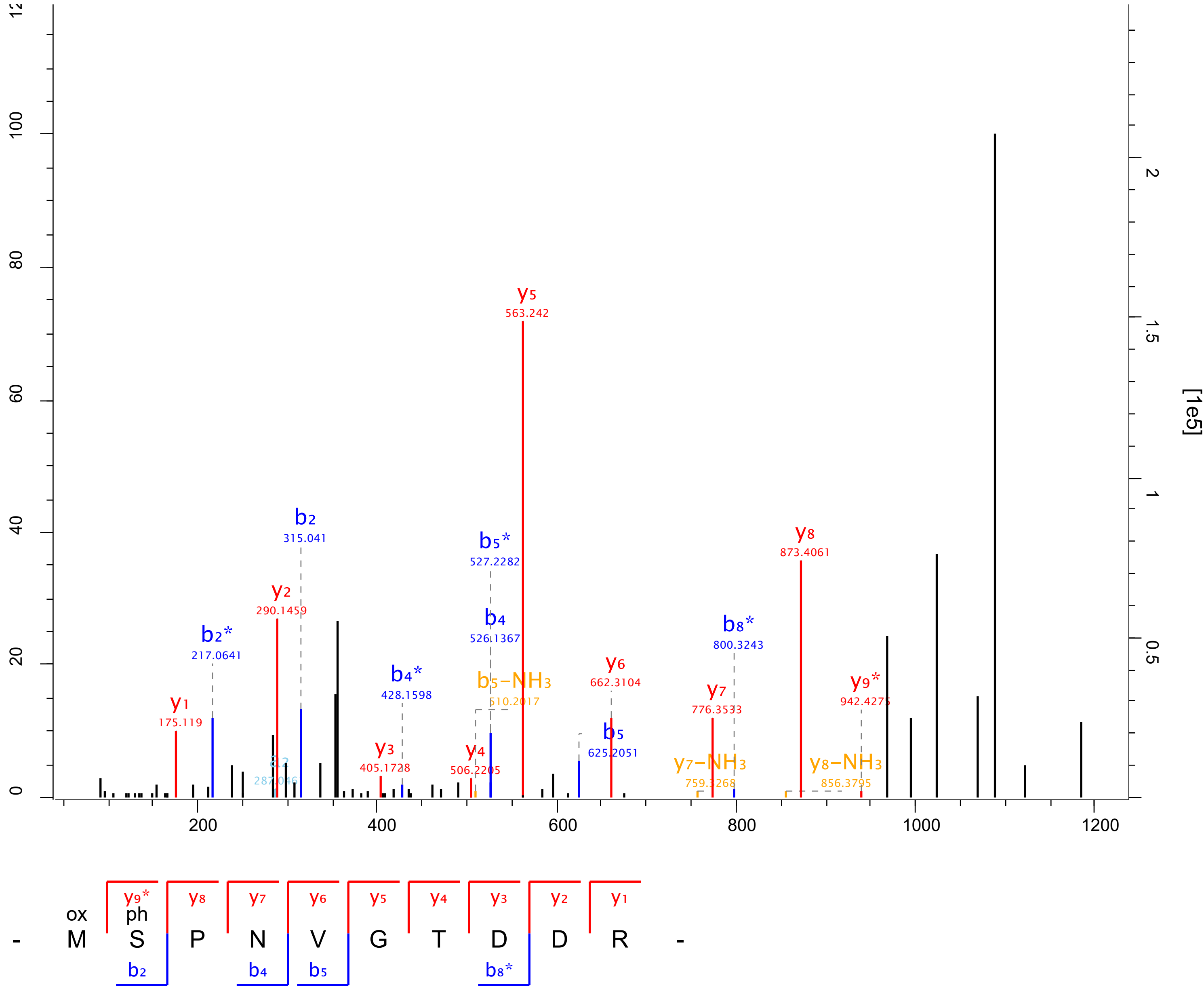

| Raw file | Scan | Method | Score | m/z |
|---|---|---|---|---|
| water_02_1 | 4535 | FTMS; HCD | 129.15 | 658.28 |

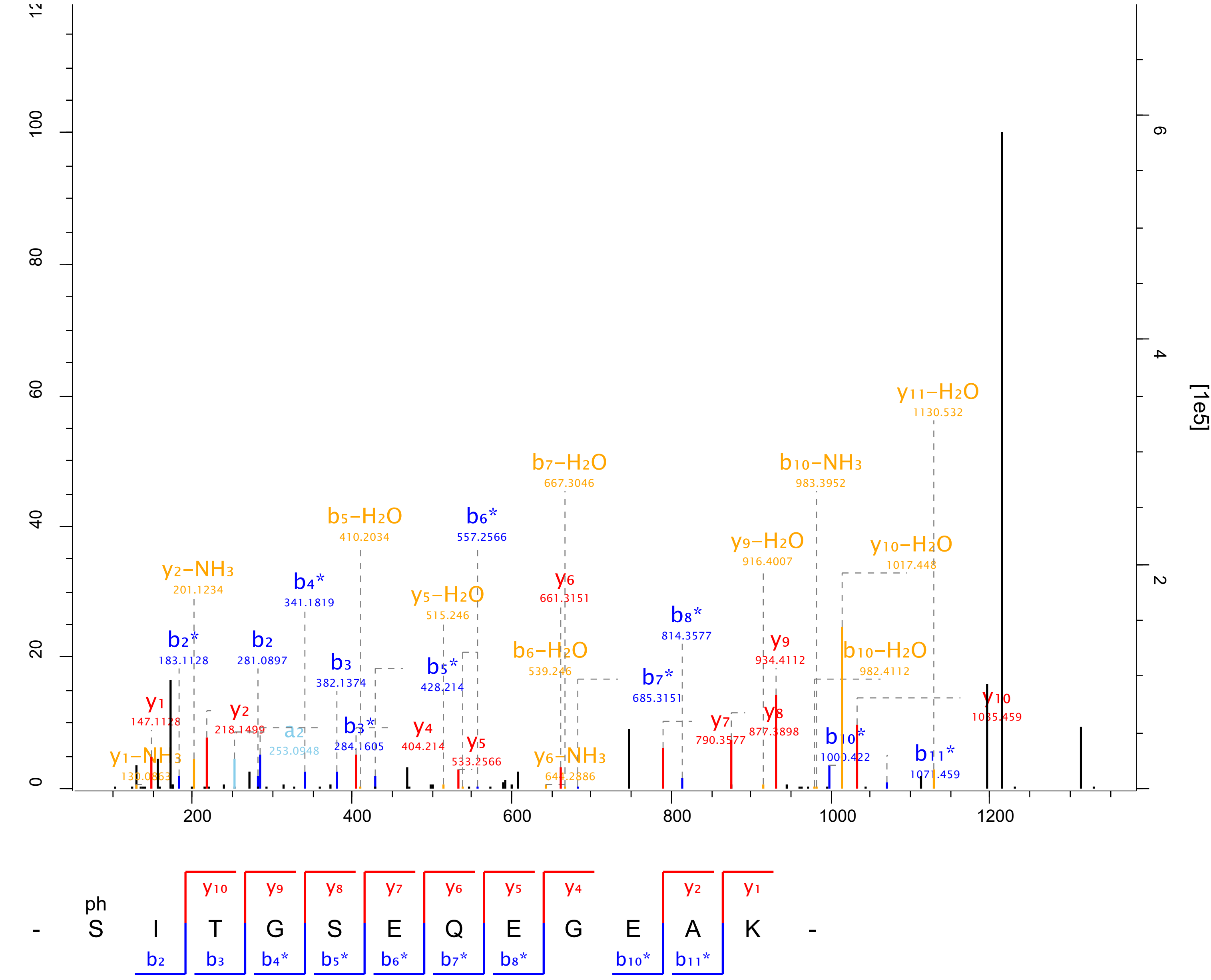

| Raw file | Scan | Method | Score | m/z |
|---|---|---|---|---|
| water_02_1 | 4537 | FTMS; HCD | 42.95 | 620.26 |

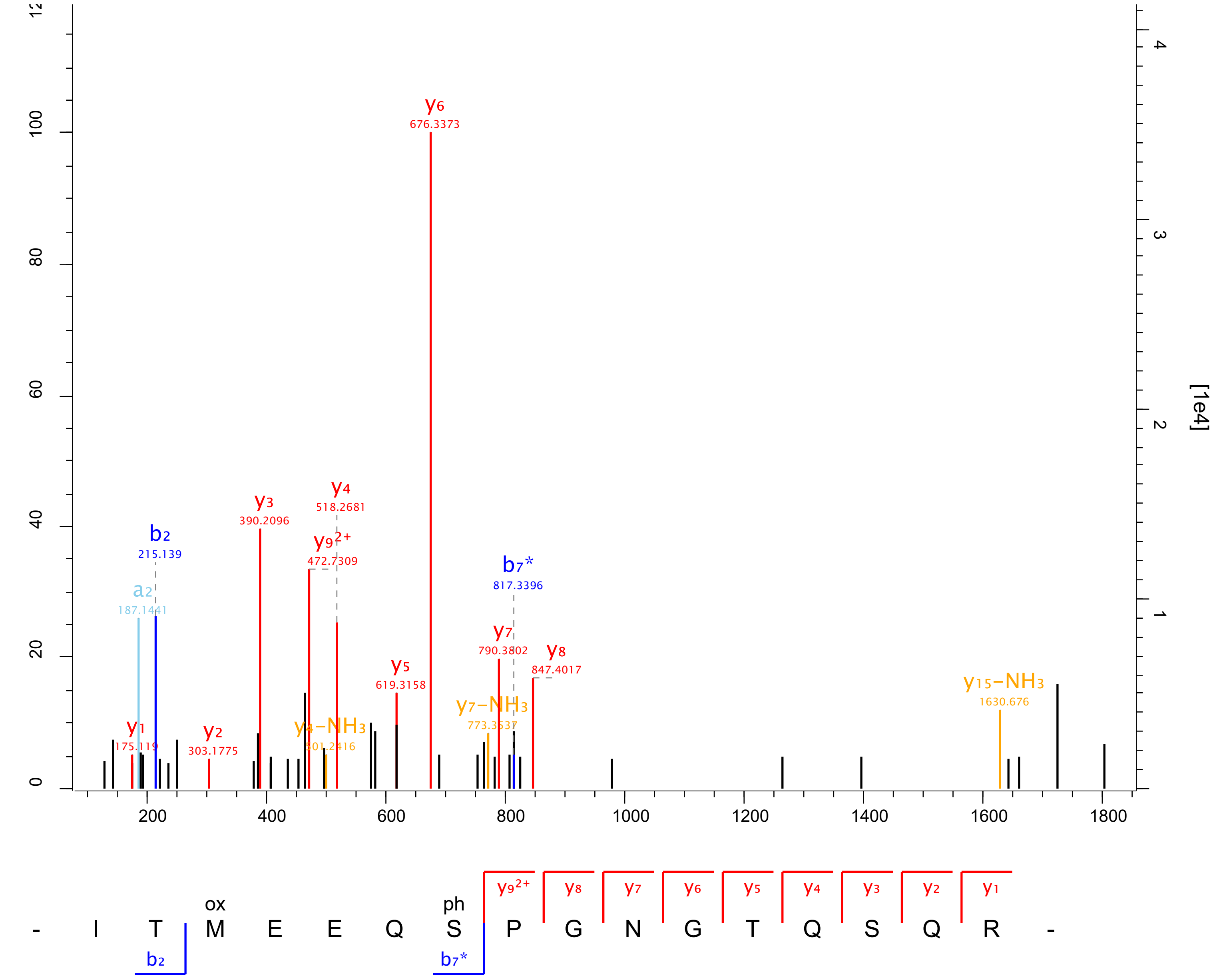

| Raw file | Scan | Method | Score | m/z |
|---|---|---|---|---|
| water_02_1 | 4598 | FTMS; HCD | 69.6 | 363.18 |

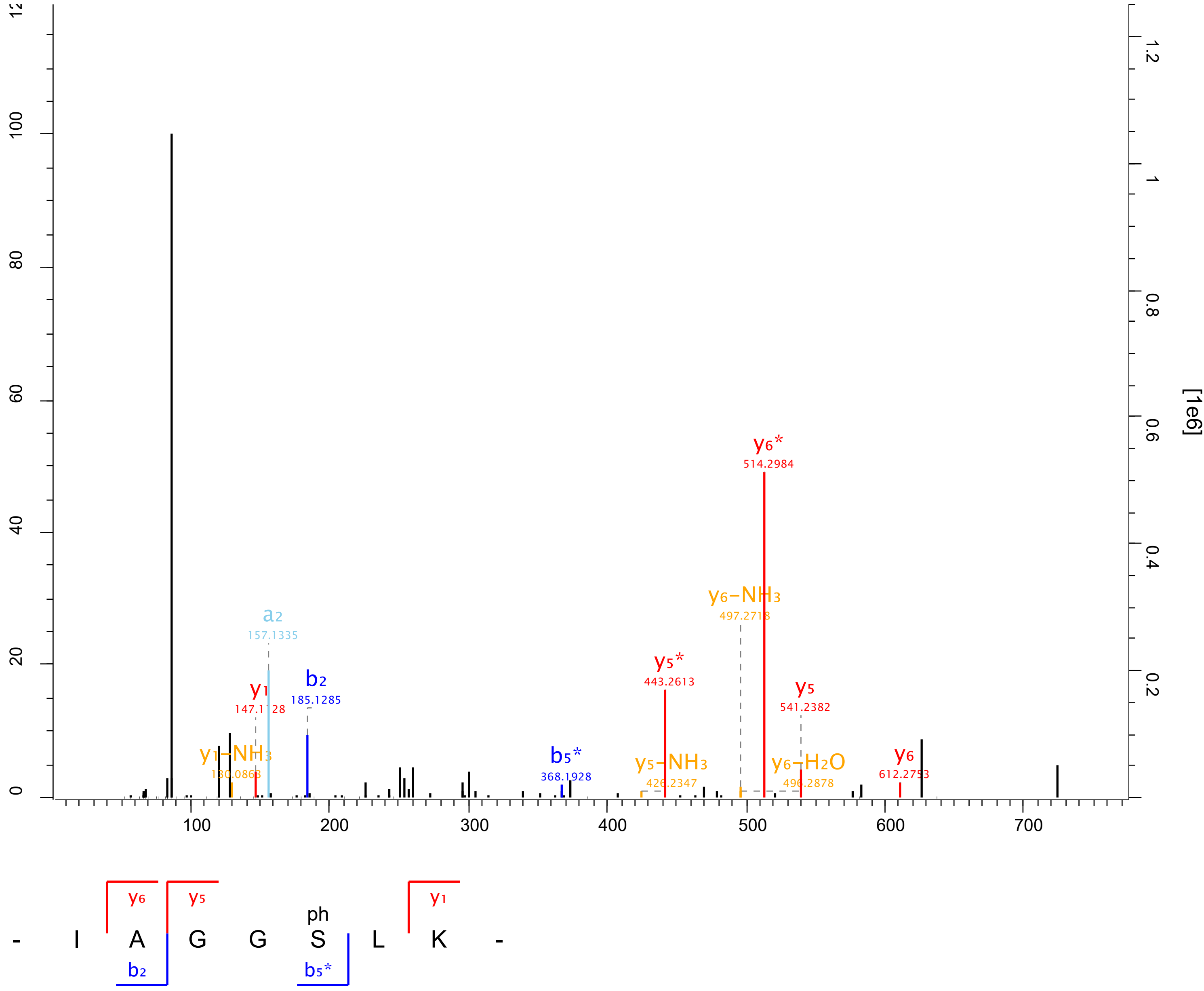

| Raw file | Scan | Method | Score | m/z |
|---|---|---|---|---|
| water_02_1 | 46185 | FTMS; HCD | 78.69 | 725.34 |

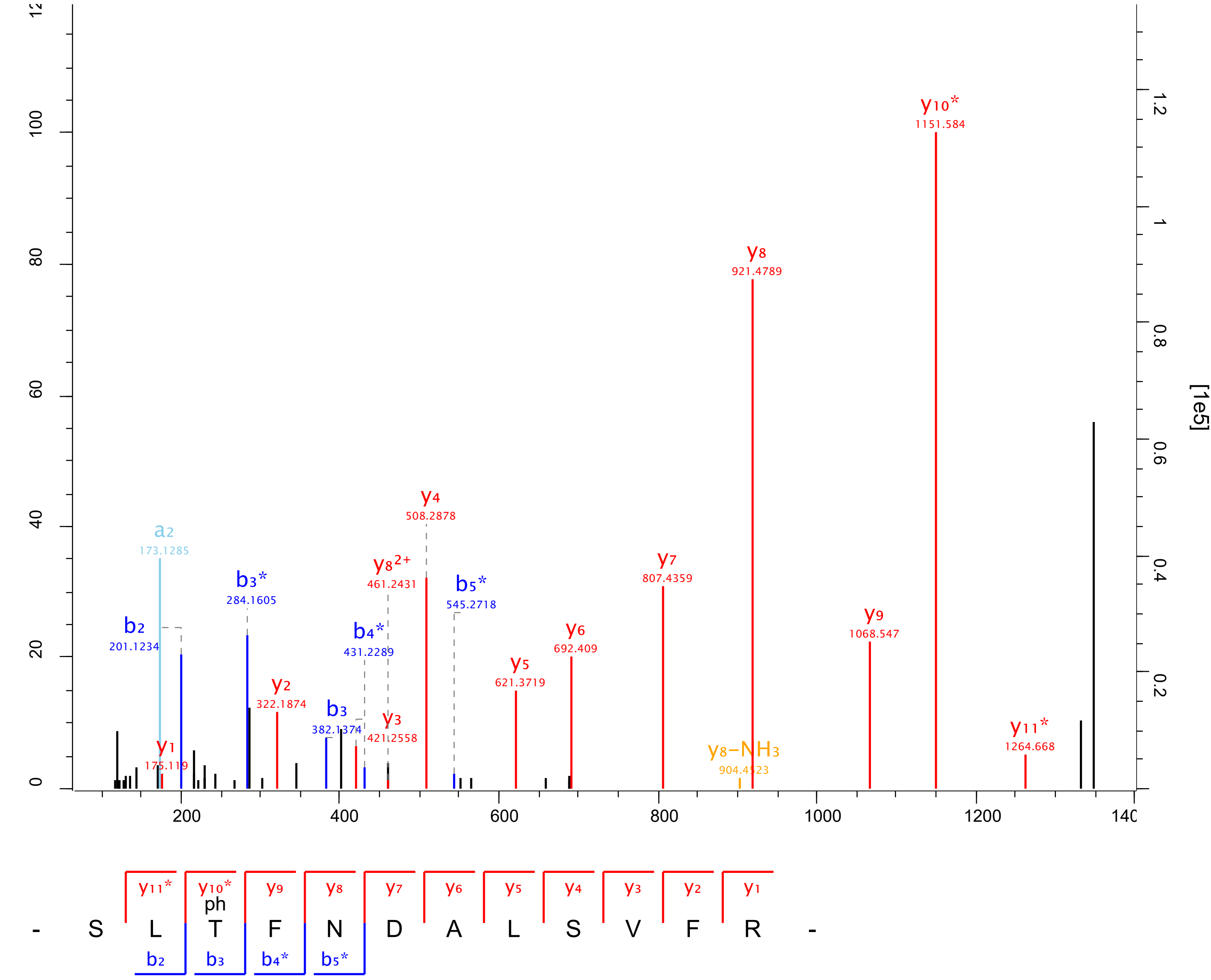

| Raw file | Scan | Method | Score | m/z |
|---|---|---|---|---|
| water_02_1 | 4636 | FTMS; HCD | 78.1 | 702.32 |

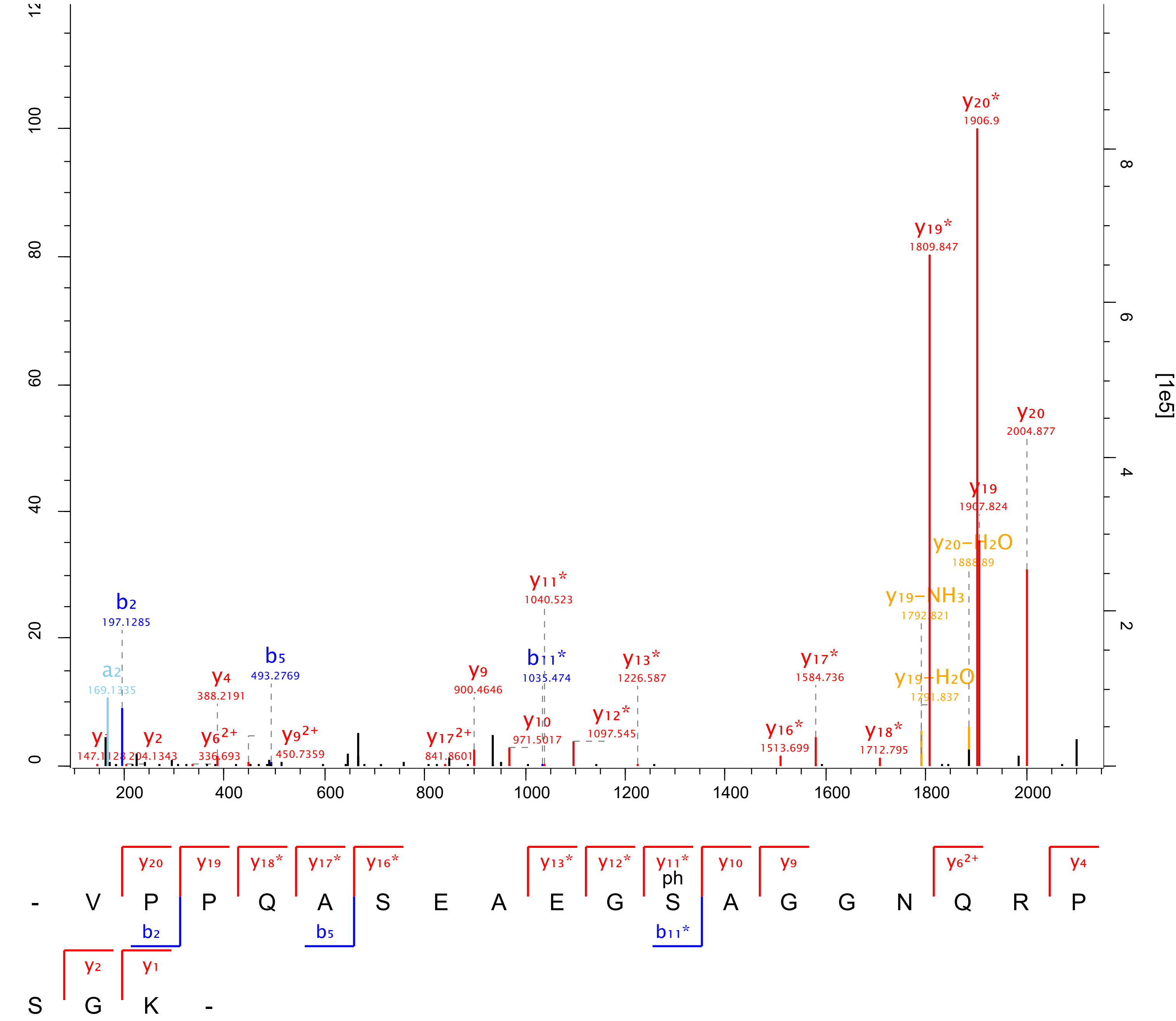

| Raw file | Scan | Method | Score | m/z |
| --- | --- | --- | --- | --- |
| water_02_1 | 46487 | FTMS; HCD | 47.33 | 1053.1 |

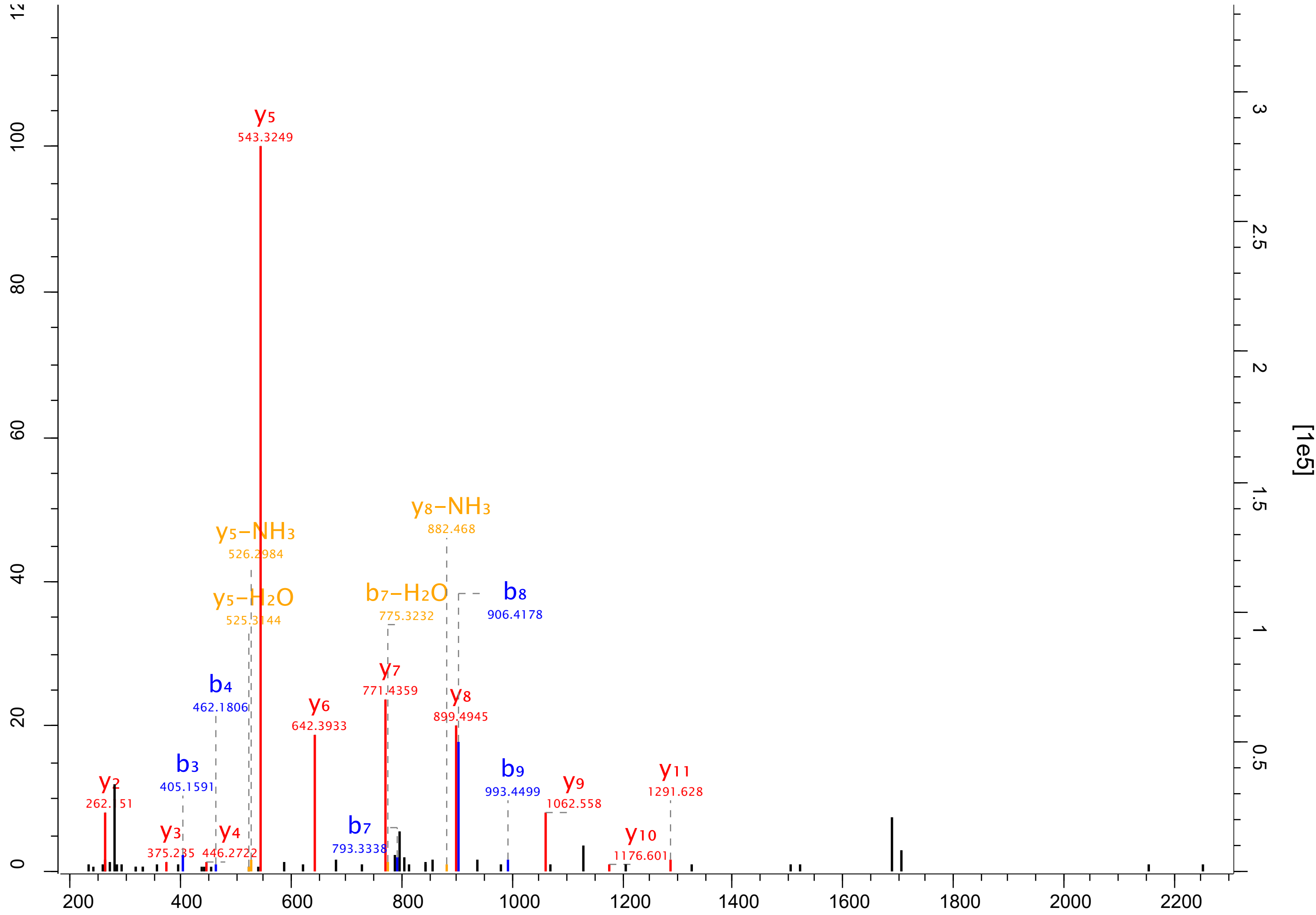


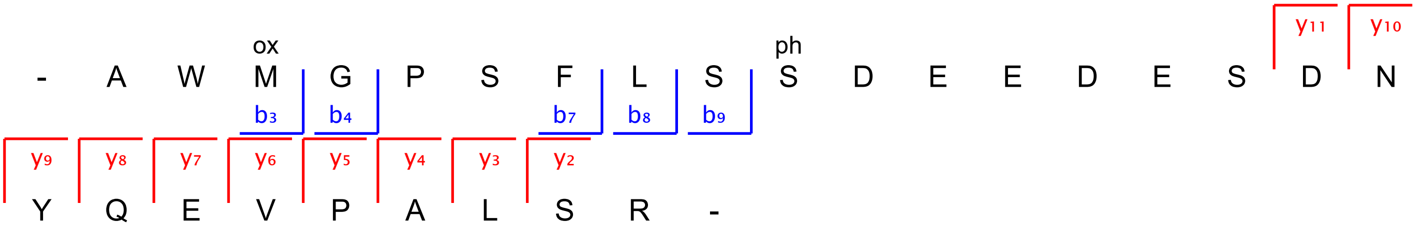

| Raw file | Scan | Method | Score | m/z |
| --- | --- | --- | --- | --- |
| water_02_1 | 46597 | FTMS; HCD | 91.66 | 789.94 |

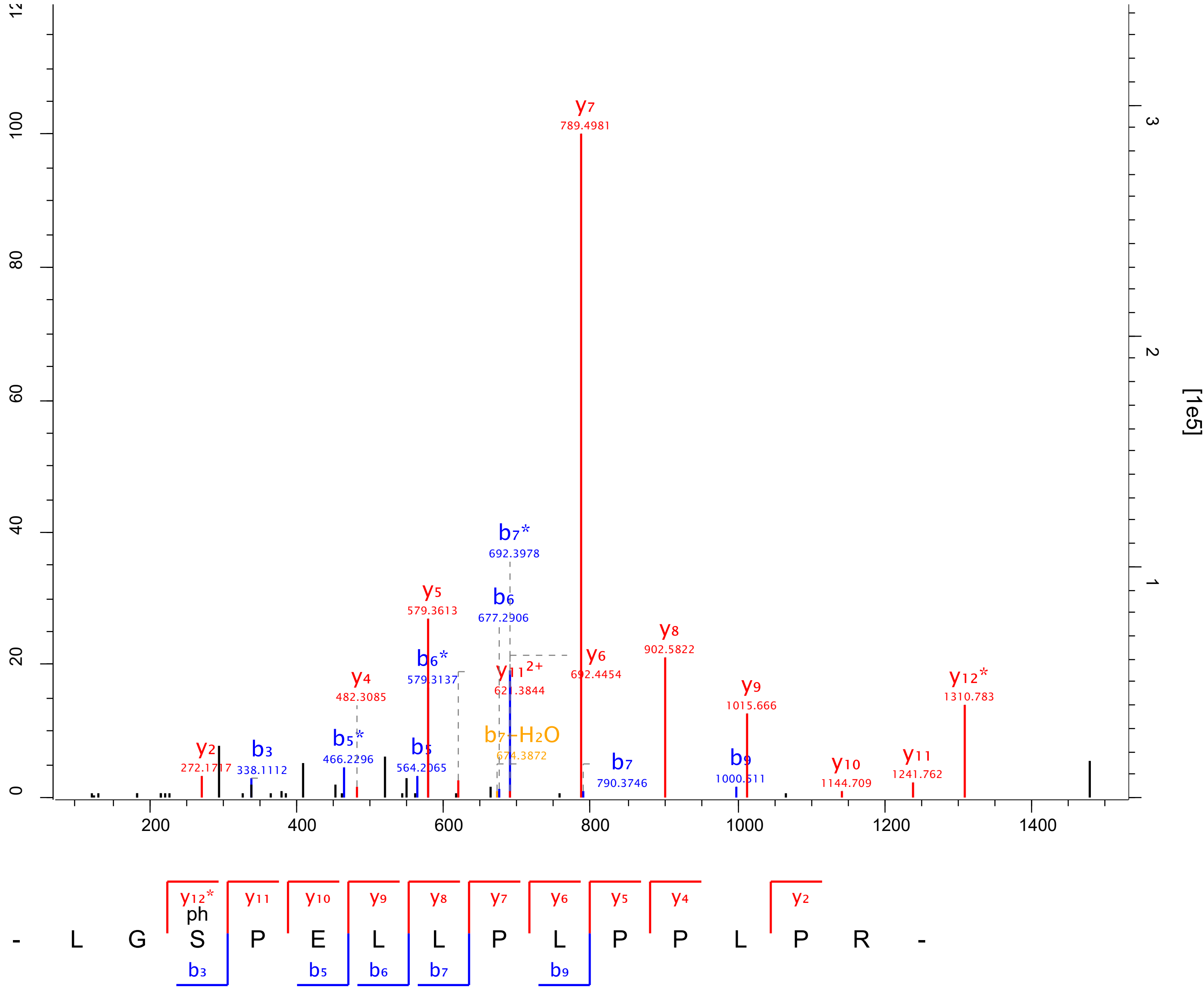

| Raw file | Scan | Method | Score | m/z |
|---|---|---|---|---|
| water_02_1 | 46693 | FTMS; HCD | 147.42 | 941.08 |

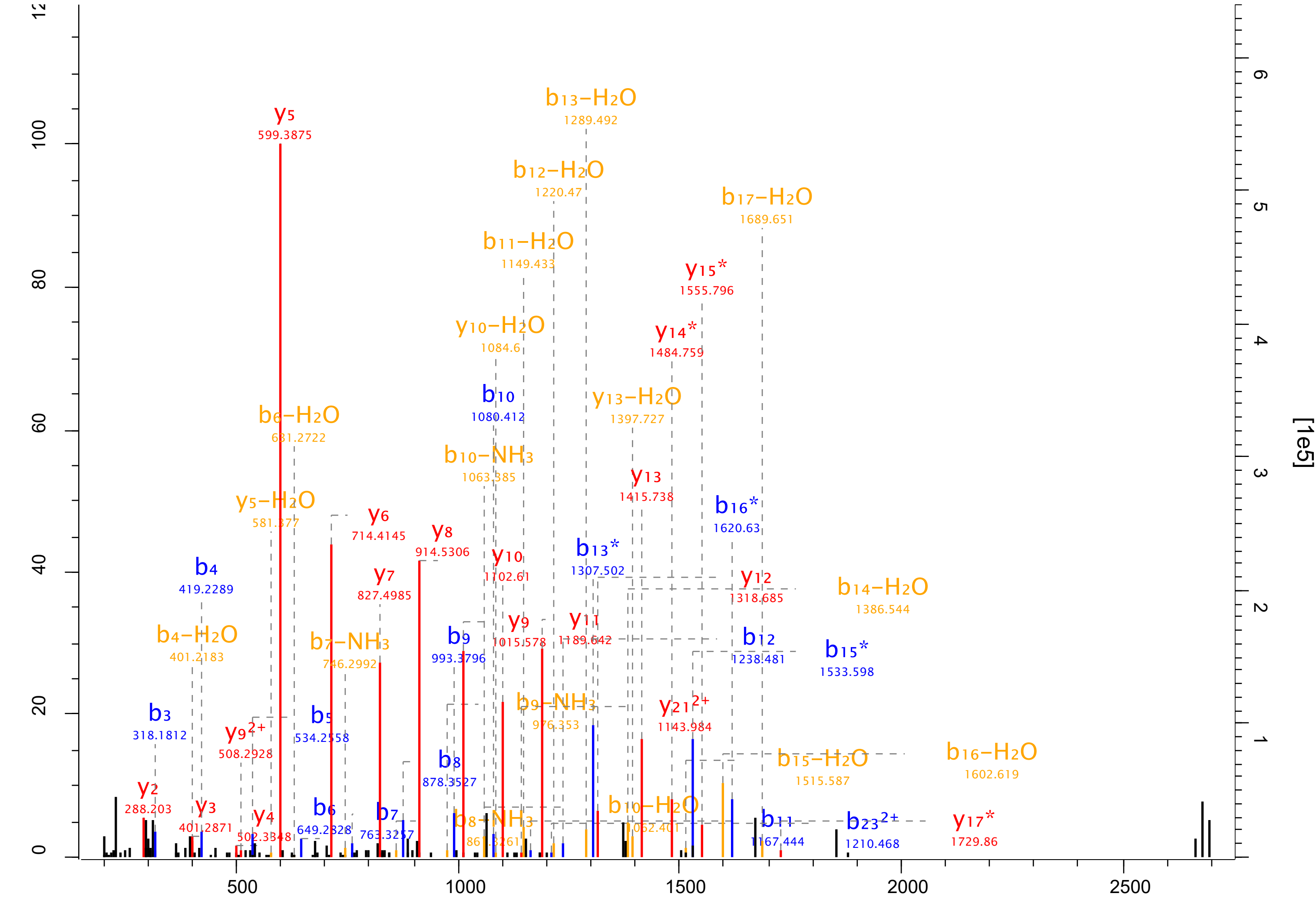


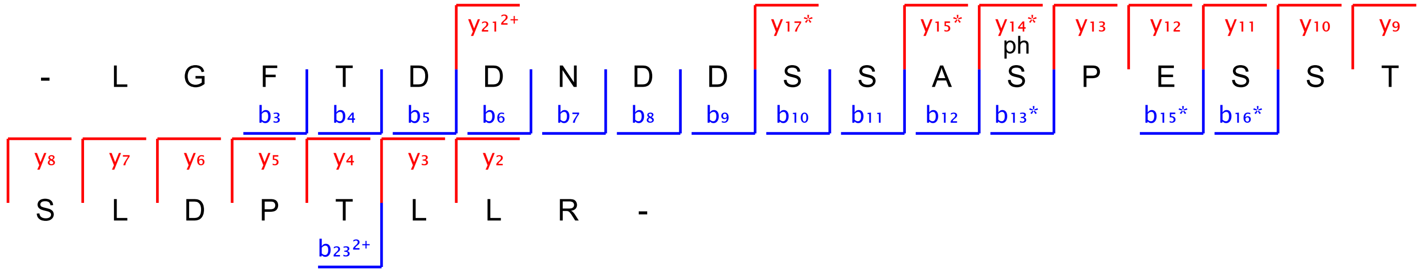

| Raw file | Scan | Method | Score | m/z |
|---|---|---|---|---|
| water_02_1 | 4709 | FTMS; HCD | 42.84 | 499.23 |

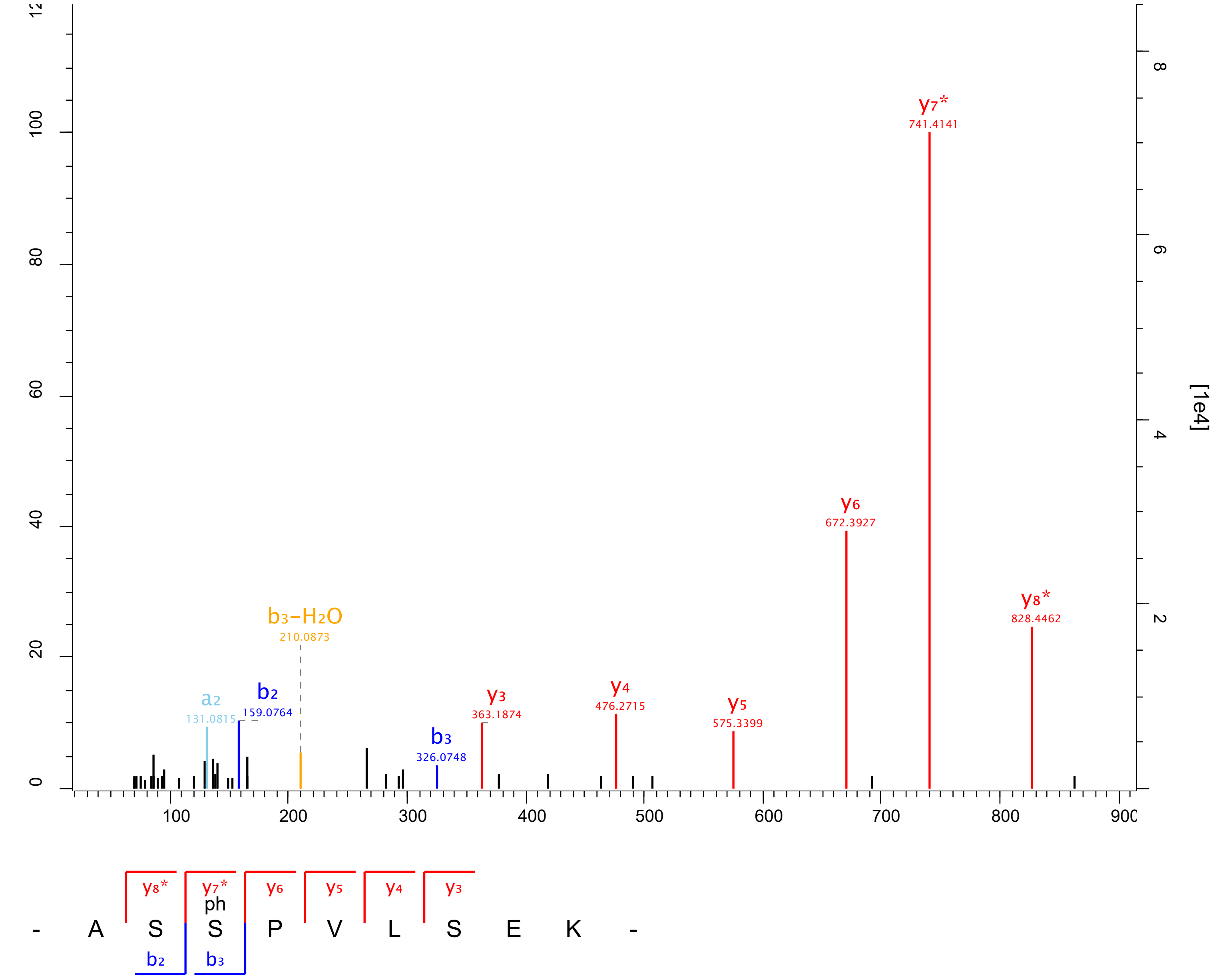

| Raw file | Scan | Method | Score | m/z |
| --- | --- | --- | --- | --- |
| water_02_1 | 4719 | FTMS; HCD | 47.71 | 437.21 |

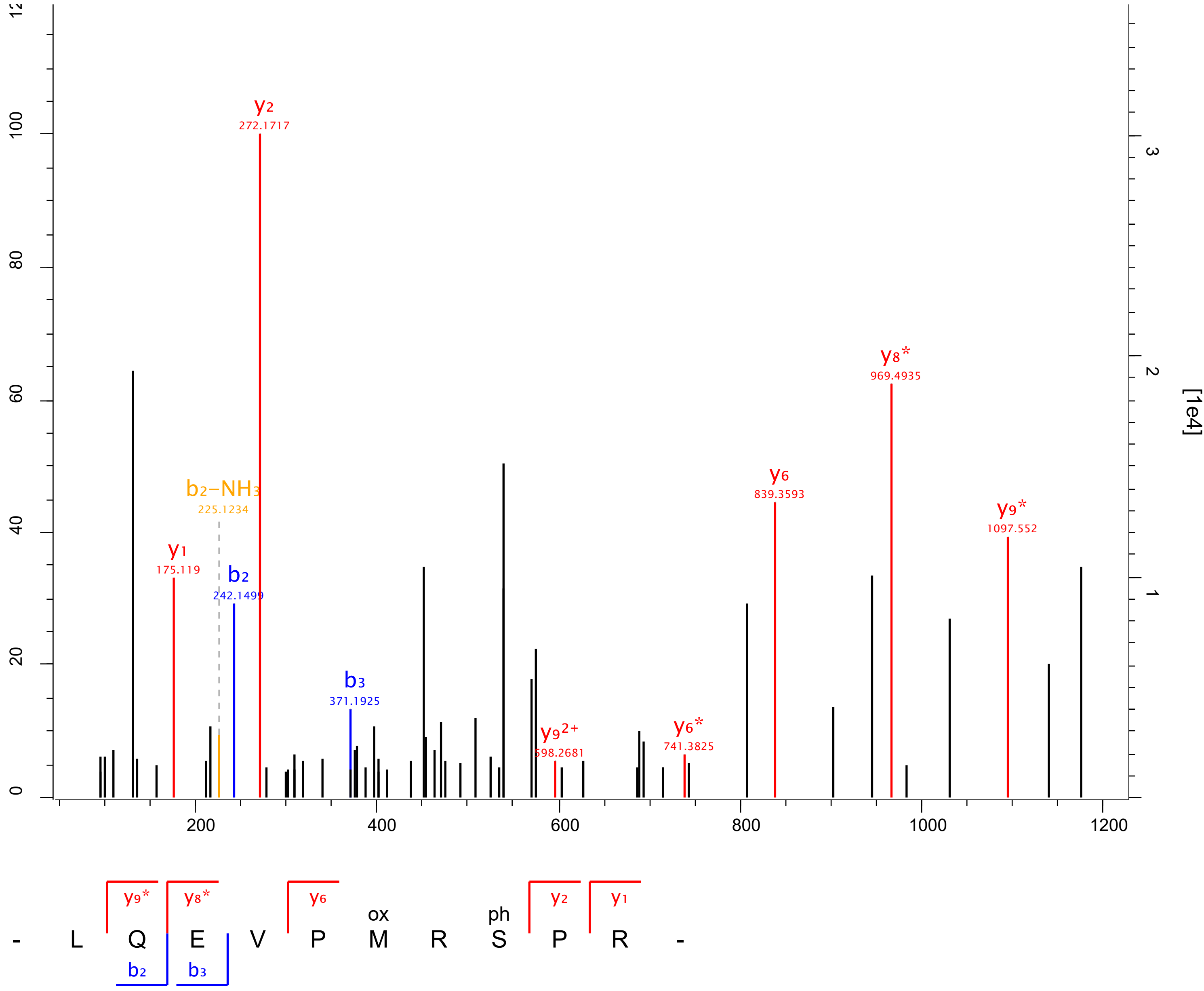

| Raw file | Scan | Method | Score | m/z |
| --- | --- | --- | --- | --- |
| water_02_1 | 47198 | FTMS; HCD | 59.37 | 768.33 |

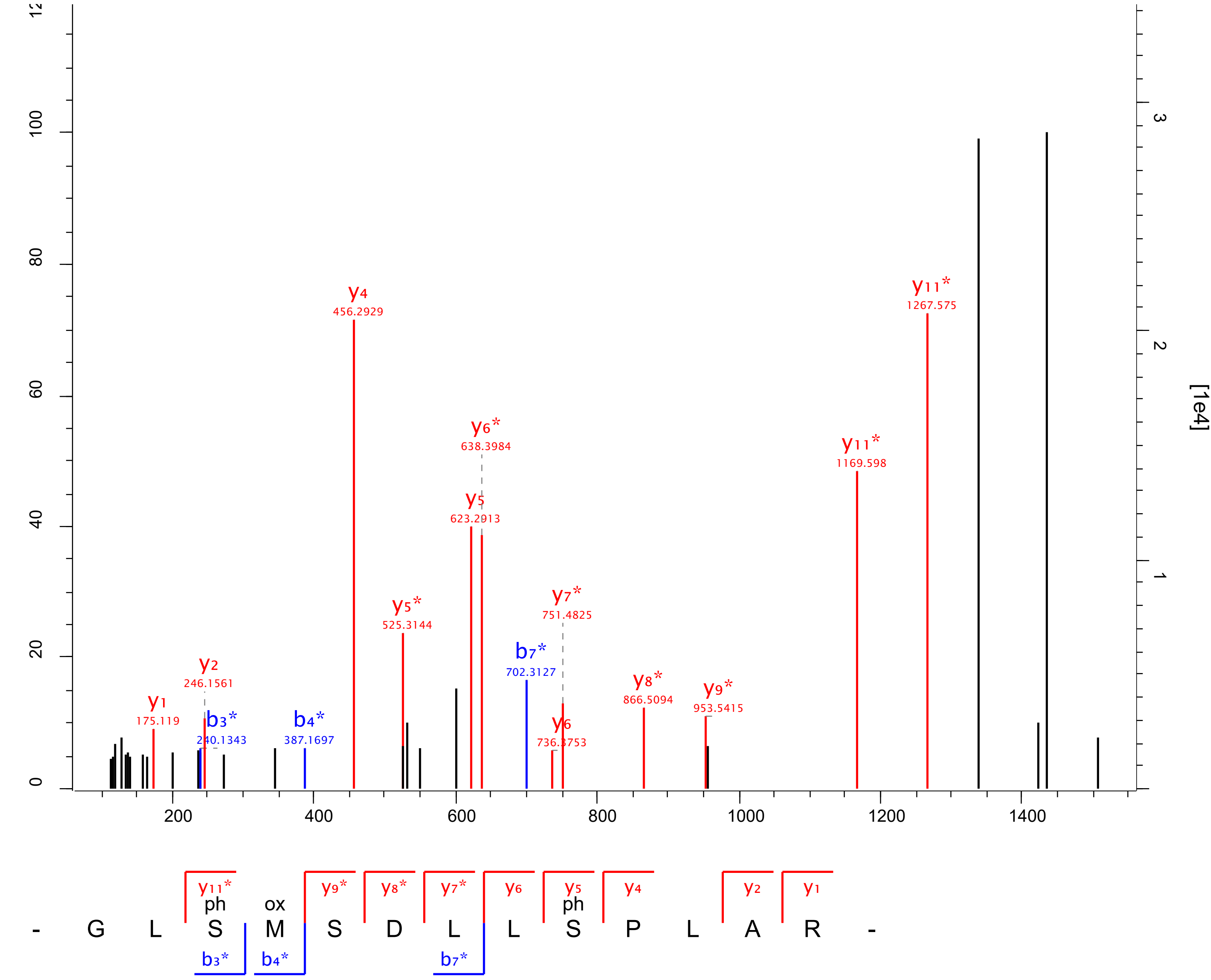

| Raw file | Scan | Method | Score | m/z |
| --- | --- | --- | --- | --- |
| water_02_1 | 47203 | FTMS; HCD | 69.73 | 1071.14 |

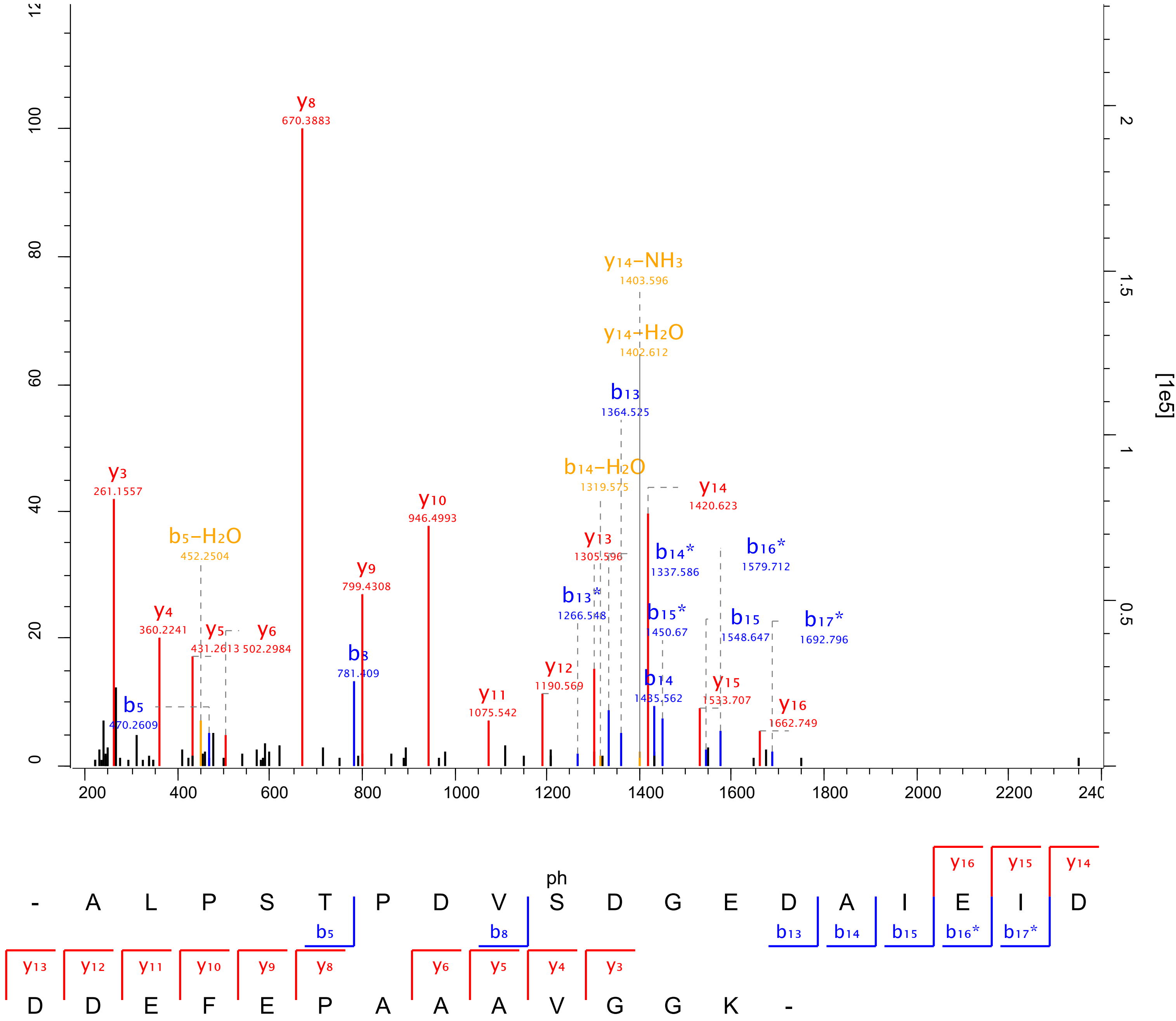

| Raw file | Scan | Method | Score | m/z |
|---|---|---|---|---|
| water_02_1 | 47229 | FTMS; HCD | 78.44 | 838.32 |

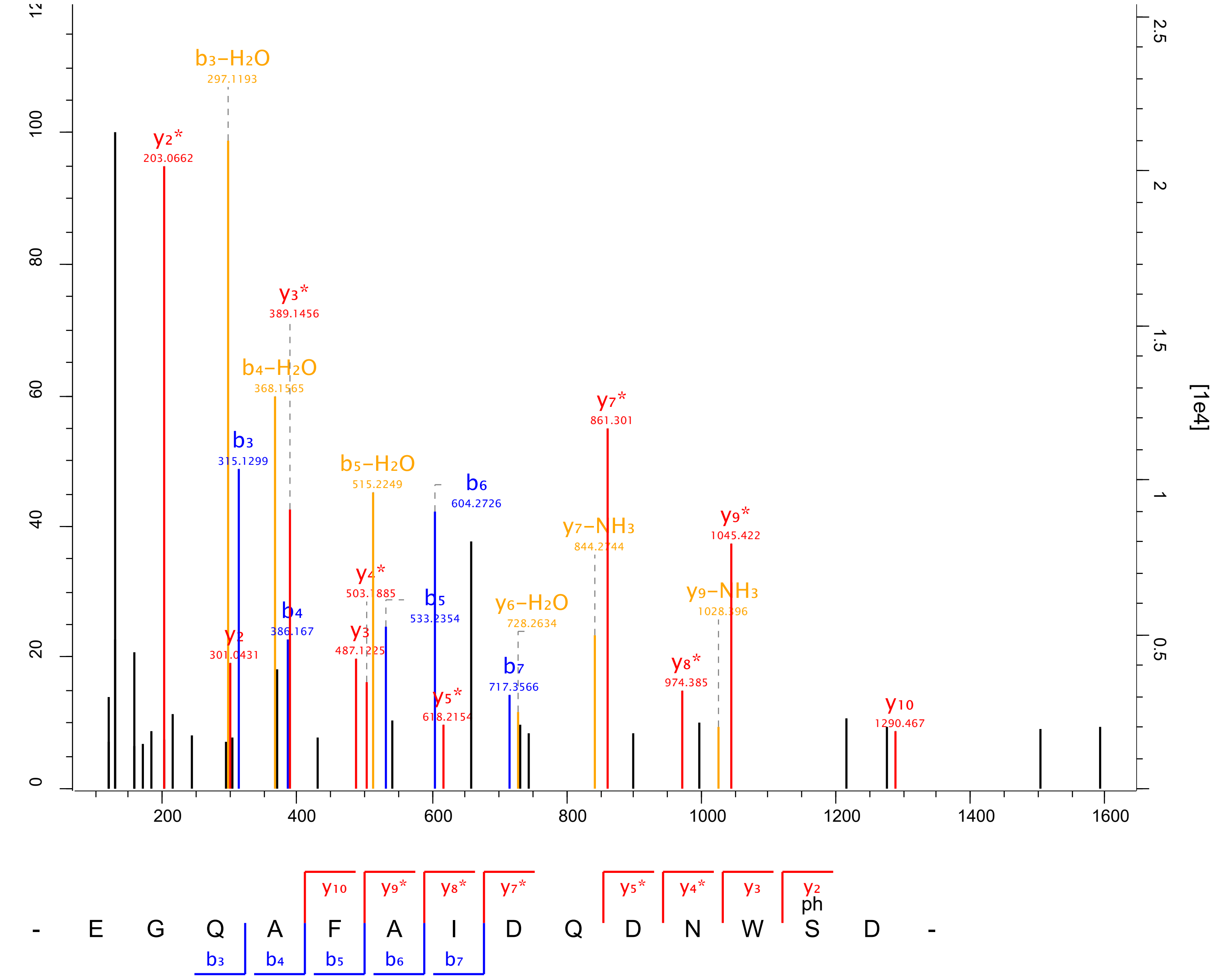

| Raw file | Scan | Method | Score | m/z |
|---|---|---|---|---|
| water_02_1 | 47448 | FTMS; HCD | 133.93 | 885.89 |

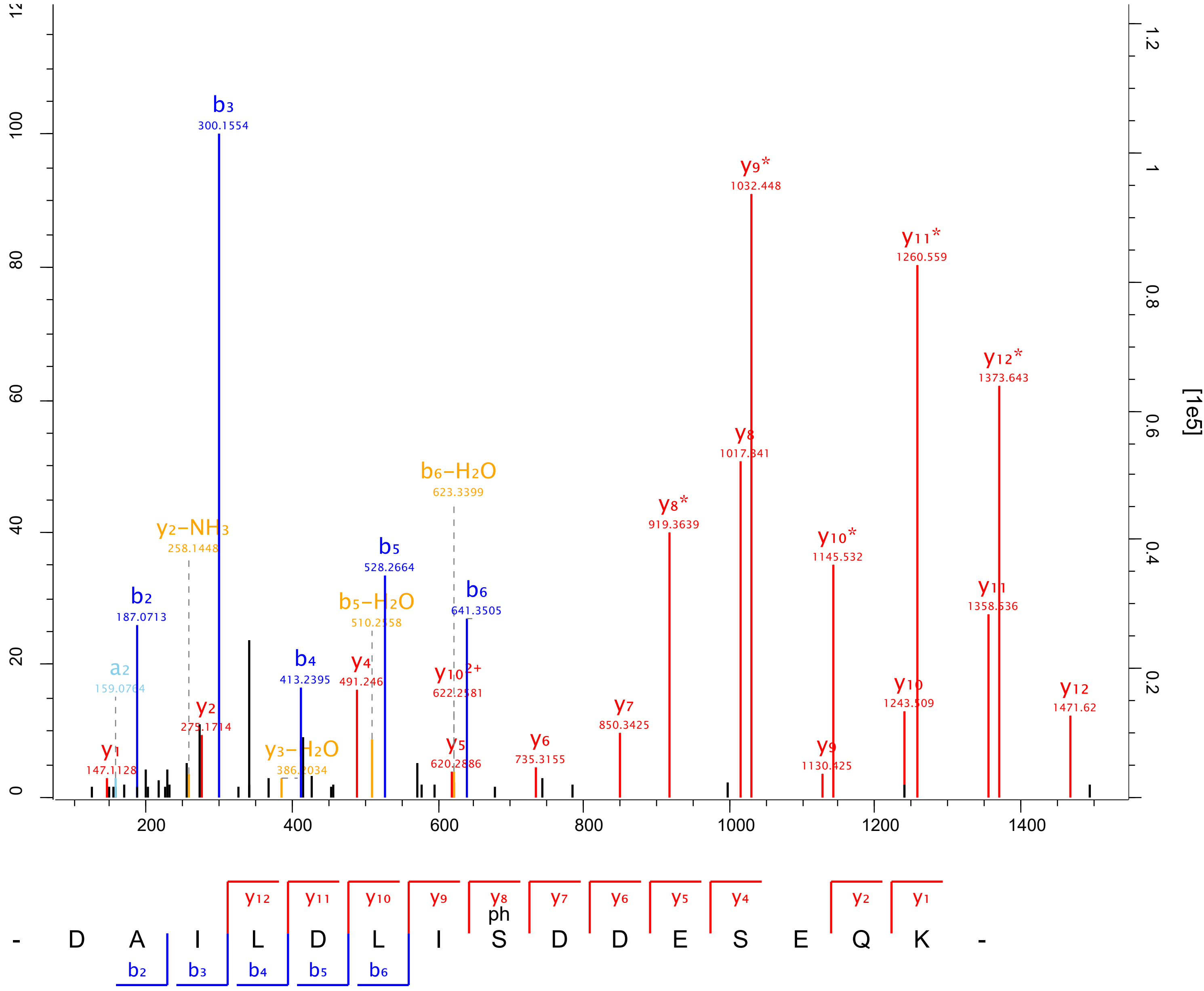

| Raw file | Scan | Method | Score | m/z |
|---|---|---|---|---|
| water_02_1 | 4755 | FTMS; HCD | 50.35 | 582.72 |

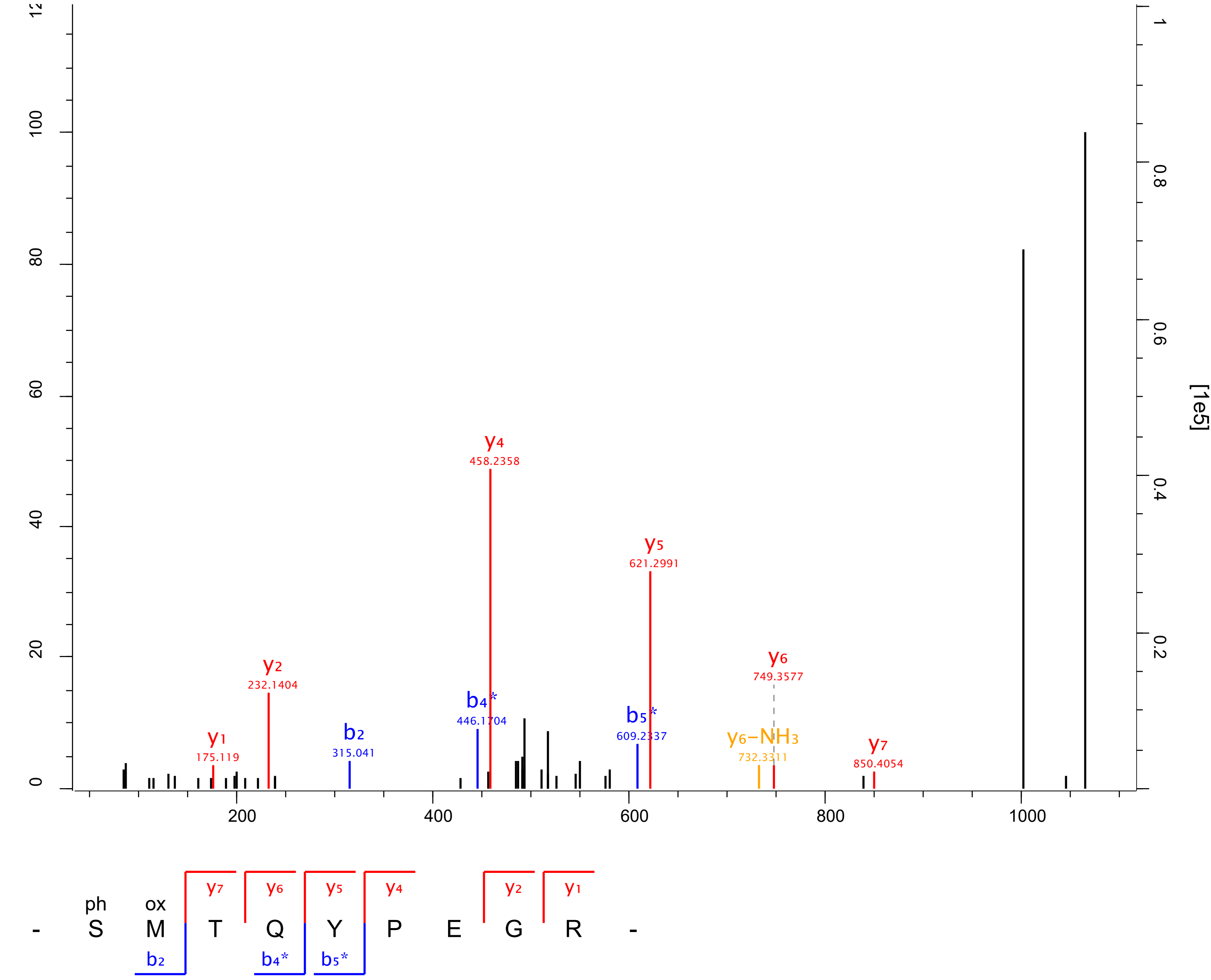

| Raw file | Scan | Method | Score | m/z |
| --- | --- | --- | --- | --- |
| water_02_1 | 4776 | FTMS; HCD | 51.03 | 472.89 |

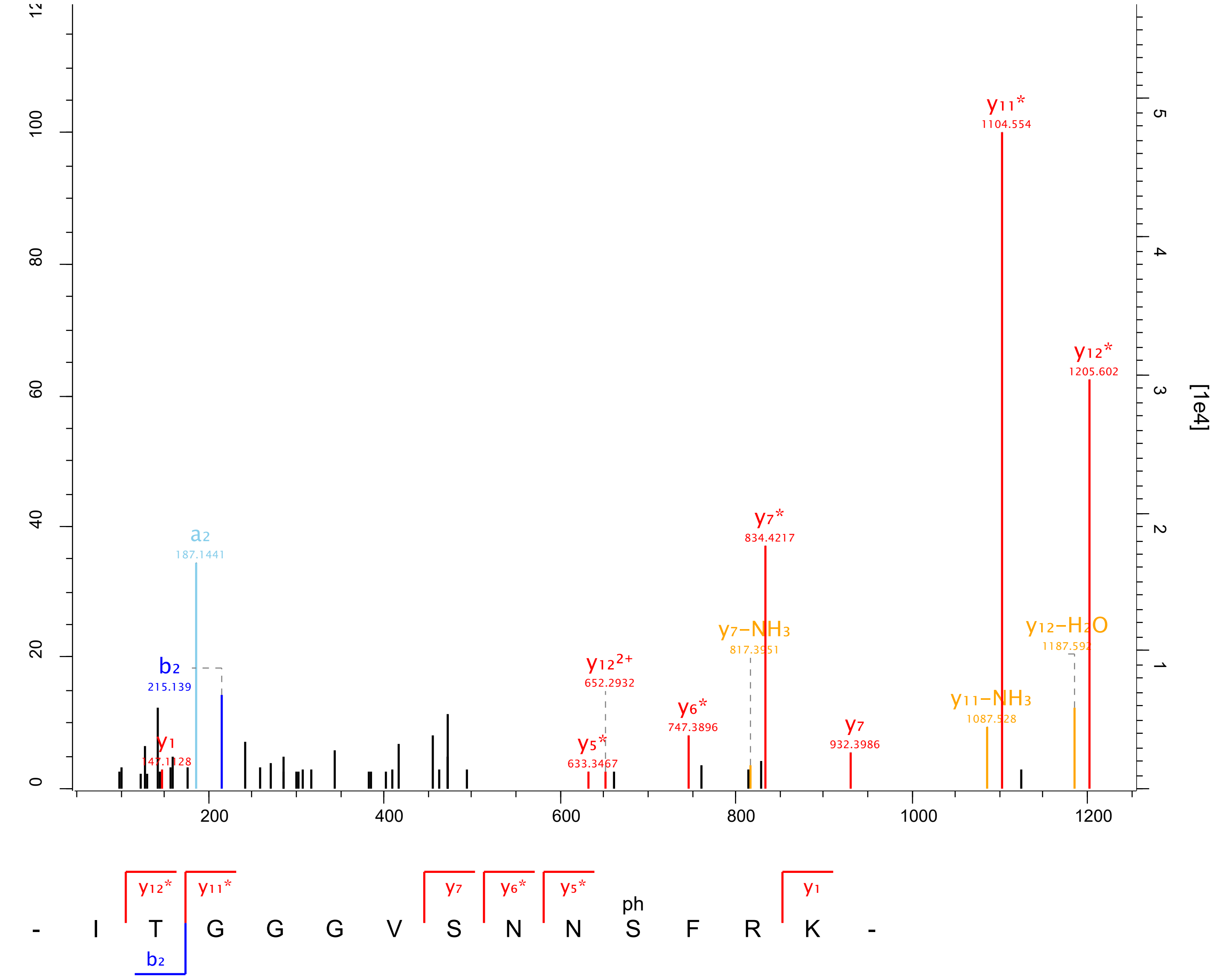

| Raw file | Scan | Method | Score | m/z |
|---|---|---|---|---|
| water_02_1 | 47802 | FTMS; HCD | 63.89 | 833.05 |

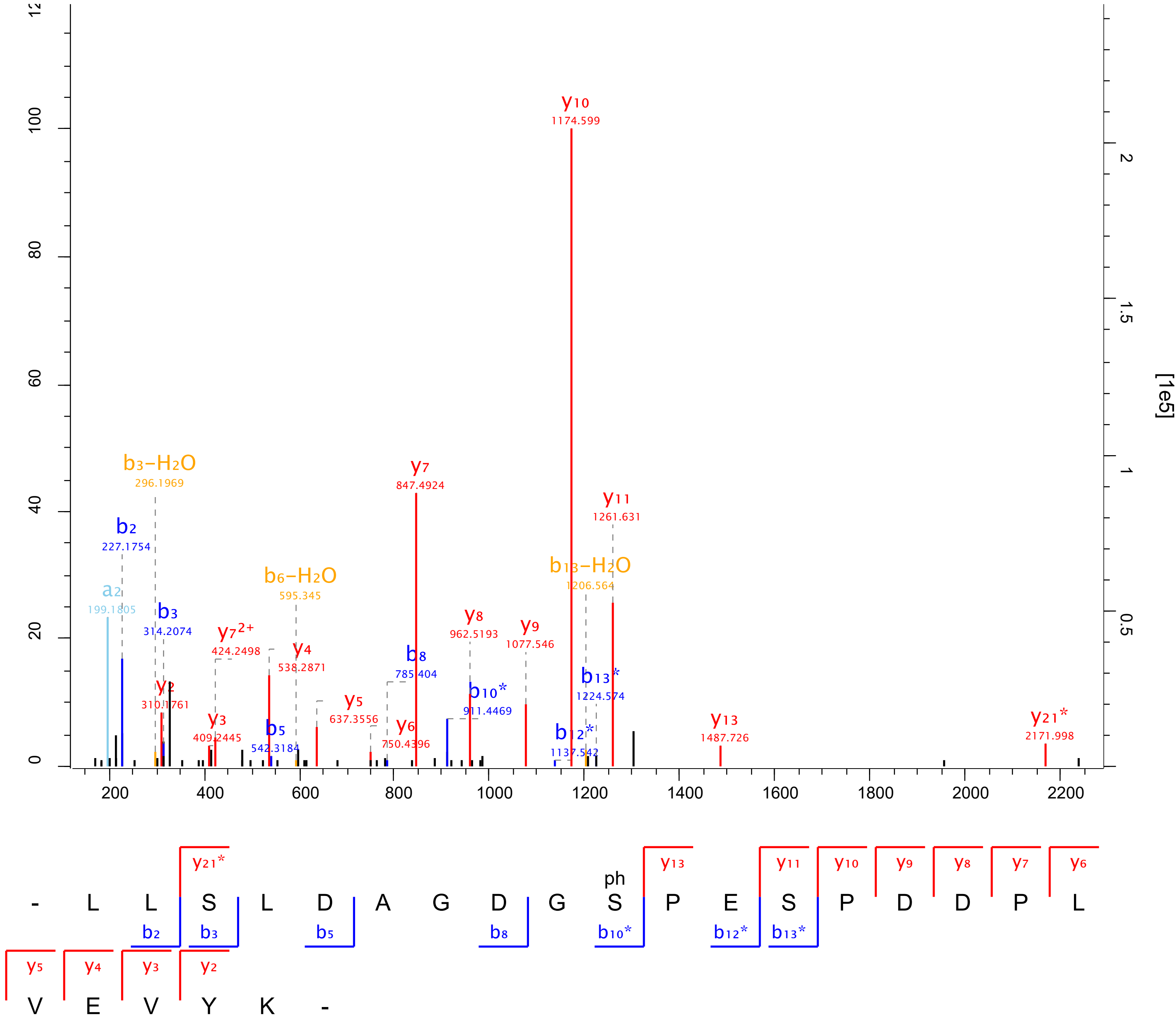

| Raw file | Scan | Method | Score | m/z |
|---|---|---|---|---|
| water_02_1 | 48238 | FTMS; HCD | 79.38 | 995.49 |

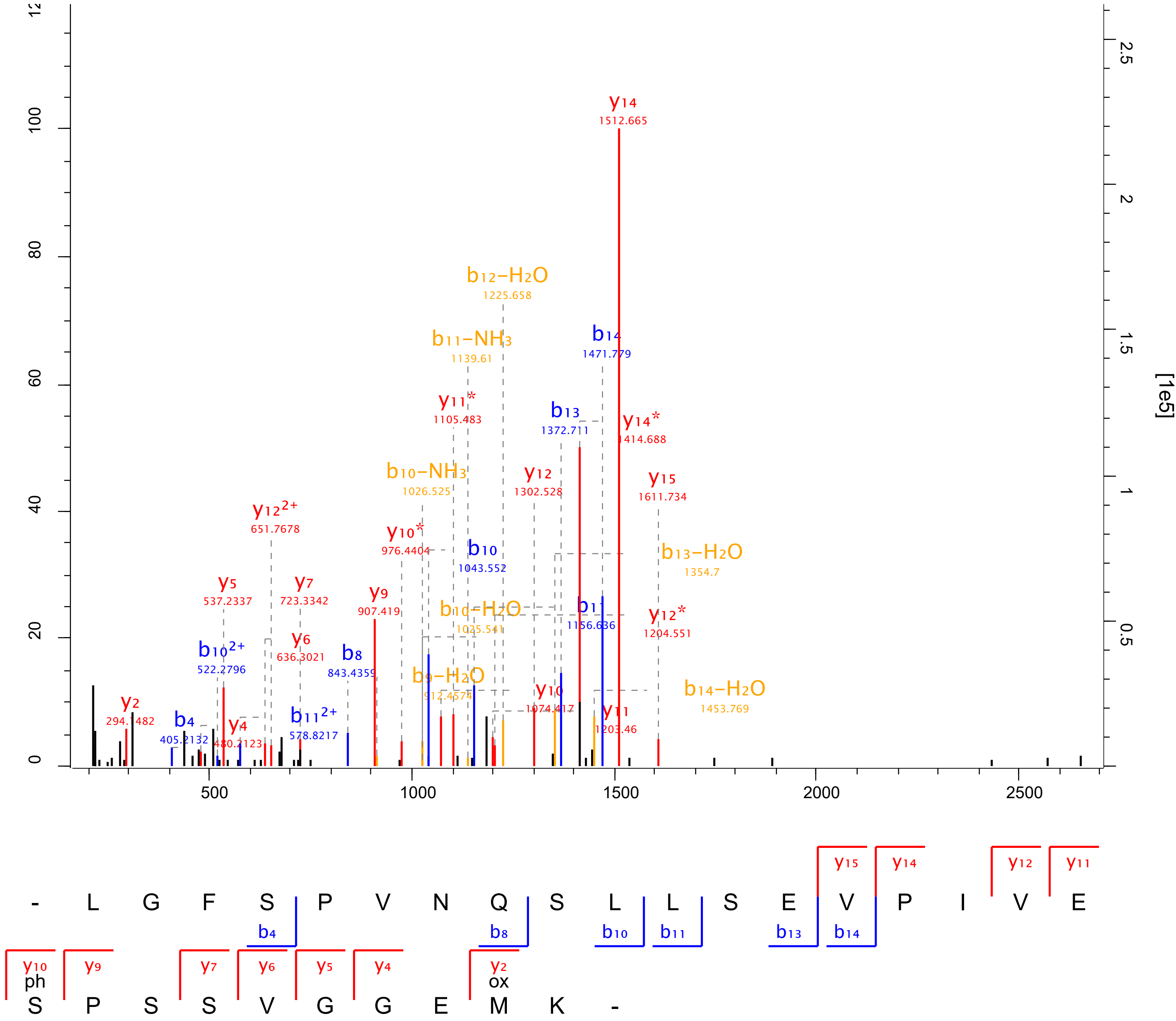

| Raw file | Scan | Method | Score | m/z |
|---|---|---|---|---|
| water_02_1 | 48515 | FTMS; HCD | 154.34 | 944.42 |

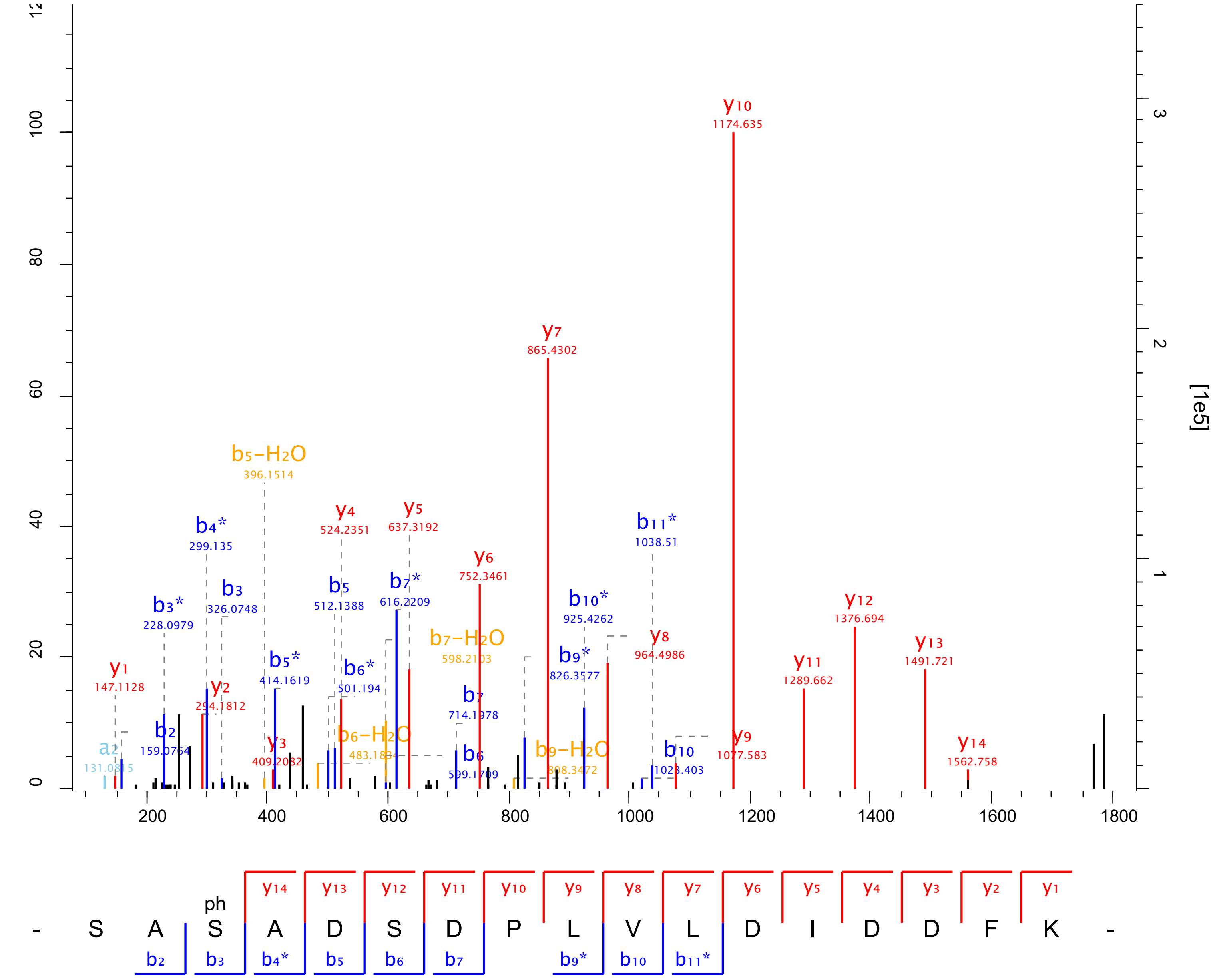

| Raw file | Scan | Method | Score | m/z |
| --- | --- | --- | --- | --- |
| water_02_1 | 4852 | FTMS; HCD | 45.99 | 653.93 |

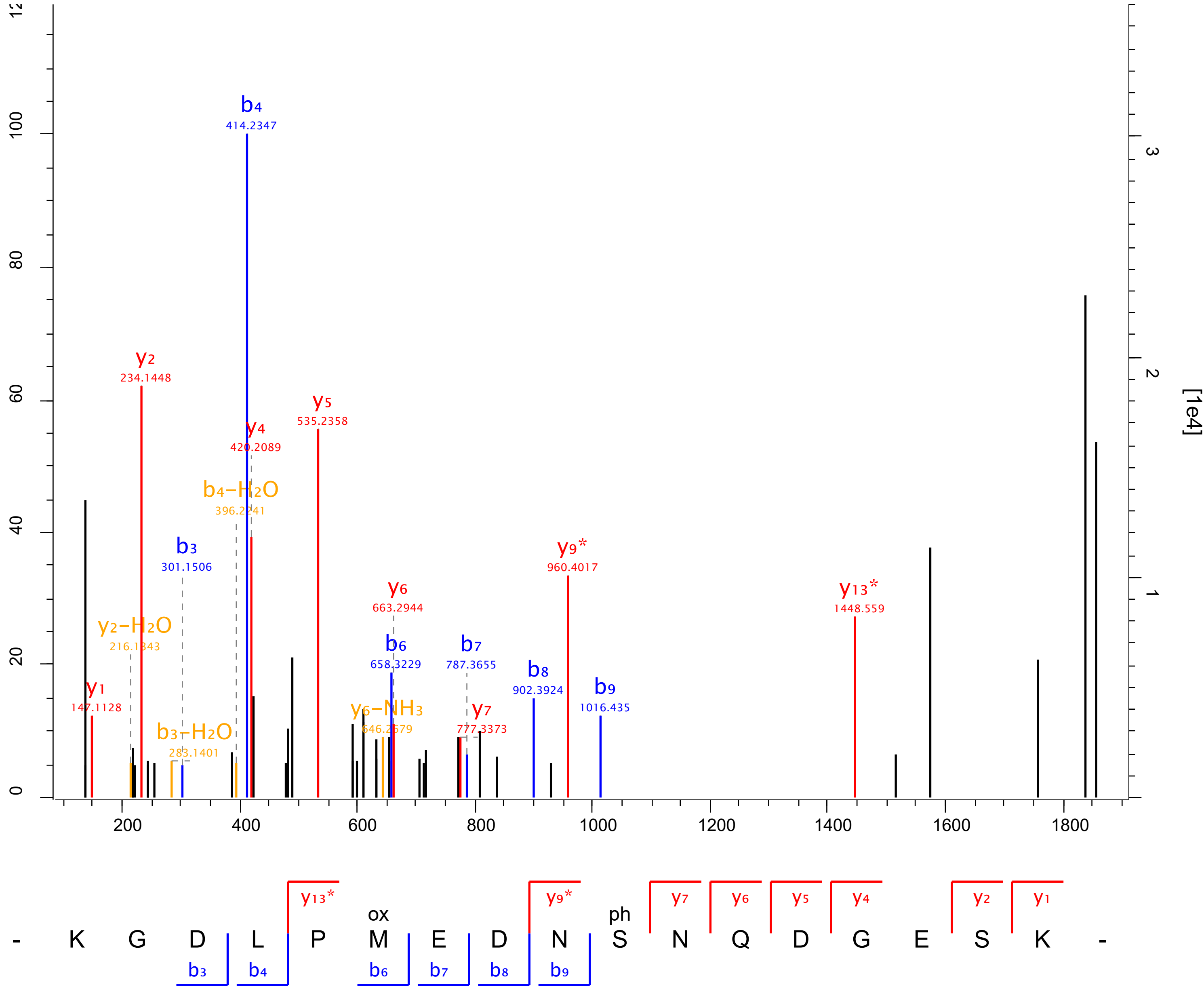

| Raw file | Scan | Method | Score | m/z |
|---|---|---|---|---|
| water_02_1 | 48535 | FTMS; HCD | 50.03 | 969.14 |

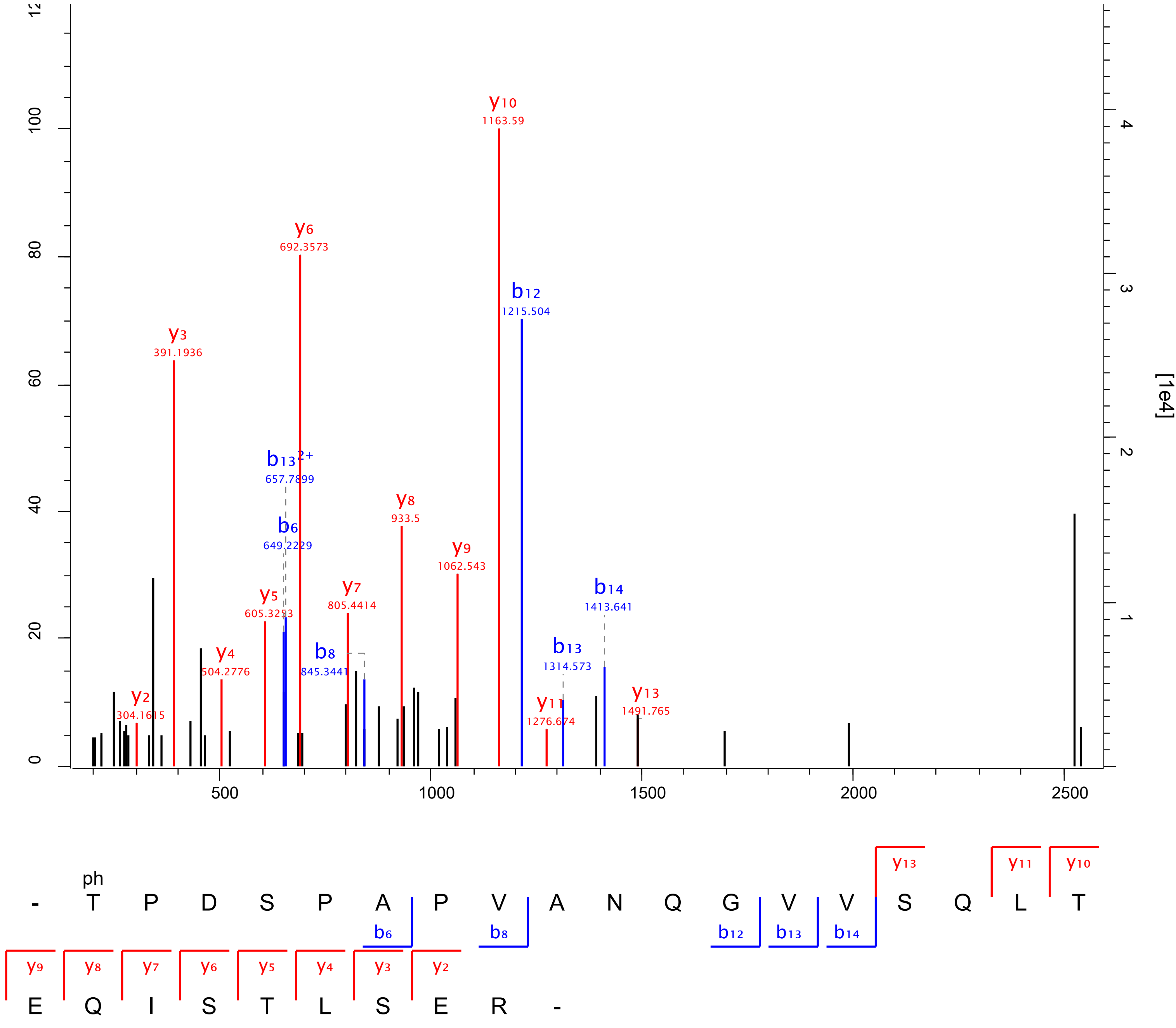

| Raw file | Scan | Method | Score | m/z |
| --- | --- | --- | --- | --- |
| water_02_1 | 48575 | FTMS; HCD | 85.09 | 924.91 |

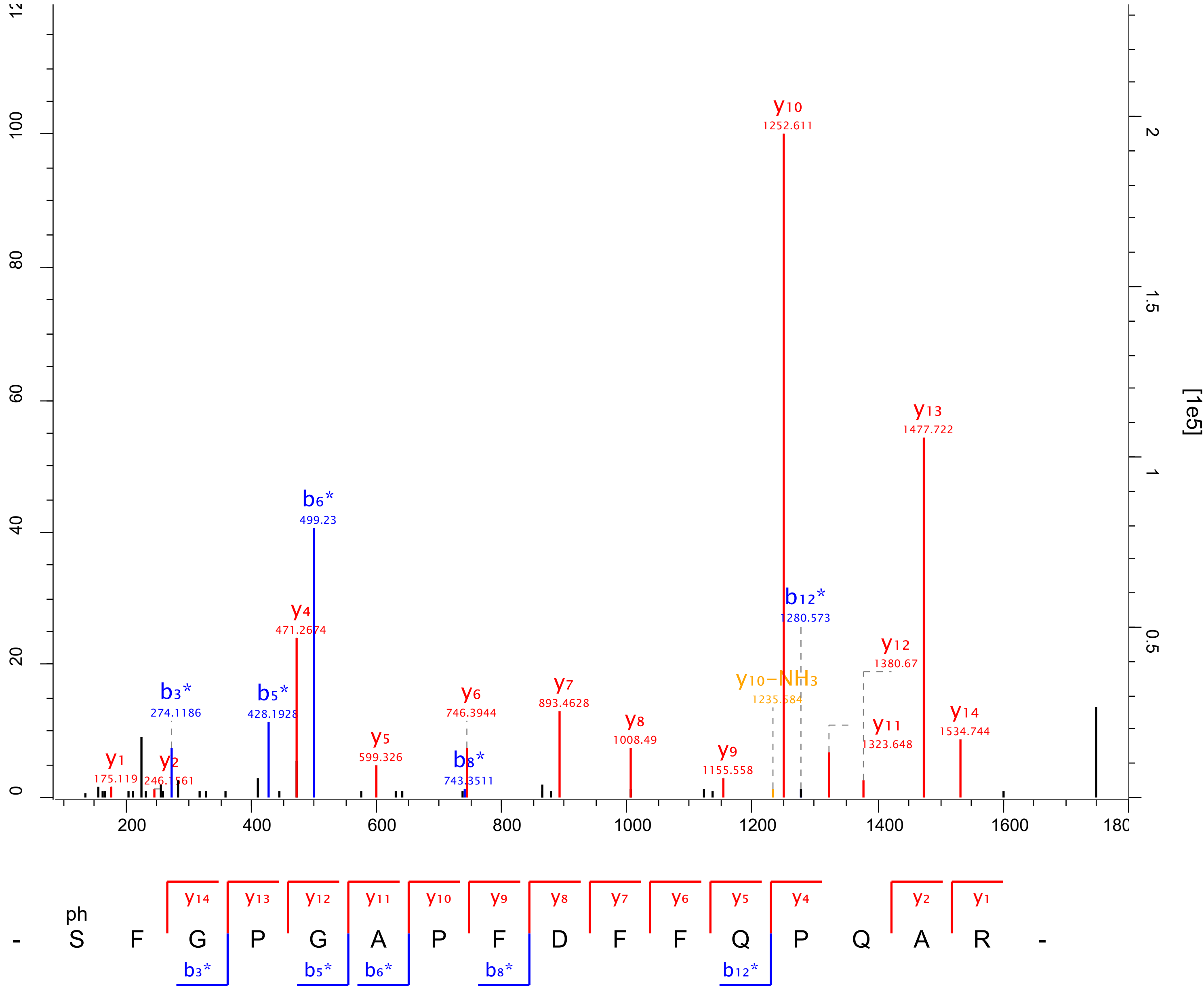

| Raw file | Scan | Method | Score | m/z |
|---|---|---|---|---|
| water_02_1 | 4867 | FTMS; HCD | 54.67 | 545.22 |

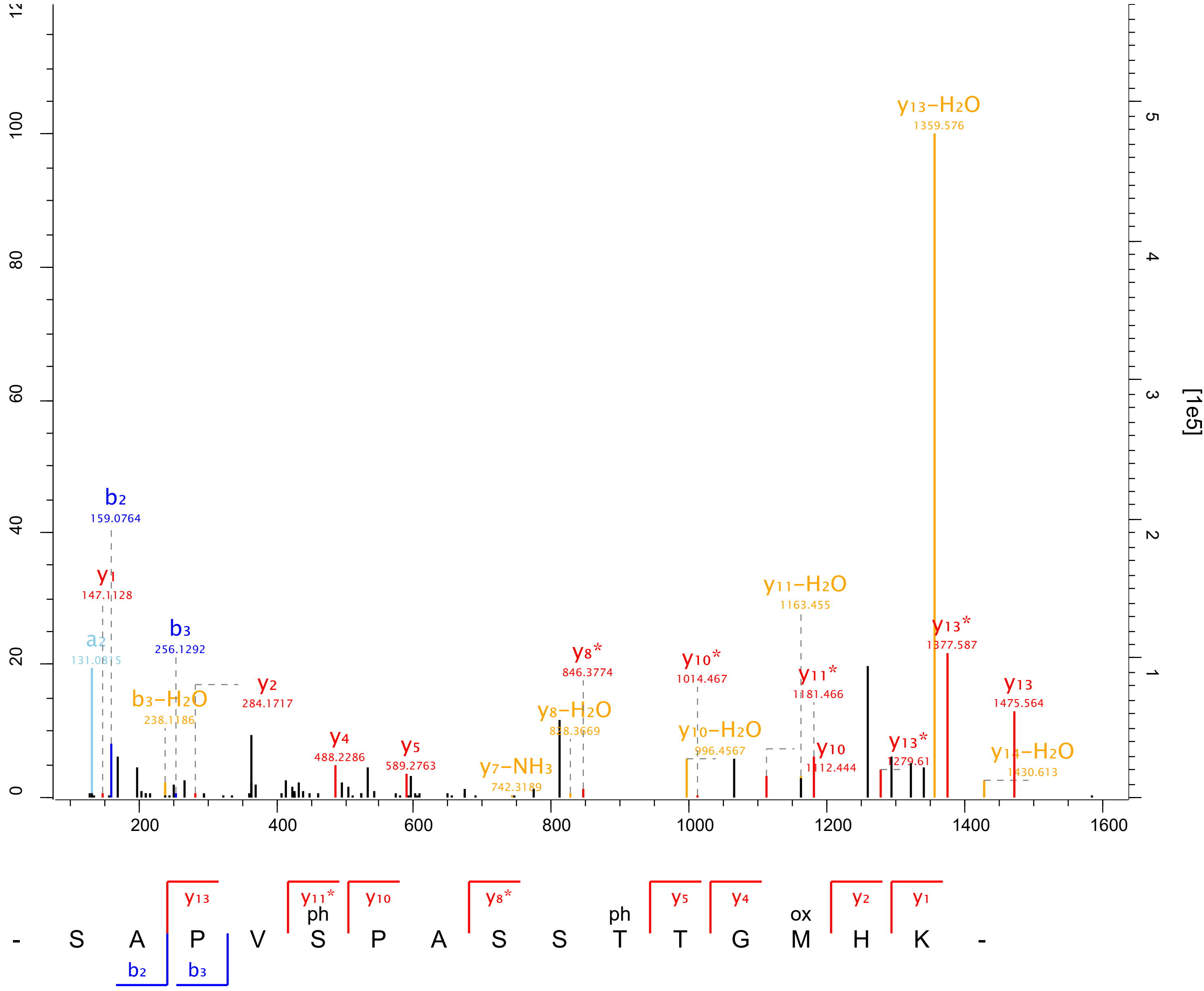

| Raw file | Scan | Method | Score | m/z |
|---|---|---|---|---|
| water_02_1 | 4879 | FTMS; HCD | 59.25 | 656.25 |

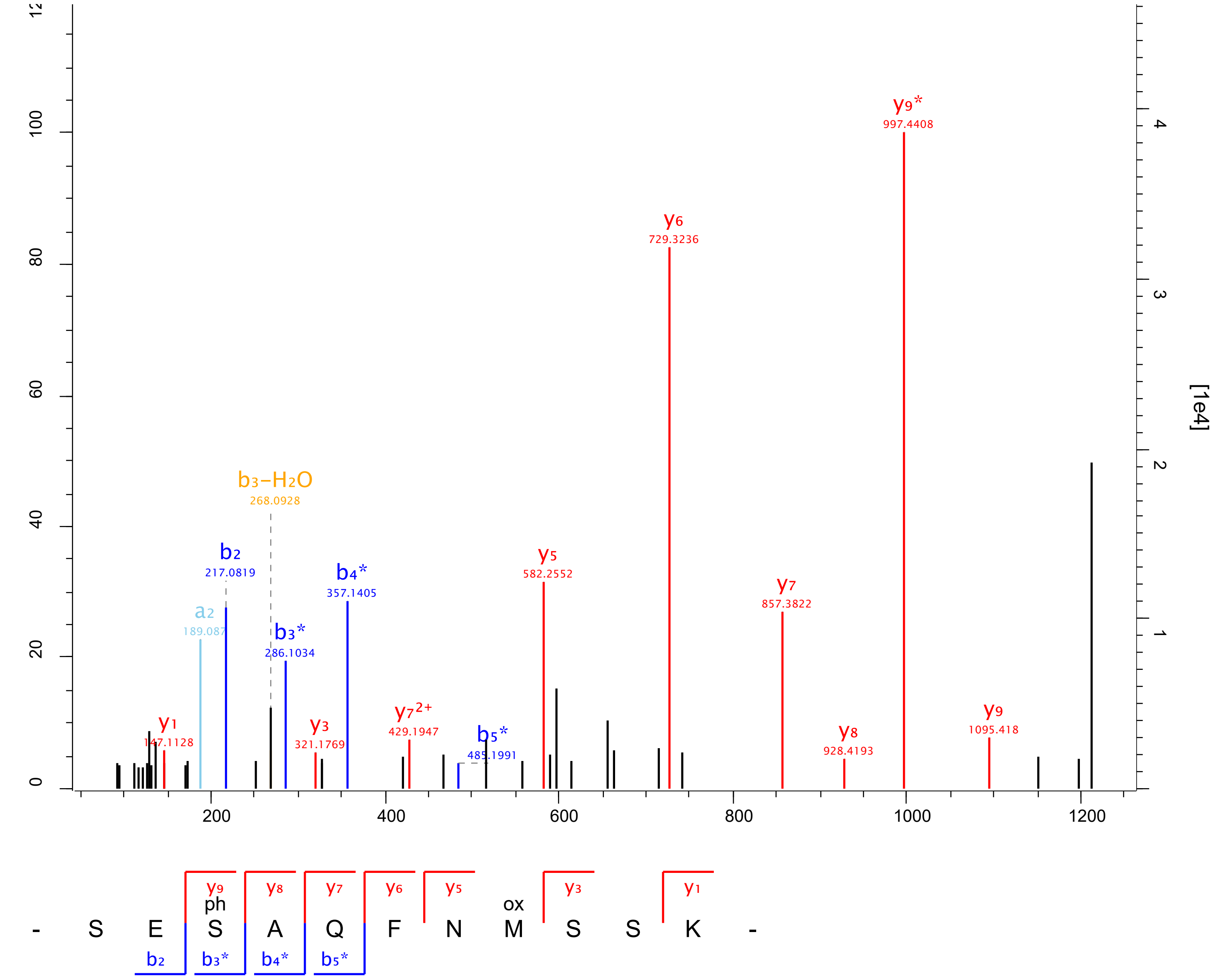

| Raw file | Scan | Method | Score | m/z |
|---|---|---|---|---|
| water_02_1 | 48857 | FTMS; HCD | 81.14 | 814.36 |

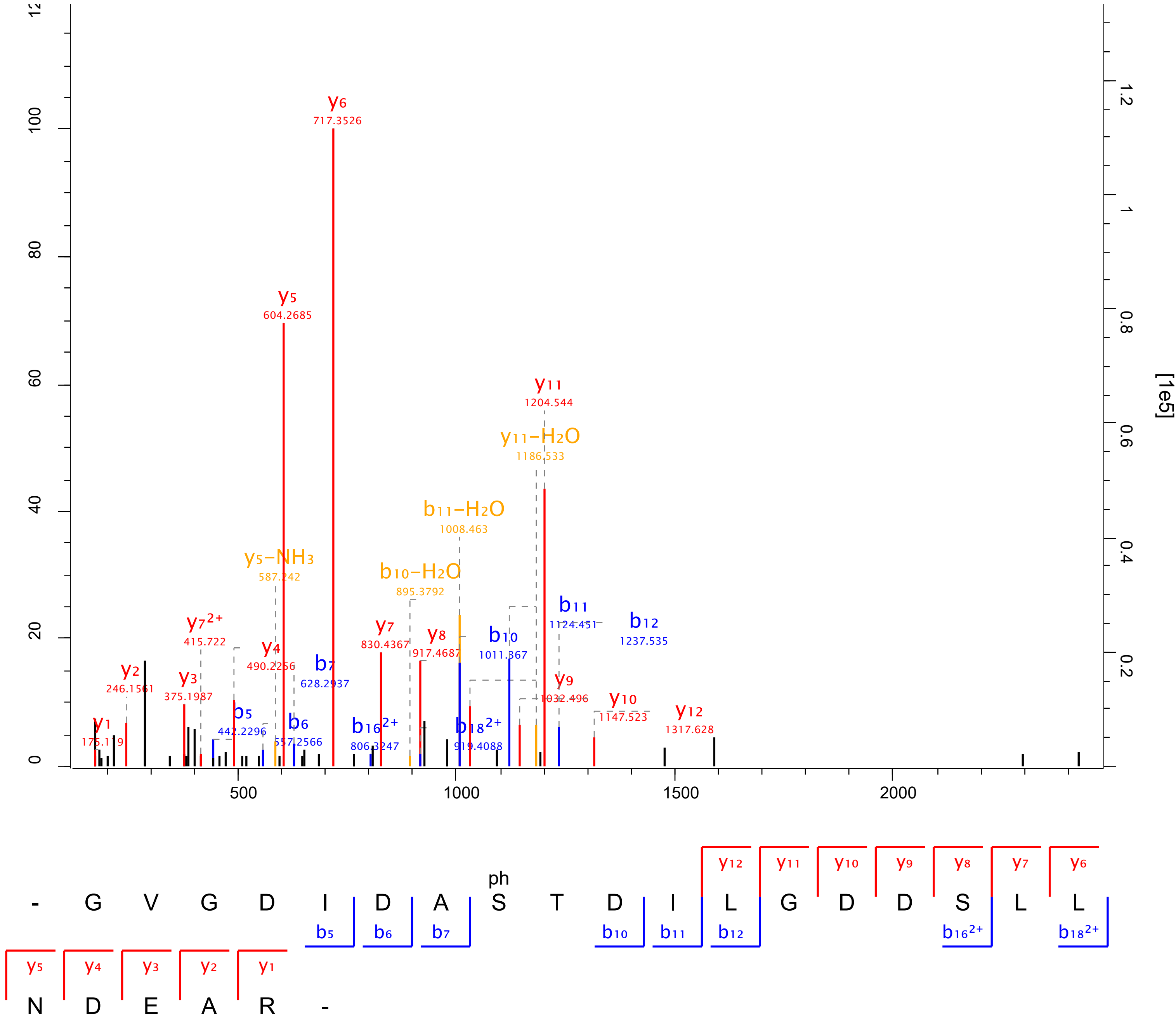

| Raw file | Scan | Method | Score | m/z |
|---|---|---|---|---|
| water_02_1 | 48897 | FTMS; HCD | 55.63 | 1121.5 |

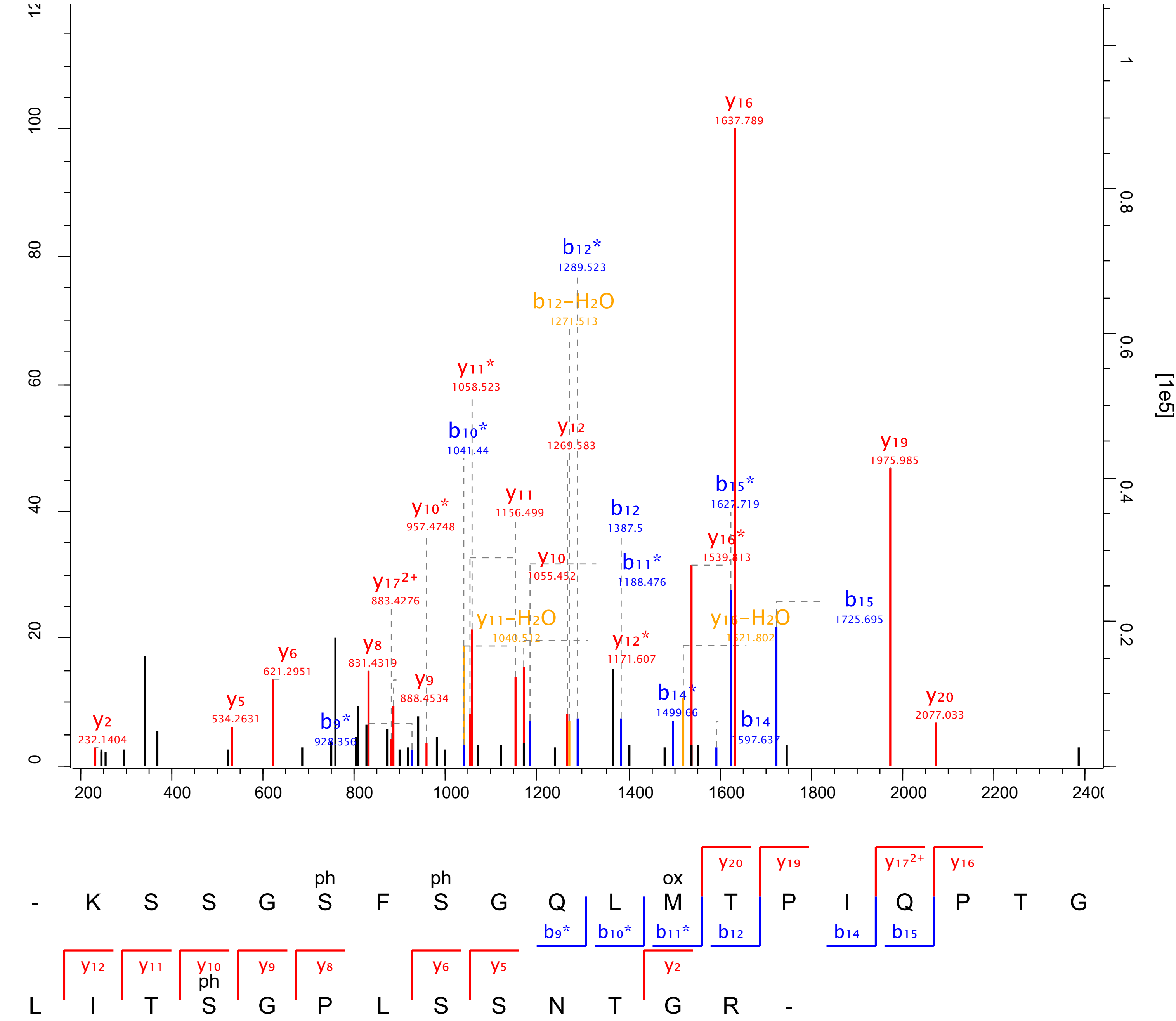

| Raw file | Scan | Method | Score | m/z |
| --- | --- | --- | --- | --- |
| water_02_1 | 48944 | FTMS; HCD | 51.64 | 974.32 |

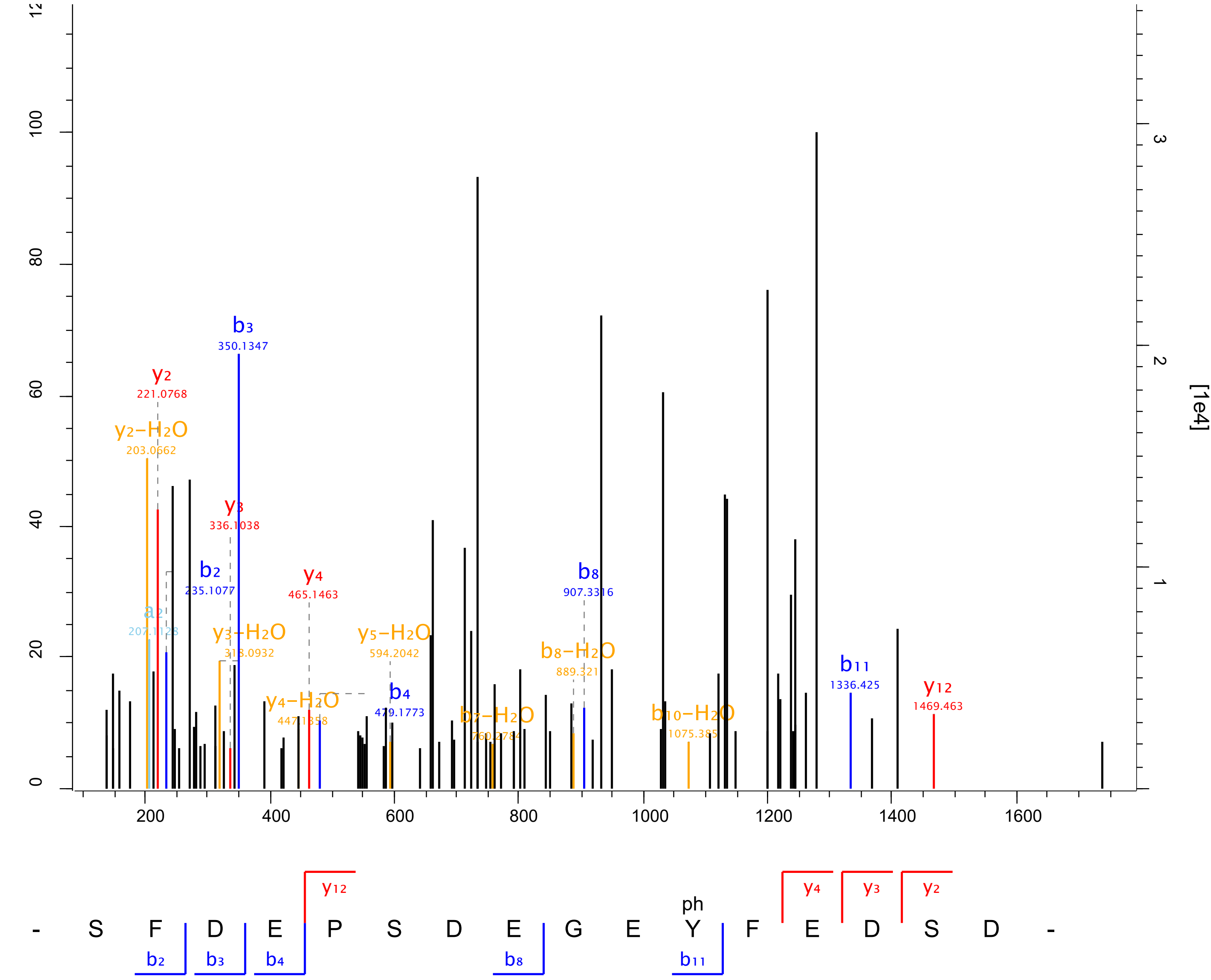

| Raw file | Scan | Method | Score | m/z |
|---|---|---|---|---|
| water_02_1 | 49253 | FTMS; HCD | 182.4 | 881.7 |

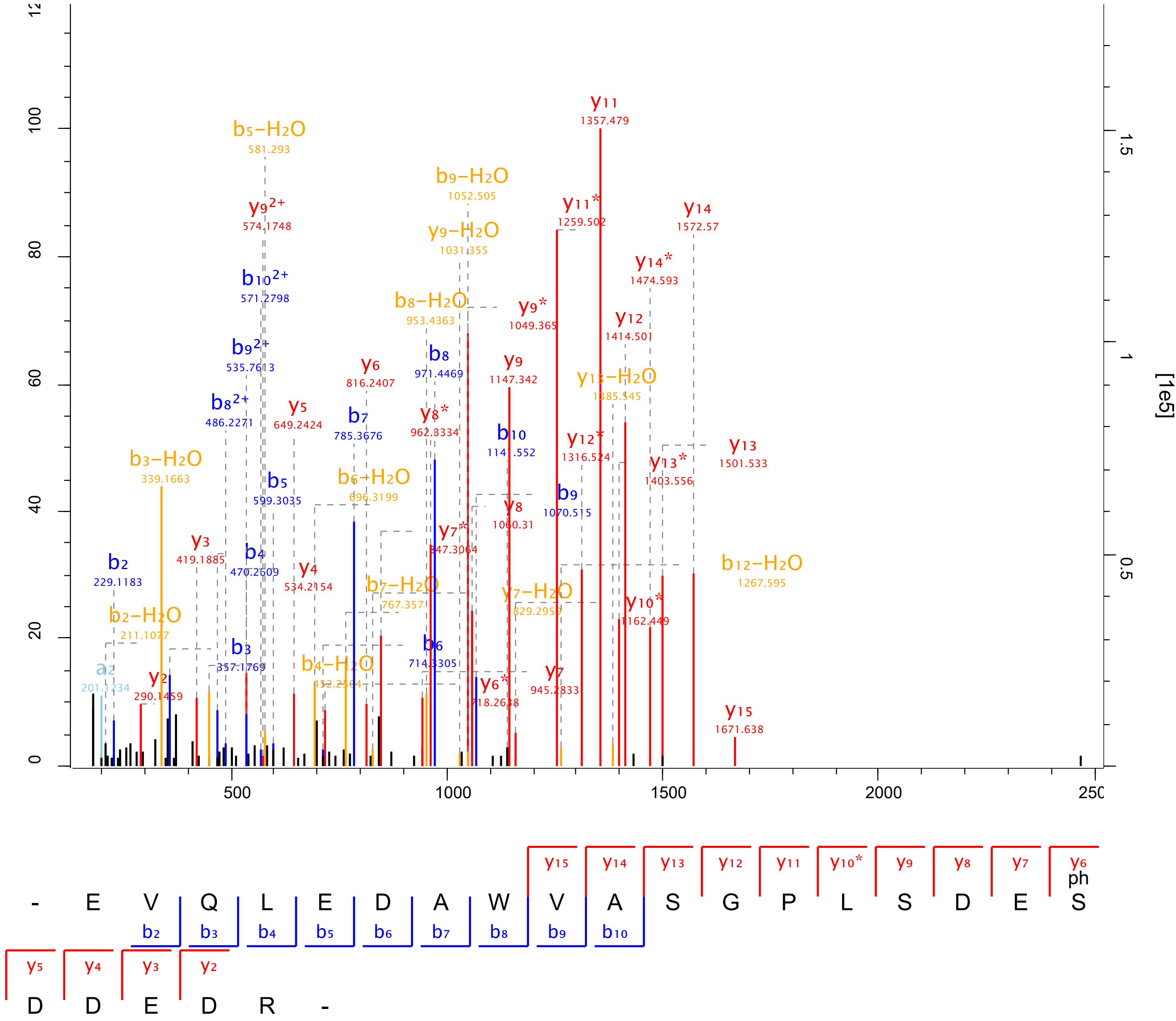

| Raw file | Scan | Method | Score | m/z |
|---|---|---|---|---|
| water_02_1 | 4930 | FTMS; HCD | 76.28 | 451.69 |

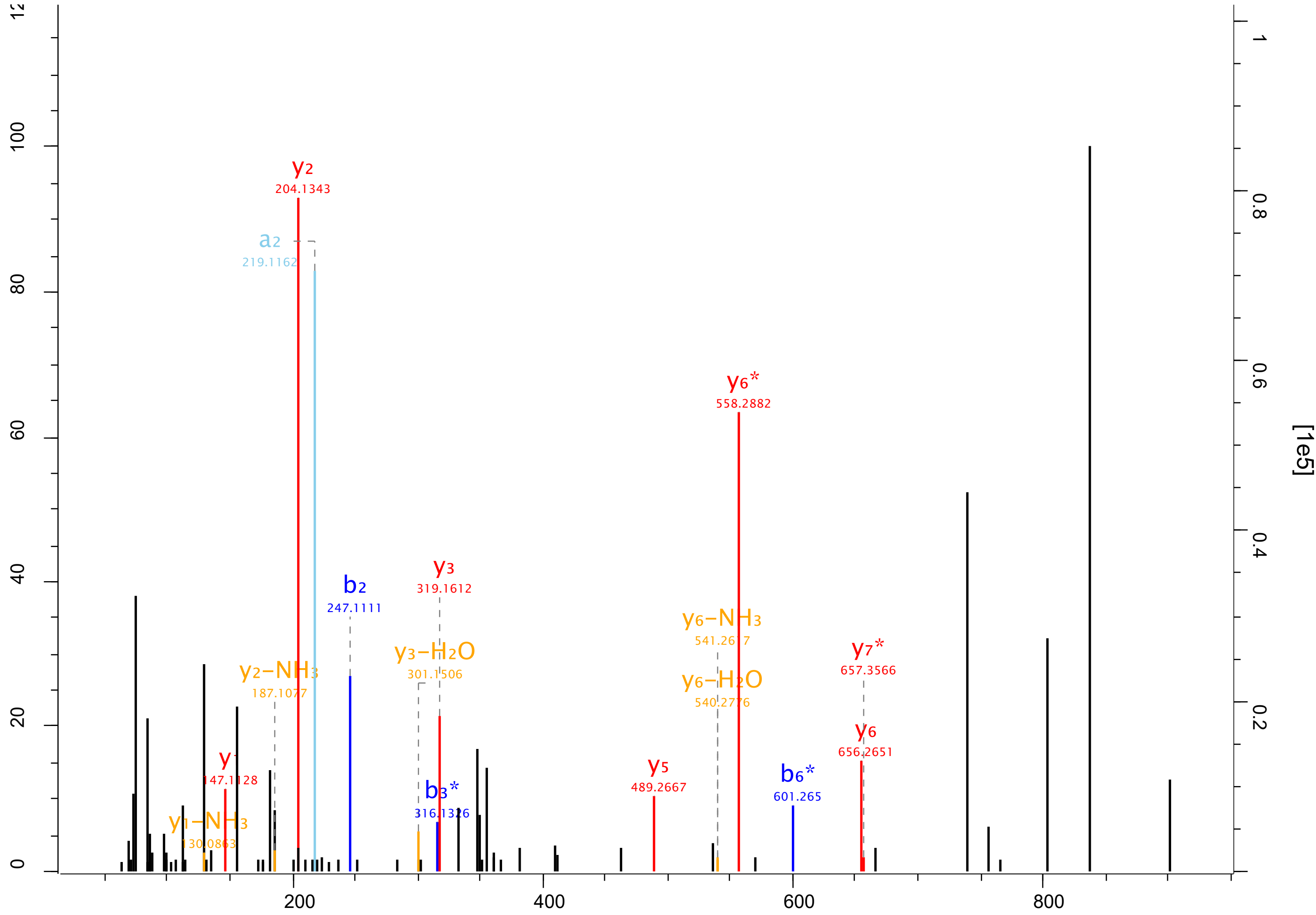


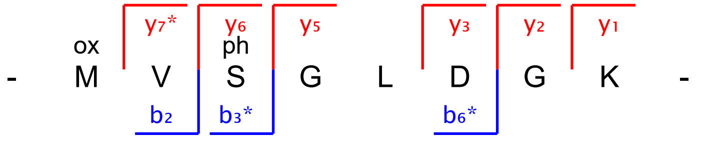

| Raw file | Scan | Method | Score | m/z |
|---|---|---|---|---|
| water_02_1 | 49535 | FTMS; HCD | 91.04 | 1052.48 |

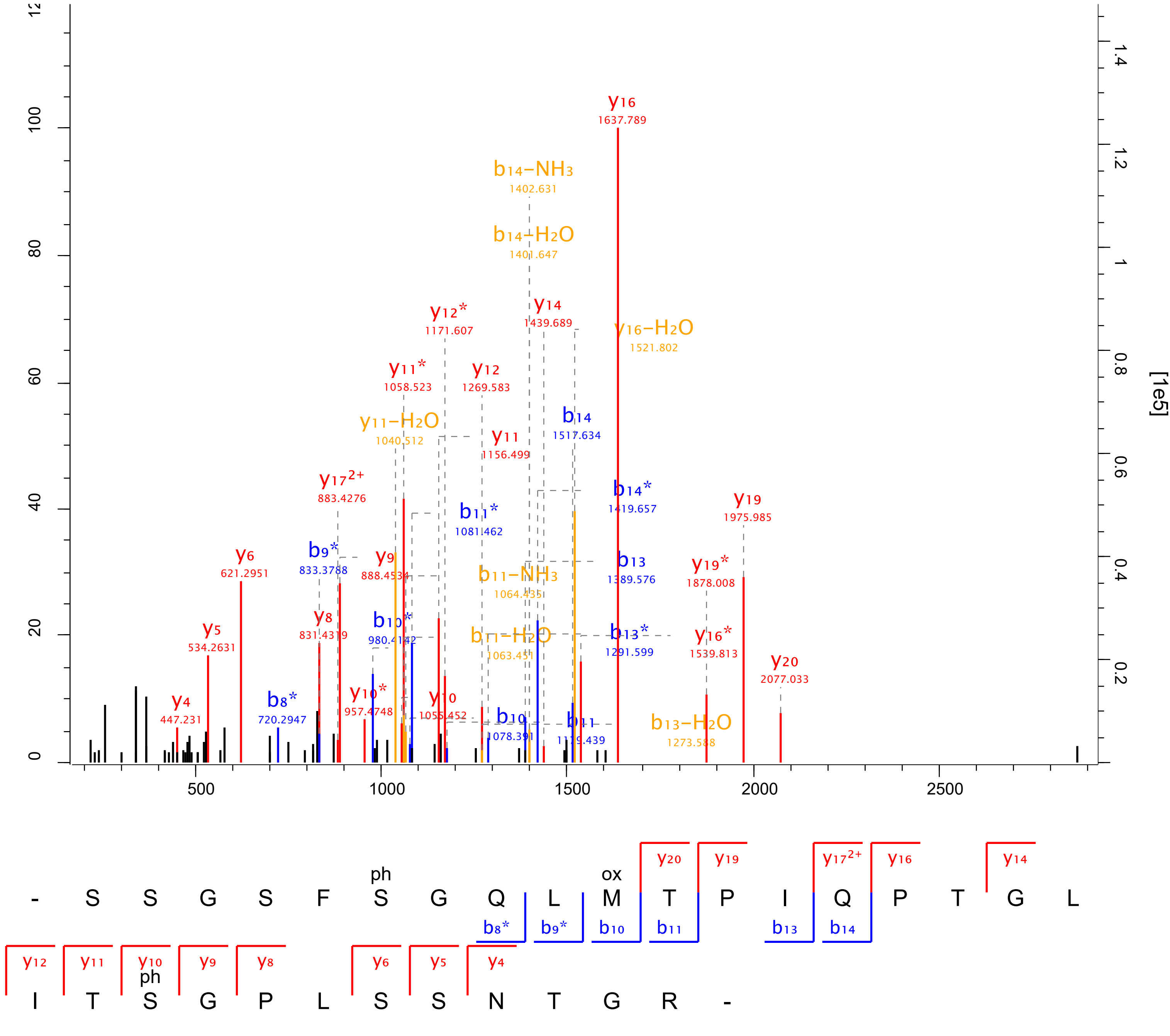

| Raw file | Scan | Method | Score | m/z |
| --- | --- | --- | --- | --- |
| water_02_1 | 4995 | FTMS; HCD | 89.08 | 500.7 |

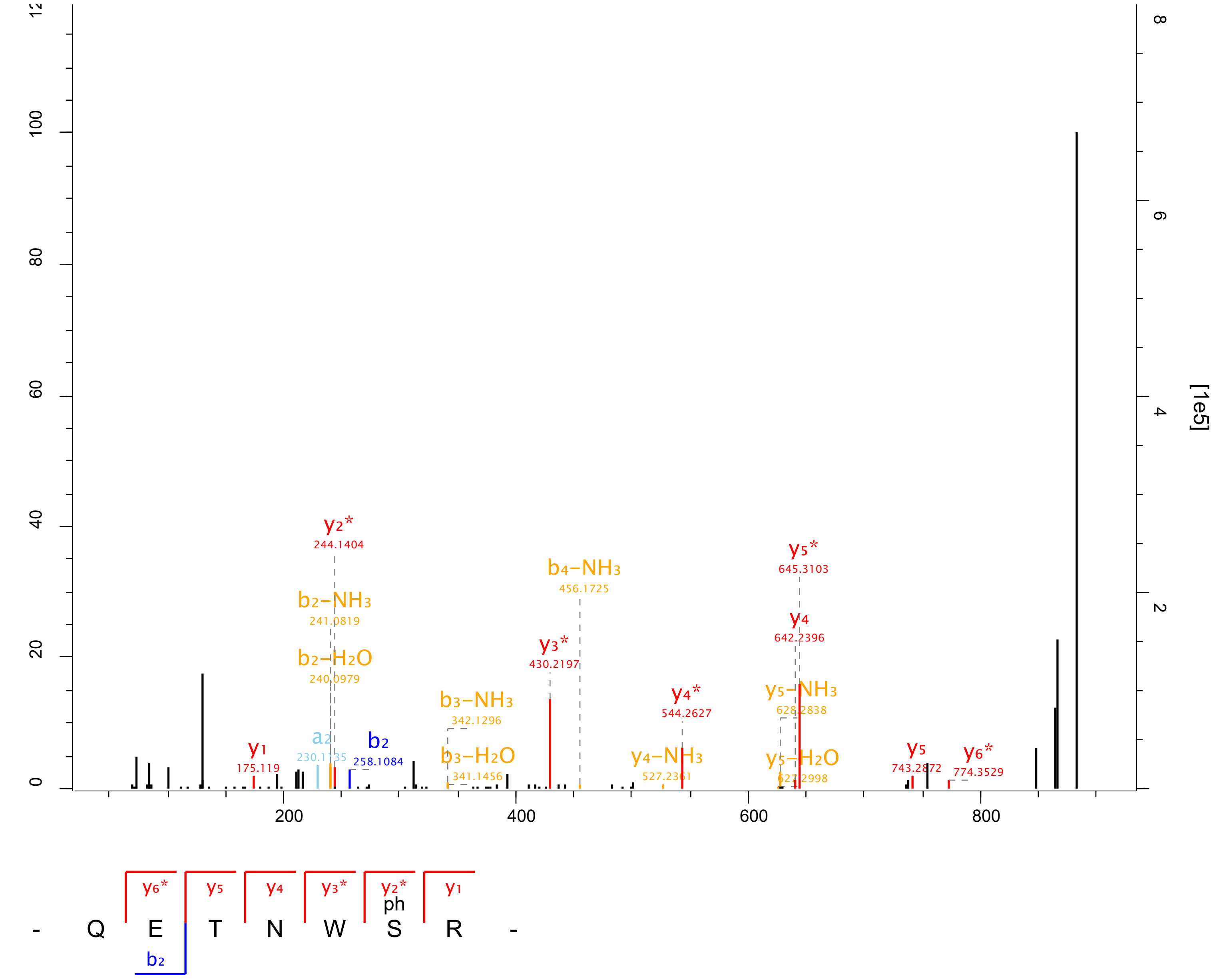

| Raw file | Scan | Method | Score | m/z |
| --- | --- | --- | --- | --- |
| water_02_1 | 50019 | FTMS; HCD | 87.88 | 989.75 |

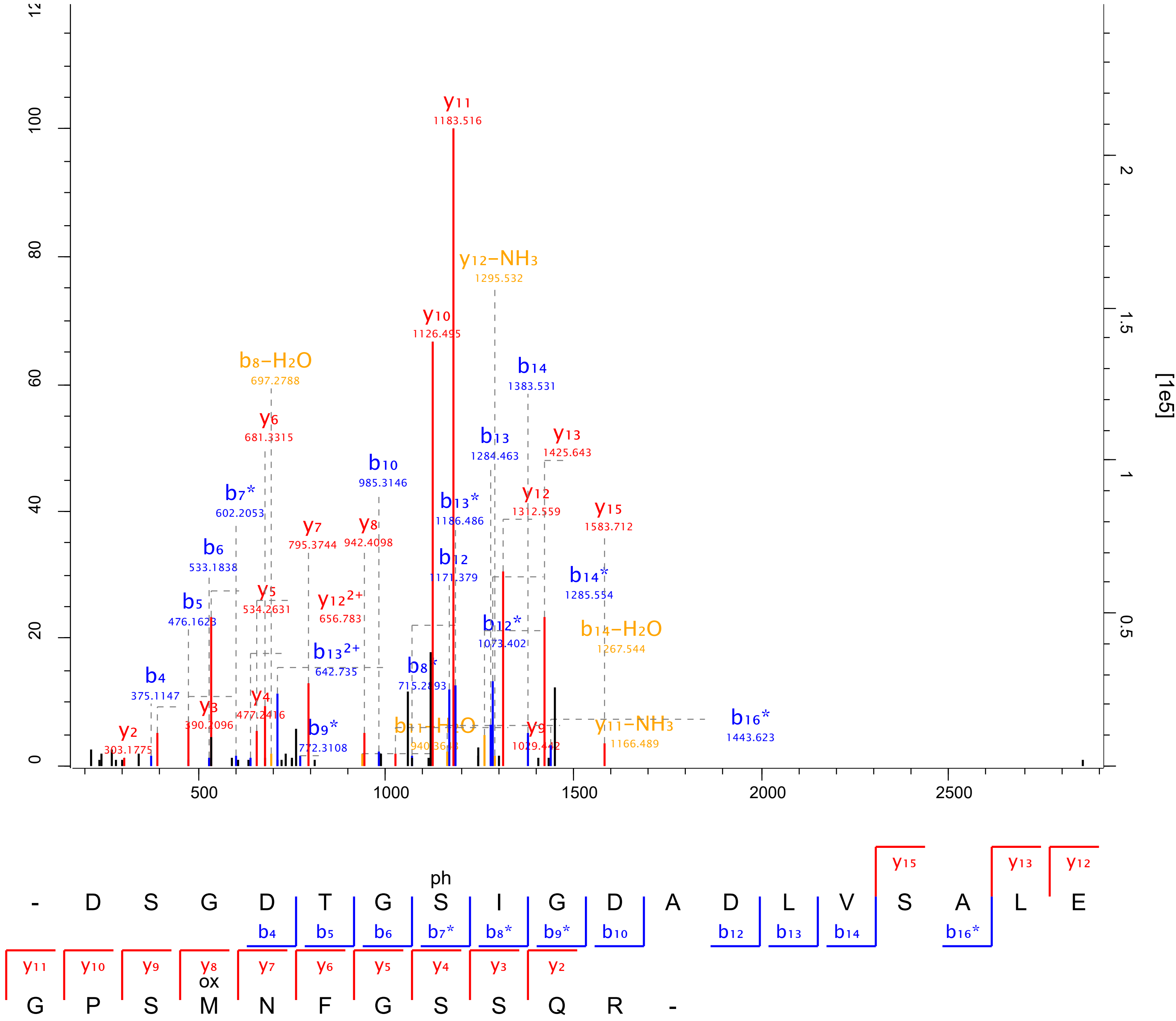

| Raw file | Scan | Method | Score | m/z |
|---|---|---|---|---|
| water_02_1 | 5020 | FTMS; HCD | 94.09 | 659.25 |

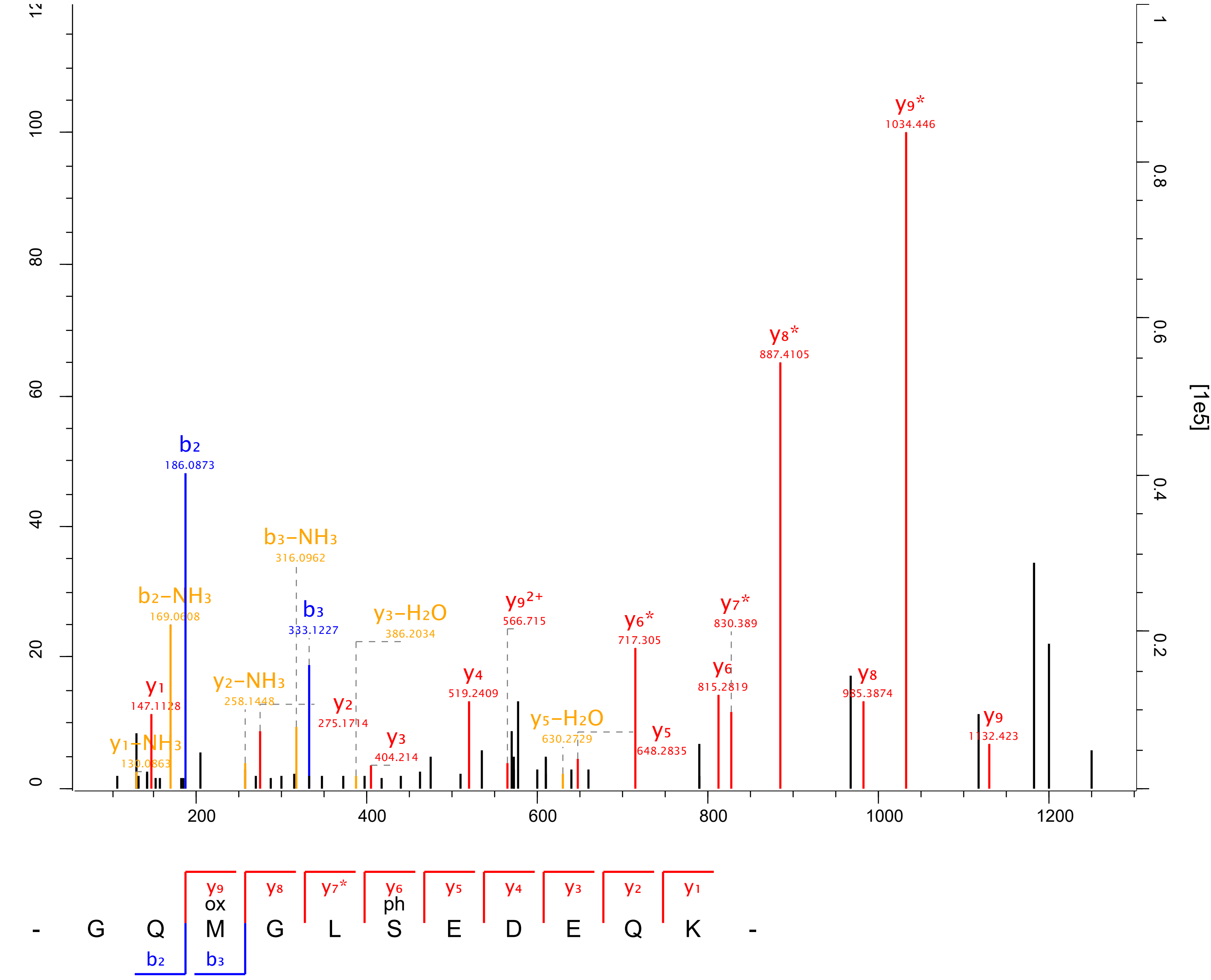

| Raw file | Scan | Method | Score | m/z |
|---|---|---|---|---|
| water_02_1 | 50632 | FTMS; HCD | 221.25 | 937.37 |

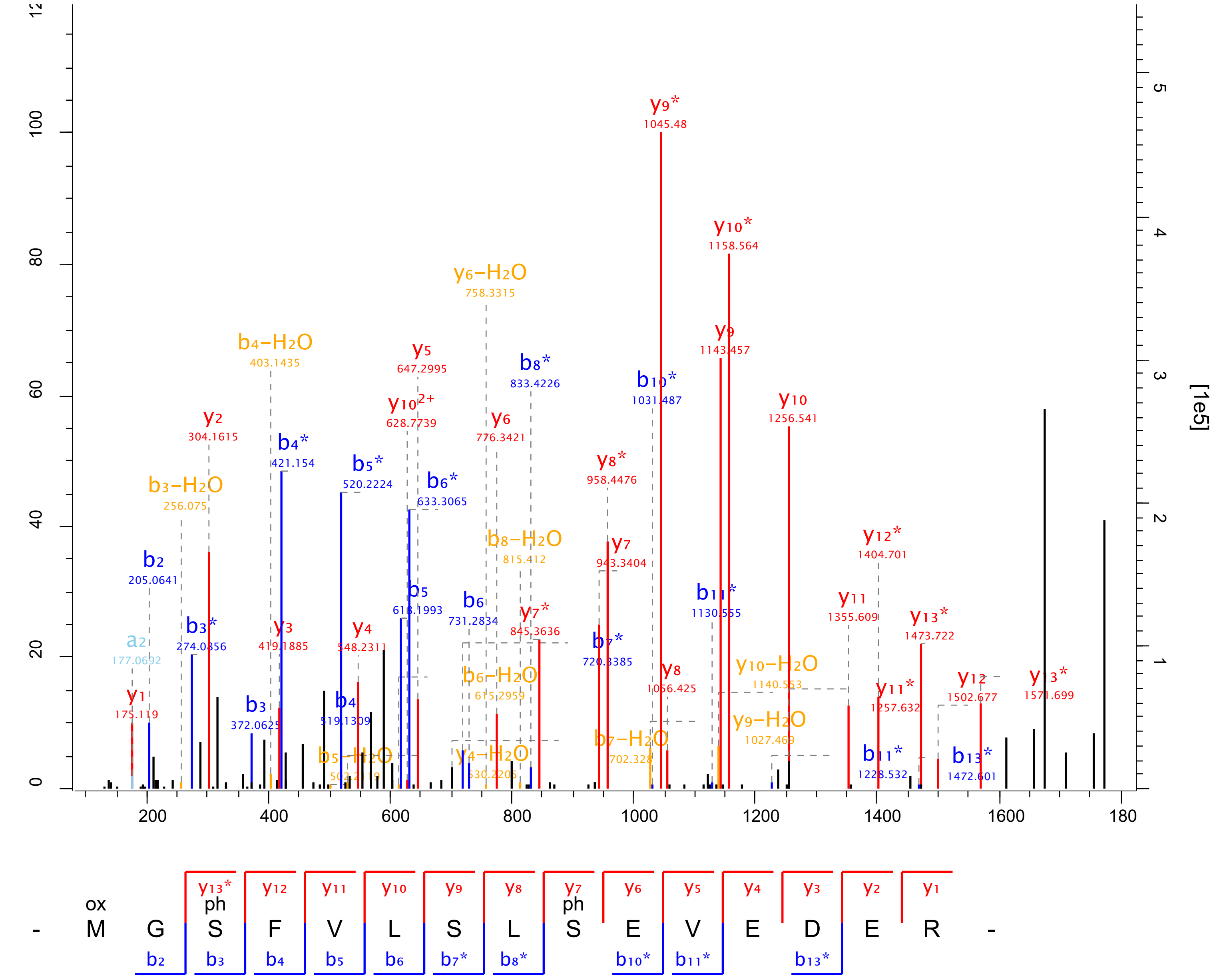

| Raw file | Scan | Method | Score | m/z |
| --- | --- | --- | --- | --- |
| water_02_1 | 50637 | FTMS; HCD | 165.34 | 848.08 |

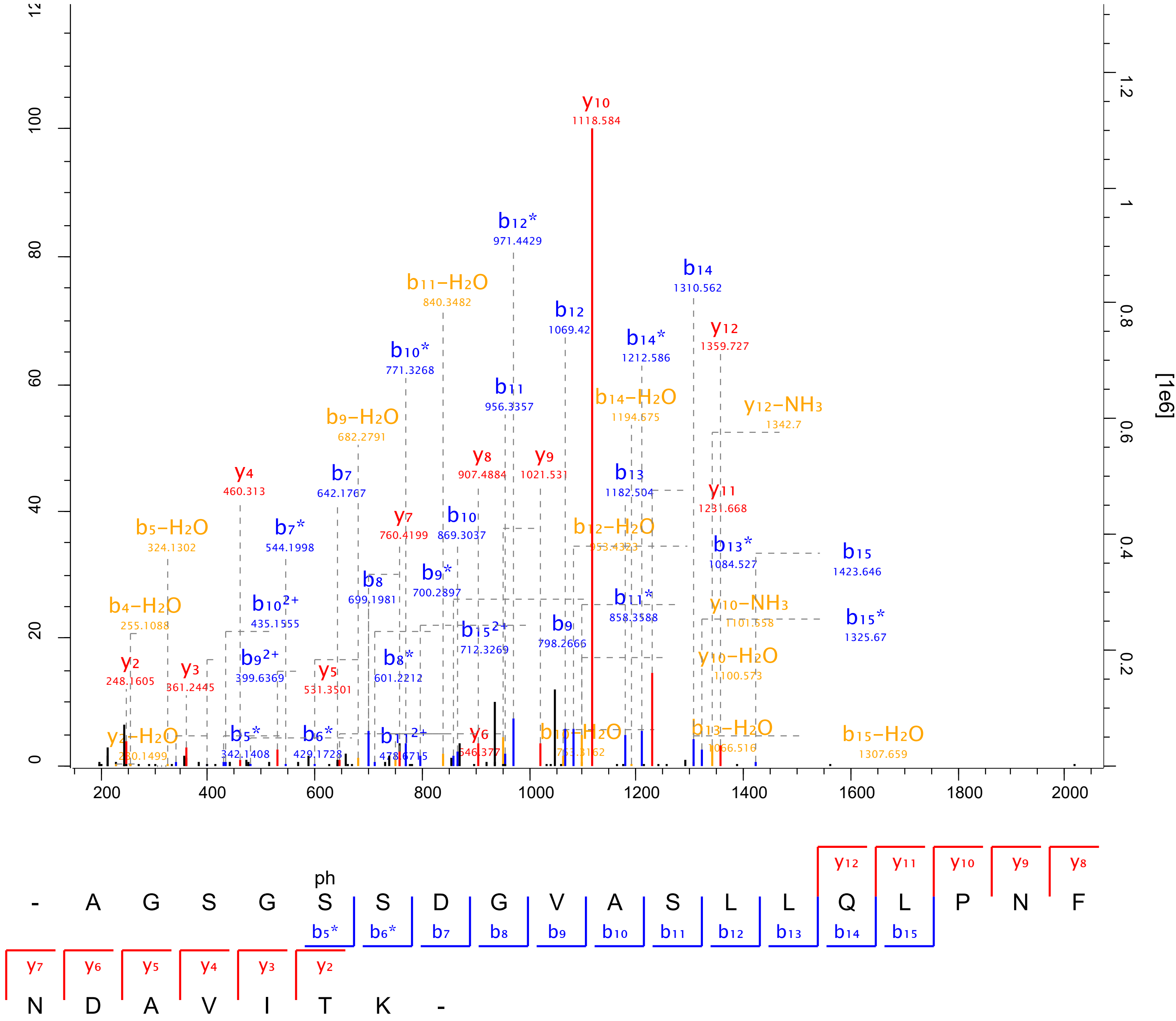

| Raw file | Scan | Method | Score | m/z |
|---|---|---|---|---|
| water_02_1 | 5133 | FTMS; HCD | 63.47 | 471.21 |

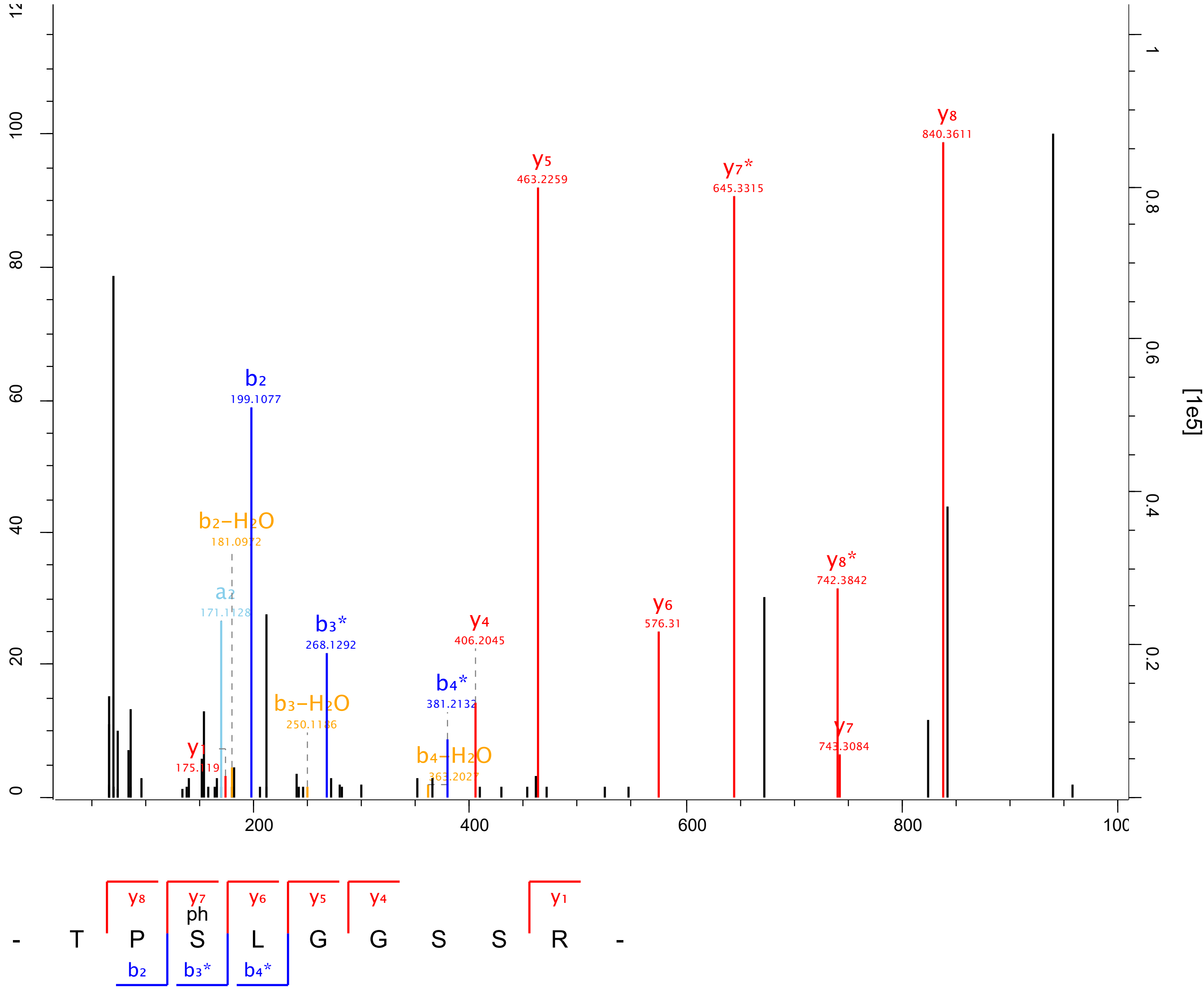

| Raw file | Scan | Method | Score | m/z |
|---|---|---|---|---|
| water_02_1 | 5144 | FTMS; HCD | 79.49 | 879.86 |

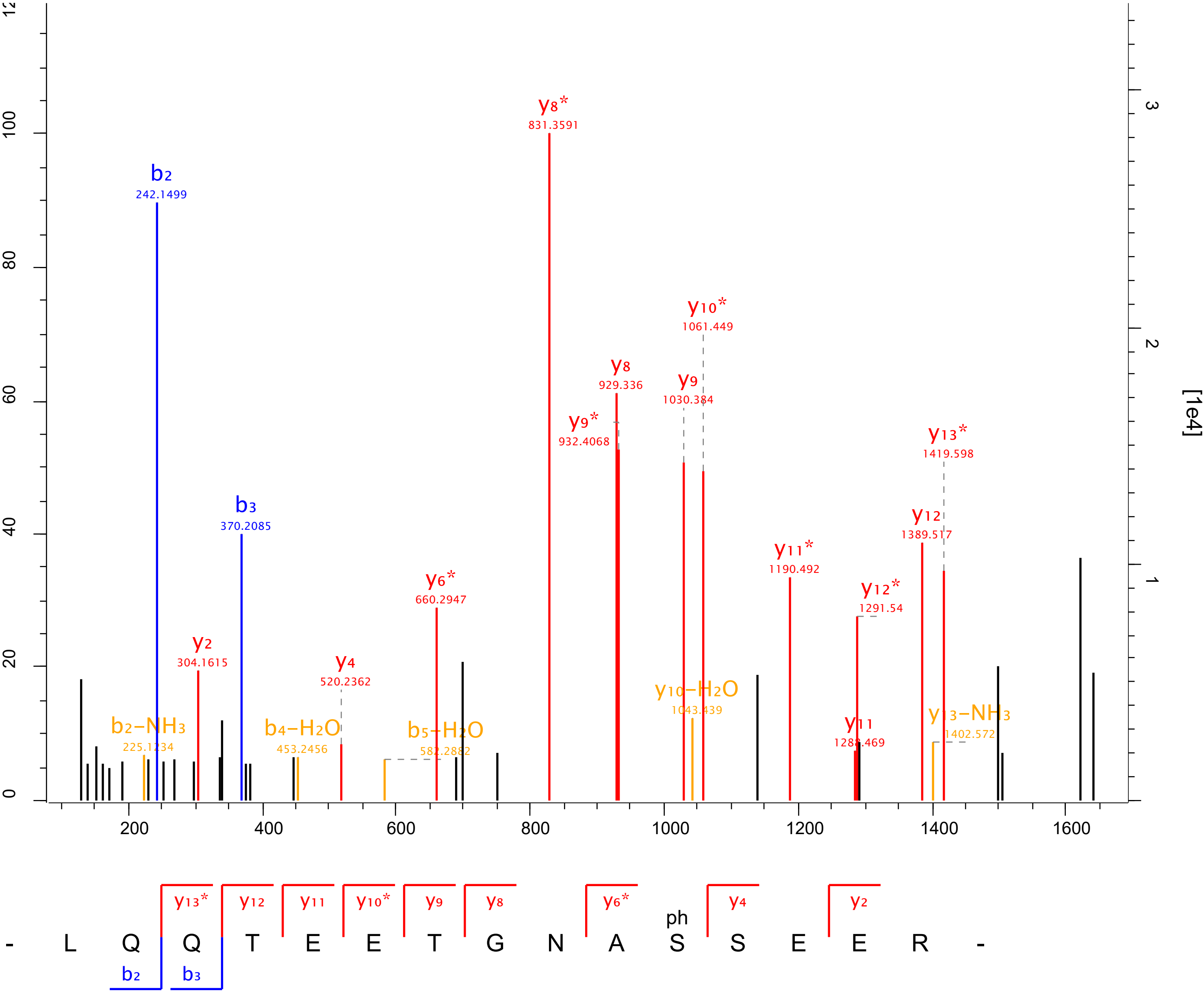

| Raw file | Scan | Method | Score | m/z |
| --- | --- | --- | --- | --- |
| water_02_1 | 5166 | FTMS; HCD | 86.74 | 684.28 |

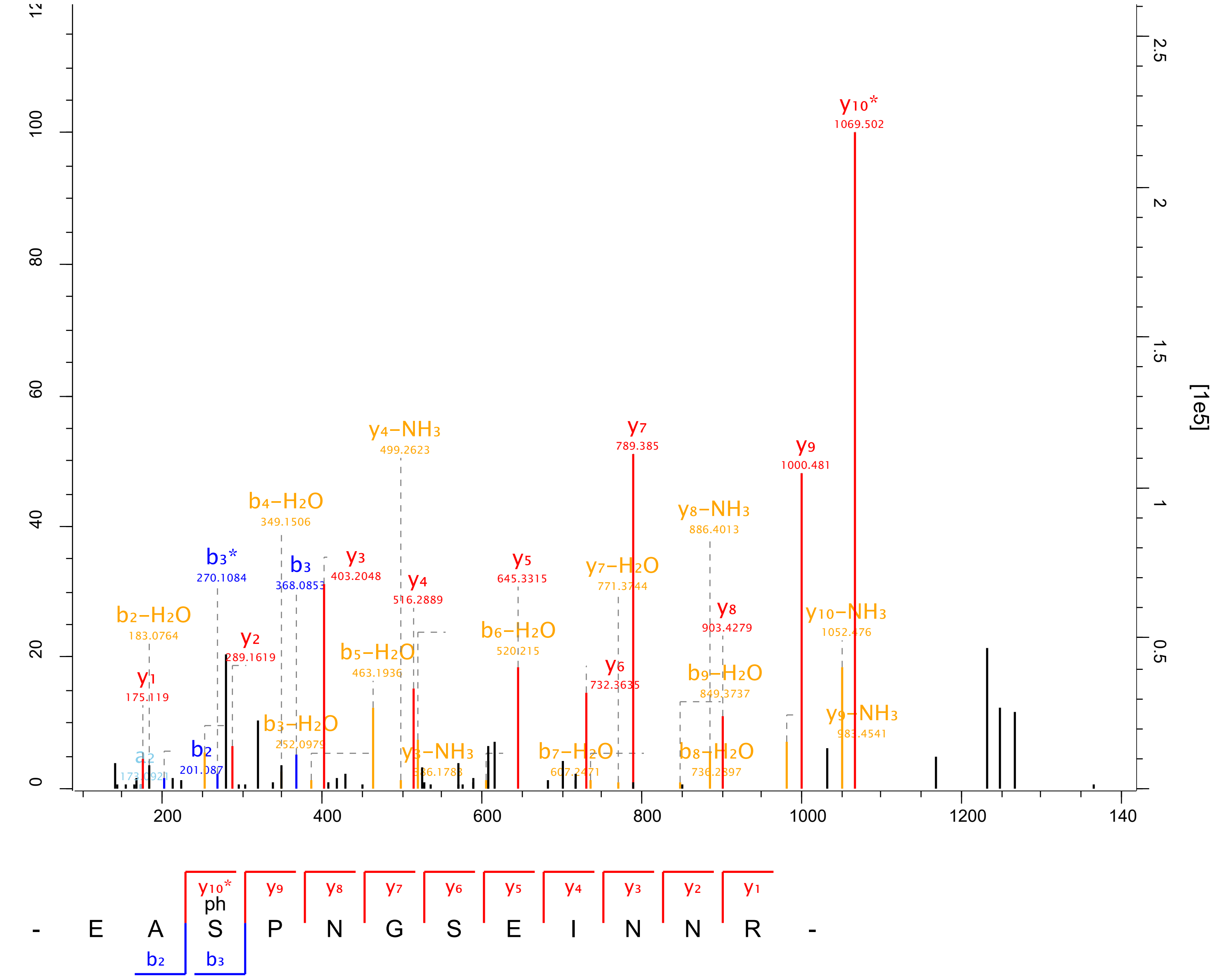

| Raw file | Scan | Method | Score | m/z |
|---|---|---|---|---|
| water_02_1 | 51703 | FTMS; HCD | 142.68 | 1097.47 |

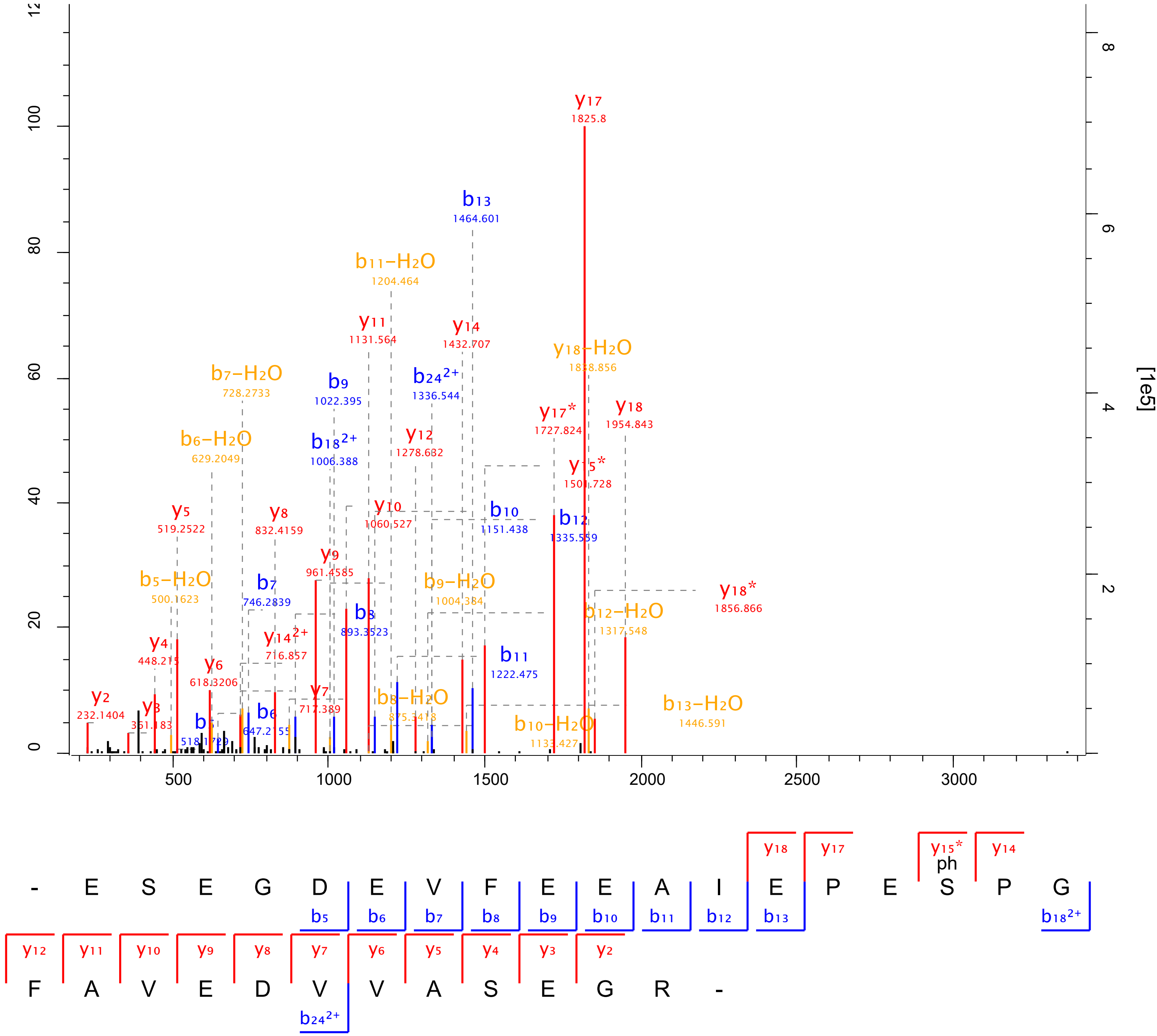

| Raw file | Scan | Method | Score | m/z |
|---|---|---|---|---|
| water_02_1 | 5197 | FTMS; HCD | 57.11 | 529.23 |

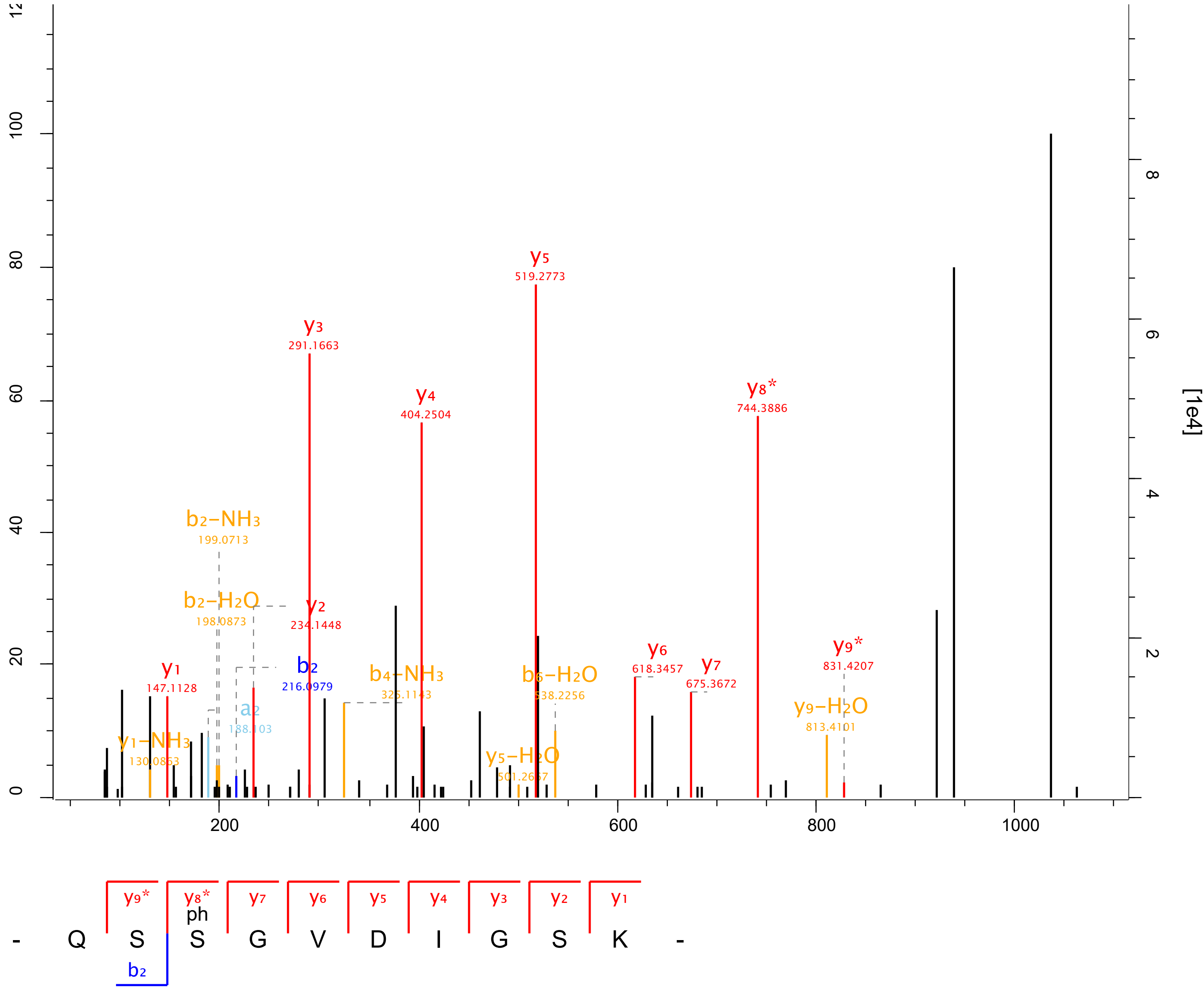

| Raw file | Scan | Method | Score | m/z |
| --- | --- | --- | --- | --- |
| water_02_1 | 5217 | FTMS; HCD | 51.29 | 536.76 |

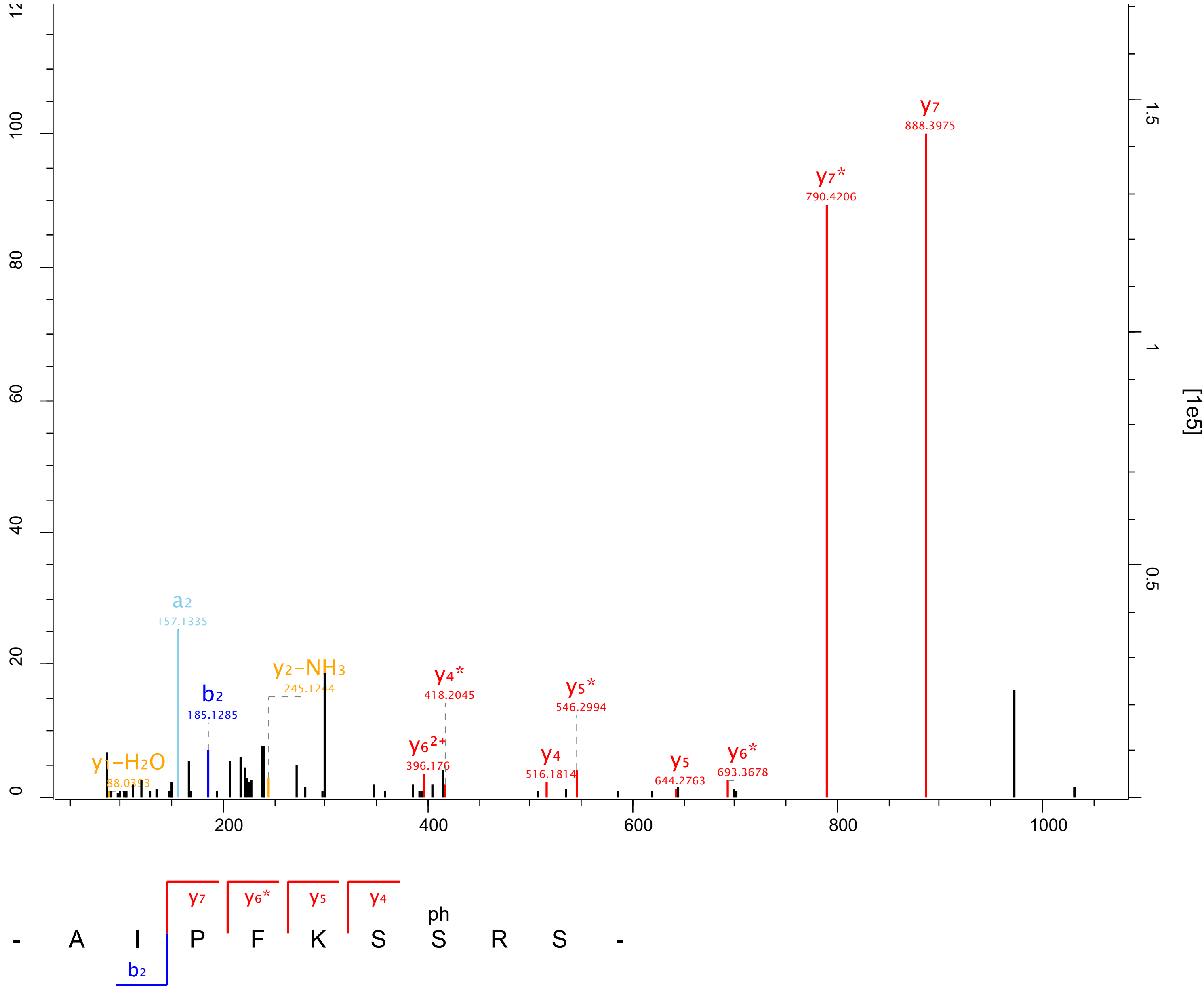

| Raw file | Scan | Method | Score | m/z |
| --- | --- | --- | --- | --- |
| water_02_1 | 5259 | FTMS; HCD | 51.47 | 628.27 |

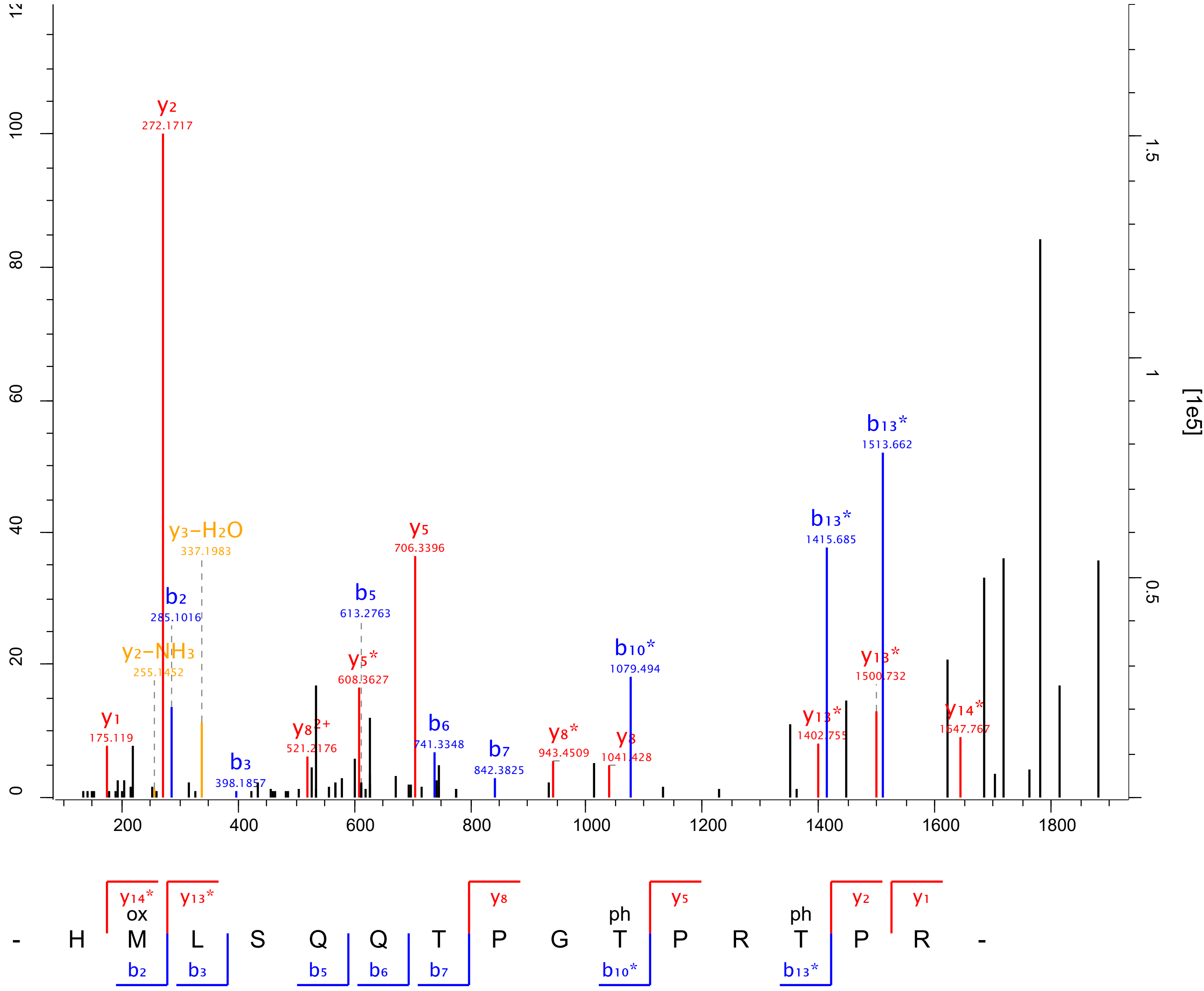

| Raw file | Scan | Method | Score | m/z |
|---|---|---|---|---|
| water_02_1 | 5262 | FTMS; HCD | 76.07 | 569.24 |

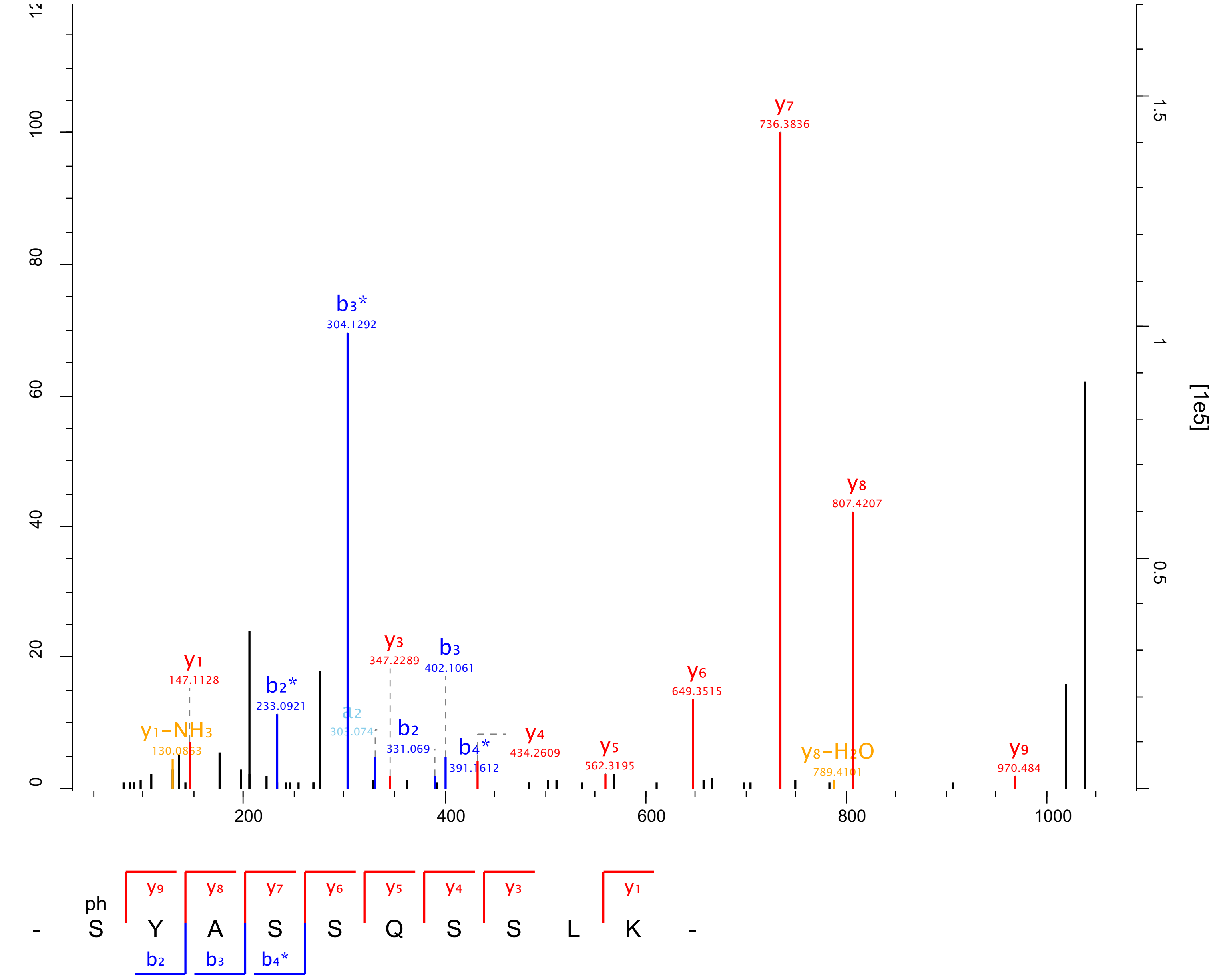

| Raw file | Scan | Method | Score | m/z |
|---|---|---|---|---|
| water_02_1 | 5313 | FTMS; HCD | 87.67 | 484.71 |

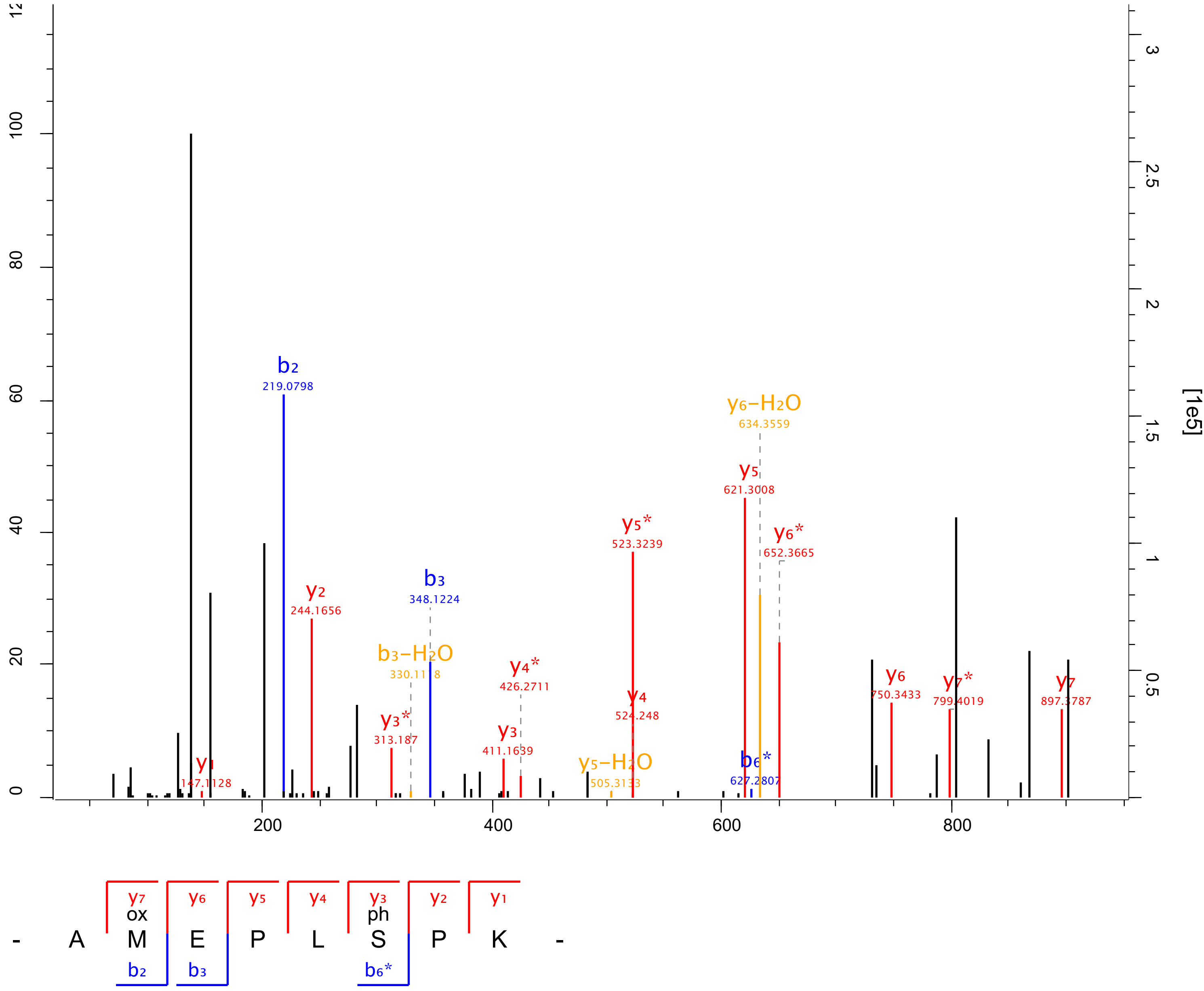

| Raw file | Scan | Method | Score | m/z |
|---|---|---|---|---|
| water_02_1 | 5441 | FTMS; HCD | 273.41 | 895.86 |

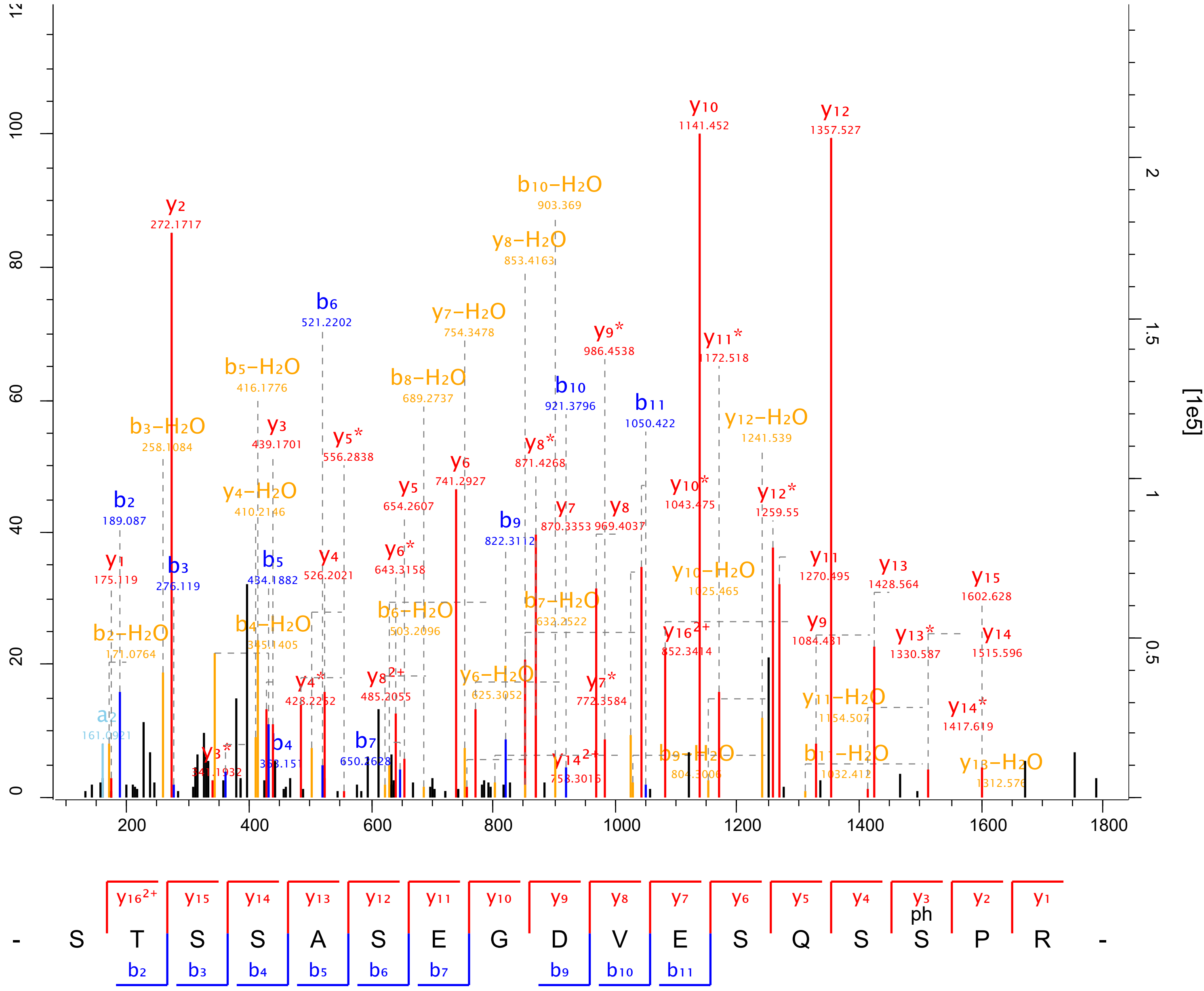

| Raw file | Scan | Method | Score | m/z |
|---|---|---|---|---|
| water_02_1 | 5466 | FTMS; HCD | 102.08 | 729.29 |

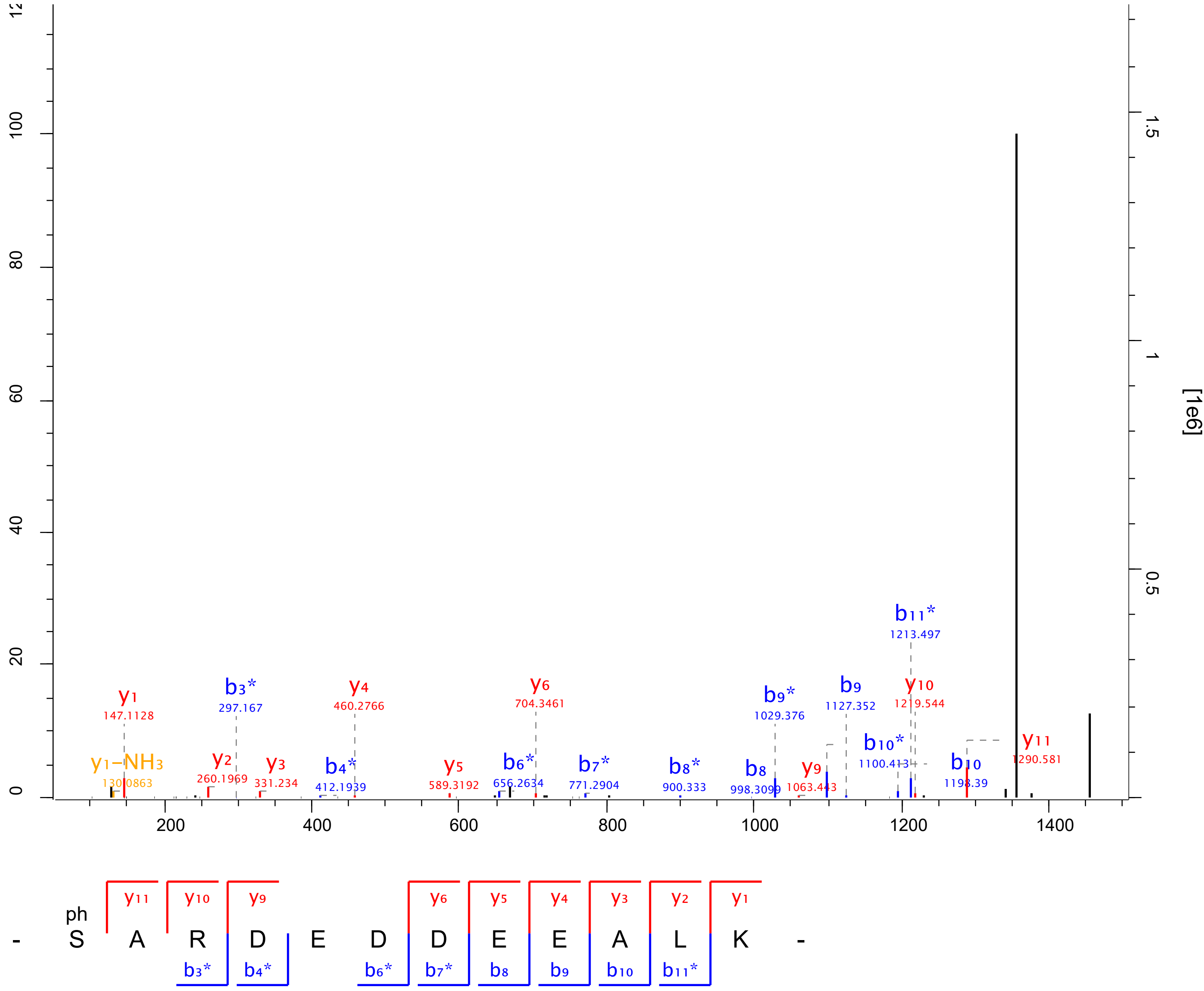

| Raw file | Scan | Method | Score | m/z |
| --- | --- | --- | --- | --- |
| water_02_1 | 5470 | FTMS; HCD | 123.02 | 497.56 |

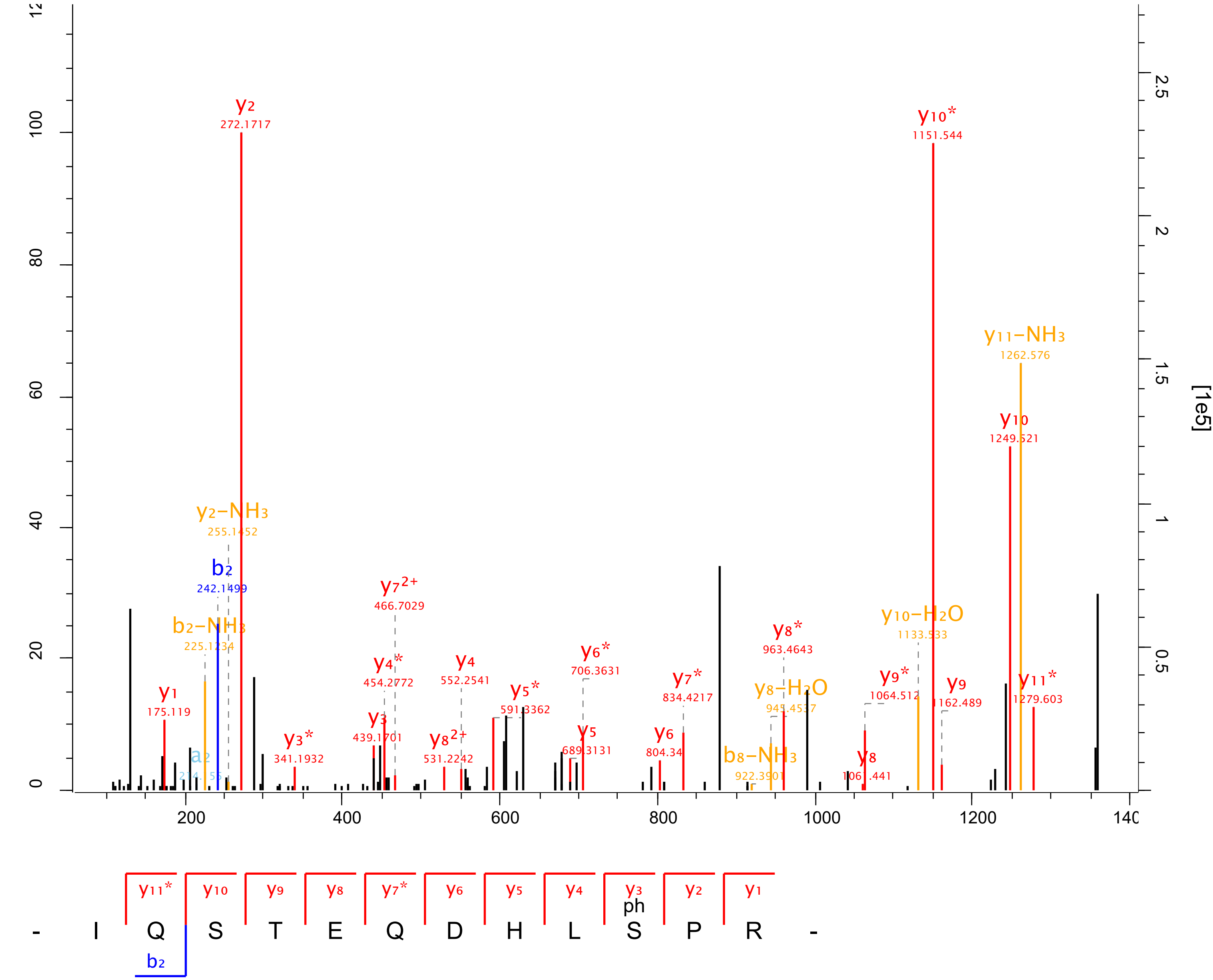

| Raw file | Scan | Method | Score | m/z |
| --- | --- | --- | --- | --- |
| water_02_1 | 5508 | FTMS; HCD | 45.28 | 409.18 |

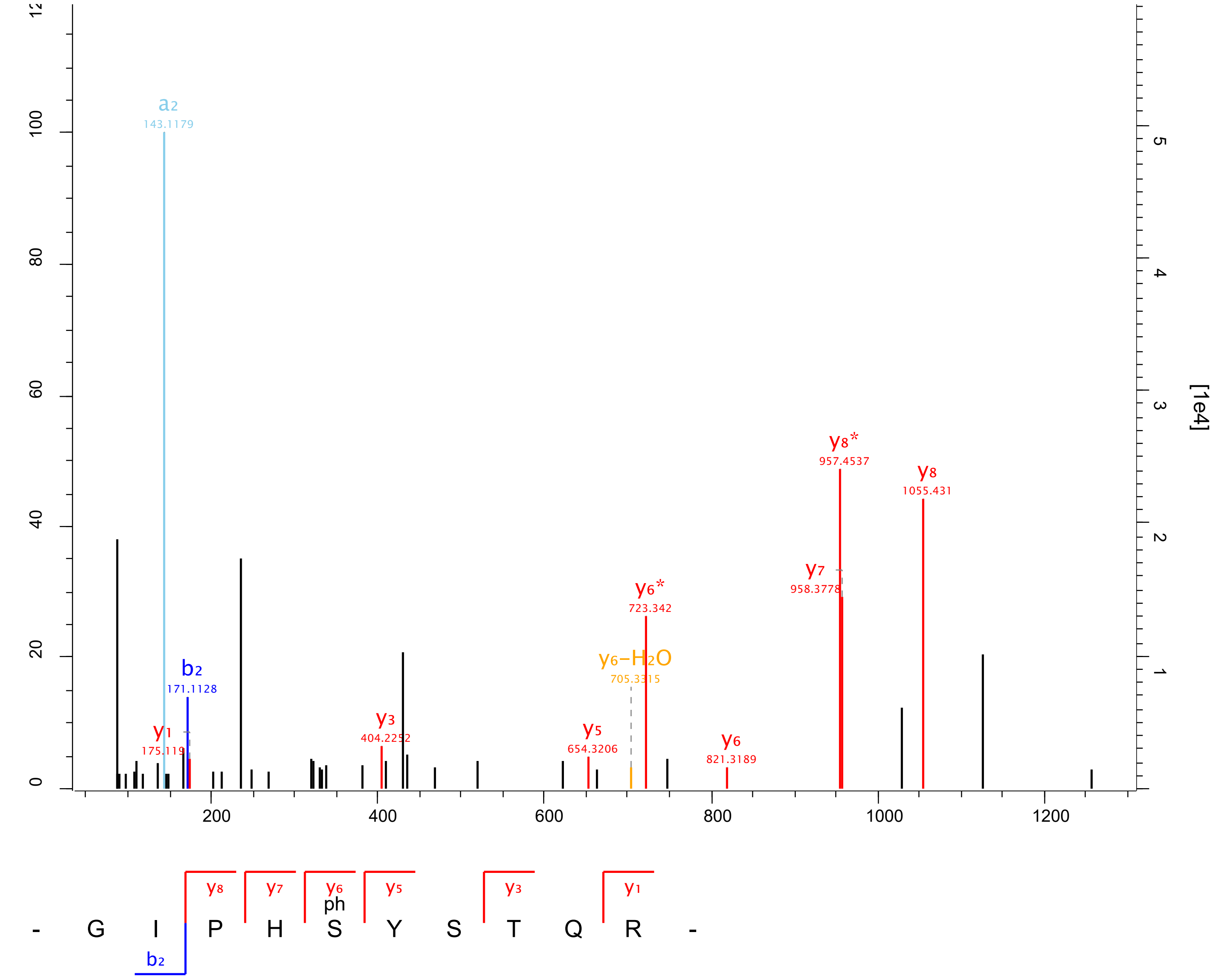

| Raw file | Scan | Method | Score | m/z |
|---|---|---|---|---|
| water_02_1 | 5538 | FTMS; HCD | 174.18 | 713.33 |

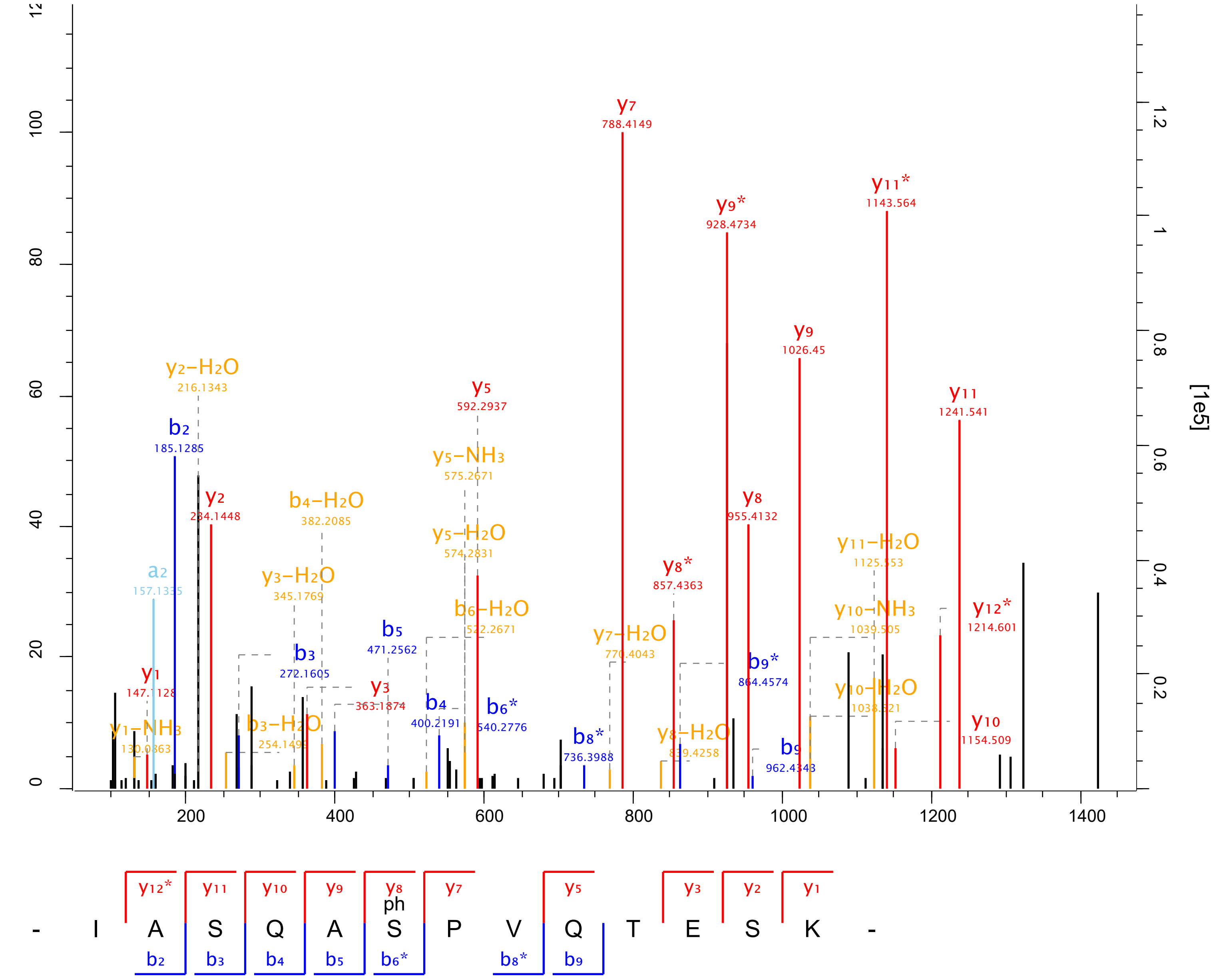

| Raw file | Scan | Method | Score | m/z |
| --- | --- | --- | --- | --- |
| water_02_1 | 5542 | FTMS; HCD | 129.76 | 525.26 |

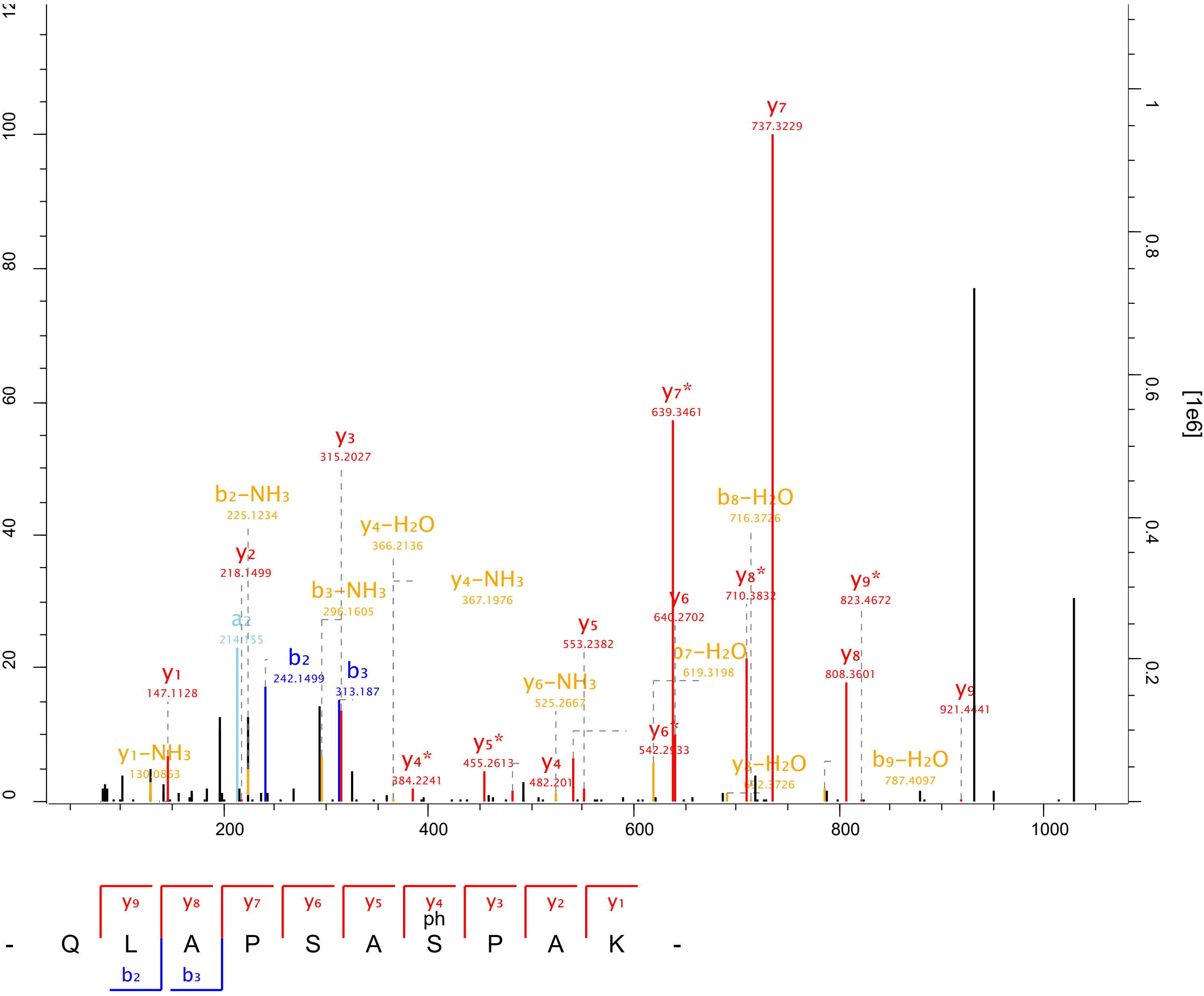

| Raw file | Scan | Method | Score | m/z |
|---|---|---|---|---|
| water_02_1 | 5559 | FTMS; HCD | 66.99 | 400.86 |

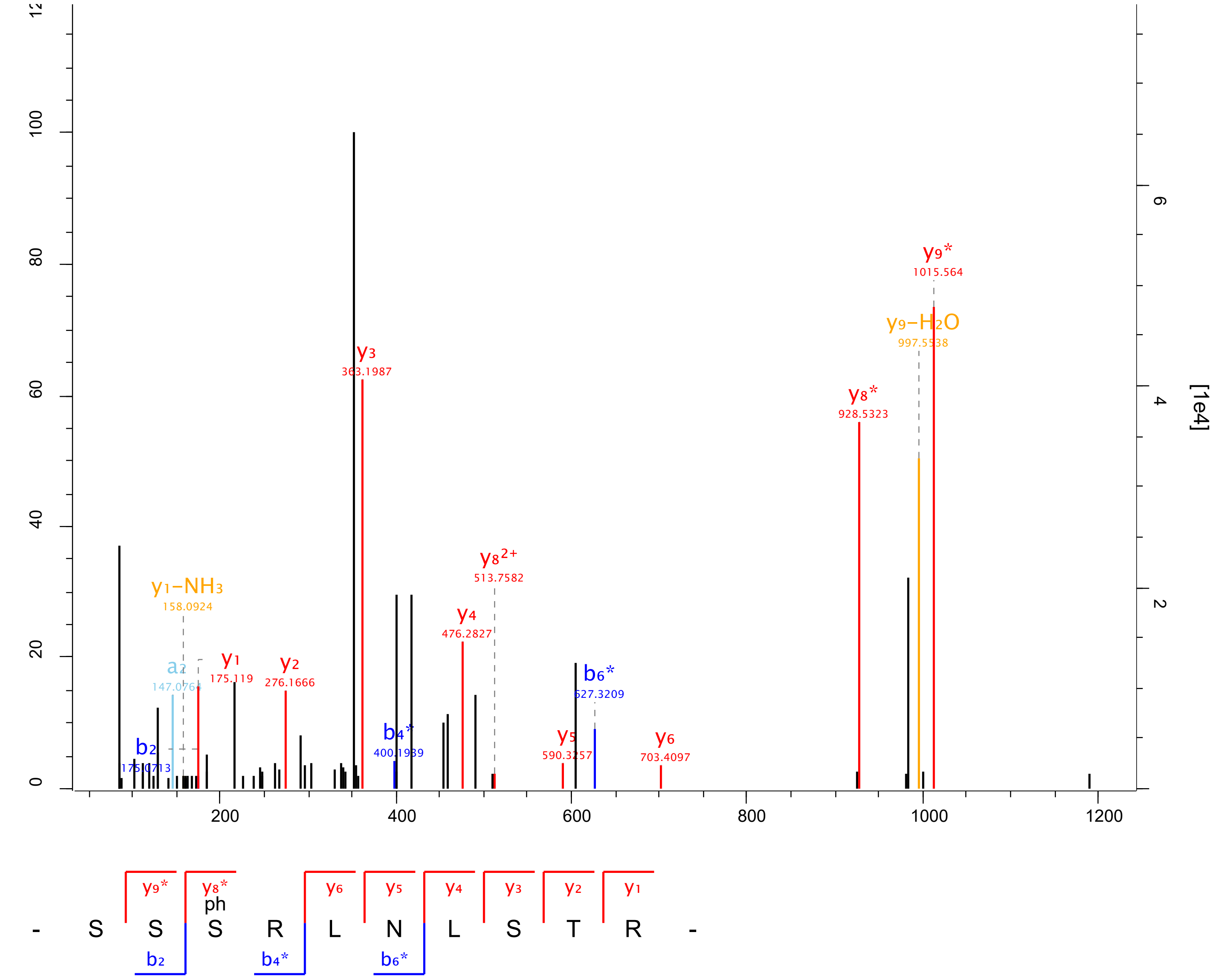

| Raw file | Scan | Method | Score | m/z |
|---|---|---|---|---|
| water_02_1 | 5563 | FTMS; HCD | 153.11 | 500.75 |

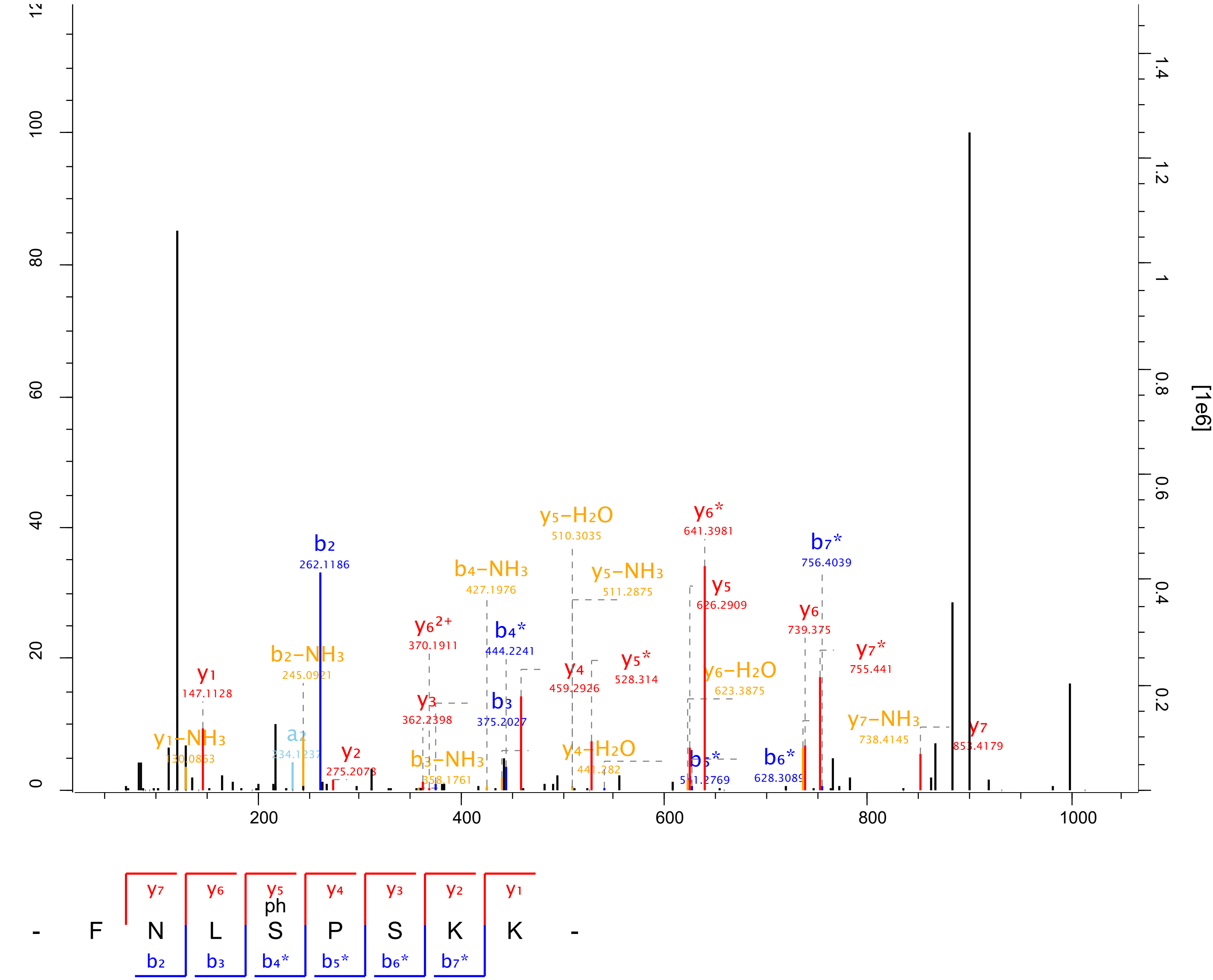

| Raw file | Scan | Method | Score | m/z |
|---|---|---|---|---|
| water_02_1 | 5572 | FTMS; HCD | 157.28 | 580.24 |

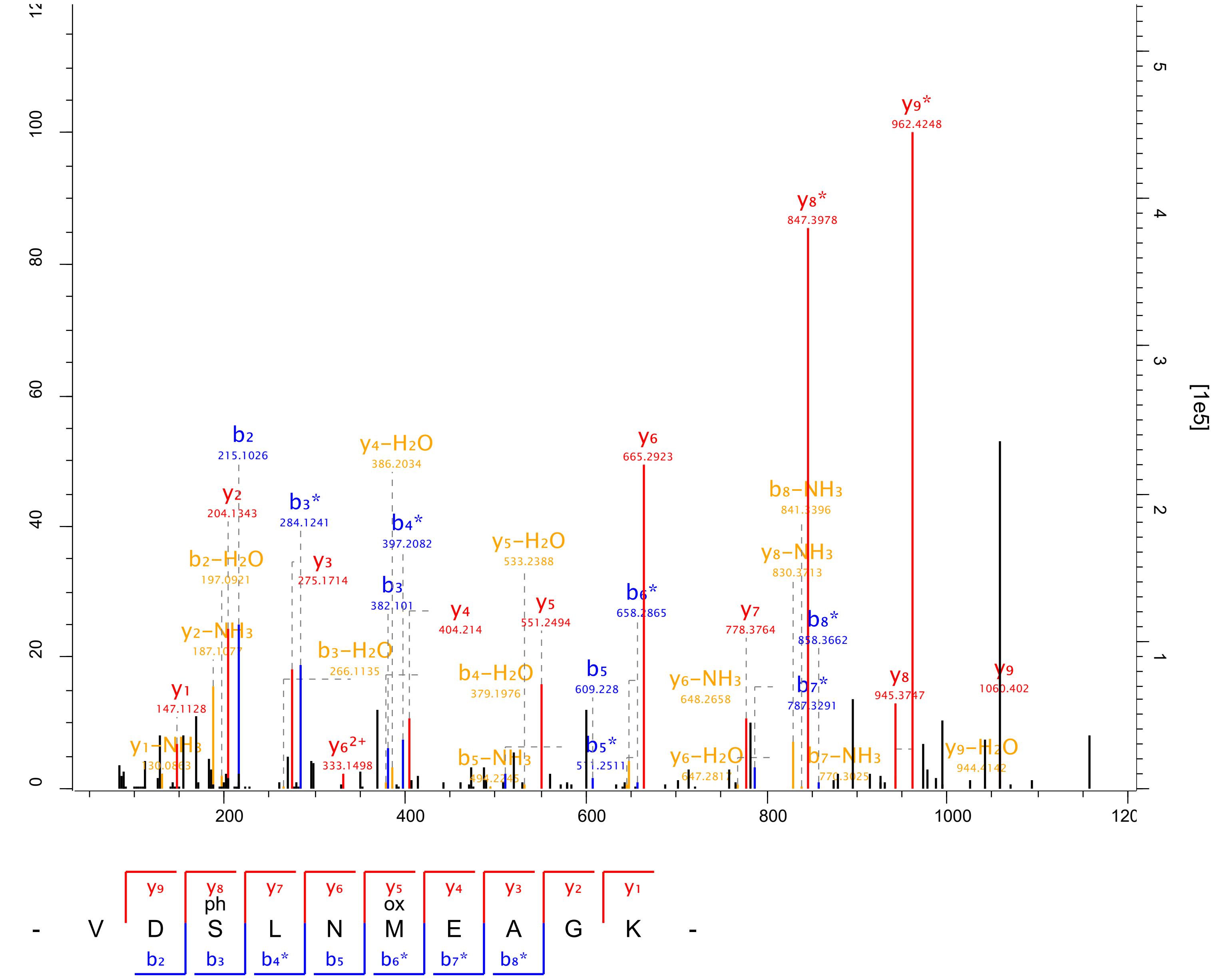

| Raw file | Scan | Method | Score | m/z |
| --- | --- | --- | --- | --- |
| water_02_1 | 5579 | FTMS; HCD | 220.98 | 695.65 |

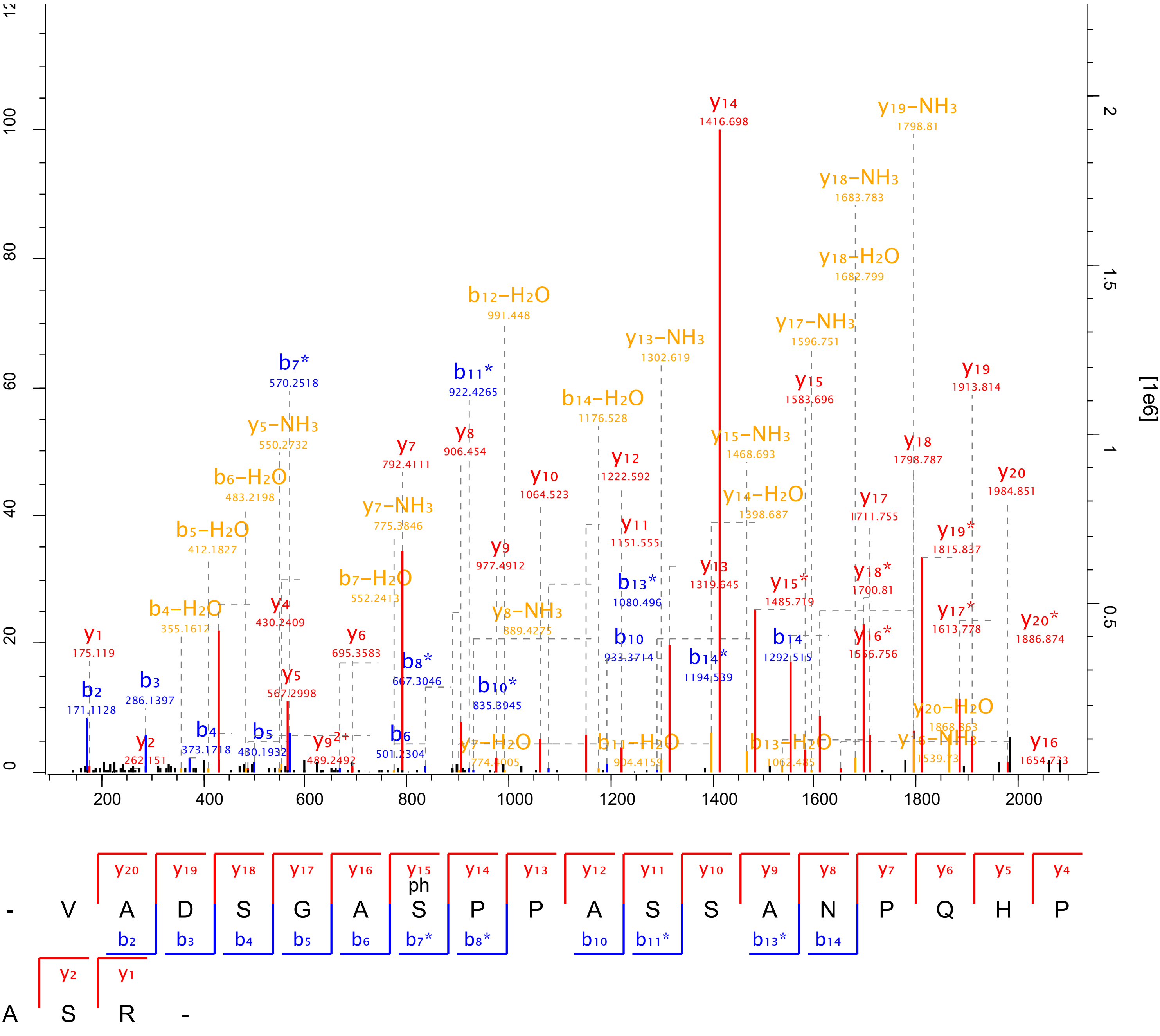

| Raw file | Scan | Method | Score | m/z |
|---|---|---|---|---|
| water_02_1 | 5662 | FTMS; HCD | 48.05 | 941.34 |

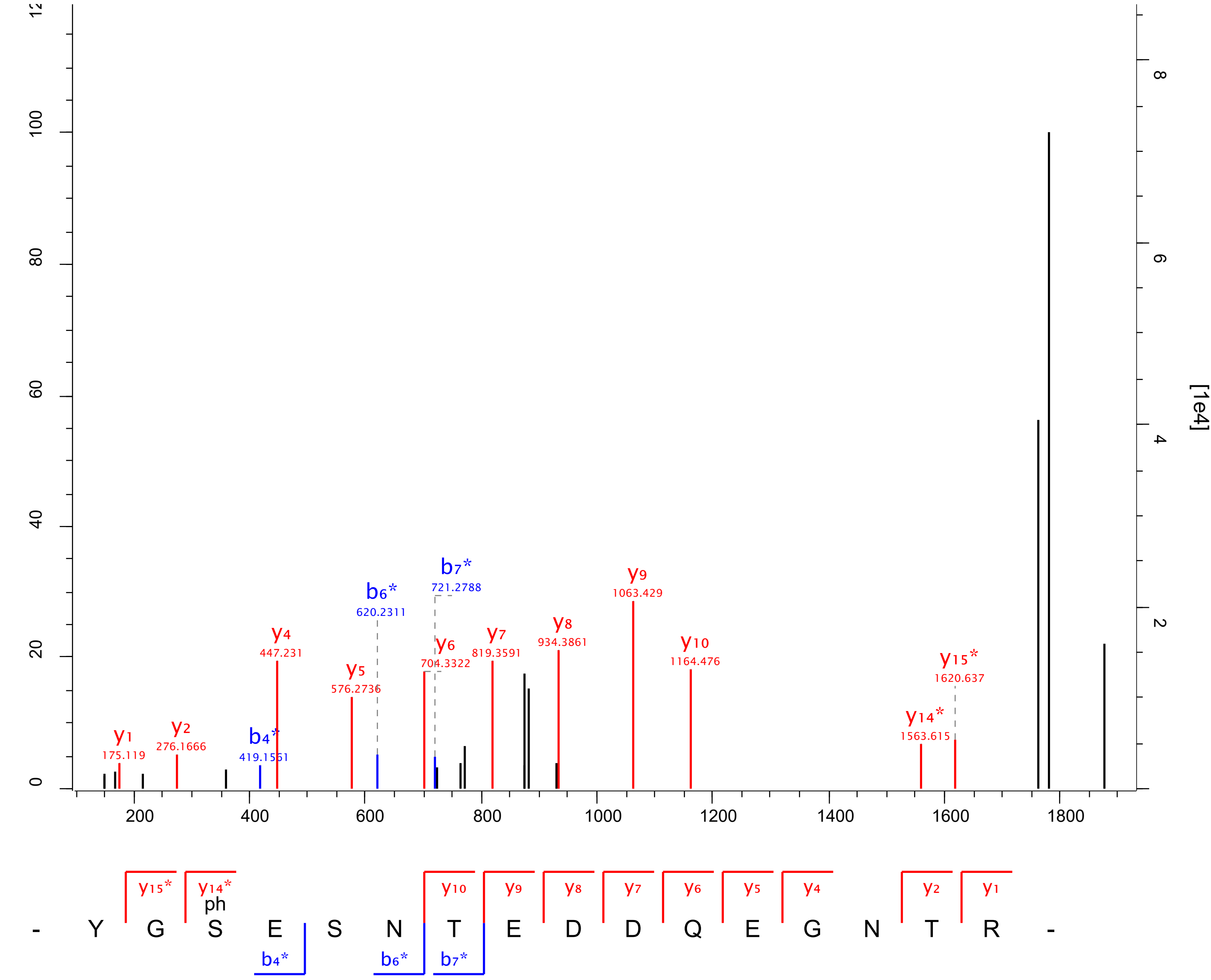

| Raw file | Scan | Method | Score | m/z |
|---|---|---|---|---|
| water_02_1 | 5774 | FTMS; HCD | 61.76 | 499.2 |

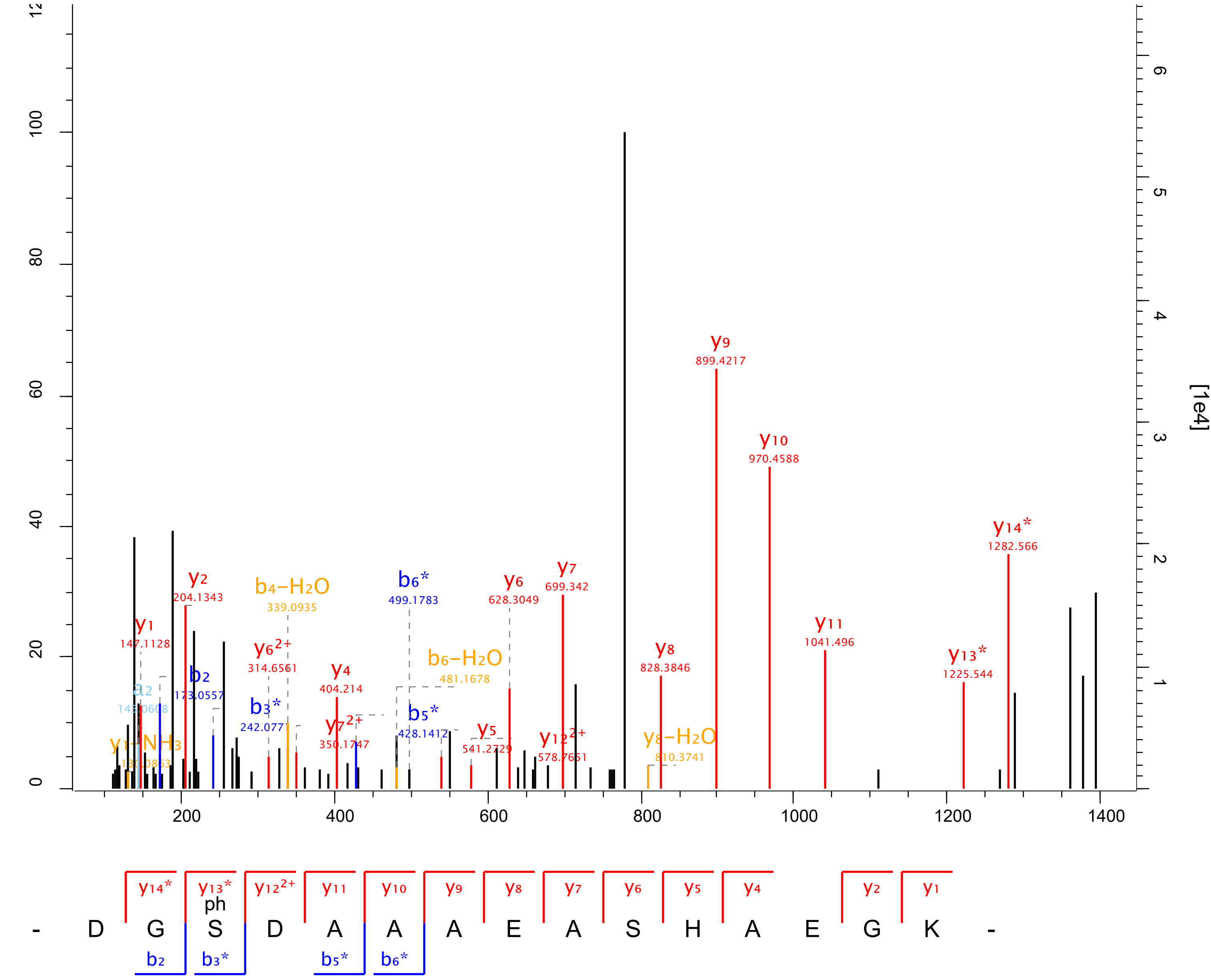

| Raw file | Scan | Method | Score | m/z |
| --- | --- | --- | --- | --- |
| water_02_1 | 5805 | FTMS; HCD | 61.44 | 775.31 |

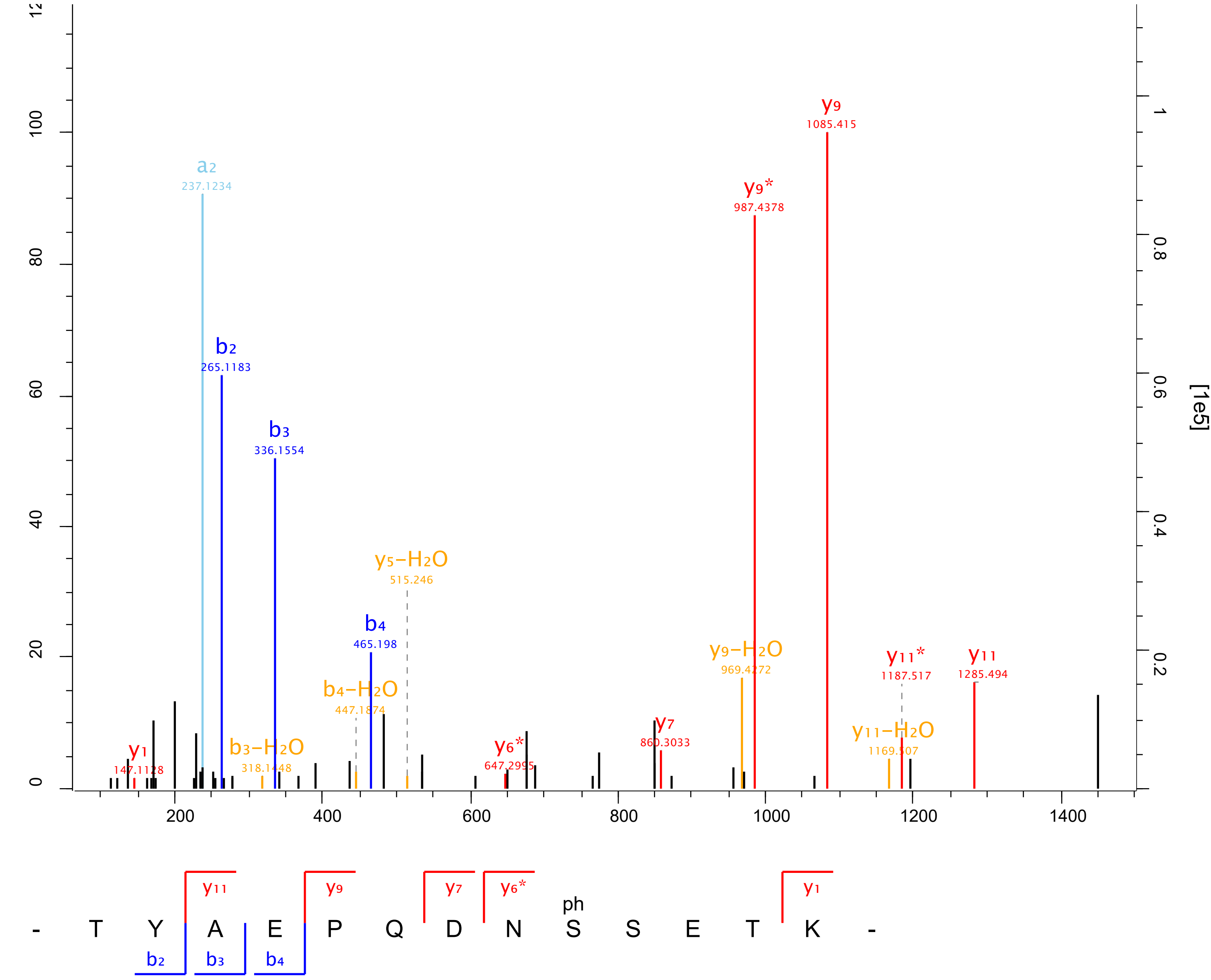

| Raw file | Scan | Method | Score | m/z |
|---|---|---|---|---|
| water_02_1 | 5823 | FTMS; HCD | 45.54 | 468.55 |

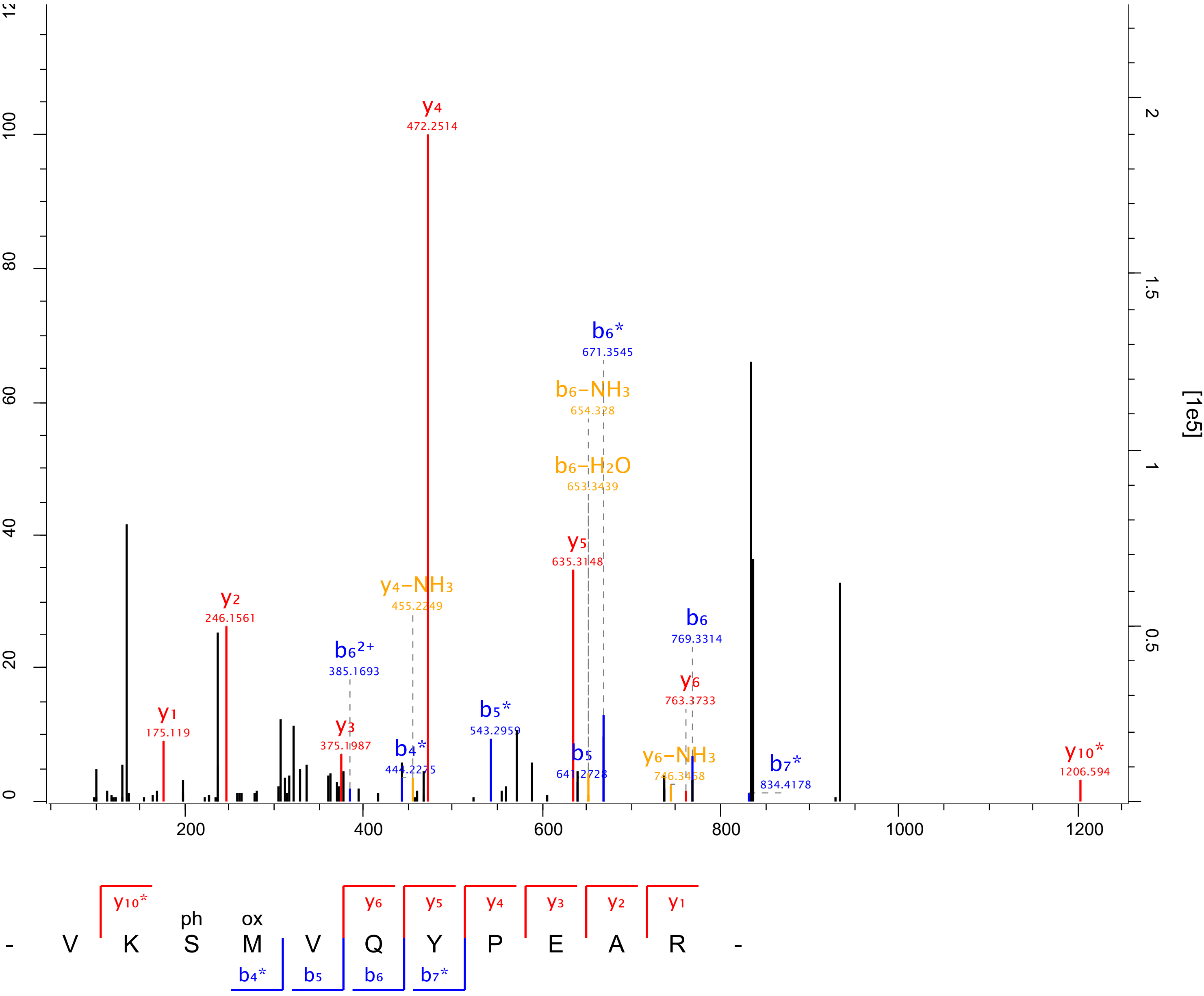

| Raw file | Scan | Method | Score | m/z |
| --- | --- | --- | --- | --- |
| water_02_1 | 5916 | FTMS; HCD | 76.22 | 437.2 |

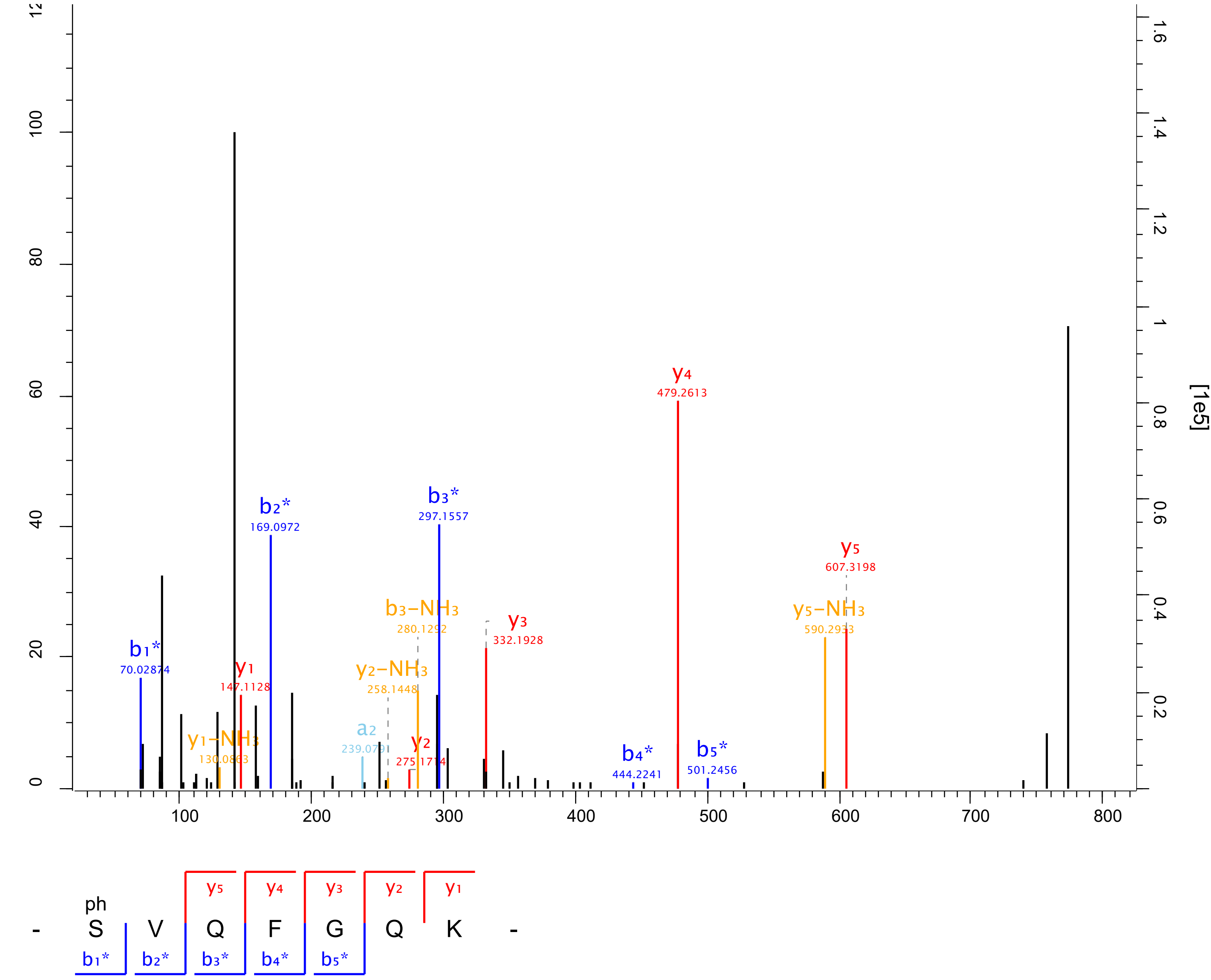

| Raw file | Scan | Method | Score | m/z |
|---|---|---|---|---|
| water_02_1 | 6109 | FTMS; HCD | 89.05 | 1070.39 |

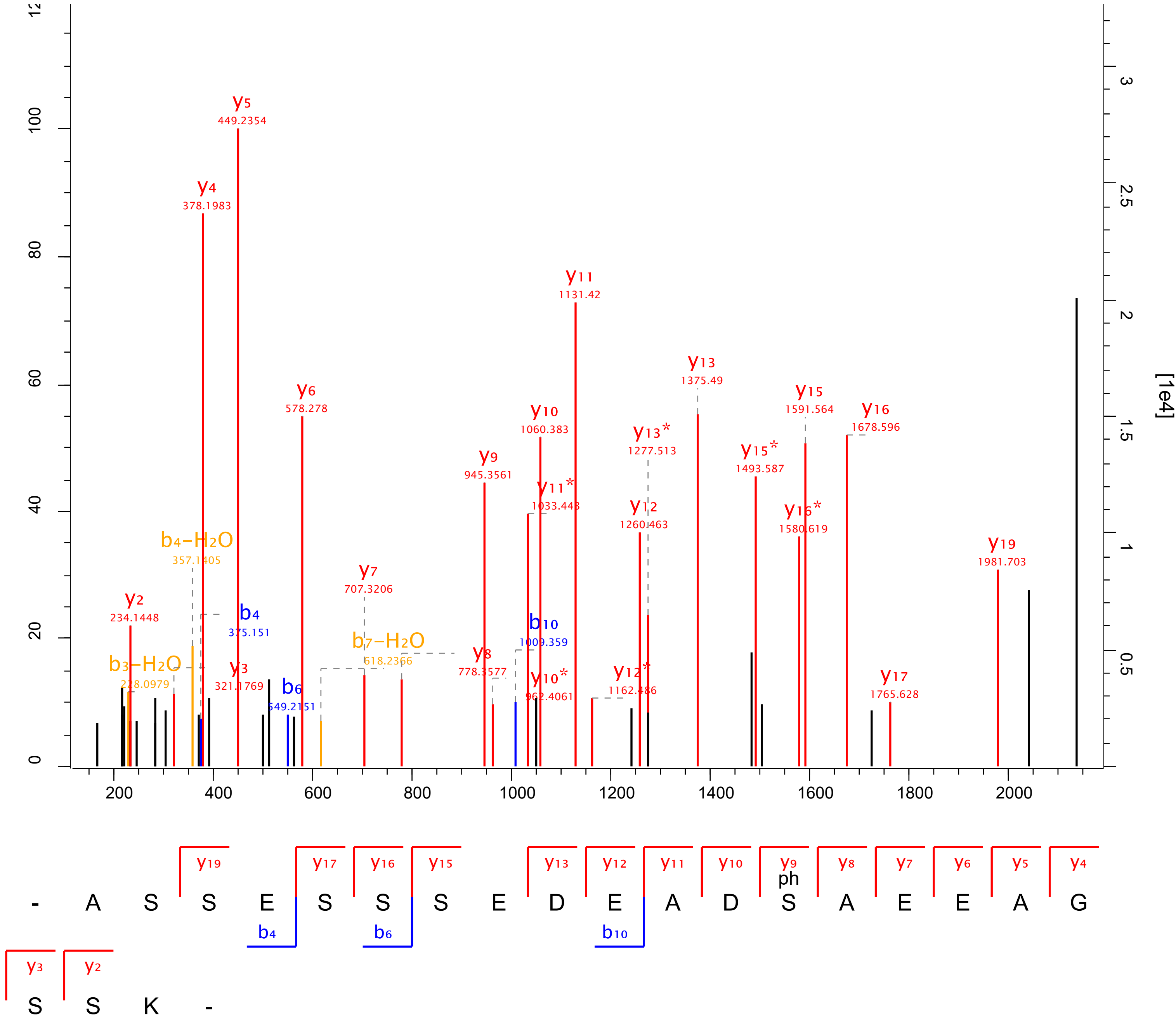

| Raw file | Scan | Method | Score | m/z |
|---|---|---|---|---|
| water_02_1 | 6128 | FTMS; HCD | 80.39 | 452.2 |

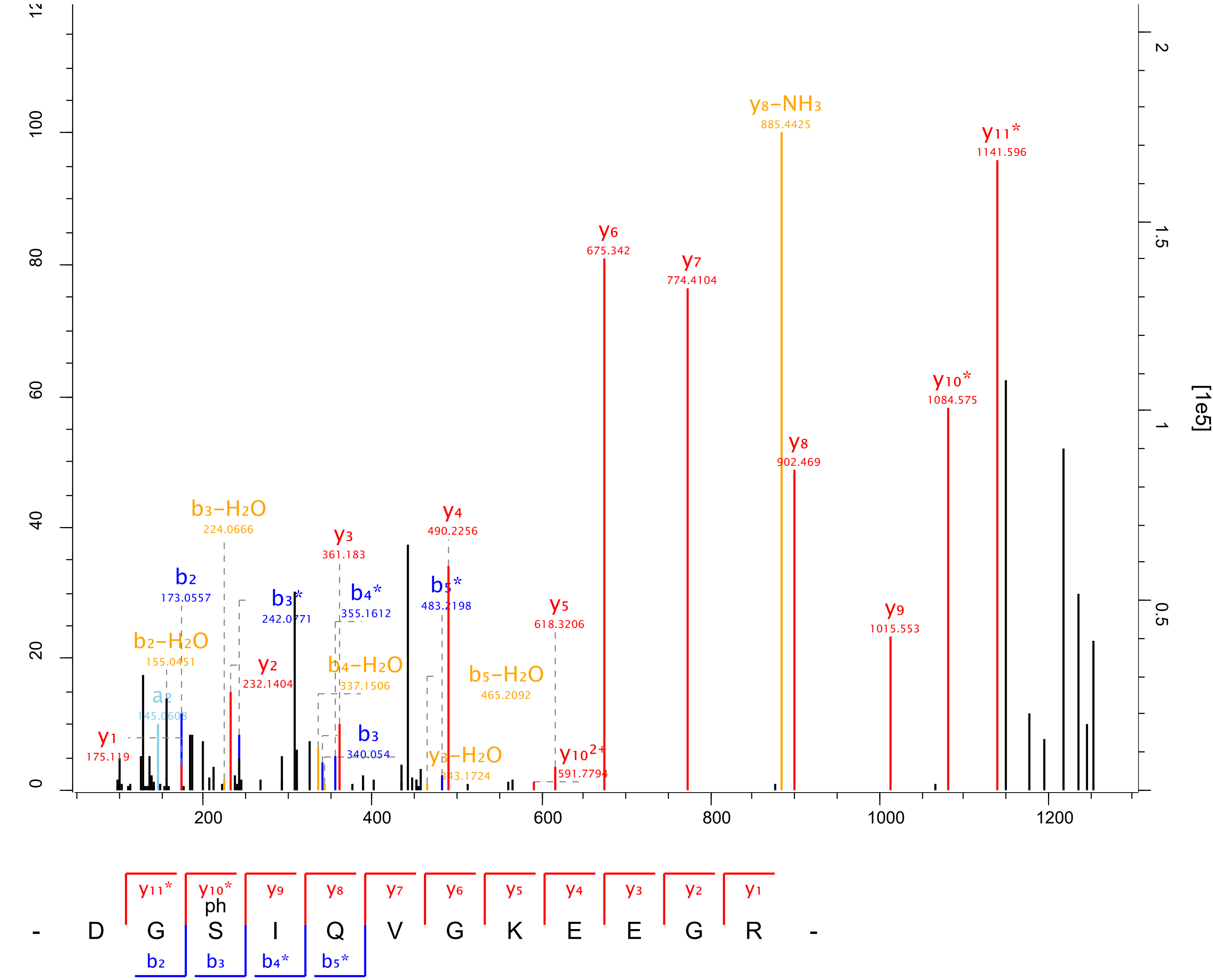

| Raw file | Scan | Method | Score | m/z |
| --- | --- | --- | --- | --- |
| water_02_1 | 6133 | FTMS; HCD | 80.96 | 585.75 |

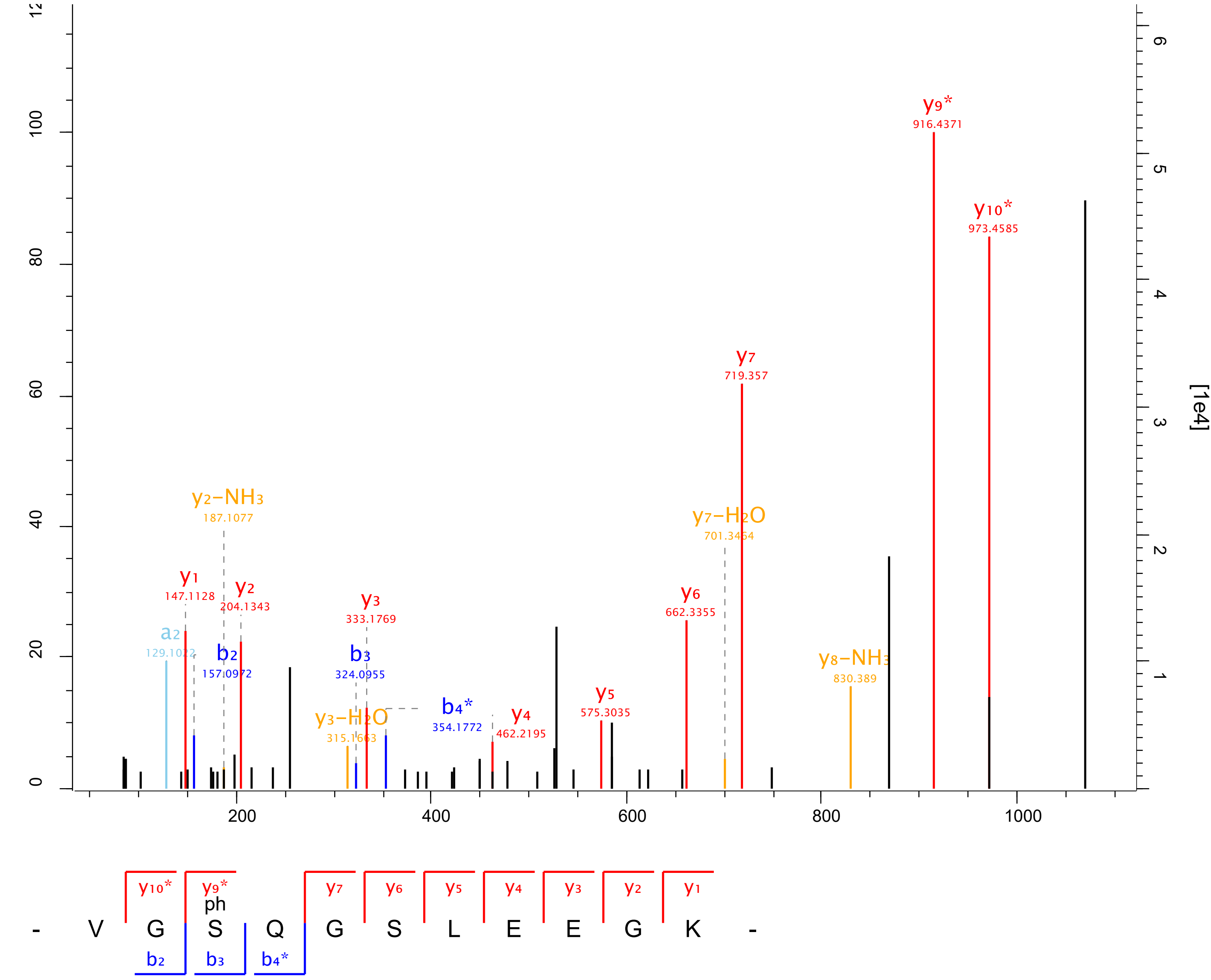

| Raw file | Scan | Method | Score | m/z |
|---|---|---|---|---|
| water_02_1 | 6182 | FTMS; HCD | 119.03 | 462.73 |

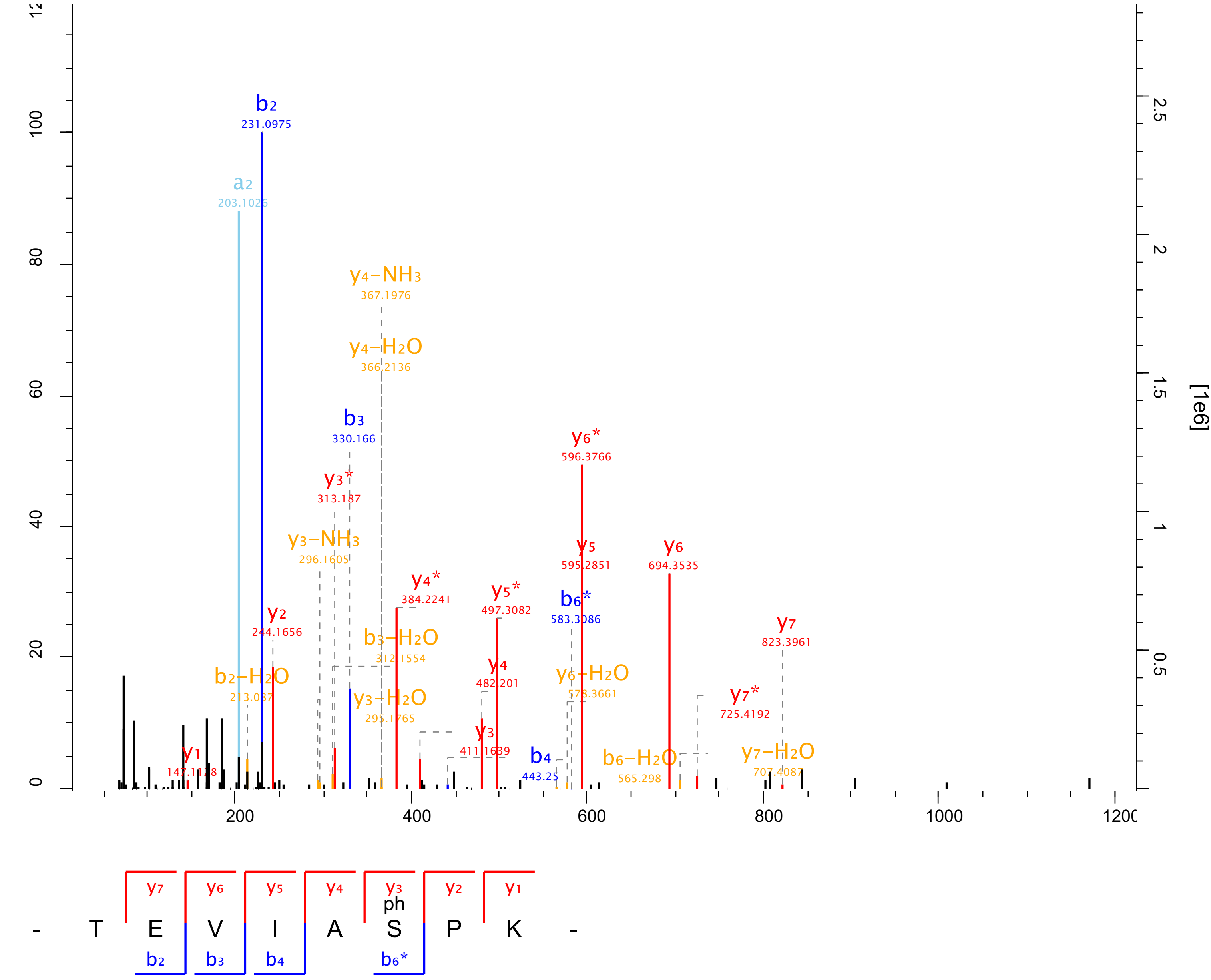

| Raw file | Scan | Method | Score | m/z |
|---|---|---|---|---|
| water_02_1 | 6192 | FTMS; HCD | 186.58 | 593.76 |

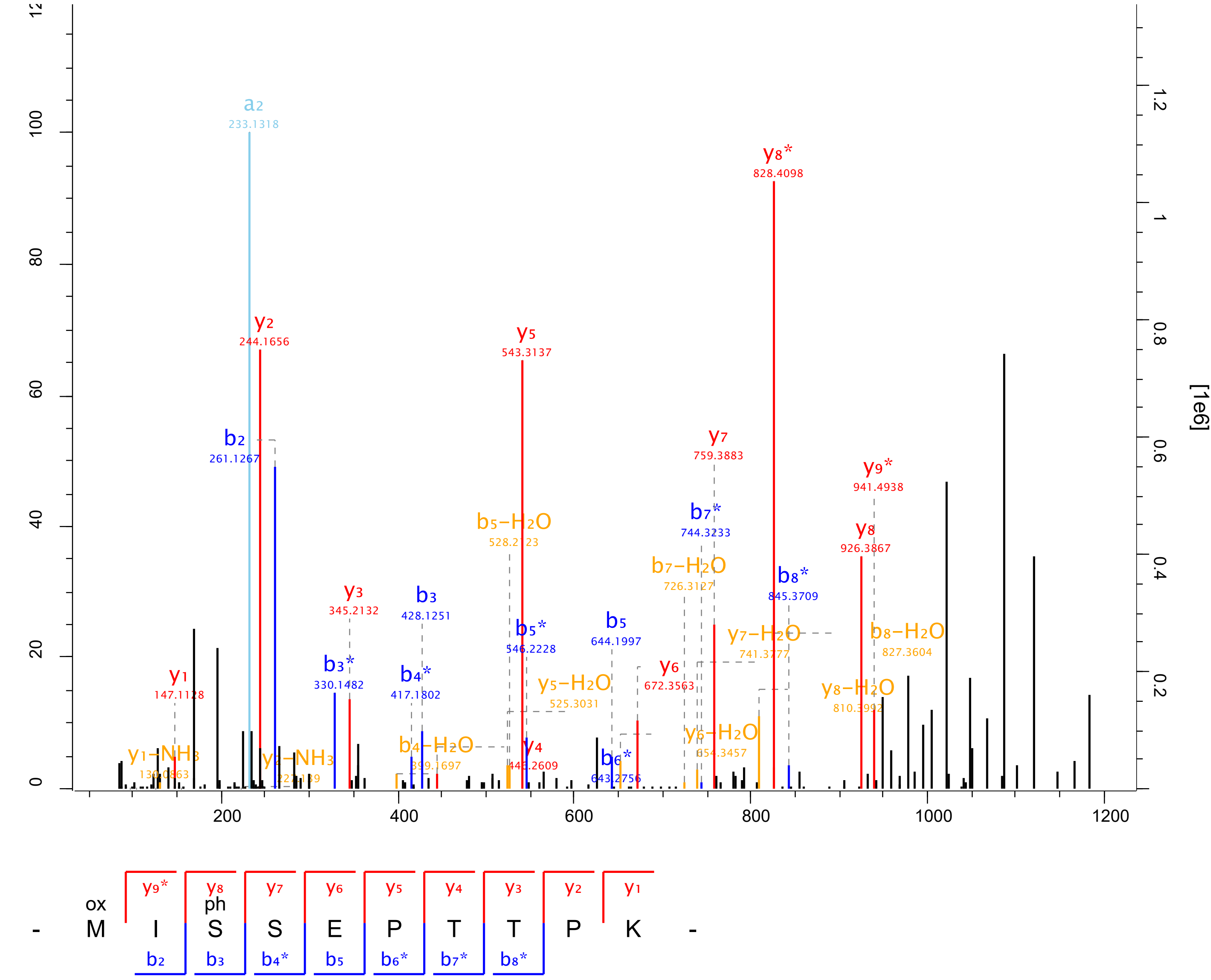

| Raw file | Scan | Method | Score | m/z |
|---|---|---|---|---|
| water_02_1 | 6223 | FTMS; HCD | 174.26 | 632.62 |

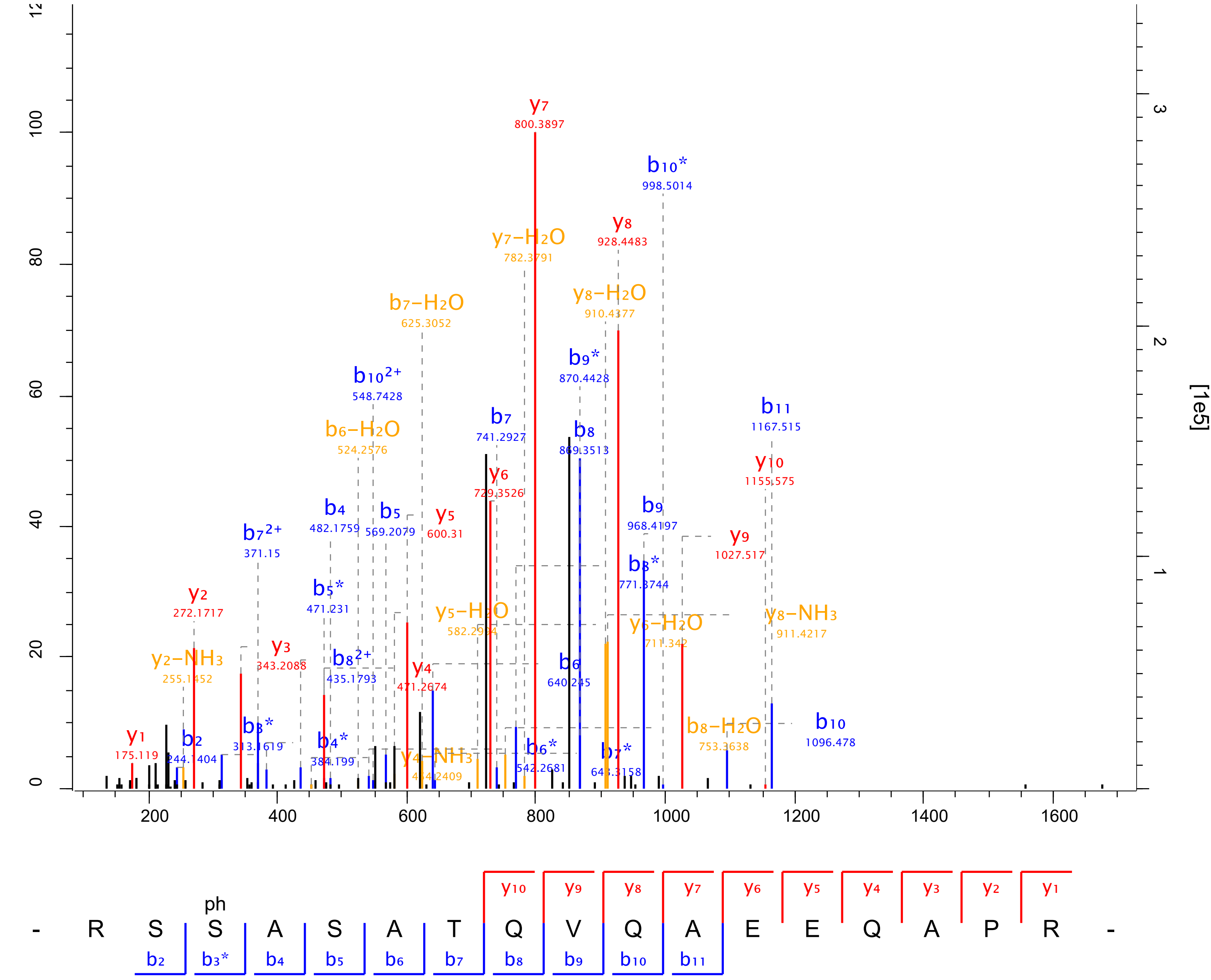

| Raw file | Scan | Method | Score | m/z |
|---|---|---|---|---|
| water_02_1 | 6288 | FTMS; HCD | 92.47 | 764.82 |

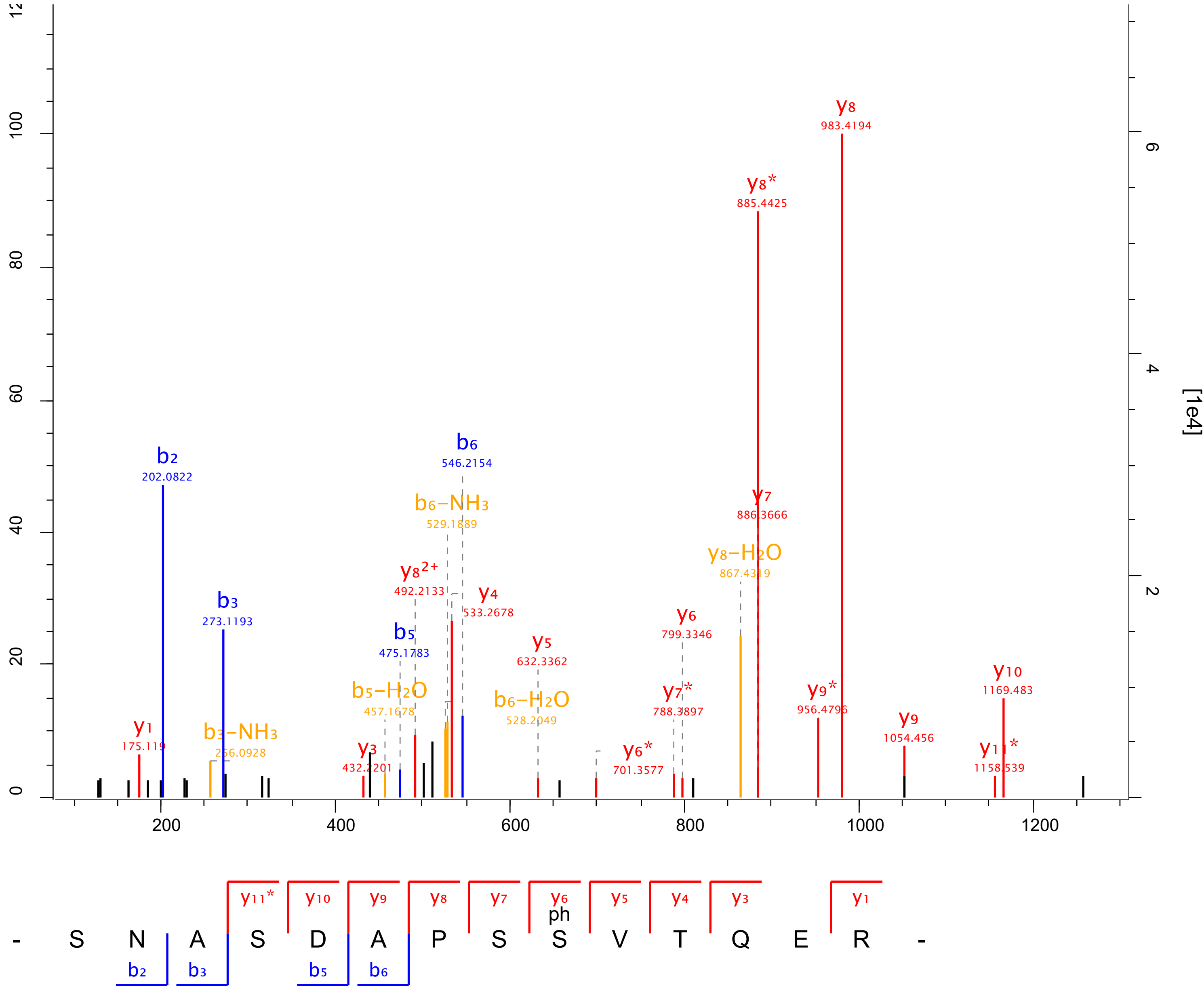

| Raw file | Scan | Method | Score | m/z |
|---|---|---|---|---|
| water_02_1 | 6342 | FTMS; HCD | 60.91 | 469.71 |

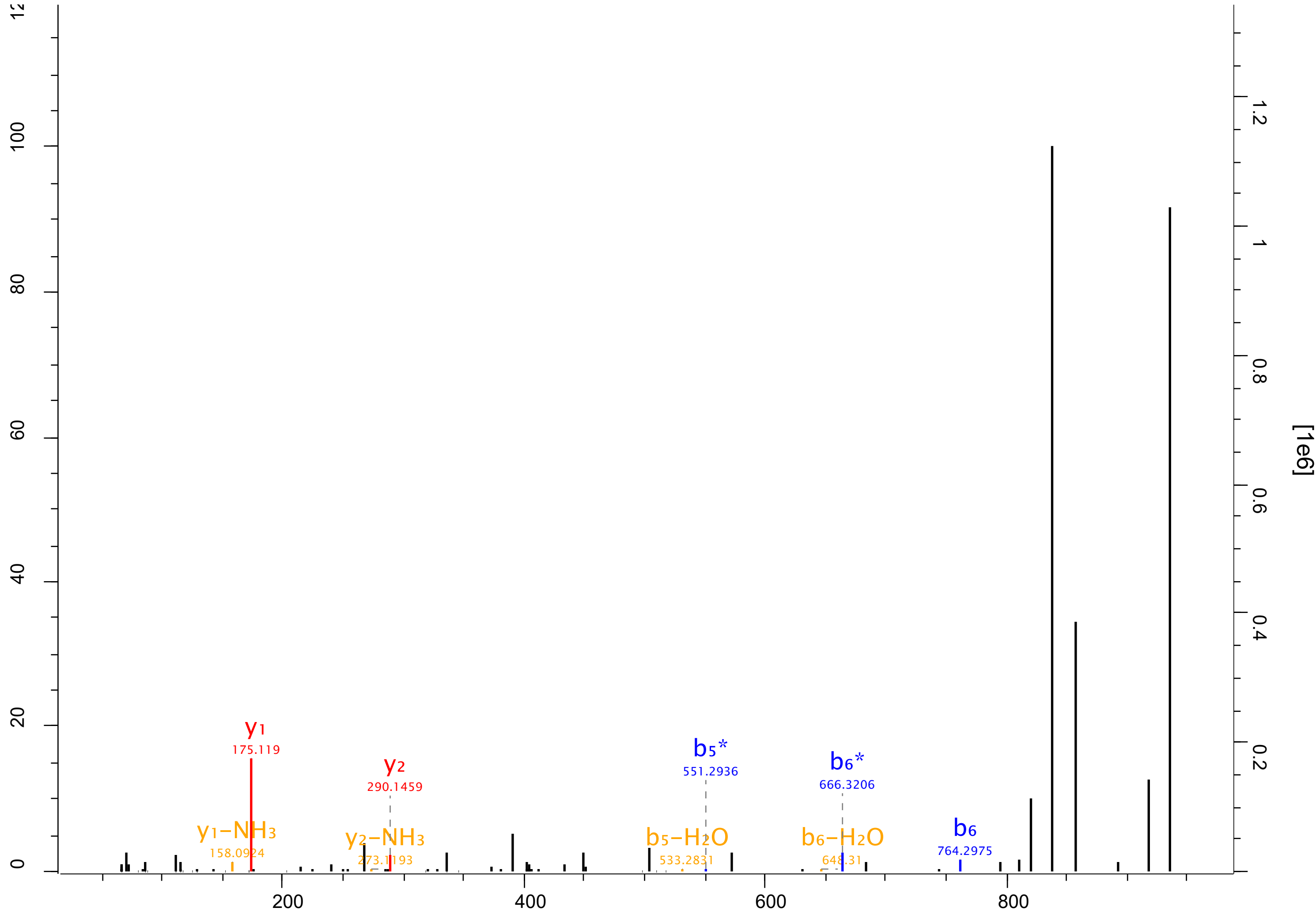


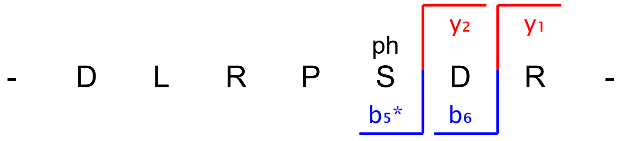

| Raw file | Scan | Method | Score | m/z |
| --- | --- | --- | --- | --- |
| water_02_1 | 6356 | FTMS; HCD | 136.43 | 785.31 |

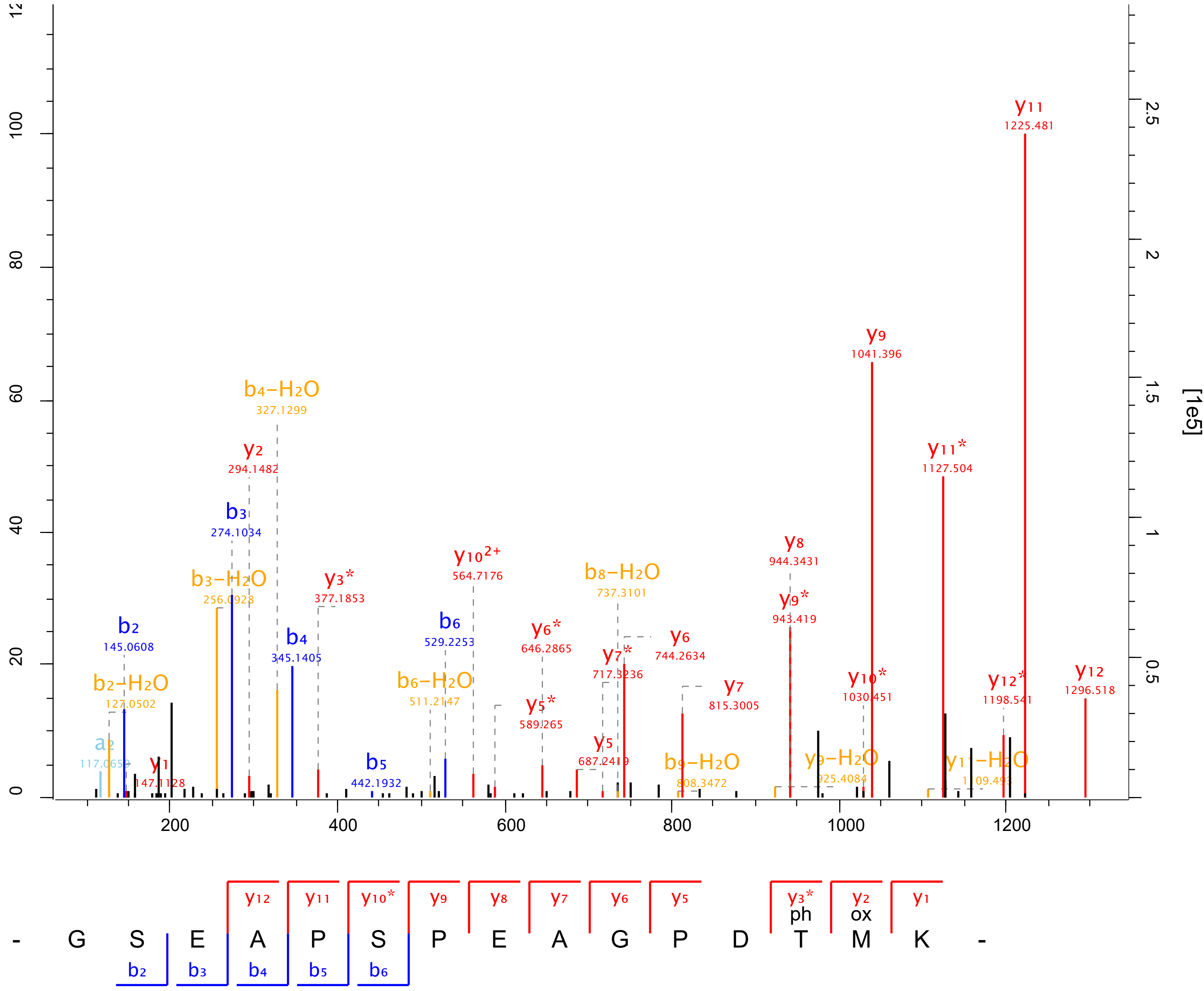

| Raw file | Scan | Method | Score | m/z |
|---|---|---|---|---|
| water_02_1 | 6477 | FTMS; HCD | 125.3 | 480.71 |

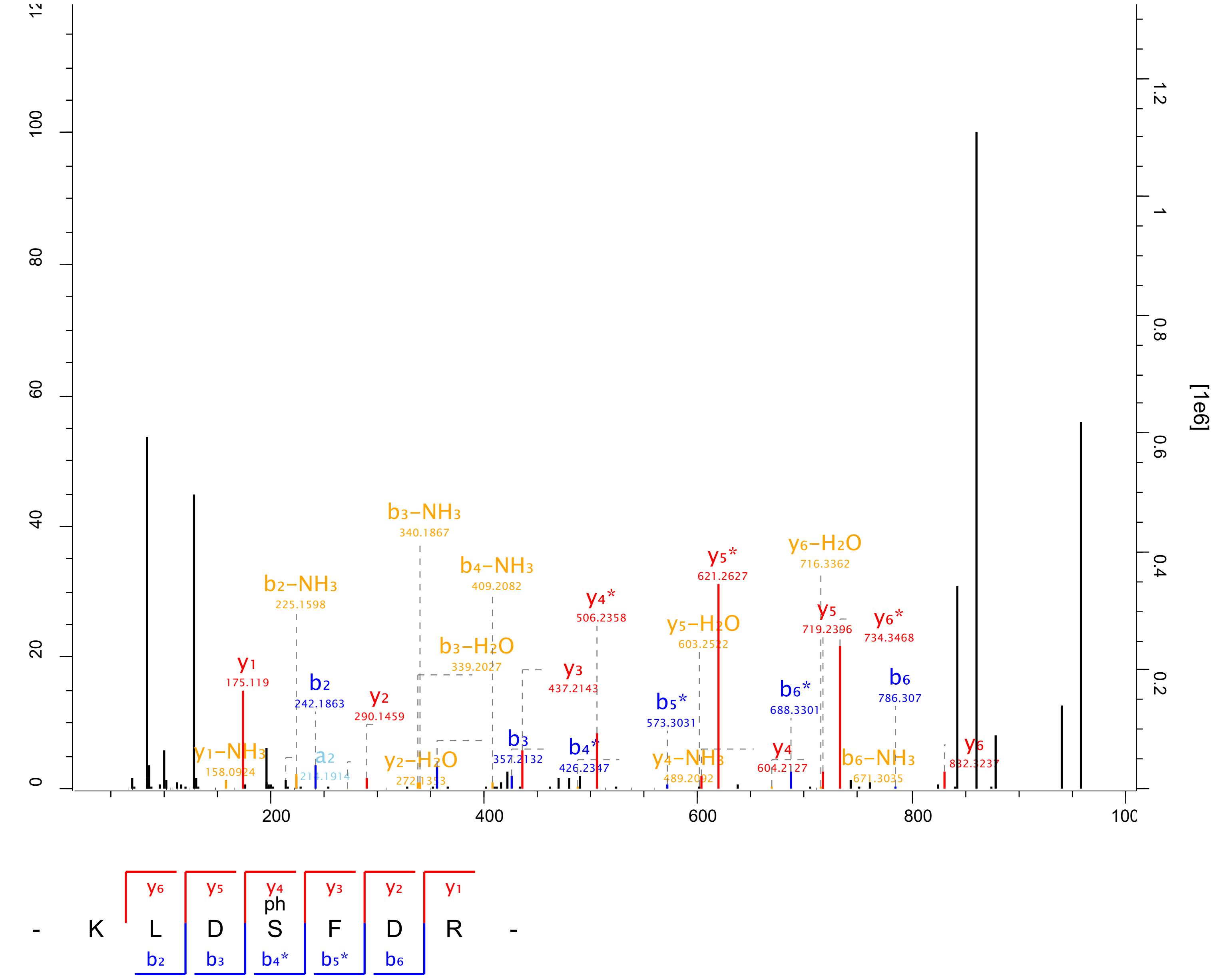

| Raw file | Scan | Method | Score | m/z |
|---|---|---|---|---|
| water_02_1 | 6508 | FTMS; HCD | 90.83 | 469.72 |

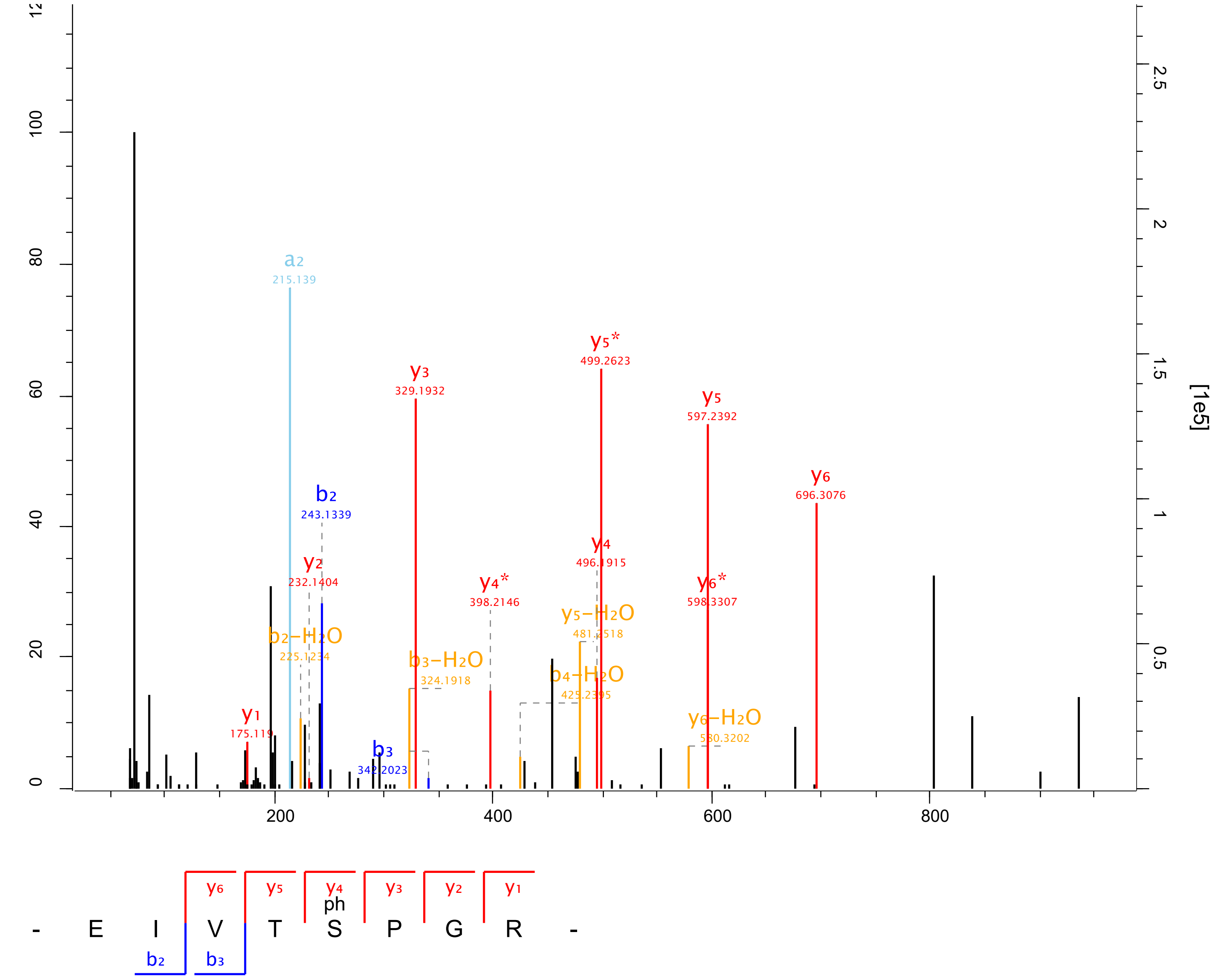

| Raw file | Scan | Method | Score | m/z |
| --- | --- | --- | --- | --- |
| water_02_1 | 6545 | FTMS; HCD | 88.57 | 580 |

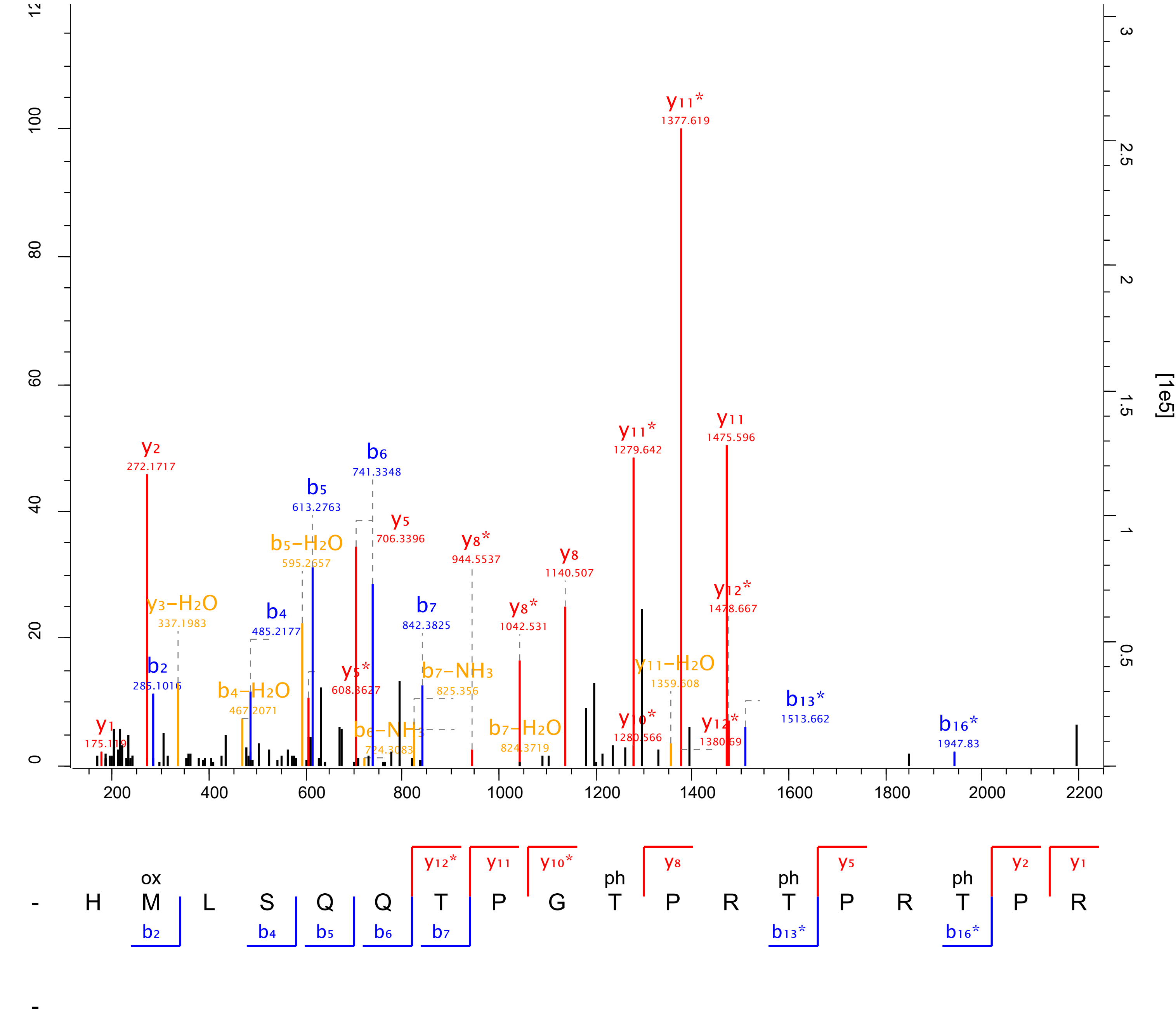

| Raw file | Scan | Method | Score | m/z |
| --- | --- | --- | --- | --- |
| water_02_1 | 6555 | FTMS; HCD | 105.06 | 565.23 |

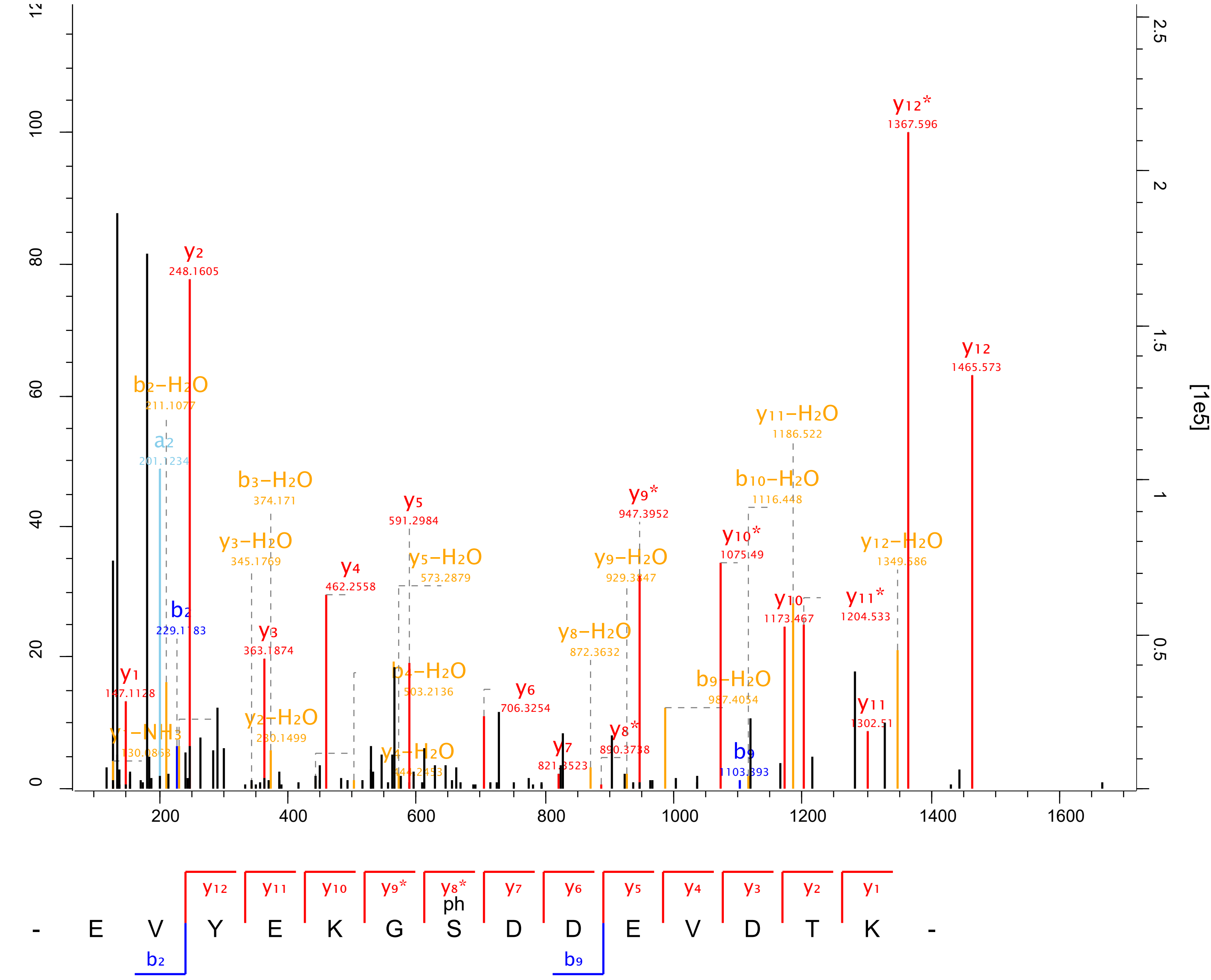

| Raw file | Scan | Method | Score | m/z |
| --- | --- | --- | --- | --- |
| water_02_1 | 6564 | FTMS; HCD | 160.65 | 661.8 |

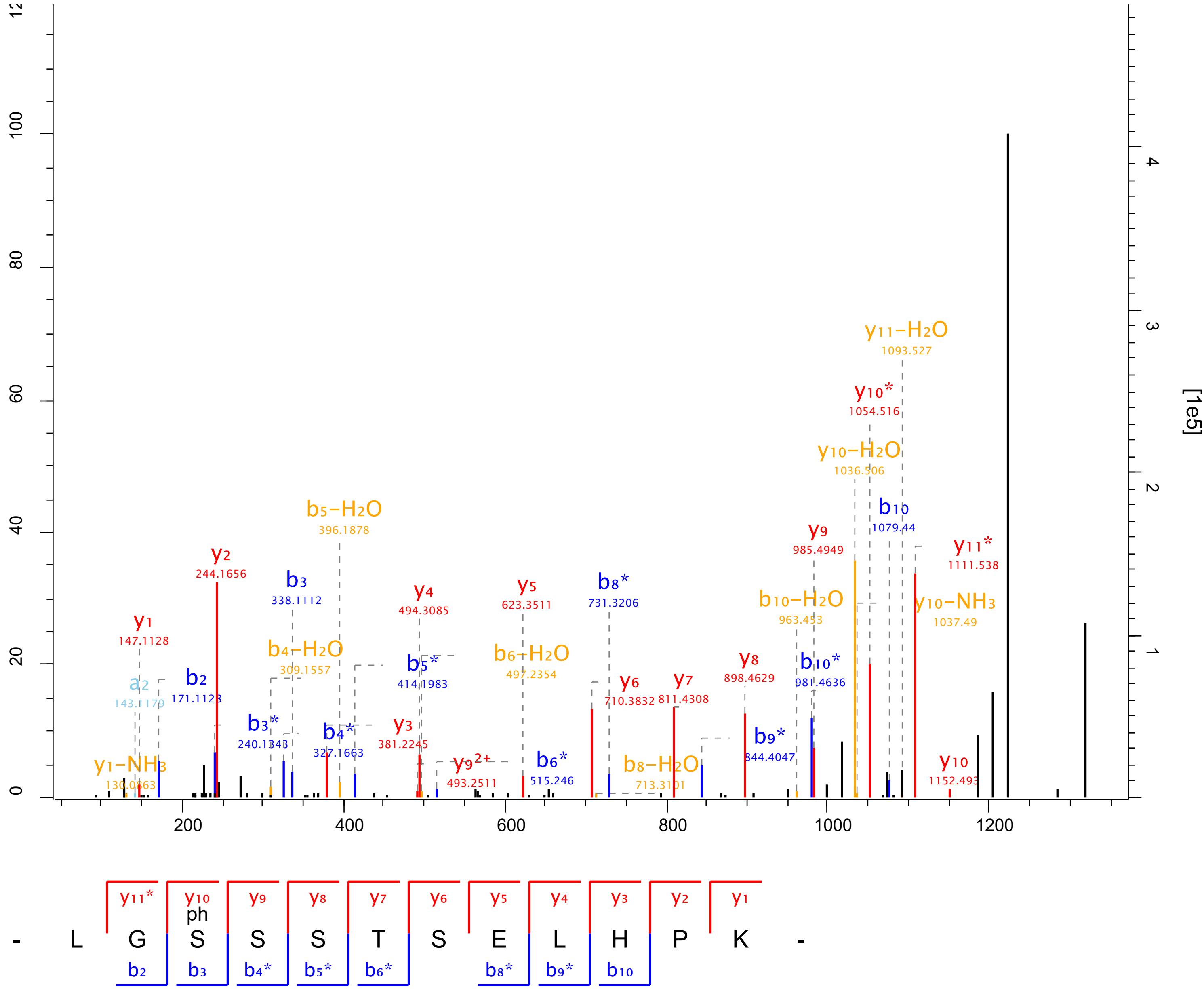

| Raw file | Scan | Method | Score | m/z |
|---|---|---|---|---|
| water_02_1 | 6581 | FTMS; HCD | 140.95 | 851.82 |

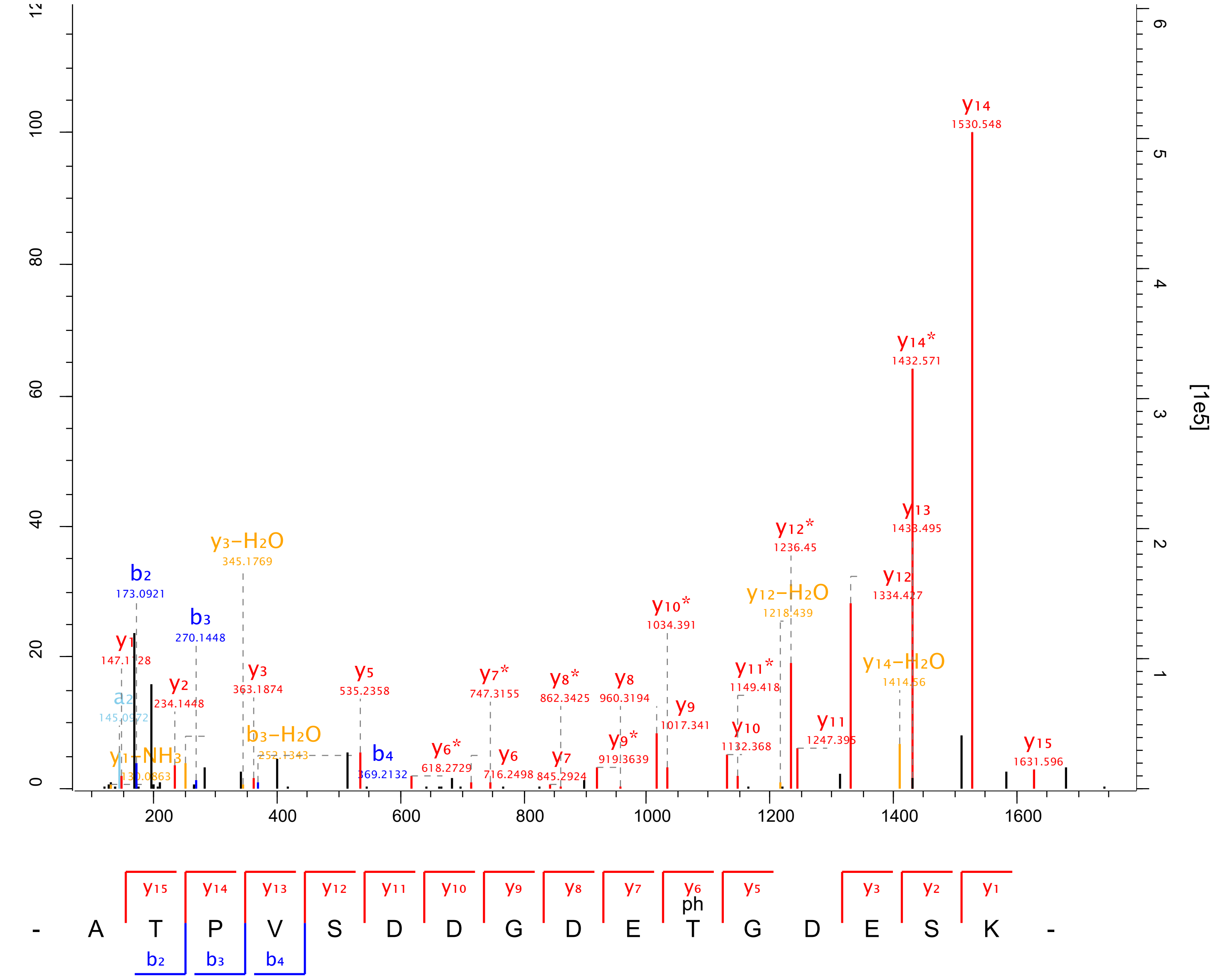

| Raw file | Scan | Method | Score | m/z |
|---|---|---|---|---|
| water_02_1 | 6607 | FTMS; HCD | 50.35 | 540.22 |

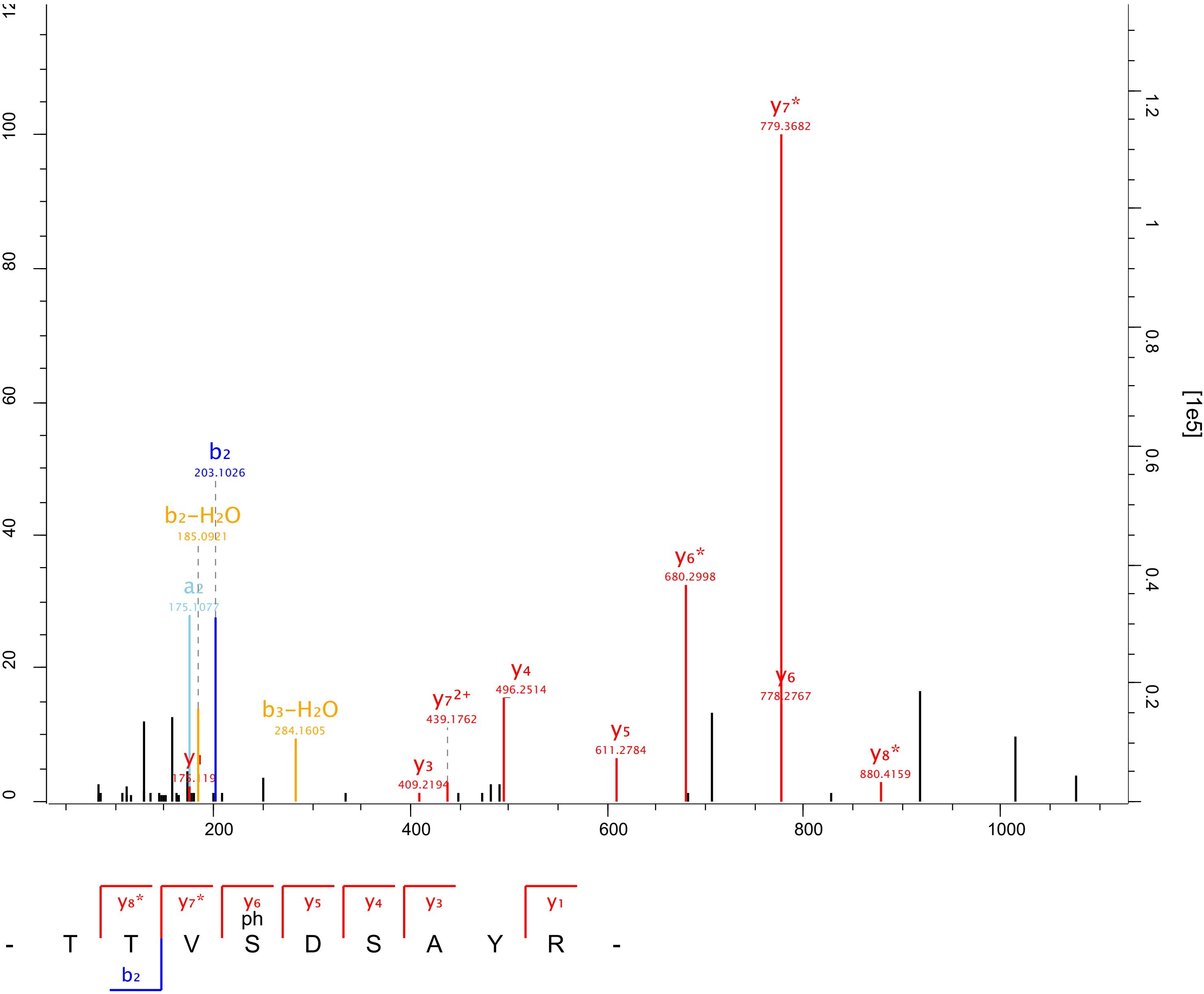

| Raw file | Scan | Method | Score | m/z |
| --- | --- | --- | --- | --- |
| water_02_1 | 6628 | FTMS; HCD | 101.48 | 628.29 |

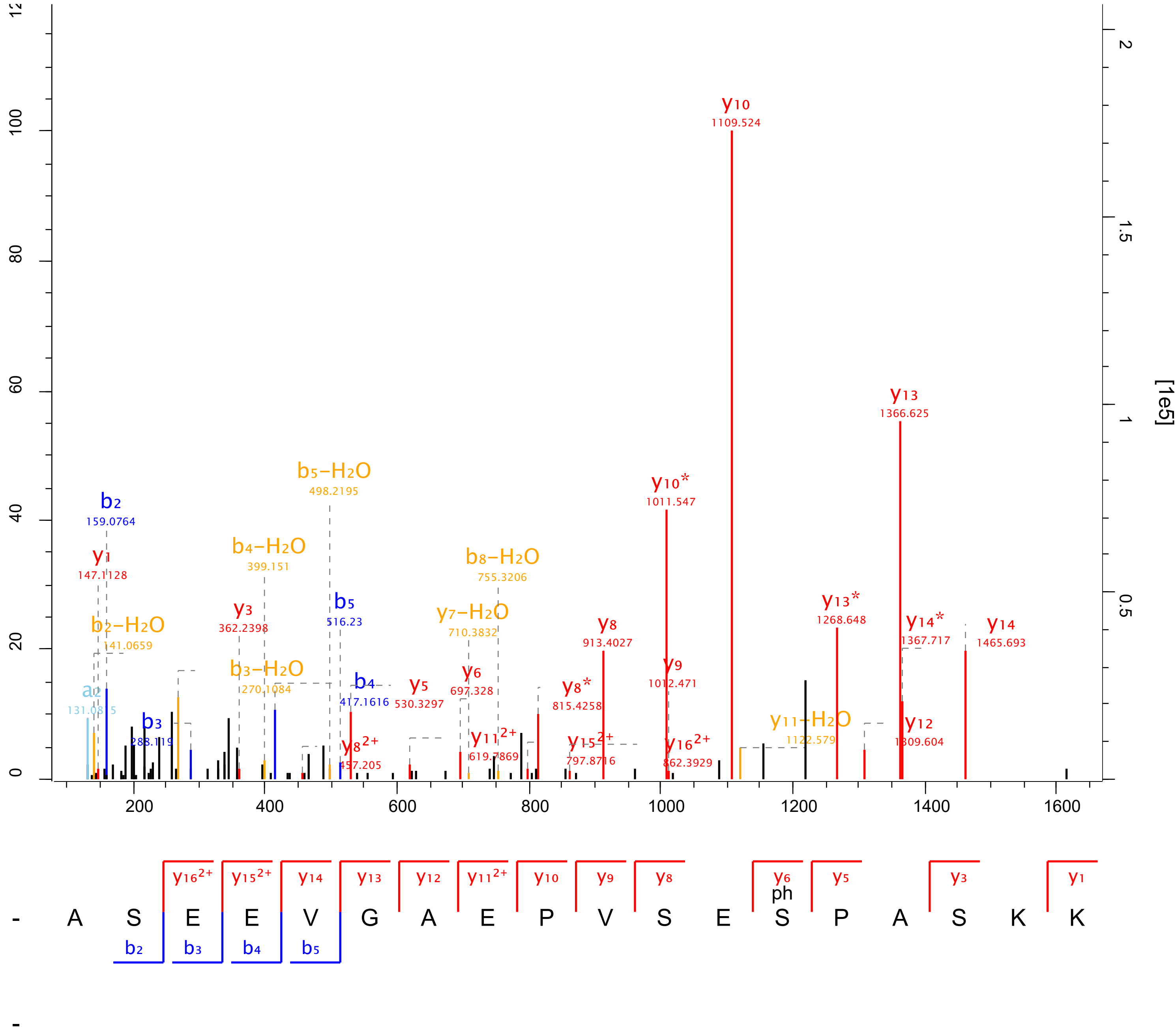

| Raw file | Scan | Method | Score | m/z |
|---|---|---|---|---|
| water_02_1 | 6632 | FTMS; HCD | 121.46 | 633.27 |

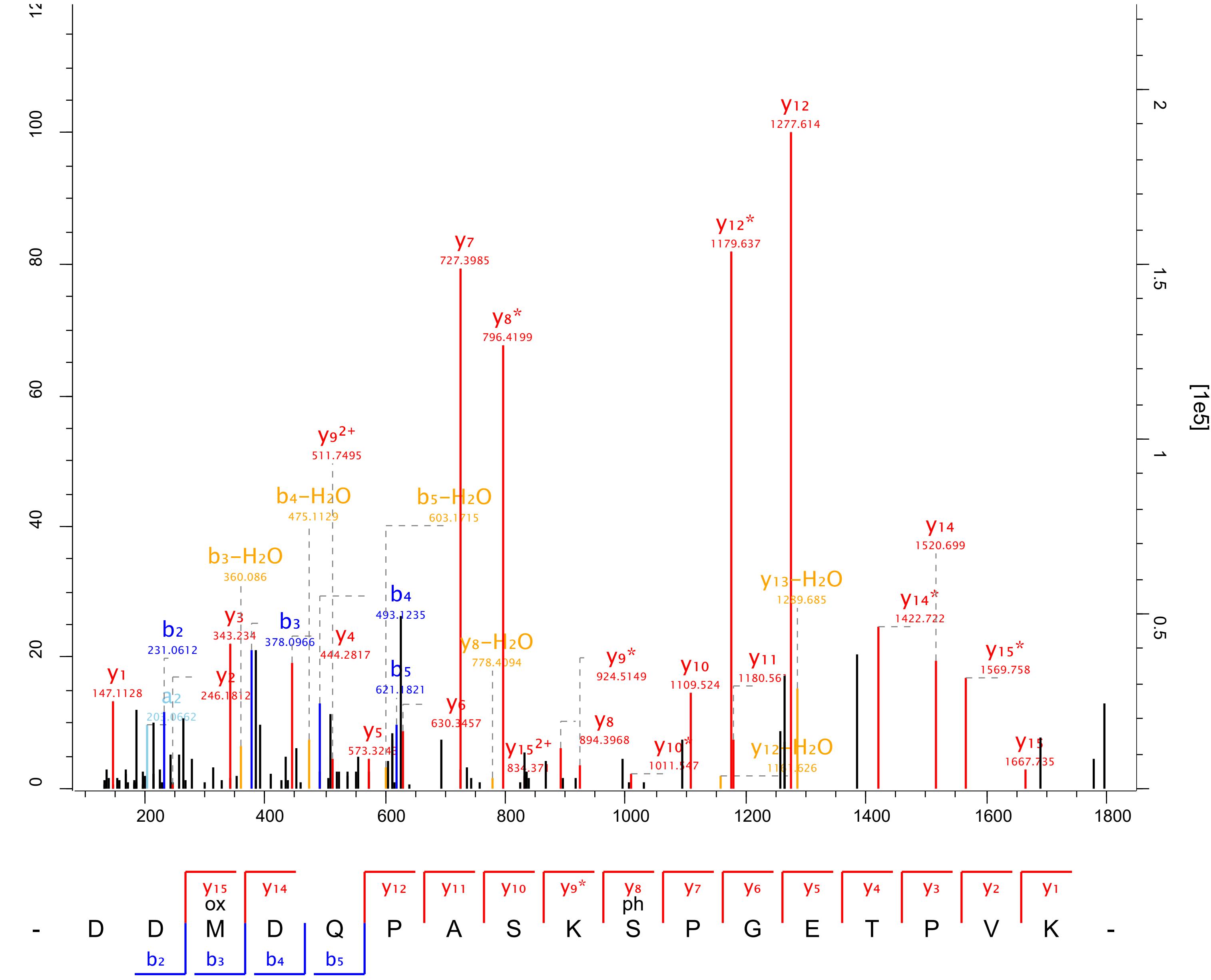

| Raw file | Scan | Method | Score | m/z |
|---|---|---|---|---|
| water_02_1 | 6688 | FTMS; HCD | 114.46 | 653.95 |

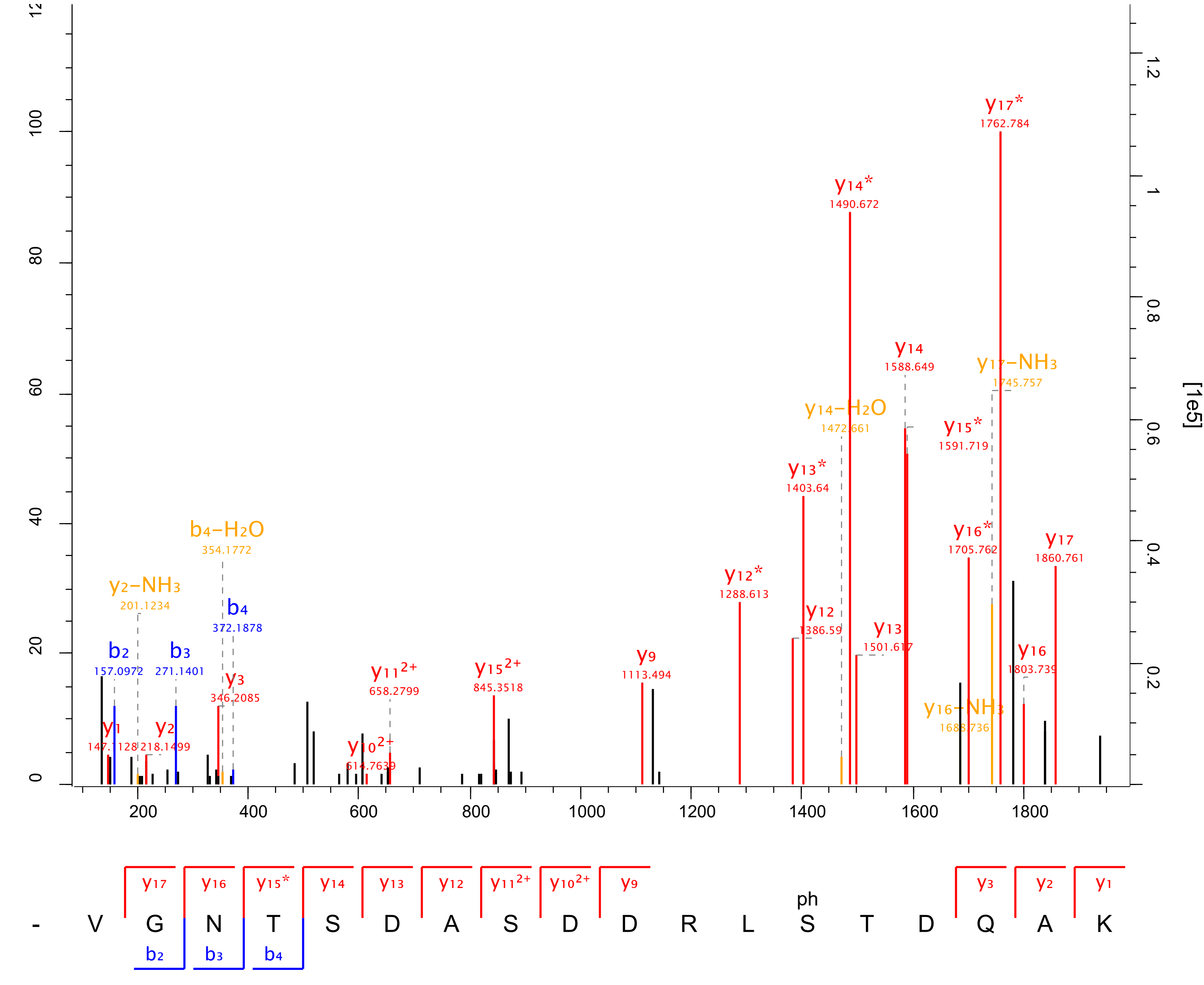

| Raw file | Scan | Method | Score | m/z |
|---|---|---|---|---|
| water_02_1 | 6712 | FTMS; HCD | 89.51 | 479.55 |

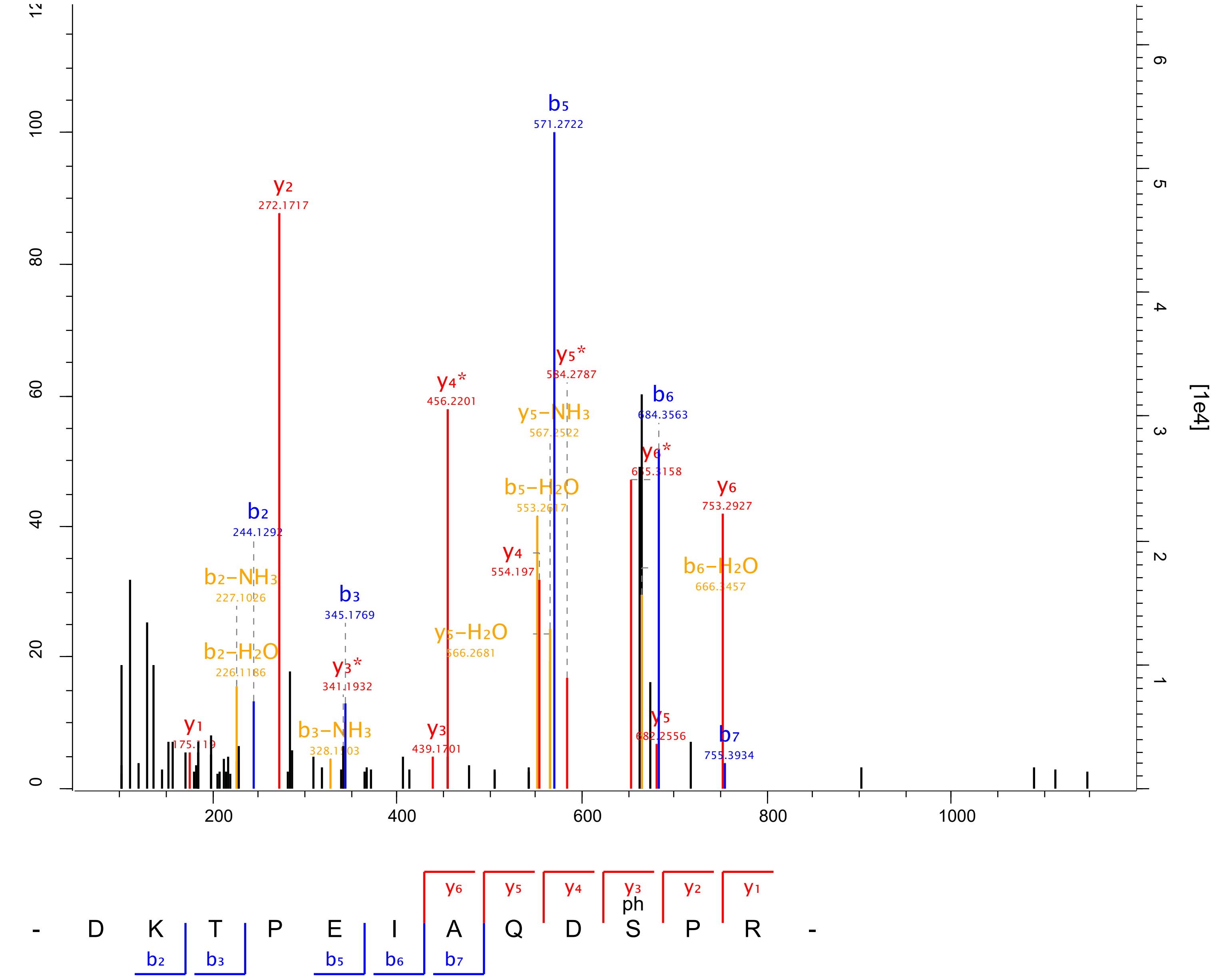

| Raw file | Scan | Method | Score | m/z |
|---|---|---|---|---|
| water_02_1 | 6758 | FTMS; HCD | 51.29 | 525.2 |

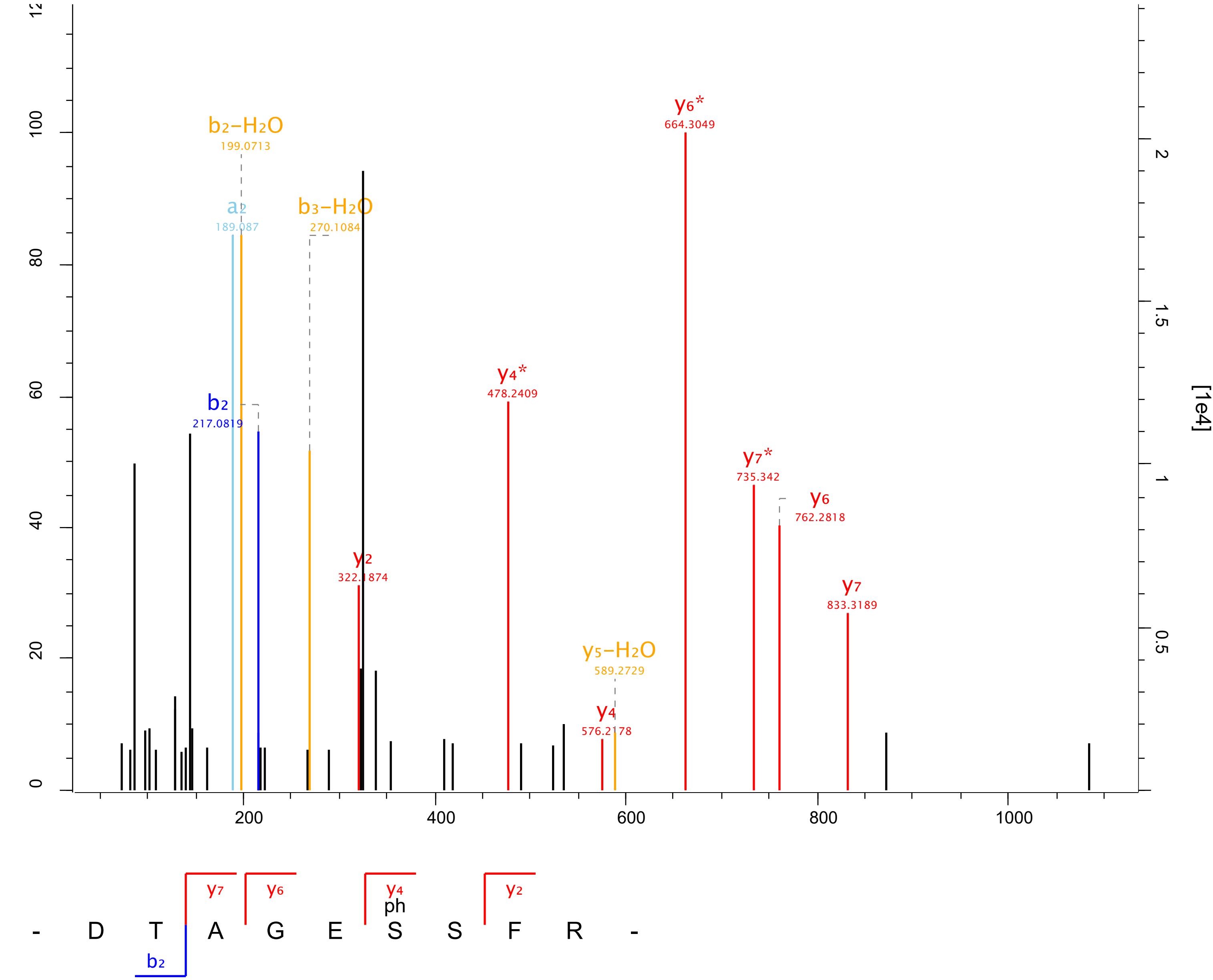

| Raw file | Scan | Method | Score | m/z |
|---|---|---|---|---|
| water_02_1 | 6792 | FTMS; HCD | 49.99 | 614.6 |

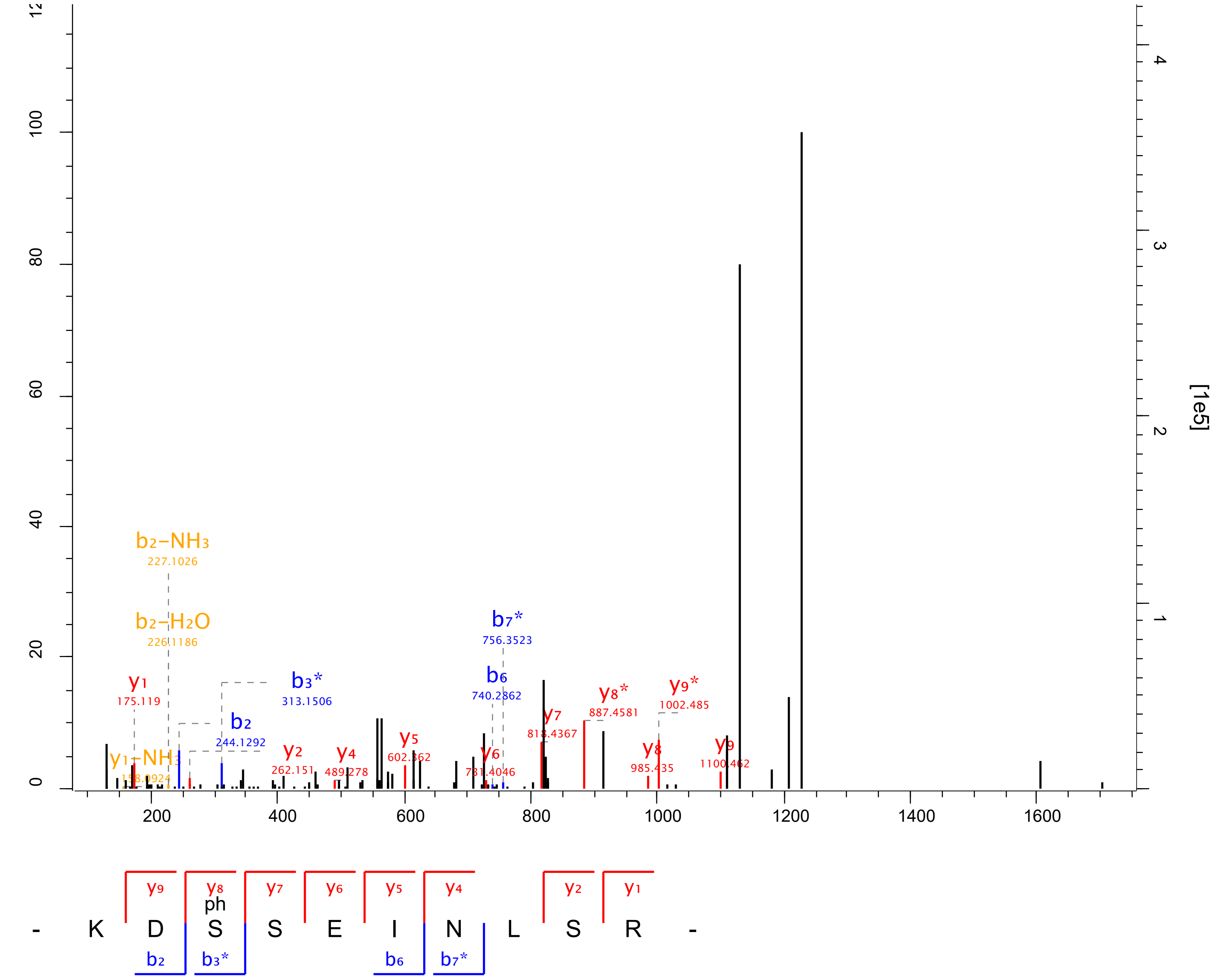

| Raw file | Scan | Method | Score | m/z |
|---|---|---|---|---|
| water_02_1 | 6817 | FTMS; HCD | 47.77 | 646.8 |

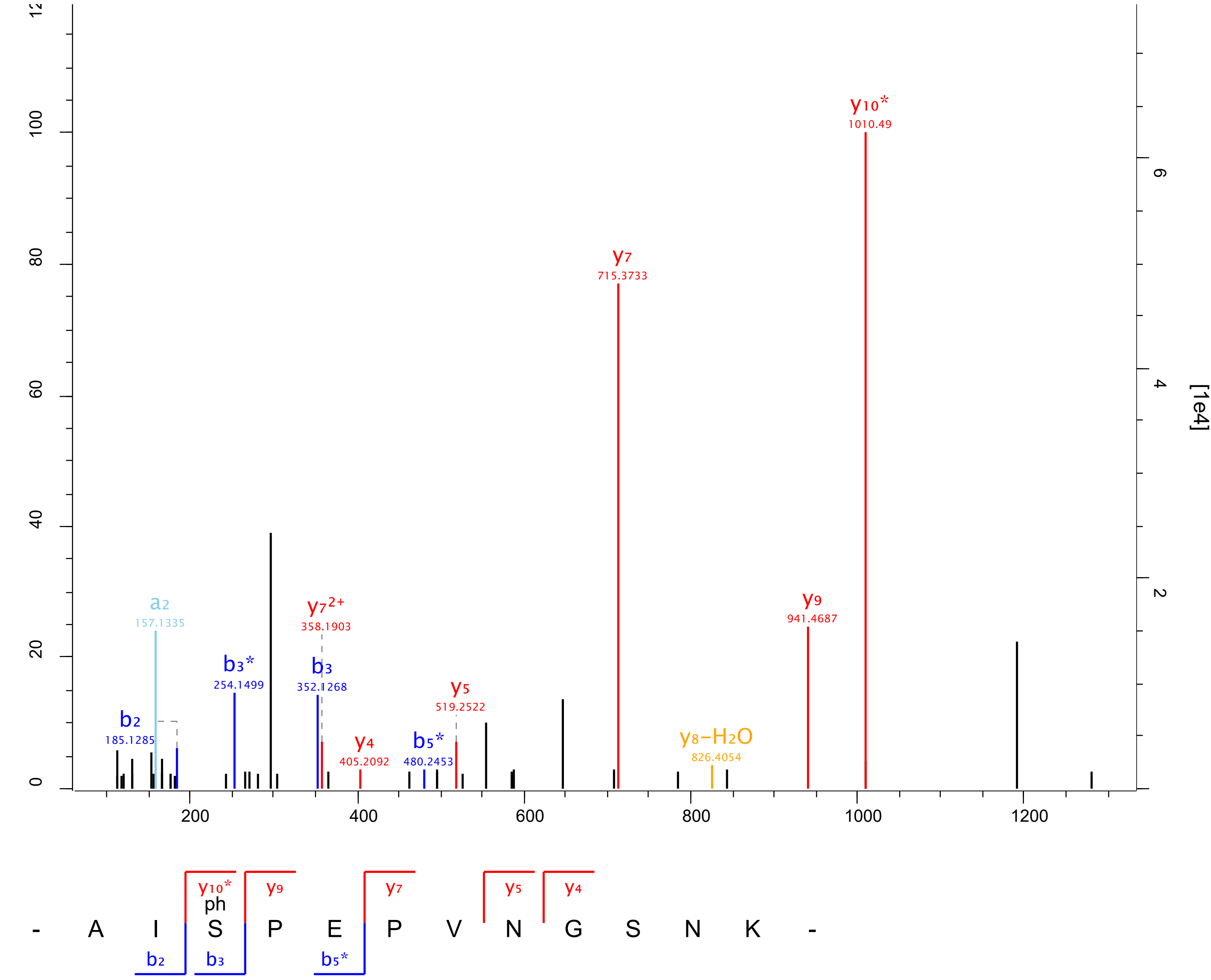

| Raw file | Scan | Method | Score | m/z |
|---|---|---|---|---|
| water_02_1 | 6820 | FTMS; HCD | 105.46 | 709.25 |

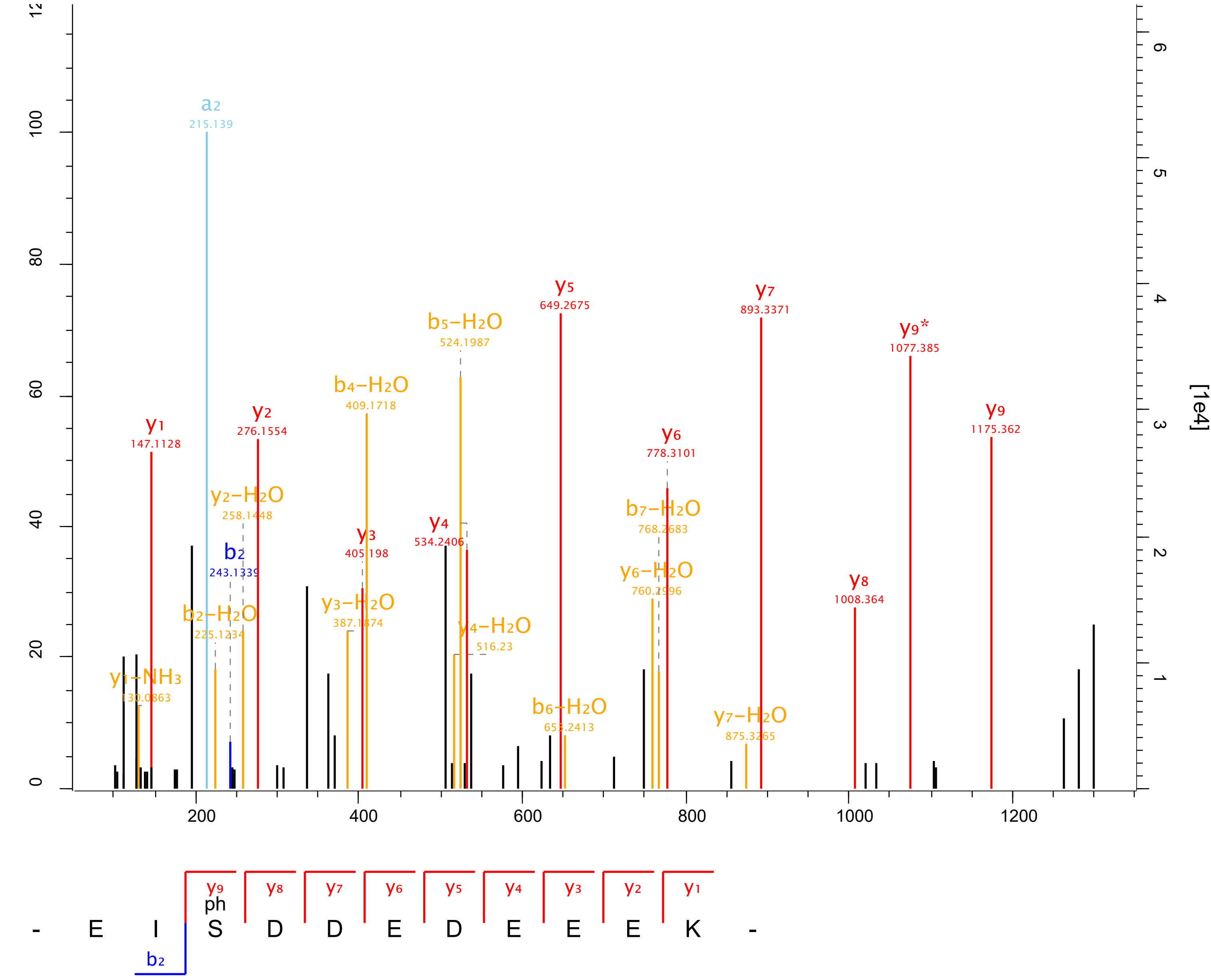

| Raw file | Scan | Method | Score | m/z |
|---|---|---|---|---|
| water_02_1 | 6936 | FTMS; HCD | 45 | 724.29 |

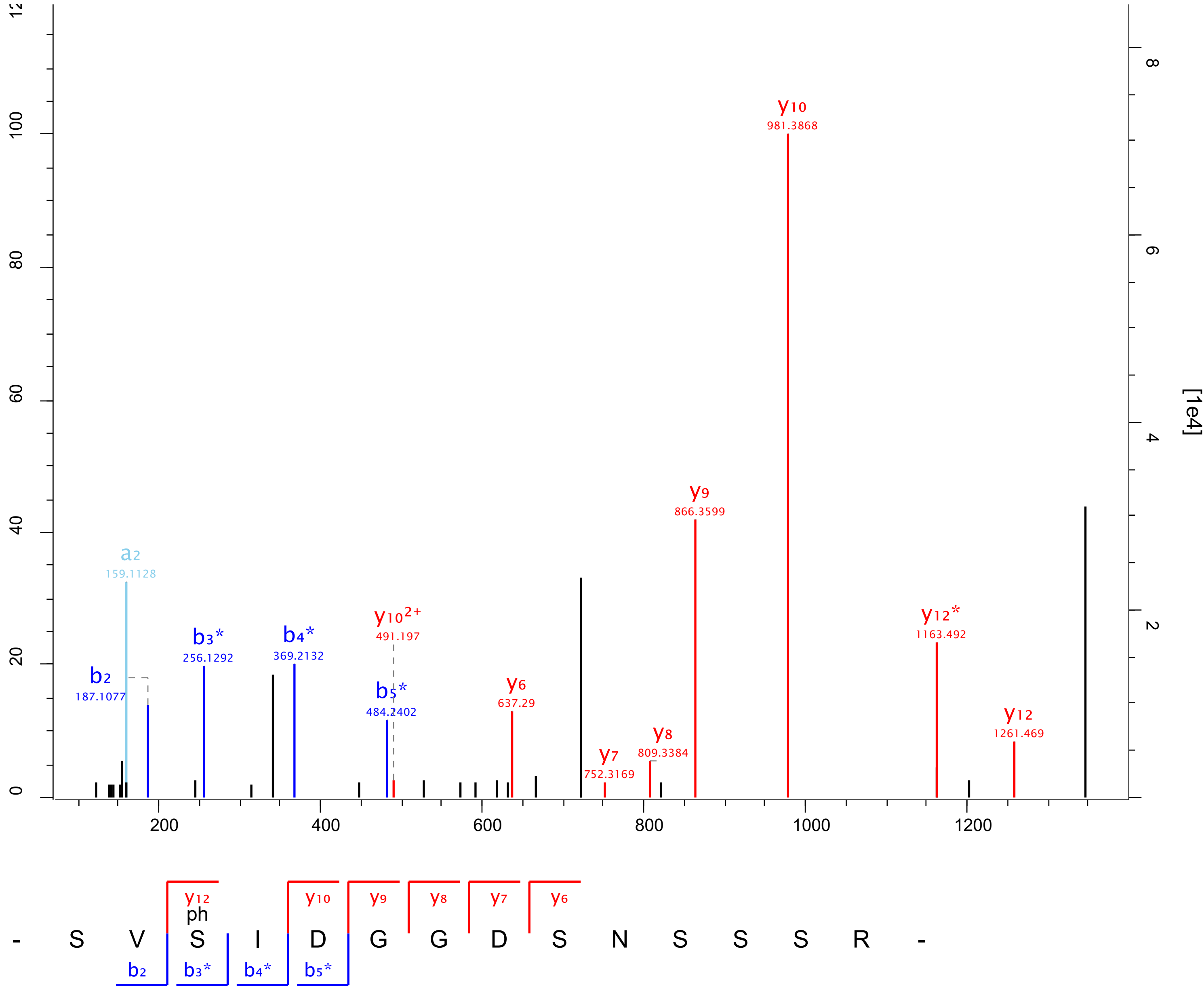

| Raw file | Scan | Method | Score | m/z |
|---|---|---|---|---|
| water_02_1 | 6944 | FTMS; HCD | 58.32 | 591.71 |

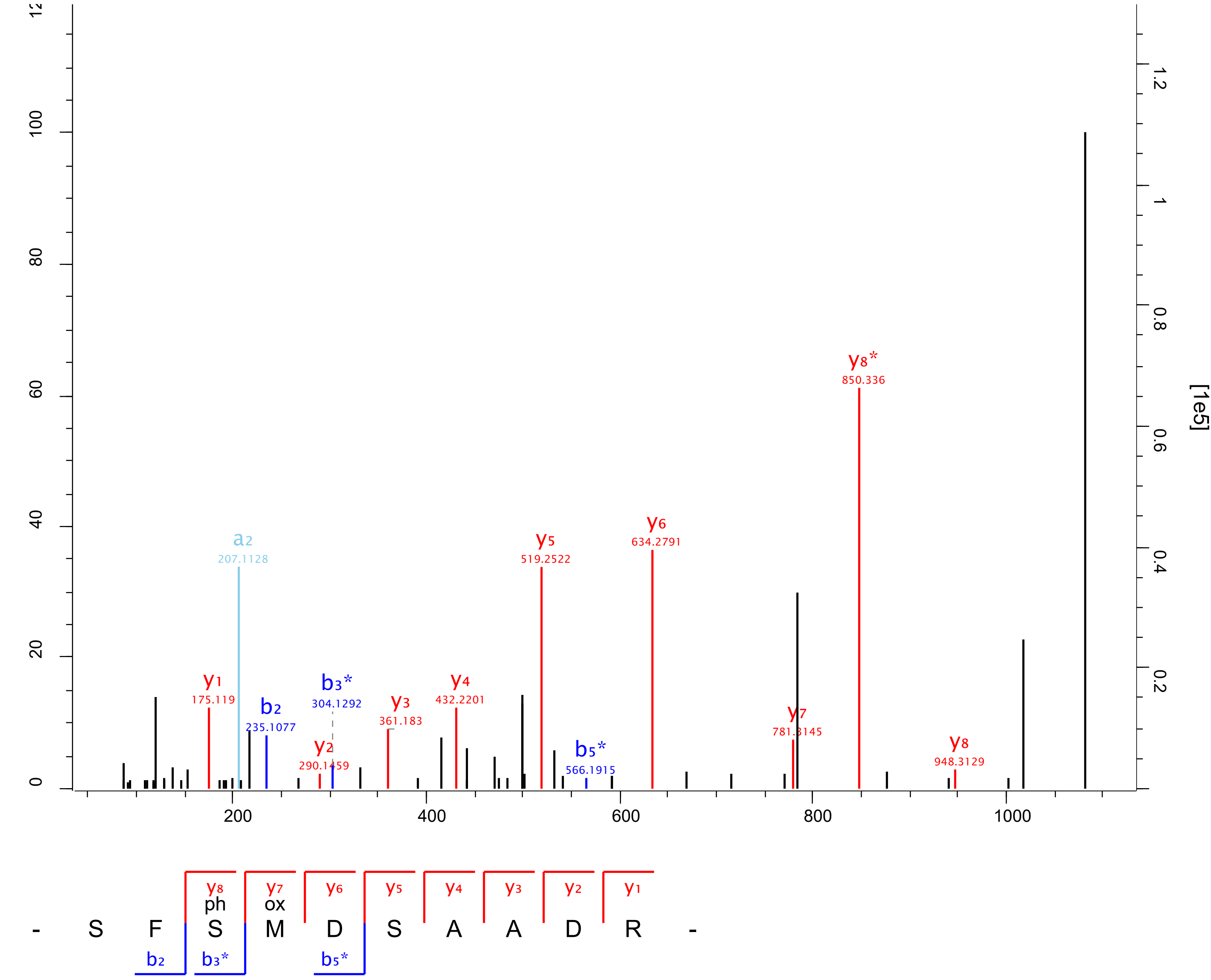

| Raw file | Scan | Method | Score | m/z |
| --- | --- | --- | --- | --- |
| water_02_1 | 6993 | FTMS; HCD | 42.6 | 526.28 |

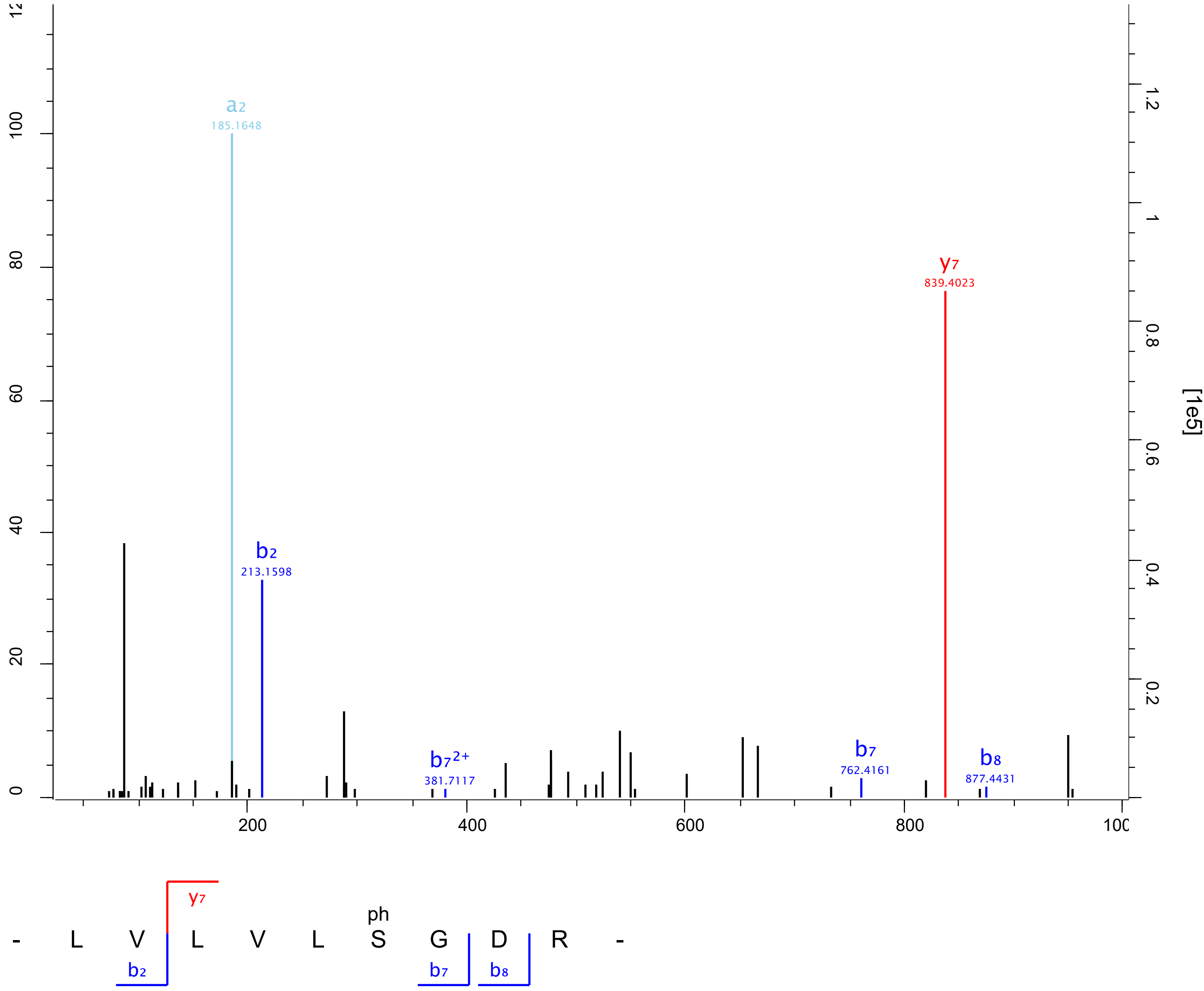

| Raw file | Scan | Method | Score | m/z |
| --- | --- | --- | --- | --- |
| water_02_1 | 7157 | FTMS; HCD | 99.93 | 658.29 |

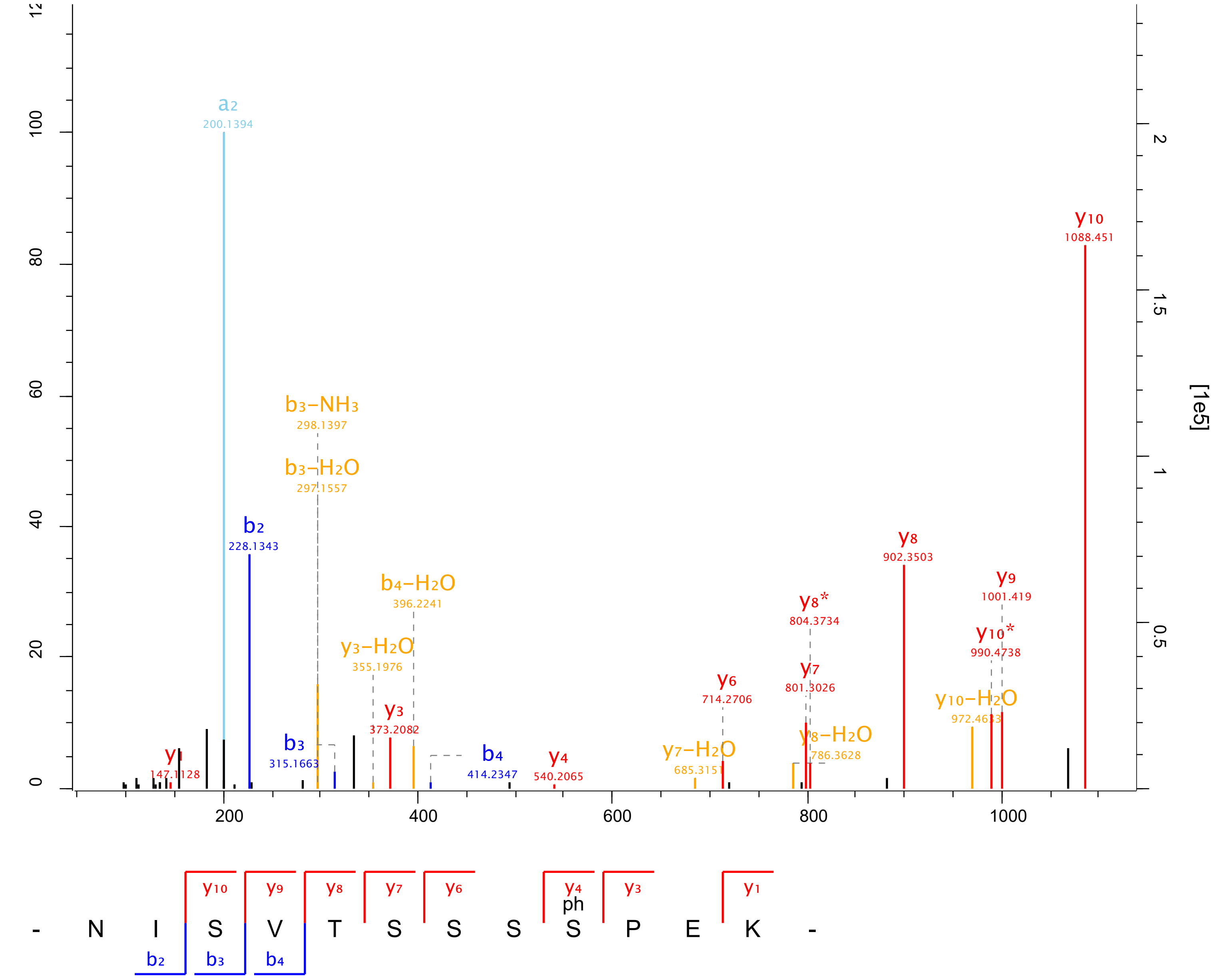

| Raw file | Scan | Method | Score | m/z |
| --- | --- | --- | --- | --- |
| water_02_1 | 7160 | FTMS; HCD | 85.91 | 573.75 |

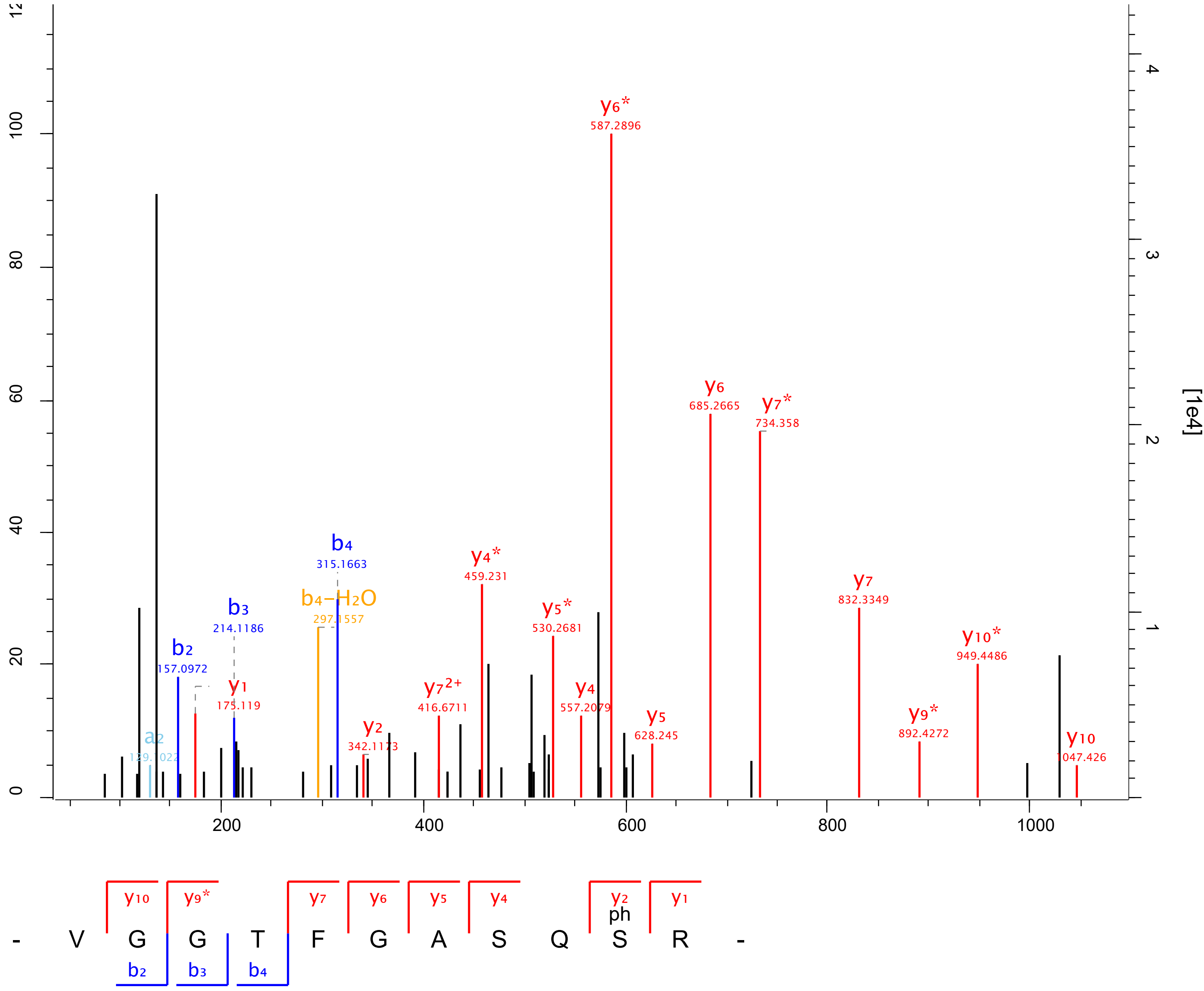

| Raw file | Scan | Method | Score | m/z |
|---|---|---|---|---|
| water_02_1 | 7180 | FTMS; HCD | 70.91 | 469.2 |

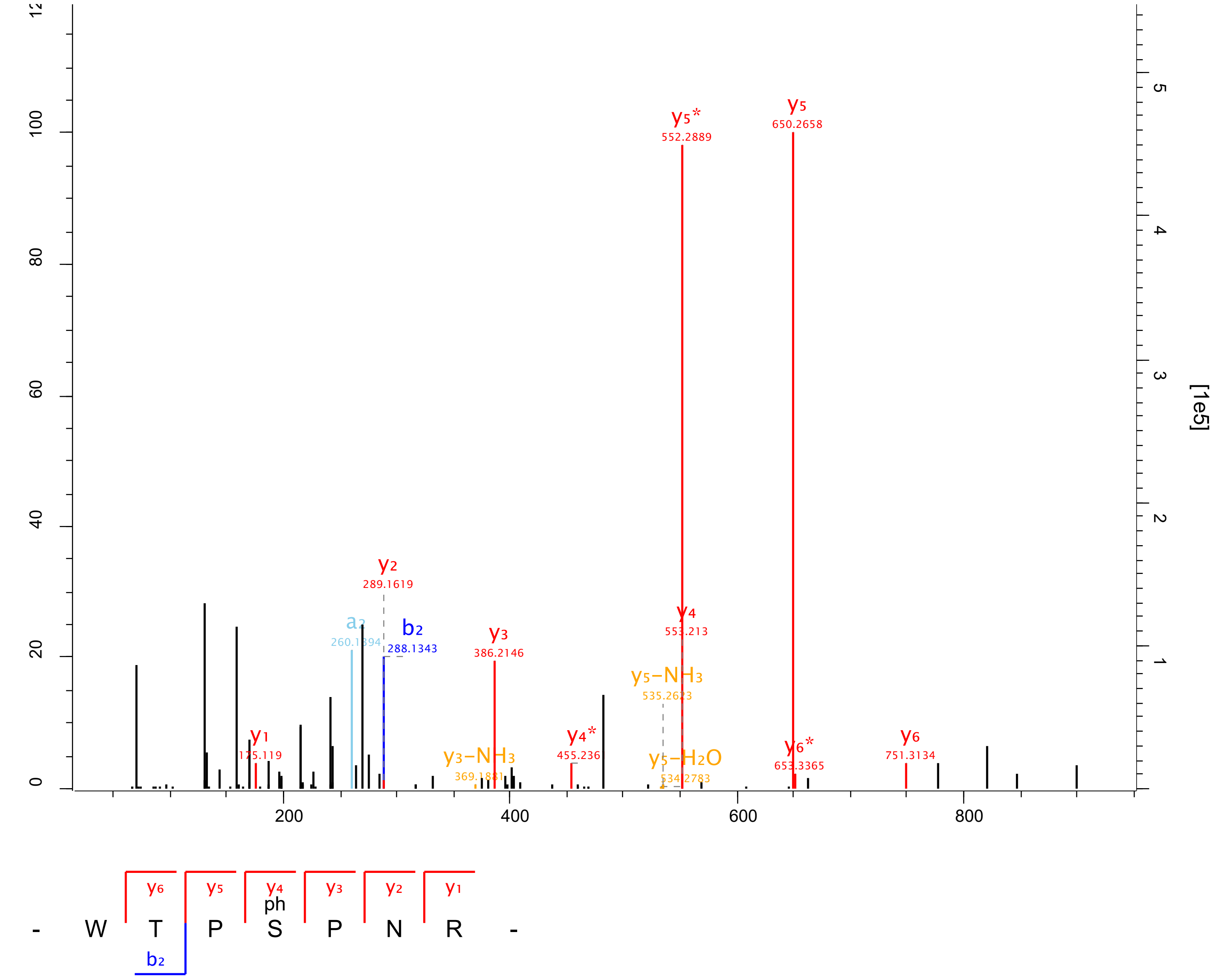

| Raw file | Scan | Method | Score | m/z |
|---|---|---|---|---|
| water_02_1 | 7185 | FTMS; HCD | 111.31 | 606.72 |

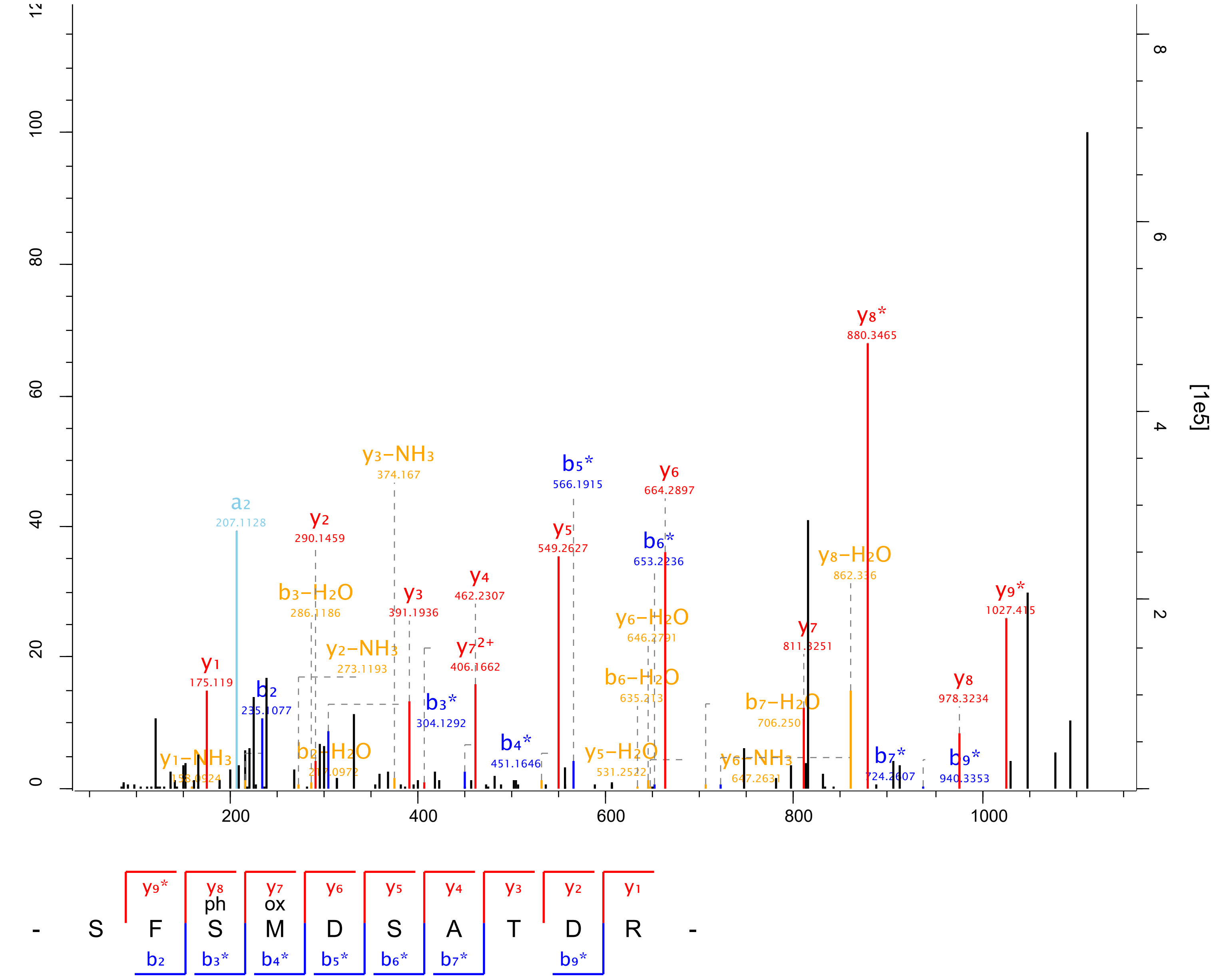

| Raw file | Scan | Method | Score | m/z |
| --- | --- | --- | --- | --- |
| water_02_1 | 7239 | FTMS; HCD | 45.99 | 752.93 |

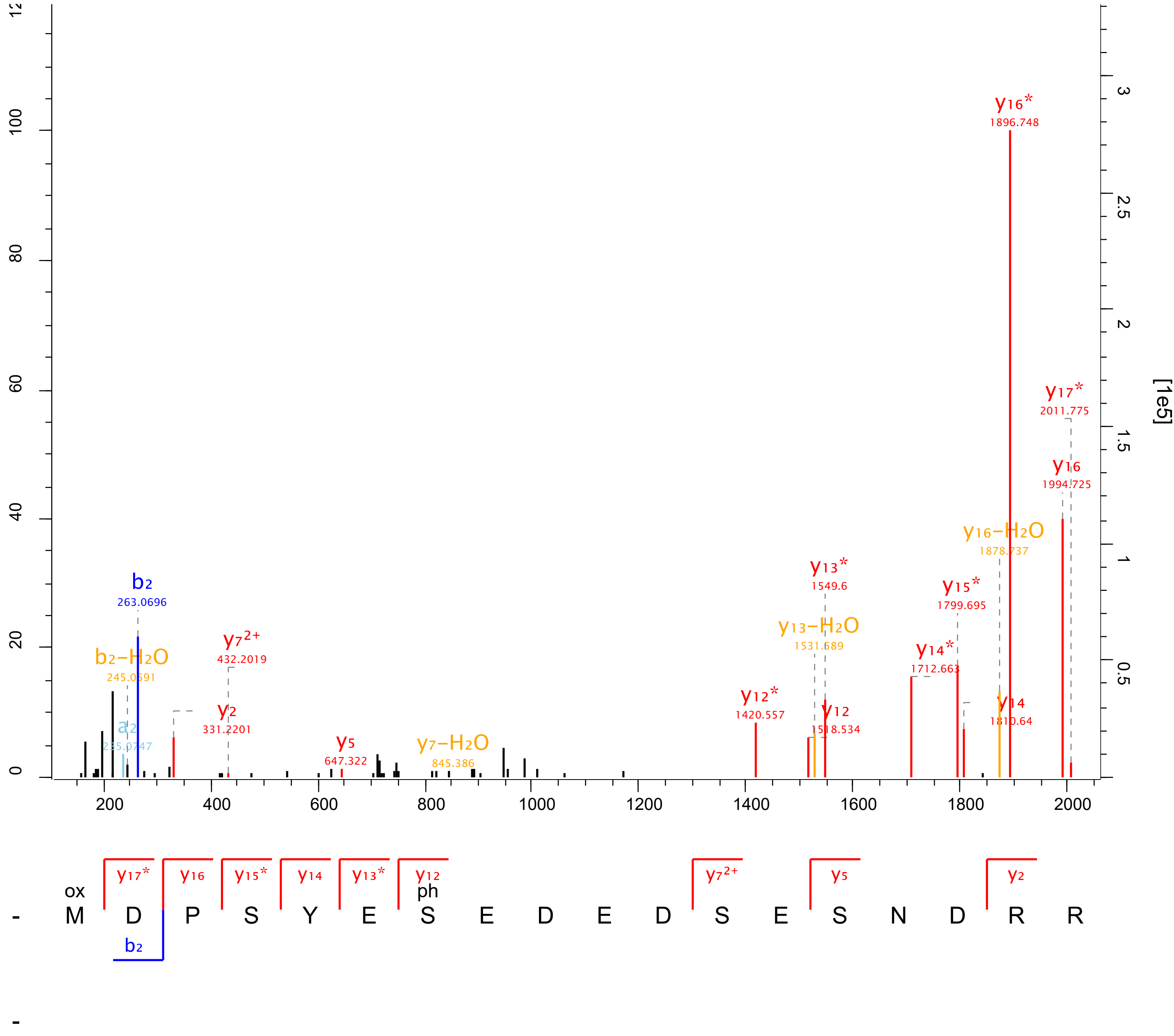

| Raw file | Scan | Method | Score | m/z |
| --- | --- | --- | --- | --- |
| water_02_1 | 7269 | FTMS; HCD | 59.91 | 526.74 |

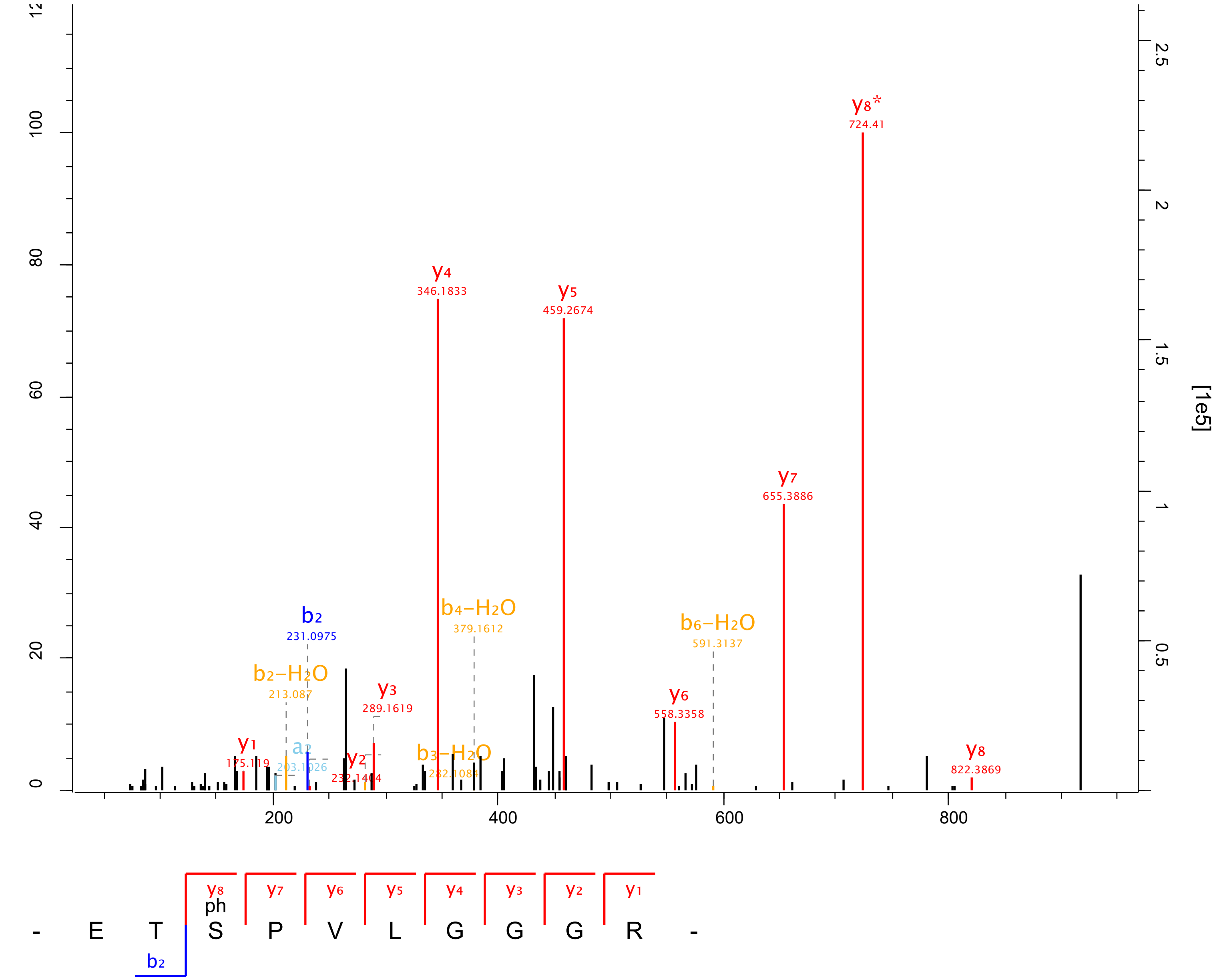

| Raw file | Scan | Method | Score | m/z |
|---|---|---|---|---|
| water_02_1 | 7275 | FTMS; HCD | 215.64 | 557.56 |

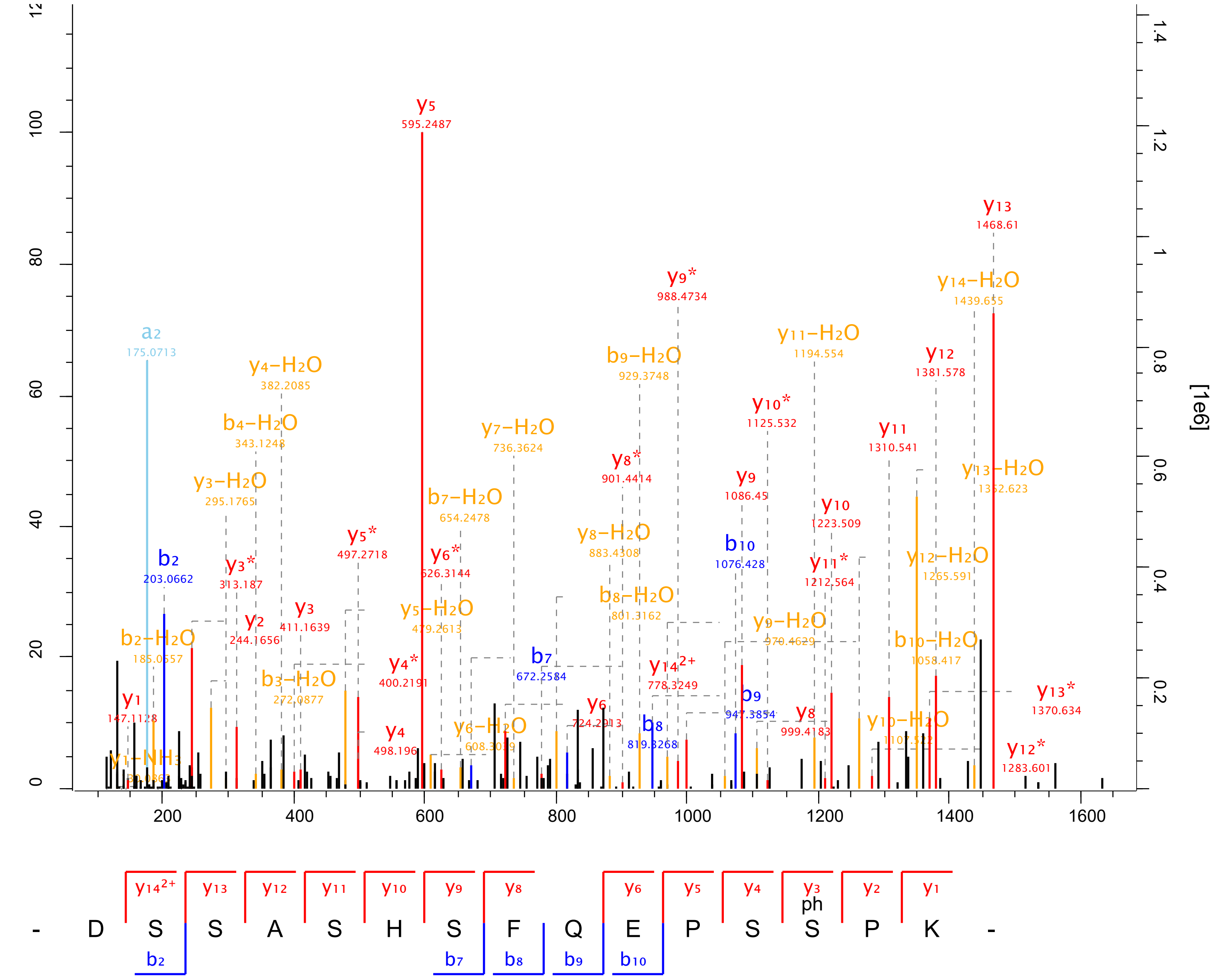

| Raw file | Scan | Method | Score | m/z |
| --- | --- | --- | --- | --- |
| water_02_1 | 7280 | FTMS; HCD | 124.57 | 824 |

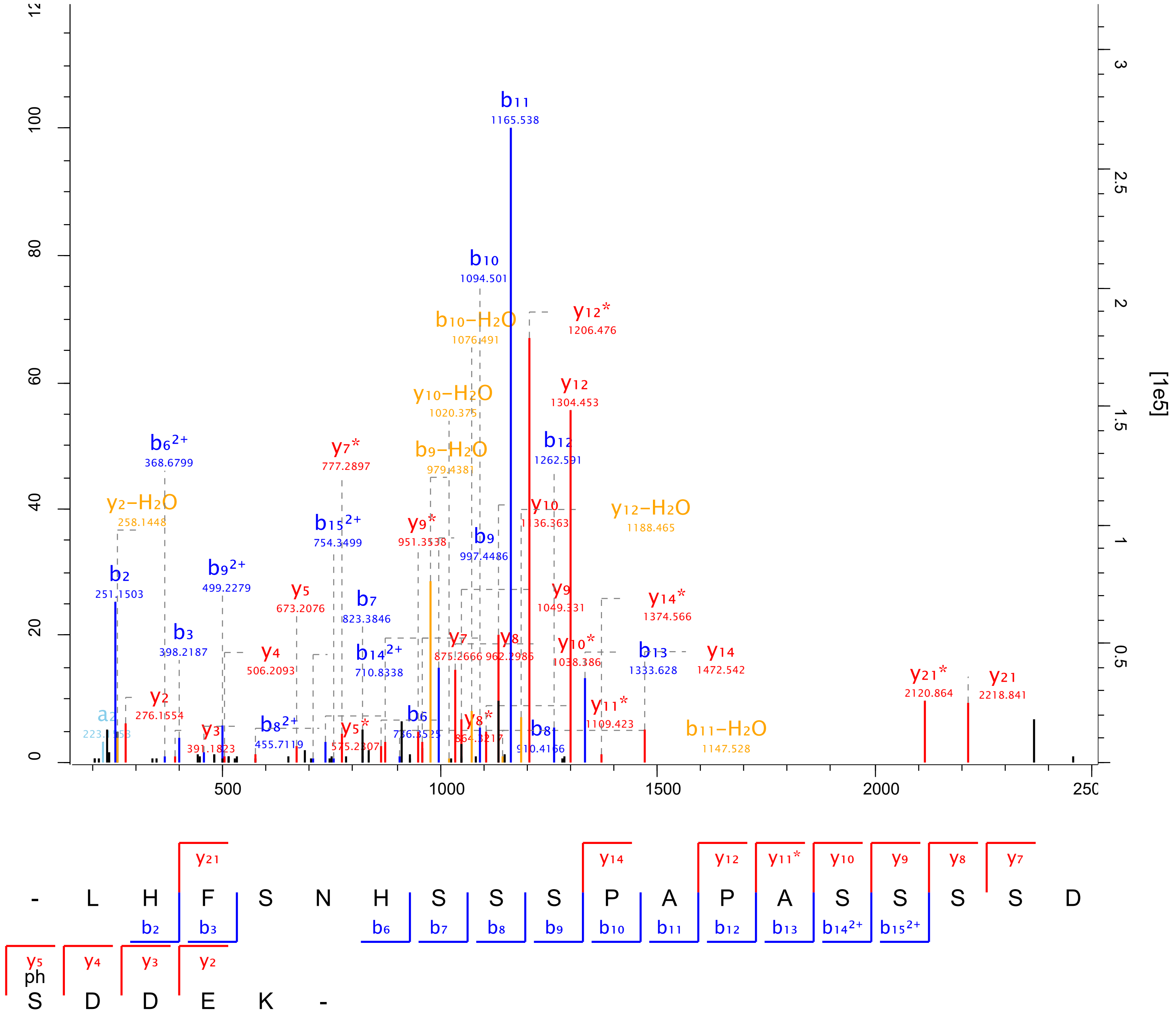

| Raw file | Scan | Method | Score | m/z |
|---|---|---|---|---|
| water_02_1 | 7287 | FTMS; HCD | 45.28 | 510.75 |

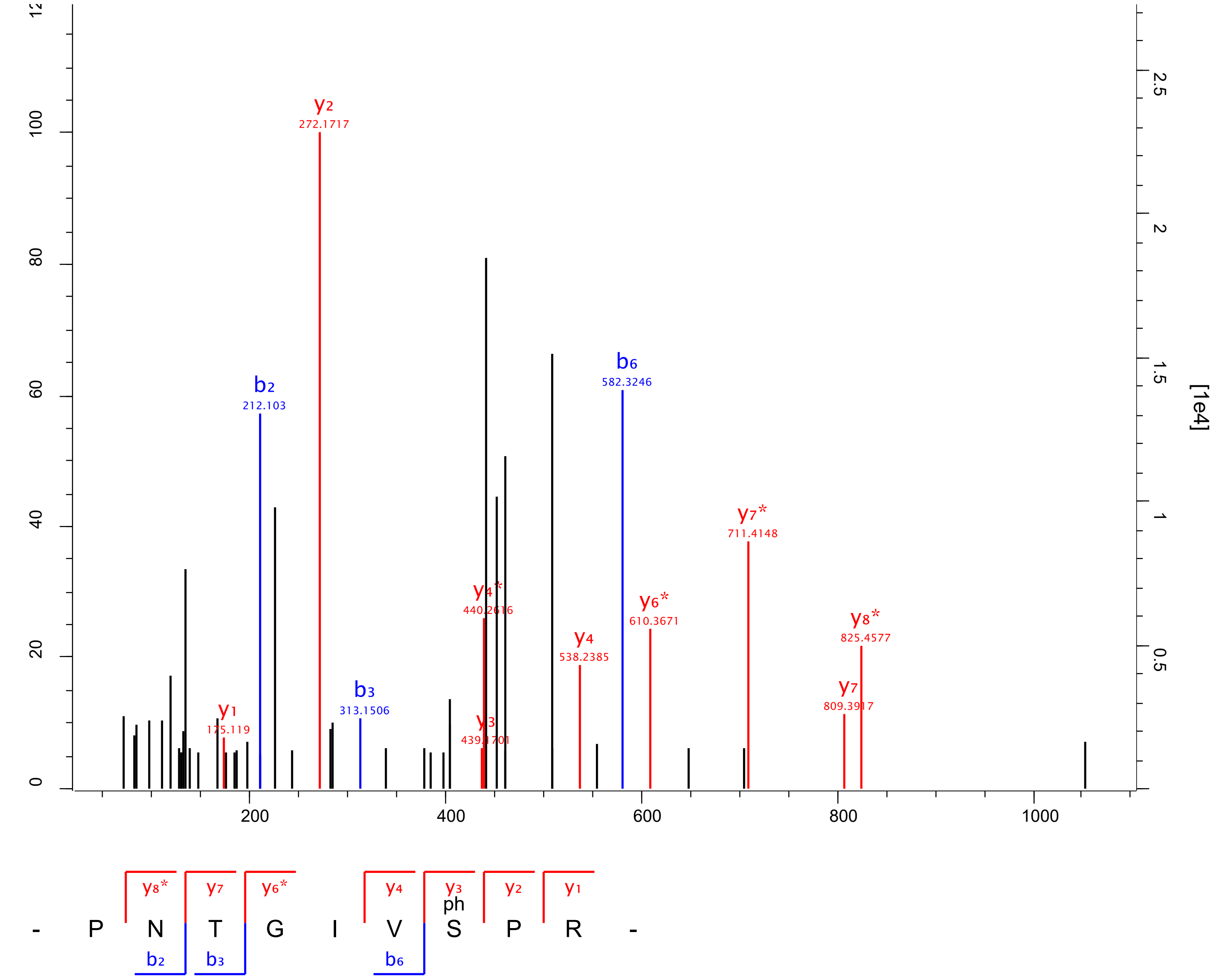

| Raw file | Scan | Method | Score | m/z |
|---|---|---|---|---|
| water_02_1 | 7355 | FTMS; HCD | 166.35 | 1011.85 |

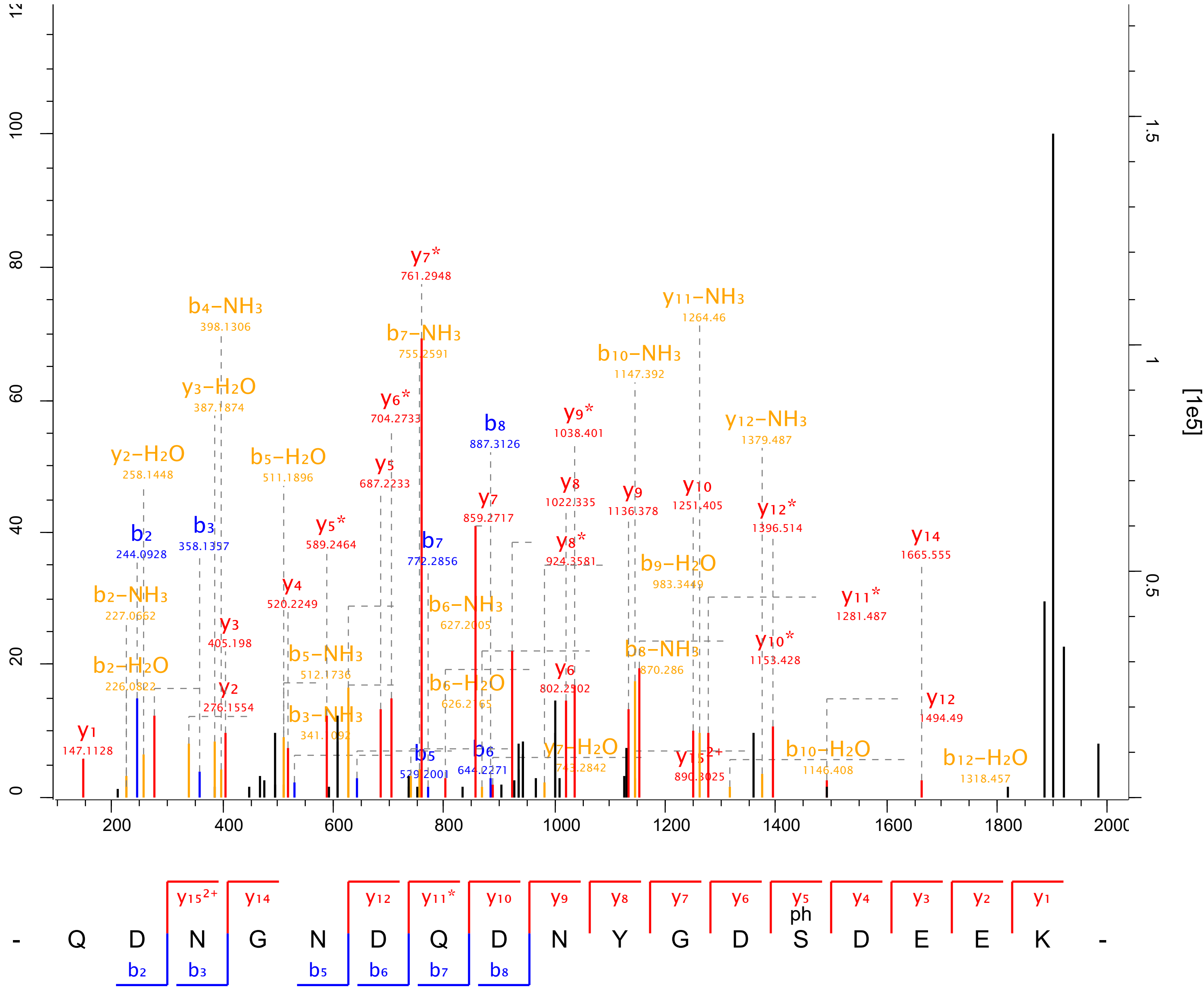

| Raw file | Scan | Method | Score | m/z |
|---|---|---|---|---|
| water_02_1 | 7403 | FTMS; HCD | 191.89 | 776.85 |

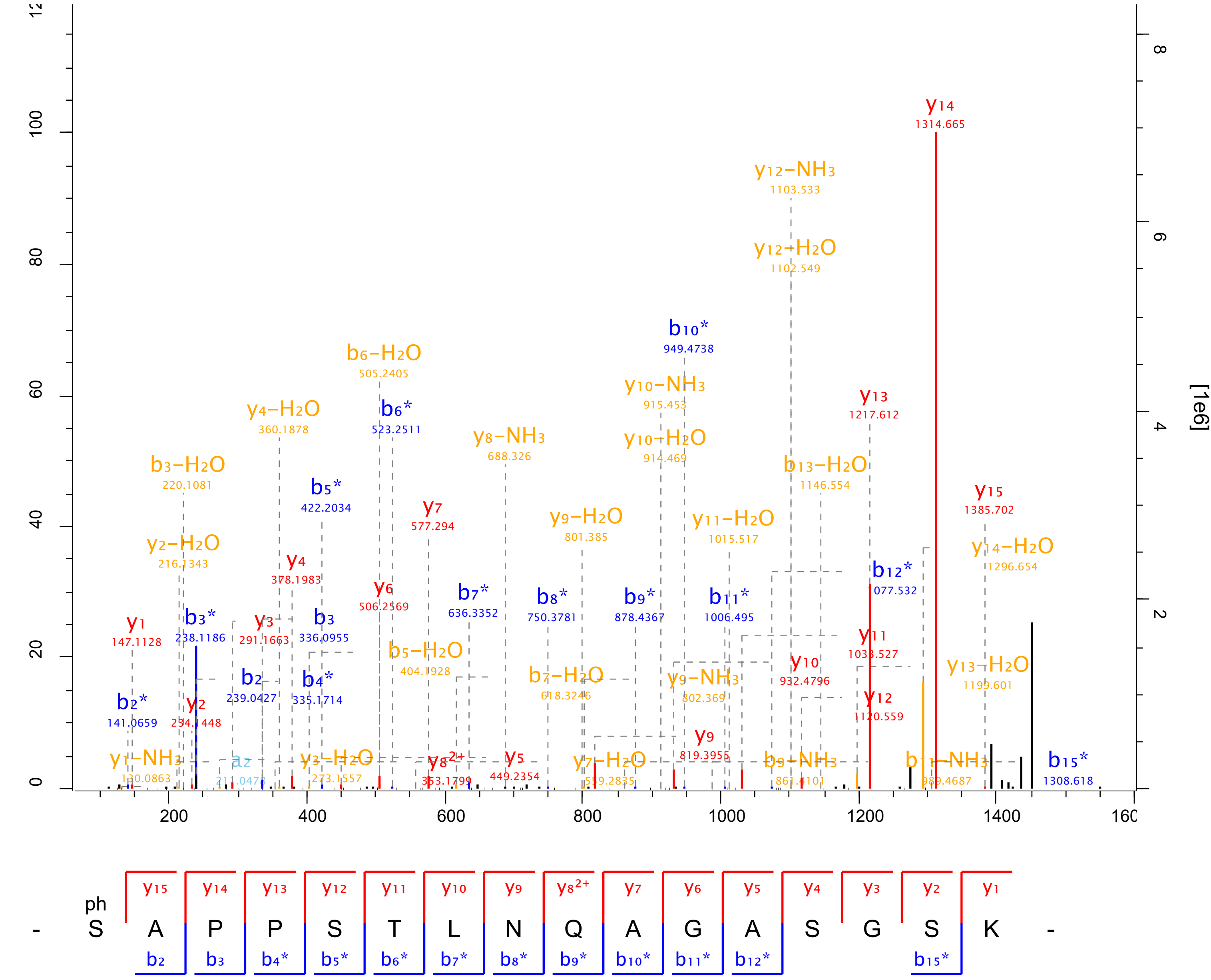

| Raw file | Scan | Method | Score | m/z |
| --- | --- | --- | --- | --- |
| water_02_1 | 7404 | FTMS; HCD | 157.47 | 572.76 |

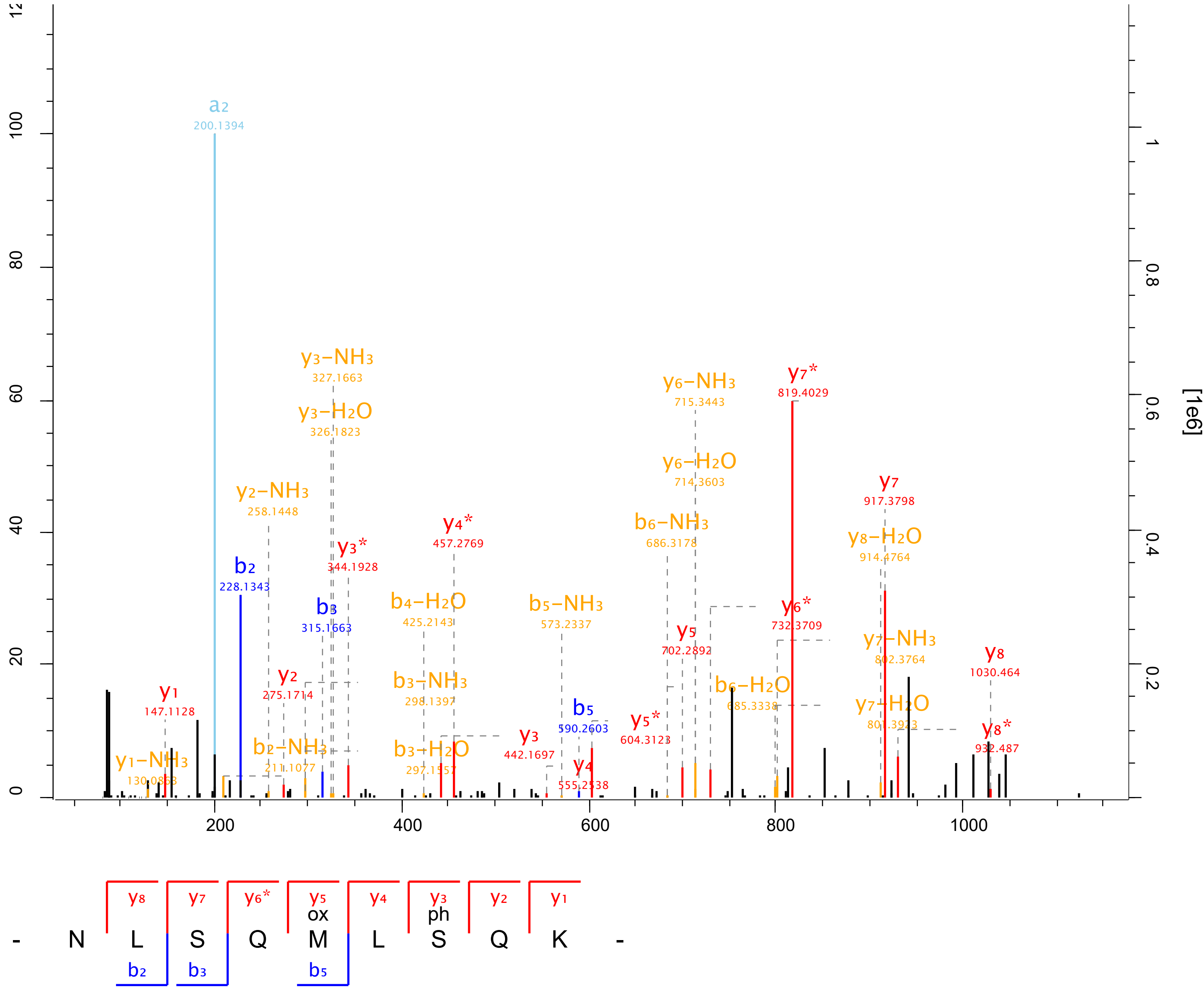

| Raw file | Scan | Method | Score | m/z |
| --- | --- | --- | --- | --- |
| water_02_1 | 7407 | FTMS; HCD | 120.55 | 585.27 |

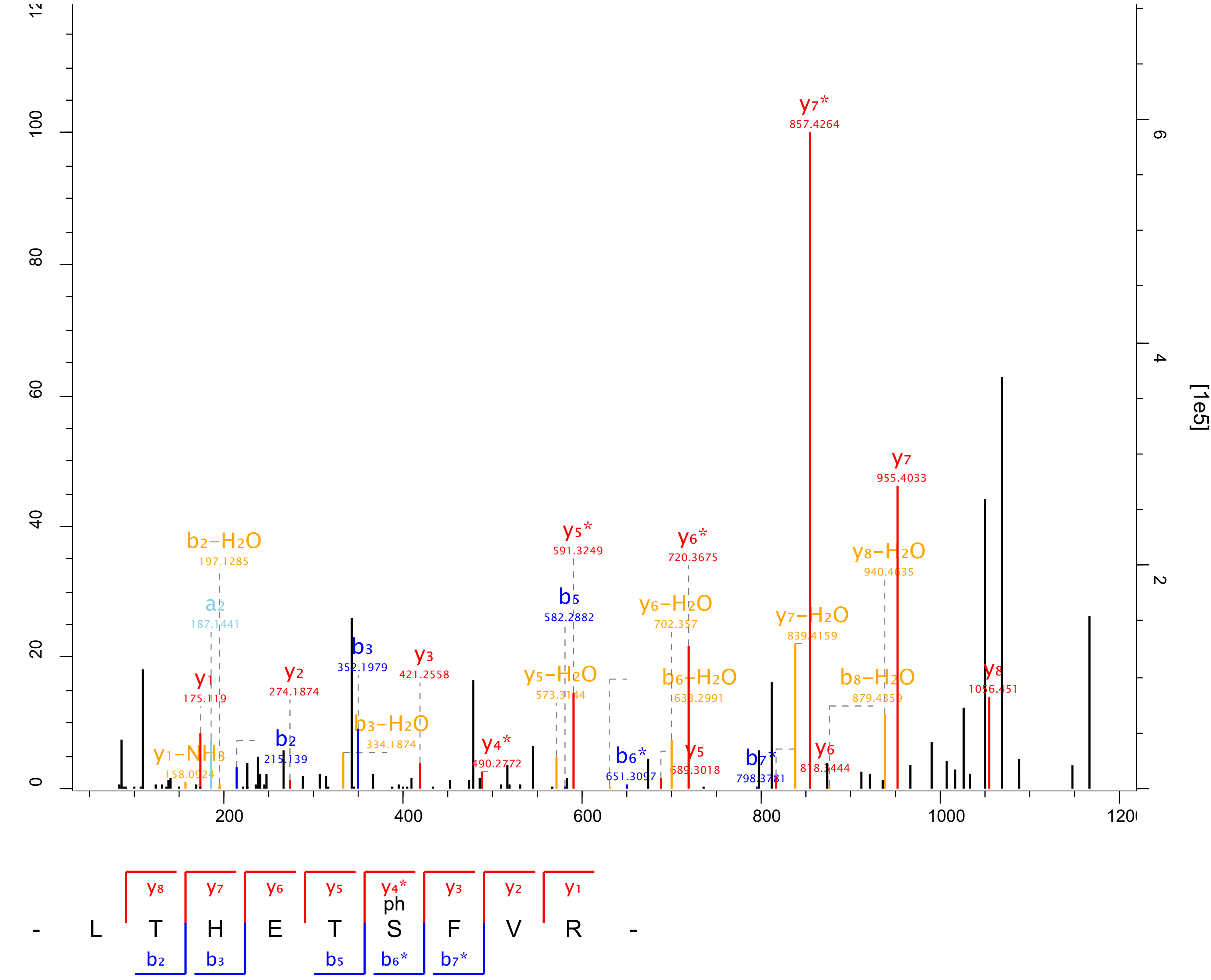

| Raw file | Scan | Method | Score | m/z |
|---|---|---|---|---|
| water_02_1 | 7520 | FTMS; HCD | 131.35 | 634.28 |

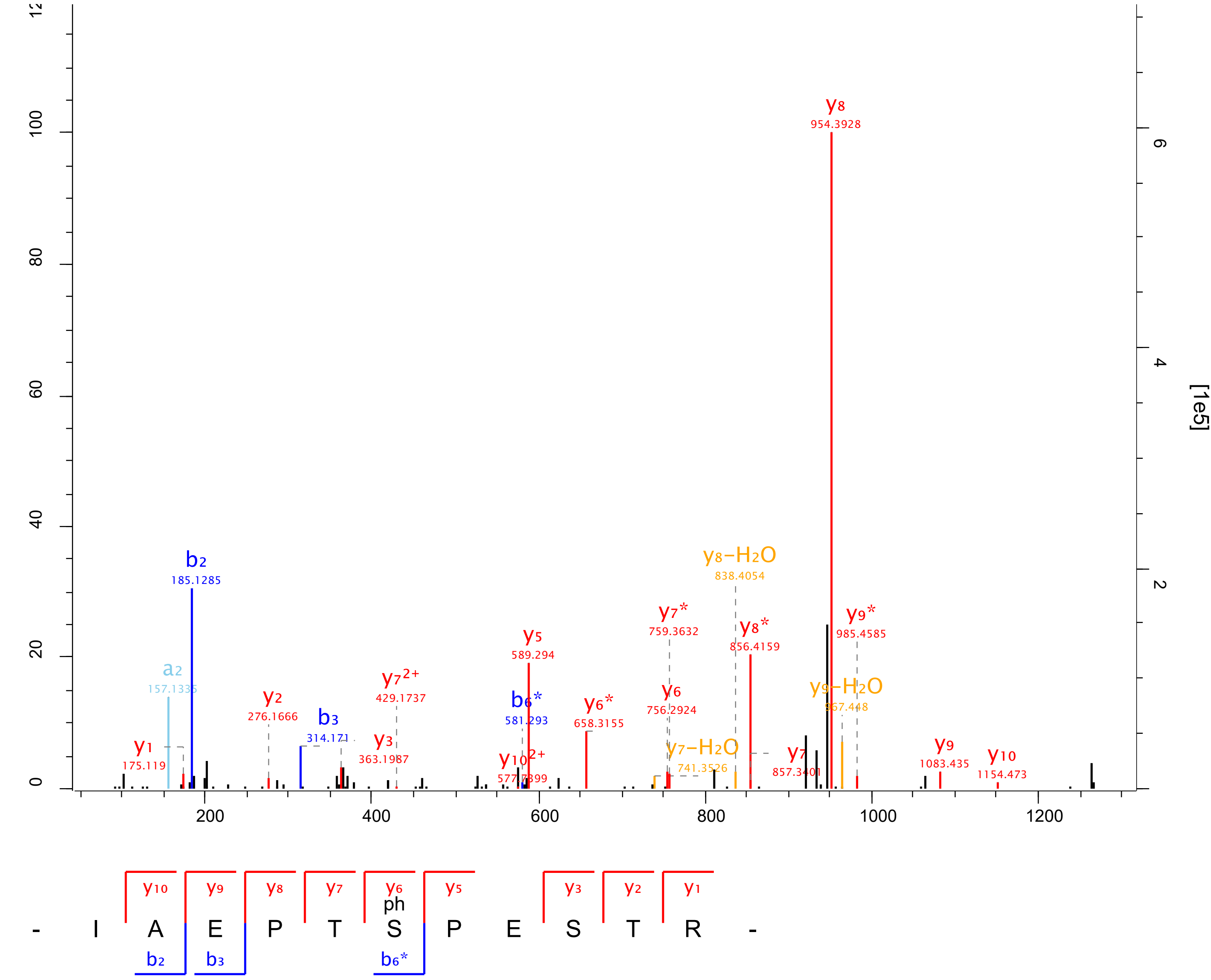

| Raw file | Scan | Method | Score | m/z |
|---|---|---|---|---|
| water_02_1 | 7659 | FTMS; HCD | 118.51 | 649.28 |

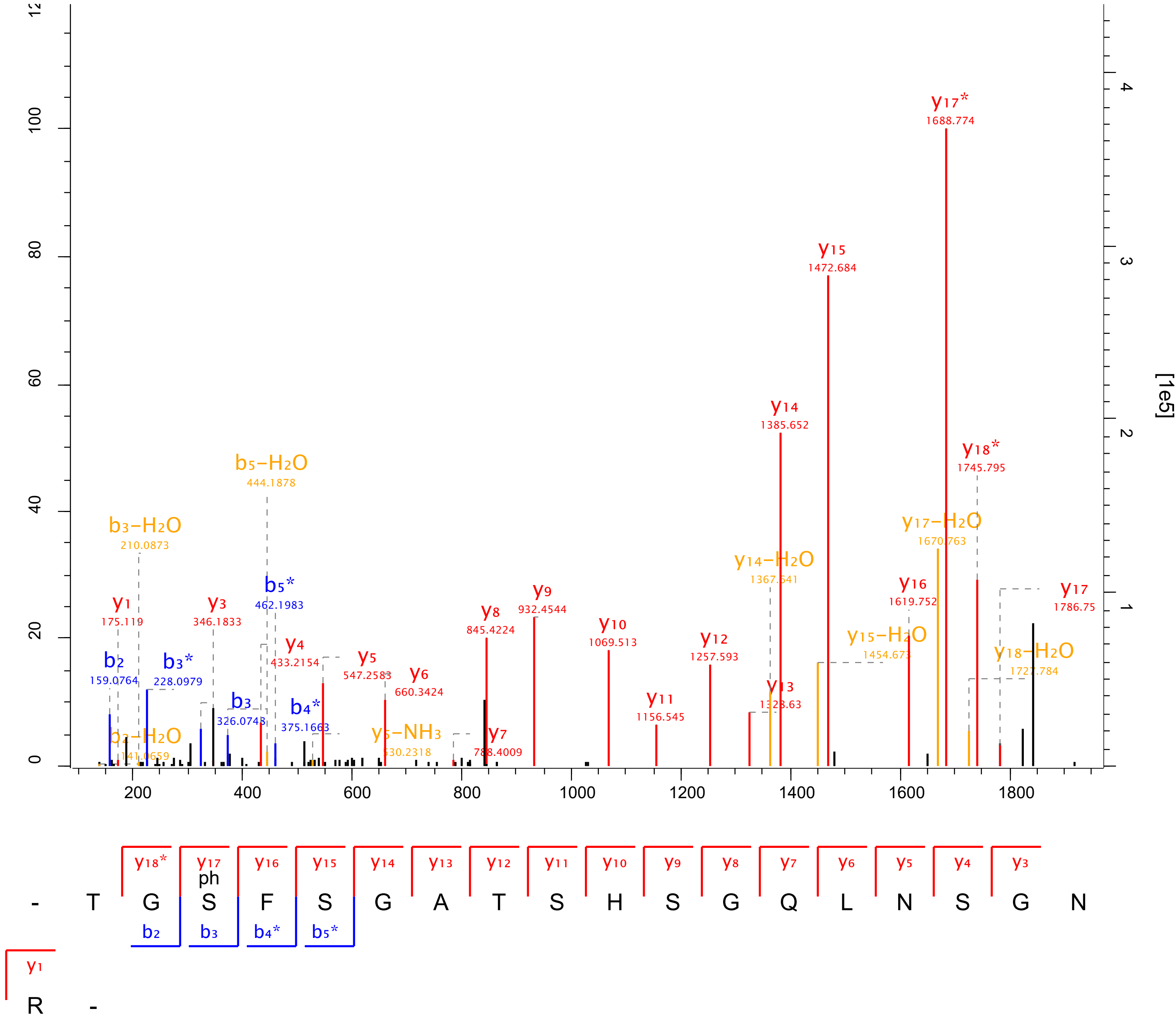

| Raw file | Scan | Method | Score | m/z |
| --- | --- | --- | --- | --- |
| water_02_1 | 7660 | FTMS; HCD | 144.97 | 783.79 |

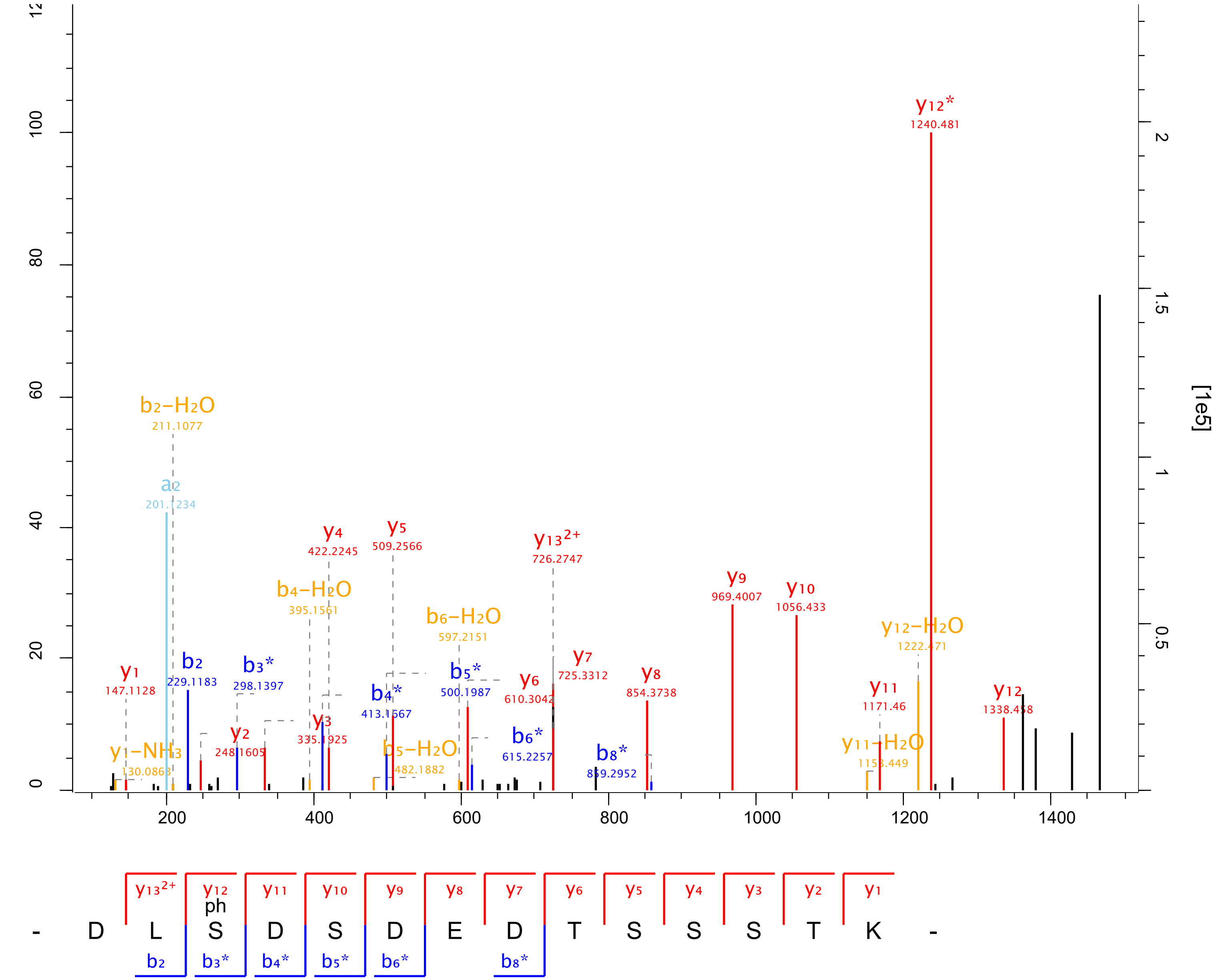

| Raw file | Scan | Method | Score | m/z |
|---|---|---|---|---|
| water_02_1 | 7818 | FTMS; HCD | 51.03 | 478.9 |

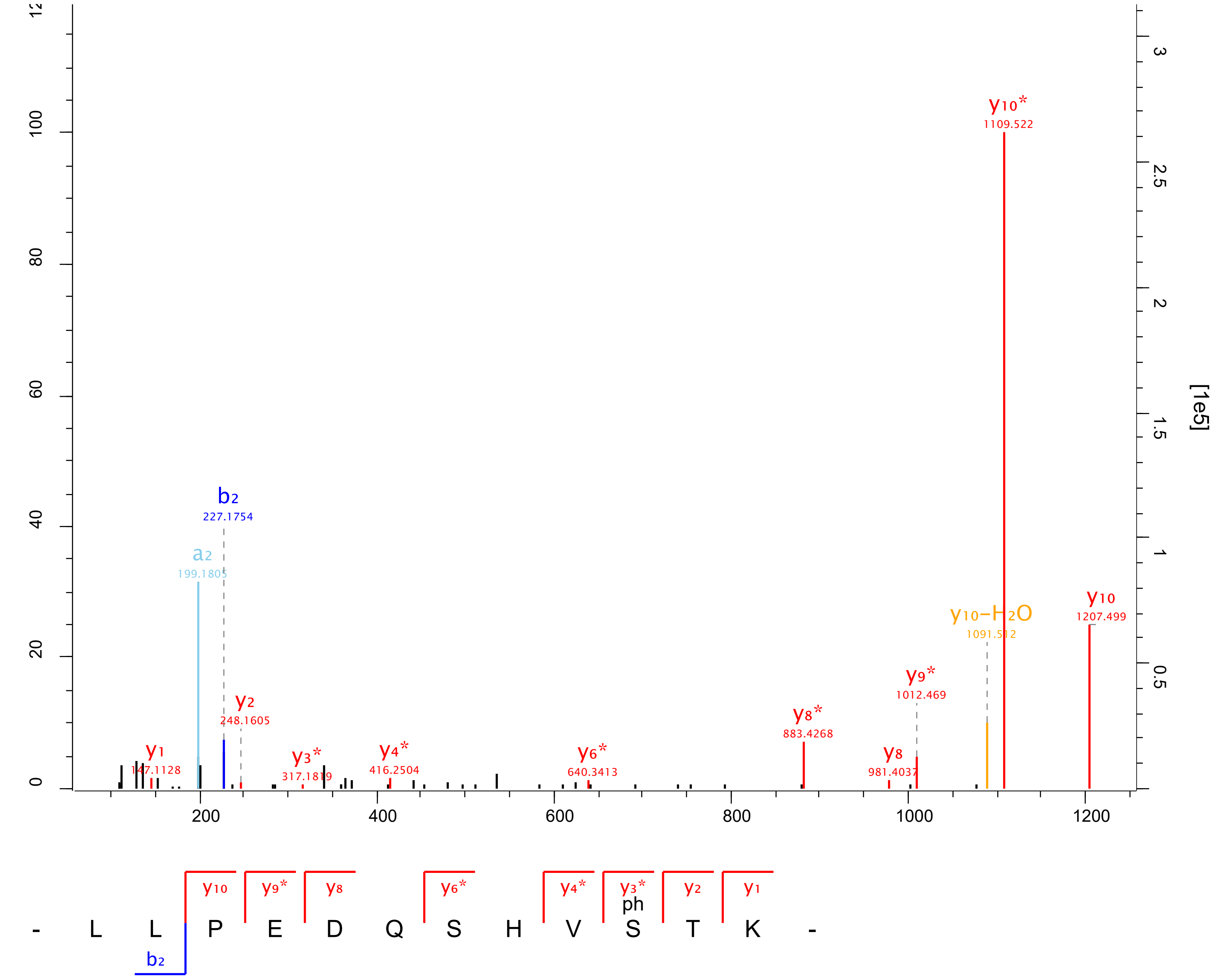

| Raw file | Scan | Method | Score | m/z |
|---|---|---|---|---|
| water_02_1 | 7820 | FTMS; HCD | 66.69 | 513.24 |

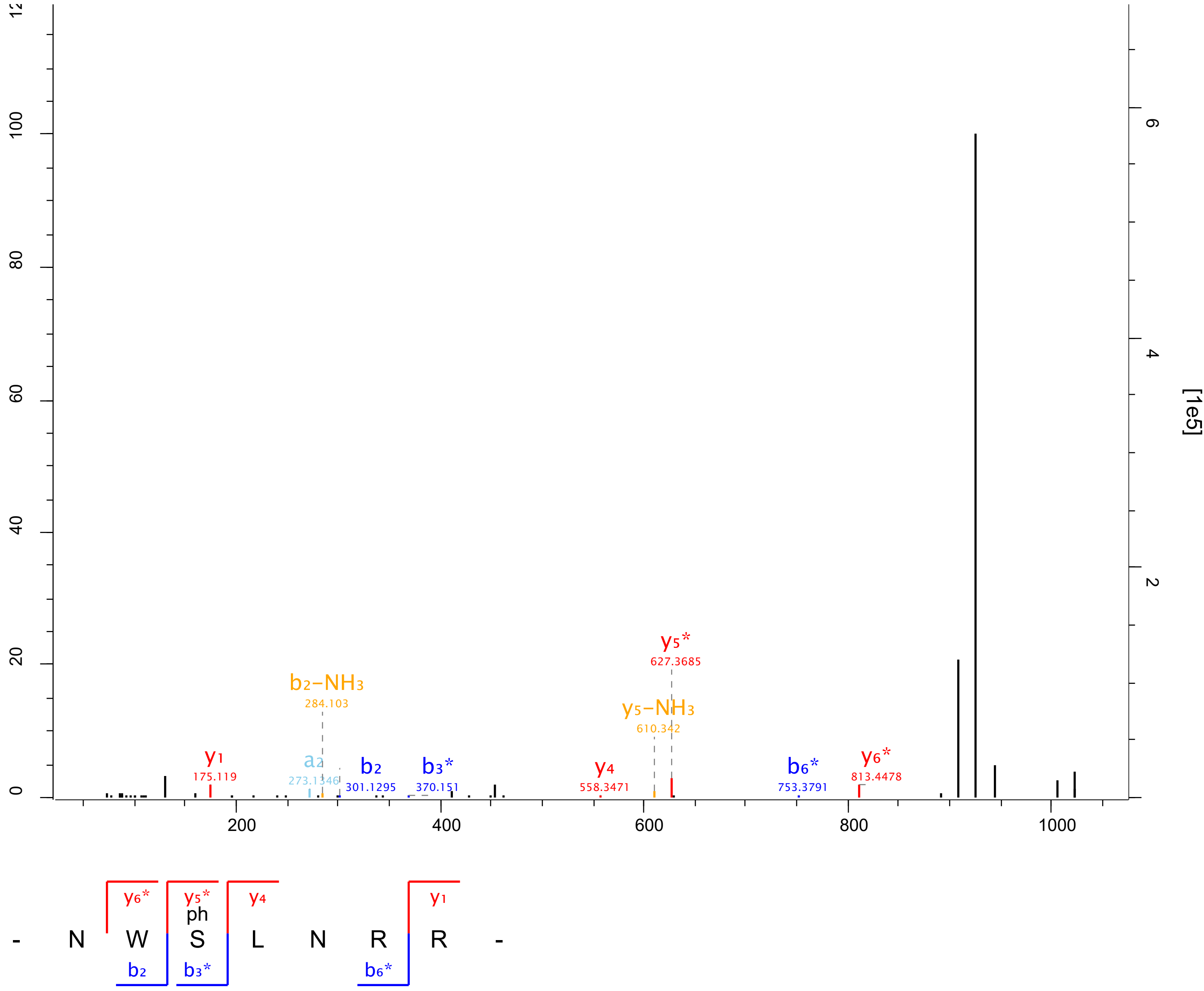

| Raw file | Scan | Method | Score | m/z |
| --- | --- | --- | --- | --- |
| water_02_1 | 7832 | FTMS; HCD | 52.53 | 913.3 |

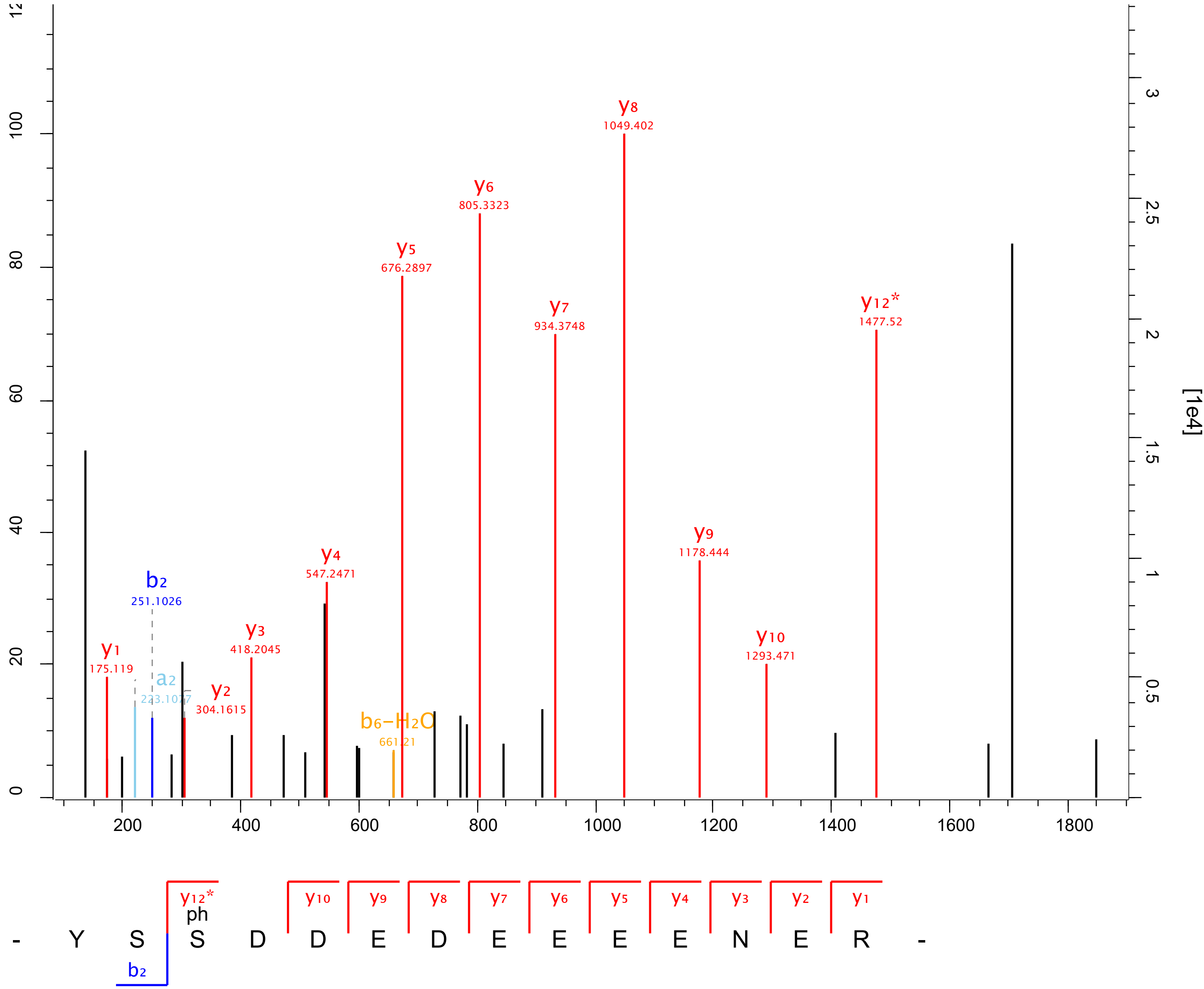

| Raw file | Scan | Method | Score | m/z |
|---|---|---|---|---|
| water_02_1 | 7860 | FTMS; HCD | 100.11 | 660.81 |

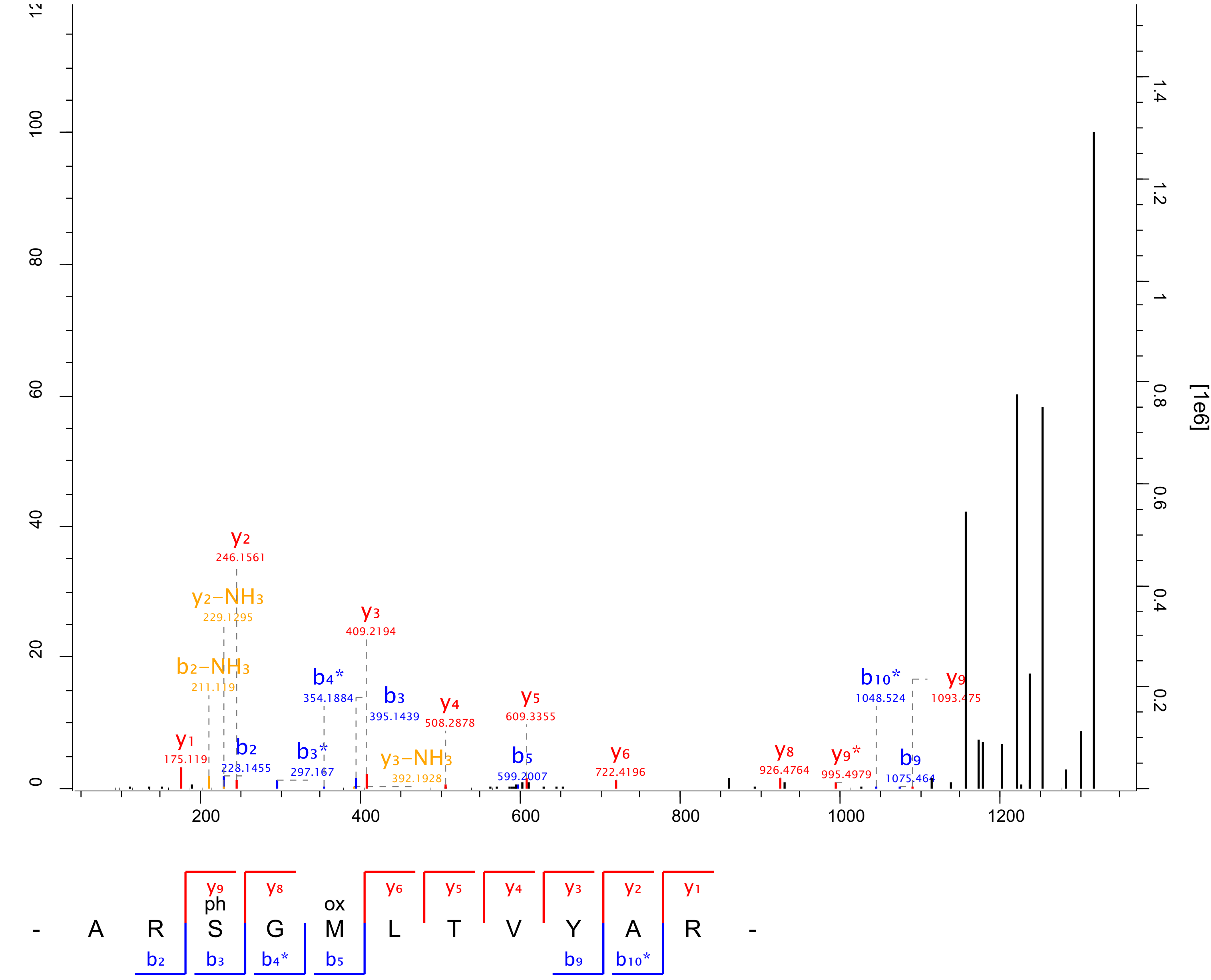

| Raw file | Scan | Method | Score | m/z |
|---|---|---|---|---|
| water_02_1 | 7872 | FTMS; HCD | 80.31 | 527.74 |

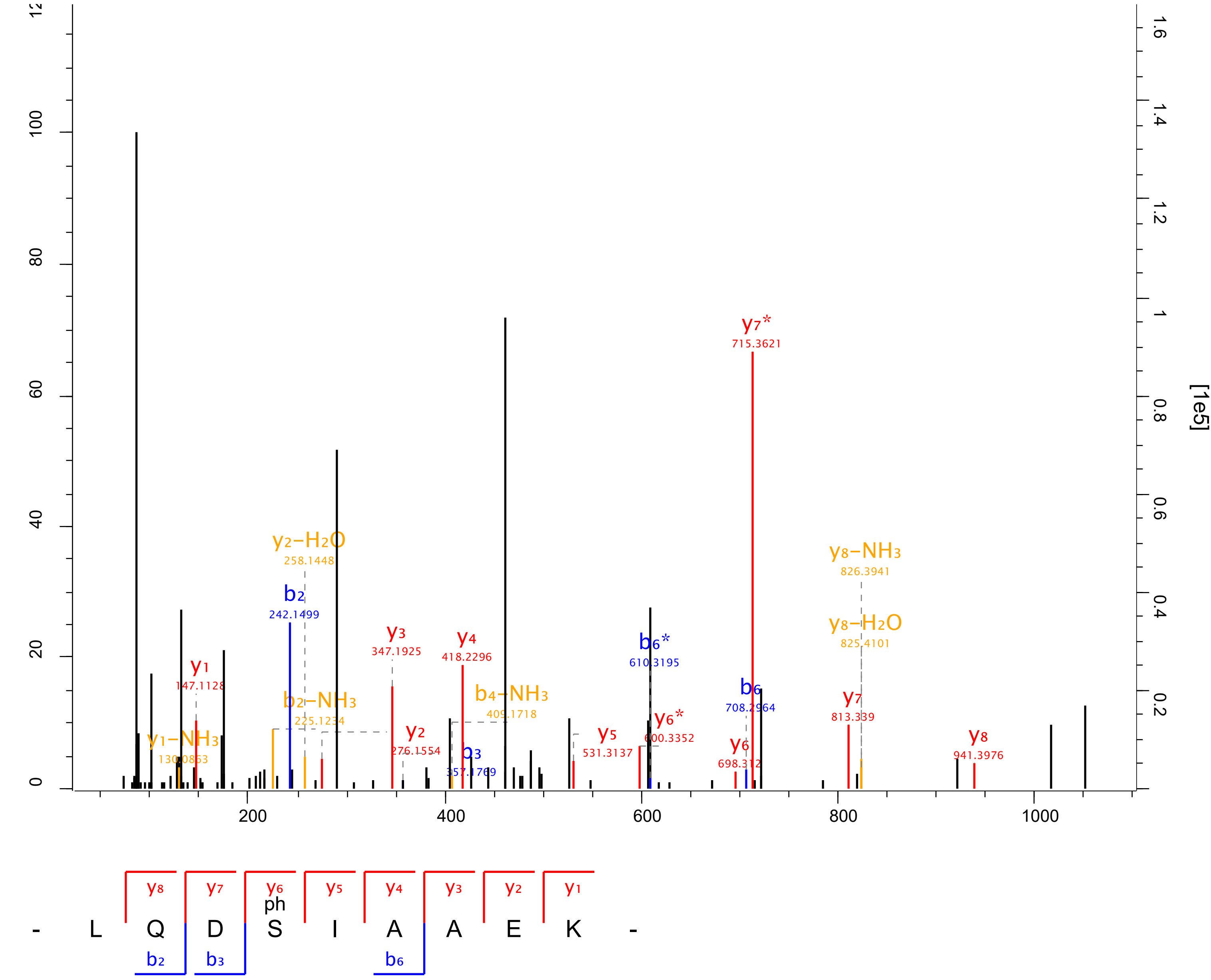

| Raw file | Scan | Method | Score | m/z |
|---|---|---|---|---|
| water_02_1 | 7876 | FTMS; HCD | 42.08 | 587.75 |

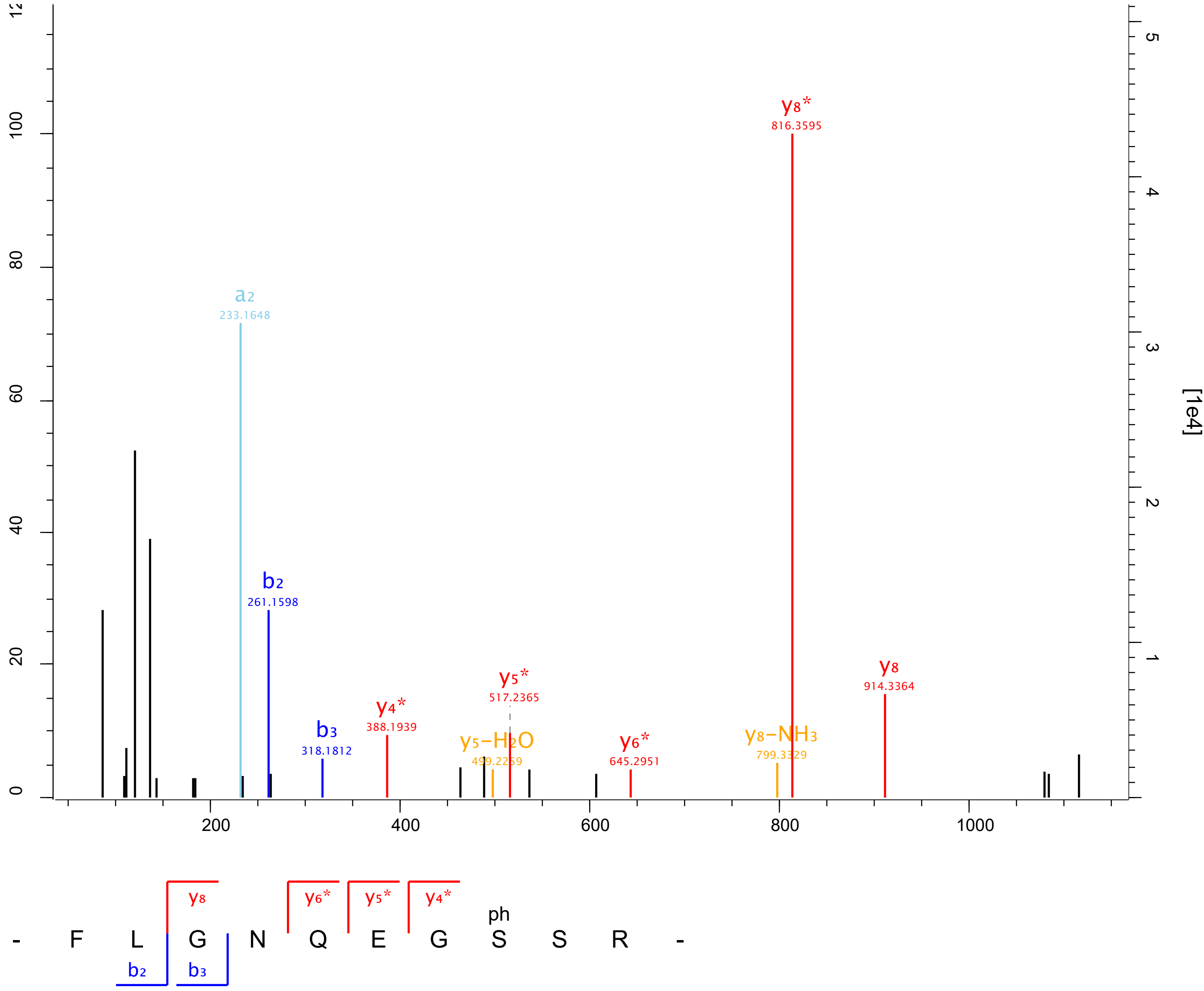

| Raw file | Scan | Method | Score | m/z |
| --- | --- | --- | --- | --- |
| water_02_1 | 8007 | FTMS; HCD | 65.65 | 481.23 |

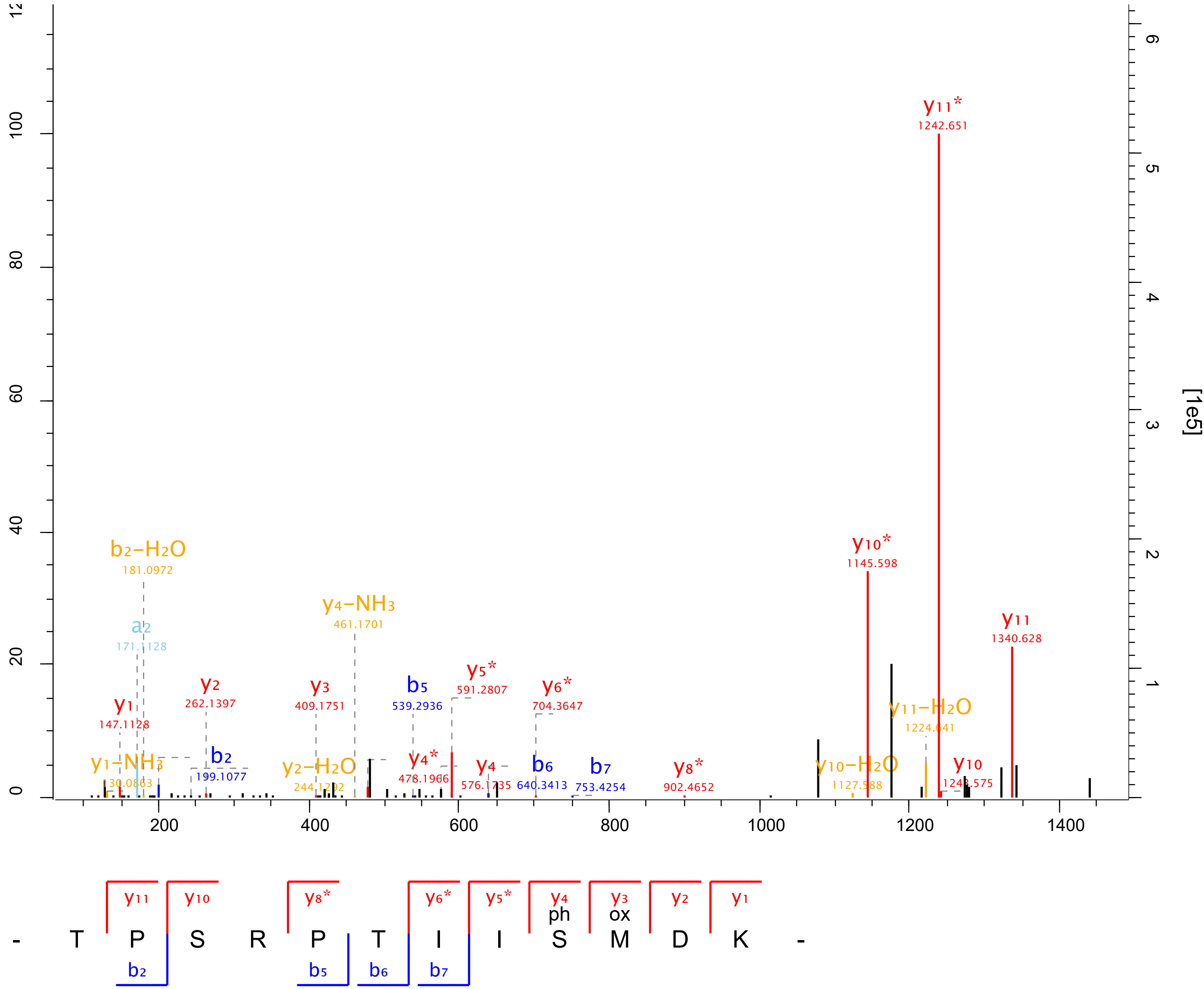

| Raw file | Scan | Method | Score | m/z |
|---|---|---|---|---|
| water_02_1 | 8095 | FTMS; HCD | 125.5 | 636.74 |

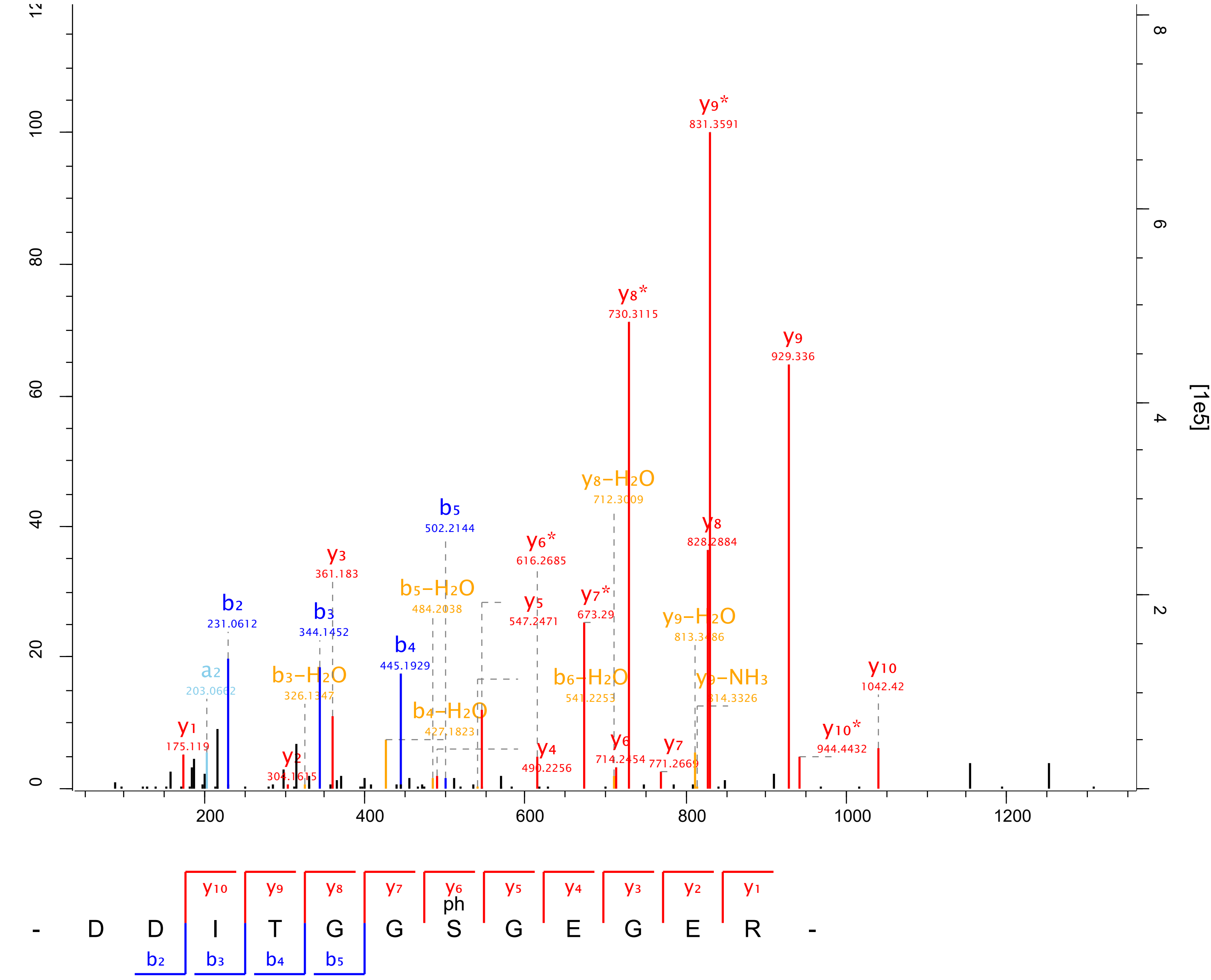

| Raw file | Scan | Method | Score | m/z |
|---|---|---|---|---|
| water_02_1 | 8123 | FTMS; HCD | 40.48 | 494.23 |

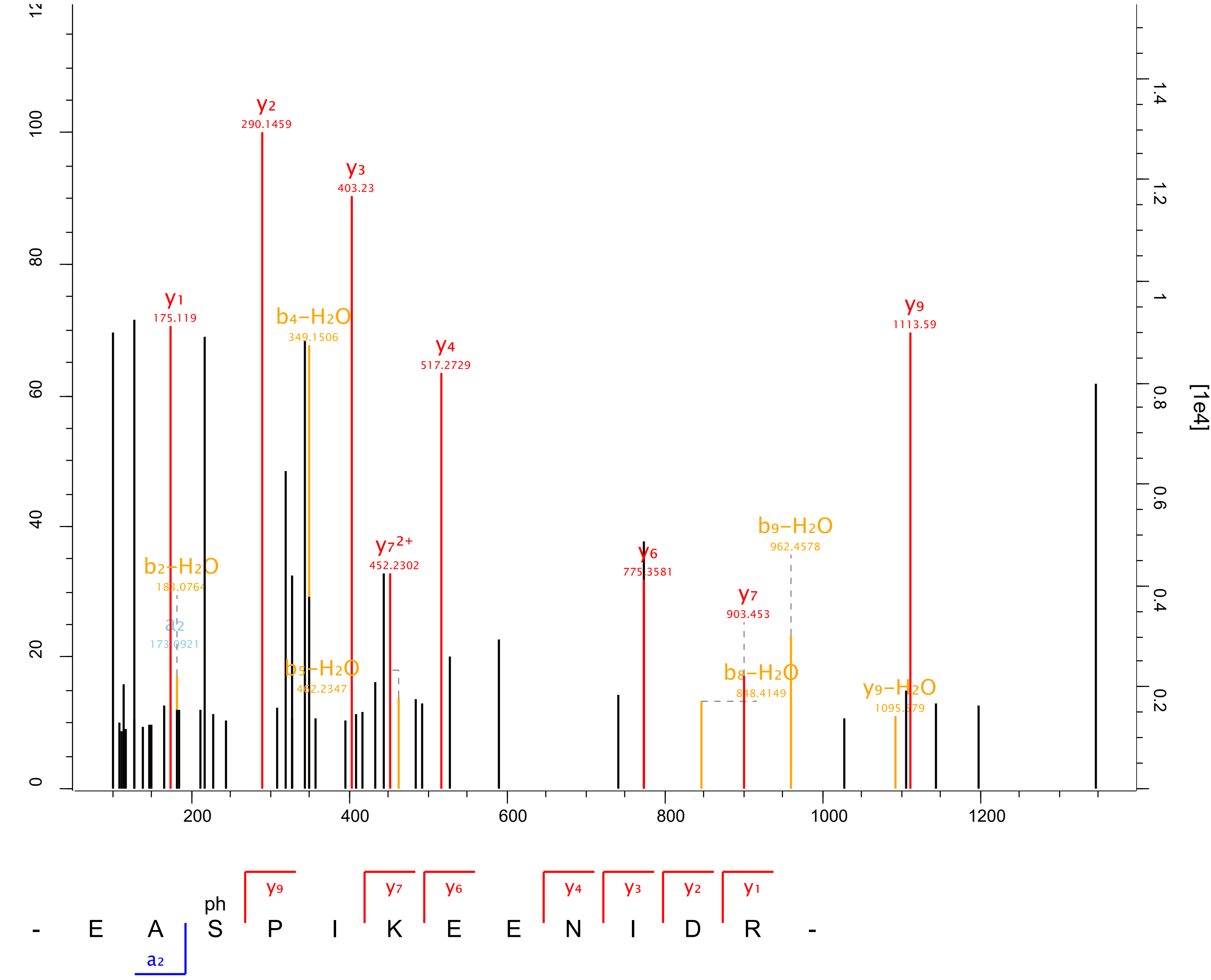

| Raw file | Scan | Method | Score | m/z |
| --- | --- | --- | --- | --- |
| water_02_1 | 8151 | FTMS; HCD | 54.49 | 515.23 |

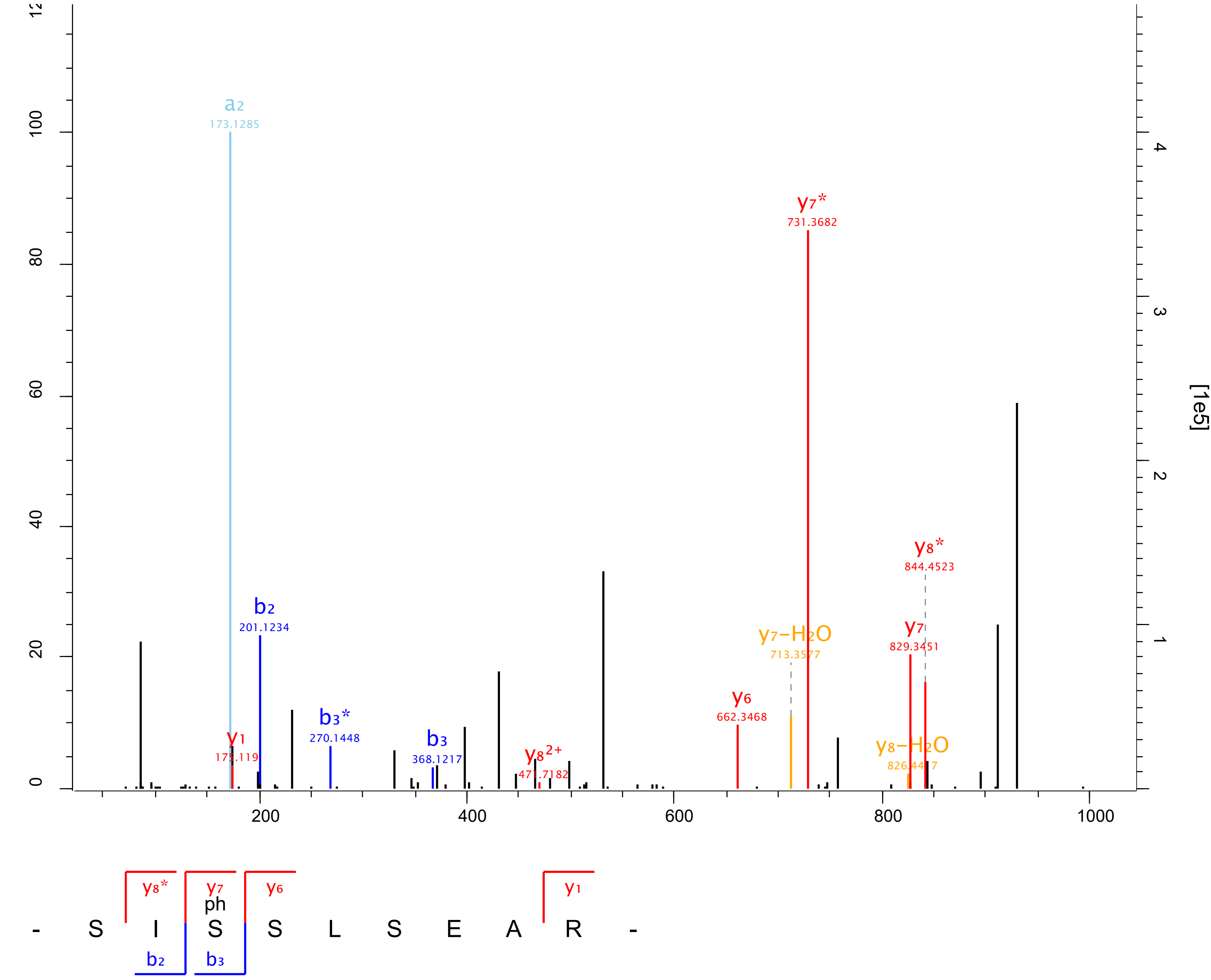

| Raw file | Scan | Method | Score | m/z |
|---|---|---|---|---|
| water_02_1 | 8197 | FTMS; HCD | 44.93 | 505.57 |

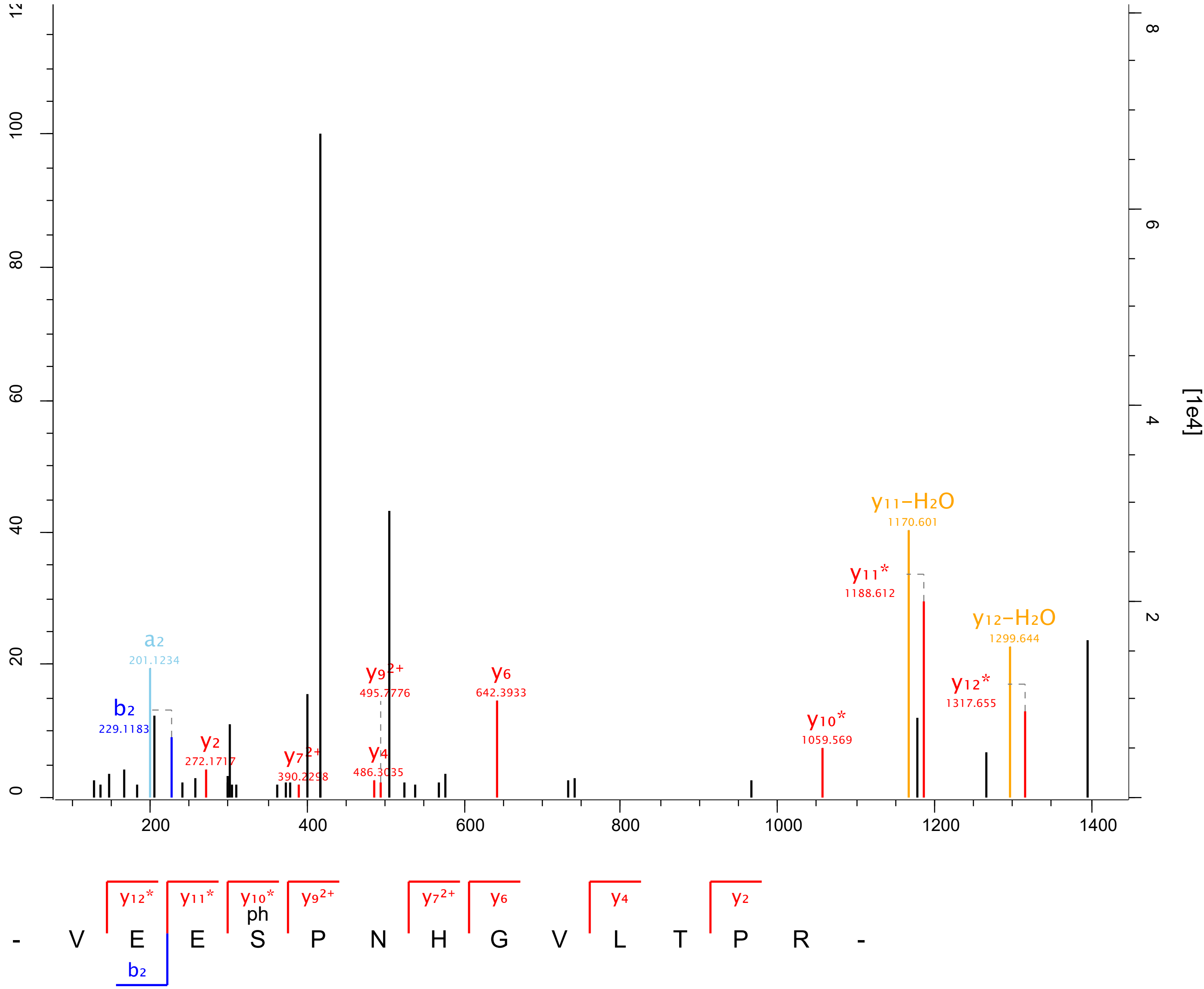

| Raw file | Scan | Method | Score | m/z |
|---|---|---|---|---|
| water_02_1 | 8281 | FTMS; HCD | 167.29 | 670.34 |

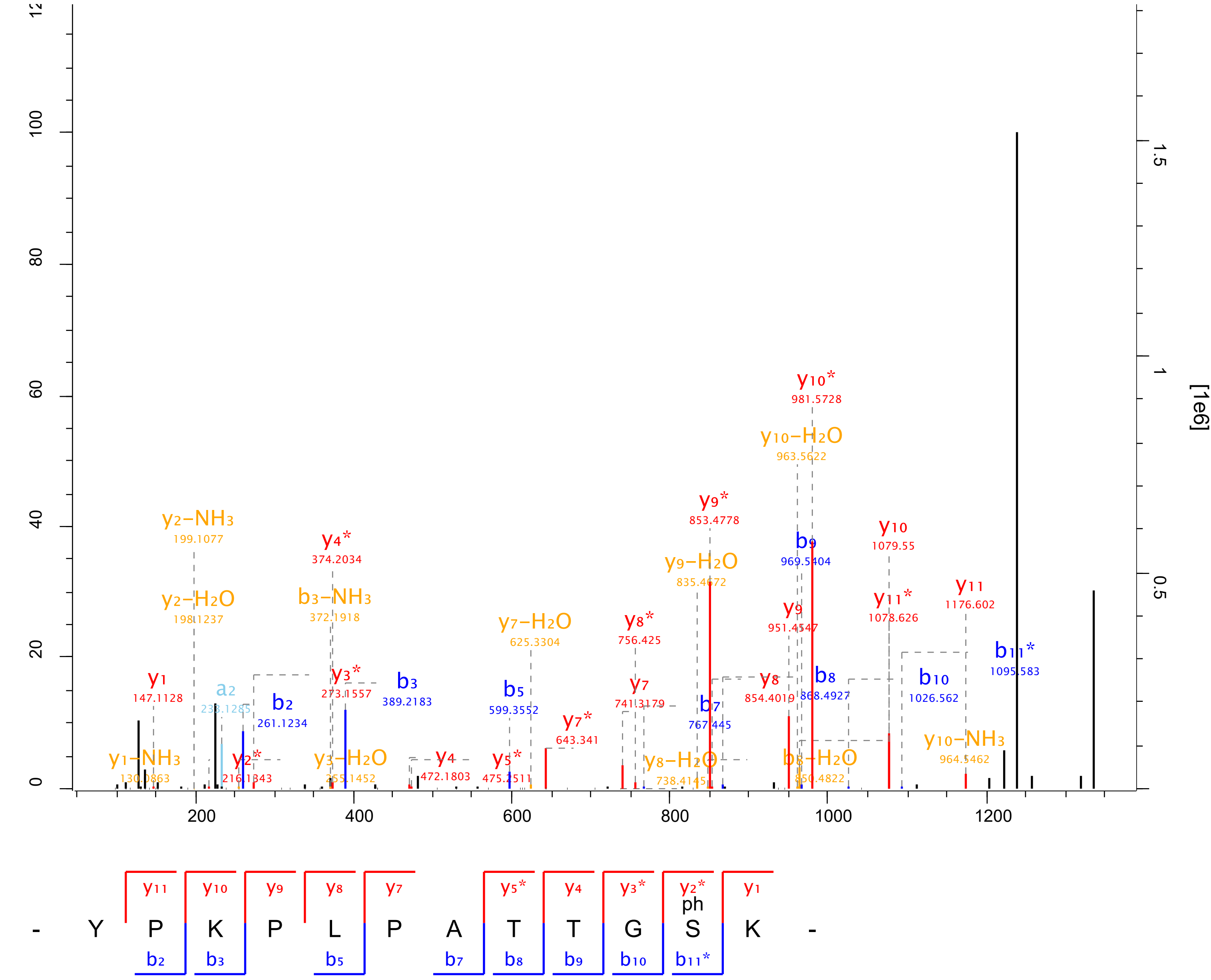

| Raw file | Scan | Method | Score | m/z |
|---|---|---|---|---|
| water_02_1 | 8294 | FTMS; HCD | 48.4 | 539.93 |

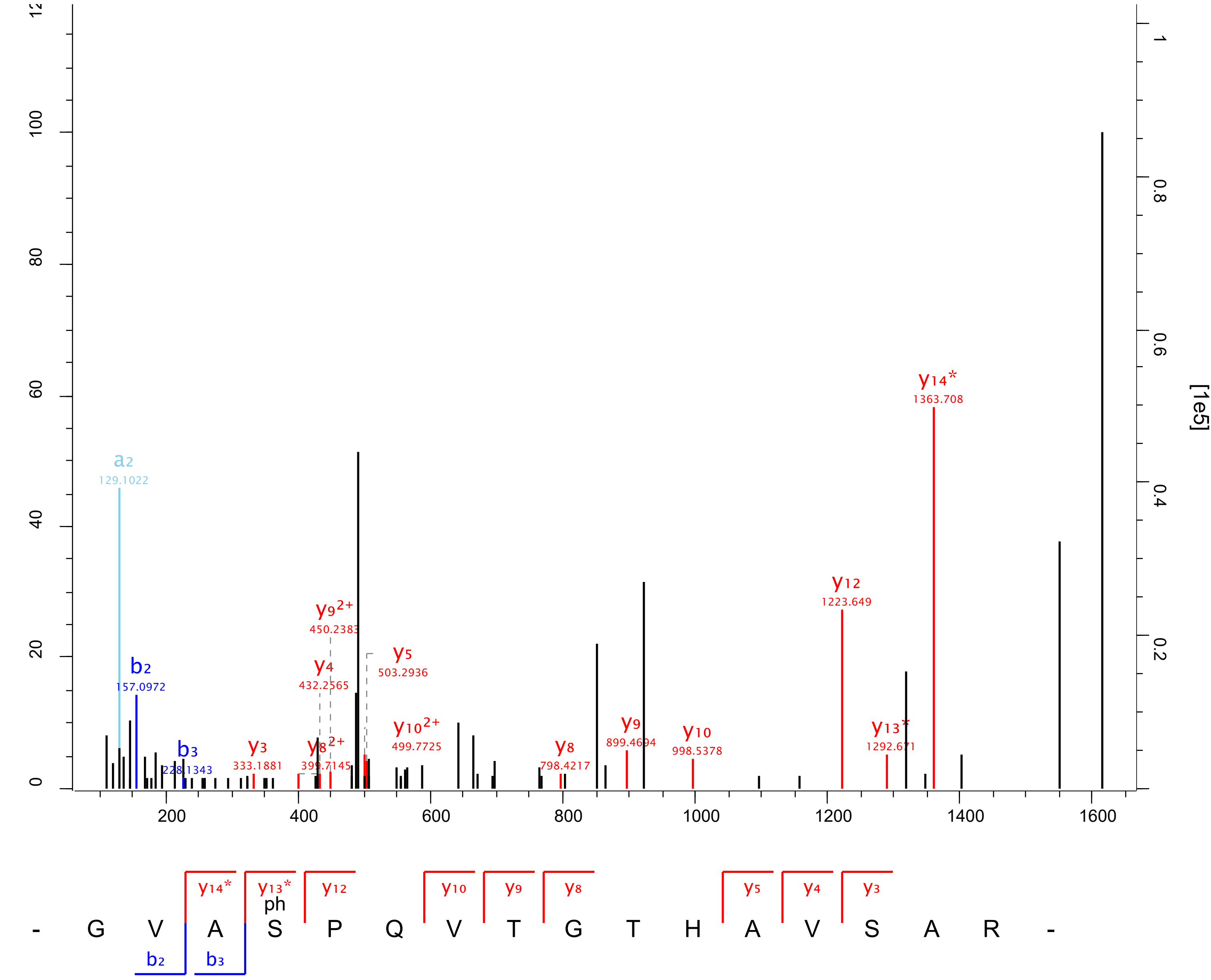

| Raw file | Scan | Method | Score | m/z |
|---|---|---|---|---|
| water_02_1 | 8321 | FTMS; HCD | 94.51 | 580.74 |

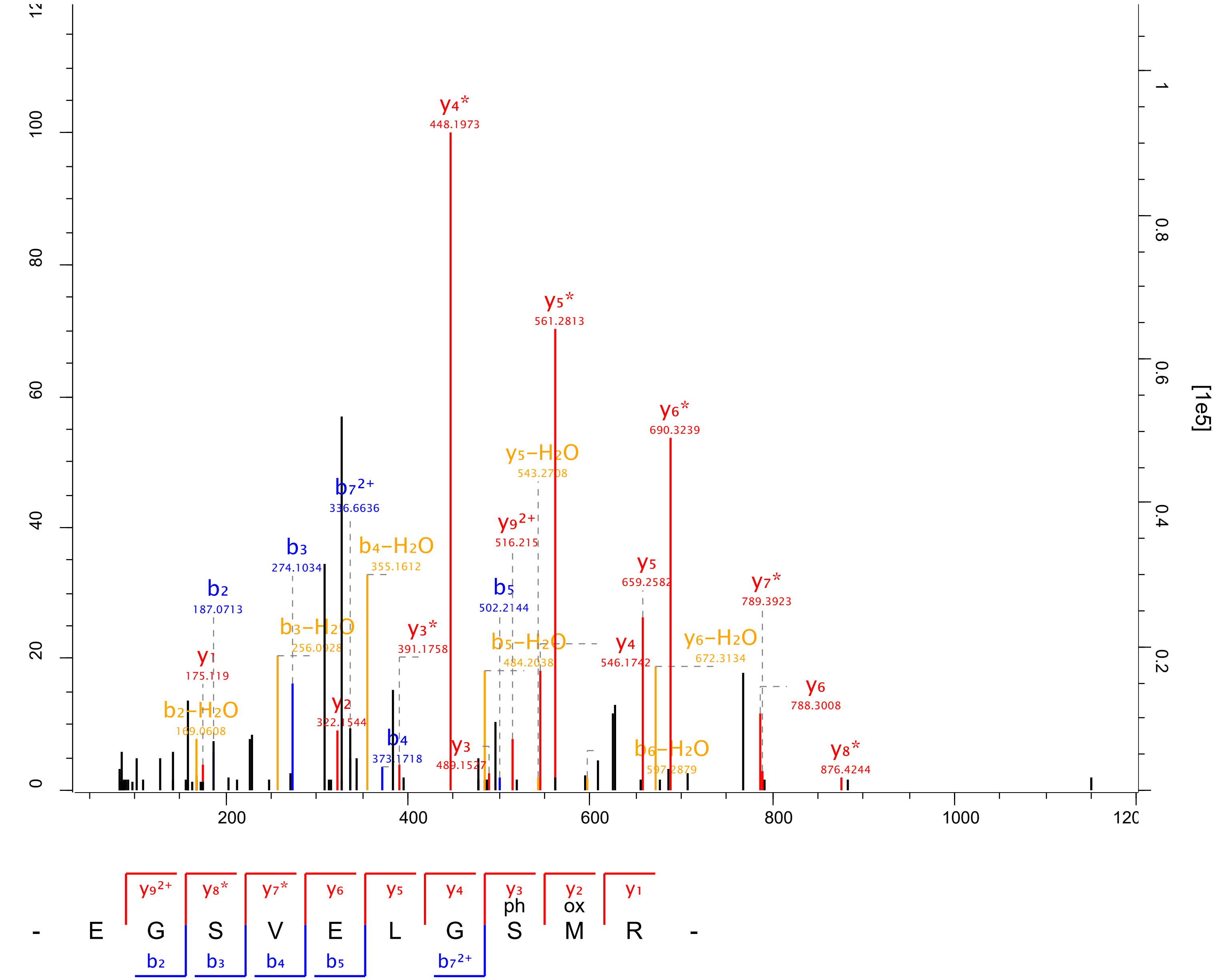

| Raw file | Scan | Method | Score | m/z |
|---|---|---|---|---|
| water_02_1 | 8331 | FTMS; HCD | 50.15 | 686.24 |

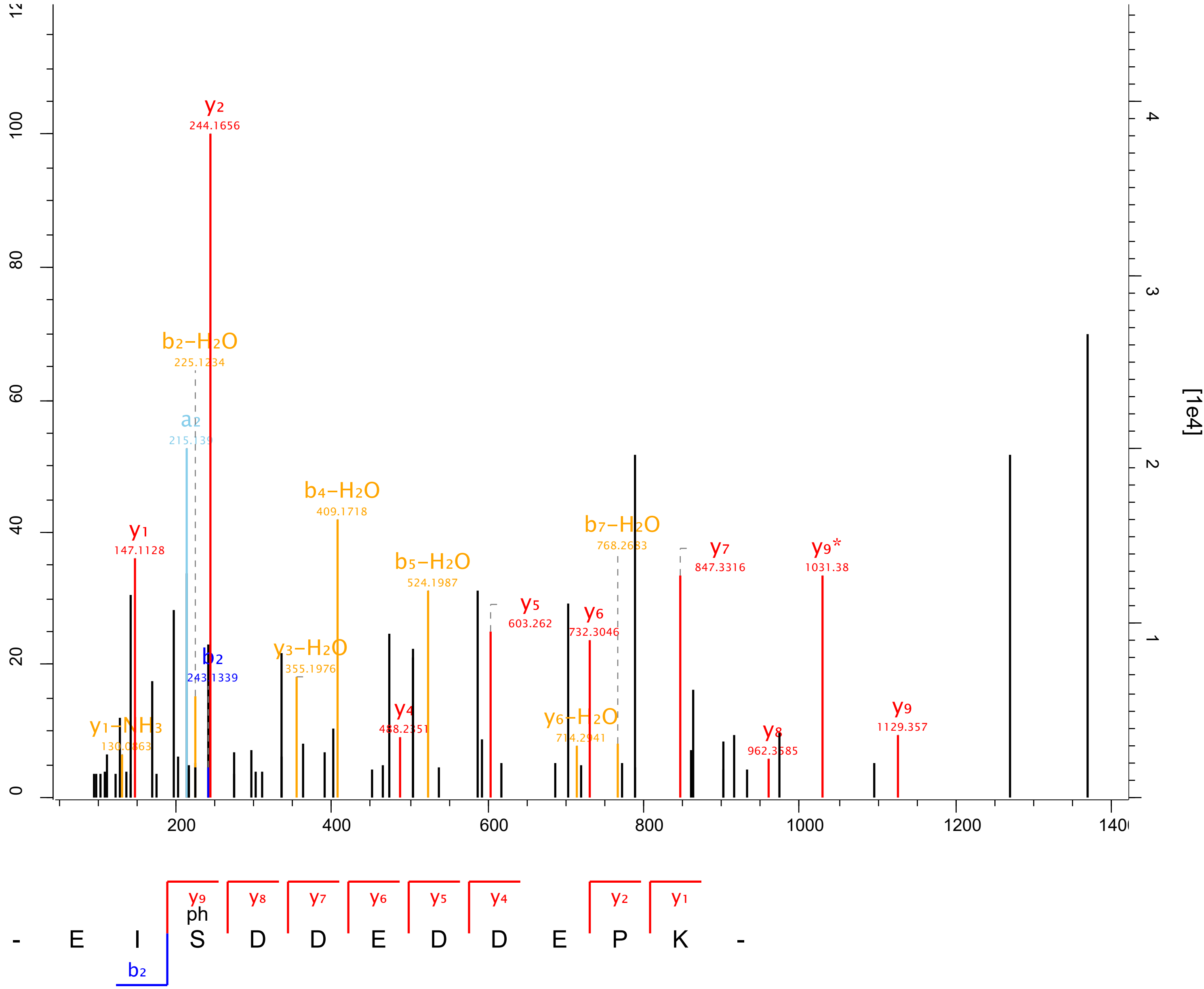

| Raw file | Scan | Method | Score | m/z |
|---|---|---|---|---|
| water_02_1 | 8336 | FTMS; HCD | 92.91 | 739.65 |

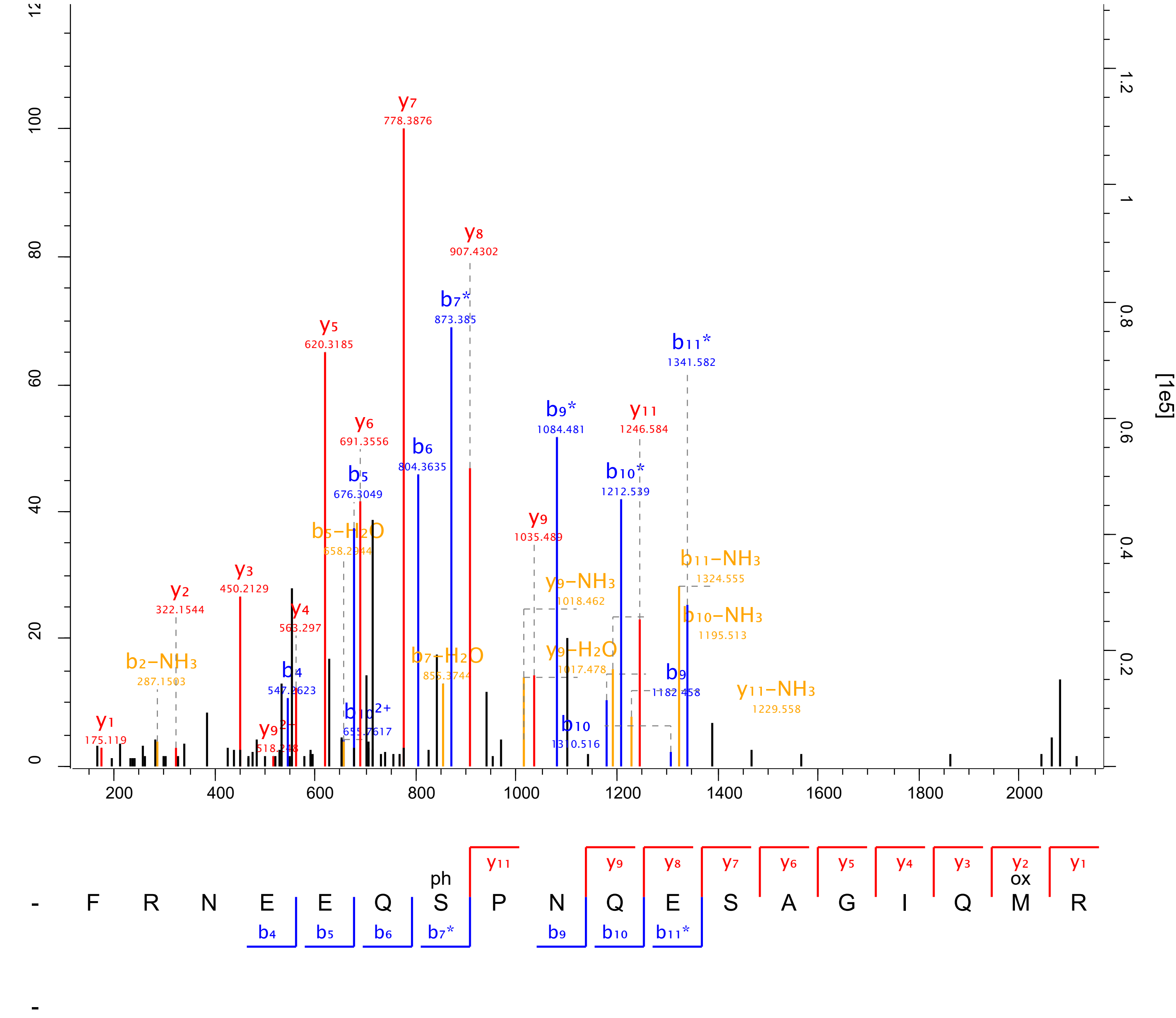

| Raw file | Scan | Method | Score | m/z |
| --- | --- | --- | --- | --- |
| water_02_1 | 8373 | FTMS; HCD | 118.48 | 798.8 |

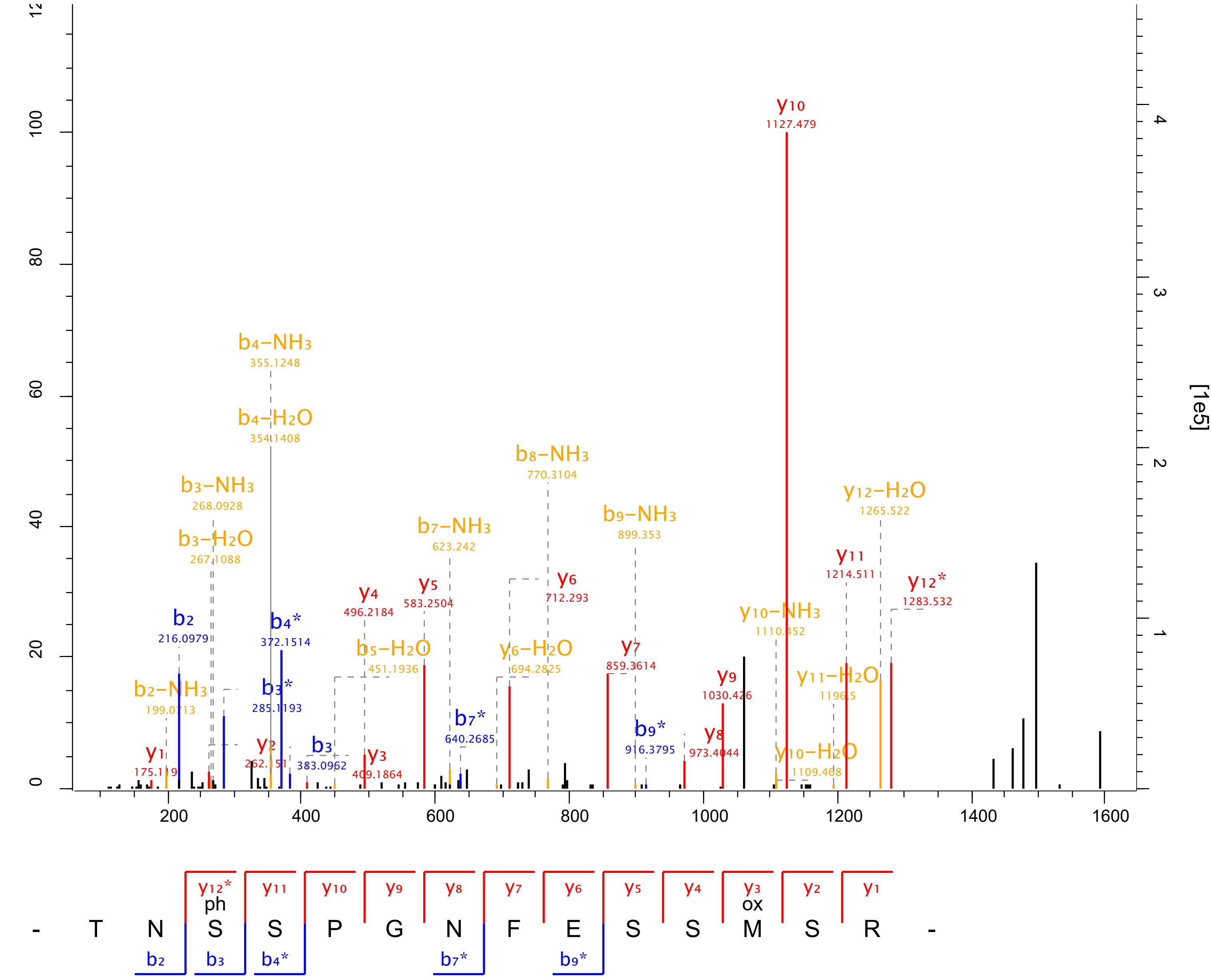

| Raw file | Scan | Method | Score | m/z |
|---|---|---|---|---|
| water_02_1 | 8381 | FTMS; HCD | 93.58 | 681.65 |

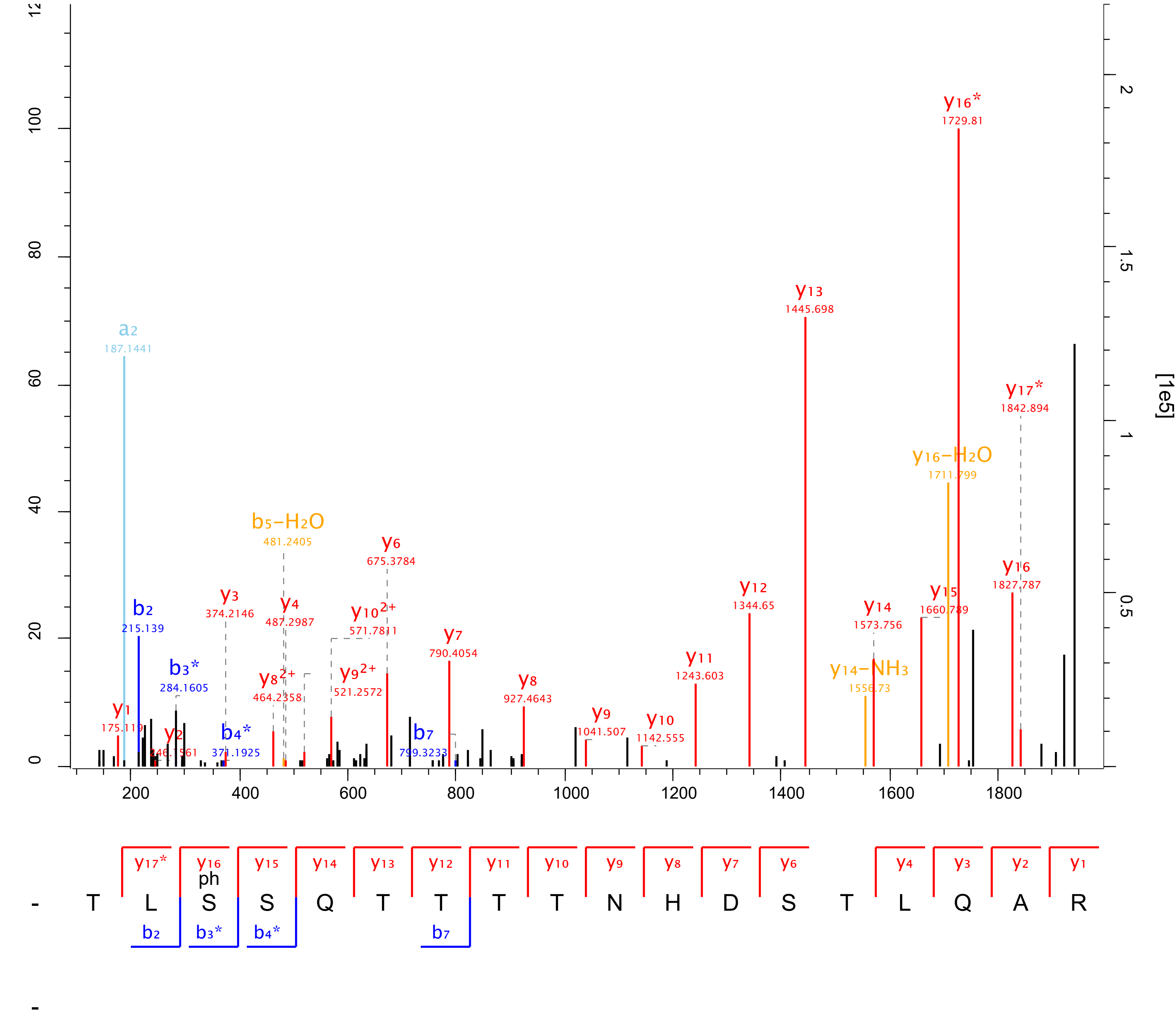

| Raw file | Scan | Method | Score | m/z |
|---|---|---|---|---|
| water_02_1 | 8432 | FTMS; HCD | 74.27 | 516.91 |

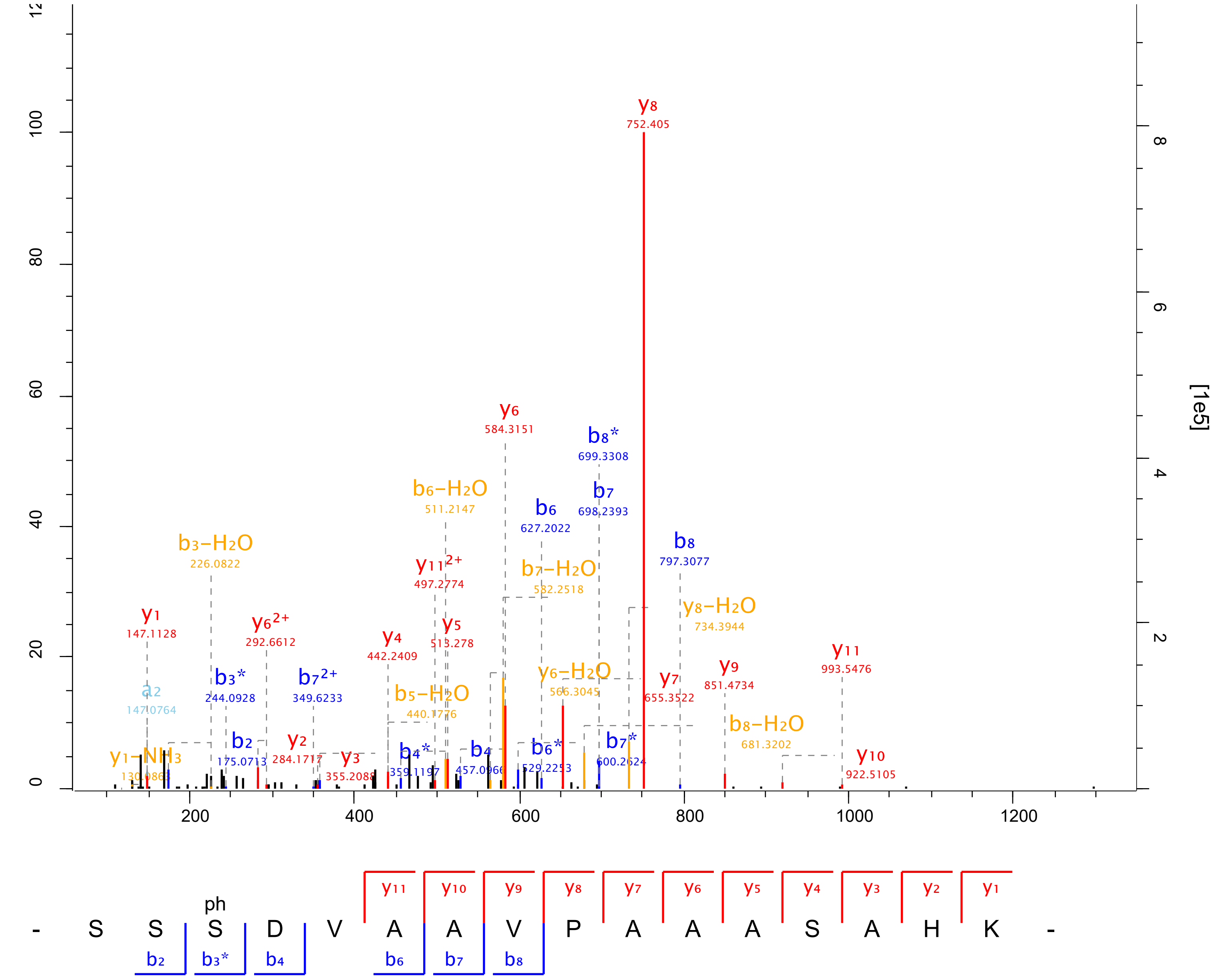

| Raw file | Scan | Method | Score | m/z |
|---|---|---|---|---|
| water_02_1 | 8528 | FTMS; HCD | 62.23 | 549.75 |

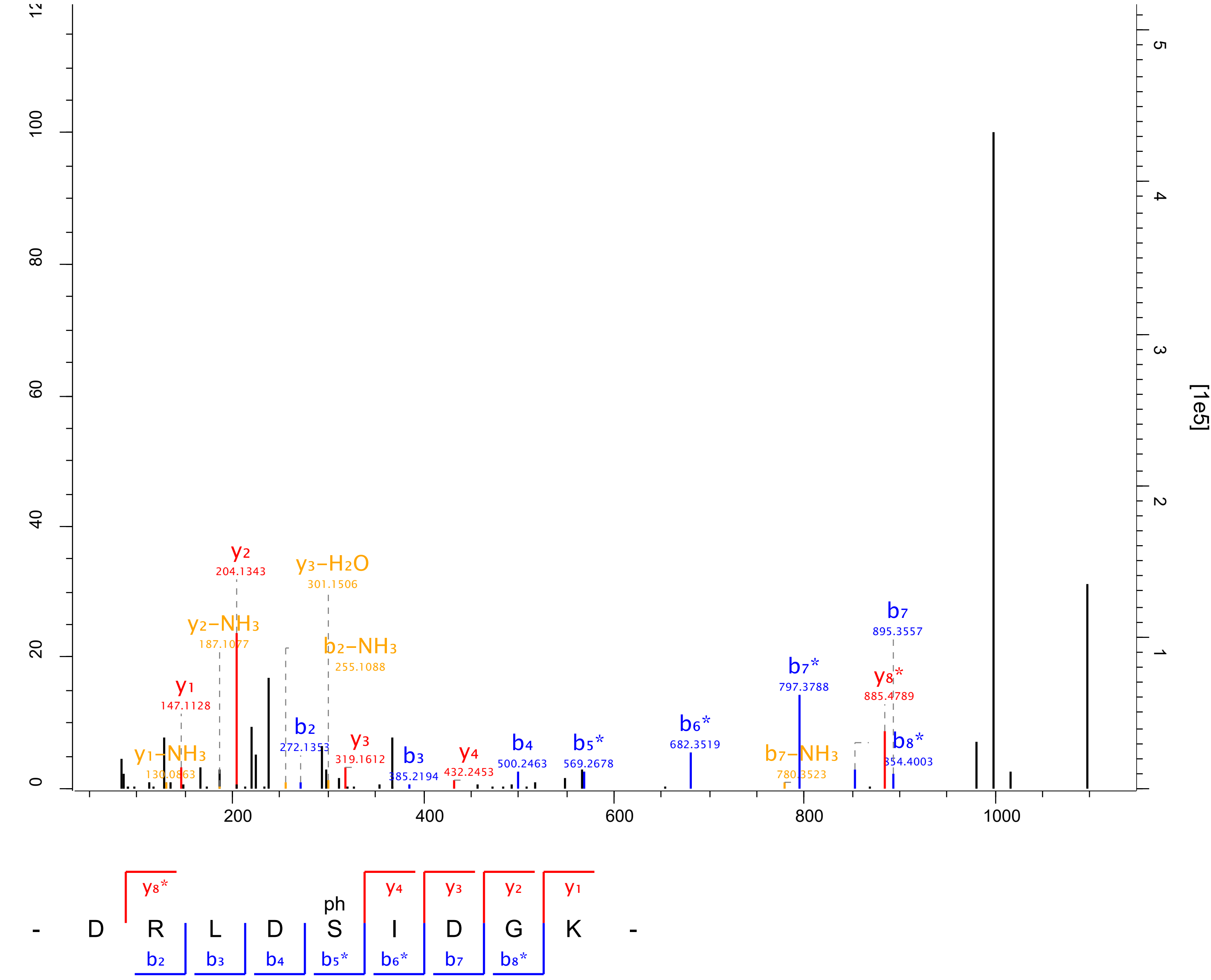

| Raw file | Scan | Method | Score | m/z |
| --- | --- | --- | --- | --- |
| water_02_1 | 8566 | FTMS; HCD | 79.84 | 480.24 |

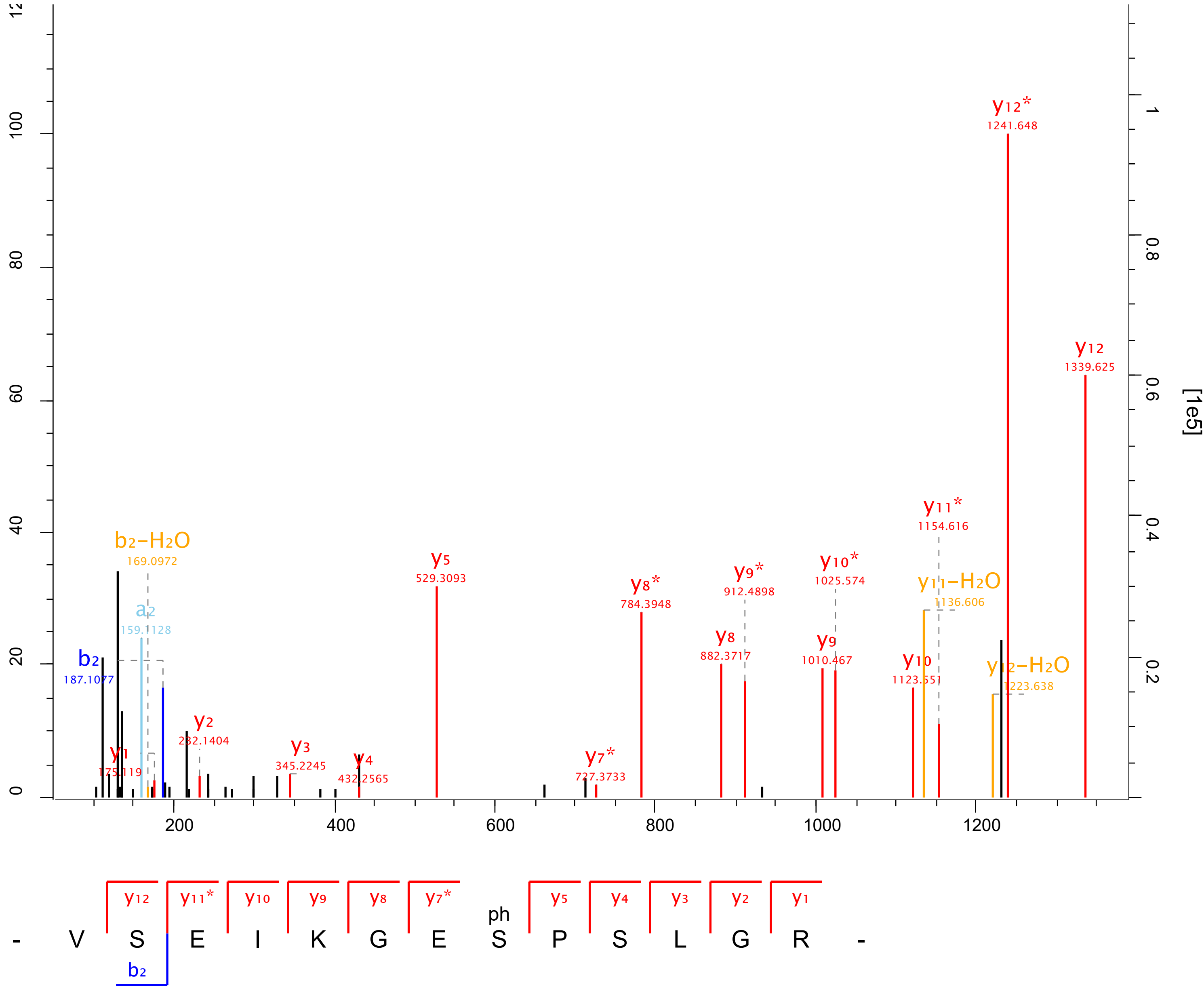

| Raw file | Scan | Method | Score | m/z |
| --- | --- | --- | --- | --- |
| water_02_1 | 8589 | FTMS; HCD | 128.05 | 572.93 |

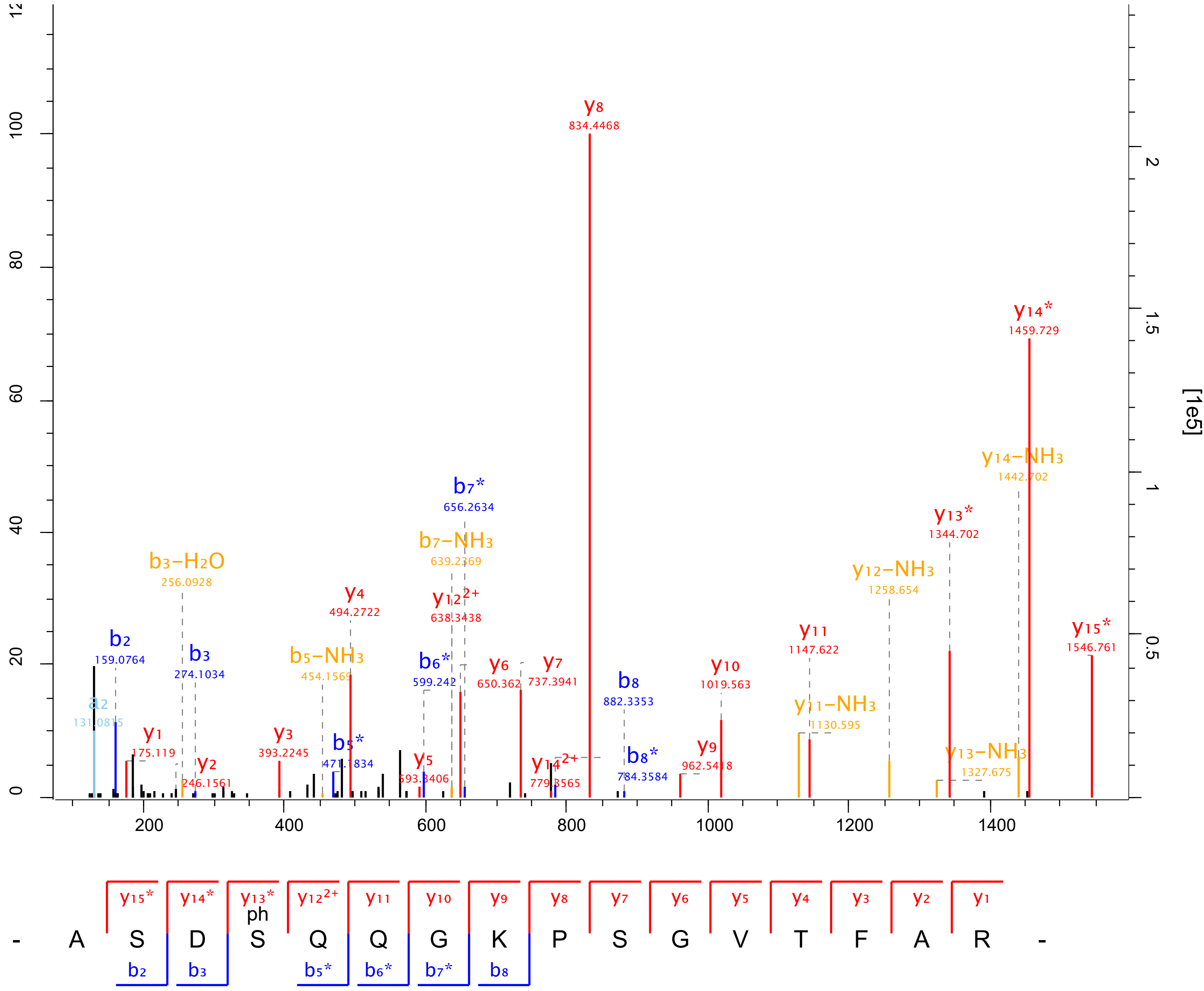

| Raw file | Scan | Method | Score | m/z |
| --- | --- | --- | --- | --- |
| water_02_1 | 8614 | FTMS; HCD | 58.32 | 649.31 |

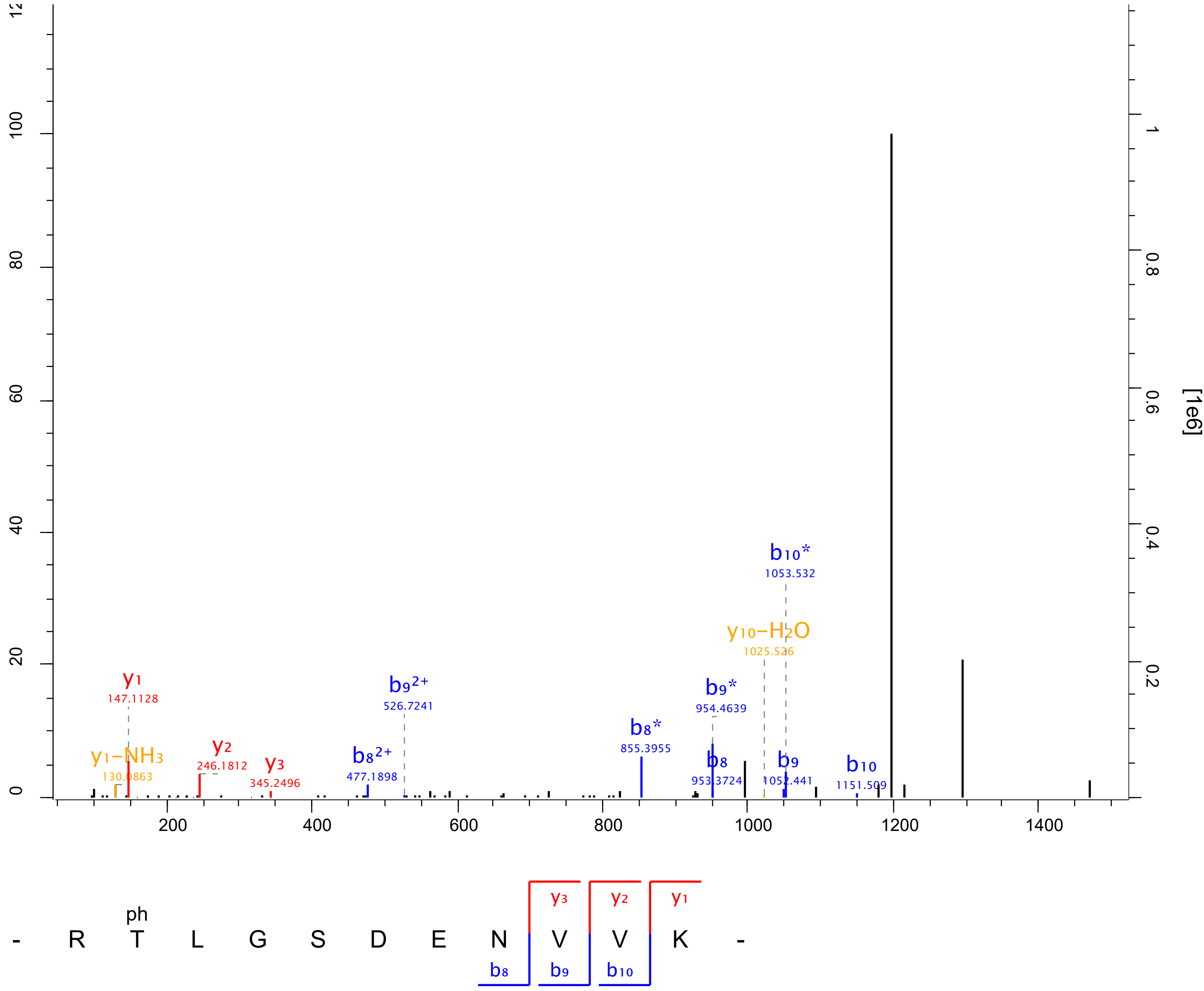

| Raw file | Scan | Method | Score | m/z |
|---|---|---|---|---|
| water_02_1 | 8628 | FTMS; HCD | 96.02 | 683.28 |

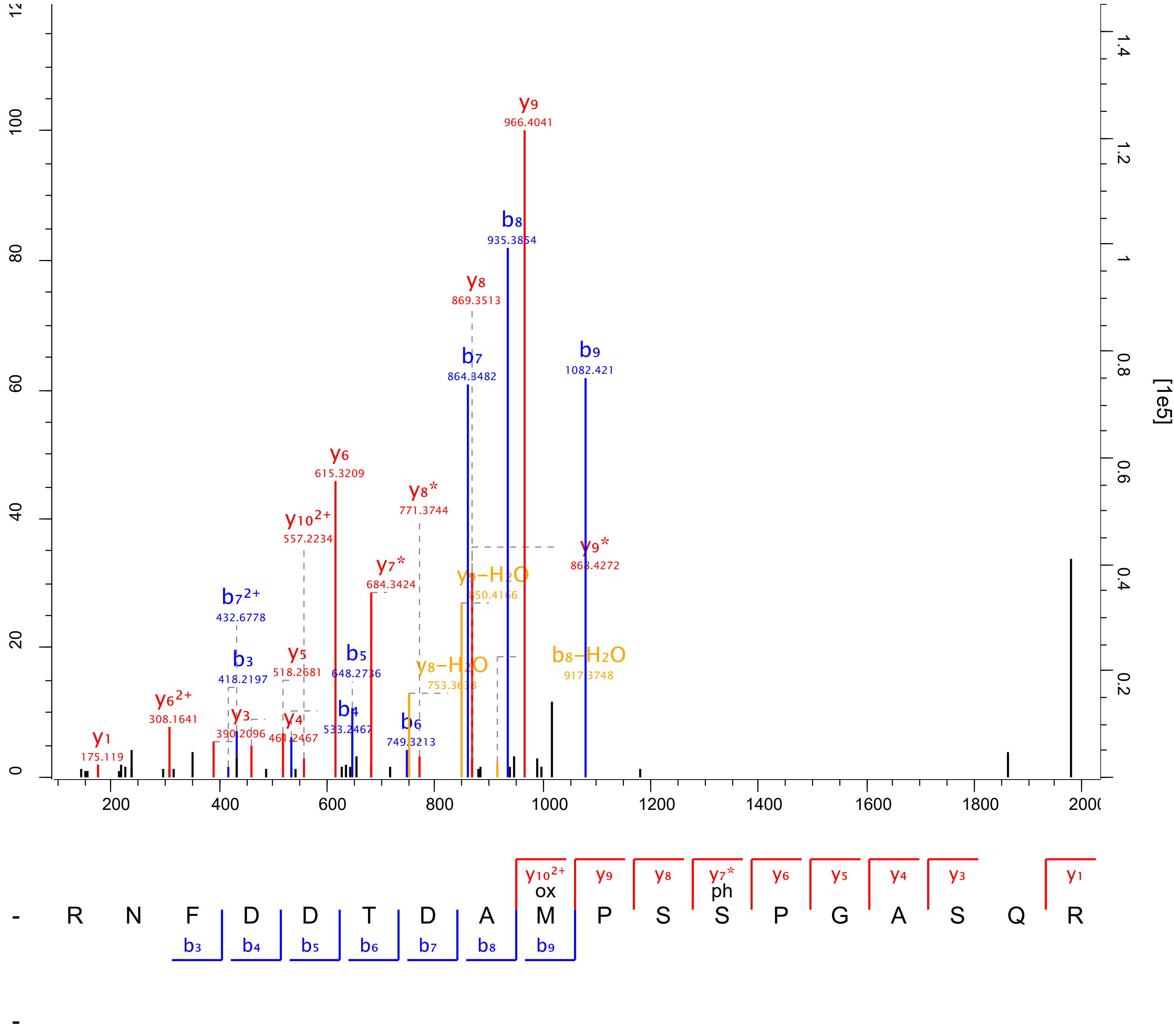

| Raw file | Scan | Method | Score | m/z |
| --- | --- | --- | --- | --- |
| water_02_1 | 8731 | FTMS; HCD | 142.76 | 588.74 |

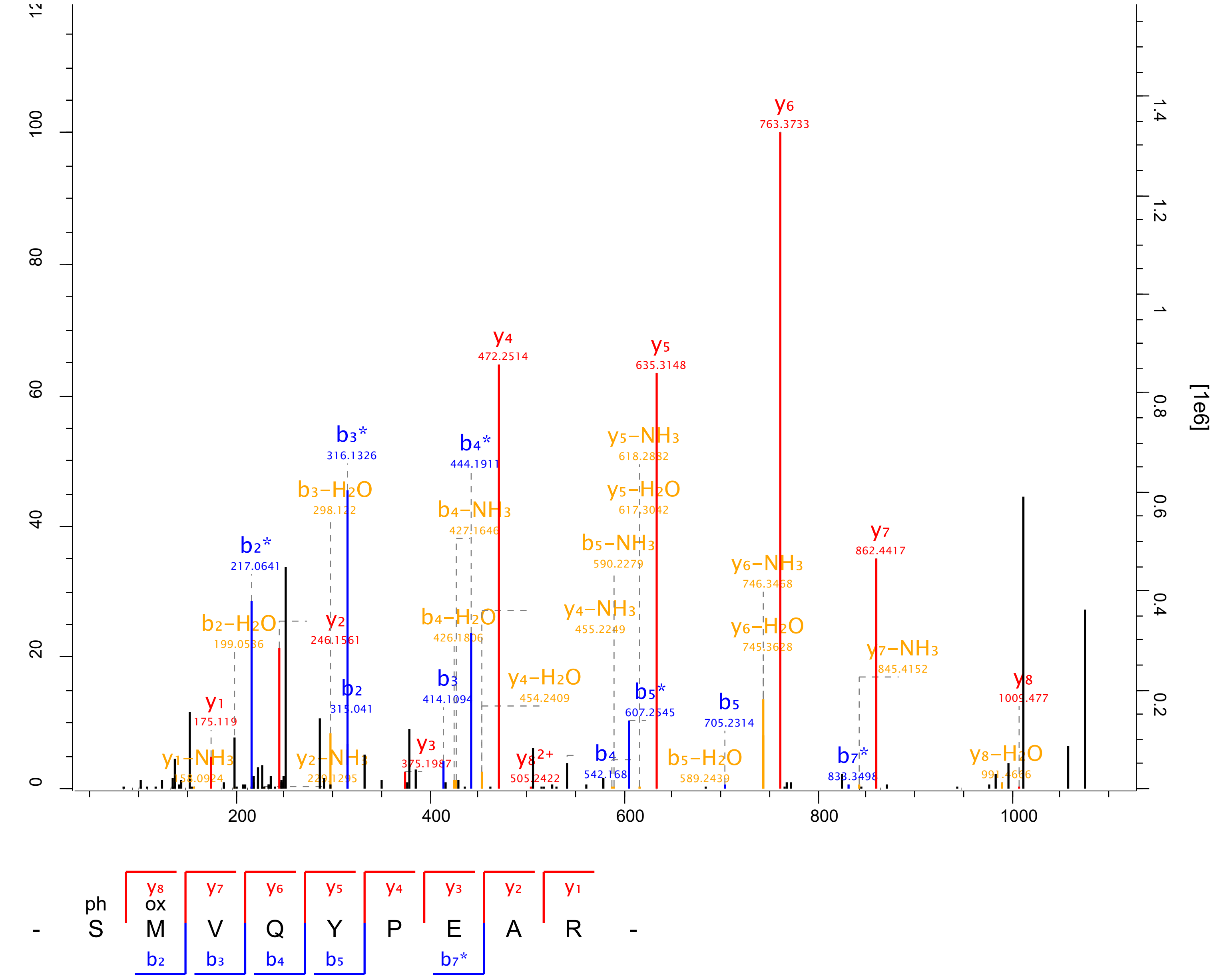

| Raw file | Scan | Method | Score | m/z |
|---|---|---|---|---|
| water_02_1 | 8732 | FTMS; HCD | 133.52 | 669.84 |

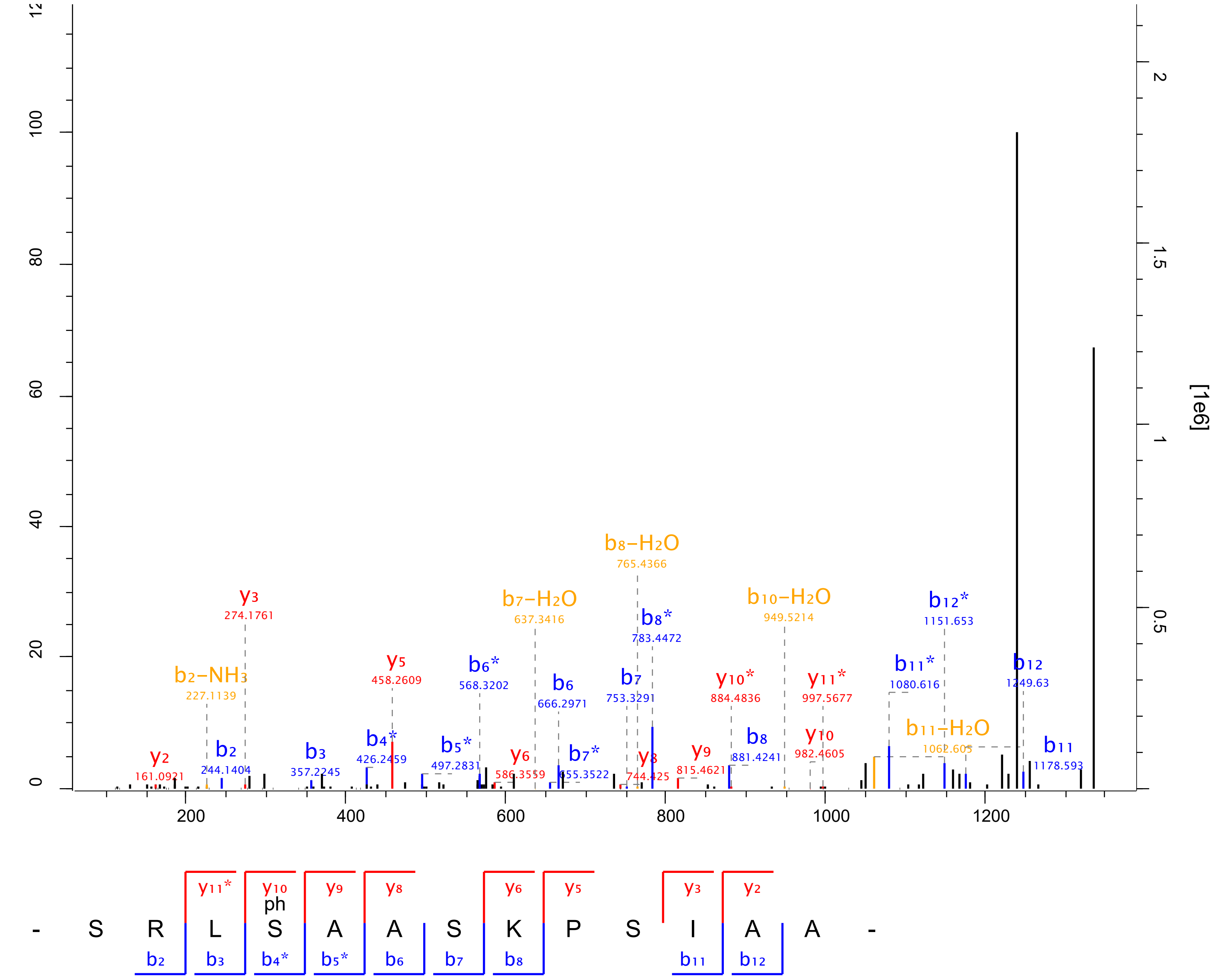

| Raw file | Scan | Method | Score | m/z |
|---|---|---|---|---|
| water_02_1 | 8742 | FTMS; HCD | 97.27 | 780.35 |

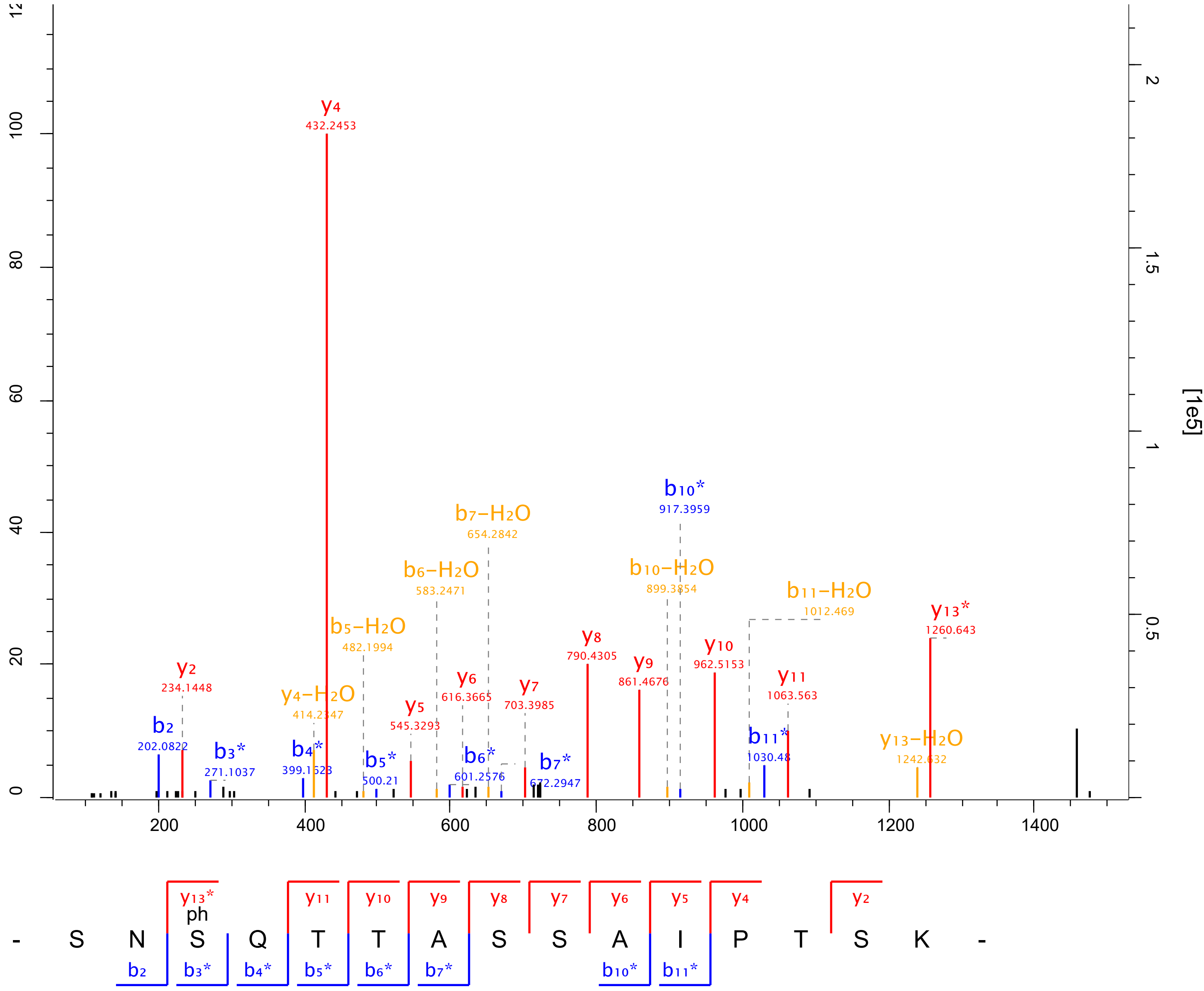

| Raw file | Scan | Method | Score | m/z |
|---|---|---|---|---|
| water_02_1 | 8801 | FTMS; HCD | 124.77 | 889.34 |

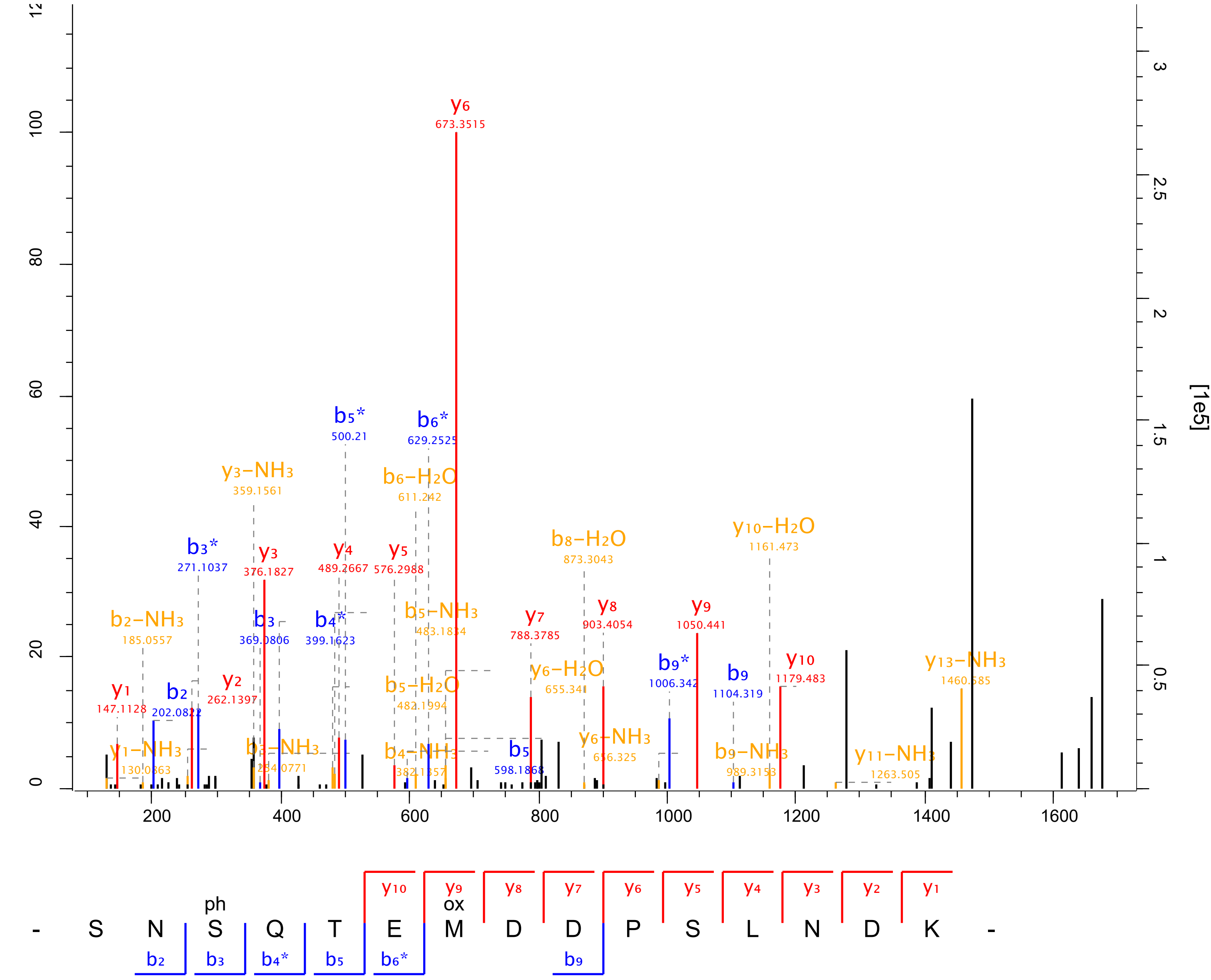

| Raw file | Scan | Method | Score | m/z |
| --- | --- | --- | --- | --- |
| water_02_1 | 8862 | FTMS; HCD | 53.75 | 493.72 |

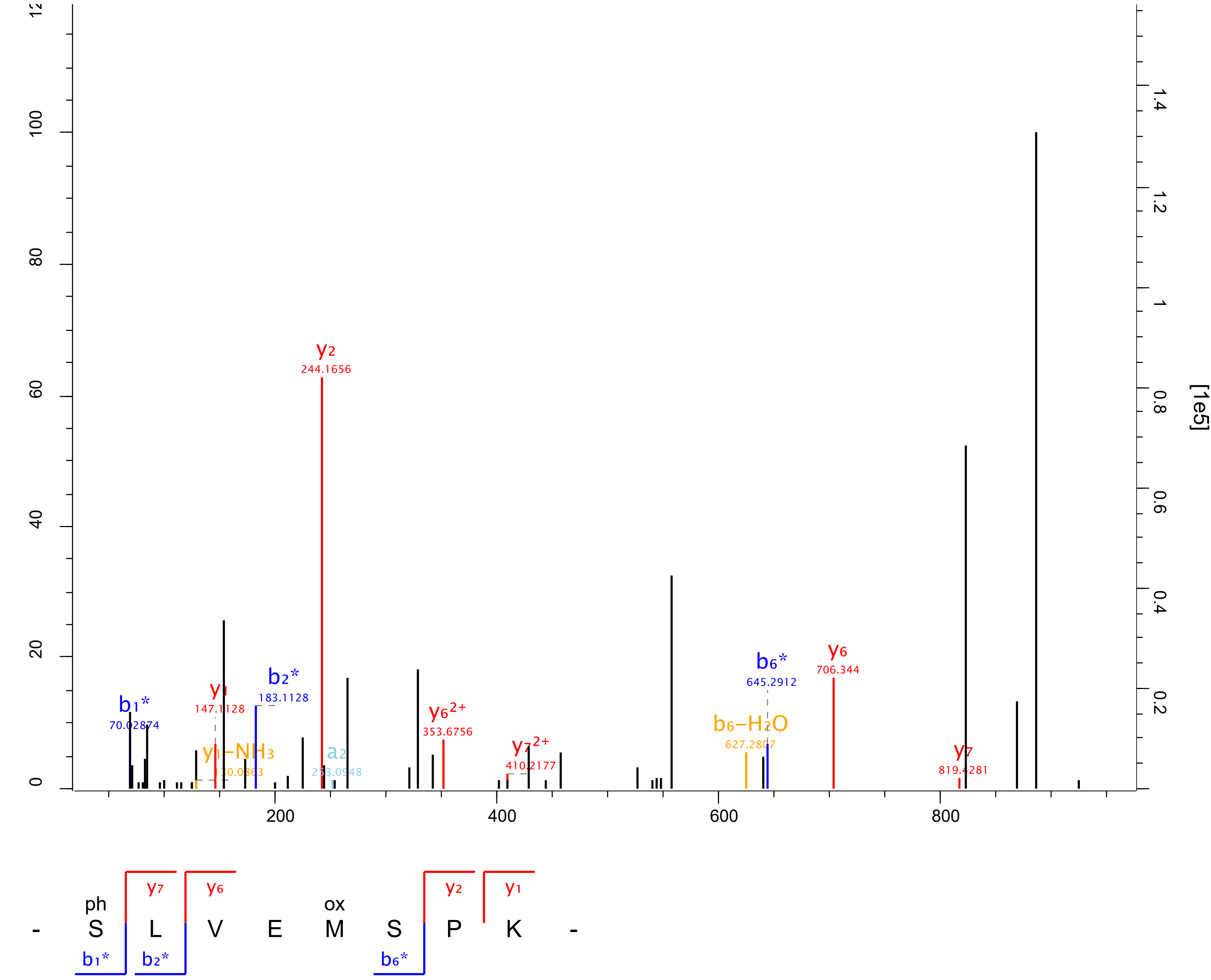

| Raw file | Scan | Method | Score | m/z |
|---|---|---|---|---|
| water_02_1 | 8997 | FTMS; HCD | 43.95 | 594.62 |

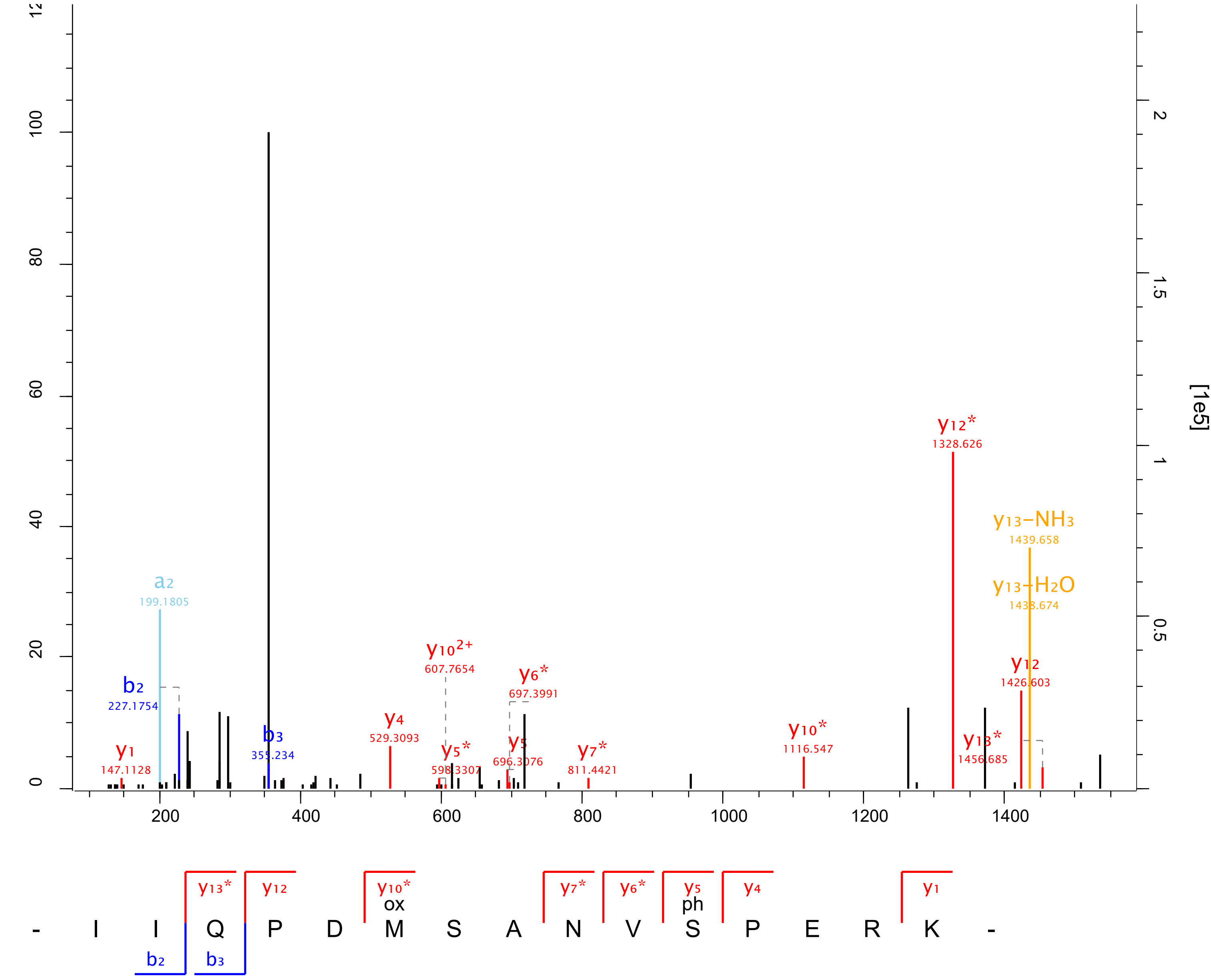

| Raw file | Scan | Method | Score | m/z |
|---|---|---|---|---|
| water_02_1 | 9005 | FTMS; HCD | 40.53 | 762.32 |

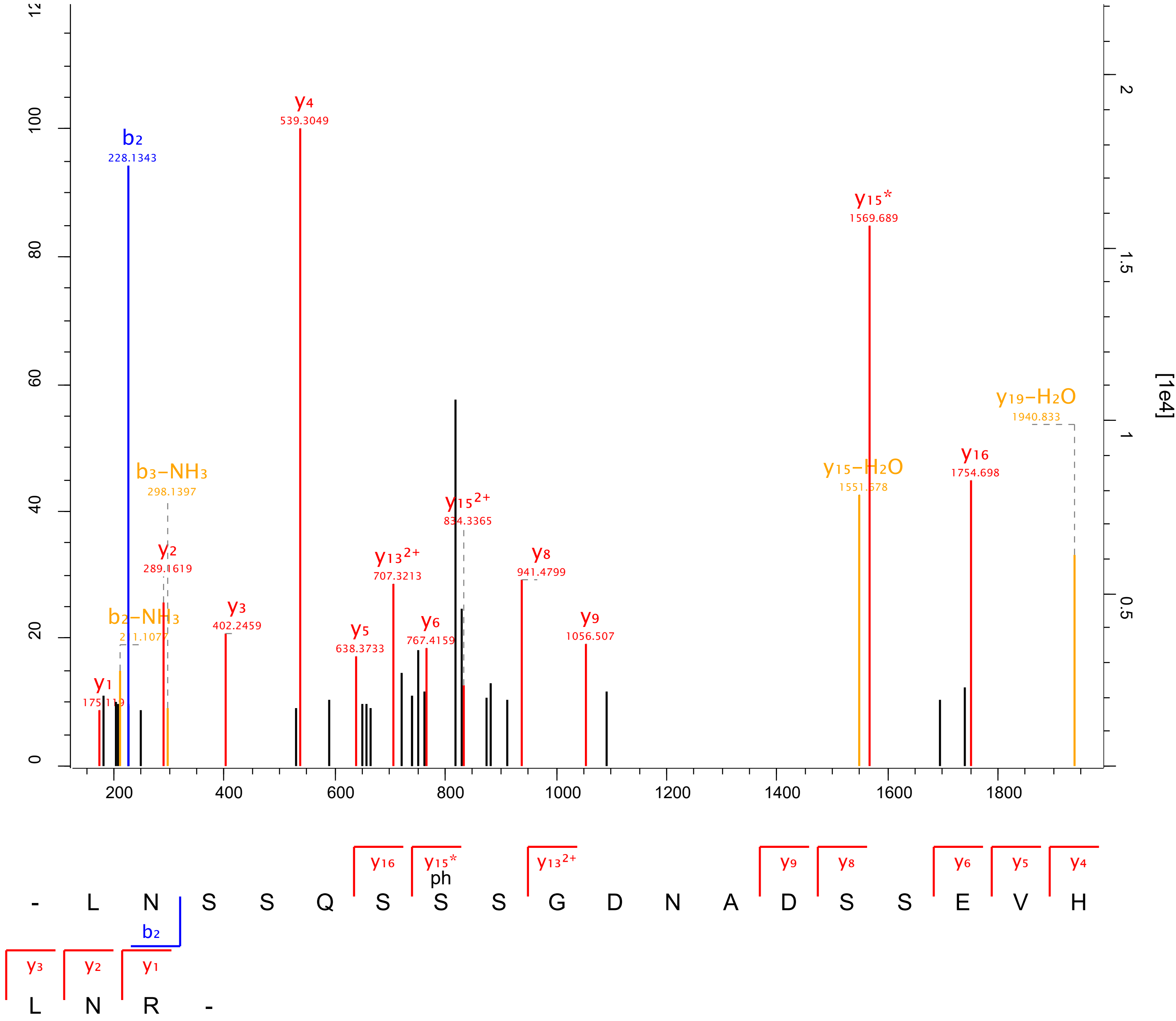

| Raw file | Scan | Method | Score | m/z |
|---|---|---|---|---|
| water_02_1 | 9025 | FTMS; HCD | 94.45 | 652.77 |

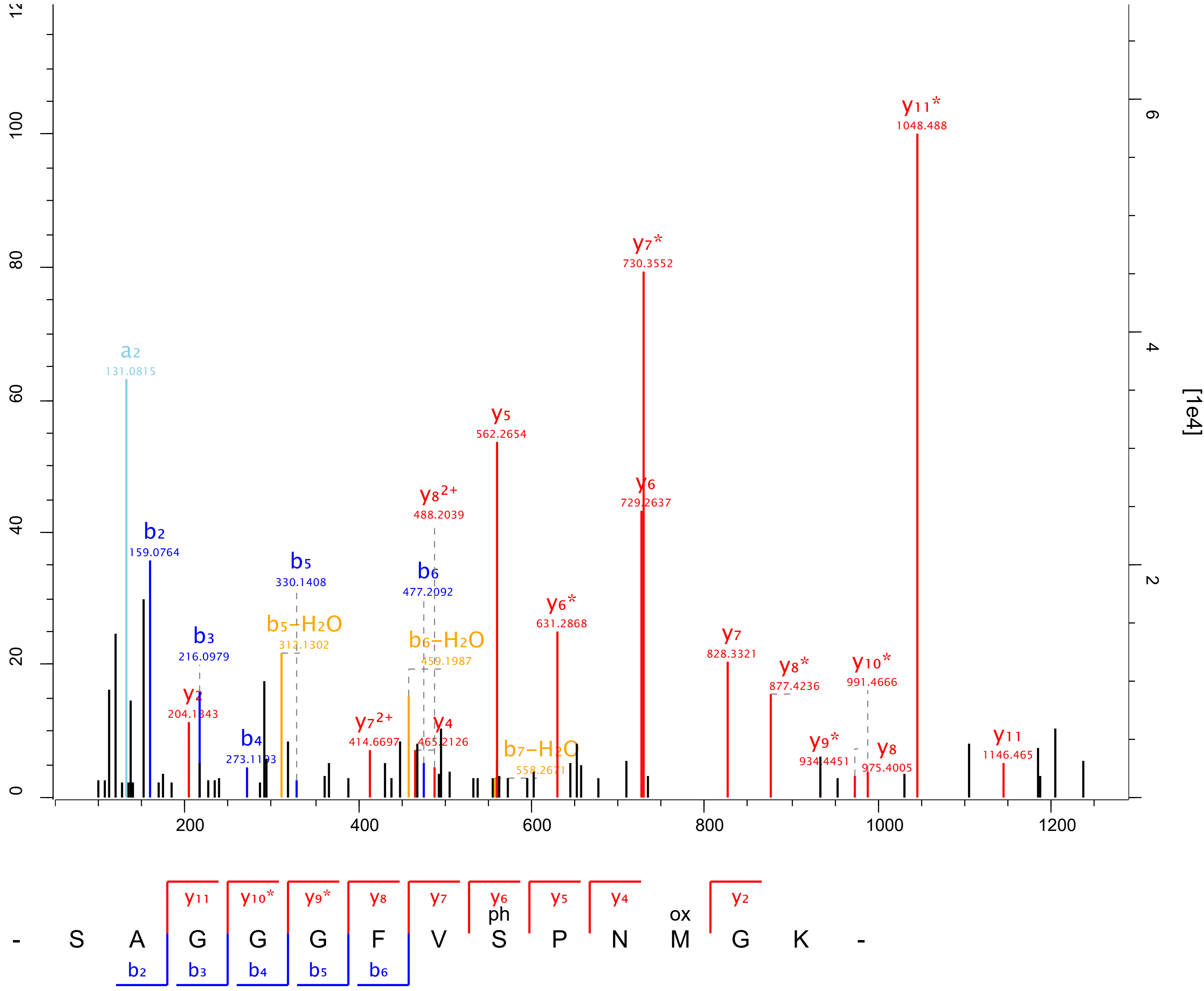

| Raw file | Scan | Method | Score | m/z |
|---|---|---|---|---|
| water_02_1 | 9078 | FTMS; HCD | 63.37 | 685.82 |

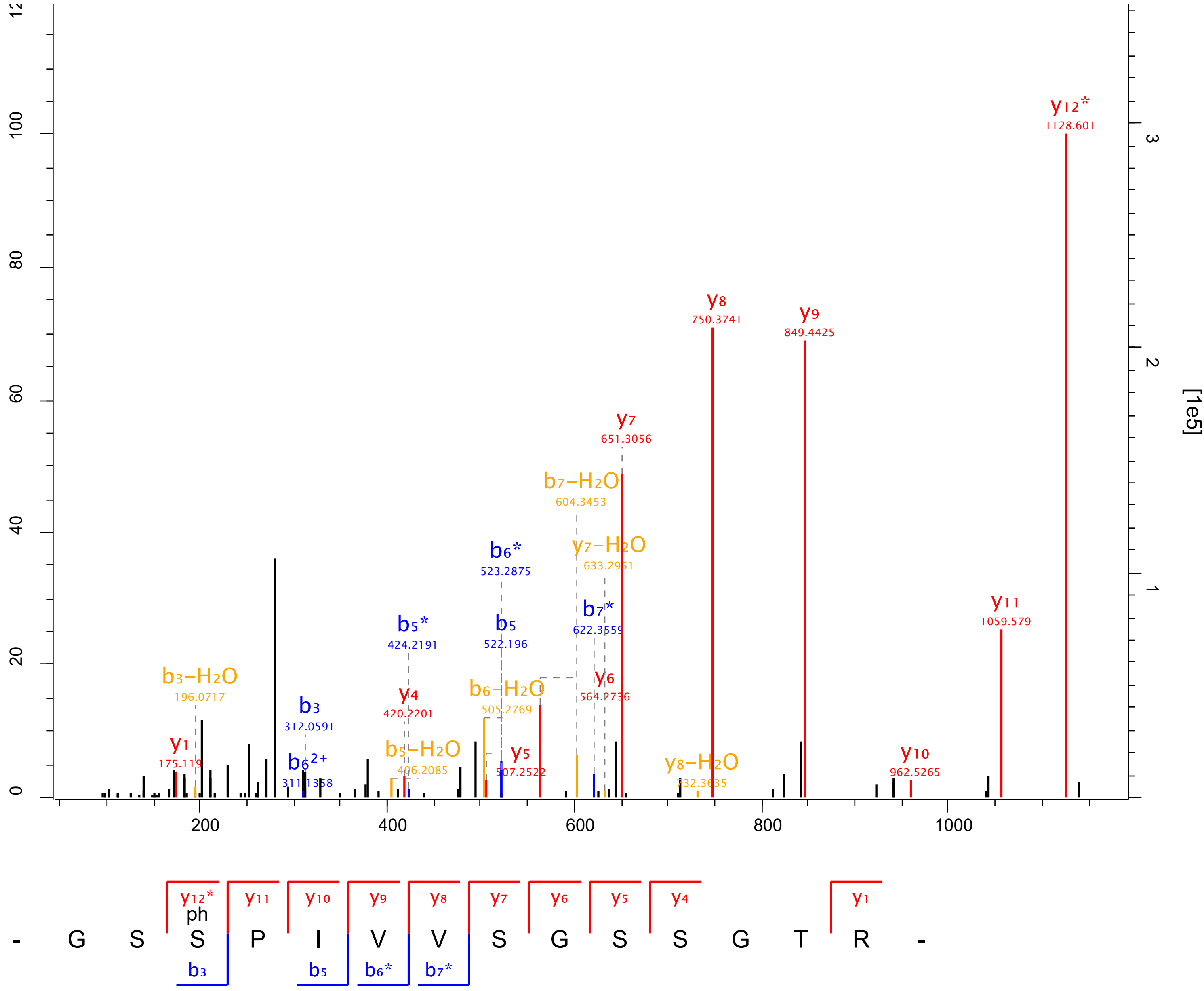

| Raw file | Scan | Method | Score | m/z |
| --- | --- | --- | --- | --- |
| water_02_1 | 9082 | FTMS; HCD | 71.67 | 856.03 |

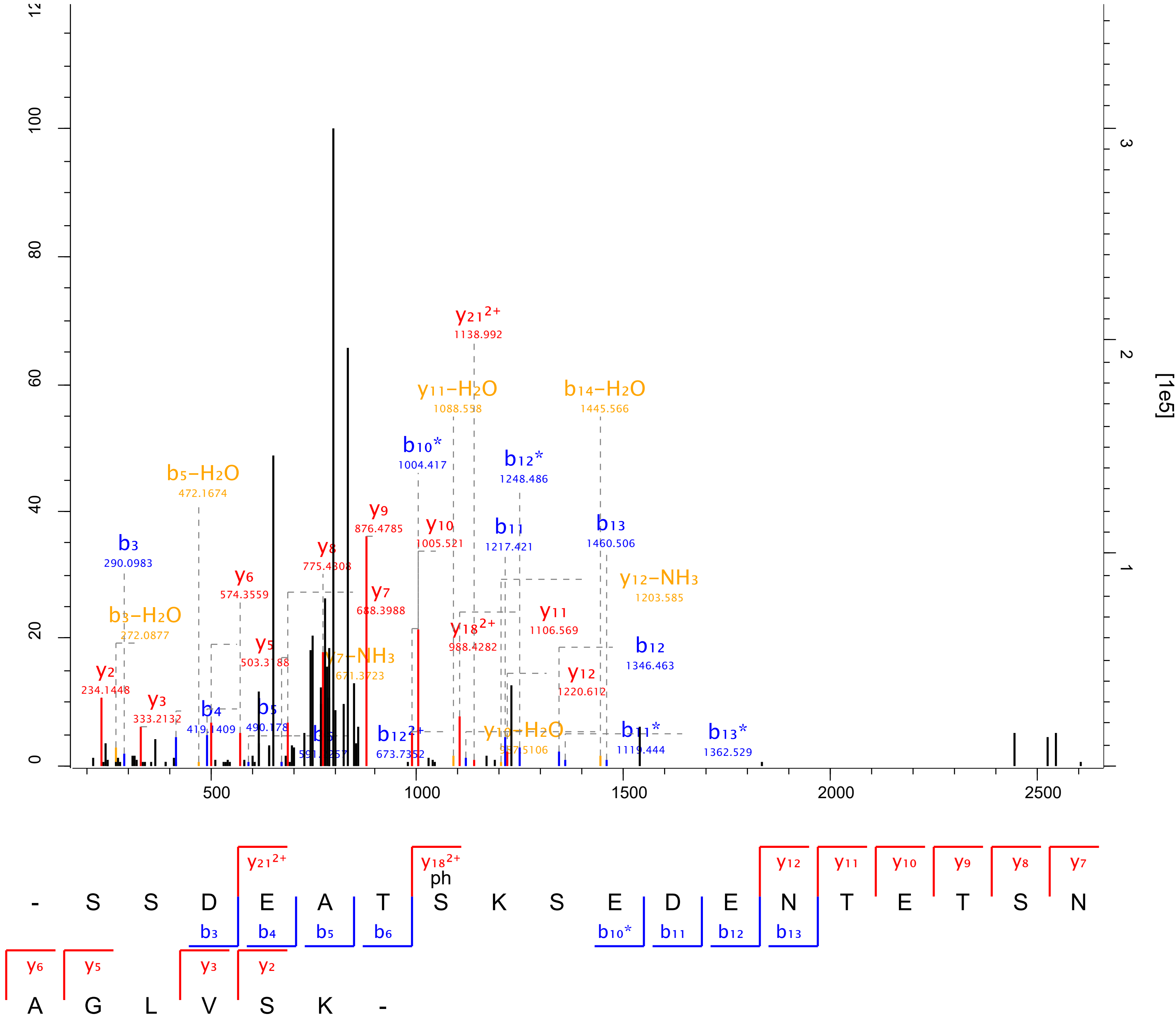

| Raw file | Scan | Method | Score | m/z |
|---|---|---|---|---|
| water_02_1 | 9102 | FTMS; HCD | 182.61 | 777.79 |

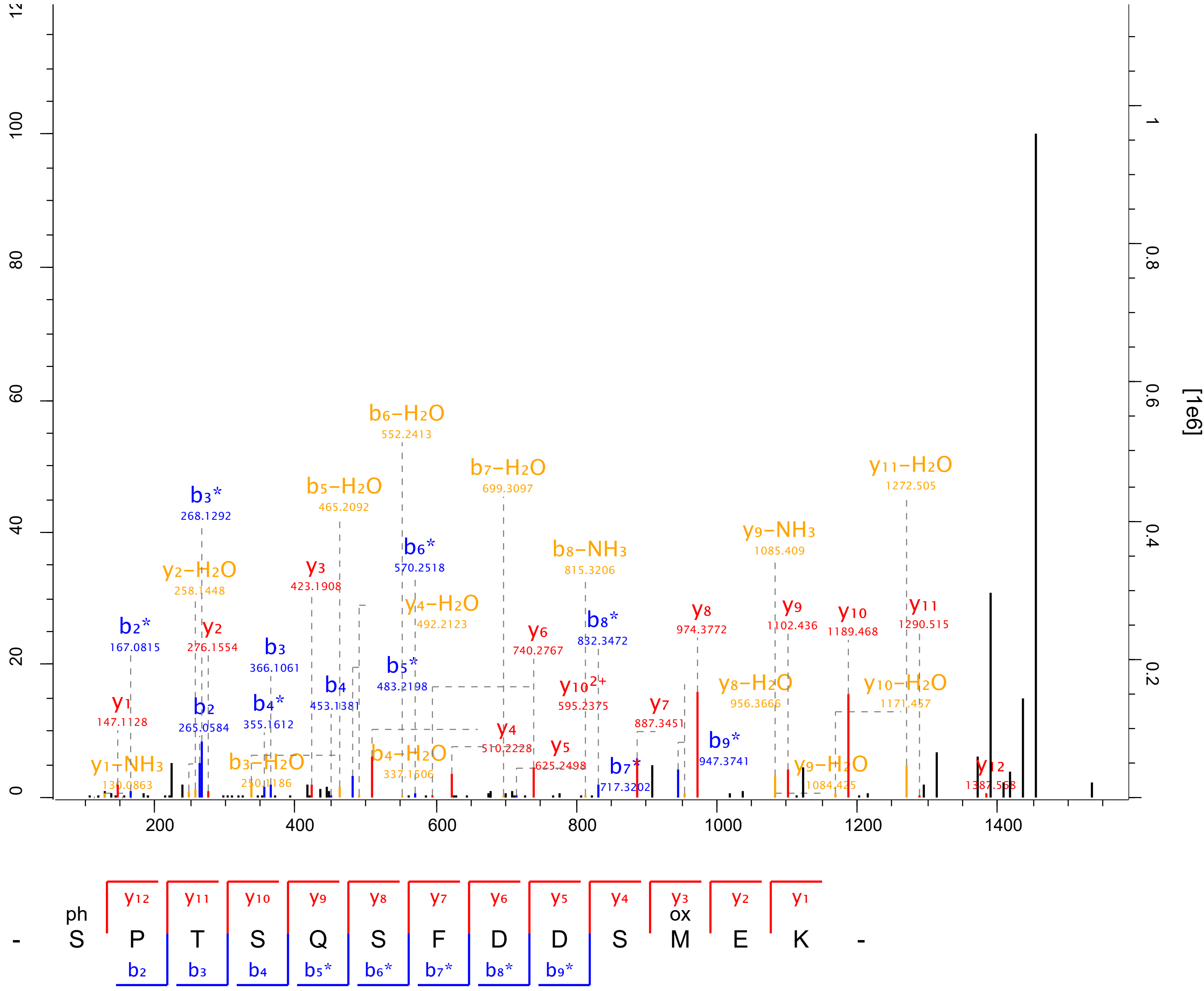

| Raw file | Scan | Method | Score | m/z |
|---|---|---|---|---|
| water_02_1 | 9113 | FTMS; HCD | 128.22 | 733.24 |

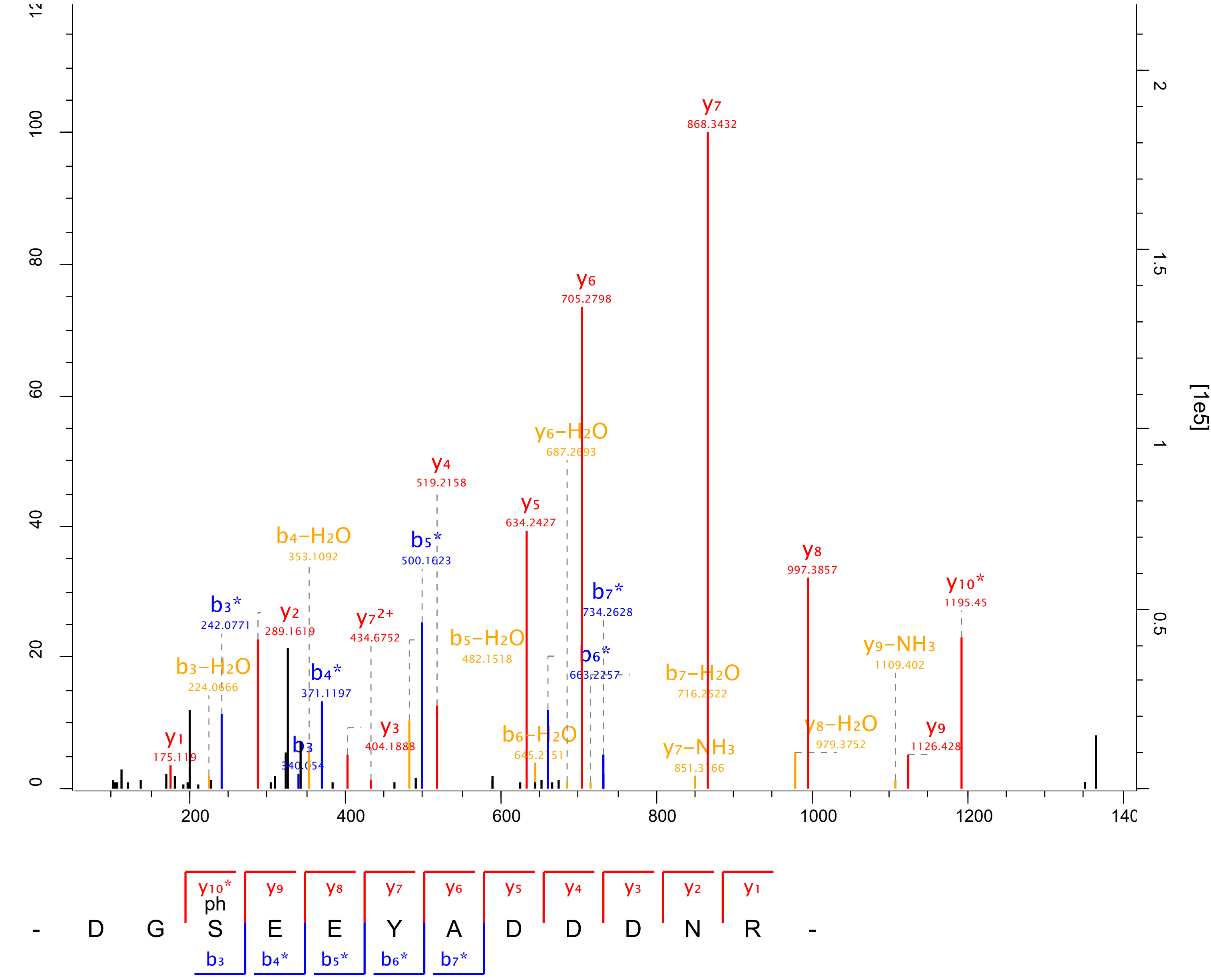

| Raw file | Scan | Method | Score | m/z |
|---|---|---|---|---|
| water_02_1 | 9116 | FTMS; HCD | 48.9 | 716.31 |

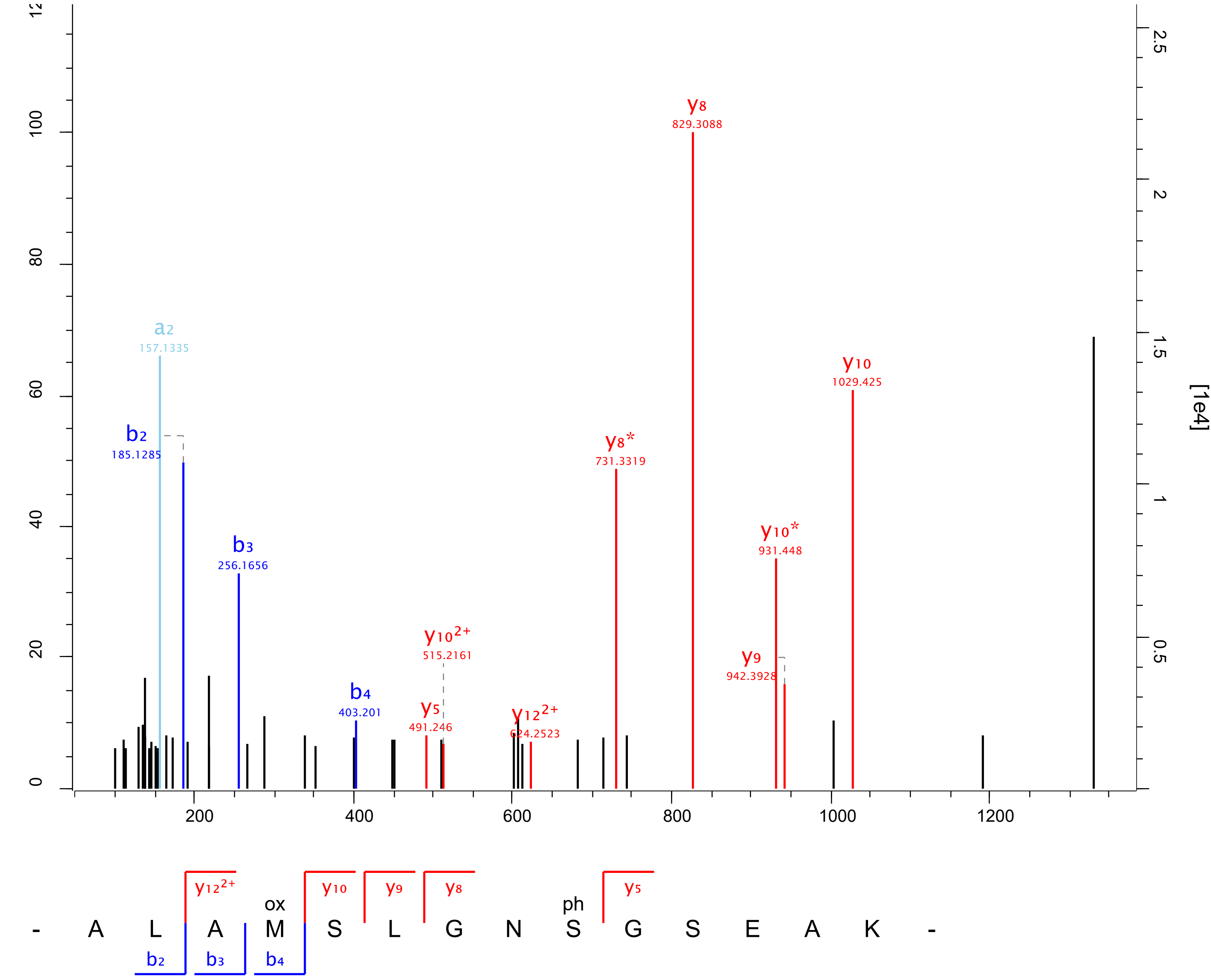

| Raw file | Scan | Method | Score | m/z |
|---|---|---|---|---|
| water_02_1 | 9149 | FTMS; HCD | 227.46 | 763.85 |

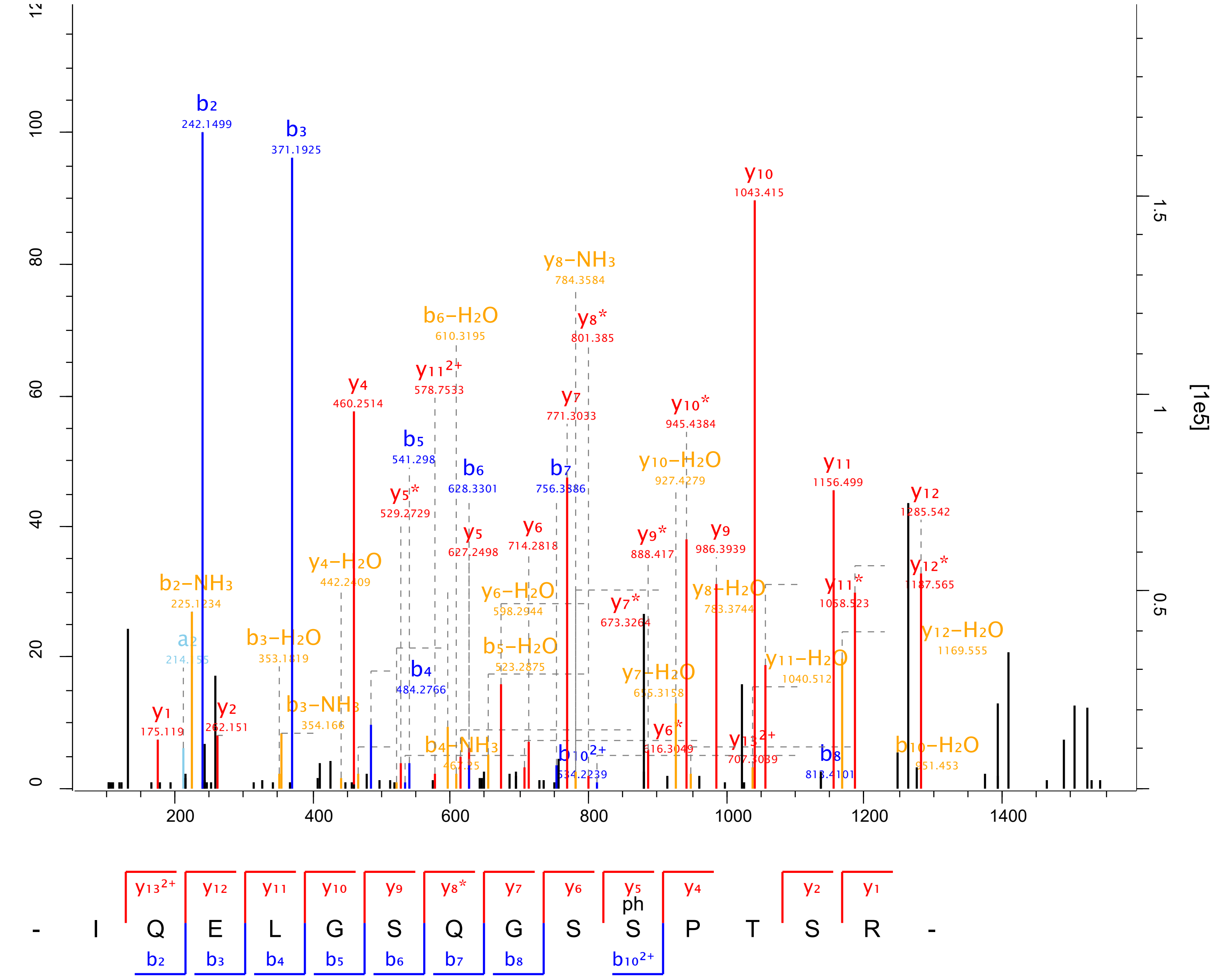

| Raw file | Scan | Method | Score | m/z |
| --- | --- | --- | --- | --- |
| water_02_1 | 9173 | FTMS; HCD | 129.11 | 557.24 |

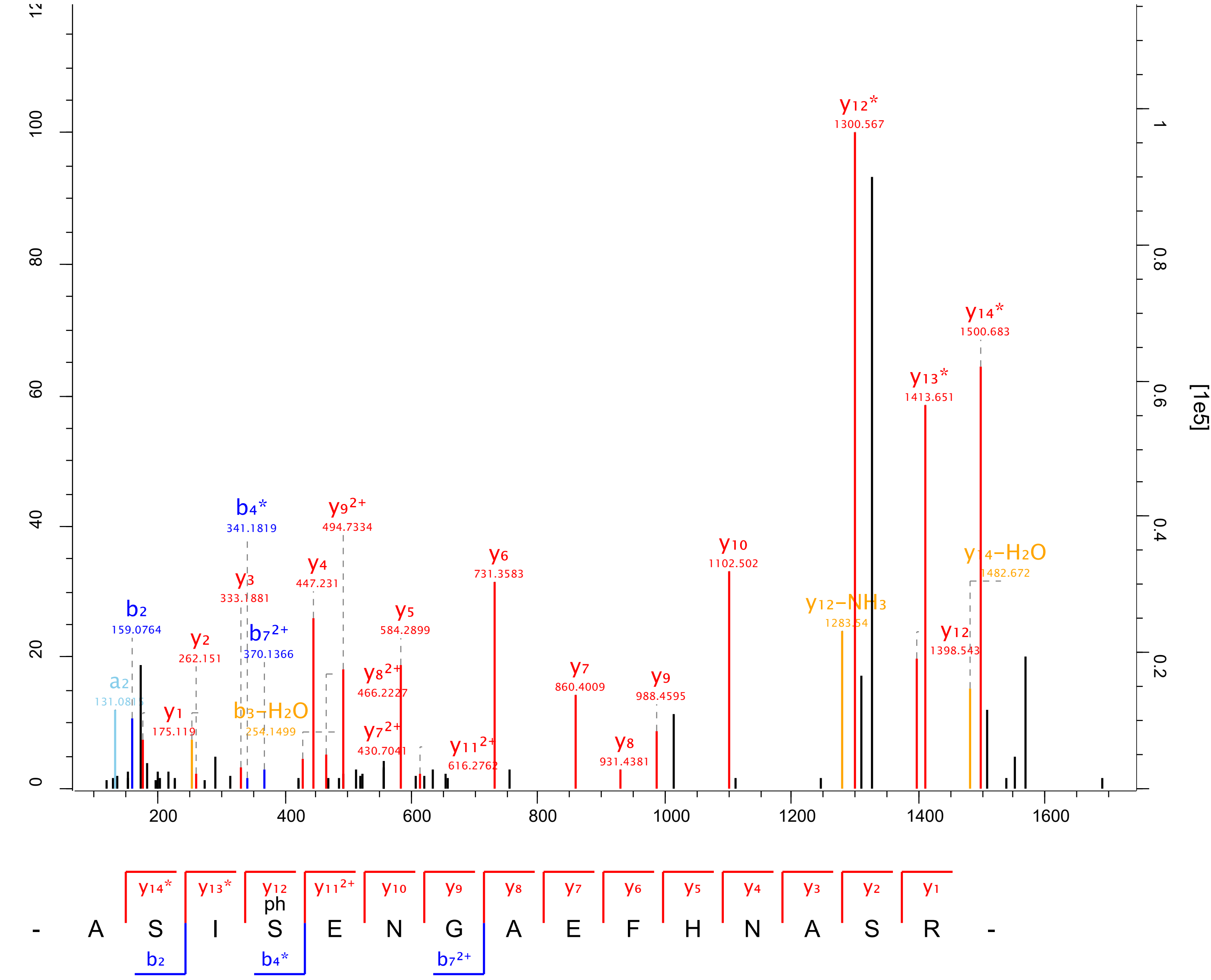

| Raw file | Scan | Method | Score | m/z |
|---|---|---|---|---|
| water_02_1 | 9236 | FTMS; HCD | 260.14 | 676.3 |

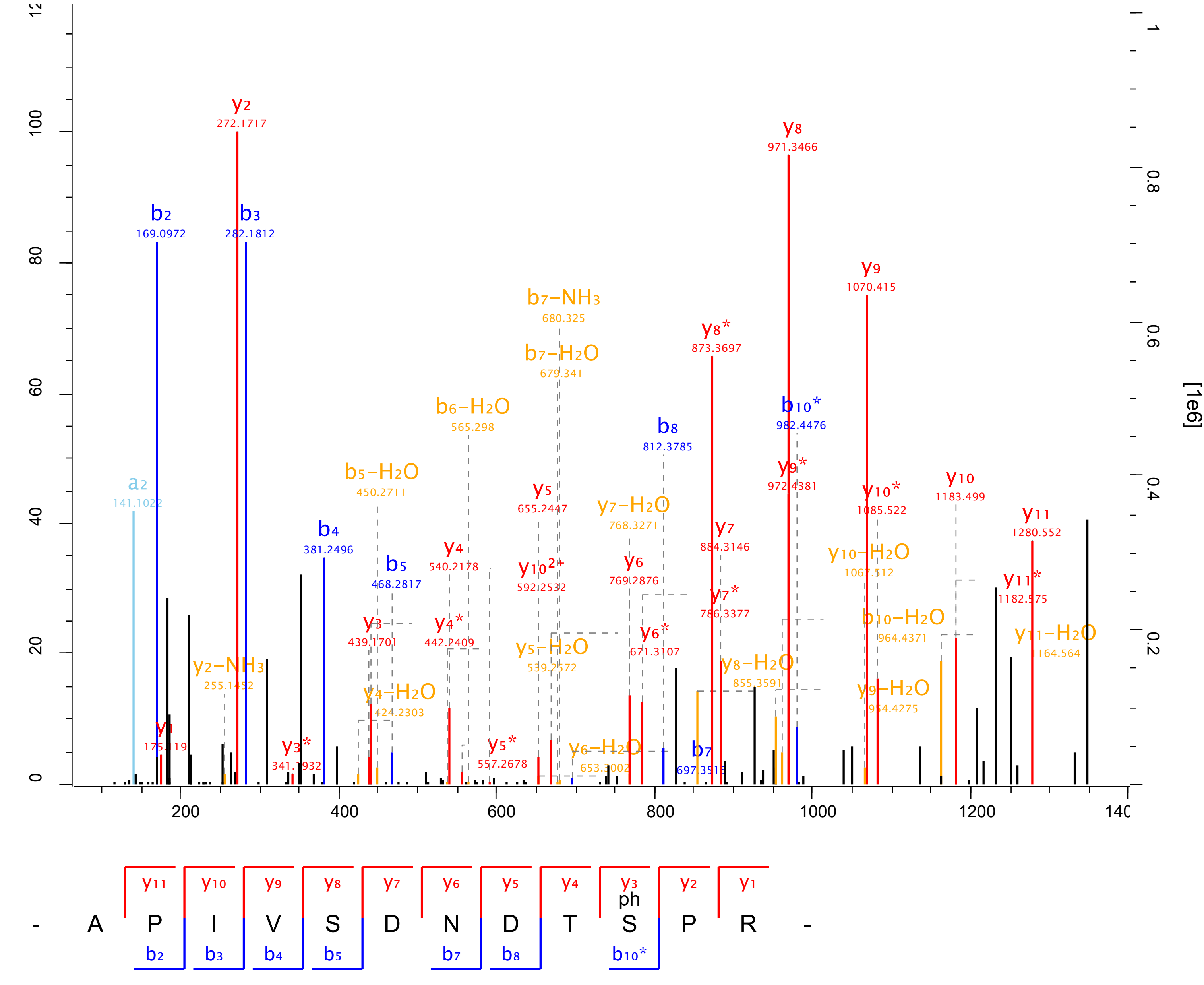

| Raw file | Scan | Method | Score | m/z |
|---|---|---|---|---|
| water_02_1 | 9275 | FTMS; HCD | 112.08 | 621.28 |

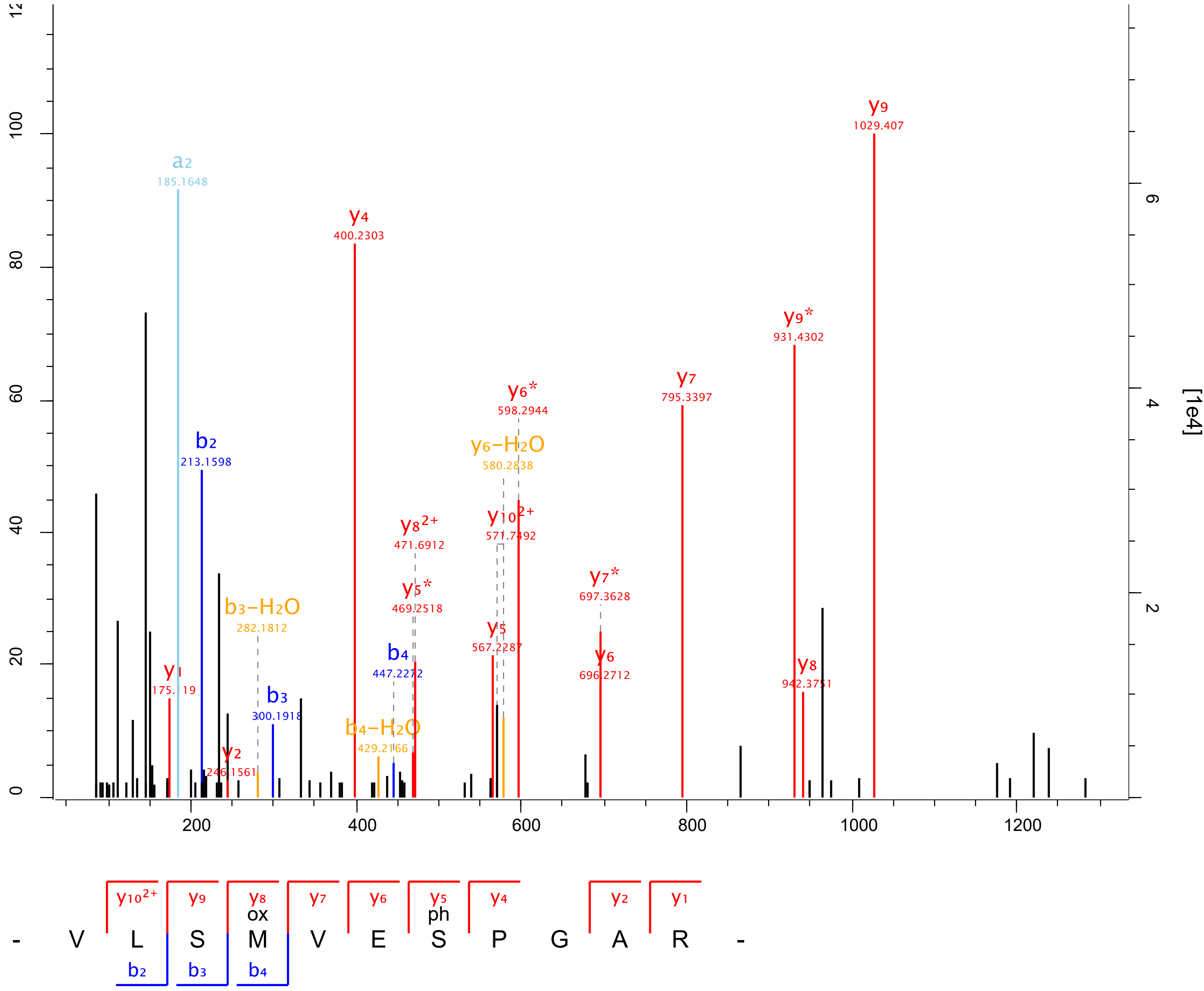

| Raw file | Scan | Method | Score | m/z |
|---|---|---|---|---|
| water_02_1 | 9297 | FTMS; HCD | 67.65 | 690.26 |

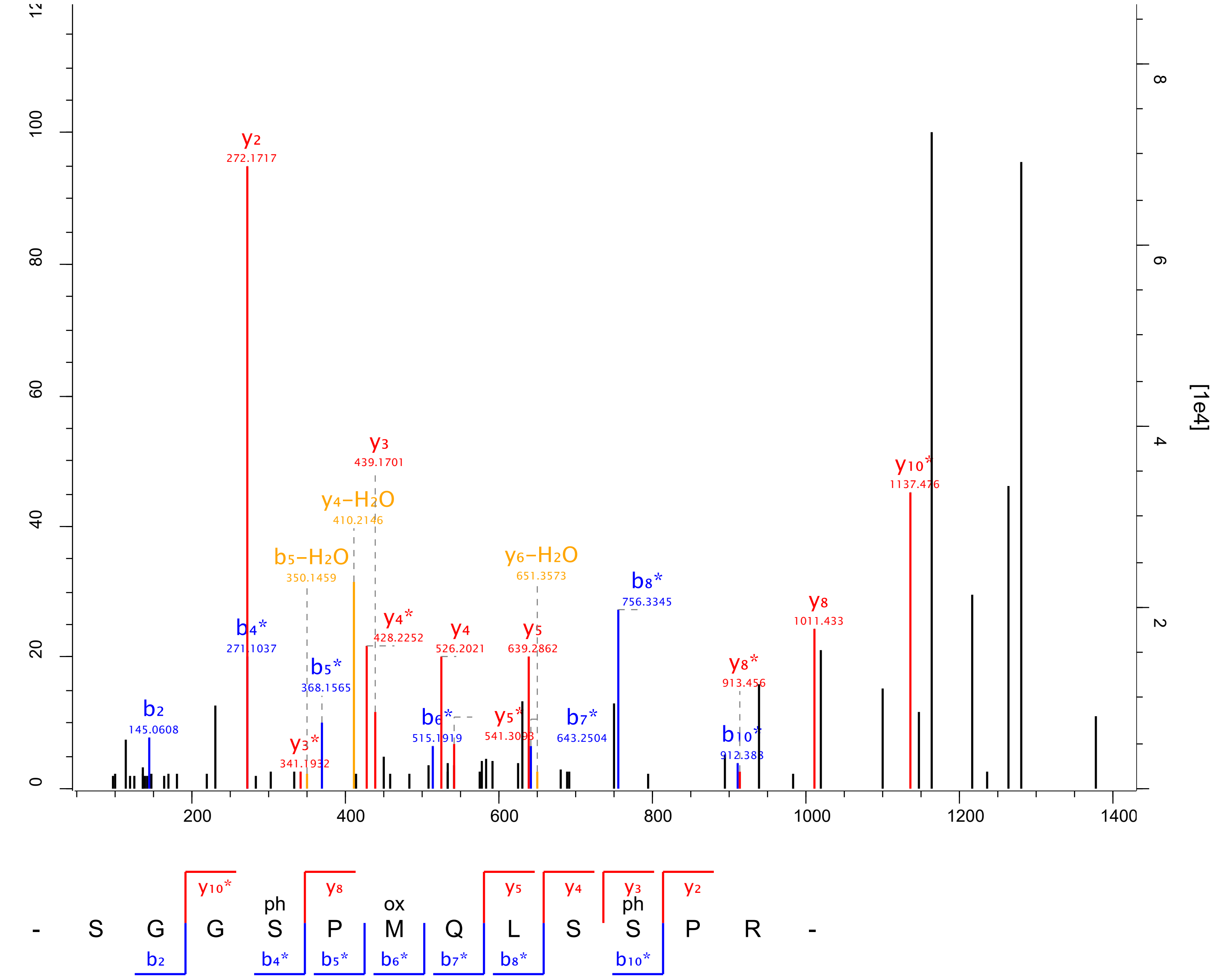

| Raw file | Scan | Method | Score | m/z |
|---|---|---|---|---|
| water_02_1 | 9375 | FTMS; HCD | 62.29 | 604.24 |

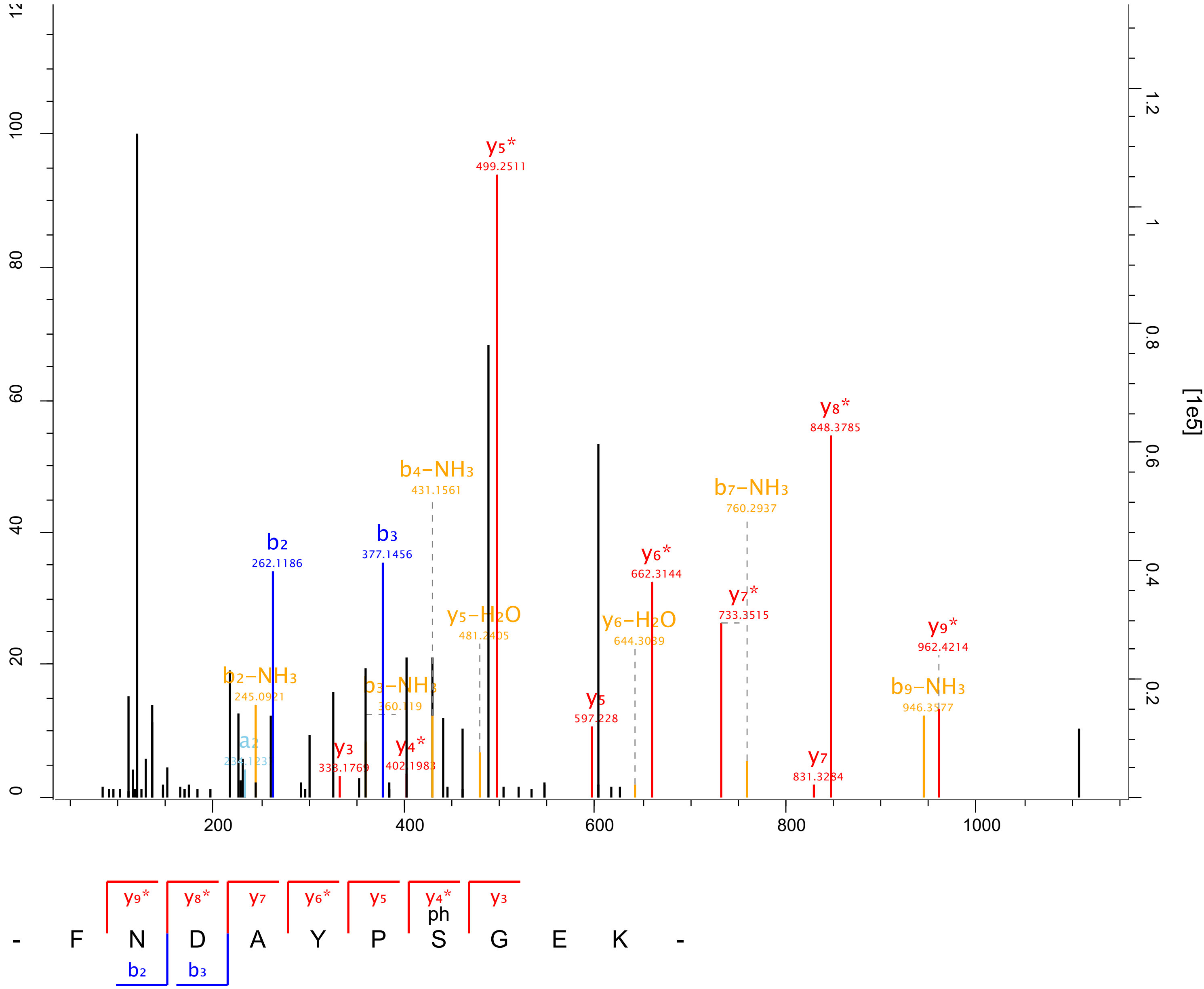

| Raw file | Scan | Method | Score | m/z |
|---|---|---|---|---|
| water_02_1 | 9382 | FTMS; HCD | 87.68 | 624.93 |

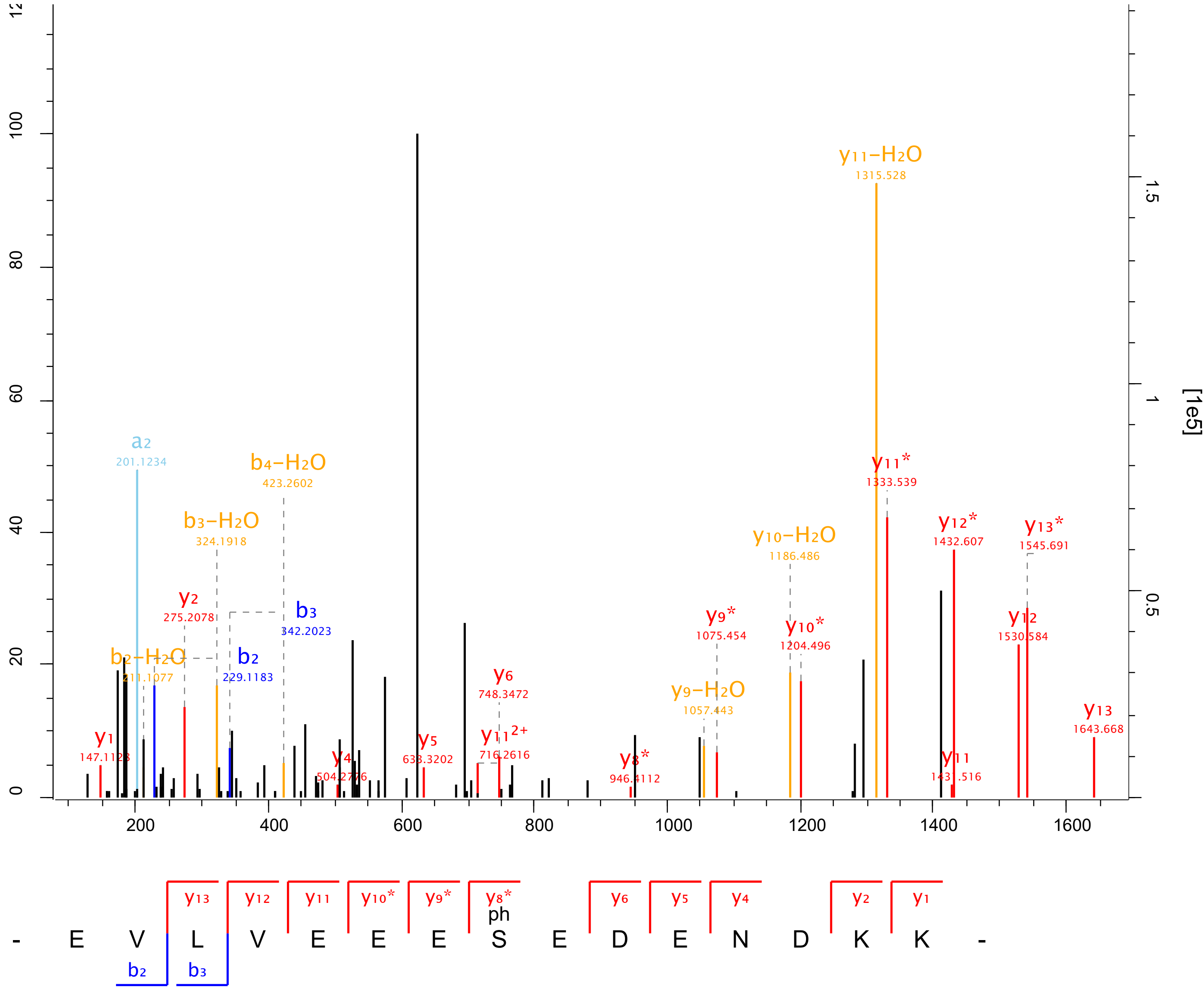

| Raw file | Scan | Method | Score | m/z |
|---|---|---|---|---|
| water_02_1 | 9409 | FTMS; HCD | 78.9 | 627.24 |

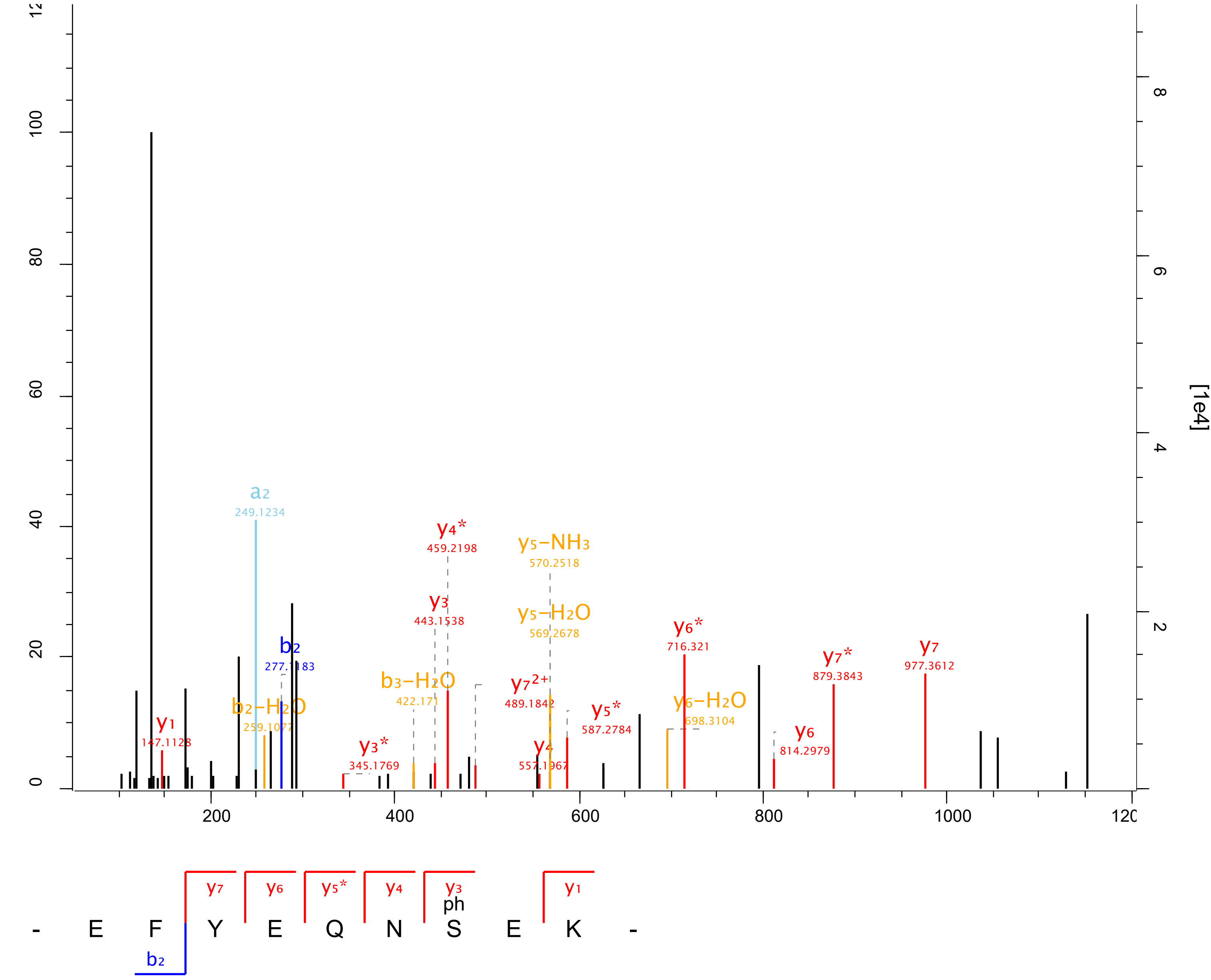

| Raw file | Scan | Method | Score | m/z |
|---|---|---|---|---|
| water_02_1 | 9448 | FTMS; HCD | 45.65 | 574.24 |

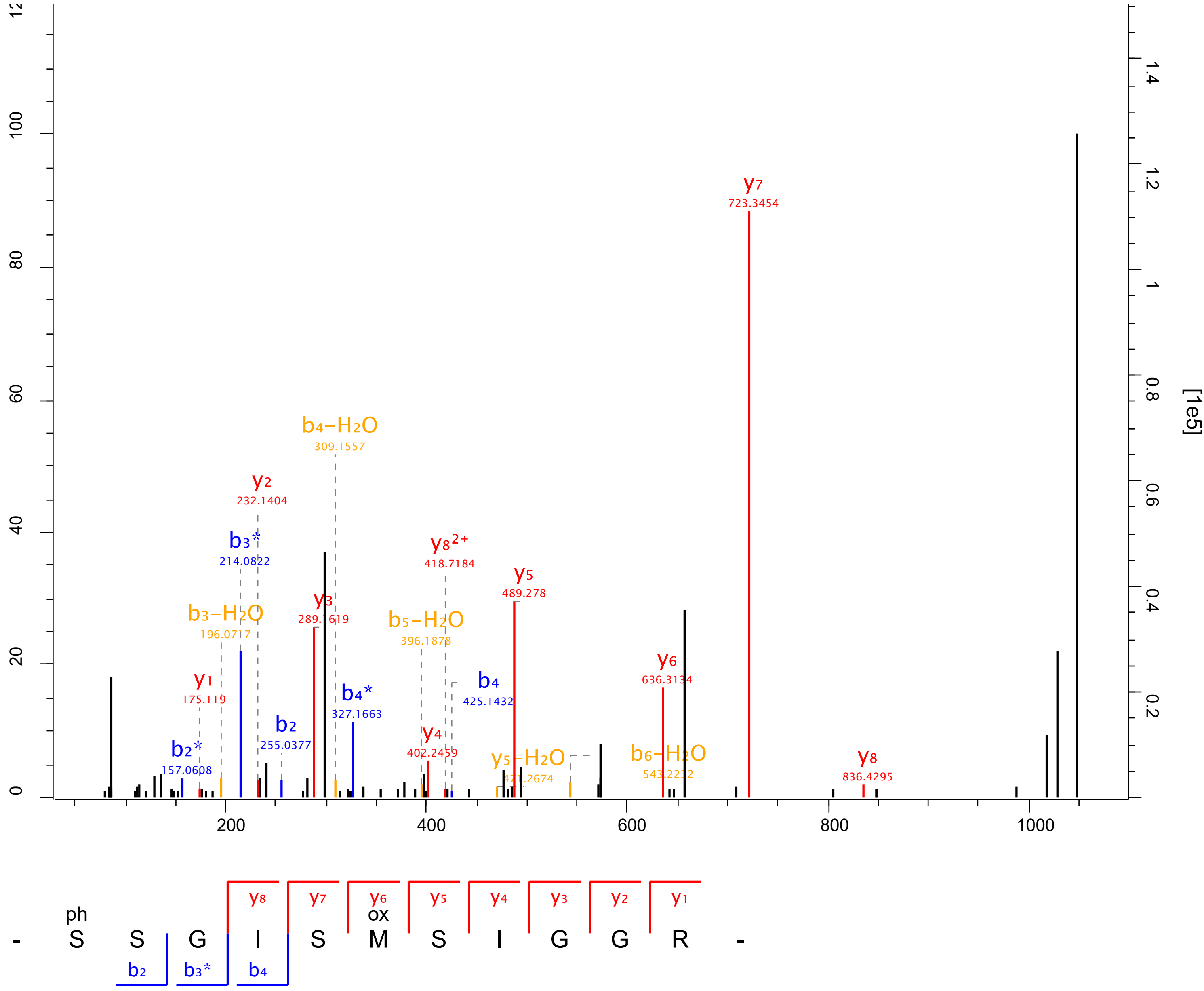

| Raw file | Scan | Method | Score | m/z |
|---|---|---|---|---|
| water_02_1 | 9487 | FTMS; HCD | 71.15 | 982.4 |

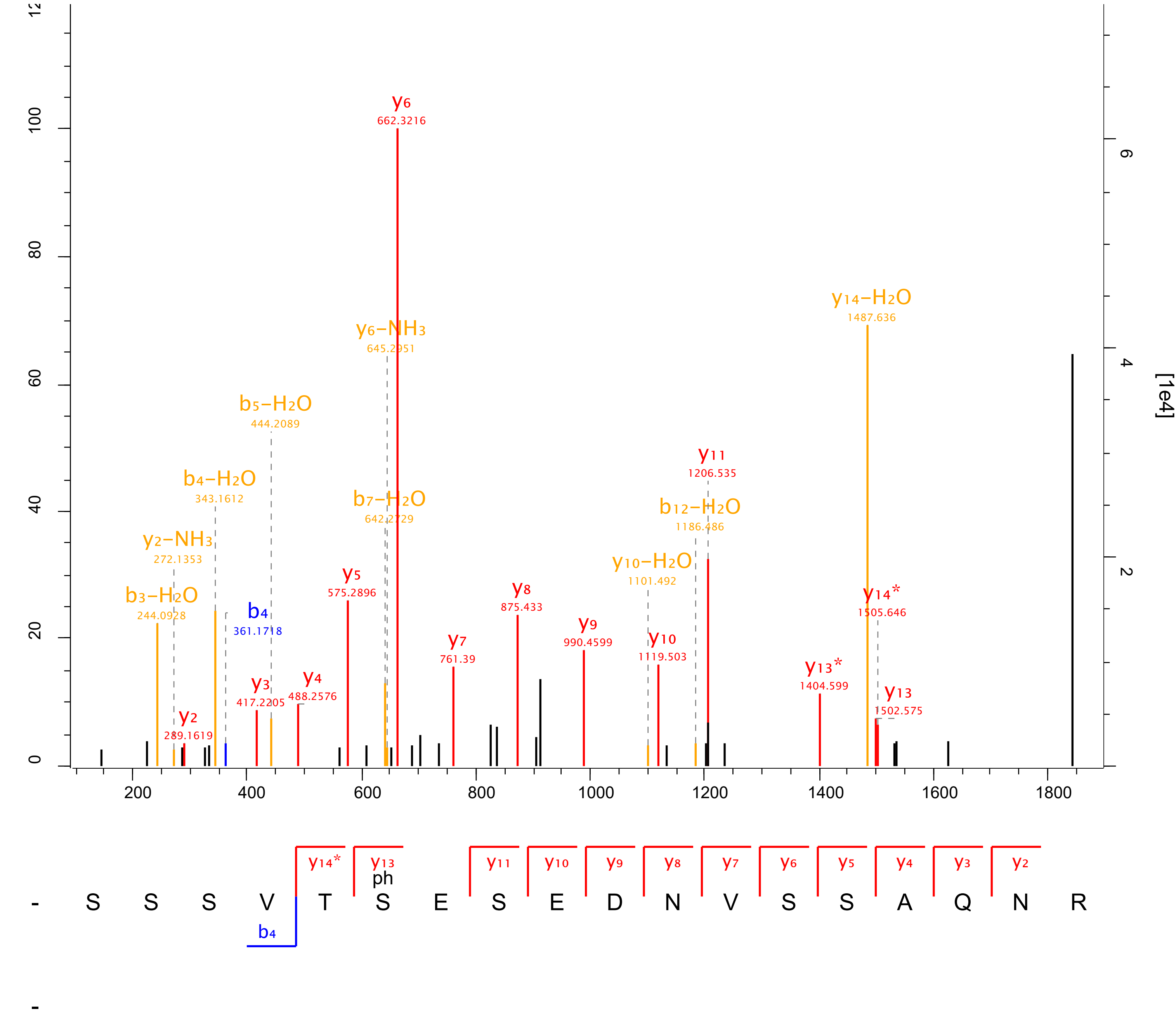

| Raw file | Scan | Method | Score | m/z |
|---|---|---|---|---|
| water_02_1 | 9595 | FTMS; HCD | 103.88 | 449.57 |

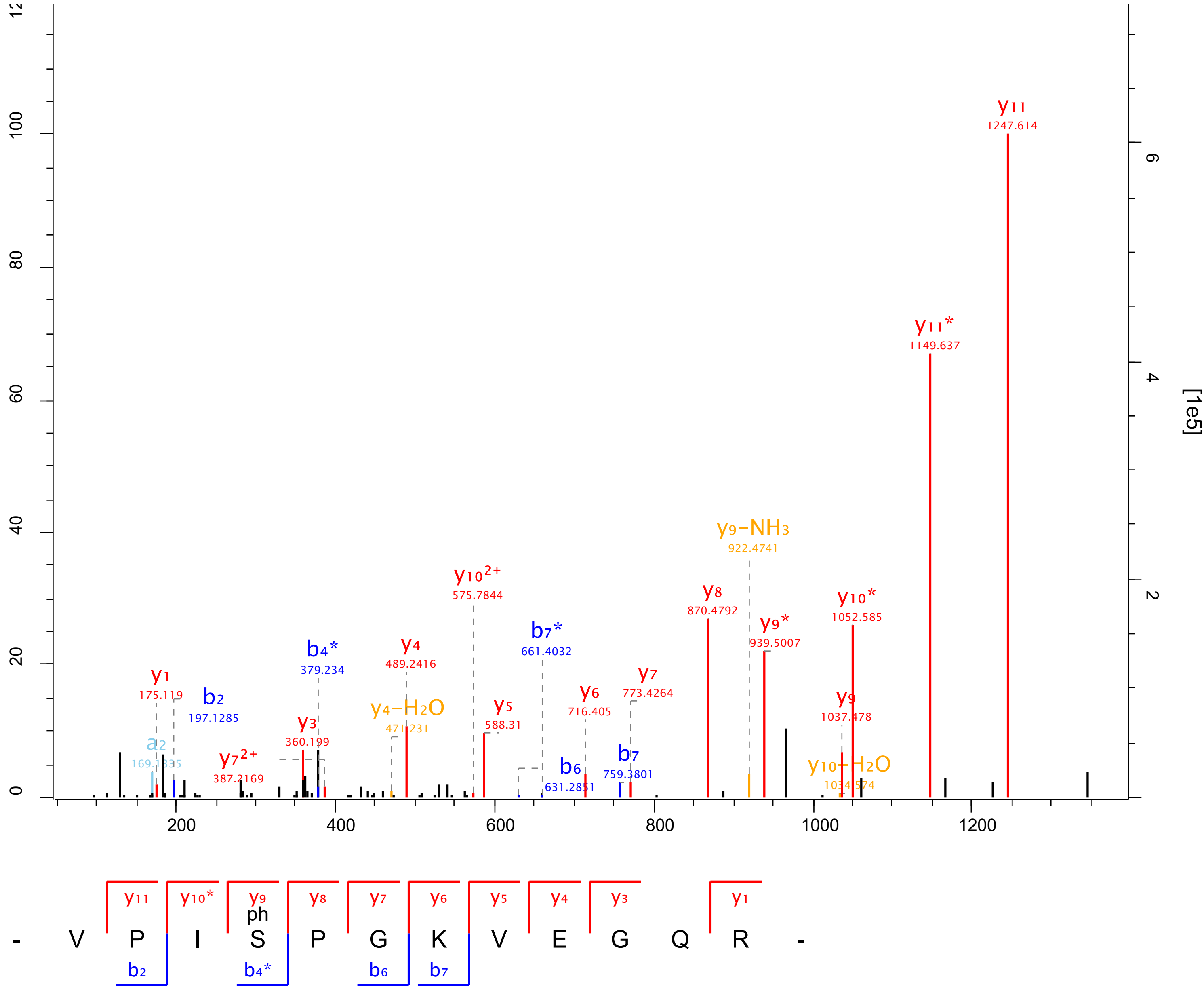

| Raw file | Scan | Method | Score | m/z |
| --- | --- | --- | --- | --- |
| water_02_1 | 9629 | FTMS; HCD | 66.98 | 707.82 |

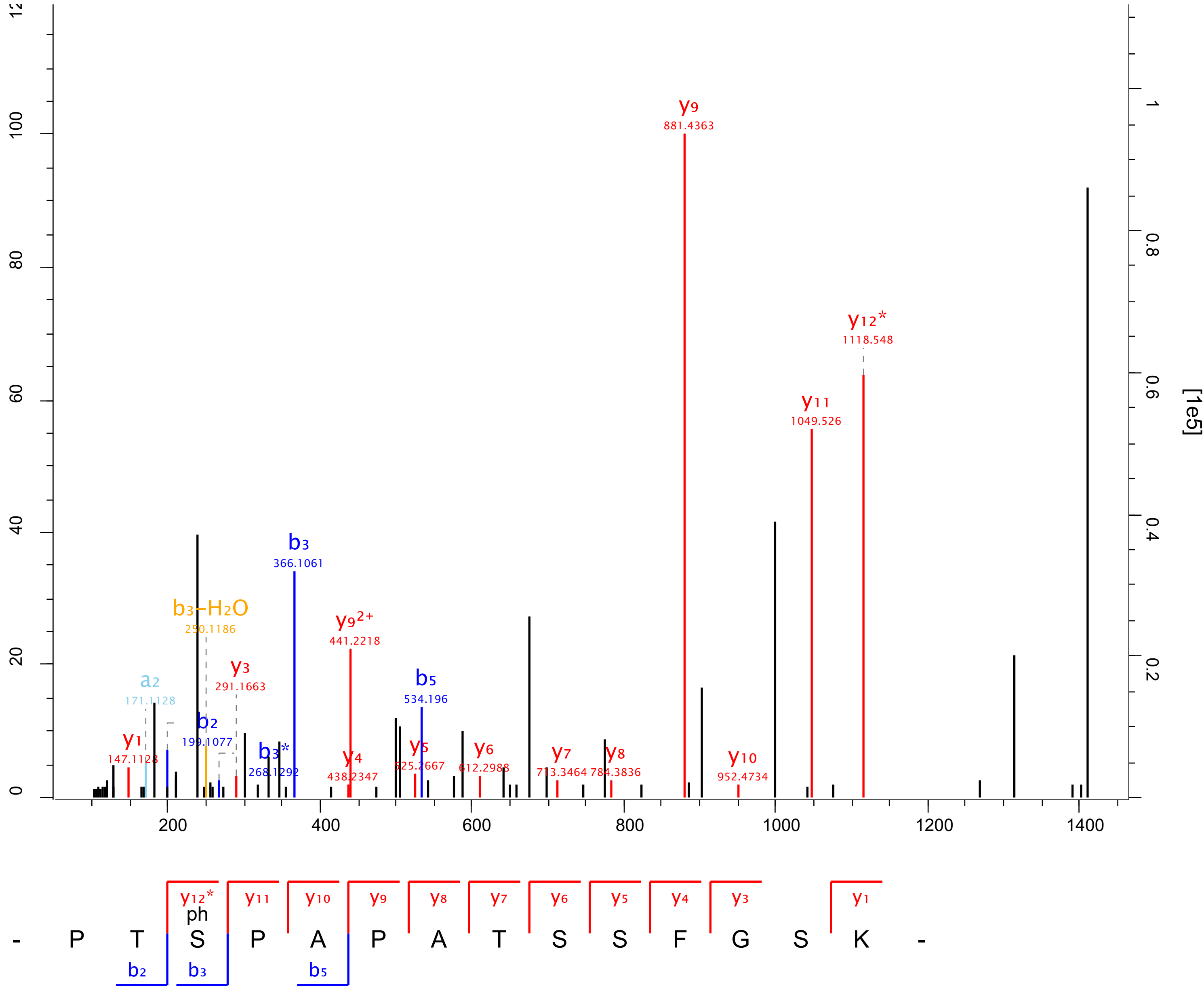

| Raw file | Scan | Method | Score | m/z |
| --- | --- | --- | --- | --- |
| water_02_1 | 9725 | FTMS; HCD | 119.76 | 853.3 |

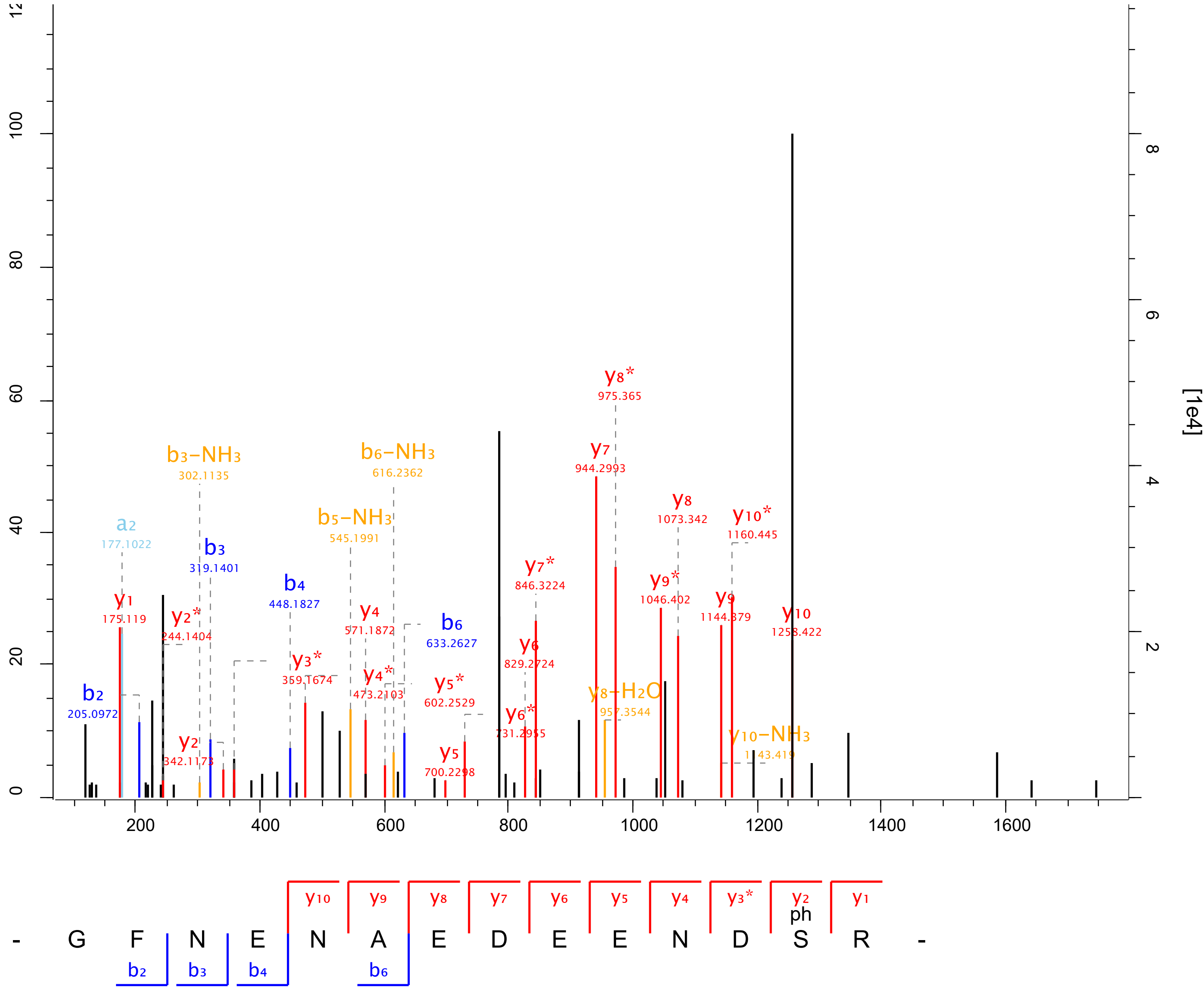

| Raw file | Scan | Method | Score | m/z |
| --- | --- | --- | --- | --- |
| water_02_1 | 9809 | FTMS; HCD | 127.52 | 561.23 |

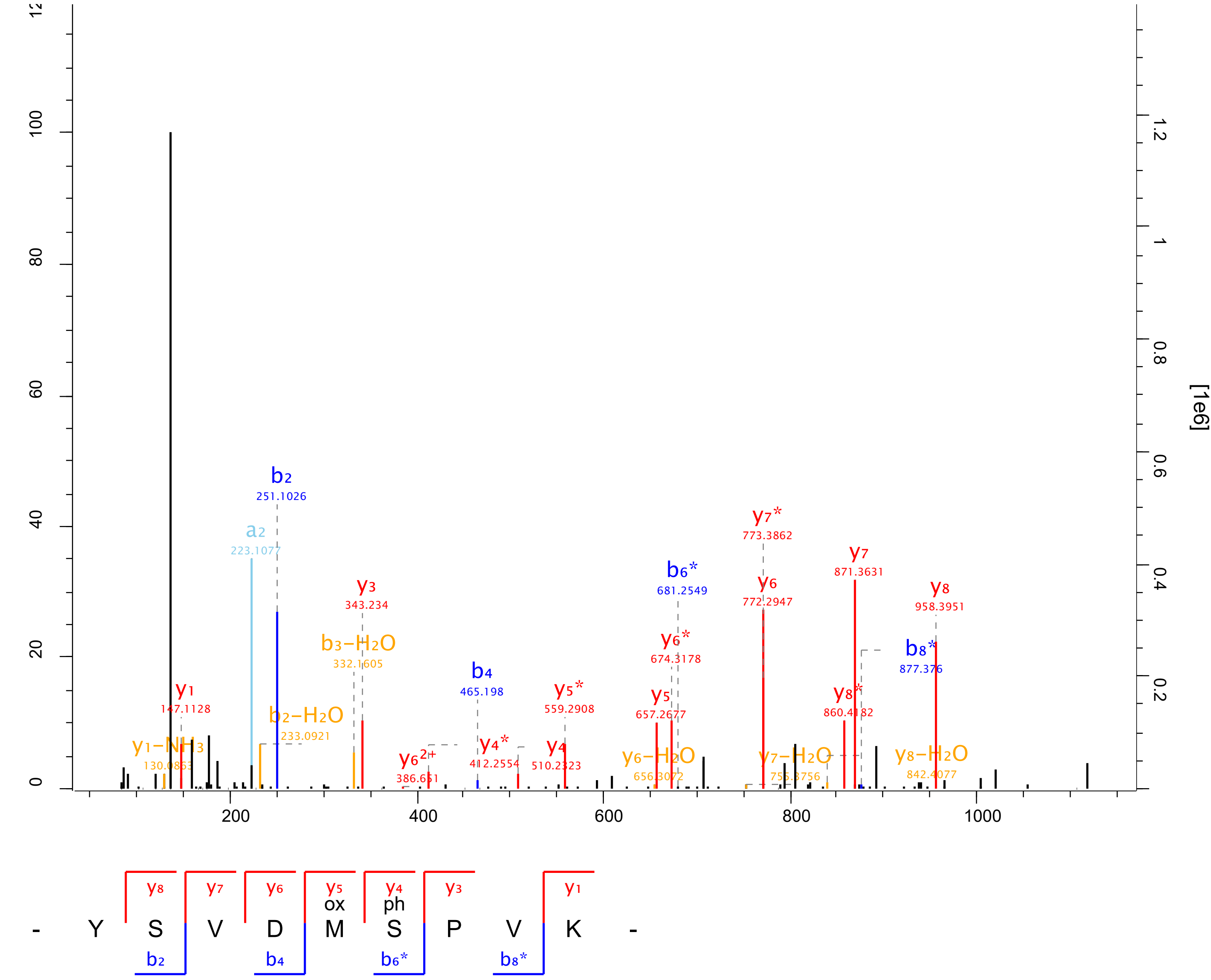

| Raw file | Scan | Method | Score | m/z |
|---|---|---|---|---|
| water_02_1 | 9876 | FTMS; HCD | 107.79 | 486.23 |

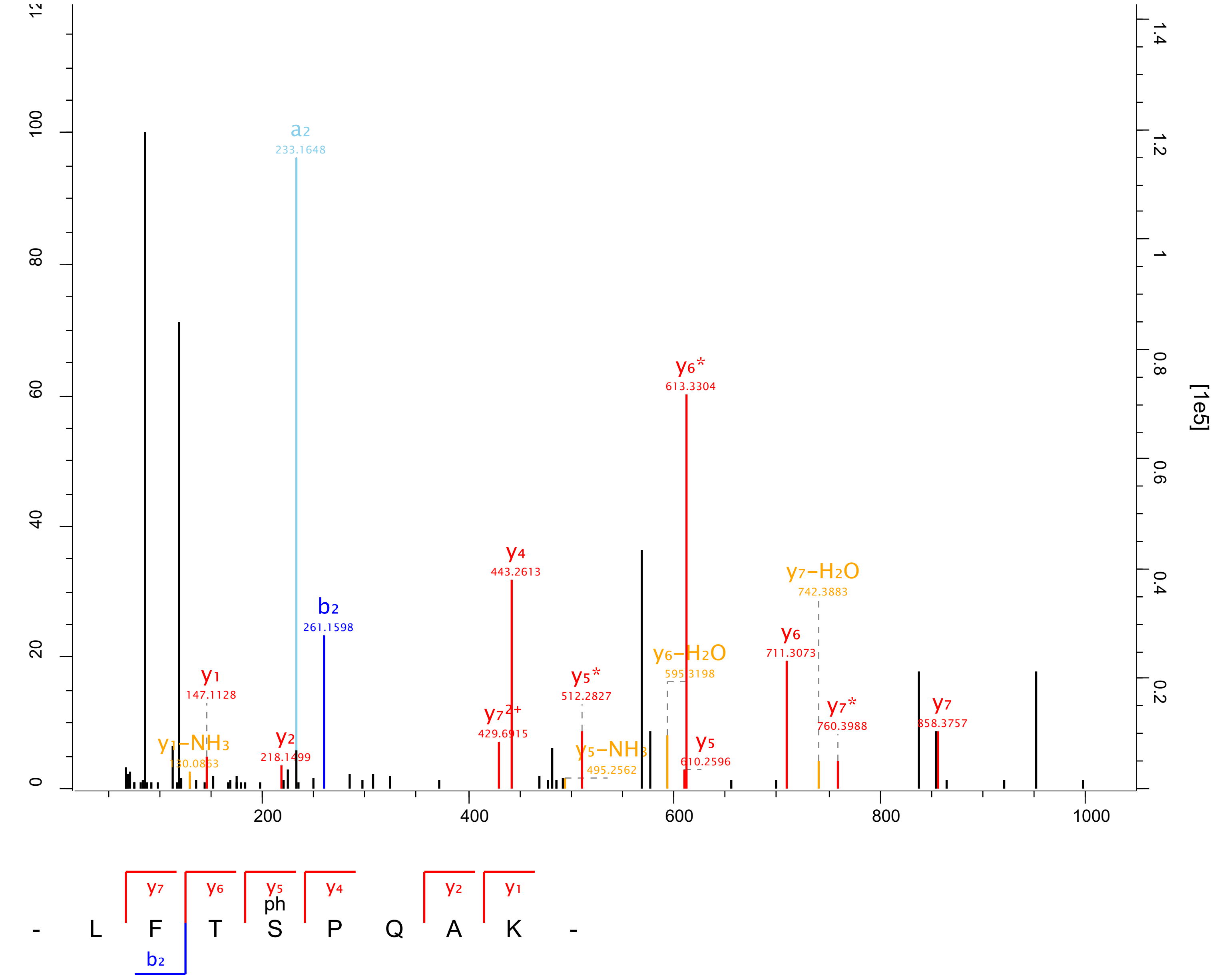

| Raw file | Scan | Method | Score | m/z |
|---|---|---|---|---|
| water_02_1 | 9893 | FTMS; HCD | 74.38 | 398.19 |

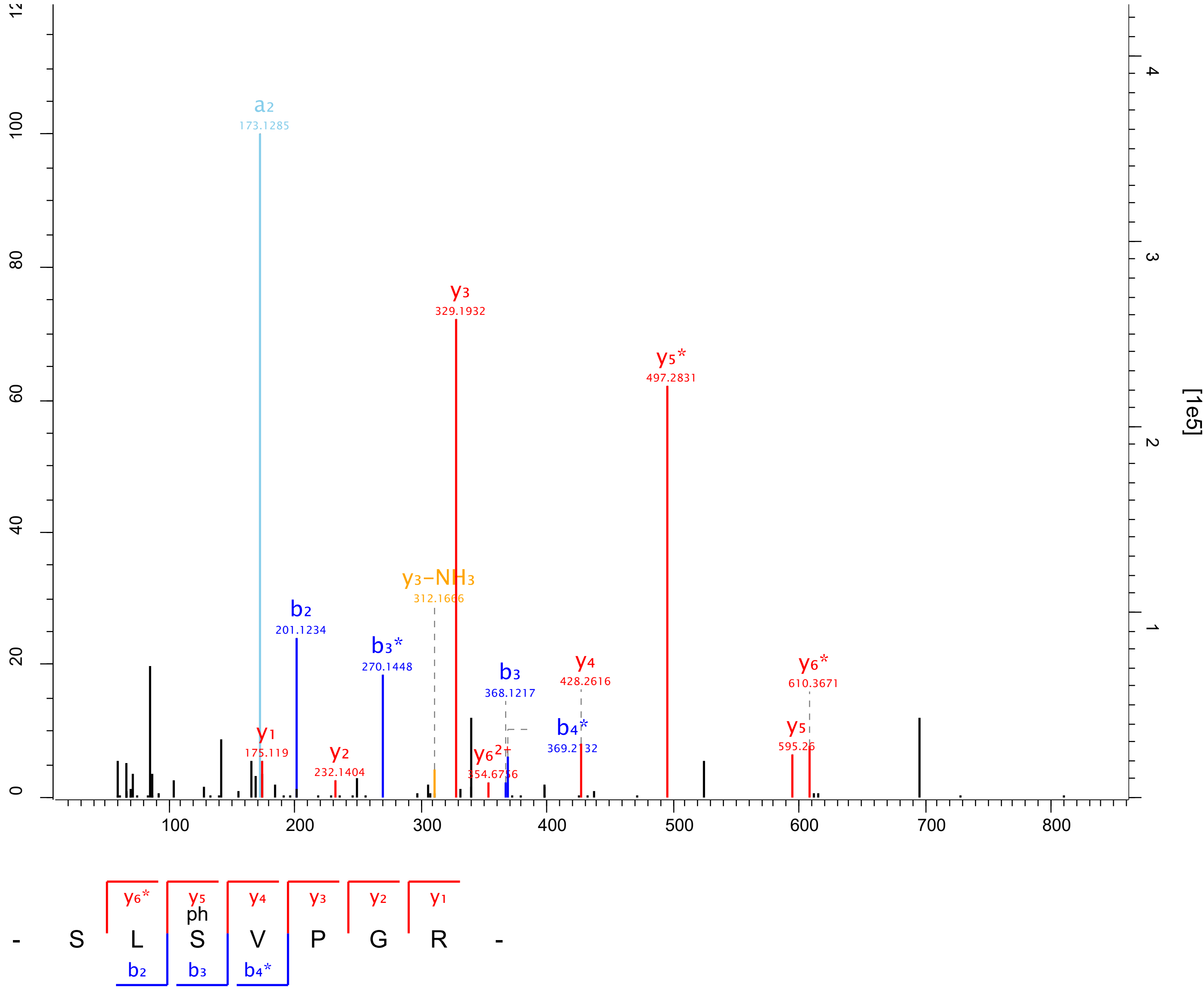

| Raw file | Scan | Method | Score | m/z |
|---|---|---|---|---|
| water_02_1 | 9941 | FTMS; HCD | 49.99 | 642.77 |

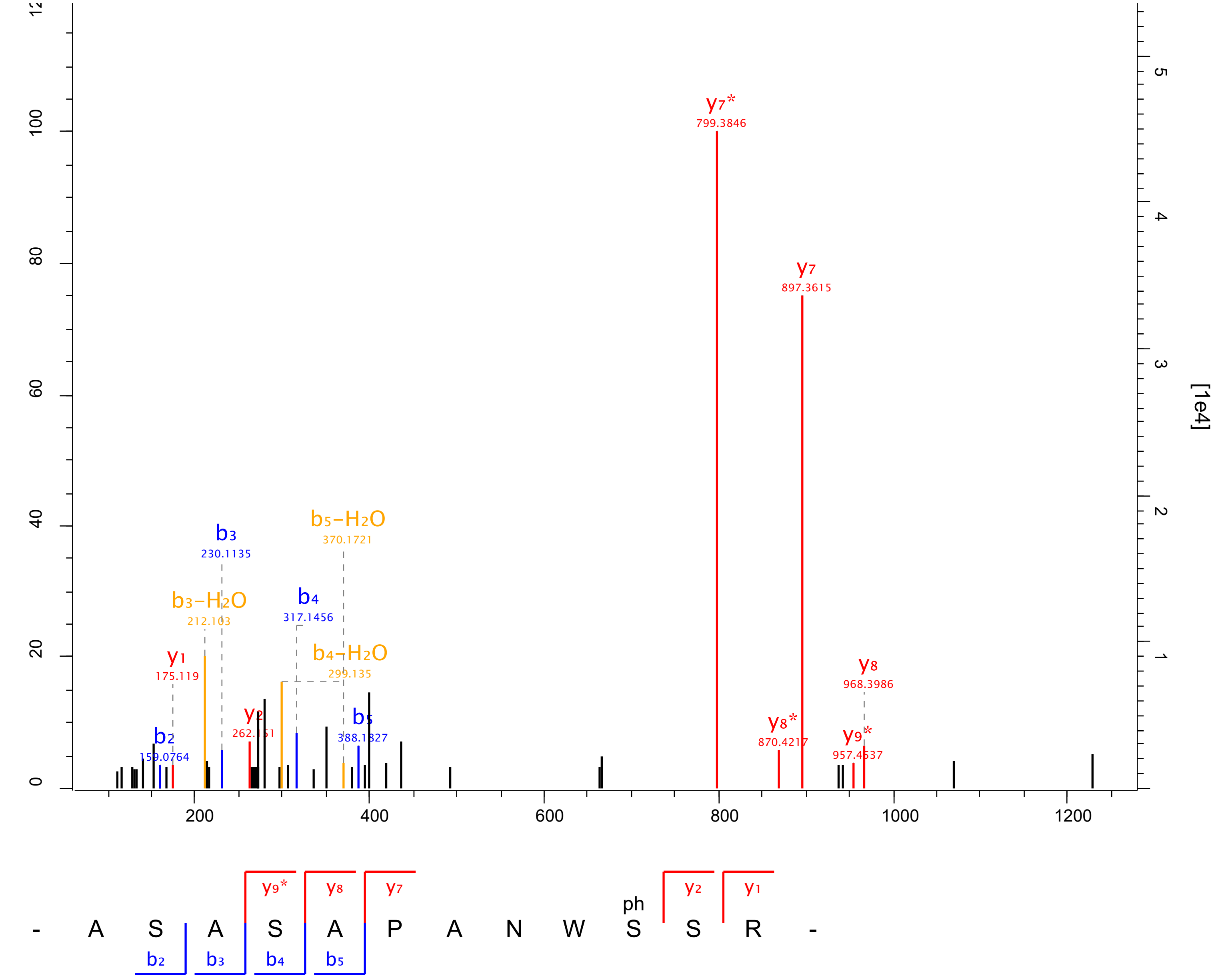

| Raw file | Scan | Method | Score | m/z |
|---|---|---|---|---|
| water_02_1 | 9959 | FTMS; HCD | 66 | 616.22 |

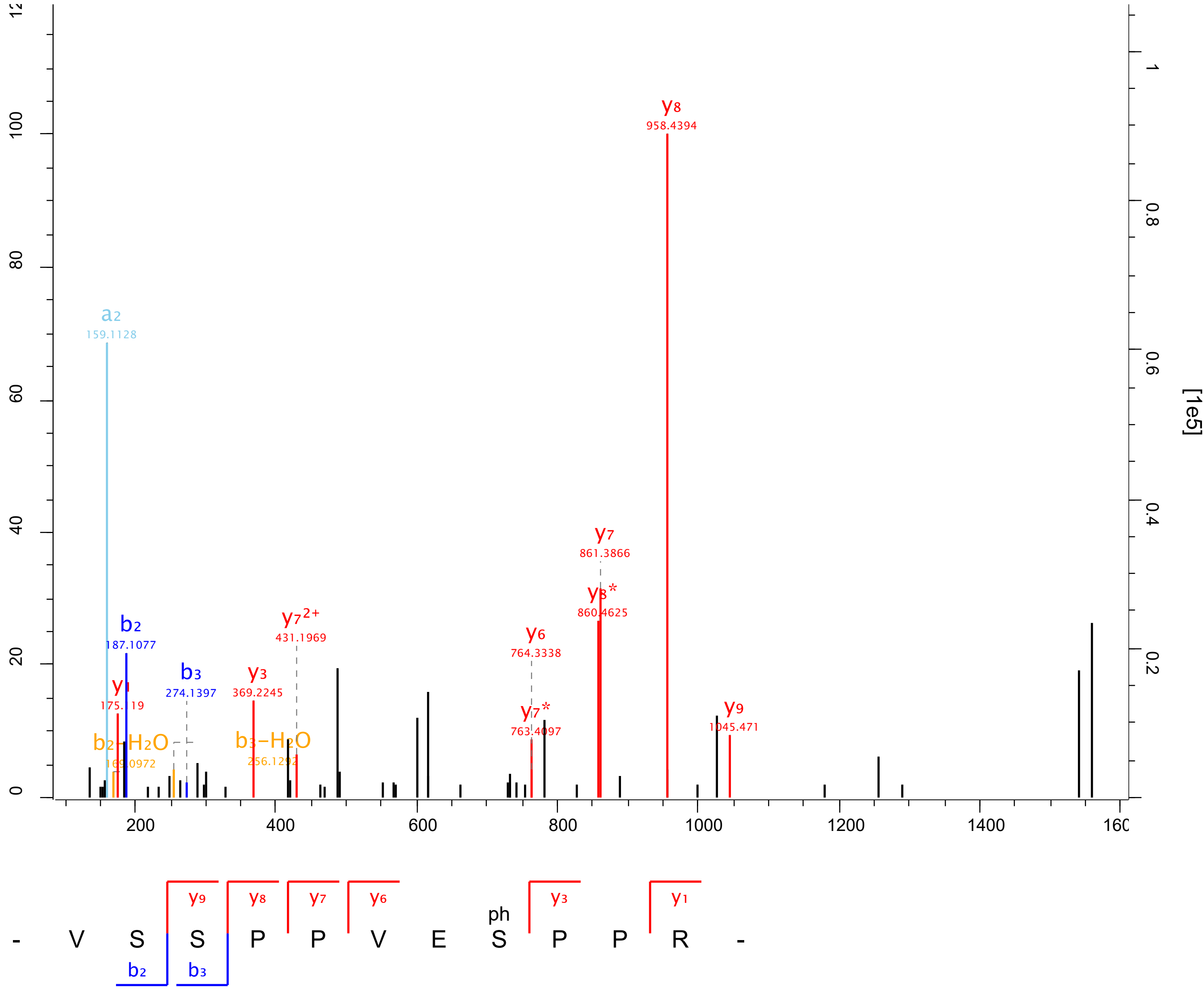

| Raw file | Scan | Method | Score | m/z |
| --- | --- | --- | --- | --- |
| water_02_2 | 10024 | FTMS; HCD | 141.65 | 563.24 |

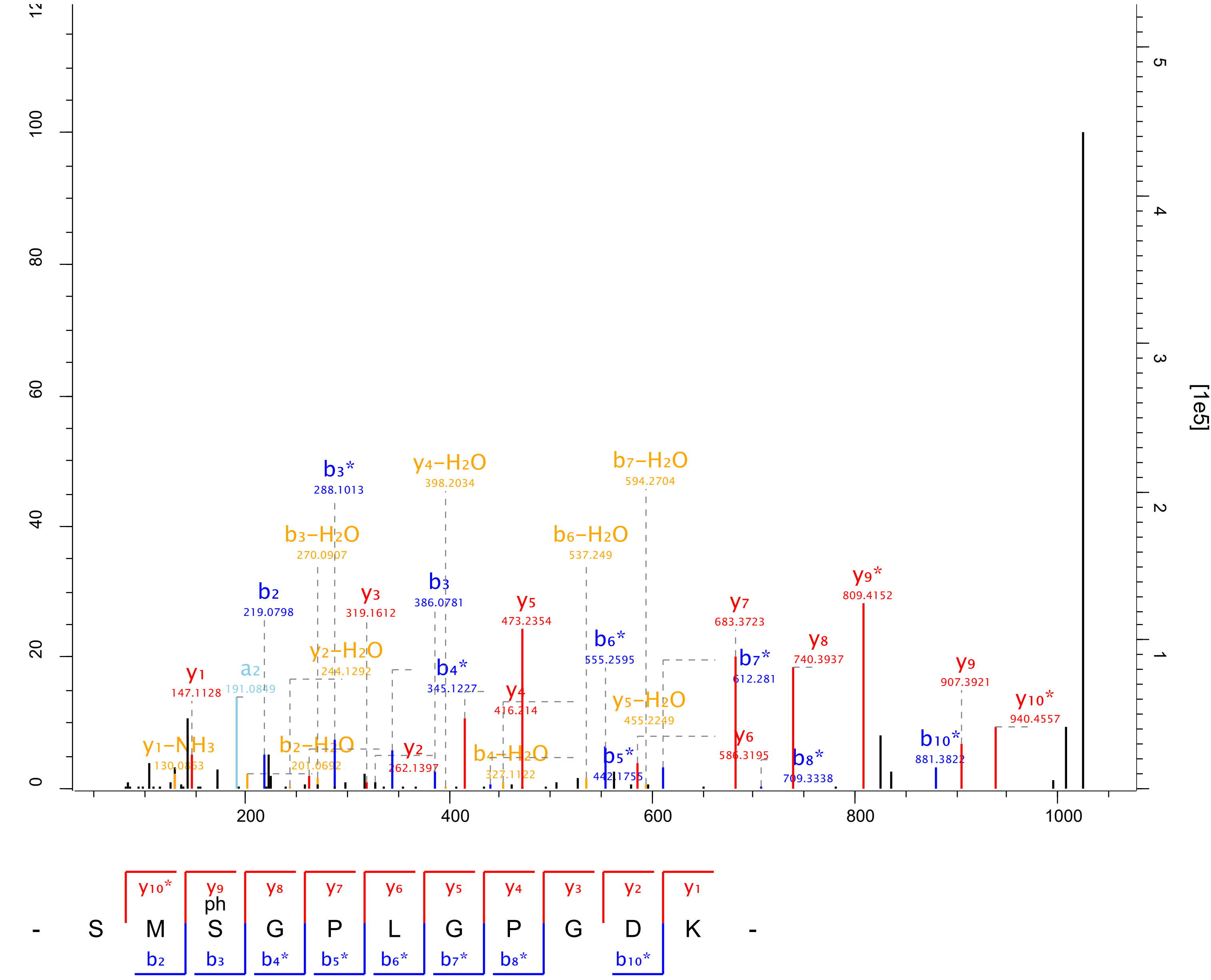

| Raw file | Scan | Method | Score | m/z |
|---|---|---|---|---|
| water_02_2 | 10759 | FTMS; HCD | 44.44 | 435.54 |

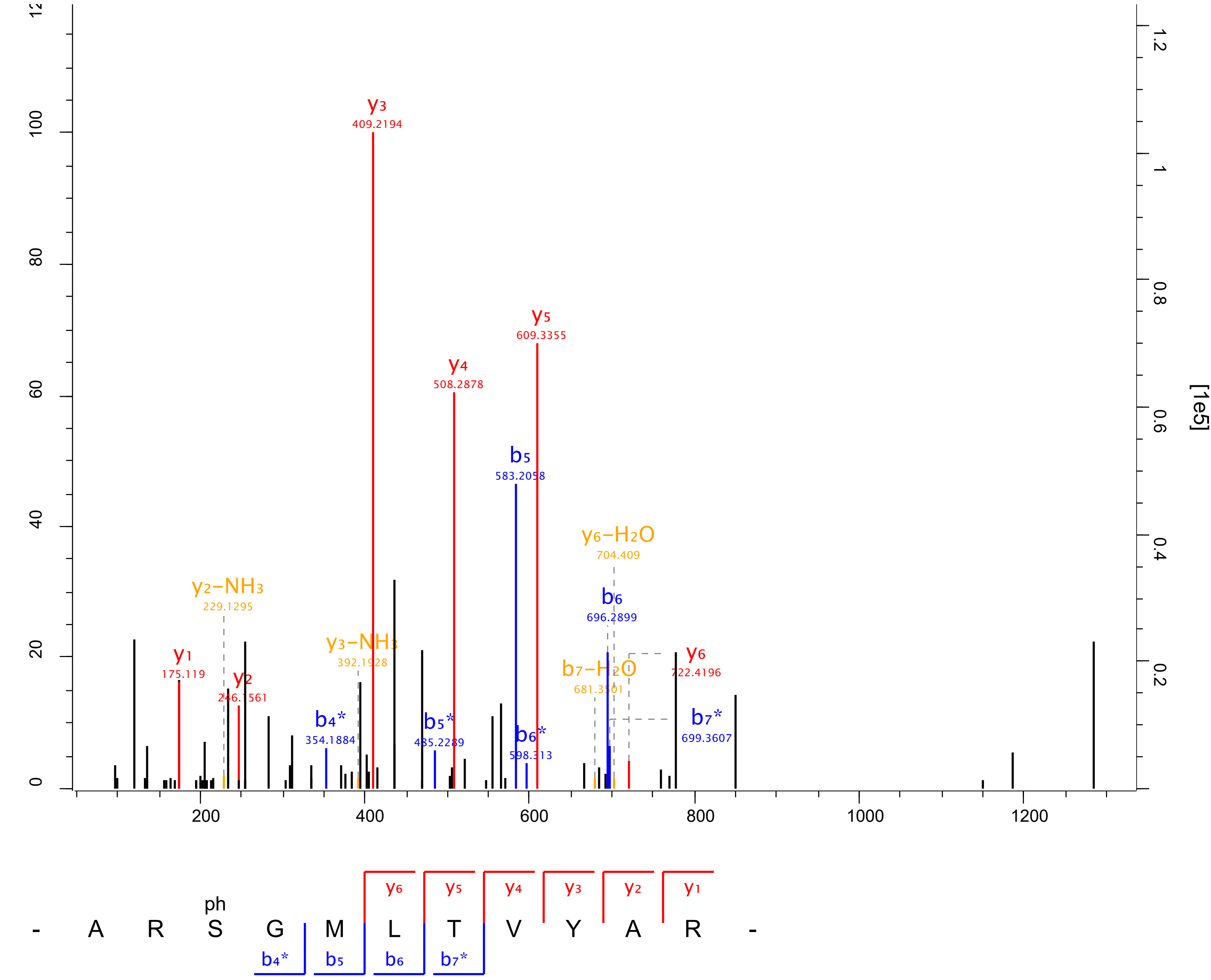

| Raw file | Scan | Method | Score | m/z |
| --- | --- | --- | --- | --- |
| water_02_2 | 12643 | FTMS; HCD | 87.43 | 620.79 |

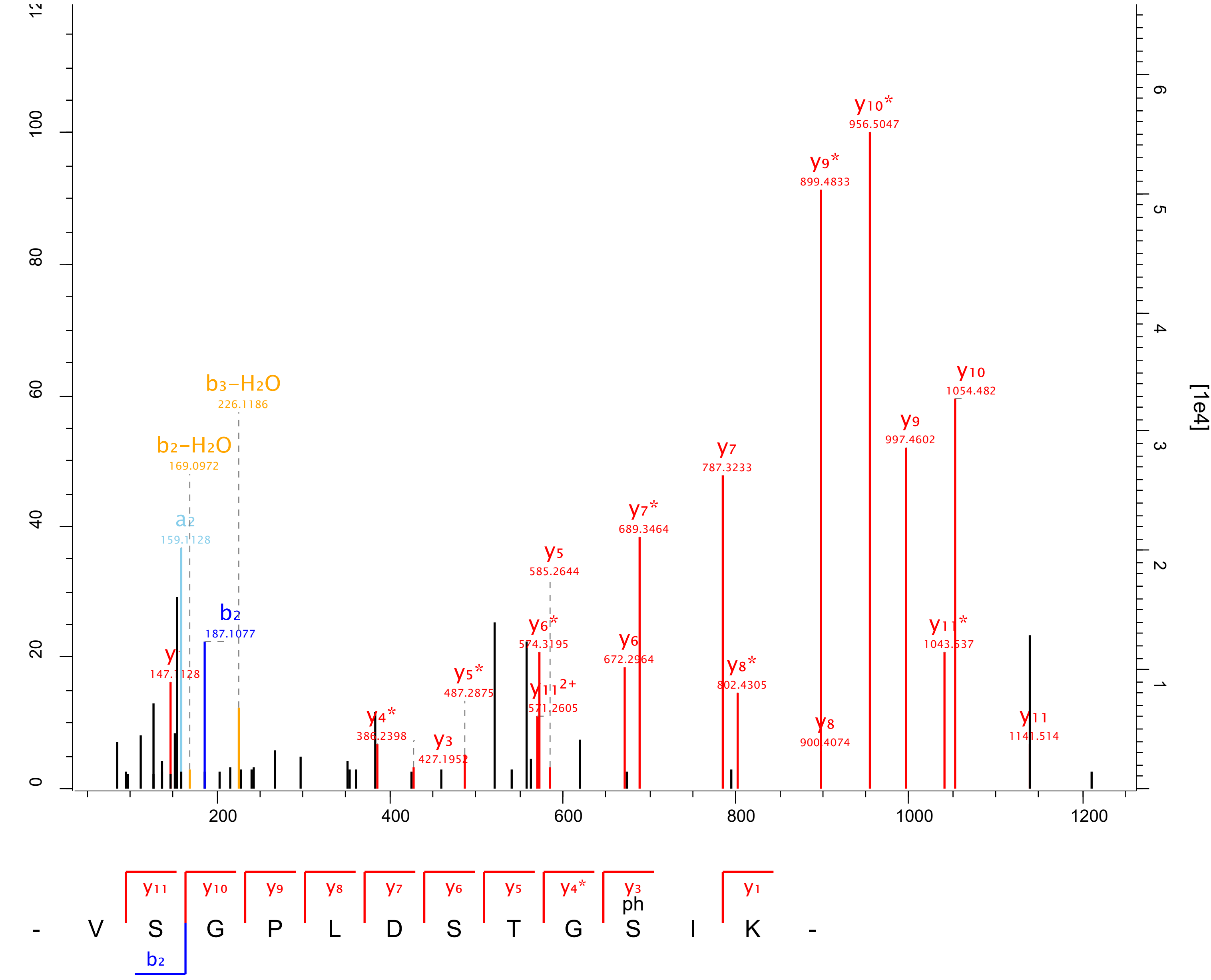

| Raw file | Scan | Method | Score | m/z |
|---|---|---|---|---|
| water_02_2 | 13072 | FTMS; HCD | 135.72 | 741.81 |

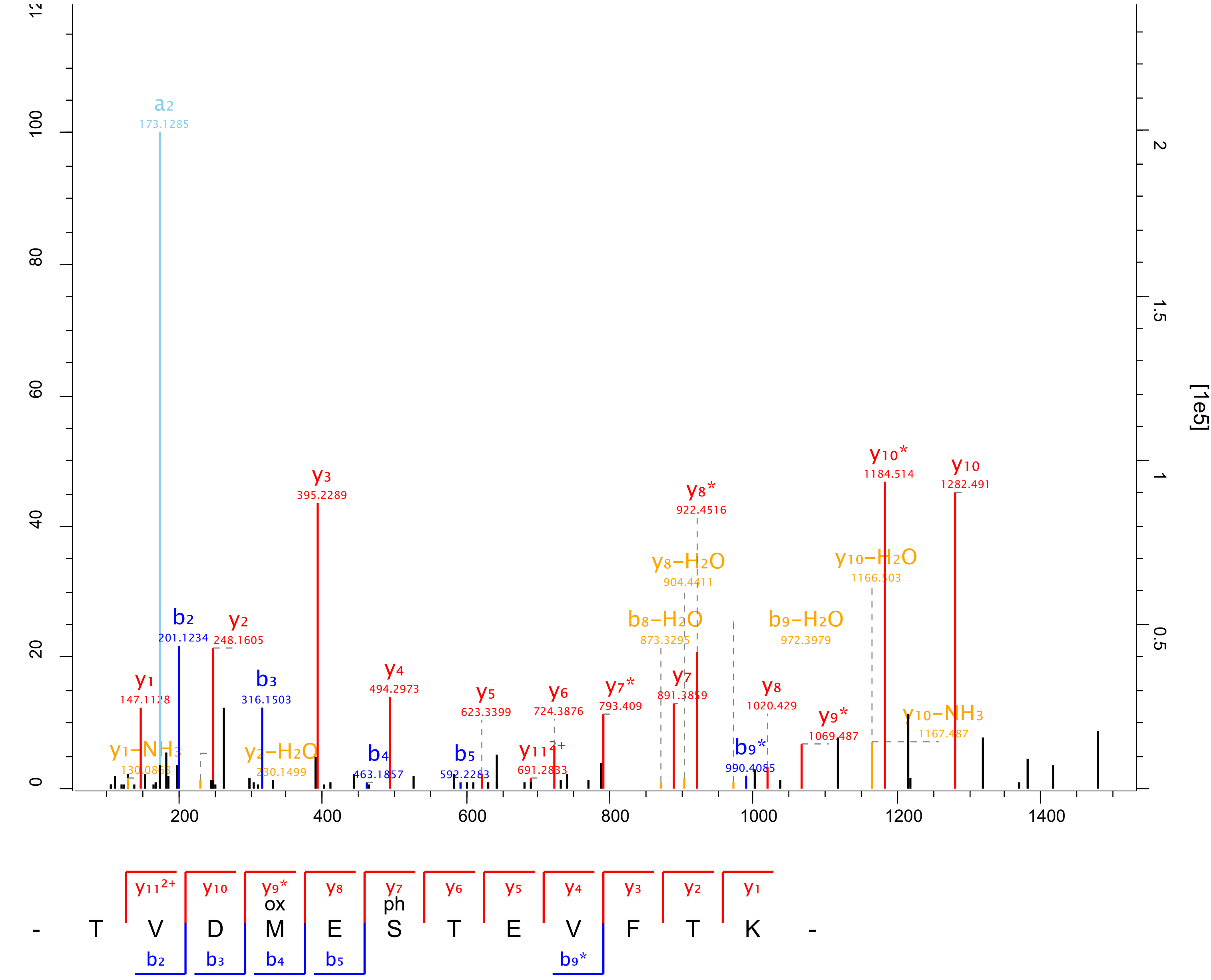

| Raw file | Scan | Method | Score | m/z |
| --- | --- | --- | --- | --- |
| water_02_2 | 14109 | FTMS; HCD | 141.1 | 661.29 |

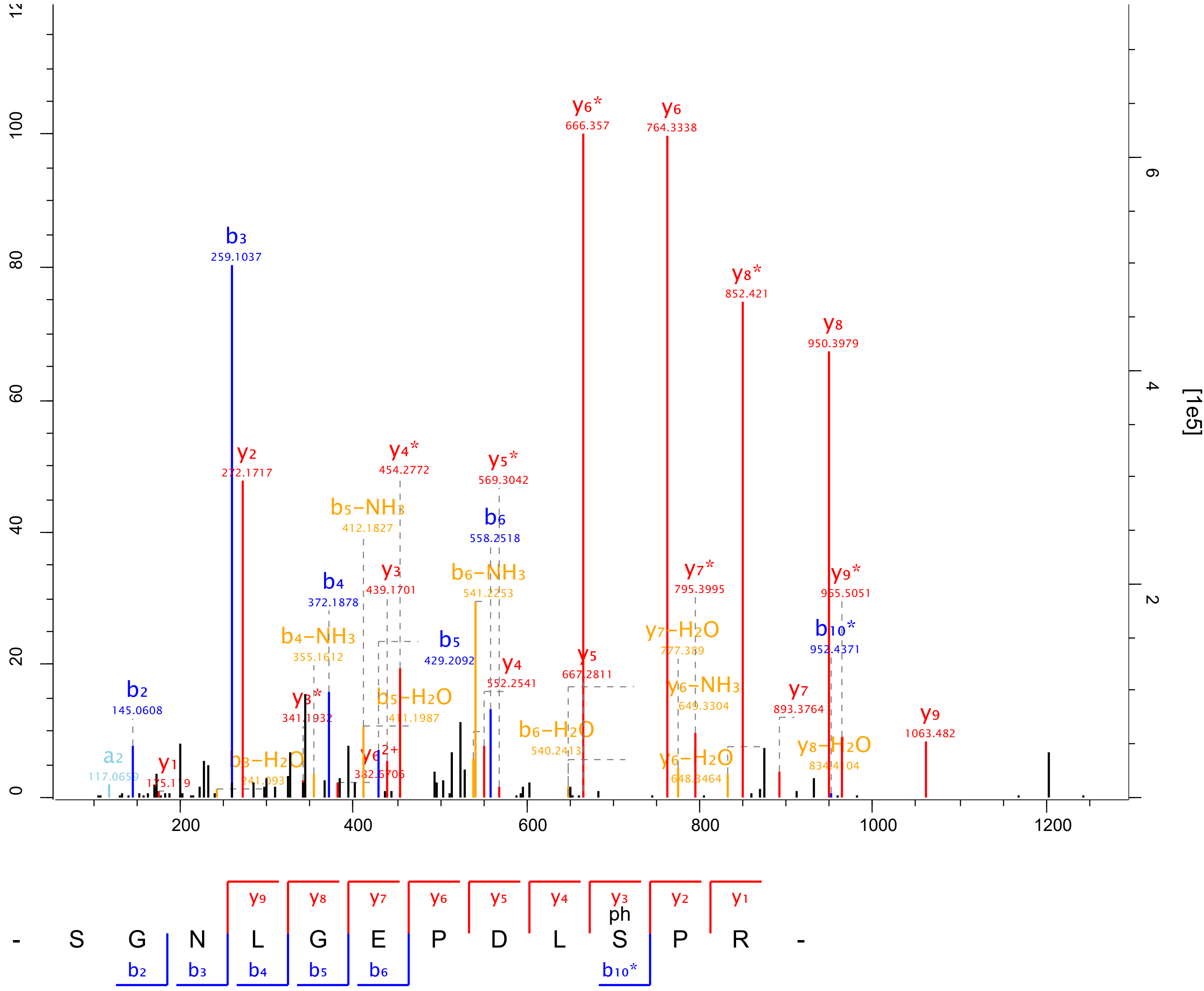

| Raw file | Scan | Method | Score | m/z |
|---|---|---|---|---|
| water_02_2 | 15054 | FTMS; HCD | 439.88 | 1031.42 |

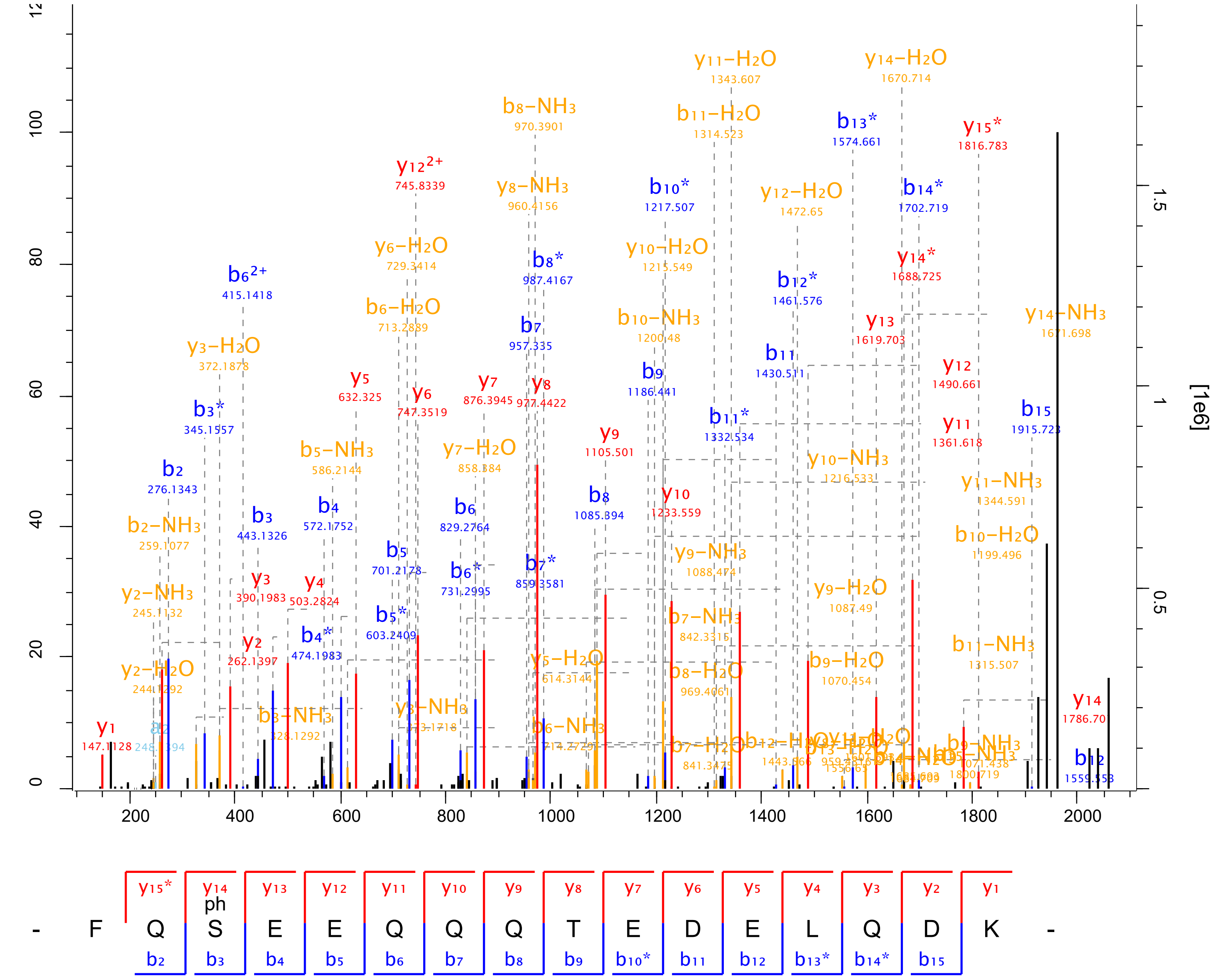

| Raw file | Scan | Method | Score | m/z |
|---|---|---|---|---|
| water_02_2 | 1531 | FTMS; HCD | 43.31 | 353.53 |

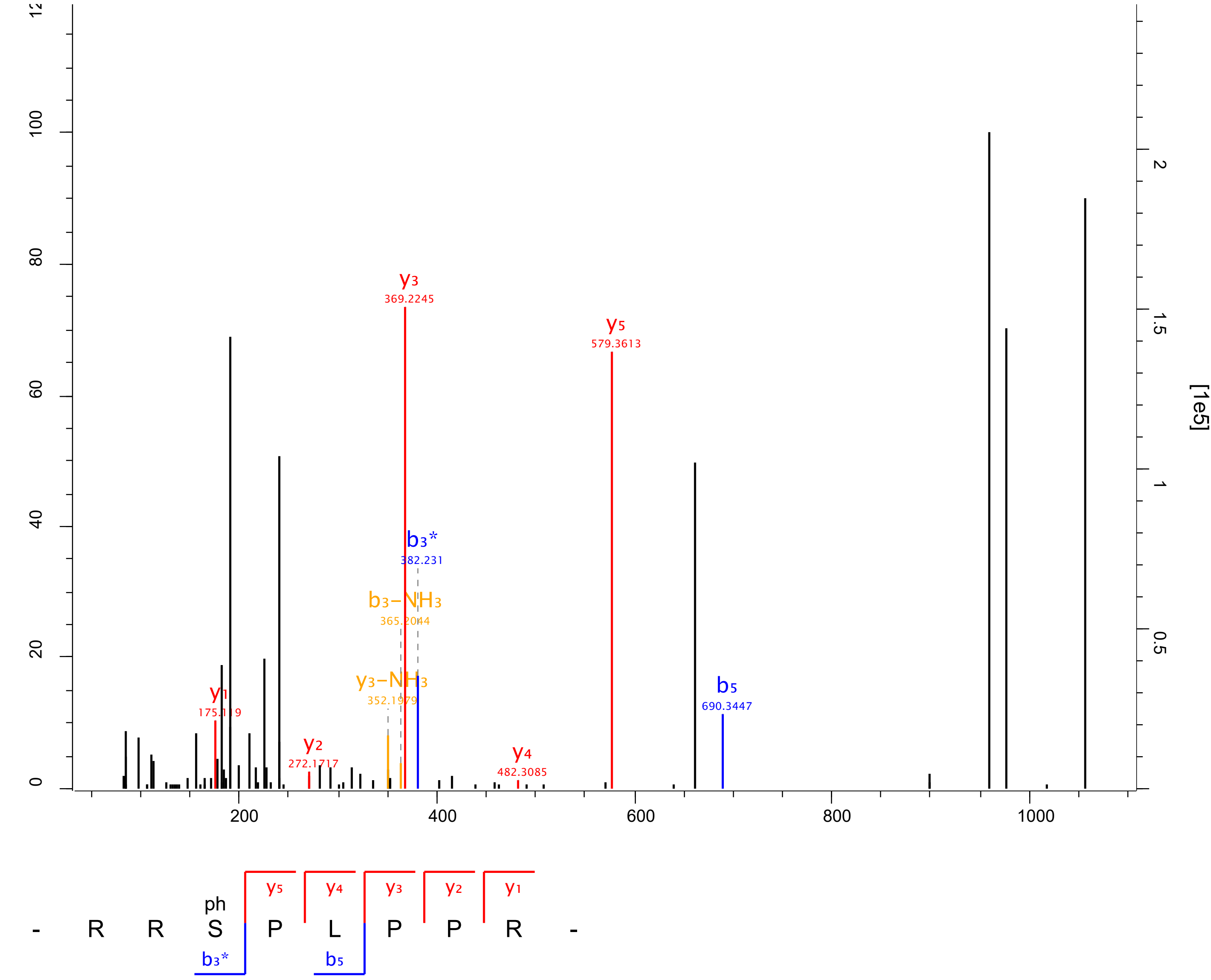

| Raw file | Scan | Method | Score | m/z |
|---|---|---|---|---|
| water_02_2 | 19956 | FTMS; HCD | 147.12 | 733.81 |

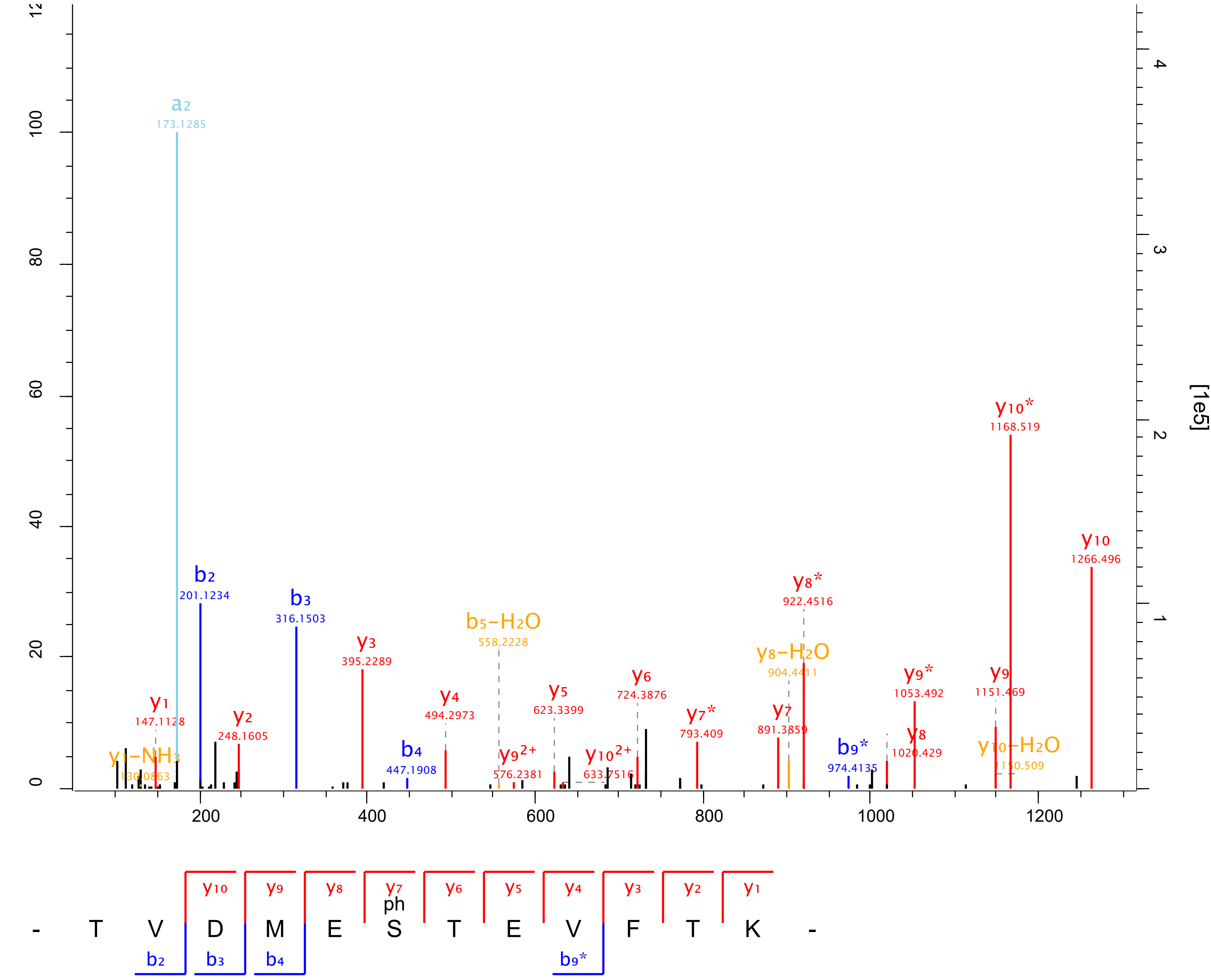

| Raw file | Scan | Method | Score | m/z |
|---|---|---|---|---|
| water_02_2 | 22285 | FTMS; HCD | 89.36 | 736.83 |

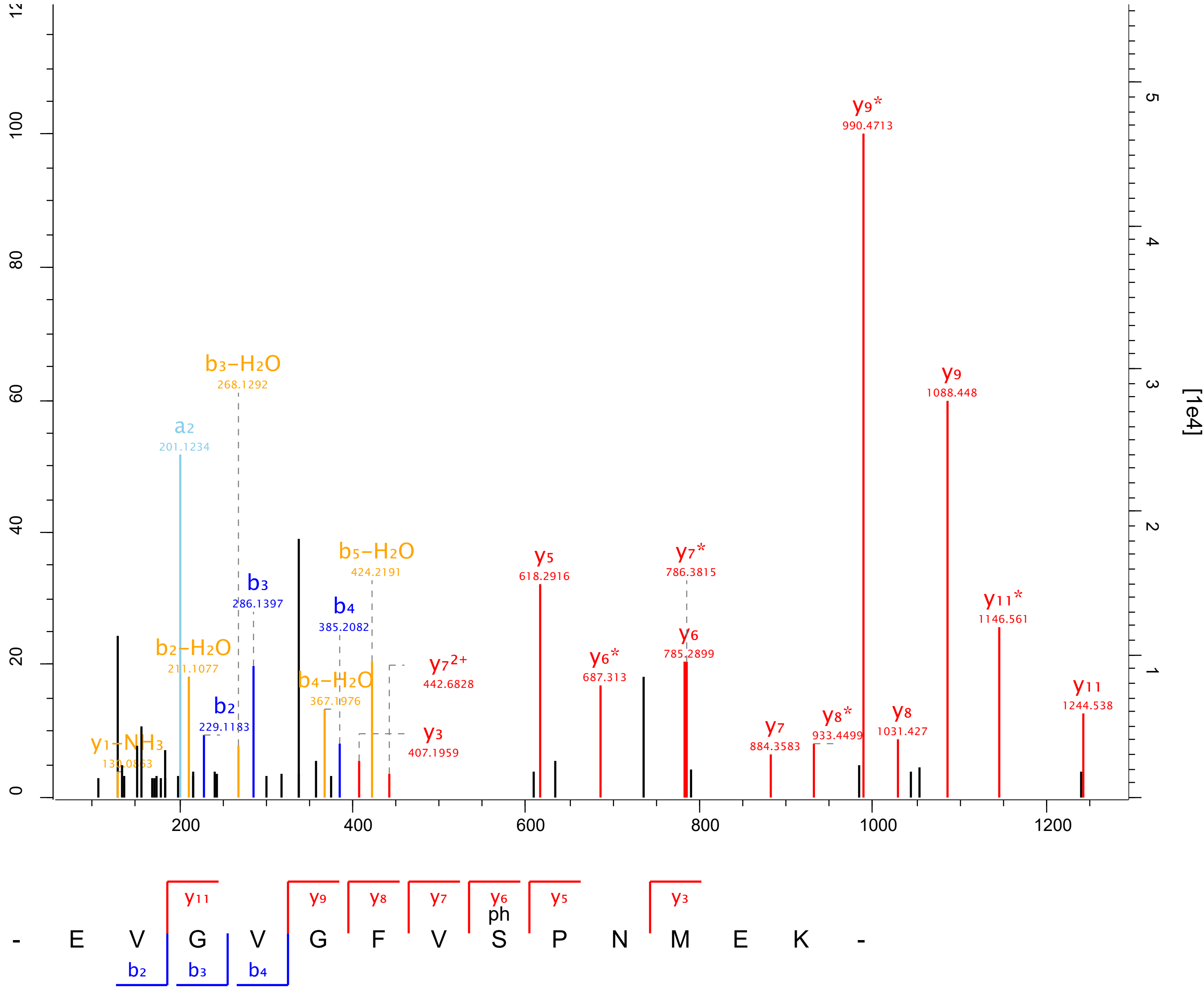

| Raw file | Scan | Method | Score | m/z |
|---|---|---|---|---|
| water_02_2 | 22582 | FTMS; HCD | 41.62 | 760.85 |

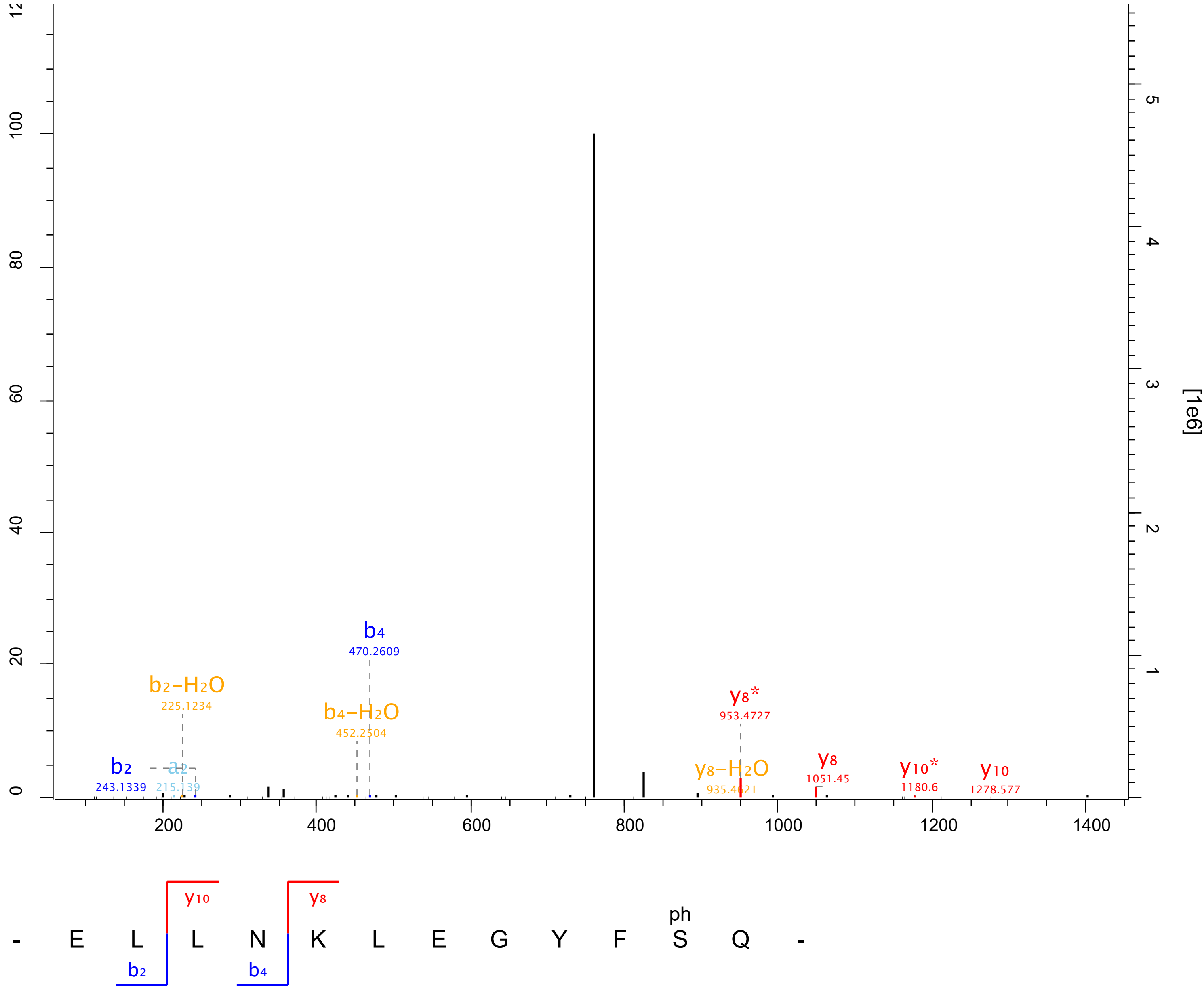

| Raw file | Scan | Method | Score | m/z |
|---|---|---|---|---|
| water_02_2 | 23450 | FTMS; HCD | 253.09 | 830.9 |

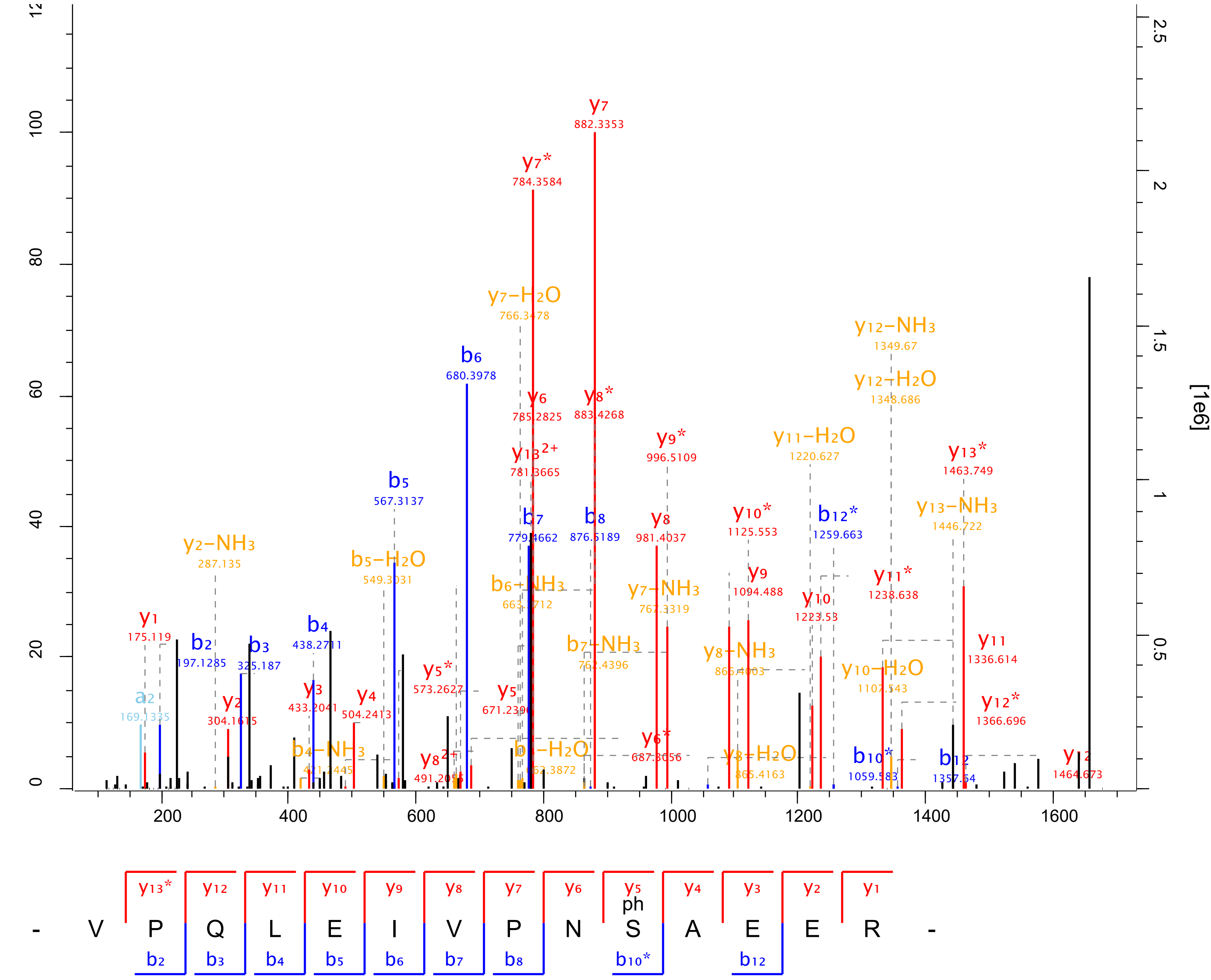

| Raw file | Scan | Method | Score | m/z |
| --- | --- | --- | --- | --- |
| water_02_2 | 24066 | FTMS; HCD | 42.63 | 806.85 |

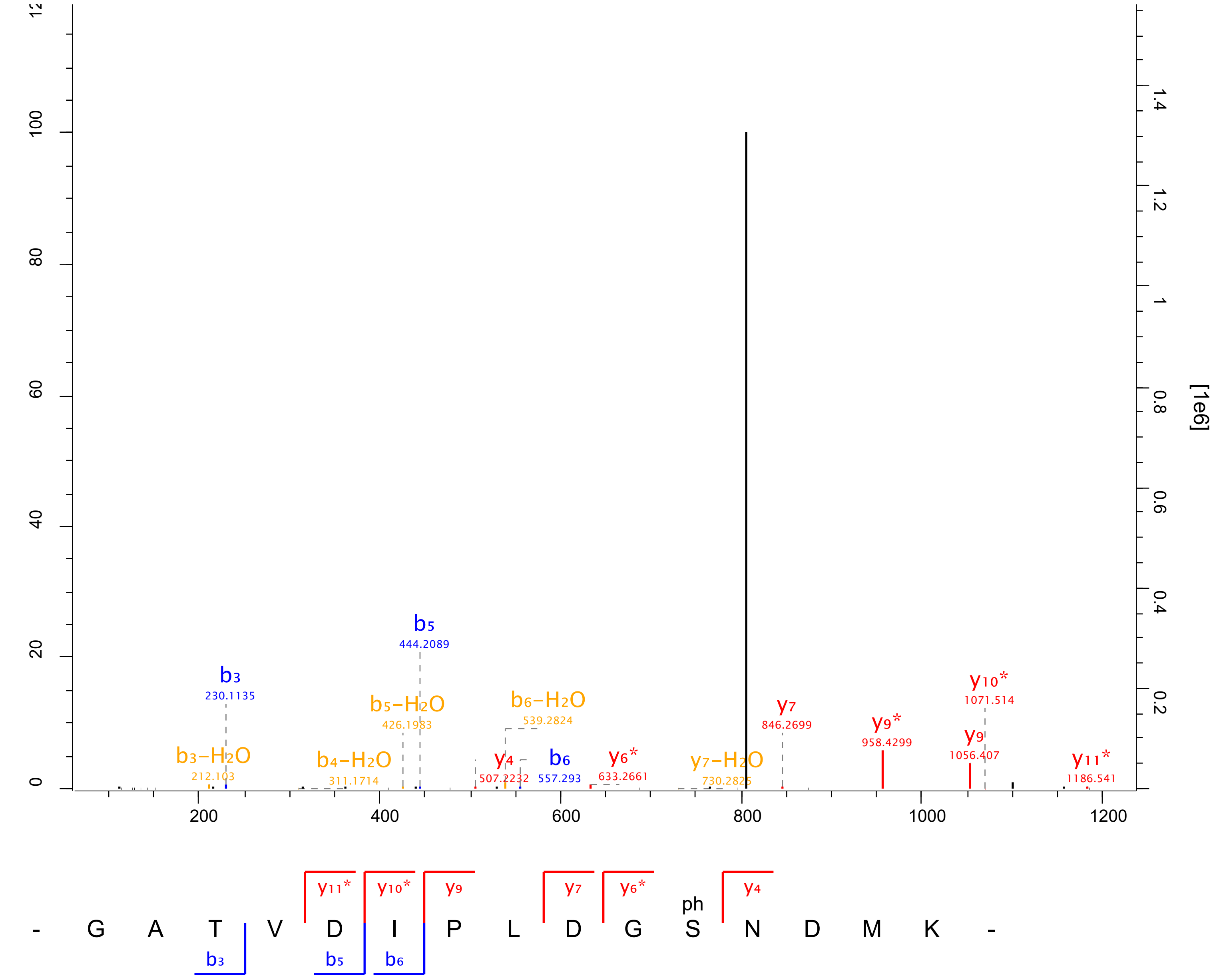

| Raw file | Scan | Method | Score | m/z |
| --- | --- | --- | --- | --- |
| water_02_2 | 24067 | FTMS; HCD | 48.9 | 760.32 |

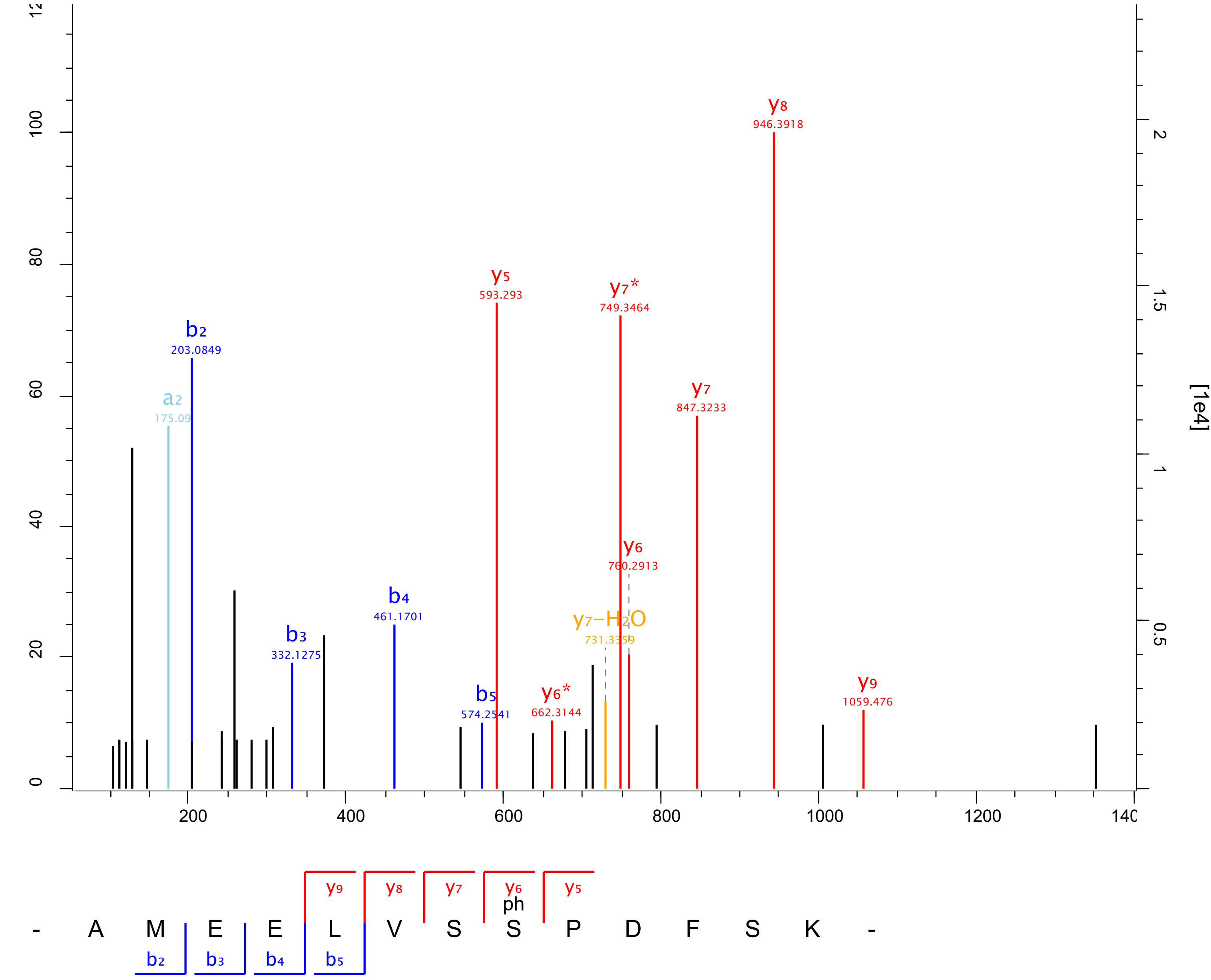

| Raw file | Scan | Method | Score | m/z |
|---|---|---|---|---|
| water_02_2 | 24300 | FTMS; HCD | 53.62 | 912.4 |

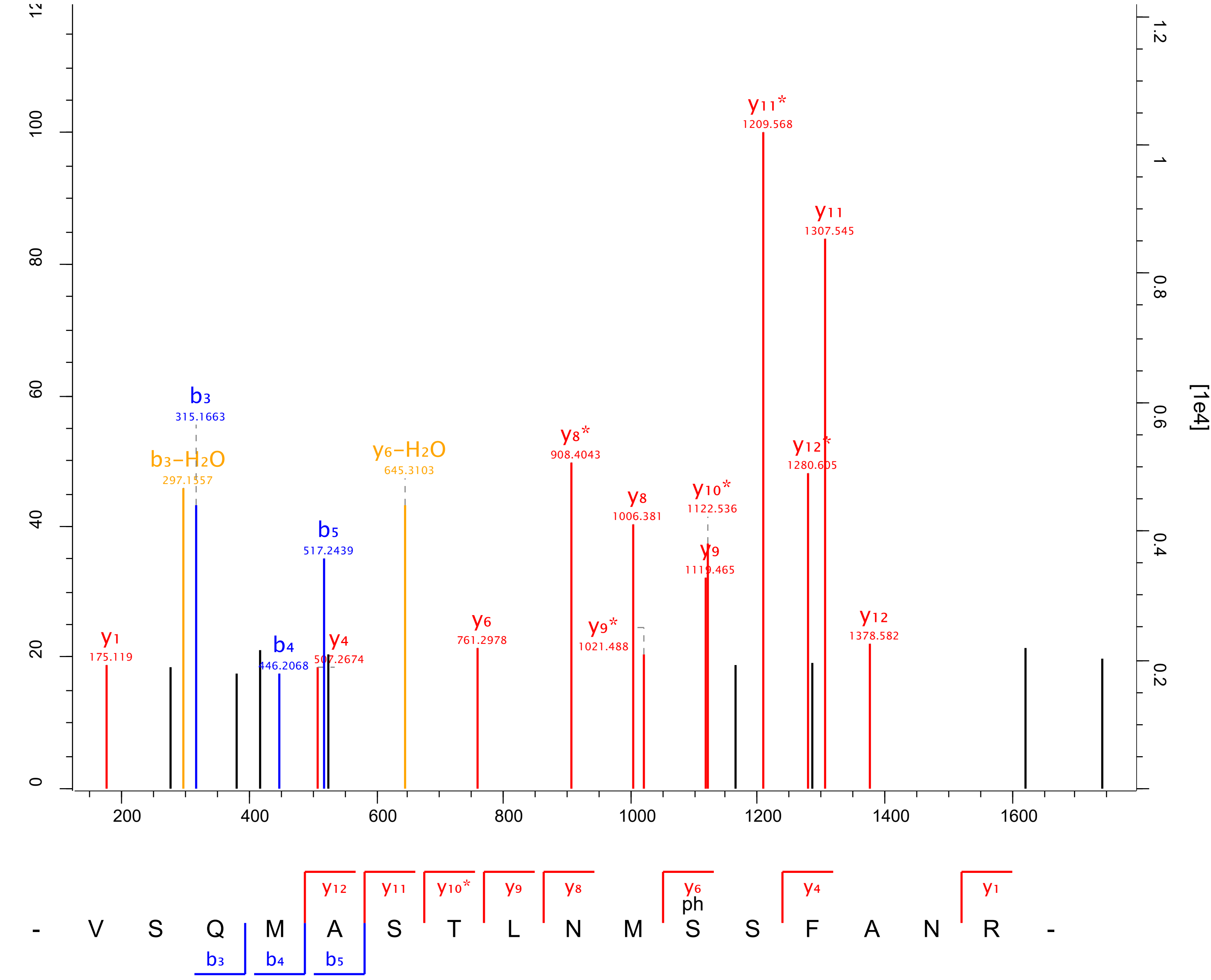

| Raw file | Scan | Method | Score | m/z |
|---|---|---|---|---|
| water_02_2 | 29007 | FTMS; HCD | 59.37 | 816.37 |

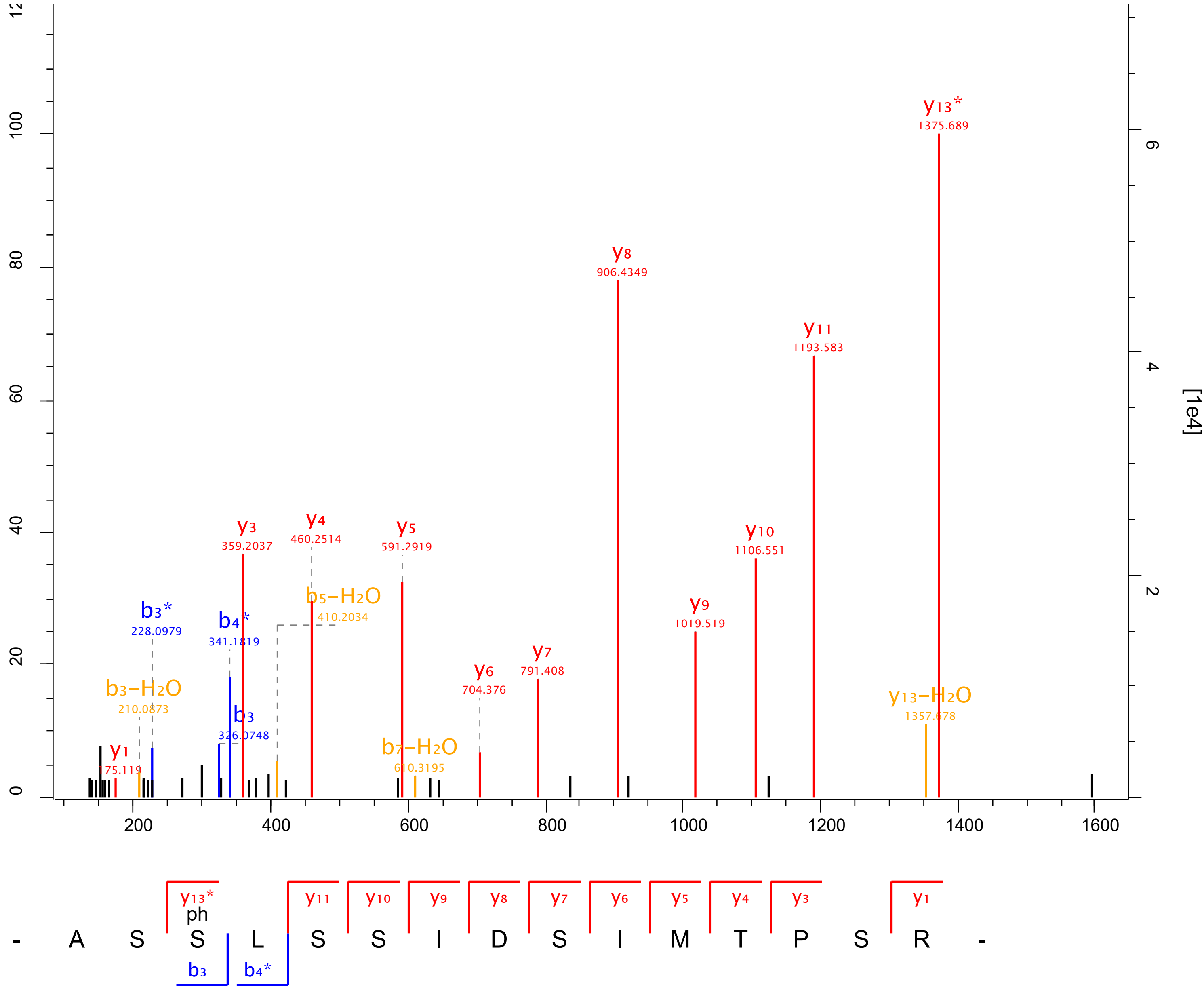

| Raw file | Scan | Method | Score | m/z |
| --- | --- | --- | --- | --- |
| water_02_2 | 30127 | FTMS; HCD | 76.73 | 686.81 |

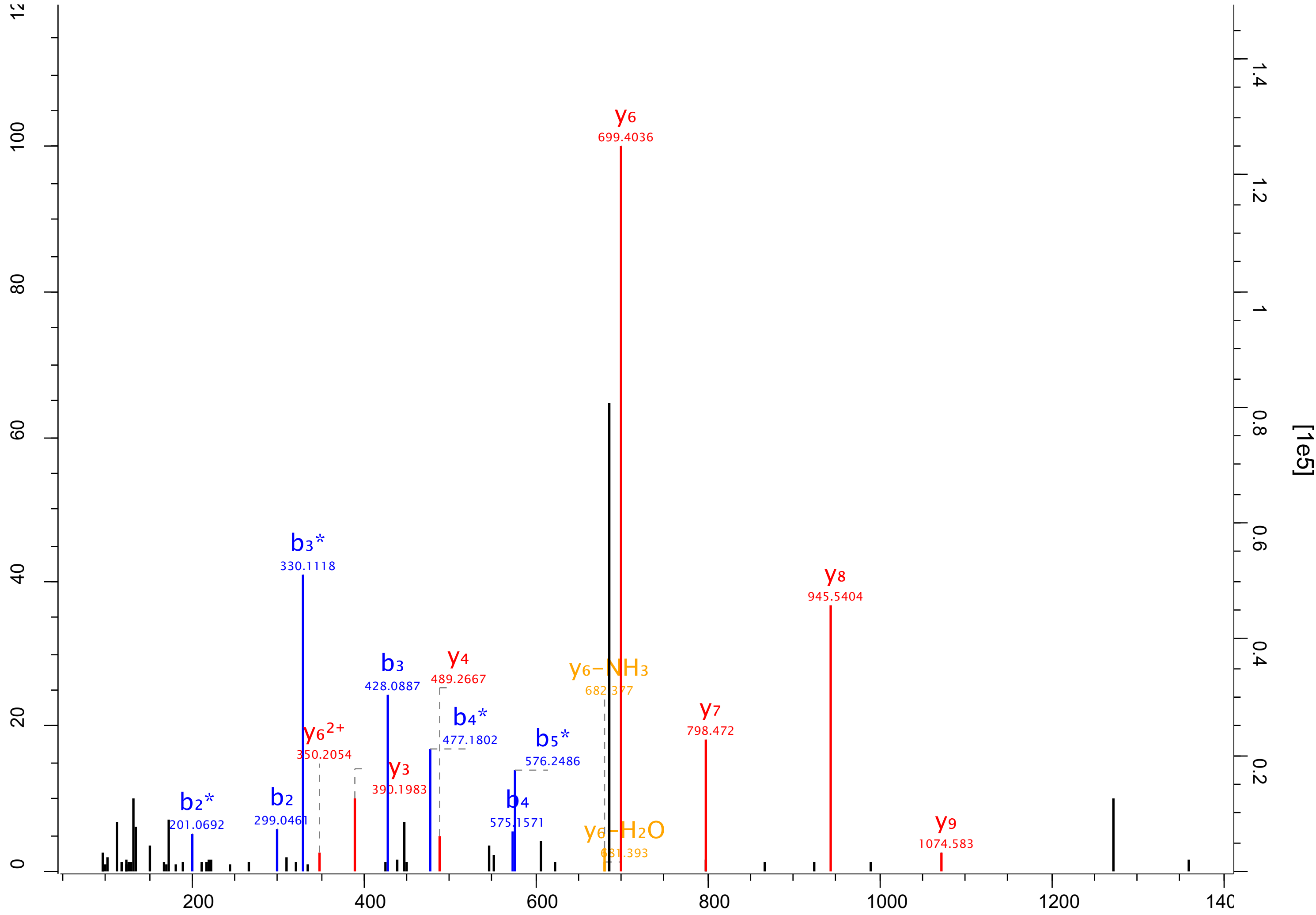


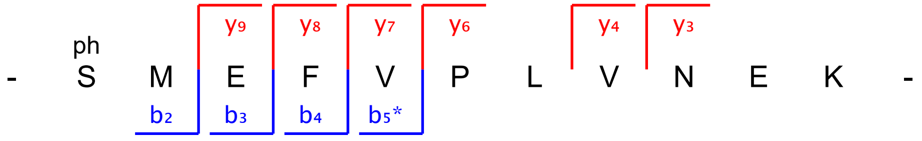

| Raw file | Scan | Method | Score | m/z |
| --- | --- | --- | --- | --- |
| water_02_2 | 3125 | FTMS; HCD | 62.59 | 557.61 |

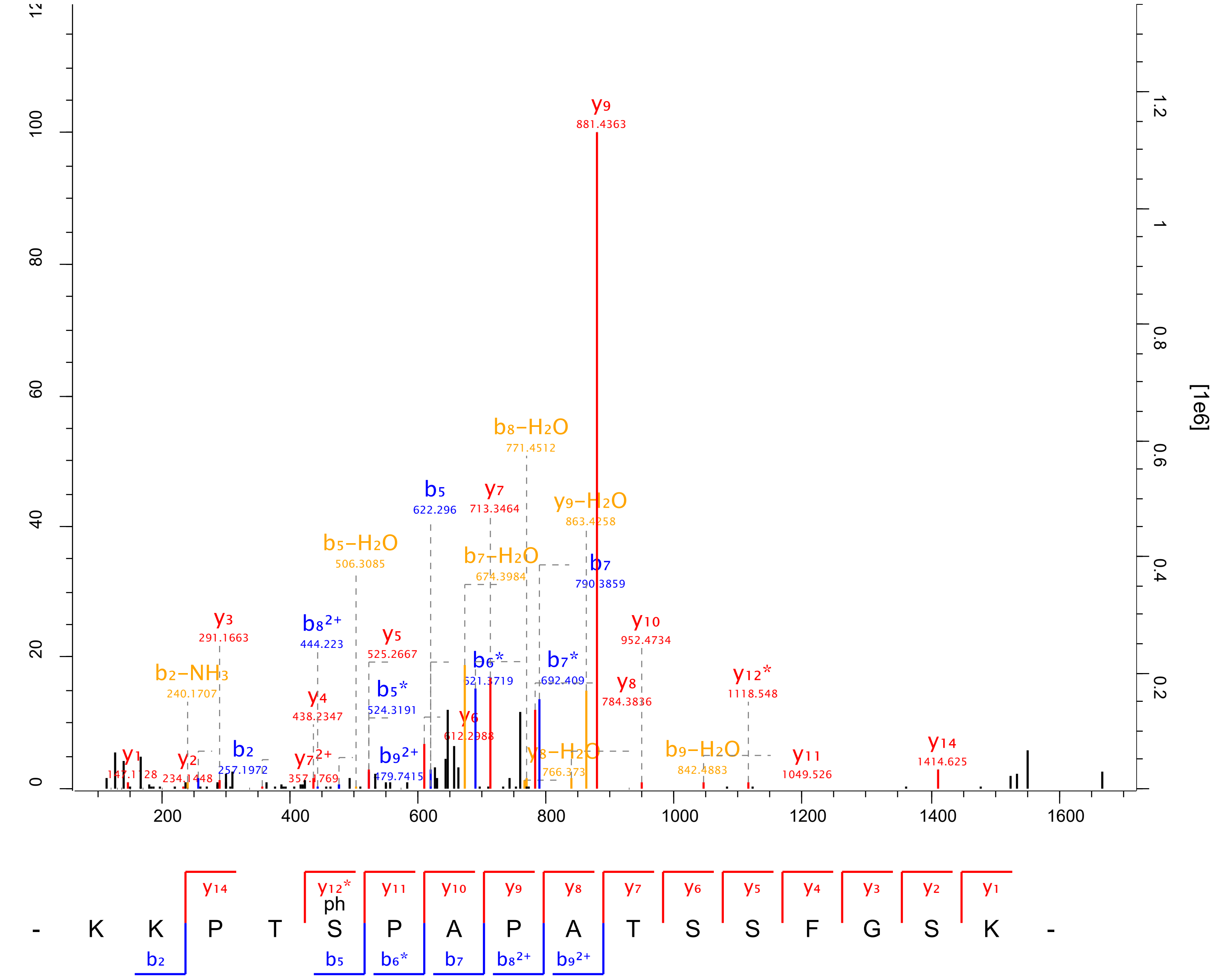

| Raw file | Scan | Method | Score | m/z |
|---|---|---|---|---|
| water_02_2 | 32723 | FTMS; HCD | 93.42 | 836.83 |

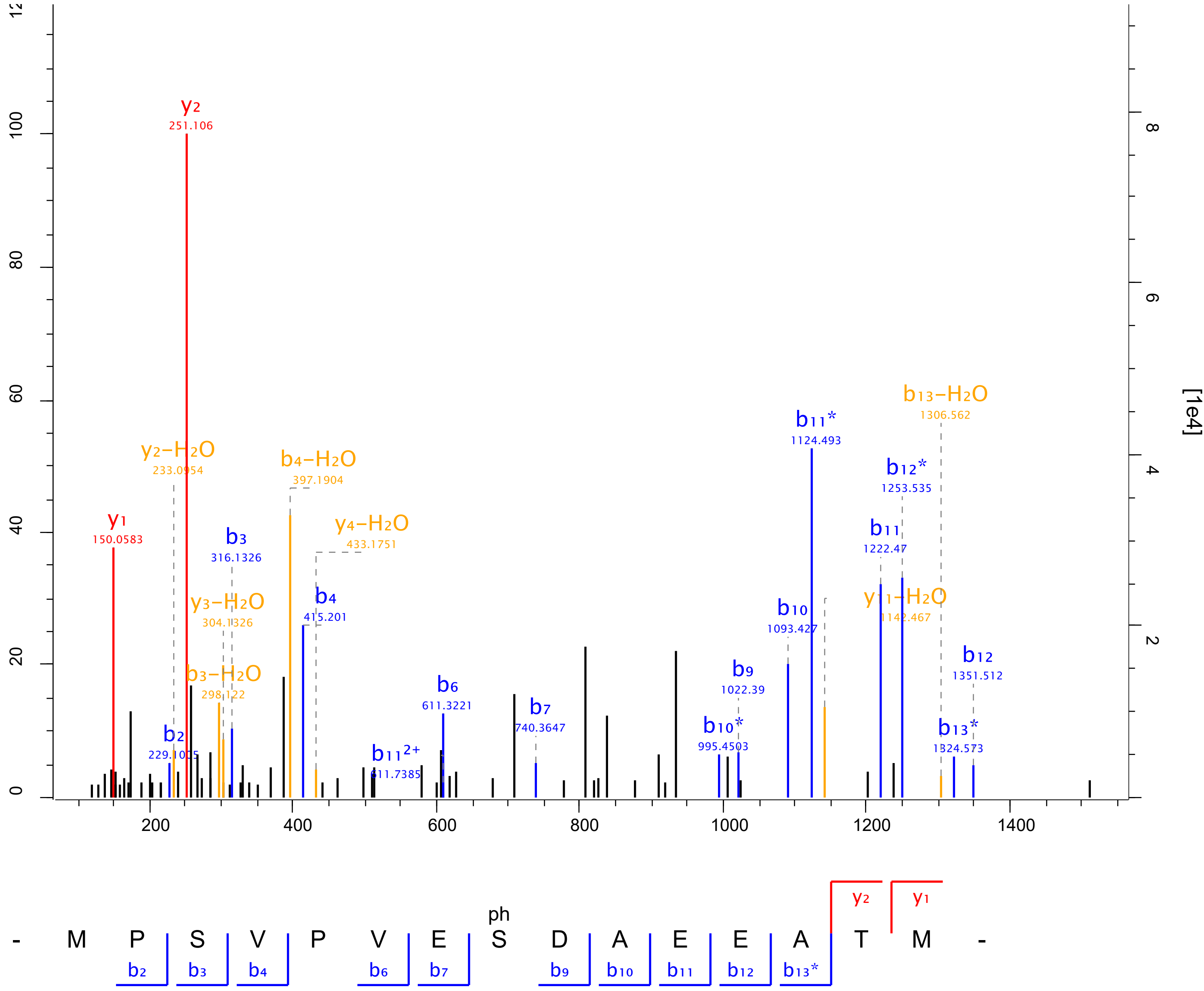

| Raw file | Scan | Method | Score | m/z |
|---|---|---|---|---|
| water_02_2 | 3647 | FTMS; HCD | 164.13 | 546.78 |

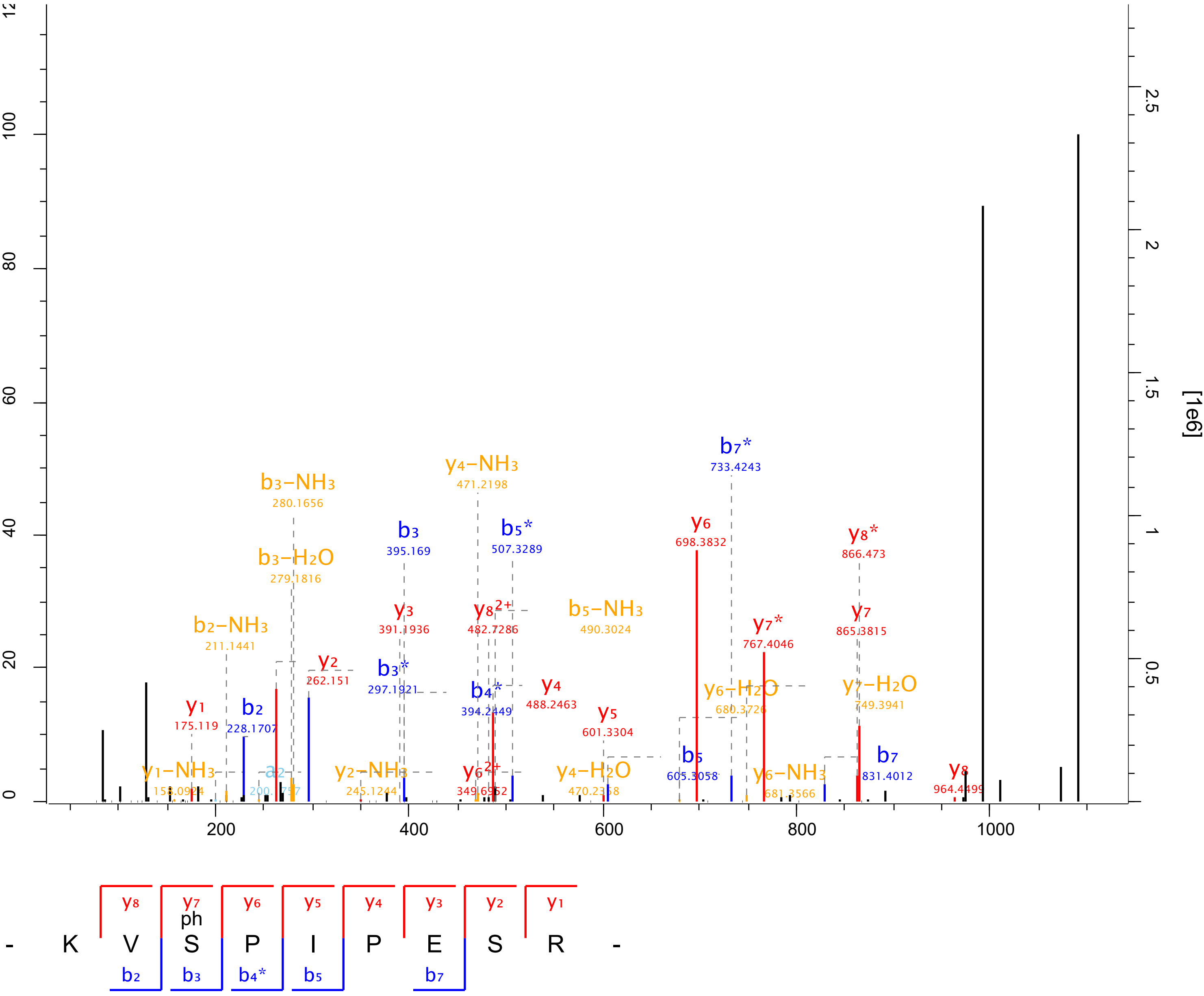

| Raw file | Scan | Method | Score | m/z |
|---|---|---|---|---|
| water_02_2 | 4797 | FTMS; HCD | 248.78 | 771.86 |

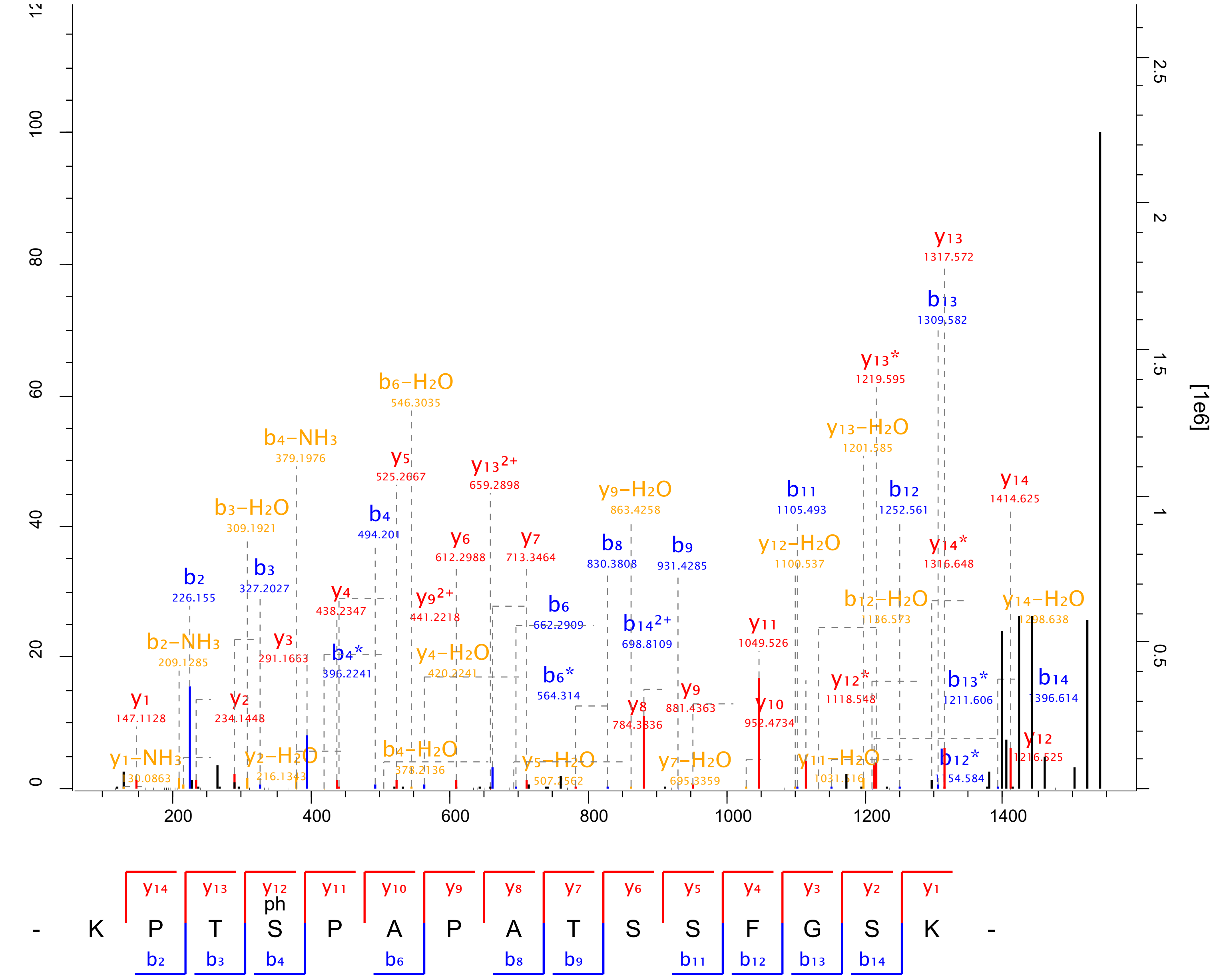

| Raw file | Scan | Method | Score | m/z |
| --- | --- | --- | --- | --- |
| water_02_2 | 5045 | FTMS; HCD | 63.62 | 477.71 |

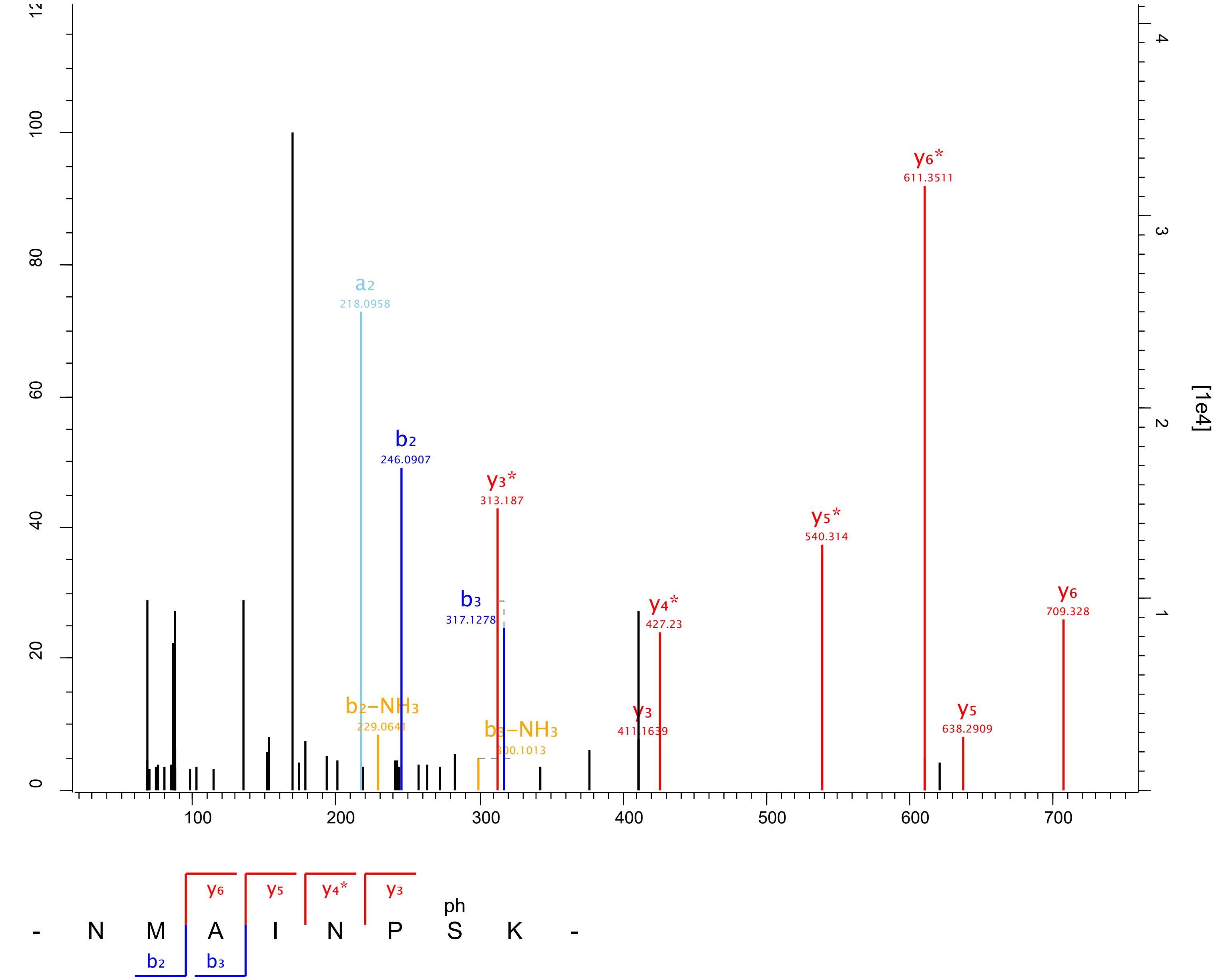

| Raw file | Scan | Method | Score | m/z |
| --- | --- | --- | --- | --- |
| water_02_2 | 5093 | FTMS; HCD | 42.34 | 585.76 |

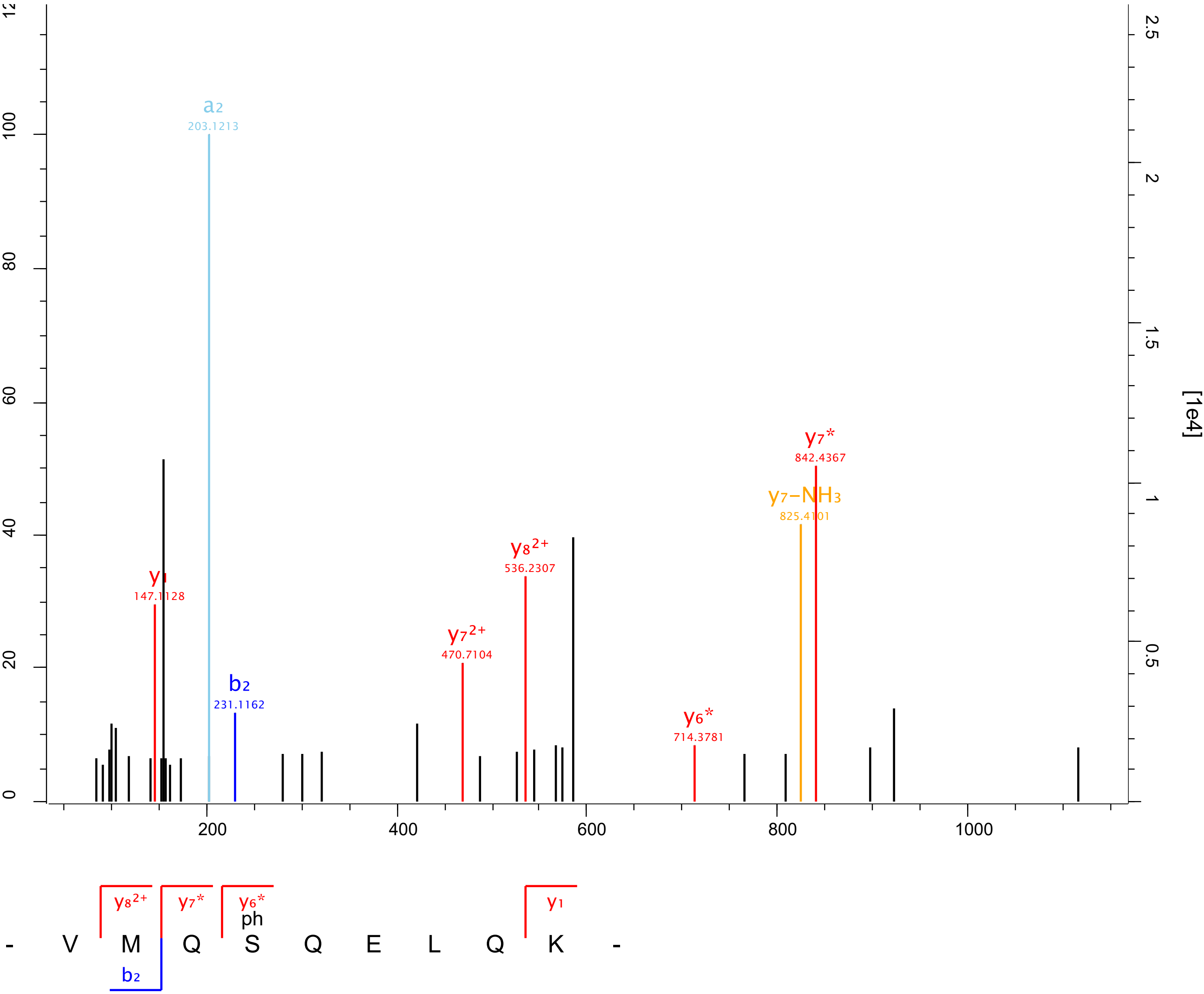

| Raw file | Scan | Method | Score | m/z |
| --- | --- | --- | --- | --- |
| water_02_2 | 5433 | FTMS; HCD | 69.92 | 478.22 |

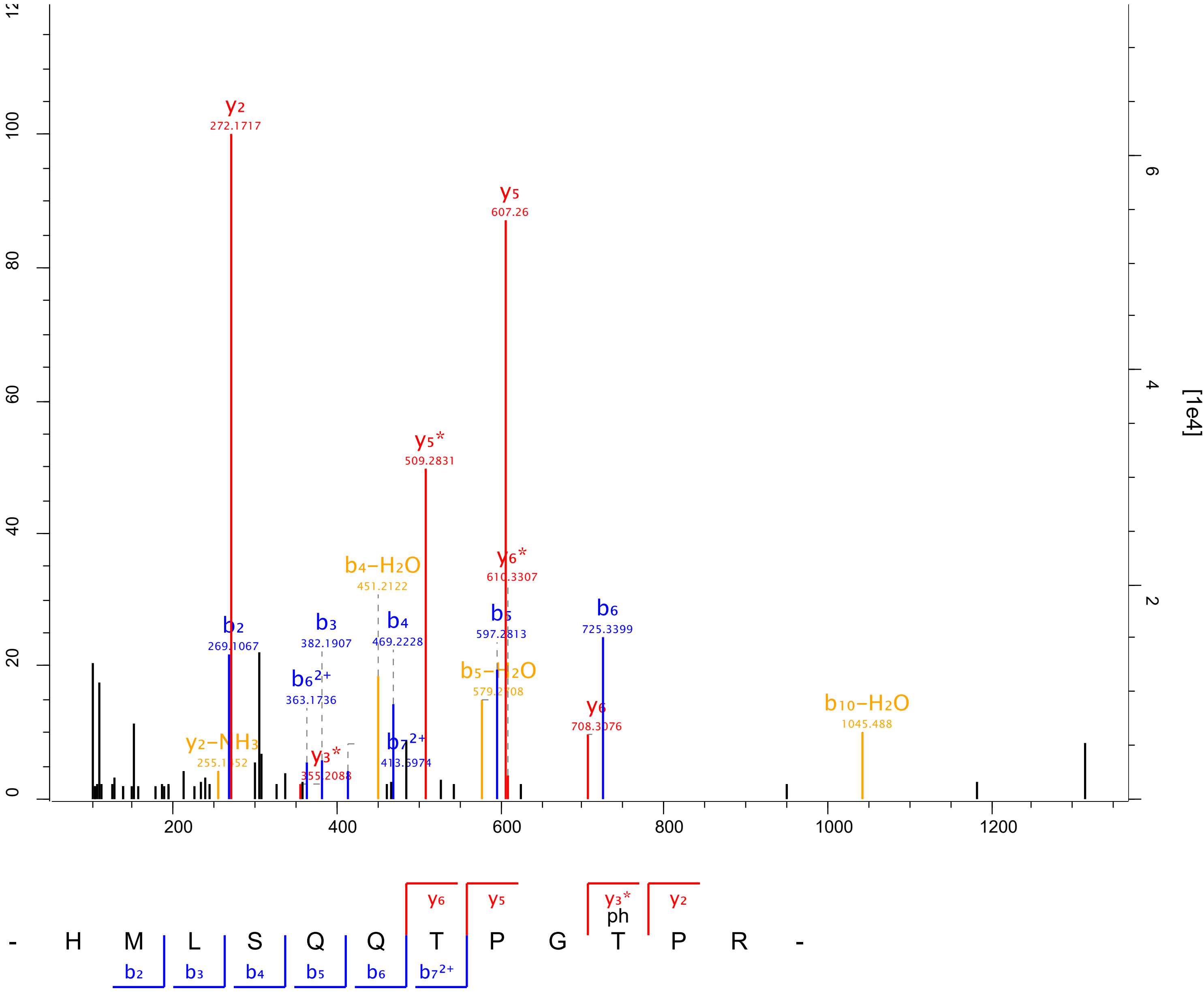

| Raw file | Scan | Method | Score | m/z |
| --- | --- | --- | --- | --- |
| water_02_2 | 5588 | FTMS; HCD | 50.9 | 570.75 |

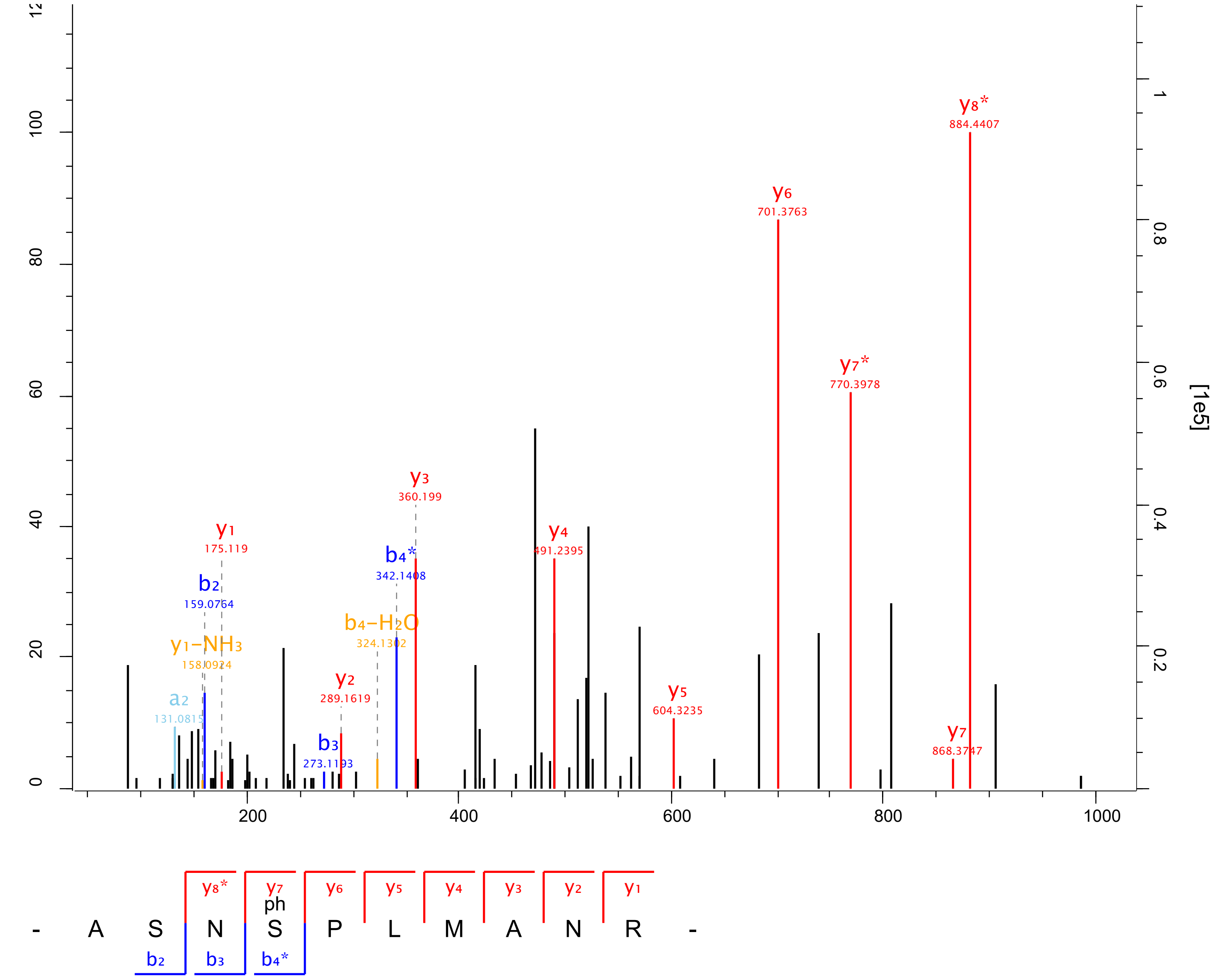

| Raw file | Scan | Method | Score | m/z |
|---|---|---|---|---|
| water_02_2 | 5844 | FTMS; HCD | 46.88 | 538.25 |

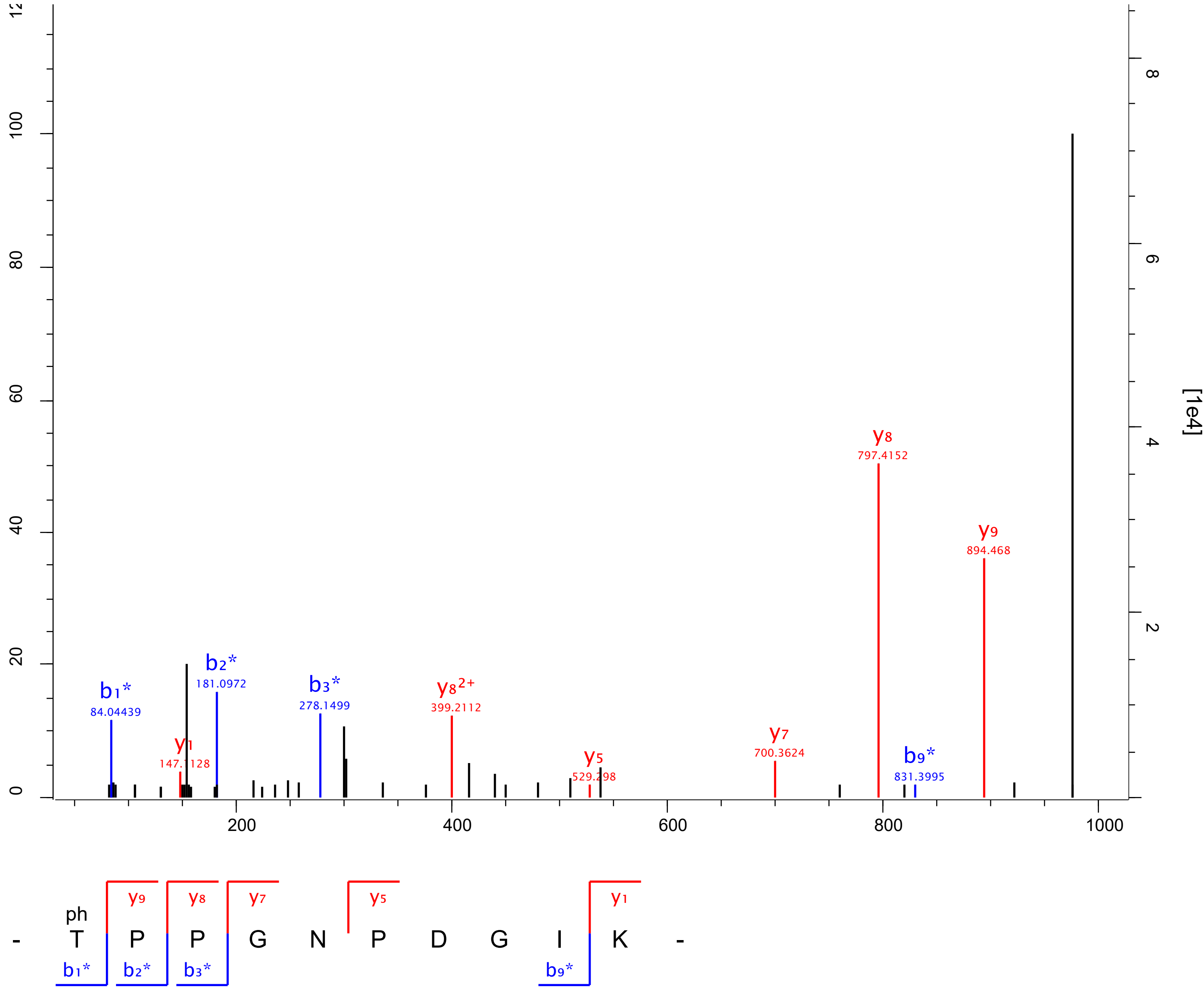

| Raw file | Scan | Method | Score | m/z |
|---|---|---|---|---|
| water_02_2 | 7538 | FTMS; HCD | 73.5 | 469.72 |

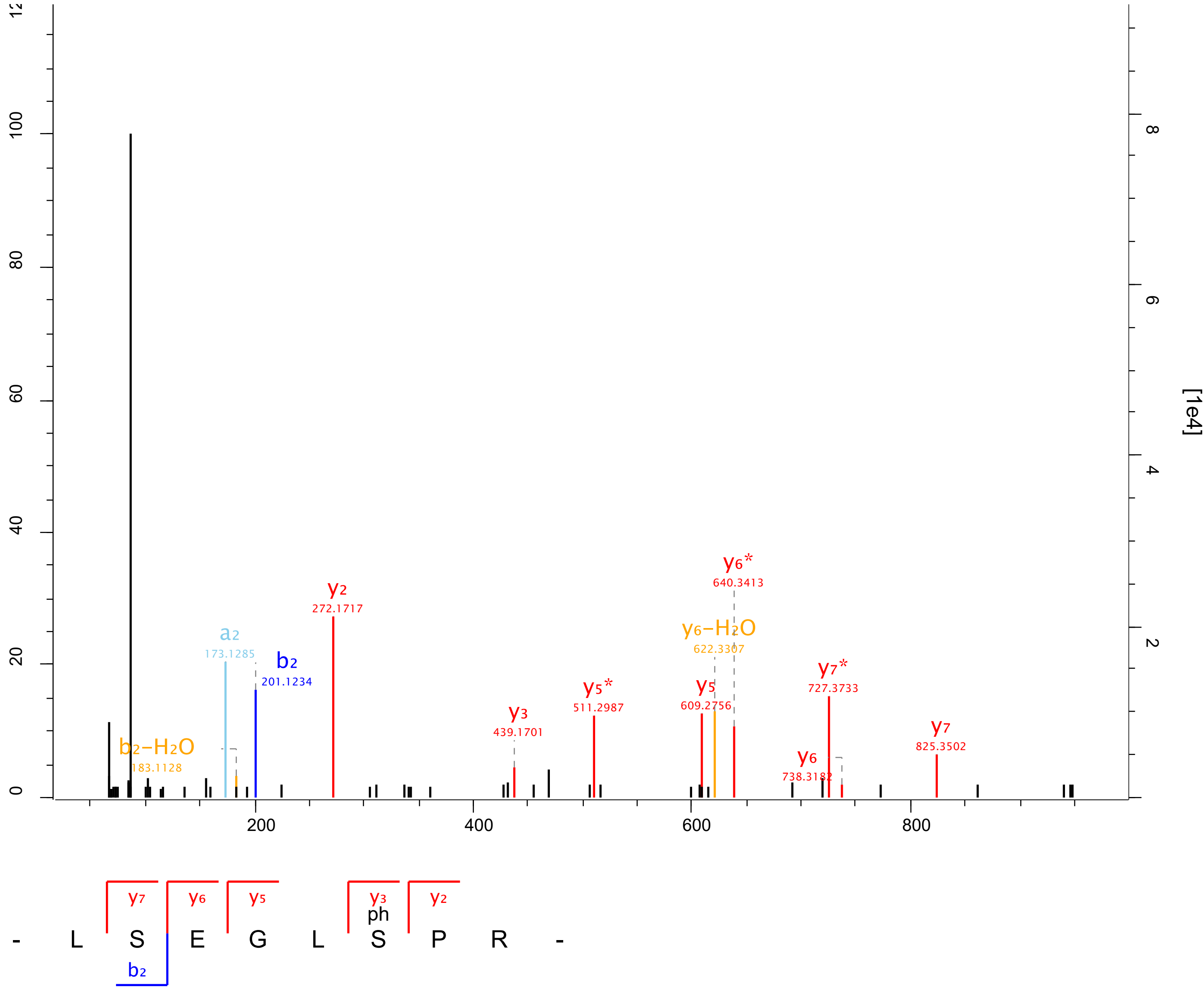

| Raw file | Scan | Method | Score | m/z |
|---|---|---|---|---|
| water_05_1 | 11055 | FTMS; HCD | 137.89 | 515.24 |

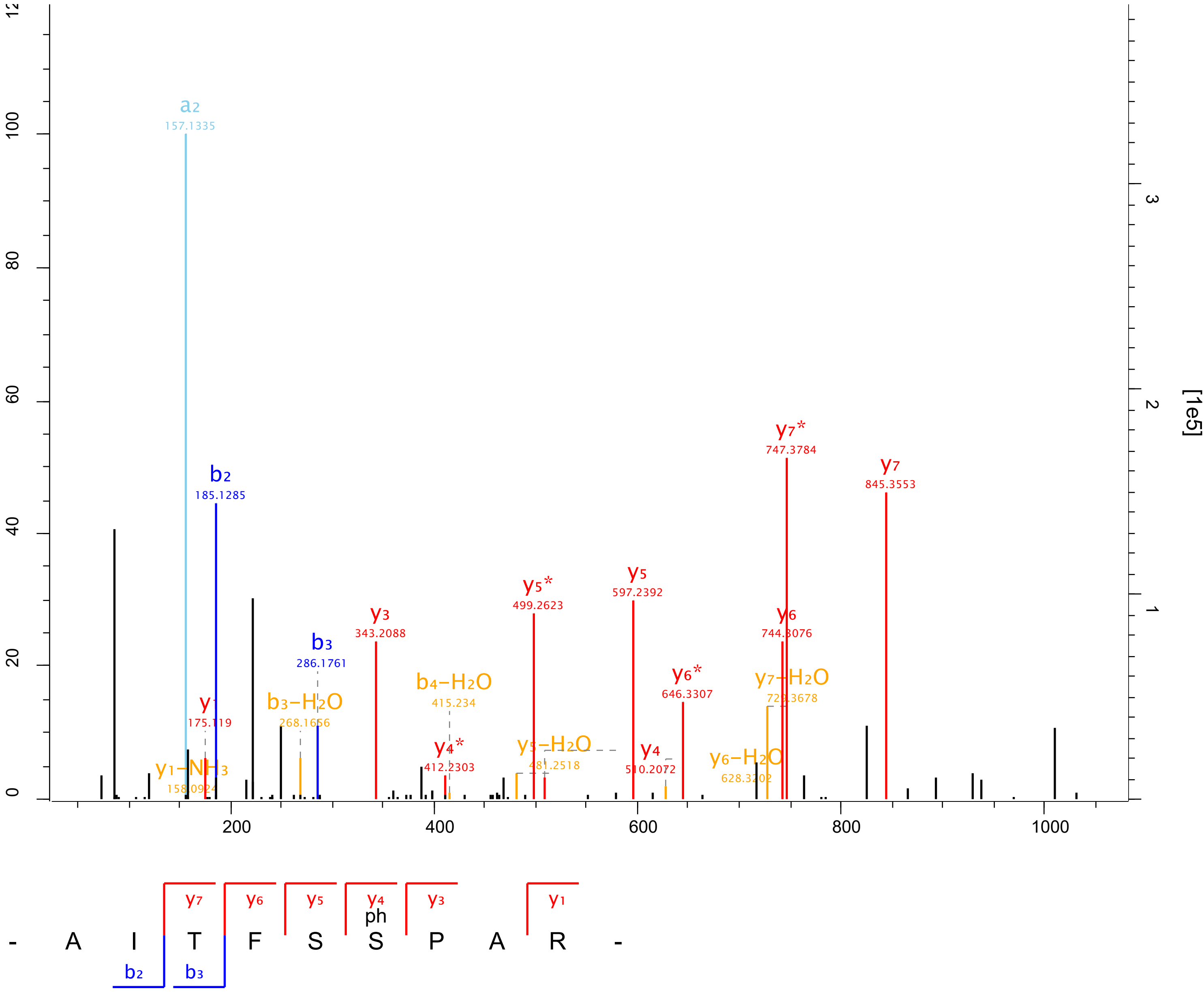

| Raw file | Scan | Method | Score | m/z |
| --- | --- | --- | --- | --- |
| water_05_1 | 16754 | FTMS; HCD | 81.53 | 1027.04 |

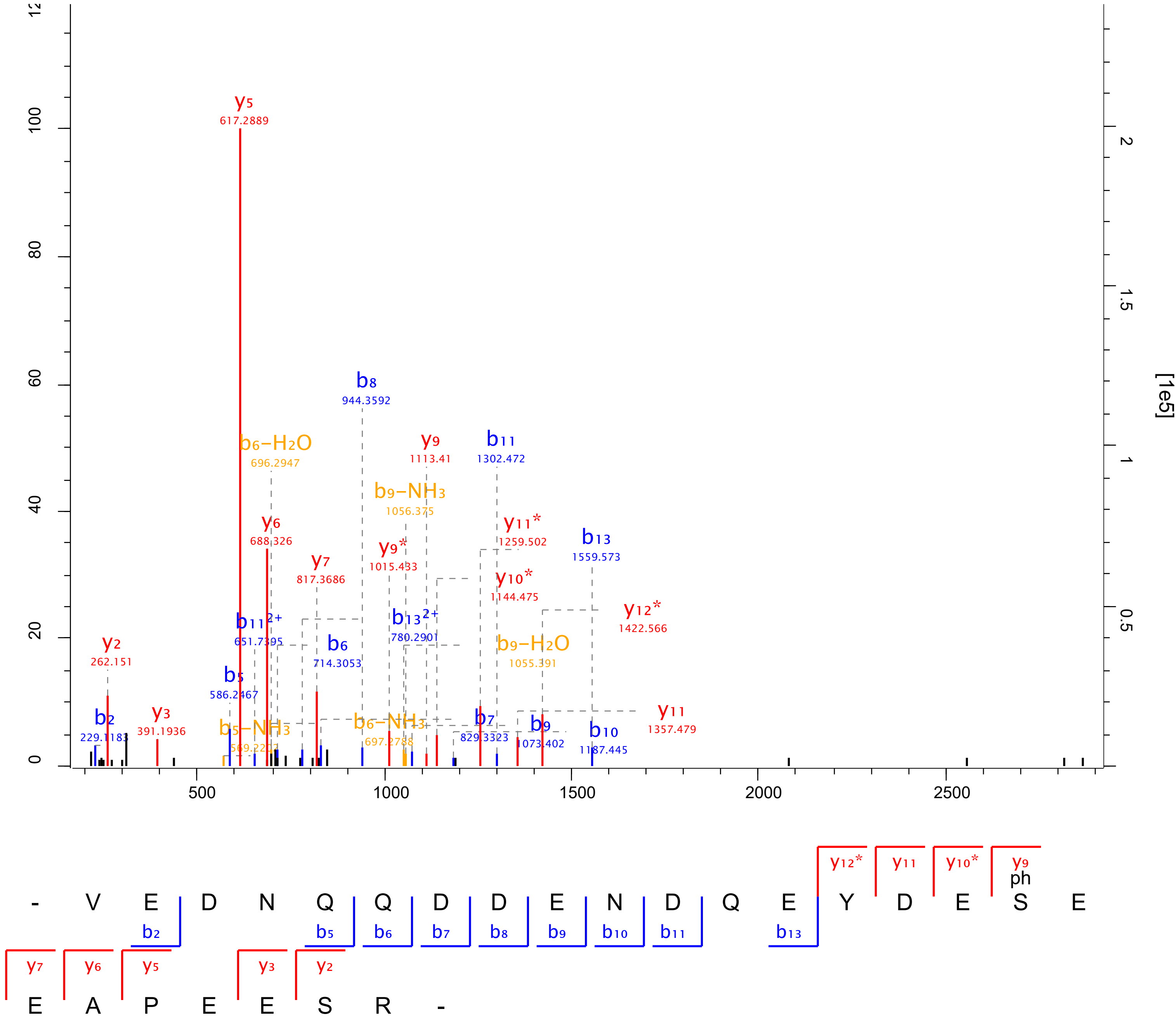

| Raw file | Scan | Method | Score | m/z |
|---|---|---|---|---|
| water_05_1 | 1992 | FTMS; HCD | 65.84 | 562.75 |

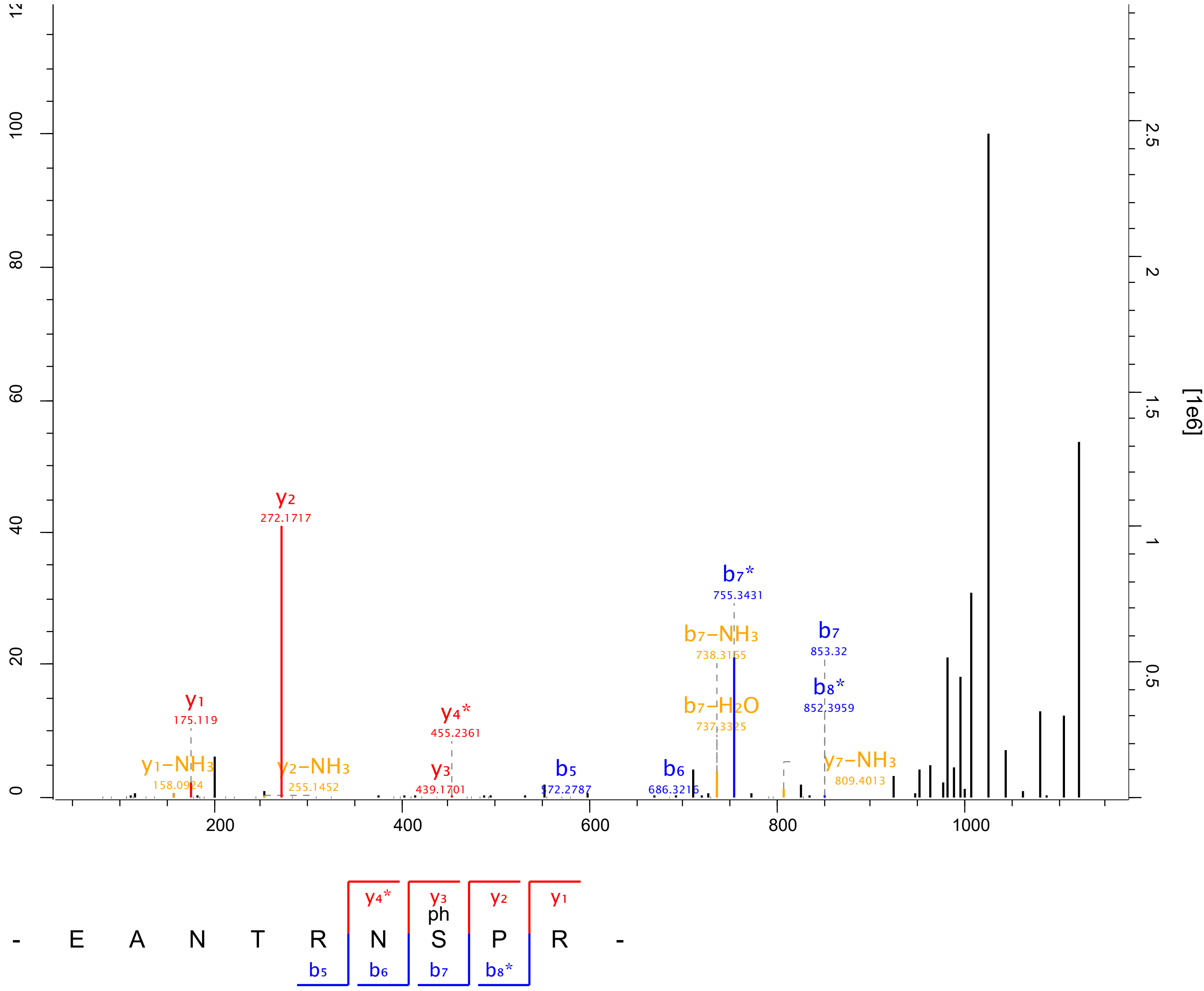

| Raw file | Scan | Method | Score | m/z |
| --- | --- | --- | --- | --- |
| water_05_1 | 2046 | FTMS; HCD | 143.58 | 592.75 |

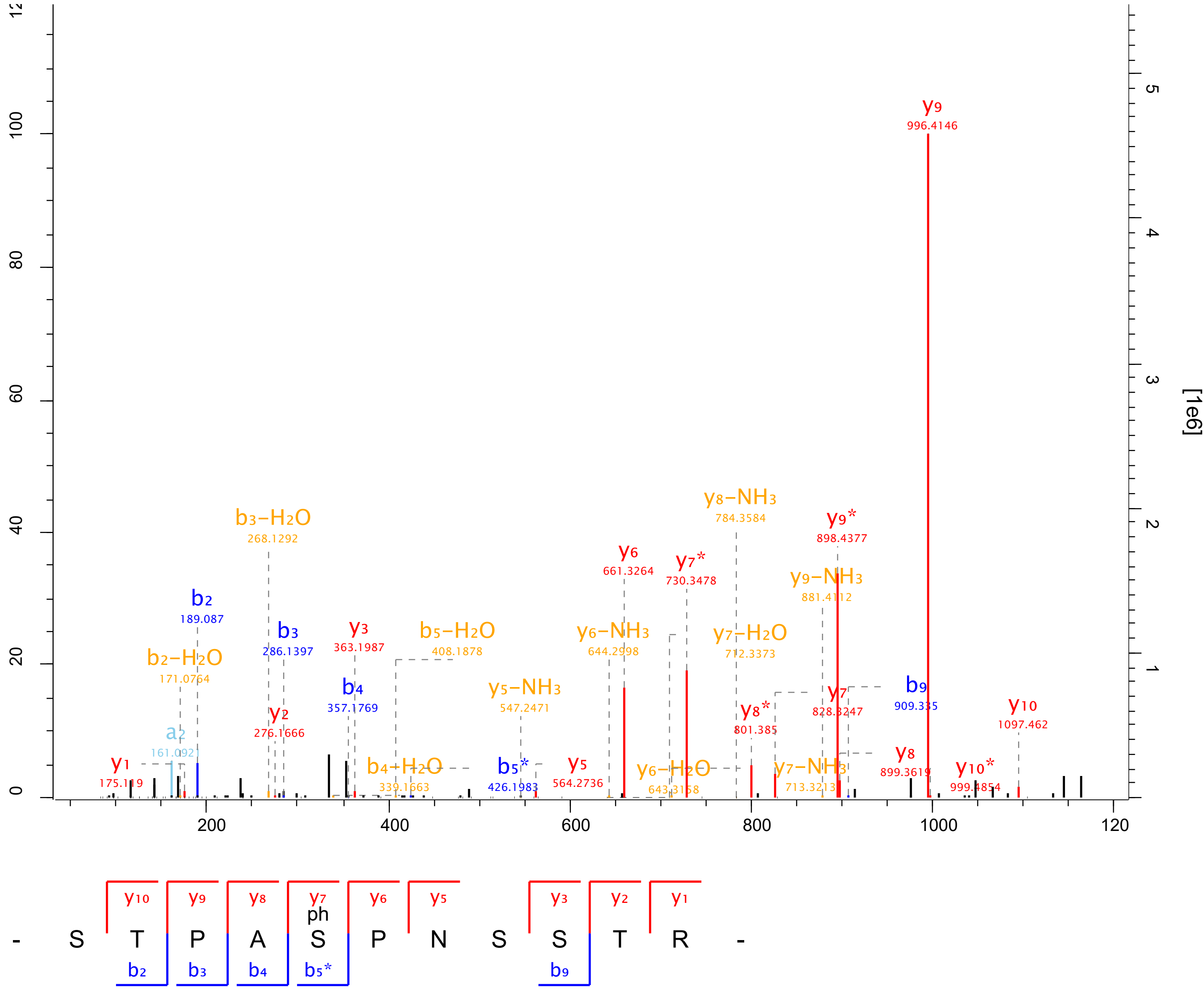

| Raw file | Scan | Method | Score | m/z |
| --- | --- | --- | --- | --- |
| water_05_1 | 3176 | FTMS; HCD | 40.48 | 556.89 |

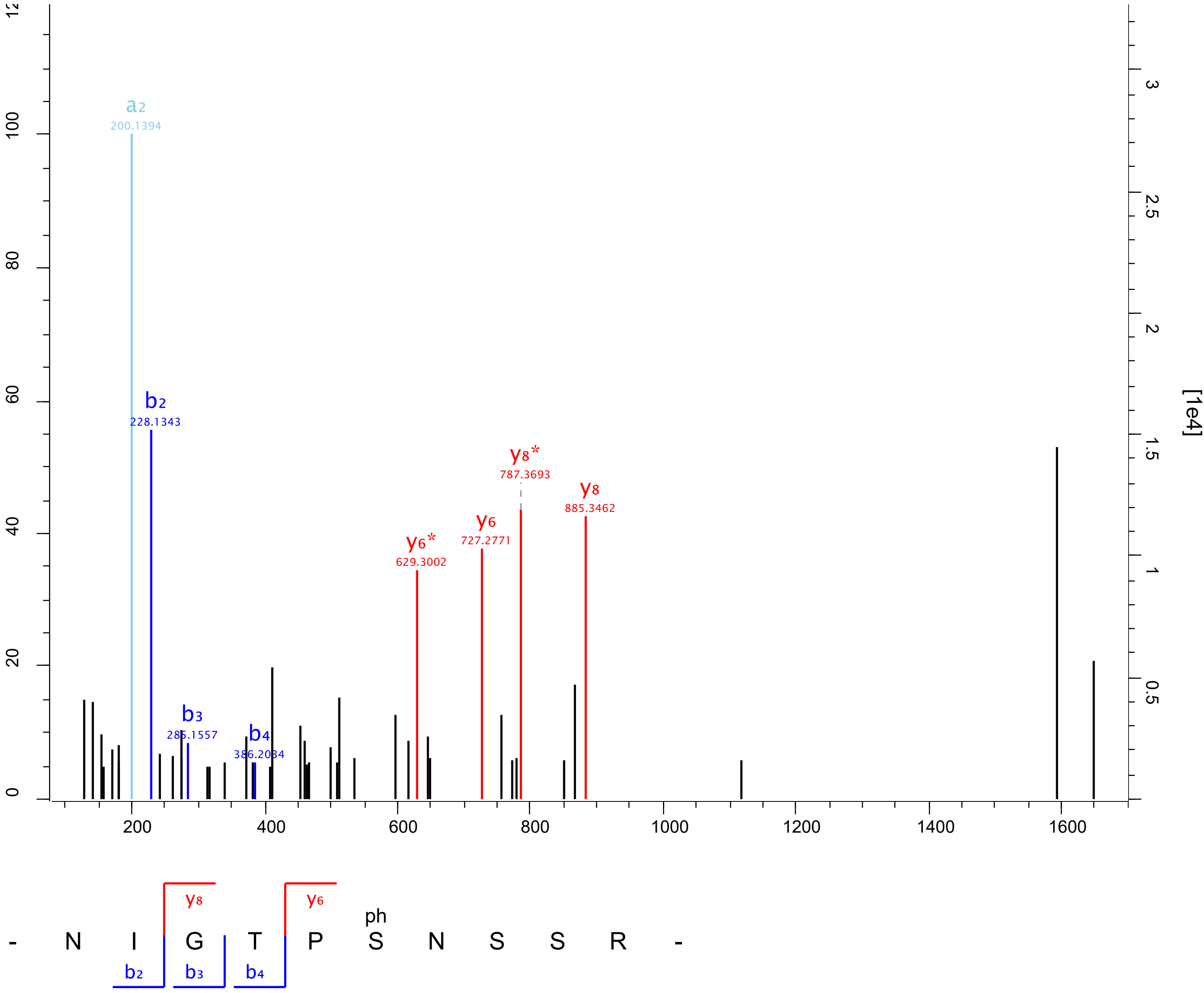

| Raw file | Scan | Method | Score | m/z |
|---|---|---|---|---|
| water_05_1 | 34844 | FTMS; HCD | 109 | 801.76 |

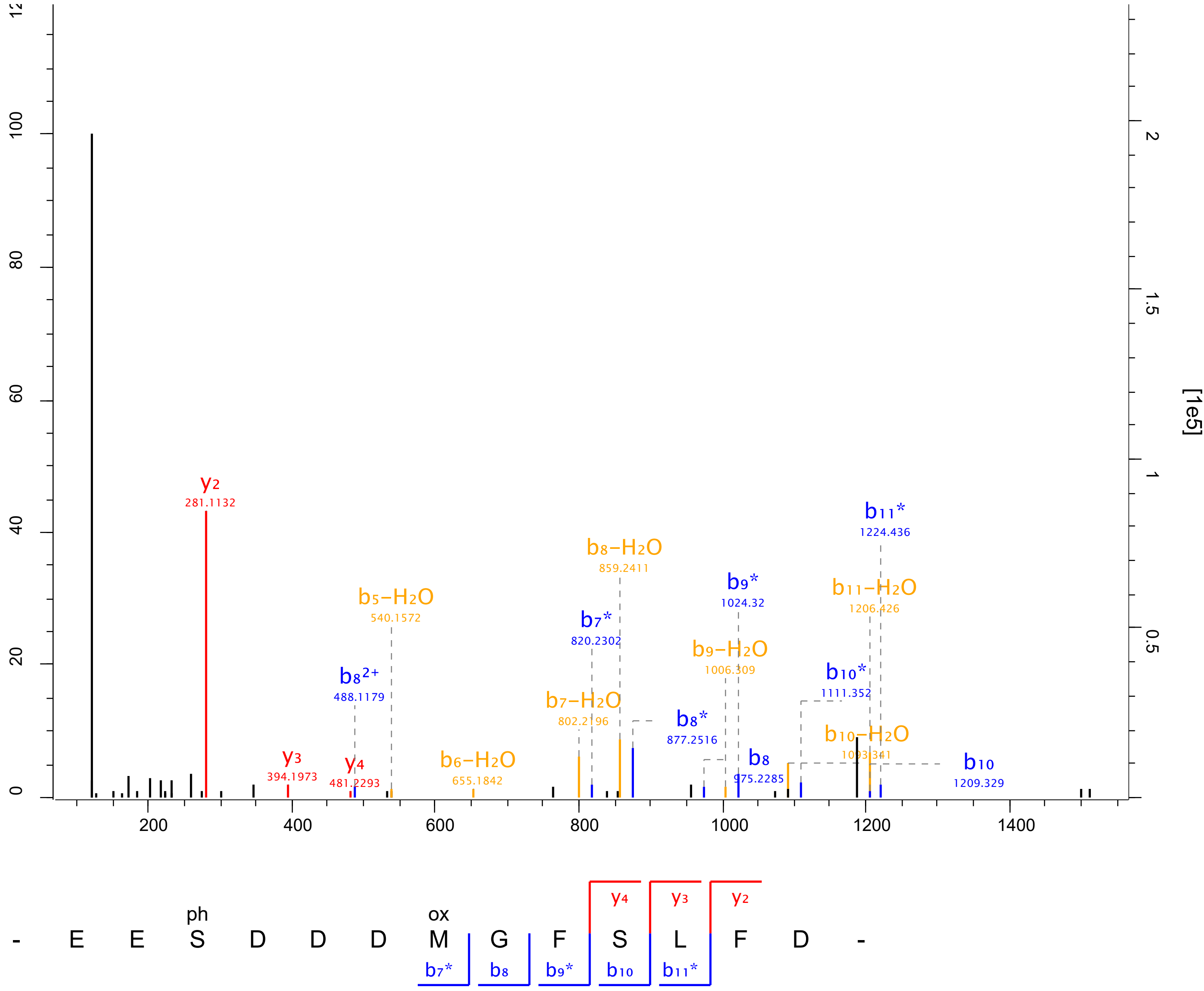

| Raw file | Scan | Method | Score | m/z |
|---|---|---|---|---|
| water_05_1 | 3934 | FTMS; HCD | 70.94 | 337.16 |

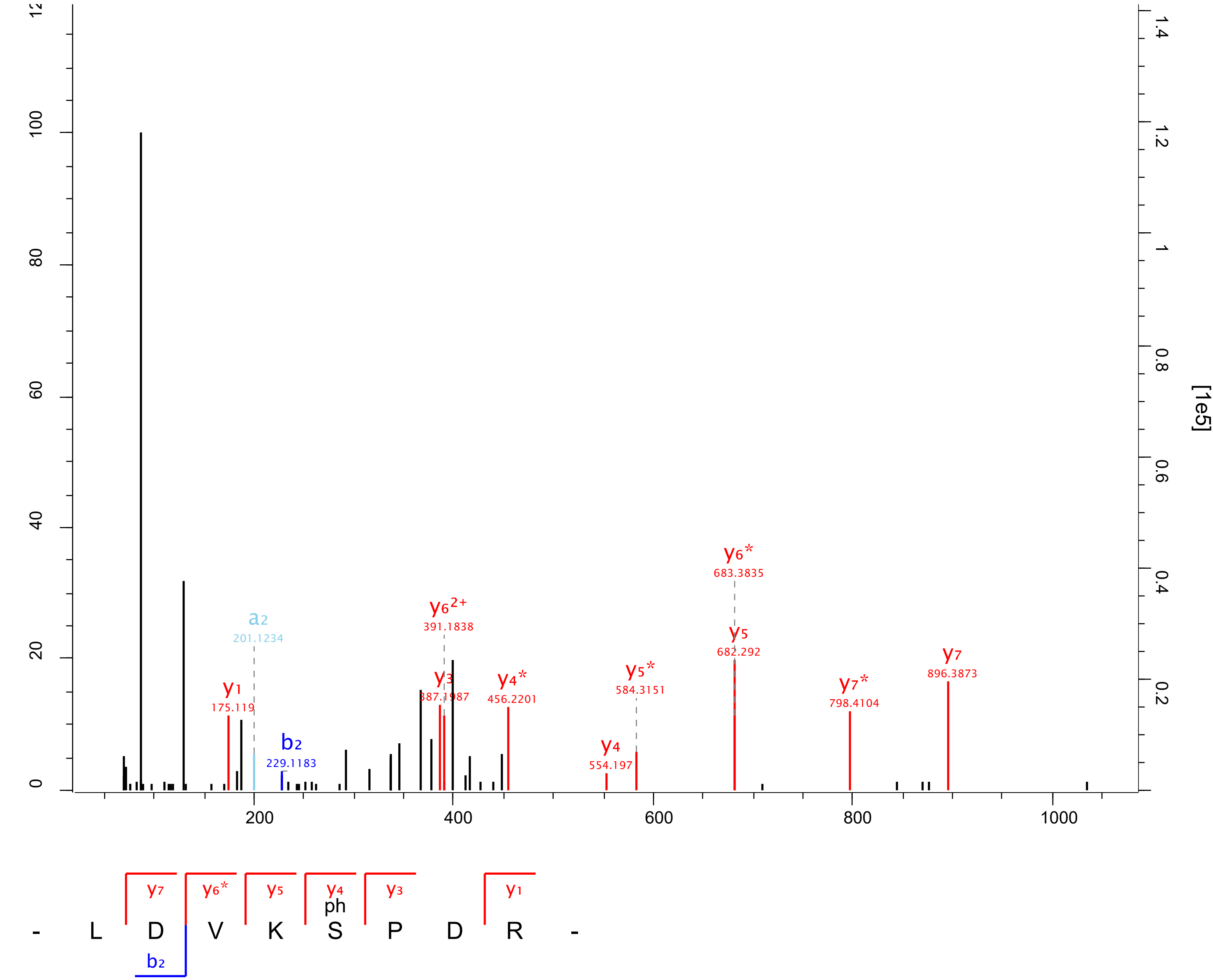

| Raw file | Scan | Method | Score | m/z |
|---|---|---|---|---|
| water_05_1 | 5103 | FTMS; HCD | 100.88 | 427.7 |

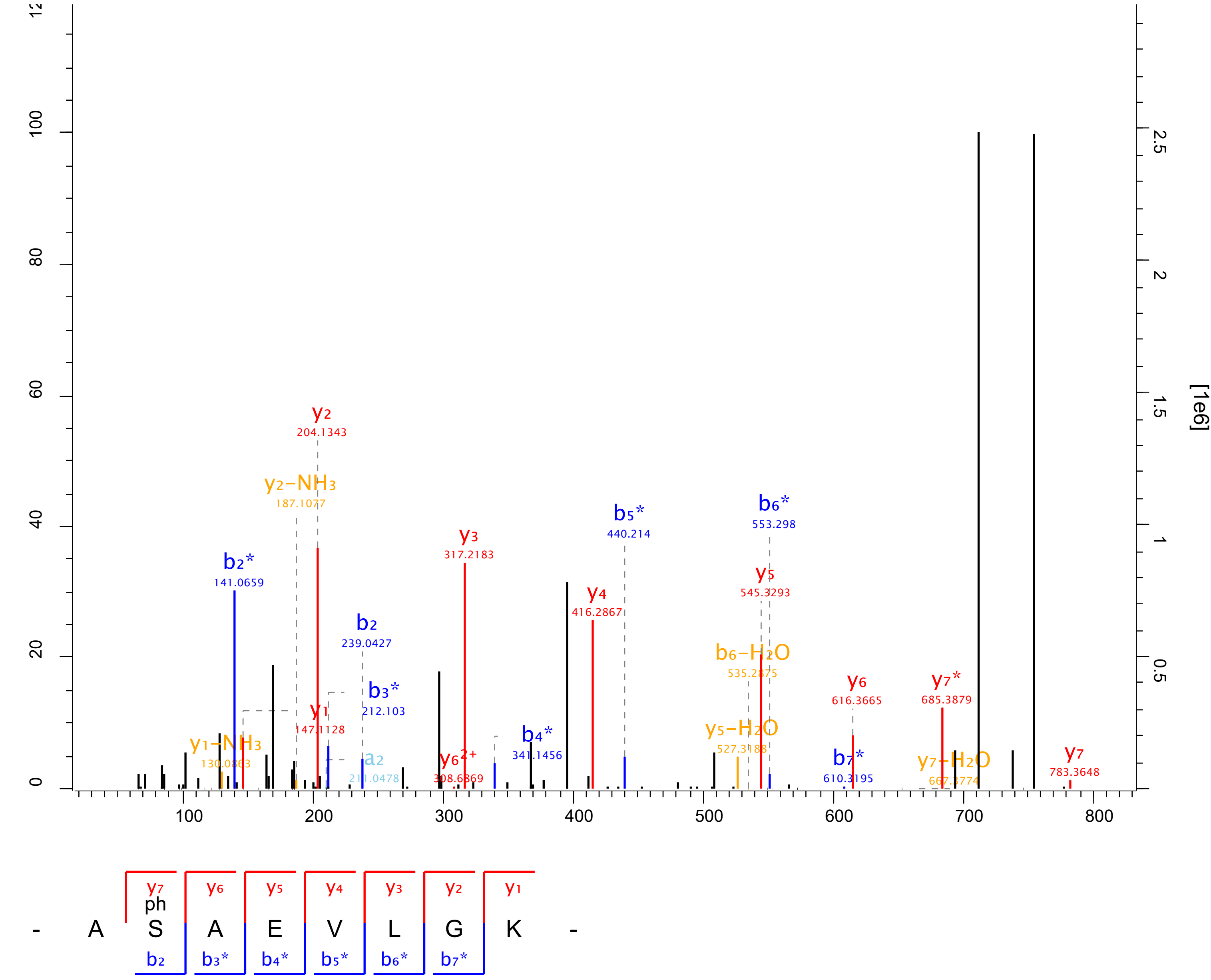

| Raw file | Scan | Method | Score | m/z |
|---|---|---|---|---|
| water_05_1 | 6272 | FTMS; HCD | 138.76 | 493.22 |

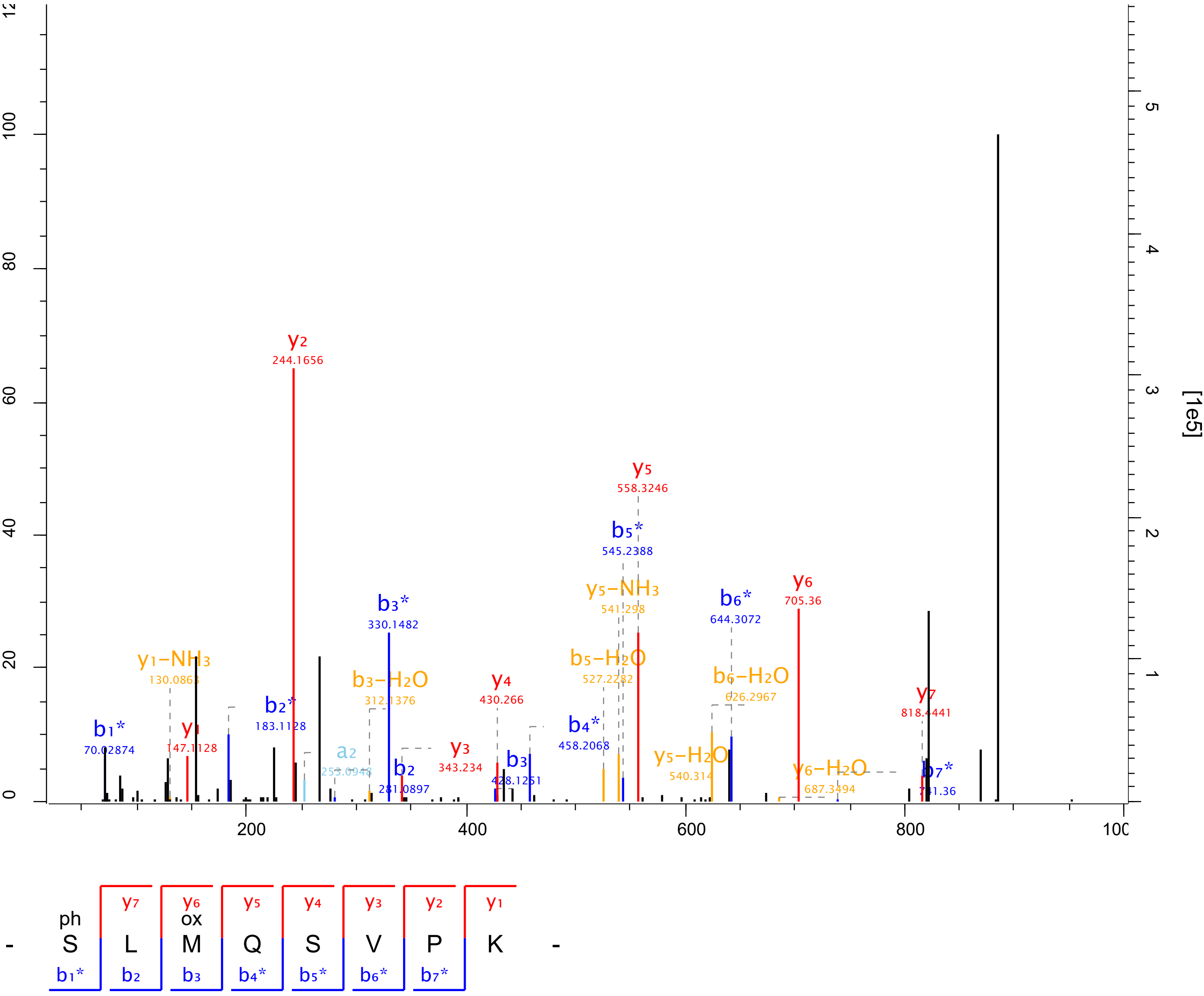

| Raw file | Scan | Method | Score | m/z |
|---|---|---|---|---|
| water_05_1 | 6411 | FTMS; HCD | 51.29 | 550.74 |

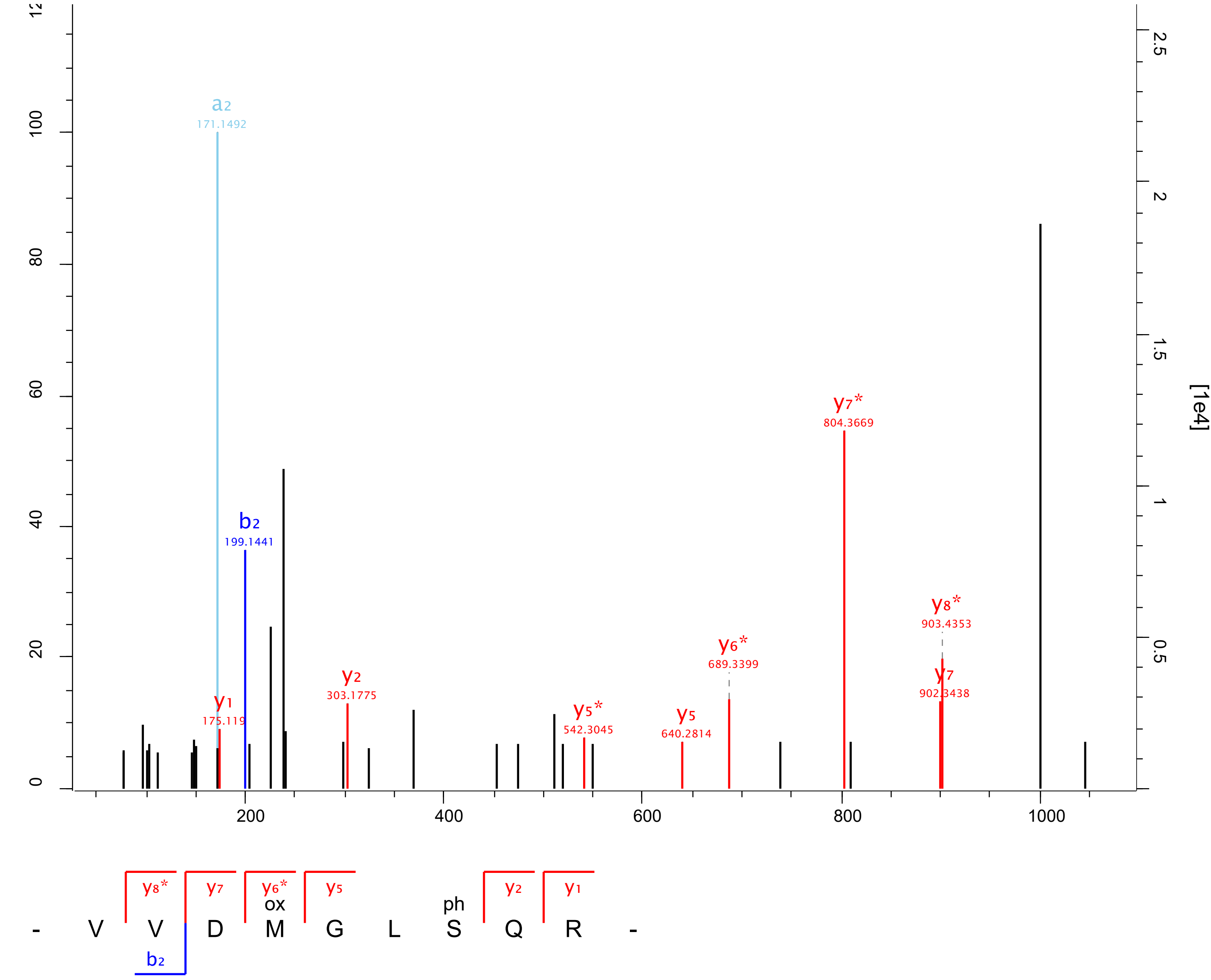

| Raw file | Scan | Method | Score | m/z |
|---|---|---|---|---|
| water_05_1 | 8444 | FTMS; HCD | 155.84 | 441.71 |

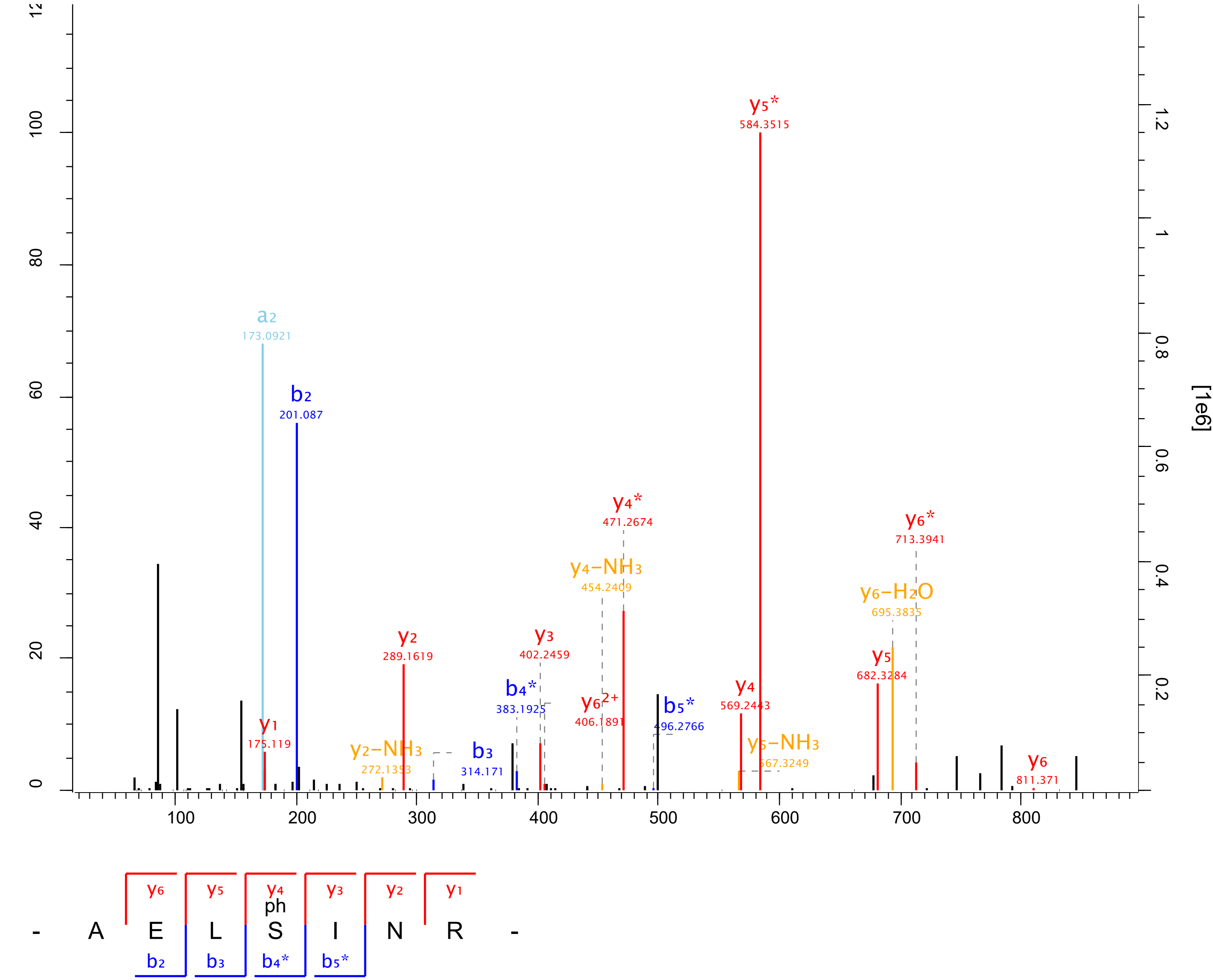

| Raw file | Scan | Method | Score | m/z |
| --- | --- | --- | --- | --- |
| water_05_1 | 9198 | FTMS; HCD | 216.03 | 560.24 |

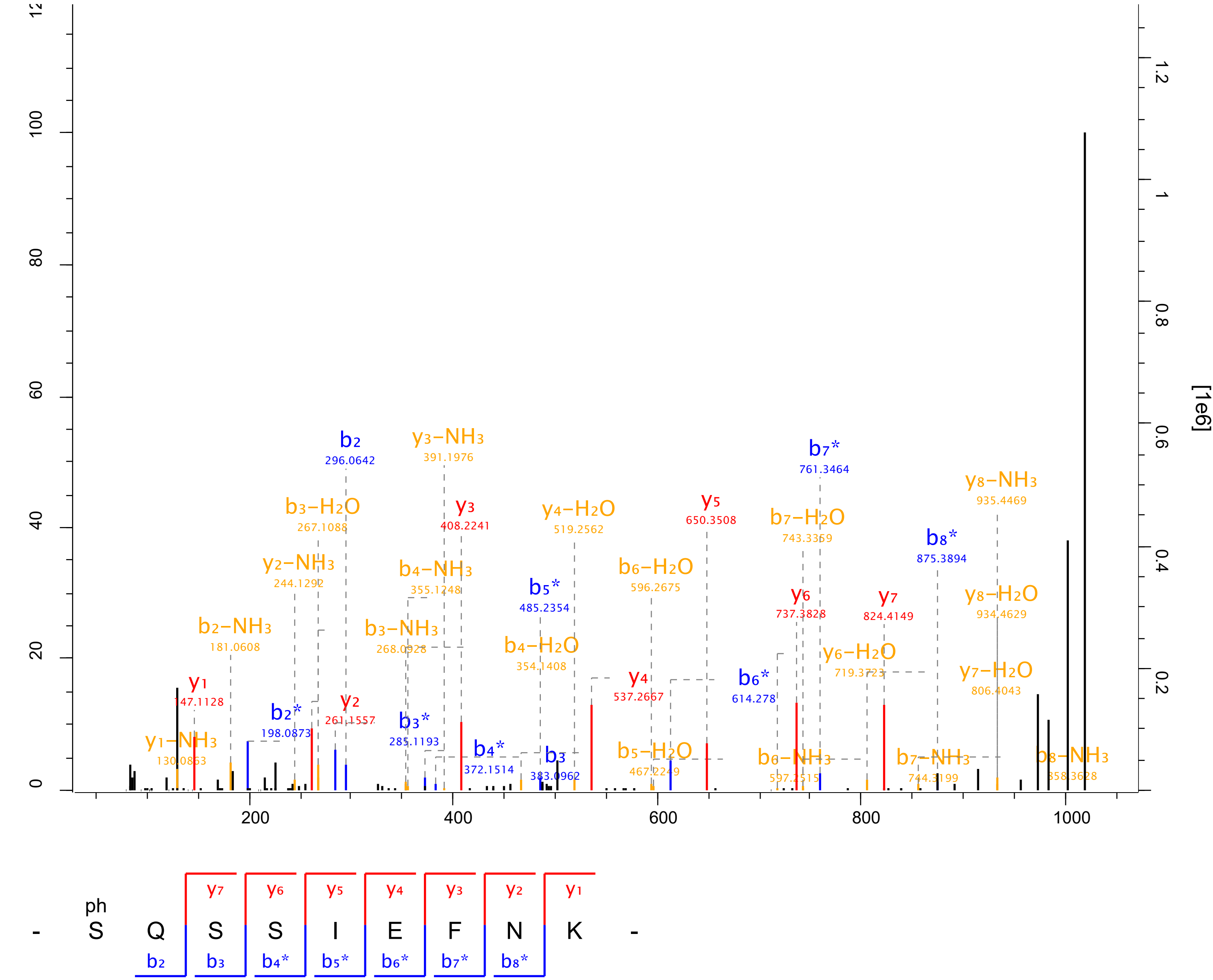

| Raw file | Scan | Method | Score | m/z |
| --- | --- | --- | --- | --- |
| water_05_2 | 14009 | FTMS; HCD | 48.53 | 707.33 |

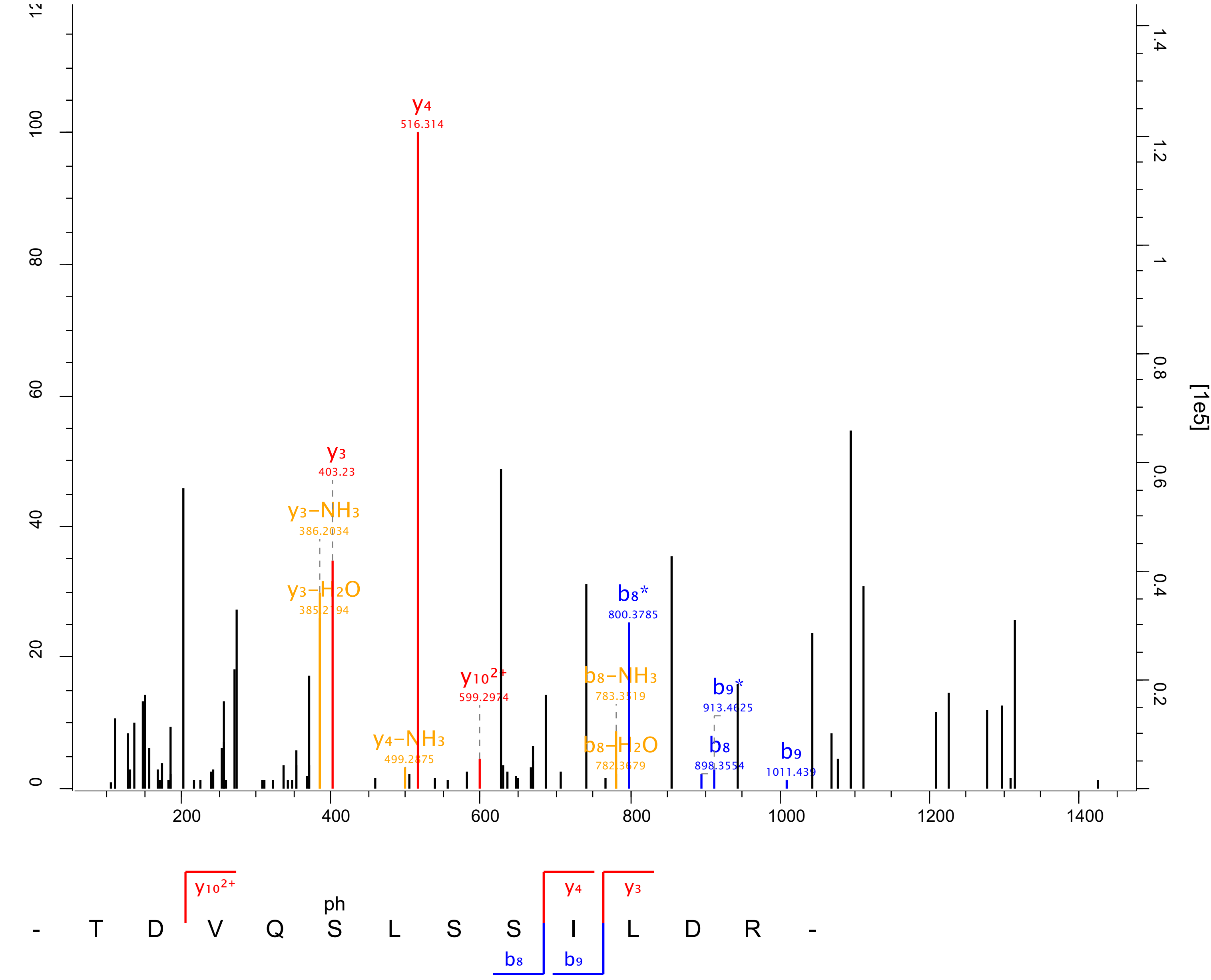

| Raw file | Scan | Method | Score | m/z |
| --- | --- | --- | --- | --- |
| water_05_2 | 18849 | FTMS; HCD | 41.24 | 646.3 |

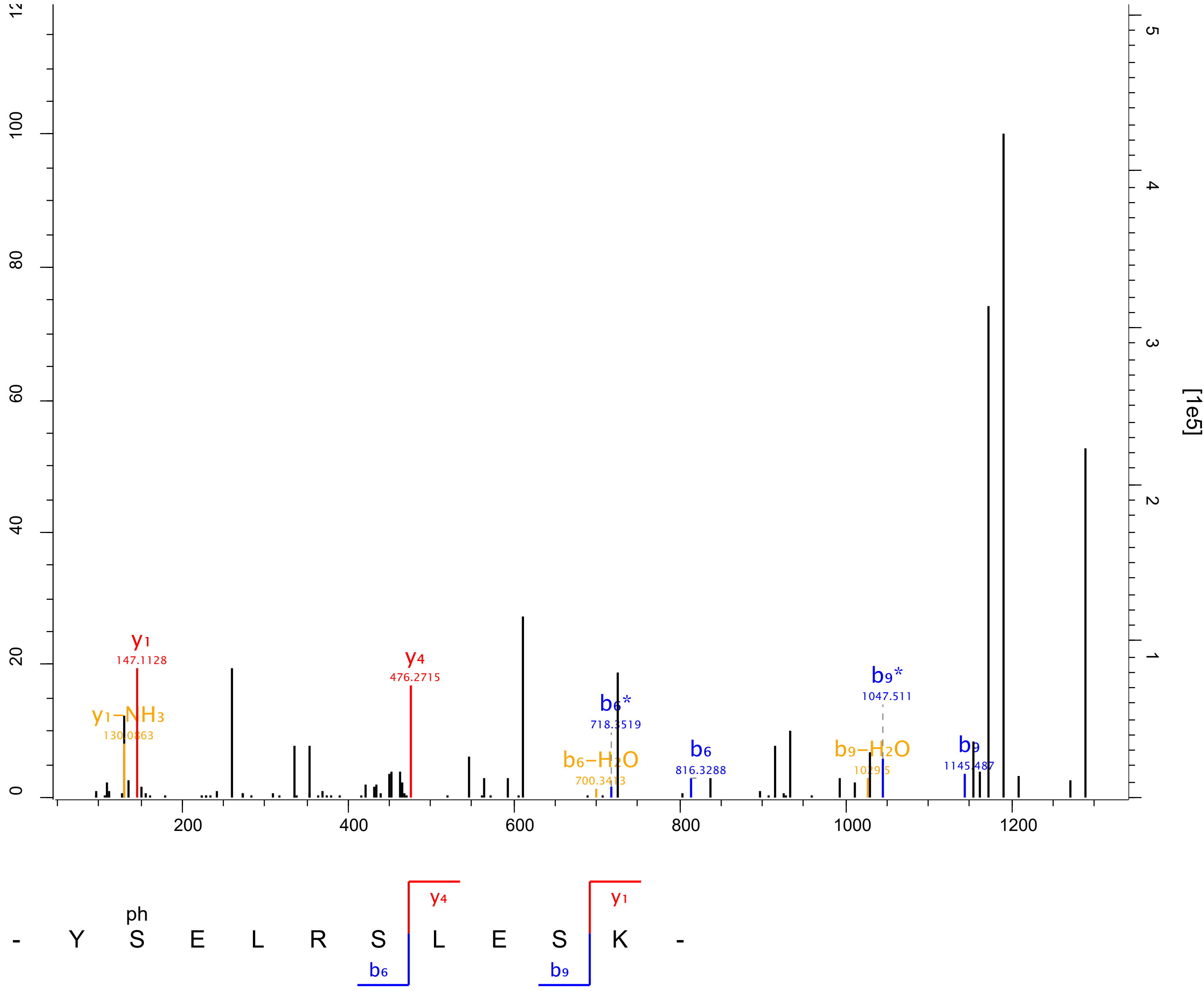

| Raw file | Scan | Method | Score | m/z |
| --- | --- | --- | --- | --- |
| water_05_2 | 5287 | FTMS; HCD | 57.59 | 409.2 |

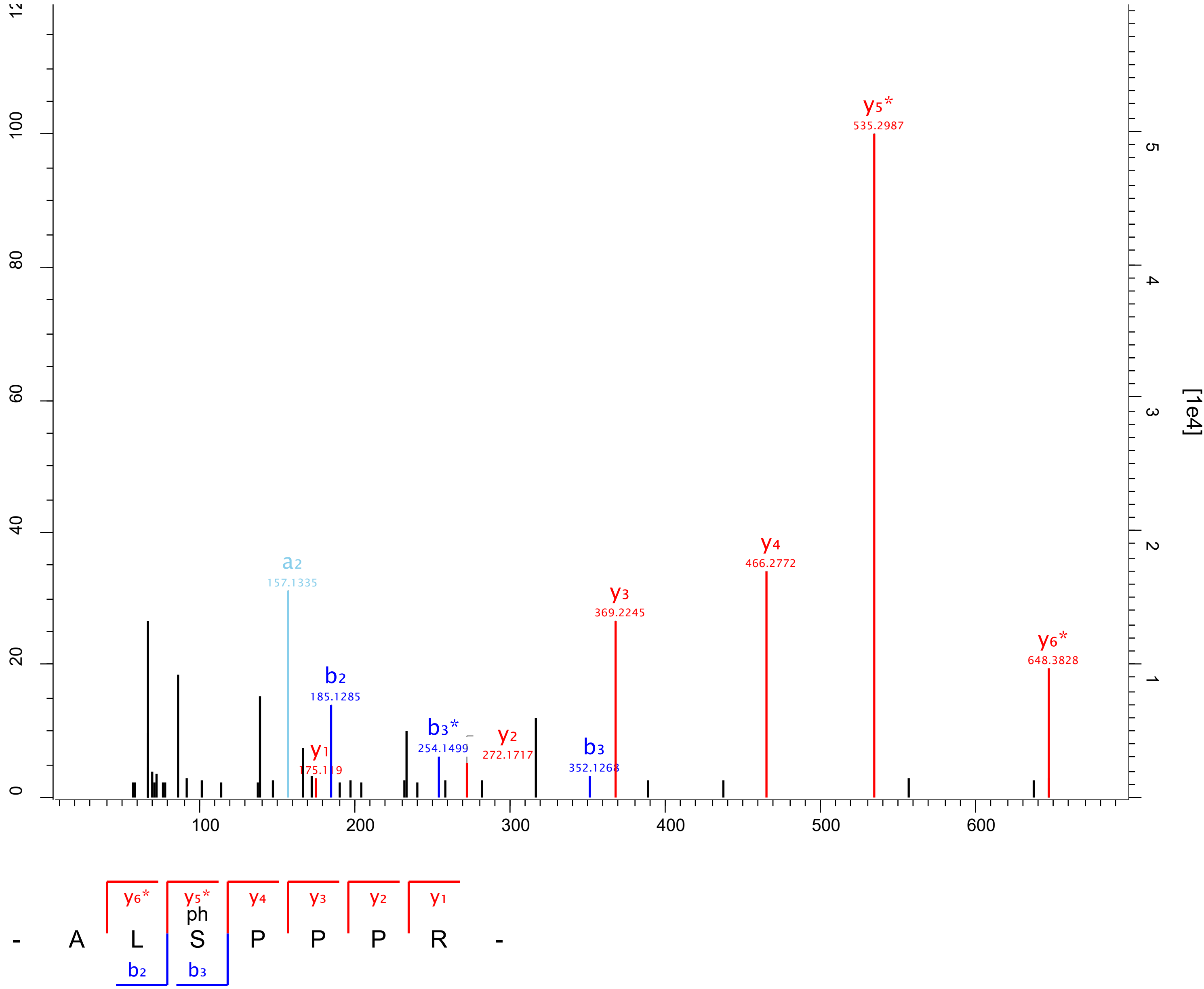

| Raw file | Scan | Method | Score | m/z |
| --- | --- | --- | --- | --- |
| water_05_2 | 7233 | FTMS; HCD | 147.62 | 478.7 |

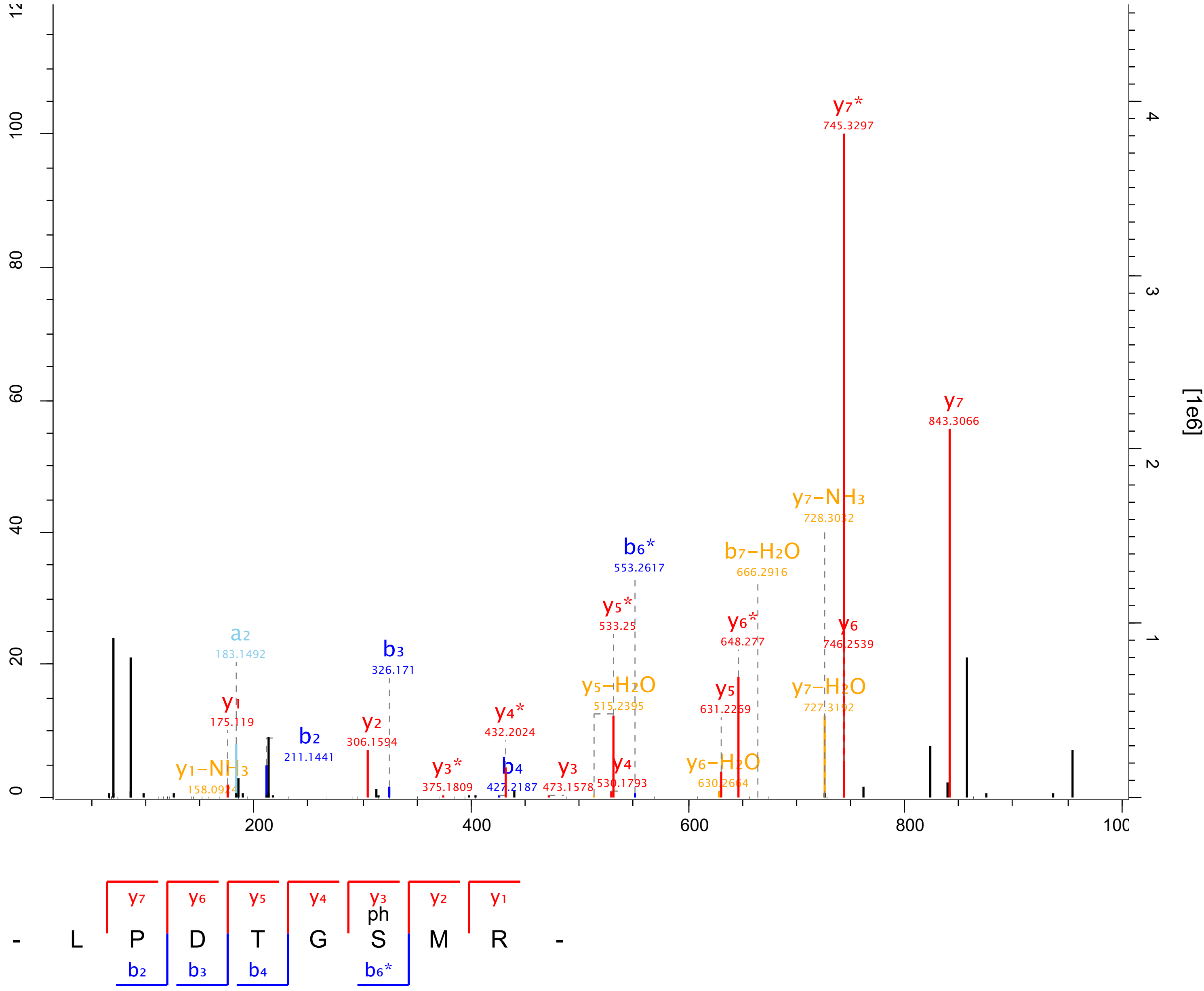

| Raw file | Scan | Method | Score | m/z |
|---|---|---|---|---|
| water_15_1 | 10271 | FTMS; HCD | 87.45 | 697.28 |

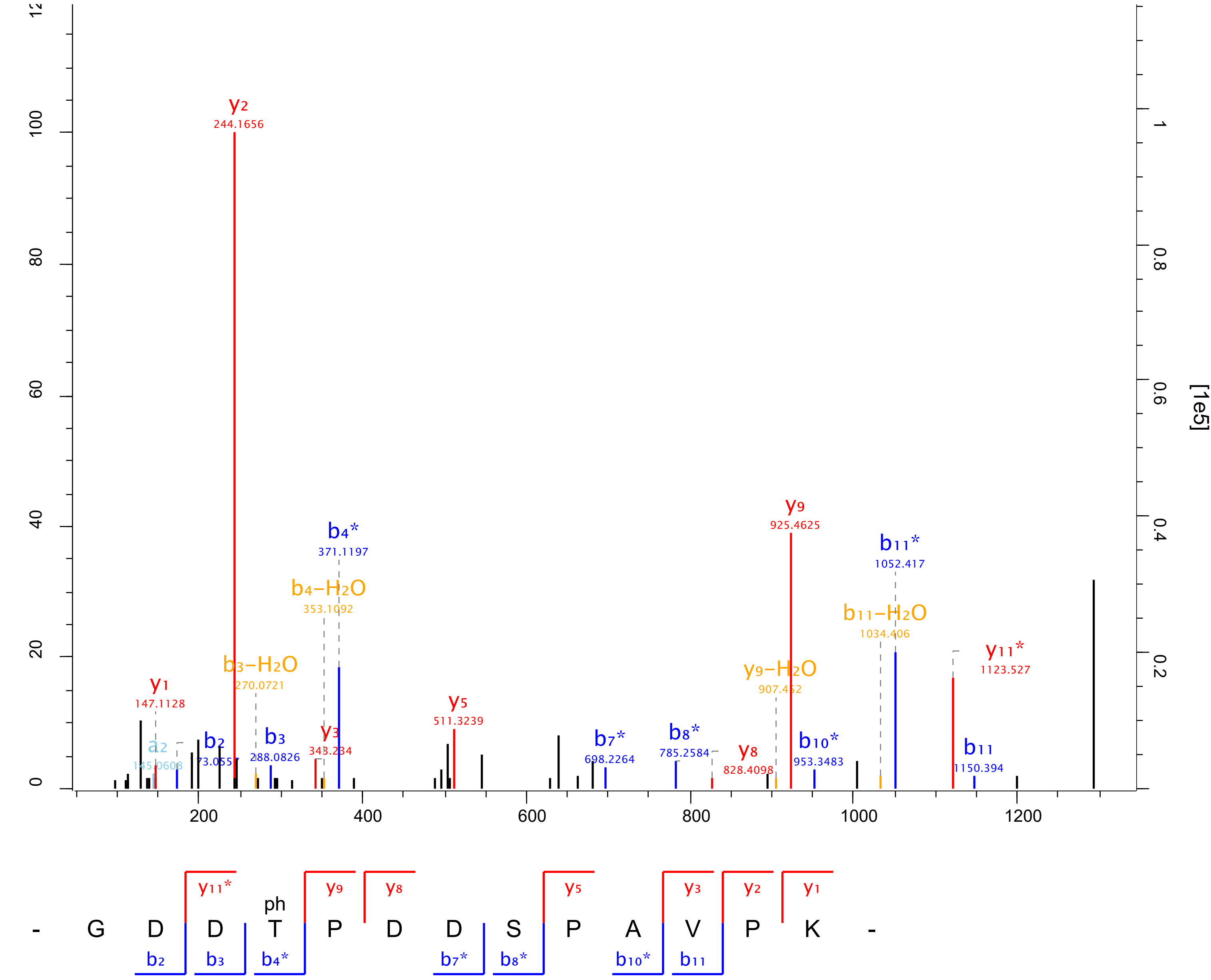

| Raw file | Scan | Method | Score | m/z |
| --- | --- | --- | --- | --- |
| water_15_1 | 10422 | FTMS; HCD | 97.27 | 556.25 |

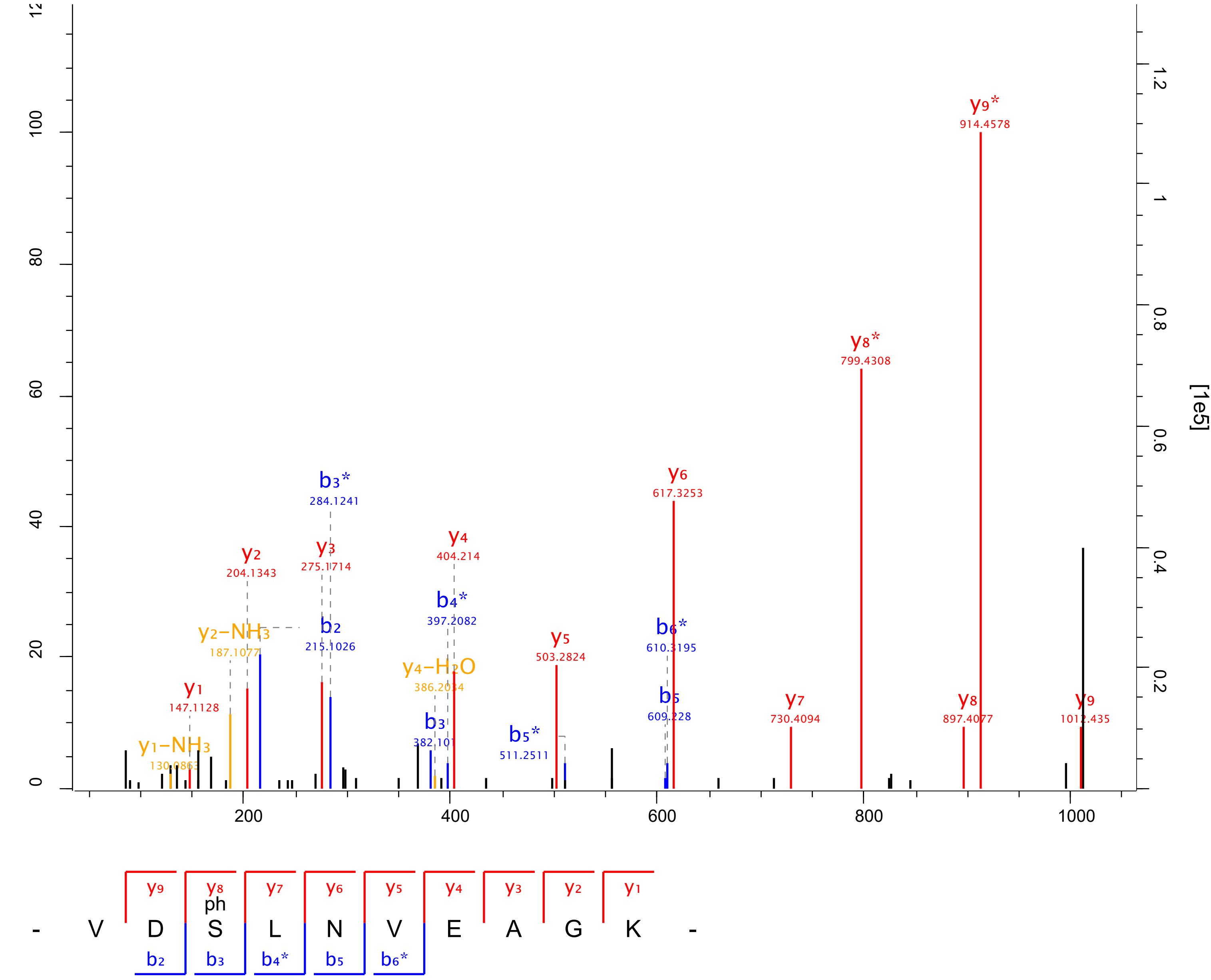

| Raw file | Scan | Method | Score | m/z |
|---|---|---|---|---|
| water_15_1 | 10717 | FTMS; HCD | 229.75 | 602.27 |

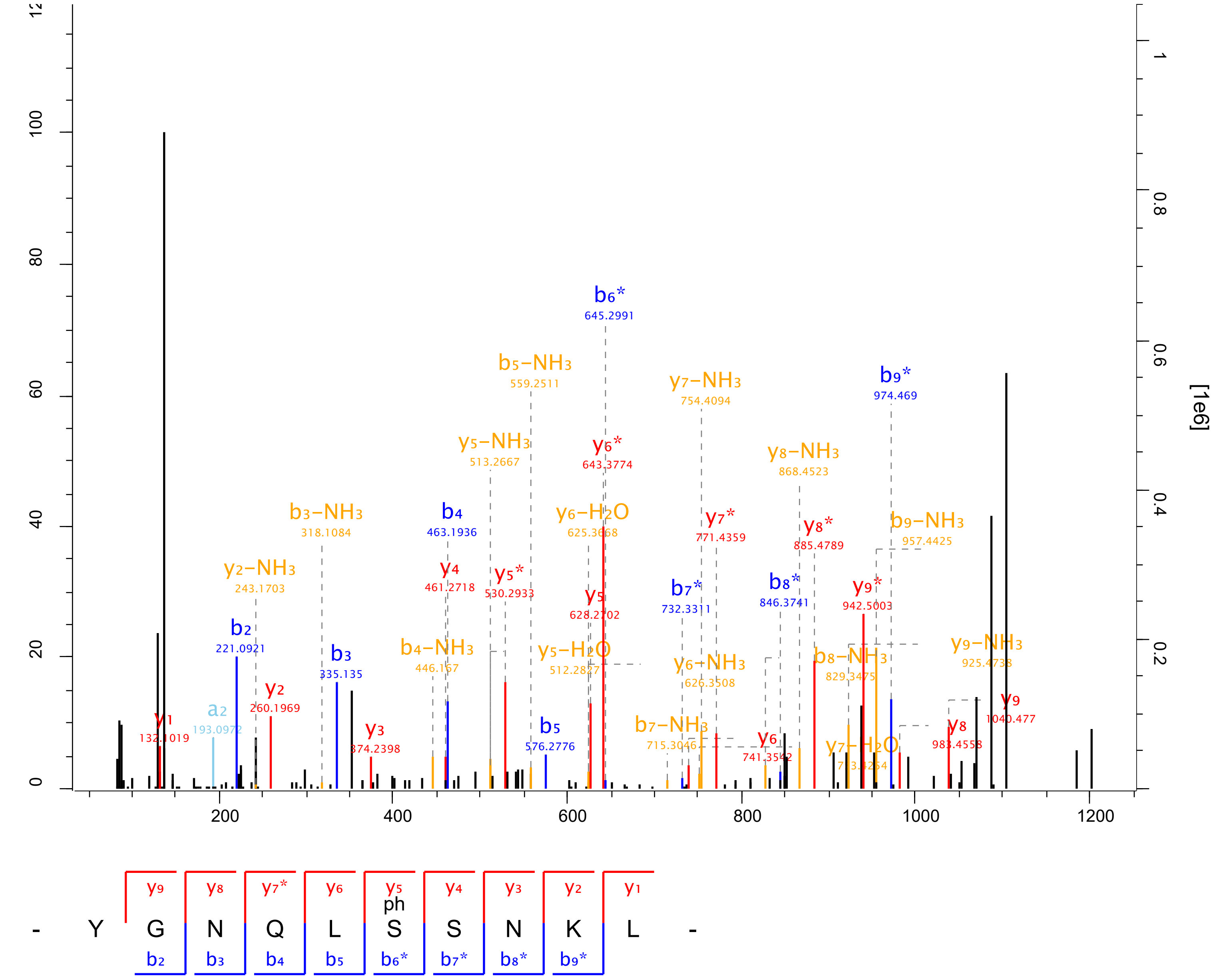

| Raw file | Scan | Method | Score | m/z |
| --- | --- | --- | --- | --- |
| water_15_1 | 10812 | FTMS; HCD | 124.62 | 730.31 |

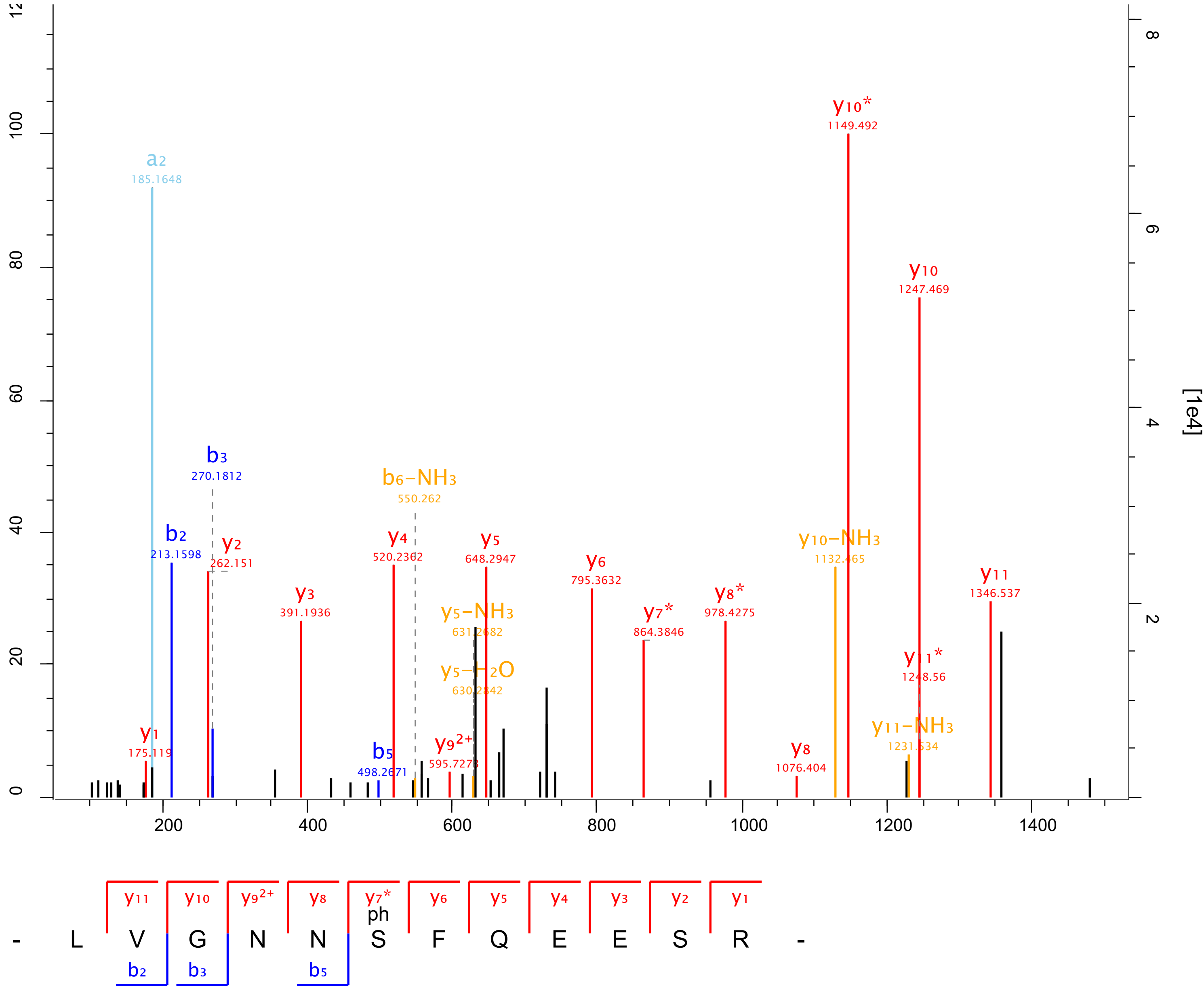

| Raw file | Scan | Method | Score | m/z |
| --- | --- | --- | --- | --- |
| water_15_1 | 10823 | FTMS; HCD | 92.86 | 586.75 |

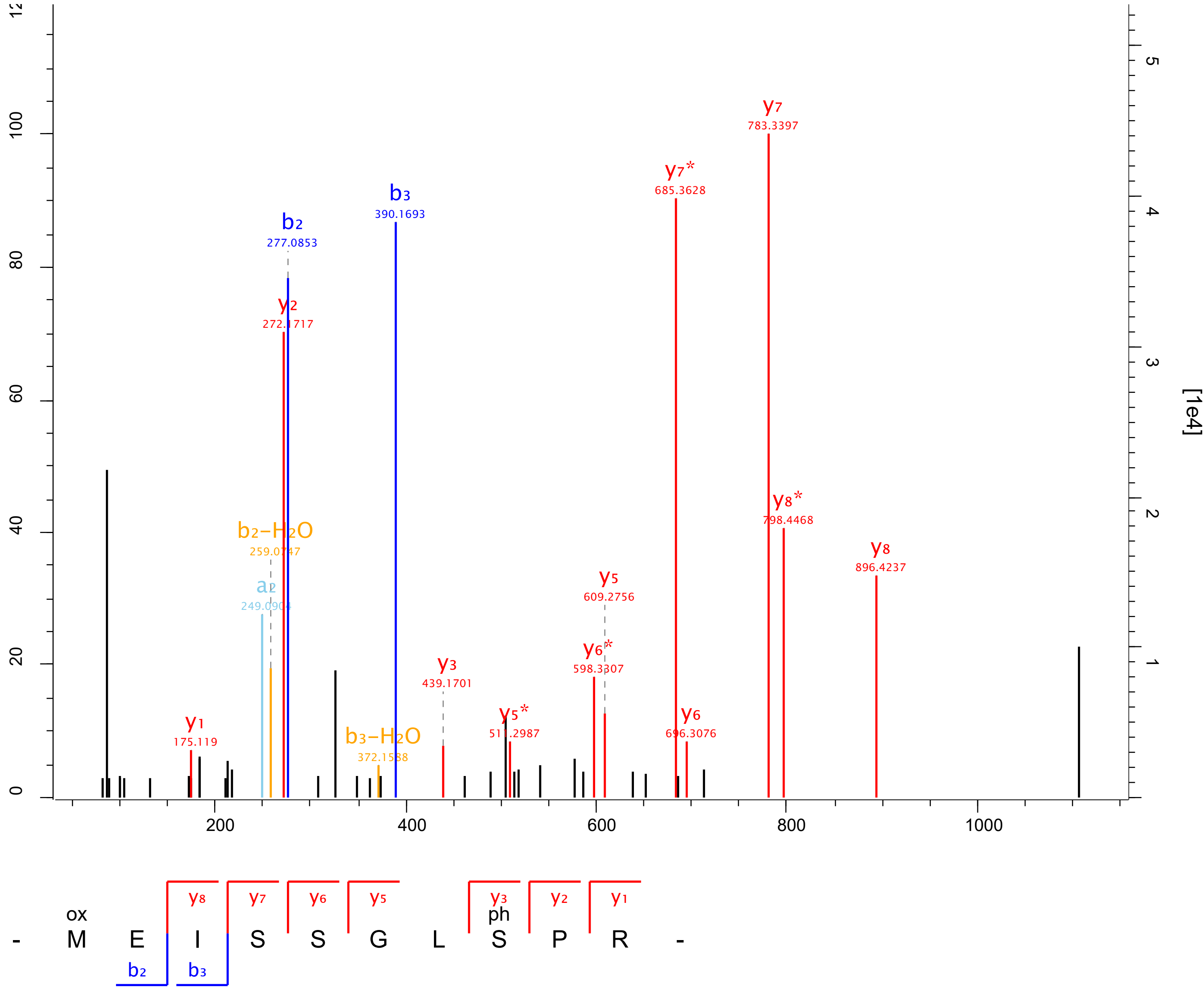

| Raw file | Scan | Method | Score | m/z |
| --- | --- | --- | --- | --- |
| water_15_1 | 10859 | FTMS; HCD | 115.92 | 531.73 |

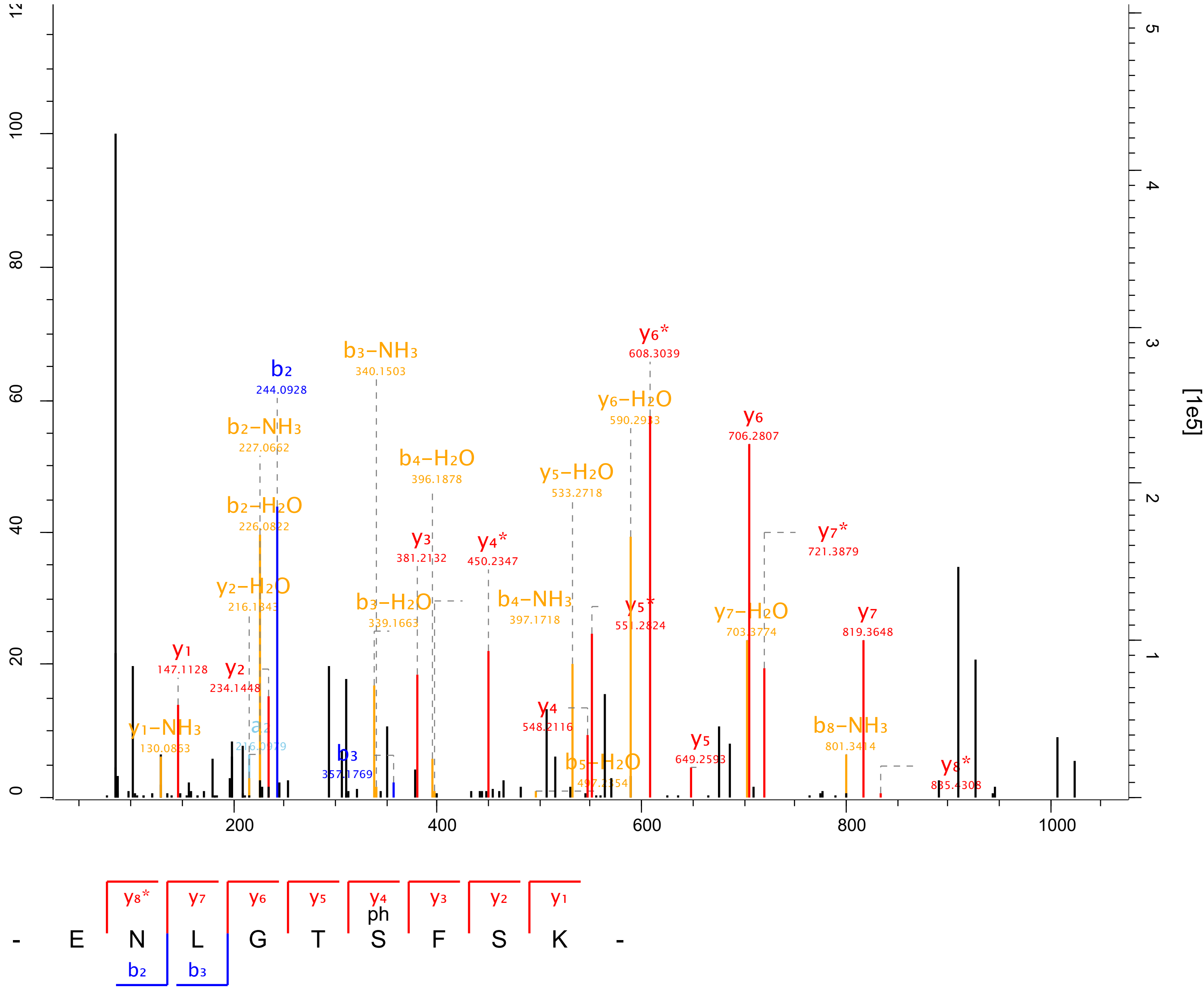

| Raw file | Scan | Method | Score | m/z |
|---|---|---|---|---|
| water_15_1 | 11133 | FTMS; HCD | 74.17 | 411.22 |

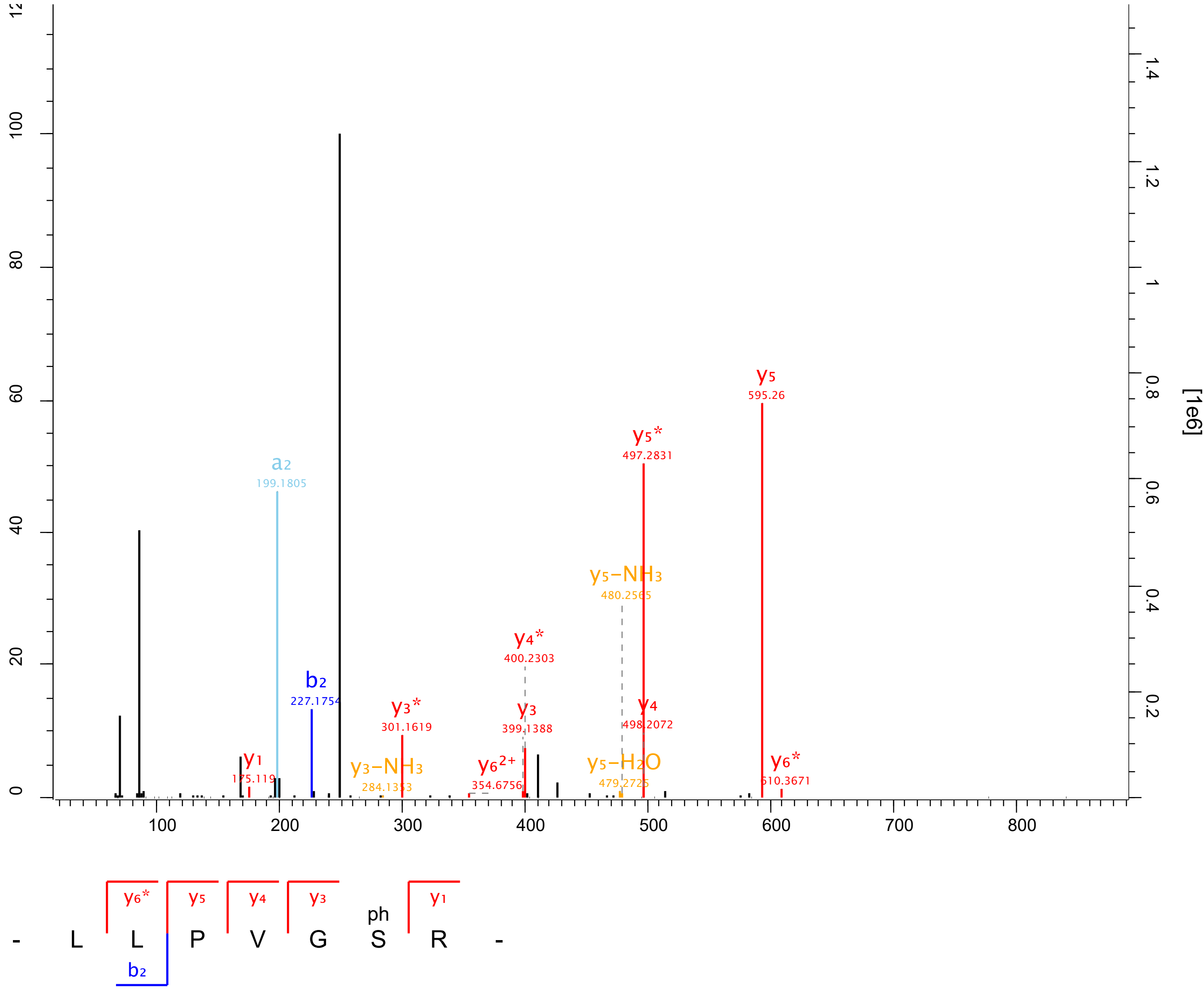

| Raw file | Scan | Method | Score | m/z |
| --- | --- | --- | --- | --- |
| water_15_1 | 11257 | FTMS; HCD | 95.2 | 677.31 |

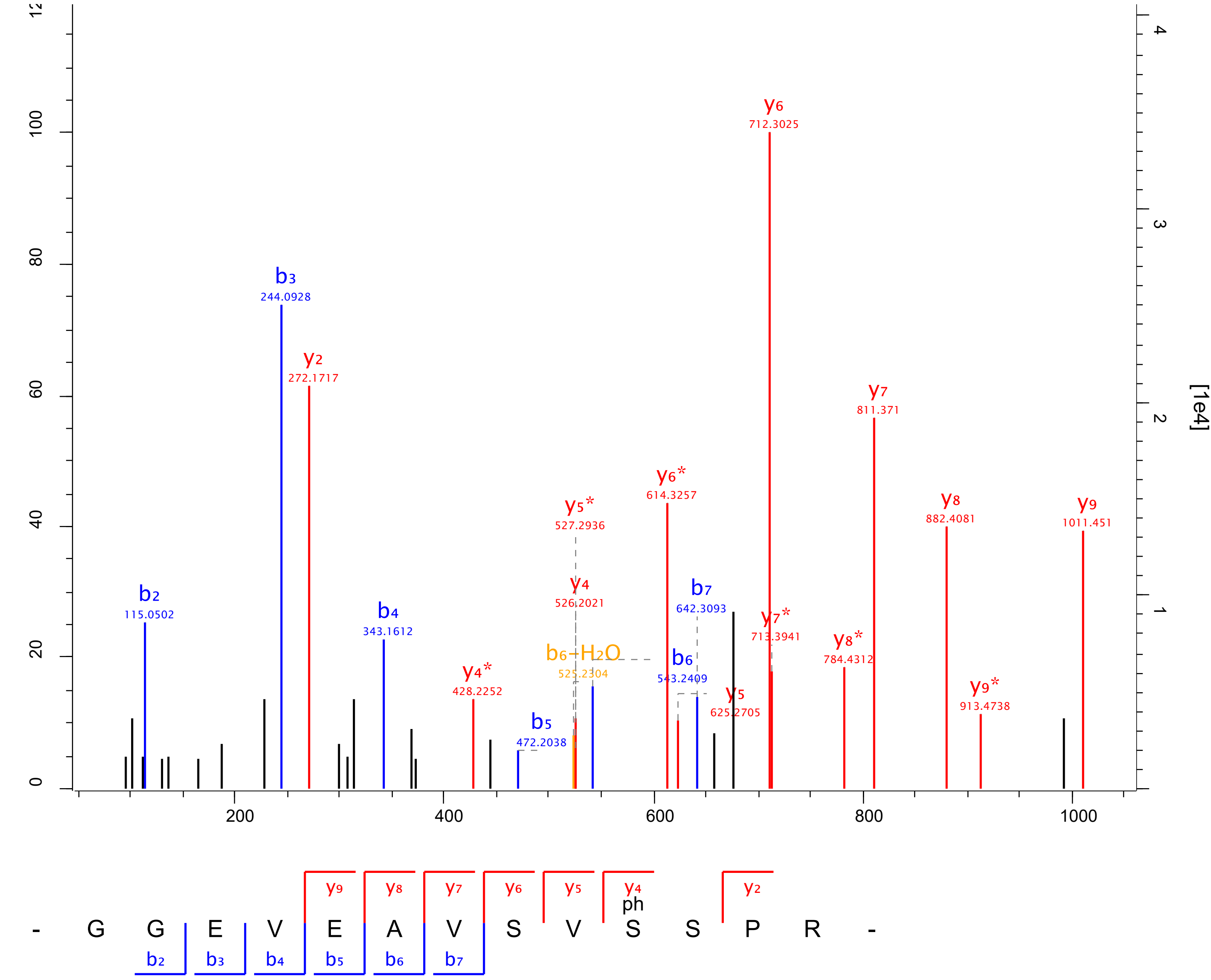

| Raw file | Scan | Method | Score | m/z |
|---|---|---|---|---|
| water_15_1 | 11298 | FTMS; HCD | 80.92 | 526.21 |

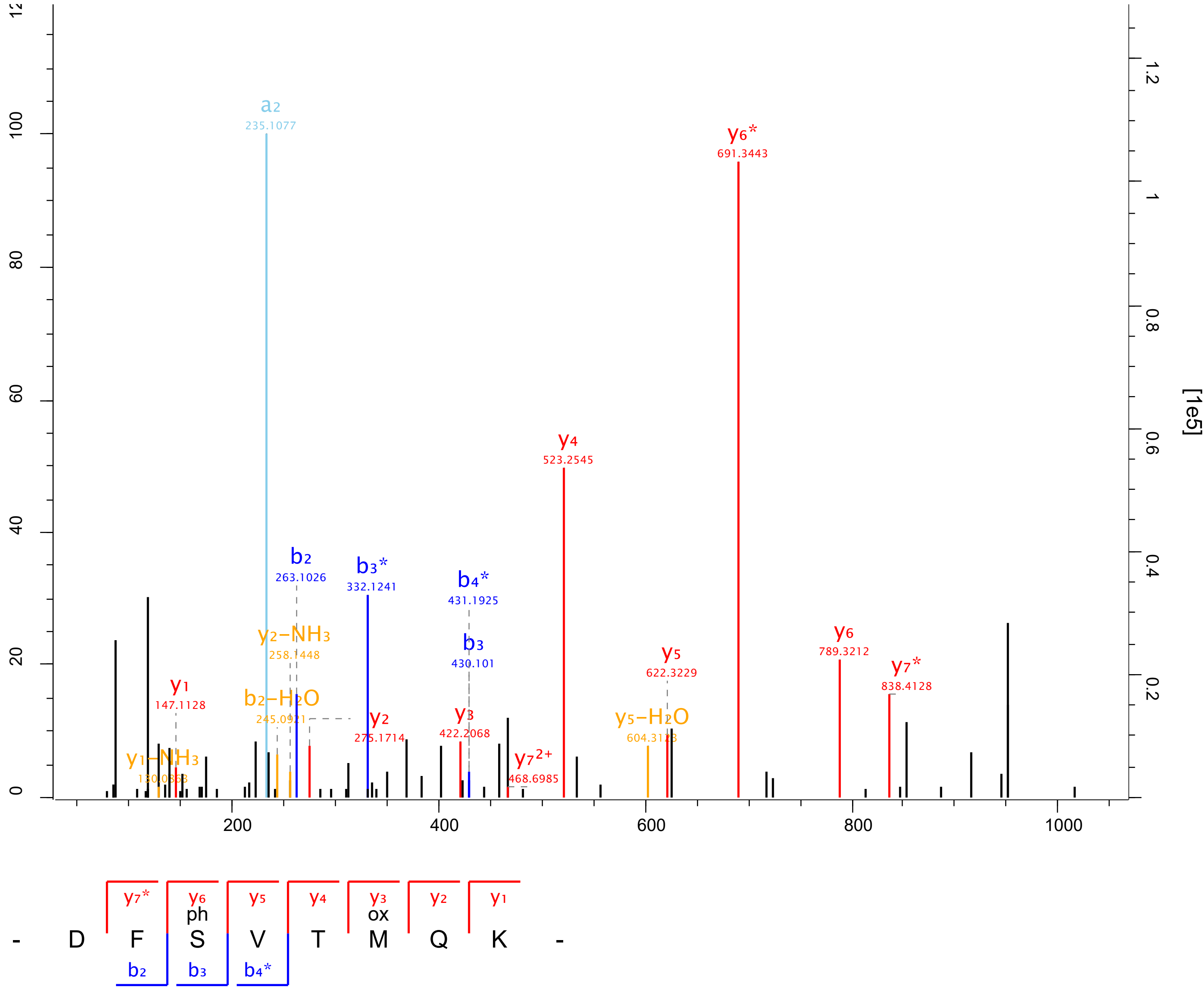

| Raw file | Scan | Method | Score | m/z |
|---|---|---|---|---|
| water_15_1 | 11385 | FTMS; HCD | 323.47 | 886.35 |

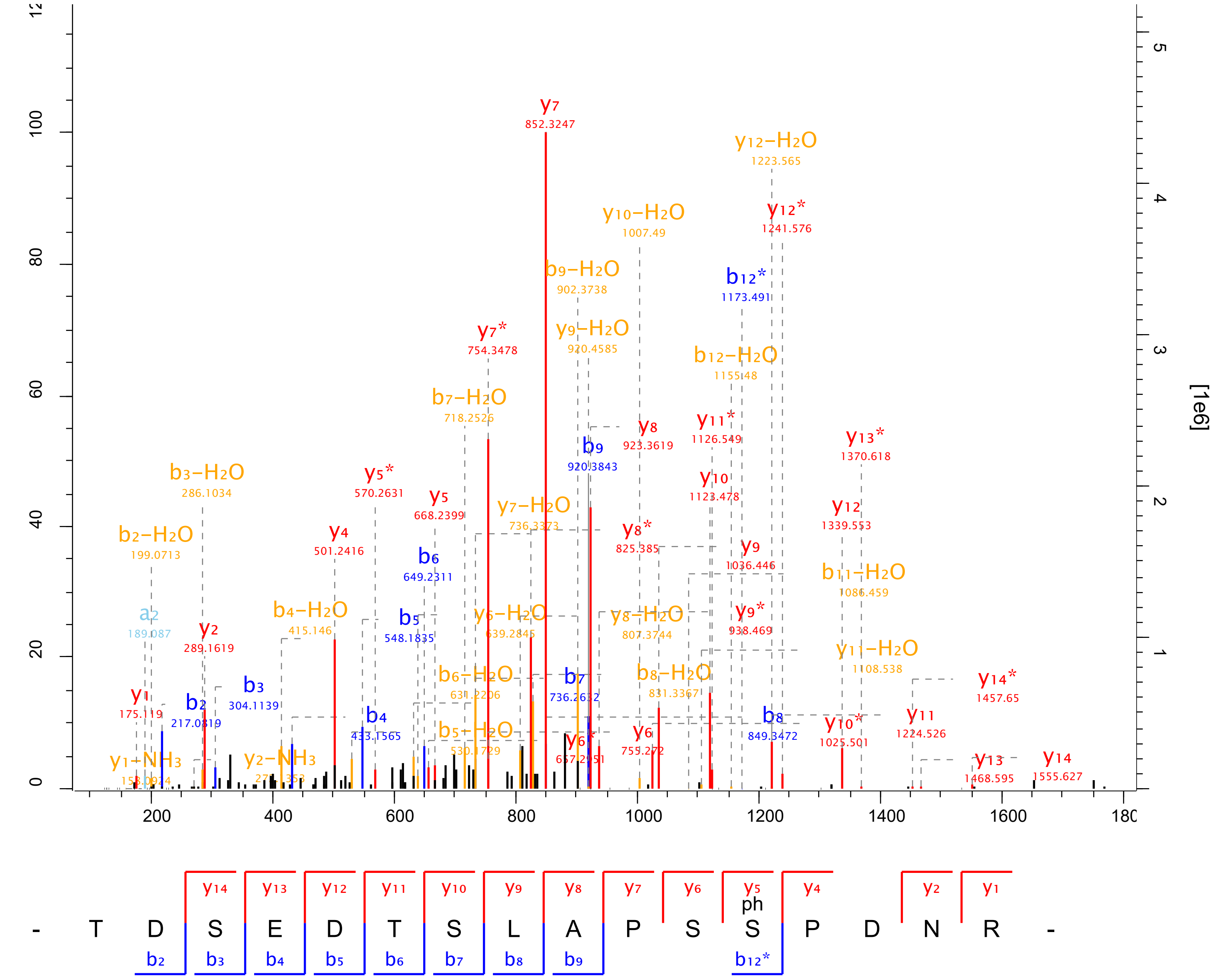

| Raw file | Scan | Method | Score | m/z |
|---|---|---|---|---|
| water_15_1 | 11475 | FTMS; HCD | 76.82 | 683.29 |

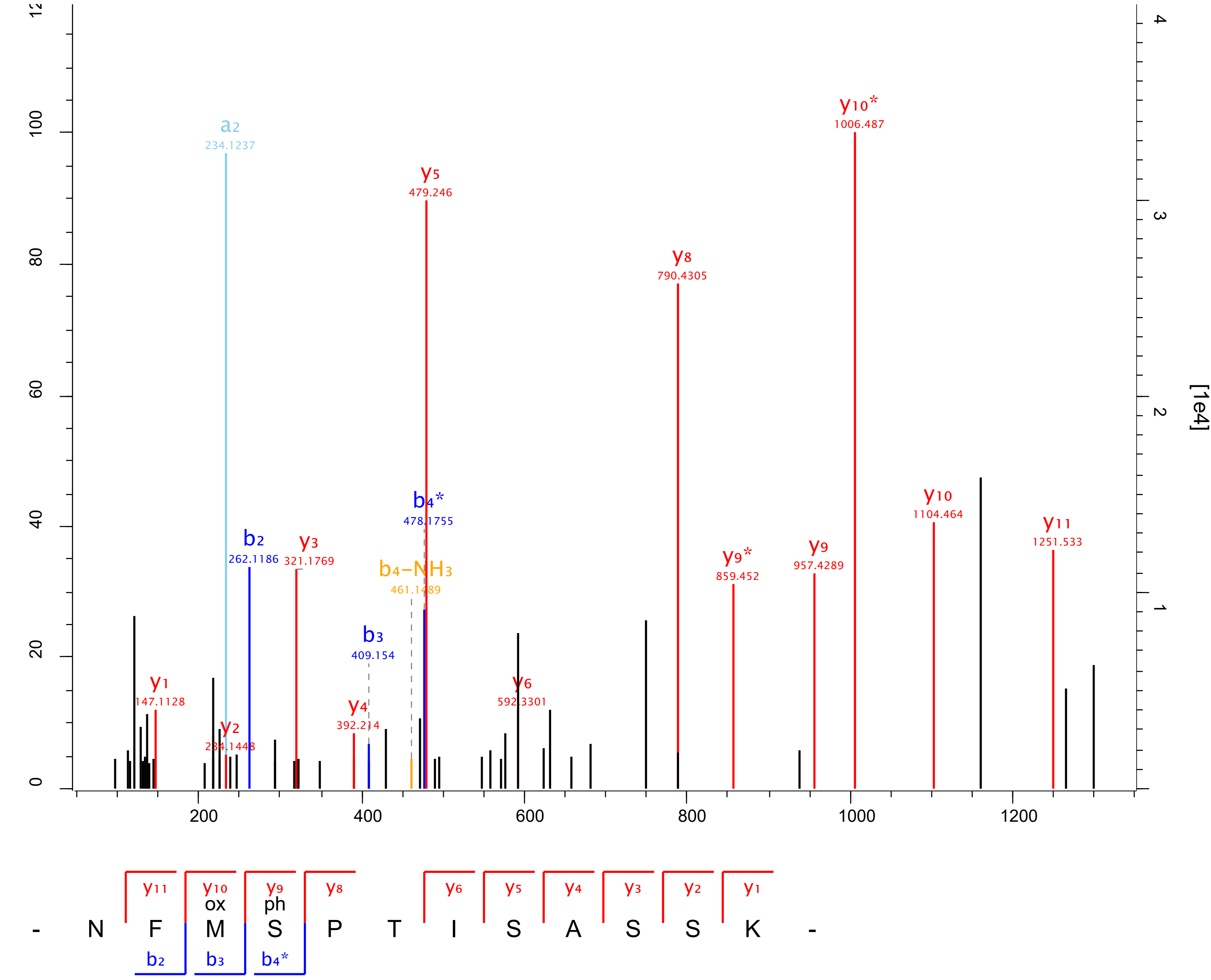

| Raw file | Scan | Method | Score | m/z |
| --- | --- | --- | --- | --- |
| water_15_1 | 11810 | FTMS; HCD | 50.83 | 773.33 |

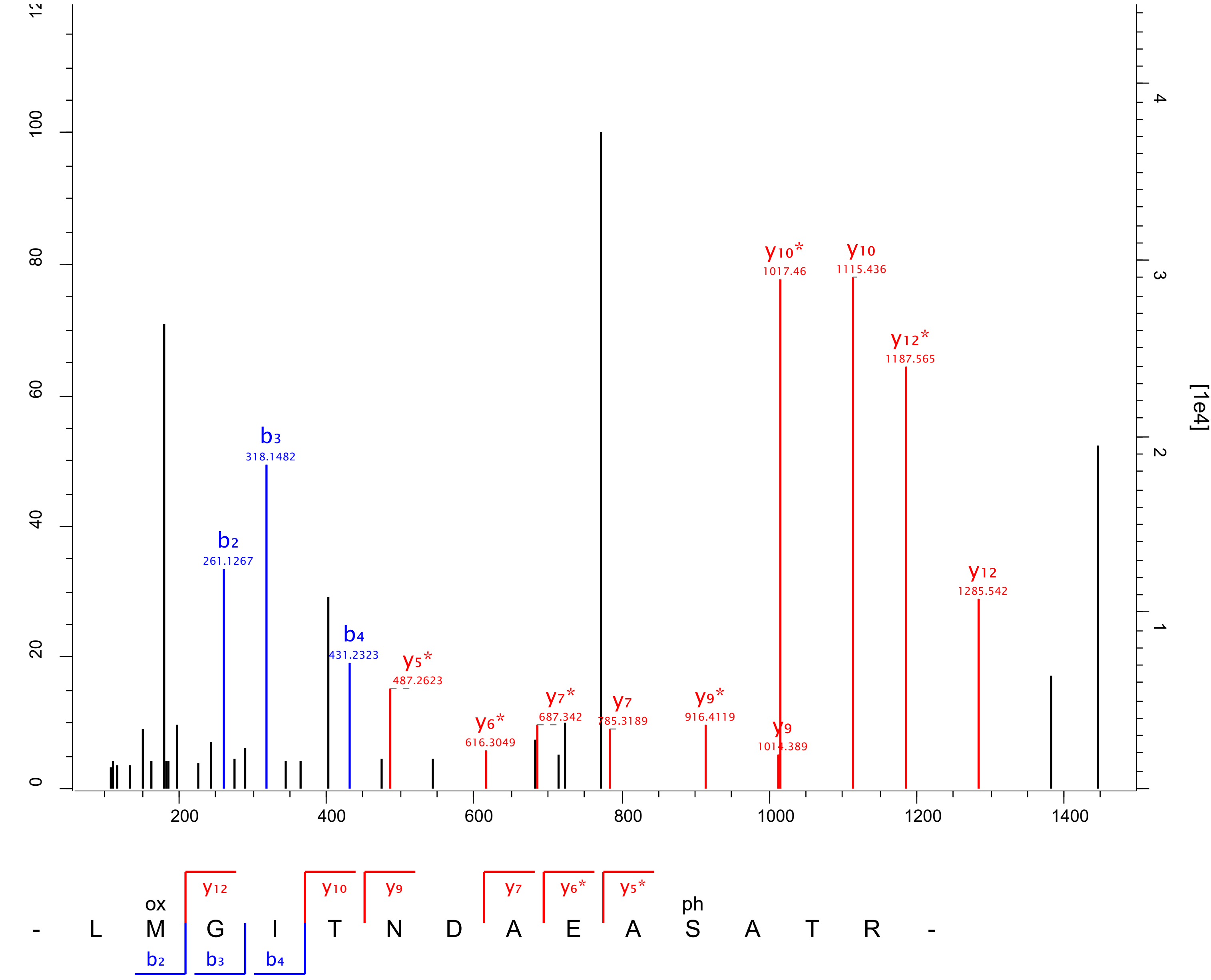

| Raw file | Scan | Method | Score | m/z |
|---|---|---|---|---|
| water_15_1 | 11871 | FTMS; HCD | 69.81 | 398.2 |

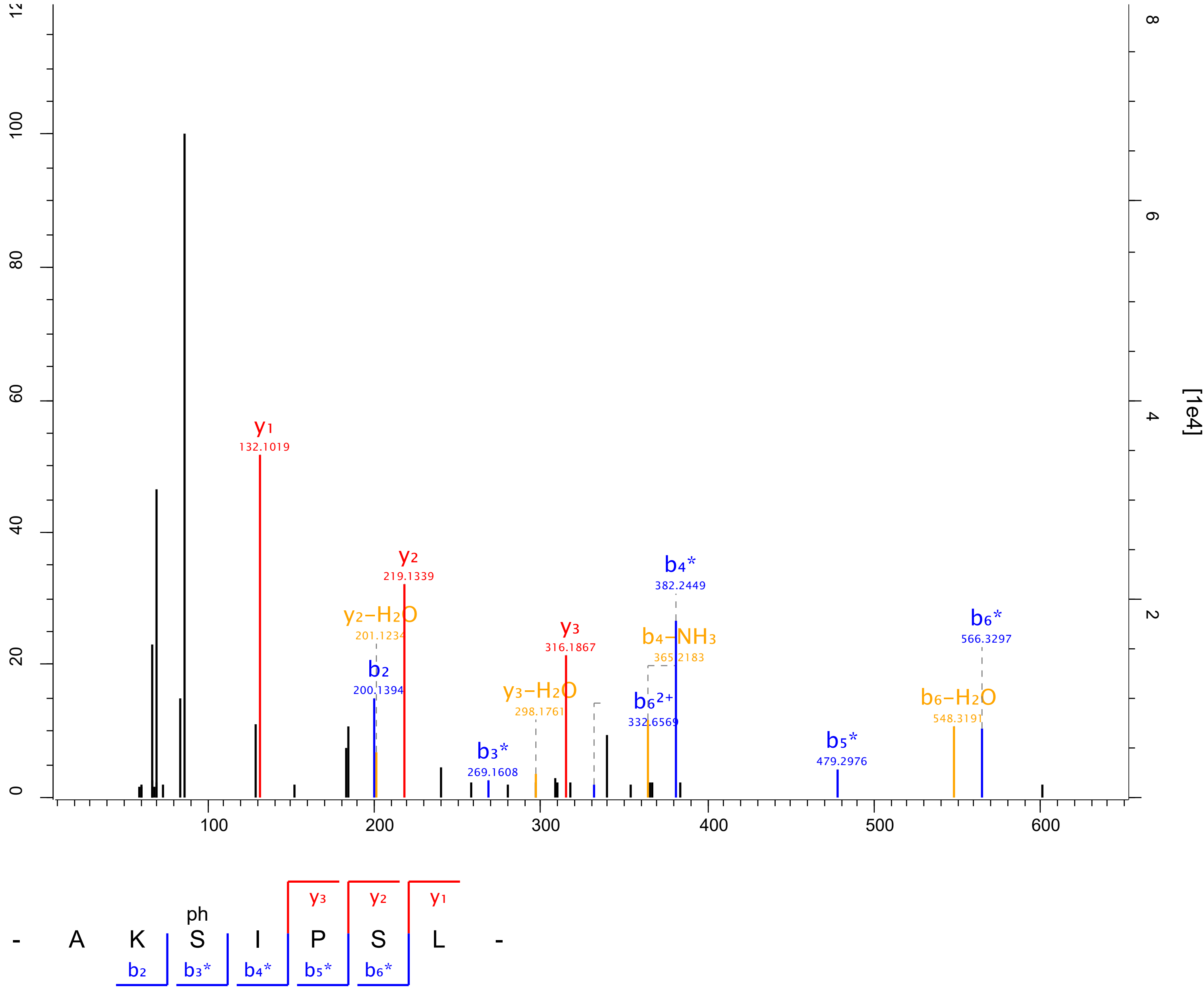

| Raw file | Scan | Method | Score | m/z |
|---|---|---|---|---|
| water_15_1 | 11879 | FTMS; HCD | 159.28 | 445.19 |

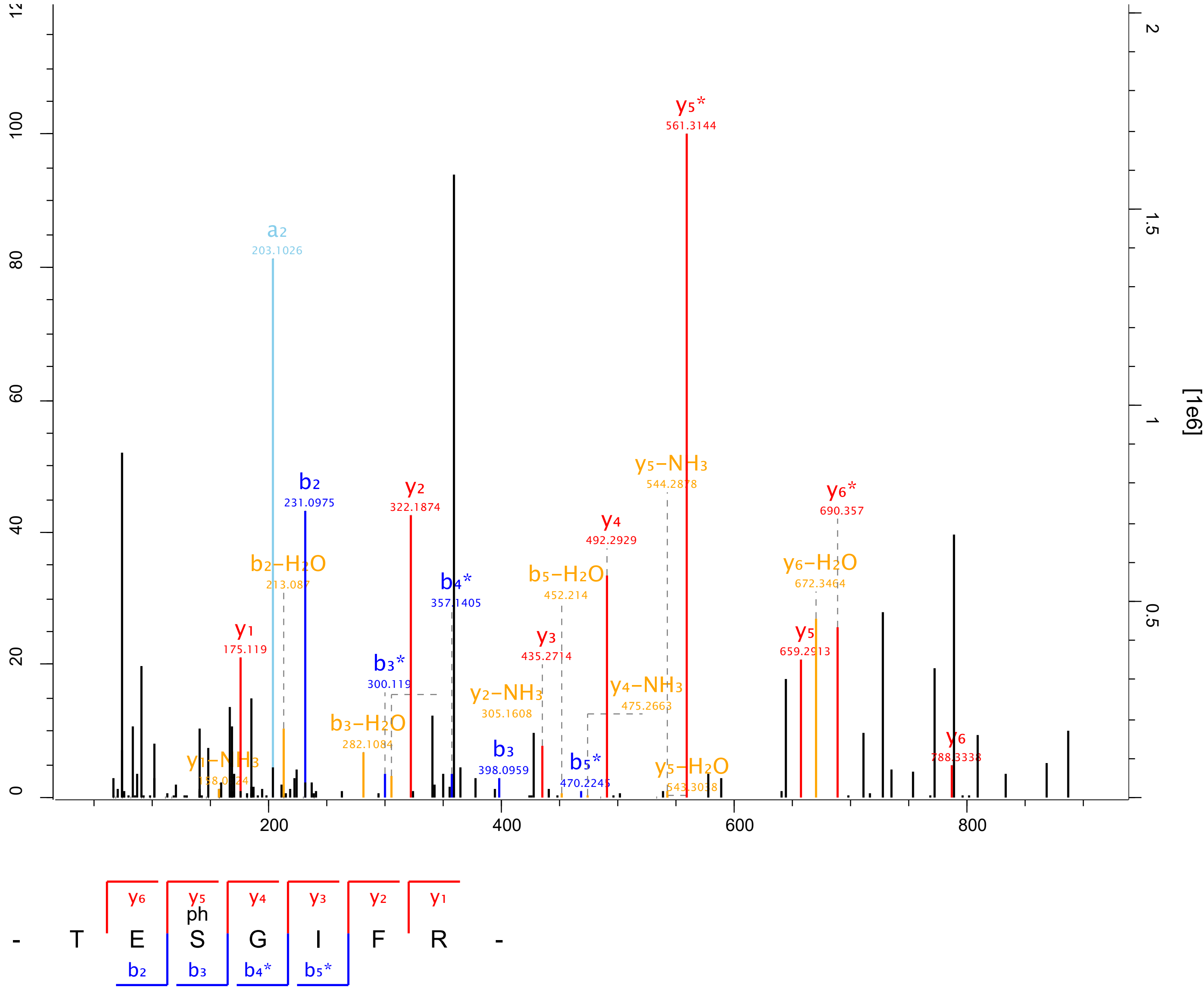

| Raw file | Scan | Method | Score | m/z |
| --- | --- | --- | --- | --- |
| water_15_1 | 12305 | FTMS; HCD | 46.88 | 560.23 |

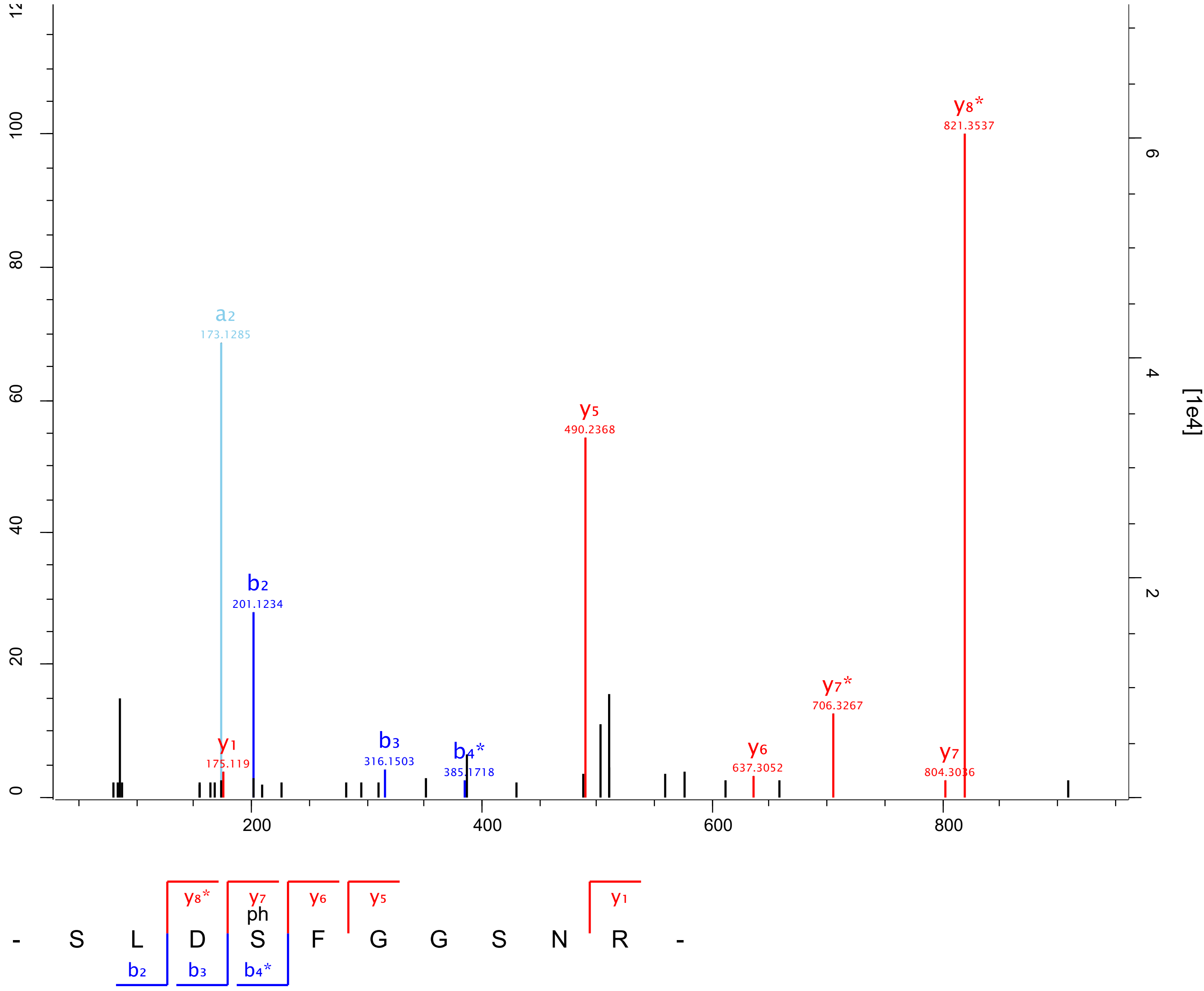

| Raw file | Scan | Method | Score | m/z |
|---|---|---|---|---|
| water_15_1 | 12472 | FTMS; HCD | 175.25 | 547.24 |

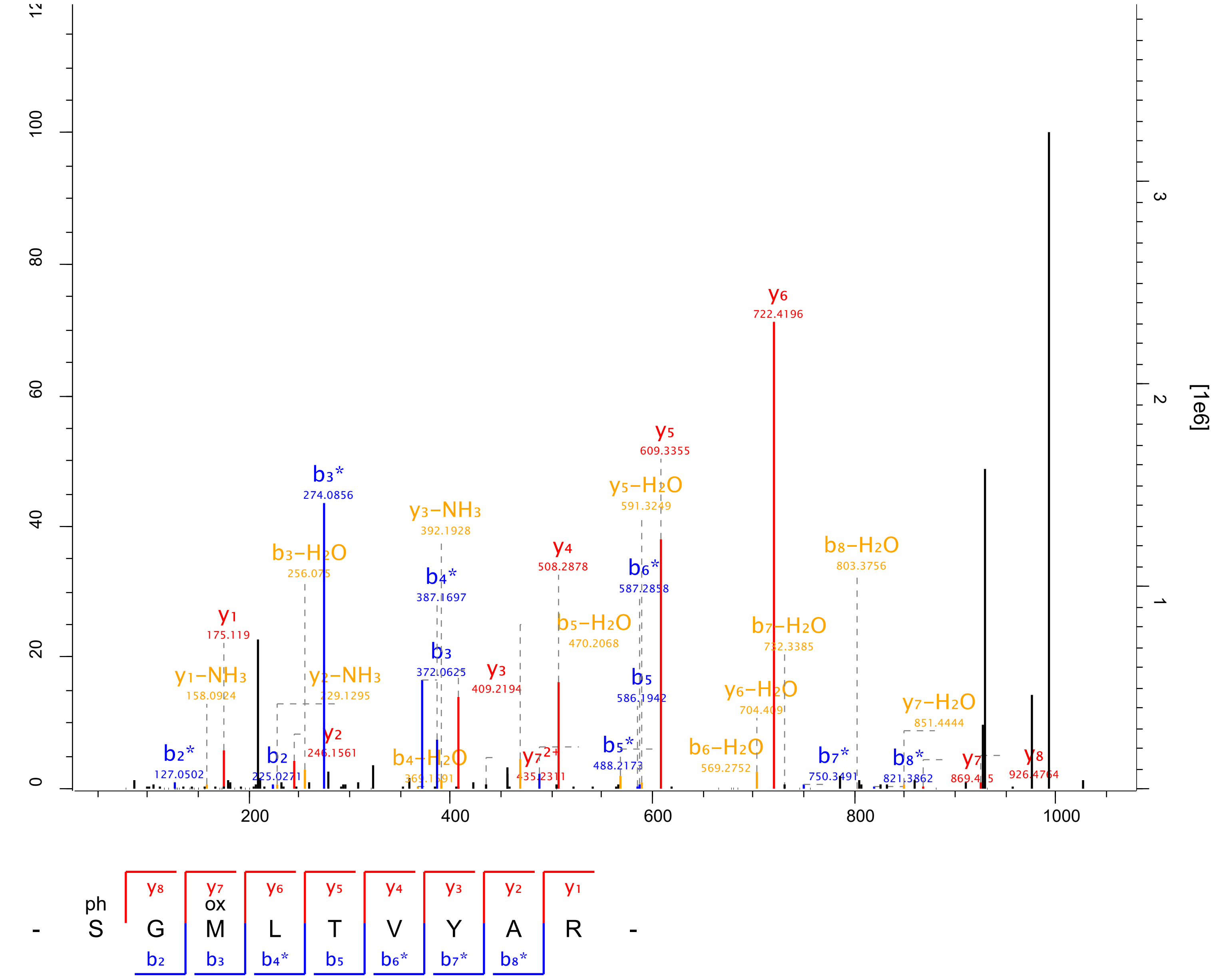

| Raw file | Scan | Method | Score | m/z |
|---|---|---|---|---|
| water_15_1 | 12844 | FTMS; HCD | 93.23 | 610.26 |

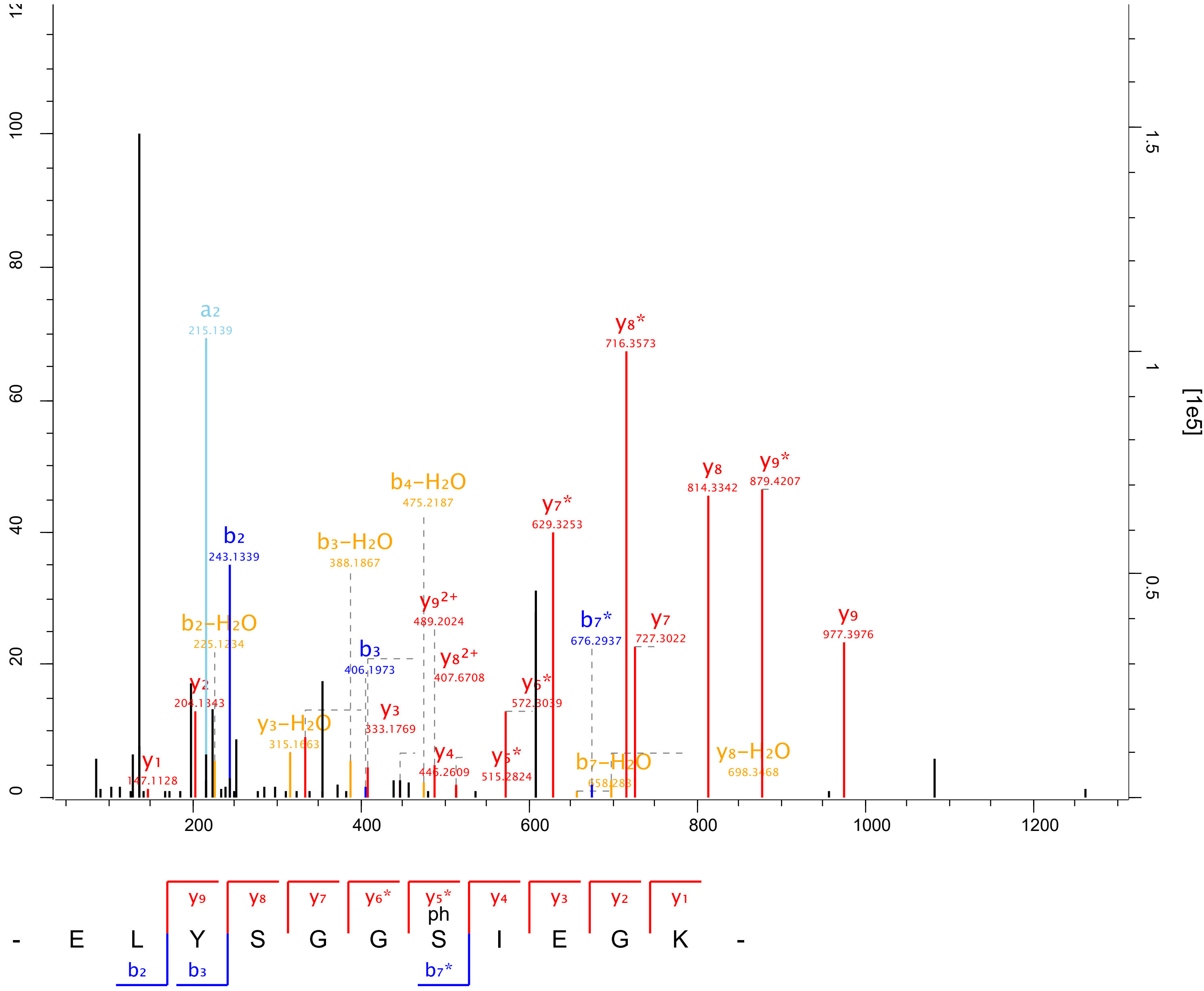

| Raw file | Scan | Method | Score | m/z |
| --- | --- | --- | --- | --- |
| water_15_1 | 13184 | FTMS; HCD | 79.09 | 561.79 |

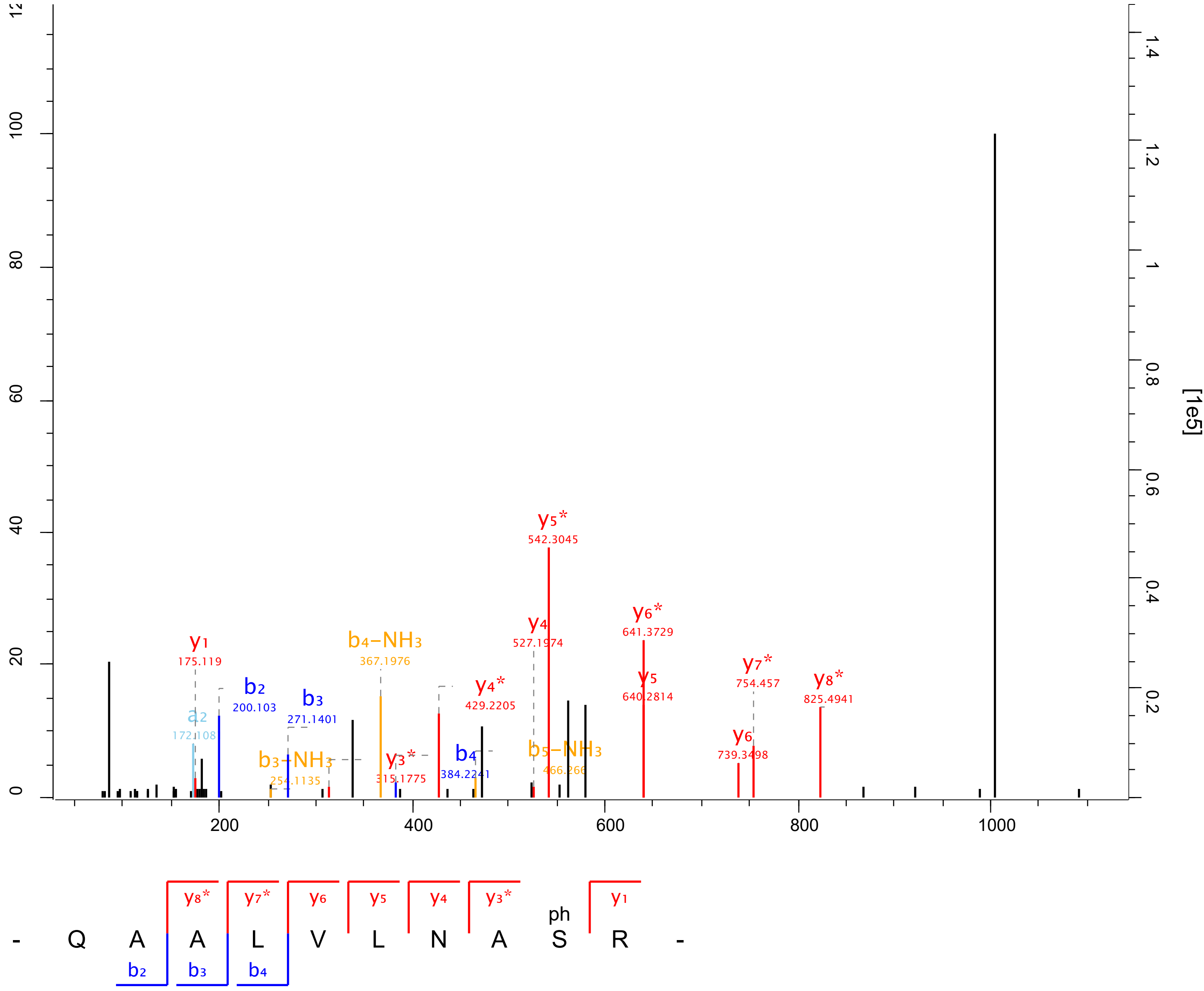

| Raw file | Scan | Method | Score | m/z |
|---|---|---|---|---|
| water_15_1 | 13313 | FTMS; HCD | 83.18 | 542.26 |

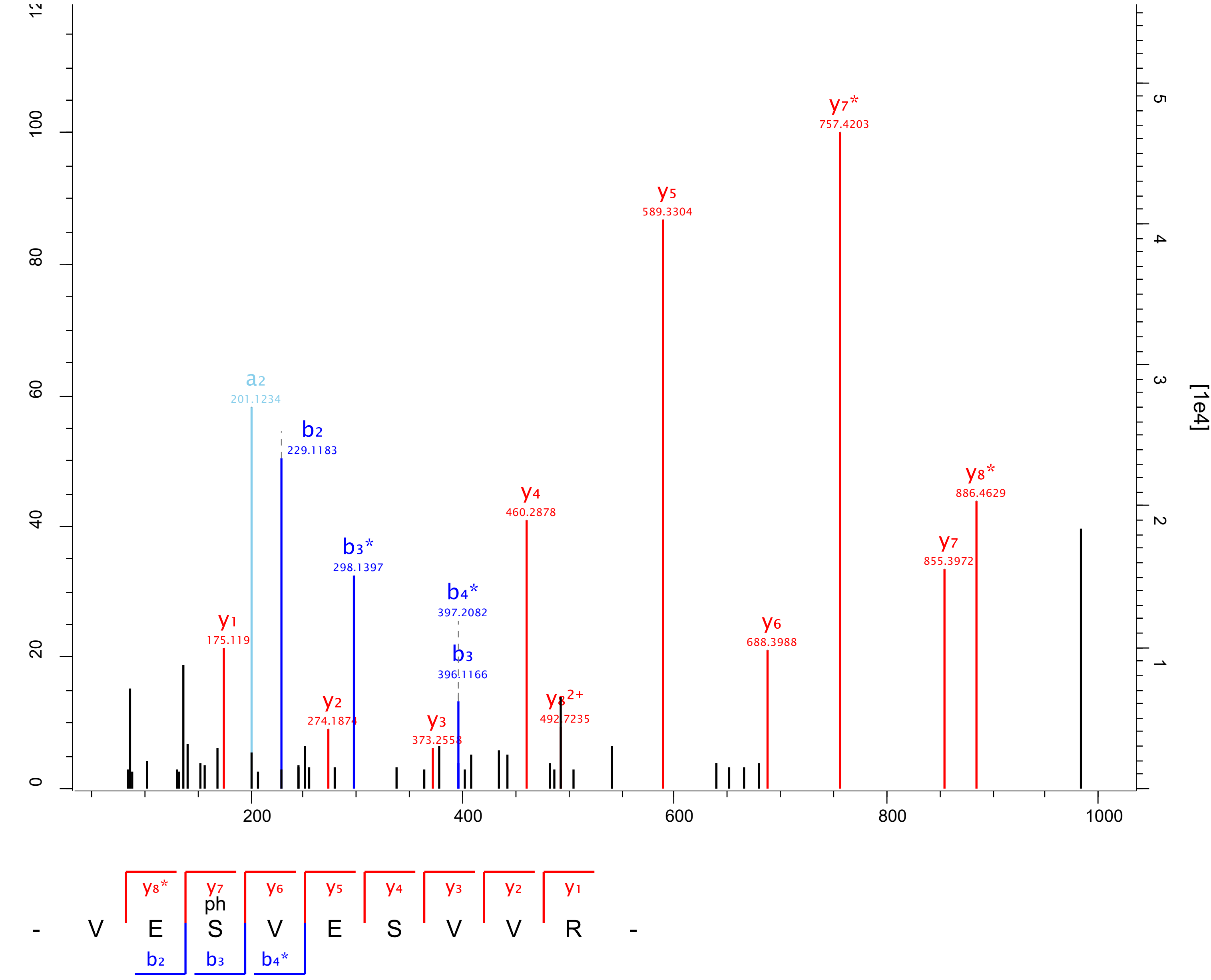

| Raw file | Scan | Method | Score | m/z |
|---|---|---|---|---|
| water_15_1 | 13782 | FTMS; HCD | 195.36 | 643.79 |

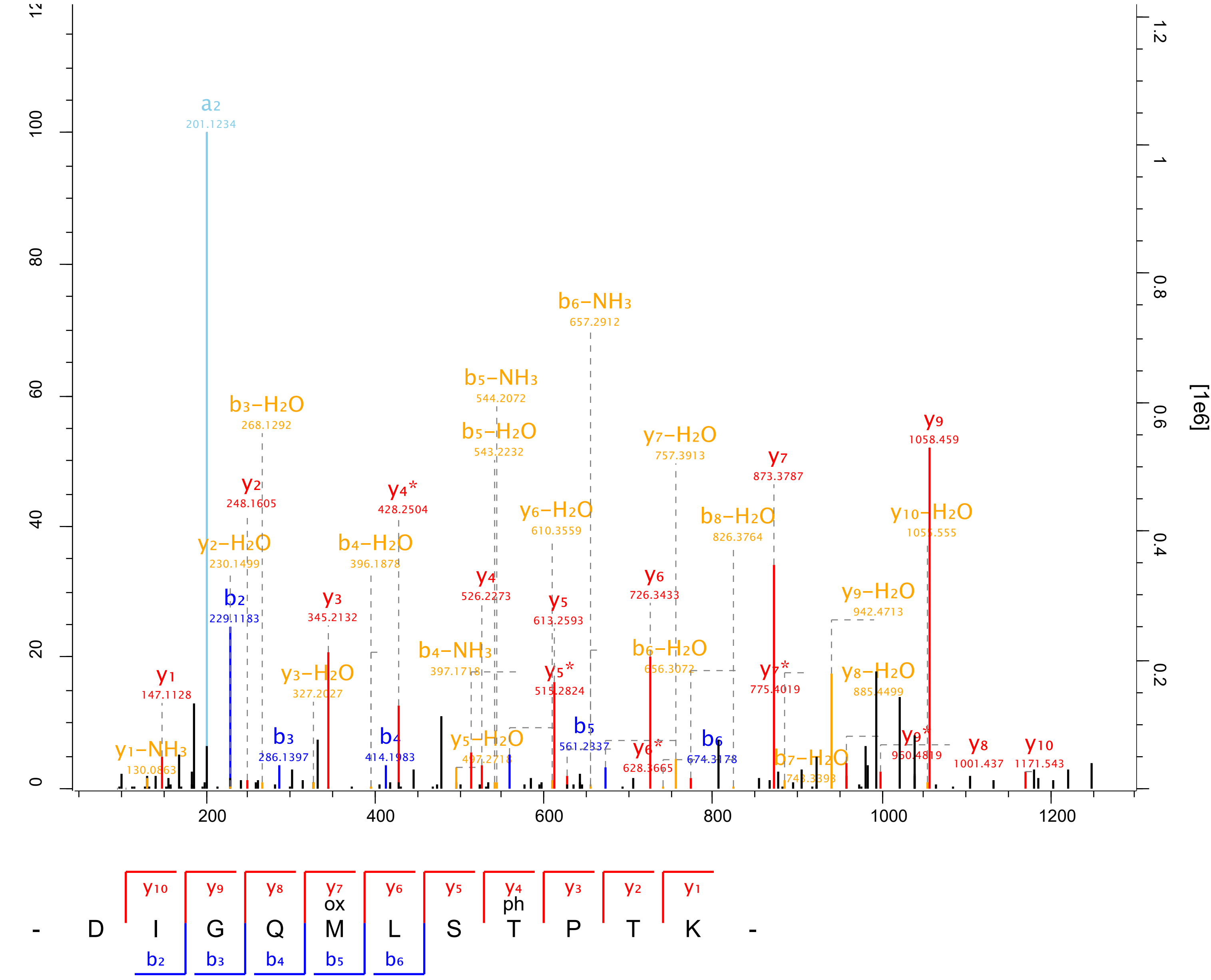

| Raw file | Scan | Method | Score | m/z |
| --- | --- | --- | --- | --- |
| water_15_1 | 14003 | FTMS; HCD | 84.36 | 562.26 |

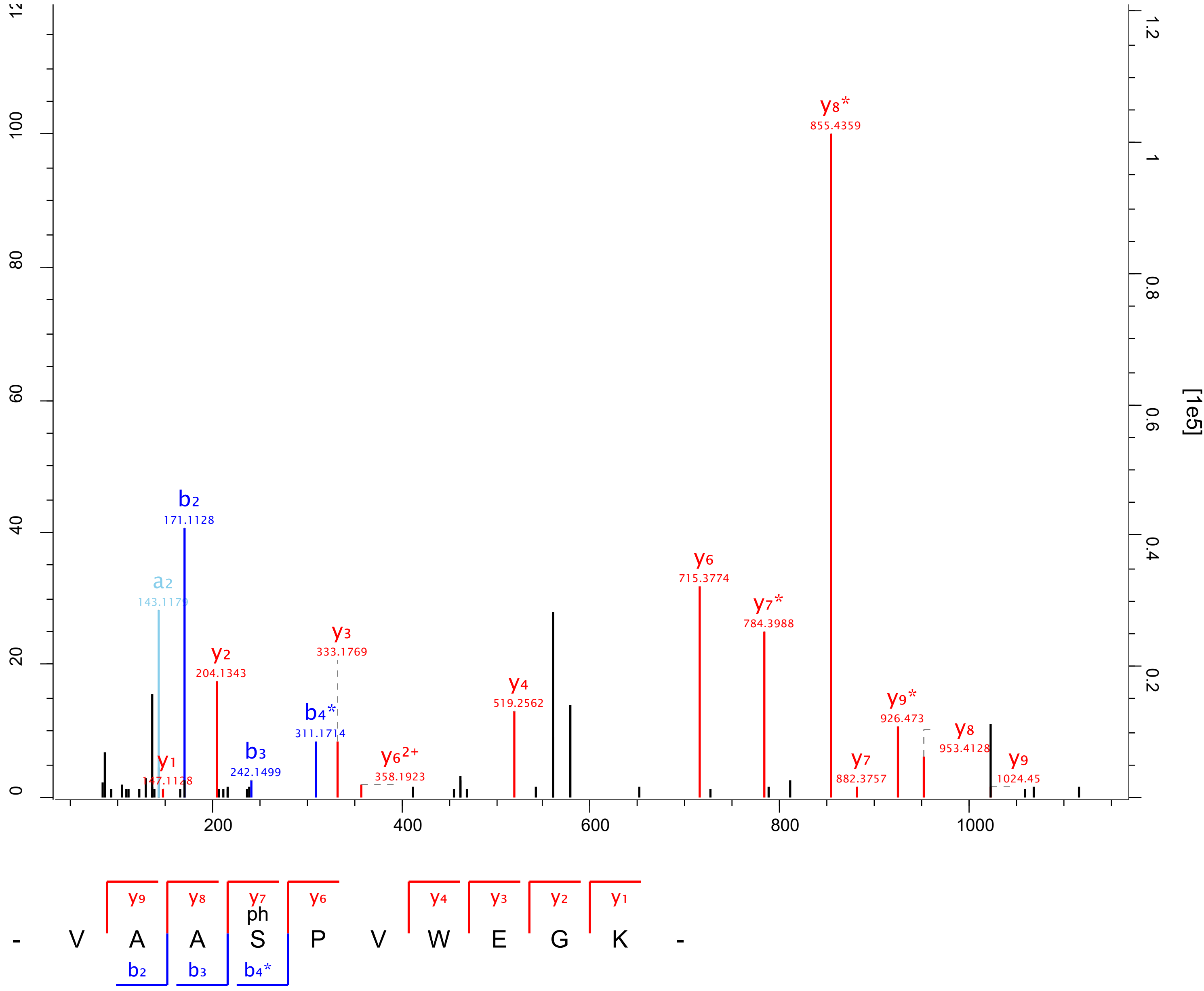

| Raw file | Scan | Method | Score | m/z |
|---|---|---|---|---|
| water_15_1 | 14538 | FTMS; HCD | 68 | 551.75 |

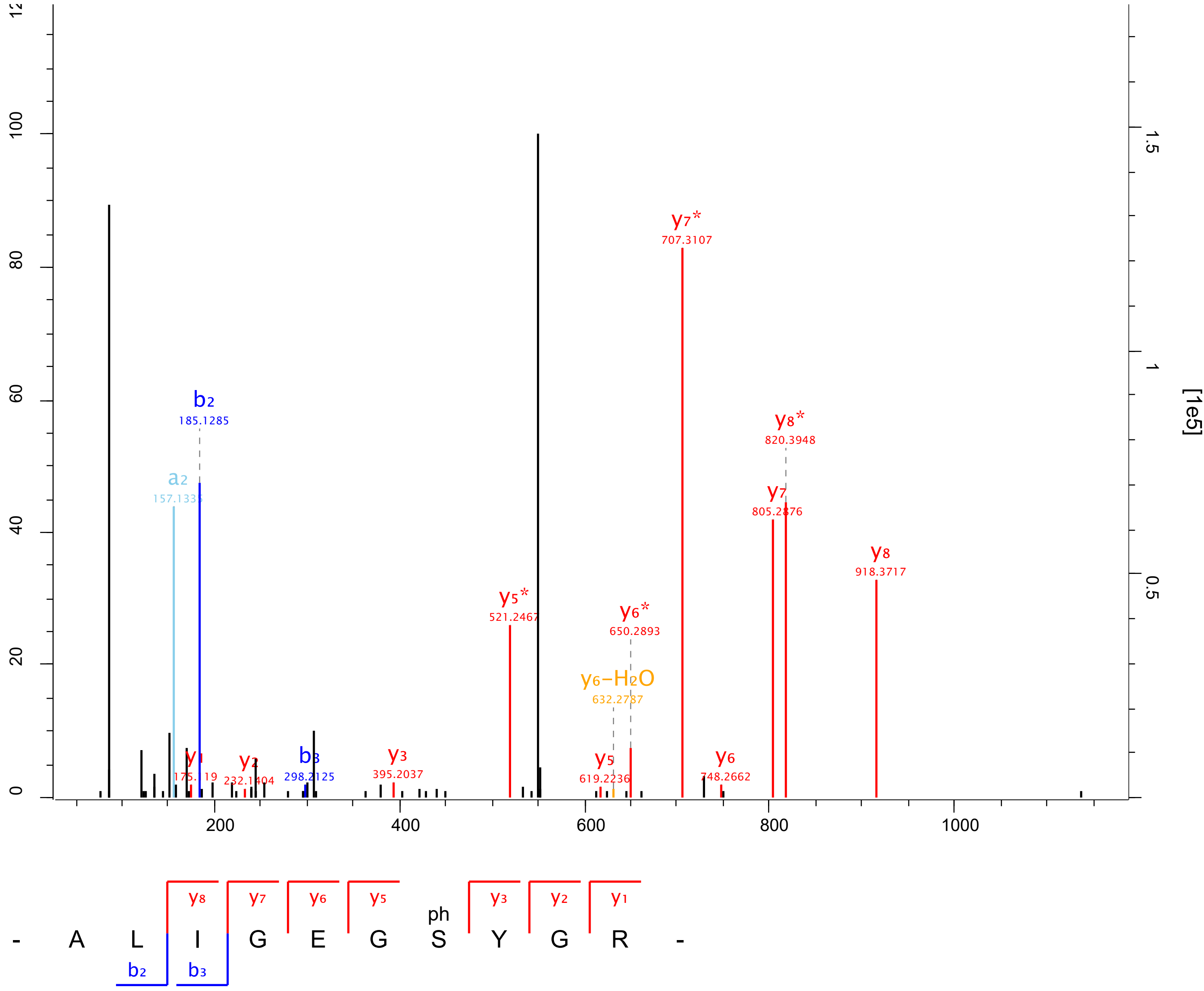

| Raw file | Scan | Method | Score | m/z |
|---|---|---|---|---|
| water_15_1 | 14609 | FTMS; HCD | 127.03 | 480.2 |

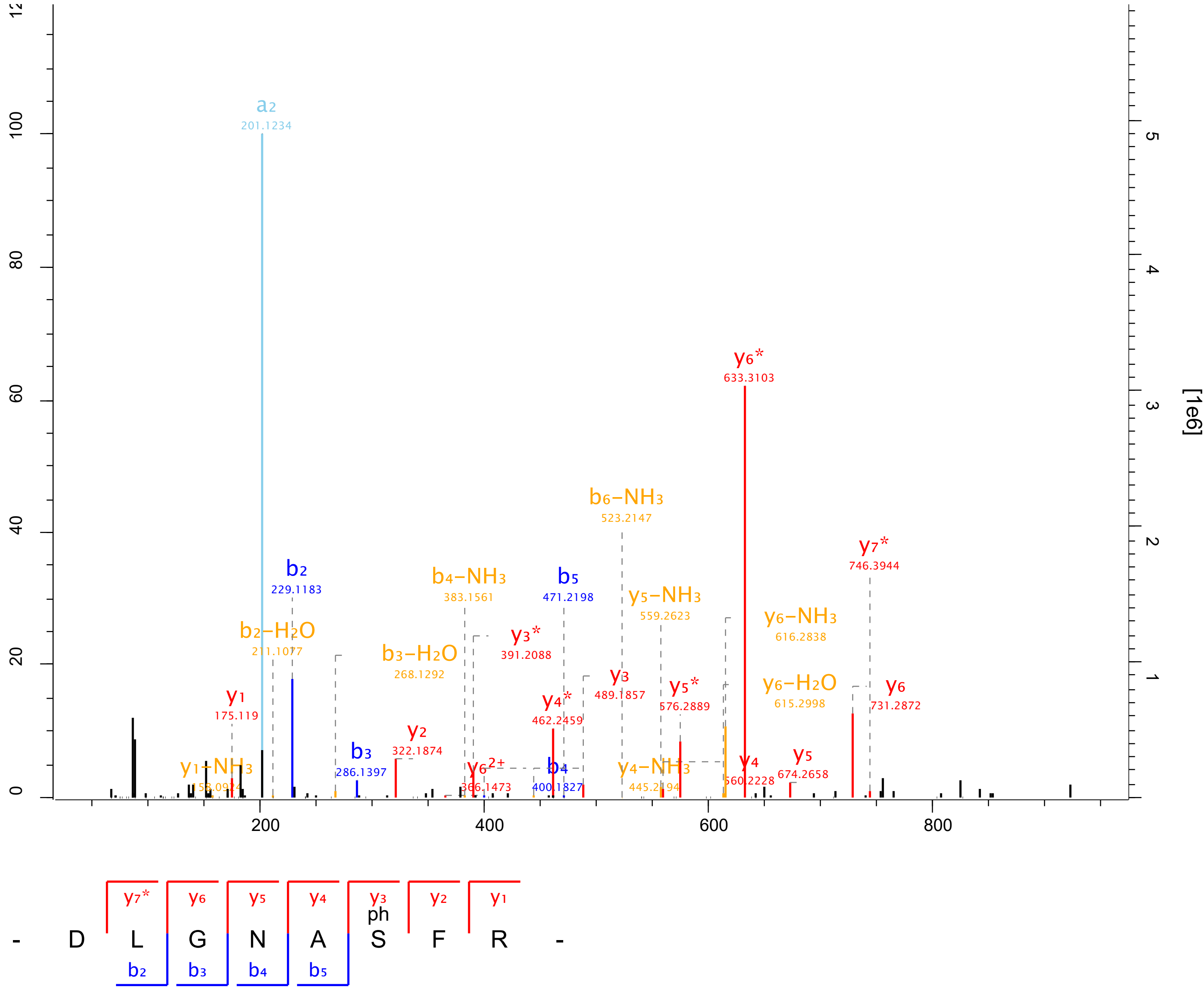

| Raw file | Scan | Method | Score | m/z |
| --- | --- | --- | --- | --- |
| water_15_1 | 15014 | FTMS; HCD | 46.32 | 681.3 |

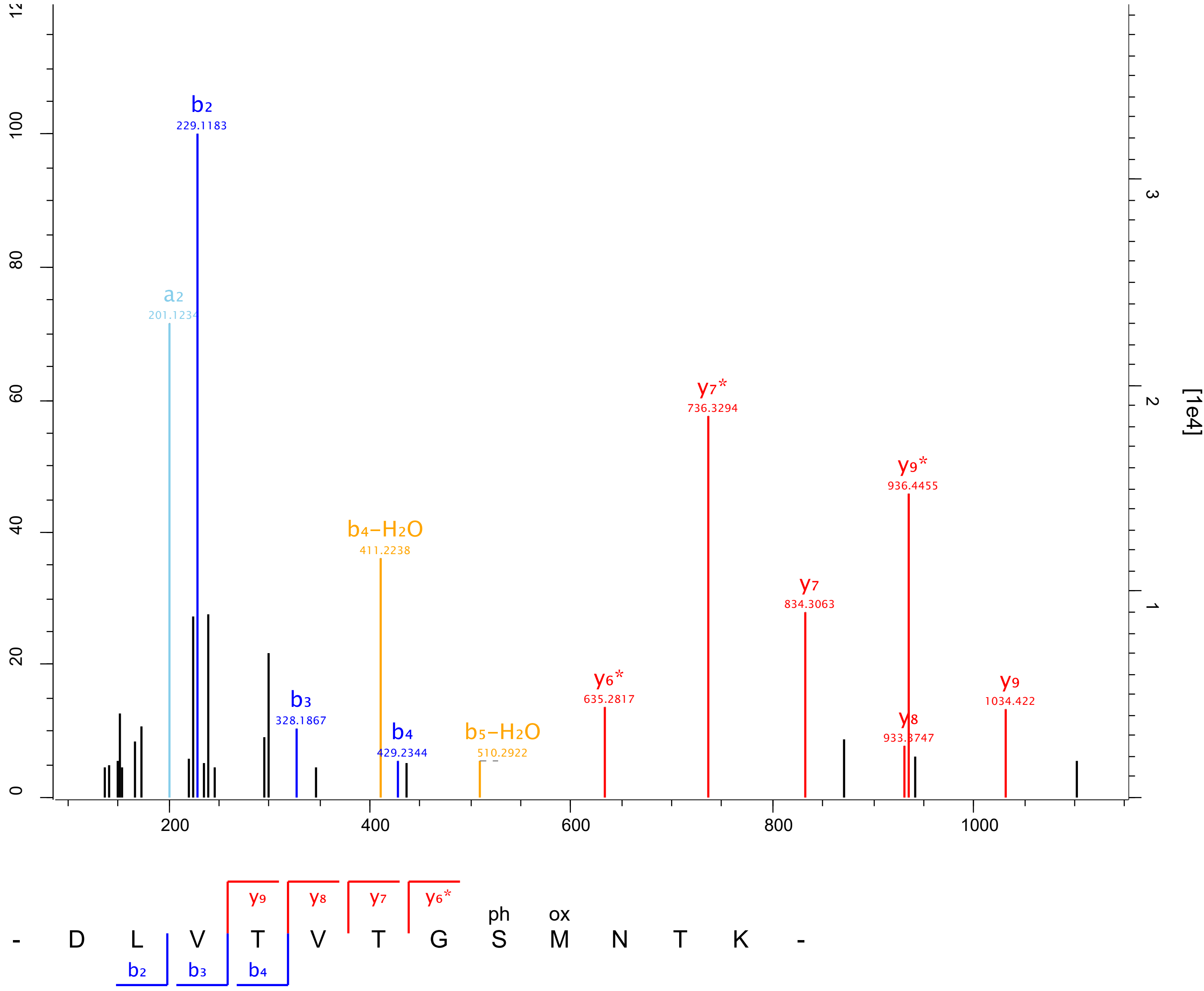

| Raw file | Scan | Method | Score | m/z |
|---|---|---|---|---|
| water_15_1 | 15175 | FTMS; HCD | 50.18 | 751.3 |

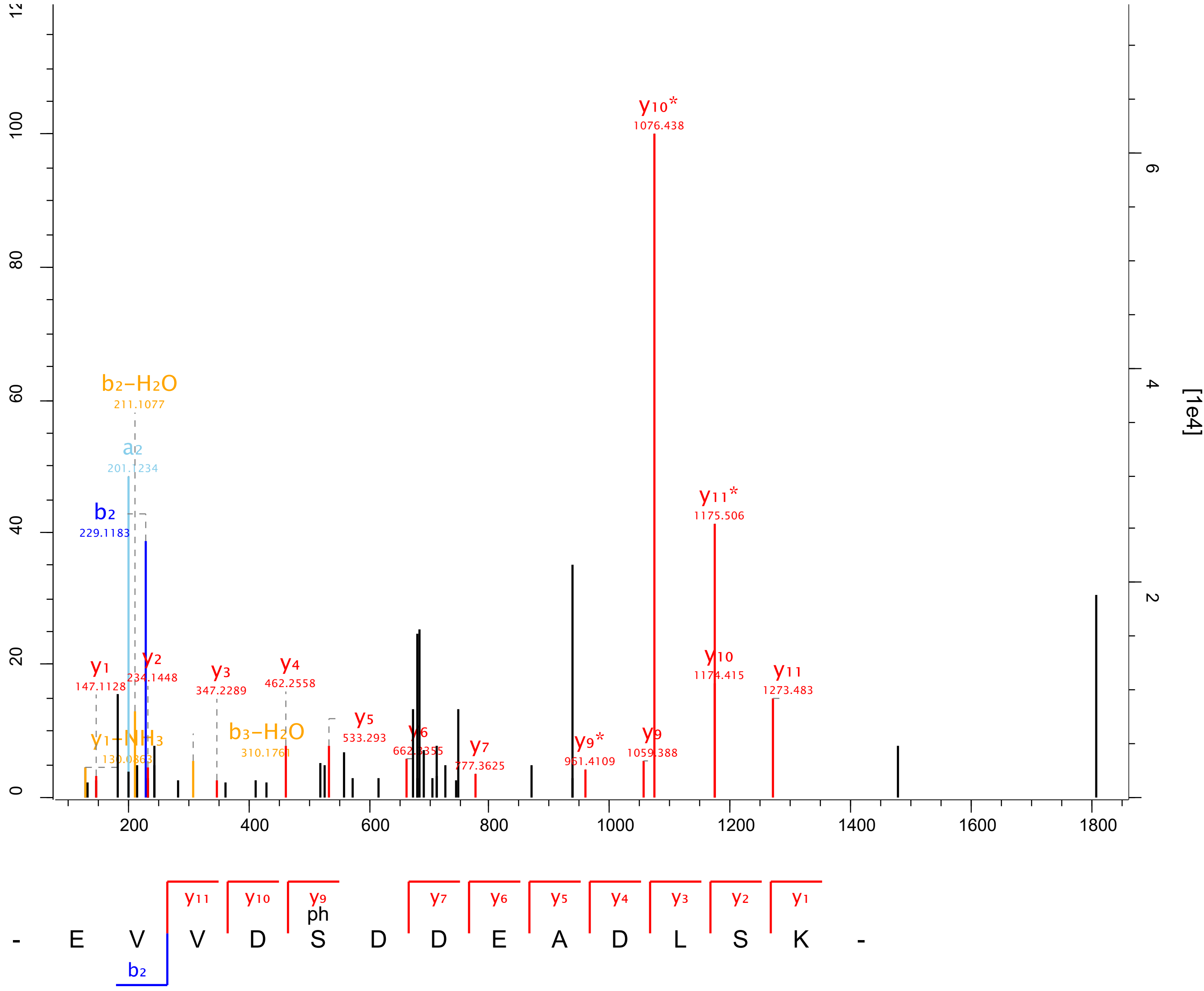

| Raw file | Scan | Method | Score | m/z |
| --- | --- | --- | --- | --- |
| water_15_1 | 15314 | FTMS; HCD | 56.12 | 761.84 |

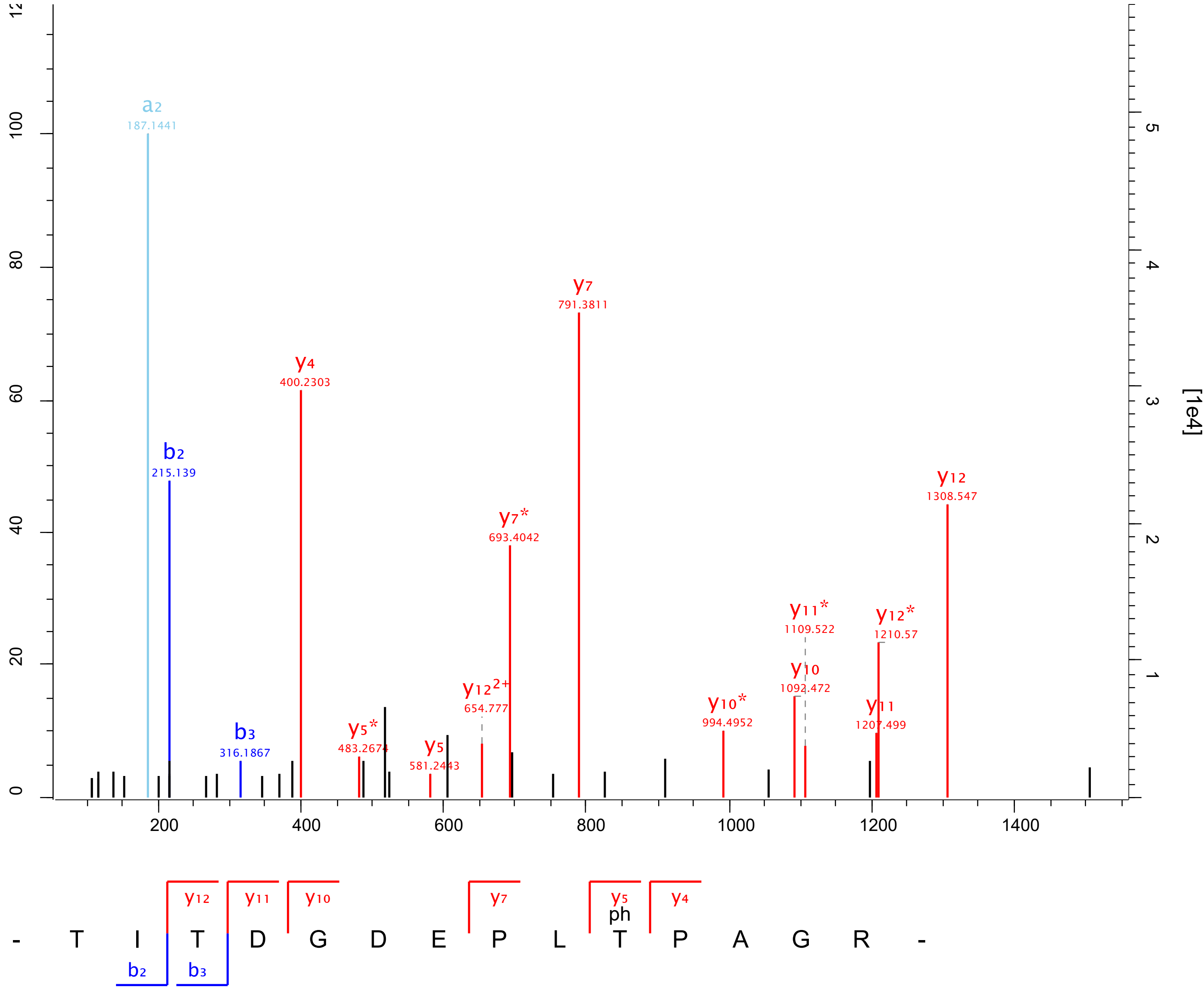

| Raw file | Scan | Method | Score | m/z |
|---|---|---|---|---|
| water_15_1 | 15636 | FTMS; HCD | 95.48 | 527.73 |

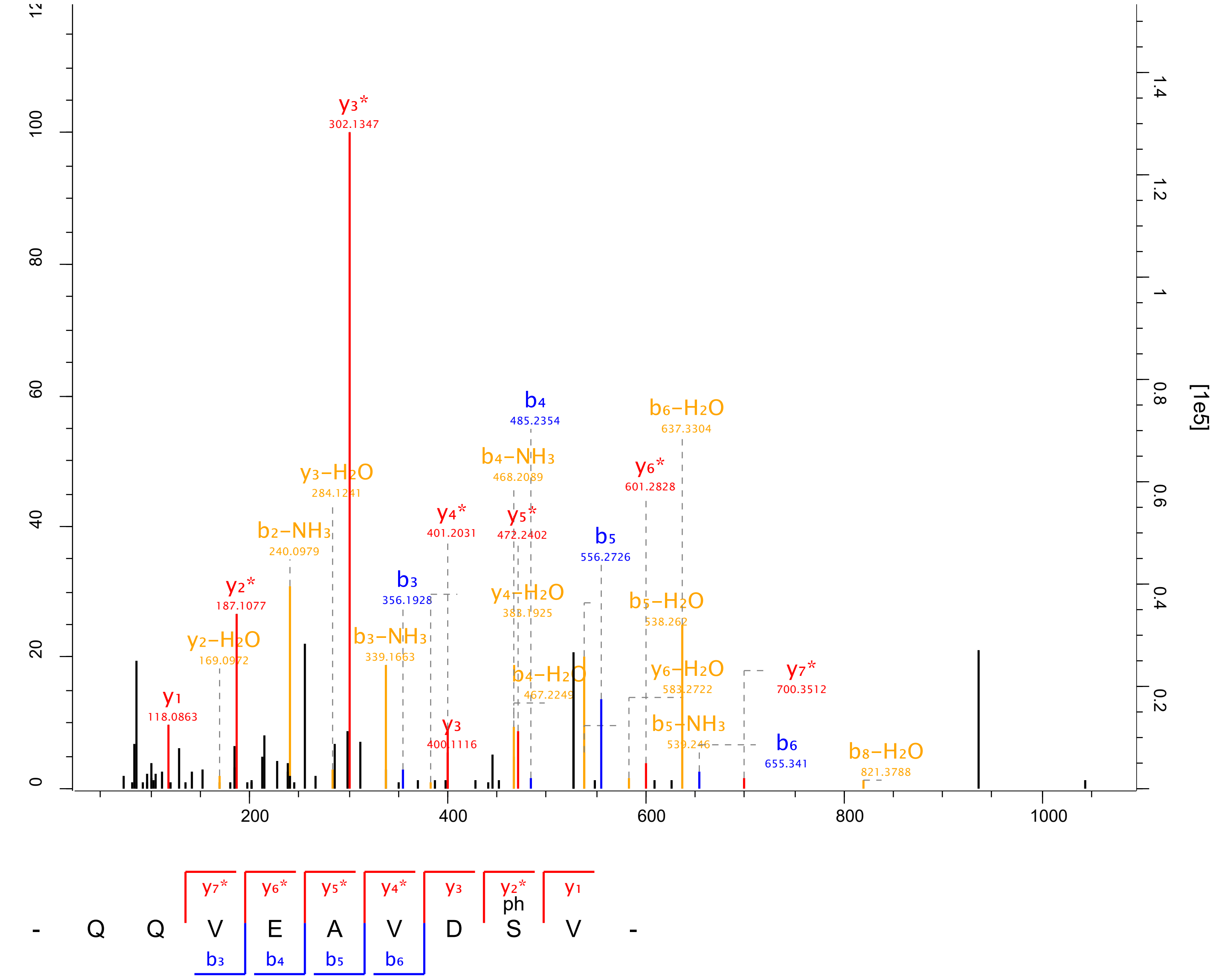

| Raw file | Scan | Method | Score | m/z |
| --- | --- | --- | --- | --- |
| water_15_1 | 1566 | FTMS; HCD | 72.27 | 421.17 |

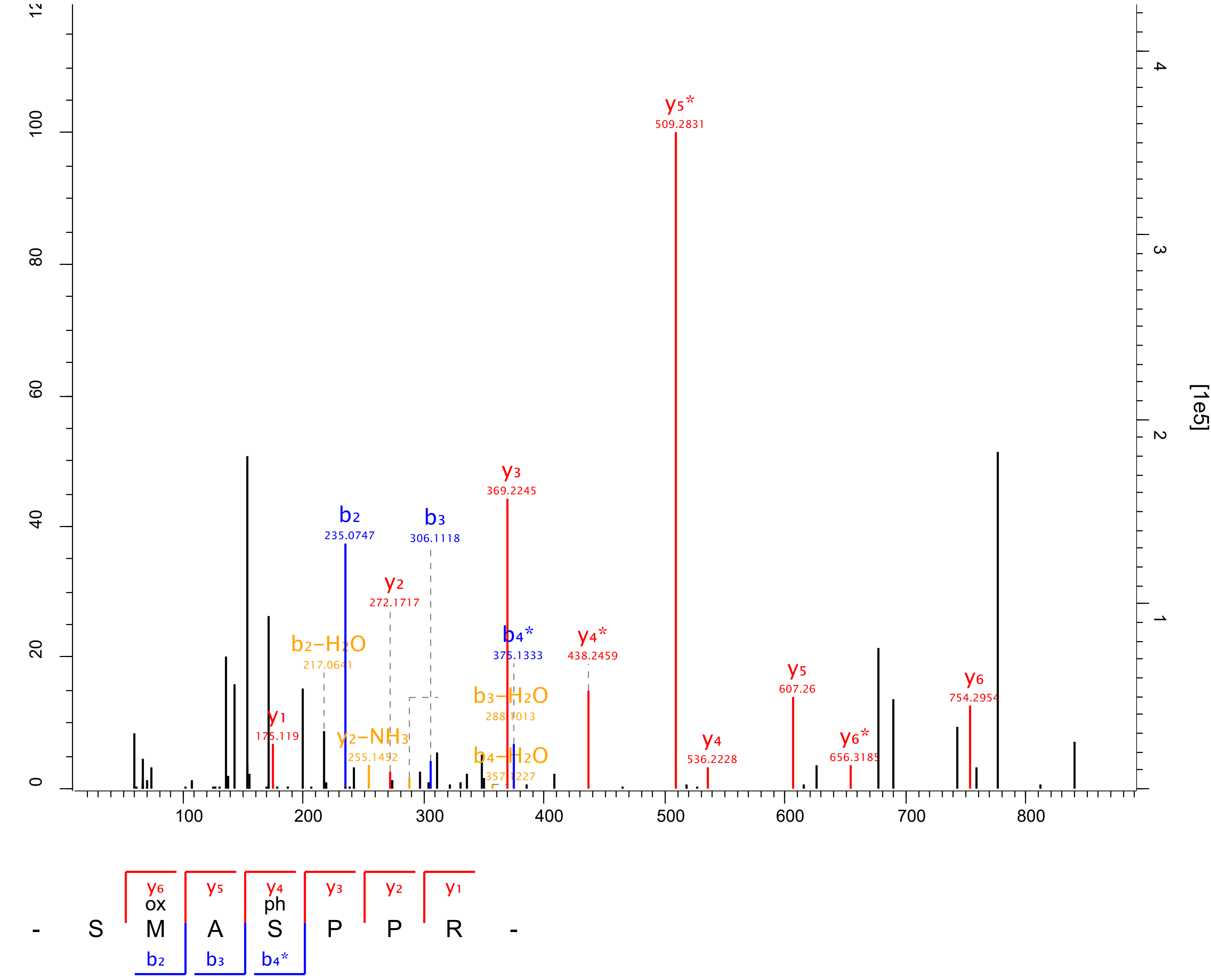

| Raw file | Scan | Method | Score | m/z |
|---|---|---|---|---|
| water_15_1 | 15934 | FTMS; HCD | 55.72 | 557.22 |

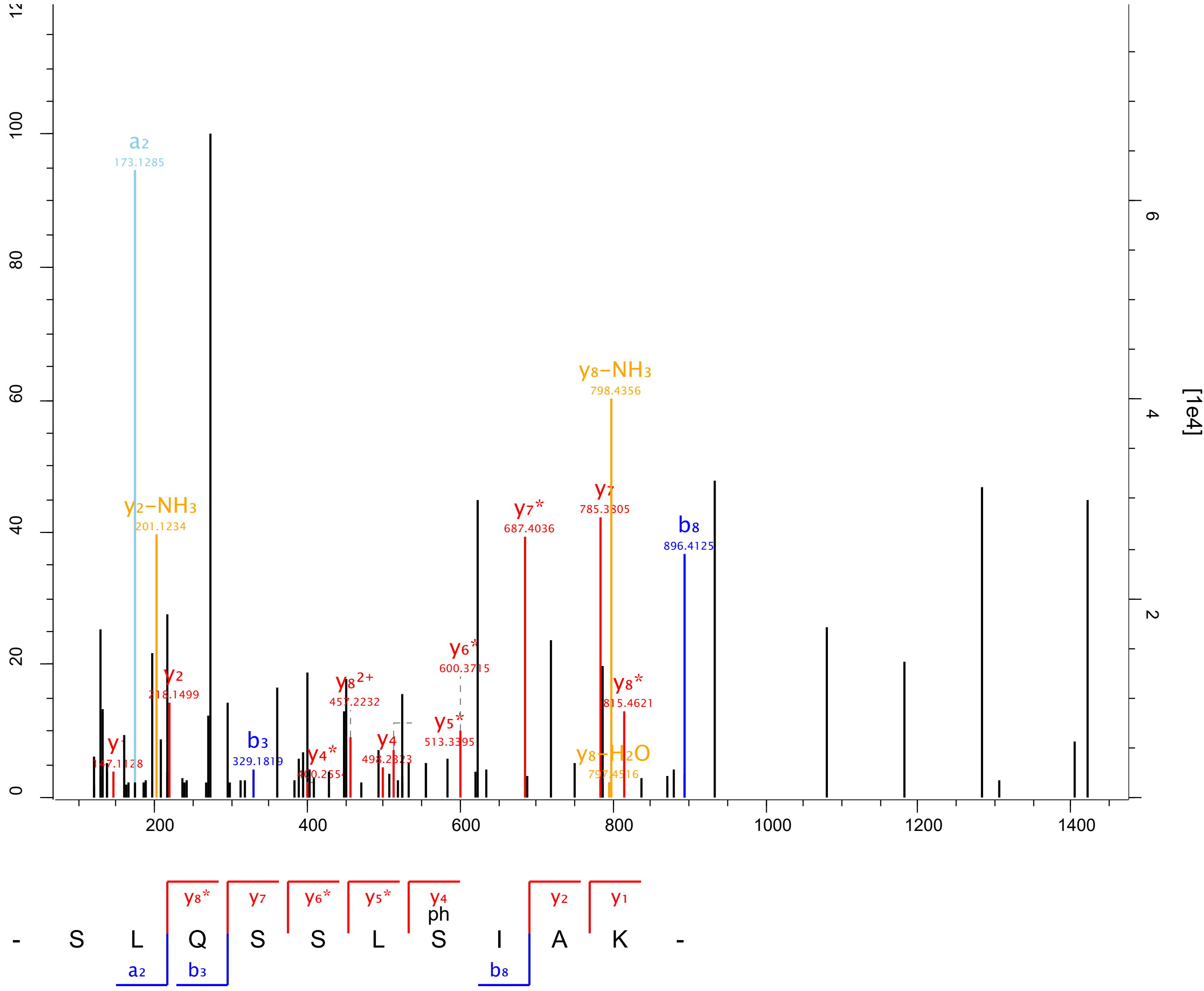

| Raw file | Scan | Method | Score | m/z |
| --- | --- | --- | --- | --- |
| water_15_1 | 15972 | FTMS; HCD | 83.09 | 605.25 |

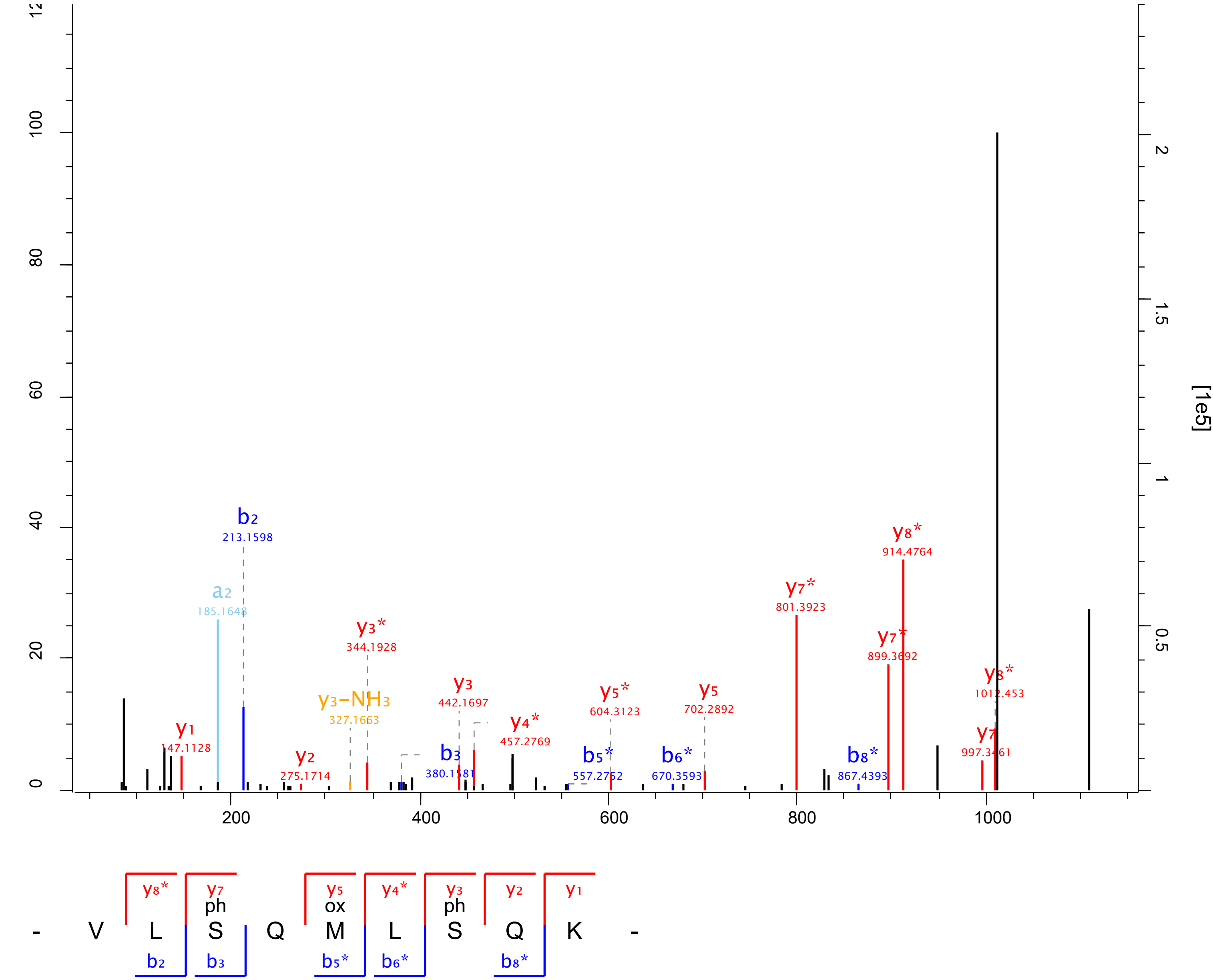

| Raw file | Scan | Method | Score | m/z |
|---|---|---|---|---|
| water_15_1 | 1612 | FTMS; HCD | 96.34 | 537.71 |

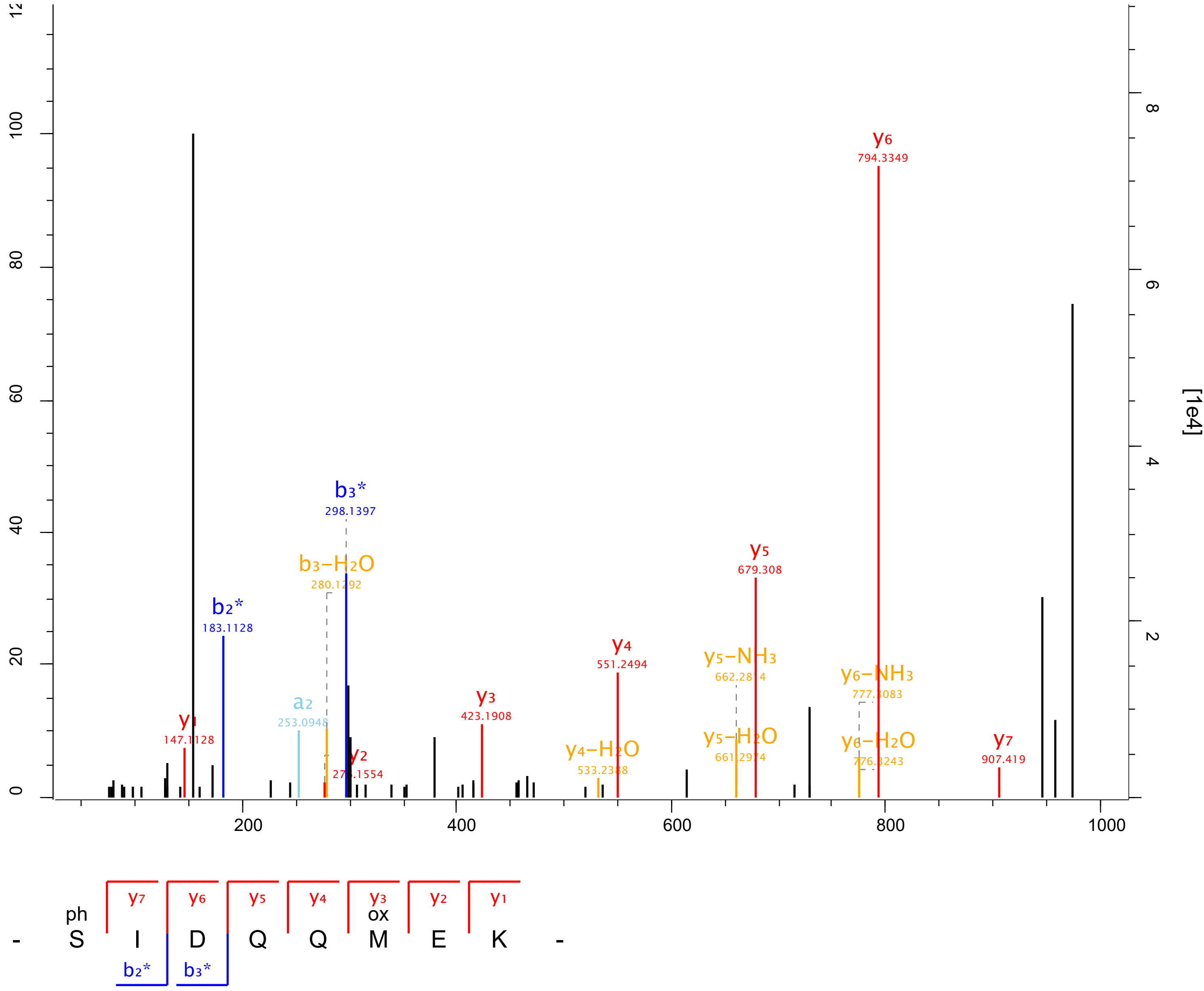

| Raw file | Scan | Method | Score | m/z |
| --- | --- | --- | --- | --- |
| water_15_1 | 16176 | FTMS; HCD | 40.86 | 611.76 |

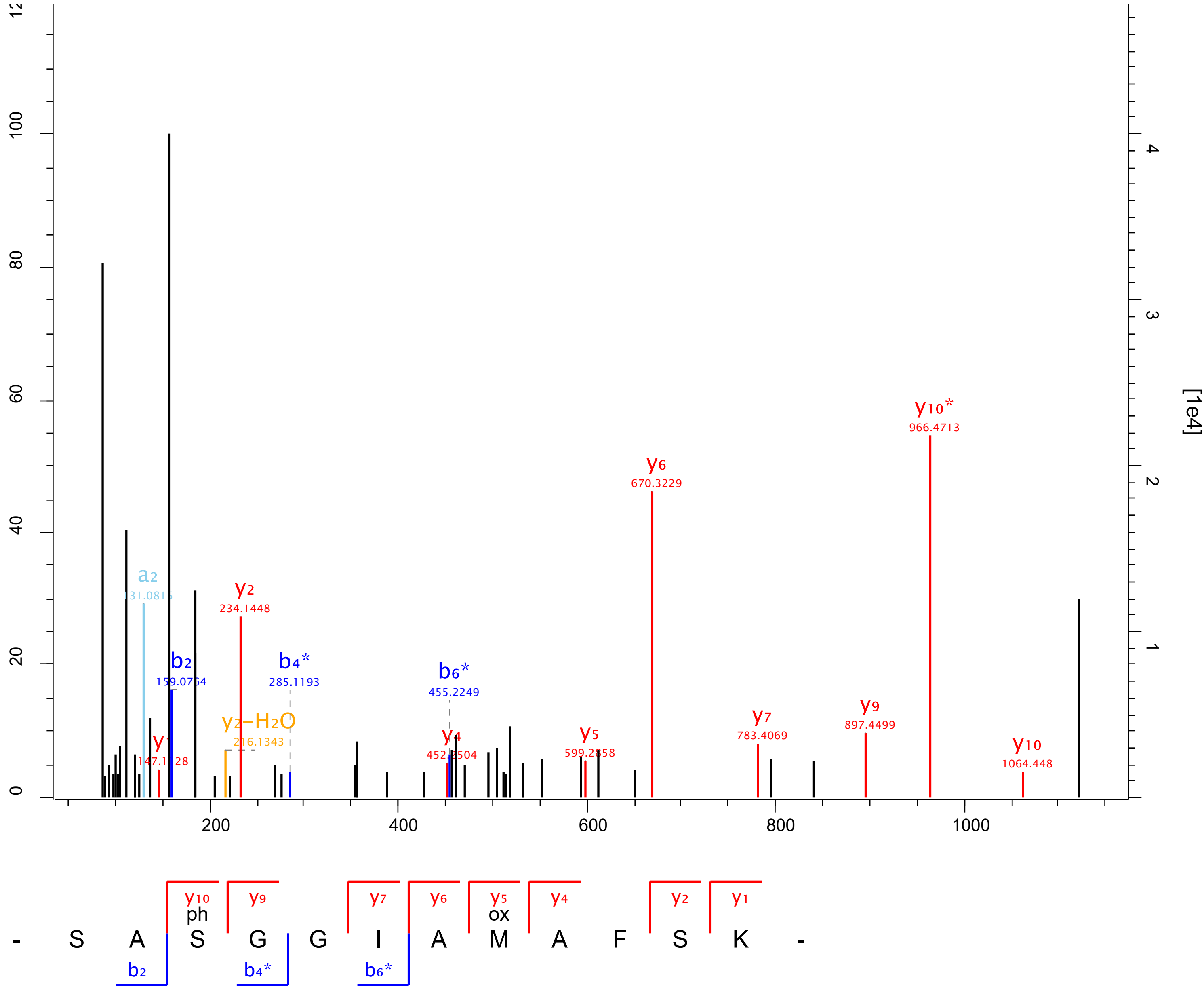

| Raw file | Scan | Method | Score | m/z |
| --- | --- | --- | --- | --- |
| water_15_1 | 16321 | FTMS; HCD | 68.58 | 720.83 |

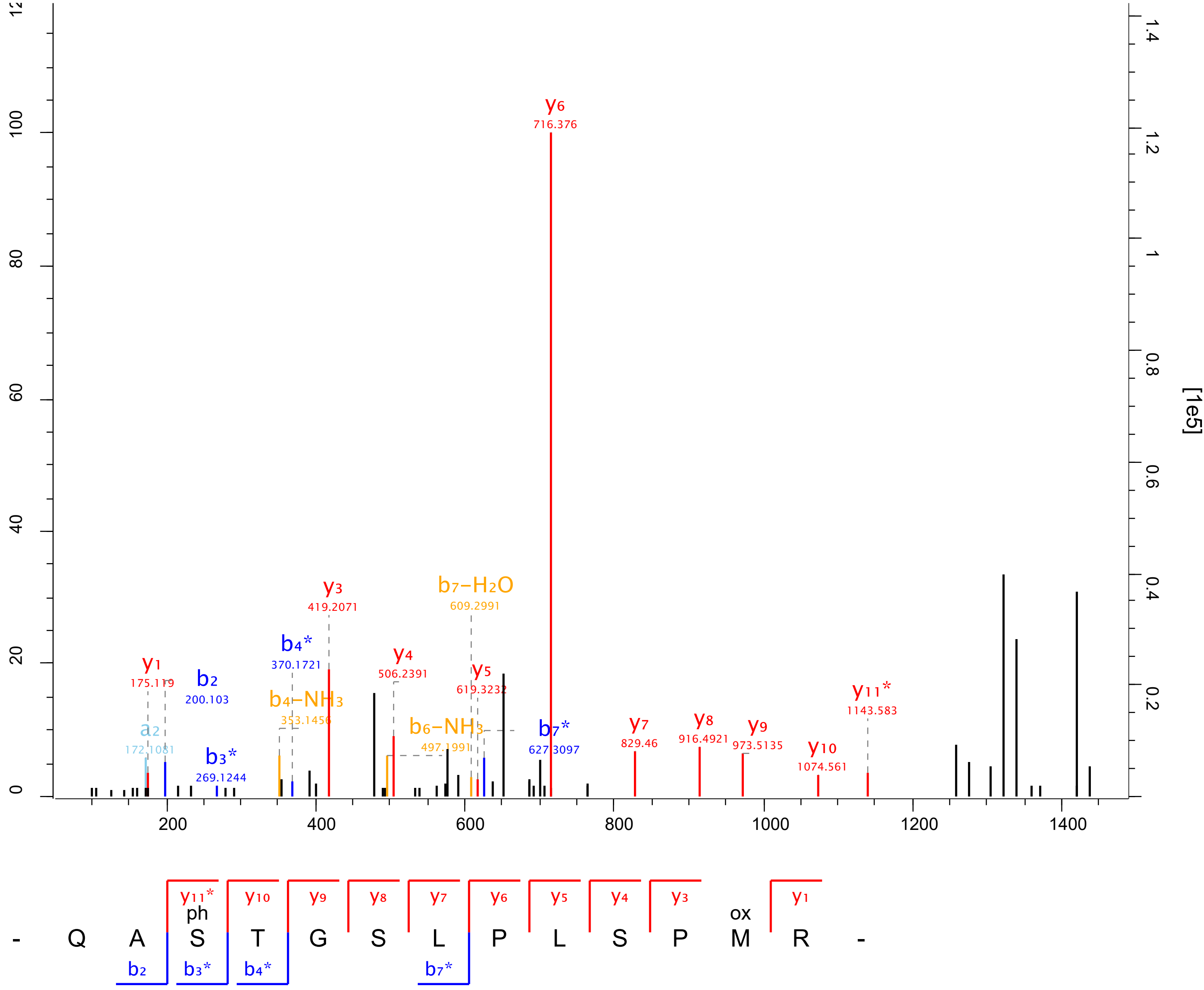

| Raw file | Scan | Method | Score | m/z |
| --- | --- | --- | --- | --- |
| water_15_1 | 16398 | FTMS; HCD | 62.29 | 606.77 |

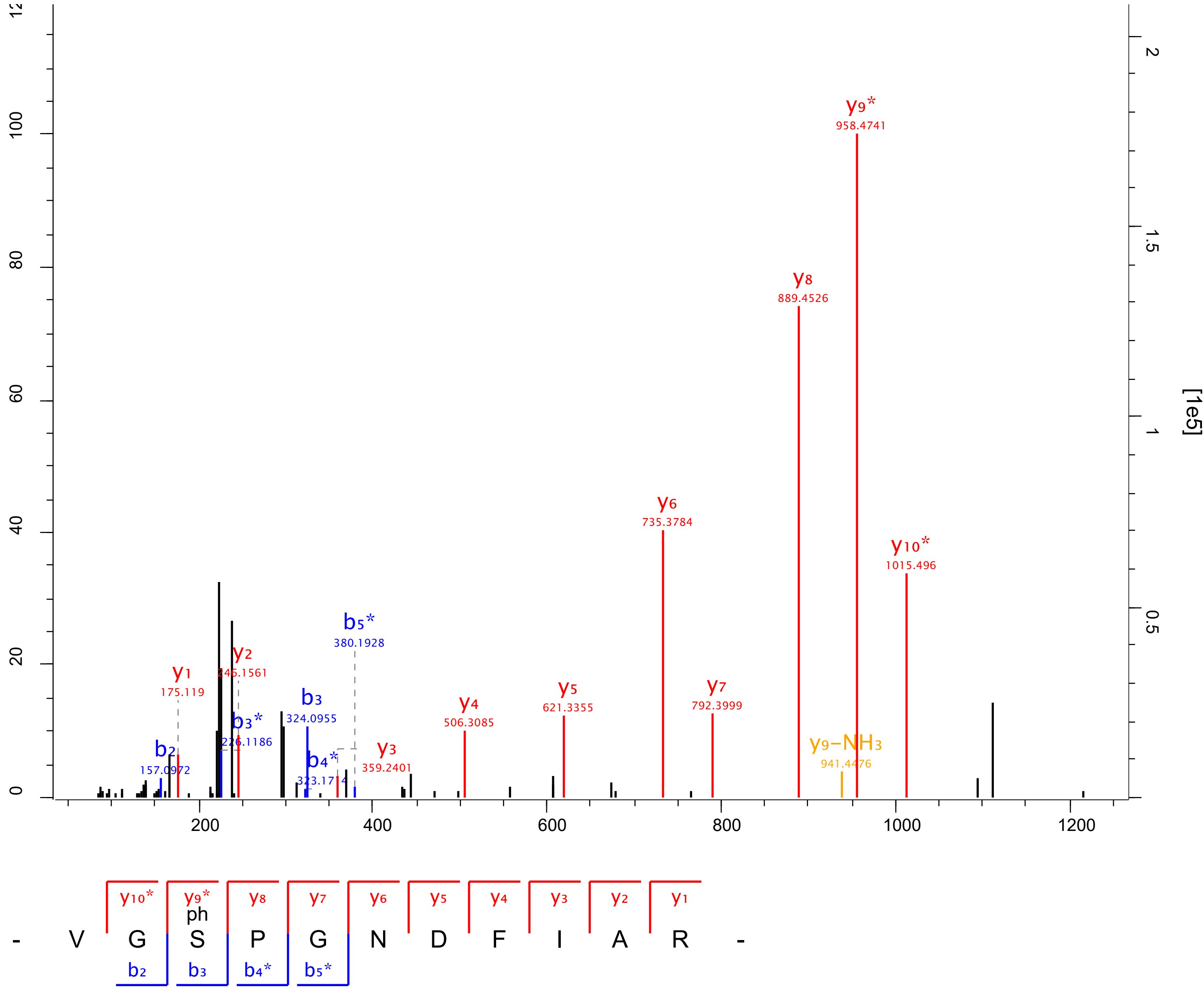

| Raw file | Scan | Method | Score | m/z |
| --- | --- | --- | --- | --- |
| water_15_1 | 16657 | FTMS; HCD | 132.62 | 769.37 |

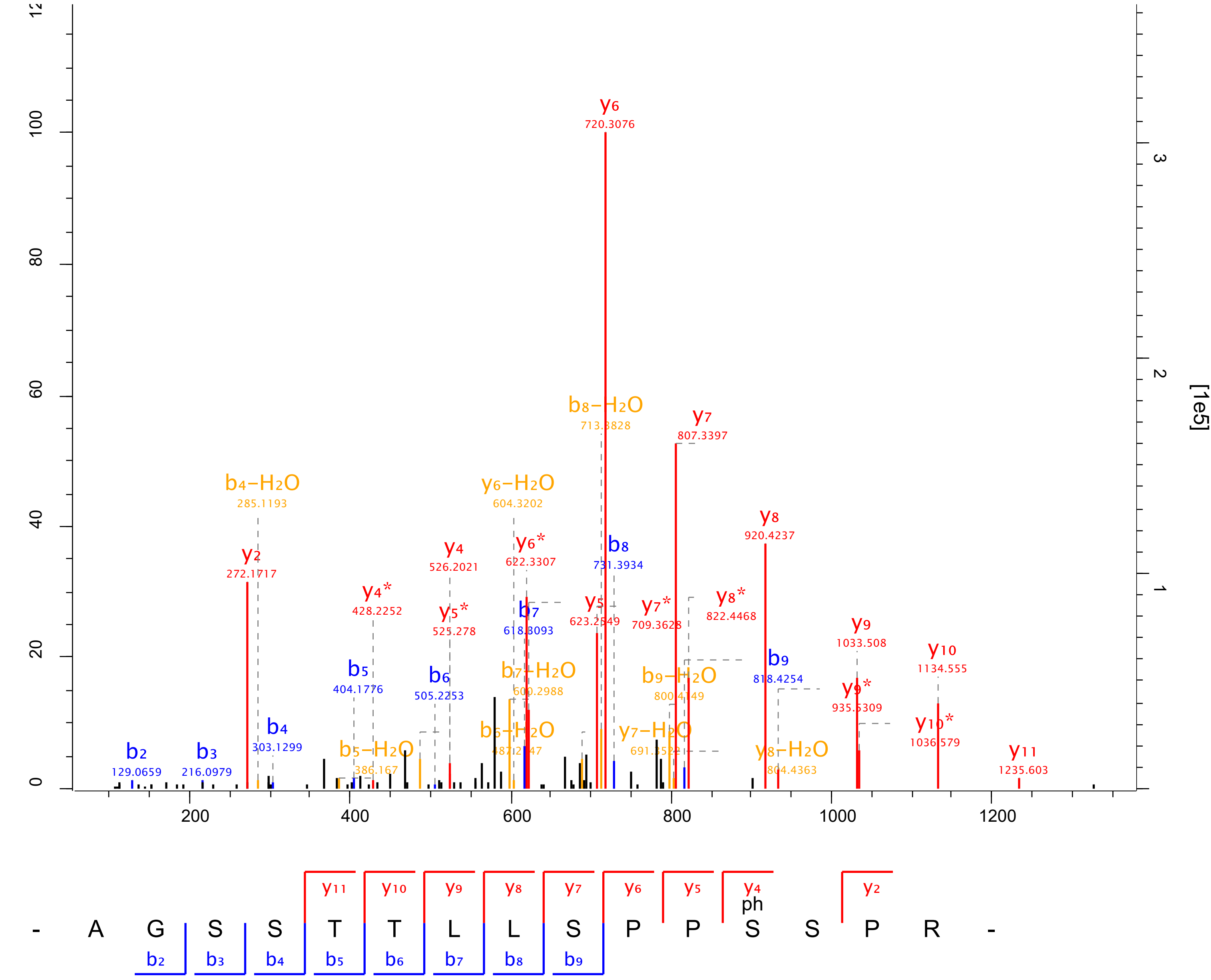

| Raw file | Scan | Method | Score | m/z |
| --- | --- | --- | --- | --- |
| water_15_1 | 16865 | FTMS; HCD | 195.77 | 707.84 |

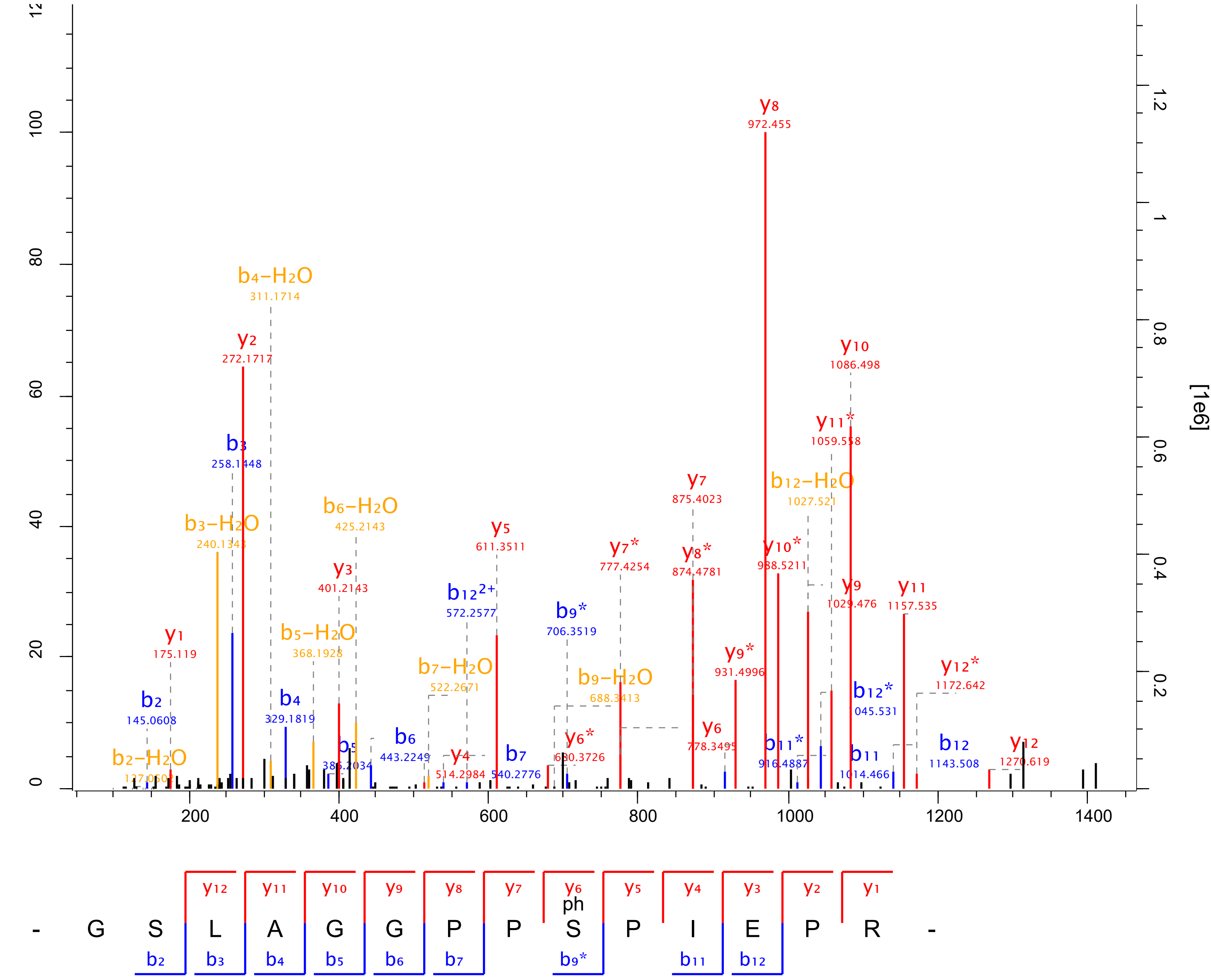

| Raw file | Scan | Method | Score | m/z |
|---|---|---|---|---|
| water_15_1 | 17396 | FTMS; HCD | 91.32 | 839.87 |

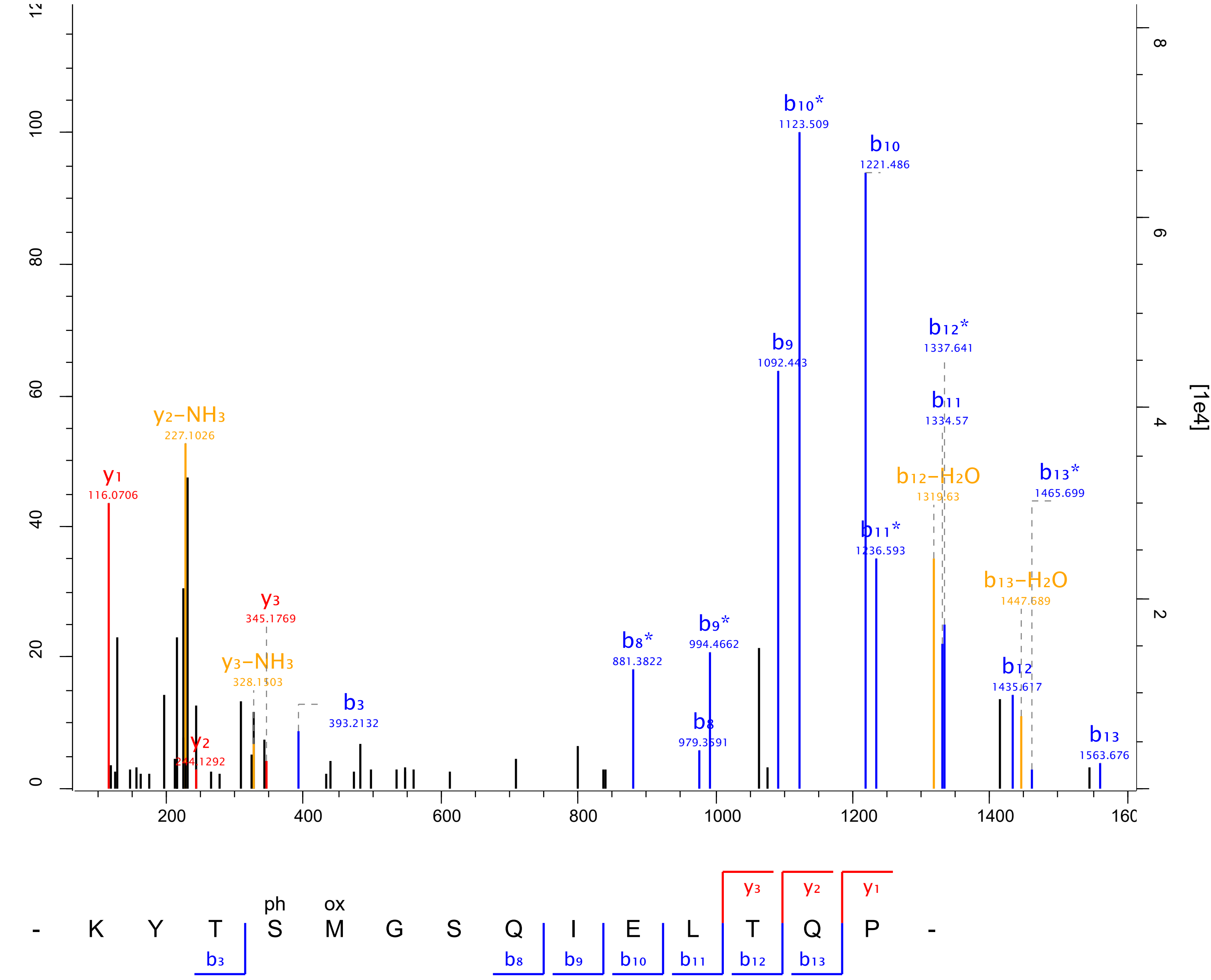

| Raw file | Scan | Method | Score | m/z |
|---|---|---|---|---|
| water_15_1 | 17759 | FTMS; HCD | 111.01 | 723.27 |

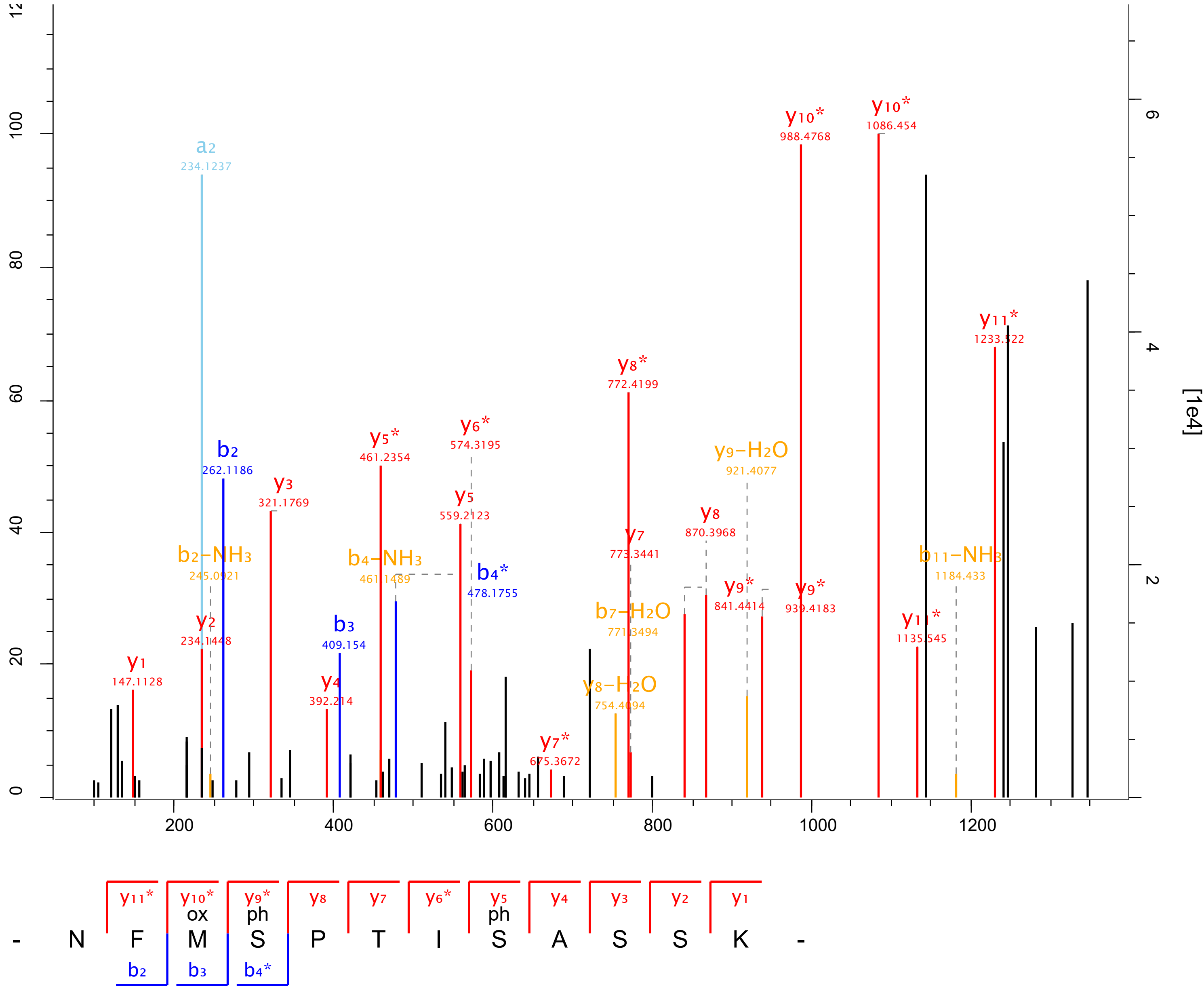

| Raw file | Scan | Method | Score | m/z |
| --- | --- | --- | --- | --- |
| water_15_1 | 19946 | FTMS; HCD | 51.09 | 715.28 |

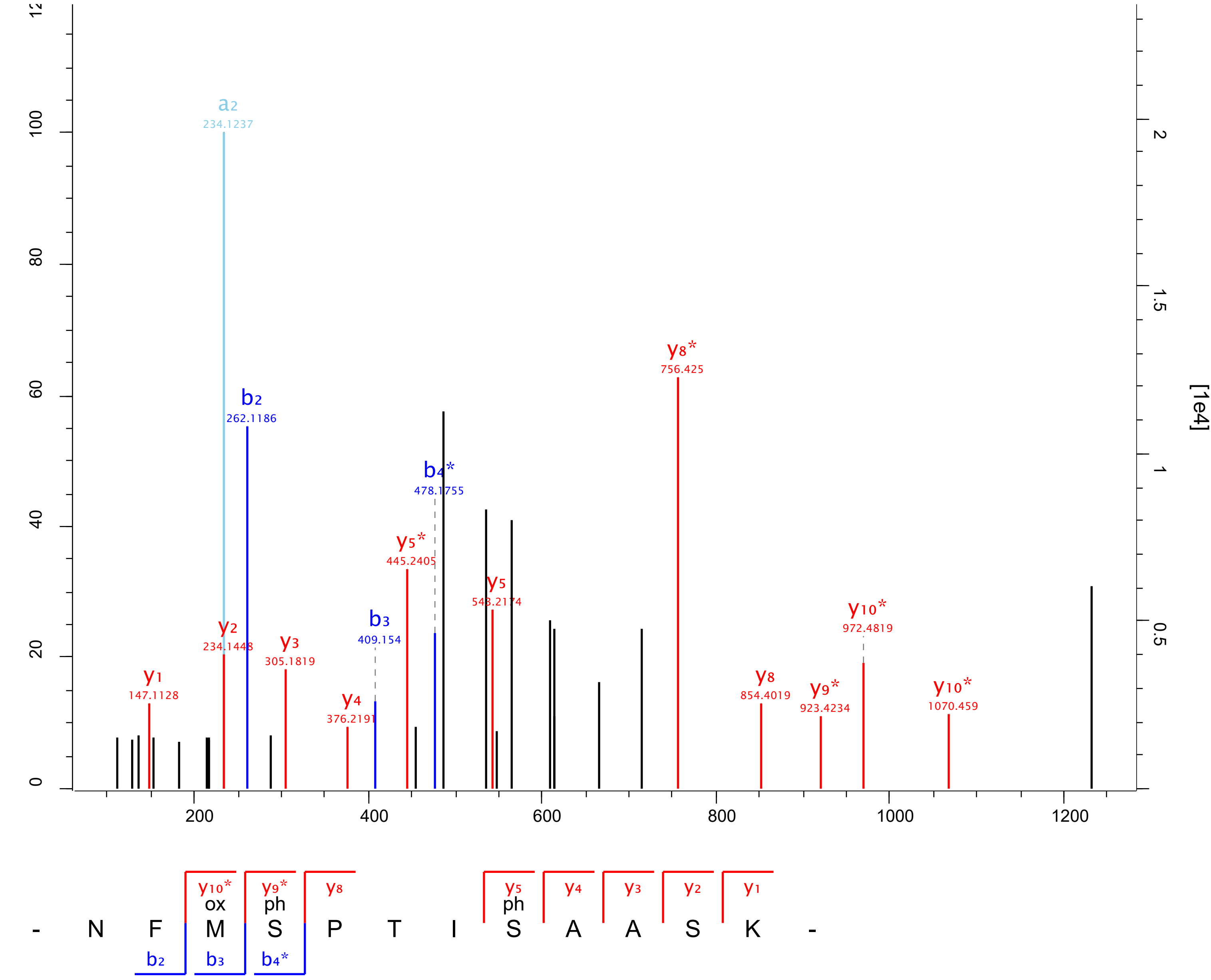

| Raw file | Scan | Method | Score | m/z |
| --- | --- | --- | --- | --- |
| water_15_1 | 19976 | FTMS; HCD | 80.24 | 572.76 |

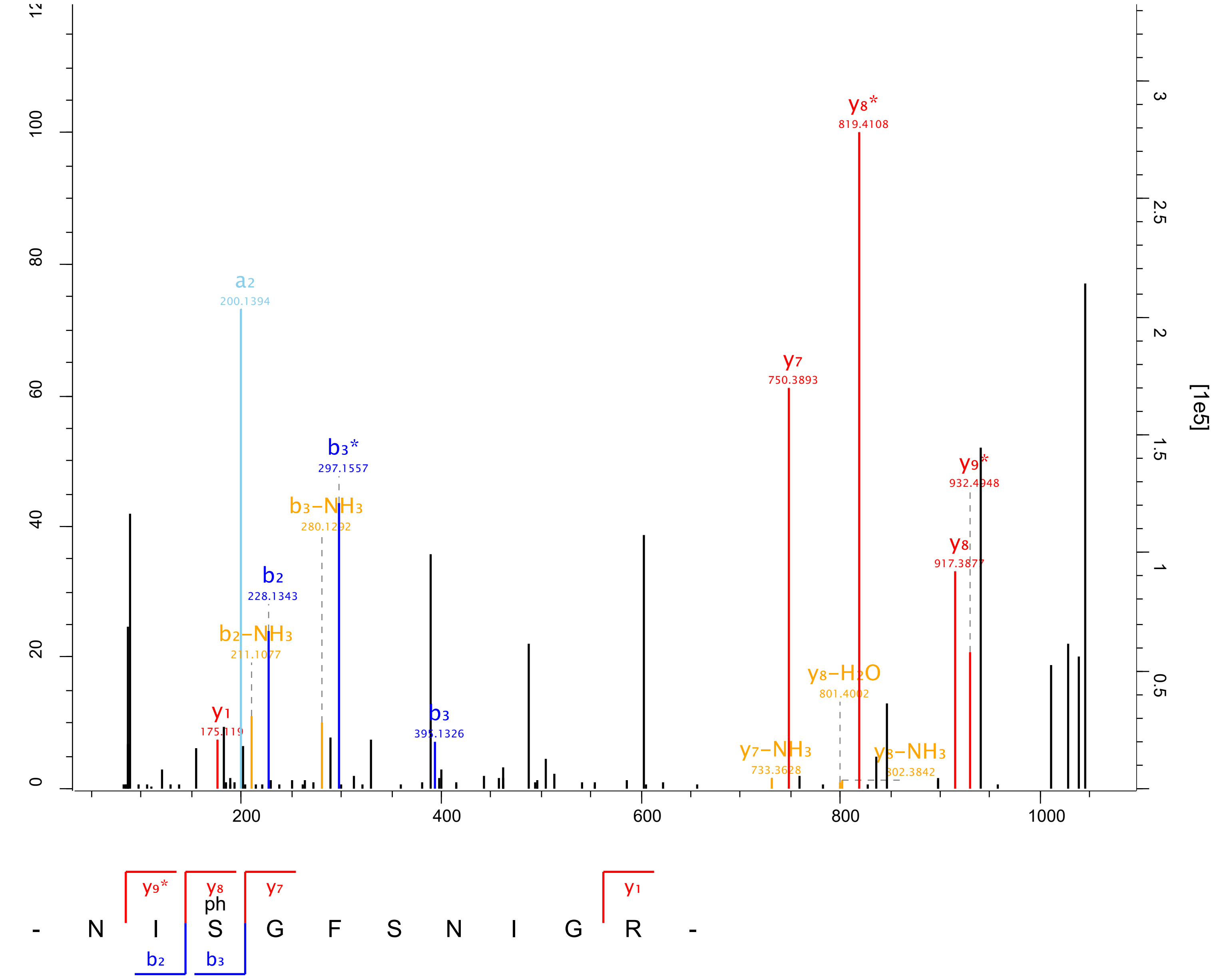

| Raw file | Scan | Method | Score | m/z |
|---|---|---|---|---|
| water_15_1 | 20601 | FTMS; HCD | 53.45 | 870.36 |

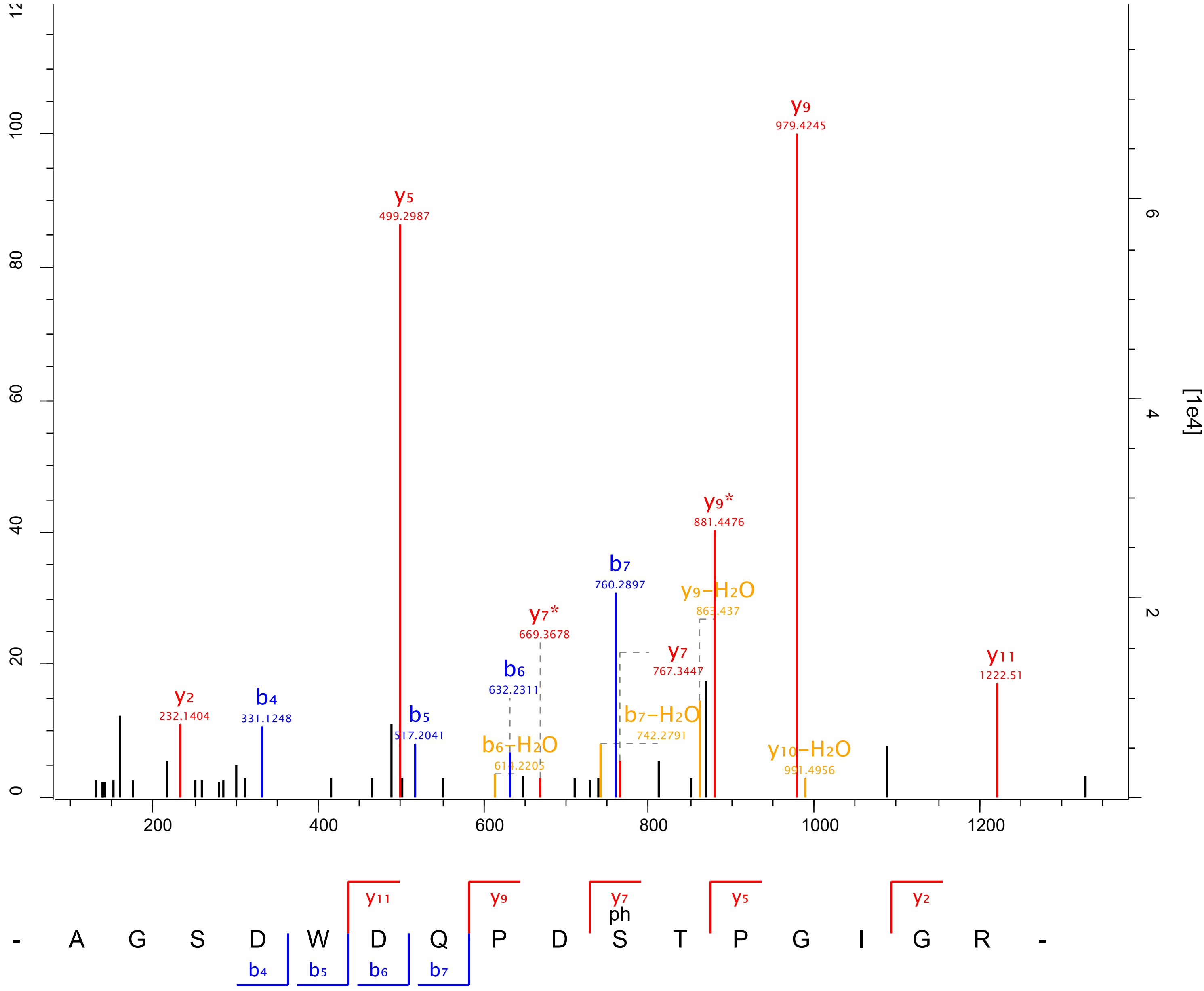

| Raw file | Scan | Method | Score | m/z |
| --- | --- | --- | --- | --- |
| water_15_1 | 20706 | FTMS; HCD | 145.31 | 537.77 |

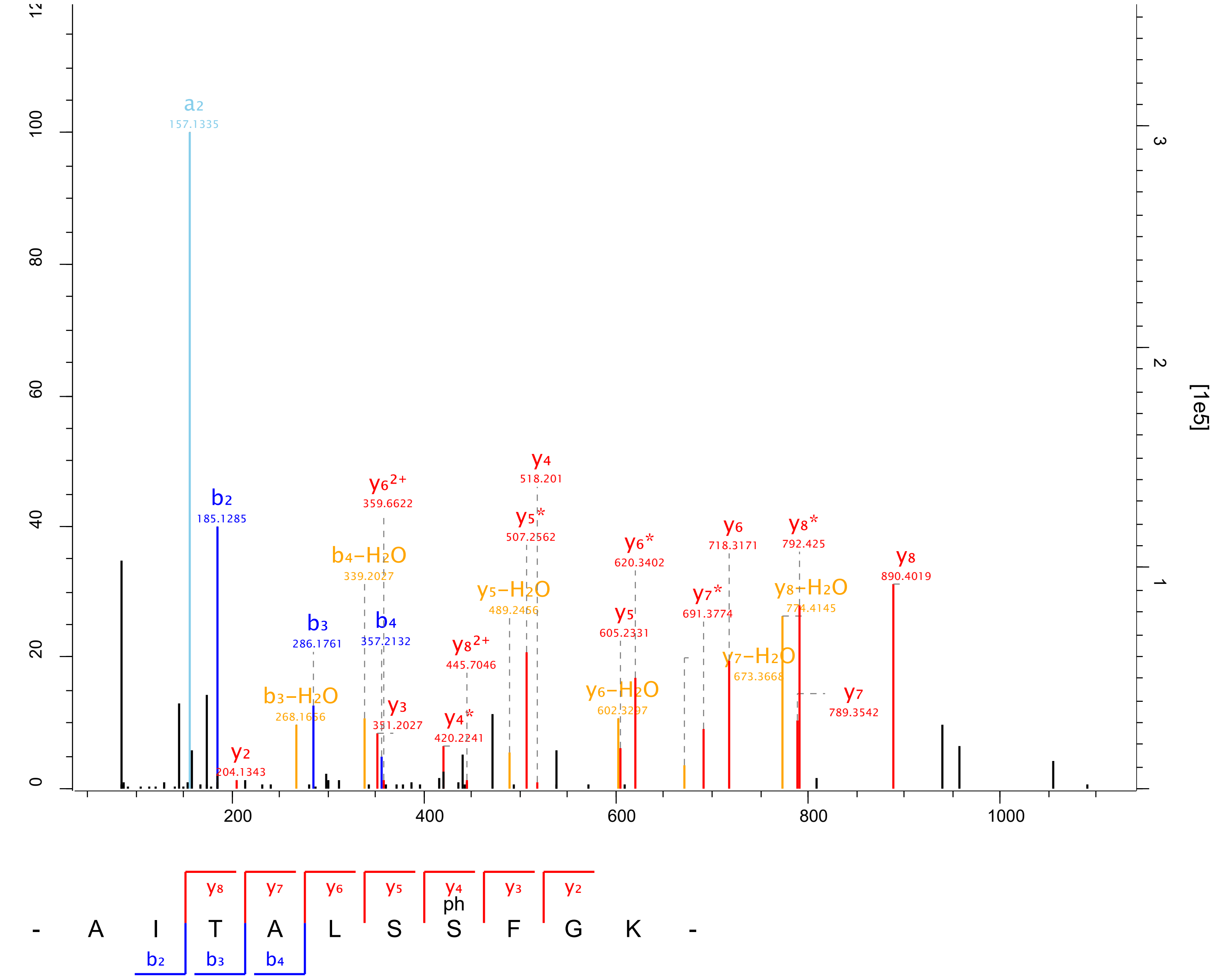

| Raw file | Scan | Method | Score | m/z |
|---|---|---|---|---|
| water_15_1 | 21086 | FTMS; HCD | 63.69 | 700.81 |

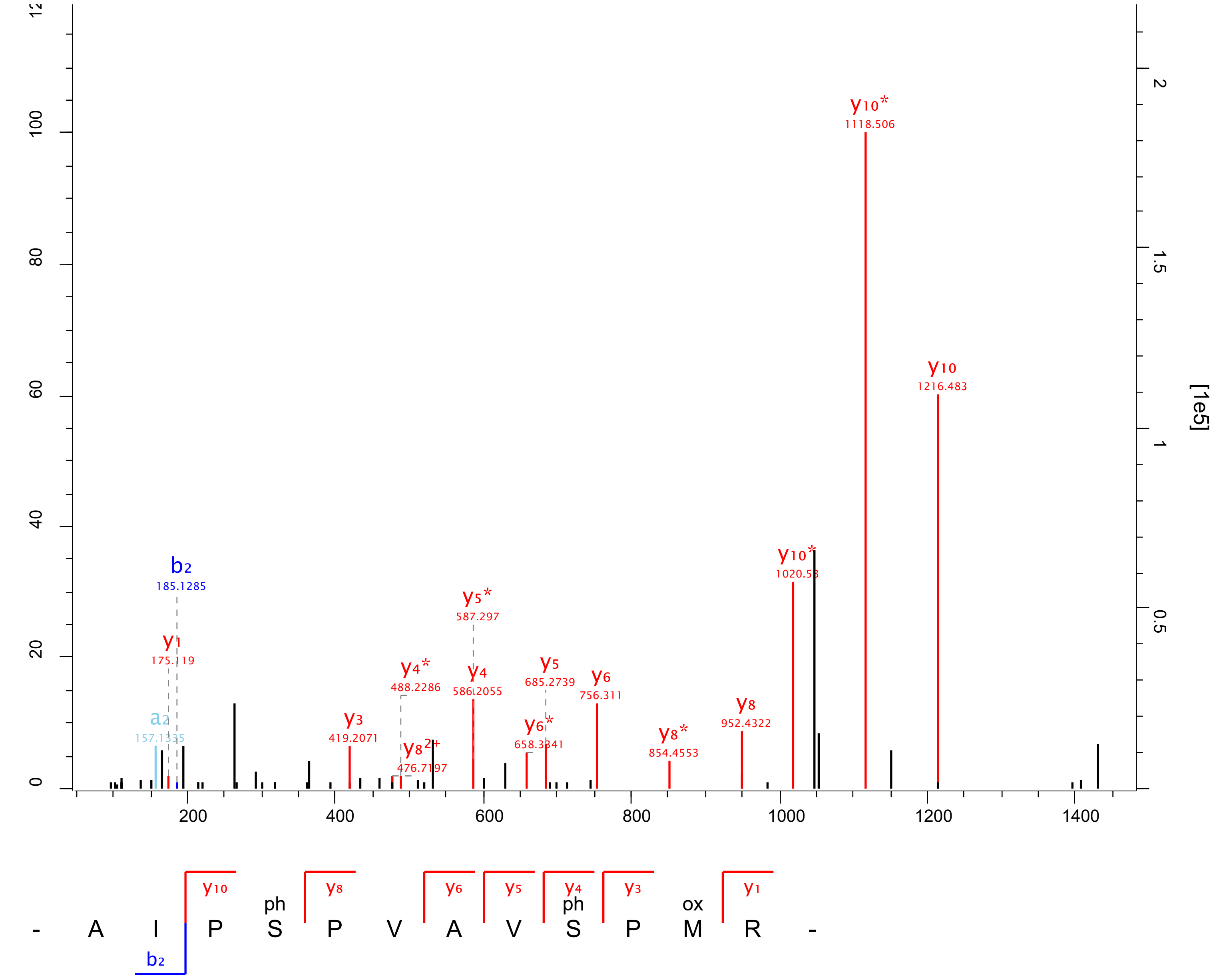

| Raw file | Scan | Method | Score | m/z |
| --- | --- | --- | --- | --- |
| water_15_1 | 21191 | FTMS; HCD | 99.94 | 674.8 |

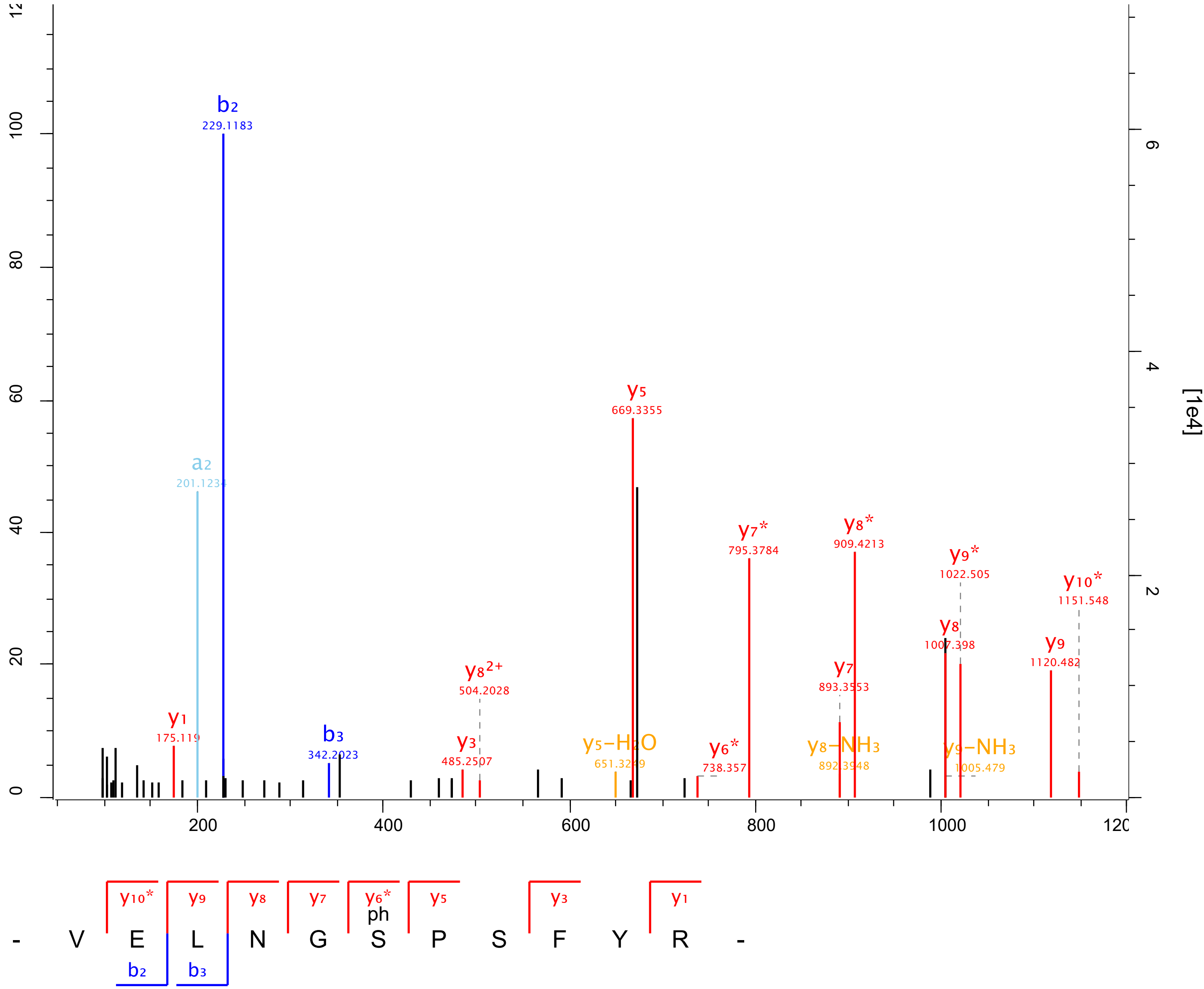

| Raw file | Scan | Method | Score | m/z |
| --- | --- | --- | --- | --- |
| water_15_1 | 21215 | FTMS; HCD | 77.22 | 732.82 |

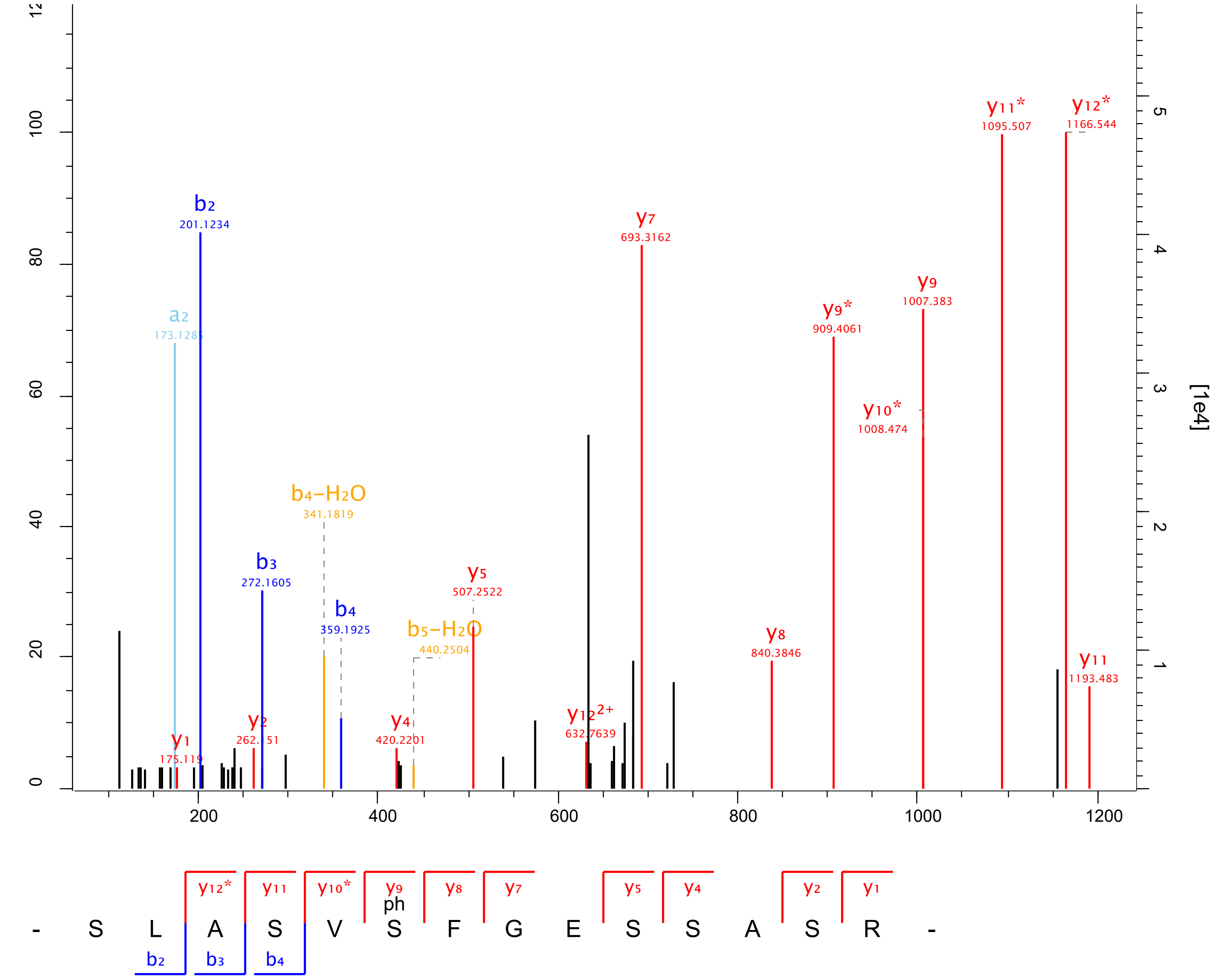

| Raw file | Scan | Method | Score | m/z |
|---|---|---|---|---|
| water_15_1 | 21318 | FTMS; HCD | 117.02 | 499.26 |

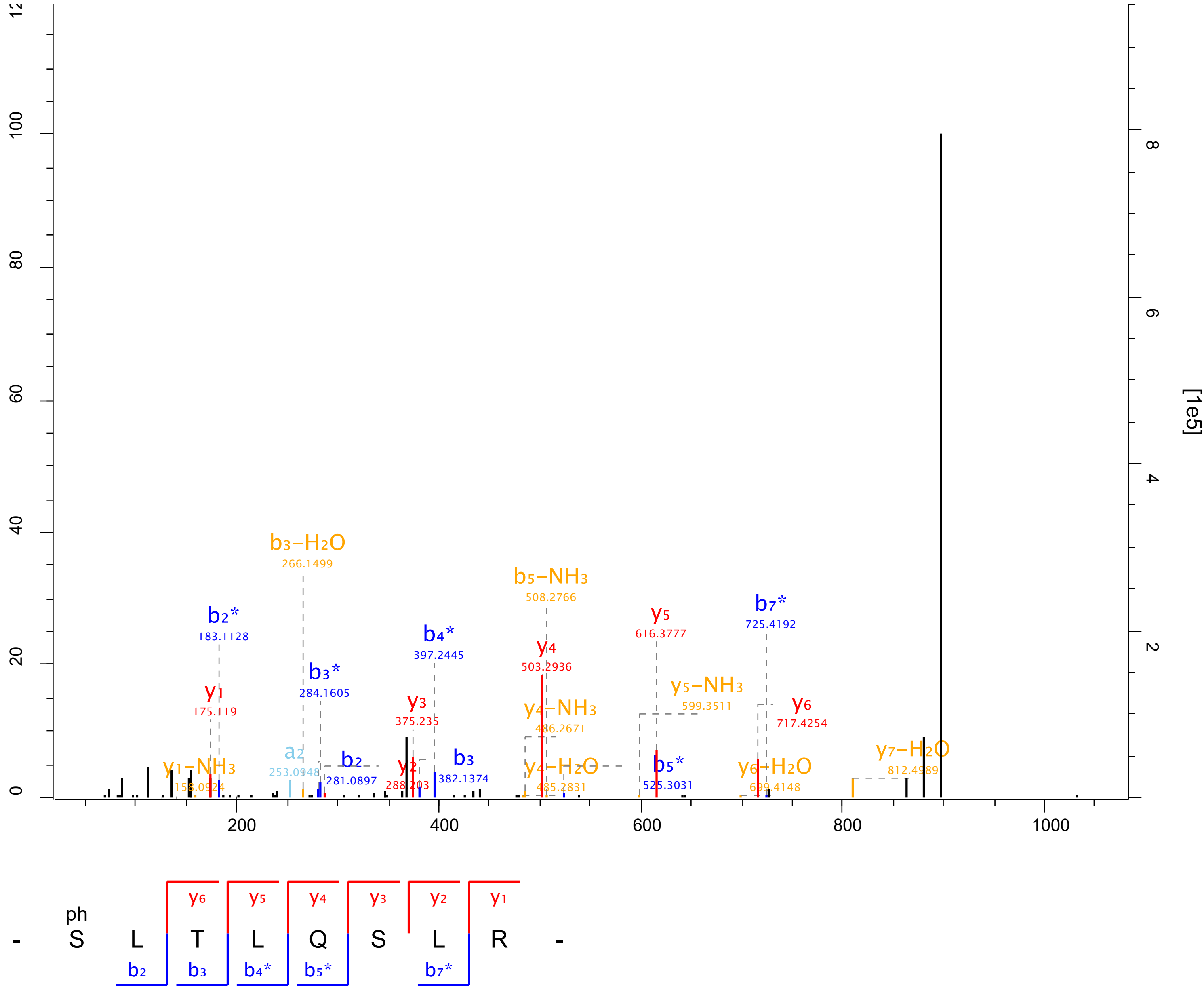

| Raw file | Scan | Method | Score | m/z |
| --- | --- | --- | --- | --- |
| water_15_1 | 21431 | FTMS; HCD | 208.87 | 782.89 |

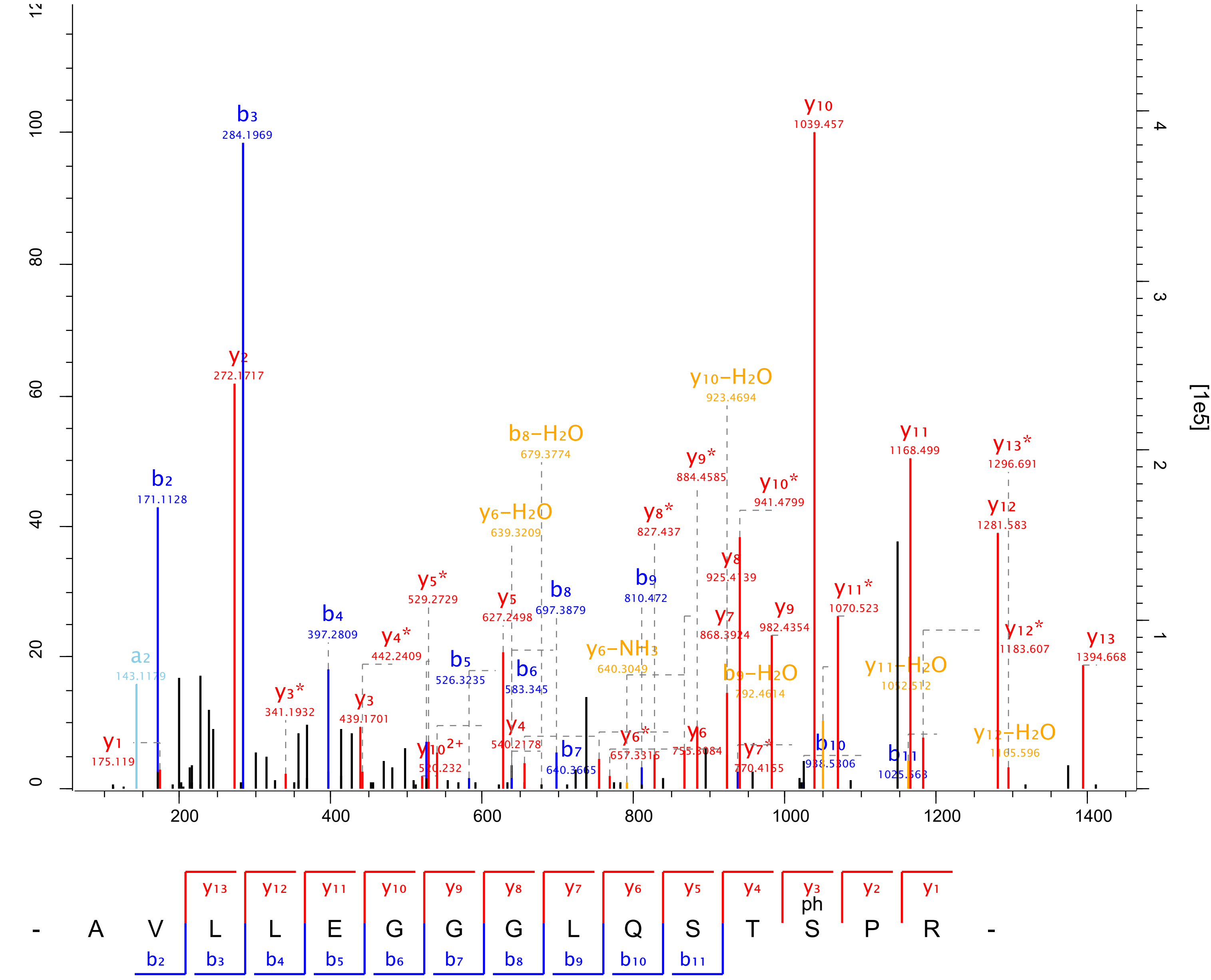

| Raw file | Scan | Method | Score | m/z |
| --- | --- | --- | --- | --- |
| water_15_1 | 21693 | FTMS; HCD | 154.49 | 799.85 |

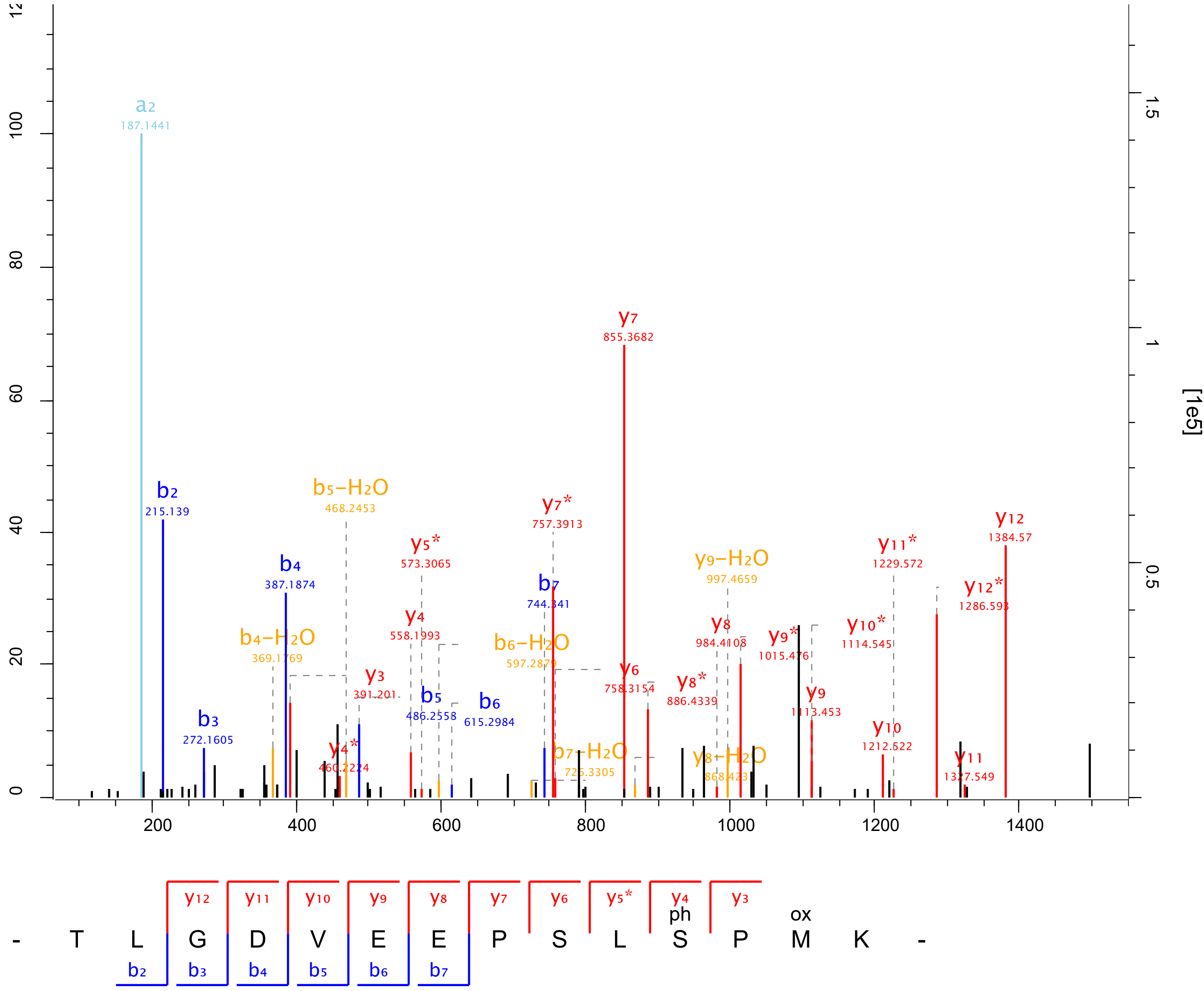

| Raw file | Scan | Method | Score | m/z |
| --- | --- | --- | --- | --- |
| water_15_1 | 21911 | FTMS; HCD | 146.3 | 713.83 |

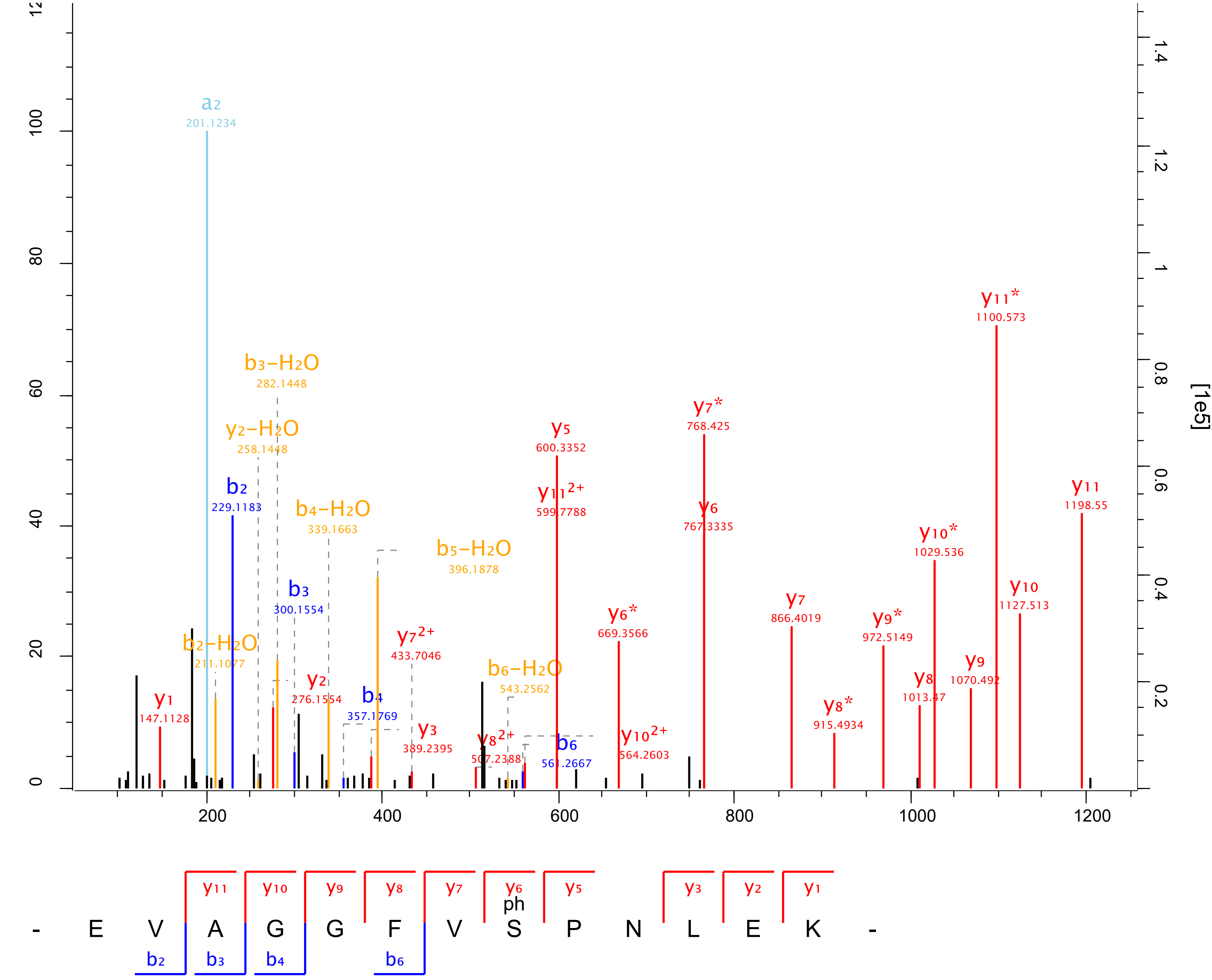

| Raw file | Scan | Method | Score | m/z |
|---|---|---|---|---|
| water_15_1 | 22520 | FTMS; HCD | 186.98 | 682.35 |

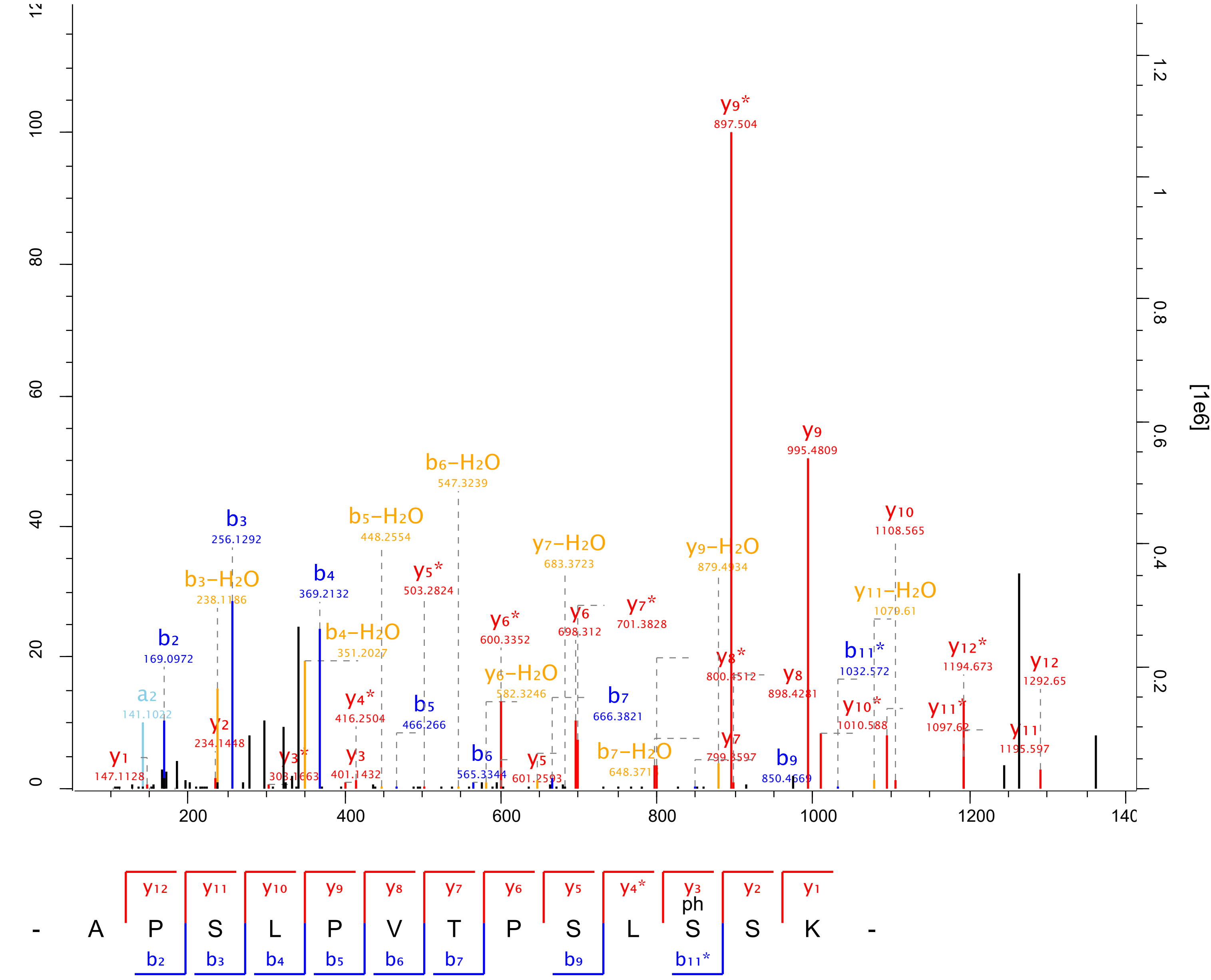

| Raw file | Scan | Method | Score | m/z |
| --- | --- | --- | --- | --- |
| water_15_1 | 23101 | FTMS; HCD | 87.35 | 934.92 |

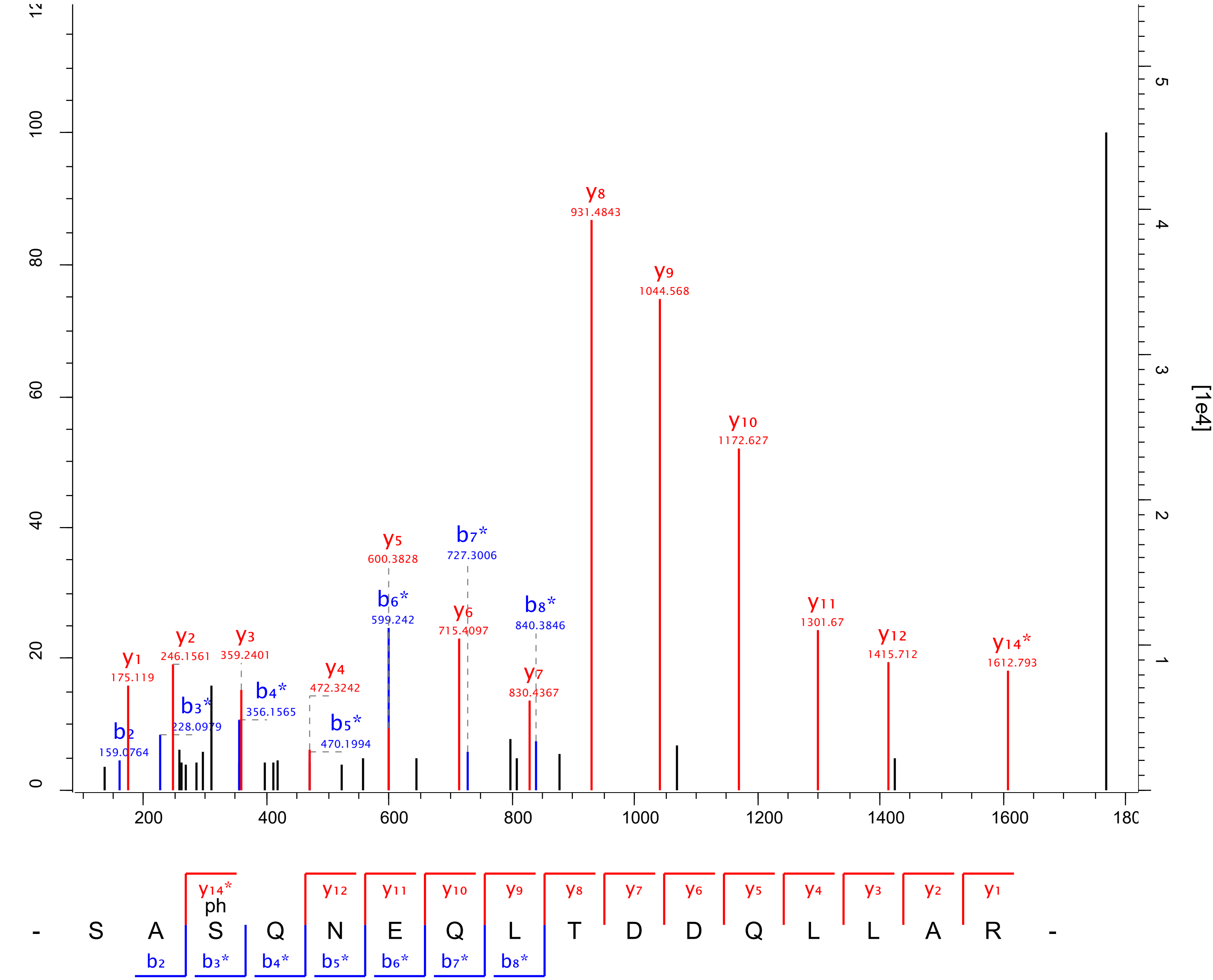

| Raw file | Scan | Method | Score | m/z |
|---|---|---|---|---|
| water_15_1 | 23500 | FTMS; HCD | 70.09 | 625.77 |

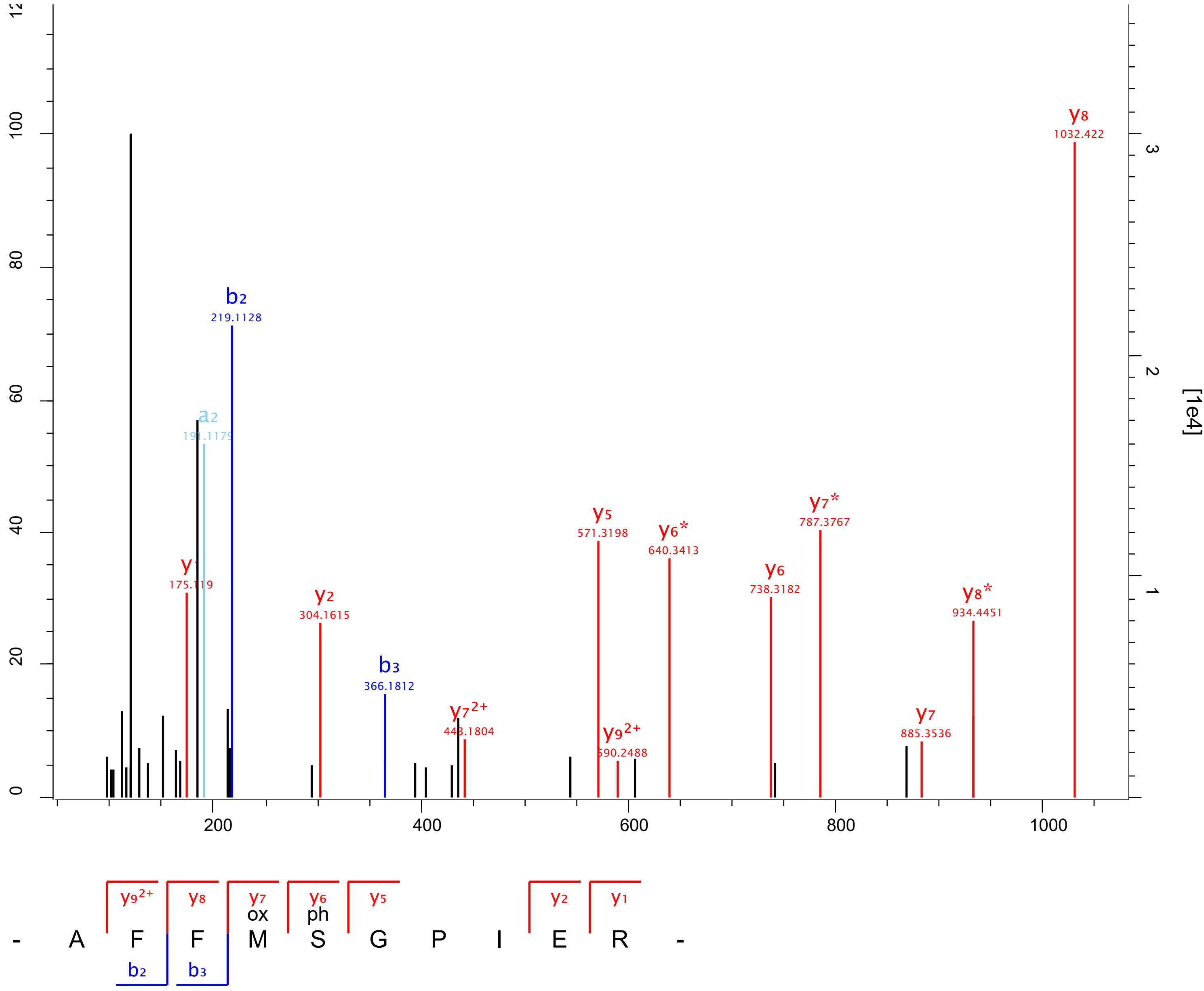

| Raw file | Scan | Method | Score | m/z |
| --- | --- | --- | --- | --- |
| water_15_1 | 23555 | FTMS; HCD | 103.84 | 774.65 |

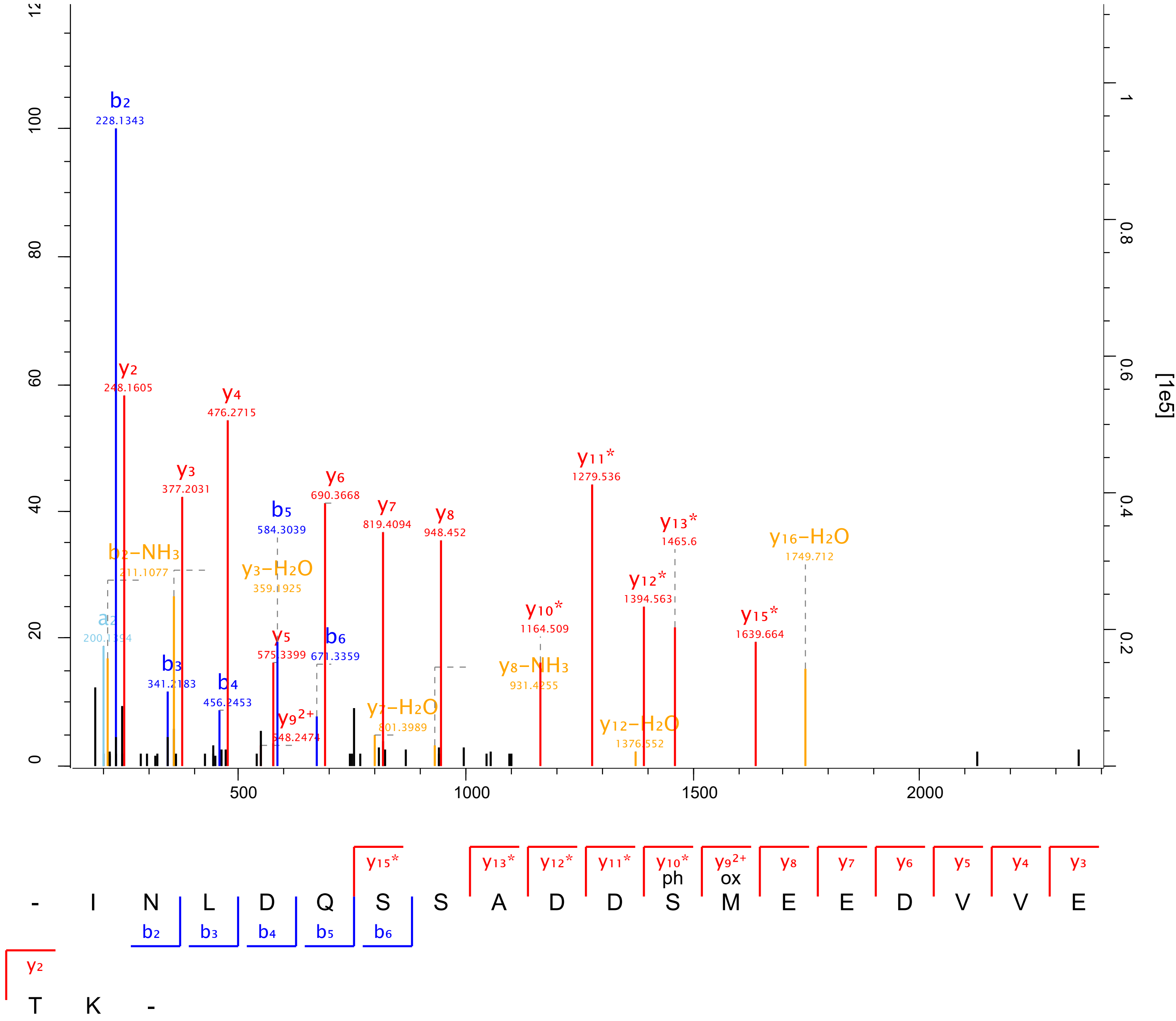

| Raw file | Scan | Method | Score | m/z |
|---|---|---|---|---|
| water_15_1 | 23827 | FTMS; HCD | 159.41 | 706.34 |

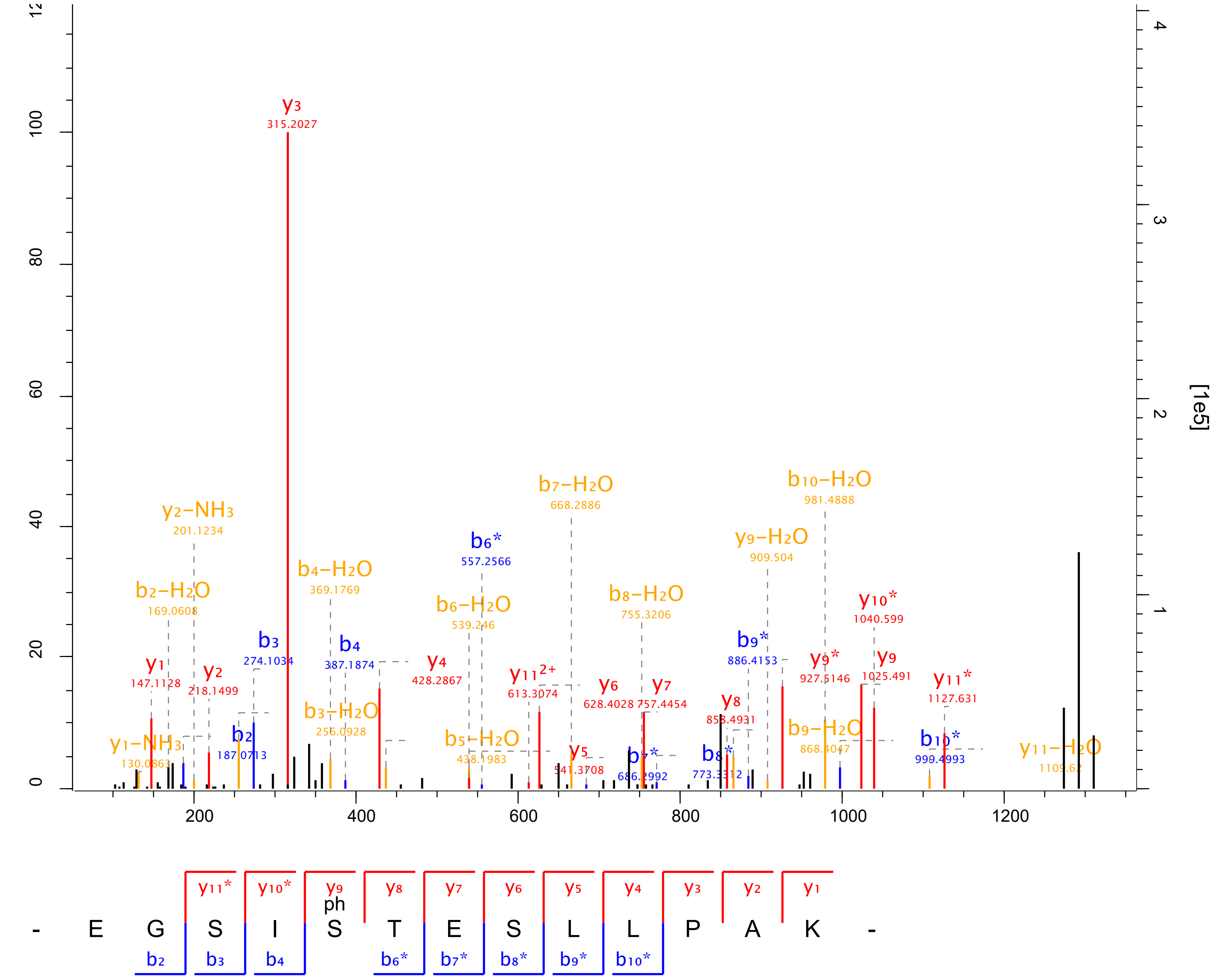

| Raw file | Scan | Method | Score | m/z |
|---|---|---|---|---|
| water_15_1 | 23926 | FTMS; HCD | 101.75 | 606.77 |

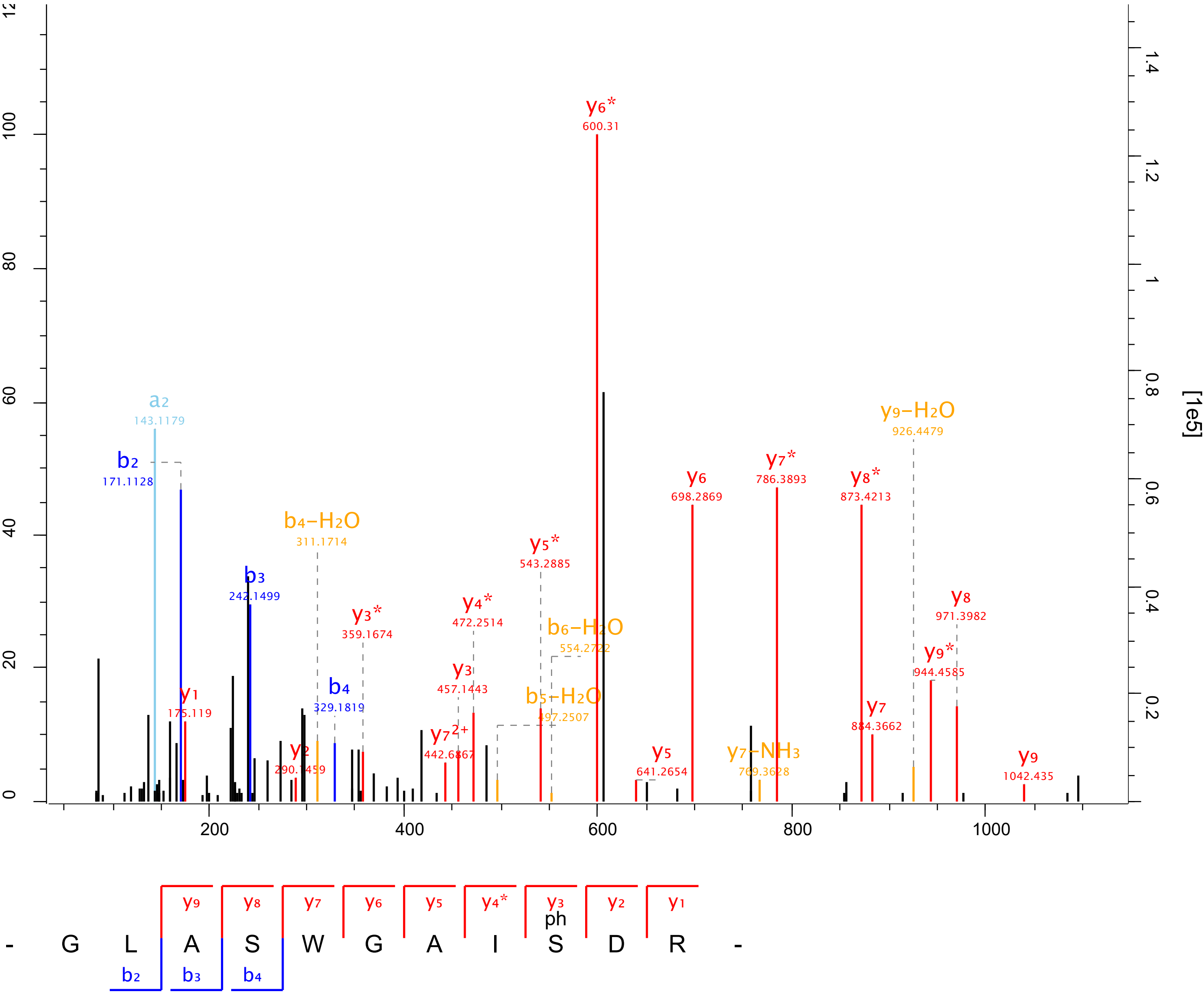

| Raw file | Scan | Method | Score | m/z |
|---|---|---|---|---|
| water_15_1 | 24141 | FTMS; HCD | 80.79 | 954.93 |

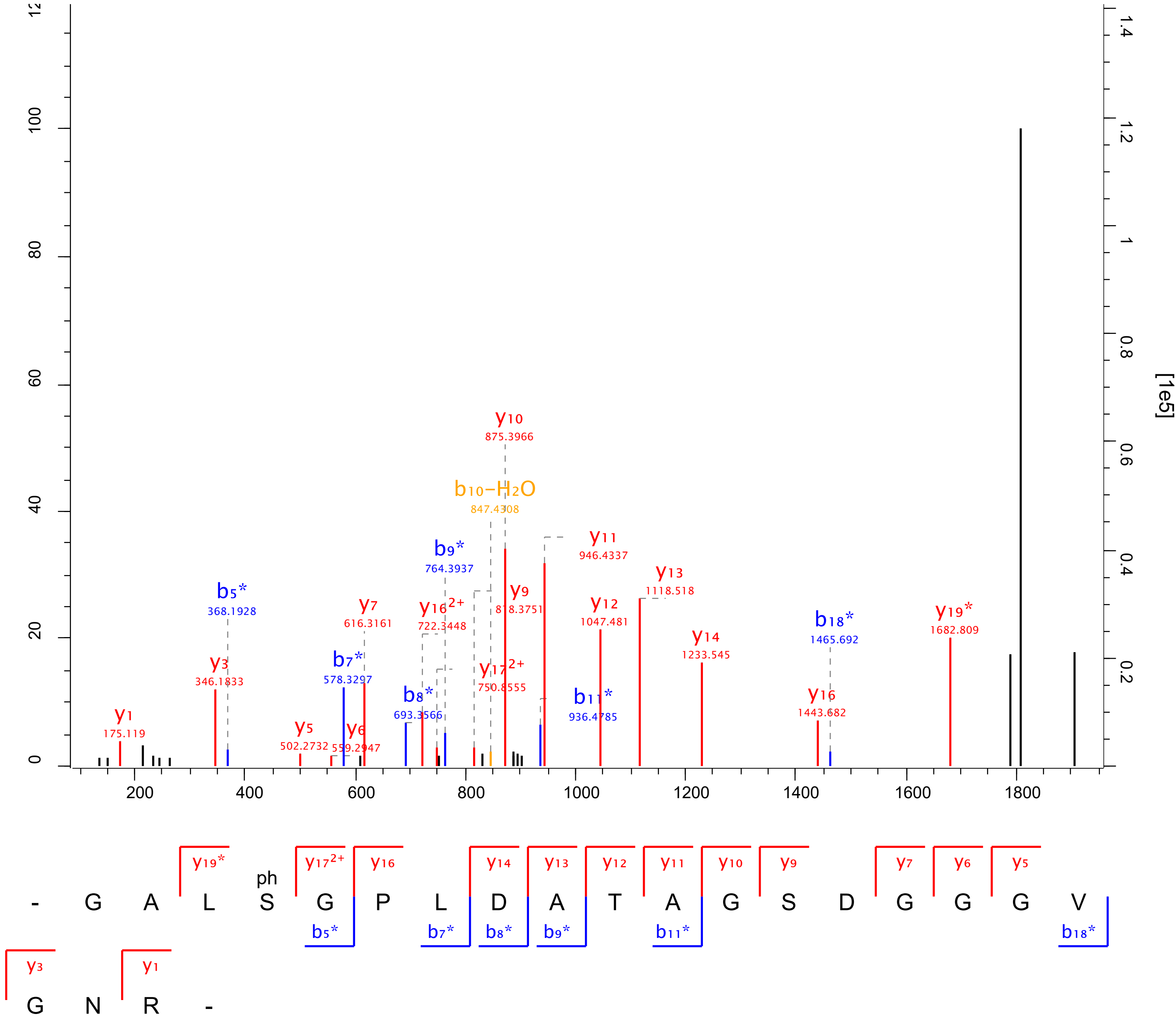

| Raw file | Scan | Method | Score | m/z |
| --- | --- | --- | --- | --- |
| water_15_1 | 24206 | FTMS; HCD | 59.73 | 737.37 |

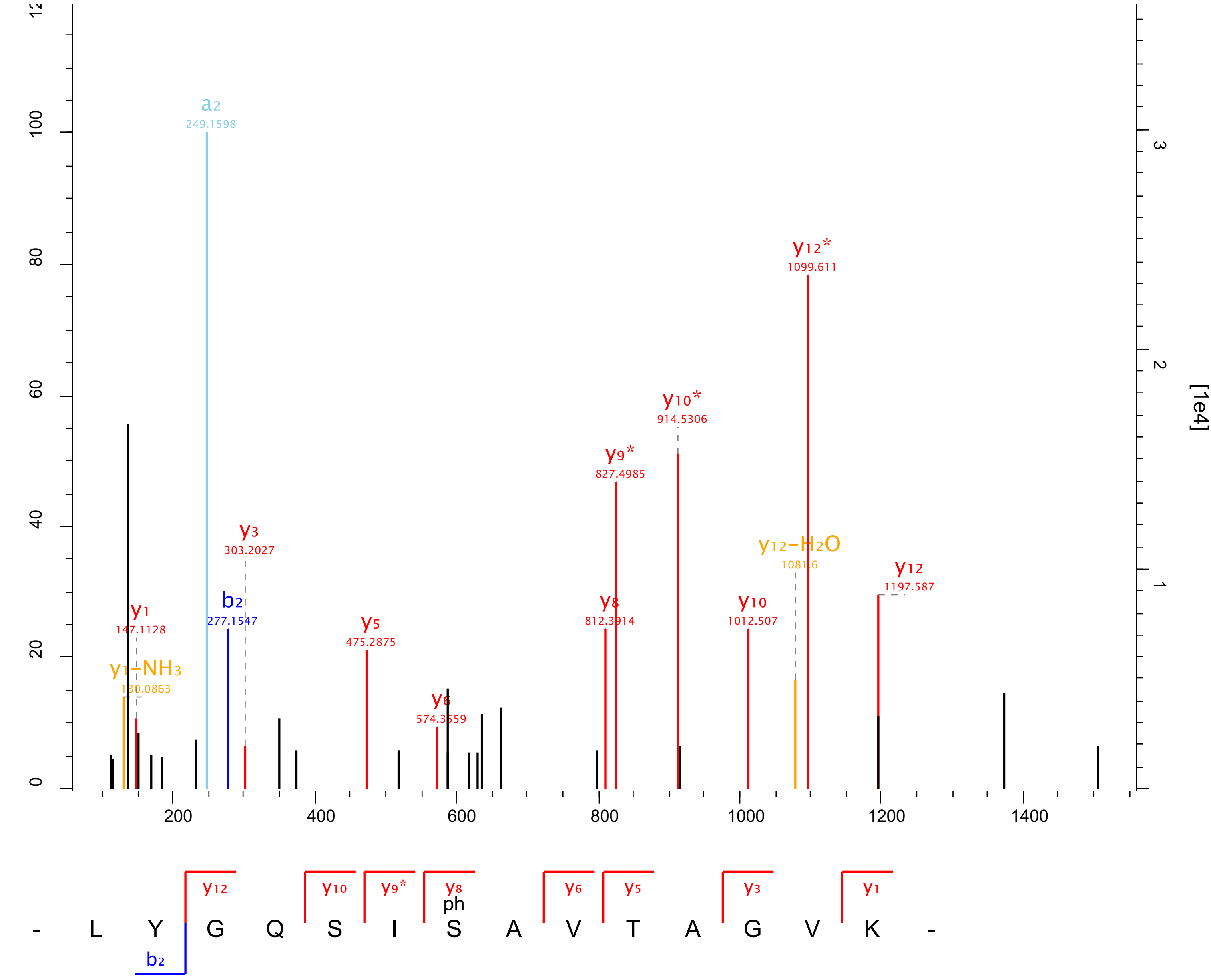

| Raw file | Scan | Method | Score | m/z |
|---|---|---|---|---|
| water_15_1 | 24367 | FTMS; HCD | 162.67 | 570.29 |

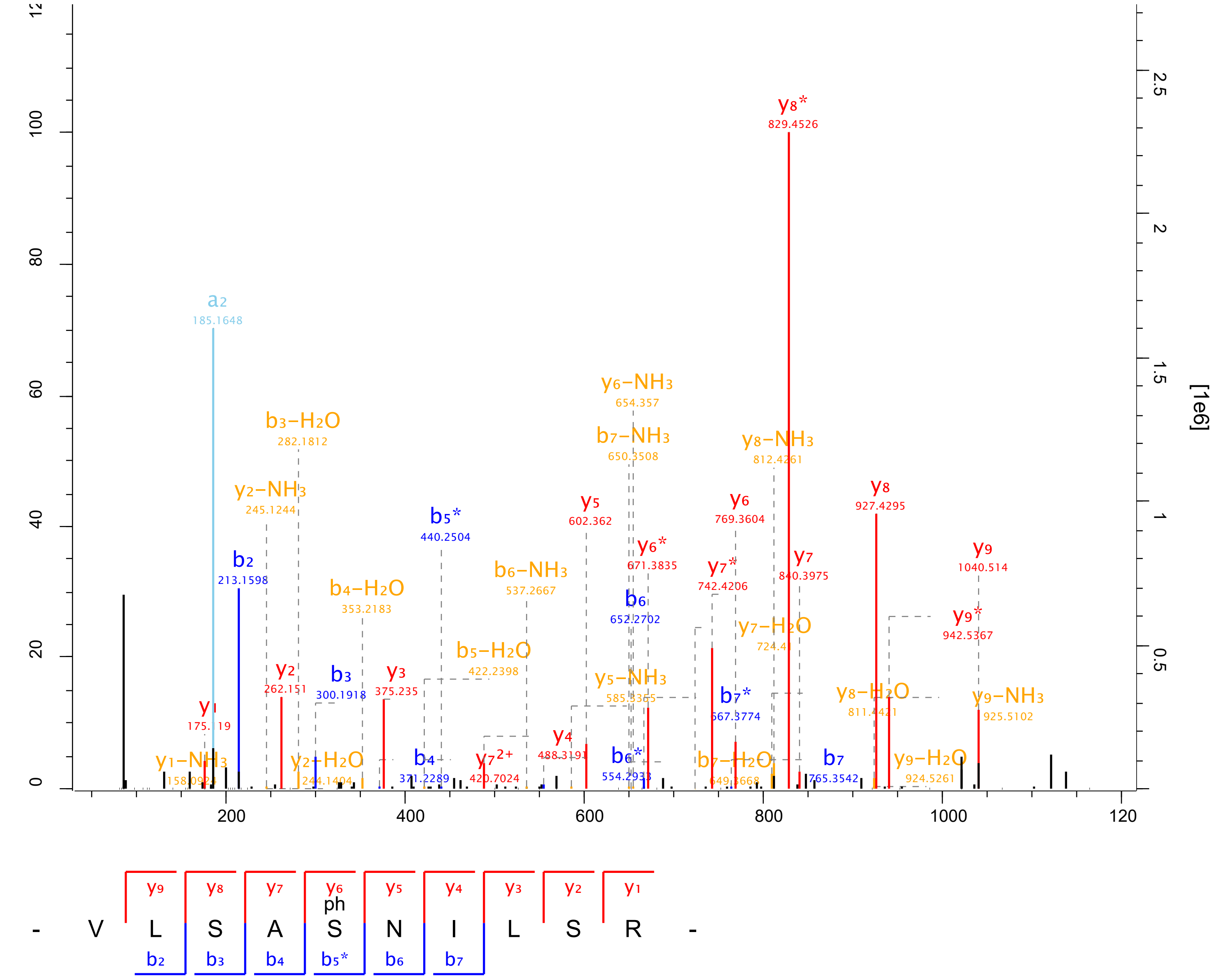

| Raw file | Scan | Method | Score | m/z |
|---|---|---|---|---|
| water_15_1 | 24644 | FTMS; HCD | 148.52 | 799.89 |

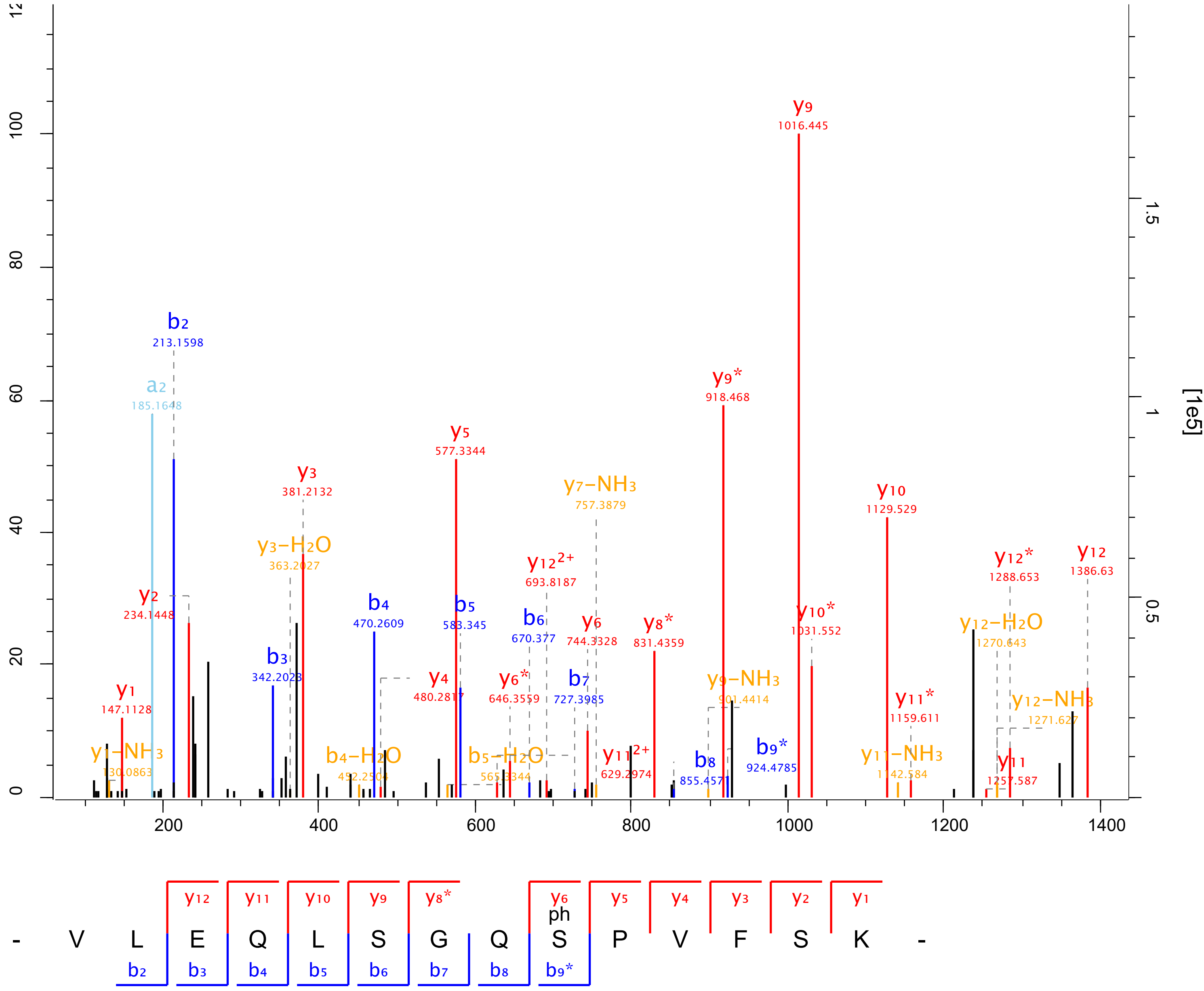

| Raw file | Scan | Method | Score | m/z |
| --- | --- | --- | --- | --- |
| water_15_1 | 24858 | FTMS; HCD | 126.71 | 499.7 |

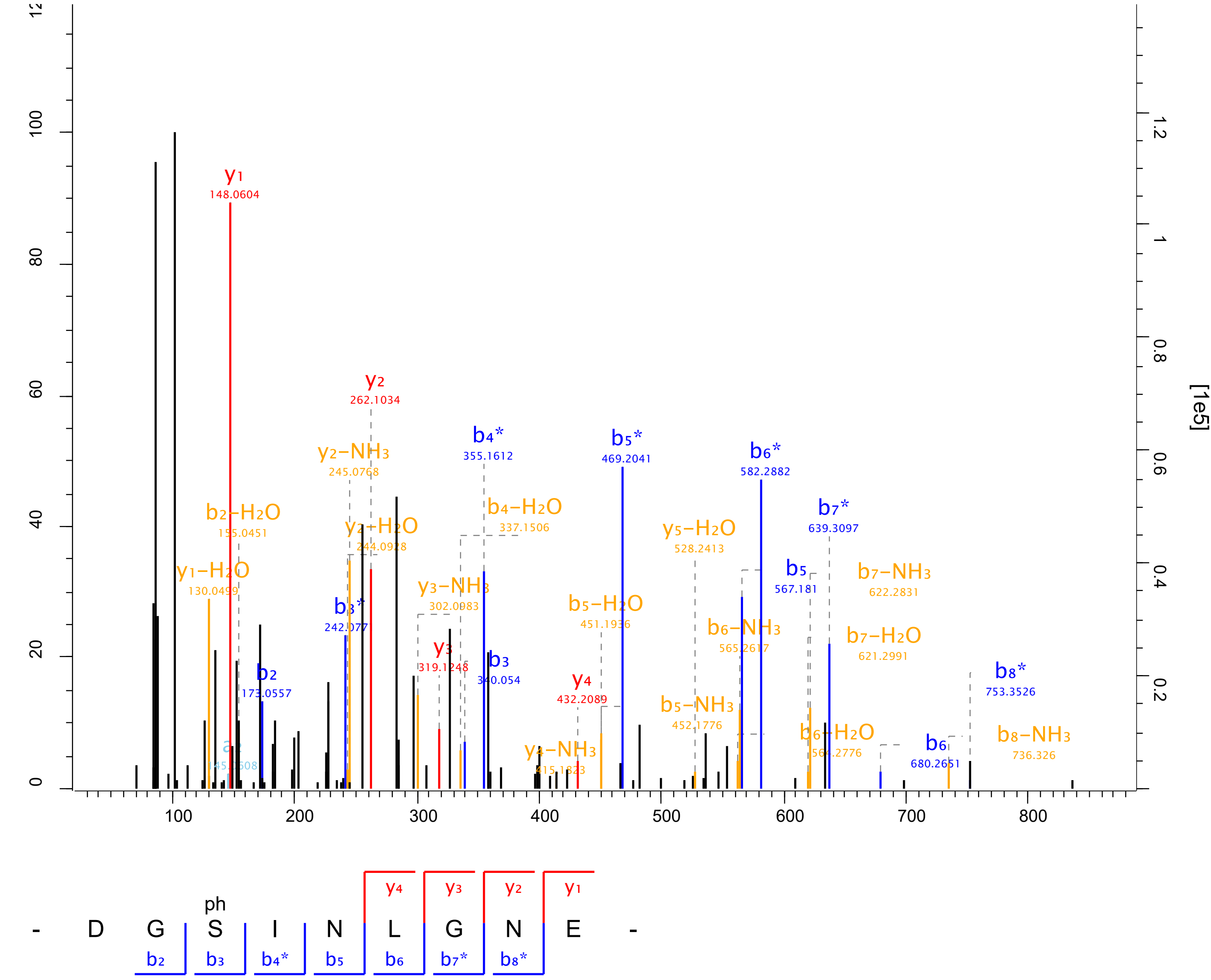

| Raw file | Scan | Method | Score | m/z |
|---|---|---|---|---|
| water_15_1 | 24903 | FTMS; HCD | 78.08 | 854.9 |

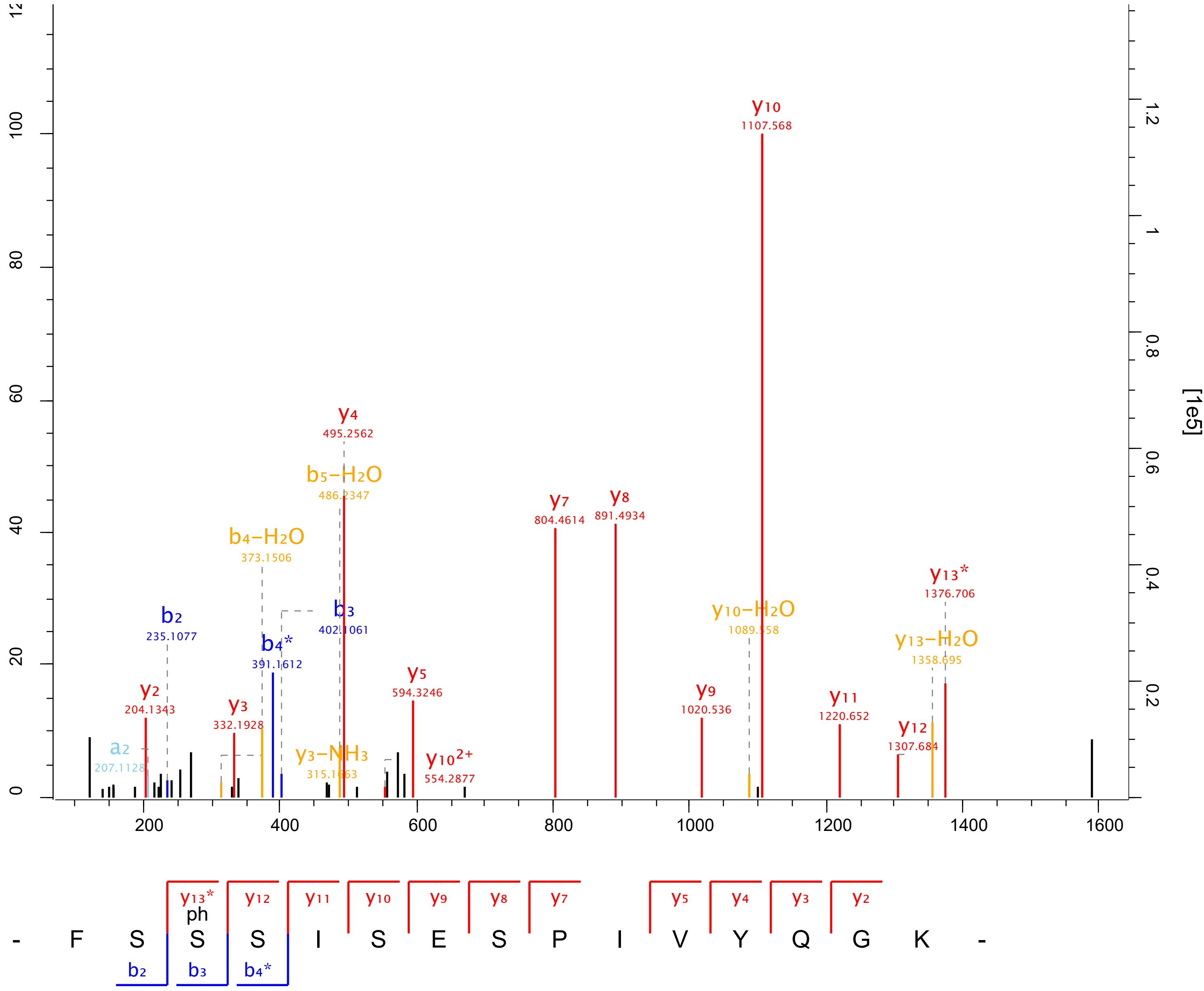

| Raw file | Scan | Method | Score | m/z |
|---|---|---|---|---|
| water_15_1 | 25316 | FTMS; HCD | 80.63 | 775.83 |

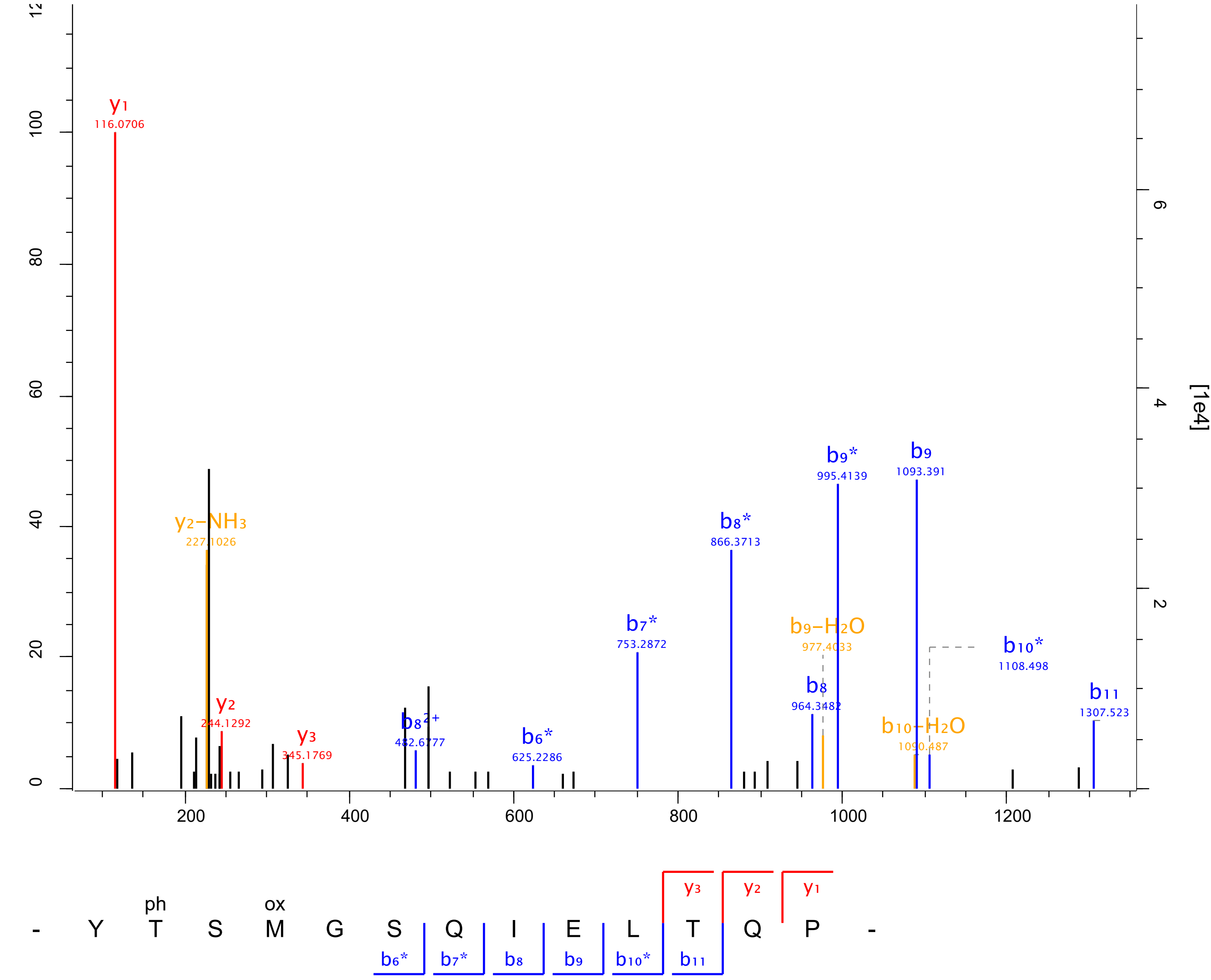

| Raw file | Scan | Method | Score | m/z |
|---|---|---|---|---|
| water_15_1 | 26250 | FTMS; HCD | 100.43 | 730.33 |

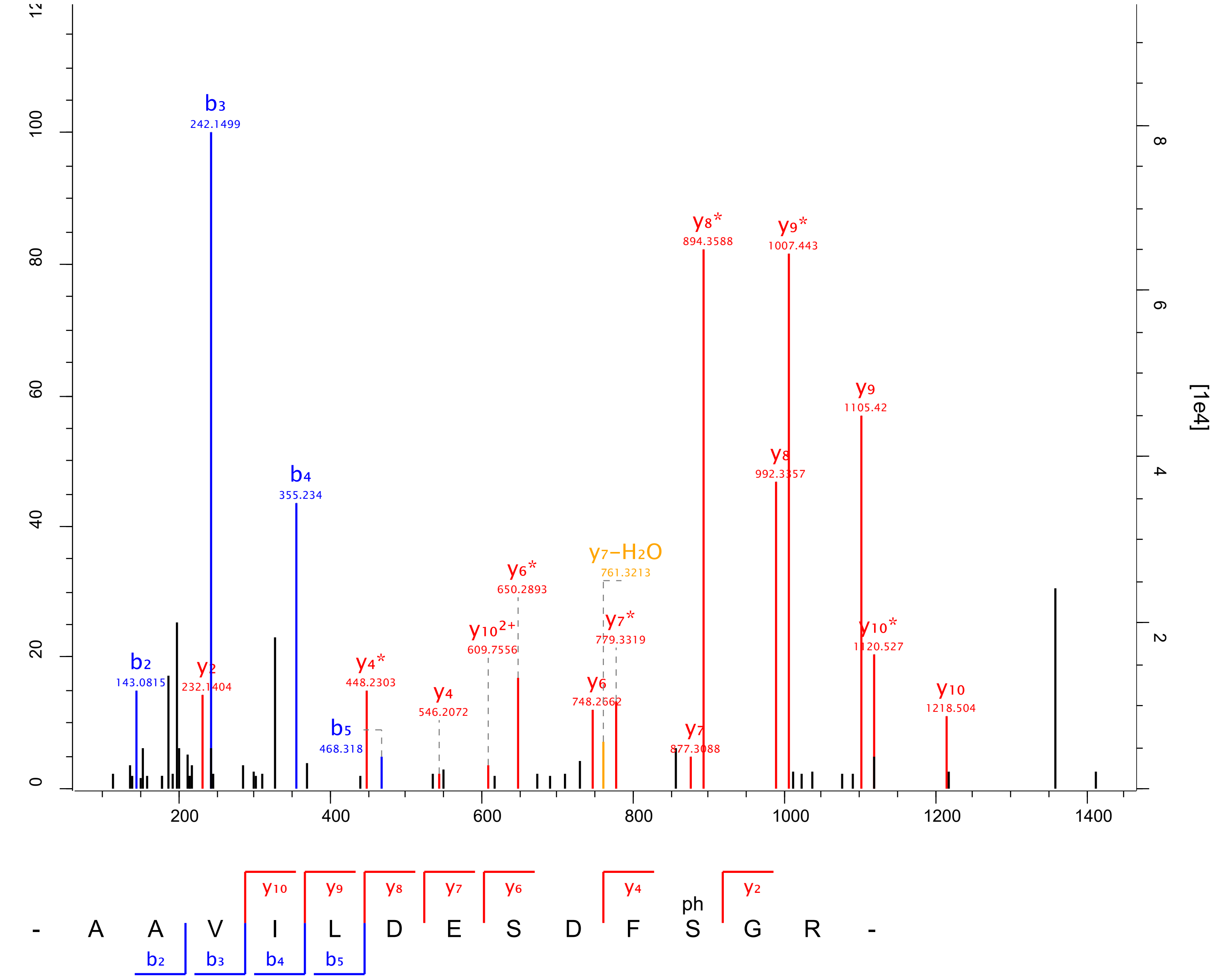

| Raw file | Scan | Method | Score | m/z |
|---|---|---|---|---|
| water_15_1 | 26549 | FTMS; HCD | 294.65 | 827.85 |

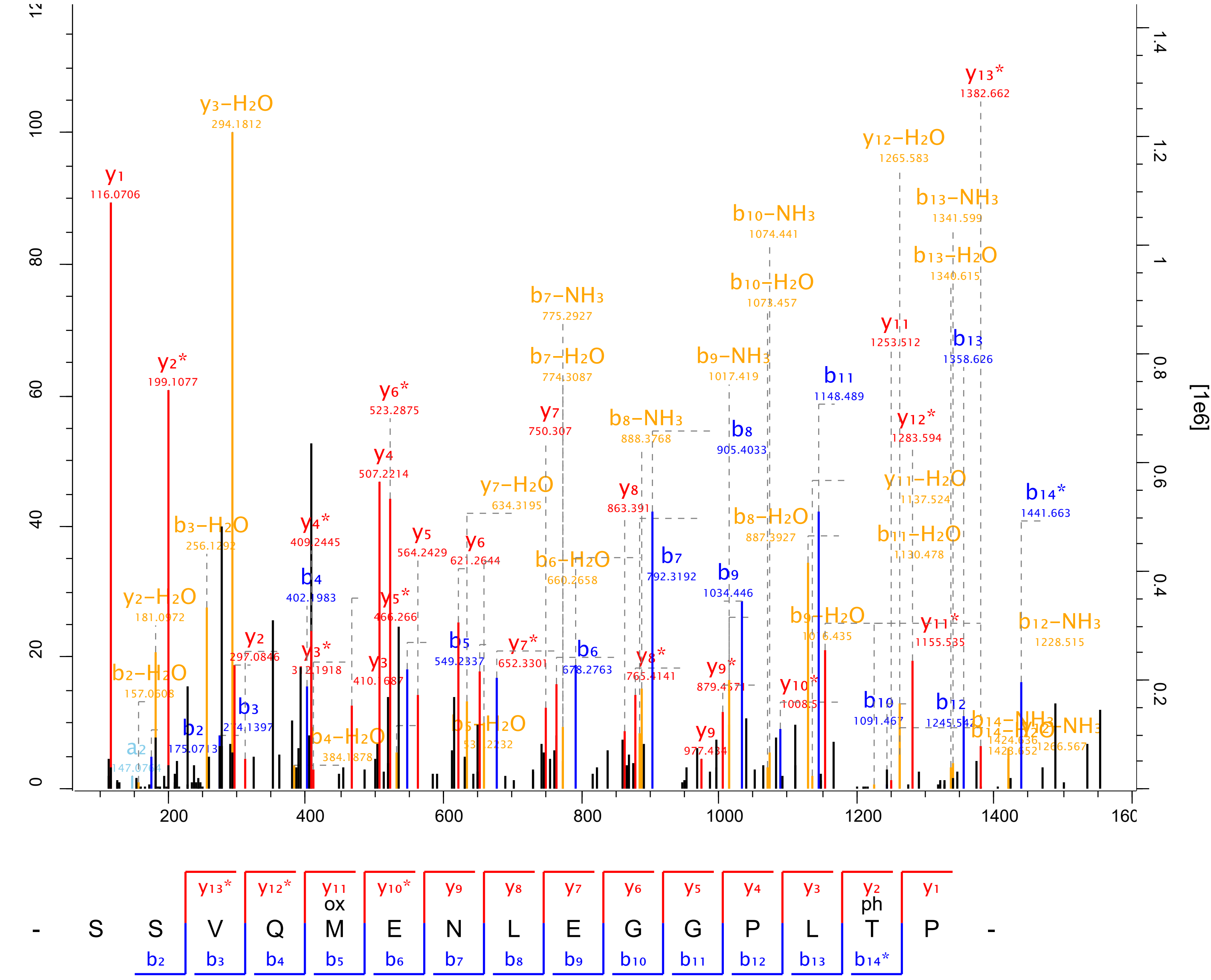

| Raw file | Scan | Method | Score | m/z |
| --- | --- | --- | --- | --- |
| water_15_1 | 26741 | FTMS; HCD | 147.7 | 692.34 |

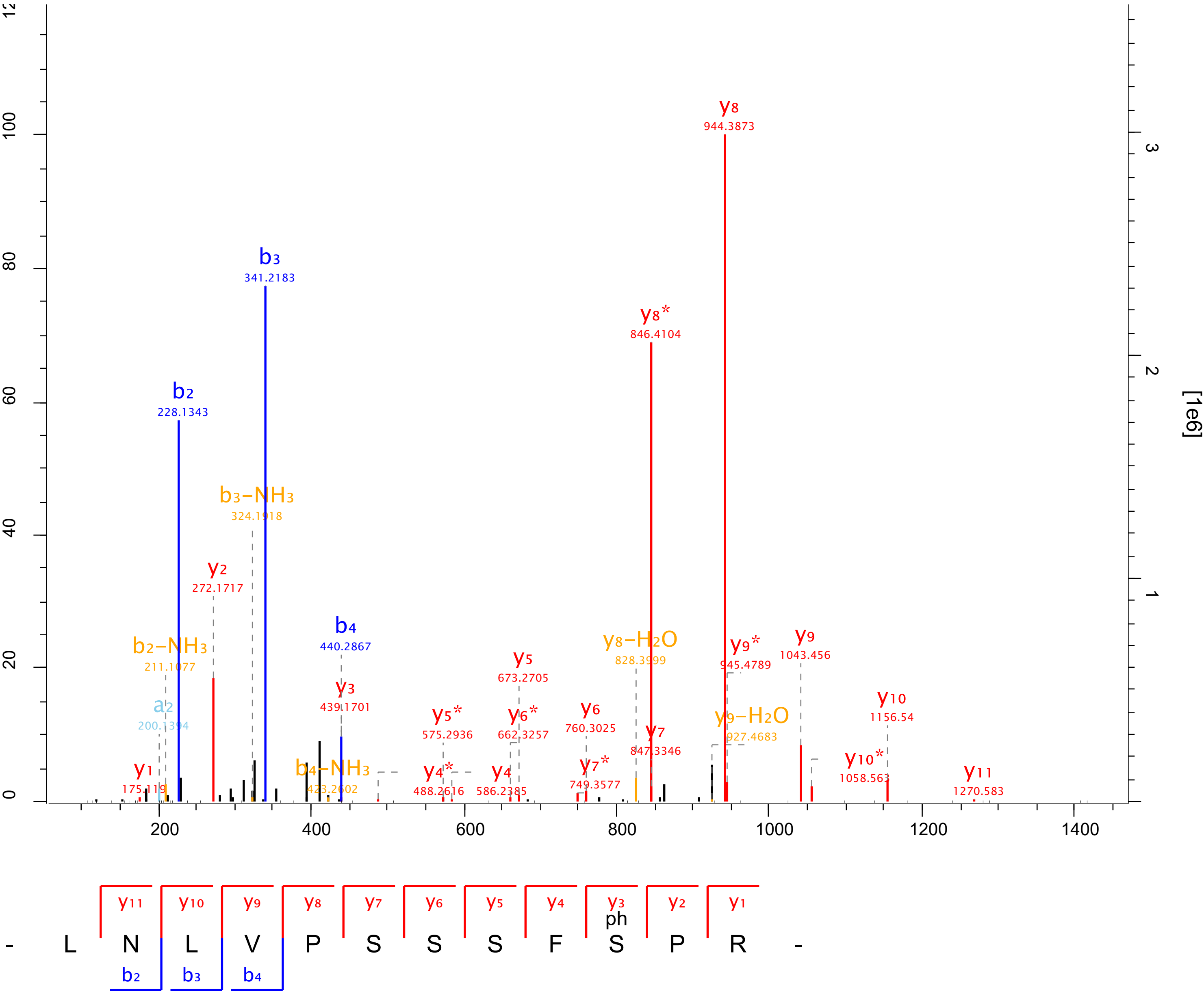

| Raw file | Scan | Method | Score | m/z |
|---|---|---|---|---|
| water_15_1 | 27973 | FTMS; HCD | 117.08 | 649.8 |

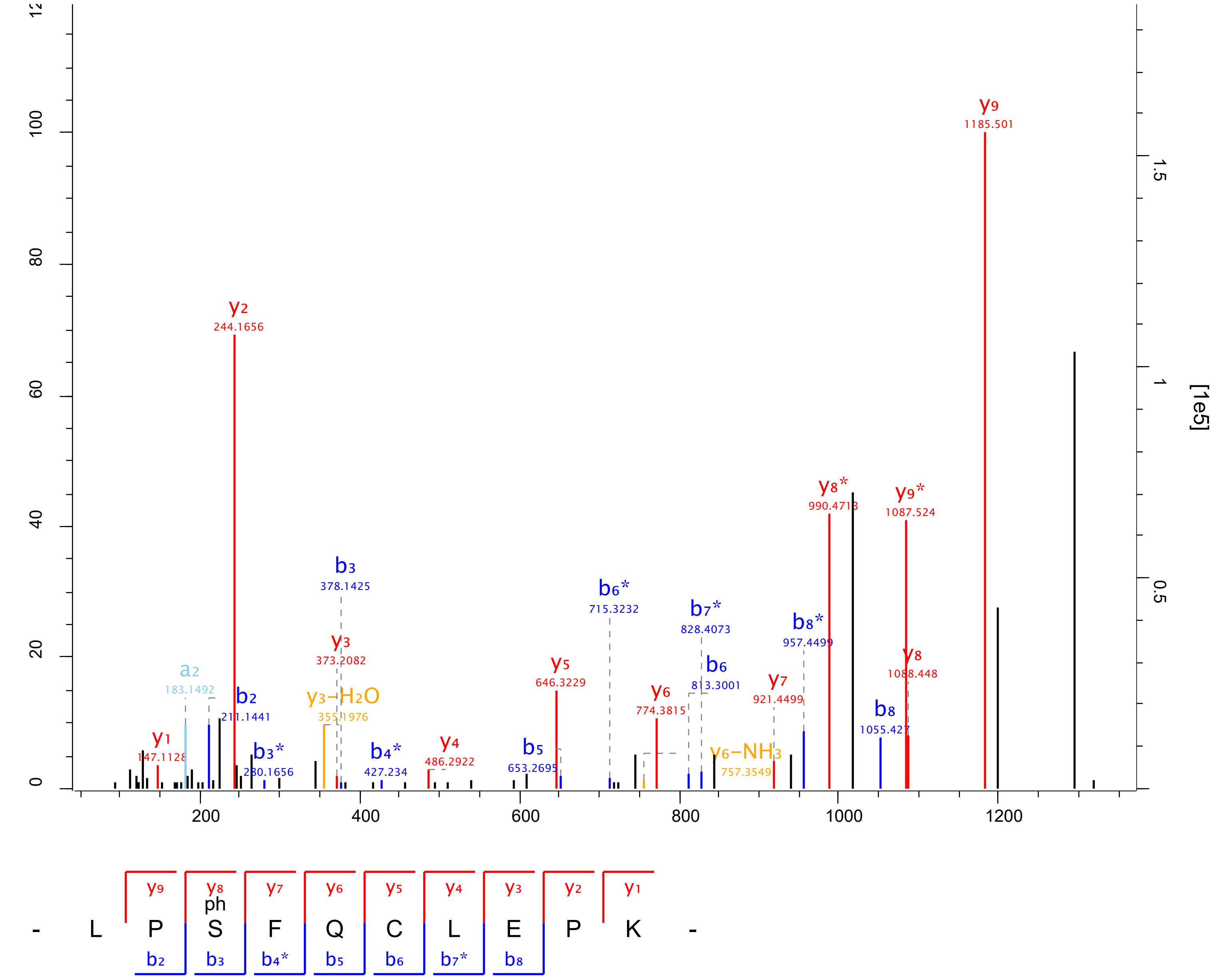

| Raw file | Scan | Method | Score | m/z |
|---|---|---|---|---|
| water_15_1 | 29114 | FTMS; HCD | 71.18 | 590.81 |

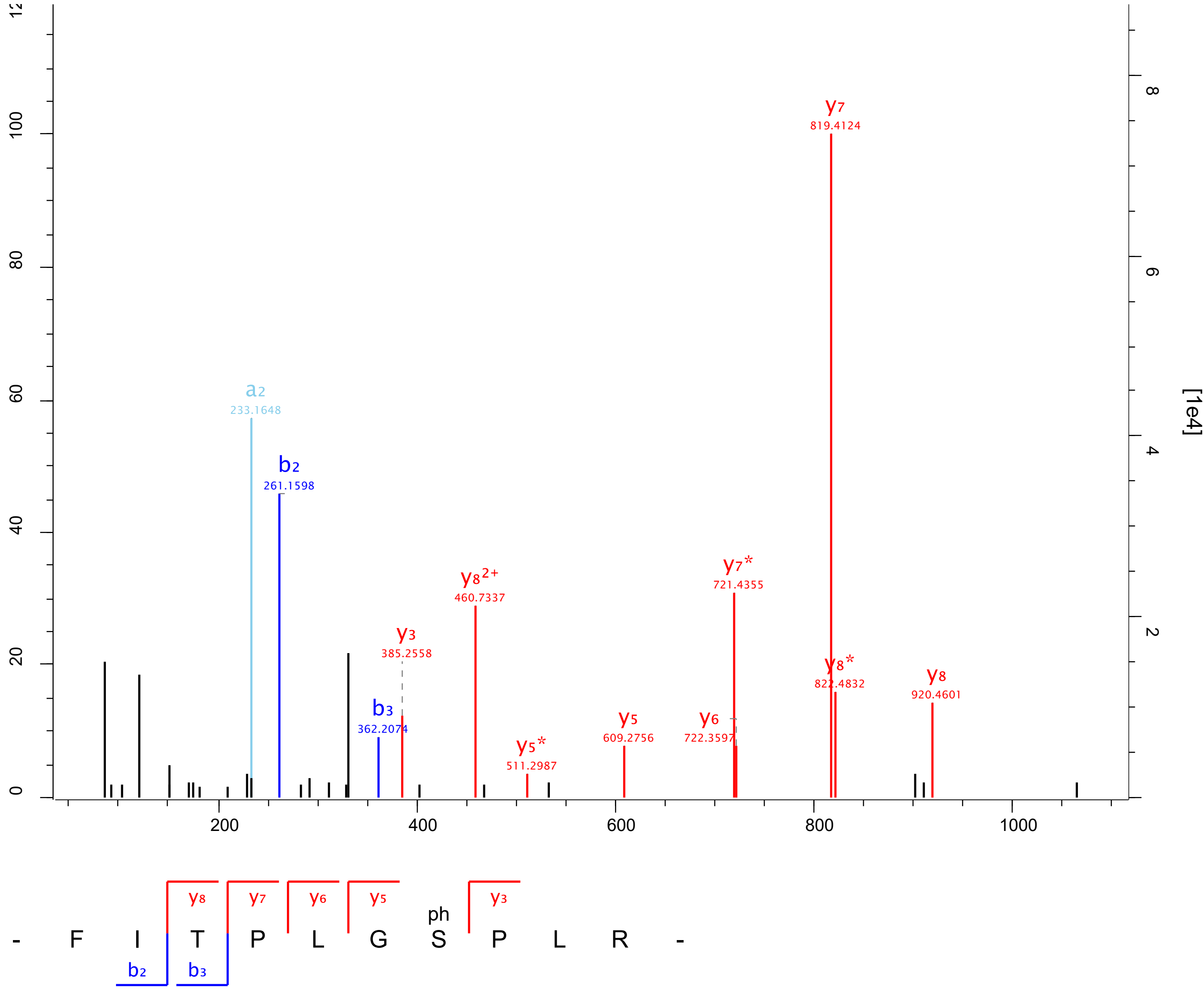

| Raw file | Scan | Method | Score | m/z |
|---|---|---|---|---|
| water_15_1 | 29131 | FTMS; HCD | 193.47 | 742.34 |

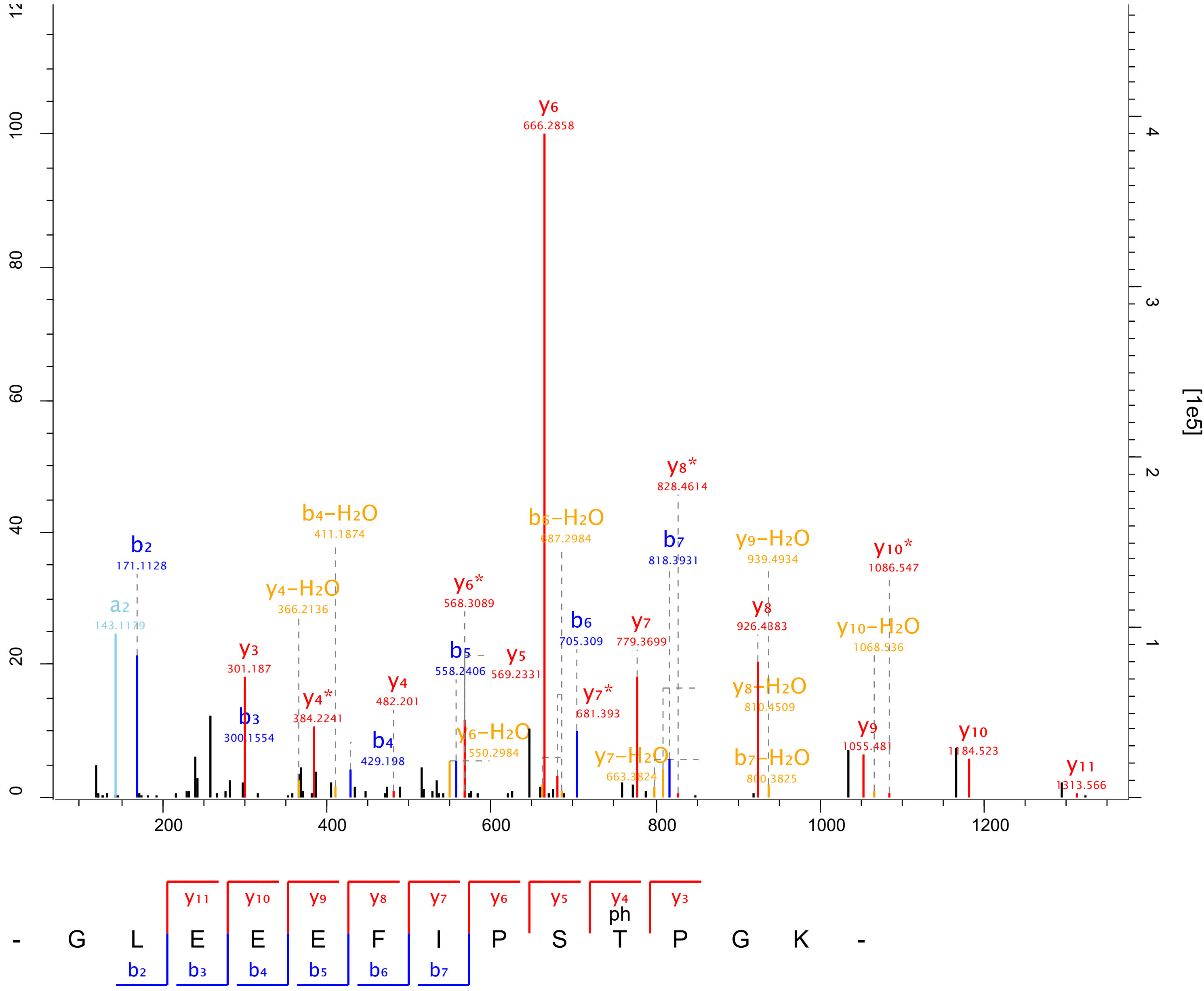

| Raw file | Scan | Method | Score | m/z |
| --- | --- | --- | --- | --- |
| water_15_1 | 2978 | FTMS; HCD | 60.91 | 506.18 |

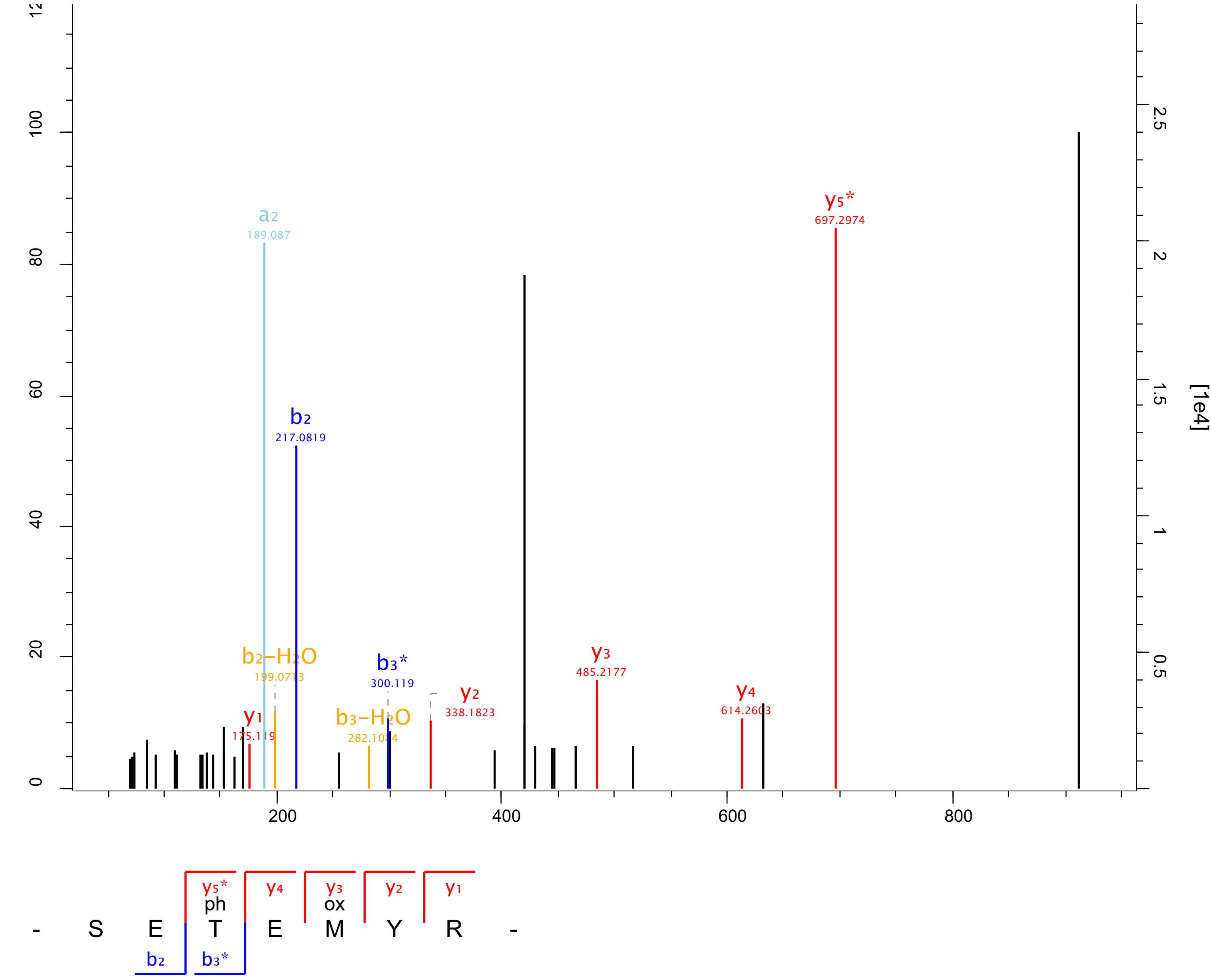

| Raw file | Scan | Method | Score | m/z |
|---|---|---|---|---|
| water_15_1 | 29837 | FTMS; HCD | 206.18 | 637.79 |

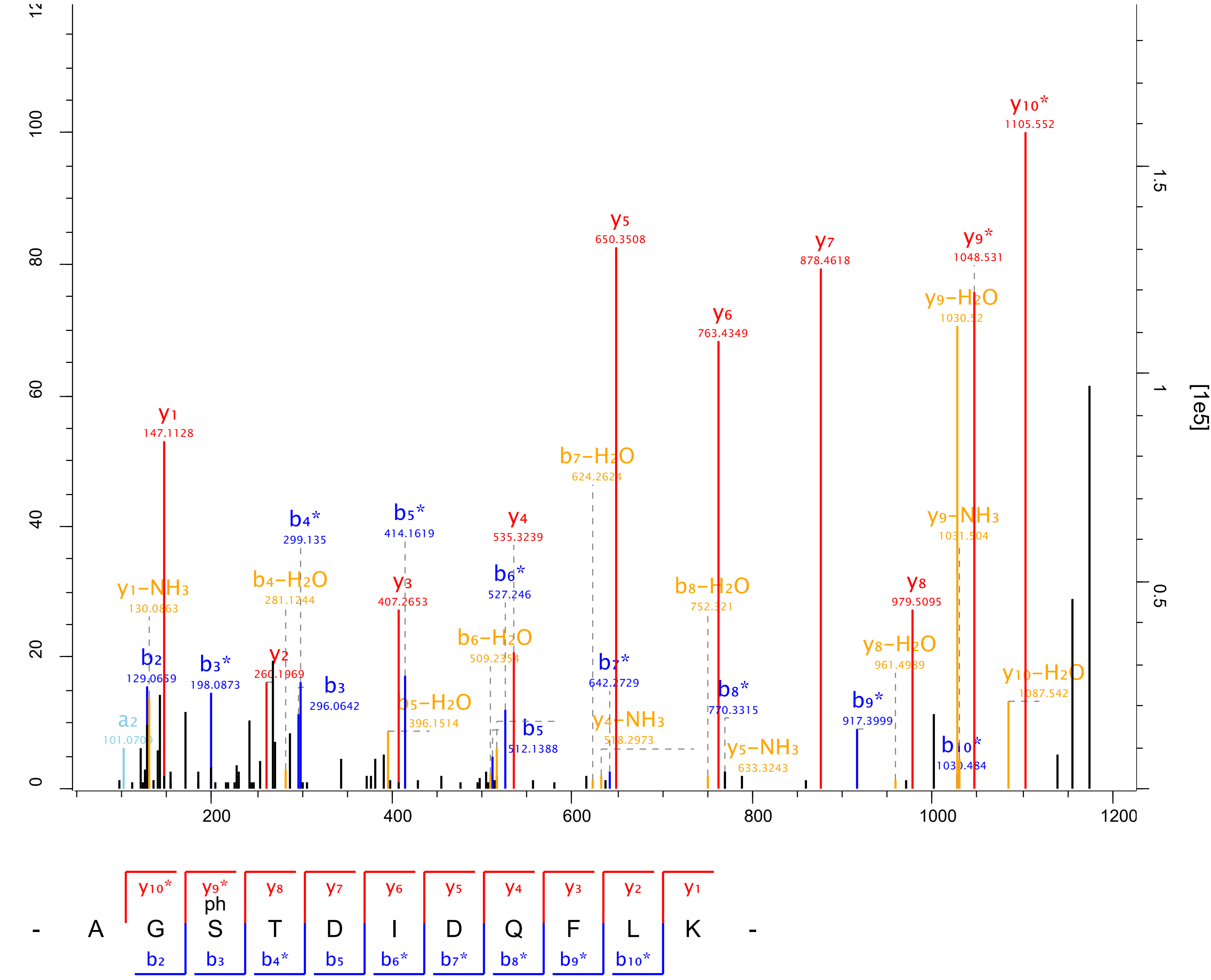

| Raw file | Scan | Method | Score | m/z |
|---|---|---|---|---|
| water_15_1 | 30131 | FTMS; HCD | 111.22 | 837.33 |

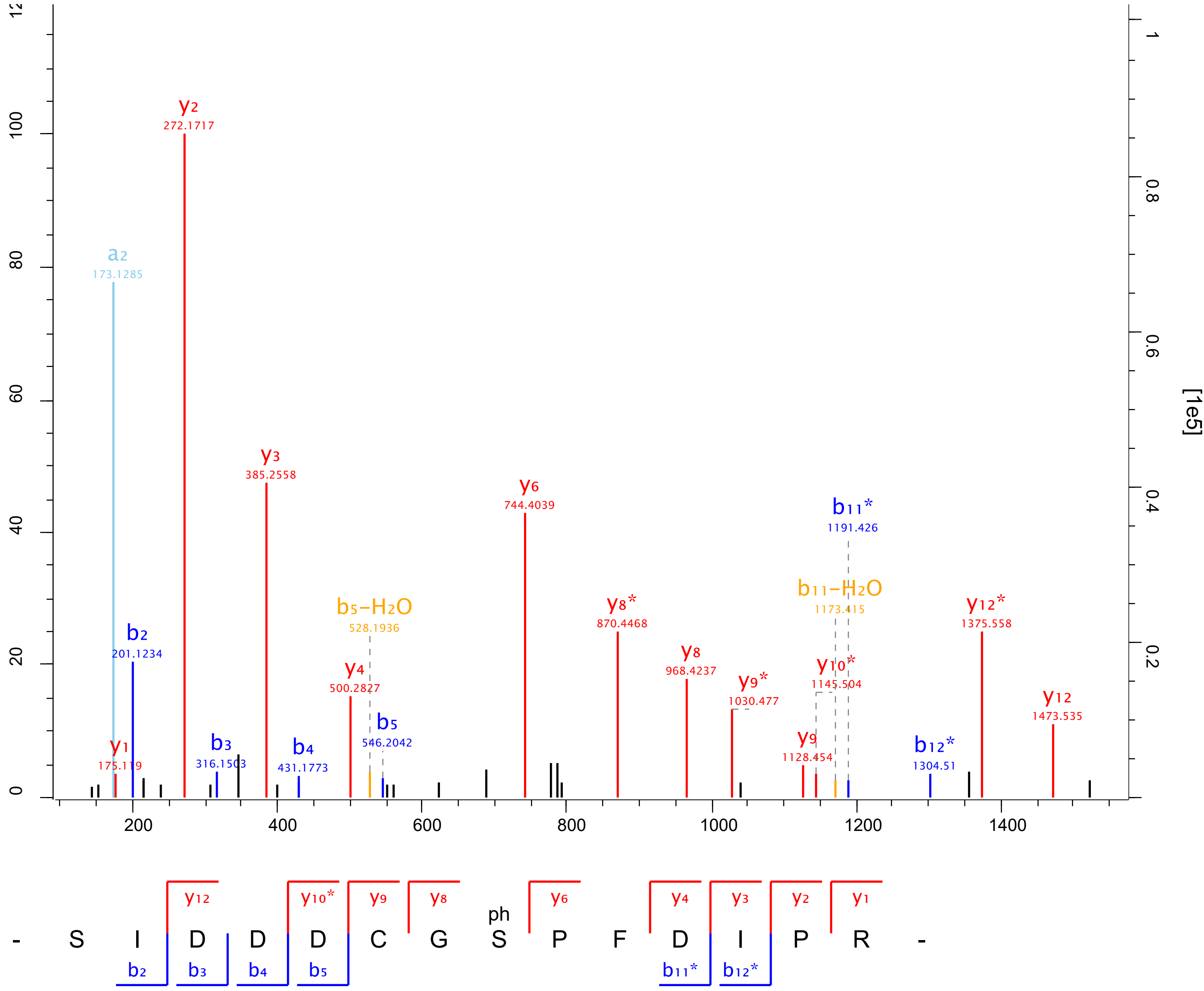

| Raw file | Scan | Method | Score | m/z |
| --- | --- | --- | --- | --- |
| water_15_1 | 30318 | FTMS; HCD | 41.24 | 592.28 |

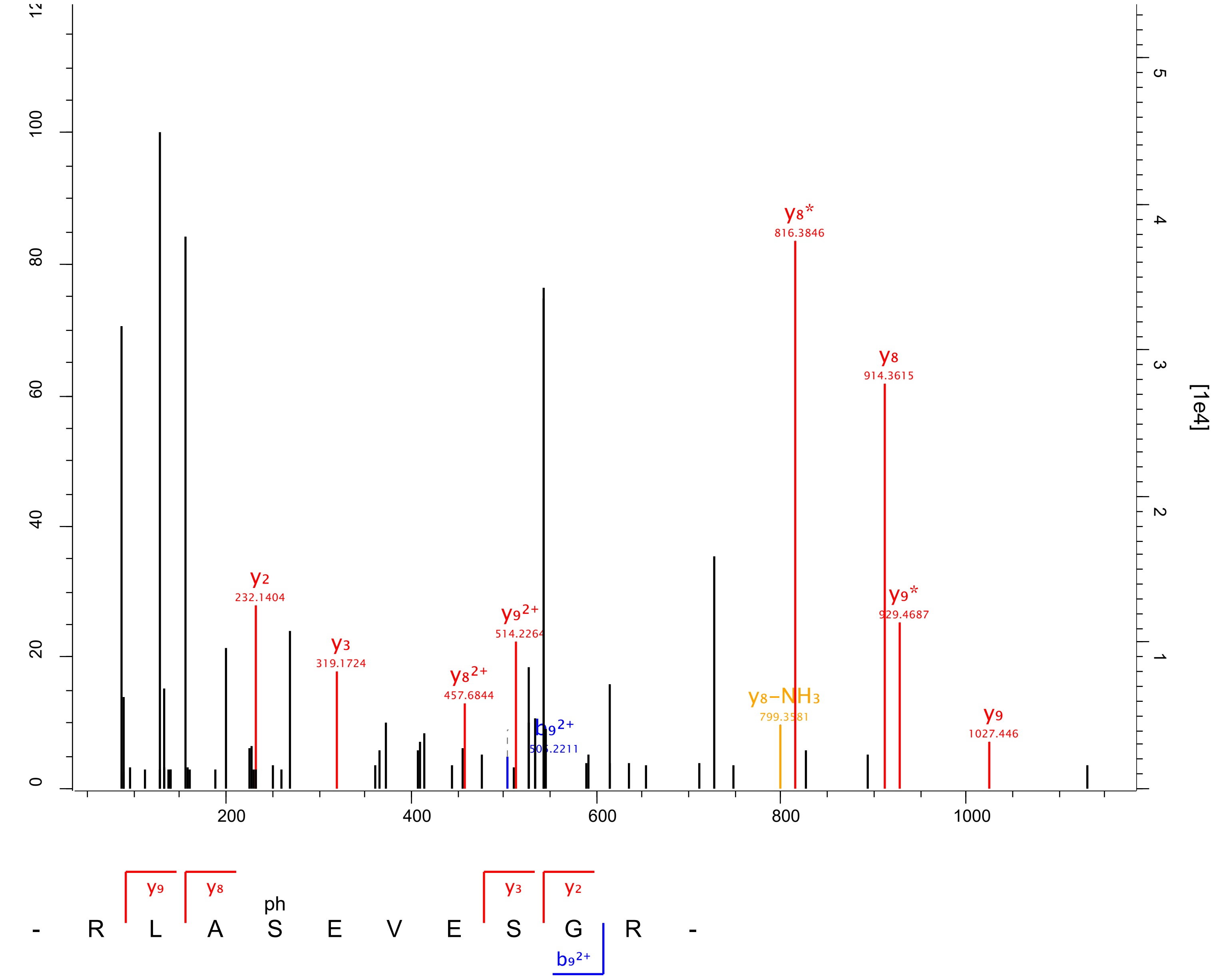

| Raw file | Scan | Method | Score | m/z |
| --- | --- | --- | --- | --- |
| water_15_1 | 30885 | FTMS; HCD | 81.45 | 700.32 |

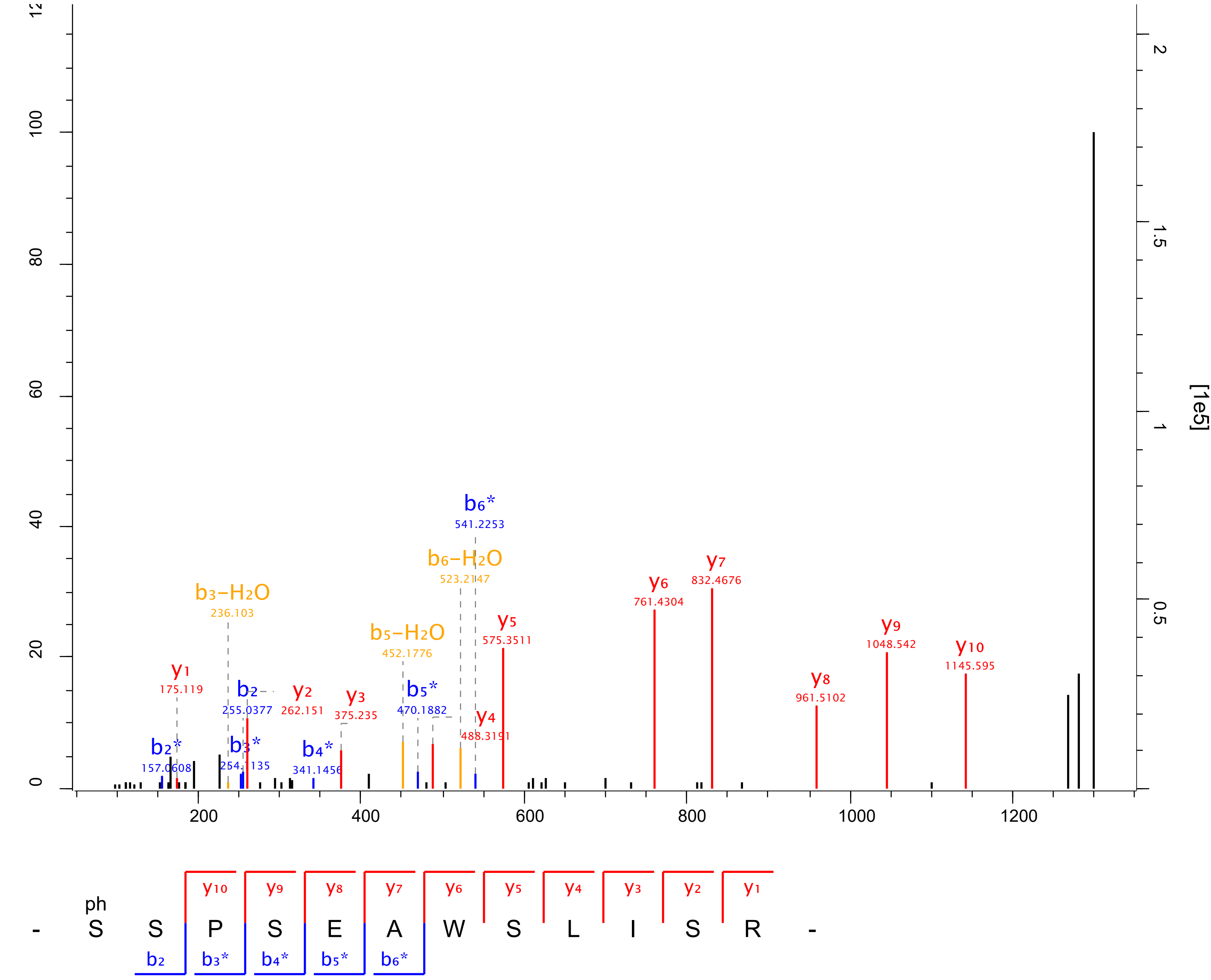

| Raw file | Scan | Method | Score | m/z |
| --- | --- | --- | --- | --- |
| water_15_1 | 32653 | FTMS; HCD | 43.59 | 759.35 |

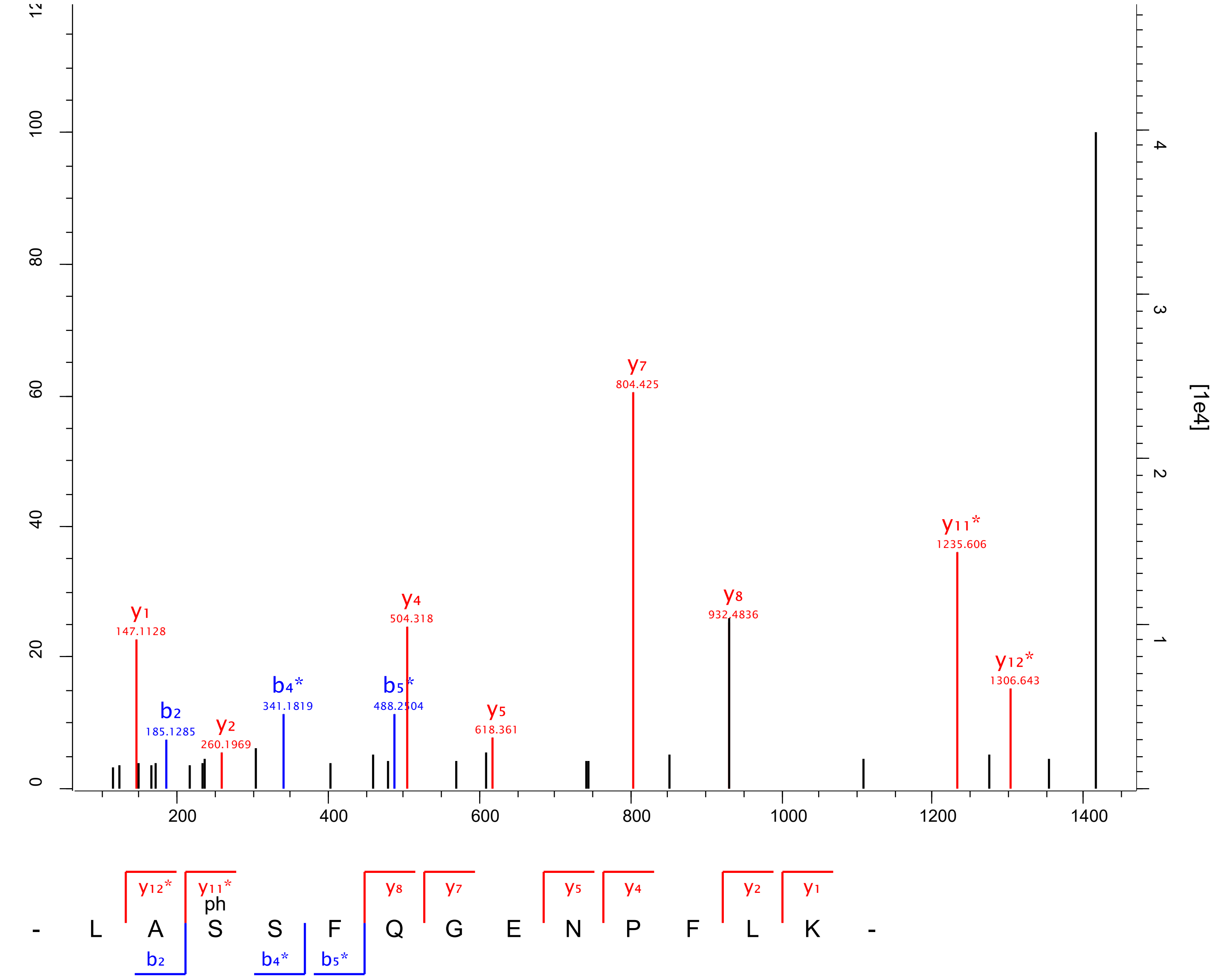

| Raw file | Scan | Method | Score | m/z |
| --- | --- | --- | --- | --- |
| water_15_1 | 32767 | FTMS; HCD | 192.78 | 742.83 |

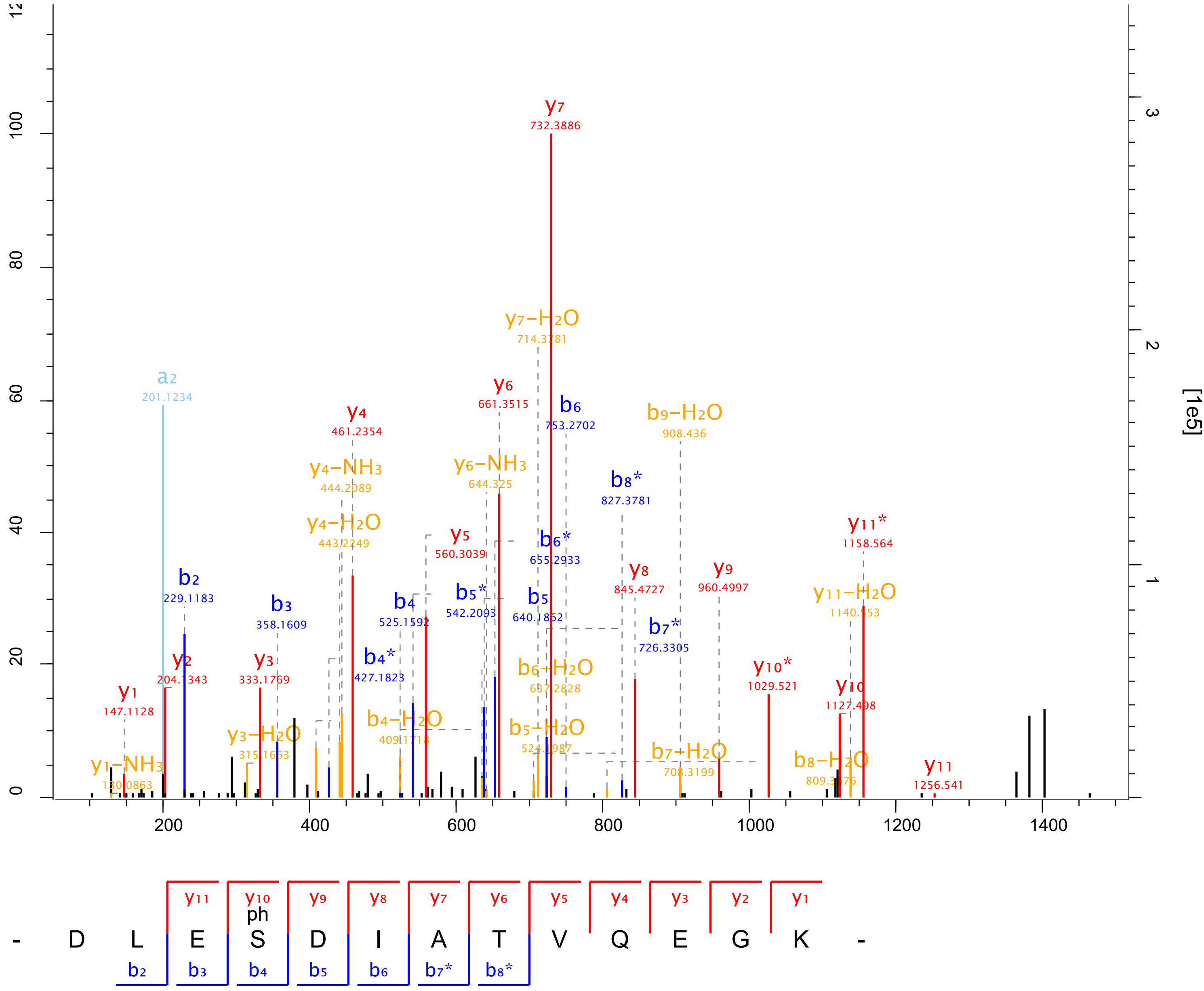

| Raw file | Scan | Method | Score | m/z |
| --- | --- | --- | --- | --- |
| water_15_1 | 3308 | FTMS; HCD | 112.94 | 536.73 |

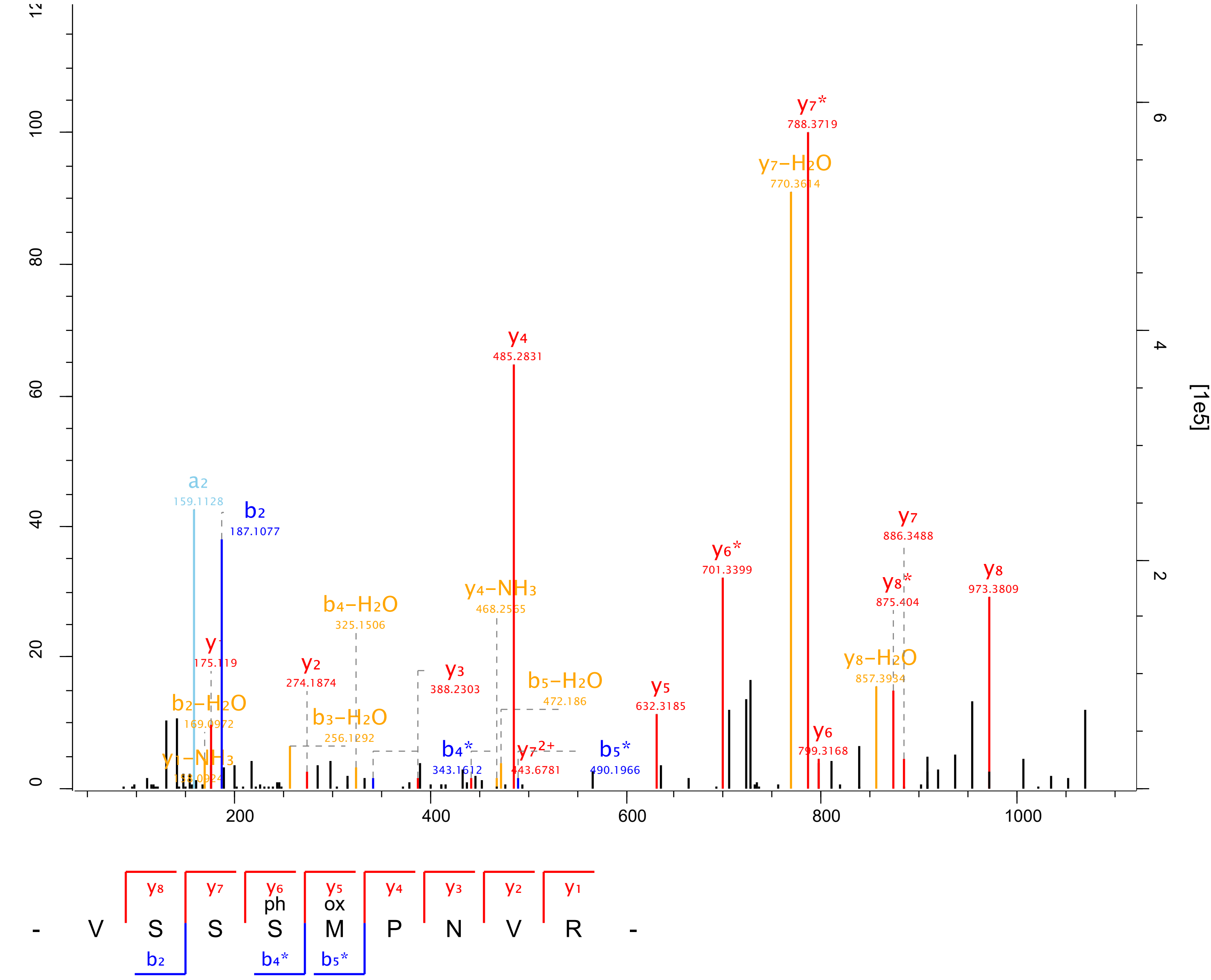

| Raw file | Scan | Method | Score | m/z |
| --- | --- | --- | --- | --- |
| water_15_1 | 3632 | FTMS; HCD | 132.93 | 578.74 |

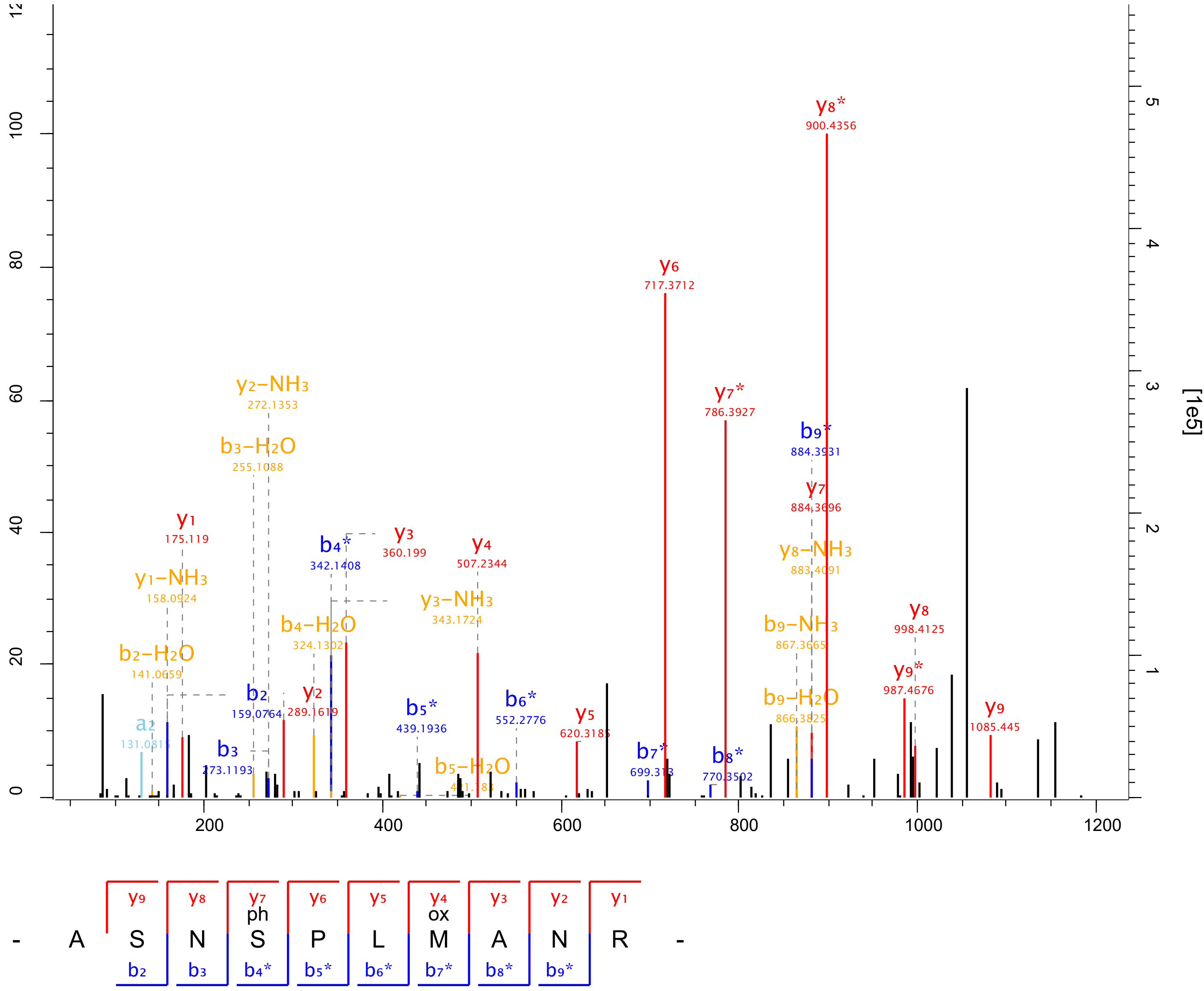

| Raw file | Scan | Method | Score | m/z |
|---|---|---|---|---|
| water_15_1 | 3654 | FTMS; HCD | 79.66 | 485.71 |

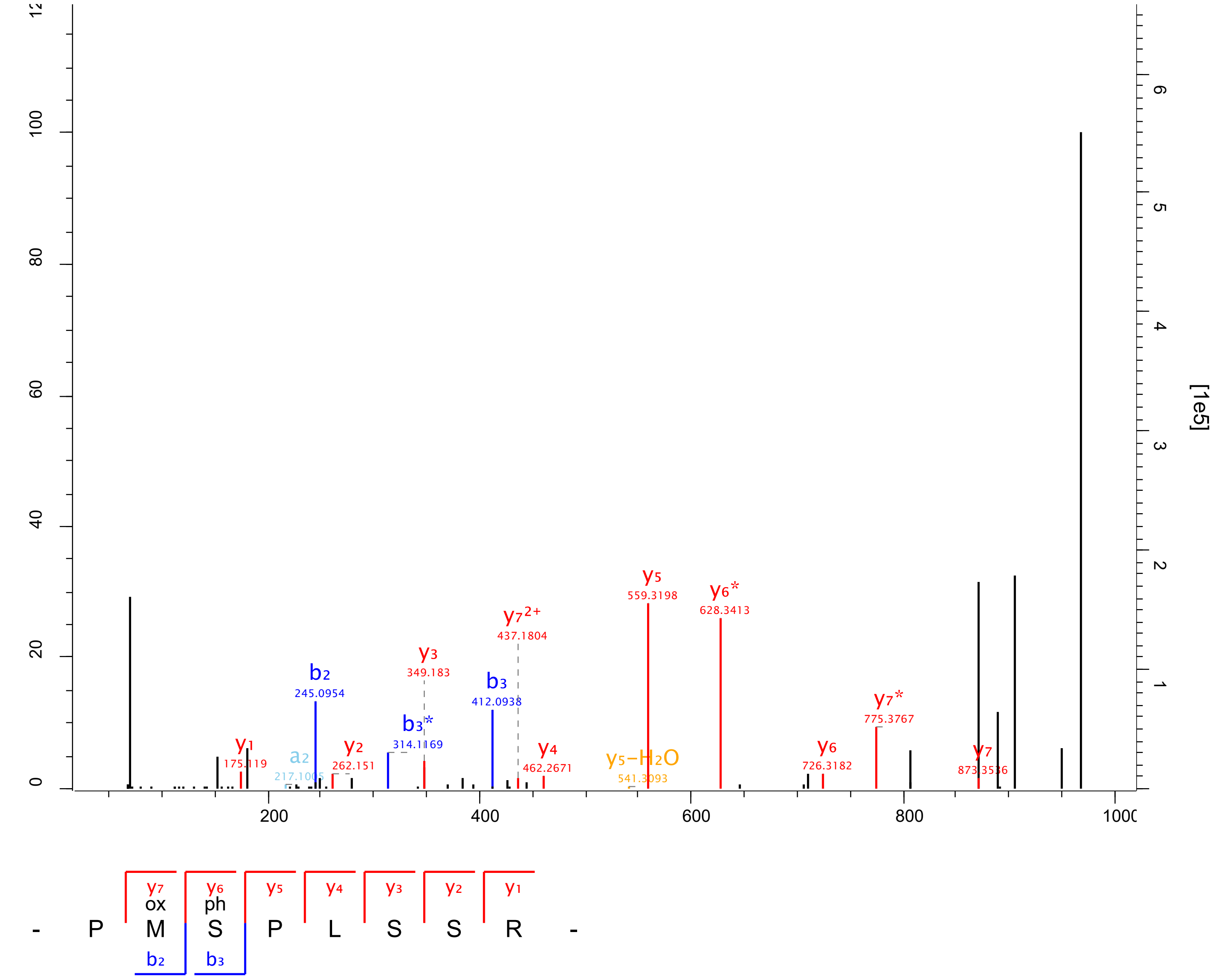

| Raw file | Scan | Method | Score | m/z |
| --- | --- | --- | --- | --- |
| water_15_1 | 38096 | FTMS; HCD | 291.89 | 993.42 |

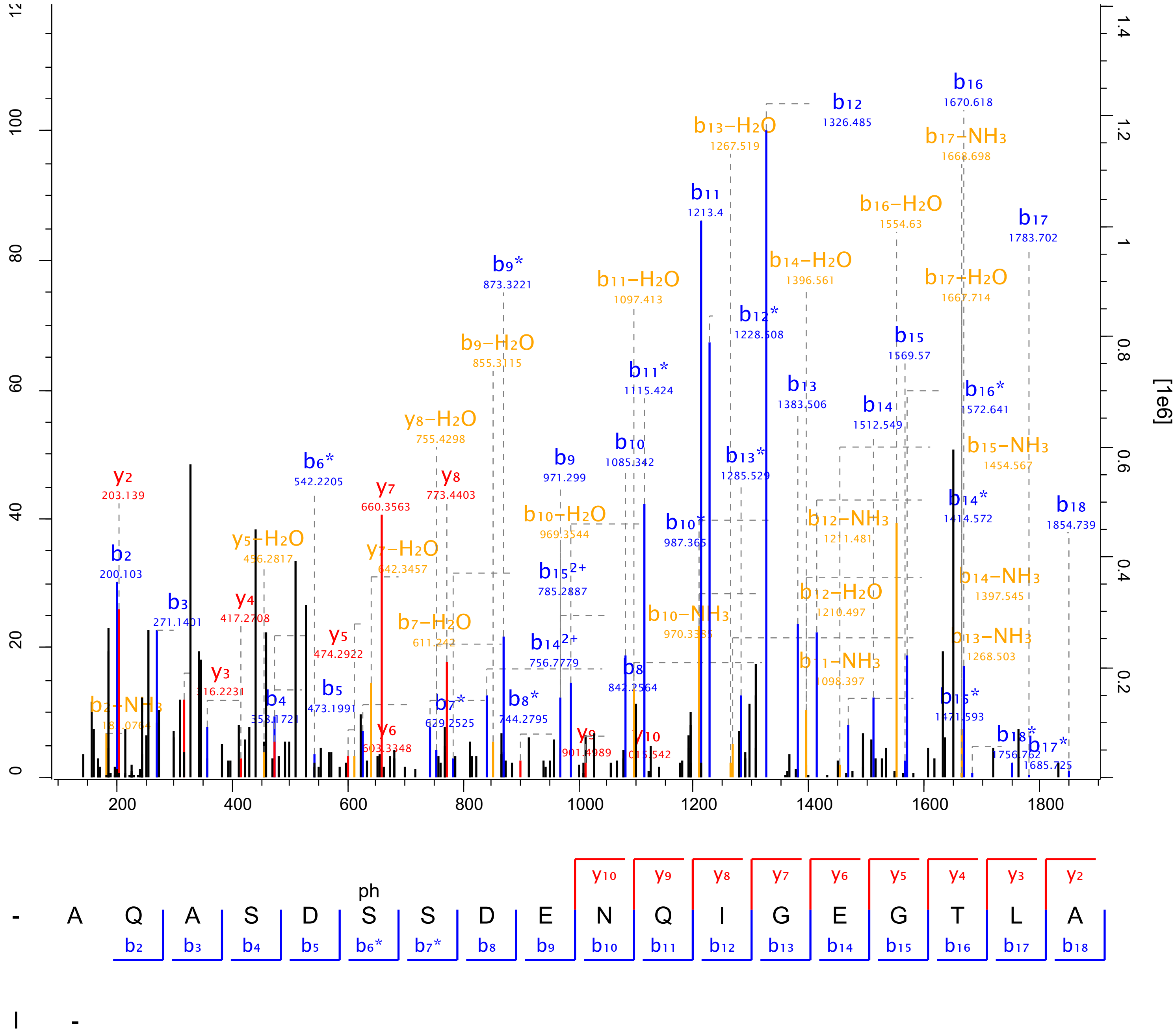

| Raw file | Scan | Method | Score | m/z |
|---|---|---|---|---|
| water_15_1 | 3877 | FTMS; HCD | 61.24 | 563.2 |

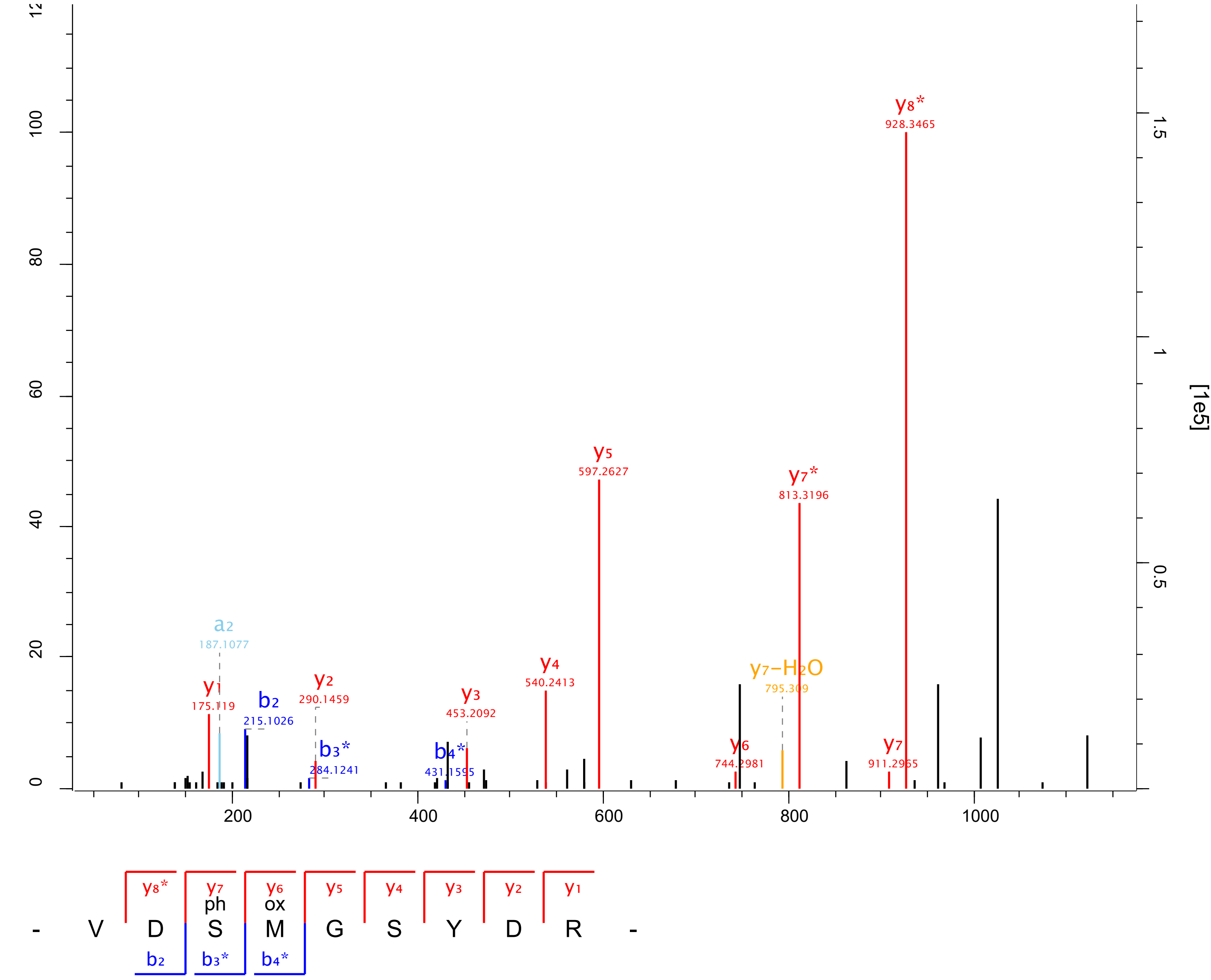

| Raw file | Scan | Method | Score | m/z |
| --- | --- | --- | --- | --- |
| water_15_1 | 3927 | FTMS; HCD | 110.08 | 461.71 |

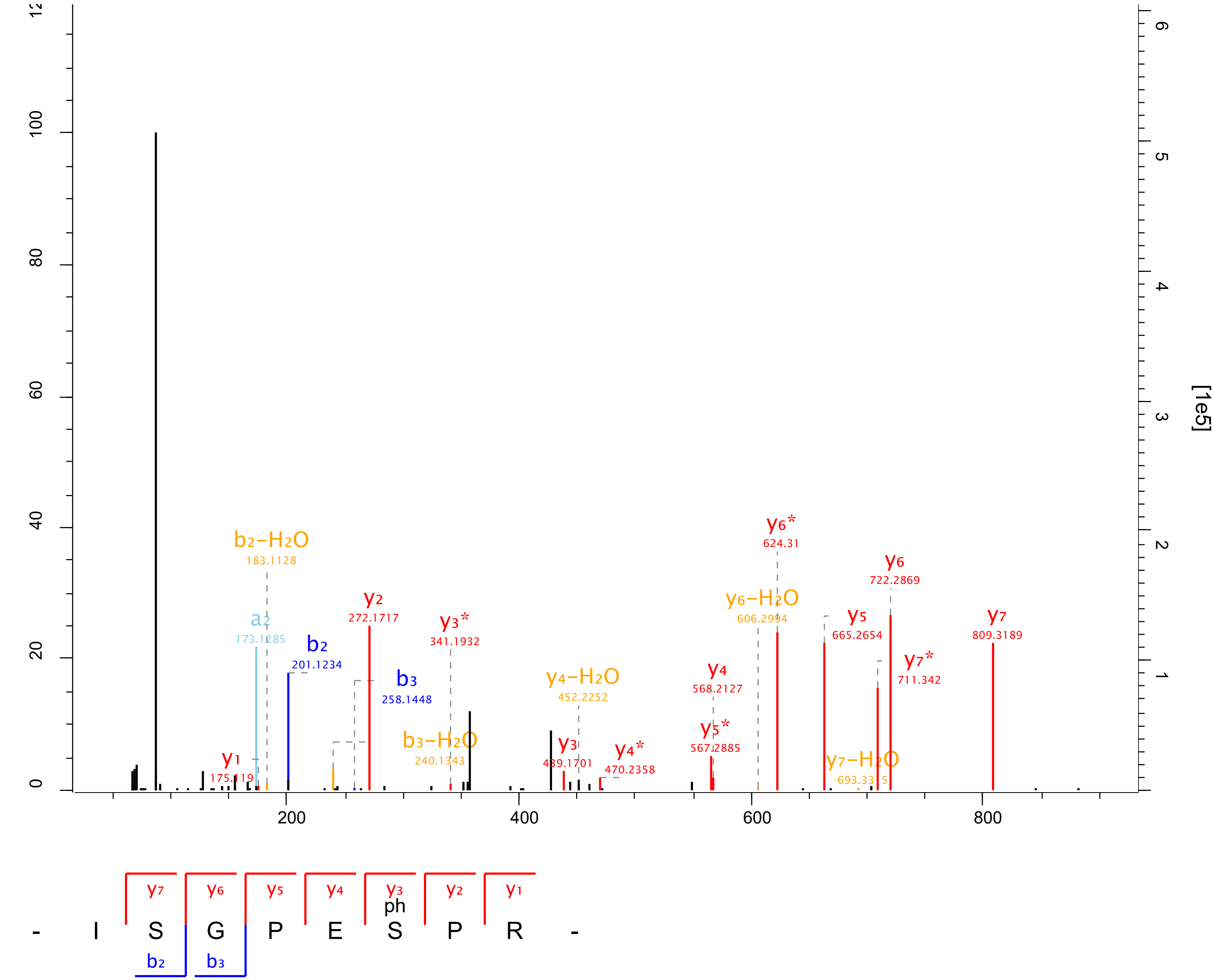

| Raw file | Scan | Method | Score | m/z |
|---|---|---|---|---|
| water_15_1 | 4090 | FTMS; HCD | 106.29 | 536.25 |

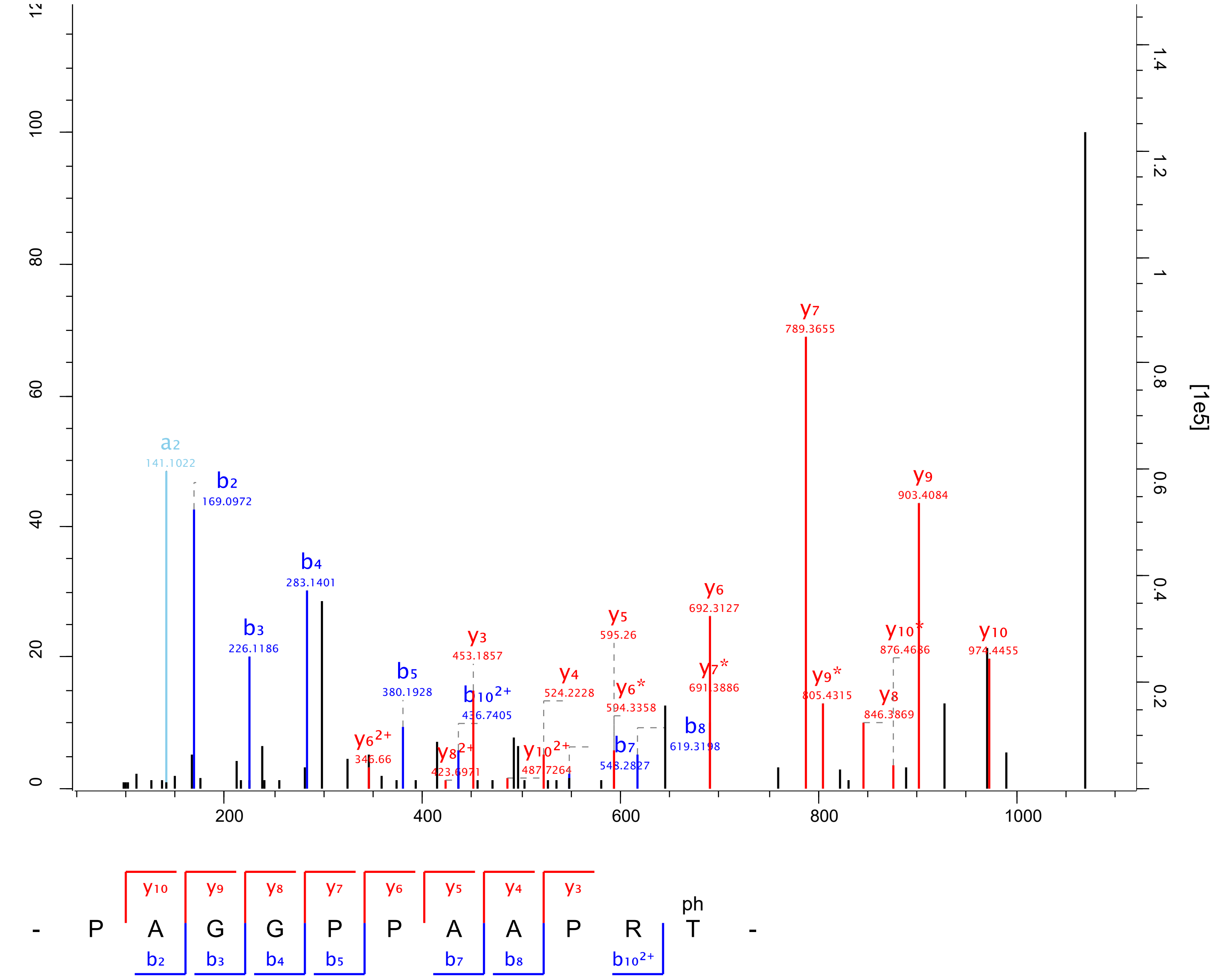

| Raw file | Scan | Method | Score | m/z |
| --- | --- | --- | --- | --- |
| water_15_1 | 41313 | FTMS; HCD | 94.63 | 1044.39 |

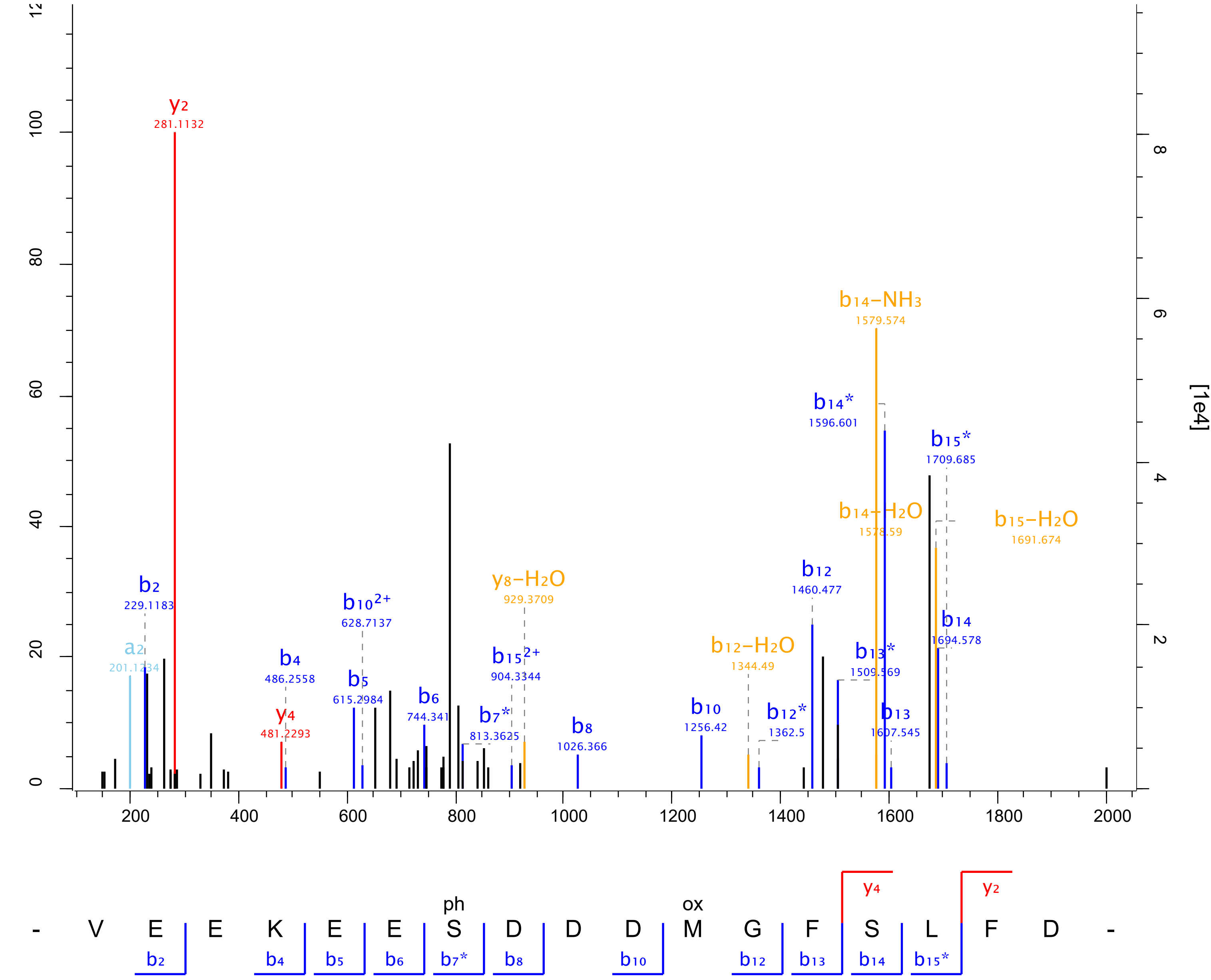

| Raw file | Scan | Method | Score | m/z |
|---|---|---|---|---|
| water_15_1 | 43117 | FTMS; HCD | 162.92 | 1043.38 |

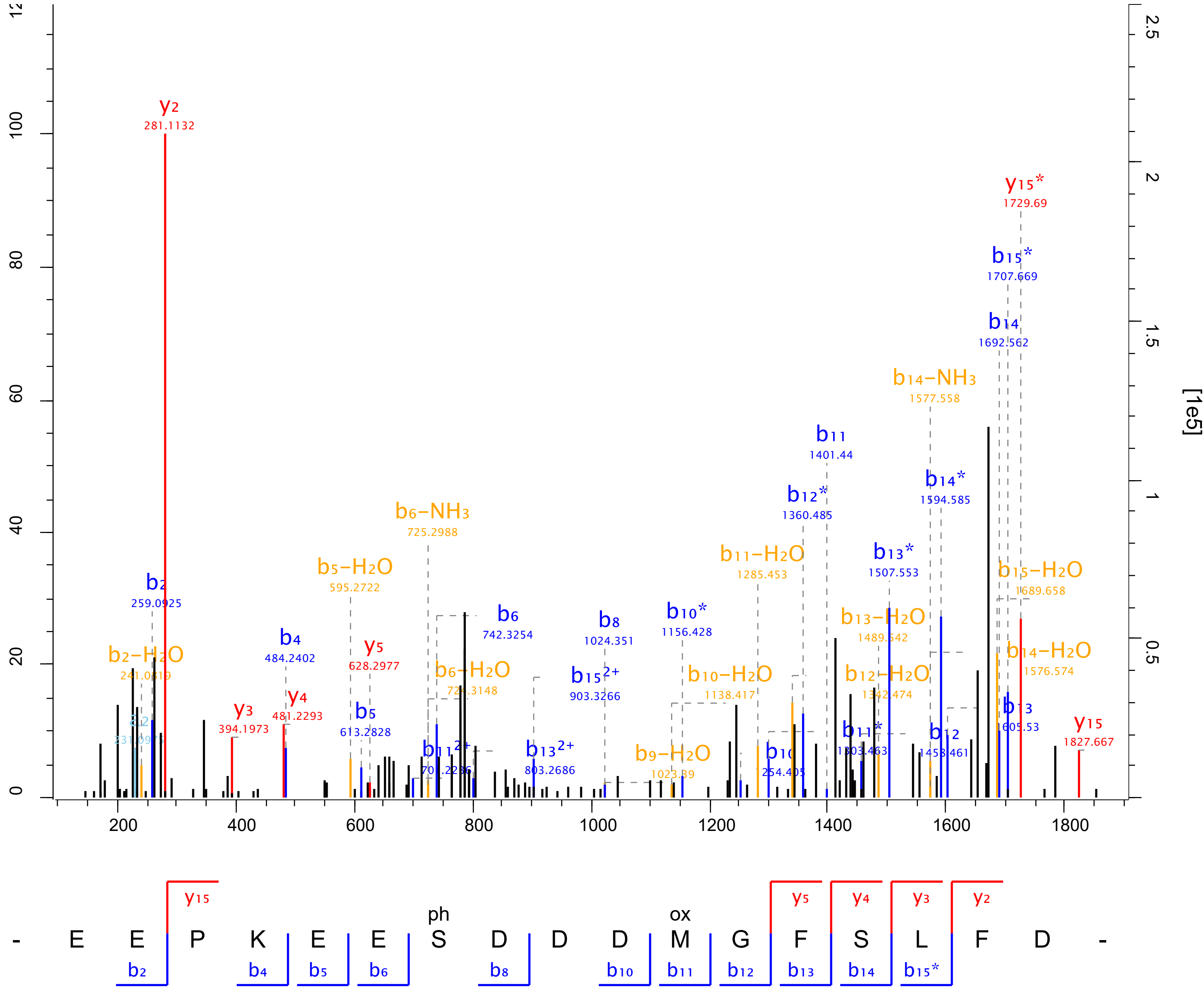

| Raw file | Scan | Method | Score | m/z |
|---|---|---|---|---|
| water_15_1 | 43401 | FTMS; HCD | 53.3 | 665.3 |

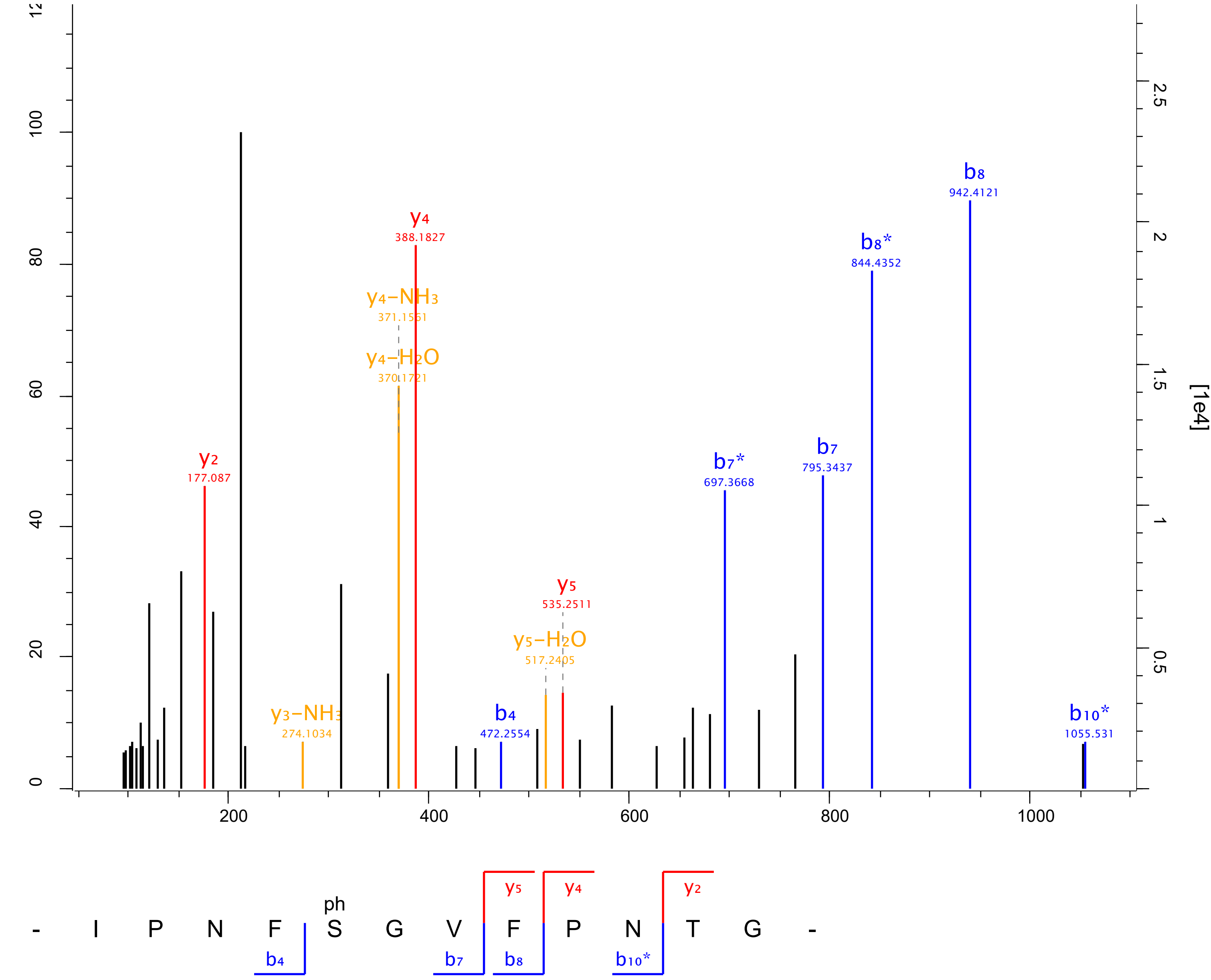

| Raw file | Scan | Method | Score | m/z |
| --- | --- | --- | --- | --- |
| water_15_1 | 4428 | FTMS; HCD | 47.3 | 492.72 |

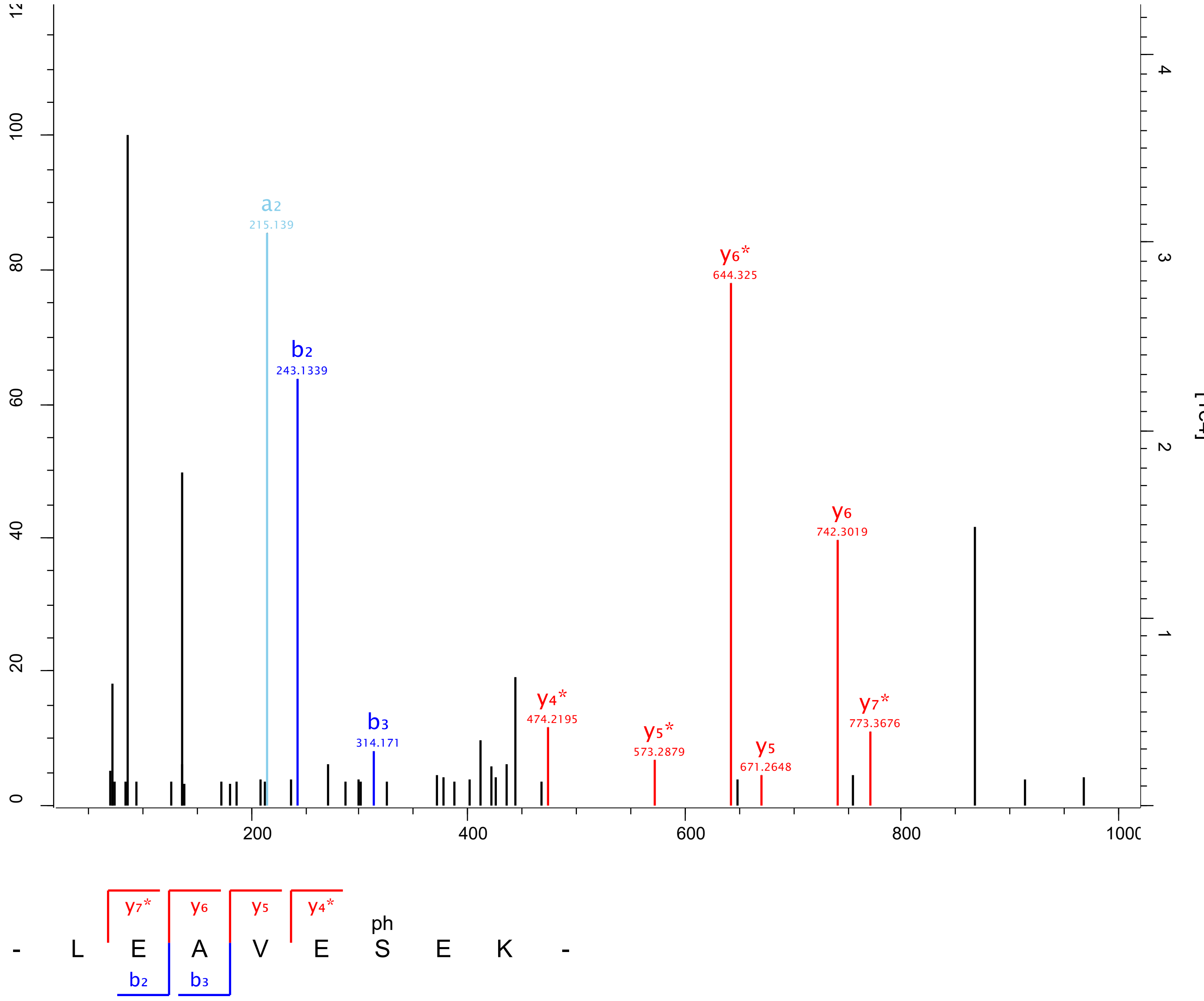

| Raw file | Scan | Method | Score | m/z |
|---|---|---|---|---|
| water_15_1 | 44643 | FTMS; HCD | 52.19 | 889.44 |

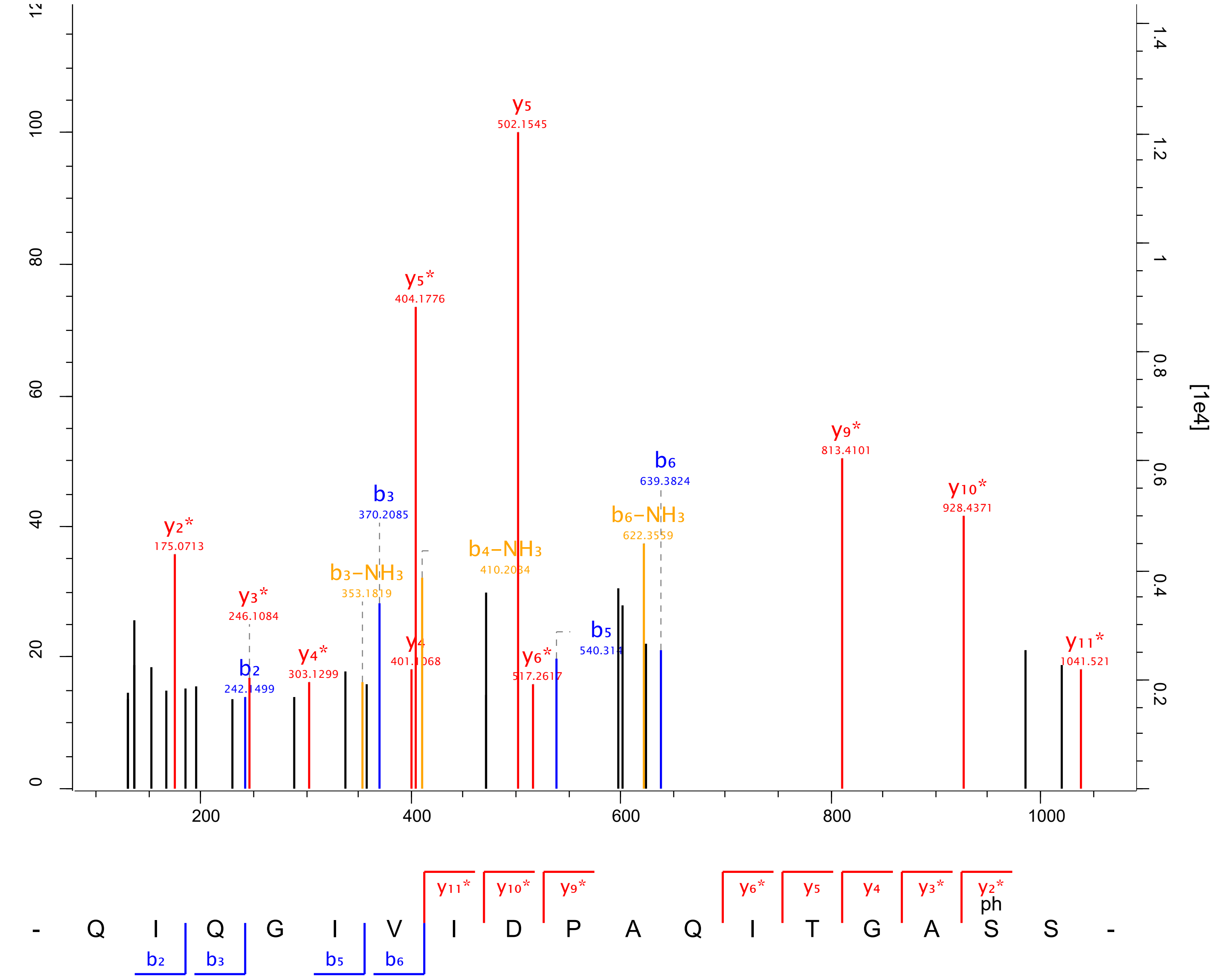

| Raw file | Scan | Method | Score | m/z |
|---|---|---|---|---|
| water_15_1 | 4654 | FTMS; HCD | 88.13 | 457.22 |

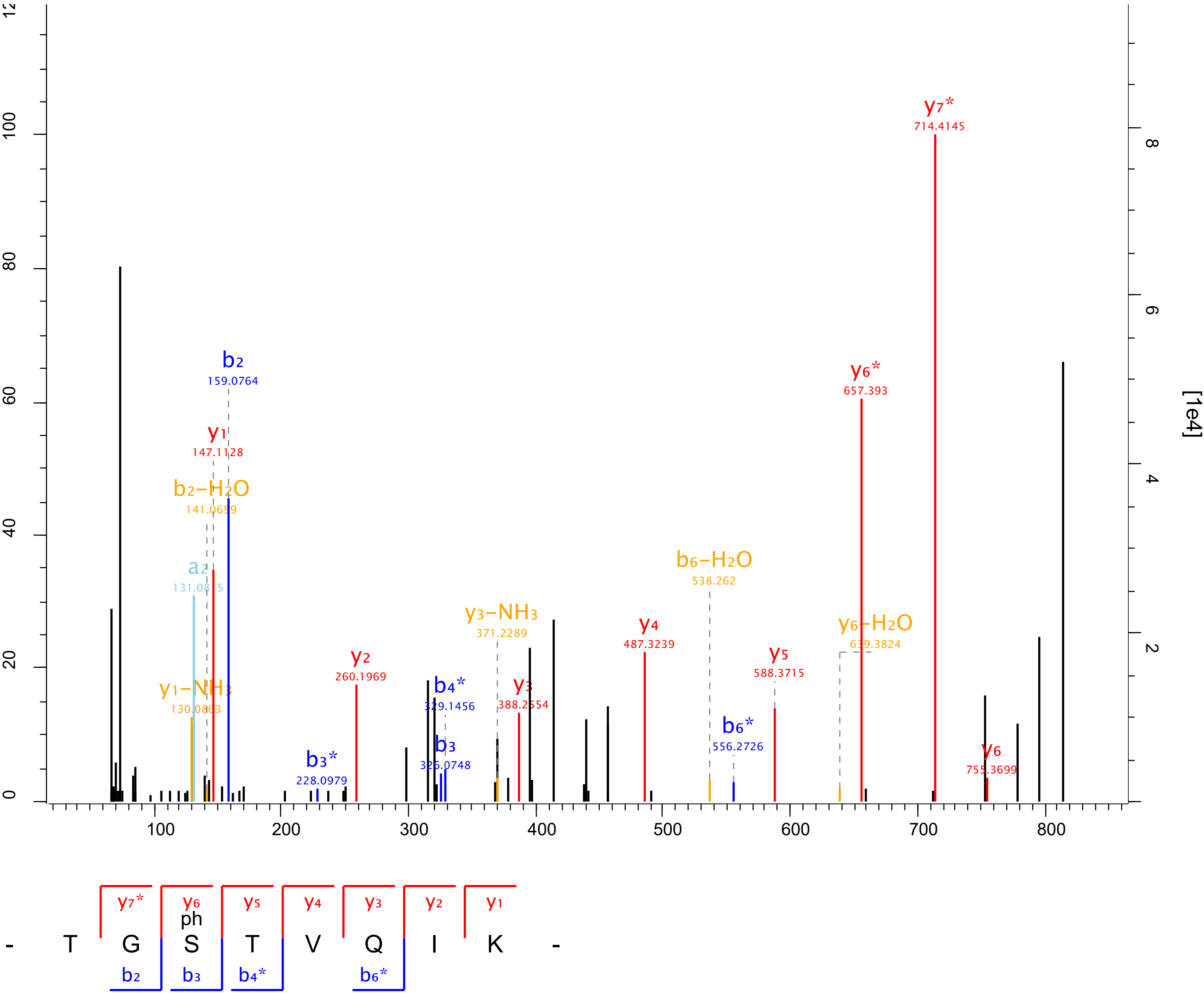

| Raw file | Scan | Method | Score | m/z |
|---|---|---|---|---|
| water_15_1 | 4906 | FTMS; HCD | 101.3 | 462.7 |

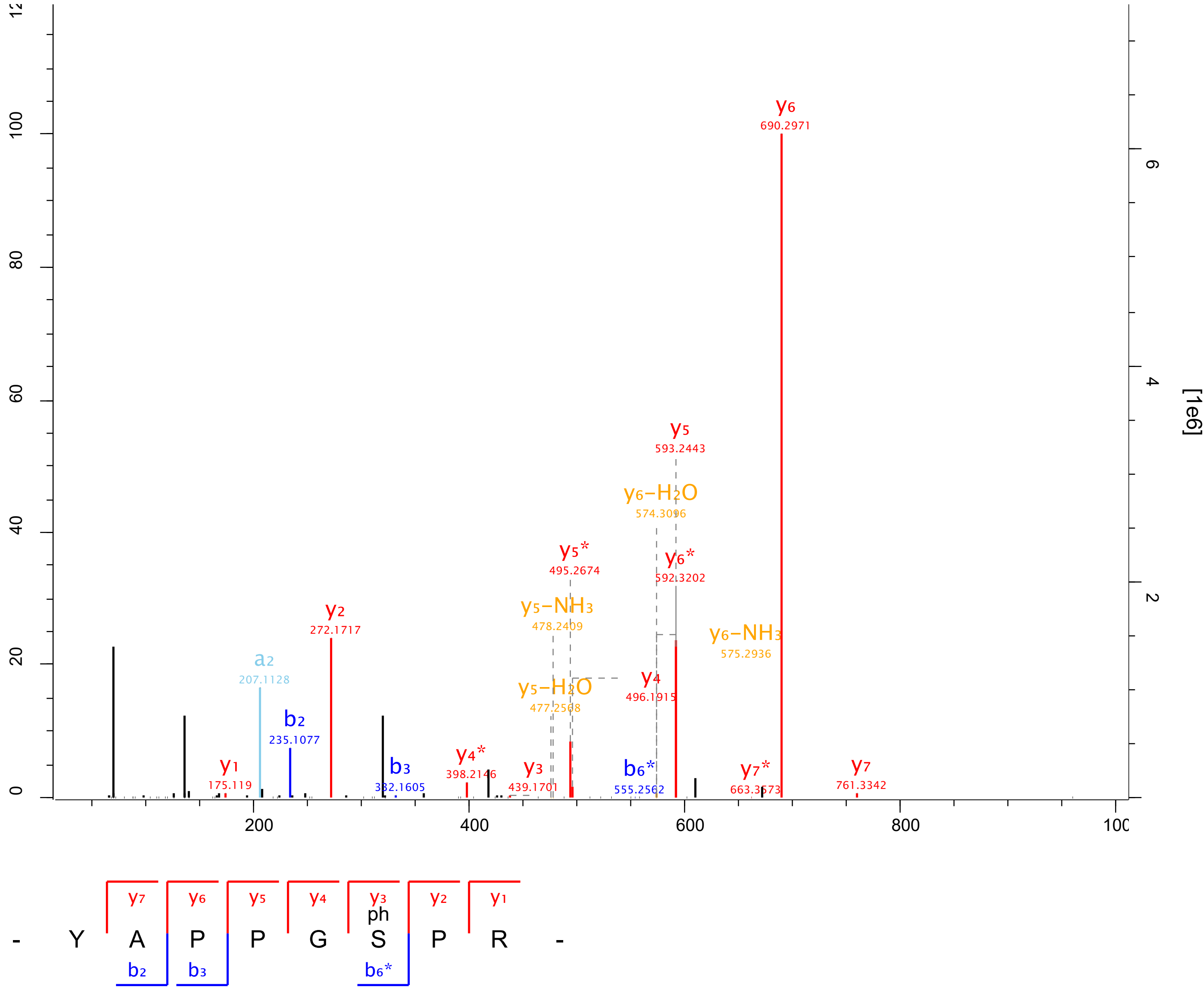

| Raw file | Scan | Method | Score | m/z |
|---|---|---|---|---|
| water_15_1 | 5062 | FTMS; HCD | 62.47 | 513.75 |

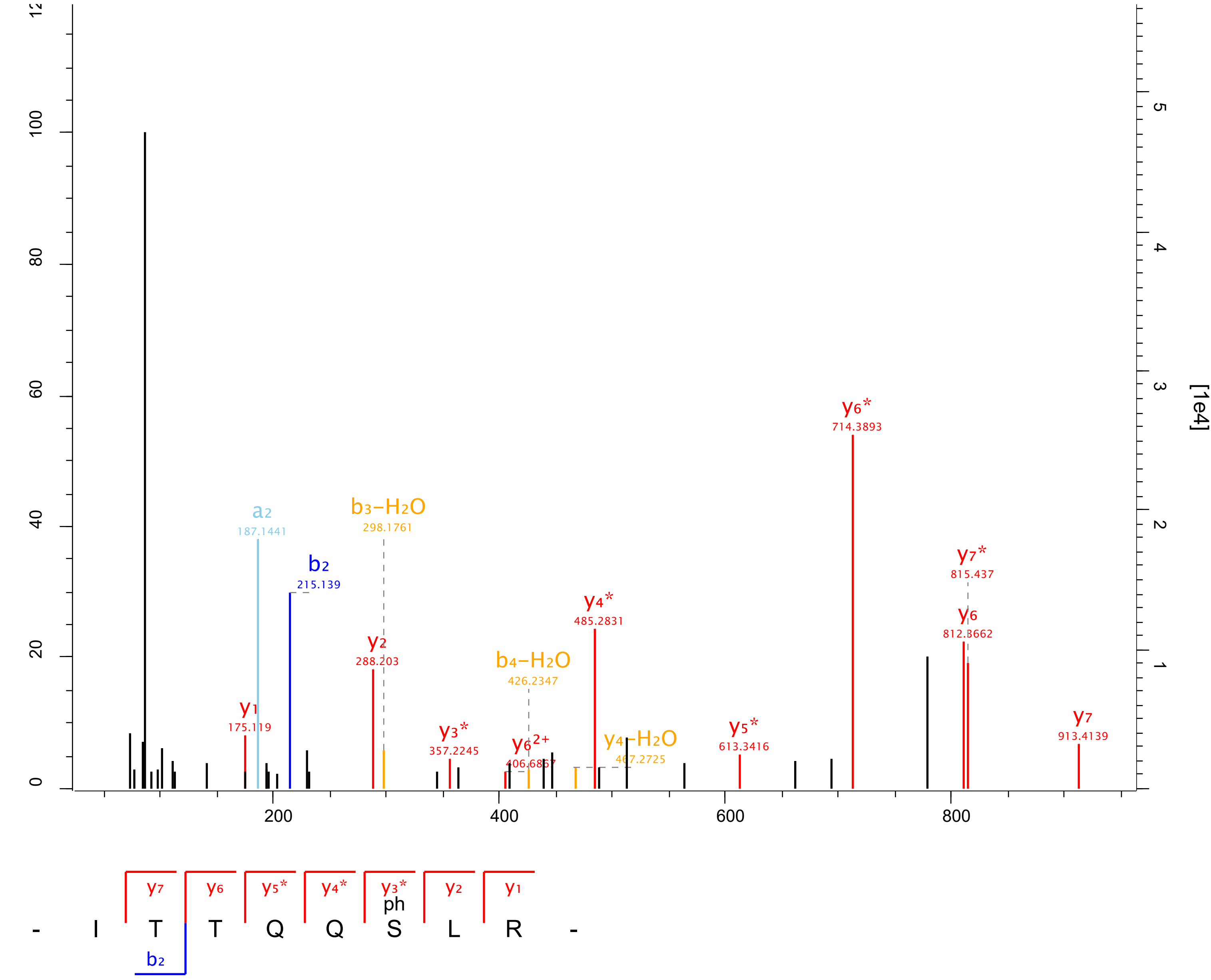

| Raw file | Scan | Method | Score | m/z |
|---|---|---|---|---|
| water_15_1 | 5210 | FTMS; HCD | 136.96 | 492.74 |

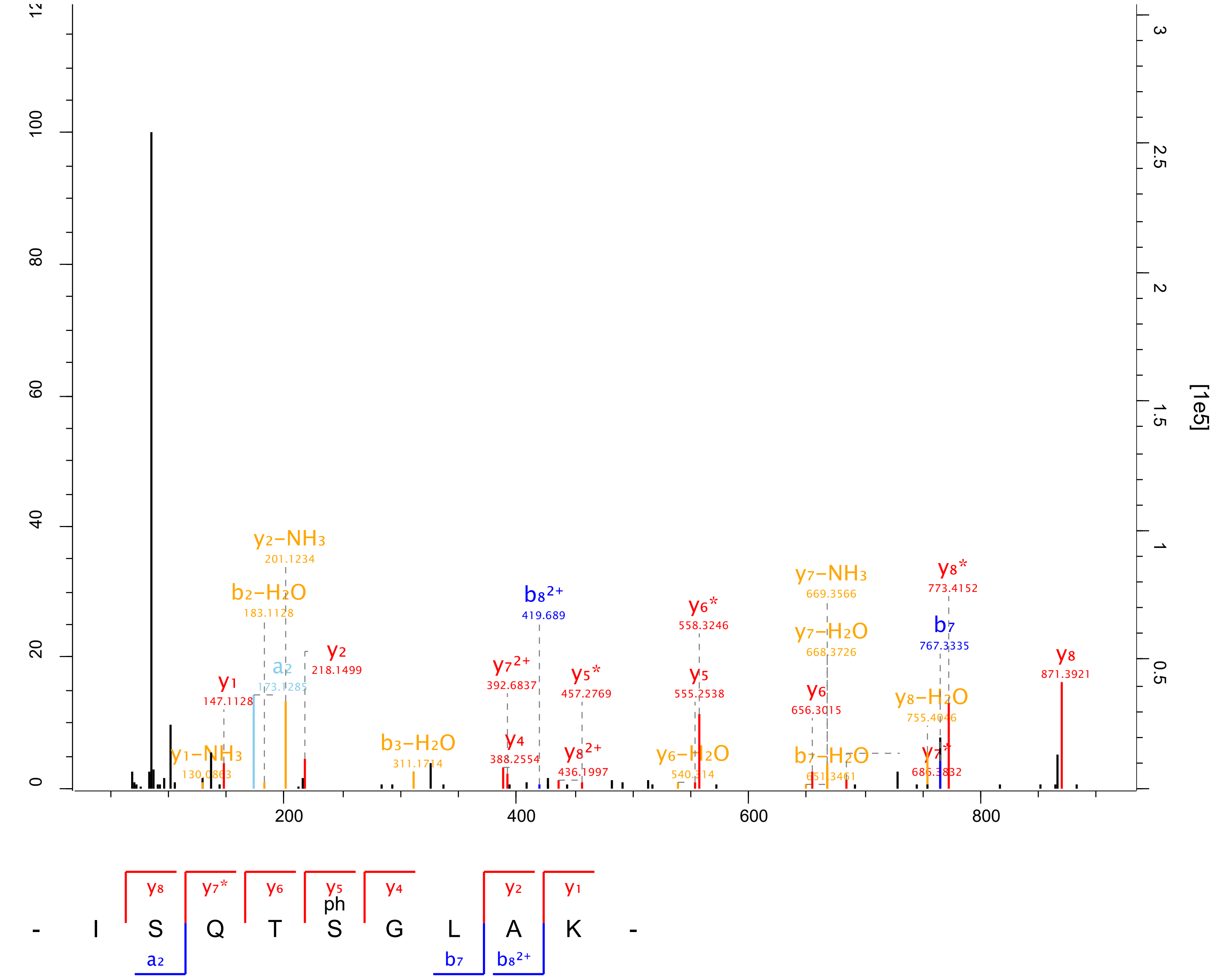

| Raw file | Scan | Method | Score | m/z |
|---|---|---|---|---|
| water_15_1 | 5647 | FTMS; HCD | 106.62 | 503.22 |

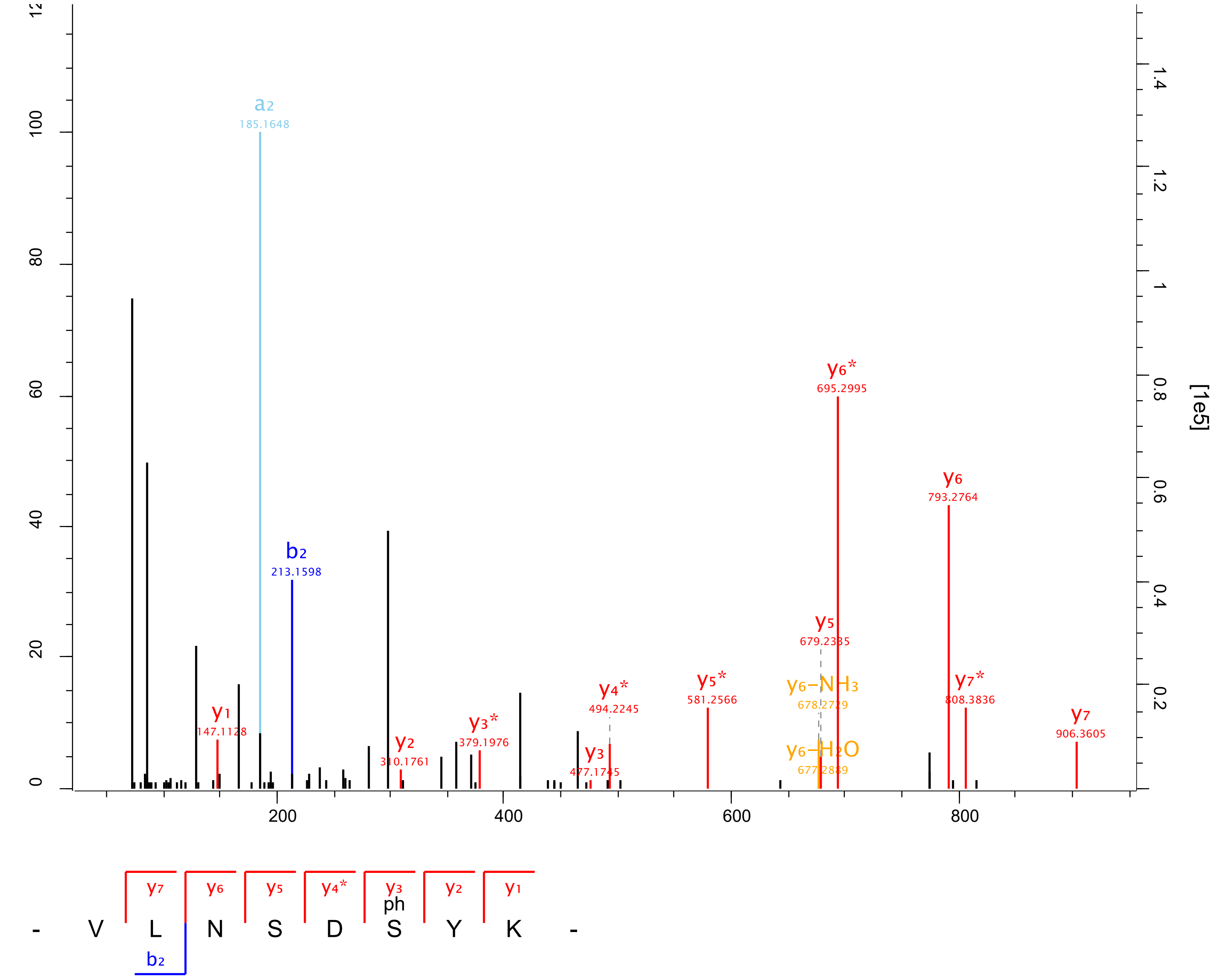

| Raw file | Scan | Method | Score | m/z |
|---|---|---|---|---|
| water_15_1 | 5858 | FTMS; HCD | 193.62 | 571.23 |

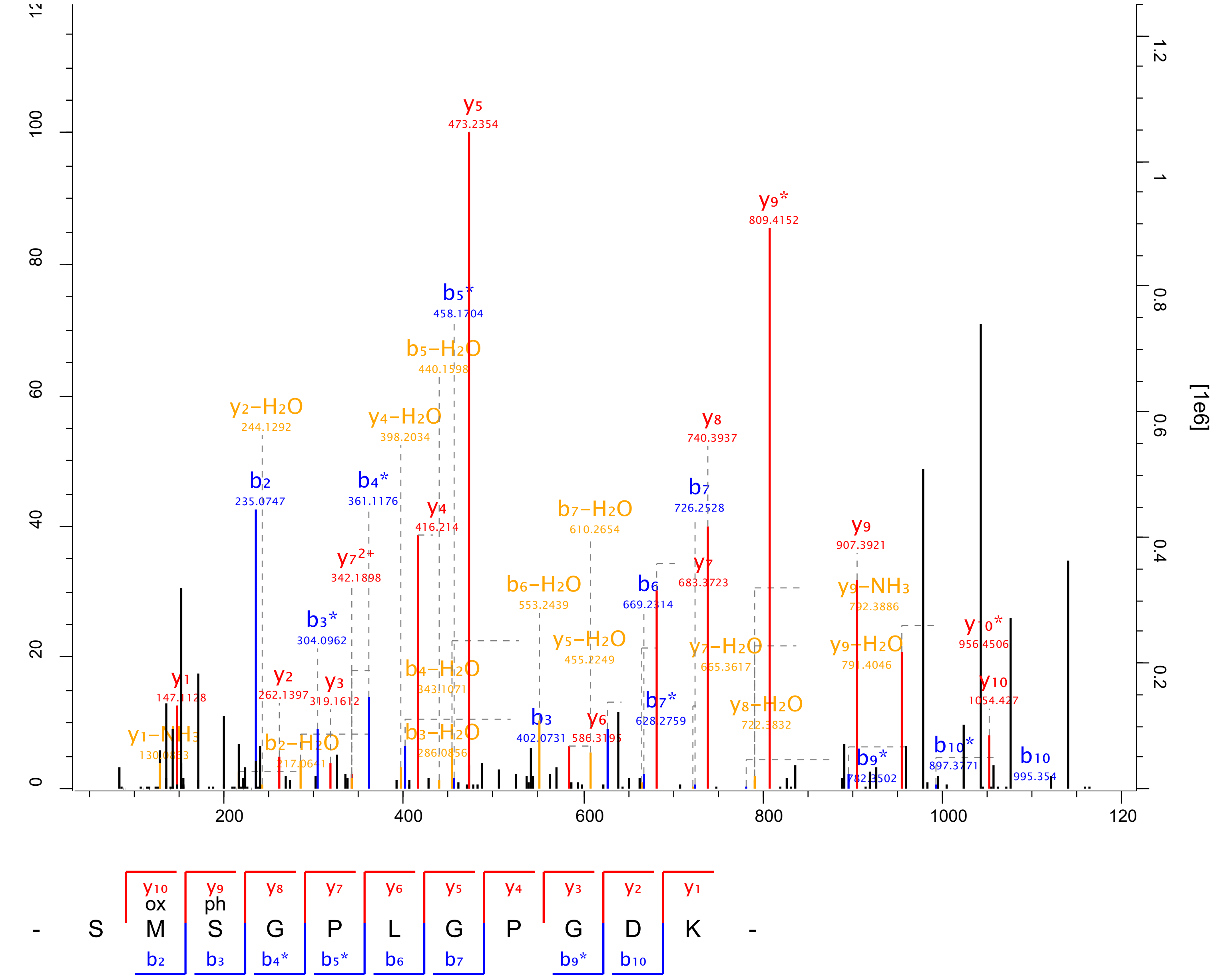

| Raw file | Scan | Method | Score | m/z |
|---|---|---|---|---|
| water_15_1 | 5909 | FTMS; HCD | 107.03 | 551.73 |

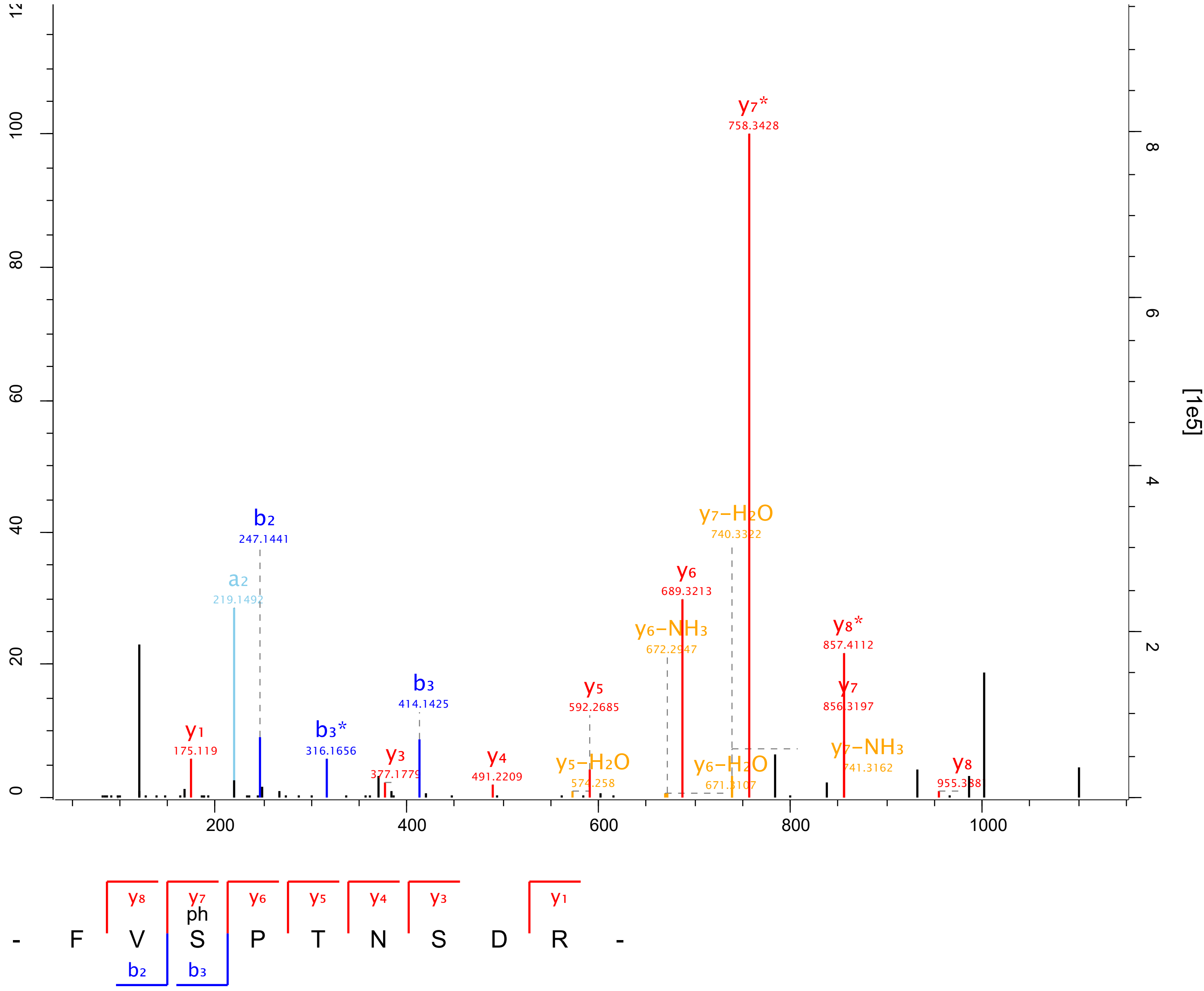

| Raw file | Scan | Method | Score | m/z |
| --- | --- | --- | --- | --- |
| water_15_1 | 5943 | FTMS; HCD | 62.09 | 638.72 |

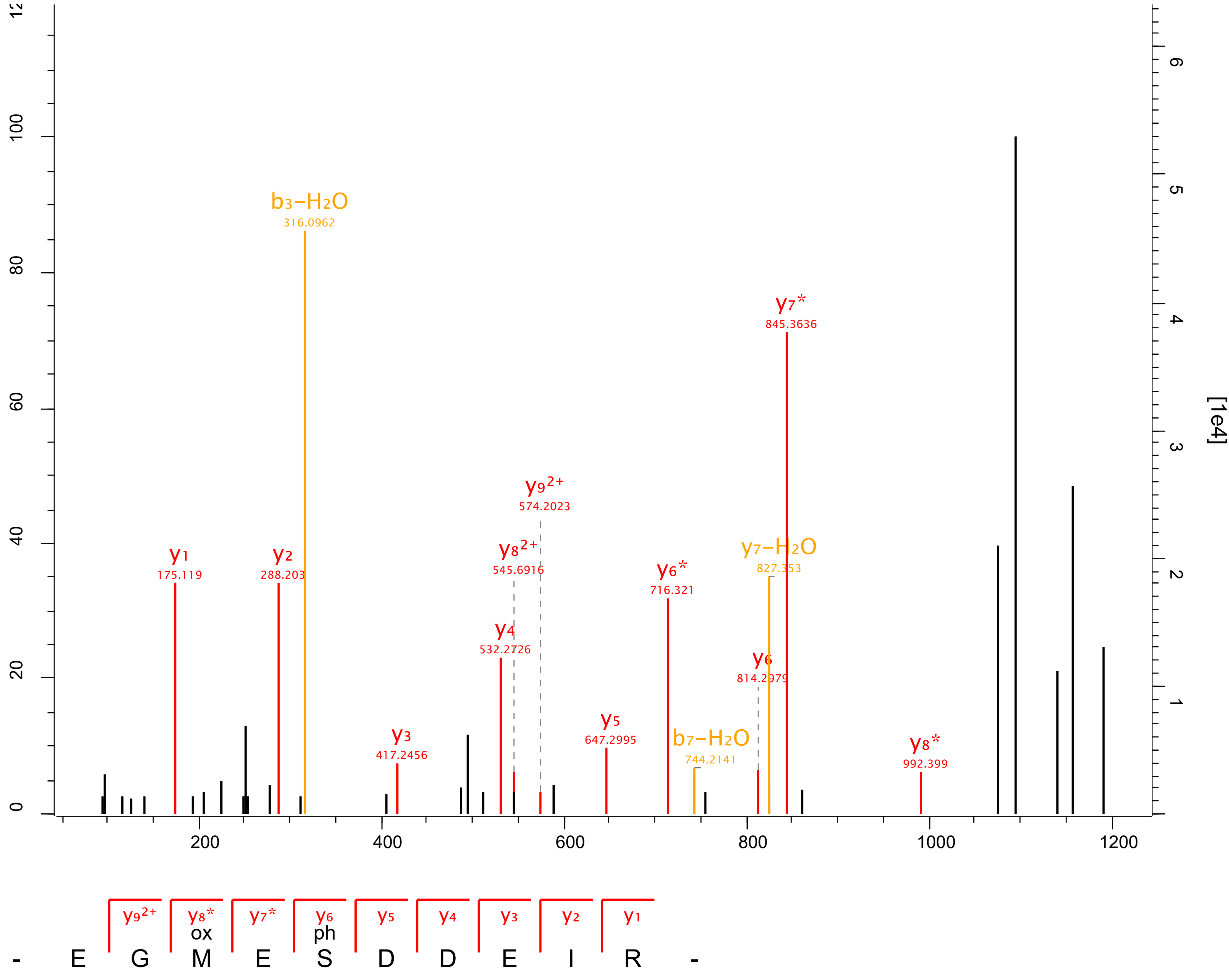

| Raw file | Scan | Method | Score | m/z |
| --- | --- | --- | --- | --- |
| water_15_1 | 6224 | FTMS; HCD | 233.43 | 663.27 |

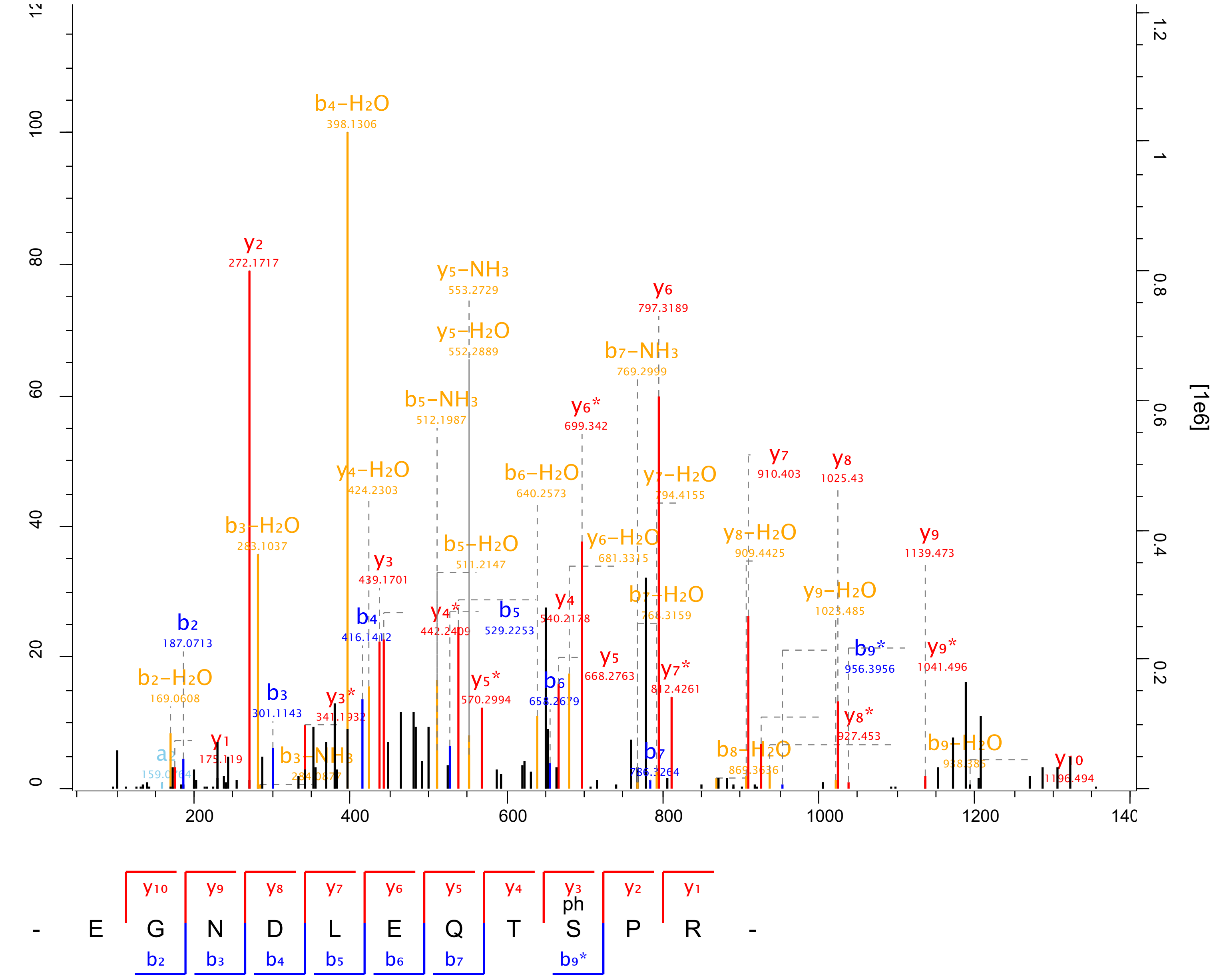

| Raw file | Scan | Method | Score | m/z |
|---|---|---|---|---|
| water_15_1 | 6606 | FTMS; HCD | 202.01 | 625.29 |

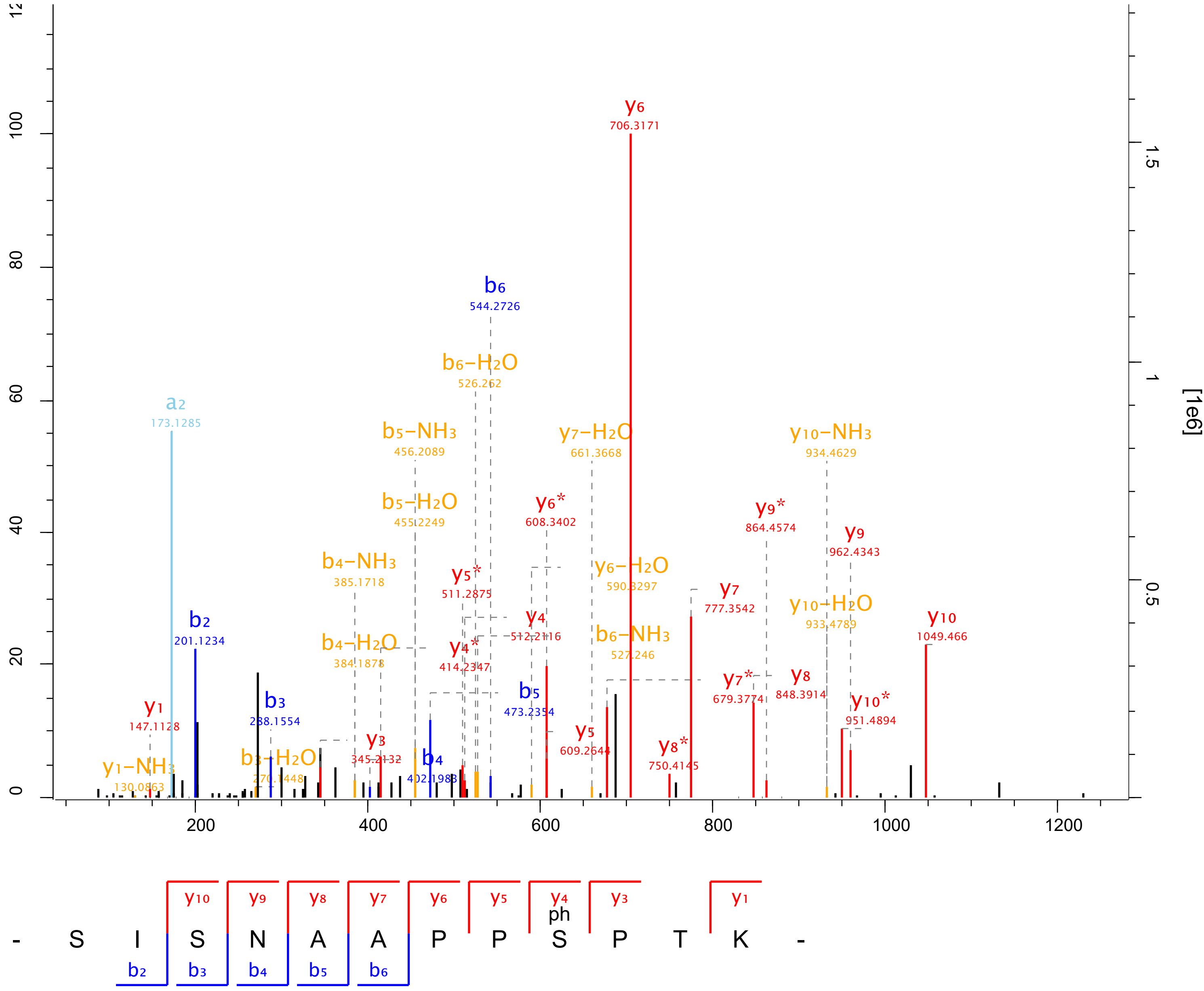

| Raw file | Scan | Method | Score | m/z |
|---|---|---|---|---|
| water_15_1 | 6737 | FTMS; HCD | 175.54 | 565.27 |

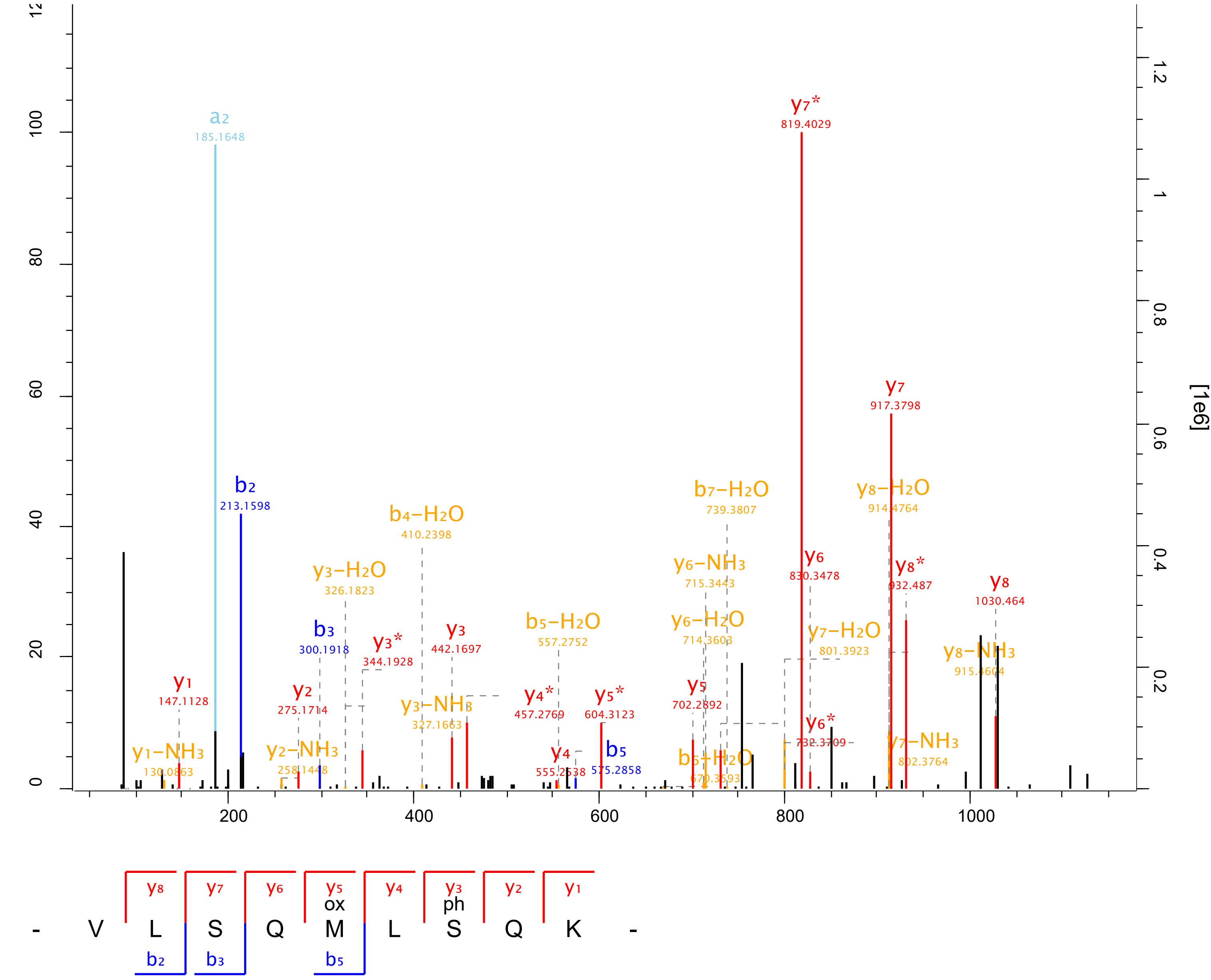

| Raw file | Scan | Method | Score | m/z |
|---|---|---|---|---|
| water_15_1 | 6950 | FTMS; HCD | 119.75 | 441.22 |

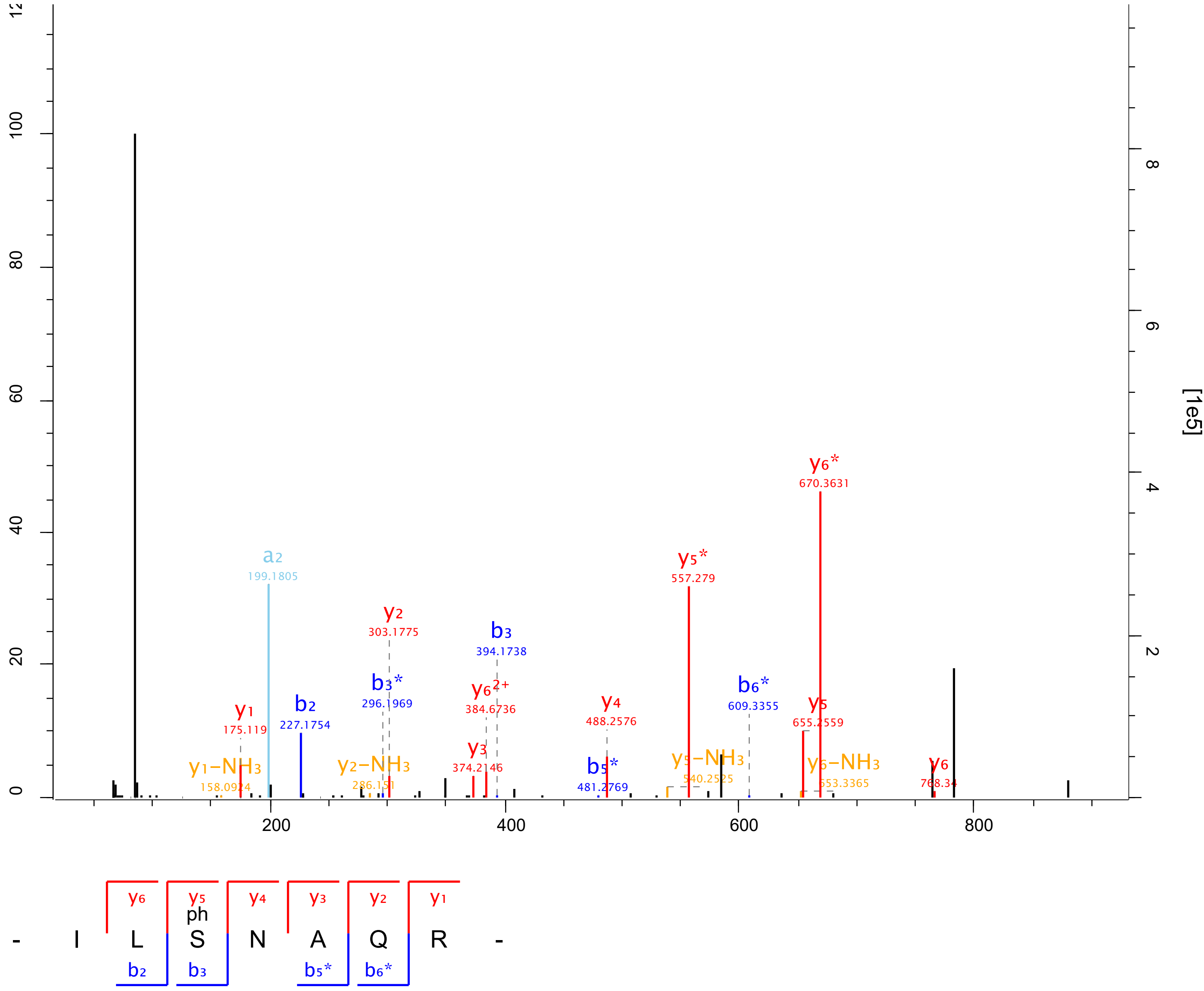

| Raw file | Scan | Method | Score | m/z |
|---|---|---|---|---|
| water_15_1 | 7326 | FTMS; HCD | 100.93 | 576.23 |

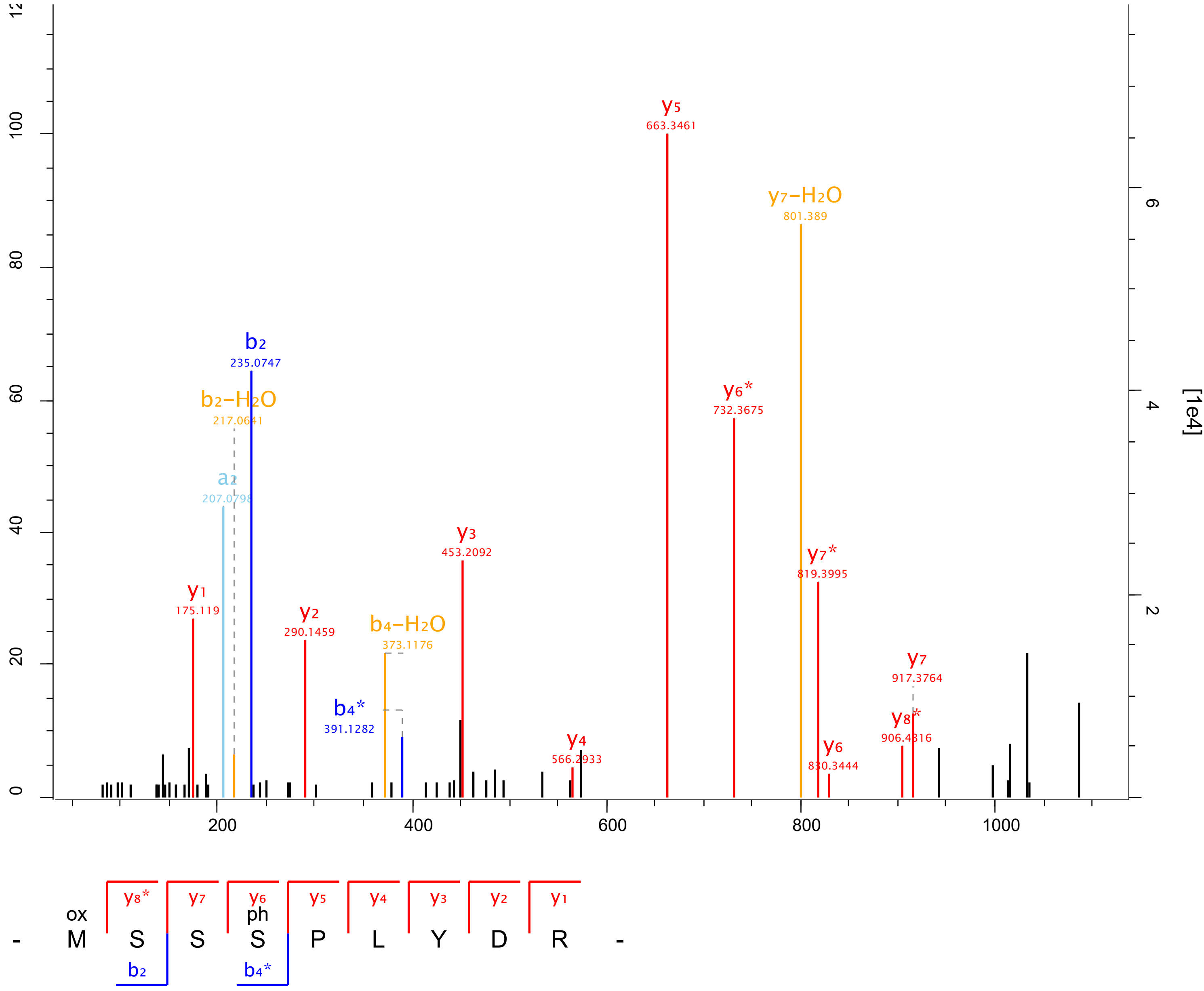

| Raw file | Scan | Method | Score | m/z |
| --- | --- | --- | --- | --- |
| water_15_1 | 7515 | FTMS; HCD | 210.91 | 622.28 |

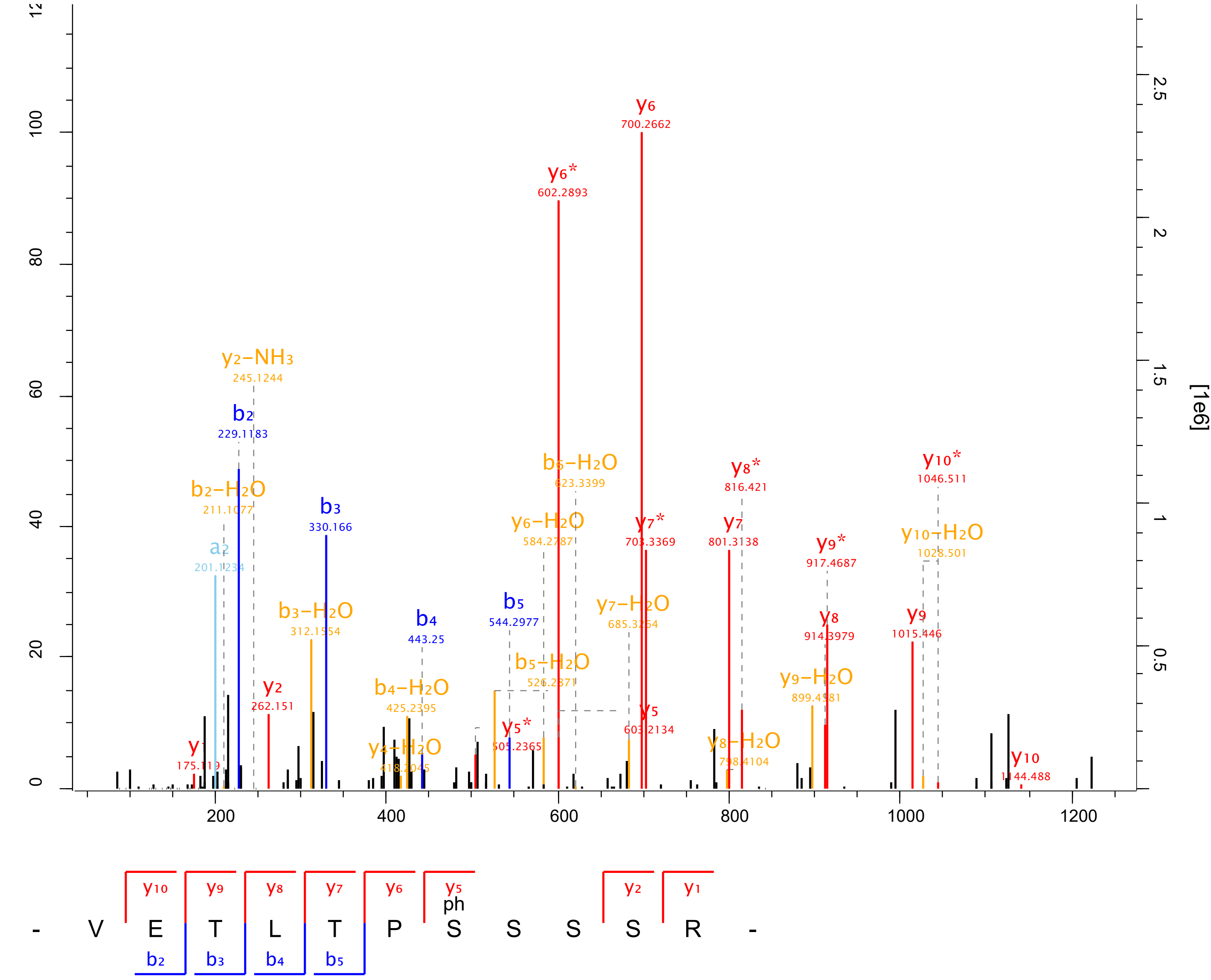

| Raw file | Scan | Method | Score | m/z |
| --- | --- | --- | --- | --- |
| water_15_1 | 7593 | FTMS; HCD | 128.87 | 430.7 |

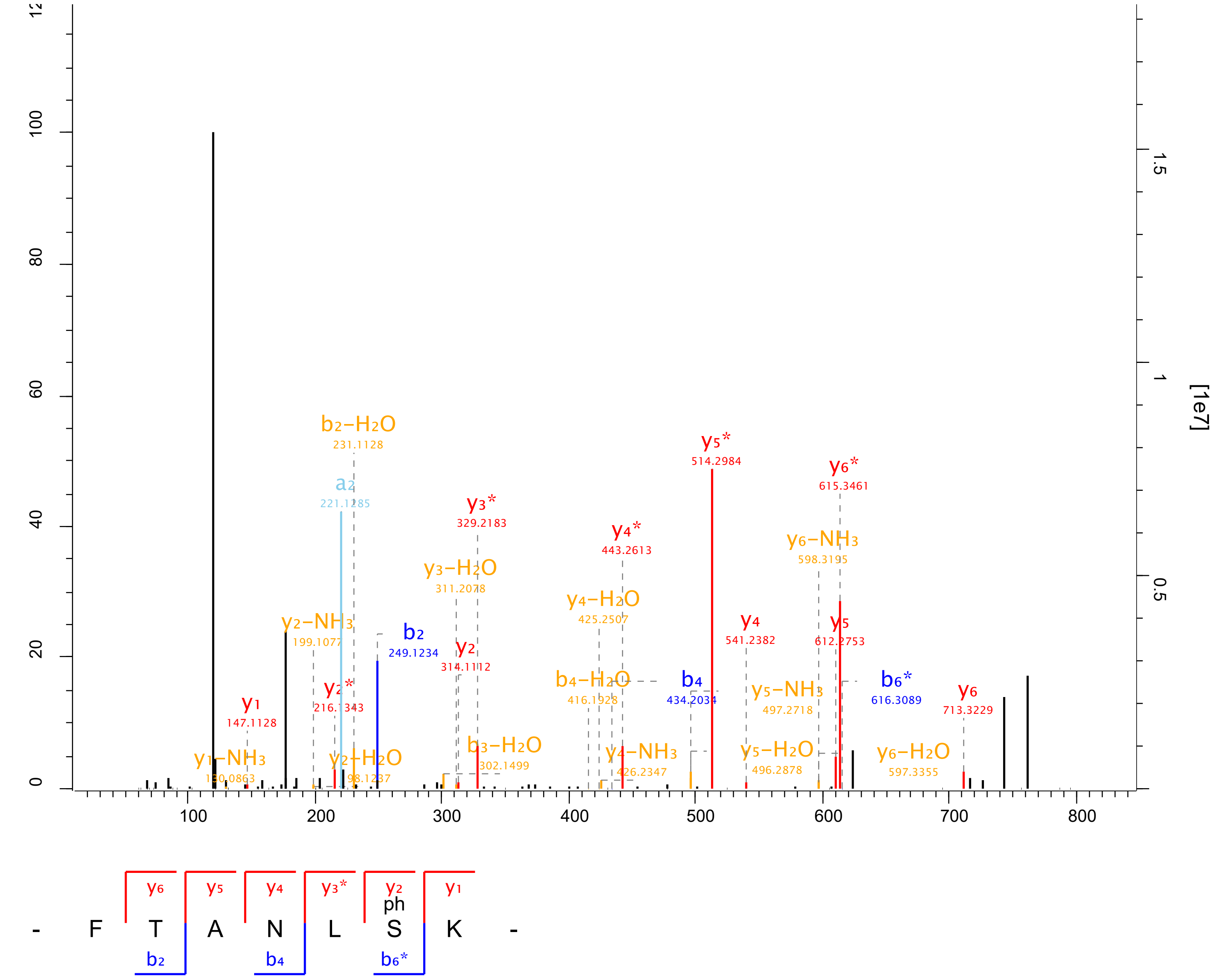

| Raw file | Scan | Method | Score | m/z |
|---|---|---|---|---|
| water_15_1 | 7974 | FTMS; HCD | 164.3 | 633.74 |

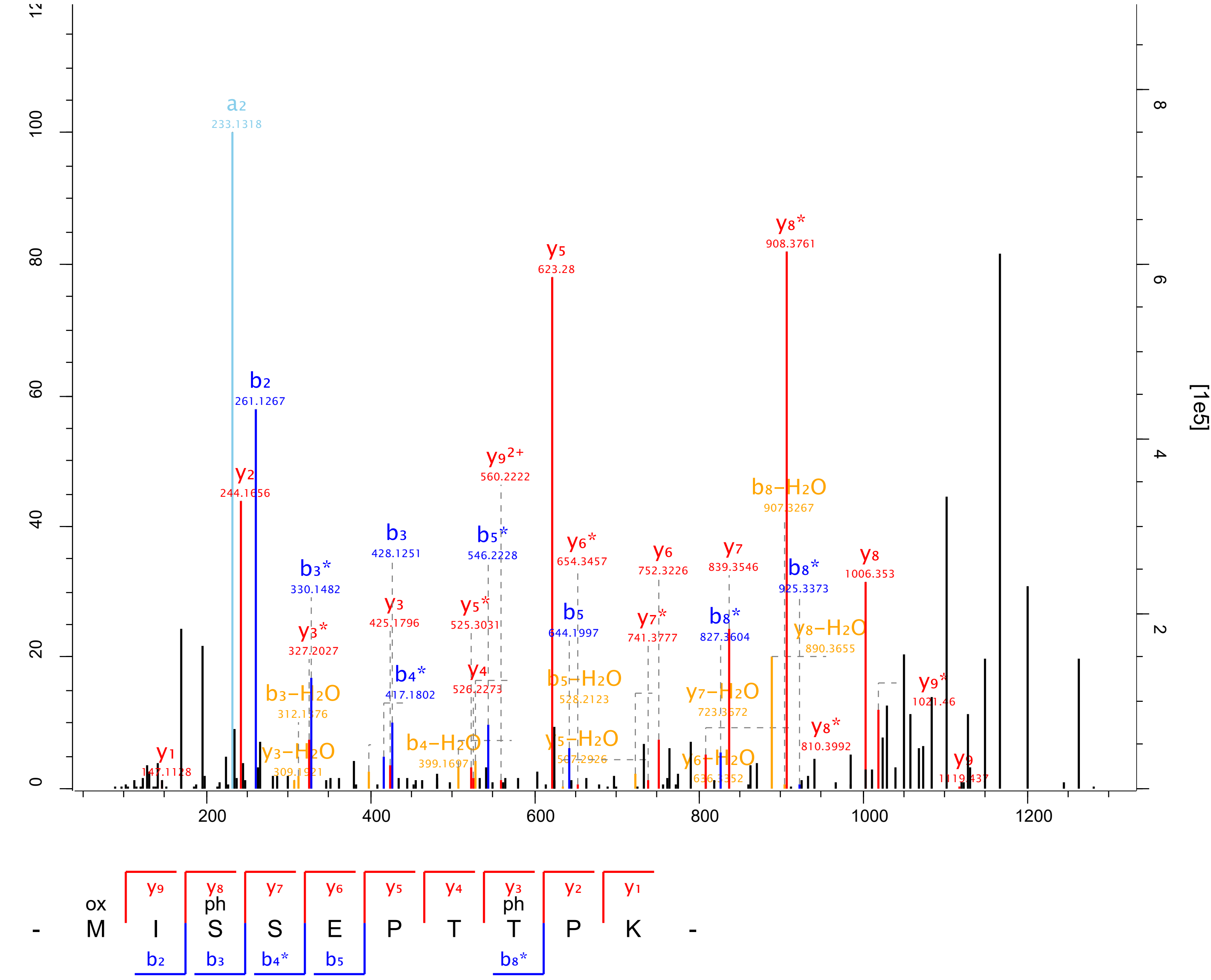

| Raw file | Scan | Method | Score | m/z |
| --- | --- | --- | --- | --- |
| water_15_1 | 8137 | FTMS; HCD | 52.58 | 512.2 |

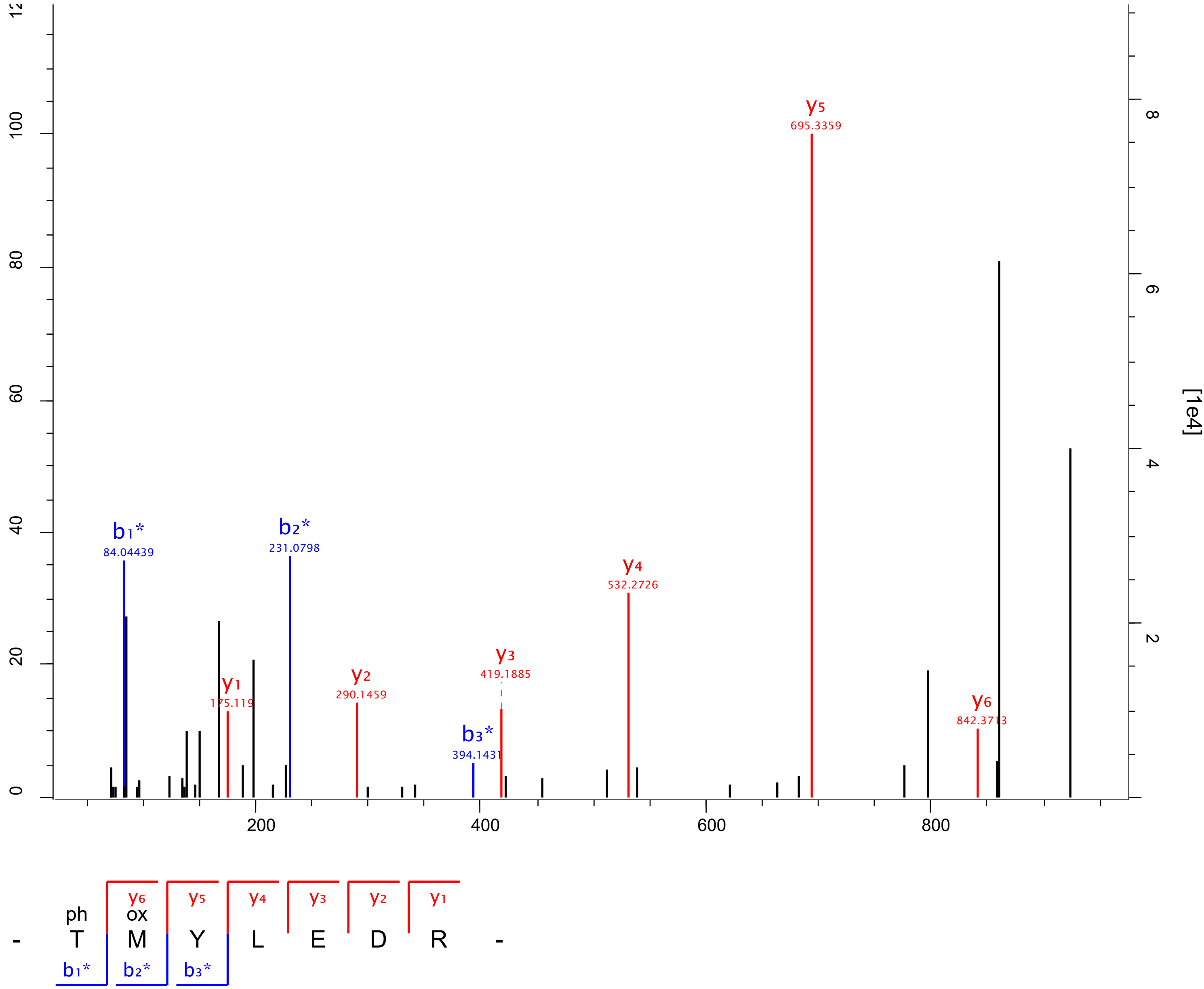

| Raw file | Scan | Method | Score | m/z |
|---|---|---|---|---|
| water_15_1 | 8245 | FTMS; HCD | 54.98 | 427.22 |

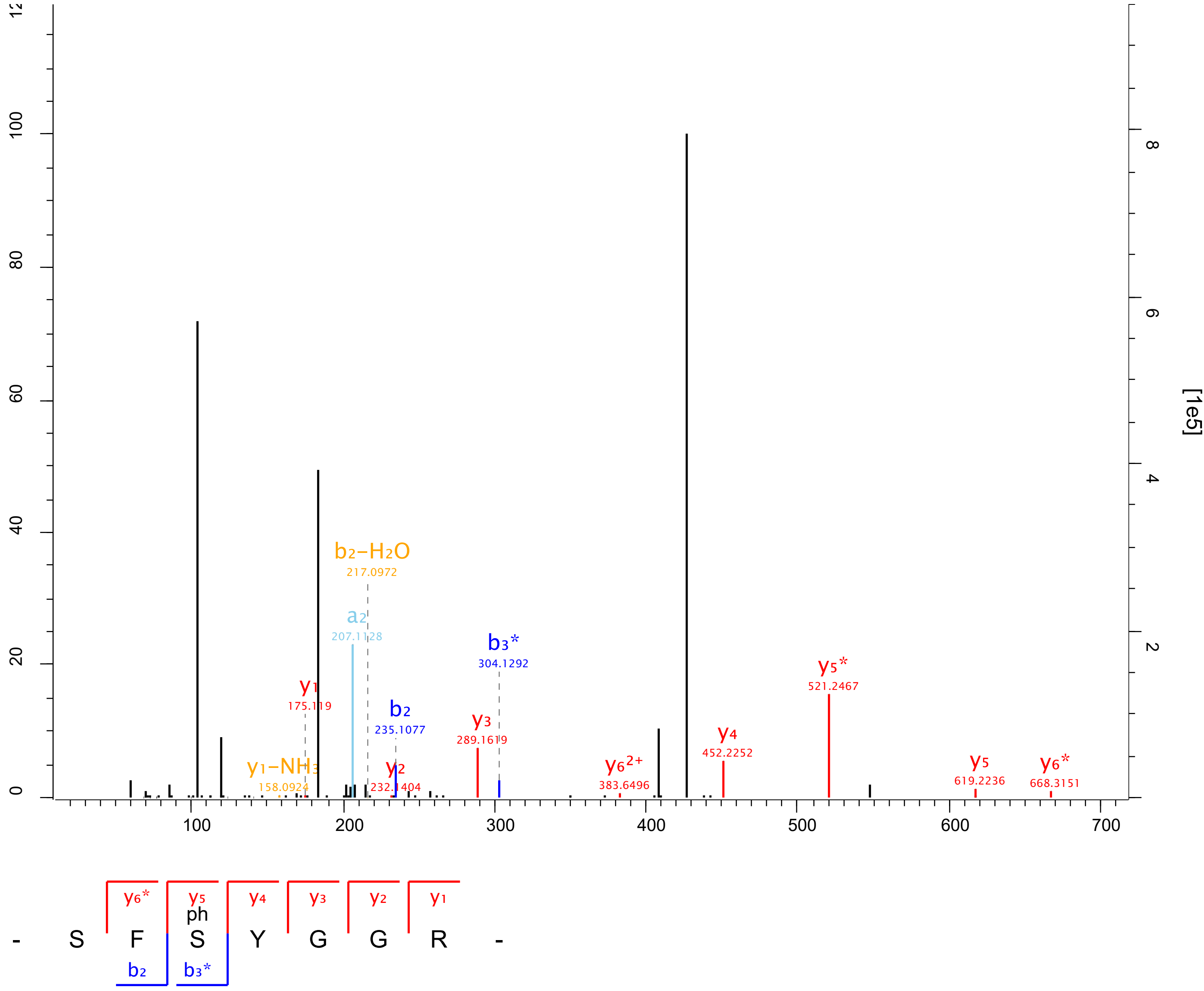

| Raw file | Scan | Method | Score | m/z |
|---|---|---|---|---|
| water_15_1 | 8279 | FTMS; HCD | 104.22 | 496.69 |

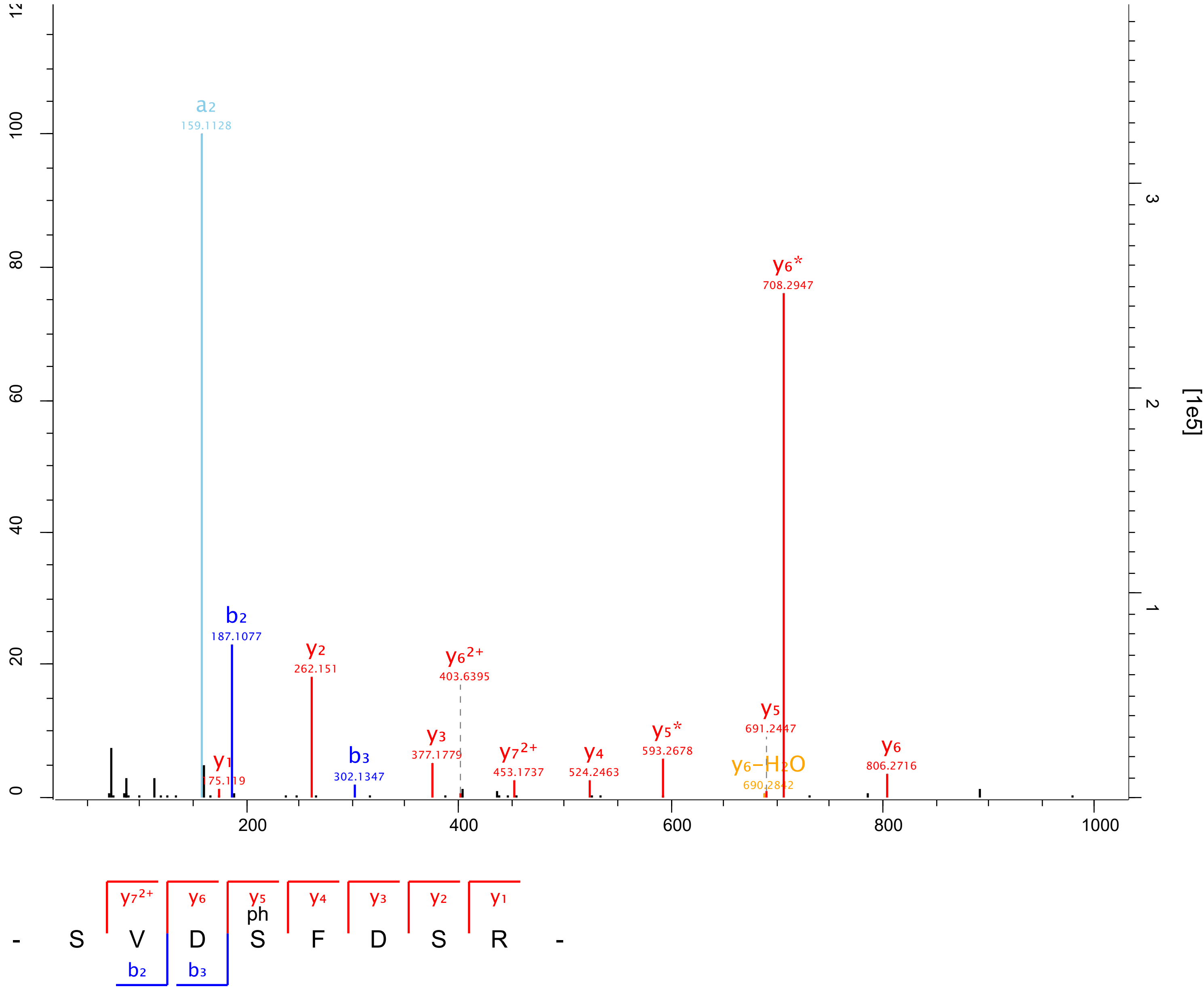

| Raw file | Scan | Method | Score | m/z |
| --- | --- | --- | --- | --- |
| water_15_1 | 8518 | FTMS; HCD | 140.82 | 650.24 |

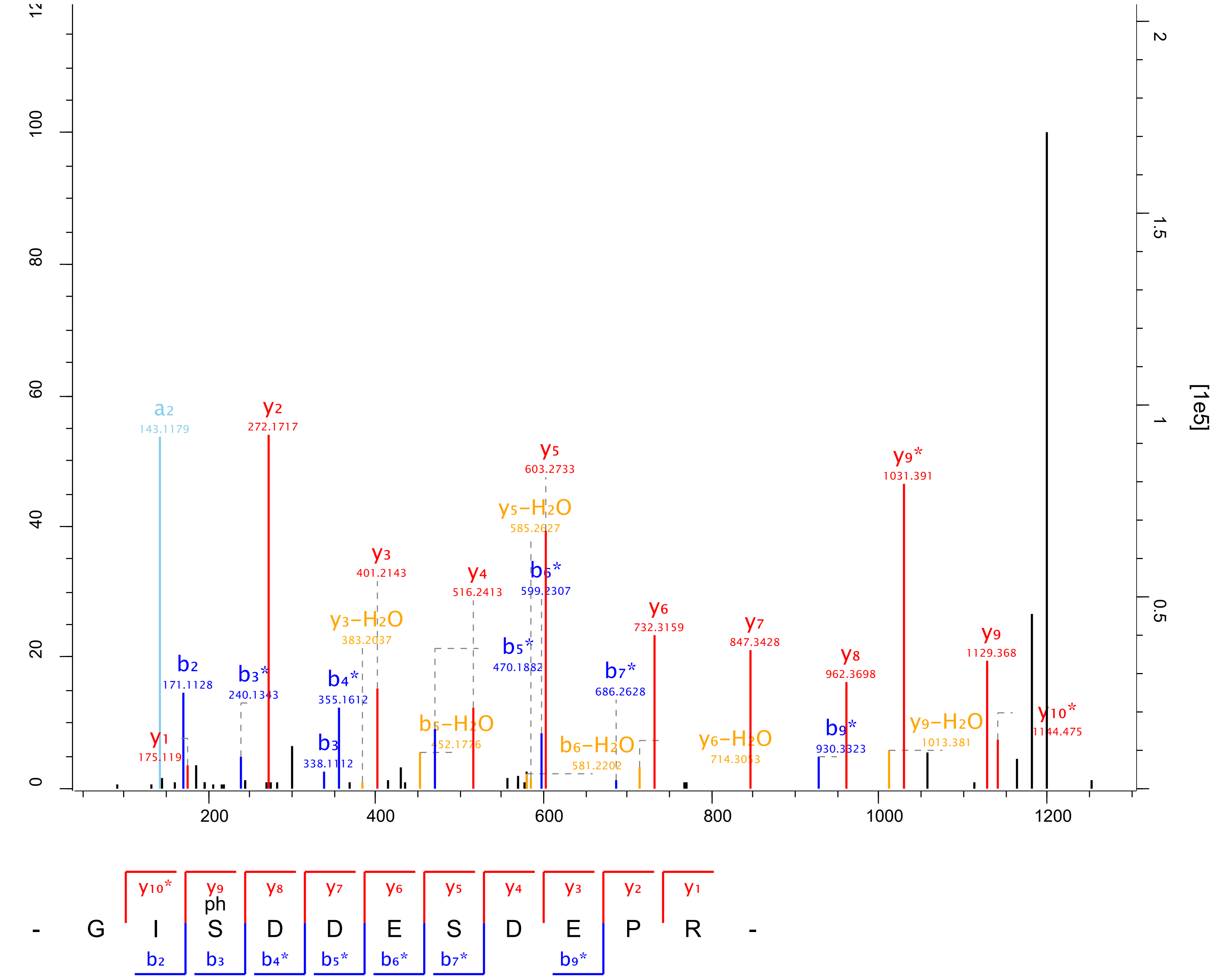

| Raw file | Scan | Method | Score | m/z |
|---|---|---|---|---|
| water_15_1 | 8585 | FTMS; HCD | 192.54 | 671.79 |

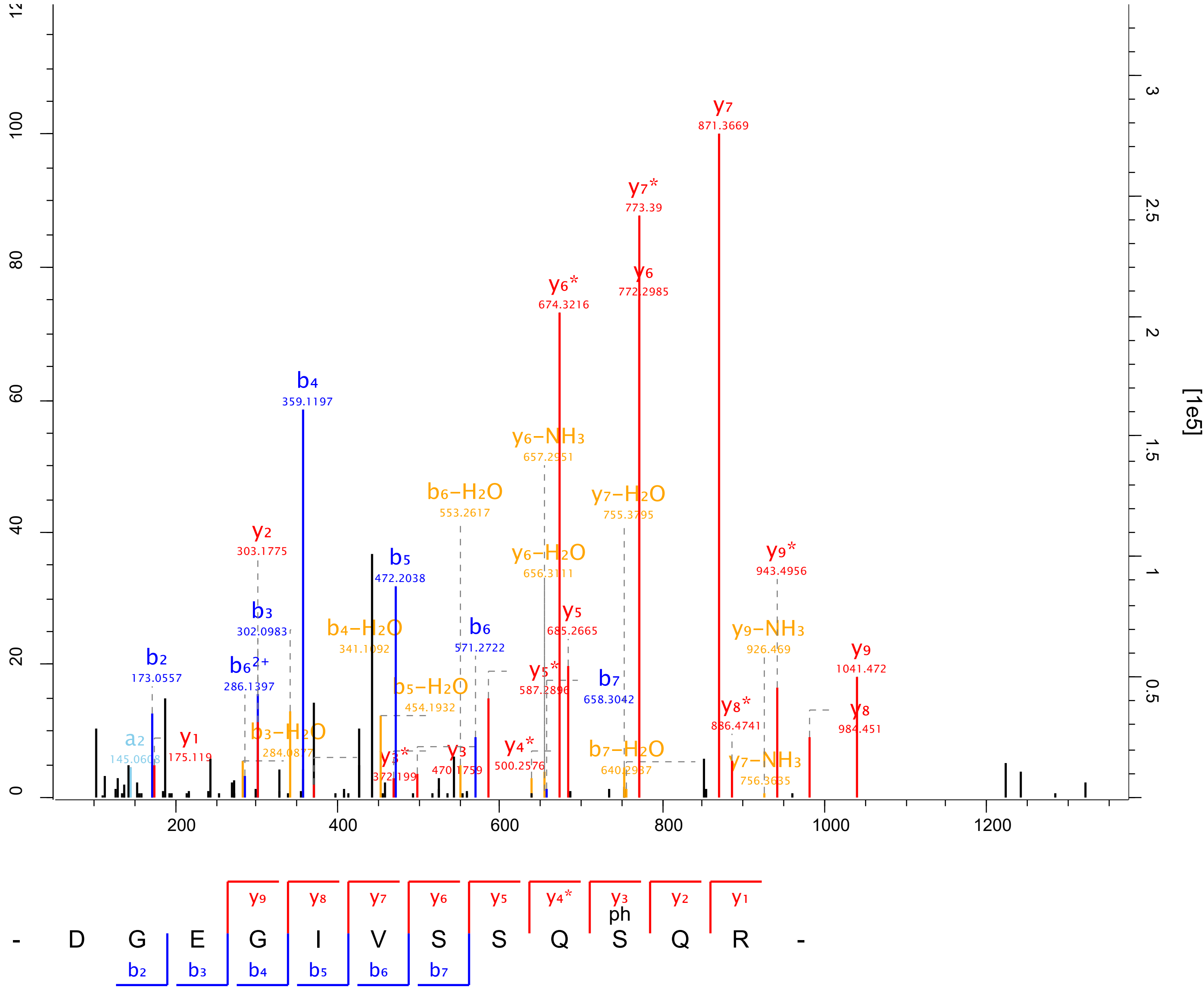

| Raw file | Scan | Method | Score | m/z |
| --- | --- | --- | --- | --- |
| water_15_1 | 8706 | FTMS; HCD | 145.09 | 562.76 |

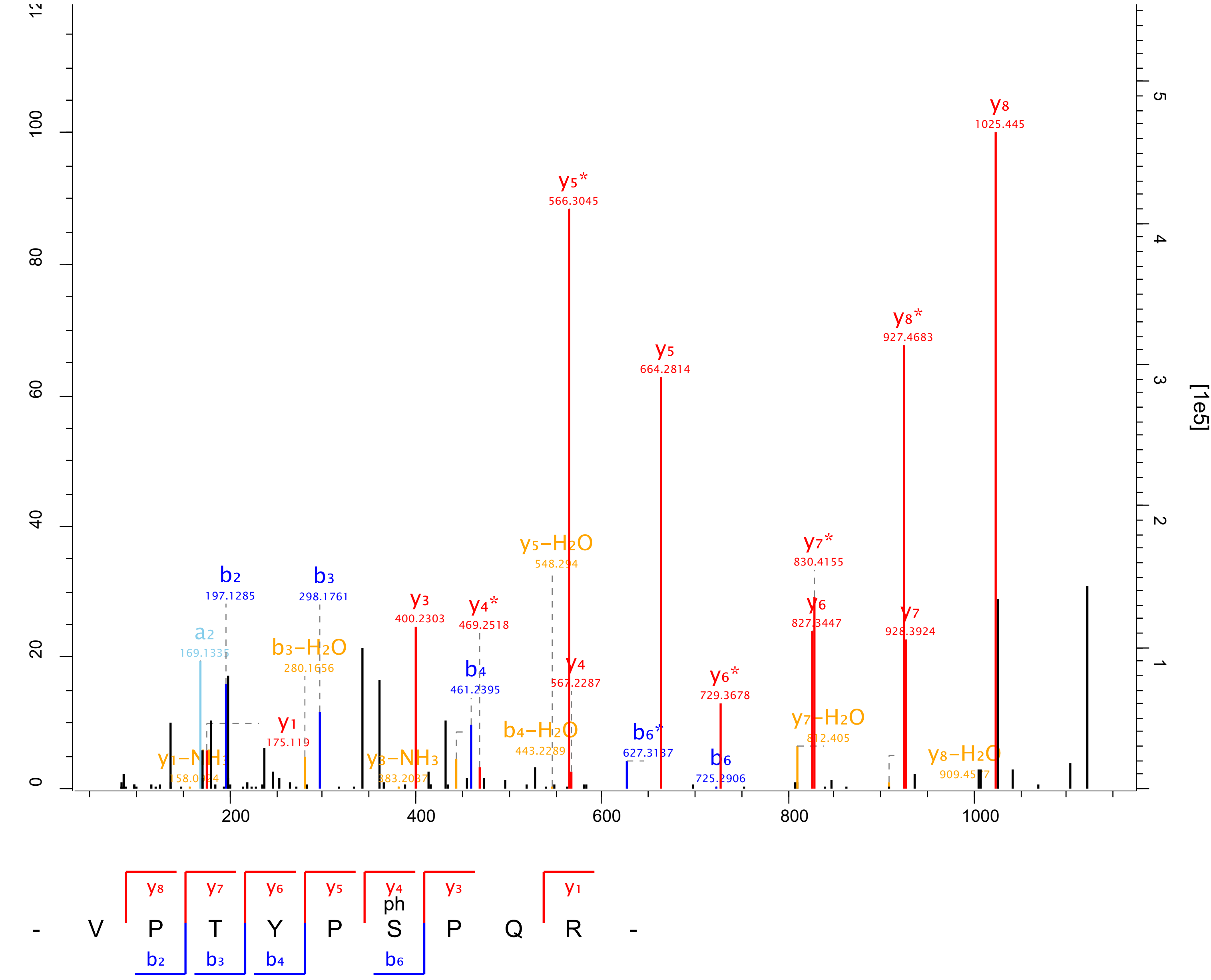

| Raw file | Scan | Method | Score | m/z |
|---|---|---|---|---|
| water_15_1 | 8707 | FTMS; HCD | 78.15 | 421.7 |

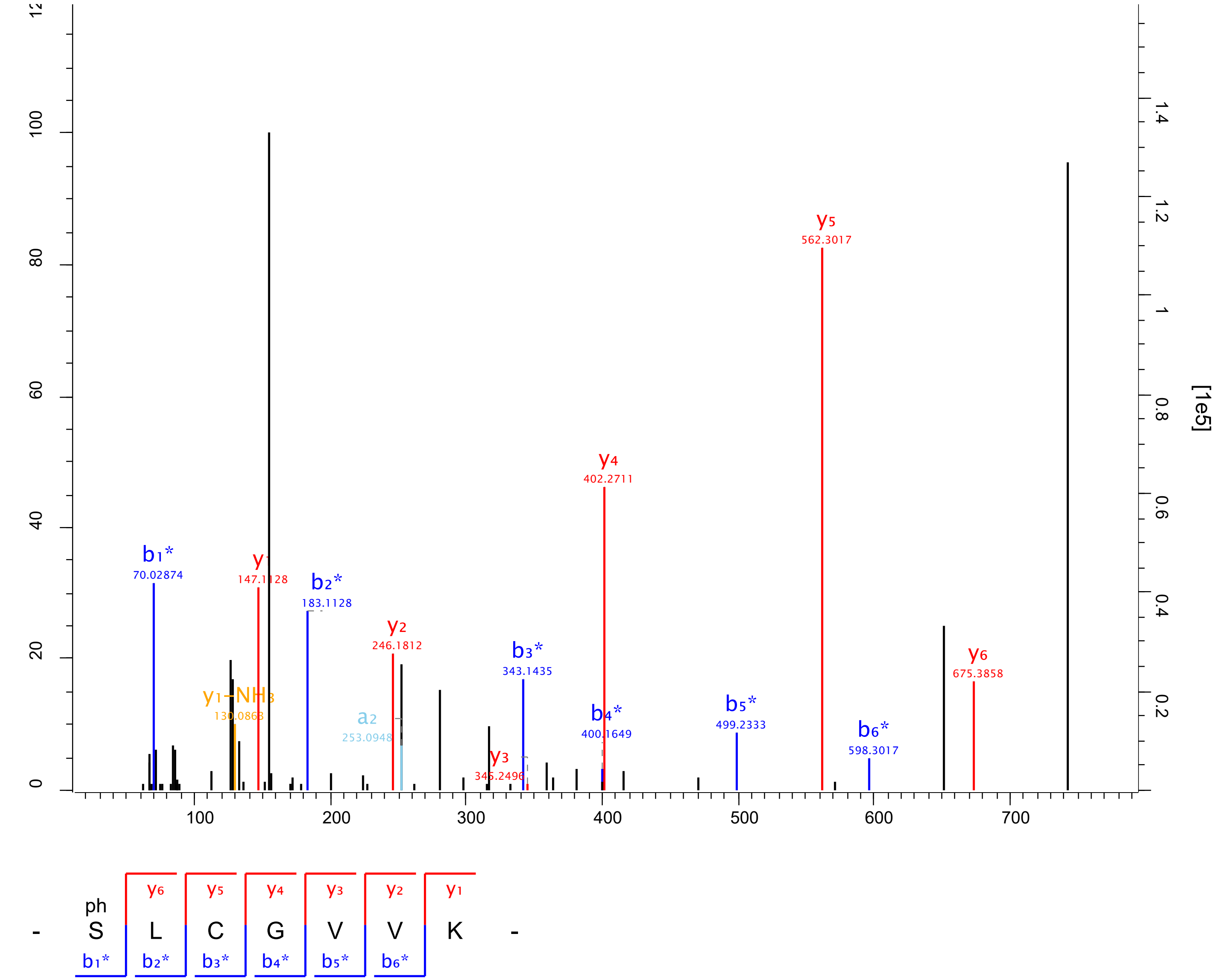

| Raw file | Scan | Method | Score | m/z |
|---|---|---|---|---|
| water_15_1 | 8778 | FTMS; HCD | 160.72 | 495.72 |

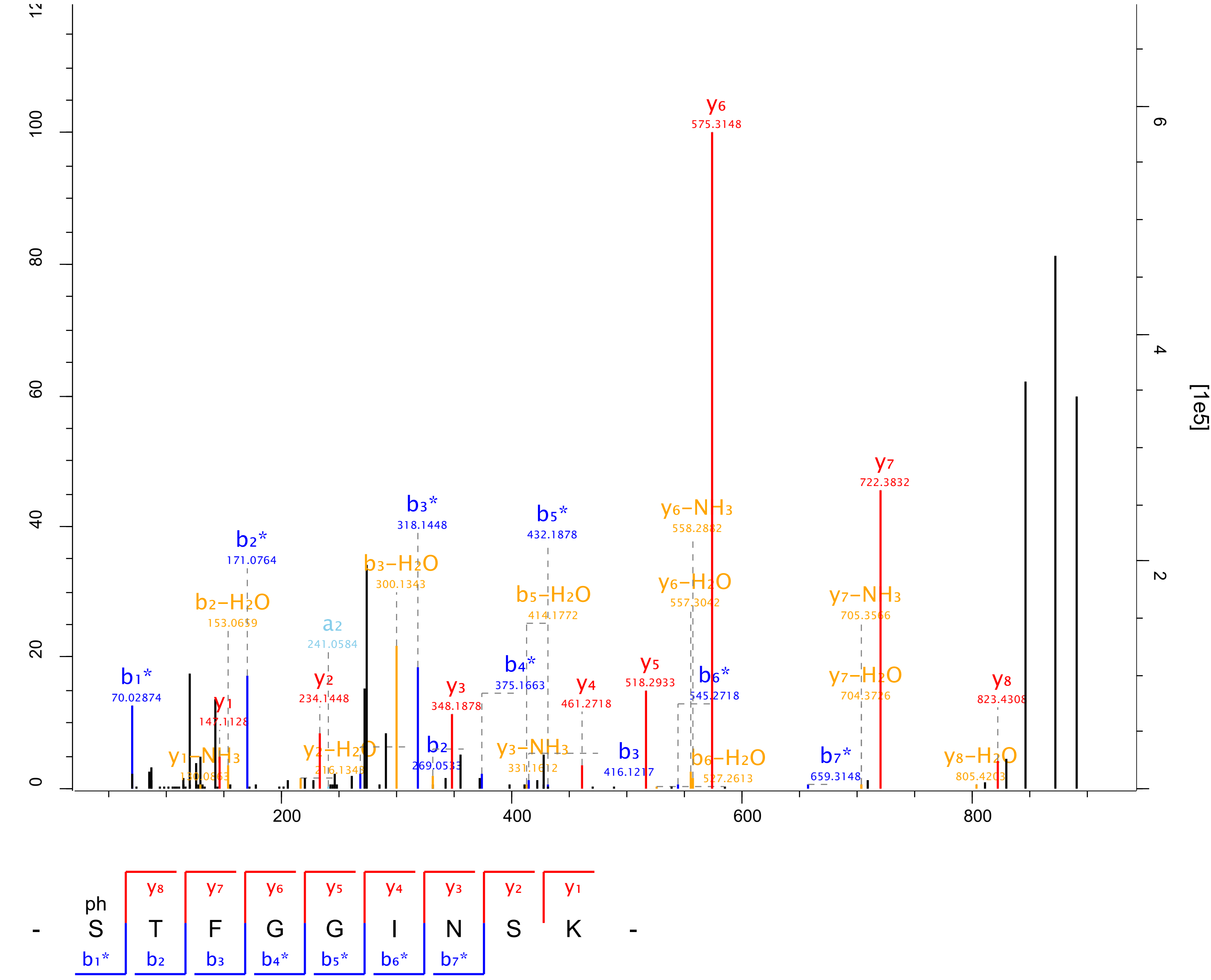

| Raw file | Scan | Method | Score | m/z |
|---|---|---|---|---|
| water_15_1 | 8917 | FTMS; HCD | 164.48 | 534.74 |

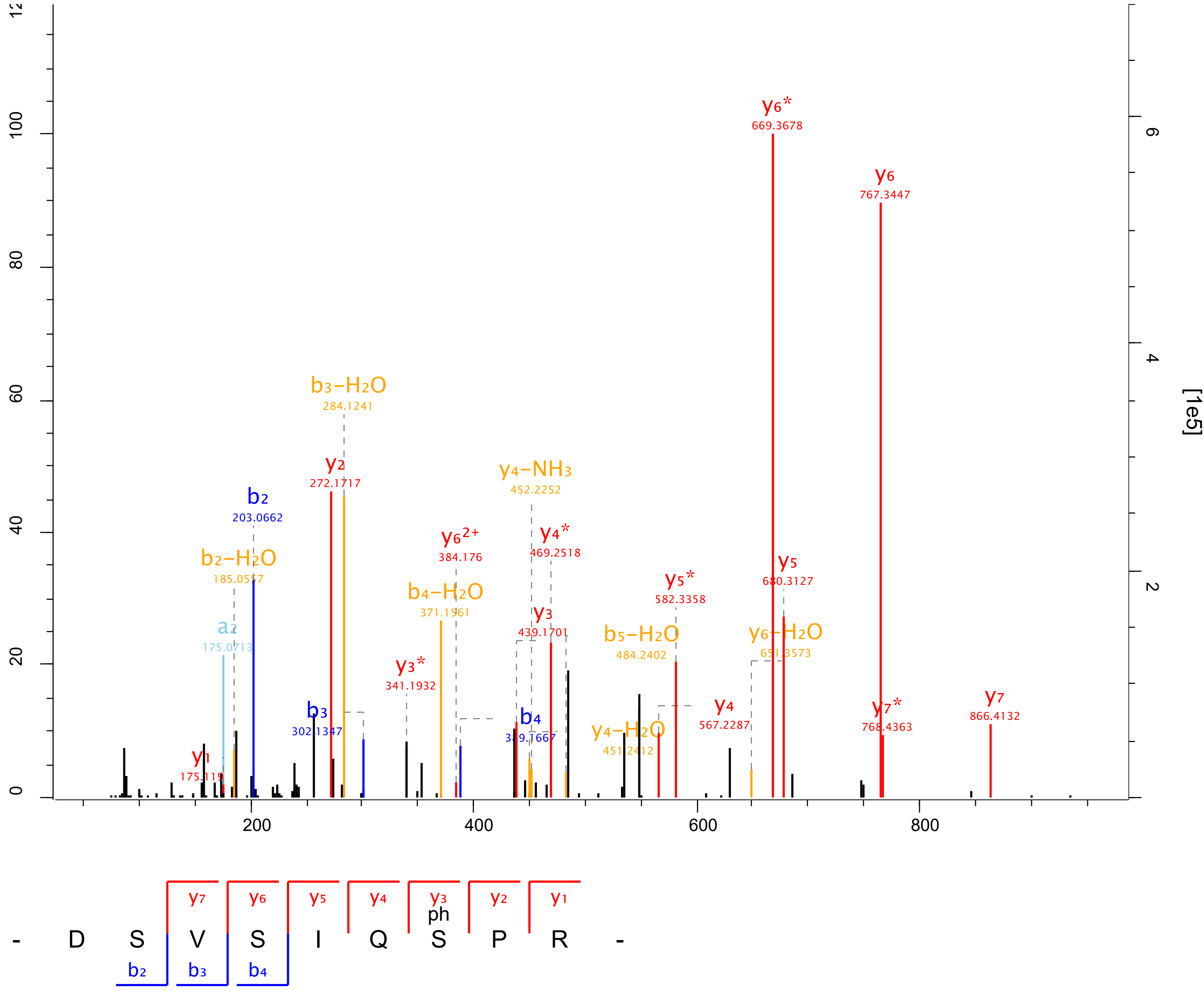

| Raw file | Scan | Method | Score | m/z |
|---|---|---|---|---|
| water_15_1 | 9315 | FTMS; HCD | 64.71 | 431.19 |

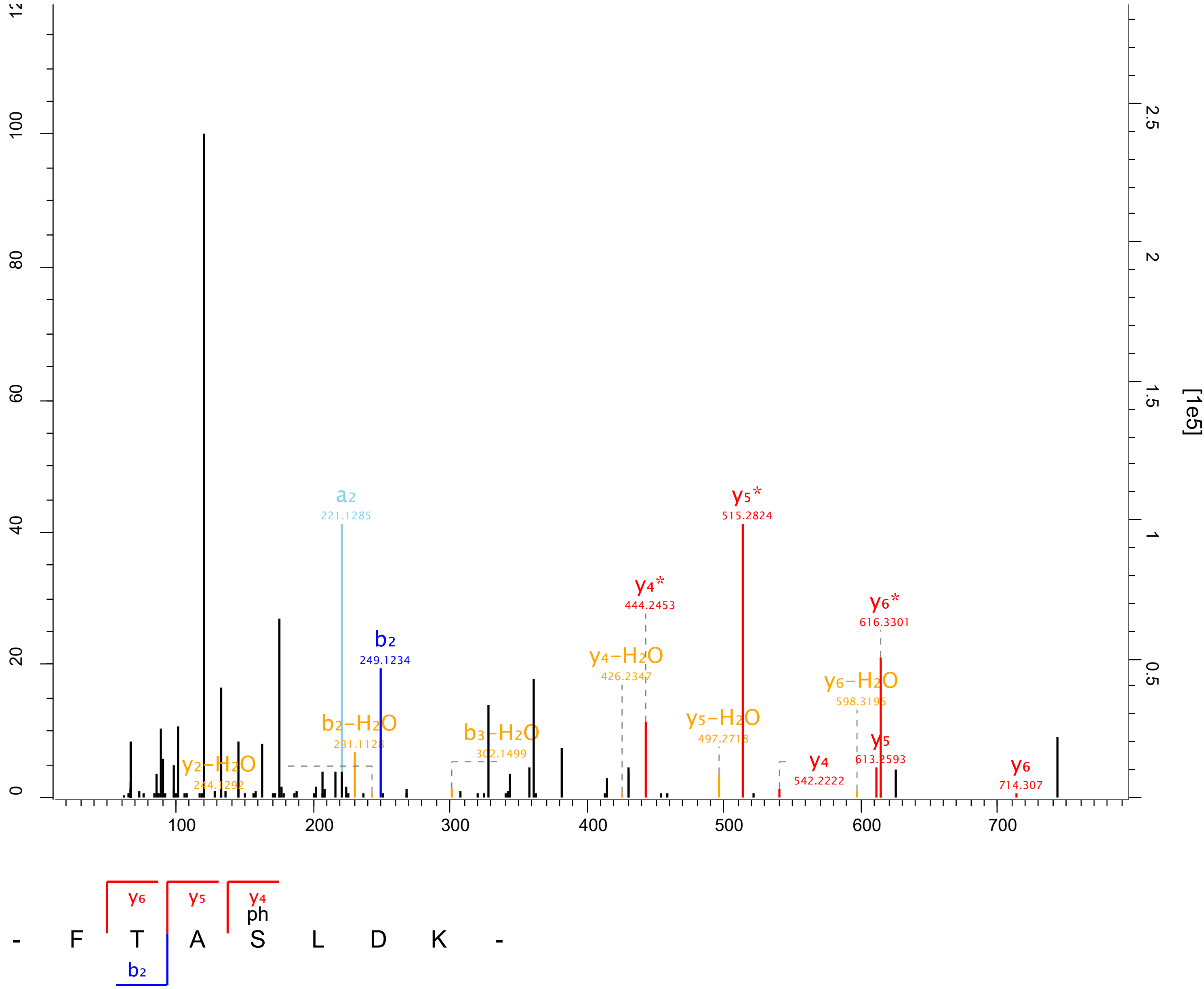

| Raw file | Scan | Method | Score | m/z |
|---|---|---|---|---|
| water_15_1 | 9811 | FTMS; HCD | 114.7 | 692.26 |

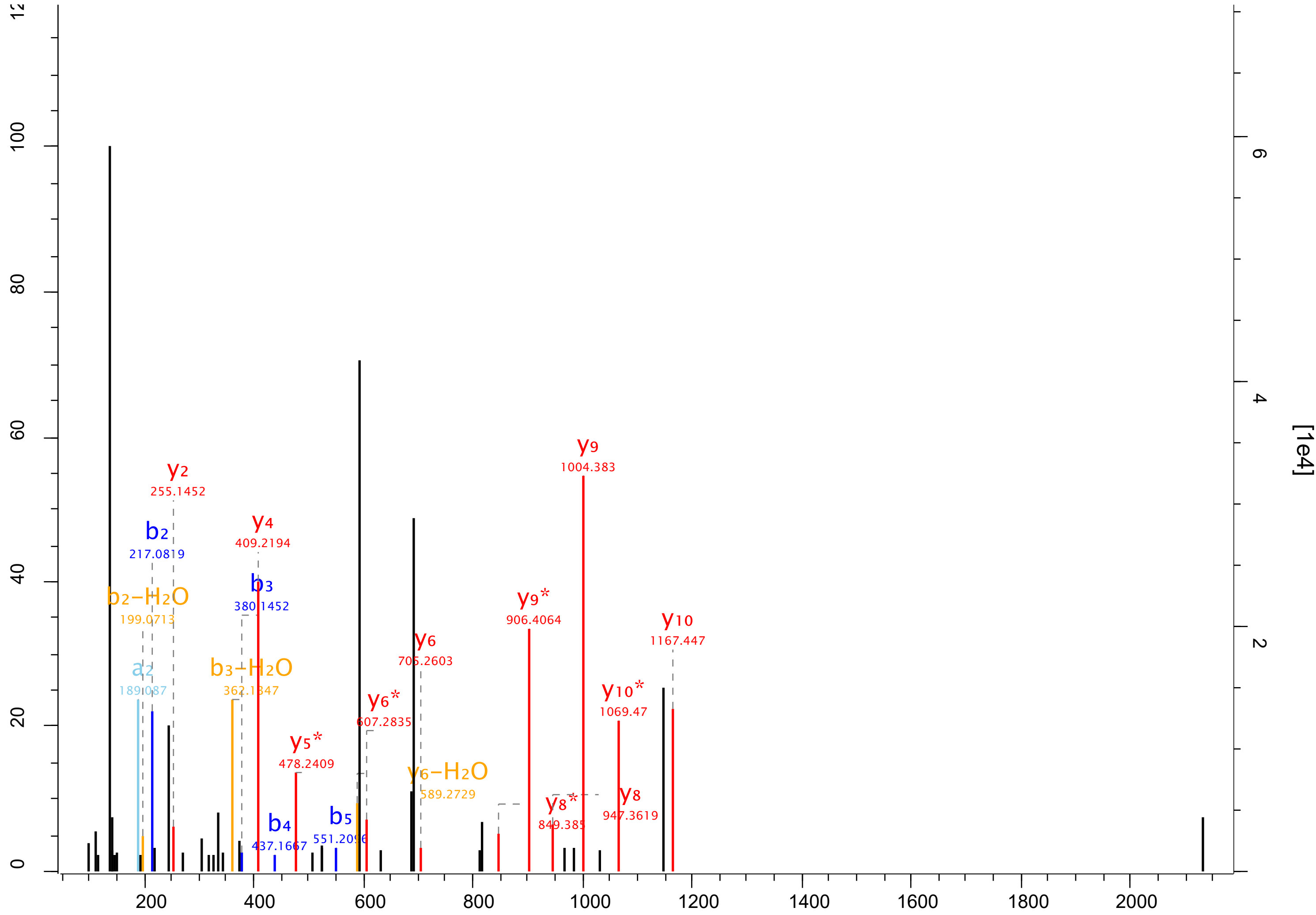


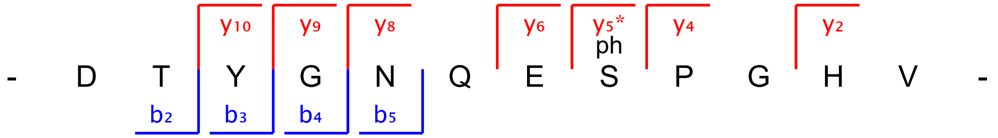

| Raw file | Scan | Method | Score | m/z |
| --- | --- | --- | --- | --- |
| water_15_1 | 9865 | FTMS; HCD | 51.45 | 420.21 |

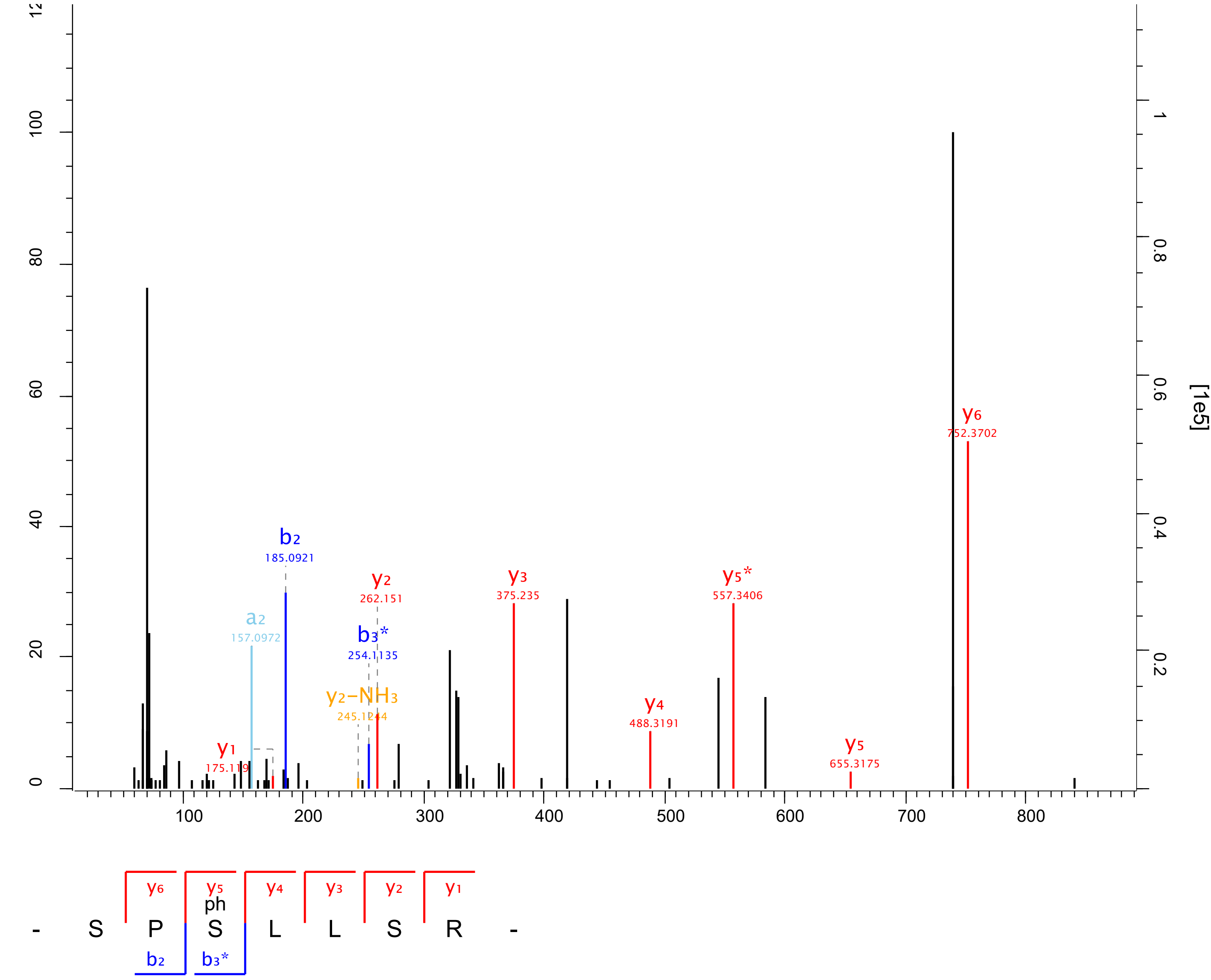

| Raw file | Scan | Method | Score | m/z |
| --- | --- | --- | --- | --- |
| water_15_1 | 9884 | FTMS; HCD | 128.74 | 548.28 |

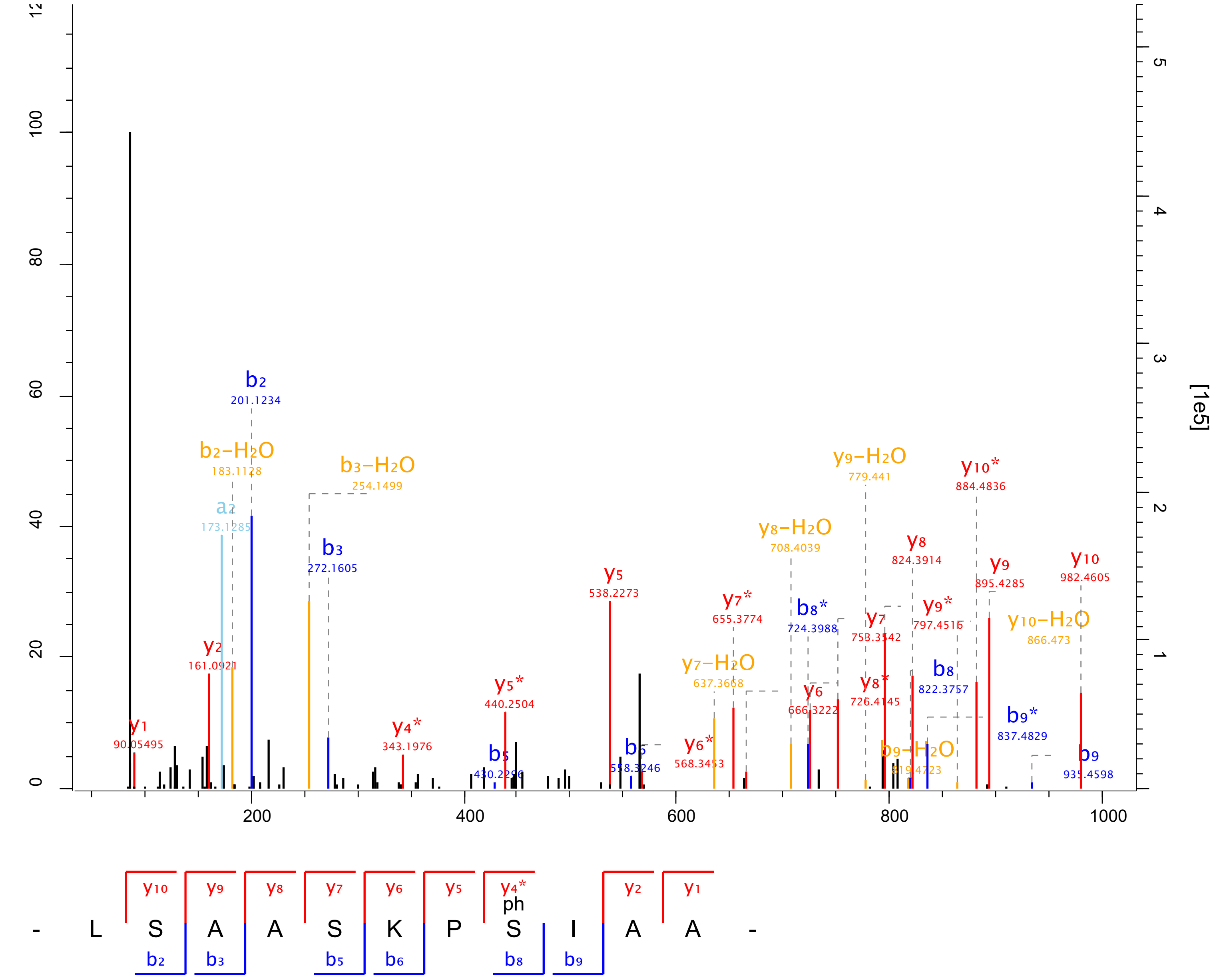

| Raw file | Scan | Method | Score | m/z |
|---|---|---|---|---|
| water_15_2 | 10064 | FTMS; HCD | 112.34 | 676.8 |

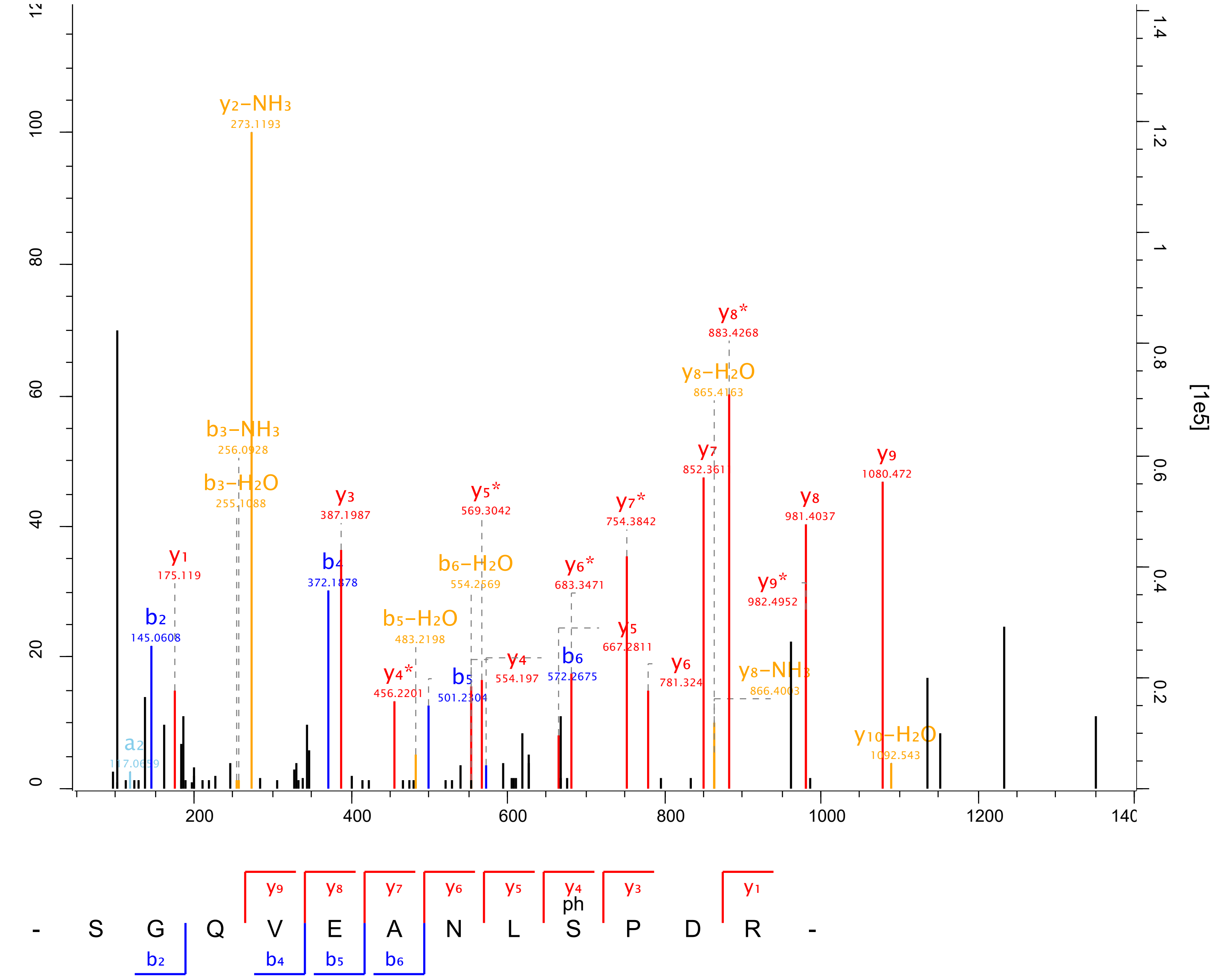

| Raw file | Scan | Method | Score | m/z |
| --- | --- | --- | --- | --- |
| water_15_2 | 10075 | FTMS; HCD | 94.57 | 753.8 |

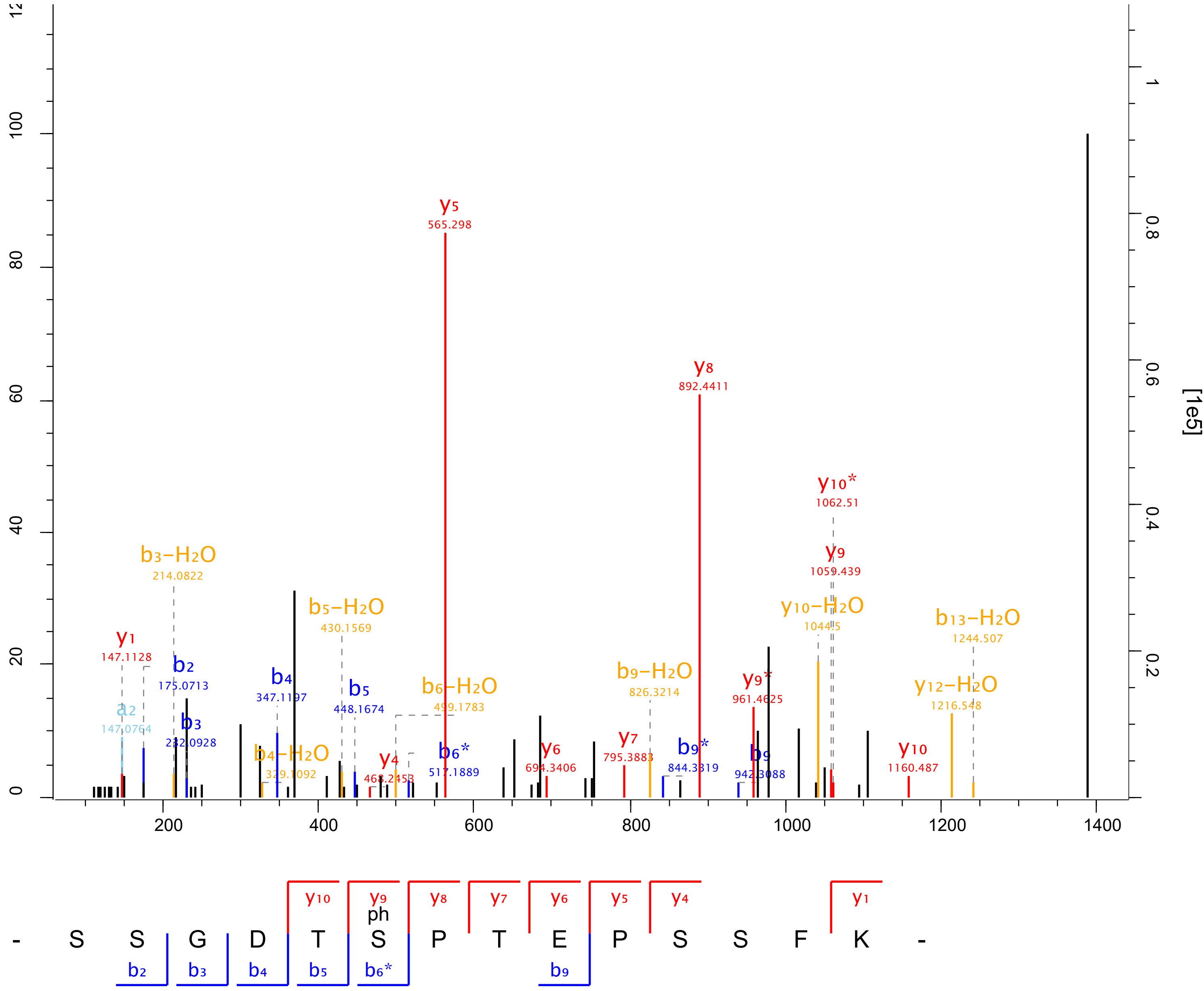

| Raw file | Scan | Method | Score | m/z |
|---|---|---|---|---|
| water_15_2 | 10087 | FTMS; HCD | 216.42 | 440.72 |

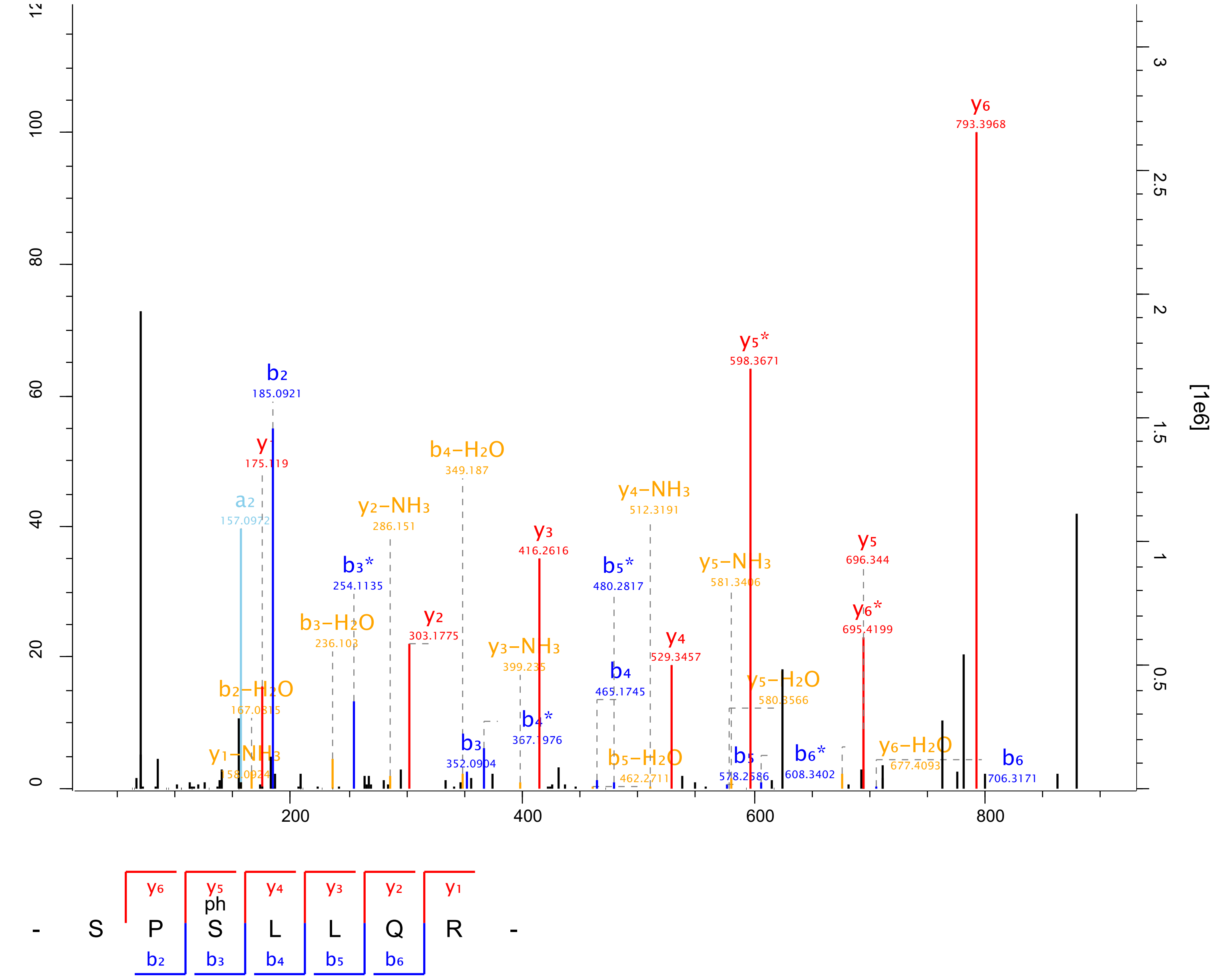

| Raw file | Scan | Method | Score | m/z |
|---|---|---|---|---|
| water_15_2 | 10101 | FTMS; HCD | 82.07 | 573.76 |

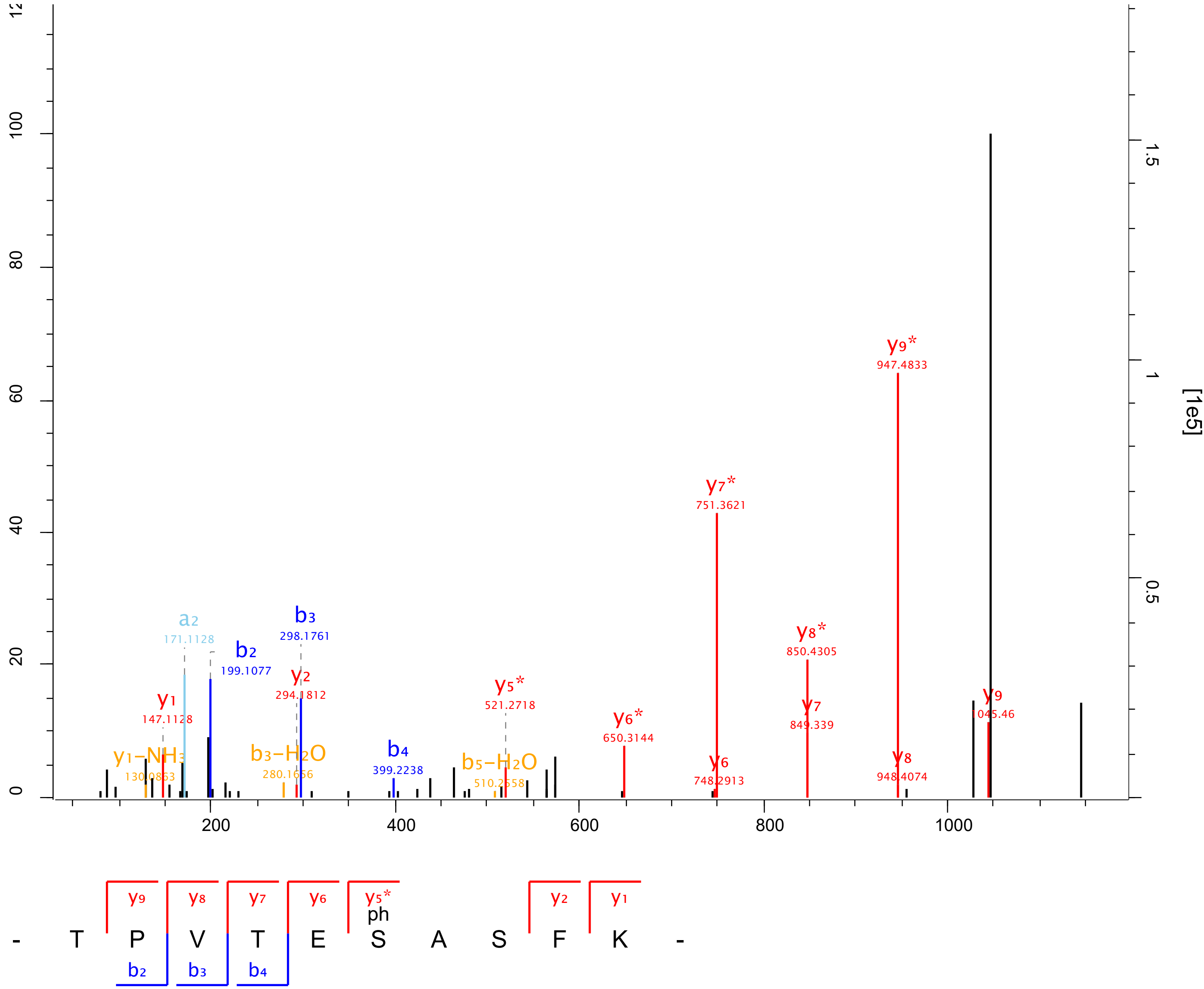

| Raw file | Scan | Method | Score | m/z |
|---|---|---|---|---|
| water_15_2 | 10177 | FTMS; HCD | 58.08 | 671.78 |

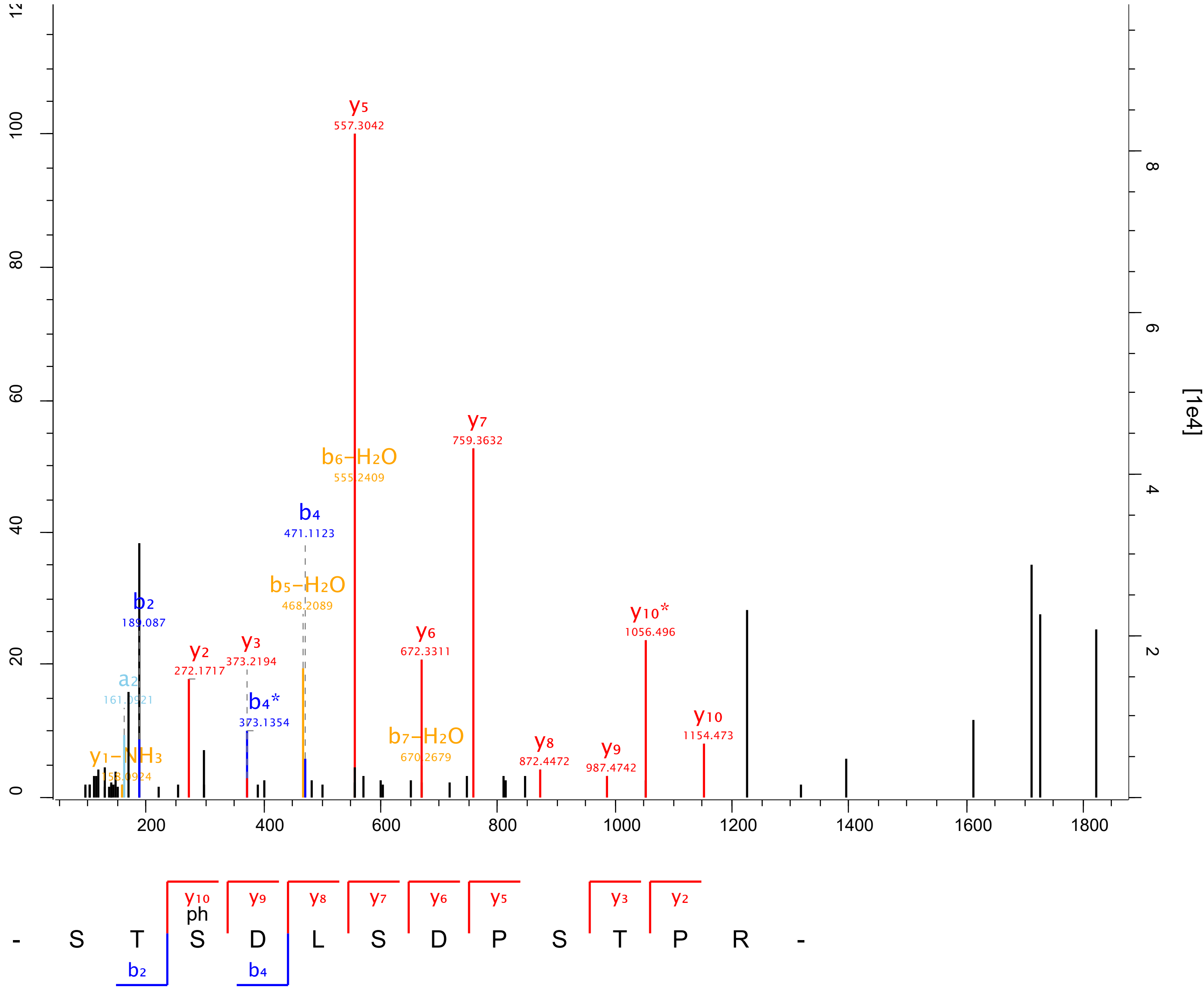

| Raw file | Scan | Method | Score | m/z |
| --- | --- | --- | --- | --- |
| water_15_2 | 10187 | FTMS; HCD | 71.45 | 427.21 |

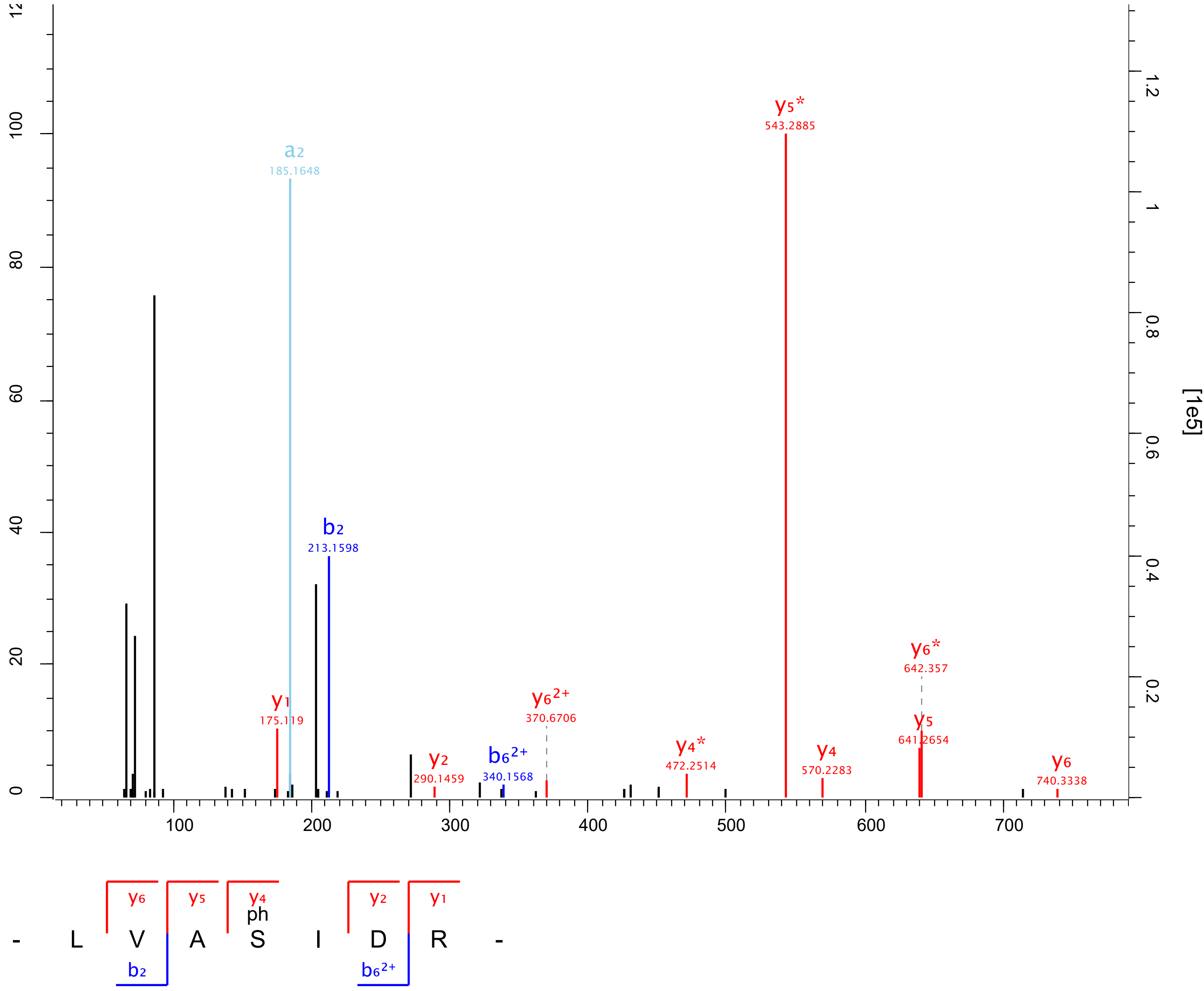

| Raw file | Scan | Method | Score | m/z |
|---|---|---|---|---|
| water_15_2 | 10277 | FTMS; HCD | 60.16 | 495.22 |

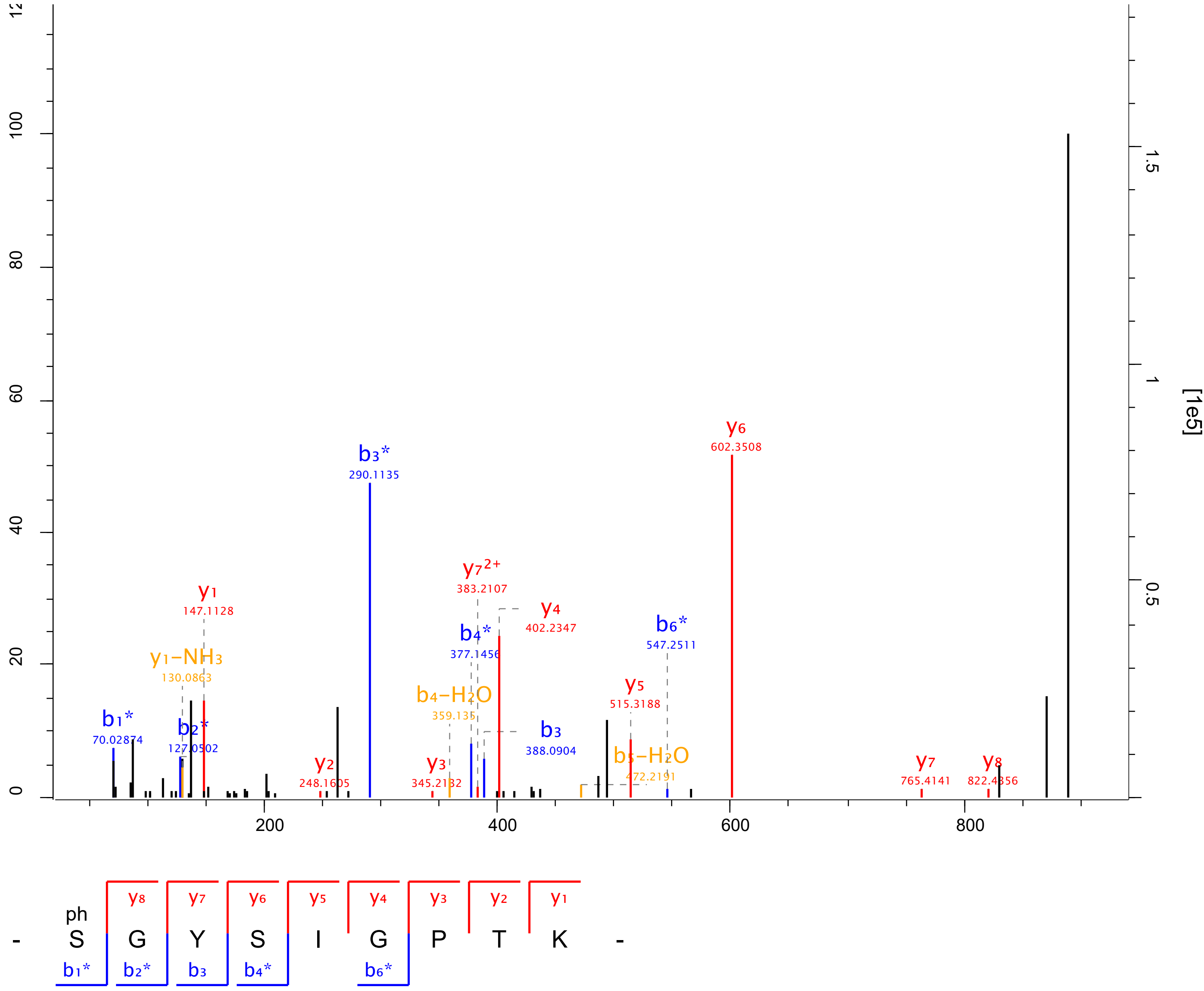

| Raw file | Scan | Method | Score | m/z |
| --- | --- | --- | --- | --- |
| water_15_2 | 10322 | FTMS; HCD | 174.47 | 609.27 |

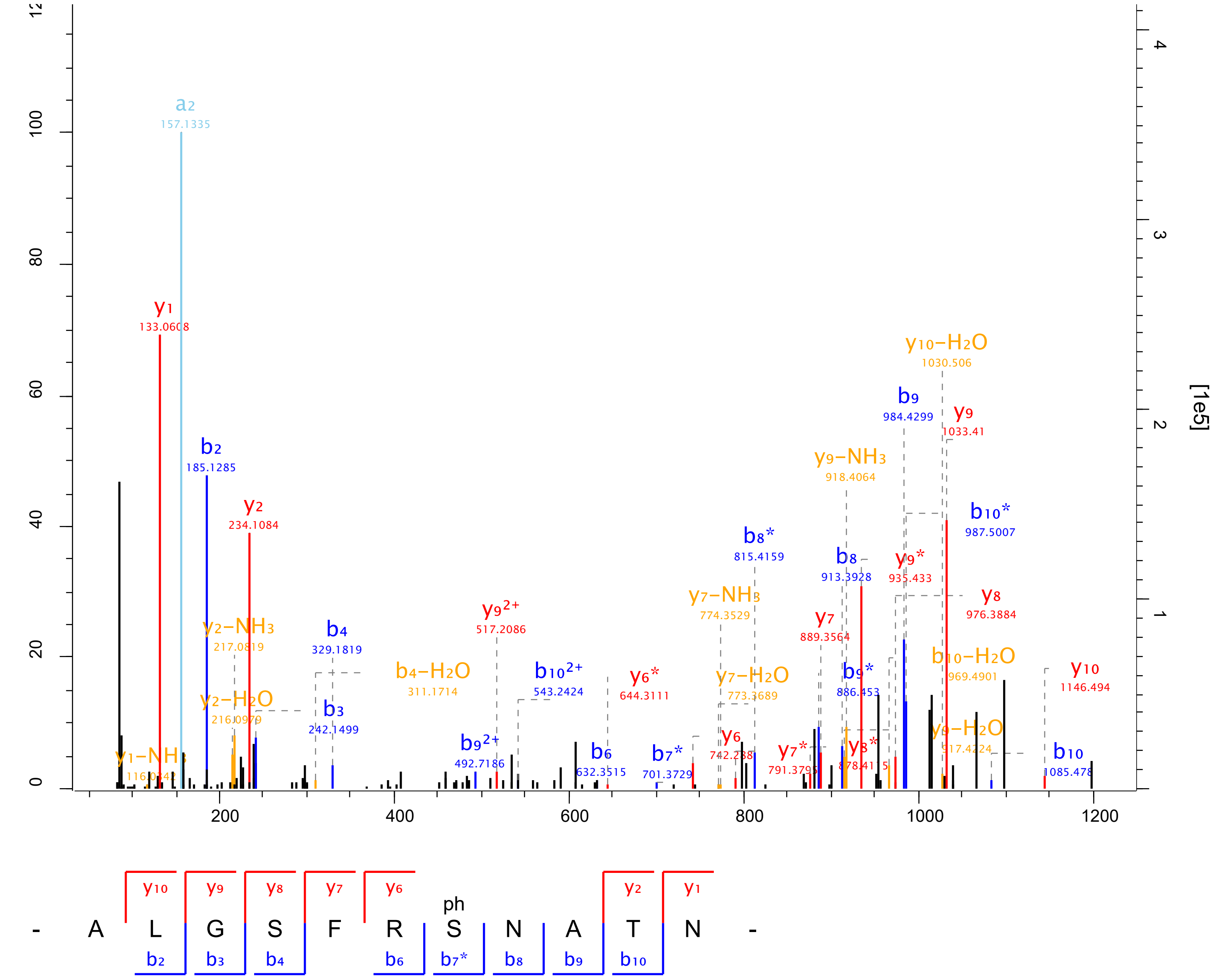

| Raw file | Scan | Method | Score | m/z |
|---|---|---|---|---|
| water_15_2 | 10353 | FTMS; HCD | 108.38 | 681.81 |

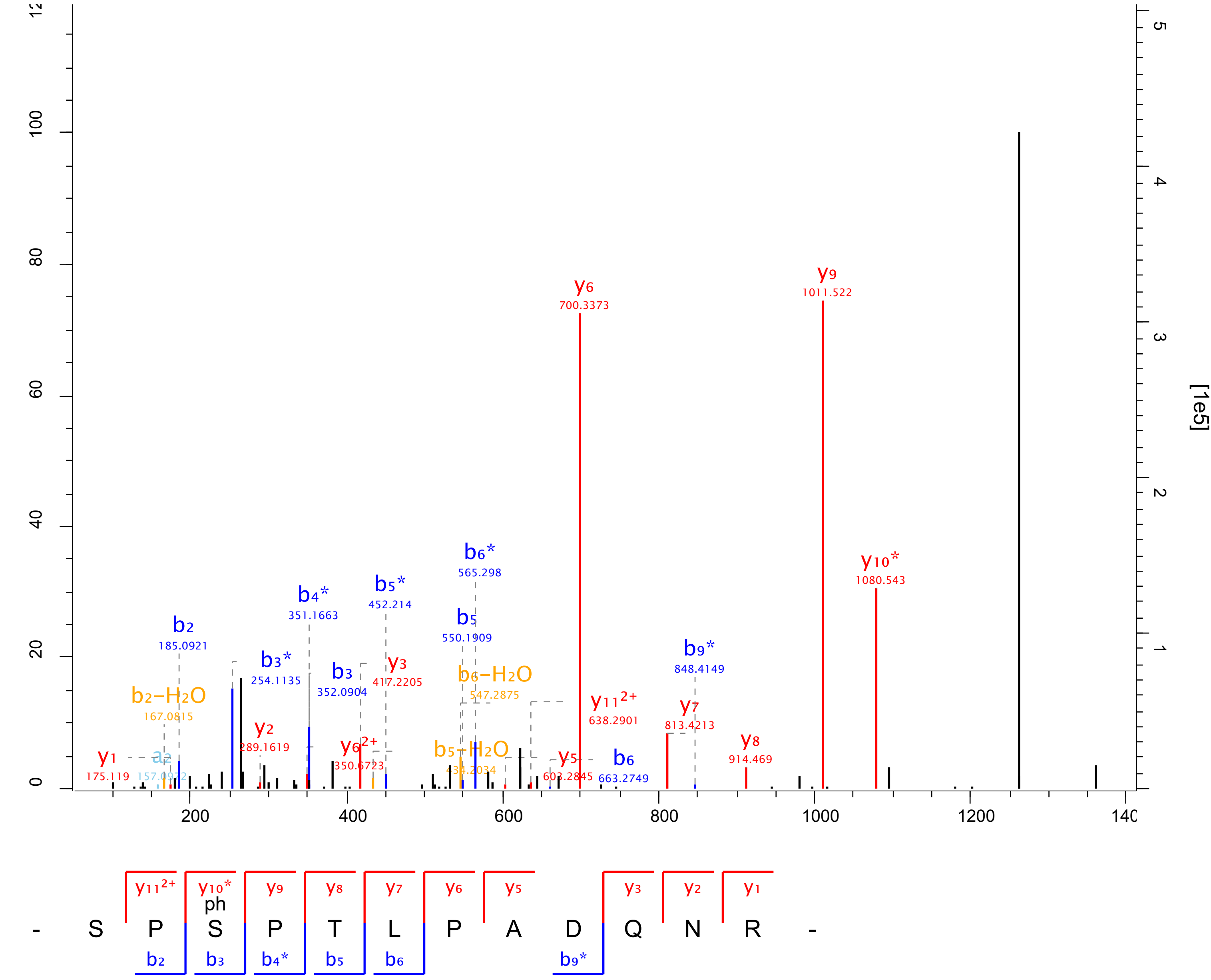

| Raw file | Scan | Method | Score | m/z |
| --- | --- | --- | --- | --- |
| water_15_2 | 10368 | FTMS; HCD | 110.74 | 753.82 |

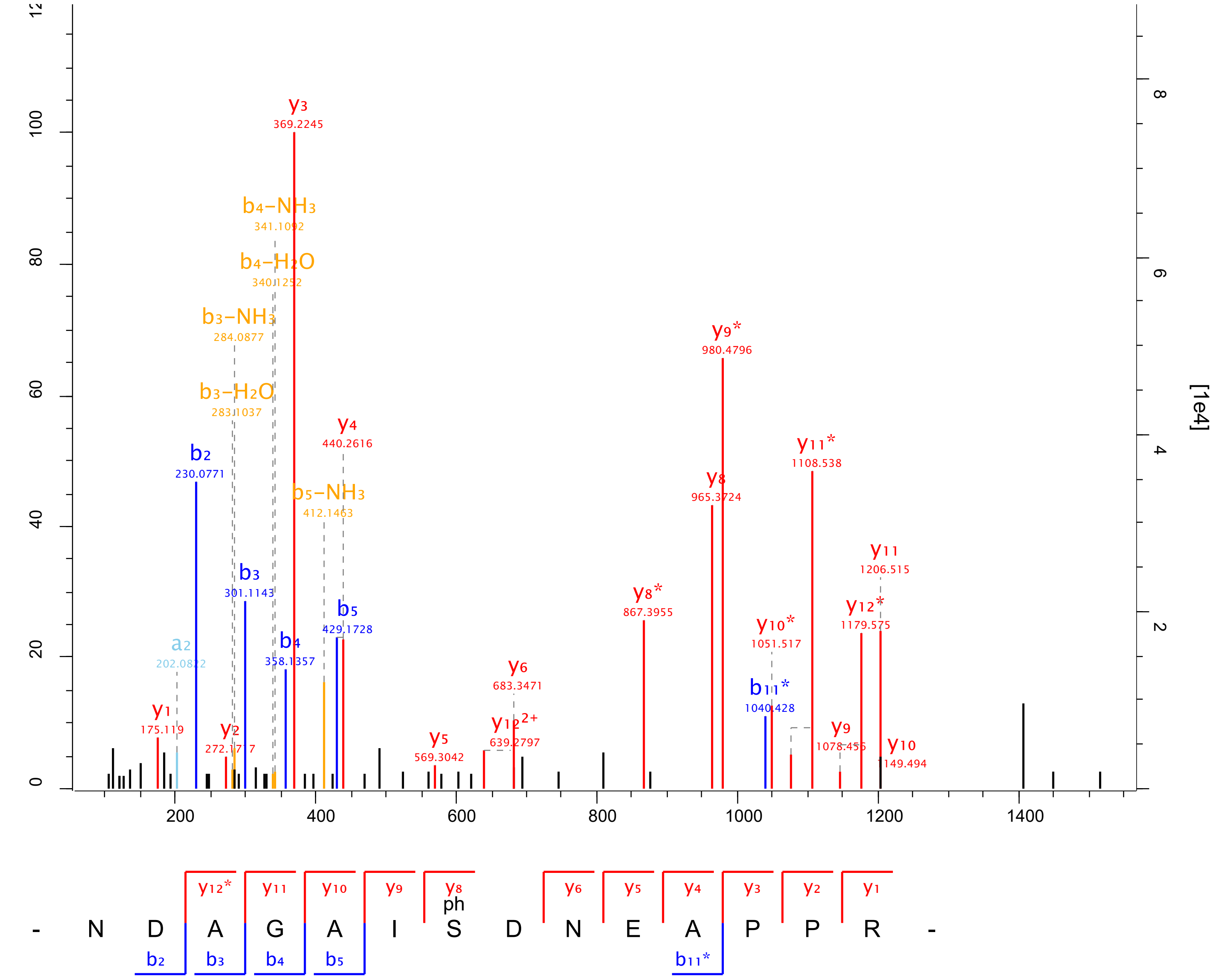

| Raw file | Scan | Method | Score | m/z |
|---|---|---|---|---|
| water_15_2 | 10395 | FTMS; HCD | 40.35 | 629.75 |

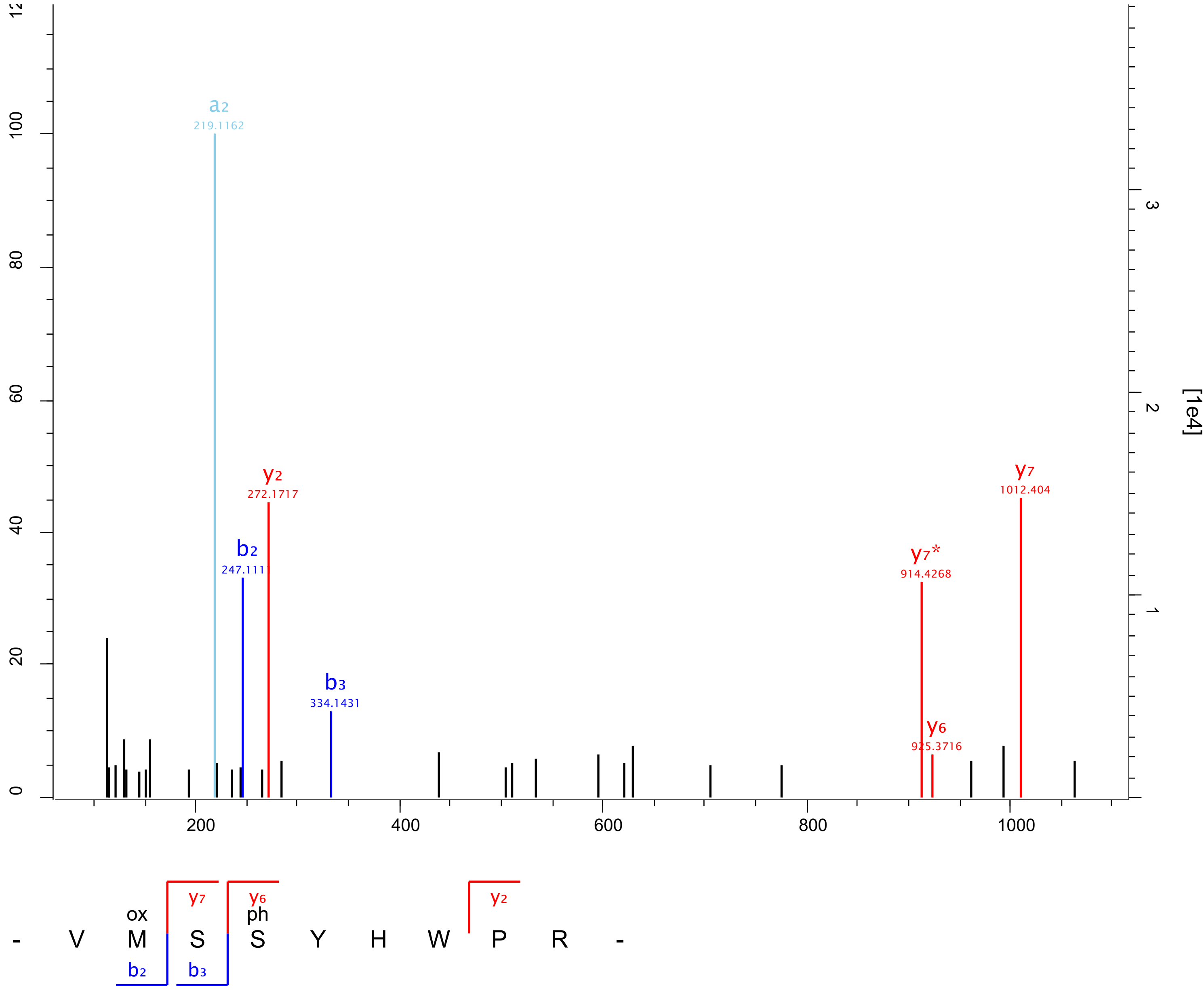

| Raw file | Scan | Method | Score | m/z |
|---|---|---|---|---|
| water_15_2 | 10440 | FTMS; HCD | 121.45 | 636.78 |

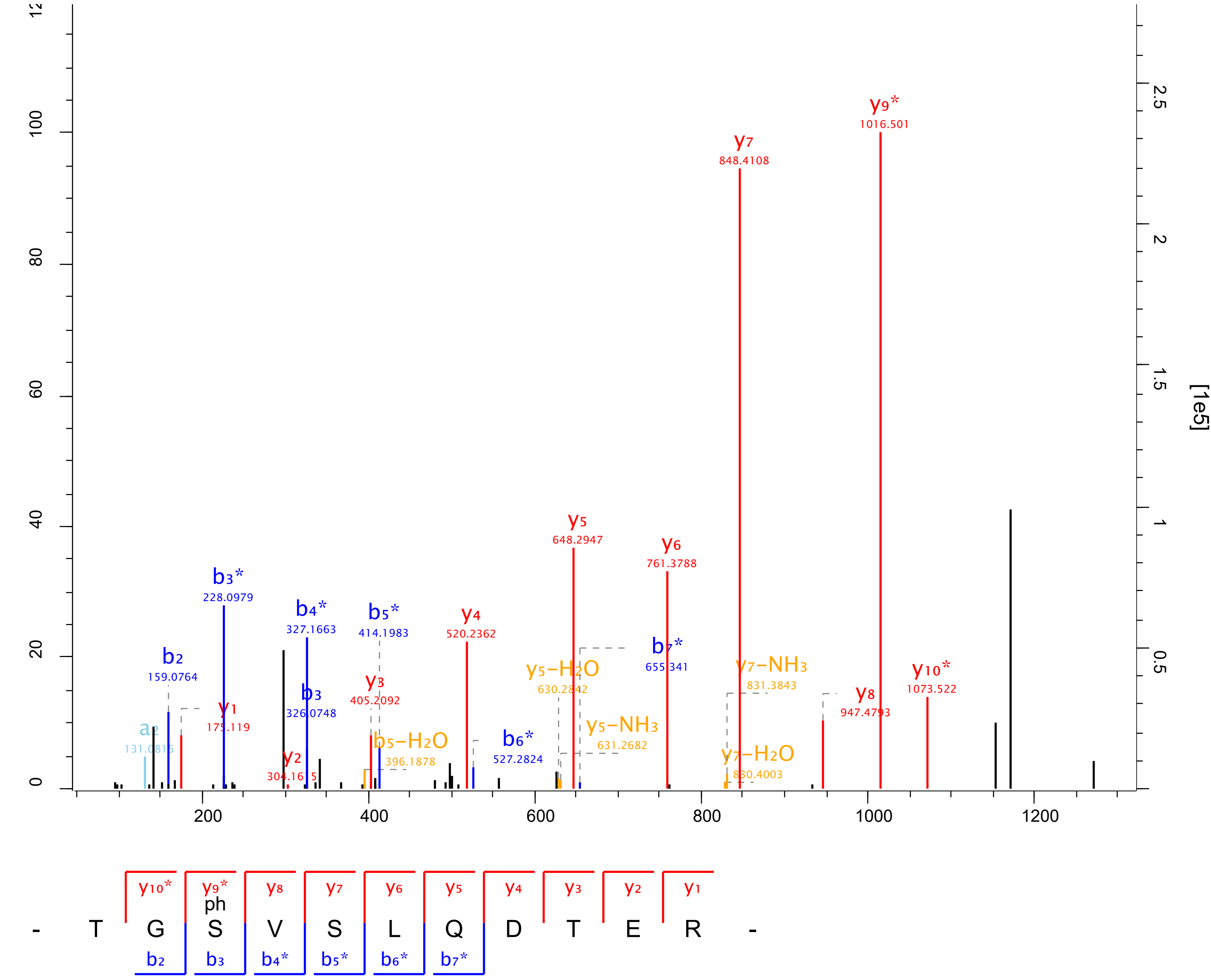

| Raw file | Scan | Method | Score | m/z |
| --- | --- | --- | --- | --- |
| water_15_2 | 10468 | FTMS; HCD | 99.75 | 524.22 |

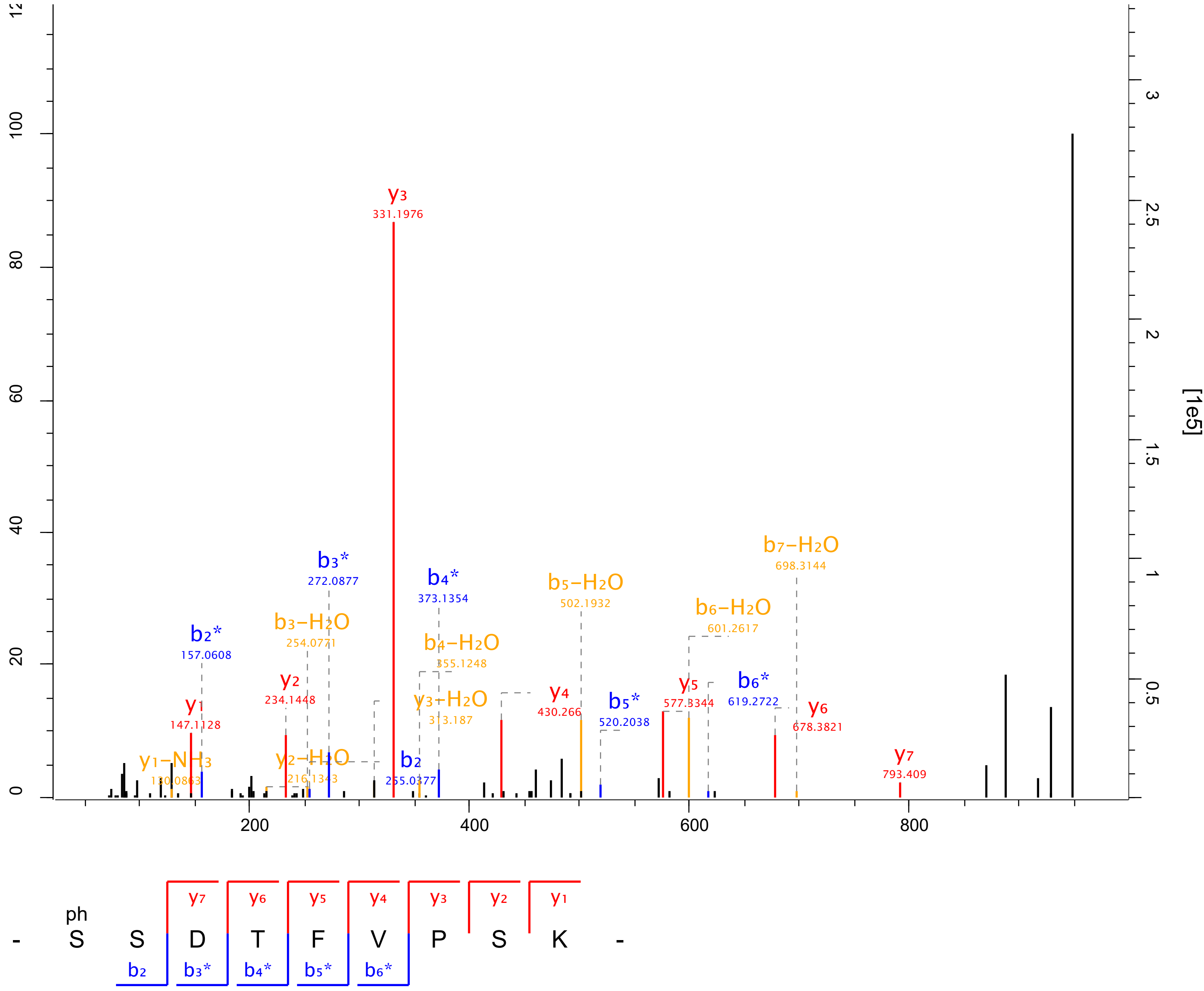

| Raw file | Scan | Method | Score | m/z |
| --- | --- | --- | --- | --- |
| water_15_2 | 10480 | FTMS; HCD | 42.79 | 723.66 |

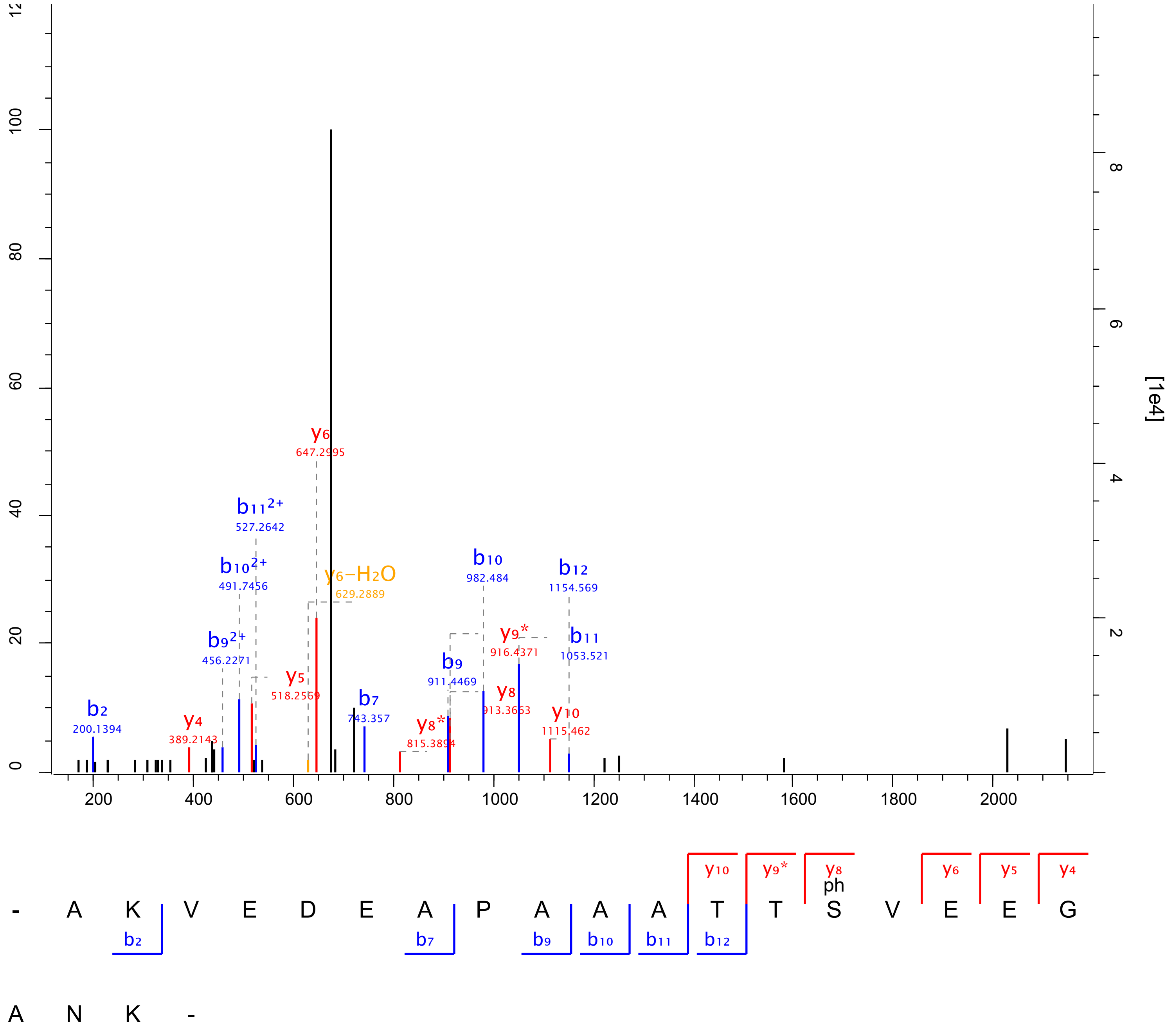

| Raw file | Scan | Method | Score | m/z |
|---|---|---|---|---|
| water_15_2 | 10637 | FTMS; HCD | 221.51 | 877.38 |

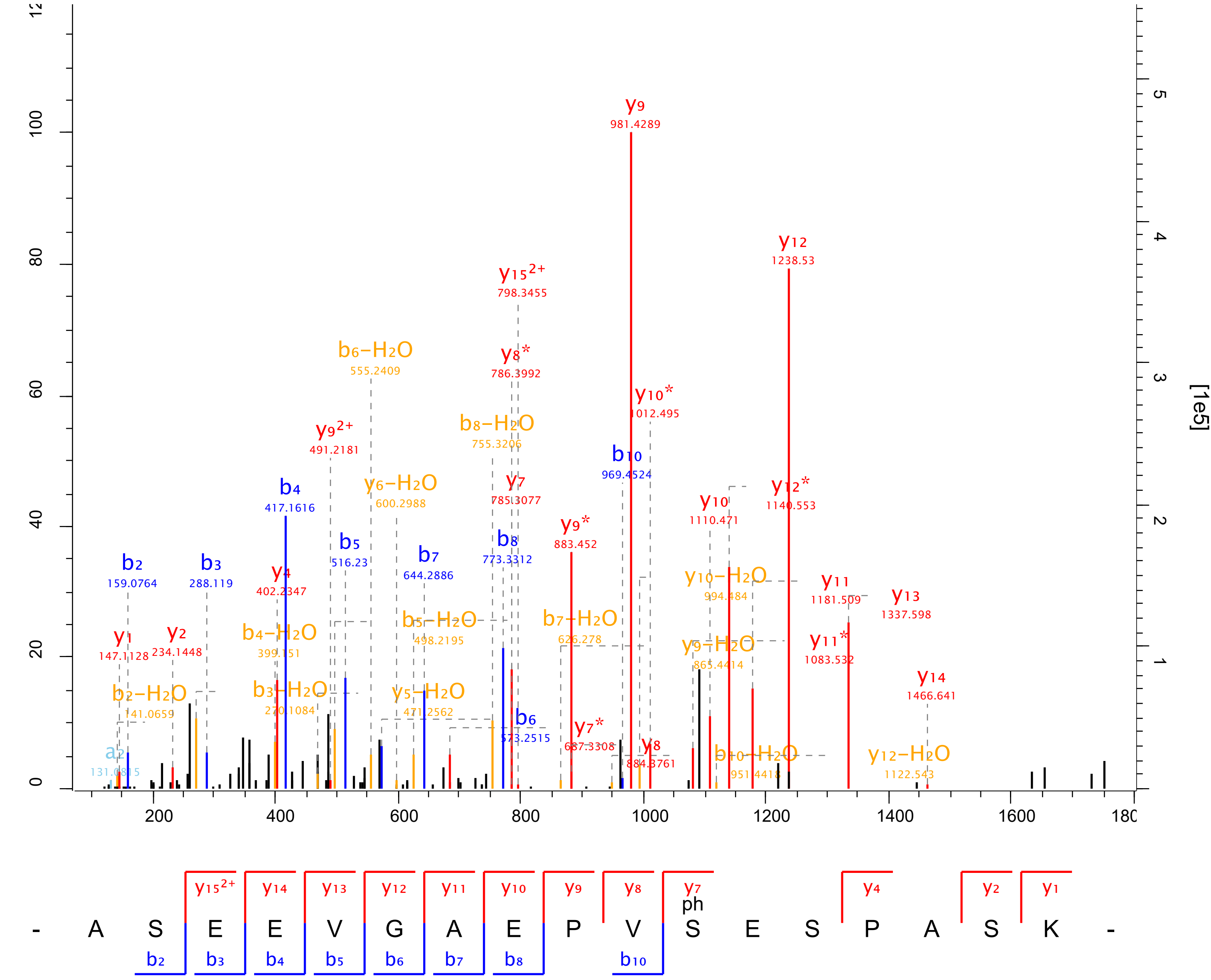

| Raw file | Scan | Method | Score | m/z |
| --- | --- | --- | --- | --- |
| water_15_2 | 10643 | FTMS; HCD | 85.74 | 788.85 |

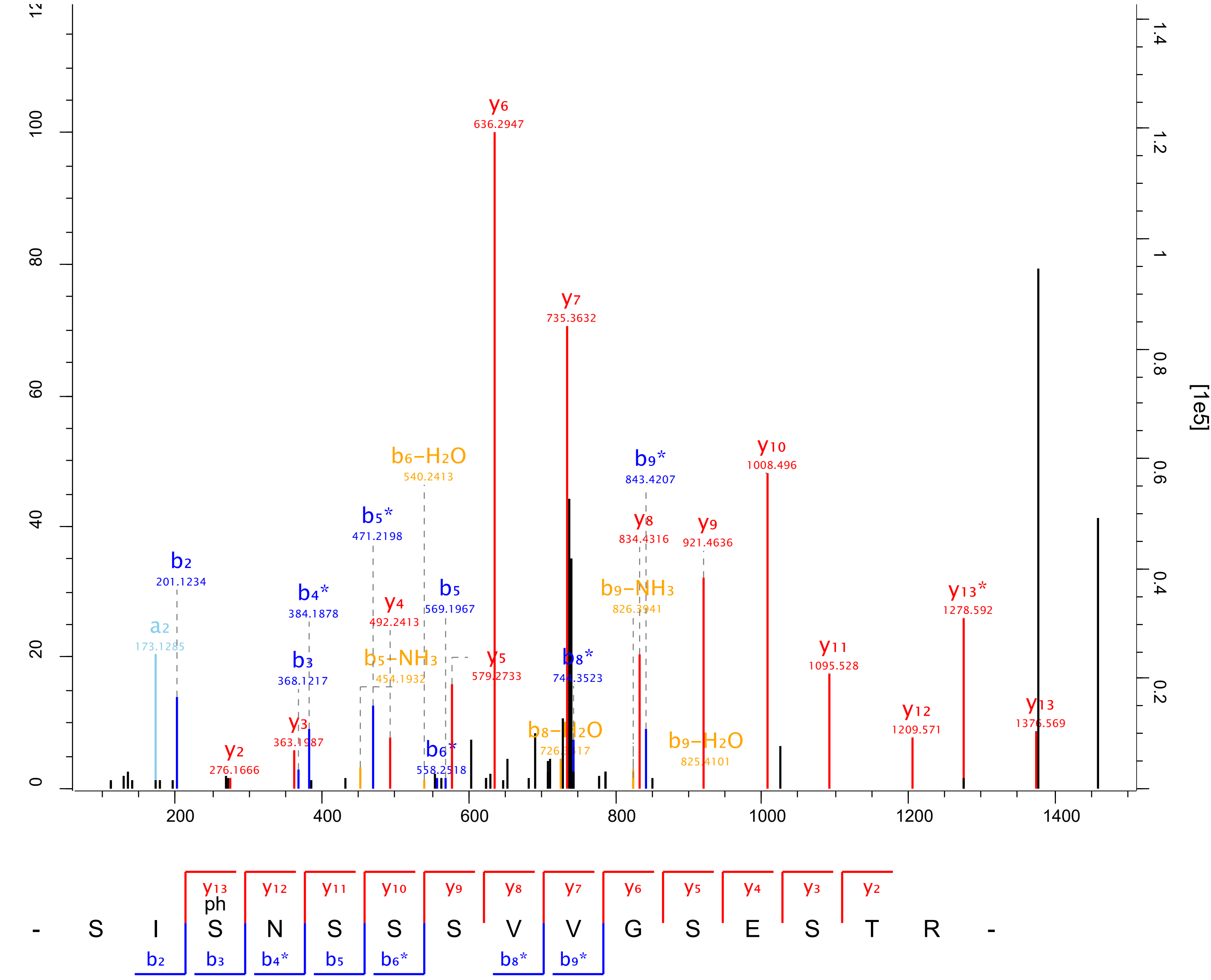

| Raw file | Scan | Method | Score | m/z |
|---|---|---|---|---|
| water_15_2 | 10651 | FTMS; HCD | 201.3 | 600.28 |

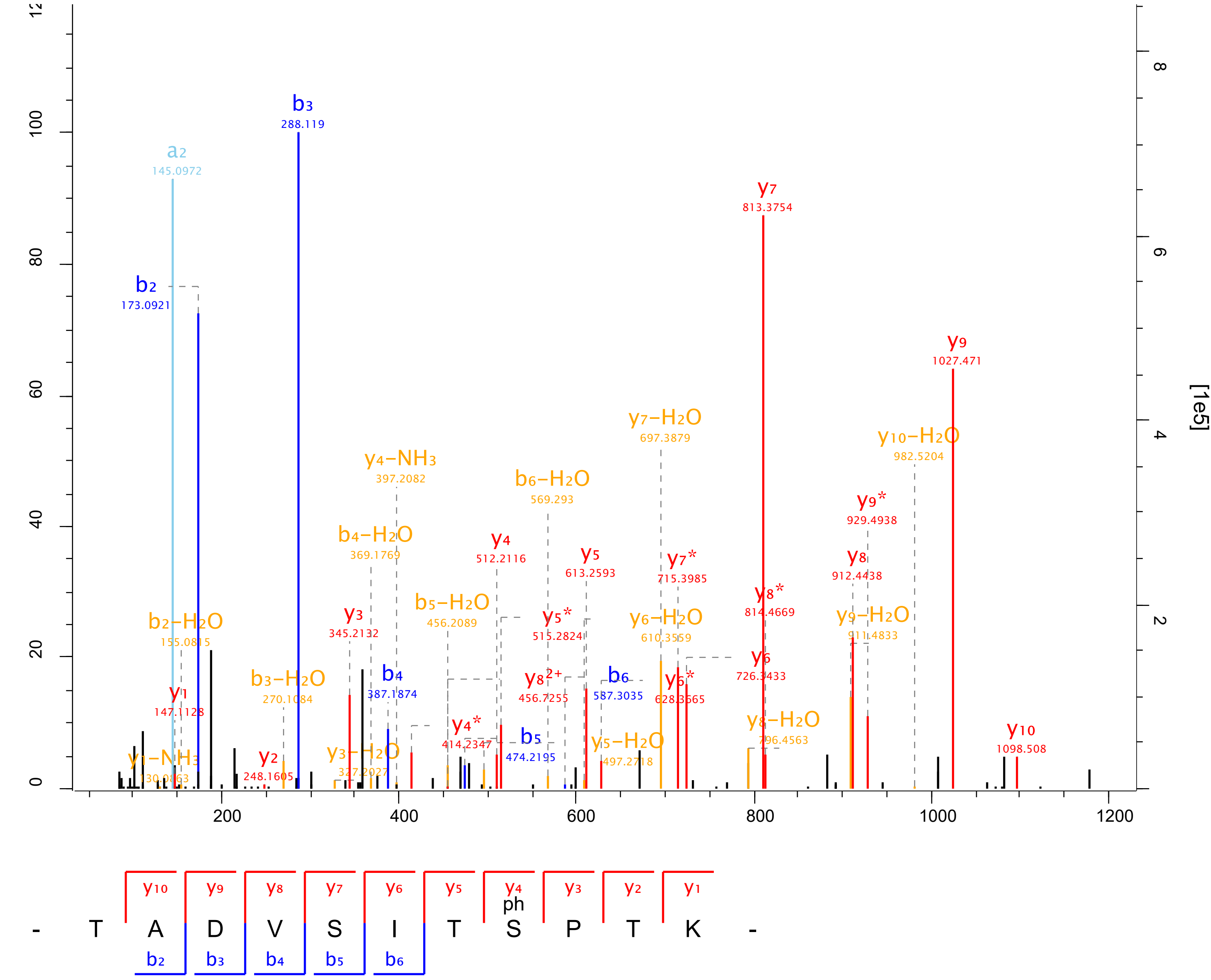

| Raw file | Scan | Method | Score | m/z |
| --- | --- | --- | --- | --- |
| water_15_2 | 10662 | FTMS; HCD | 60.26 | 647.31 |

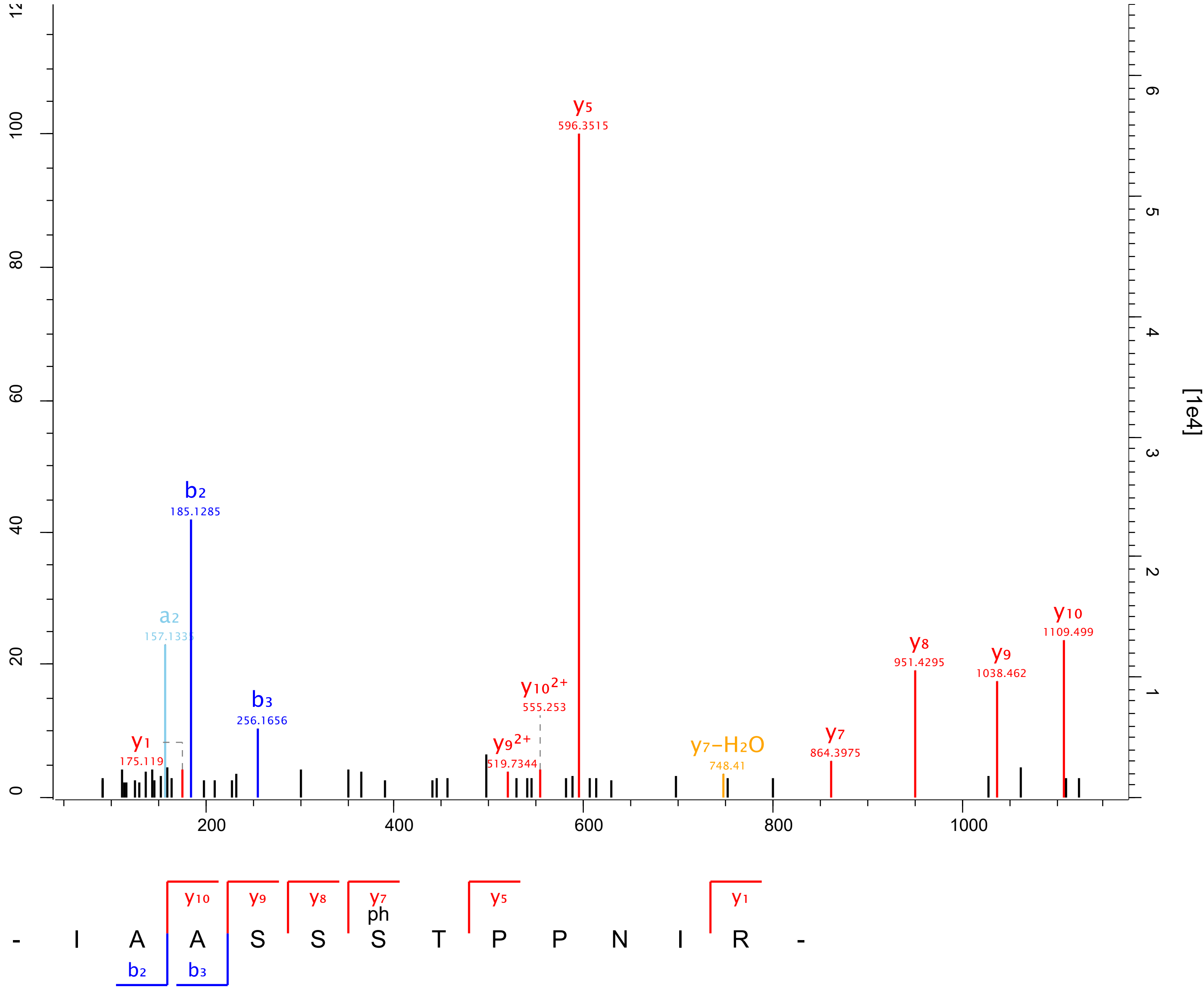

| Raw file | Scan | Method | Score | m/z |
|---|---|---|---|---|
| water_15_2 | 10666 | FTMS; HCD | 42.95 | 668.26 |

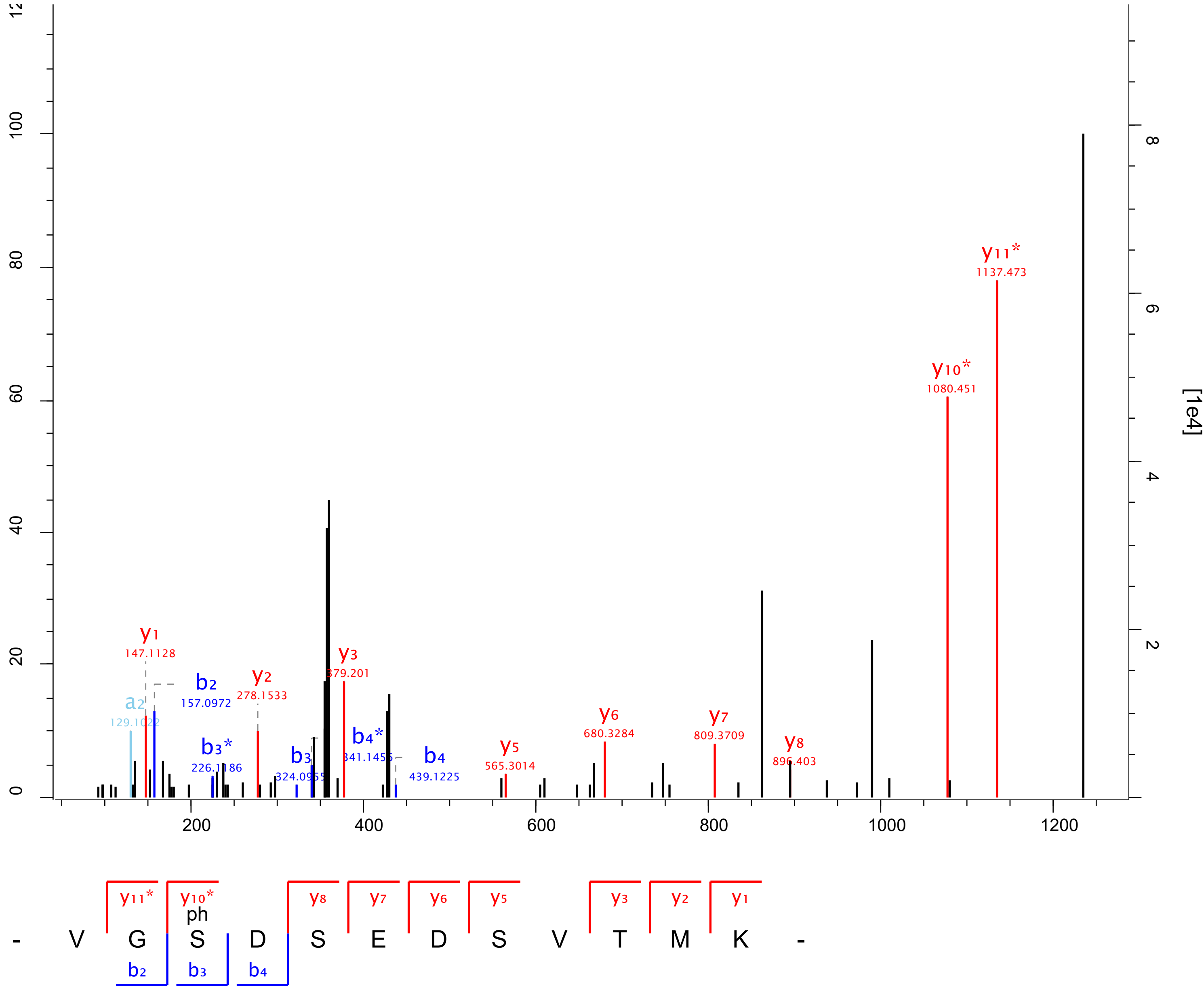

| Raw file | Scan | Method | Score | m/z |
| --- | --- | --- | --- | --- |
| water_15_2 | 10677 | FTMS; HCD | 63.47 | 520.23 |

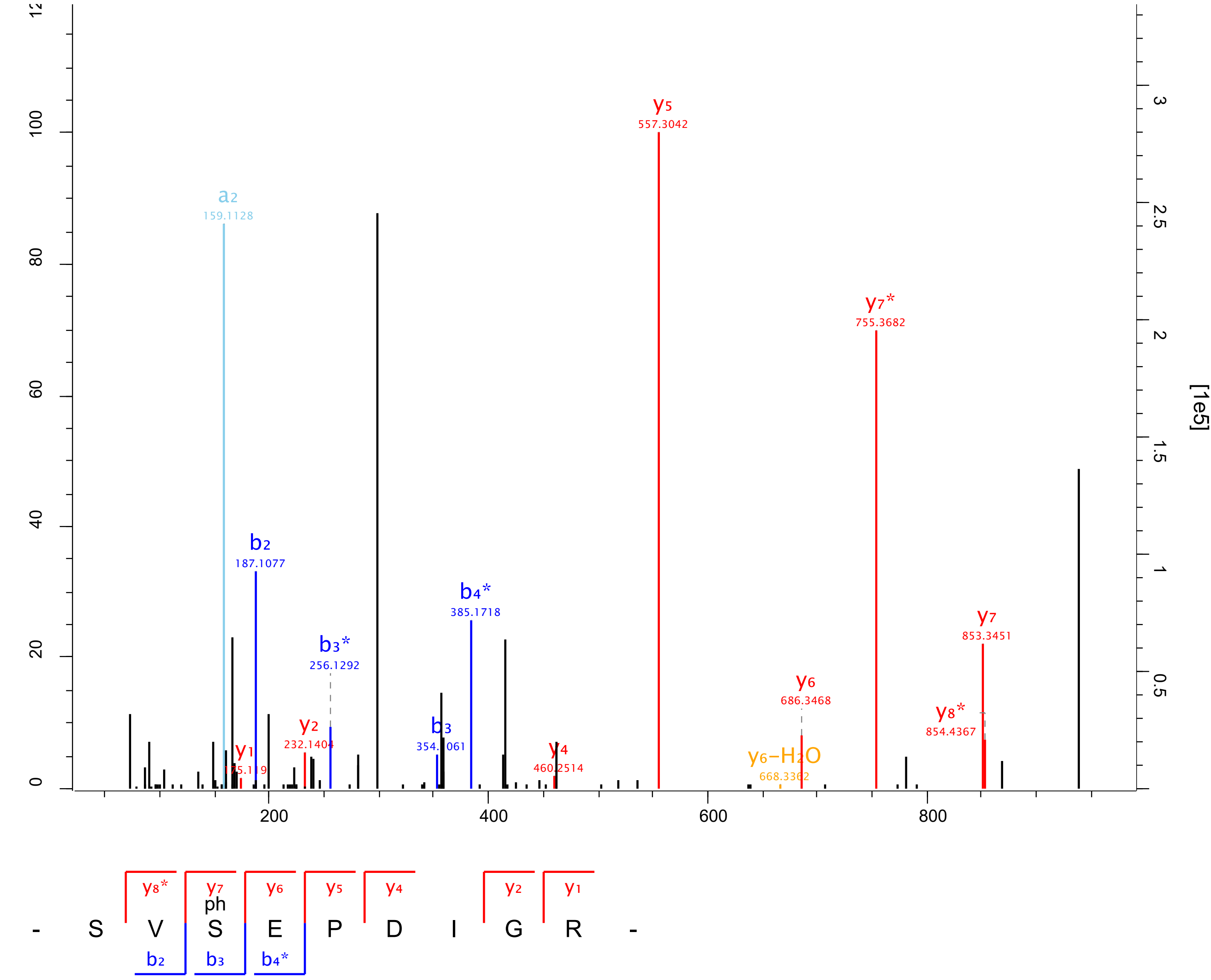

| Raw file | Scan | Method | Score | m/z |
|---|---|---|---|---|
| water_15_2 | 10689 | FTMS; HCD | 160.16 | 585.73 |

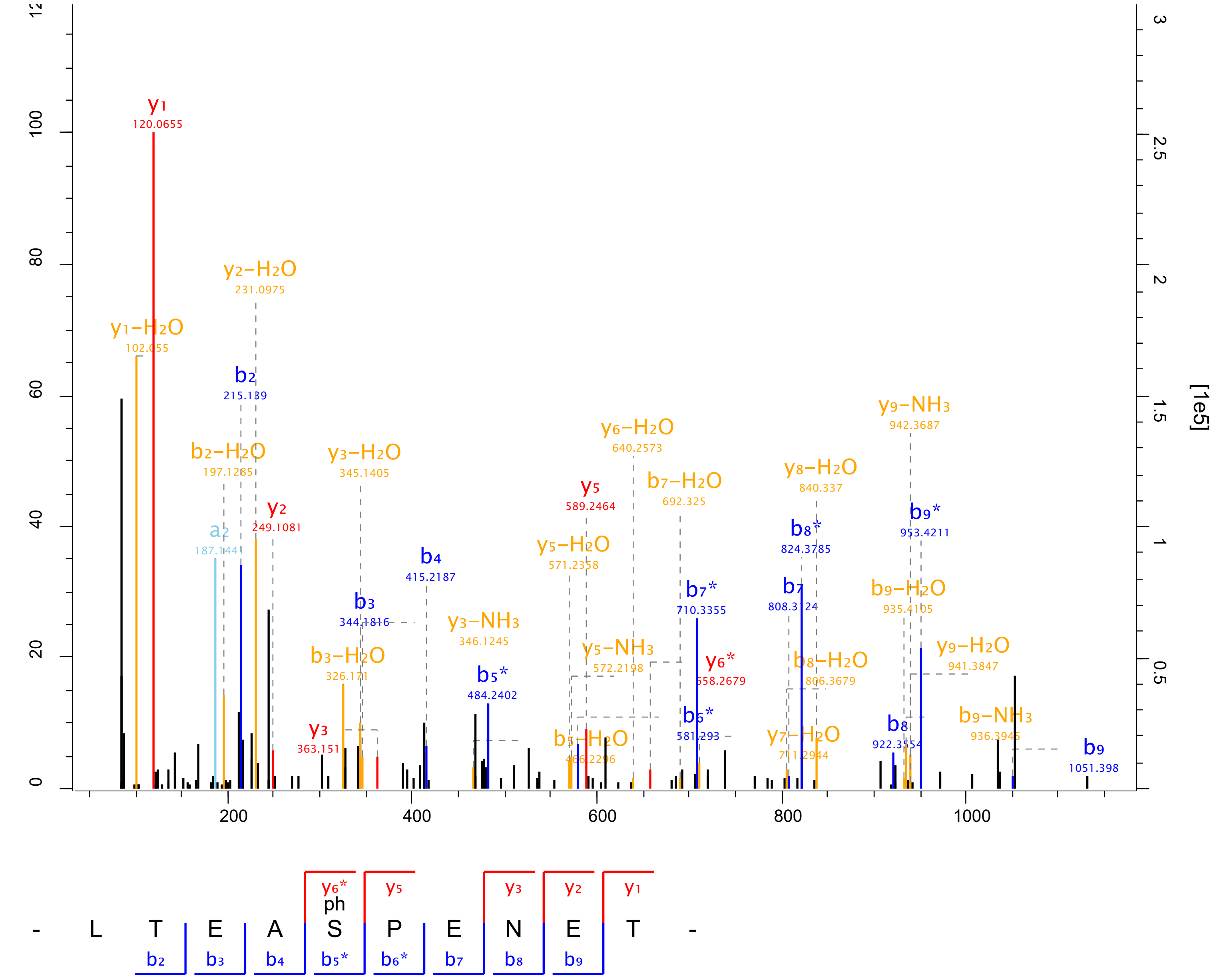

| Raw file | Scan | Method | Score | m/z |
| --- | --- | --- | --- | --- |
| water_15_2 | 10731 | FTMS; HCD | 60.16 | 572.25 |

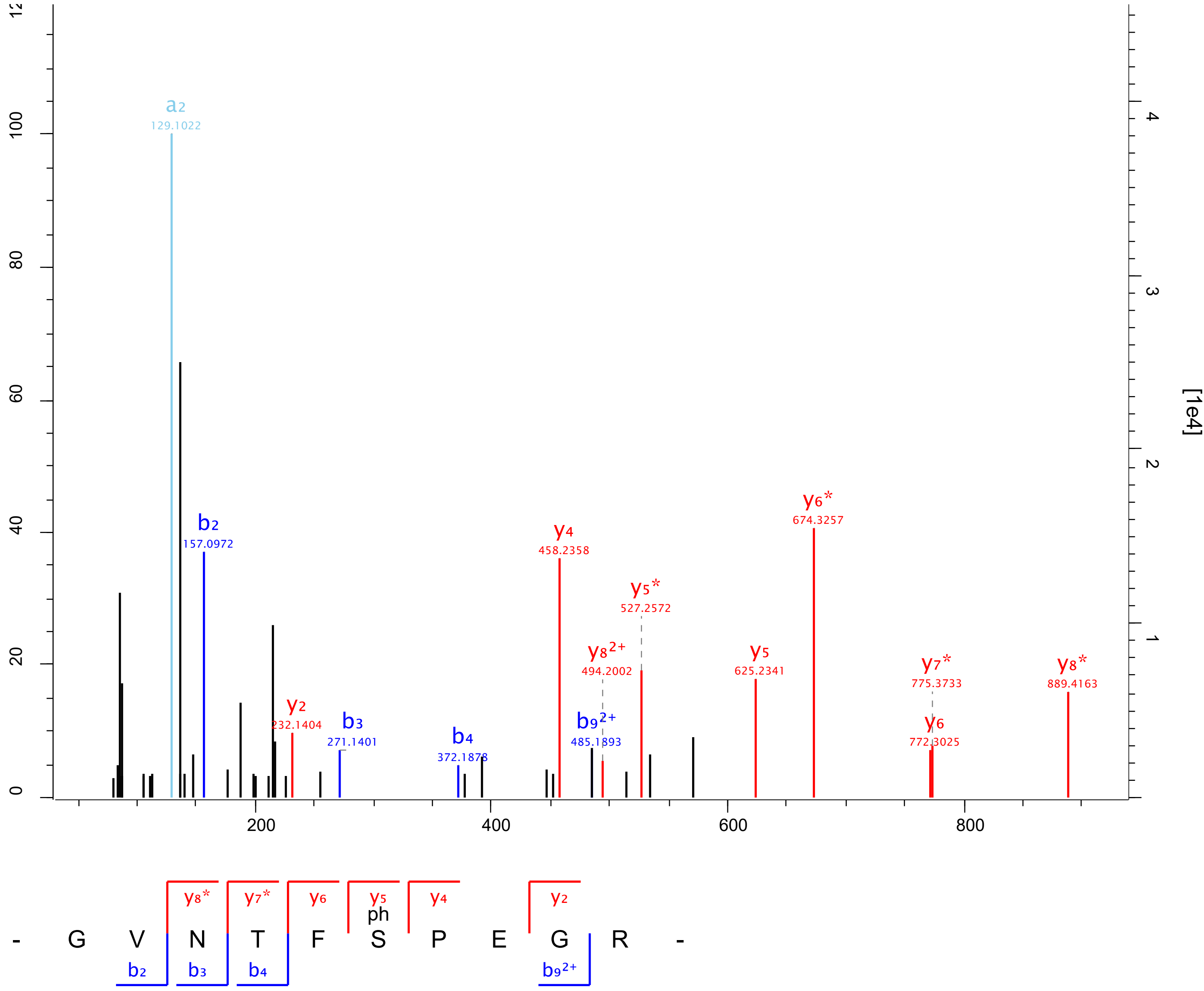

| Raw file | Scan | Method | Score | m/z |
|---|---|---|---|---|
| water_15_2 | 10780 | FTMS; HCD | 65.18 | 544.73 |

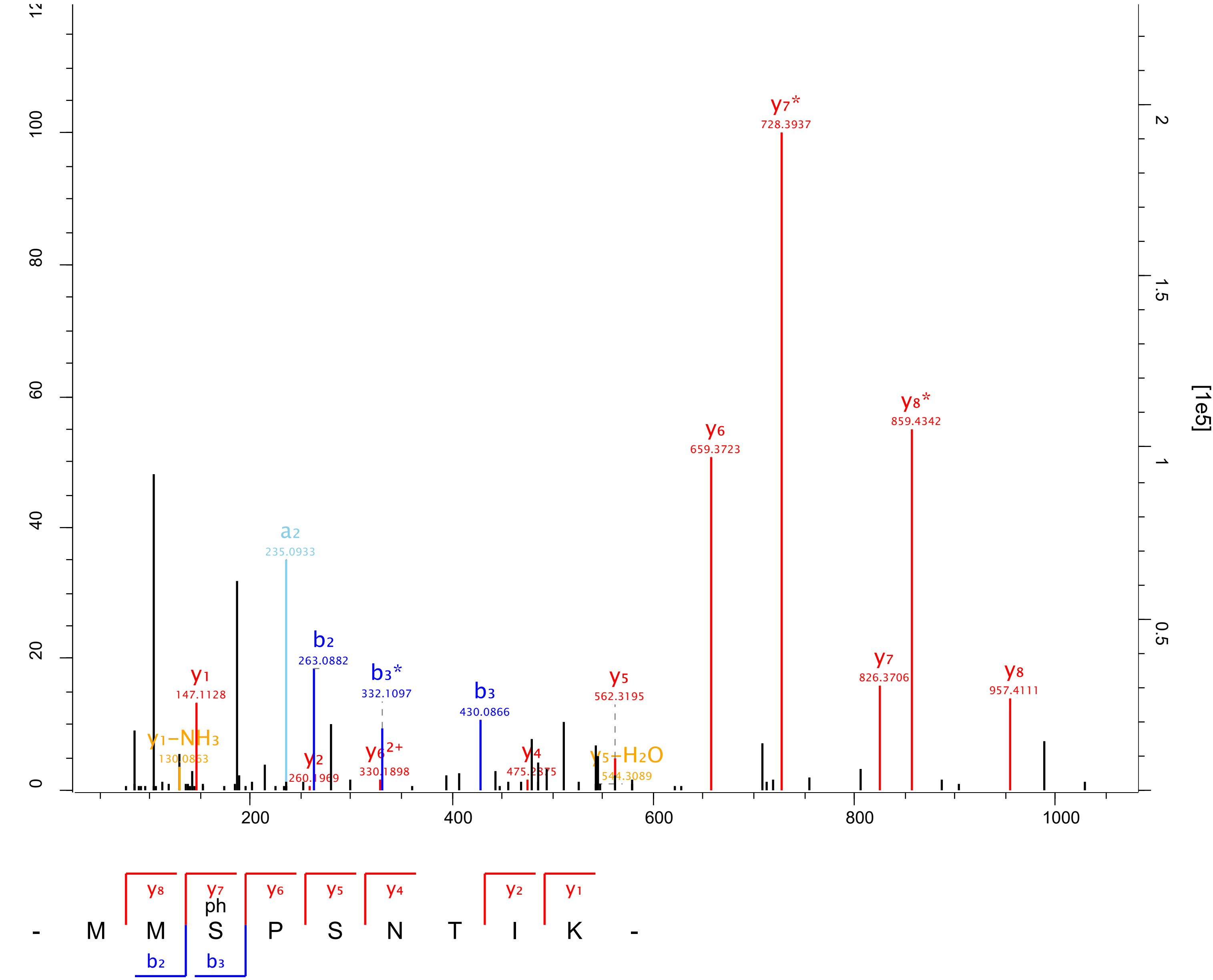

| Raw file | Scan | Method | Score | m/z |
|---|---|---|---|---|
| water_15_2 | 10791 | FTMS; HCD | 164.53 | 681.78 |

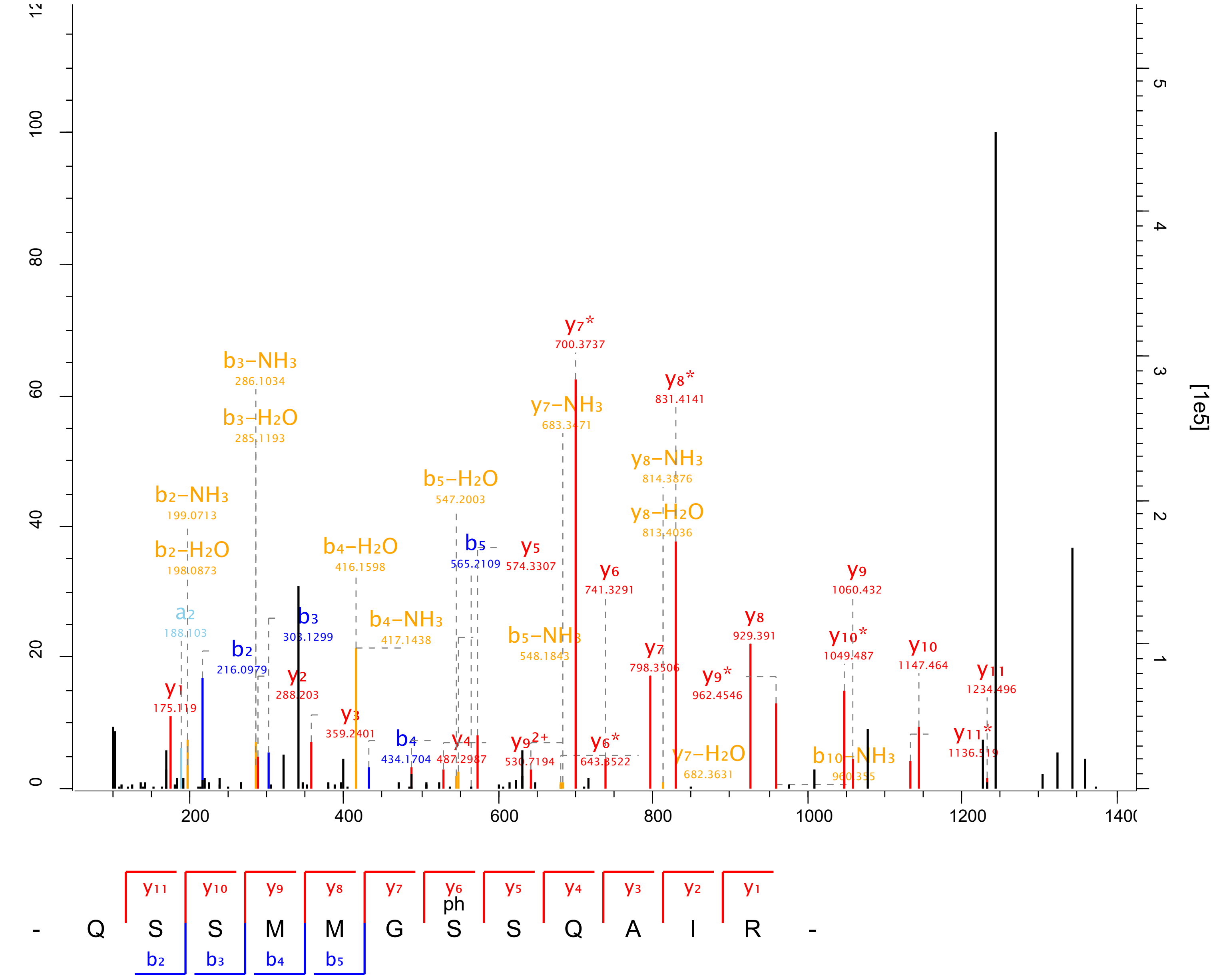

| Raw file | Scan | Method | Score | m/z |
| --- | --- | --- | --- | --- |
| water_15_2 | 10822 | FTMS; HCD | 47.3 | 478.71 |

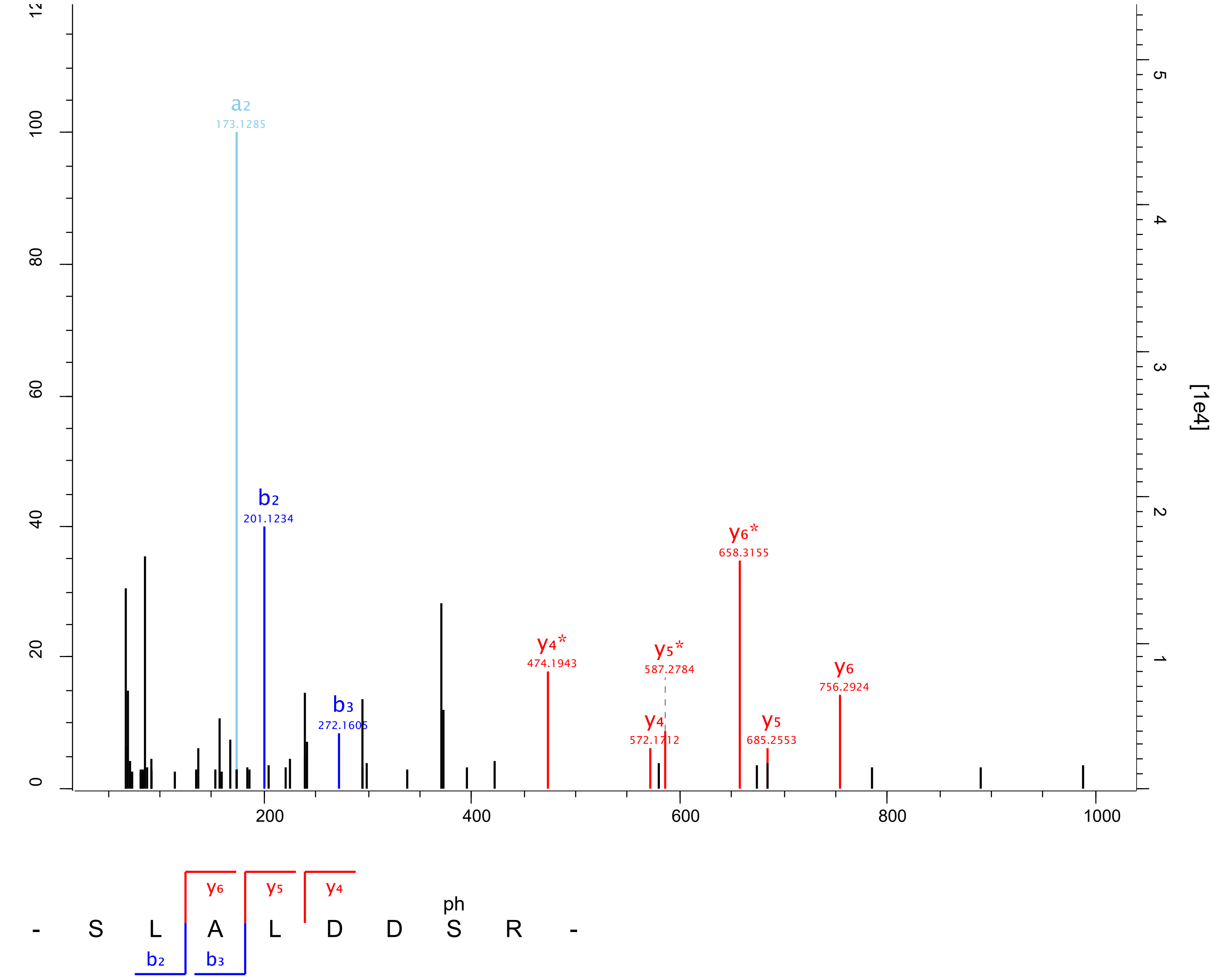

| Raw file | Scan | Method | Score | m/z |
|---|---|---|---|---|
| water_15_2 | 10825 | FTMS; HCD | 121.21 | 627.93 |

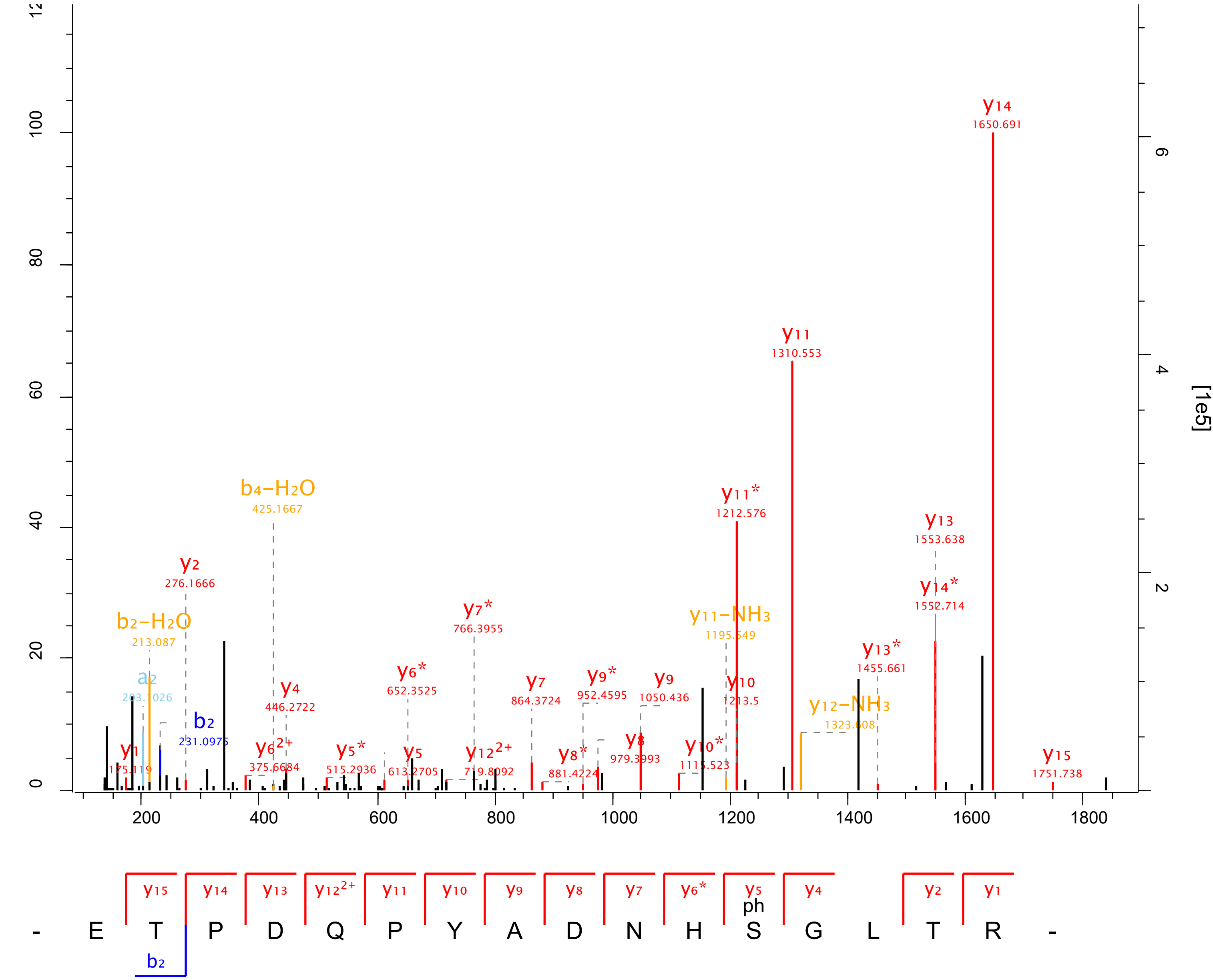

| Raw file | Scan | Method | Score | m/z |
|---|---|---|---|---|
| water_15_2 | 10826 | FTMS; HCD | 71.18 | 524.21 |

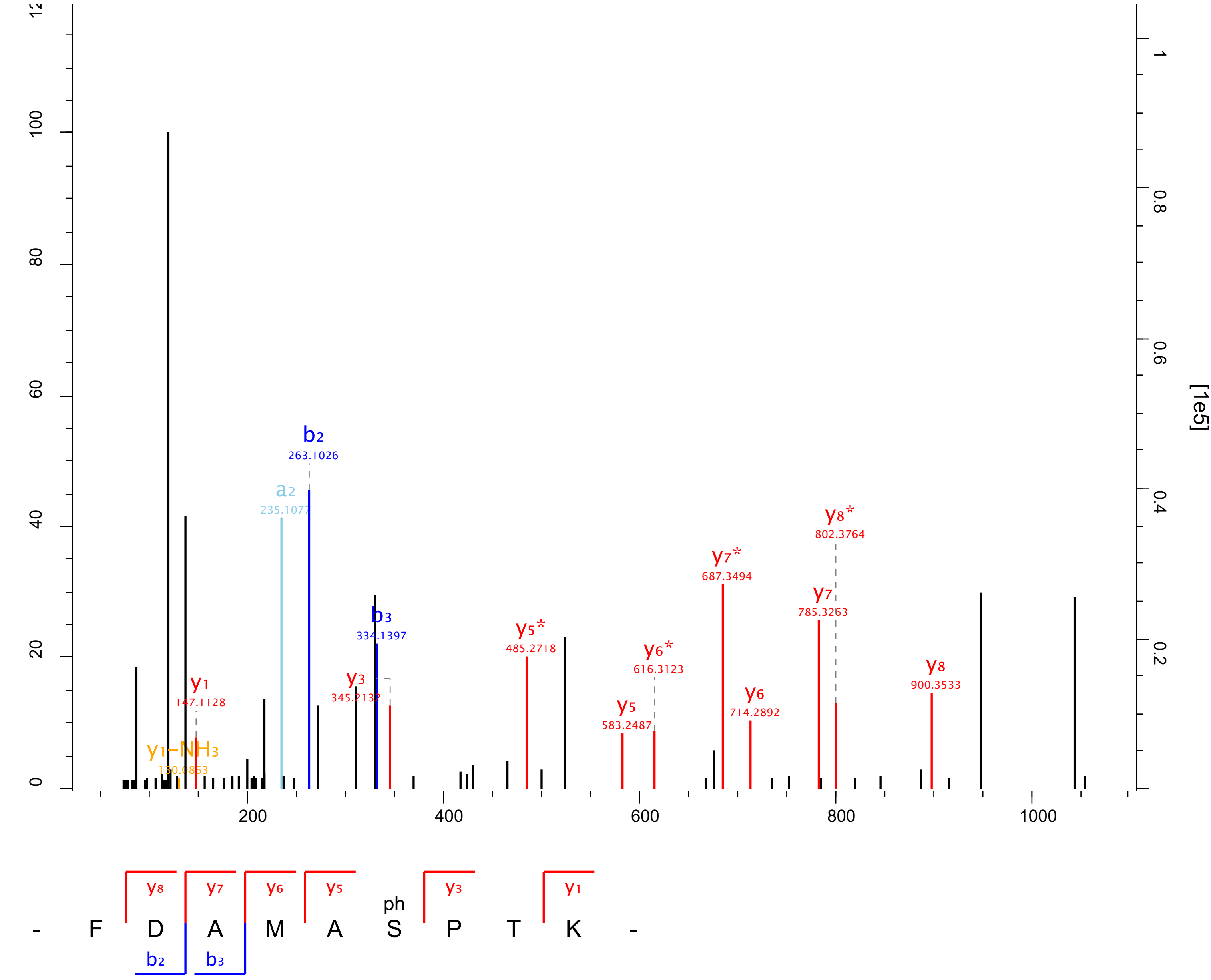

| Raw file | Scan | Method | Score | m/z |
|---|---|---|---|---|
| water_15_2 | 10849 | FTMS; HCD | 78.14 | 487.7 |

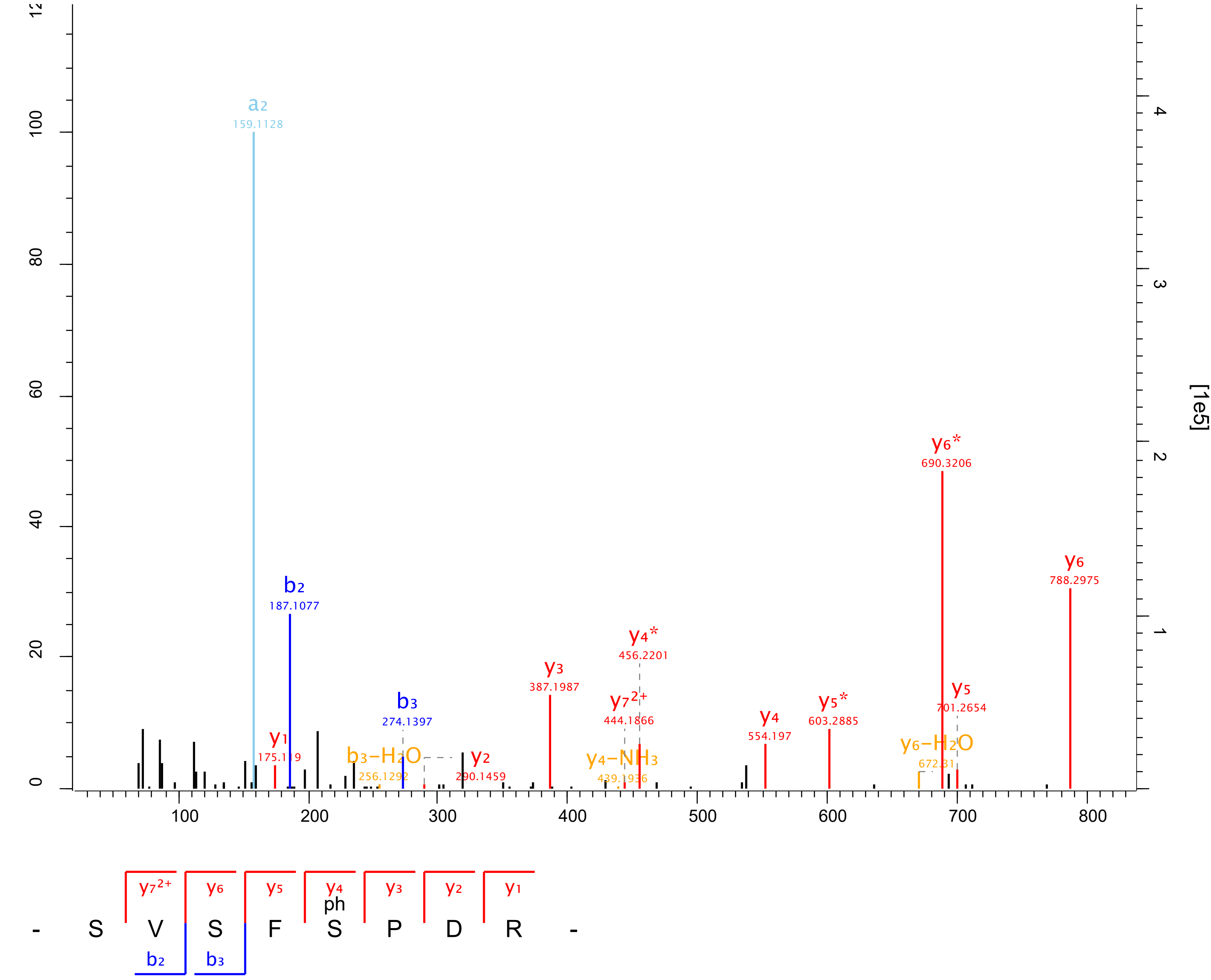

| Raw file | Scan | Method | Score | m/z |
|---|---|---|---|---|
| water_15_2 | 10855 | FTMS; HCD | 85.91 | 665.79 |

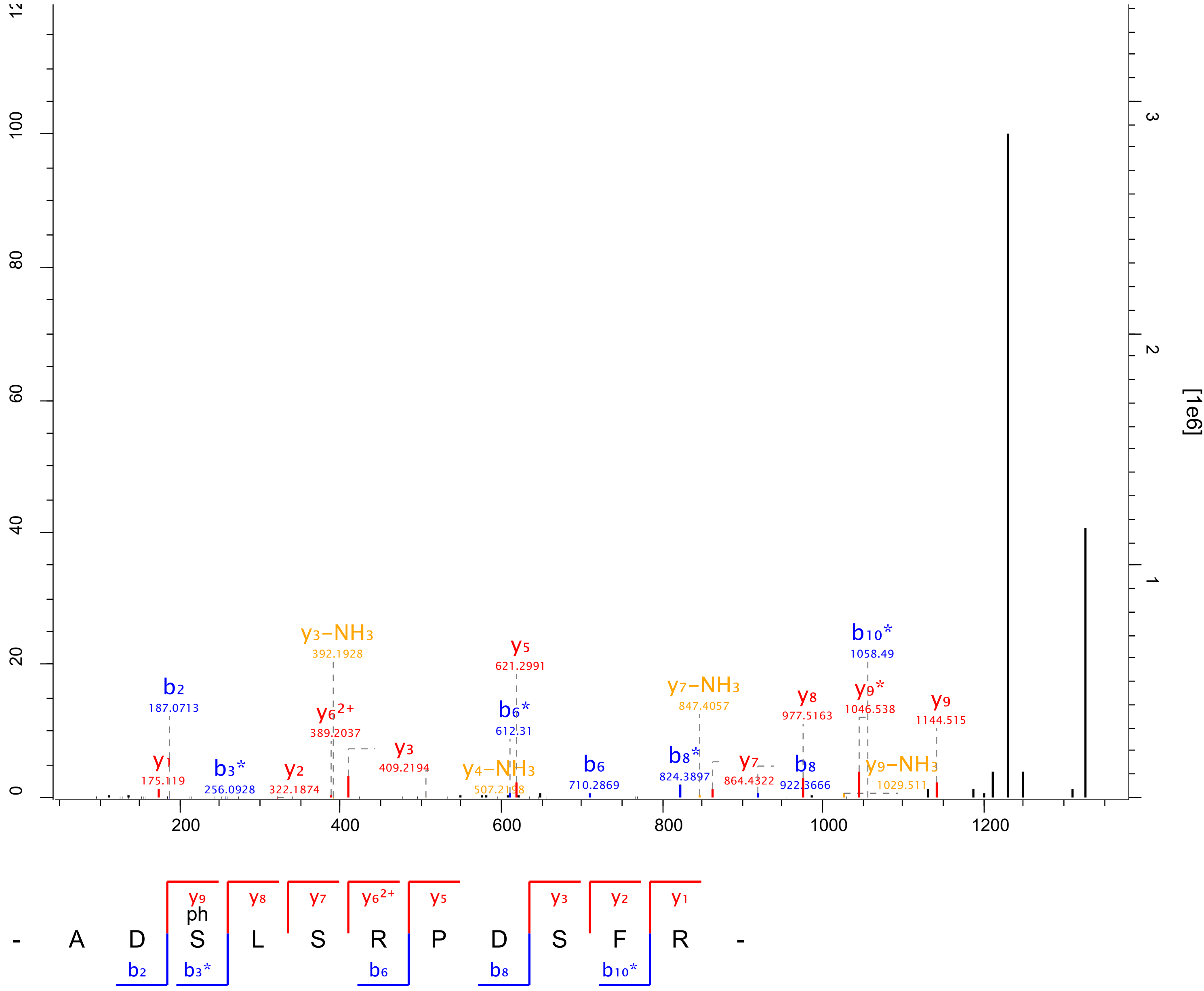

| Raw file | Scan | Method | Score | m/z |
|---|---|---|---|---|
| water_15_2 | 10891 | FTMS; HCD | 51.03 | 495.92 |

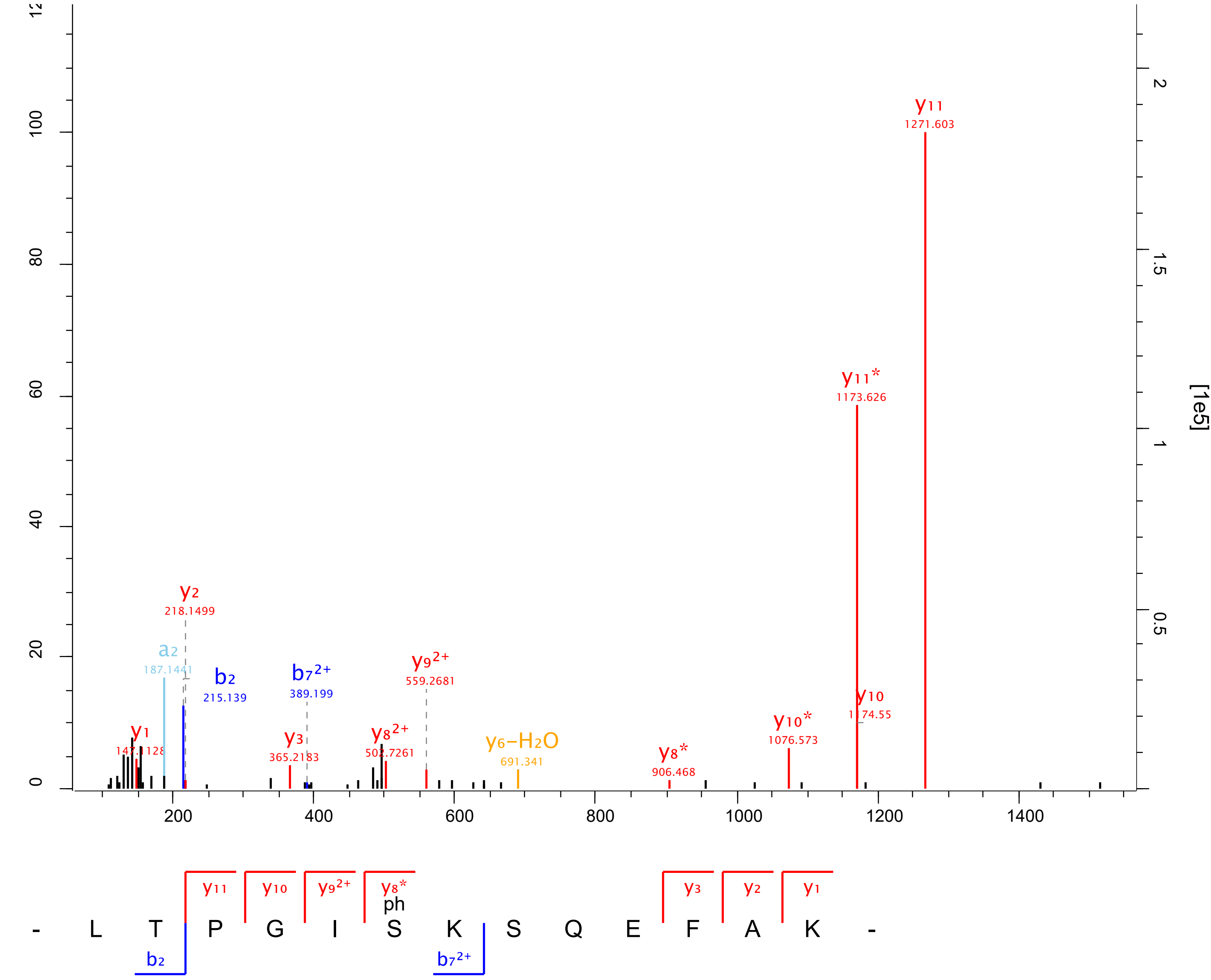

| Raw file | Scan | Method | Score | m/z |
|---|---|---|---|---|
| water_15_2 | 10951 | FTMS; HCD | 245.22 | 829.35 |

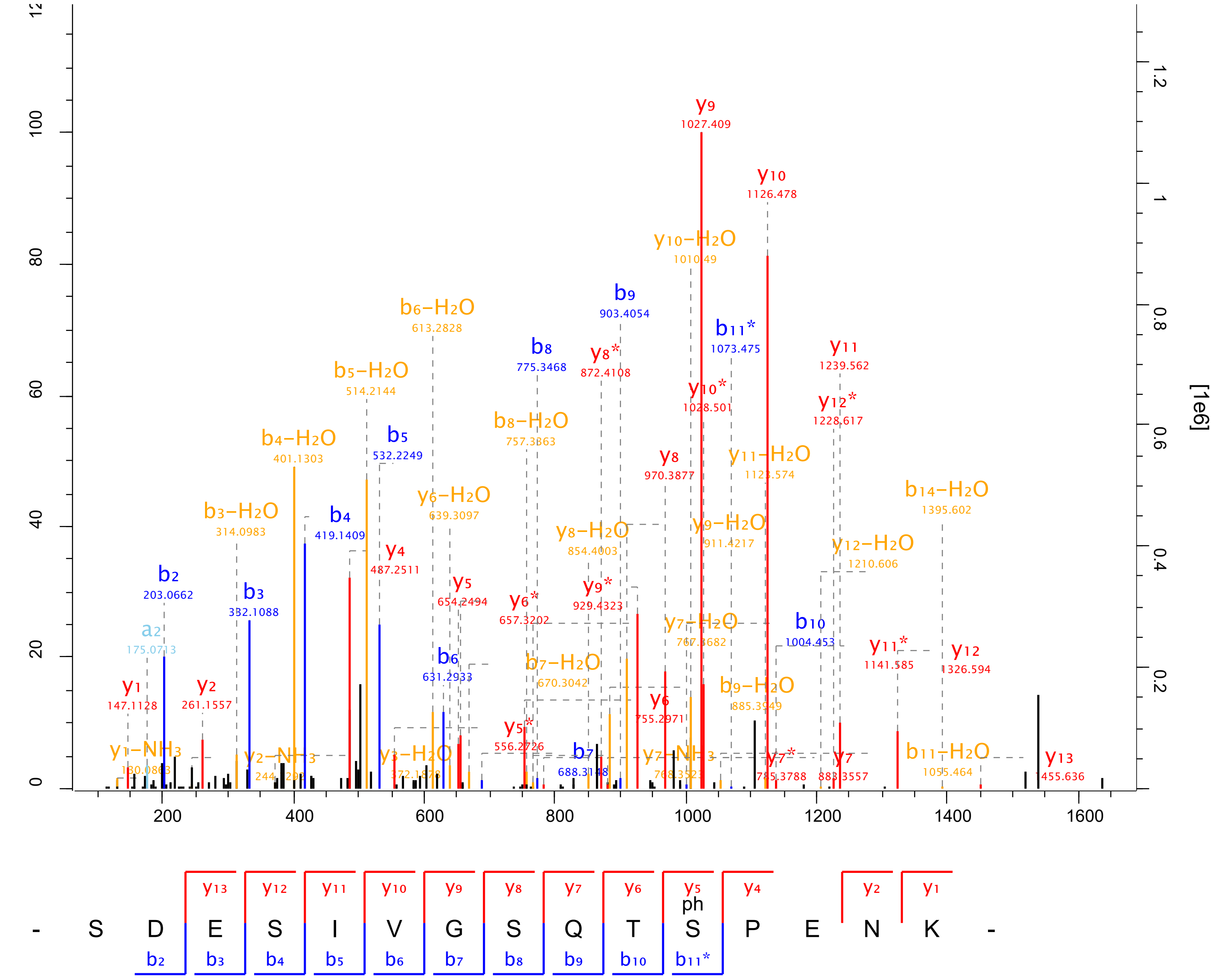

| Raw file | Scan | Method | Score | m/z |
| --- | --- | --- | --- | --- |
| water_15_2 | 11100 | FTMS; HCD | 118.72 | 777.31 |

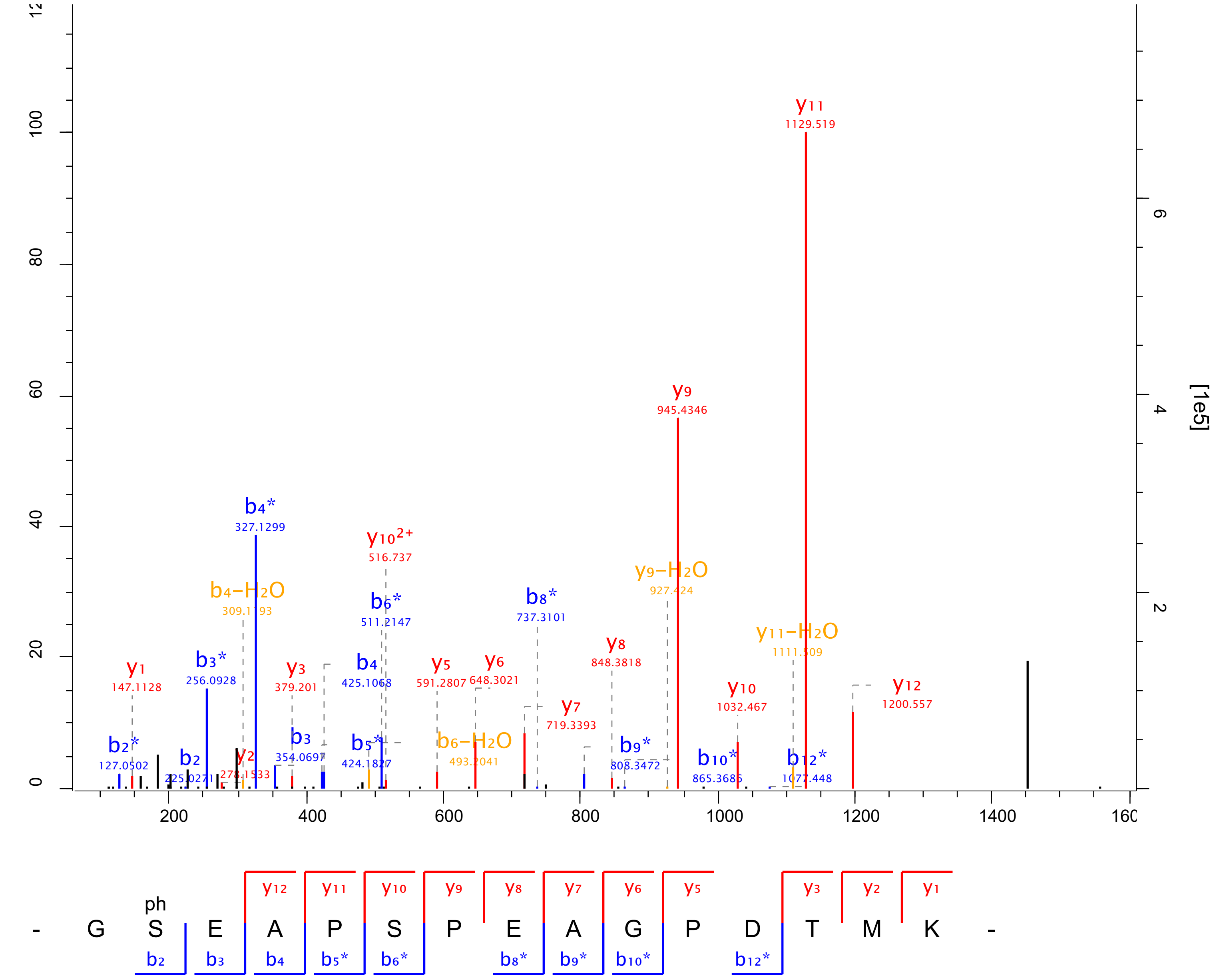

| Raw file | Scan | Method | Score | m/z |
| --- | --- | --- | --- | --- |
| water_15_2 | 11105 | FTMS; HCD | 142.91 | 672.29 |

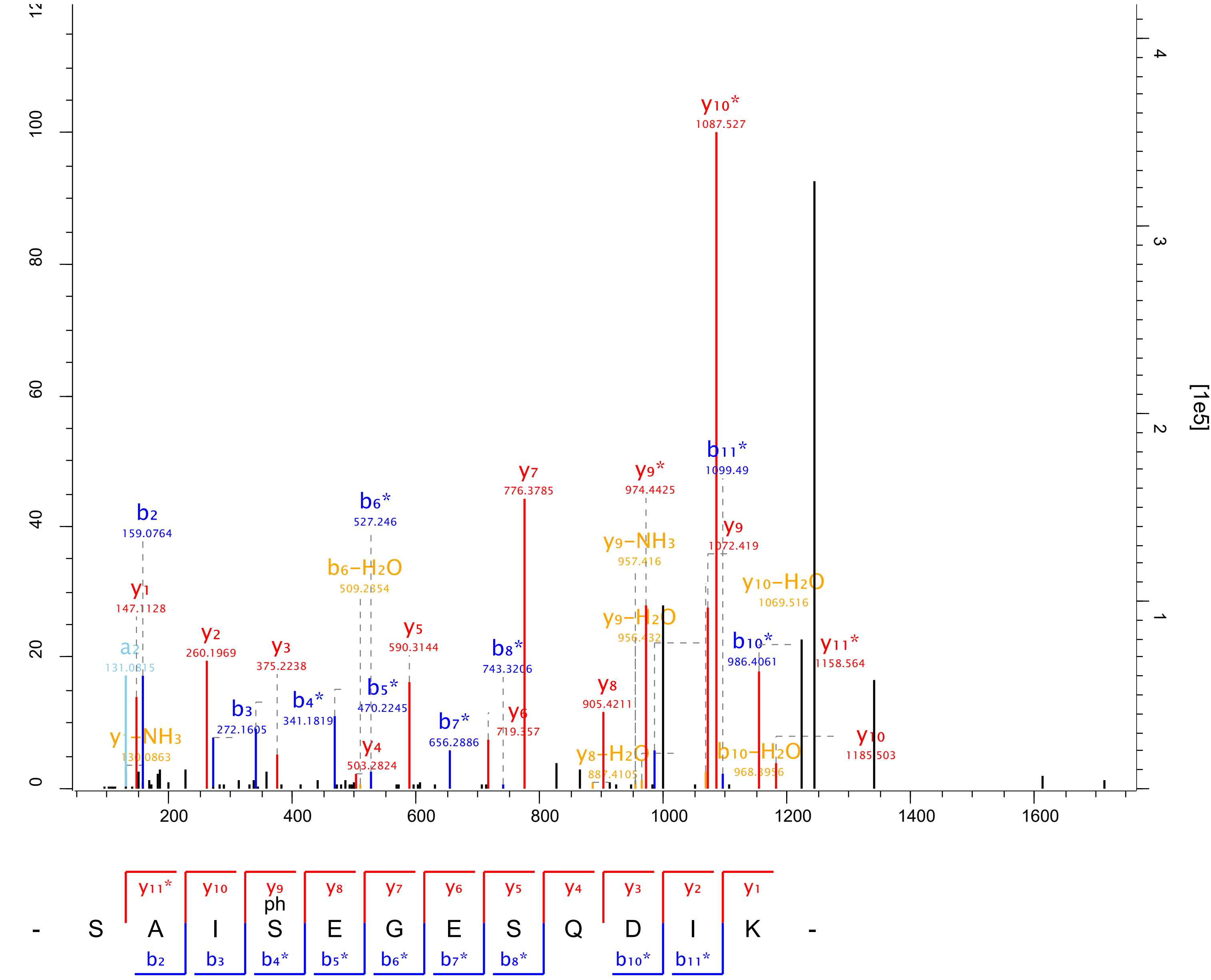

| Raw file | Scan | Method | Score | m/z |
|---|---|---|---|---|
| water_15_2 | 11187 | FTMS; HCD | 44.85 | 493.22 |

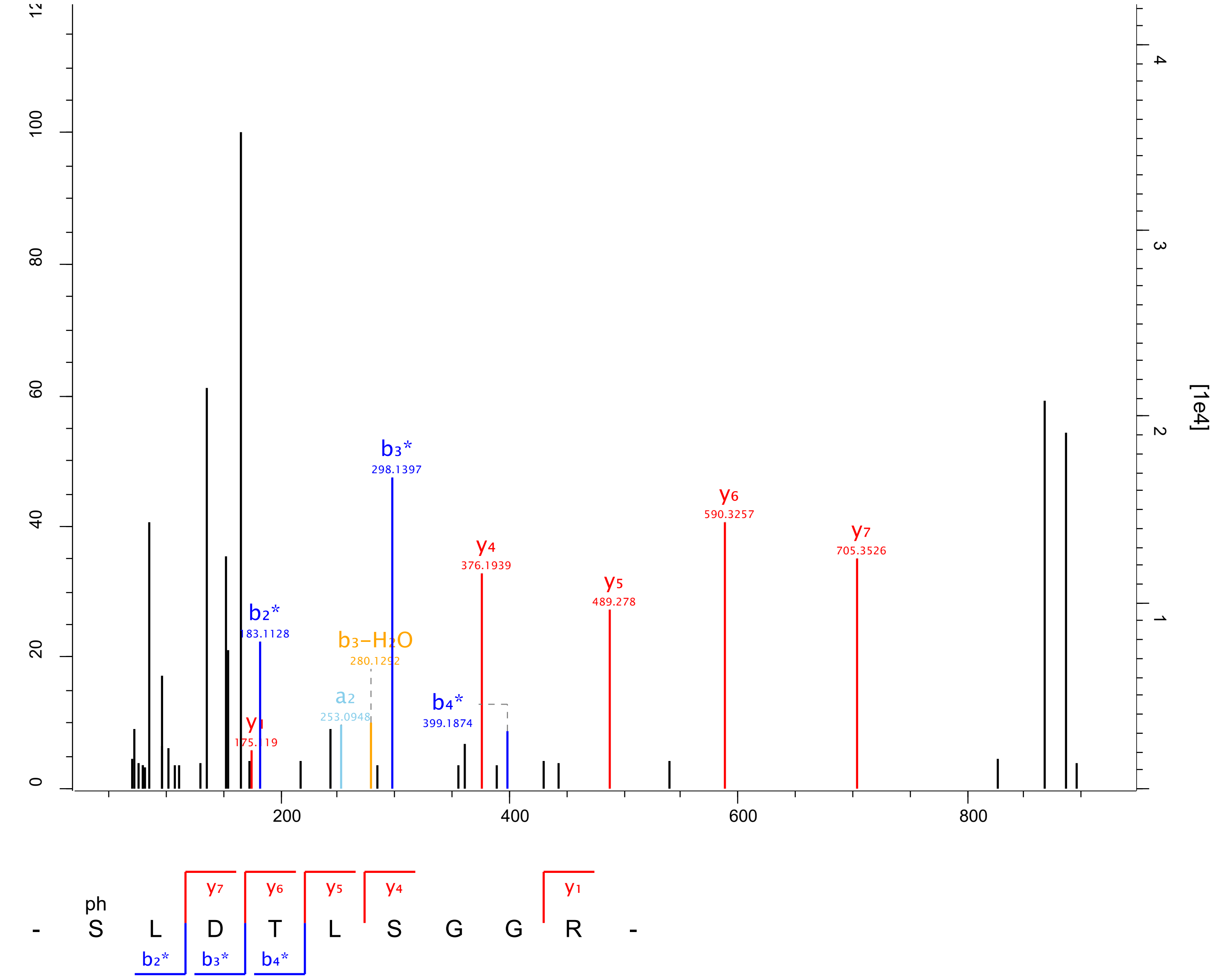

| Raw file | Scan | Method | Score | m/z |
| --- | --- | --- | --- | --- |
| water_15_2 | 11232 | FTMS; HCD | 52.18 | 832.89 |

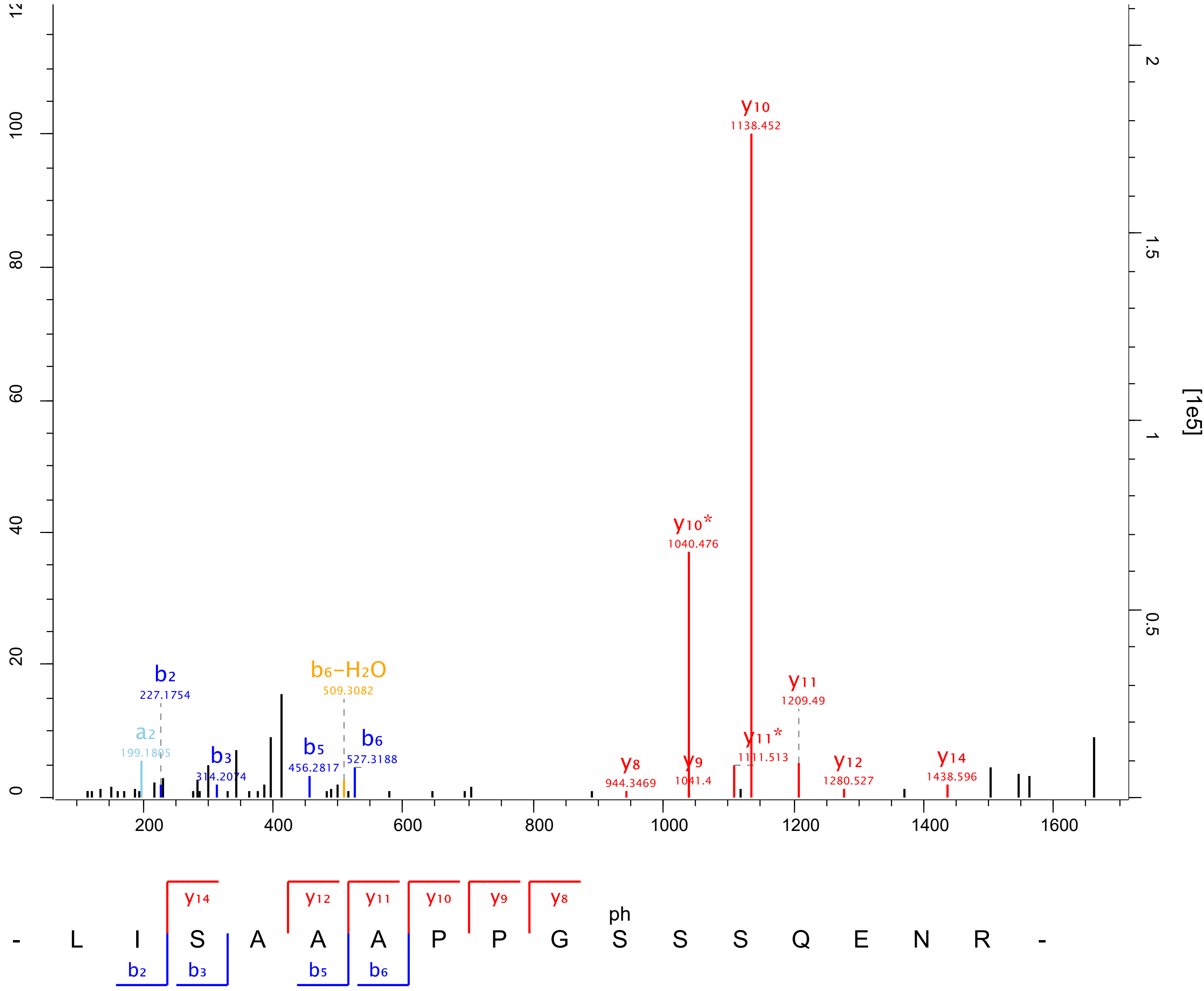

| Raw file | Scan | Method | Score | m/z |
|---|---|---|---|---|
| water_15_2 | 11266 | FTMS; HCD | 55.26 | 604.28 |

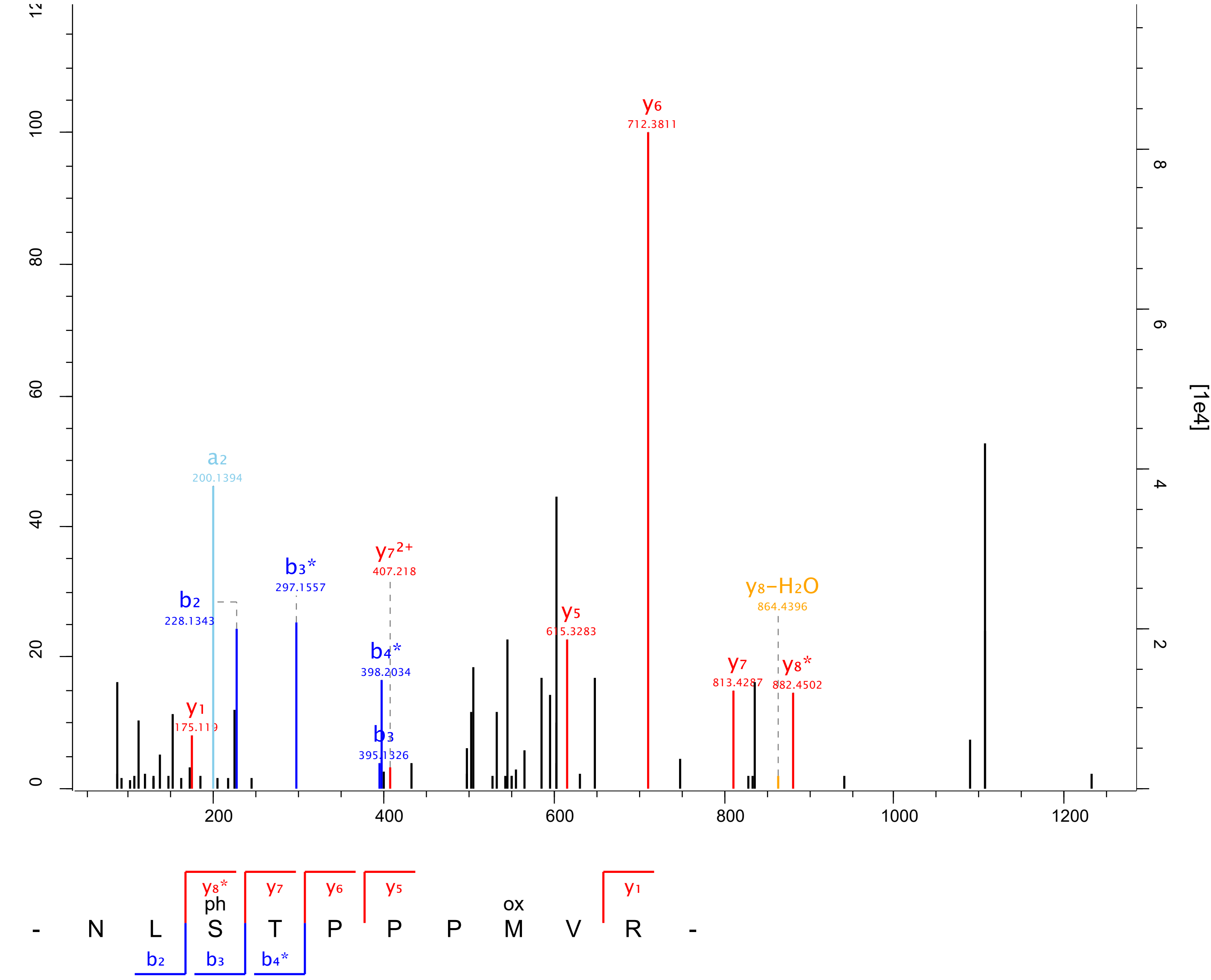

| Raw file | Scan | Method | Score | m/z |
|---|---|---|---|---|
| water_15_2 | 11286 | FTMS; HCD | 66.81 | 687.75 |

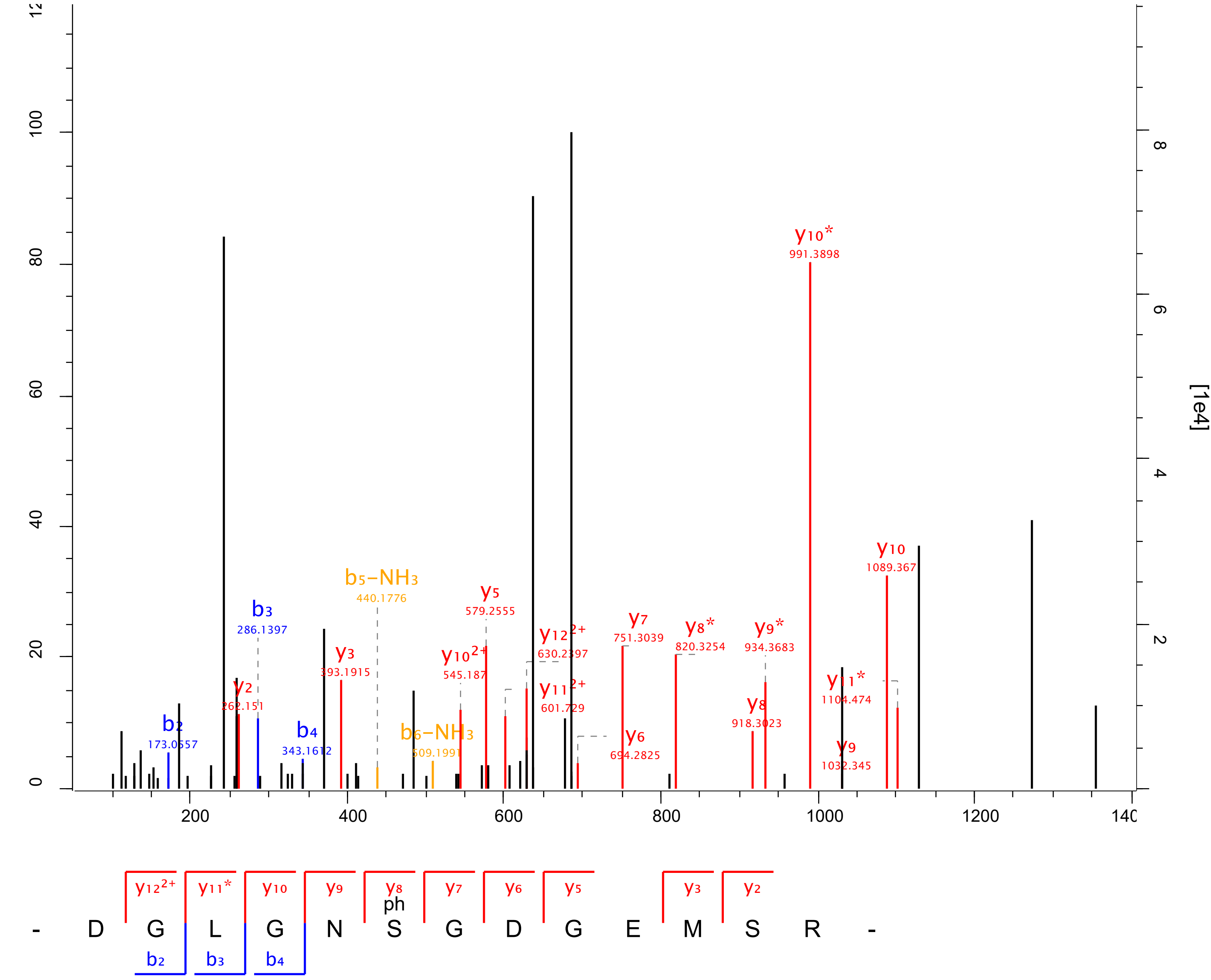

| Raw file | Scan | Method | Score | m/z |
| --- | --- | --- | --- | --- |
| water_15_2 | 11317 | FTMS; HCD | 74.78 | 613.93 |

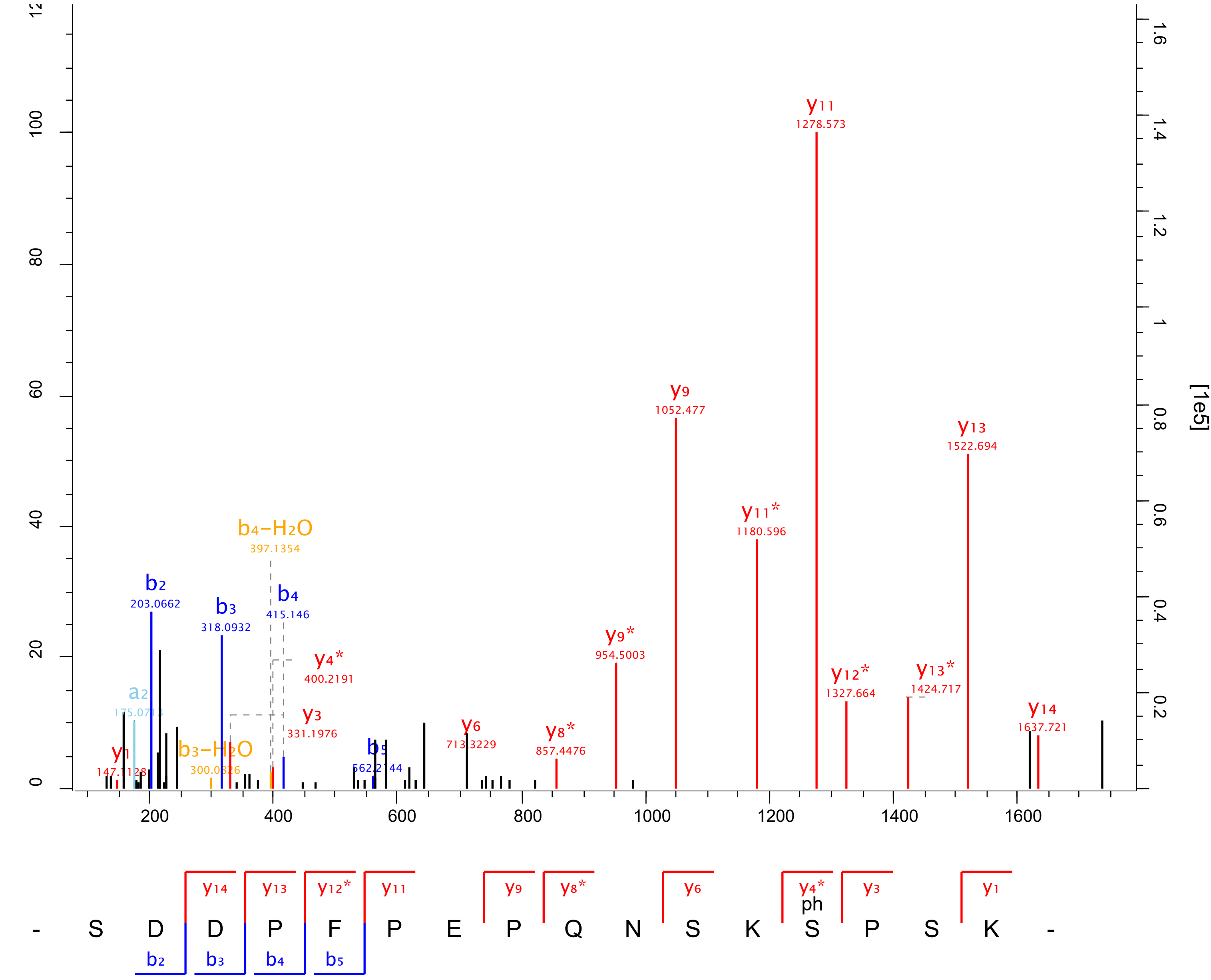

| Raw file | Scan | Method | Score | m/z |
|---|---|---|---|---|
| water_15_2 | 11326 | FTMS; HCD | 78.81 | 589.76 |

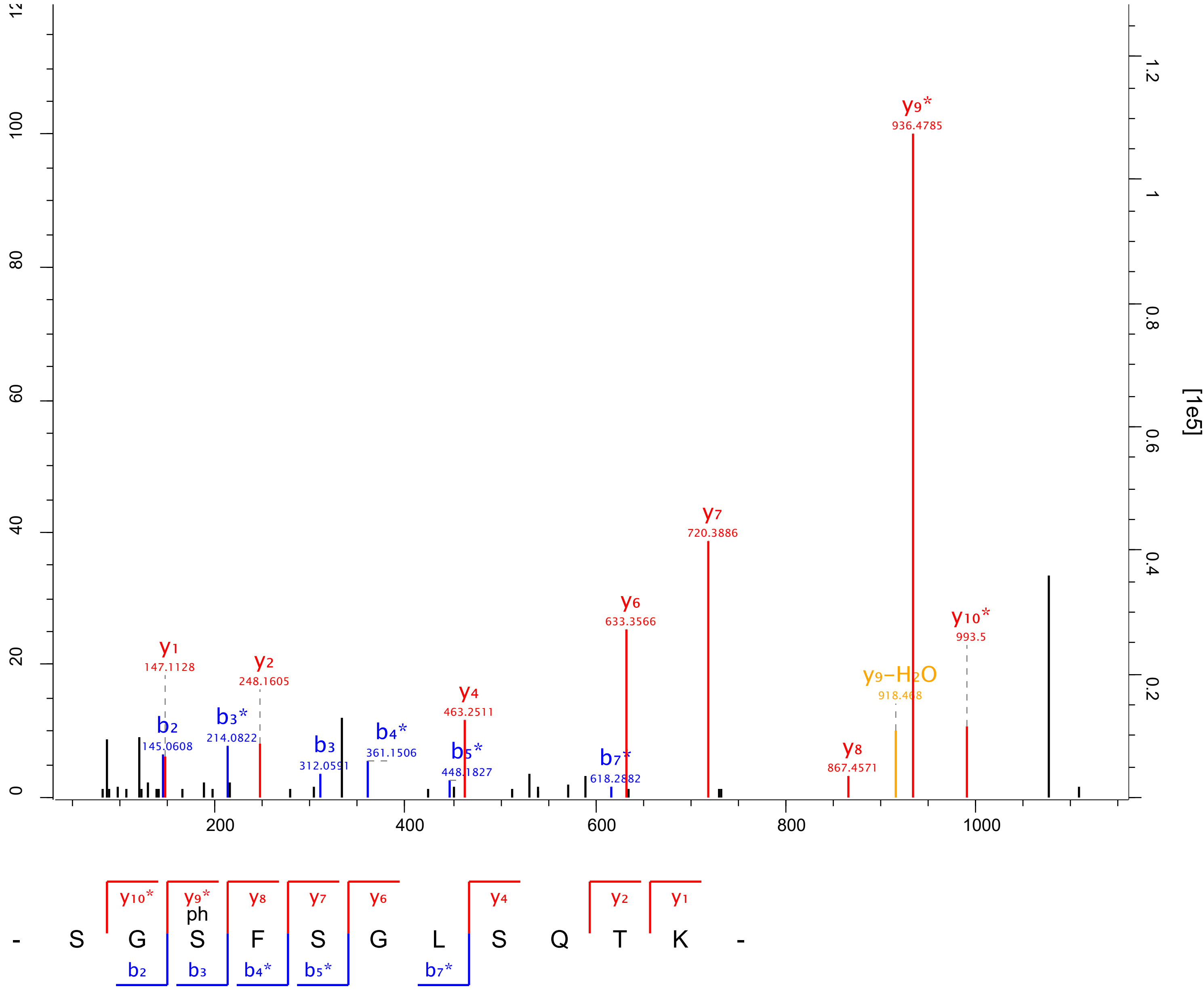

| Raw file | Scan | Method | Score | m/z |
| --- | --- | --- | --- | --- |
| water_15_2 | 11334 | FTMS; HCD | 42.34 | 553.27 |

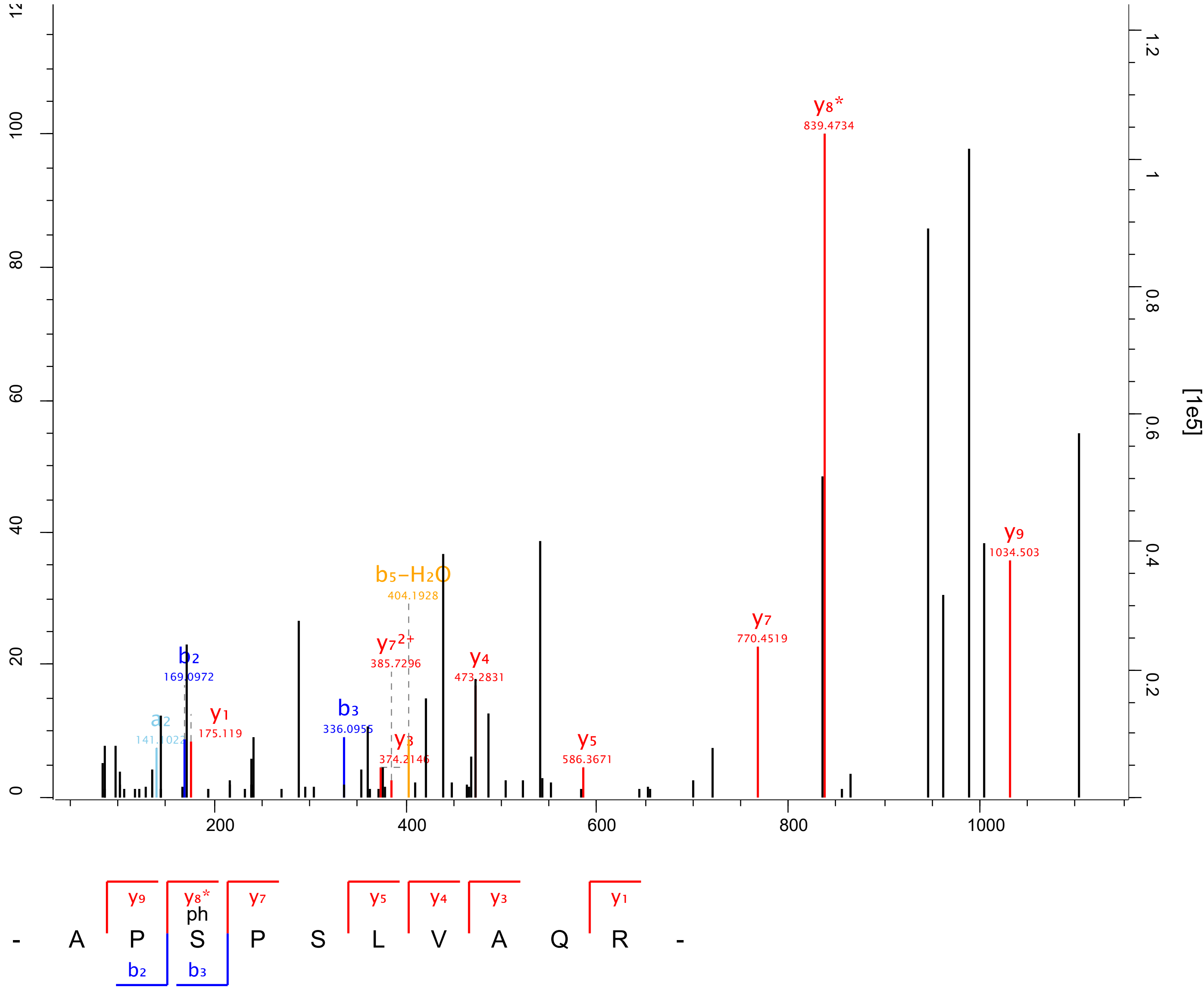

| Raw file | Scan | Method | Score | m/z |
|---|---|---|---|---|
| water_15_2 | 11365 | FTMS; HCD | 62.09 | 507.23 |

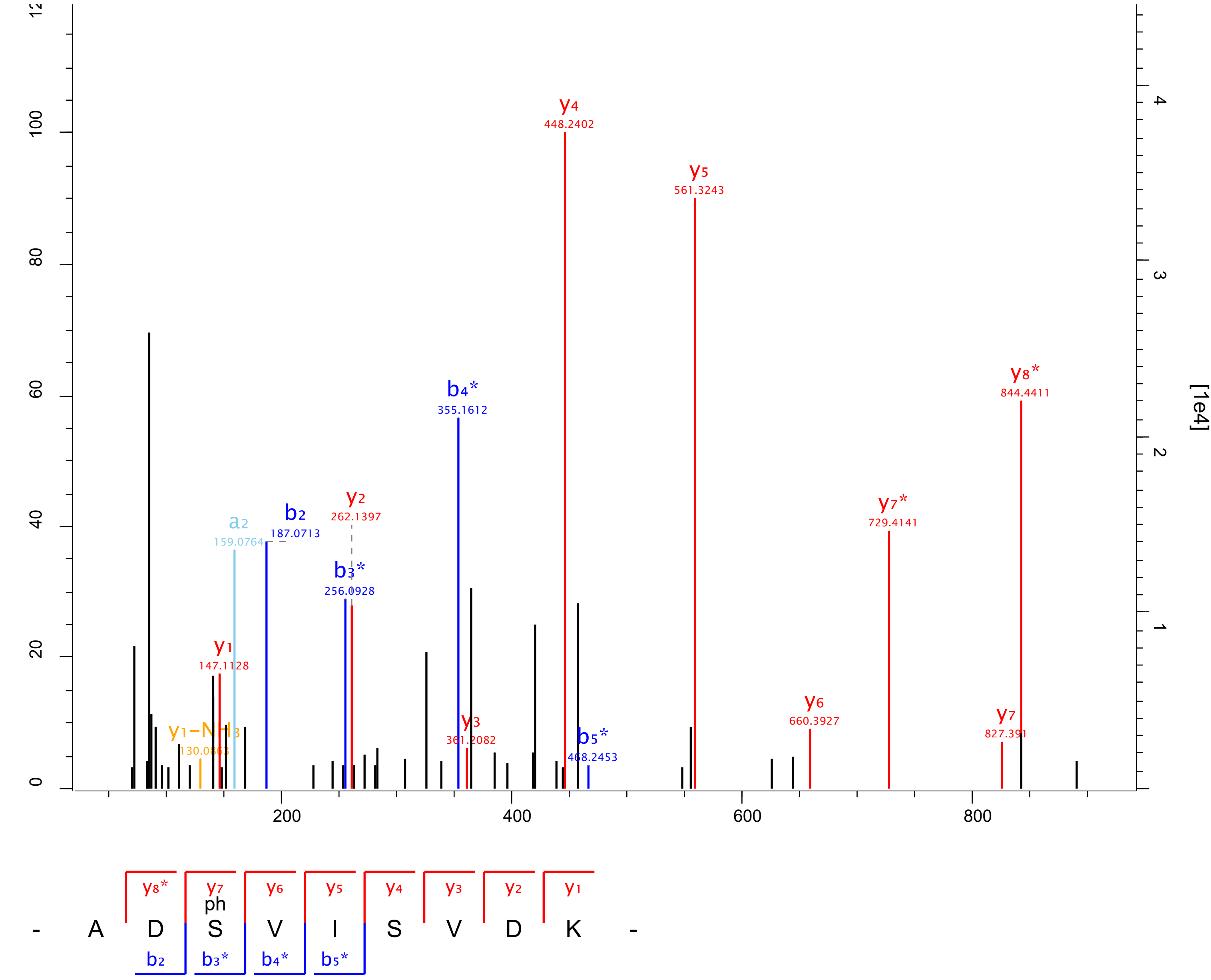

| Raw file | Scan | Method | Score | m/z |
| --- | --- | --- | --- | --- |
| water_15_2 | 11470 | FTMS; HCD | 67.33 | 455.72 |

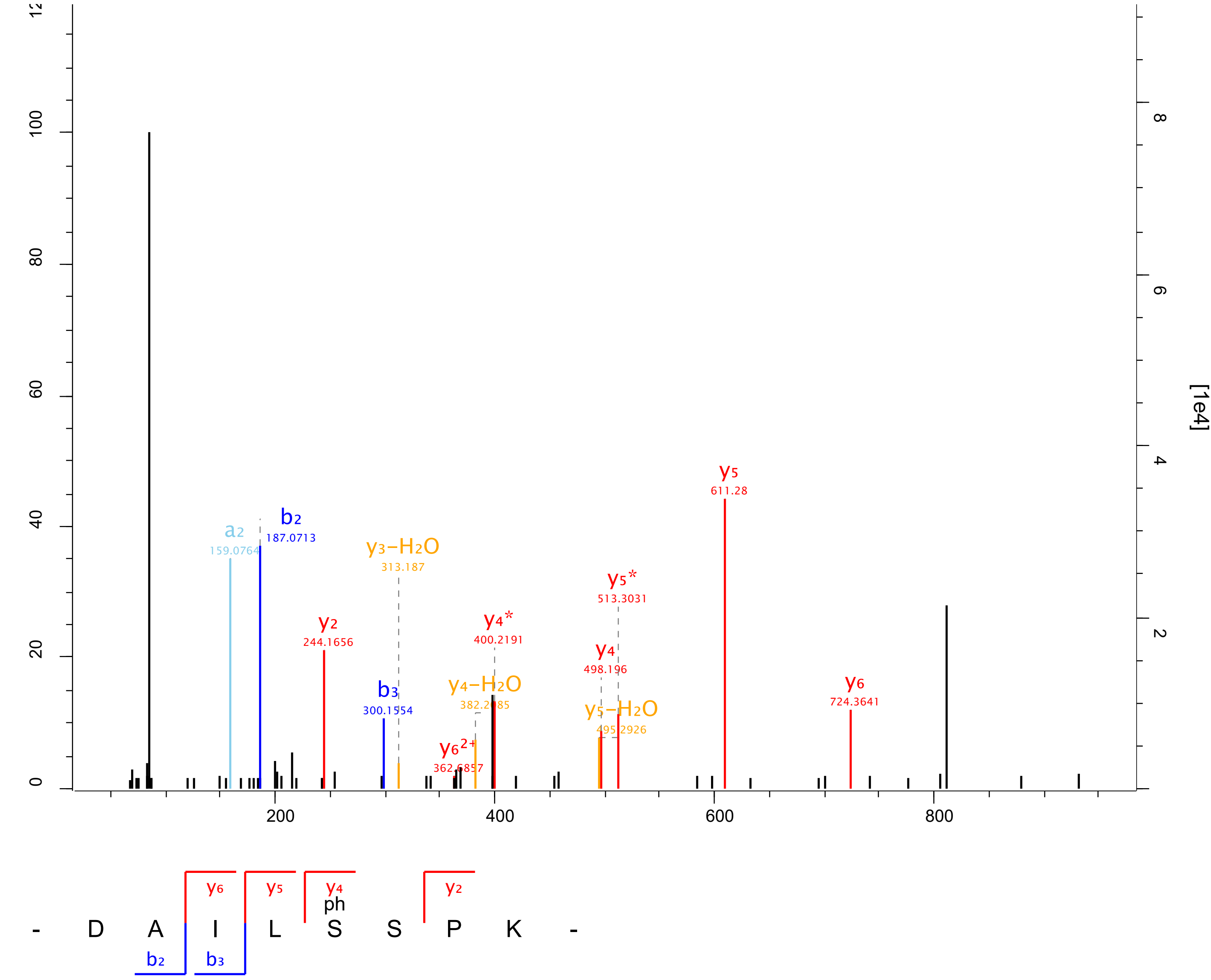

| Raw file | Scan | Method | Score | m/z |
| --- | --- | --- | --- | --- |
| water_15_2 | 11473 | FTMS; HCD | 101.3 | 603.74 |

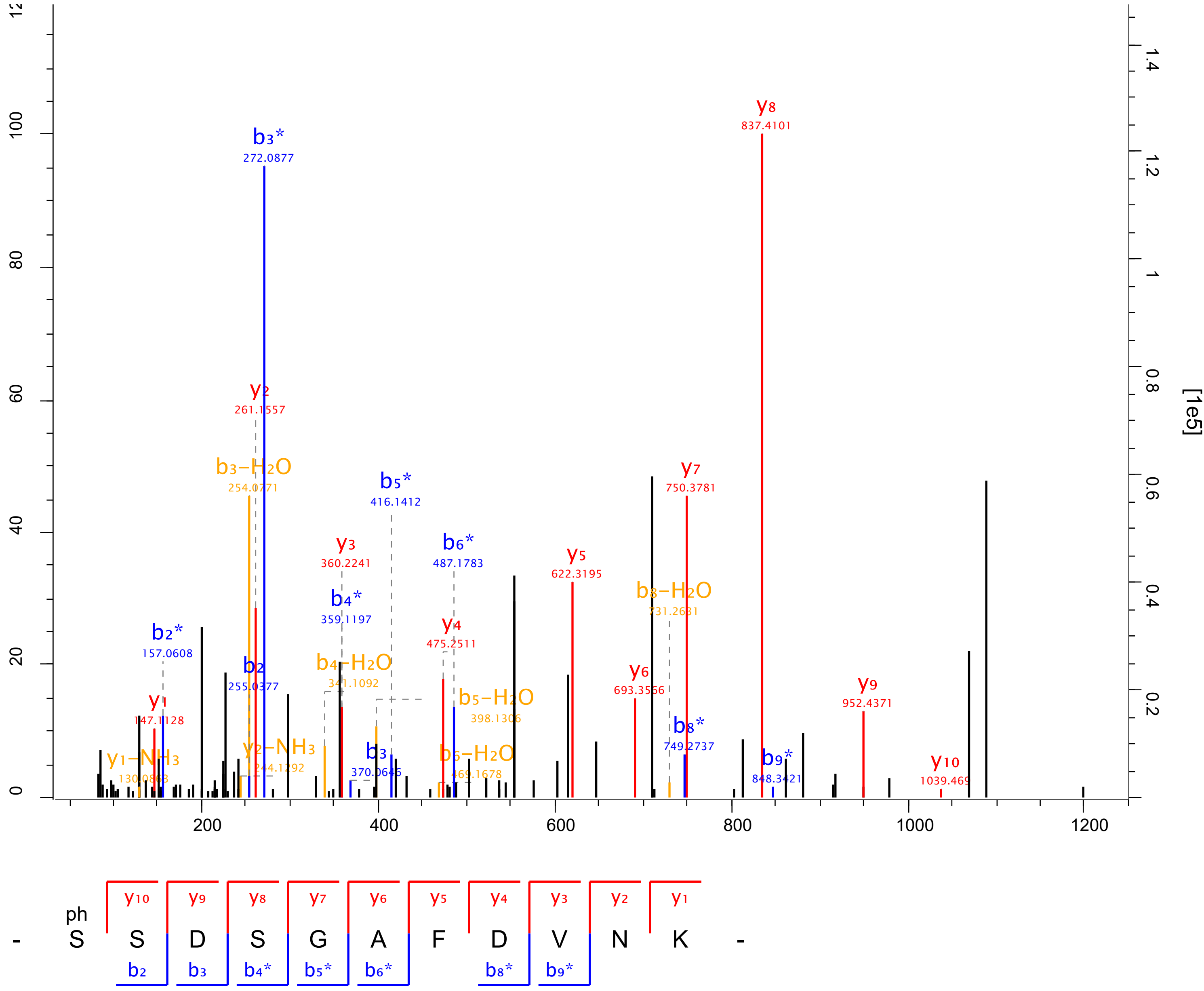

| Raw file | Scan | Method | Score | m/z |
| --- | --- | --- | --- | --- |
| water_15_2 | 11511 | FTMS; HCD | 95.42 | 510.72 |

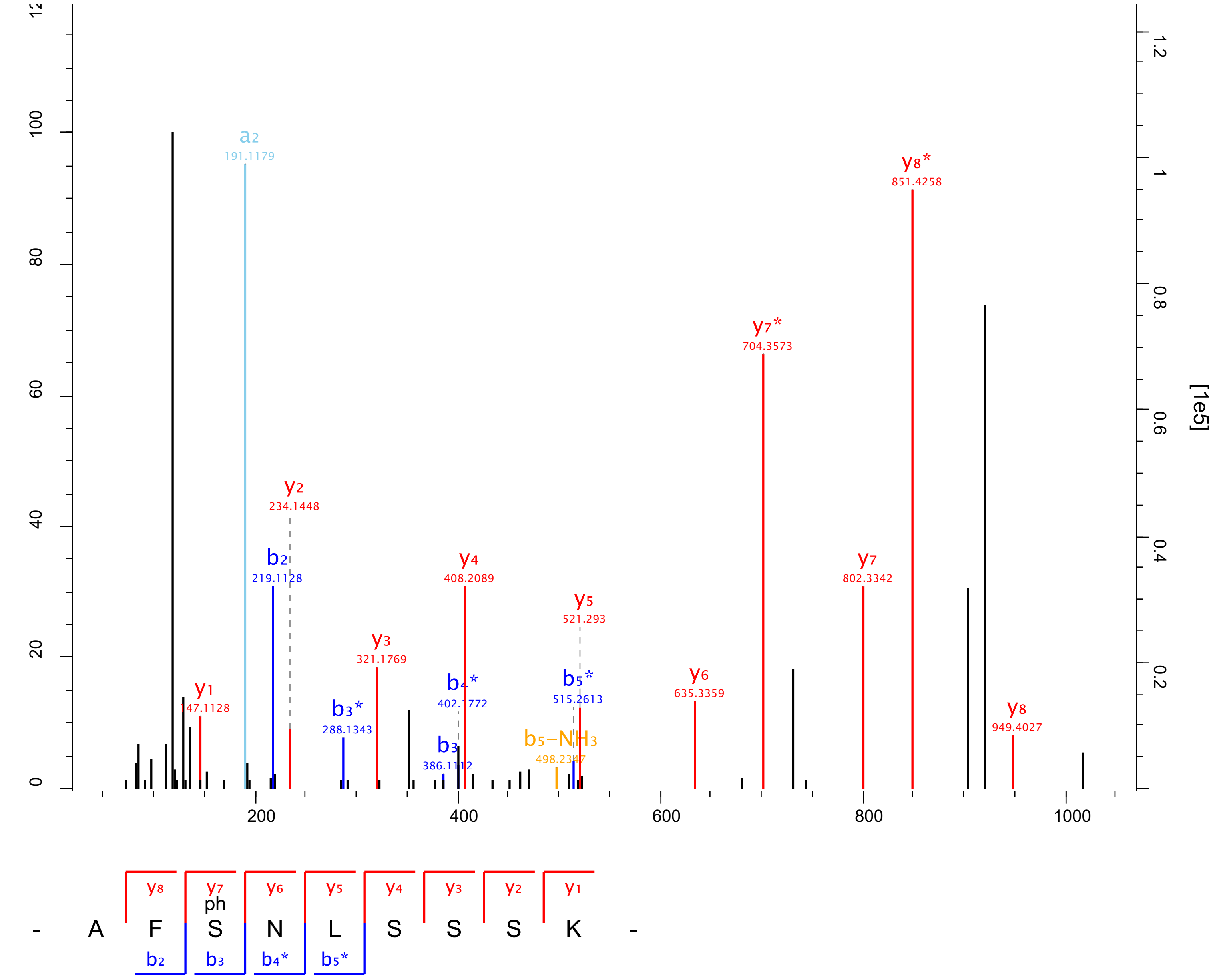

| Raw file | Scan | Method | Score | m/z |
|---|---|---|---|---|
| water_15_2 | 11593 | FTMS; HCD | 235.78 | 756.34 |

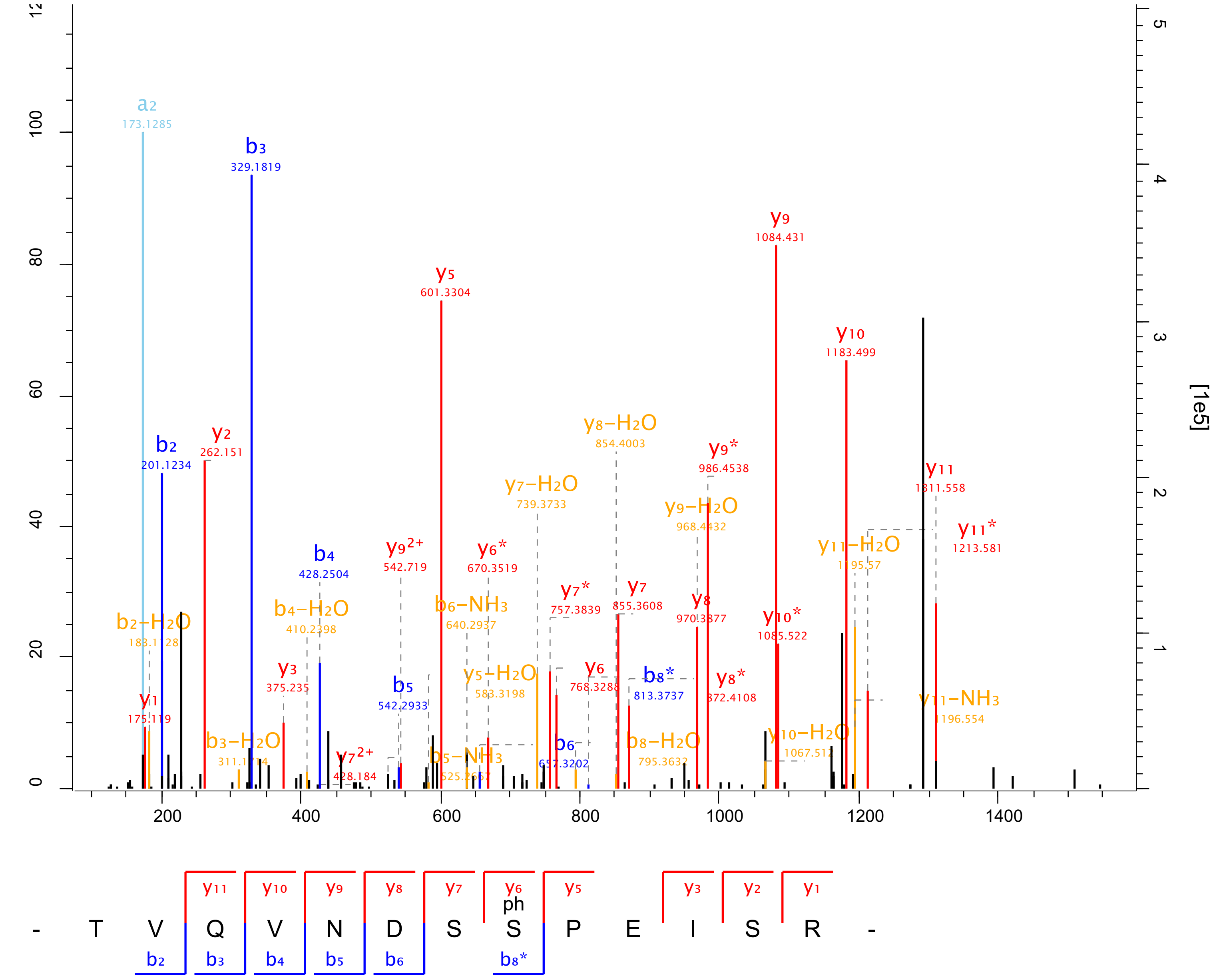

| Raw file | Scan | Method | Score | m/z |
| --- | --- | --- | --- | --- |
| water_15_2 | 11659 | FTMS; HCD | 55.04 | 512.25 |

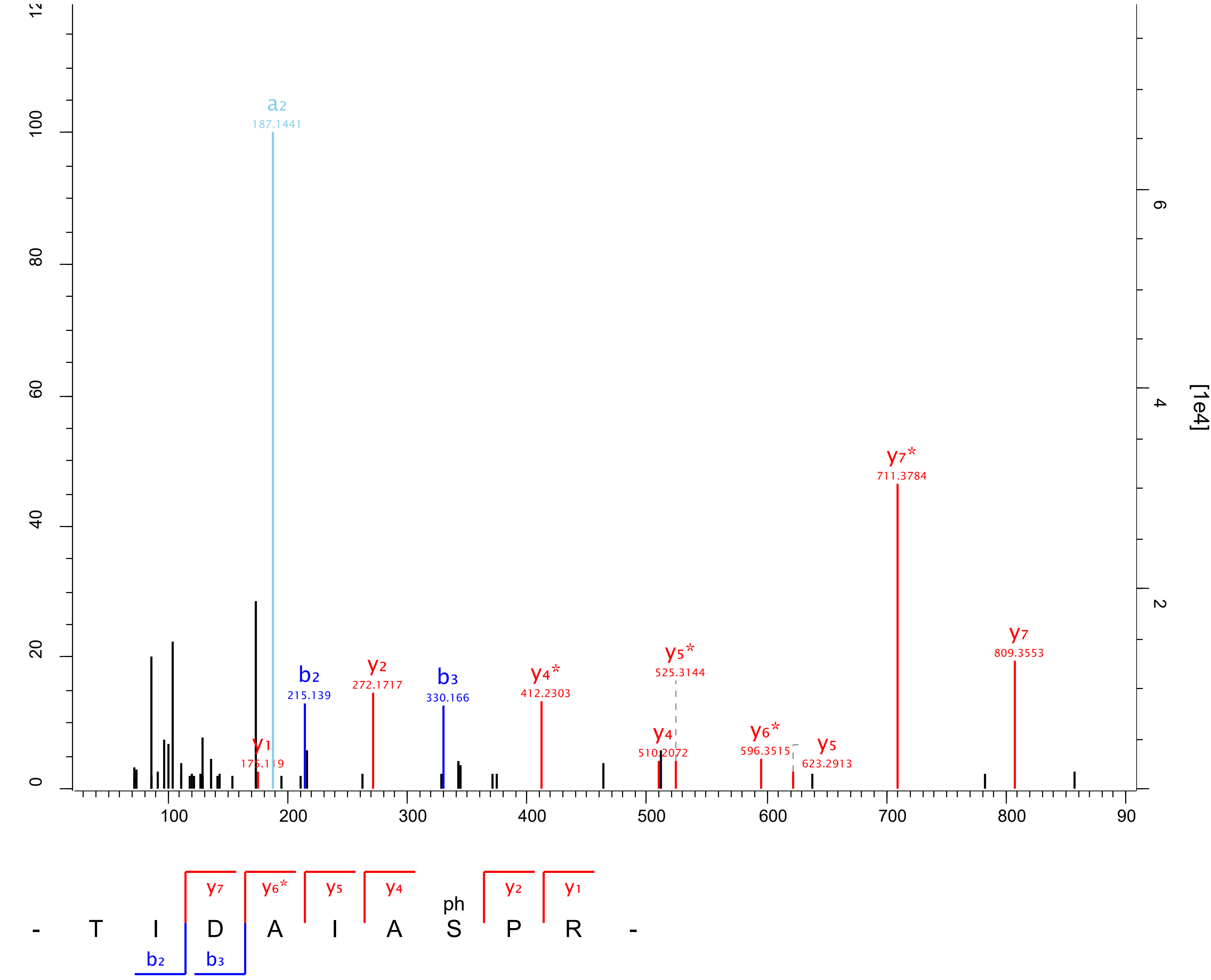

| Raw file | Scan | Method | Score | m/z |
|---|---|---|---|---|
| water_15_2 | 11742 | FTMS; HCD | 169.6 | 1002.9 |

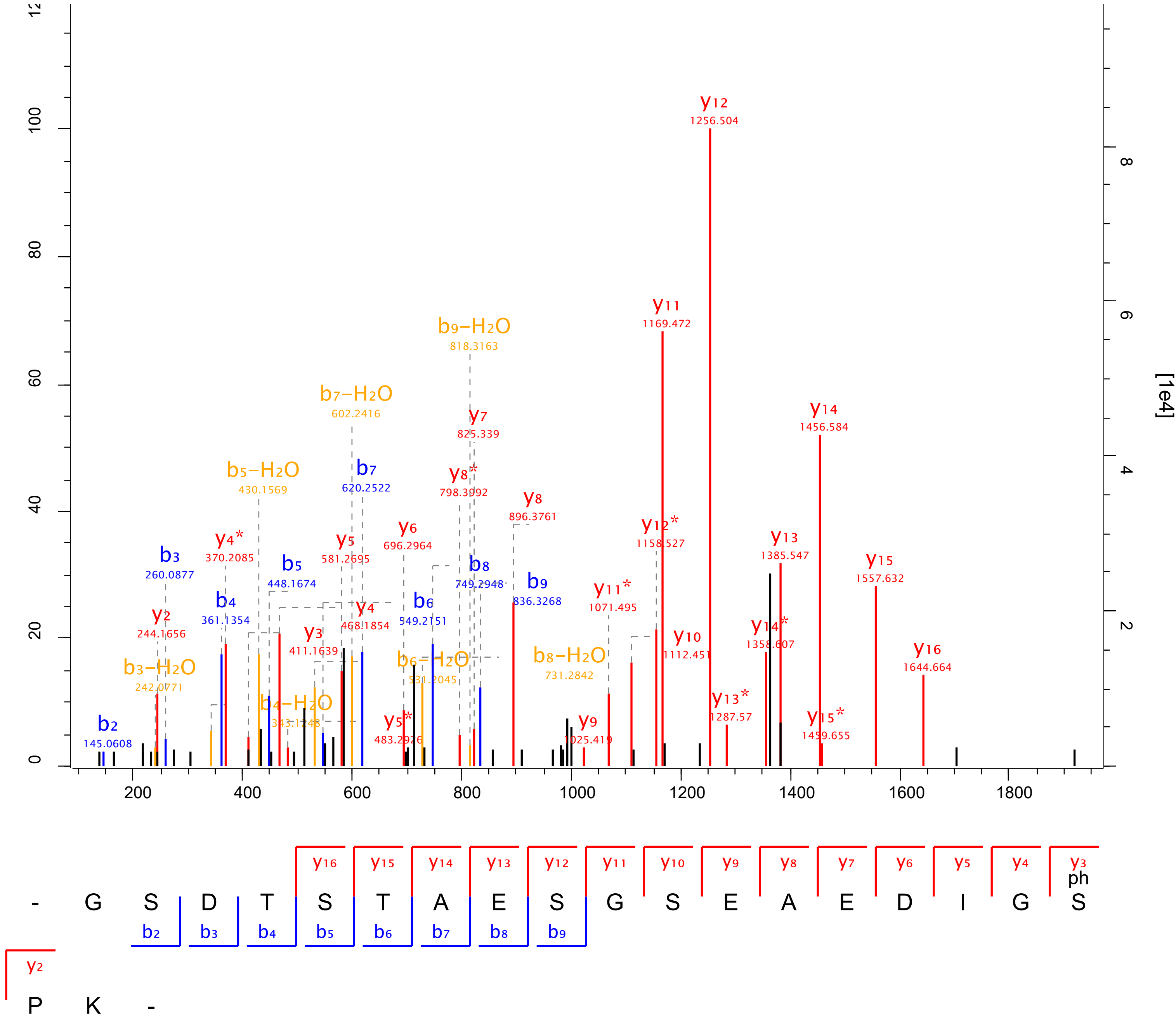

| Raw file | Scan | Method | Score | m/z |
| --- | --- | --- | --- | --- |
| water_15_2 | 11785 | FTMS; HCD | 124.51 | 635.27 |

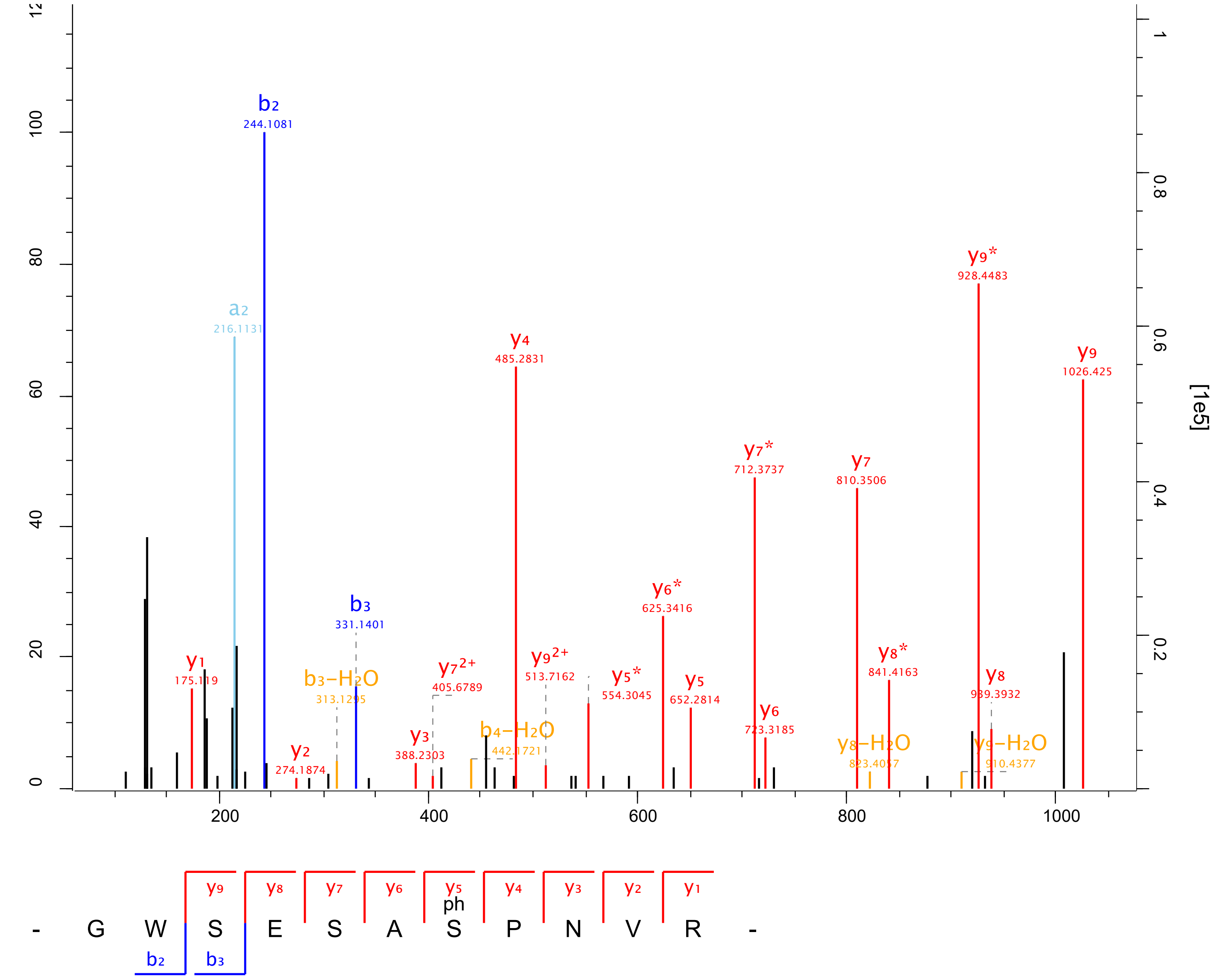

| Raw file | Scan | Method | Score | m/z |
|---|---|---|---|---|
| water_15_2 | 11805 | FTMS; HCD | 52.87 | 809.35 |

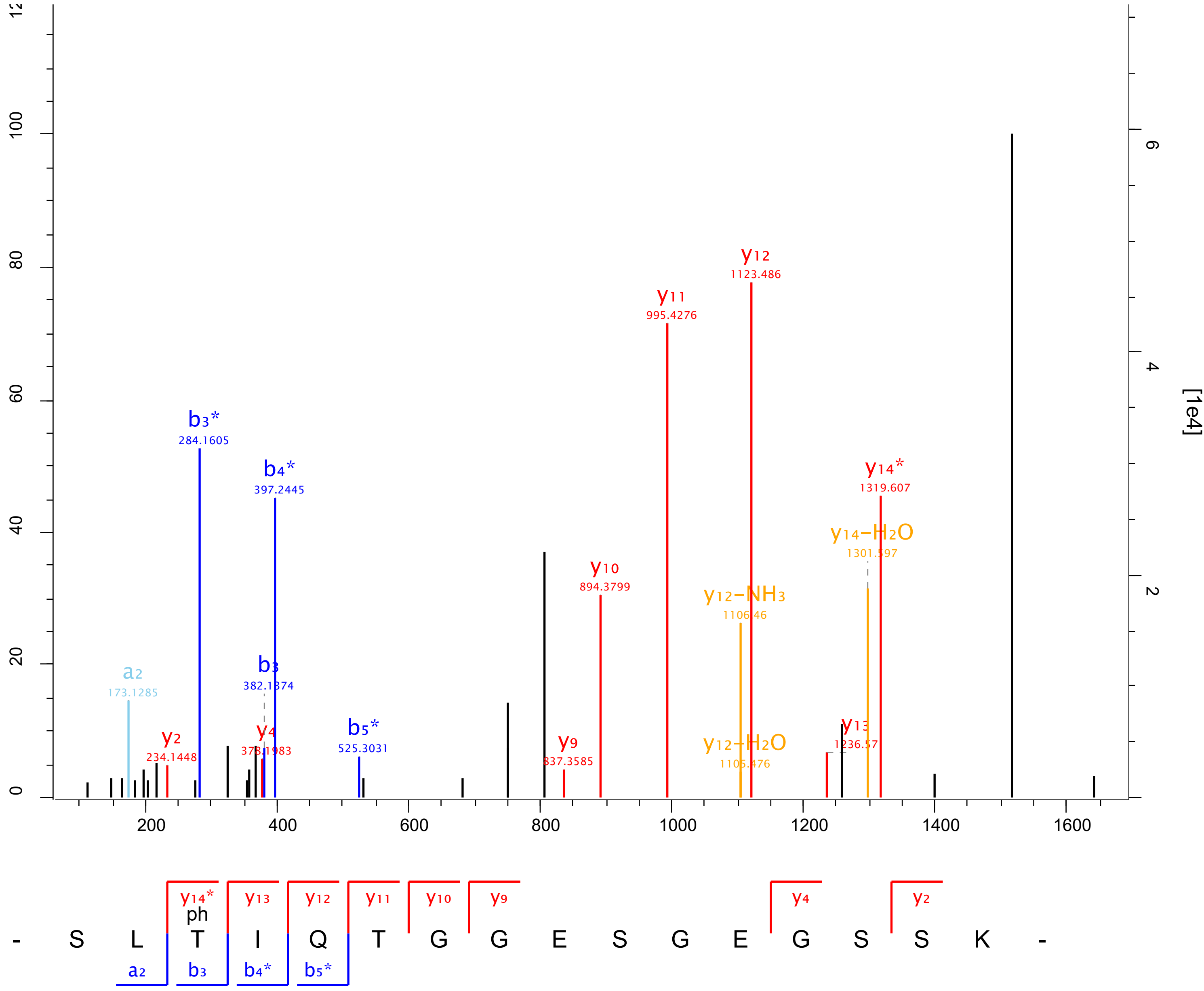

| Raw file | Scan | Method | Score | m/z |
| --- | --- | --- | --- | --- |
| water_15_2 | 11827 | FTMS; HCD | 58.69 | 898.33 |

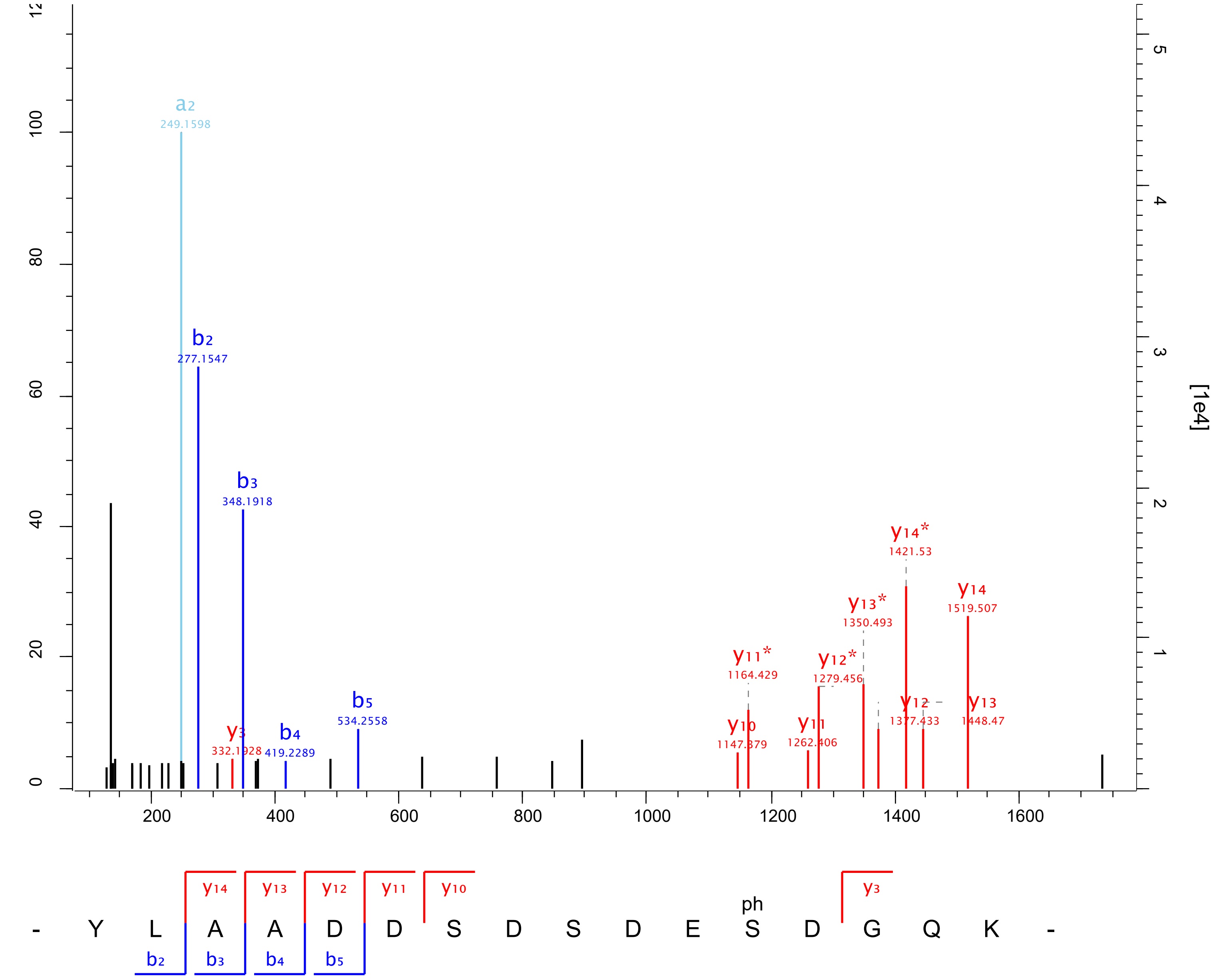

| Raw file | Scan | Method | Score | m/z |
|---|---|---|---|---|
| water_15_2 | 11832 | FTMS; HCD | 106.57 | 849.39 |

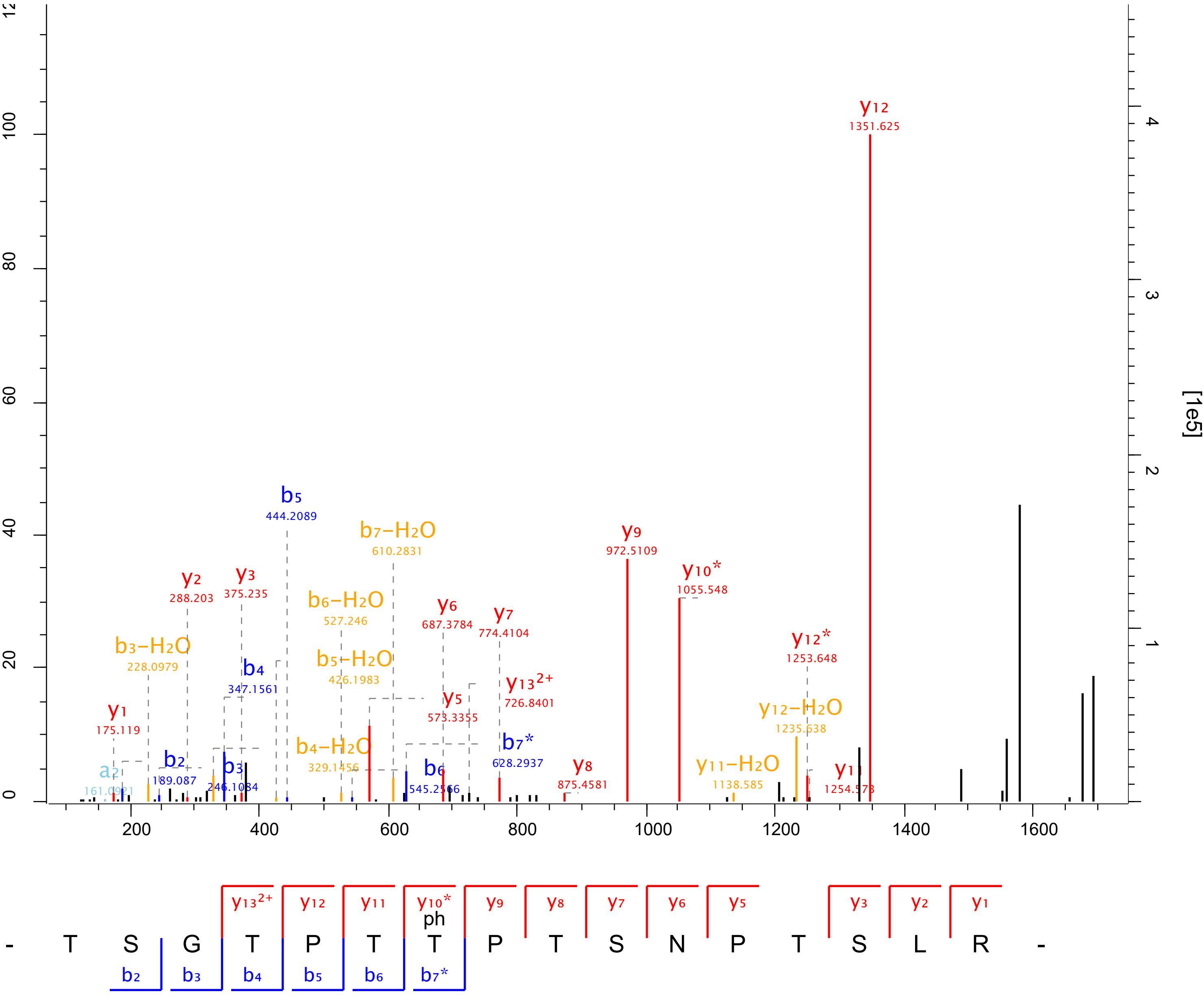

| Raw file | Scan | Method | Score | m/z |
|---|---|---|---|---|
| water_15_2 | 11883 | FTMS; HCD | 98.94 | 572.24 |

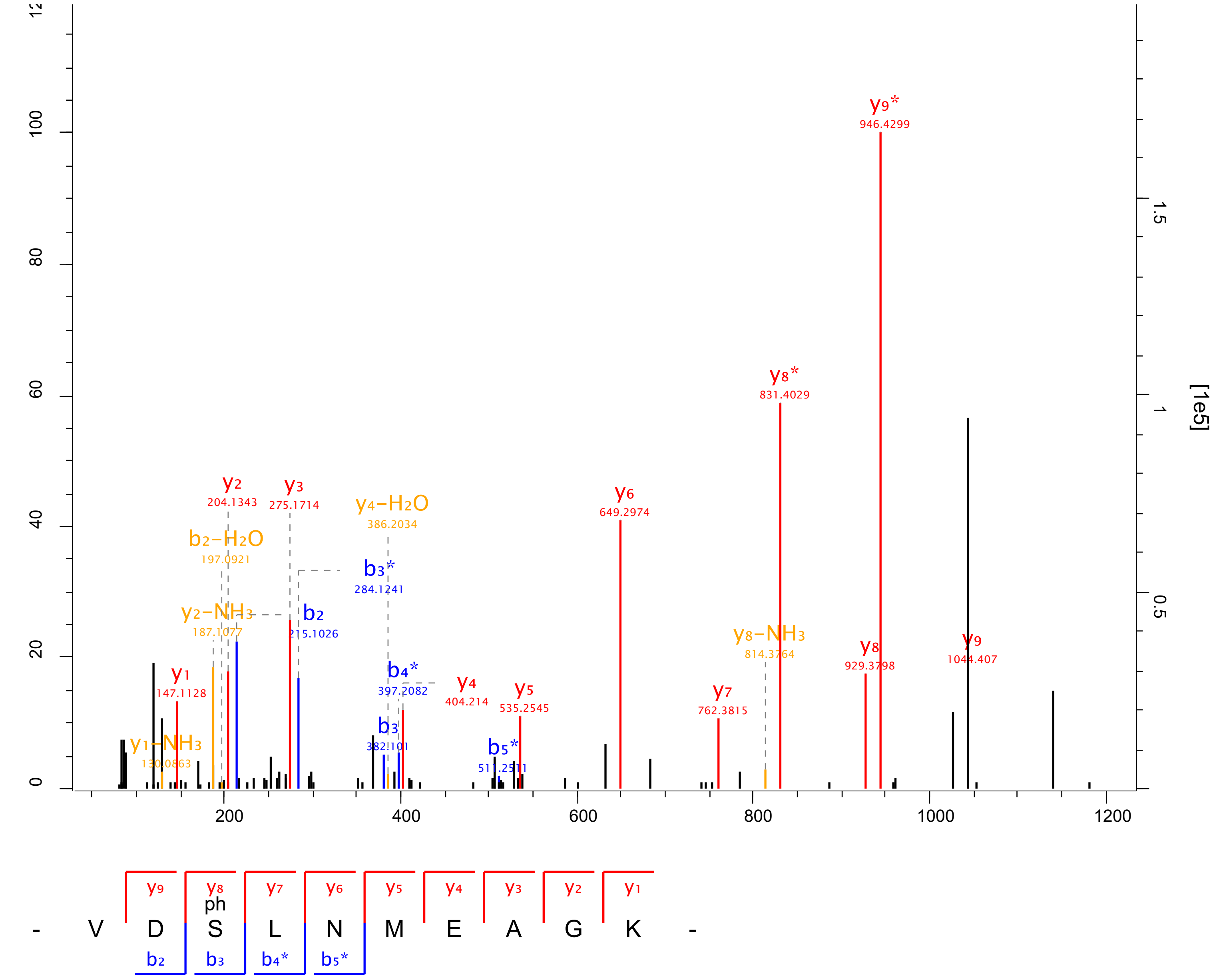

| Raw file | Scan | Method | Score | m/z |
|---|---|---|---|---|
| water_15_2 | 11894 | FTMS; HCD | 41.7 | 711.83 |

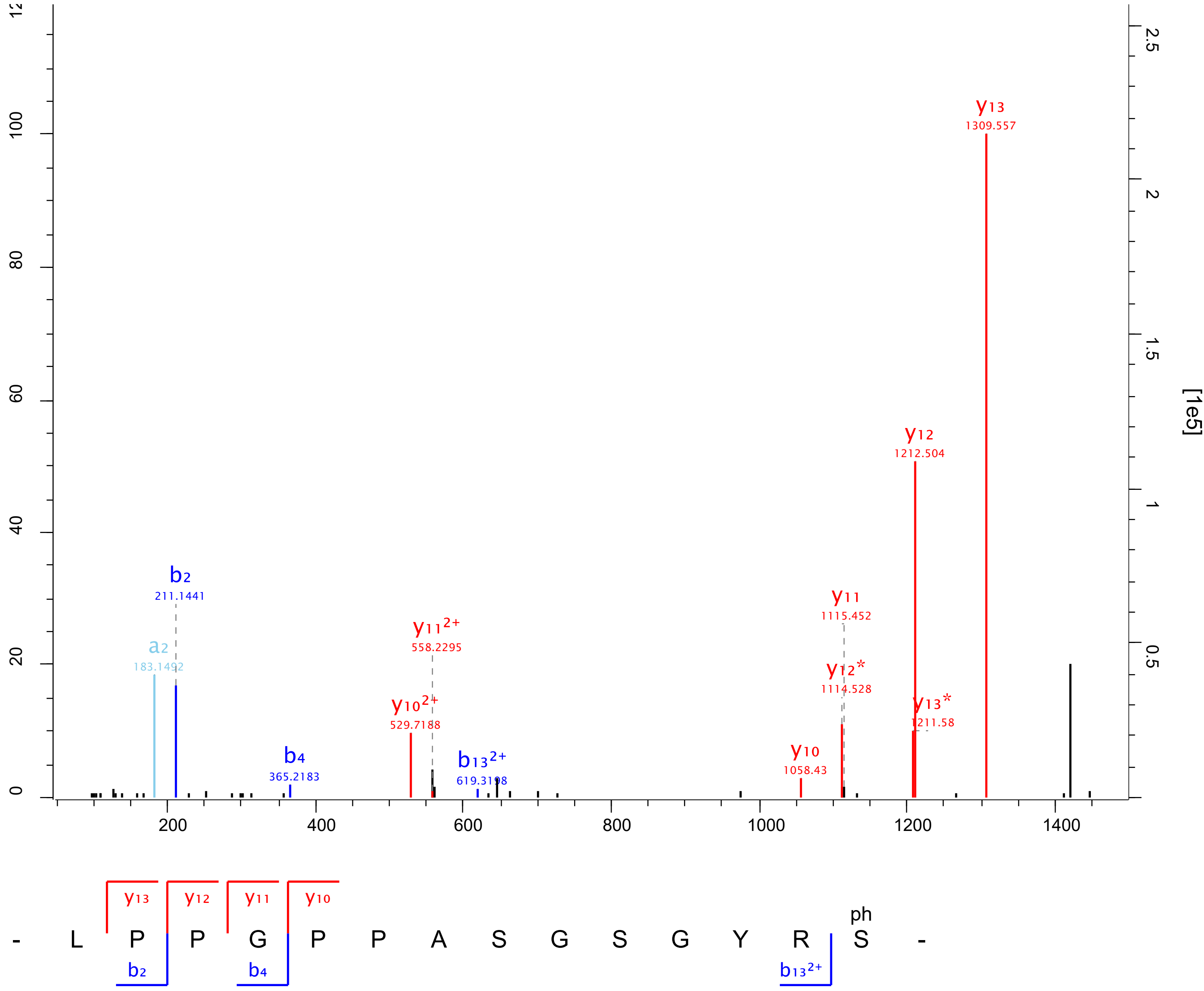

| Raw file | Scan | Method | Score | m/z |
|---|---|---|---|---|
| water_15_2 | 11925 | FTMS; HCD | 281.67 | 763.82 |

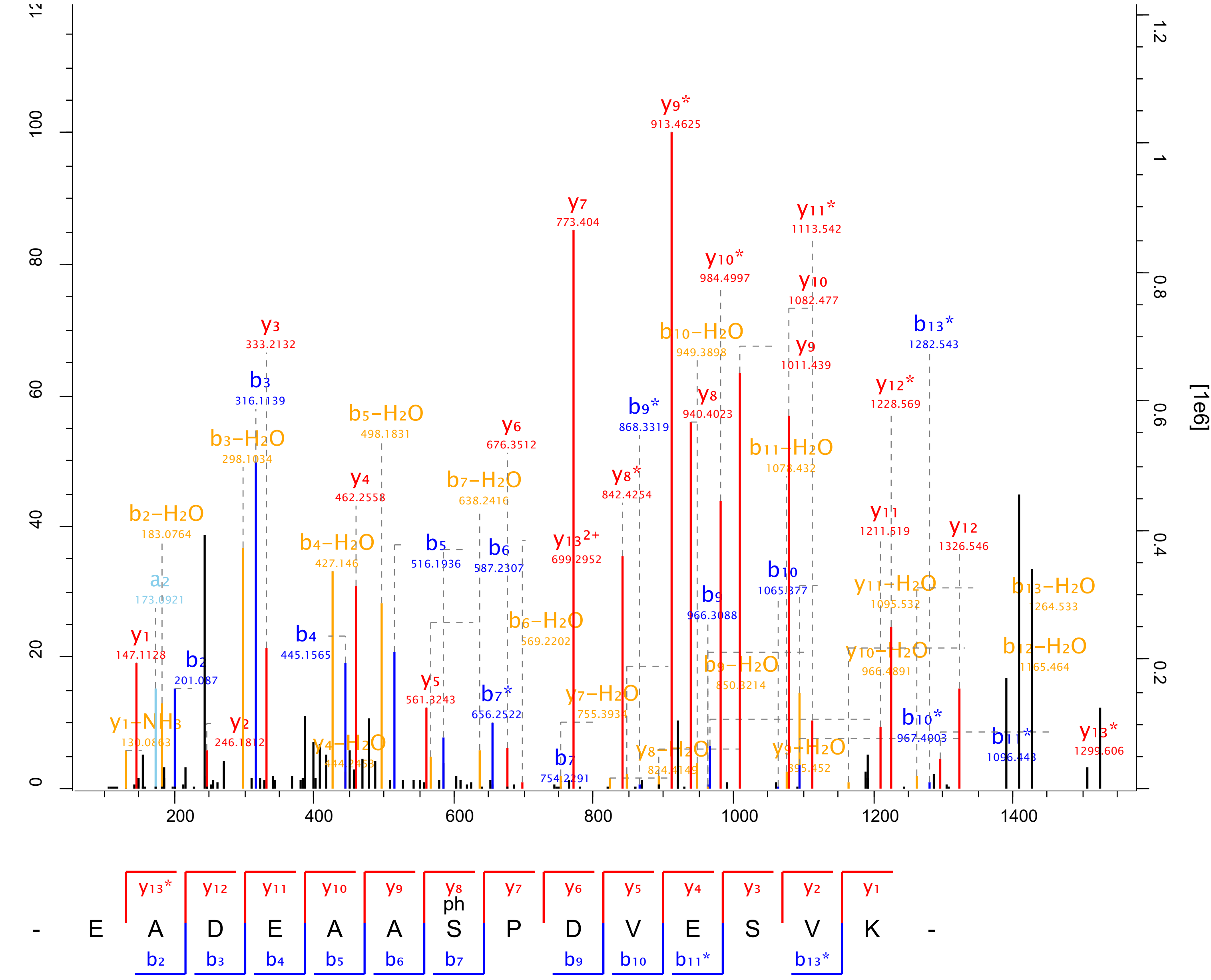

| Raw file | Scan | Method | Score | m/z |
|---|---|---|---|---|
| water_15_2 | 11956 | FTMS; HCD | 79.99 | 450.2 |

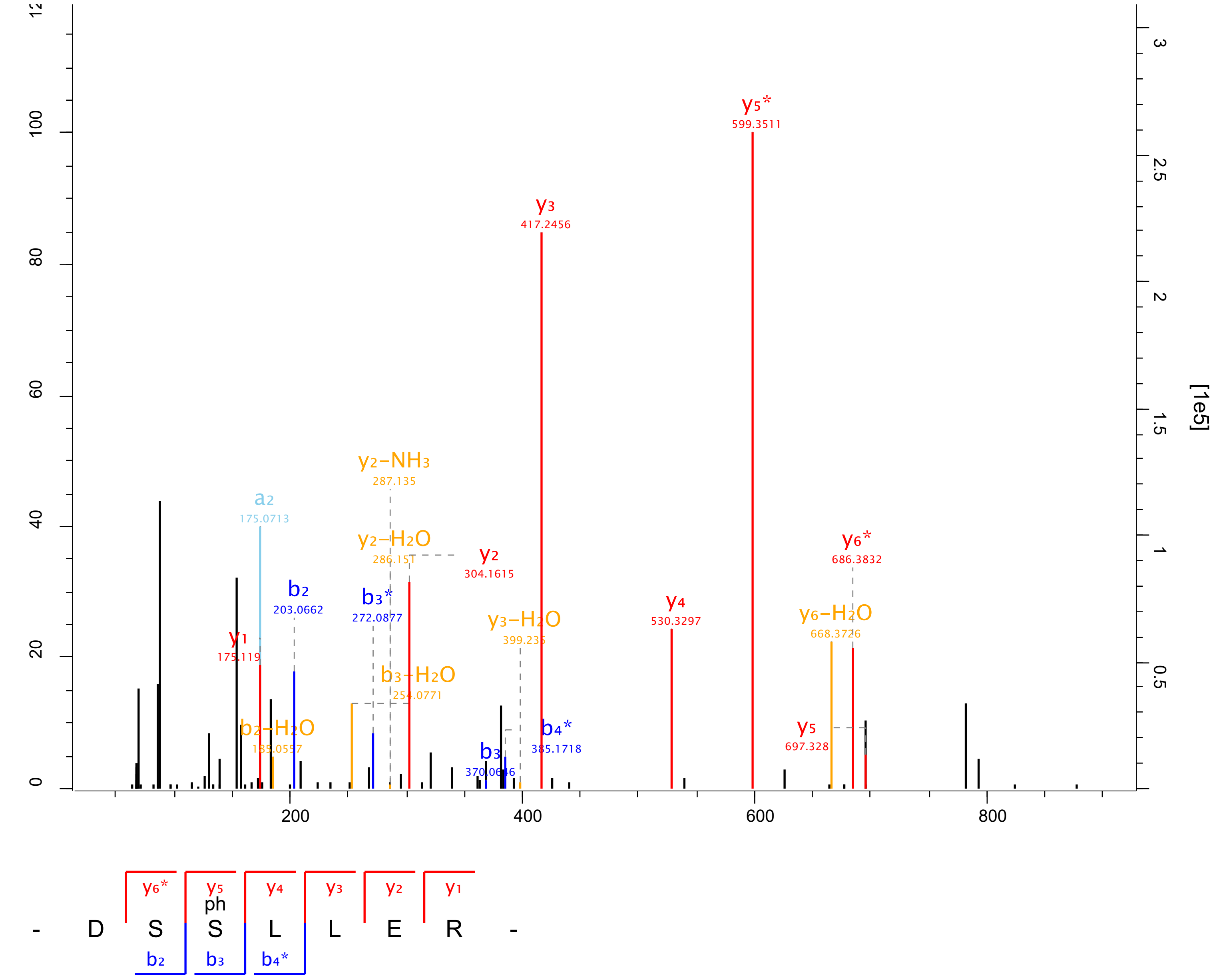

| Raw file | Scan | Method | Score | m/z |
| --- | --- | --- | --- | --- |
| water_15_2 | 12019 | FTMS; HCD | 209.11 | 603.29 |

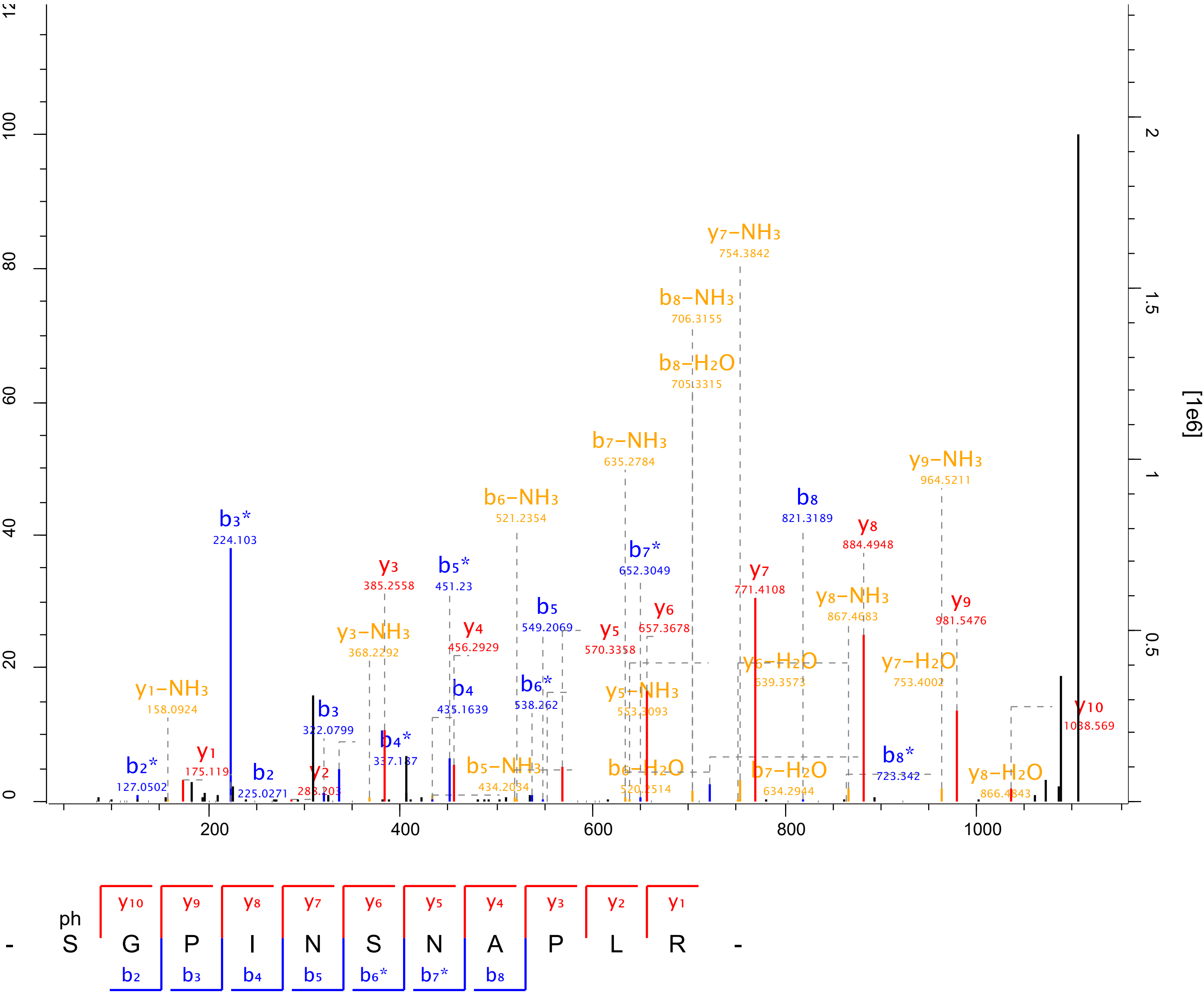

| Raw file | Scan | Method | Score | m/z |
|---|---|---|---|---|
| water_15_2 | 12124 | FTMS; HCD | 147.26 | 556.77 |

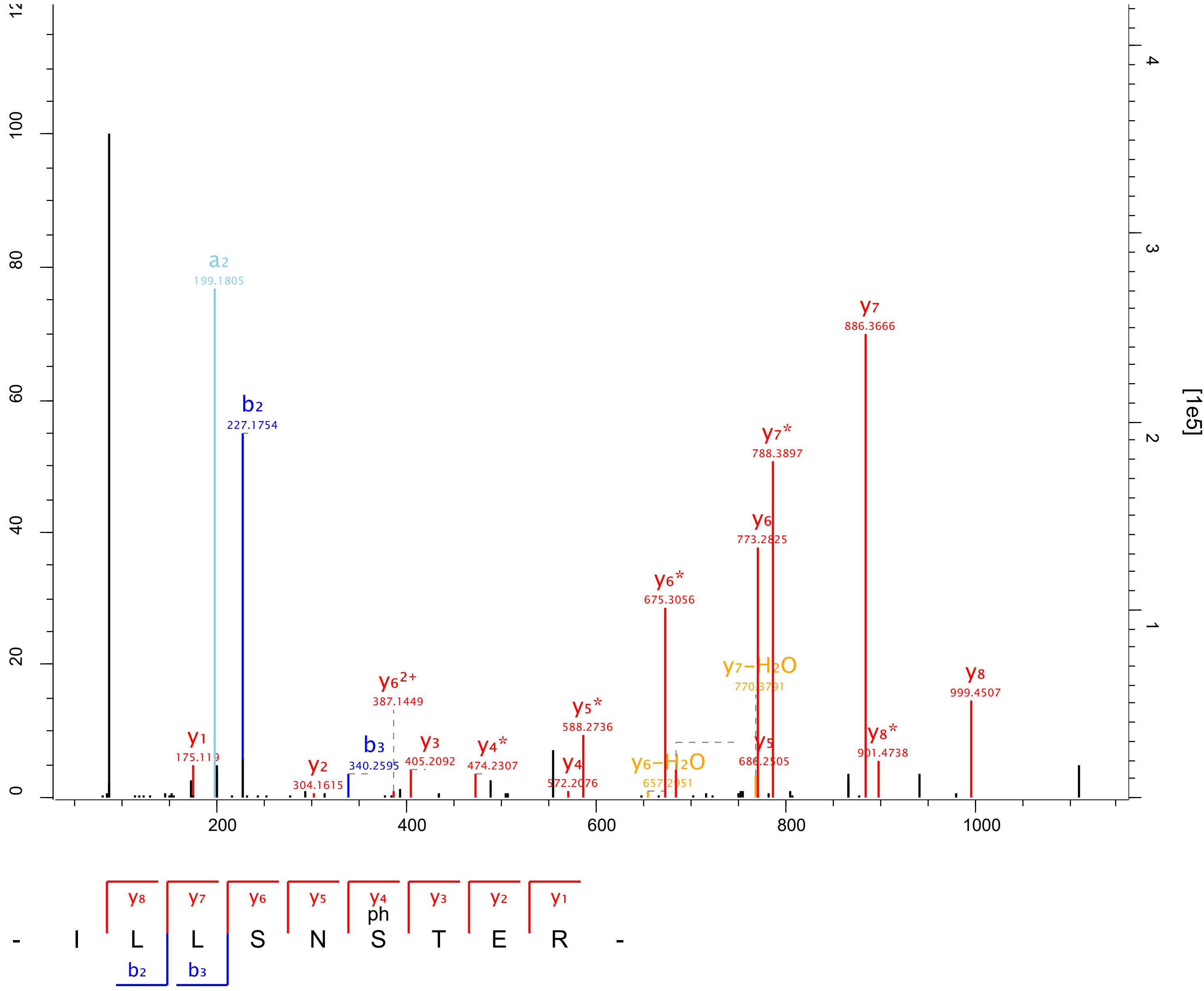

| Raw file | Scan | Method | Score | m/z |
|---|---|---|---|---|
| water_15_2 | 12225 | FTMS; HCD | 61.34 | 418.7 |

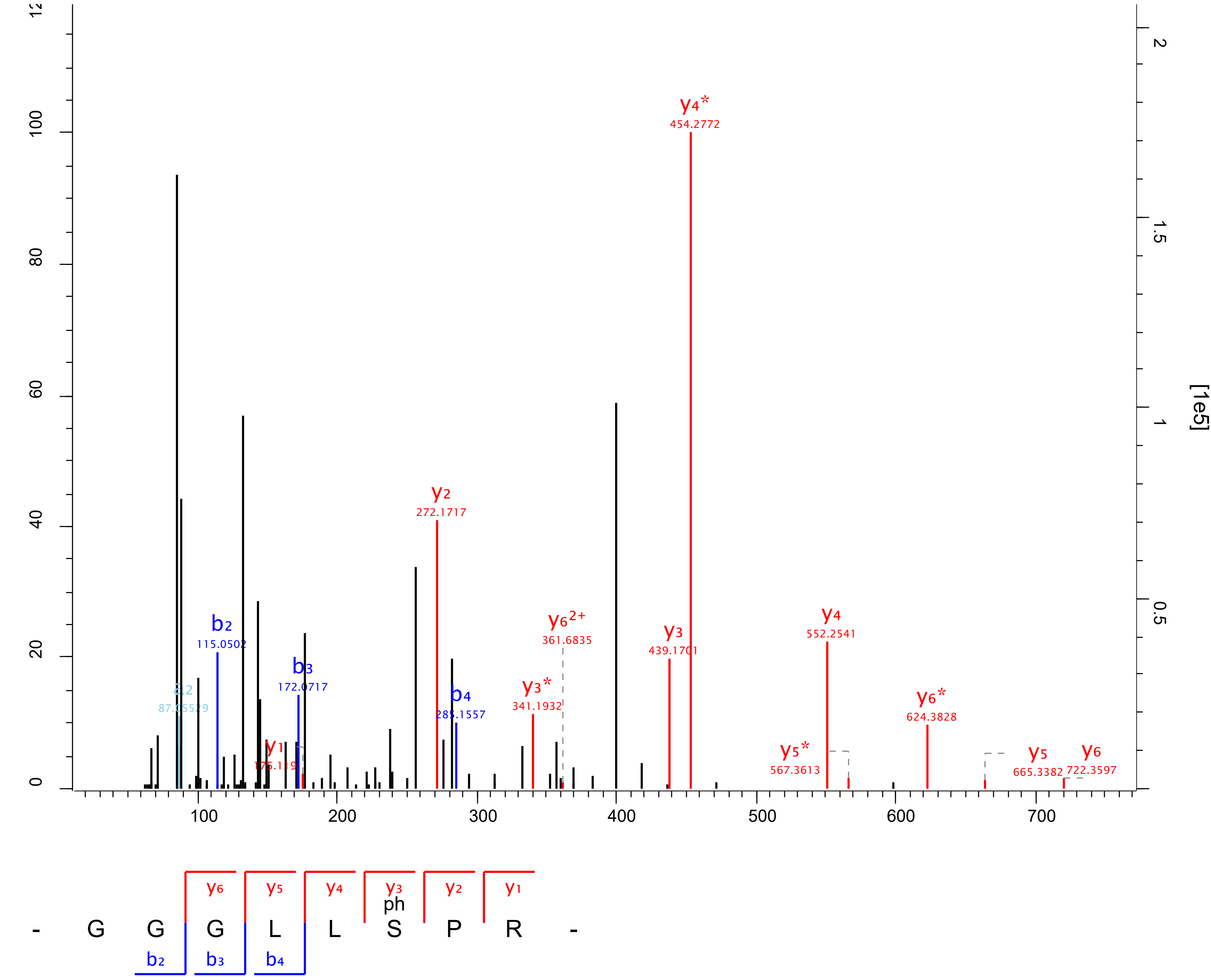

| Raw file | Scan | Method | Score | m/z |
|---|---|---|---|---|
| water_15_2 | 12276 | FTMS; HCD | 66.02 | 555.25 |

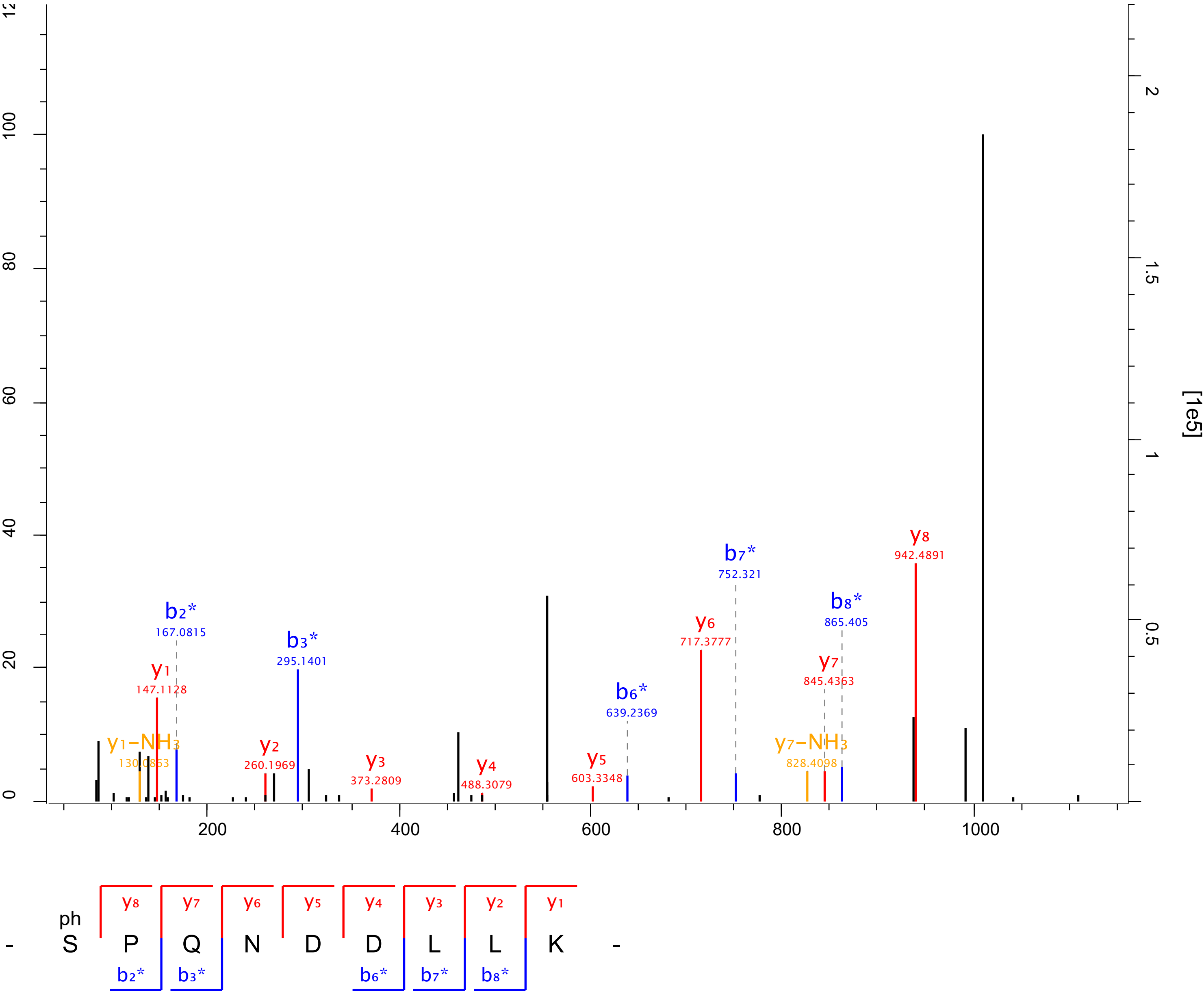

| Raw file | Scan | Method | Score | m/z |
| --- | --- | --- | --- | --- |
| water_15_2 | 12334 | FTMS; HCD | 58.7 | 508.74 |

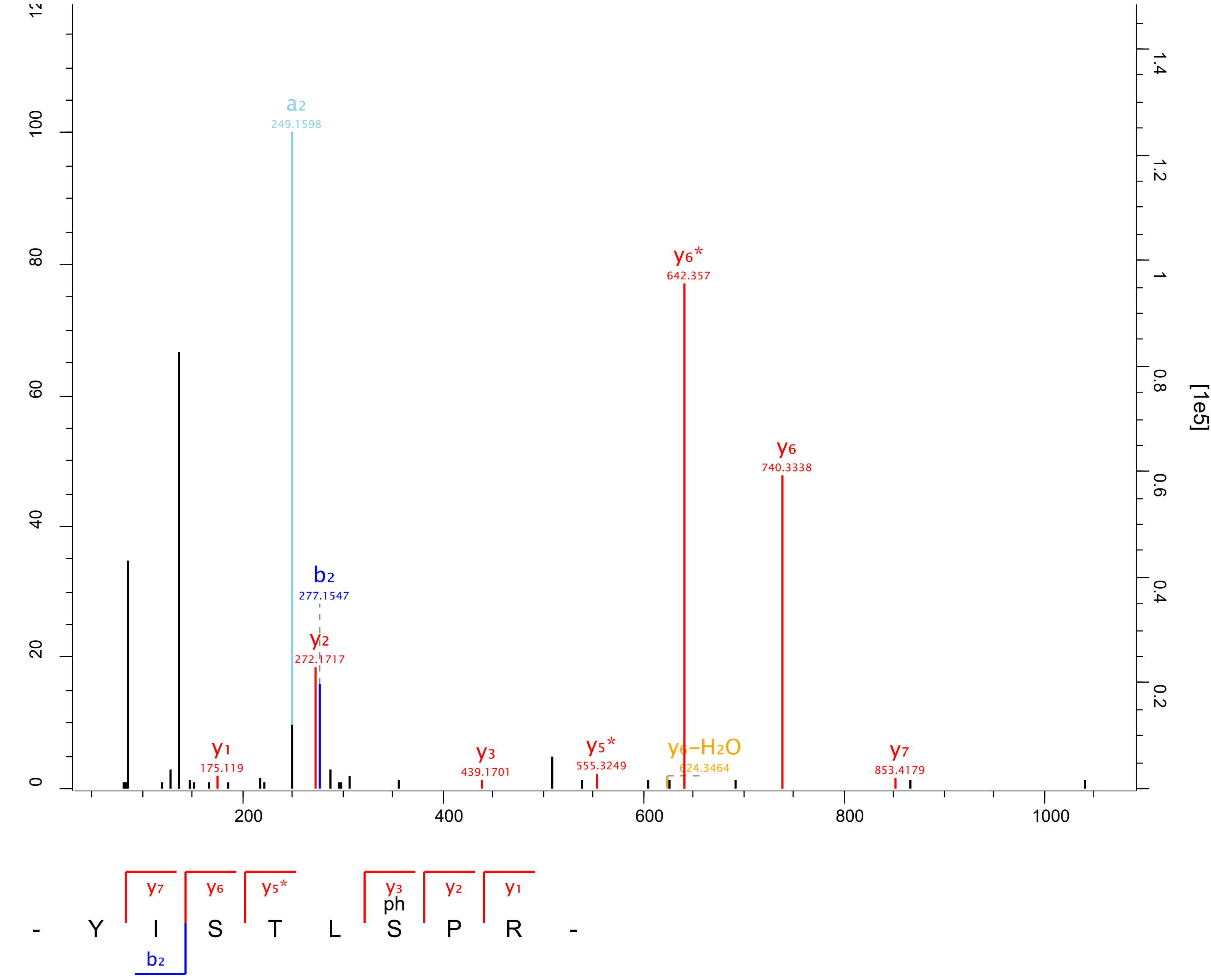

| Raw file | Scan | Method | Score | m/z |
|---|---|---|---|---|
| water_15_2 | 12350 | FTMS; HCD | 100.93 | 568.23 |

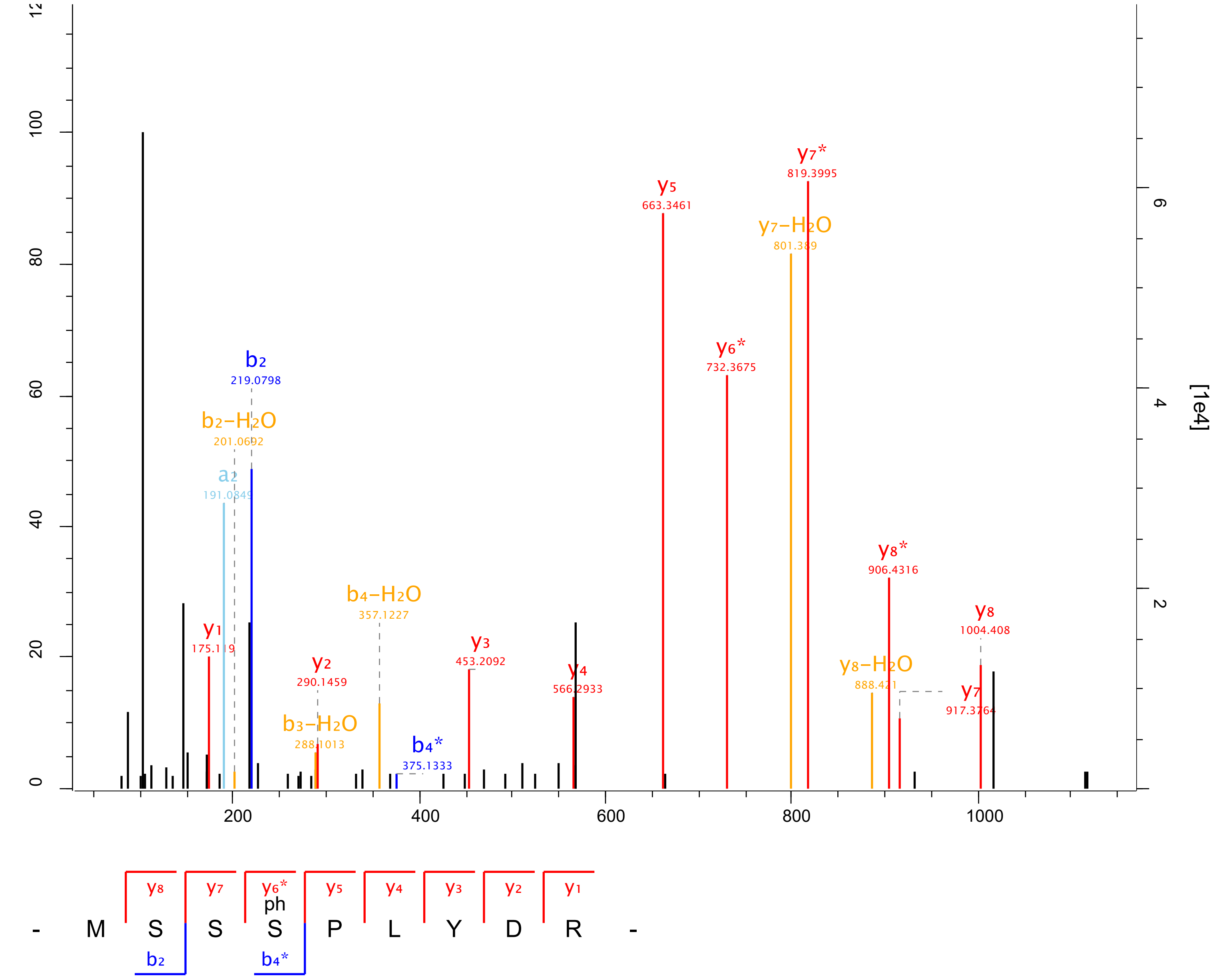

| Raw file | Scan | Method | Score | m/z |
|---|---|---|---|---|
| water_15_2 | 12408 | FTMS; HCD | 77.73 | 487.7 |

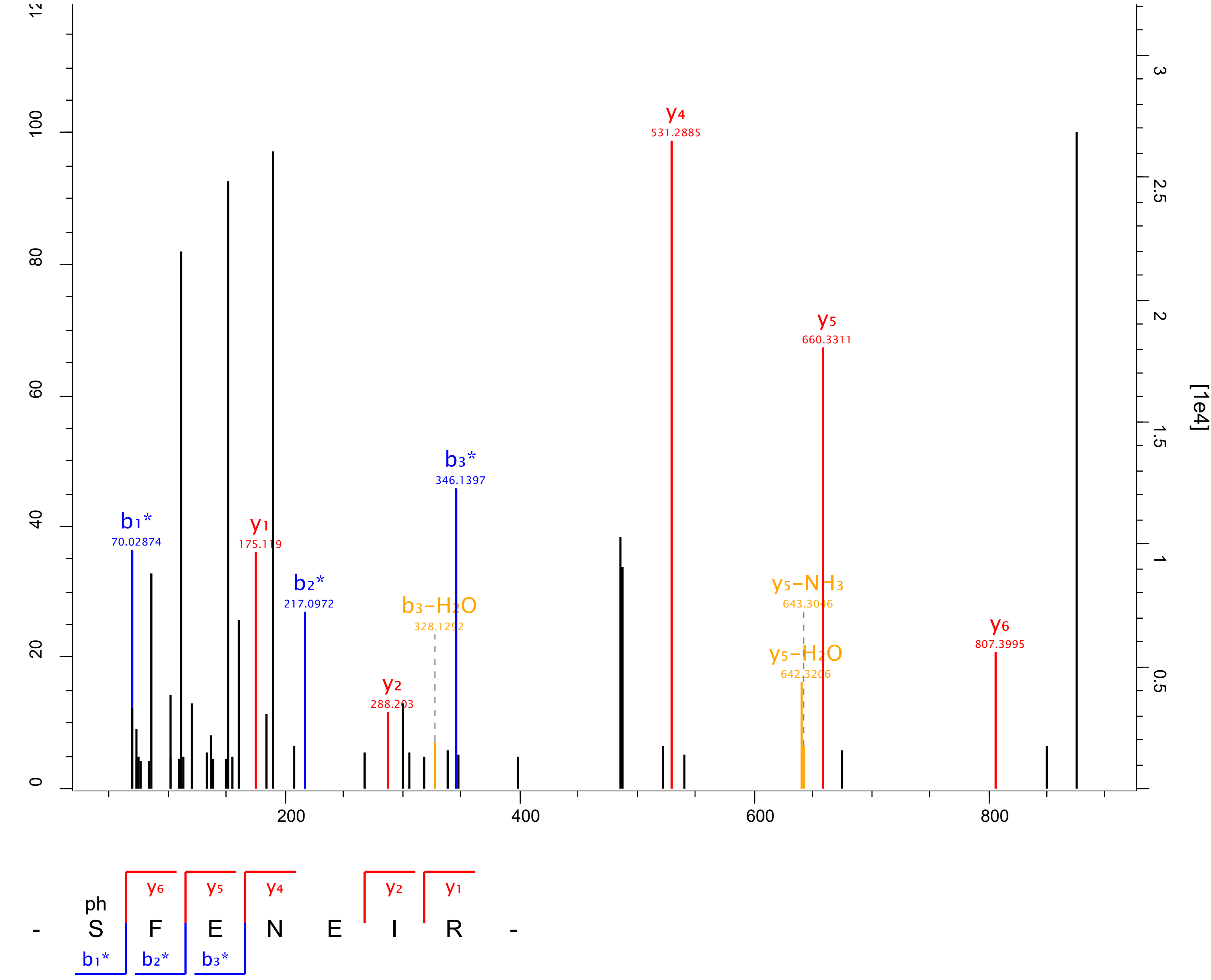

| Raw file | Scan | Method | Score | m/z |
|---|---|---|---|---|
| water_15_2 | 12462 | FTMS; HCD | 92.94 | 898.91 |

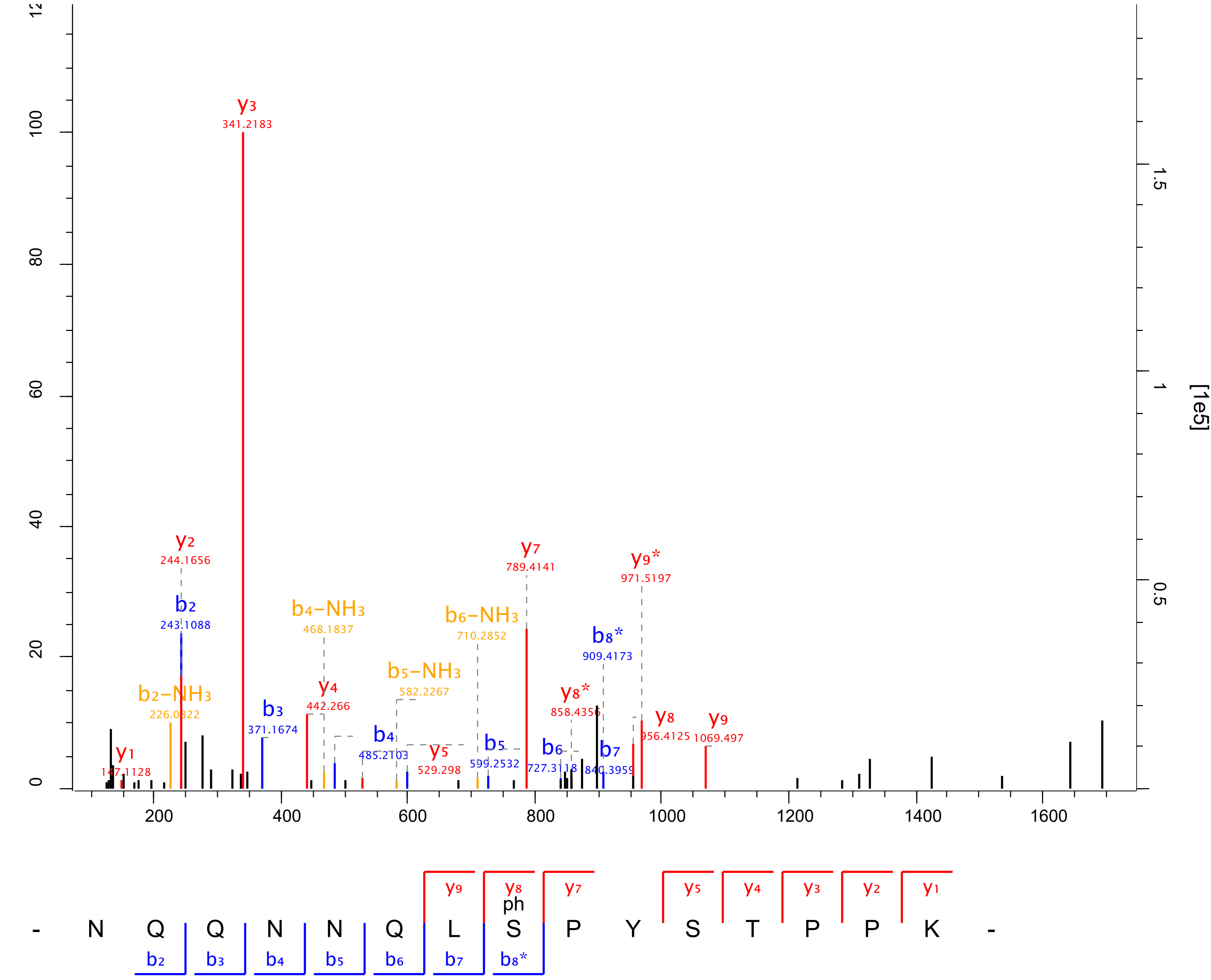

| Raw file | Scan | Method | Score | m/z |
|---|---|---|---|---|
| water_15_2 | 12503 | FTMS; HCD | 88.31 | 686.31 |

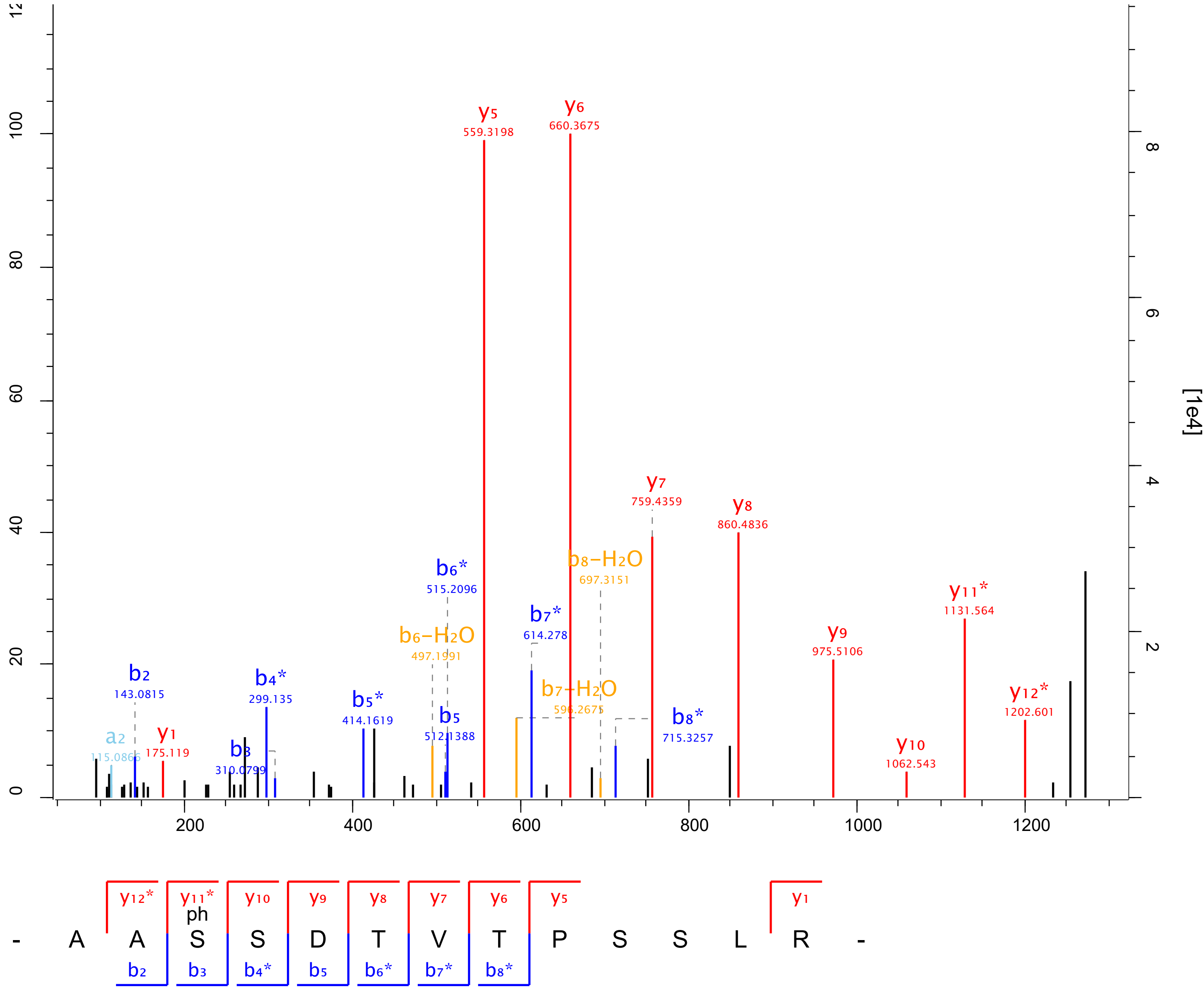

| Raw file | Scan | Method | Score | m/z |
|---|---|---|---|---|
| water_15_2 | 12529 | FTMS; HCD | 98.9 | 435.7 |

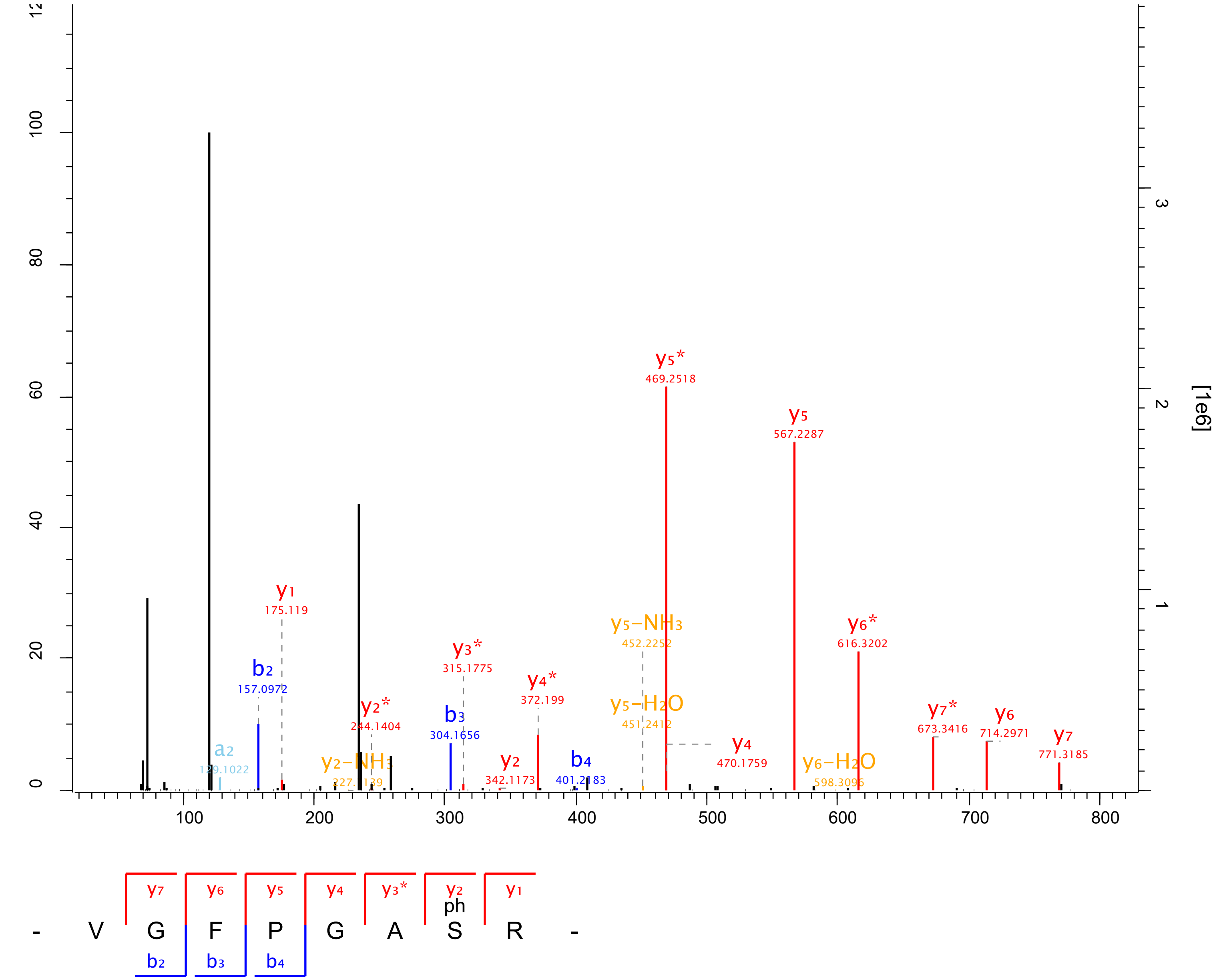

| Raw file | Scan | Method | Score | m/z |
|---|---|---|---|---|
| water_15_2 | 12781 | FTMS; HCD | 187.62 | 549.76 |

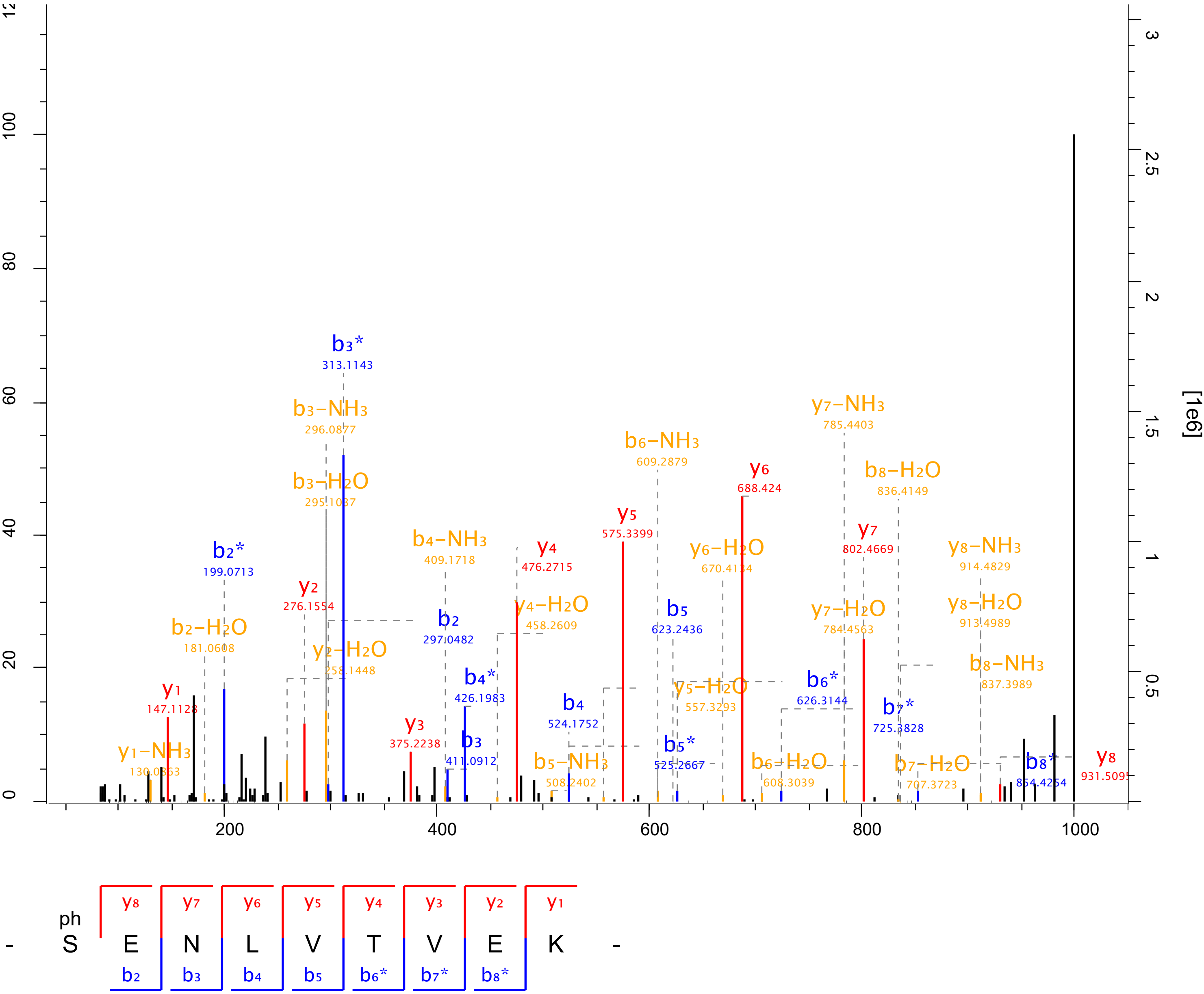

| Raw file | Scan | Method | Score | m/z |
| --- | --- | --- | --- | --- |
| water_15_2 | 12806 | FTMS; HCD | 75.02 | 774.83 |

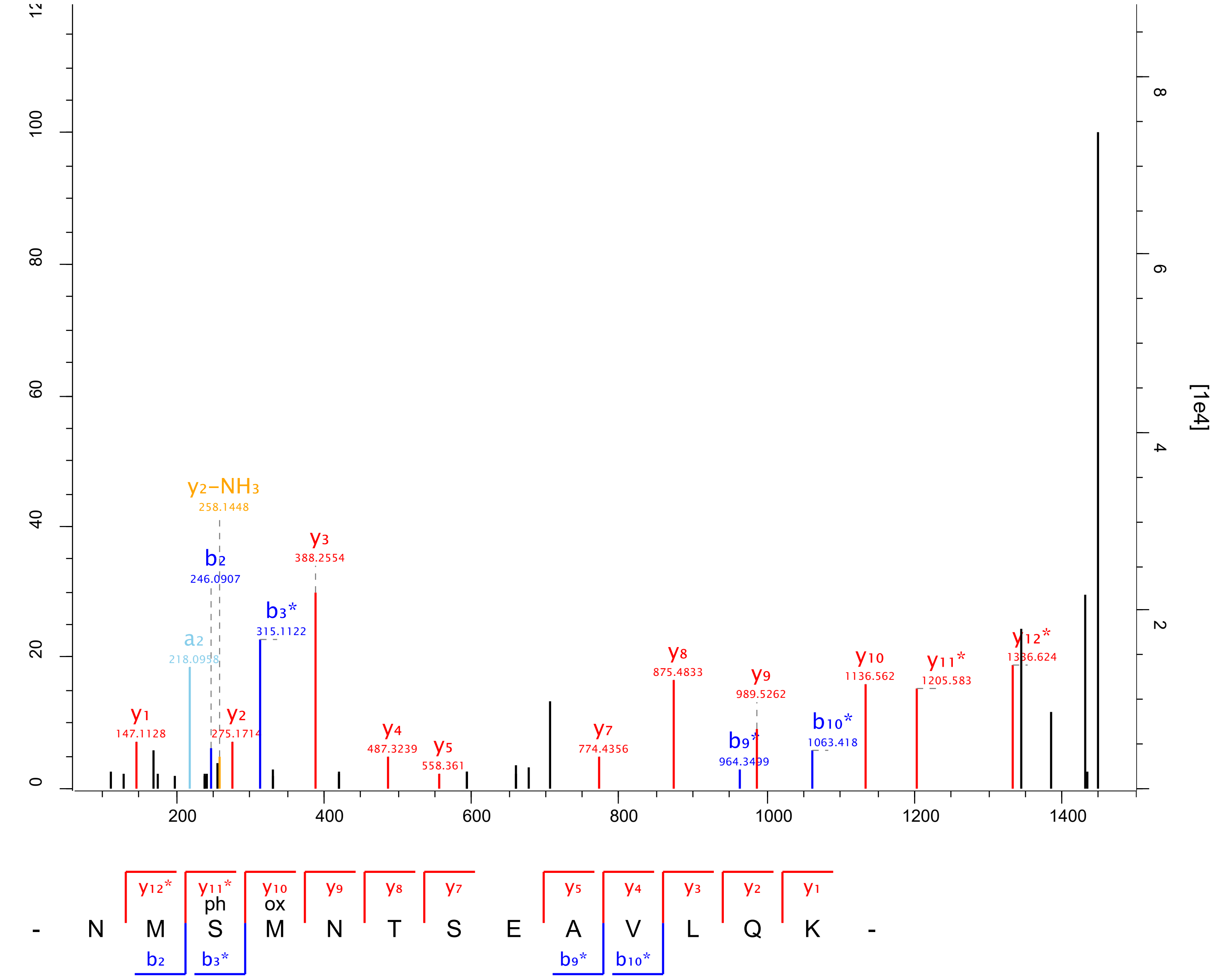

| Raw file | Scan | Method | Score | m/z |
|---|---|---|---|---|
| water_15_2 | 12829 | FTMS; HCD | 88.08 | 579.76 |

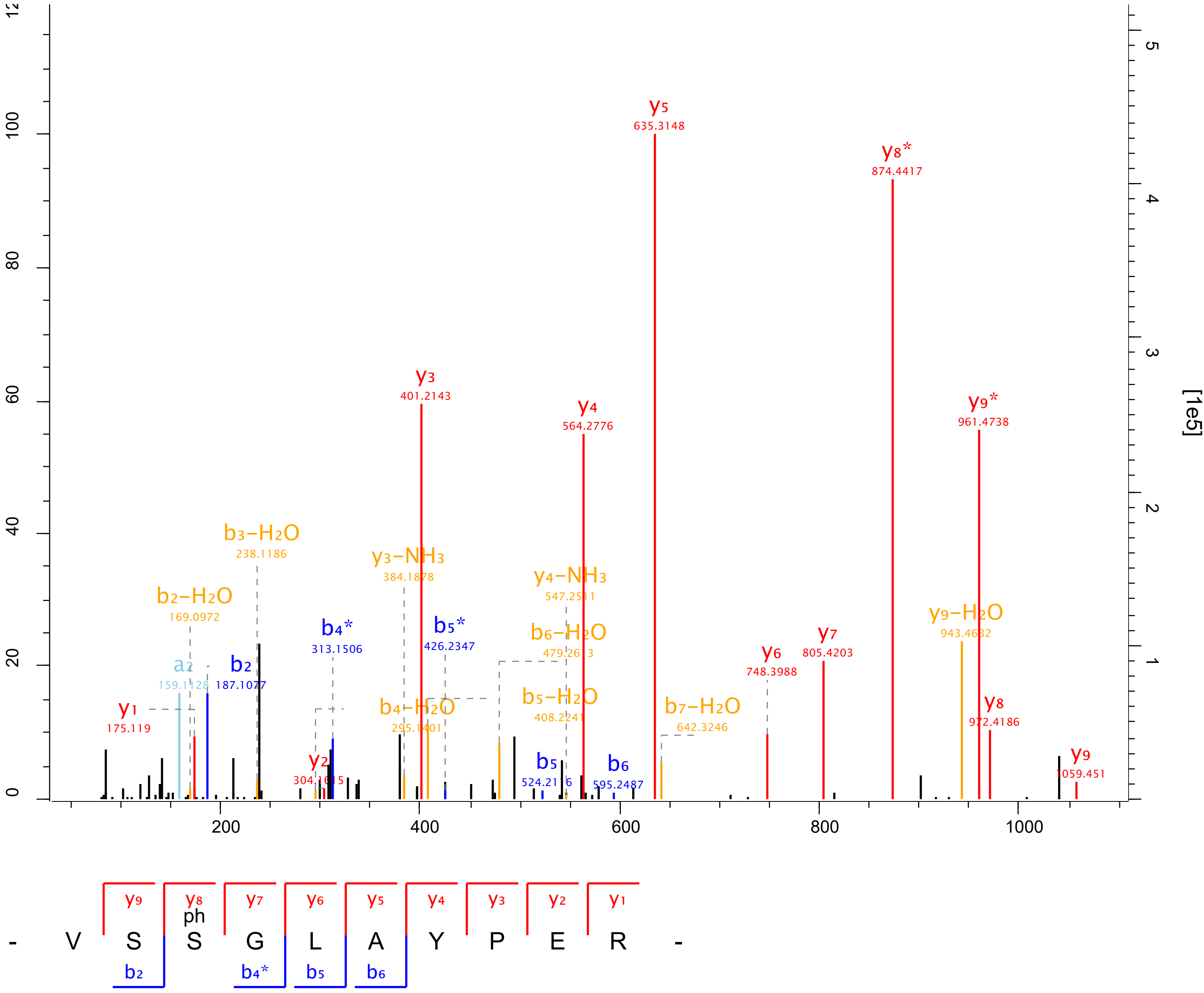

| Raw file | Scan | Method | Score | m/z |
| --- | --- | --- | --- | --- |
| water_15_2 | 12863 | FTMS; HCD | 72.89 | 587.76 |

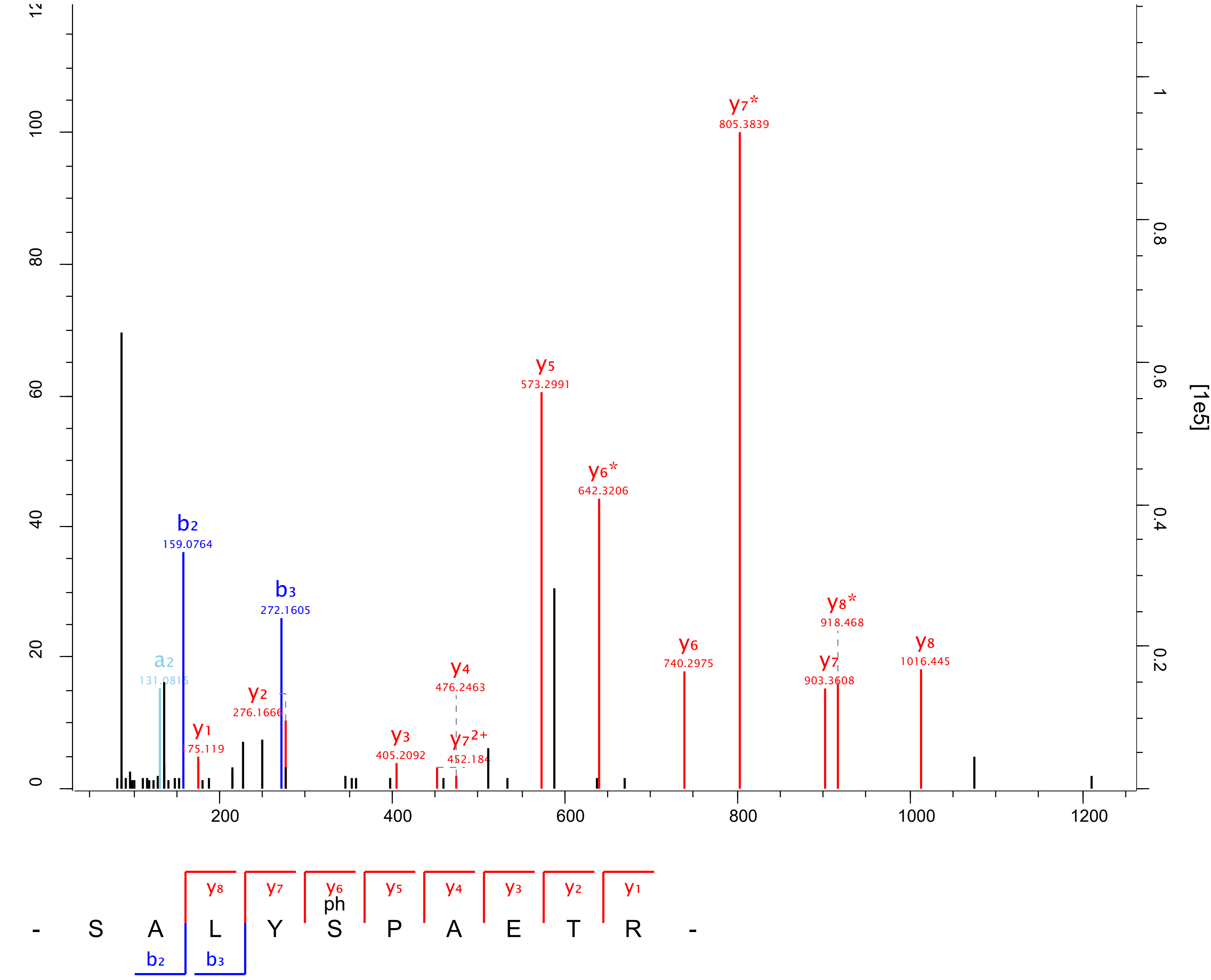

| Raw file | Scan | Method | Score | m/z |
| --- | --- | --- | --- | --- |
| water_15_2 | 12864 | FTMS; HCD | 54.04 | 948.72 |

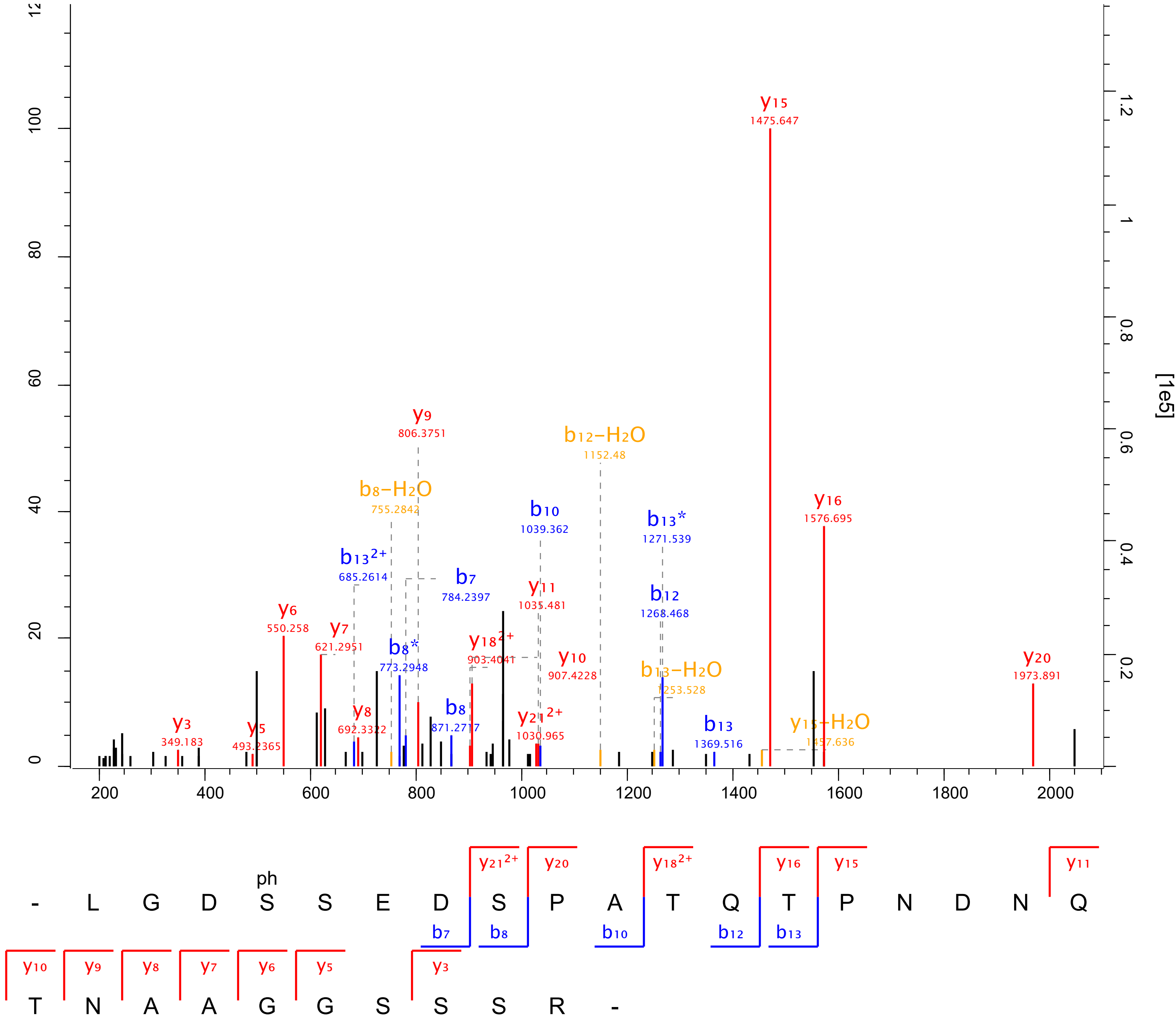

| Raw file | Scan | Method | Score | m/z |
| --- | --- | --- | --- | --- |
| water_15_2 | 12912 | FTMS; HCD | 62.29 | 577.27 |

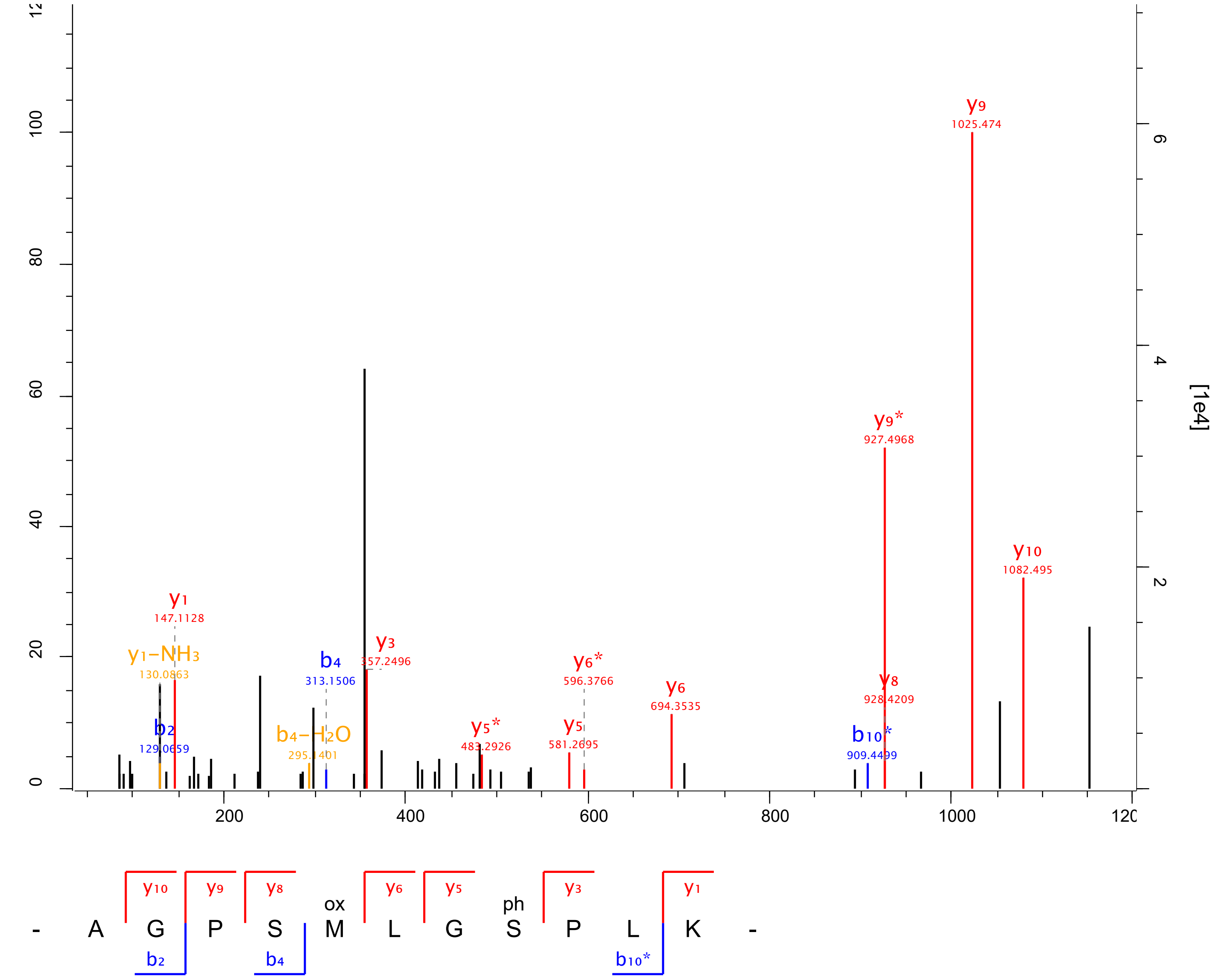

| Raw file | Scan | Method | Score | m/z |
|---|---|---|---|---|
| water_15_2 | 12928 | FTMS; HCD | 276.63 | 698.33 |

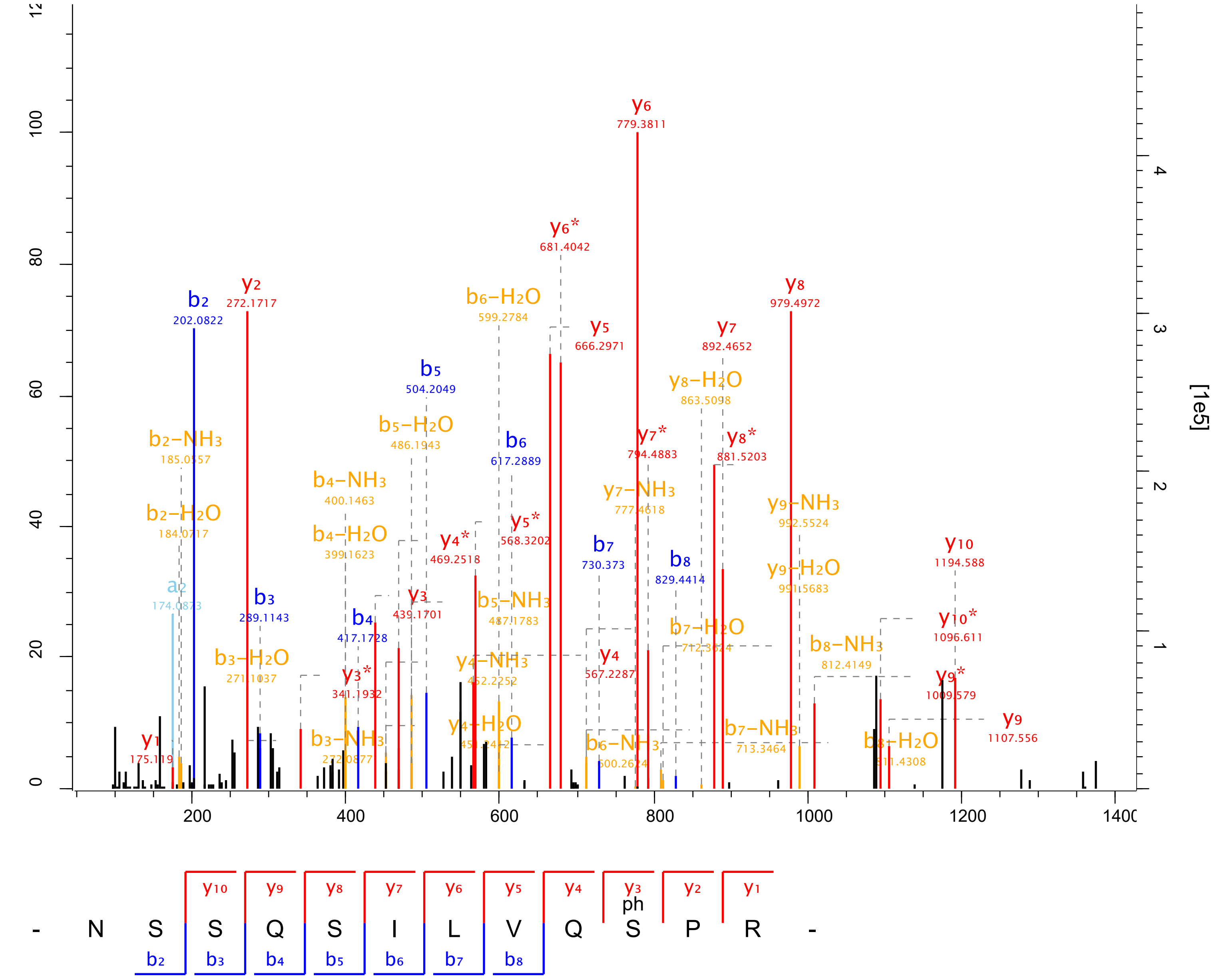

| Raw file | Scan | Method | Score | m/z |
| --- | --- | --- | --- | --- |
| water_15_2 | 12977 | FTMS; HCD | 56.88 | 656.95 |

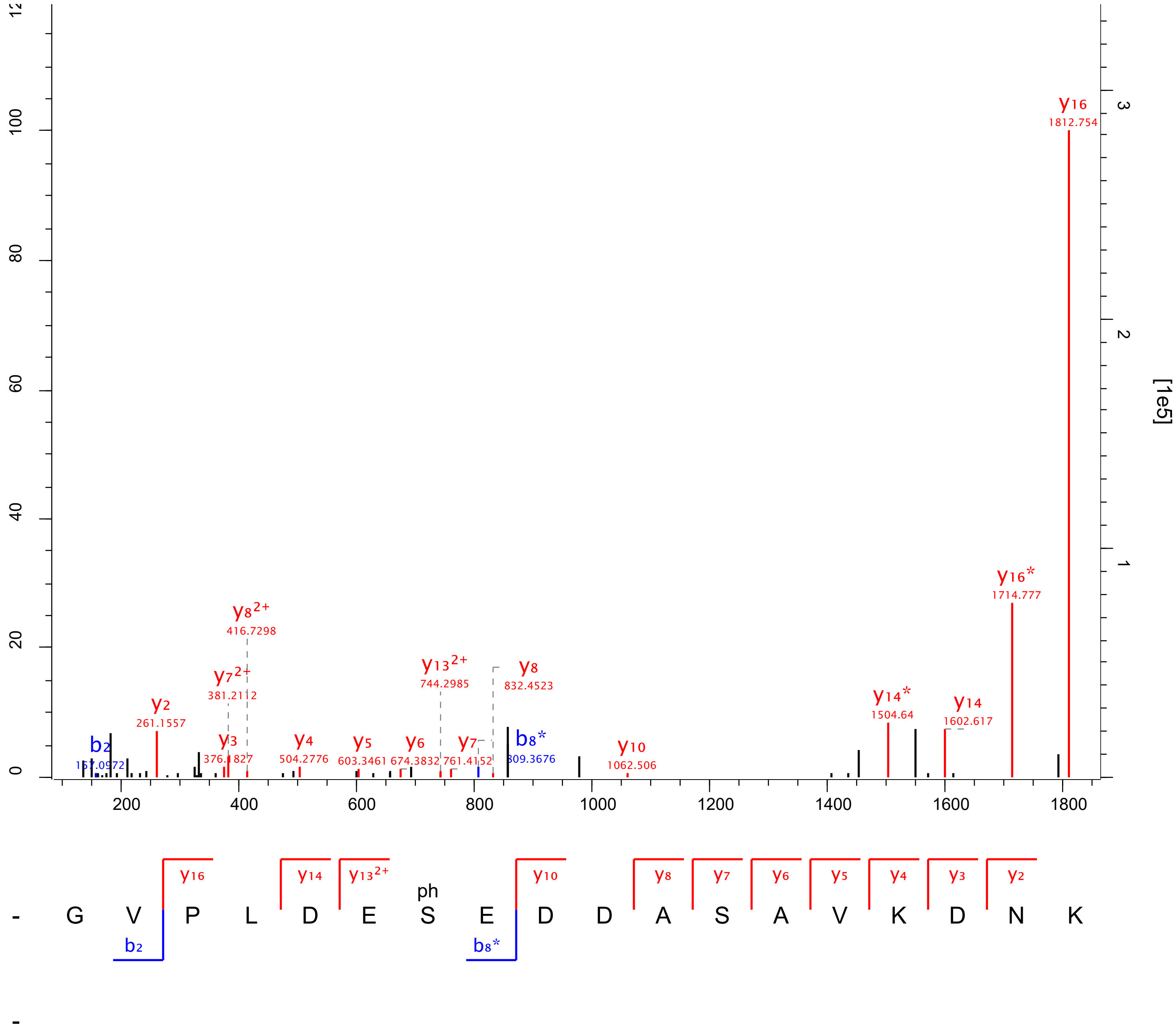

| Raw file | Scan | Method | Score | m/z |
| --- | --- | --- | --- | --- |
| water_15_2 | 13099 | FTMS; HCD | 104.95 | 610.79 |

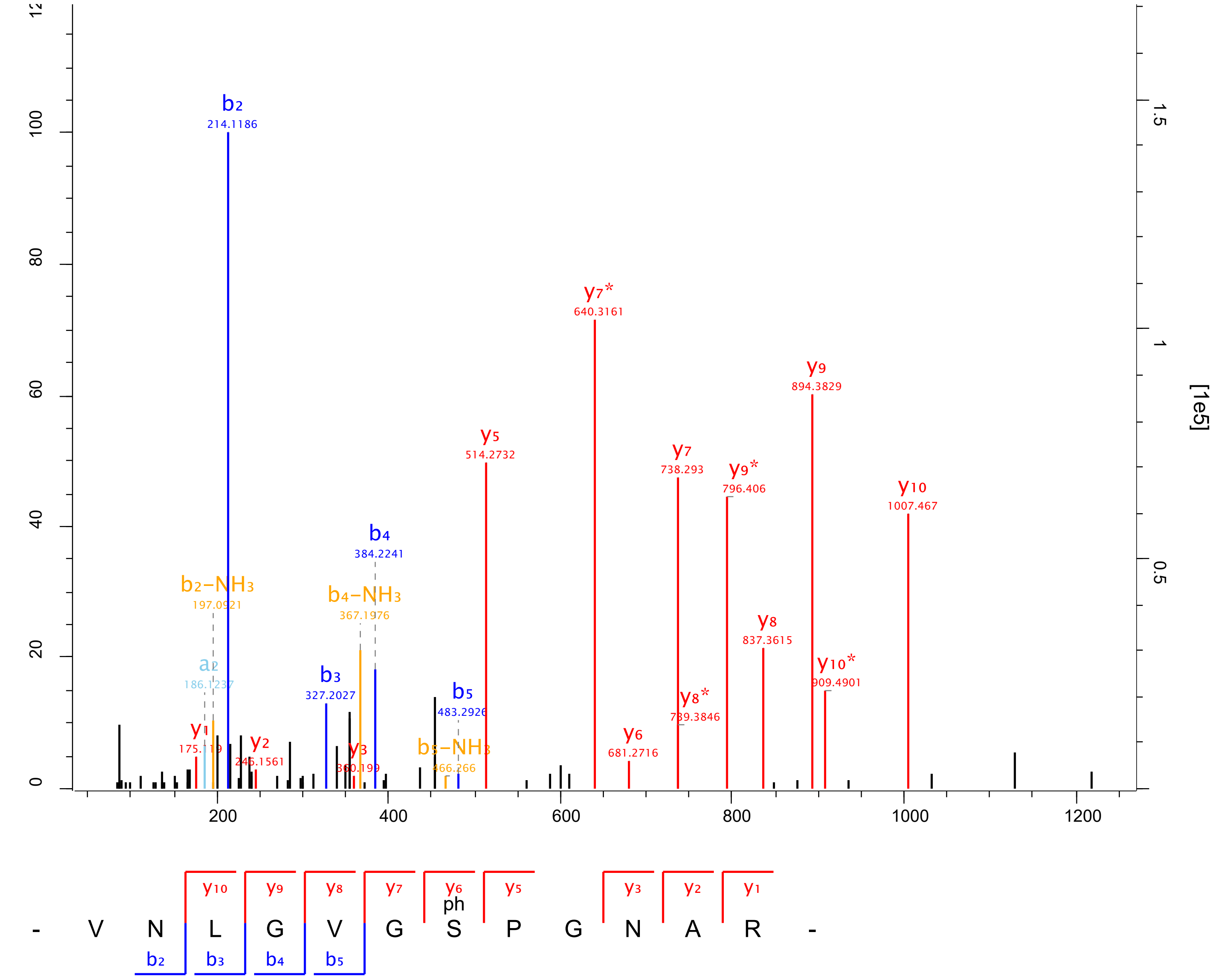

| Raw file | Scan | Method | Score | m/z |
|---|---|---|---|---|
| water_15_2 | 13192 | FTMS; HCD | 147.31 | 598.28 |

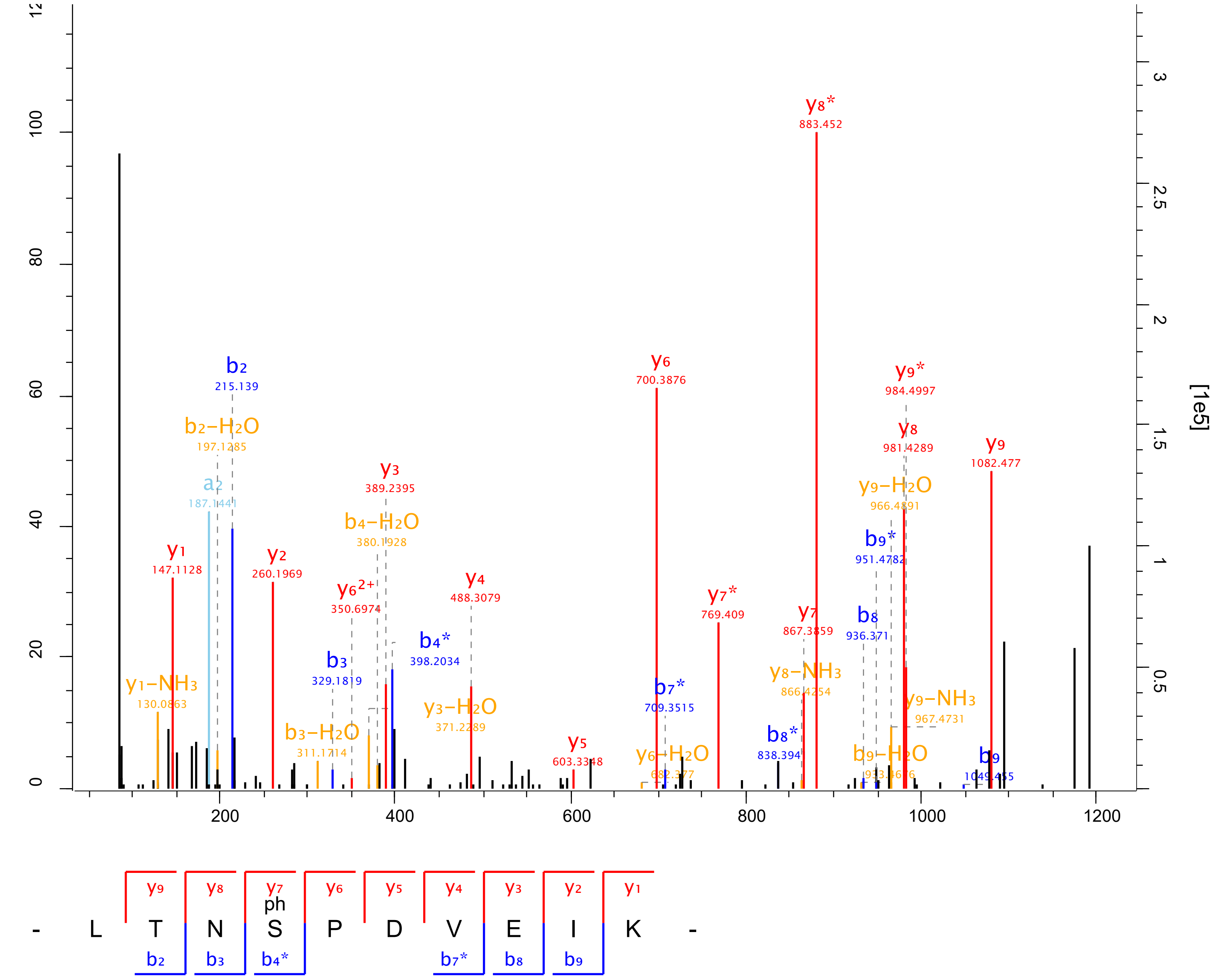

| Raw file | Scan | Method | Score | m/z |
|---|---|---|---|---|
| water_15_2 | 13198 | FTMS; HCD | 64.82 | 454.24 |

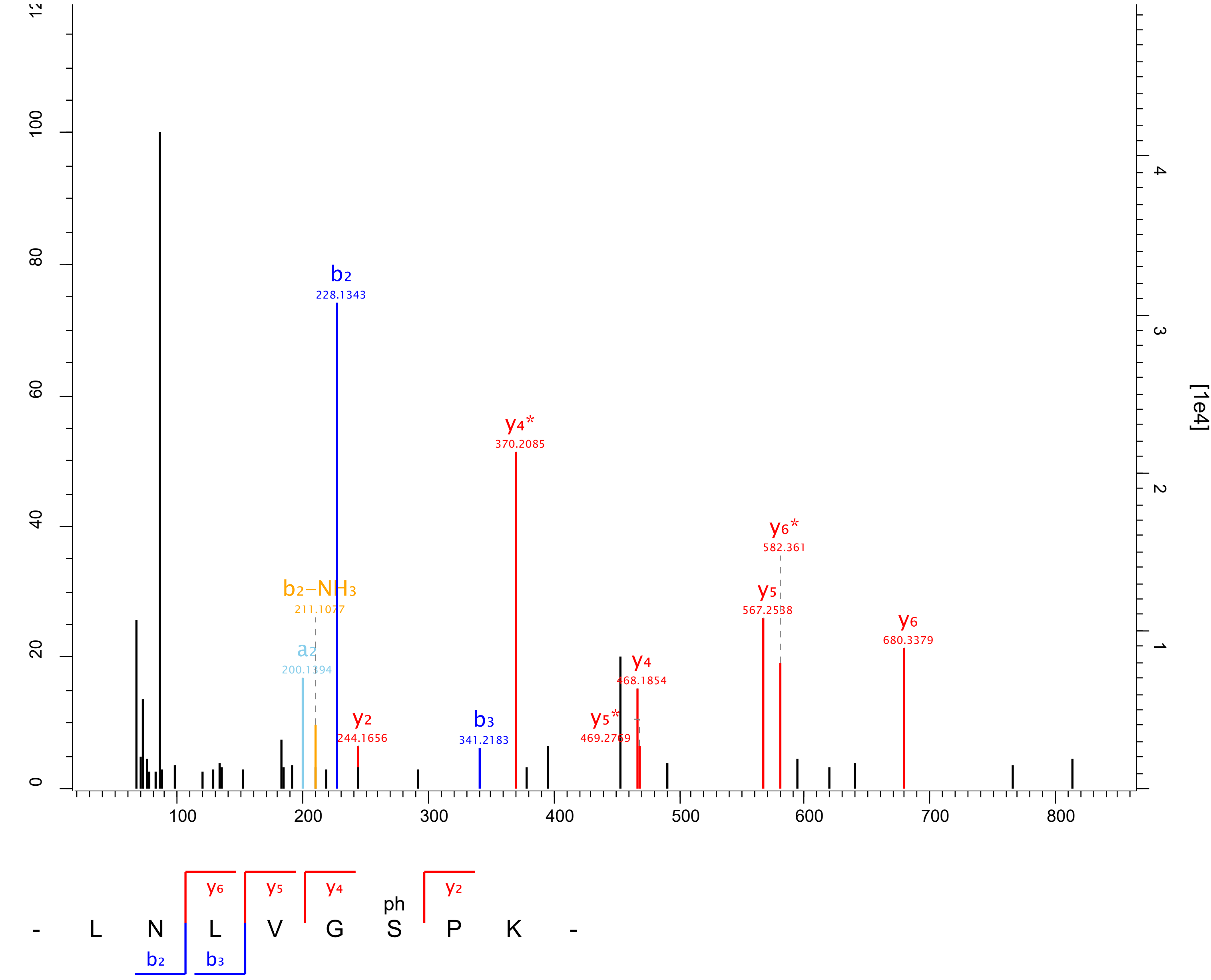

| Raw file | Scan | Method | Score | m/z |
| --- | --- | --- | --- | --- |
| water_15_2 | 13232 | FTMS; HCD | 125.22 | 547.27 |

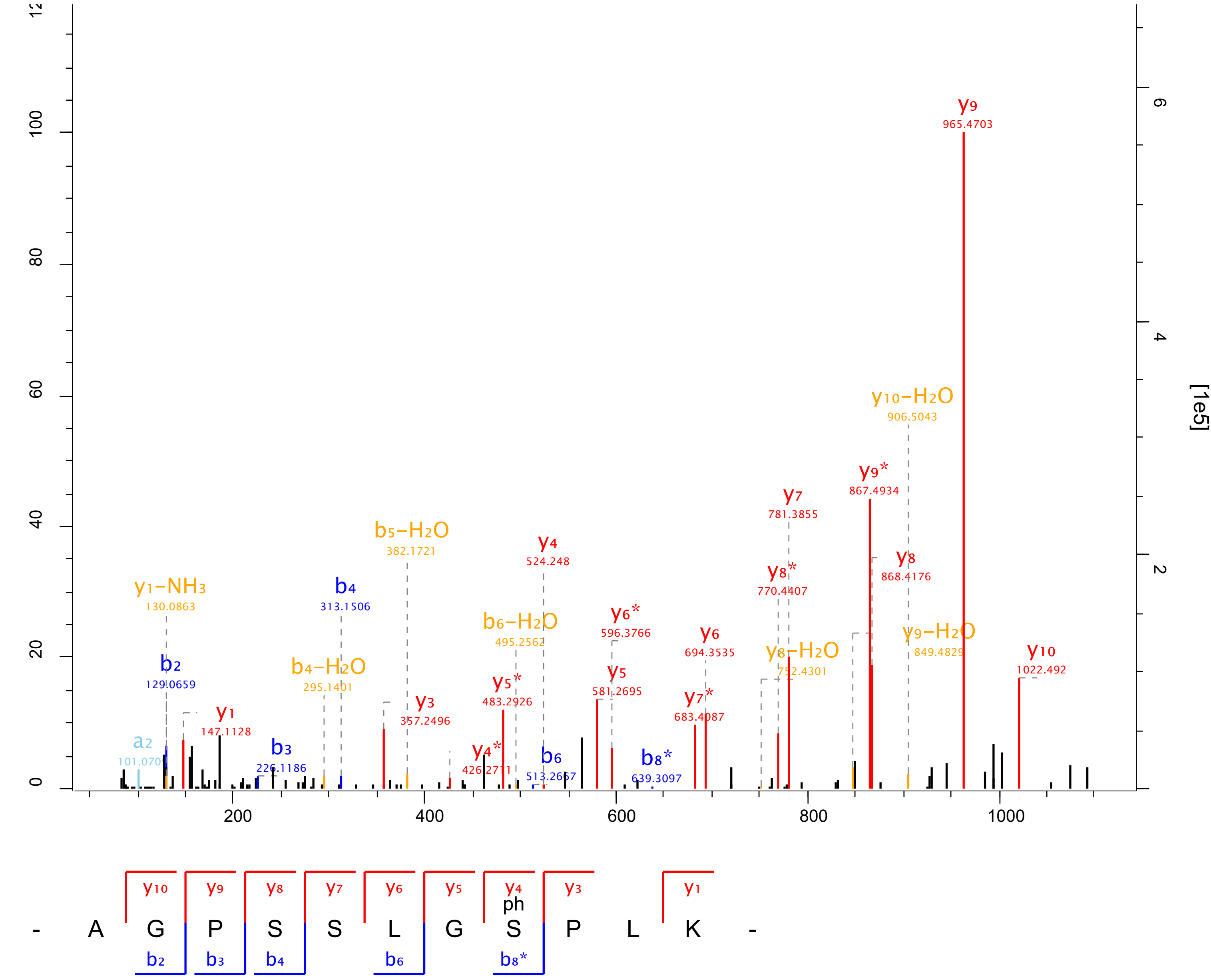

| Raw file | Scan | Method | Score | m/z |
|---|---|---|---|---|
| water_15_2 | 13243 | FTMS; HCD | 131.09 | 554.22 |

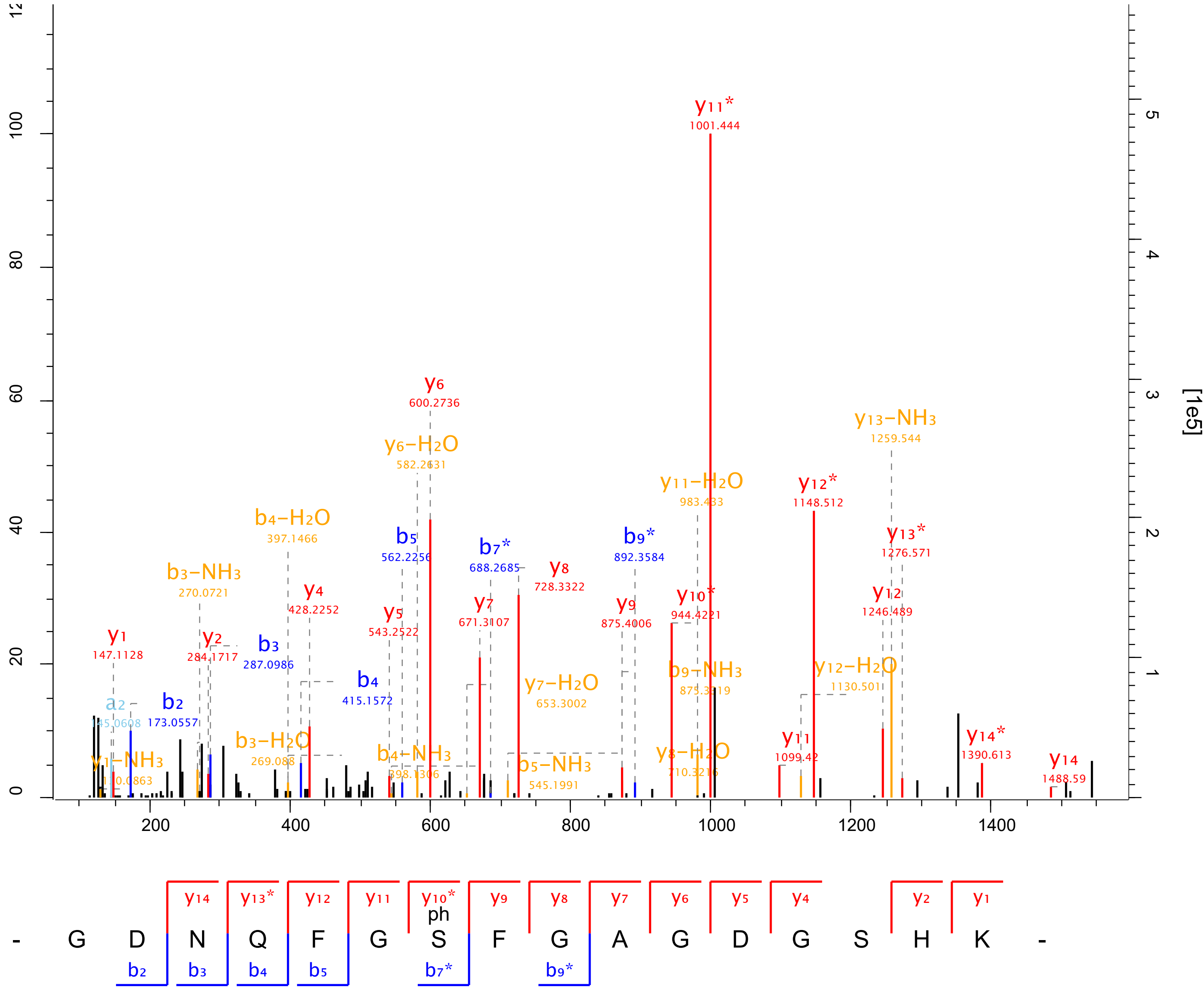

| Raw file | Scan | Method | Score | m/z |
| --- | --- | --- | --- | --- |
| water_15_2 | 13248 | FTMS; HCD | 87.43 | 816.8 |

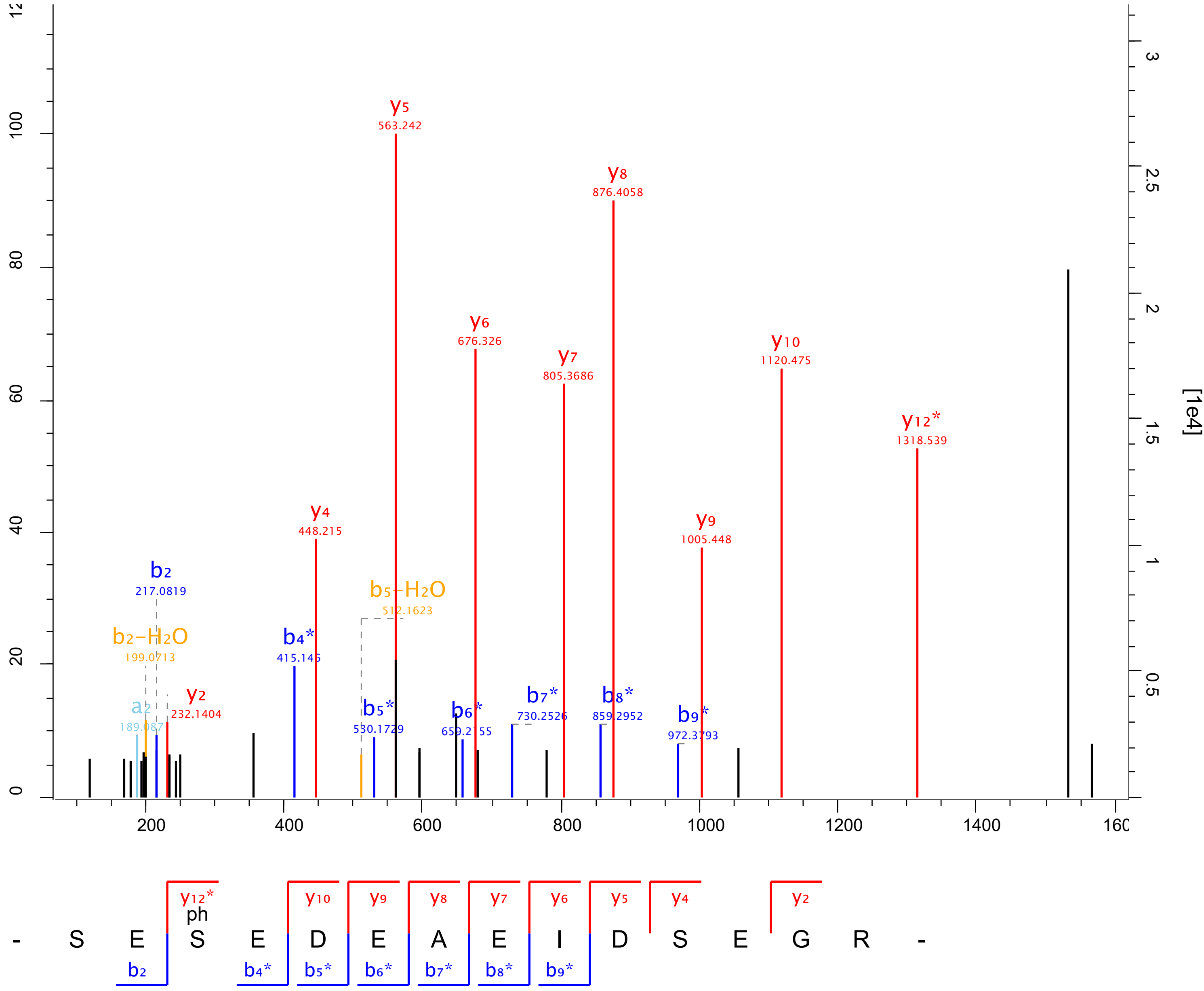

| Raw file | Scan | Method | Score | m/z |
| --- | --- | --- | --- | --- |
| water_15_2 | 13335 | FTMS; HCD | 227 | 728.82 |

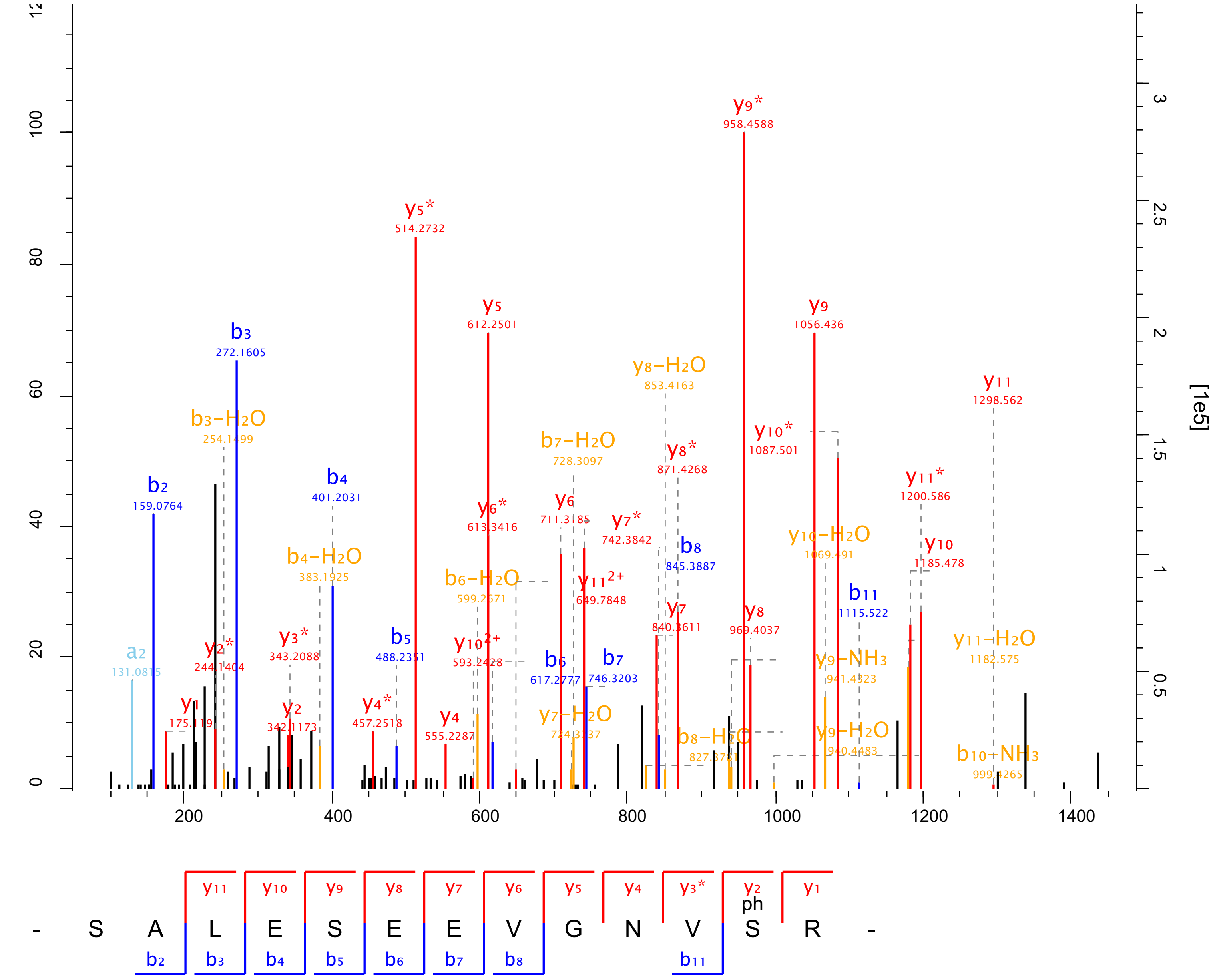

| Raw file | Scan | Method | Score | m/z |
| --- | --- | --- | --- | --- |
| water_15_2 | 13350 | FTMS; HCD | 88.34 | 524.23 |

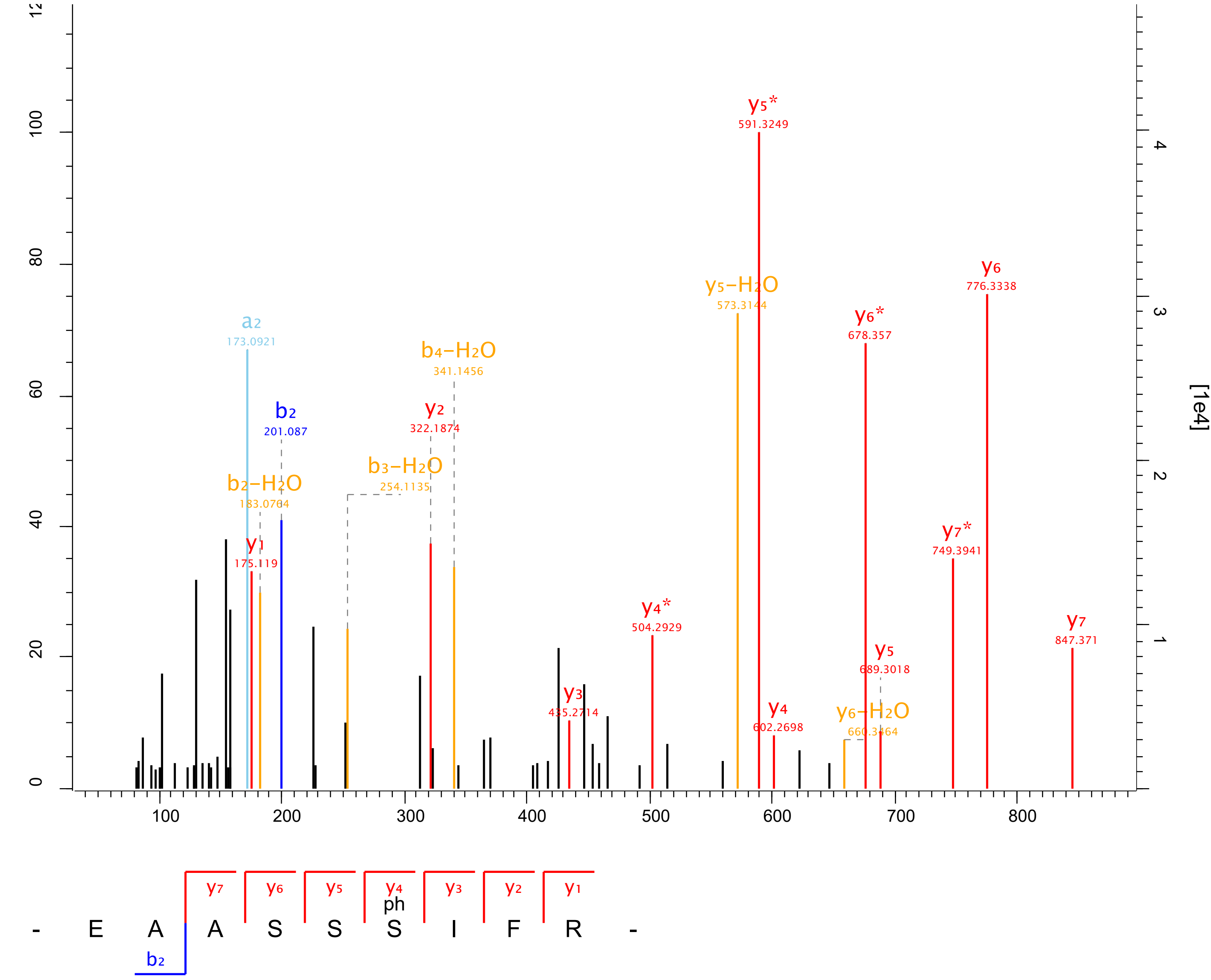

| Raw file | Scan | Method | Score | m/z |
|---|---|---|---|---|
| water_15_2 | 13356 | FTMS; HCD | 95.93 | 780.69 |

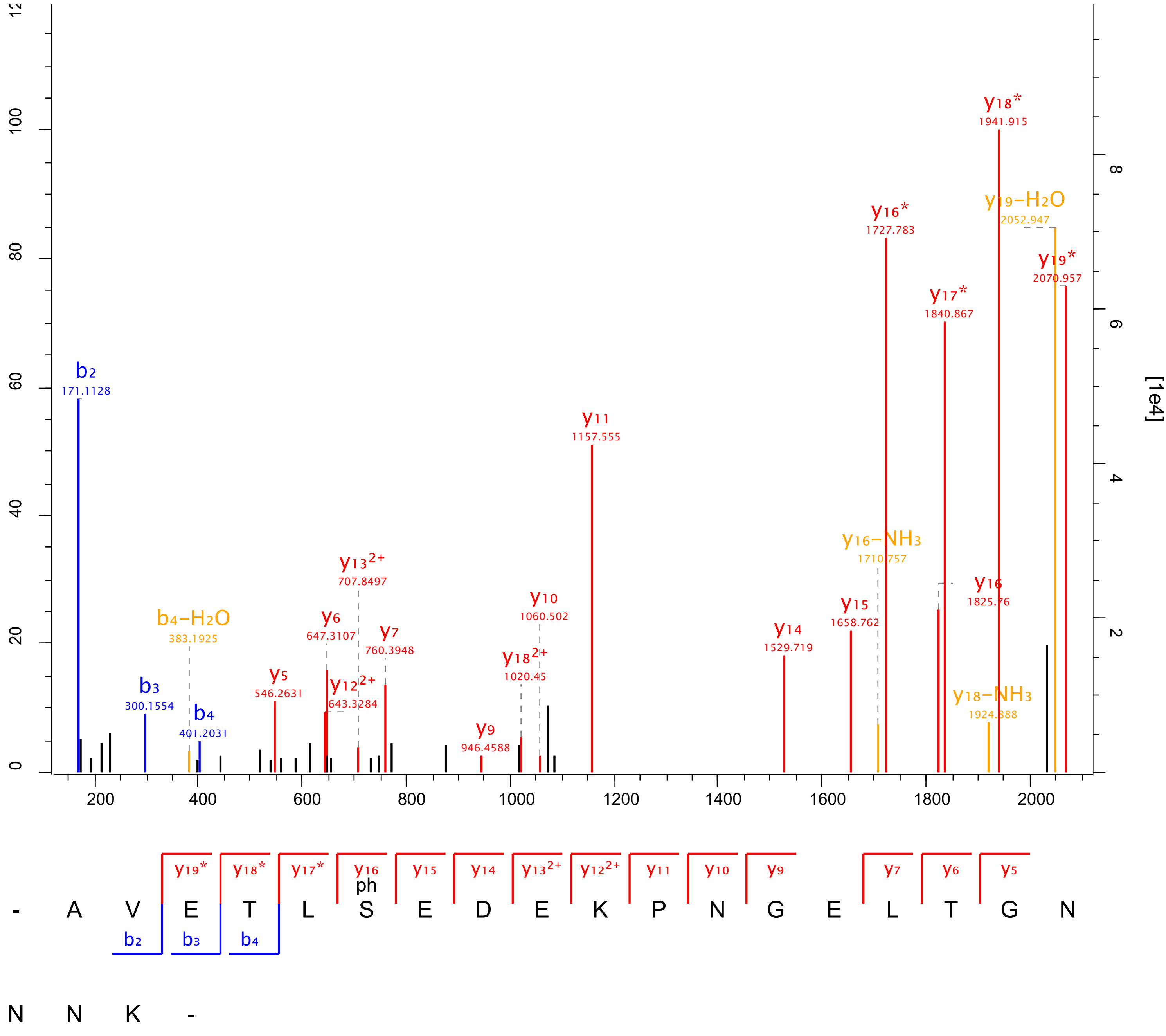

| Raw file | Scan | Method | Score | m/z |
| --- | --- | --- | --- | --- |
| water_15_2 | 13403 | FTMS; HCD | 106.29 | 556.75 |

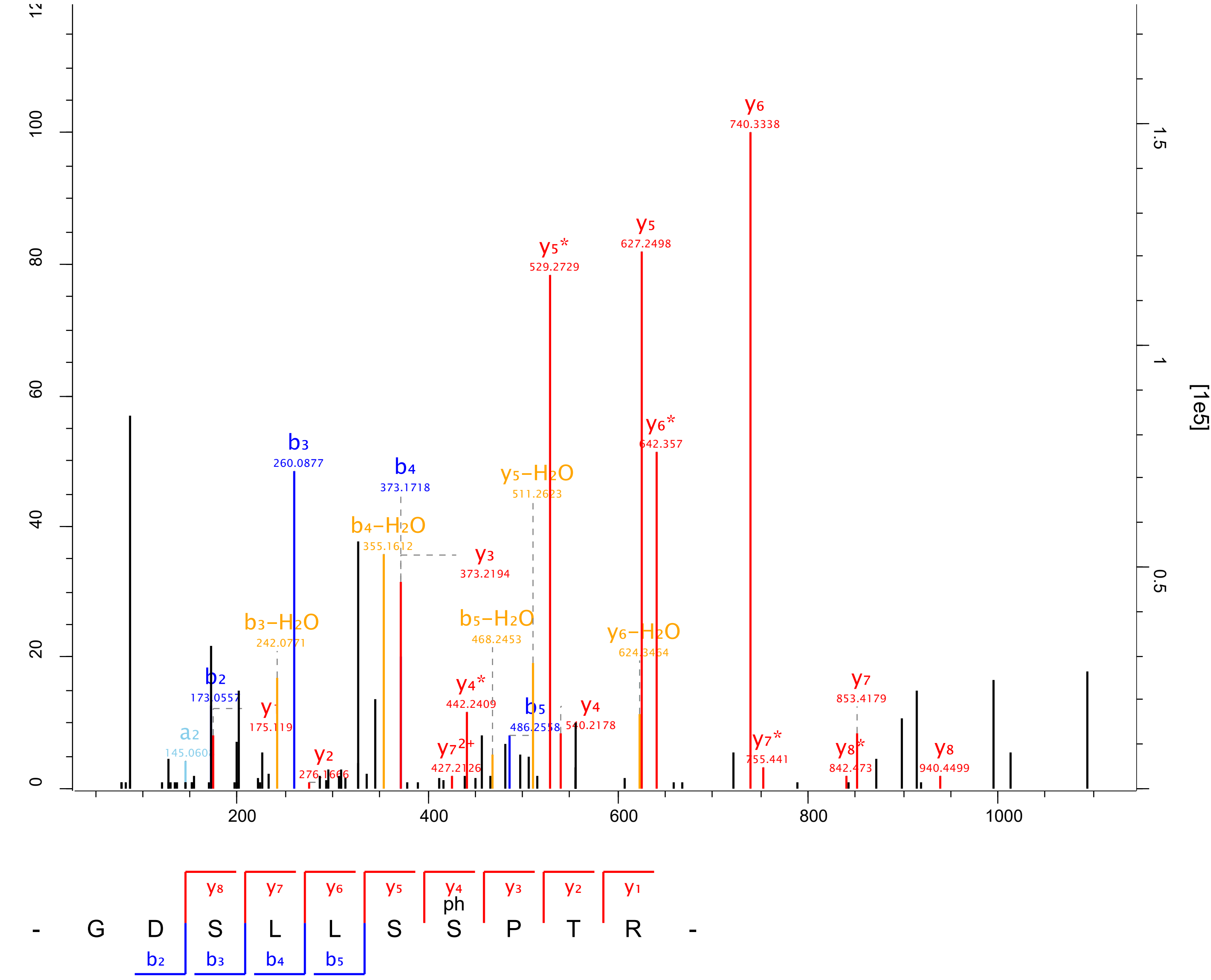

| Raw file | Scan | Method | Score | m/z |
| --- | --- | --- | --- | --- |
| water_15_2 | 13407 | FTMS; HCD | 140.96 | 669.78 |

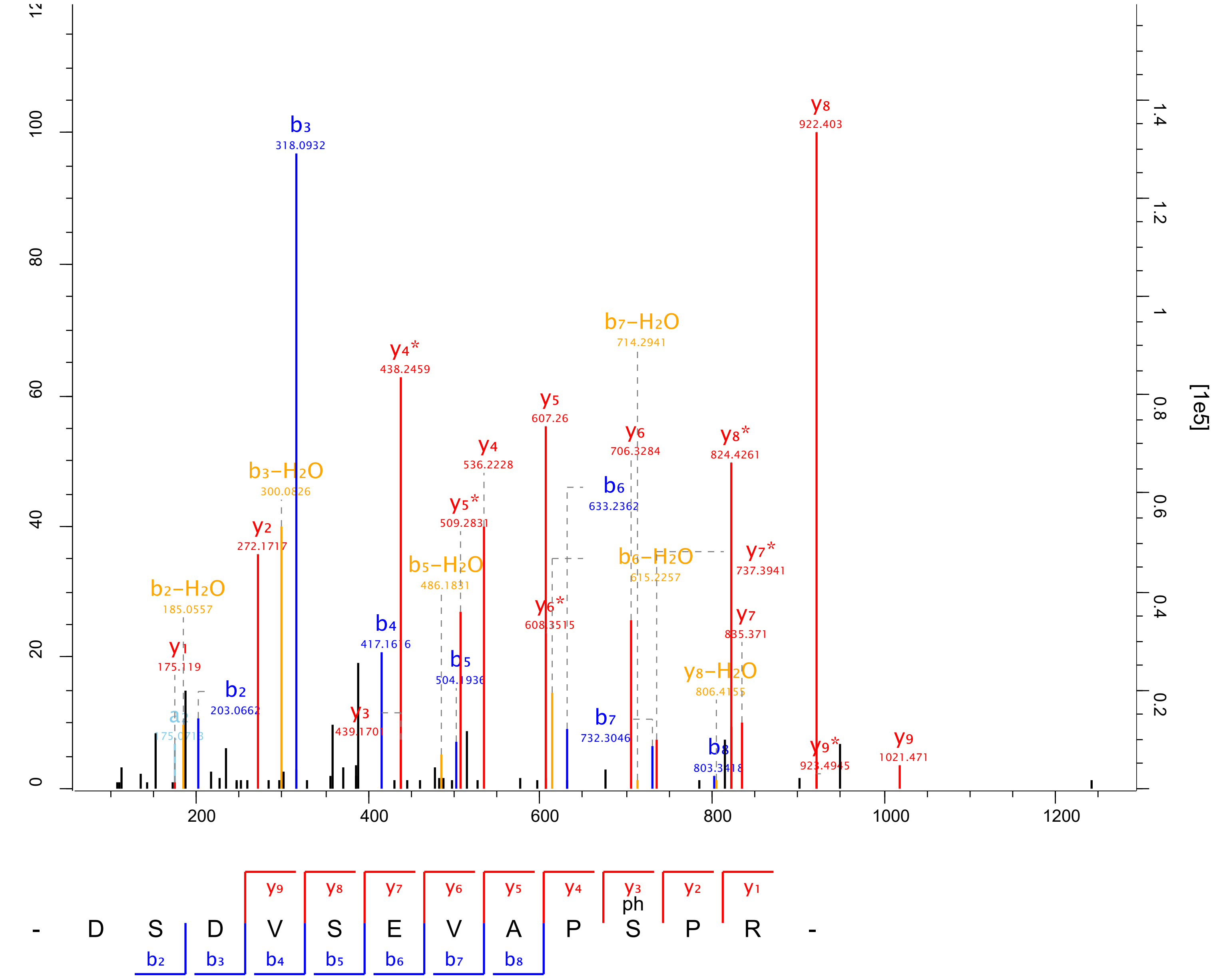

| Raw file | Scan | Method | Score | m/z |
|---|---|---|---|---|
| water_15_2 | 13425 | FTMS; HCD | 101.39 | 580.74 |

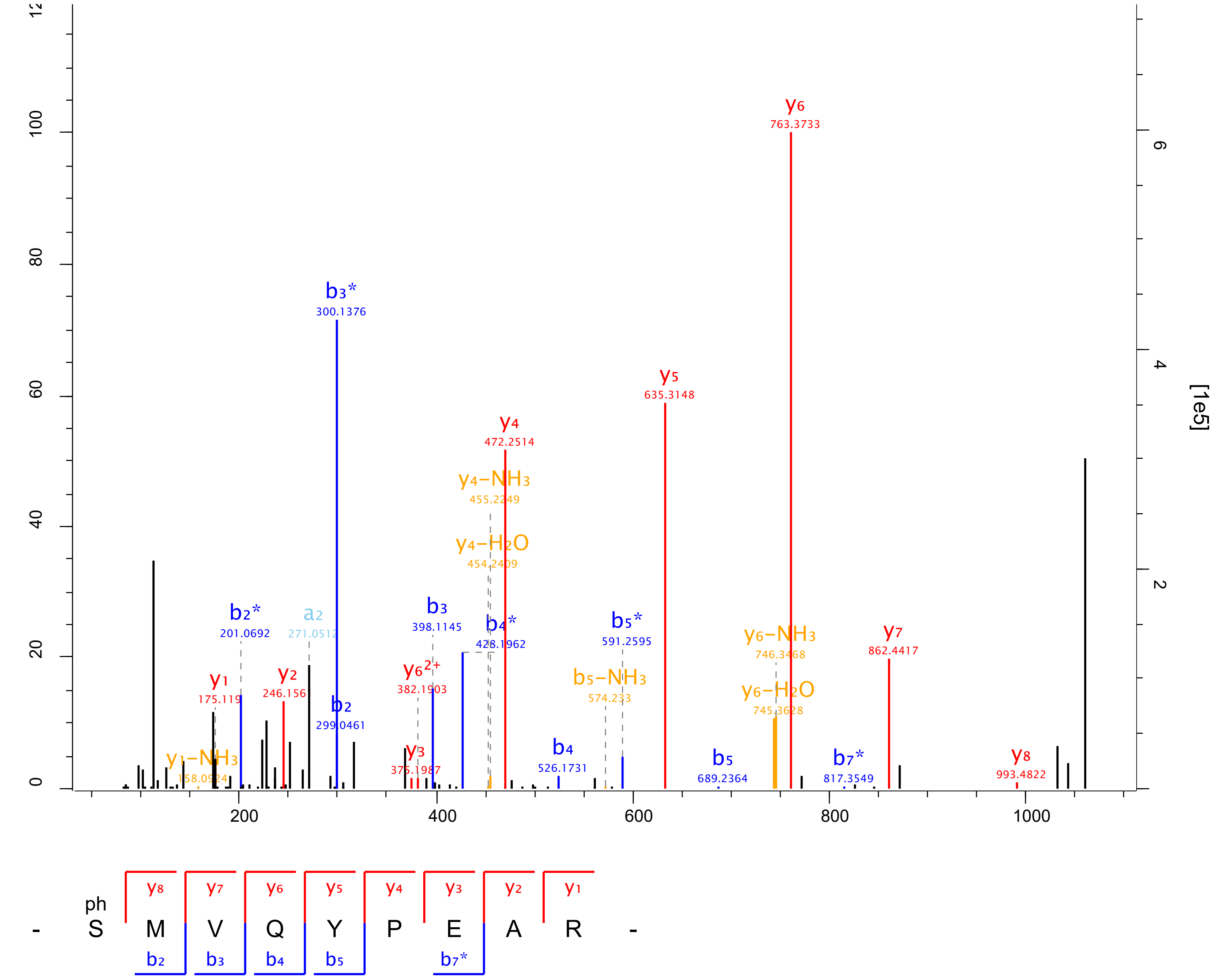

| Raw file | Scan | Method | Score | m/z |
| --- | --- | --- | --- | --- |
| water_15_2 | 13455 | FTMS; HCD | 84.82 | 565.73 |

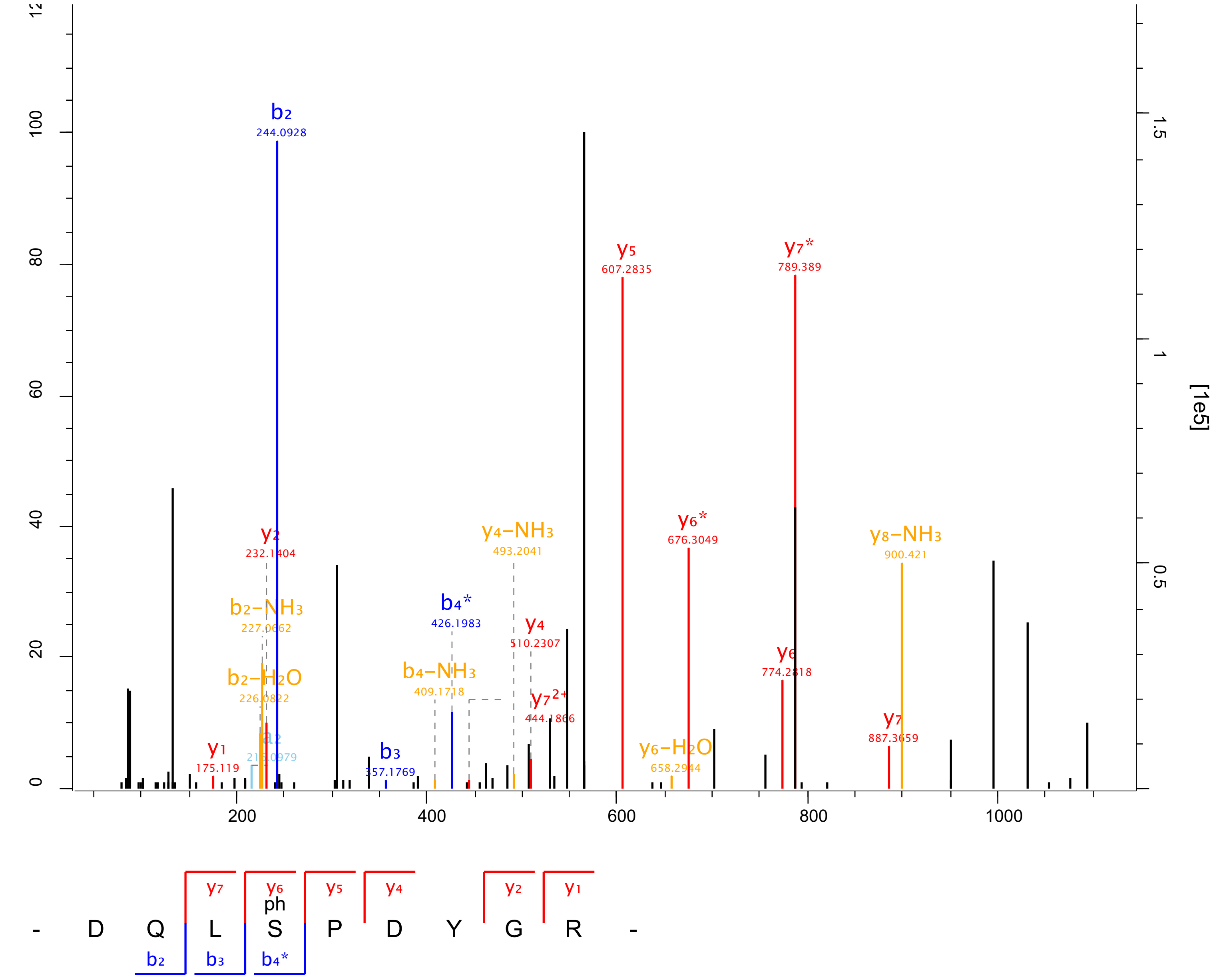

| Raw file | Scan | Method | Score | m/z |
|---|---|---|---|---|
| water_15_2 | 13463 | FTMS; HCD | 162.82 | 1012.43 |

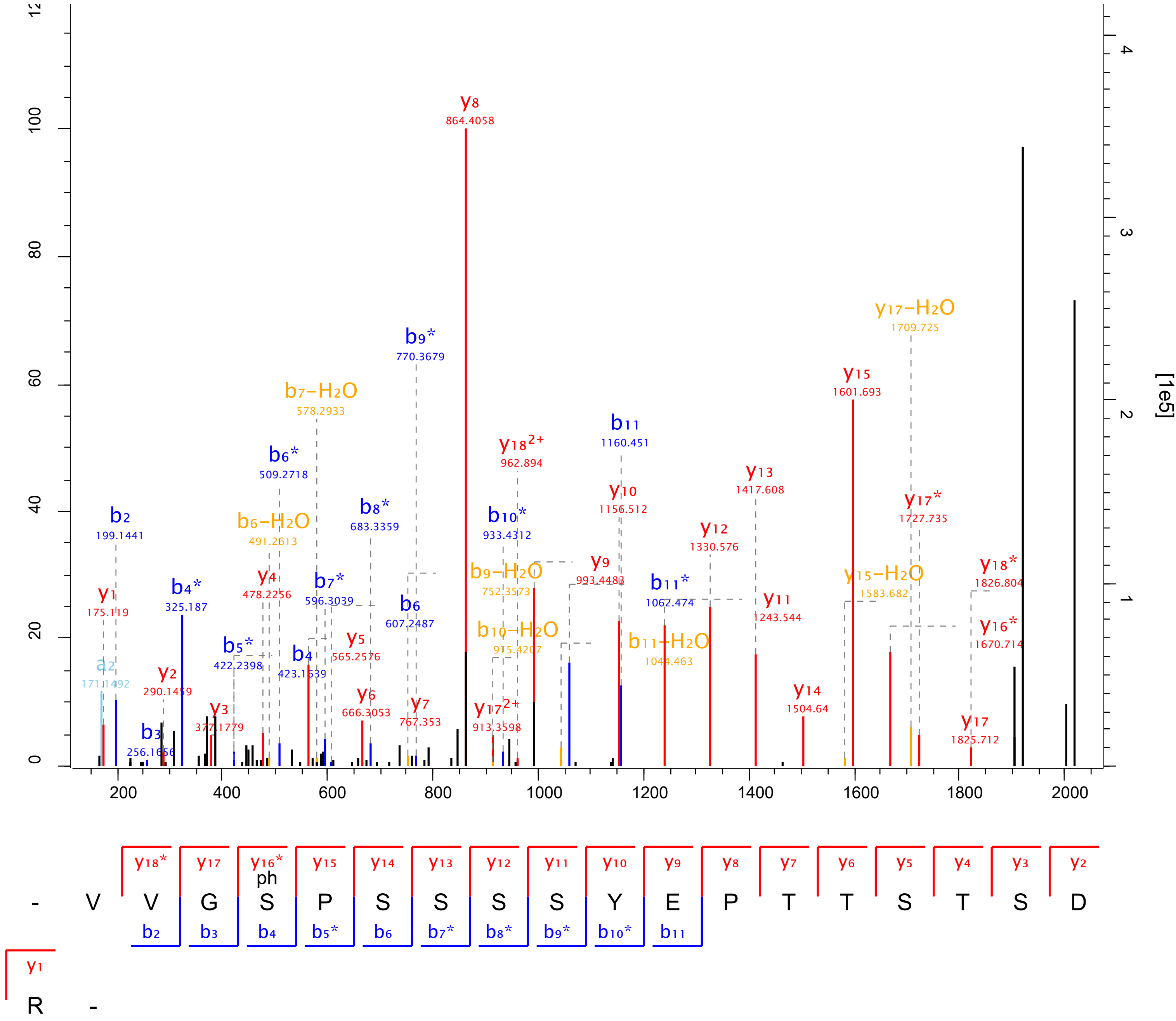

| Raw file | Scan | Method | Score | m/z |
|---|---|---|---|---|
| water_15_2 | 13516 | FTMS; HCD | 45.35 | 949.39 |

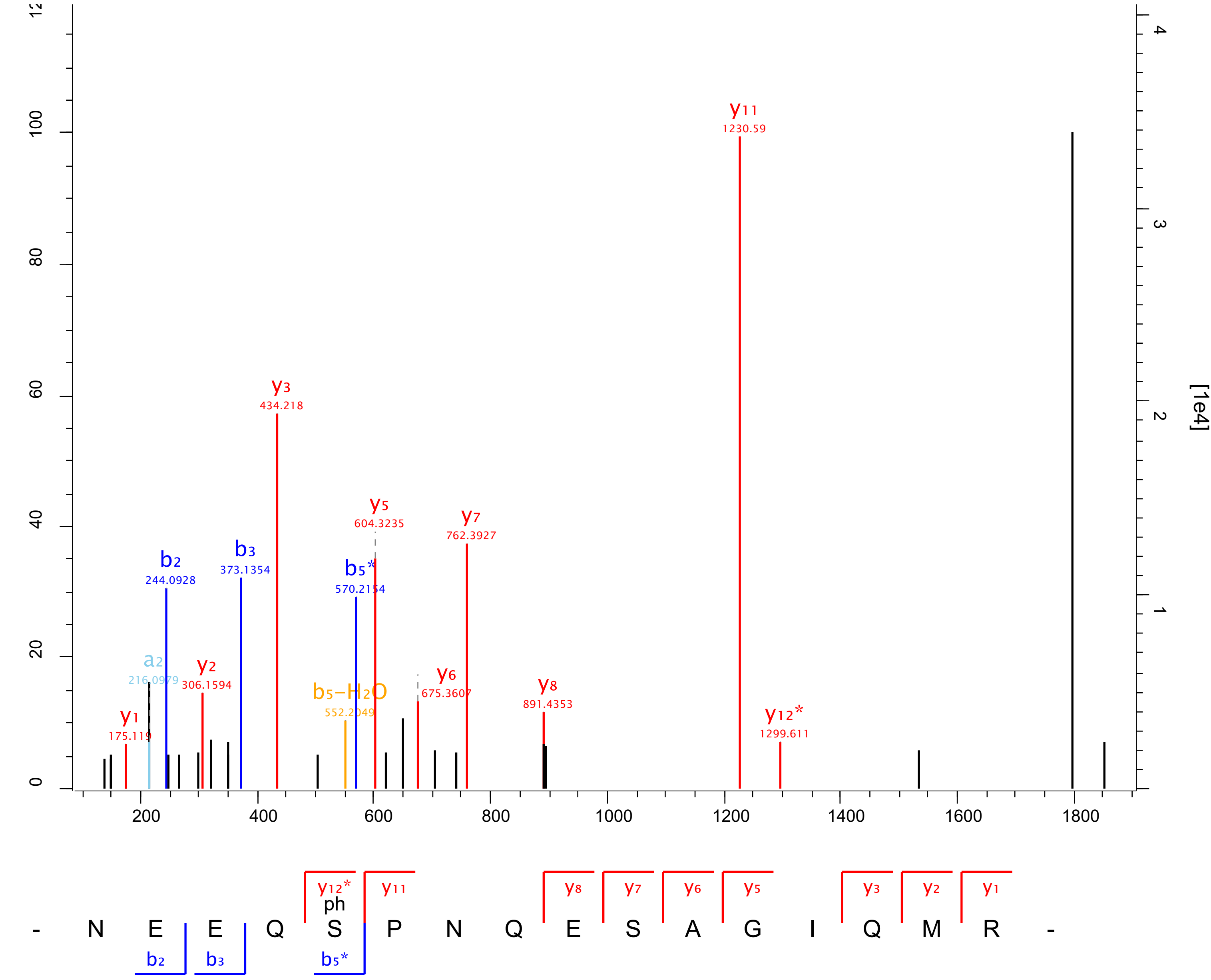

| Raw file | Scan | Method | Score | m/z |
| --- | --- | --- | --- | --- |
| water_15_2 | 13552 | FTMS; HCD | 83.37 | 760.8 |

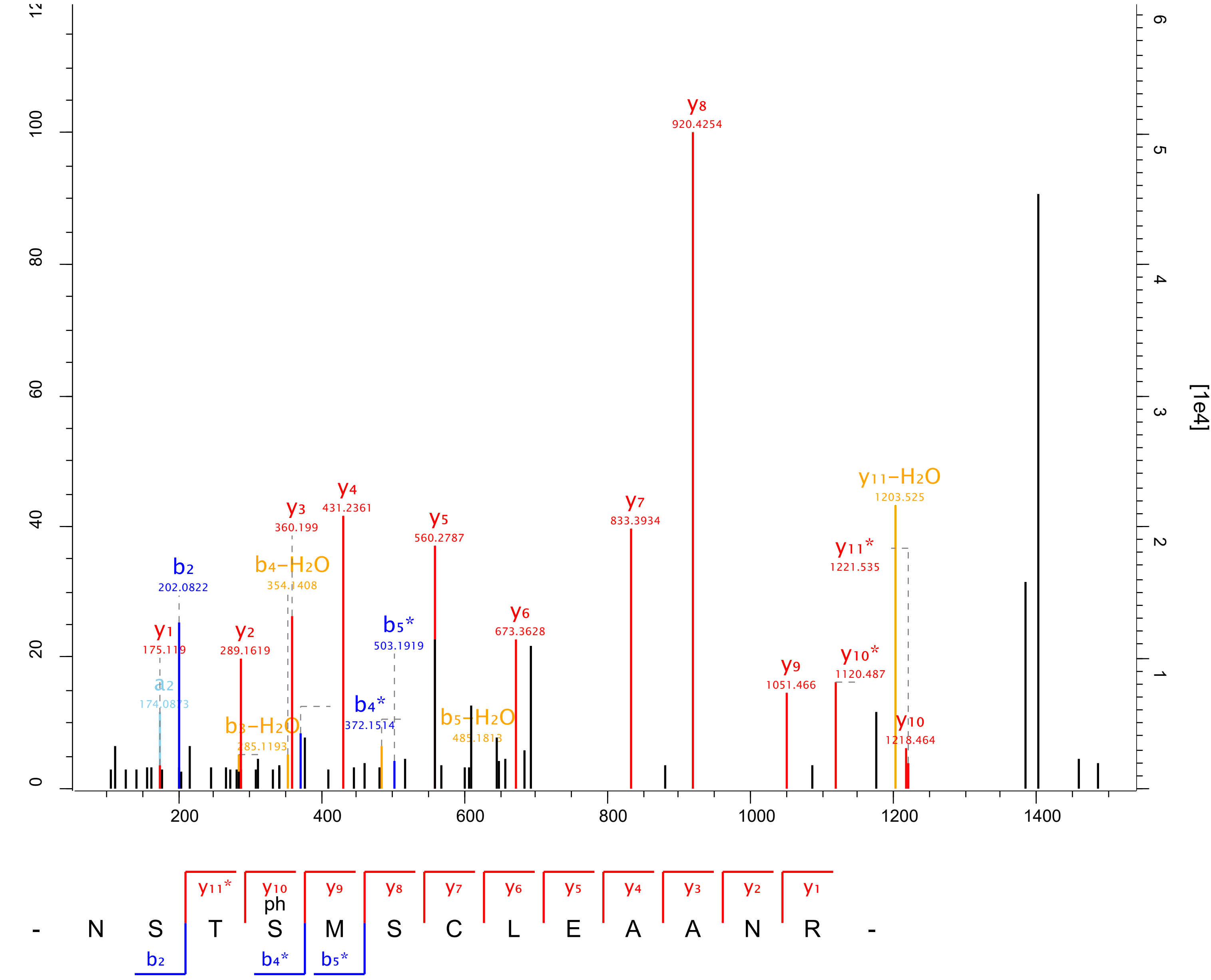

| Raw file | Scan | Method | Score | m/z |
| --- | --- | --- | --- | --- |
| water_15_2 | 13649 | FTMS; HCD | 83.4 | 670.81 |

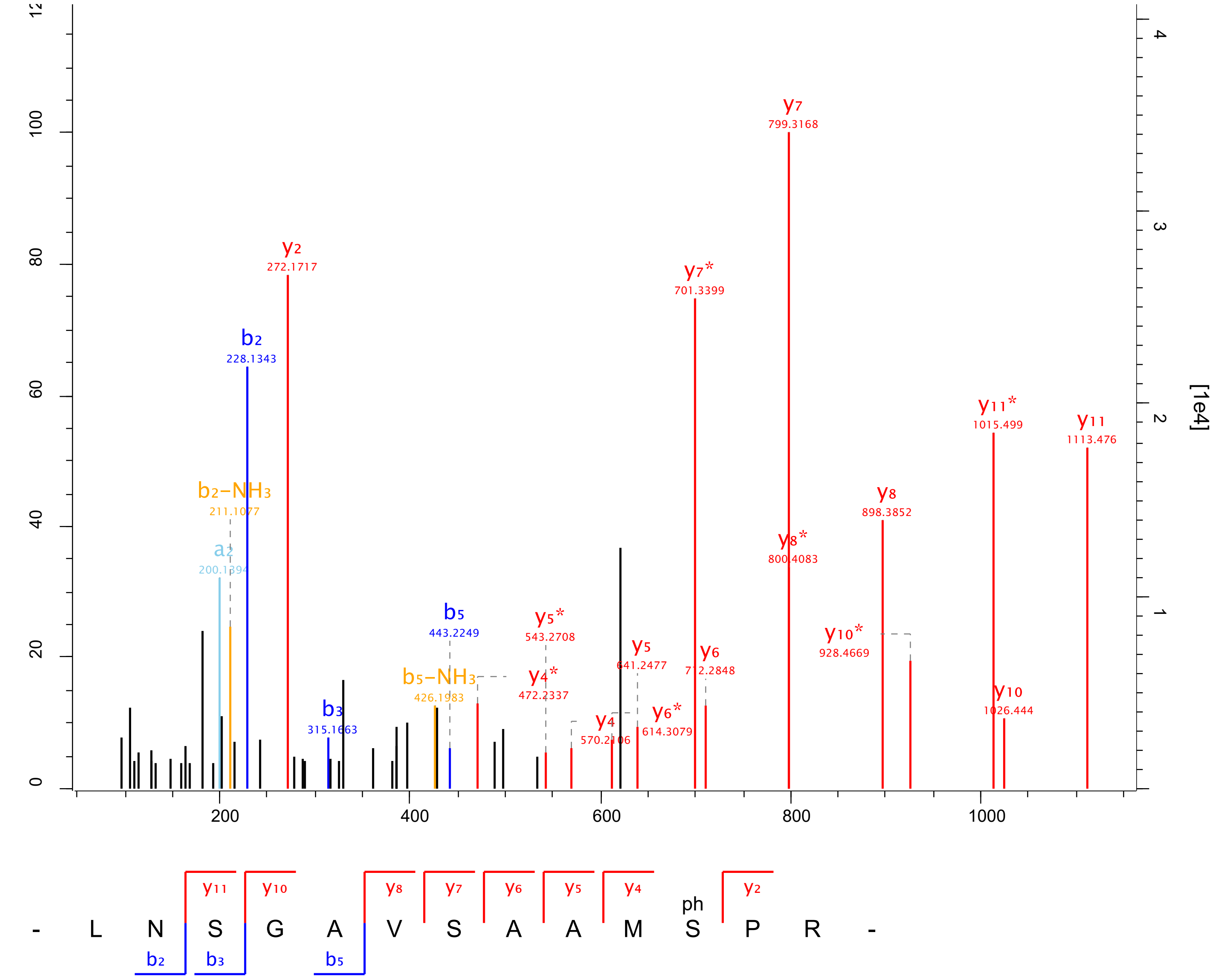

| Raw file | Scan | Method | Score | m/z |
| --- | --- | --- | --- | --- |
| water_15_2 | 13701 | FTMS; HCD | 69.36 | 880.84 |

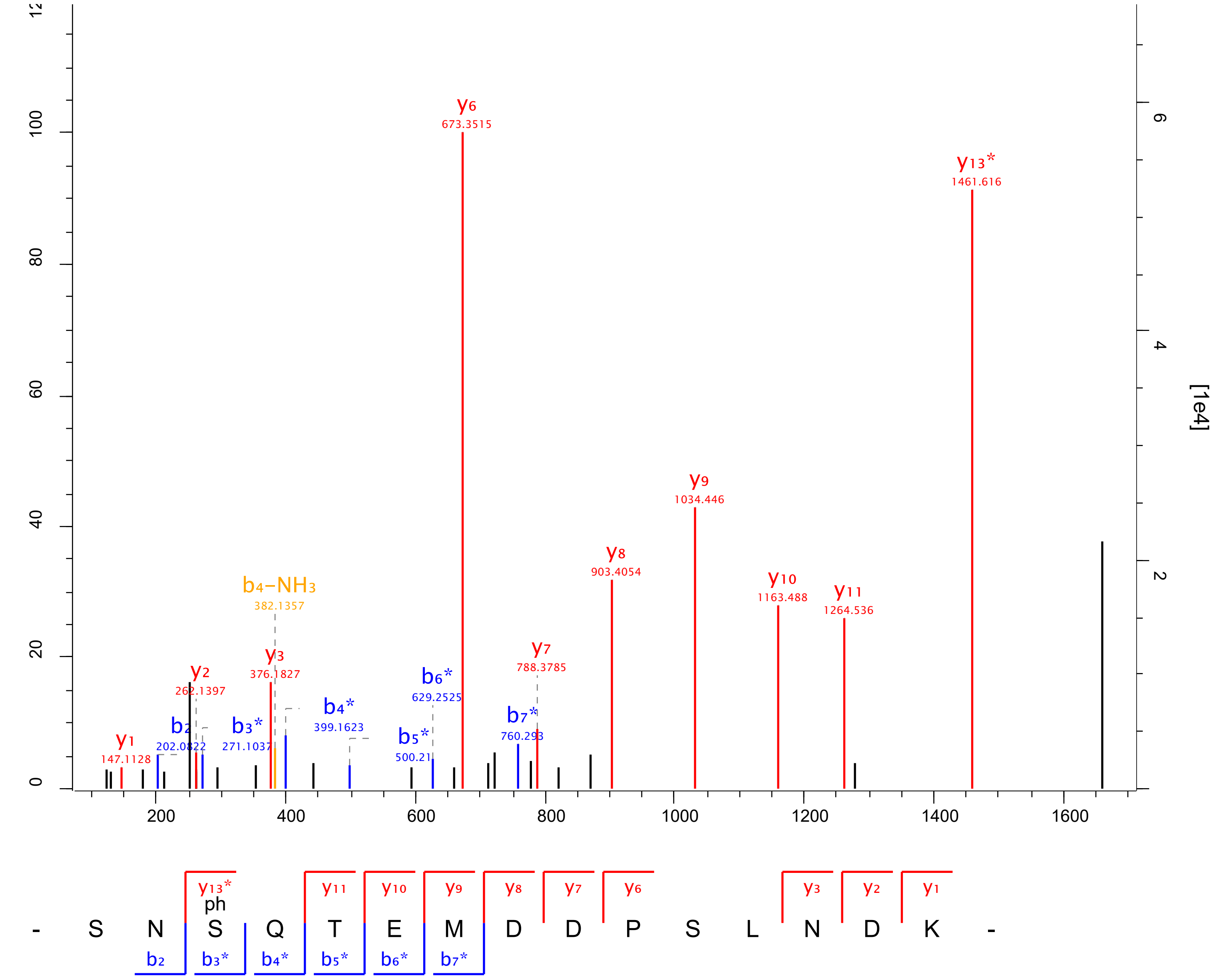

| Raw file | Scan | Method | Score | m/z |
|---|---|---|---|---|
| water_15_2 | 13703 | FTMS; HCD | 192.91 | 651.28 |

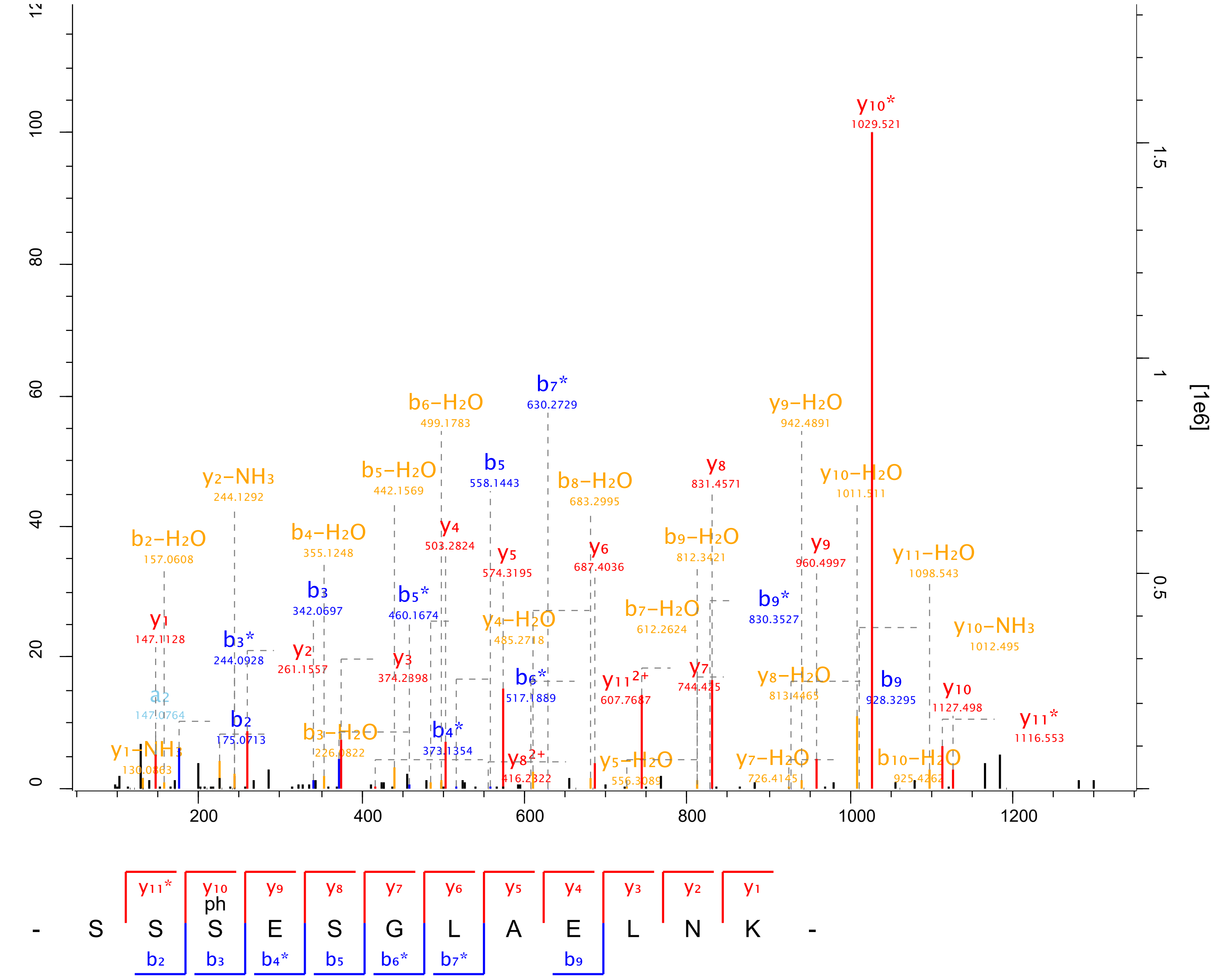

| Raw file | Scan | Method | Score | m/z |
| --- | --- | --- | --- | --- |
| water_15_2 | 13710 | FTMS; HCD | 239.87 | 725.29 |

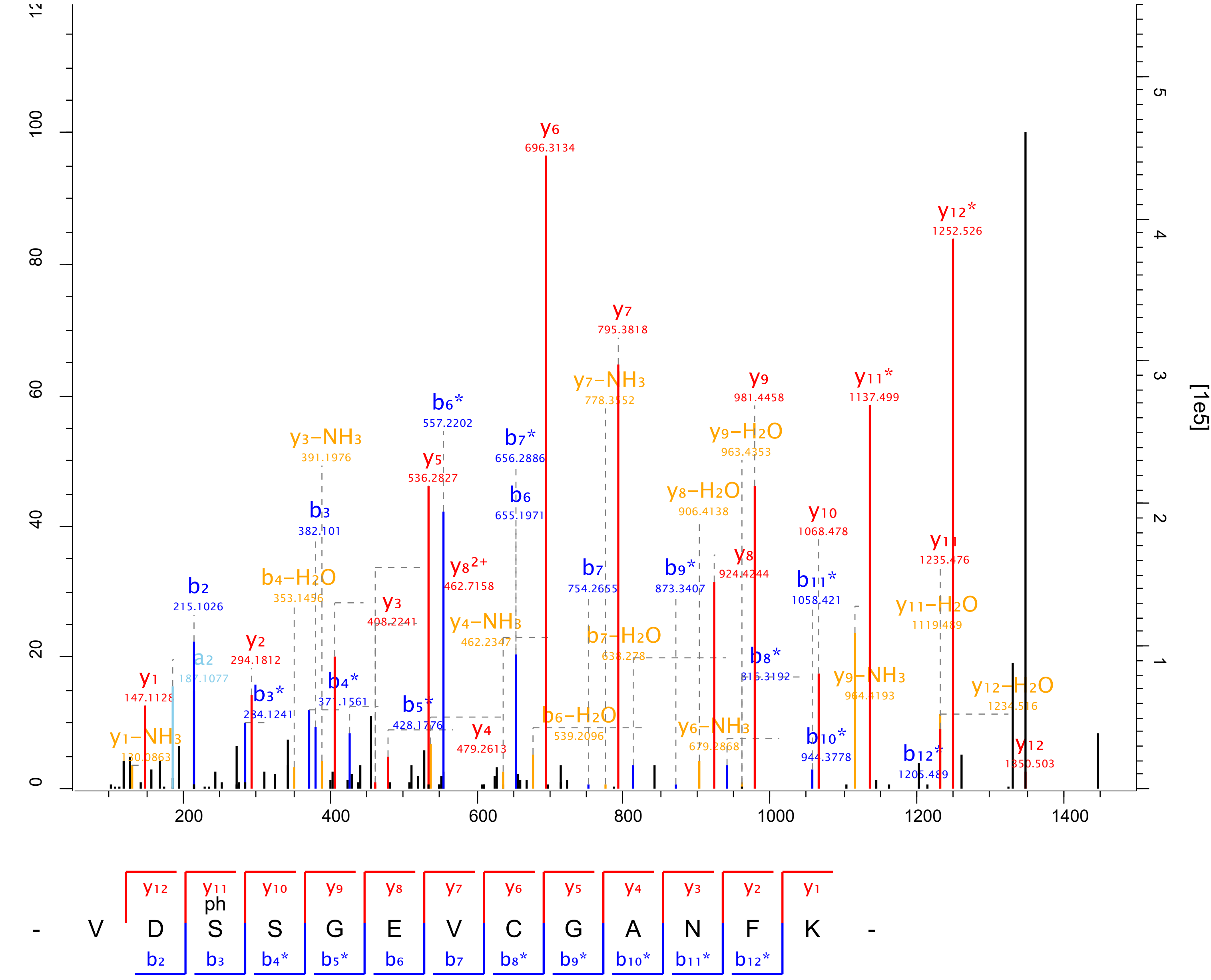

| Raw file | Scan | Method | Score | m/z |
| --- | --- | --- | --- | --- |
| water_15_2 | 13847 | FTMS; HCD | 107.5 | 670.3 |

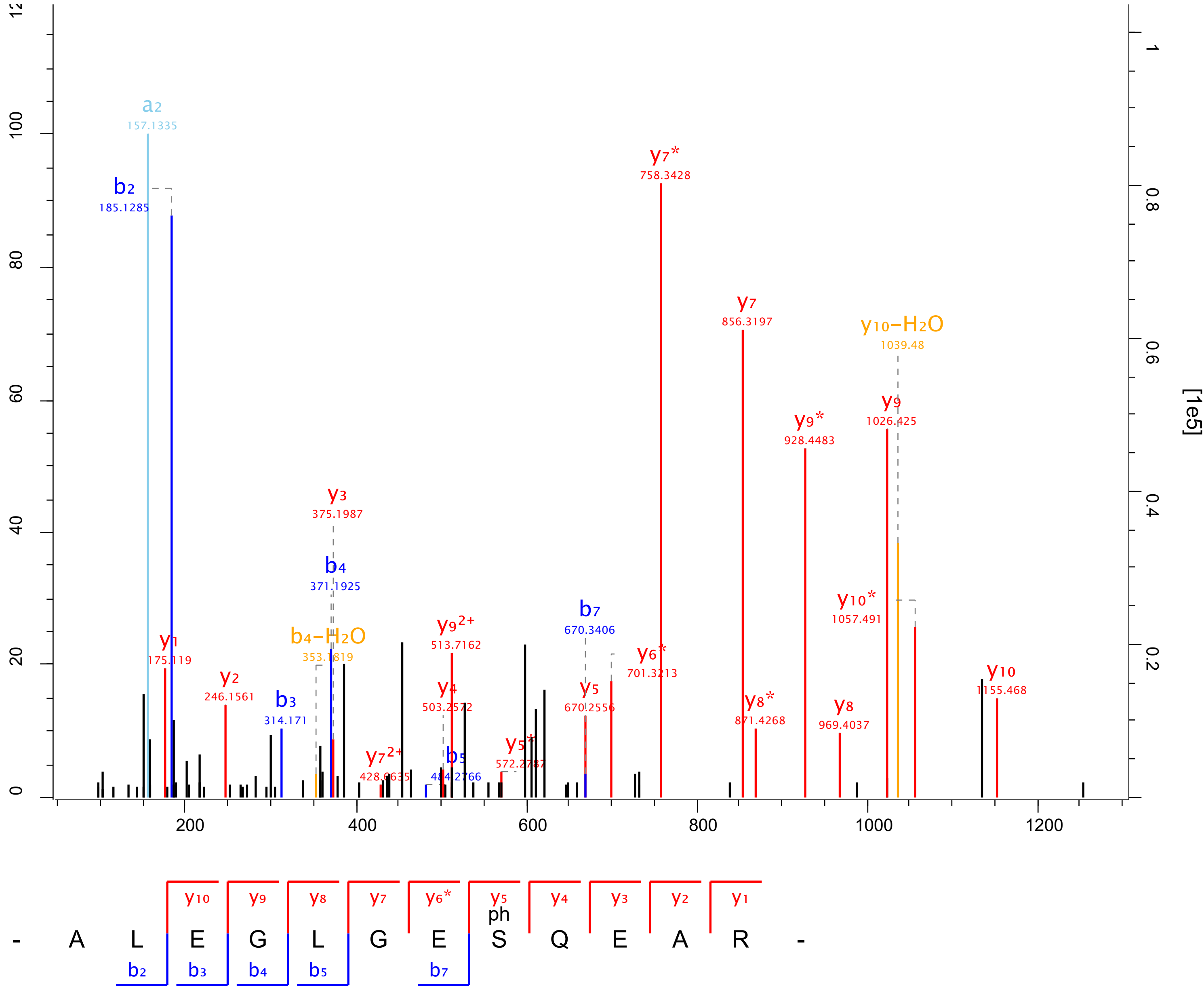

| Raw file | Scan | Method | Score | m/z |
|---|---|---|---|---|
| water_15_2 | 13861 | FTMS; HCD | 89.23 | 601.27 |

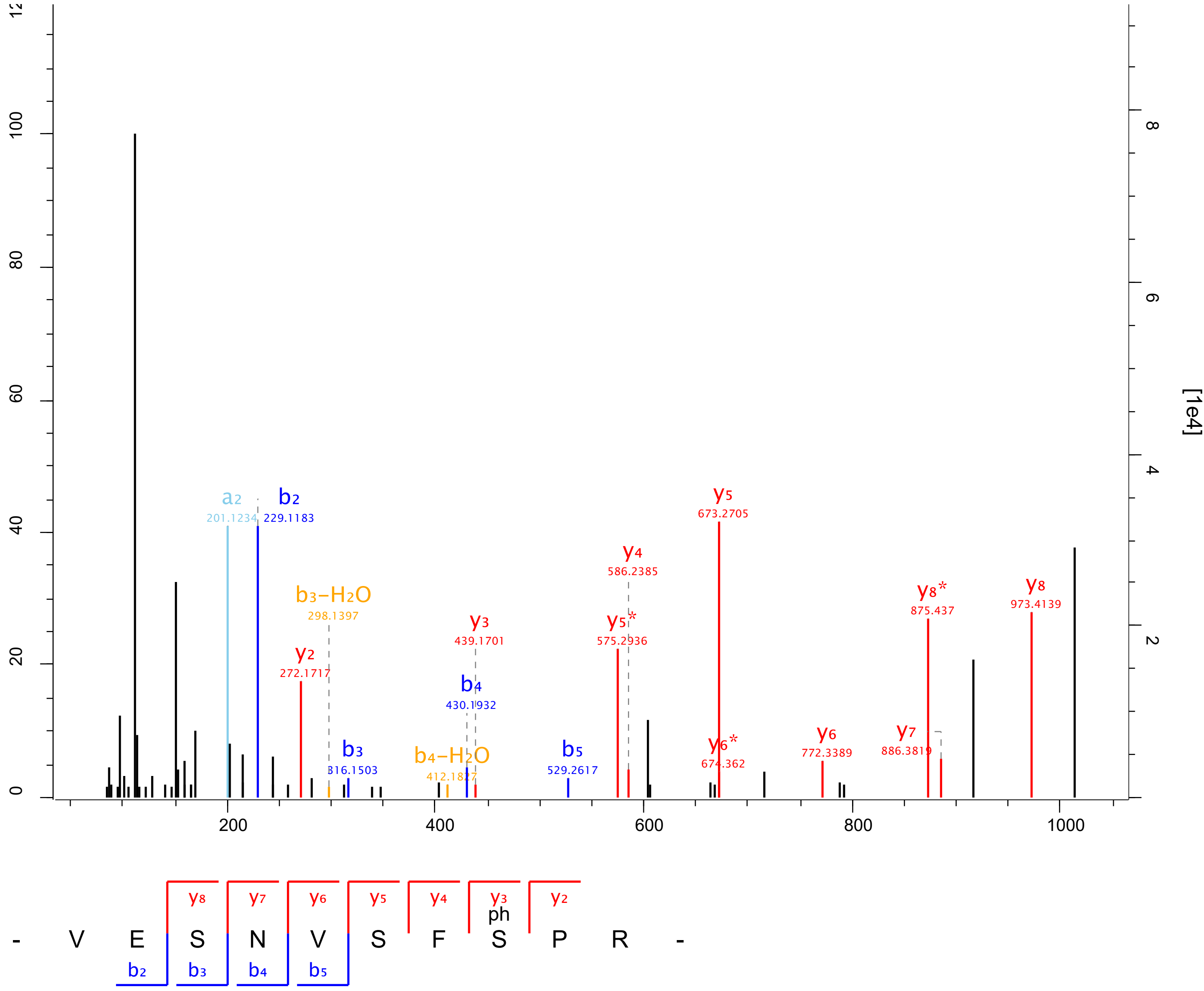

| Raw file | Scan | Method | Score | m/z |
| --- | --- | --- | --- | --- |
| water_15_2 | 13928 | FTMS; HCD | 310.57 | 682.77 |

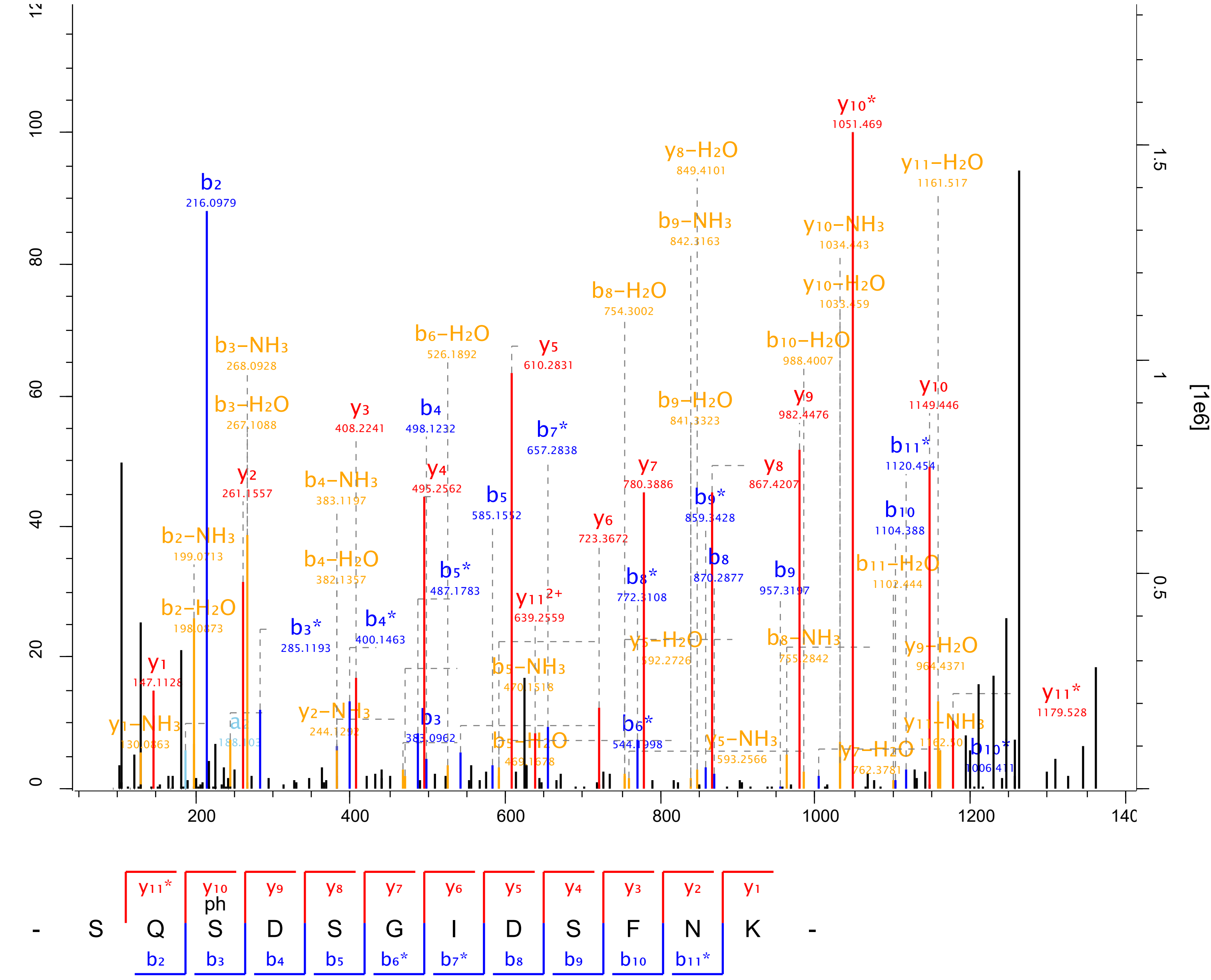

| Raw file | Scan | Method | Score | m/z |
|---|---|---|---|---|
| water_15_2 | 14009 | FTMS; HCD | 65.18 | 491.24 |

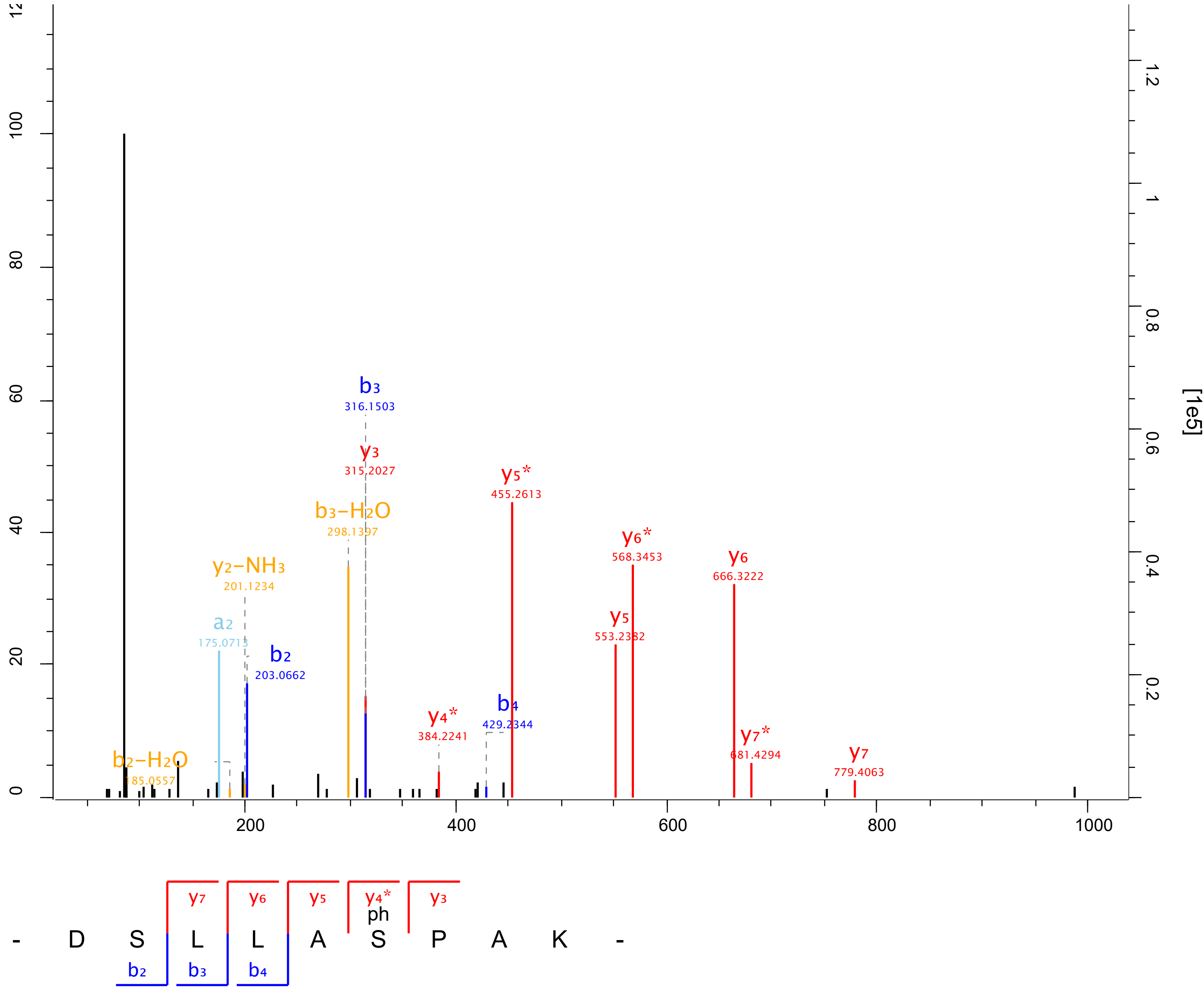

| Raw file | Scan | Method | Score | m/z |
|---|---|---|---|---|
| water_15_2 | 14038 | FTMS; HCD | 96.14 | 876.85 |

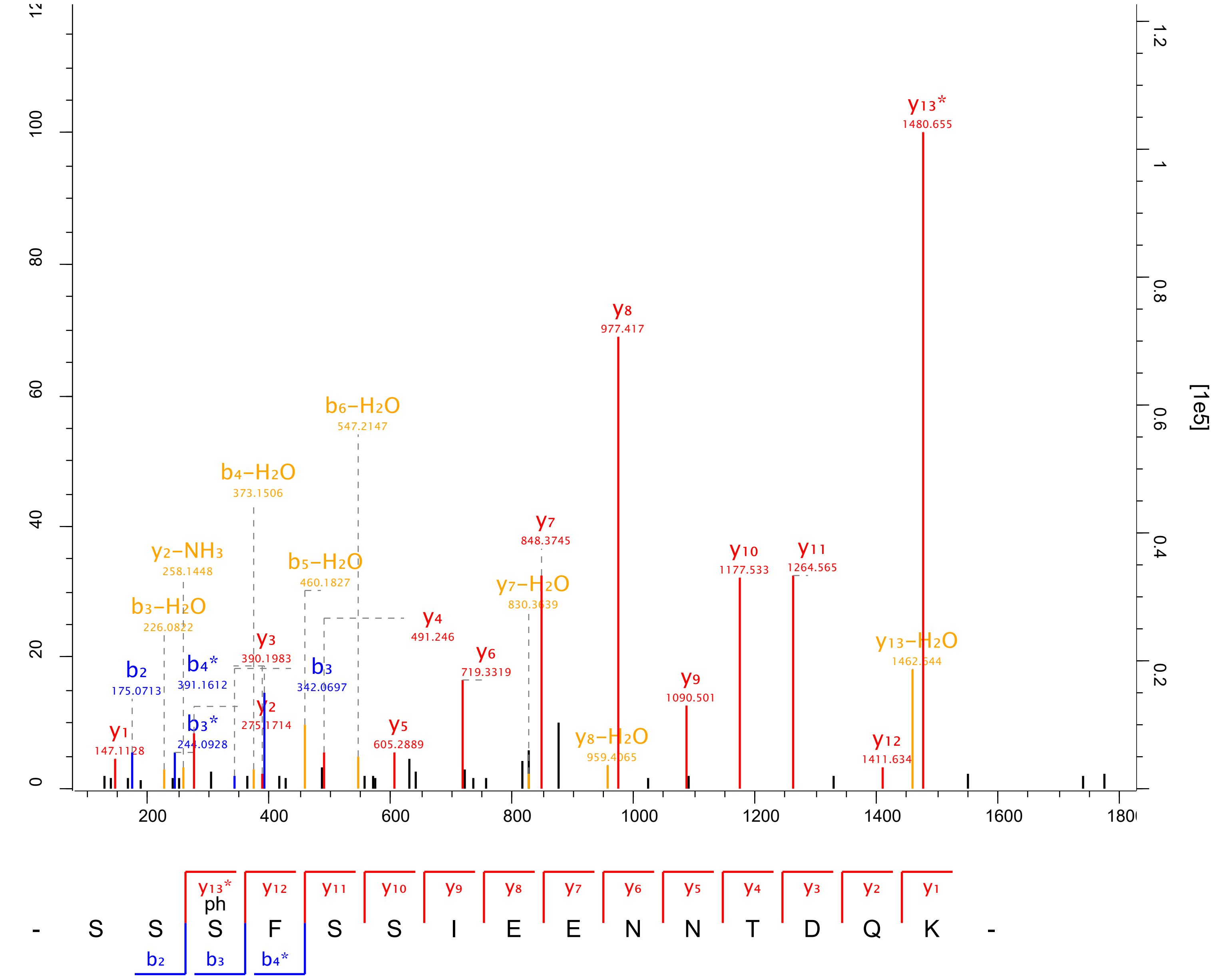

| Raw file | Scan | Method | Score | m/z |
| --- | --- | --- | --- | --- |
| water_15_2 | 14206 | FTMS; HCD | 43.24 | 685.67 |

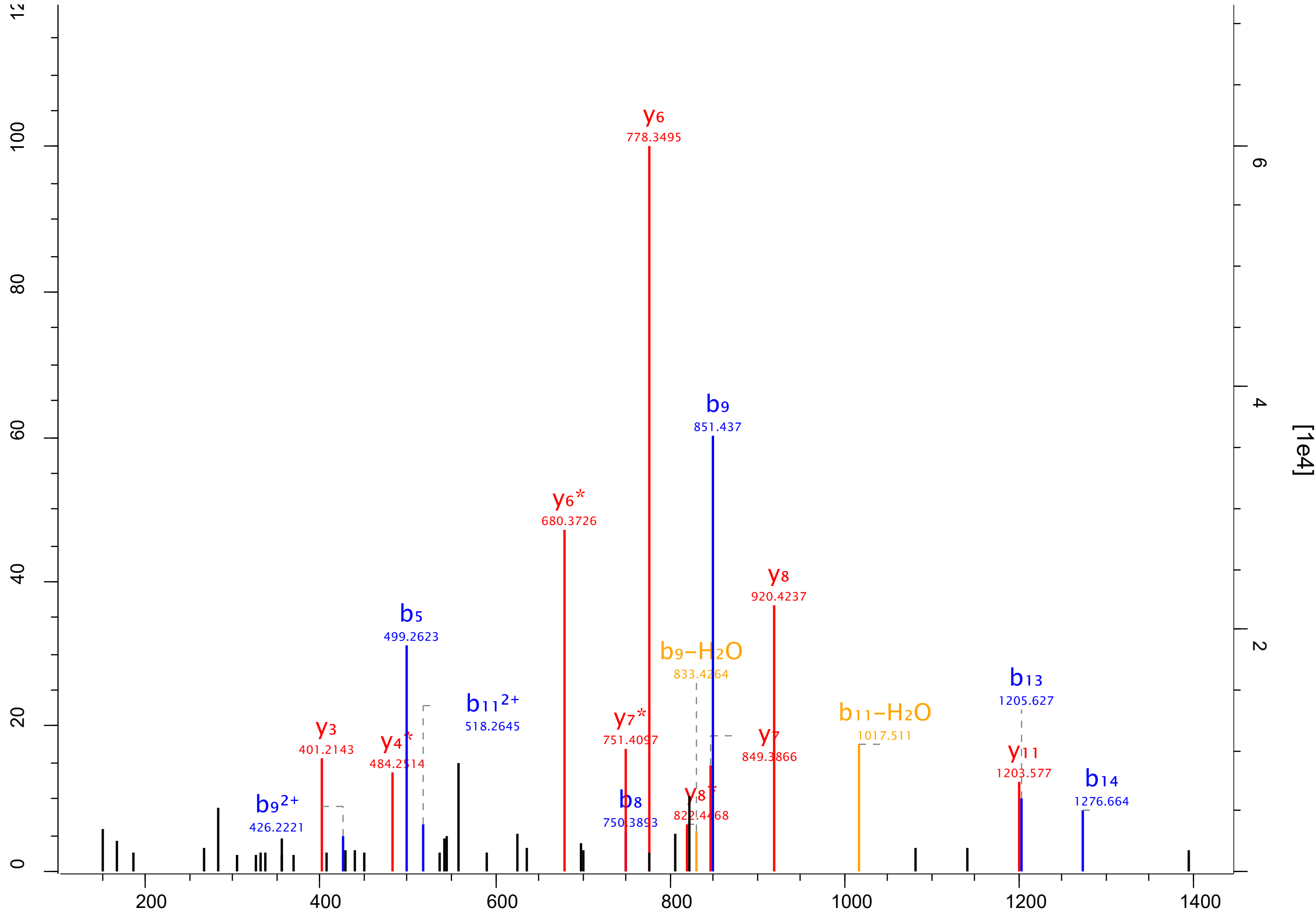


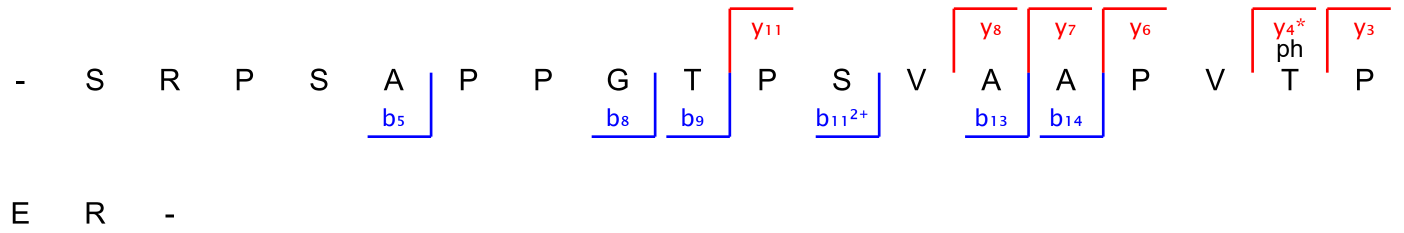

| Raw file | Scan | Method | Score | m/z |
|---|---|---|---|---|
| water_15_2 | 14216 | FTMS; HCD | 72.7 | 584.79 |

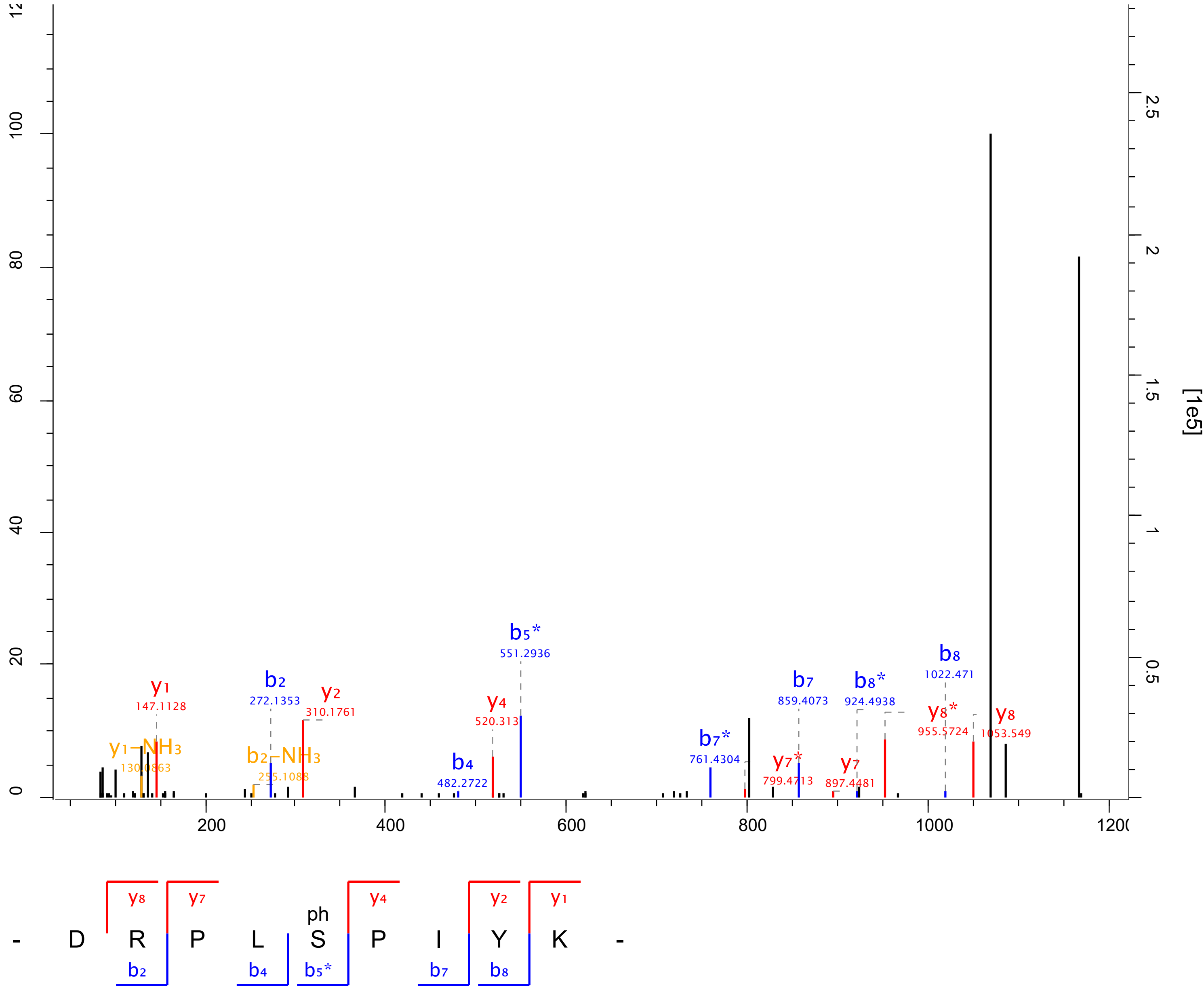

| Raw file | Scan | Method | Score | m/z |
|---|---|---|---|---|
| water_15_2 | 14248 | FTMS; HCD | 54.22 | 936.91 |

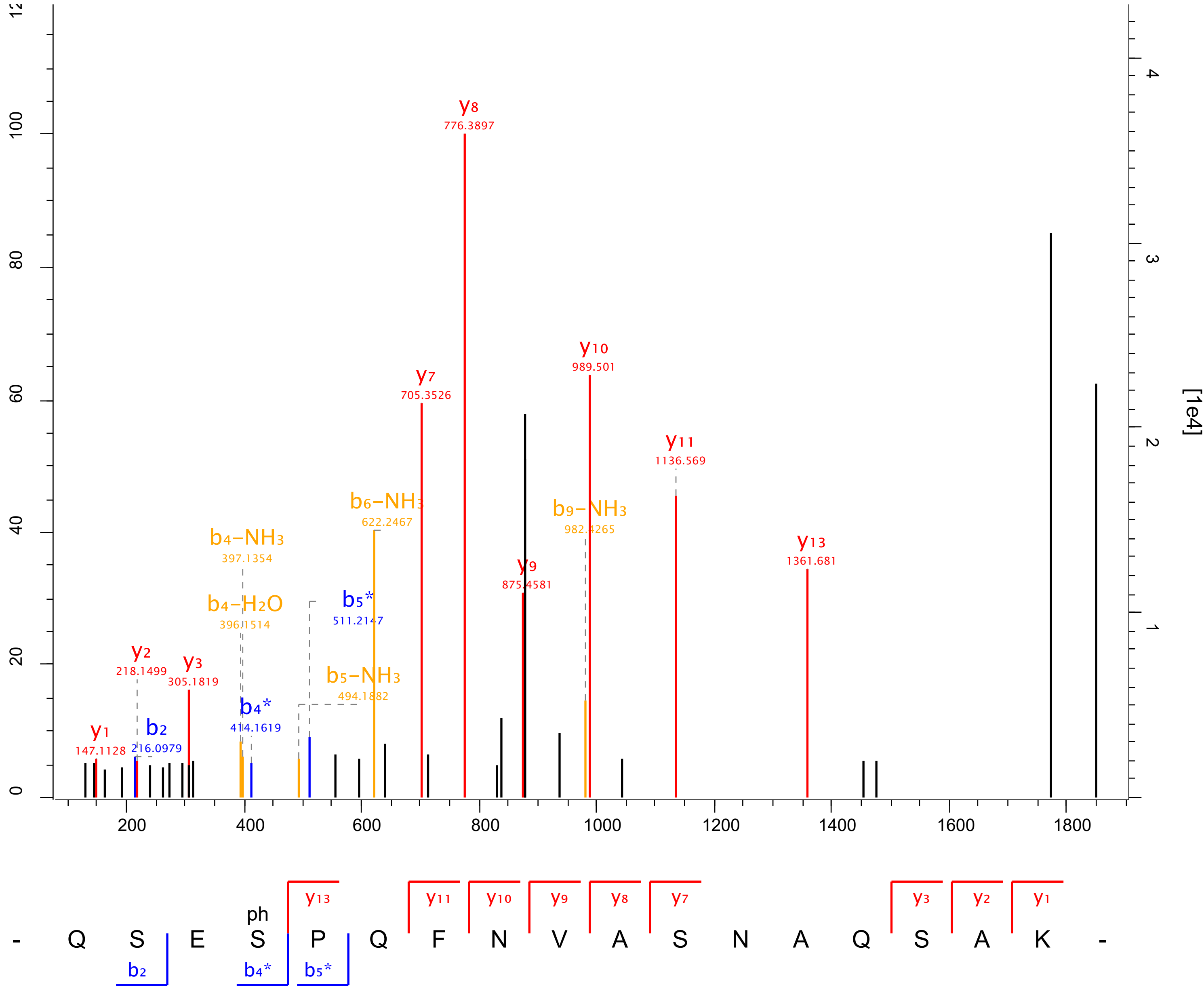

| Raw file | Scan | Method | Score | m/z |
|---|---|---|---|---|
| water_15_2 | 14310 | FTMS; HCD | 124.67 | 485.23 |

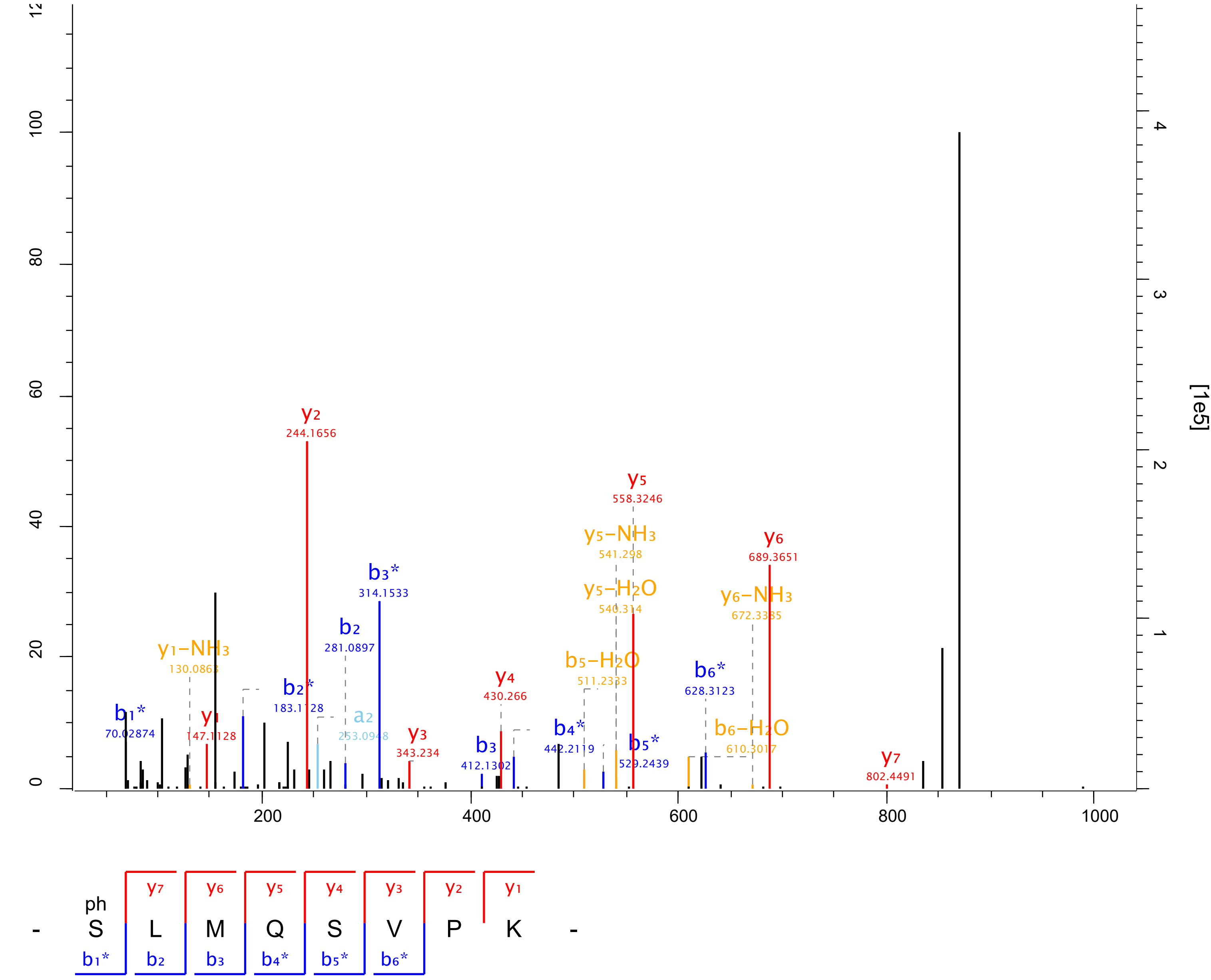

| Raw file | Scan | Method | Score | m/z |
|---|---|---|---|---|
| water_15_2 | 14334 | FTMS; HCD | 53.24 | 512.26 |

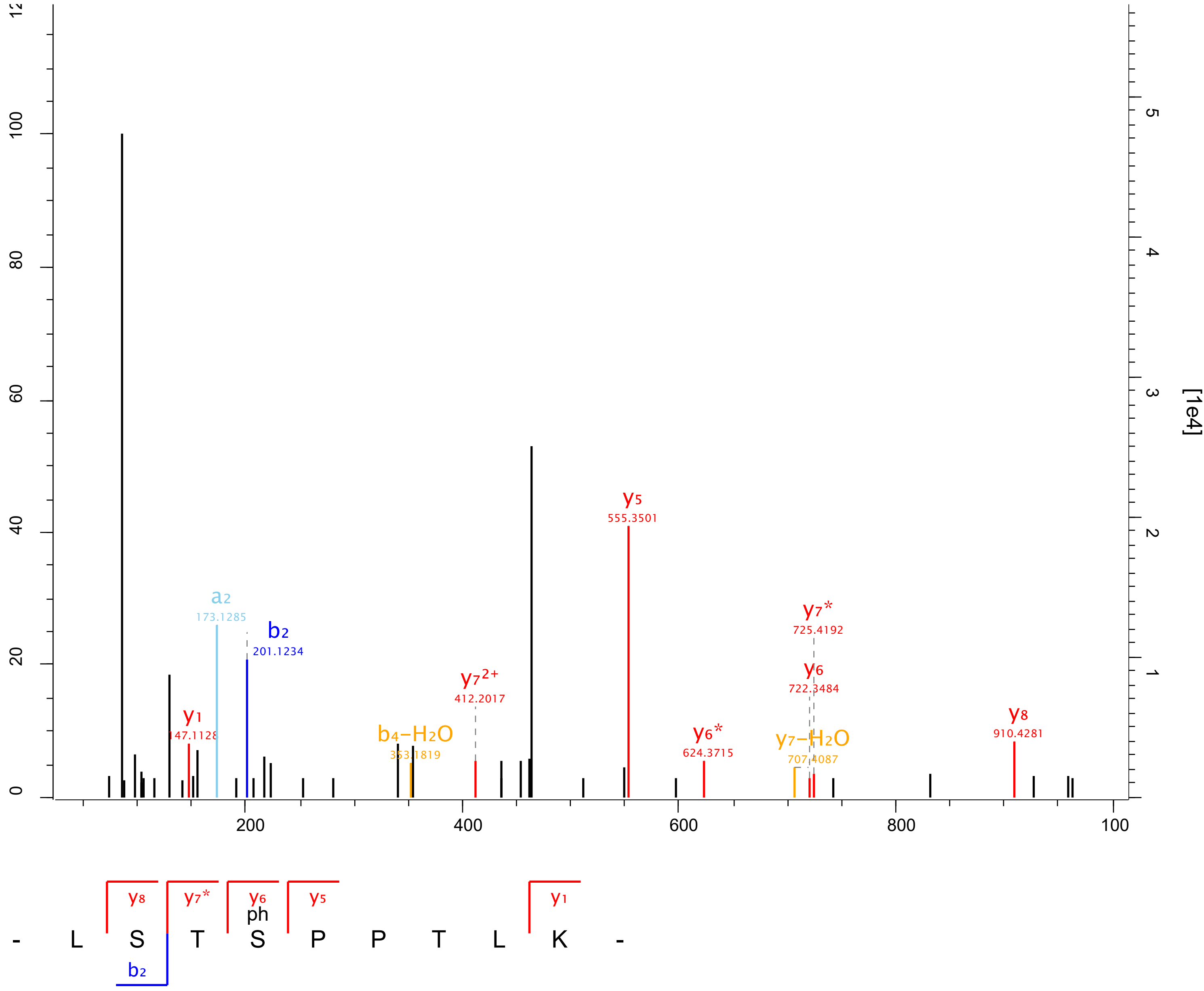

| Raw file | Scan | Method | Score | m/z |
| --- | --- | --- | --- | --- |
| water_15_2 | 14380 | FTMS; HCD | 264.52 | 840.79 |

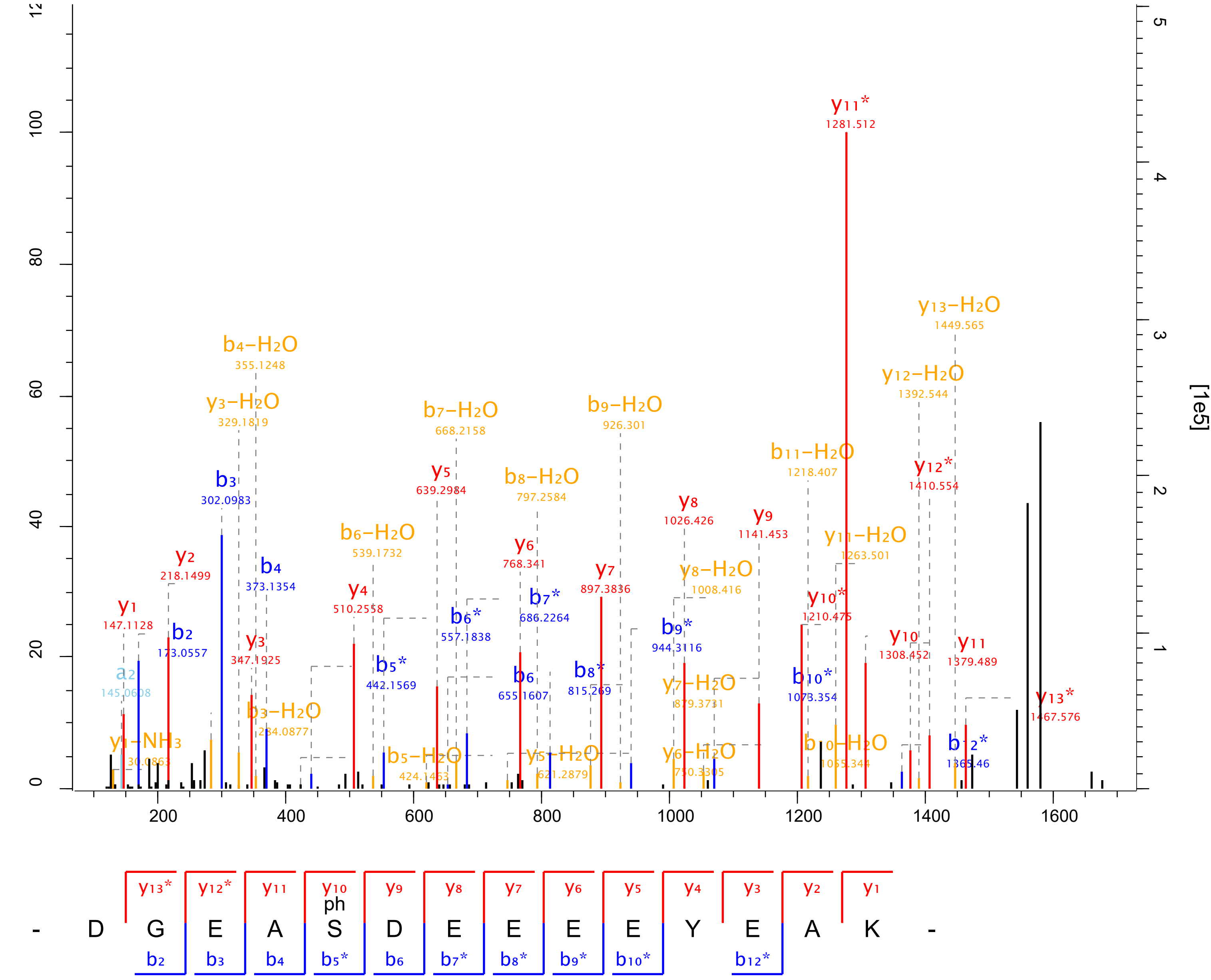

| Raw file | Scan | Method | Score | m/z |
|---|---|---|---|---|
| water_15_2 | 14397 | FTMS; HCD | 114.7 | 676.3 |

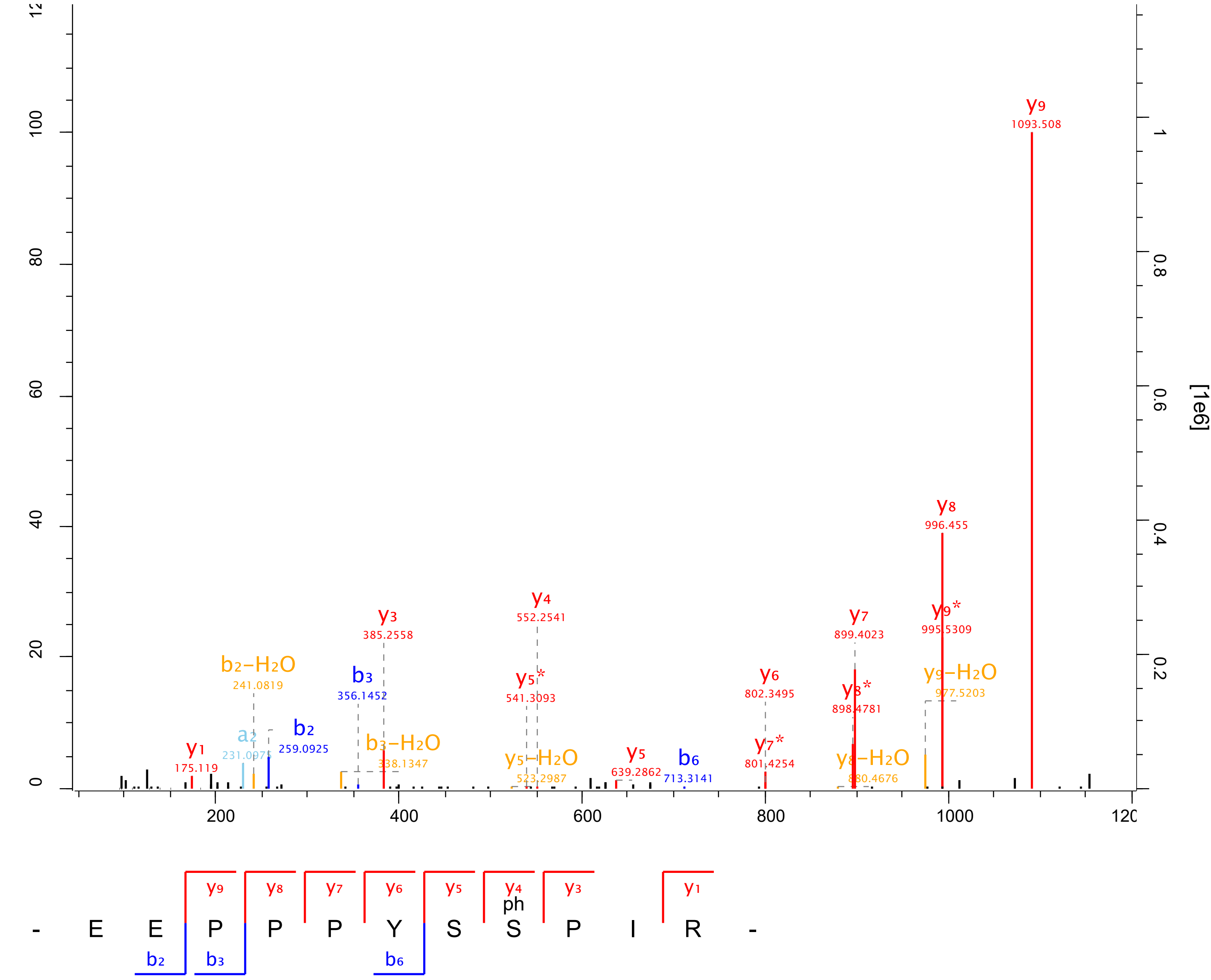

| Raw file | Scan | Method | Score | m/z |
|---|---|---|---|---|
| water_15_2 | 14443 | FTMS; HCD | 45.83 | 515.22 |

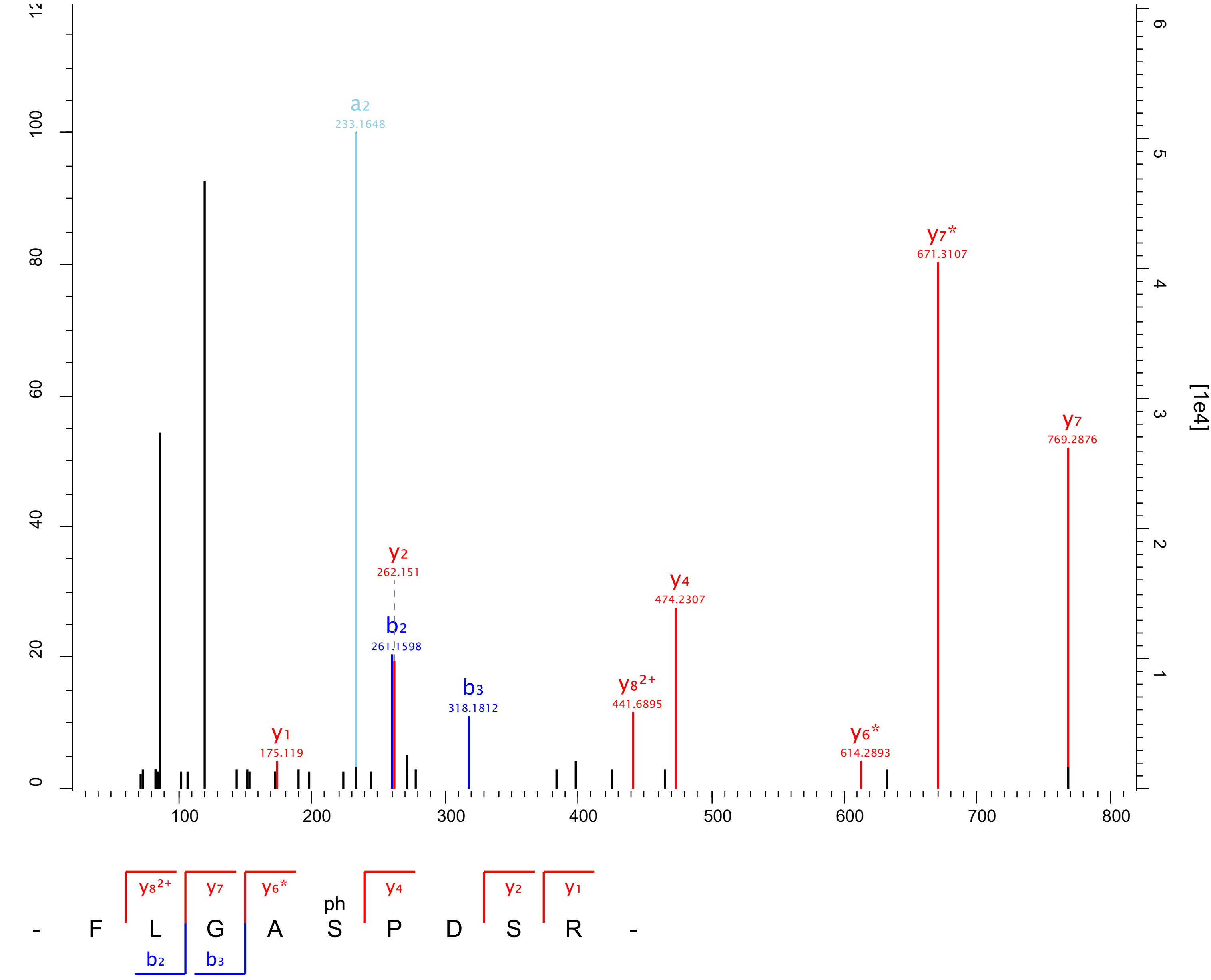

| Raw file | Scan | Method | Score | m/z |
|---|---|---|---|---|
| water_15_2 | 14465 | FTMS; HCD | 66.27 | 568.21 |

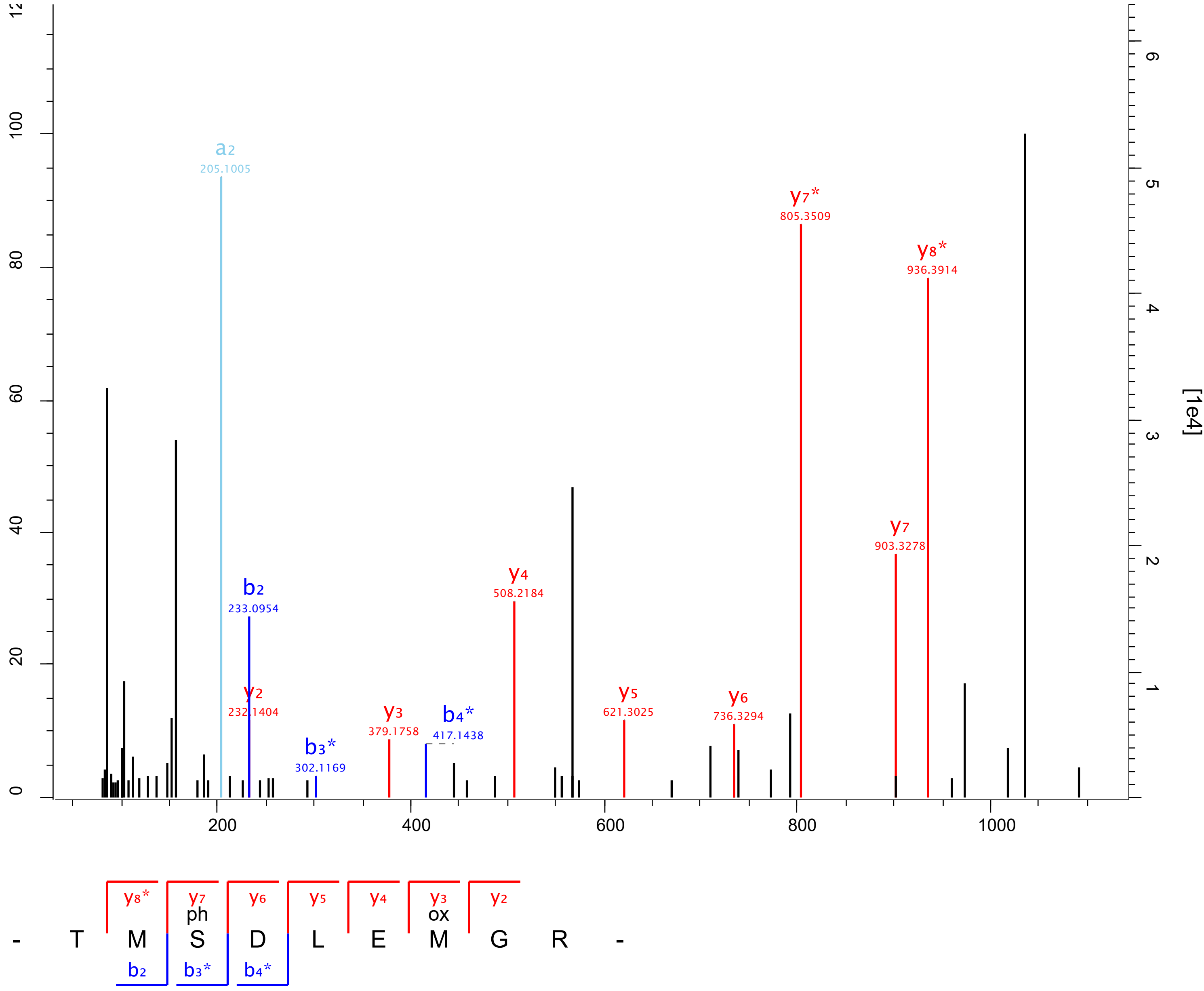

| Raw file | Scan | Method | Score | m/z |
| --- | --- | --- | --- | --- |
| water_15_2 | 14532 | FTMS; HCD | 54.02 | 572.74 |

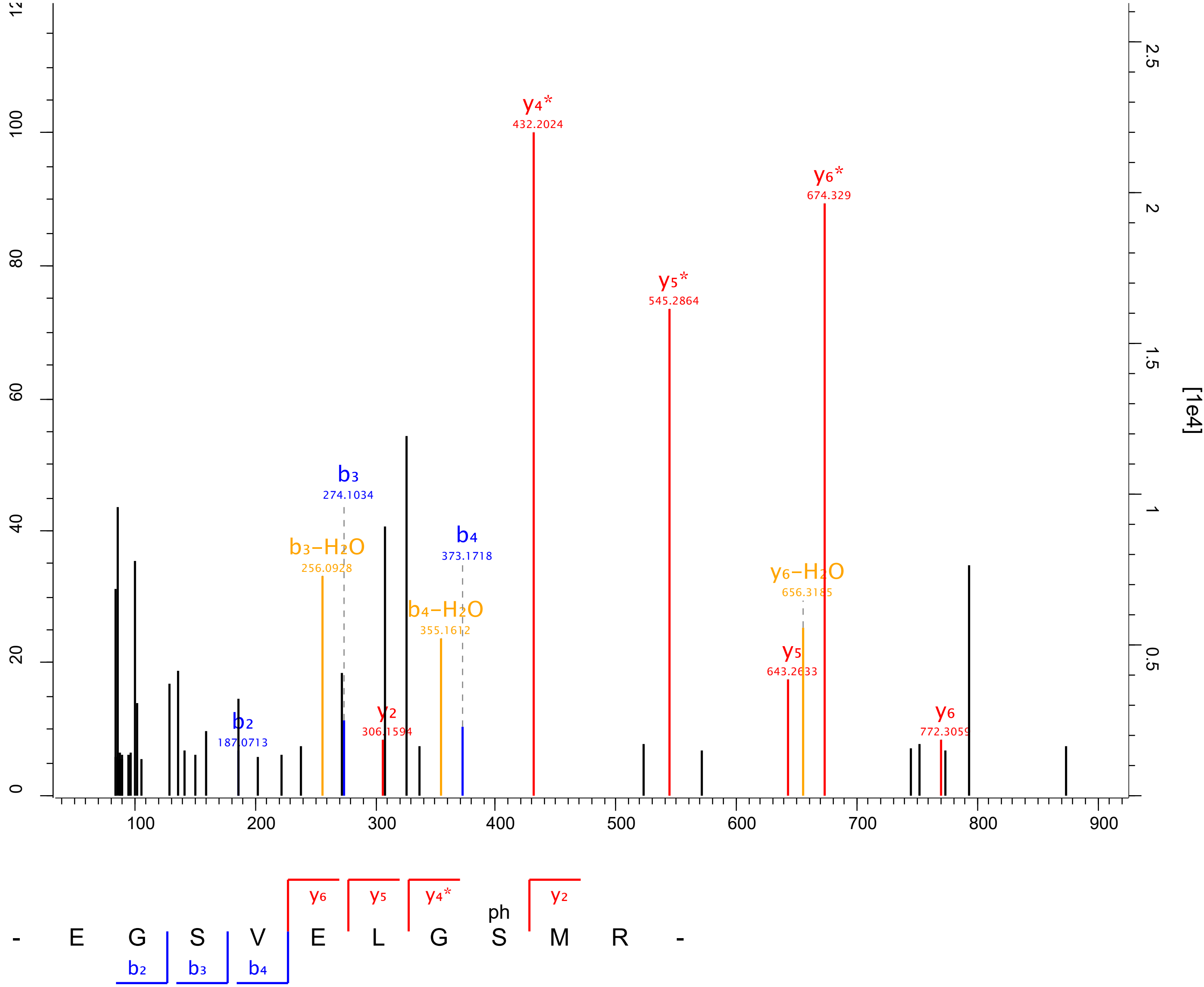

| Raw file | Scan | Method | Score | m/z |
|---|---|---|---|---|
| water_15_2 | 14551 | FTMS; HCD | 91.31 | 528.73 |

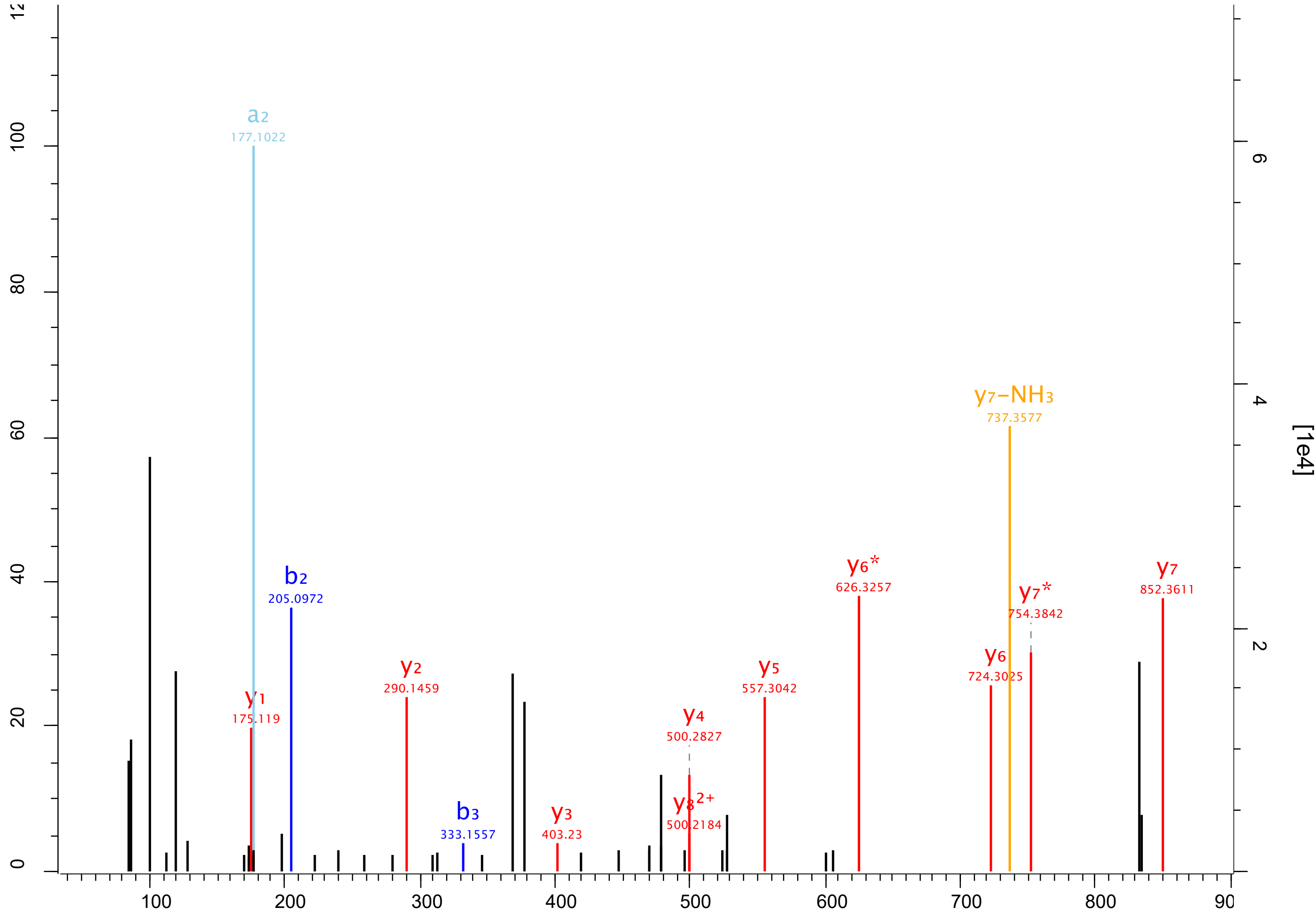


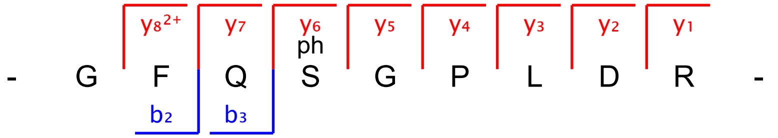

| Raw file | Scan | Method | Score | m/z |
|---|---|---|---|---|
| water_15_2 | 14560 | FTMS; HCD | 74.14 | 695.31 |

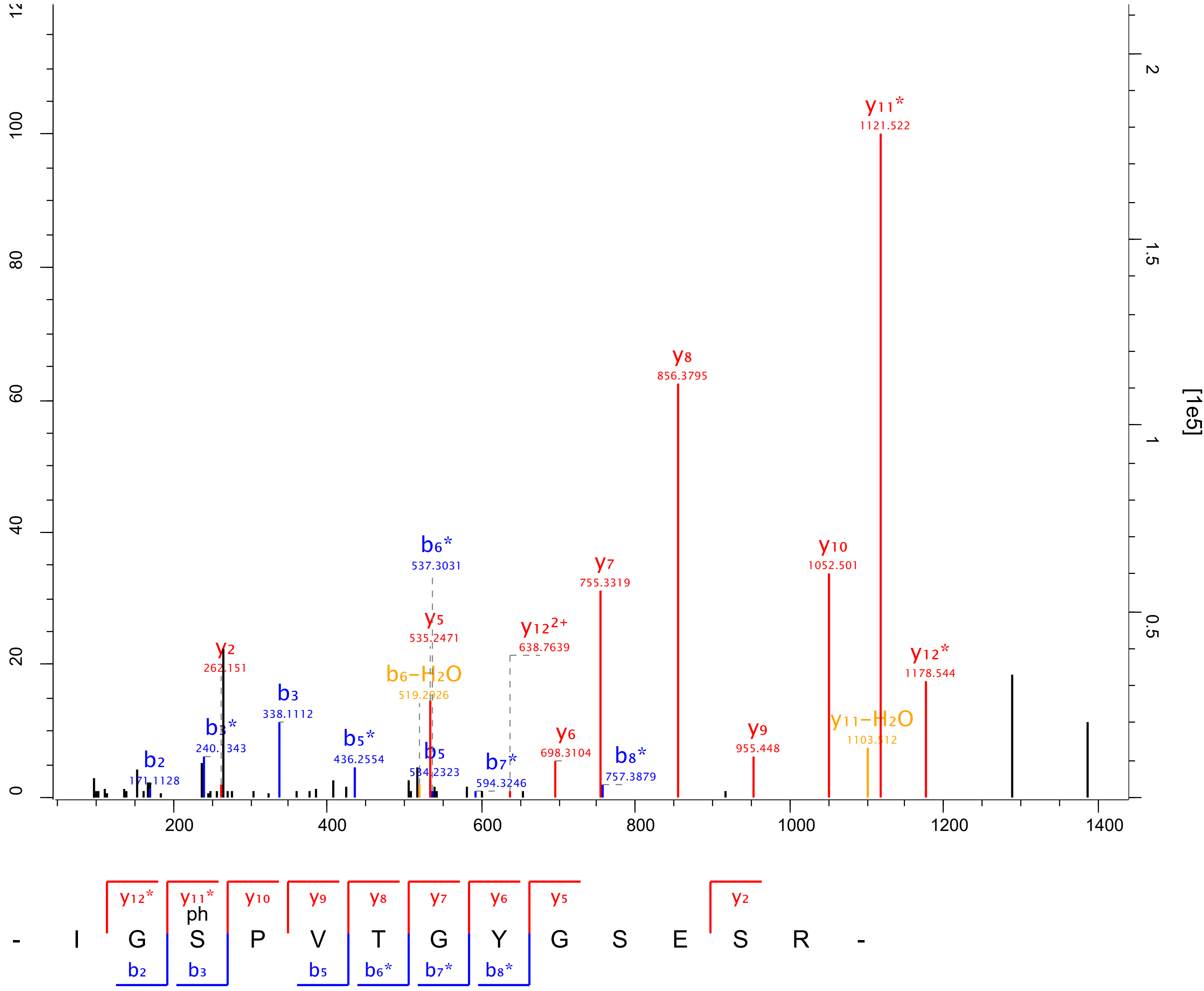

| Raw file | Scan | Method | Score | m/z |
|---|---|---|---|---|
| water_15_2 | 14573 | FTMS; HCD | 97.08 | 769.79 |

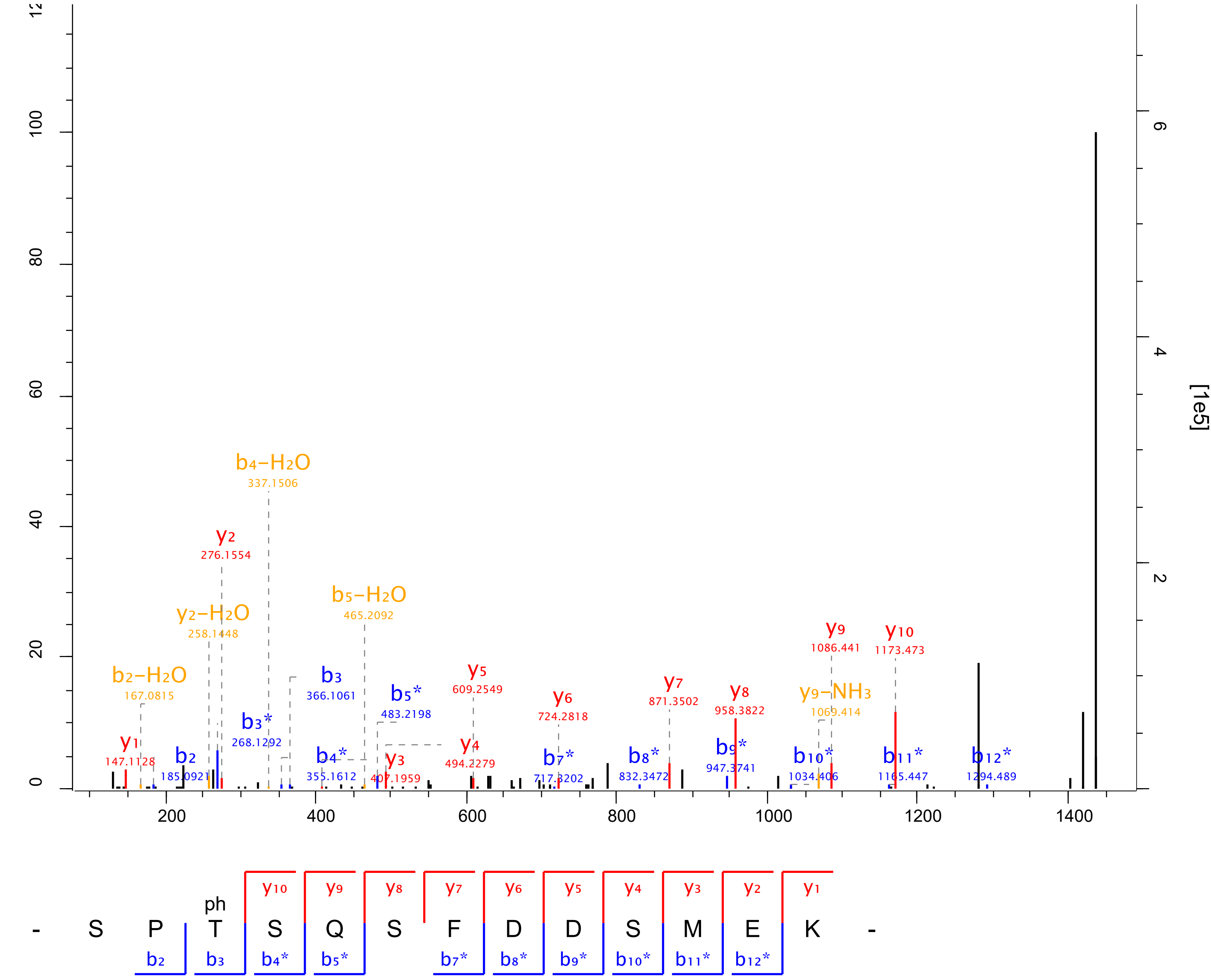

| Raw file | Scan | Method | Score | m/z |
|---|---|---|---|---|
| water_15_2 | 14623 | FTMS; HCD | 88.31 | 529.9 |

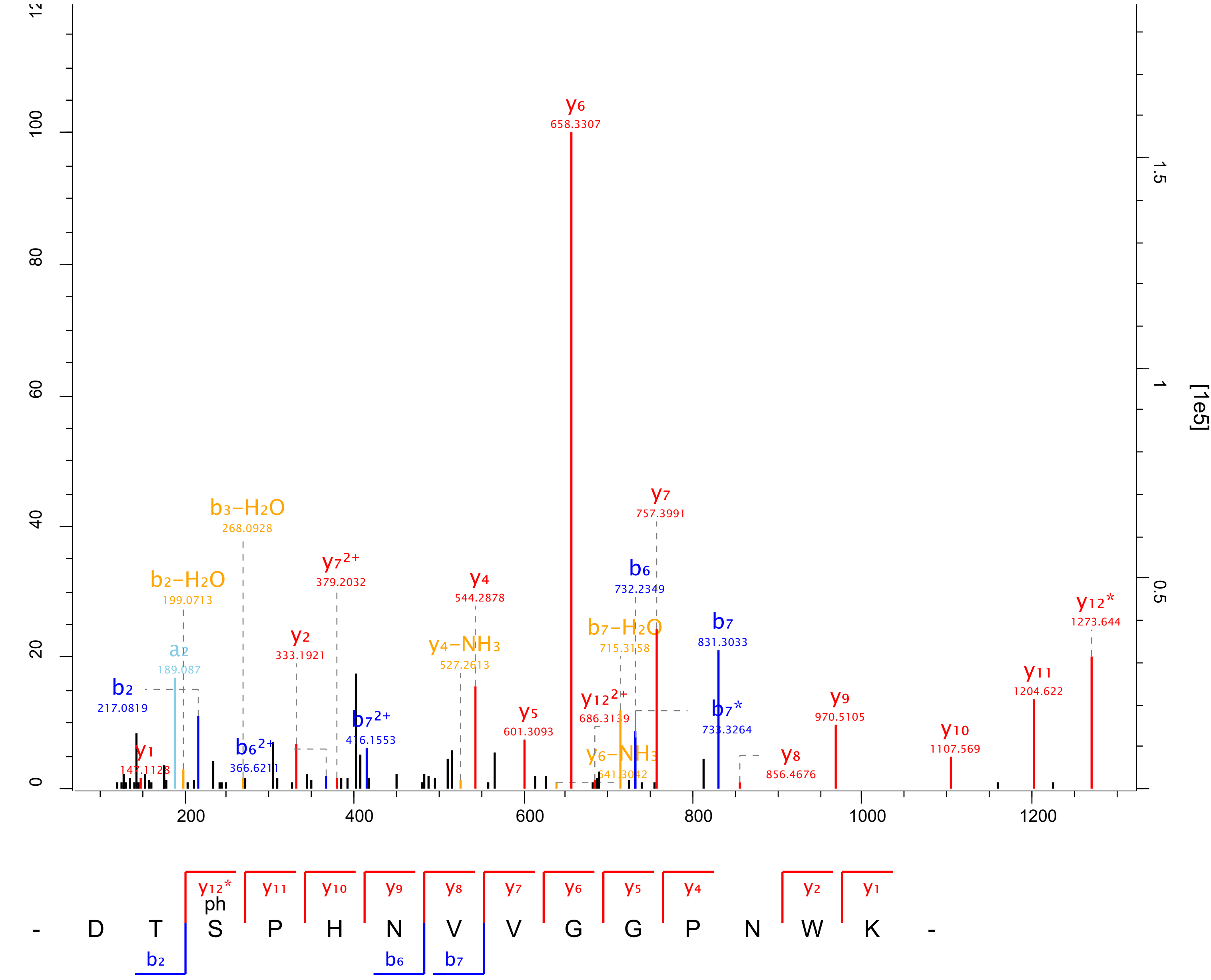

| Raw file | Scan | Method | Score | m/z |
| --- | --- | --- | --- | --- |
| water_15_2 | 14674 | FTMS; HCD | 78.92 | 790.81 |

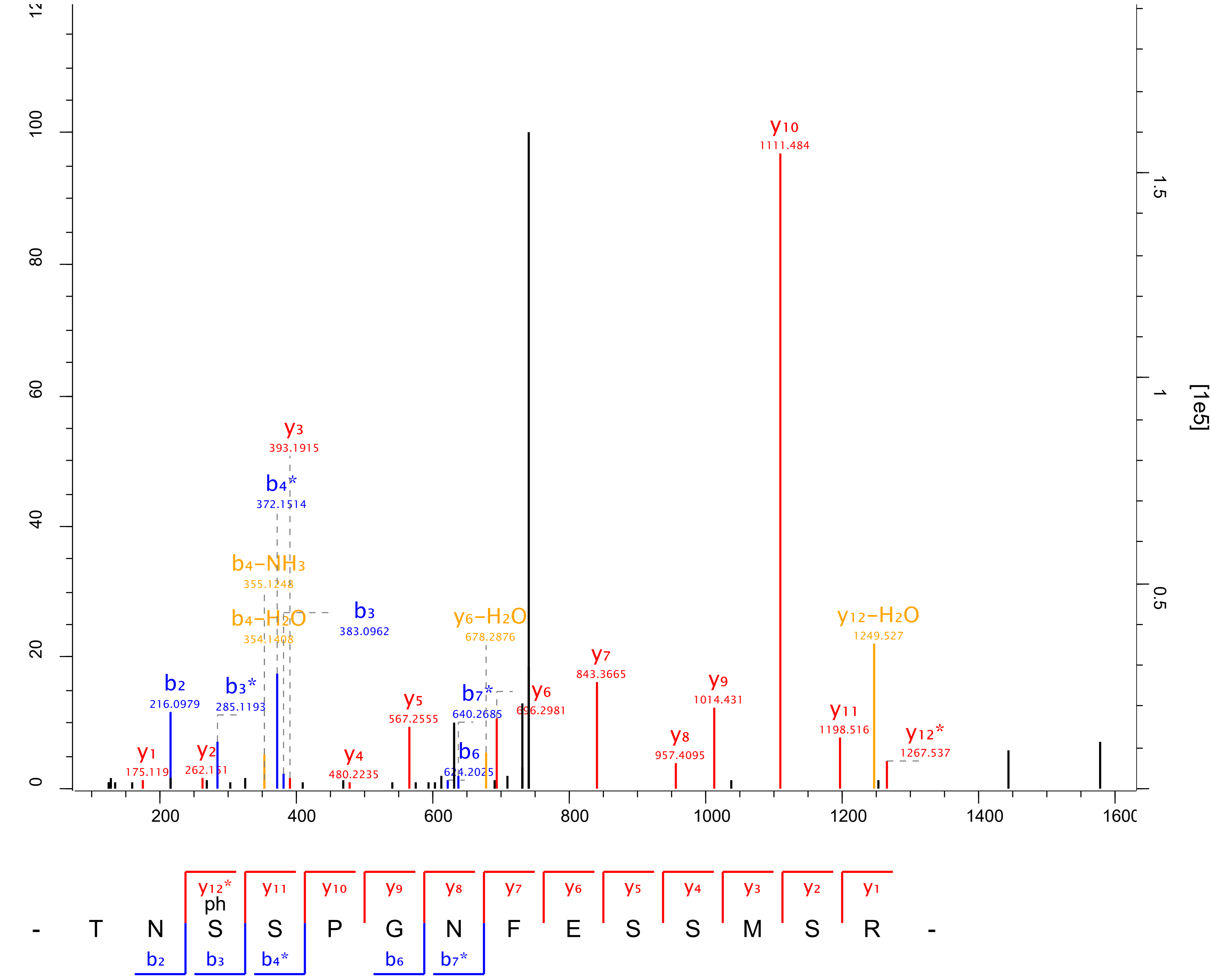

| Raw file | Scan | Method | Score | m/z |
|---|---|---|---|---|
| water_15_2 | 14676 | FTMS; HCD | 142.45 | 620.24 |

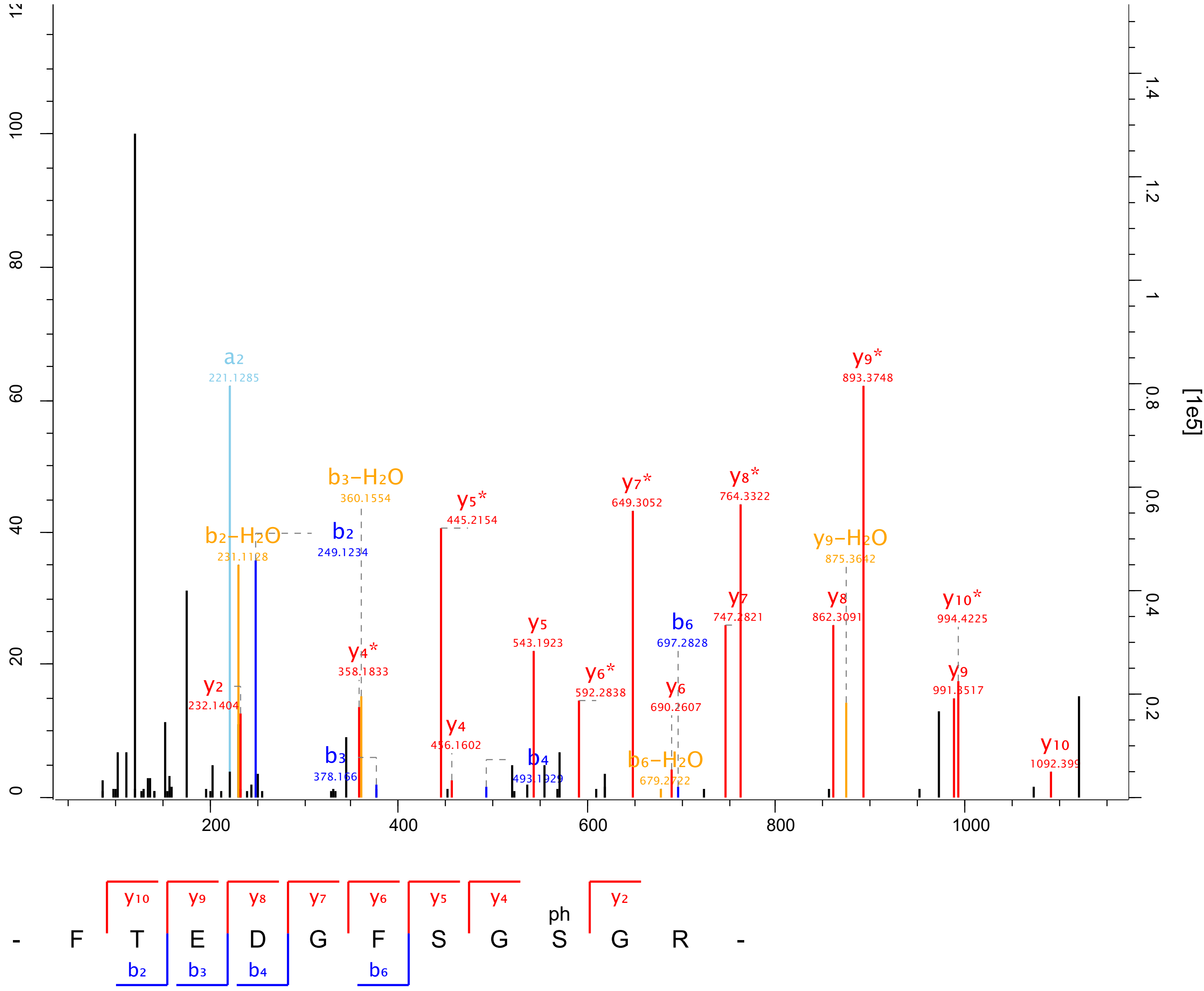

| Raw file | Scan | Method | Score | m/z |
|---|---|---|---|---|
| water_15_2 | 14679 | FTMS; HCD | 83.76 | 811.87 |

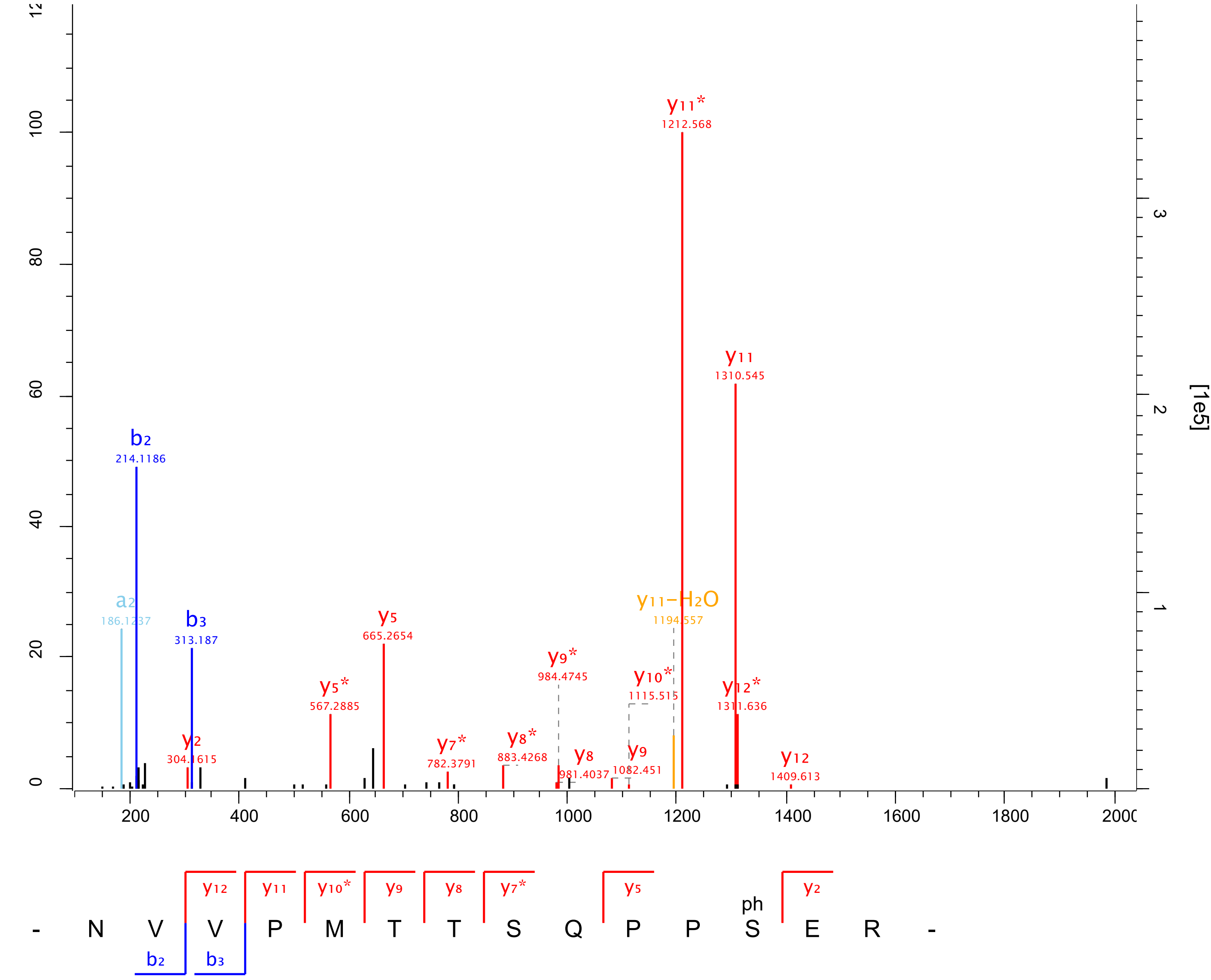

| Raw file | Scan | Method | Score | m/z |
| --- | --- | --- | --- | --- |
| water_15_2 | 14702 | FTMS; HCD | 94.69 | 587.76 |

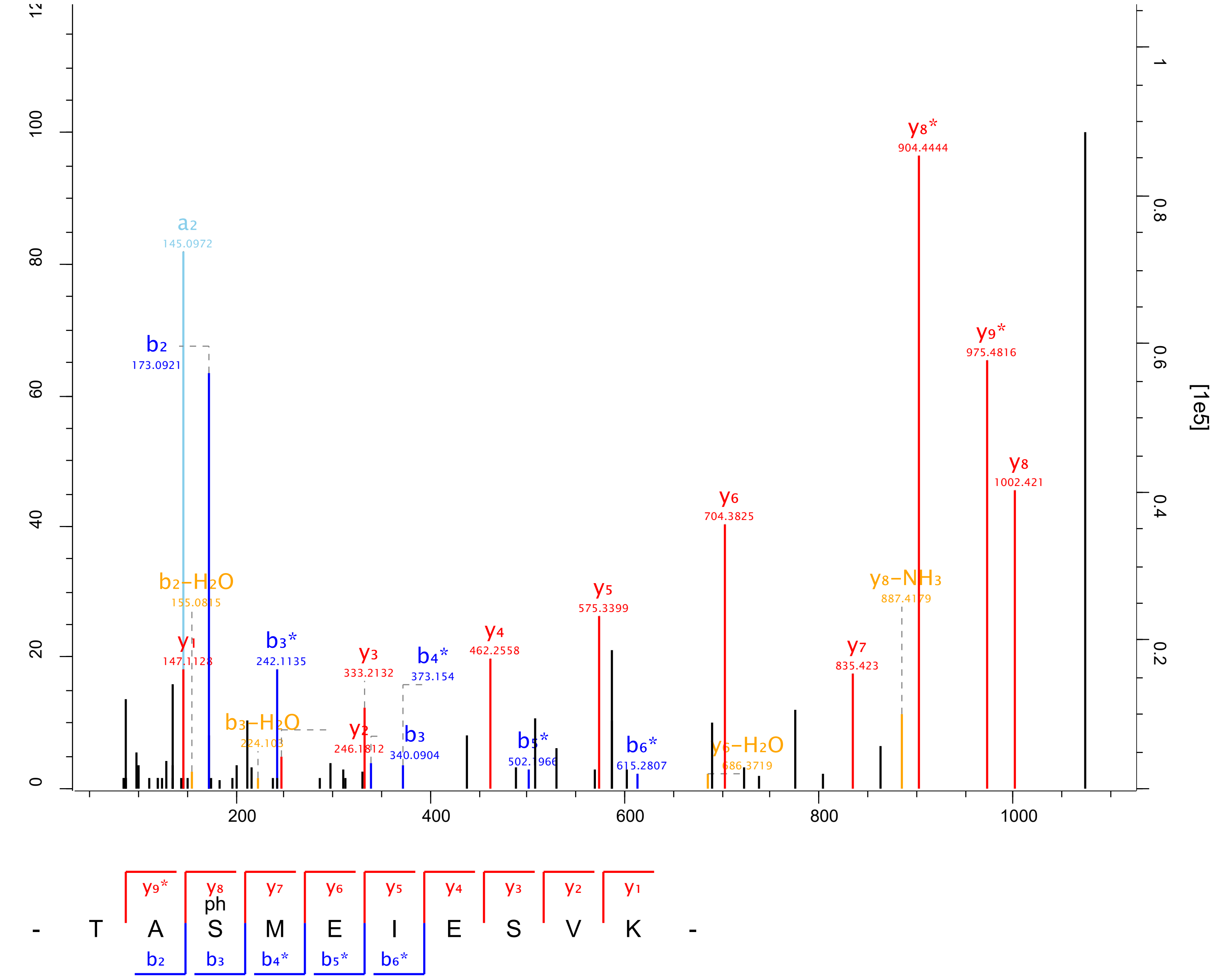

| Raw file | Scan | Method | Score | m/z |
| --- | --- | --- | --- | --- |
| water_15_2 | 14705 | FTMS; HCD | 61.48 | 681.28 |

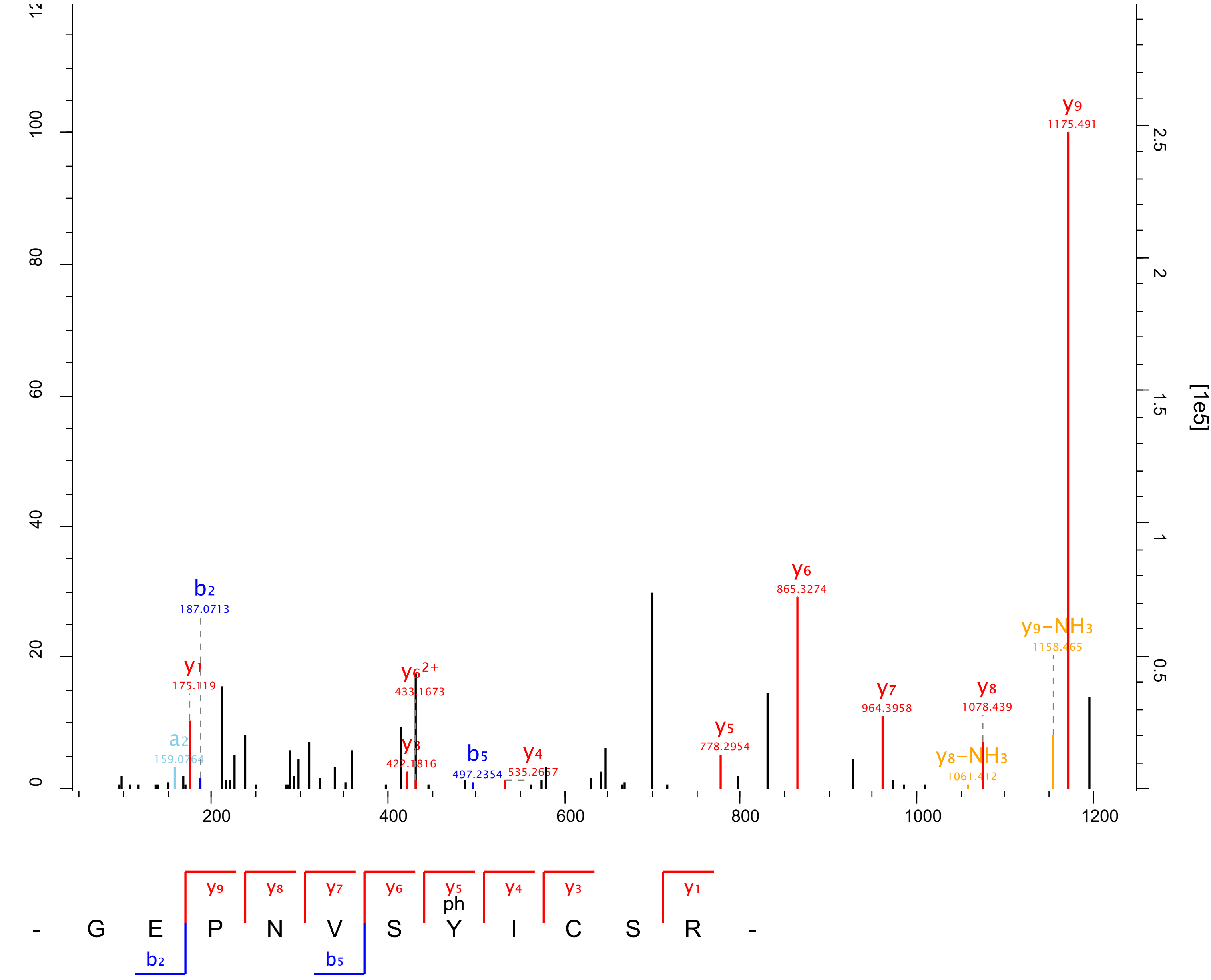

| Raw file | Scan | Method | Score | m/z |
|---|---|---|---|---|
| water_15_2 | 14732 | FTMS; HCD | 76.73 | 598.72 |

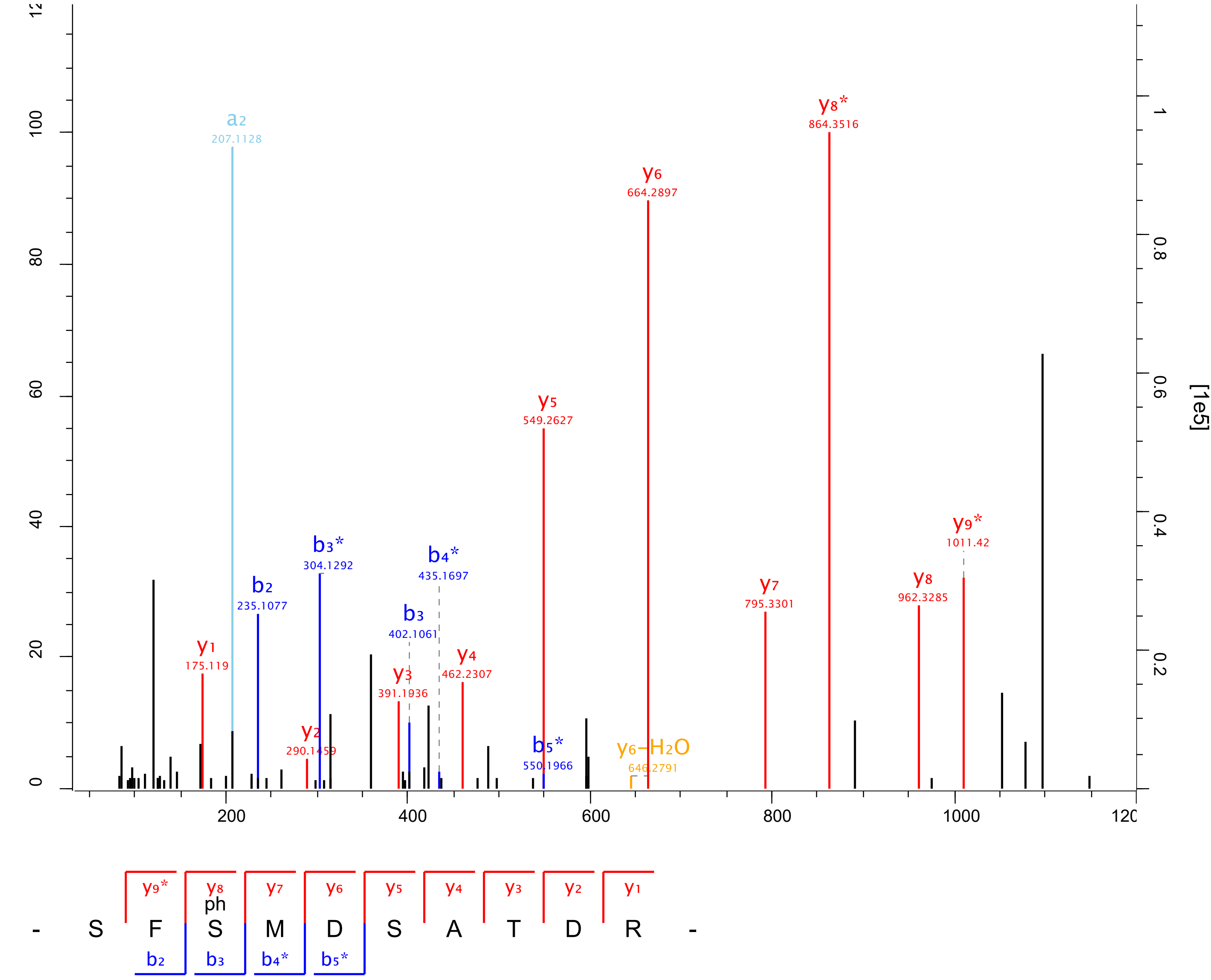

| Raw file | Scan | Method | Score | m/z |
|---|---|---|---|---|
| water_15_2 | 14837 | FTMS; HCD | 58.32 | 563.76 |

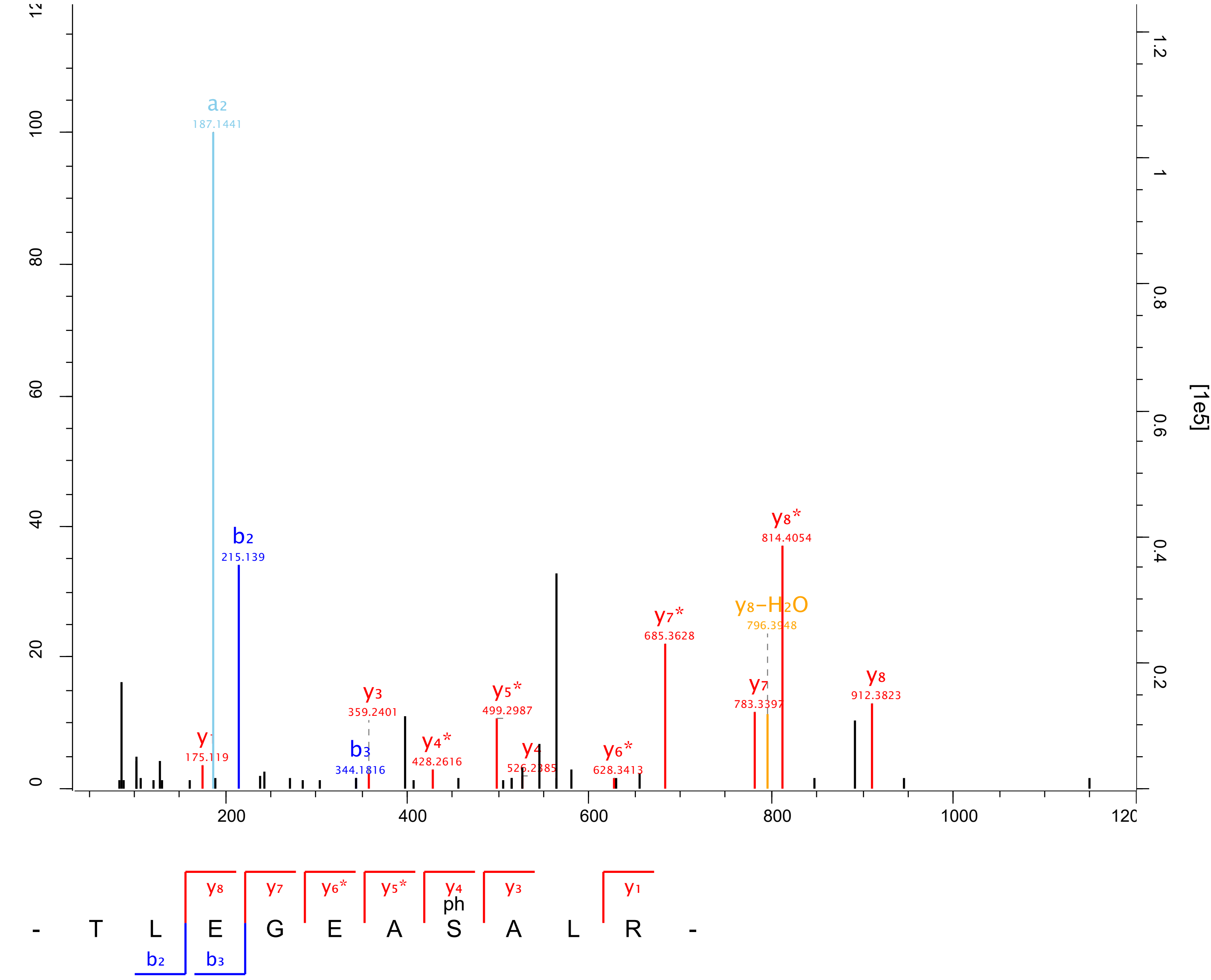

| Raw file | Scan | Method | Score | m/z |
|---|---|---|---|---|
| water_15_2 | 14887 | FTMS; HCD | 218.14 | 782.84 |

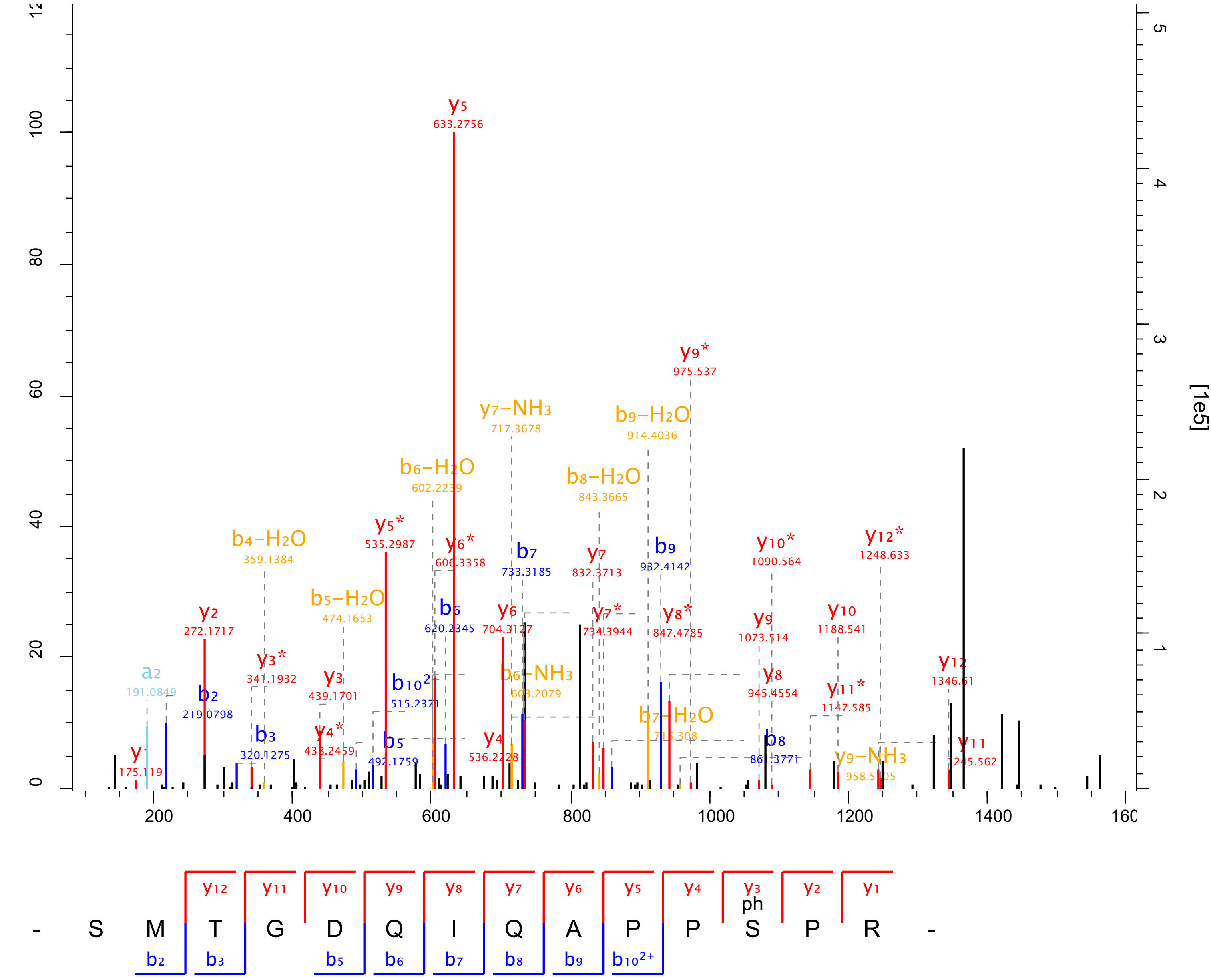

| Raw file | Scan | Method | Score | m/z |
|---|---|---|---|---|
| water_15_2 | 14930 | FTMS; HCD | 80.75 | 633.81 |

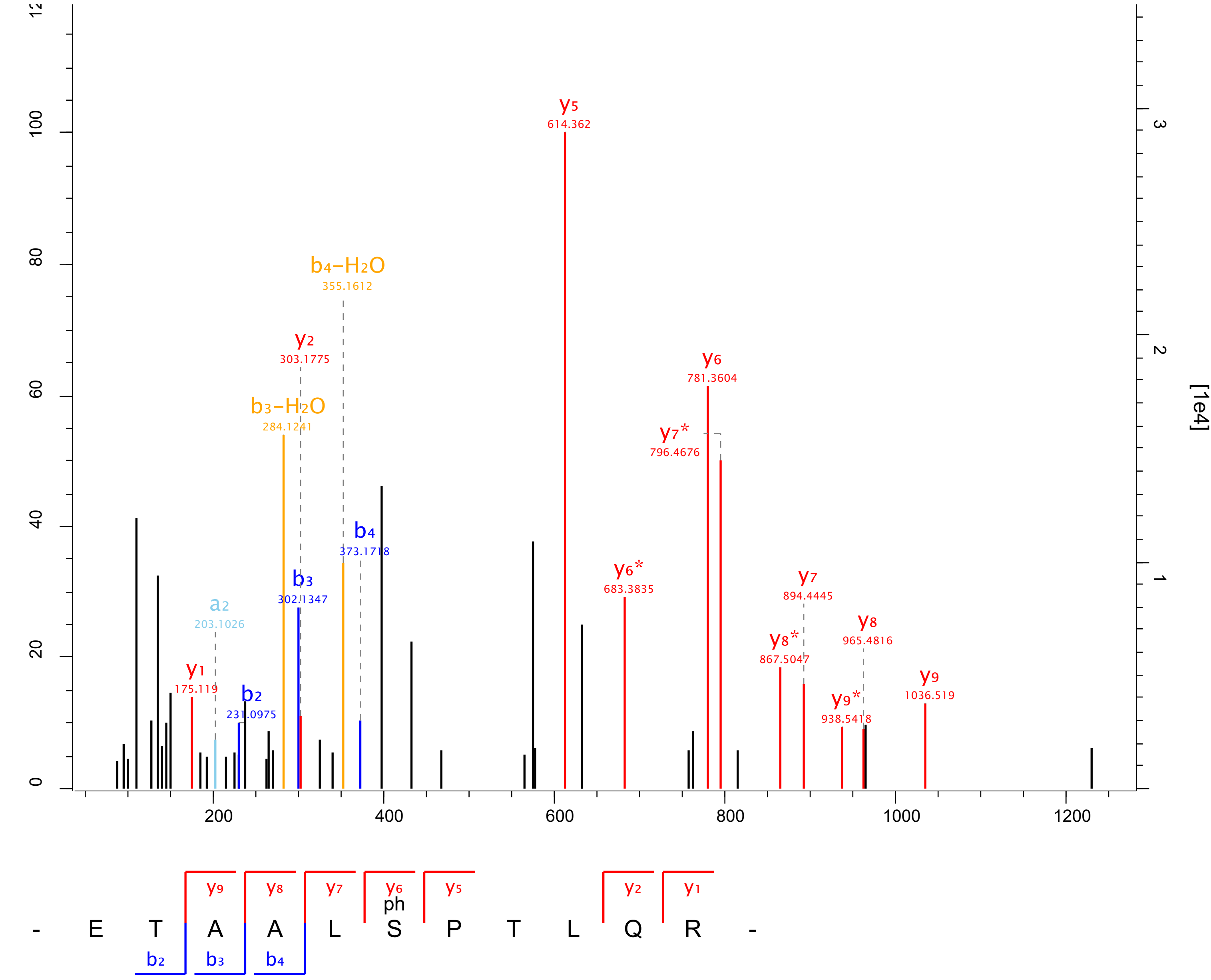

| Raw file | Scan | Method | Score | m/z |
| --- | --- | --- | --- | --- |
| water_15_2 | 14937 | FTMS; HCD | 131.83 | 511.22 |

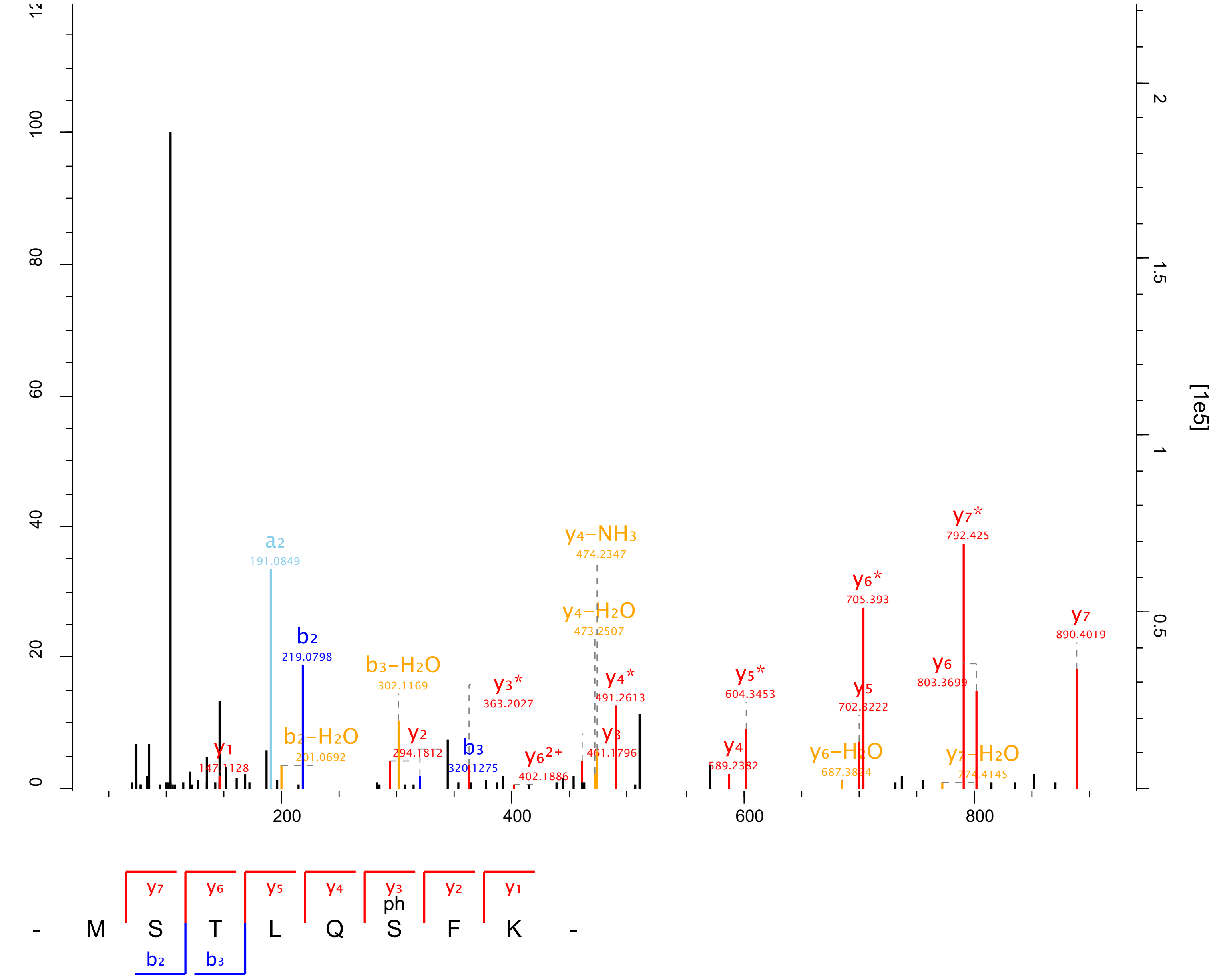

| Raw file | Scan | Method | Score | m/z |
| --- | --- | --- | --- | --- |
| water_15_2 | 14974 | FTMS; HCD | 114.46 | 958.86 |

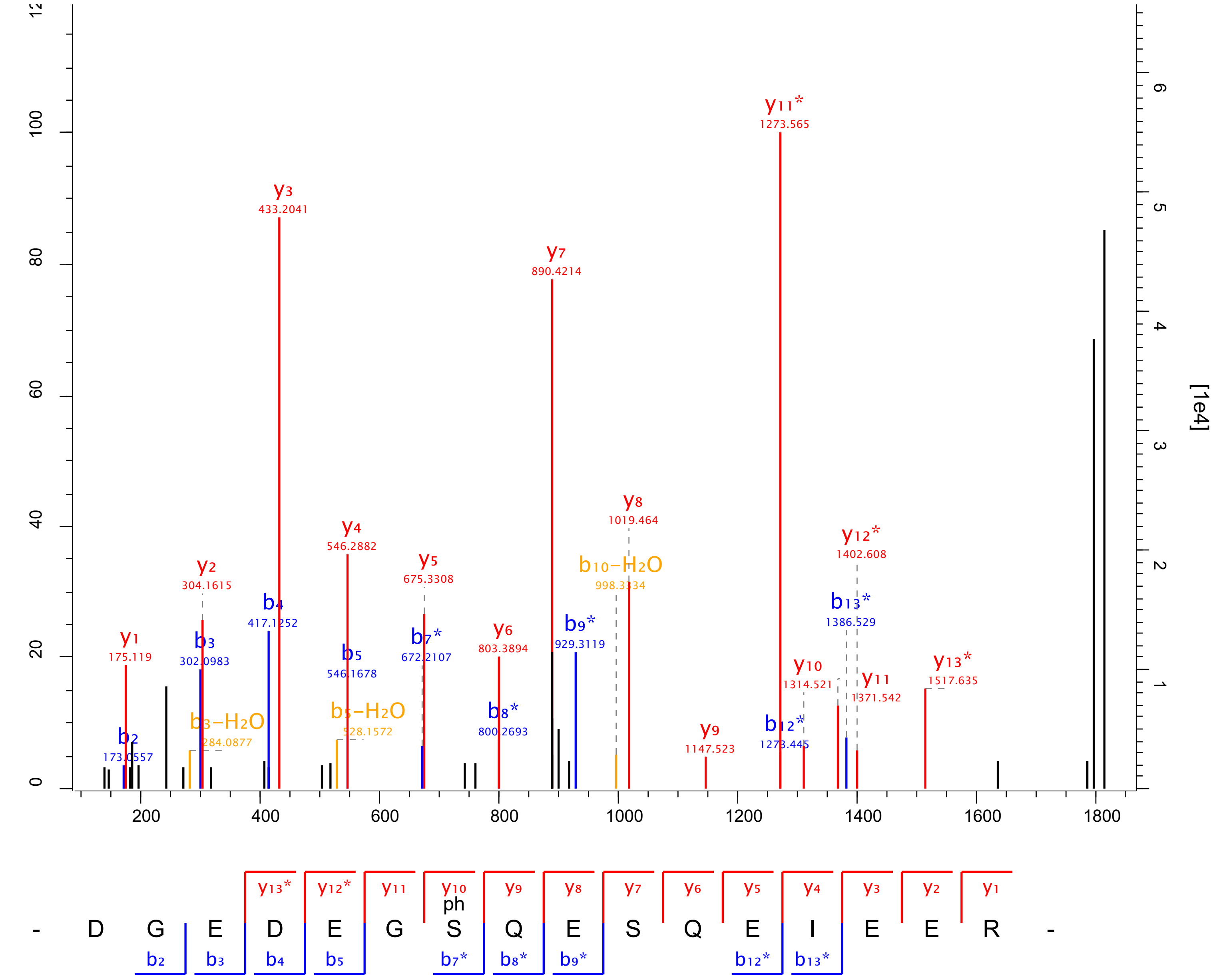

| Raw file | Scan | Method | Score | m/z |
| --- | --- | --- | --- | --- |
| water_15_2 | 14980 | FTMS; HCD | 117.86 | 675.28 |

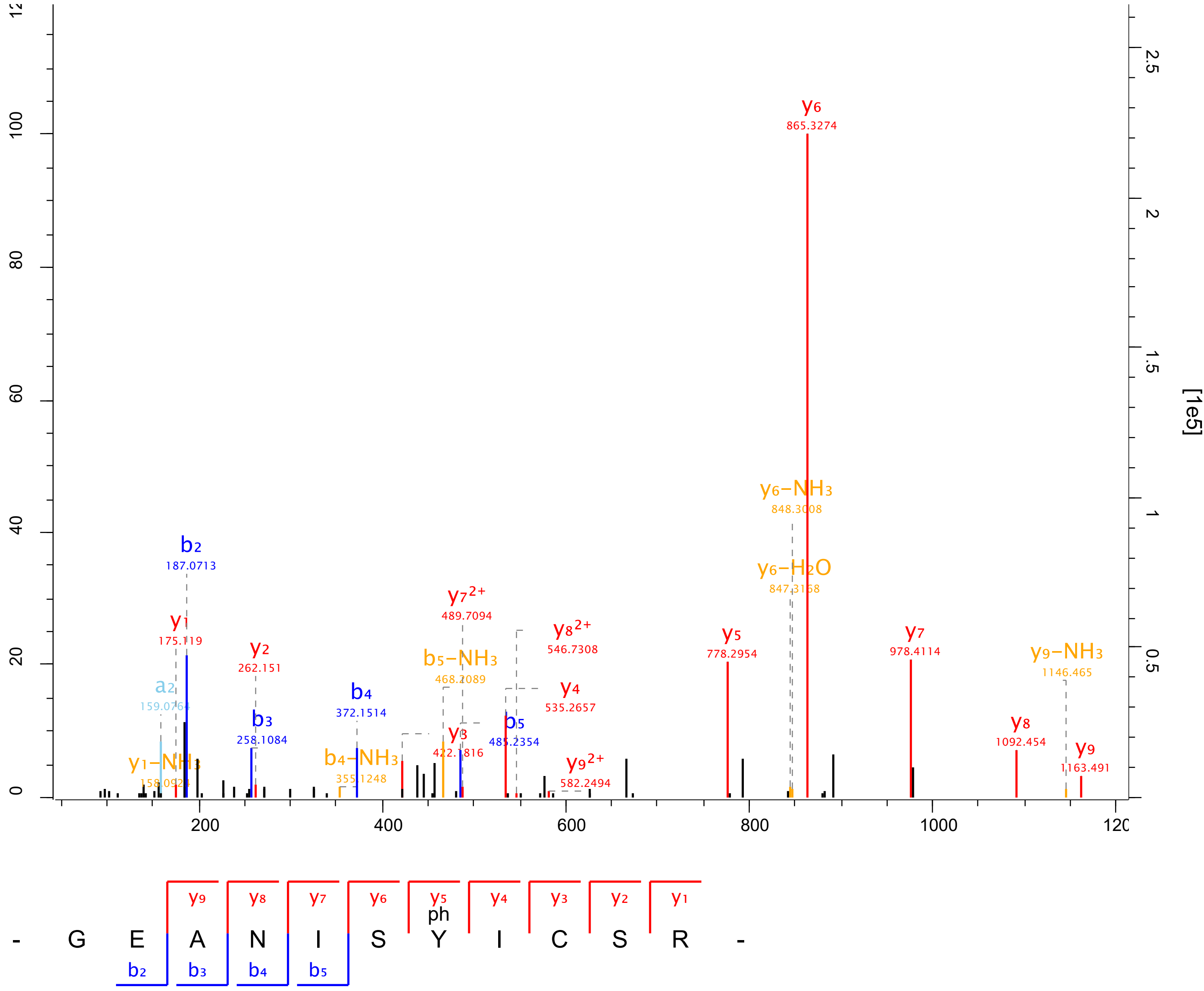

| Raw file | Scan | Method | Score | m/z |
| --- | --- | --- | --- | --- |
| water_15_2 | 15100 | FTMS; HCD | 70.2 | 759.82 |

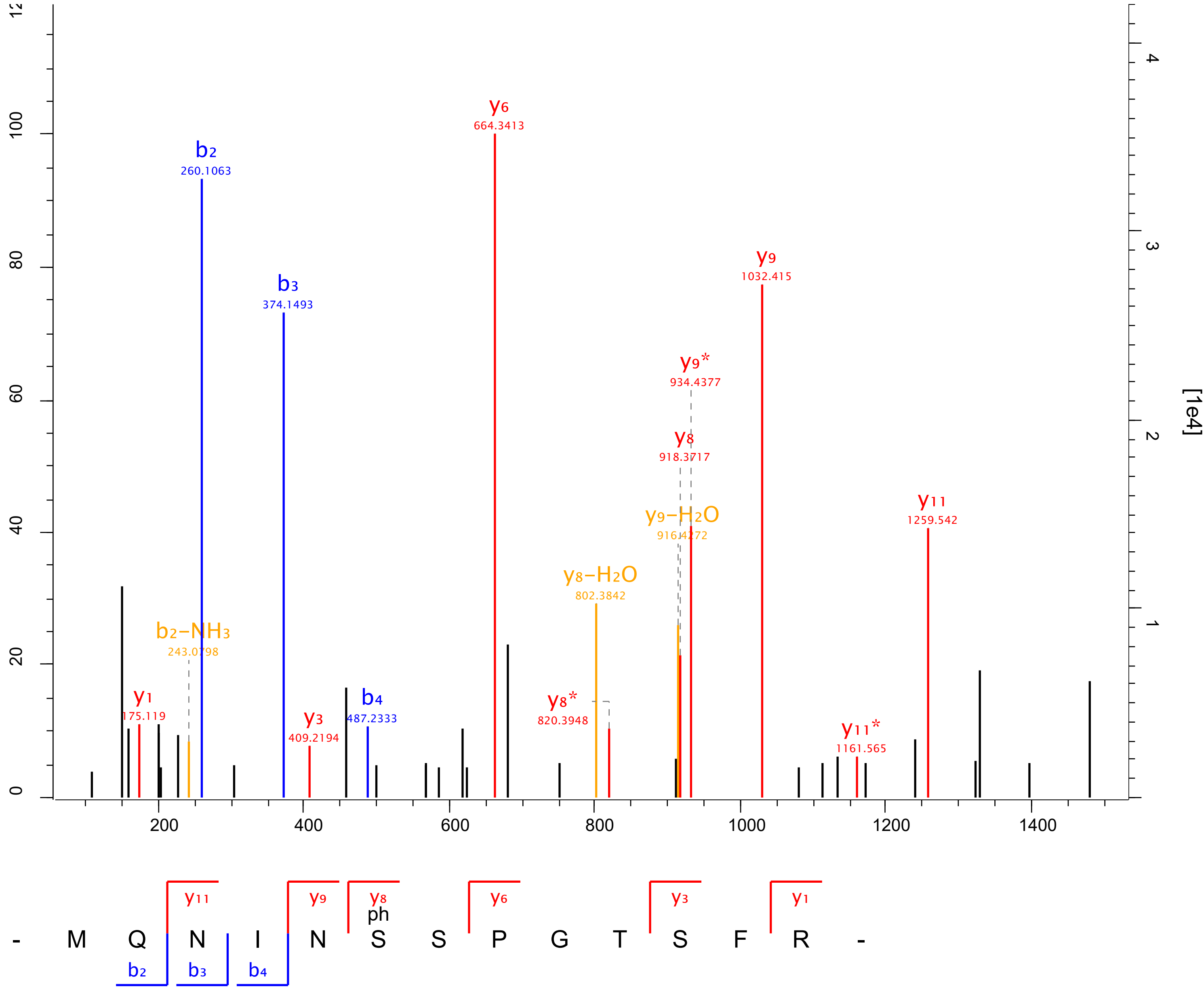

| Raw file | Scan | Method | Score | m/z |
|---|---|---|---|---|
| water_15_2 | 15181 | FTMS; HCD | 122.34 | 557.27 |

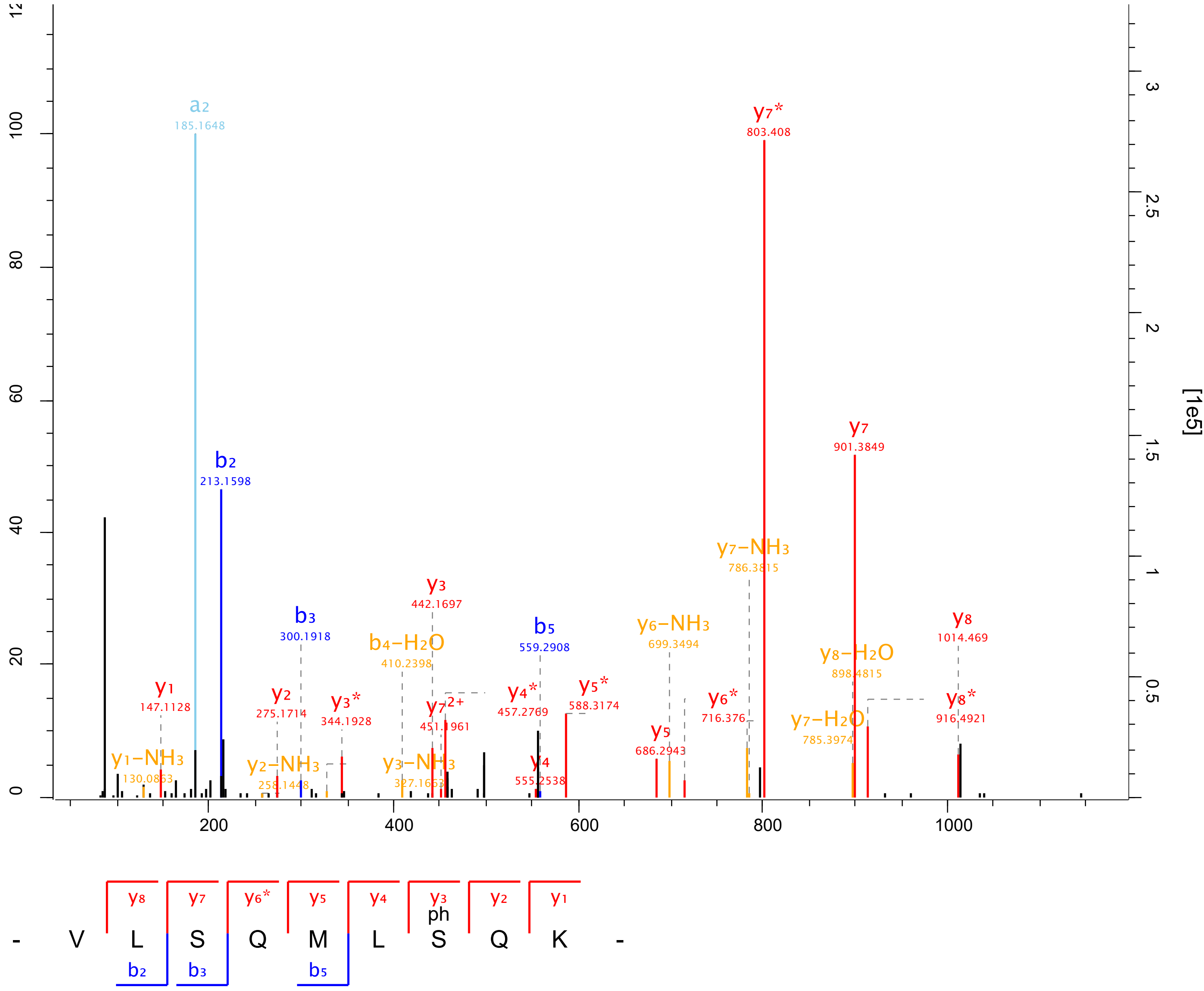

| Raw file | Scan | Method | Score | m/z |
|---|---|---|---|---|
| water_15_2 | 15226 | FTMS; HCD | 43.76 | 615.27 |

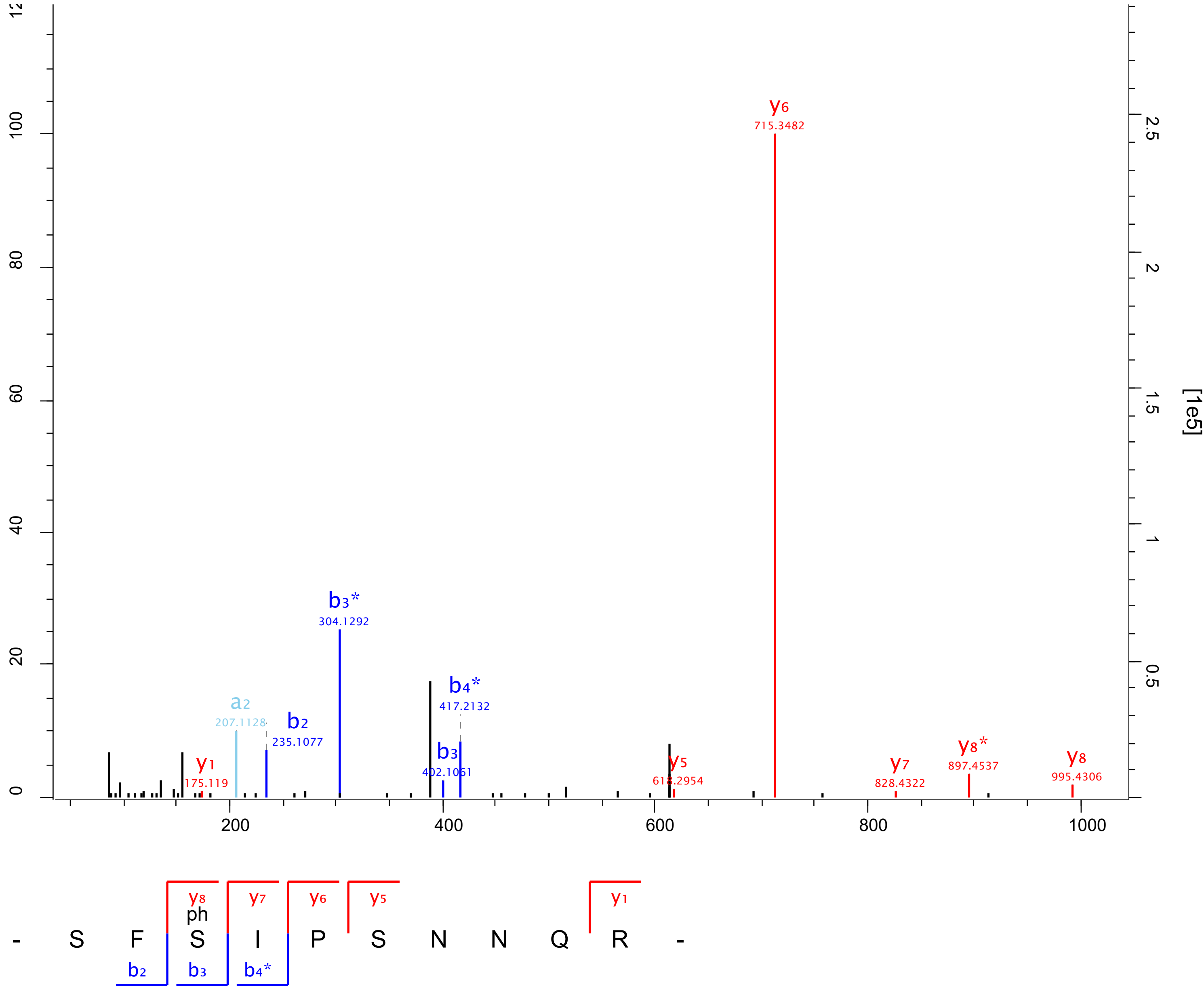

| Raw file | Scan | Method | Score | m/z |
|---|---|---|---|---|
| water_15_2 | 15240 | FTMS; HCD | 50.35 | 571.27 |

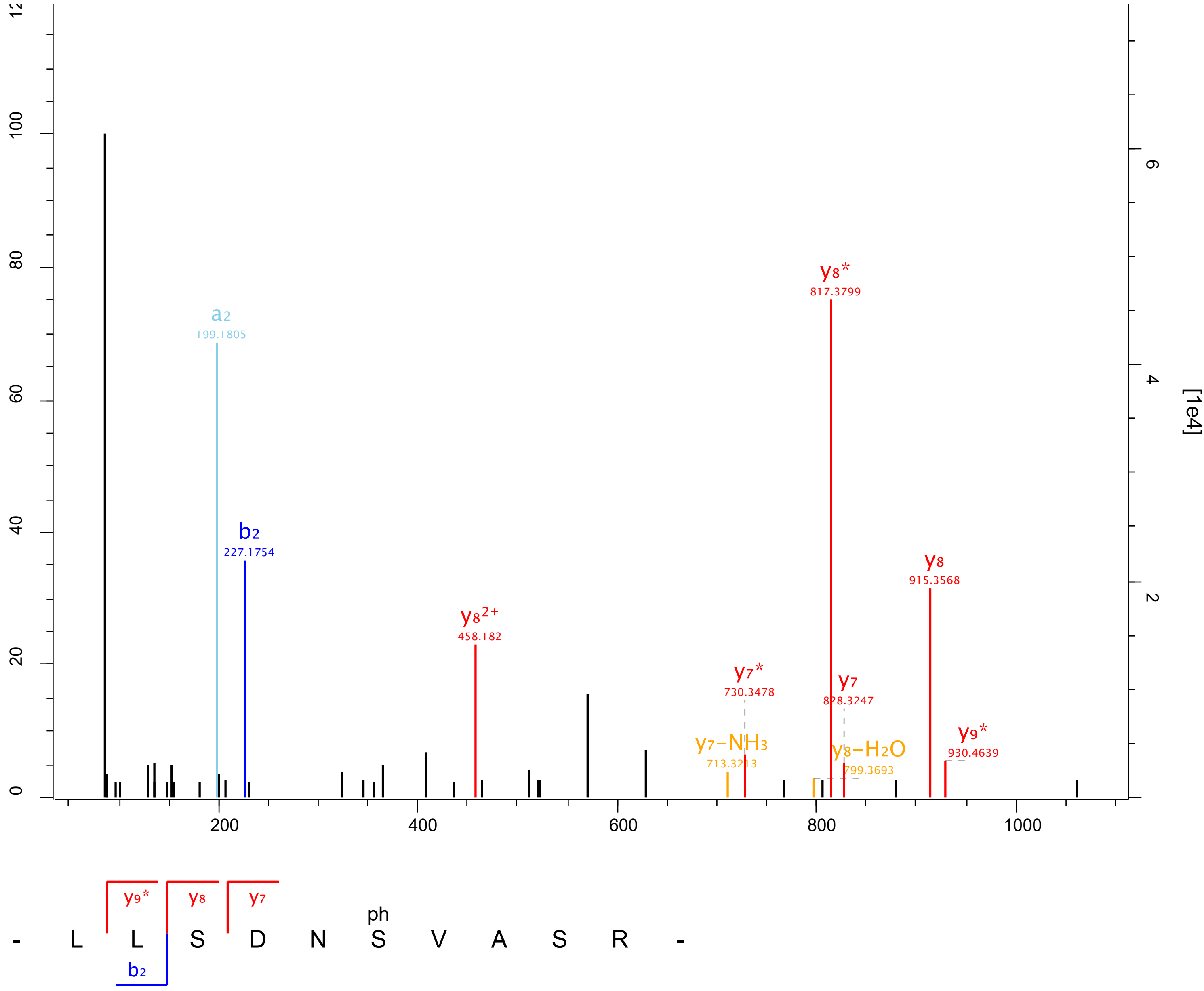

| Raw file | Scan | Method | Score | m/z |
|---|---|---|---|---|
| water_15_2 | 15278 | FTMS; HCD | 62.29 | 530.92 |

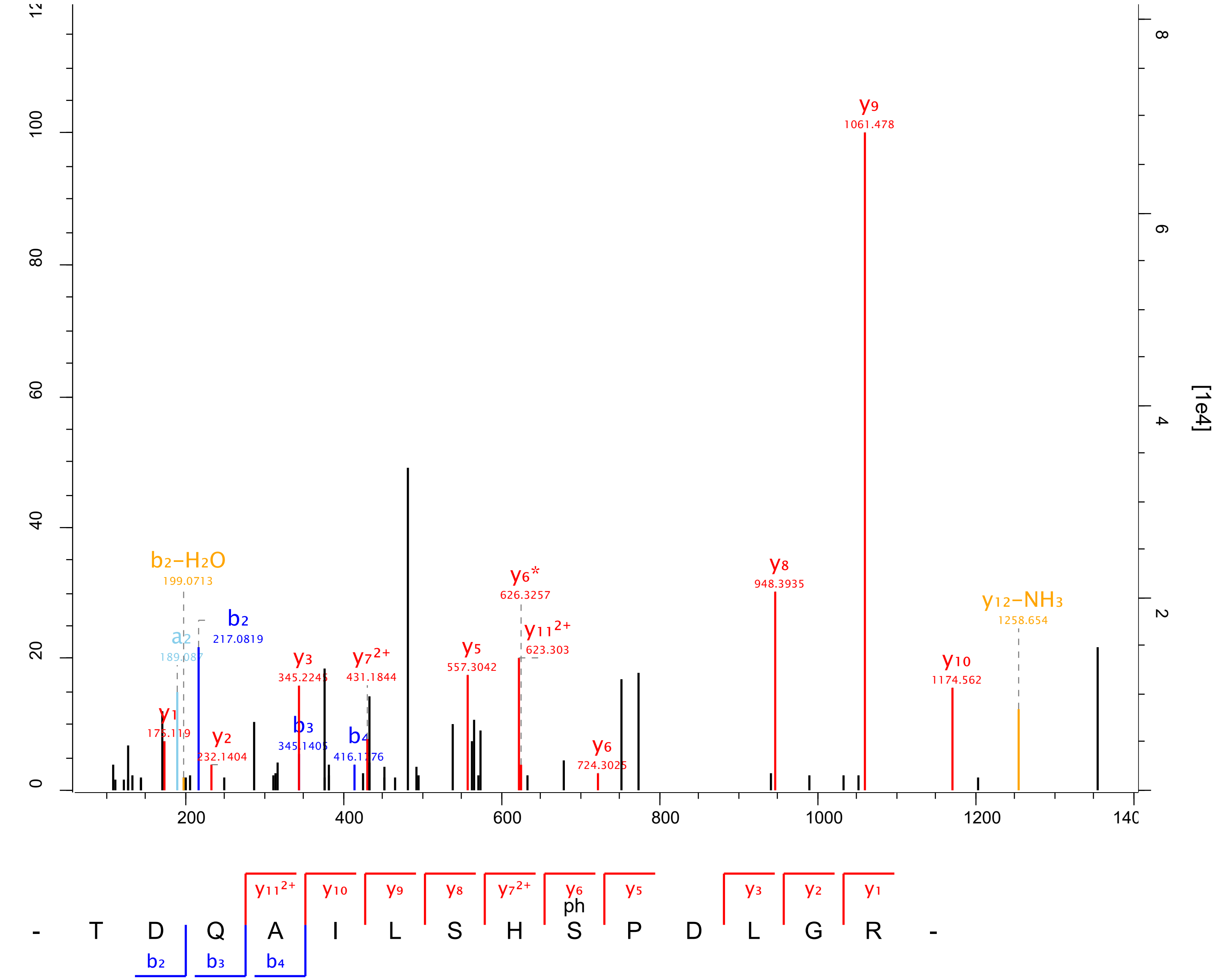

| Raw file | Scan | Method | Score | m/z |
|---|---|---|---|---|
| water_15_2 | 15286 | FTMS; HCD | 141.55 | 686.29 |

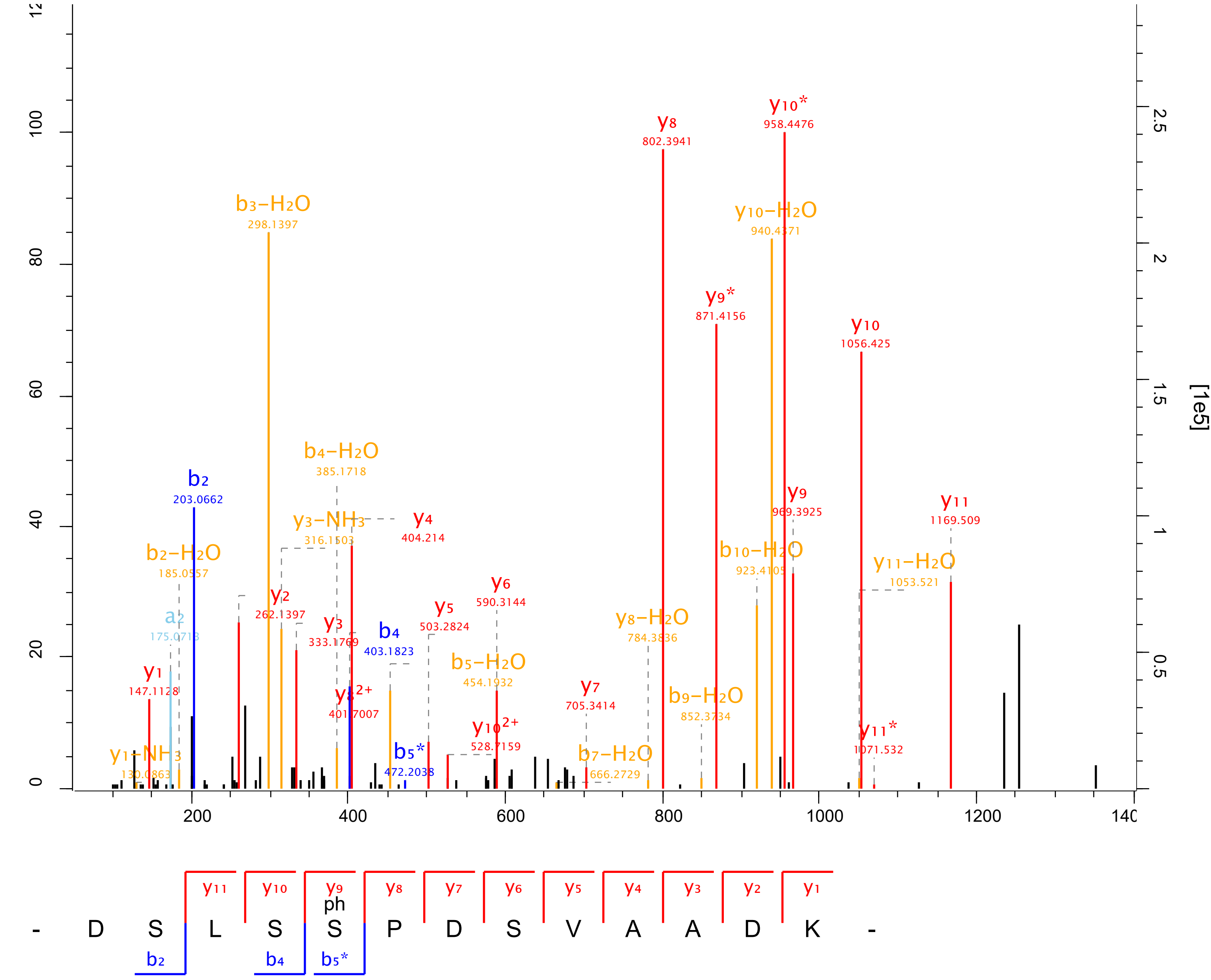

| Raw file | Scan | Method | Score | m/z |
|---|---|---|---|---|
| water_15_2 | 15287 | FTMS; HCD | 167.84 | 841.34 |

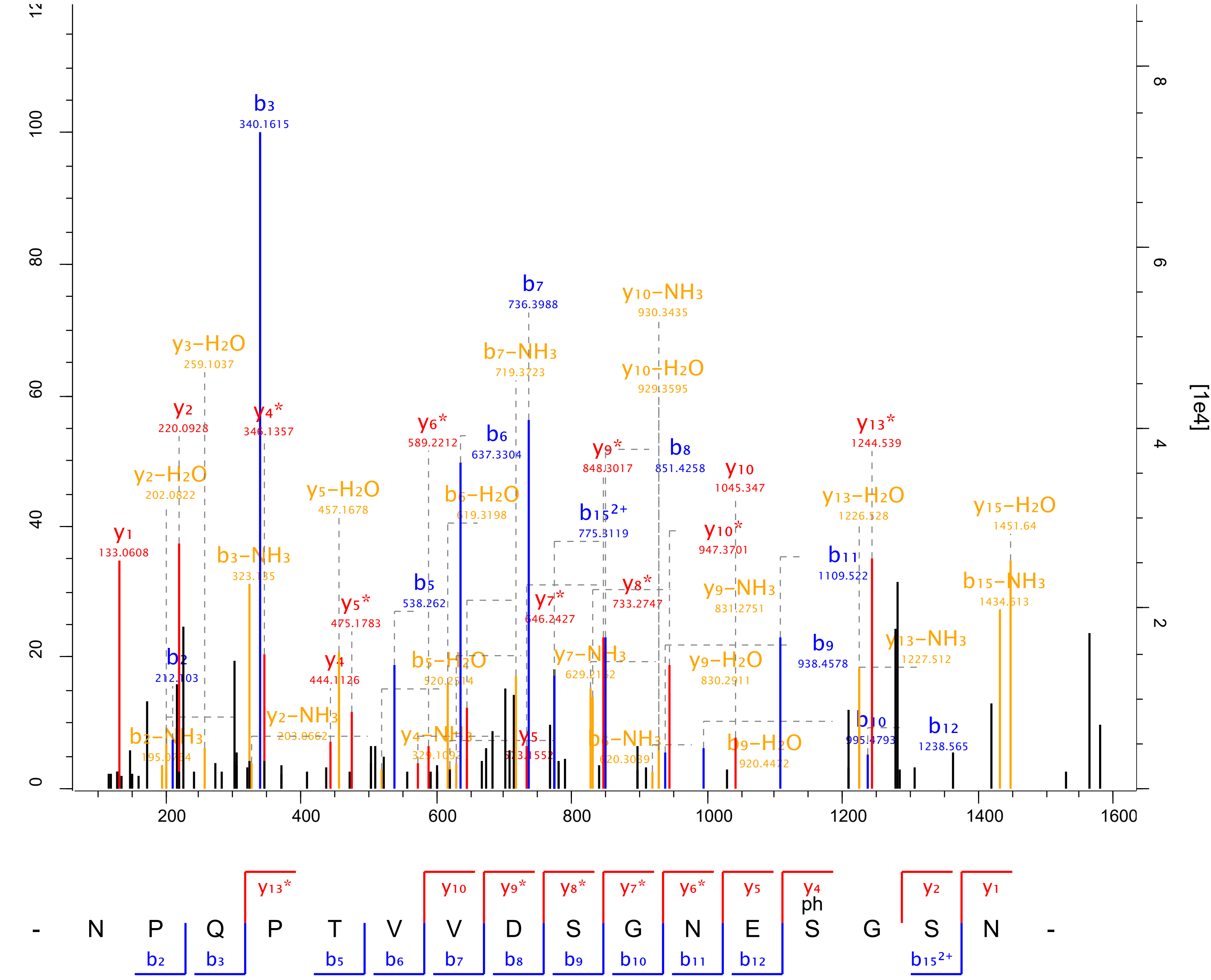

| Raw file | Scan | Method | Score | m/z |
|---|---|---|---|---|
| water_15_2 | 15291 | FTMS; HCD | 176.04 | 729.82 |

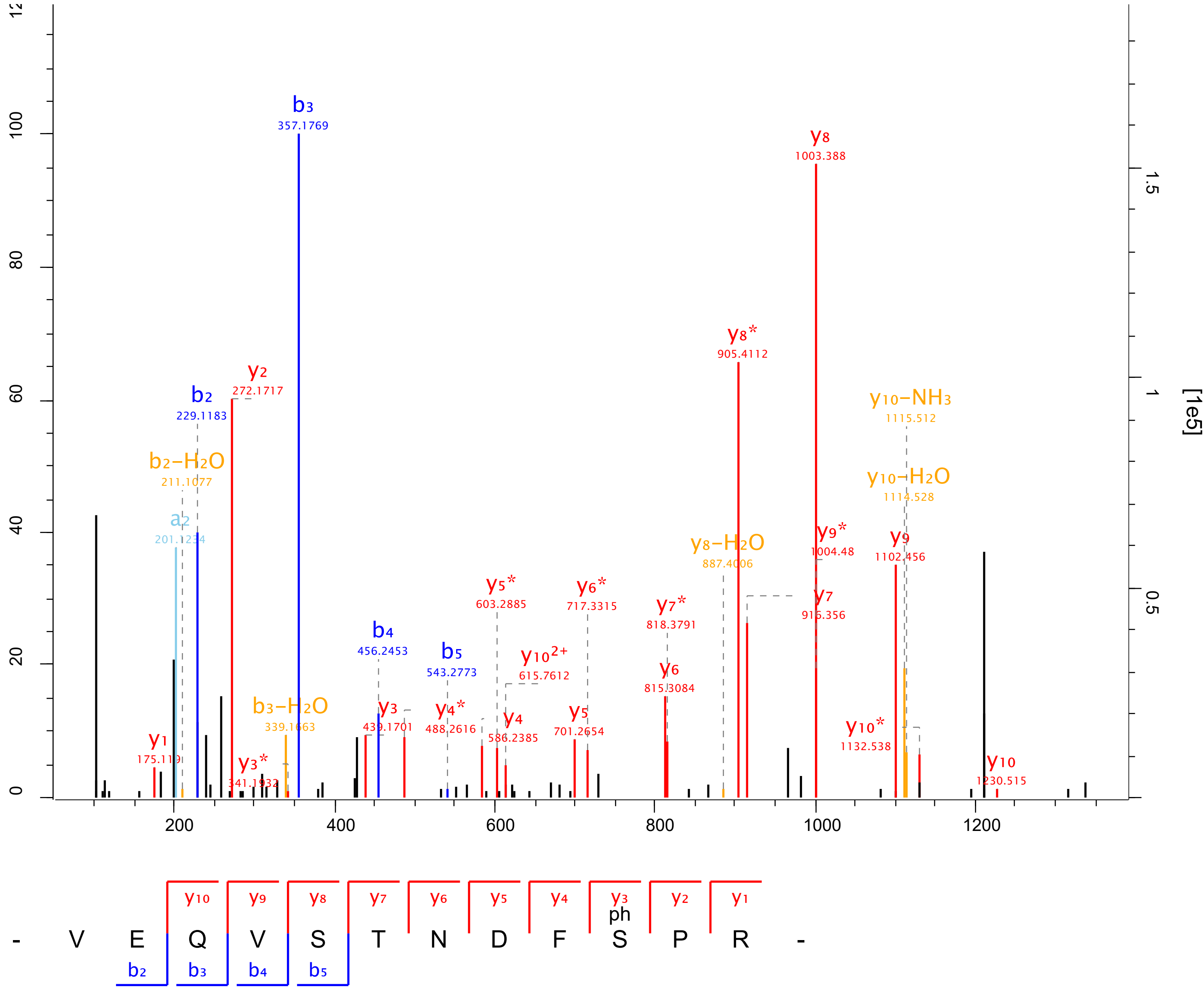

| Raw file | Scan | Method | Score | m/z |
|---|---|---|---|---|
| water_15_2 | 15326 | FTMS; HCD | 53.45 | 421.88 |

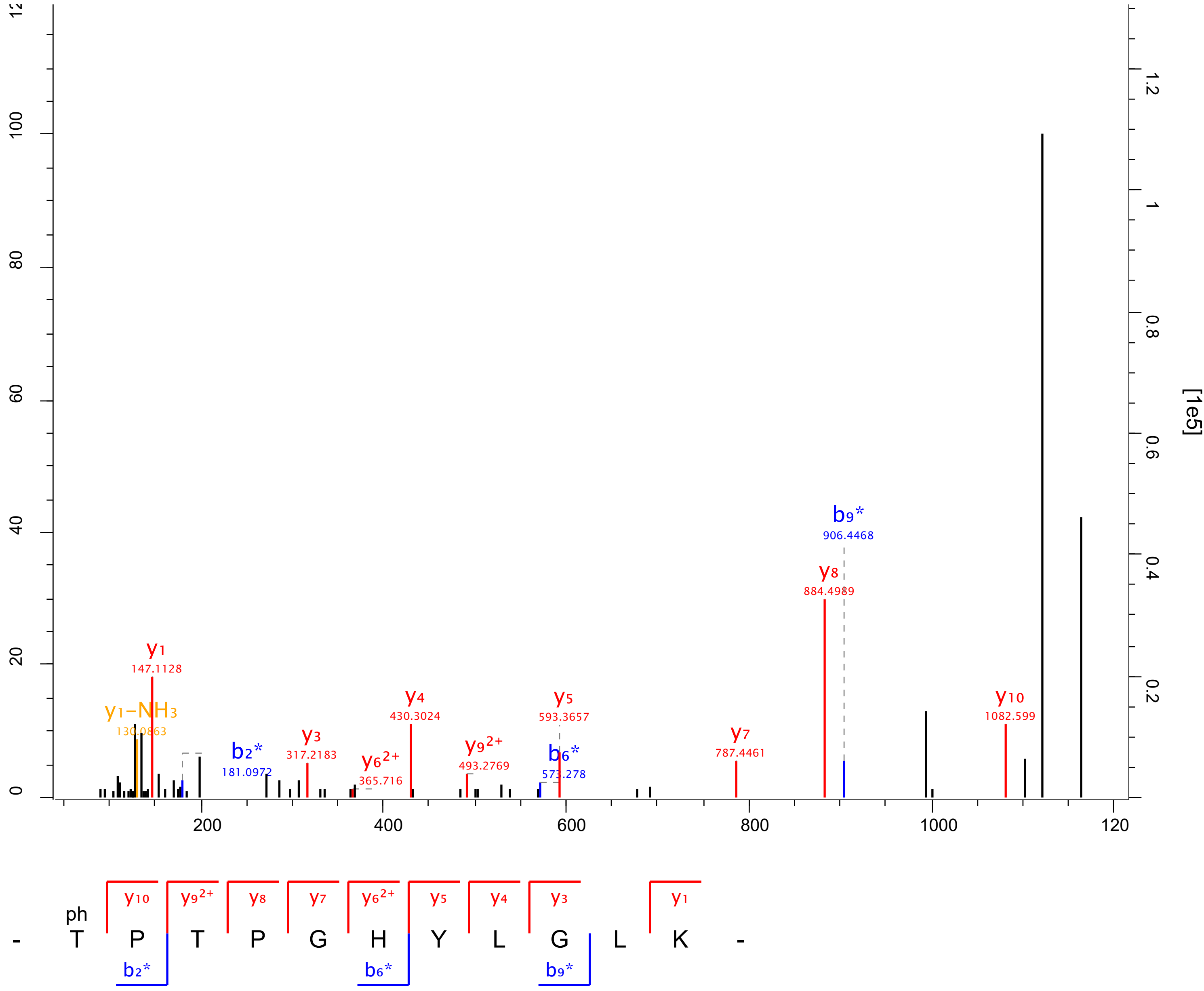

| Raw file | Scan | Method | Score | m/z |
|---|---|---|---|---|
| water_15_2 | 15339 | FTMS; HCD | 103.88 | 762.33 |

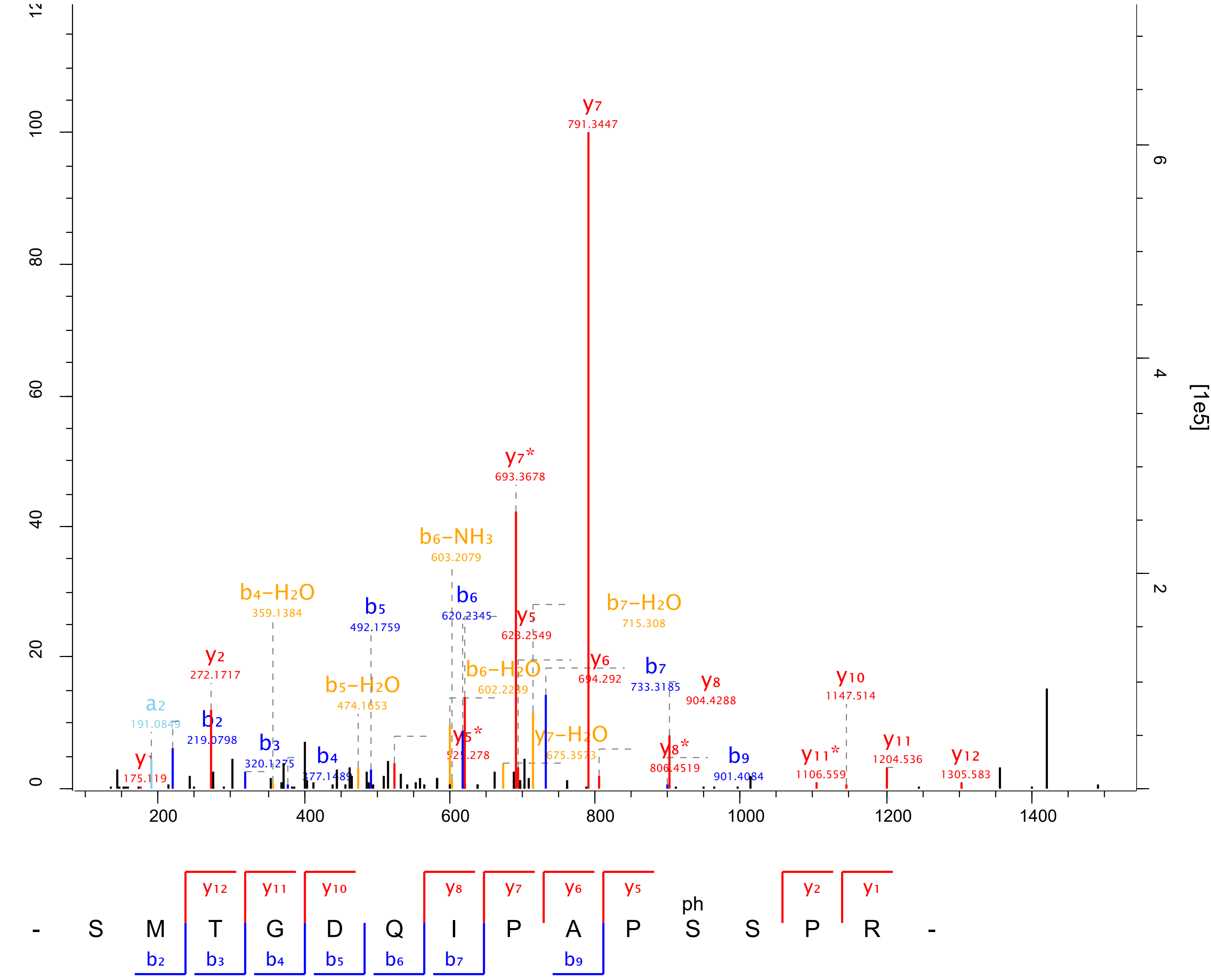

| Raw file | Scan | Method | Score | m/z |
|---|---|---|---|---|
| water_15_2 | 15344 | FTMS; HCD | 110.12 | 644.77 |

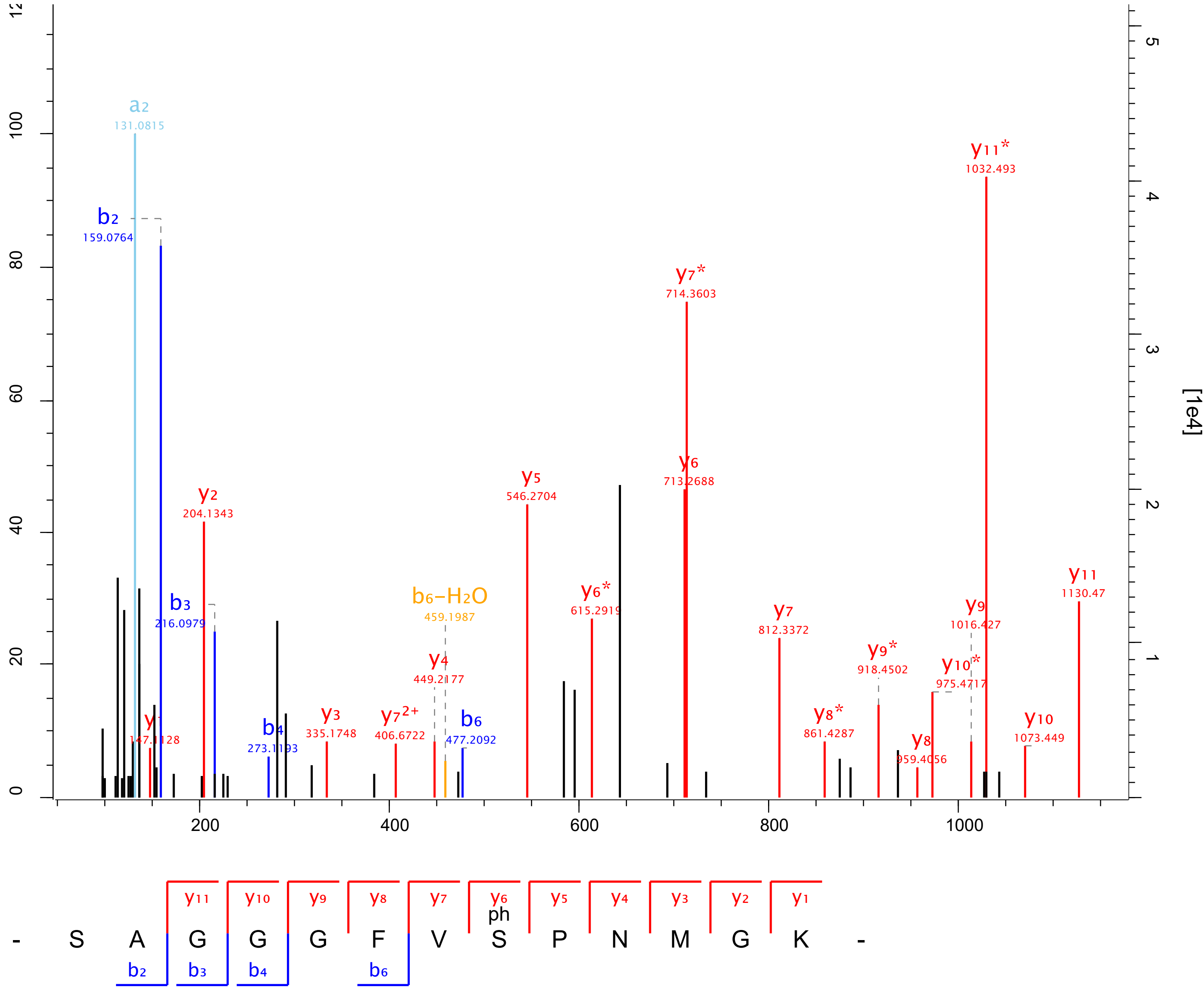

| Raw file | Scan | Method | Score | m/z |
| --- | --- | --- | --- | --- |
| water_15_2 | 15346 | FTMS; HCD | 59.2 | 514.24 |

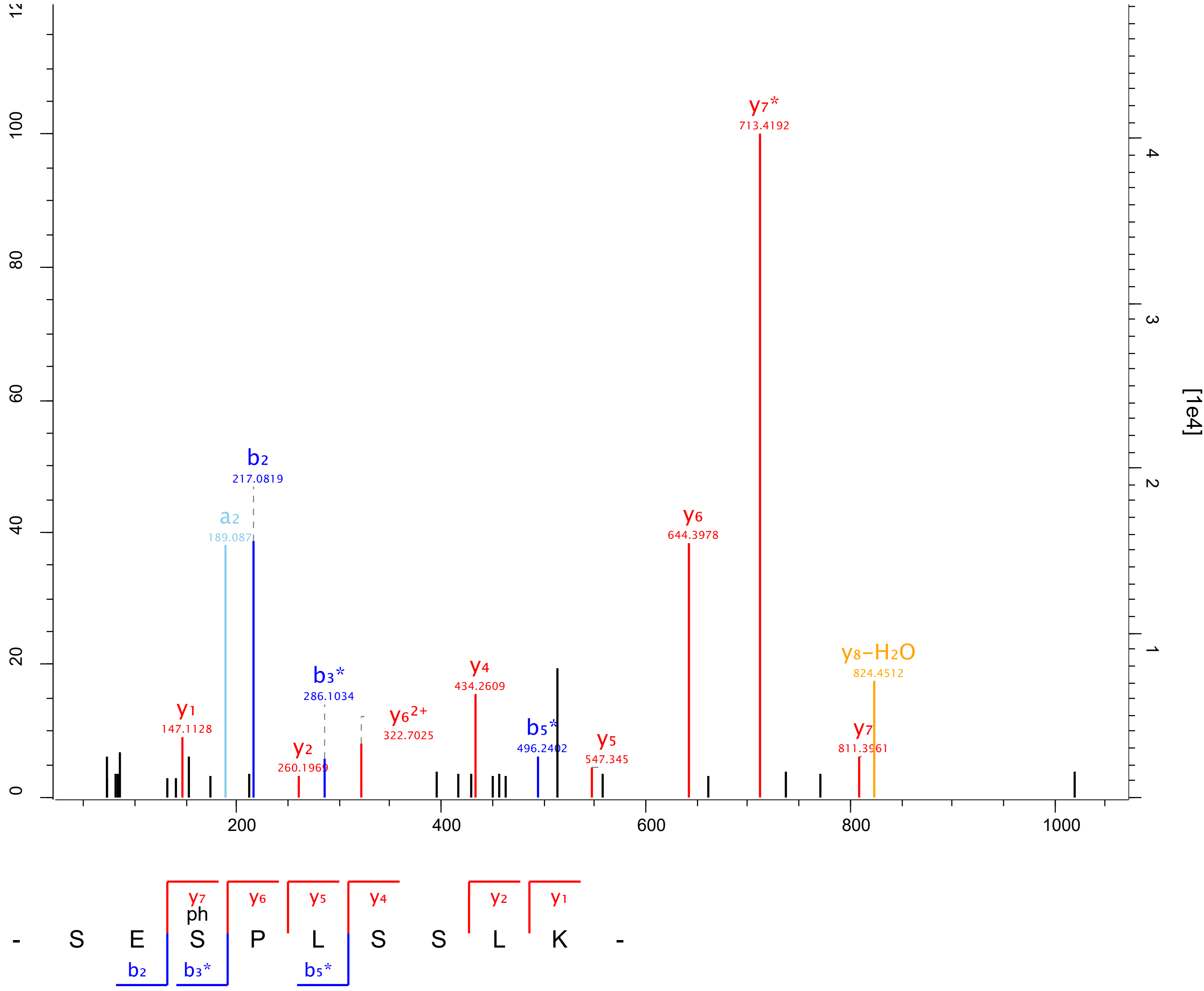

| Raw file | Scan | Method | Score | m/z |
|---|---|---|---|---|
| water_15_2 | 15348 | FTMS; HCD | 96.76 | 568.75 |

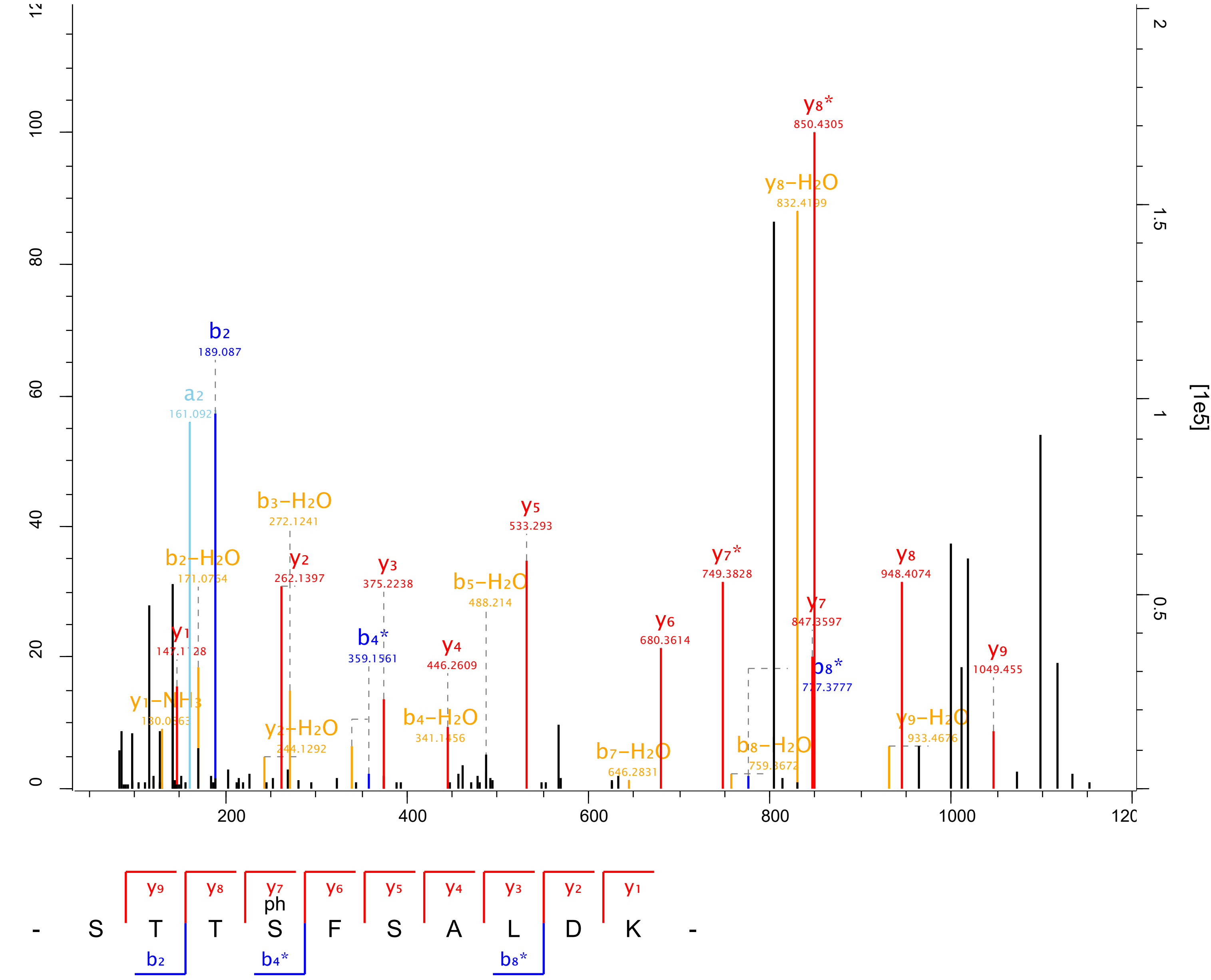

| Raw file | Scan | Method | Score | m/z |
| --- | --- | --- | --- | --- |
| water_15_2 | 15375 | FTMS; HCD | 101.42 | 727.82 |

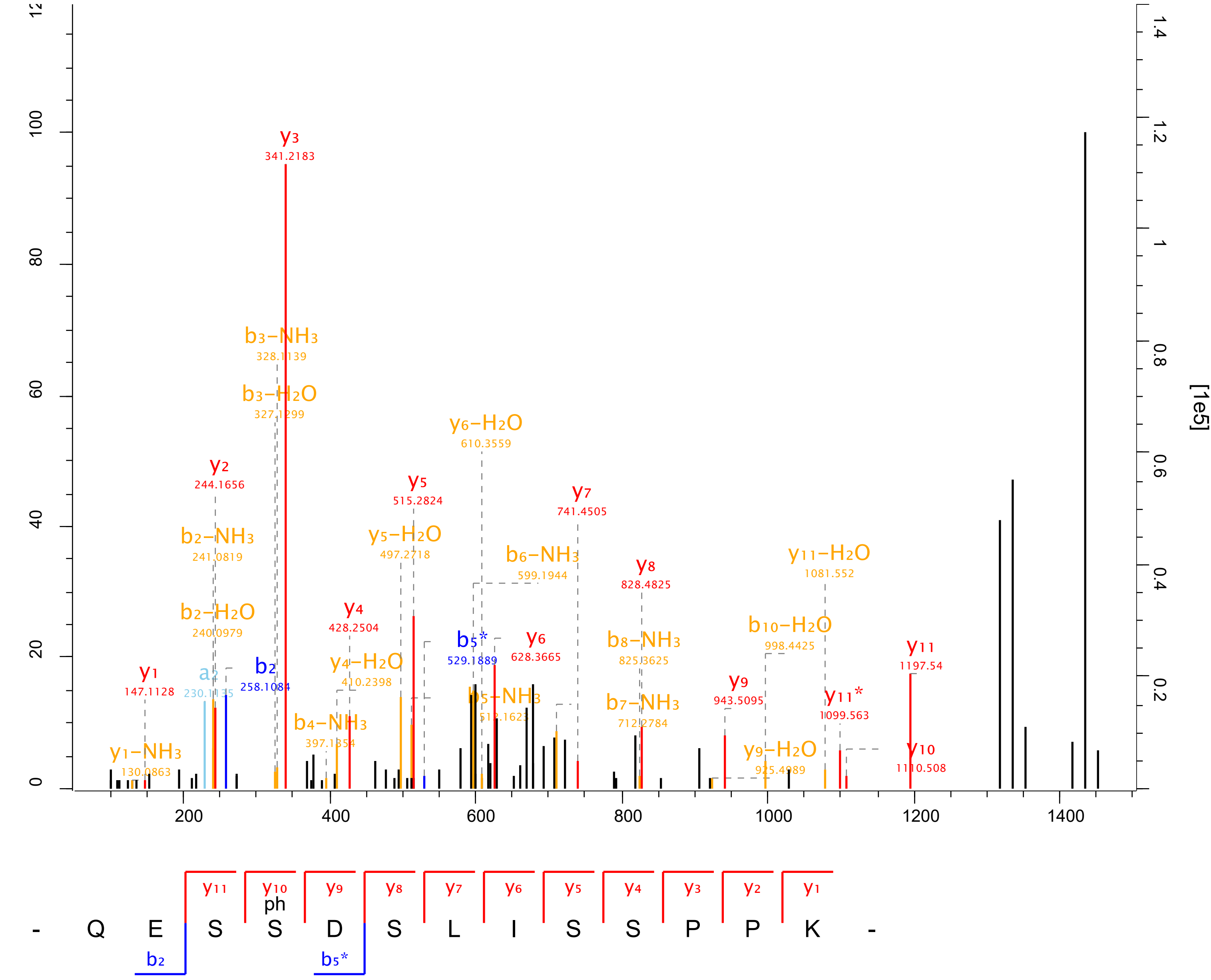

| Raw file | Scan | Method | Score | m/z |
|---|---|---|---|---|
| water_15_2 | 15440 | FTMS; HCD | 72.68 | 540.76 |

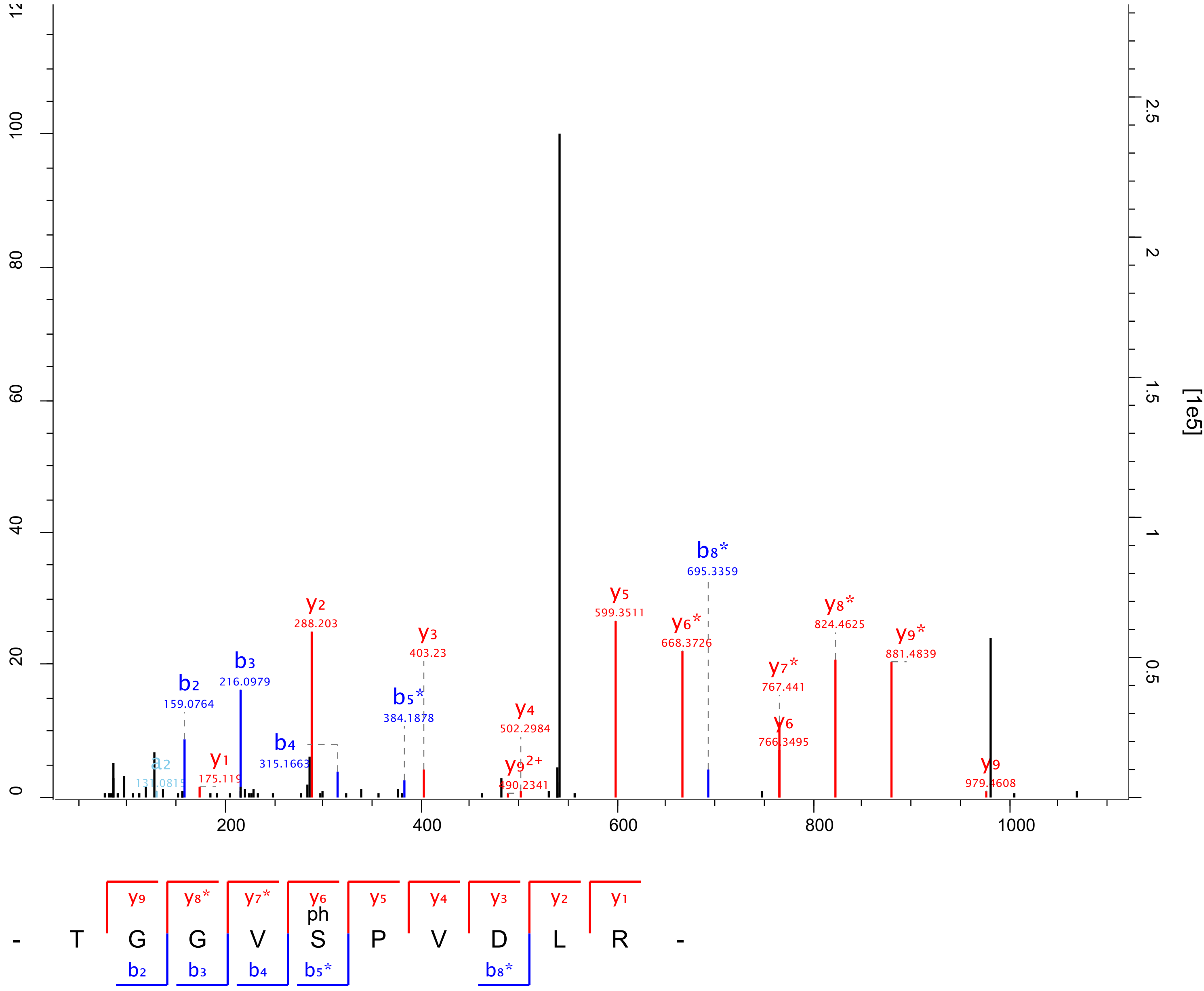

| Raw file | Scan | Method | Score | m/z |
|---|---|---|---|---|
| water_15_2 | 15509 | FTMS; HCD | 155.56 | 692.81 |

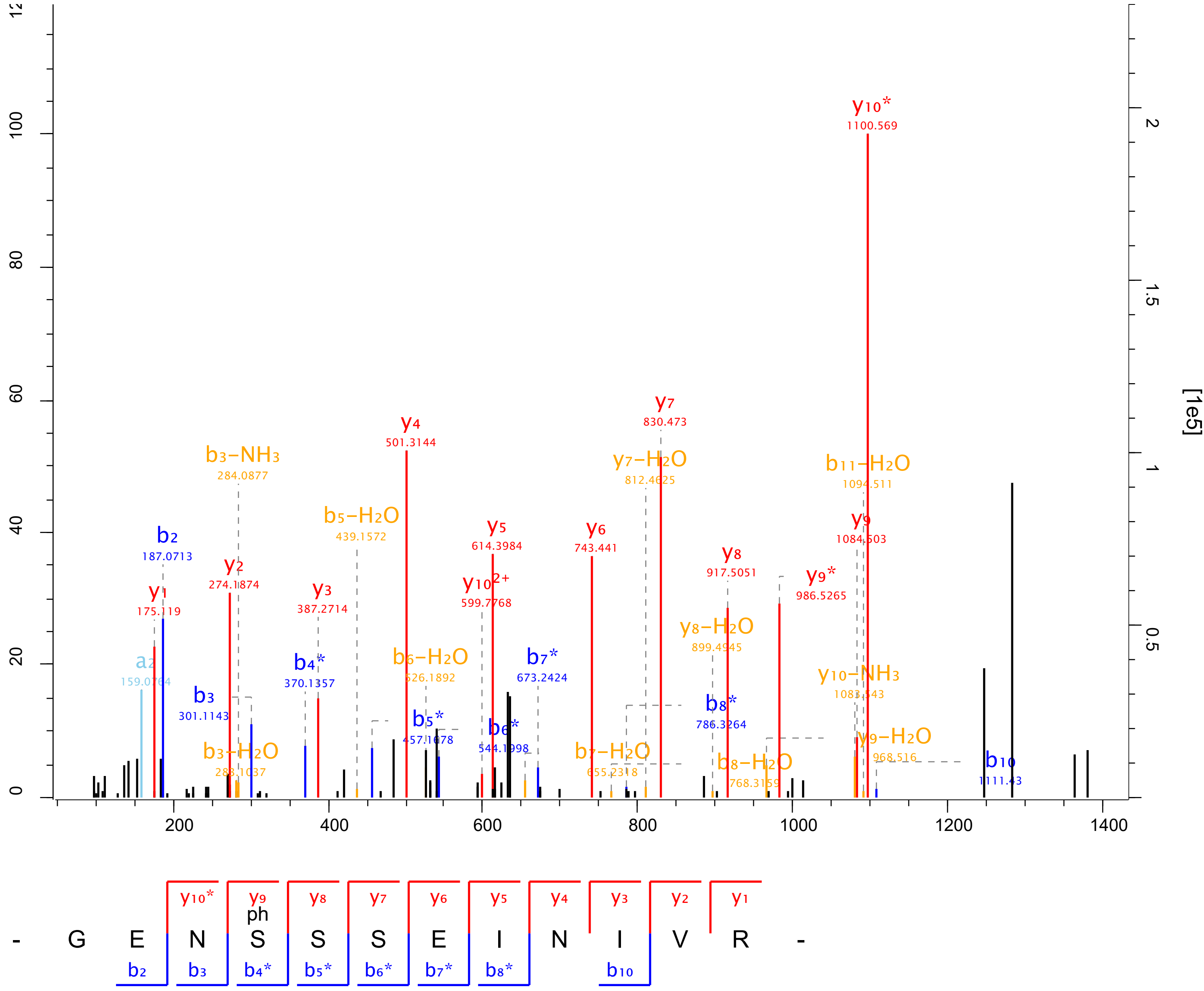

| Raw file | Scan | Method | Score | m/z |
|---|---|---|---|---|
| water_15_2 | 1560 | FTMS; HCD | 41.62 | 430.88 |

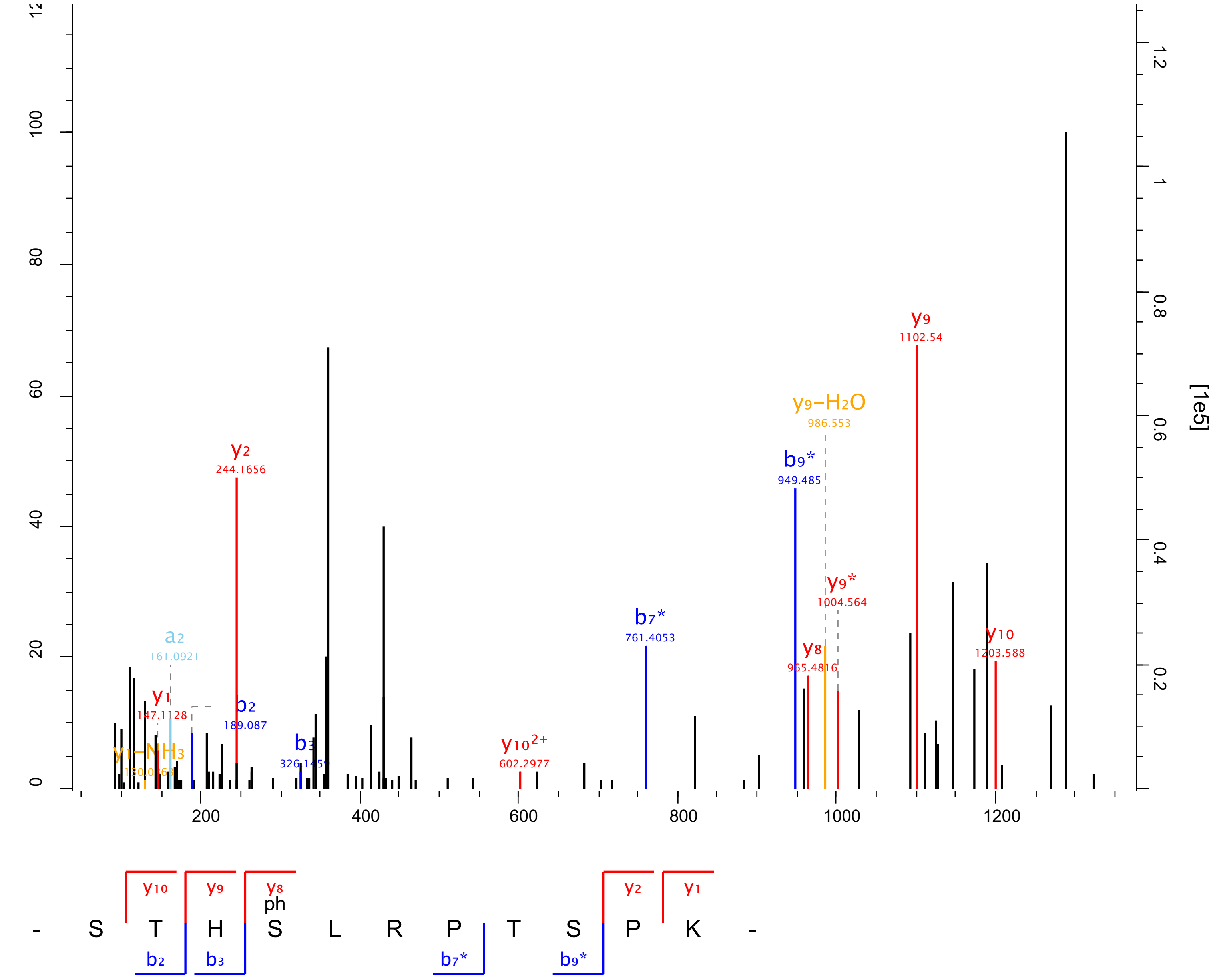

| Raw file | Scan | Method | Score | m/z |
|---|---|---|---|---|
| water_15_2 | 15617 | FTMS; HCD | 88.5 | 598.77 |

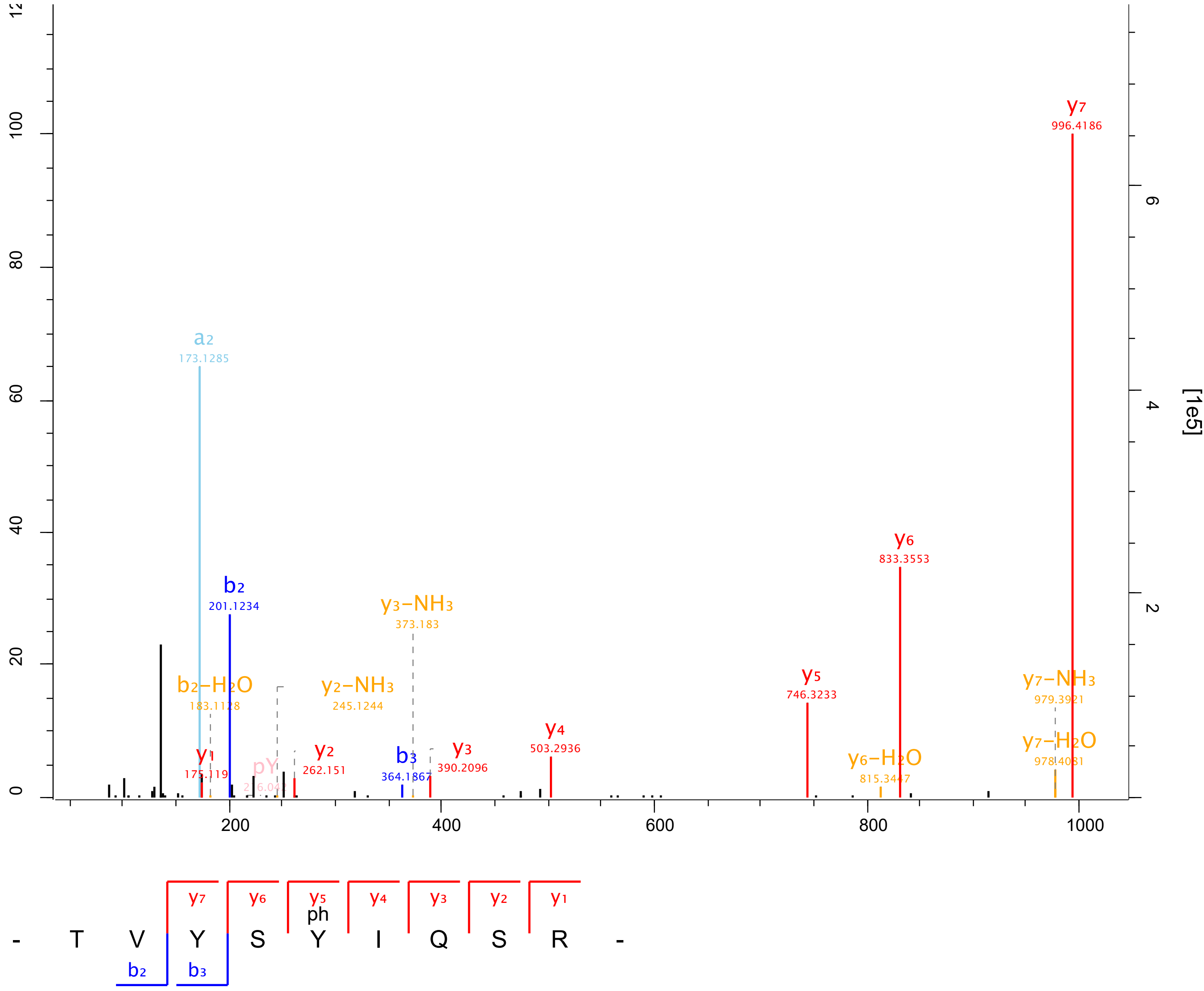

| Raw file | Scan | Method | Score | m/z |
|---|---|---|---|---|
| water_15_2 | 15677 | FTMS; HCD | 133.23 | 727.82 |

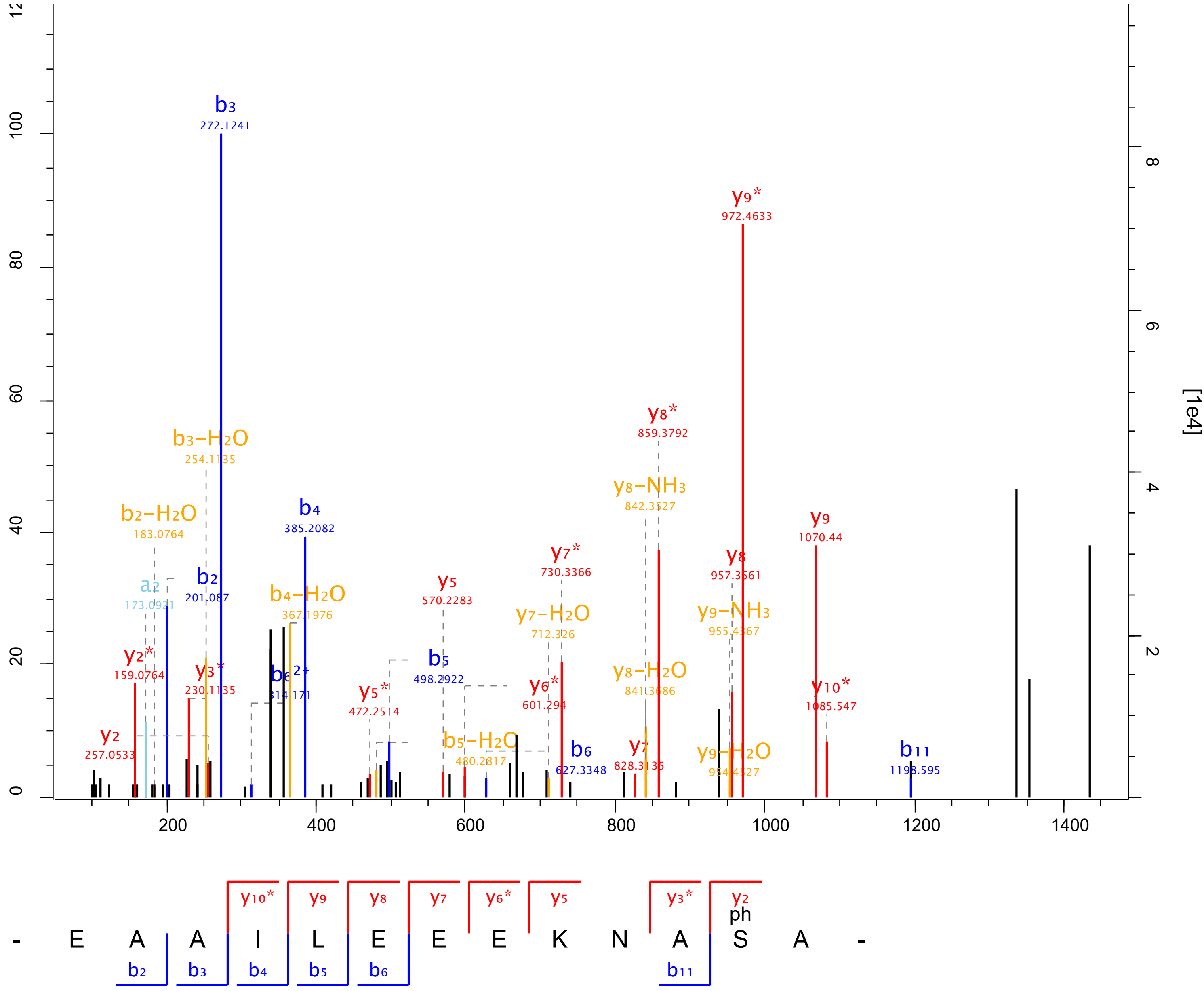

| Raw file | Scan | Method | Score | m/z |
|---|---|---|---|---|
| water_15_2 | 15802 | FTMS; HCD | 65.37 | 672.79 |

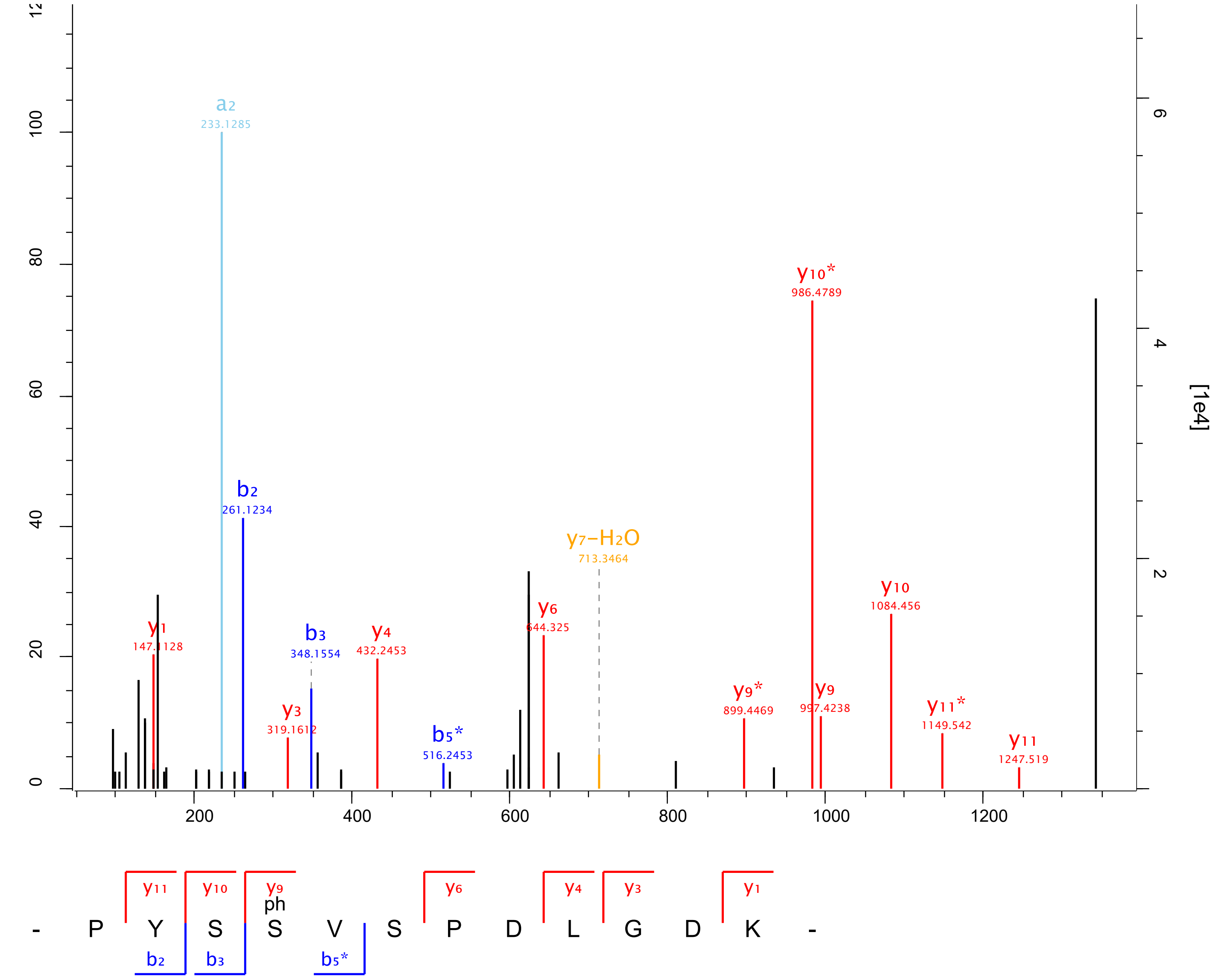

| Raw file | Scan | Method | Score | m/z |
|---|---|---|---|---|
| water_15_2 | 15814 | FTMS; HCD | 76.34 | 759.83 |

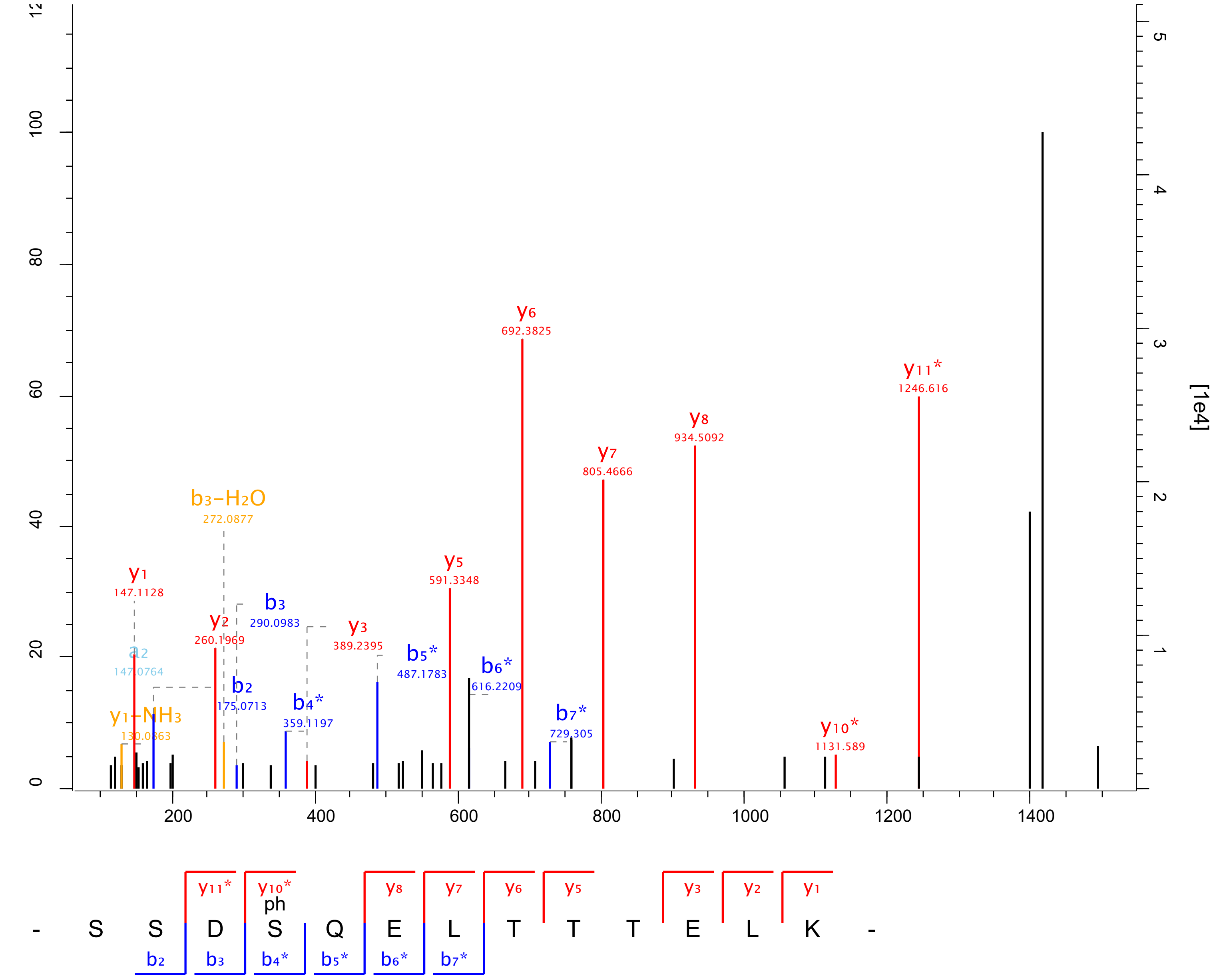

| Raw file | Scan | Method | Score | m/z |
|---|---|---|---|---|
| water_15_2 | 1585 | FTMS; HCD | 65.04 | 415.86 |

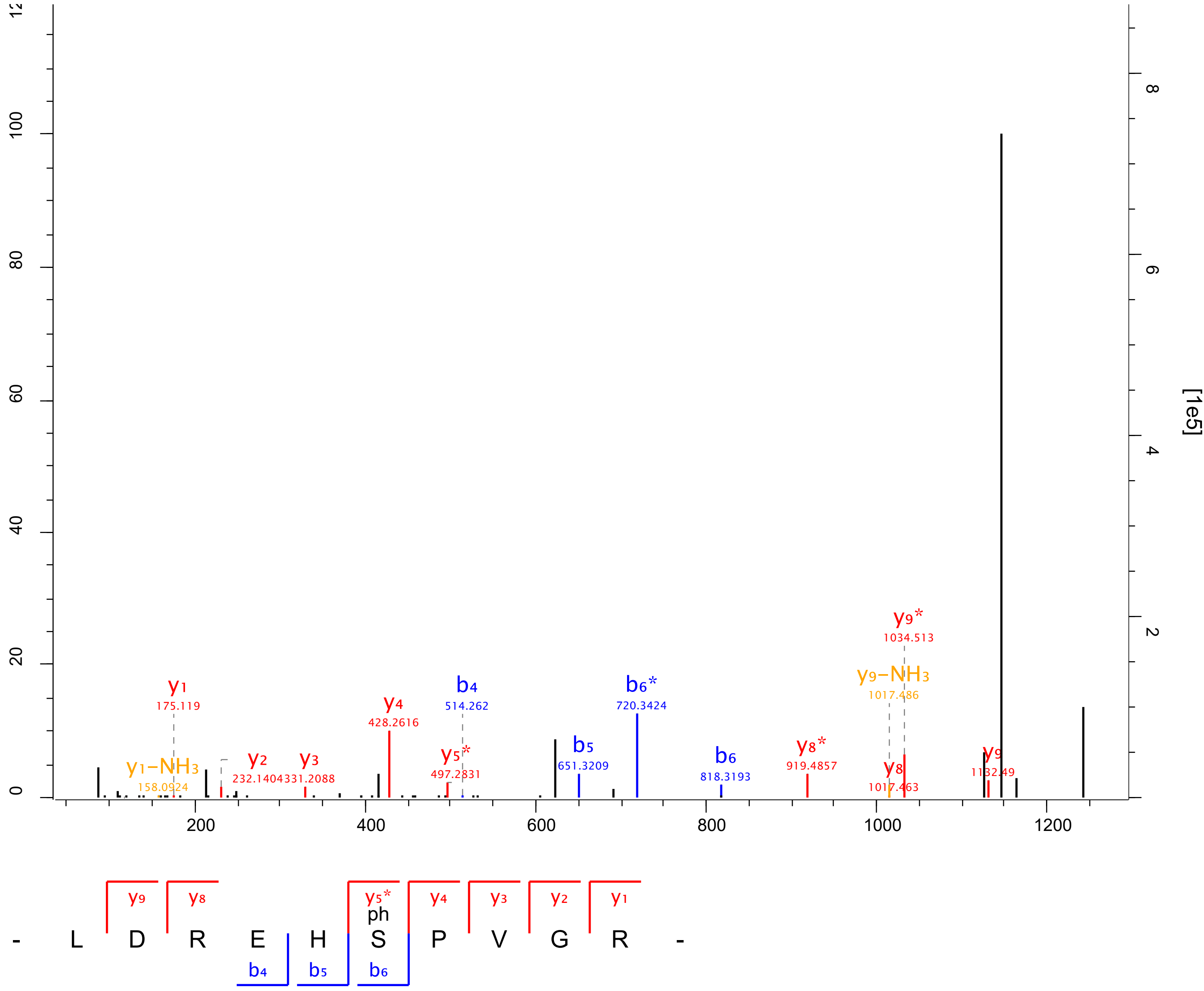

| Raw file | Scan | Method | Score | m/z |
| --- | --- | --- | --- | --- |
| water_15_2 | 15866 | FTMS; HCD | 54.26 | 482.73 |

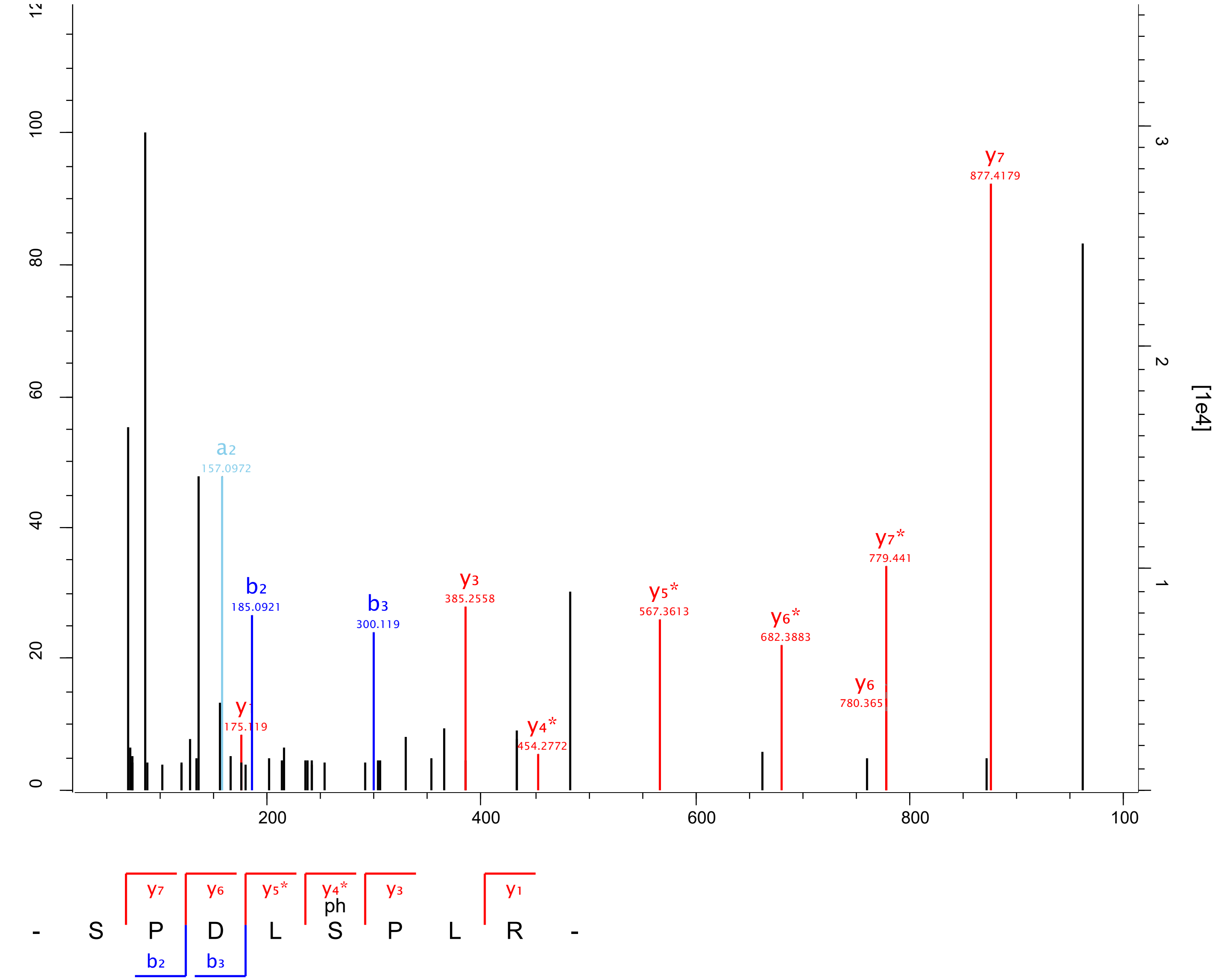

| Raw file | Scan | Method | Score | m/z |
| --- | --- | --- | --- | --- |
| water_15_2 | 15927 | FTMS; HCD | 91.81 | 642.8 |

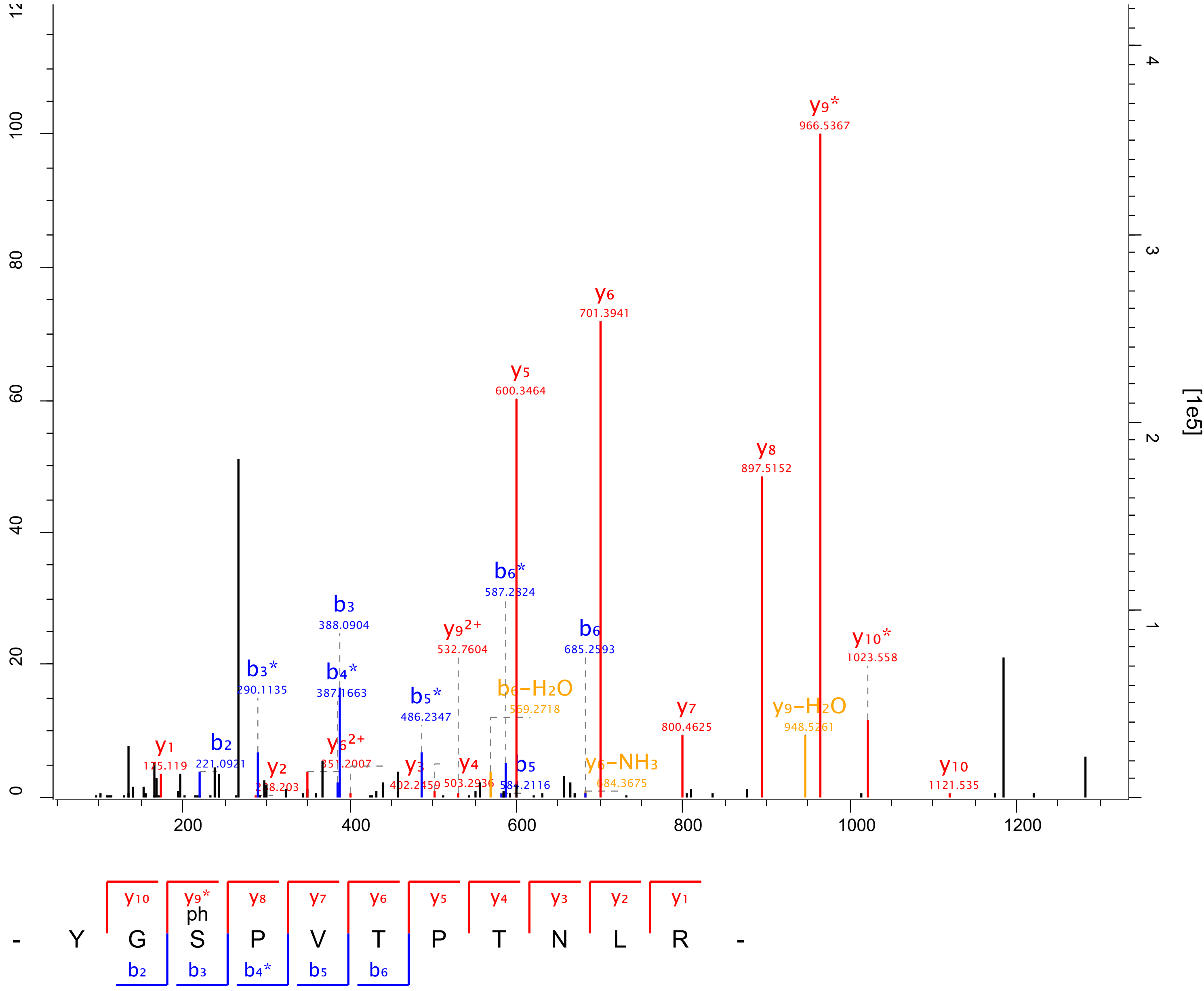

| Raw file | Scan | Method | Score | m/z |
| --- | --- | --- | --- | --- |
| water_15_2 | 15976 | FTMS; HCD | 85.06 | 979.91 |

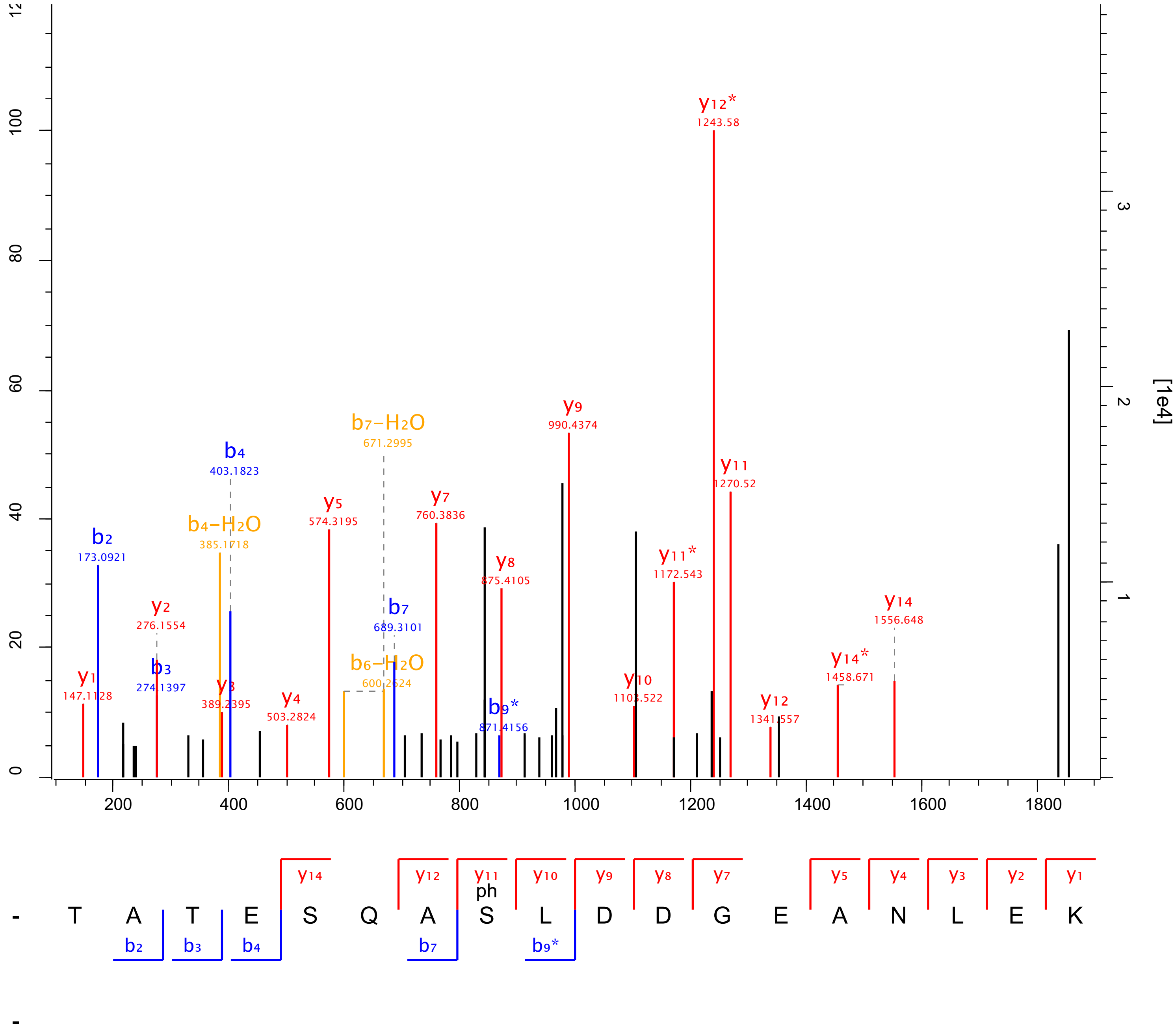

| Raw file | Scan | Method | Score | m/z |
| --- | --- | --- | --- | --- |
| water_15_2 | 16011 | FTMS; HCD | 64.71 | 437.2 |

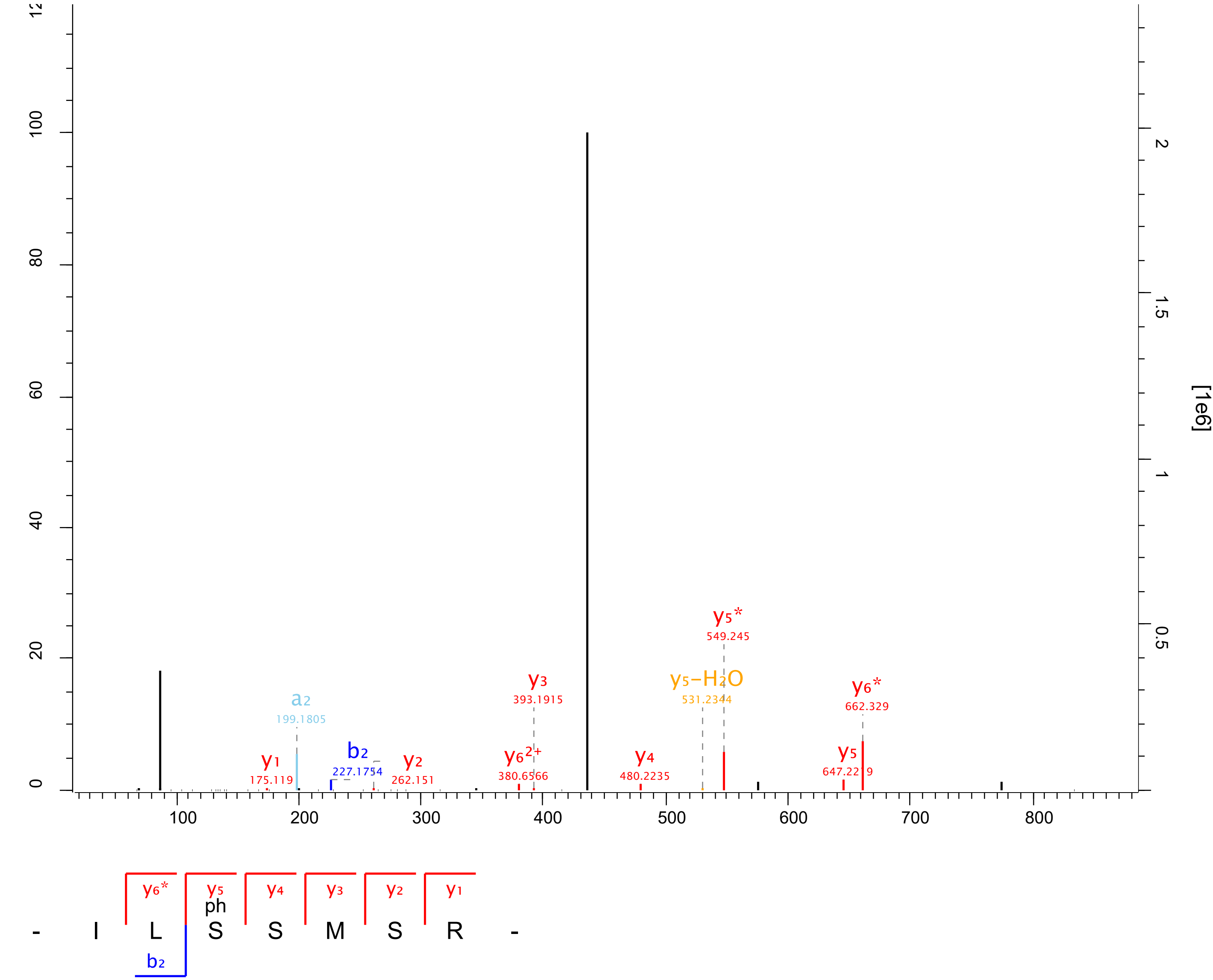

| Raw file | Scan | Method | Score | m/z |
|---|---|---|---|---|
| water_15_2 | 16106 | FTMS; HCD | 67.23 | 635.26 |

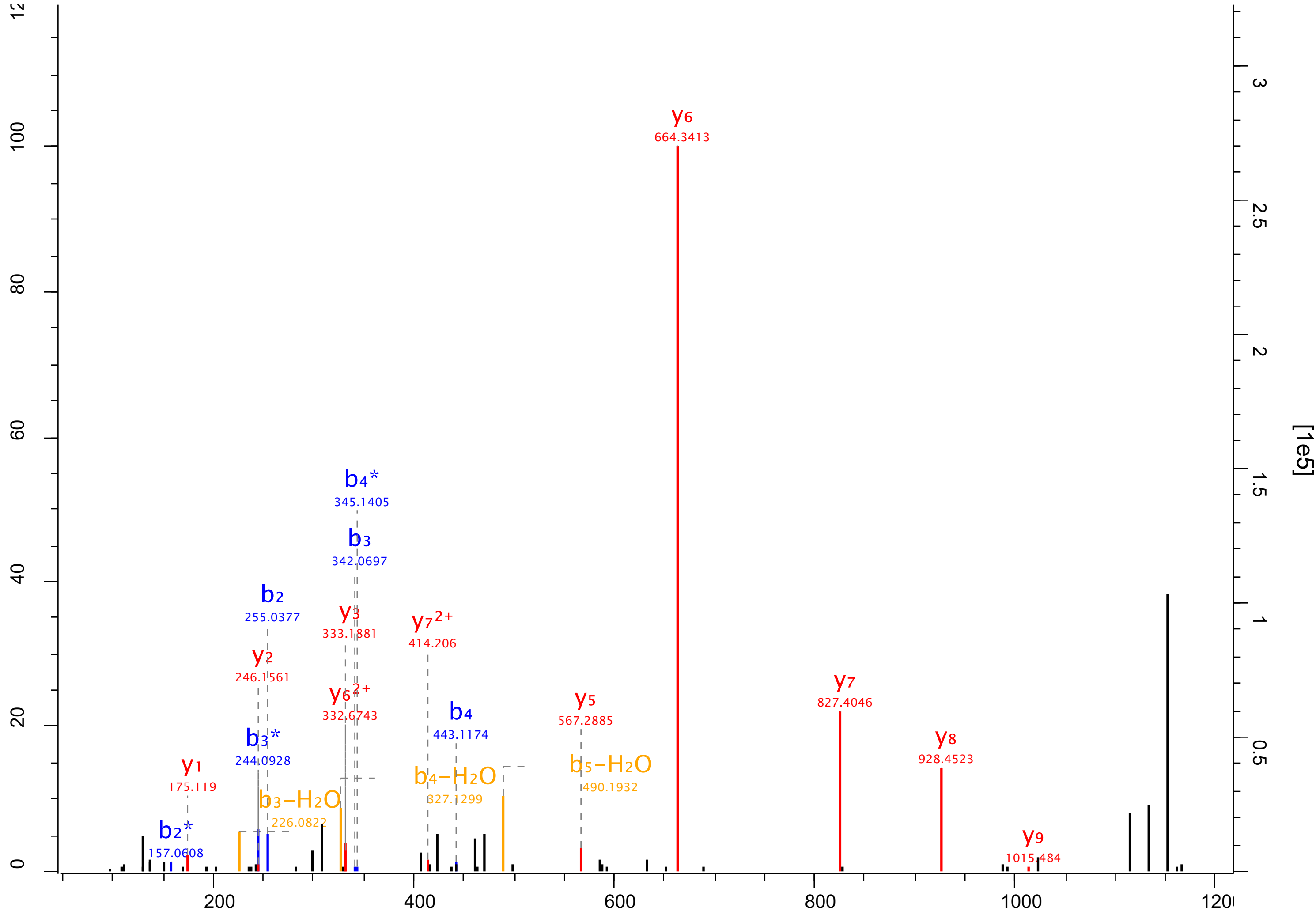


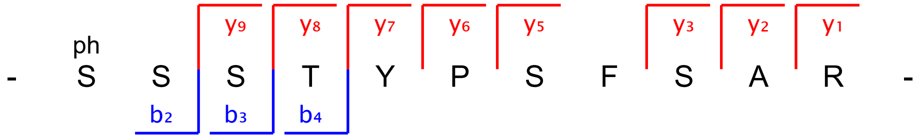

| Raw file | Scan | Method | Score | m/z |
|---|---|---|---|---|
| water_15_2 | 16181 | FTMS; HCD | 160.57 | 913.9 |

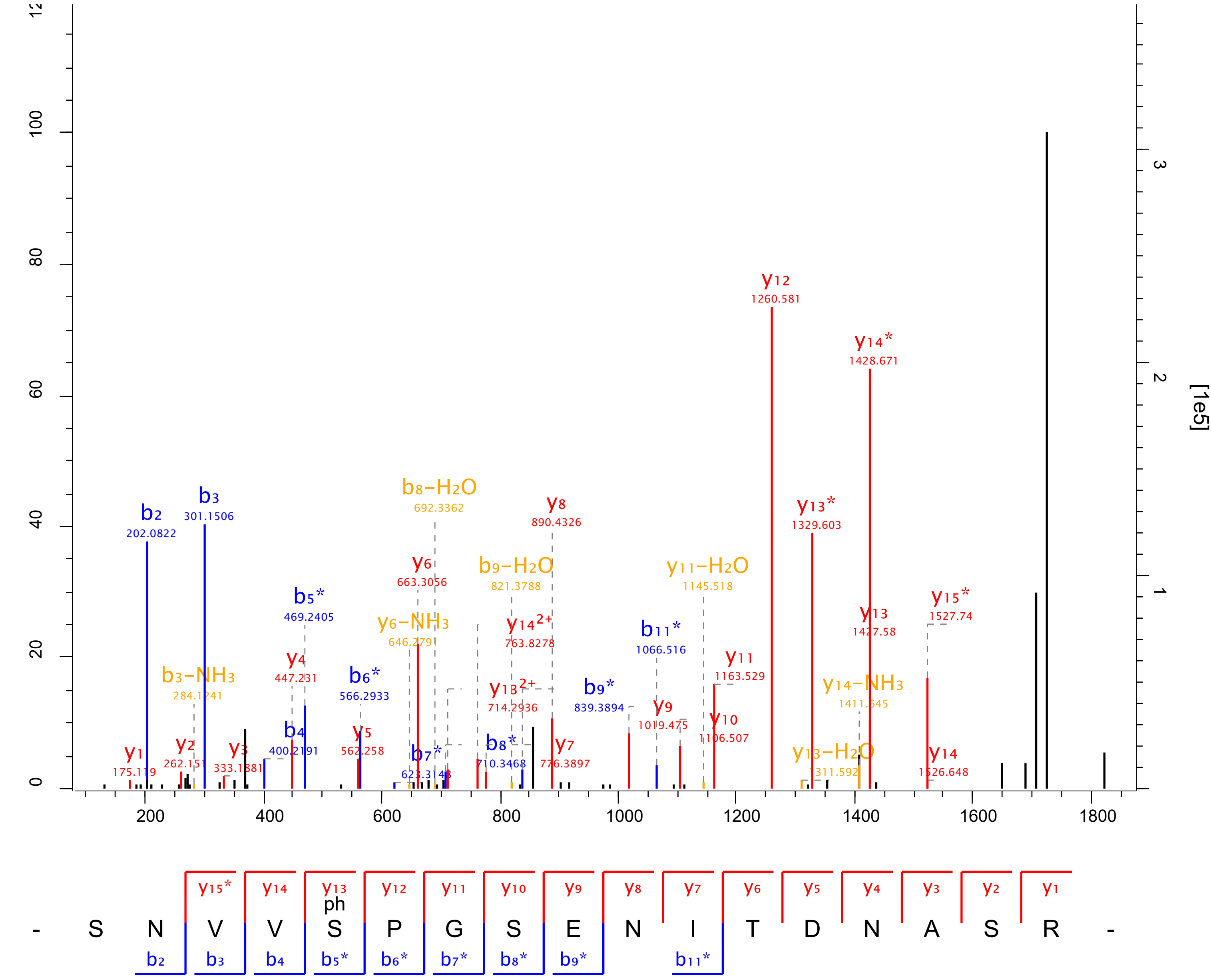

| Raw file | Scan | Method | Score | m/z |
| --- | --- | --- | --- | --- |
| water_15_2 | 16224 | FTMS; HCD | 142.97 | 576.26 |

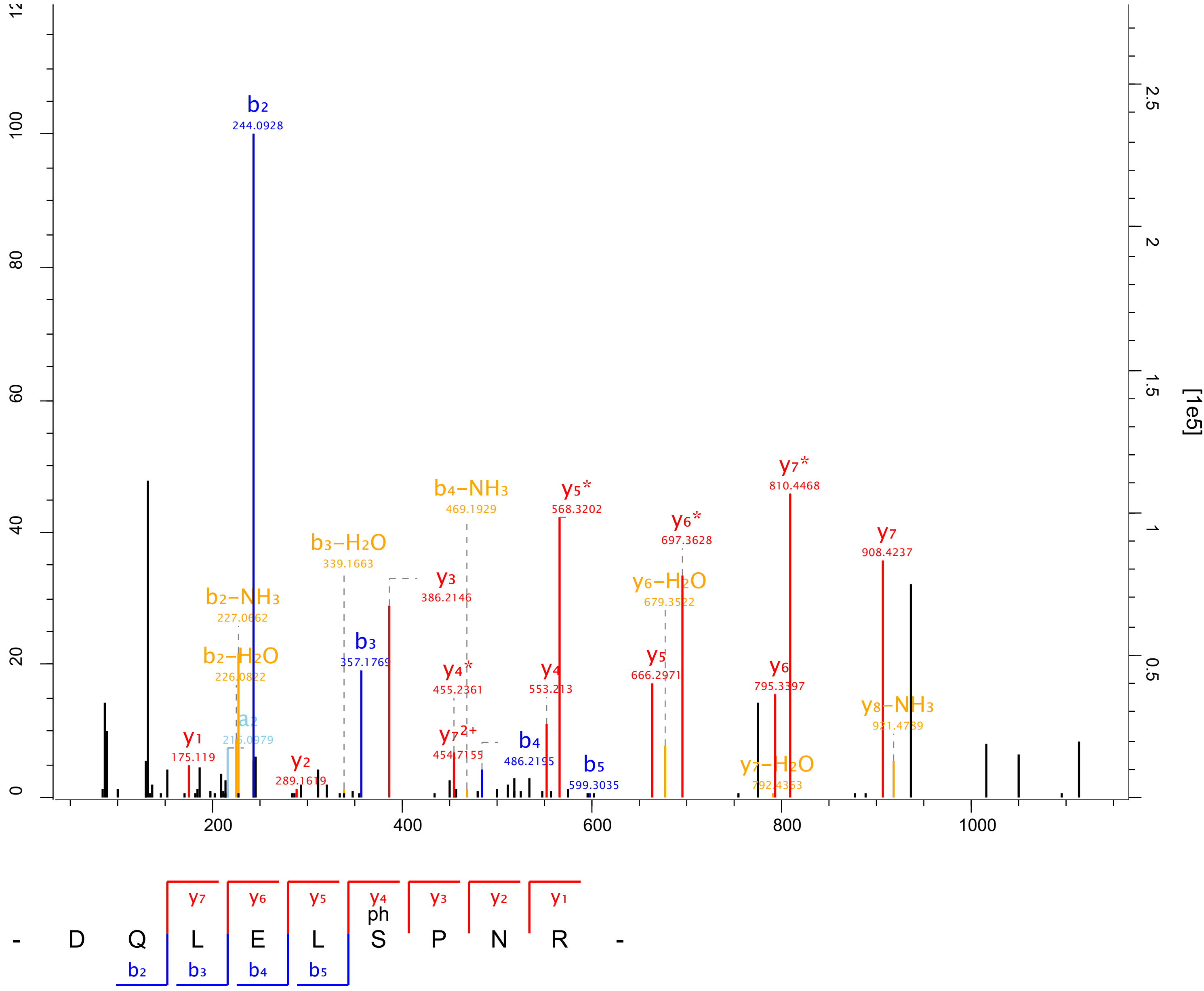

| Raw file | Scan | Method | Score | m/z |
| --- | --- | --- | --- | --- |
| water_15_2 | 16234 | FTMS; HCD | 98.94 | 620.77 |

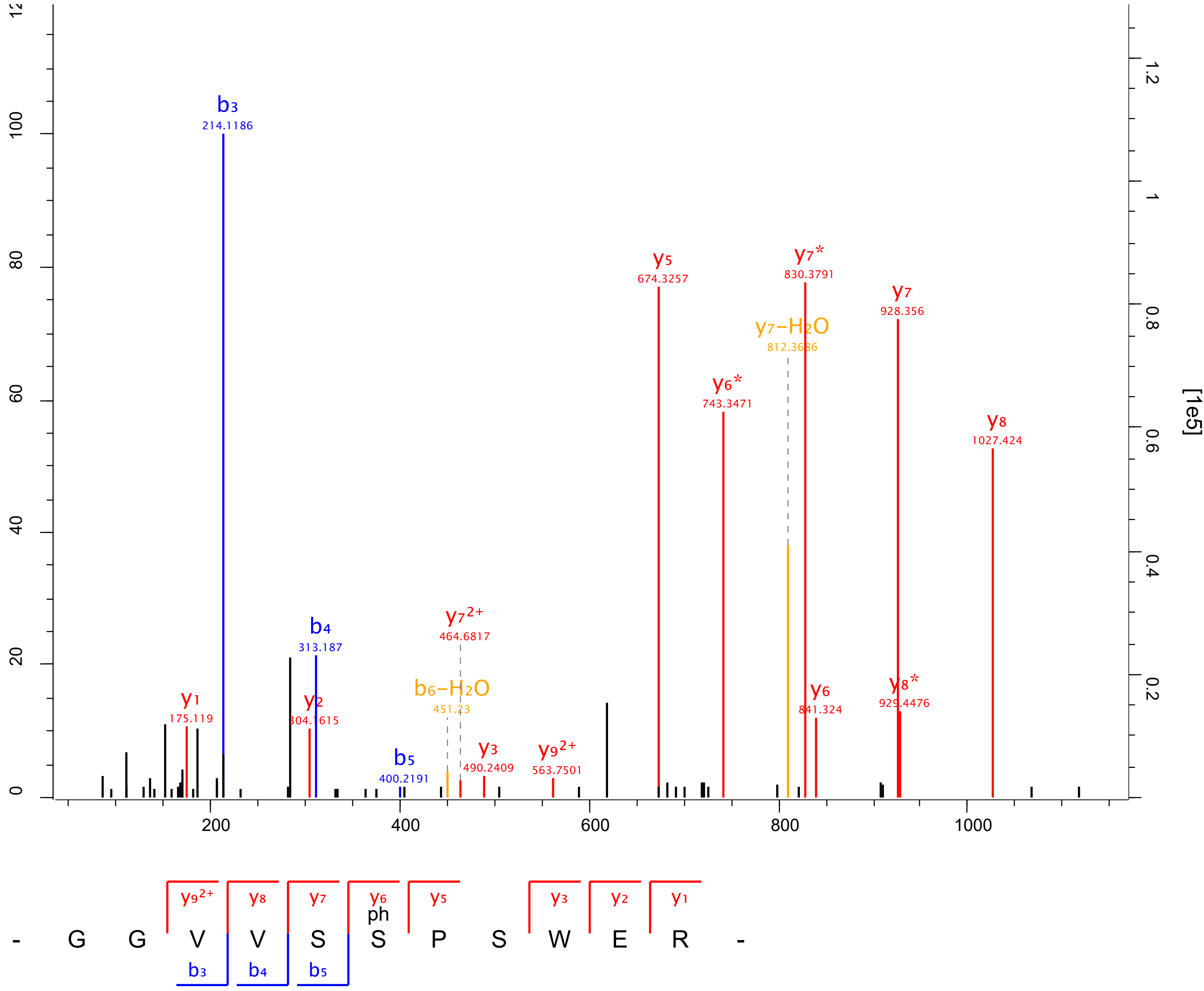

| Raw file | Scan | Method | Score | m/z |
| --- | --- | --- | --- | --- |
| water_15_2 | 16281 | FTMS; HCD | 97.27 | 613.29 |

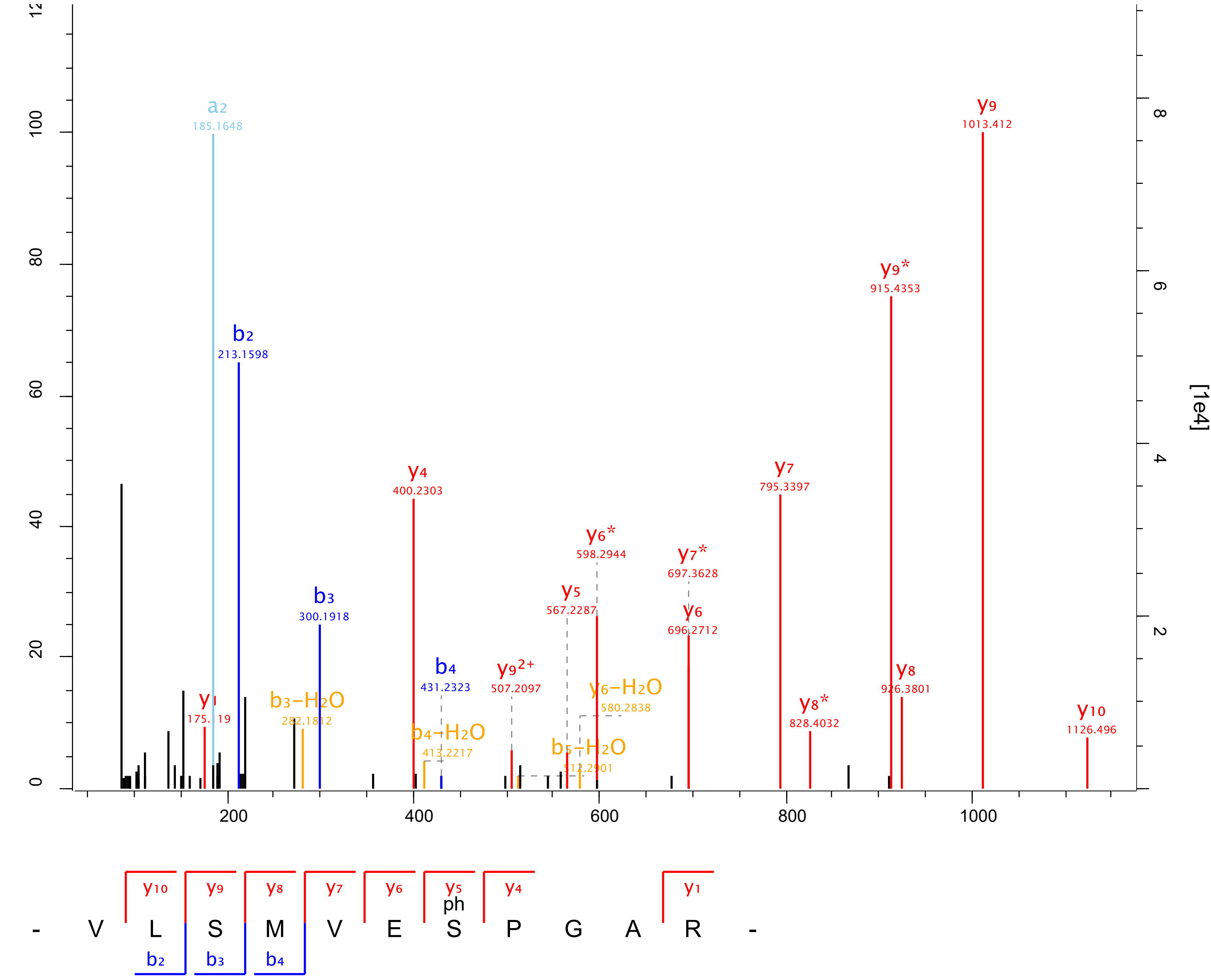

| Raw file | Scan | Method | Score | m/z |
|---|---|---|---|---|
| water_15_2 | 16301 | FTMS; HCD | 143.97 | 700.81 |

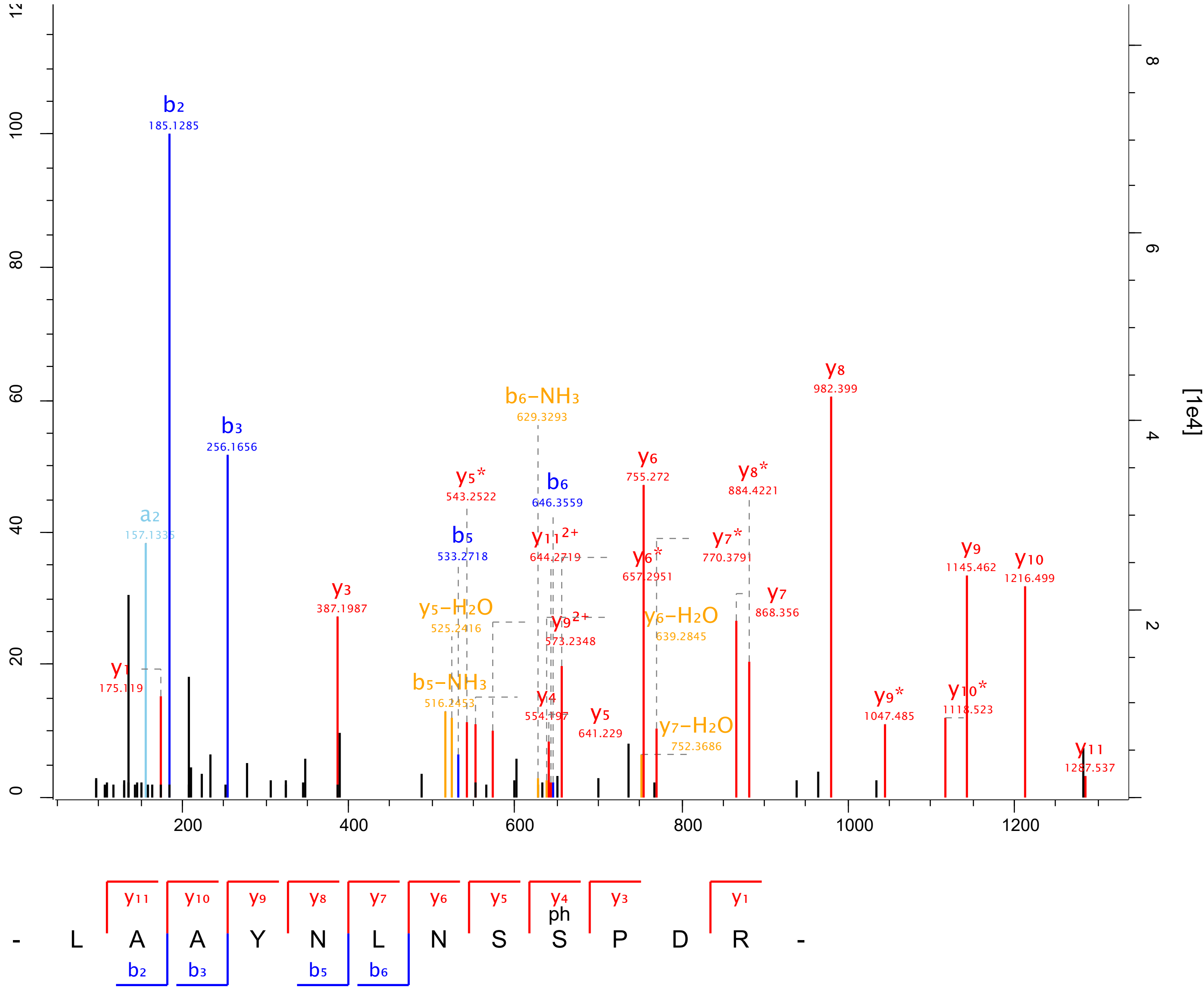

| Raw file | Scan | Method | Score | m/z |
|---|---|---|---|---|
| water_15_2 | 16351 | FTMS; HCD | 41.62 | 626.78 |

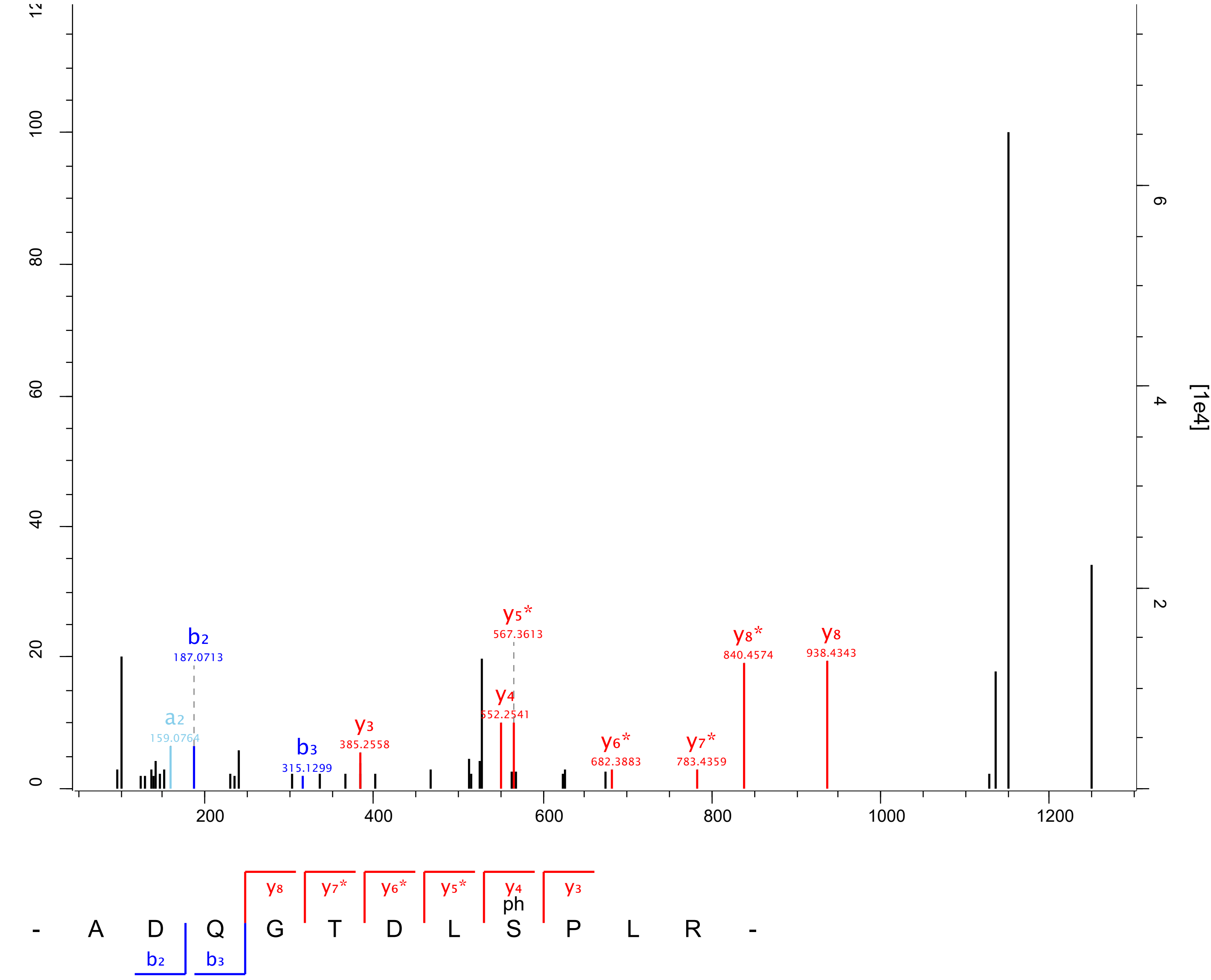

| Raw file | Scan | Method | Score | m/z |
|---|---|---|---|---|
| water_15_2 | 16356 | FTMS; HCD | 69.72 | 483.73 |

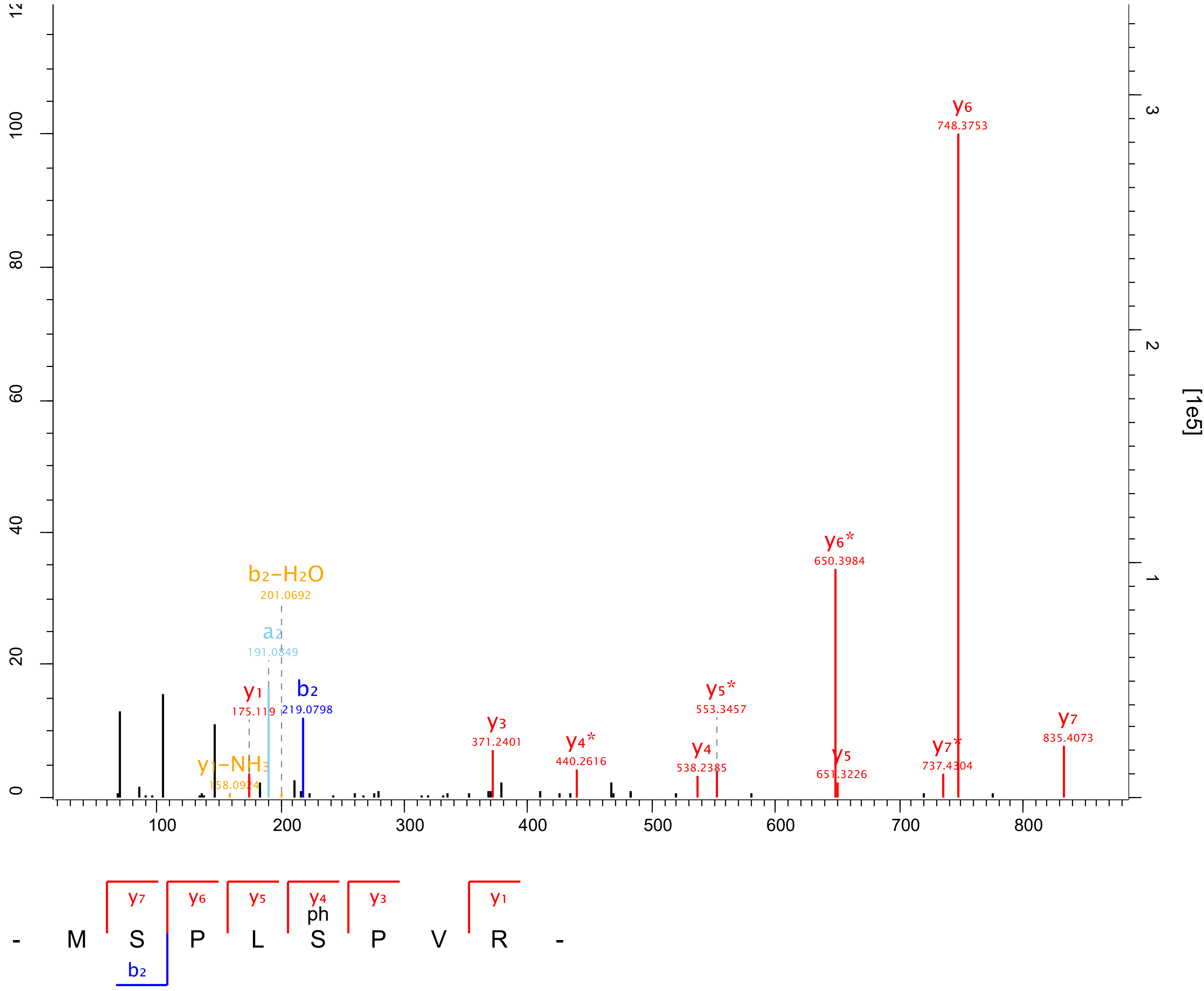

| Raw file | Scan | Method | Score | m/z |
|---|---|---|---|---|
| water_15_2 | 16367 | FTMS; HCD | 116.51 | 580.75 |

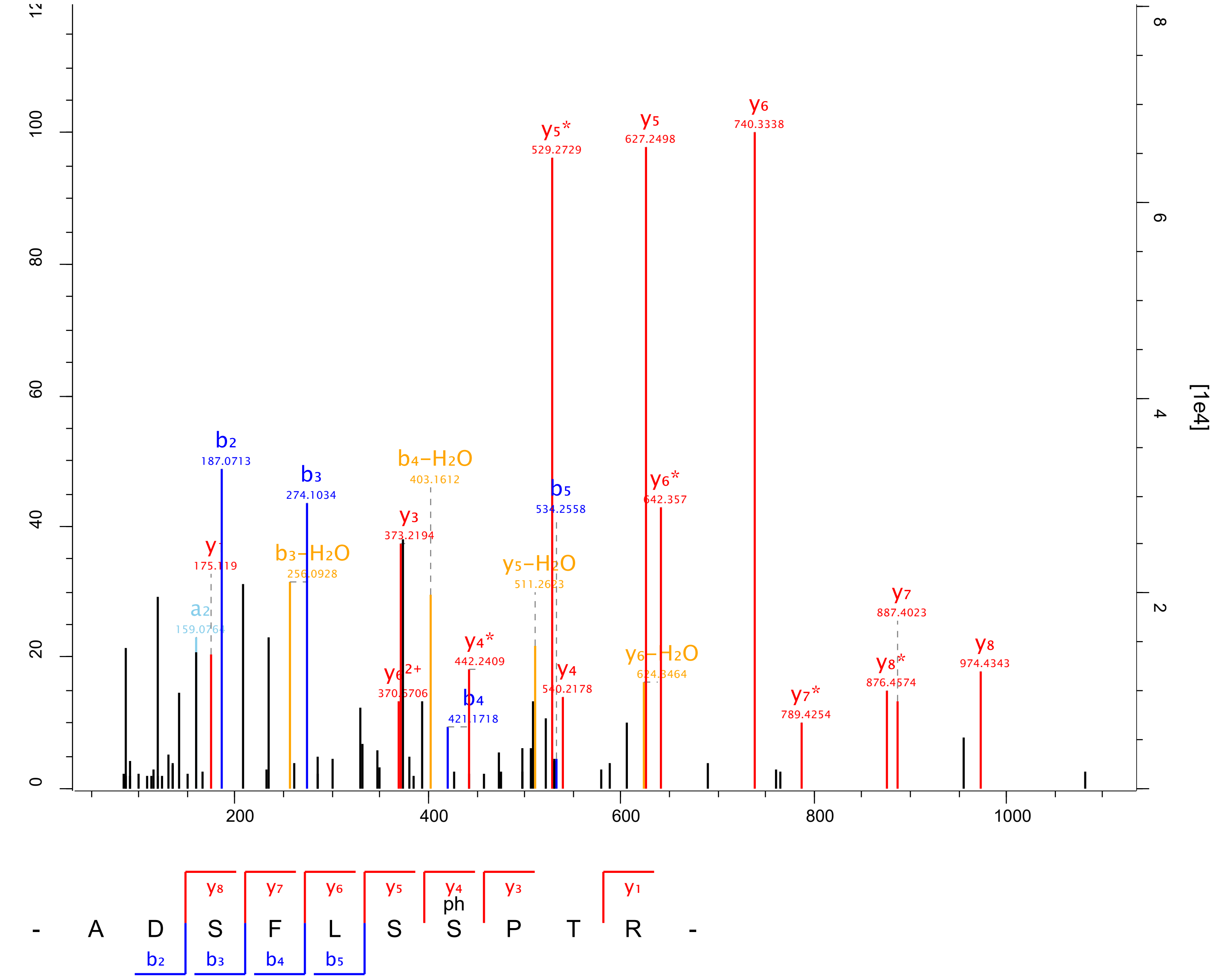

| Raw file | Scan | Method | Score | m/z |
| --- | --- | --- | --- | --- |
| water_15_2 | 16373 | FTMS; HCD | 65.23 | 735.83 |

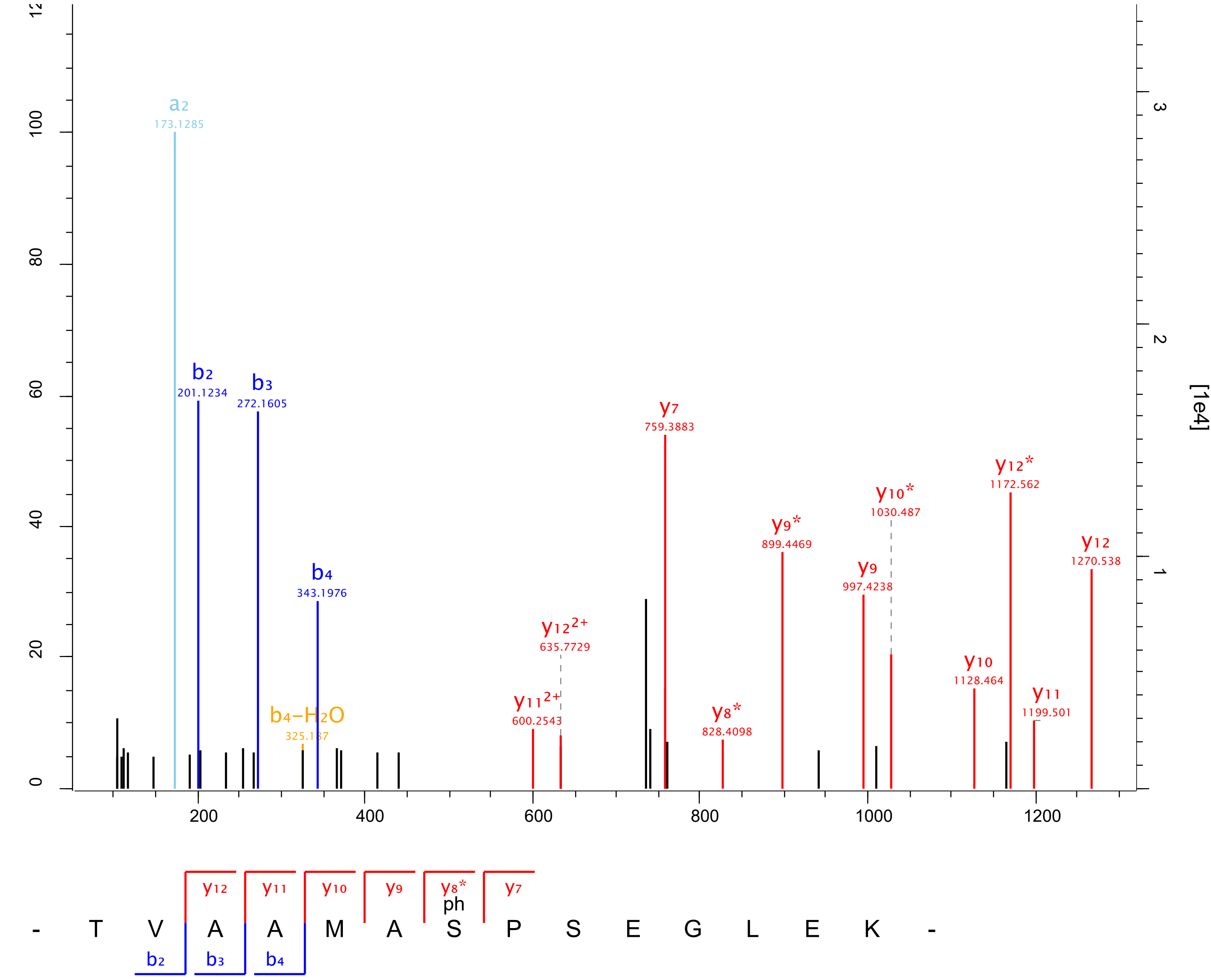

| Raw file | Scan | Method | Score | m/z |
|---|---|---|---|---|
| water_15_2 | 16383 | FTMS; HCD | 62.34 | 500.25 |

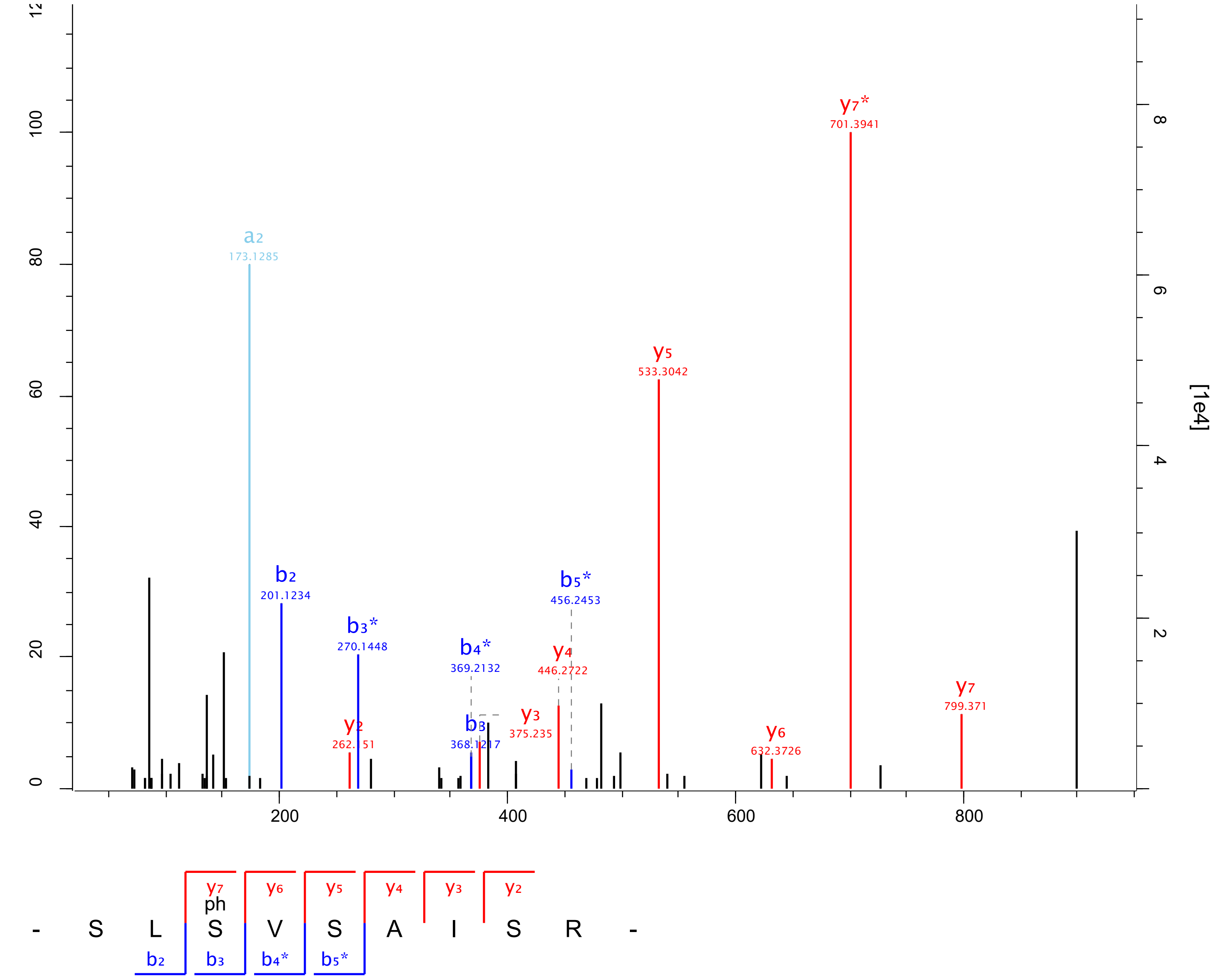

| Raw file | Scan | Method | Score | m/z |
|---|---|---|---|---|
| water_15_2 | 1640 | FTMS; HCD | 162.79 | 531.53 |

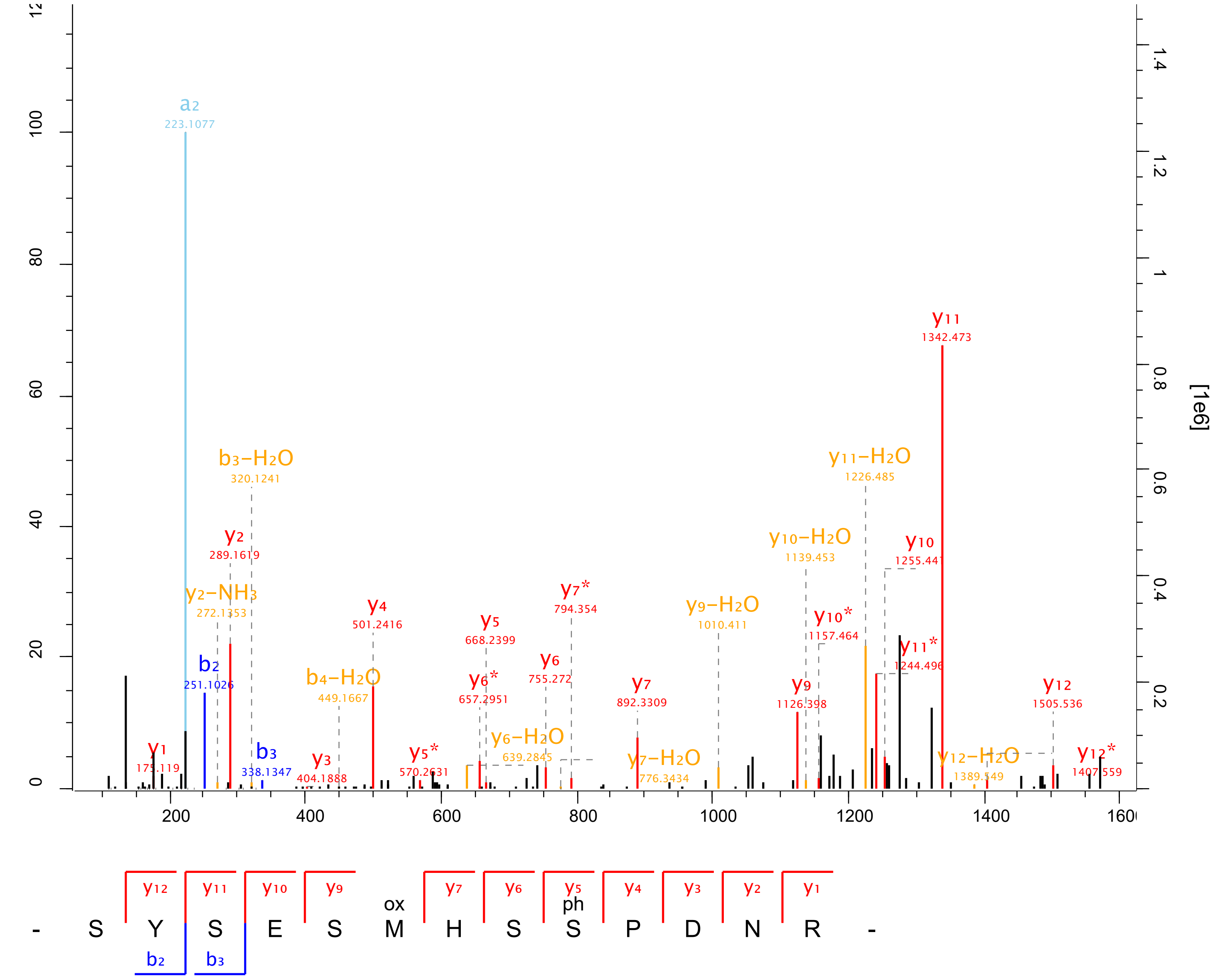

| Raw file | Scan | Method | Score | m/z |
| --- | --- | --- | --- | --- |
| water_15_2 | 16464 | FTMS; HCD | 43.69 | 631.83 |

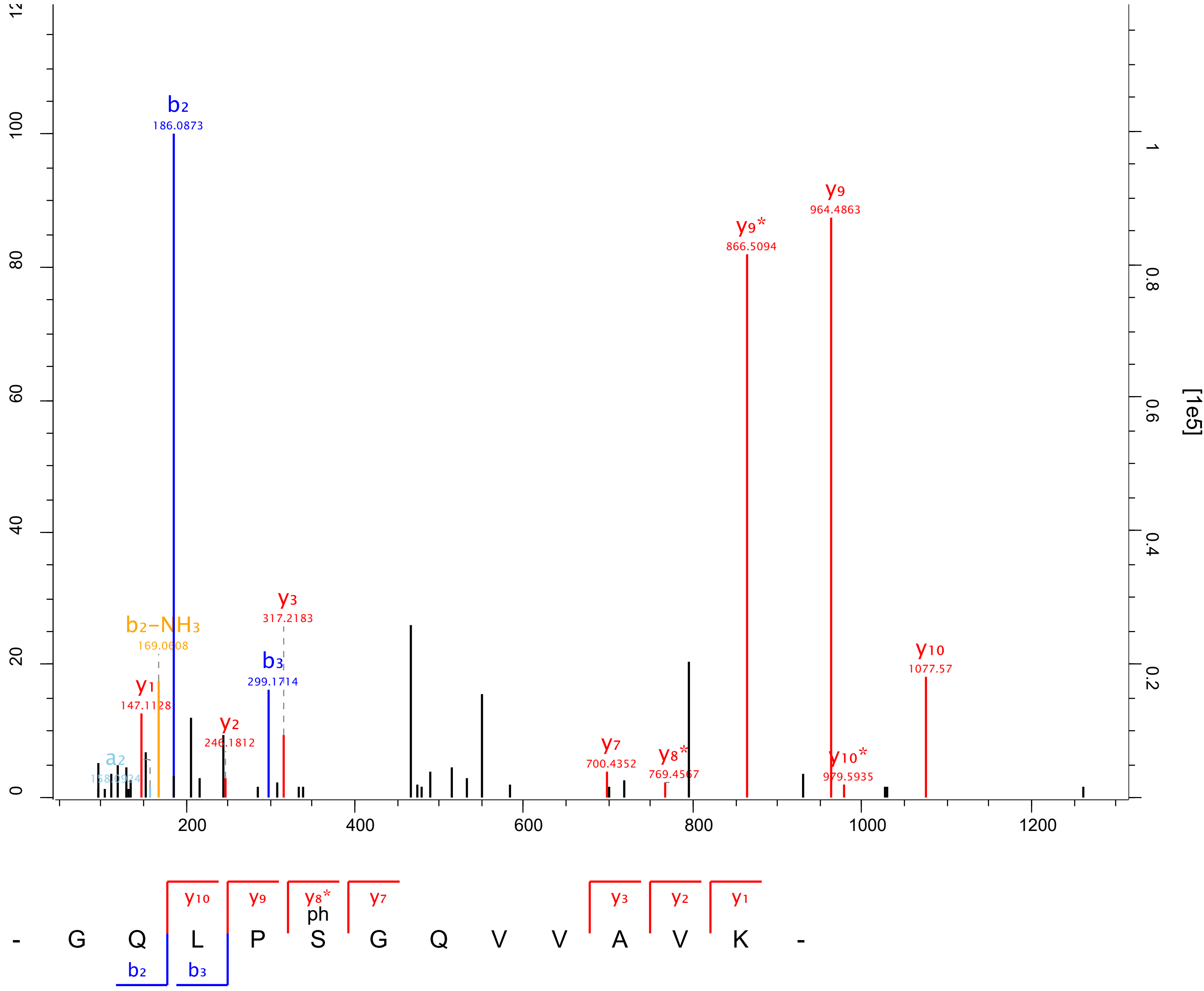

| Raw file | Scan | Method | Score | m/z |
|---|---|---|---|---|
| water_15_2 | 16479 | FTMS; HCD | 52.58 | 598.27 |

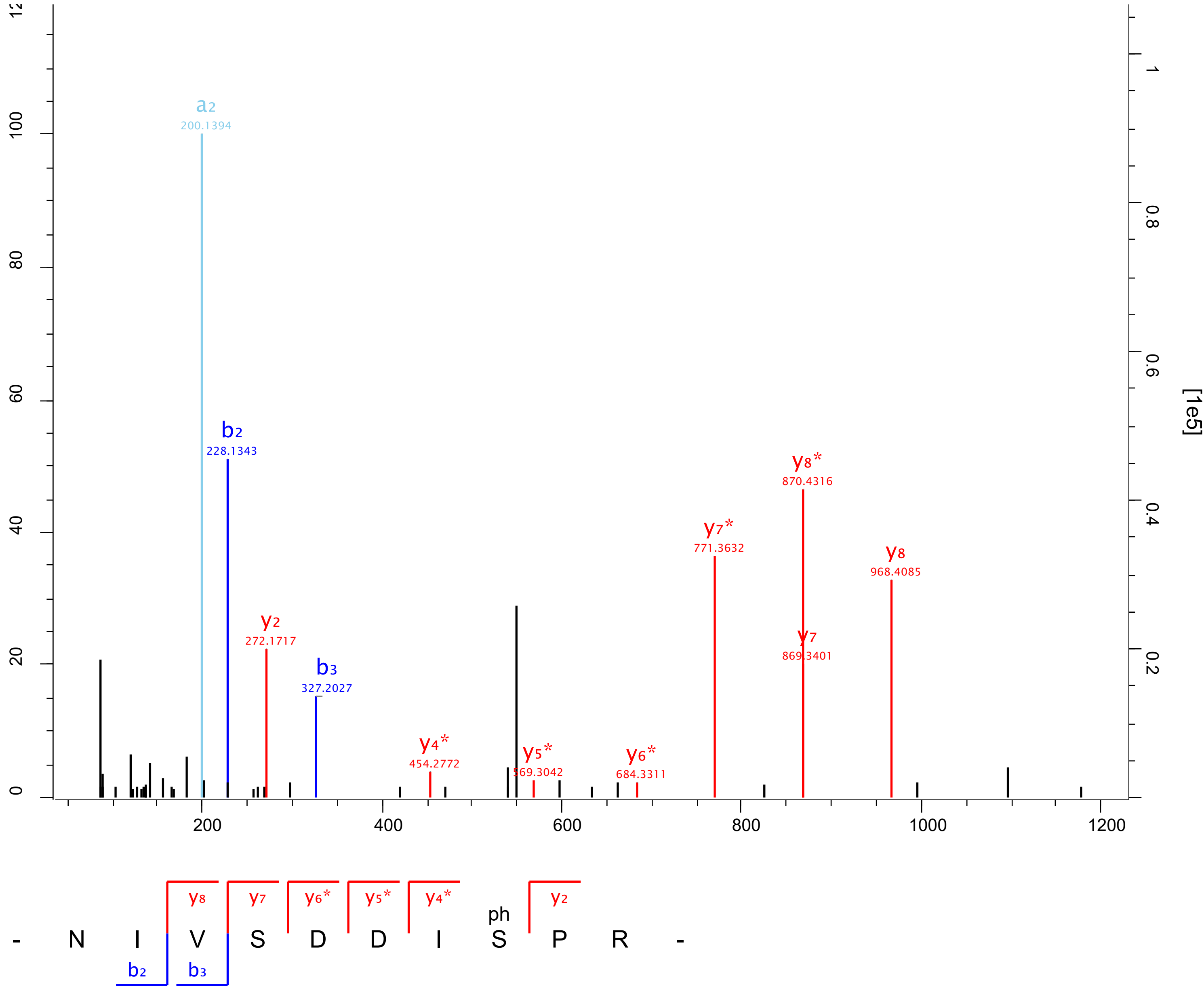

| Raw file | Scan | Method | Score | m/z |
|---|---|---|---|---|
| water_15_2 | 16510 | FTMS; HCD | 84.76 | 695.34 |

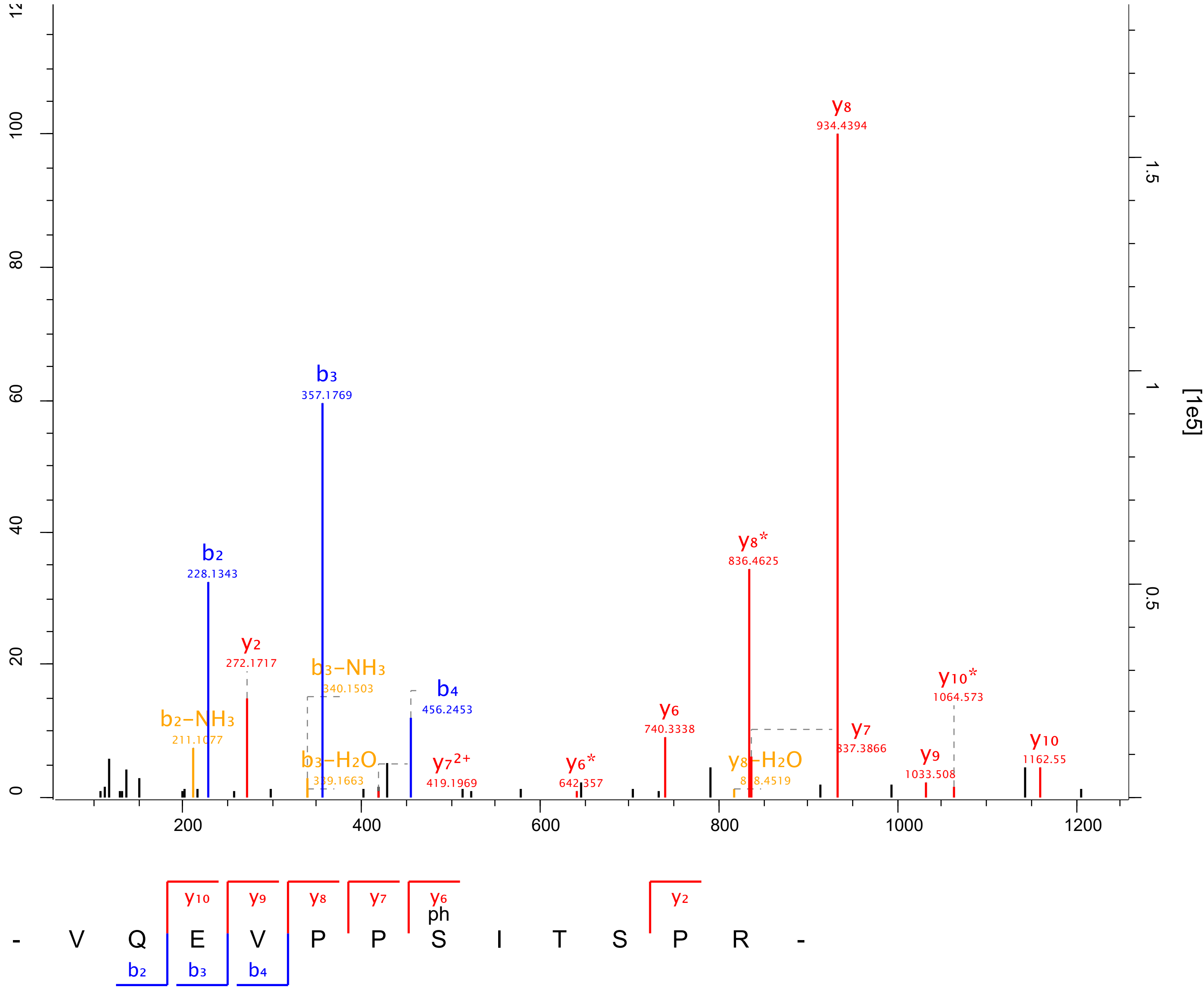

| Raw file | Scan | Method | Score | m/z |
|---|---|---|---|---|
| water_15_2 | 16577 | FTMS; HCD | 81.66 | 681.31 |

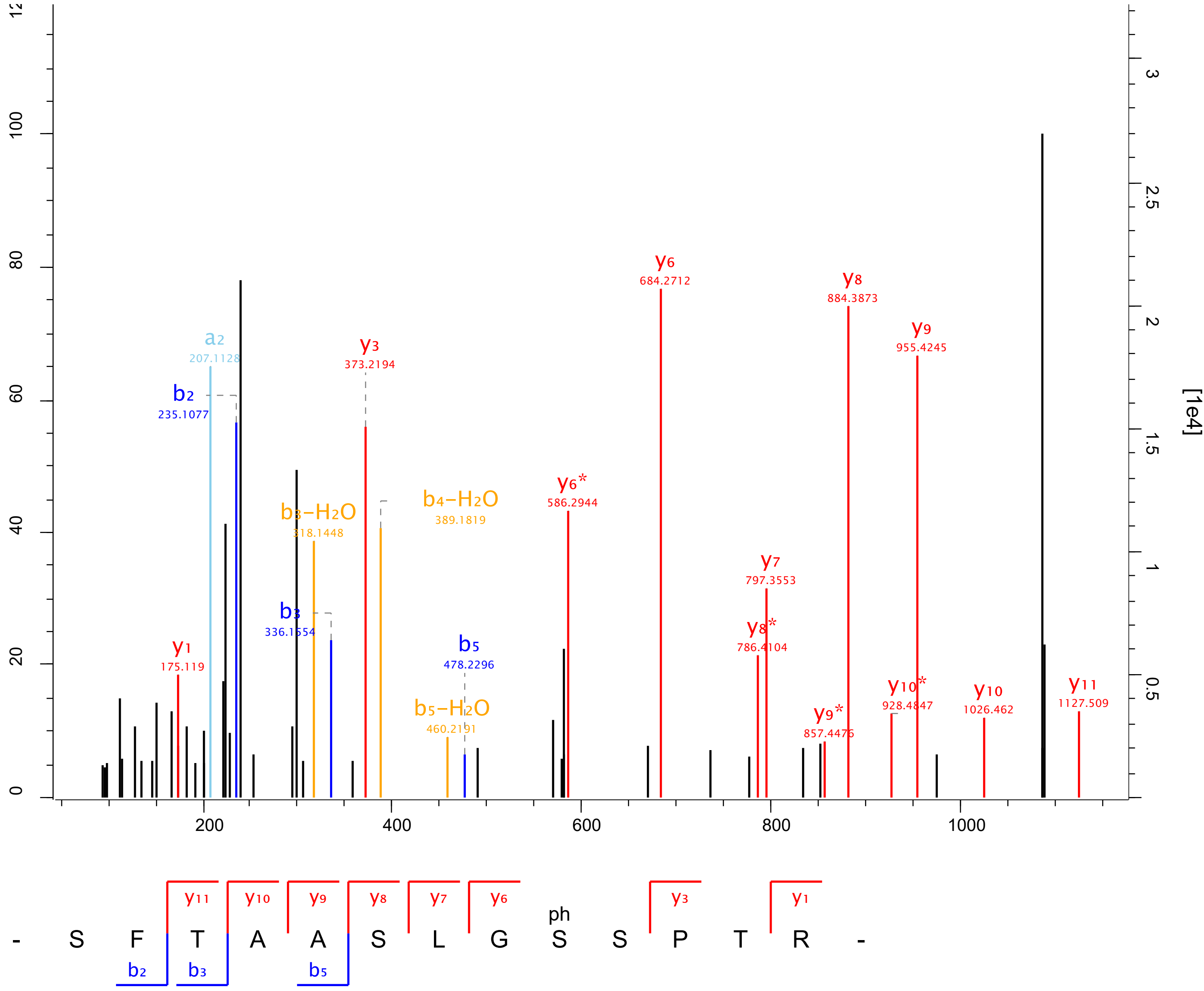

| Raw file | Scan | Method | Score | m/z |
|---|---|---|---|---|
| water_15_2 | 16651 | FTMS; HCD | 60.9 | 707.33 |

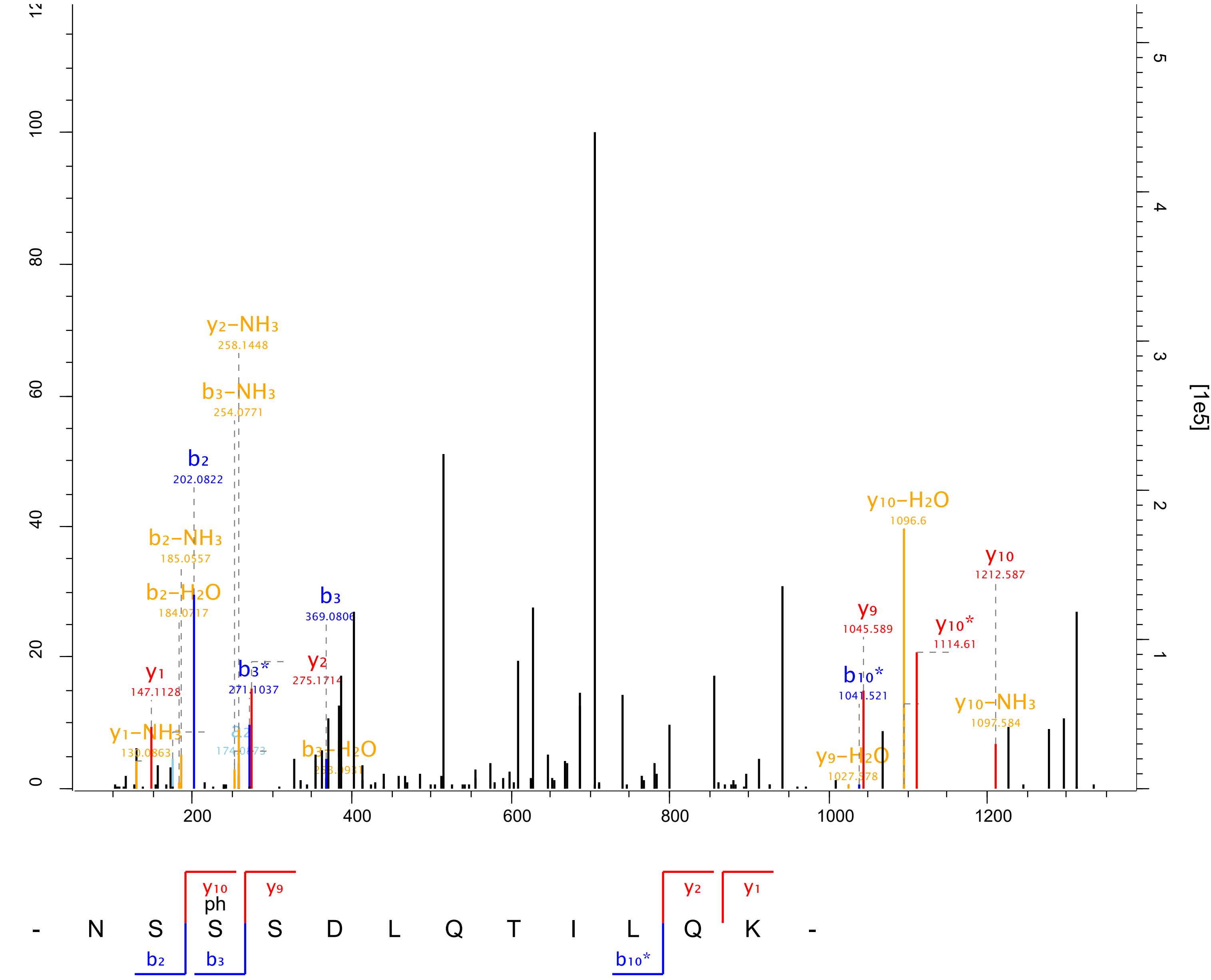

| Raw file | Scan | Method | Score | m/z |
|---|---|---|---|---|
| water_15_2 | 16813 | FTMS; HCD | 95.66 | 872.35 |

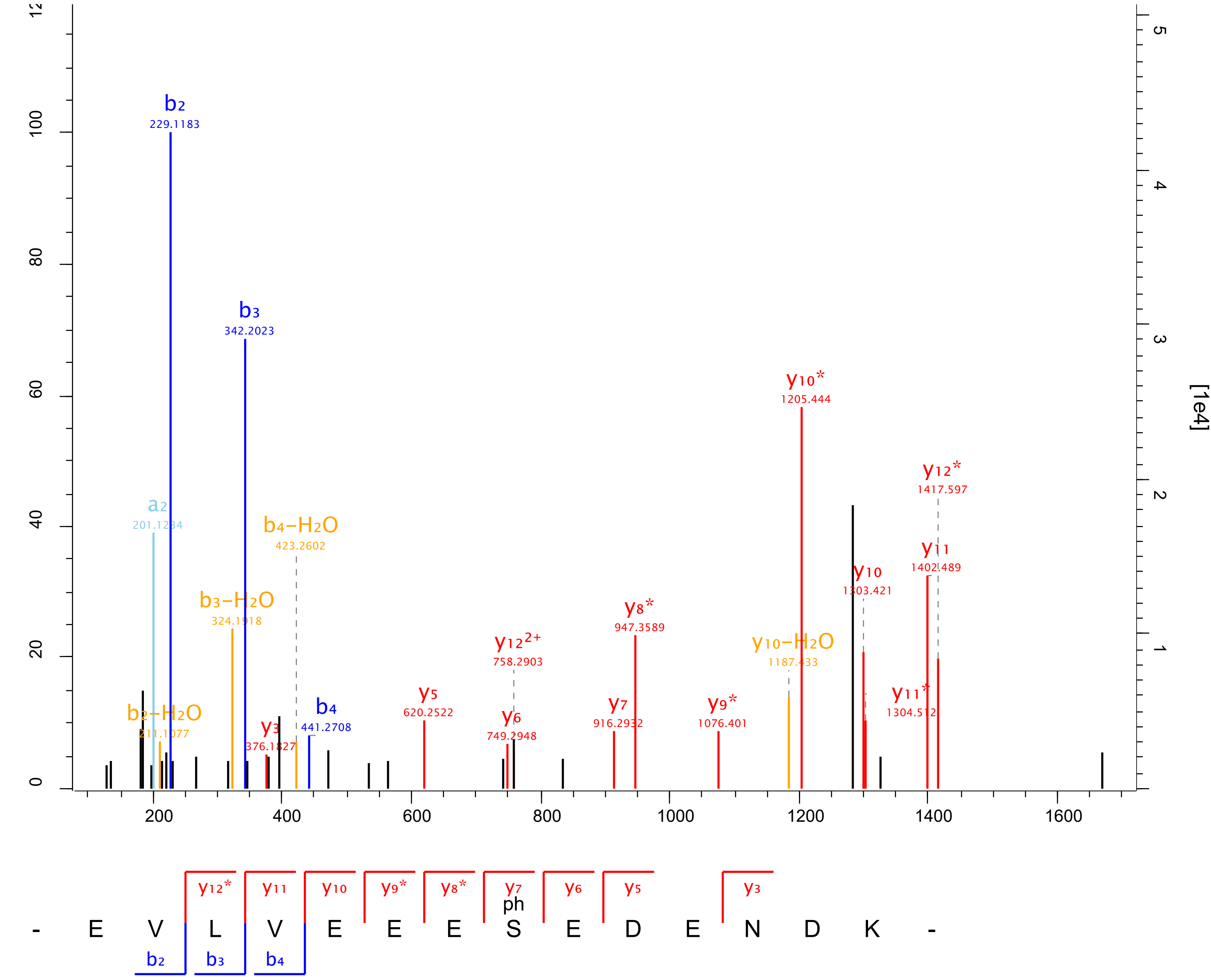

| Raw file | Scan | Method | Score | m/z |
| --- | --- | --- | --- | --- |
| water_15_2 | 16846 | FTMS; HCD | 209.89 | 705.82 |

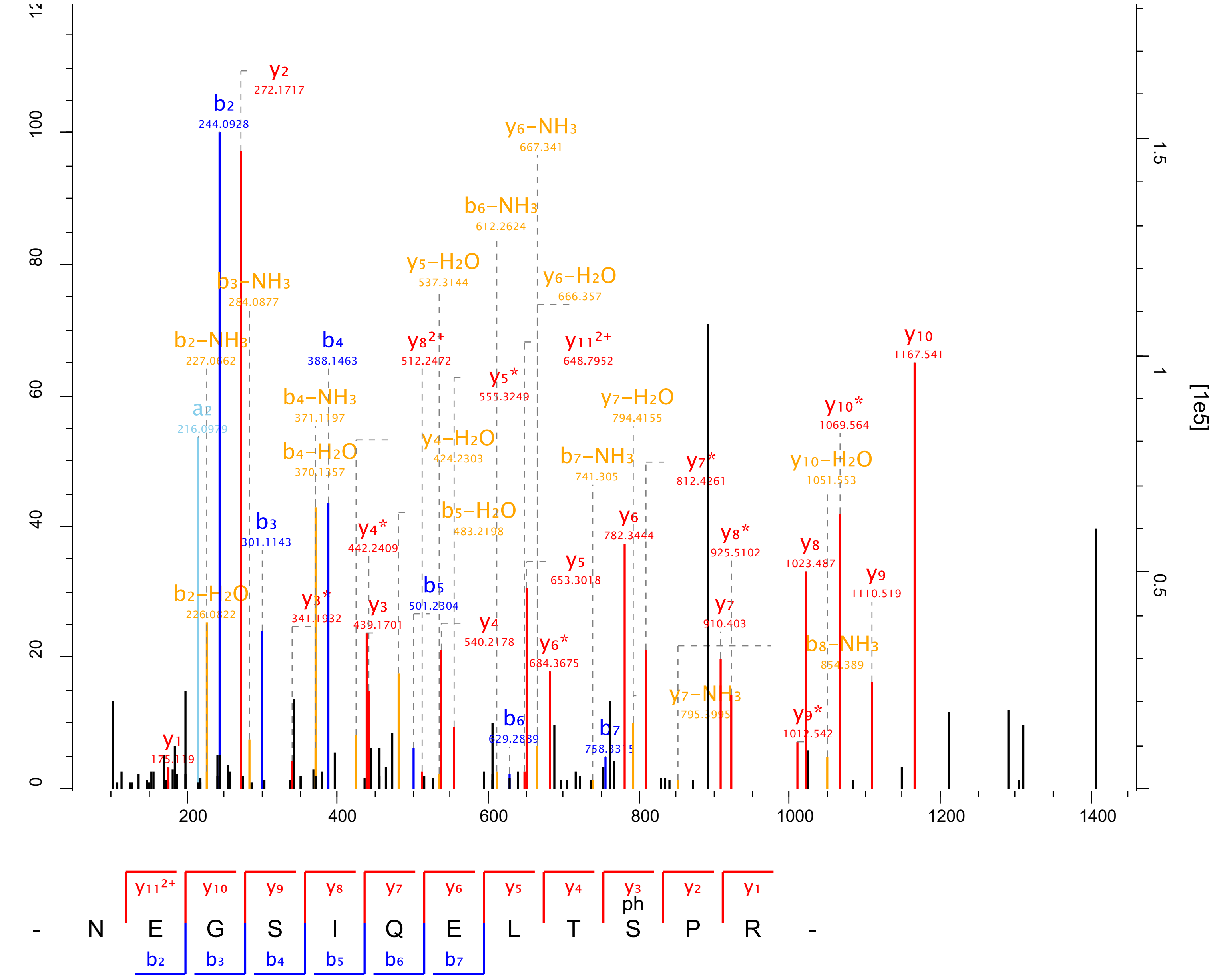

| Raw file | Scan | Method | Score | m/z |
|---|---|---|---|---|
| water_15_2 | 16870 | FTMS; HCD | 84.76 | 657.74 |

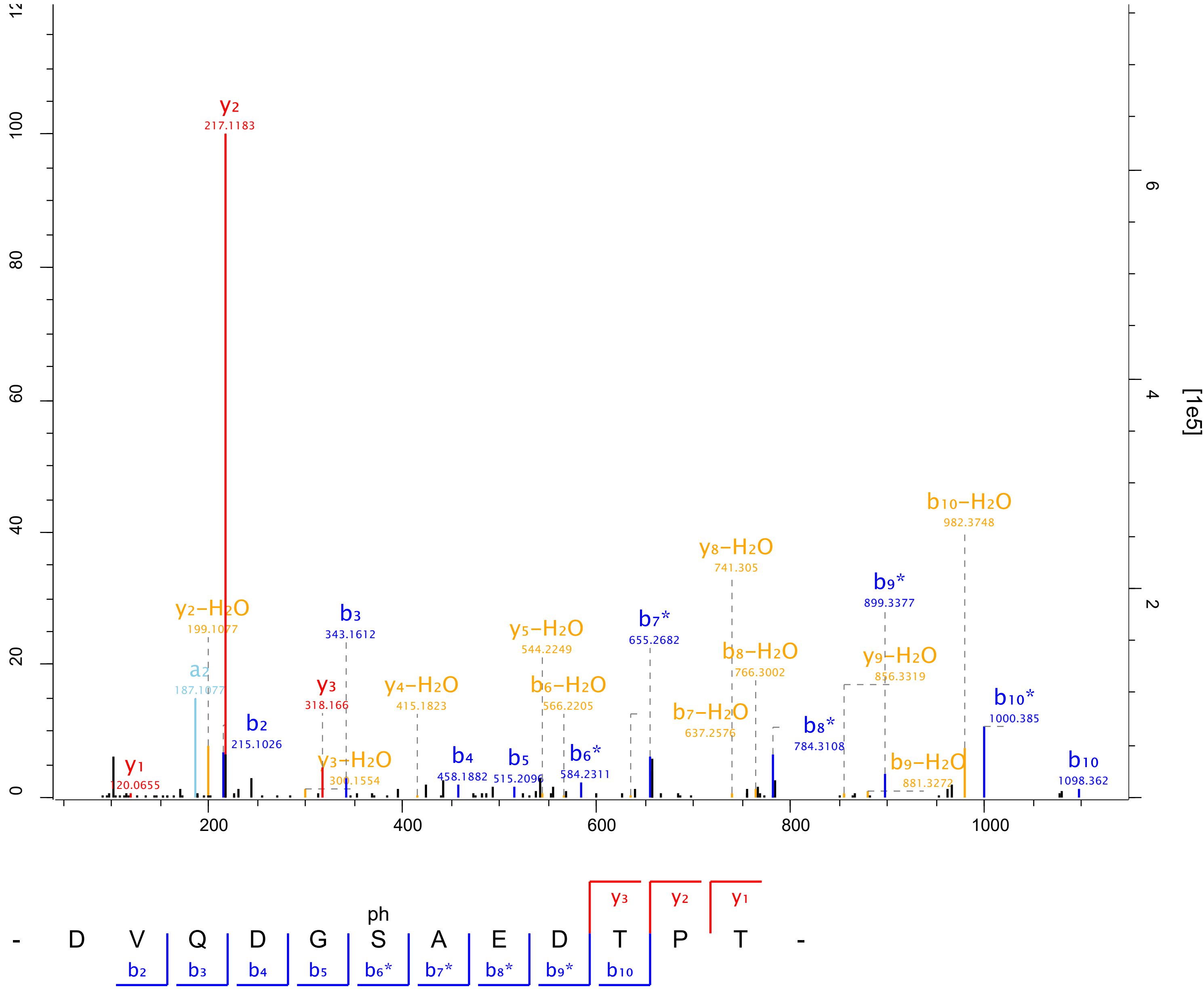

| Raw file | Scan | Method | Score | m/z |
| --- | --- | --- | --- | --- |
| water_15_2 | 16908 | FTMS; HCD | 102.73 | 557.27 |

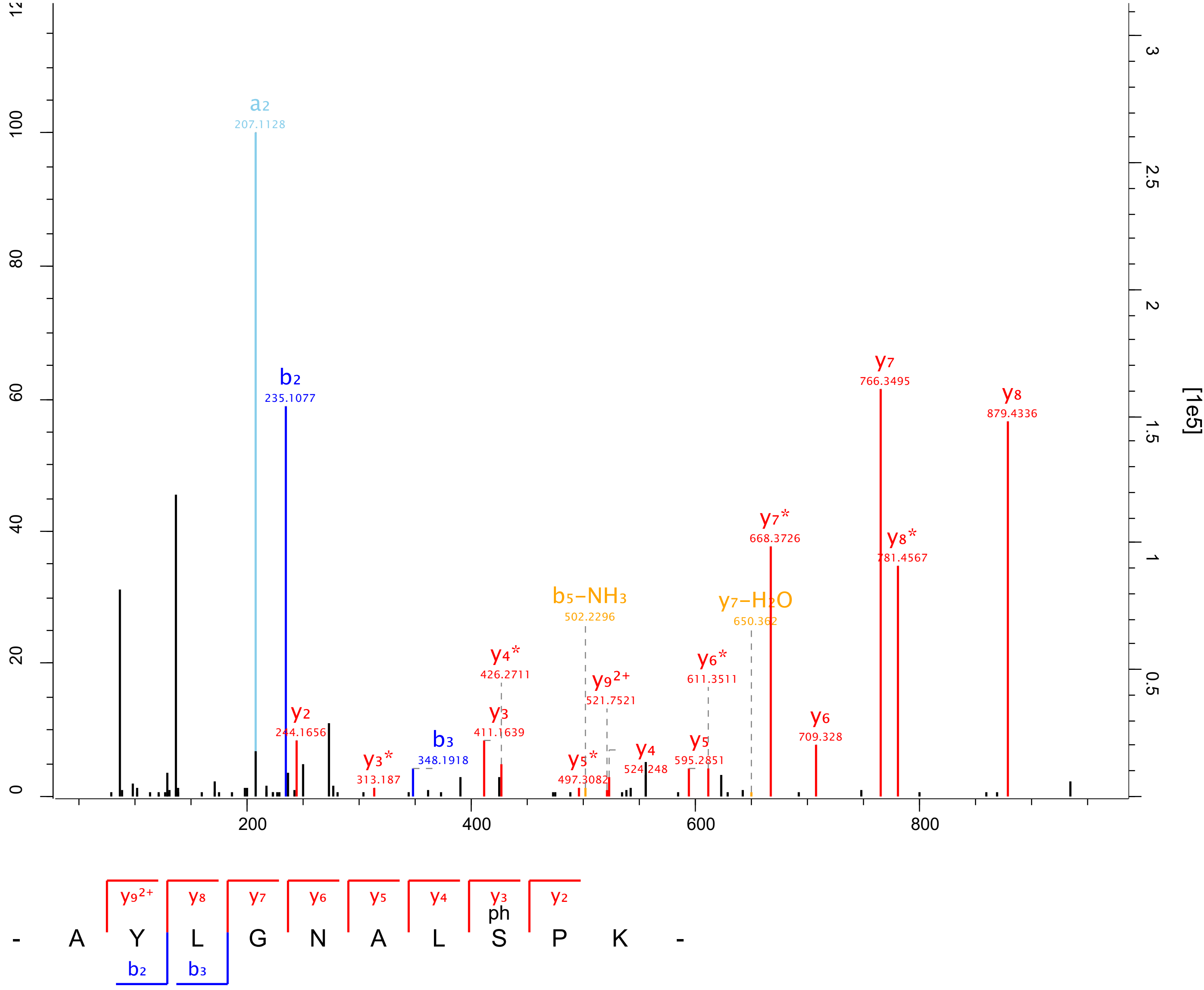

| Raw file | Scan | Method | Score | m/z |
|---|---|---|---|---|
| water_15_2 | 17022 | FTMS; HCD | 45.28 | 553.73 |

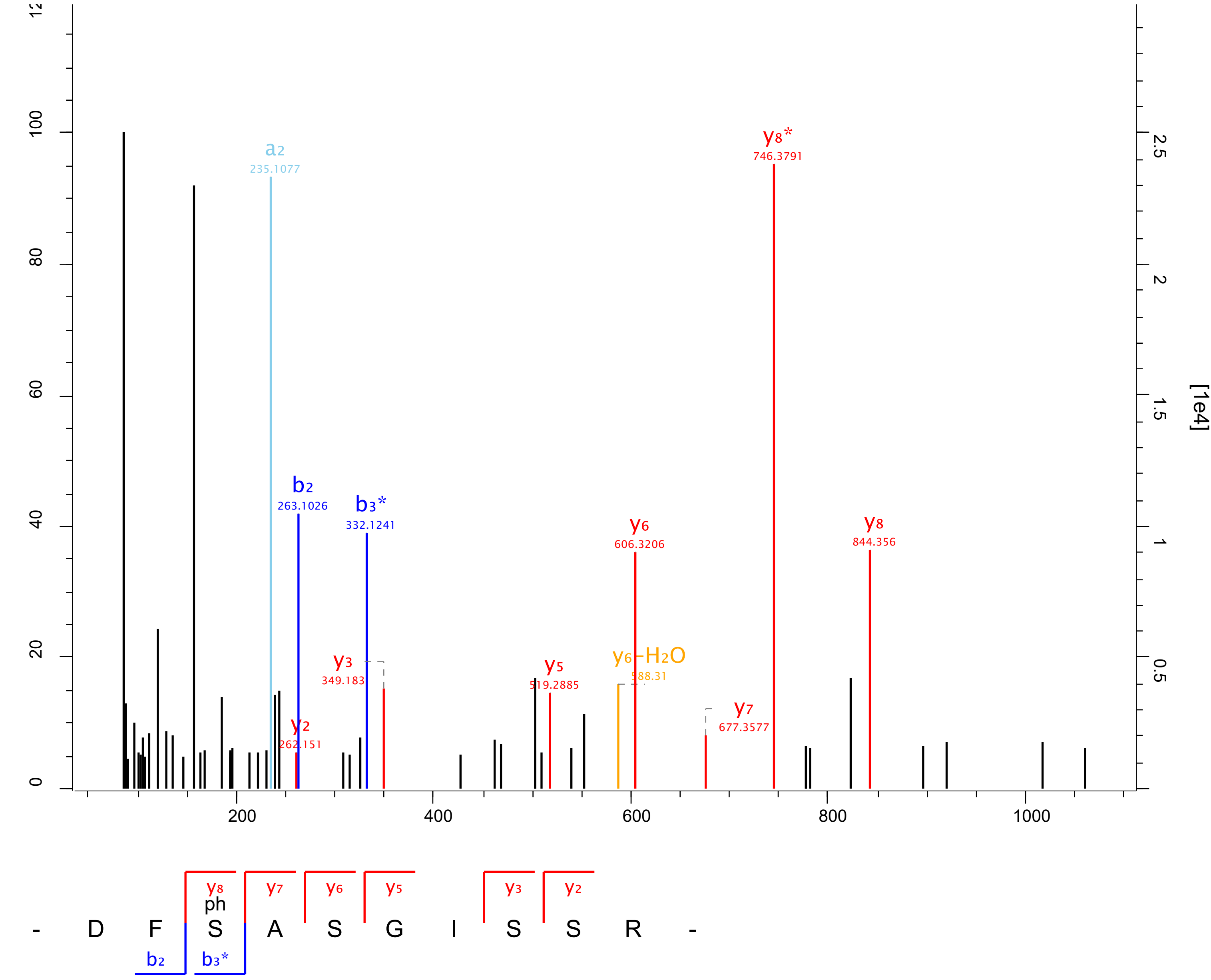

| Raw file | Scan | Method | Score | m/z |
|---|---|---|---|---|
| water_15_2 | 17029 | FTMS; HCD | 76.94 | 481.7 |

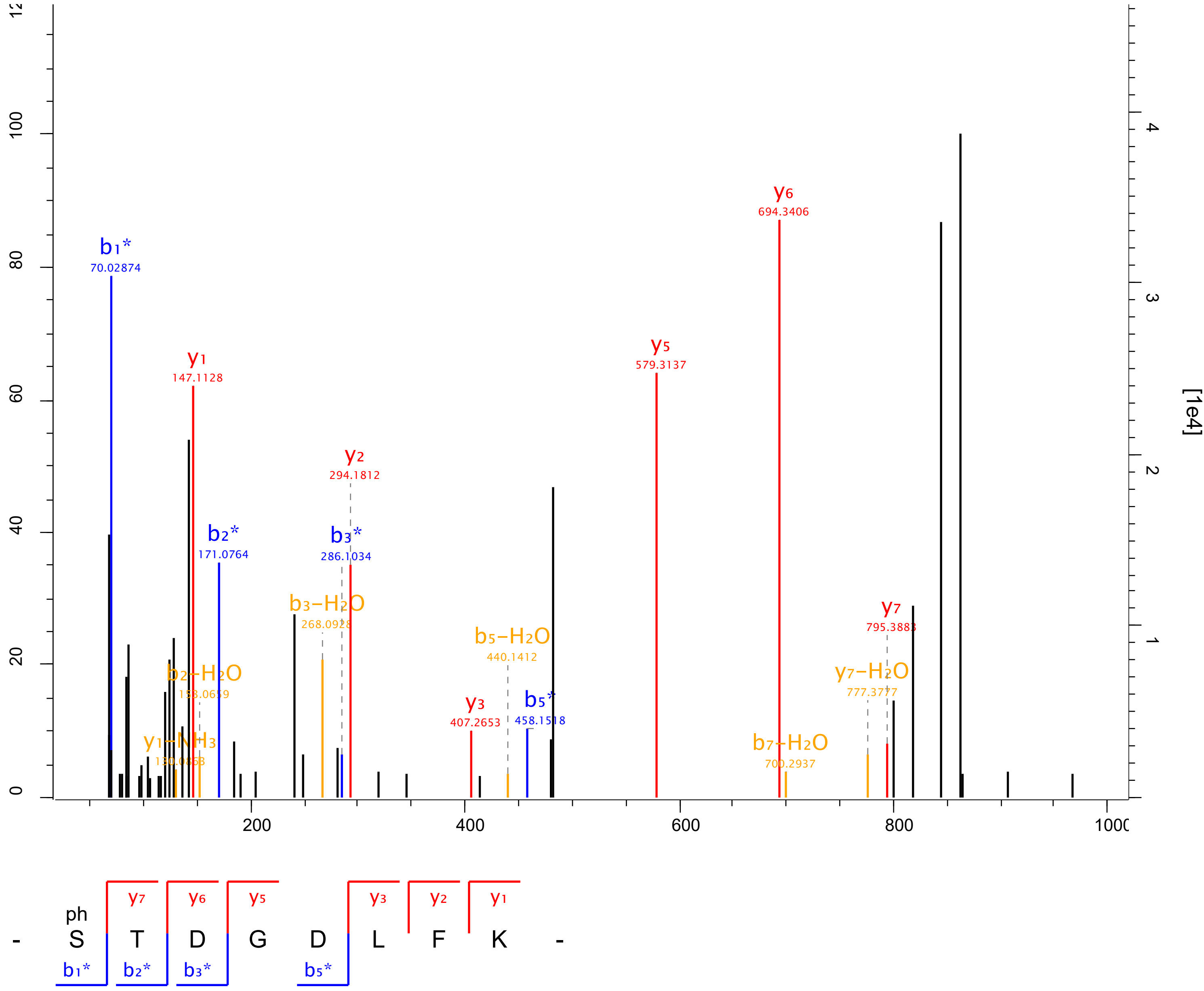

| Raw file | Scan | Method | Score | m/z |
|---|---|---|---|---|
| water_15_2 | 1704 | FTMS; HCD | 77.18 | 530.75 |

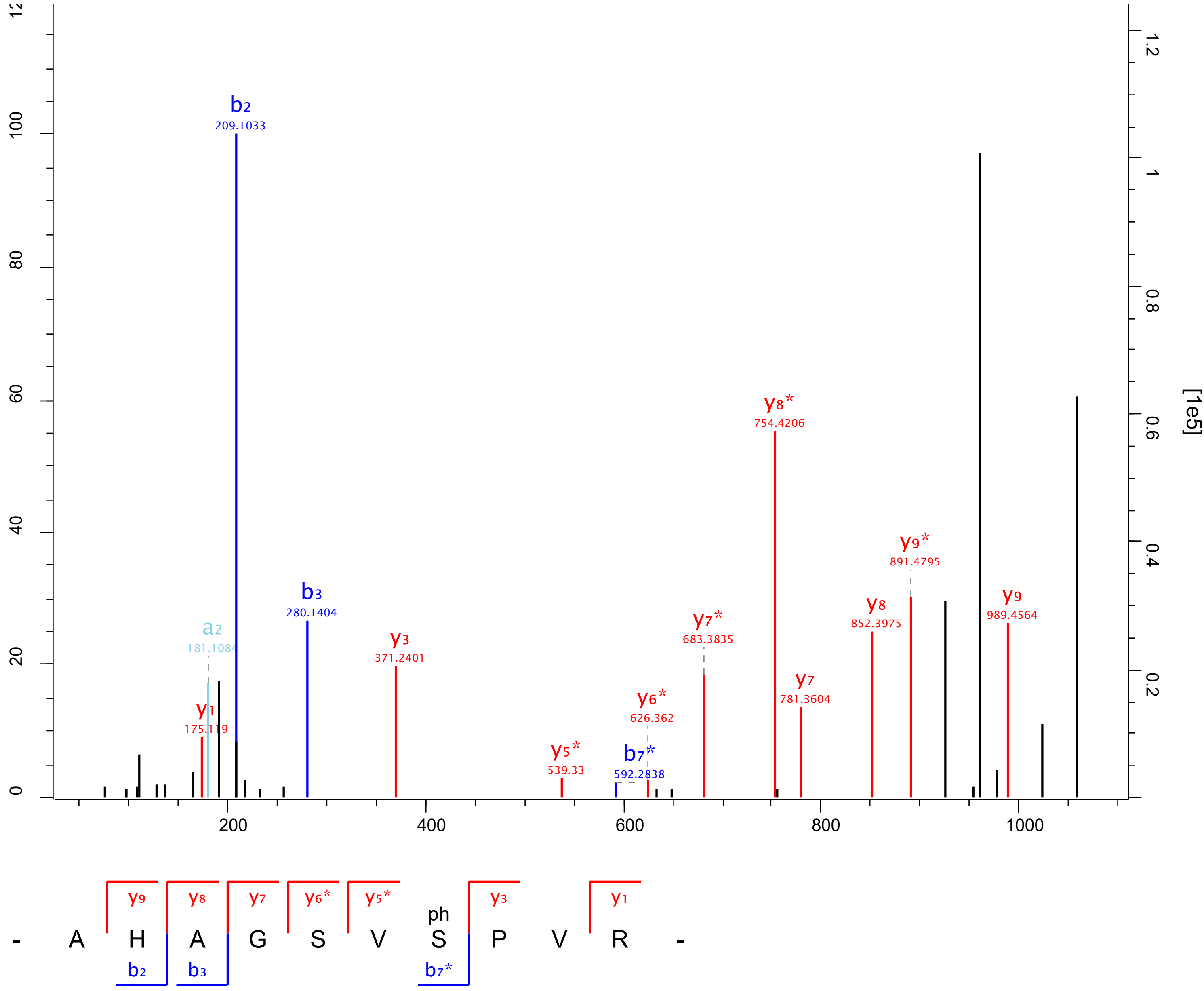

| Raw file | Scan | Method | Score | m/z |
|---|---|---|---|---|
| water_15_2 | 17063 | FTMS; HCD | 69.72 | 589.26 |

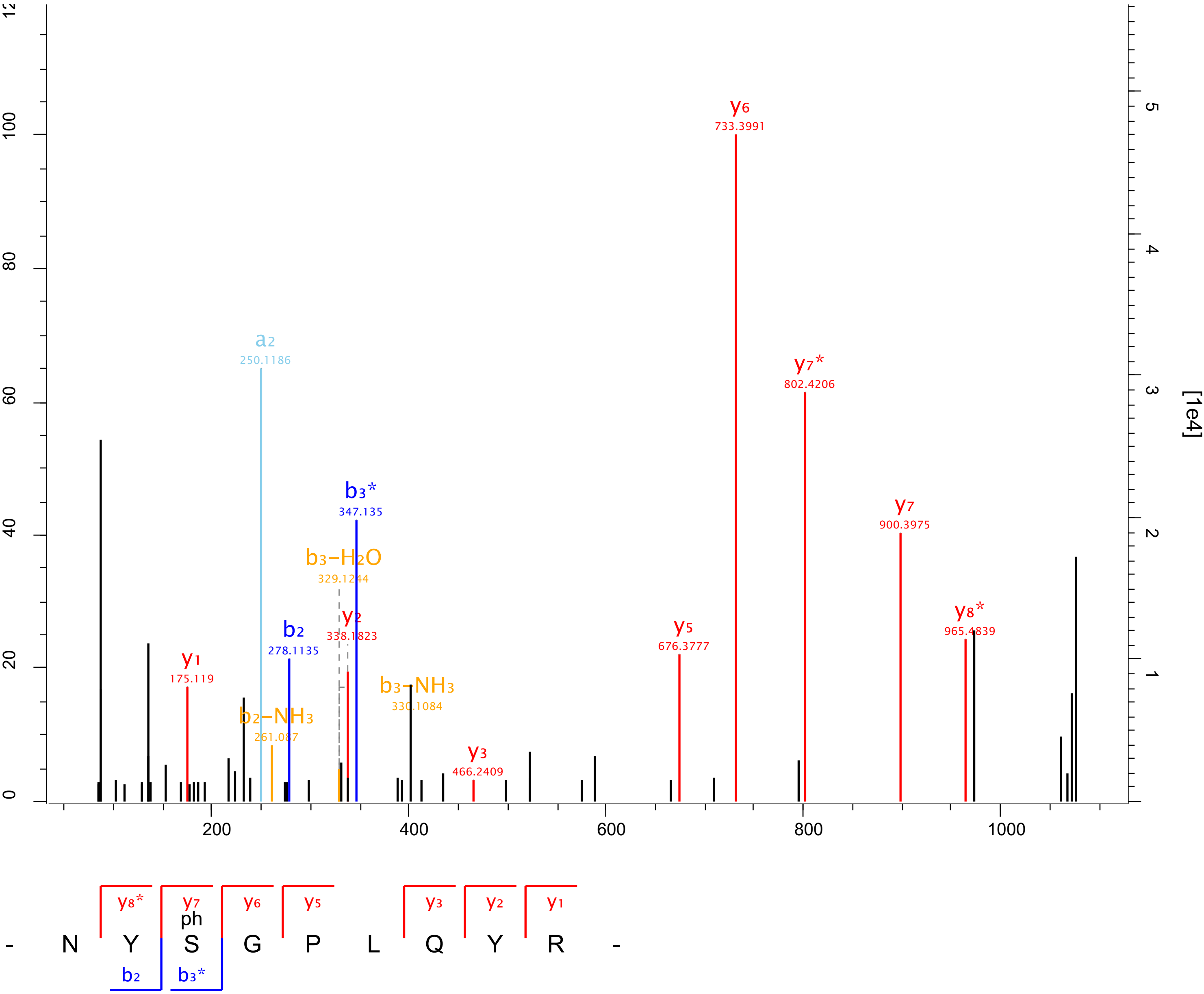

| Raw file | Scan | Method | Score | m/z |
|---|---|---|---|---|
| water_15_2 | 17067 | FTMS; HCD | 205.65 | 761.88 |

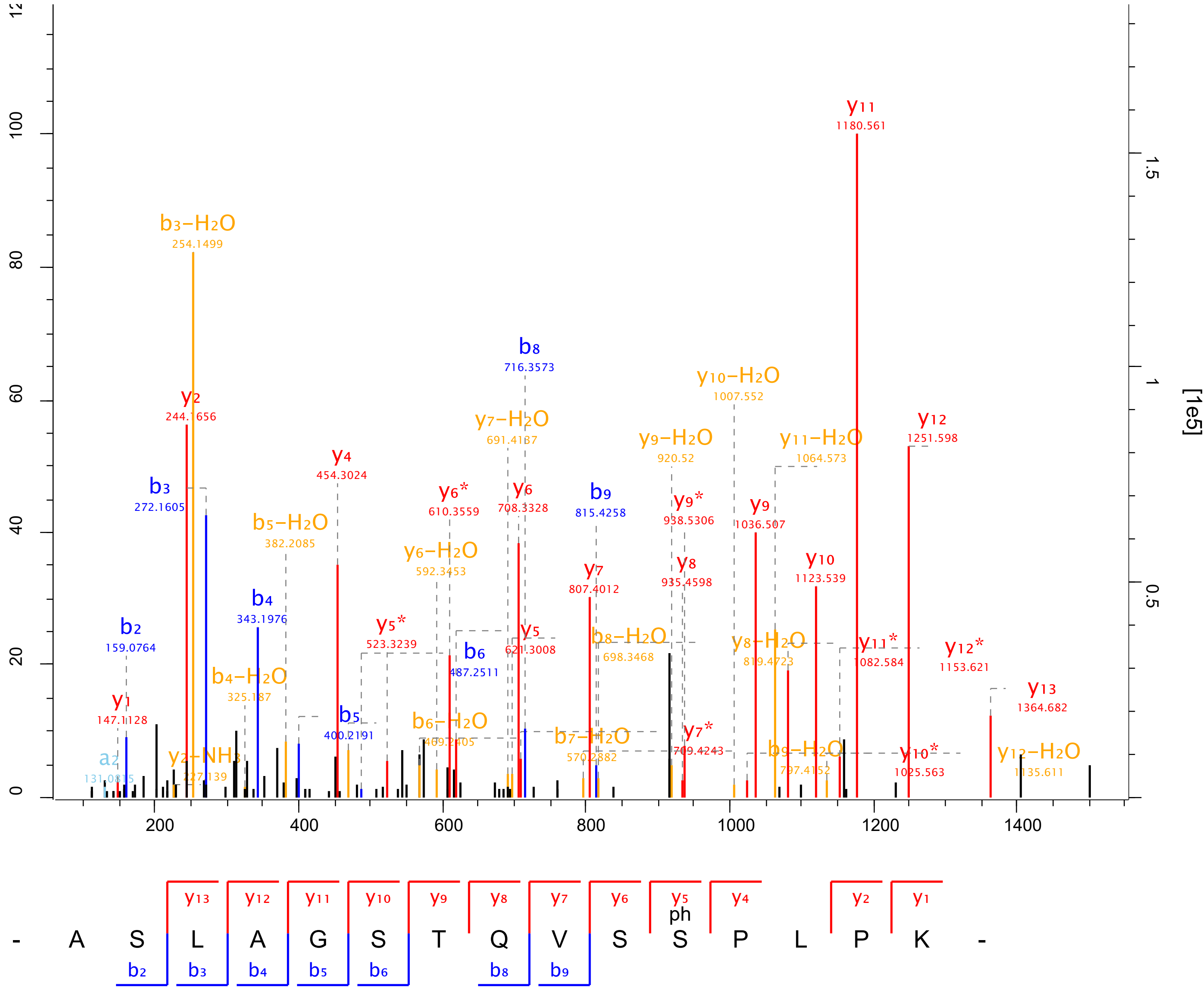

| Raw file | Scan | Method | Score | m/z |
|---|---|---|---|---|
| water_15_2 | 17178 | FTMS; HCD | 47.08 | 685.31 |

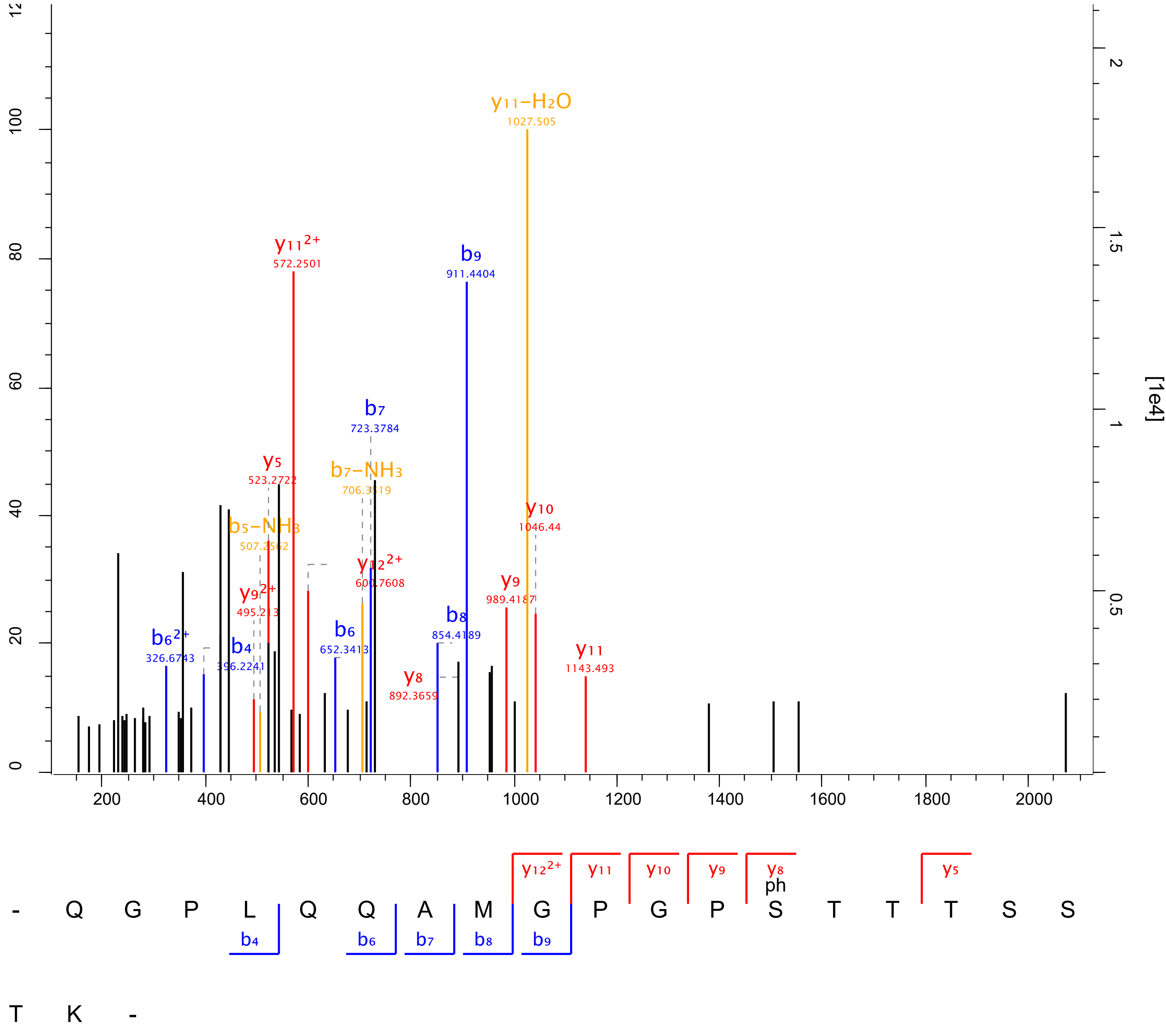

| Raw file | Scan | Method | Score | m/z |
|---|---|---|---|---|
| water_15_2 | 17237 | FTMS; HCD | 117.79 | 553.24 |

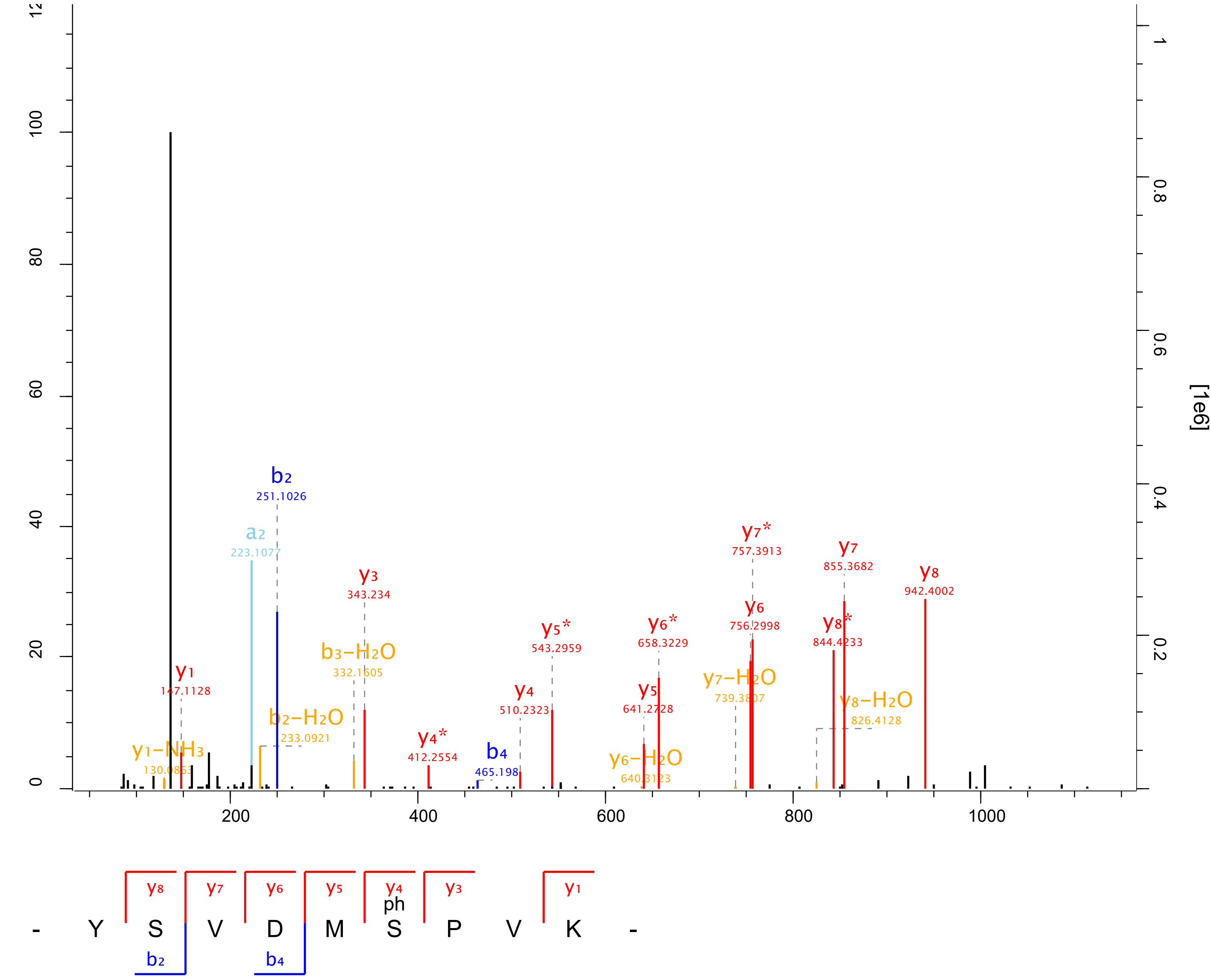

| Raw file | Scan | Method | Score | m/z |
| --- | --- | --- | --- | --- |
| water_15_2 | 17240 | FTMS; HCD | 40.97 | 840.87 |

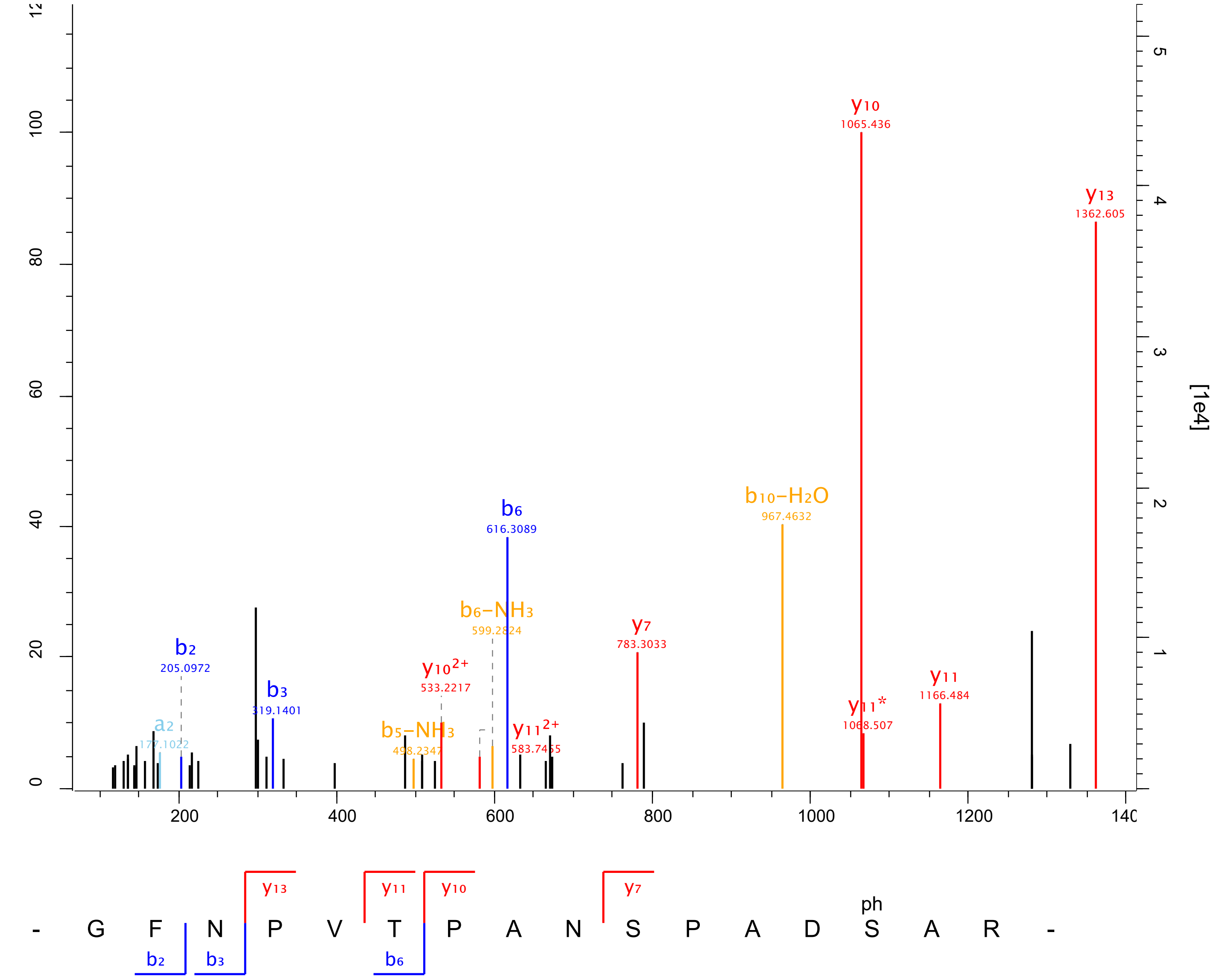

| Raw file | Scan | Method | Score | m/z |
|---|---|---|---|---|
| water_15_2 | 17299 | FTMS; HCD | 65.22 | 593.75 |

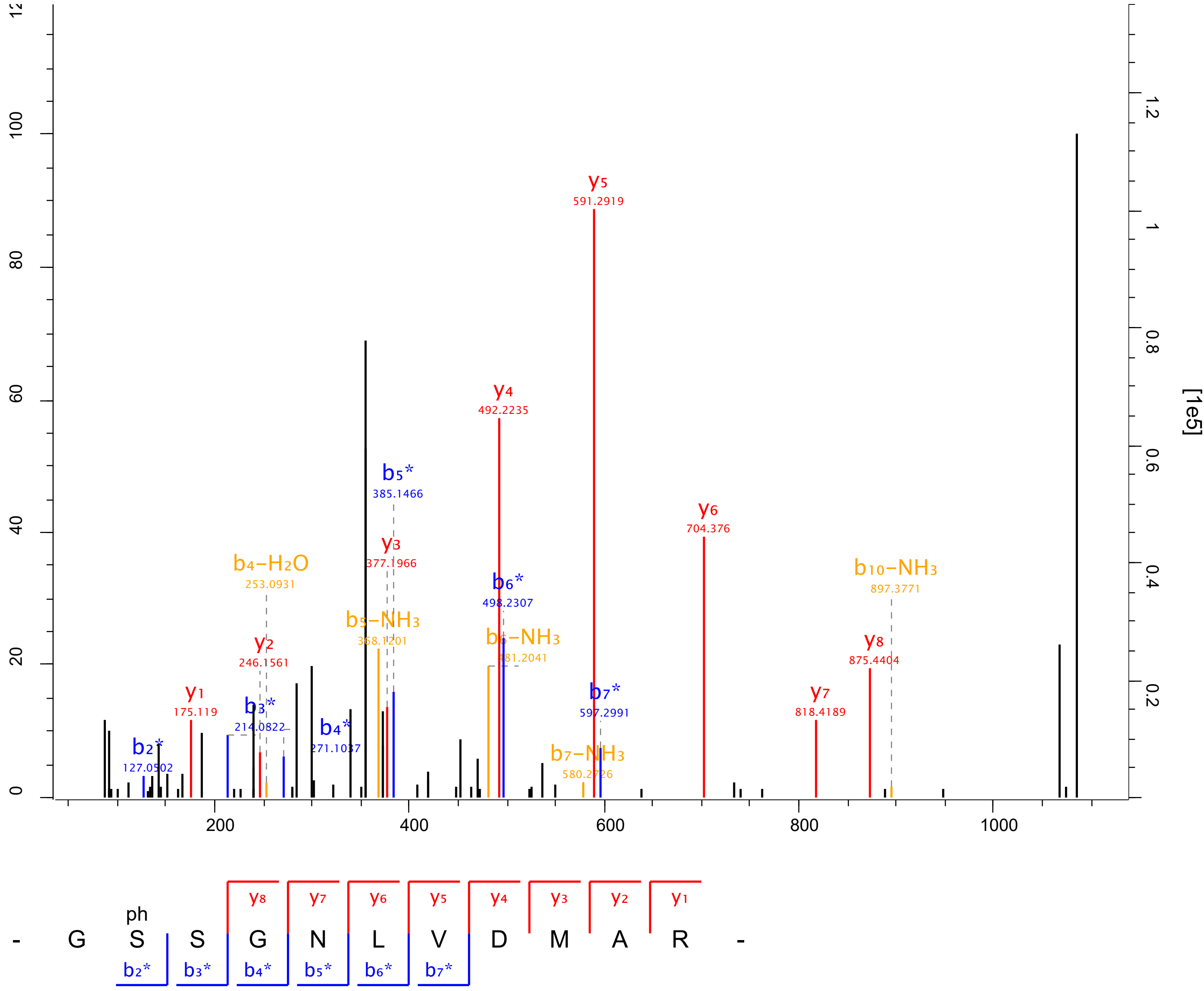

| Raw file | Scan | Method | Score | m/z |
|---|---|---|---|---|
| water_15_2 | 17311 | FTMS; HCD | 70.89 | 603.75 |

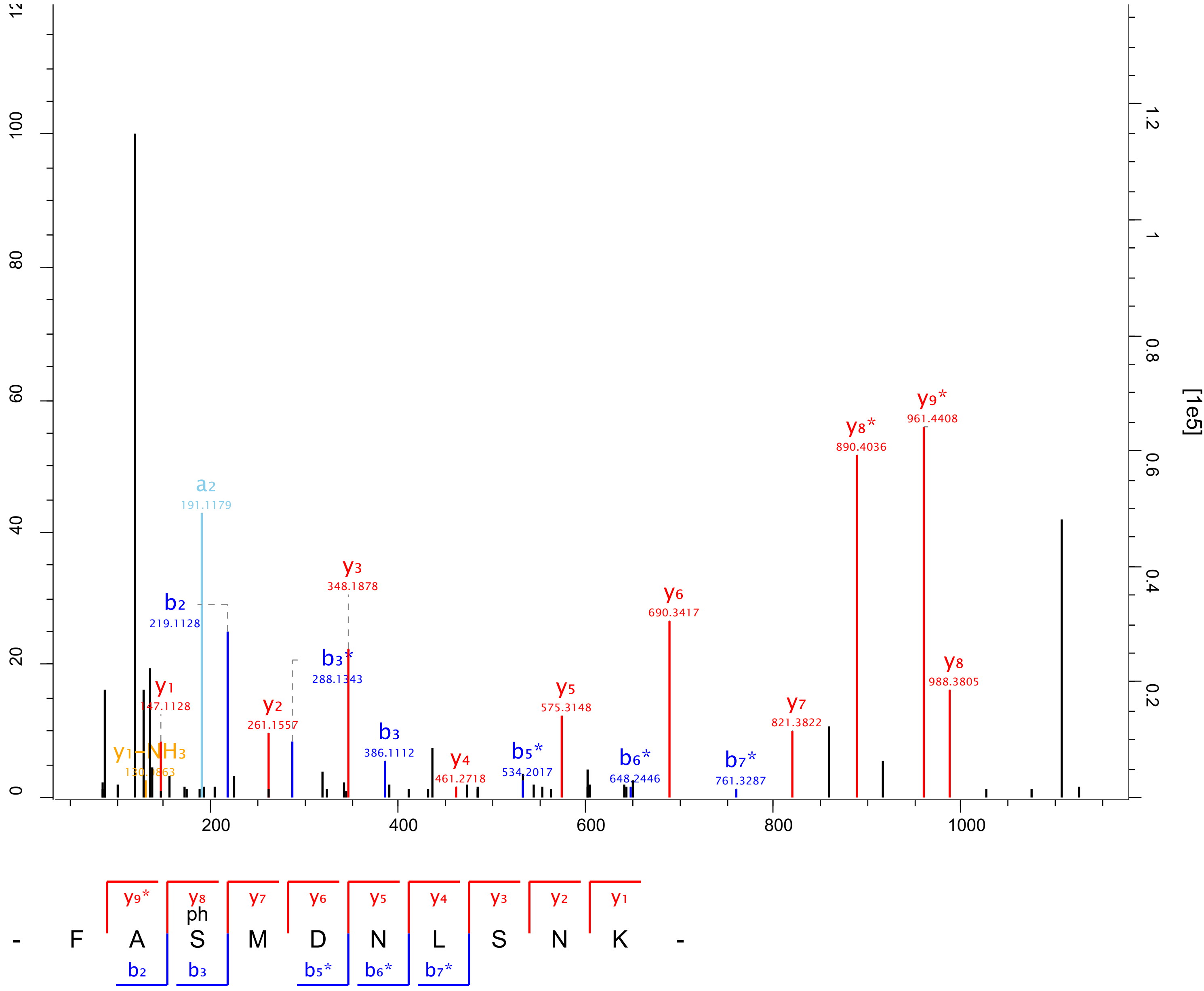

| Raw file | Scan | Method | Score | m/z |
|---|---|---|---|---|
| water_15_2 | 17413 | FTMS; HCD | 58.89 | 657.78 |

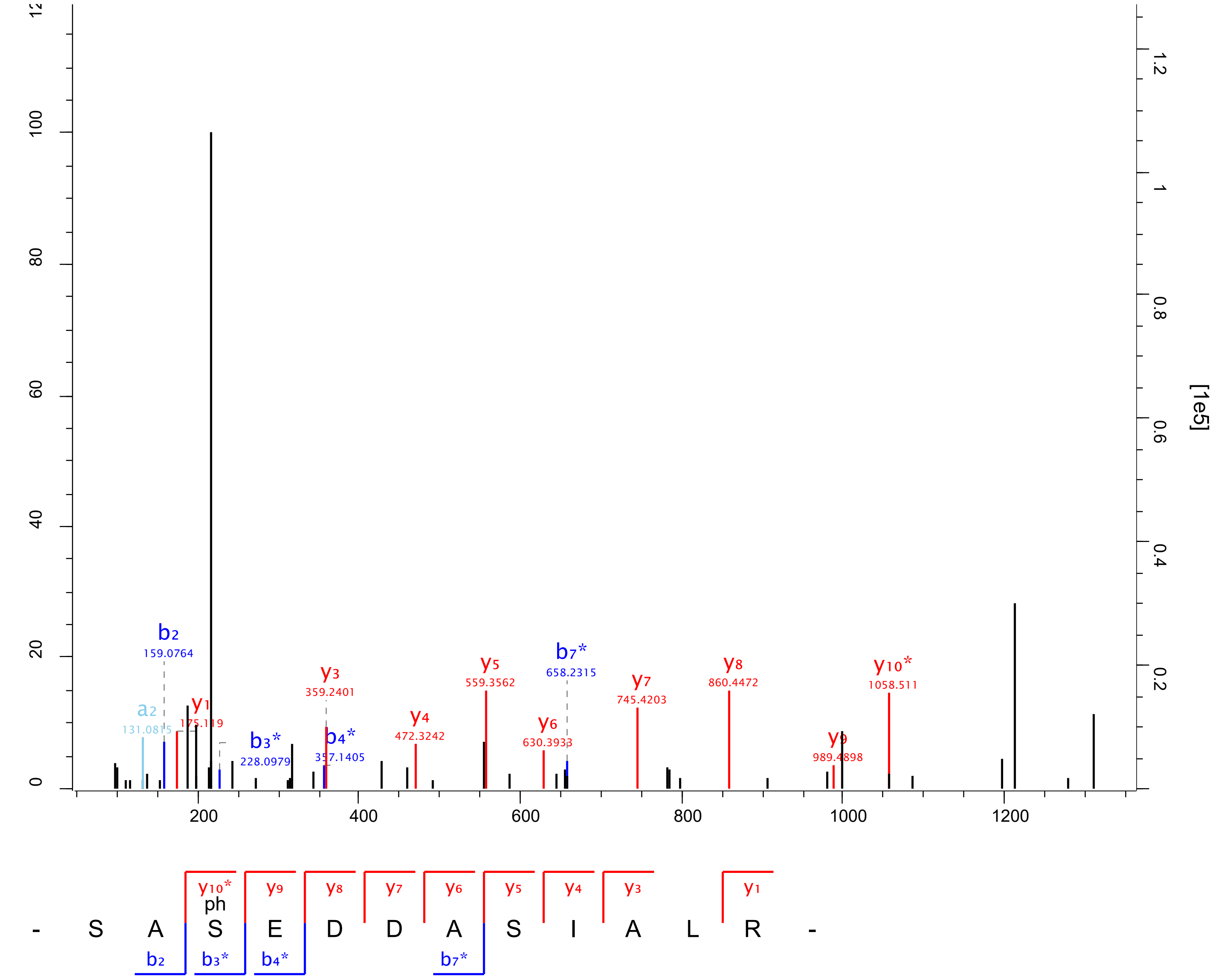

| Raw file | Scan | Method | Score | m/z |
| --- | --- | --- | --- | --- |
| water_15_2 | 17498 | FTMS; HCD | 59.23 | 596.28 |

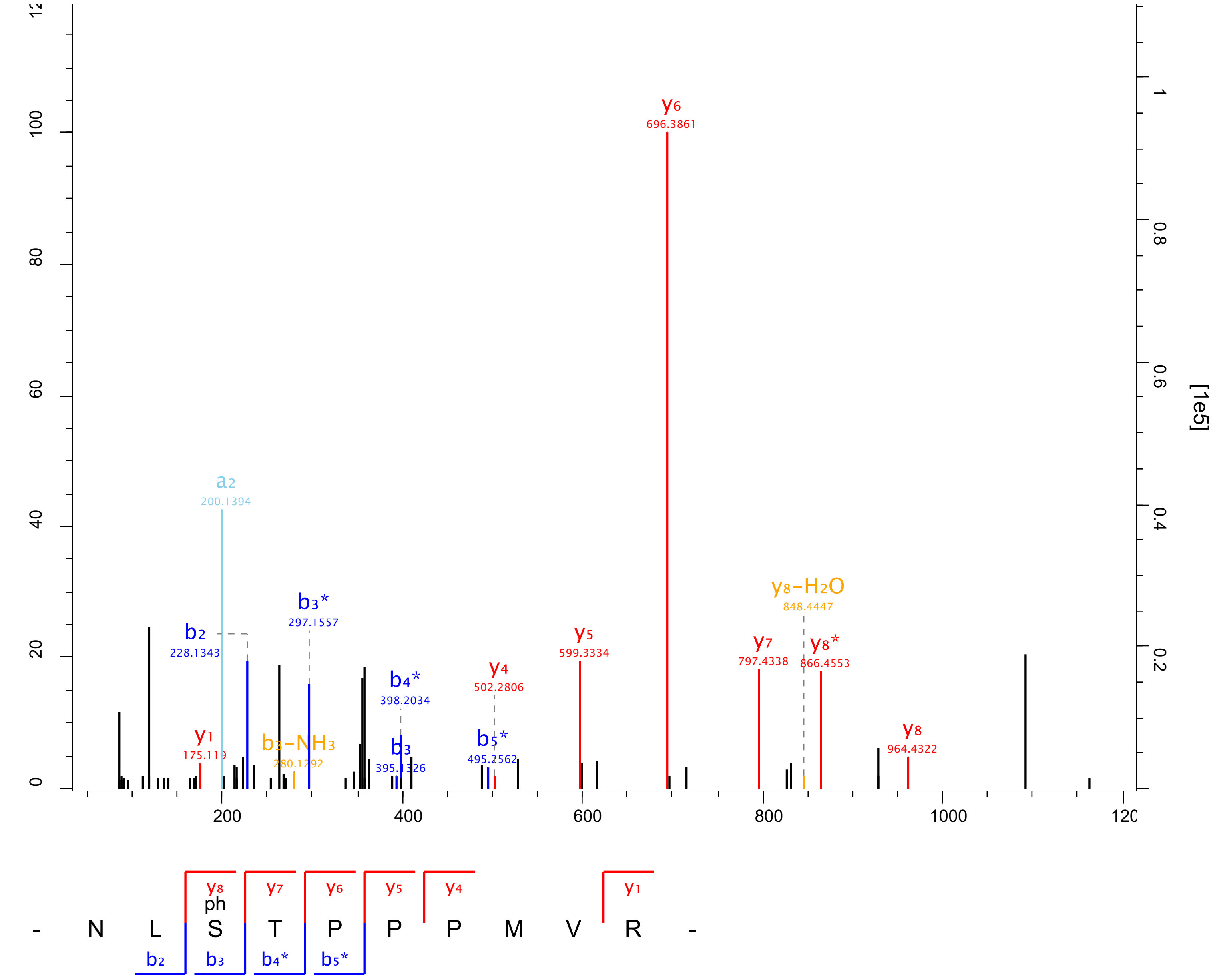

| Raw file | Scan | Method | Score | m/z |
|---|---|---|---|---|
| water_15_2 | 17515 | FTMS; HCD | 57.29 | 740.33 |

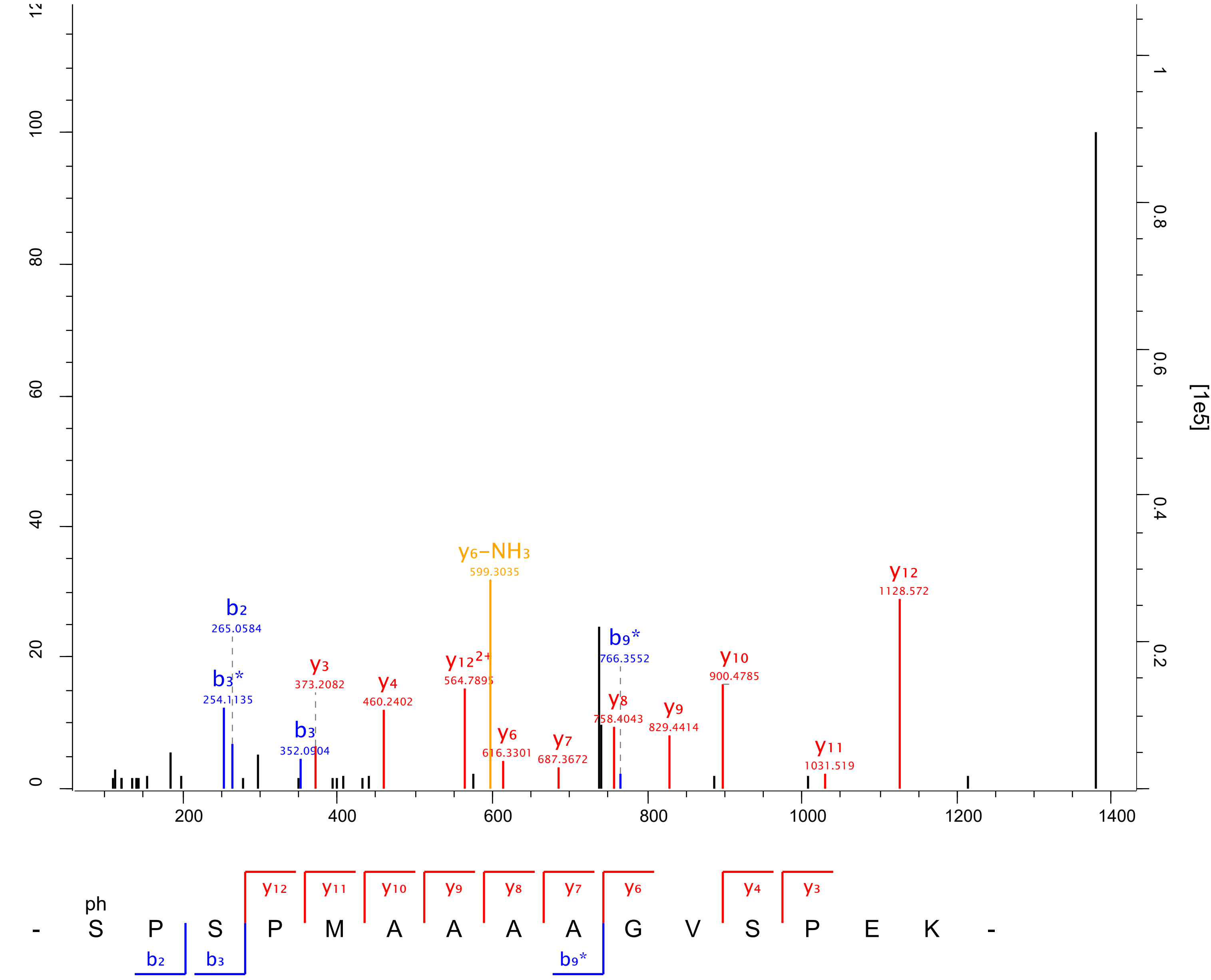

| Raw file | Scan | Method | Score | m/z |
|---|---|---|---|---|
| water_15_2 | 17644 | FTMS; HCD | 111.79 | 567.76 |

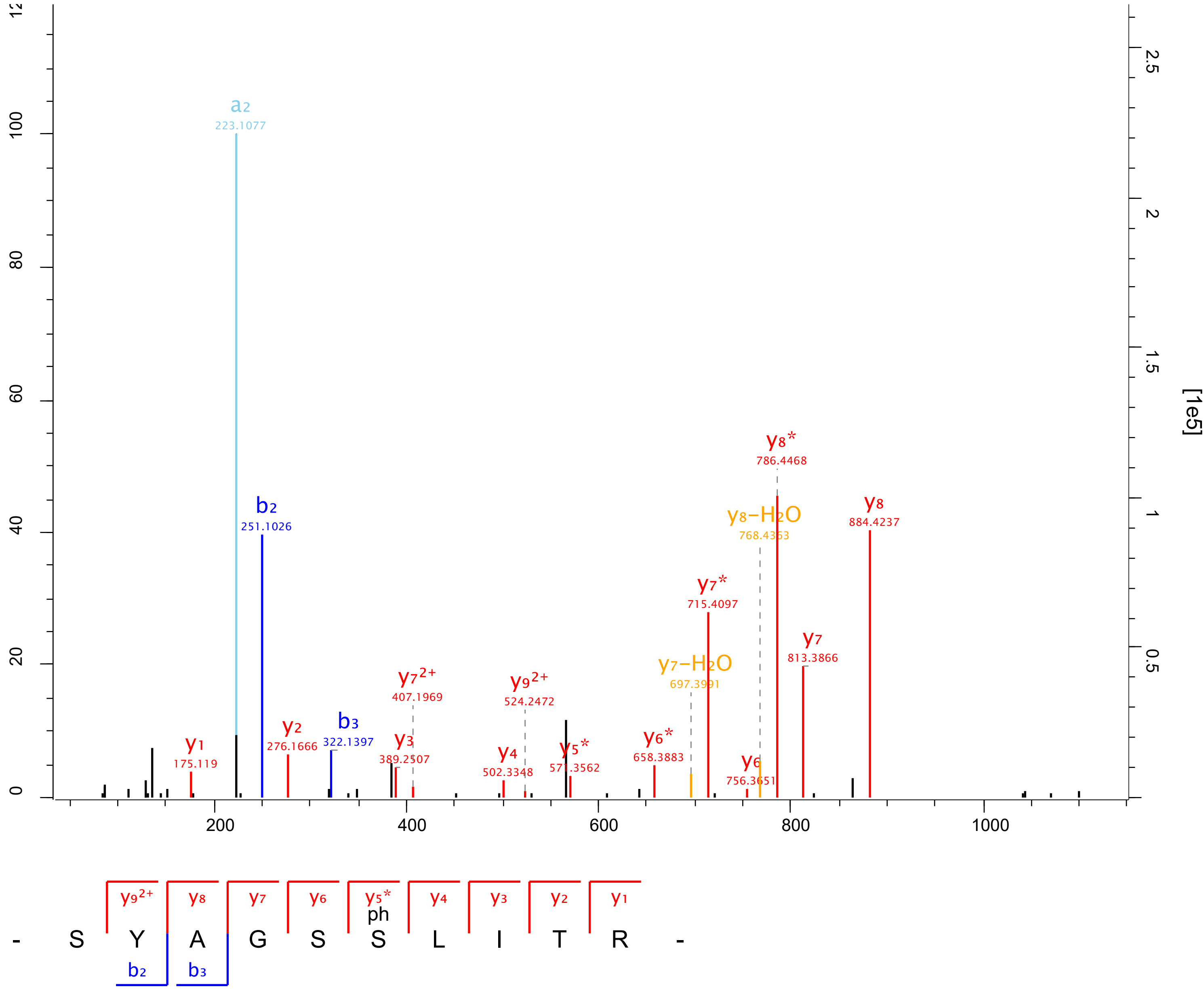

| Raw file | Scan | Method | Score | m/z |
|---|---|---|---|---|
| water_15_2 | 17692 | FTMS; HCD | 88.32 | 760.27 |

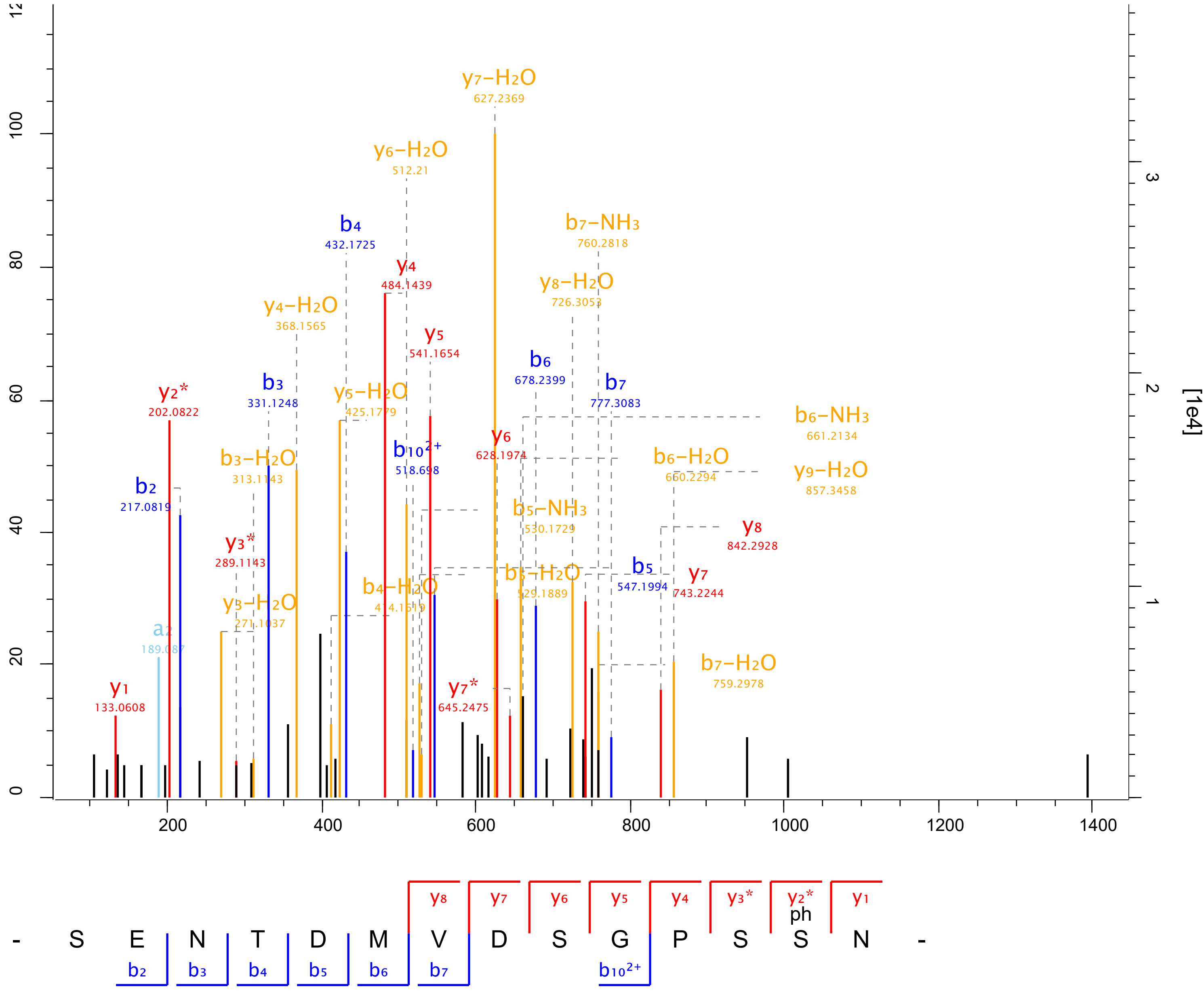

| Raw file | Scan | Method | Score | m/z |
|---|---|---|---|---|
| water_15_2 | 17871 | FTMS; HCD | 105.5 | 544.57 |

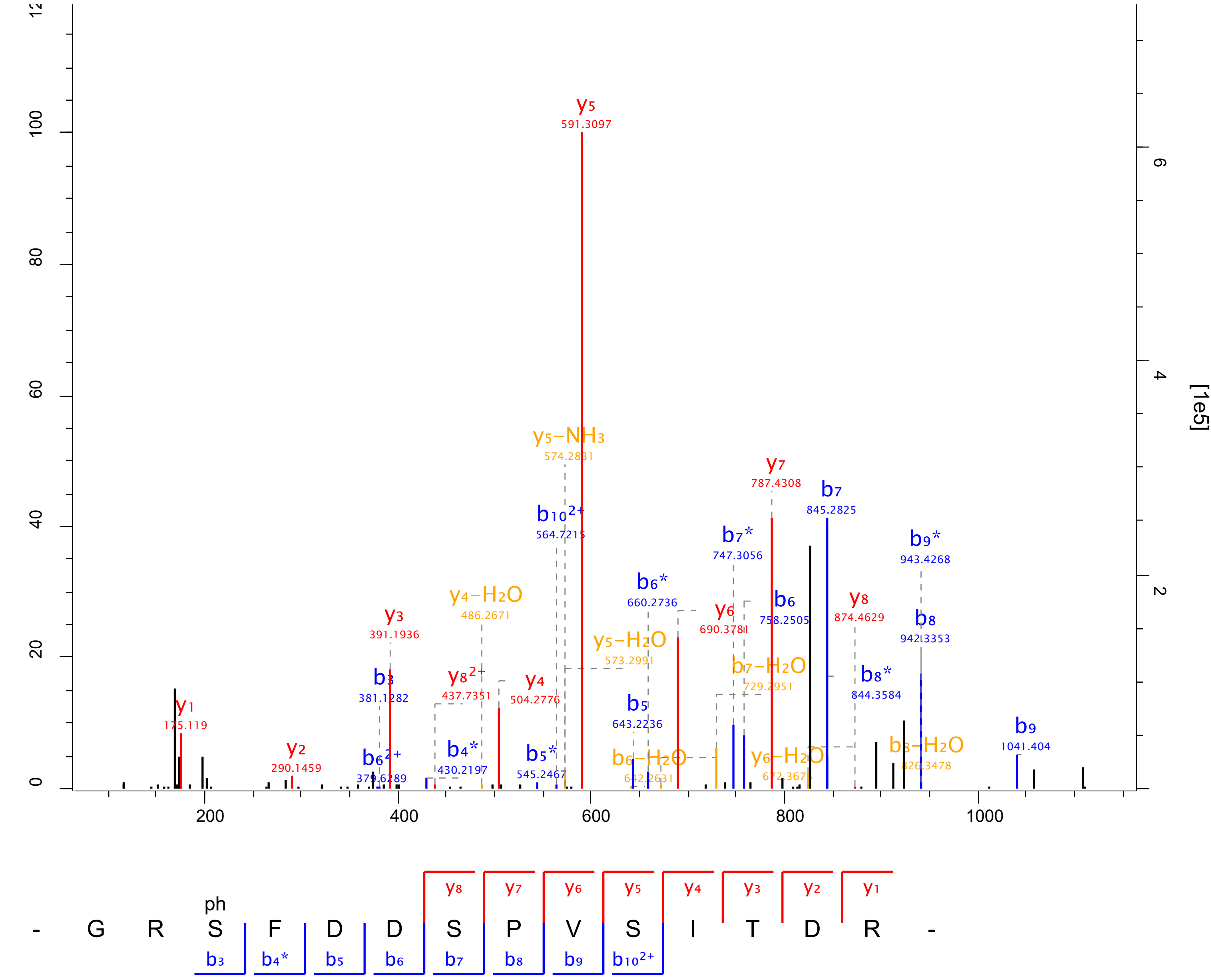

| Raw file | Scan | Method | Score | m/z |
|---|---|---|---|---|
| water_15_2 | 18038 | FTMS; HCD | 45.62 | 702.8 |

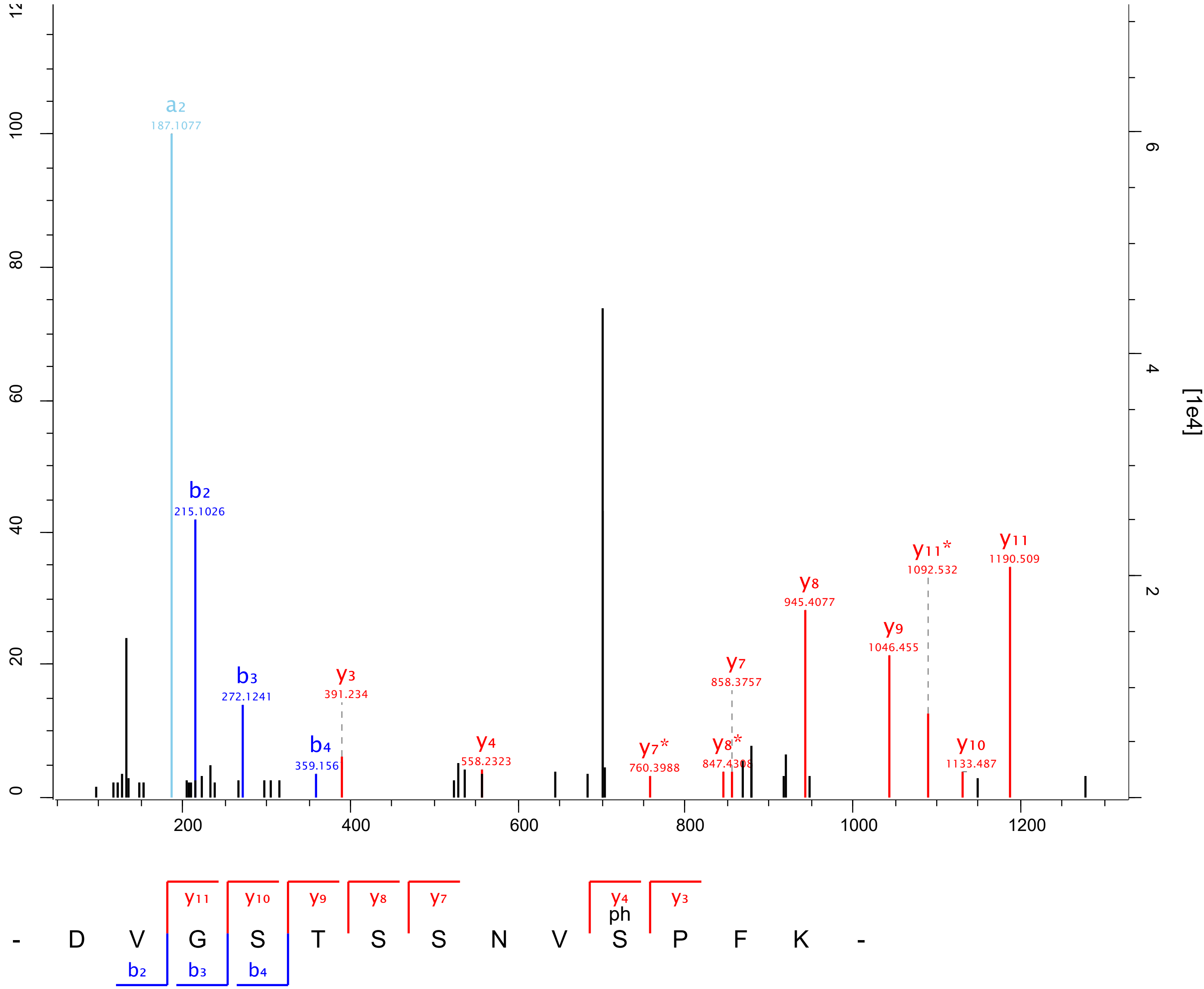

| Raw file | Scan | Method | Score | m/z |
|---|---|---|---|---|
| water_15_2 | 18124 | FTMS; HCD | 135.94 | 574.77 |

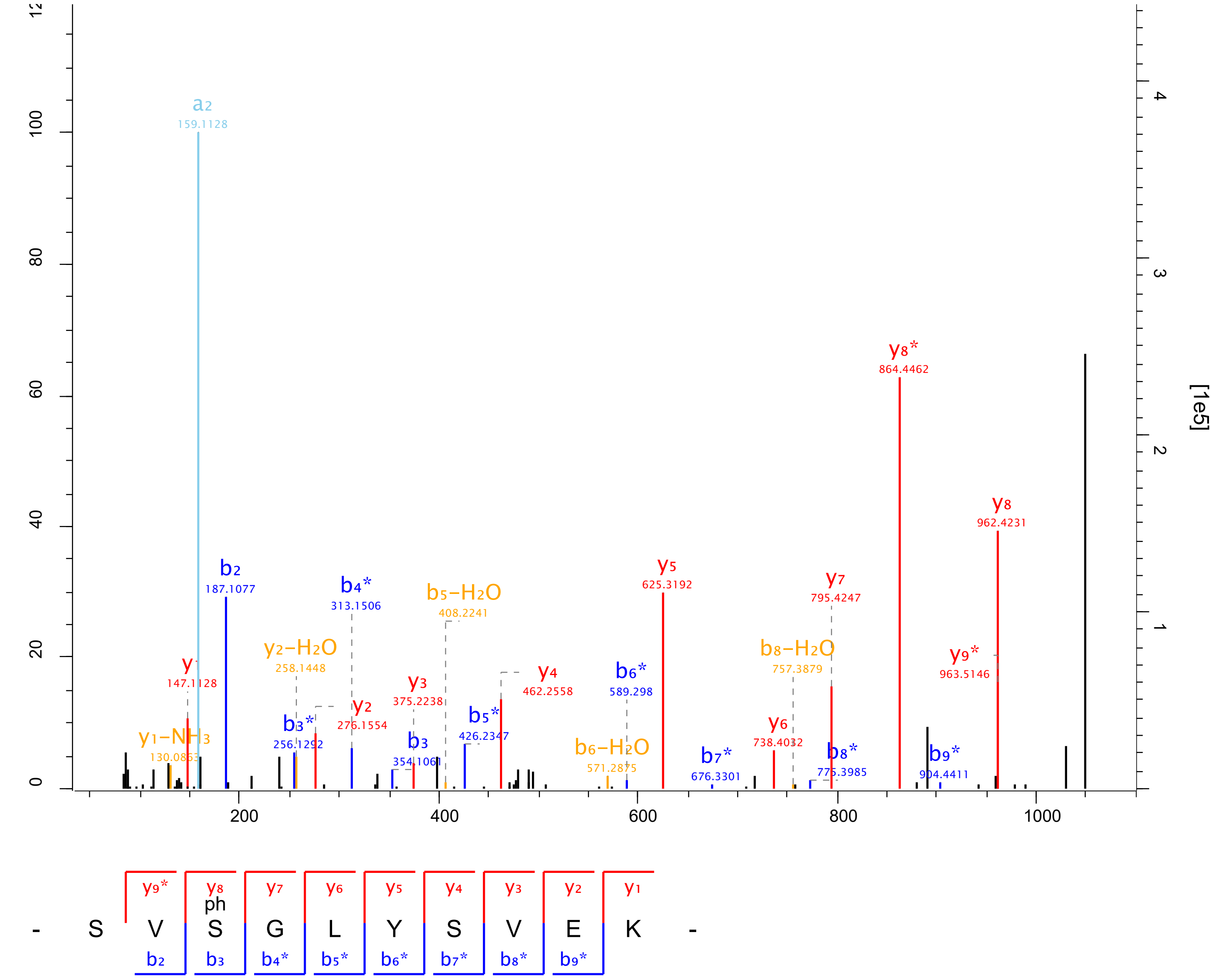

| Raw file | Scan | Method | Score | m/z |
|---|---|---|---|---|
| water_15_2 | 18153 | FTMS; HCD | 52.81 | 852.84 |

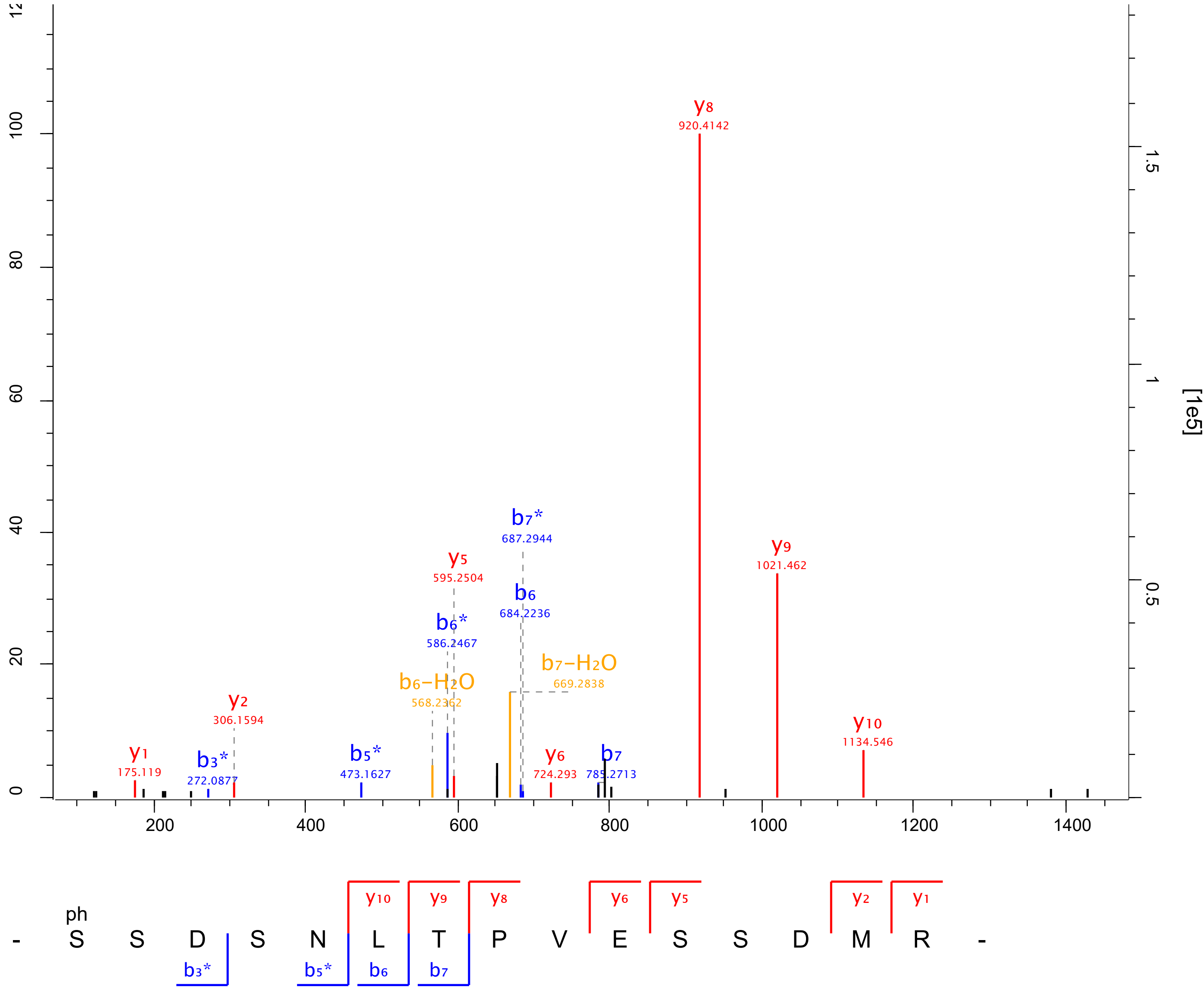

| Raw file | Scan | Method | Score | m/z |
| --- | --- | --- | --- | --- |
| water_15_2 | 18226 | FTMS; HCD | 47.51 | 698.79 |

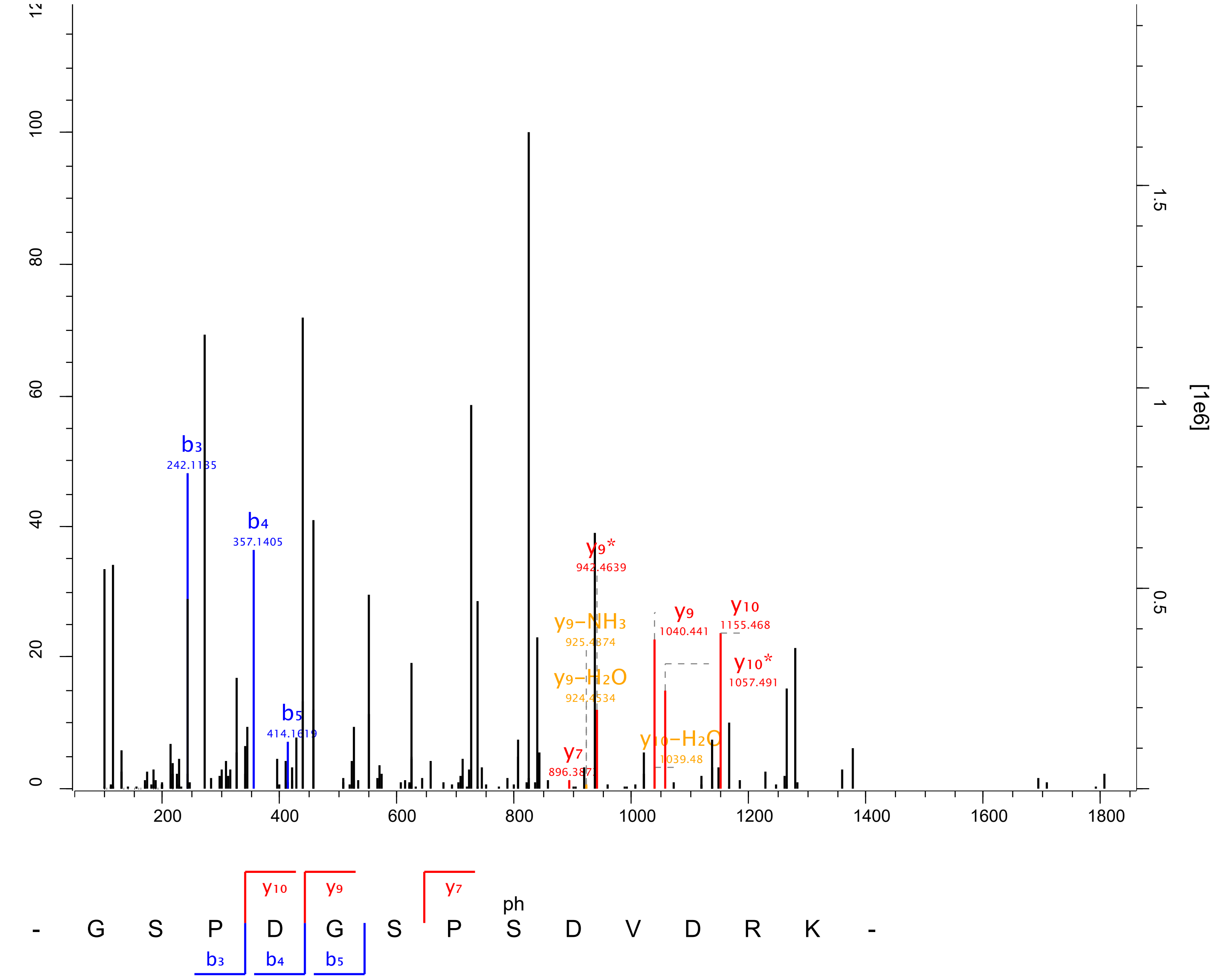

| Raw file | Scan | Method | Score | m/z |
| --- | --- | --- | --- | --- |
| water_15_2 | 18434 | FTMS; HCD | 179.5 | 635.28 |

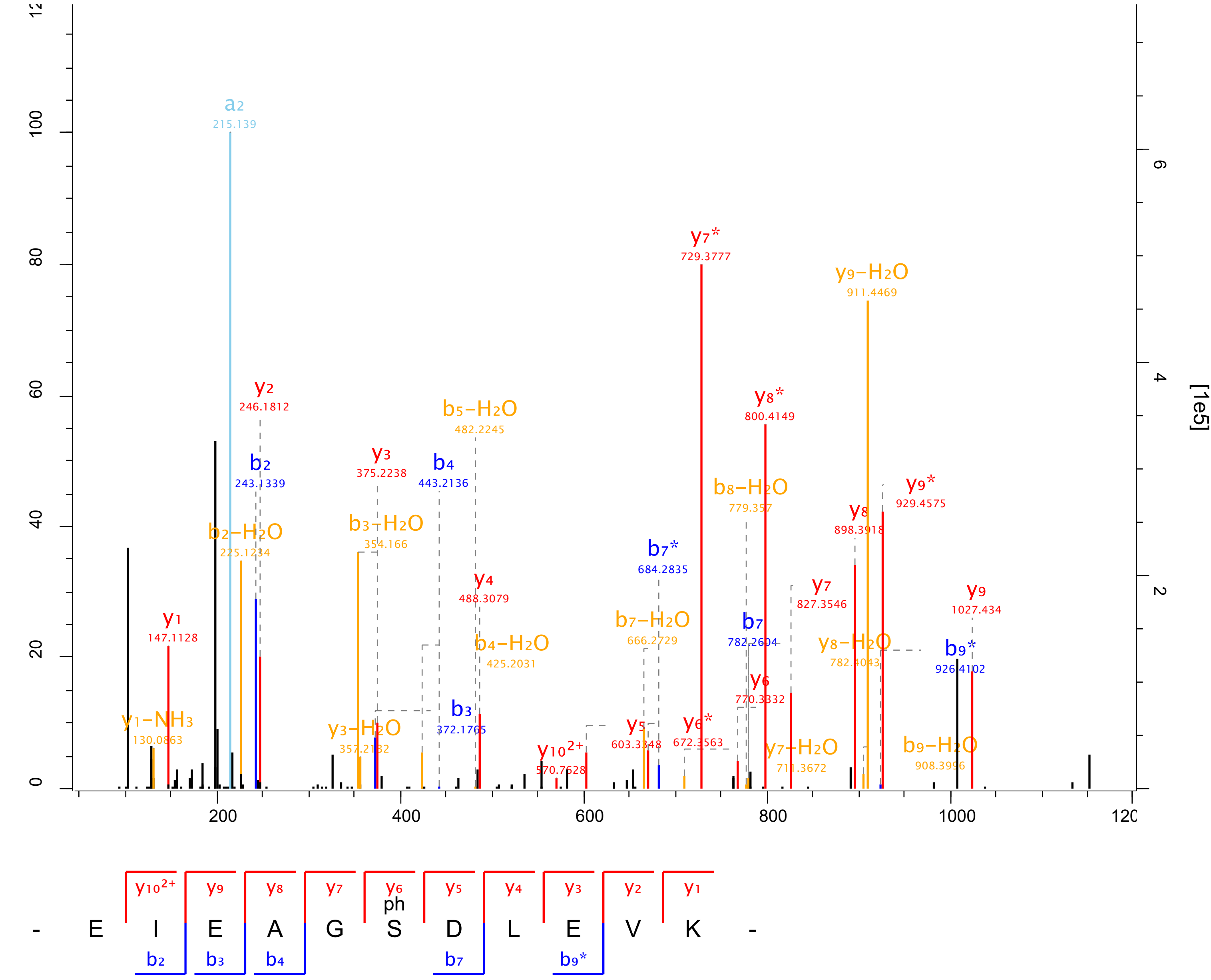

| Raw file | Scan | Method | Score | m/z |
|---|---|---|---|---|
| water_15_2 | 18441 | FTMS; HCD | 193.72 | 883.87 |

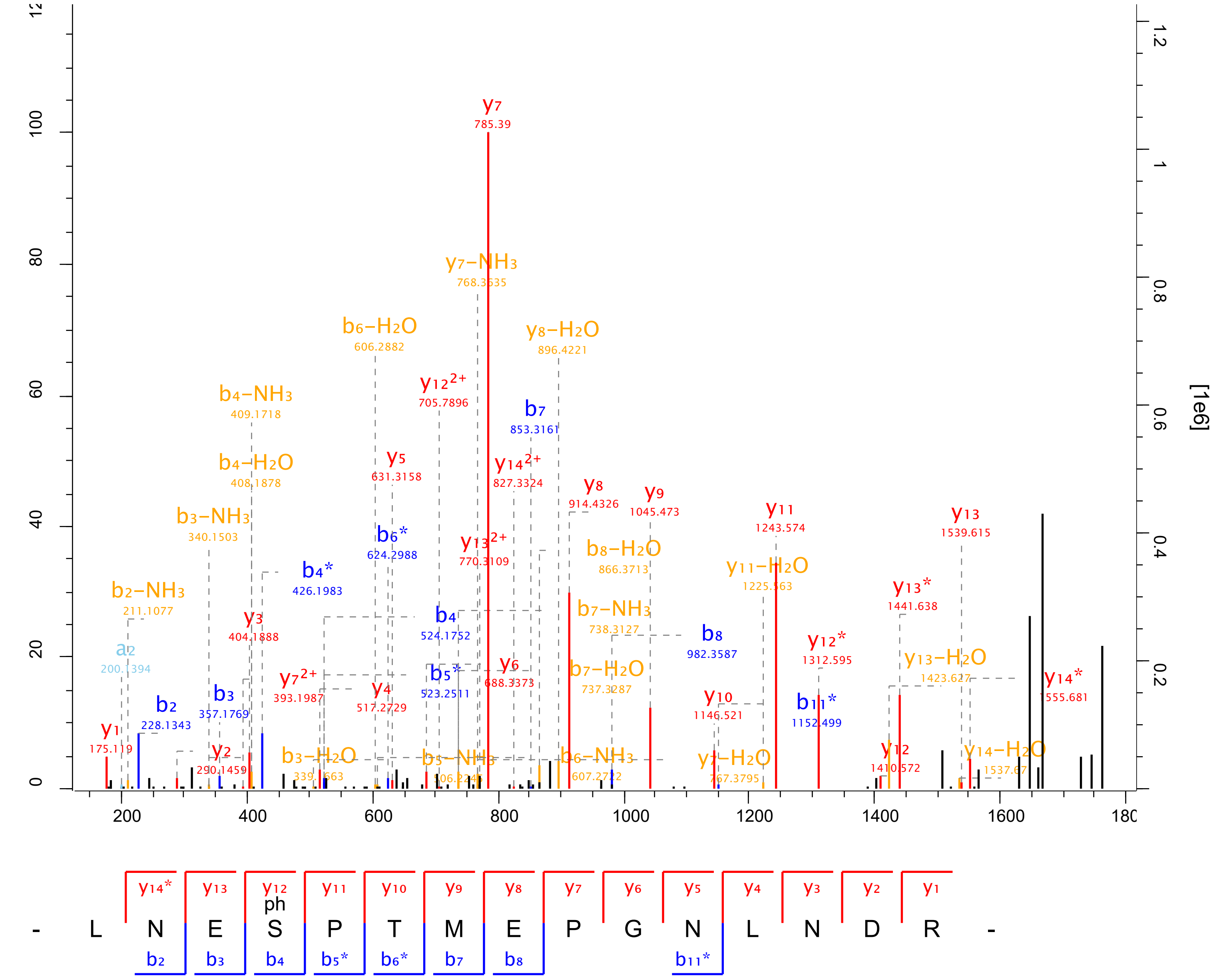

| Raw file | Scan | Method | Score | m/z |
| --- | --- | --- | --- | --- |
| water_15_2 | 18504 | FTMS; HCD | 62.09 | 525.79 |

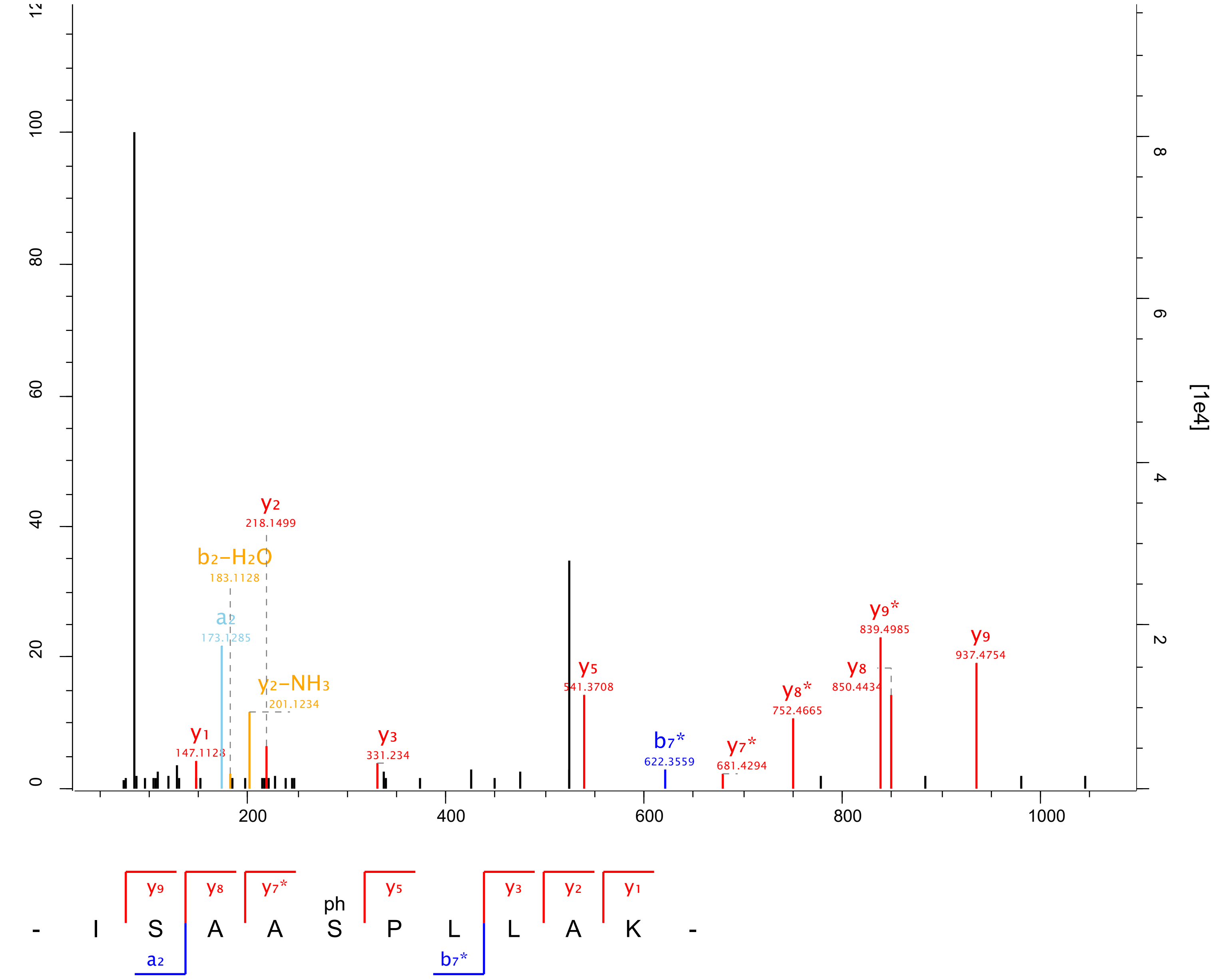

| Raw file | Scan | Method | Score | m/z |
| --- | --- | --- | --- | --- |
| water_15_2 | 18585 | FTMS; HCD | 108.32 | 607.74 |

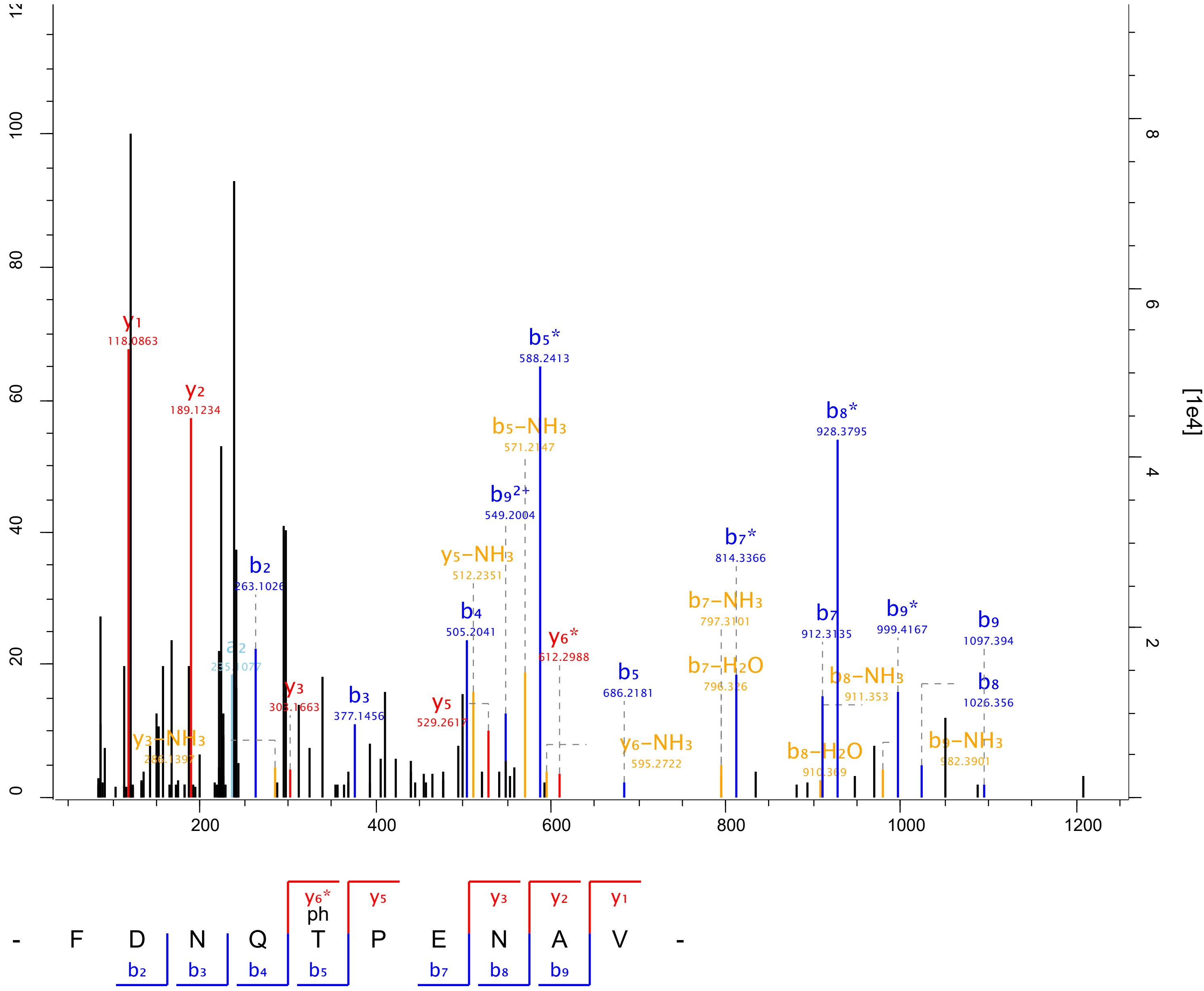

| Raw file | Scan | Method | Score | m/z |
| --- | --- | --- | --- | --- |
| water_15_2 | 18710 | FTMS; HCD | 58.37 | 732.96 |

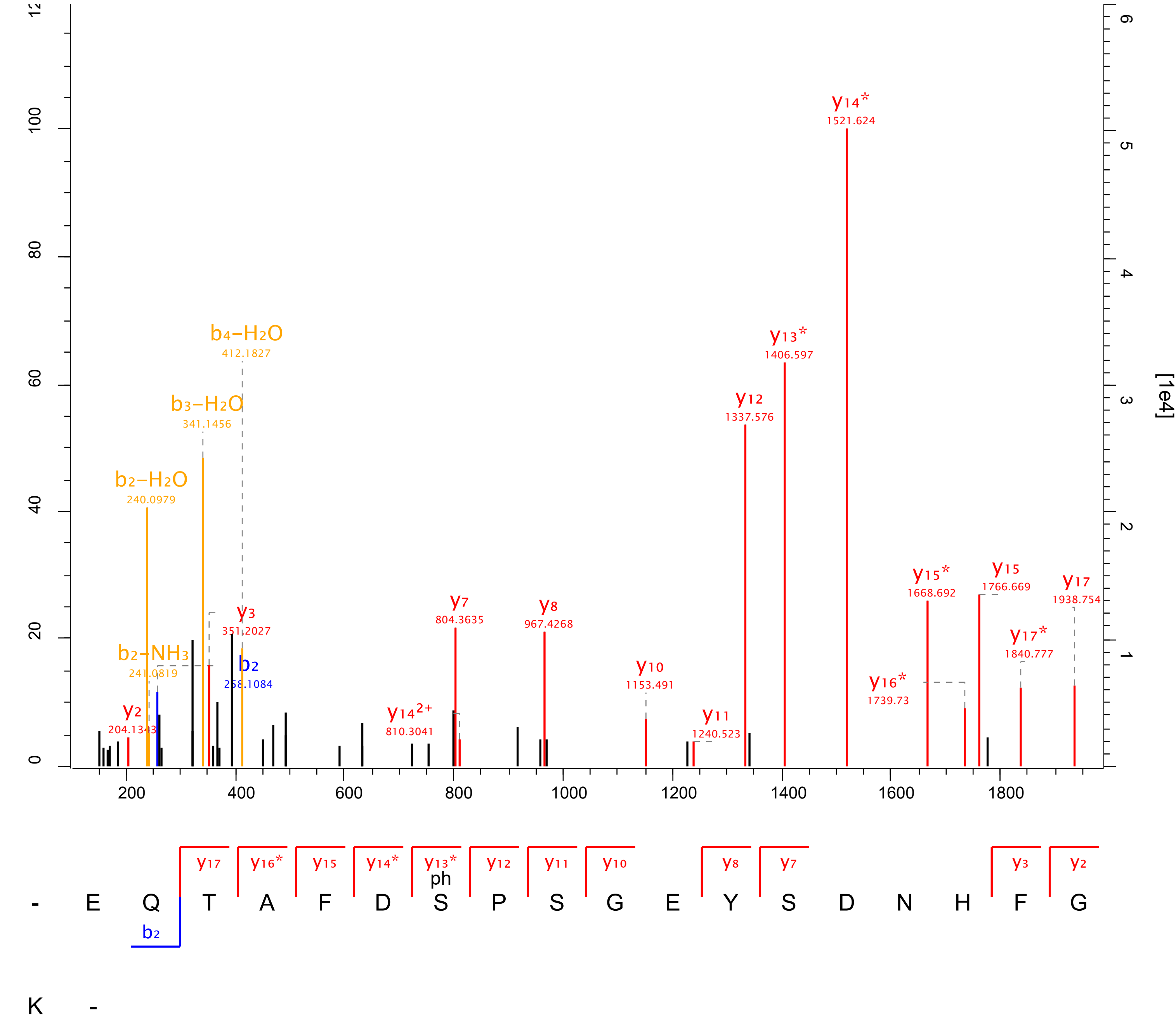

| Raw file | Scan | Method | Score | m/z |
|---|---|---|---|---|
| water_15_2 | 18768 | FTMS; HCD | 138.46 | 787.34 |

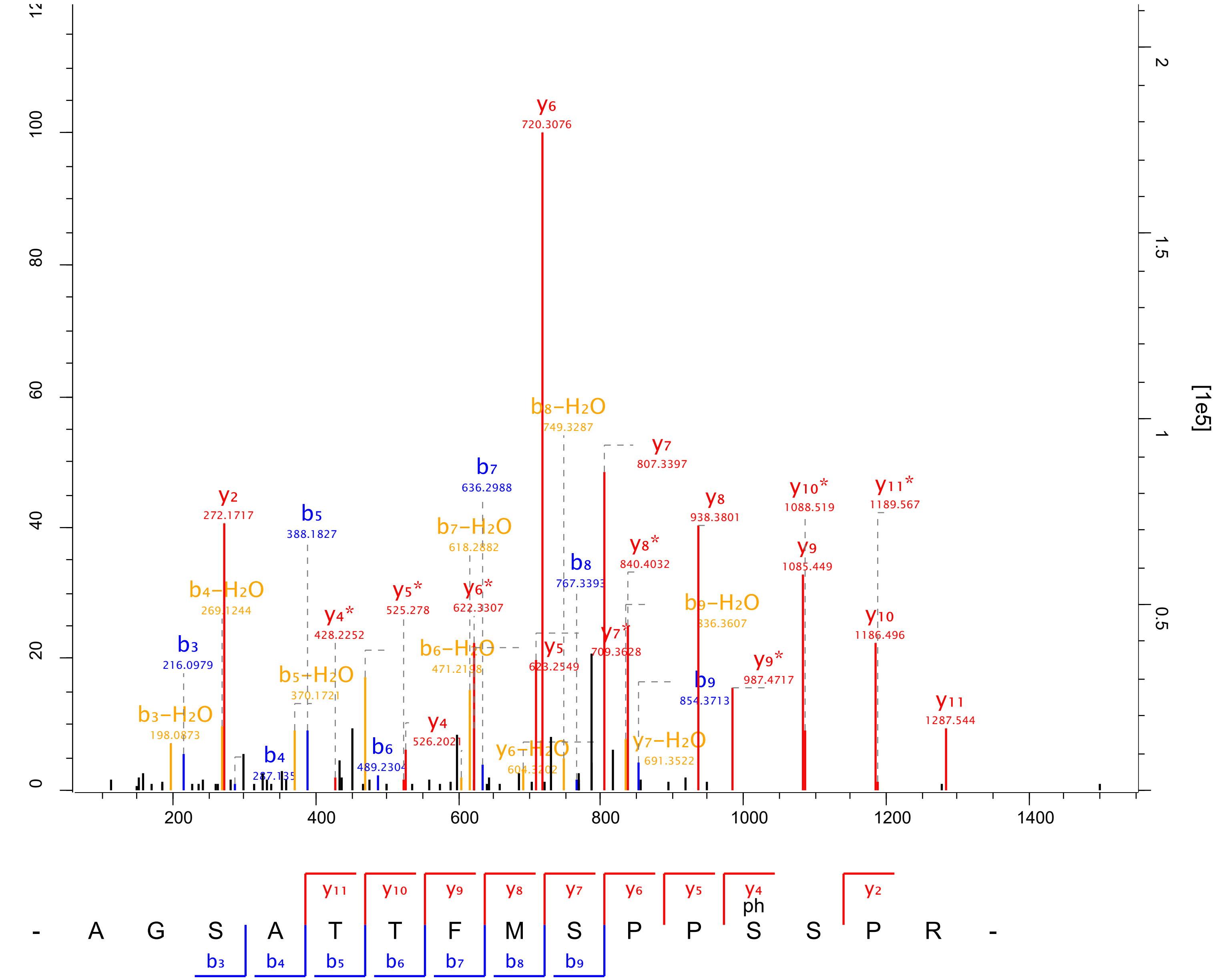

| Raw file | Scan | Method | Score | m/z |
|---|---|---|---|---|
| water_15_2 | 18811 | FTMS; HCD | 53.76 | 492.71 |

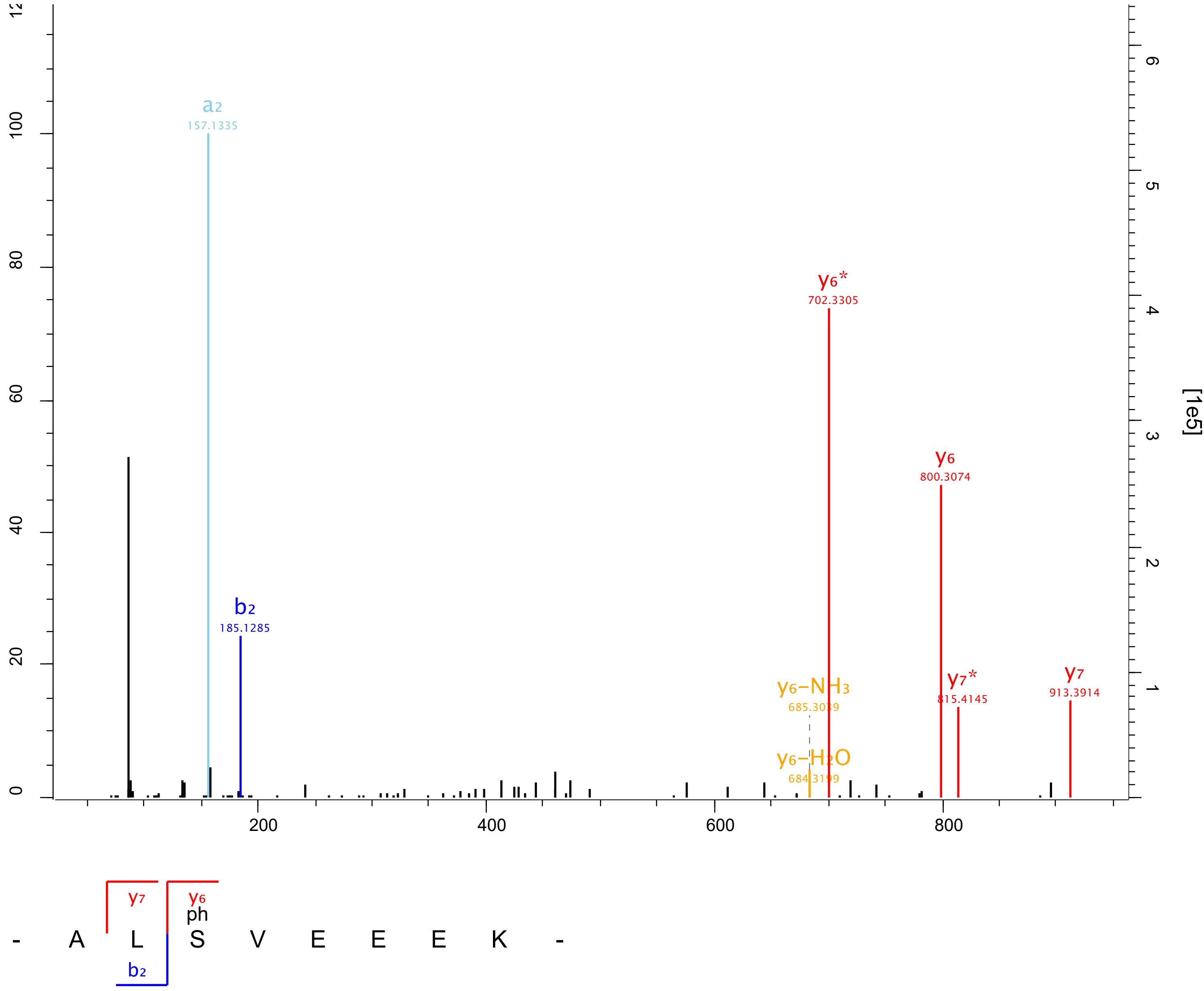

| Raw file | Scan | Method | Score | m/z |
| --- | --- | --- | --- | --- |
| water_15_2 | 18853 | FTMS; HCD | 121.44 | 822.87 |

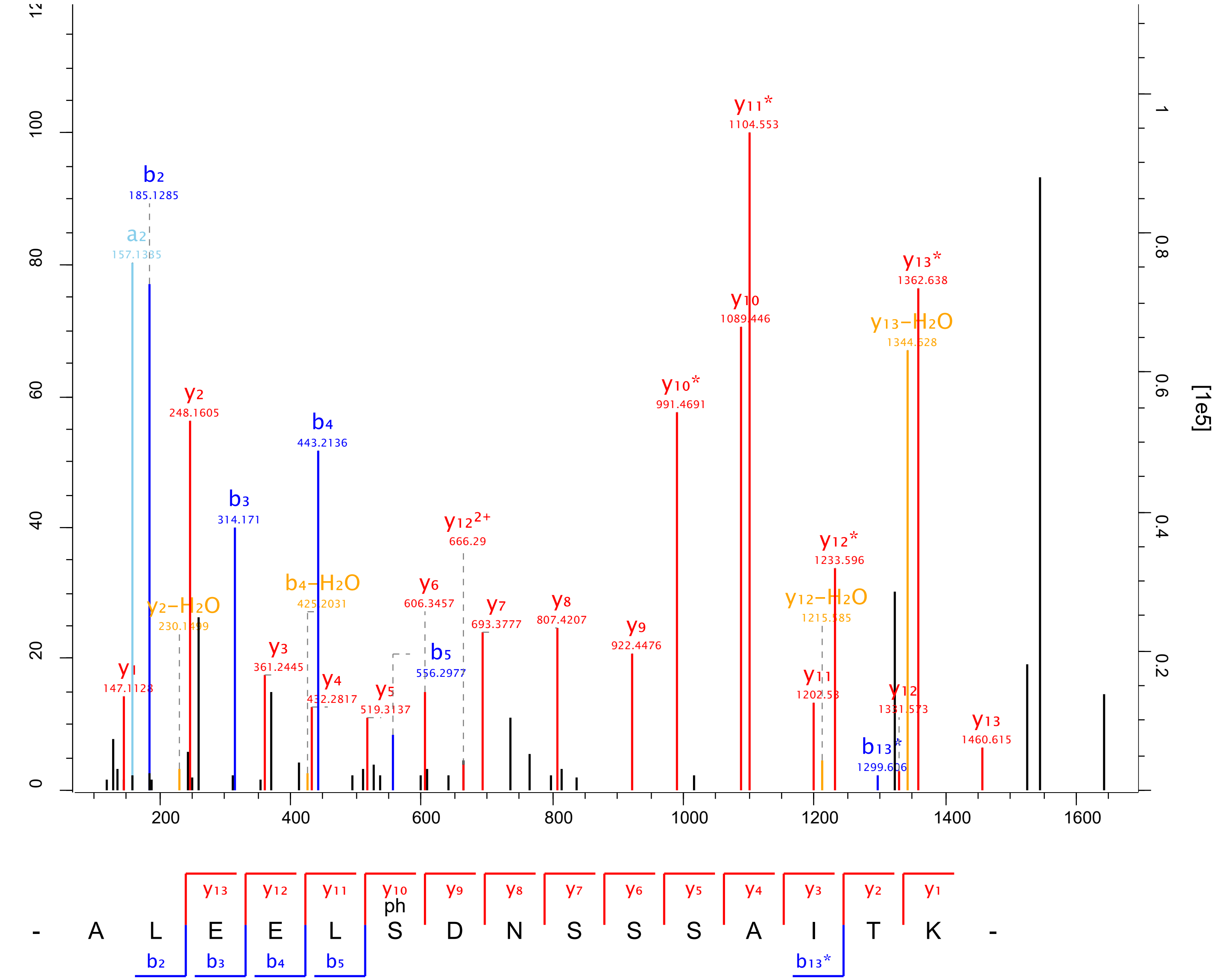

| Raw file | Scan | Method | Score | m/z |
|---|---|---|---|---|
| water_15_2 | 1892 | FTMS; HCD | 139.86 | 427.53 |

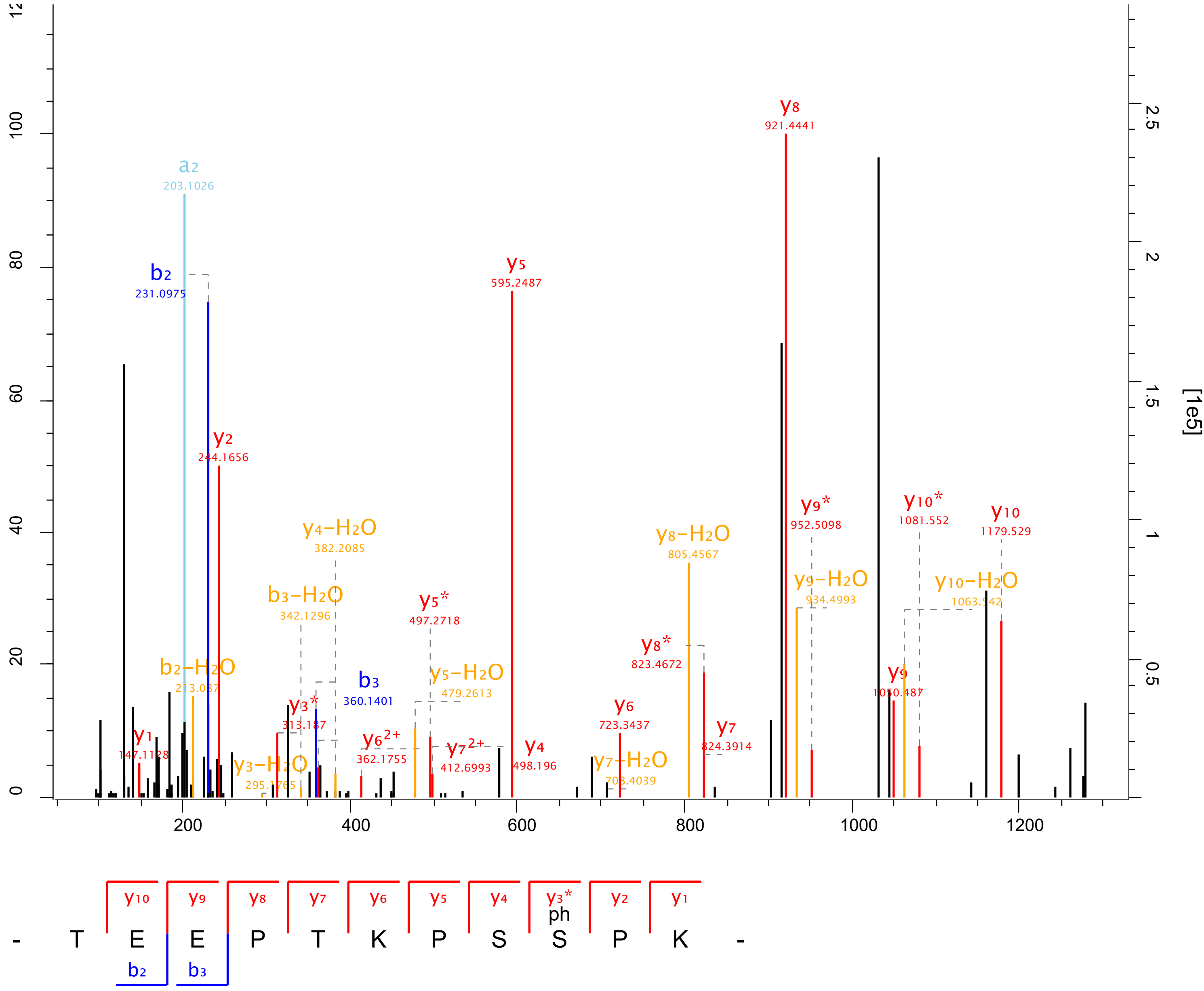

| Raw file | Scan | Method | Score | m/z |
| --- | --- | --- | --- | --- |
| water_15_2 | 19168 | FTMS; HCD | 116.03 | 688.29 |

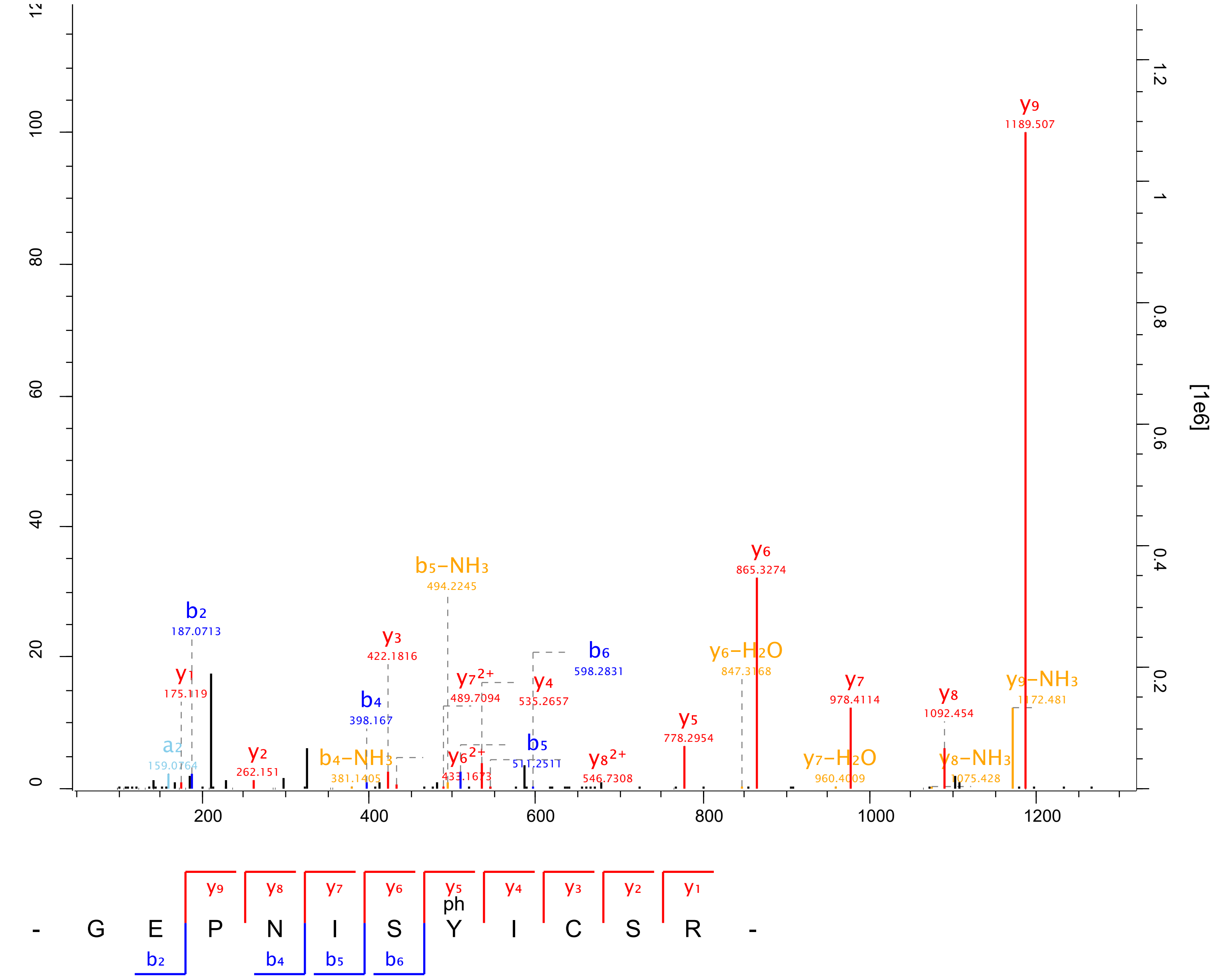

| Raw file | Scan | Method | Score | m/z |
|---|---|---|---|---|
| water_15_2 | 19187 | FTMS; HCD | 199.09 | 857.85 |

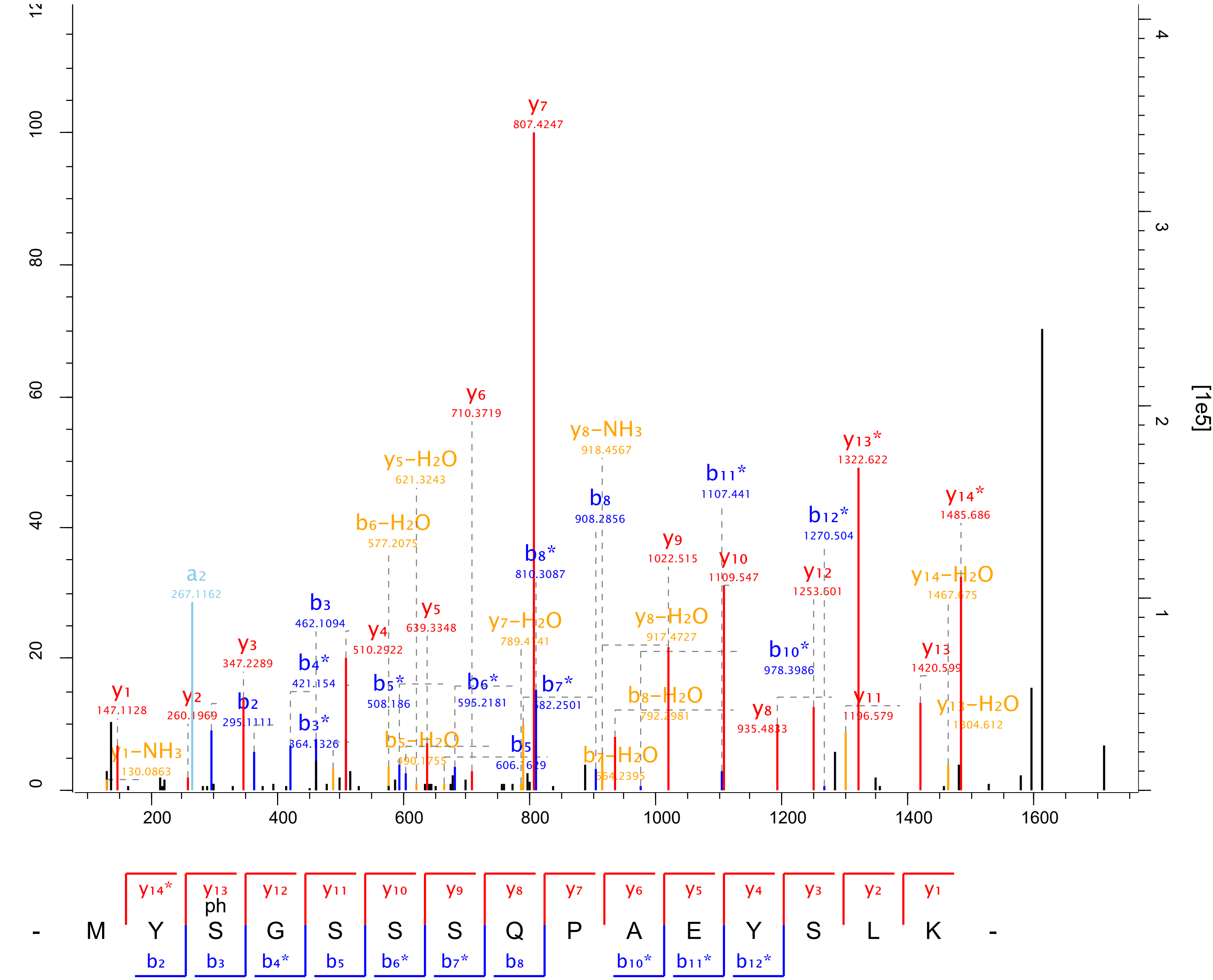

| Raw file | Scan | Method | Score | m/z |
| --- | --- | --- | --- | --- |
| water_15_2 | 19415 | FTMS; HCD | 111.77 | 806.88 |

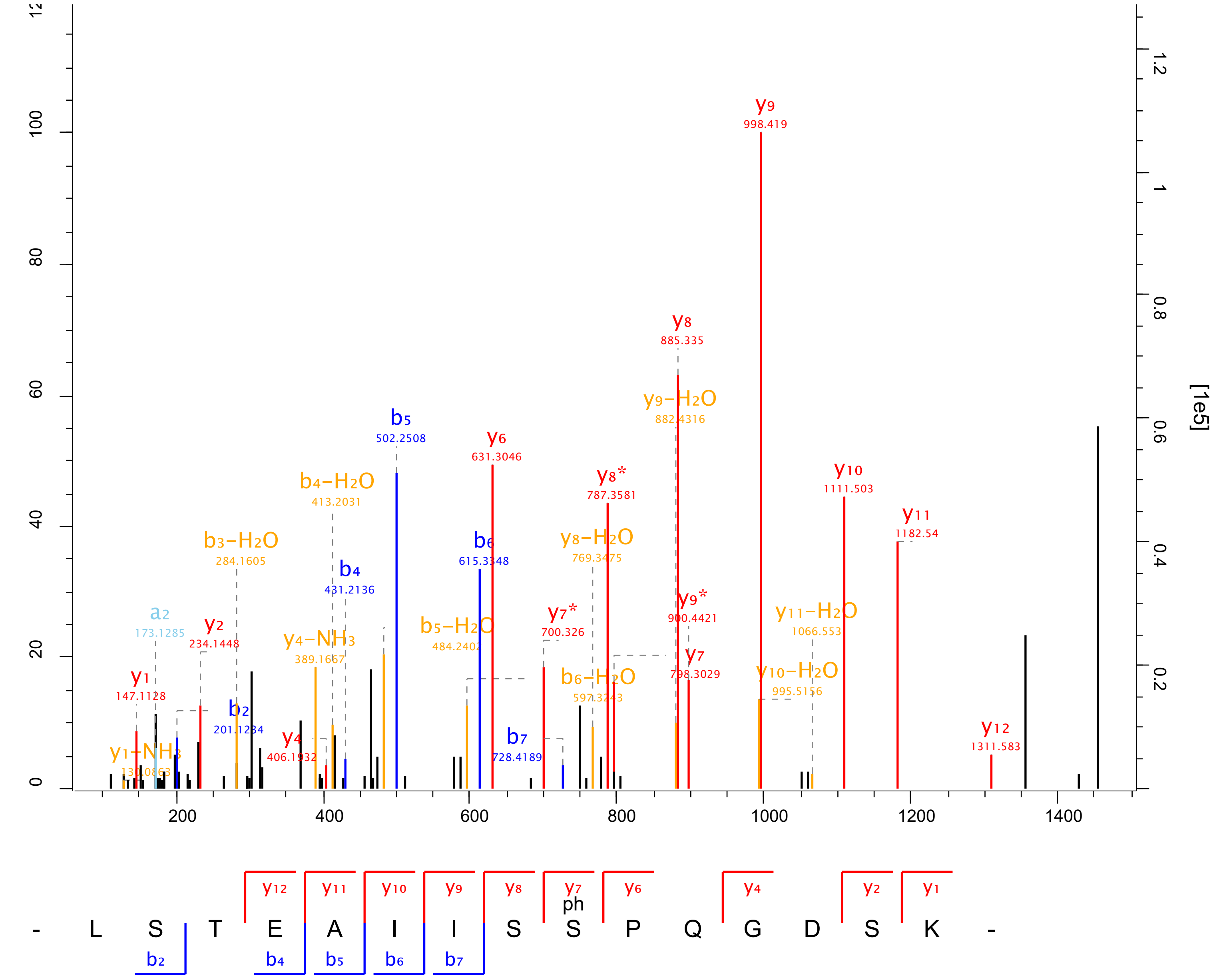

| Raw file | Scan | Method | Score | m/z |
|---|---|---|---|---|
| water_15_2 | 19474 | FTMS; HCD | 41.7 | 705.78 |

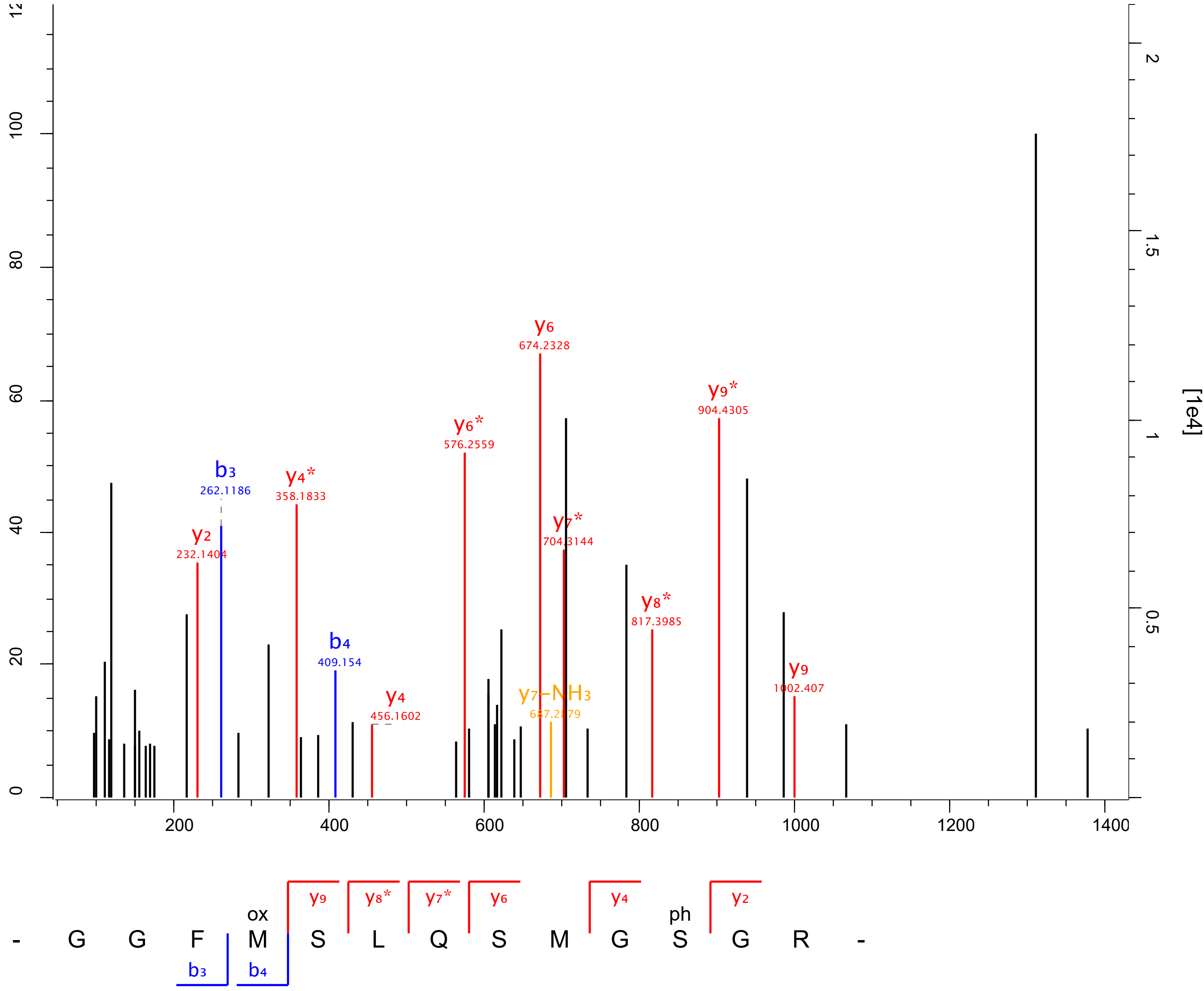

| Raw file | Scan | Method | Score | m/z |
| --- | --- | --- | --- | --- |
| water_15_2 | 19523 | FTMS; HCD | 93.32 | 506.73 |

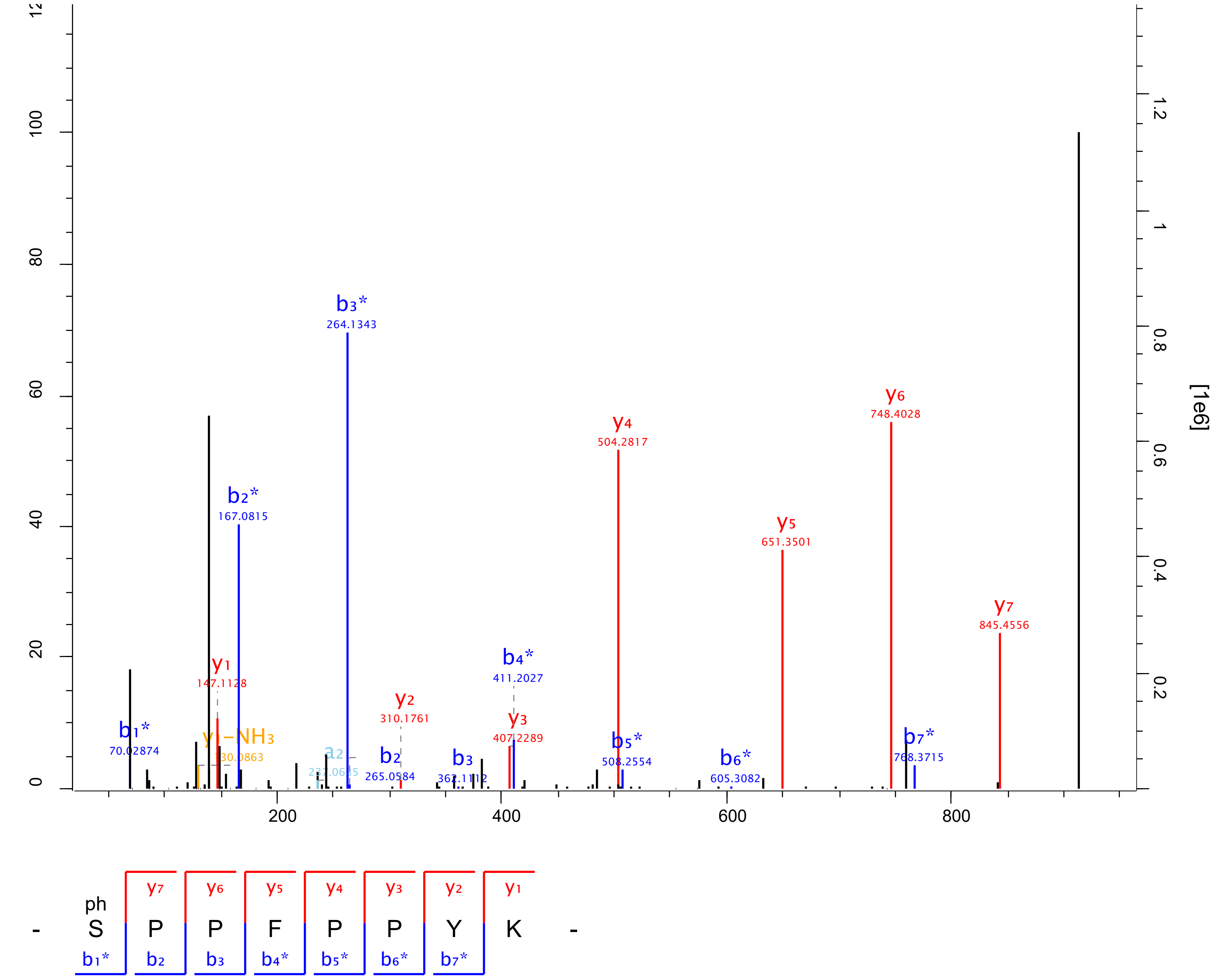

| Raw file | Scan | Method | Score | m/z |
| --- | --- | --- | --- | --- |
| water_15_2 | 19558 | FTMS; HCD | 68.48 | 768.31 |

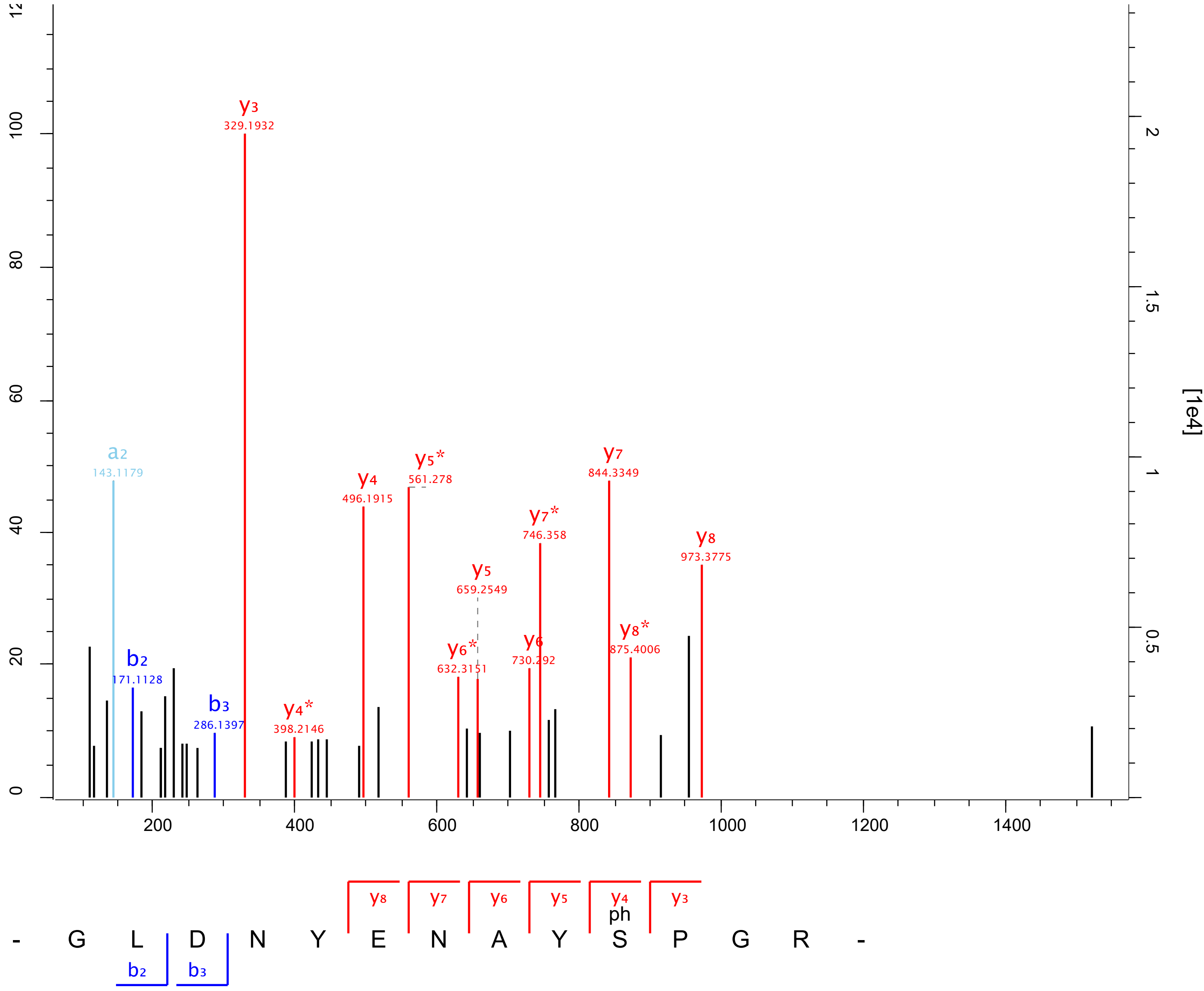

| Raw file | Scan | Method | Score | m/z |
| --- | --- | --- | --- | --- |
| water_15_2 | 19767 | FTMS; HCD | 61.96 | 766.83 |

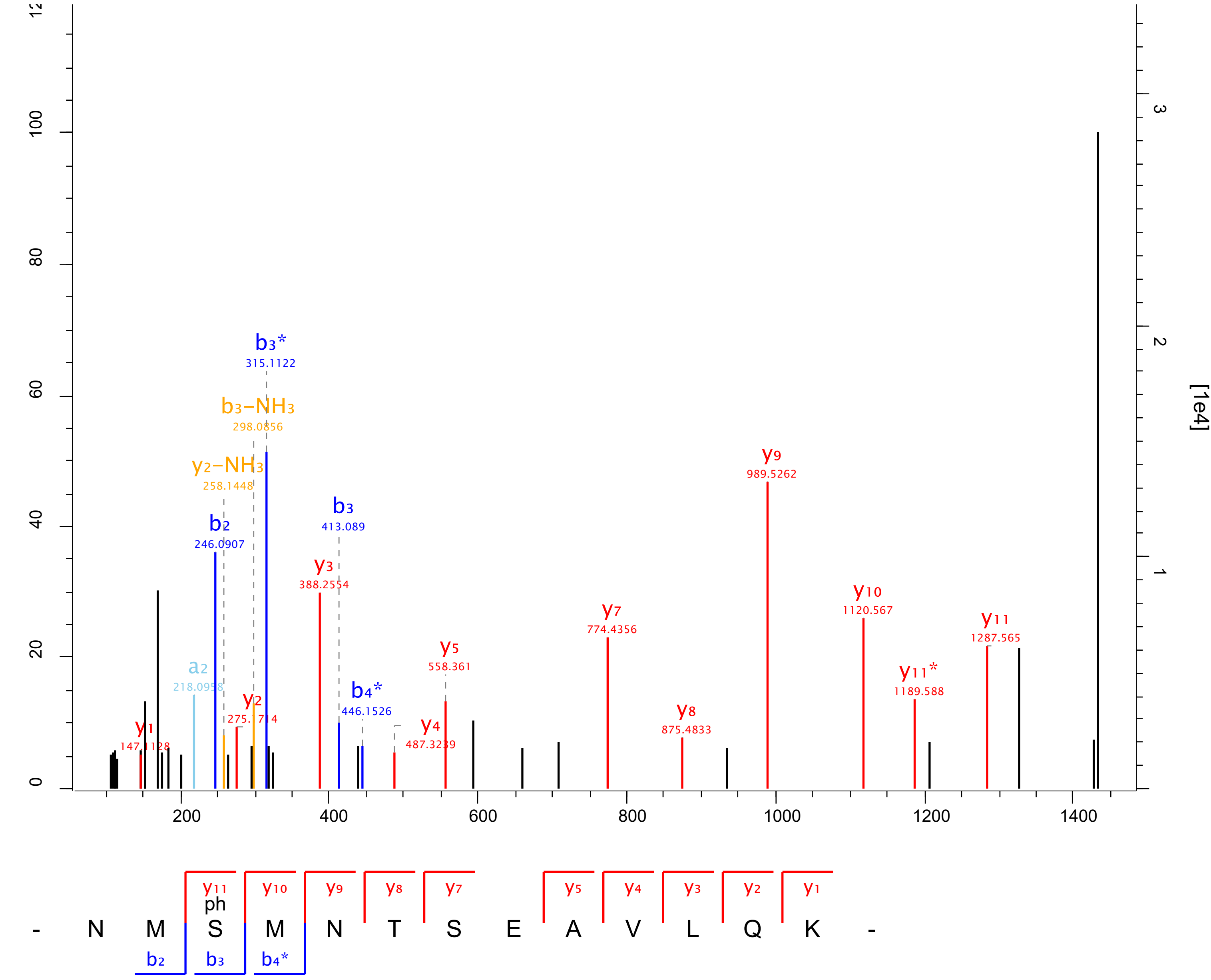

| Raw file | Scan | Method | Score | m/z |
| --- | --- | --- | --- | --- |
| water_15_2 | 19987 | FTMS; HCD | 202.62 | 938.36 |

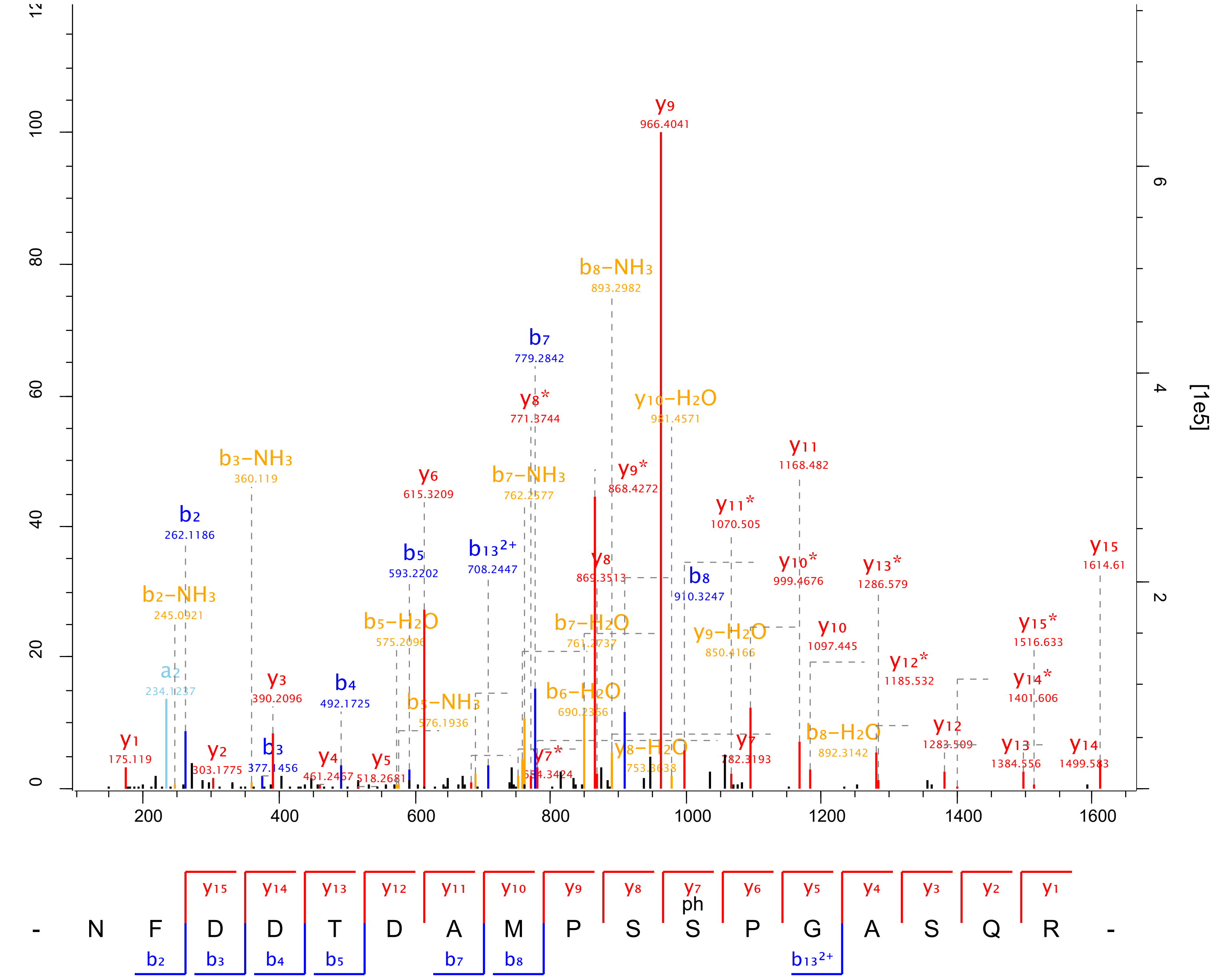

| Raw file | Scan | Method | Score | m/z |
|---|---|---|---|---|
| water_15_2 | 20051 | FTMS; HCD | 113.13 | 1080.92 |

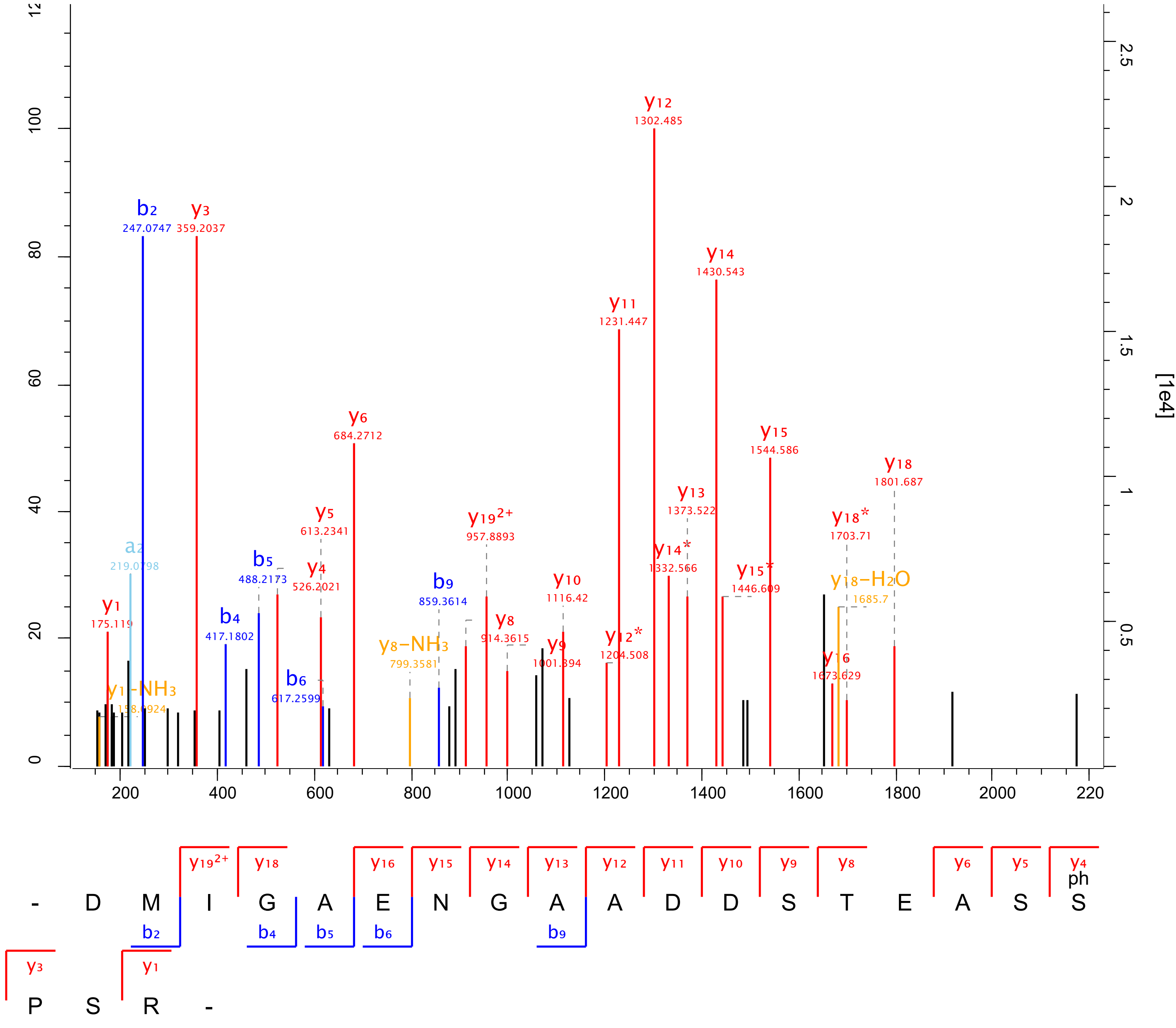

| Raw file | Scan | Method | Score | m/z |
| --- | --- | --- | --- | --- |
| water_15_2 | 20081 | FTMS; HCD | 57.29 | 569.27 |

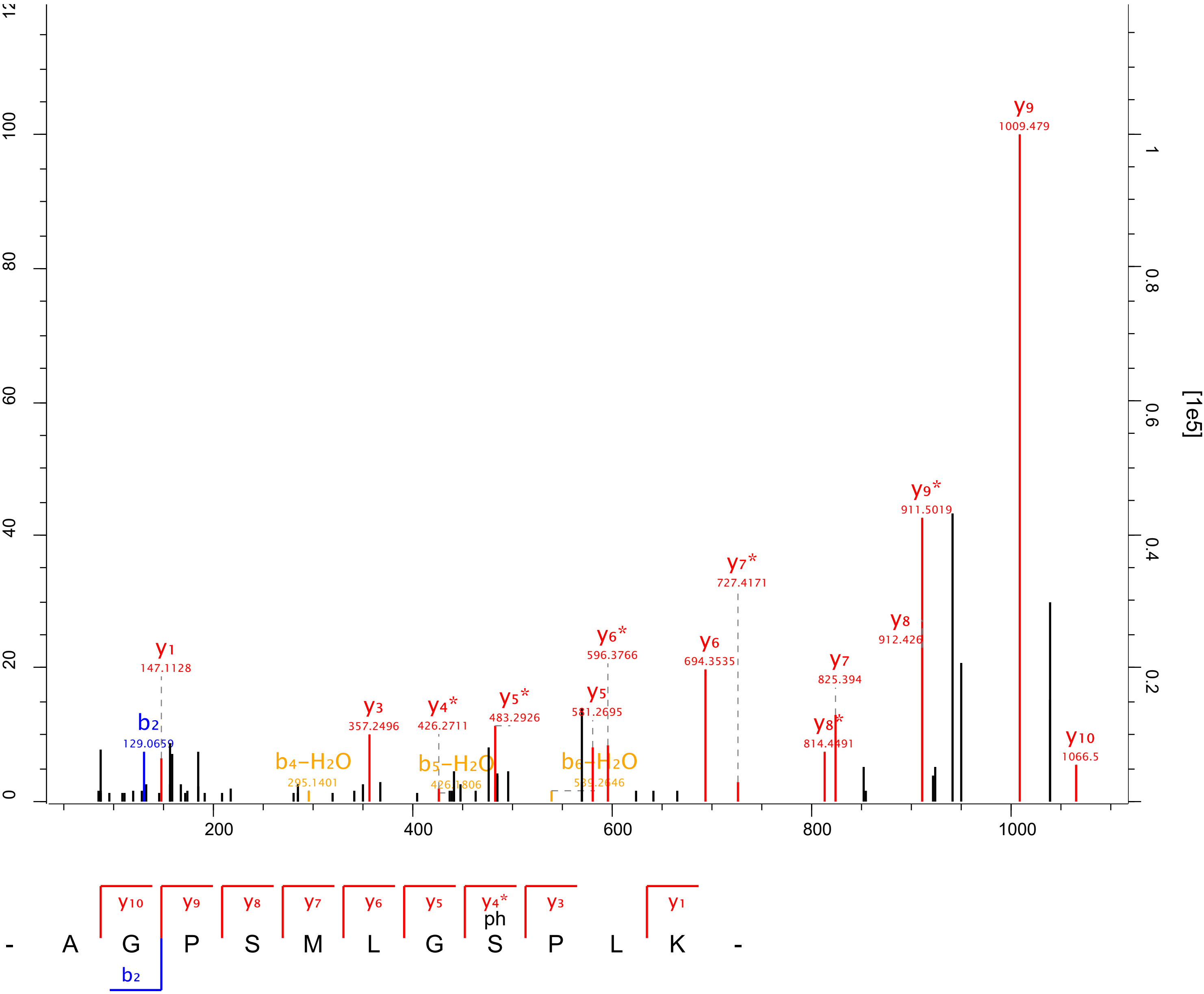

| Raw file | Scan | Method | Score | m/z |
|---|---|---|---|---|
| water_15_2 | 20123 | FTMS; HCD | 201.64 | 831.38 |

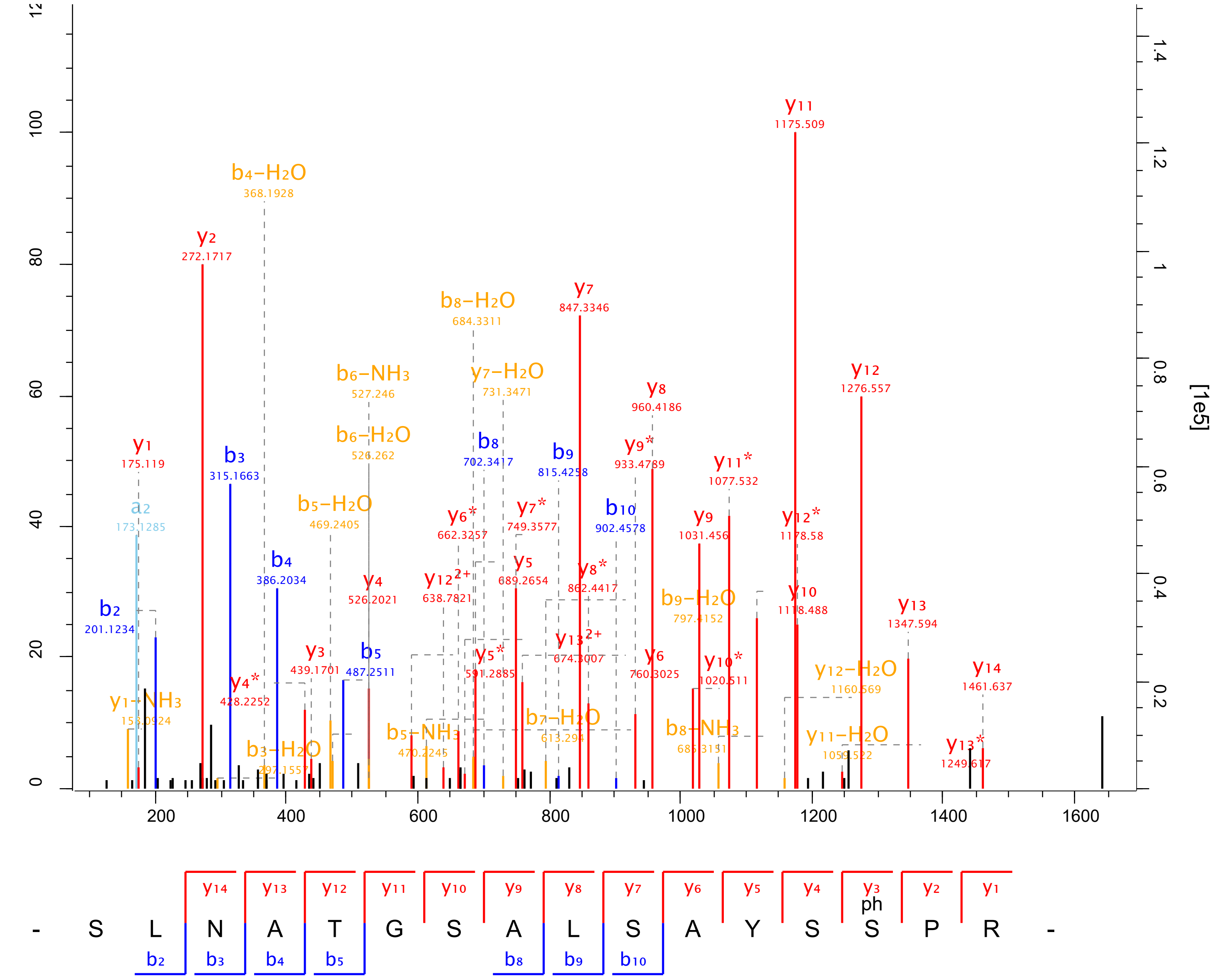

| Raw file | Scan | Method | Score | m/z |
|---|---|---|---|---|
| water_15_2 | 20137 | FTMS; HCD | 66.54 | 966.46 |

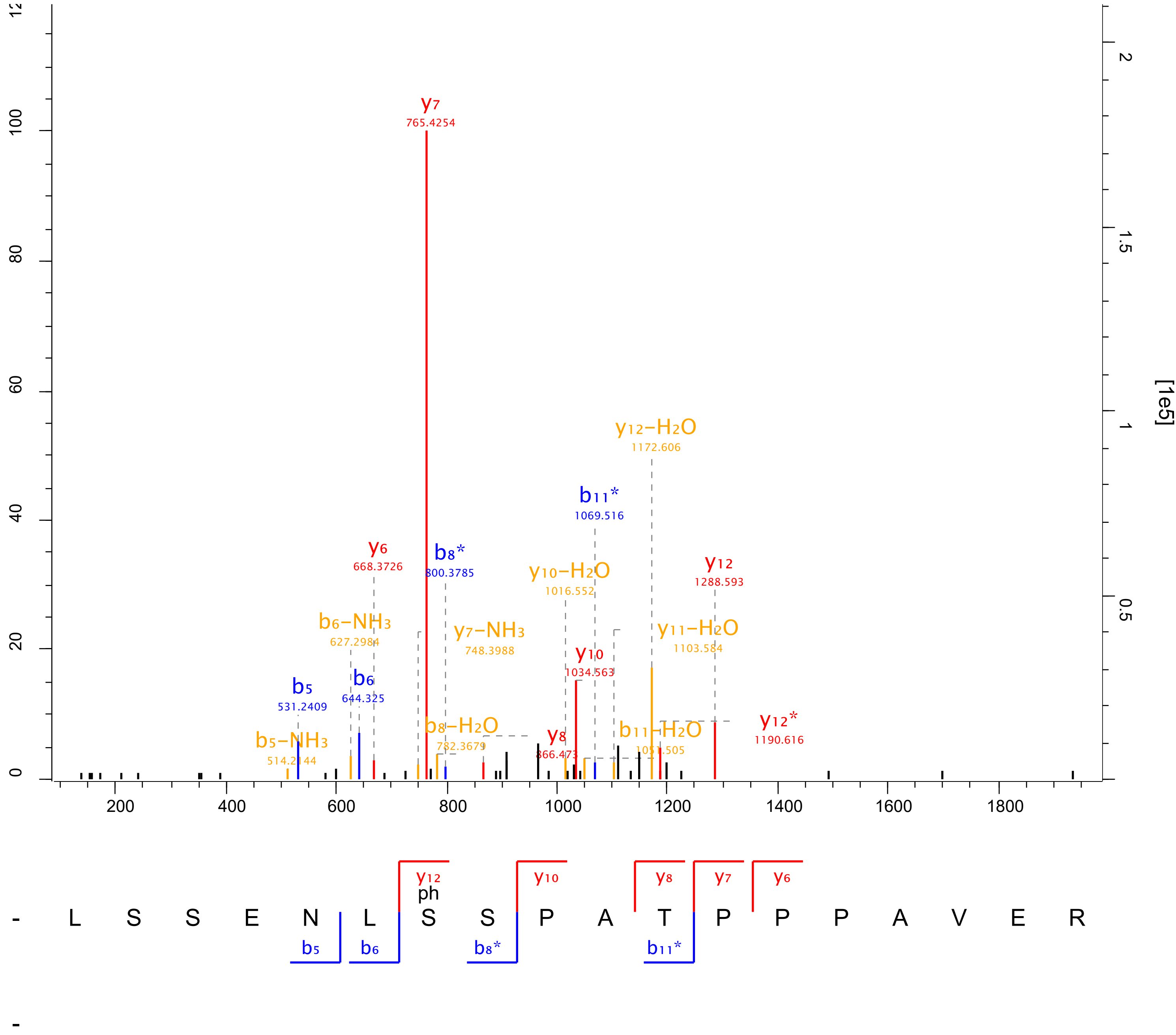

| Raw file | Scan | Method | Score | m/z |
|---|---|---|---|---|
| water_15_2 | 20222 | FTMS; HCD | 78.34 | 524.72 |

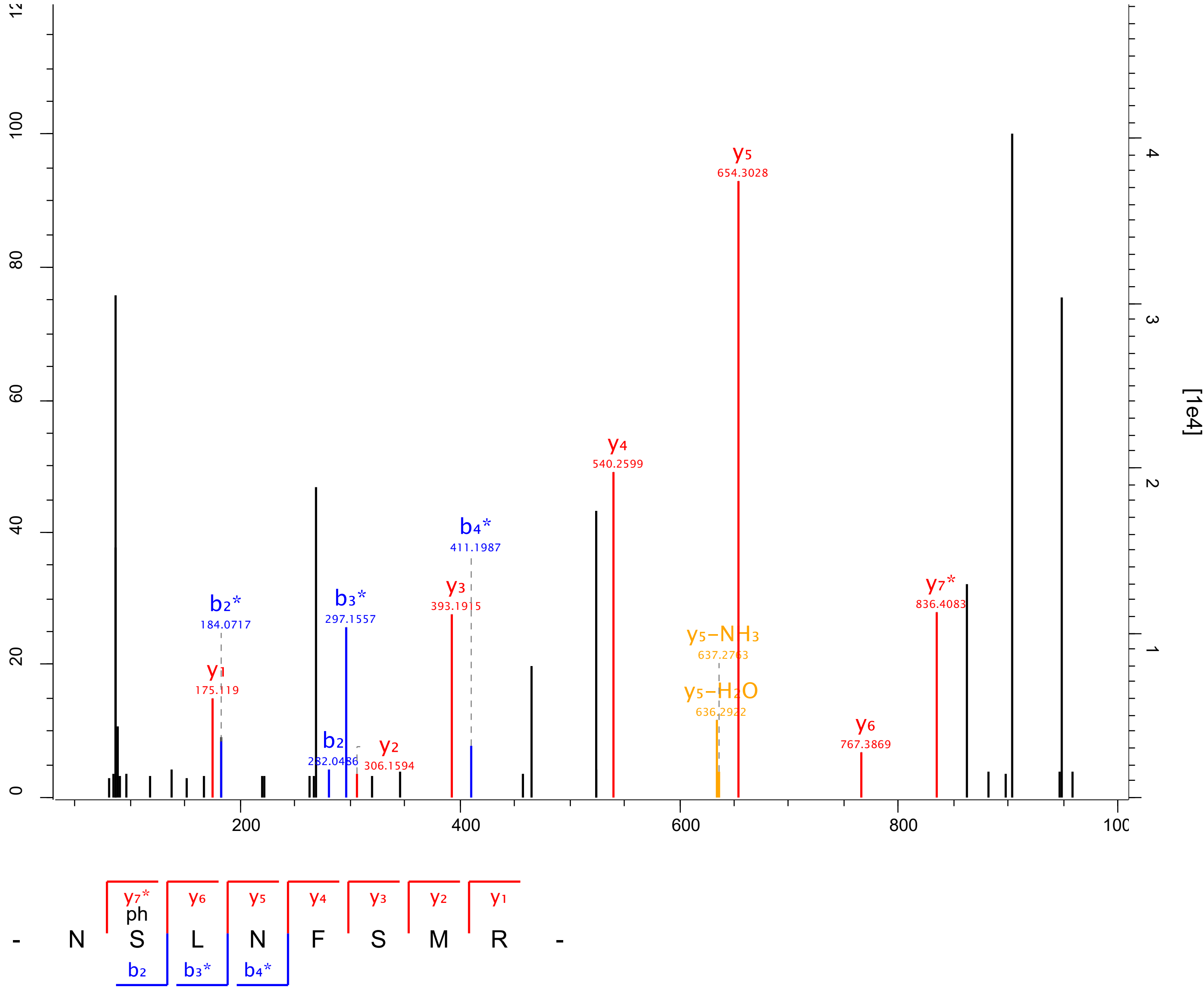

| Raw file | Scan | Method | Score | m/z |
|---|---|---|---|---|
| water_15_2 | 20264 | FTMS; HCD | 69.72 | 501.23 |

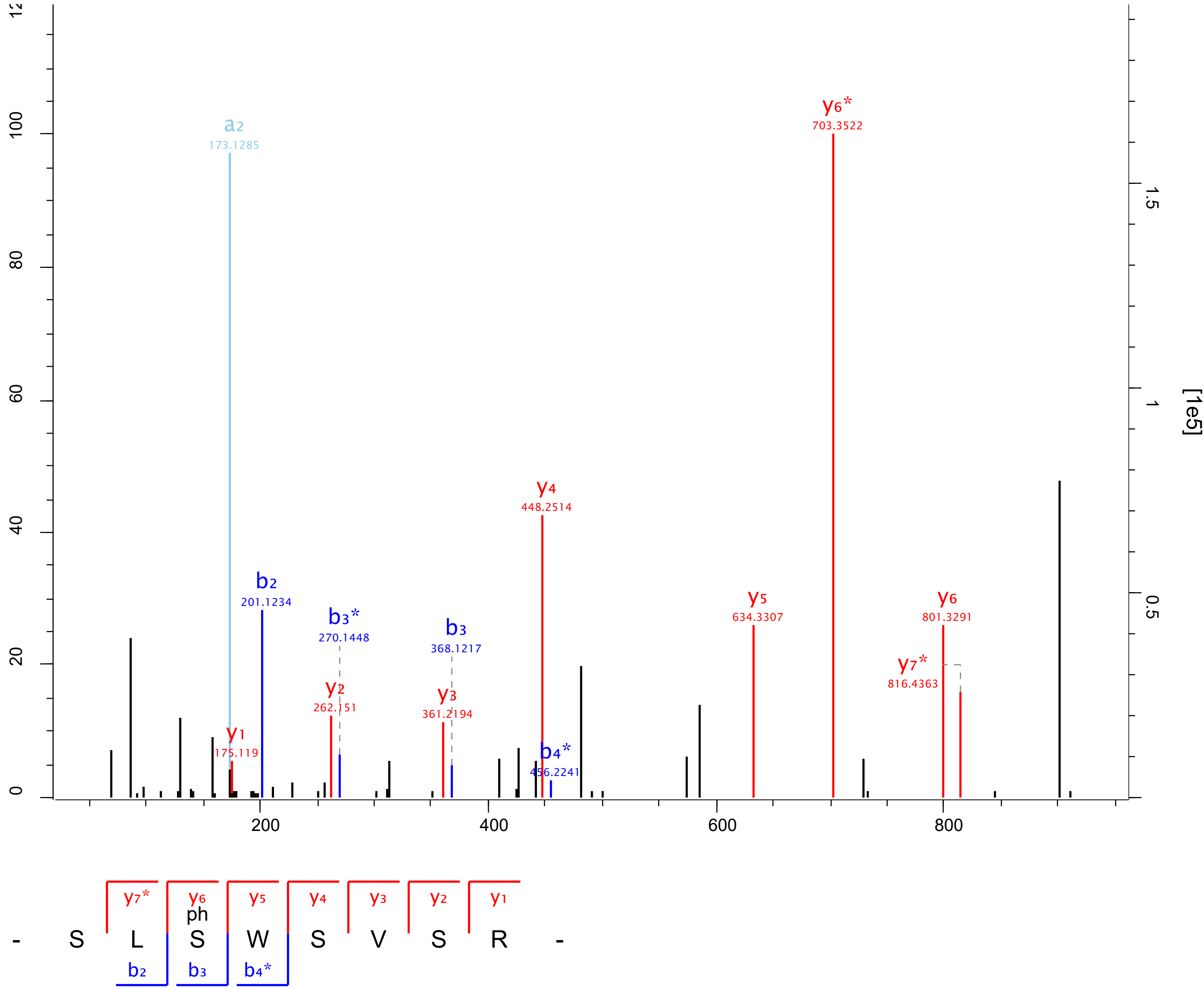

| Raw file | Scan | Method | Score | m/z |
| --- | --- | --- | --- | --- |
| water_15_2 | 20394 | FTMS; HCD | 42.12 | 850.4 |

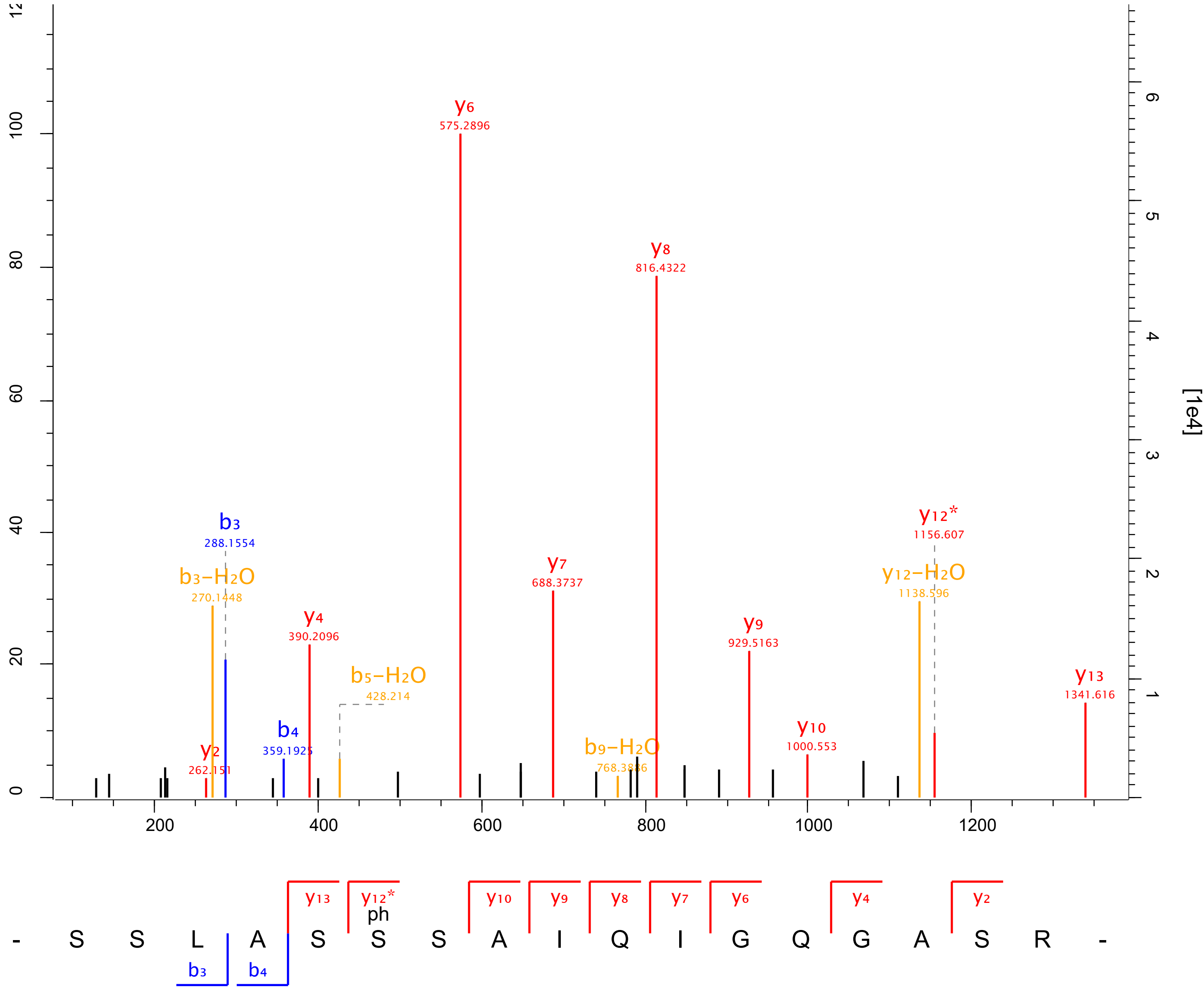

| Raw file | Scan | Method | Score | m/z |
|---|---|---|---|---|
| water_15_2 | 20479 | FTMS; HCD | 96.77 | 734.28 |

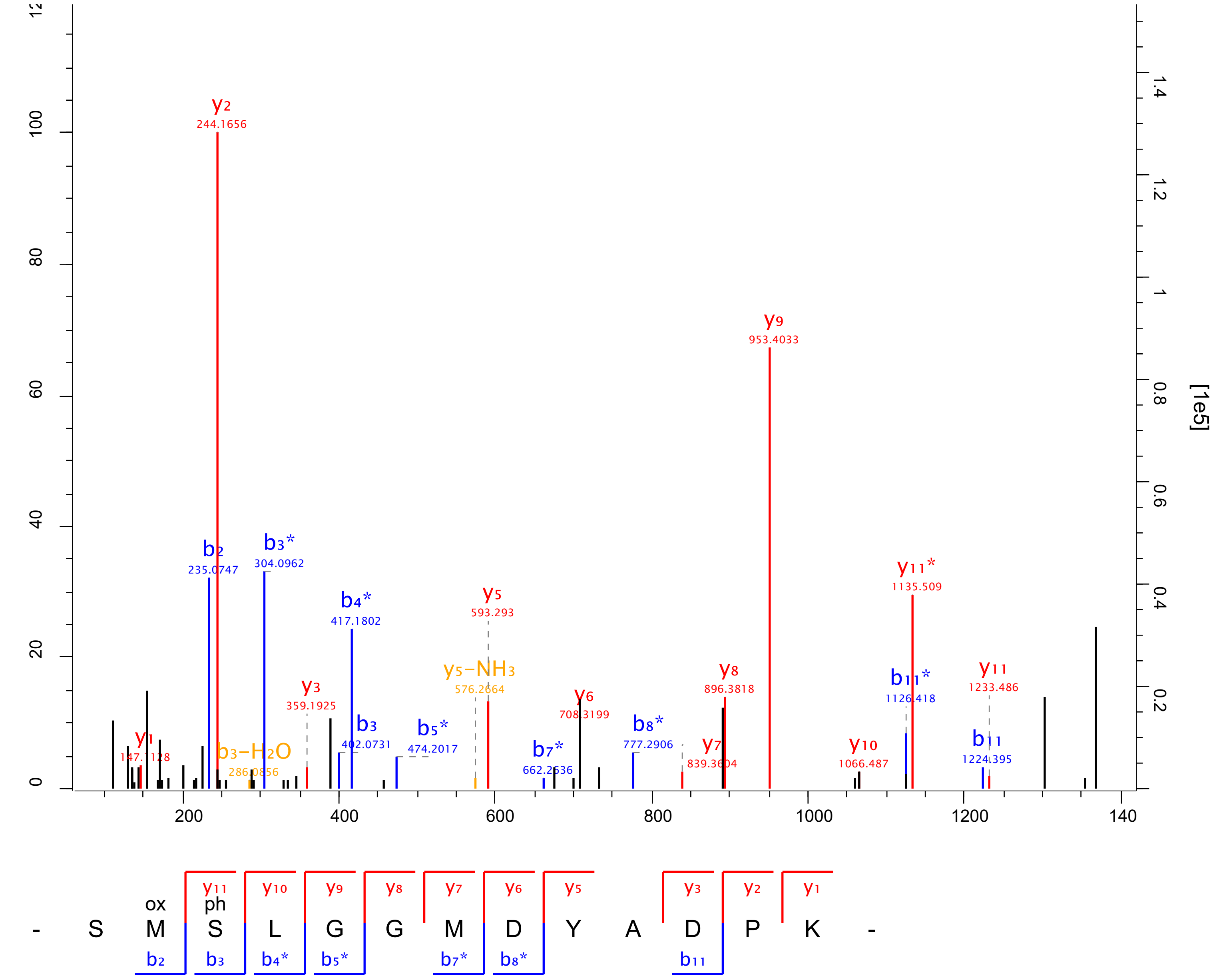

| Raw file | Scan | Method | Score | m/z |
|---|---|---|---|---|
| water_15_2 | 20490 | FTMS; HCD | 46.06 | 698.57 |

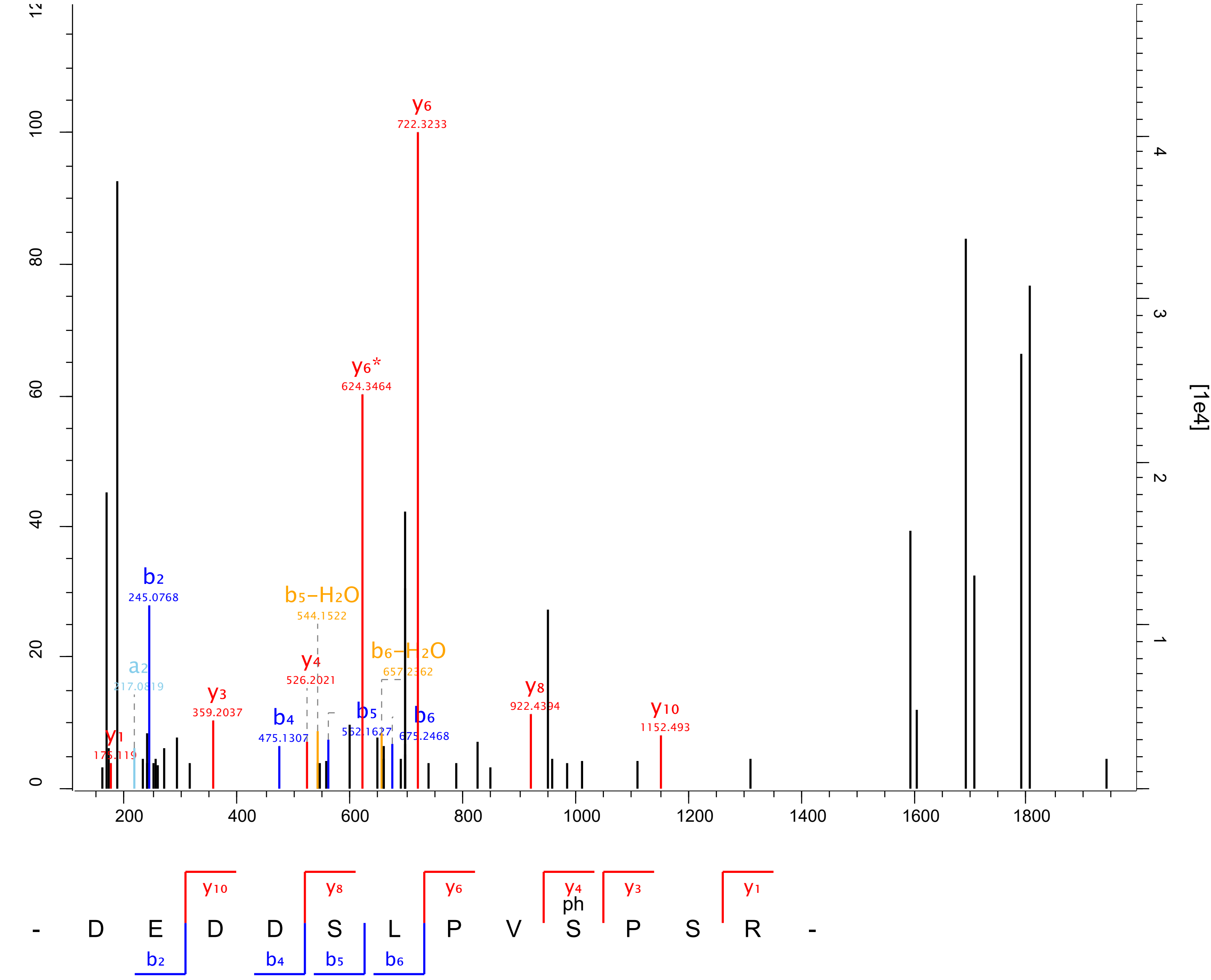

| Raw file | Scan | Method | Score | m/z |
| --- | --- | --- | --- | --- |
| water_15_2 | 20505 | FTMS; HCD | 105.28 | 796.35 |

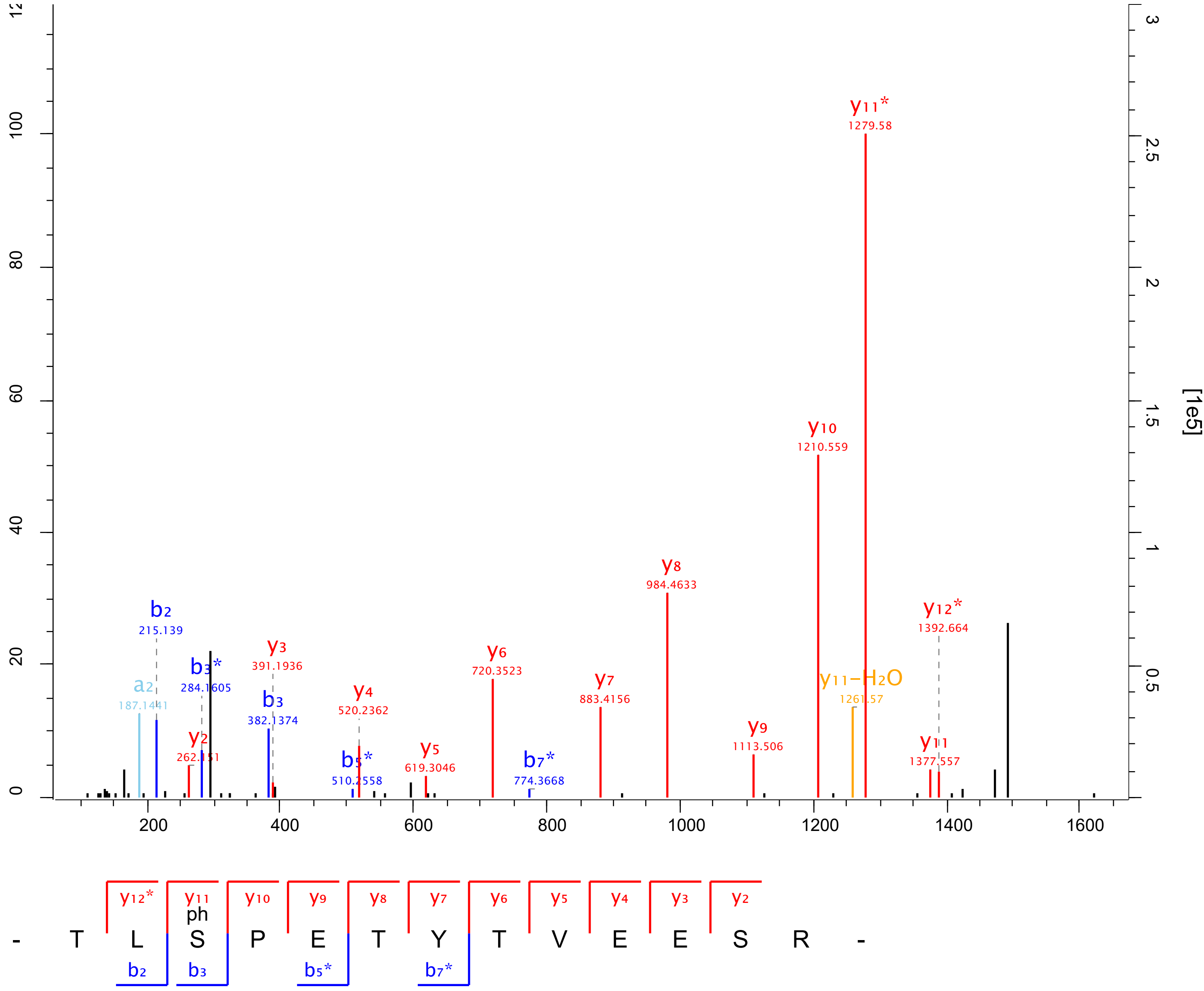

| Raw file | Scan | Method | Score | m/z |
| --- | --- | --- | --- | --- |
| water_15_2 | 20558 | FTMS; HCD | 54.9 | 663.79 |

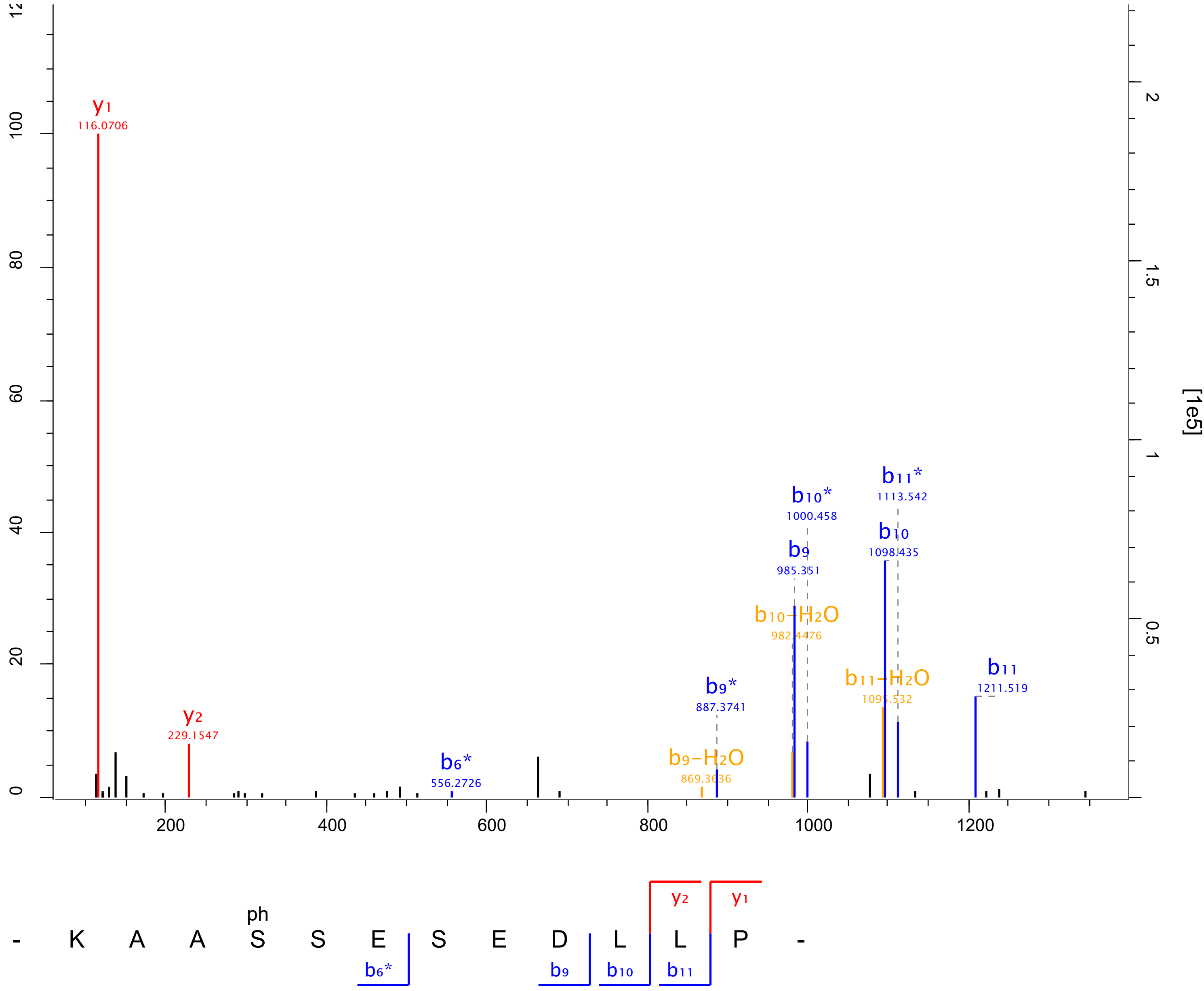

| Raw file | Scan | Method | Score | m/z |
| --- | --- | --- | --- | --- |
| water_15_2 | 20676 | FTMS; HCD | 88.6 | 480.73 |

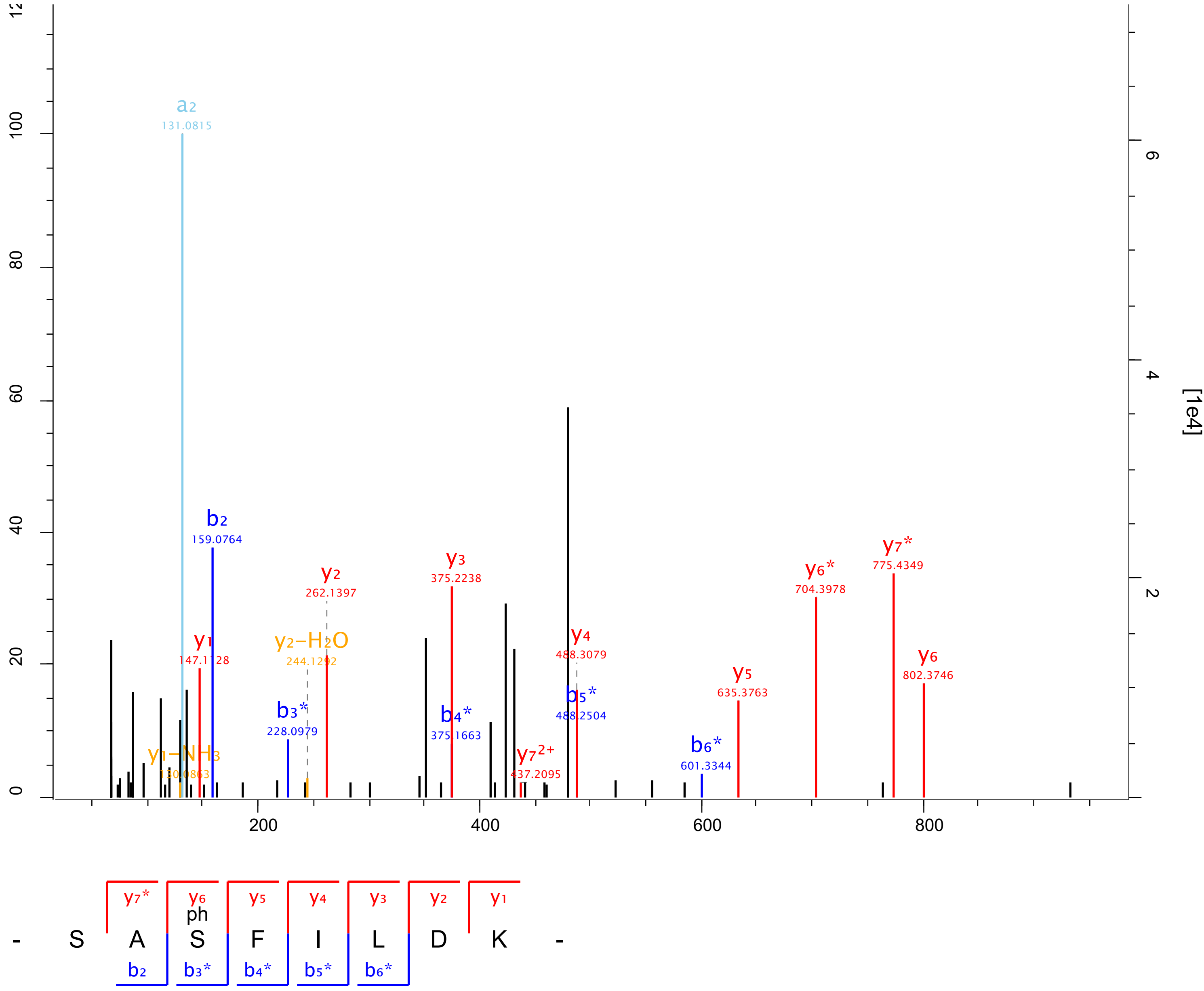

| Raw file | Scan | Method | Score | m/z |
|---|---|---|---|---|
| water_15_2 | 20715 | FTMS; HCD | 48.57 | 560.68 |

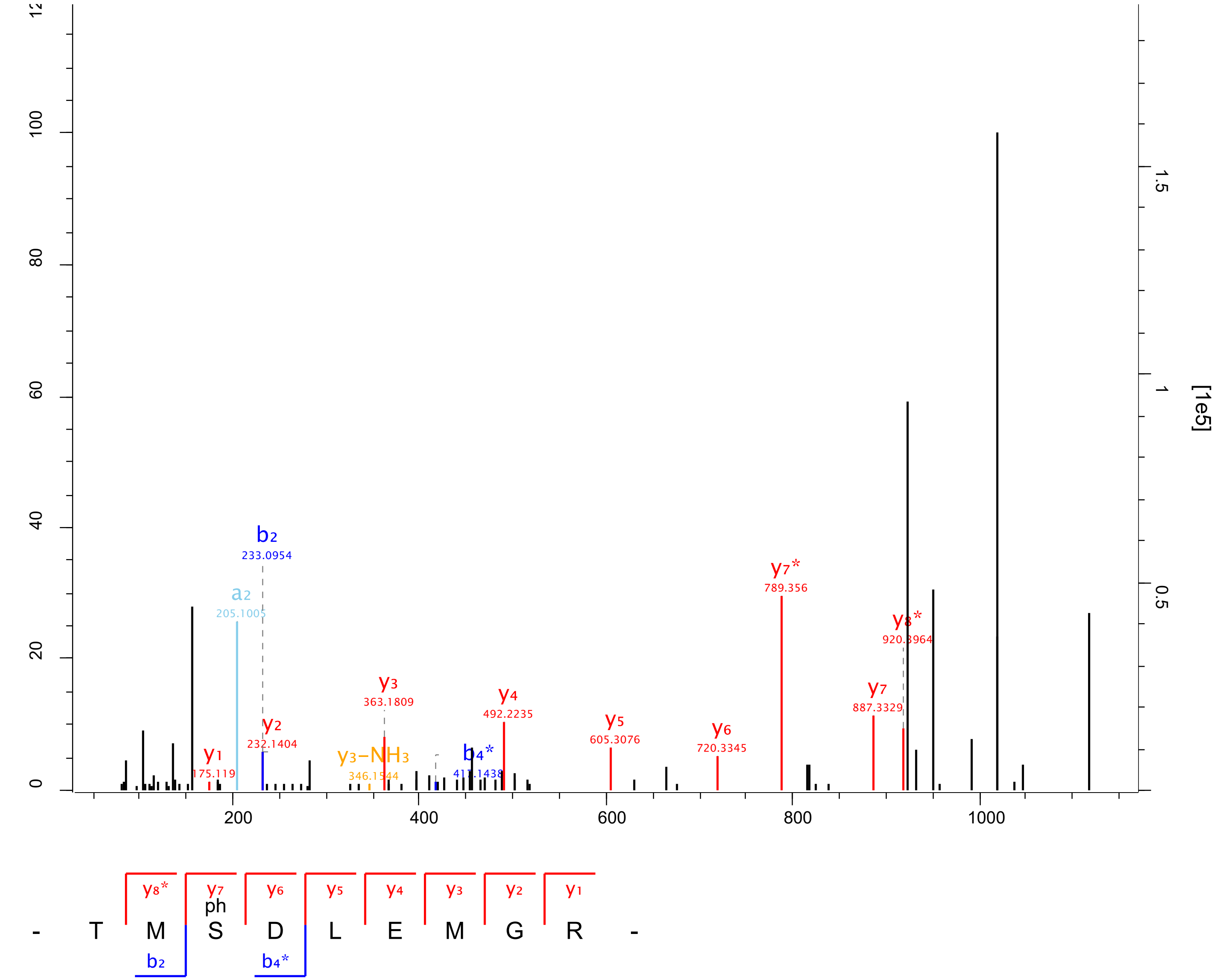

| Raw file | Scan | Method | Score | m/z |
|---|---|---|---|---|
| water_15_2 | 20722 | FTMS; HCD | 157.54 | 810.89 |

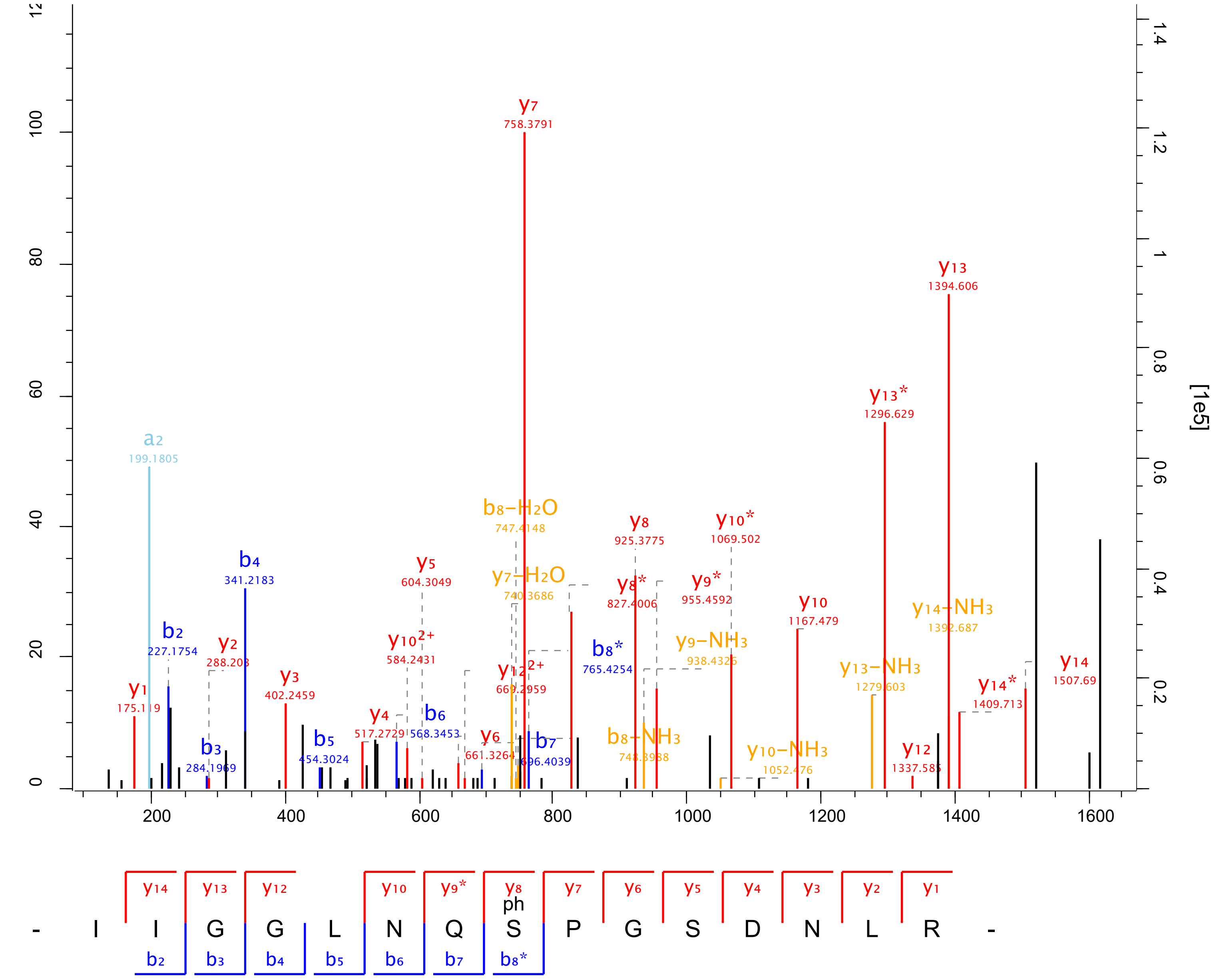

| Raw file | Scan | Method | Score | m/z |
|---|---|---|---|---|
| water_15_2 | 21000 | FTMS; HCD | 78.69 | 595.27 |

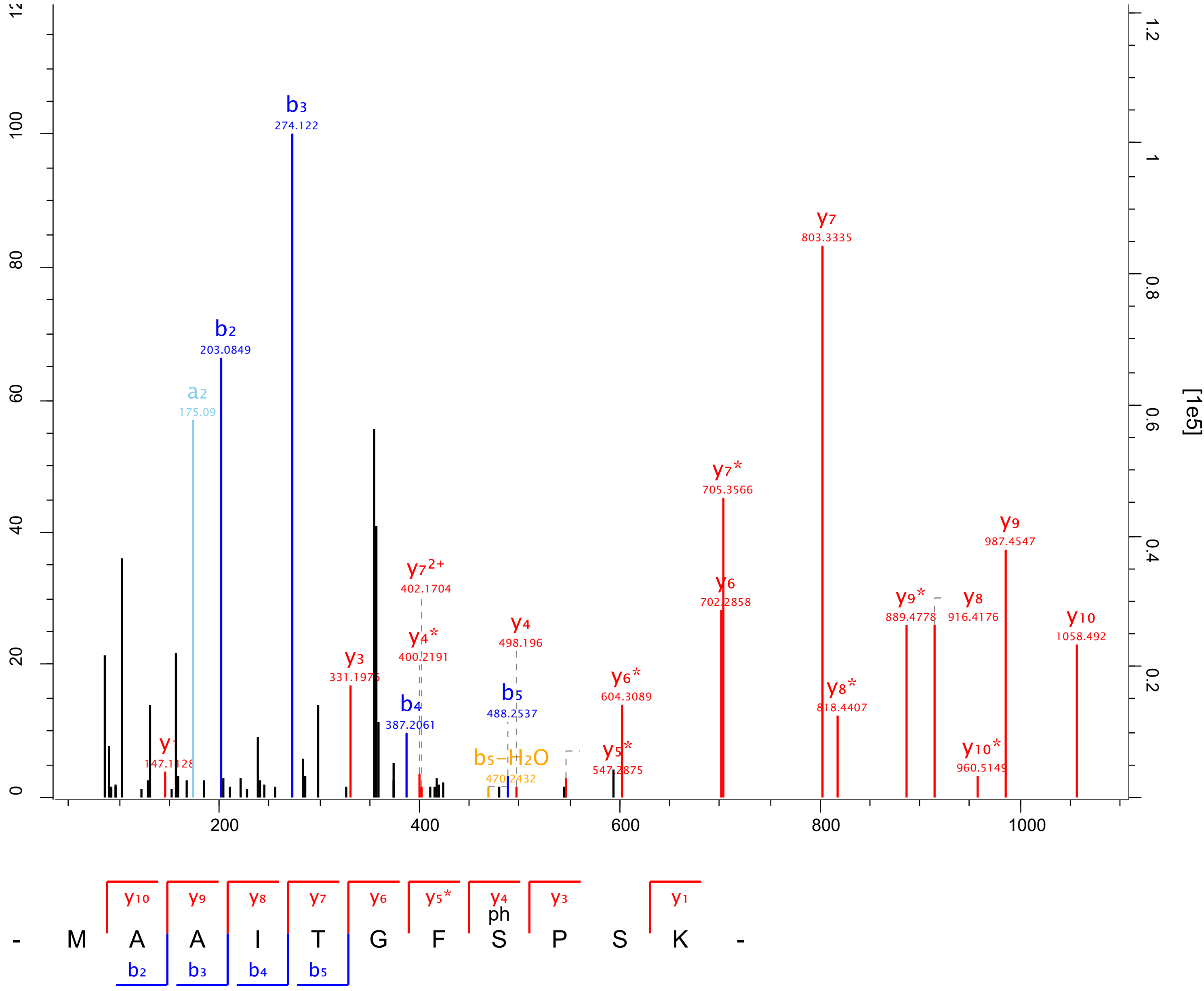

| Raw file | Scan | Method | Score | m/z |
|---|---|---|---|---|
| water_15_2 | 21165 | FTMS; HCD | 43.77 | 469.74 |

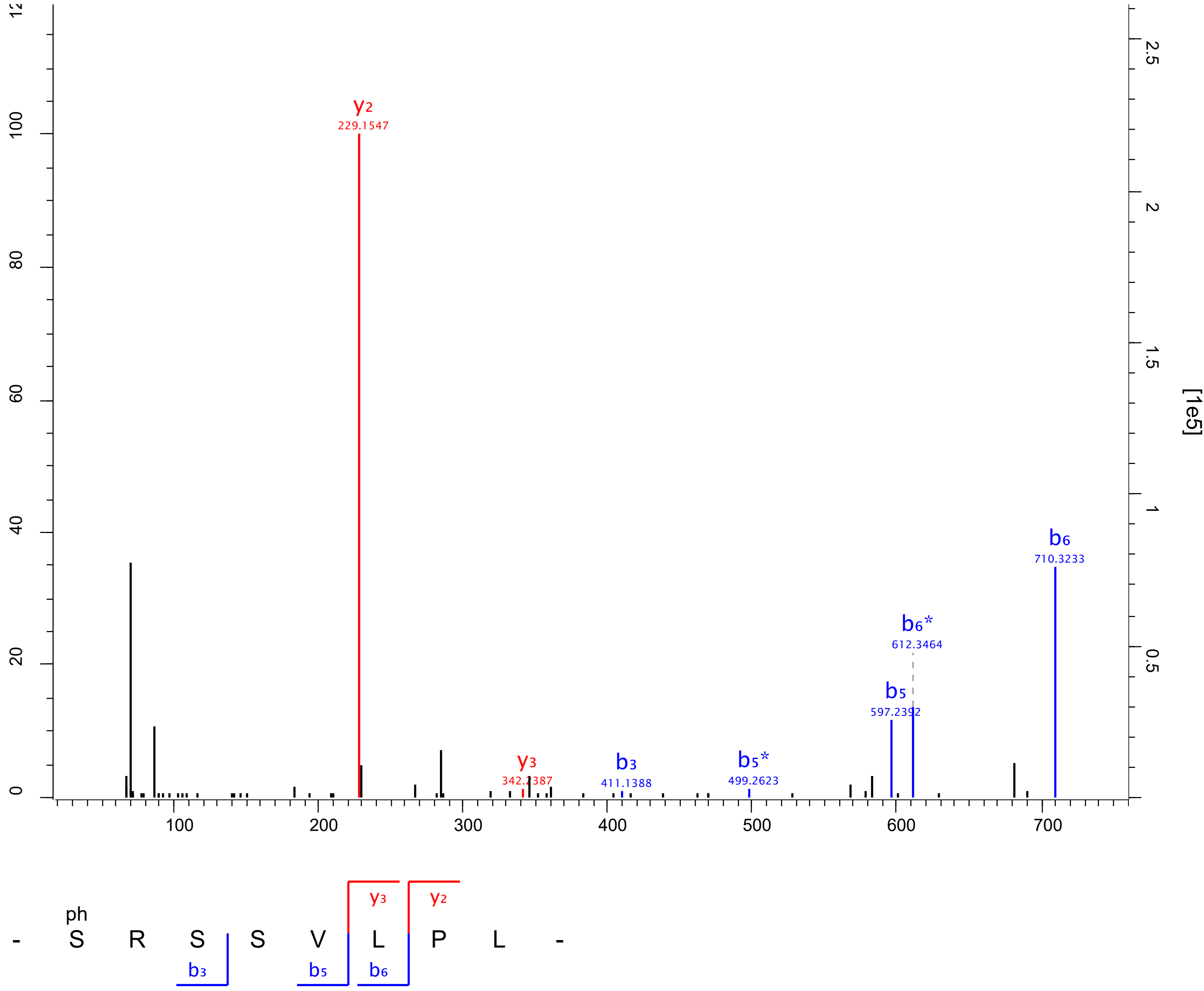

| Raw file | Scan | Method | Score | m/z |
|---|---|---|---|---|
| water_15_2 | 21283 | FTMS; HCD | 44.54 | 529.24 |

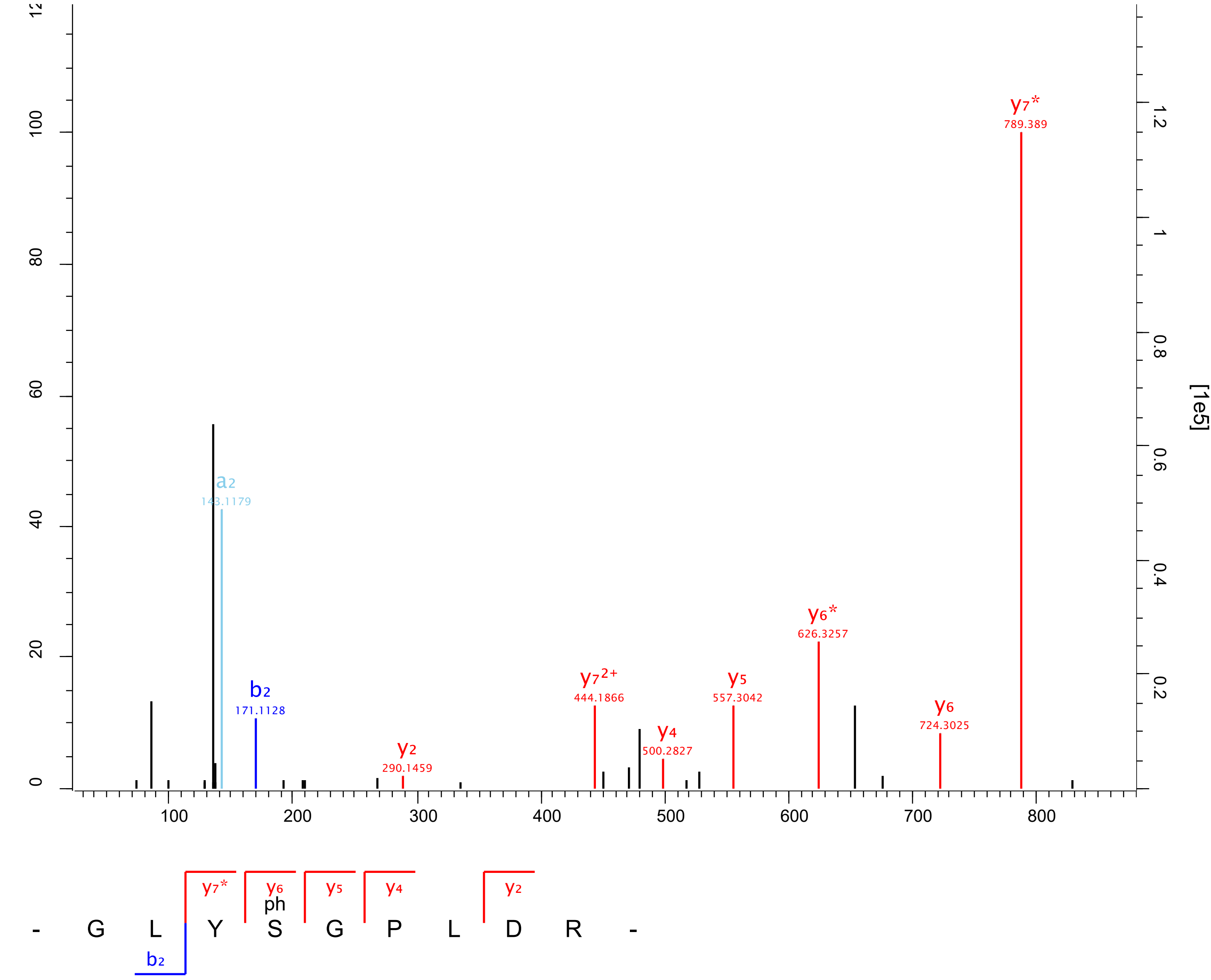

| Raw file | Scan | Method | Score | m/z |
|---|---|---|---|---|
| water_15_2 | 21352 | FTMS; HCD | 137.78 | 539.24 |

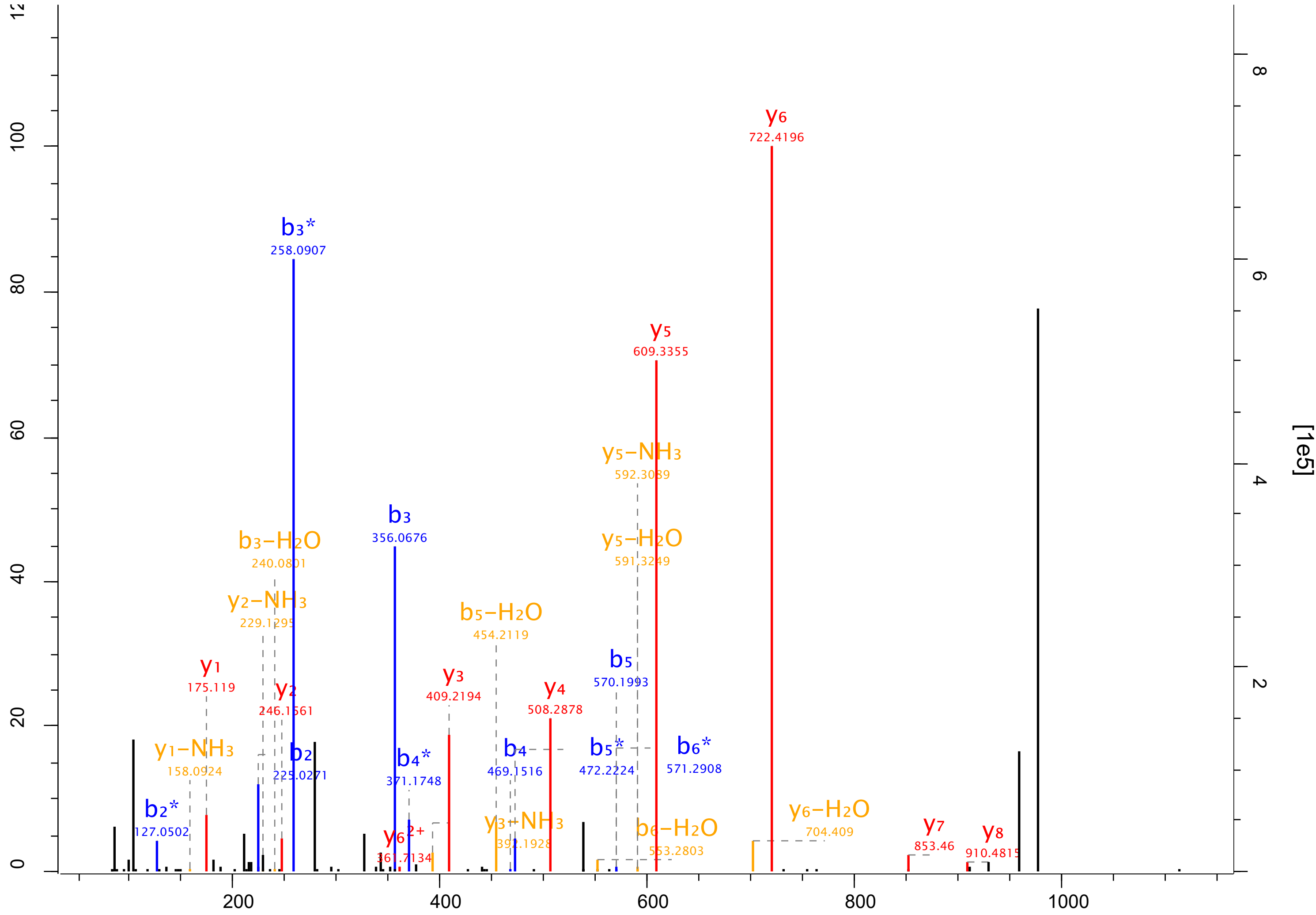


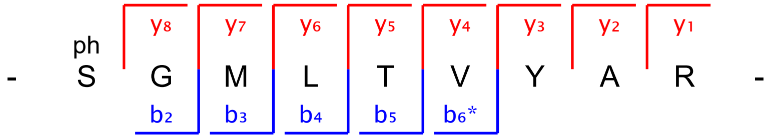

| Raw file | Scan | Method | Score | m/z |
|---|---|---|---|---|
| water_15_2 | 21369 | FTMS; HCD | 50.09 | 681.8 |

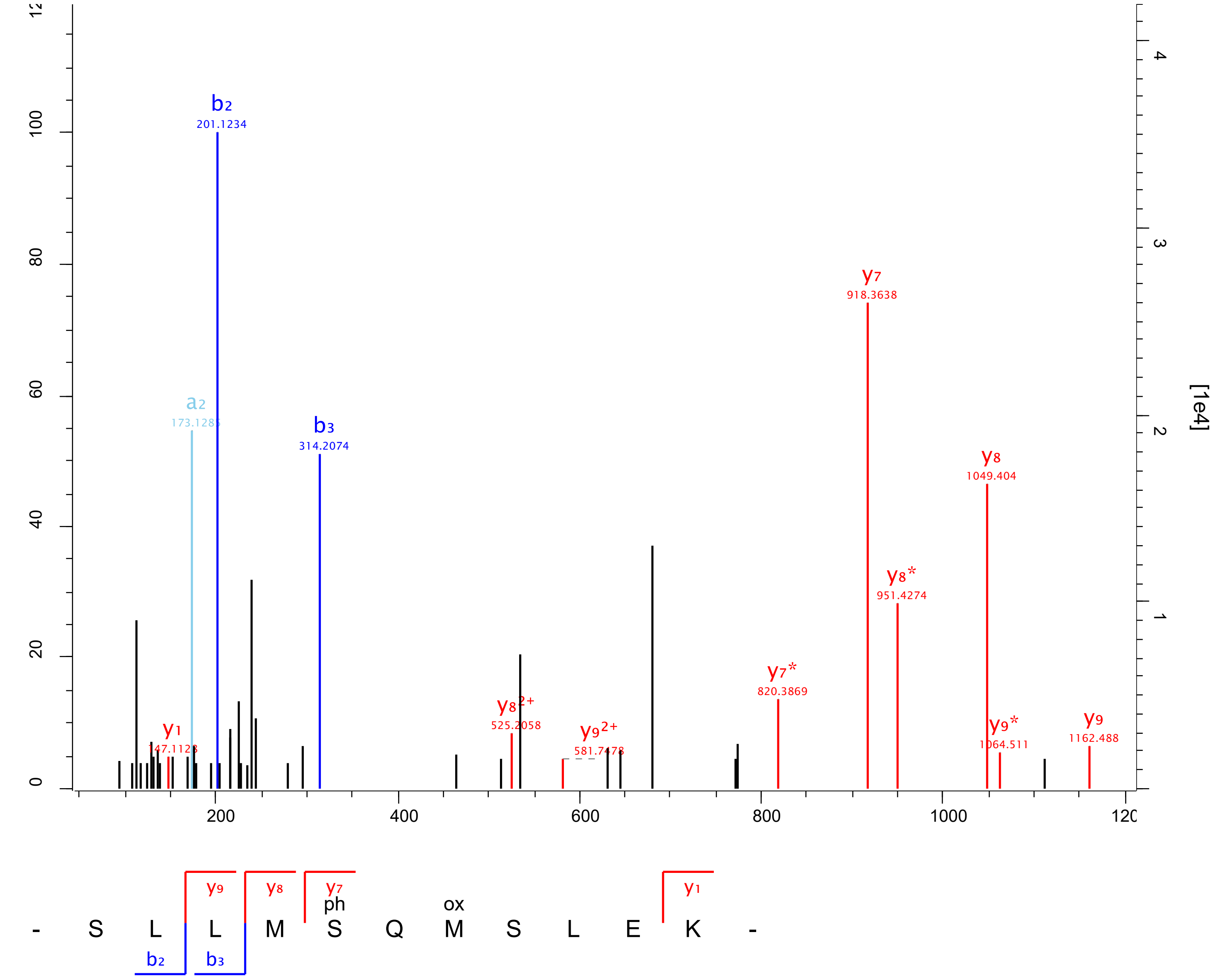

| Raw file | Scan | Method | Score | m/z |
|---|---|---|---|---|
| water_15_2 | 2142 | FTMS; HCD | 54.86 | 410.2 |

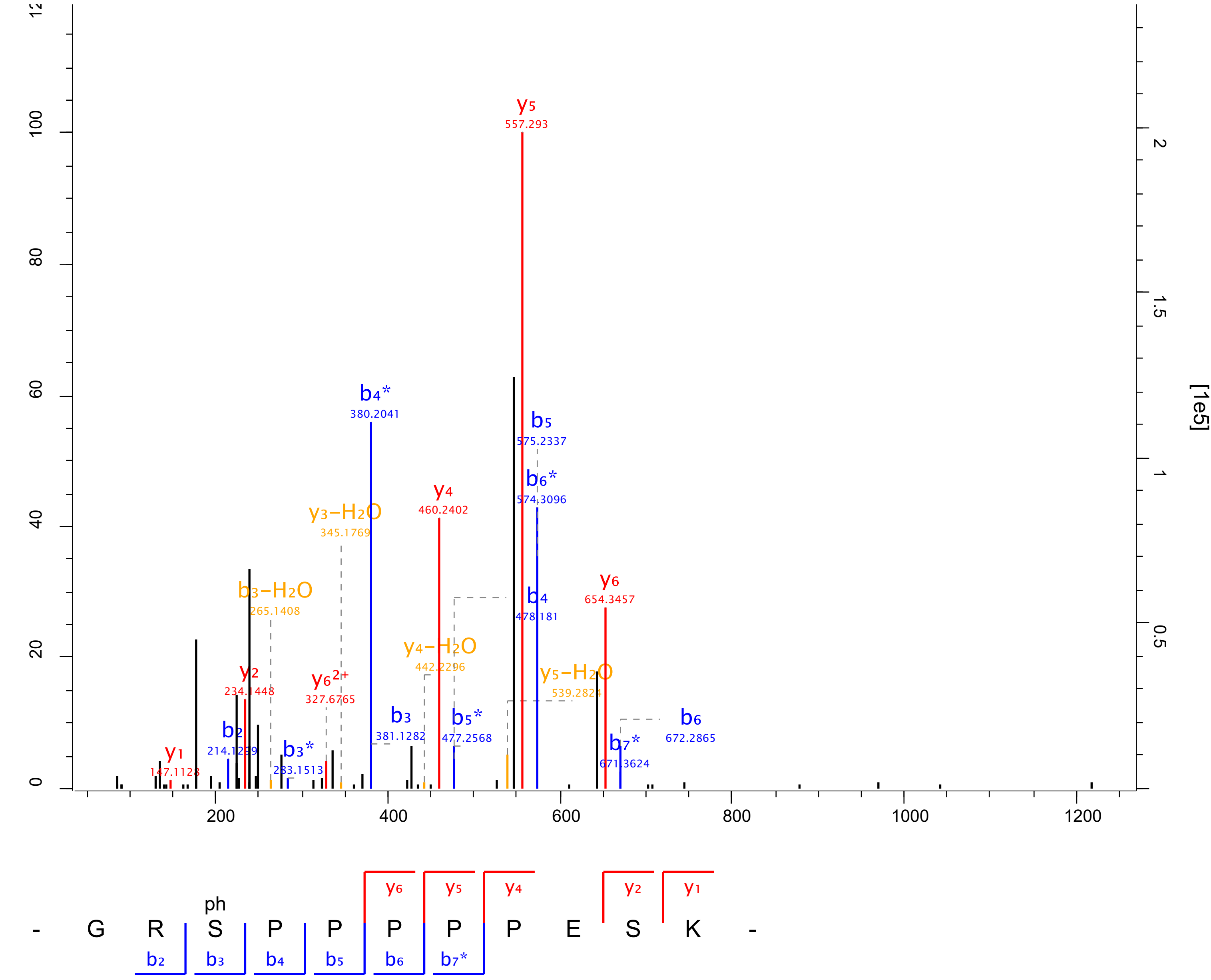

| Raw file | Scan | Method | Score | m/z |
| --- | --- | --- | --- | --- |
| water_15_2 | 21584 | FTMS; HCD | 71.6 | 751.84 |

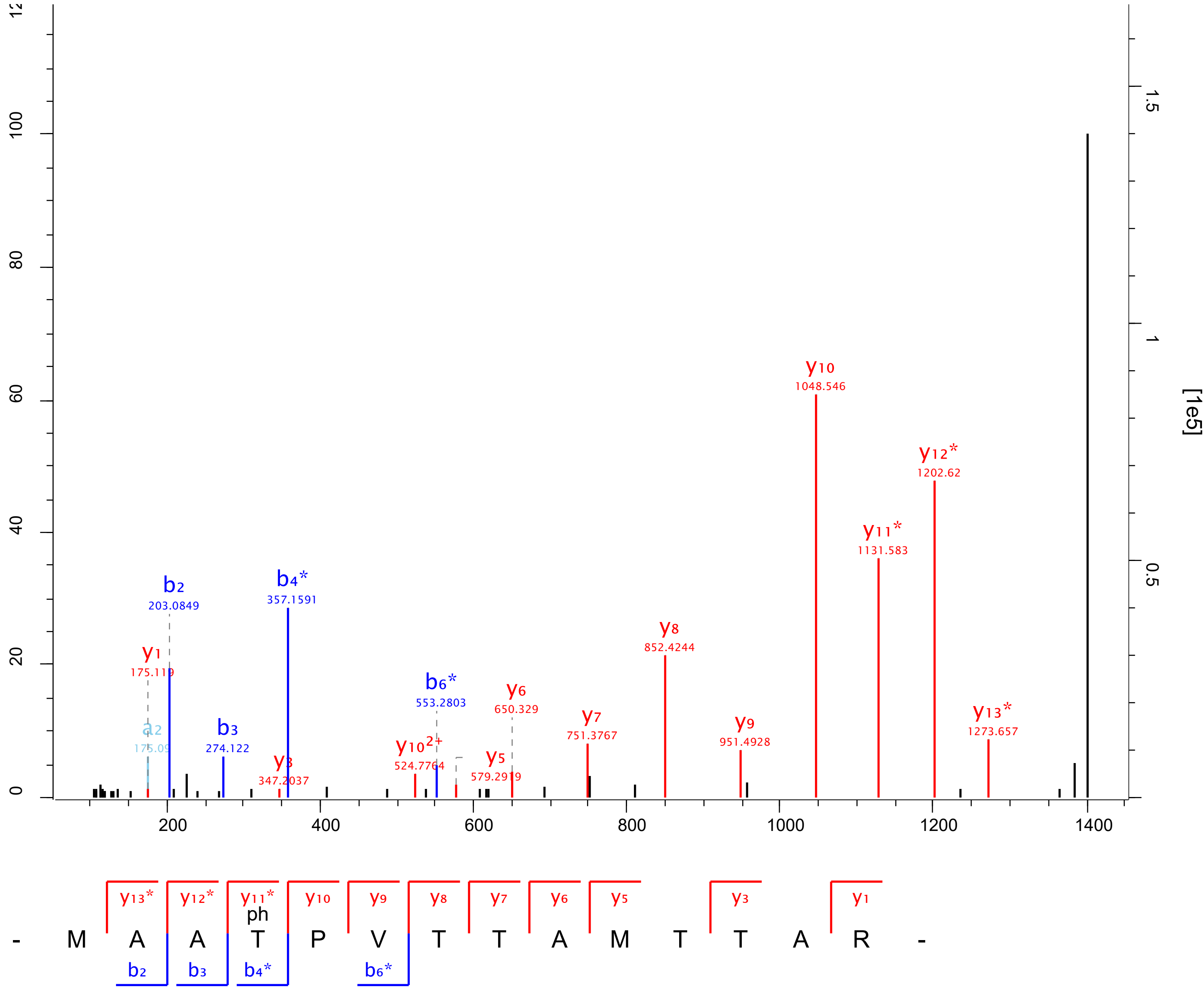

| Raw file | Scan | Method | Score | m/z |
|---|---|---|---|---|
| water_15_2 | 21622 | FTMS; HCD | 122.39 | 937.41 |

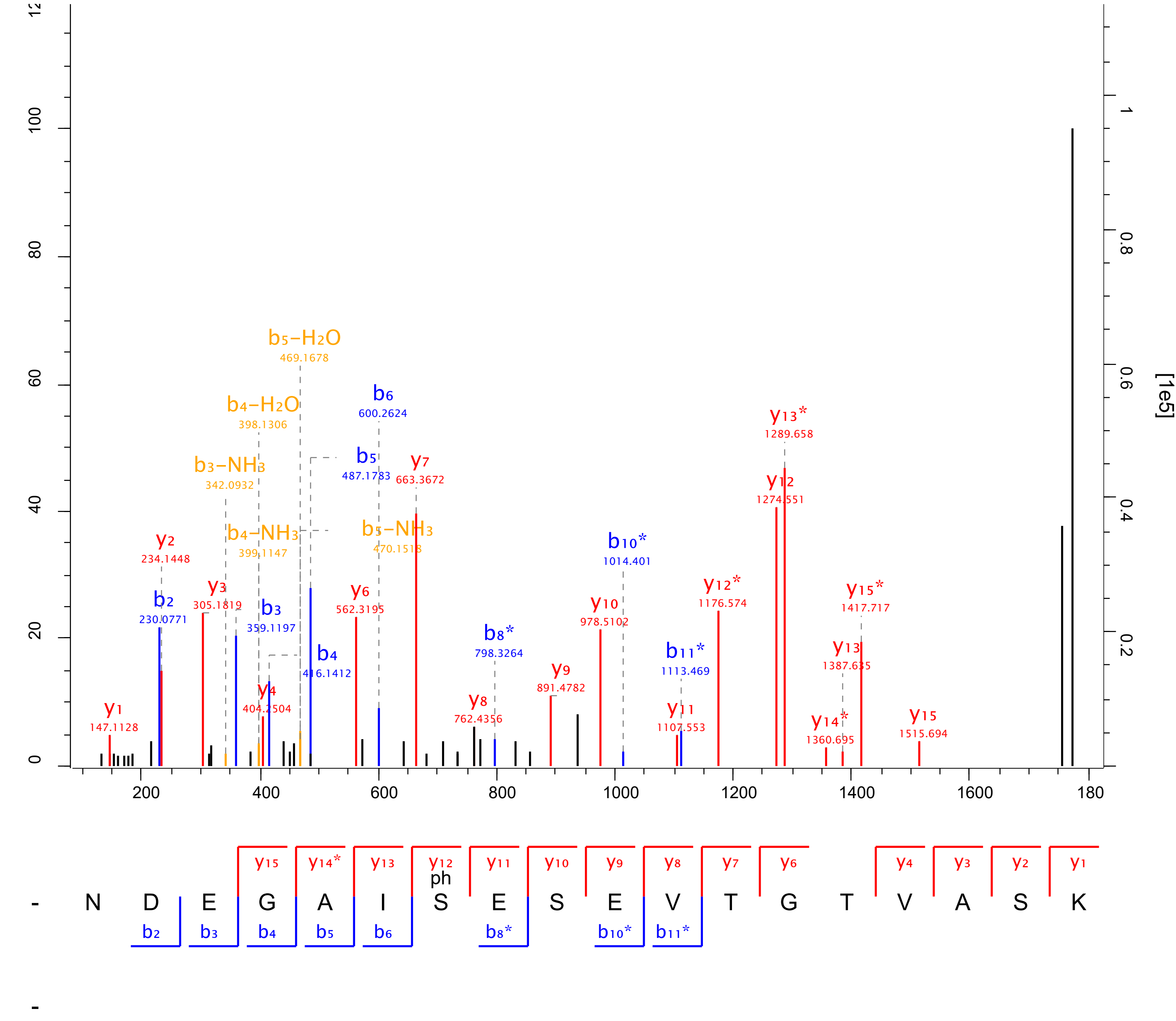

| Raw file | Scan | Method | Score | m/z |
|---|---|---|---|---|
| water_15_2 | 21732 | FTMS; HCD | 109.66 | 636.31 |

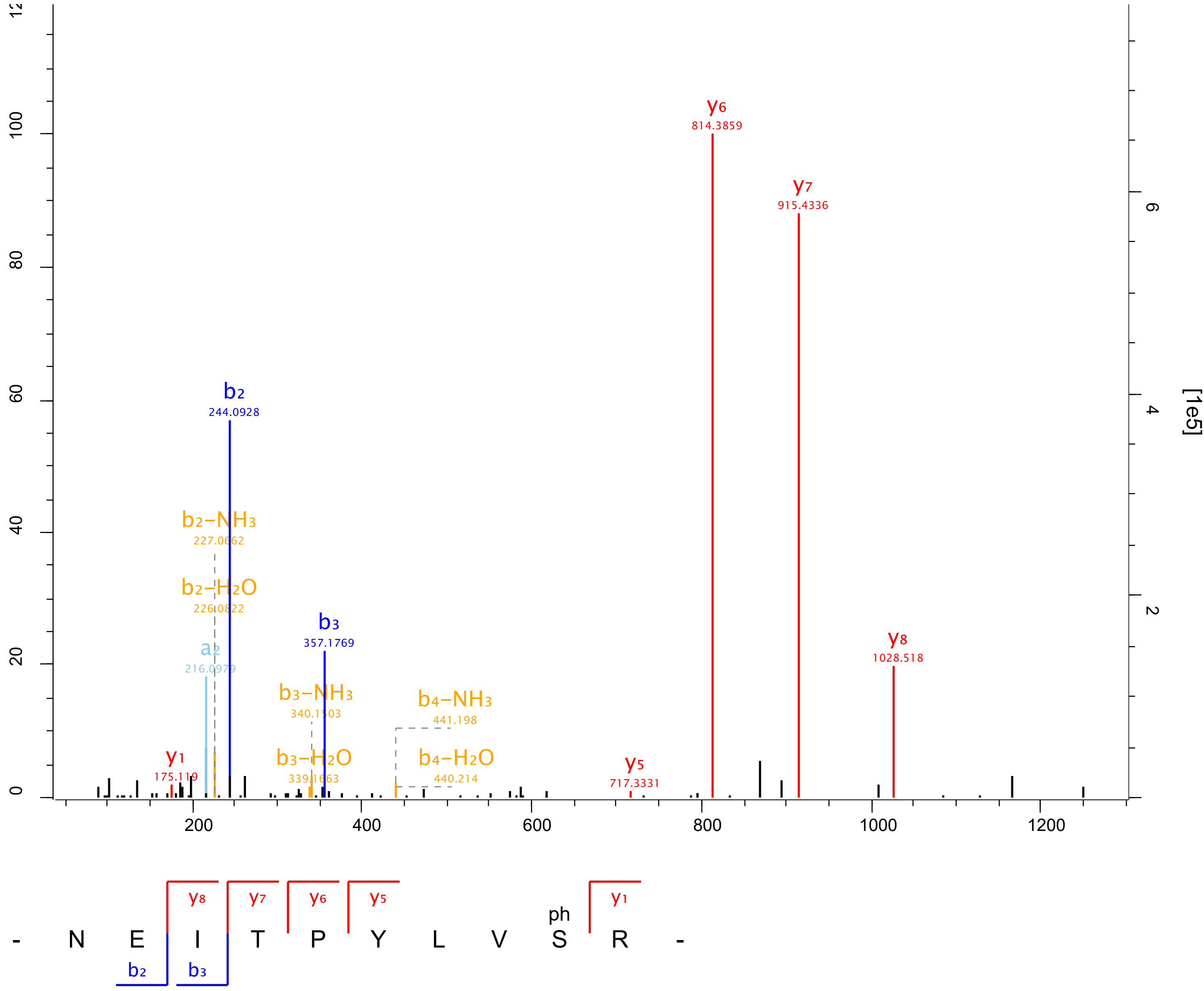

| Raw file | Scan | Method | Score | m/z |
| --- | --- | --- | --- | --- |
| water_15_2 | 21816 | FTMS; HCD | 239.22 | 969.92 |

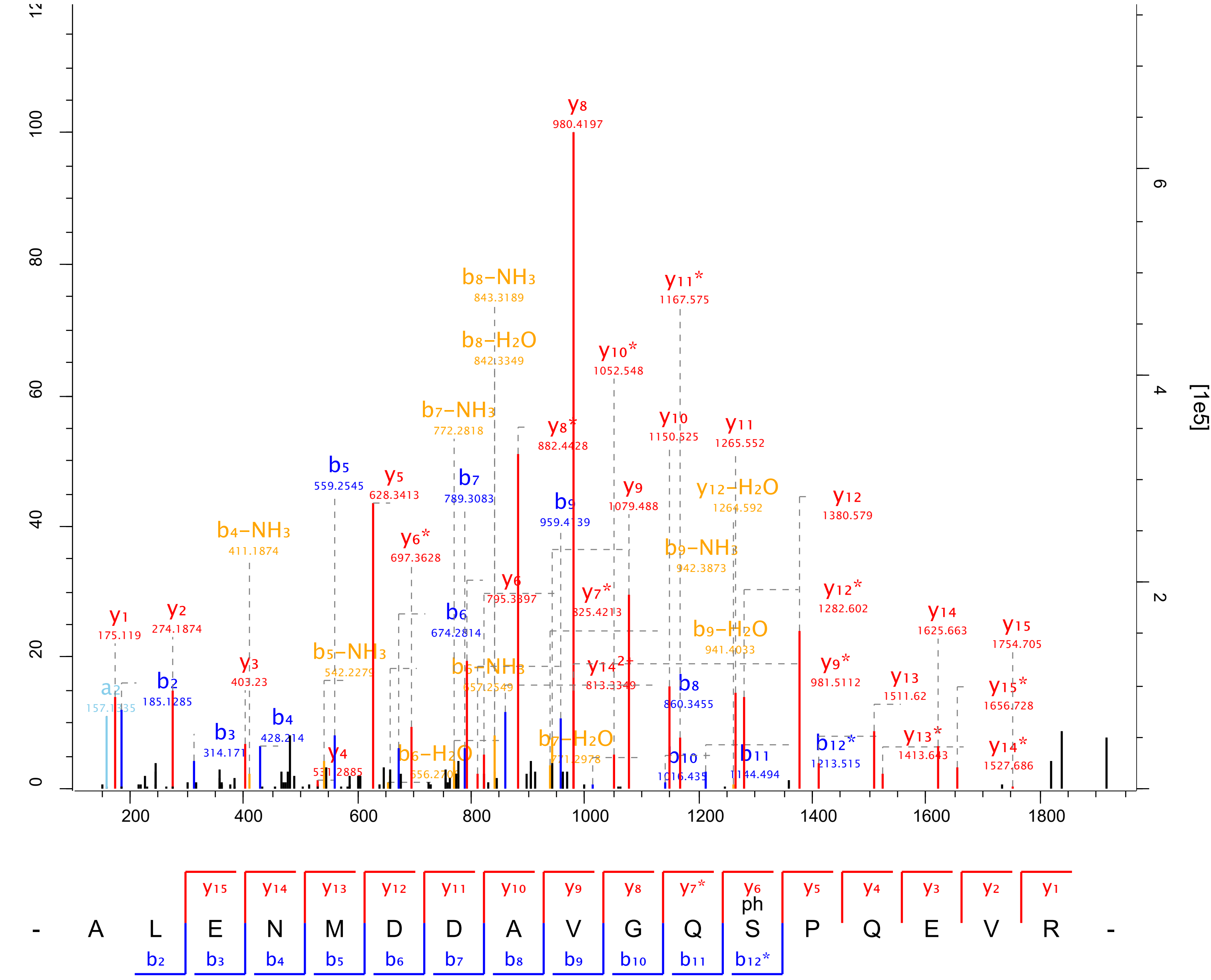

| Raw file | Scan | Method | Score | m/z |
|---|---|---|---|---|
| water_15_2 | 21879 | FTMS; HCD | 80.74 | 748.85 |

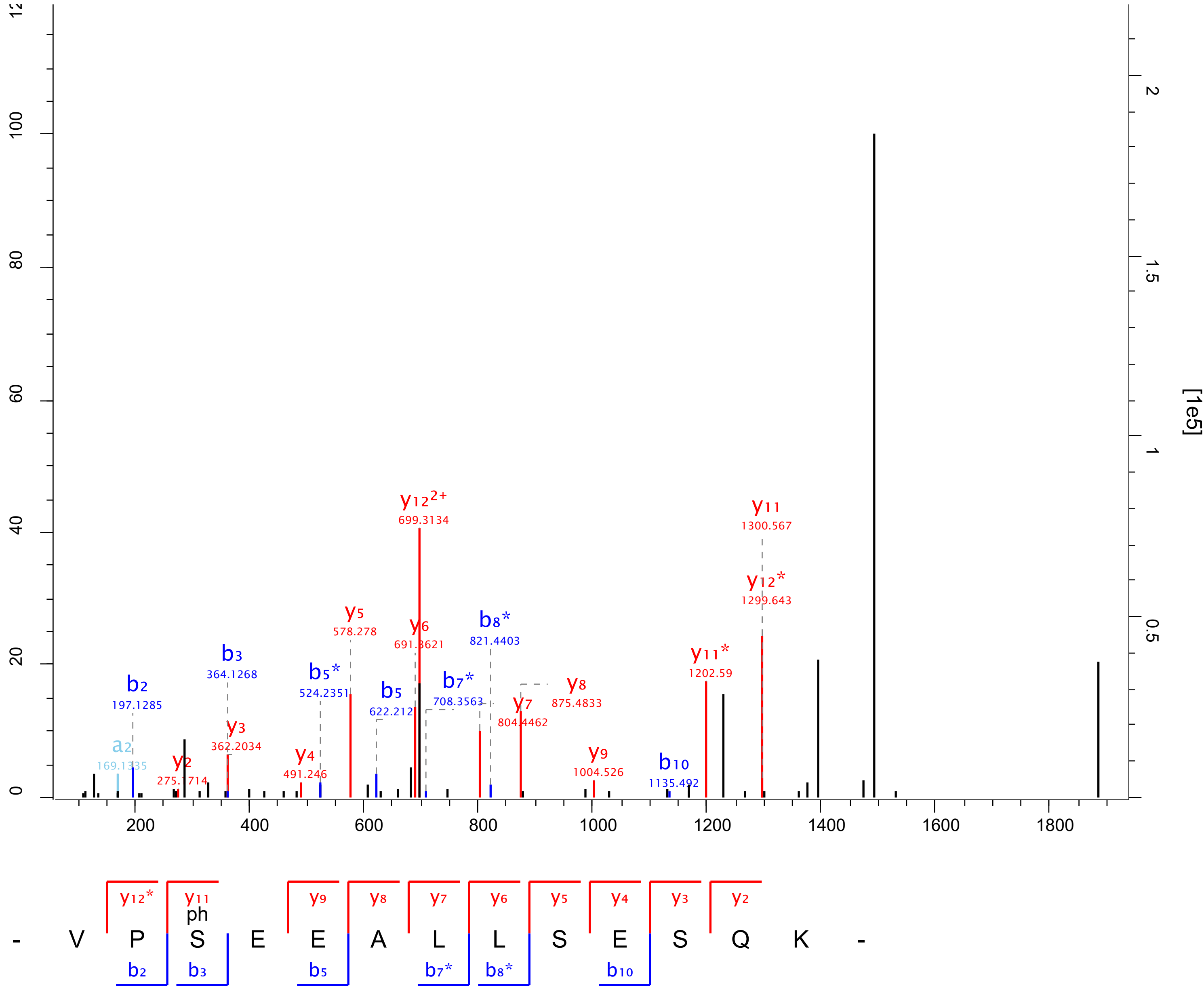

| Raw file | Scan | Method | Score | m/z |
|---|---|---|---|---|
| water_15_2 | 22060 | FTMS; HCD | 98.34 | 581.76 |

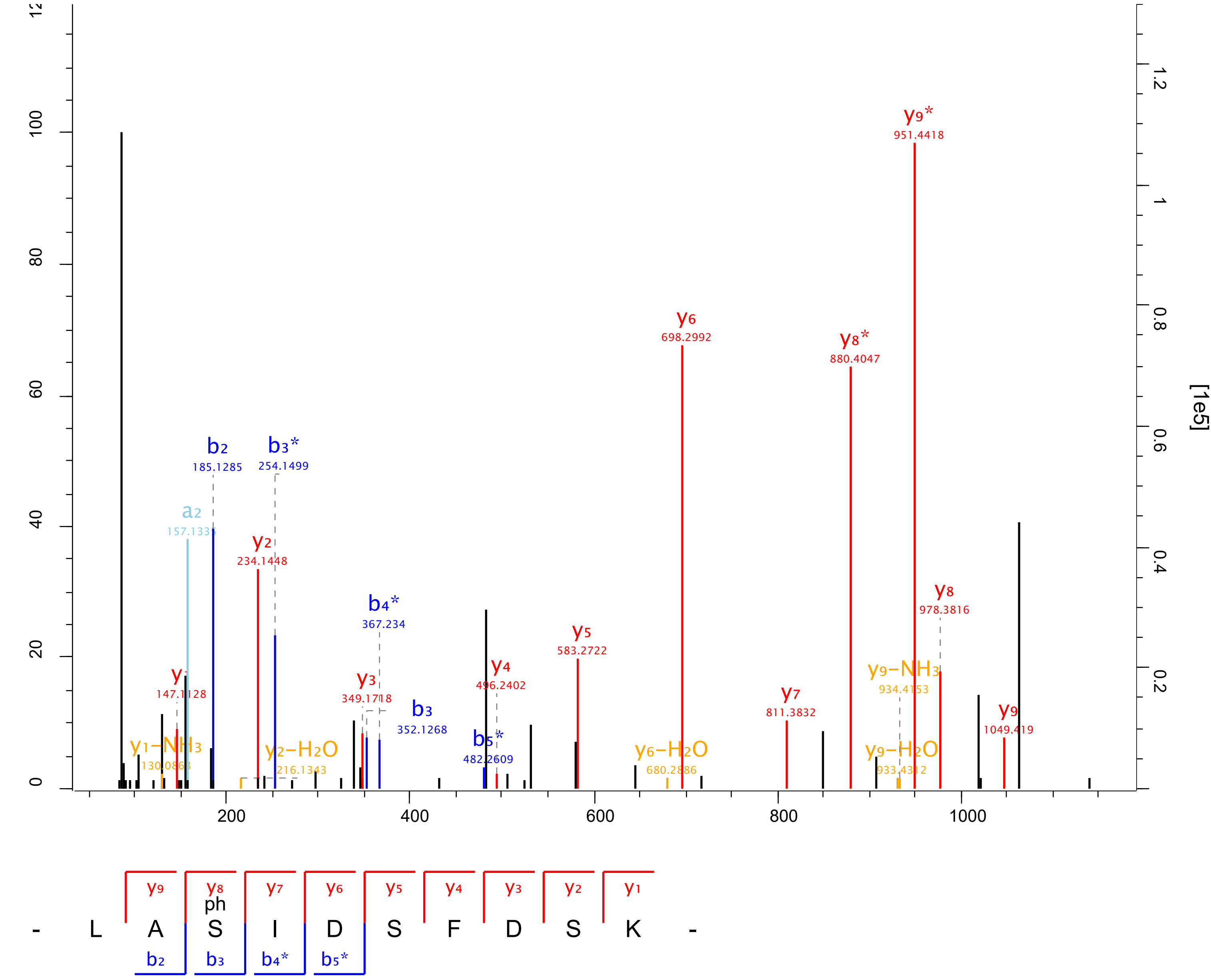

| Raw file | Scan | Method | Score | m/z |
| --- | --- | --- | --- | --- |
| water_15_2 | 22091 | FTMS; HCD | 42.95 | 739.85 |

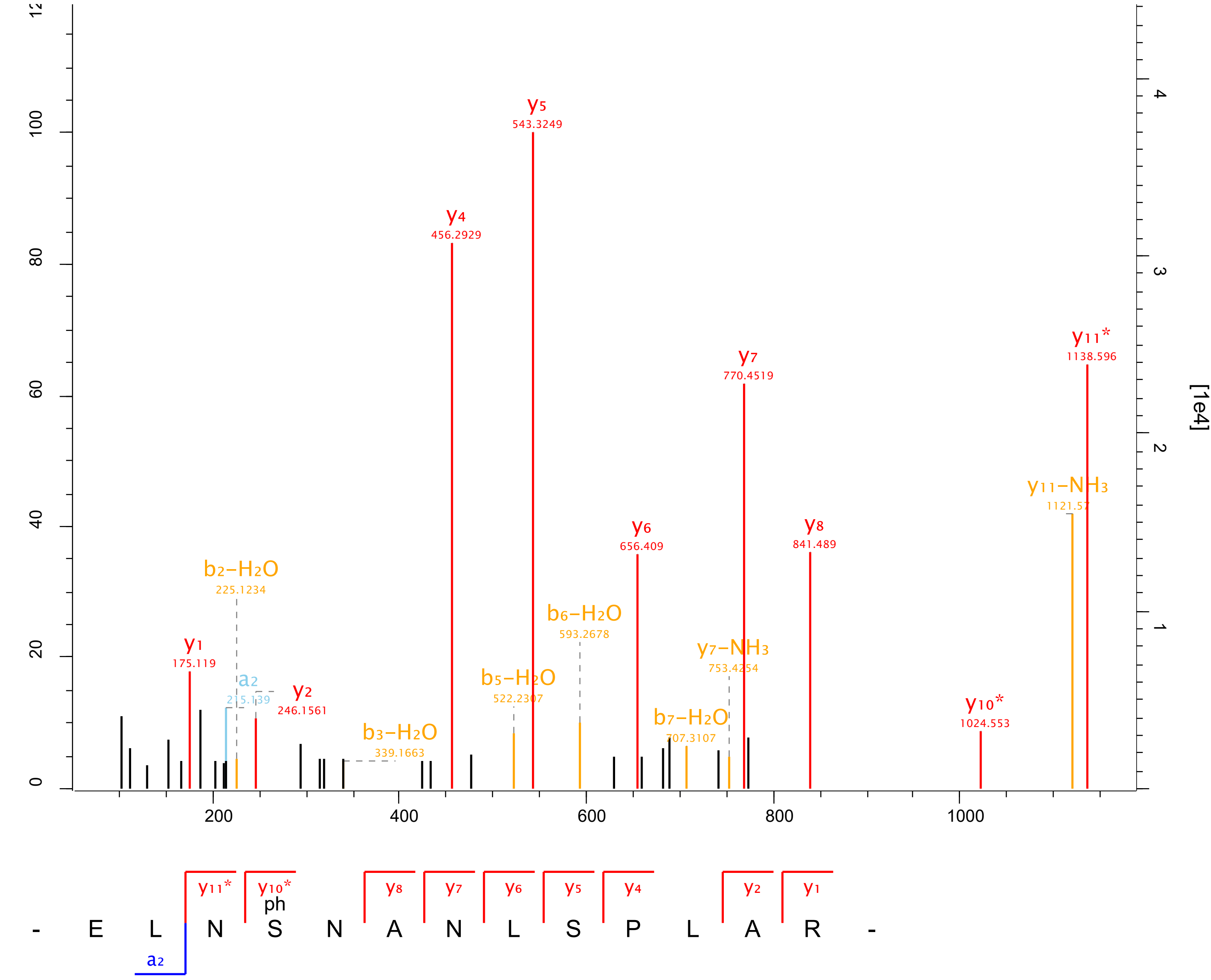

| Raw file | Scan | Method | Score | m/z |
| --- | --- | --- | --- | --- |
| water_15_2 | 22197 | FTMS; HCD | 40.49 | 756.33 |

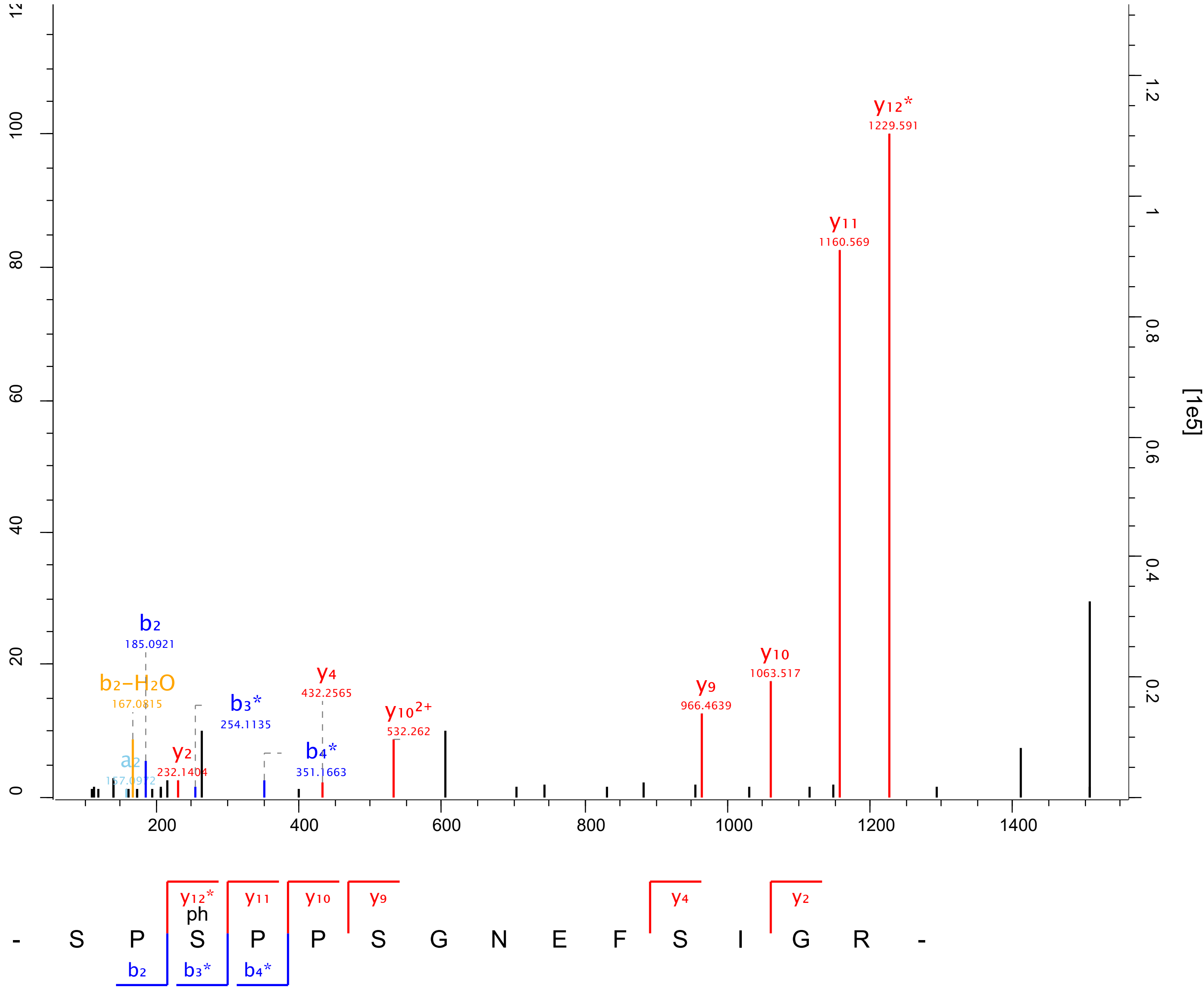

| Raw file | Scan | Method | Score | m/z |
|---|---|---|---|---|
| water_15_2 | 22324 | FTMS; HCD | 70.09 | 585.79 |

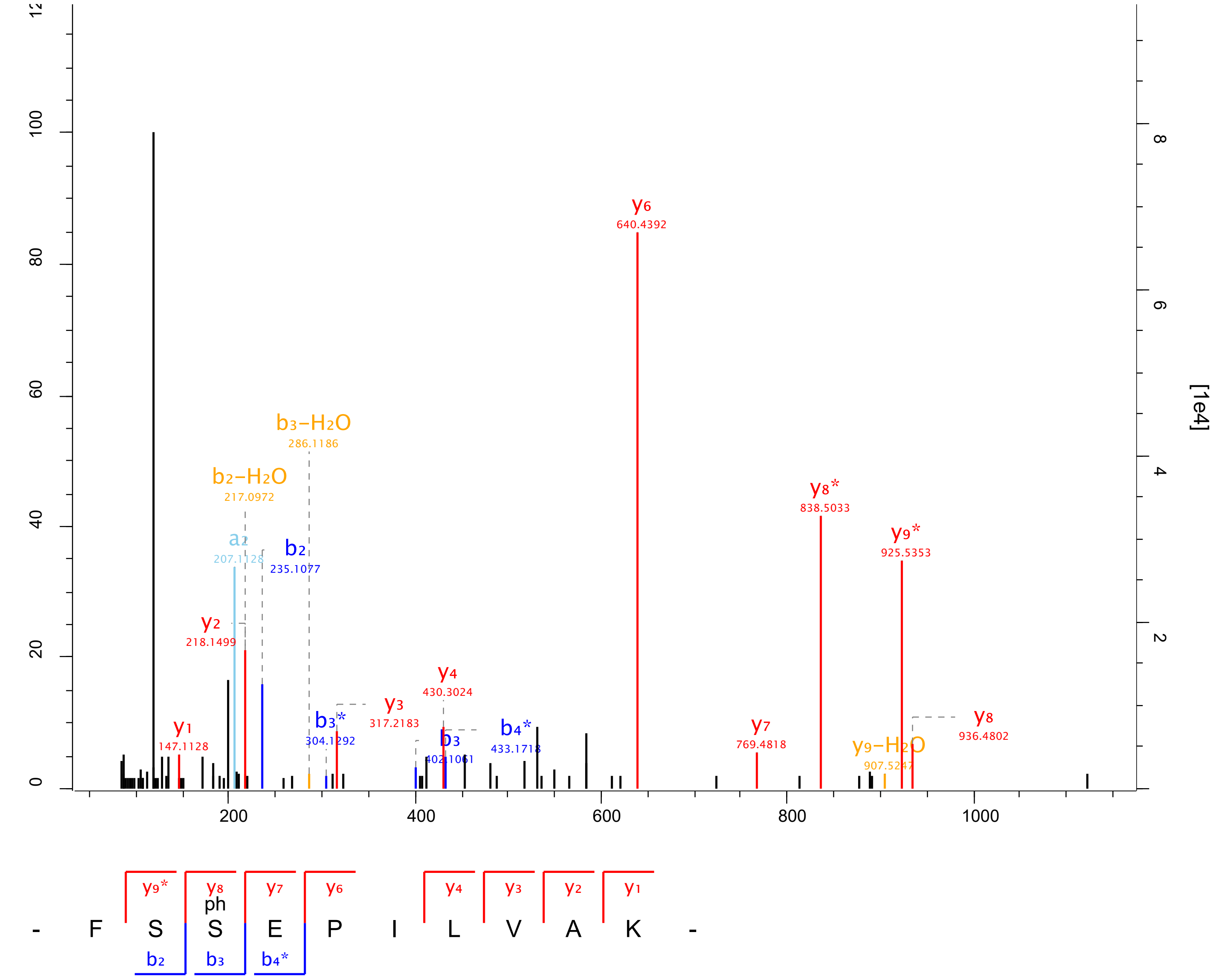

| Raw file | Scan | Method | Score | m/z |
| --- | --- | --- | --- | --- |
| water_15_2 | 22346 | FTMS; HCD | 114.96 | 697.86 |

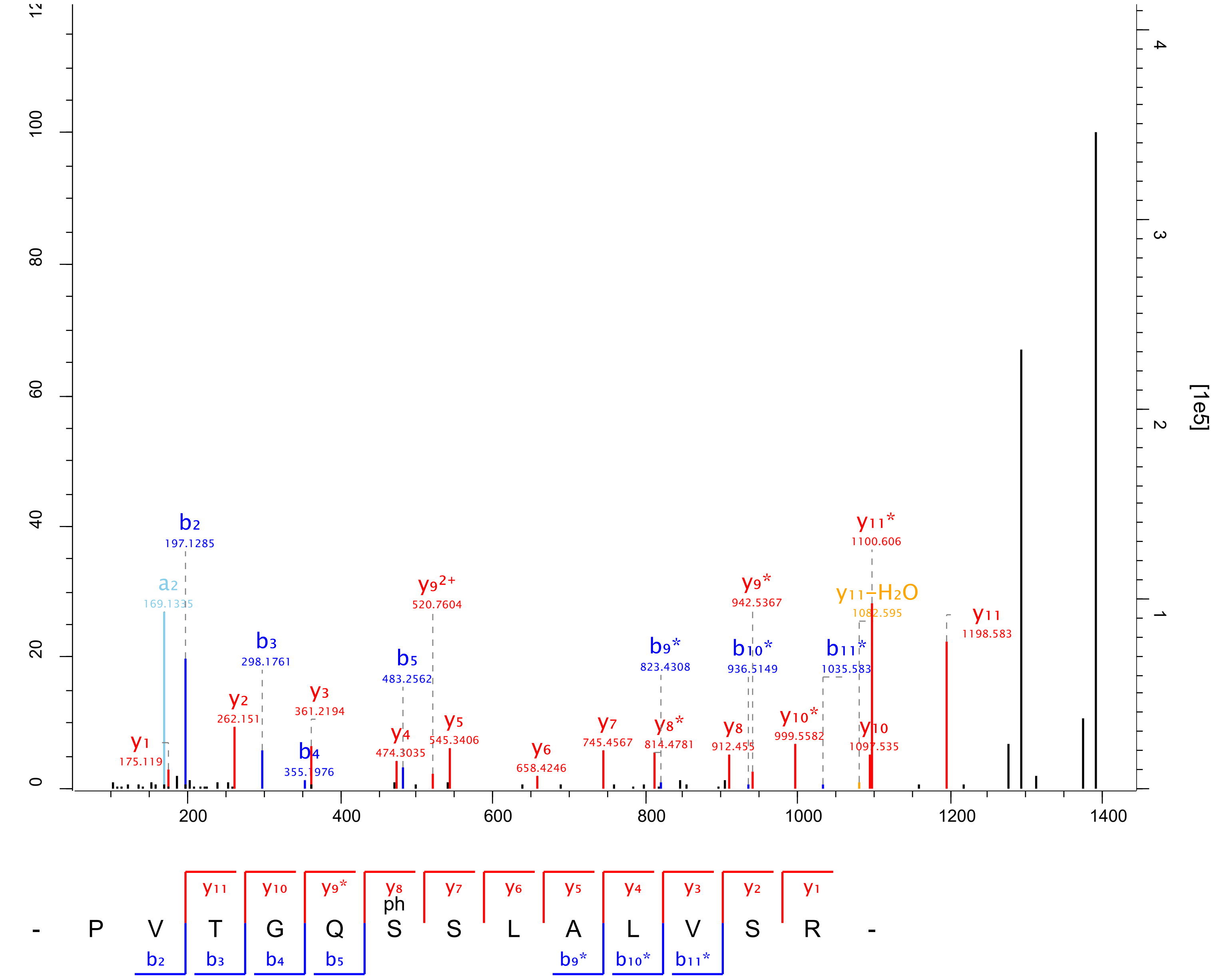

| Raw file | Scan | Method | Score | m/z |
| --- | --- | --- | --- | --- |
| water_15_2 | 22404 | FTMS; HCD | 53.62 | 920.39 |

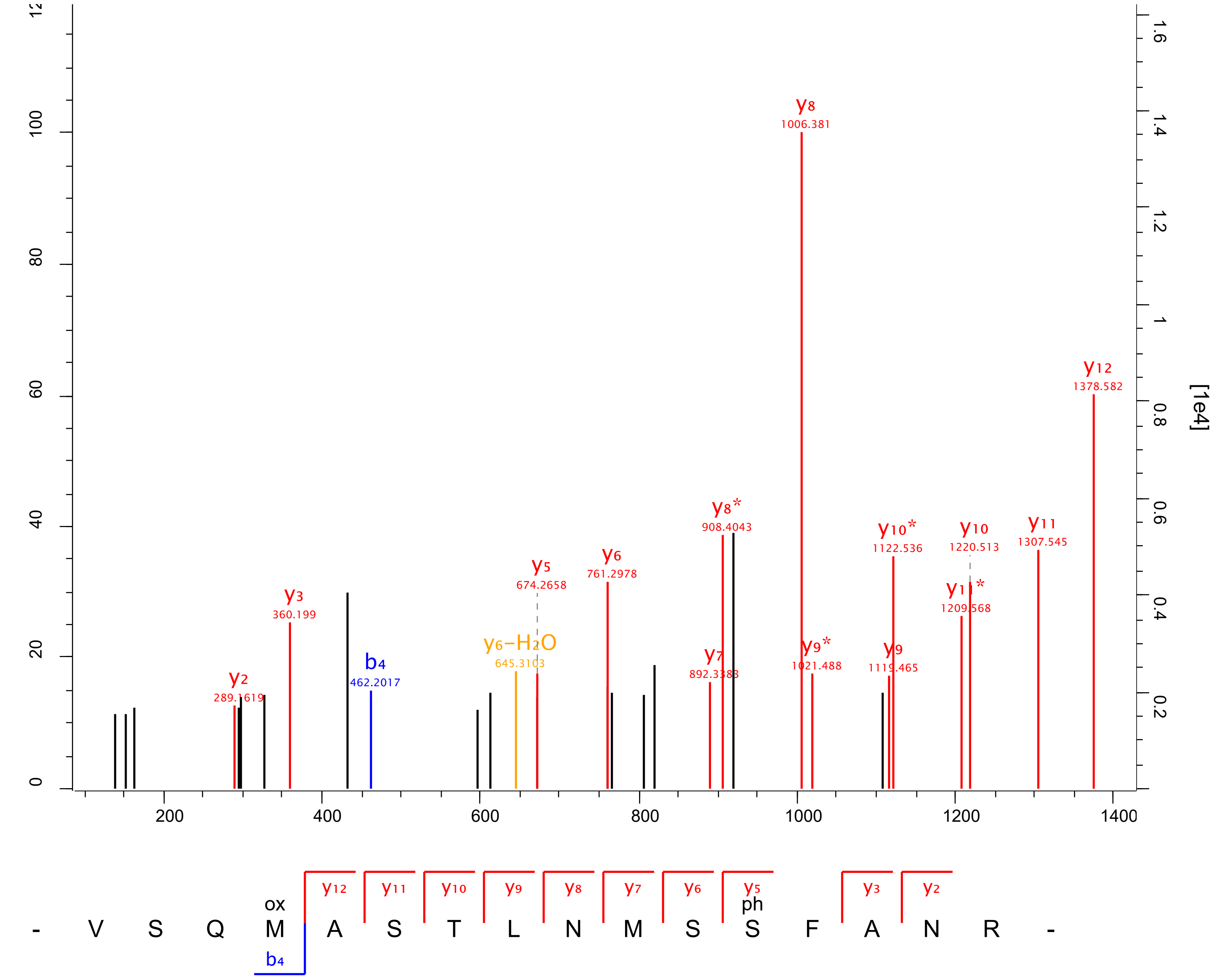

| Raw file | Scan | Method | Score | m/z |
|---|---|---|---|---|
| water_15_2 | 22420 | FTMS; HCD | 43.5 | 481.26 |

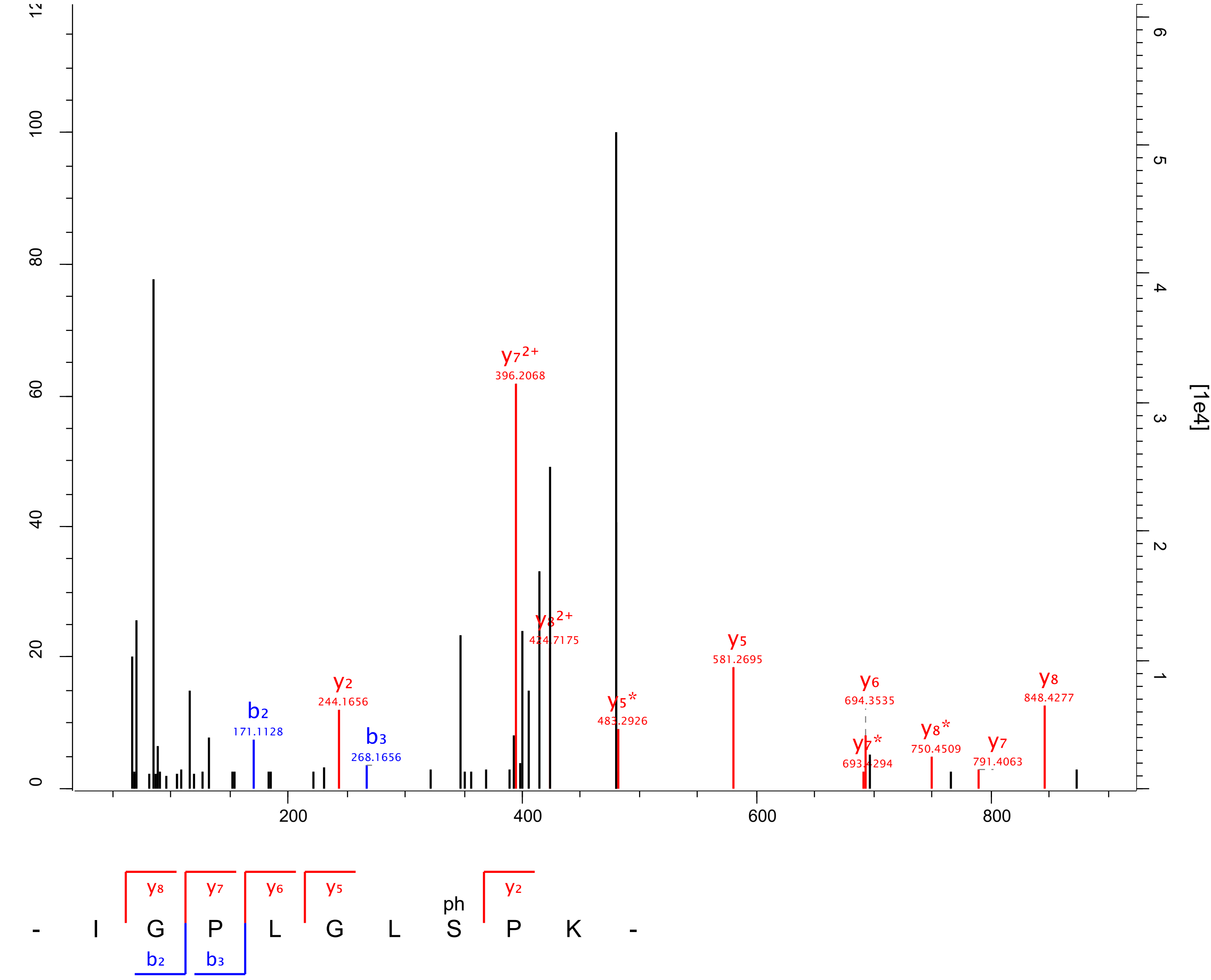

| Raw file | Scan | Method | Score | m/z |
|---|---|---|---|---|
| water_15_2 | 22447 | FTMS; HCD | 204.13 | 627.3 |

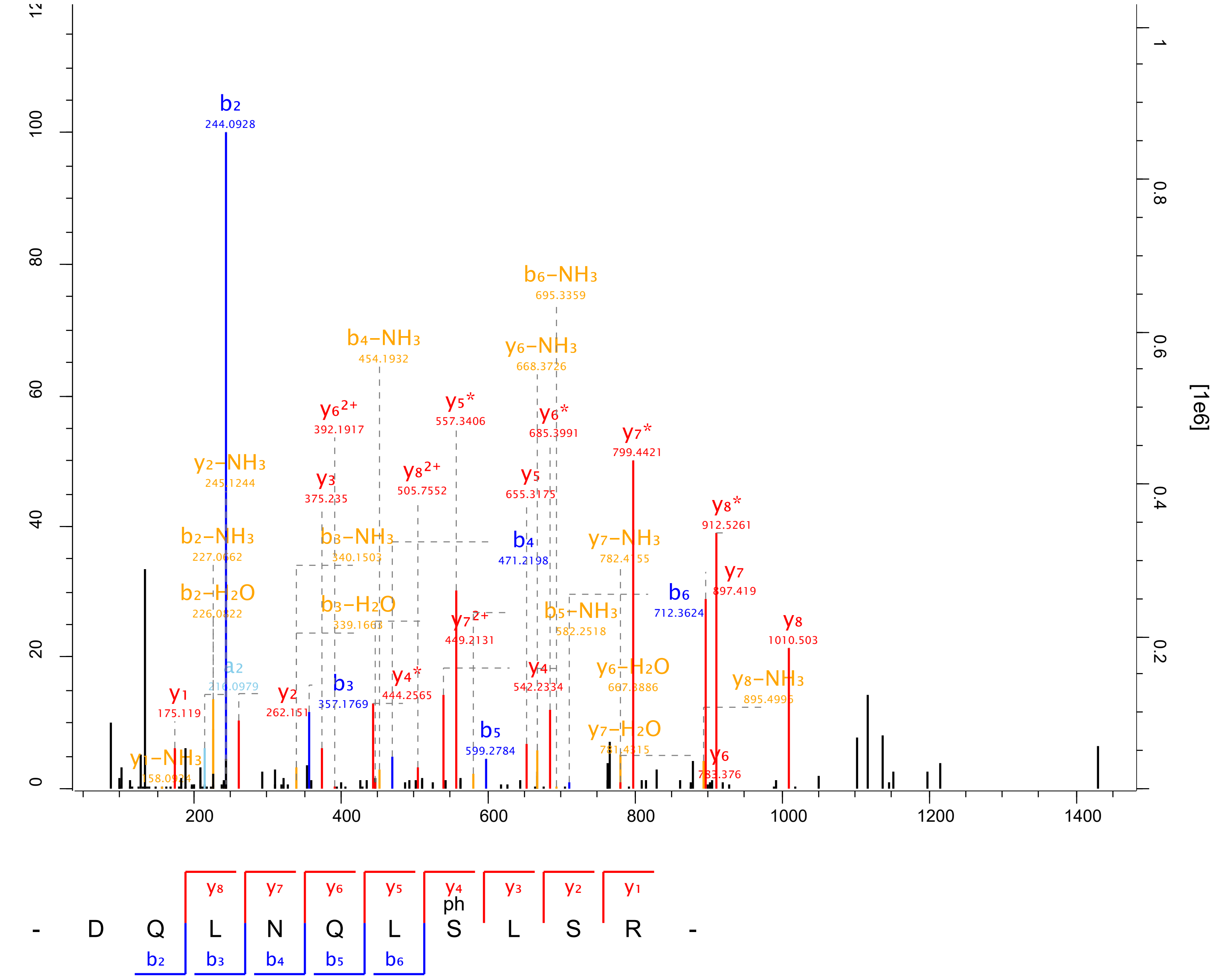

| Raw file | Scan | Method | Score | m/z |
| --- | --- | --- | --- | --- |
| water_15_2 | 22593 | FTMS; HCD | 156.49 | 752.82 |

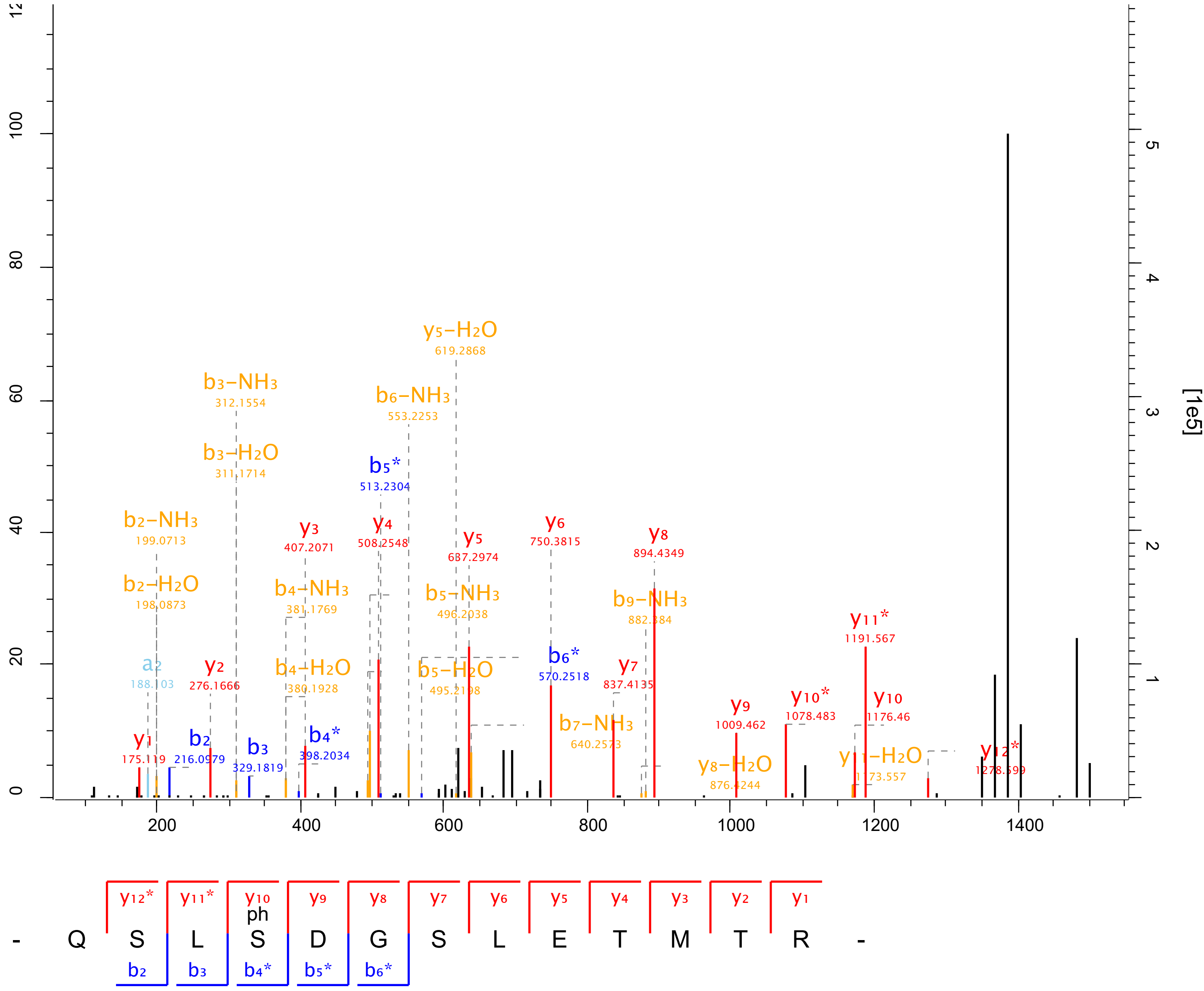

| Raw file | Scan | Method | Score | m/z |
| --- | --- | --- | --- | --- |
| water_15_2 | 22624 | FTMS; HCD | 44.34 | 790.72 |

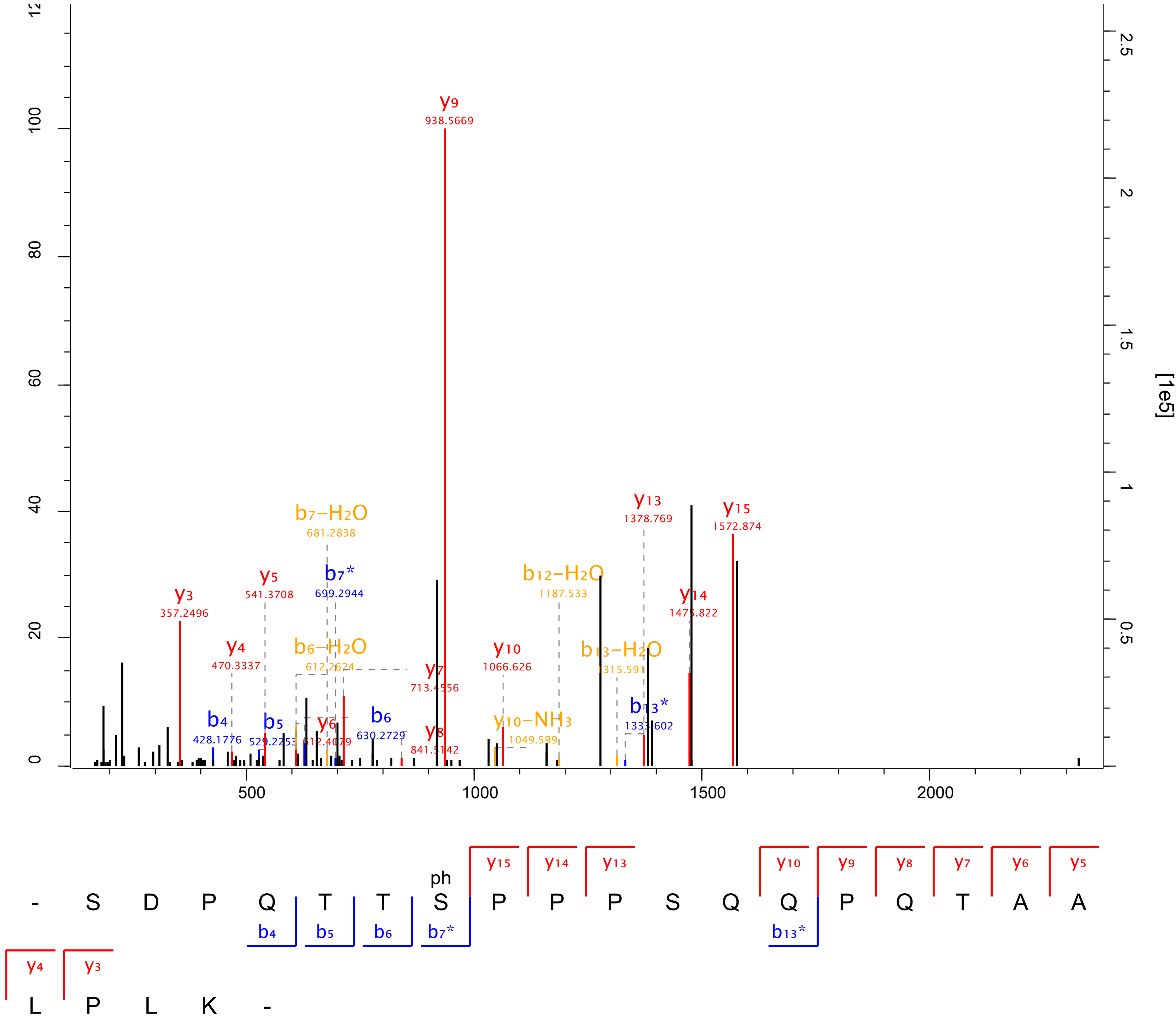

| Raw file | Scan | Method | Score | m/z |
|---|---|---|---|---|
| water_15_2 | 22659 | FTMS; HCD | 97.24 | 767.3 |

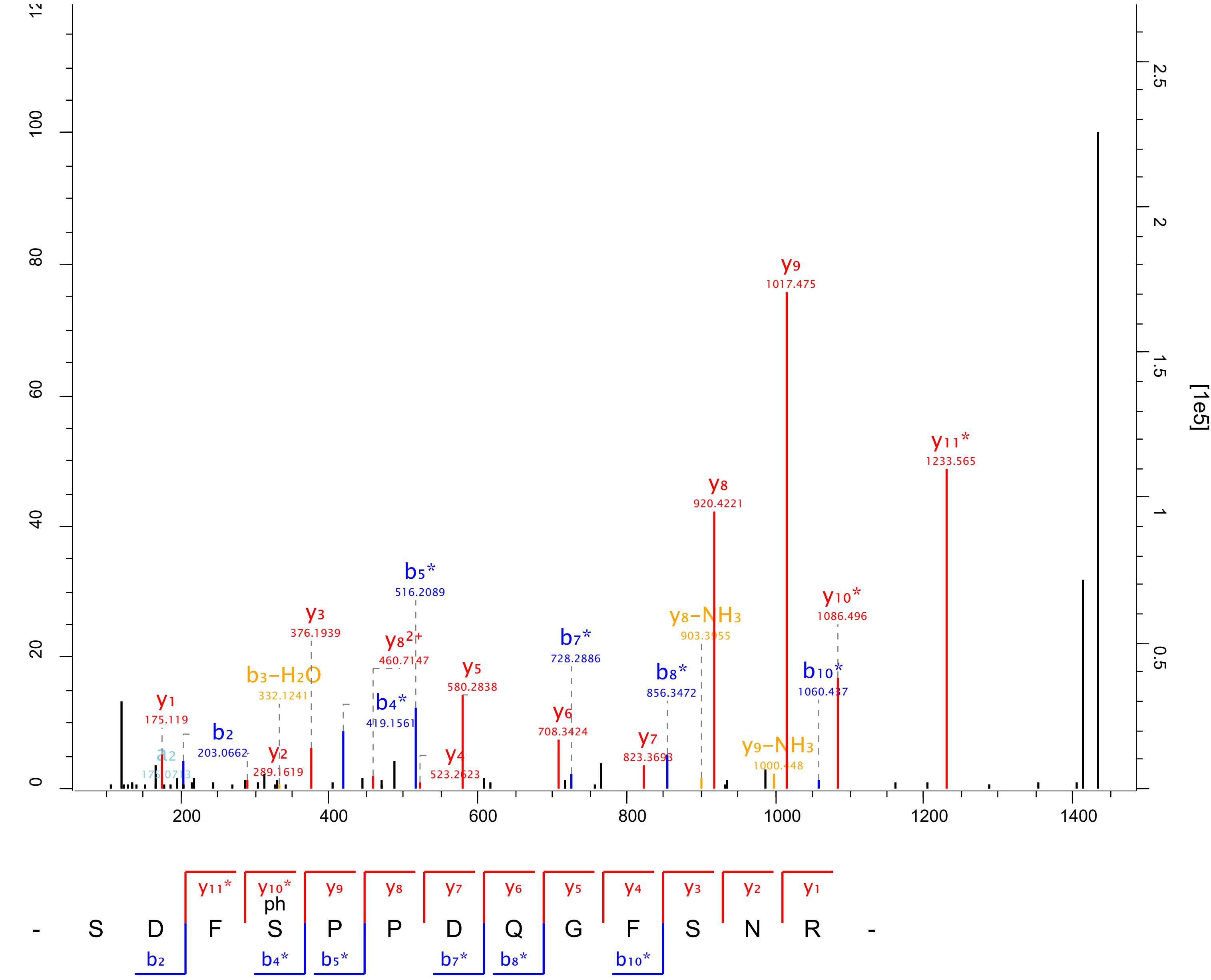

| Raw file | Scan | Method | Score | m/z |
|---|---|---|---|---|
| water_15_2 | 22680 | FTMS; HCD | 145.72 | 917.37 |

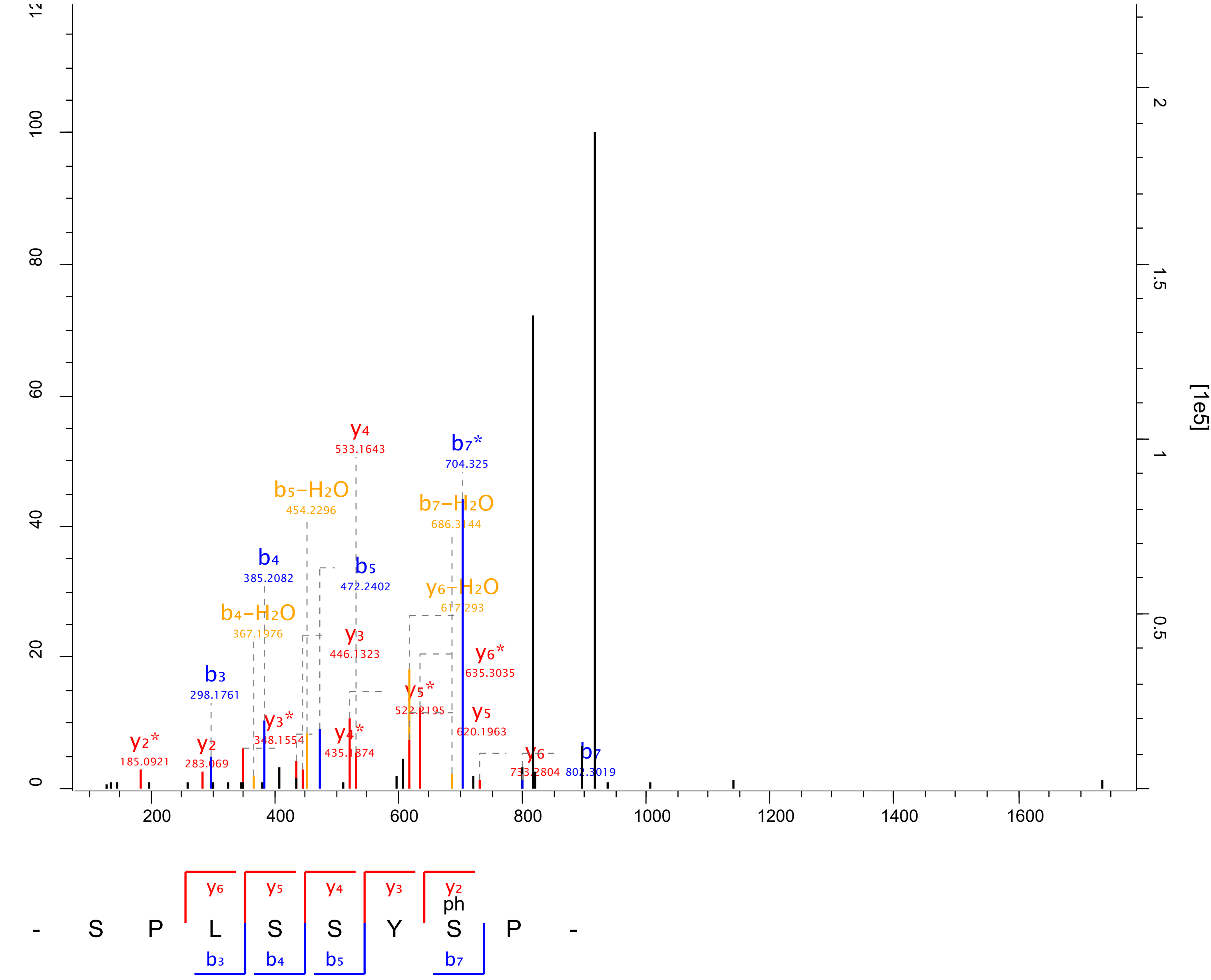

| Raw file | Scan | Method | Score | m/z |
|---|---|---|---|---|
| water_15_2 | 22683 | FTMS; HCD | 98.54 | 880.9 |

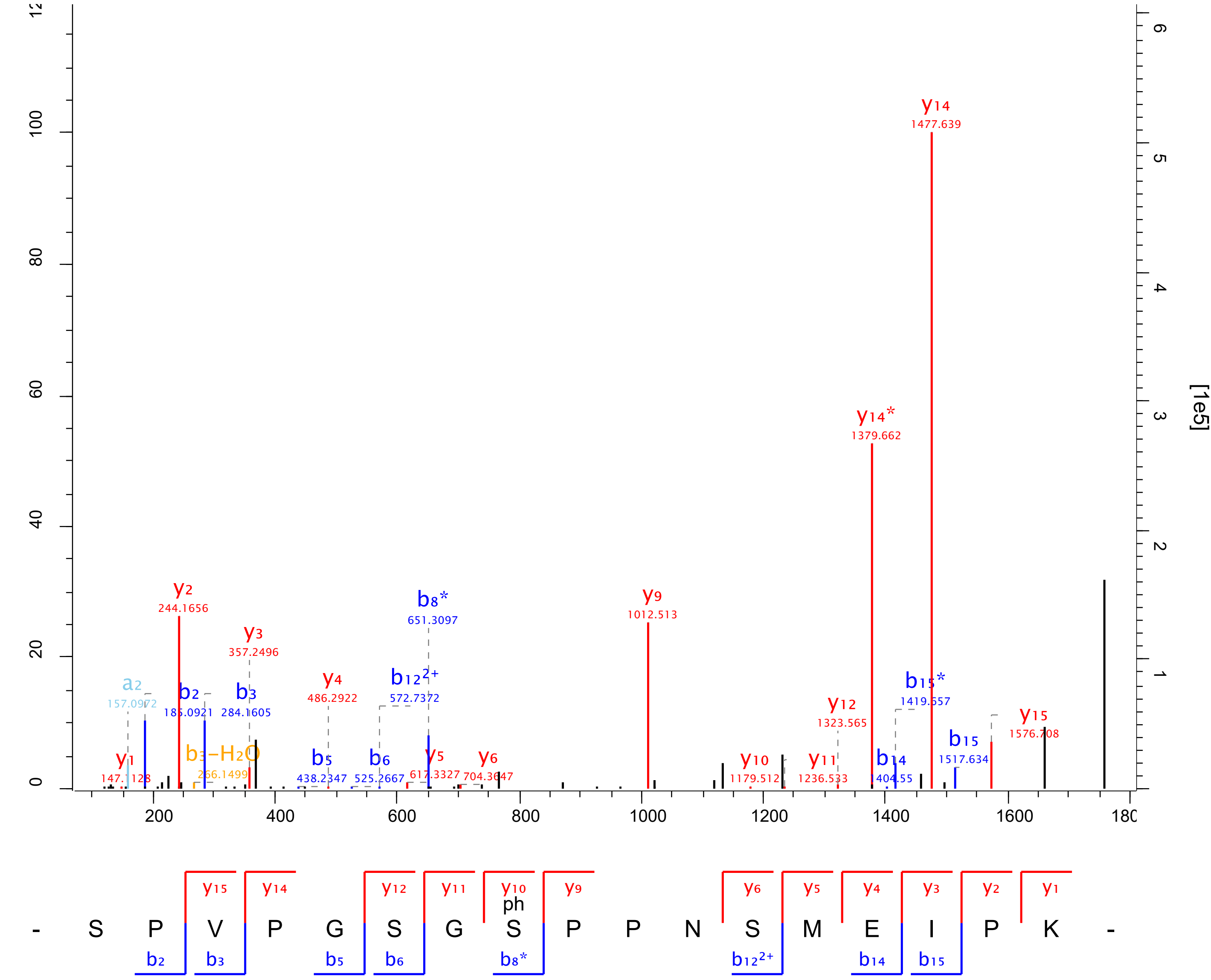

| Raw file | Scan | Method | Score | m/z |
|---|---|---|---|---|
| water_15_2 | 22808 | FTMS; HCD | 144.27 | 819.87 |

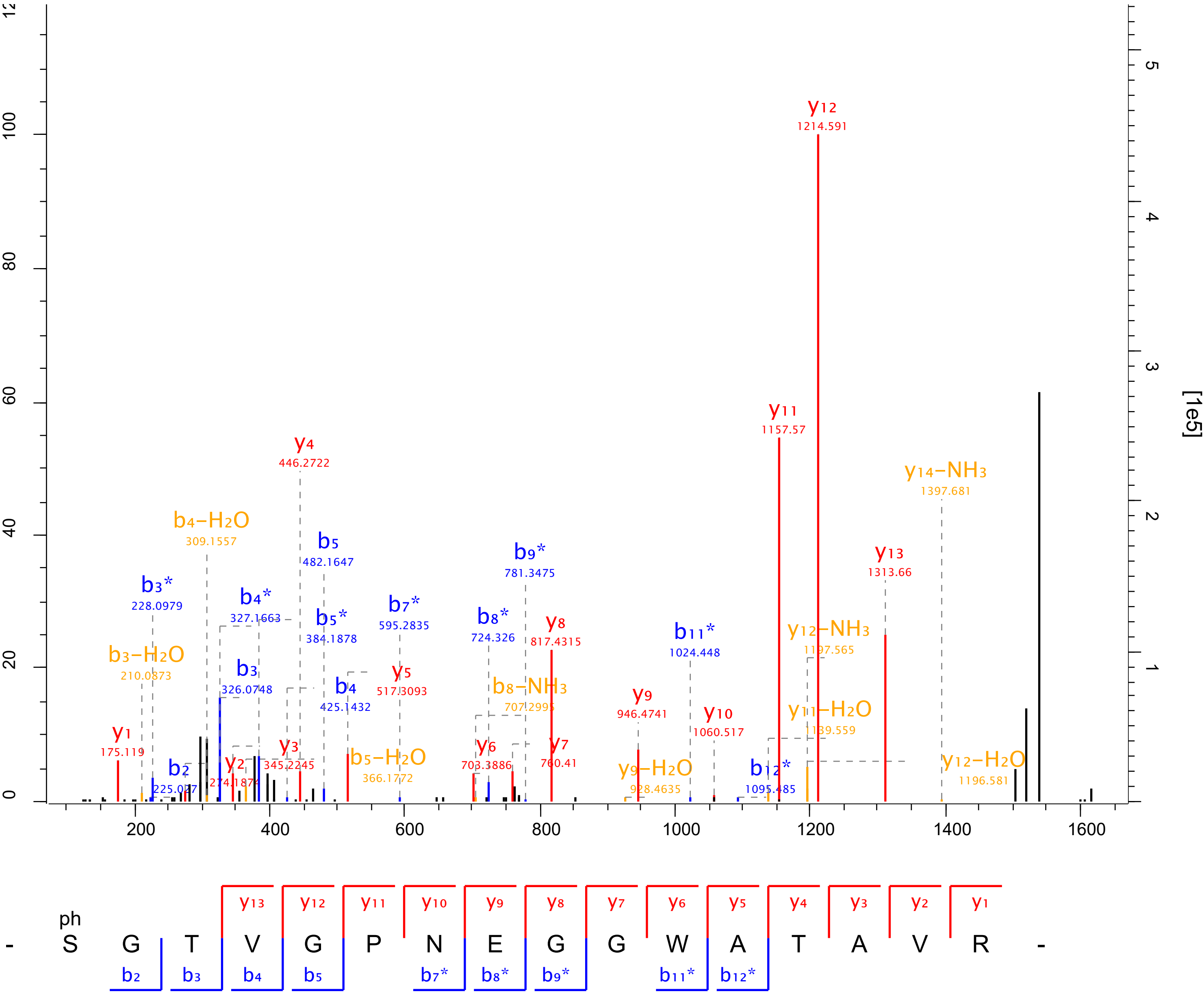

| Raw file | Scan | Method | Score | m/z |
|---|---|---|---|---|
| water_15_2 | 22833 | FTMS; HCD | 230.23 | 713.84 |

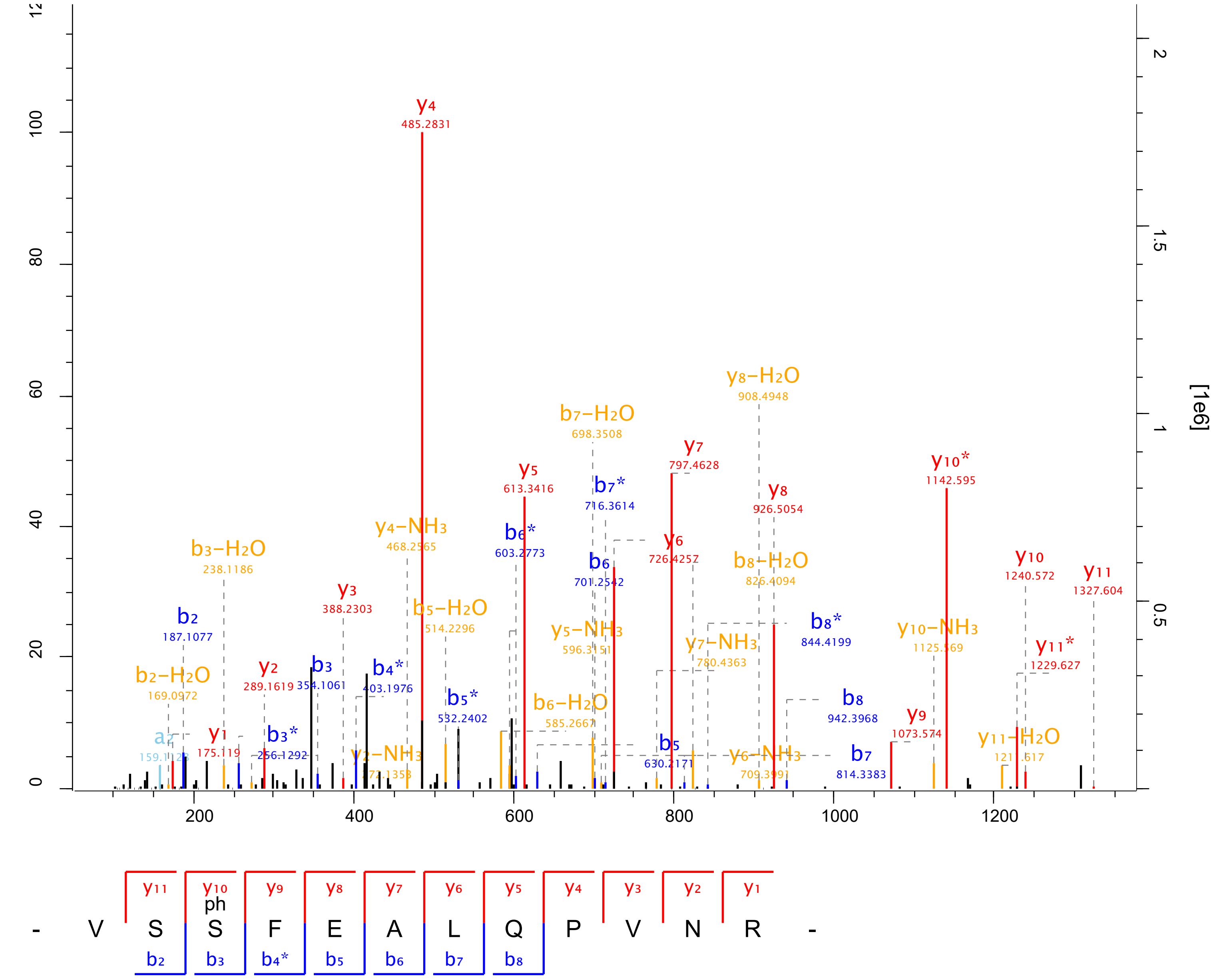

| Raw file | Scan | Method | Score | m/z |
|---|---|---|---|---|
| water_15_2 | 23088 | FTMS; HCD | 252.55 | 919.35 |

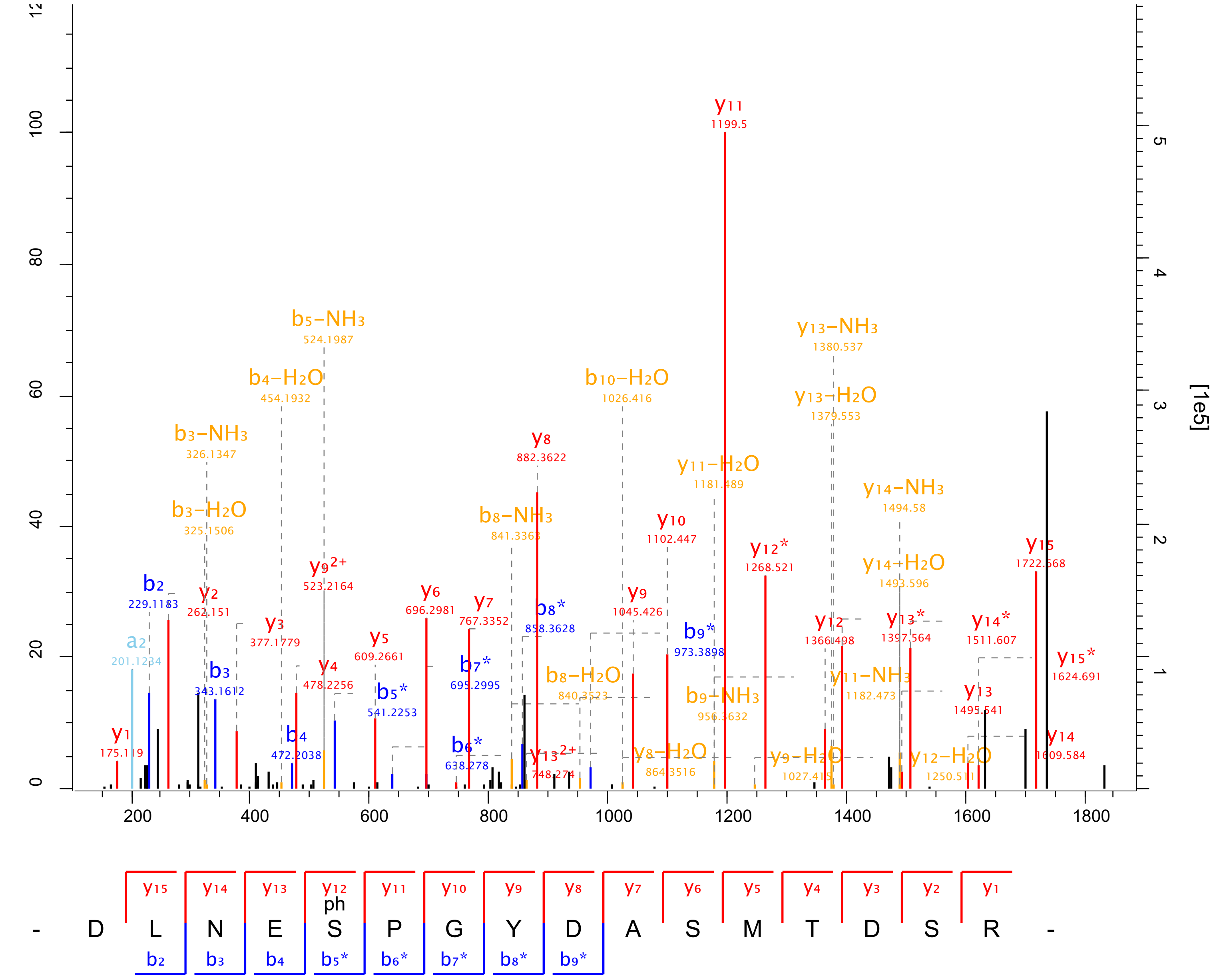

| Raw file | Scan | Method | Score | m/z |
| --- | --- | --- | --- | --- |
| water_15_2 | 23170 | FTMS; HCD | 40.74 | 708.66 |

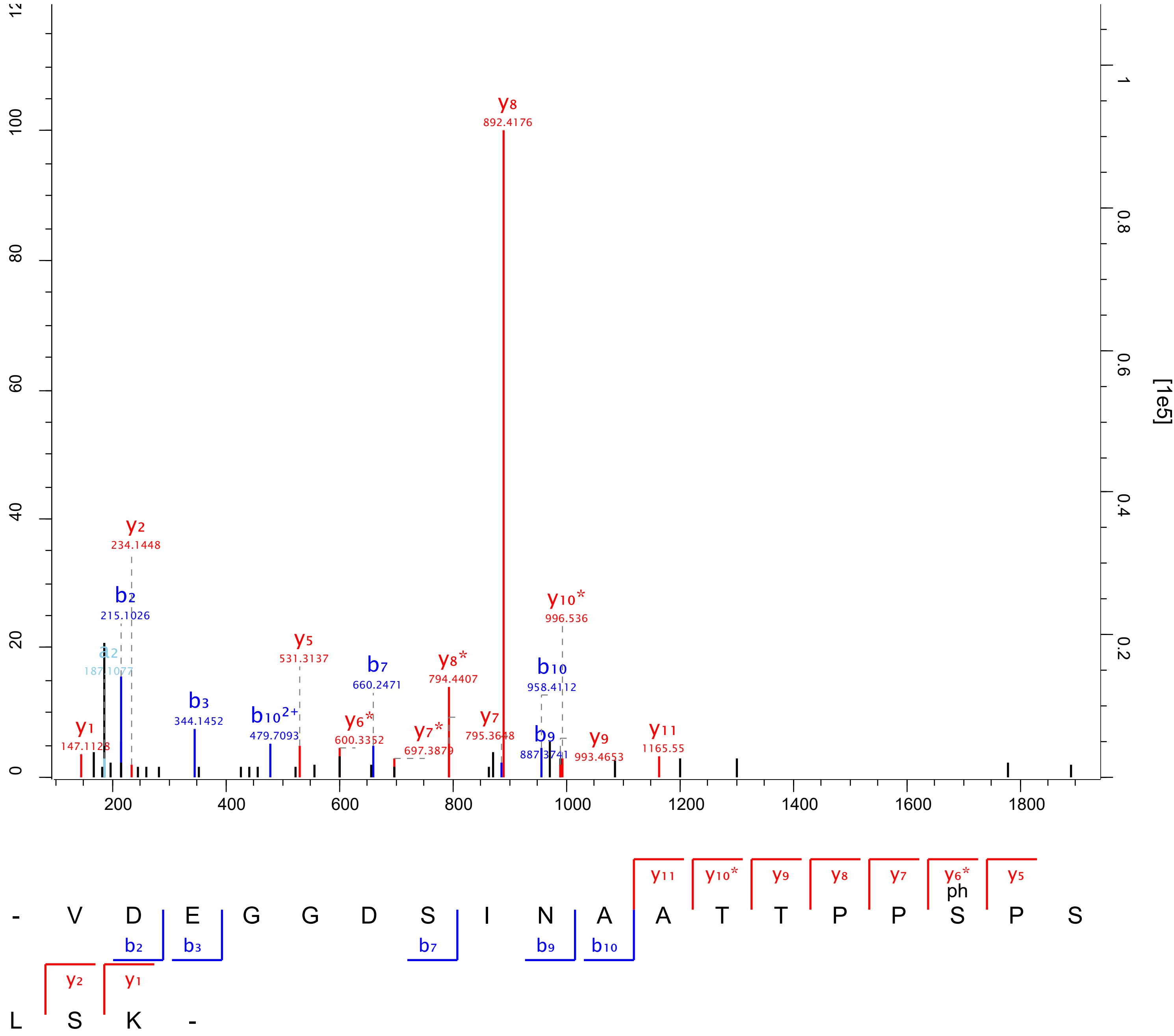

| Raw file | Scan | Method | Score | m/z |
| --- | --- | --- | --- | --- |
| water_15_2 | 23289 | FTMS; HCD | 98.28 | 715.3 |

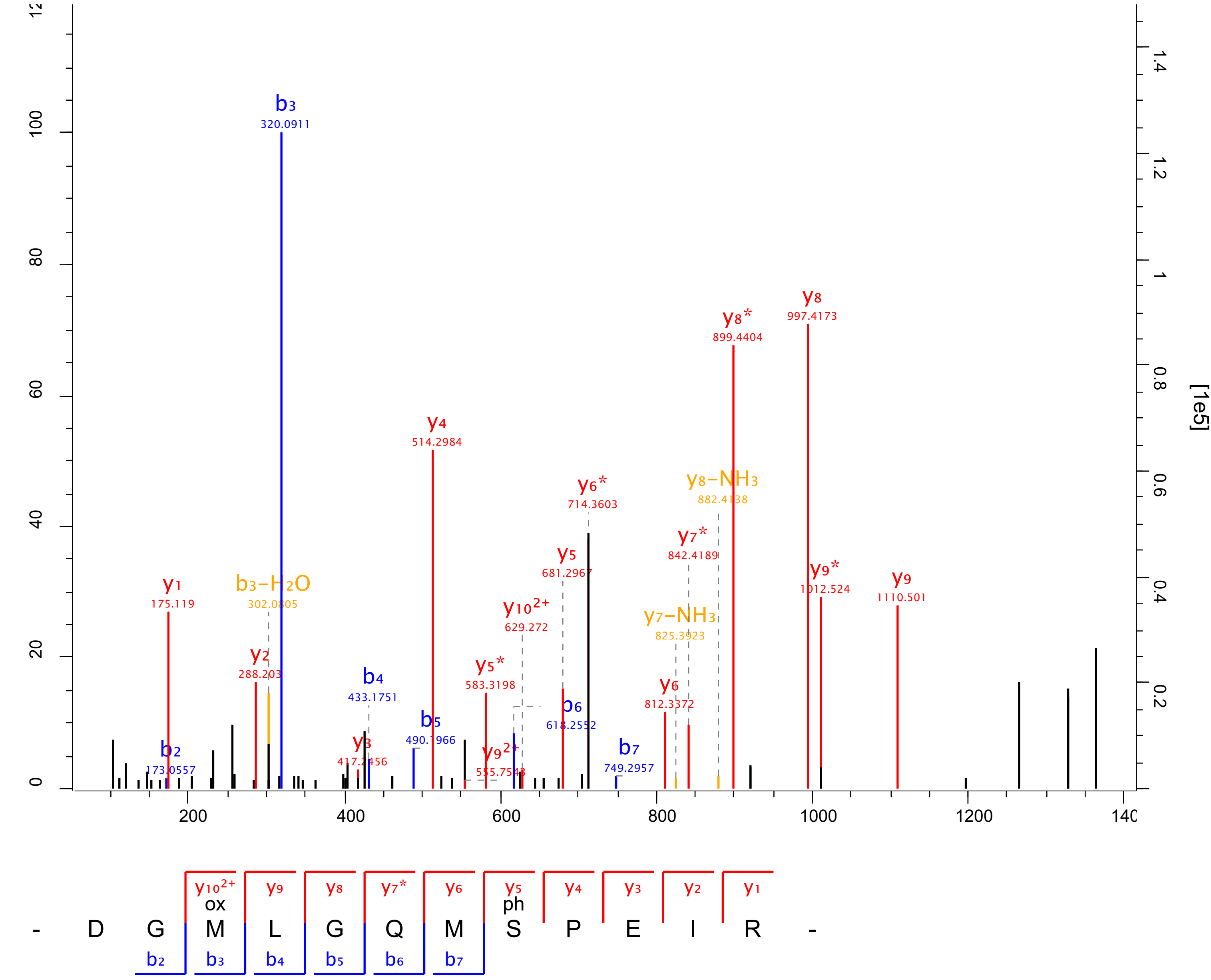

| Raw file | Scan | Method | Score | m/z |
| --- | --- | --- | --- | --- |
| water_15_2 | 23332 | FTMS; HCD | 56.57 | 696.28 |

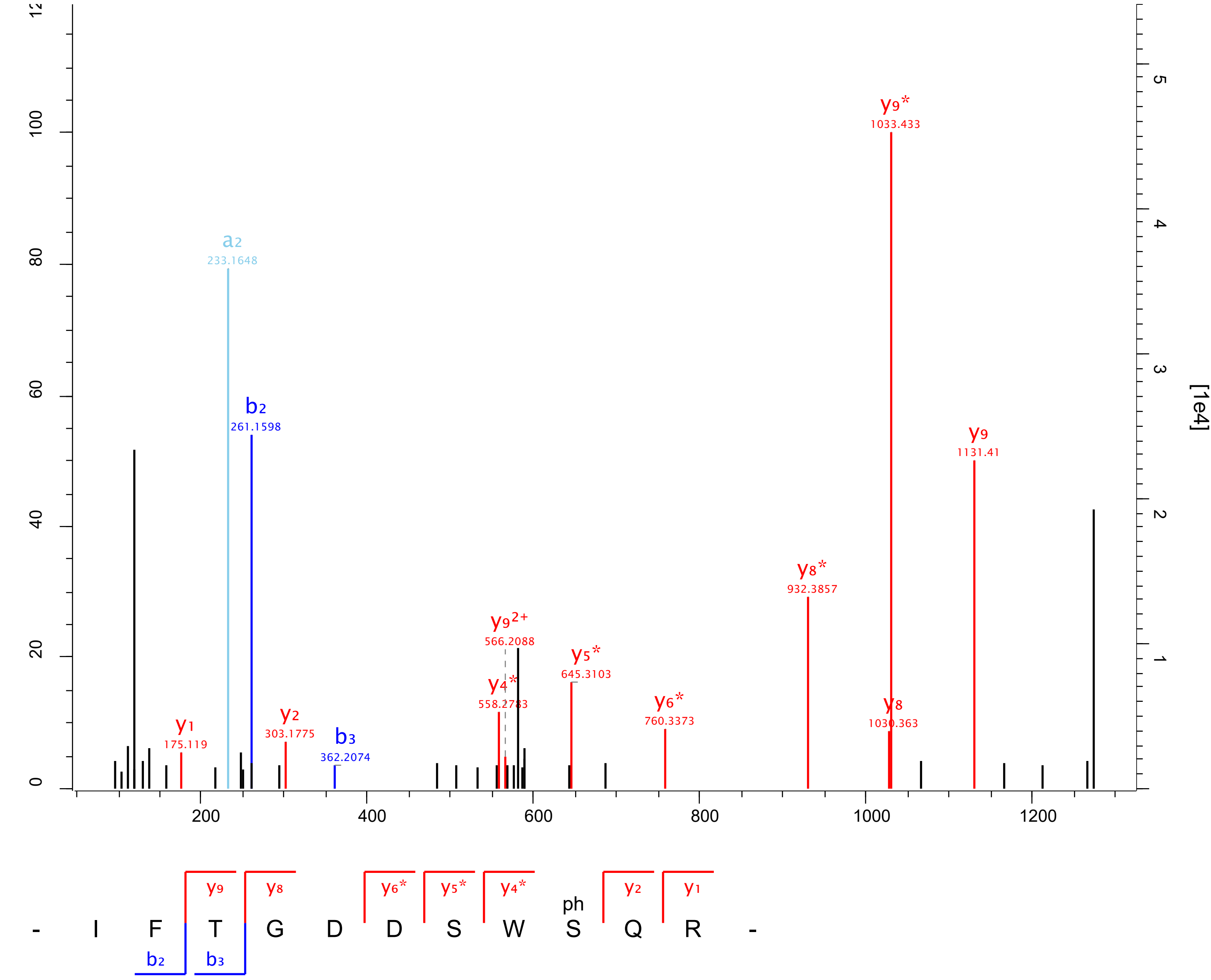

| Raw file | Scan | Method | Score | m/z |
|---|---|---|---|---|
| water_15_2 | 23378 | FTMS; HCD | 101.01 | 682.3 |

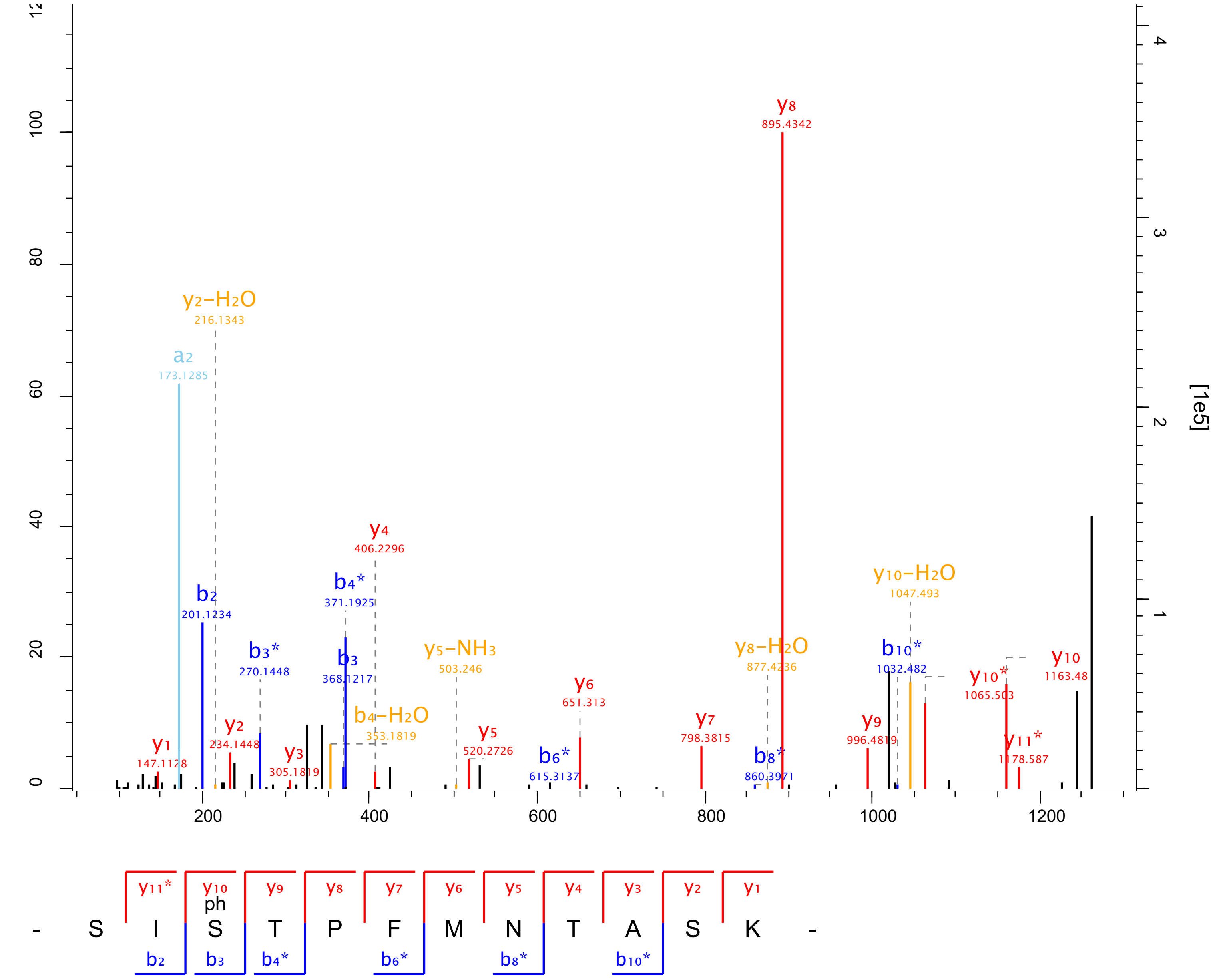

| Raw file | Scan | Method | Score | m/z |
|---|---|---|---|---|
| water_15_2 | 23389 | FTMS; HCD | 77.69 | 962.46 |

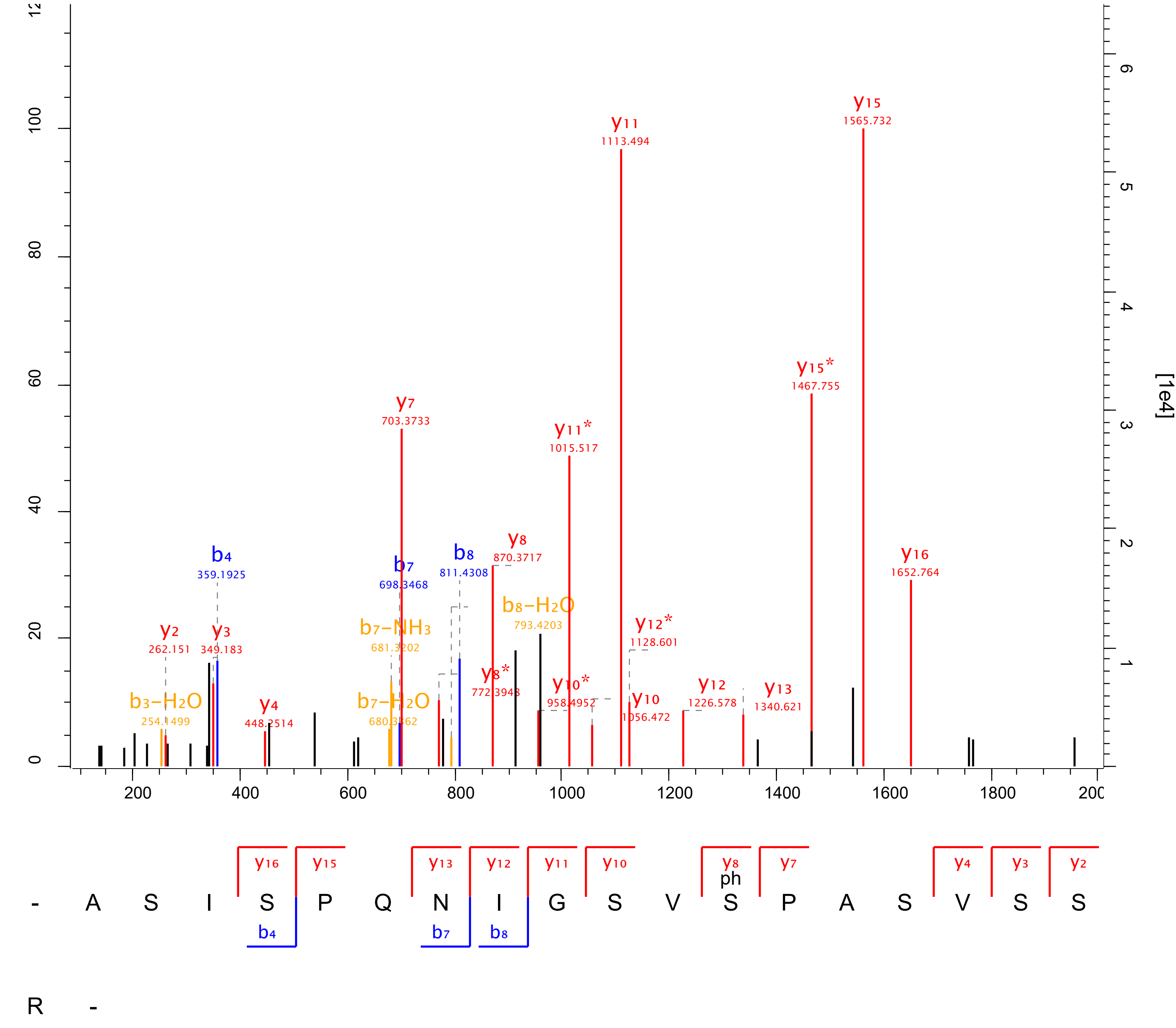

| Raw file | Scan | Method | Score | m/z |
| --- | --- | --- | --- | --- |
| water_15_2 | 23413 | FTMS; HCD | 52.17 | 650.3 |

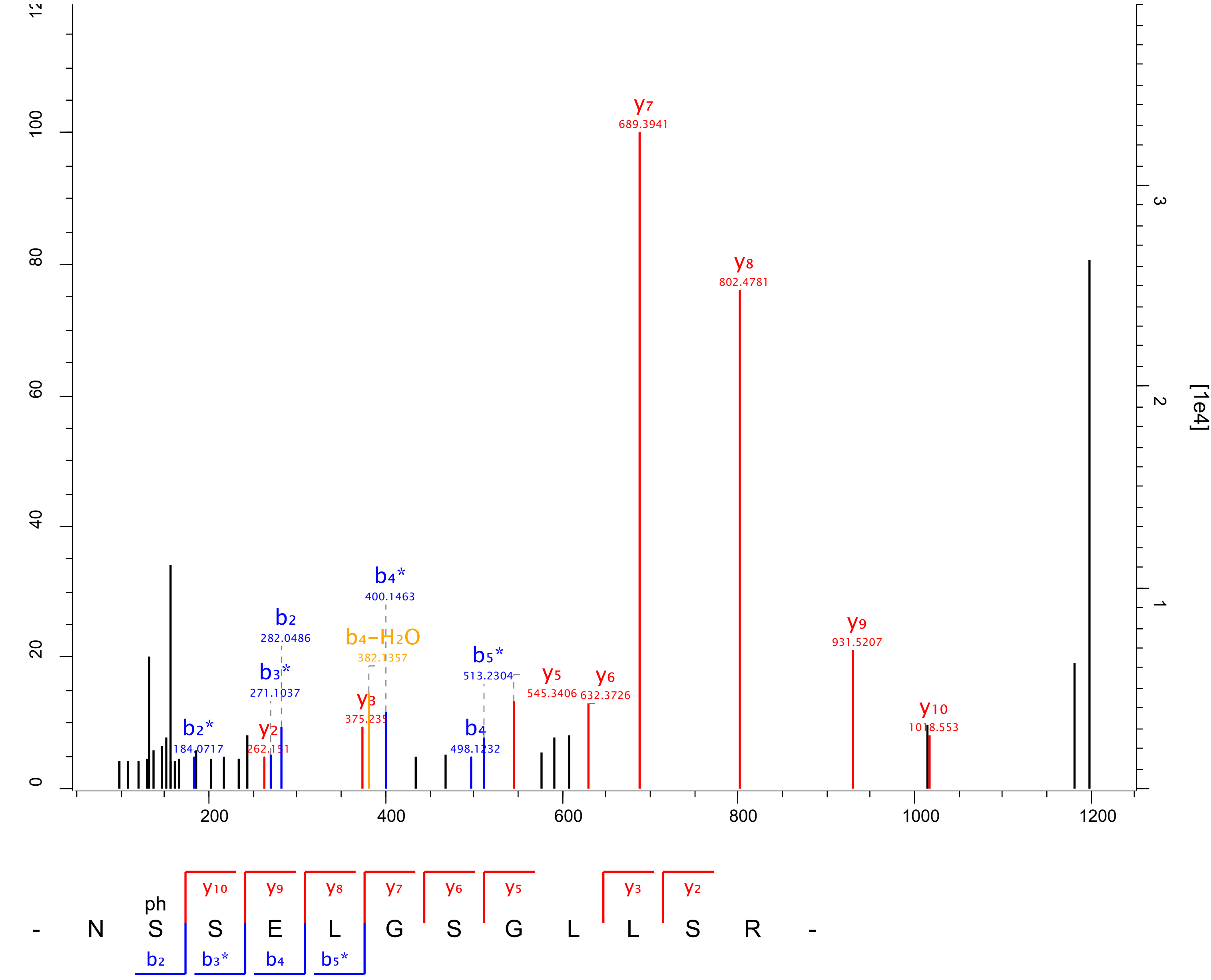

| Raw file | Scan | Method | Score | m/z |
|---|---|---|---|---|
| water_15_2 | 23416 | FTMS; HCD | 42.38 | 972.91 |

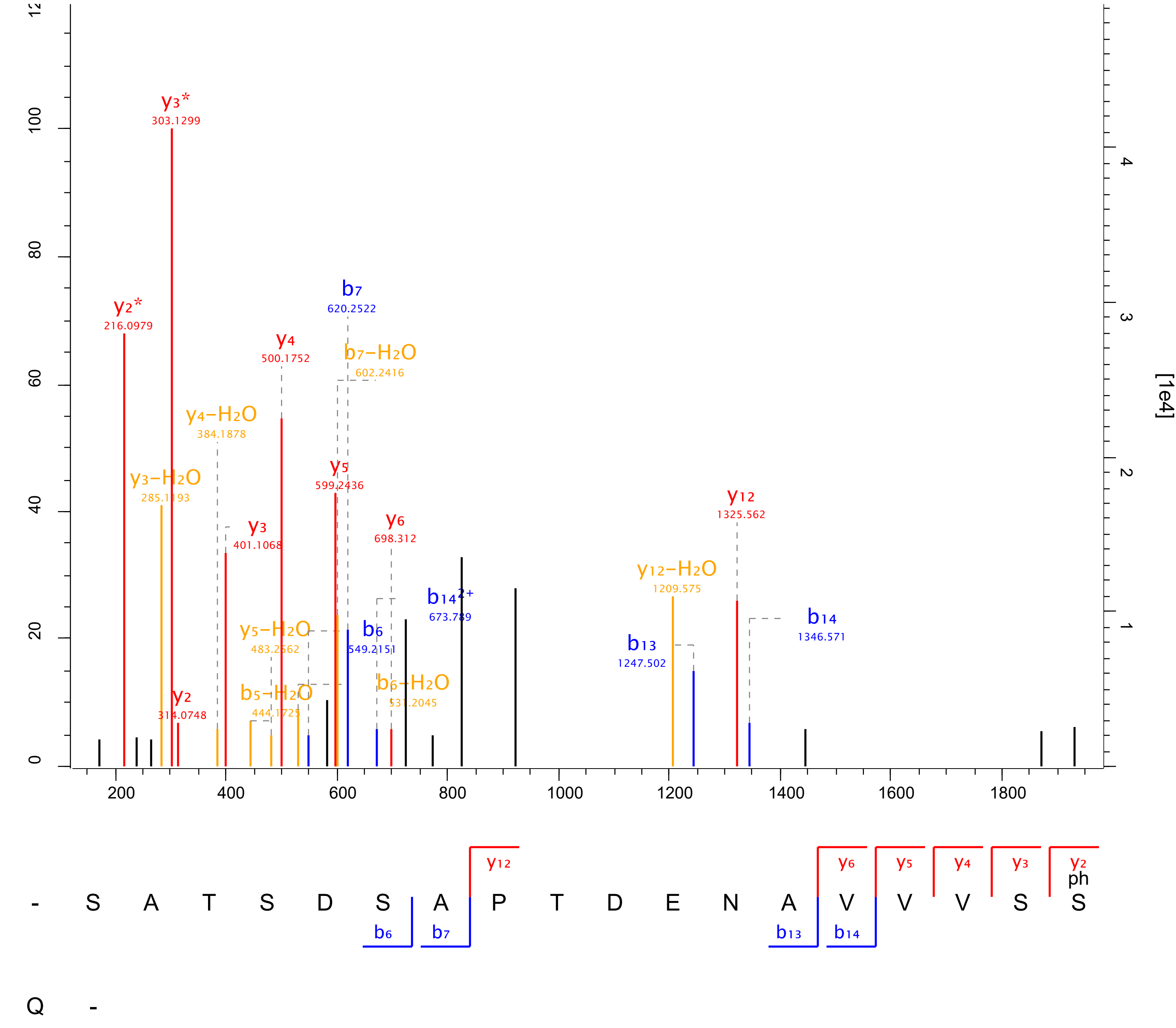

| Raw file | Scan | Method | Score | m/z |
|---|---|---|---|---|
| water_15_2 | 23582 | FTMS; HCD | 254.87 | 920.4 |

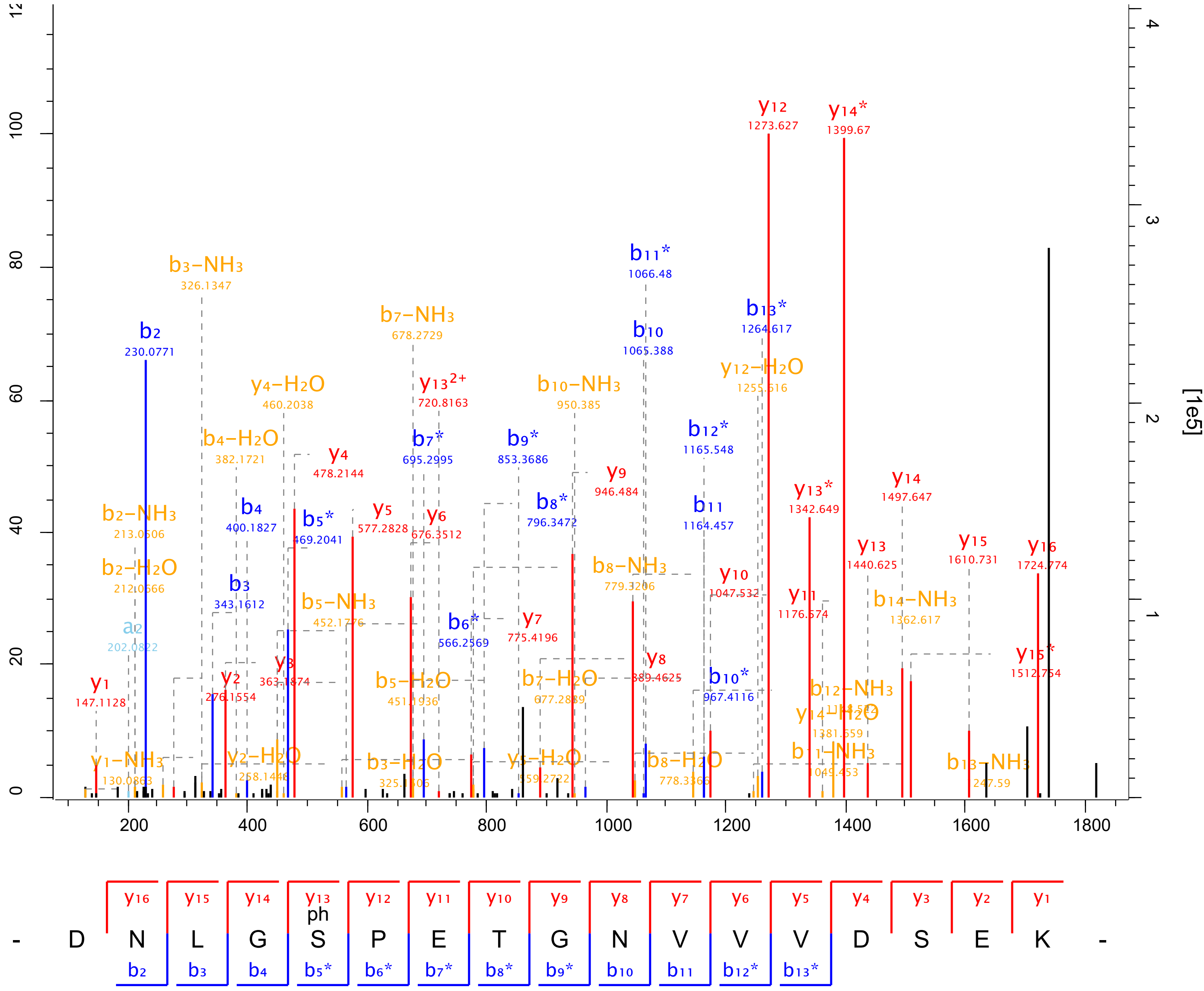

| Raw file | Scan | Method | Score | m/z |
|---|---|---|---|---|
| water_15_2 | 23612 | FTMS; HCD | 67.23 | 671.27 |

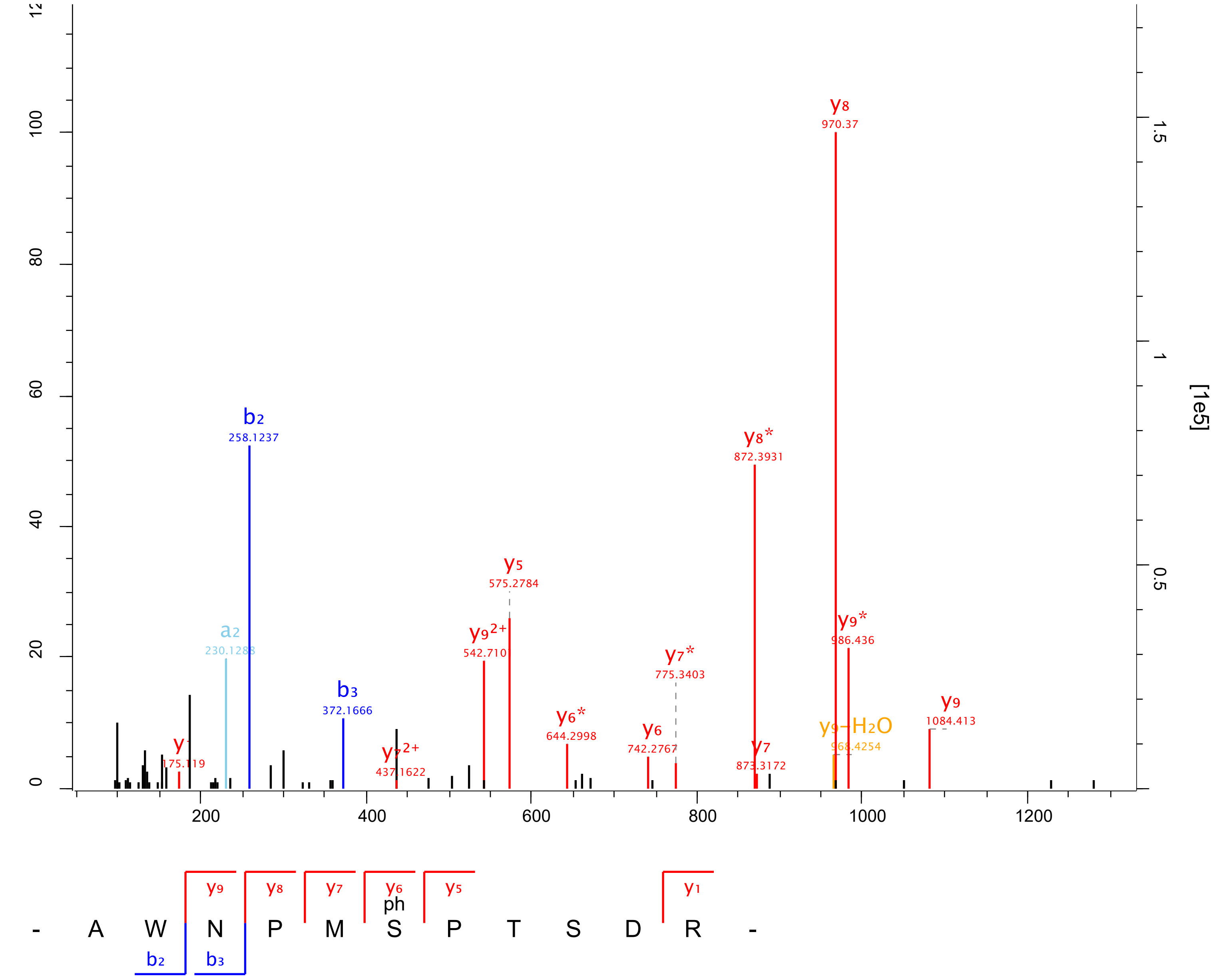

| Raw file | Scan | Method | Score | m/z |
|---|---|---|---|---|
| water_15_2 | 23735 | FTMS; HCD | 66.99 | 693.8 |

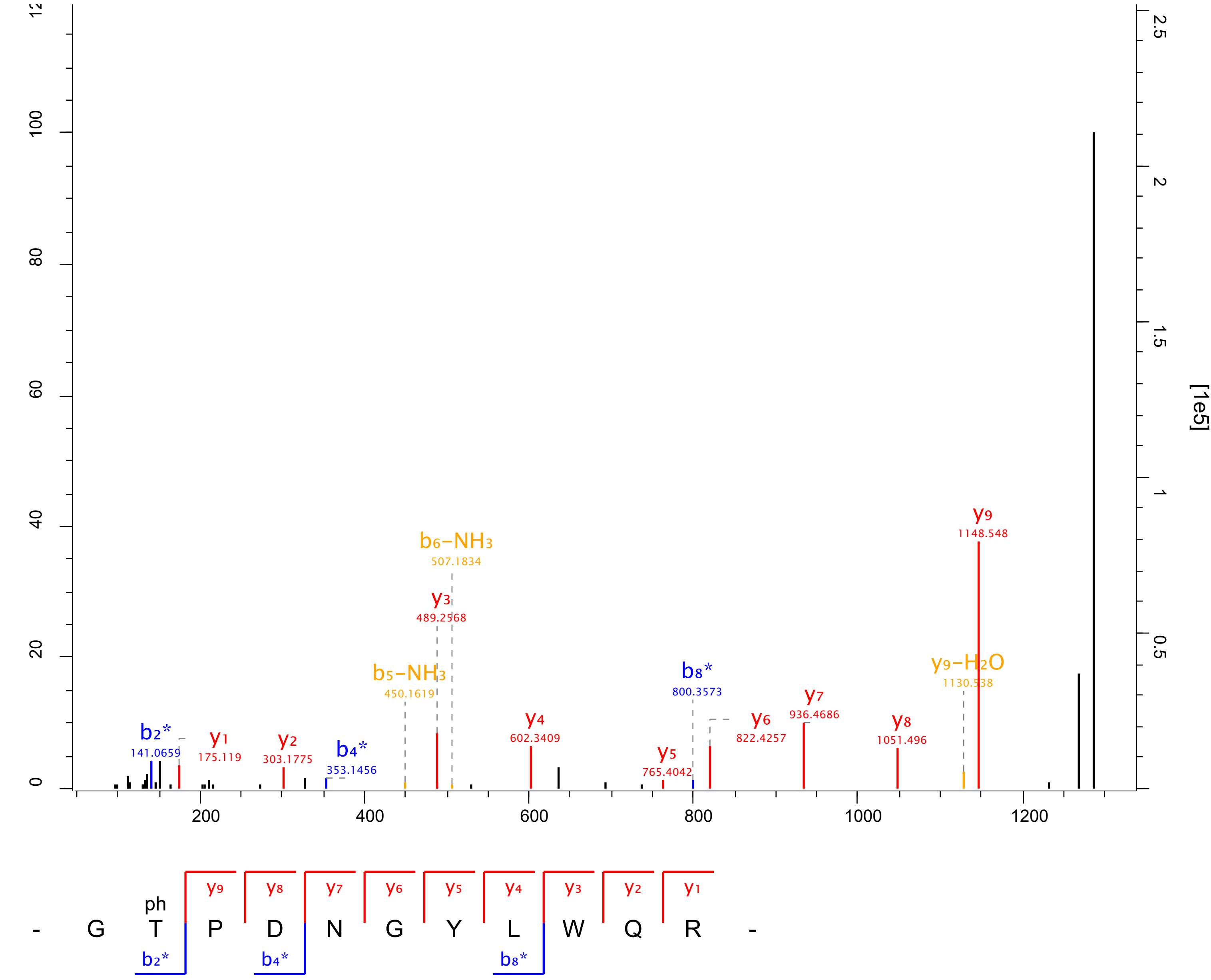

| Raw file | Scan | Method | Score | m/z |
|---|---|---|---|---|
| water_15_2 | 23774 | FTMS; HCD | 154.34 | 748.83 |

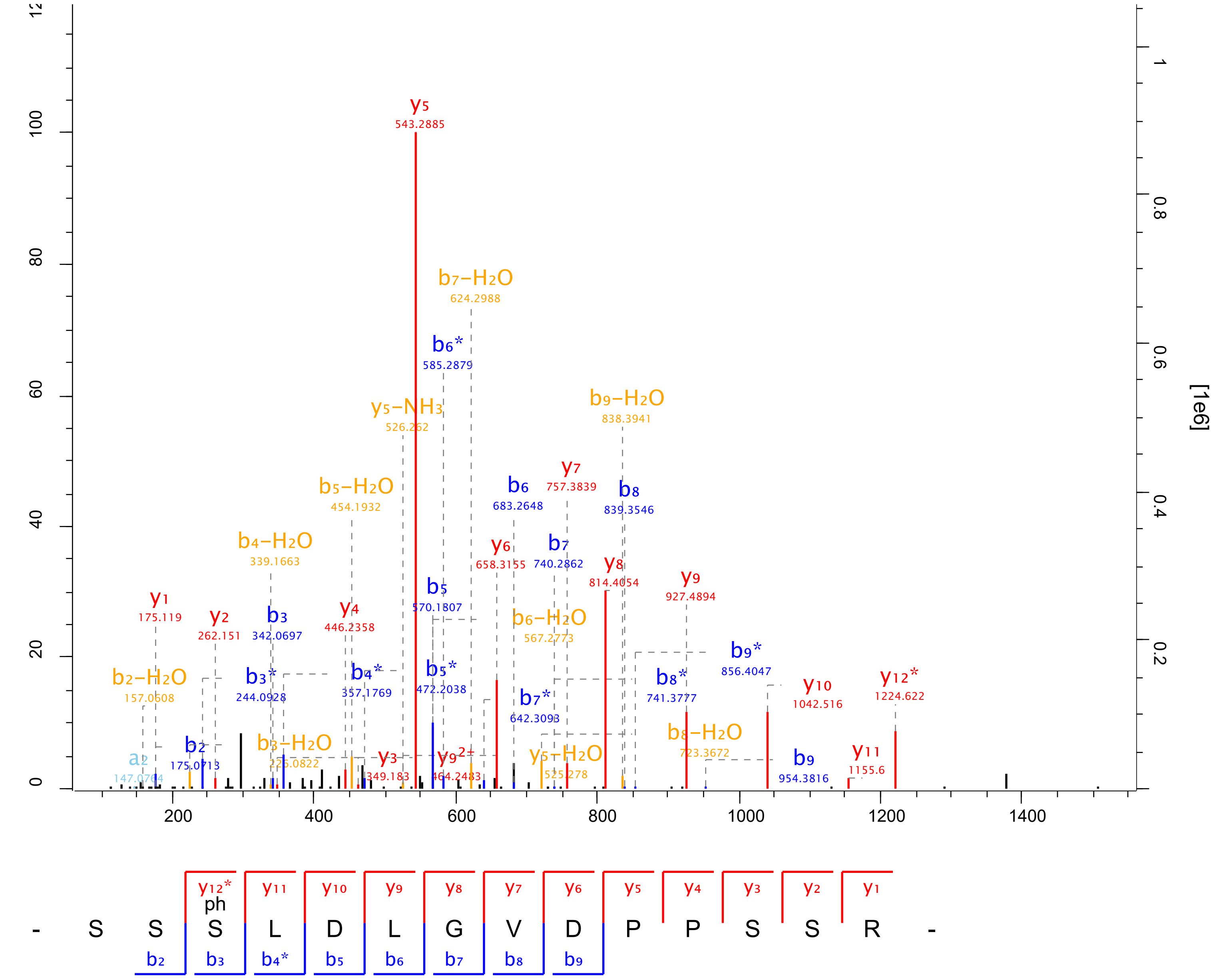

| Raw file | Scan | Method | Score | m/z |
| --- | --- | --- | --- | --- |
| water_15_2 | 23889 | FTMS; HCD | 77.75 | 730.31 |

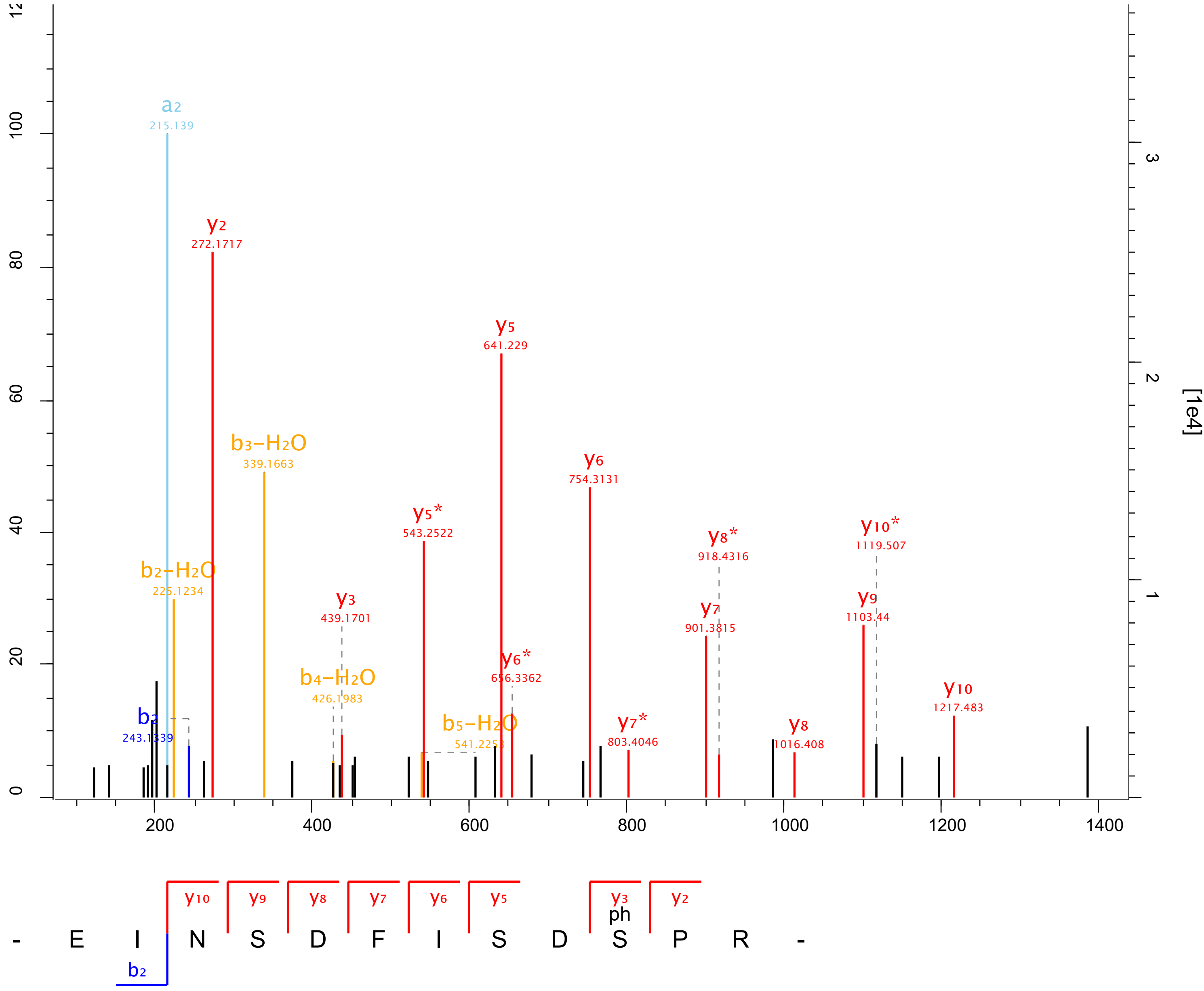

| Raw file | Scan | Method | Score | m/z |
|---|---|---|---|---|
| water_15_2 | 24268 | FTMS; HCD | 67.23 | 692.81 |

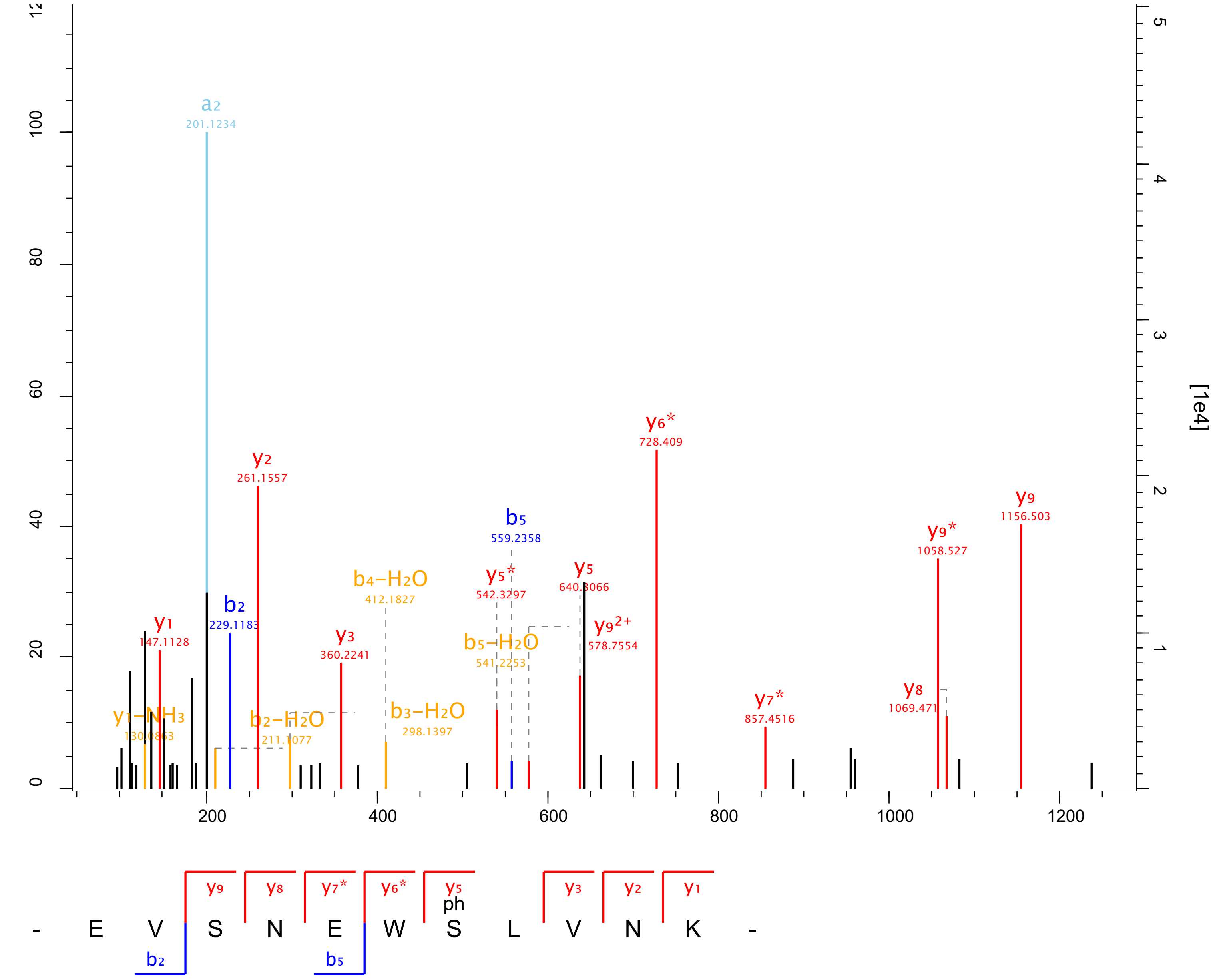

| Raw file | Scan | Method | Score | m/z |
| --- | --- | --- | --- | --- |
| water_15_2 | 24417 | FTMS; HCD | 123.4 | 729.85 |

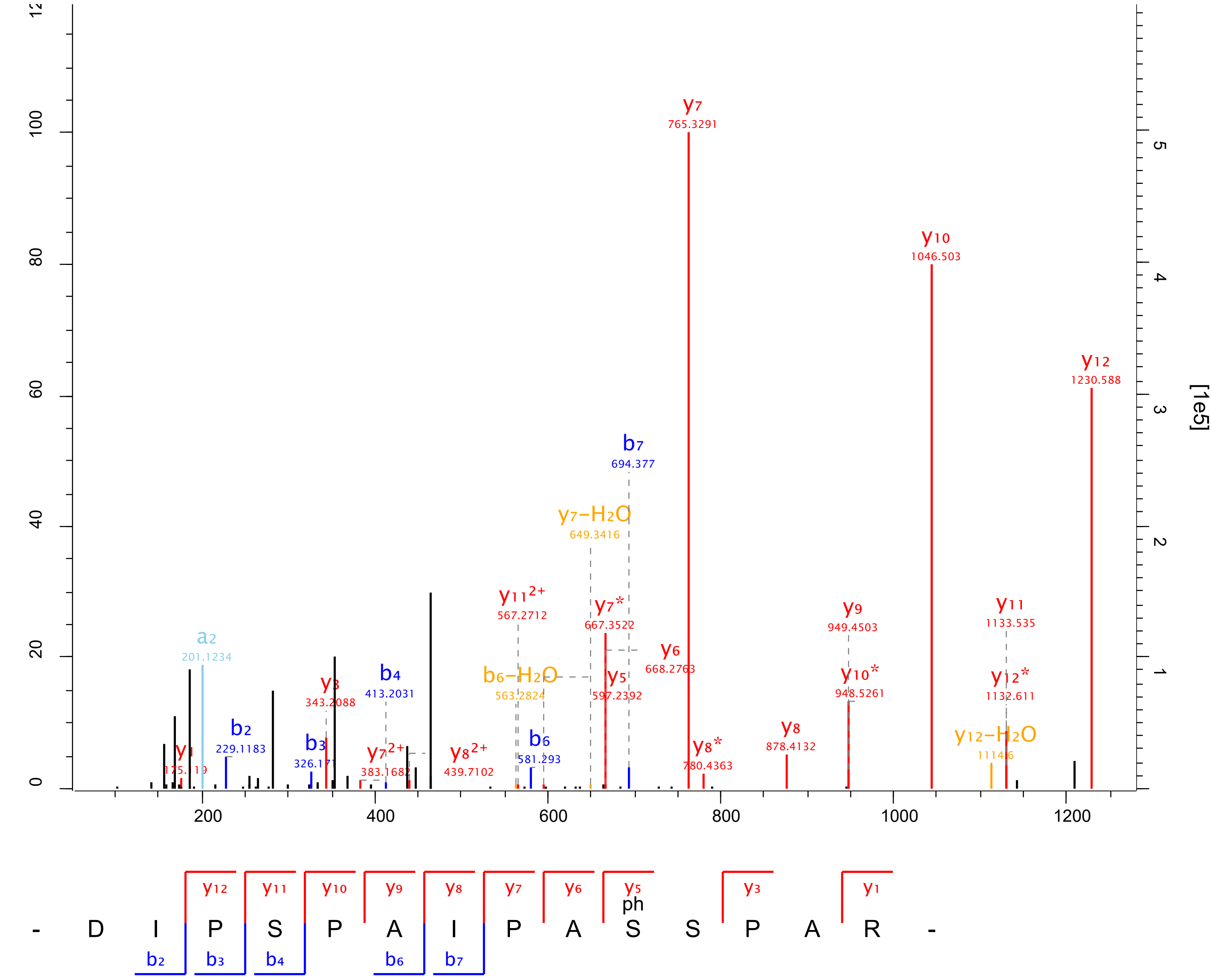

| Raw file | Scan | Method | Score | m/z |
| --- | --- | --- | --- | --- |
| water_15_2 | 24526 | FTMS; HCD | 63.3 | 871.4 |

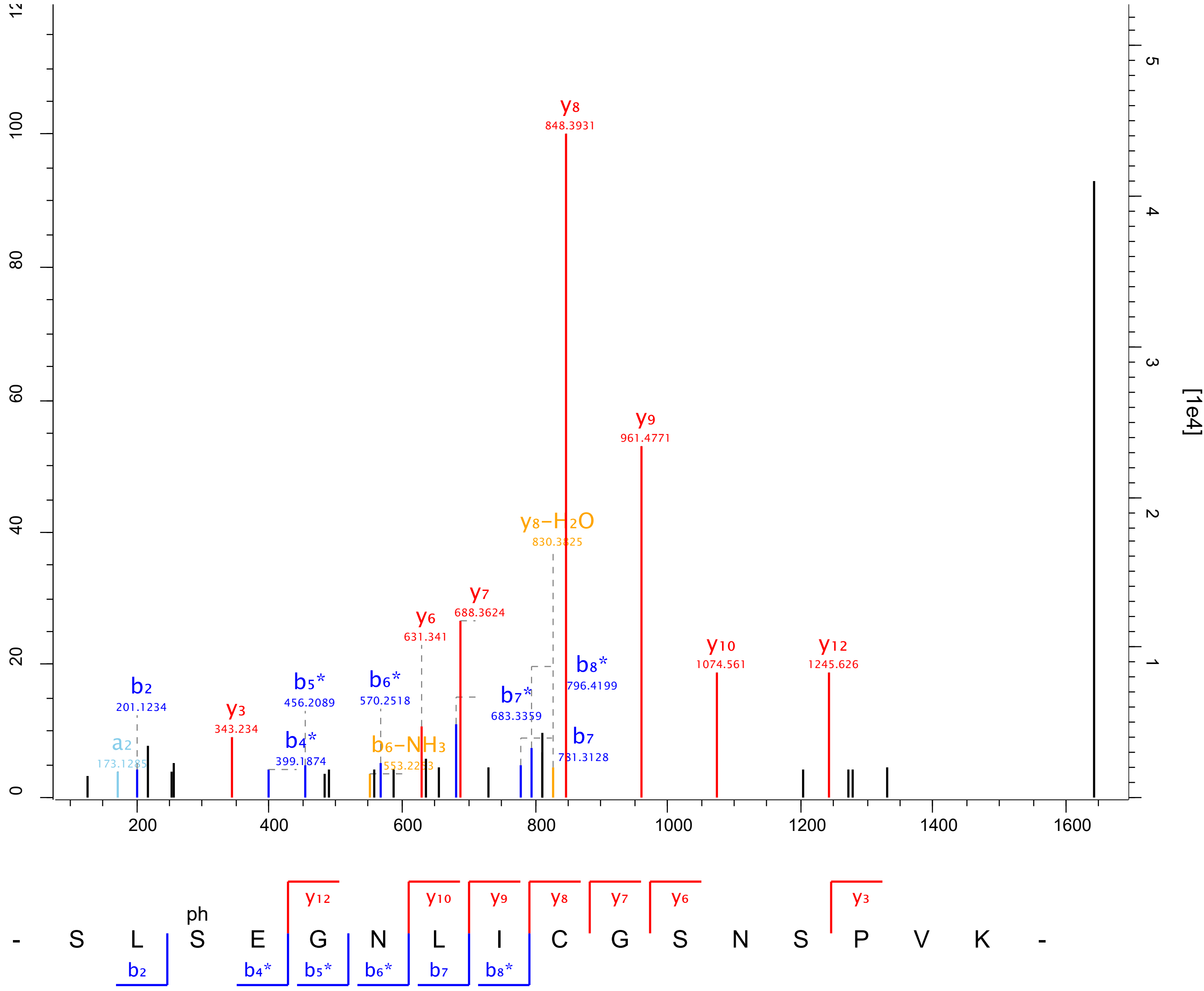

| Raw file | Scan | Method | Score | m/z |
| --- | --- | --- | --- | --- |
| water_15_2 | 2462 | FTMS; HCD | 71.18 | 554.24 |

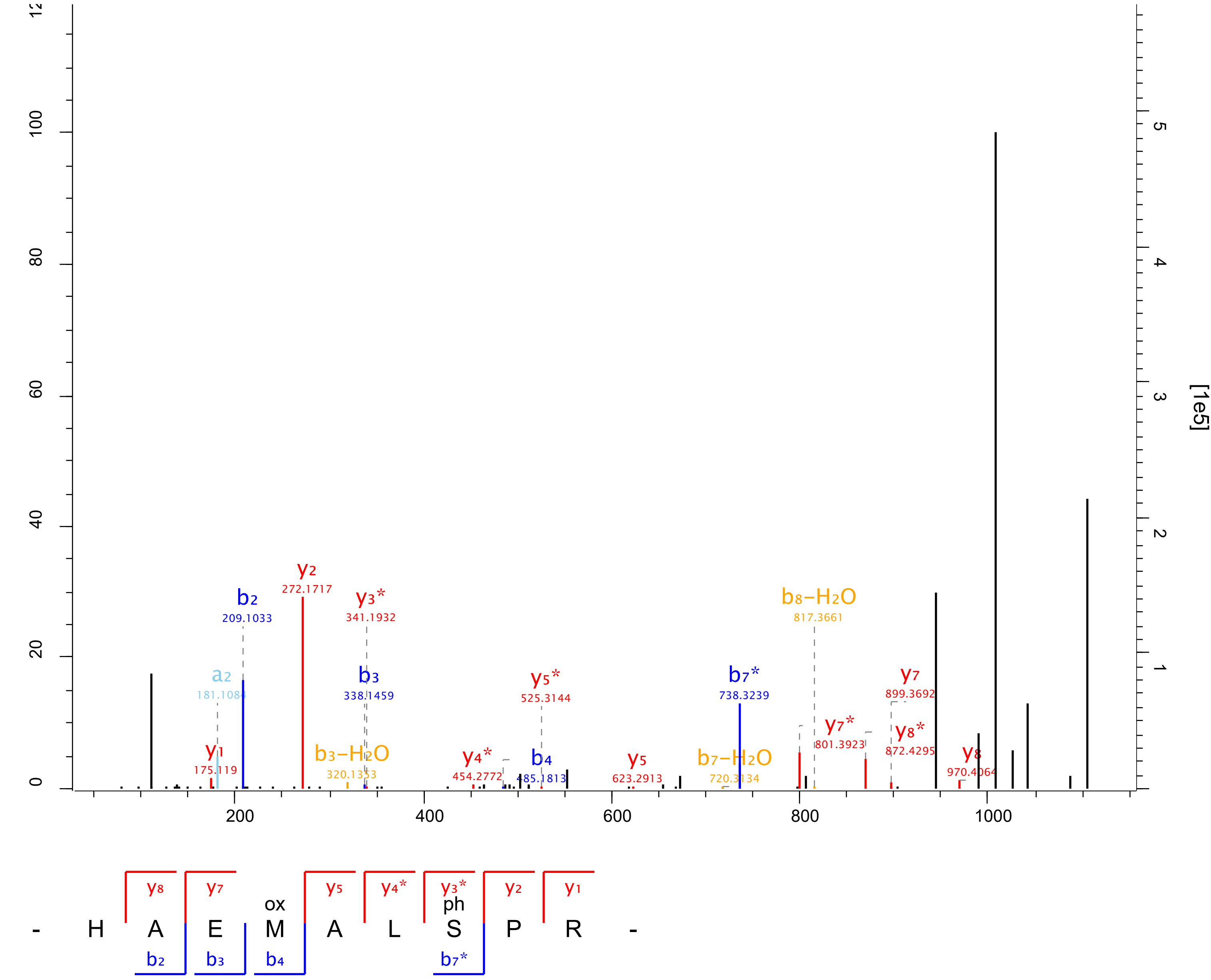

| Raw file | Scan | Method | Score | m/z |
| --- | --- | --- | --- | --- |
| water_15_2 | 24837 | FTMS; HCD | 121.56 | 576.23 |

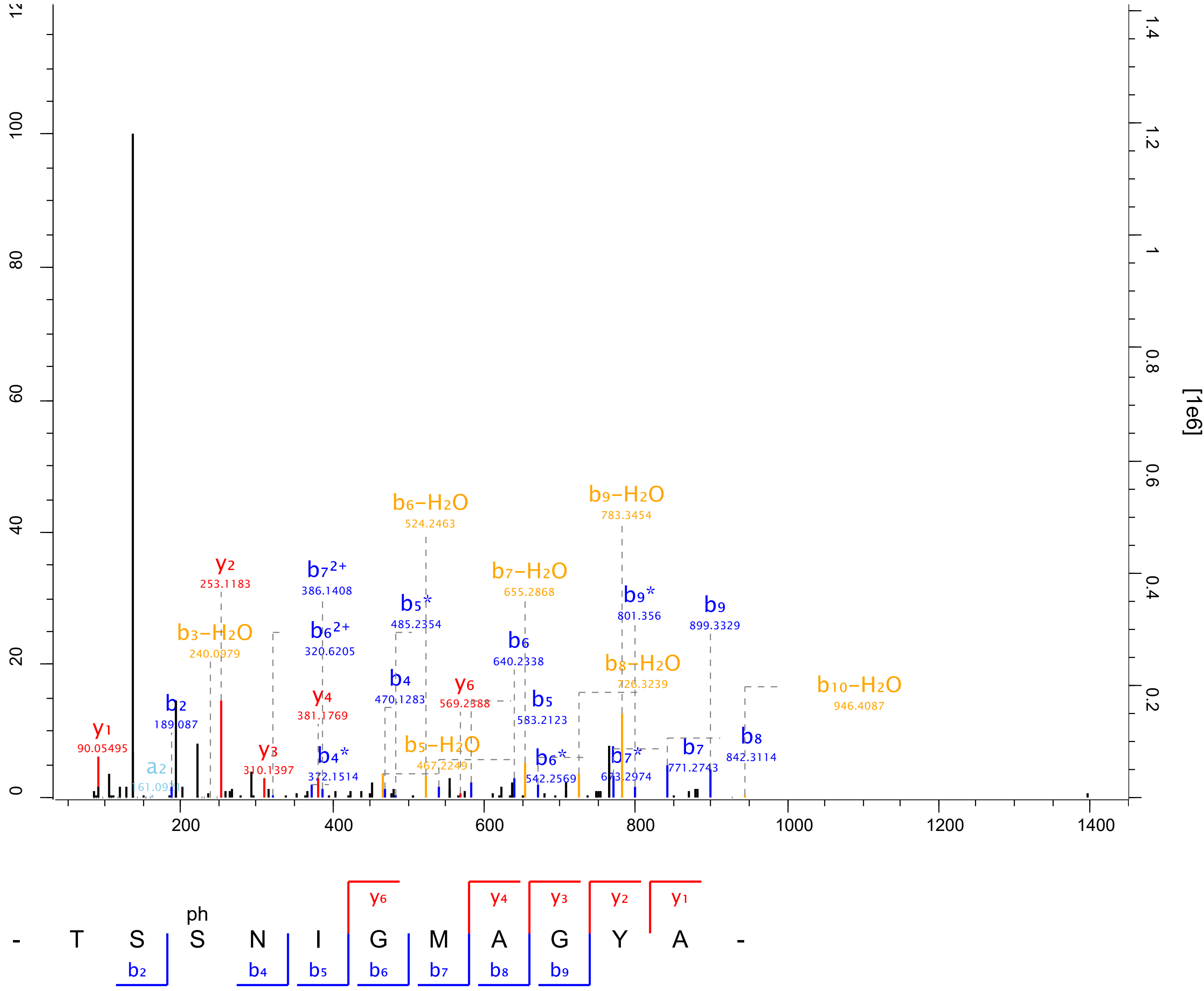

| Raw file | Scan | Method | Score | m/z |
|---|---|---|---|---|
| water_15_2 | 24980 | FTMS; HCD | 135.02 | 635.79 |

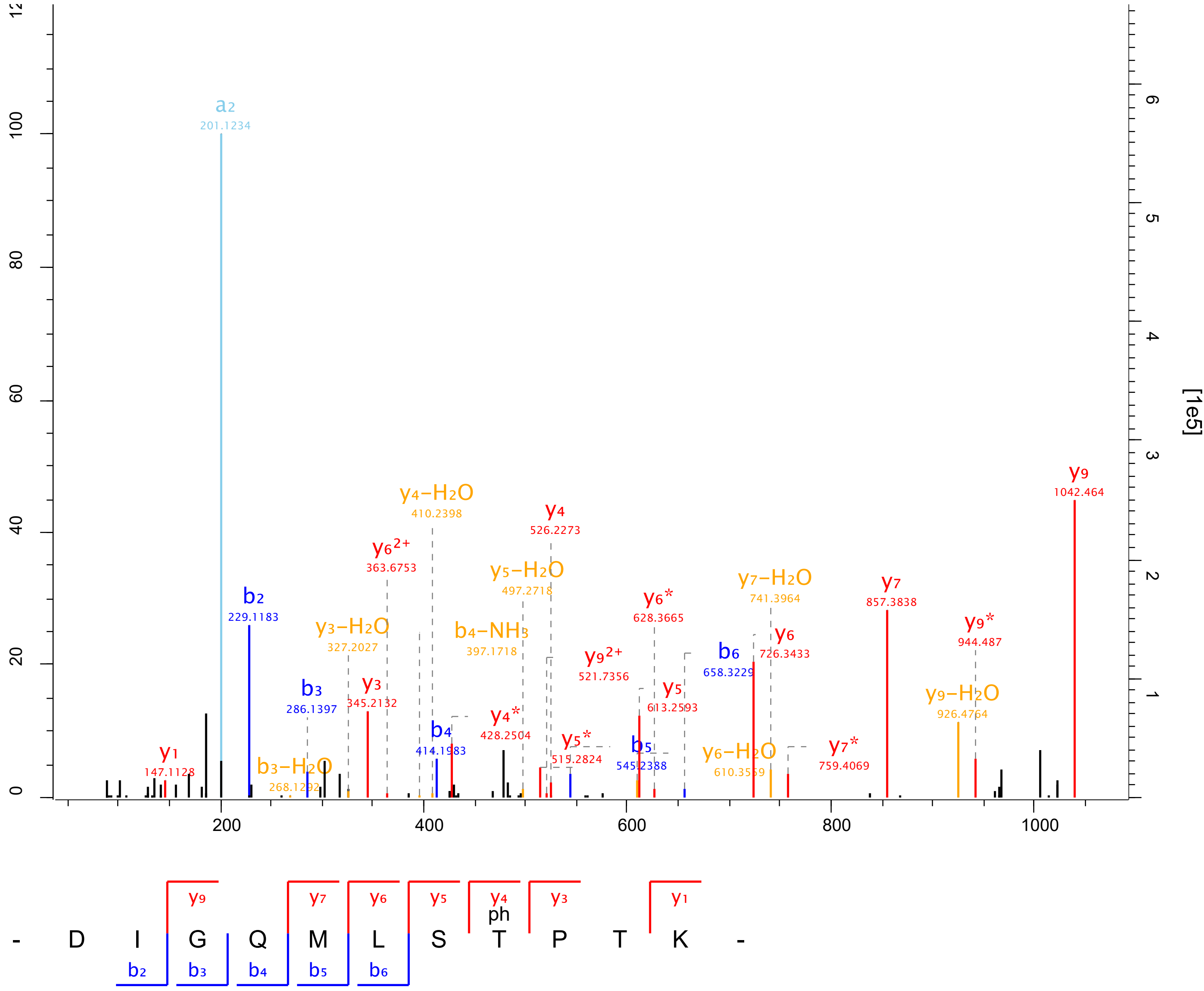

| Raw file | Scan | Method | Score | m/z |
| --- | --- | --- | --- | --- |
| water_15_2 | 25014 | FTMS; HCD | 80.63 | 755.79 |

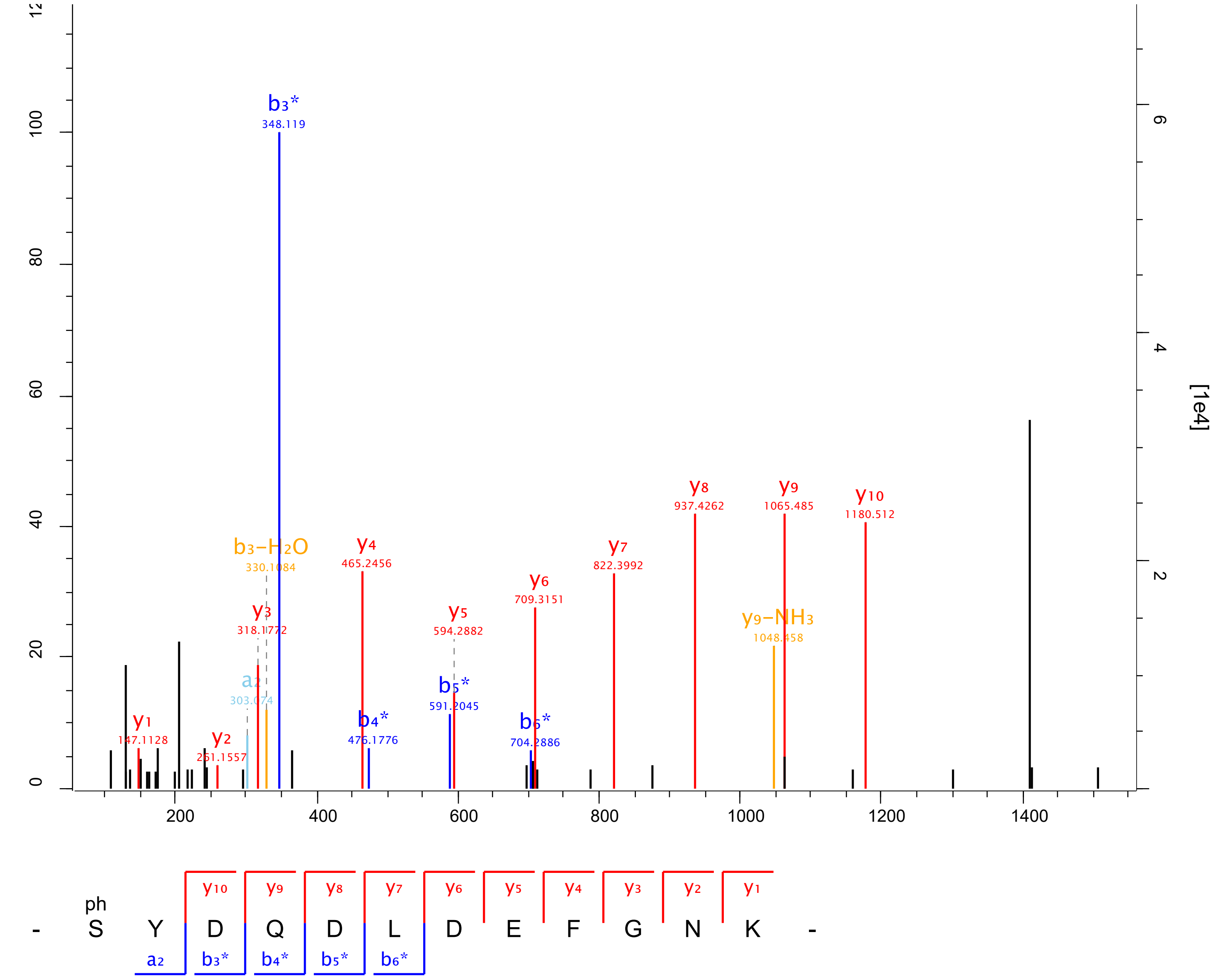

| Raw file | Scan | Method | Score | m/z |
| --- | --- | --- | --- | --- |
| water_15_2 | 25046 | FTMS; HCD | 139.49 | 799.32 |

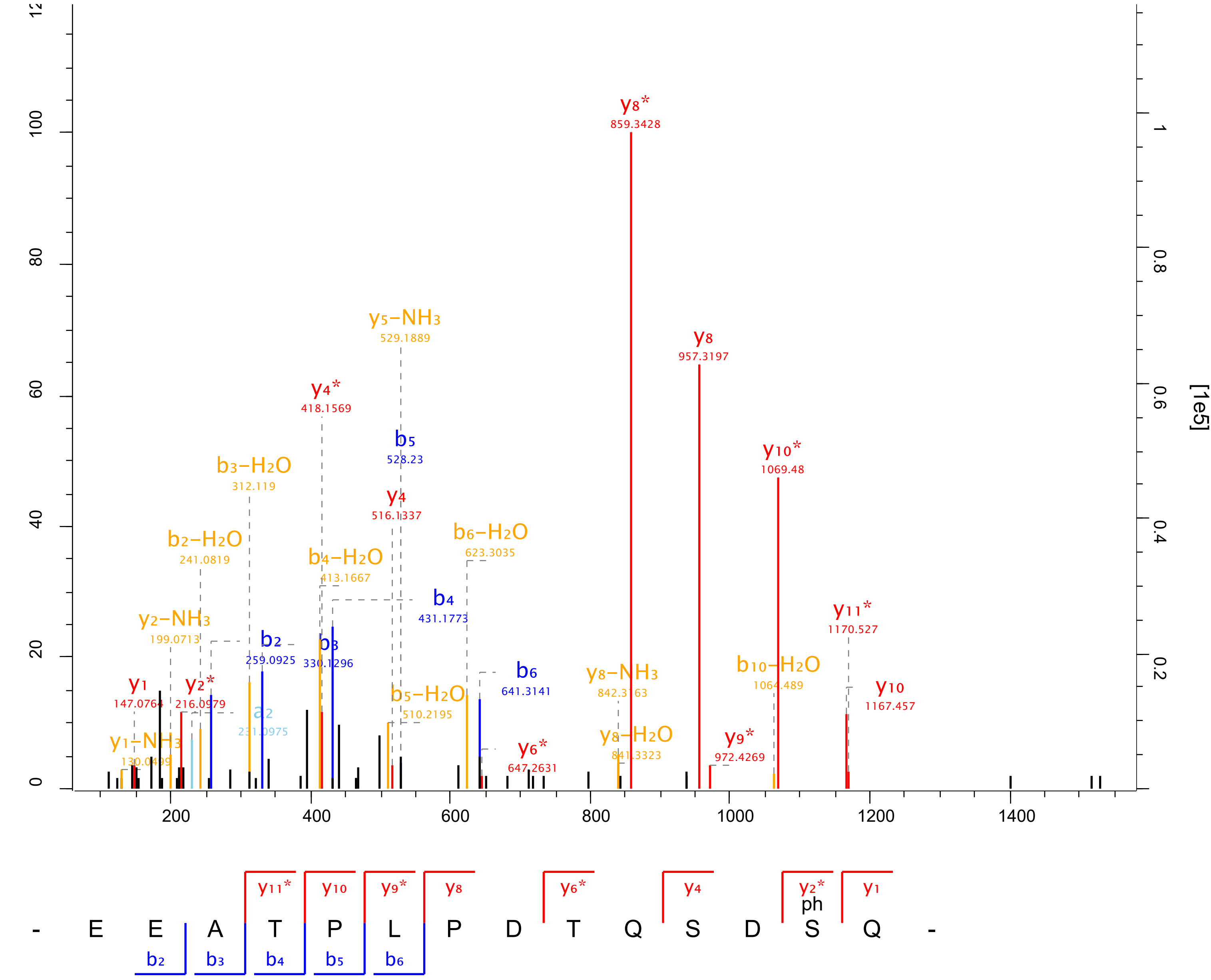

| Raw file | Scan | Method | Score | m/z |
|---|---|---|---|---|
| water_15_2 | 25091 | FTMS; HCD | 45.6 | 497.72 |

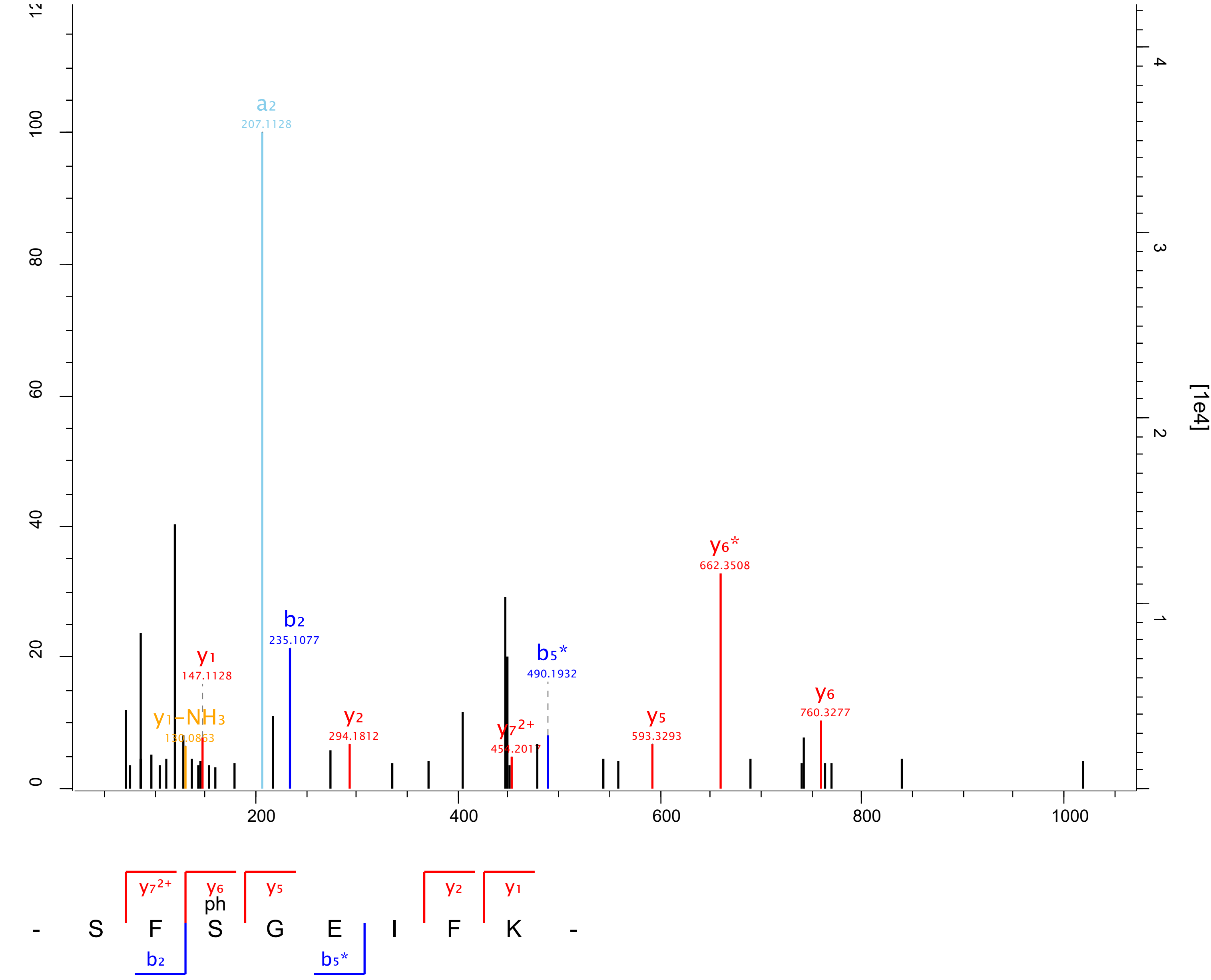

| Raw file | Scan | Method | Score | m/z |
|---|---|---|---|---|
| water_15_2 | 25094 | FTMS; HCD | 105.51 | 1049.93 |

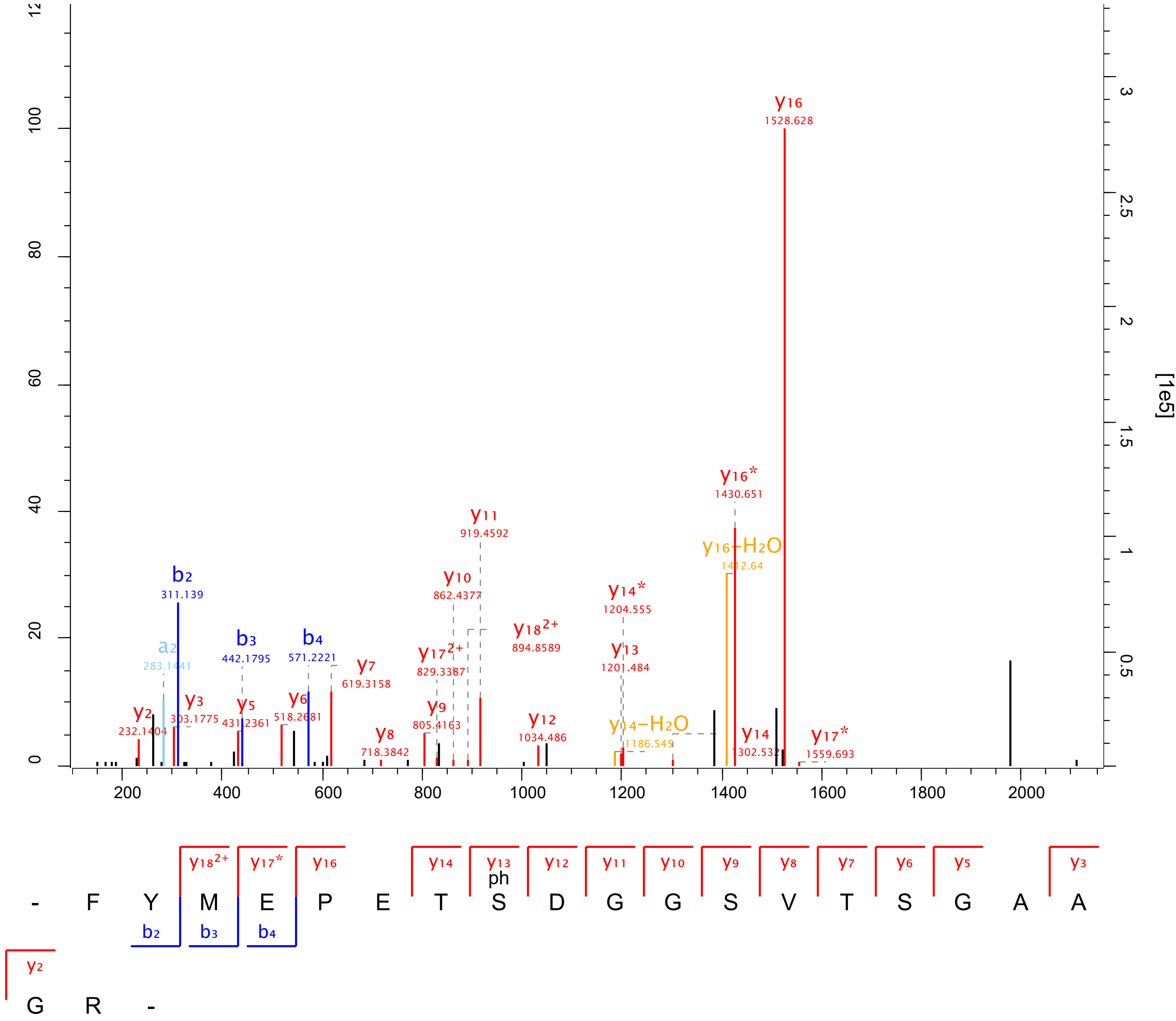

| Raw file | Scan | Method | Score | m/z |
|---|---|---|---|---|
| water_15_2 | 2514 | FTMS; HCD | 166.04 | 625.81 |

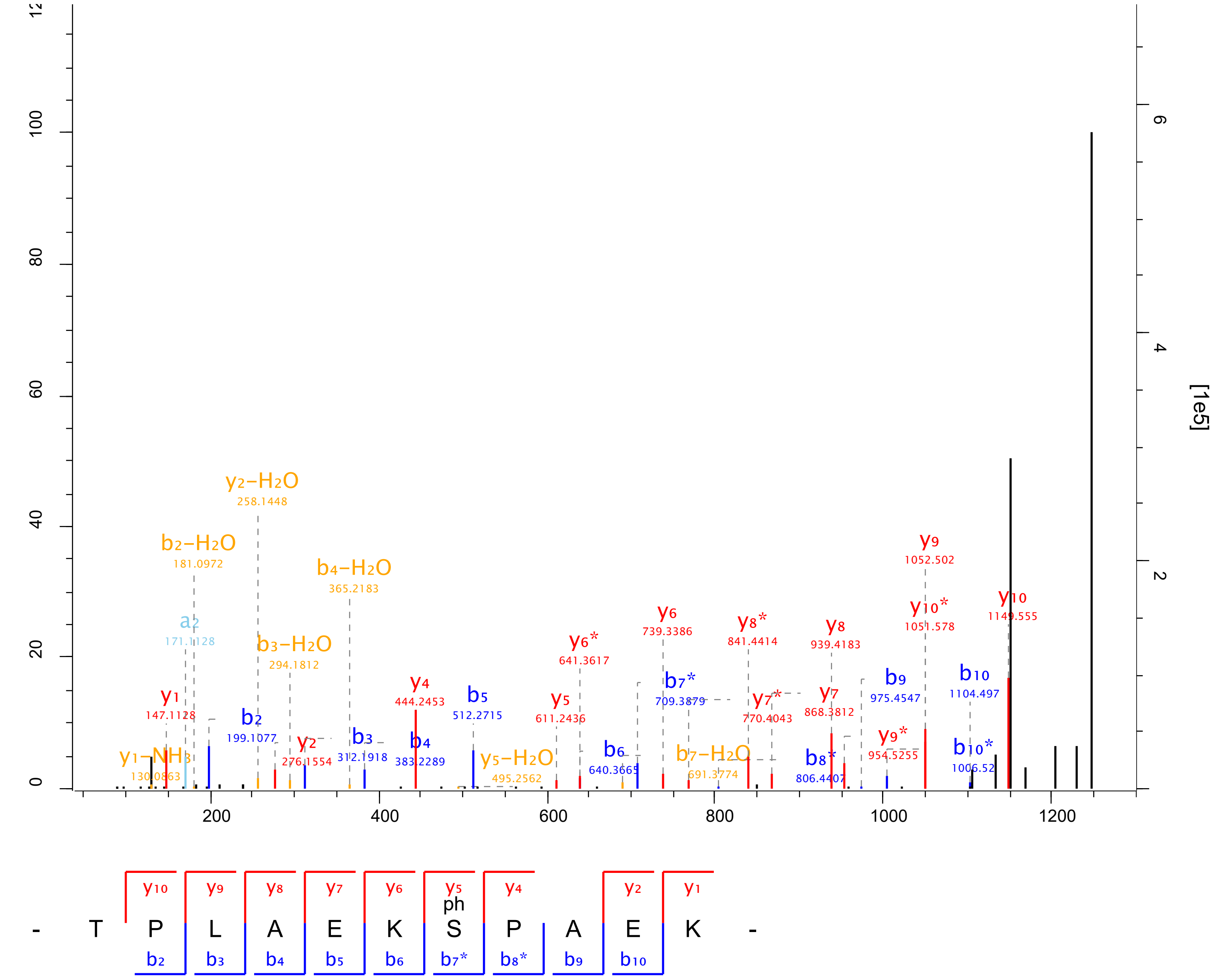

| Raw file | Scan | Method | Score | m/z |
|---|---|---|---|---|
| water_15_2 | 25247 | FTMS; HCD | 46.42 | 941.93 |

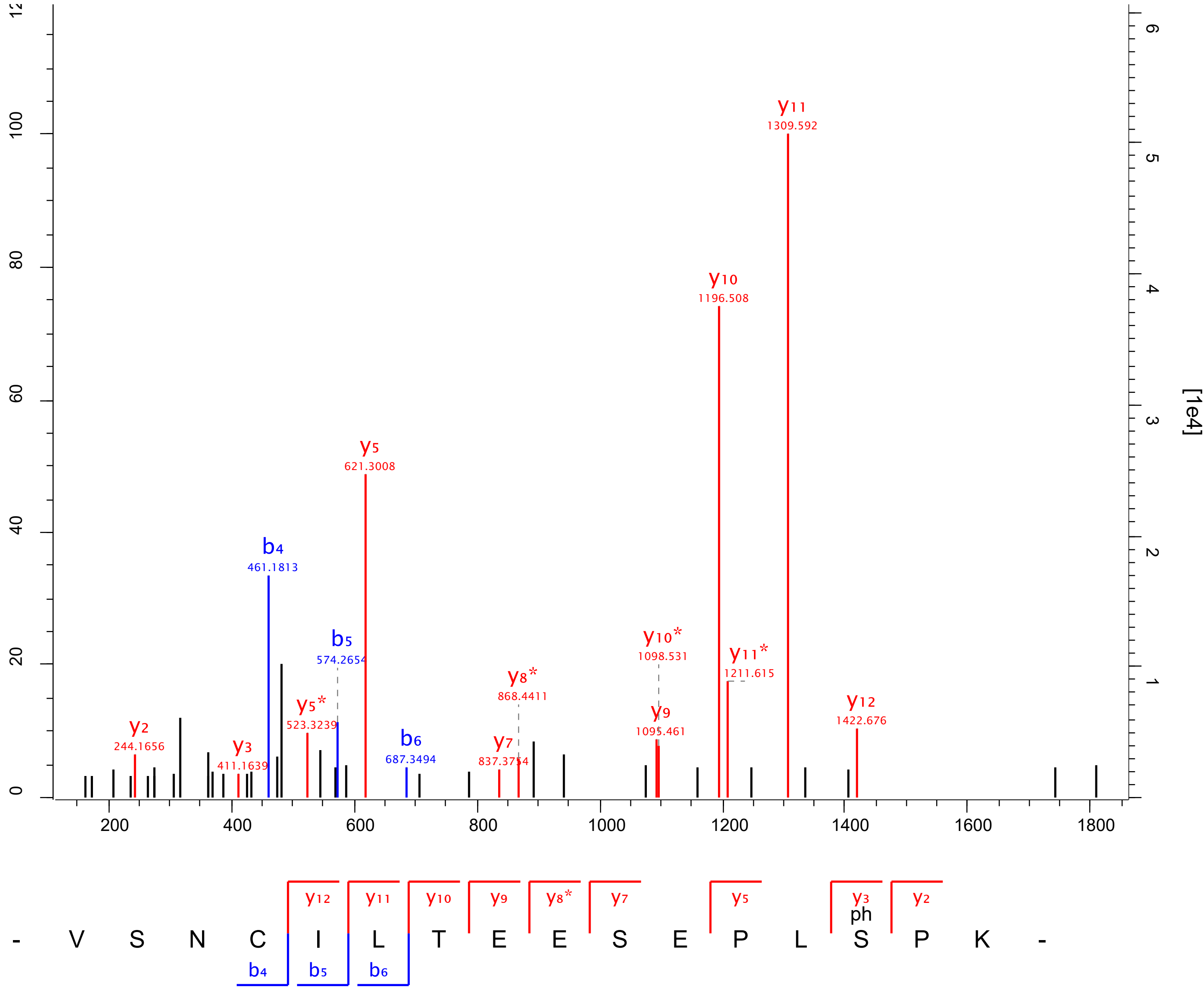

| Raw file | Scan | Method | Score | m/z |
|---|---|---|---|---|
| water_15_2 | 25322 | FTMS; HCD | 143.25 | 643.29 |

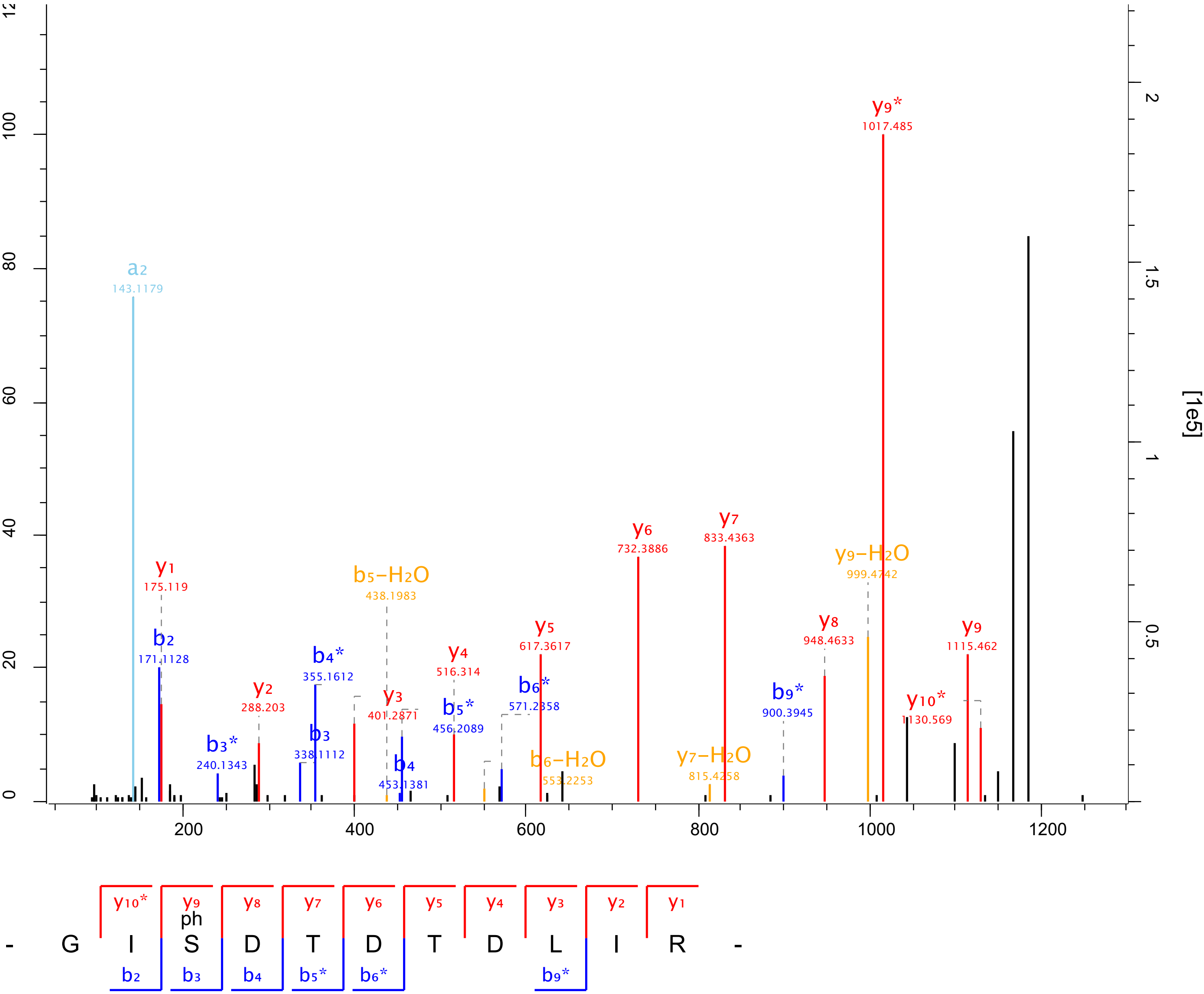

| Raw file | Scan | Method | Score | m/z |
|---|---|---|---|---|
| water_15_2 | 25554 | FTMS; HCD | 65.86 | 871.9 |

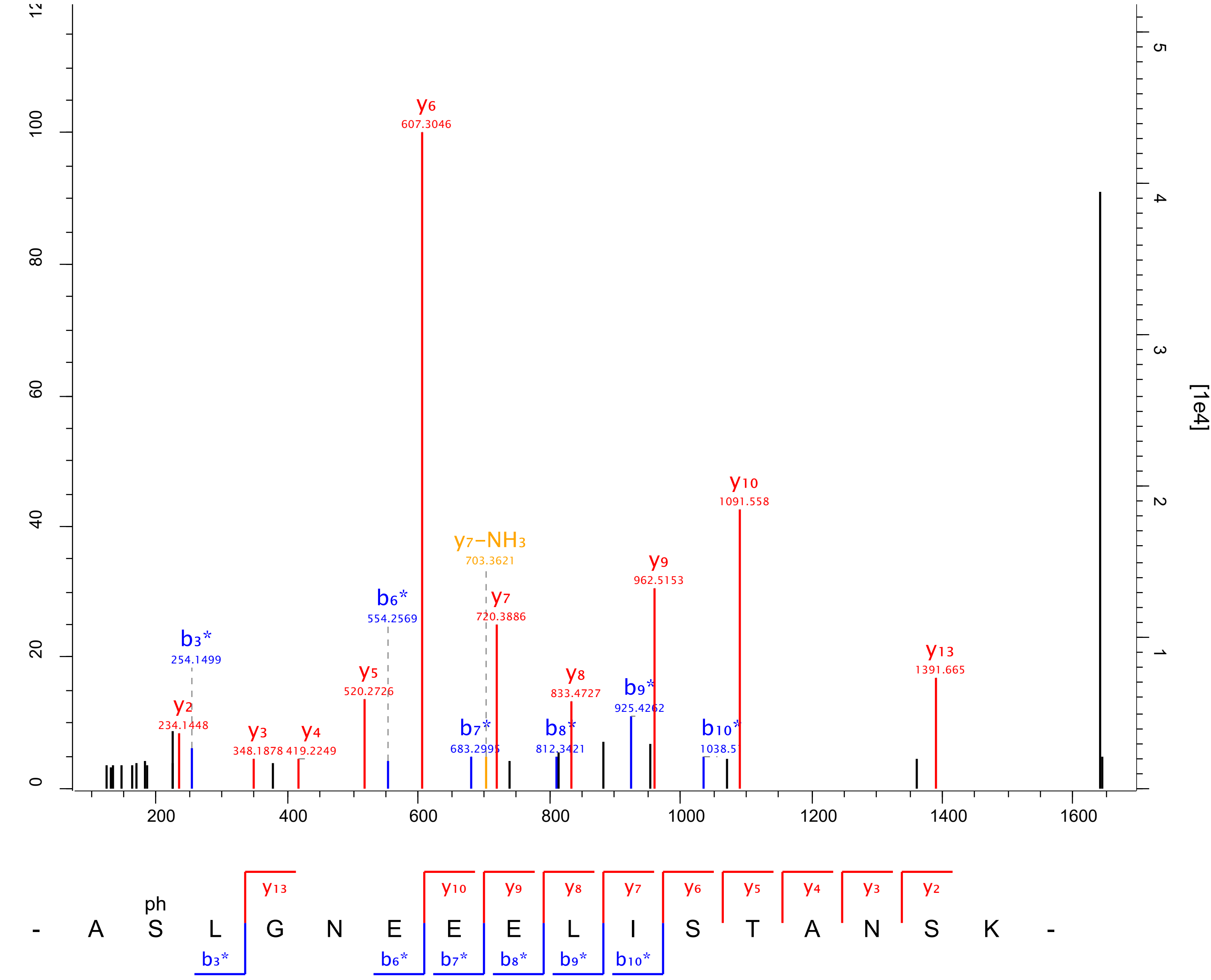

| Raw file | Scan | Method | Score | m/z |
|---|---|---|---|---|
| water_15_2 | 2575 | FTMS; HCD | 97.08 | 499.24 |

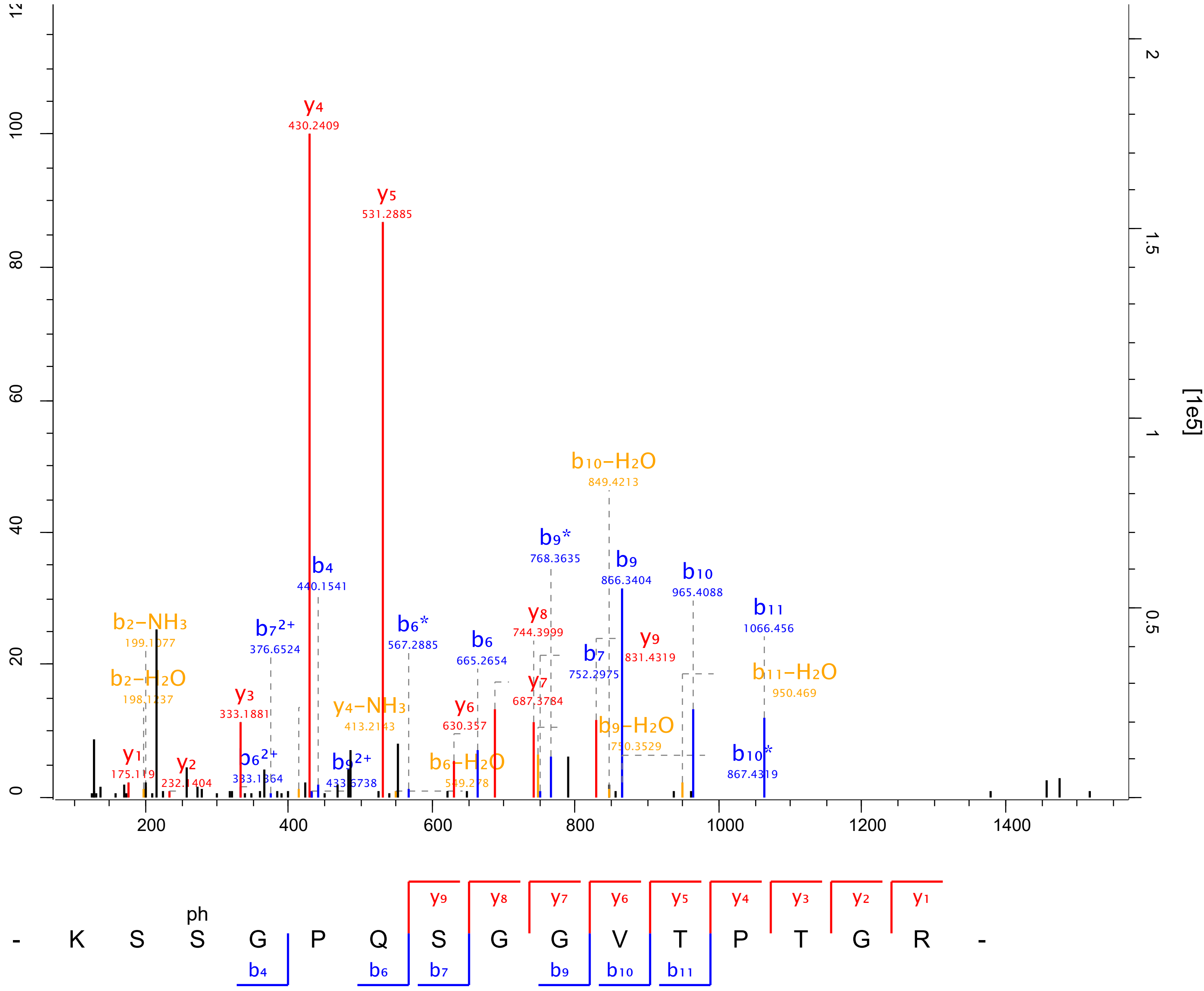

| Raw file | Scan | Method | Score | m/z |
| --- | --- | --- | --- | --- |
| water_15_2 | 25855 | FTMS; HCD | 124.2 | 896.89 |

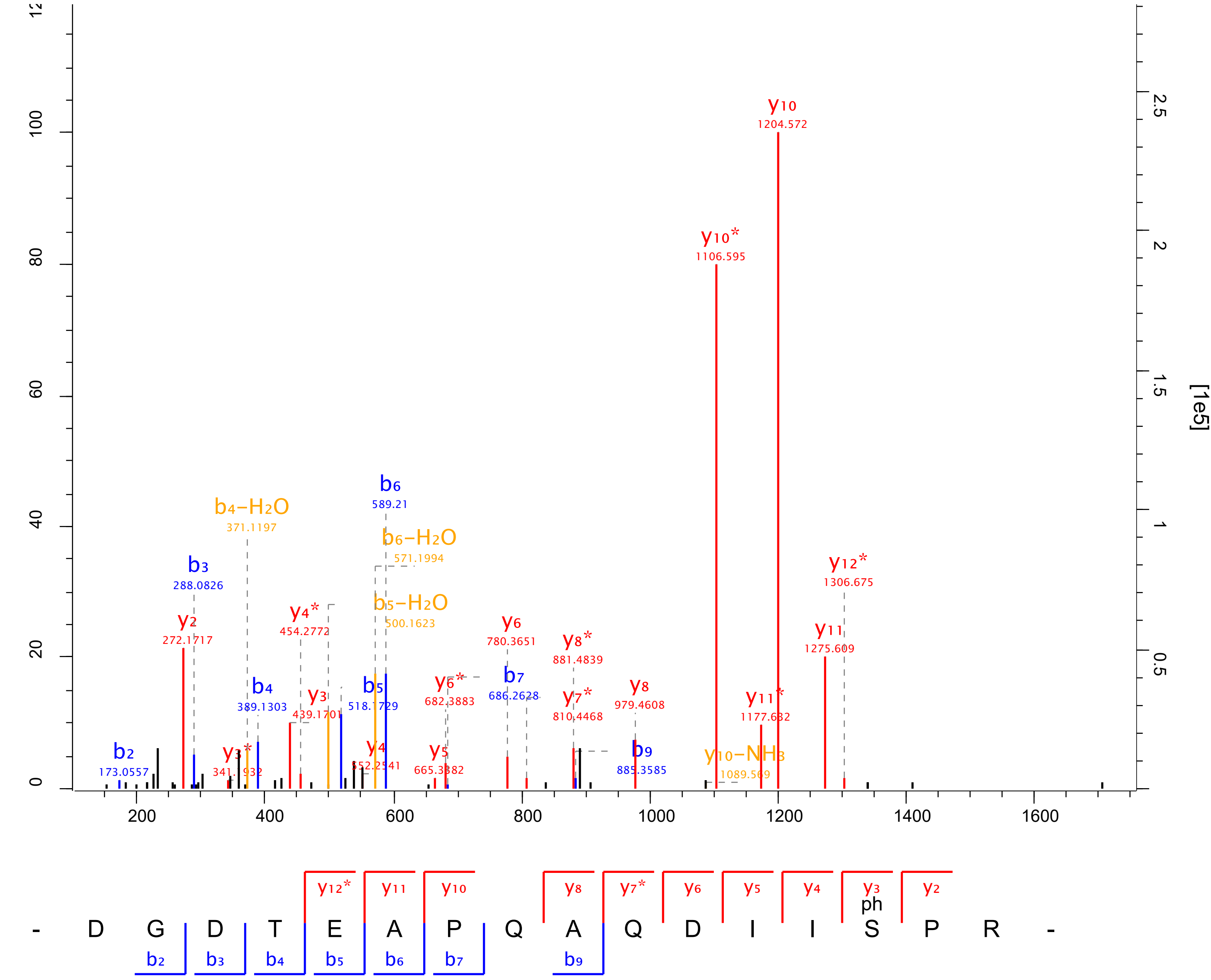

| Raw file | Scan | Method | Score | m/z |
|---|---|---|---|---|
| water_15_2 | 26030 | FTMS; HCD | 160.8 | 686.79 |

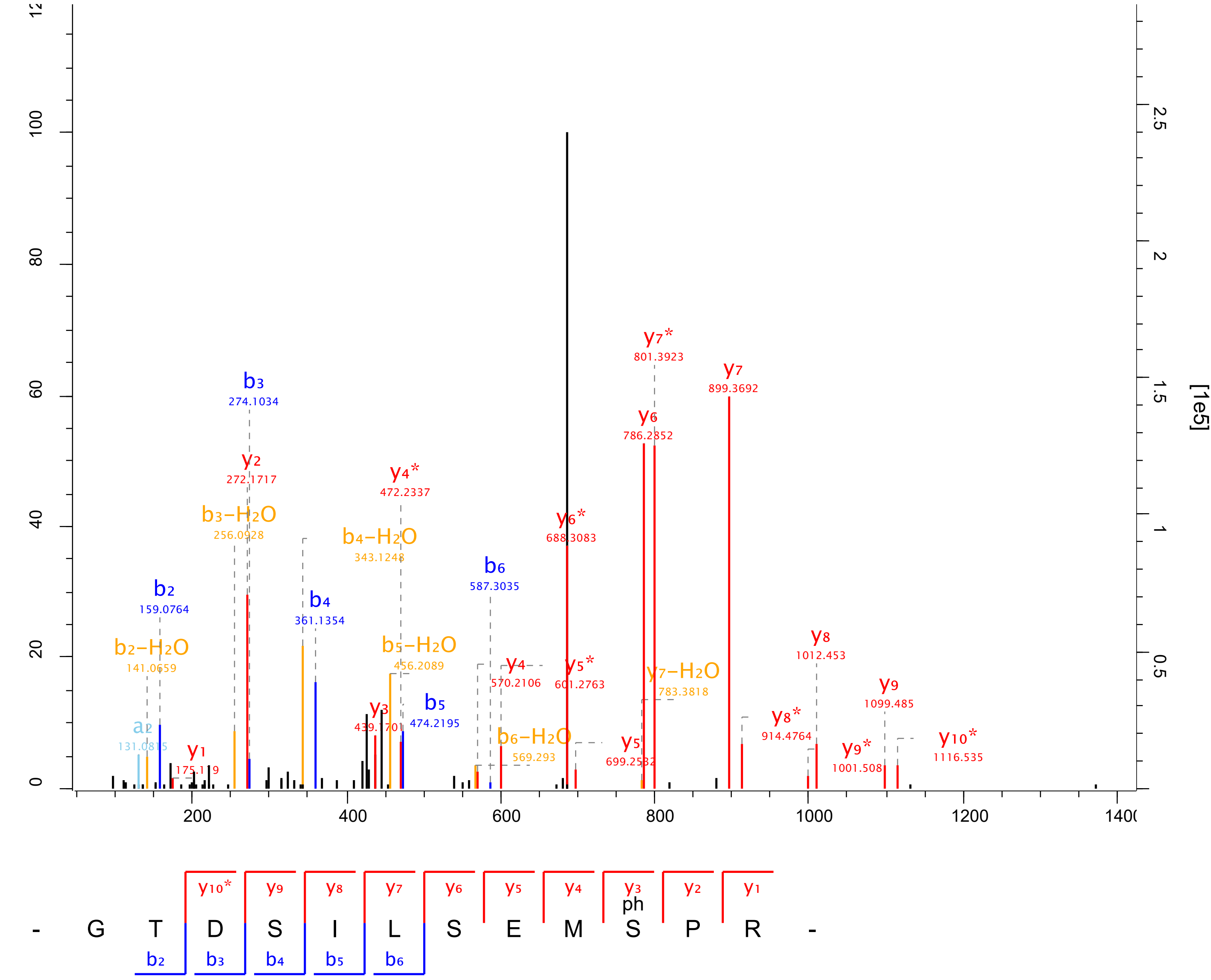

| Raw file | Scan | Method | Score | m/z |
|---|---|---|---|---|
| water_15_2 | 26033 | FTMS; HCD | 74.92 | 485.7 |

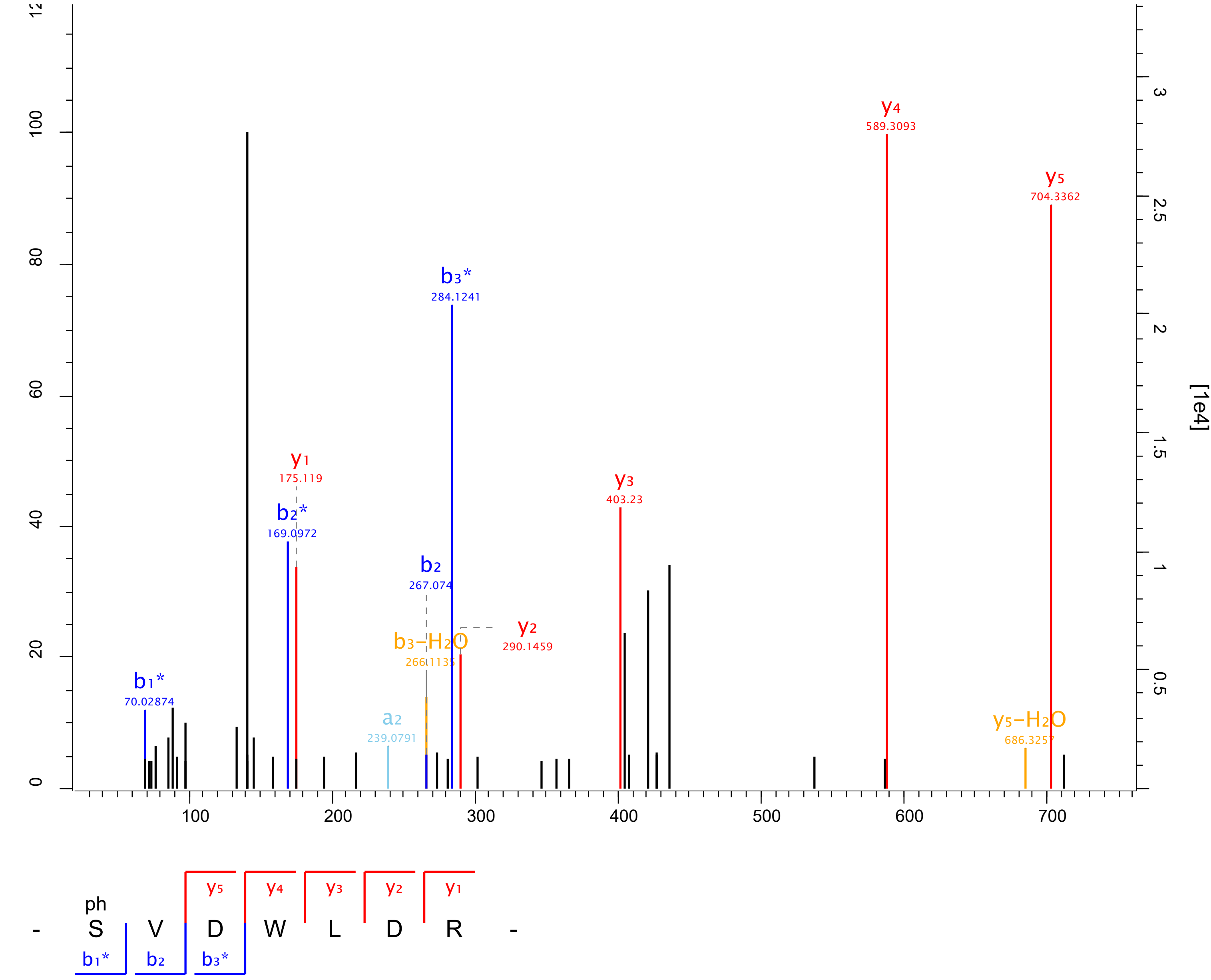

| Raw file | Scan | Method | Score | m/z |
|---|---|---|---|---|
| water_15_2 | 26374 | FTMS; HCD | 42.08 | 587.26 |

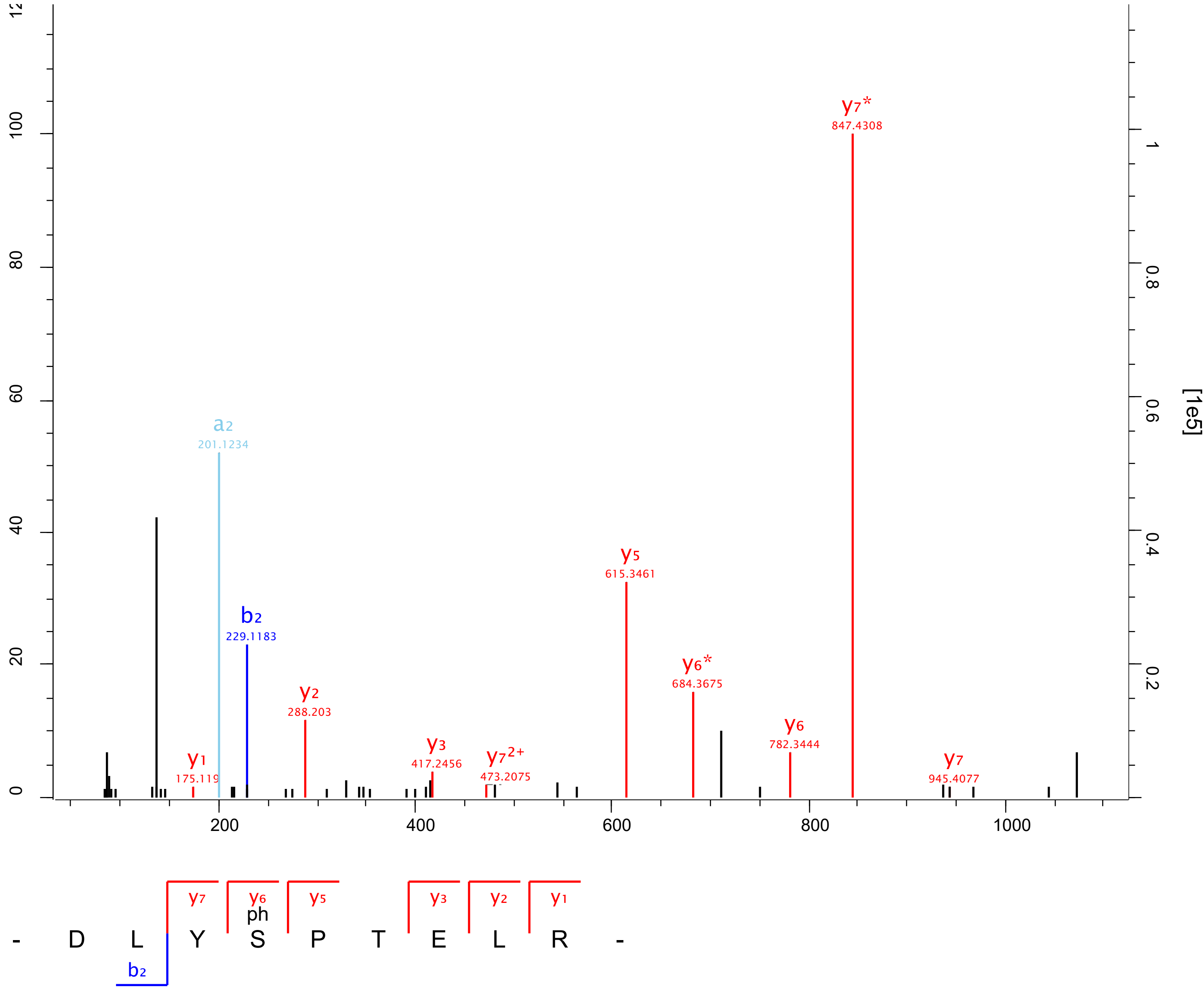

| Raw file | Scan | Method | Score | m/z |
|---|---|---|---|---|
| water_15_2 | 26406 | FTMS; HCD | 102.08 | 795.35 |

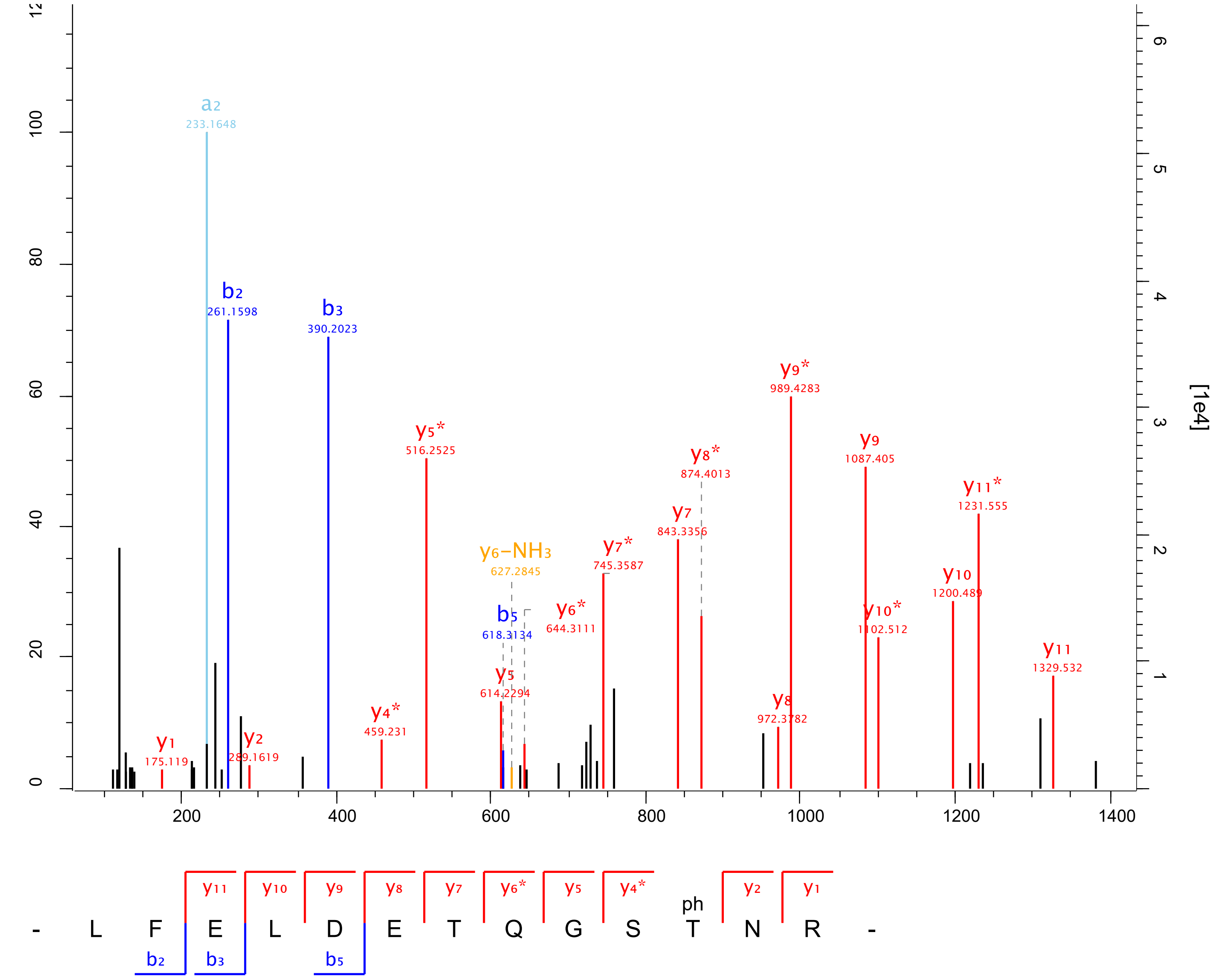

| Raw file | Scan | Method | Score | m/z |
| --- | --- | --- | --- | --- |
| water_15_2 | 26531 | FTMS; HCD | 171.76 | 709.8 |

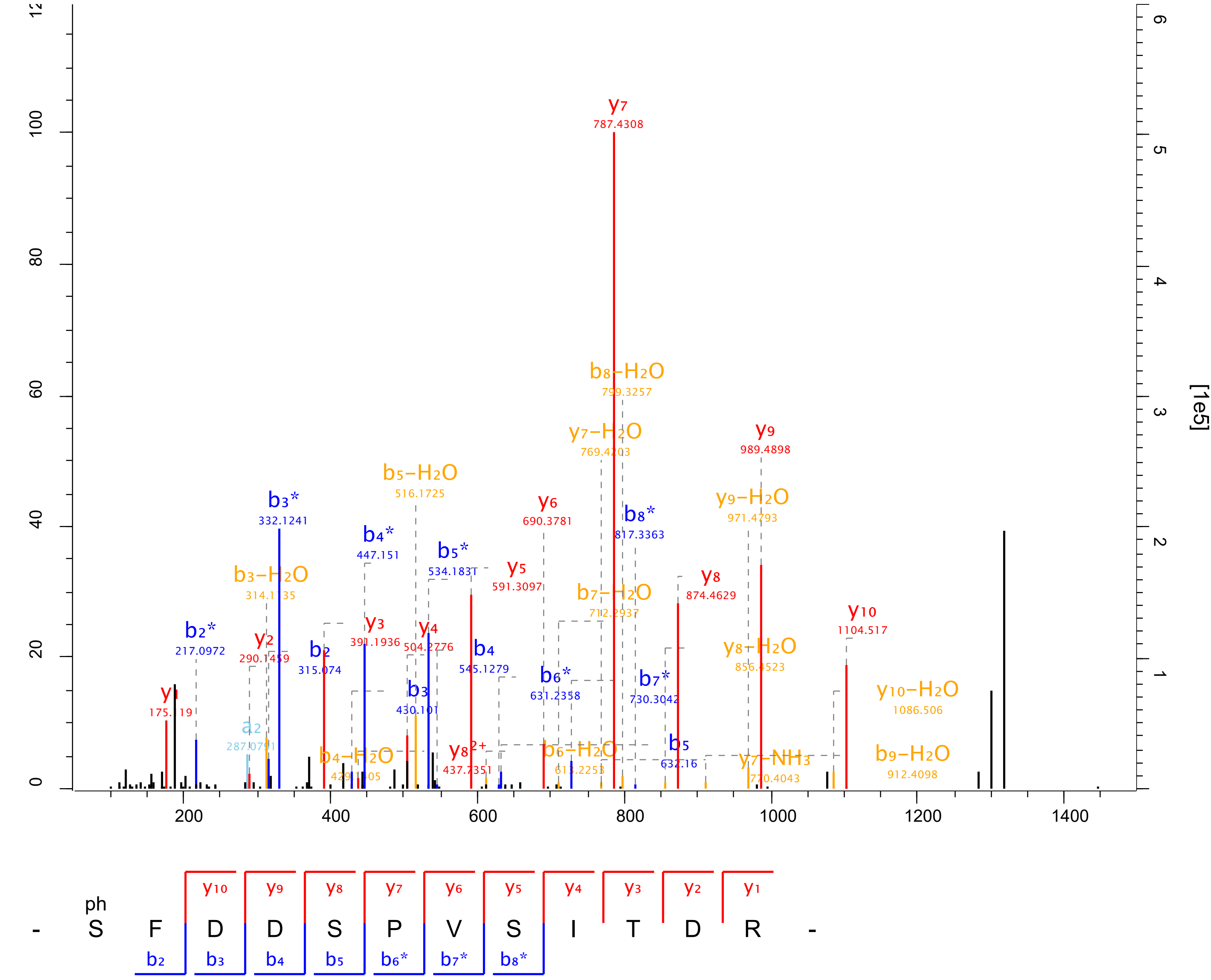

| Raw file | Scan | Method | Score | m/z |
|---|---|---|---|---|
| water_15_2 | 26746 | FTMS; HCD | 115.91 | 781.31 |

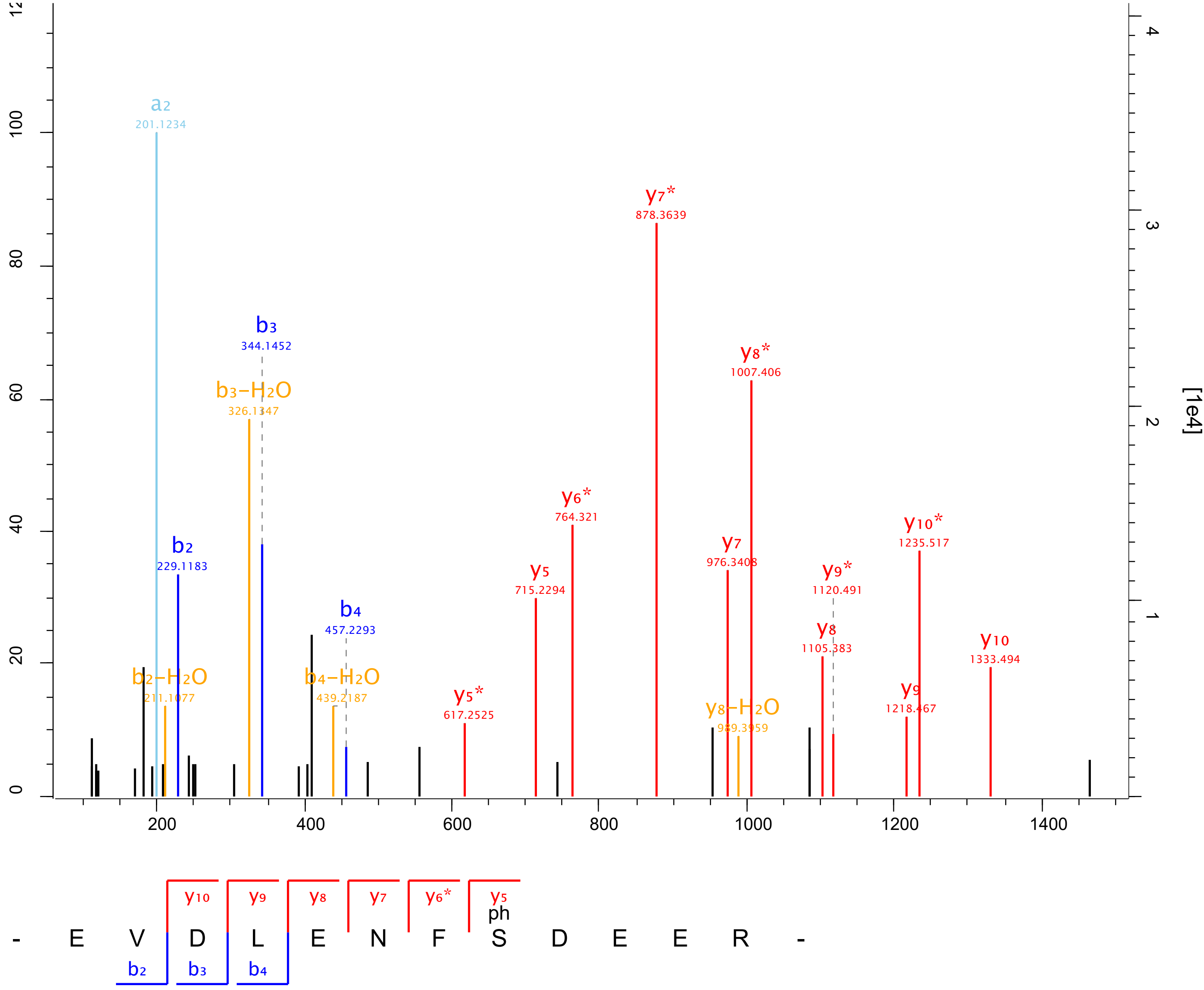

| Raw file | Scan | Method | Score | m/z |
| --- | --- | --- | --- | --- |
| water_15_2 | 27088 | FTMS; HCD | 59.2 | 525.78 |

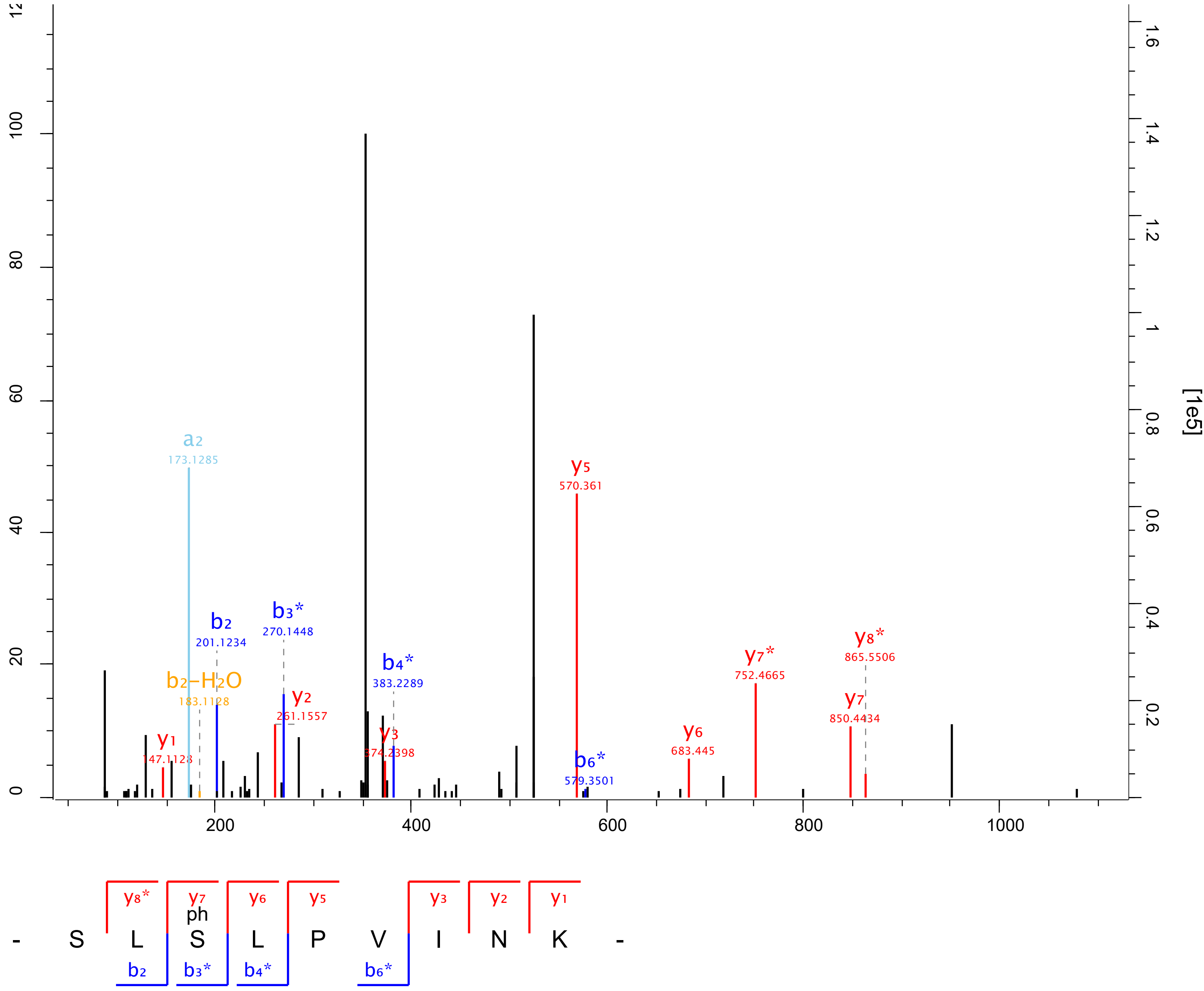

| Raw file | Scan | Method | Score | m/z |
|---|---|---|---|---|
| water_15_2 | 27149 | FTMS; HCD | 87.43 | 726.28 |

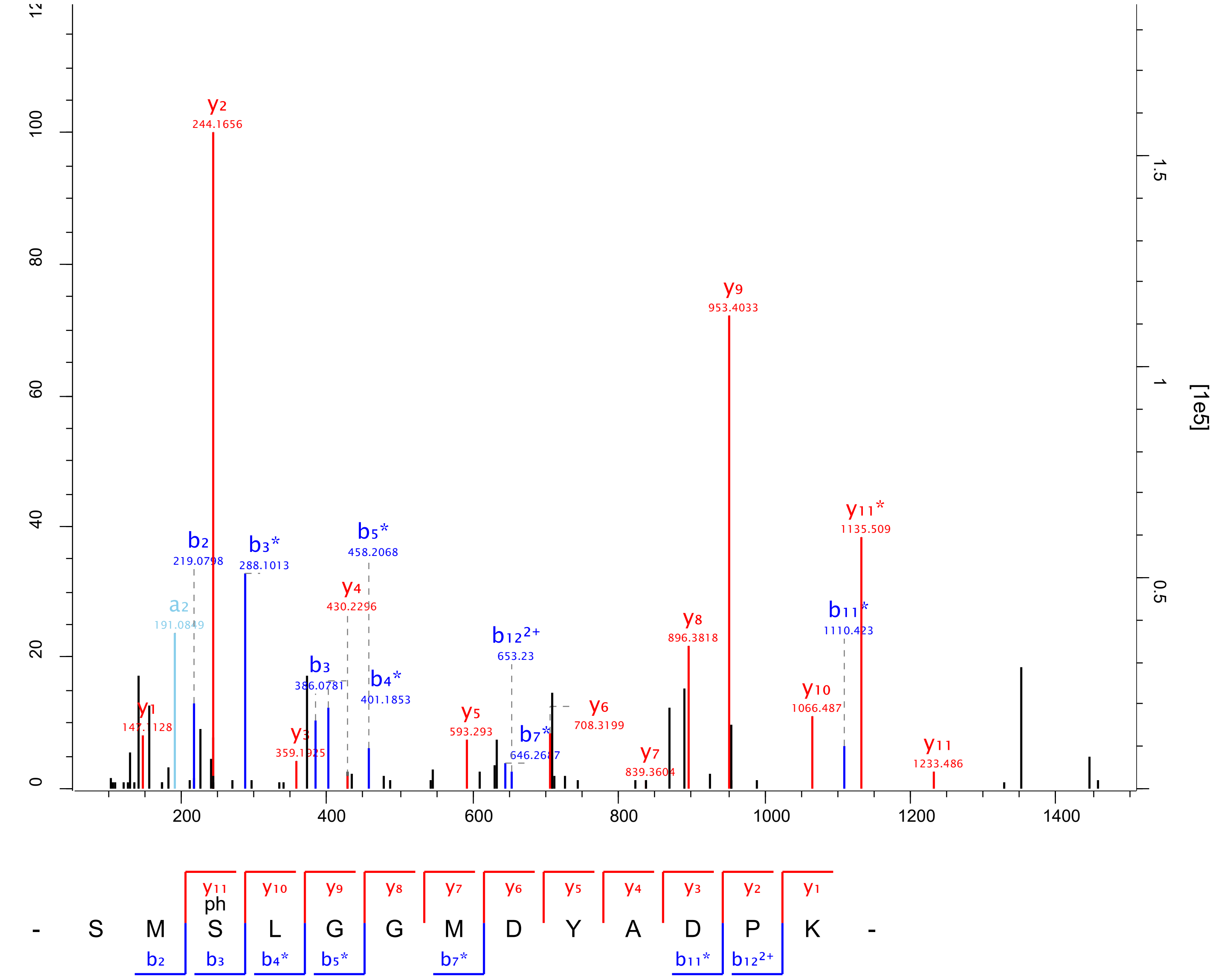

| Raw file | Scan | Method | Score | m/z |
| --- | --- | --- | --- | --- |
| water_15_2 | 27515 | FTMS; HCD | 141.86 | 887.88 |

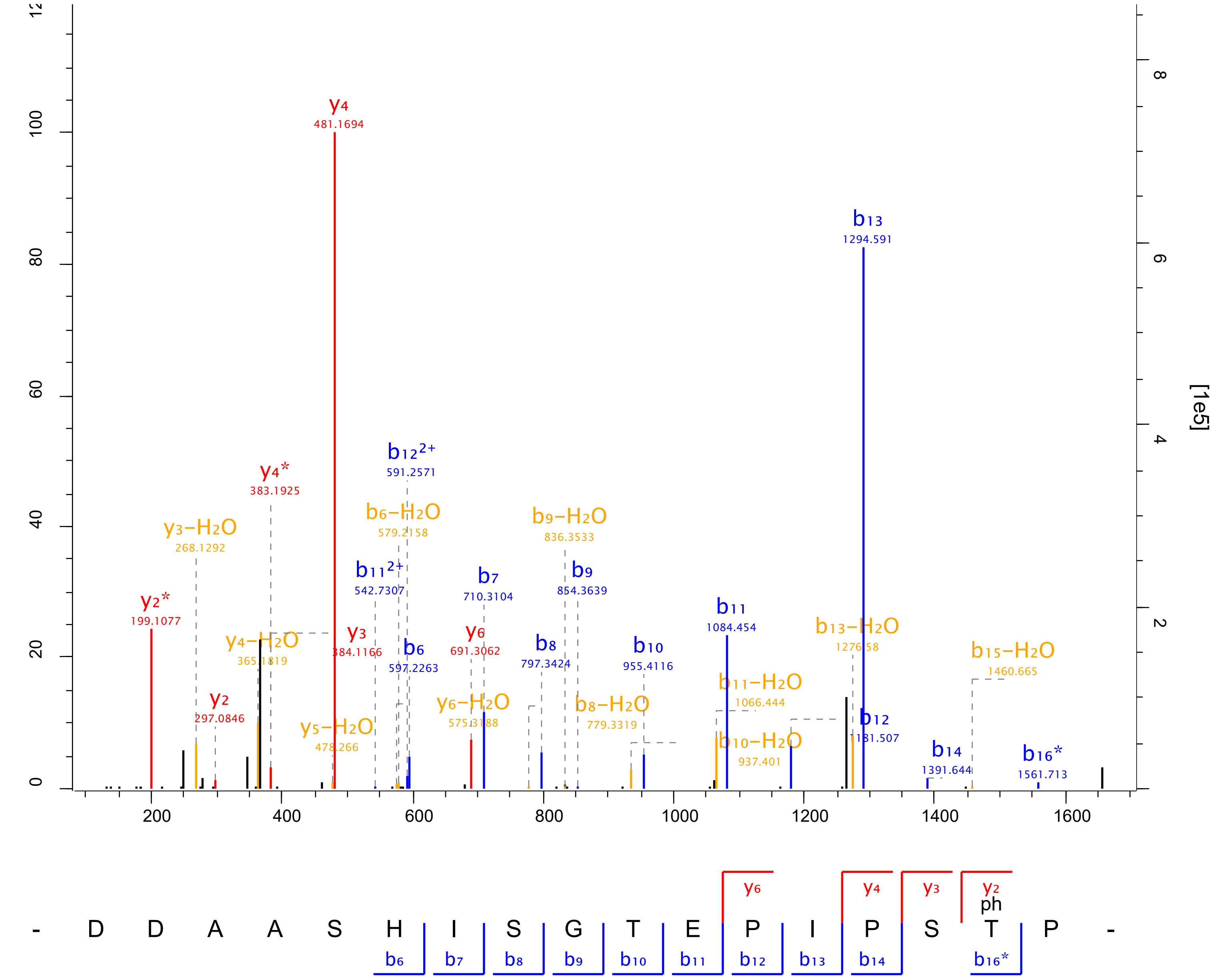

| Raw file | Scan | Method | Score | m/z |
|---|---|---|---|---|
| water_15_2 | 27682 | FTMS; HCD | 144.3 | 760.87 |

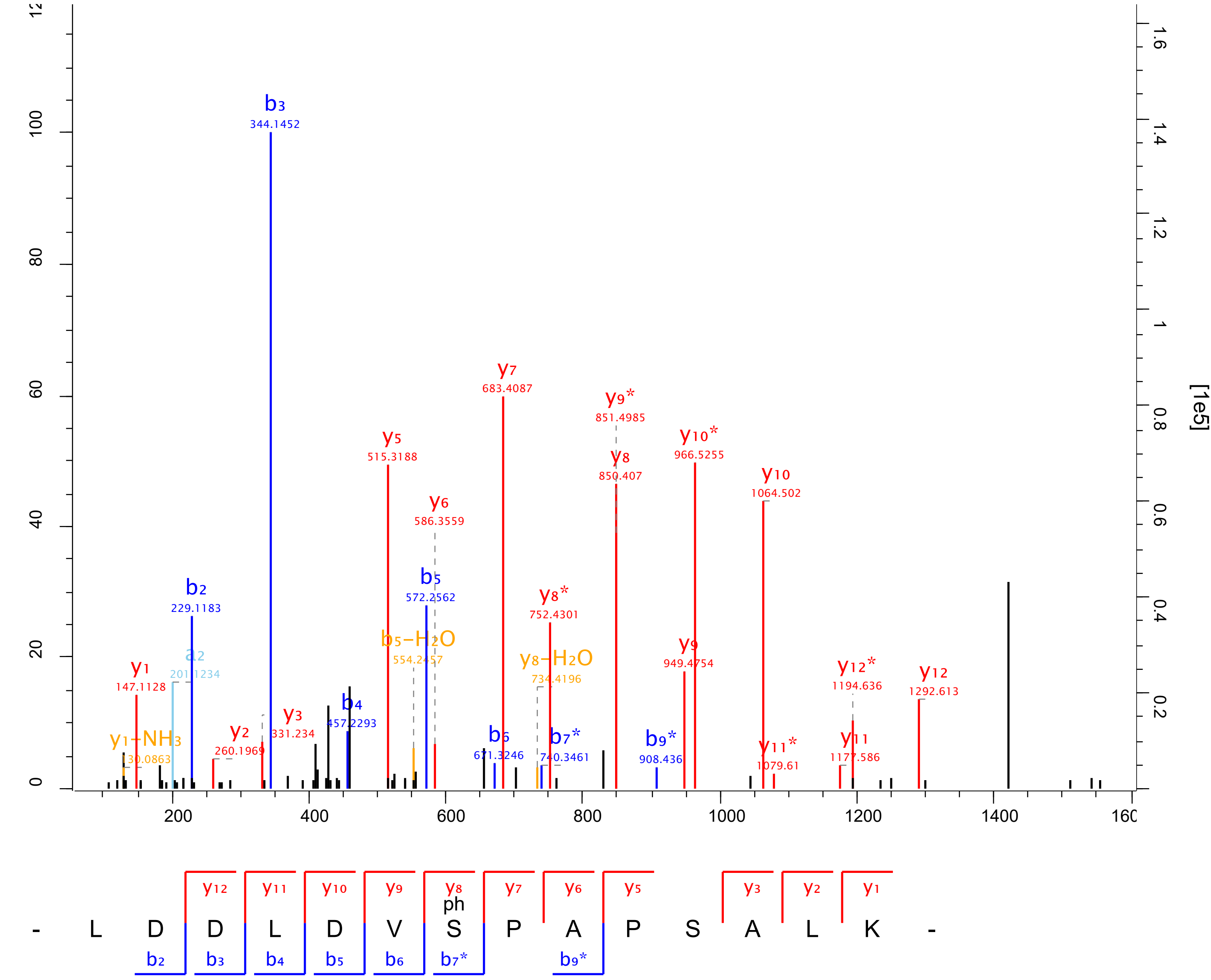

| Raw file | Scan | Method | Score | m/z |
|---|---|---|---|---|
| water_15_2 | 27687 | FTMS; HCD | 107.93 | 632.79 |

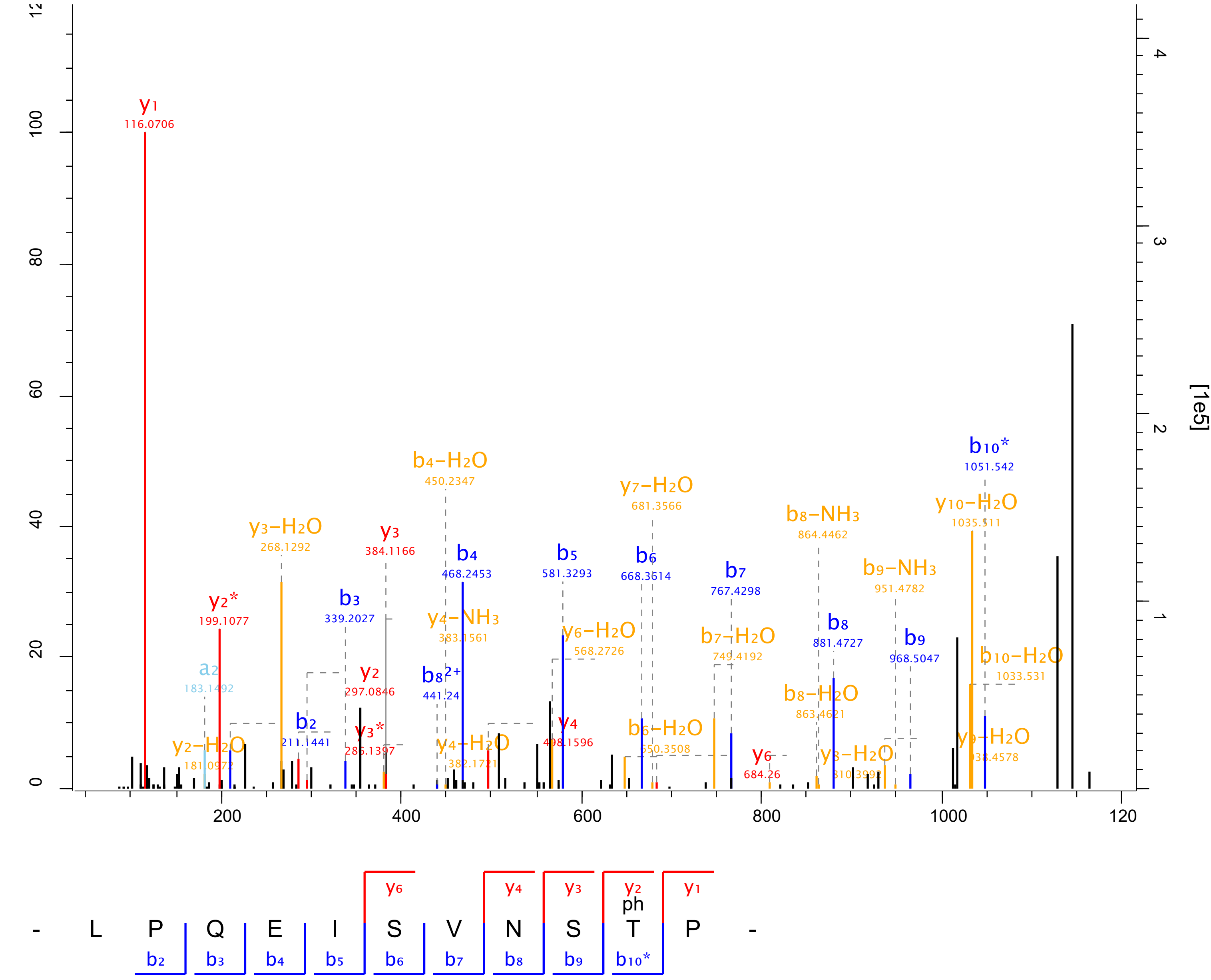

| Raw file | Scan | Method | Score | m/z |
| --- | --- | --- | --- | --- |
| water_15_2 | 28152 | FTMS; HCD | 61.23 | 549.23 |

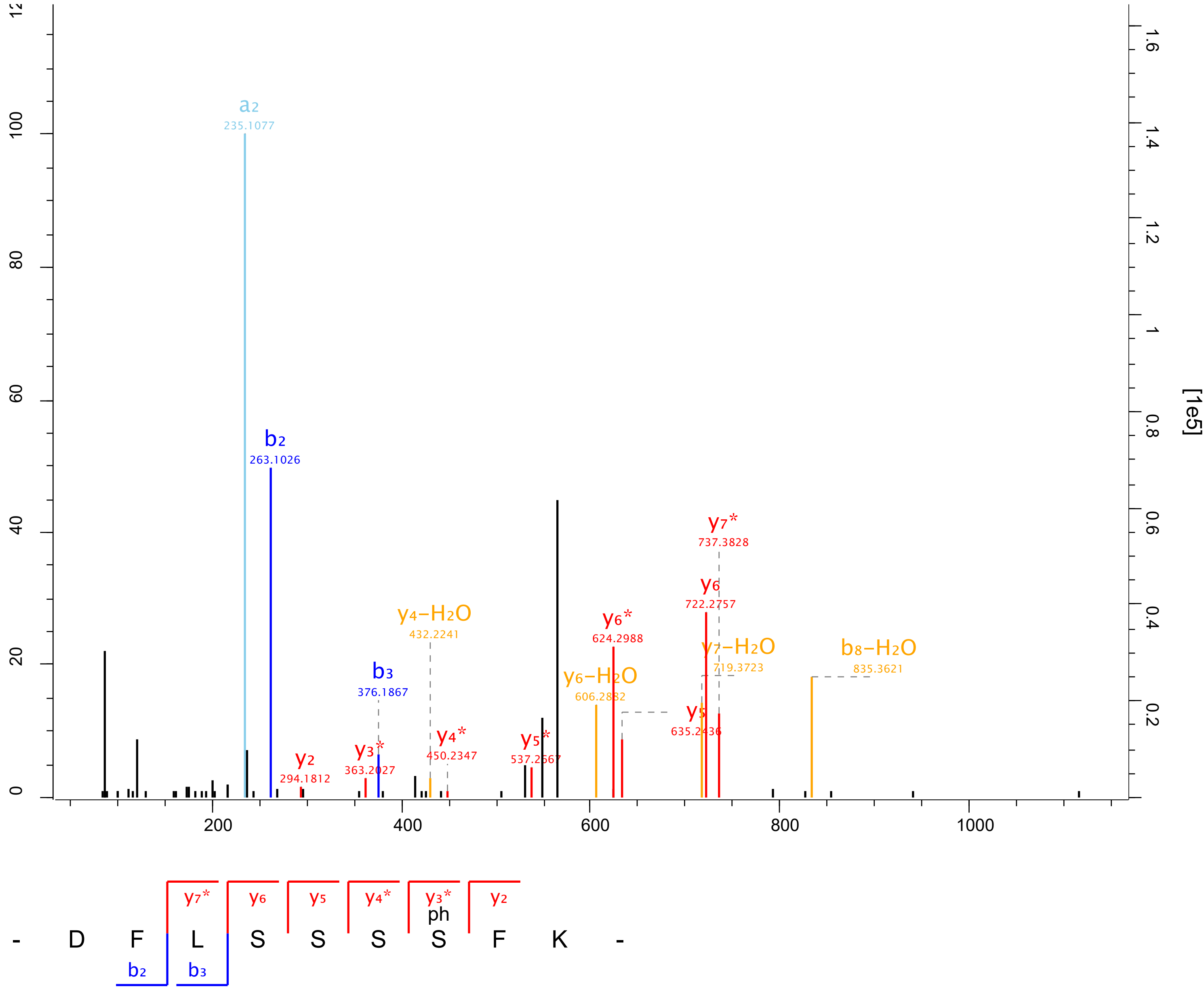

| Raw file | Scan | Method | Score | m/z |
| --- | --- | --- | --- | --- |
| water_15_2 | 2835 | FTMS; HCD | 42.72 | 588.79 |

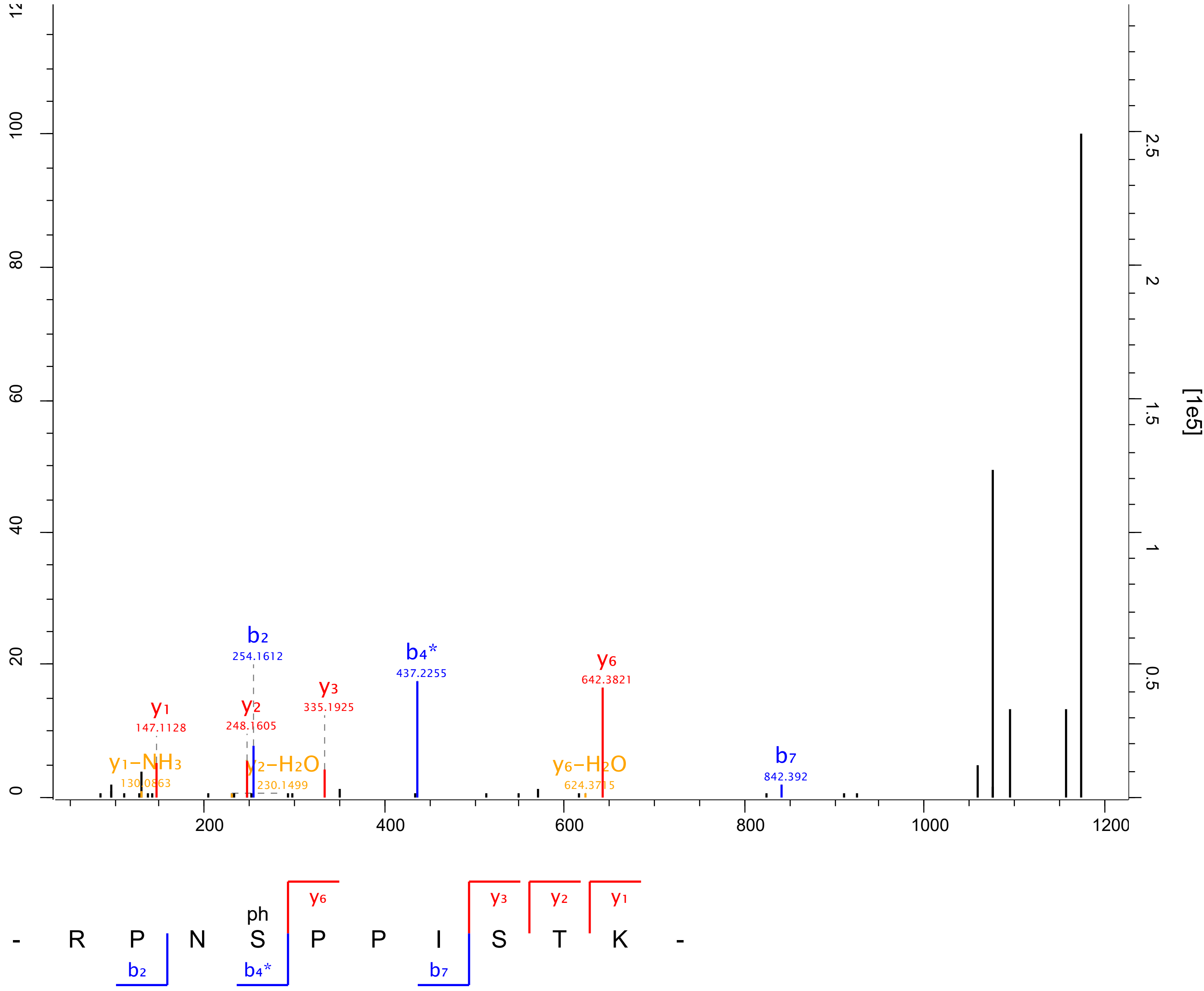

| Raw file | Scan | Method | Score | m/z |
| --- | --- | --- | --- | --- |
| water_15_2 | 28502 | FTMS; HCD | 118.31 | 791.86 |

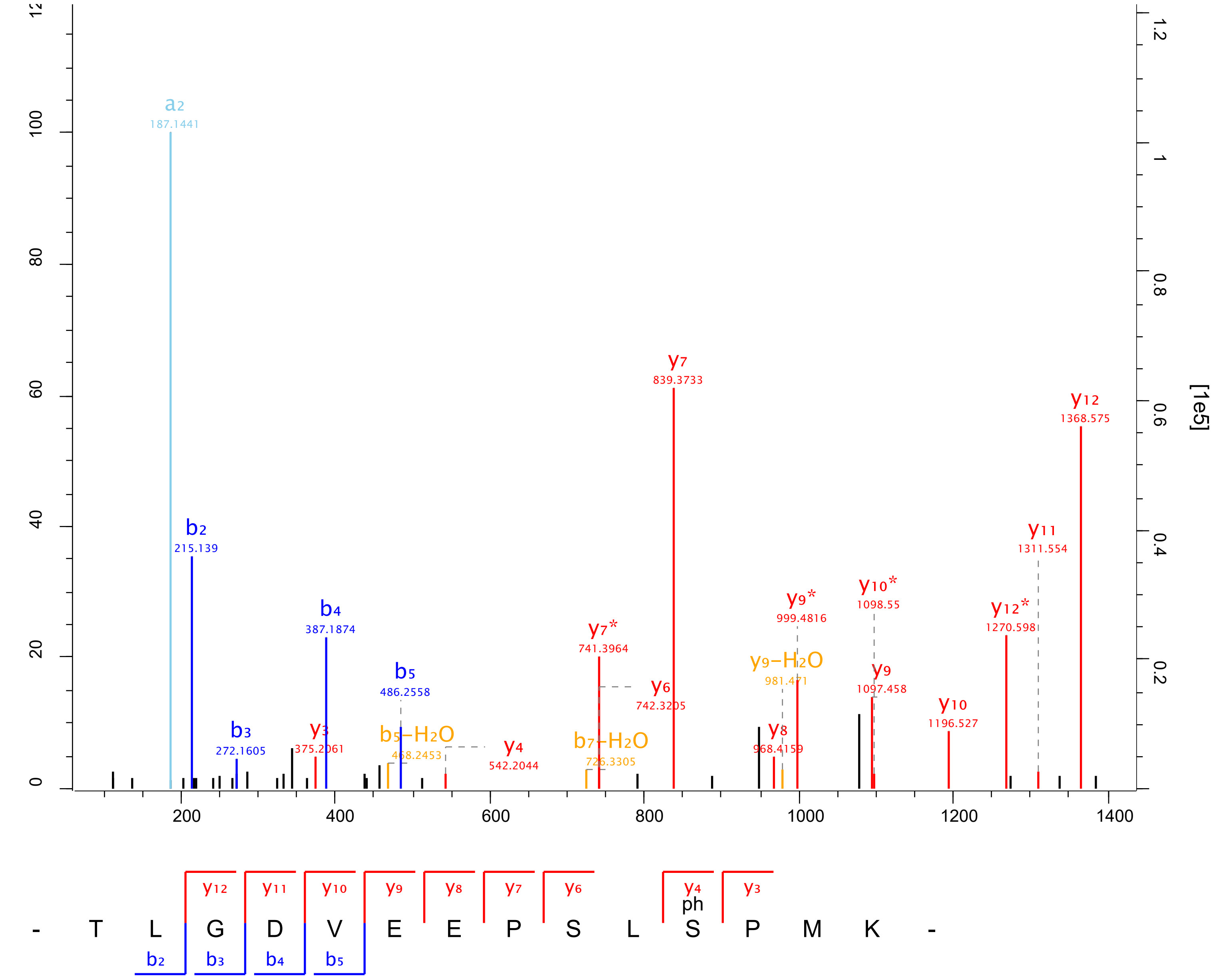

| Raw file | Scan | Method | Score | m/z |
|---|---|---|---|---|
| water_15_2 | 28512 | FTMS; HCD | 104.84 | 779.89 |

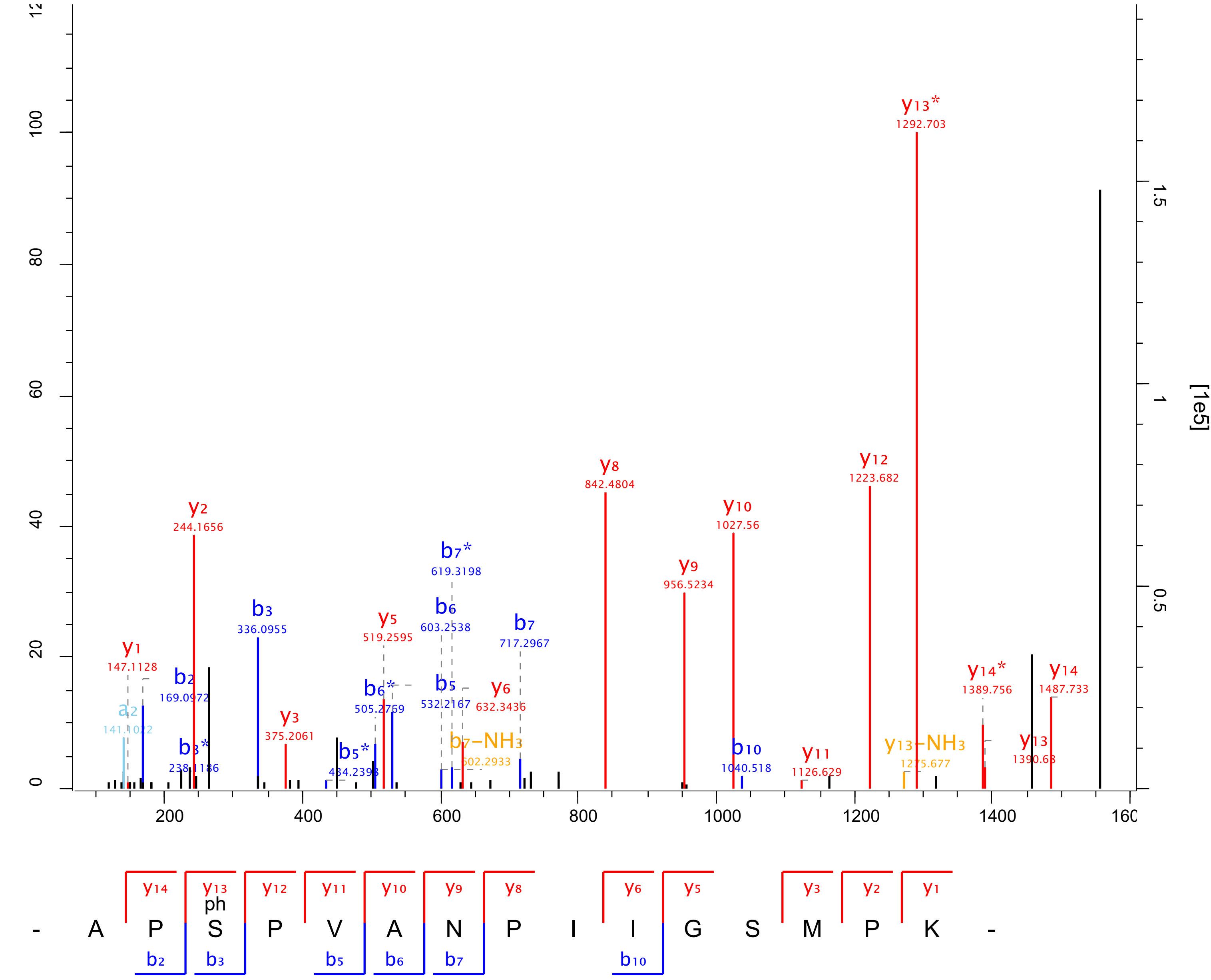

| Raw file | Scan | Method | Score | m/z |
| --- | --- | --- | --- | --- |
| water_15_2 | 28959 | FTMS; HCD | 201.04 | 642.29 |

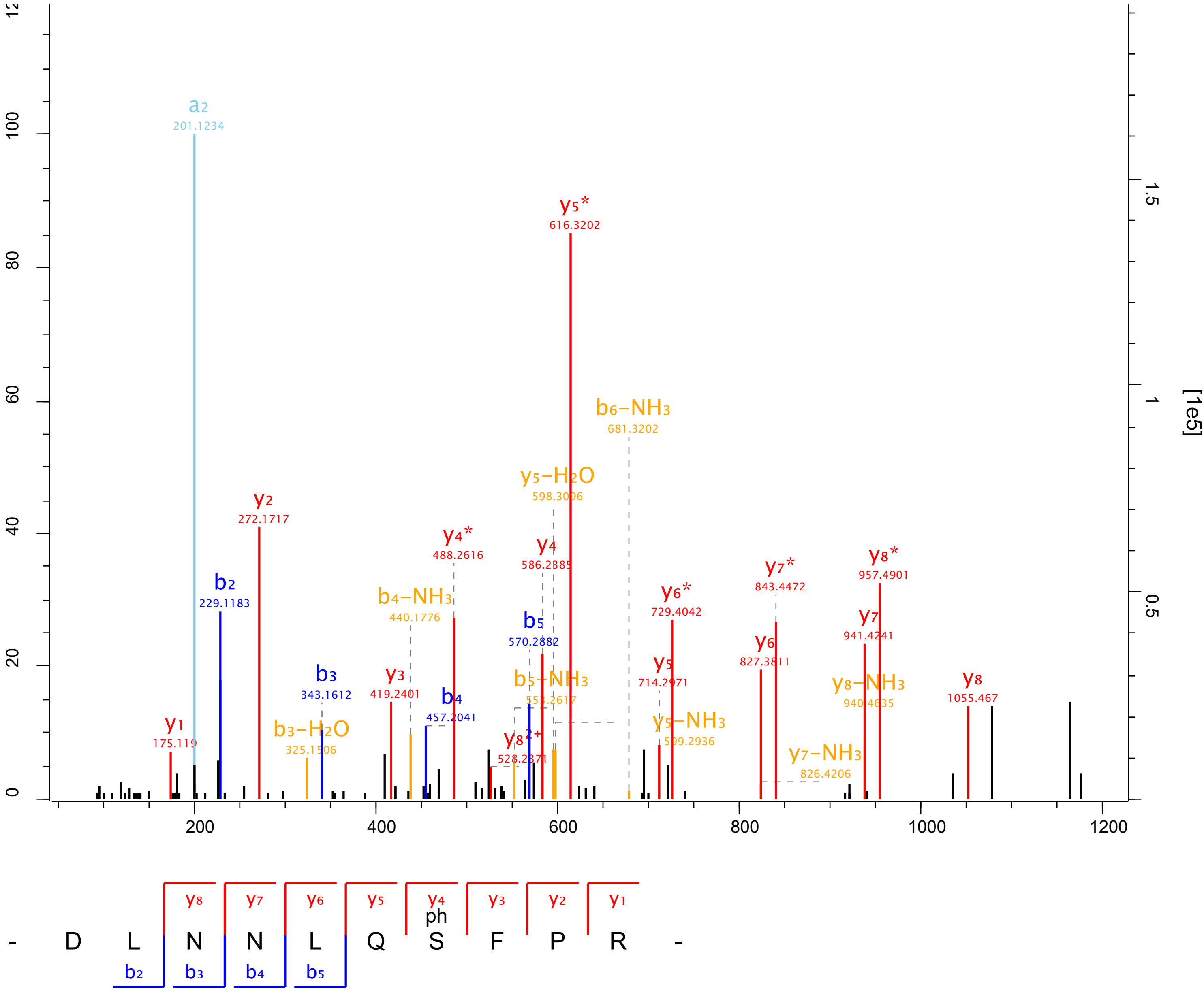

| Raw file | Scan | Method | Score | m/z |
|---|---|---|---|---|
| water_15_2 | 2923 | FTMS; HCD | 81.97 | 441.23 |

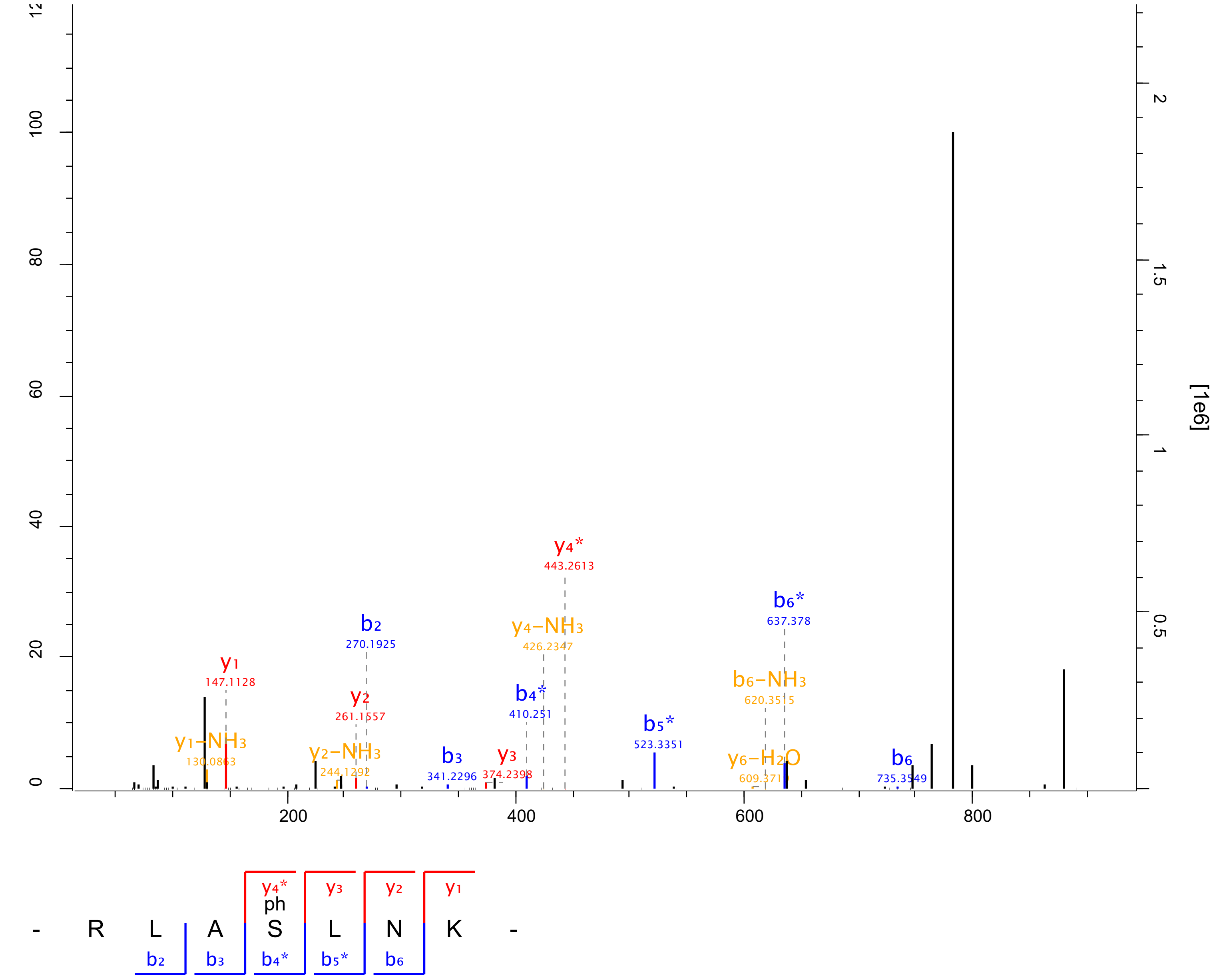

| Raw file | Scan | Method | Score | m/z |
|---|---|---|---|---|
| water_15_2 | 29244 | FTMS; HCD | 179.35 | 582.75 |

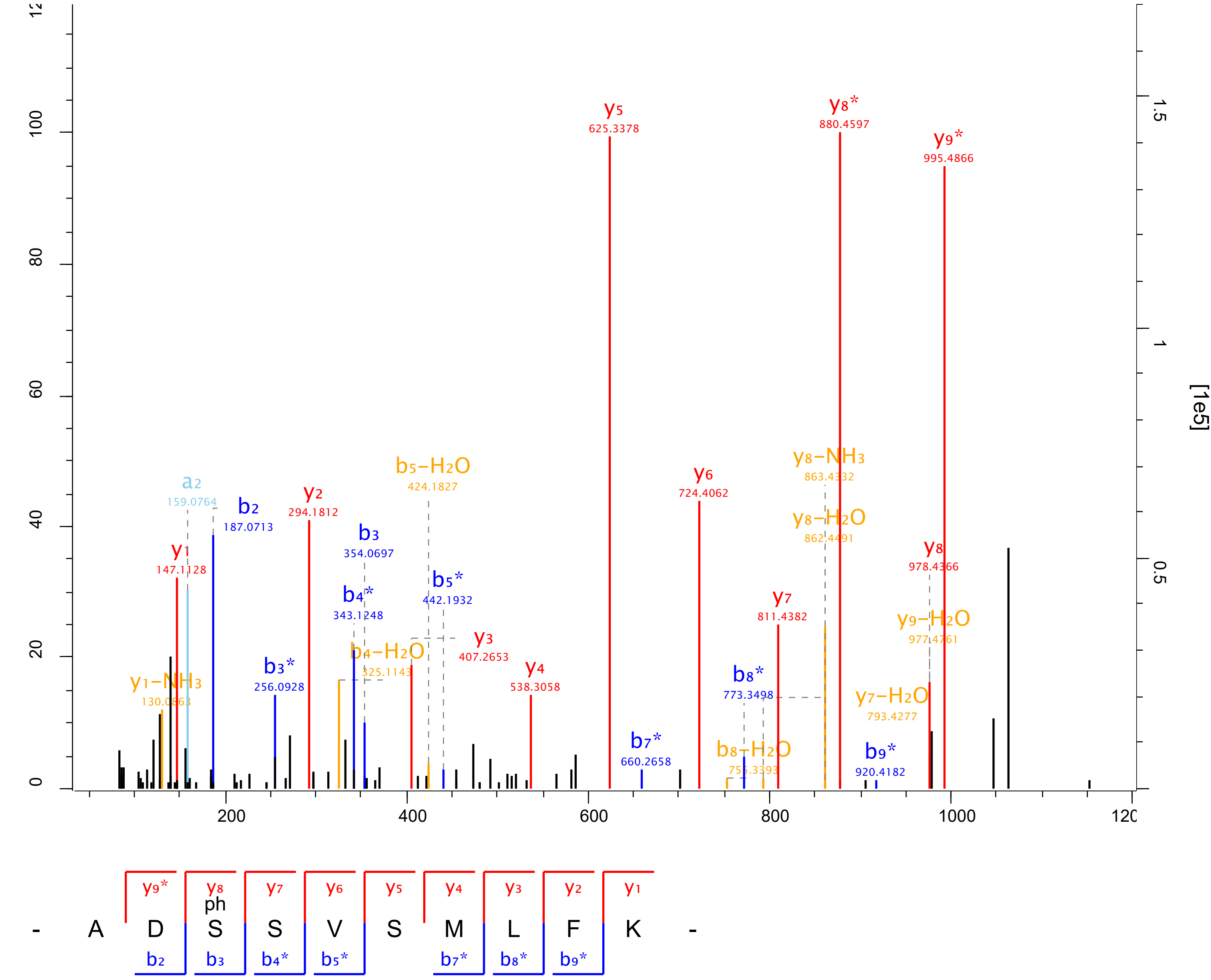

| Raw file | Scan | Method | Score | m/z |
| --- | --- | --- | --- | --- |
| water_15_2 | 29697 | FTMS; HCD | 48.9 | 760.33 |

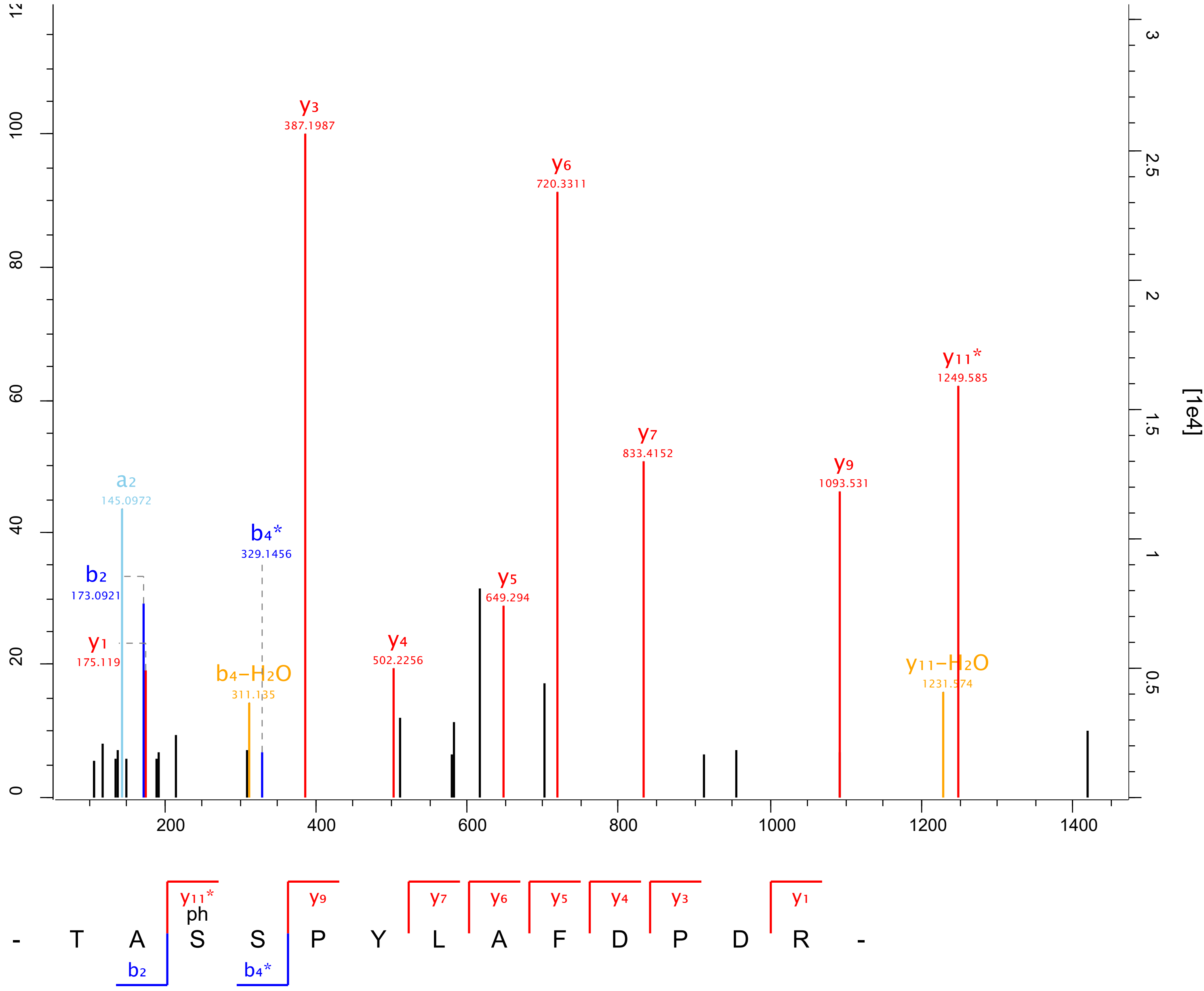

| Raw file | Scan | Method | Score | m/z |
|---|---|---|---|---|
| water_15_2 | 29772 | FTMS; HCD | 132.84 | 719.82 |

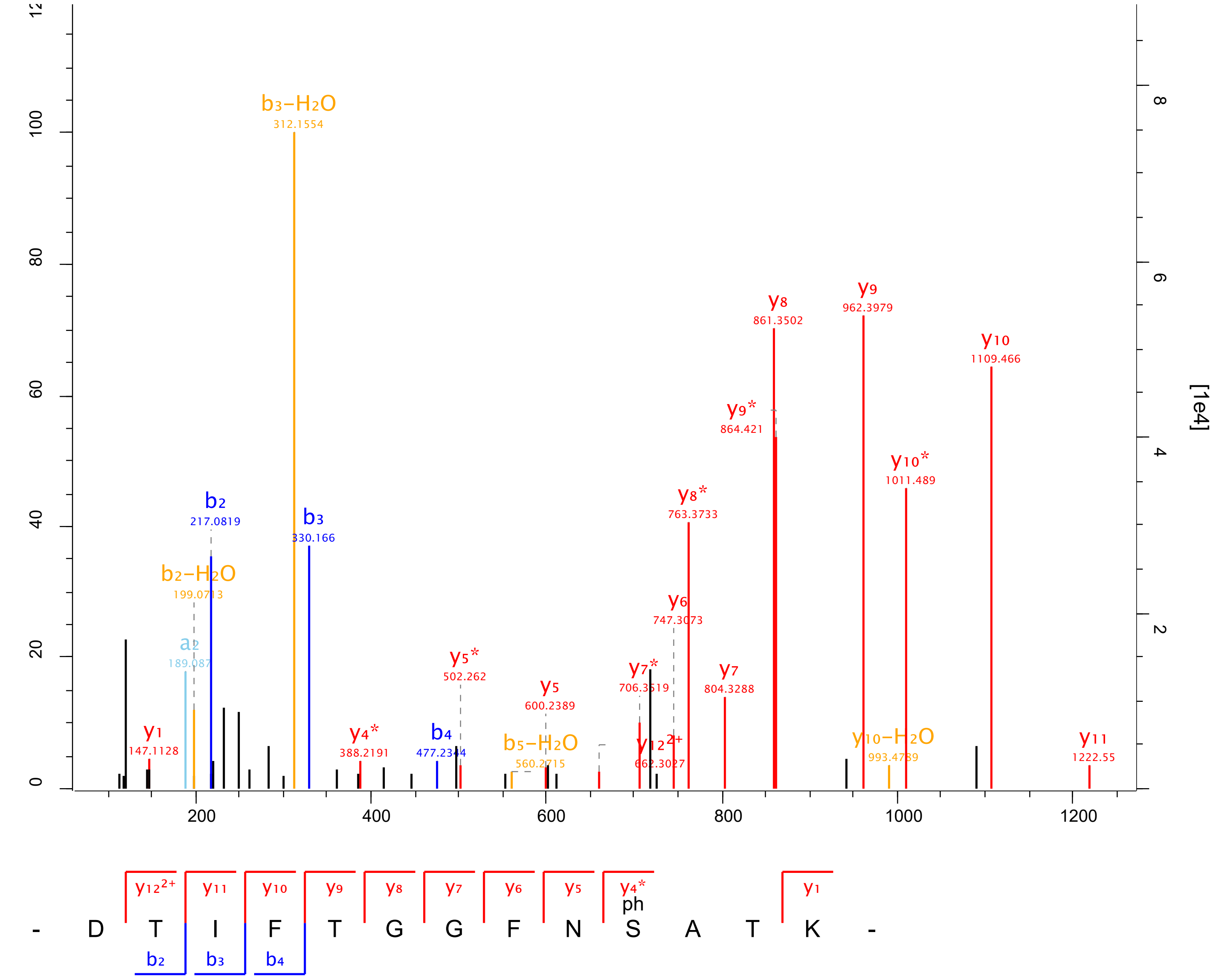

| Raw file | Scan | Method | Score | m/z |
|---|---|---|---|---|
| water_15_2 | 30932 | FTMS; HCD | 100.43 | 707.3 |

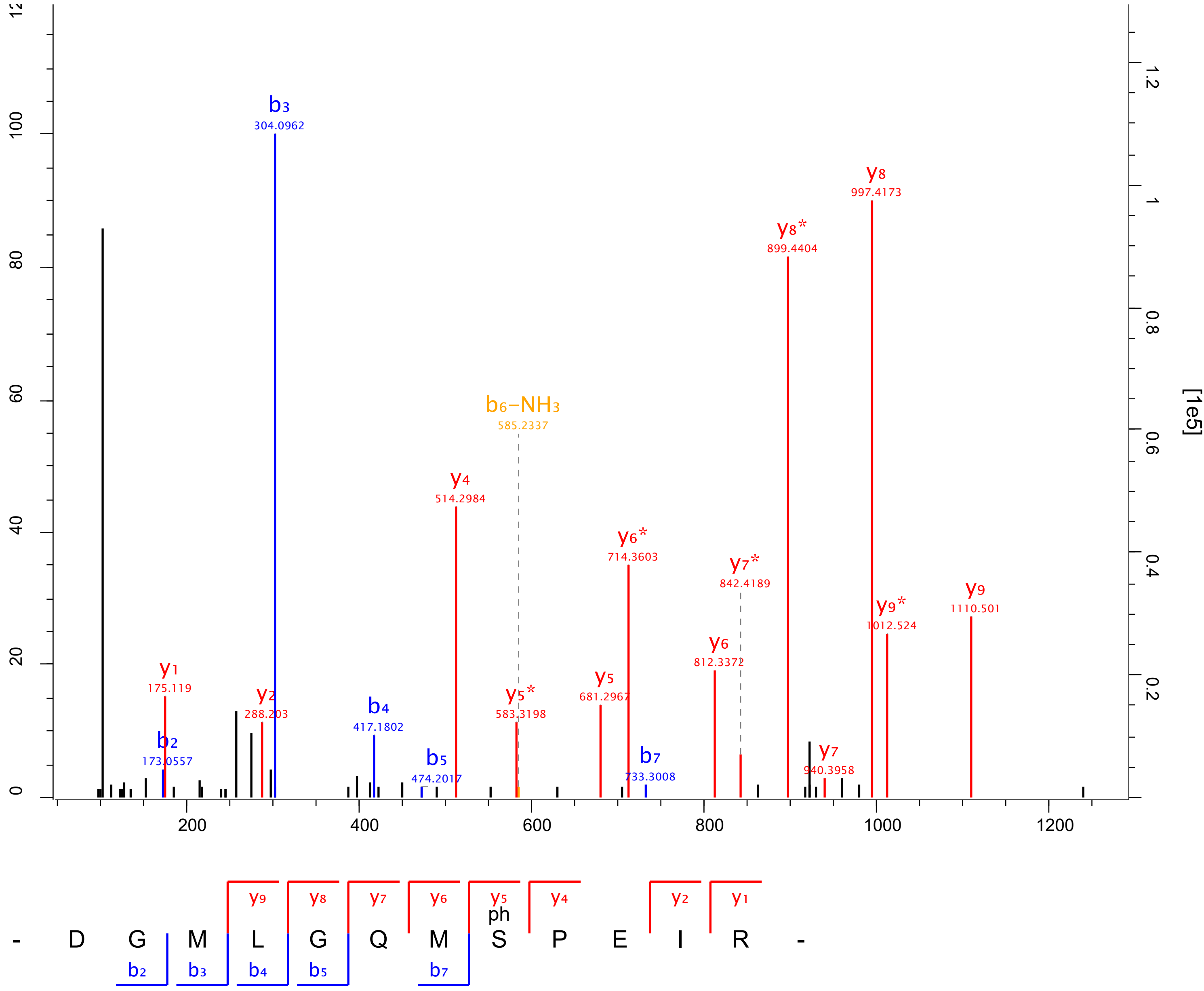

| Raw file | Scan | Method | Score | m/z |
|---|---|---|---|---|
| water_15_2 | 30991 | FTMS; HCD | 121.95 | 701.34 |

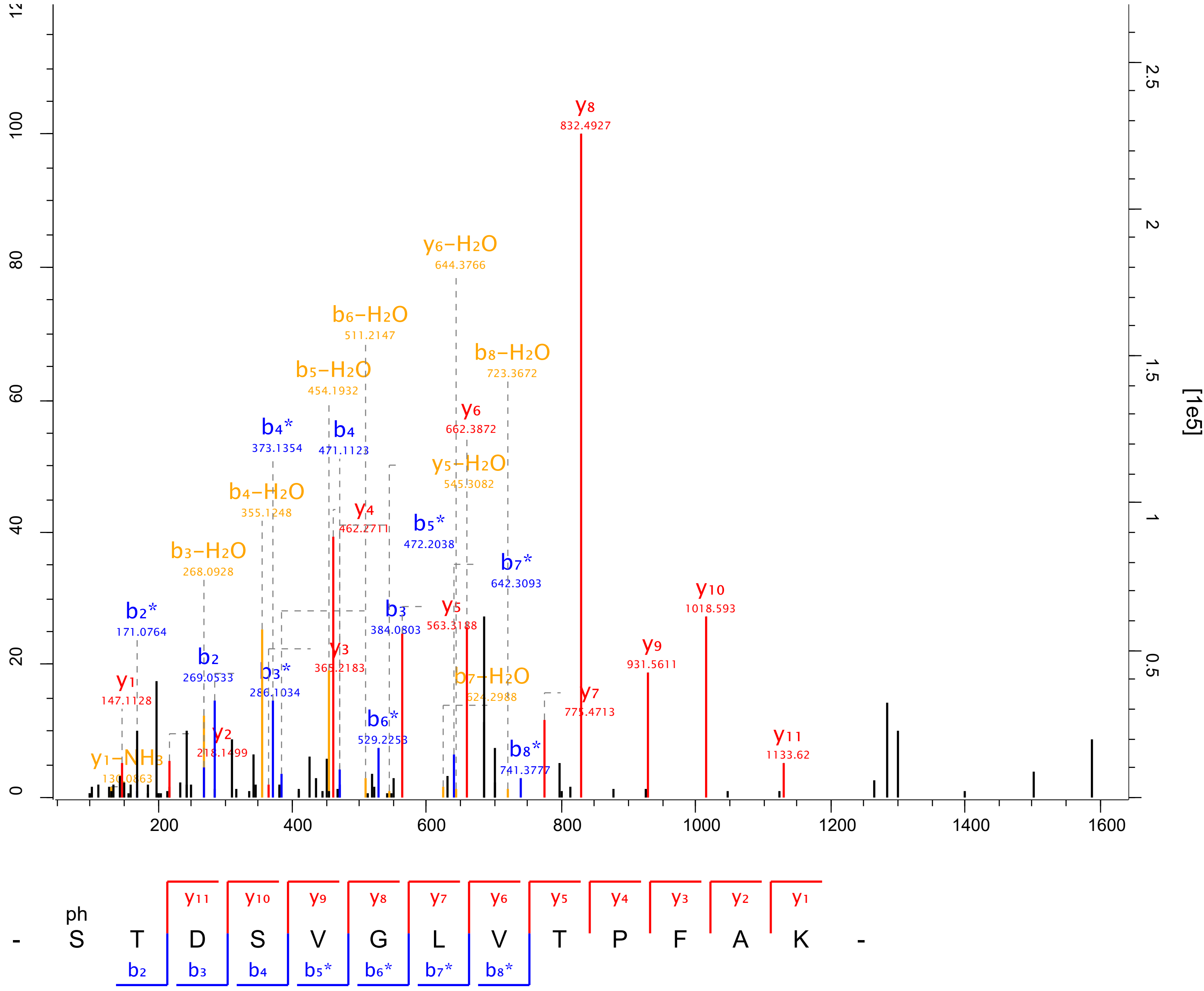

| Raw file | Scan | Method | Score | m/z |
|---|---|---|---|---|
| water_15_2 | 31201 | FTMS; HCD | 231.62 | 818.37 |

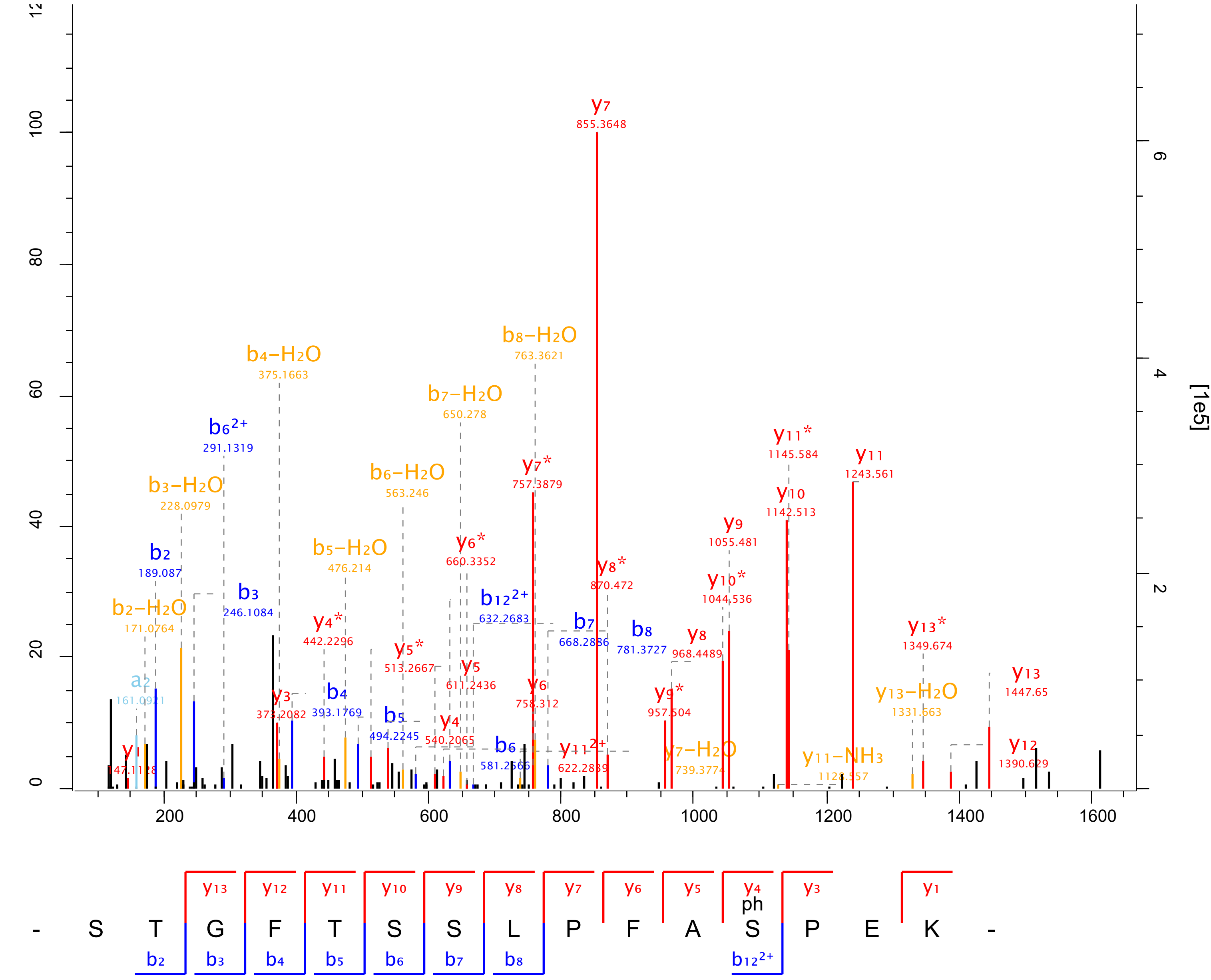

| Raw file | Scan | Method | Score | m/z |
|---|---|---|---|---|
| water_15_2 | 3145 | FTMS; HCD | 58.89 | 430.18 |

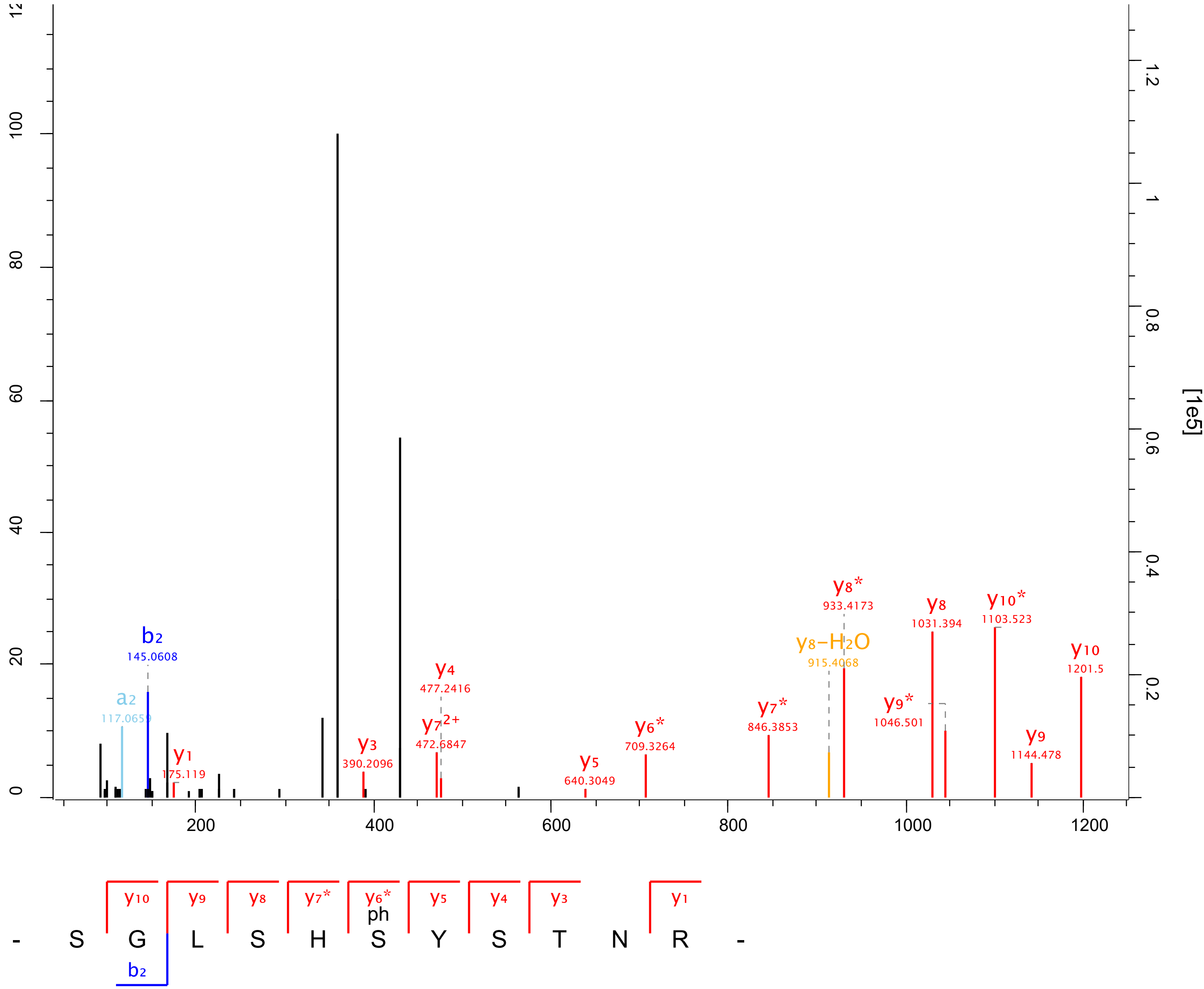

| Raw file | Scan | Method | Score | m/z |
|---|---|---|---|---|
| water_15_2 | 32515 | FTMS; HCD | 344.4 | 868.69 |

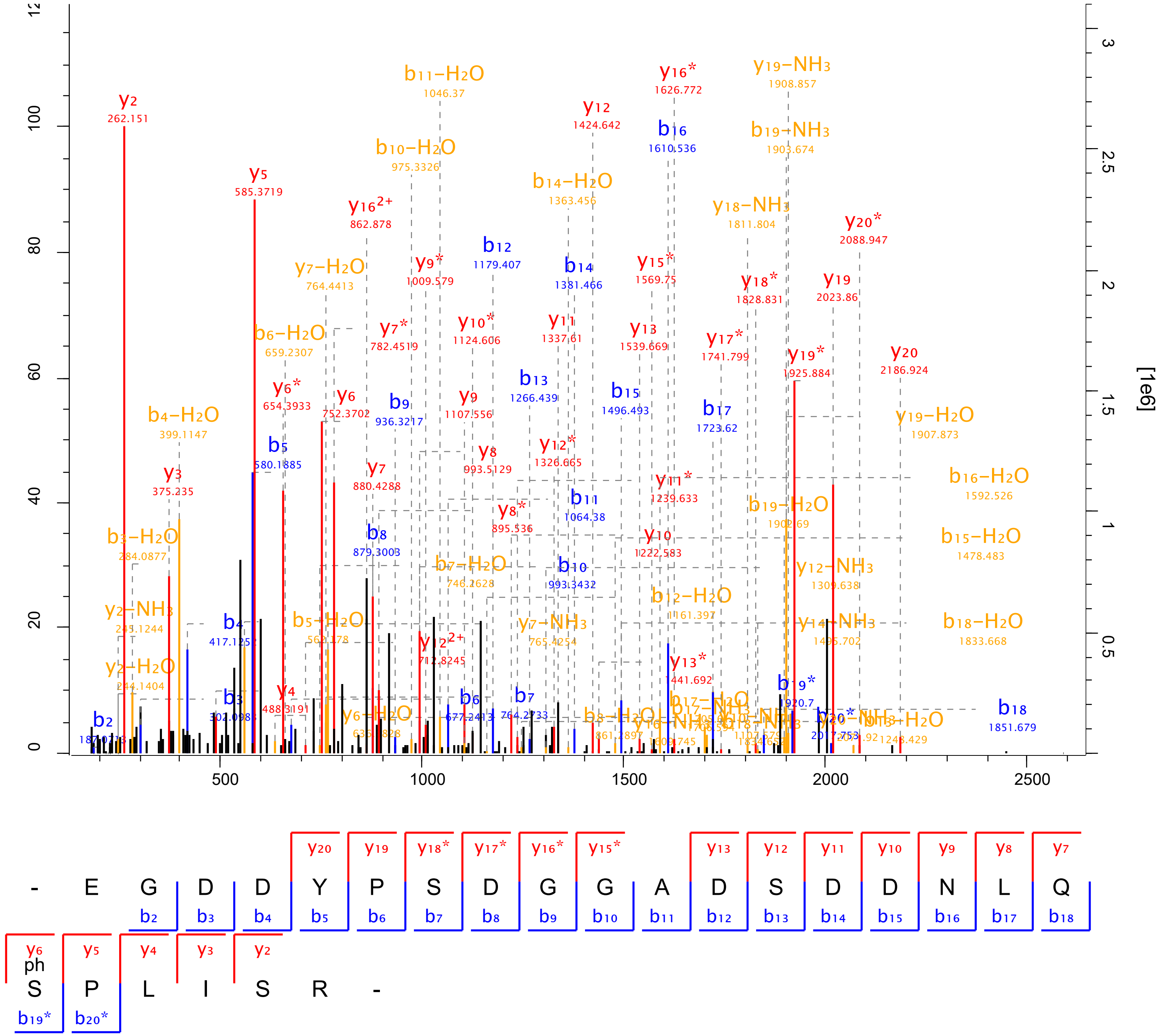

| Raw file | Scan | Method | Score | m/z |
|---|---|---|---|---|
| water_15_2 | 32905 | FTMS; HCD | 128.86 | 469.22 |

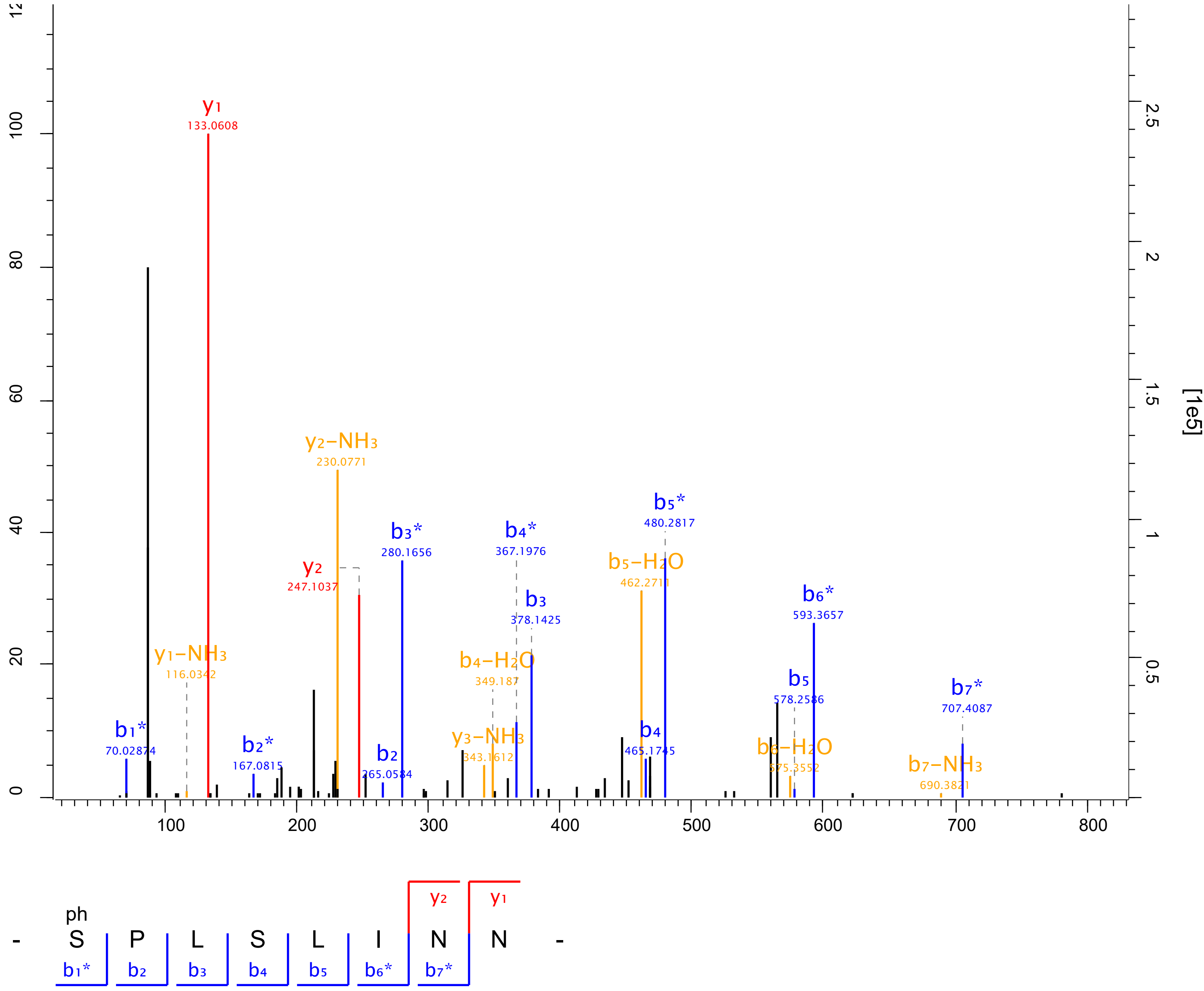

| Raw file | Scan | Method | Score | m/z |
|---|---|---|---|---|
| water_15_2 | 3328 | FTMS; HCD | 42.12 | 649.61 |

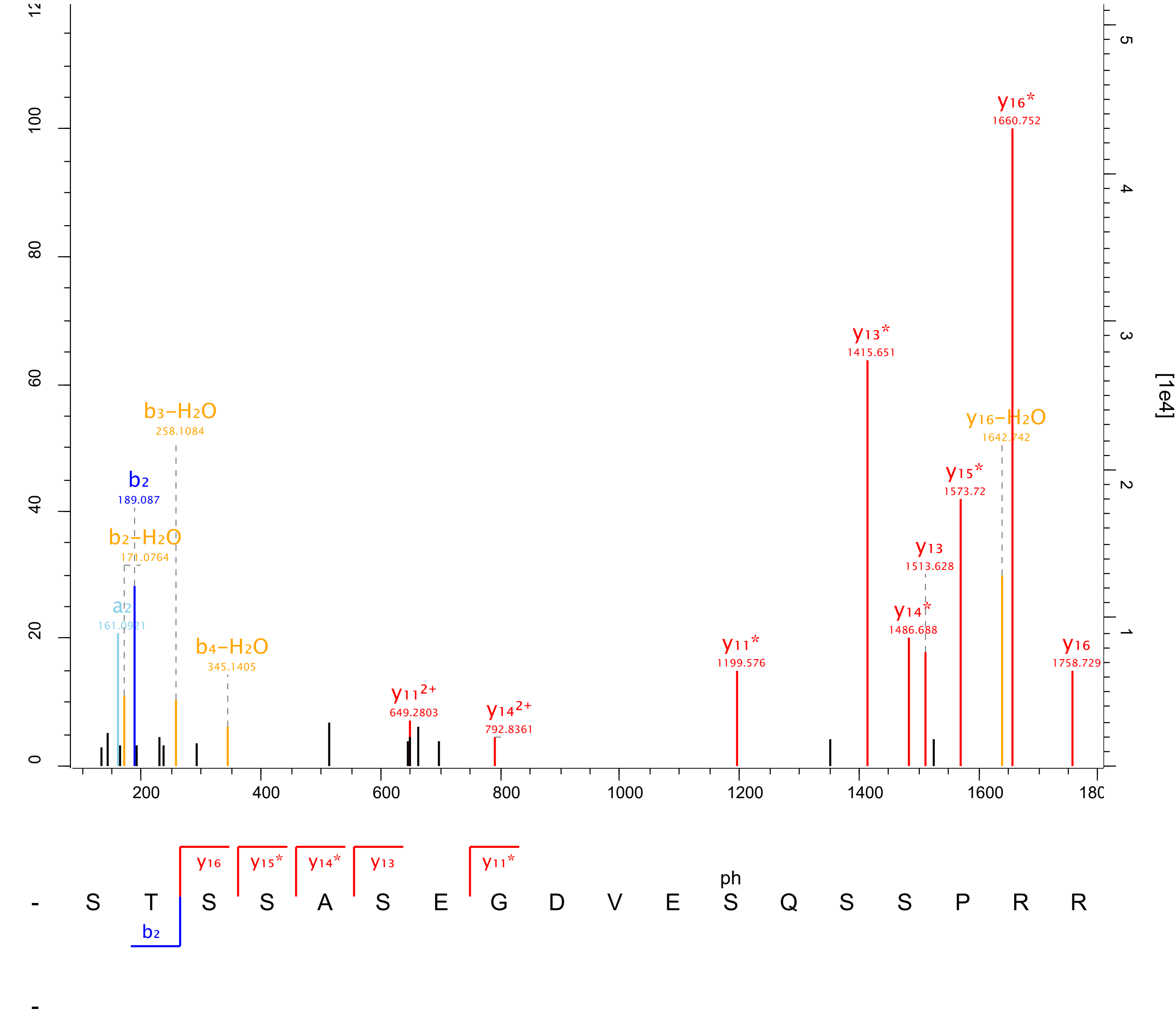

| Raw file | Scan | Method | Score | m/z |
| --- | --- | --- | --- | --- |
| water_15_2 | 3372 | FTMS; HCD | 174.37 | 611.77 |

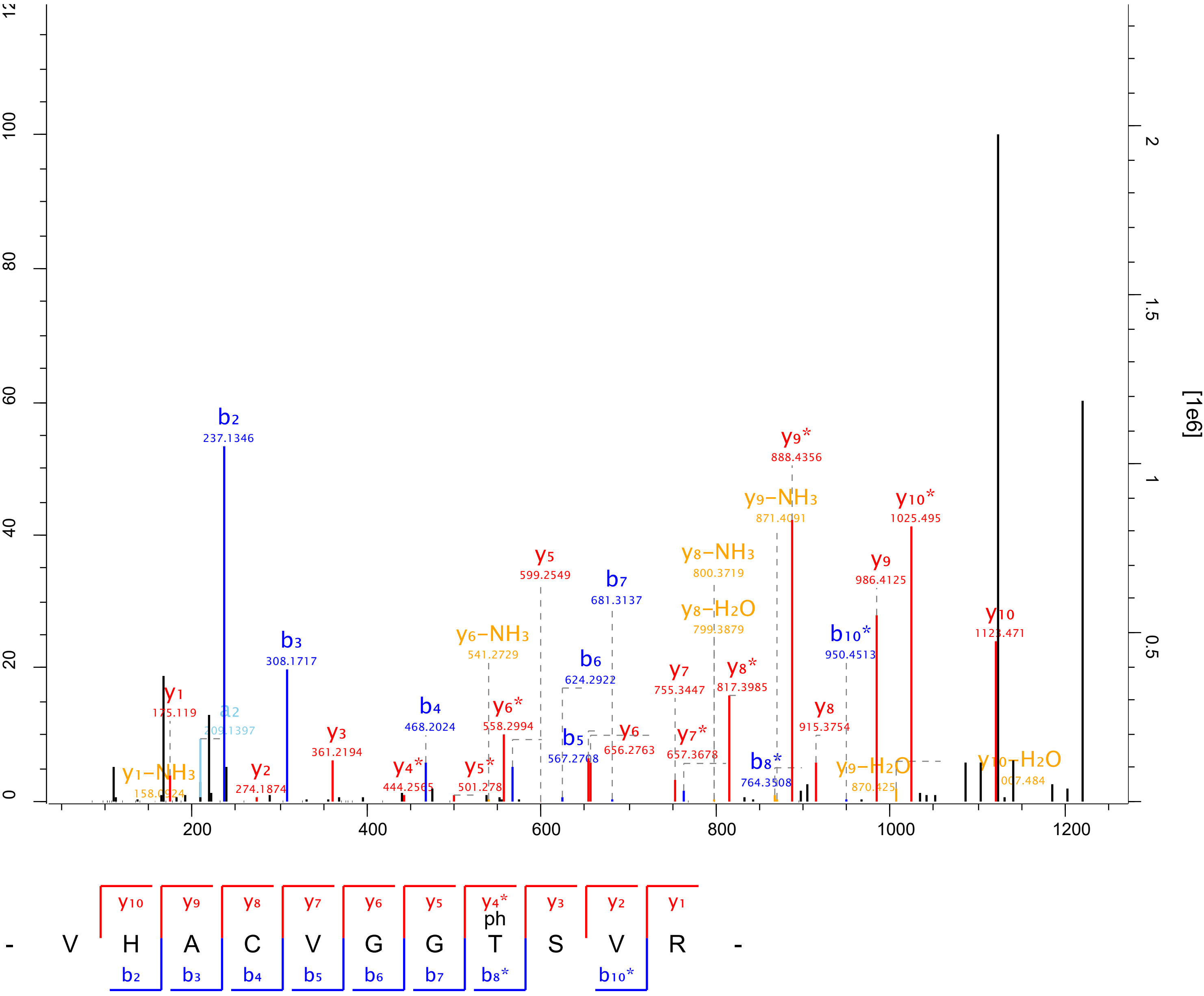

| Raw file | Scan | Method | Score | m/z |
| --- | --- | --- | --- | --- |
| water_15_2 | 33811 | FTMS; HCD | 53.03 | 738.62 |

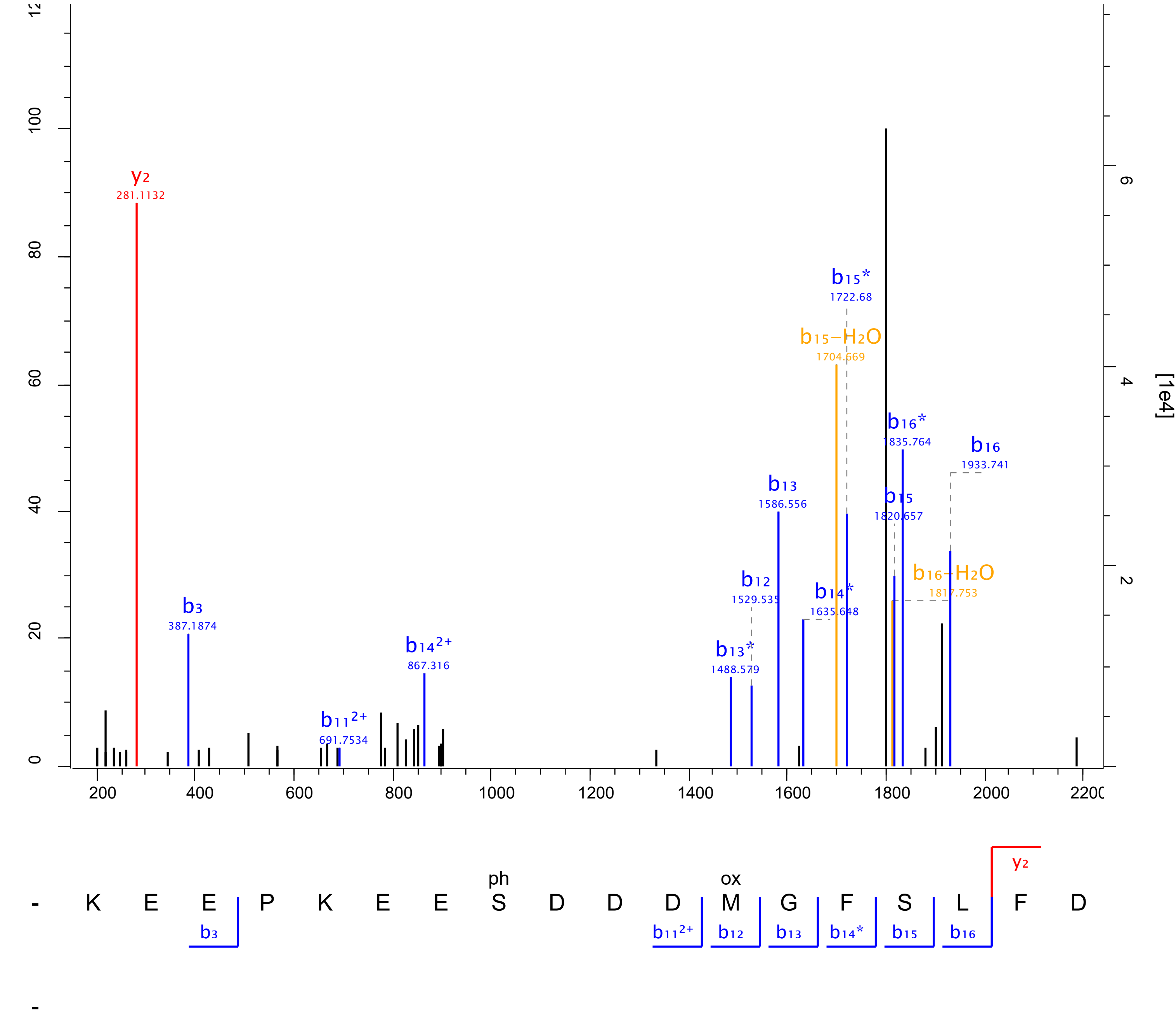

| Raw file | Scan | Method | Score | m/z |
|---|---|---|---|---|
| water_15_2 | 34054 | FTMS; HCD | 41.91 | 1052.45 |

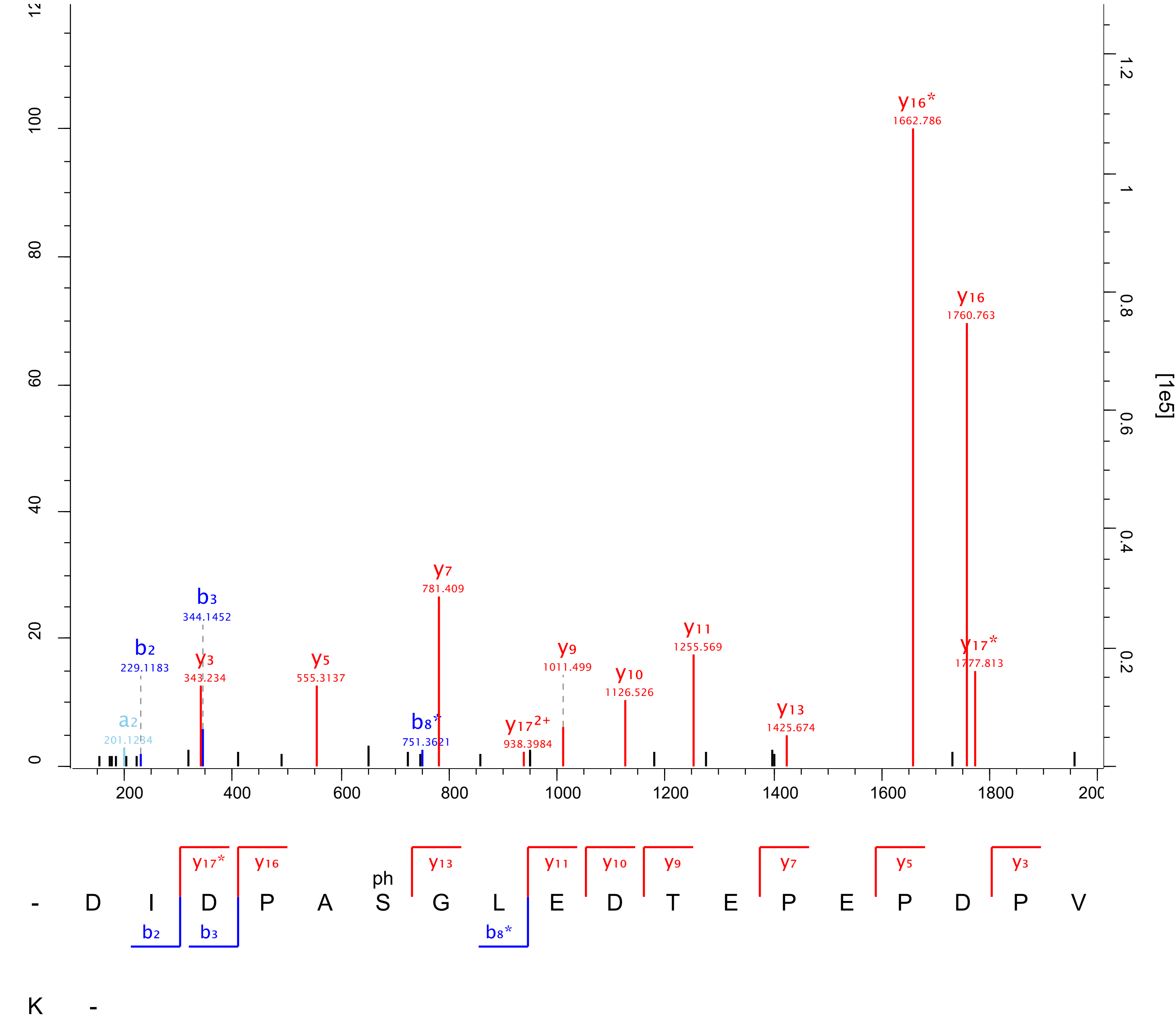

| Raw file | Scan | Method | Score | m/z |
|---|---|---|---|---|
| water_15_2 | 3421 | FTMS; HCD | 46.32 | 470.54 |

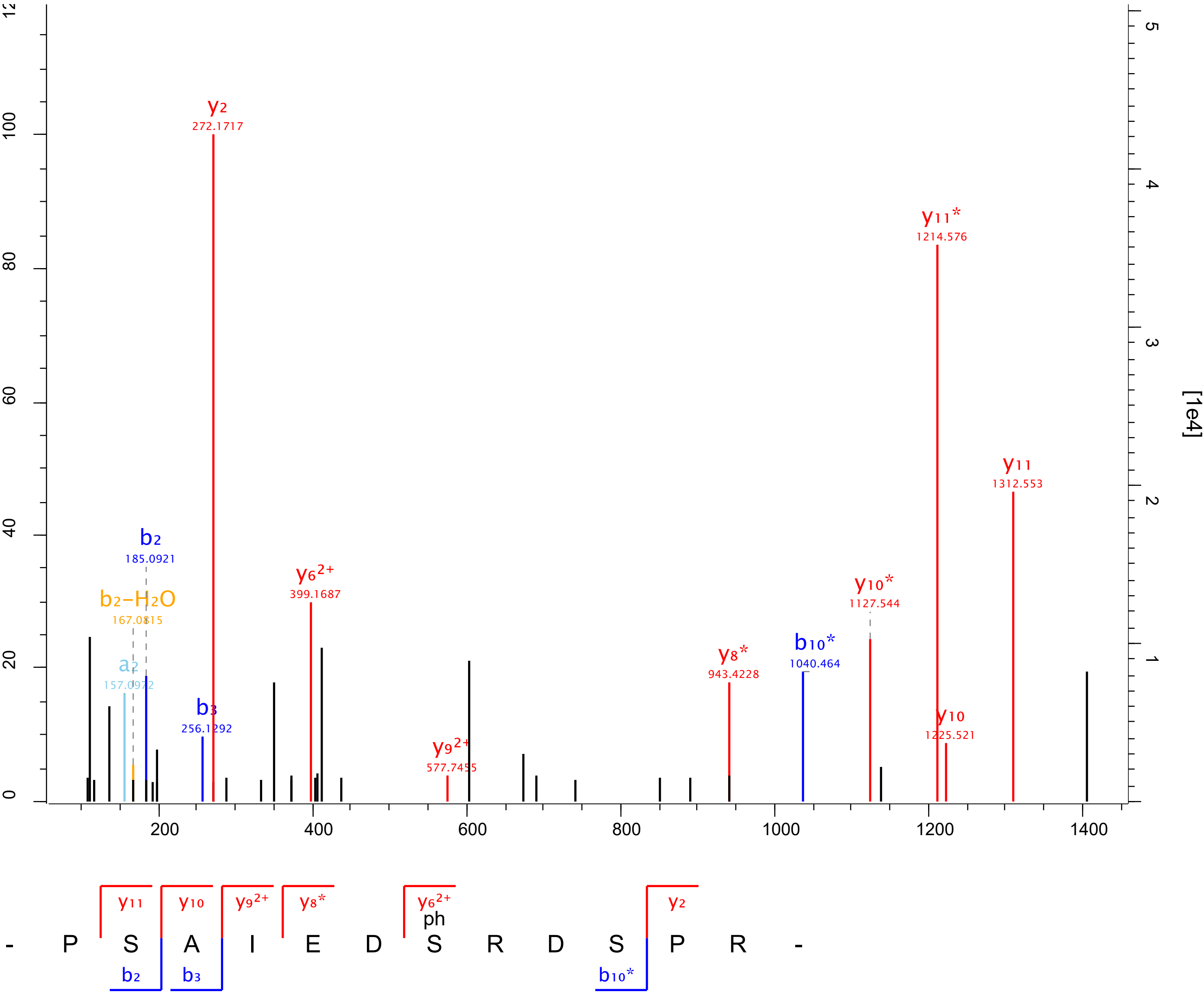

| Raw file | Scan | Method | Score | m/z |
| --- | --- | --- | --- | --- |
| water_15_2 | 3473 | FTMS; HCD | 109.79 | 641.79 |

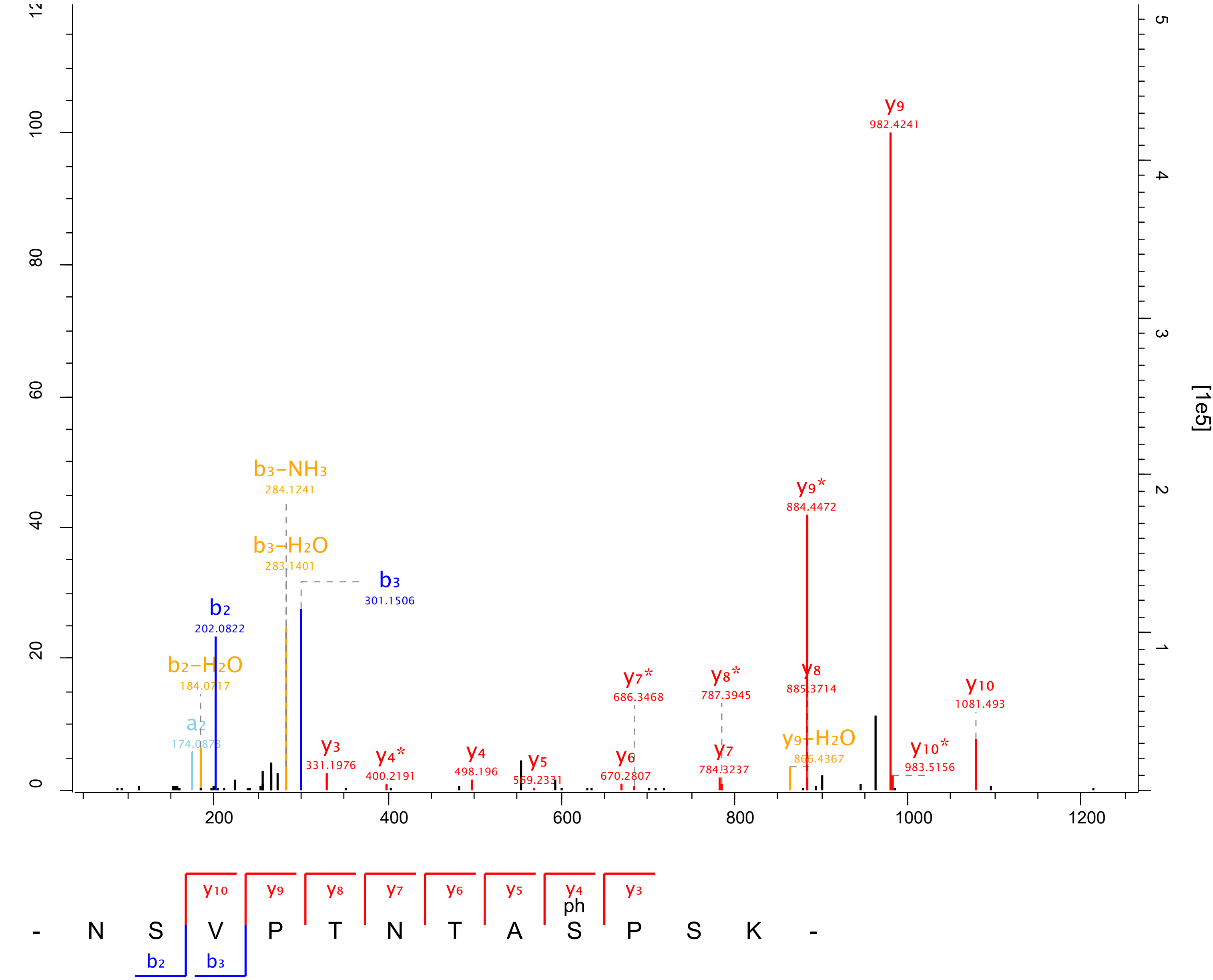

| Raw file | Scan | Method | Score | m/z |
| --- | --- | --- | --- | --- |
| water_15_2 | 3485 | FTMS; HCD | 51.88 | 870.37 |

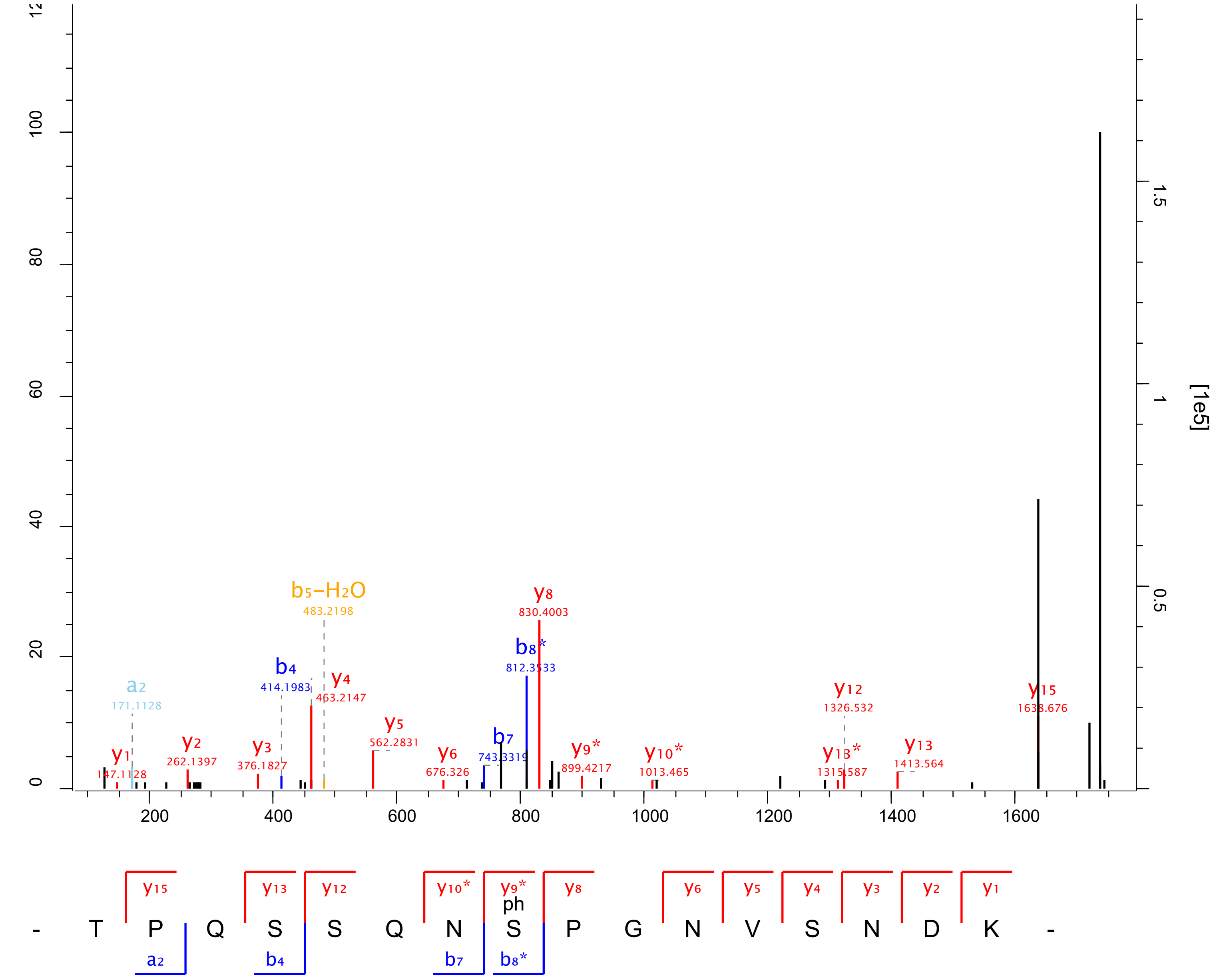

| Raw file | Scan | Method | Score | m/z |
|---|---|---|---|---|
| water_15_2 | 3517 | FTMS; HCD | 47.77 | 604.31 |

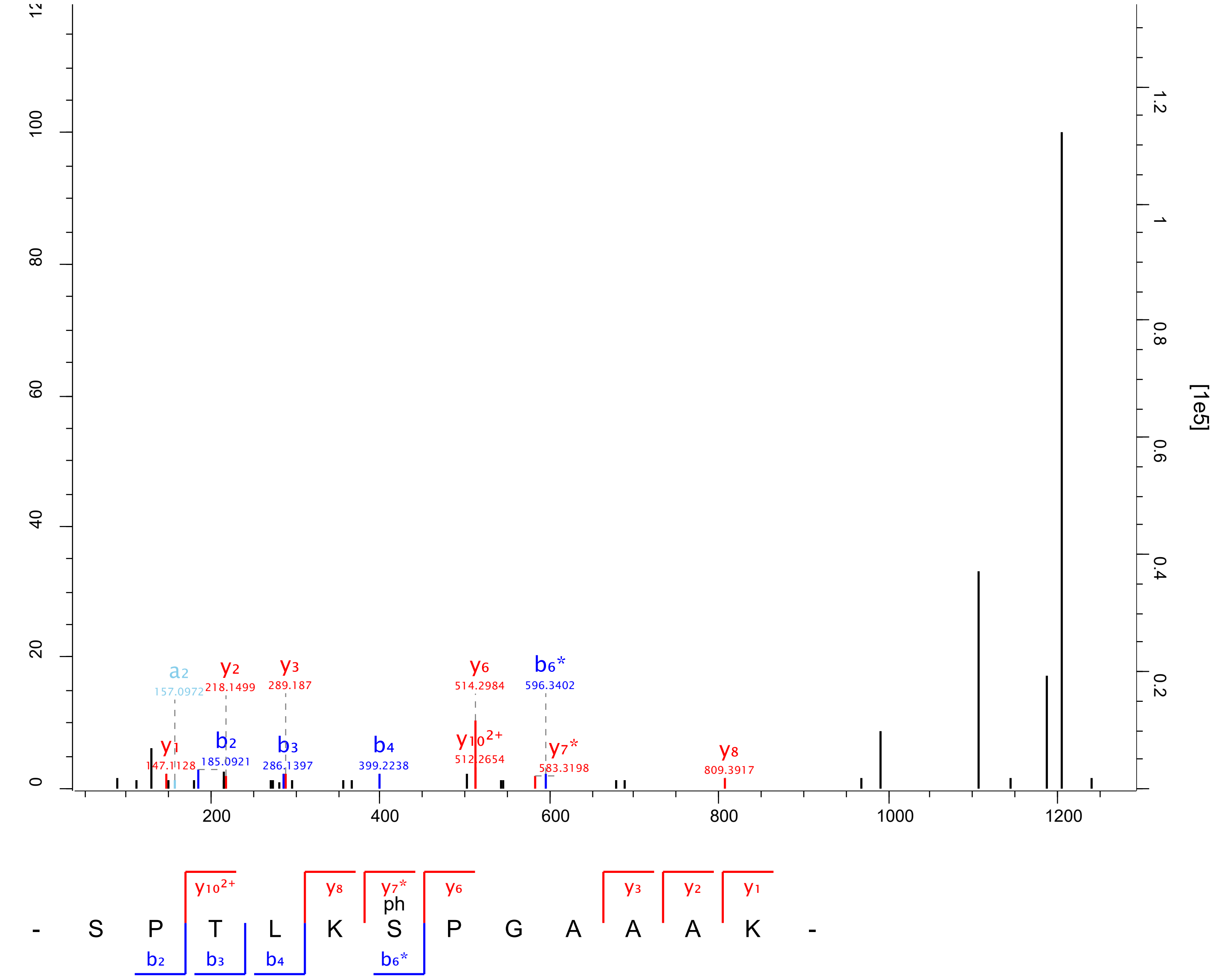

| Raw file | Scan | Method | Score | m/z |
| --- | --- | --- | --- | --- |
| water_15_2 | 35728 | FTMS; HCD | 300.28 | 819.86 |

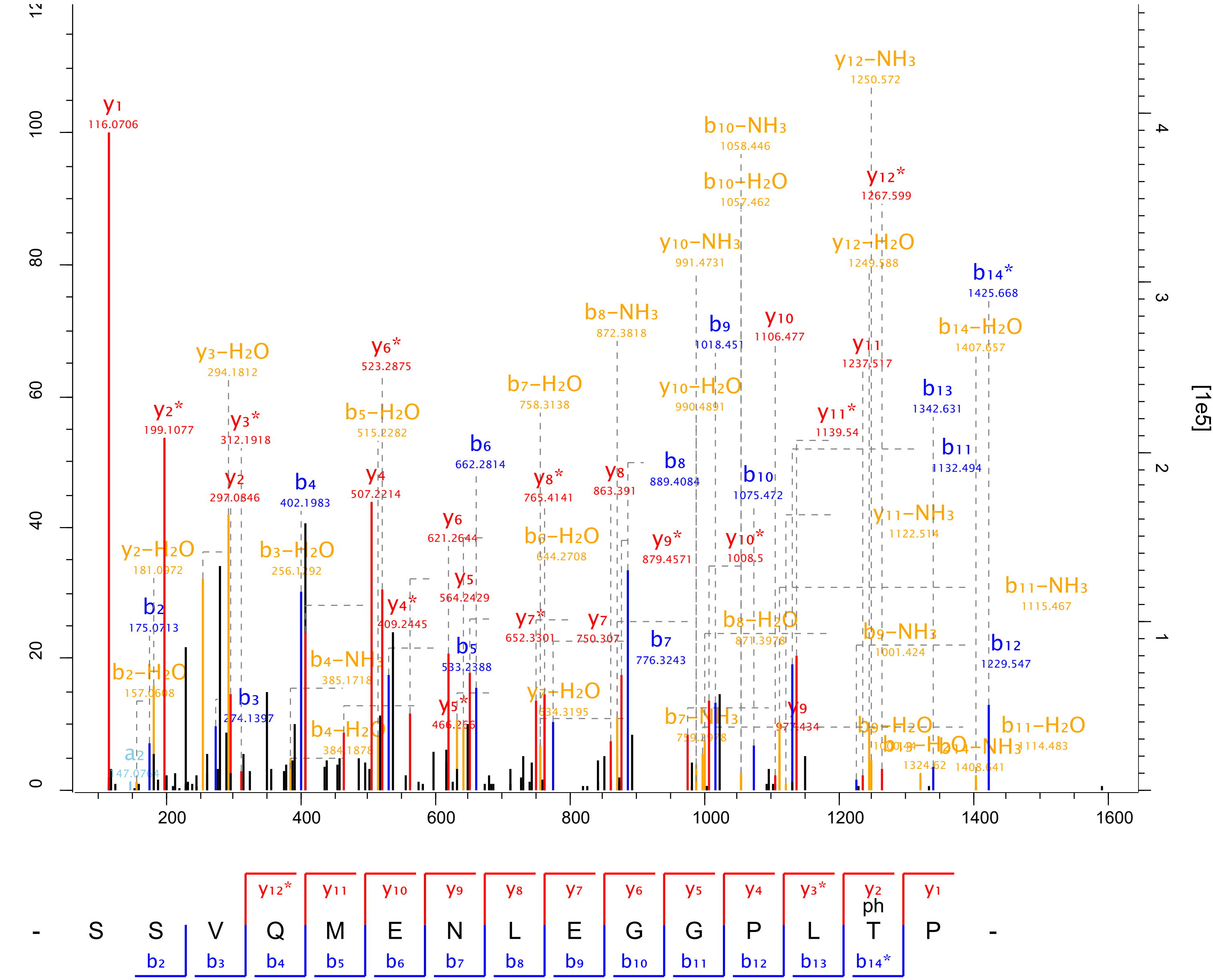

| Raw file | Scan | Method | Score | m/z |
|---|---|---|---|---|
| water_15_2 | 3610 | FTMS; HCD | 65.23 | 526.2 |

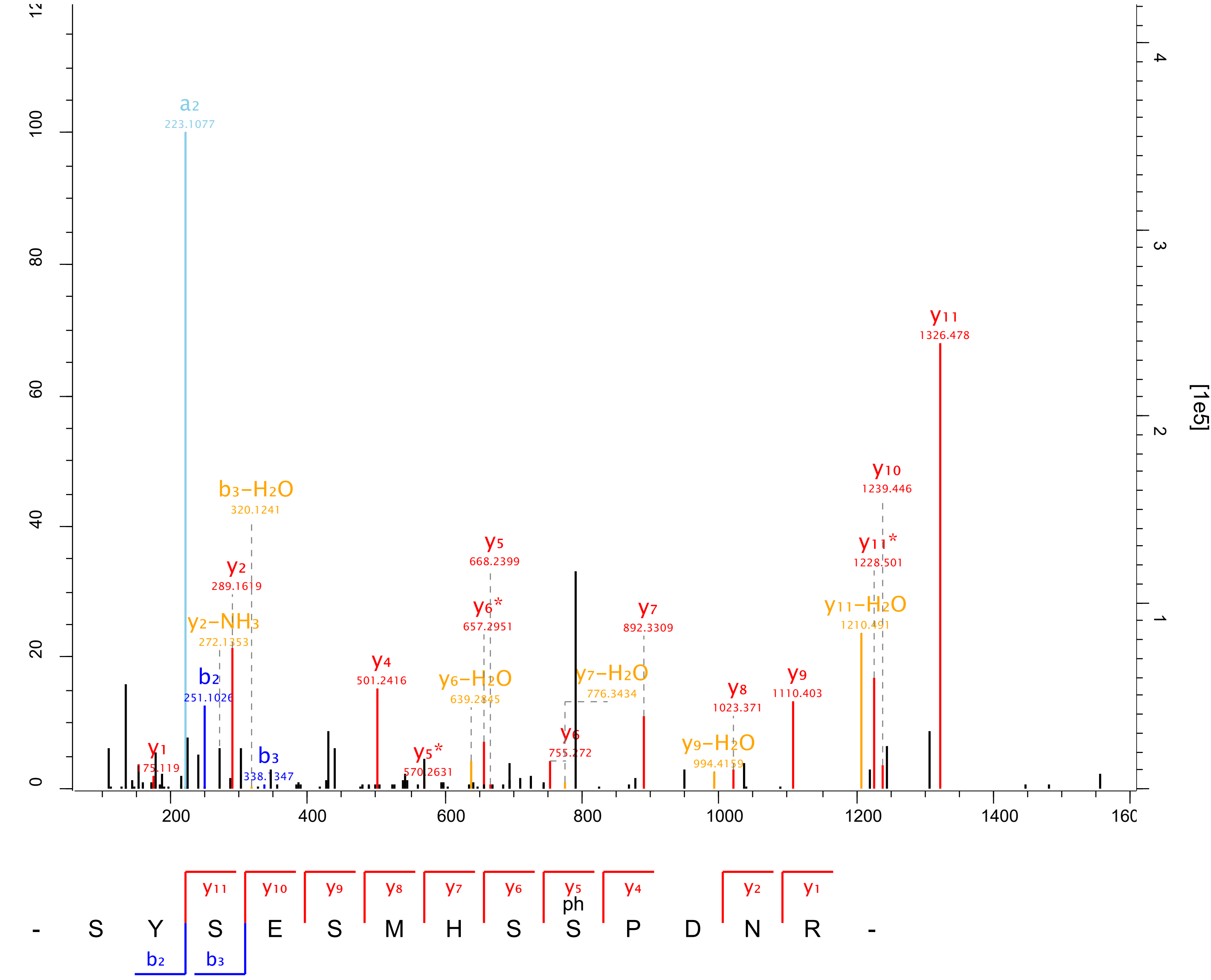

| Raw file | Scan | Method | Score | m/z |
|---|---|---|---|---|
| water_15_2 | 3660 | FTMS; HCD | 81.88 | 980.36 |

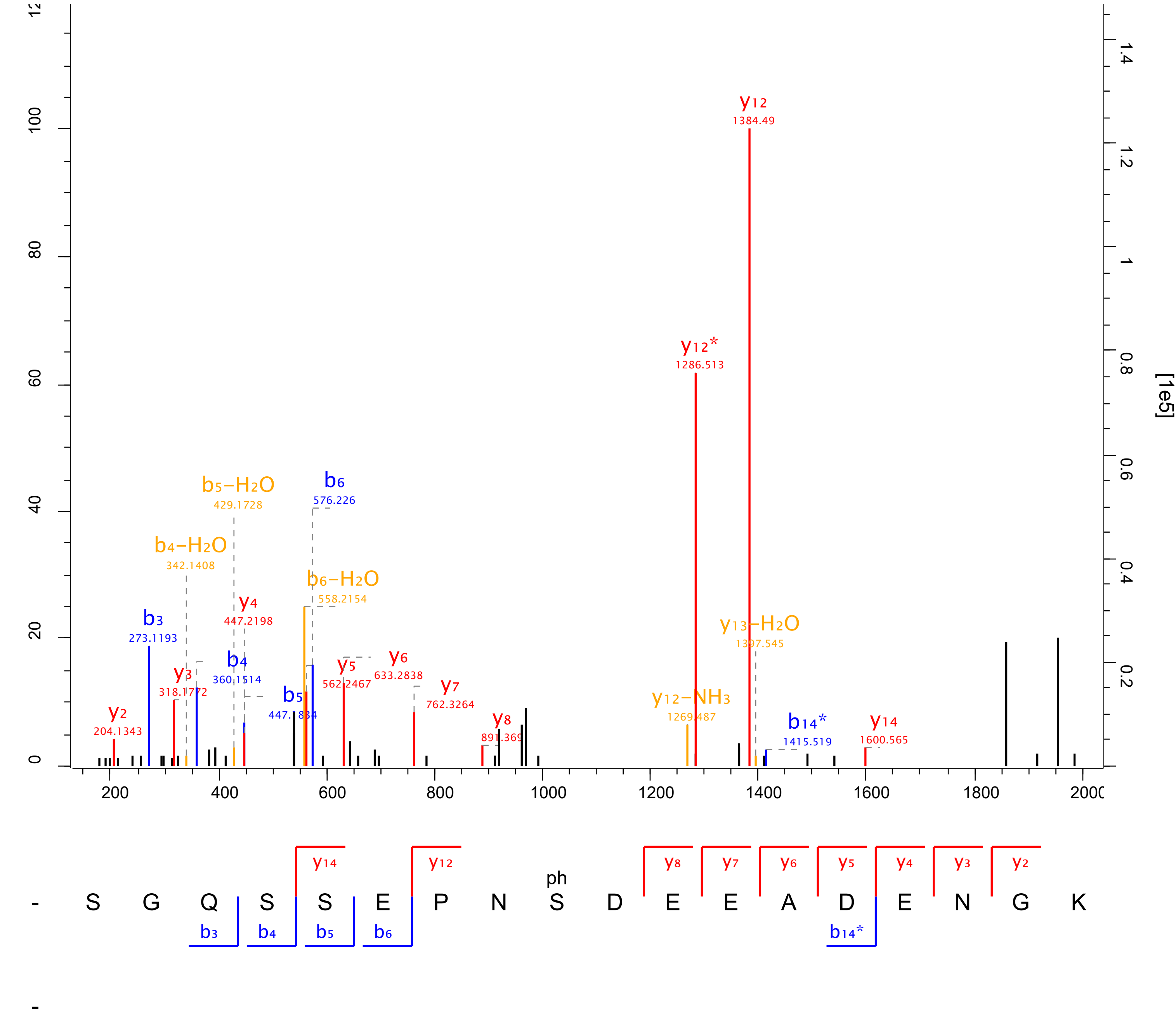

| Raw file | Scan | Method | Score | m/z |
| --- | --- | --- | --- | --- |
| water_15_2 | 3693 | FTMS; HCD | 60.55 | 465.89 |

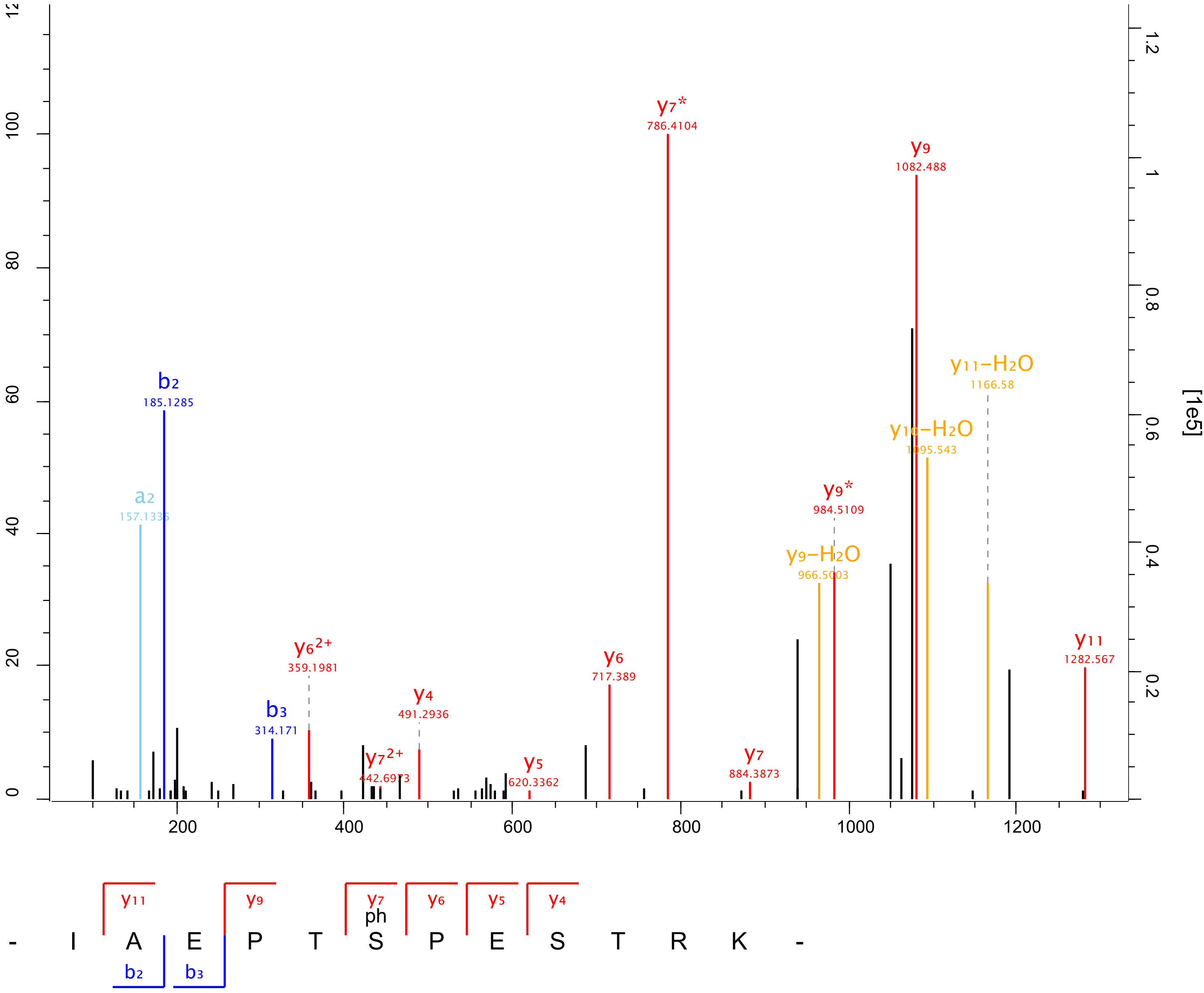

| Raw file | Scan | Method | Score | m/z |
|---|---|---|---|---|
| water_15_2 | 37369 | FTMS; HCD | 71.88 | 620.3 |

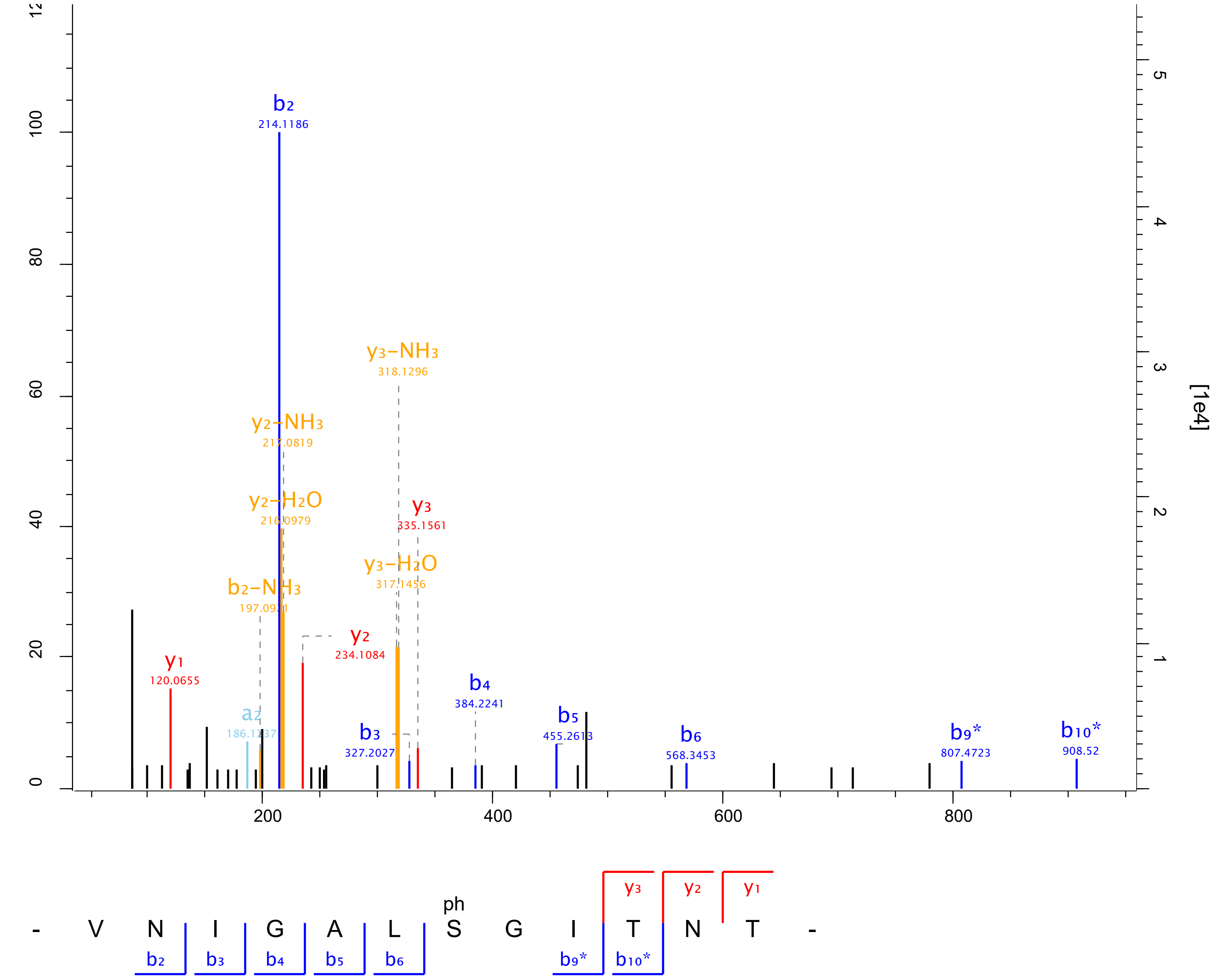

| Raw file | Scan | Method | Score | m/z |
|---|---|---|---|---|
| water_15_2 | 3742 | FTMS; HCD | 53.56 | 481.56 |

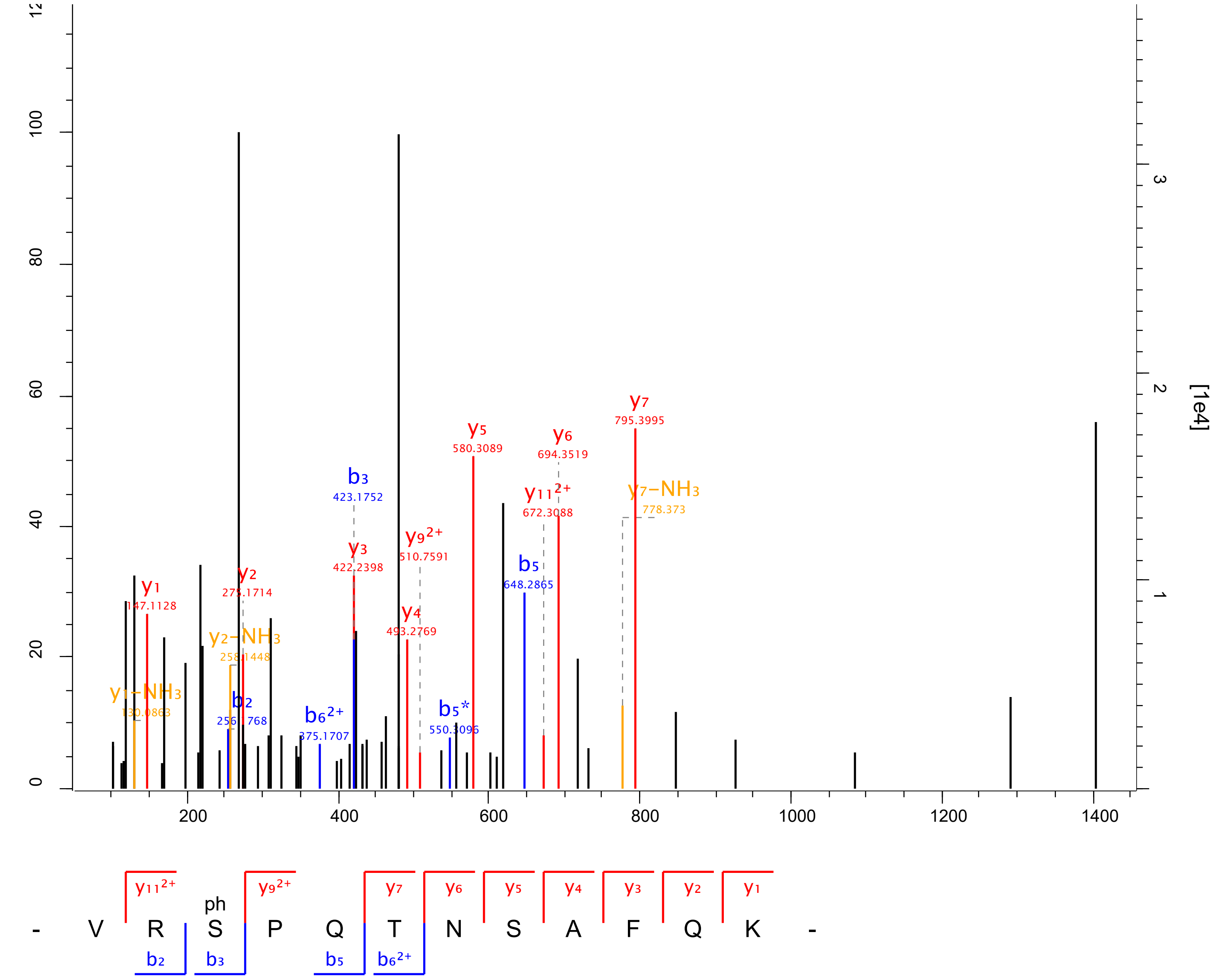

| Raw file | Scan | Method | Score | m/z |
|---|---|---|---|---|
| water_15_2 | 37886 | FTMS; HCD | 139.98 | 508.71 |

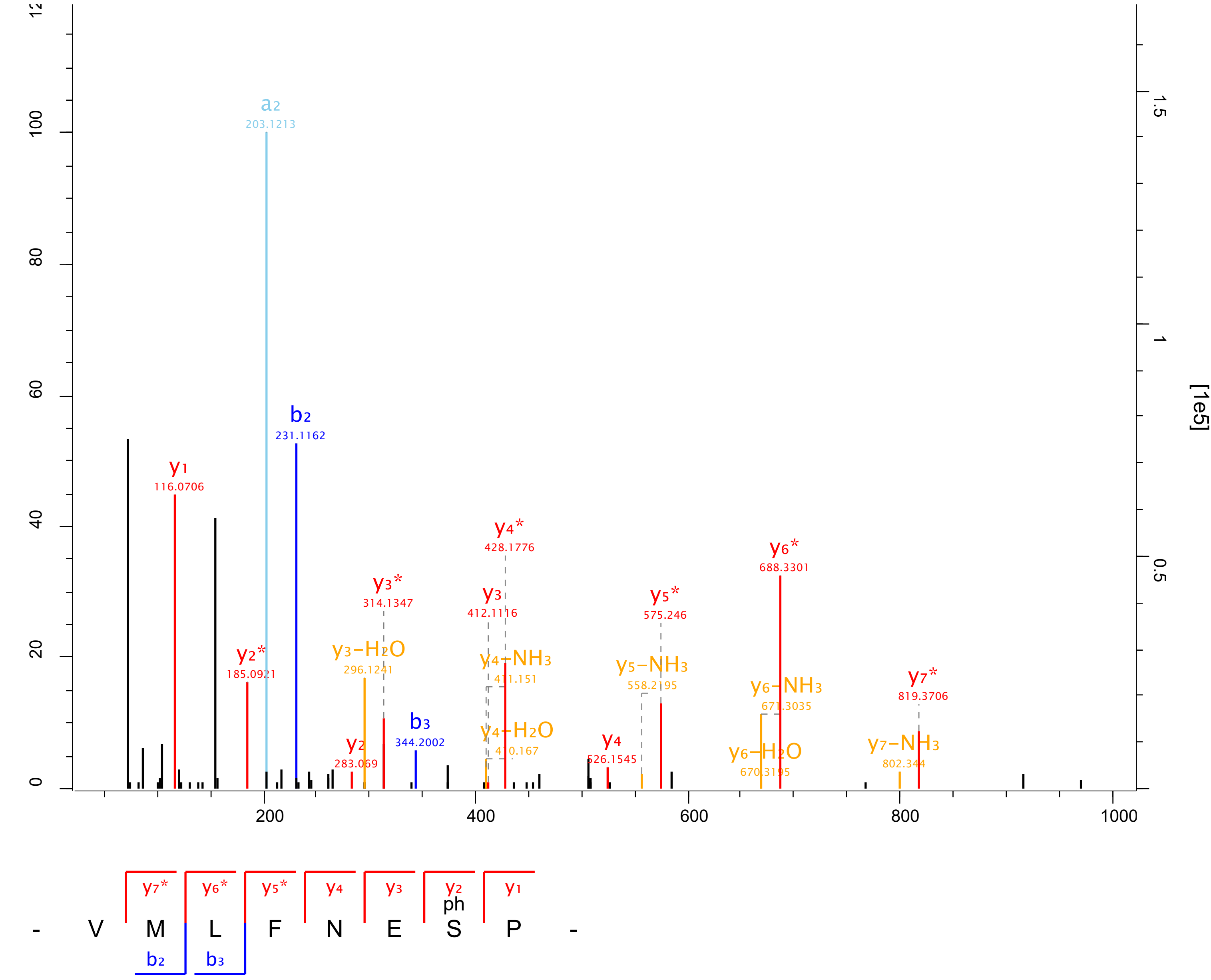

| Raw file | Scan | Method | Score | m/z |
| --- | --- | --- | --- | --- |
| water_15_2 | 3844 | FTMS; HCD | 47.55 | 464.88 |

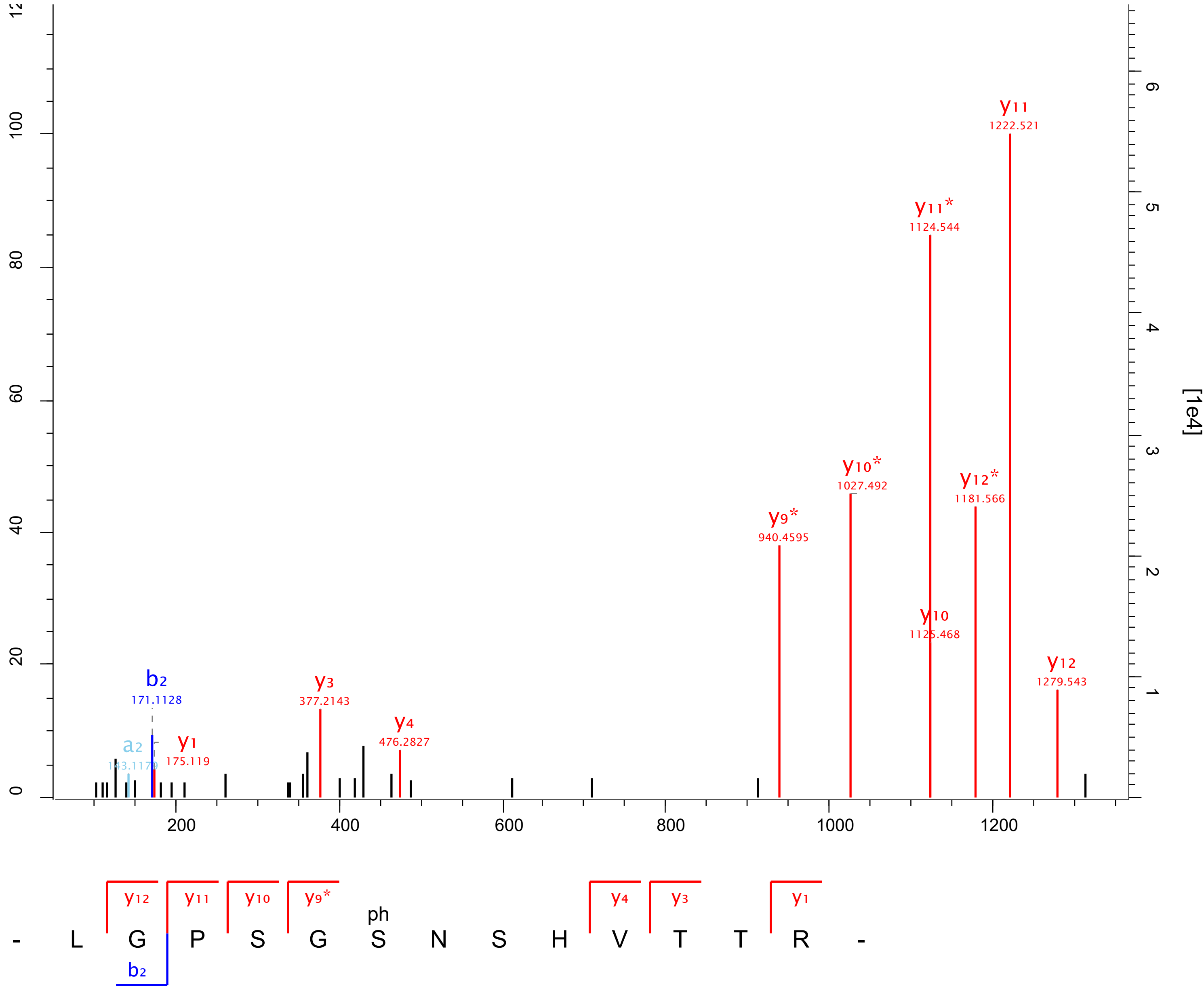

| Raw file | Scan | Method | Score | m/z |
|---|---|---|---|---|
| water_15_2 | 3953 | FTMS; HCD | 73.44 | 381.49 |

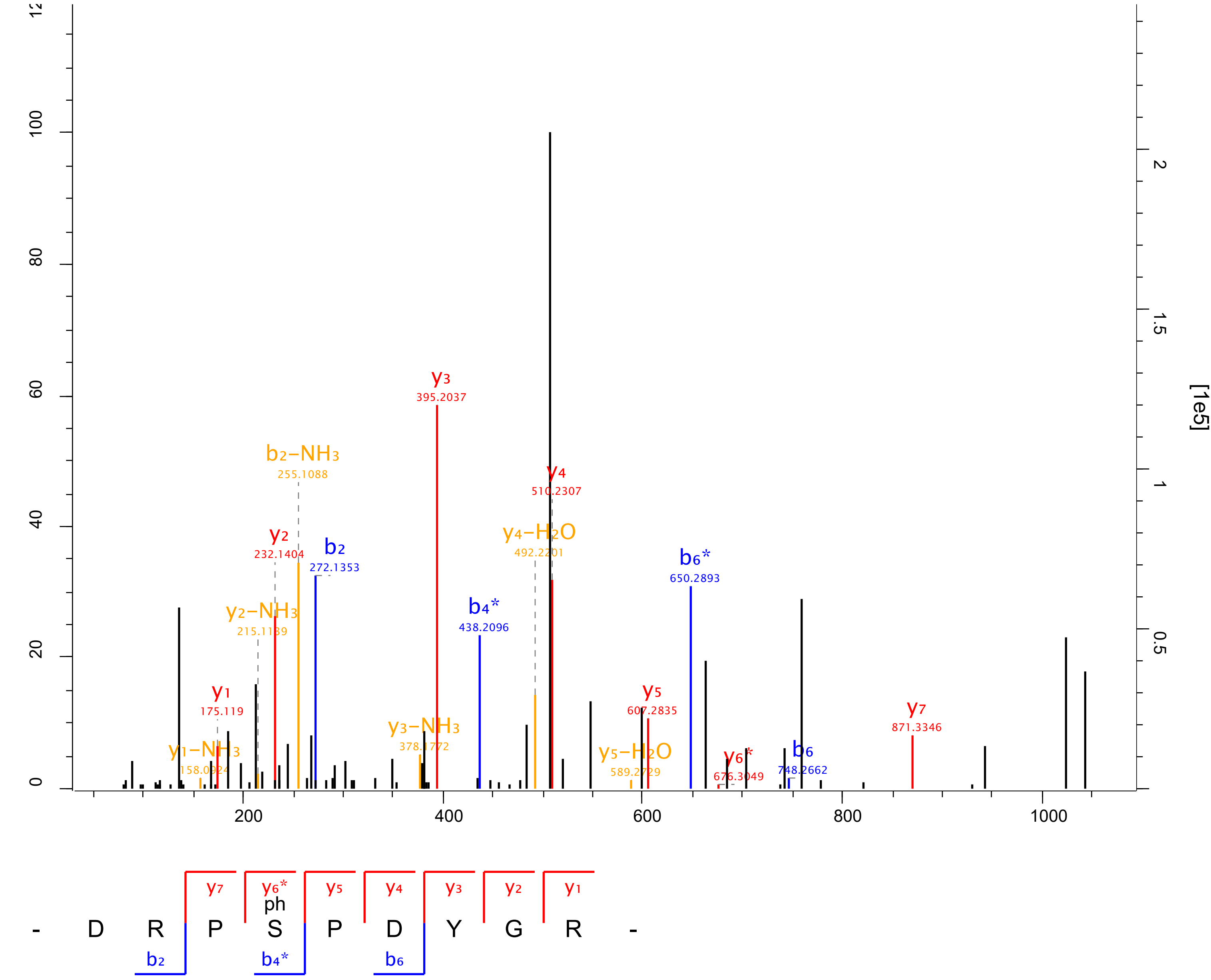

| Raw file | Scan | Method | Score | m/z |
| --- | --- | --- | --- | --- |
| water_15_2 | 4047 | FTMS; HCD | 92.38 | 672.3 |

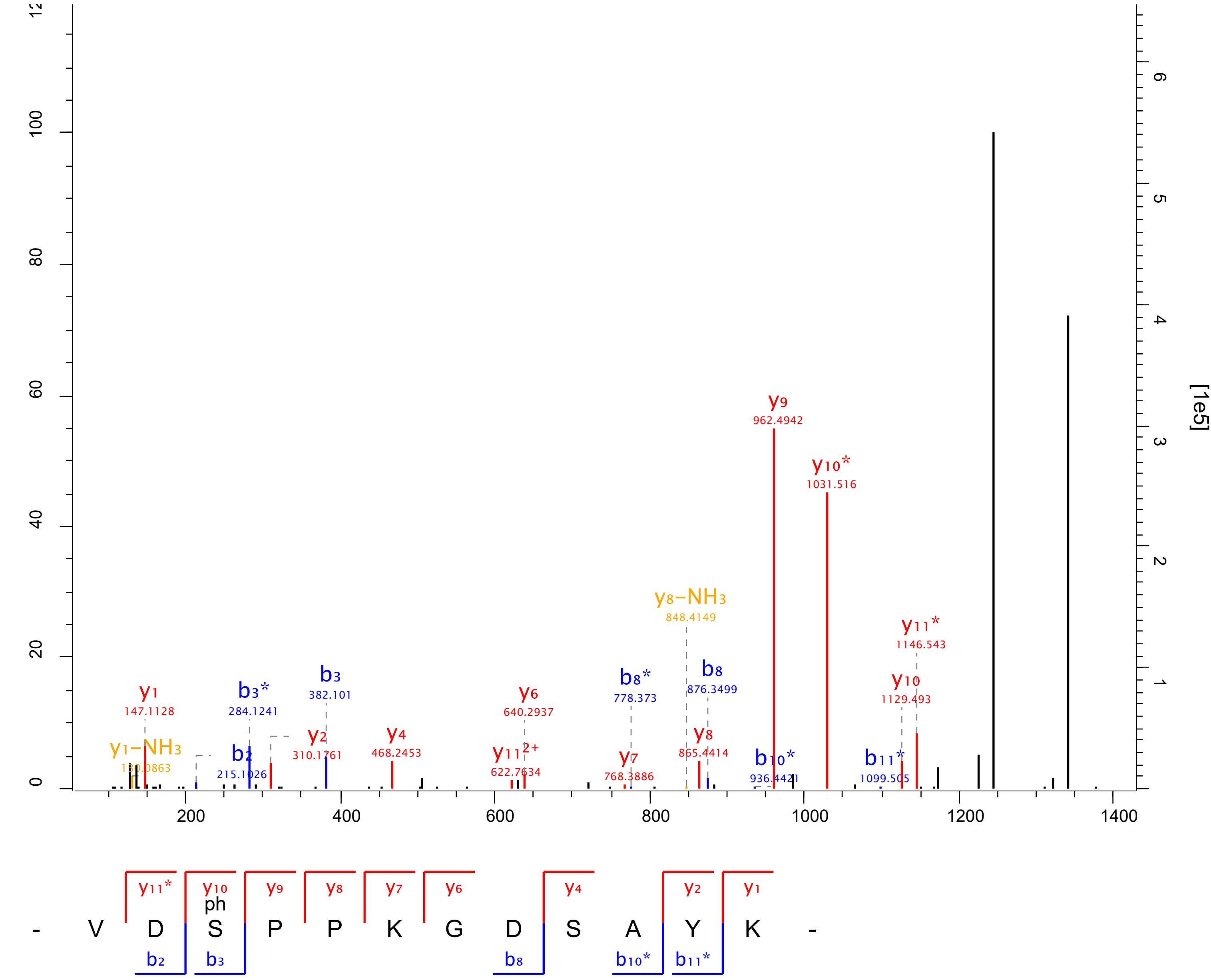

| Raw file | Scan | Method | Score | m/z |
| --- | --- | --- | --- | --- |
| water_15_2 | 4056 | FTMS; HCD | 113.93 | 693.64 |

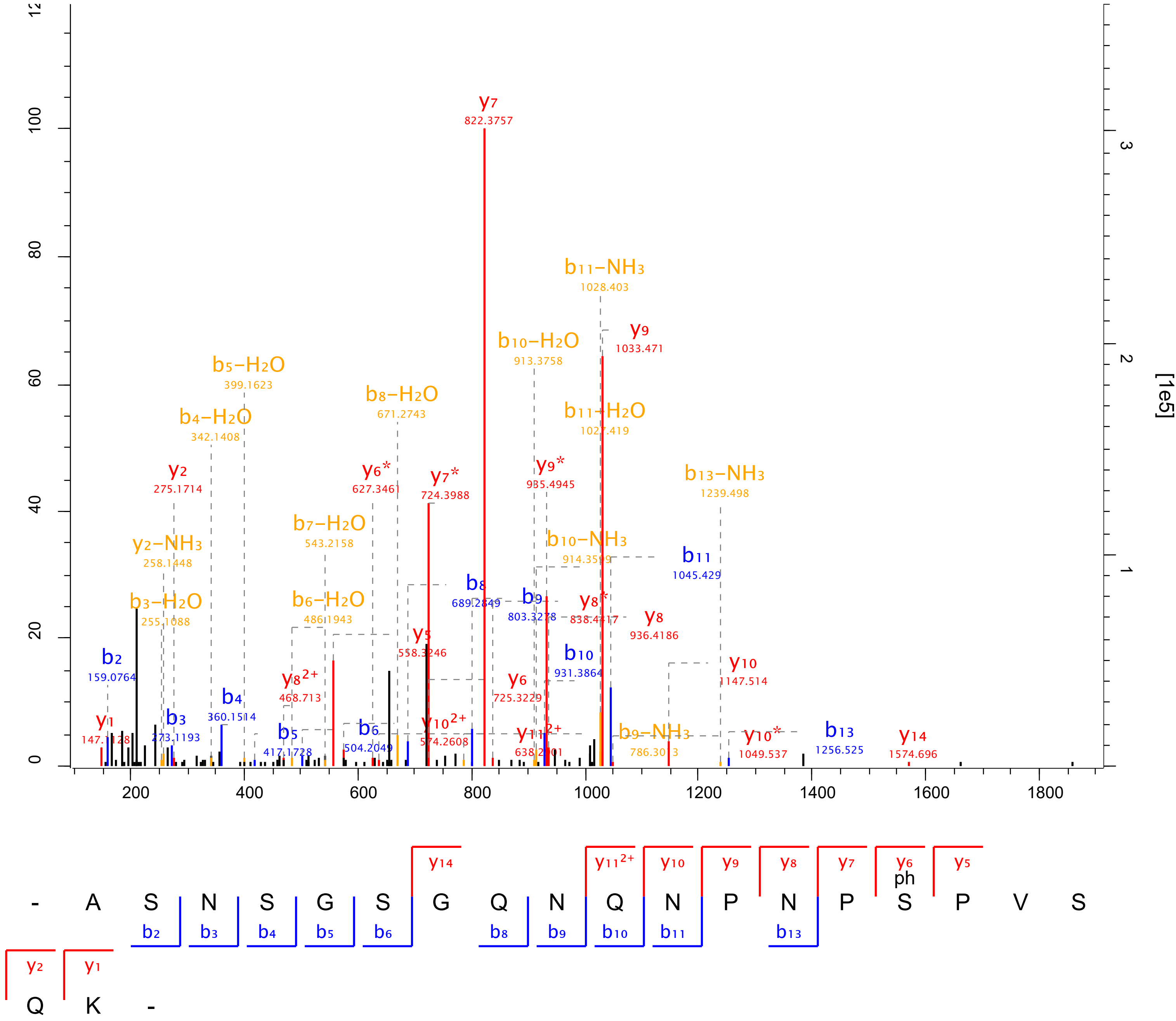

| Raw file | Scan | Method | Score | m/z |
|---|---|---|---|---|
| water_15_2 | 4296 | FTMS; HCD | 180.44 | 886.85 |

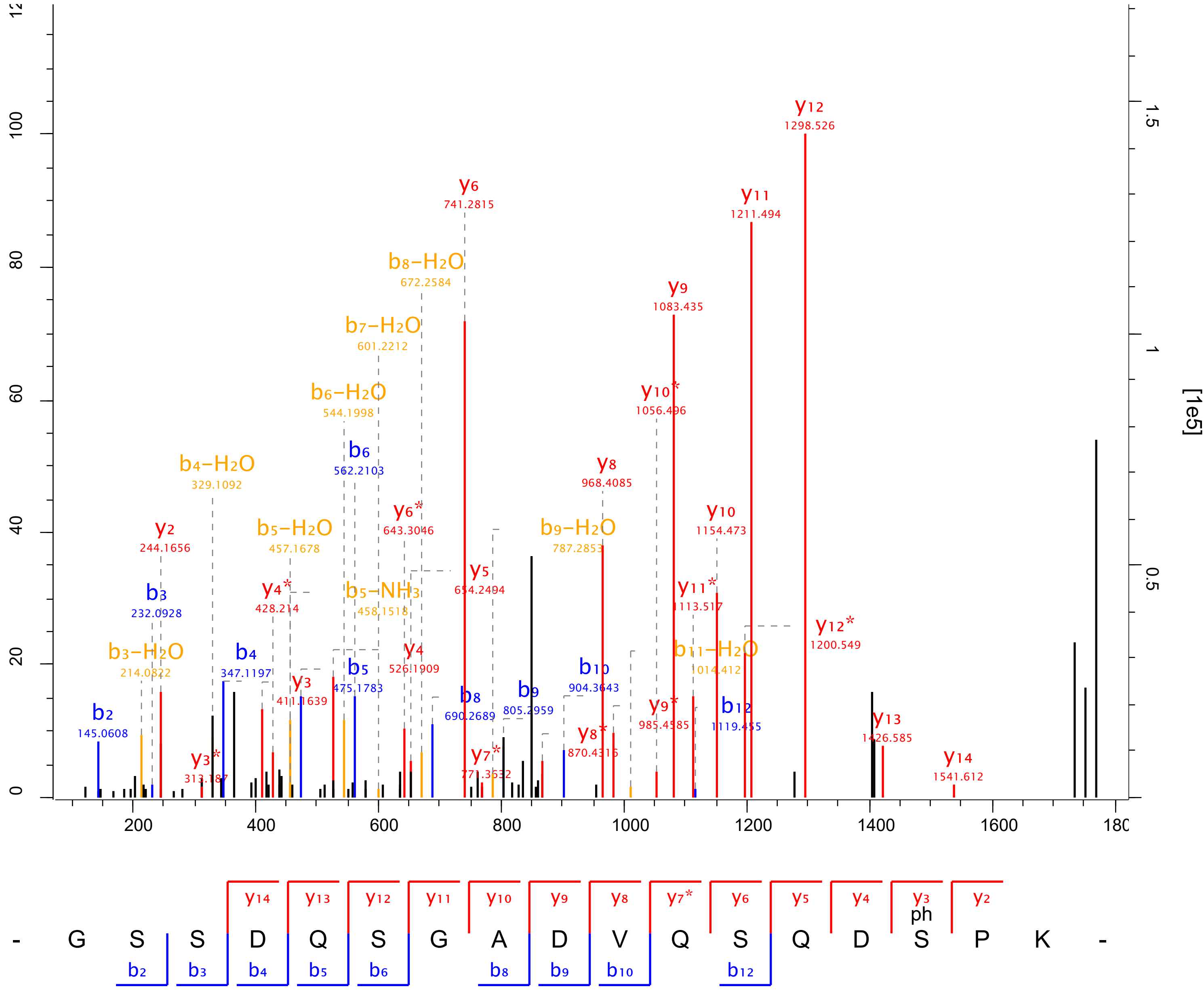

| Raw file | Scan | Method | Score | m/z |
|---|---|---|---|---|
| water_15_2 | 4379 | FTMS; HCD | 42.24 | 469.55 |

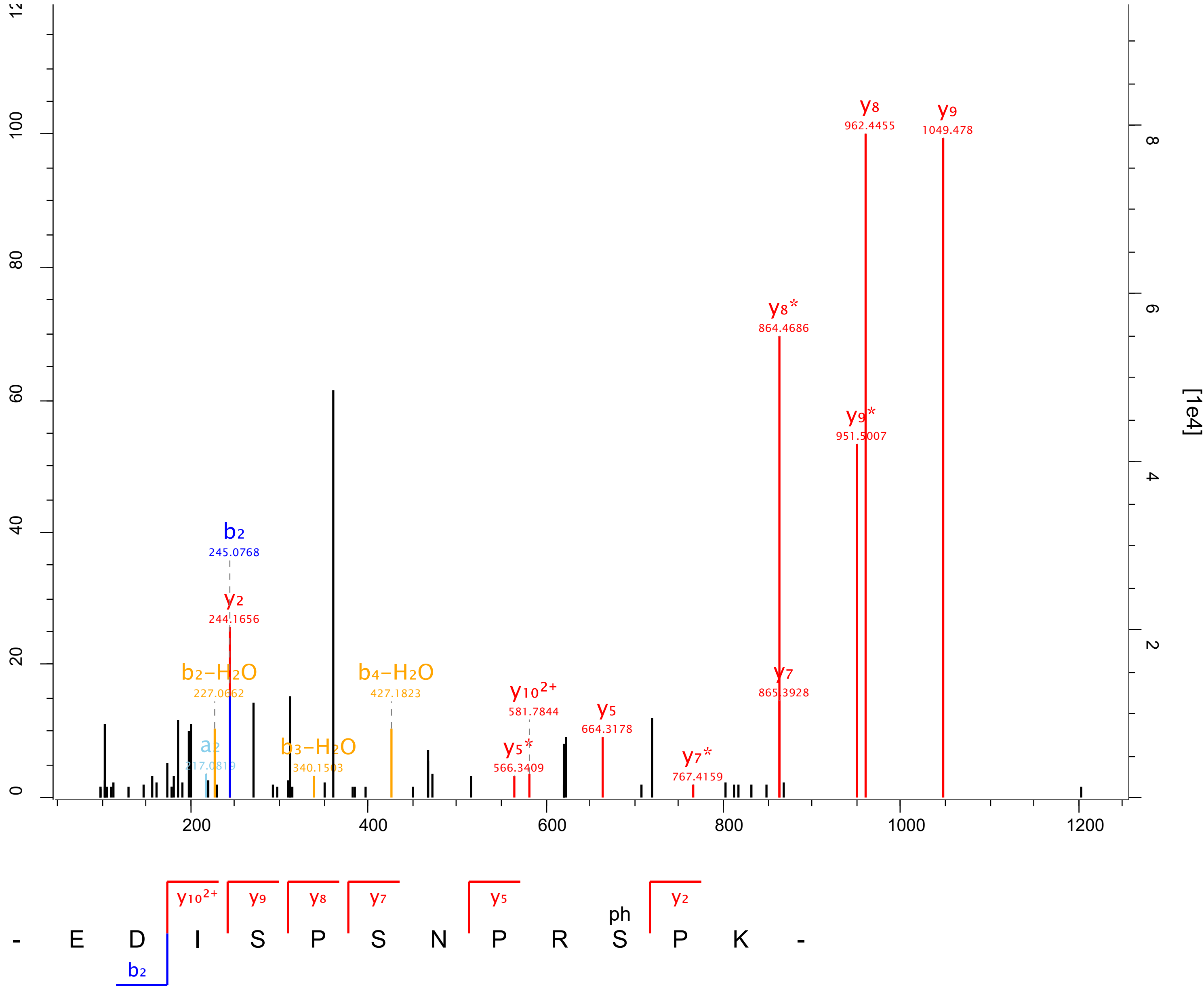

| Raw file | Scan | Method | Score | m/z |
|---|---|---|---|---|
| water_15_2 | 4393 | FTMS; HCD | 53.45 | 599.74 |

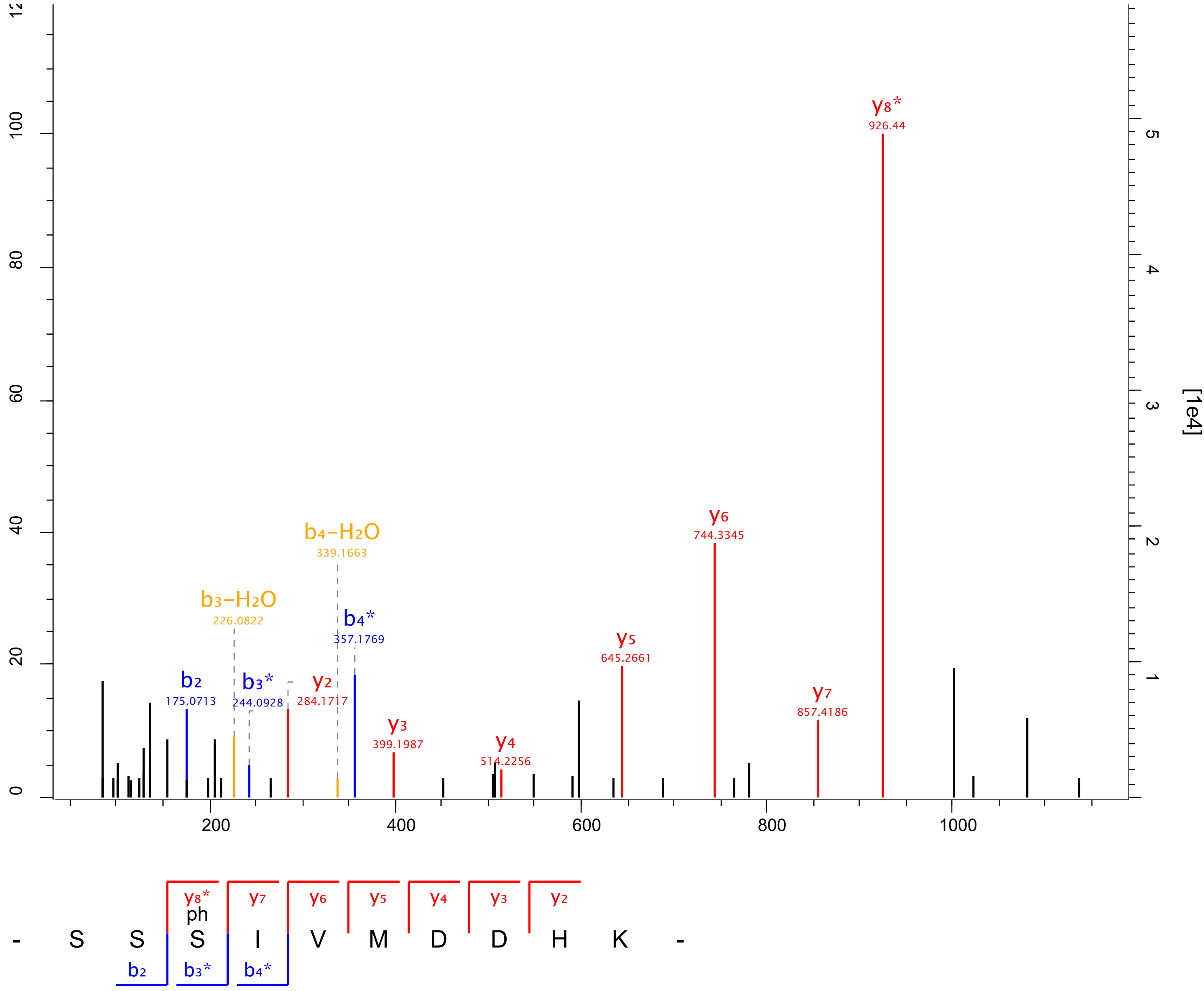

| Raw file | Scan | Method | Score | m/z |
|---|---|---|---|---|
| water_15_2 | 4409 | FTMS; HCD | 175.57 | 507.72 |

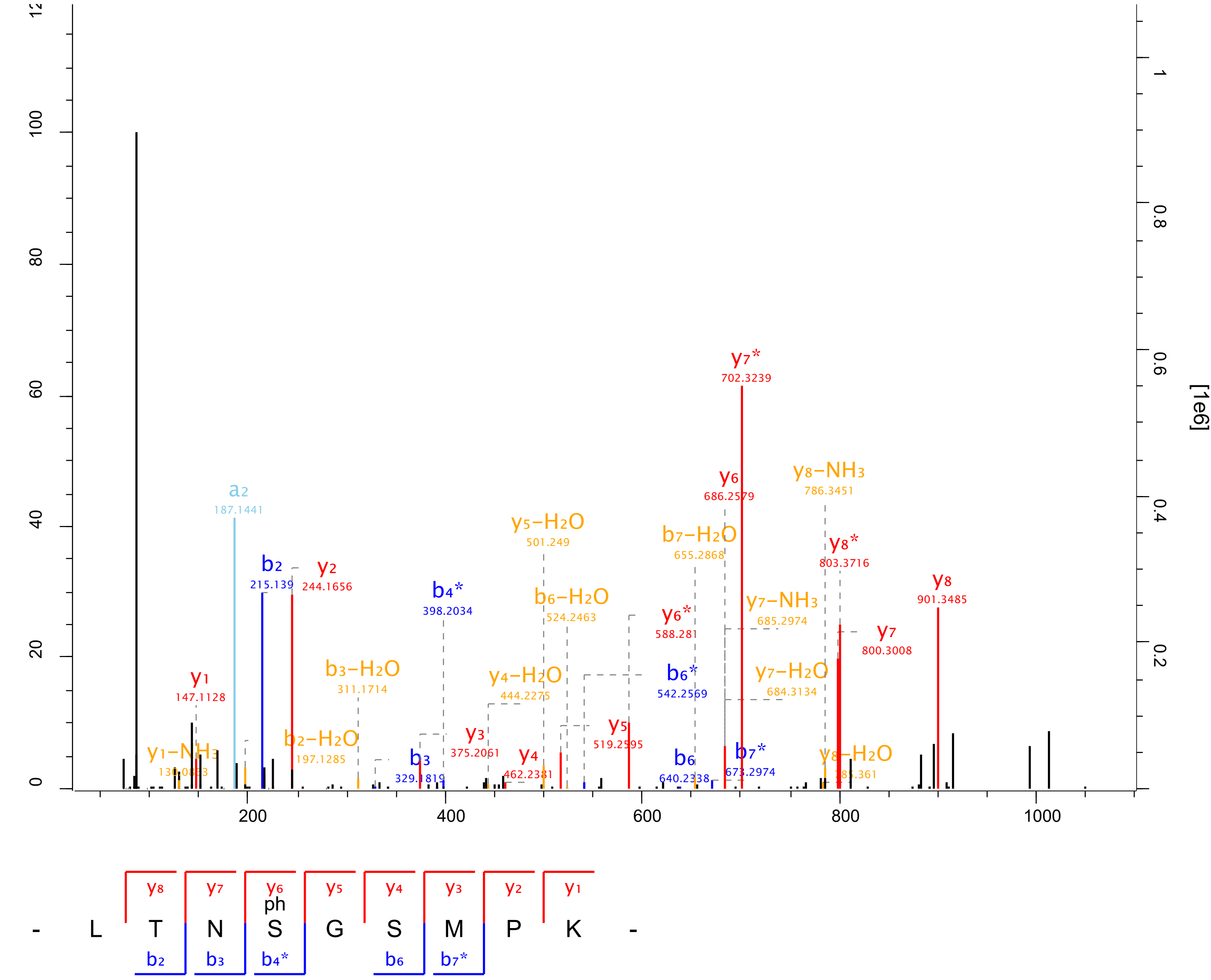

| Raw file | Scan | Method | Score | m/z |
| --- | --- | --- | --- | --- |
| water_15_2 | 4462 | FTMS; HCD | 44.43 | 604.77 |

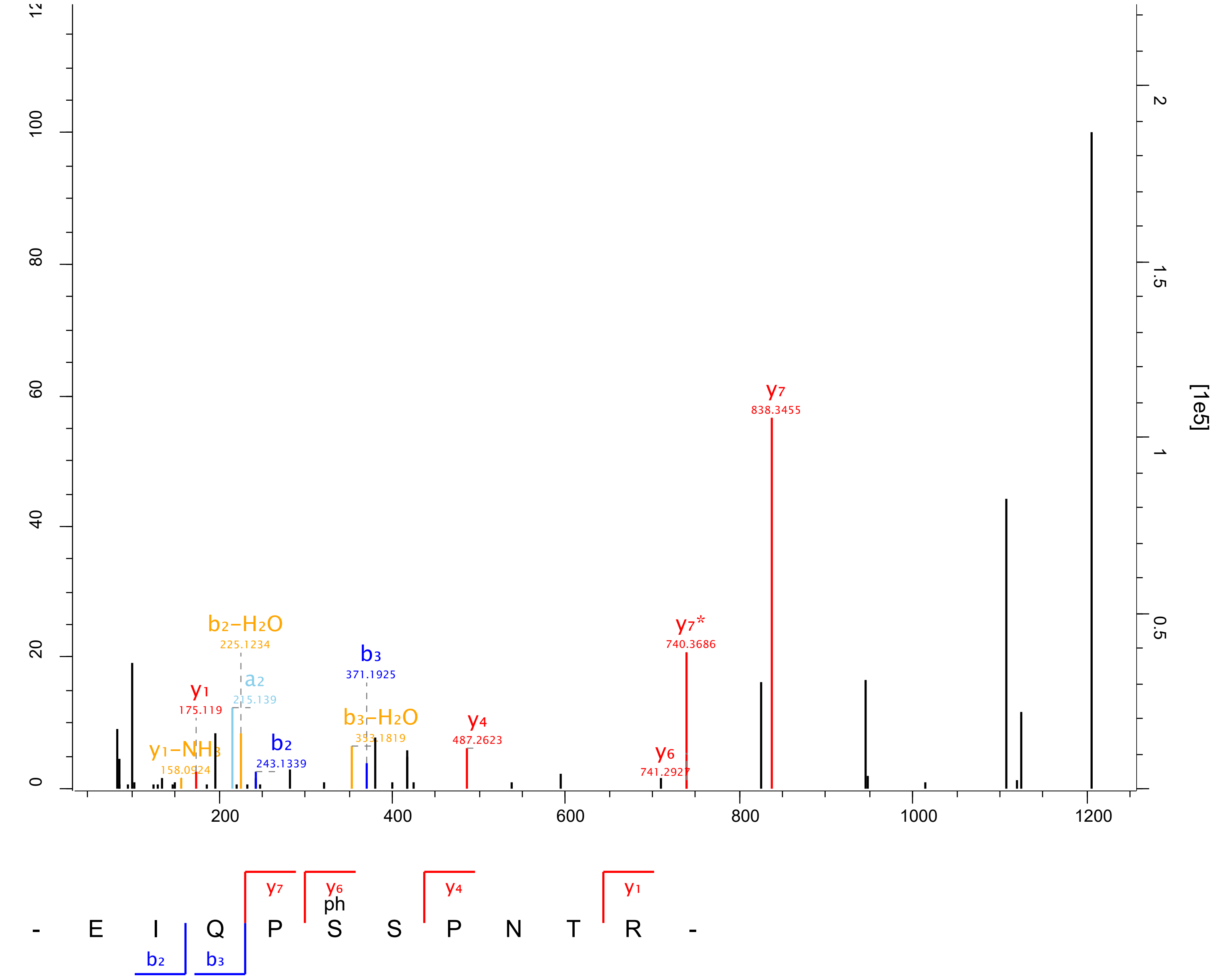

| Raw file | Scan | Method | Score | m/z |
|---|---|---|---|---|
| water_15_2 | 4473 | FTMS; HCD | 118.71 | 677.78 |

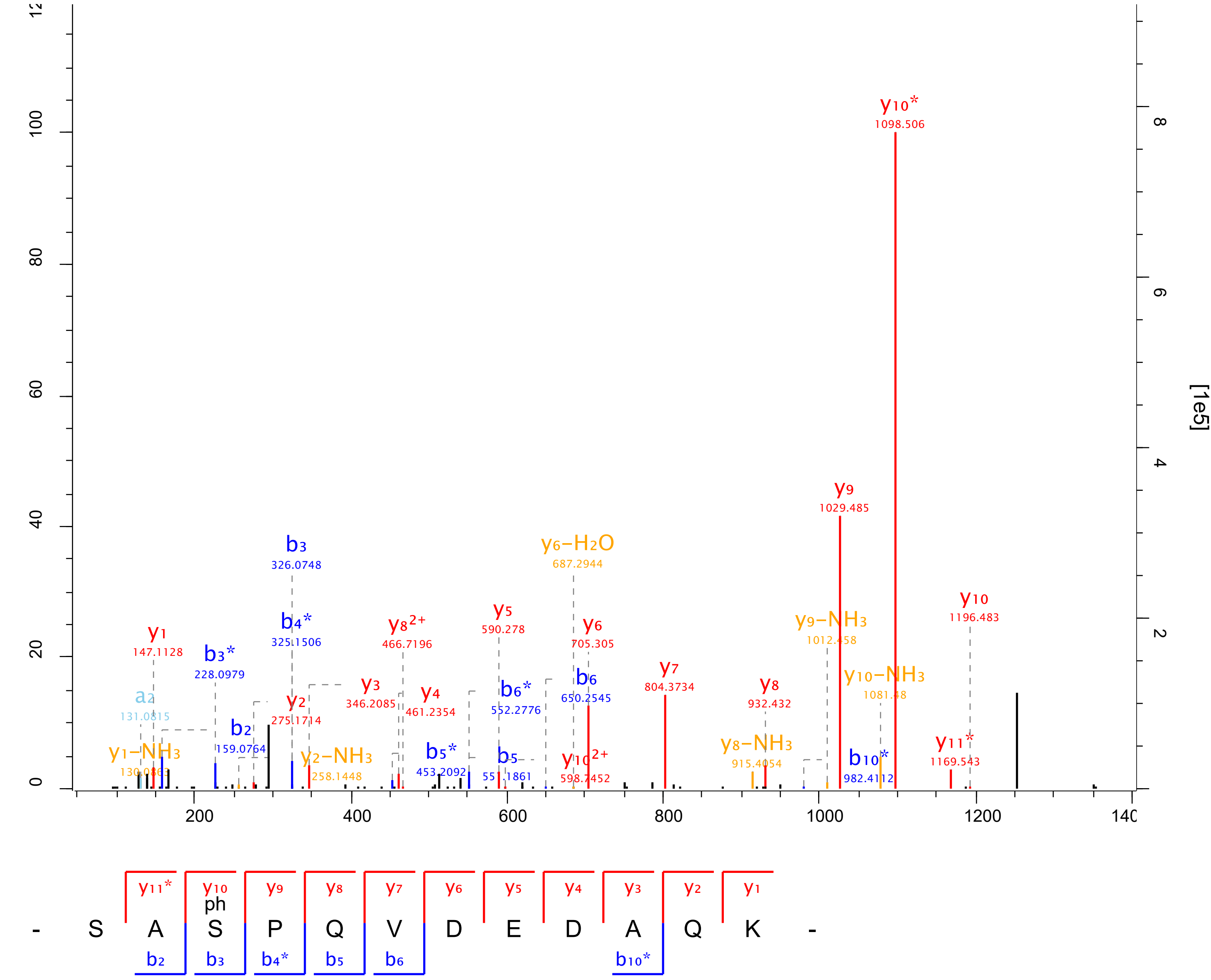

| Raw file | Scan | Method | Score | m/z |
|---|---|---|---|---|
| water_15_2 | 4506 | FTMS; HCD | 43.79 | 623.92 |

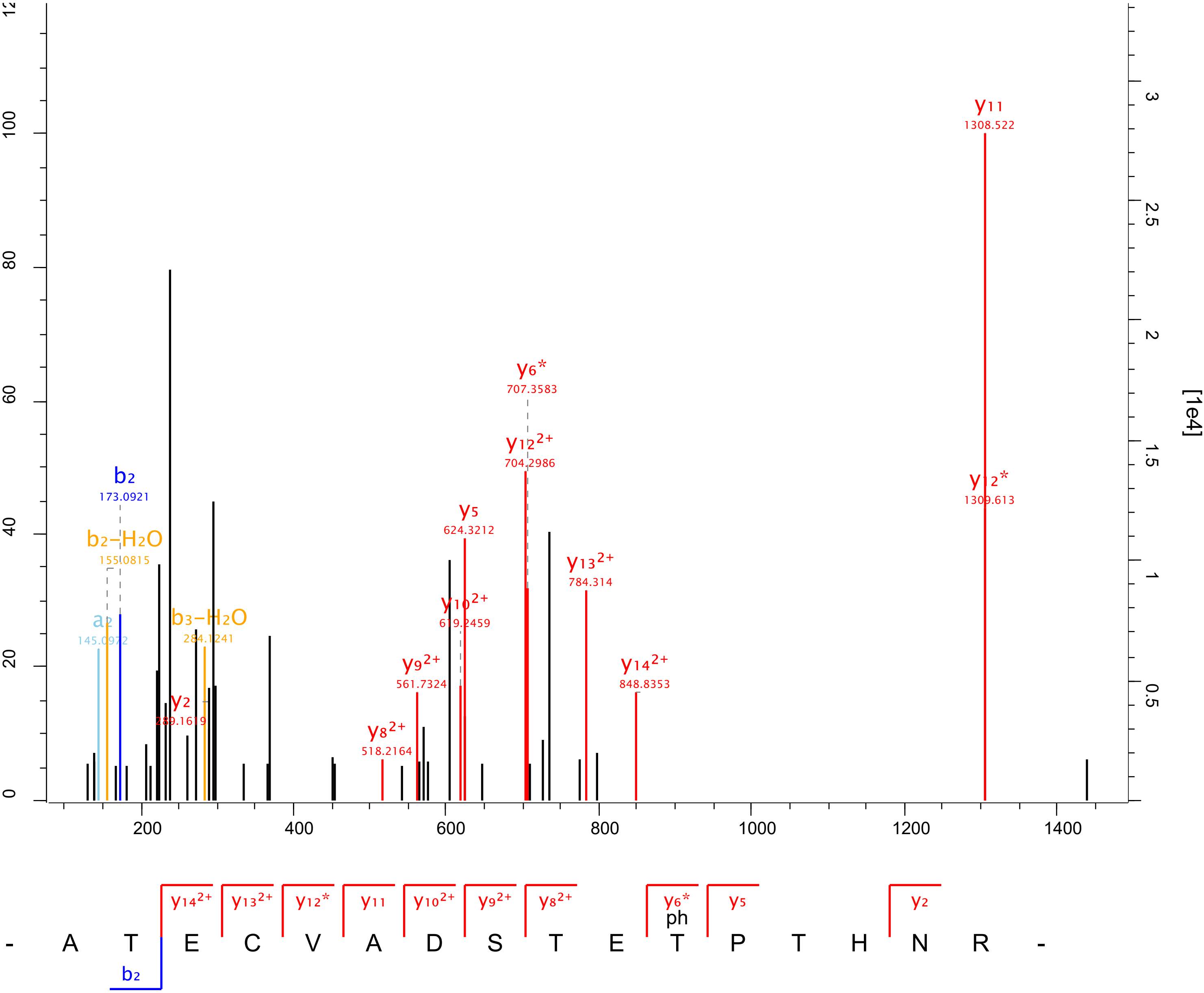

| Raw file | Scan | Method | Score | m/z |
|---|---|---|---|---|
| water_15_2 | 4531 | FTMS; HCD | 40.49 | 481.24 |

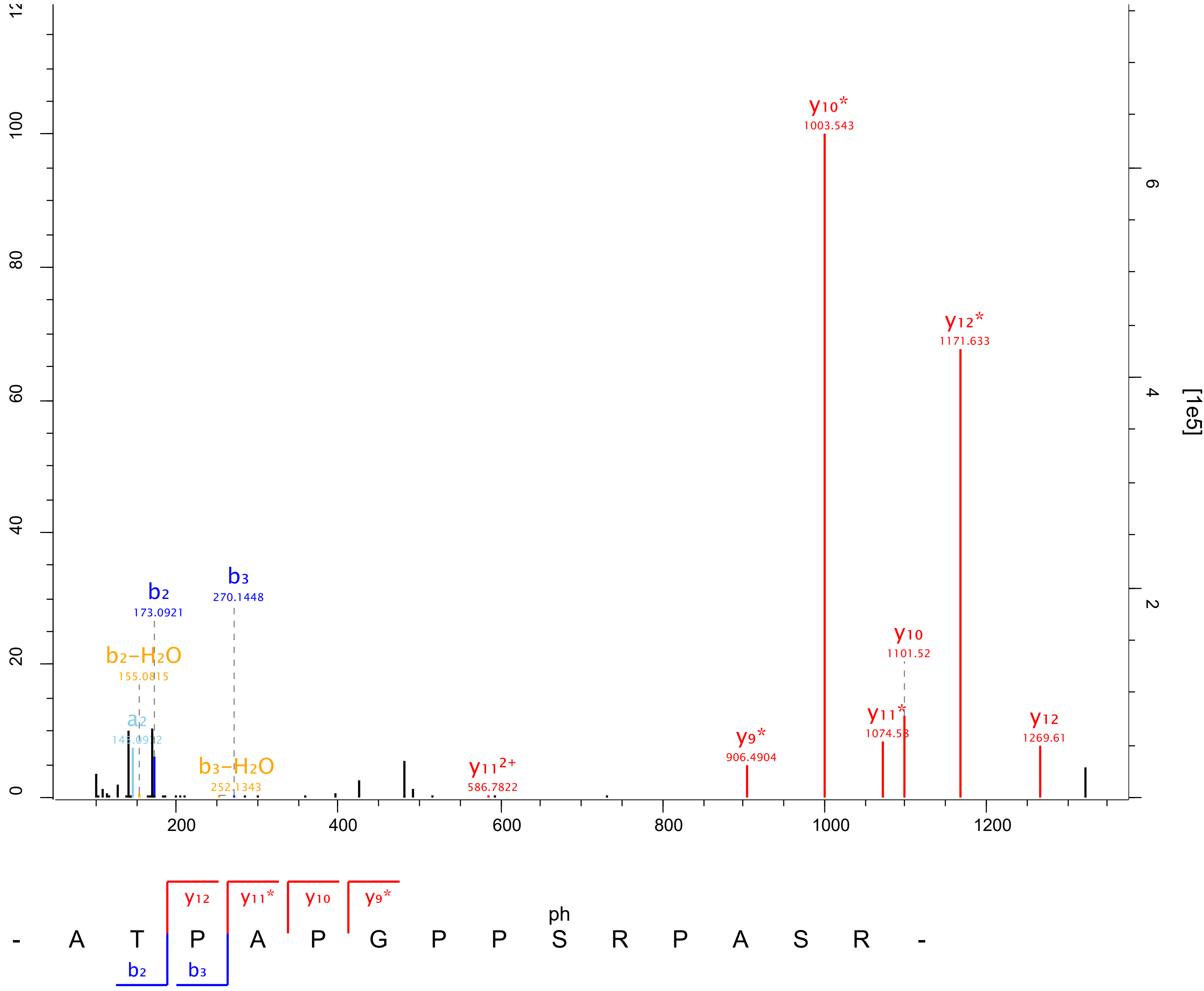

| Raw file | Scan | Method | Score | m/z |
|---|---|---|---|---|
| water_15_2 | 4572 | FTMS; HCD | 45.62 | 469.57 |

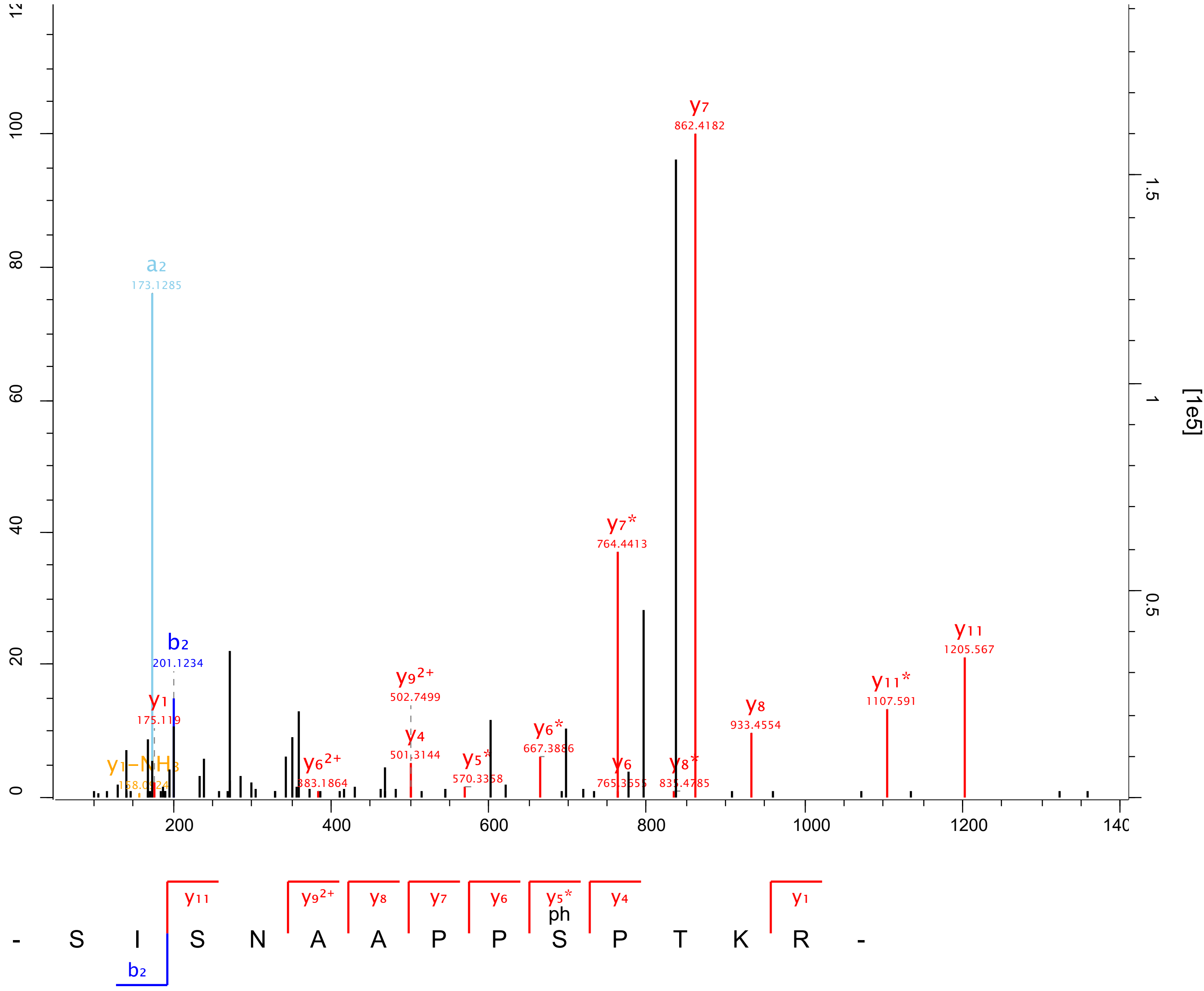

| Raw file | Scan | Method | Score | m/z |
|---|---|---|---|---|
| water_15_2 | 4617 | FTMS; HCD | 115.68 | 628.28 |

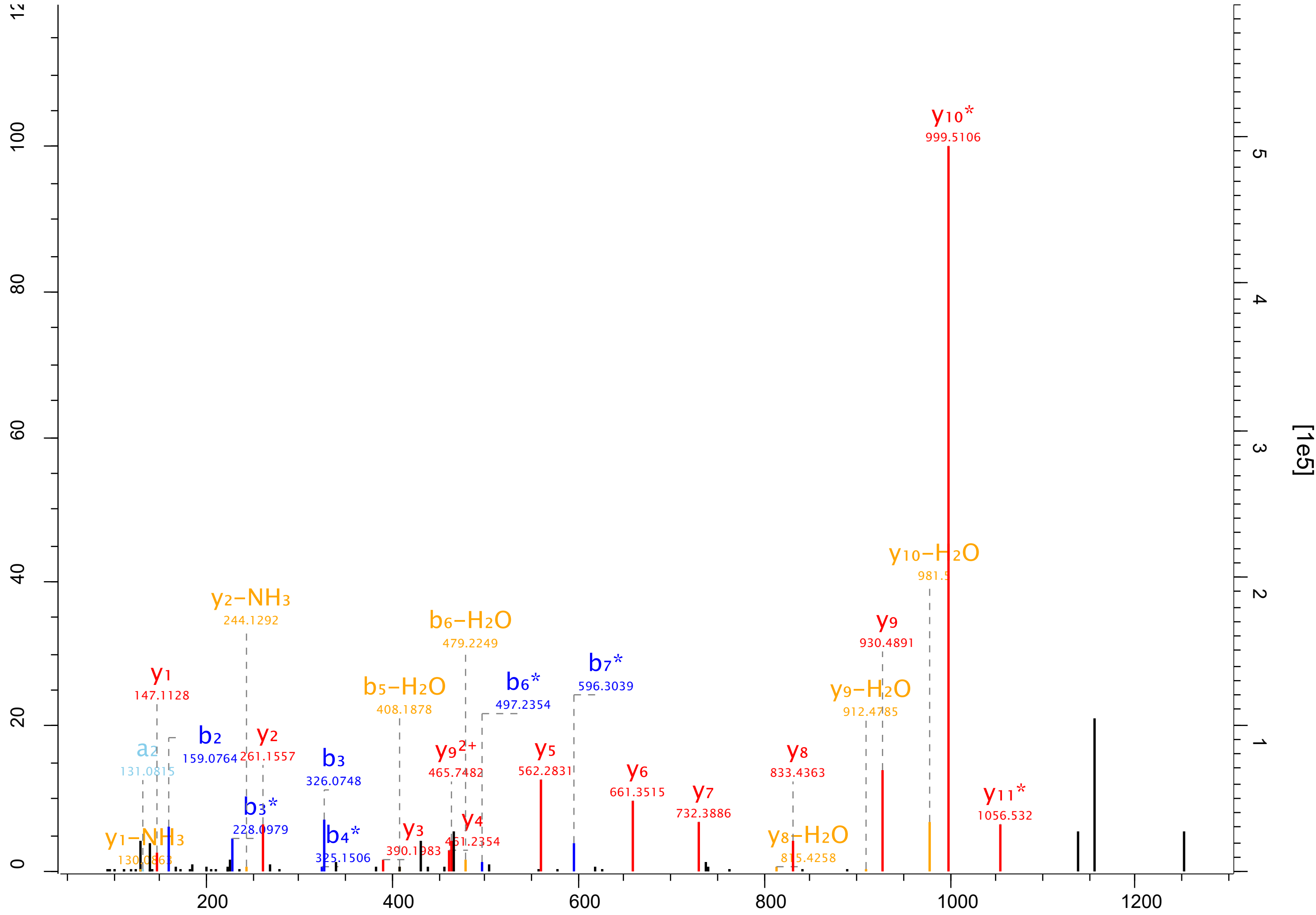


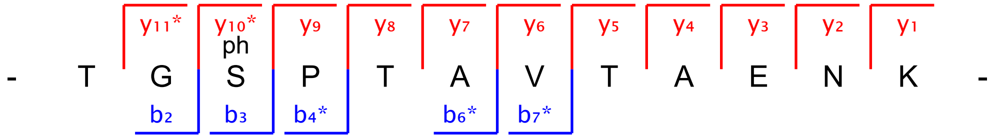

| Raw file | Scan | Method | Score | m/z |
|---|---|---|---|---|
| water_15_2 | 4680 | FTMS; HCD | 57.43 | 617.79 |

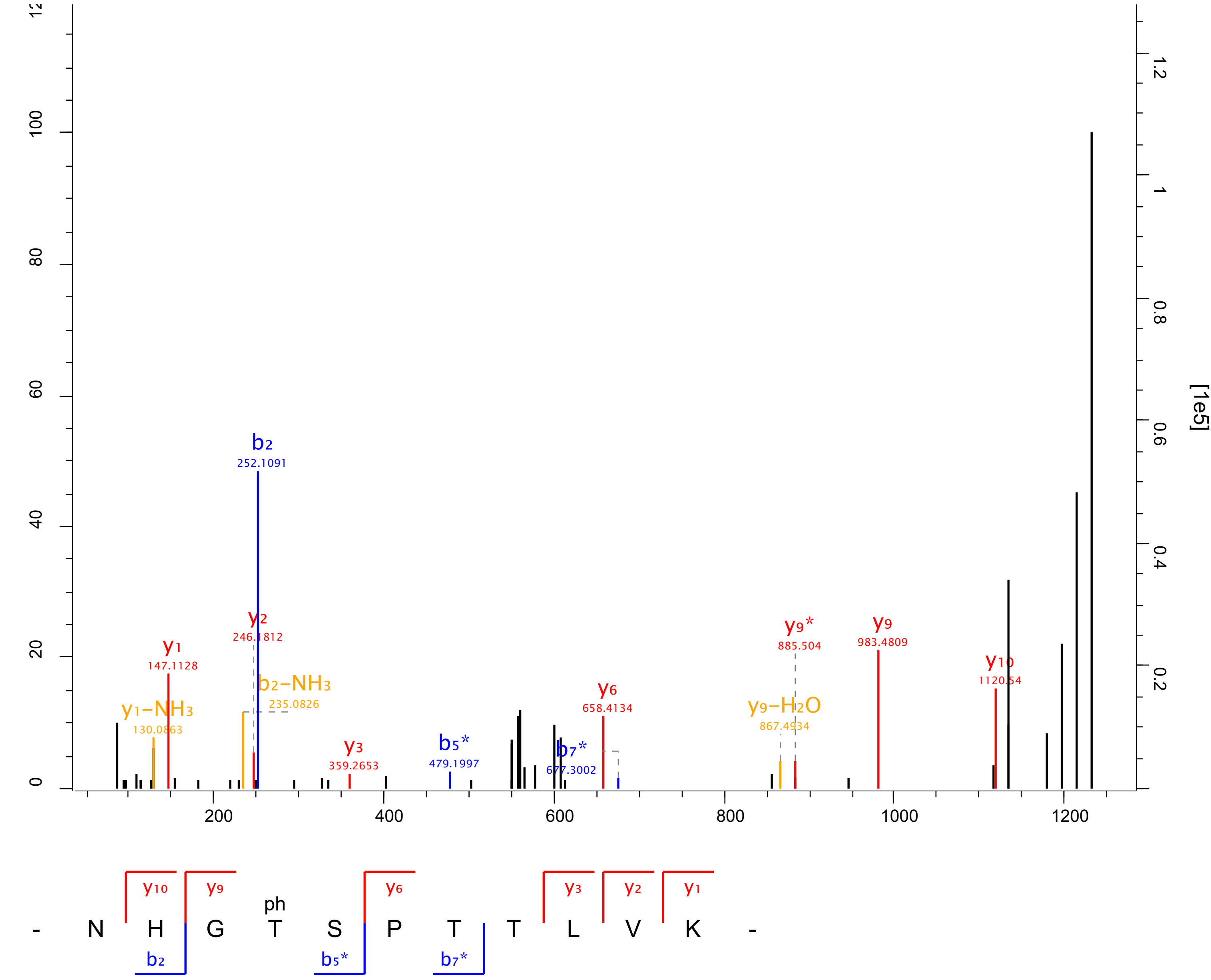

| Raw file | Scan | Method | Score | m/z |
|---|---|---|---|---|
| water_15_2 | 4753 | FTMS; HCD | 50.35 | 484.23 |

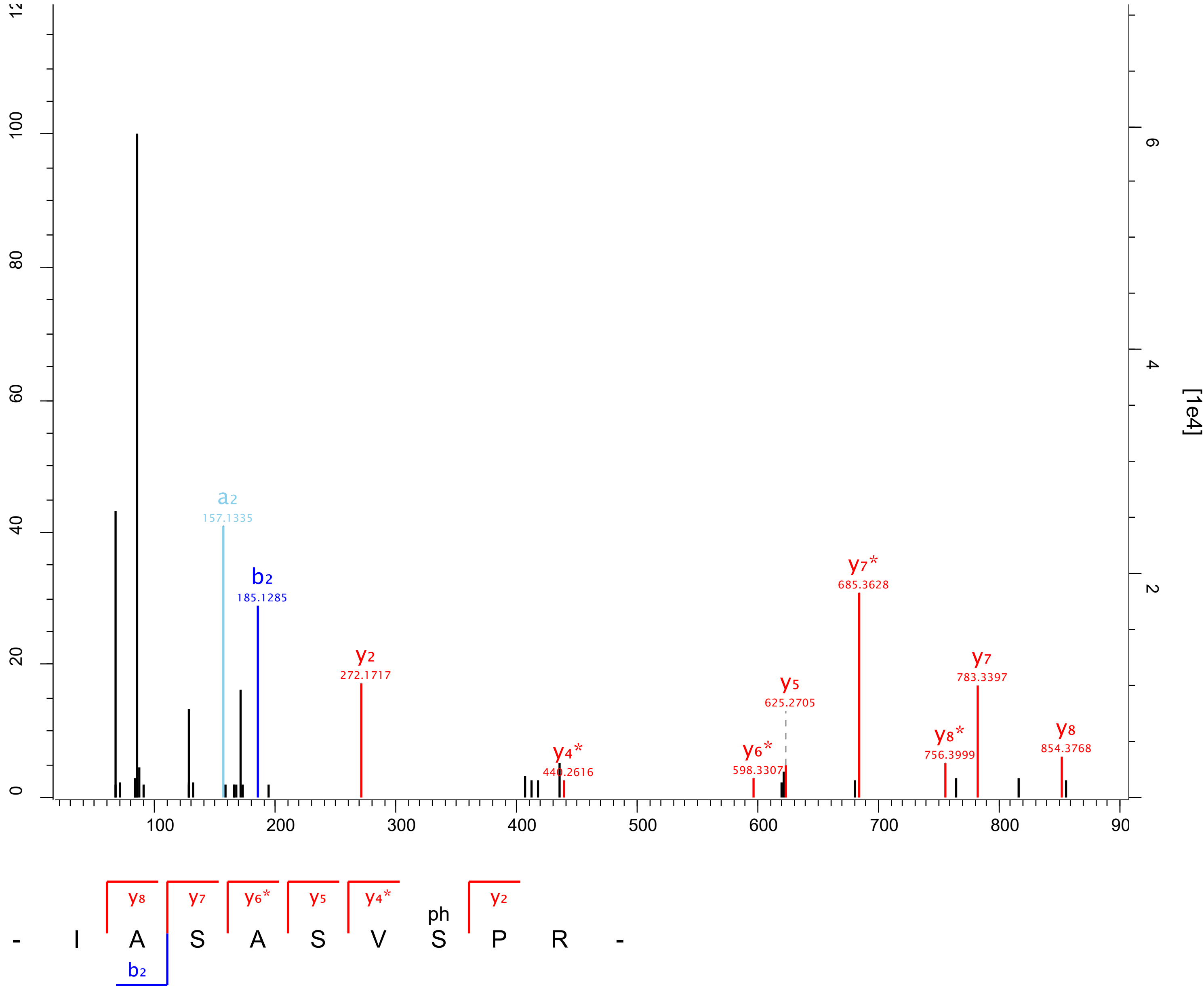

| Raw file | Scan | Method | Score | m/z |
|---|---|---|---|---|
| water_15_2 | 4823 | FTMS; HCD | 87.45 | 627.28 |

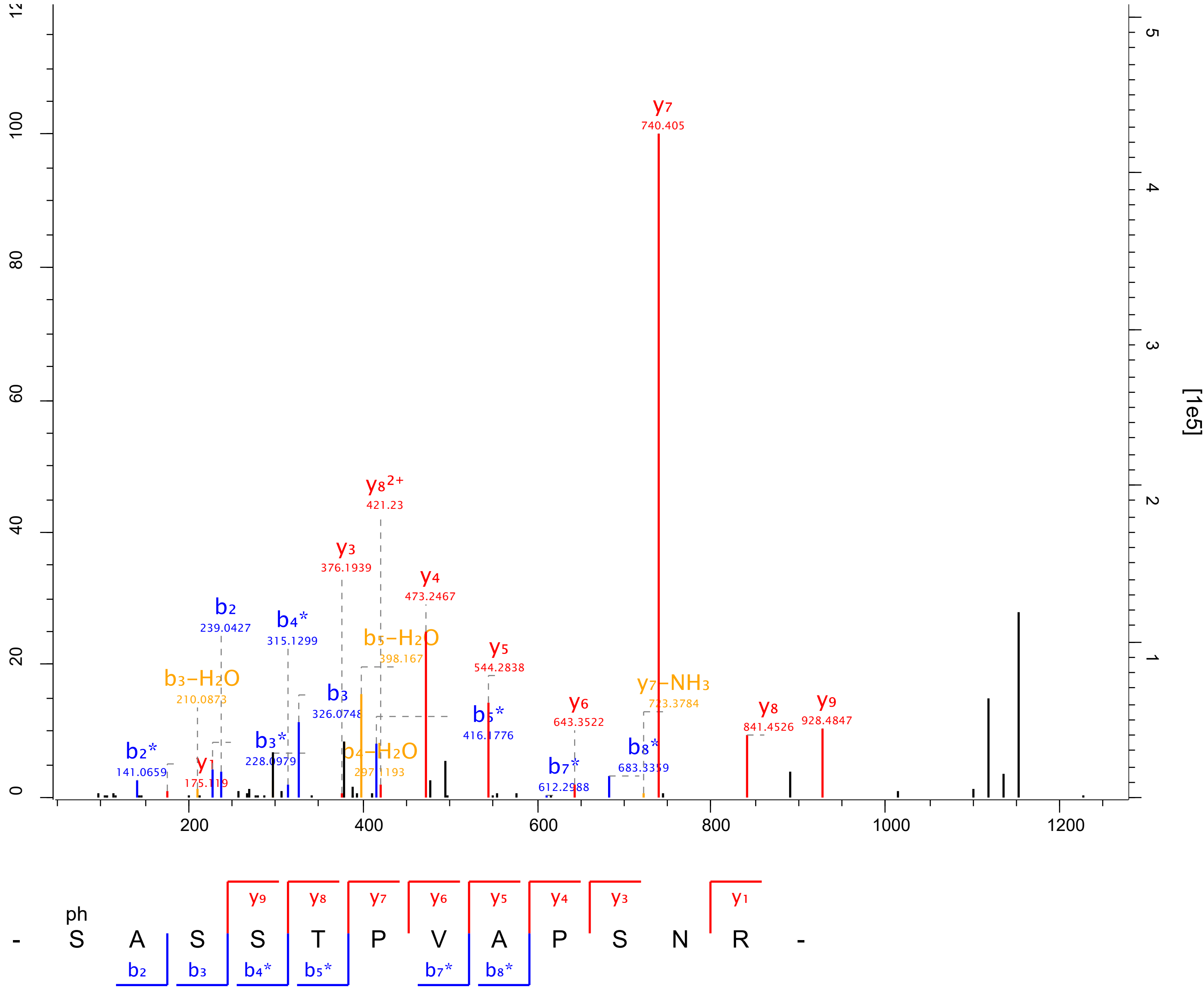

| Raw file | Scan | Method | Score | m/z |
|---|---|---|---|---|
| water_15_2 | 4888 | FTMS; HCD | 51.03 | 665.79 |

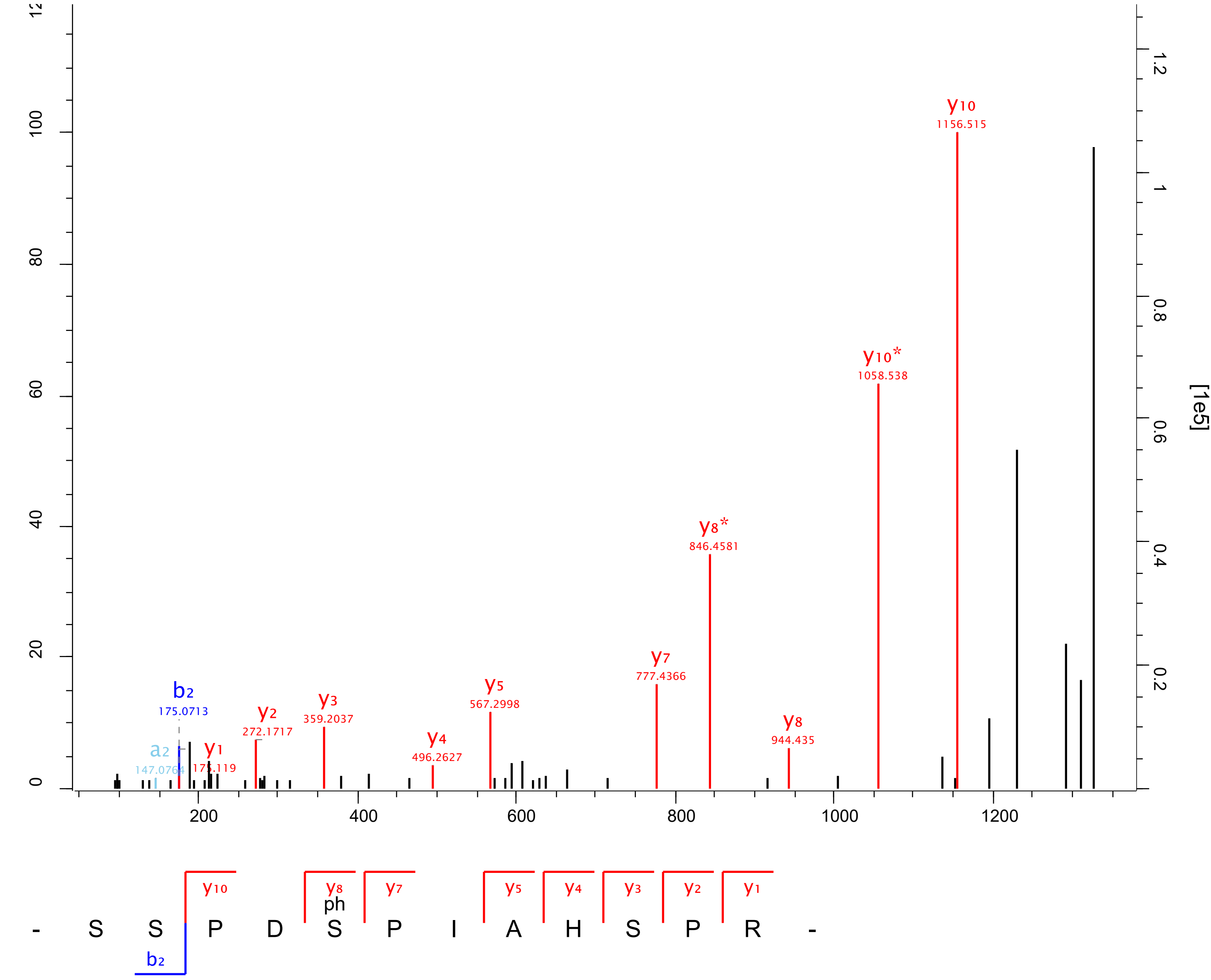

| Raw file | Scan | Method | Score | m/z |
| --- | --- | --- | --- | --- |
| water_15_2 | 4931 | FTMS; HCD | 64.24 | 721.31 |

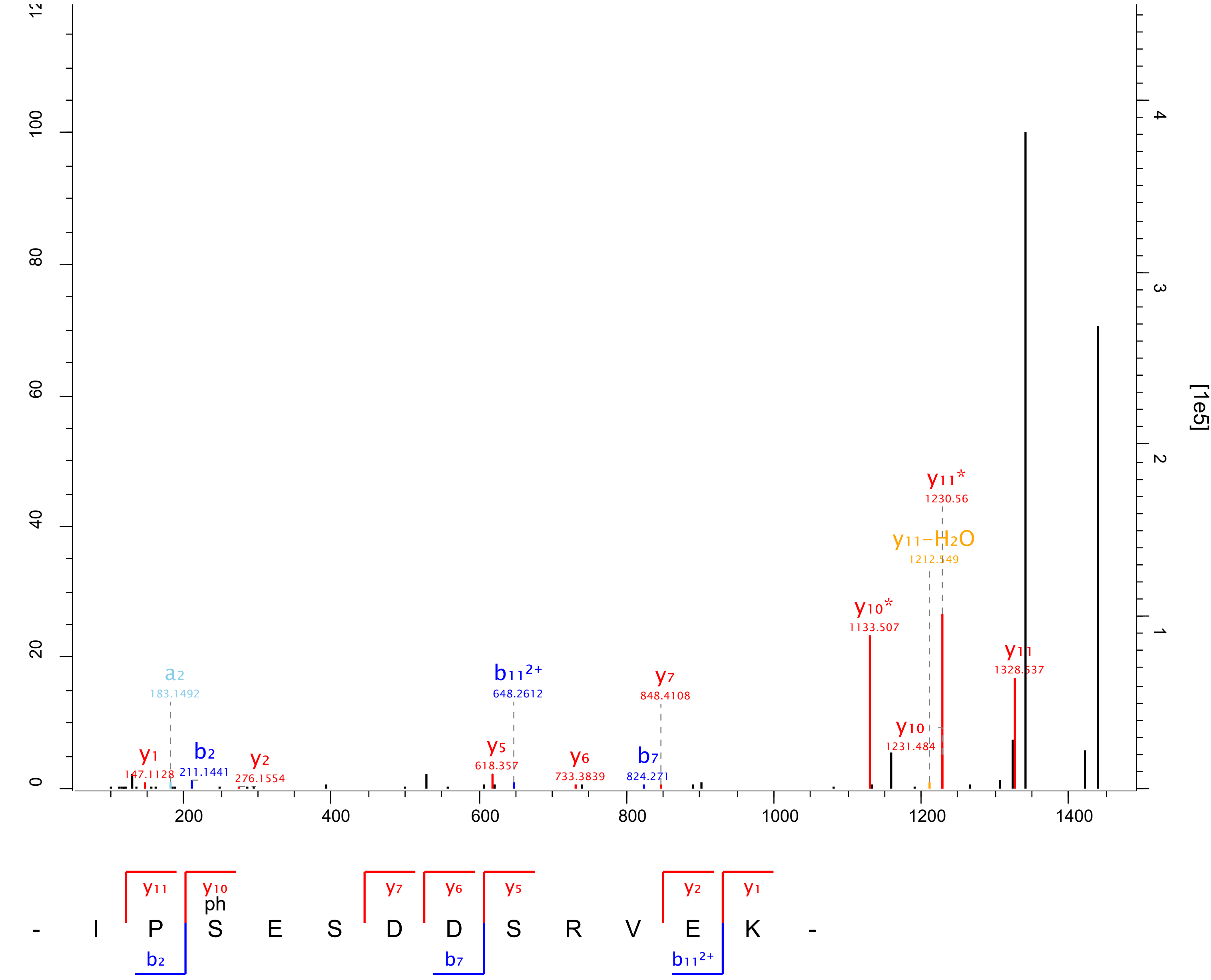

| Raw file | Scan | Method | Score | m/z |
|---|---|---|---|---|
| water_15_2 | 4943 | FTMS; HCD | 49.3 | 555.76 |

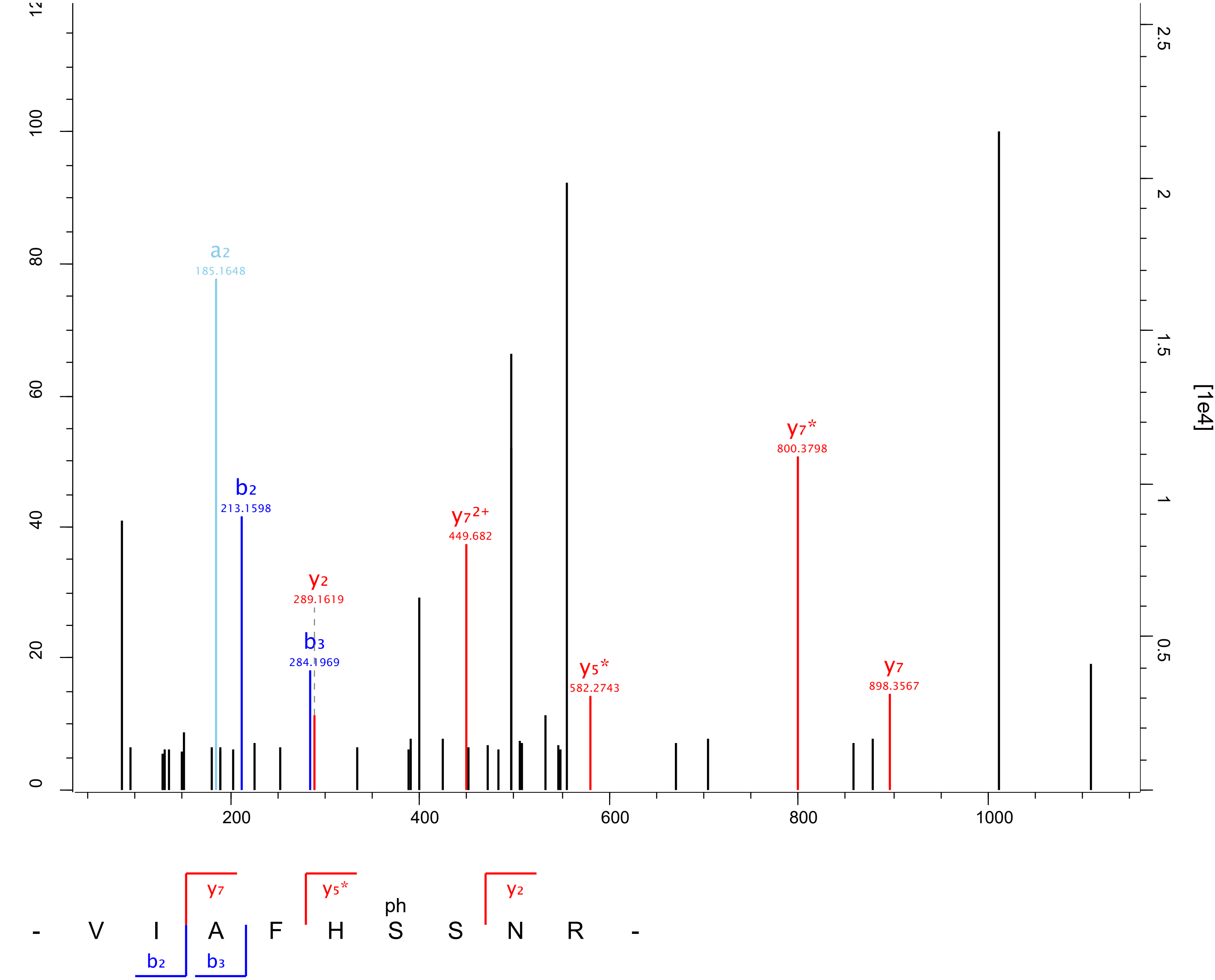

| Raw file | Scan | Method | Score | m/z |
| --- | --- | --- | --- | --- |
| water_15_2 | 5041 | FTMS; HCD | 83.76 | 600.59 |

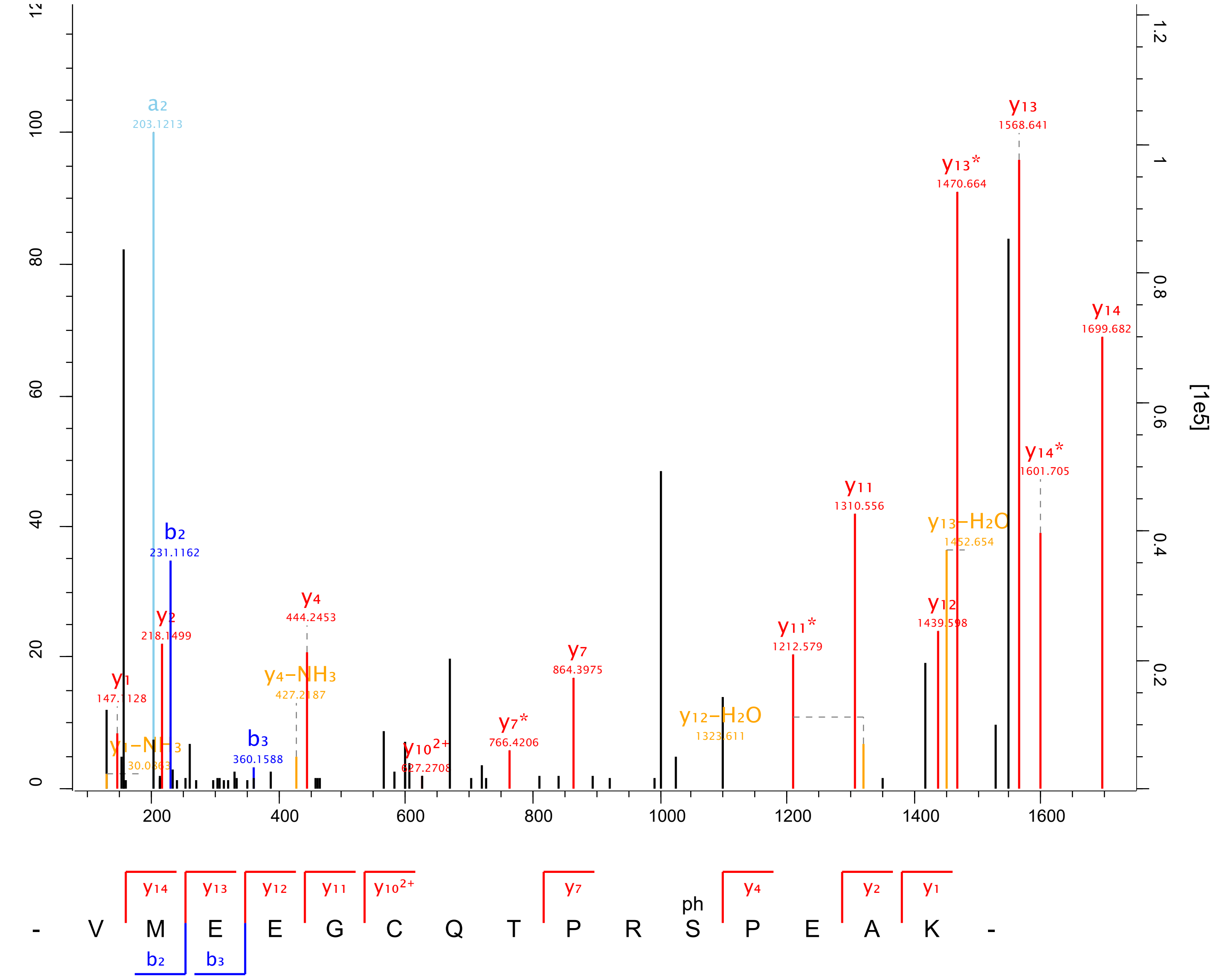

| Raw file | Scan | Method | Score | m/z |
| --- | --- | --- | --- | --- |
| water_15_2 | 5151 | FTMS; HCD | 108.57 | 450.2 |

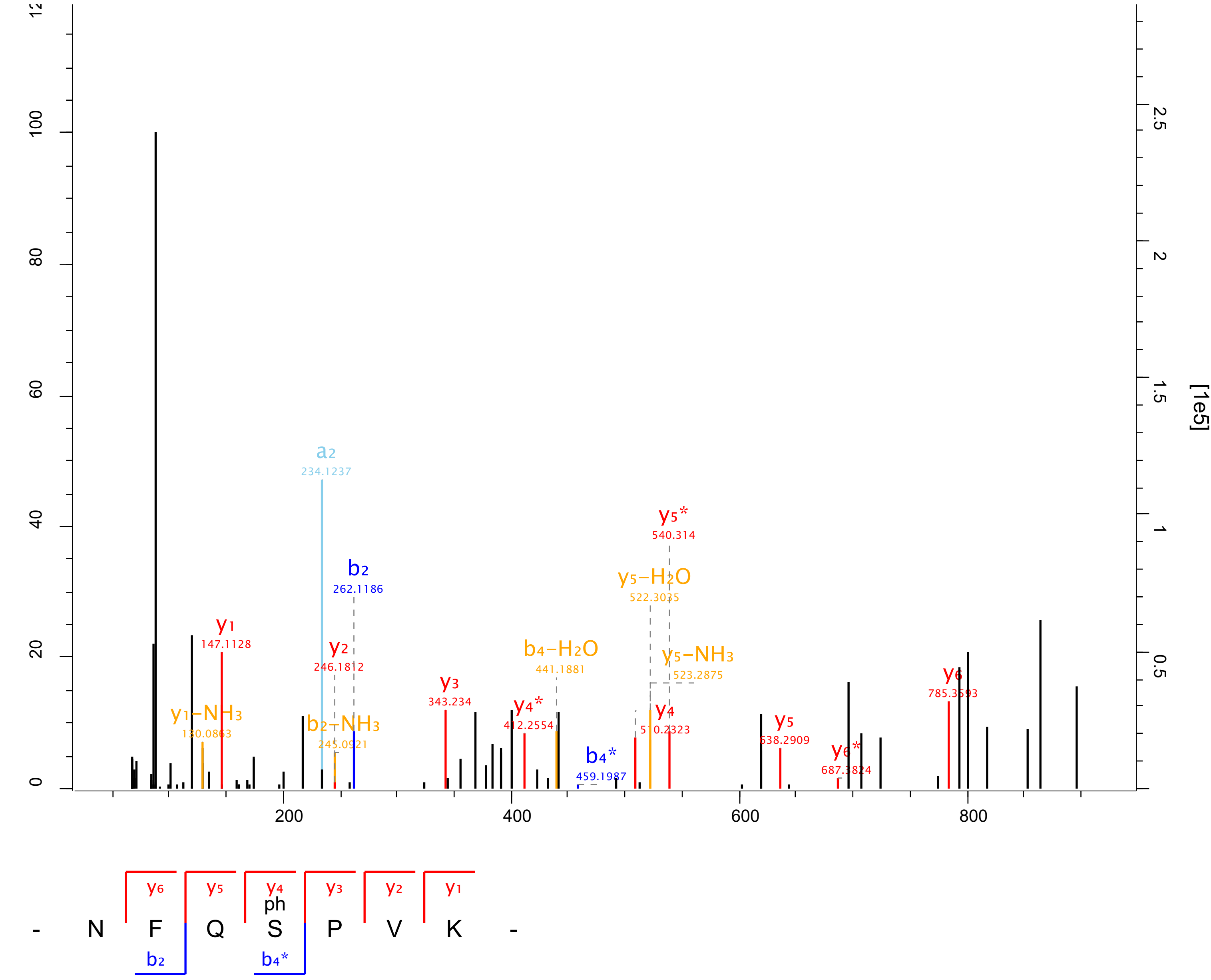

| Raw file | Scan | Method | Score | m/z |
| --- | --- | --- | --- | --- |
| water_15_2 | 5189 | FTMS; HCD | 148.52 | 578.24 |

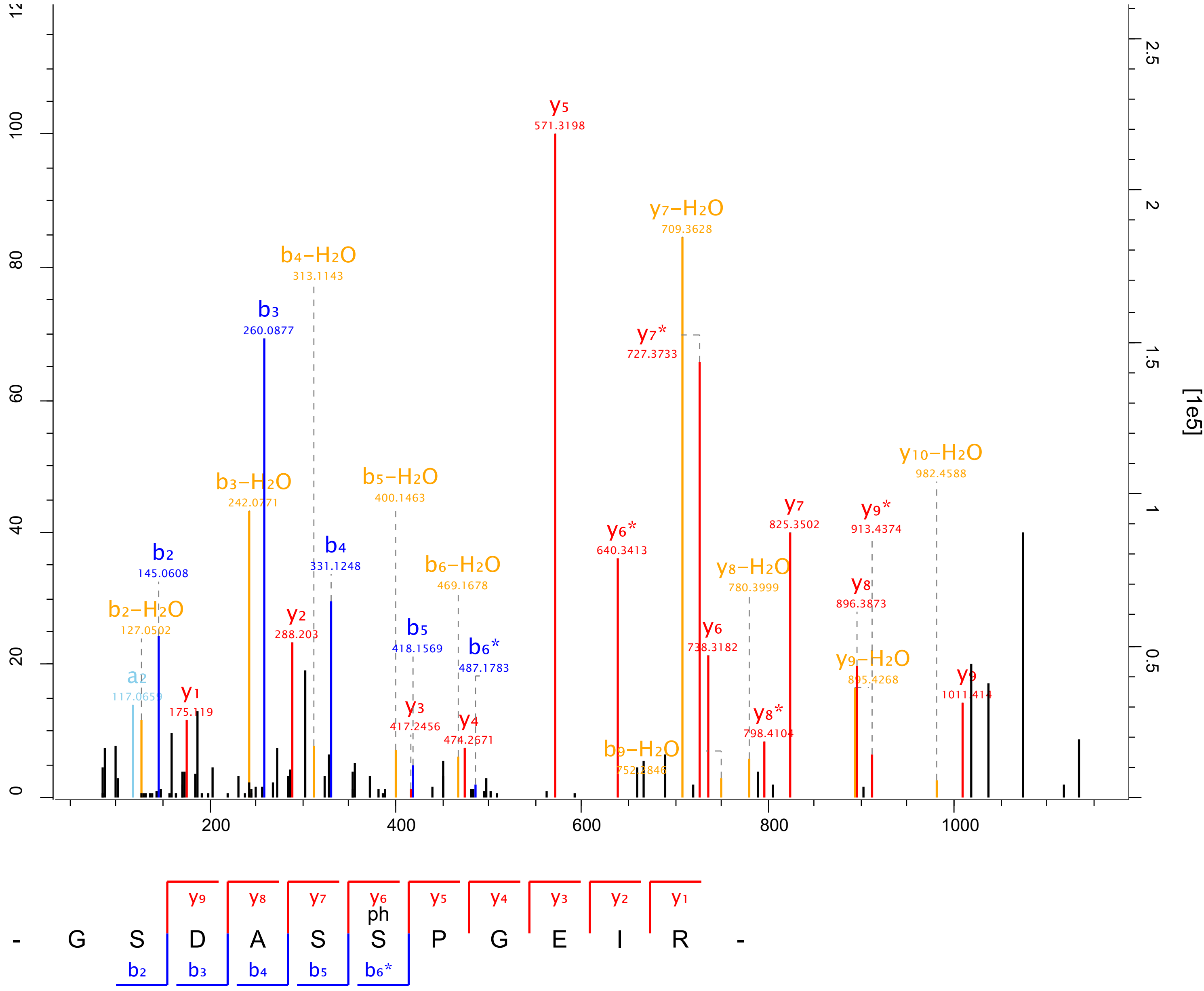

| Raw file | Scan | Method | Score | m/z |
|---|---|---|---|---|
| water_15_2 | 5205 | FTMS; HCD | 131.96 | 591.76 |

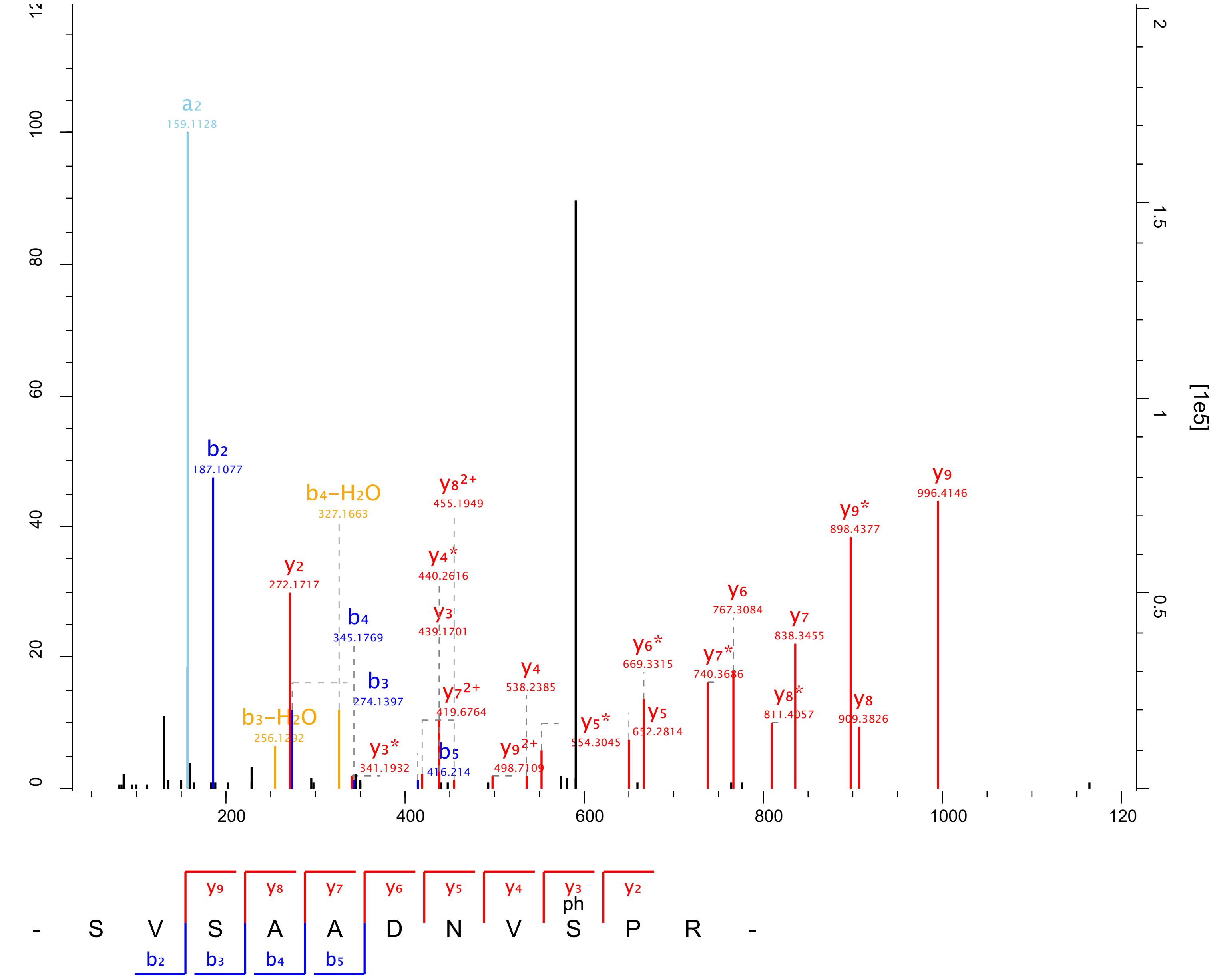

| Raw file | Scan | Method | Score | m/z |
|---|---|---|---|---|
| water_15_2 | 5301 | FTMS; HCD | 82.77 | 541.25 |

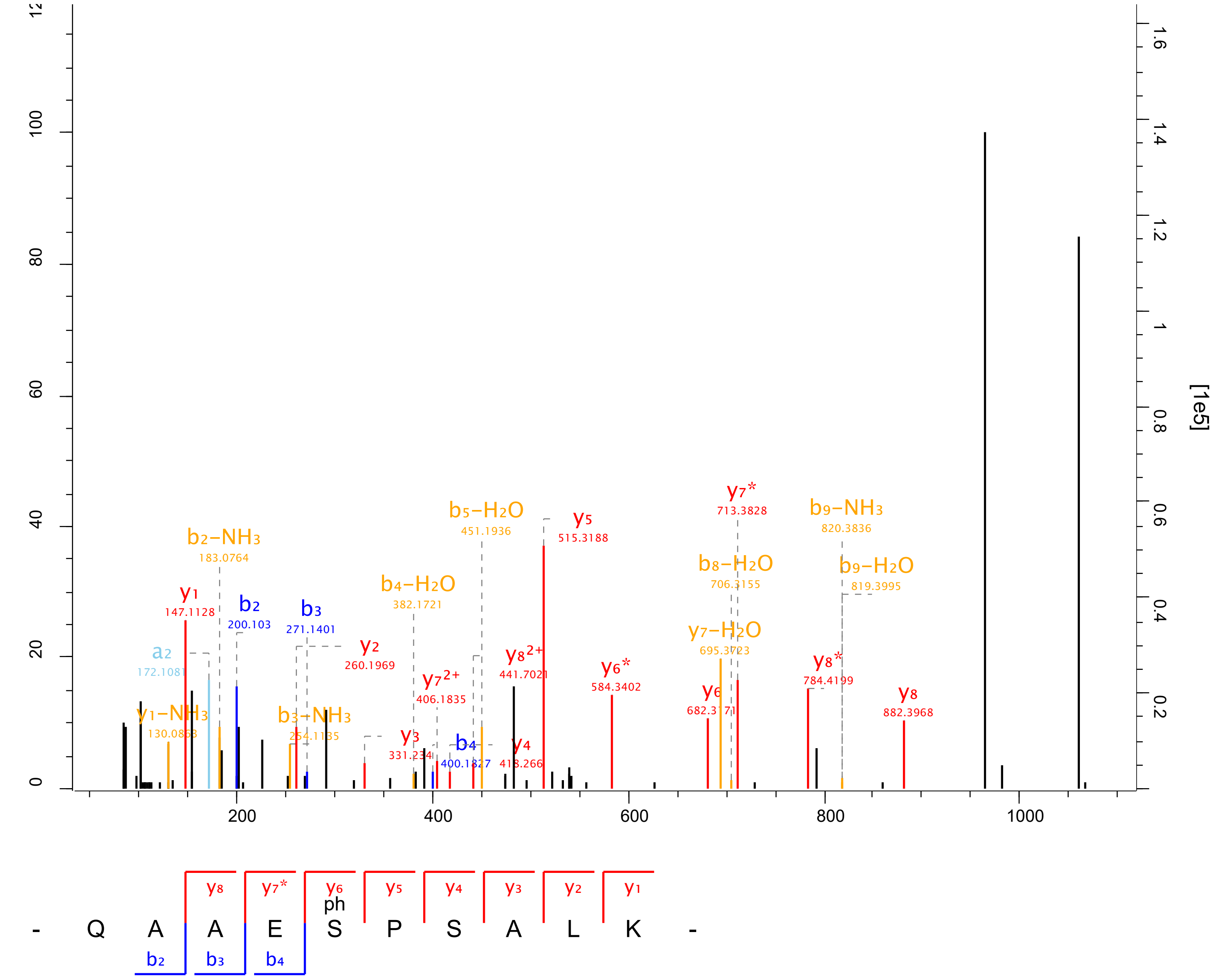

| Raw file | Scan | Method | Score | m/z |
|---|---|---|---|---|
| water_15_2 | 5337 | FTMS; HCD | 118.21 | 512.54 |

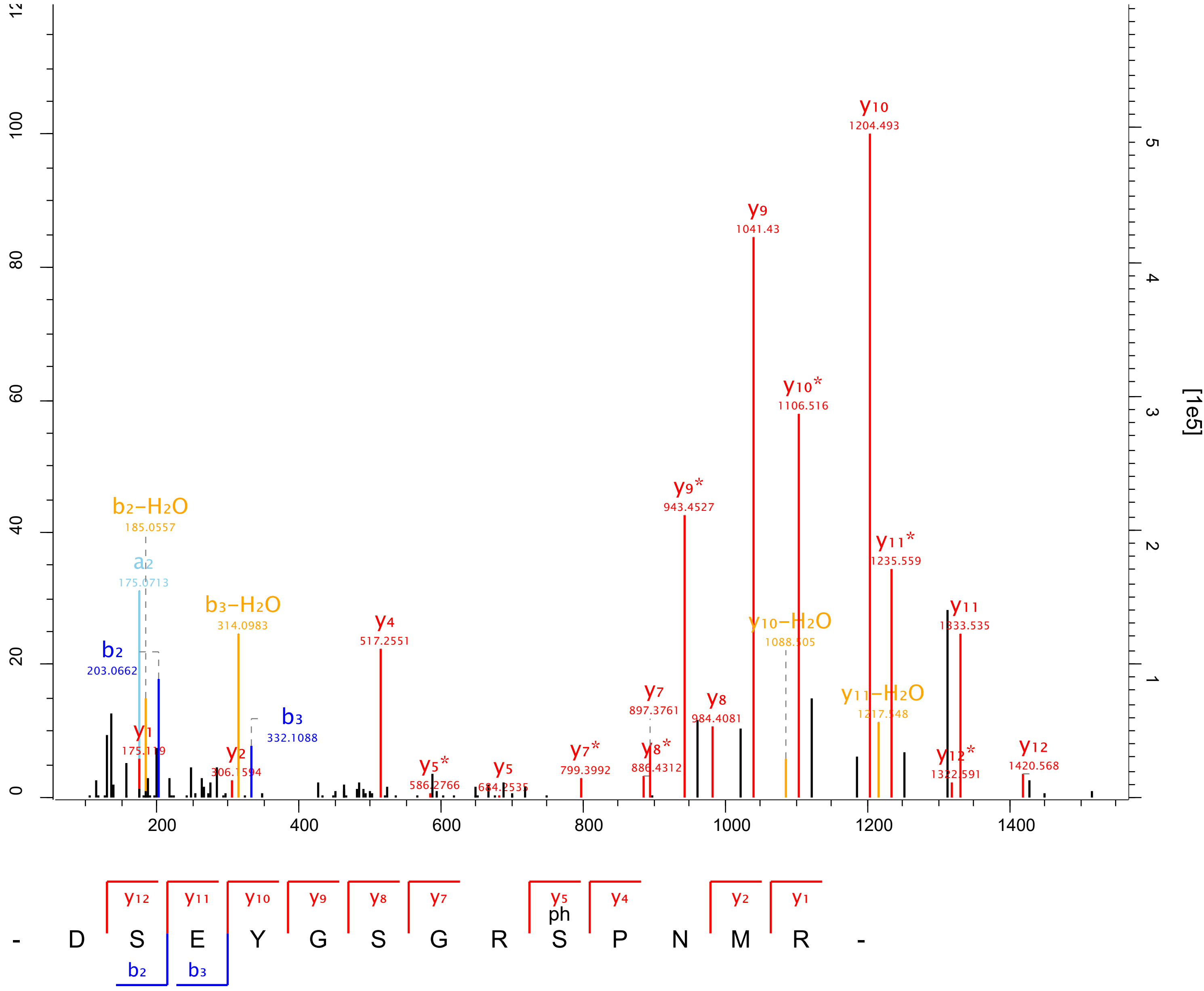

| Raw file | Scan | Method | Score | m/z |
| --- | --- | --- | --- | --- |
| water_15_2 | 5349 | FTMS; HCD | 187.01 | 635.76 |

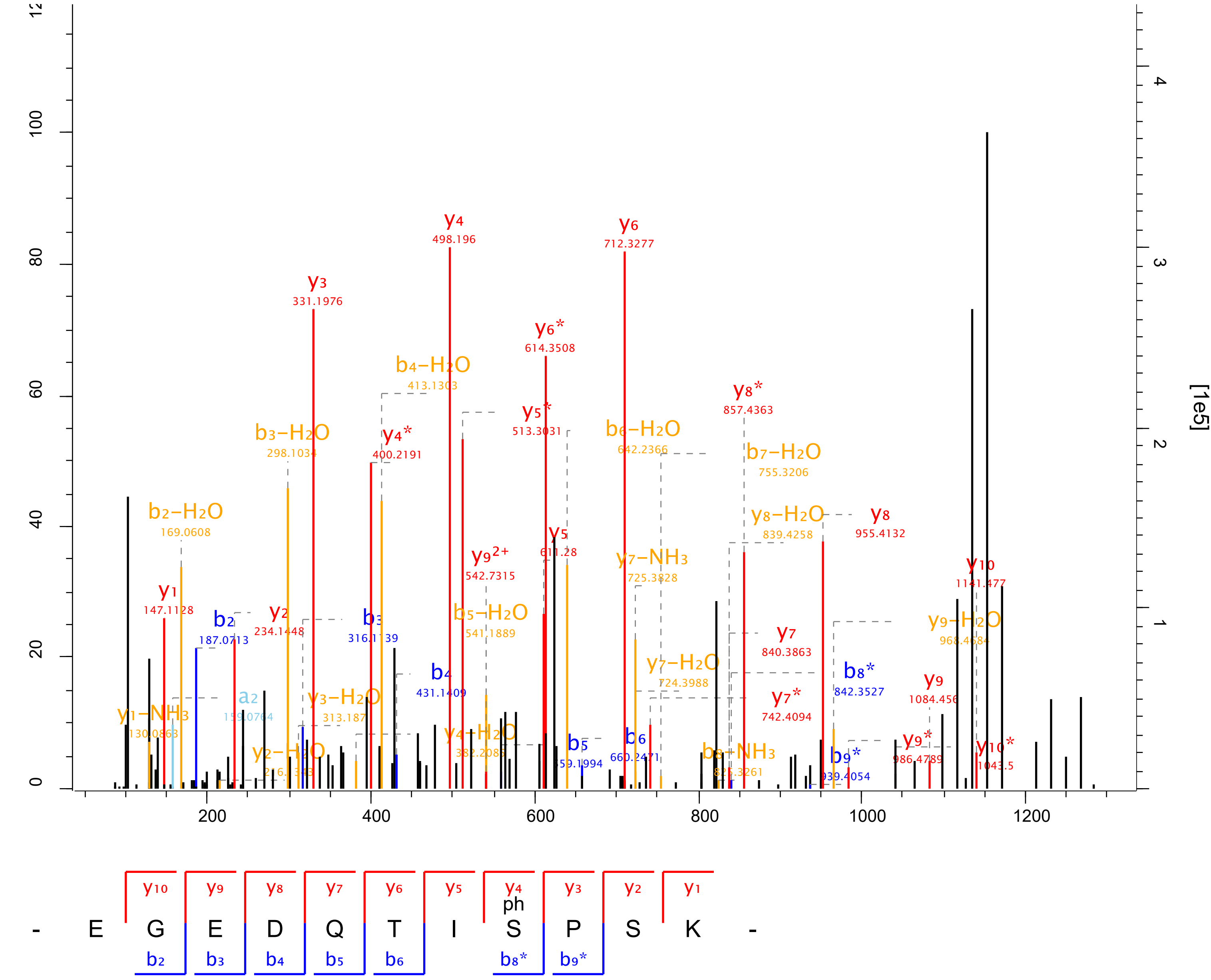

| Raw file | Scan | Method | Score | m/z |
|---|---|---|---|---|
| water_15_2 | 5496 | FTMS; HCD | 213.59 | 529.23 |

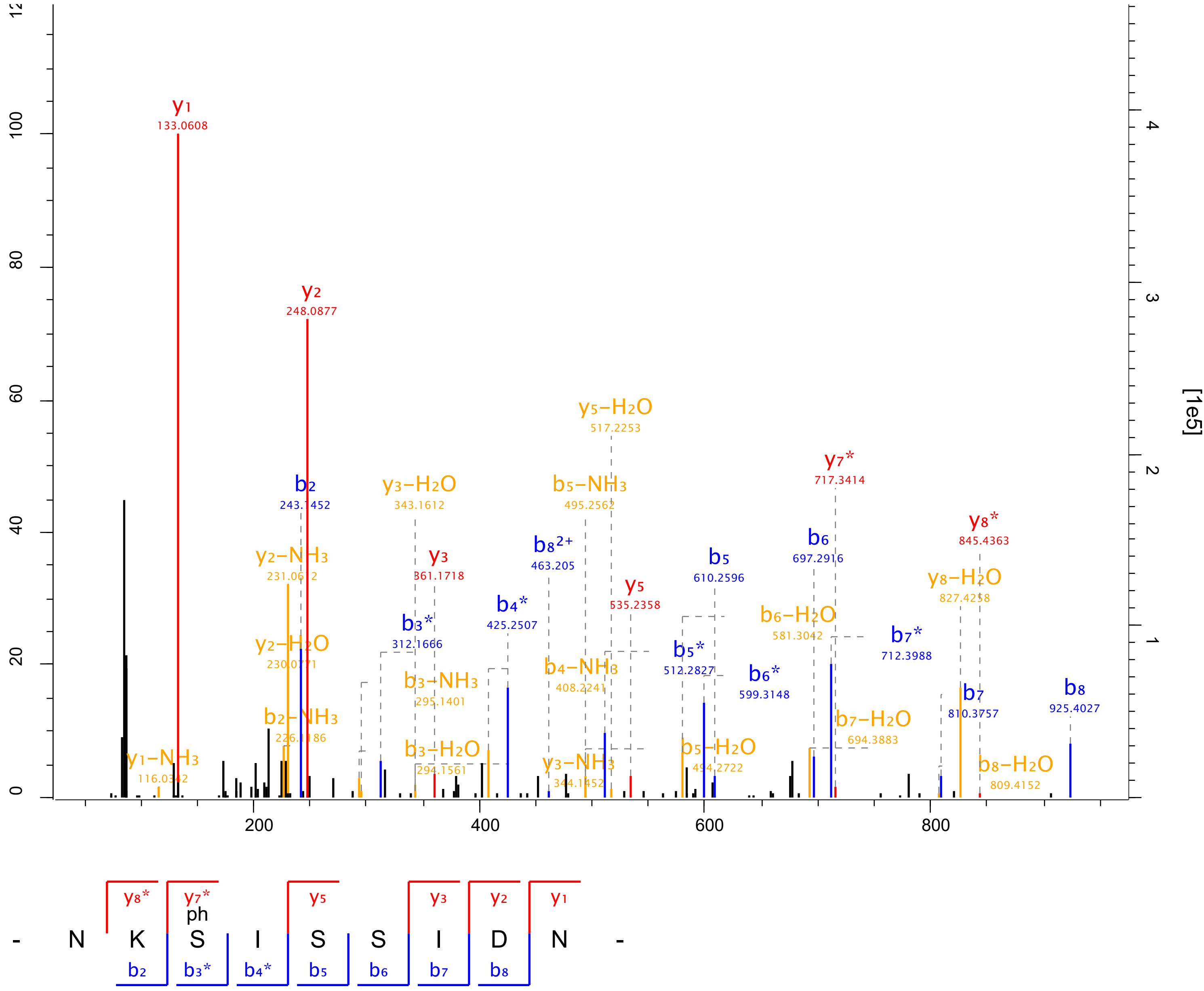

| Raw file | Scan | Method | Score | m/z |
|---|---|---|---|---|
| water_15_2 | 5497 | FTMS; HCD | 83.69 | 593.75 |

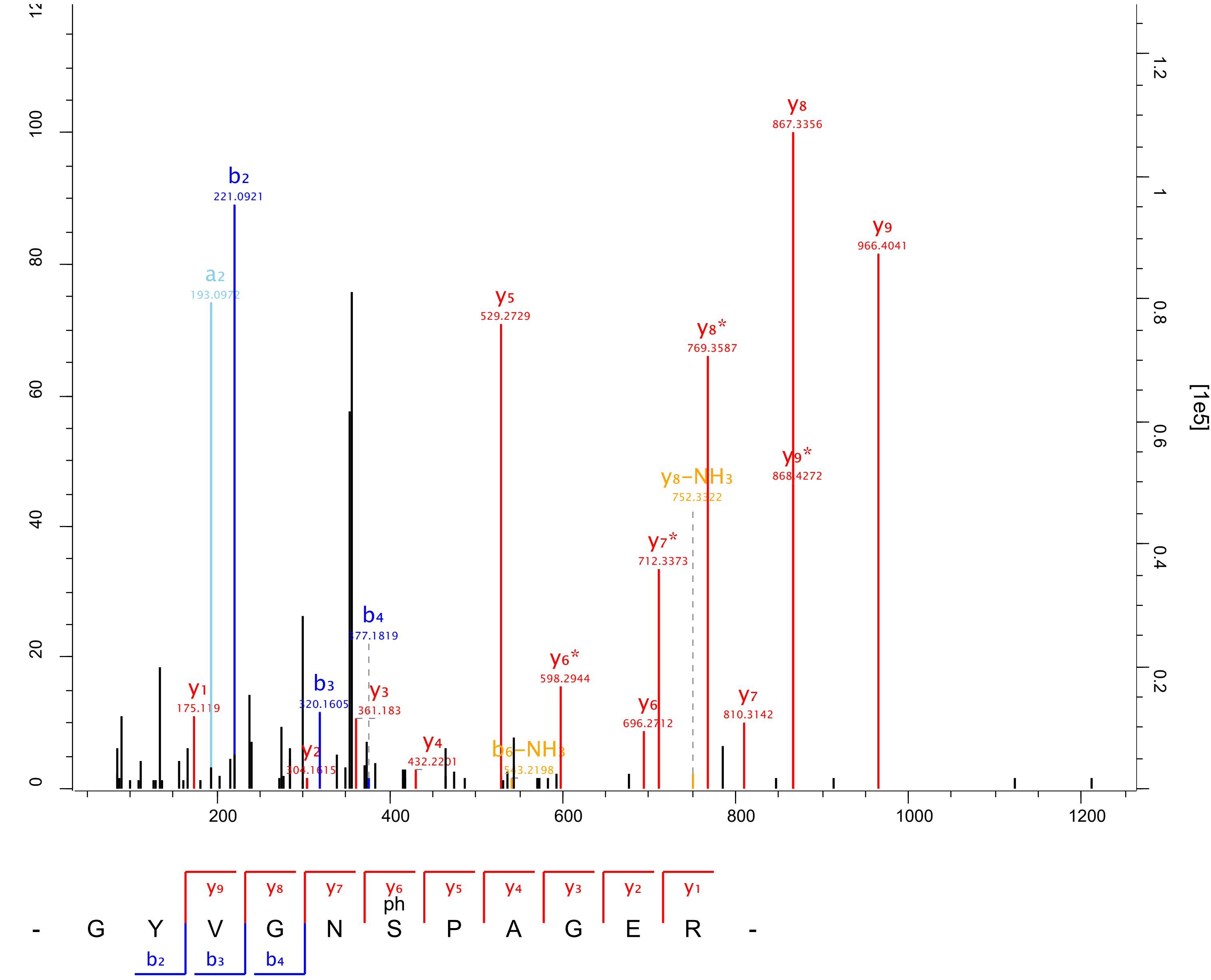

| Raw file | Scan | Method | Score | m/z |
| --- | --- | --- | --- | --- |
| water_15_2 | 5528 | FTMS; HCD | 143.99 | 523.89 |

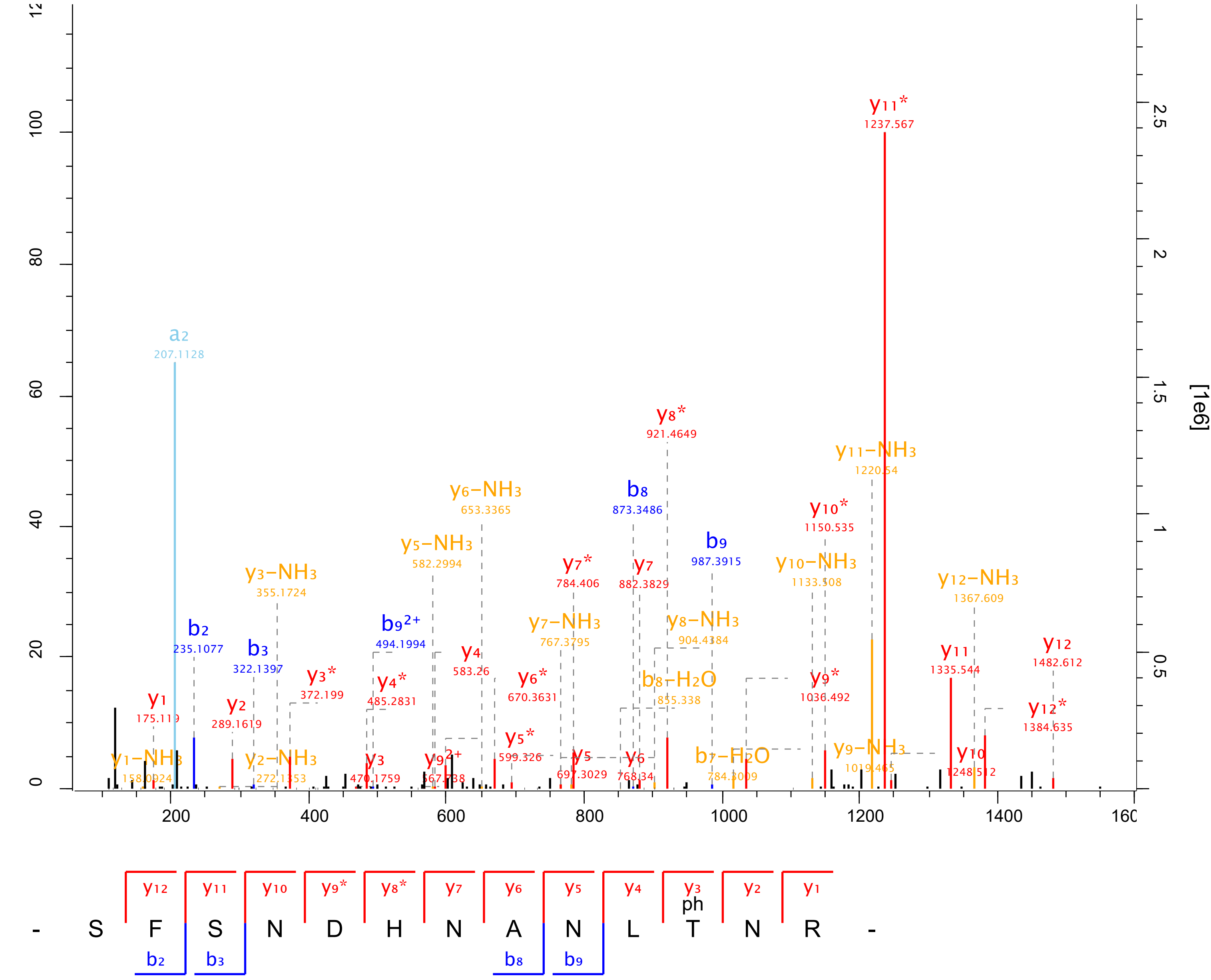

| Raw file | Scan | Method | Score | m/z |
|---|---|---|---|---|
| water_15_2 | 5558 | FTMS; HCD | 52.58 | 506.74 |

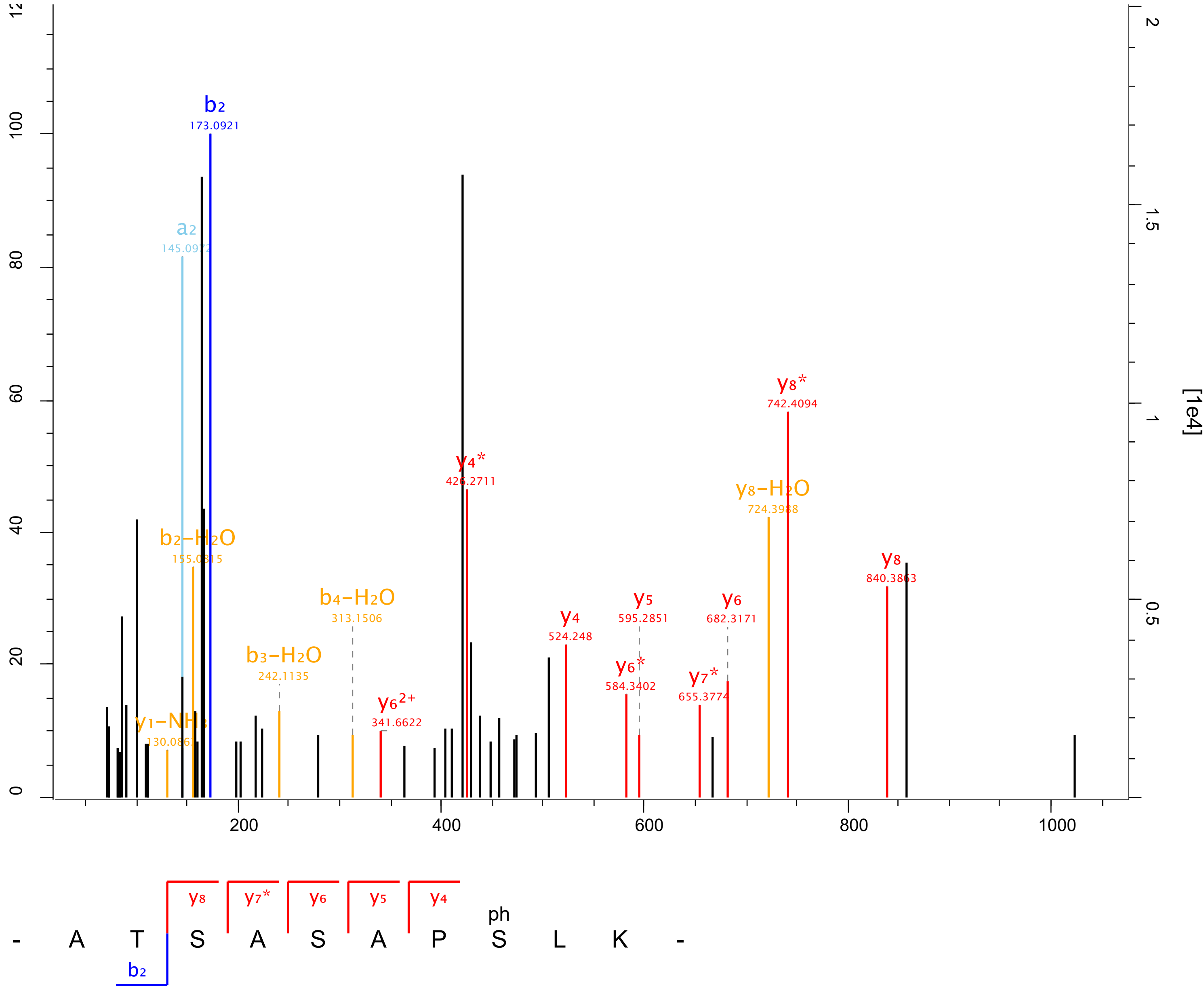

| Raw file | Scan | Method | Score | m/z |
|---|---|---|---|---|
| water_15_2 | 5562 | FTMS; HCD | 229.41 | 656.6 |

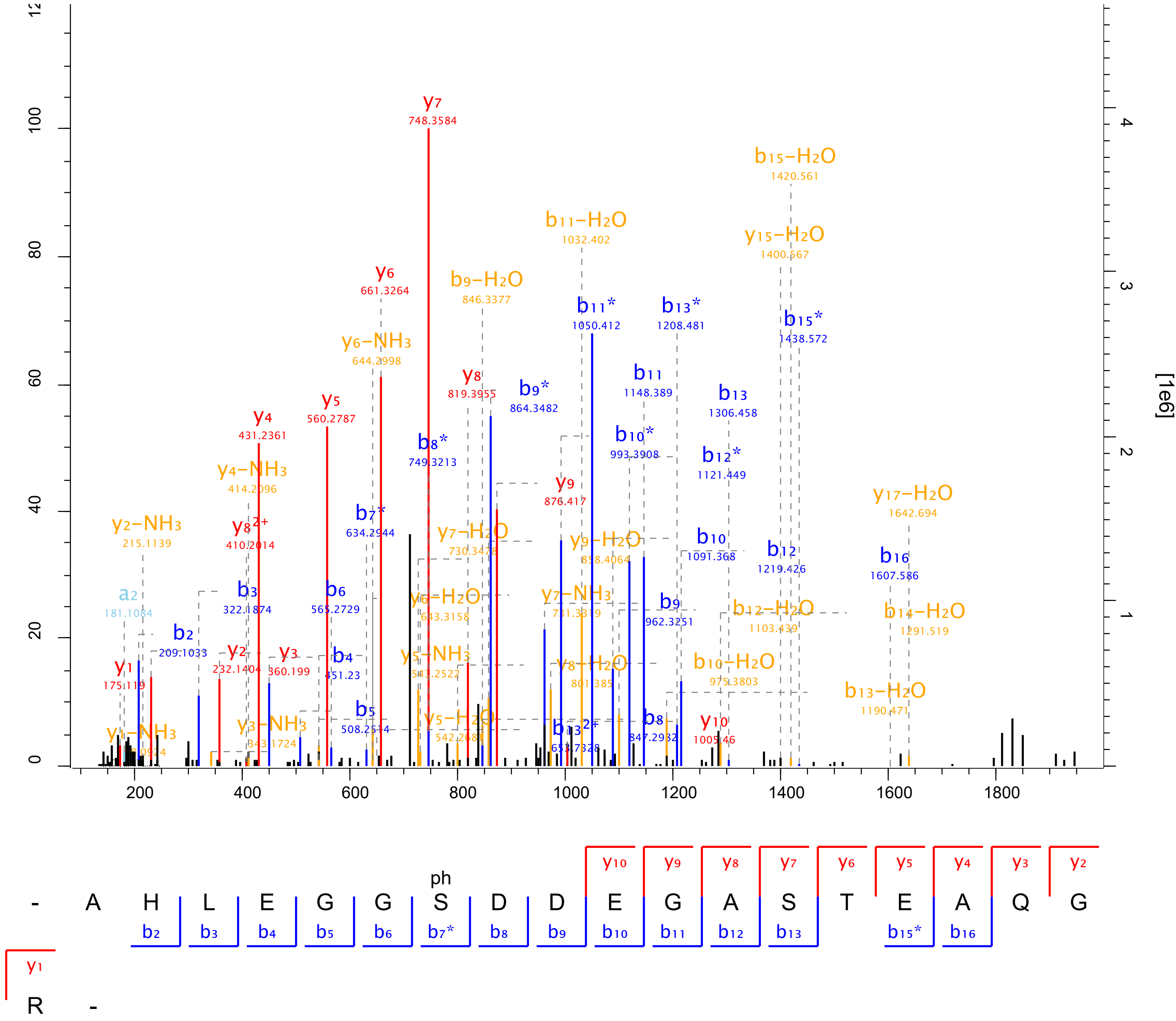

| Raw file | Scan | Method | Score | m/z |
| --- | --- | --- | --- | --- |
| water_15_2 | 5654 | FTMS; HCD | 44.56 | 522.72 |

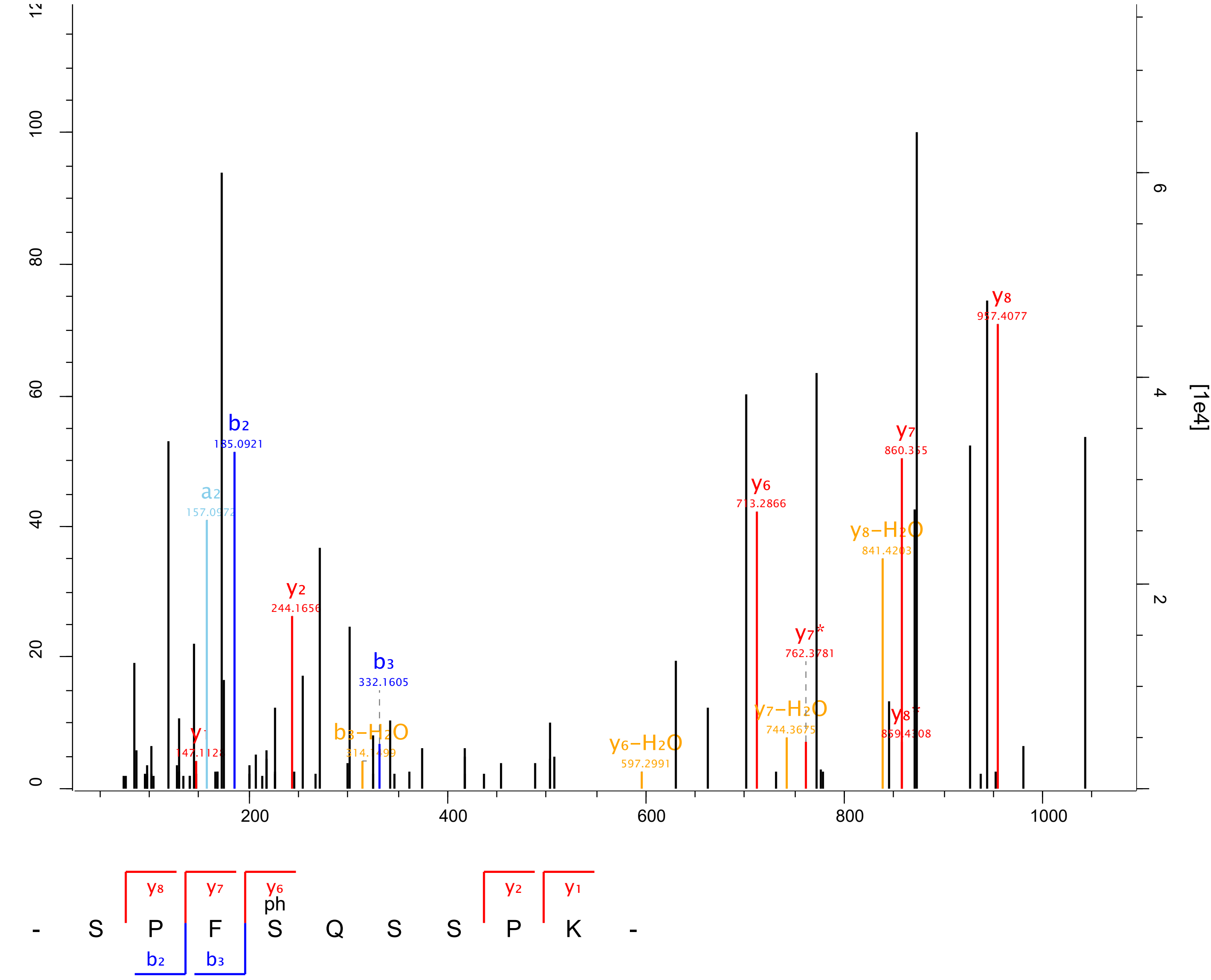

| Raw file | Scan | Method | Score | m/z |
| --- | --- | --- | --- | --- |
| water_15_2 | 5664 | FTMS; HCD | 139.49 | 689.78 |

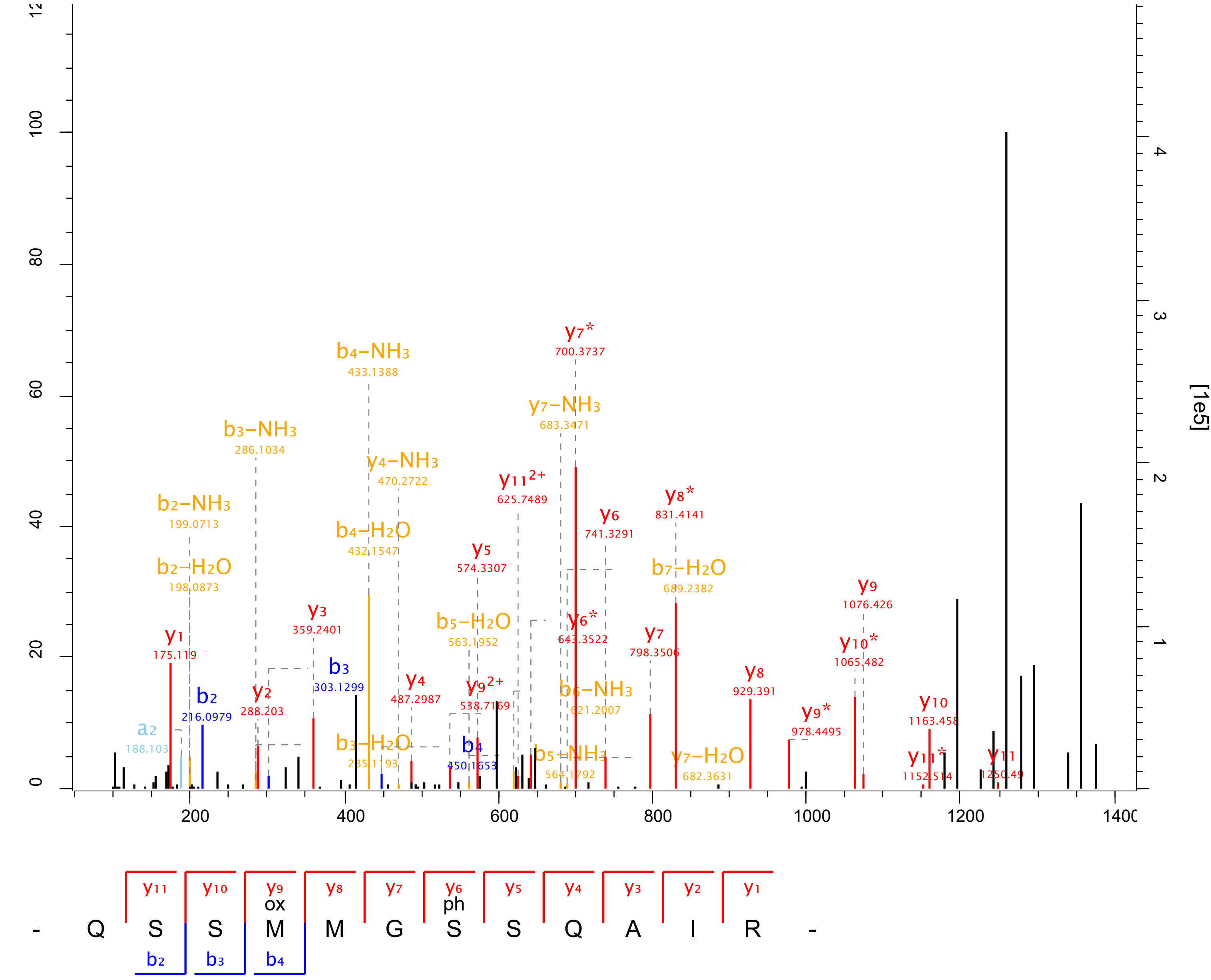

| Raw file | Scan | Method | Score | m/z |
|---|---|---|---|---|
| water_15_2 | 5665 | FTMS; HCD | 125.45 | 584.55 |

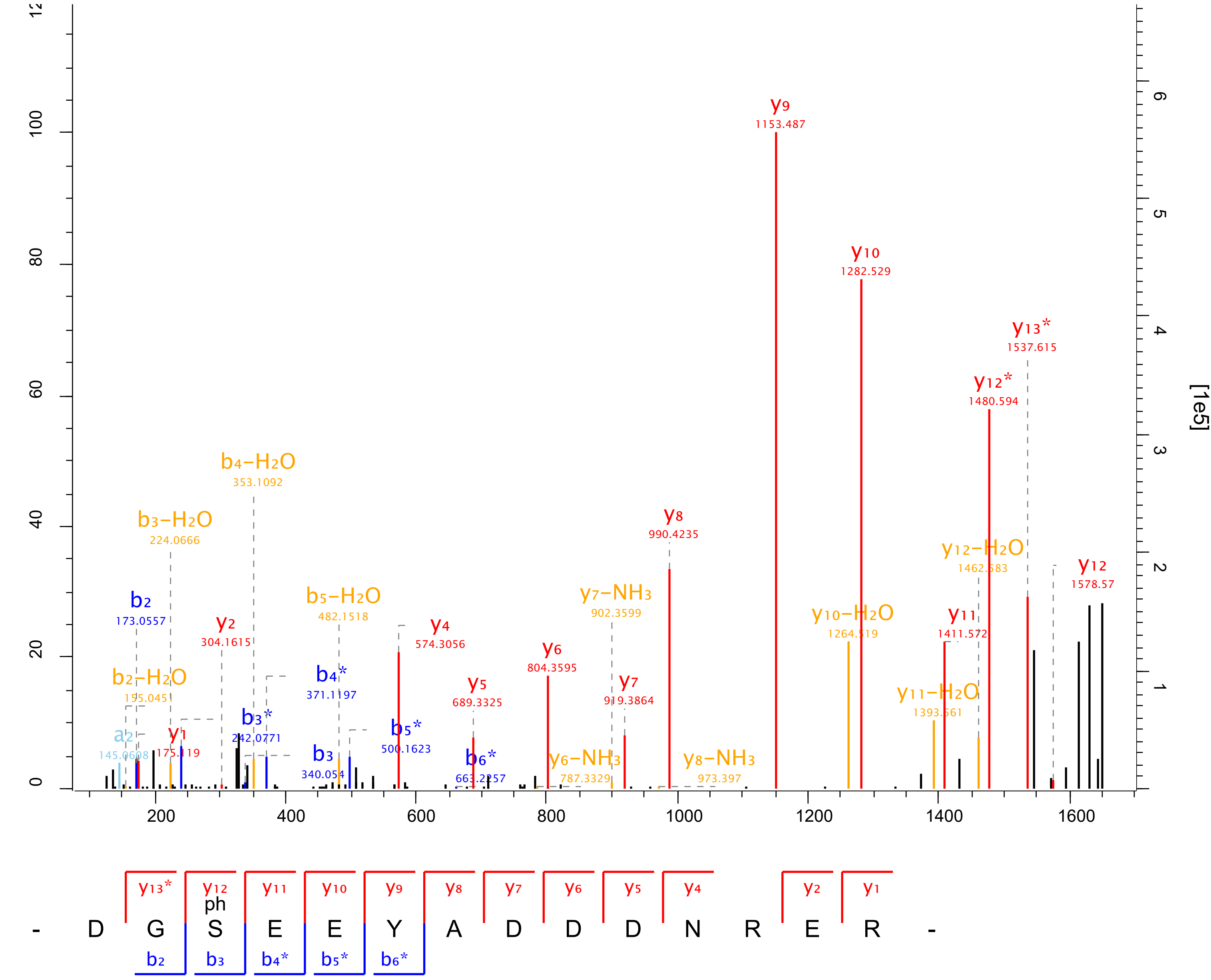

| Raw file | Scan | Method | Score | m/z |
|---|---|---|---|---|
| water_15_2 | 5672 | FTMS; HCD | 83.69 | 636.27 |

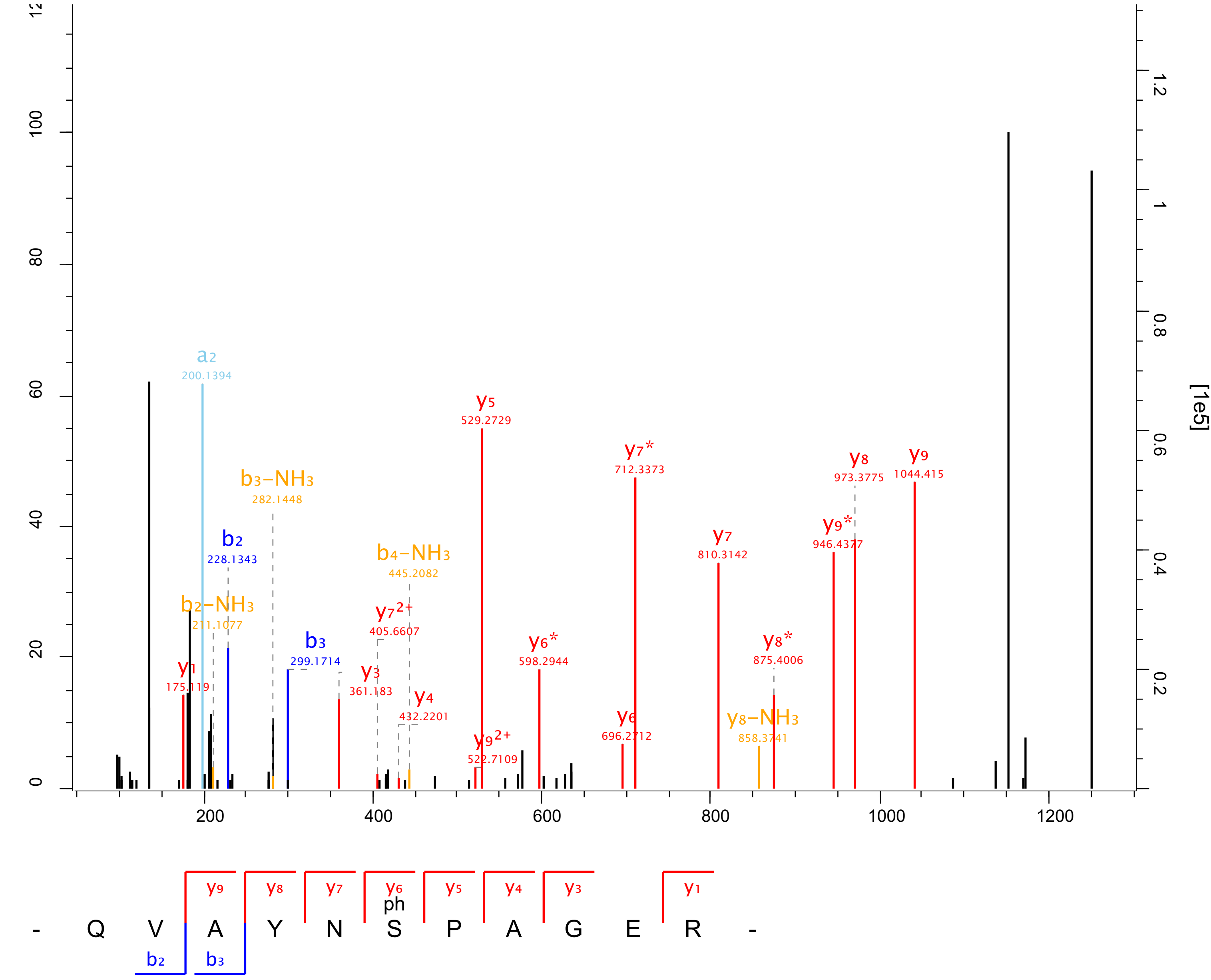

| Raw file | Scan | Method | Score | m/z |
|---|---|---|---|---|
| water_15_2 | 5684 | FTMS; HCD | 105.96 | 626.25 |

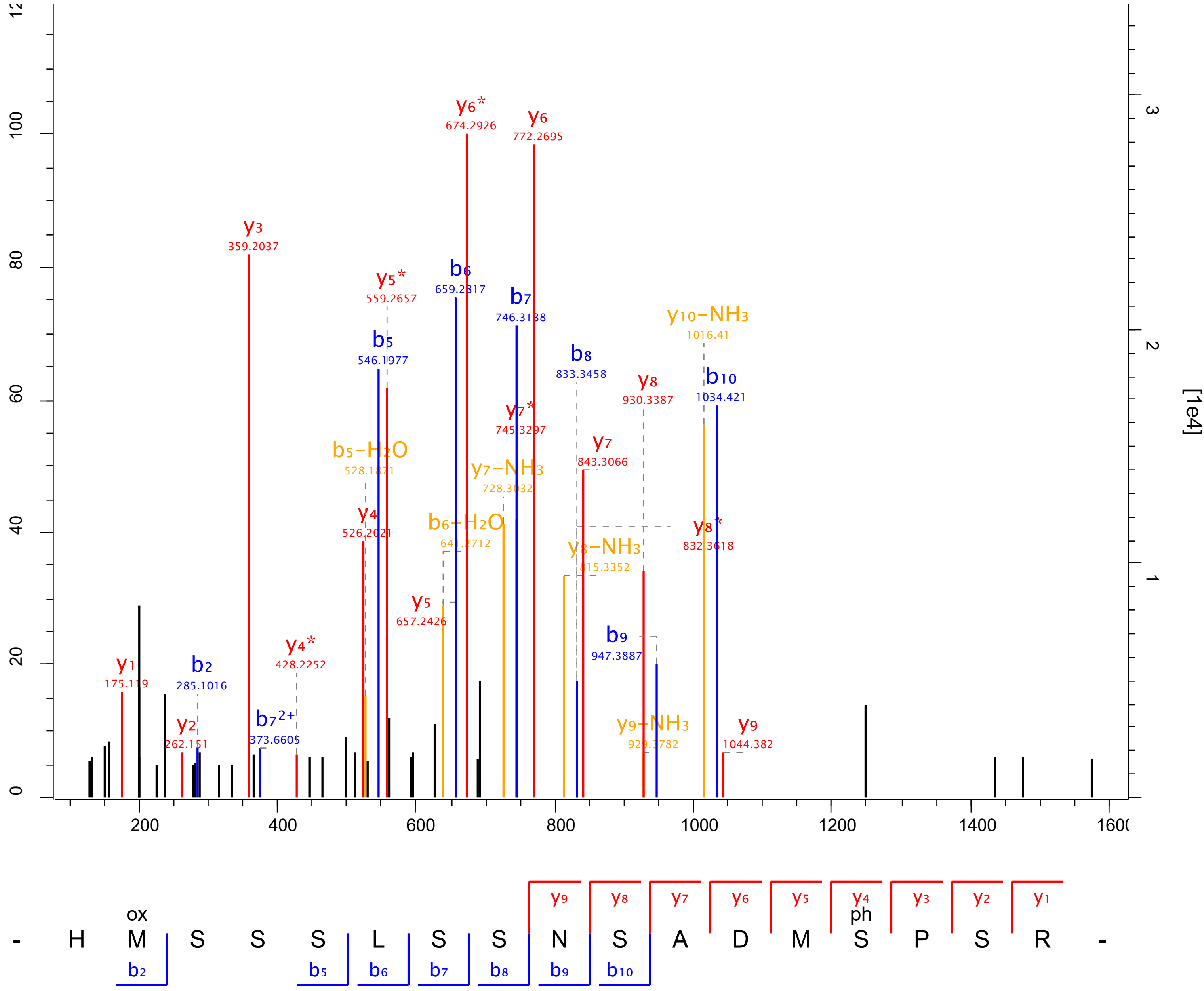

| Raw file | Scan | Method | Score | m/z |
|---|---|---|---|---|
| water_15_2 | 5738 | FTMS; HCD | 190.46 | 504.88 |

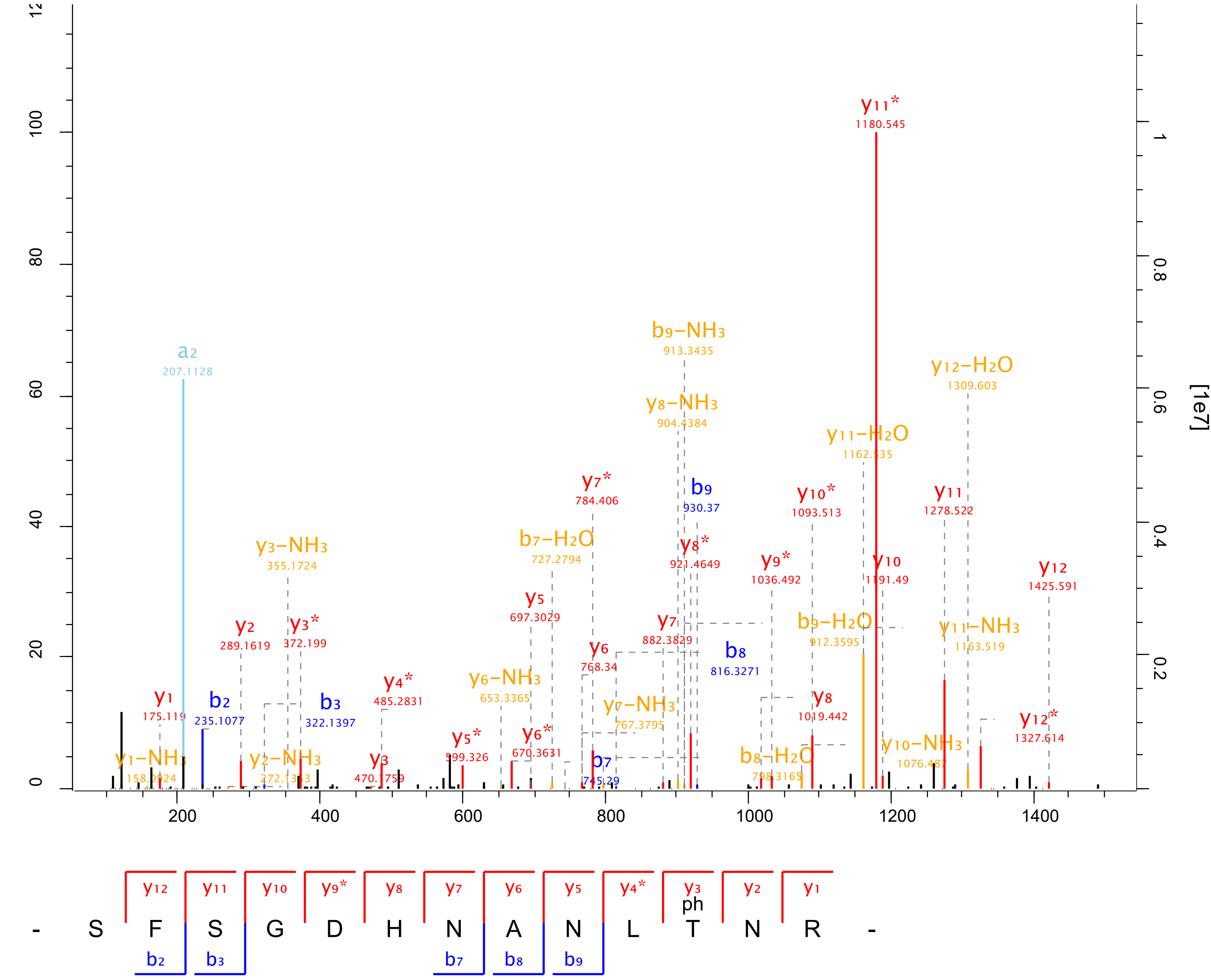

| Raw file | Scan | Method | Score | m/z |
| --- | --- | --- | --- | --- |
| water_15_2 | 5761 | FTMS; HCD | 97.46 | 434.87 |

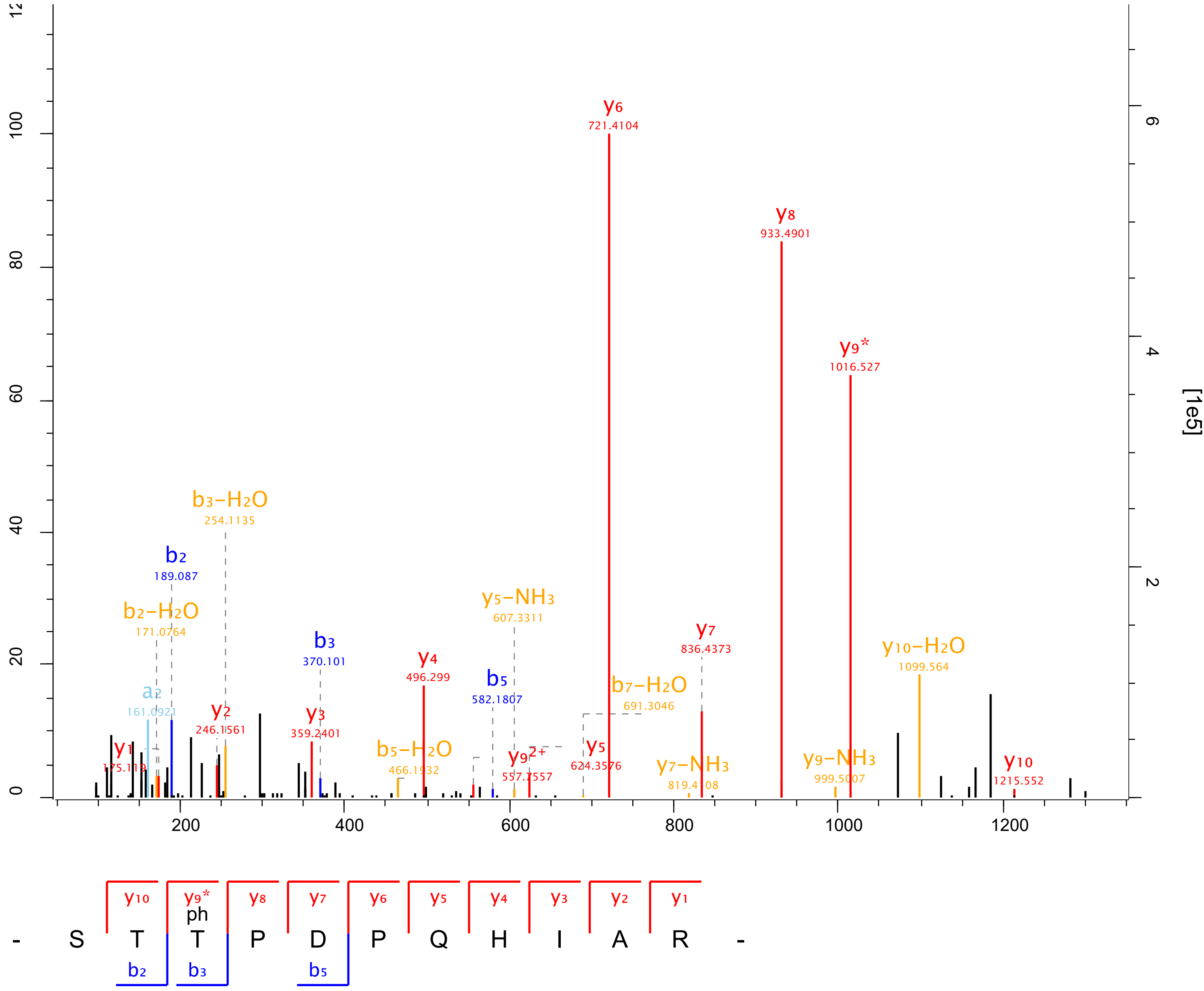

| Raw file | Scan | Method | Score | m/z |
| --- | --- | --- | --- | --- |
| water_15_2 | 5793 | FTMS; HCD | 131.54 | 622.62 |

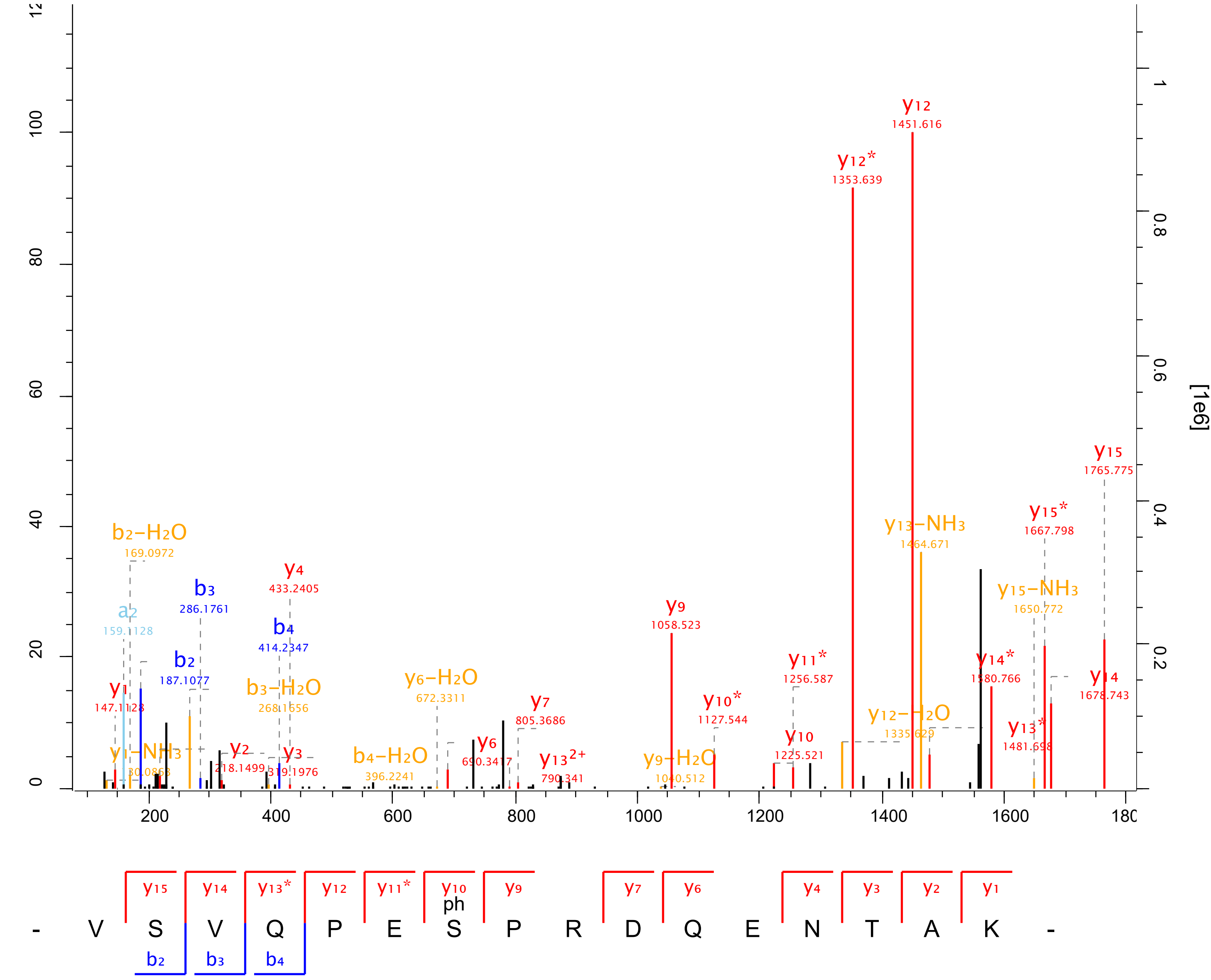

| Raw file | Scan | Method | Score | m/z |
|---|---|---|---|---|
| water_15_2 | 5814 | FTMS; HCD | 228.9 | 551.56 |

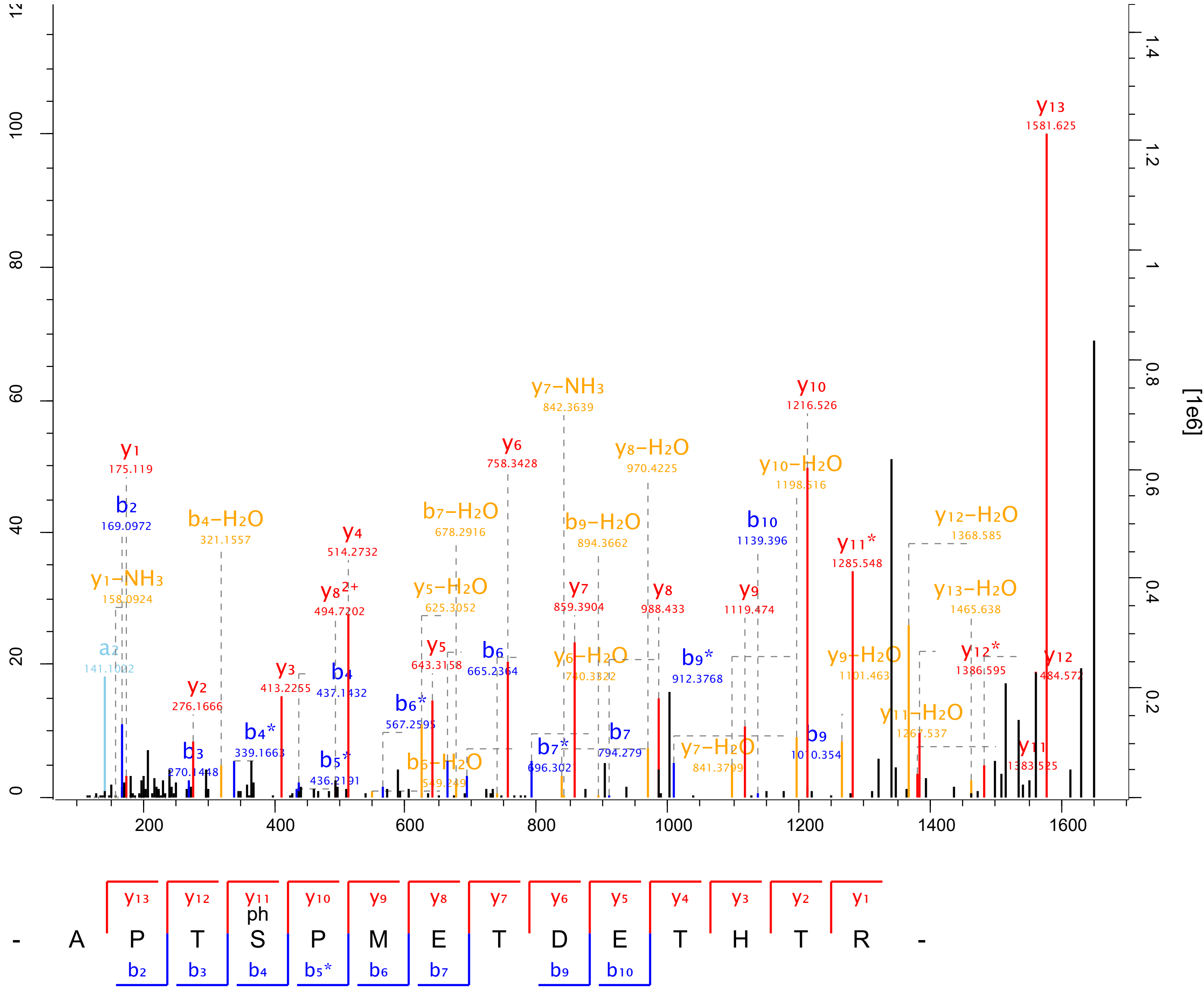

| Raw file | Scan | Method | Score | m/z |
|---|---|---|---|---|
| water_15_2 | 5845 | FTMS; HCD | 90.83 | 529.72 |

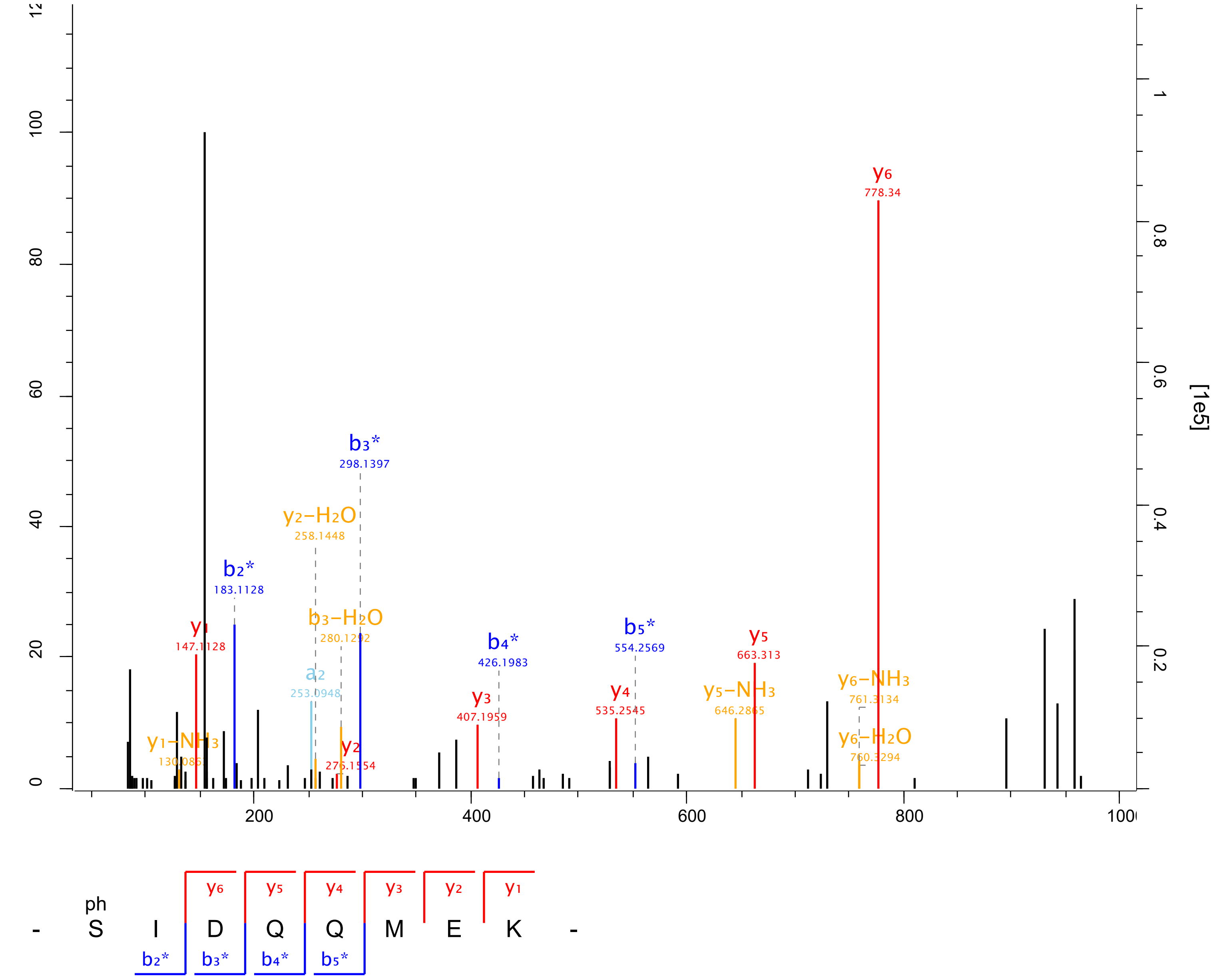

| Raw file | Scan | Method | Score | m/z |
| --- | --- | --- | --- | --- |
| water_15_2 | 5884 | FTMS; HCD | 50.83 | 789.3 |

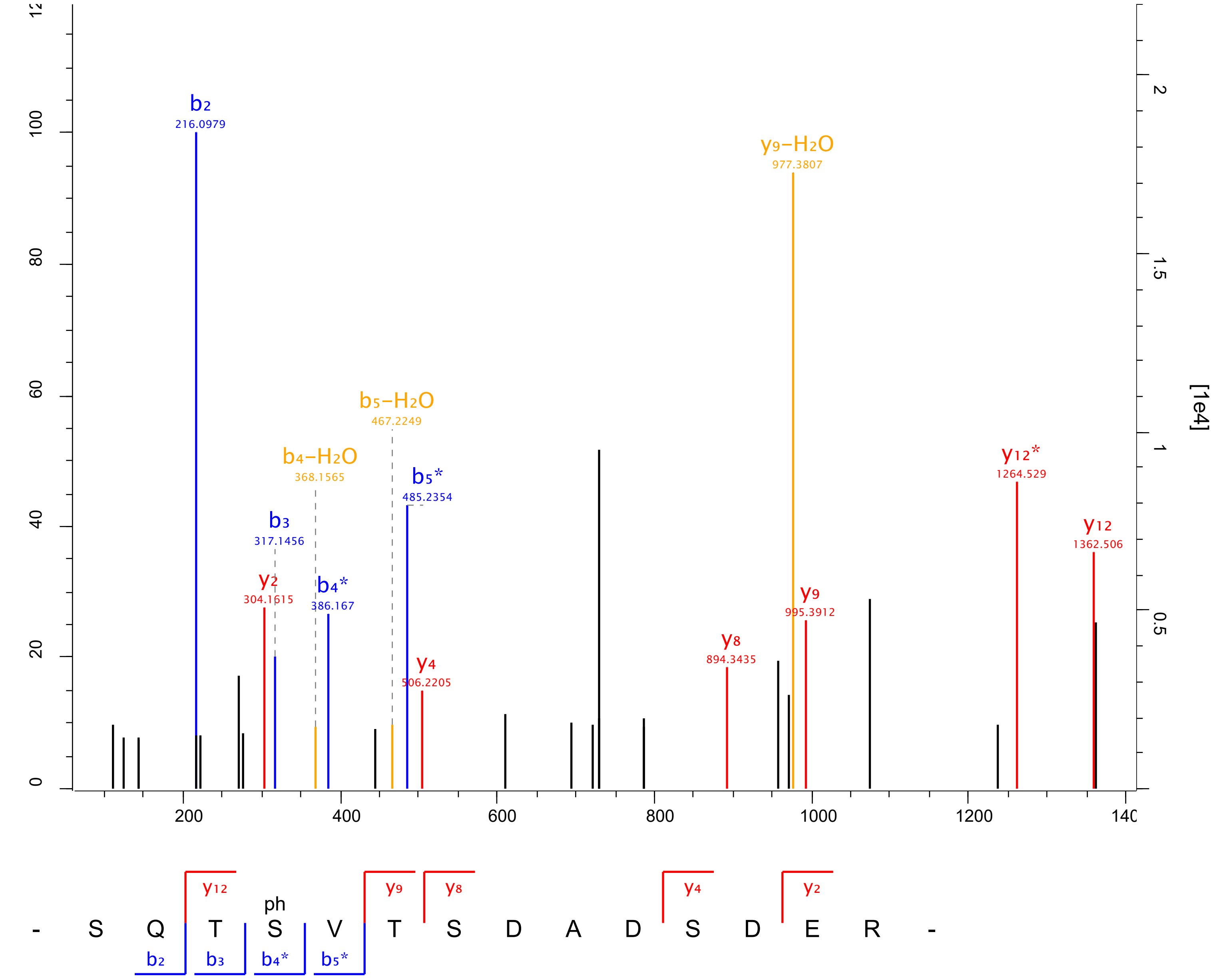

| Raw file | Scan | Method | Score | m/z |
| --- | --- | --- | --- | --- |
| water_15_2 | 5887 | FTMS; HCD | 90.61 | 574.73 |

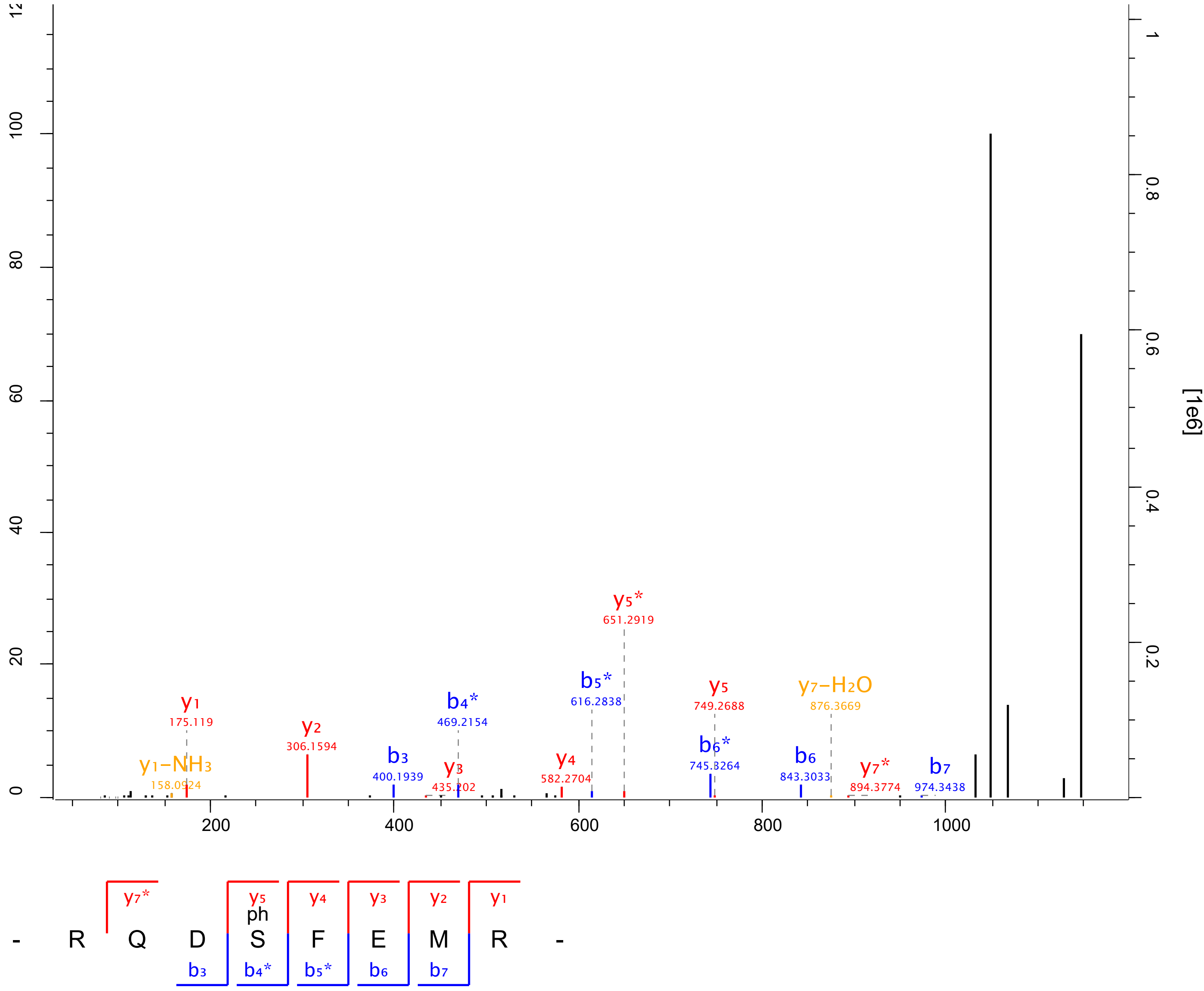

| Raw file | Scan | Method | Score | m/z |
|---|---|---|---|---|
| water_15_2 | 5950 | FTMS; HCD | 129.82 | 495.21 |

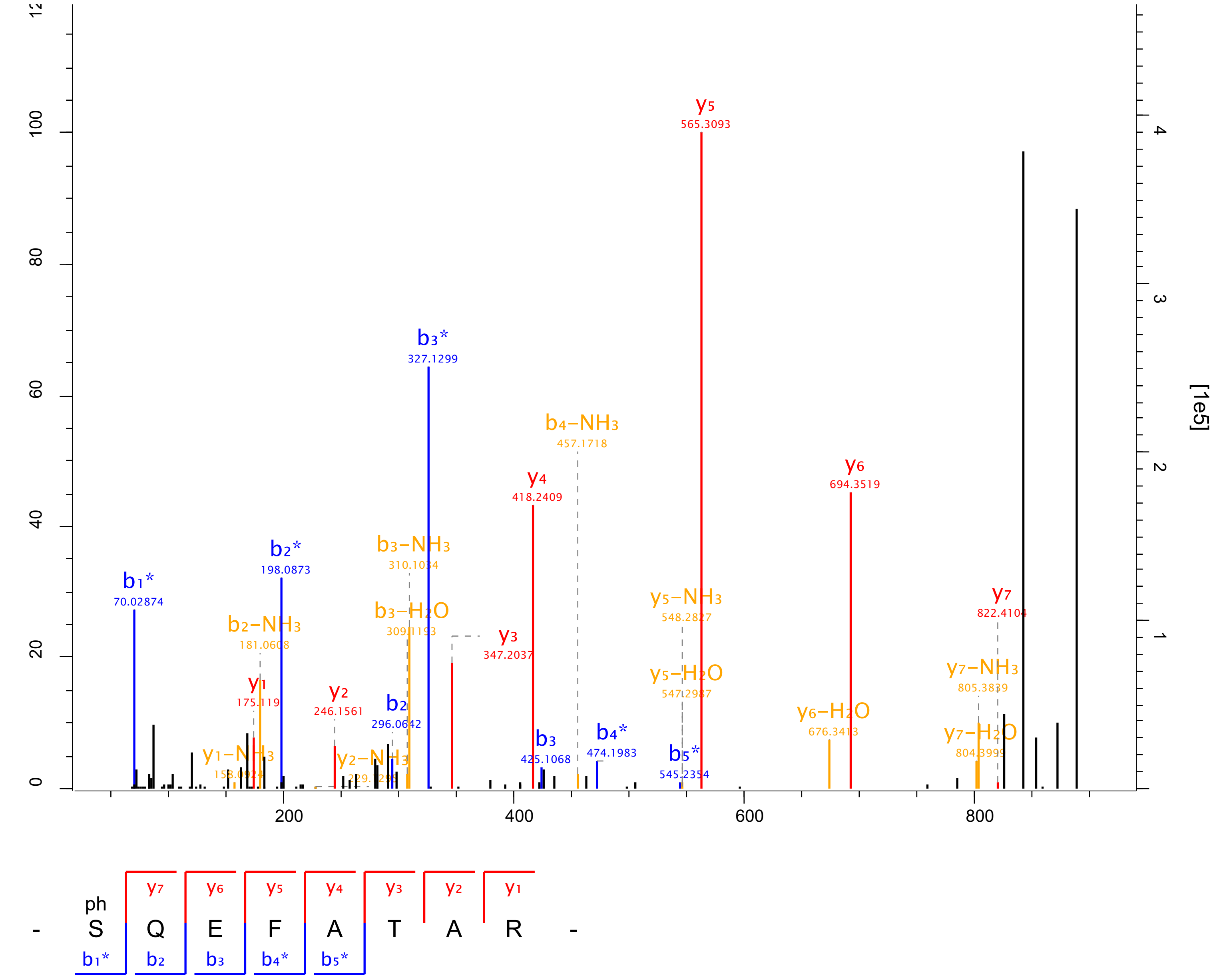

| Raw file | Scan | Method | Score | m/z |
|---|---|---|---|---|
| water_15_2 | 5978 | FTMS; HCD | 140.14 | 539.75 |

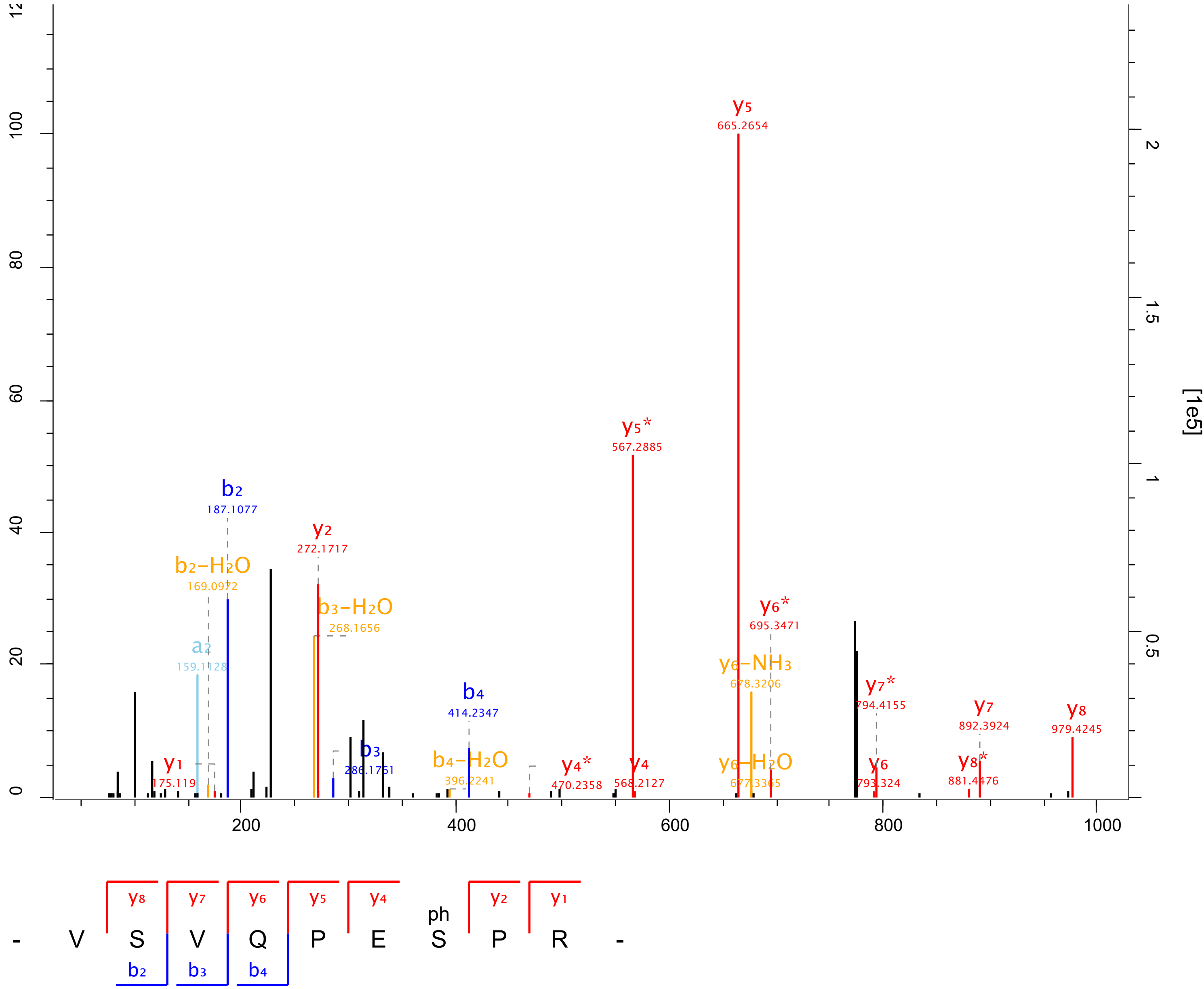

| Raw file | Scan | Method | Score | m/z |
| --- | --- | --- | --- | --- |
| water_15_2 | 5979 | FTMS; HCD | 103.9 | 651.95 |

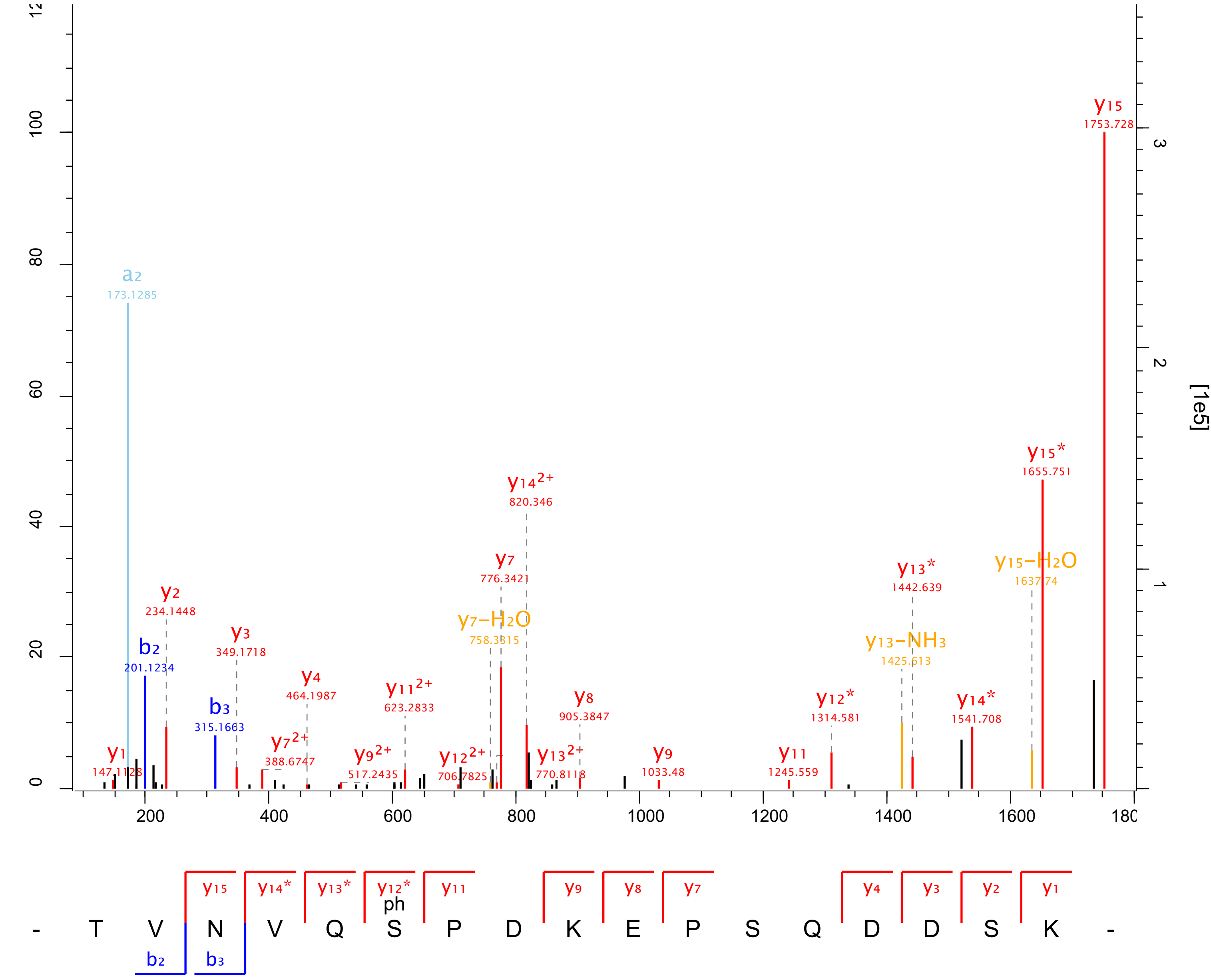

| Raw file | Scan | Method | Score | m/z |
| --- | --- | --- | --- | --- |
| water_15_2 | 5983 | FTMS; HCD | 161.49 | 548.28 |

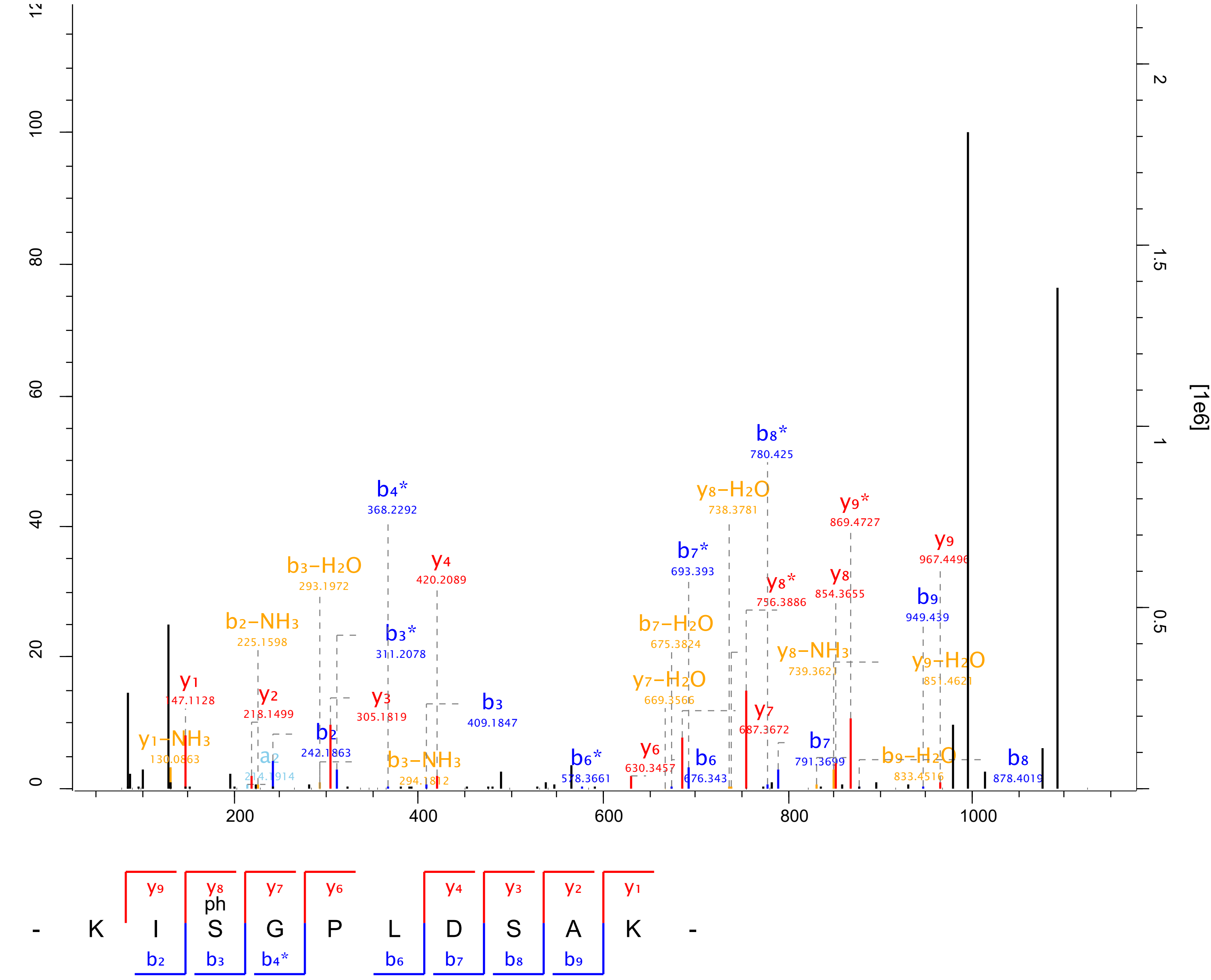

| Raw file | Scan | Method | Score | m/z |
|---|---|---|---|---|
| water_15_2 | 5997 | FTMS; HCD | 66.39 | 602.77 |

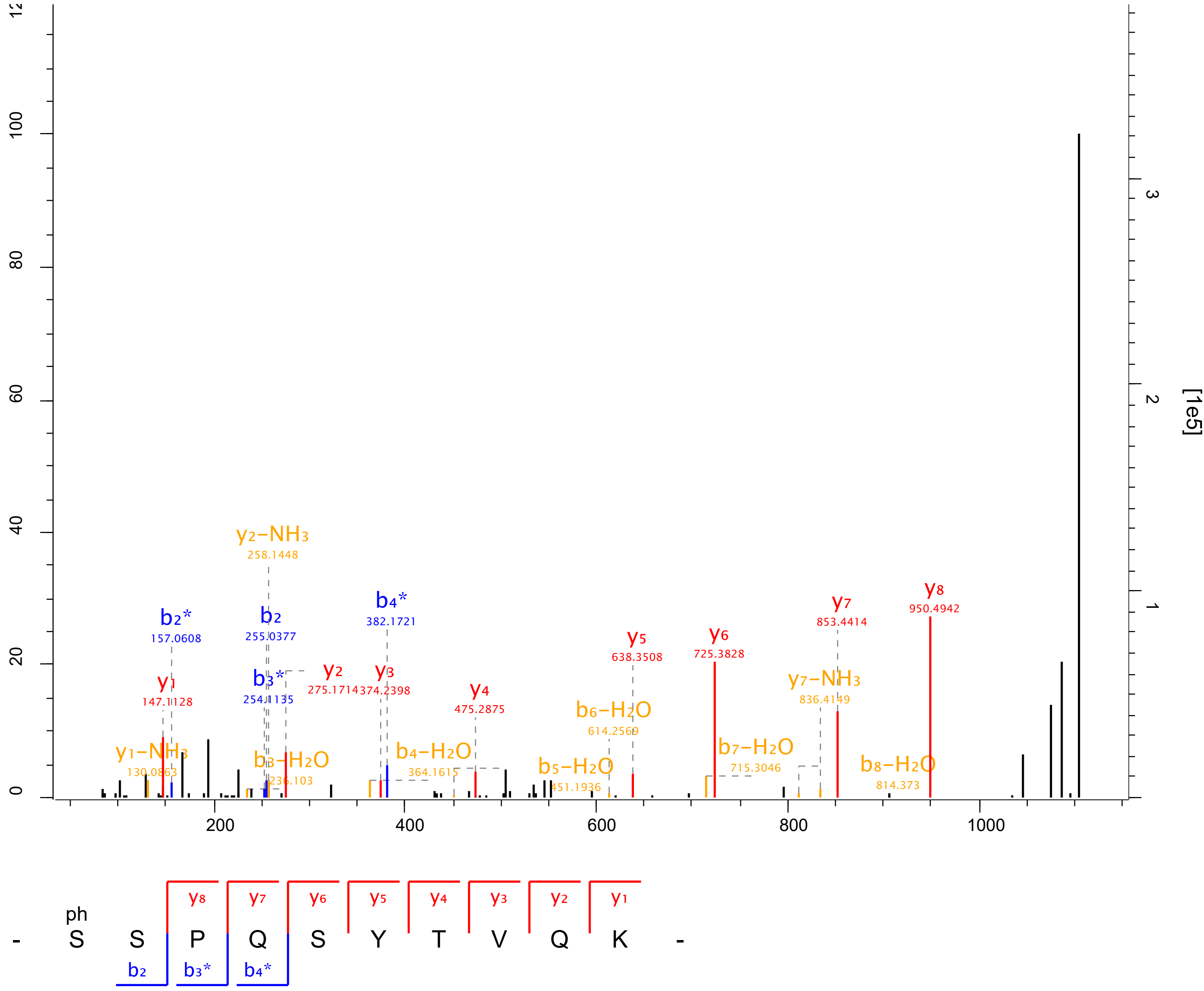

| Raw file | Scan | Method | Score | m/z |
|---|---|---|---|---|
| water_15_2 | 6009 | FTMS; HCD | 156.13 | 777.32 |

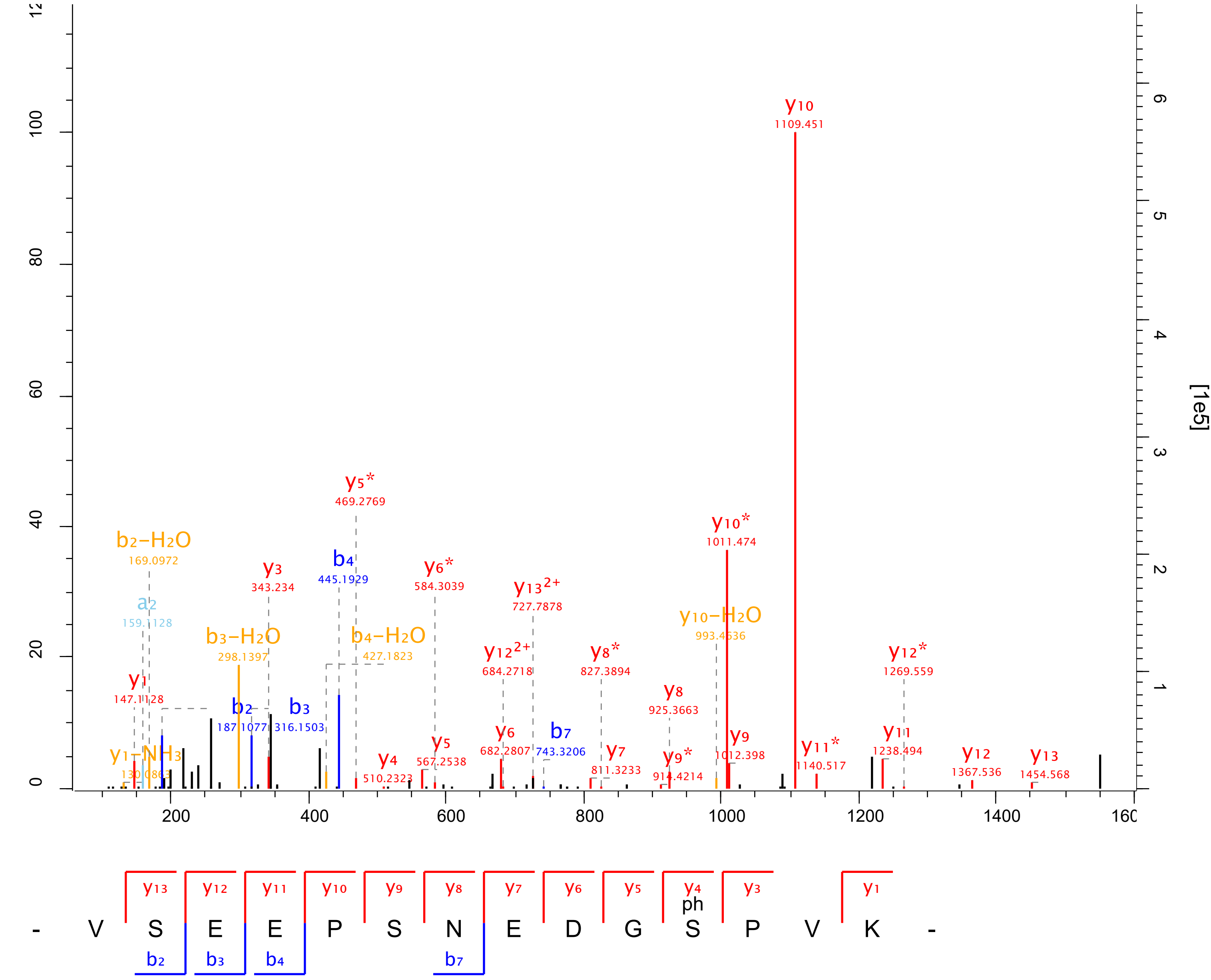

| Raw file | Scan | Method | Score | m/z |
| --- | --- | --- | --- | --- |
| water_15_2 | 6049 | FTMS; HCD | 54.9 | 629.76 |

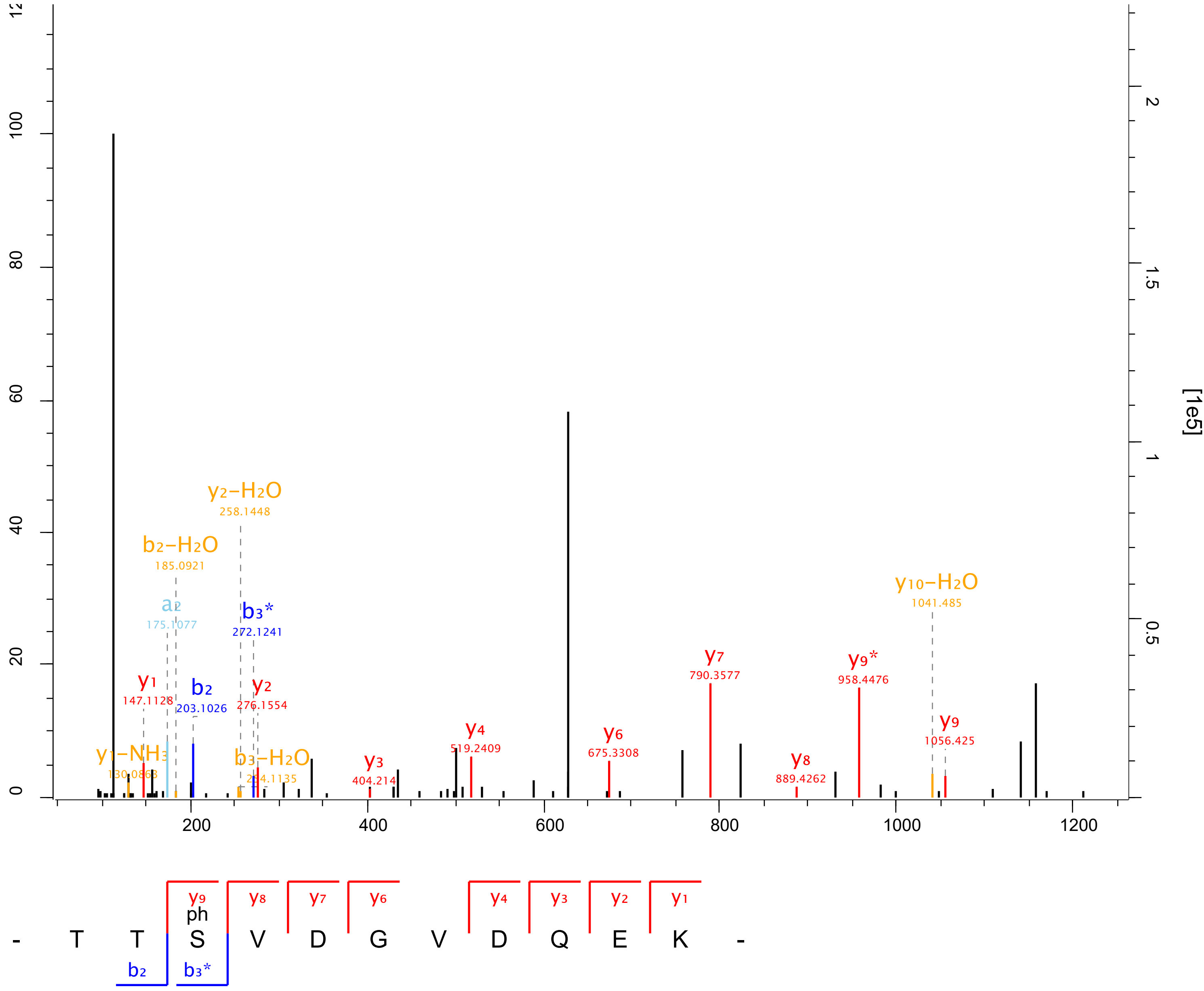

| Raw file | Scan | Method | Score | m/z |
|---|---|---|---|---|
| water_15_2 | 6062 | FTMS; HCD | 55.35 | 495.21 |

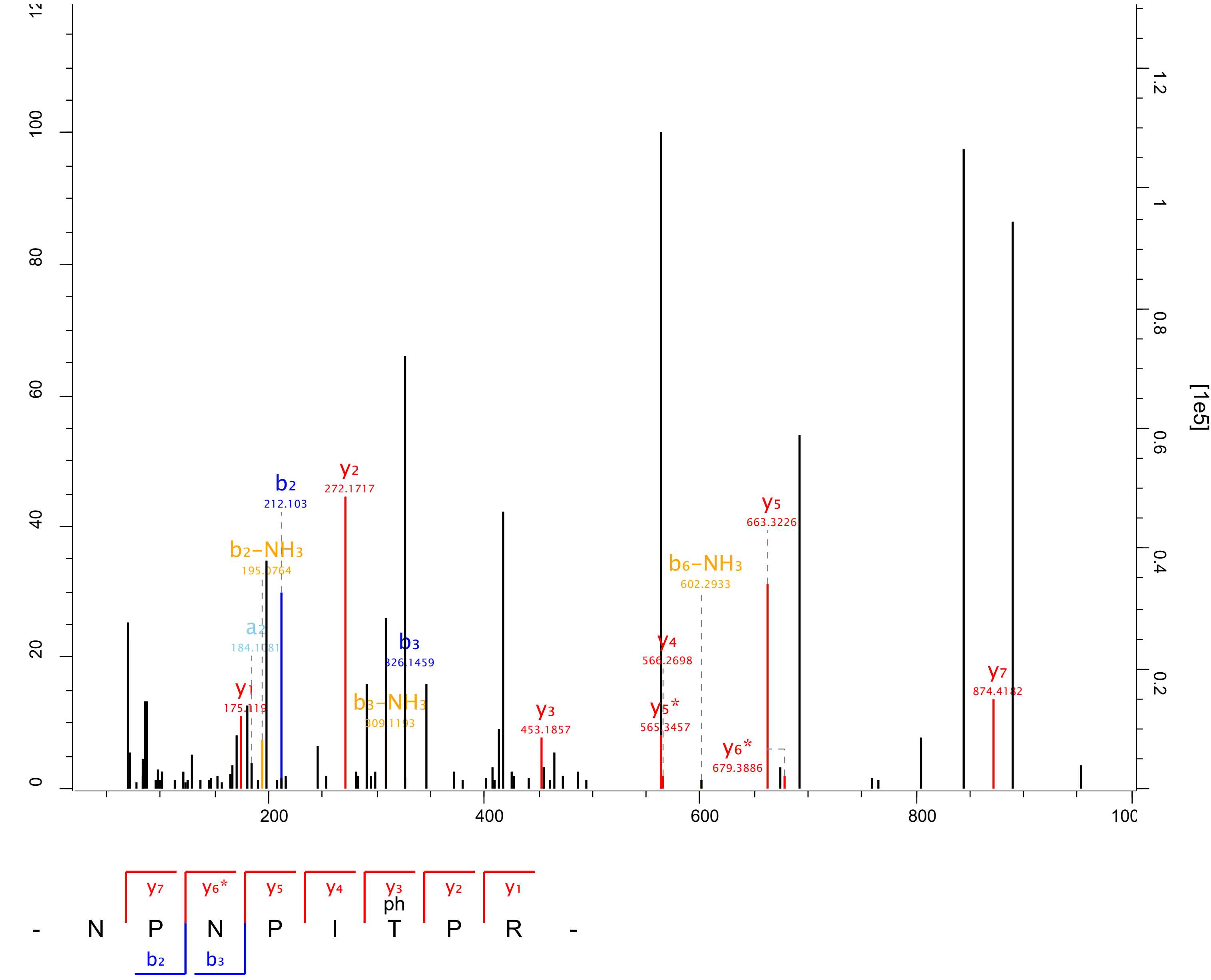

| Raw file | Scan | Method | Score | m/z |
| --- | --- | --- | --- | --- |
| water_15_2 | 6137 | FTMS; HCD | 49.71 | 762.35 |

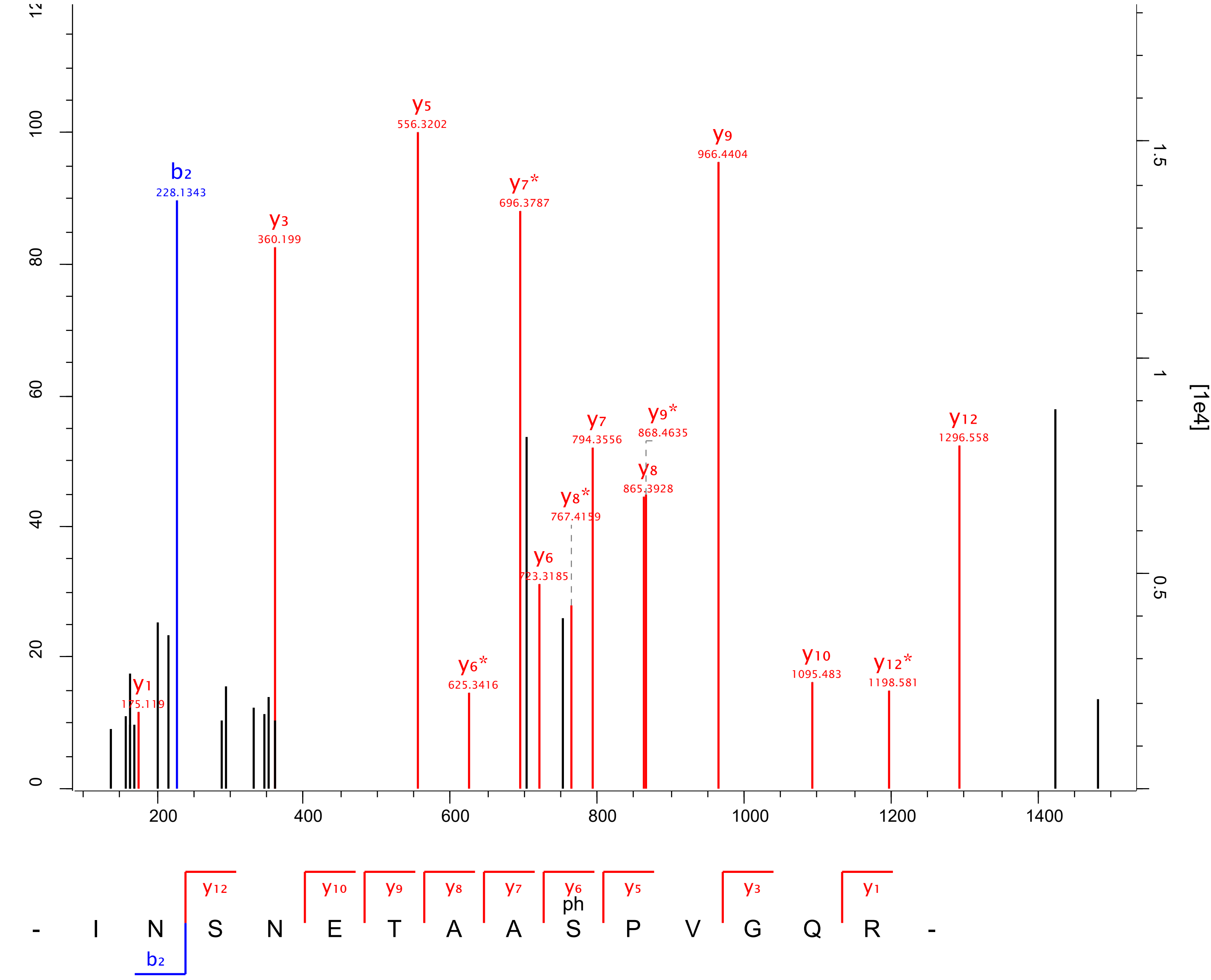

| Raw file | Scan | Method | Score | m/z |
|---|---|---|---|---|
| water_15_2 | 6231 | FTMS; HCD | 168.1 | 588.26 |

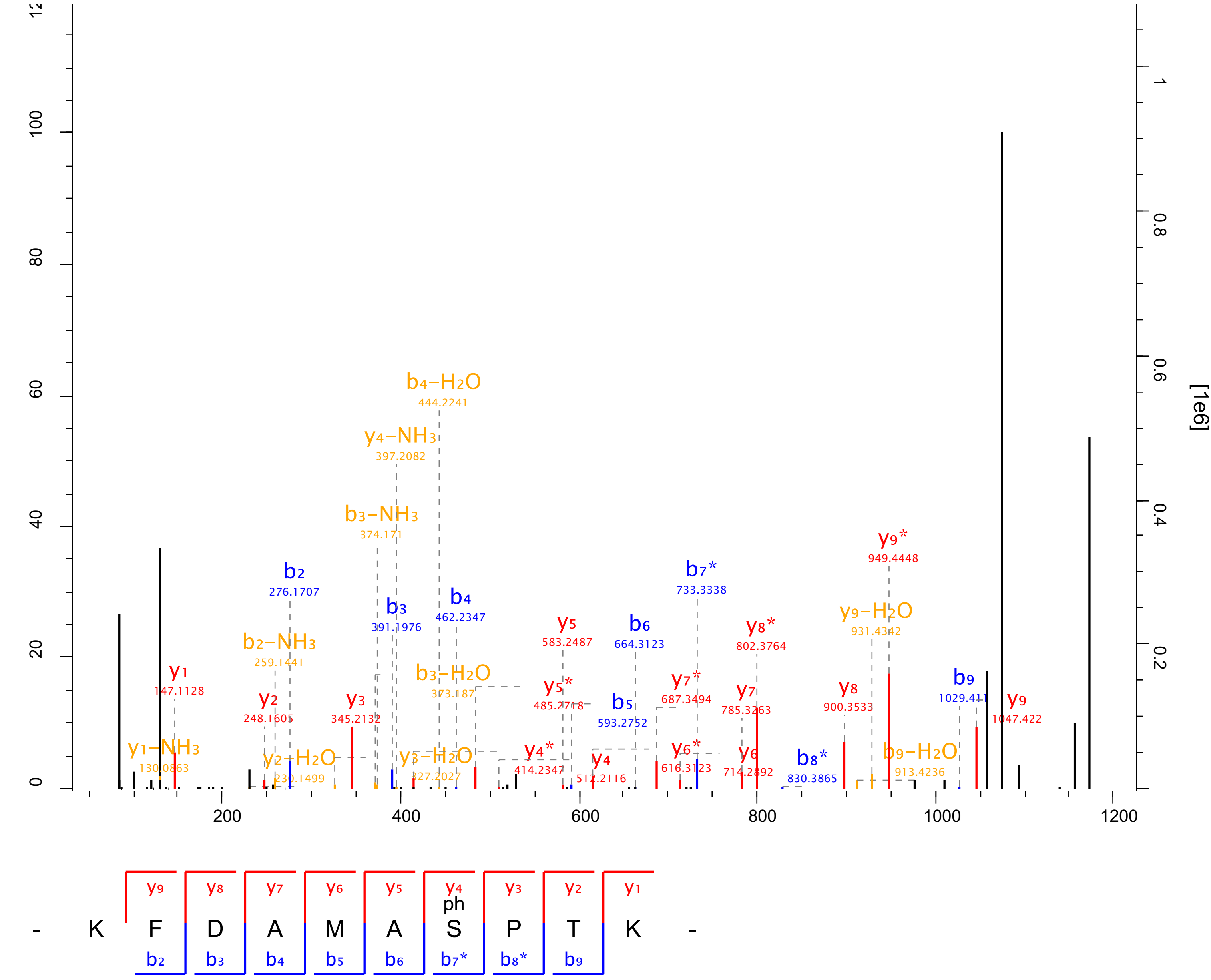

| Raw file | Scan | Method | Score | m/z |
|---|---|---|---|---|
| water_15_2 | 6232 | FTMS; HCD | 98.16 | 546.24 |

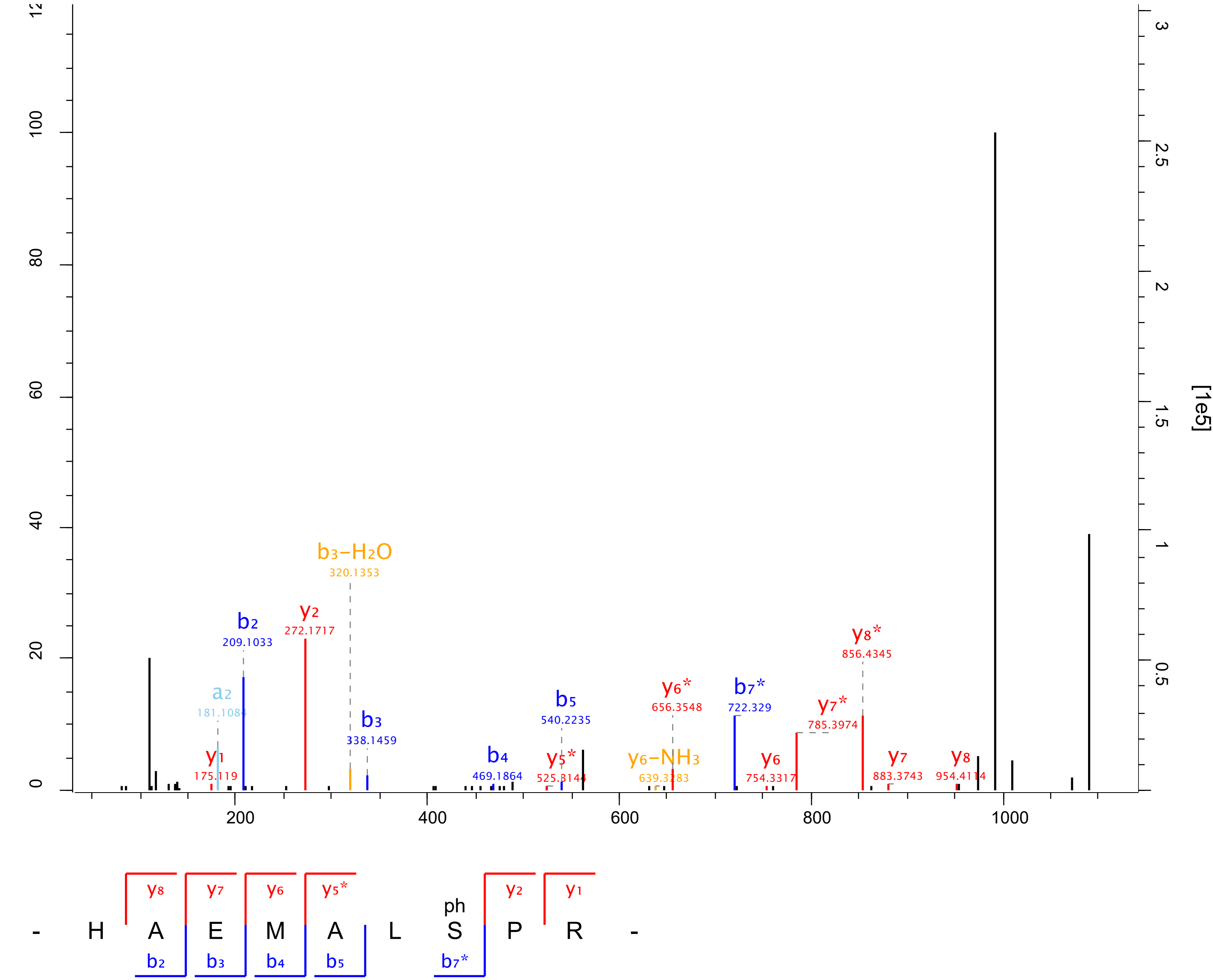

| Raw file | Scan | Method | Score | m/z |
| --- | --- | --- | --- | --- |
| water_15_2 | 6321 | FTMS; HCD | 98.59 | 608.24 |

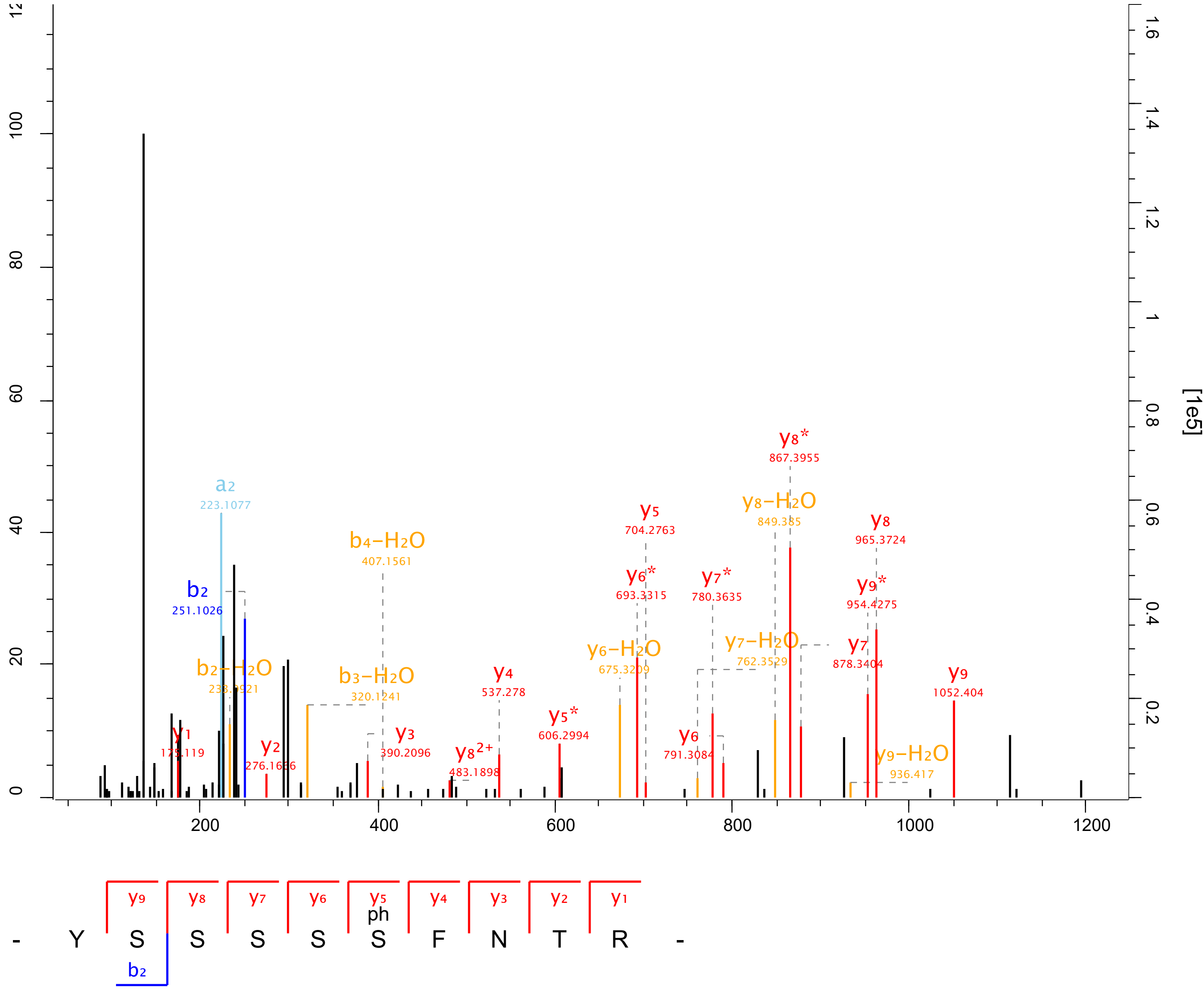

| Raw file | Scan | Method | Score | m/z |
|---|---|---|---|---|
| water_15_2 | 6388 | FTMS; HCD | 59.04 | 448.71 |

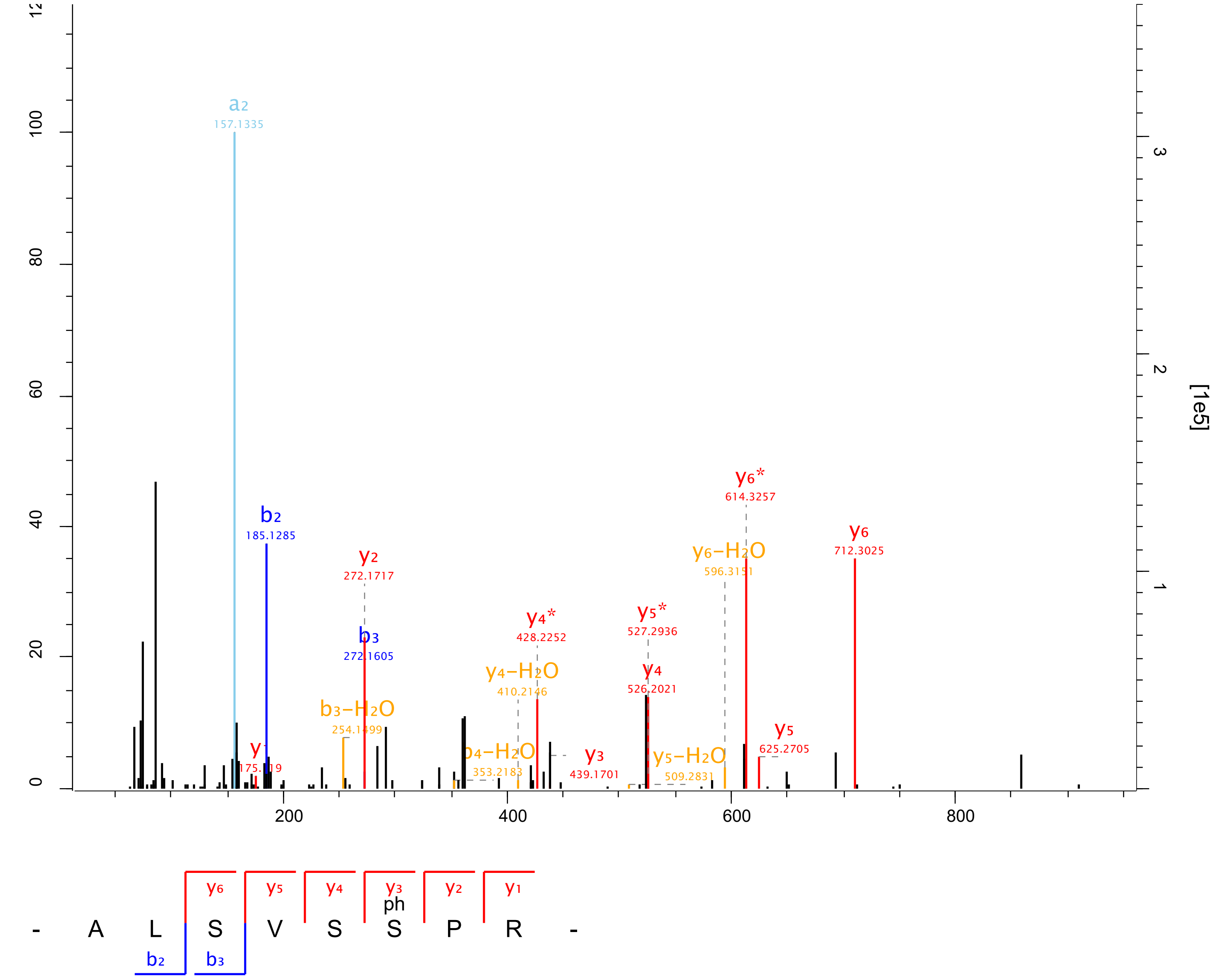

| Raw file | Scan | Method | Score | m/z |
| --- | --- | --- | --- | --- |
| water_15_2 | 6410 | FTMS; HCD | 52.58 | 609.78 |

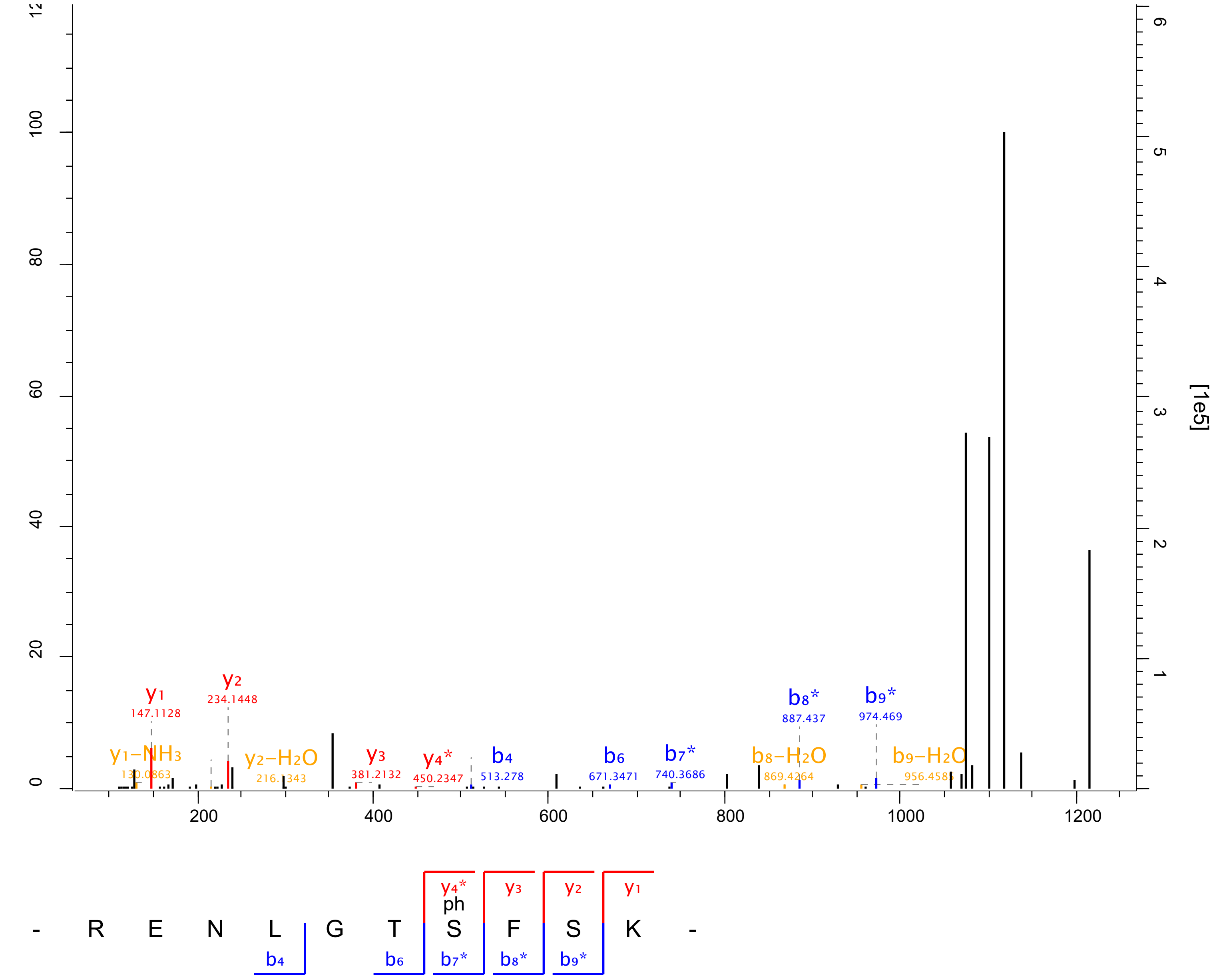

| Raw file | Scan | Method | Score | m/z |
|---|---|---|---|---|
| water_15_2 | 6426 | FTMS; HCD | 134.76 | 671.8 |

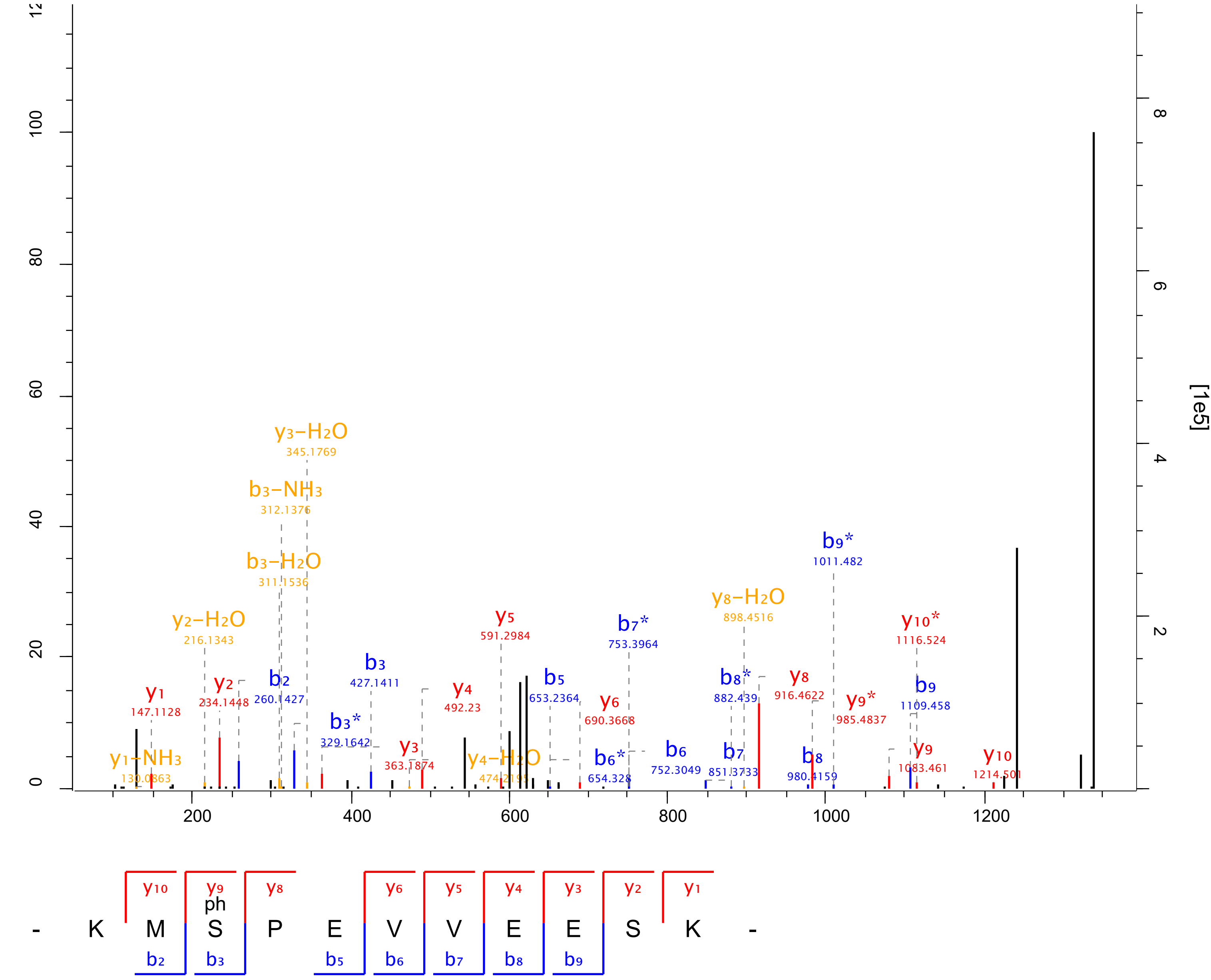

| Raw file | Scan | Method | Score | m/z |
| --- | --- | --- | --- | --- |
| water_15_2 | 6459 | FTMS; HCD | 45.61 | 553.24 |

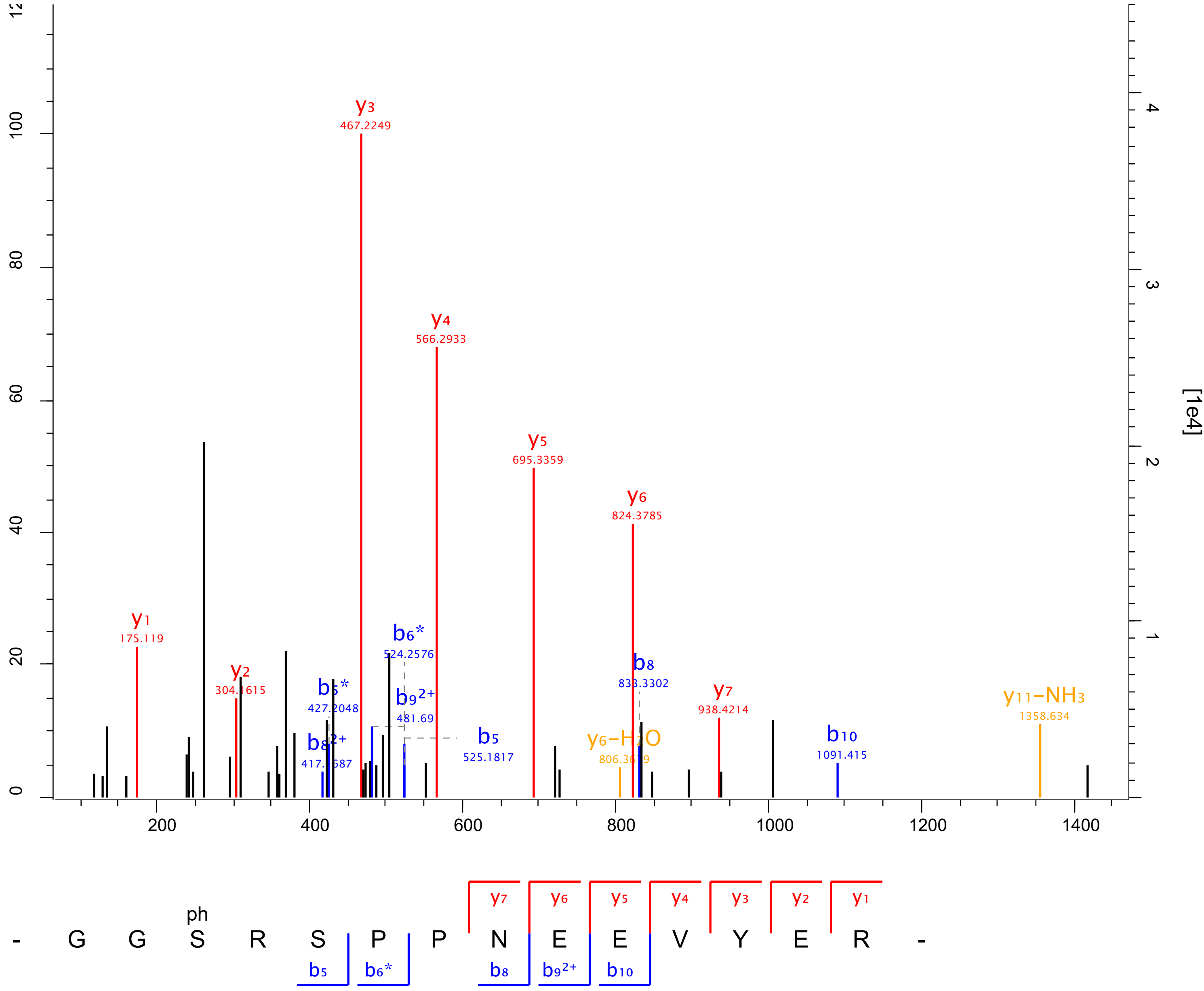

| Raw file | Scan | Method | Score | m/z |
|---|---|---|---|---|
| water_15_2 | 6496 | FTMS; HCD | 47.71 | 509.23 |

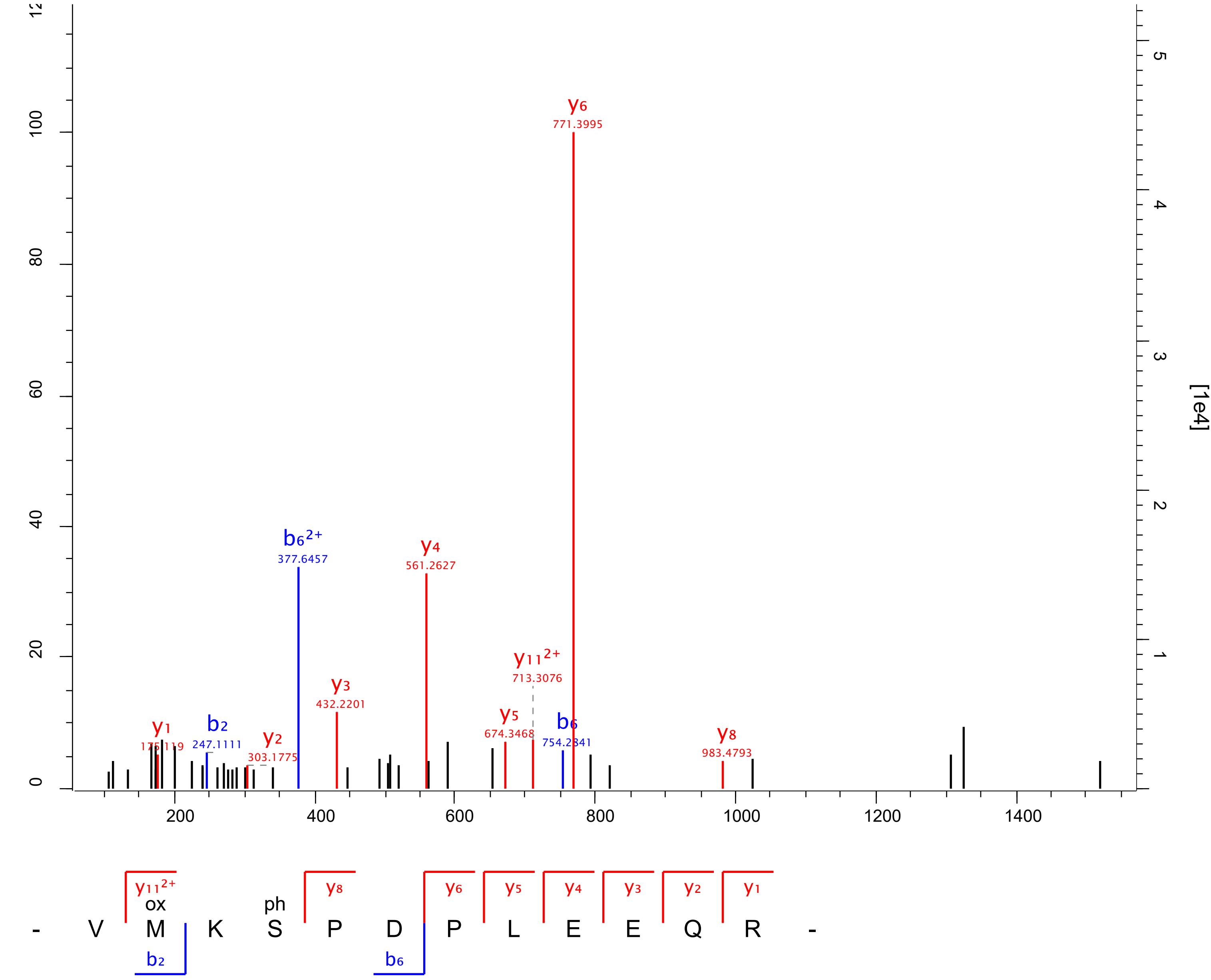

| Raw file | Scan | Method | Score | m/z |
| --- | --- | --- | --- | --- |
| water_15_2 | 6621 | FTMS; HCD | 40.49 | 508.59 |

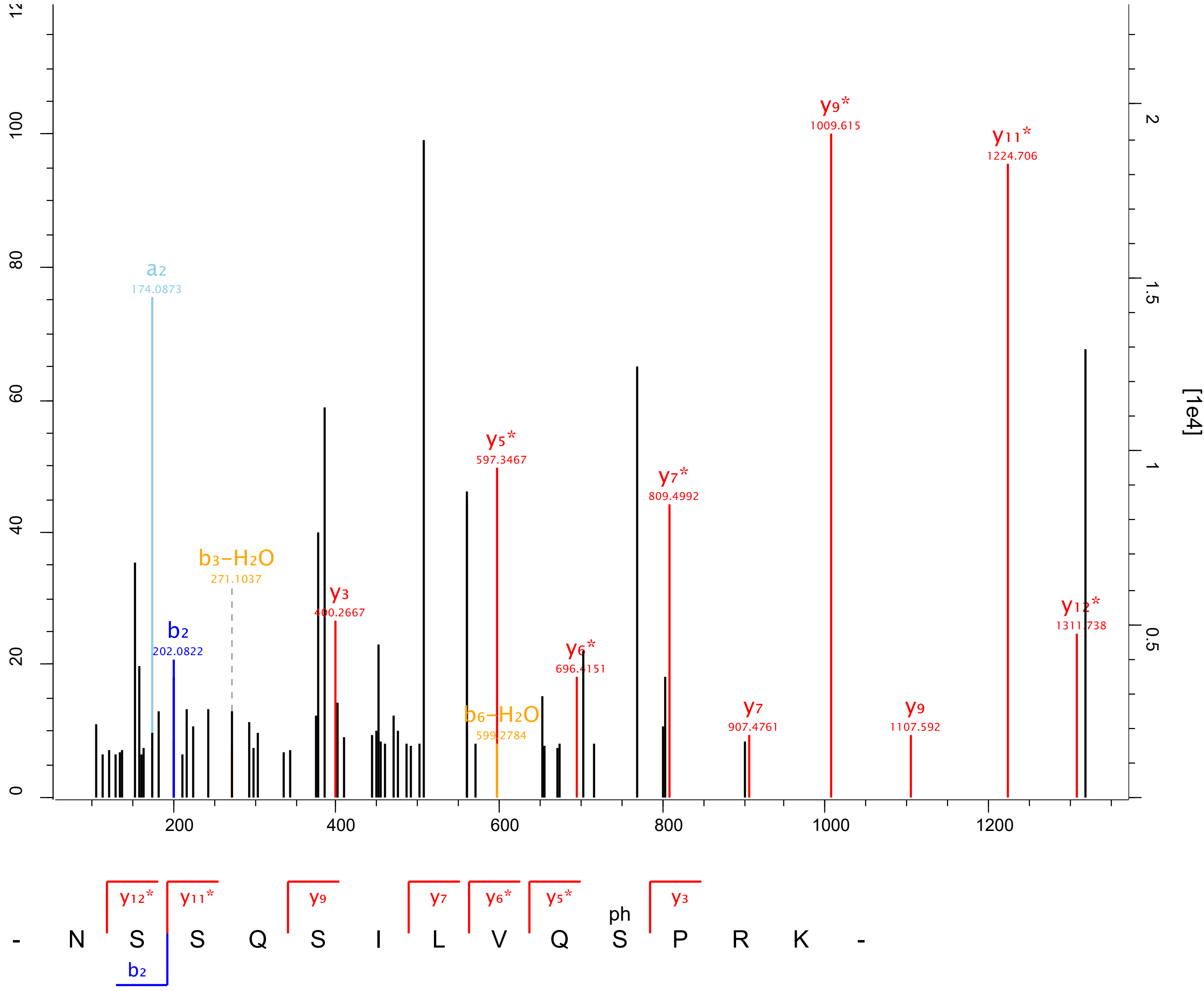

| Raw file | Scan | Method | Score | m/z |
| --- | --- | --- | --- | --- |
| water_15_2 | 6745 | FTMS; HCD | 76.17 | 419.2 |

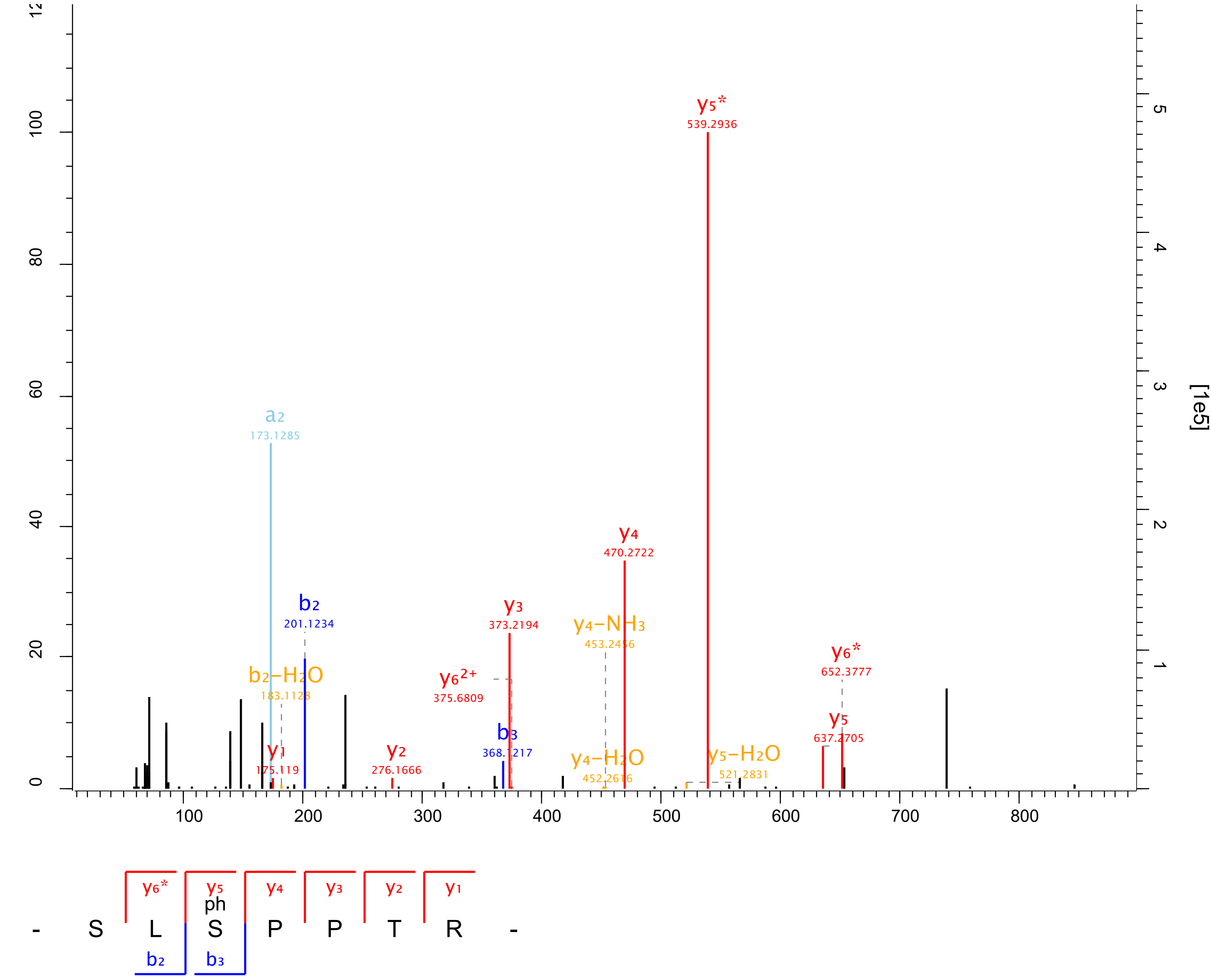

| Raw file | Scan | Method | Score | m/z |
|---|---|---|---|---|
| water_15_2 | 6811 | FTMS; HCD | 74.56 | 616.62 |

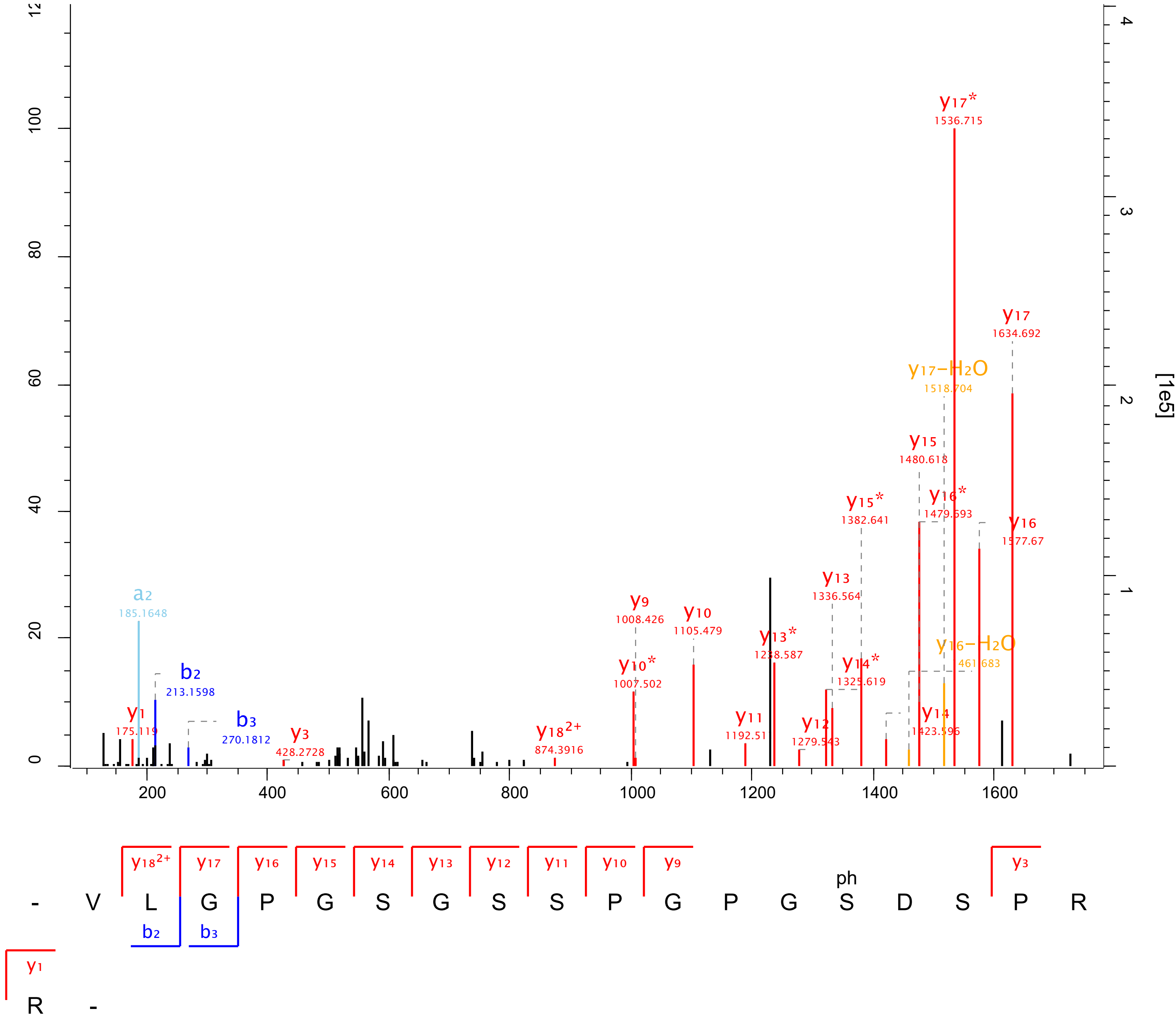

| Raw file | Scan | Method | Score | m/z |
|---|---|---|---|---|
| water_15_2 | 6828 | FTMS; HCD | 335.48 | 827.86 |

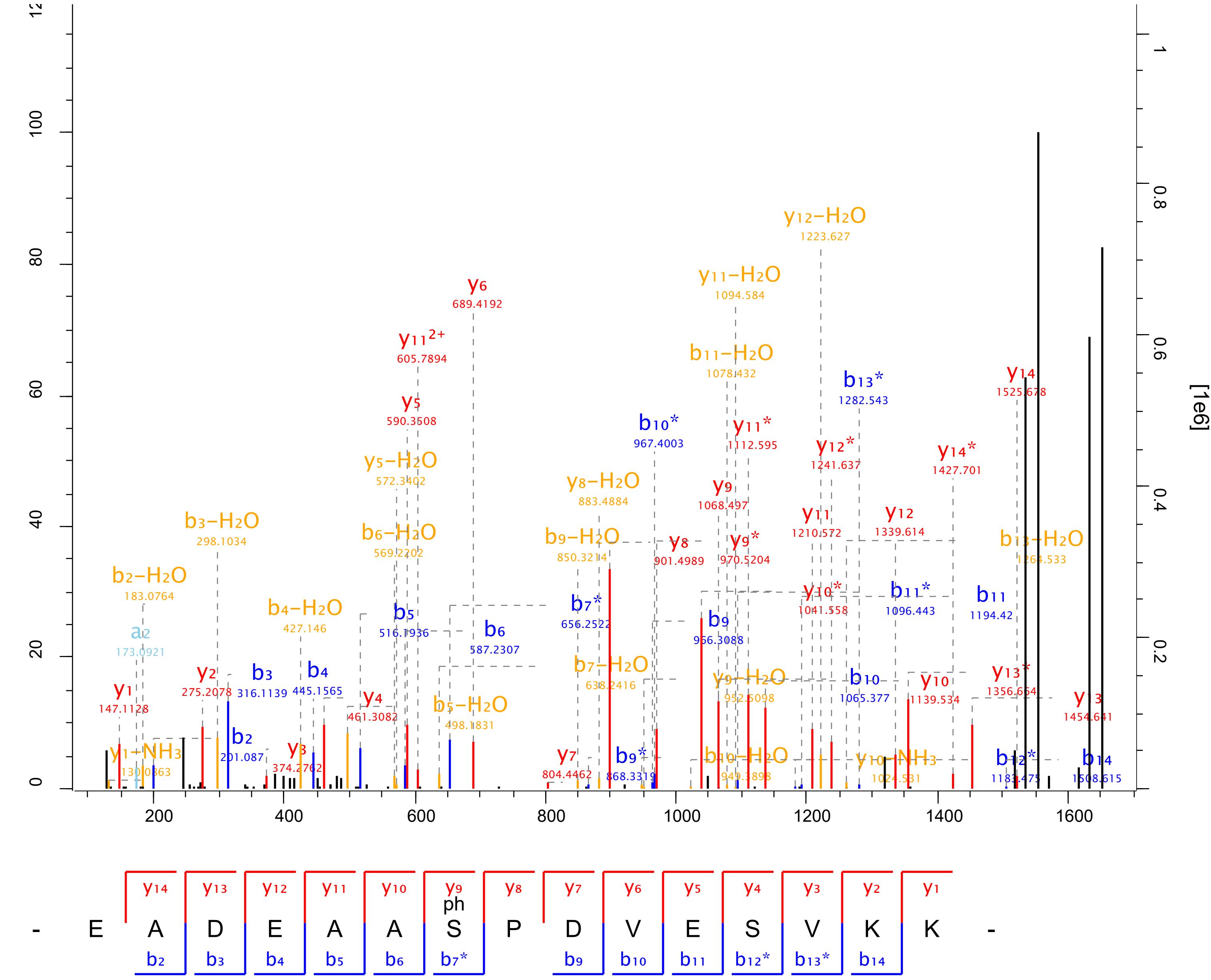

| Raw file | Scan | Method | Score | m/z |
|---|---|---|---|---|
| water_15_2 | 6862 | FTMS; HCD | 55.85 | 756.32 |

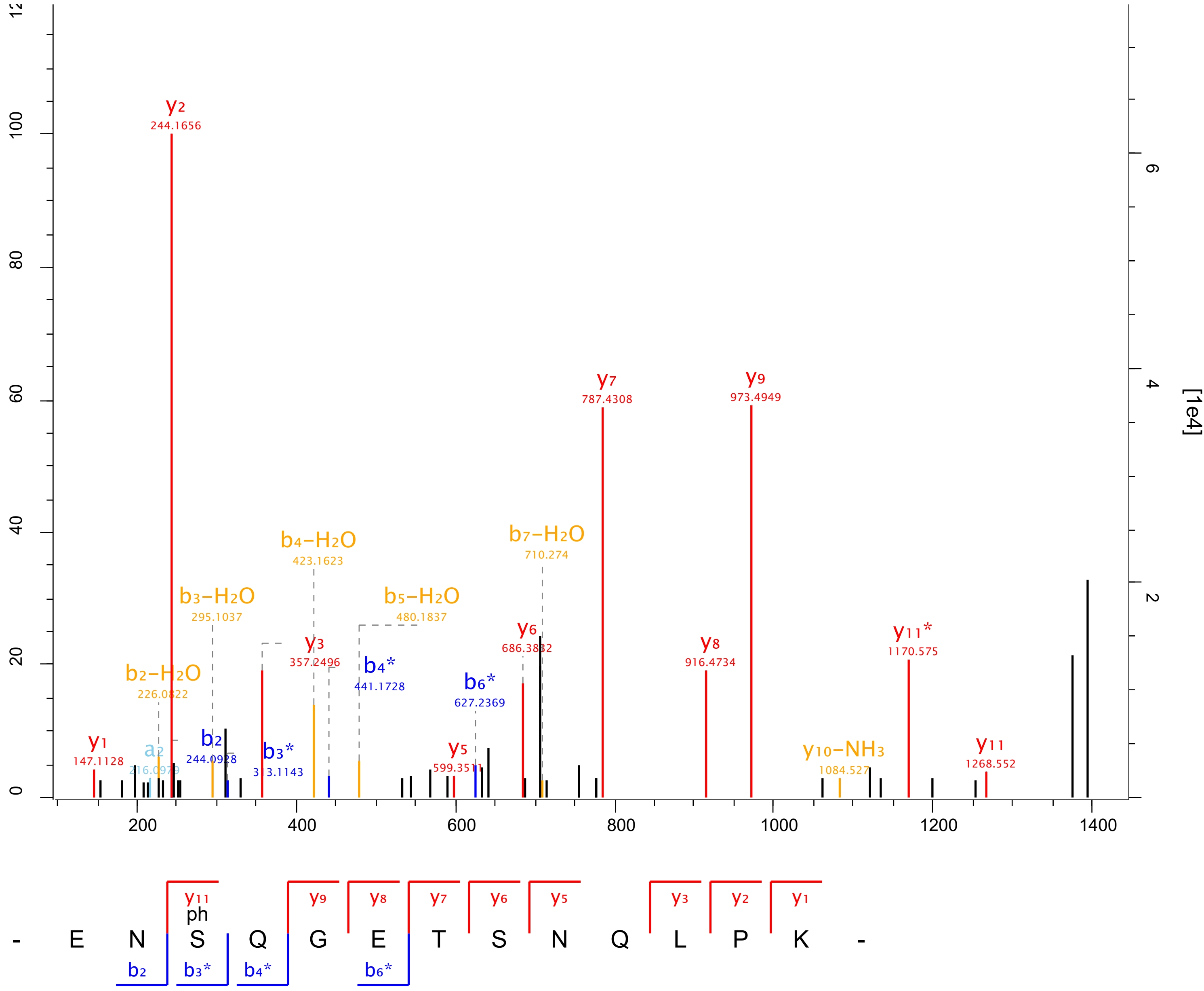

| Raw file | Scan | Method | Score | m/z |
| --- | --- | --- | --- | --- |
| water_15_2 | 6880 | FTMS; HCD | 50.26 | 613.94 |

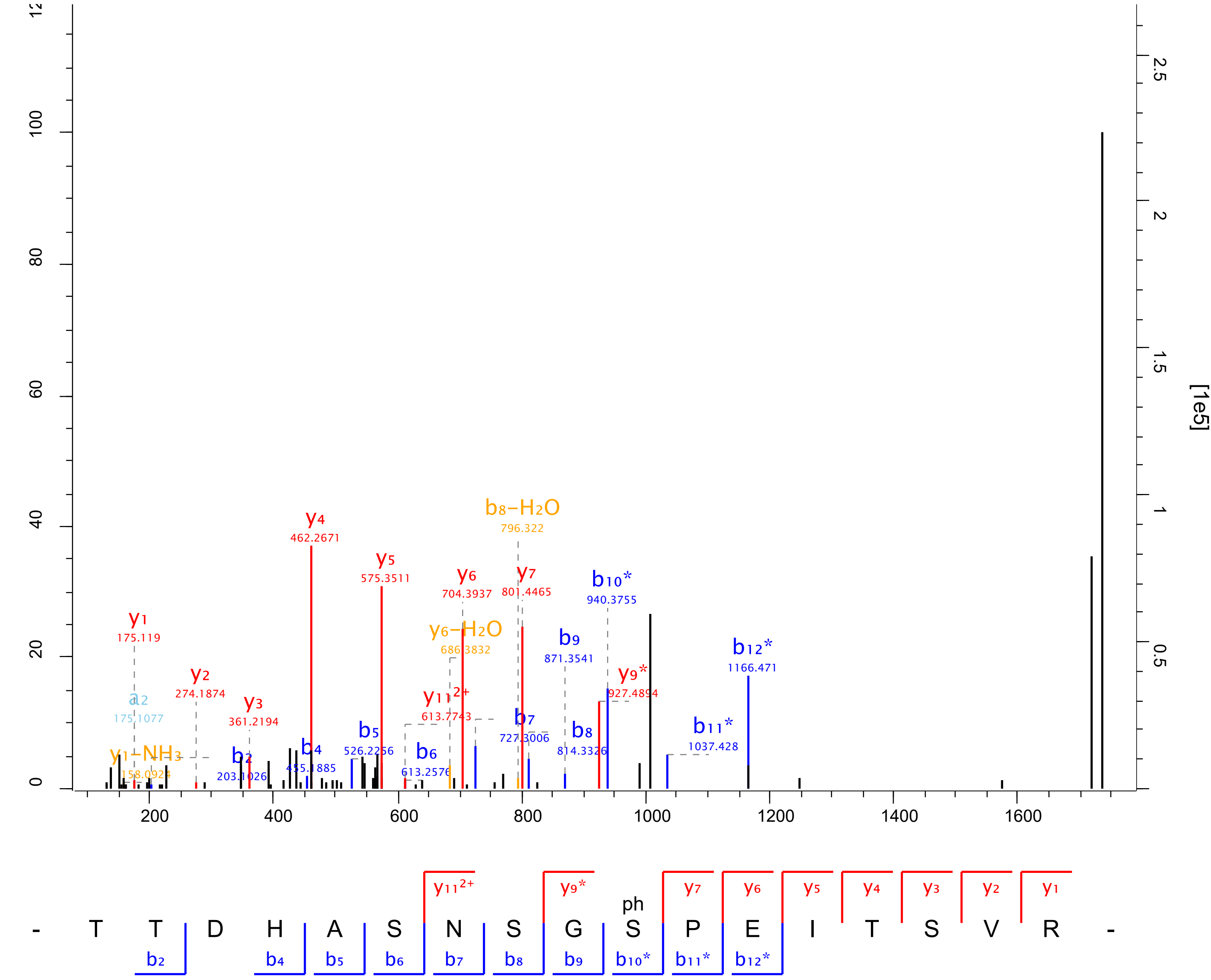

| Raw file | Scan | Method | Score | m/z |
| --- | --- | --- | --- | --- |
| water_15_2 | 6904 | FTMS; HCD | 48.53 | 648.81 |

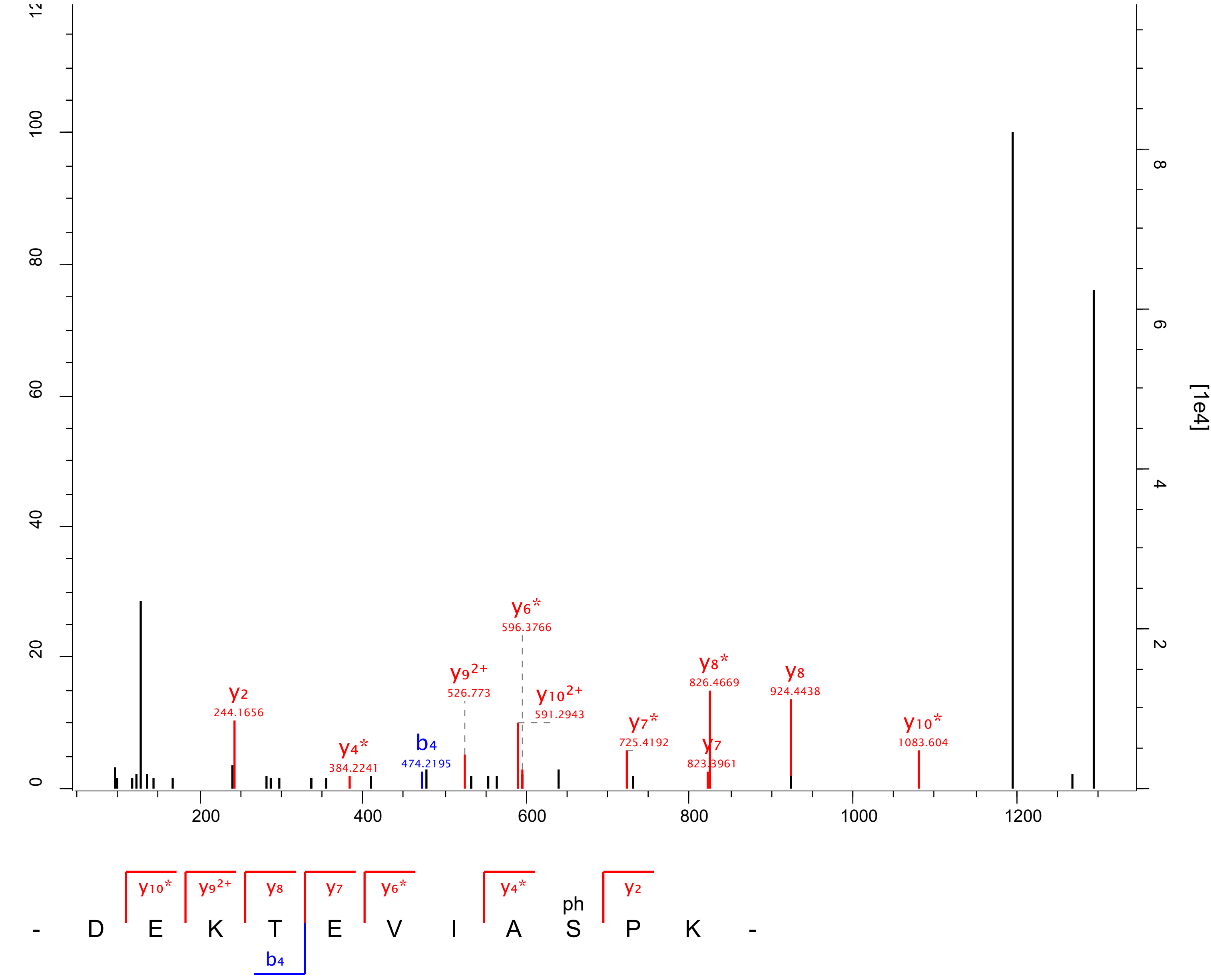

| Raw file | Scan | Method | Score | m/z |
| --- | --- | --- | --- | --- |
| water_15_2 | 6978 | FTMS; HCD | 109 | 716.26 |

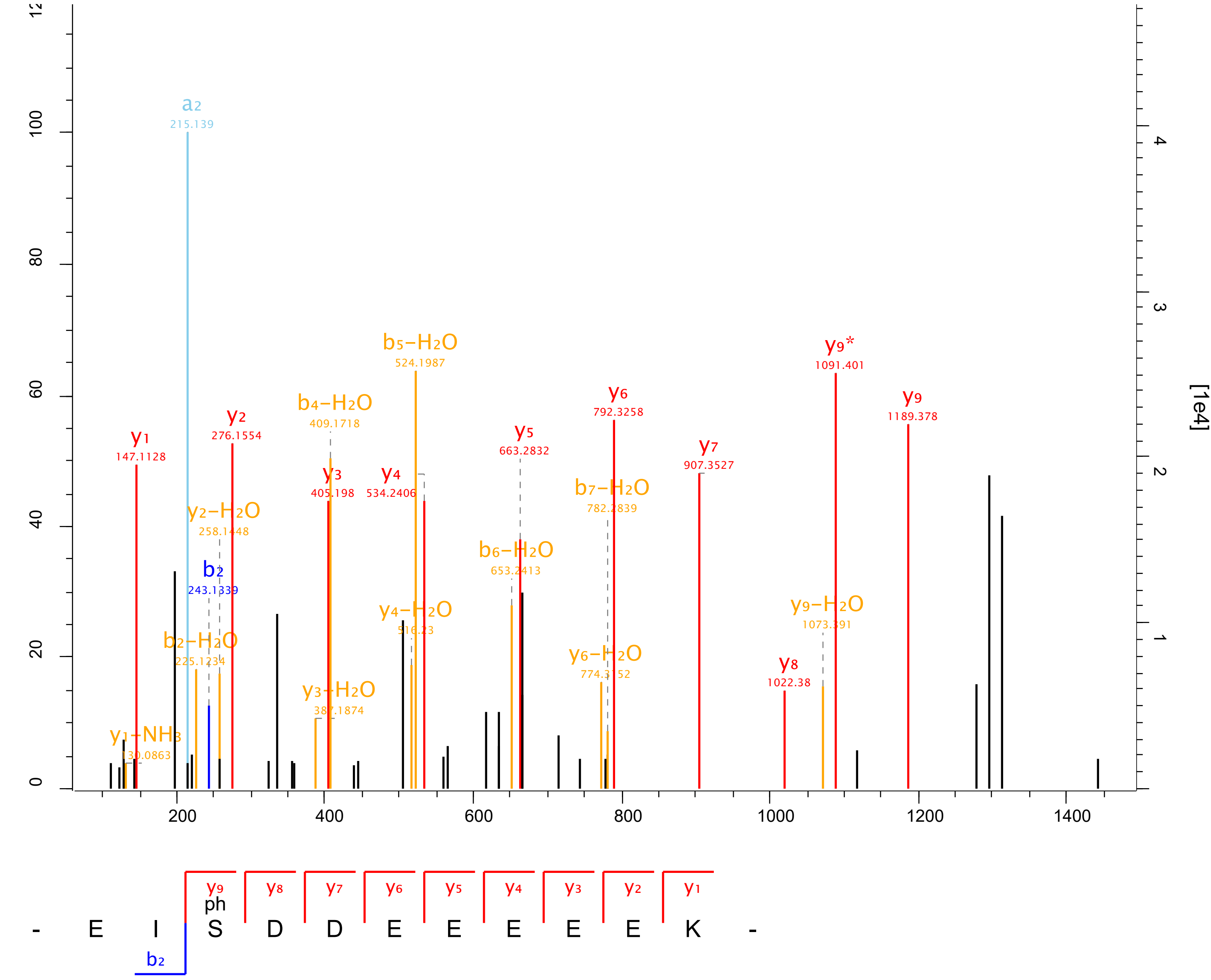

| Raw file | Scan | Method | Score | m/z |
|---|---|---|---|---|
| water_15_2 | 7067 | FTMS; HCD | 60.16 | 535.72 |

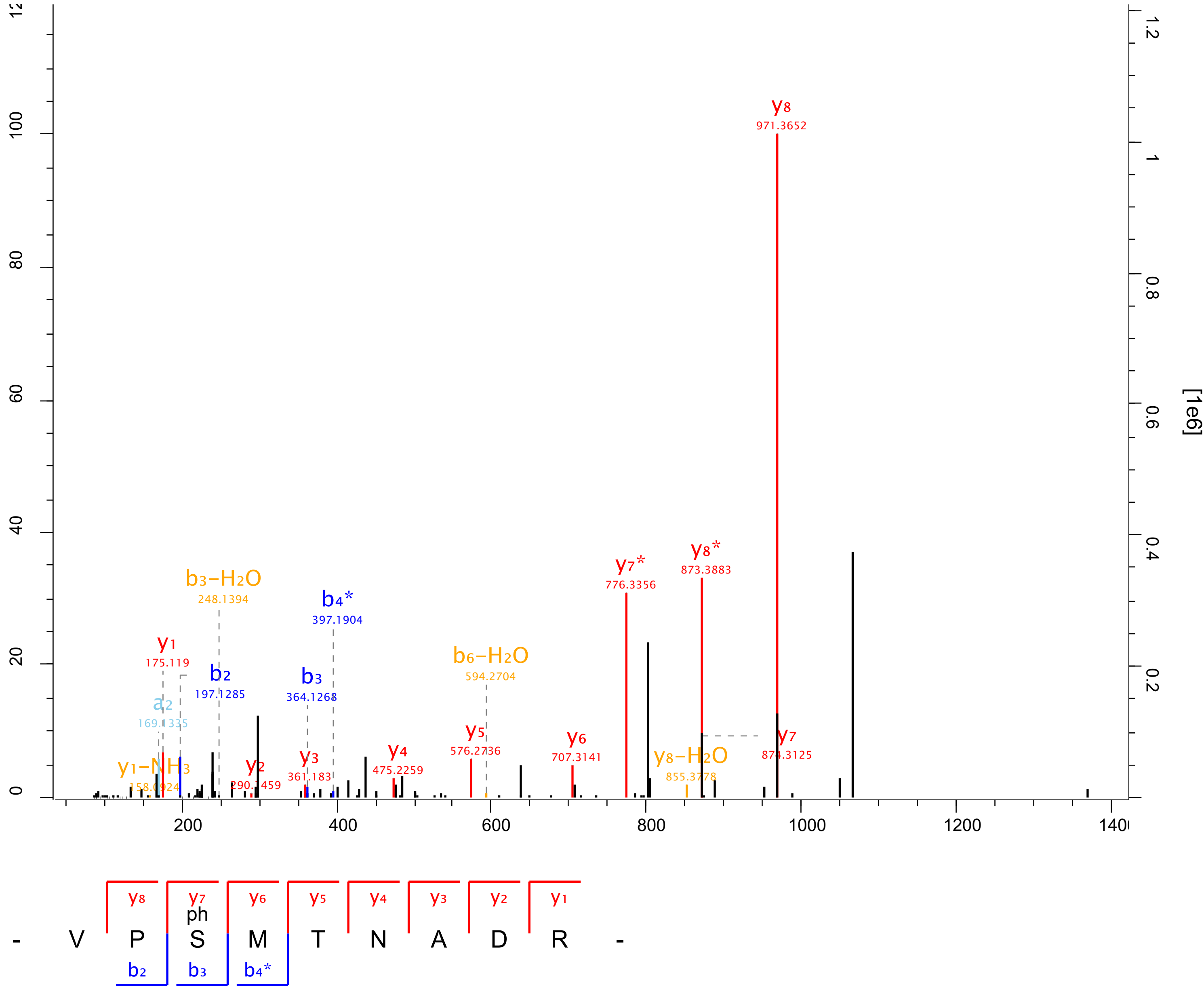

| Raw file | Scan | Method | Score | m/z |
|---|---|---|---|---|
| water_15_2 | 7195 | FTMS; HCD | 110.55 | 597.26 |

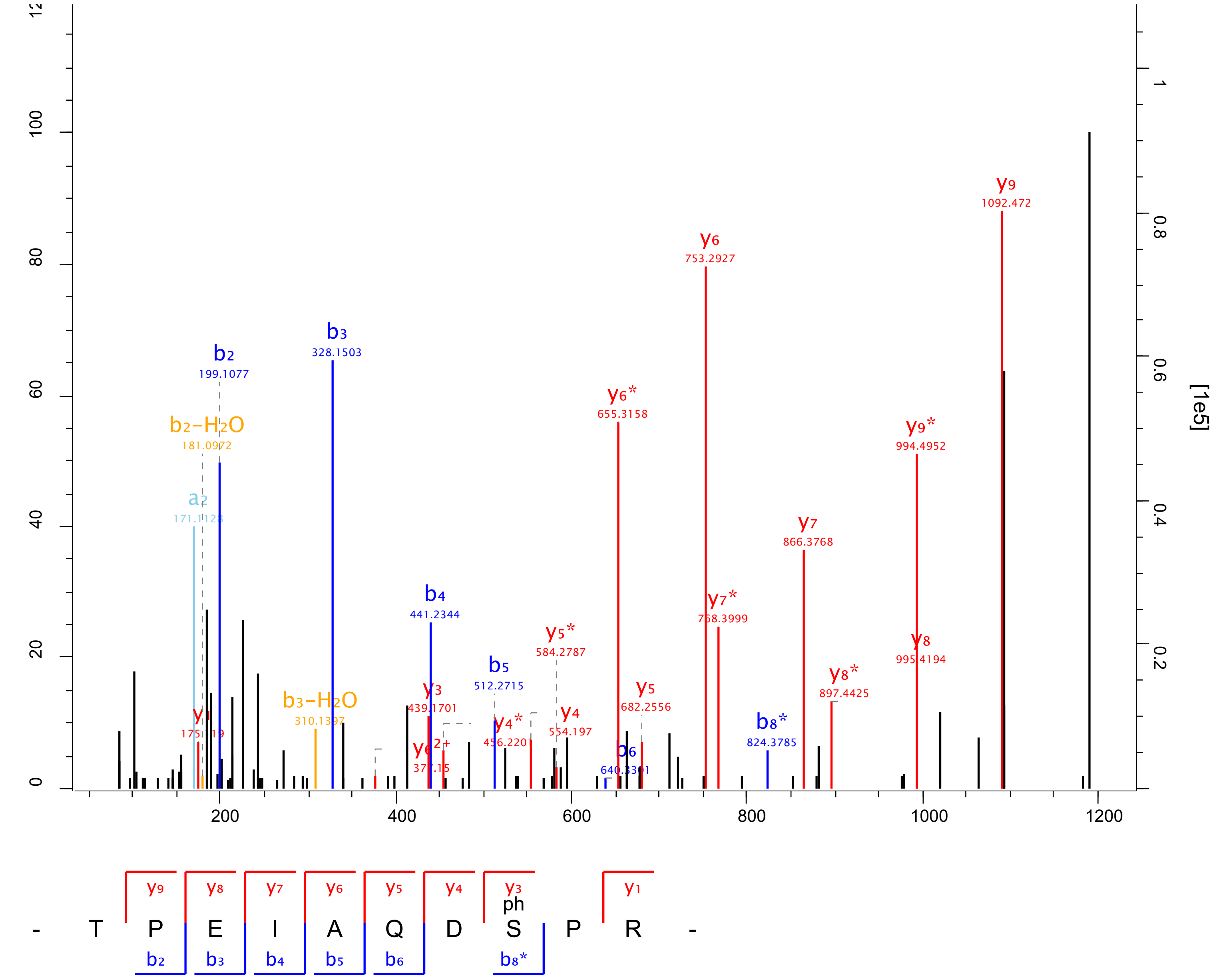

| Raw file | Scan | Method | Score | m/z |
| --- | --- | --- | --- | --- |
| water_15_2 | 7234 | FTMS; HCD | 64.22 | 634.28 |

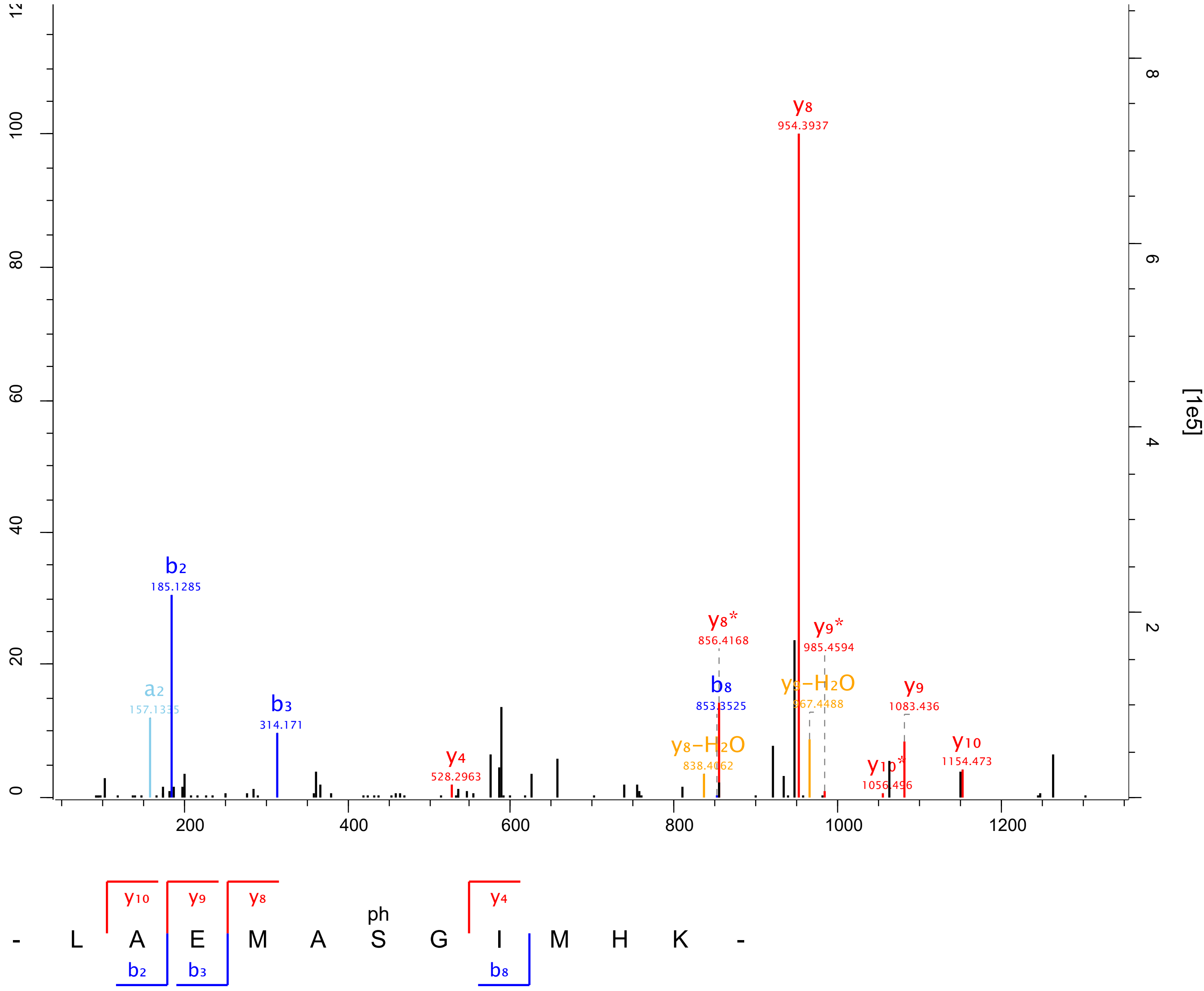

| Raw file | Scan | Method | Score | m/z |
|---|---|---|---|---|
| water_15_2 | 7237 | FTMS; HCD | 73.88 | 622.75 |

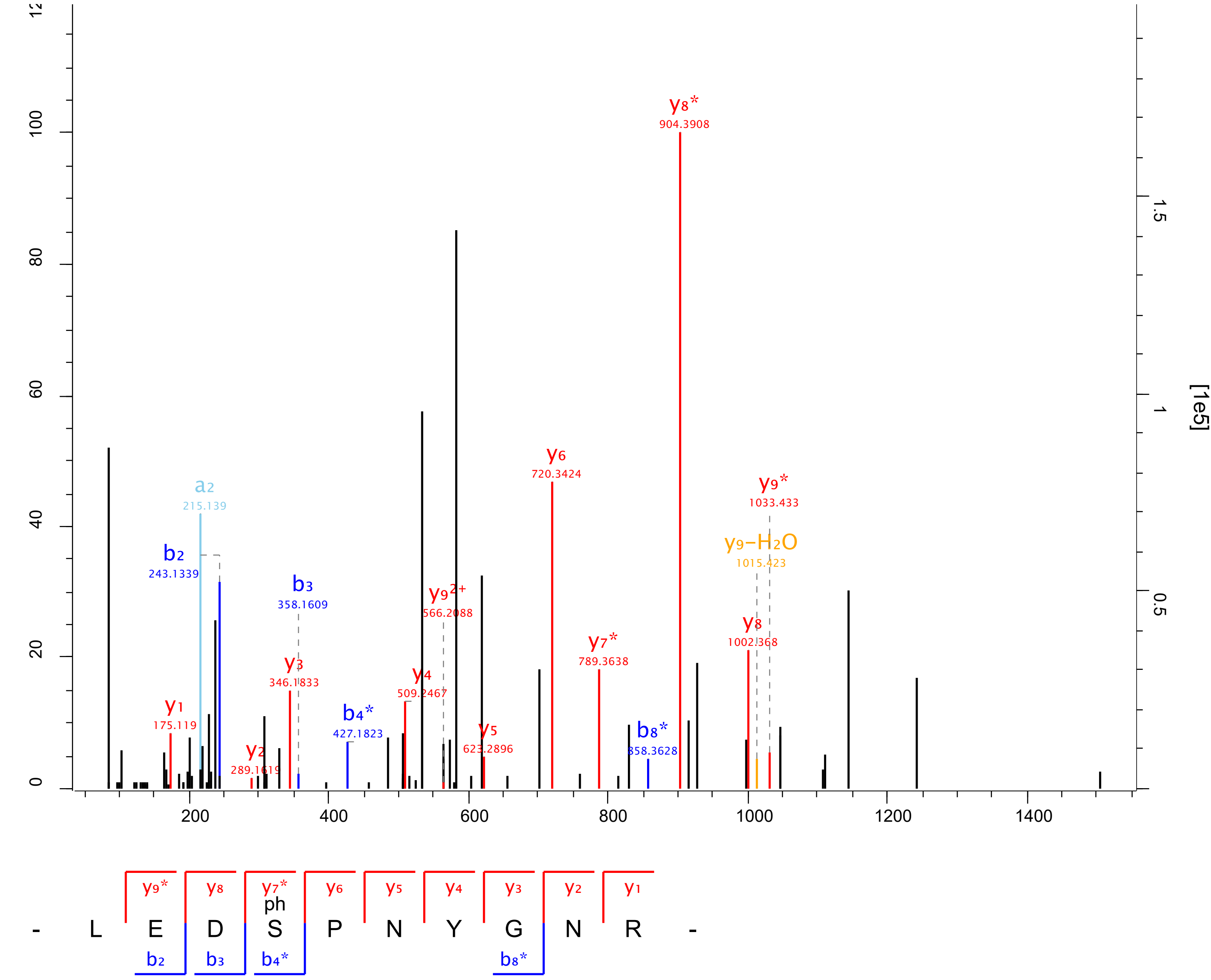

| Raw file | Scan | Method | Score | m/z |
| --- | --- | --- | --- | --- |
| water_15_2 | 7241 | FTMS; HCD | 66.27 | 530.74 |

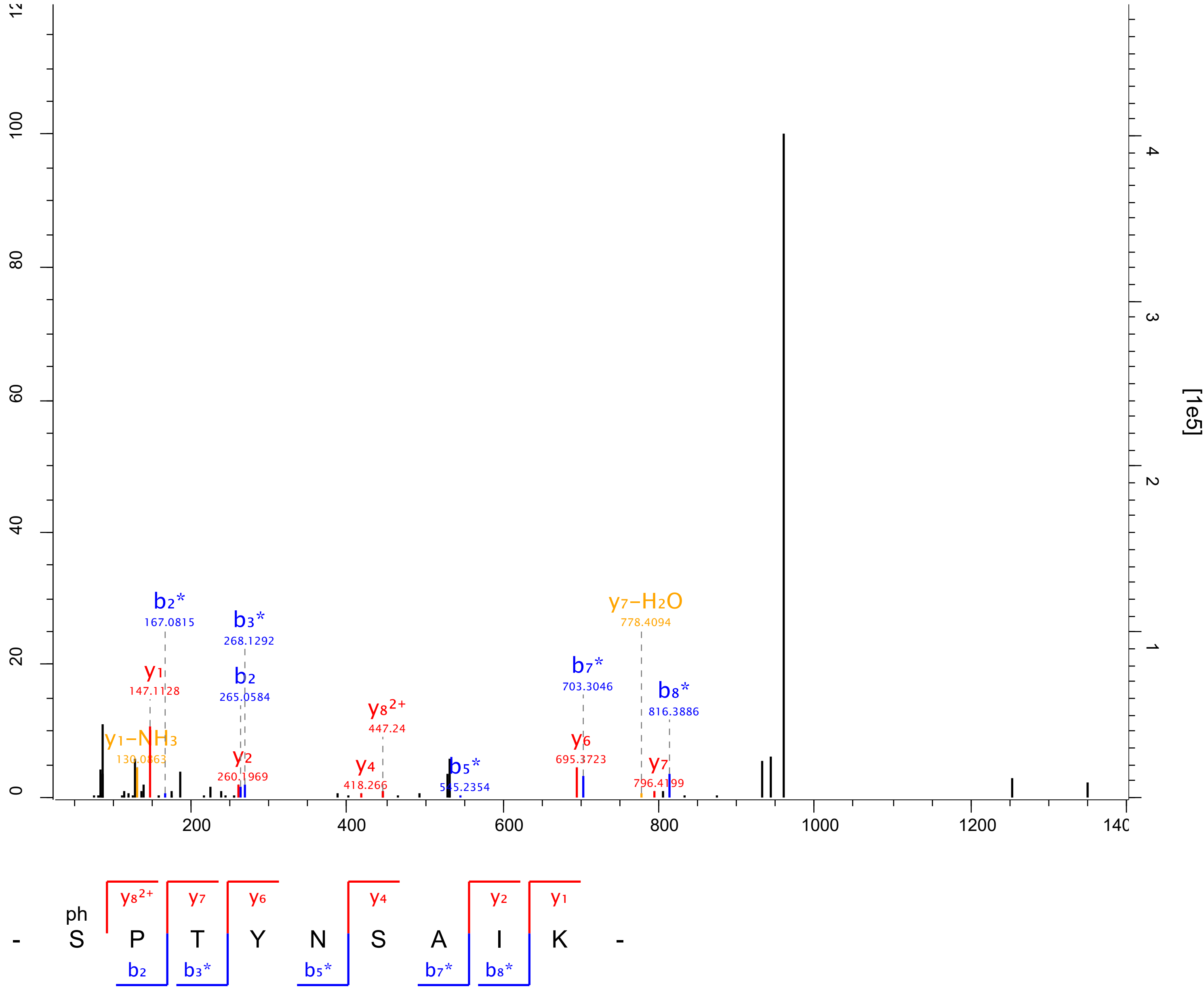

| Raw file | Scan | Method | Score | m/z |
|---|---|---|---|---|
| water_15_2 | 7271 | FTMS; HCD | 112.02 | 576.73 |

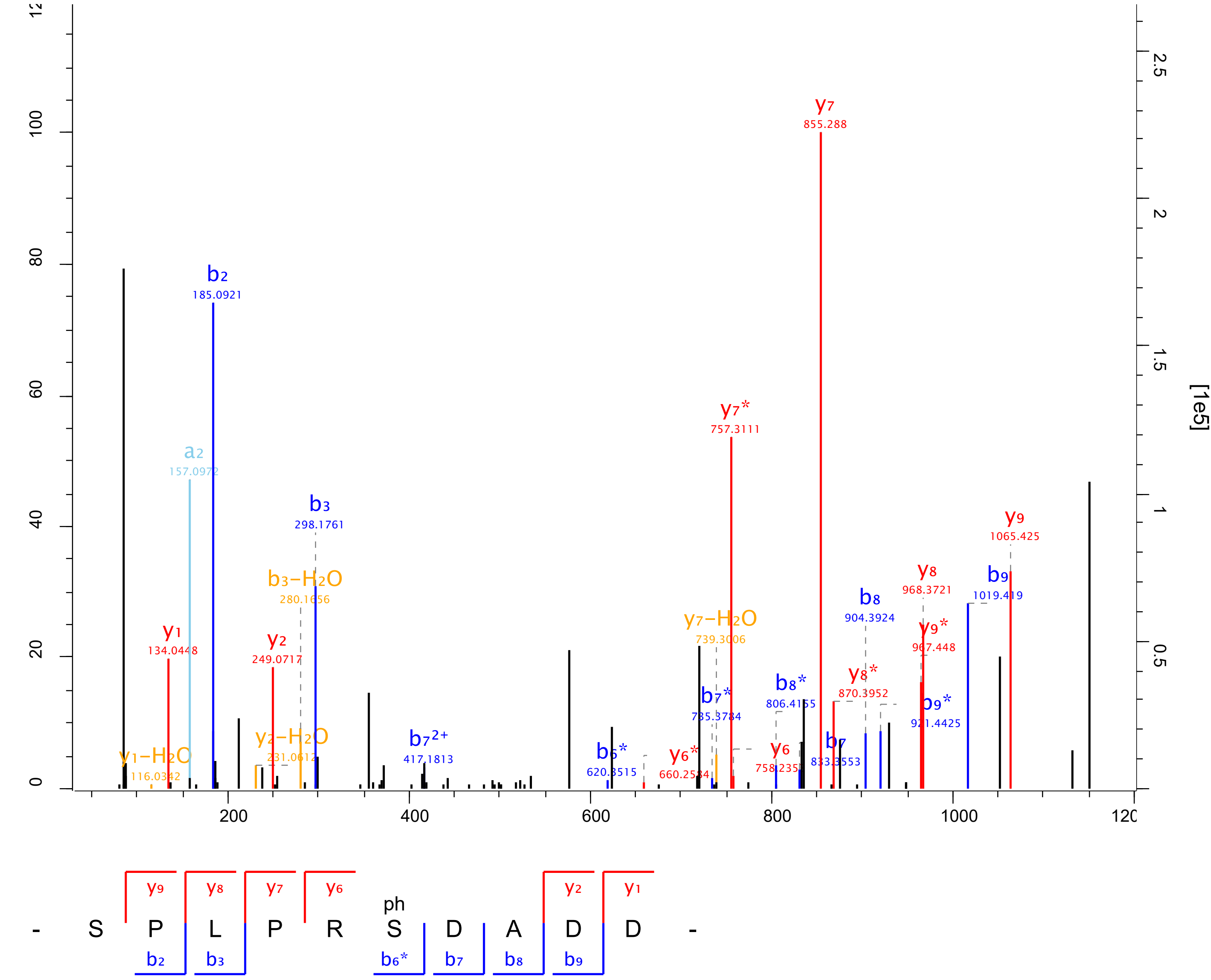

| Raw file | Scan | Method | Score | m/z |
|---|---|---|---|---|
| water_15_2 | 7292 | FTMS; HCD | 54.9 | 612.3 |

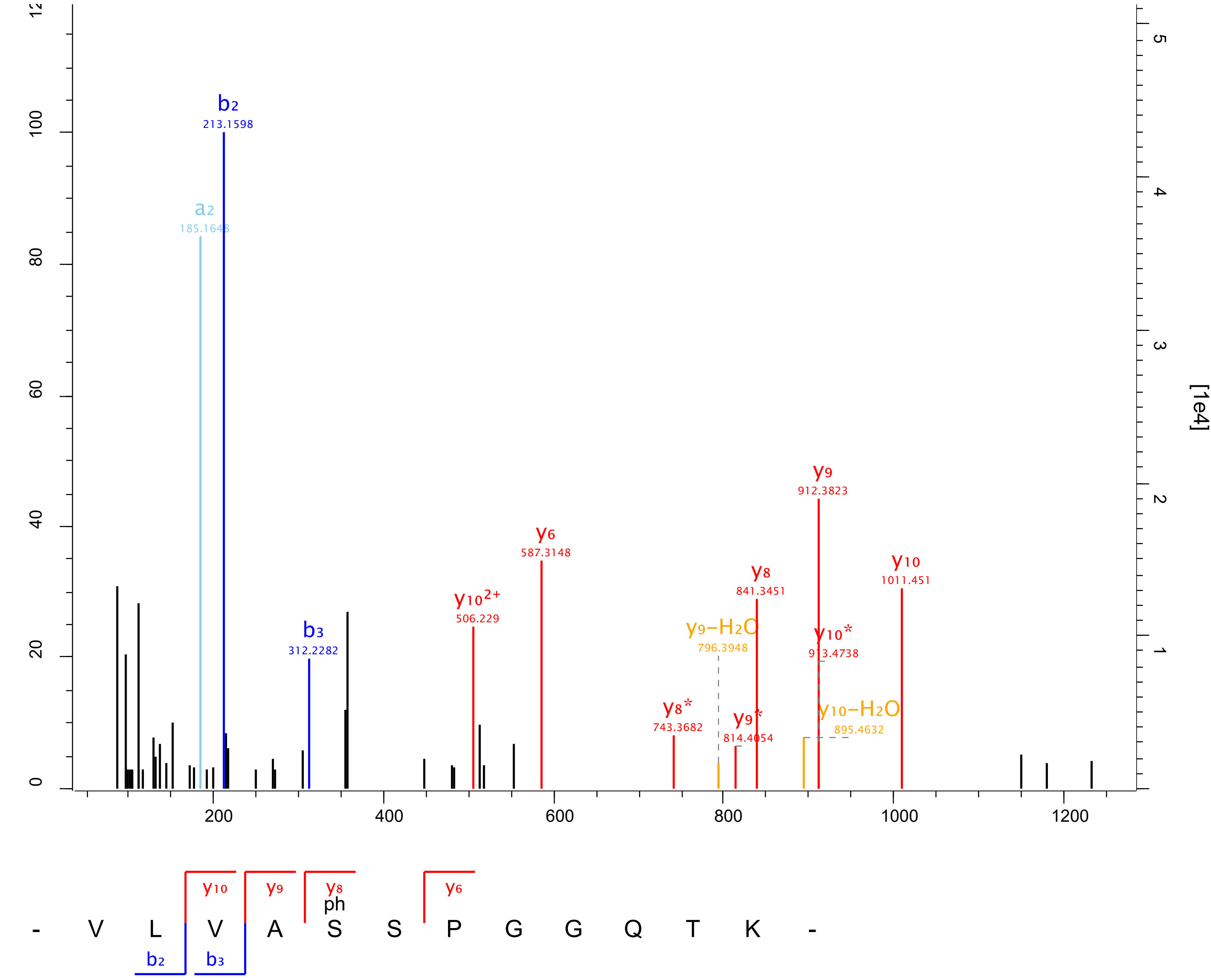

| Raw file | Scan | Method | Score | m/z |
|---|---|---|---|---|
| water_15_2 | 7436 | FTMS; HCD | 172.09 | 726.79 |

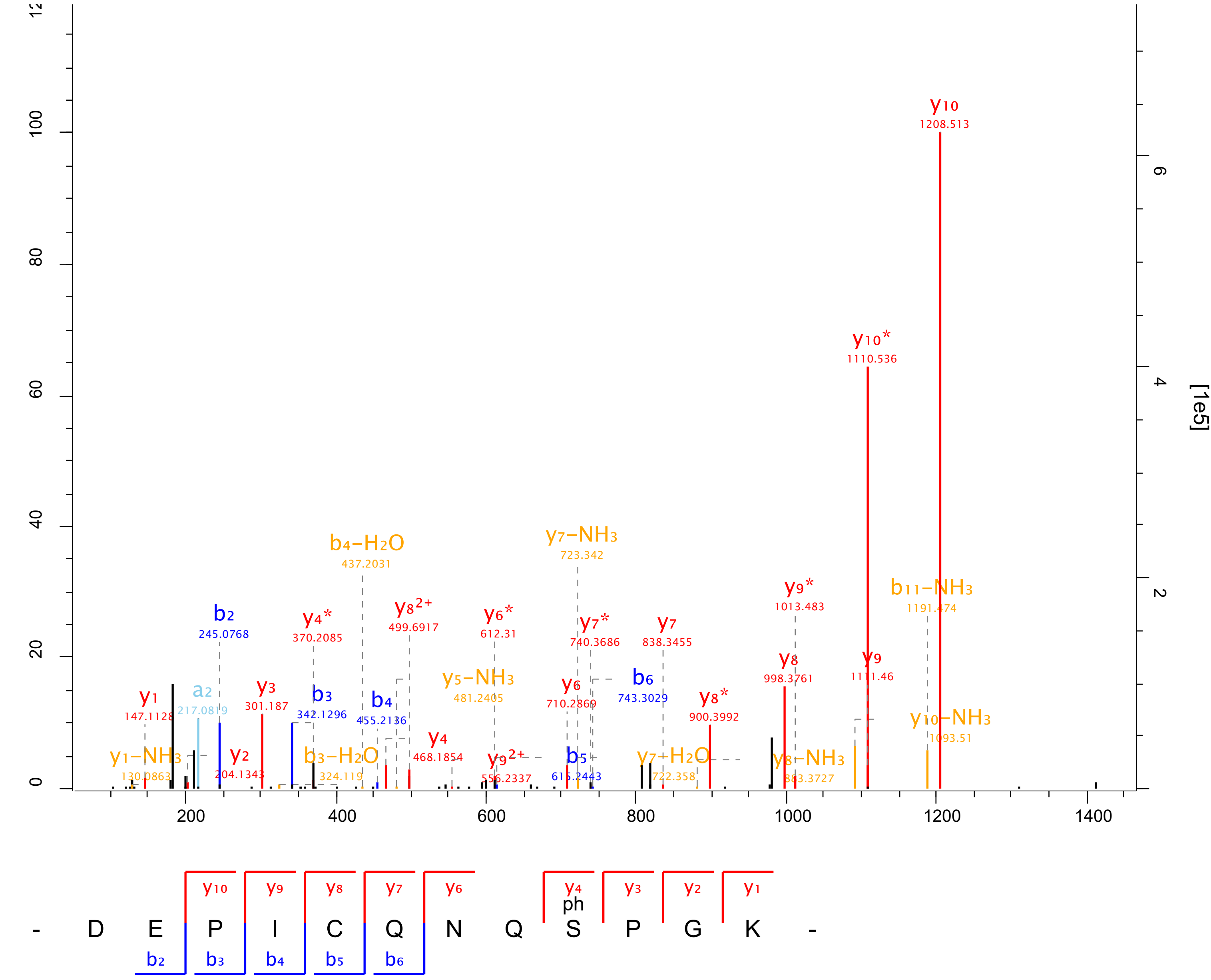

| Raw file | Scan | Method | Score | m/z |
| --- | --- | --- | --- | --- |
| water_15_2 | 7437 | FTMS; HCD | 58.7 | 429.18 |

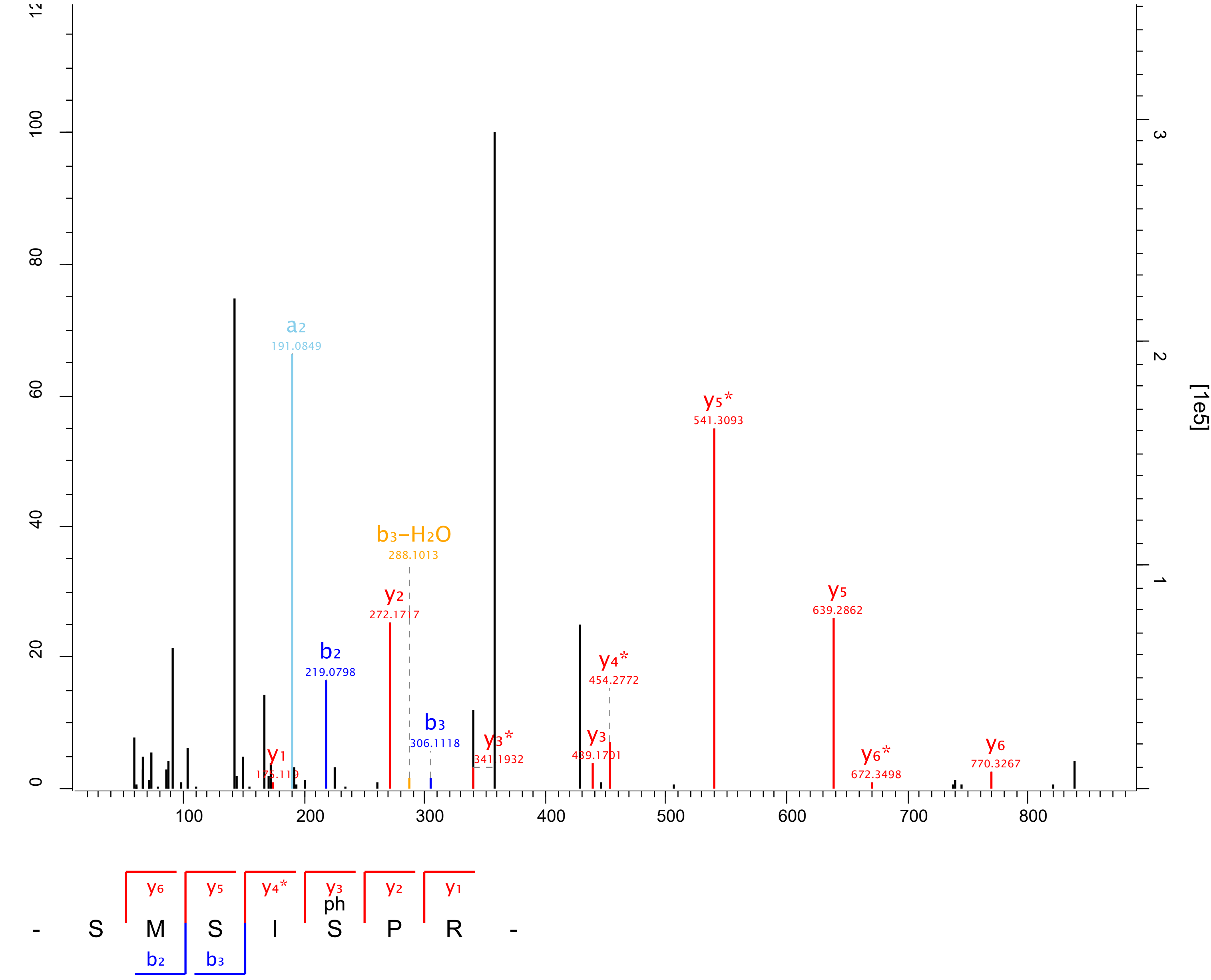

| Raw file | Scan | Method | Score | m/z |
|---|---|---|---|---|
| water_15_2 | 7447 | FTMS; HCD | 124.16 | 628.28 |

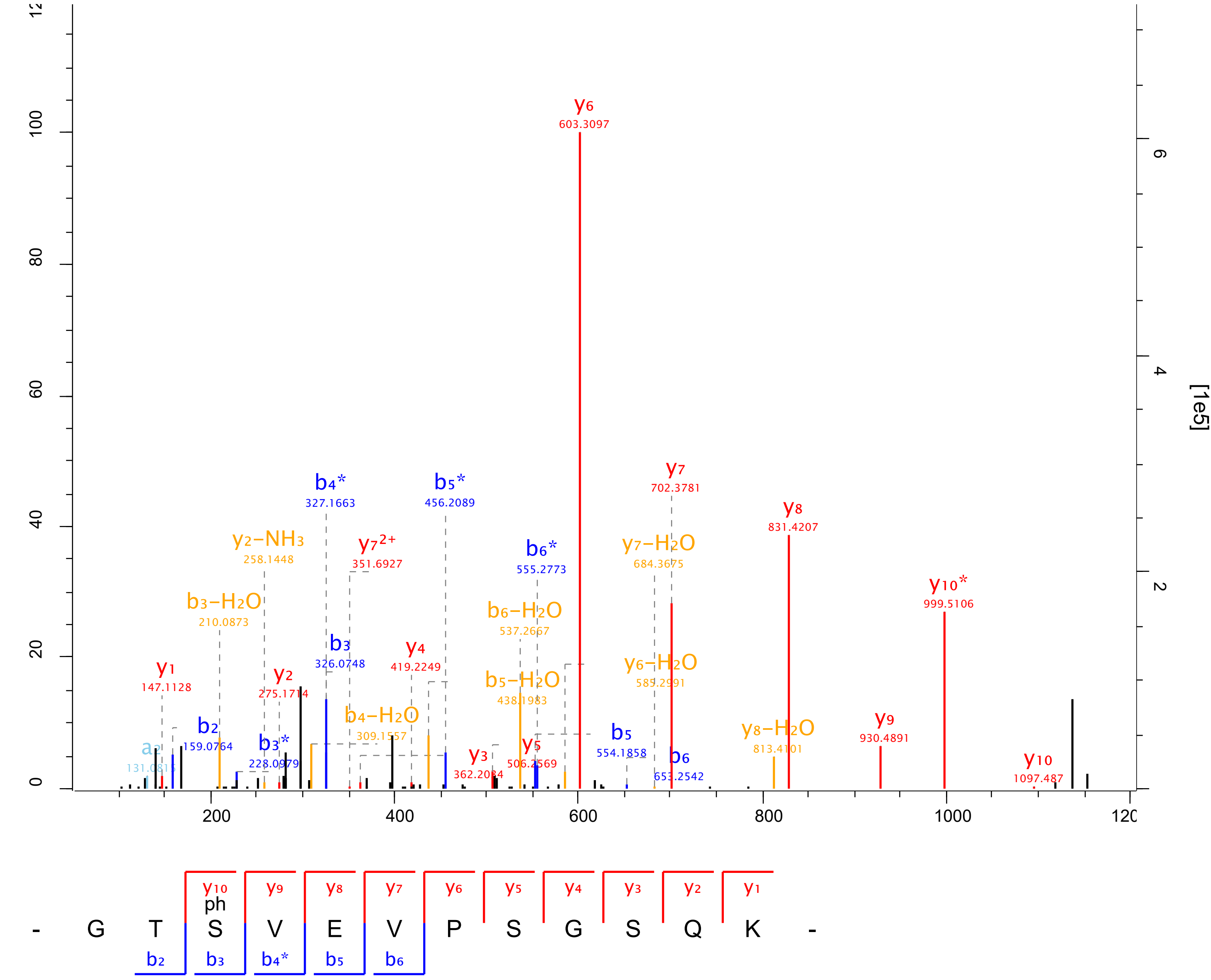

| Raw file | Scan | Method | Score | m/z |
| --- | --- | --- | --- | --- |
| water_15_2 | 7453 | FTMS; HCD | 99.28 | 498.23 |

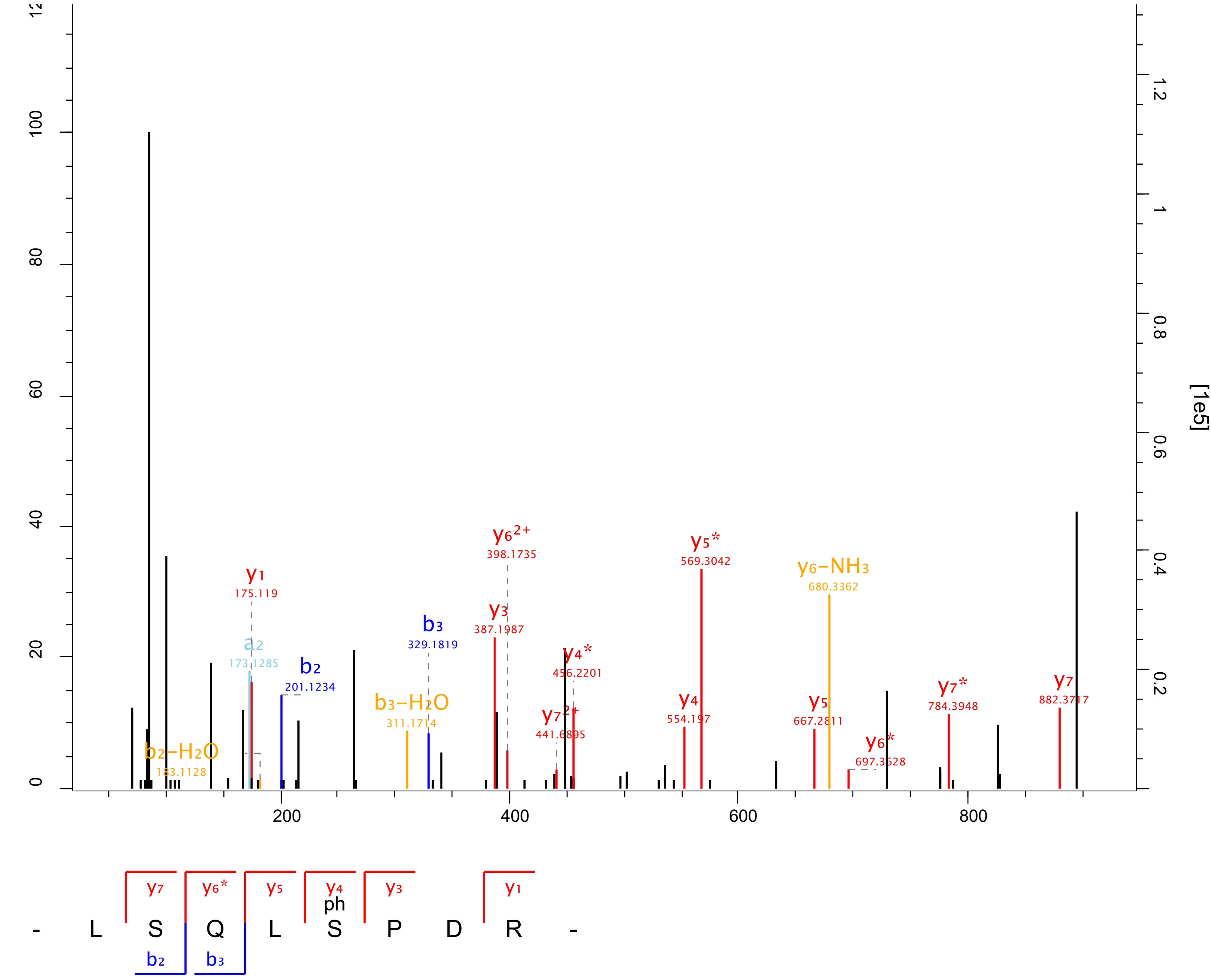

| Raw file | Scan | Method | Score | m/z |
|---|---|---|---|---|
| water_15_2 | 7543 | FTMS; HCD | 193.56 | 735.8 |

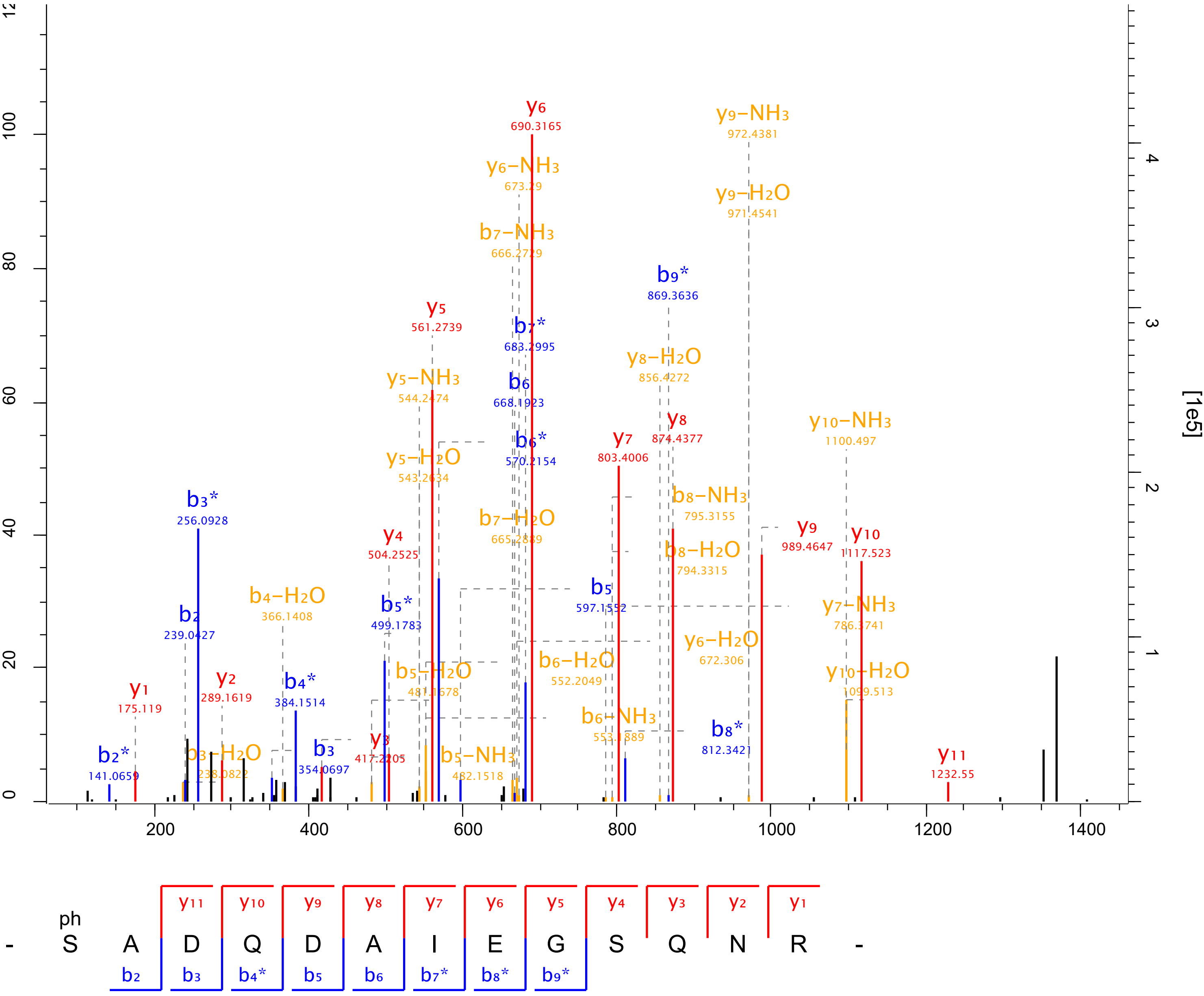

| Raw file | Scan | Method | Score | m/z |
| --- | --- | --- | --- | --- |
| water_15_2 | 7596 | FTMS; HCD | 49.45 | 579.75 |

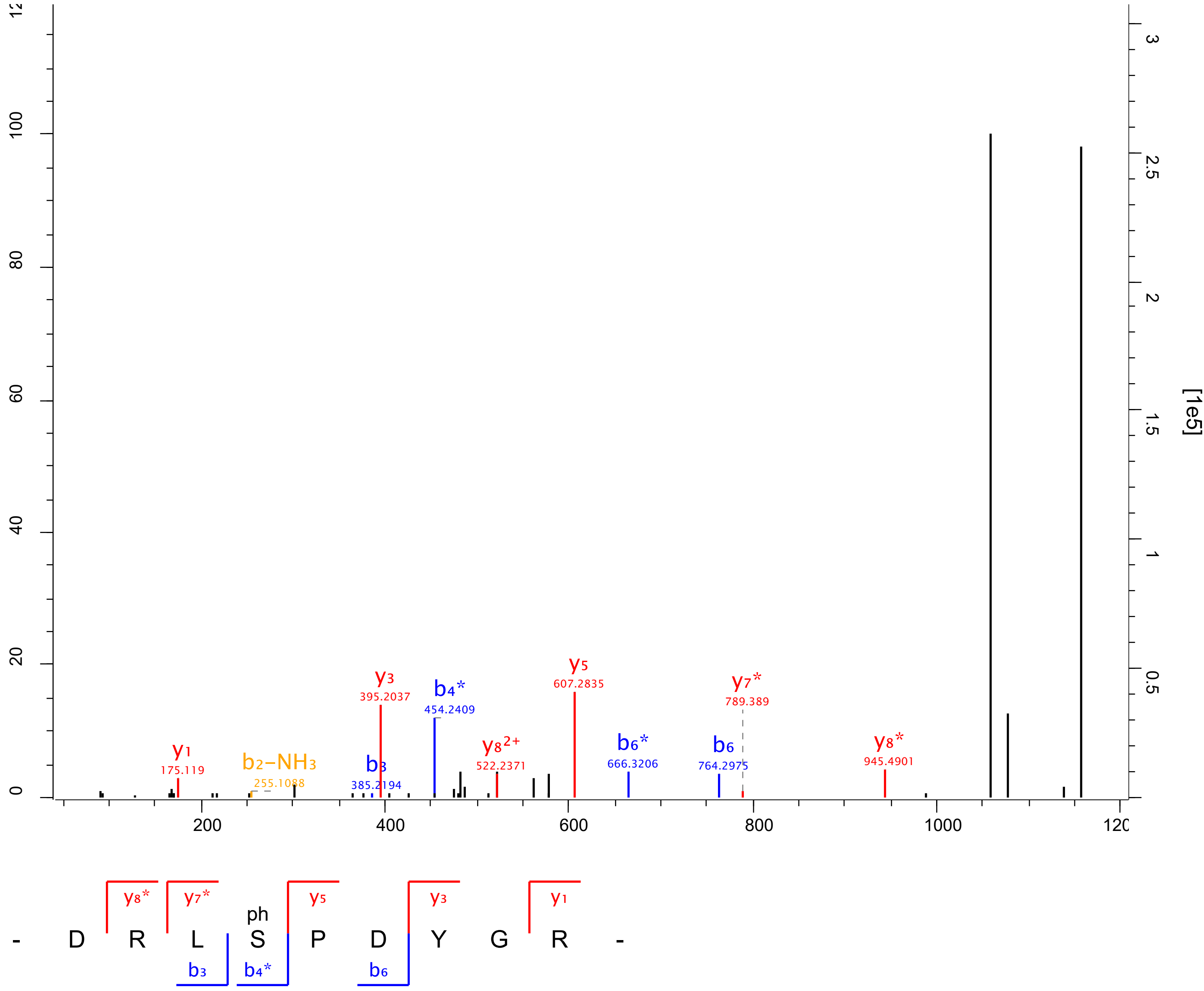

| Raw file | Scan | Method | Score | m/z |
|---|---|---|---|---|
| water_15_2 | 7702 | FTMS; HCD | 165.58 | 703.82 |

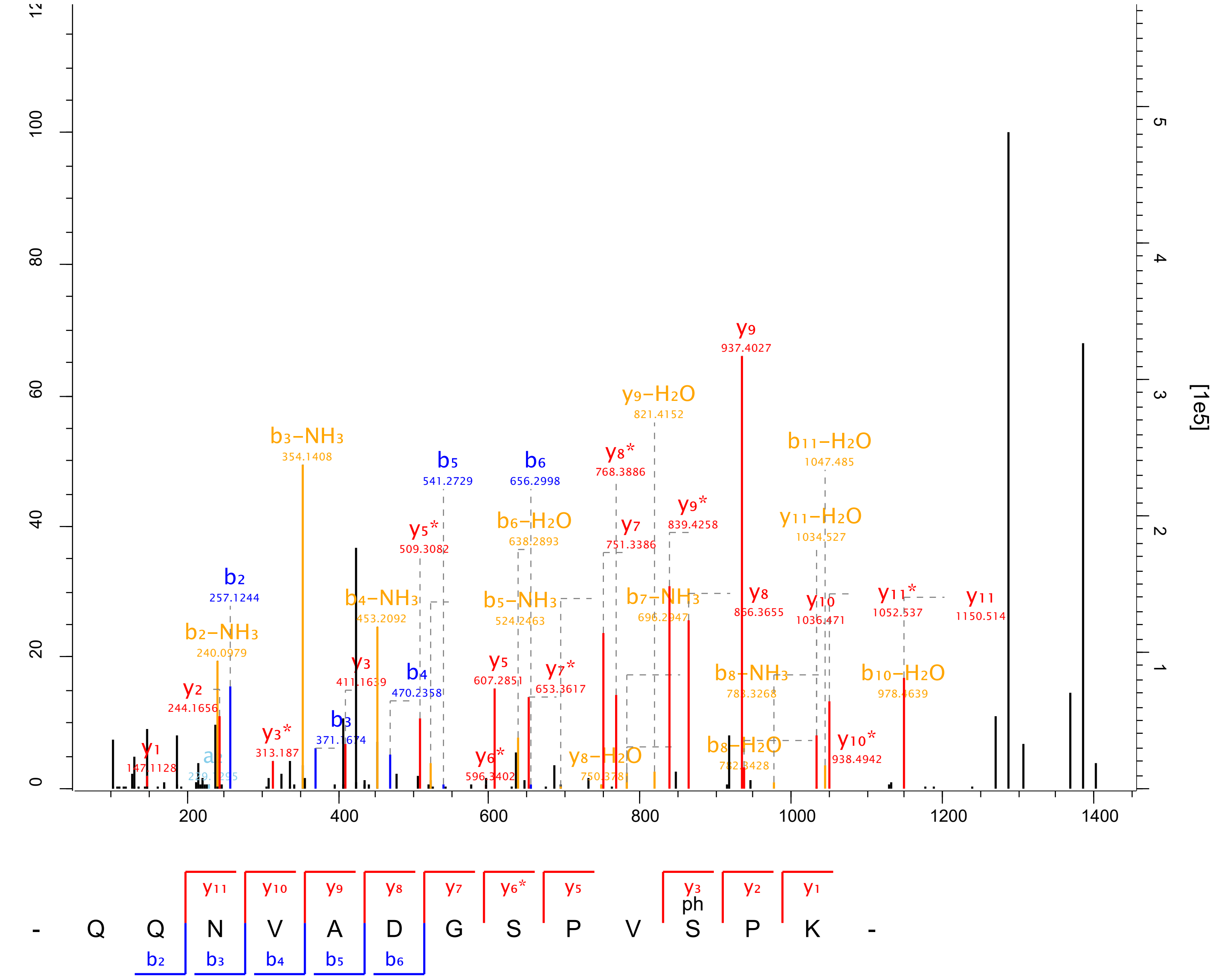

| Raw file | Scan | Method | Score | m/z |
|---|---|---|---|---|
| water_15_2 | 7751 | FTMS; HCD | 181.94 | 655.77 |

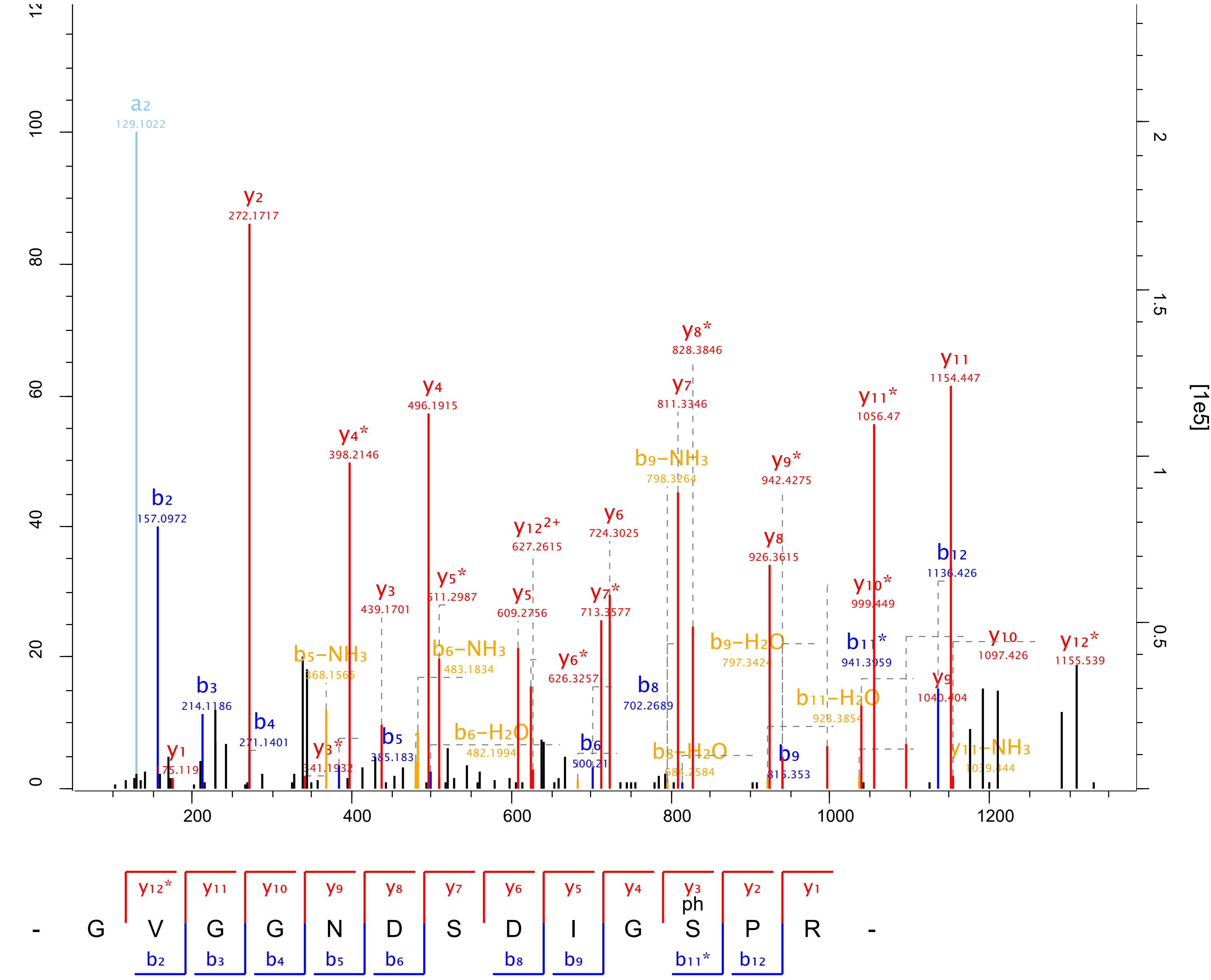

| Raw file | Scan | Method | Score | m/z |
| --- | --- | --- | --- | --- |
| water_15_2 | 7769 | FTMS; HCD | 44.25 | 671.82 |

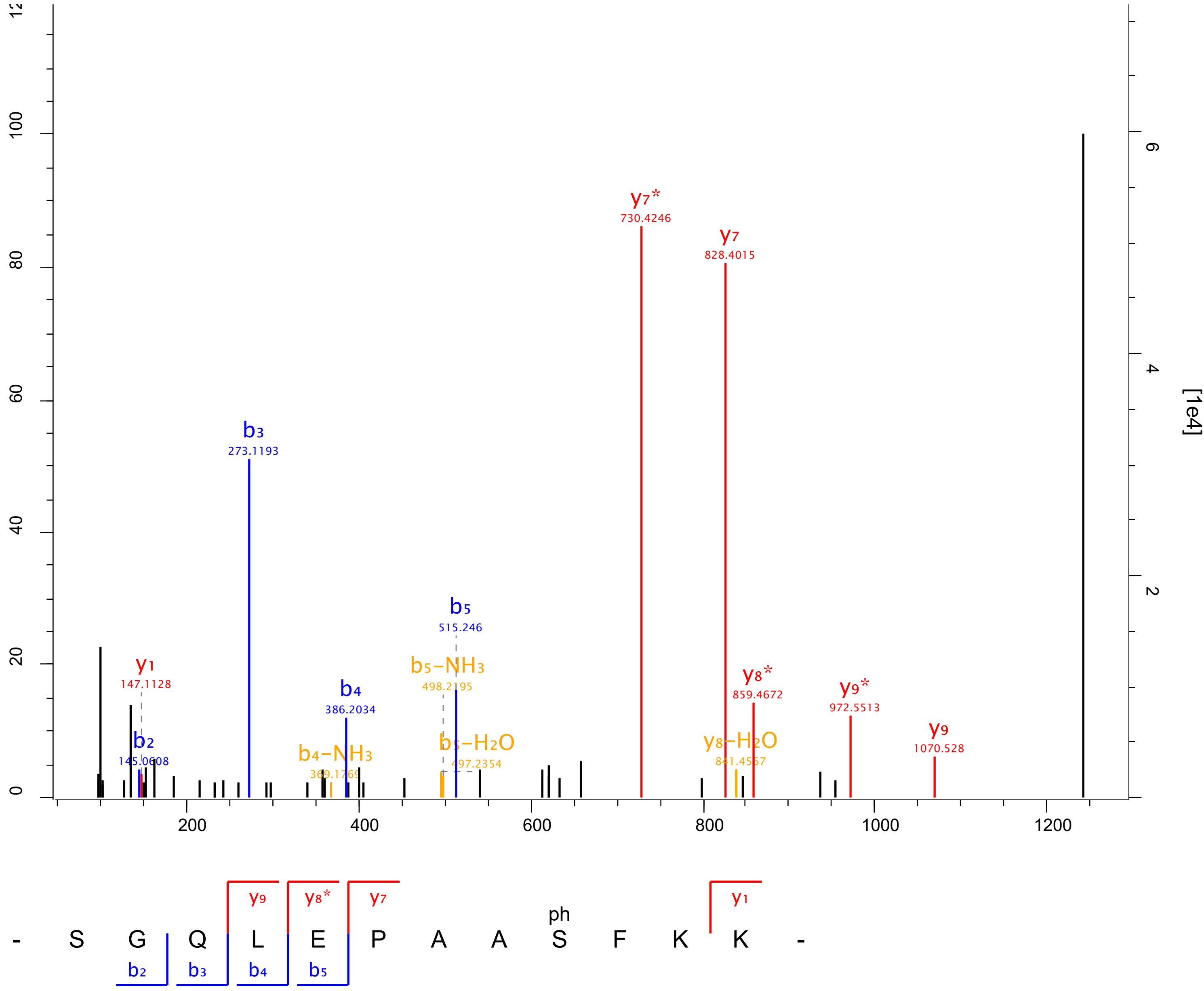

| Raw file | Scan | Method | Score | m/z |
| --- | --- | --- | --- | --- |
| water_15_2 | 7786 | FTMS; HCD | 157.07 | 796.86 |

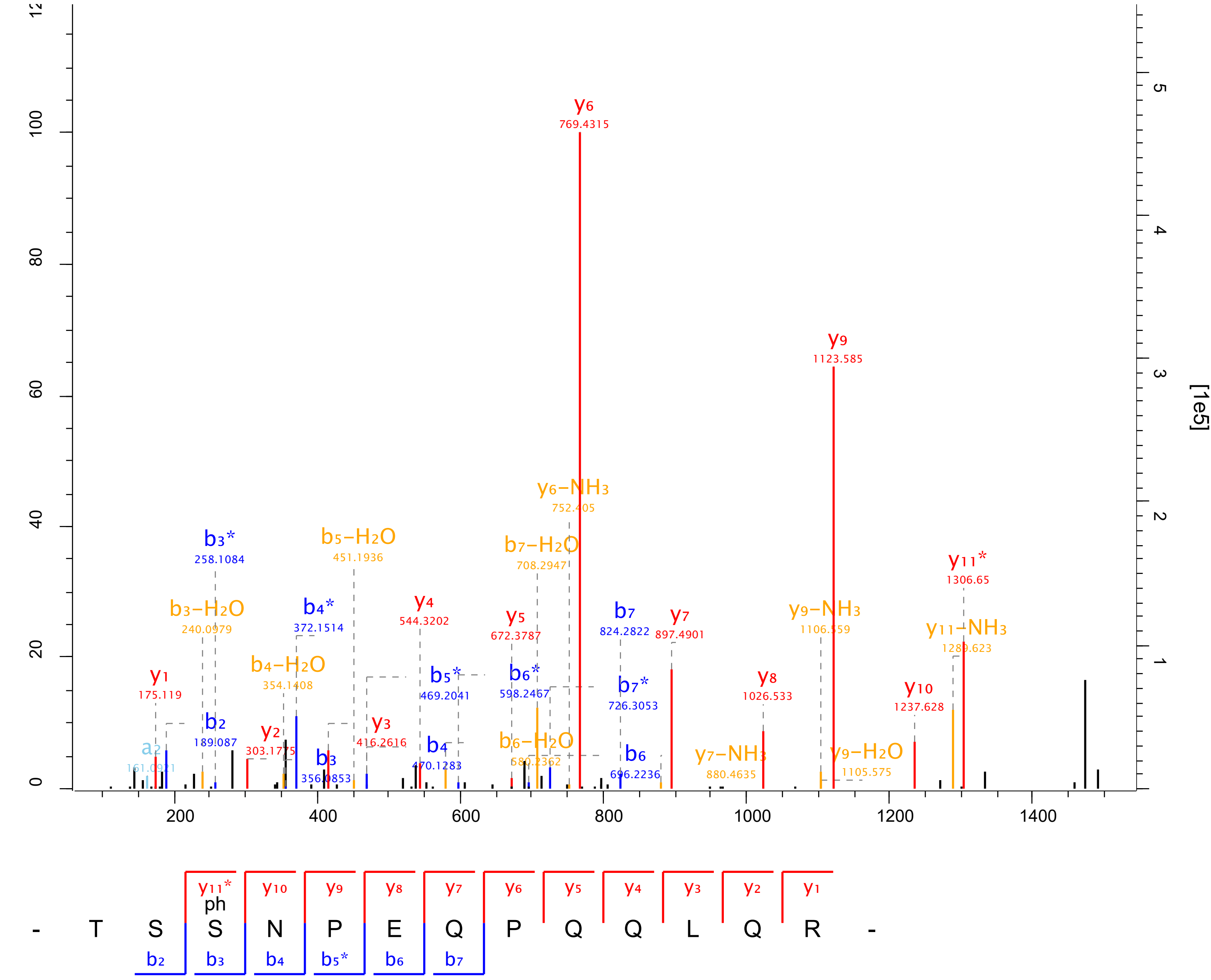

| Raw file | Scan | Method | Score | m/z |
| --- | --- | --- | --- | --- |
| water_15_2 | 7835 | FTMS; HCD | 78.69 | 640.29 |

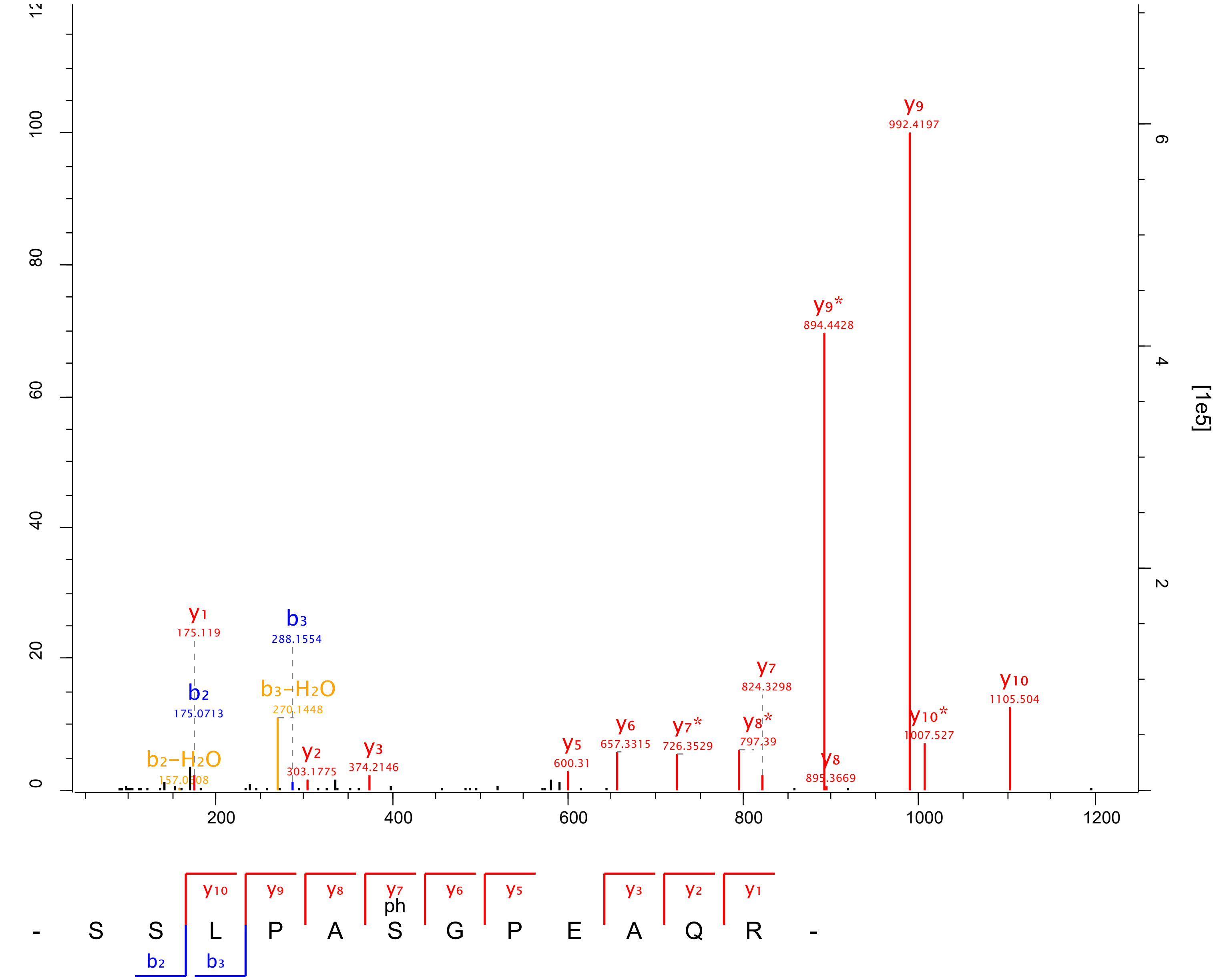

| Raw file | Scan | Method | Score | m/z |
|---|---|---|---|---|
| water_15_2 | 7861 | FTMS; HCD | 83.35 | 586.23 |

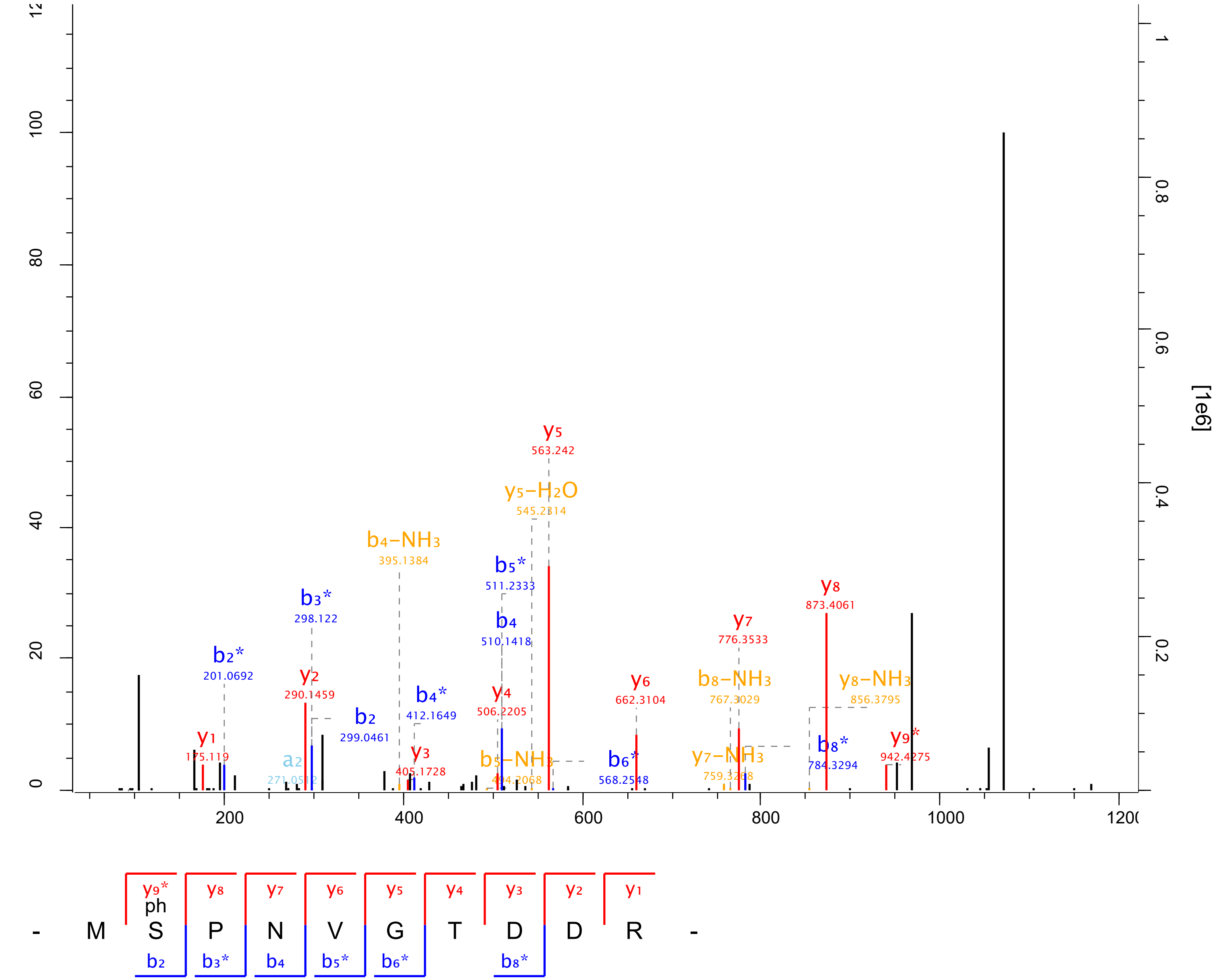

| Raw file | Scan | Method | Score | m/z |
|---|---|---|---|---|
| water_15_2 | 7918 | FTMS; HCD | 235.75 | 743.82 |

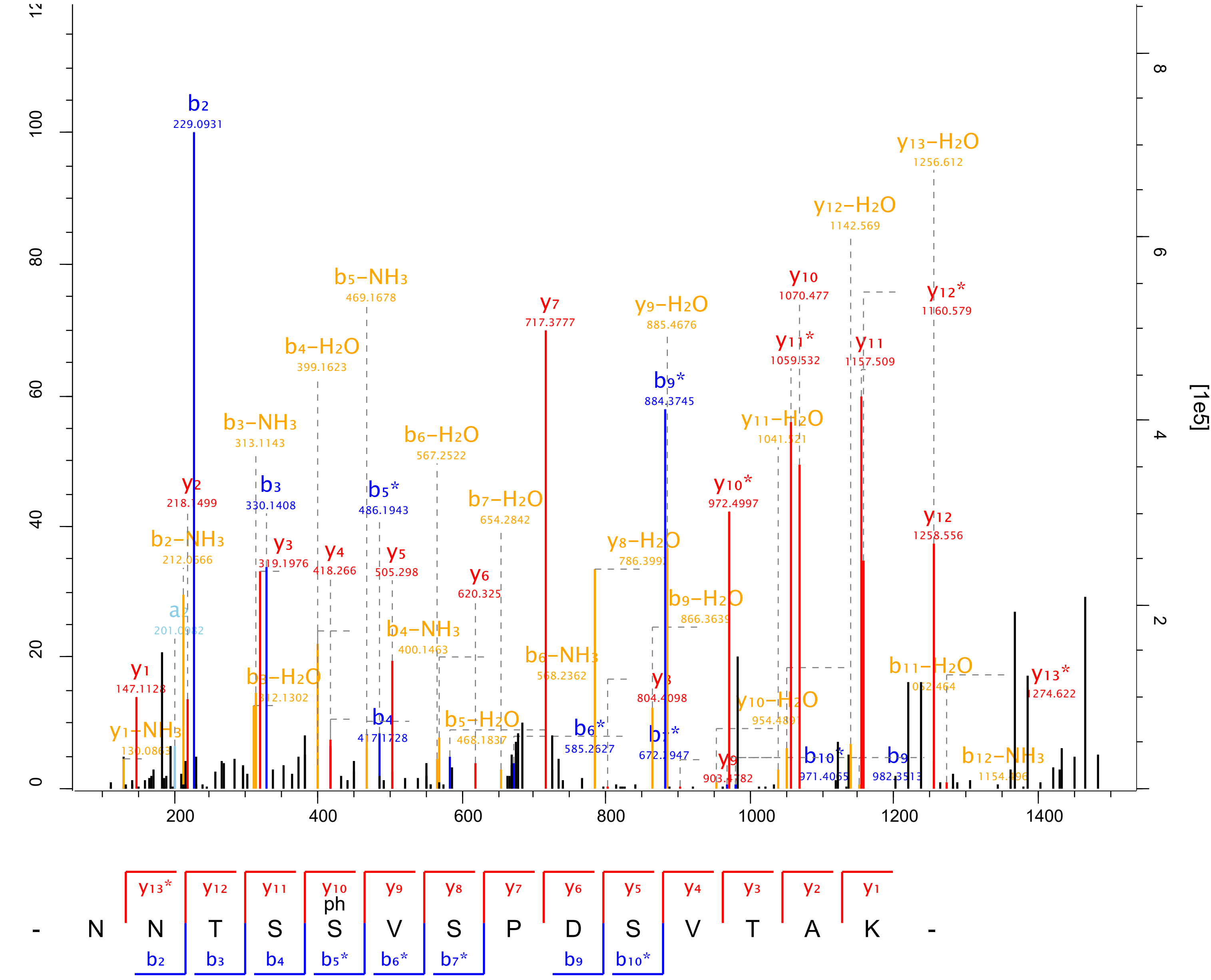

| Raw file | Scan | Method | Score | m/z |
| --- | --- | --- | --- | --- |
| water_15_2 | 7934 | FTMS; HCD | 237.46 | 680.29 |

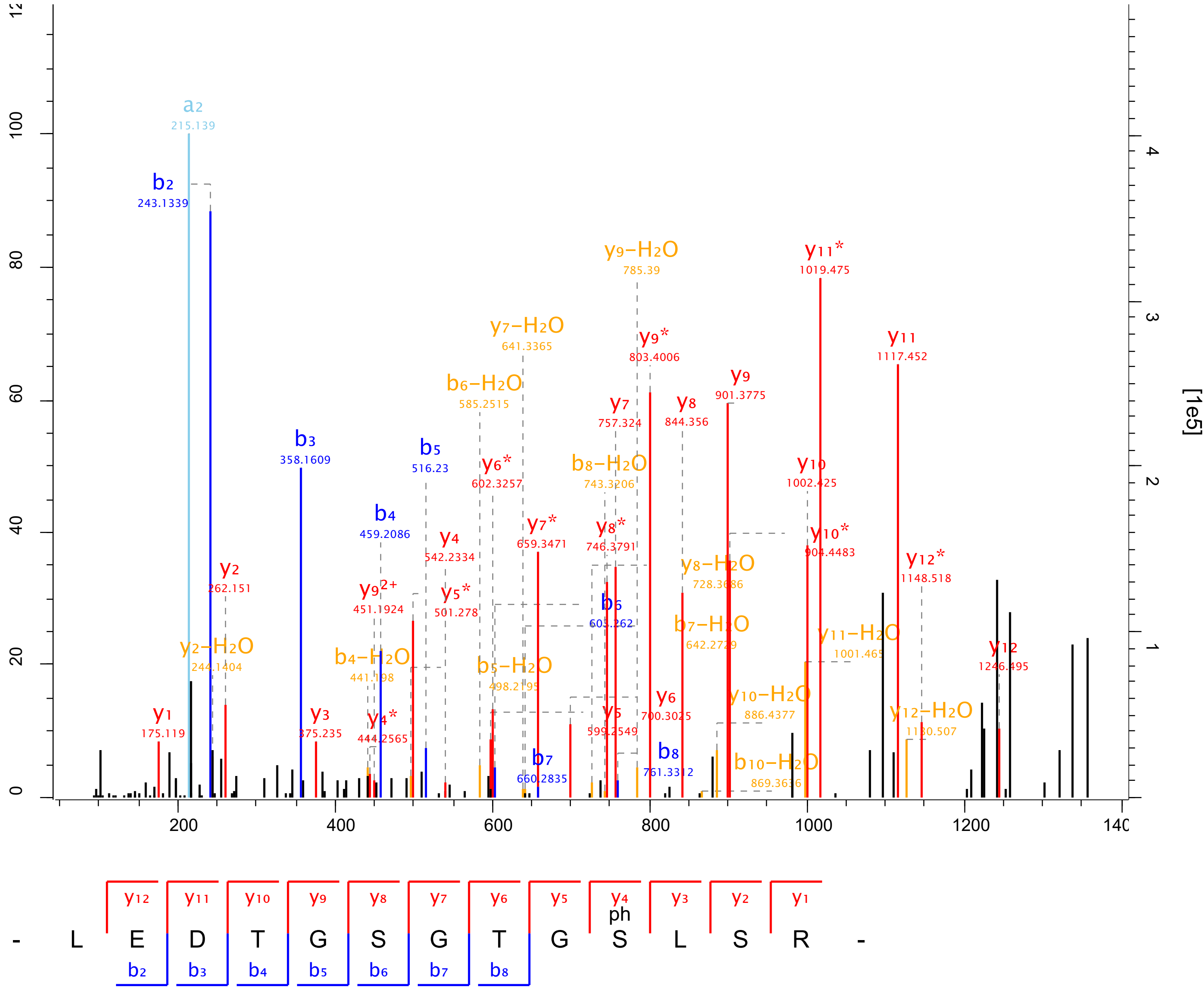

| Raw file | Scan | Method | Score | m/z |
| --- | --- | --- | --- | --- |
| water_15_2 | 8087 | FTMS; HCD | 47.55 | 728.33 |

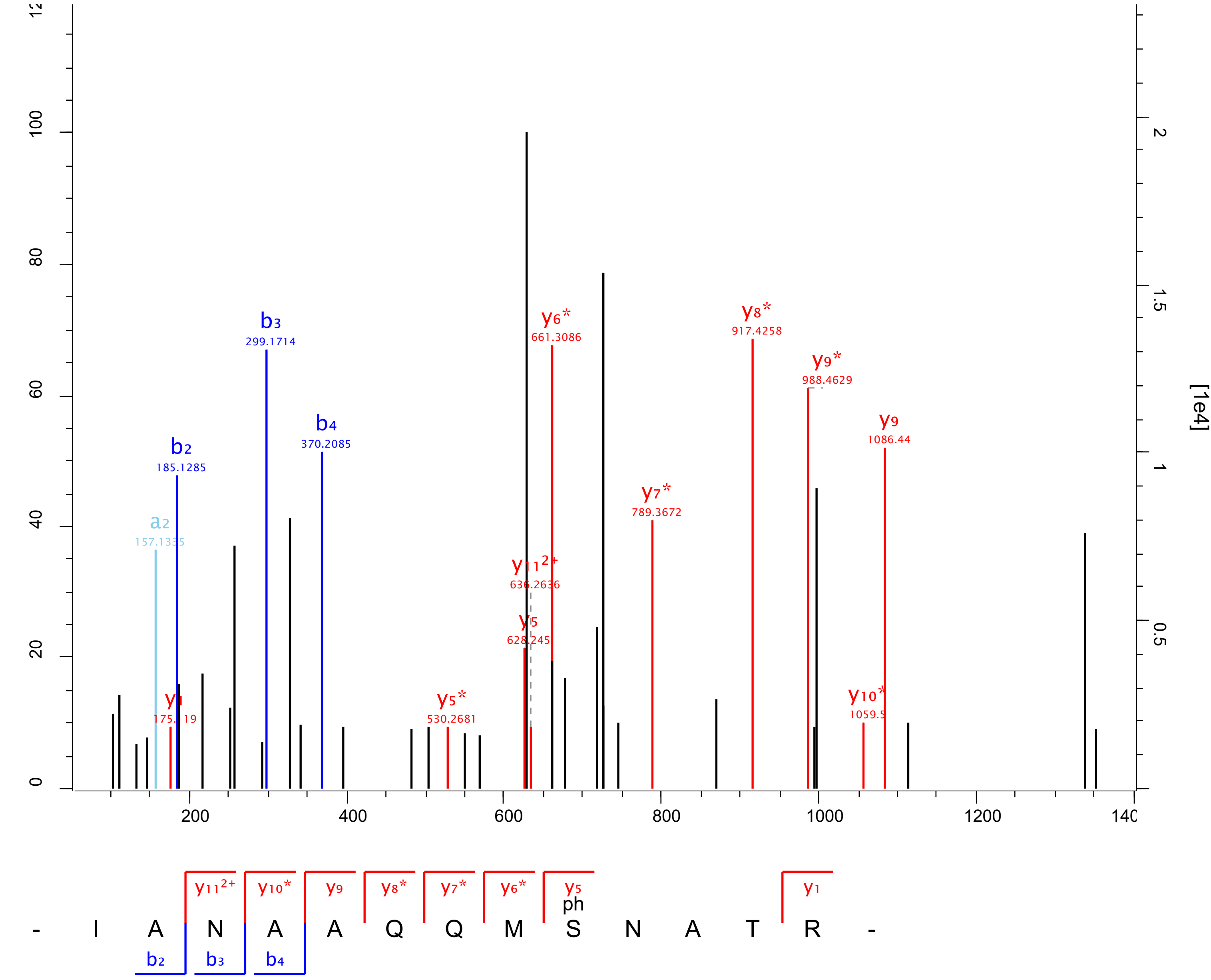

| Raw file | Scan | Method | Score | m/z |
|---|---|---|---|---|
| water_15_2 | 8096 | FTMS; HCD | 291.21 | 826.89 |

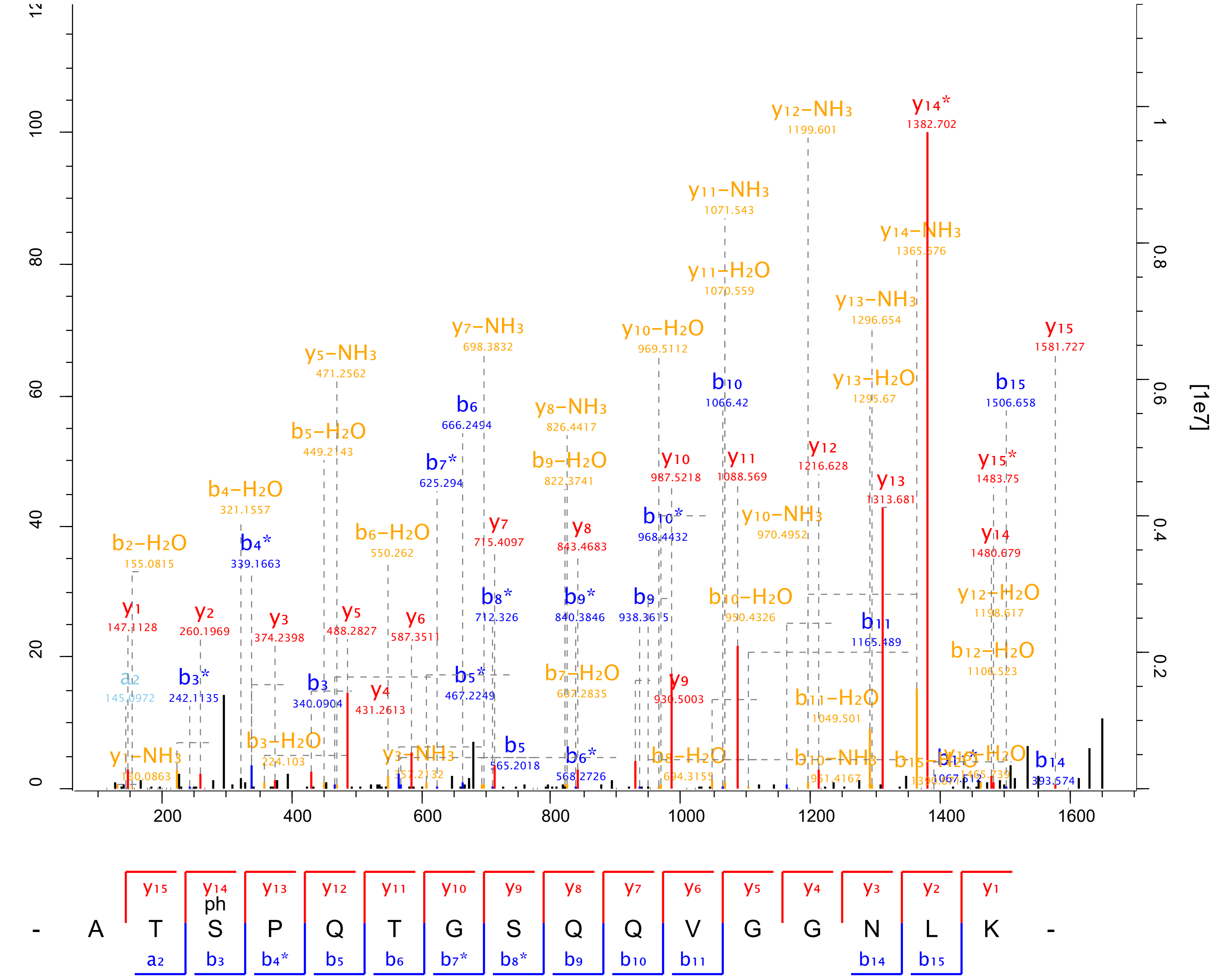

| Raw file | Scan | Method | Score | m/z |
| --- | --- | --- | --- | --- |
| water_15_2 | 8100 | FTMS; HCD | 42.32 | 633.78 |

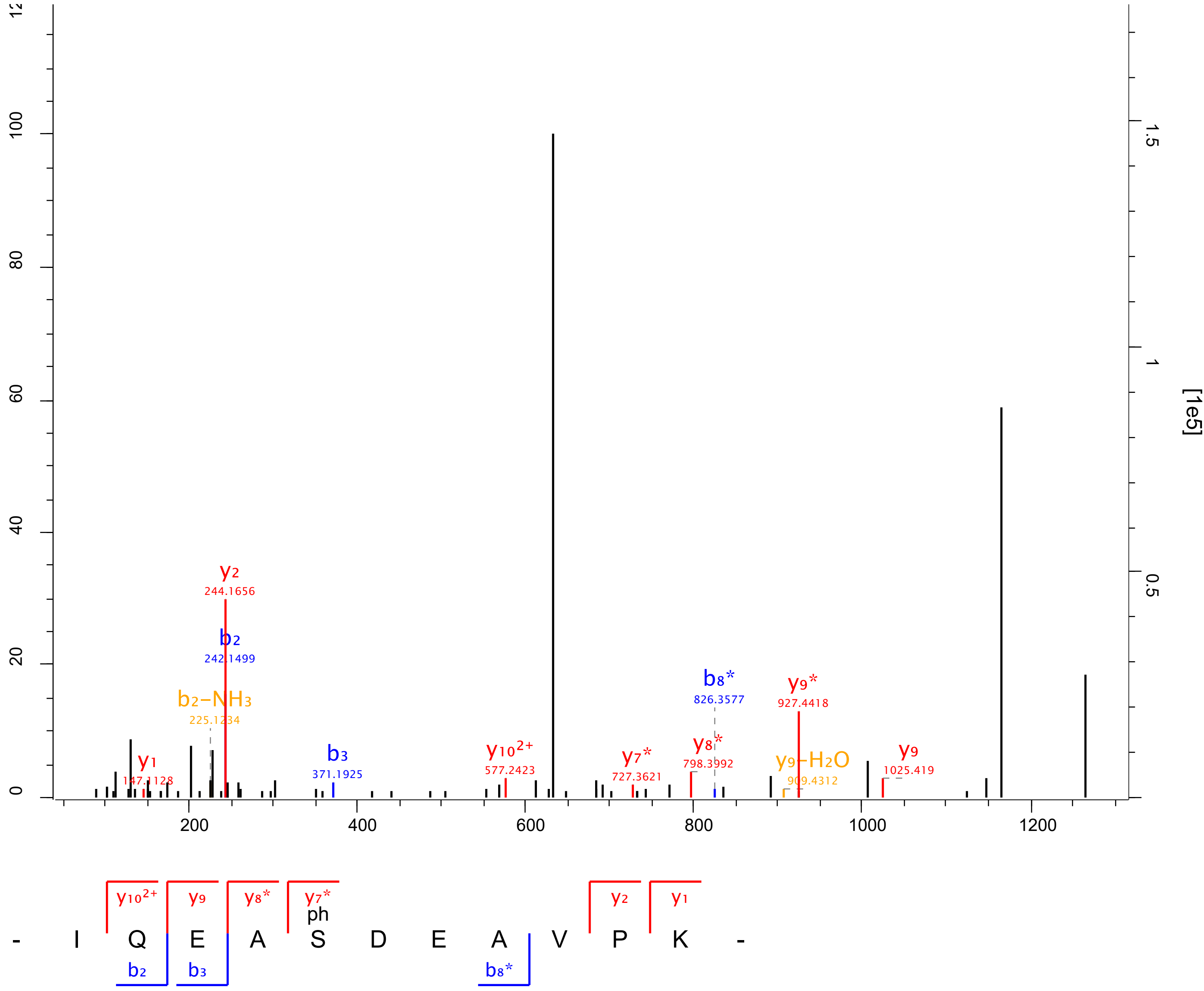

| Raw file | Scan | Method | Score | m/z |
| --- | --- | --- | --- | --- |
| water_15_2 | 8180 | FTMS; HCD | 60.16 | 578.26 |

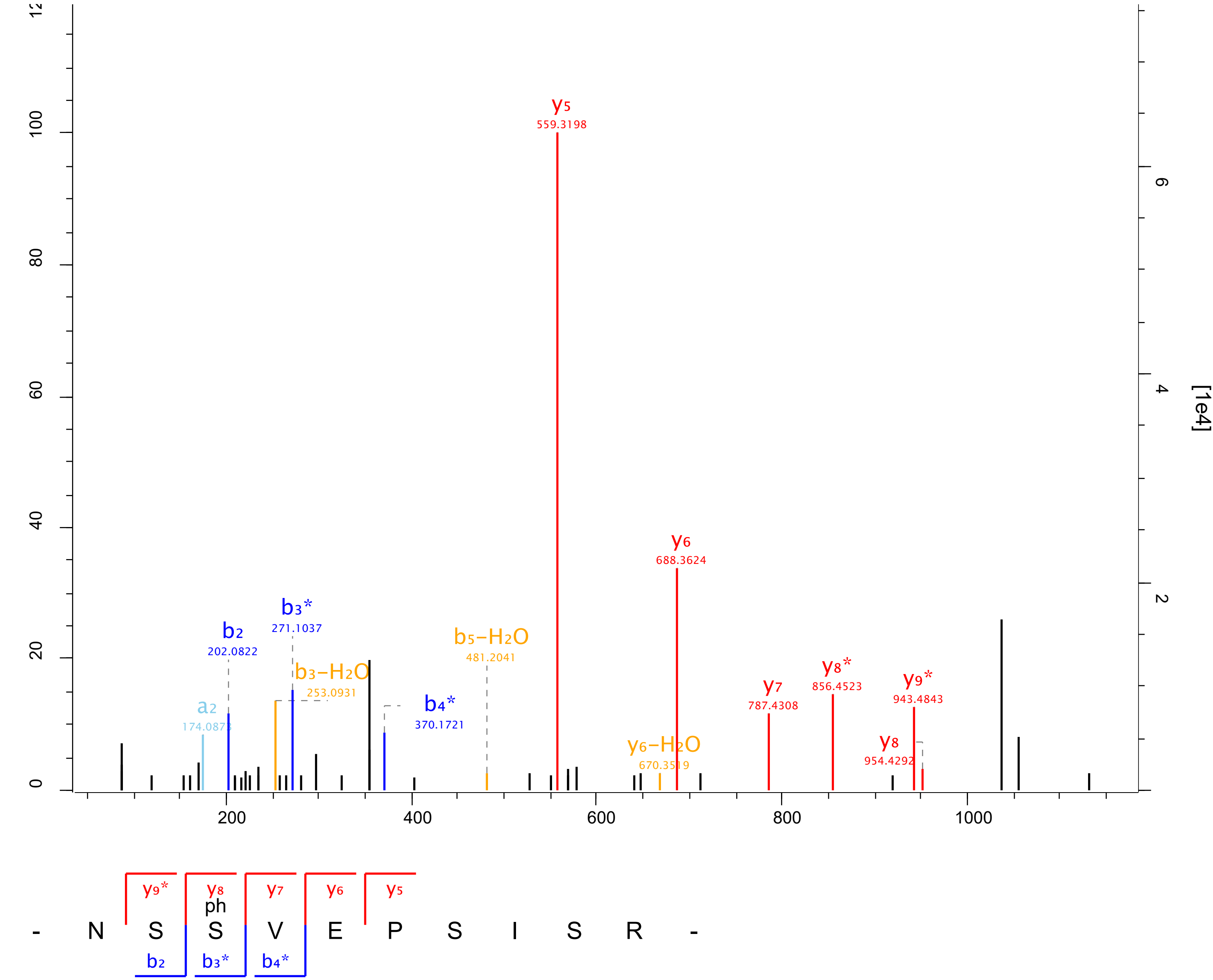

| Raw file | Scan | Method | Score | m/z |
|---|---|---|---|---|
| water_15_2 | 8304 | FTMS; HCD | 40.69 | 1029.87 |

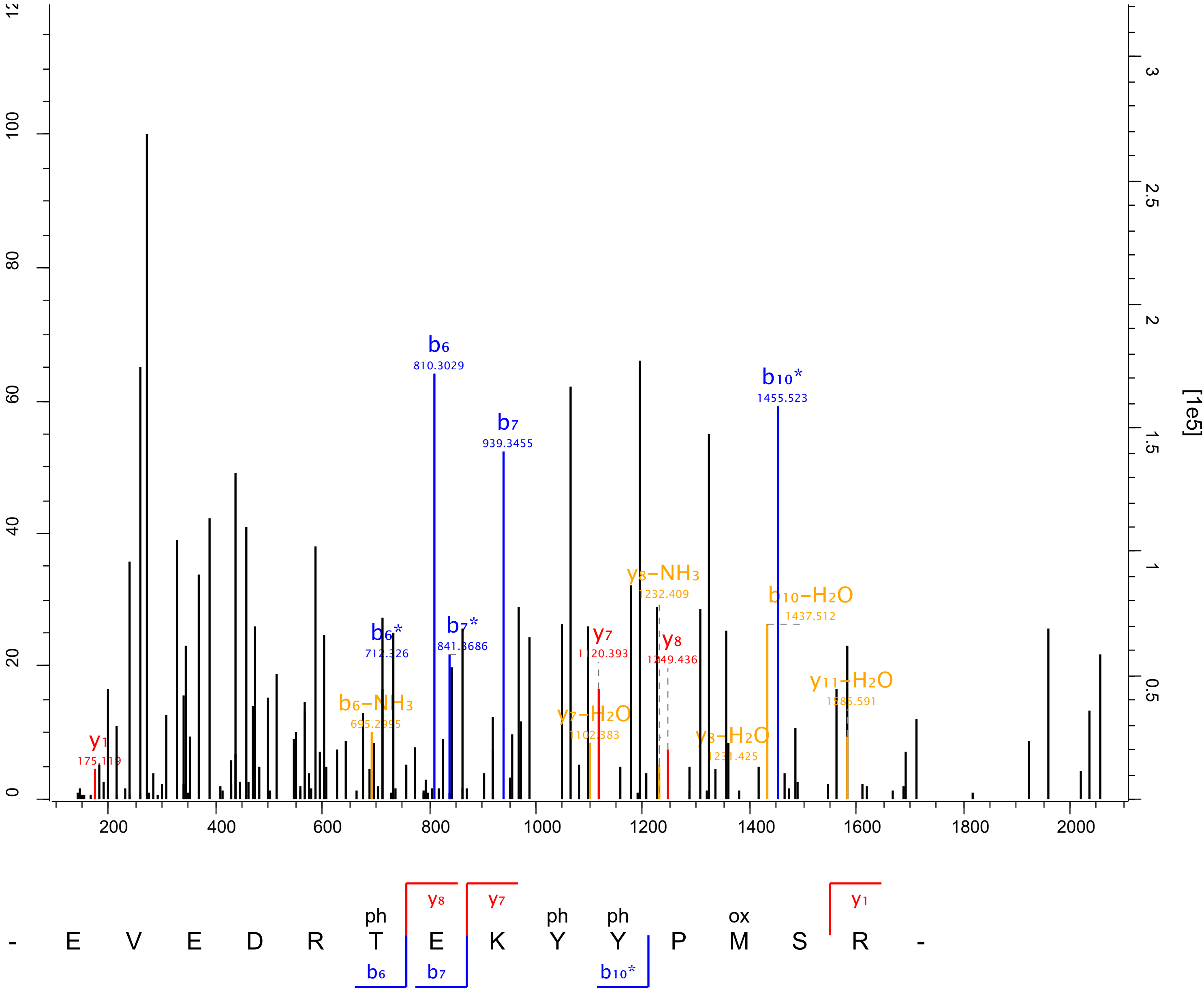

| Raw file | Scan | Method | Score | m/z |
|---|---|---|---|---|
| water_15_2 | 8451 | FTMS; HCD | 43.21 | 561.23 |

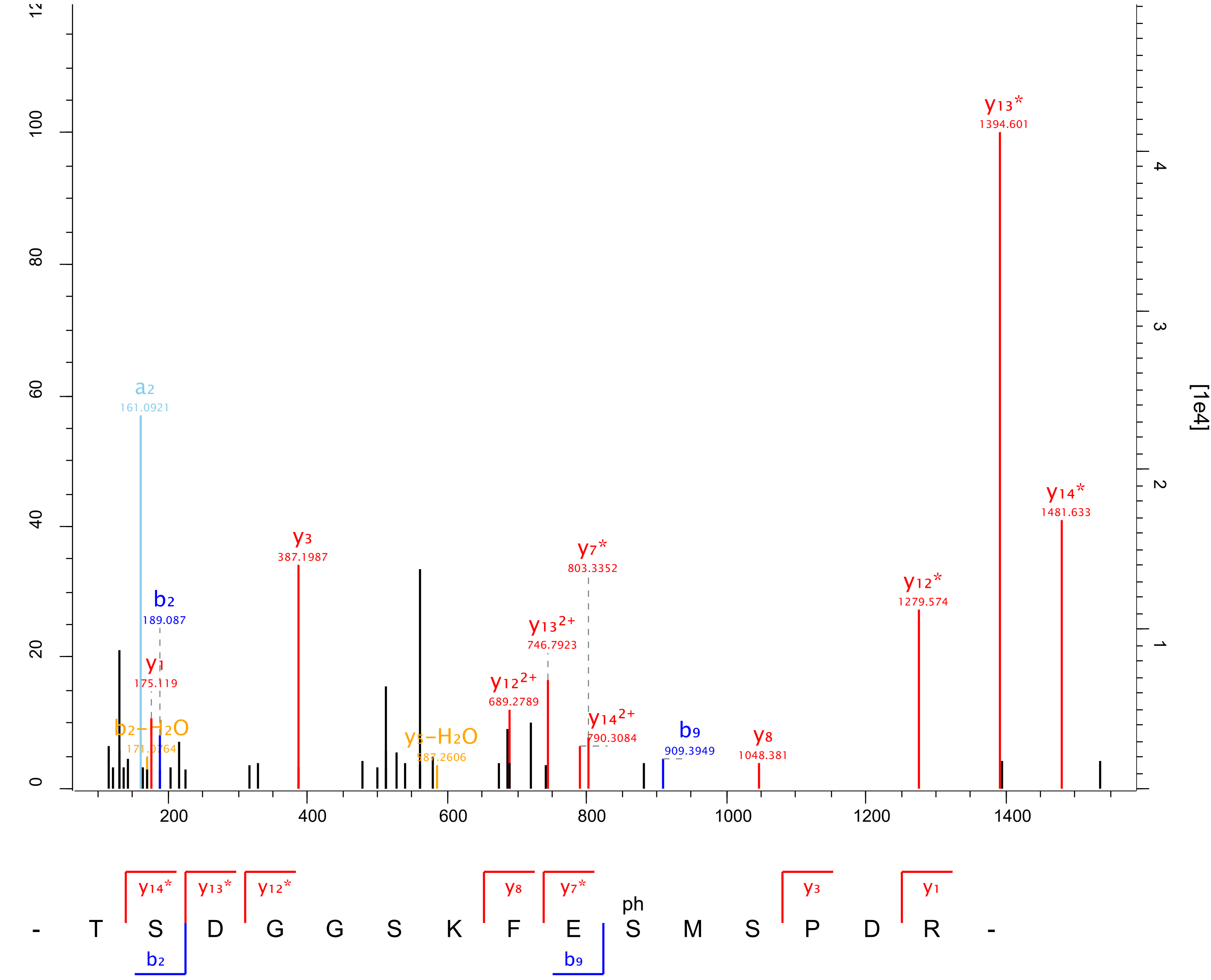

| Raw file | Scan | Method | Score | m/z |
|---|---|---|---|---|
| water_15_2 | 8483 | FTMS; HCD | 140.22 | 734.78 |

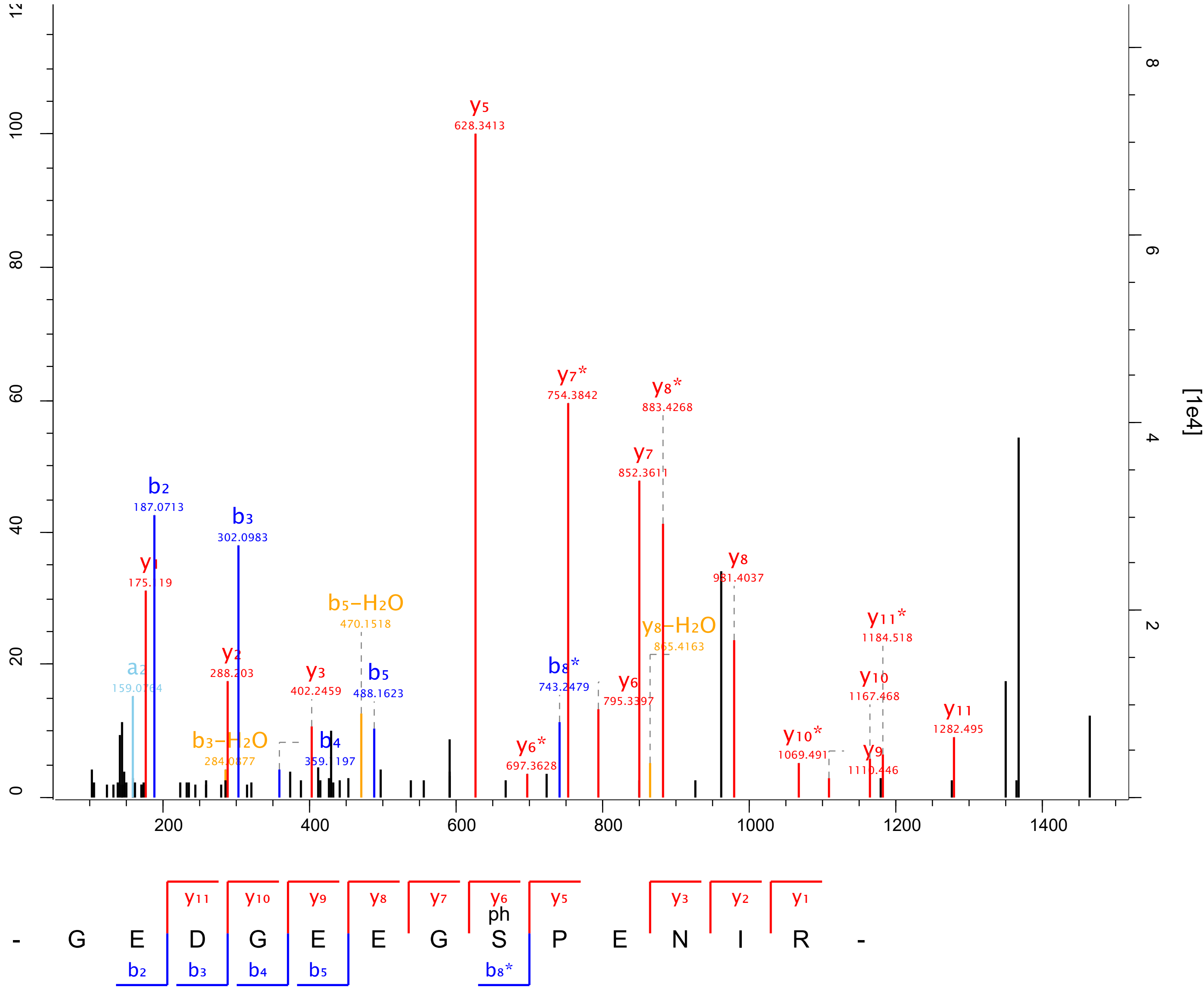

| Raw file | Scan | Method | Score | m/z |
| --- | --- | --- | --- | --- |
| water_15_2 | 8547 | FTMS; HCD | 227.88 | 750.81 |

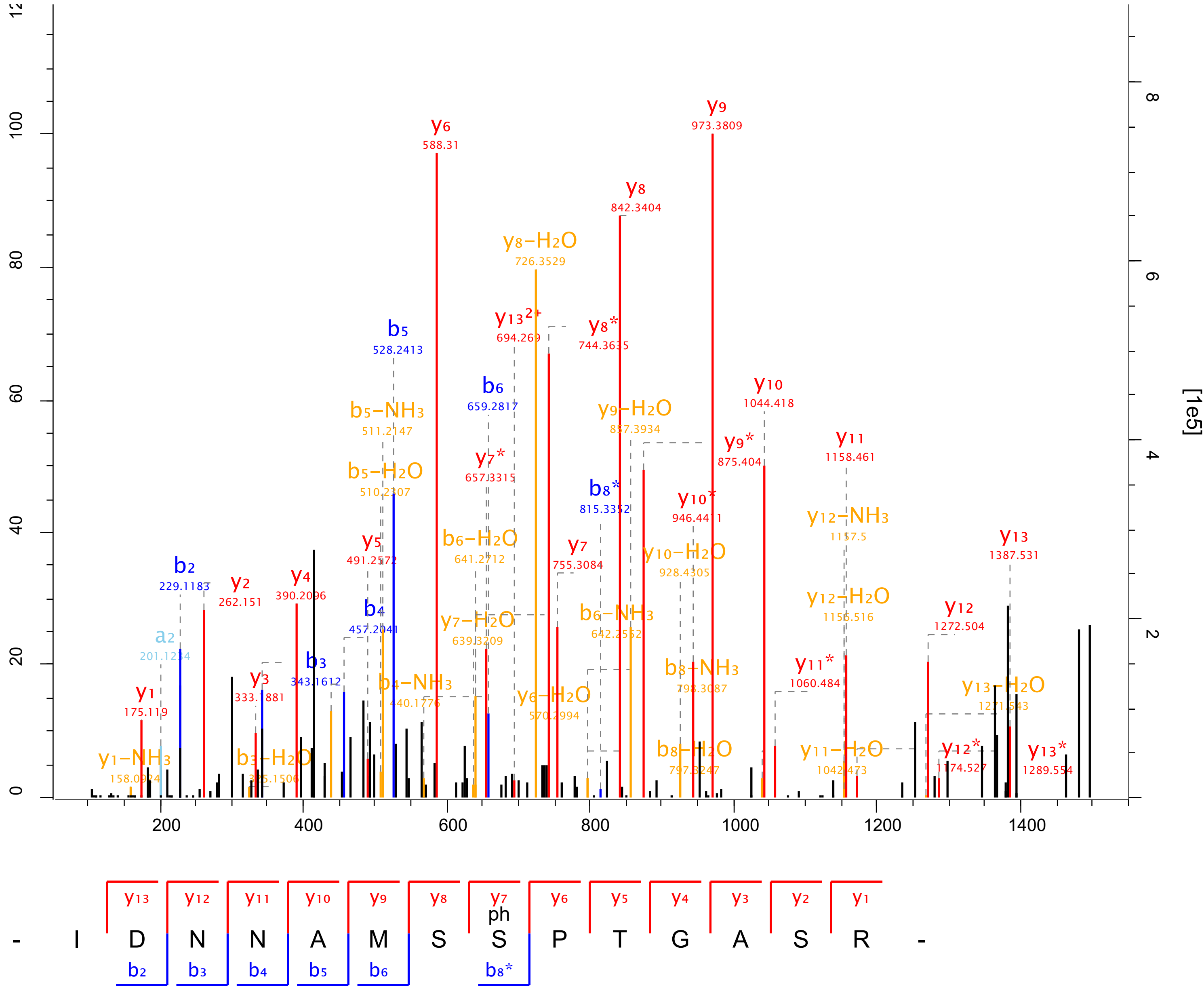

| Raw file | Scan | Method | Score | m/z |
| --- | --- | --- | --- | --- |
| water_15_2 | 8548 | FTMS; HCD | 157.68 | 663.79 |

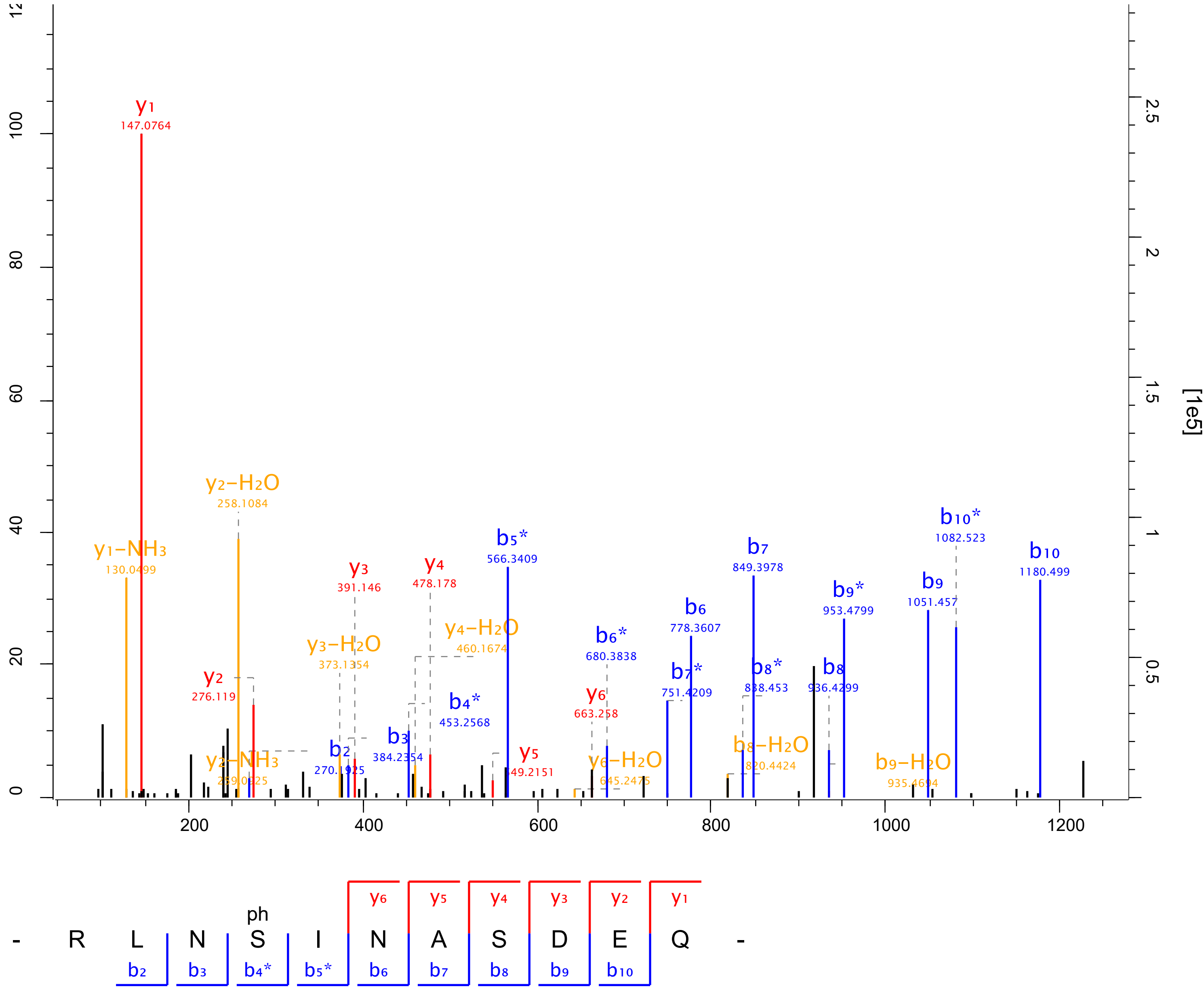

| Raw file | Scan | Method | Score | m/z |
|---|---|---|---|---|
| water_15_2 | 8555 | FTMS; HCD | 129.38 | 688.28 |

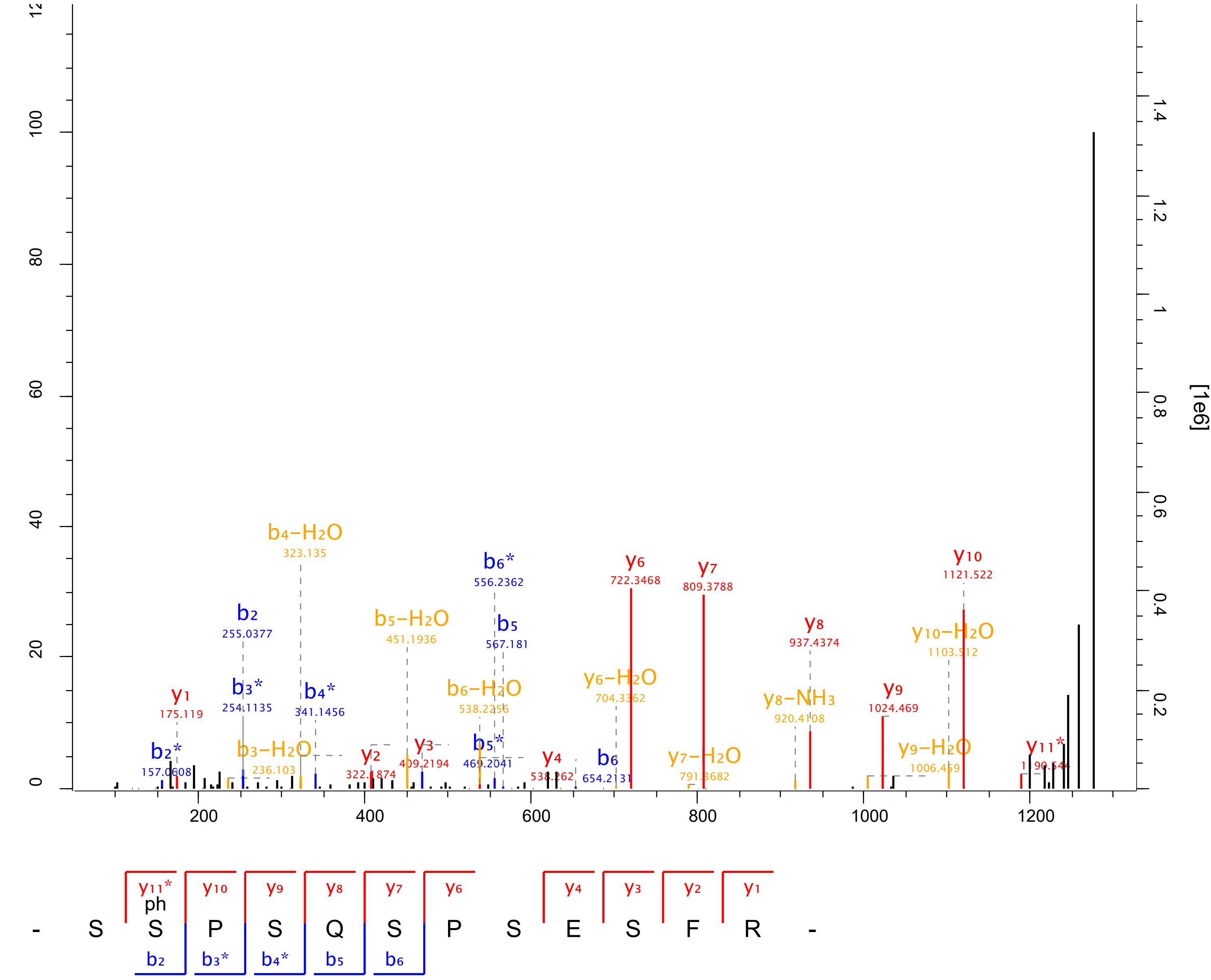

| Raw file | Scan | Method | Score | m/z |
|---|---|---|---|---|
| water_15_2 | 8676 | FTMS; HCD | 163.33 | 630.77 |

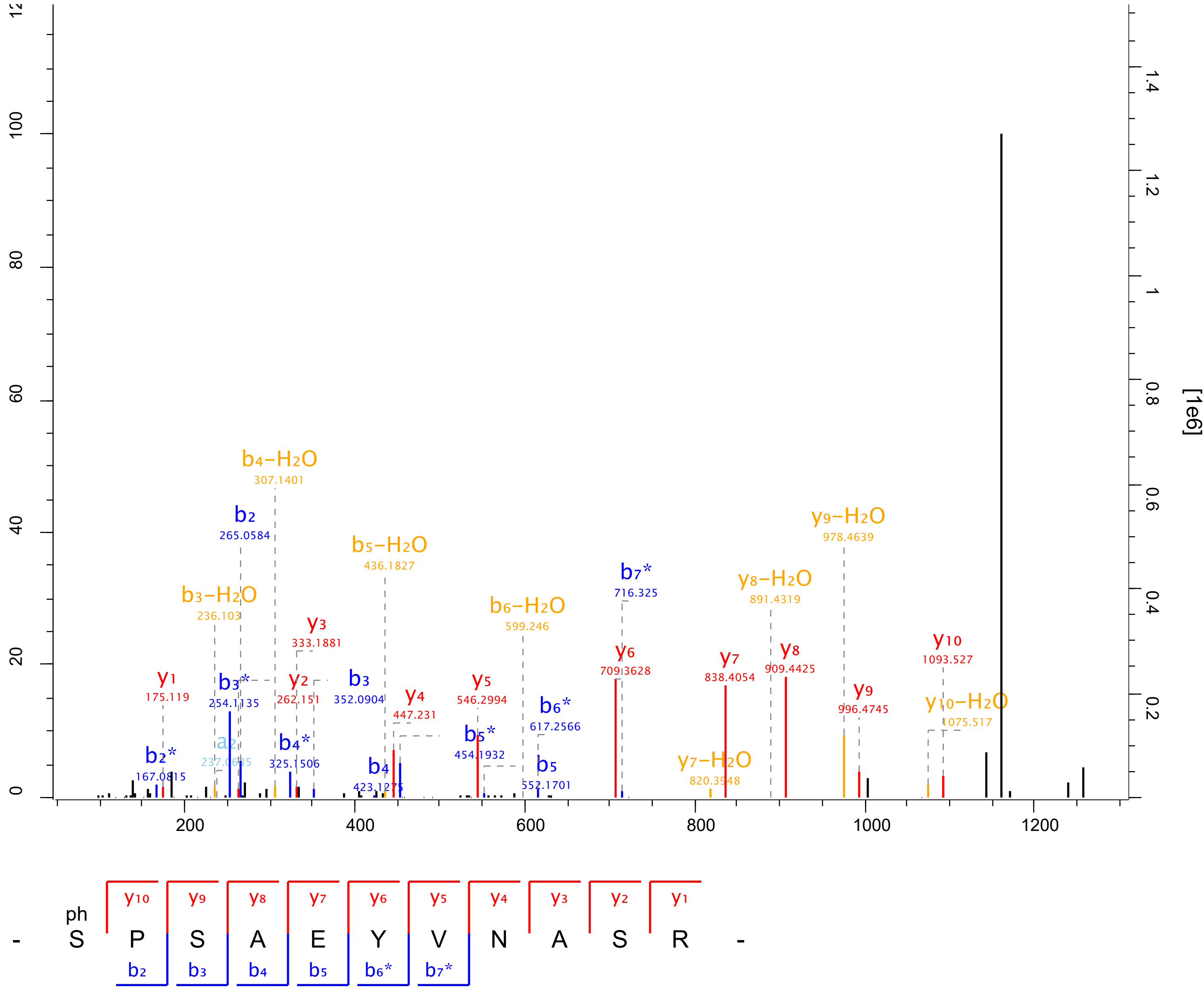

| Raw file | Scan | Method | Score | m/z |
|---|---|---|---|---|
| water_15_2 | 8706 | FTMS; HCD | 96.23 | 528.73 |

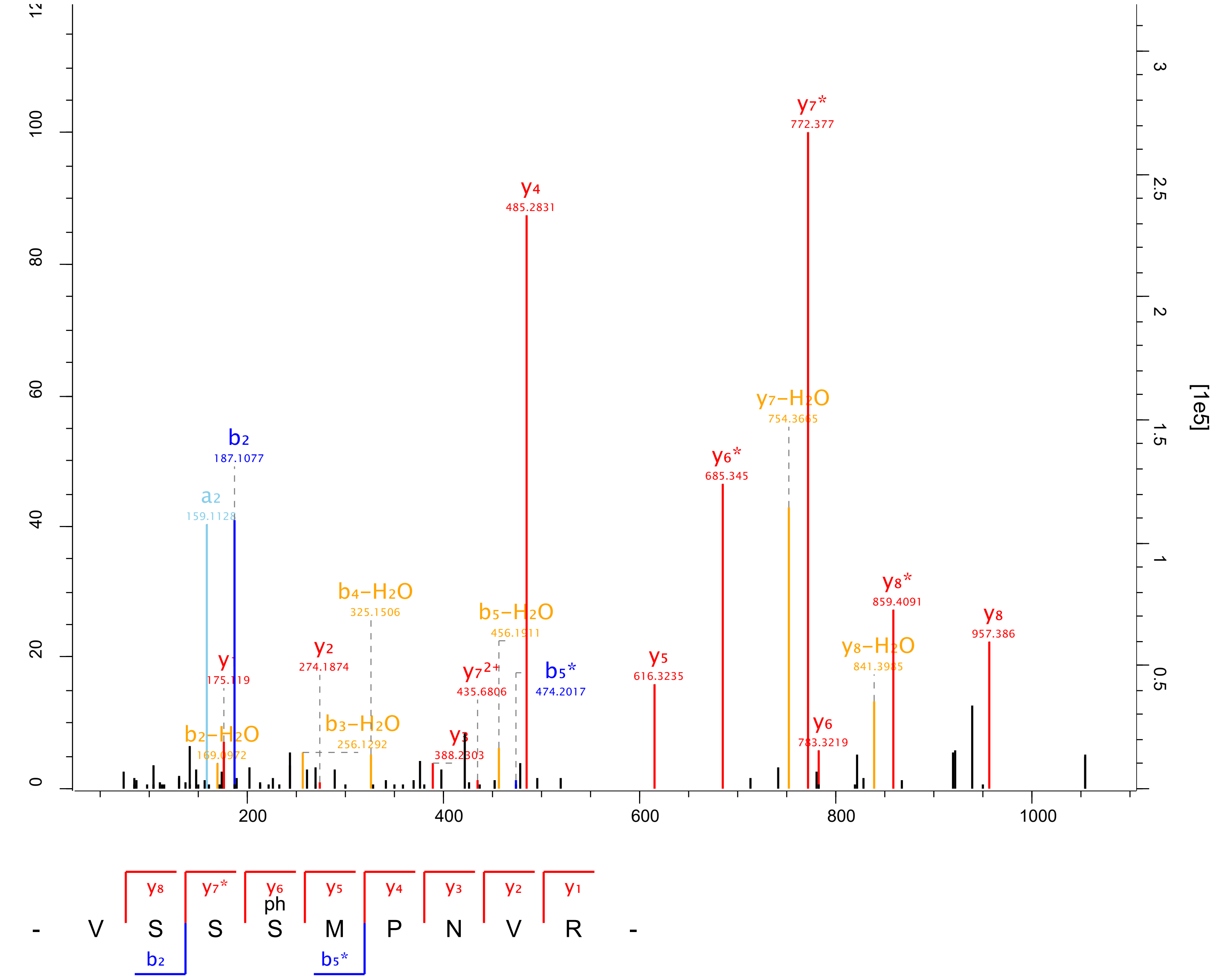

| Raw file | Scan | Method | Score | m/z |
|---|---|---|---|---|
| water_15_2 | 8711 | FTMS; HCD | 114.46 | 578.58 |

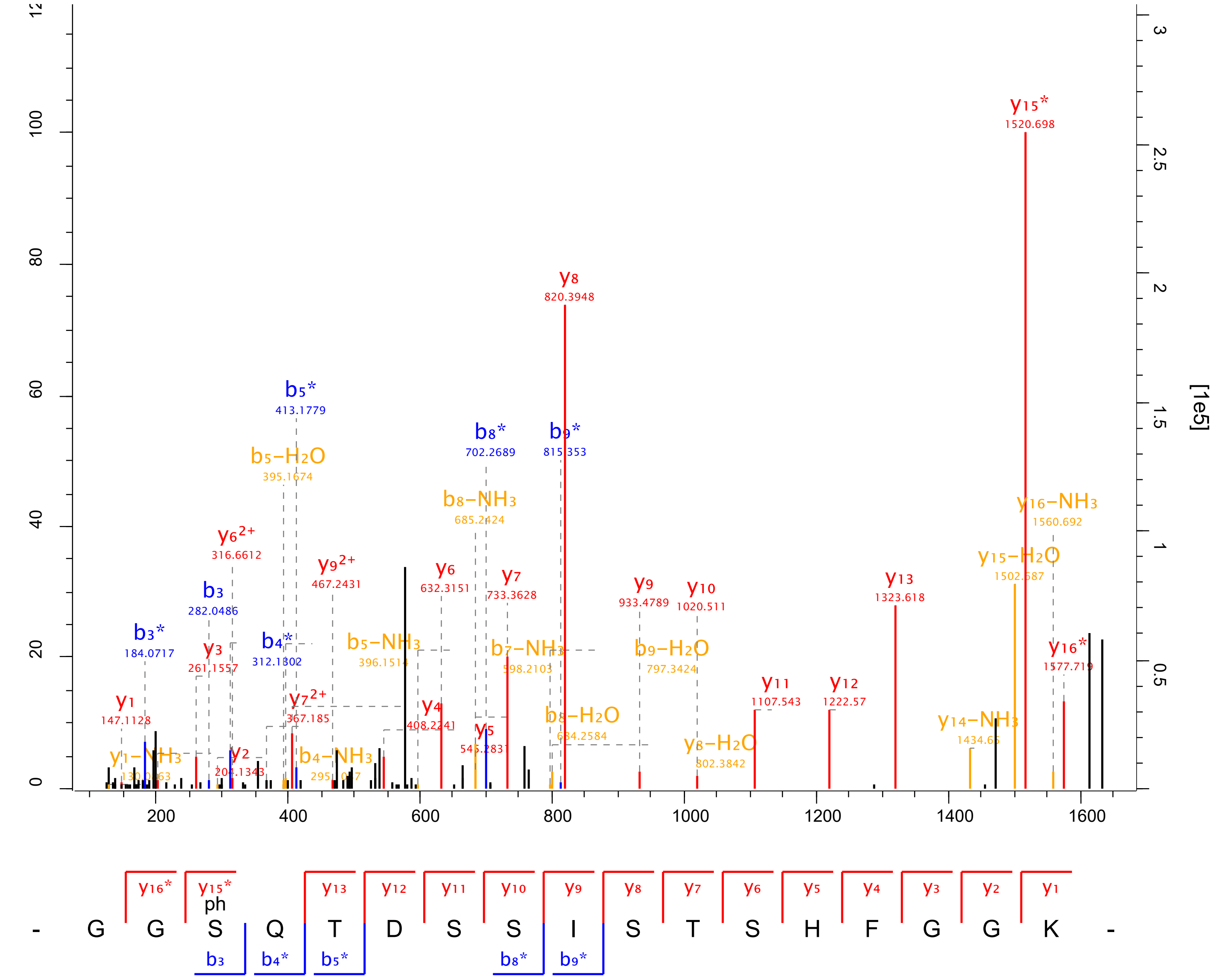

| Raw file | Scan | Method | Score | m/z |
| --- | --- | --- | --- | --- |
| water_15_2 | 8718 | FTMS; HCD | 42.63 | 559.26 |

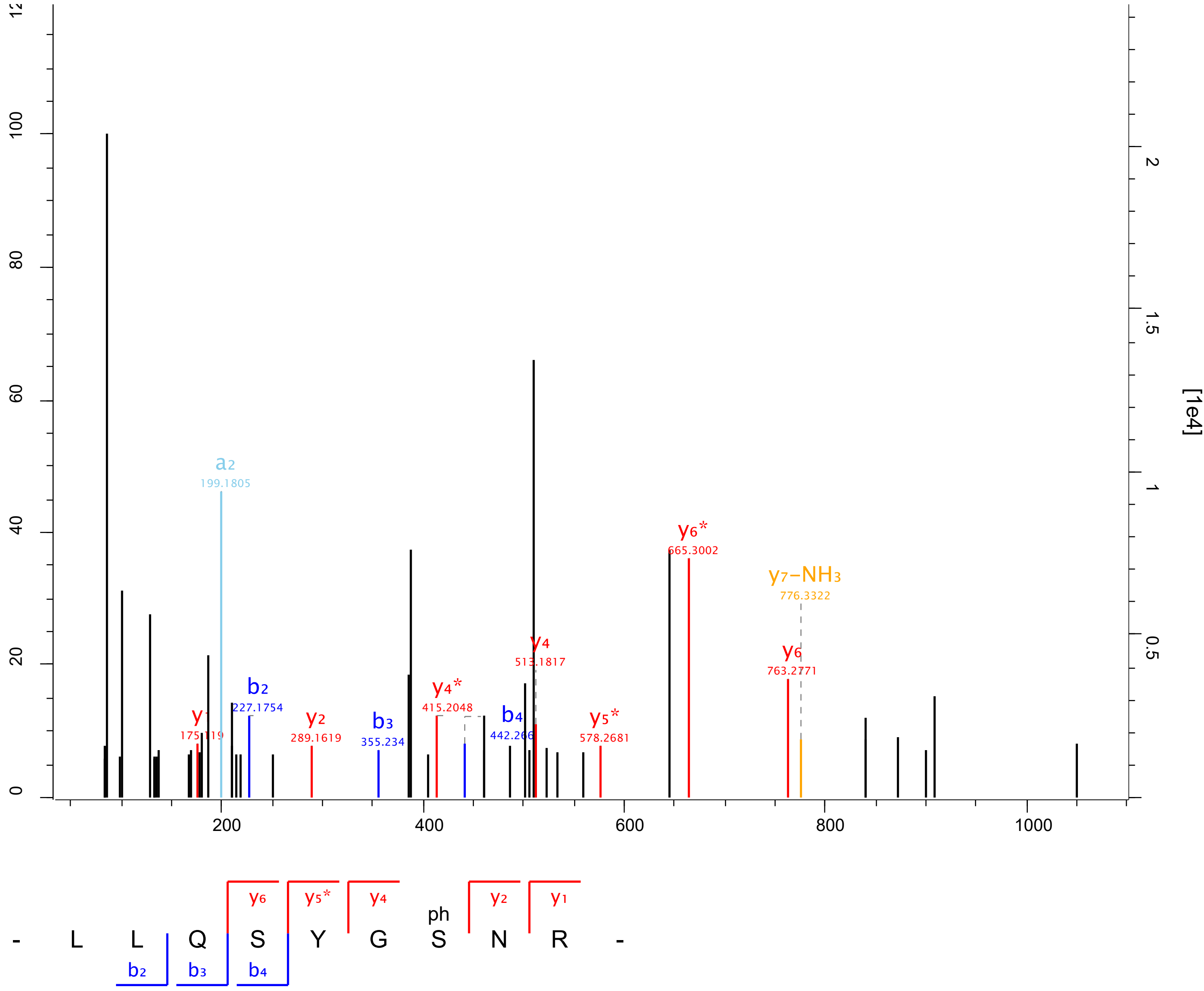

| Raw file | Scan | Method | Score | m/z |
| --- | --- | --- | --- | --- |
| water_15_2 | 8730 | FTMS; HCD | 143.31 | 521.73 |

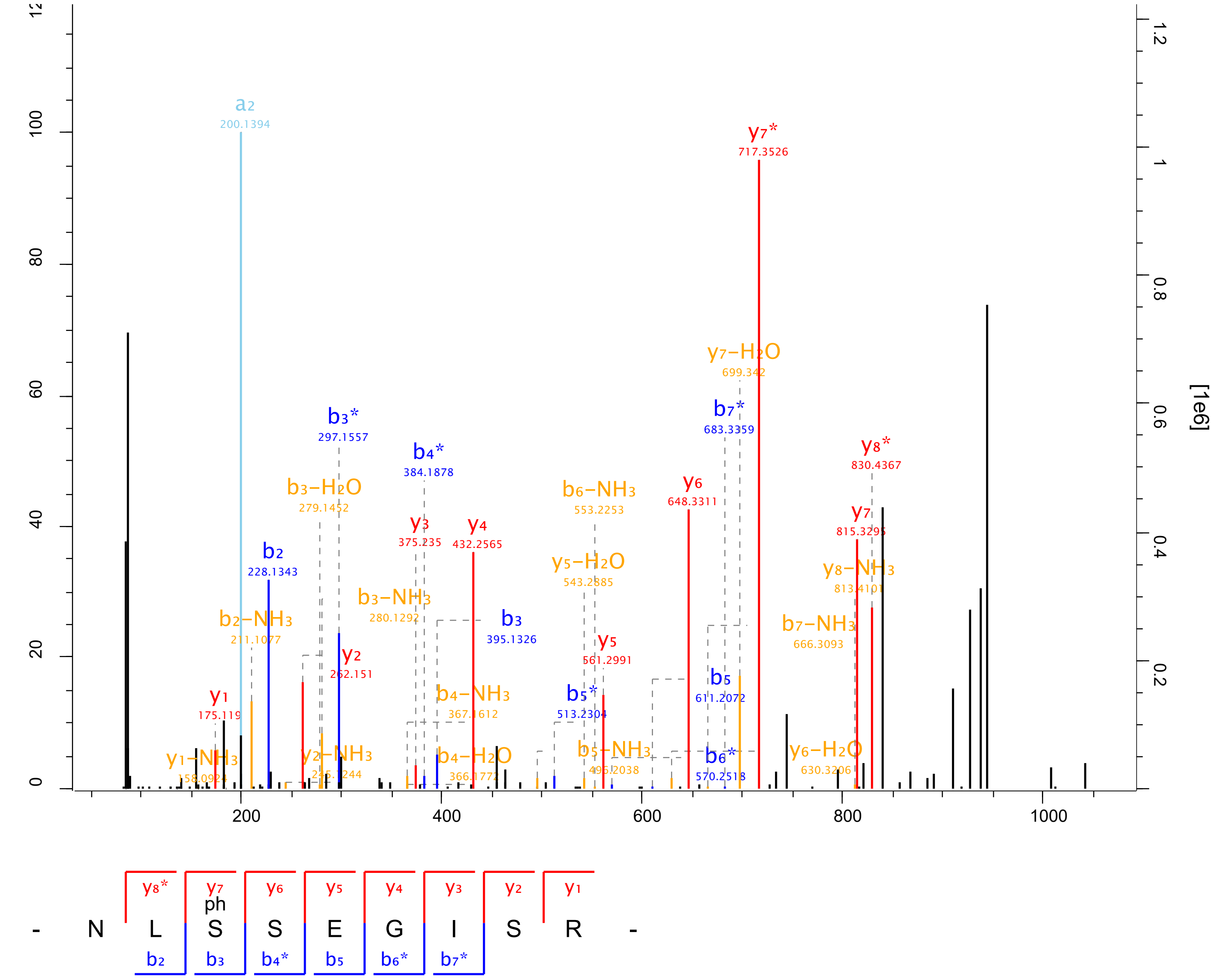

| Raw file | Scan | Method | Score | m/z |
|---|---|---|---|---|
| water_15_2 | 8744 | FTMS; HCD | 186.56 | 684.79 |

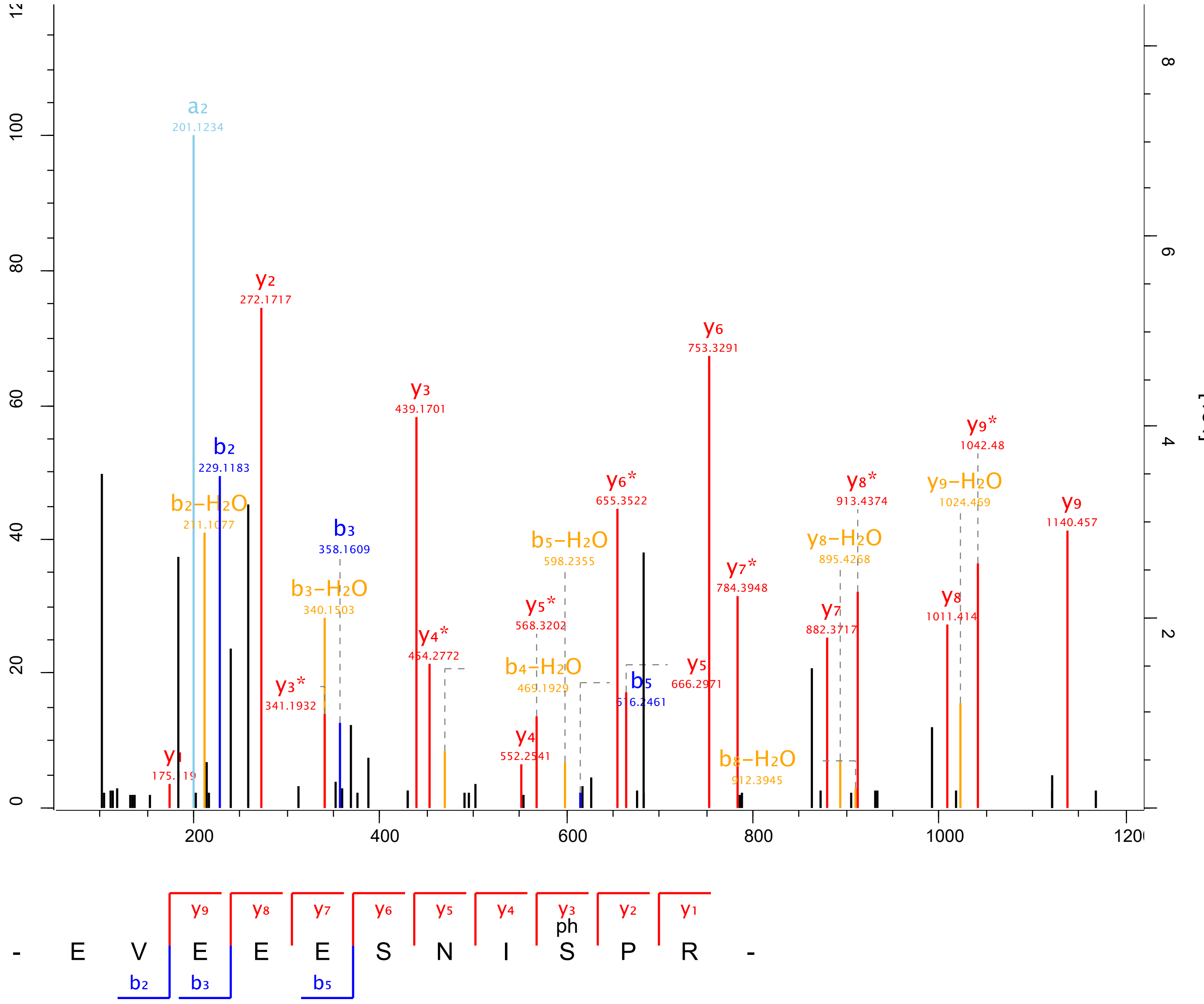

| Raw file | Scan | Method | Score | m/z |
|---|---|---|---|---|
| water_15_2 | 8785 | FTMS; HCD | 115.56 | 707.77 |

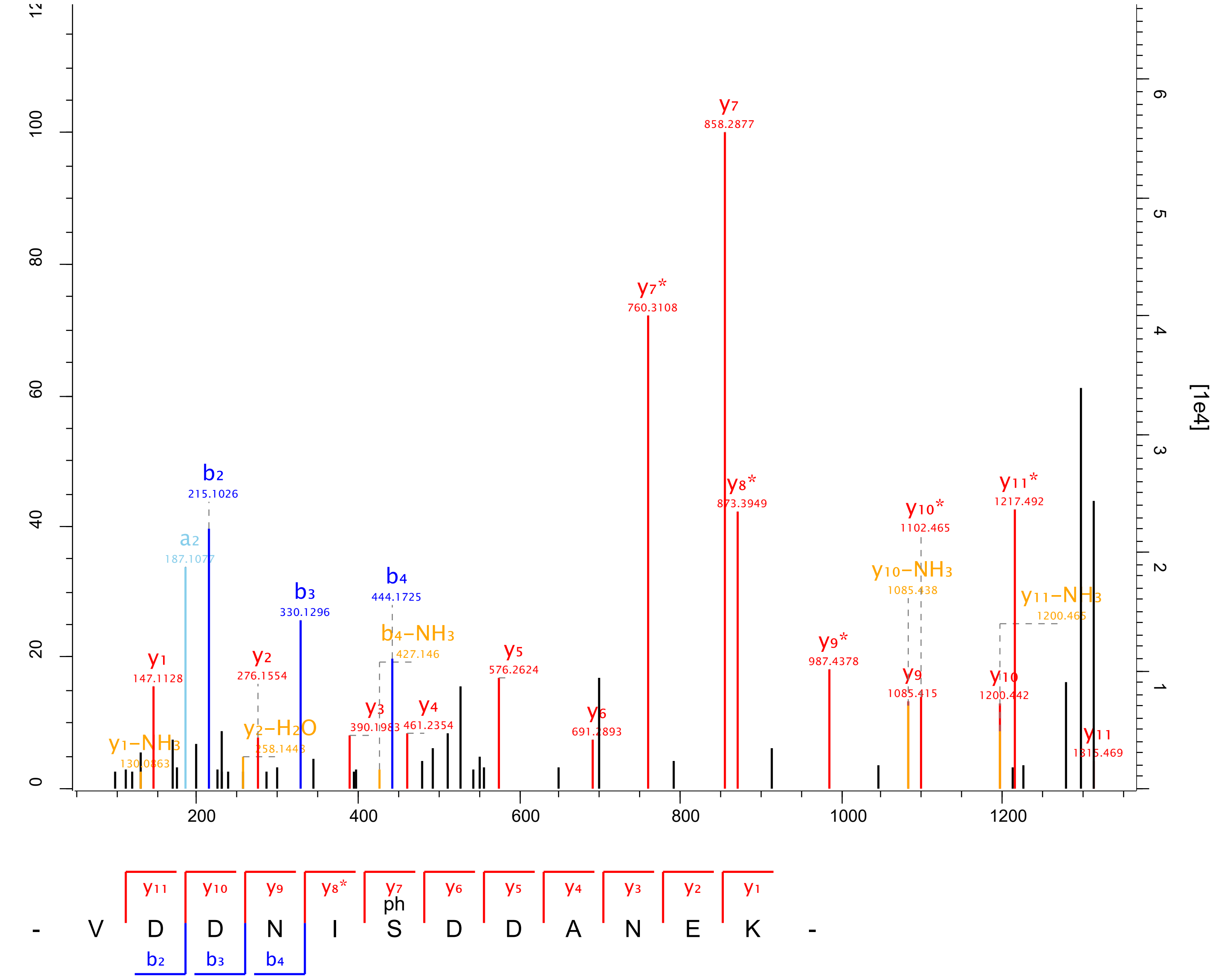

| Raw file | Scan | Method | Score | m/z |
|---|---|---|---|---|
| water_15_2 | 8787 | FTMS; HCD | 62.09 | 542.76 |

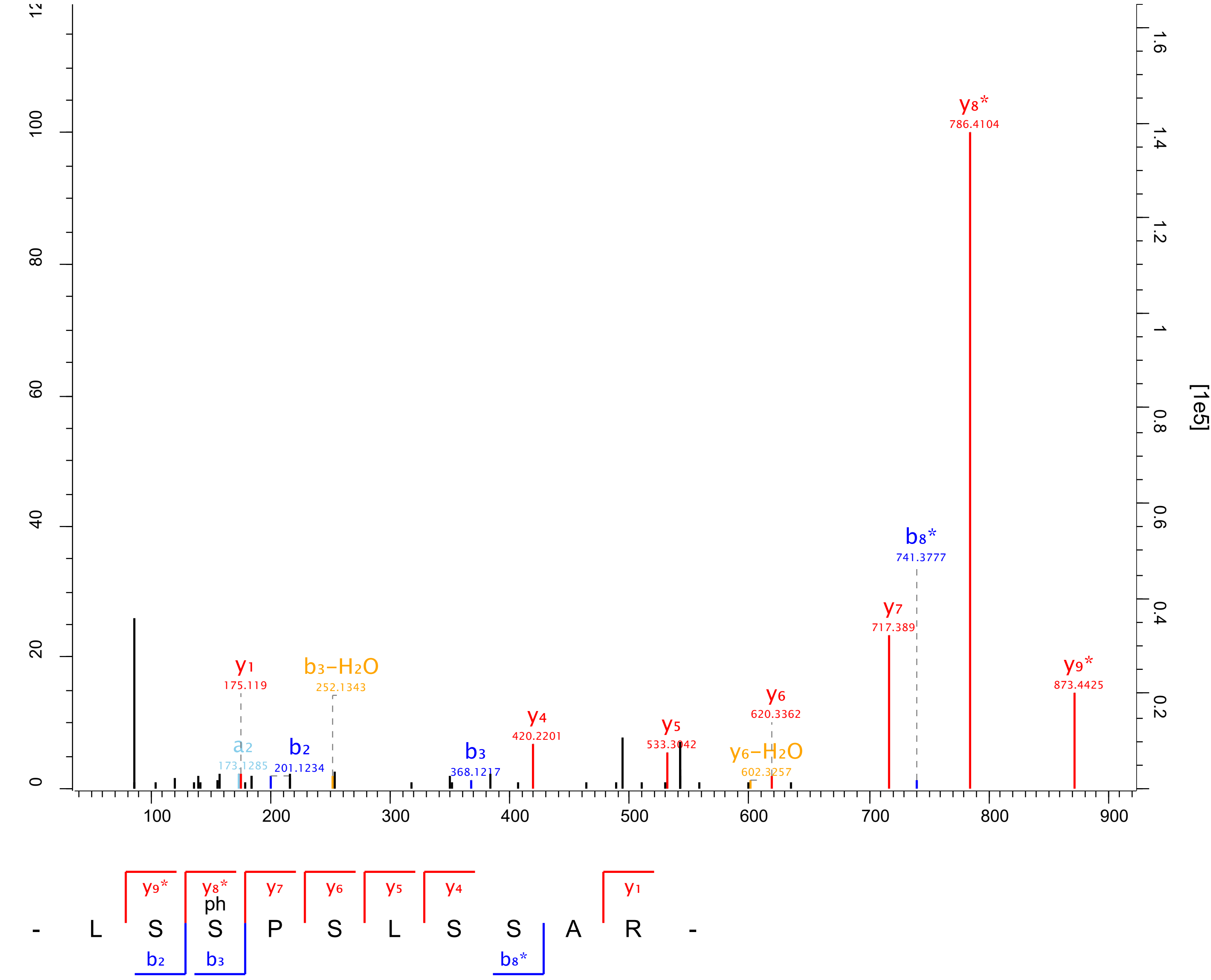

| Raw file | Scan | Method | Score | m/z |
| --- | --- | --- | --- | --- |
| water_15_2 | 8800 | FTMS; HCD | 47.71 | 415.88 |

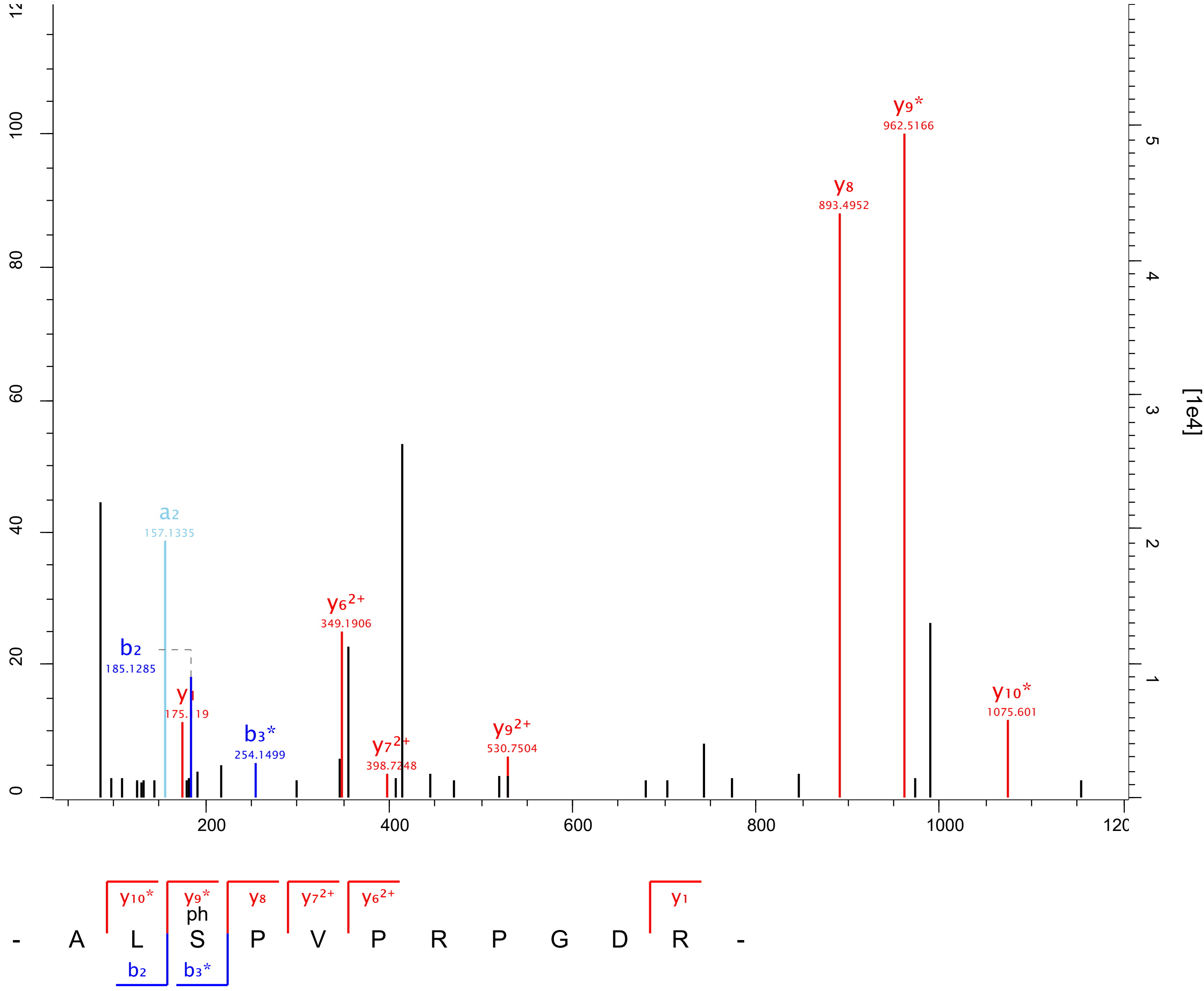

| Raw file | Scan | Method | Score | m/z |
| --- | --- | --- | --- | --- |
| water_15_2 | 8830 | FTMS; HCD | 197.38 | 570.75 |

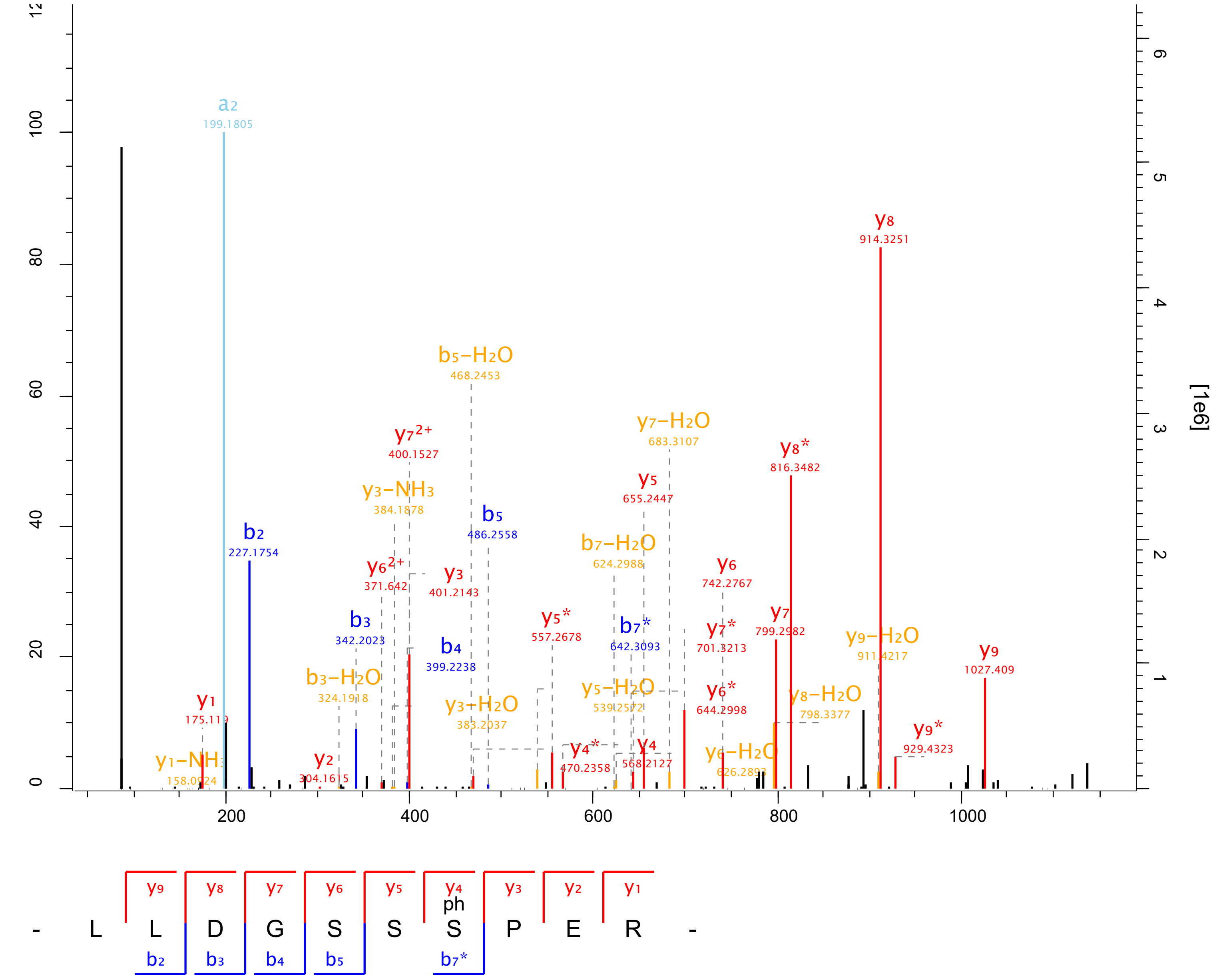

| Raw file | Scan | Method | Score | m/z |
| --- | --- | --- | --- | --- |
| water_15_2 | 8869 | FTMS; HCD | 165.38 | 1038.91 |

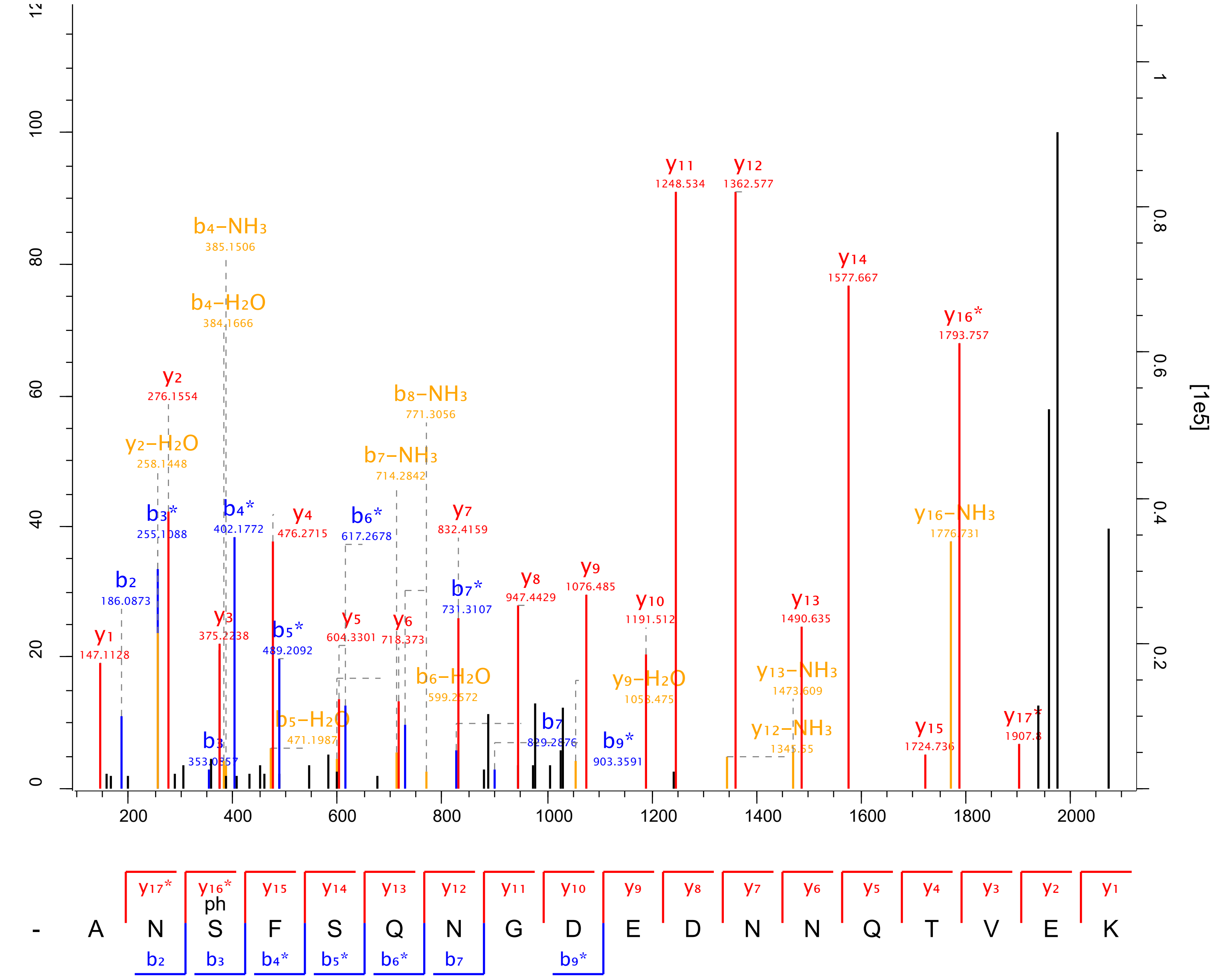

| Raw file | Scan | Method | Score | m/z |
| --- | --- | --- | --- | --- |
| water_15_2 | 8956 | FTMS; HCD | 41.24 | 536.74 |

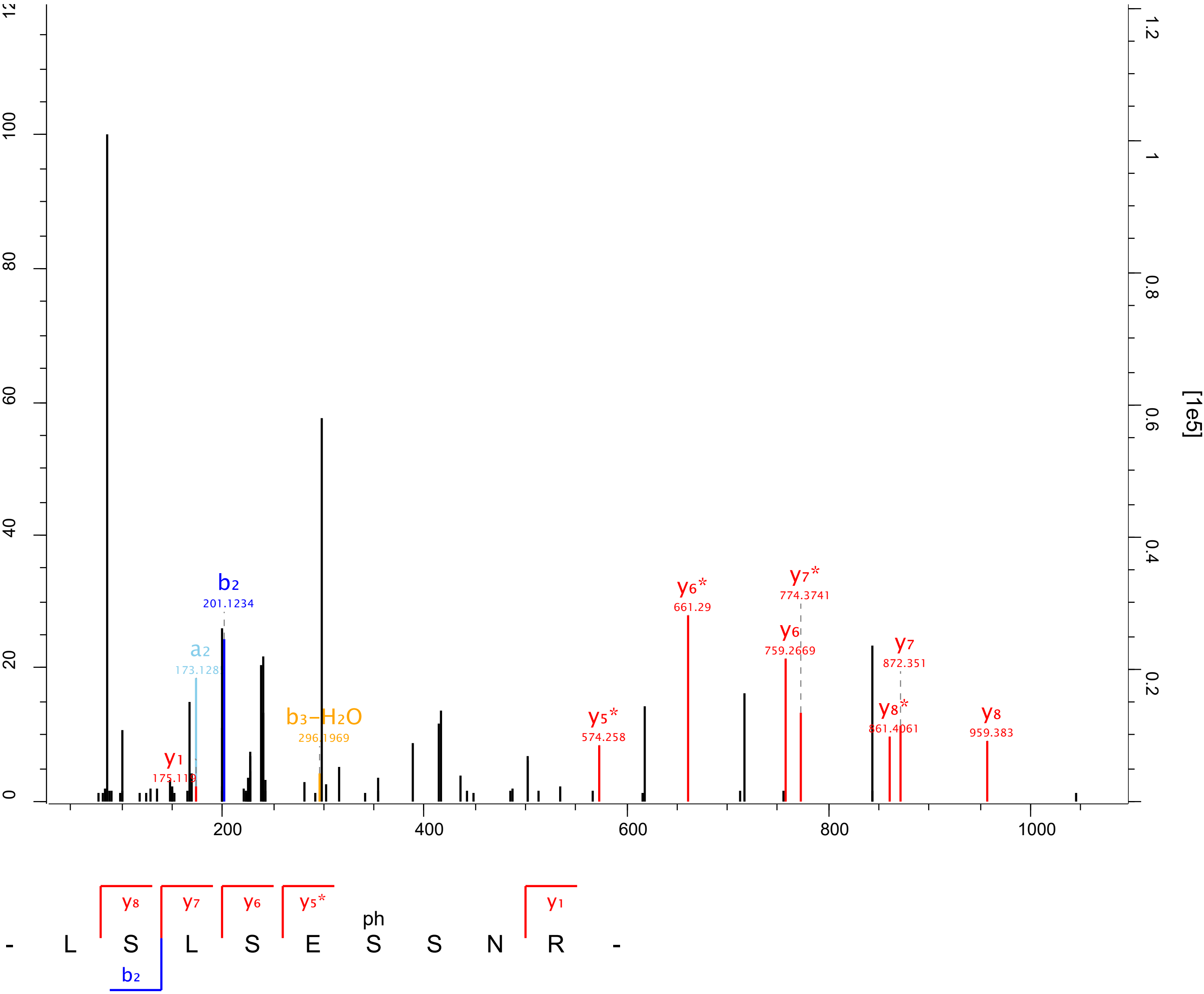

| Raw file | Scan | Method | Score | m/z |
|---|---|---|---|---|
| water_15_2 | 8967 | FTMS; HCD | 59.23 | 514.72 |

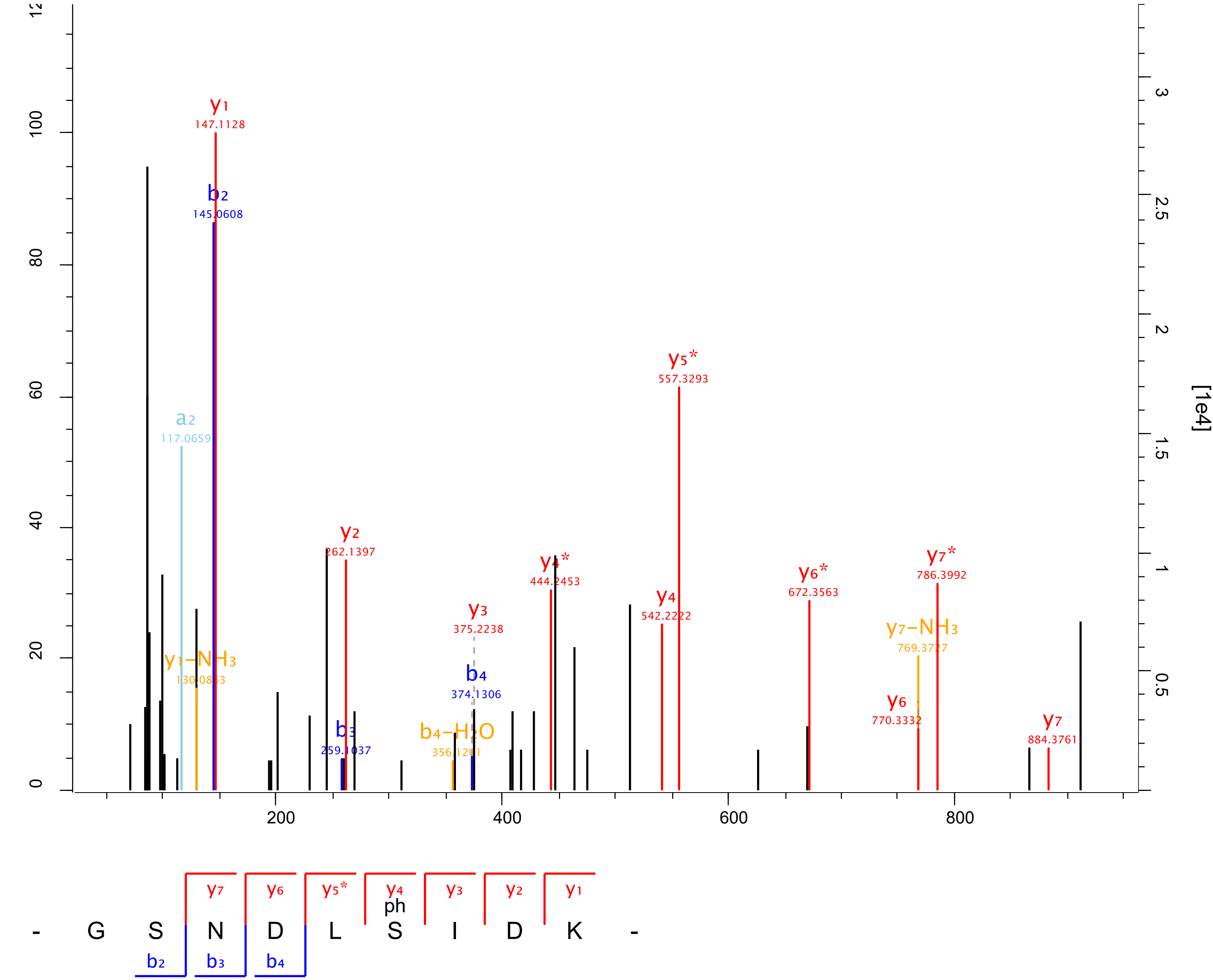

| Raw file | Scan | Method | Score | m/z |
|---|---|---|---|---|
| water_15_2 | 9001 | FTMS; HCD | 54.28 | 693.29 |

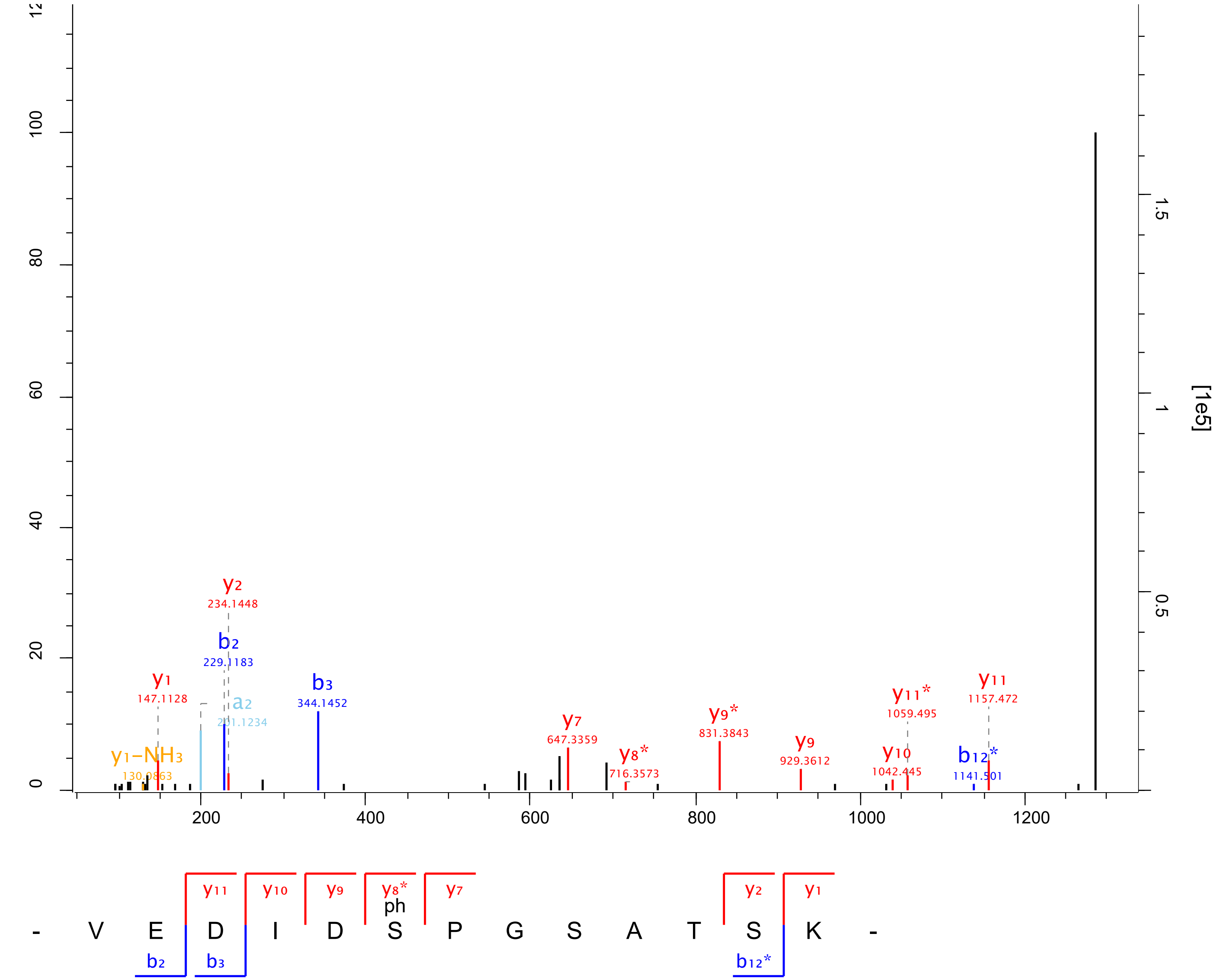

| Raw file | Scan | Method | Score | m/z |
| --- | --- | --- | --- | --- |
| water_15_2 | 9023 | FTMS; HCD | 92.38 | 484.87 |

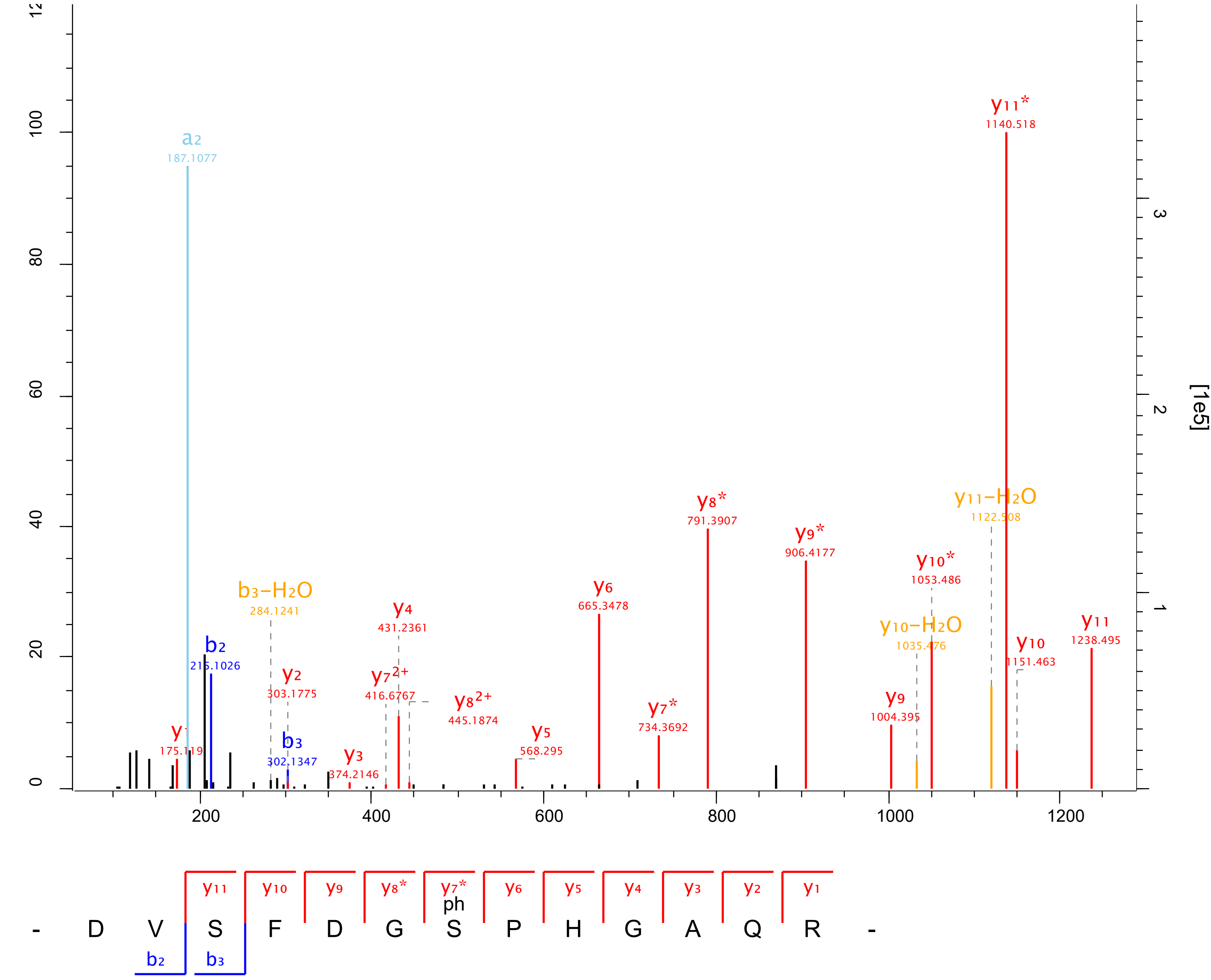

| Raw file | Scan | Method | Score | m/z |
|---|---|---|---|---|
| water_15_2 | 9114 | FTMS; HCD | 158.44 | 786.34 |

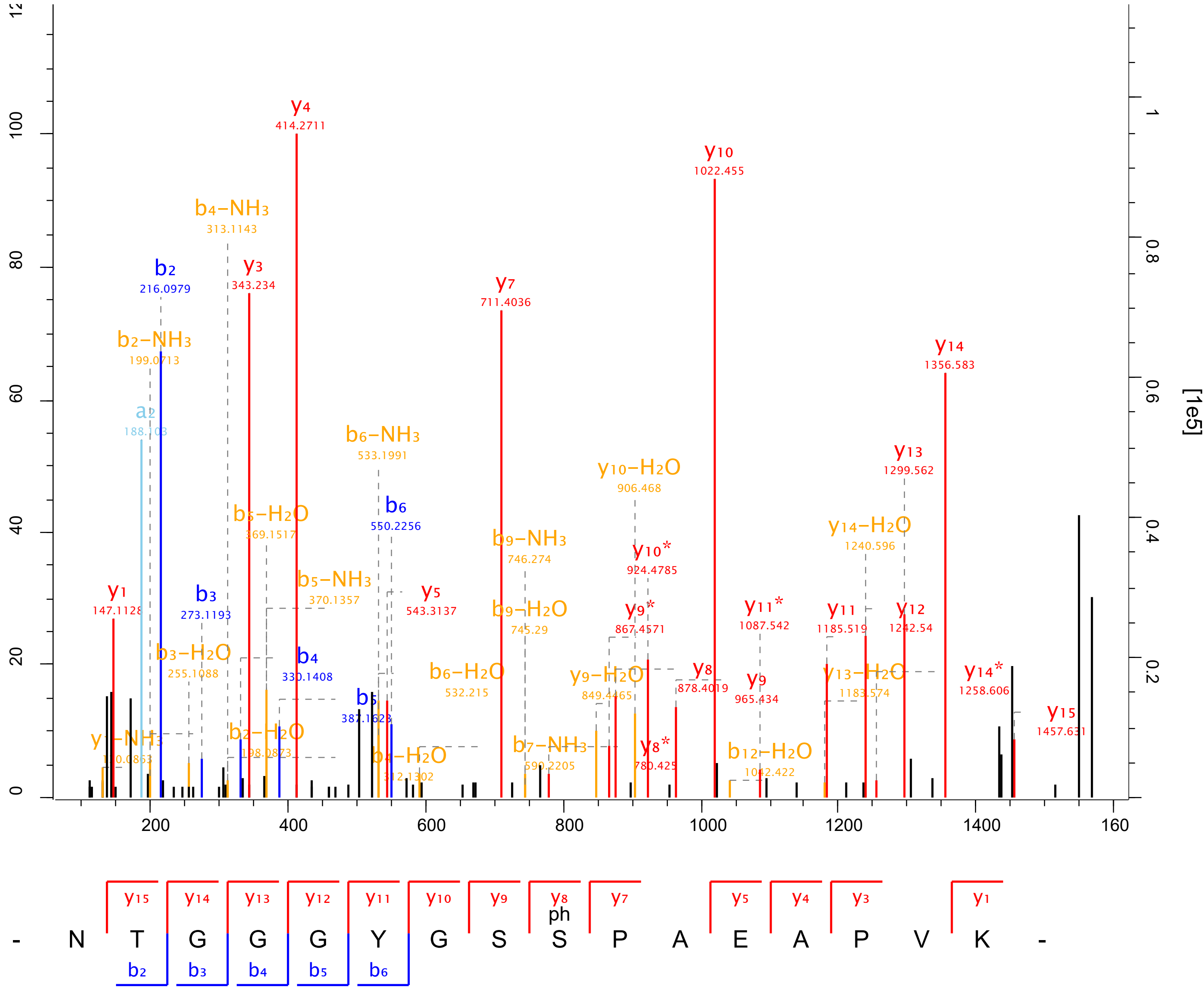

| Raw file | Scan | Method | Score | m/z |
| --- | --- | --- | --- | --- |
| water_15_2 | 9125 | FTMS; HCD | 61.28 | 627.94 |

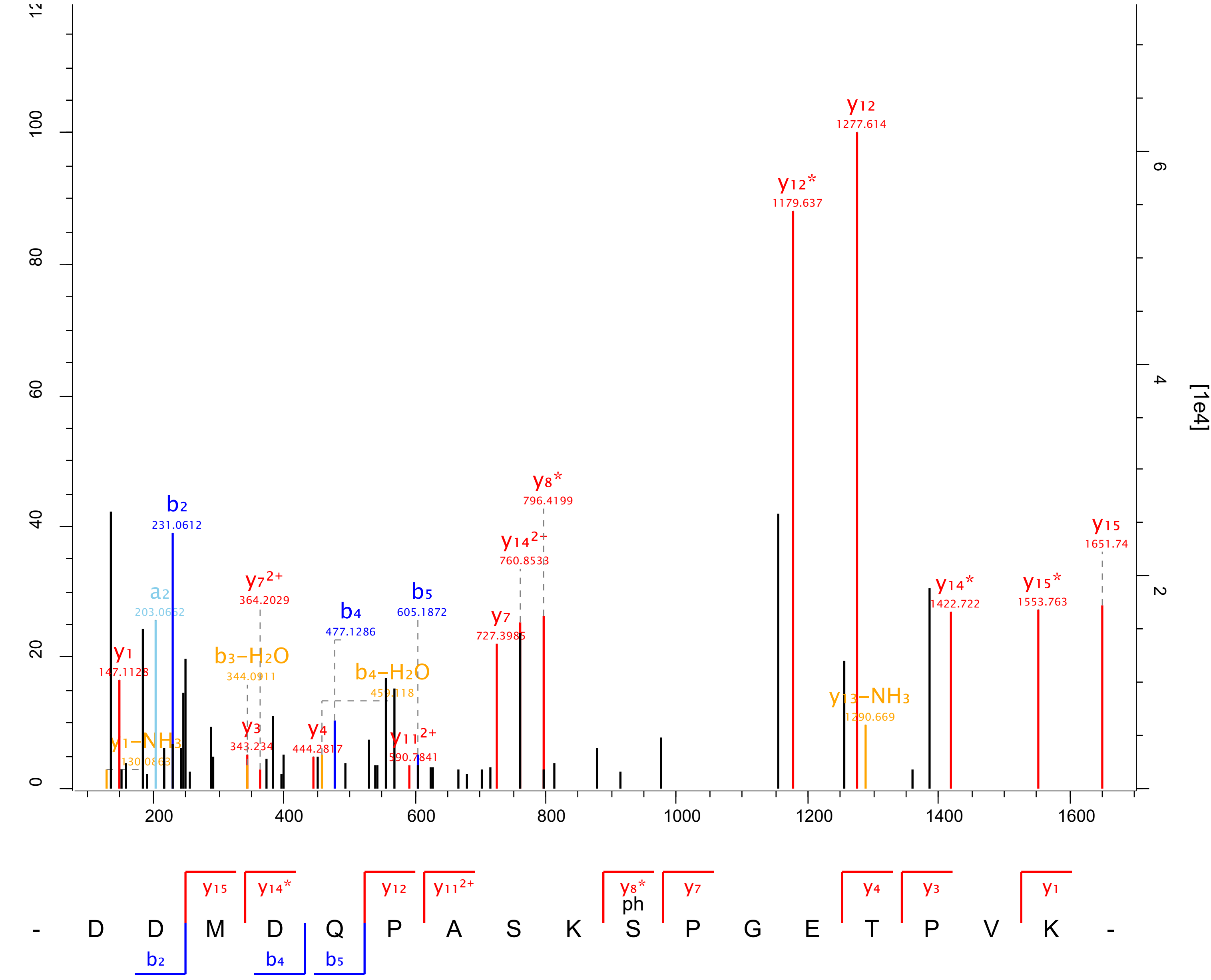

| Raw file | Scan | Method | Score | m/z |
|---|---|---|---|---|
| water_15_2 | 9159 | FTMS; HCD | 174.22 | 650.93 |

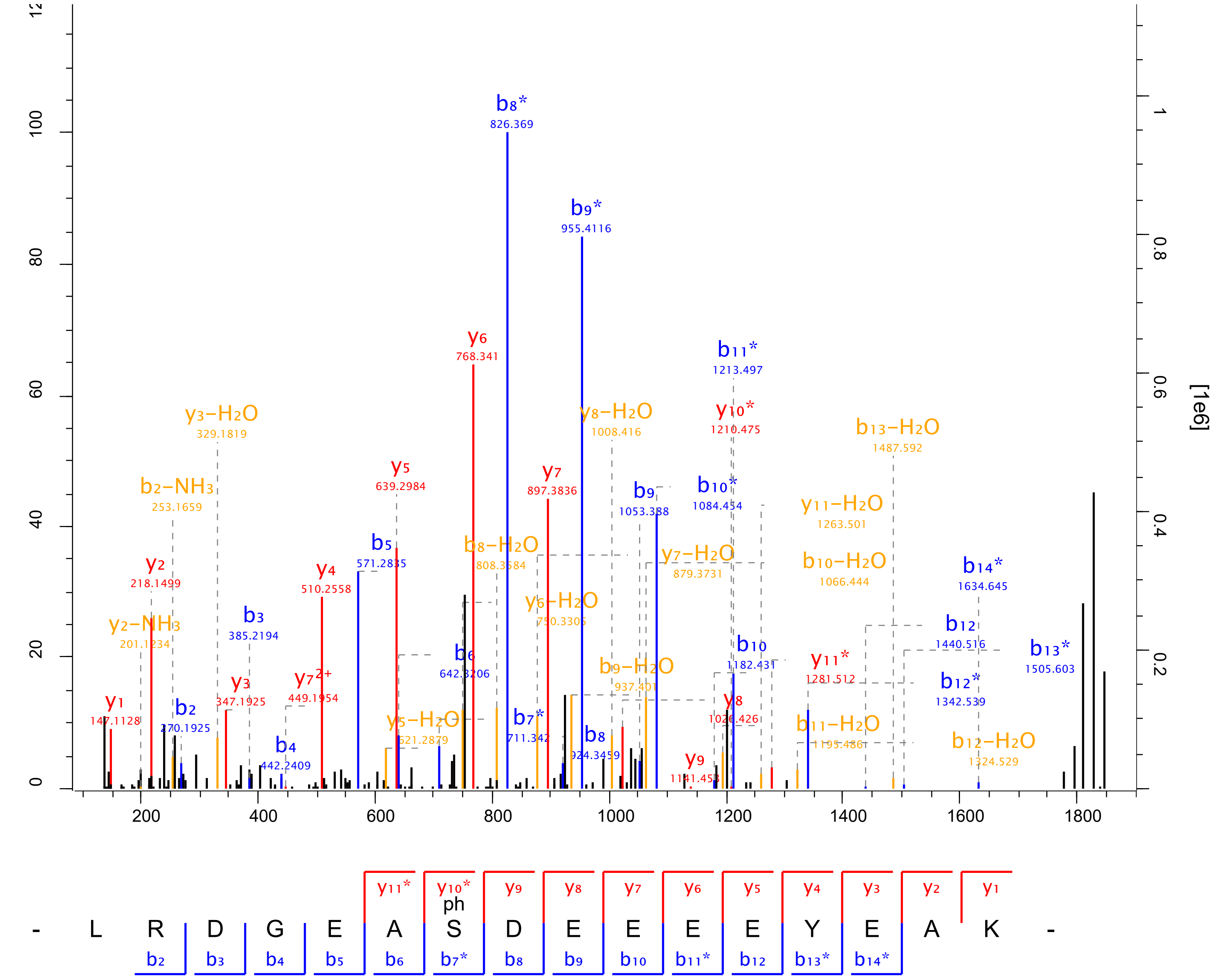

| Raw file | Scan | Method | Score | m/z |
|---|---|---|---|---|
| water_15_2 | 9213 | FTMS; HCD | 184.44 | 566.25 |

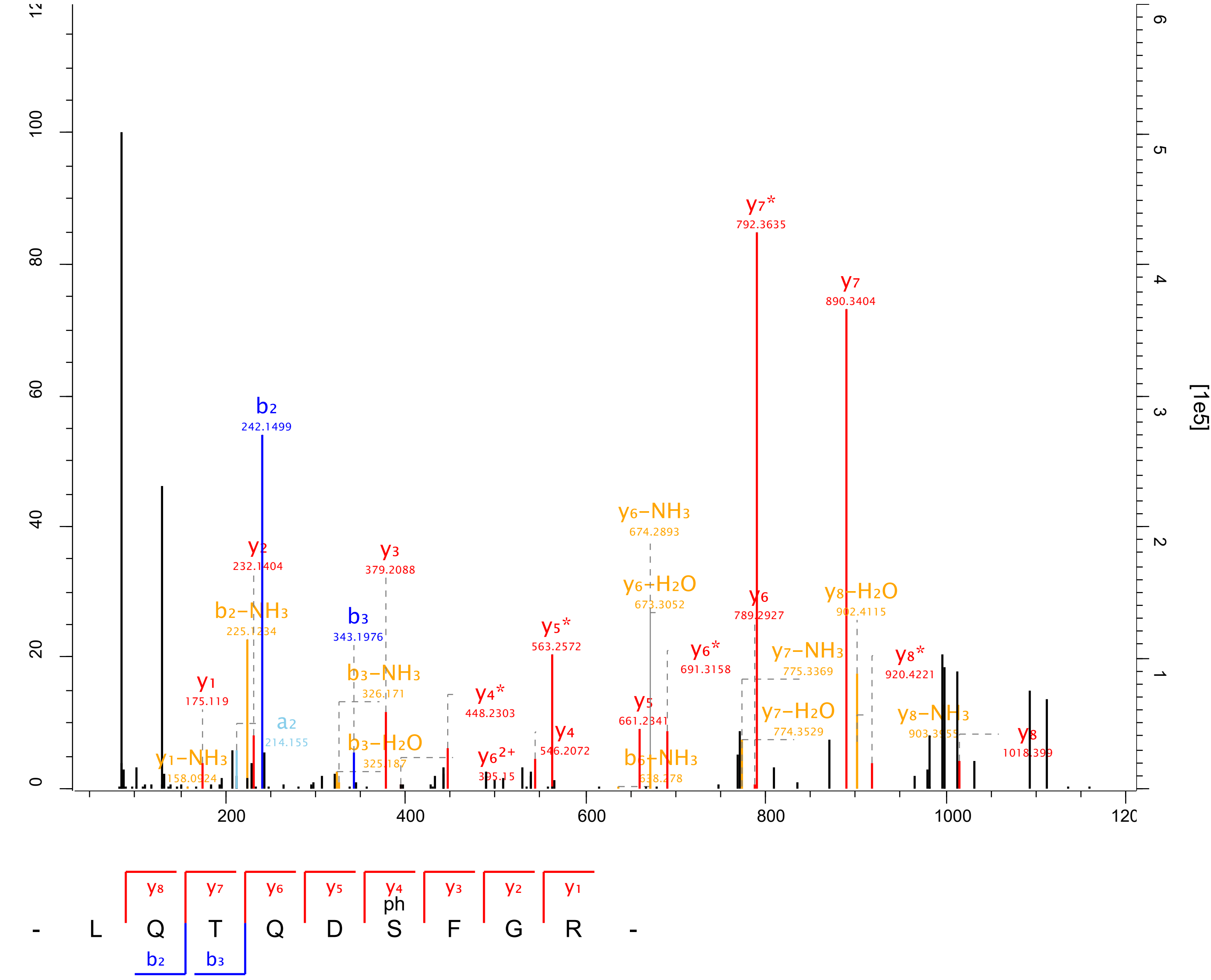

| Raw file | Scan | Method | Score | m/z |
|---|---|---|---|---|
| water_15_2 | 9224 | FTMS; HCD | 81.62 | 508.22 |

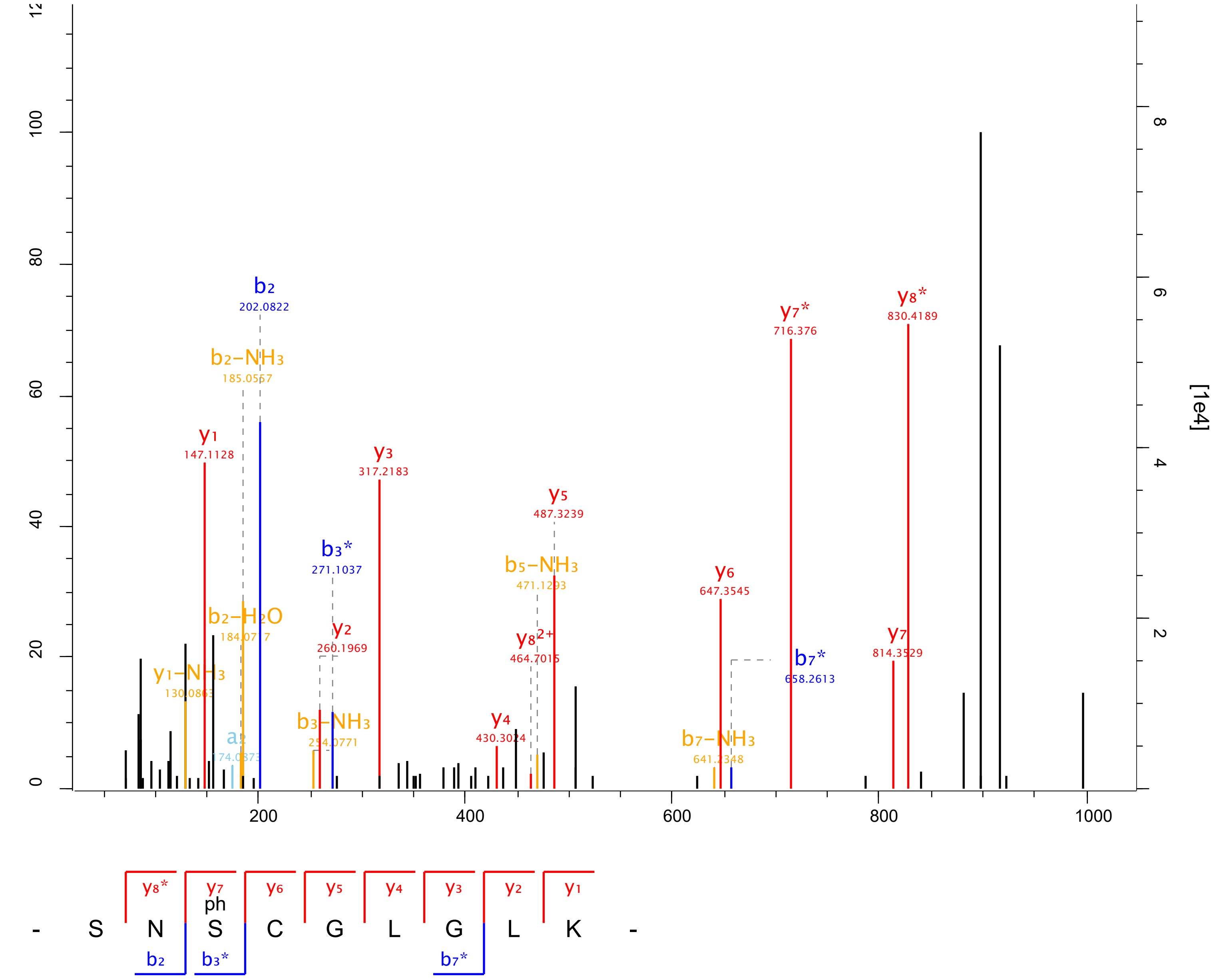

| Raw file | Scan | Method | Score | m/z |
|---|---|---|---|---|
| water_15_2 | 9229 | FTMS; HCD | 95.26 | 601.75 |

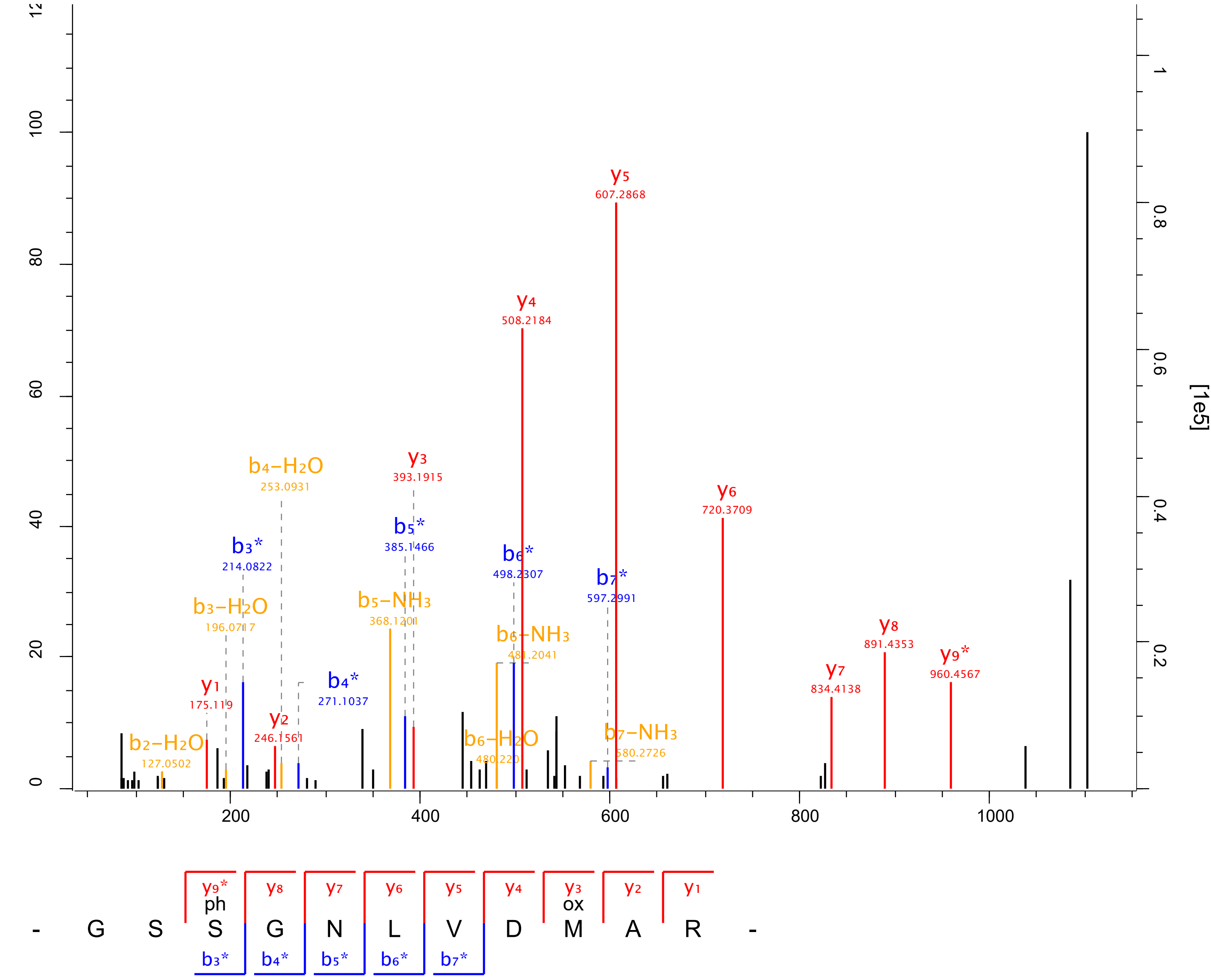

| Raw file | Scan | Method | Score | m/z |
| --- | --- | --- | --- | --- |
| water_15_2 | 9279 | FTMS; HCD | 120.06 | 524.69 |

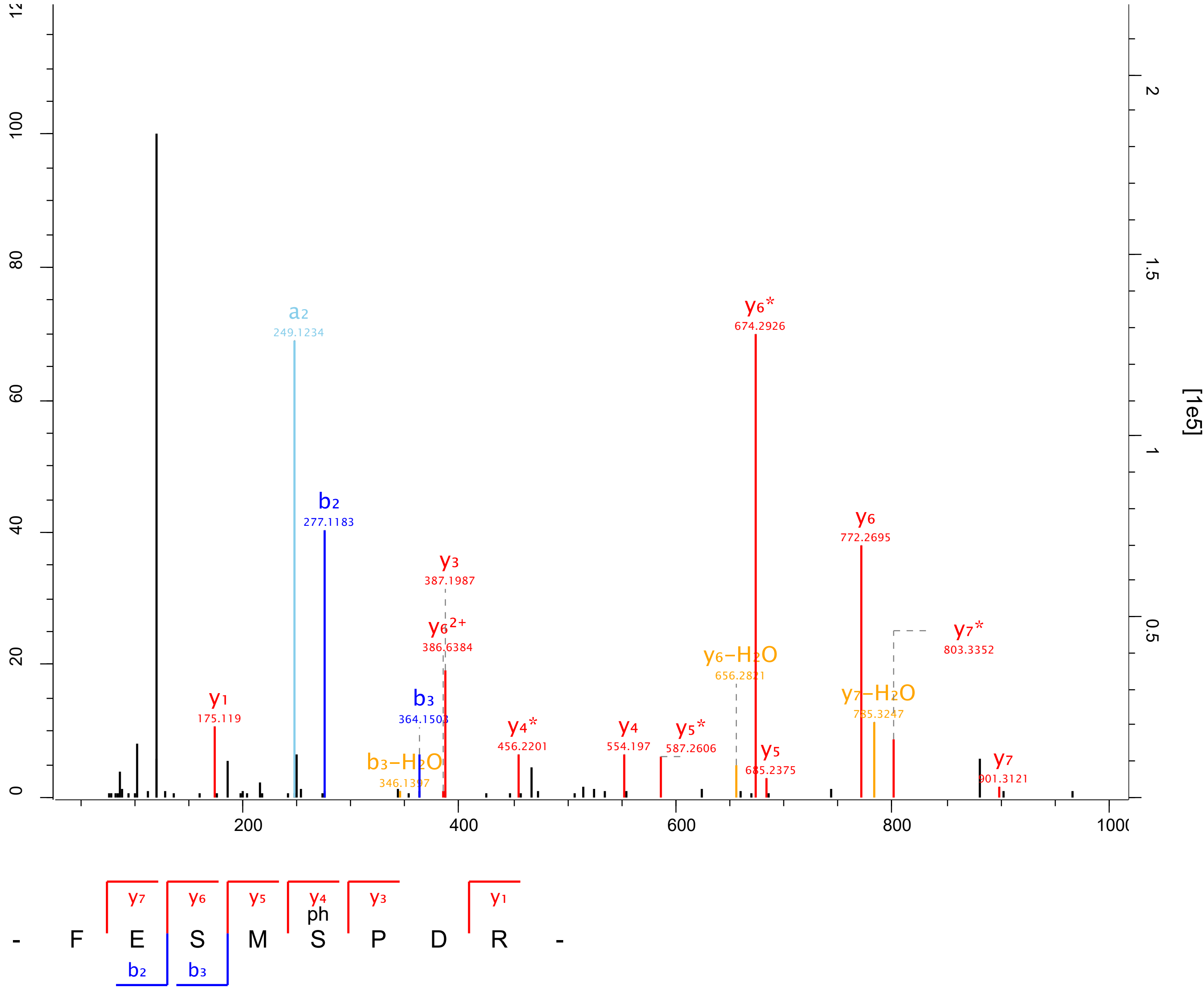

| Raw file | Scan | Method | Score | m/z |
|---|---|---|---|---|
| water_15_2 | 9292 | FTMS; HCD | 166.7 | 585.76 |

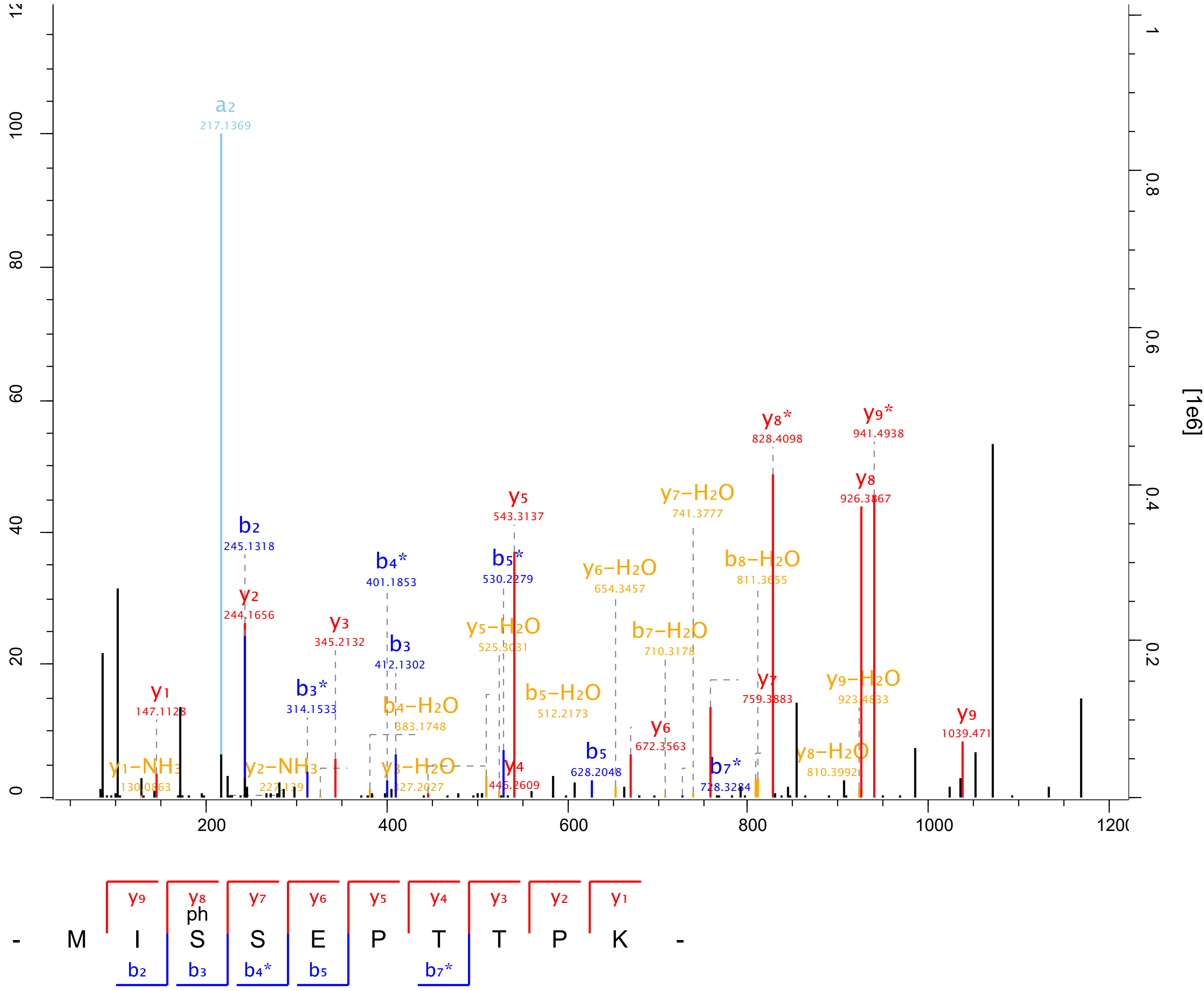

| Raw file | Scan | Method | Score | m/z |
|---|---|---|---|---|
| water_15_2 | 9329 | FTMS; HCD | 70.03 | 418.22 |

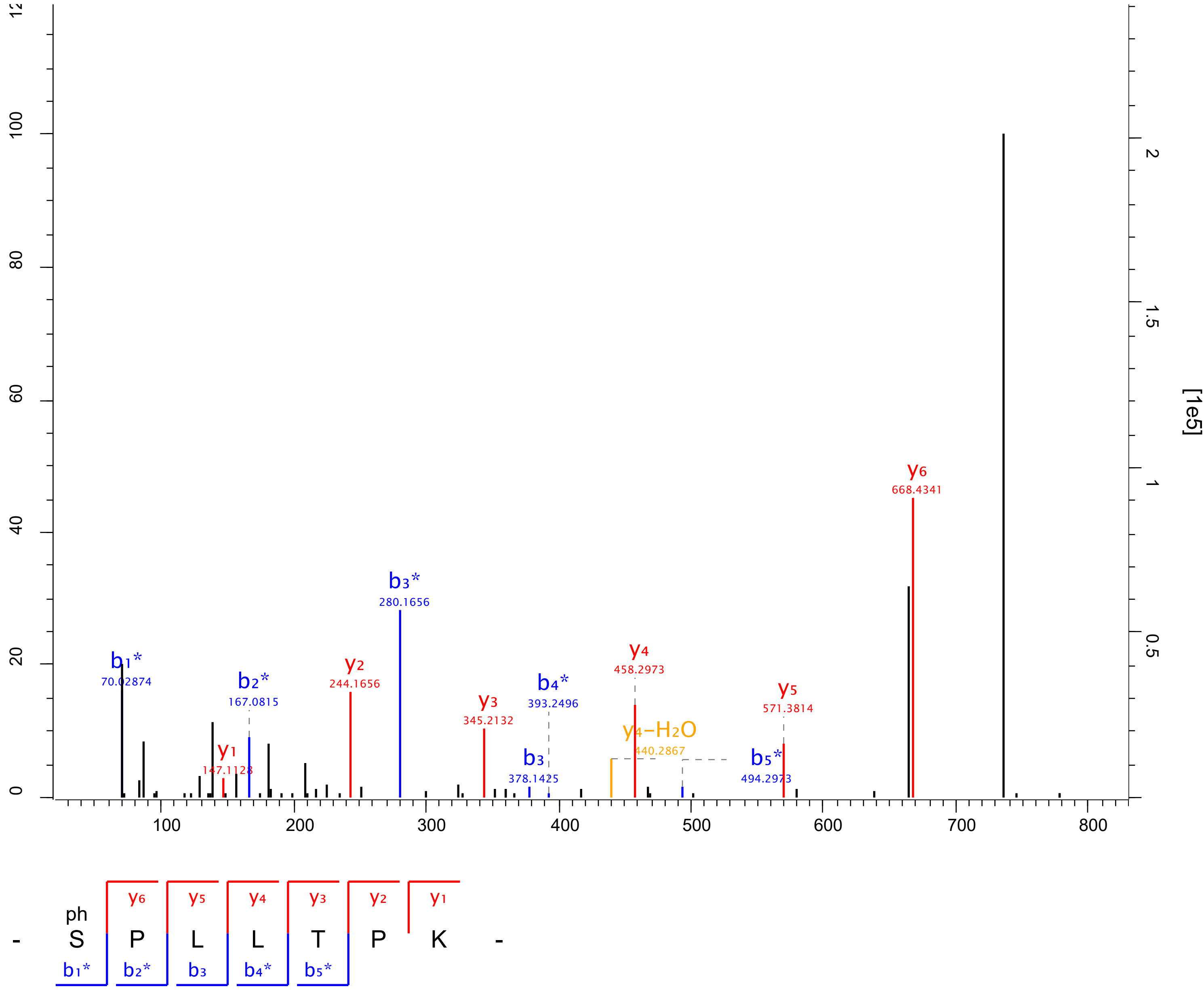

| Raw file | Scan | Method | Score | m/z |
| --- | --- | --- | --- | --- |
| water_15_2 | 9349 | FTMS; HCD | 295.03 | 812.85 |

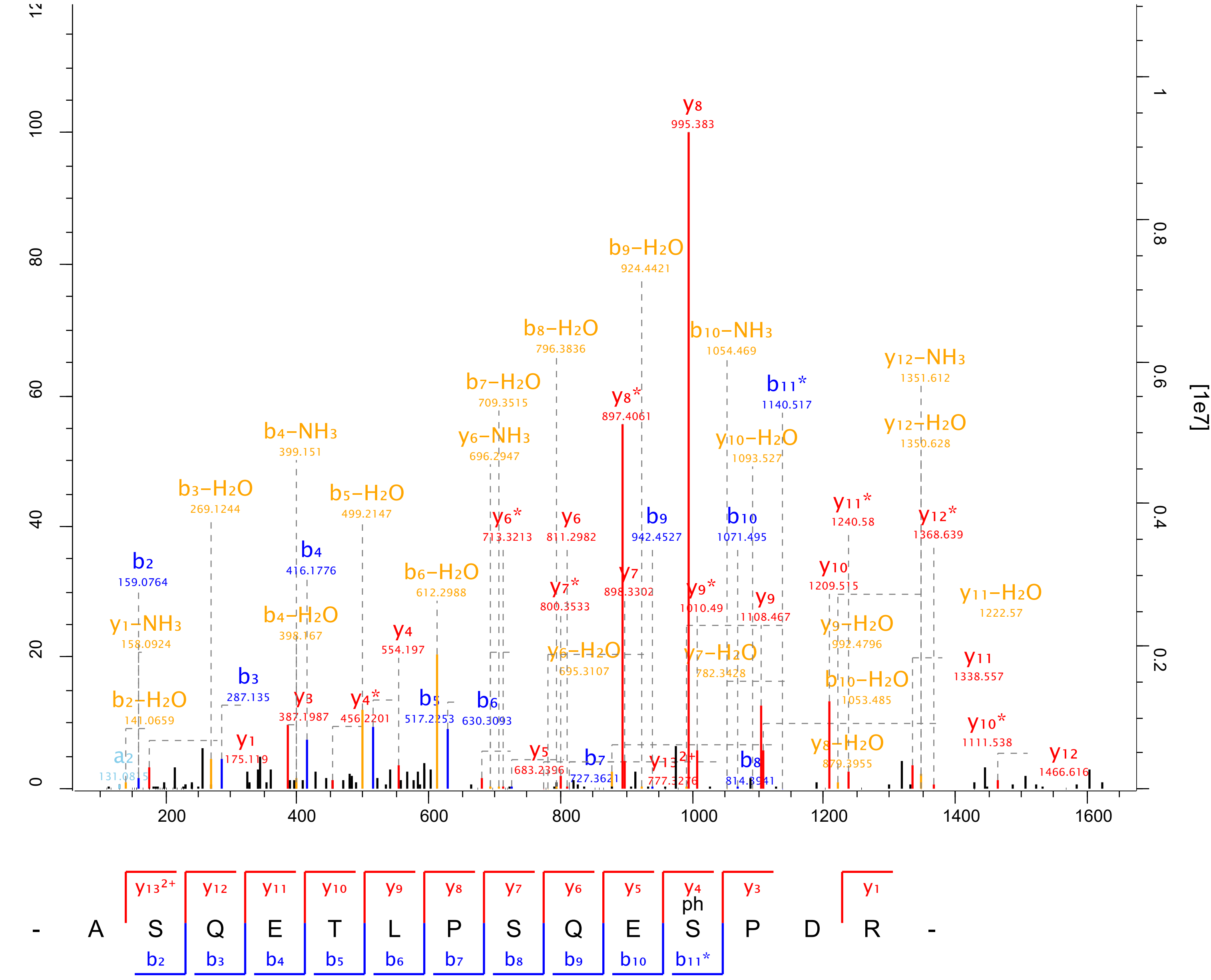

| Raw file | Scan | Method | Score | m/z |
| --- | --- | --- | --- | --- |
| water_15_2 | 9365 | FTMS; HCD | 90.05 | 465.72 |

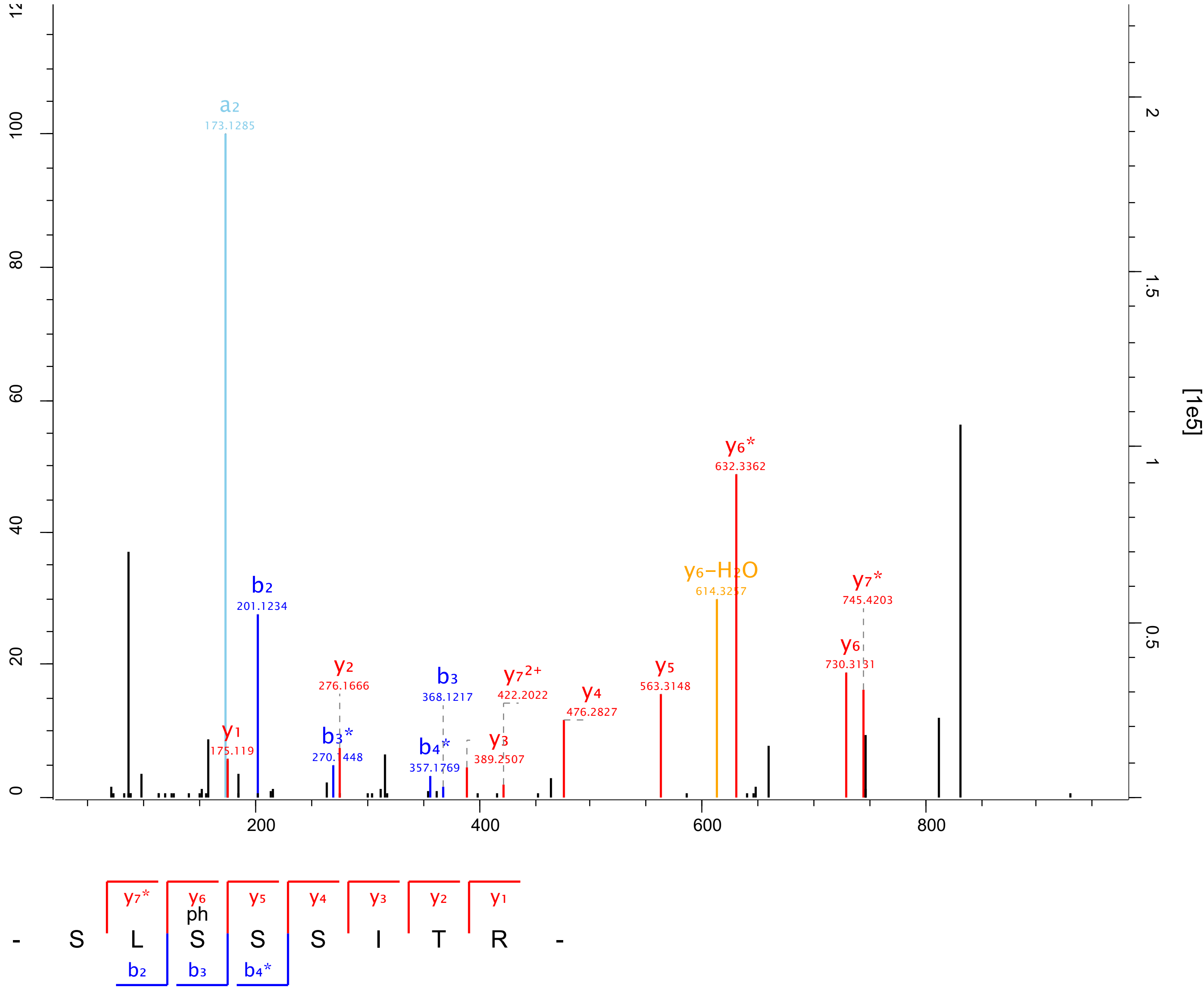

| Raw file | Scan | Method | Score | m/z |
| --- | --- | --- | --- | --- |
| water_15_2 | 9396 | FTMS; HCD | 104.82 | 427.72 |

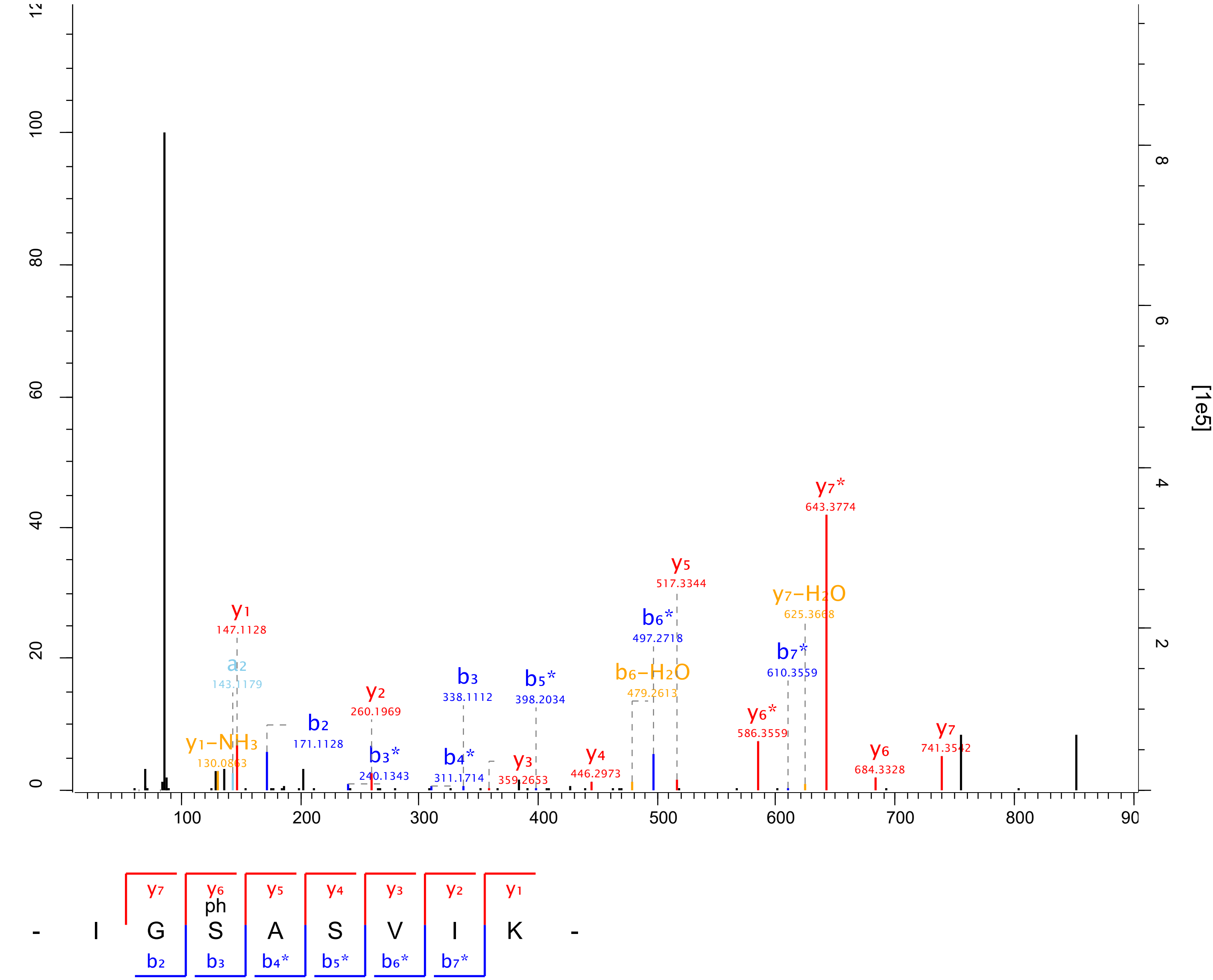

| Raw file | Scan | Method | Score | m/z |
|---|---|---|---|---|
| water_15_2 | 9479 | FTMS; HCD | 51.07 | 516.71 |

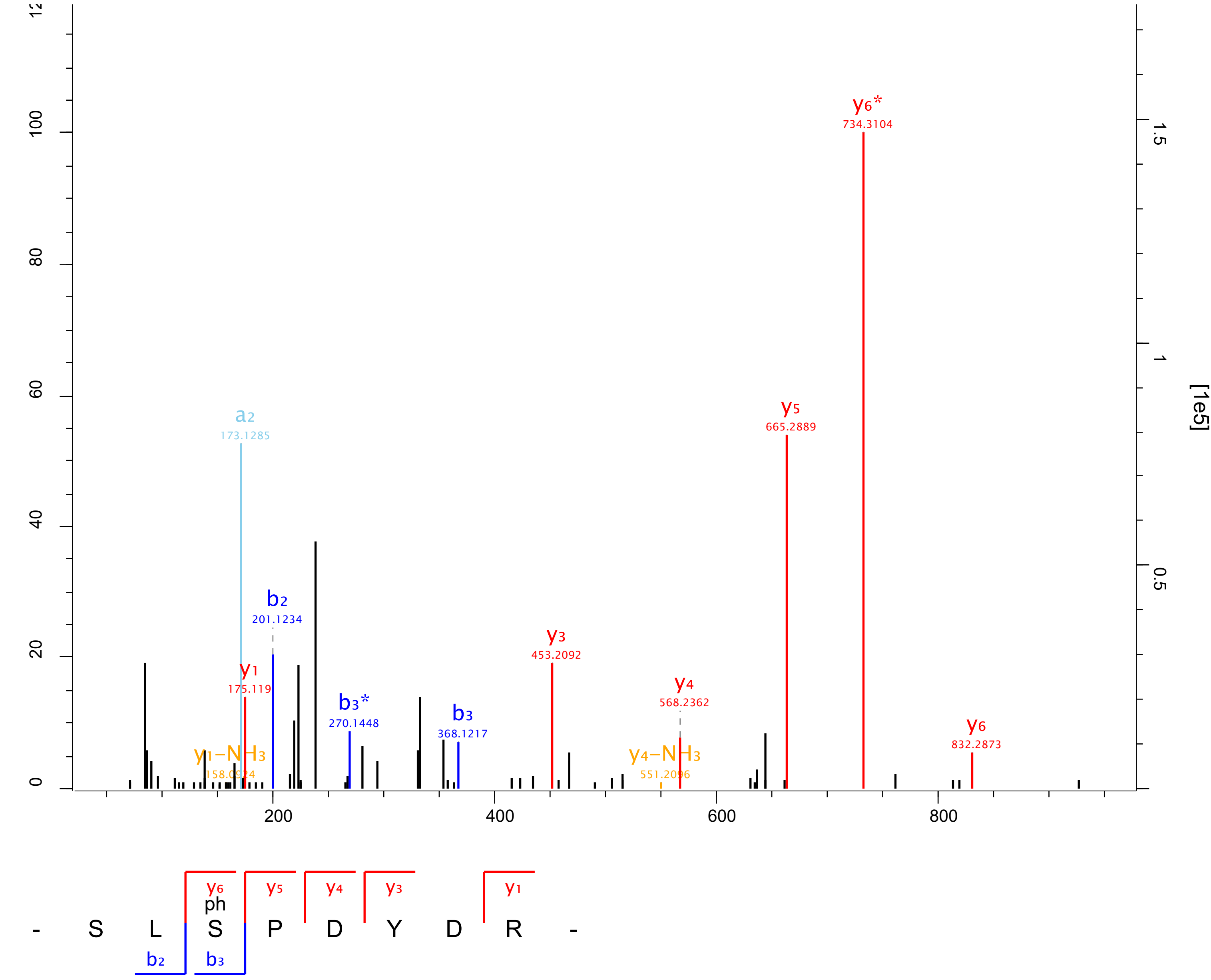

| Raw file | Scan | Method | Score | m/z |
|---|---|---|---|---|
| water_15_2 | 9483 | FTMS; HCD | 176.29 | 713.83 |

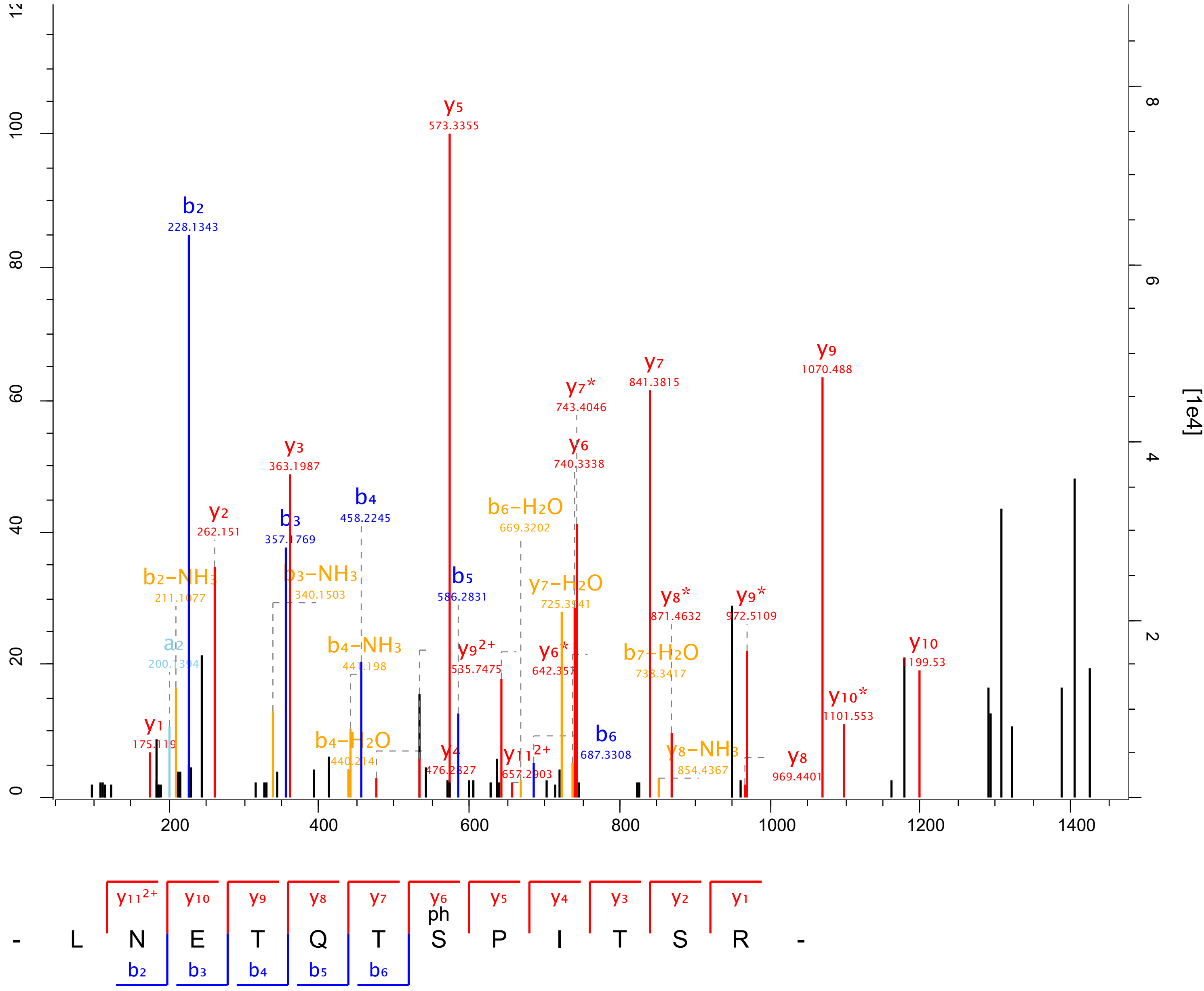

| Raw file | Scan | Method | Score | m/z |
| --- | --- | --- | --- | --- |
| water_15_2 | 9512 | FTMS; HCD | 76.07 | 540.75 |

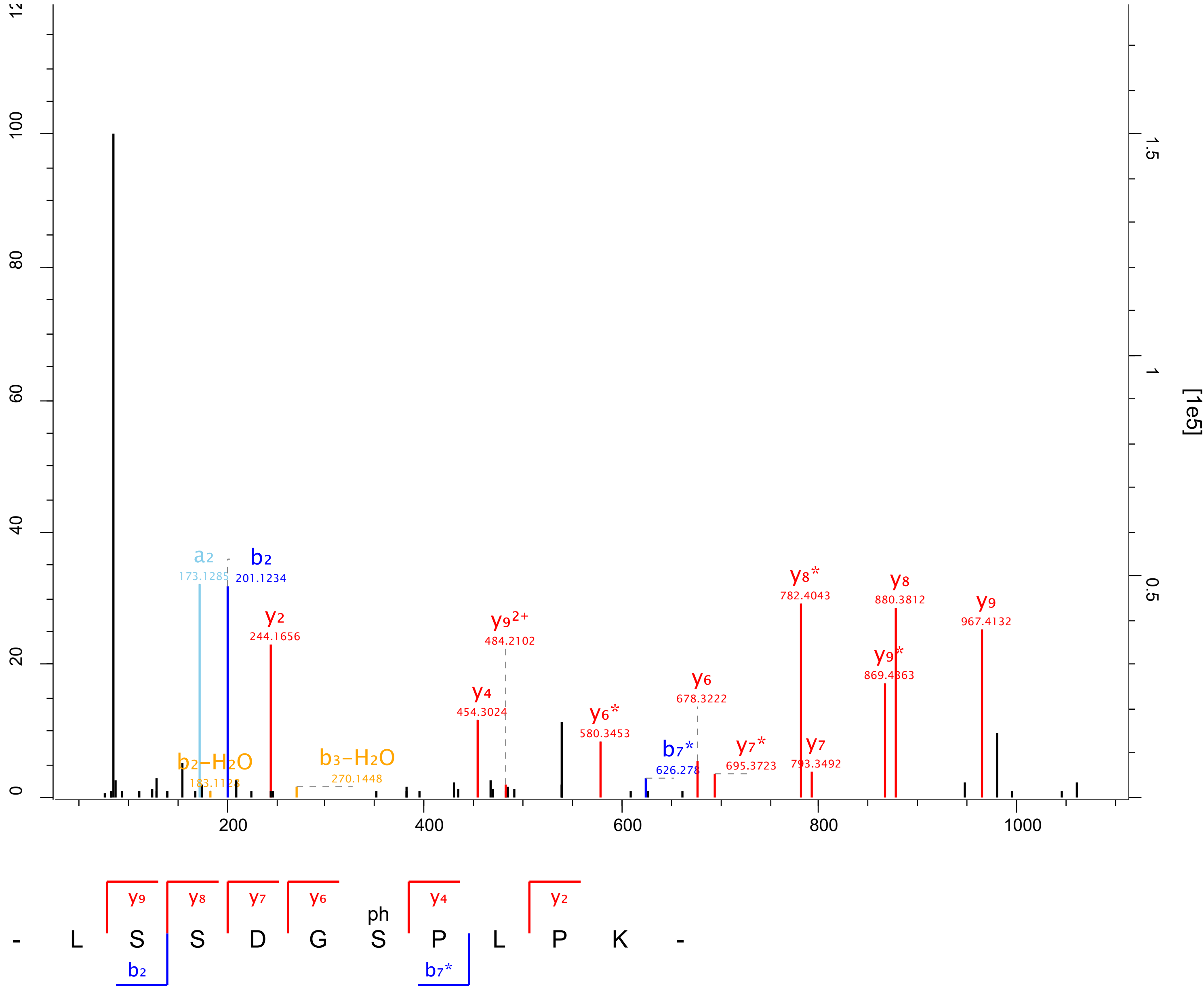

| Raw file | Scan | Method | Score | m/z |
| --- | --- | --- | --- | --- |
| water_15_2 | 9529 | FTMS; HCD | 98.65 | 645.8 |

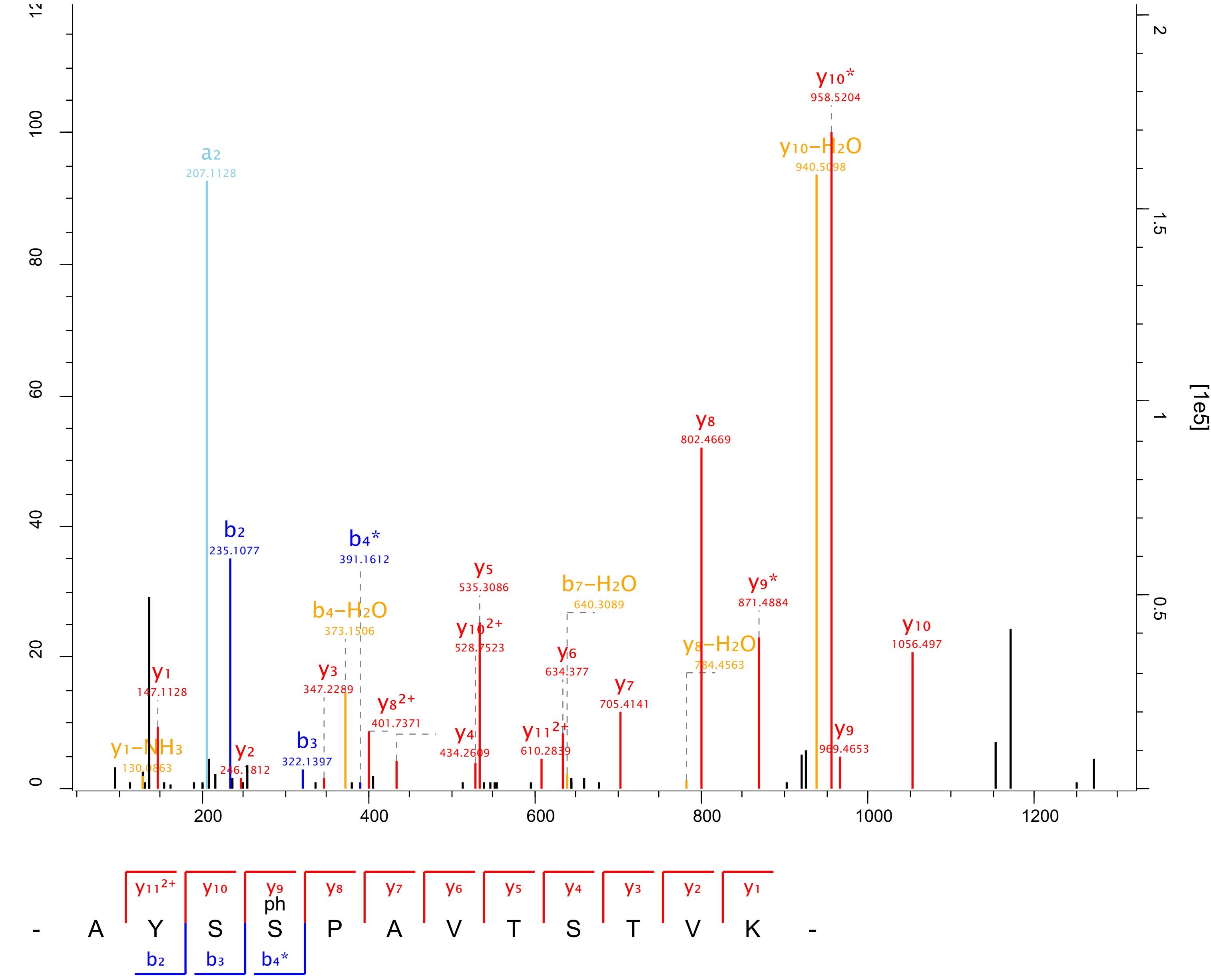

| Raw file | Scan | Method | Score | m/z |
|---|---|---|---|---|
| water_15_2 | 9550 | FTMS; HCD | 62.06 | 830.82 |

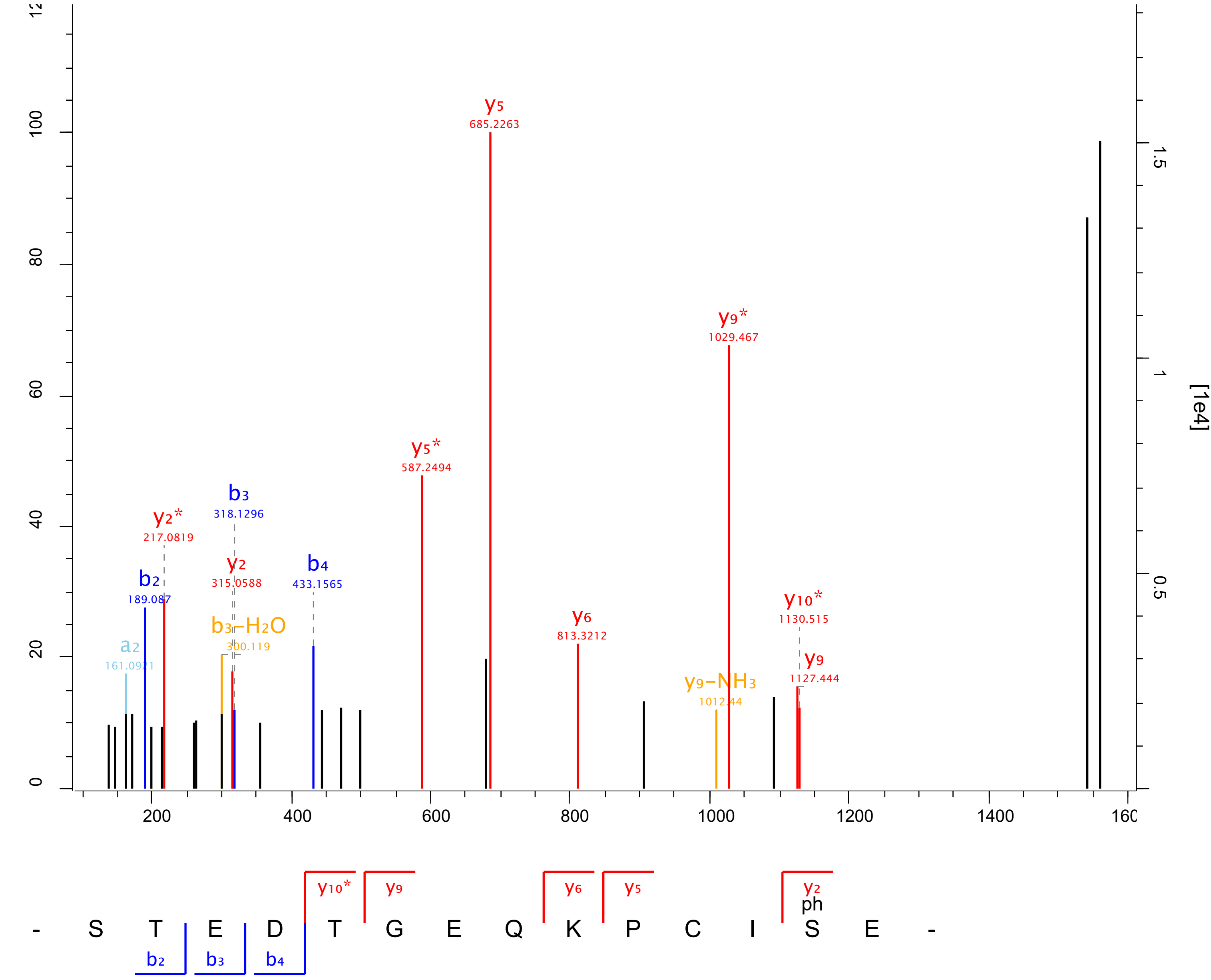

| Raw file | Scan | Method | Score | m/z |
| --- | --- | --- | --- | --- |
| water_15_2 | 9581 | FTMS; HCD | 127.83 | 672.3 |

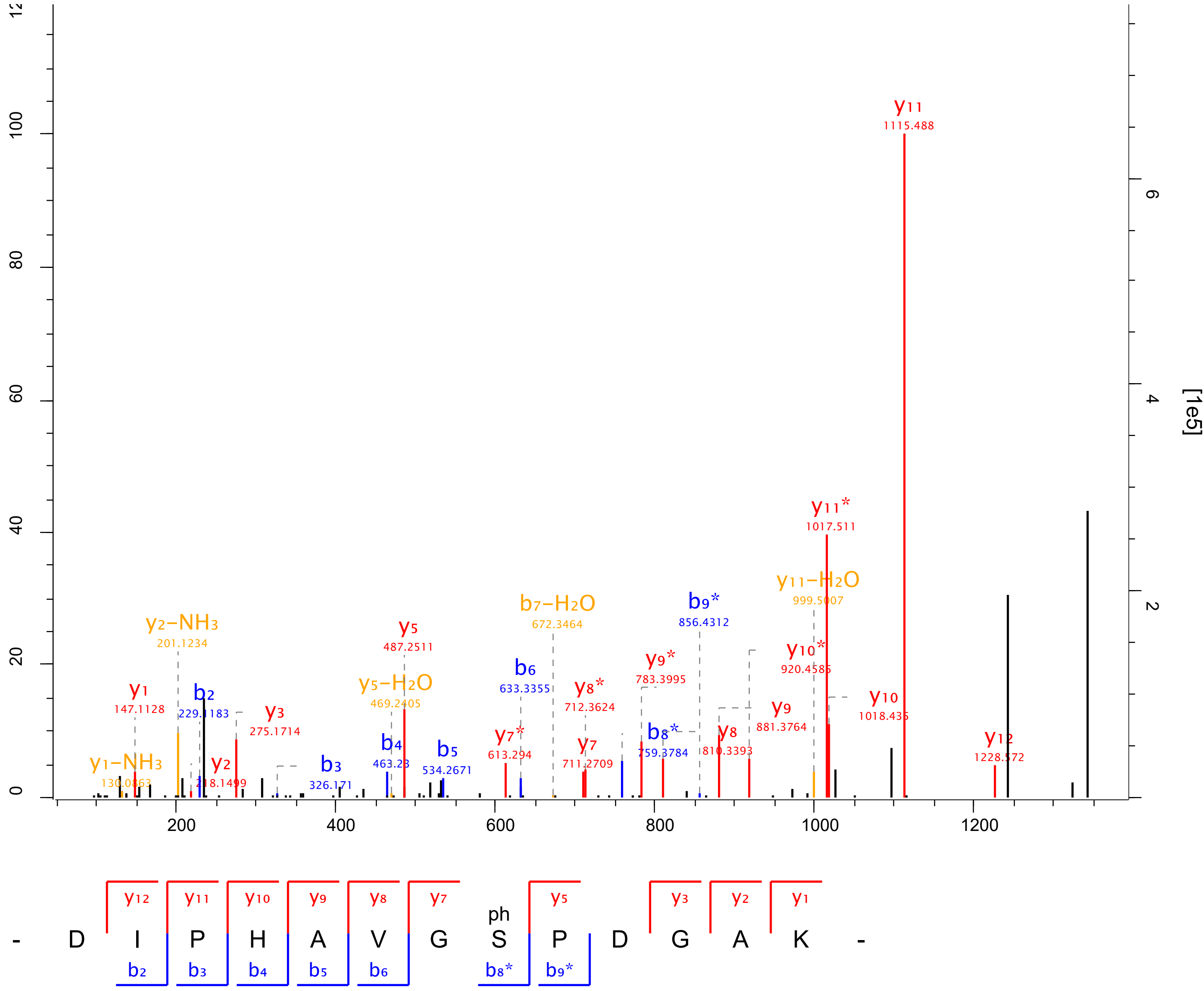

| Raw file | Scan | Method | Score | m/z |
|---|---|---|---|---|
| water_15_2 | 9595 | FTMS; HCD | 68.17 | 620.3 |

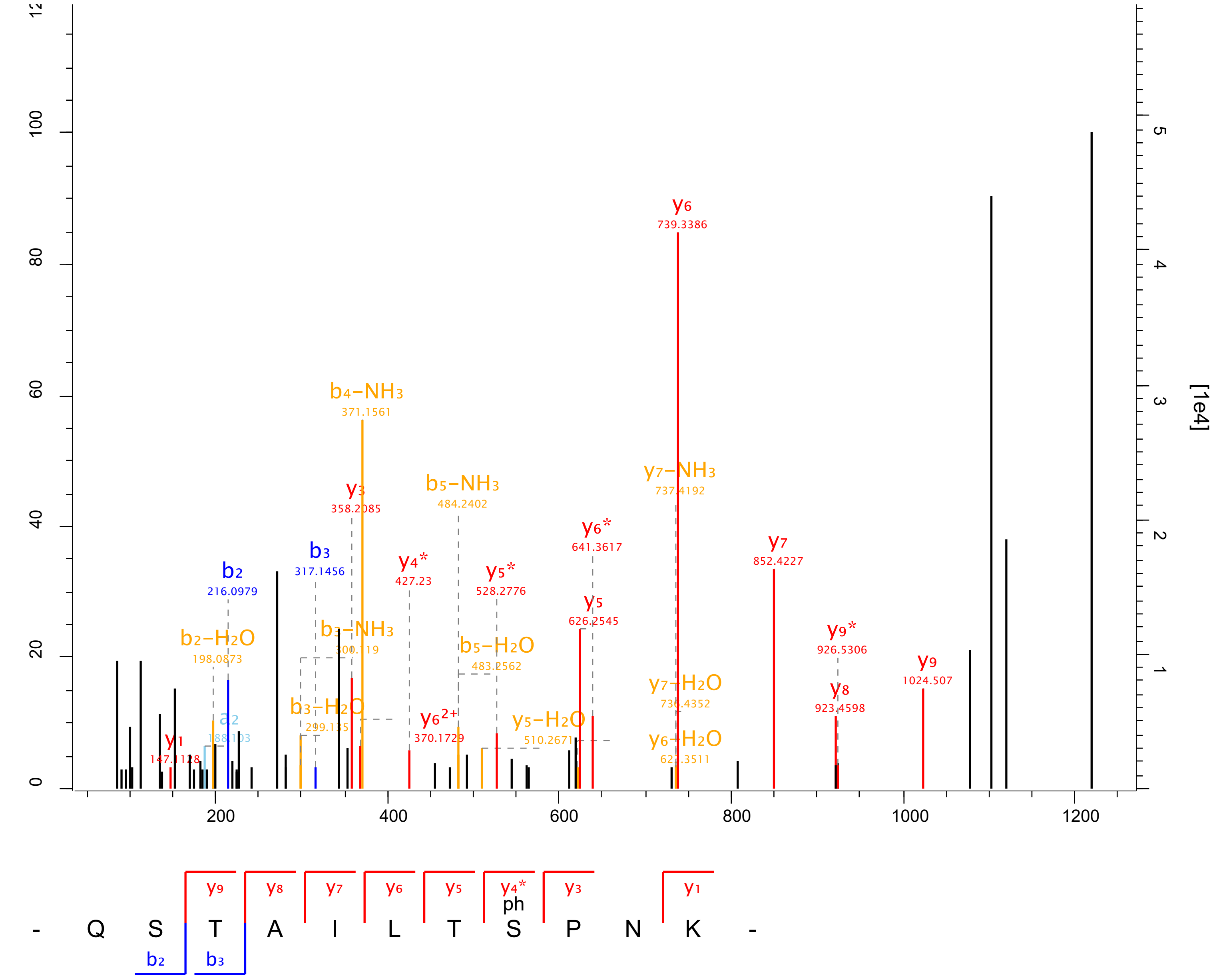

| Raw file | Scan | Method | Score | m/z |
| --- | --- | --- | --- | --- |
| water_15_2 | 9632 | FTMS; HCD | 59.5 | 502.58 |

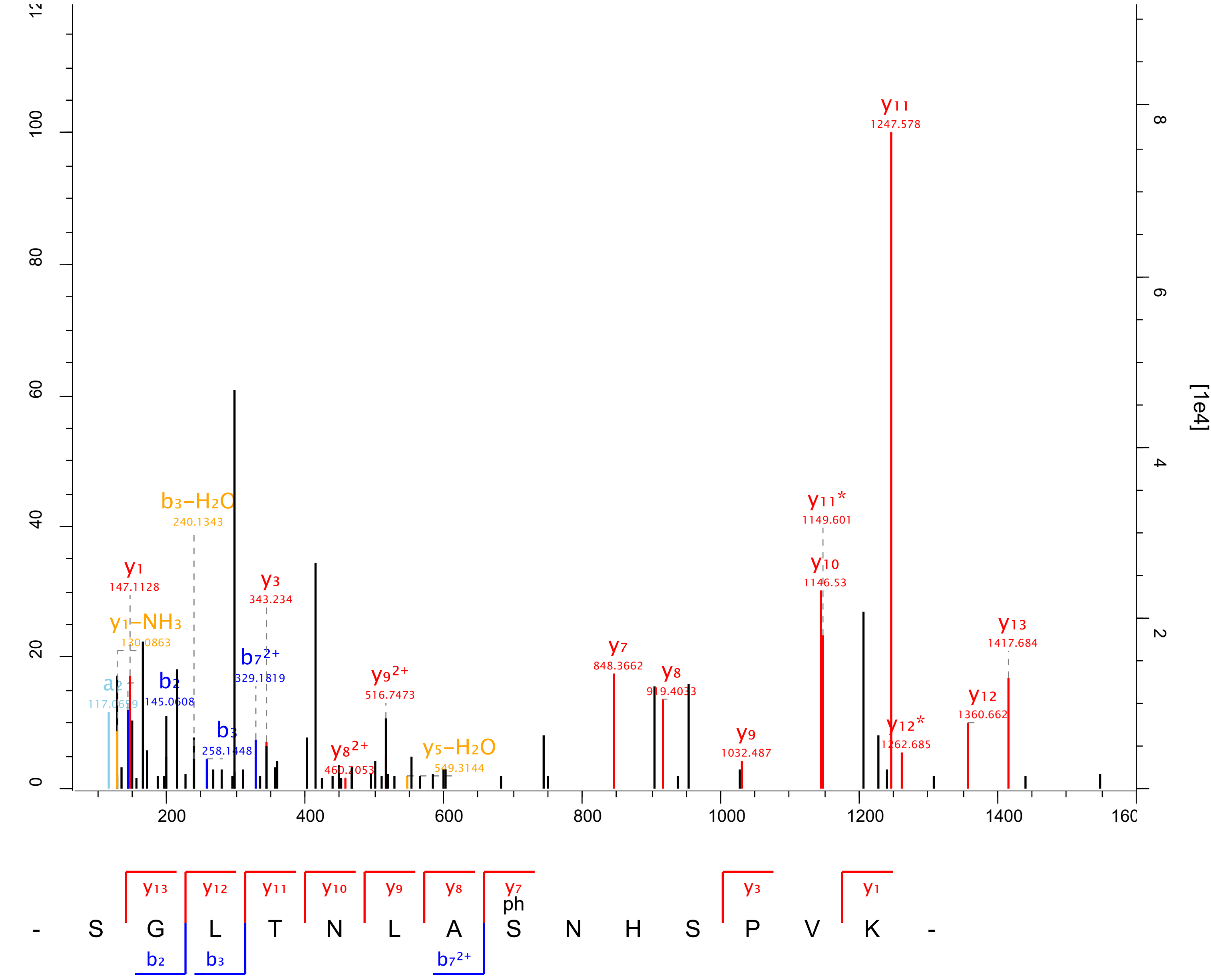

| Raw file | Scan | Method | Score | m/z |
| --- | --- | --- | --- | --- |
| water_15_2 | 9643 | FTMS; HCD | 79.64 | 611.74 |

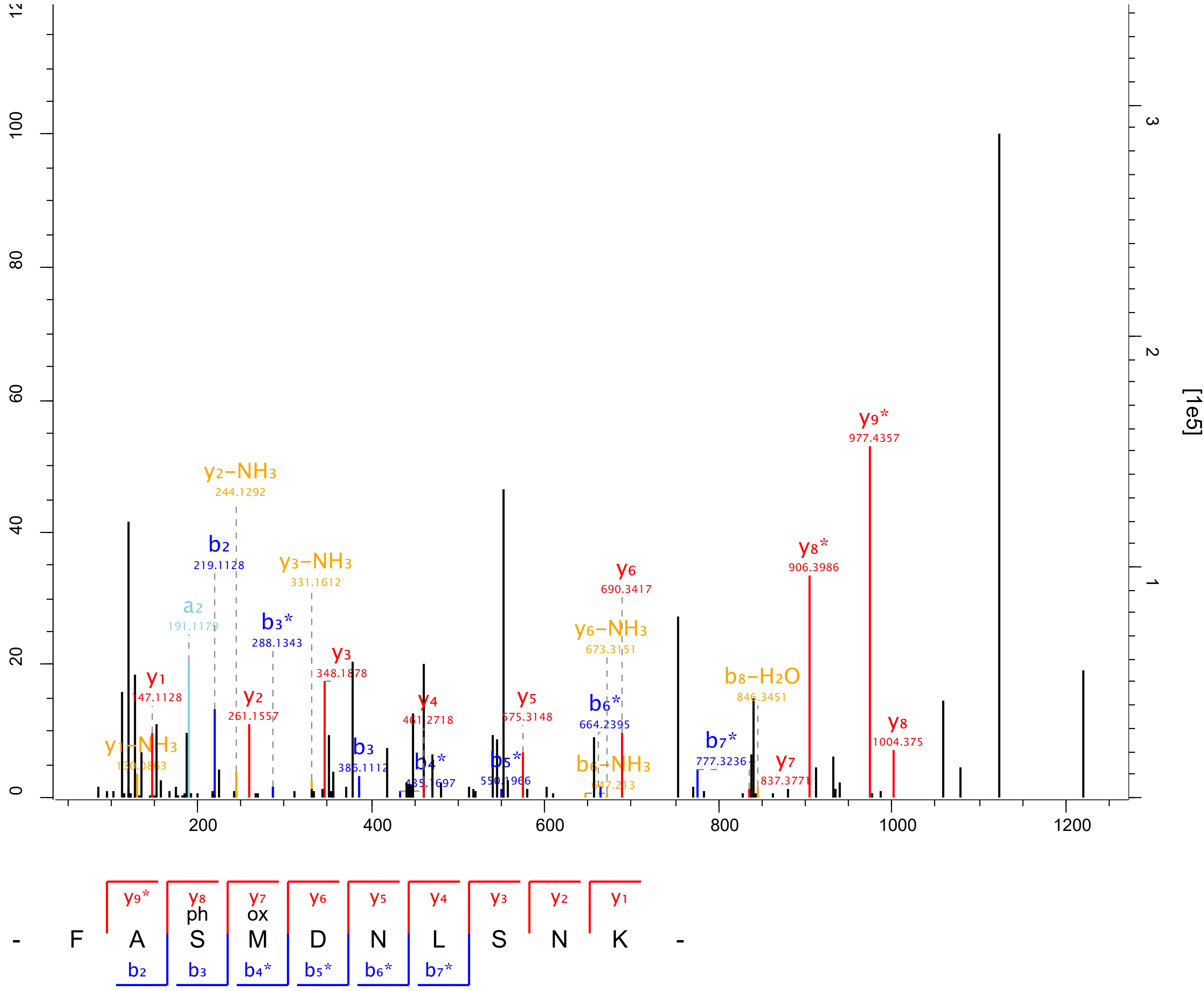

| Raw file | Scan | Method | Score | m/z |
| --- | --- | --- | --- | --- |
| water_15_2 | 9656 | FTMS; HCD | 96.06 | 560.78 |

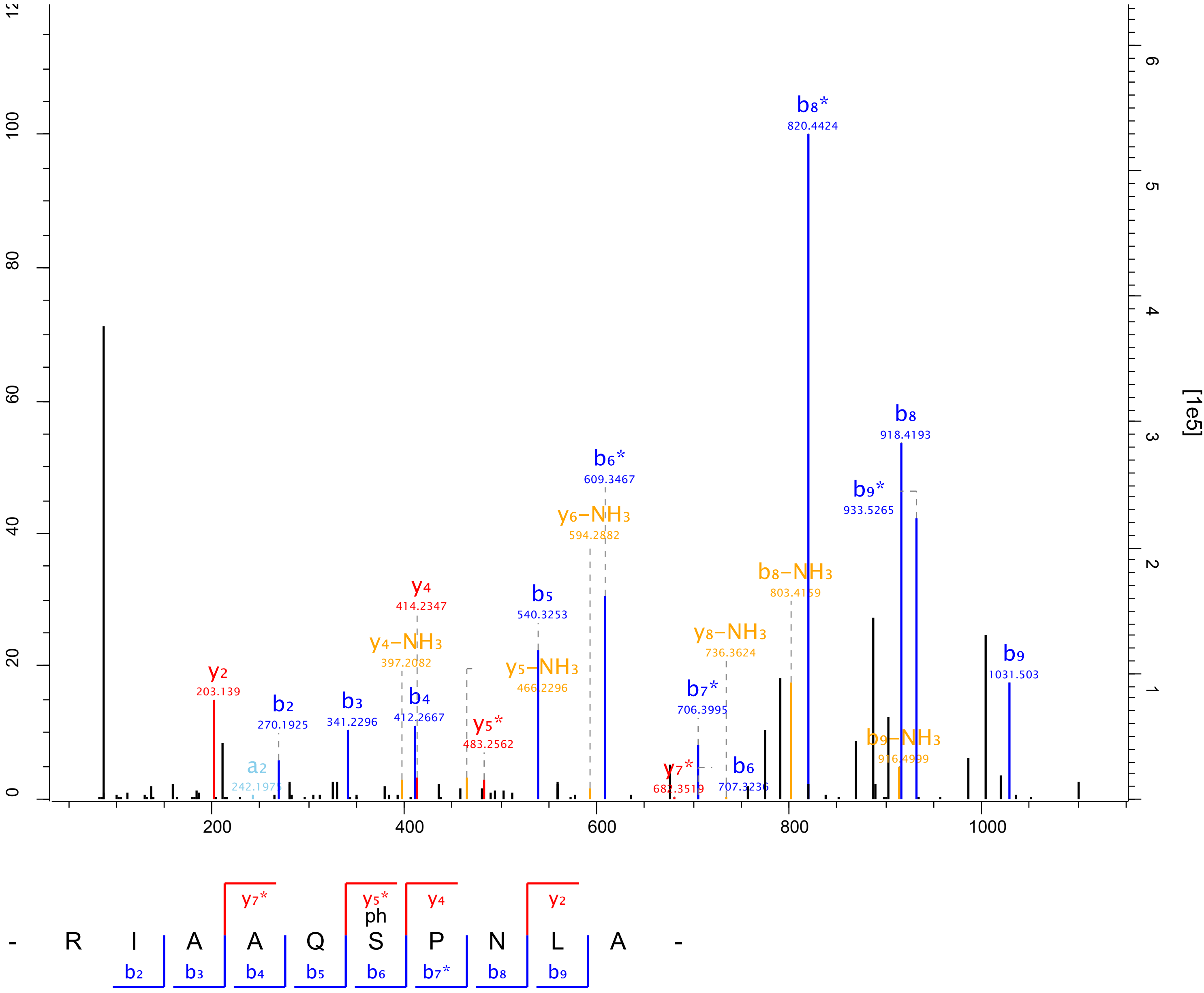

| Raw file | Scan | Method | Score | m/z |
|---|---|---|---|---|
| water_15_2 | 9684 | FTMS; HCD | 82.42 | 483.74 |

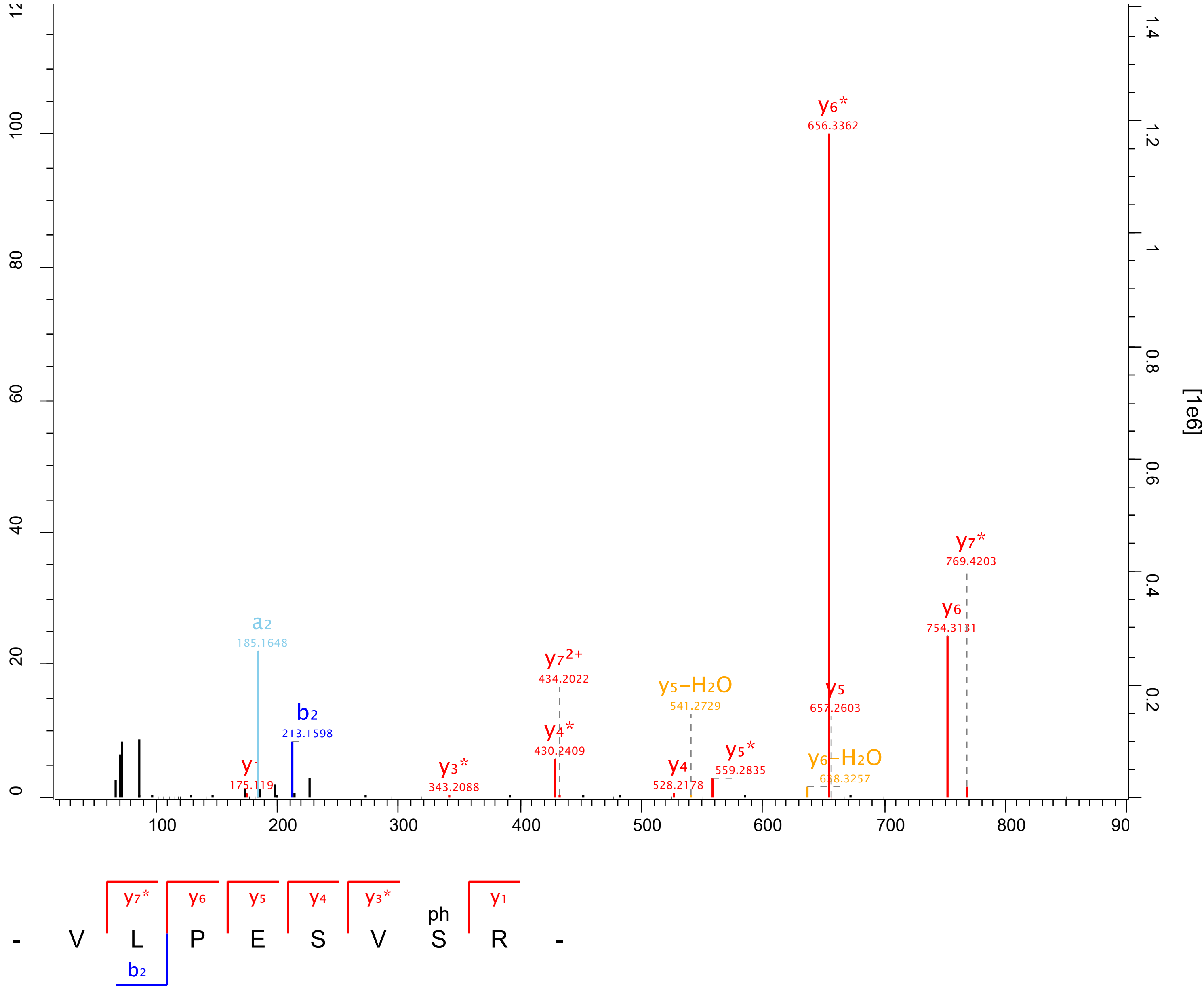

| Raw file | Scan | Method | Score | m/z |
| --- | --- | --- | --- | --- |
| water_15_2 | 9685 | FTMS; HCD | 107.17 | 538.76 |

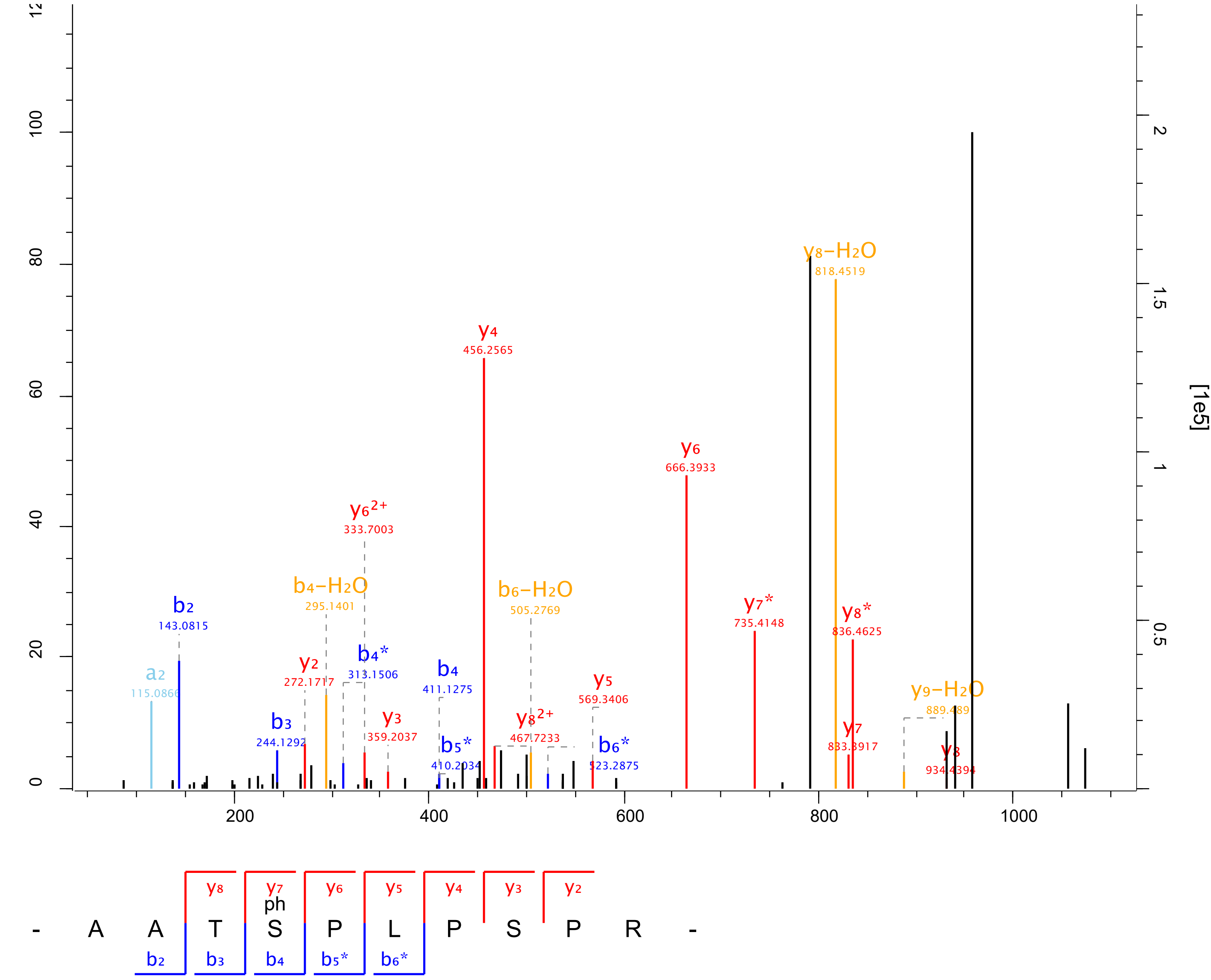

| Raw file | Scan | Method | Score | m/z |
|---|---|---|---|---|
| water_15_2 | 9694 | FTMS; HCD | 46.07 | 394.2 |

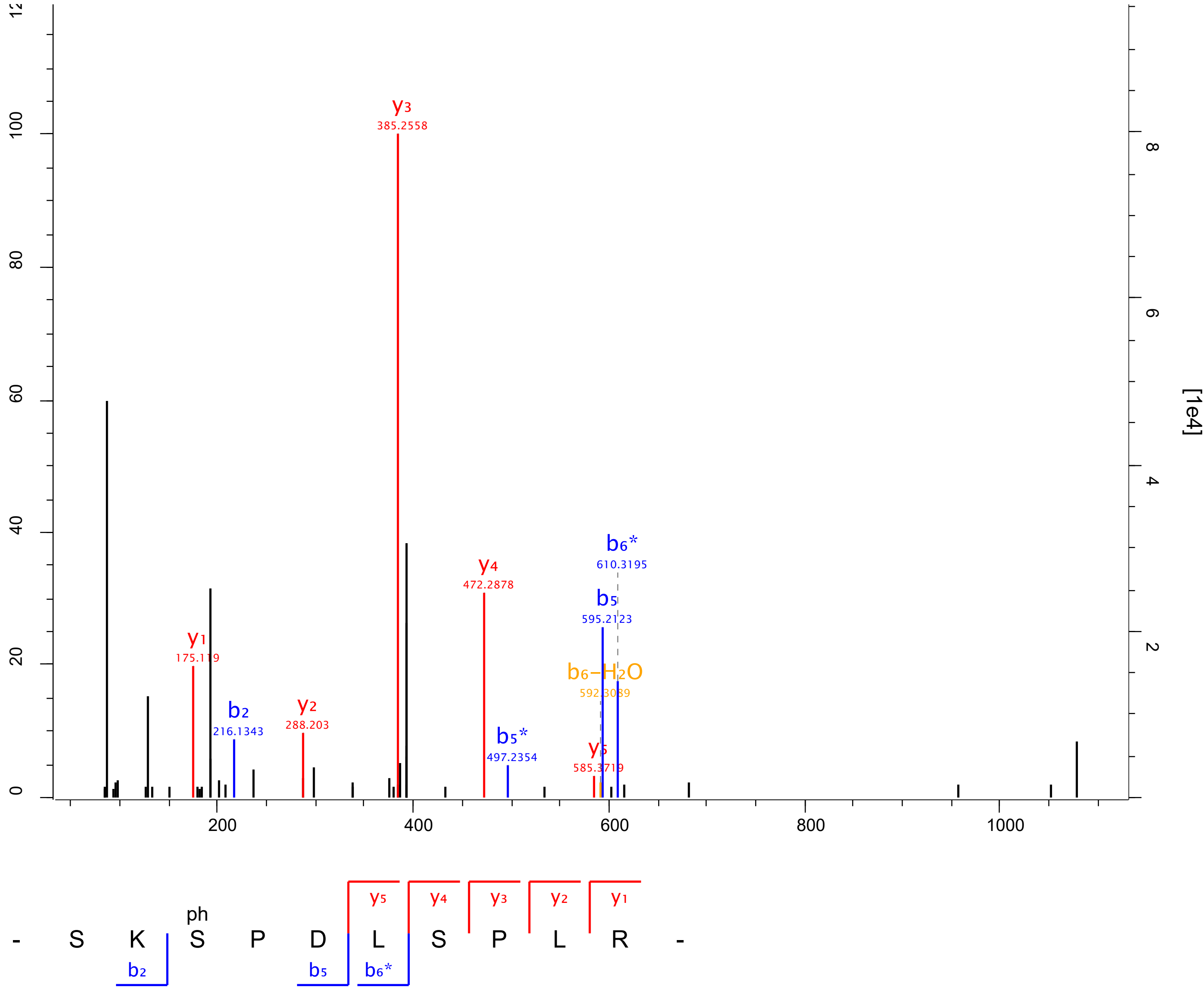

| Raw file | Scan | Method | Score | m/z |
| --- | --- | --- | --- | --- |
| water_15_2 | 9780 | FTMS; HCD | 141.13 | 557.76 |

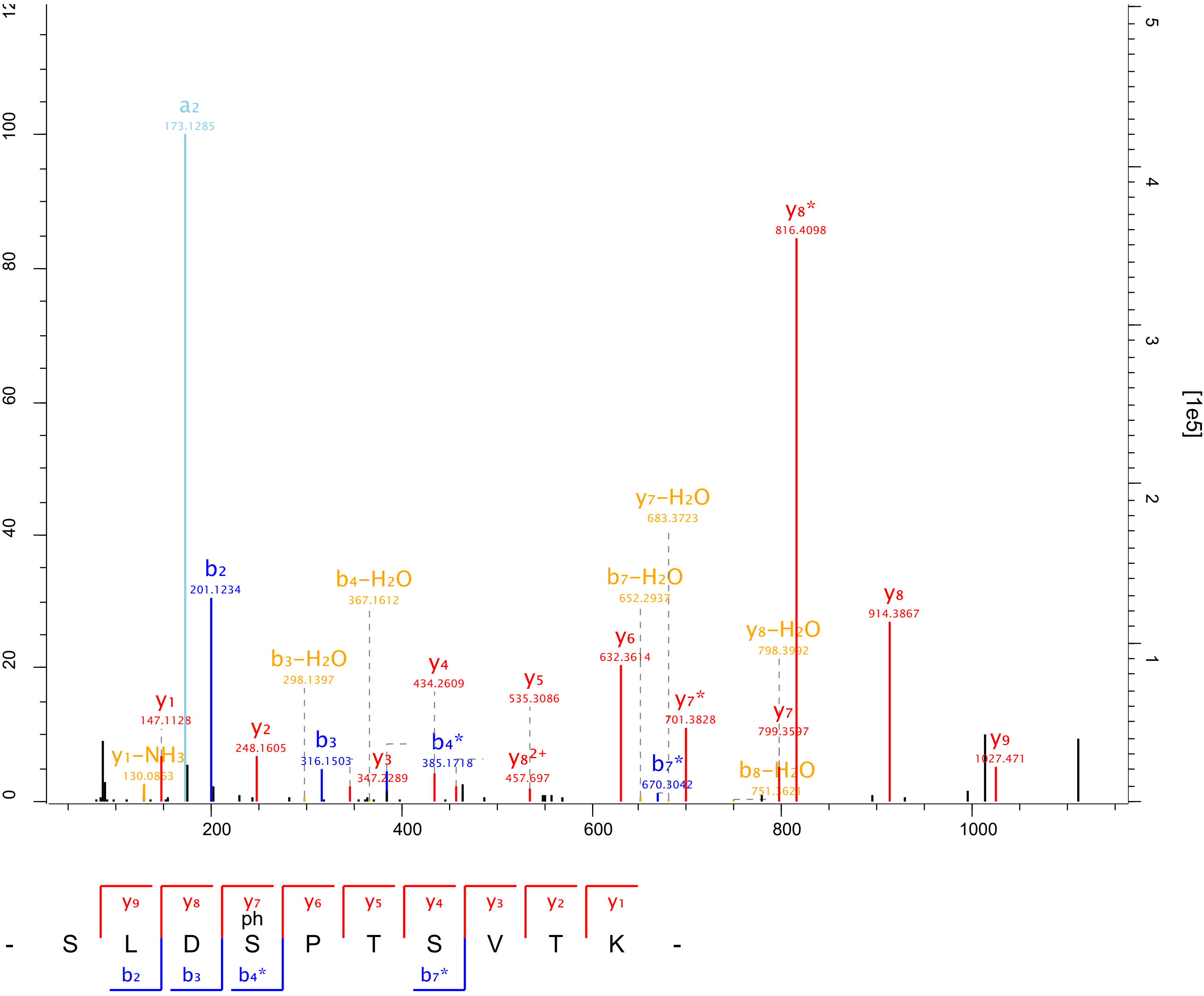

| Raw file | Scan | Method | Score | m/z |
| --- | --- | --- | --- | --- |
| water_15_2 | 9787 | FTMS; HCD | 137.4 | 591.28 |

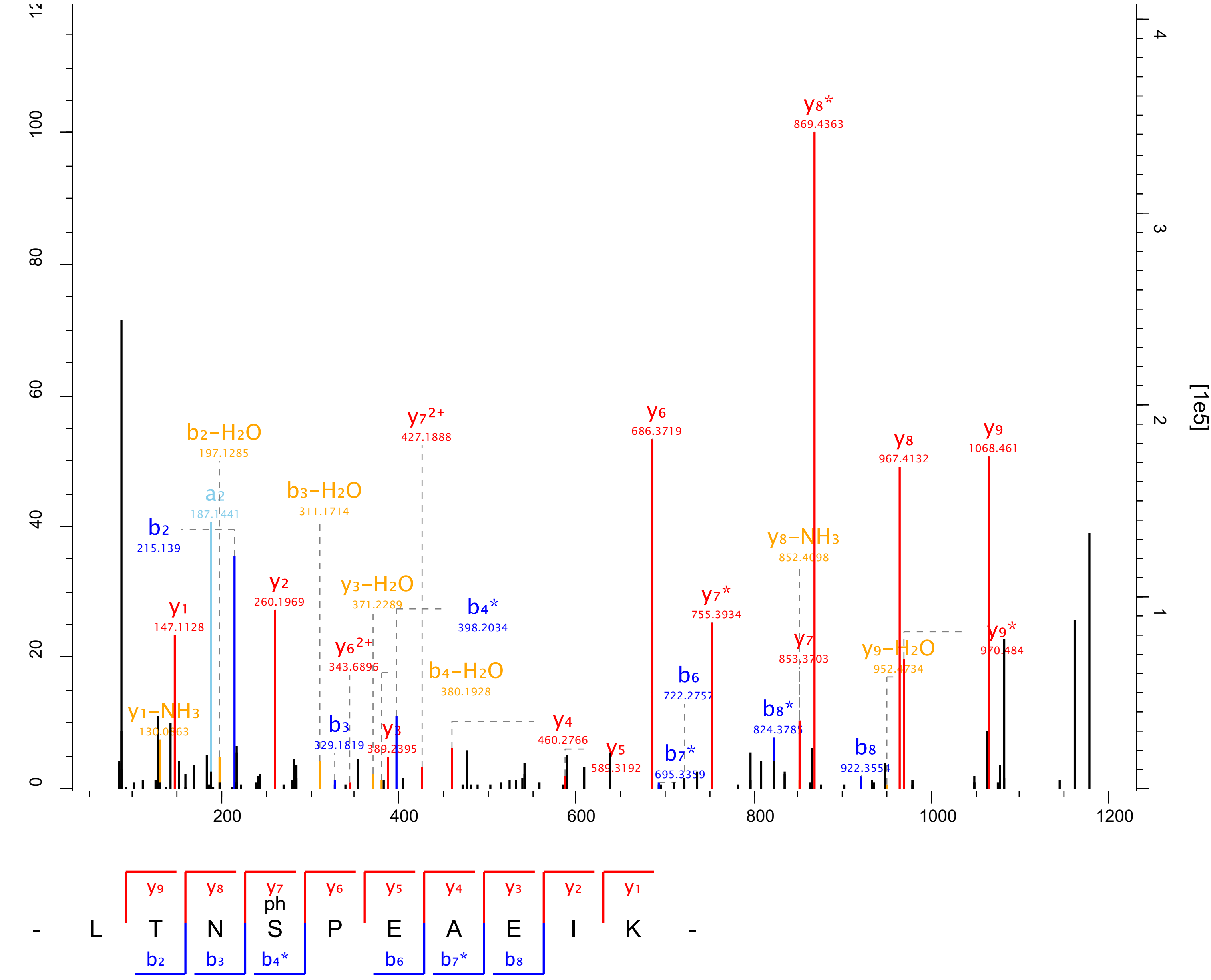

| Raw file | Scan | Method | Score | m/z |
| --- | --- | --- | --- | --- |
| water_15_2 | 9793 | FTMS; HCD | 131.4 | 637.27 |

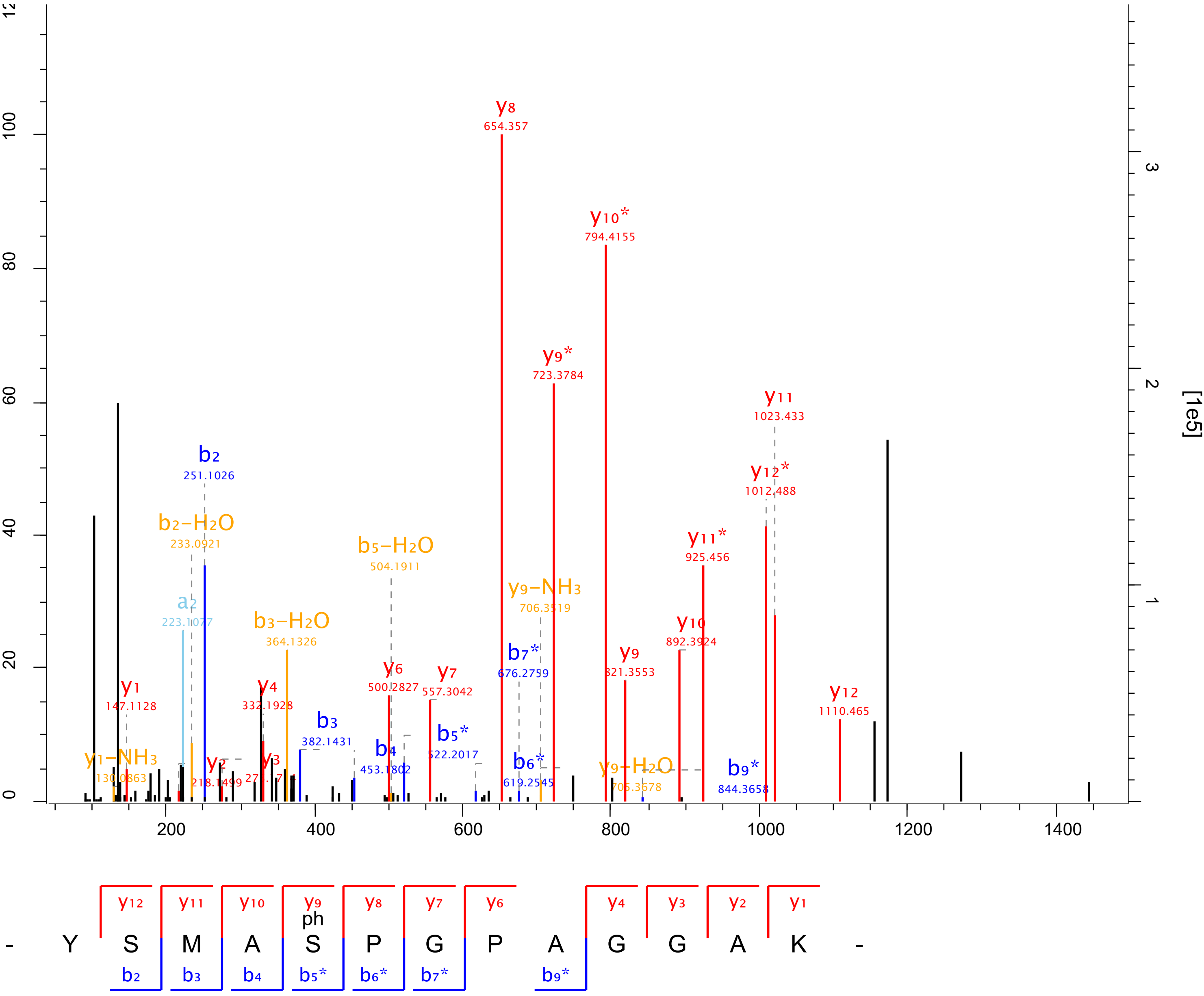

| Raw file | Scan | Method | Score | m/z |
|---|---|---|---|---|
| water_15_2 | 9818 | FTMS; HCD | 94.53 | 627.79 |

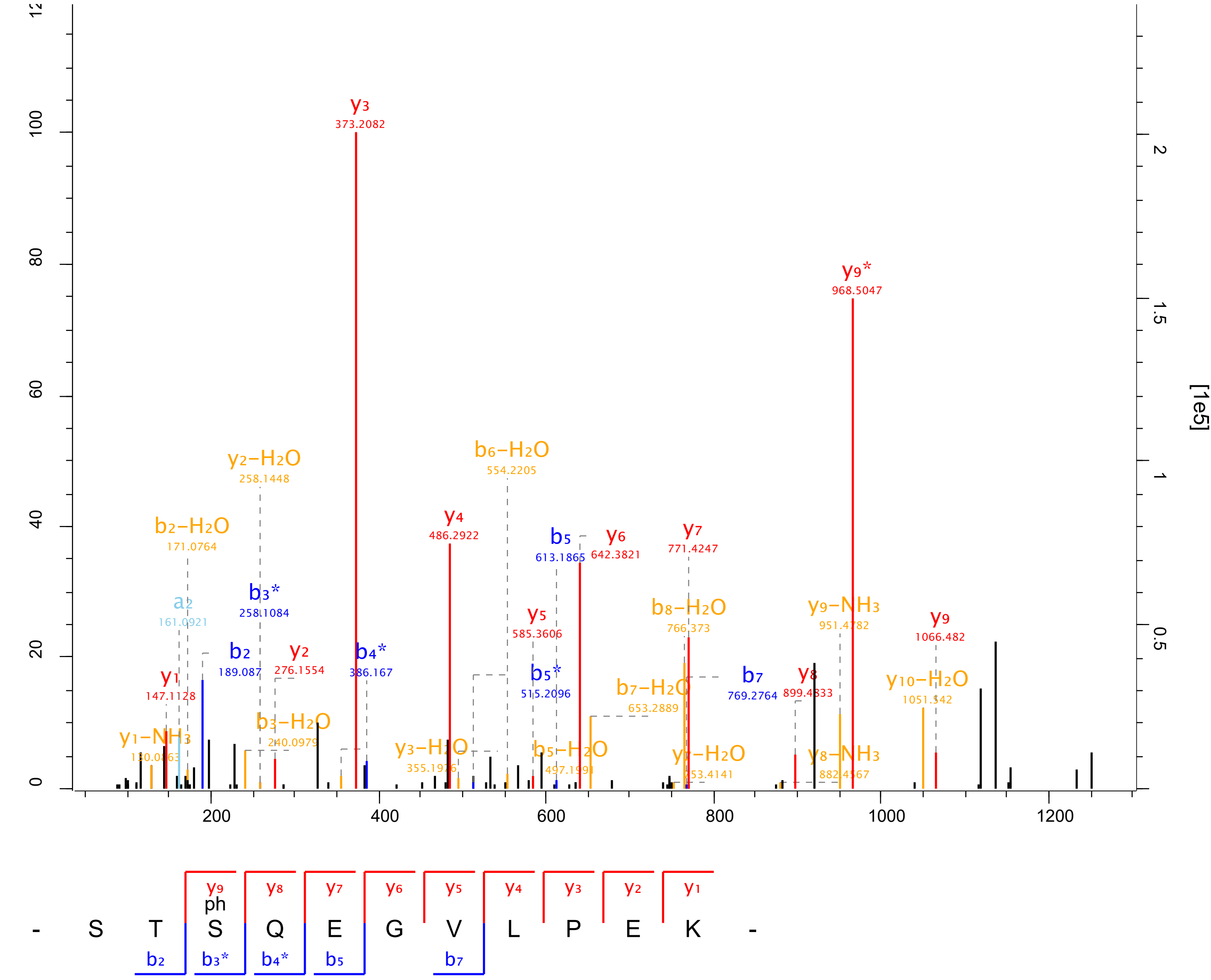

| Raw file | Scan | Method | Score | m/z |
|---|---|---|---|---|
| water_15_2 | 9922 | FTMS; HCD | 101.05 | 679.8 |

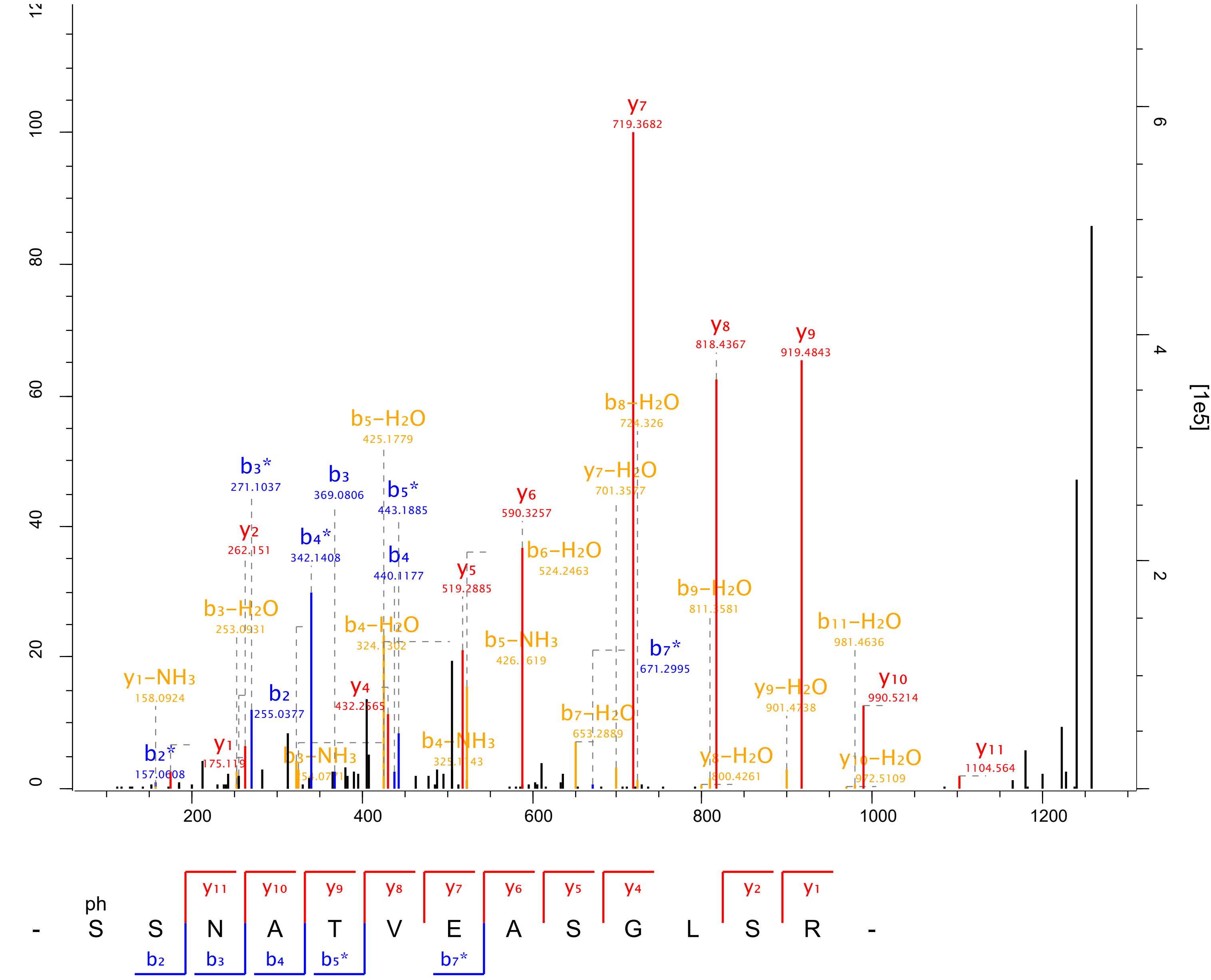

| Raw file | Scan | Method | Score | m/z |
|---|---|---|---|---|
| water_15_2 | 9927 | FTMS; HCD | 165.3 | 684.84 |

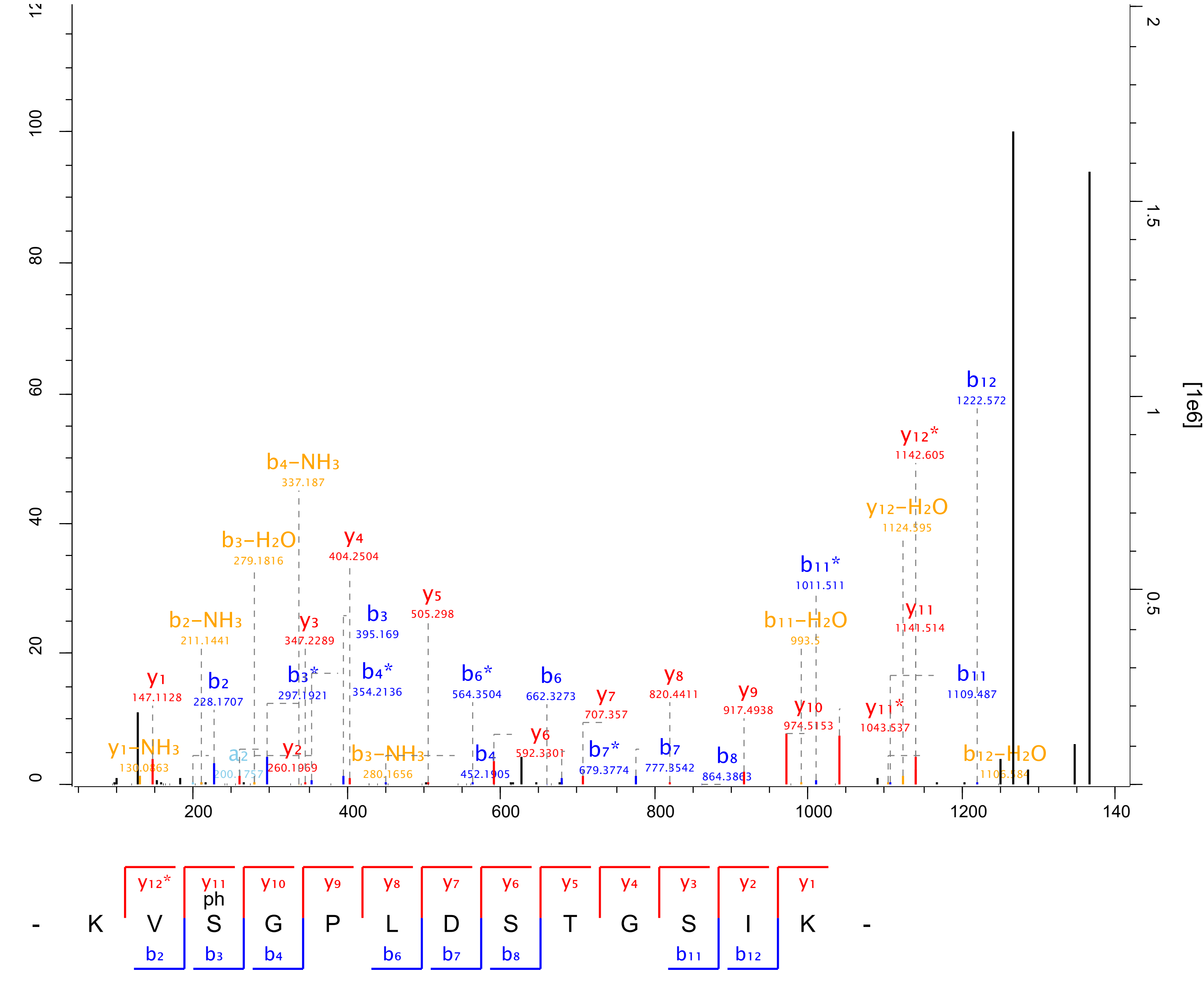

| Raw file | Scan | Method | Score | m/z |
| --- | --- | --- | --- | --- |
| water_15_2 | 9949 | FTMS; HCD | 40.94 | 572.76 |

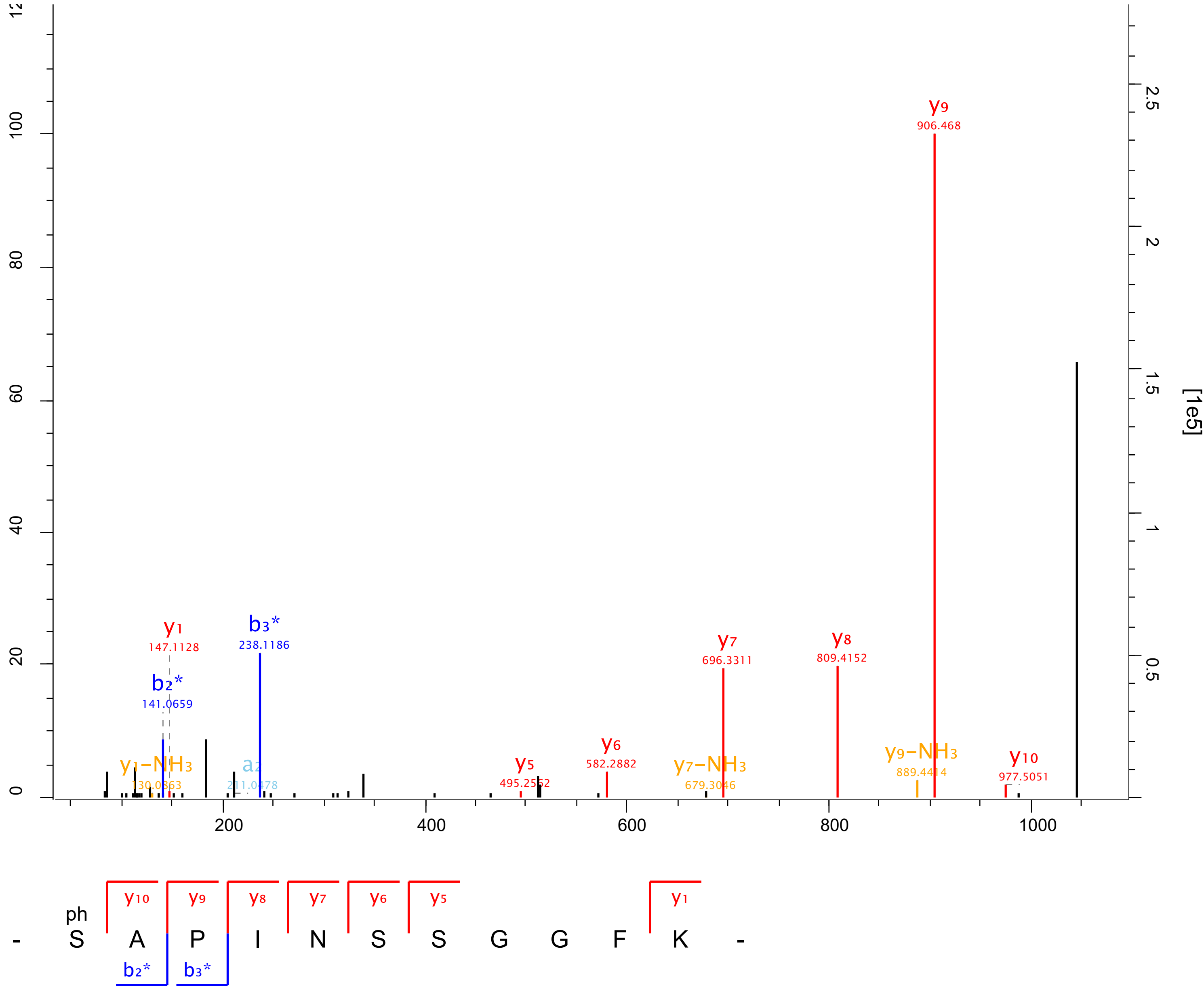

| Raw file | Scan | Method | Score | m/z |
|---|---|---|---|---|
| water_15_2 | 9971 | FTMS; HCD | 94.47 | 476.71 |

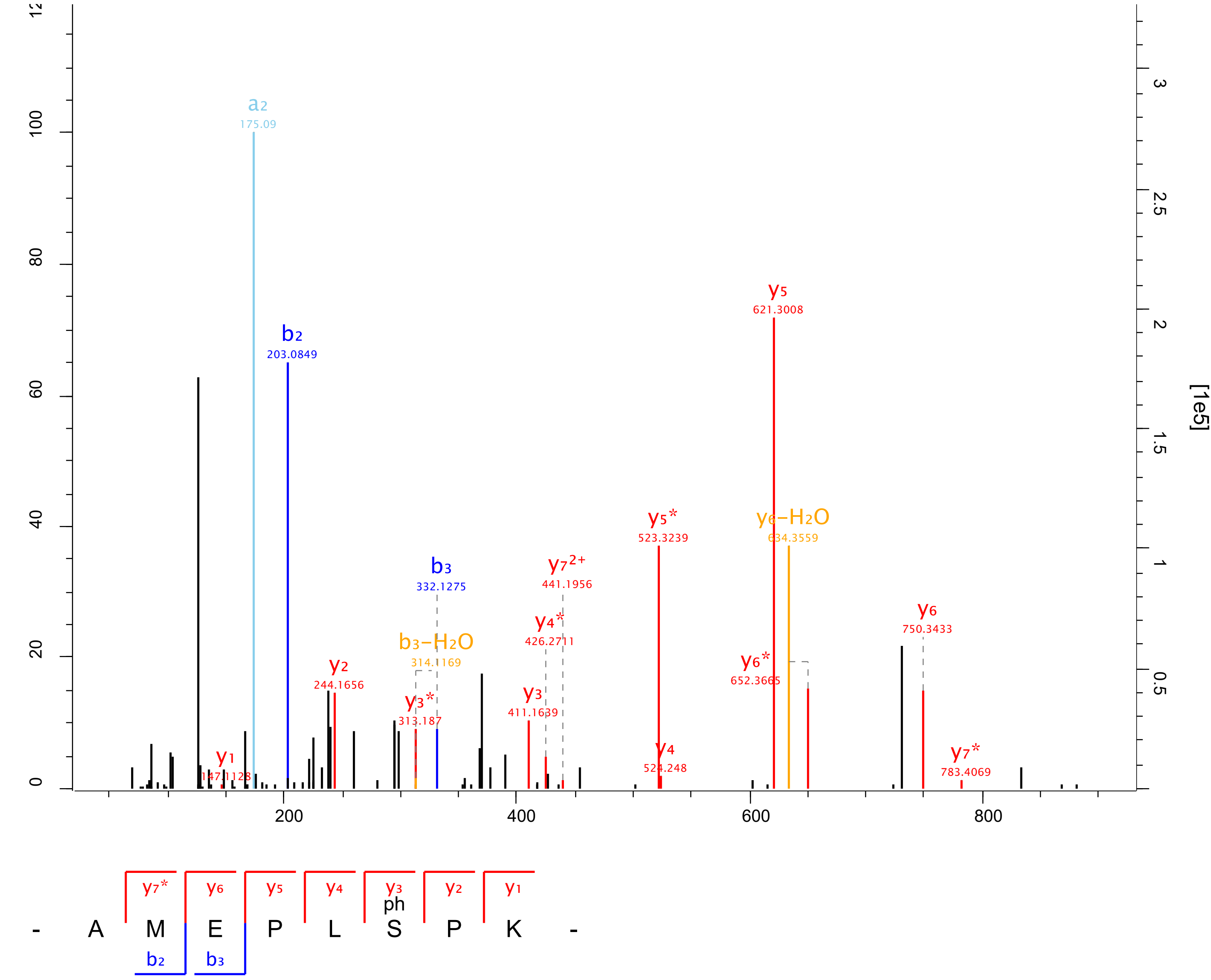

| Raw file | Scan | Method | Score | m/z |
|---|---|---|---|---|
| water_15_2 | 9995 | FTMS; HCD | 70.89 | 620.26 |

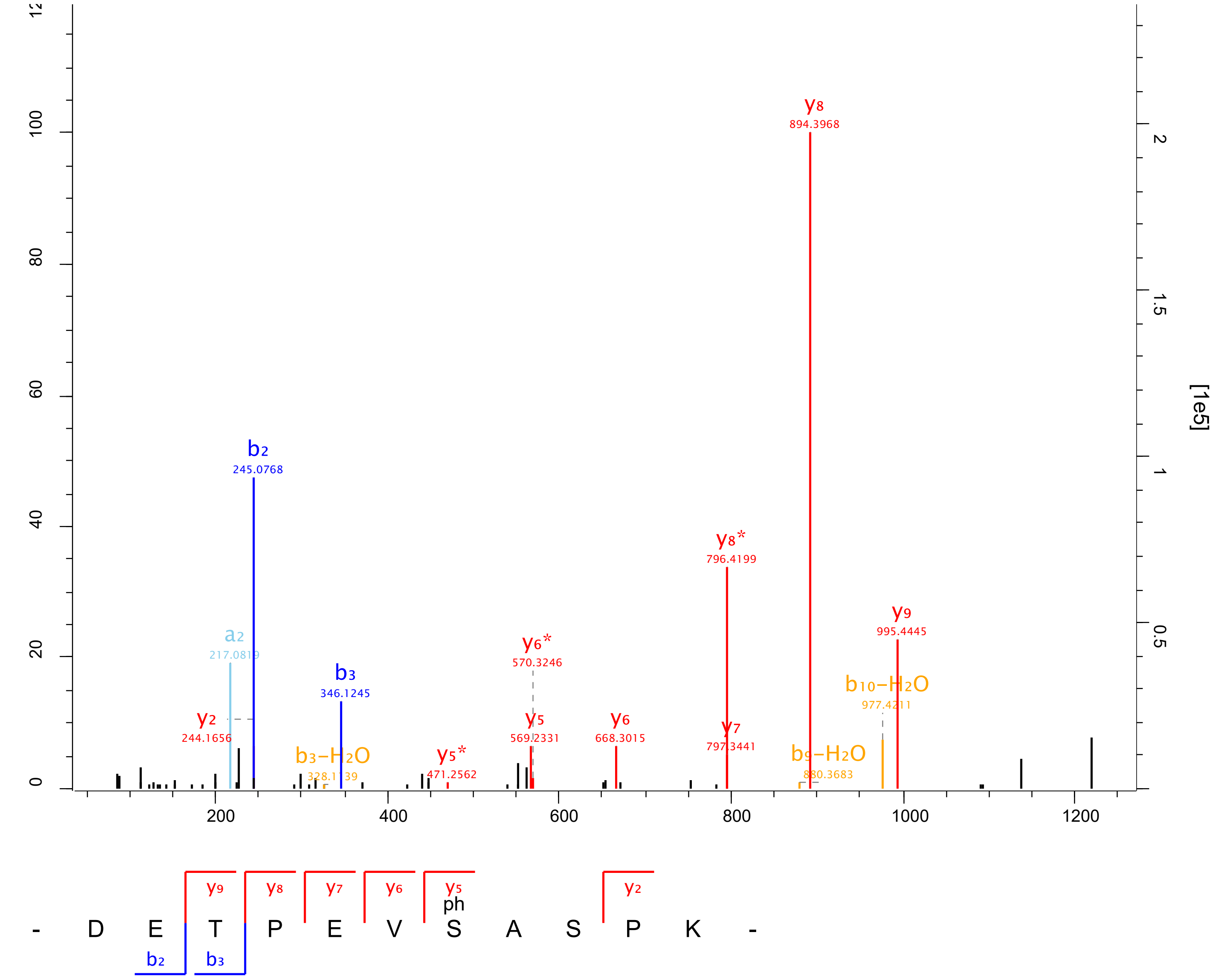

| Raw file | Scan | Method | Score | m/z |
| --- | --- | --- | --- | --- |
| water_15_3 | 12791 | FTMS; HCD | 57.86 | 457.71 |

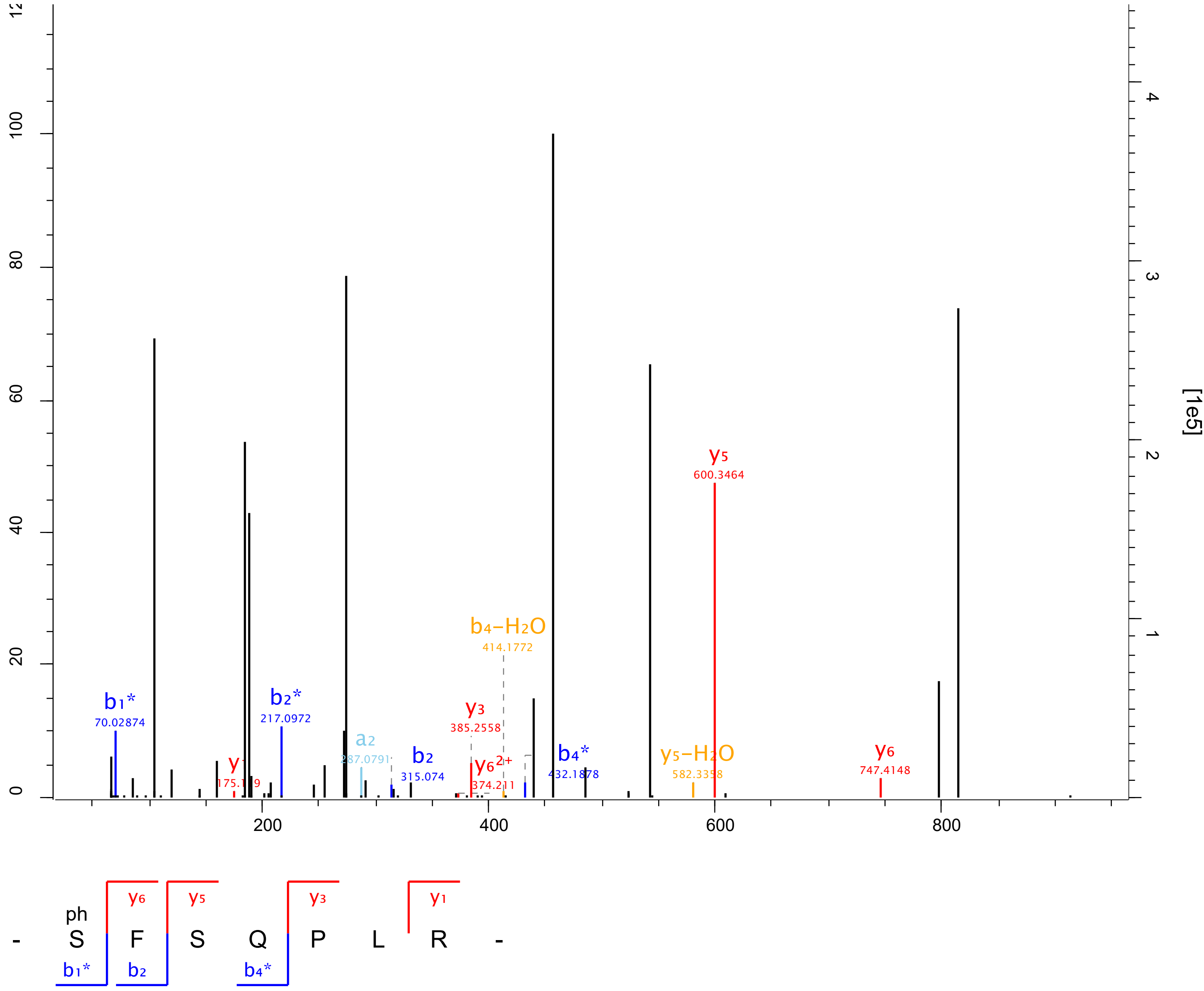

| Raw file | Scan | Method | Score | m/z |
|---|---|---|---|---|
| water_15_3 | 14420 | FTMS; HCD | 176.29 | 750.33 |

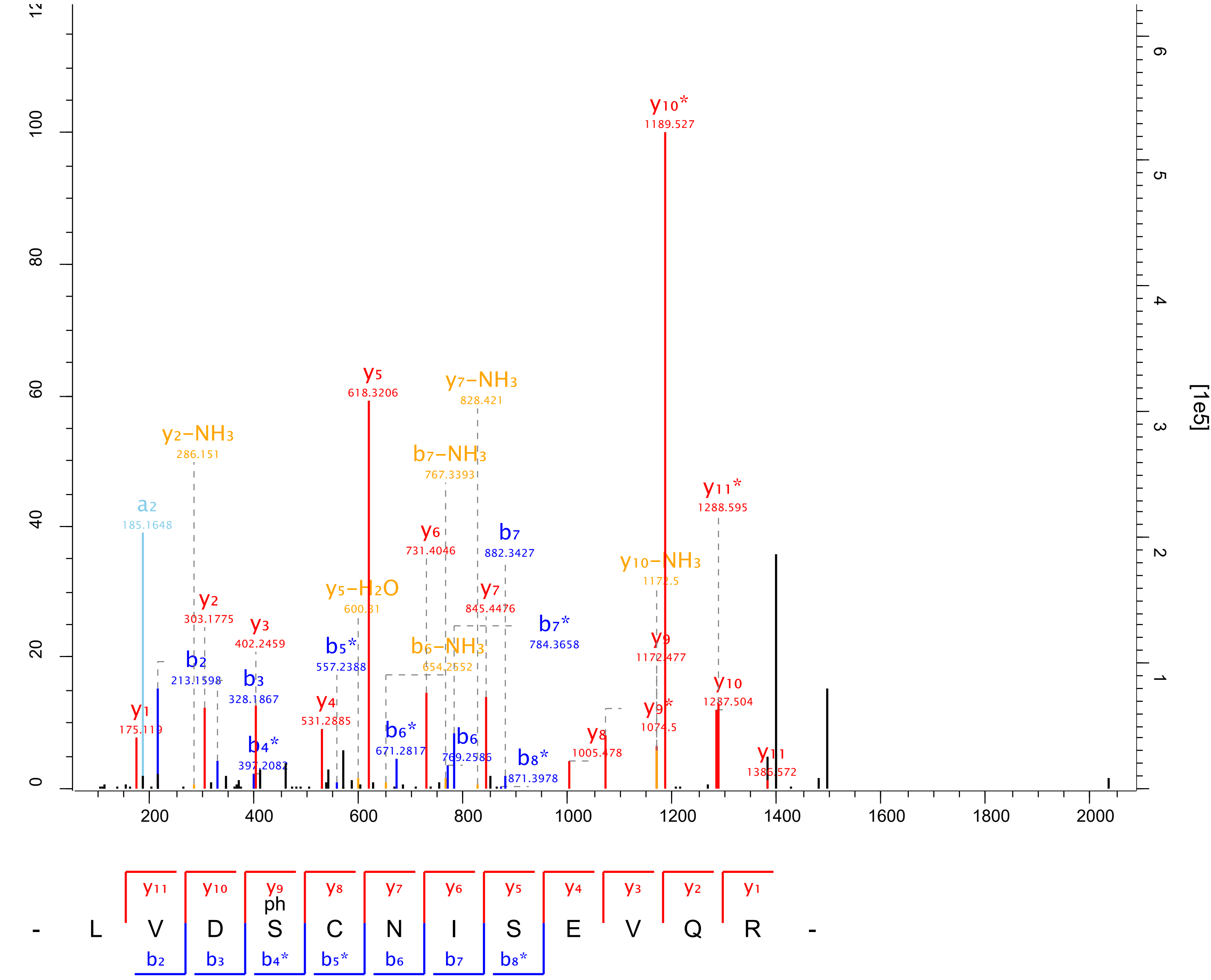

| Raw file | Scan | Method | Score | m/z |
|---|---|---|---|---|
| water_15_3 | 17435 | FTMS; HCD | 87.18 | 475.76 |

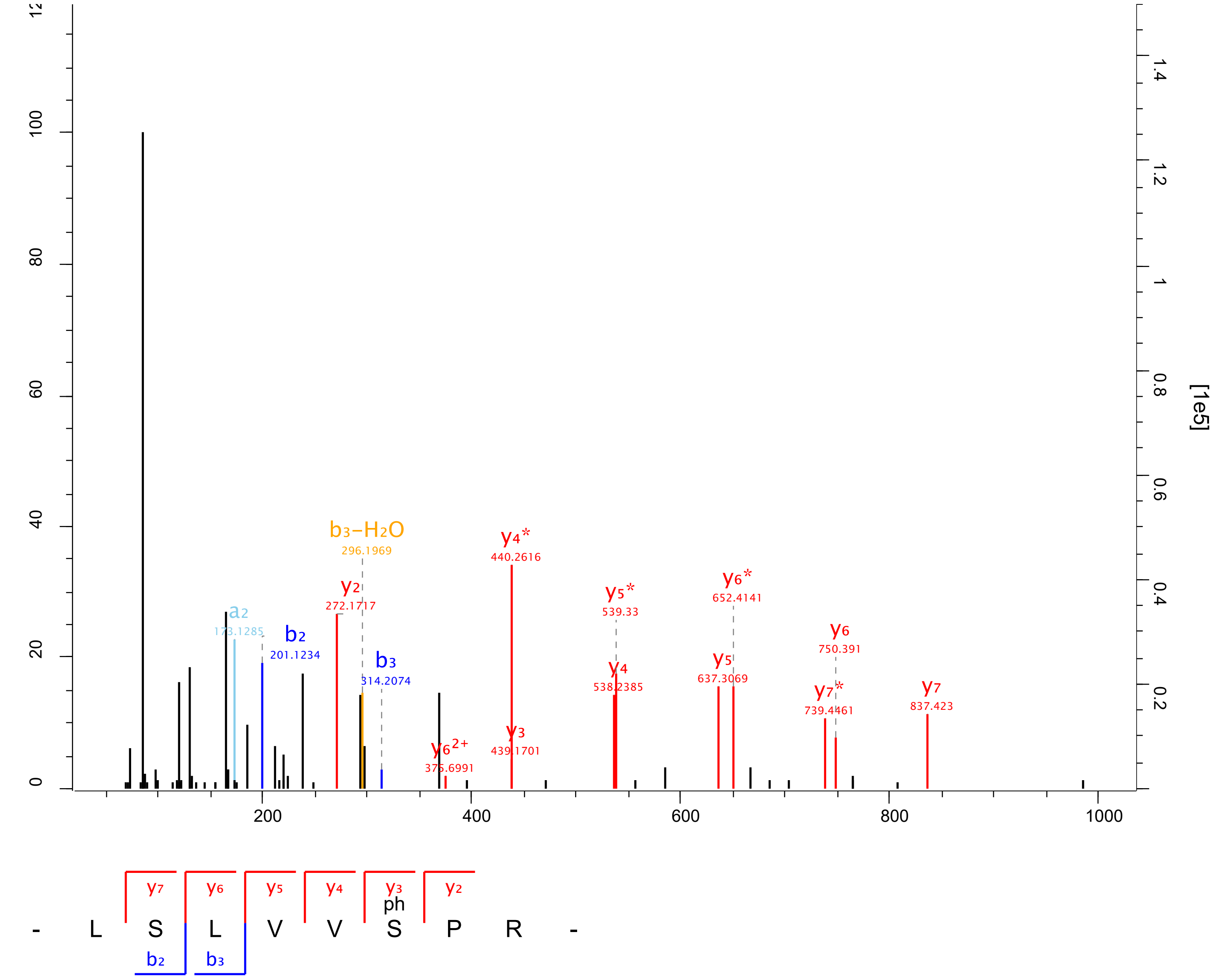

| Raw file | Scan | Method | Score | m/z |
|---|---|---|---|---|
| water_15_3 | 3958 | FTMS; HCD | 156.72 | 645.27 |

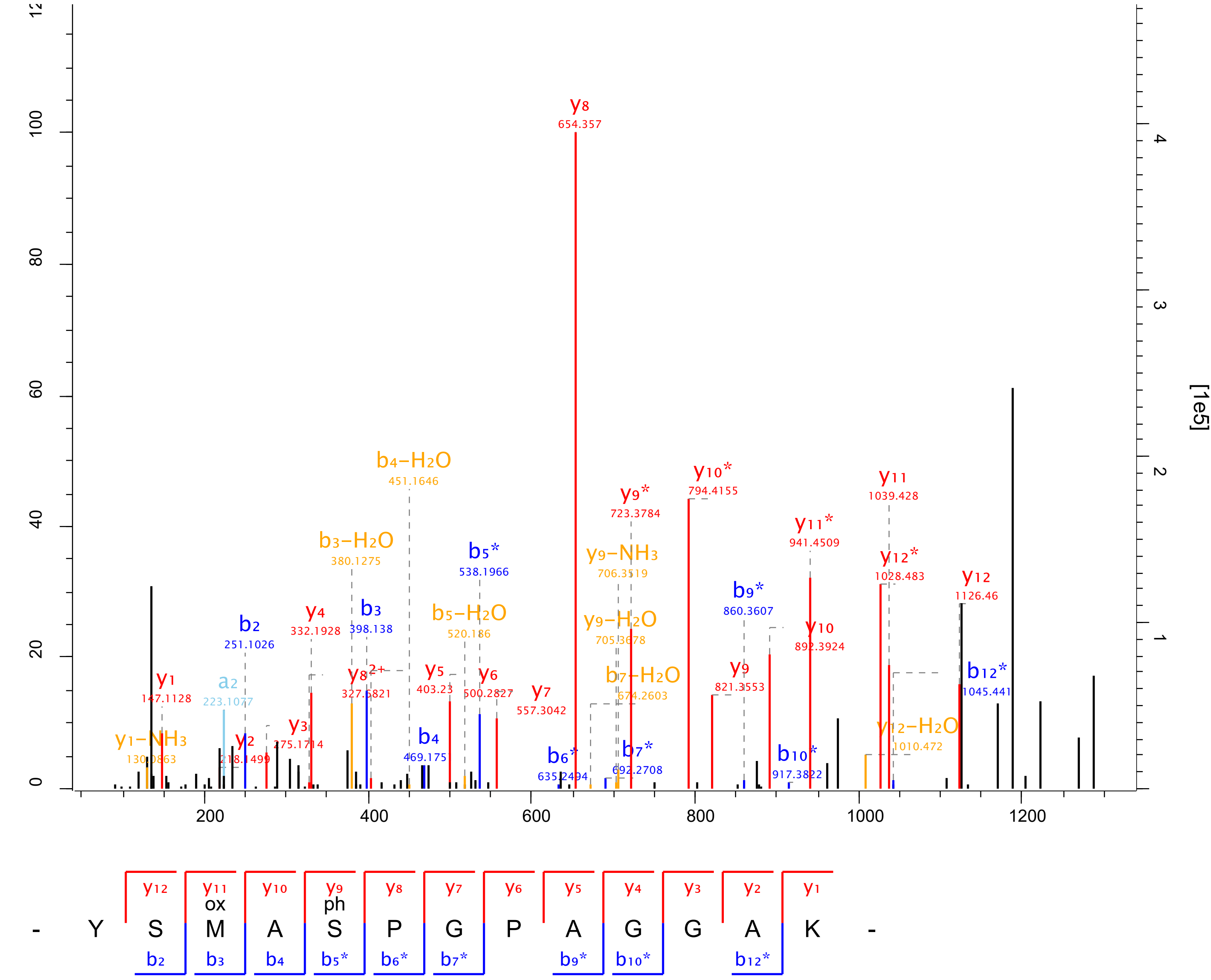

| Raw file | Scan | Method | Score | m/z |
|---|---|---|---|---|
| water_15_3 | 4513 | FTMS; HCD | 42.34 | 492.23 |

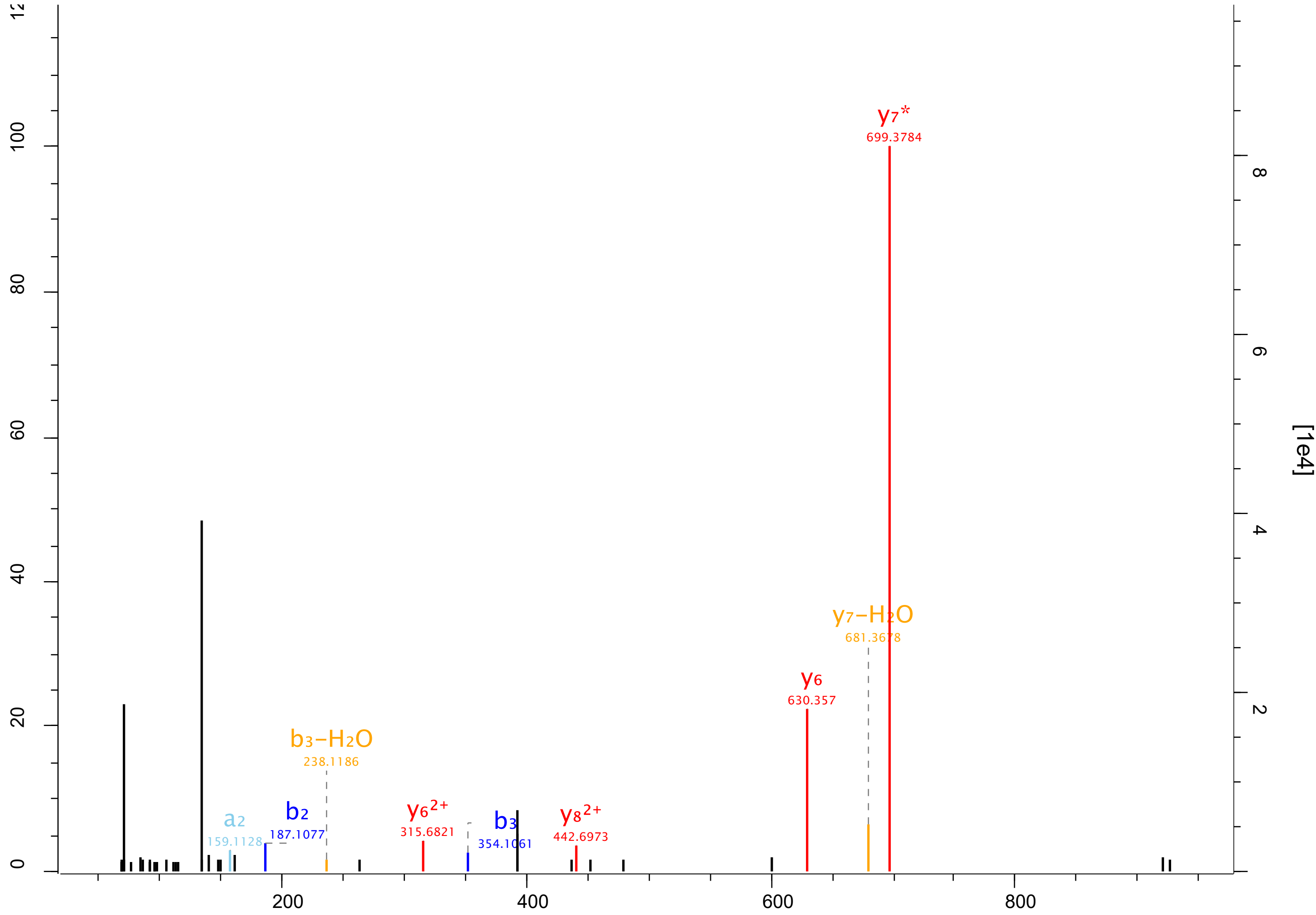


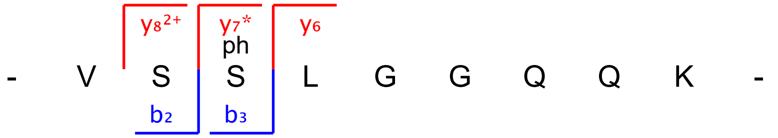

| Raw file | Scan | Method | Score | m/z |
|---|---|---|---|---|
| water_15_3 | 6852 | FTMS; HCD | 73.23 | 410.21 |

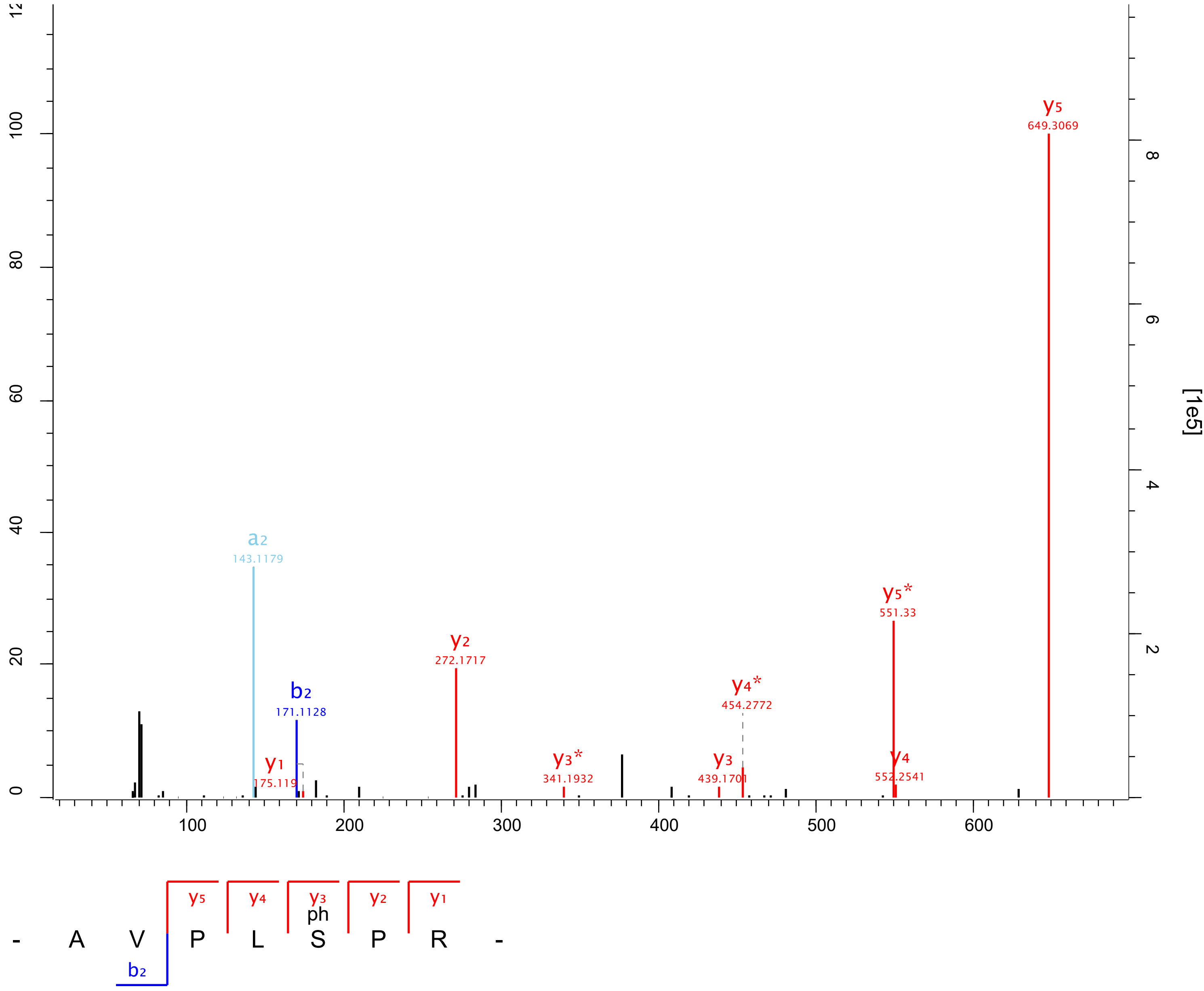

| Raw file | Scan | Method | Score | m/z |
|---|---|---|---|---|
| water_15_3 | 7223 | FTMS; HCD | 66.27 | 410.21 |

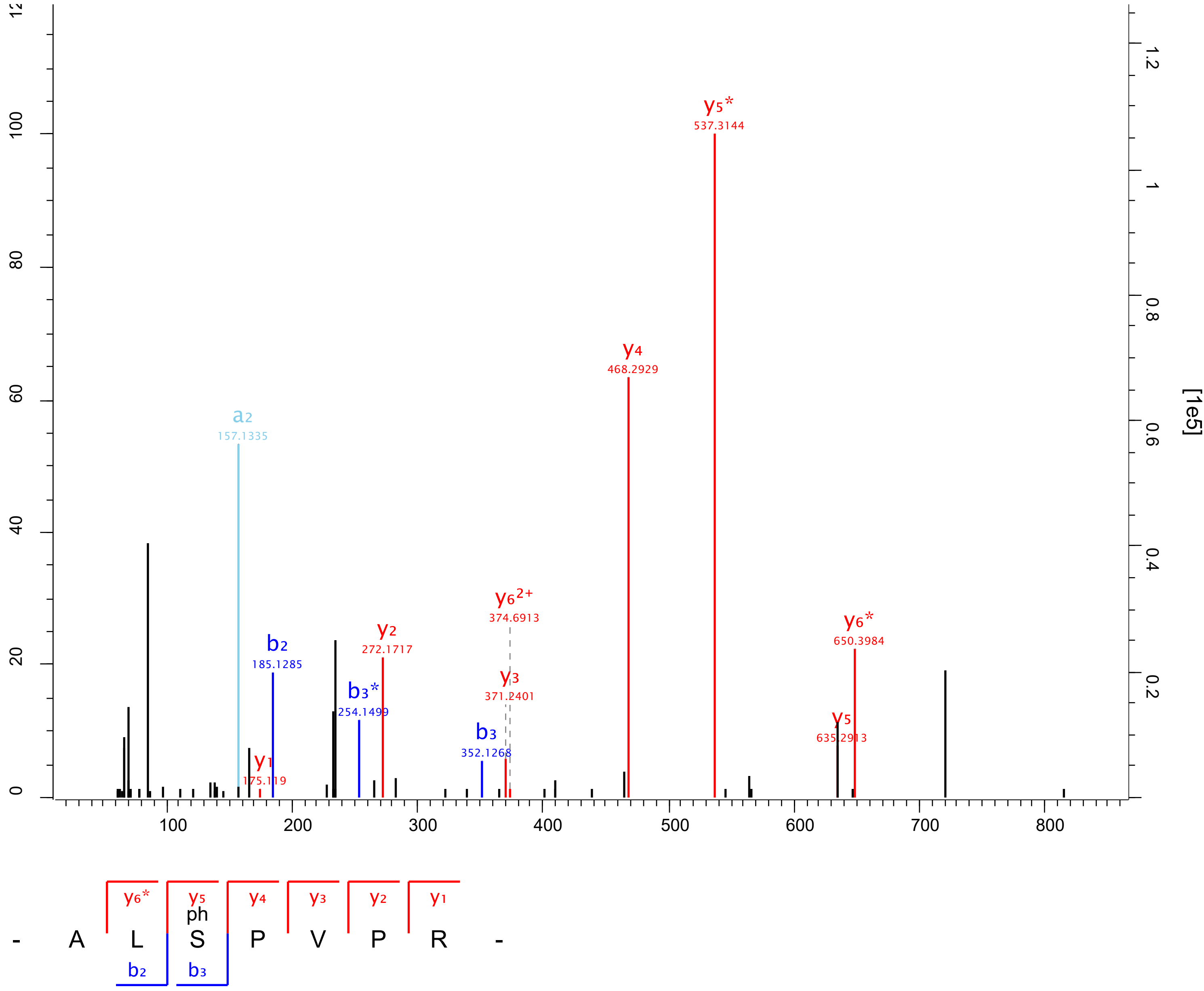

| Raw file | Scan | Method | Score | m/z |
| --- | --- | --- | --- | --- |
| water_15_3 | 8553 | FTMS; HCD | 108.56 | 517.72 |

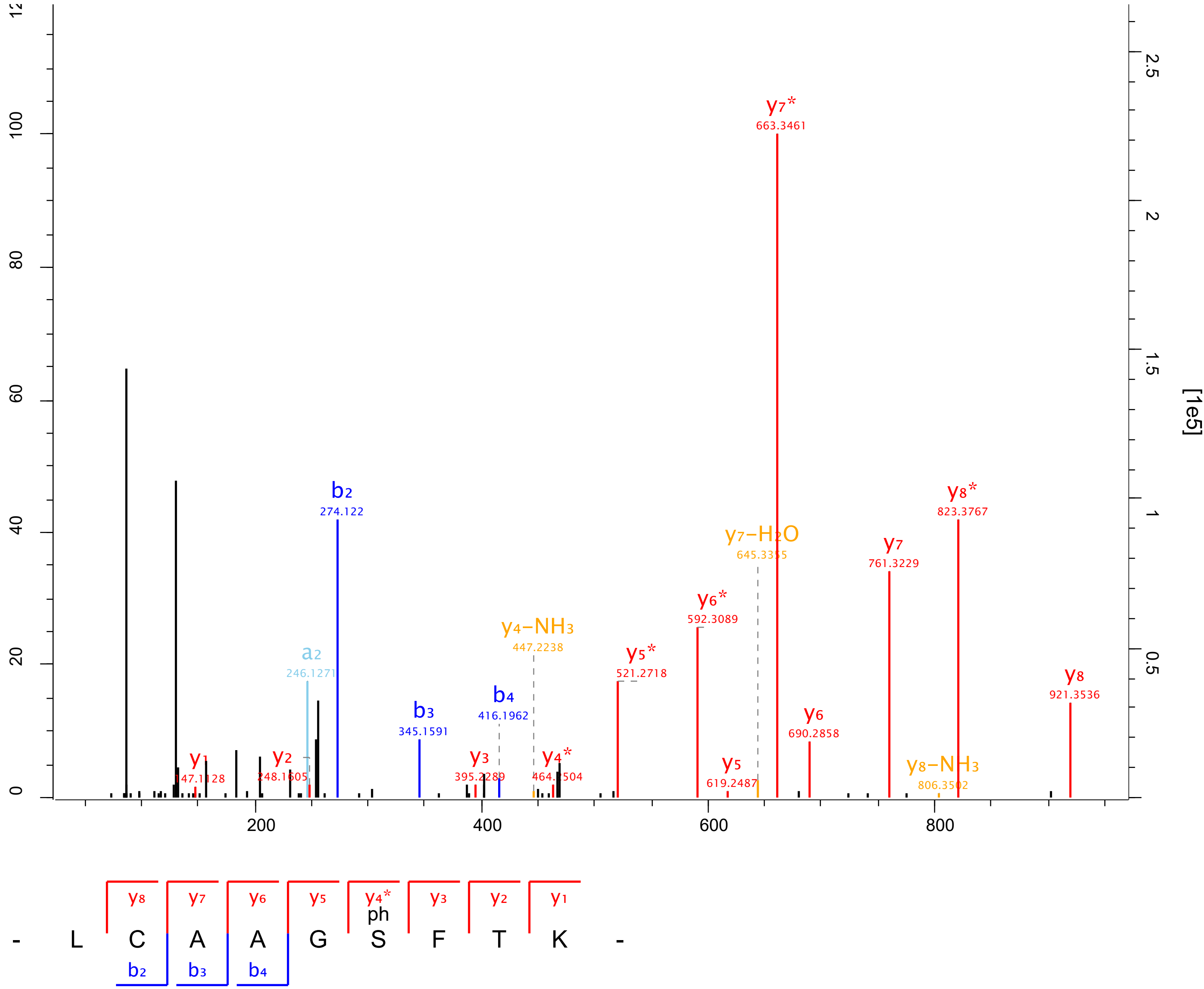

| Raw file | Scan | Method | Score | m/z |
|---|---|---|---|---|
| water_15_3 | 8922 | FTMS; HCD | 105.52 | 454.22 |

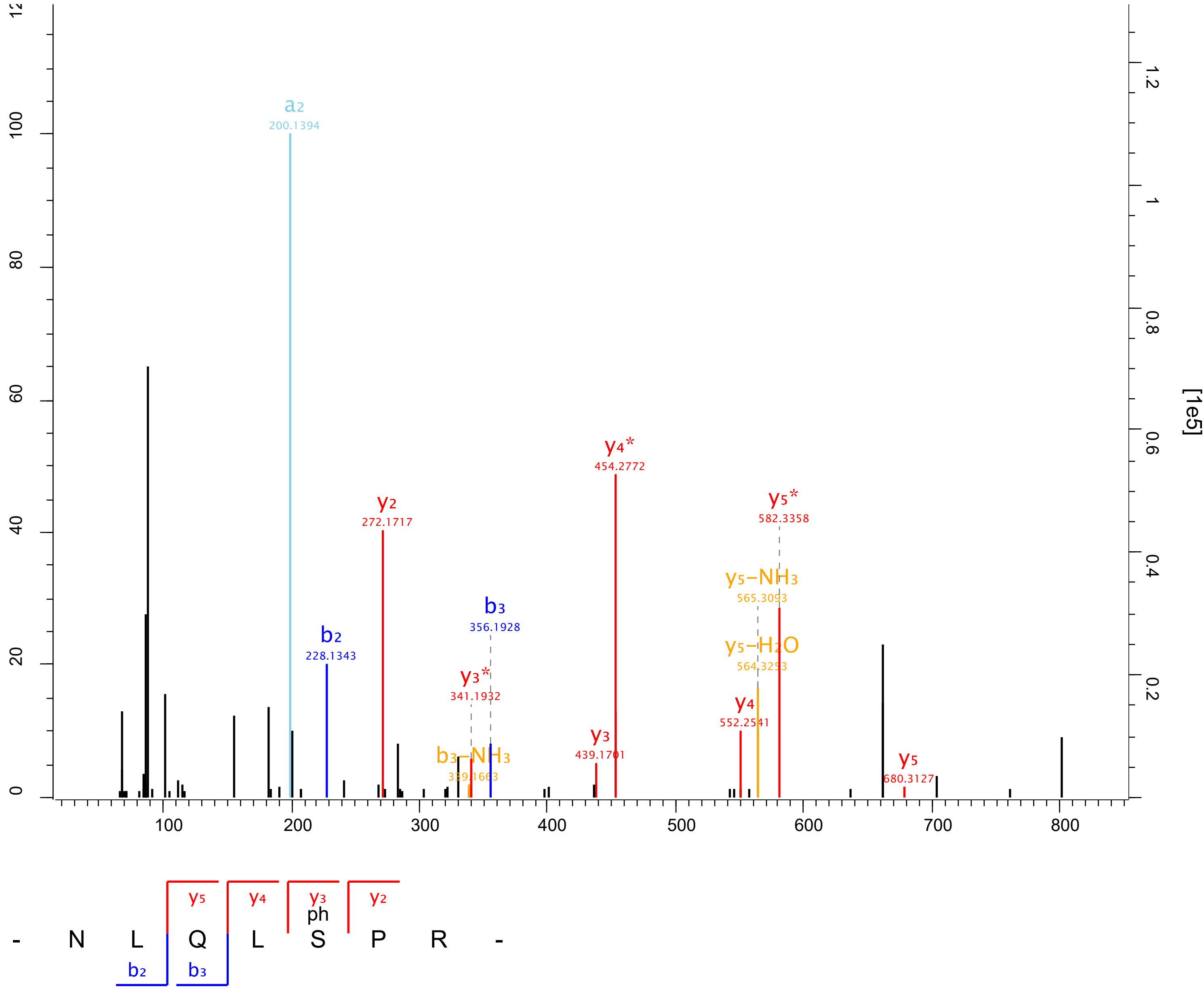

| Raw file | Scan | Method | Score | m/z |
|---|---|---|---|---|
| water_15_3 | 9288 | FTMS; HCD | 49.3 | 457.74 |

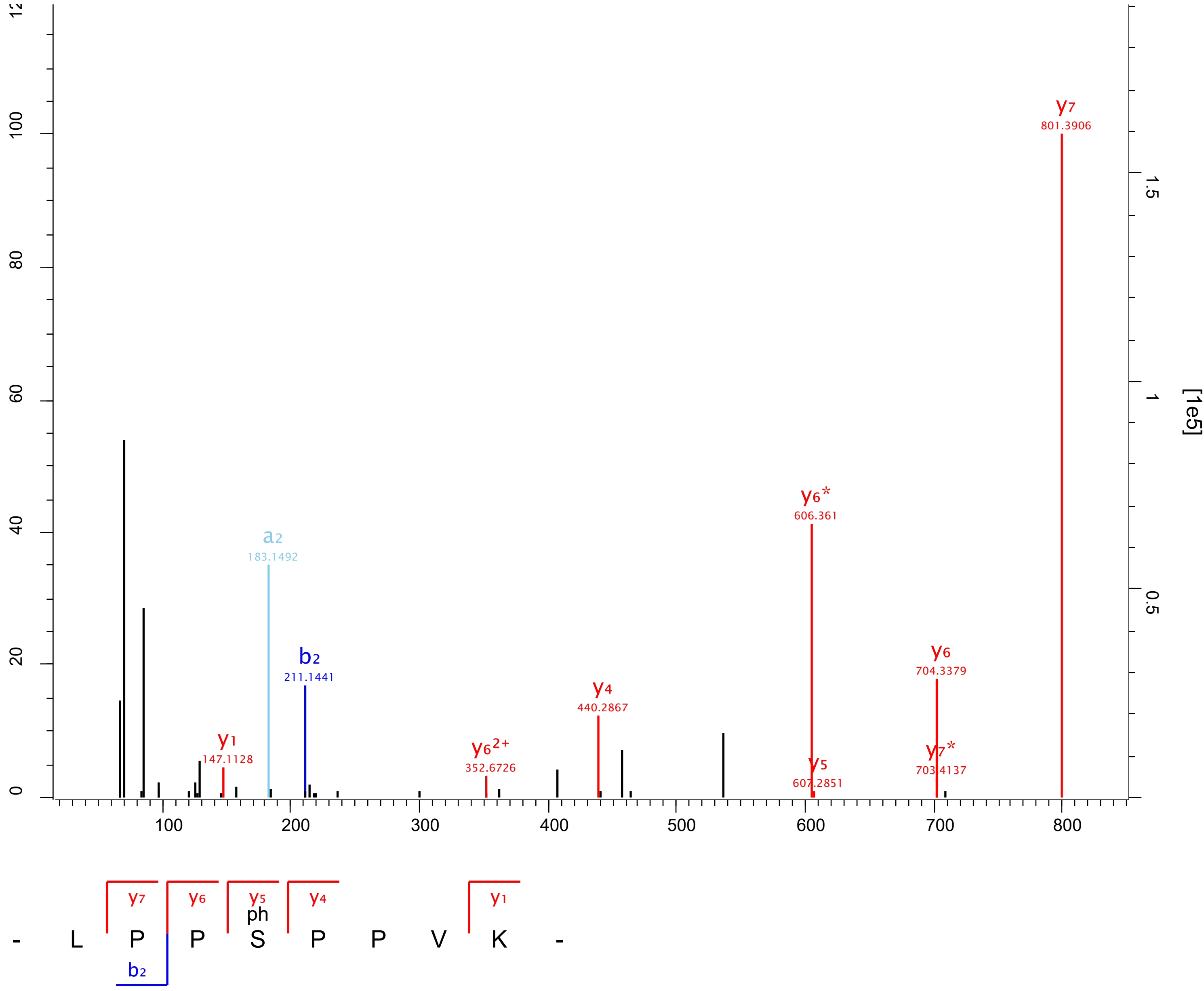

| Raw file | Scan | Method | Score | m/z |
|---|---|---|---|---|
| water_30_2 | 19626 | FTMS; HCD | 227.67 | 788.88 |

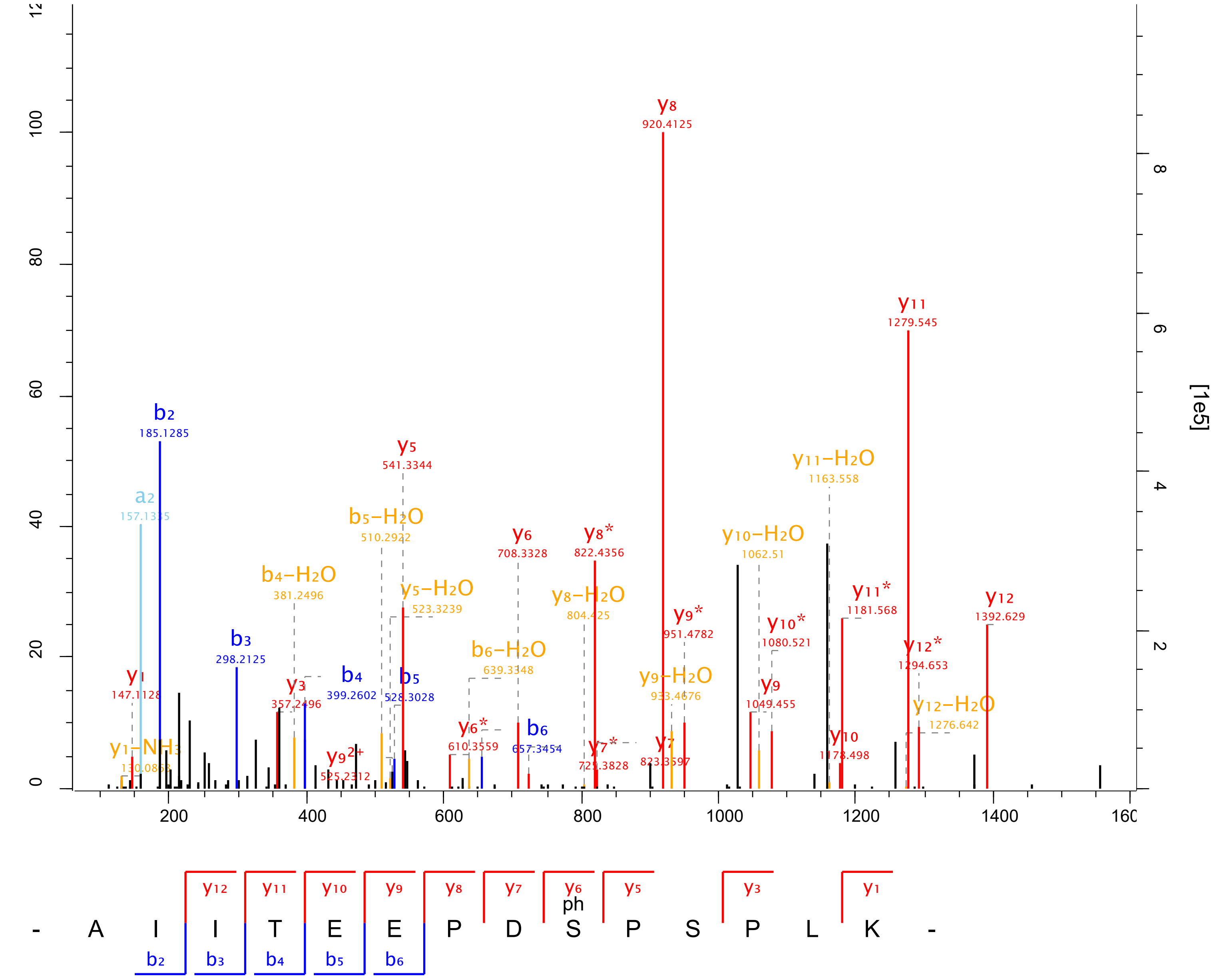

| Raw file | Scan | Method | Score | m/z |
|---|---|---|---|---|
| water_30_2 | 29313 | FTMS; HCD | 69.48 | 799.37 |

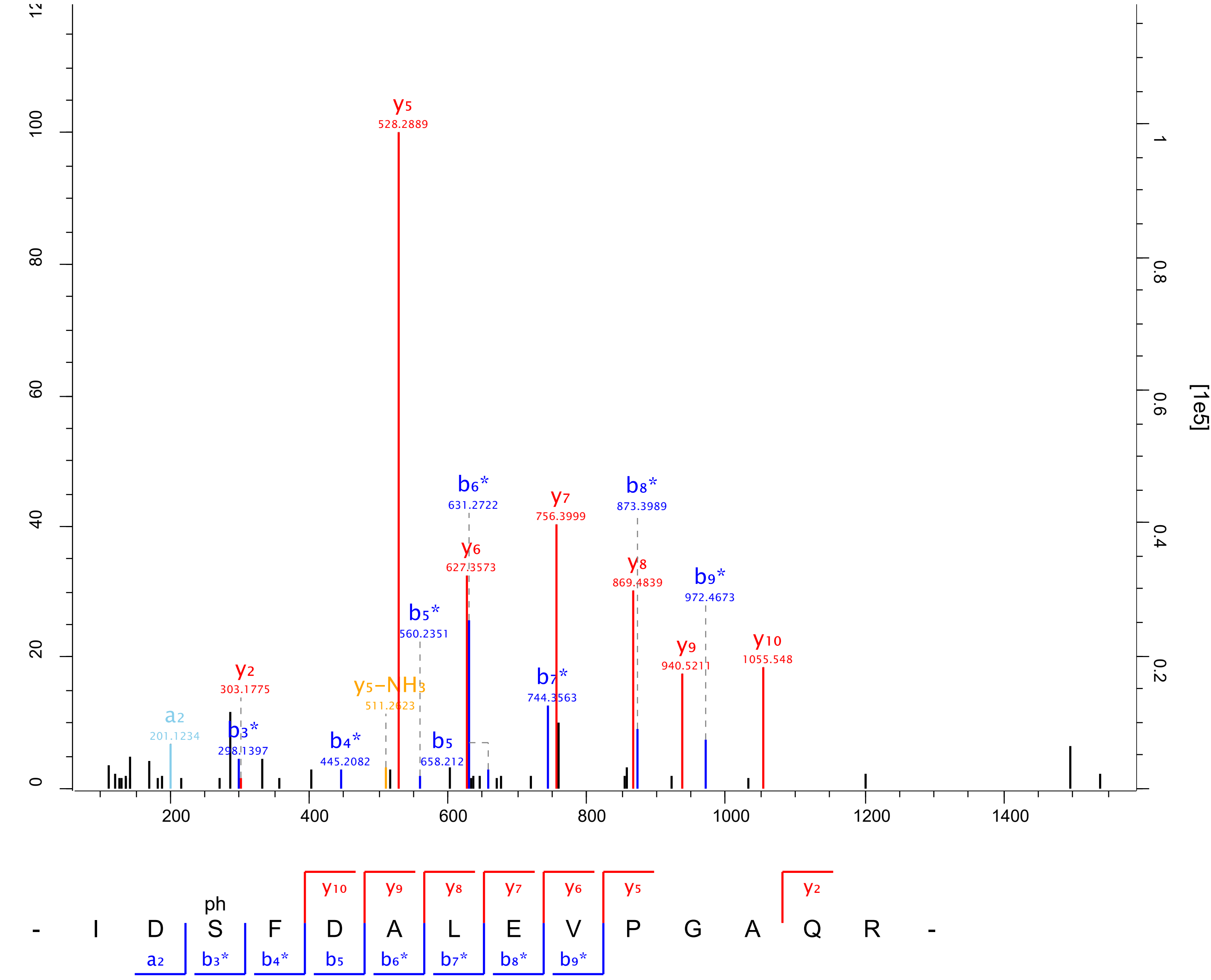

| Raw file | Scan | Method | Score | m/z |
|---|---|---|---|---|
| water_30_2 | 3488 | FTMS; HCD | 62.46 | 482.19 |

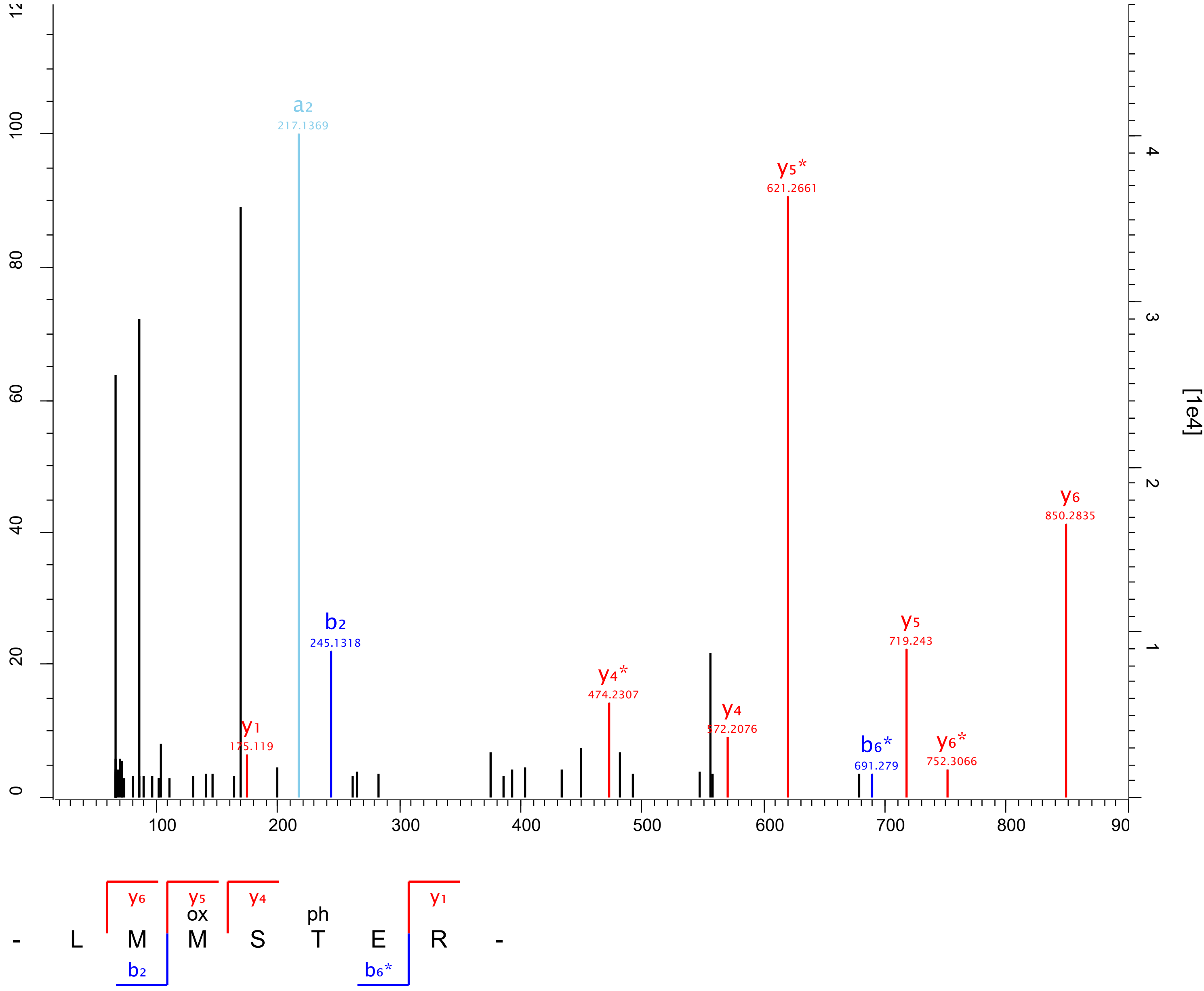

| Raw file | Scan | Method | Score | m/z |
|---|---|---|---|---|
| water_30_2 | 4157 | FTMS; HCD | 60.16 | 605.78 |

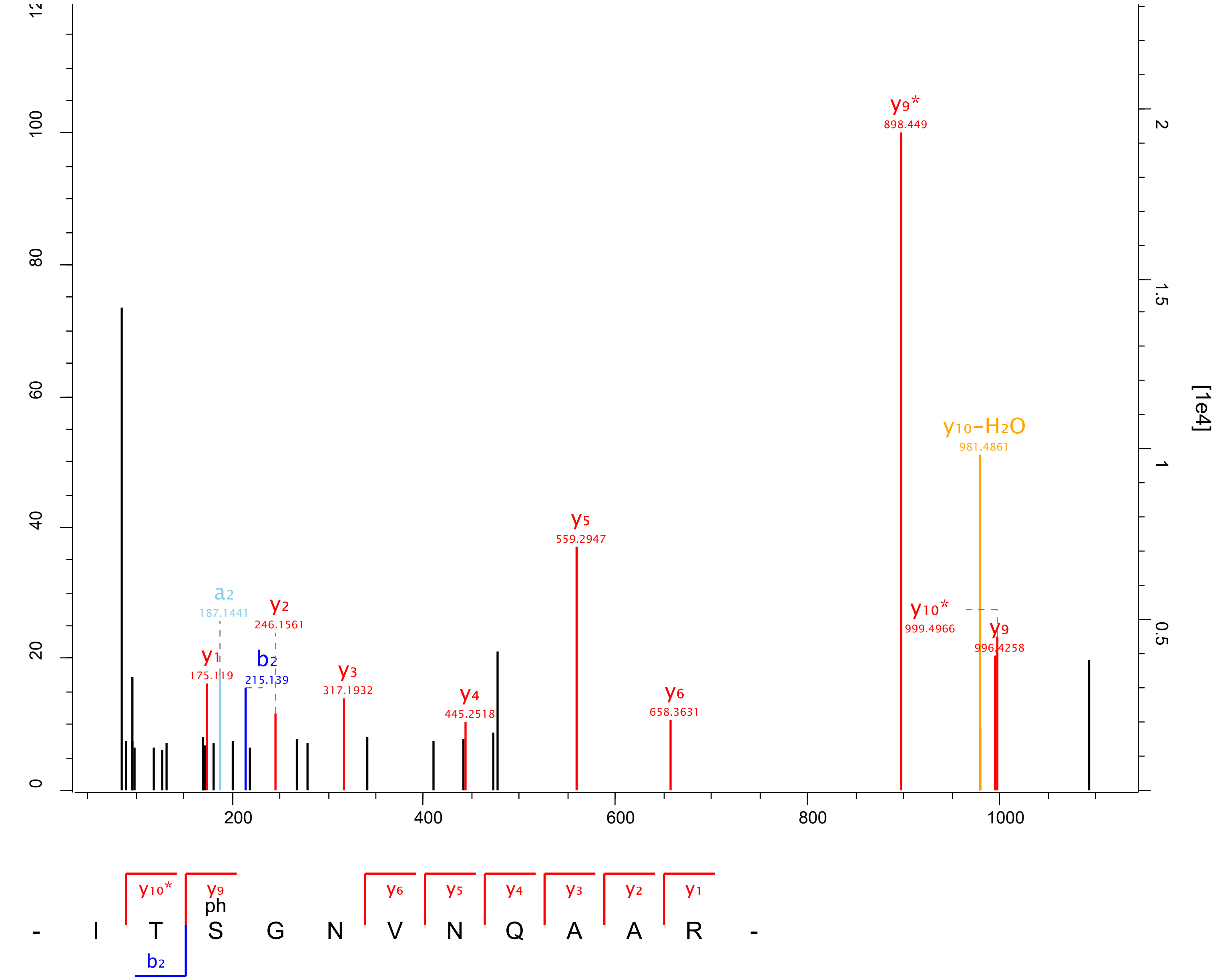

| Raw file | Scan | Method | Score | m/z |
| --- | --- | --- | --- | --- |
| water_30_2 | 5804 | FTMS; HCD | 42.98 | 529.74 |

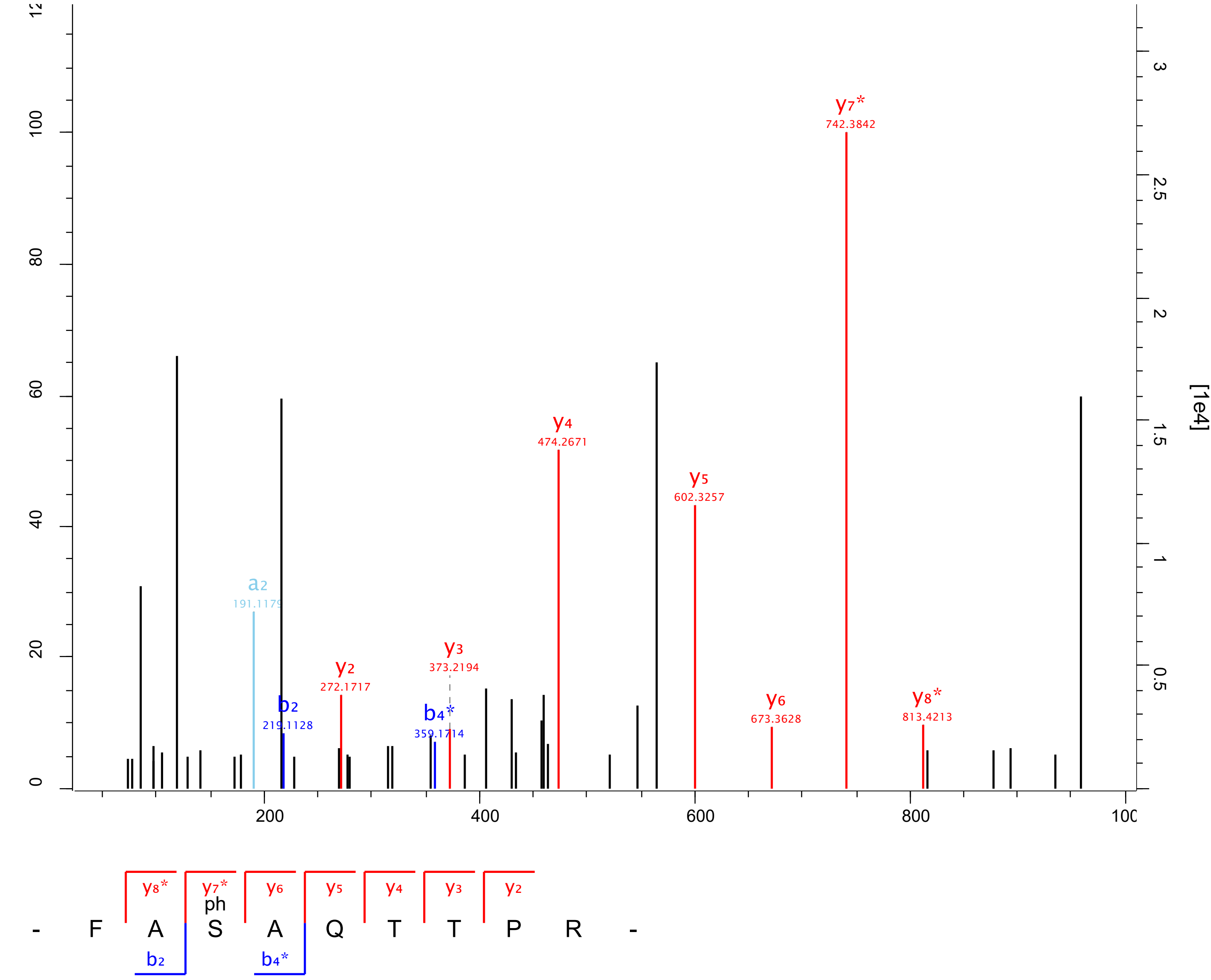

| Raw file | Scan | Method | Score | m/z |
|---|---|---|---|---|
| water_30_2 | 8064 | FTMS; HCD | 174.68 | 637.78 |

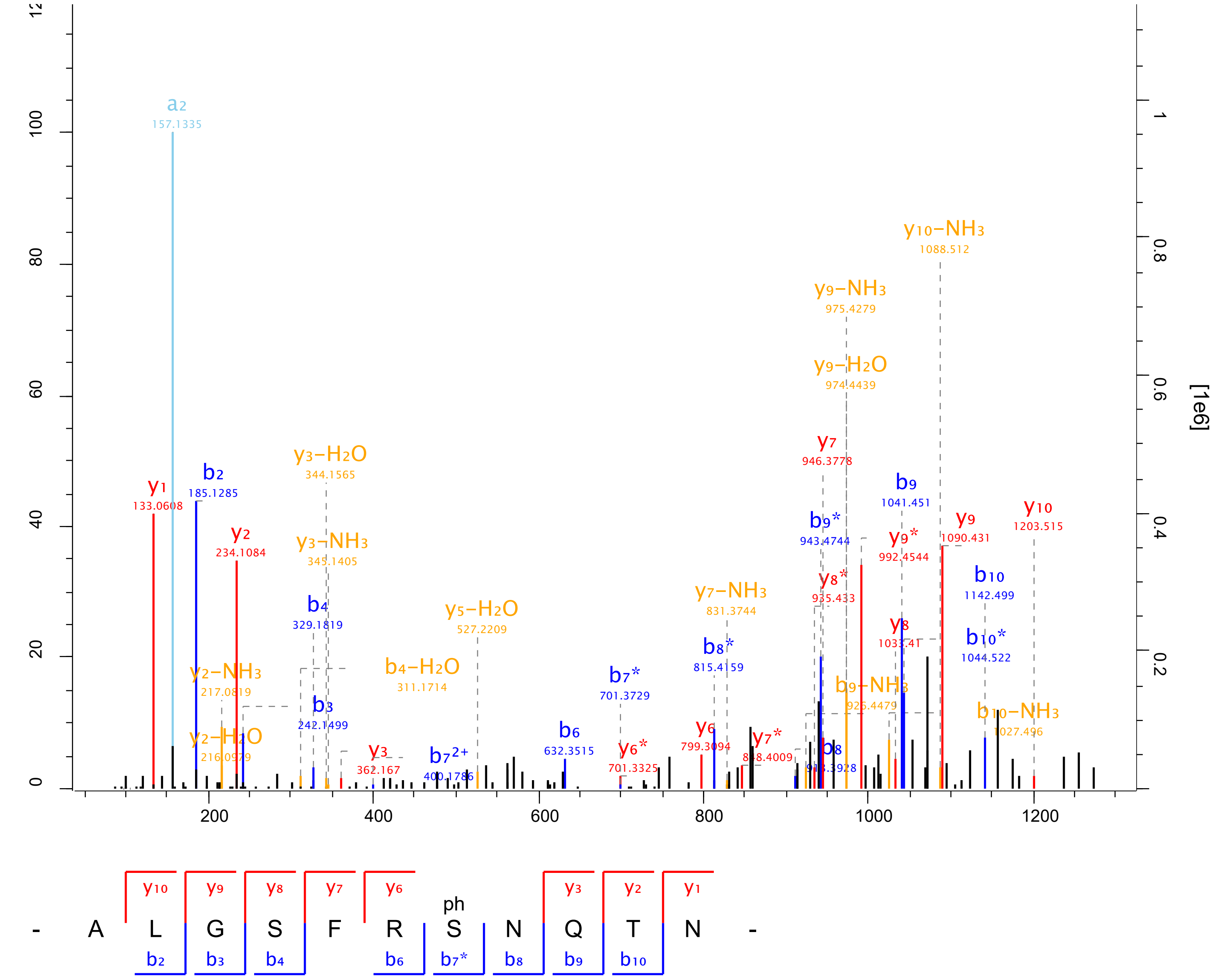

| Raw file | Scan | Method | Score | m/z |
|---|---|---|---|---|
| water_30_3 | 11158 | FTMS; HCD | 45.83 | 671.32 |

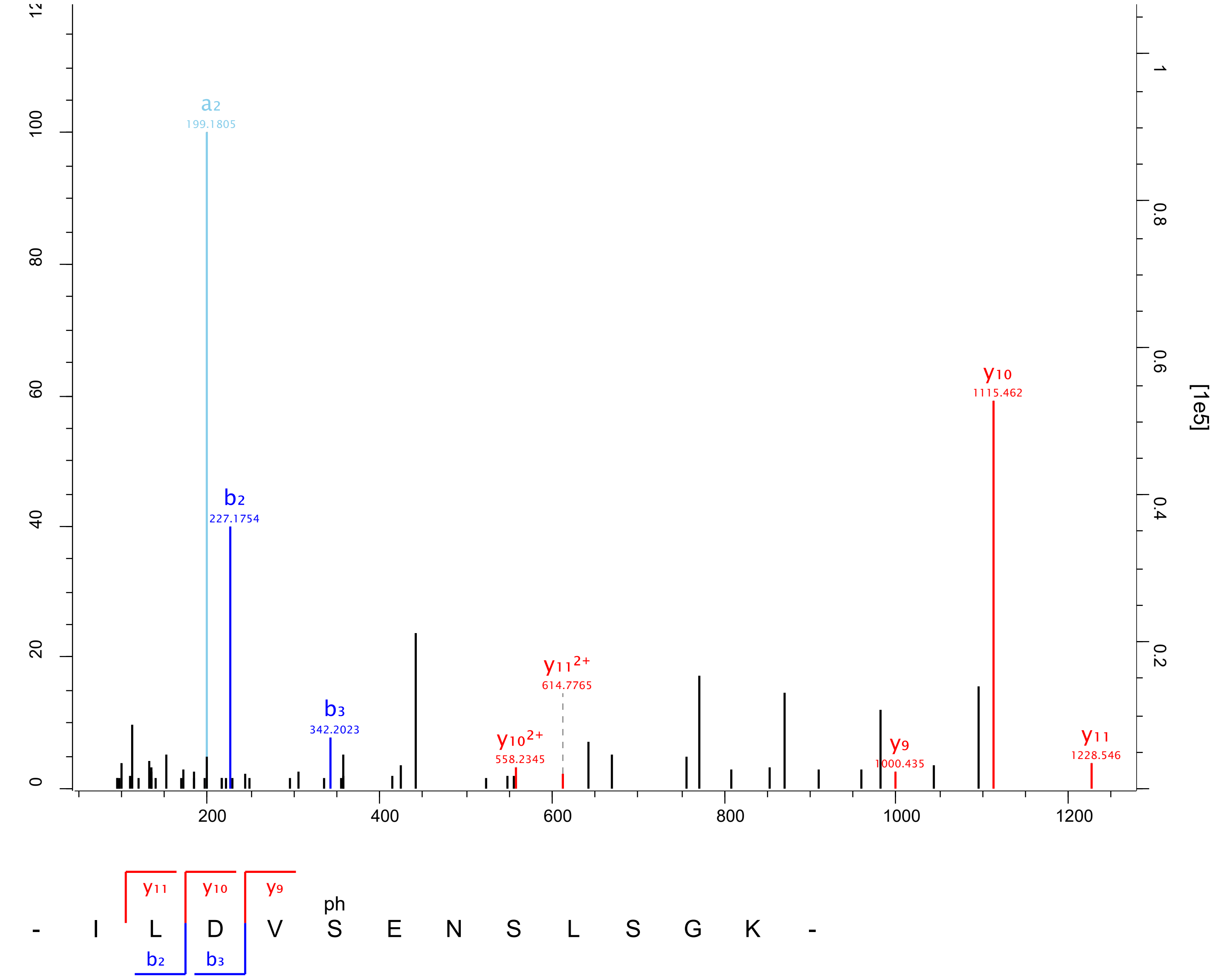

| Raw file | Scan | Method | Score | m/z |
| --- | --- | --- | --- | --- |
| water_30_3 | 15639 | FTMS; HCD | 80.69 | 564.76 |

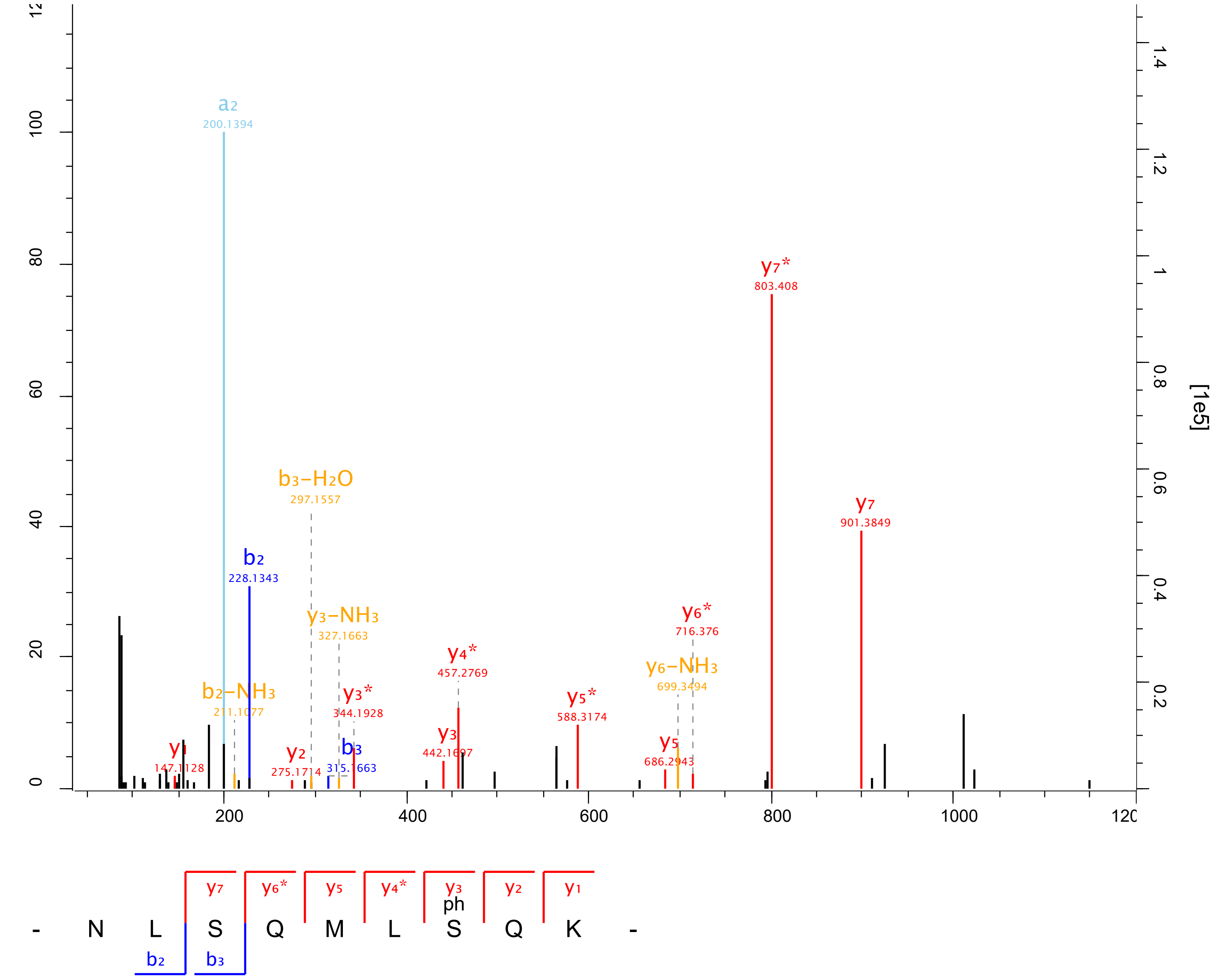
